# Adaptive introgression as a driver of local adaptation to climate in European white oaks

**DOI:** 10.1101/584847

**Authors:** Thibault Leroy, Jean-Marc Louvet, Céline Lalanne, Grégoire Le Provost, Karine Labadie, Jean-Marc Aury, Sylvain Delzon, Christophe Plomion, Antoine Kremer

## Abstract

Latitudinal and elevational gradients provide valuable experimental settings for studies of the potential impact of global warming on forest tree species. The availability of long-term phenological surveys in common garden experiments for traits associated with climate, such as bud flushing, for sessile oaks (Quercus petraea), provides an ideal opportunity to investigate this impact.

We sequenced 18 sessile oak populations, and used available sequencing data for three other closely related European white oak species (Q. pyrenaica, Q. pubescens, Q. robur), to explore the evolutionary processes responsible for shaping the genetic variation across latitudinal and elevational gradients in extant sessile oaks. We used phenotypic surveys in common garden experiments and climatic data for the population of origin, to perform genome-wide scans for population differentiation, genotype-environment (GEA) and genotype-phenotype associations (GPA).

The inferred historical relationships between *Q. petraea* populations suggest that interspecific gene flow occurred between *Q. robur* and *Q. petraea* populations from cooler or wetter areas. A genome-wide scan of differentiation between *Q. petraea* populations identified SNPs displaying strong interspecific relative divergence between these two species. These SNPs followed genetic clines along climatic or phenotypic gradients, providing further support for the likely contribution of introgression to the adaptive divergence of *Q. petraea* populations.

Overall, the results of this study indicate that adaptive introgression of *Q. robur* alleles has occurred. We discuss the results of this study in the framework of the post-glacial colonization scenario, in which introgression and diversifying selection have been proposed as essential drivers of *Q. petraea* microevolution.

## Introduction

Evolutionary biologists are becoming increasingly fascinated by the tracking of adaptive genetic changes, as our understanding of paleoecology and genomics (Shapiro & Hofreiter, 2014) and climate reconstructions at various spatial and temporal scales (Mauri *et al.*, 2015) improve. Ultimately, assembling data relating to historical and genetic changes will increase our understanding of how, when and how rapidly evolutionary shifts have enhanced adaptation. The recent report that introgression with Neanderthals or Denisovans increased the adaptation of modern Eurasian humans provides an emblematic example of a major evolutionary shift supported by historical and genomic evidence (Dannemann & Racimo, 2018). Adaptive shifts have also been predicted in non-model plants and animals that have repeatedly witnessed large-scale environmental changes over larger temporal scales due to quaternary climatic oscillations (Dynesius & Jansson, 2000). Forest trees are relevant target species for explorations of evolutionary changes, as their past history and distribution can be reconstructed easily, due to the availability of large quantities of fossil remains (Brewer *et al.*, 2017; Wagner *et al.*, 2018). Furthermore, understanding how trees adapt to changing environments has become a major topic of interest in theoretical and applied ecological genetics and genomics. On the one hand, there are concerns that trees may not cope with the velocity of ongoing climatic change, and on the other, land managers and foresters are seeking methods of adaptive management (Lindner *et al.*, 2010). Adaptation can be triggered by new alleles originating from new mutations, neutral standing genetic variation or adaptive introgression. Adaptation can be investigated empirically by assessing adaptive divergence in common garden experiments, or by tracking genetic changes over successive generations. The common garden approach has been widely used for long-lived species, such as trees, and has revealed the existence of high levels of adaptive divergence between extant populations distributed along large geographic gradients (Alberto *et al.*, 2013). Moreover, adaptive divergence is maintained between tree populations, despite extensive gene flow potentially causing pollen swamping effects that might constrain adaptation (Savolainen *et al.*, 2007). In many European tree species, pollen swamping results from intra- or inter-specific gene flow, as many congeneric species live together in the same stands and interspecific hybridization is widespread. By contrast to the pollen swamping effect, it has also been suggested that gene flow may actually enhance adaptation, by increasing selection response (Kremer & Le Corre, 2012; Yeaman, 2015). It has also recently been suggested that interspecific gene flow may enhance adaptation by facilitating adaptive introgression (Suarez-Gonzalez *et al.*, 2018). Here, we explore the genomic footprints of adaptation variation in sessile oak (*Quercus petraea* (Matt.) Liebl.), a widespread species in Europe, and address the potential contribution of interspecific gene flow to local adaptation.

The distribution range of sessile oak extends from Spain to Southern Scandinavian and this species mostly grows in forests also containing the closely related pedunculate oak (*Q. robur*), another white oak species with a range that extends farther north and east, to the Ural Mountains (Leroy *et al,.* 2018). At more southern latitudes, sessile oak also shares habitats with pubescent (*Q. pubescens*) and Pyrenean (*Q. pyrenaica*) oaks. Hybridization between these four European white oak species has been reported in sympatric stands (Curtu *et al.*, 2007; Lepais & Gerber, 2011). Furthermore hybridization is thought to have played a major role in the expansion of *Q. petraea* populations during postglacial recolonization, as suggested by the widespread sharing of chloroplast genomes between *Q. petraea* and *Q. robur* in sympatric stands (Petit *al.*, 2002). Interestingly, demographic inferences based on approximate Bayesian computation simulations support scenarios in which these four European white oak species came into secondary contact, at the onset of the last glacial period (Leroy *et al.*, 2017; 2018). A number of questions relating to this scenario remain unanswered. How much did introgression actually contribute to todays’ distribution of *Q. petraea?* Did introgression facilitate the adaptation of *Q. petraea* or was it a purely neutral process? Which adaptive alleles were introgressed, and from which other white oak species? We tackle these questions here by exploring imprints of historical population splits and admixture events through studies of whole-genome sequences from *Q. petraea* populations. We also performed genome scans to search for genomic footprints of divergence between *Q. petraea* populations, with either phenotypic data from common garden experiments or climatic data for the populations of origin. To this end, we sequenced the genome of 18 *Q. petraea* populations distributed along latitudinal and altitudinal gradients in Europe, using a pooled sequencing (pool-seq) strategy, and performed whole-genome scans for differentiation, and for genotype-environment (GEA) and genotype-phenotype associations (GPA) for phenological traits associated with climate. In previous studies, the date of leaf unfolding was found to display clinal variation according to the spring temperature of the populations of origin (Alberto *et al.*, 2013; Firmat *et* al., 2017). Here, we examine whether similar patterns of clinal variation can be identified at the level of the genome. We first explore whether the genome of the 18 *Q. petraea* populations studied displays any signs of introgression of genes from the other closely related European white oak species (*Q. robur, Q. pubescens, Q. pyrenaica*), and the geographic variation of introgression.

## Materials and methods

### Sampling and sequencing

We sampled eight *Q. petraea* populations (with up to 20 individuals per pool, Table 1) from lowlands to middle elevations in the Pyrenees (up to 1,600 meters) in South-West France. These eight populations are distributed along two neighboring elevational transects (the Ossau & Luz valleys) and are stands of natural origin (elevation of 100 m to 1600 m). We also collected data for 10 *Q. petraea* populations (with up to 25 individuals per pool, Table 1) growing in a large common garden experiment in Sillégny, Eastern France, corresponding to a total of 116 sessile oak provenances in Europe (Saenz-Romero *et al.*, 2017).

**Table 1.** Geographic and climatic data for the Q. petraea populations studied. Date of leaf unfolding expressed as standardized values for common gardens (see text). Negative values indicate early flushing, and positive values, late flushing

| Code | Location | Elevation (m) | Latitude | Longitude | Temperature | Precipitation<br>(mm/year) | Leaf<br>Unfolding | Sample Size |
| --- | --- | --- | --- | --- | --- | --- | --- | --- |
| **Elevational gradient (French Pyrenees)** |  |  |  |  |  |  |  |  |
| L1 | Laveyron, Luz Valley, France | 131 | 43.75 | -0.22 | 12.33 | 901 | -1.333 | 20 |
| L8 | Chèze, Luz Valley, France | 803 | 42.92 | -0.03 | 9.27 | 914 | 0.817 | 20 |
| L12 | Gèdre, Luz Valley, France | 1235 | 42.78 | 0.02 | 7.12 | 1016 | 1.011 | 20 |
| L16 | Péguères, Luz Valley, France | 1630 | 42.87 | -0.12 | 6.58 | 982 | 1.724 | 18 |
| O1 | Josbaig, Ossau Valley, France | 259 | 43.22 | -0.73 | 12.24 | 979 | -1.309 | 20 |
| O8 | Le Hourcq, Ossau Valley, France | 841 | 42.90 | -0.43 | 9.16 | 933 | -0.324 | 20 |
| O12 | Gabas, Ossau Valley, France | 1194 | 42.88 | -0.42 | 7.35 | 1031 | 0.036 | 20 |
| O16 | Artouste, Ossau Valley, France | 1614 | 42.88 | -0.40 | 5.16 | 1164 | 0.427 | 10 |
| **Latitudinal gradient** |  |  |  |  |  |  |  |  |
| 9 | Saint Sauvant, France | 155 | 46.38 | 0.12 | 11.78 | 786 | -0.166 | 25 |
| 97 | Grésigne, France | 310 | 44.04 | 1.75 | 12.05 | 791 | -1.139 | 25 |
| 124 | Killarney, Ireland | 50 | 52.01 | -9.50 | 9.96 | 1362 | 4.084 | 25 |
| 204 | Bézanges, France | 275 | 48.76 | 6.49 | 9.50 | 751 | 0.371 | 25 |
| 217 | Bercé, France | 165 | 47.81 | 0.39 | 10.65 | 698 | 0.434 | 25 |
| 218 | Longchamp, France | 235 | 47.26 | 5.31 | 10.59 | 801 | -0.920 | 22 |
| 219 | Tronçais, France | 245 | 46.68 | 2.83 | 10.63 | 742 | 1.350 | 25 |
| 233 | Vachères, France | 650 | 43.98 | 5.63 | 10.22 | 797 | -1.532 | 25 |
| 253 | Göhrde, Germany | 85 | 53.10 | 10.86 | 8.30 | 635 | 0.953 | 25 |
| 256 | Lappwald, Germany | 180 | 52.26 | 10.99 | 8.50 | 597 | 0.650 | 25 |

The selected populations were sampled along a latitudinal gradient extending from southern France to northern Germany (latitude 44° to 53°). We also included a reference population of the four main white oak species in Europe *(Q. petraea, Q. robur, Q. pubescens, Q. pyrenaica*) corresponding to four oak stands located in South-West France analyzed in a companion paper (Leroy et al., 2018). The *Q. petraea* reference population of Leroy et al. (2018) corresponded to a pool of 13 individuals from the same low-elevation population of the Luz valley (L1: Laveyron in Table 1). We used the two populations as pseudoreplicates, to check the accuracy of our pool-sequencing strategy.

DNA was extracted from individual trees with a CTAB DNA extraction protocol (Doyle and Doyle, 1987) (latitudinal gradient,) or with the Invisorb Spin Plant Mini Kit (Startec Molecular; elevational gradient) according to the manufacturer’s instructions (Startec Molecular, GmbH, Berlin, Germany). DNA yields were evaluated with a NanoDrop 1000 spectrophotometer (NanoDrop Technologies, Wilmington, DE, USA) (elevational gradient) or with an Infinite F200 (Tecan Group Ltd., Männedorf, Switzerland), and DNA samples were mixed in equimolar amounts to obtain a single pool for each population. After trimming, the number of 2×100 bp paired-end reads retained for analyses ranged from 668 to 998 million reads per pool, corresponding to a rough estimate of sequencing coverage or 180 to 270 x, assuming an oak genome size of 740 Mb.

### Climatic and phenological data

Monthly mean climate data were obtained from WORLDCLIM (Hijmans *et al.*, 2005) for the period 1950-2000. We downscaled these values to a spatial resolution of 100 m using locally weighted regressions and a finer resolution DEM (100 m) before point overlay and accounted for topography as previously described (Zimmermann *et al.*, 2009; Dullinger *et al.*, 2012). We then calculated yearly average temperature and annual precipitations sums (Table1).

The date of leaf unfolding was recorded separately in two different common garden experiments. Saplings of populations sampled along the elevational gradient were transplanted to a common garden experiment in Toulenne in South-West France in spring 2007, and phenological observations were conducted over seven successive years (2009 to 2015) (See Firmat *et al.*, 2017 for further details). Similarly, saplings of populations sampled along the latitudinal gradient were installed in a common garden located in the North-East of France in Sillegny in 1989 and 1993, and phenological observations were conducted in 2015 (See Firmat *et al.*, 2017 and Torres-Ruiz *et al.*, 2019 for further details). Population means were calculated in each common garden and standardized values for each common garden were used to study genotype-phenotype associations across the two common gardens (Table 1).

### Mapping and SNP calling

We used the pipeline described by Leroy et al. (2018), see also https://github.com/ThibaultLeroyFr/GenomeScansByABC/tree/master/SNP_calling_filtering) to identify reliable SNPs. In brief, we used bowtie2 v2.1.0 to map sequencing data onto the v2.3 oak haplome assembly (Plomion *et al.*, 2018), removed duplicates with picard v.1.106 (http://broadinstitute.github.io/picard/) and then used *snp-frequency-diff.pl* from the Popoolation2 suite (Koffler *et al.*, 2011) to select biallelic SNPs with at least 10 alternative alleles. Positions with a mean coverage of less than 50X for any of the 18 populations and sites in the top 2% in terms of coverage per population were ignored. SNPs with a MAF lower than 0.02 were filtered out to exclude most Illumina sequencing errors.

### Population splits and gene flow inferences

We used genome-wide allele frequency data derived from allele counts to build a population tree with Treemix v. 1.12 (Pickrell & Pritchard, 2012) and to test for the presence of gene flow between populations. Given that *Q. petraea* populations are known to be frequently connected by gene flow with closely related species (Lepais & Gerber, 2011; Leroy *et al.*, 2017), we also evaluated the influence of interspecific gene flow on tree topology. We performed an analysis with a set of 1,757,476 intragenic SNPs commonly detected in the 18 previously described sessile oak populations, and in the reference populations of four white oak species (including *Q. petraea*). Data from an outgroup (*Q. suber*, cork oak) described by Leroy *et* al. (2018) were included in the analysis. Variable numbers of migration nodes (*m*), ranging from 0 to 13, were evaluated. For each fixed value of *m*, we performed 1000 replicated Treemix analyses with a python script from Michael G. Harvey (https://github.com/mgharvey/misc_python/blob/master/bin/TreeMix/treemix_tree_with_bootstraps.py). We then used SumTrees.py from the DendroPy suite to generate a consensus tree with all the bootstrap values (Sukumaran & Holder, 2010). We used the total variation explained for each replicate in the R companion script of Treemix (*plotting_funcs.R*) as a judgment criterion for evaluation of the best number of migration nodes. We also evaluated the robustness of the migration nodes, by counting the number of replicated analyses supporting the migration node.

We used the *f3*-statistic calculated with threepop v.0.1 from the TreeMix suite (-k 1000) to determine whether a focal population (X) was the product of admixture between two populations (Y and Z). Significant negative values of f3(X;Y,Z) were considered to indicate admixture (for details see Reich *et al.* 2009; Schaefer *et al.*, 2016). *F3*-statistics were calculated for all possible three-population combinations, but we report only *f3* (X;Y,*Q.robur*) values, given the known particular contribution of *Q. robur* to the evolution of the populations investigated (see the results section). No significant negative values were obtained for either *f3*(X;Y,*Q.pyrenaica*) or *f3*(X;Y,*Q.pubescens*).

### Genome scans

Genome-wide scans were performed with the BayPass v. 2.1 software package (Gautier, 2015). BayPass takes confounding demographic effects into account by estimating the covariance matrix of allele frequency between populations. The core model reports locus XtX, which is analogous to *FST* but explicitly corrected for this covariance matrix (Gunther & Coop, 2013). For each SNP, BayPass can also compare models integrating population-specific covariables (here, temperature, precipitation and date of leaf unfolding), by including a regression coefficient for the cline along the covariate gradient in the base model. All the covariates used here were scaled as described by Gautier (2015). Direct comparisons of the likelihood of models (Bayes Factors, BF) including and not including particular covariates were used to evaluate the GEA (genotype-environment association), defined as the association between changes in allele frequency and each population-specific climate variable (temperature, precipitation) and the GPA (genotype-phenotype association), defined as the association between changes in allele frequency and mean date of leaf unfolding. The identification of outliers (XtX or BF) was based on a calibration procedure using pseudo-observed datasets (PODS), as discussed by Gautier (2015). For this step, we used the BayPass_utils.R script as a source for the simulation of 100,000 SNPs. We then ran BayPass again, to generate quantile values for XtX and BF based on these PODS. Conservative and very conservative quantile values were used (“minor outlier”: 0.999 for XtX and 0.9999 for BF; “main outliers” 0.99999 for both XtX and BF) as thresholds to identify outliers. We then generated clusters of neighboring SNPs, by bulking SNPs separated by less than 10 kb. We investigated only clusters of associated SNPs containing at least two outliers, including a “main” outlier, to exclude the random associations that would be expected to occur in such a large dataset. We also investigated clusters with no “main” outlier, provided that the cluster concerned contained at least five “minor” outliers.

### Manual gene annotations

We first performed manual gene annotations based on protein blast searches (*p* values < 1.10^−5^) against the *Arabidopsis* proteome. Gene functions were identified by manual inspection of all the best hits per candidate gene per region. In this study, functional annotation was achieved by performing extensive manual literature searches for each gene, using the strategy described by Leroy *et al.* (2018) rather than automatic approaches based on gene ontology (GO)-oriented methods. This approach was preferred to ensure that we obtained accurate and detailed information about gene function, supported by checked and traceable references.

## Results

### Population history and splits

We used TreeMix to infer the evolutionary history of European white oaks, focusing, in particular, on the most likely sequence of population splits. This software infers the relationships between populations as a bifurcating tree. Interestingly, a strict drift model can explain most of the variance in relatedness between populations (mean over 1000 replicates=0.926; median=0.950; Fig. 1A). Looking at the *Q. petraea* populations (Figs. 1B & S2), the consensus tree over the 1000 replicates supports basal splits and longer branches for populations at higher elevations than for lowland populations. The only exception was population #124 (Killarney, Ireland), which was at the base of all *Q. petraea* populations. In comparisons with all other *Q. petraea* populations and the *Q. robur* reference population, population #124 also presented significant negative values of the *f3* statistic (Fig. S3), suggesting that this population is probably an admixture between *Q. petraea* and *Q. robur*. Several significant negative *f3* values were also obtained for the L16 population, for comparisons with several lowland populations (#9, #97 and #233; Fig. S3).

**Fig. 1:**
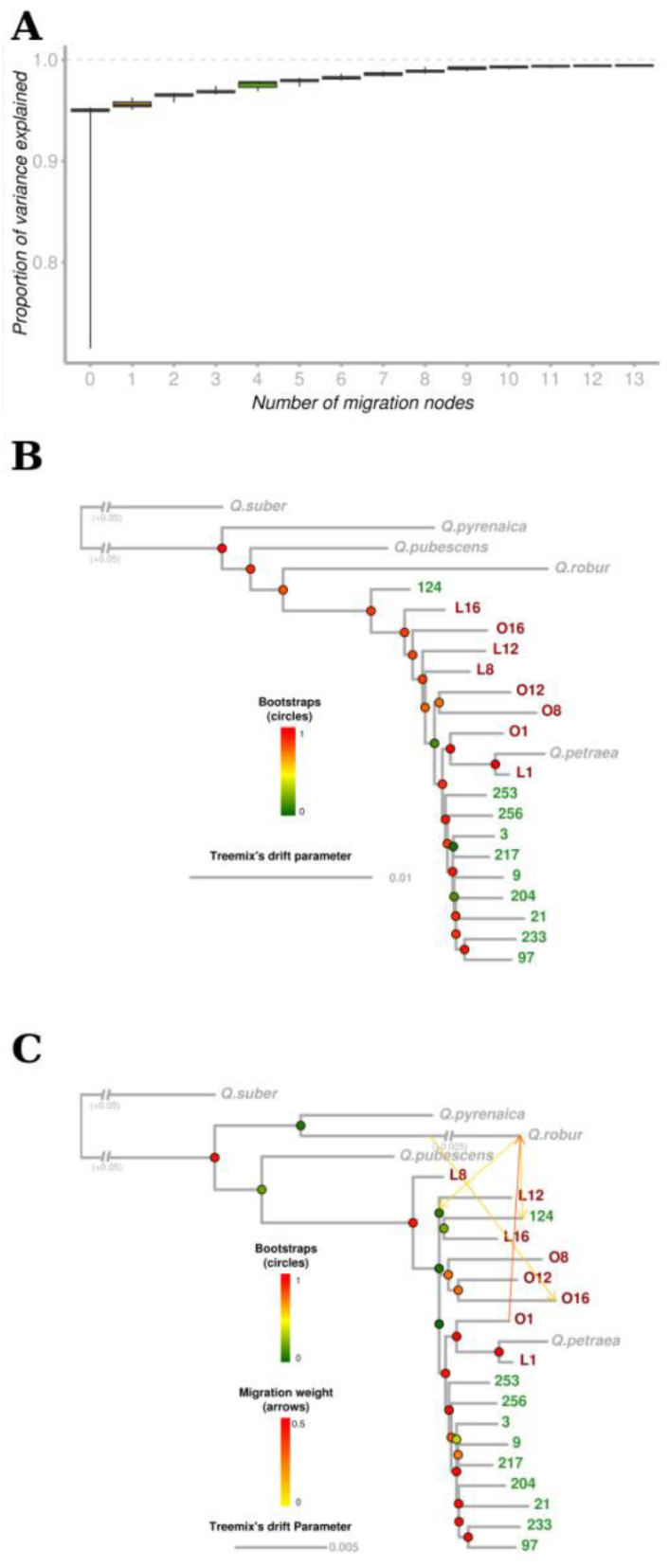
Inference of splits and migration nodes from 1,757,476 genic SNPs in Treemix. A) Boxplot of the proportion of the variance explained over 1000 replicates for 0 to 13 migration nodes. B) Consensus Treemix tree under a strict drift model (m=0). Bootstrap values are shown in circles. C) Consensus TreeMix tree for m=4. Migration events correspond to the events inferred for the best case (inference with the highest likelihood among the 1000 replicates). Bootstrap percentages and migration weights are indicated by the corresponding color scales.

We then sequentially added migration events to the tree (Fig. S2). As expected, the variance in relatedness between populations was better explained as the number of migration events increased. At *m*=1, the proportion of the variance explained increased (mean=0.955; Fig. 1A), in two thirds of replicated simulation,s due to a migration node between *Q. pubescens* and the L8 population for *m*=1 (661/1000). Otherwise, TreeMix inferred (219/1000) a migration event between *Q. robur* and population #124. The inferred tree remained inconsistent with the expected species tree (Leroy *et* al., 2017; Leroy *et al.*, 2018), but this first node helped to explain most of variance generated by the relative positions of *Q. pubescens* and *Q. robur*. At *m*=4 (Fig. 1C), the proportion of the variance explained again increased substantially (mean=0.976) and the bootstrap replicate maximizing the likelihood was, for the first time, consistent with the expected species tree.

Again, TreeMix inferred a migration event between *Q. robur* and the Irish population (#124). It also provided support for candidate introgression events from *Q. robur* for populations at the highest elevation (i.e. from *Q. robur* to O16, and from *Q. robur* to the ancestral population of the modern L12, L16 and #124 populations, Fig. 1C).

### Genome scans

For the identification of SNPs potentially subject to selection, and therefore displaying higher levels of differentiation than would be expected under a hypothesis of neutrality, we calculated the XtX statistic, an *F*_*st*_-like statistic explicitly accounting for population history (Gunther & Coop, 2013). We identified 761,554 SNPs deviating from neutral expectations (2.05% of all SNPs investigated, presenting XtX values above the 0.999 quantile threshold and referred to hereafter as “minor outliers”), including 107,764 for which the evidence was strong (0.29%, with XtX values above the 0.99999 quantile threshold referred to hereafter as “main outliers”, Table S1). These SNPs are distributed over all the chromosomes (outer circle, Fig. 2). Interestingly, XtX outliers were found to be strongly enriched in SNPs that were highly differentiated between species, particularly between *Q. robur* and *Q. petraea* (Fig. 3). More broadly, we observed a strong correlation between the intraspecific XtX value estimated for the 18 populations and the interspecific *FST* between *Q. robur* and *Q. petraea* (*P-value* < 2.2e^−16^, R^2^=0.247, Fig. S4).

**Fig. 2.**
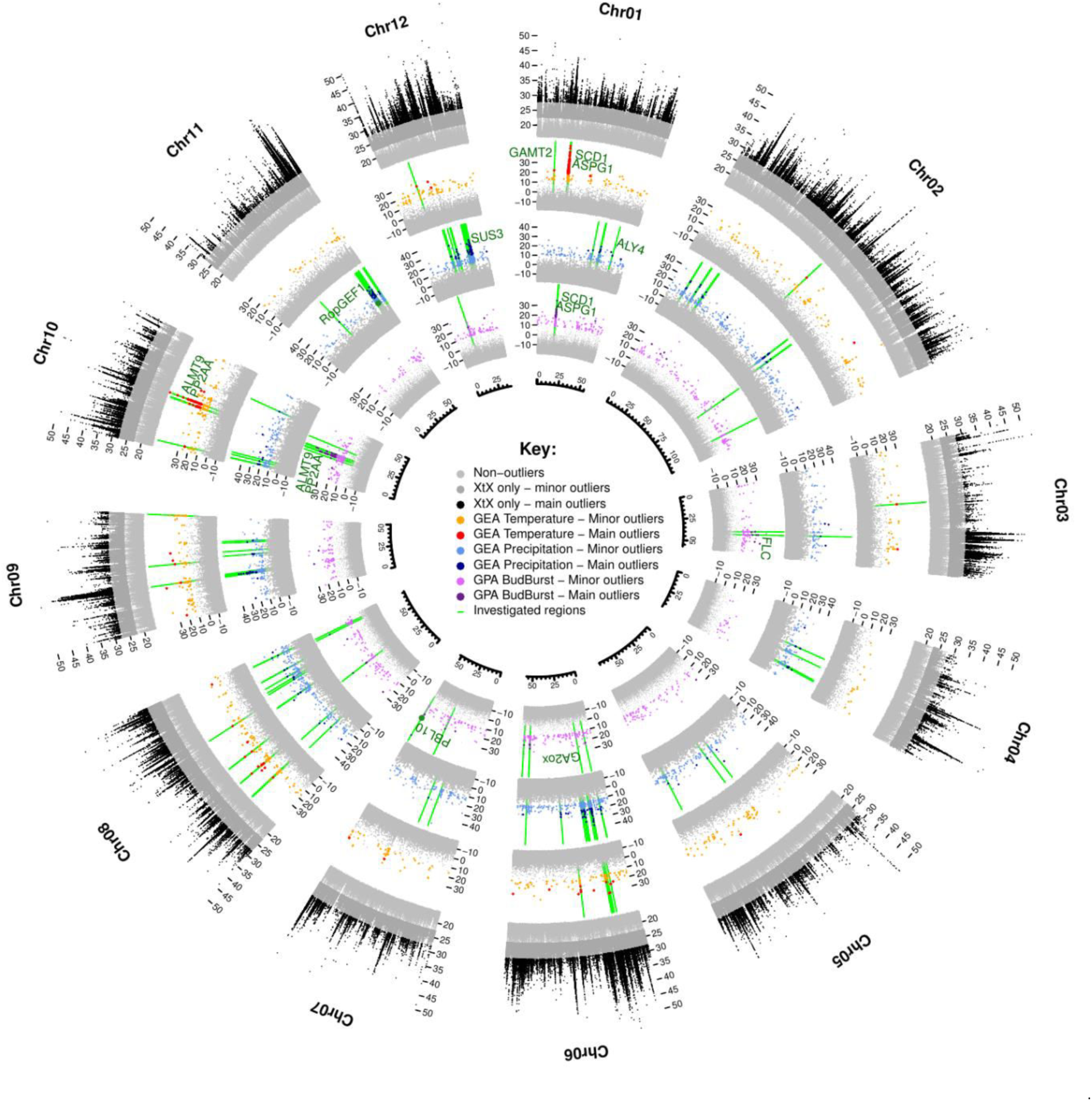
Circular summary of the genome-wide scans for divergence, GEA and GPA associations. From external to internal: divergence (XtX) and associations between temperature, precipitation and the timing of leaf unfolding and allele frequencies. Colors highlight the significance of the SNPs (light= “minor”, dark= “main”) evaluated with the calibration procedure described by Gautier, 2015. Regions investigated for the gene annotation step are shown in green, and the genes discussed in the manuscript are indicated.

**Fig. 3.**
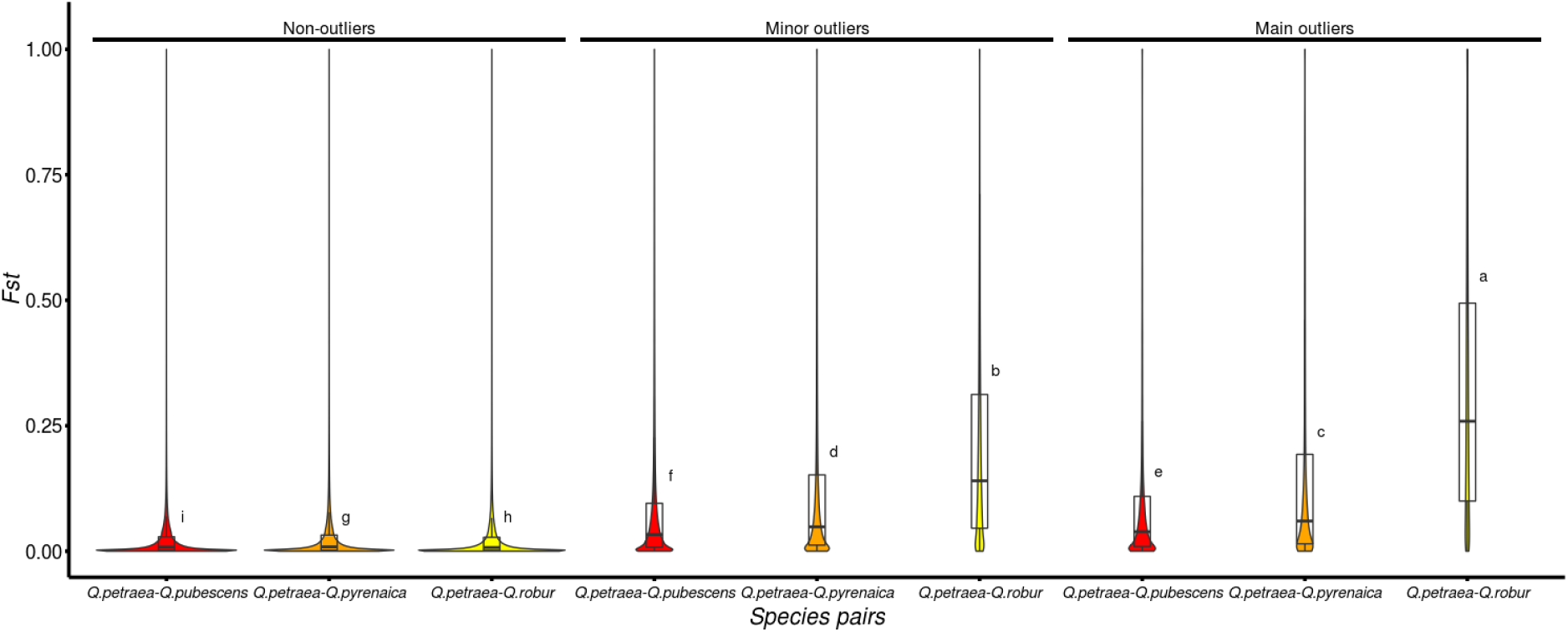
Interspecific Fst between the Q. petraea reference pool and reference populations for three other oak species (for details see Leroy et al., 2018) at XtX outlier loci. Tukey’s honestly significant difference (HSD) criterion at a significance level of 0.05 is reported. (a, b and h: Q. petraea-Q. robur differentiation; c,d and g: Q. petraea-Q. pyrenaica differentiation; e,f and i: Q. petraea-Q. pubescens differentiation)

Among the outlying SNPs detected with the XtX statistic, we identified GEA- and GPA-associated SNPs covarying with mean annual temperature, precipitation or of the date of leaf unfolding recorded in common gardens (Figs. 2 and S5-S16). In total, we identified 5,682 SNPs associated with at least one covariate. More precisely, we identified 1,331 SNPs as outliers (including 216 main outliers) associated with temperature, 2,932 as outliers (277 main outliers) associated with precipitation and 1,572 as outliers (125 main outliers) associated with leaf unfolding (Table S1). For the 153 SNPs involved in two different associations with the three covariates, the largest proportion of SNPs (143/153, 93%) was significantly associated with both temperature and leaf unfolding. The remaining set of common SNPs was found to be associated with both temperature and precipitation (6/153, 4%) or with both precipitation and leaf unfolding (4/153, 3%).

The significantly associated SNPs were highly enriched in SNPs strongly differentiated between *Q. petraea* and *Q. robur*, especially leaf unfolding- and temperature-associated SNPs (Fig. 4). We applied a binning procedure to group covariable-associated SNPs in close vicinity within the genome (less than 10 kb apart). The SNPs were located in 780, 1617 and 1033 independent genomic regions for temperature, precipitation and leaf unfolding, respectively (Table S1). No genomic regions matching the two GEA and GPA associations were detected, as expected given the near-independence of the two climate covariables (Fig. S1C). In the vast majority of cases (87.1%, 76.0% and 87.0% for temperature, precipitation and leaf unfolding, respectively), the genomic region consisted of a single associated SNP. Manual gene annotations were performed only for genes located in the 201 genomic regions containing at least two associated SNPs (see the Materials and Methods for details). These regions accounted for a total of 2,016 associated SNPs (mean=10.2 associated SNPs/region, range: 2-137). These 201 regions included 13 that partially overlapped for leaf unfolding and temperature (Fig. 2, on chromosomes 1, 8 and 10 (see also Figs. S5, S12 and S14), and on scaffold Sc0000849), three that partially overlapped for precipitation and temperature (Fig. 2, on chromosomes 6, 9 and 10; see also Figs. S10; S13 and S14) and two that partially overlapped for precipitation and leaf unfolding (Fig. 2, on chromosomes 3 and 6; see also Figs. S7 and S10).

**Fig. 4.**
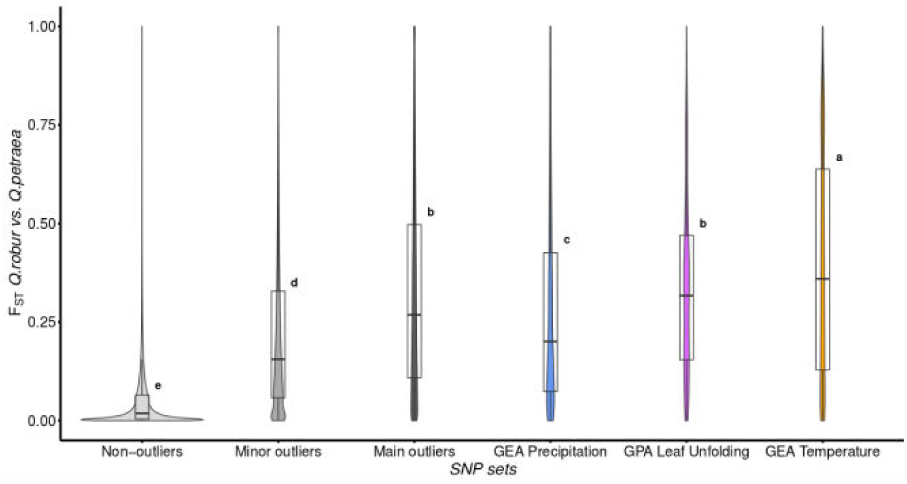
Variation in interspecific Fst between Q. robur and Q. petraea for non-outlier and outlier SNPs, including sets of GEA- and GPA-associated polymorphisms. Tukey’s honestly significant difference (HSD) criterion at a significance level of 0.05 is reported. SNPs for the GEA and GPA sets correspond to all associated SNPs (i.e. both the minor and main categories).

### Gene annotations

Manual gene annotations were performed for the genes located within or close to these 201 regions (within 5 kb on either side) to exclude border effects. This led to the identification of 167 unique candidate genes. We found regulators of various growth and development processes in plants. As expected, we identified a set of genes involved in stomatal responses to water stress (e.g. ALY4, ALMT9, PBL10, PP2AA, RopGEF1, SUS3, SCD1). Concordantly, our genome scans also revealed some temperature-associated genes acting as regulators of the production of gibberellins (e.g. GAMT2, ASPG1 or GA2ox). These genes also play important roles in various developmental processes, including seed dormancy (Shen *et al.*, 2018).

We investigated the processes at work further, by generating ecological clines for each associated SNP (Sup File 1). Most of the associated SNPs followed complex clines along the ecological gradients (*i.e.* non-linear), highlighting the advantages of model implementation in Bayenv or BayPass rather than the use of linear or logistic model-derived methods (De Mita *et al.*, 2013), but some SNPs displayed an almost linear change in allele frequency along the ecological gradient (Fig. 5). The clines were generally consistent with a continuum between the *Q. petraea* and *Q. robur* reference pools, with higher levels of *Q. robur*-like alleles in populations living in cooler and/or wetter environments, suggesting that interspecific introgression from *Q. robur* is an important source of adaptive variation for *Q. petraea* populations. Typical examples include the precipitation-associated SNPs and leaf unfolding-associated SNPs located within two important genes controlling stomatal responses, the ROPGEF1 and PBL10 (=APK1b) genes (Fig. 5 A and B respectively; Li & Liu, 2012; Elhaddad *et al.*, 2014)

**Fig. 5.**
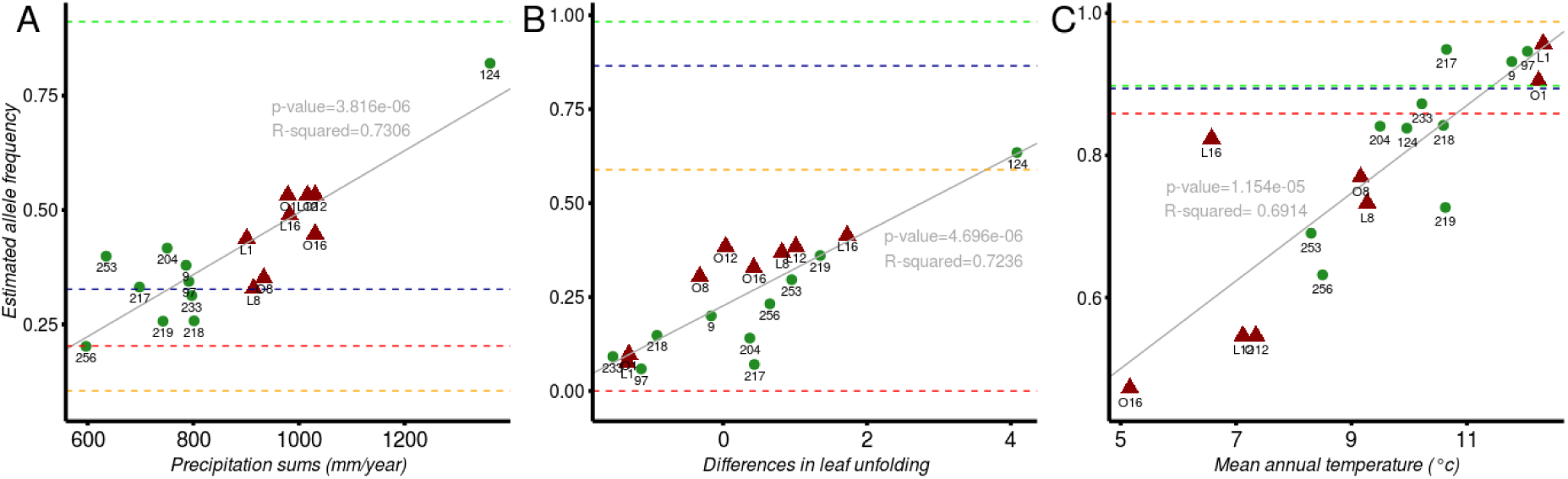
Near-linear genetic clines along ecological gradients. The clines shown are for the three SNPs detected by BayPass: the precipitation-associated SNP Chr11:47276412 (A), the leaf unfolding-associated SNP Chr07:51928762 (B) and the temperature-associated SNP Chr07:28987726 (C). Best linear regressions for each associated SNP are shown in gray. Red, blue, green and orange dotted lines correspond to allele frequencies estimated in the Q. petraea, Q. pubescens, Q. robur and Q. pyrenaica reference populations, respectively. Clines for all other associated SNPs are shown in Supplementary File 1.

## Discussion

We assembled data for phenotypic surveys in common garden experiments, climatic data for the origin of the population, and whole-genome sequences, to assess the genetic divergence of the timing of leaf unfolding between extant populations of *Quercus petraea* at the genomic level. We drew inferences about population history by searching for footprints of population splits and admixture, and conducted genome scans to explore the establishment of genetic divergence during the early Holocene in extant populations. We found that introgression from *Q. robur* had made a major contribution to population divergence of some locally adapted *Q. petraea* populations. This finding was supported by three major outcomes. First, we found that admixture events occurred mainly in *Q. petraea* populations from cooler and wetter climates. Second the SNPs displaying the highest level of genetic differentiation between *Q. petraea* populations were also highly differentiated between *Q. petraea* and *Q. robur*. Third, some of the genes contributing to phenological divergence in *Q. petraea* displayed clinal variation consistent with the geographic variation of introgression. Our results also confirmed the temperature-driven genetic cline correlated to the variation of leaf unfolding repeatedly reported in oaks (Ducousso *et al.*, 1996; Vitasse *et al.*, 2009; Alberto *et al.*, 2011; Firmat *et al.*, 2017). Finally, our findings also highlight population divergences due to precipitation gradients.

### *Historical introgression from* Q. robur *into* Q. petraea

The gene flow events inferred by *TreeMix* suggest that admixture events, mostly involving *Q. robur*, underlie the genetic differentiation of current *Q. petraea* populations over wide geographic gradients. Contemporary hybridization between other closely related white oaks and *Q. petraea* has been reported in empirical contemporary gene flow studies (Curtu *et al.*, 2007; Salvini *et al.*, 2009; Lepais & Gerber, 2011), but we found evidence only for an evolutionary footprint of admixture with *Q. robur* in our sampled populations of *Q. petraea*. Today, *Q. petraea* and *Q. robur* are frequently found in sympatric stands across Europe, due to their common pattern of post-glacial colonization dynamics. As suggested by the widespread sharing of chloroplast haplotypes between these two closely related species when present in the same stands (Petit *et al.*, 2002), hybridization followed by recurrent backcrossing between *Q. petraea* and *Q. robur* has been an essential mechanism in *Q. petraea* expansion. Similarly, hybridization and subsequent backcrossing between the pioneer (resident) species *Q. robur* and the late successional (invading) species *Q. petraea* made it possible for *Q. petraea* to migrate with *Q. robur* (Petit *et al.*, 2004; Guichoux *et al.*, 2013). Most of the admixture events inferred from the *TreeMix* model support directional introgression from *Q. robur* to *Q. petraea* (Figure 1), as predicted by theory in the case of this invasion scenario (Currat *et al.*, 2008).

The ultimate outcome of hybridization followed by unidirectional backcrossing is the restoration of *Q. petraea* within *Q. robur* stands. As suggested by artificial backcross experiments performed with breeding populations of other closely related European white oak species (Diskin *et al.*, 2006), the genome of the invading species (here *Q. petraea*) should be regenerated in a limited number of generations, ultimately maintaining a limited imprint of the introgression from *Q. robur*. However, our results show that genes introgressed from *Q. robur* are maintained in *Q. petraea* populations, particularly those from northern latitudes or higher elevations. Previous studies based on Bayesian clustering methods have already highlighted the occurrence of more admixture at higher elevations (Alberto *et al.*, 2010). Introgressed genes may still be present if admixture occurred recently, if there is a demographic imbalance between the invader and resident populations during initial contact (Currat *et al.*, 2008), or if these populations are subject to directional selection. Studies of pollen remains have indicated that temperate deciduous oaks have been present in sampled areas since about 10,000 years BP in Northern Germany (Alberto *et al.*, 2010; Giesecke, 2016) and the Pyrénées (Jalut *et al.*, 1992; Reille & Lowe, 1993) and since about 8,000 years BP in Ireland (Kelleher *et al.*, 2004). This timescale would be long enough for completion of the regeneration process and the eradication of introgression. We cannot rule out the possibility that demographic imbalance between these two species facilitated the introgression and maintenance of *Q. robur* alleles in *Q. petraea*, but the detection of introgression principally at higher elevations and latitudes suggests that some introgressed alleles were most probably maintained by selection, increasing the degree of differentiation between *Q. petraea* populations (Fig. 4). Interestingly, our results also show that genes highly differentiated between *Q. petraea* populations are also highly differentiated between *Q. petraea* and *Q. robur* (Fig. 4 and Fig. 5), suggesting their involvement in either adaptive divergence or reproductive barriers between the two species (Leroy *et al.*, 2018). These two species display subtle differences in soil preferences when present in the same forest (Timbal & Aussenac, 1996; Eaton *et al.*, 2016), but they also have different climate responses, as suggested by their allopatric (temperature- and precipitation-dependent) distributions at the edge of their ranges. *Q. robur* extends farther north (up to Finland) and east (up to the Ural Mountains) (see Leroy *et al*., 2018), and has a higher frequency in wetter climates (Eaton *et al.*, 2016). We suspect that the introgression of *Q. robur* alleles for genes involved in these differential responses may have contributed to the expansion of *Q. petraea* populations to higher elevations in the Pyrenees and wetter climates in Ireland.

### Genomic and genetic clines shaped by introgression

Association studies identified additional elements relating to the probable contribution of introgression to the adaptive divergence of *Q. petraea* populations. First, we recovered the clinal phenotypic gradient of leaf unfolding with temperature variations (Fig. S1B) as observed in previous common garden experiments (Ducousso *et* al., 1996; Vitasse *et al.*, 2009; Alberto *et al.*, 2011; Firmat *et al.*, 2017), with populations from cooler climates (higher latitudes or elevations) flushing later than populations from warmer climates. Second, we found that the genes displaying clinal variations of allelic frequency along a temperature gradient were enriched in genes differentiated between the *Q. petraea* and *Q. robur* species. This raises questions about whether the genes introgressed from *Q. robur* also contributed to the later flushing of *Q. petraea* at higher elevations. No significant interspecific differences in leaf unfolding are observed between these two closely related species in common garden experiments or *in situ* (Kleinschmit, 1993; Jensen & Hansen, 2008; Wilkinson *et al.*, 2017), although there is a slight trend towards earlier flushing in *Q. petraea*. (Kleinschmit, 1993). However, *Q. robur* is well known to display ecotypic differentiation, with the recognition of late-flushing populations that have been attributed subspecies status as *Q. robur* var. *tardiflora*, mostly in Eastern Europe, and flush almost a month later than *Q. robur* (Wesolowski & Rowinski, 2008; Utkina & Rubtsov, 2017). Similar extremely late-flushing populations have also been reported in Western Europe for *Q. robur* (Riedacker, 1968), but no such phenological differentiation has been reported for *Q. petraea*.

In conclusion, we found that adaptive introgression from *Q. robur* to *Q. petraea* contributed to the overall clinal variation for some key adaptive traits, including leaf unfolding-related traits in particular. In addition to shedding light on the evolutionary processes at work, this study identified several key oak genes involved in the stomatal behavior as involved in adaptive introgression, providing support for the general view that stomatal regulation makes a major contribution to the maintenance of homeohydry in land plants (Brodribb & McAdam, 2017). Other leaf unfolding-associated genes would be expected to have key roles, with pleiotropic effects. A typical example is provided by FLC (Flowering Locus T). FLC antagonizes the gibberellin (GA) pathway and downregulates flowering (Deng *et al.*, 2011; Li *et* al., 2015). The effects of FLC are known to be suppressed by vernalization, suggesting that FLC plays a major role in seasonal and developmental timing.

It has been suggested that diversifying selection driven by either herbivory or frost occurrence could account for the observed genetic cline of leaf unfolding (Dantec *et al.*, 2015). Non selective factors, such as assortative mating combined with gene flow (e.g. preferential pollination between late-flushing trees at lower latitudes, with earlier flushing trees at higher latitudes) have also been shown to contribute to the observed phenological clines (Soularue & Kremer, 2012; 2014).

## Supporting information

SupInfo

SupFile1

## Acknowledgments

This research was funded by the European Research Council under the European Union’s Seventh Framework Programme (TREEPEACE project, FP/2014-2019: ERC Grant Agreement no. 339728) and by the French ANR (GENOAK project, 11-BSV6-009-021). We thank the Genotoul Bioinformatics Platform Toulouse Midi-Pyrenees (Bioinfo Genotoul) and the Biogenouest BiRD core facility (Université de Nantes) for providing computing and storage resources. We thank the staff of the Experimental Units of INRA Nancy (UEFL, Unité Expérimentale Forestière de Lorraine) and INRA Toulenne (UE 0393, INRA, Domaine des Jarres, 33210-Toulenne) for their contribution during field phenological assessments and sampling. We also thank Jorge A. P. Paiva for providing access to *Q. suber* data, Nick Zimmerman for providing climate data, Mathieu Gautier for providing advice on how to make the best use of BayPass, and Quentin Rougemont for fruitful discussions. TL also thanks the project coordinator Benoit Nabholz (BirdIslandGenomic project, ANR-14-CE02-0002) for his support and feedback.

