## Supplementary material for "Adaptive introgression as a driver of local adaptation to climate in European white oaks": SupInfo

**Table S1 Number of SNPs used or outliers identified in this study**

| SNP sets | SNP counts |
|---|---|
| **Total number of SNPs (18 *Q. petraea* populations)** | **37,062,111** |
| *SNPs common to the 4 species dataset (Leroy et al.,. 2018)* | *19,806,572* |
| *Intragenic SNPs used for Treemix* | *1,757,476* |
| **All XtX outlier SNPs** | **761,554** |
| *Main XtX outlier SNPs* | *107,764* |
| **Temperature-associated SNPs** | **1,331** |
| *Main GEA outlier SNPs* | *216* |
| *Temperature-associated regions after the binning procedure* | *780* |
| *Regions investigated for annotation* | *34* |
| **Precipitation-associated SNPs** | **2,932** |
| *Main GEA-associated outlier SNPs* | *277* |
| *Precipitation-associated regions after the binning procedure* | *1,617* |
| *Regions investigated for annotation* | *132* |
| **Leaf unfolding-associated SNPs** | **1,572** |
| *Main GPA outlier SNPs* | *125* |
| *Leaf unfolding-associated regions after the binning procedure* | *1033* |
| *Regions investigated for annotation* | *35* |
| **Total number of associated SNPs** | **5835 (5682 unique SNPs)** |
| **Total number of regions investigated** | **201 (incl. 184 non-overlapping)** |

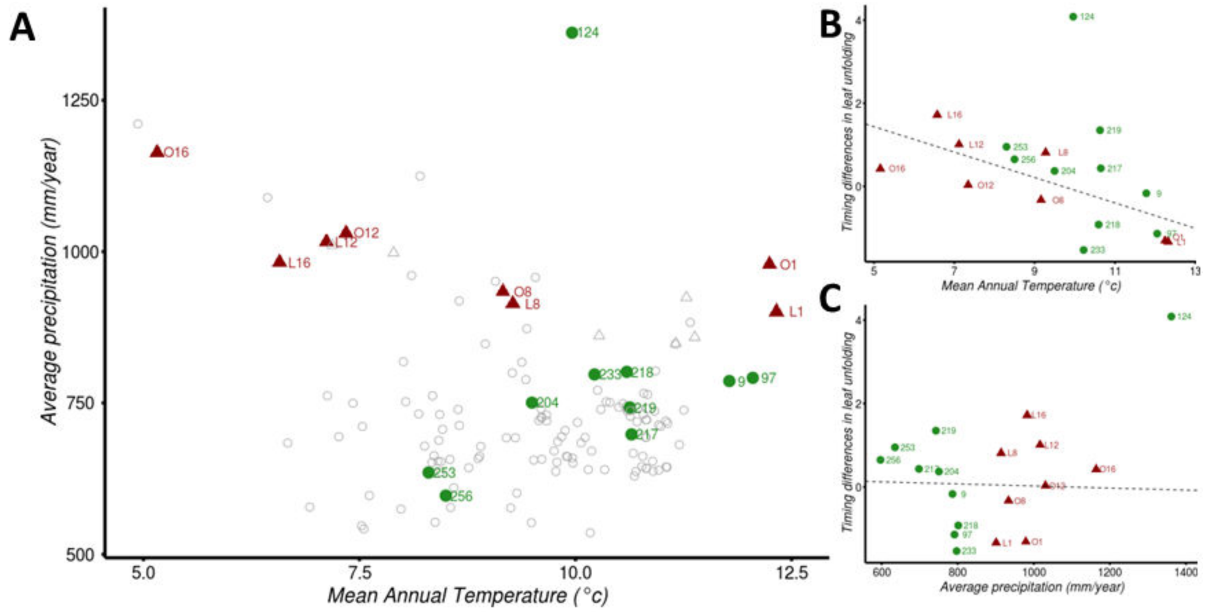


**Fig. S1 Summary of the climatic and phenological data for the *Quercus petraea* populations studied.** A) Biplot of mean annual temperature and total precipitation for the sites of origin of the populations studied. Green circles correspond to populations from the latitudinal gradients, and red triangles correspond to populations from the elevational gradients. Empty circles correspond to other populations present in the common gardens (Saenz-Romero *et al.*, 2017), indicating that the sample of the population studied corresponds to a representative sample of the climatic space of *Q. petraea*. B) Significant relationship ($R^2$=37%, $p$=0.0056) between the standardized timing of leaf unfolding and mean annual temperature, after exclusion of the climatic outlier population (124, Killarney). C) Non-significant relationship ($p$=0.8862) between the standardized timing differences in leaf unfolding and the total mean precipitation per year, after exclusion of the climatic outlier population (124, Killarney). Numbers correspond to population labels, as indicated in table 1.

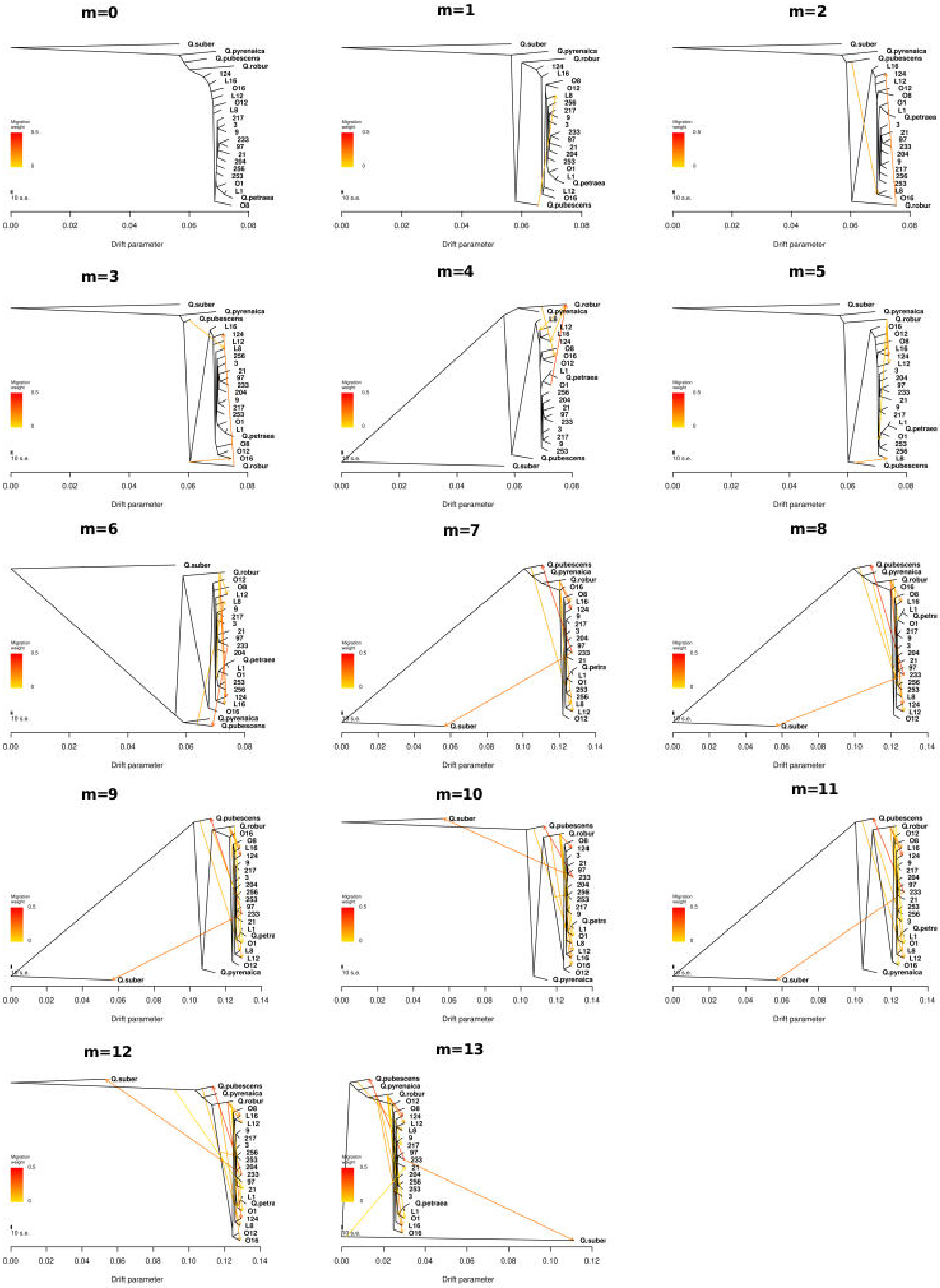


**Fig. S2:** Best-fit trees for various numbers of migration edges (*m*). The best-fit tree corresponds to the bootstrap replicate with the highest likelihood among the 1000 replicates.

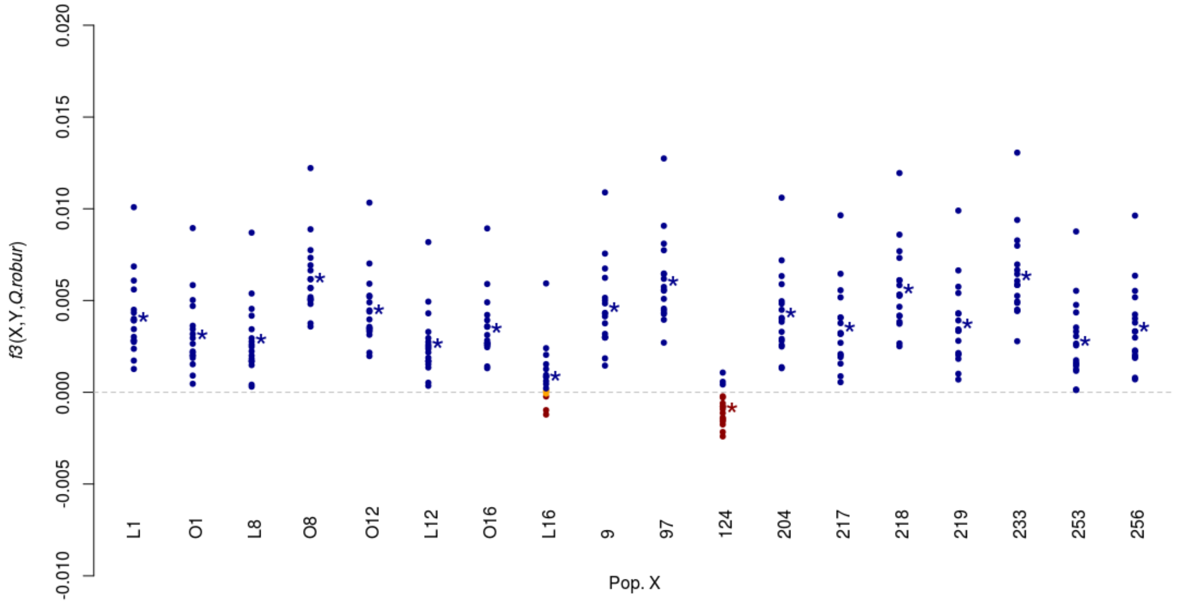


**Fig. S3 *F3*-statistics as tests for admixture between *Q. petraea* and *Q. robur* populations *[i.e. f3(X;Y,Q.robur)]*.** Mean *F3* values are shown for each focal population (X). Negative mean values indicate probable admixture between another *Q. petraea* population (Y) and *Q. robur*. Significant negative and positive values are shown in red and blue, respectively. Non-significant non-zero values (plus or minus one standard error) are shown in orange. Stars indicate the mean of the mean values over the 17 *Q. petraea* populations (*i.e.* all populations other than the focal population).

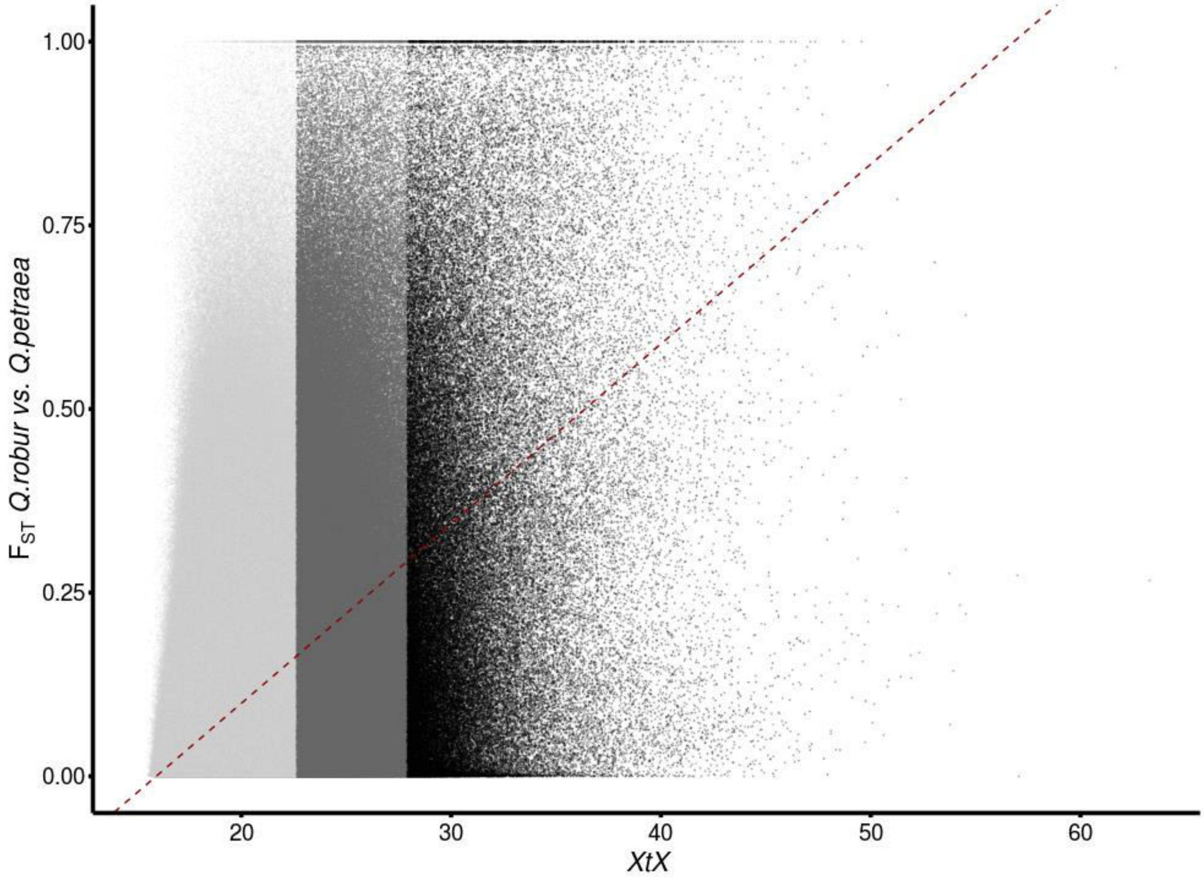


**Fig. S4: Intra- (XtX) and interspecific (Fst) differentiation between the *Q. robur* and *Q. petraea* reference populations.** Of the 19.8 million shared SNPs, XtX, calculated with our panel of 18 populations, is strongly correlated with the interspecific $F_{ST}$ calculated between *Q. robur* and *Q. petraea* by Leroy et al. (2018). Linear regression is shown in dark red (slope=0.0245, *p*-value < 2.2 e-16, $R^2$=0.247).

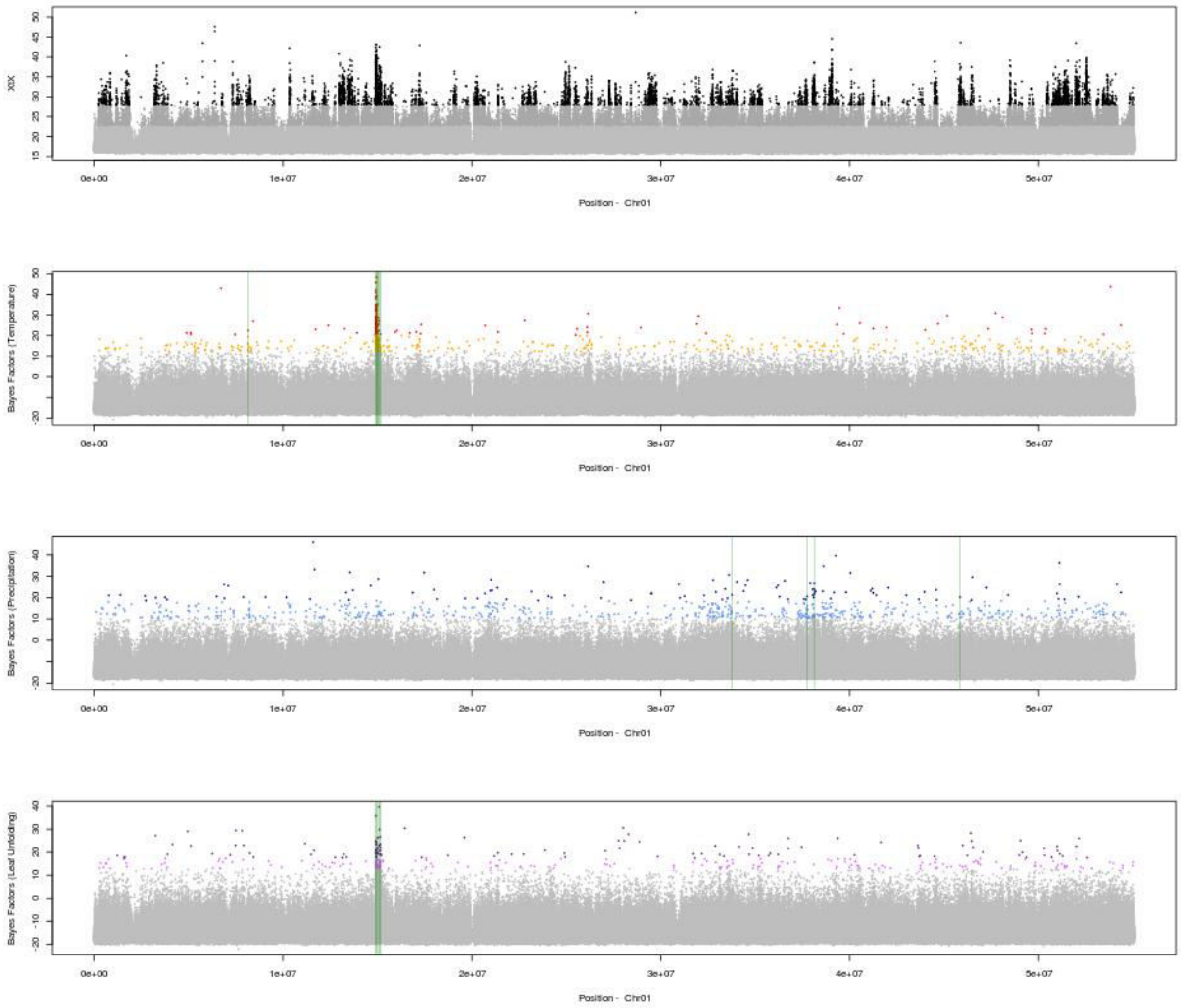


**Fig. S5: Detailed information concerning the genome-wide scans for divergence, GEA and GPA associations for SNPs on chromosome 1.** From top to bottom: relative divergence (XtX) and Bayes factors for associations between temperature, precipitation and the timing of leaf unfolding and allele frequencies, as calculated with BayPass. Colors highlight the significance of the SNPs (light = “minor”, dark = “major”) evaluated with the calibration procedure described by Gautier (2015). Regions investigated for the gene annotation step are shown in green.

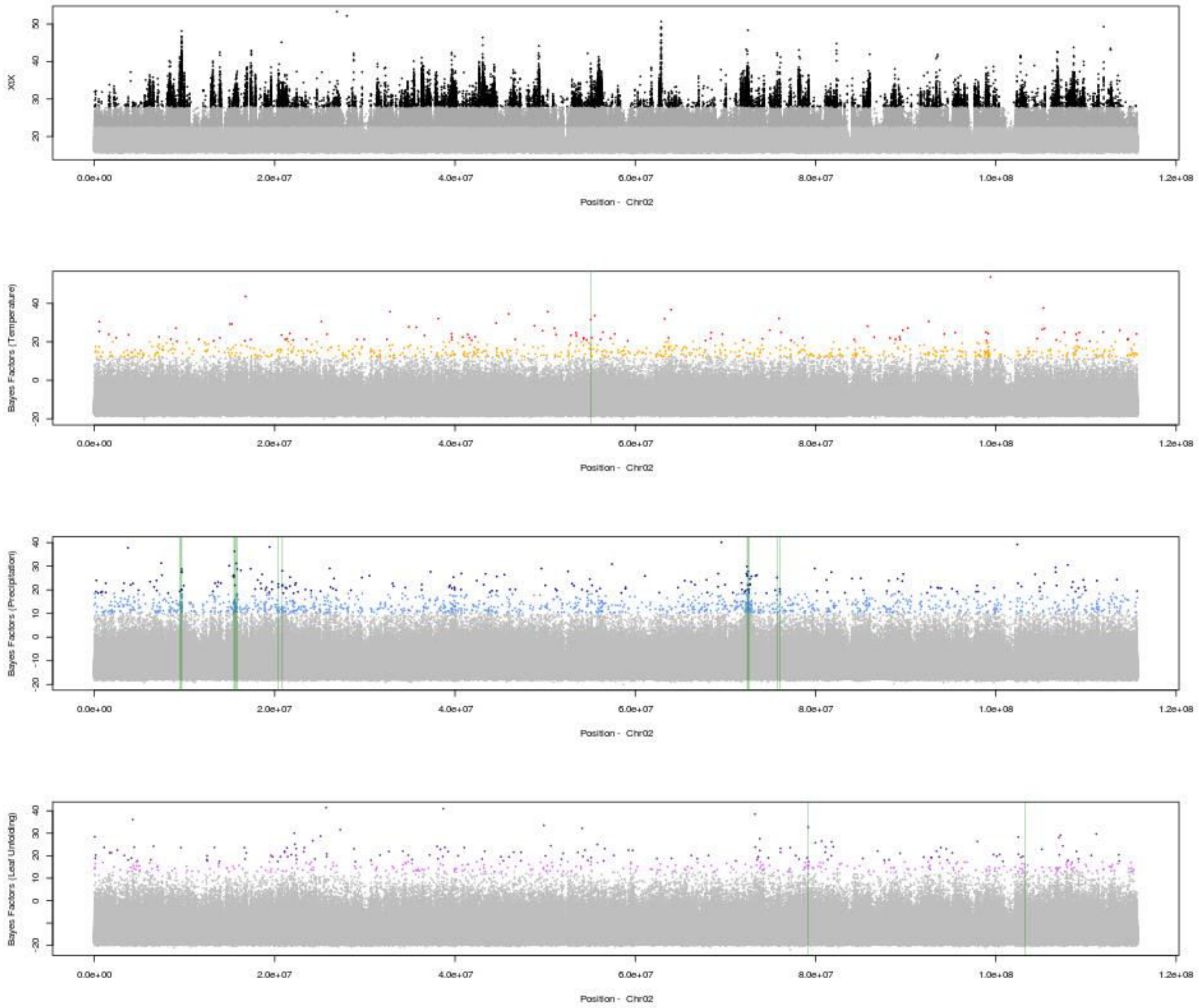


**Fig. S6: Detailed information concerning the genome-wide scans for divergence, GEA and GPA associations for SNPs on chromosome 2.** From top to bottom: relative divergence (XtX) and Bayes factors for associations between temperature, precipitation and the timing of leaf unfolding and allele frequencies, as calculated with BayPass. Colors highlight the significance of the SNPs (light = “minor”, dark = “major”) evaluated with the calibration procedure described by Gautier (2015). Regions investigated for the gene annotation step are shown in green.

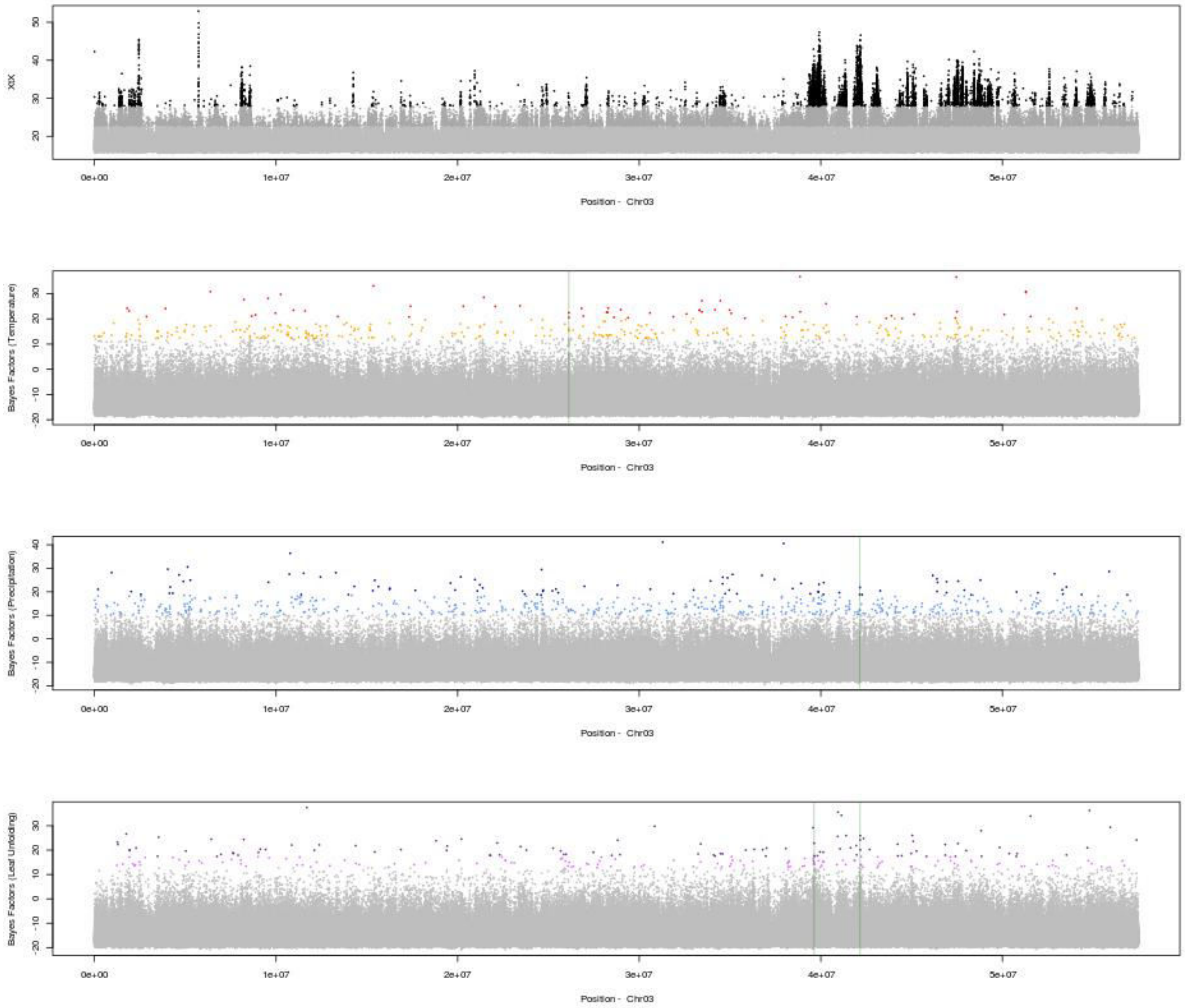


**Fig. S7: Detailed information concerning the genome-wide scans for divergence, GEA and GPA associations for SNPs on chromosome 3.** From top to bottom: relative divergence (XtX) and Bayes factors for associations between temperature, precipitation and the timing of leaf unfolding and allele frequencies, as calculated with BayPass. Colors highlight the significance of the SNPs (light = “minor”, dark = “major”) evaluated with the calibration procedure described by Gautier (2015). Regions investigated for the gene annotation step are shown in green.

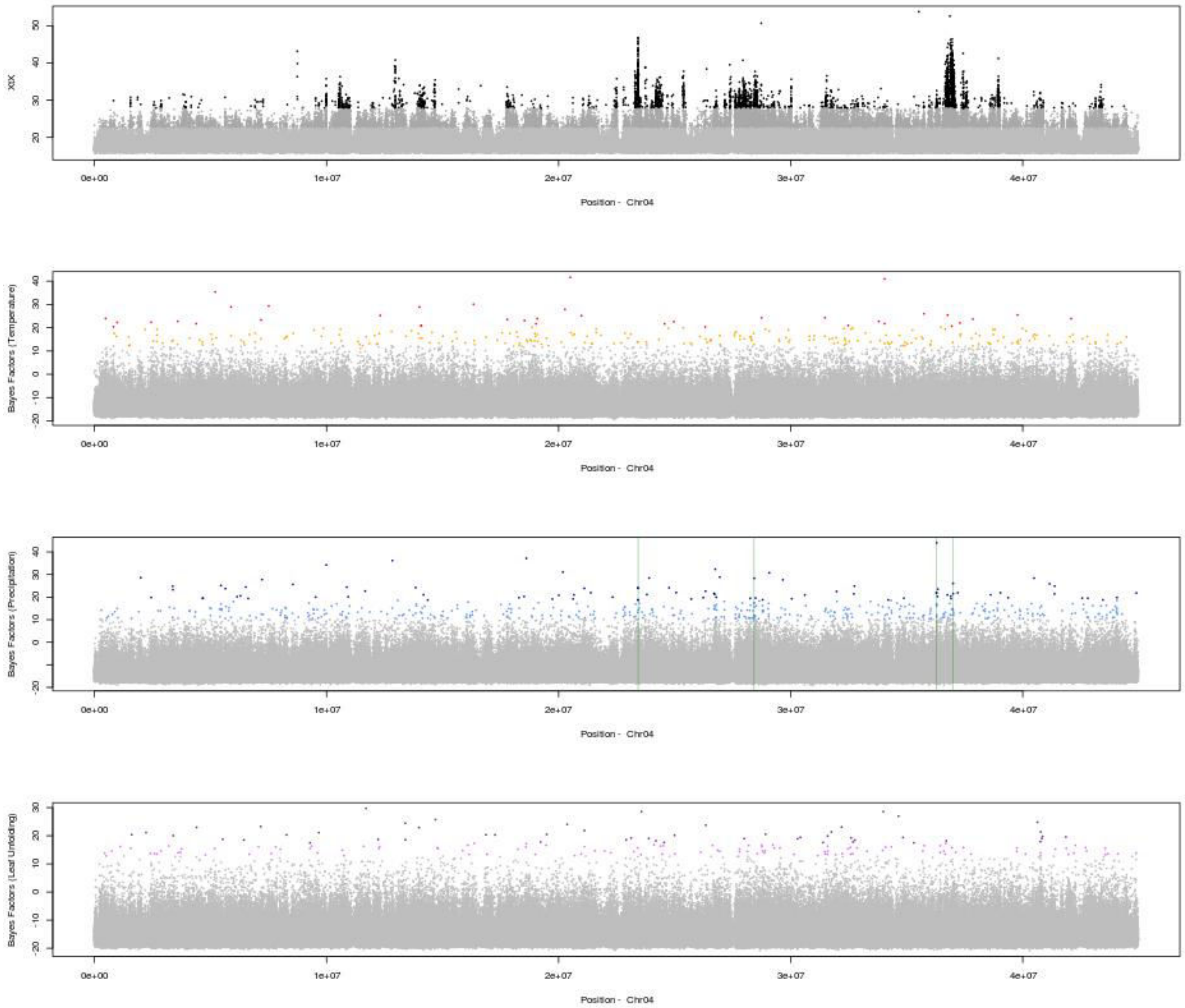


**Fig. S8: Detailed information concerning the genome-wide scans for divergence, GEA and GPA associations for SNPs on chromosome 4.** From top to bottom: relative divergence (XtX) and Bayes factors for associations between temperature, precipitation and the timing of leaf unfolding and allele frequencies, as calculated with BayPass. Colors highlight the significance of the SNPs (light = "minor", dark = "major") evaluated with the calibration procedure described by Gautier (2015). Regions investigated for the gene annotation step are shown in green.

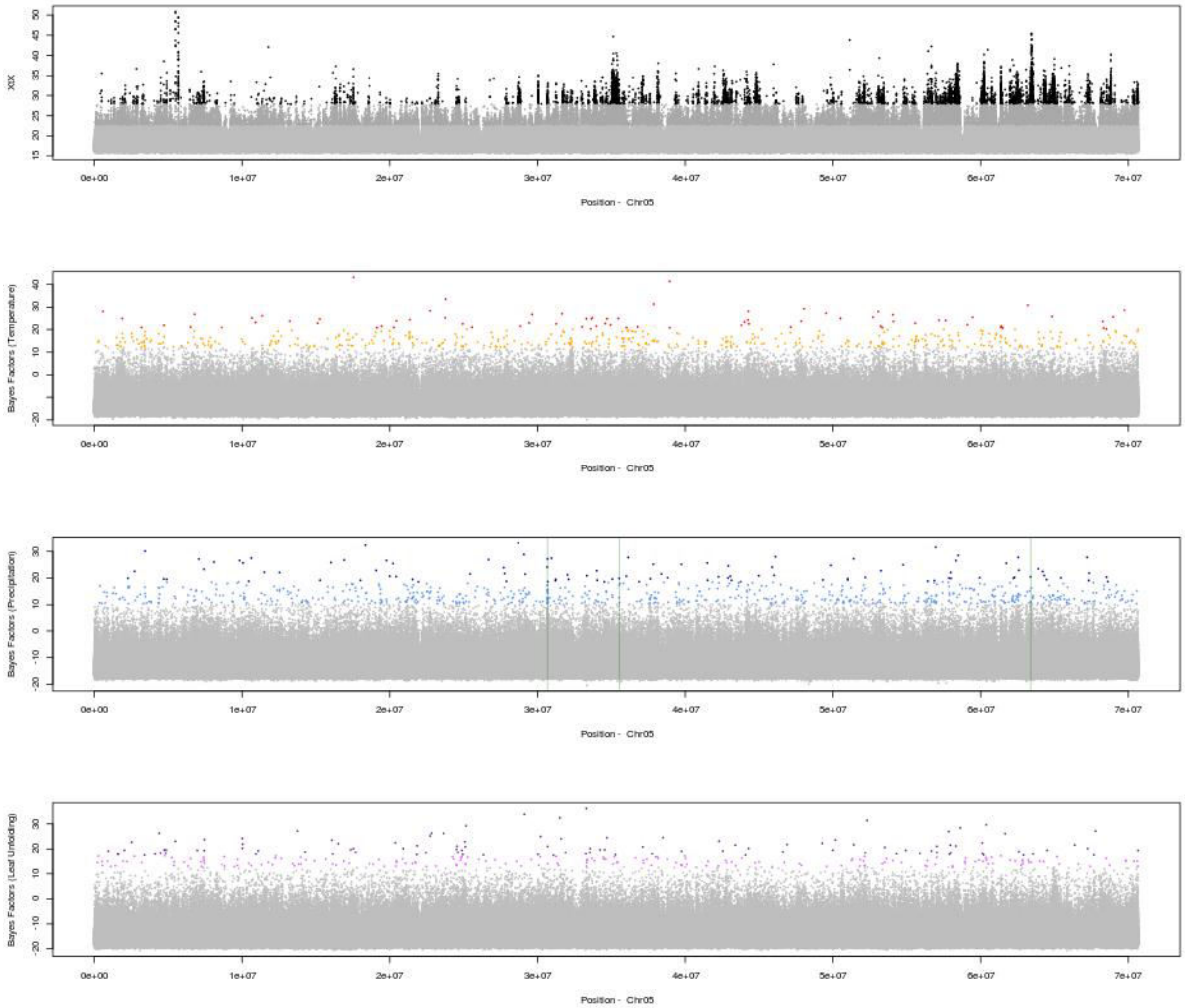


**Fig. S9: Detailed information concerning the genome-wide scans for divergence, GEA and GPA associations for SNPs on chromosome 5.** From top to bottom: relative divergence (XtX) and Bayes factors for associations between temperature, precipitation and the timing of leaf unfolding and allele frequencies, as calculated with BayPass. Colors highlight the significance of the SNPs (light = “minor”, dark = “major”) evaluated with the calibration procedure described by Gautier (2015). Regions investigated for the gene annotation step are shown in green.

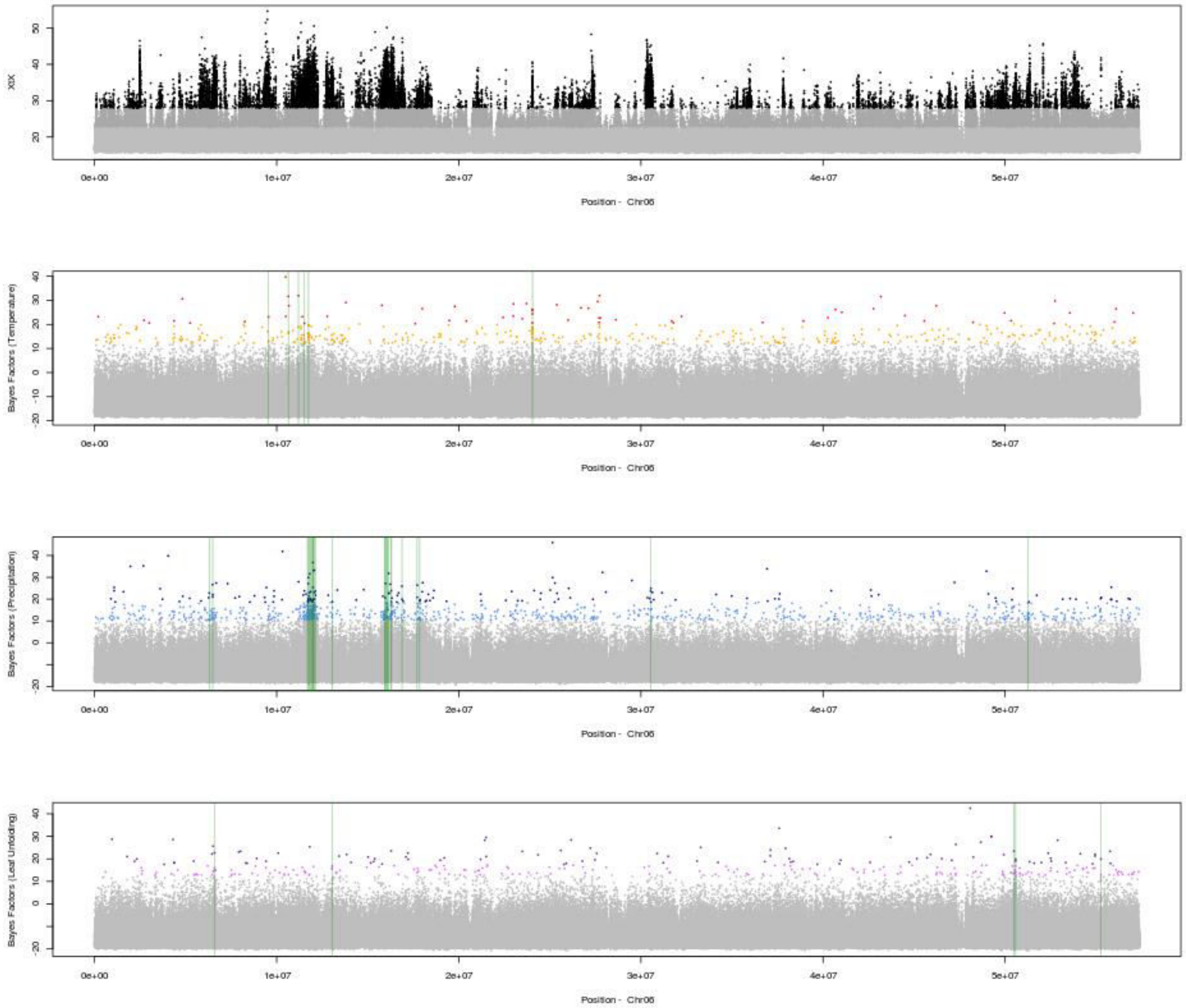


**Fig. S10: Detailed information concerning the genome-wide scans for divergence, GEA and GPA associations for SNPs on chromosome 6.** From top to bottom: relative divergence (XtX) and Bayes factors for associations between temperature, precipitation and the timing of leaf unfolding and allele frequencies, as calculated with BayPass. Colors highlight the significance of the SNPs (light = “minor”, dark = “major”) evaluated with the calibration procedure described by Gautier (2015). Regions investigated for the gene annotation step are shown in green.

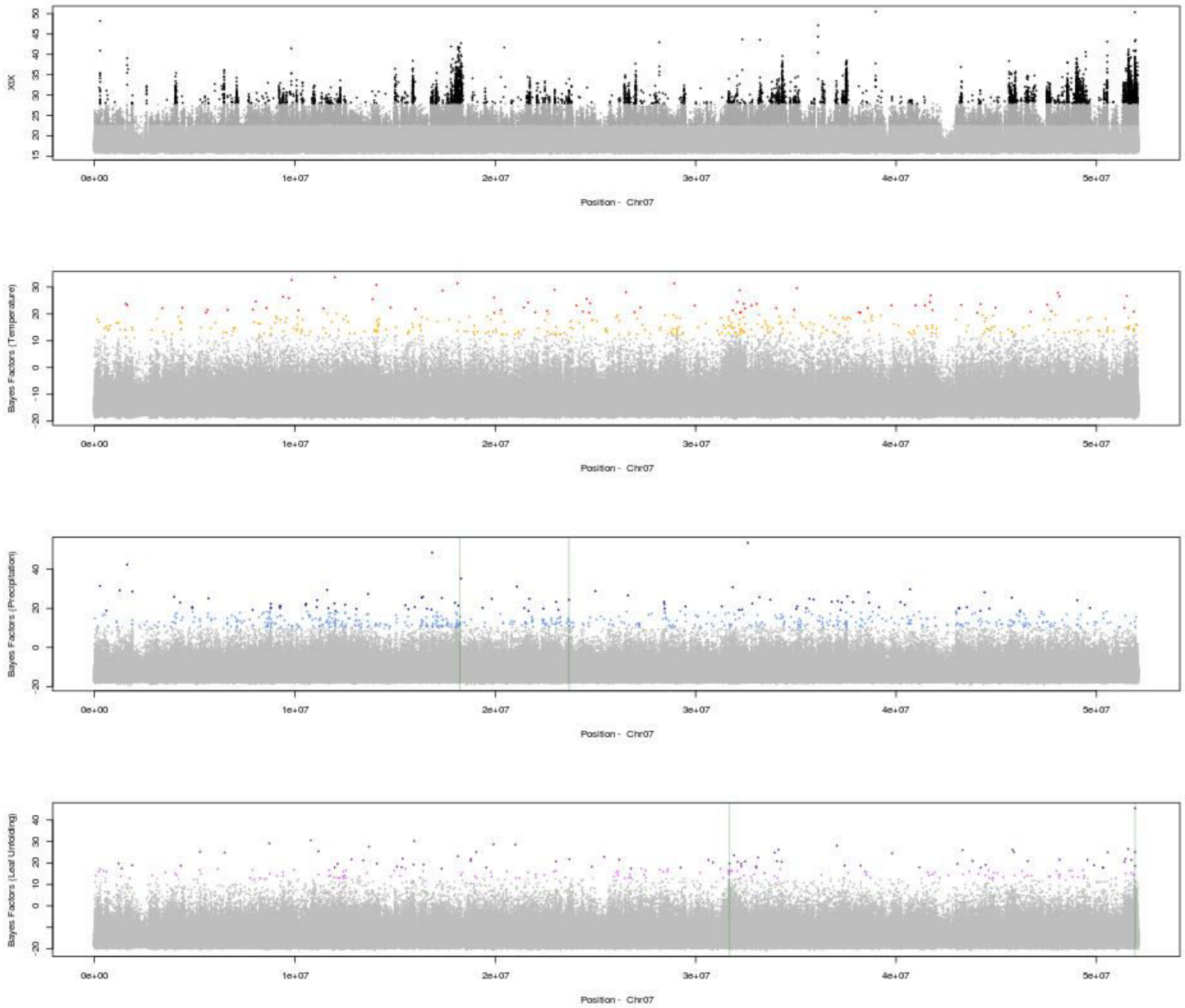


**Fig. S11: Detailed information concerning the genome-wide scans for divergence, GEA and GPA associations for SNPs on chromosome 7.** From top to bottom: relative divergence (XtX) and Bayes factors for associations between temperature, precipitation and the timing of leaf unfolding and allele frequencies, as calculated with BayPass. Colors highlight the significance of the SNPs (light = "minor", dark = "major") evaluated with the calibration procedure described by Gautier (2015). Regions investigated for the gene annotation step are shown in green.

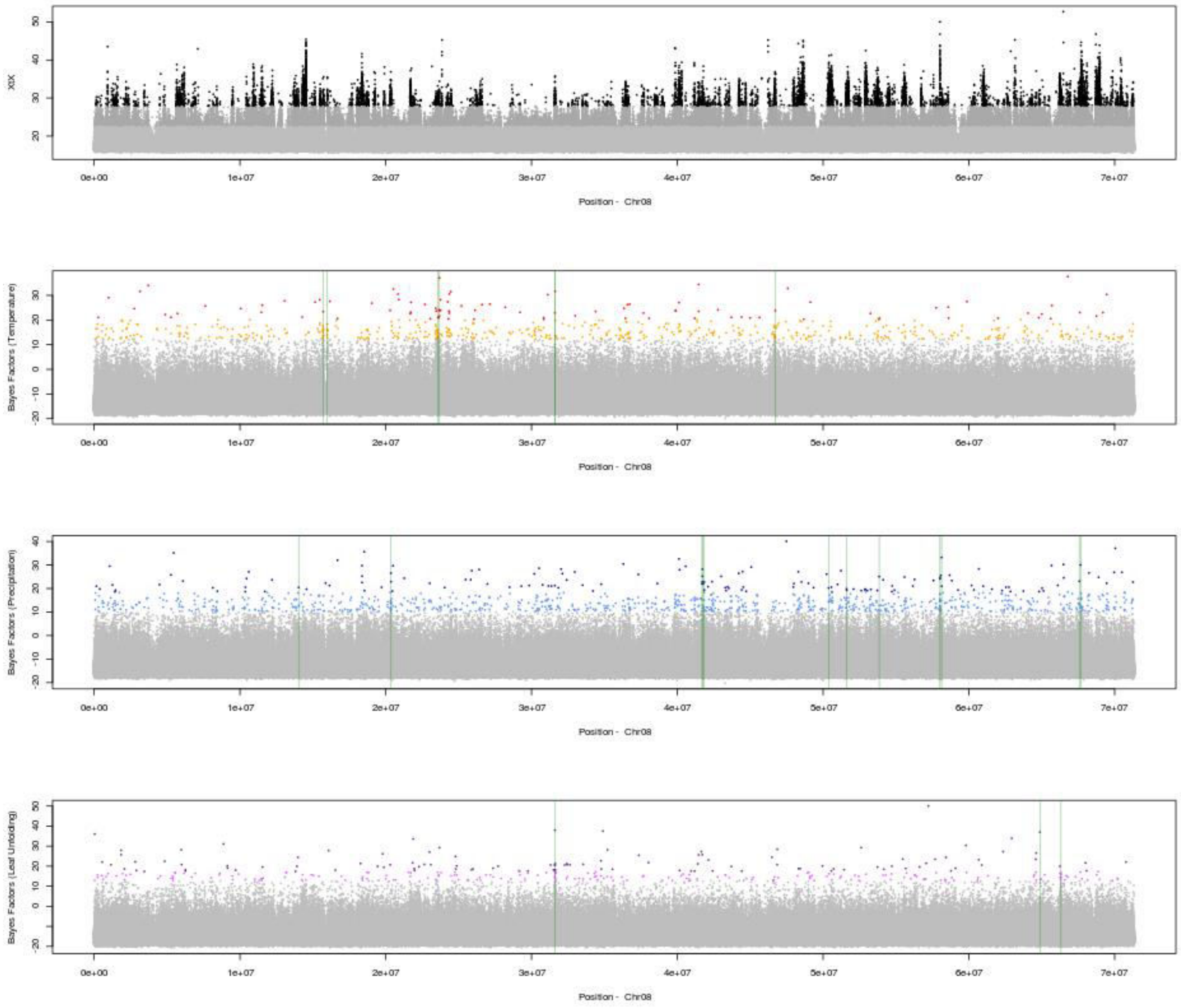


**Fig. S12: Detailed information concerning the genome-wide scans for divergence, GEA and GPA associations for SNPs on chromosome 8.** From top to bottom: relative divergence (XtX) and Bayes factors for associations between temperature, precipitation and the timing of leaf unfolding and allele frequencies, as calculated with BayPass. Colors highlight the significance of the SNPs (light = “minor”, dark = “major”) evaluated with the calibration procedure described by Gautier (2015). Regions investigated for the gene annotation step are shown in green.

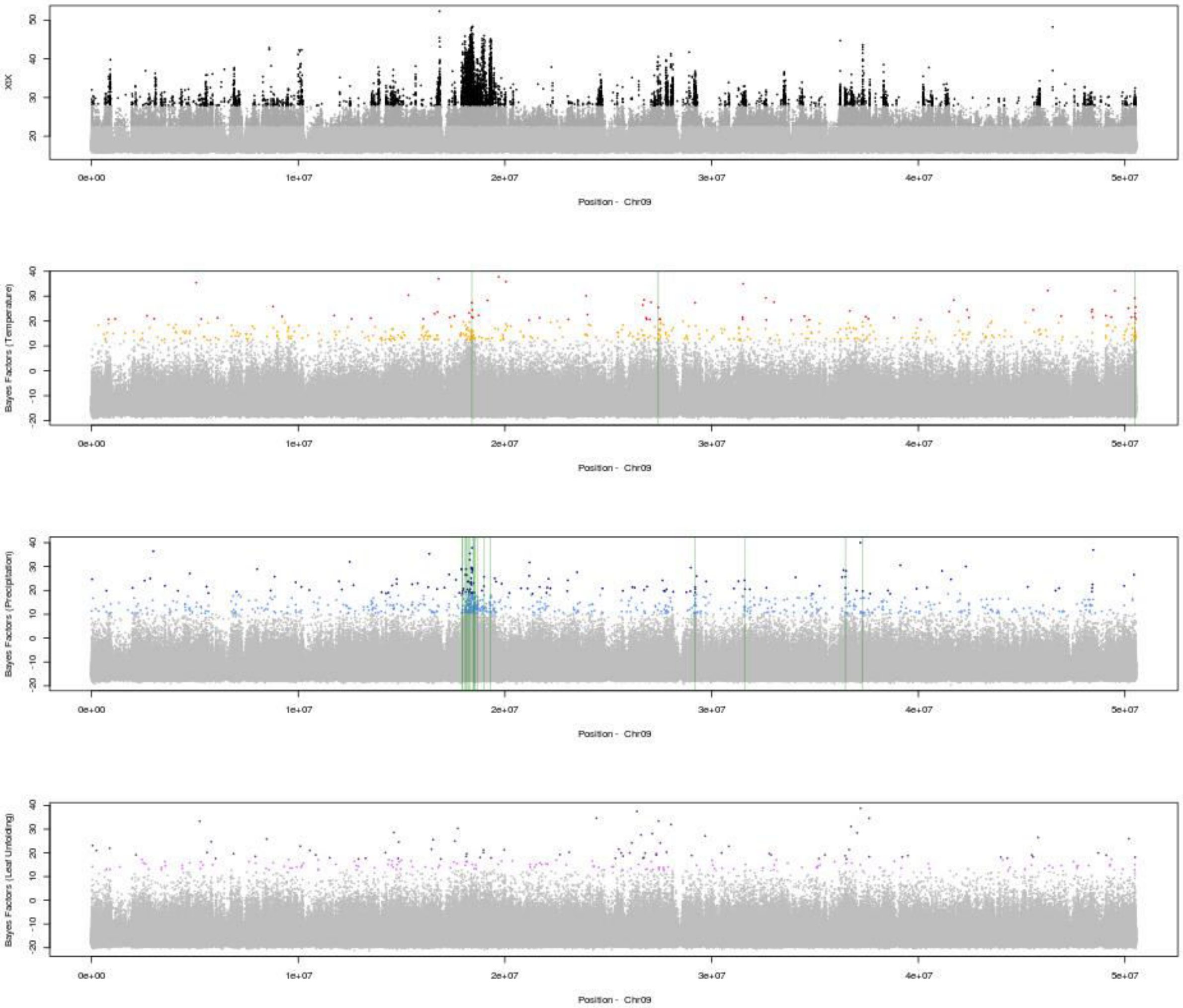


**Fig. S13: Detailed information concerning the genome-wide scans for divergence, GEA and GPA associations for SNPs on chromosome 9.** From top to bottom: relative divergence (XtX) and Bayes factors for associations between temperature, precipitation and the timing of leaf unfolding and allele frequencies, as calculated with BayPass. Colors highlight the significance of the SNPs (light = “minor”, dark = “major”) evaluated with the calibration procedure described by Gautier (2015). Regions investigated for the gene annotation step are shown in green.

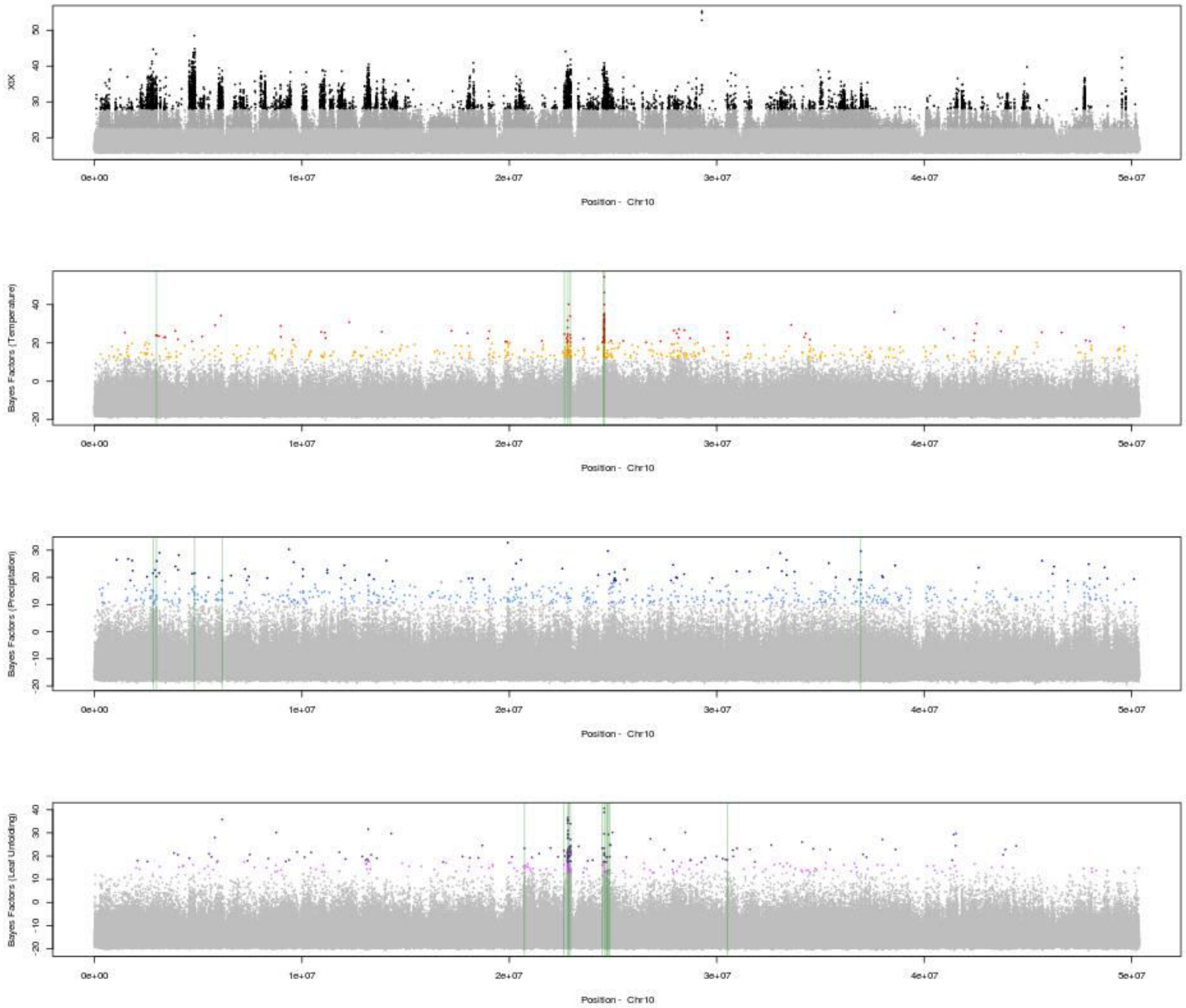


**Fig. S14: Detailed information concerning the genome-wide scans for divergence, GEA and GPA associations for SNPs on chromosome 10.** From top to bottom: relative divergence (XtX) and Bayes factors for associations between temperature, precipitation and the timing of leaf unfolding and allele frequencies, as calculated with BayPass. Colors highlight the significance of the SNPs (light = “minor”, dark = “major”) evaluated with the calibration procedure described by Gautier (2015). Regions investigated for the gene annotation step are shown in green.

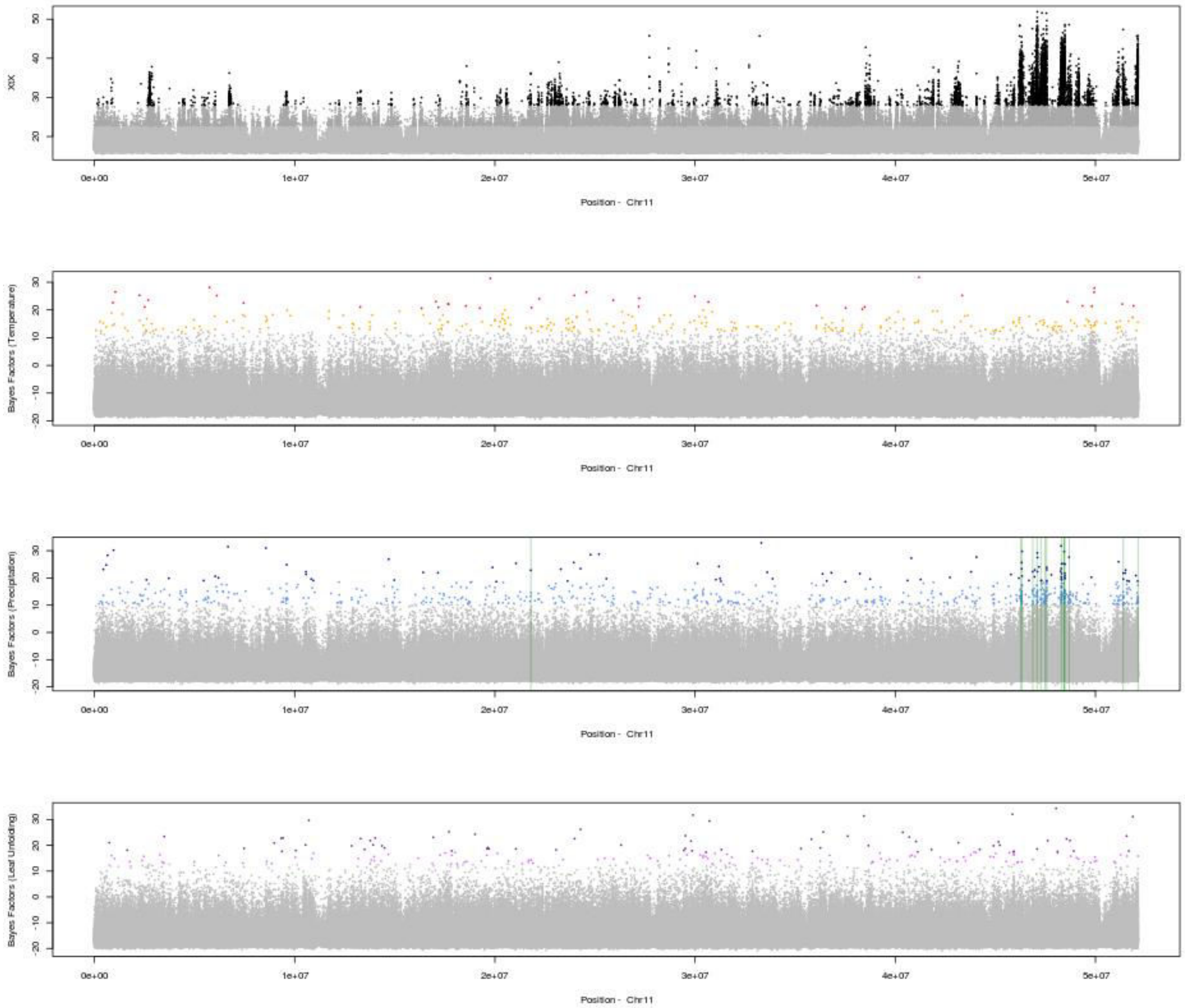


**Fig. S15: Detailed information concerning the genome-wide scans for divergence, GEA and GPA associations for SNPs on chromosome 11.** From top to bottom: relative divergence (XtX) and Bayes factors for associations between temperature, precipitation and the timing of leaf unfolding and allele frequencies, as calculated with BayPass. Colors highlight the significance of the SNPs (light = "minor", dark = "major") evaluated with the calibration procedure described by Gautier (2015). Regions investigated for the gene annotation step are shown in green.

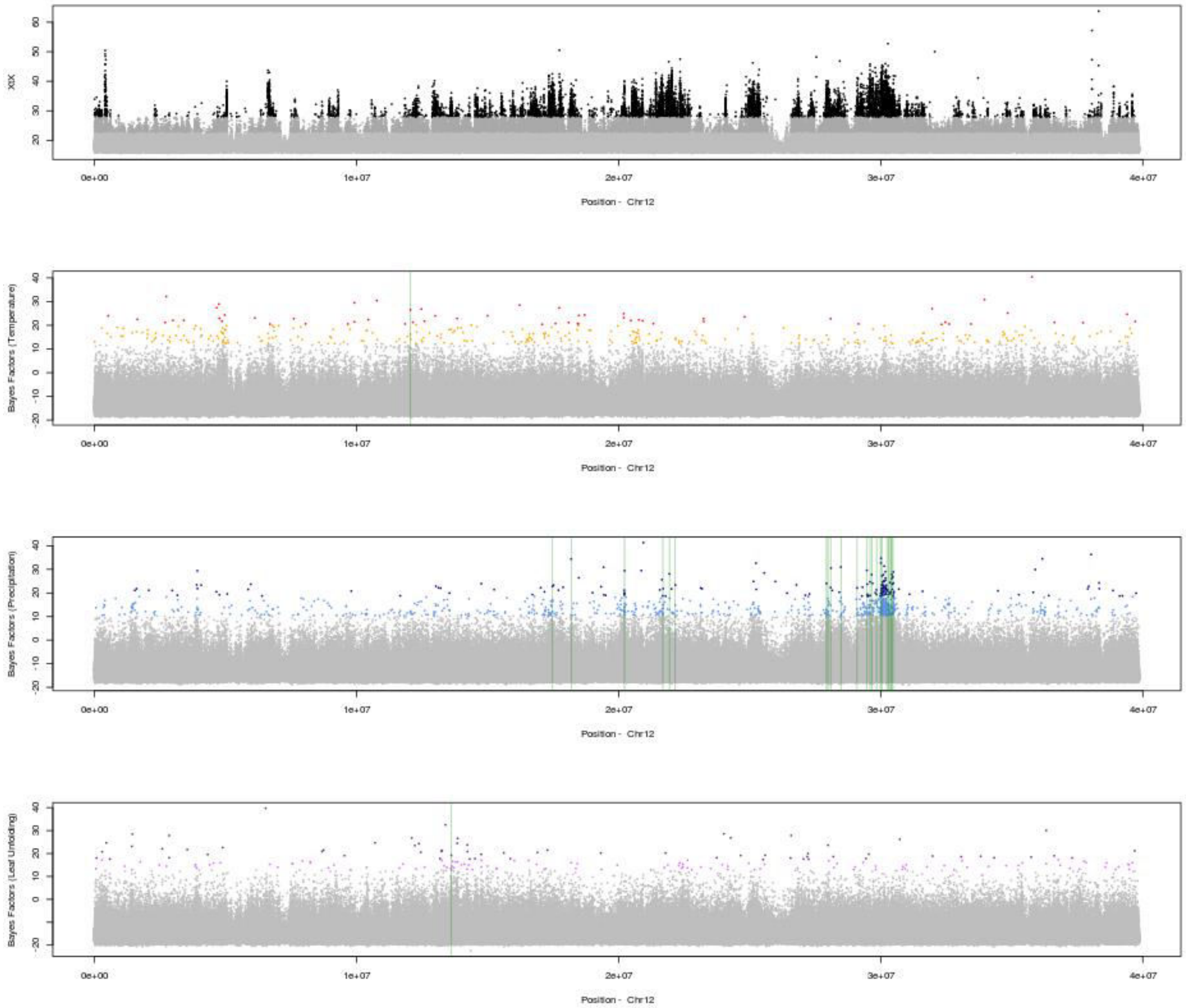


**Fig. S16: Detailed information concerning the genome-wide scans for divergence, GEA and GPA associations for SNPs on chromosome 12.** From top to bottom: relative divergence (XtX) and Bayes factors for associations between temperature, precipitation and the timing of leaf unfolding and allele frequencies, as calculated with BayPass. Colors highlight the significance of the SNPs (light = "minor", dark = "major") evaluated with the calibration procedure described by Gautier (2015). Regions investigated for the gene annotation step are shown in green.
