## Supplementary material for "Adaptive introgression as a driver of local adaptation to climate in European white oaks": SupFile1

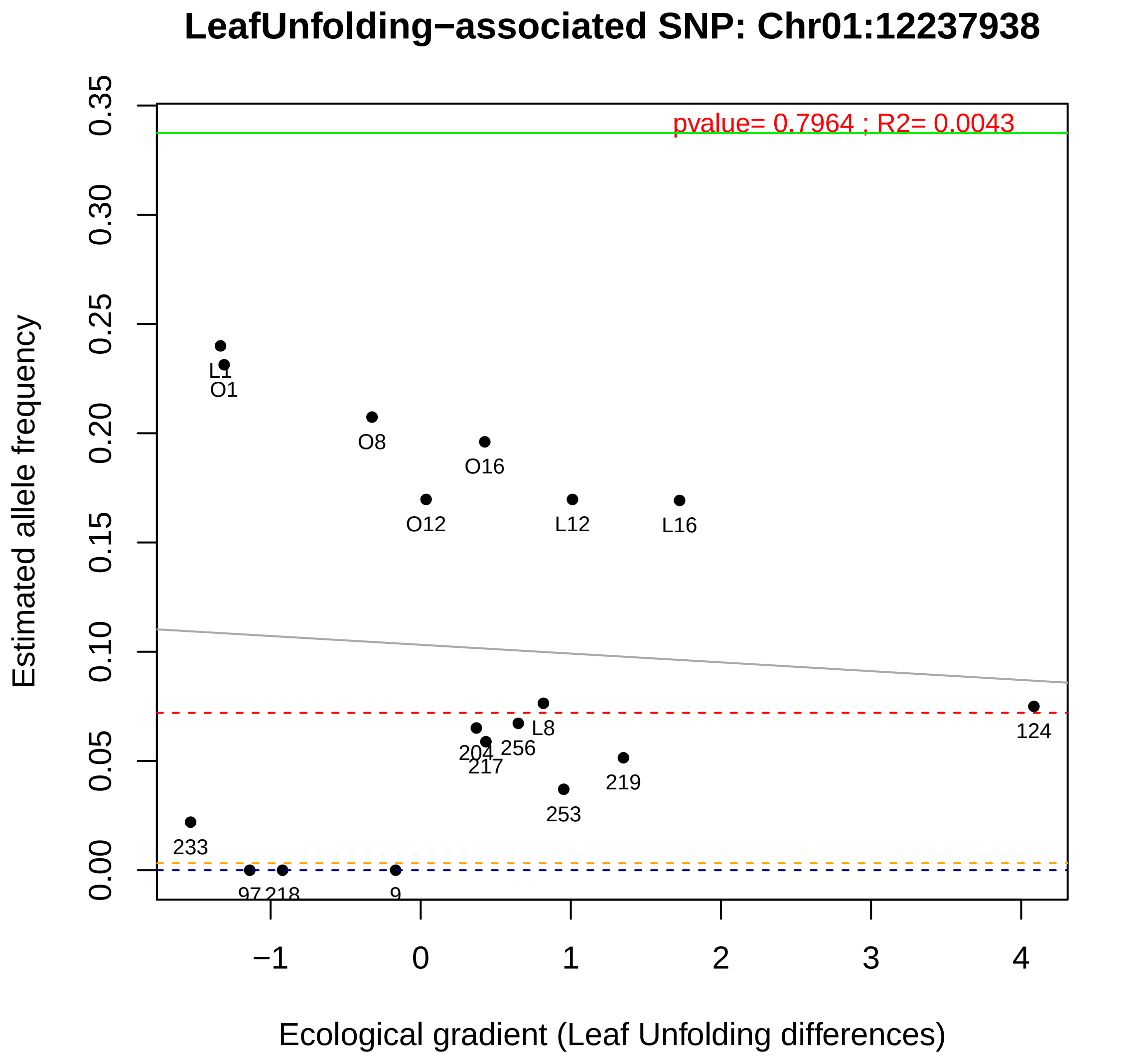

LeafUnfolding−associated SNP: Chr01:12237938
pvalue= 0.7964 ; R2= 0.0043
Estimated allele frequency
0.00
0.05
0.10
0.15
0.20
0.25
0.30
0.35
L1
O1
O8
O16
O12
L12
L16
L8
124
204
217
256
219
253
233
97
218
9
−1
0
1
2
3
4
Ecological gradient (Leaf Unfolding differences)

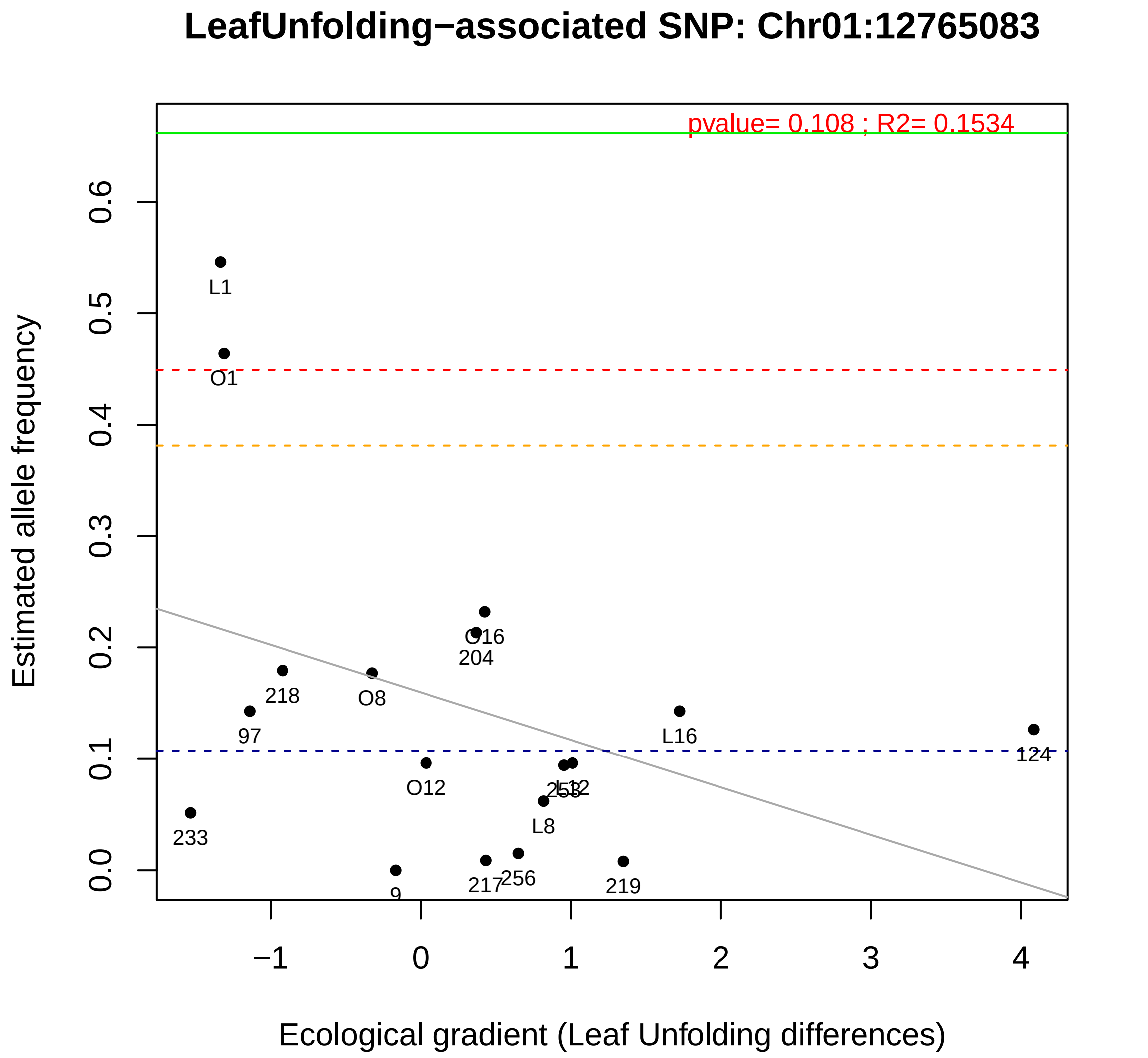
LeafUnfolding−associated SNP: Chr01:12765083
pvalue= 0.108 ; R2= 0.1534
Estimated allele frequency
0.0
0.1
0.2
0.3
0.4
0.5
0.6
−1
0
1
2
3
4
Ecological gradient (Leaf Unfolding differences)
L1
O1
O16
204
218
O8
97
L16
124
O12
233
L8
217
256
219
9

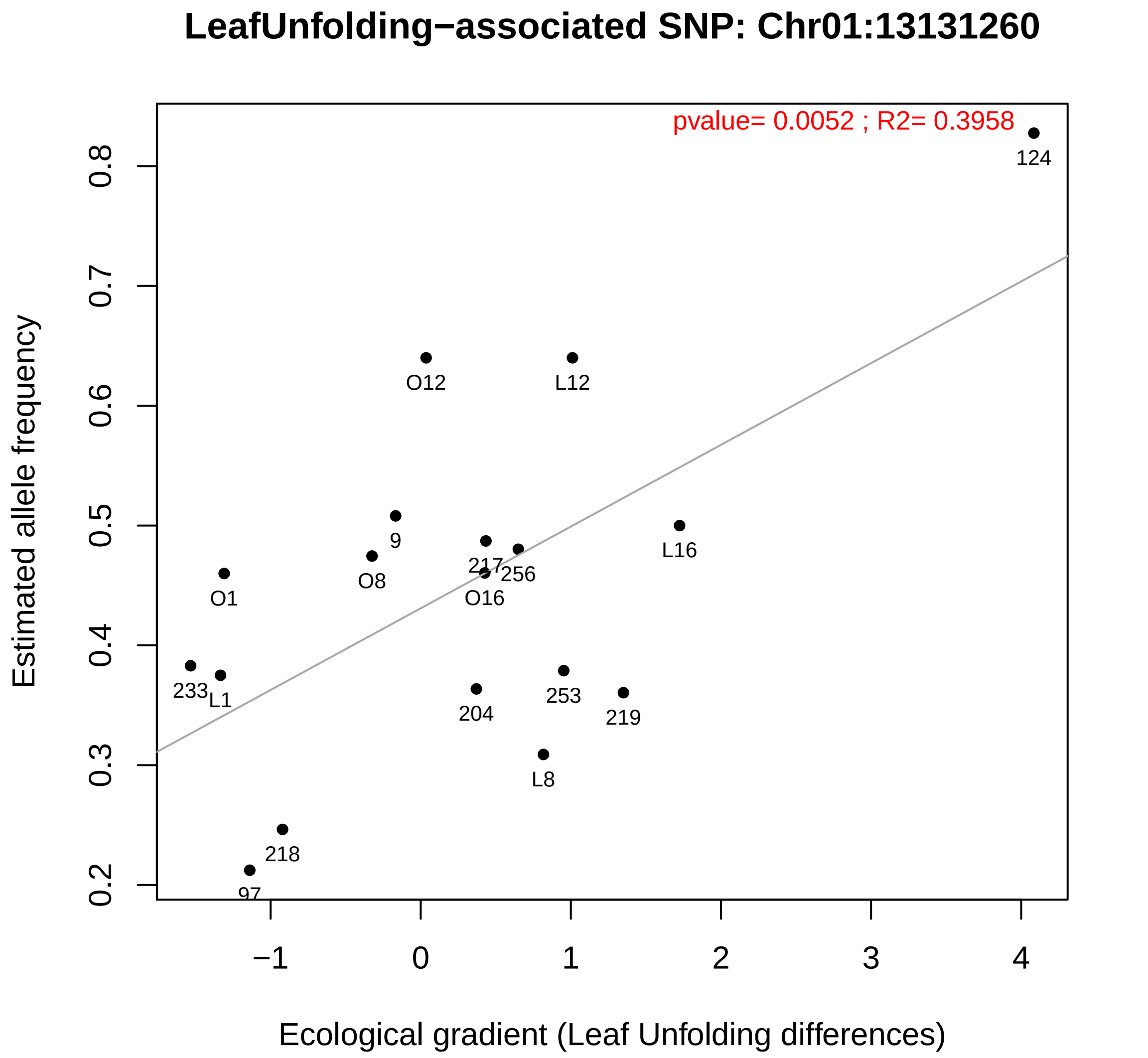

LeafUnfolding−associated SNP: Chr01:13131260
pvalue= 0.0052 ; R2= 0.3958
124
O12
L12
9
O8
217
256
O16
L16
O1
233
L1
204
253
219
L8
218
97
0.2
0.3
0.4
0.5
0.6
0.7
0.8
−1
0
1
2
3
4
Estimated allele frequency
Ecological gradient (Leaf Unfolding differences)

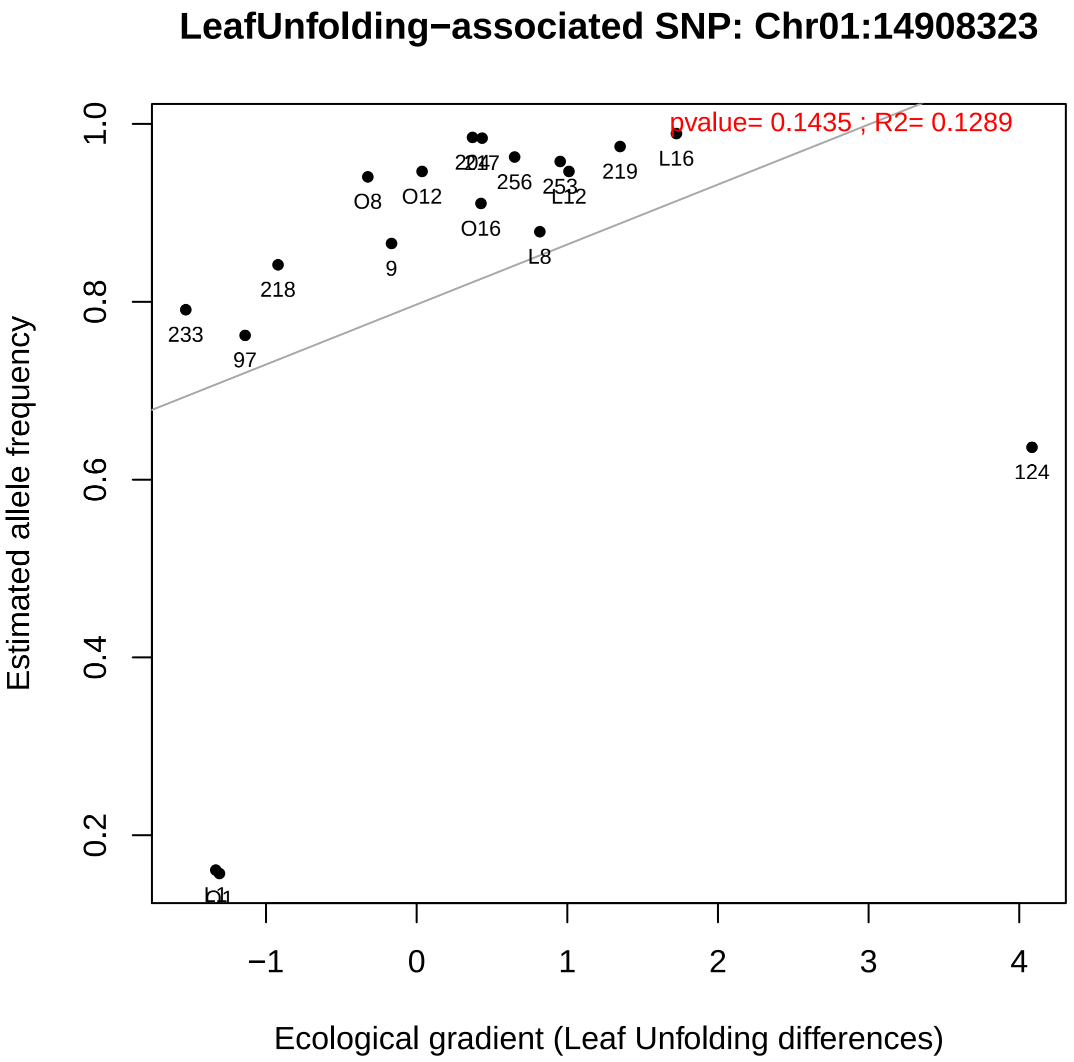
LeafUnfolding−associated SNP: Chr01:14908323
pvalue= 0.1435 ; R2= 0.1289
Estimated allele frequency
Ecological gradient (Leaf Unfolding differences)
0.2
0.4
0.6
0.8
1.0
−1
0
1
2
3
4
233
97
218
O8
9
O12
O16
256
253
L12
L8
219
L16
124

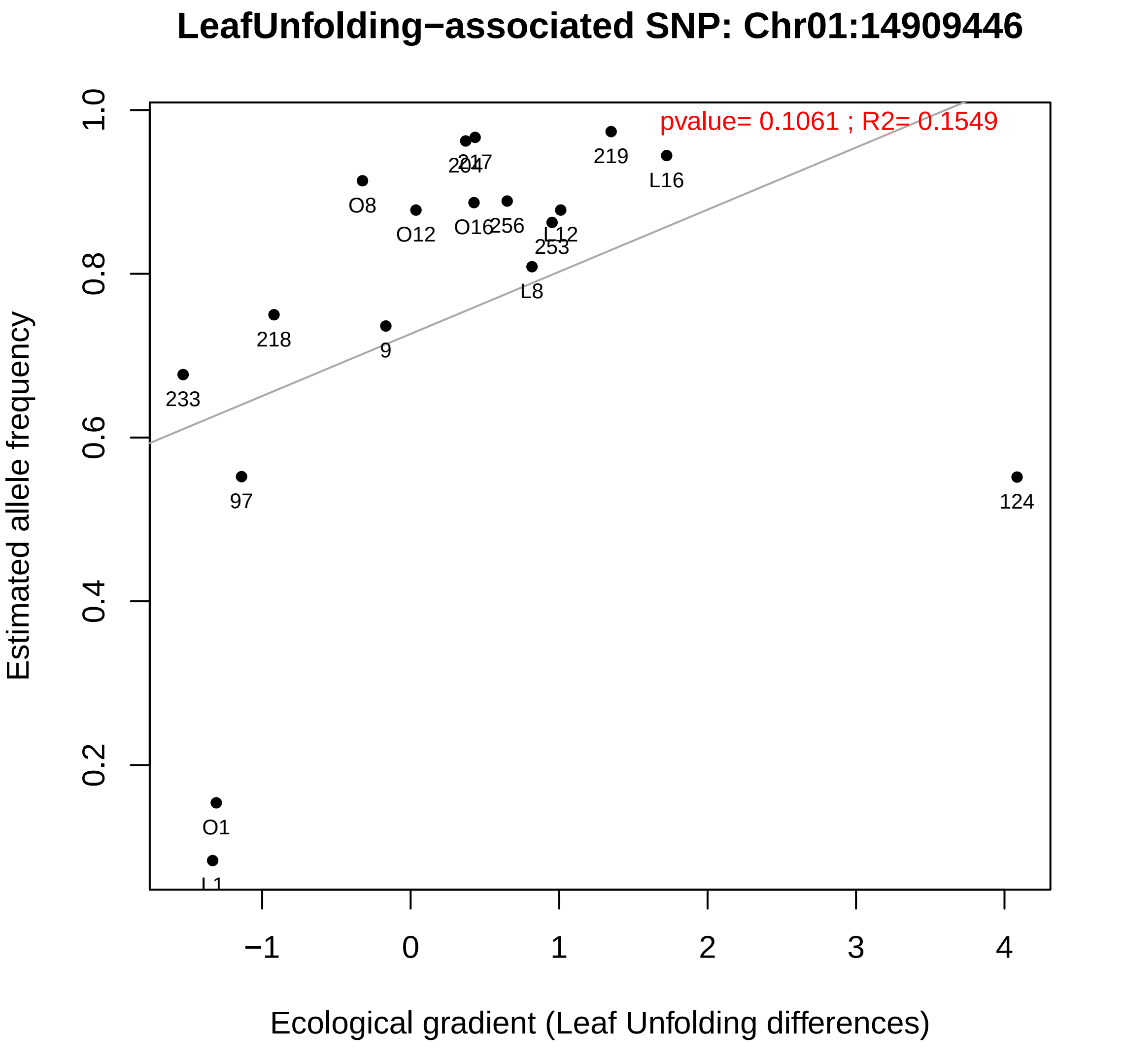
LeafUnfolding−associated SNP: Chr01:14909446
pvalue= 0.1061 ; R2= 0.1549
Estimated allele frequency
Ecological gradient (Leaf Unfolding differences)
0.2
0.4
0.6
0.8
1.0
−1
0
1
2
3
4
219
L16
O8
O12
O16
256
L12
253
L8
218
9
233
97
124
O1

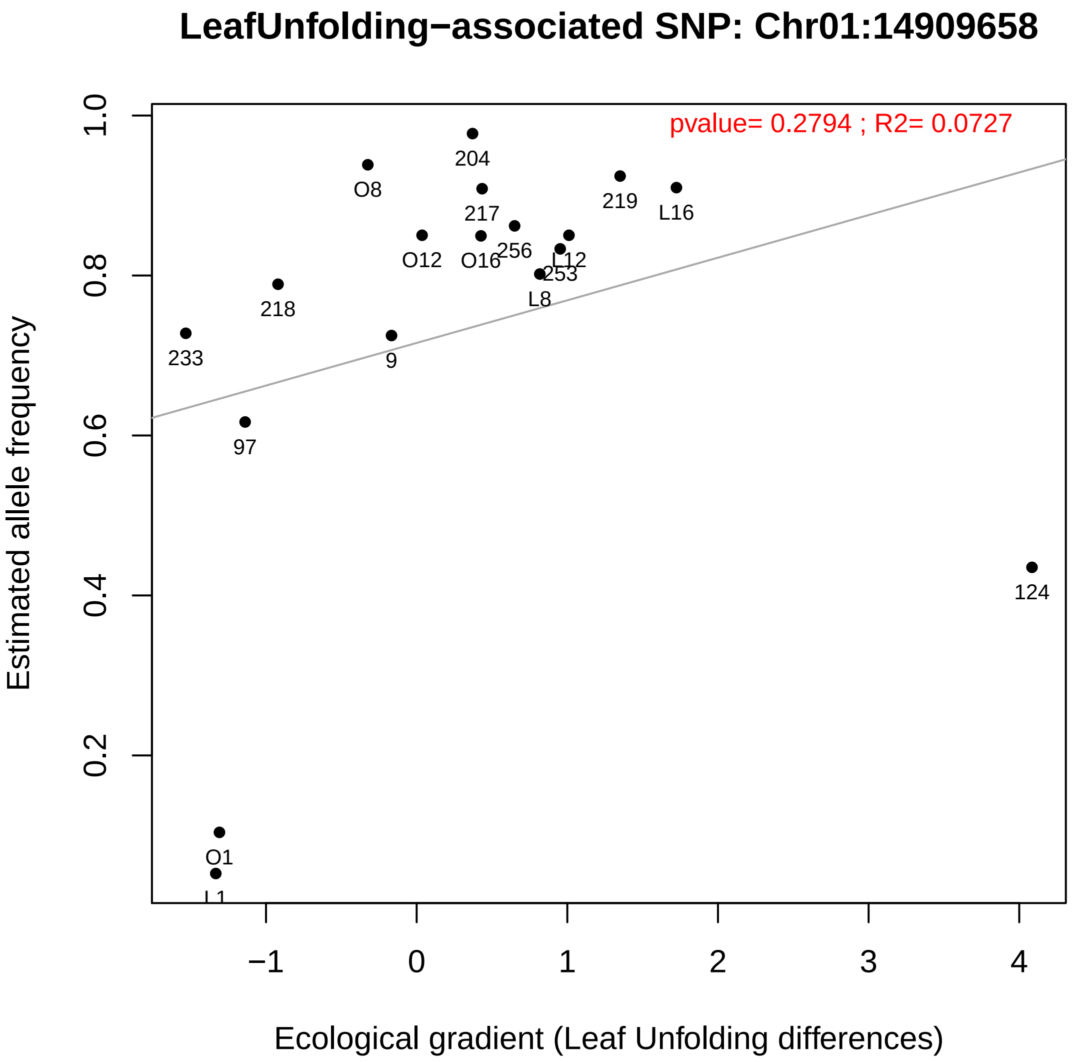
LeafUnfolding−associated SNP: Chr01:14909658
pvalue= 0.2794 ; R2= 0.0727
Estimated allele frequency
Ecological gradient (Leaf Unfolding differences)
0.2
0.4
0.6
0.8
1.0
−1
0
1
2
3
4
204
O8
217
219
L16
256
O12
O16
L12
253
L8
218
233
9
97
124
O1
L1

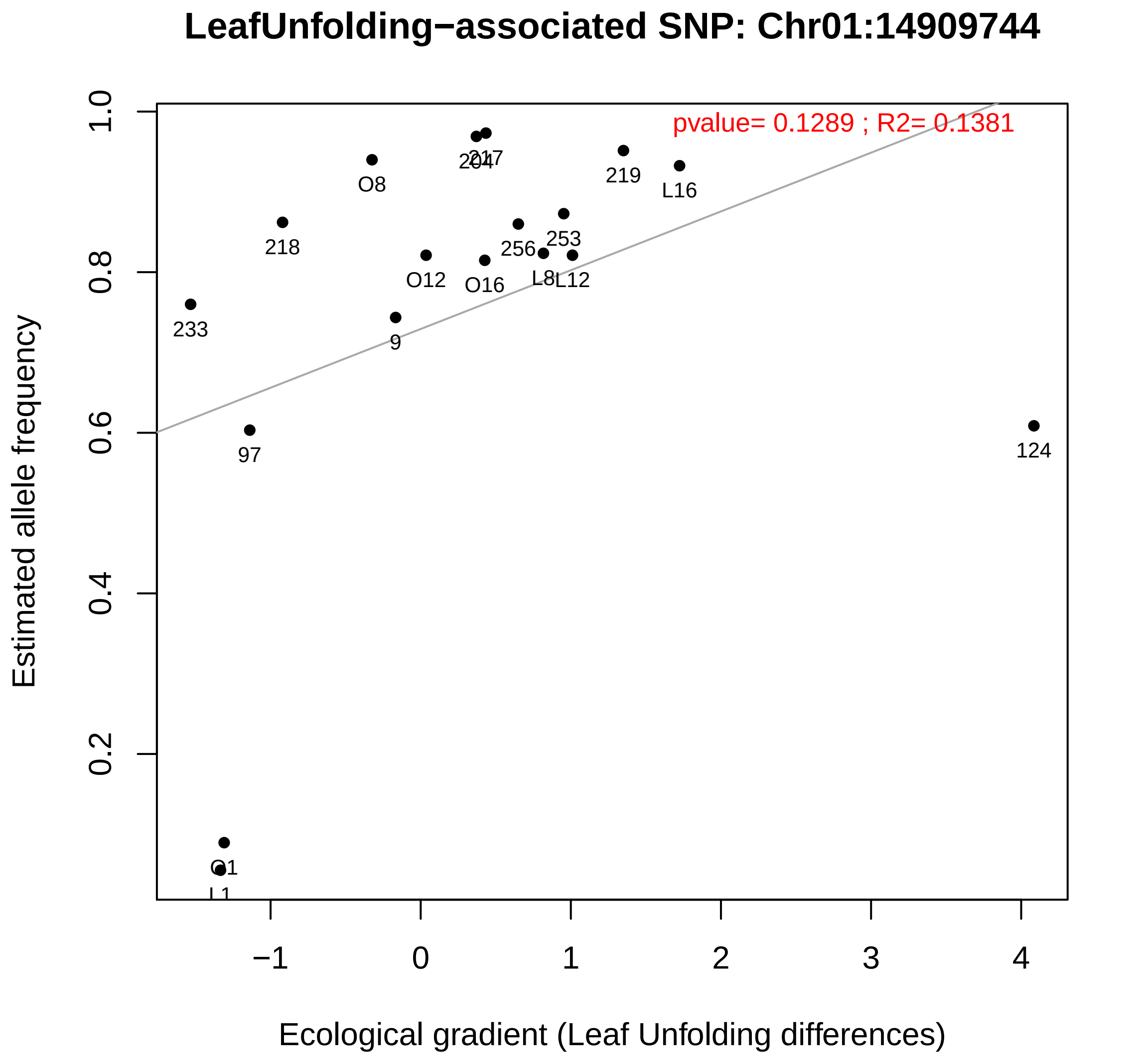
LeafUnfolding−associated SNP: Chr01:14909744
pvalue= 0.1289 ; R2= 0.1381
Estimated allele frequency
Ecological gradient (Leaf Unfolding differences)
1.0
0.8
0.6
0.4
0.2
−1
0
1
2
3
4
204
217
O8
219
L16
218
253
256
O12
O16
L8
L12
233
9
97
124
O1
L1

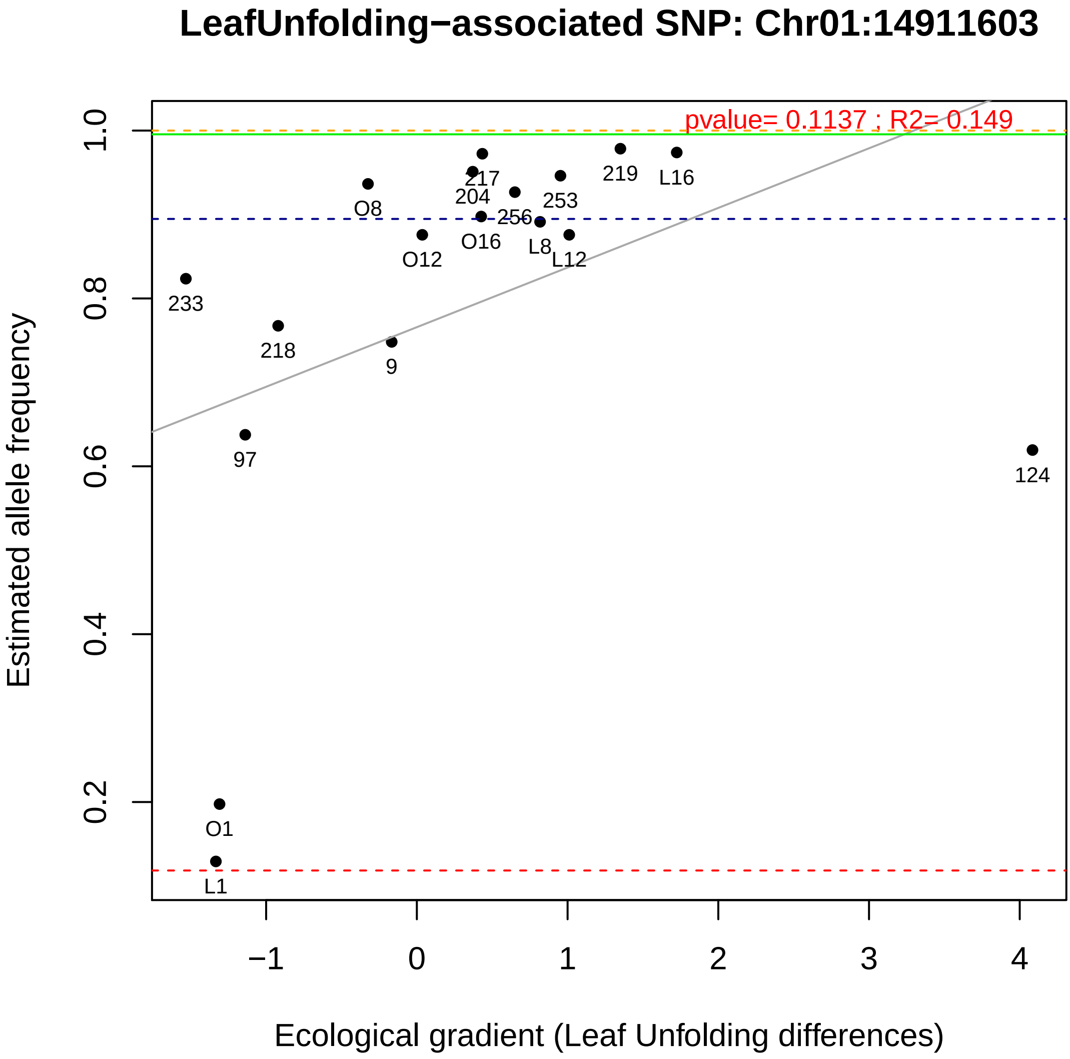
LeafUnfolding−associated SNP: Chr01:14911603
pvalue= 0.1137 ; R2= 0.149
Estimated allele frequency
1.0
0.8
0.6
0.4
0.2
−1
0
1
2
3
4
Ecological gradient (Leaf Unfolding differences)
233
218
97
O1
L1
O8
9
O12
204
217
O16
256
L8
253
L12
219
L16
124

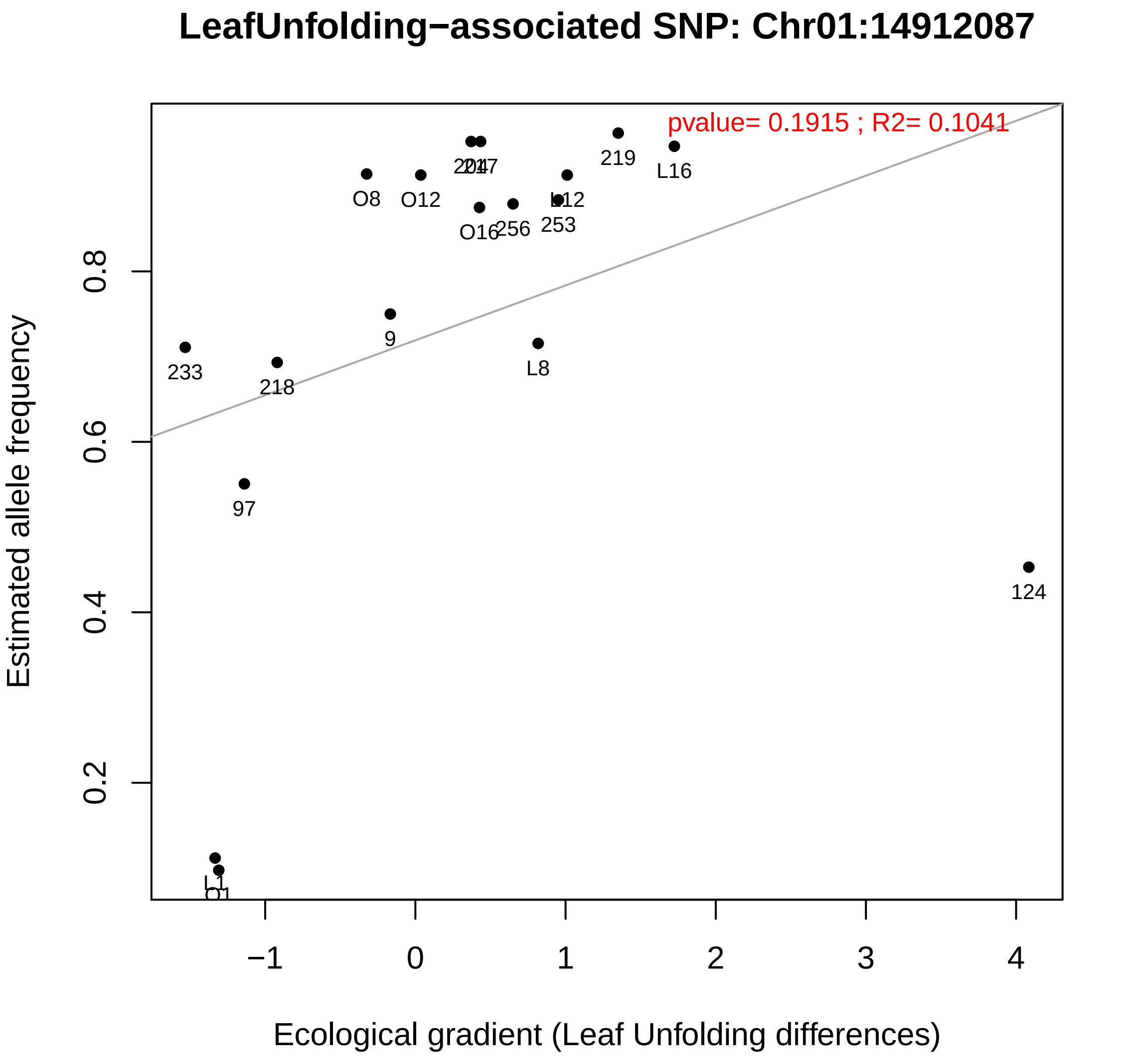
LeafUnfolding−associated SNP: Chr01:14912087
pvalue= 0.1915 ; R2= 0.1041
Estimated allele frequency
Ecological gradient (Leaf Unfolding differences)
0.2
0.4
0.6
0.8
−1
0
1
2
3
4
219
L16
O8
O12
O16
256
253
L12
9
233
218
L8
97
124
L1
O1

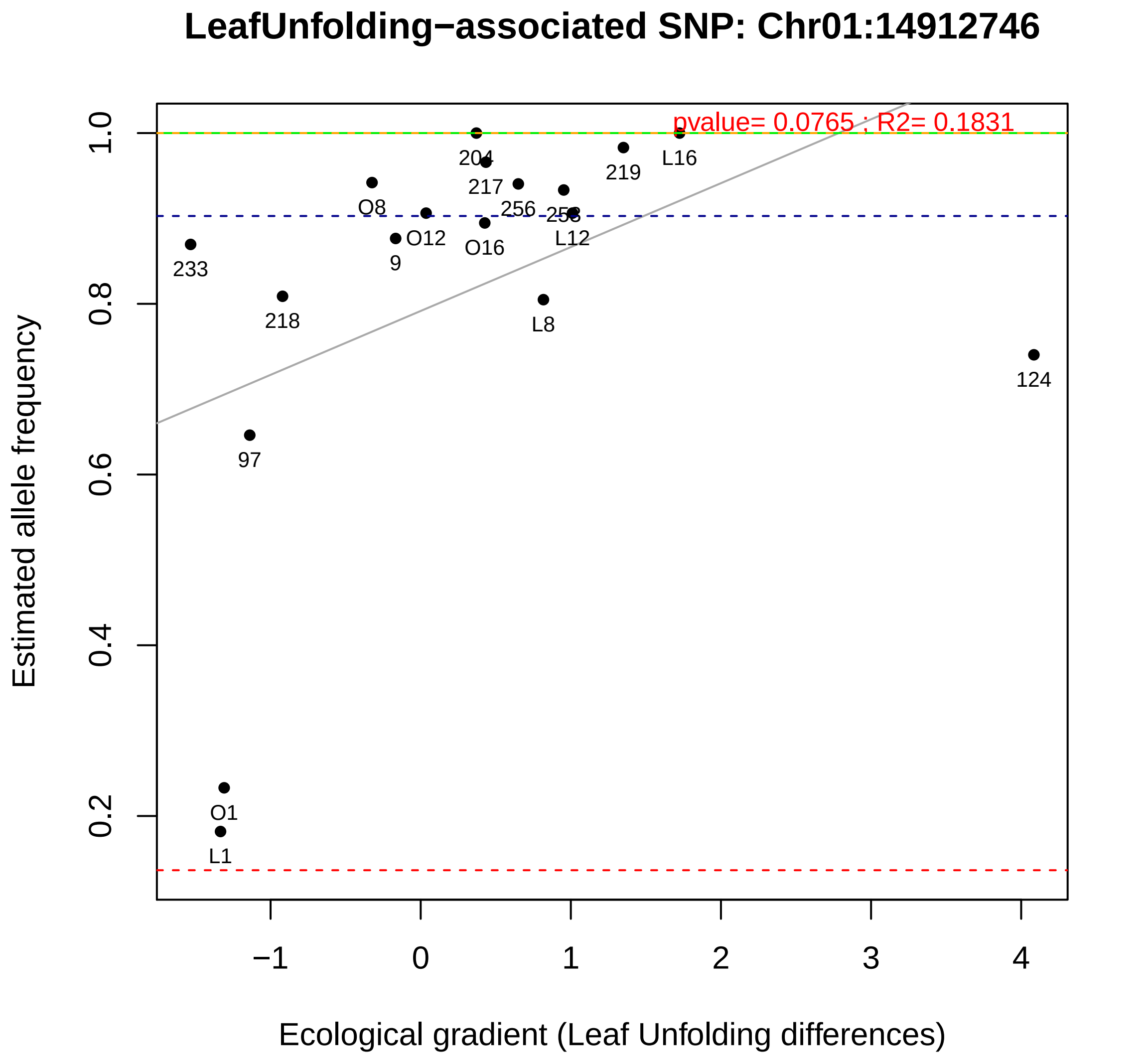

LeafUnfolding−associated SNP: Chr01:14912746
pvalue= 0.0765 ; R2= 0.1831
Estimated allele frequency
1.0
0.8
0.6
0.4
0.2
−1
0
1
2
3
4
Ecological gradient (Leaf Unfolding differences)
204
217
256
219
L16
O8
O12
9
O16
L12
233
218
L8
124
97
O1
L1

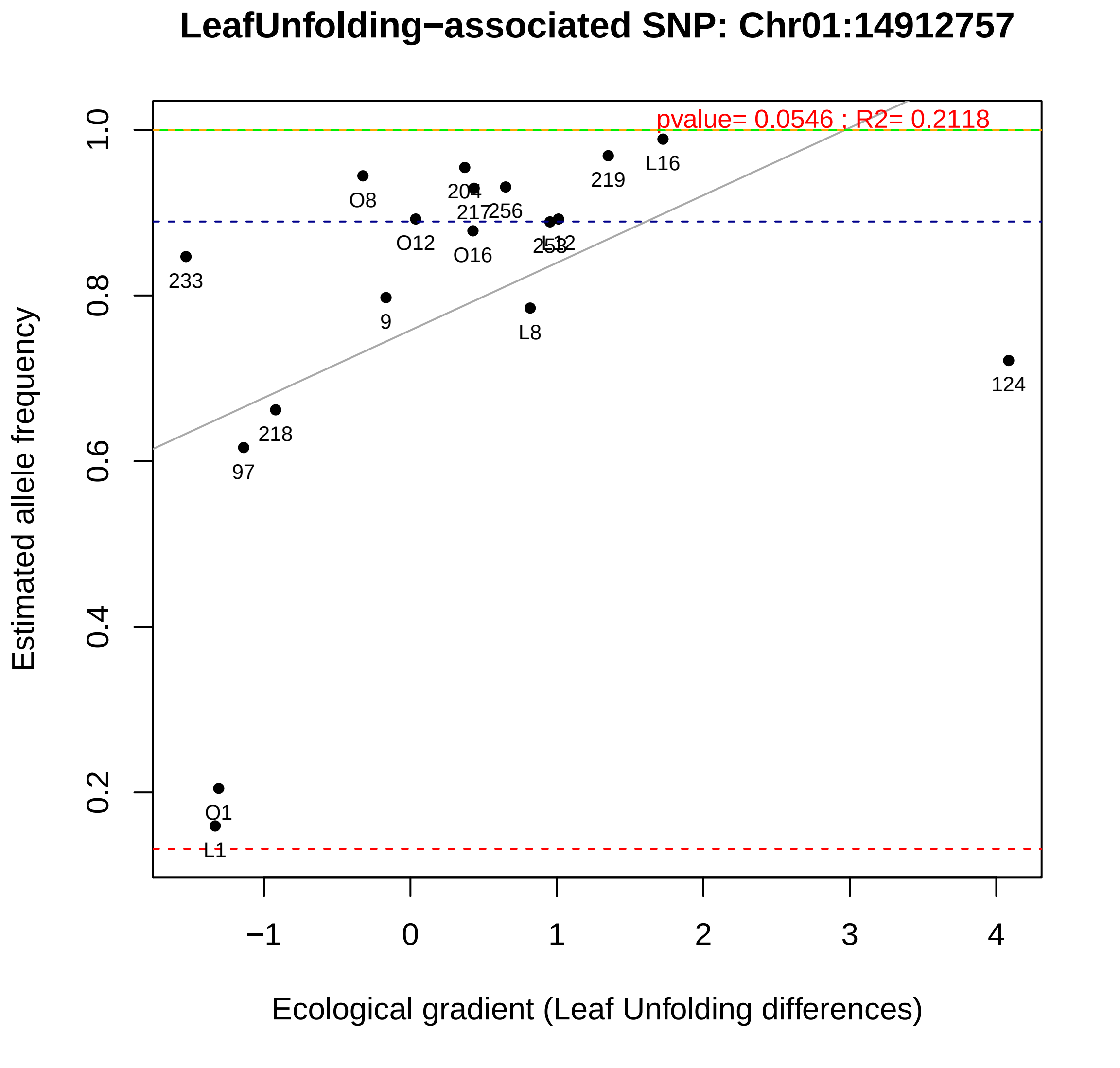

LeafUnfolding−associated SNP: Chr01:14912757
pvalue= 0.0546 ; R2= 0.2118
Estimated allele frequency
1.0
0.8
0.6
0.4
0.2
−1
0
1
2
3
4
Ecological gradient (Leaf Unfolding differences)
233
O8
204
217
256
O12
O16
219
L16
9
L8
124
218
97
O1
L1

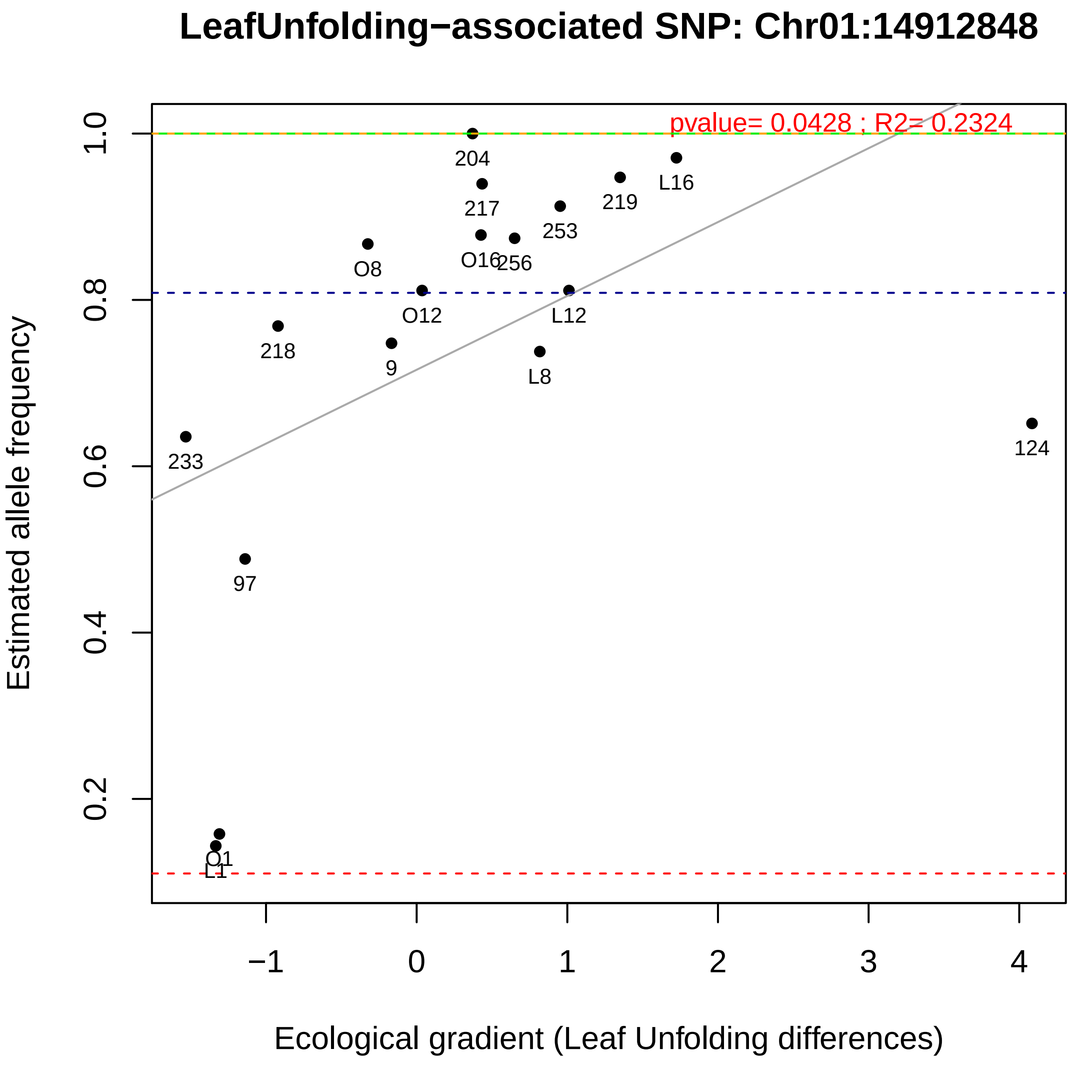
LeafUnfolding−associated SNP: Chr01:14912848
pvalue= 0.0428 ; R2= 0.2324
Estimated allele frequency
Ecological gradient (Leaf Unfolding differences)
0.2
0.4
0.6
0.8
1.0
−1
0
1
2
3
4
204
217
O16
256
253
219
L16
O8
O12
L12
218
9
L8
233
124
97
O1
L1

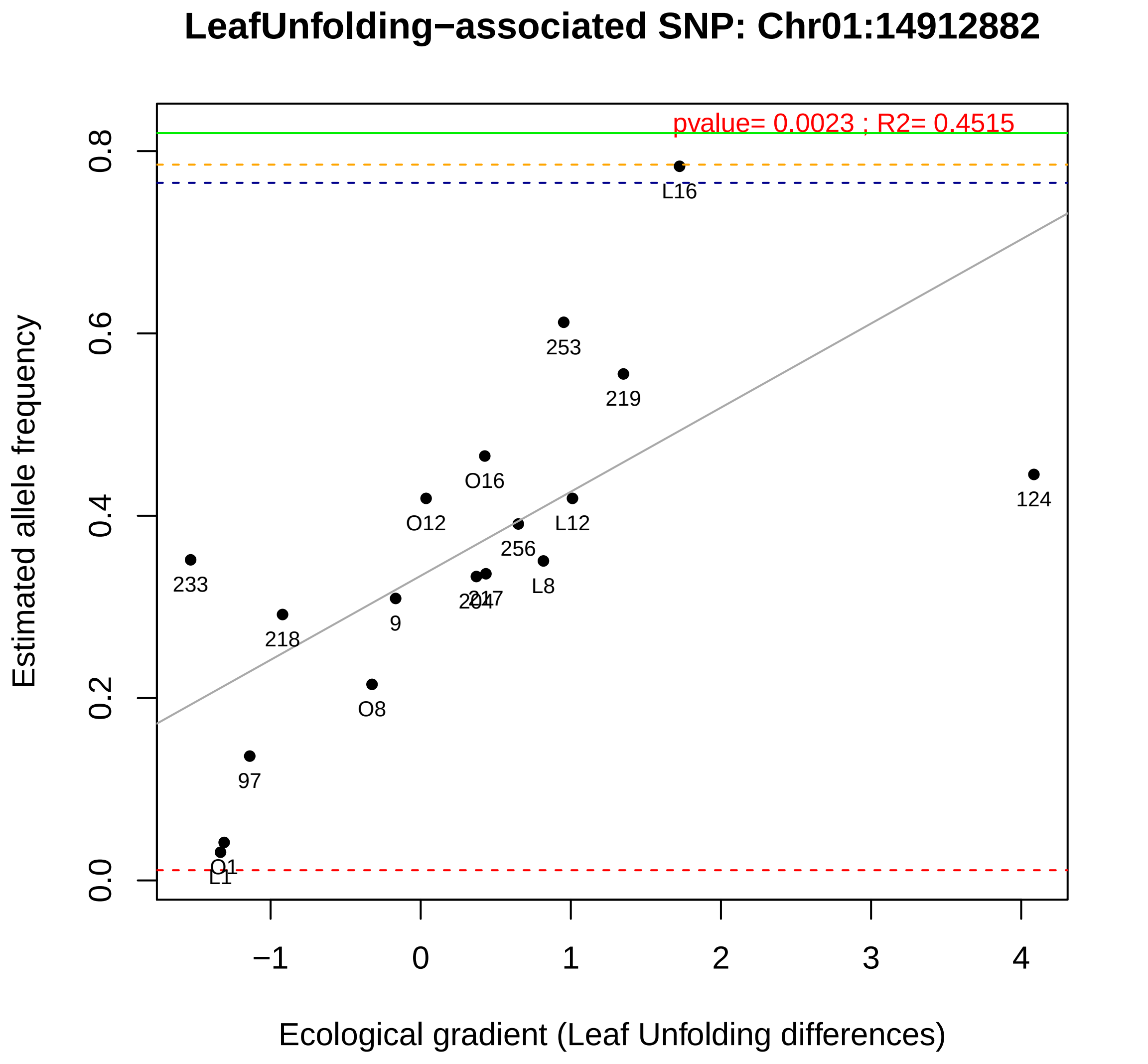
LeafUnfolding−associated SNP: Chr01:14912882
pvalue= 0.0023 ; R2= 0.4515
Estimated allele frequency
Ecological gradient (Leaf Unfolding differences)
0.0
0.2
0.4
0.6
0.8
−1
0
1
2
3
4
L16
253
219
O16
124
O12
L12
256
233
L8
217
204
9
218
O8
97
O1
L1

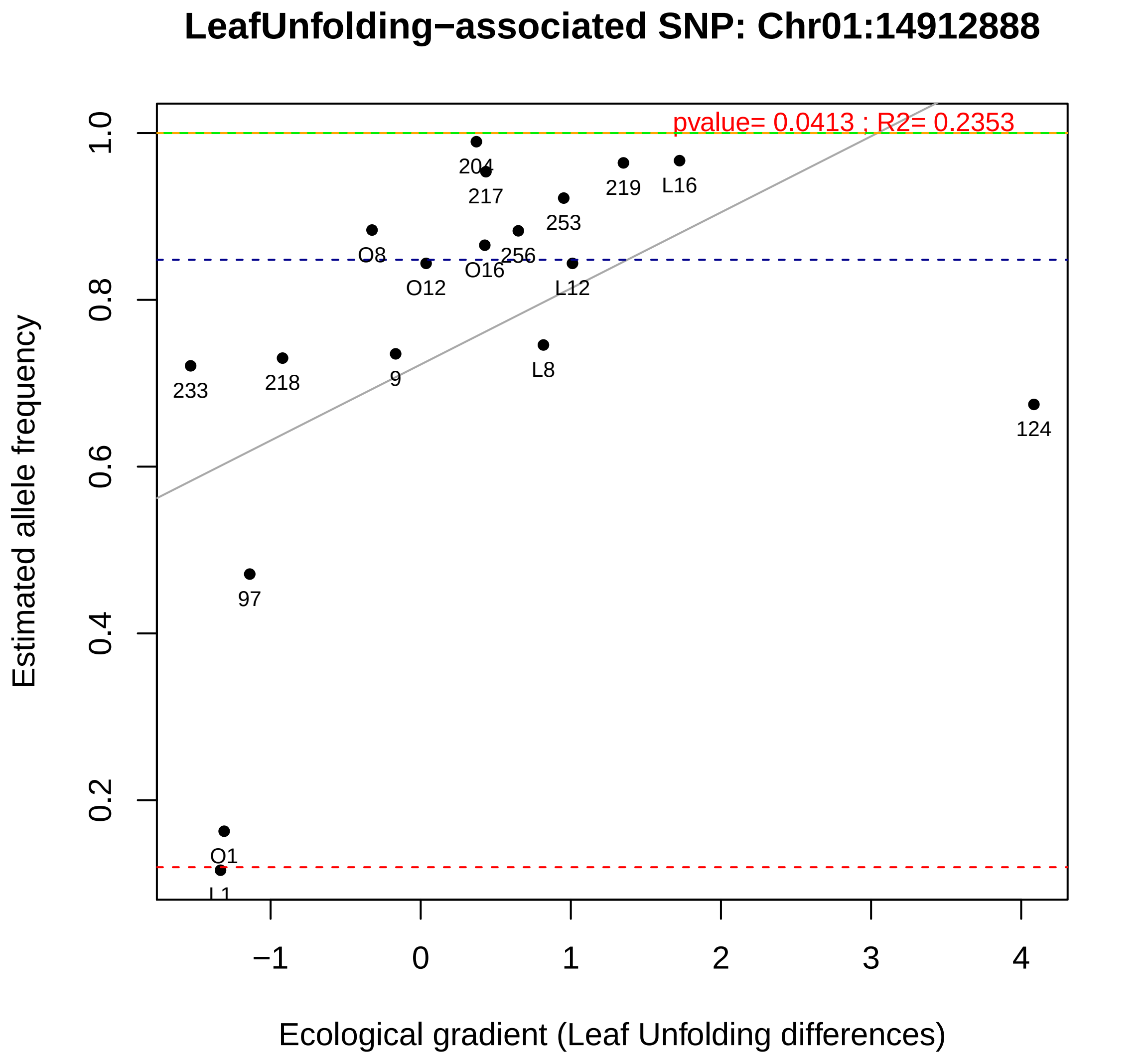

LeafUnfolding−associated SNP: Chr01:14912888
pvalue= 0.0413 ; R2= 0.2353
Estimated allele frequency
Ecological gradient (Leaf Unfolding differences)
0.2
0.4
0.6
0.8
1.0
−1
0
1
2
3
4
204
217
219
L16
253
O8
256
O16
O12
L12
233
218
9
L8
124
97
O1
L1

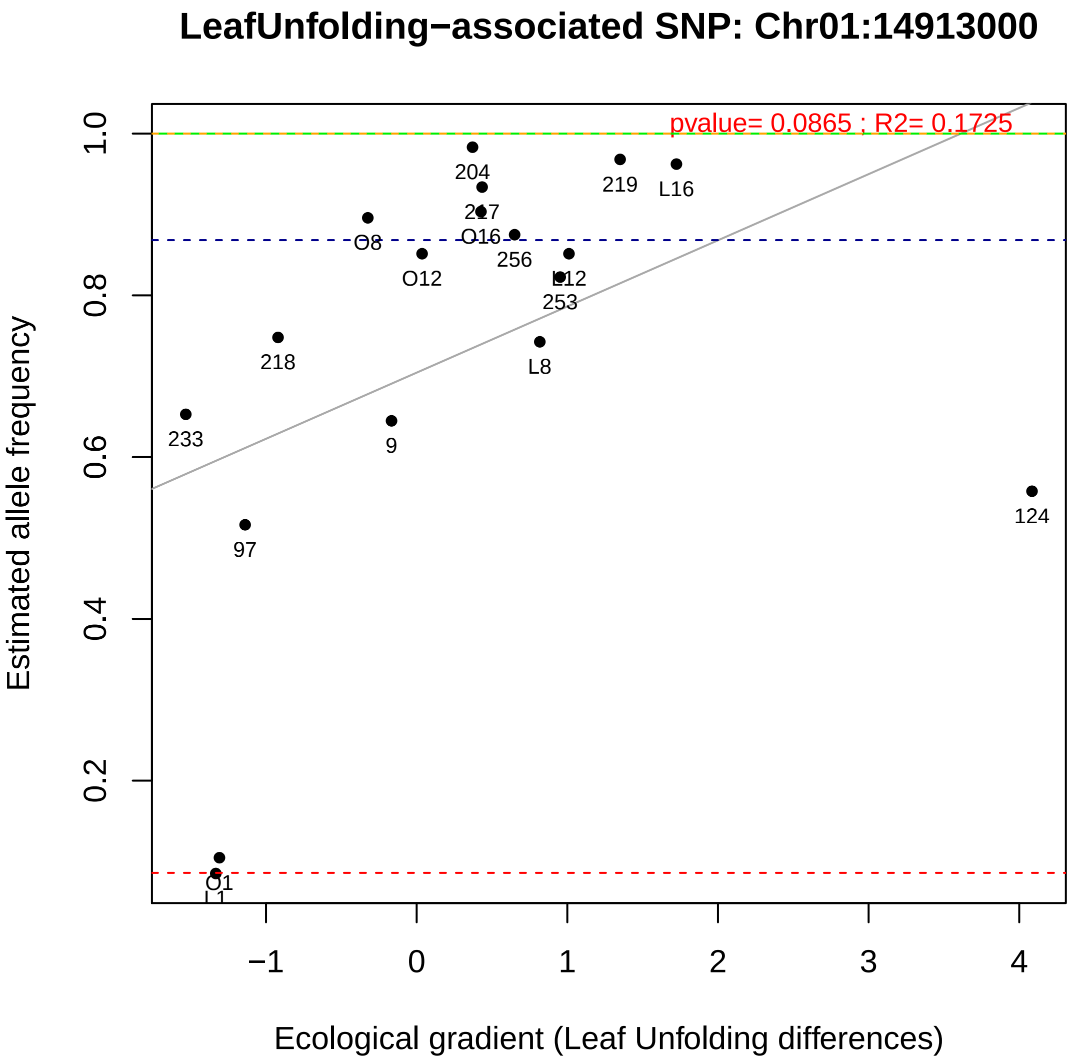

LeafUnfolding−associated SNP: Chr01:14913000
pvalue= 0.0865 ; R2= 0.1725
Estimated allele frequency
Ecological gradient (Leaf Unfolding differences)
0.2
0.4
0.6
0.8
1.0
−1
0
1
2
3
4
204
217
O16
219
L16
O8
O12
256
L12
253
218
L8
233
9
124
97
O1

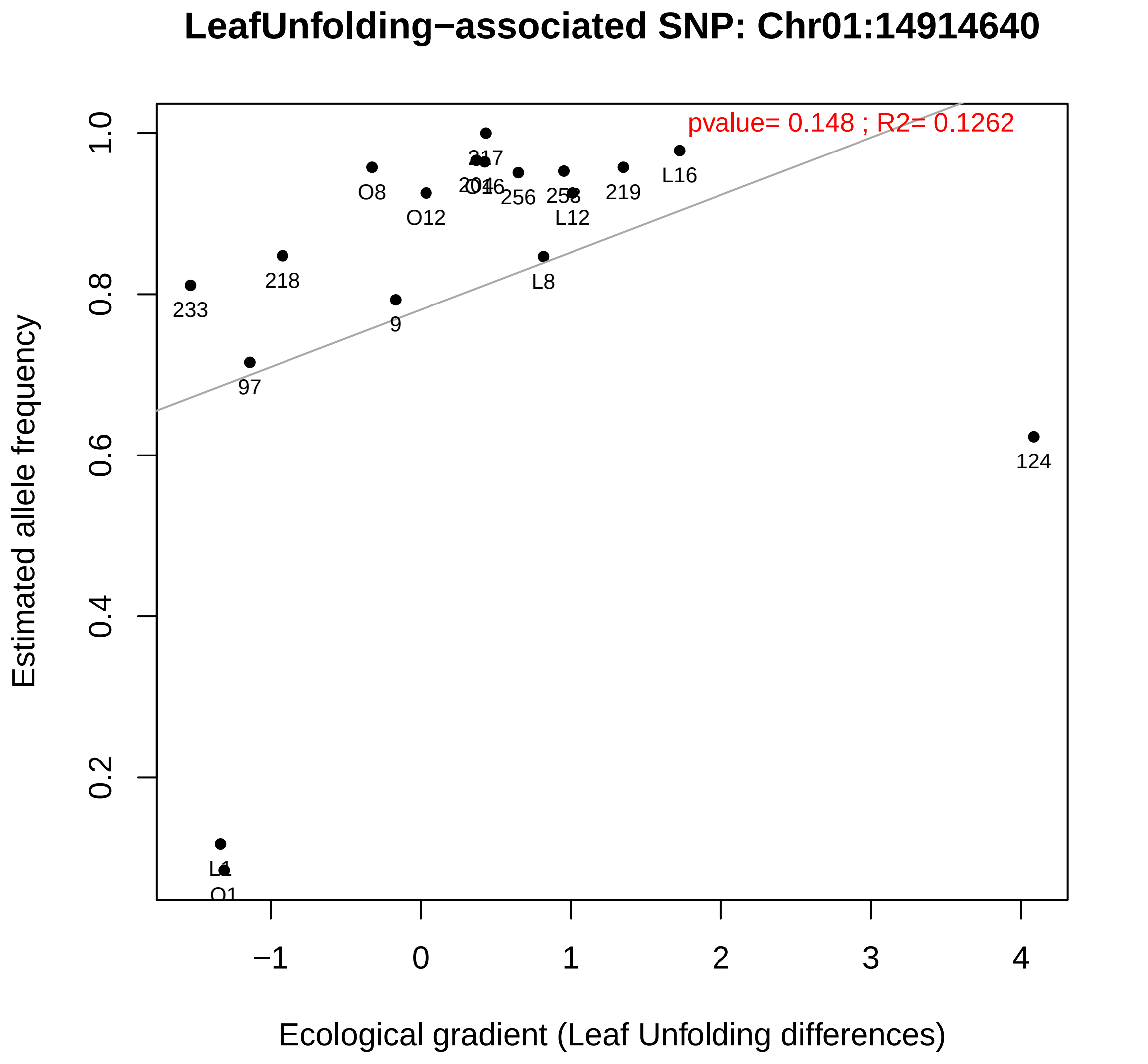
LeafUnfolding−associated SNP: Chr01:14914640
pvalue= 0.148 ; R2= 0.1262
Estimated allele frequency
Ecological gradient (Leaf Unfolding differences)
0.2
0.4
0.6
0.8
1.0
−1
0
1
2
3
4
217
O8
256
219
L16
O12
L12
218
L8
233
9
97
124
L1
O1

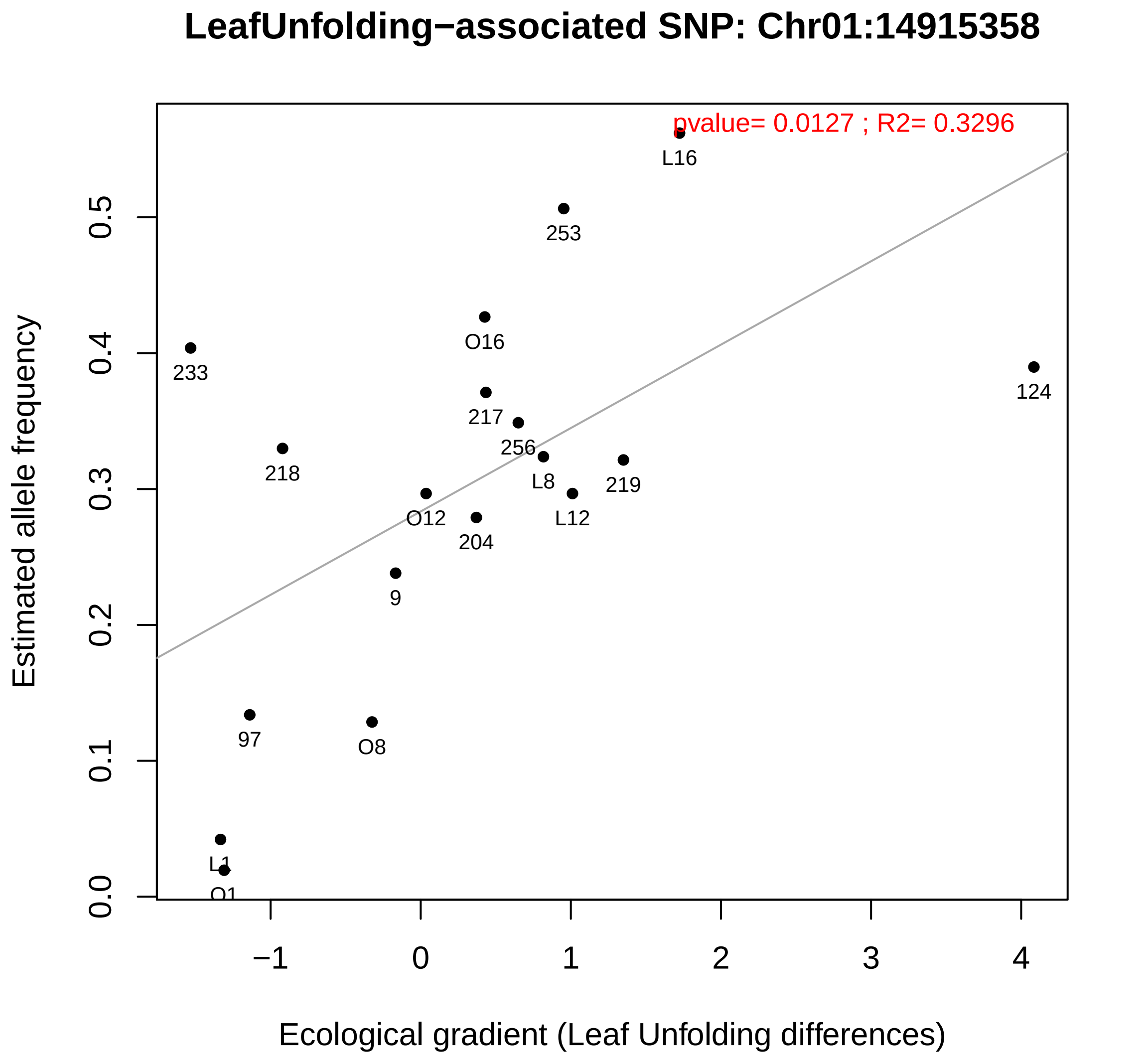
LeafUnfolding−associated SNP: Chr01:14915358
pvalue= 0.0127 ; R2= 0.3296
Estimated allele frequency
Ecological gradient (Leaf Unfolding differences)
0.0
0.1
0.2
0.3
0.4
0.5
−1
0
1
2
3
4
L16
253
O16
233
124
217
256
218
L8
219
O12
L12
204
9
97
O8
L1
O1

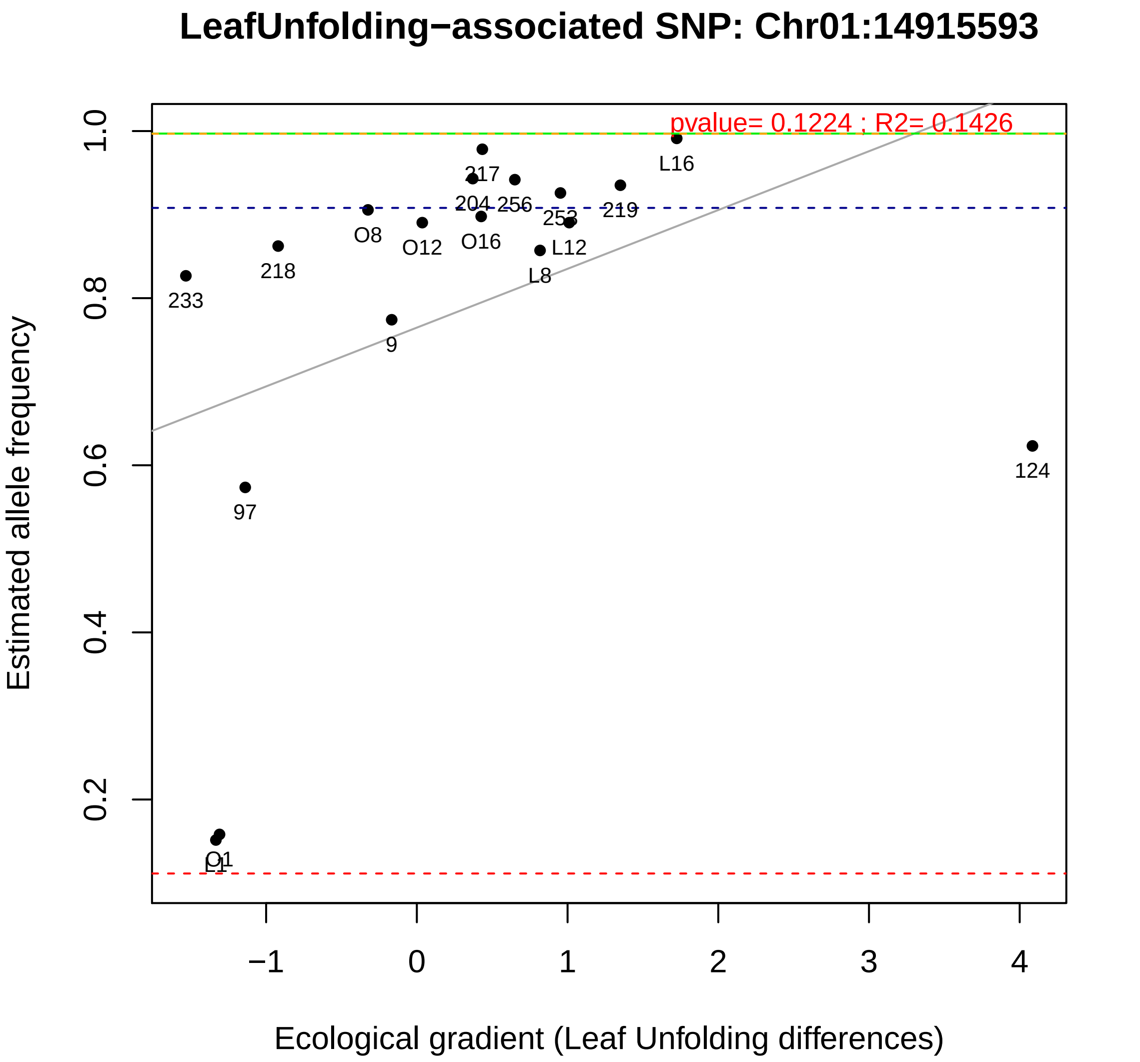
LeafUnfolding−associated SNP: Chr01:14915593
pvalue= 0.1224 ; R2= 0.1426
Estimated allele frequency
Ecological gradient (Leaf Unfolding differences)
1.0
0.8
0.6
0.4
0.2
−1
0
1
2
3
4
L16
217
204
256
253
219
O8
O12
O16
L12
L8
218
233
9
124
97
O1
L1

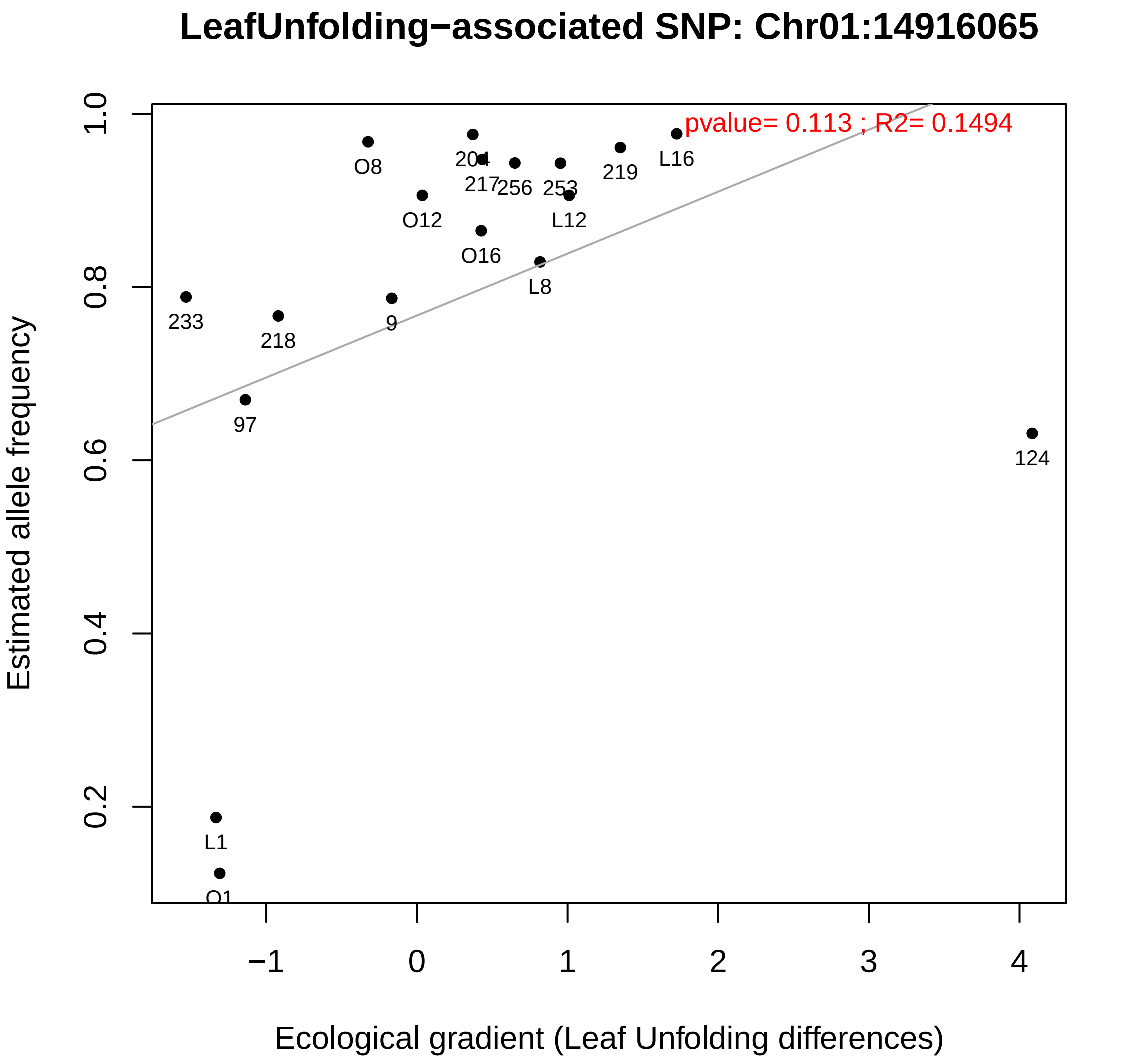

LeafUnfolding−associated SNP: Chr01:14916065
pvalue= 0.113 ; R2= 0.1494
Estimated allele frequency
0.2
0.4
0.6
0.8
1.0
−1
0
1
2
3
4
Ecological gradient (Leaf Unfolding differences)
O8
204
217
256
253
219
L16
O12
L12
O16
L8
233
218
9
97
124
L1
O1

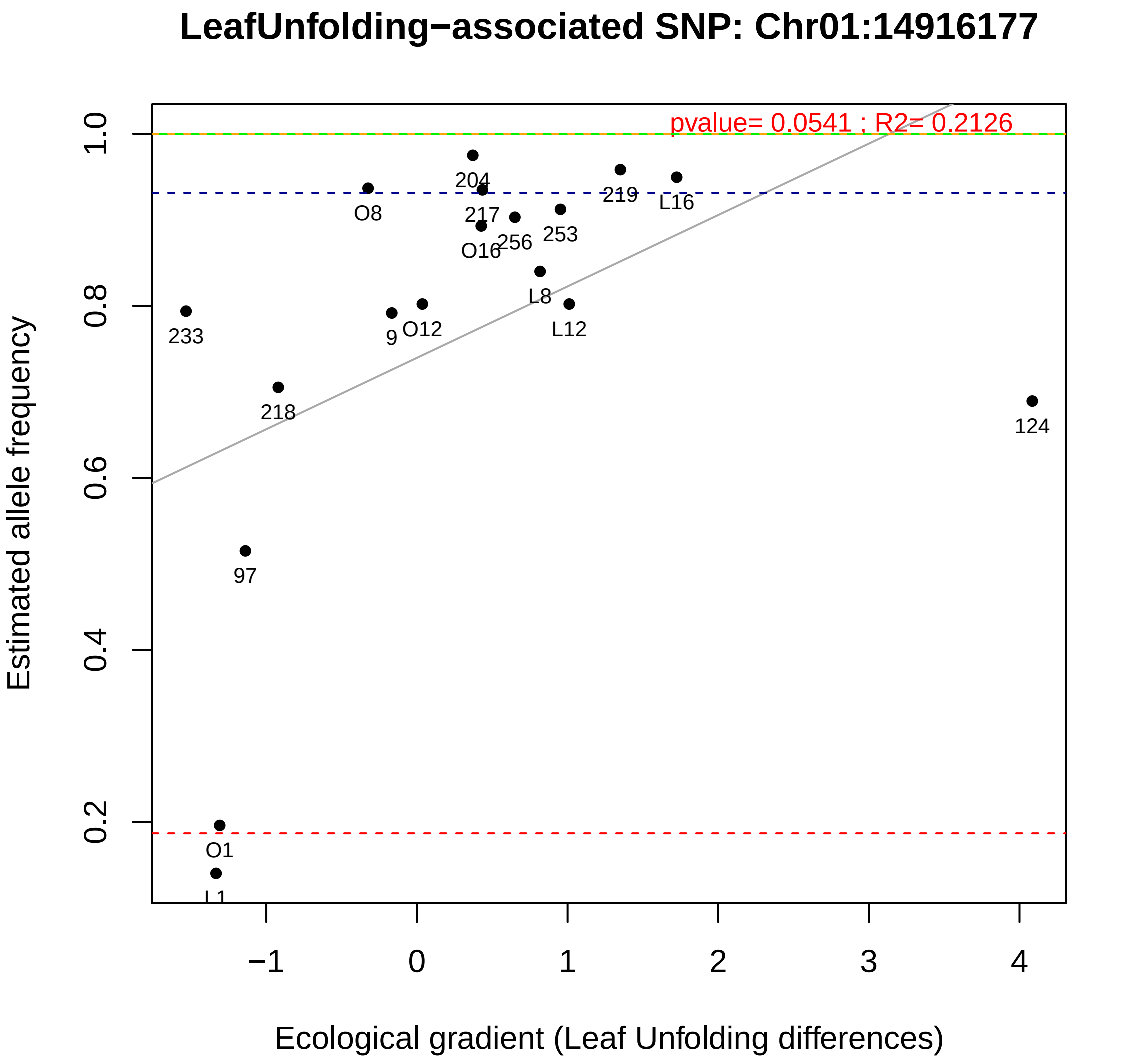
LeafUnfolding−associated SNP: Chr01:14916177
pvalue= 0.0541 ; R2= 0.2126
Estimated allele frequency
Ecological gradient (Leaf Unfolding differences)
0.2
0.4
0.6
0.8
1.0
−1
0
1
2
3
4
204
O8
217
219
L16
253
256
O16
L8
233
9
O12
L12
218
124
97
O1

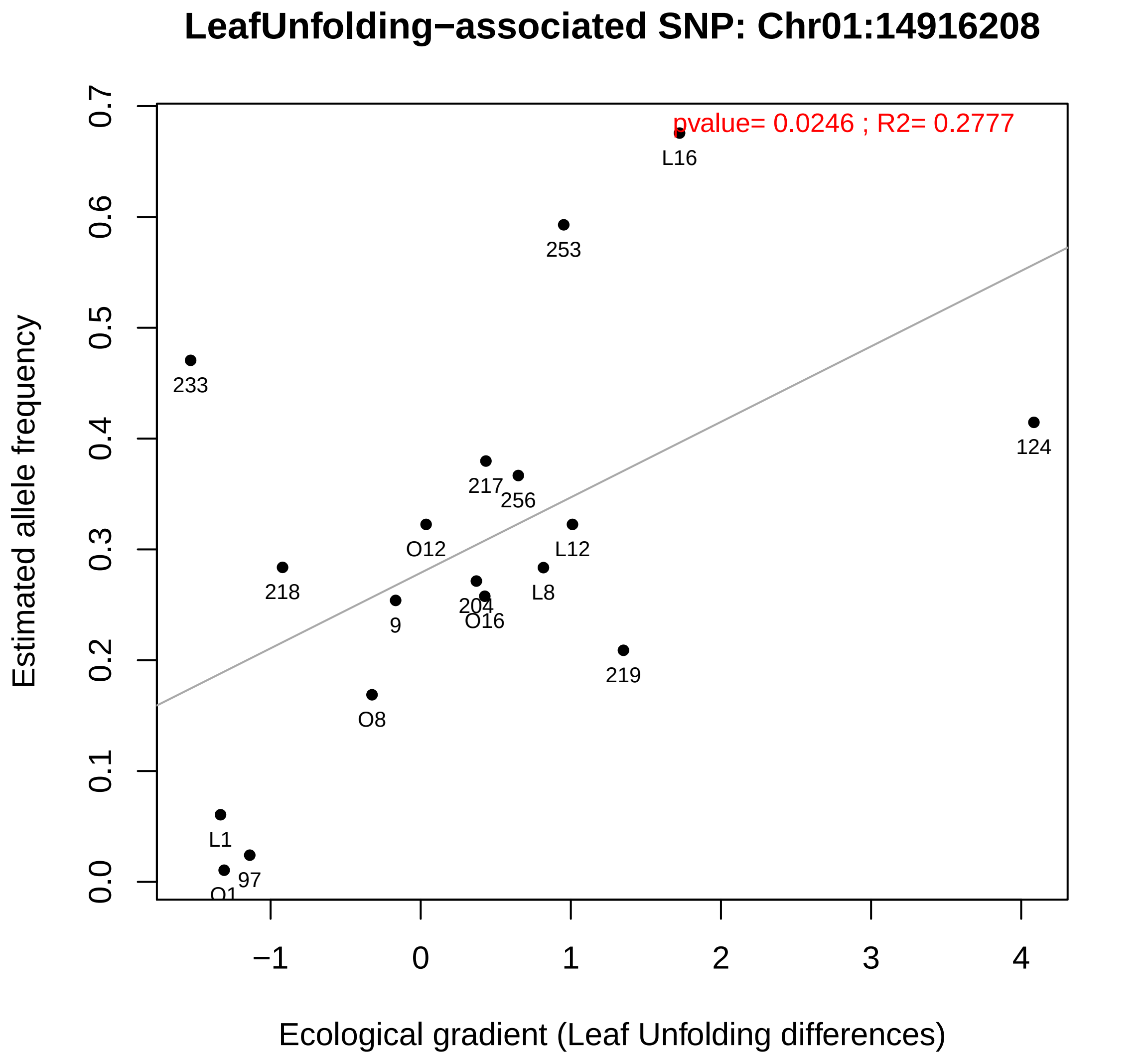

LeafUnfolding−associated SNP: Chr01:14916208
pvalue= 0.0246 ; R2= 0.2777
Estimated allele frequency
Ecological gradient (Leaf Unfolding differences)
0.0
0.1
0.2
0.3
0.4
0.5
0.6
0.7
−1
0
1
2
3
4
L16
253
233
124
217
256
O12
L12
218
9
204
O16
L8
219
O8
L1
97
O1

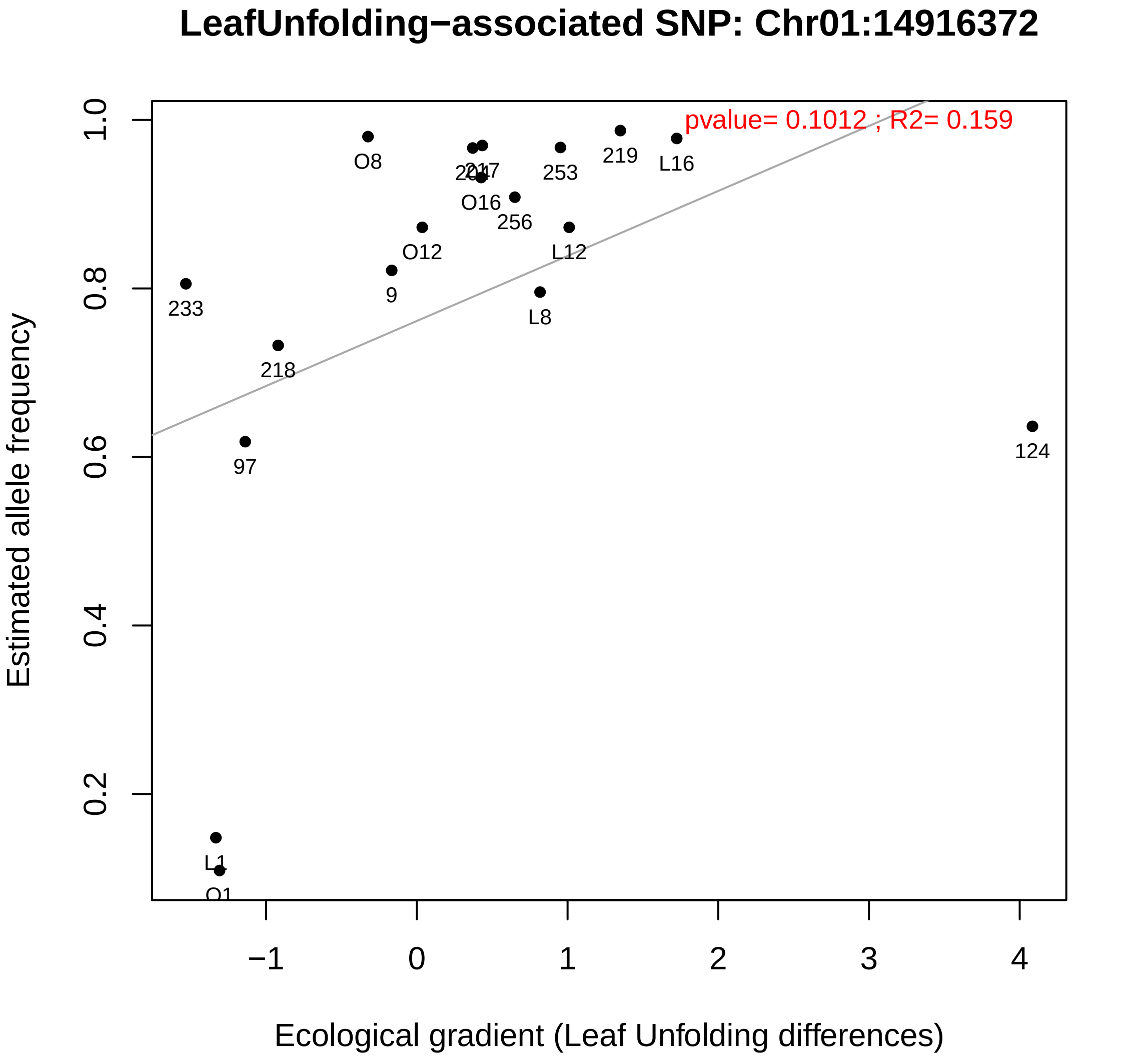

LeafUnfolding−associated SNP: Chr01:14916372
pvalue= 0.1012 ; R2= 0.159
Estimated allele frequency
Ecological gradient (Leaf Unfolding differences)
0.2
0.4
0.6
0.8
1.0
−1
0
1
2
3
4
O8
219
L16
253
O16
256
O12
L12
233
9
L8
218
97
124
L1
O1

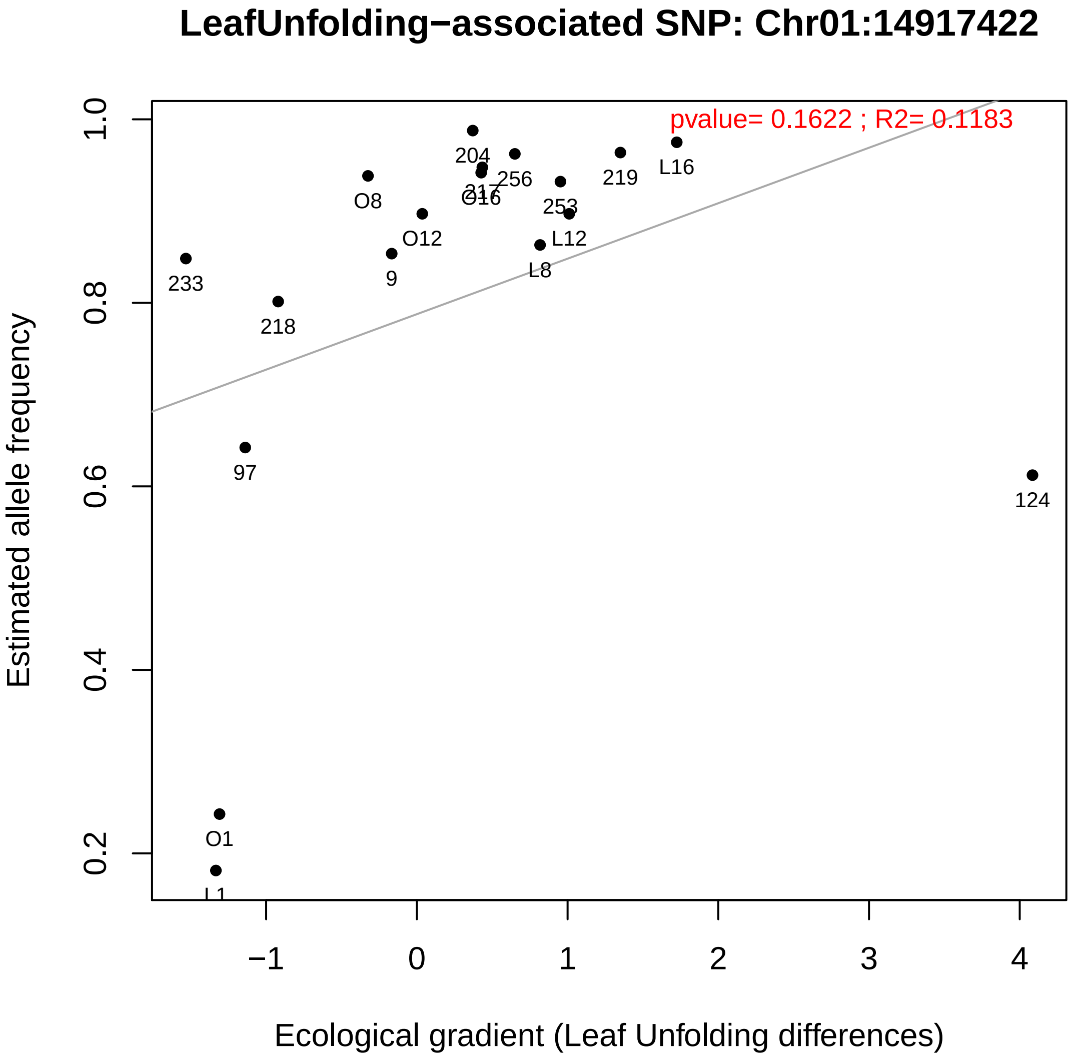
LeafUnfolding−associated SNP: Chr01:14917422
pvalue= 0.1622 ; R2= 0.1183
Estimated allele frequency
Ecological gradient (Leaf Unfolding differences)
0.2
0.4
0.6
0.8
1.0
−1
0
1
2
3
4
204
217
O16
256
O8
253
219
L16
O12
L12
233
9
L8
218
97
124
O1
L1

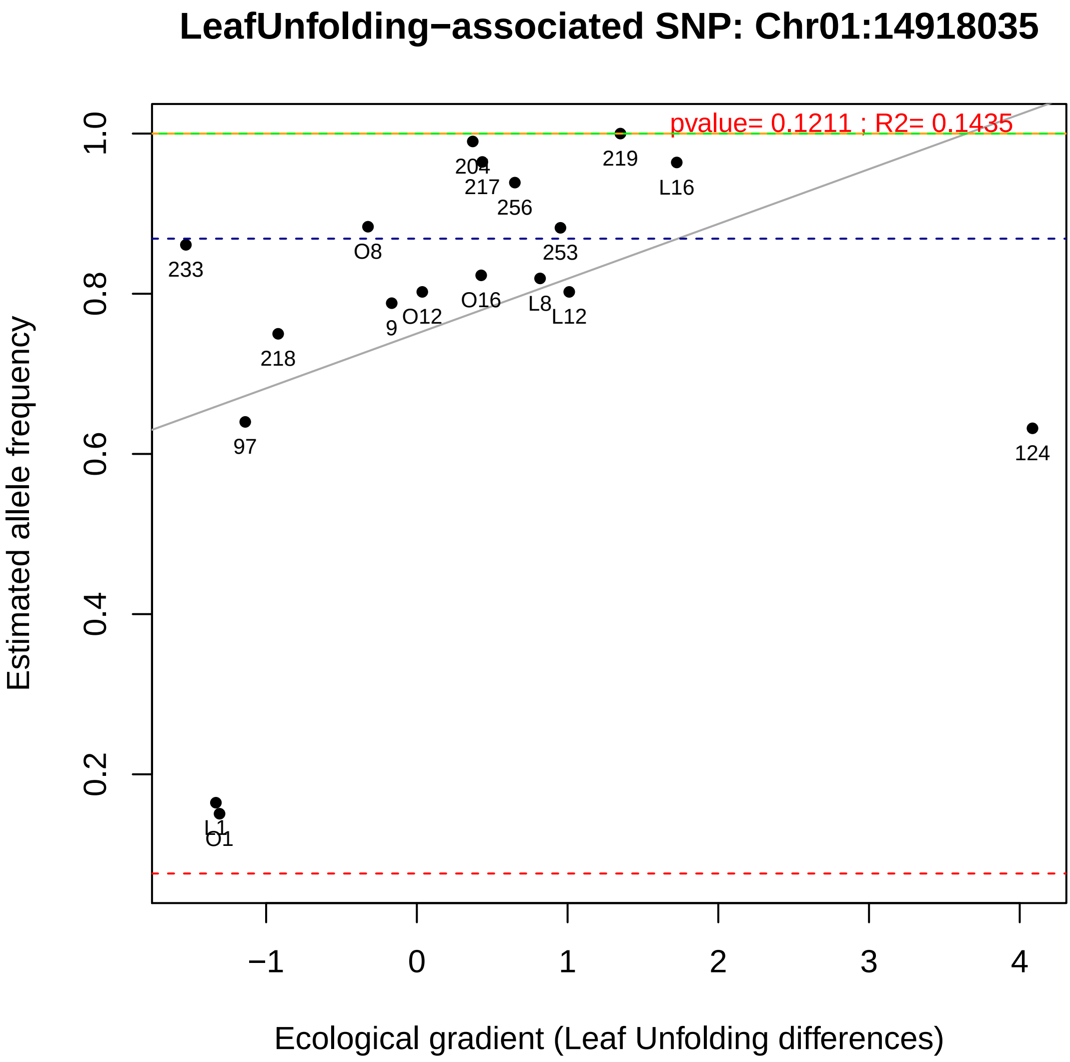

LeafUnfolding−associated SNP: Chr01:14918035
pvalue= 0.1211 ; R2= 0.1435
Estimated allele frequency
Ecological gradient (Leaf Unfolding differences)
0.2
0.4
0.6
0.8
1.0
−1
0
1
2
3
4
233
O8
204
217
256
219
L16
253
O16
L8
L12
9
O12
218
97
124
L1
O1

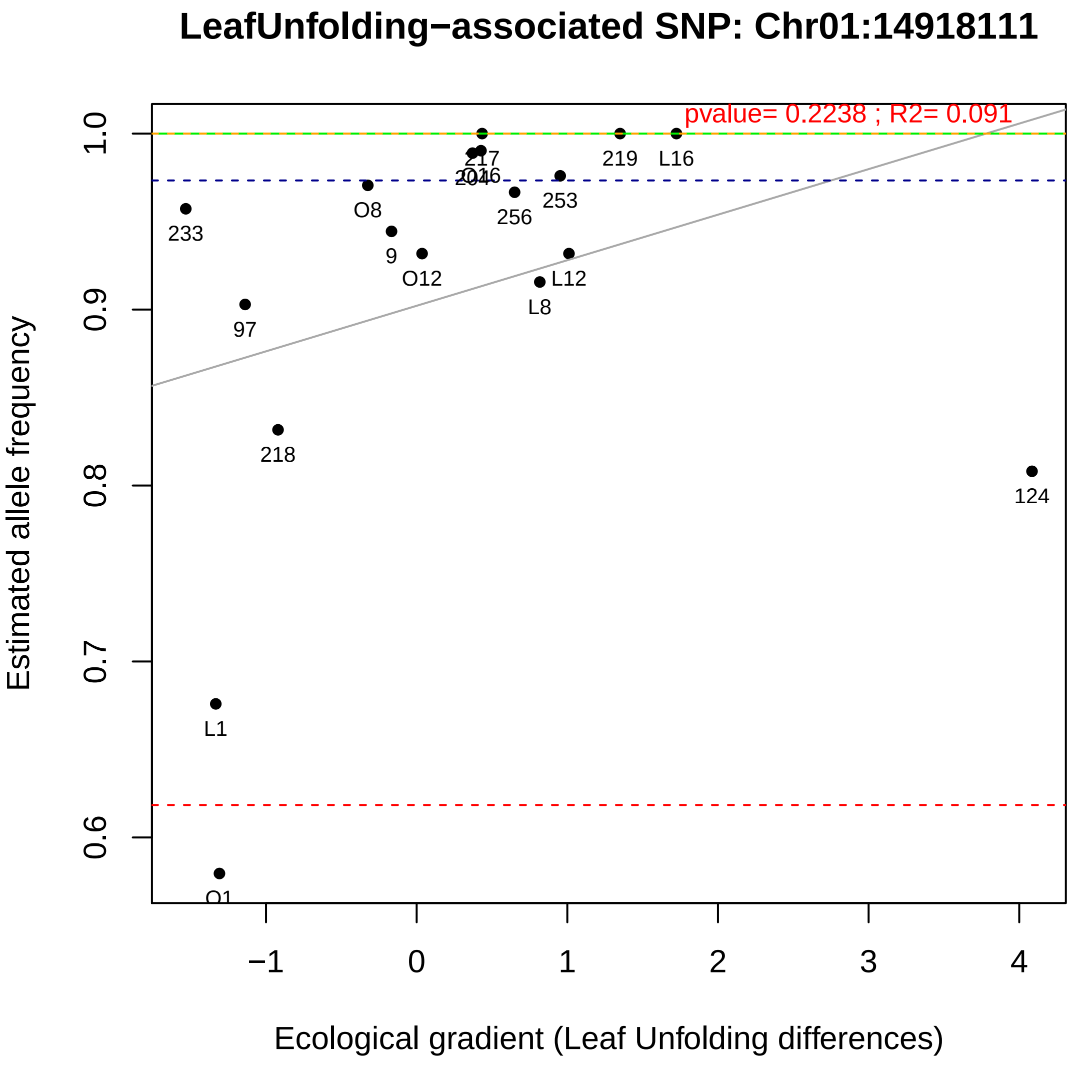

LeafUnfolding−associated SNP: Chr01:14918111
pvalue= 0.2238 ; R2= 0.091
Estimated allele frequency
Ecological gradient (Leaf Unfolding differences)
0.6
0.7
0.8
0.9
1.0
−1
0
1
2
3
4
217
219
L16
253
256
233
O8
9
O12
L12
L8
97
218
124
L1

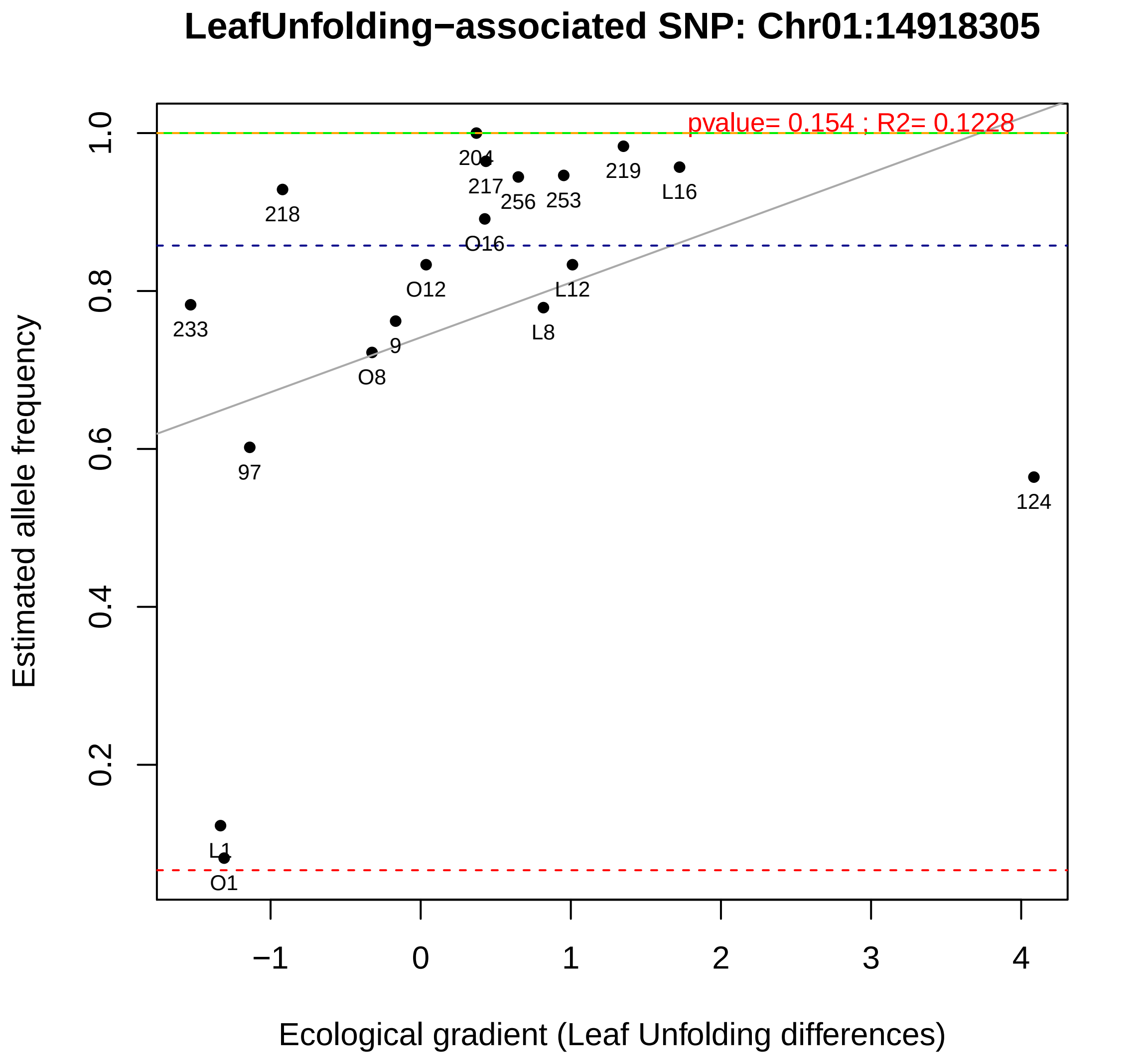
LeafUnfolding−associated SNP: Chr01:14918305
pvalue= 0.154 ; R2= 0.1228
Estimated allele frequency
Ecological gradient (Leaf Unfolding differences)
1.0
0.8
0.6
0.4
0.2
−1
0
1
2
3
4
204
217
256
253
219
L16
218
O16
O12
L12
233
9
L8
O8
97
124
L1
O1

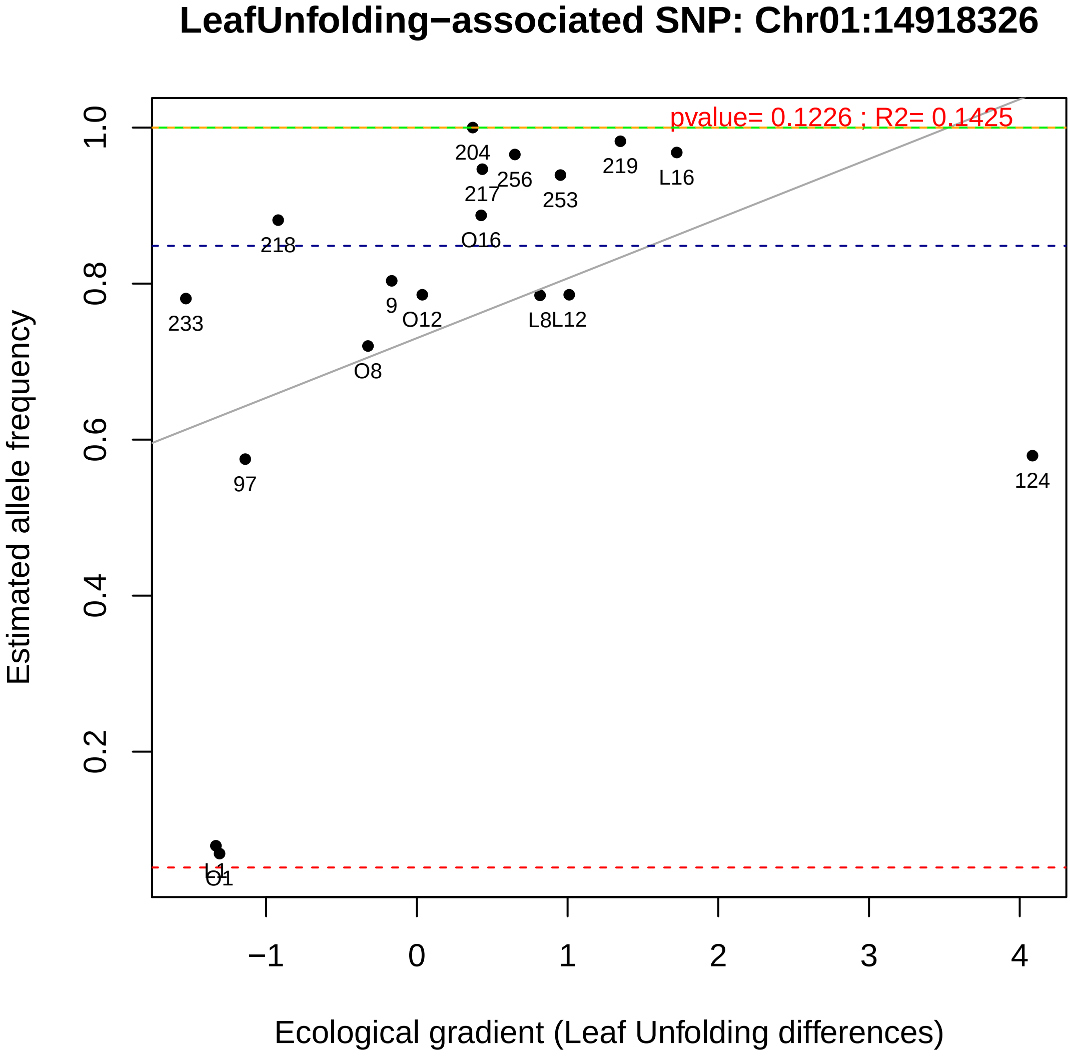

LeafUnfolding−associated SNP: Chr01:14918326
pvalue= 0.1226 ; R2= 0.1425
204
256
217
253
219
L16
218
O16
233
9
O12
L8
L12
O8
97
124
L1
O1
1.0
0.8
0.6
0.4
0.2
−1
0
1
2
3
4
Estimated allele frequency
Ecological gradient (Leaf Unfolding differences)

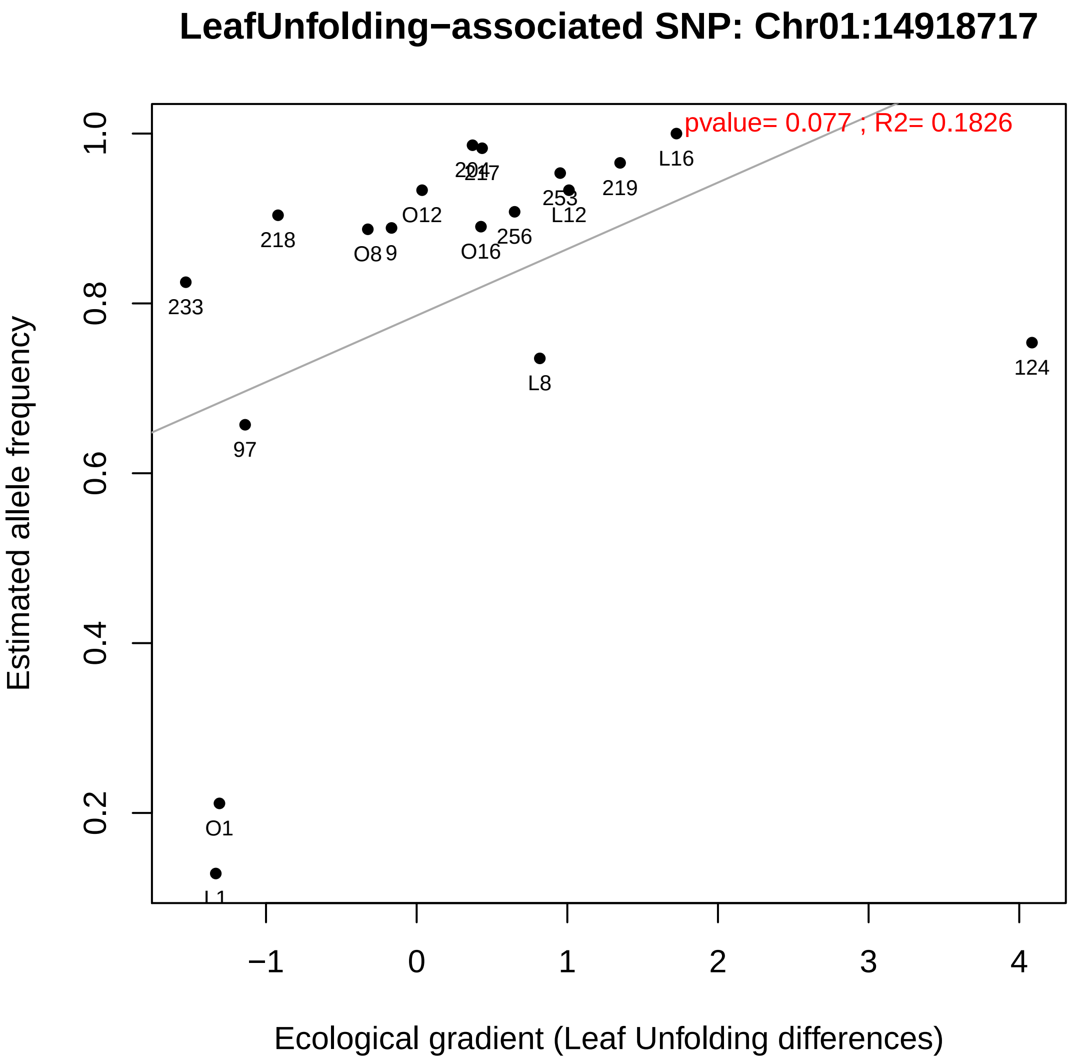
LeafUnfolding−associated SNP: Chr01:14918717
pvalue= 0.077 ; R2= 0.1826
Estimated allele frequency
Ecological gradient (Leaf Unfolding differences)
1.0
0.8
0.6
0.4
0.2
−1
0
1
2
3
4
L16
219
253
L12
O12
256
218
O8
9
O16
233
L8
124
97
O1

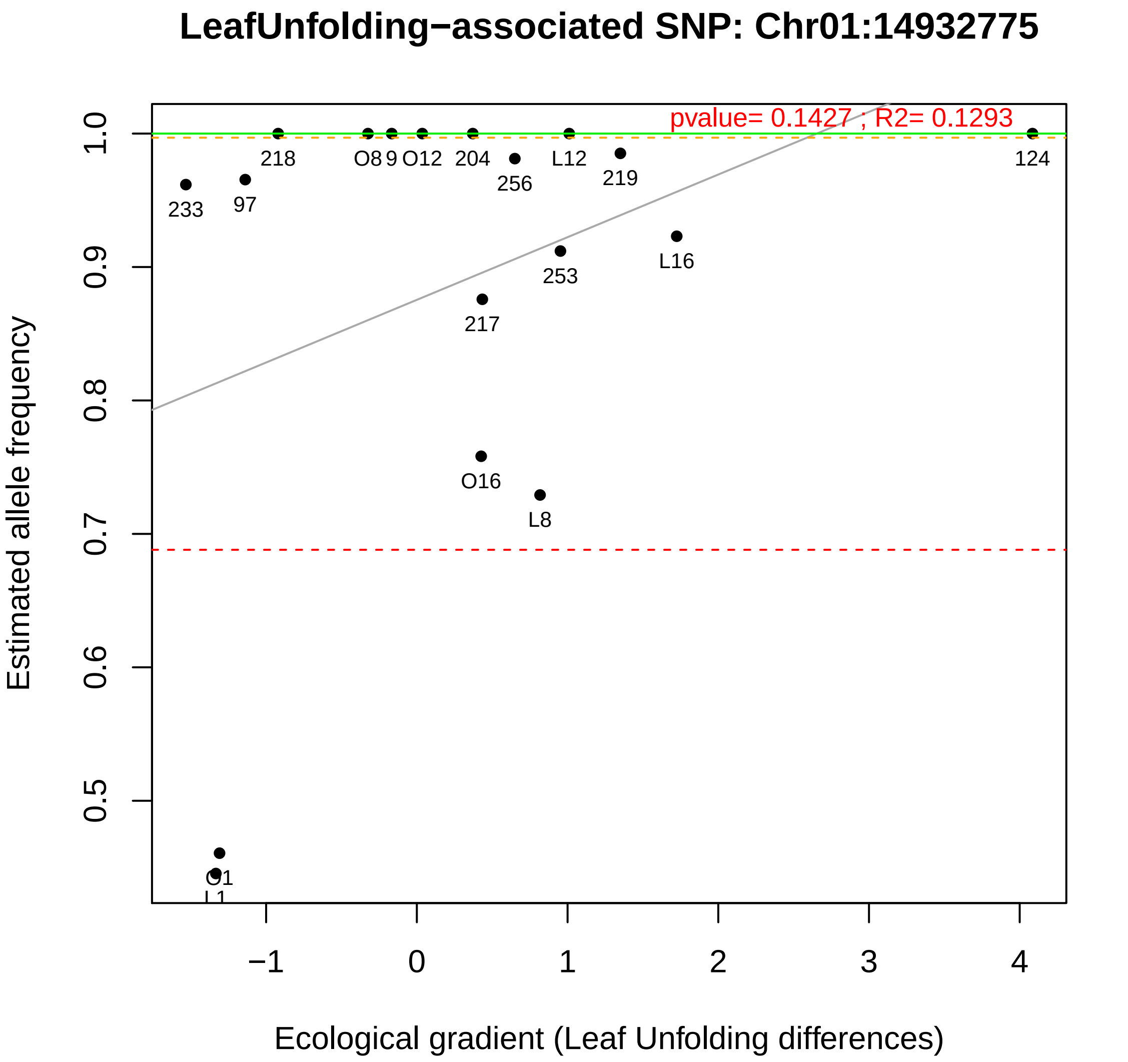
LeafUnfolding−associated SNP: Chr01:14932775
pvalue= 0.1427 ; R2= 0.1293
Estimated allele frequency
1.0
0.9
0.8
0.7
0.6
0.5
−1
0
1
2
3
4
Ecological gradient (Leaf Unfolding differences)
218
O8
9
O12
204
256
L12
219
124
233
97
253
L16
217
O16
L8
O1
L1

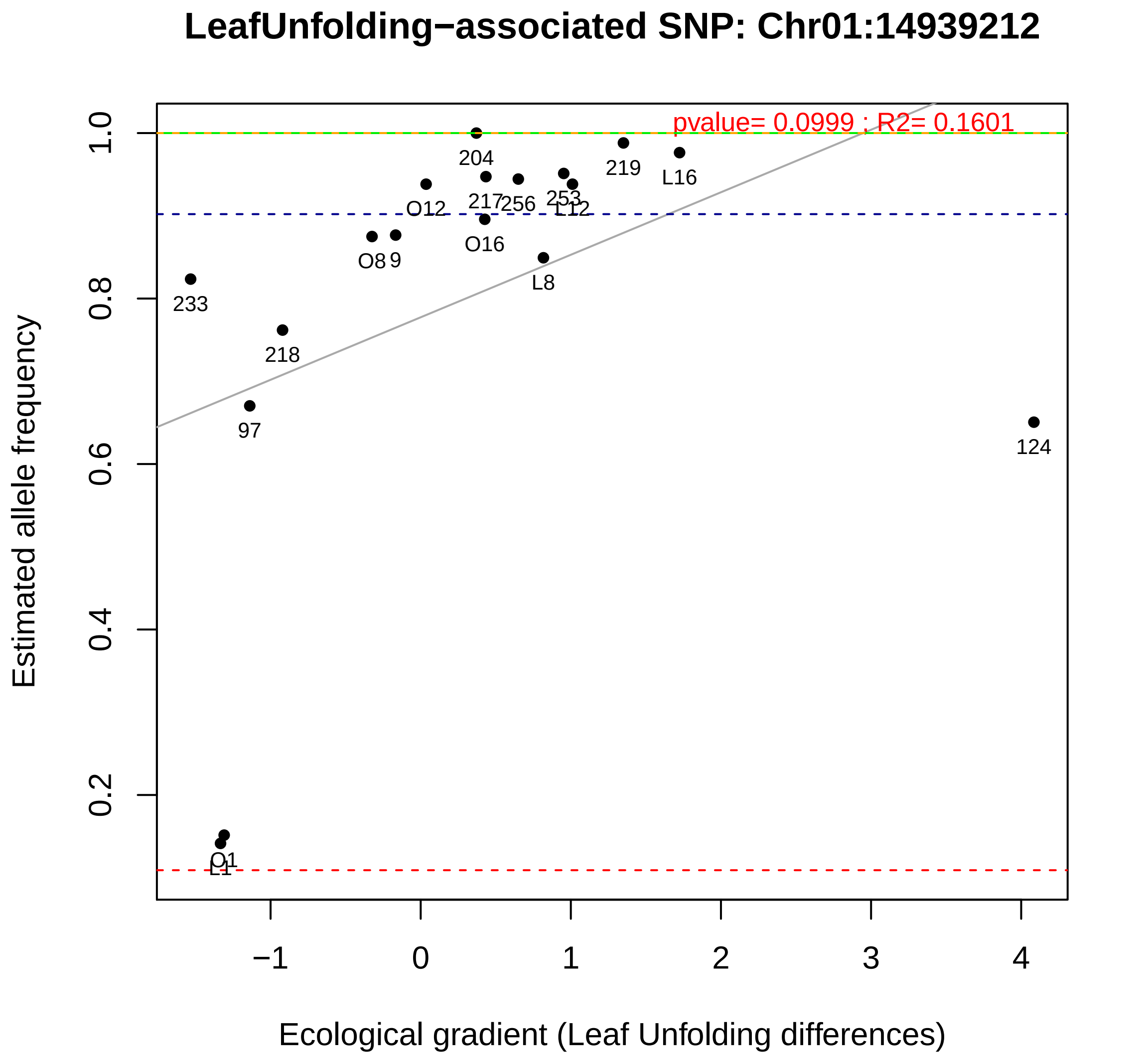

LeafUnfolding−associated SNP: Chr01:14939212
pvalue= 0.0999 ; R2= 0.1601
Estimated allele frequency
Ecological gradient (Leaf Unfolding differences)
0.2
0.4
0.6
0.8
1.0
−1
0
1
2
3
4
204
219
L16
O12
217
256
253
L12
O16
O8
9
L8
233
218
97
124
O1
L1

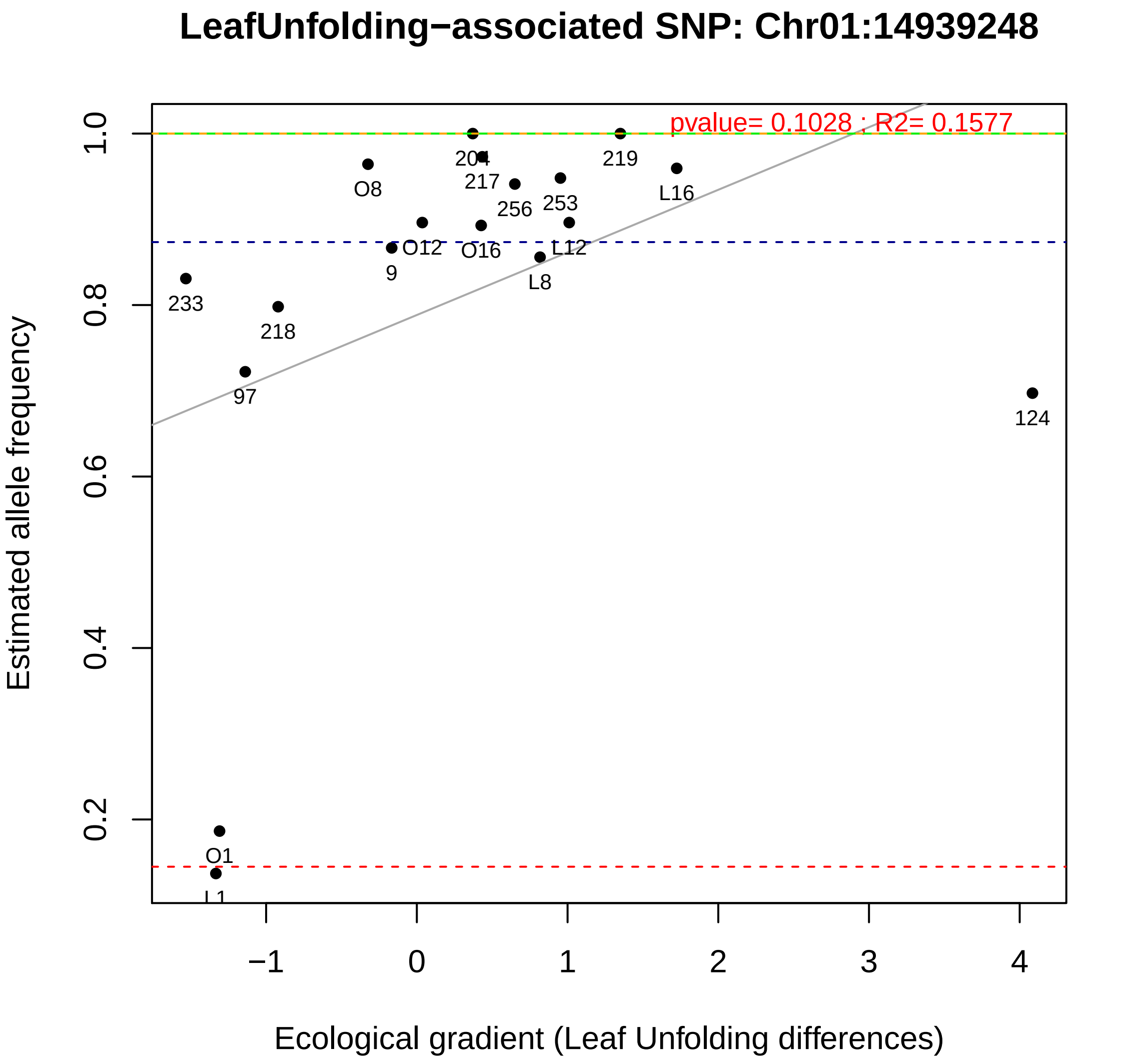
LeafUnfolding−associated SNP: Chr01:14939248
pvalue= 0.1028 ; R2= 0.1577
Estimated allele frequency
Ecological gradient (Leaf Unfolding differences)
1.0
0.8
0.6
0.4
0.2
−1
0
1
2
3
4
204
217
219
O8
256
253
L16
O12
O16
L12
9
L8
233
218
97
124
O1
L1

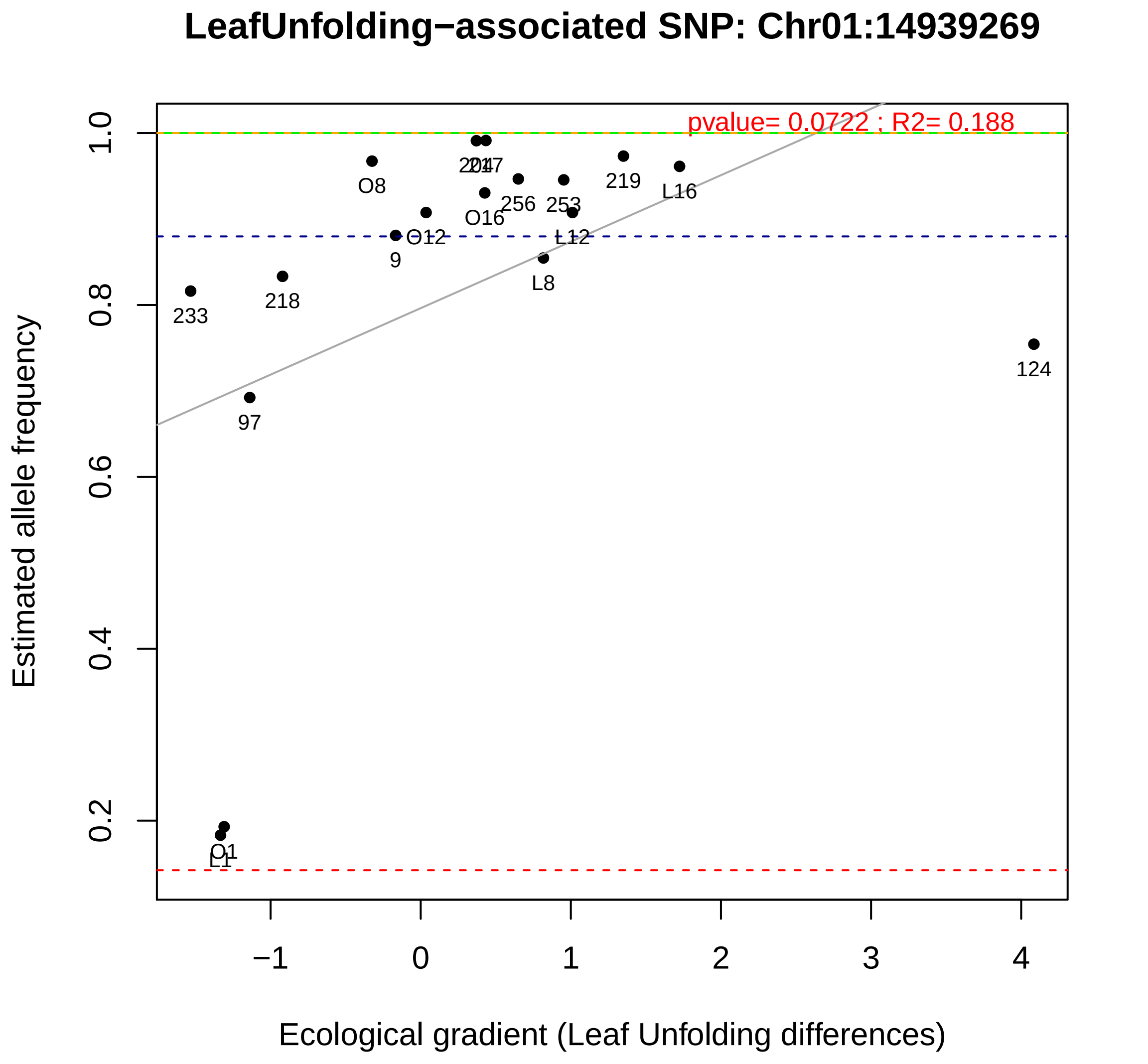

LeafUnfolding−associated SNP: Chr01:14939269
pvalue= 0.0722 ; R2= 0.188
Estimated allele frequency
Ecological gradient (Leaf Unfolding differences)
0.2
0.4
0.6
0.8
1.0
−1
0
1
2
3
4
O8
219
L16
256
253
O16
O12
L12
9
L8
233
218
97
124
O1
L1

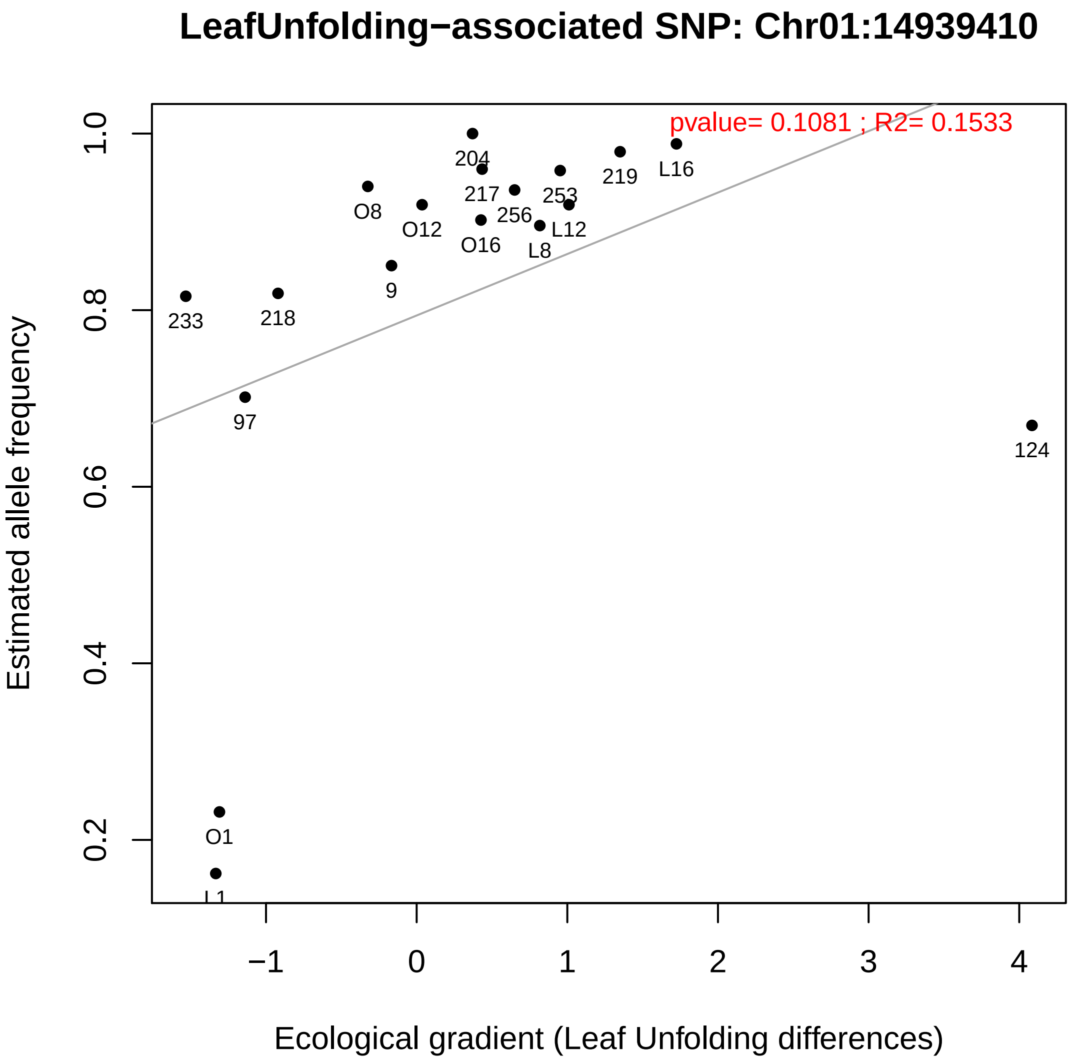
LeafUnfolding−associated SNP: Chr01:14939410
pvalue= 0.1081 ; R2= 0.1533
Estimated allele frequency
Ecological gradient (Leaf Unfolding differences)
0.2
0.4
0.6
0.8
1.0
−1
0
1
2
3
4
204
217
256
253
219
L16
O8
O12
O16
L8
L12
9
233
218
97
124
O1

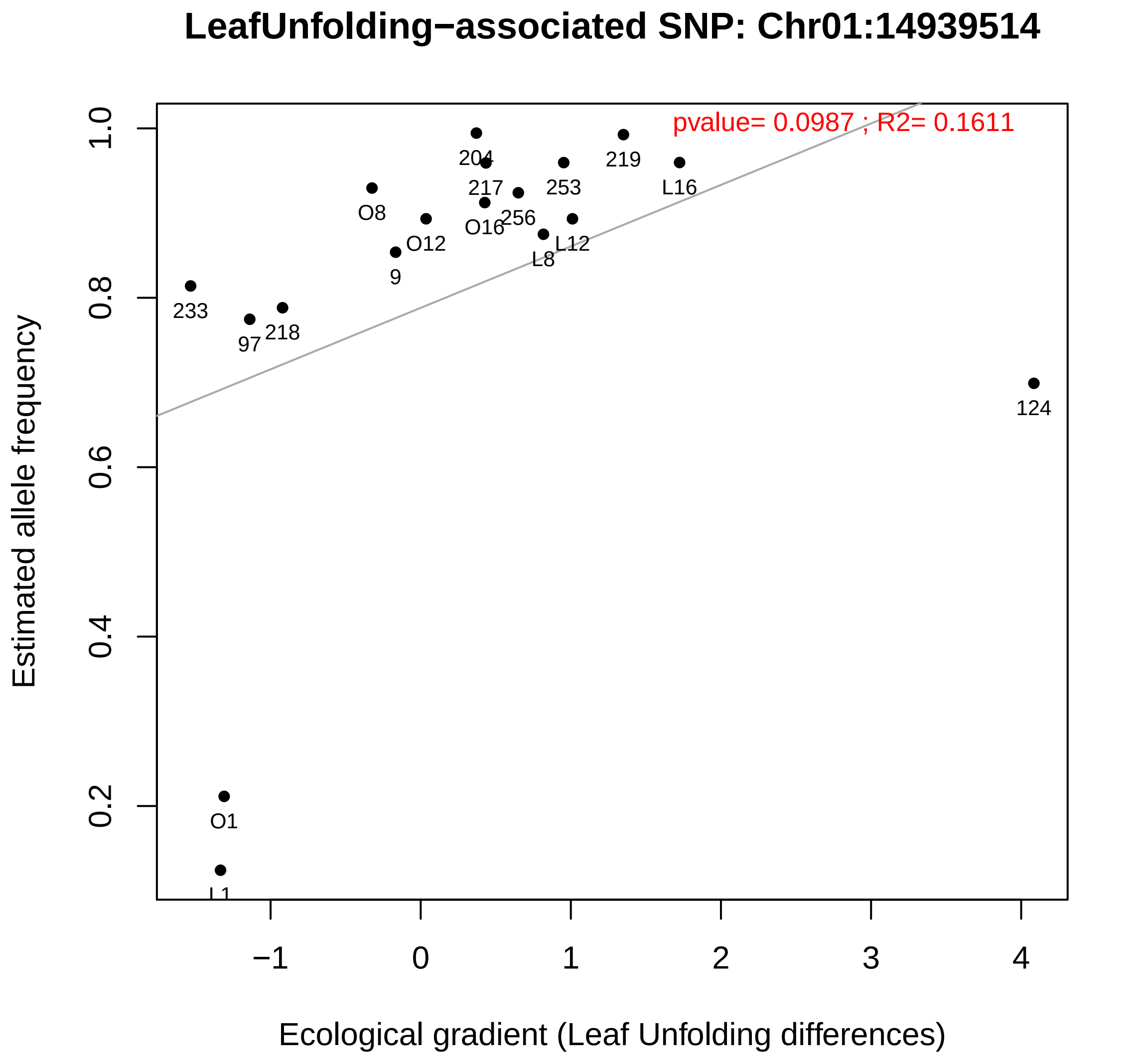

LeafUnfolding−associated SNP: Chr01:14939514
pvalue= 0.0987 ; R2= 0.1611
Estimated allele frequency
Ecological gradient (Leaf Unfolding differences)
1.0
0.8
0.6
0.4
0.2
−1
0
1
2
3
4
204
217
219
253
L16
O8
256
O16
O12
L12
L8
9
233
218
97
124
O1
L1

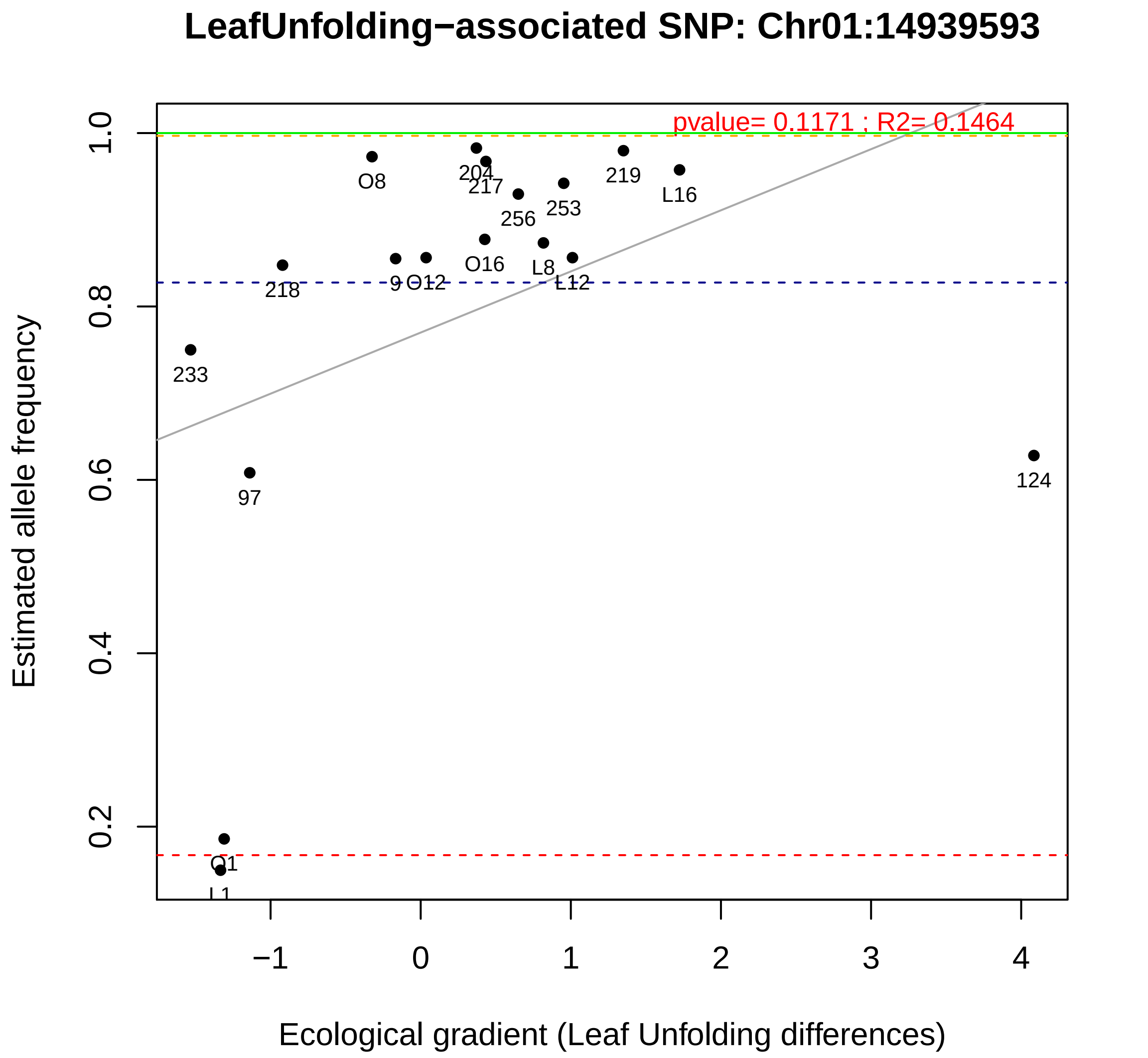

LeafUnfolding−associated SNP: Chr01:14939593
pvalue= 0.1171 ; R2= 0.1464
Estimated allele frequency
0.2
0.4
0.6
0.8
1.0
−1
0
1
2
3
4
Ecological gradient (Leaf Unfolding differences)
O8
204
217
219
L16
253
256
O16
L8
L12
218
9
O12
233
97
124
O1
L1

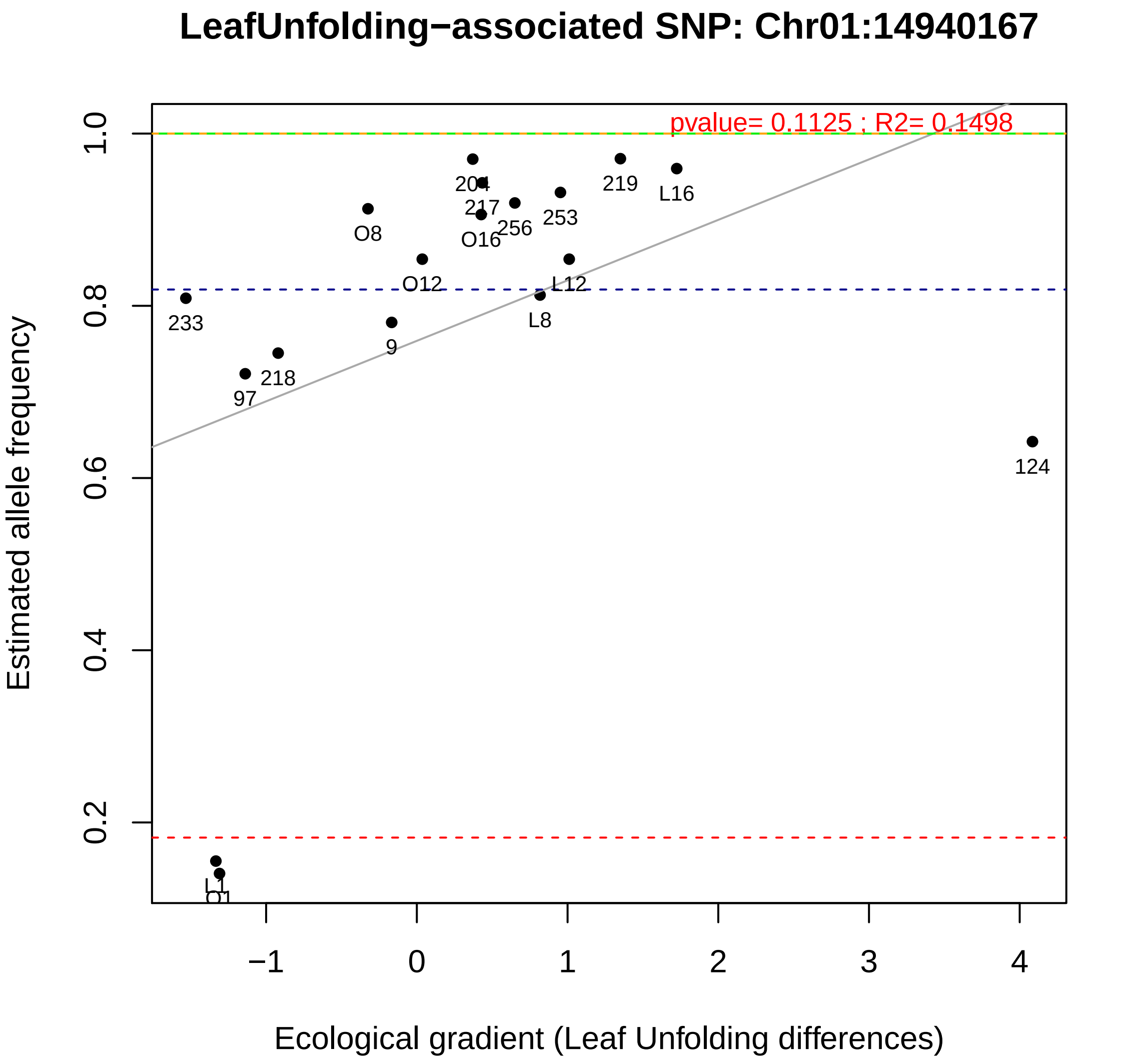

LeafUnfolding−associated SNP: Chr01:14940167
pvalue= 0.1125 ; R2= 0.1498
Estimated allele frequency
Ecological gradient (Leaf Unfolding differences)
1.0
0.8
0.6
0.4
0.2
−1
0
1
2
3
4
233
97
218
O8
9
O12
204
217
O16
256
253
L8
L12
219
L16
124
L1
O1

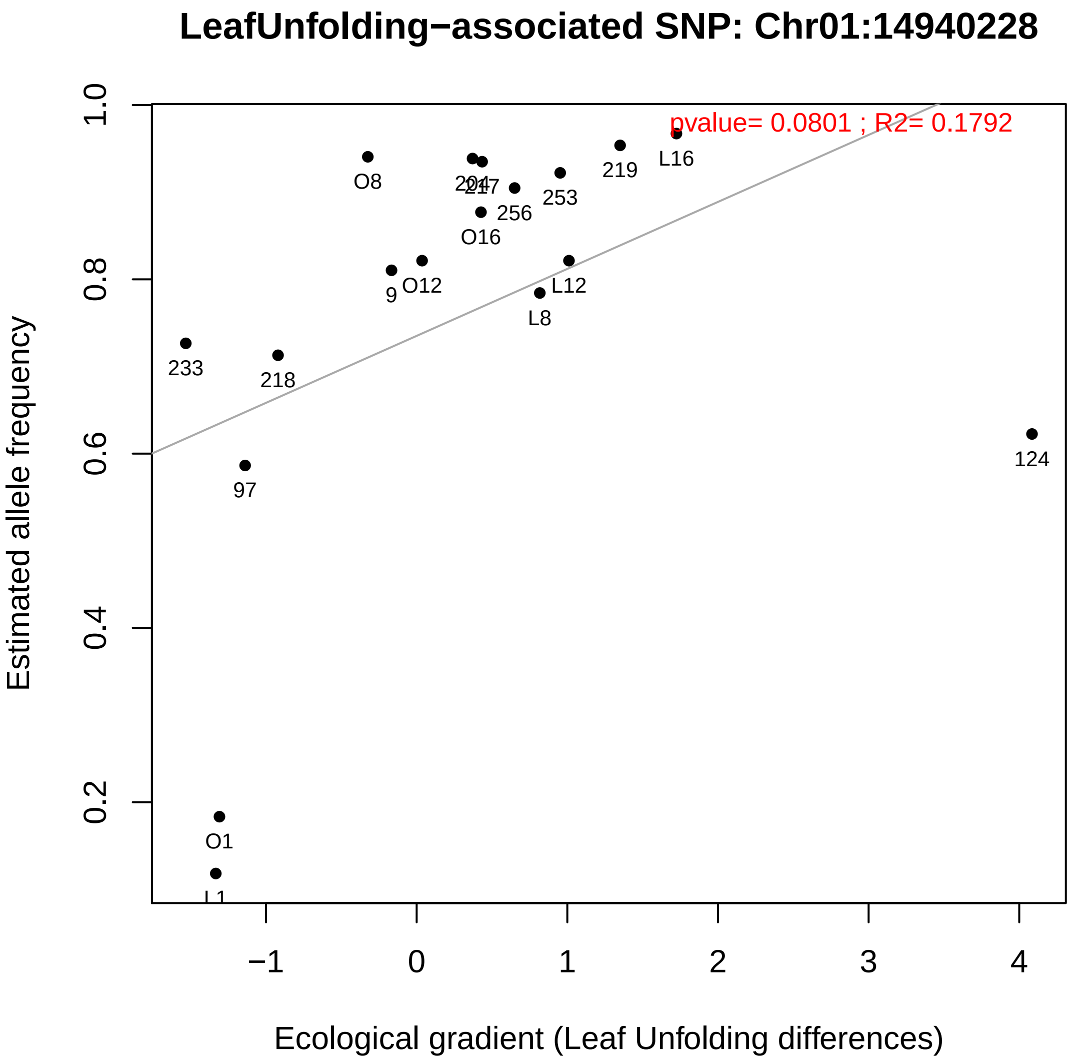

LeafUnfolding−associated SNP: Chr01:14940228
pvalue= 0.0801 ; R2= 0.1792
Estimated allele frequency
Ecological gradient (Leaf Unfolding differences)
0.2
0.4
0.6
0.8
1.0
−1
0
1
2
3
4
O8
204
217
256
O16
253
219
L16
9
O12
L12
L8
233
218
97
124
O1

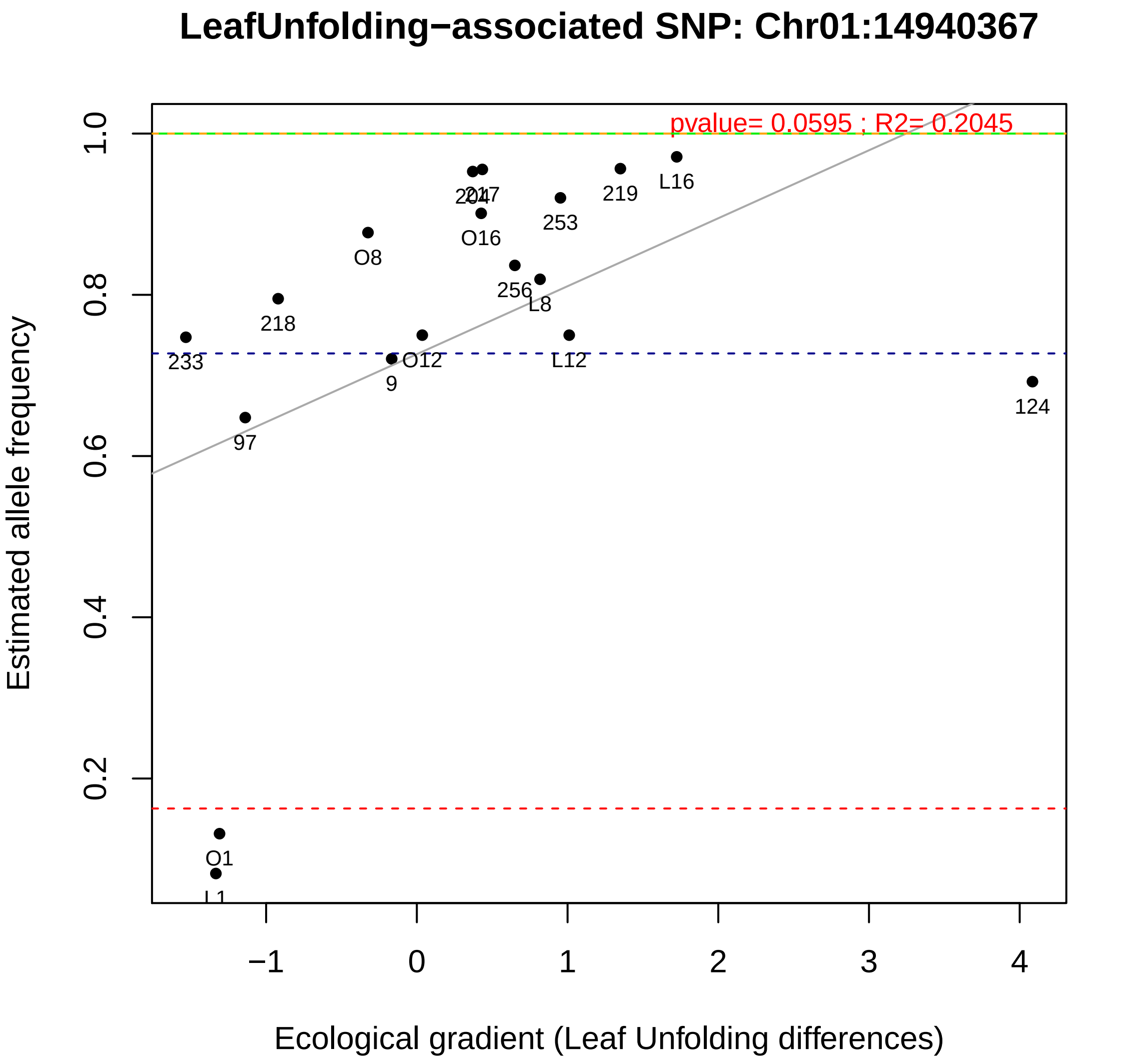
LeafUnfolding−associated SNP: Chr01:14940367
pvalue= 0.0595 ; R2= 0.2045
Estimated allele frequency
Ecological gradient (Leaf Unfolding differences)
1.0
0.8
0.6
0.4
0.2
−1
0
1
2
3
4
233
97
218
O8
9
O12
204
217
O16
256
L8
253
L12
219
L16
124
O1
L1

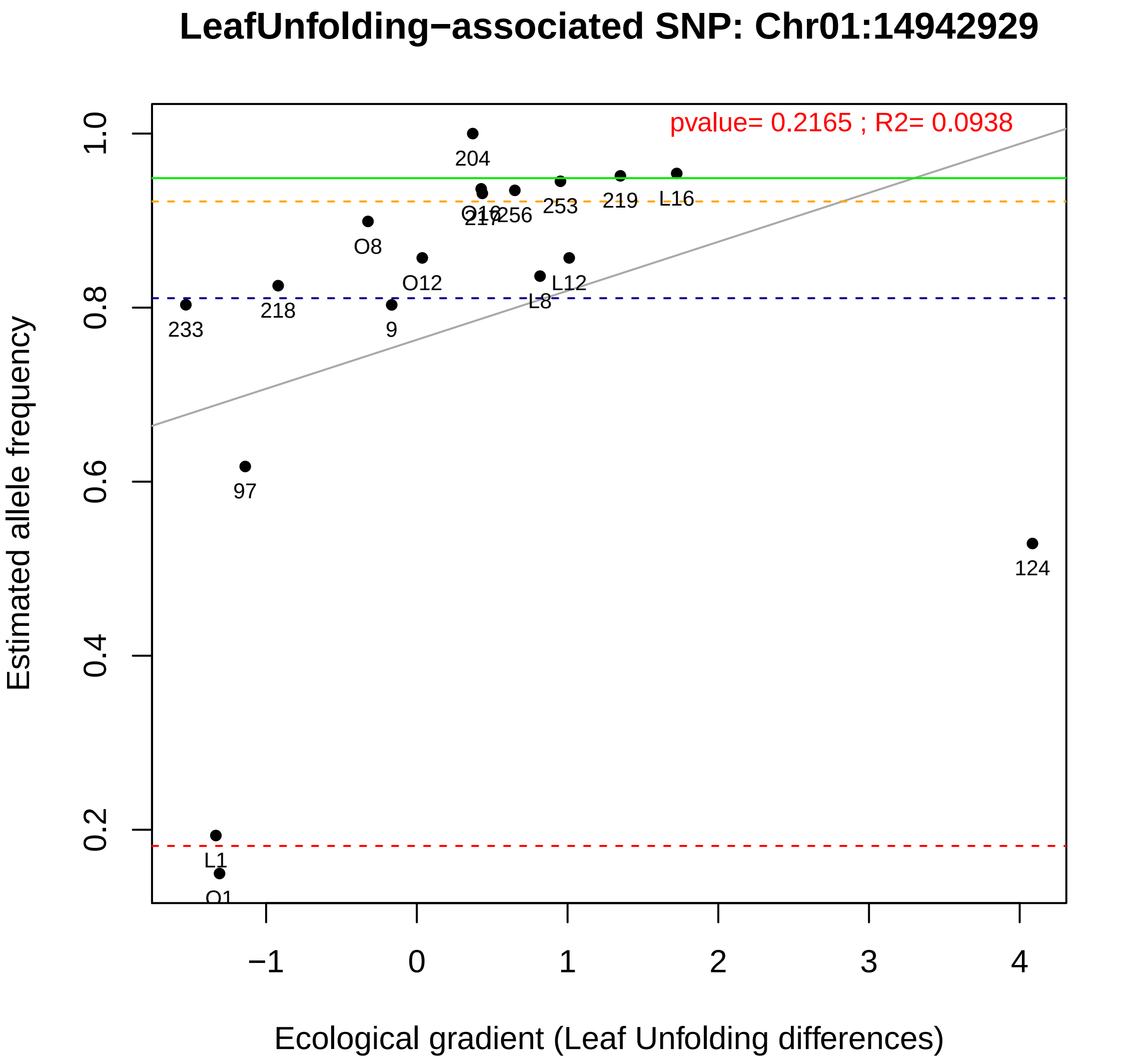

LeafUnfolding−associated SNP: Chr01:14942929
pvalue= 0.2165 ; R2= 0.0938
Estimated allele frequency
Ecological gradient (Leaf Unfolding differences)
1.0
0.8
0.6
0.4
0.2
−1
0
1
2
3
4
204
O16
217
256
253
219
L16
O8
O12
L12
L8
218
233
9
97
124
L1
O1

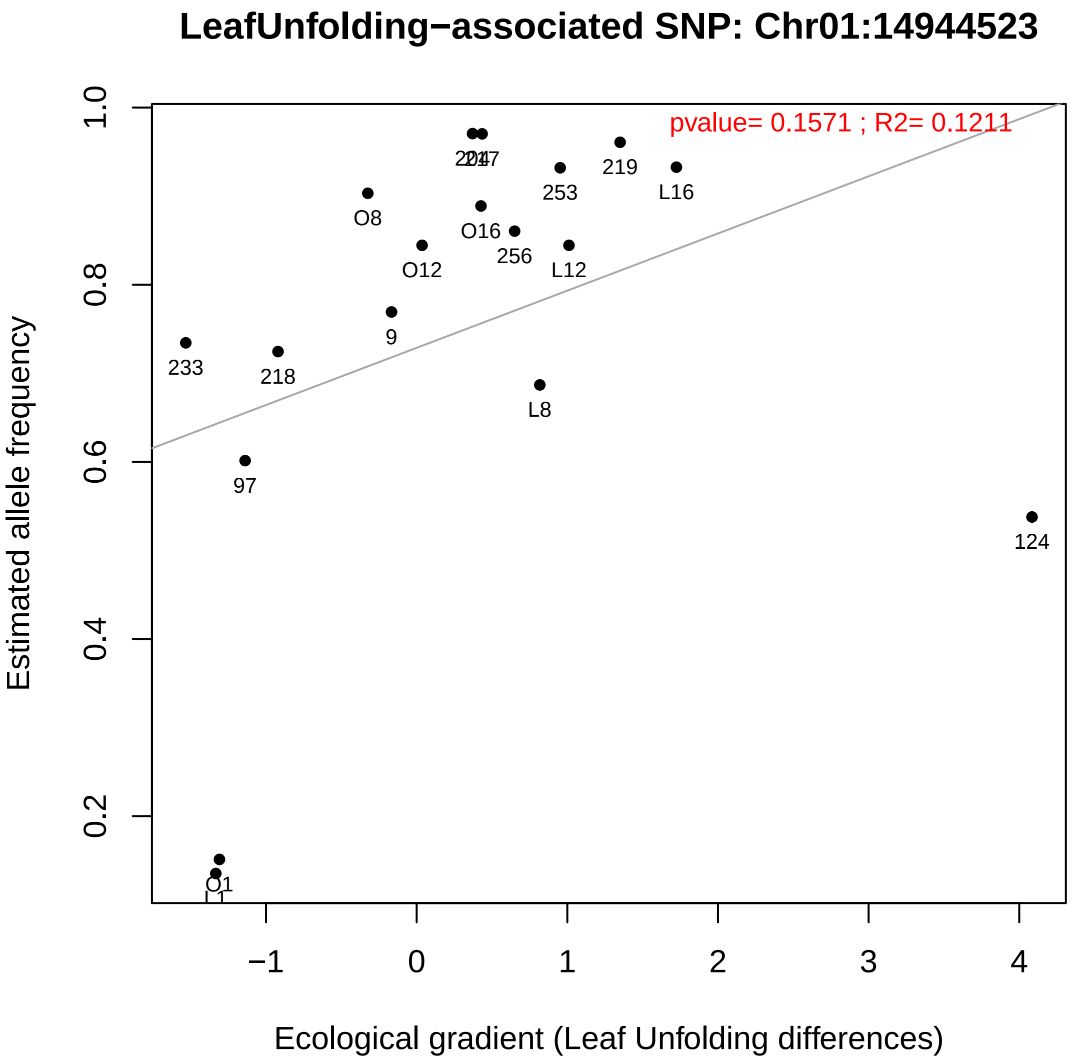
LeafUnfolding−associated SNP: Chr01:14944523
pvalue= 0.1571 ; R2= 0.1211
Estimated allele frequency
Ecological gradient (Leaf Unfolding differences)
0.2
0.4
0.6
0.8
1.0
−1
0
1
2
3
4
219
253
L16
O8
O16
256
O12
L12
9
233
218
L8
97
124
O1

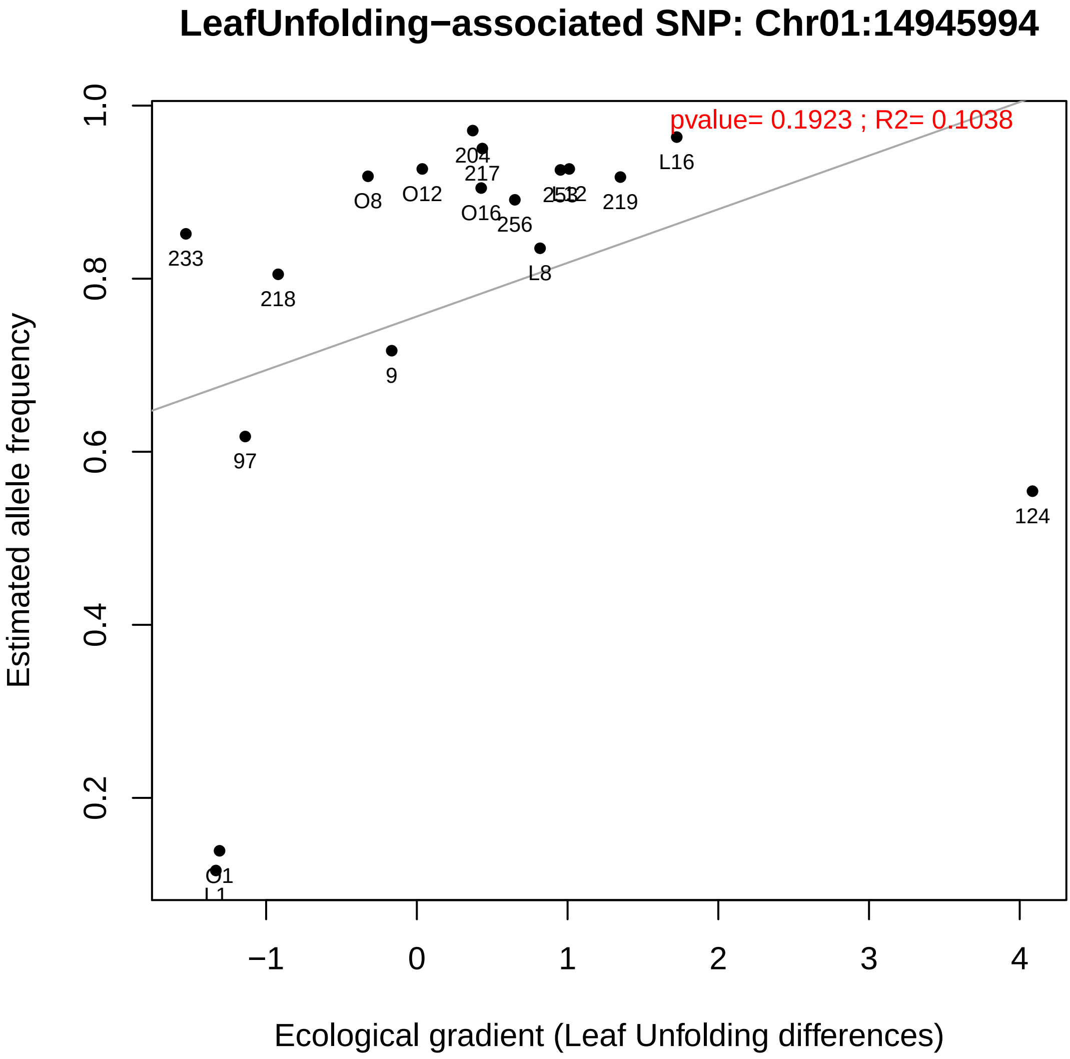

LeafUnfolding−associated SNP: Chr01:14945994
pvalue= 0.1923 ; R2= 0.1038
Estimated allele frequency
Ecological gradient (Leaf Unfolding differences)
0.2
0.4
0.6
0.8
1.0
−1
0
1
2
3
4
233
218
97
O1
L1
9
O8
O12
204
217
O16
256
L8
L12
219
L16
124

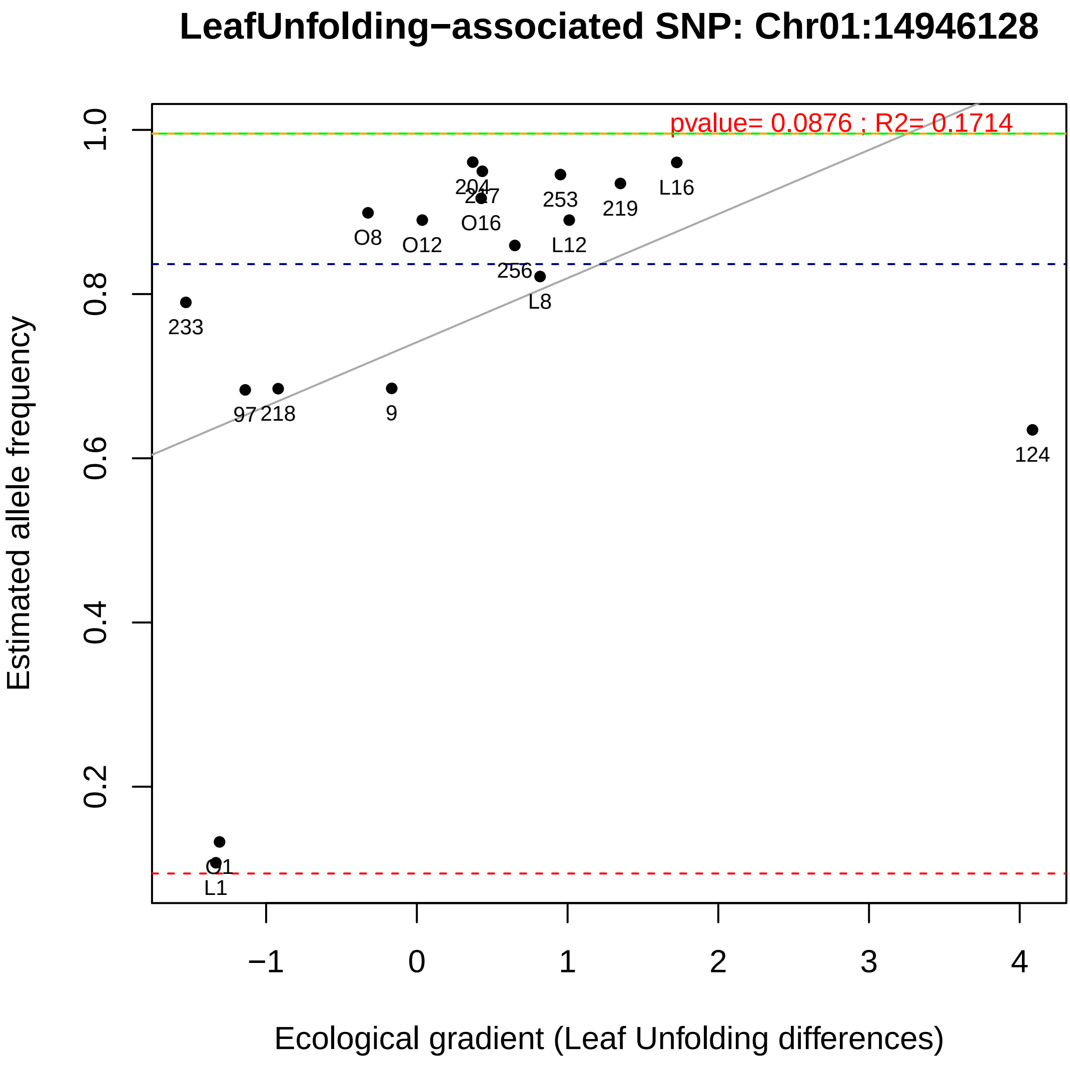
LeafUnfolding−associated SNP: Chr01:14946128
pvalue= 0.0876 ; R2= 0.1714
Estimated allele frequency
Ecological gradient (Leaf Unfolding differences)
0.2
0.4
0.6
0.8
1.0
−1
0
1
2
3
4
204
217
O16
253
219
L16
O8
O12
256
L12
L8
233
97
218
9
124
O1
L1

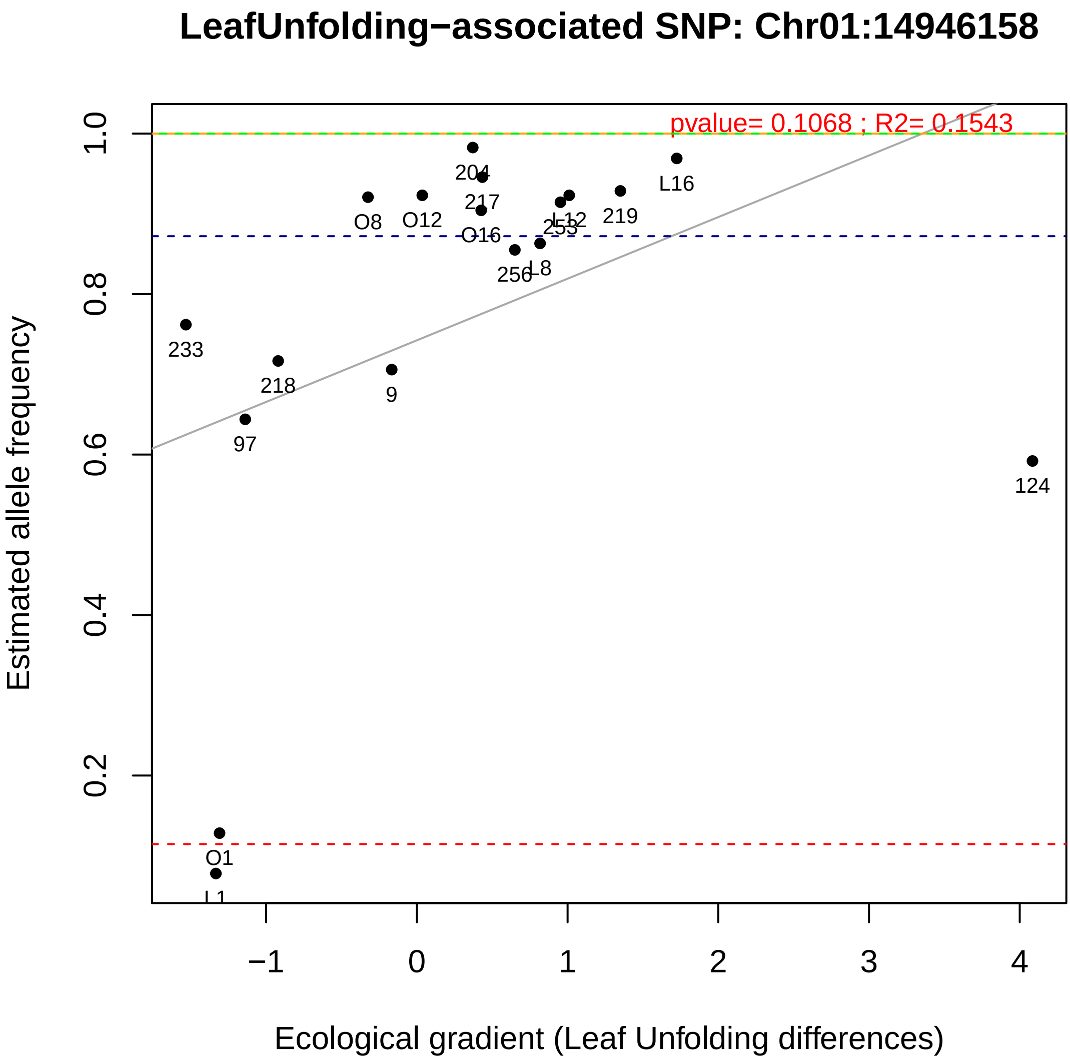
LeafUnfolding−associated SNP: Chr01:14946158
pvalue= 0.1068 ; R2= 0.1543
Estimated allele frequency
Ecological gradient (Leaf Unfolding differences)
0.2
0.4
0.6
0.8
1.0
−1
0
1
2
3
4
204
217
O8
O12
O16
L12
253
219
L16
256
L8
233
218
9
97
124
O1
L1

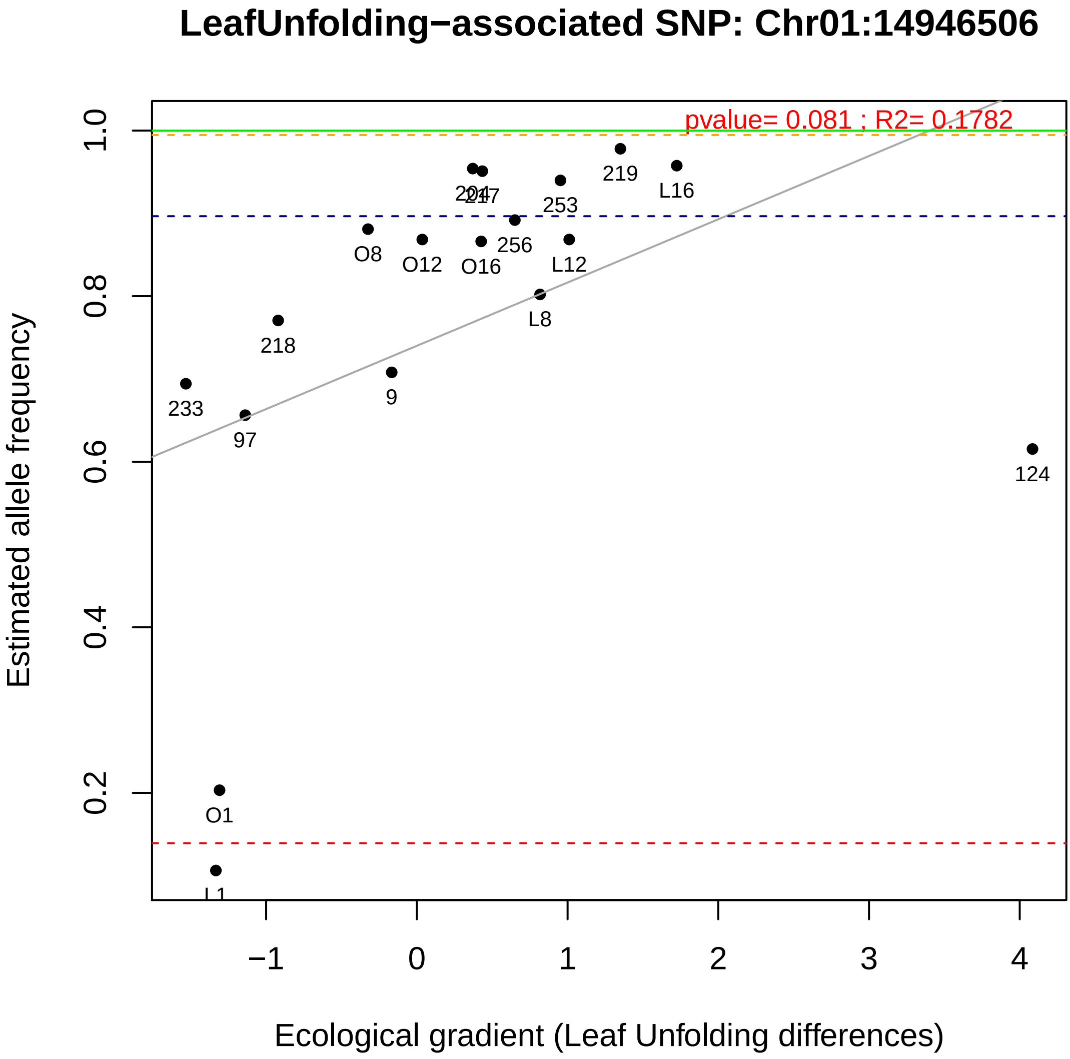
LeafUnfolding−associated SNP: Chr01:14946506
pvalue= 0.081 ; R2= 0.1782
Estimated allele frequency
Ecological gradient (Leaf Unfolding differences)
1.0
0.8
0.6
0.4
0.2
−1
0
1
2
3
4
219
L16
204
217
253
256
O8
O12
O16
L12
L8
218
233
97
9
124
O1
L1

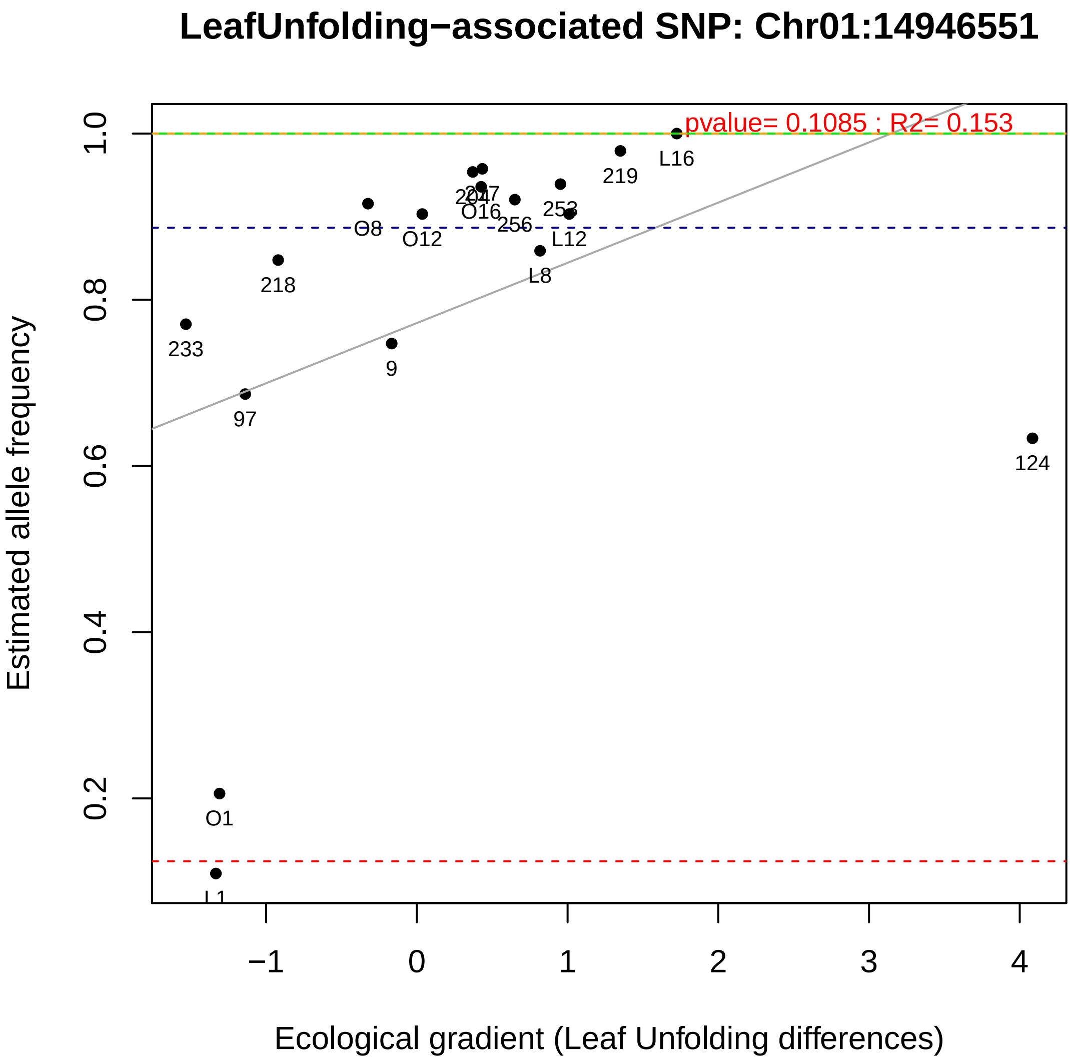
LeafUnfolding−associated SNP: Chr01:14946551
pvalue= 0.1085 ; R2= 0.153
Estimated allele frequency
Ecological gradient (Leaf Unfolding differences)
1.0
0.8
0.6
0.4
0.2
−1
0
1
2
3
4
L16
219
253
L12
256
O16
O8
O12
218
L8
233
9
97
124
O1

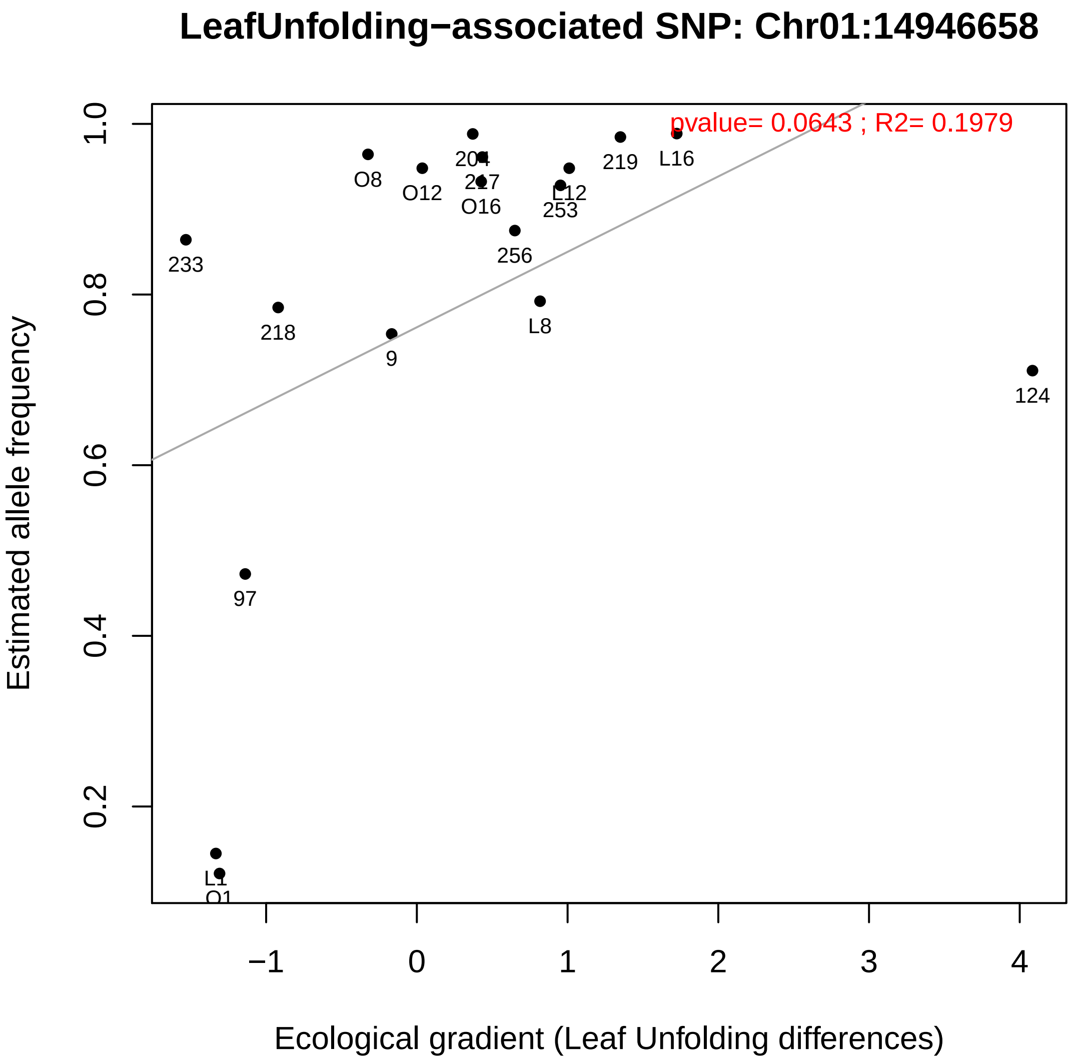
LeafUnfolding−associated SNP: Chr01:14946658
pvalue= 0.0643 ; R2= 0.1979
Estimated allele frequency
Ecological gradient (Leaf Unfolding differences)
0.2
0.4
0.6
0.8
1.0
−1
0
1
2
3
4
233
O8
O12
204
217
O16
256
L12
253
219
L16
218
9
L8
124
97
L1
O1

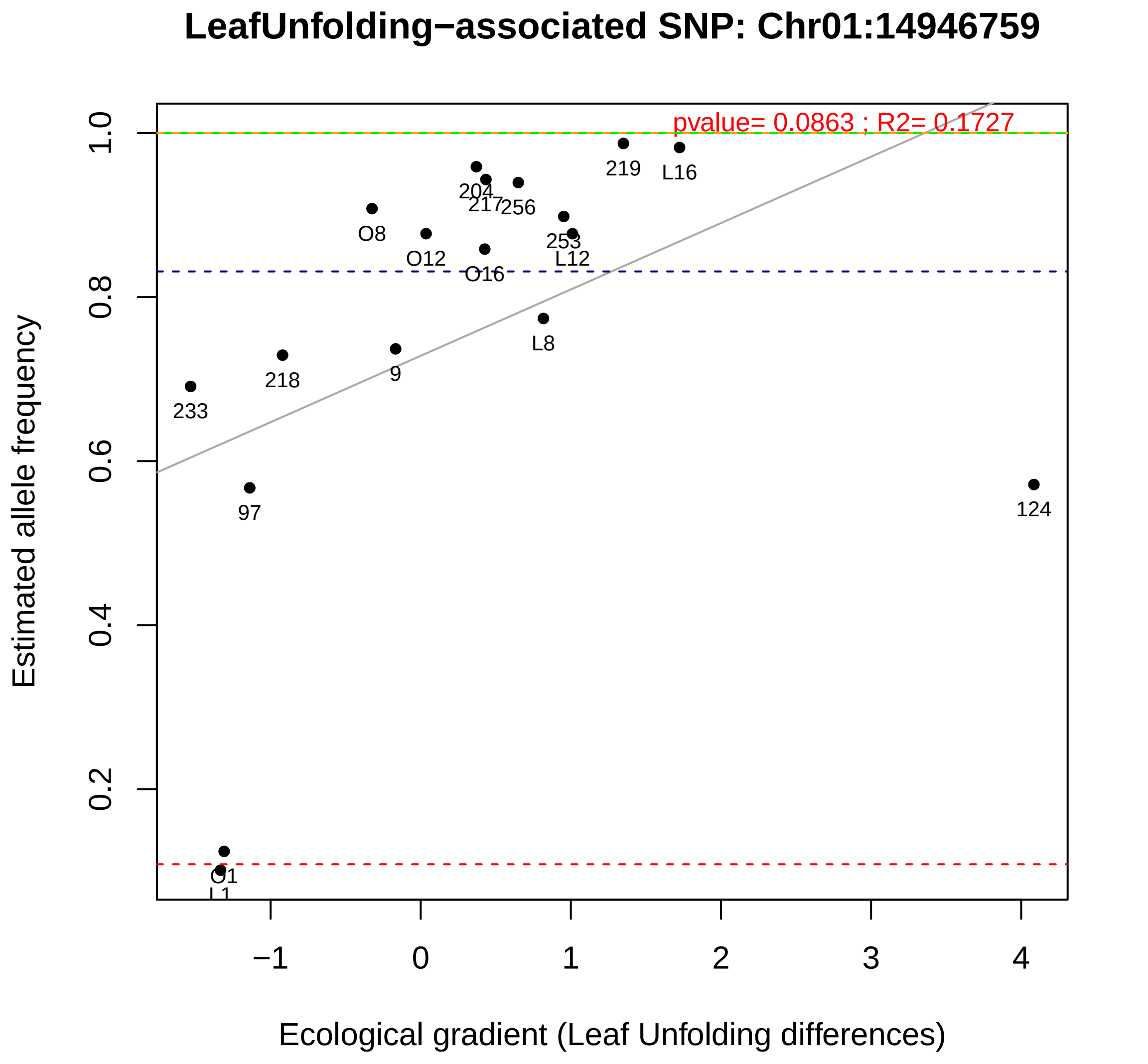
LeafUnfolding−associated SNP: Chr01:14946759
pvalue= 0.0863 ; R2= 0.1727
Estimated allele frequency
0.2
0.4
0.6
0.8
1.0
Ecological gradient (Leaf Unfolding differences)
−1
0
1
2
3
4
204
217
256
219
L16
O8
O12
O16
253
L12
L8
9
218
233
97
124
O1
L1

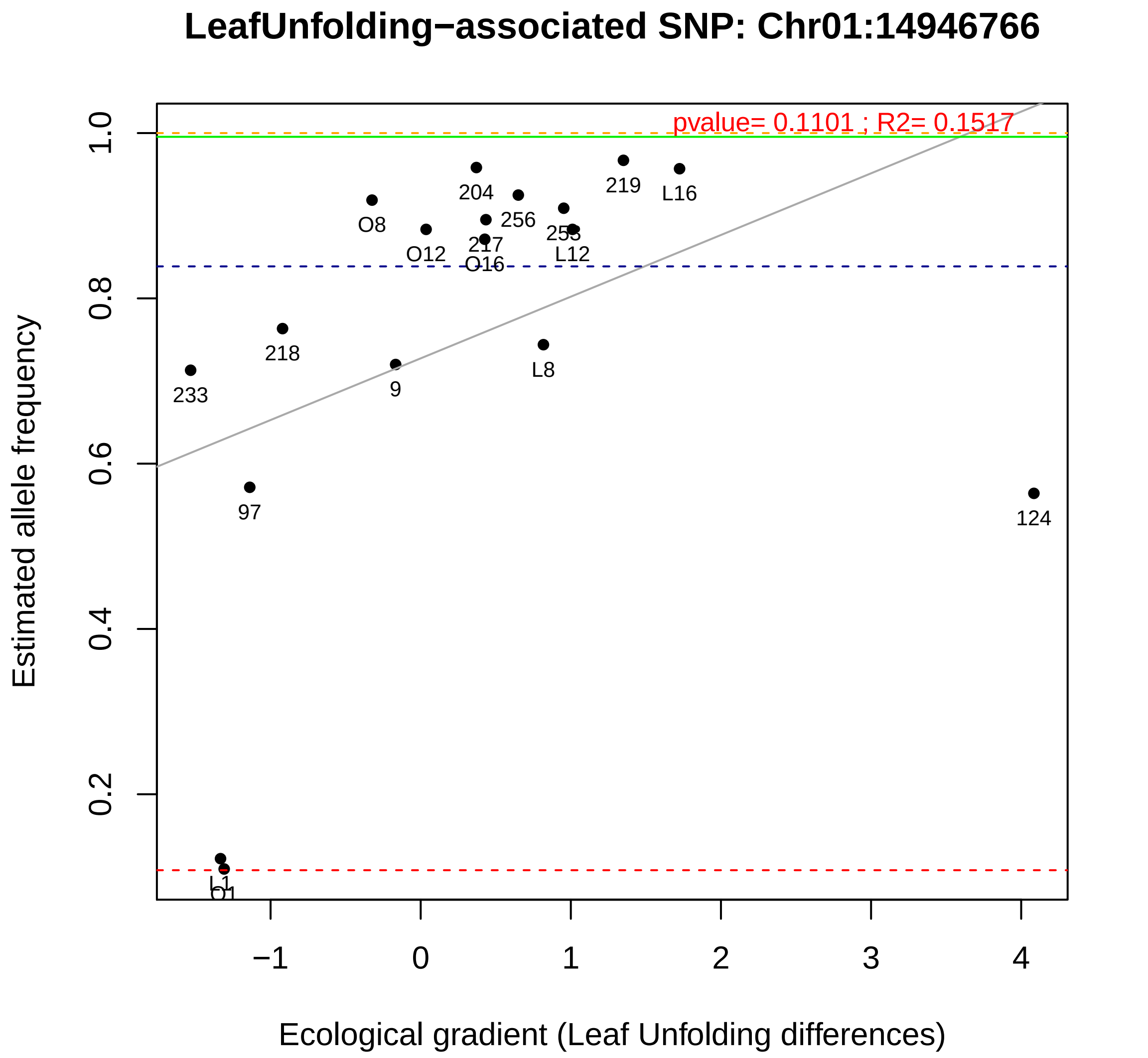
LeafUnfolding−associated SNP: Chr01:14946766
pvalue= 0.1101 ; R2= 0.1517
Estimated allele frequency
0.2
0.4
0.6
0.8
1.0
−1
0
1
2
3
4
Ecological gradient (Leaf Unfolding differences)
204
219
L16
O8
256
O12
217
O16
253
L12
218
233
9
L8
97
124
L1
O1

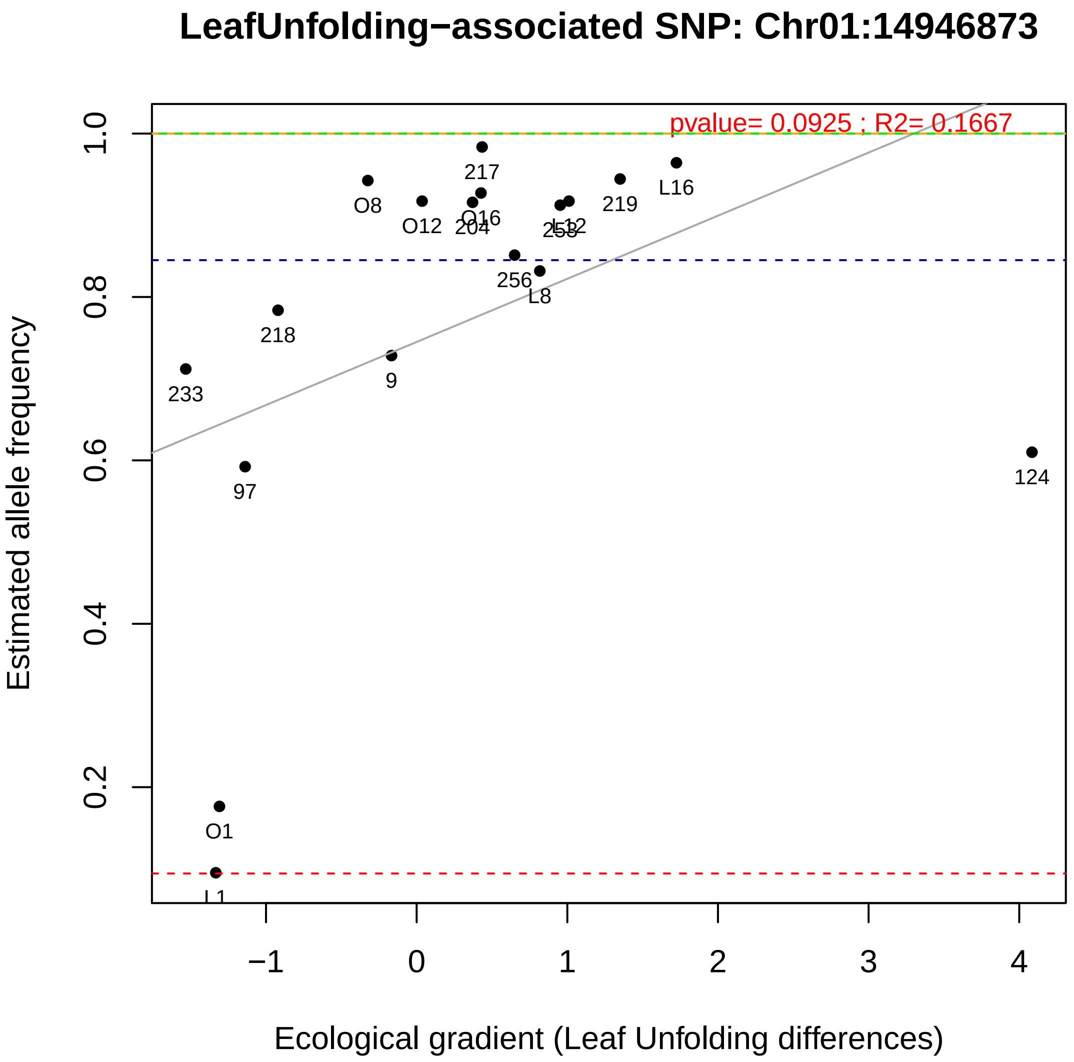

LeafUnfolding−associated SNP: Chr01:14946873
pvalue= 0.0925 ; R2= 0.1667
Estimated allele frequency
Ecological gradient (Leaf Unfolding differences)
1.0
0.8
0.6
0.4
0.2
−1
0
1
2
3
4
217
O8
O12
O16
204
L12
253
219
L16
256
L8
218
233
9
97
124
O1
L1

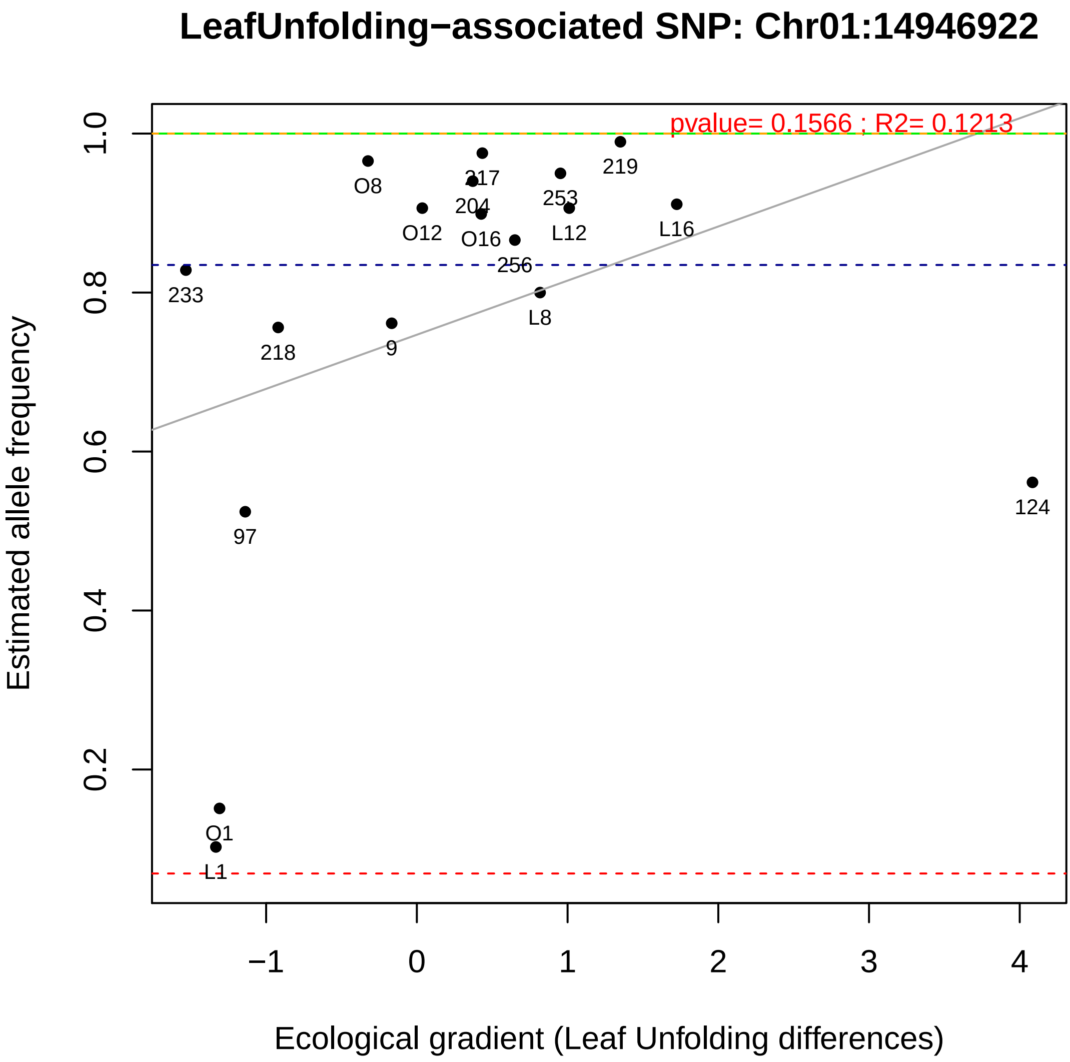
LeafUnfolding−associated SNP: Chr01:14946922
pvalue= 0.1566 ; R2= 0.1213
Estimated allele frequency
0.2
0.4
0.6
0.8
1.0
Ecological gradient (Leaf Unfolding differences)
−1
0
1
2
3
4
O8
217
204
O12
O16
256
253
L12
219
L16
233
218
9
L8
97
124
O1
L1

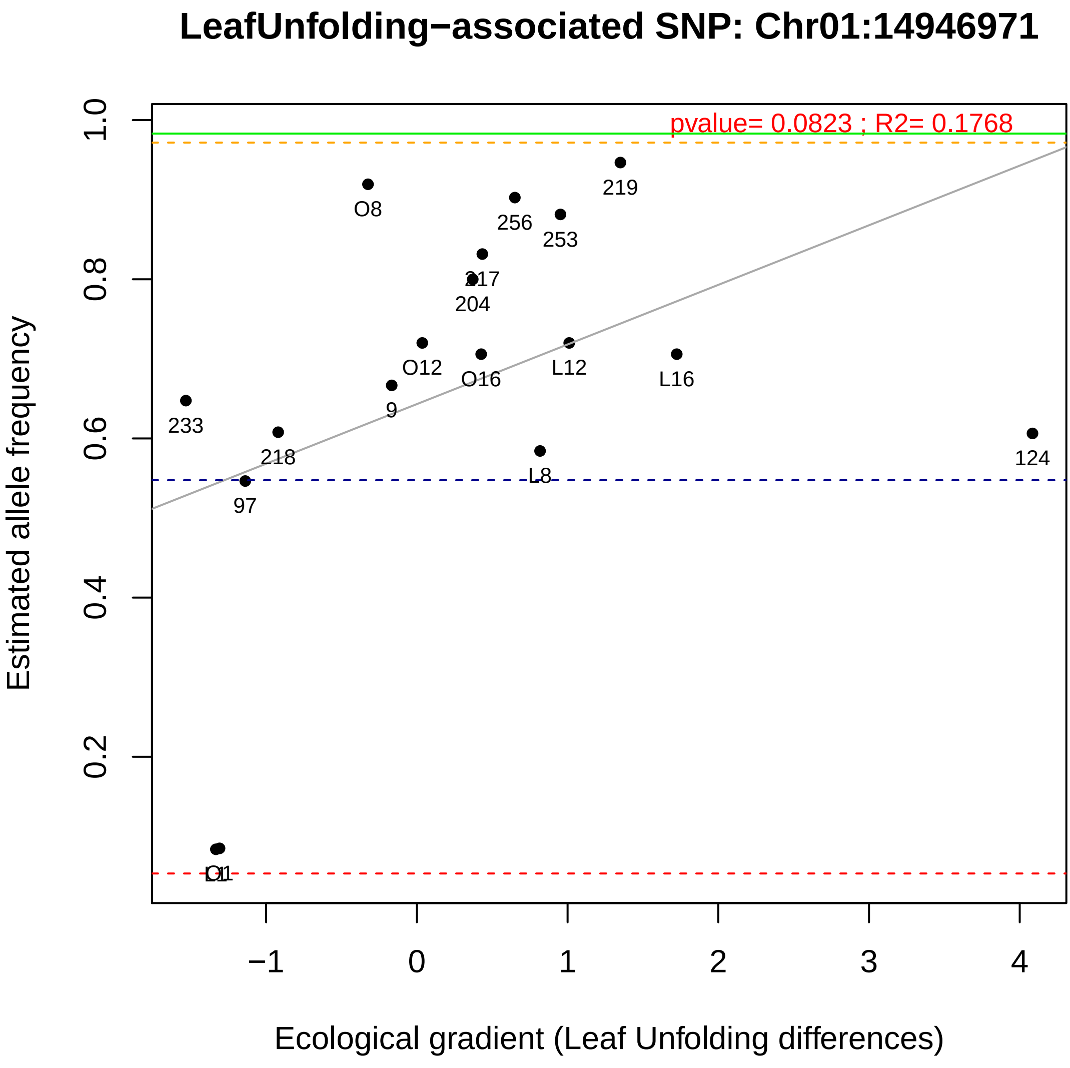

LeafUnfolding−associated SNP: Chr01:14946971
pvalue= 0.0823 ; R2= 0.1768
Estimated allele frequency
Ecological gradient (Leaf Unfolding differences)
0.2
0.4
0.6
0.8
1.0
−1
0
1
2
3
4
O8
256
253
219
217
204
O12
O16
L12
L16
9
233
218
97
L8
124

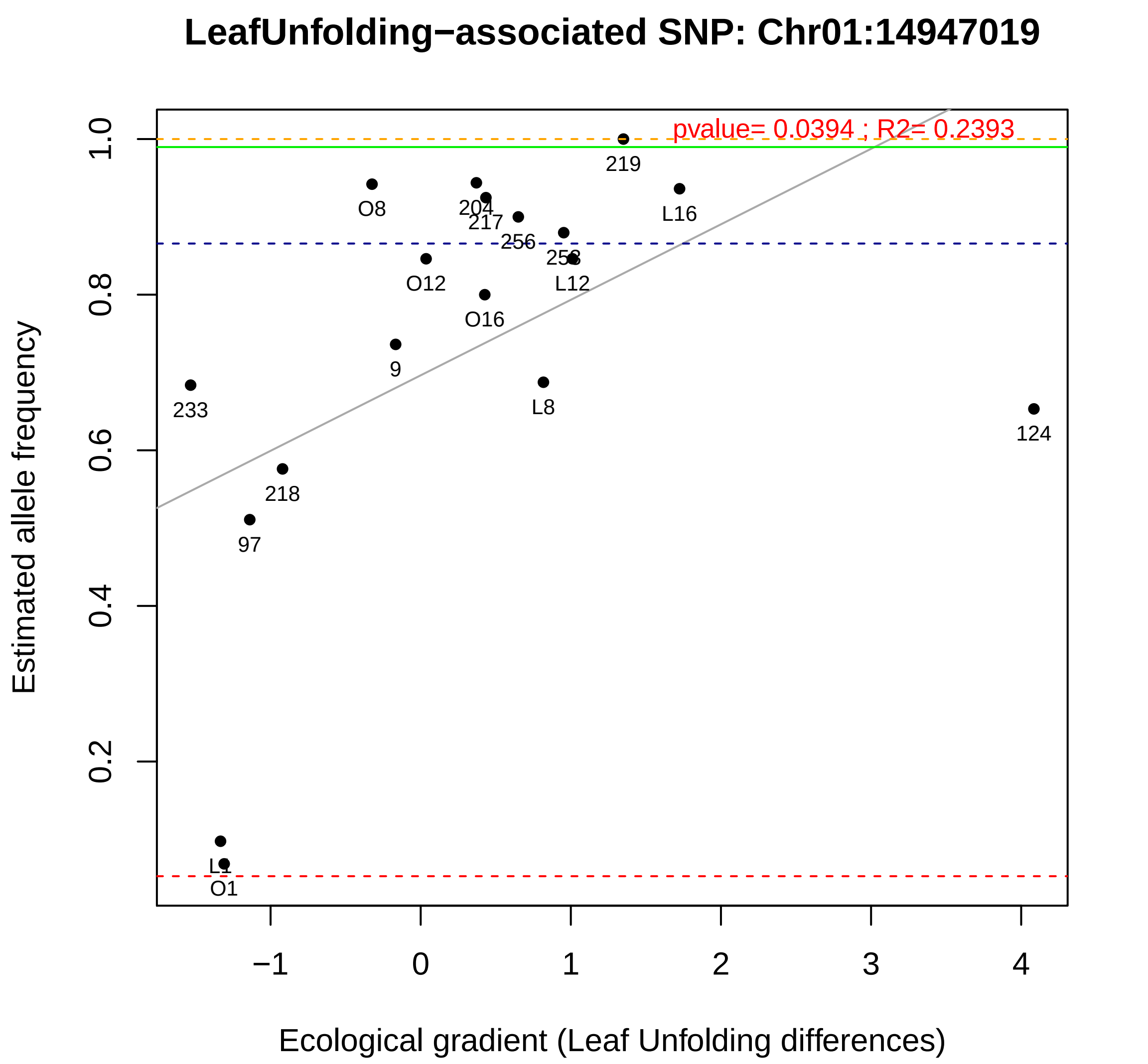
LeafUnfolding−associated SNP: Chr01:14947019
pvalue= 0.0394 ; R2= 0.2393
Estimated allele frequency
Ecological gradient (Leaf Unfolding differences)
0.2
0.4
0.6
0.8
1.0
−1
0
1
2
3
4
219
L16
O8
204
217
256
O12
L12
O16
9
L8
233
124
218
97
L1
O1

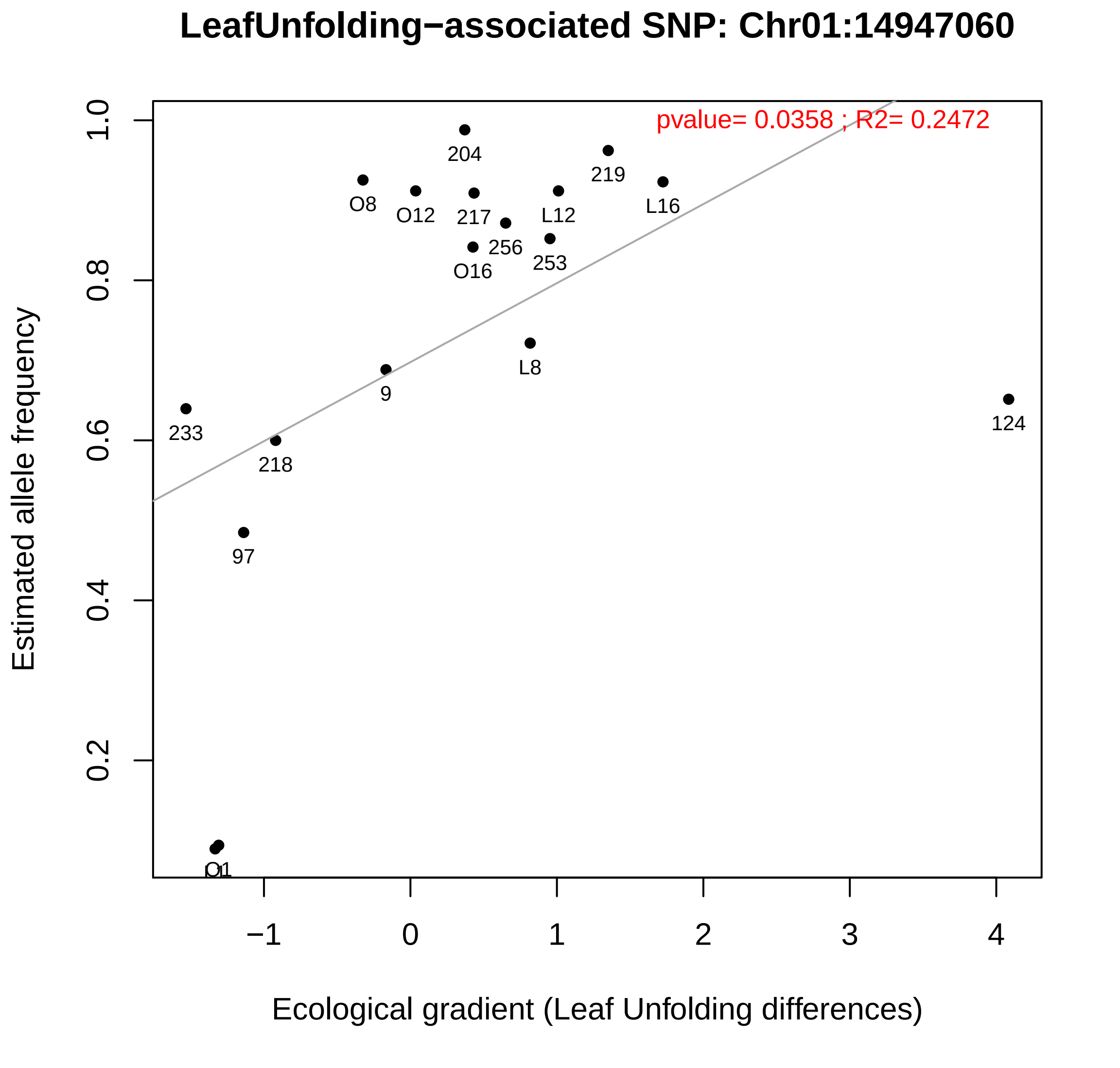
LeafUnfolding−associated SNP: Chr01:14947060
pvalue= 0.0358 ; R2= 0.2472
Estimated allele frequency
Ecological gradient (Leaf Unfolding differences)
1.0
0.8
0.6
0.4
0.2
−1
0
1
2
3
4
204
219
O8
O12
217
L12
L16
256
253
O16
L8
9
233
218
124
97
O1

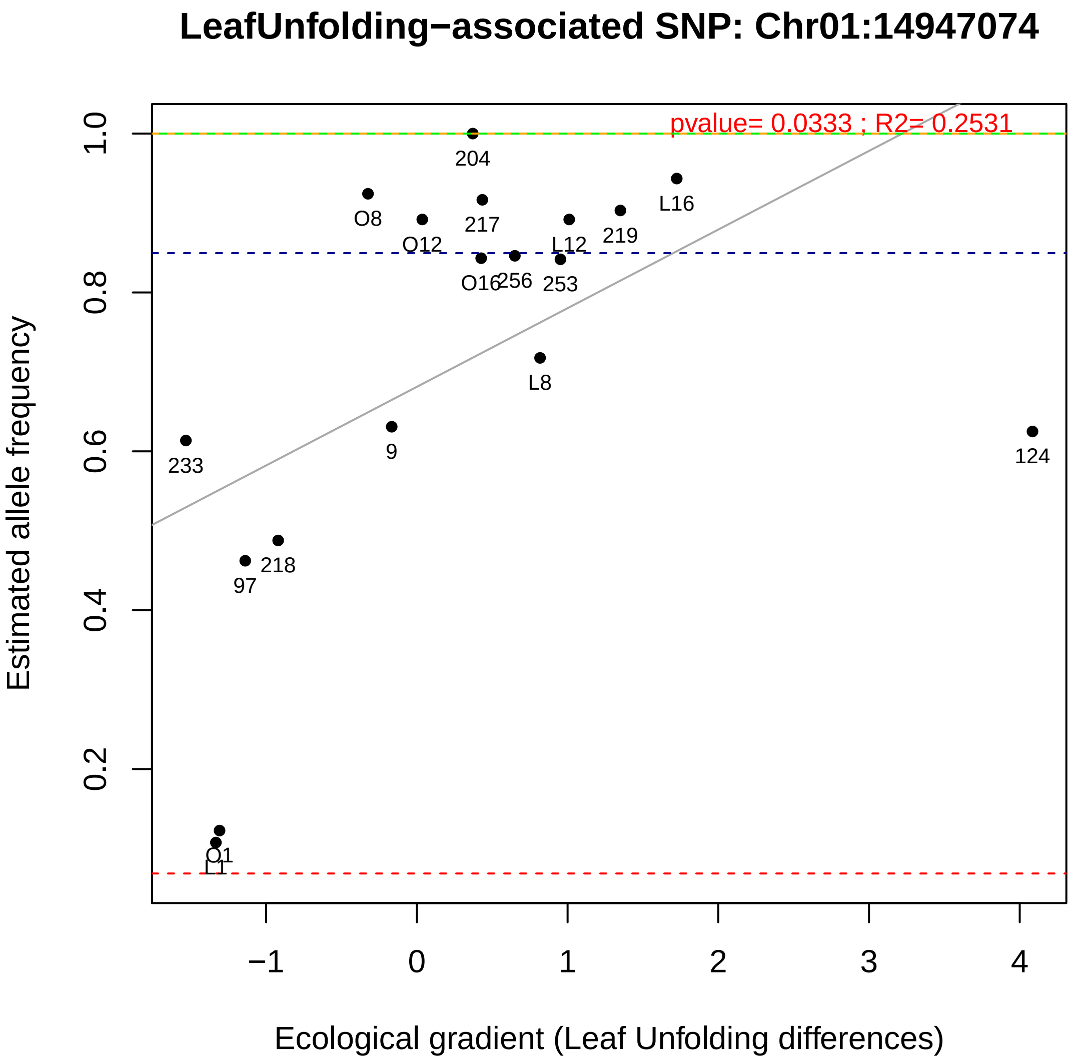

LeafUnfolding−associated SNP: Chr01:14947074
pvalue= 0.0333 ; R2= 0.2531
Estimated allele frequency
Ecological gradient (Leaf Unfolding differences)
1.0
0.8
0.6
0.4
0.2
−1
0
1
2
3
4
204
O8
217
O12
L16
219
L12
O16
256
253
L8
233
9
124
218
97
O1
L1

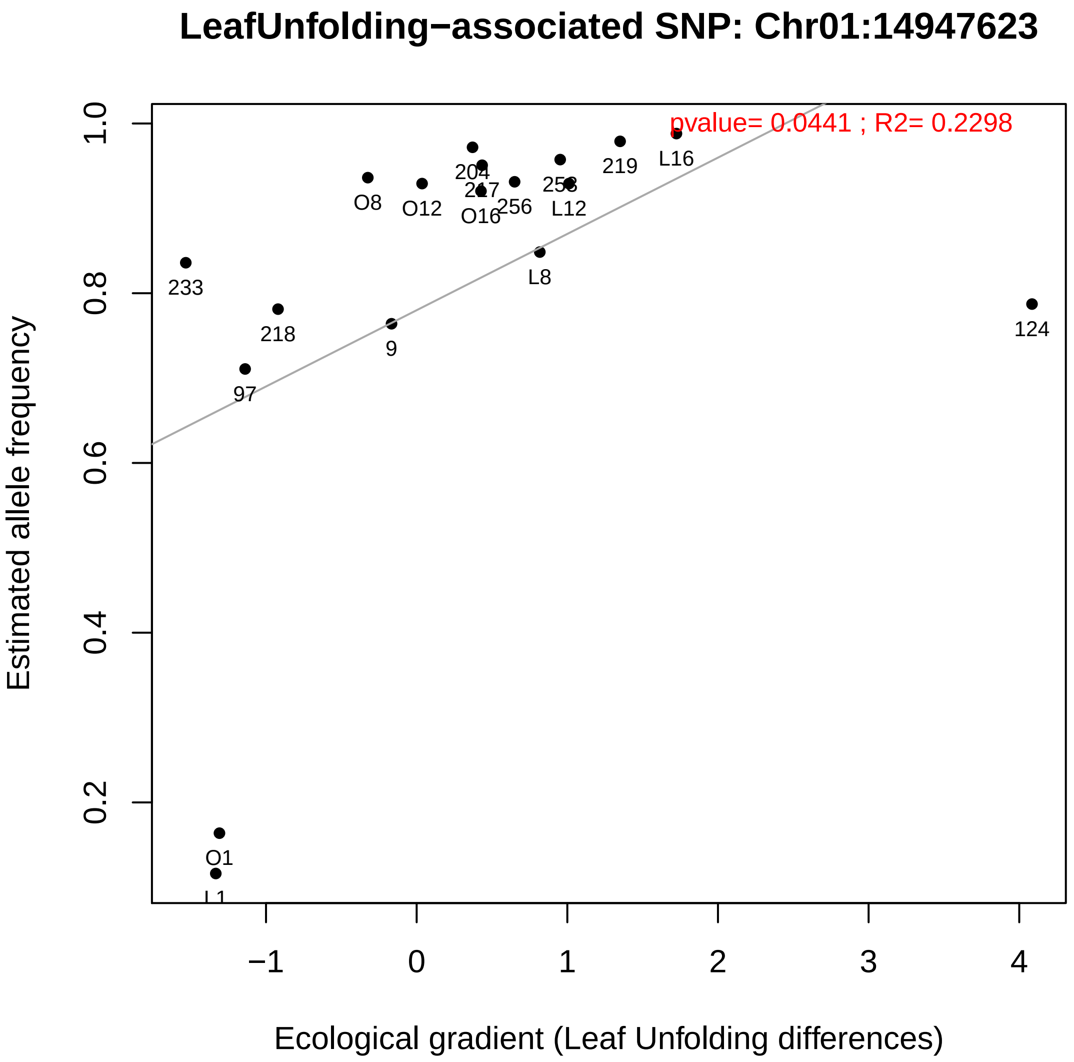
LeafUnfolding−associated SNP: Chr01:14947623
pvalue= 0.0441 ; R2= 0.2298
Estimated allele frequency
Ecological gradient (Leaf Unfolding differences)
0.2
0.4
0.6
0.8
1.0
−1
0
1
2
3
4
233
218
97
9
O8
O12
204
217
O16
256
253
L12
L8
219
L16
124
O1

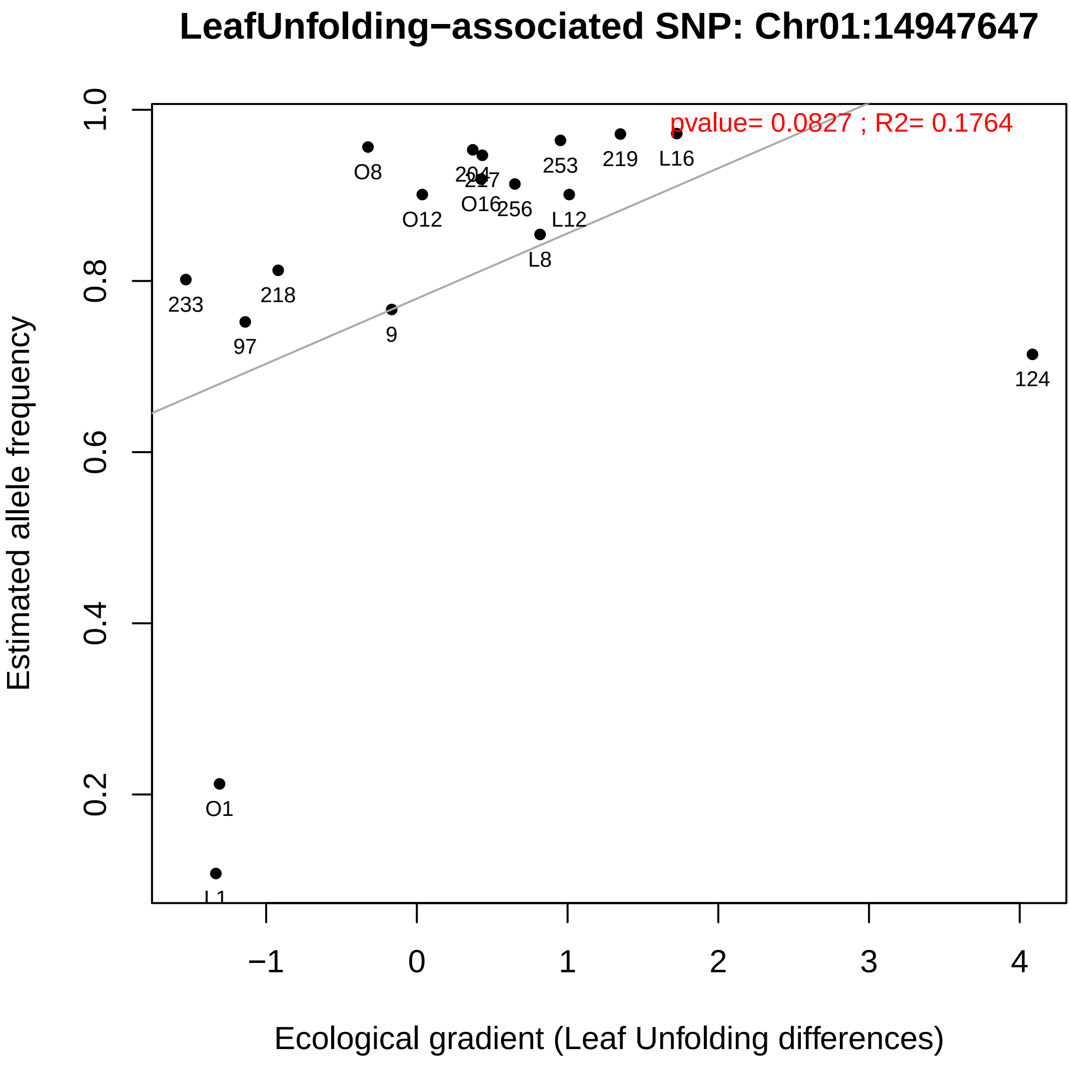

LeafUnfolding−associated SNP: Chr01:14947647
pvalue= 0.0827 ; R2= 0.1764
Estimated allele frequency
Ecological gradient (Leaf Unfolding differences)
1.0
0.8
0.6
0.4
0.2
−1
0
1
2
3
4
O8
204
217
O16
256
253
219
L16
O12
L12
L8
233
218
97
9
124
O1

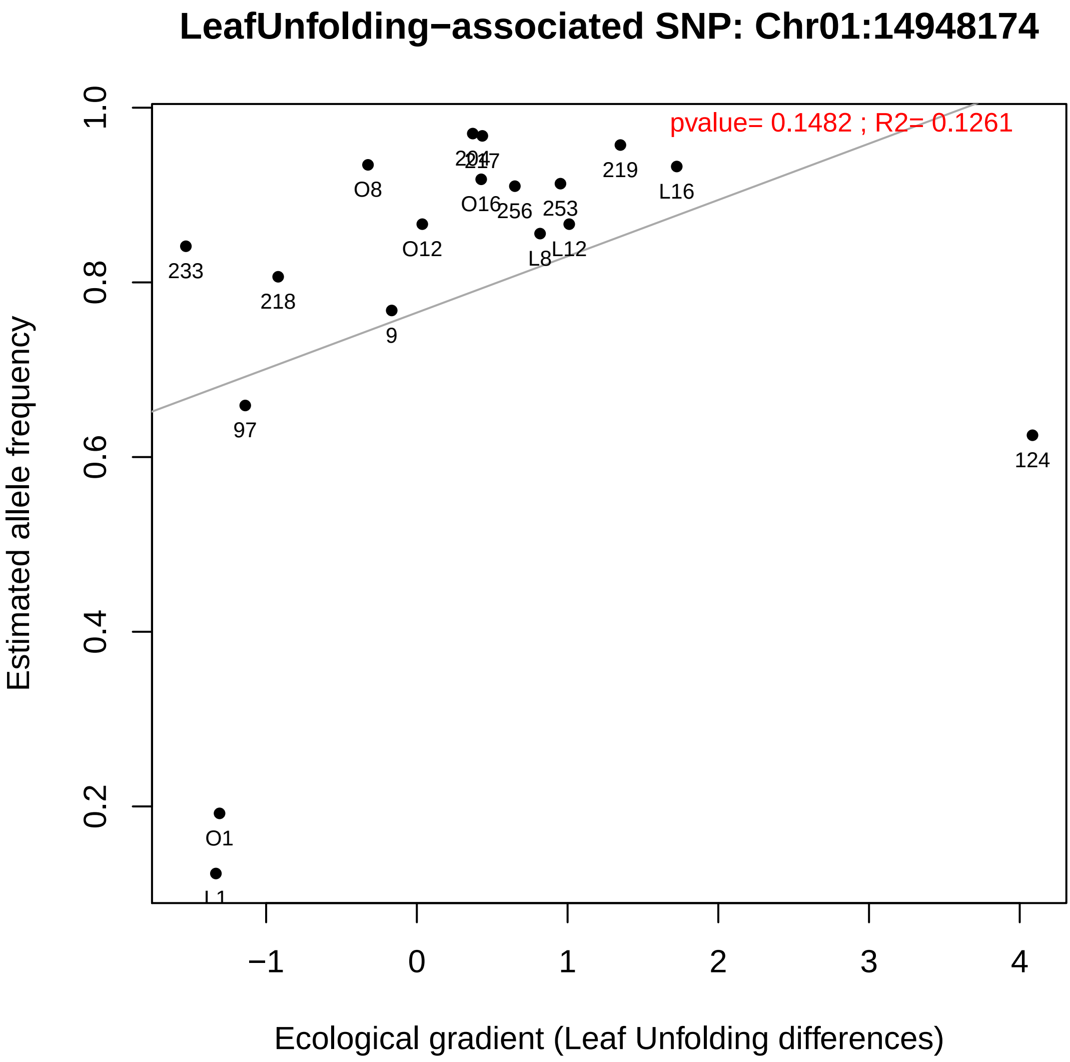
LeafUnfolding−associated SNP: Chr01:14948174
pvalue= 0.1482 ; R2= 0.1261
Estimated allele frequency
Ecological gradient (Leaf Unfolding differences)
0.2
0.4
0.6
0.8
1.0
−1
0
1
2
3
4
233
218
97
O1
O8
9
O12
204
217
O16
256
253
L8
L12
219
L16
124

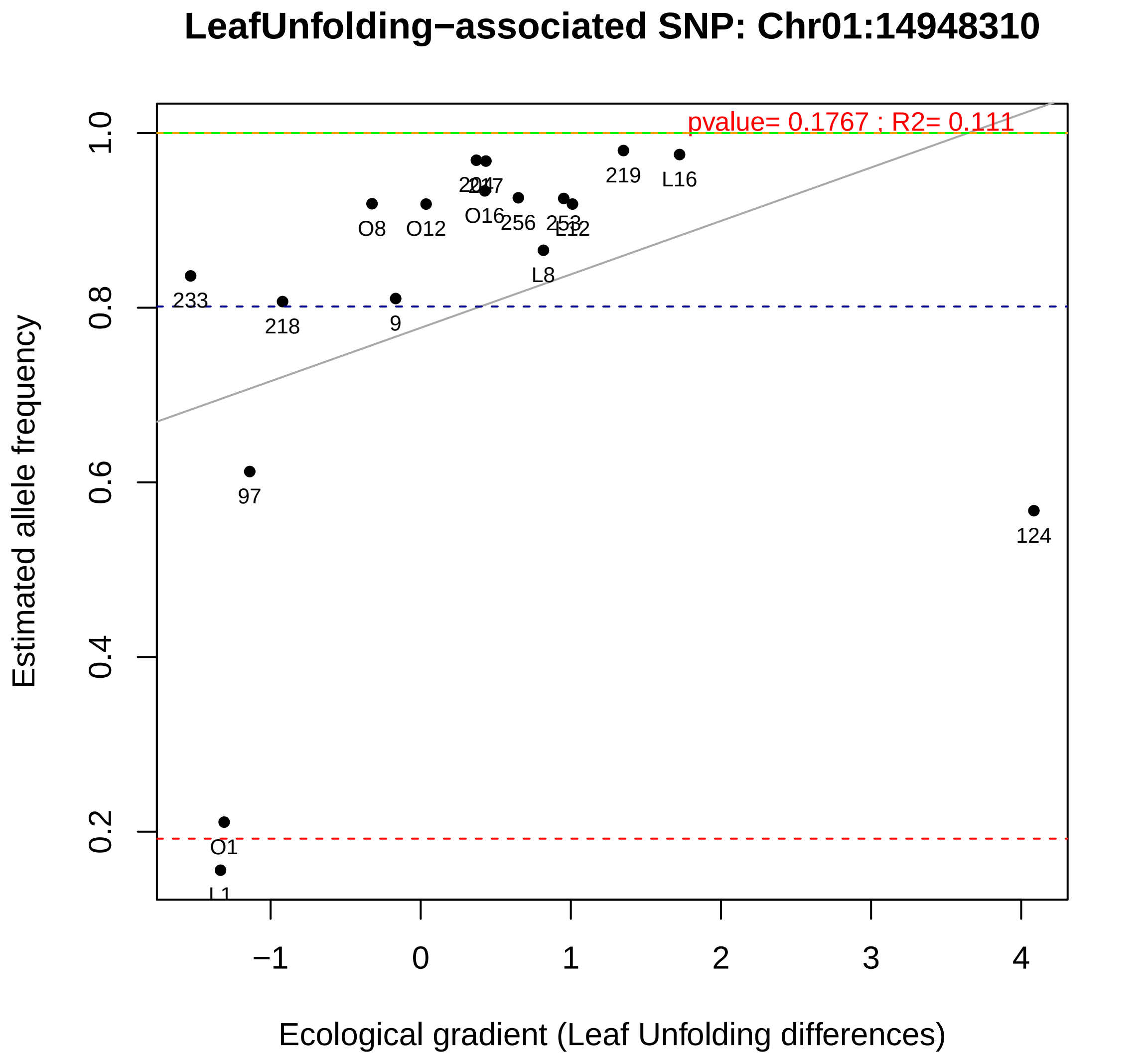
LeafUnfolding−associated SNP: Chr01:14948310
pvalue= 0.1767 ; R2= 0.111
Estimated allele frequency
Ecological gradient (Leaf Unfolding differences)
0.2
0.4
0.6
0.8
1.0
−1
0
1
2
3
4
233
218
9
O8
O12
O16
256
219
L16
L8
97
124
O1

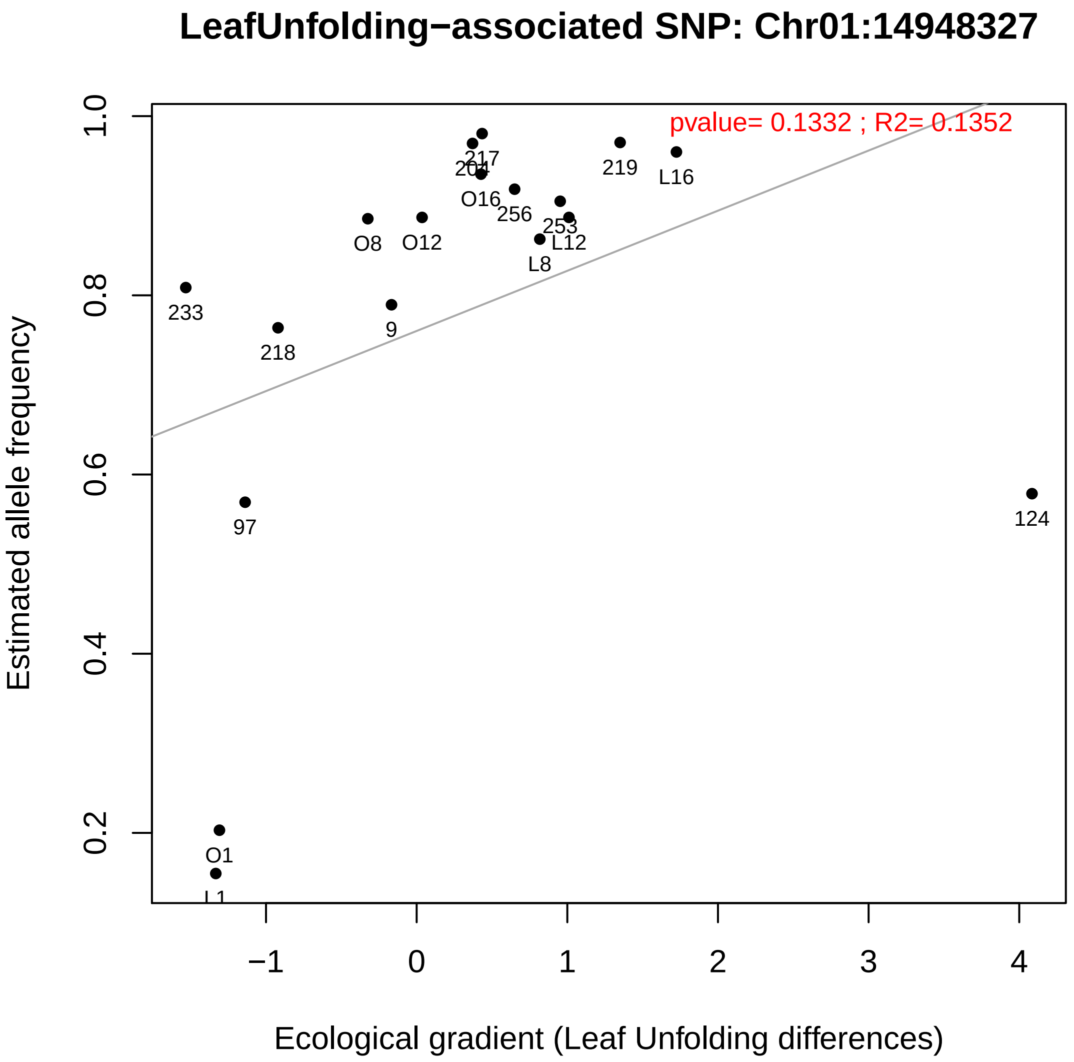
LeafUnfolding−associated SNP: Chr01:14948327
pvalue= 0.1332 ; R2= 0.1352
Estimated allele frequency
Ecological gradient (Leaf Unfolding differences)
0.2
0.4
0.6
0.8
1.0
−1
0
1
2
3
4
217
204
219
L16
O16
256
253
L12
L8
O8
O12
233
218
9
97
124
O1
L1

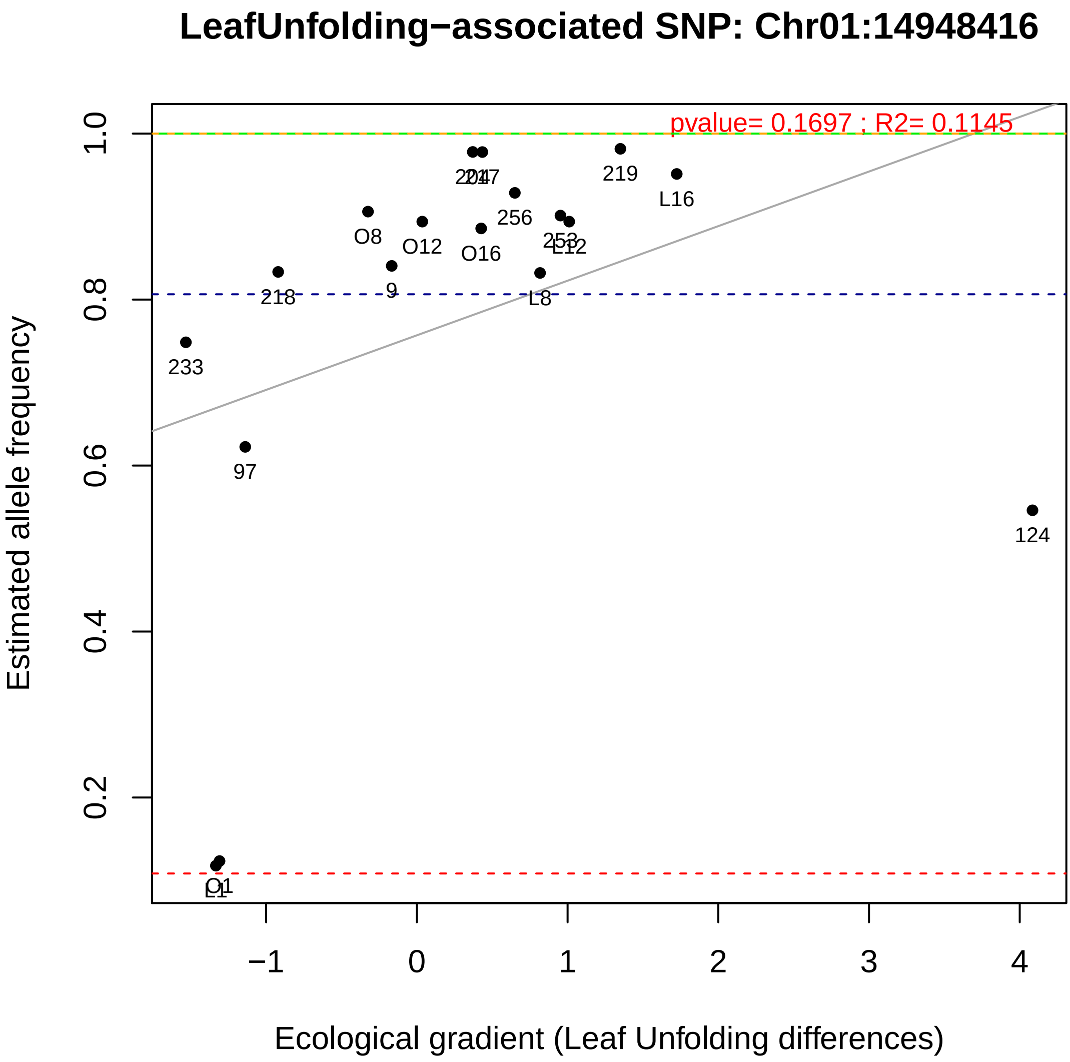

LeafUnfolding−associated SNP: Chr01:14948416
pvalue= 0.1697 ; R2= 0.1145
Estimated allele frequency
Ecological gradient (Leaf Unfolding differences)
0.2
0.4
0.6
0.8
1.0
−1
0
1
2
3
4
219
L16
256
O8
O12
O16
9
218
L8
233
97
124

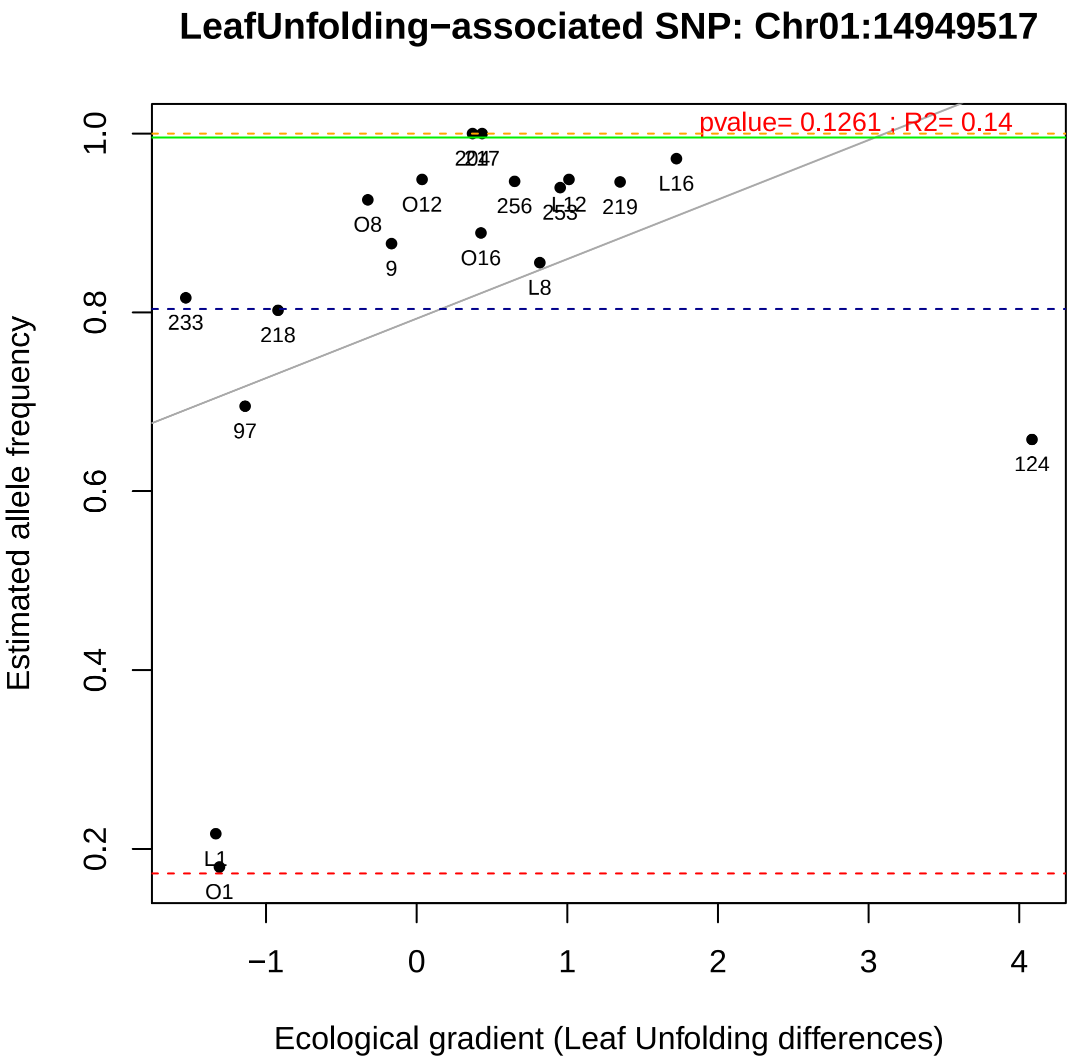

LeafUnfolding−associated SNP: Chr01:14949517
pvalue= 0.1261 ; R2= 0.14
Estimated allele frequency
Ecological gradient (Leaf Unfolding differences)
0.2
0.4
0.6
0.8
1.0
−1
0
1
2
3
4
233
218
97
L1
O1
O8
9
O12
O16
256
L8
253
L12
219
L16
124

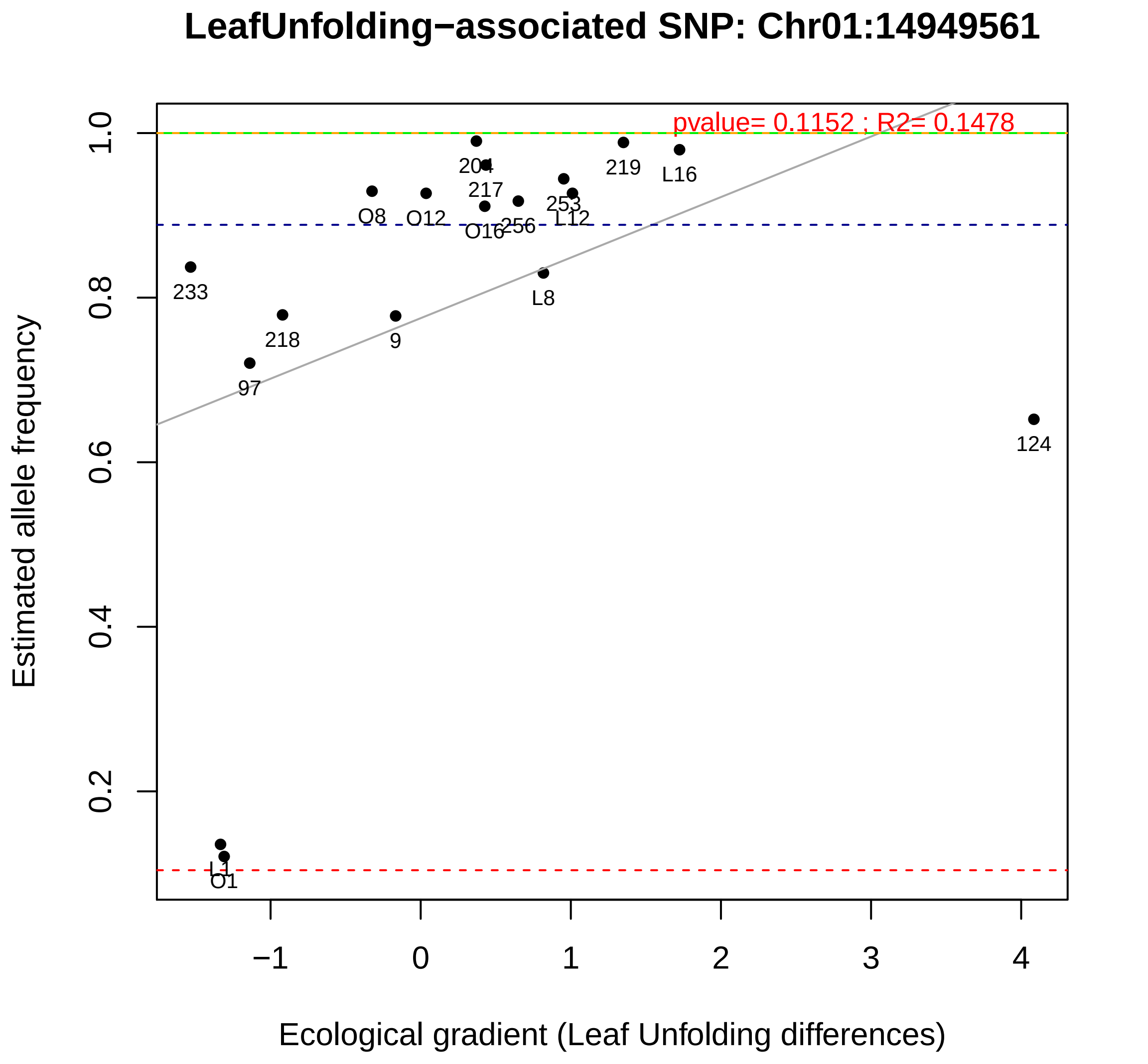
LeafUnfolding−associated SNP: Chr01:14949561
pvalue= 0.1152 ; R2= 0.1478
Estimated allele frequency
0.2
0.4
0.6
0.8
1.0
−1
0
1
2
3
4
Ecological gradient (Leaf Unfolding differences)
233
218
97
9
O8
O12
204
217
O16
256
253
L12
L8
219
L16
124
L1
O1

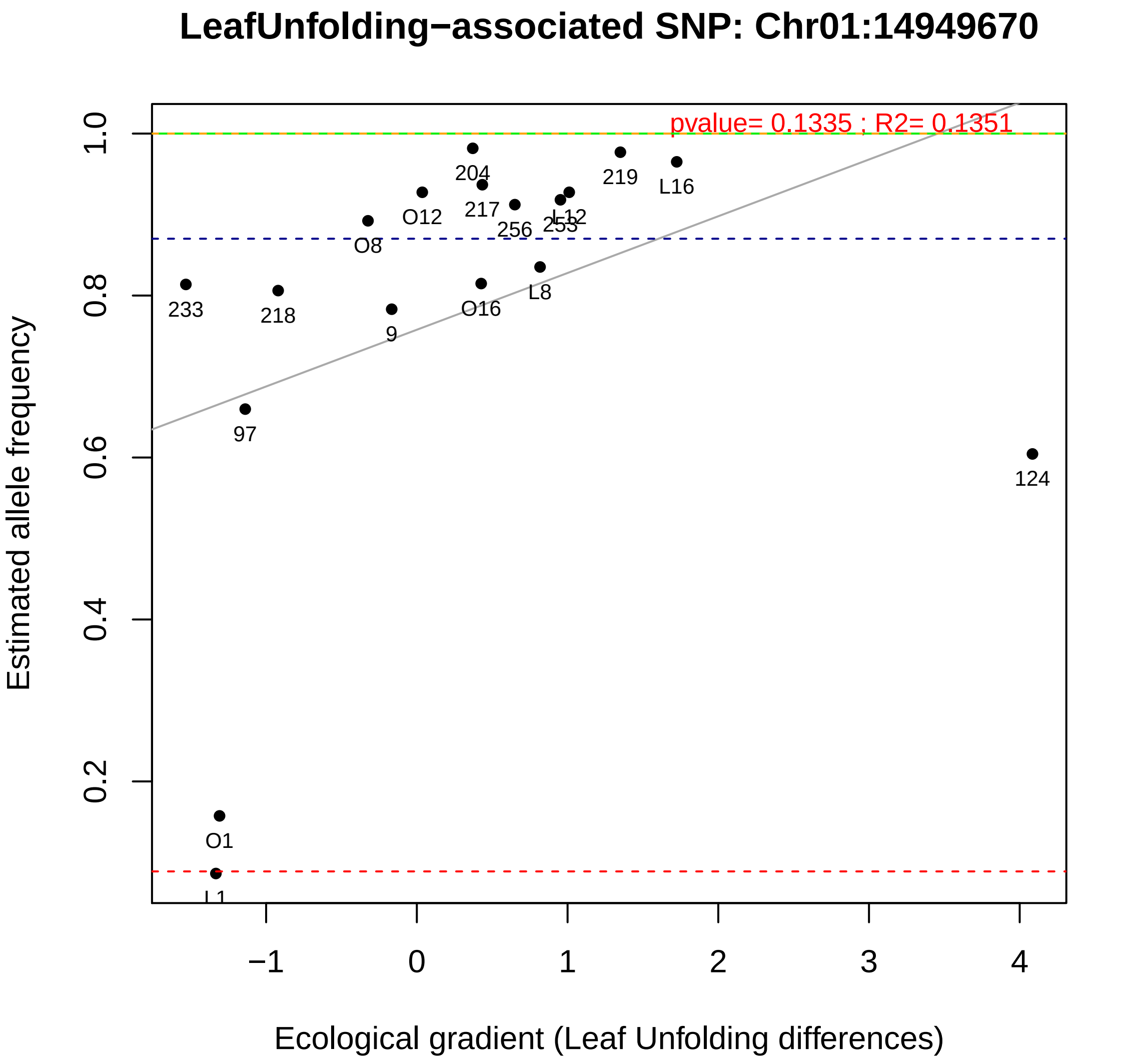
LeafUnfolding−associated SNP: Chr01:14949670
pvalue= 0.1335 ; R2= 0.1351
Estimated allele frequency
Ecological gradient (Leaf Unfolding differences)
1.0
0.8
0.6
0.4
0.2
−1
0
1
2
3
4
204
219
L16
217
O12
256
253
L12
O8
233
218
9
O16
L8
97
124
O1
L1

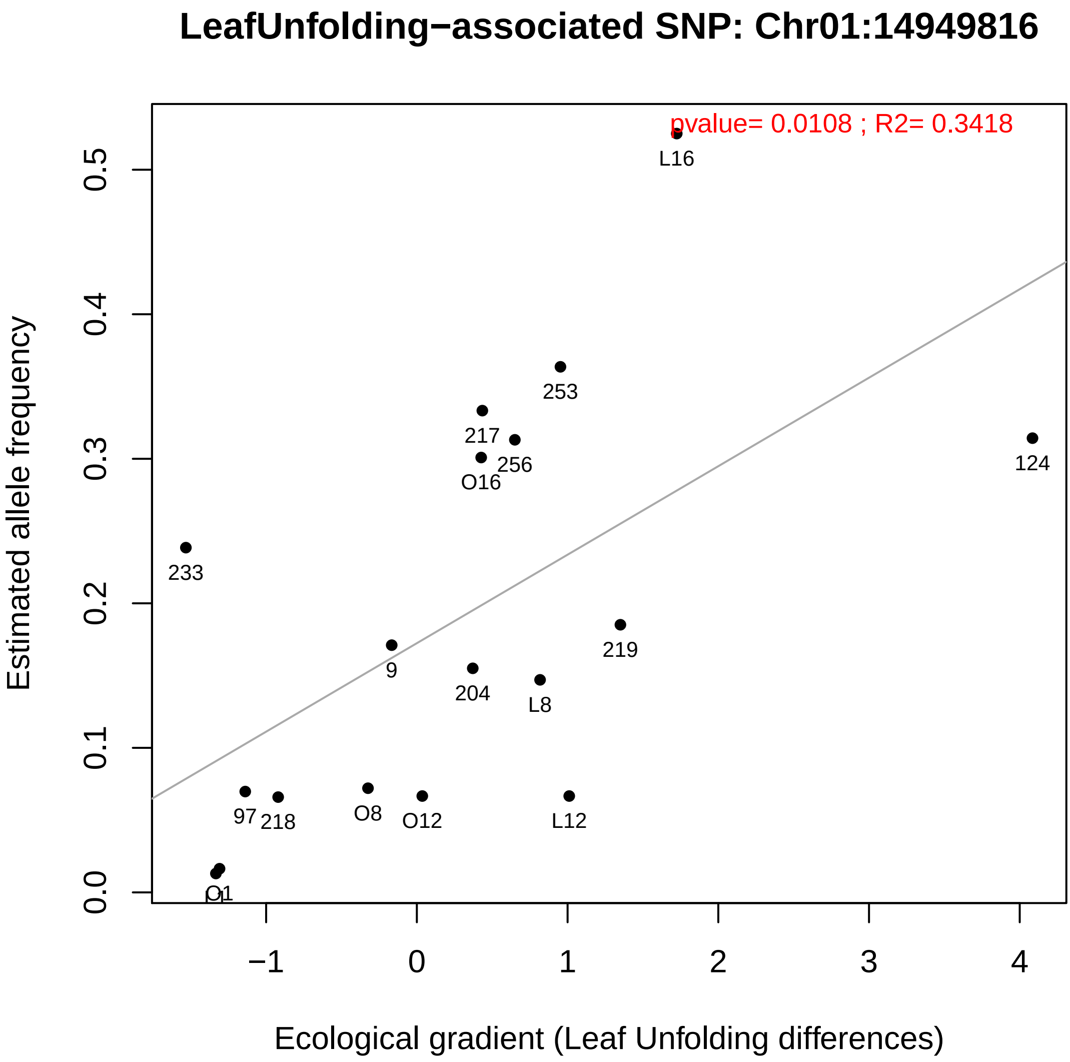

LeafUnfolding−associated SNP: Chr01:14949816
pvalue= 0.0108 ; R2= 0.3418
Estimated allele frequency
Ecological gradient (Leaf Unfolding differences)
0.0
0.1
0.2
0.3
0.4
0.5
−1
0
1
2
3
4
L16
253
217
256
O16
124
233
9
219
204
L8
97
218
O8
O12
L12
O1

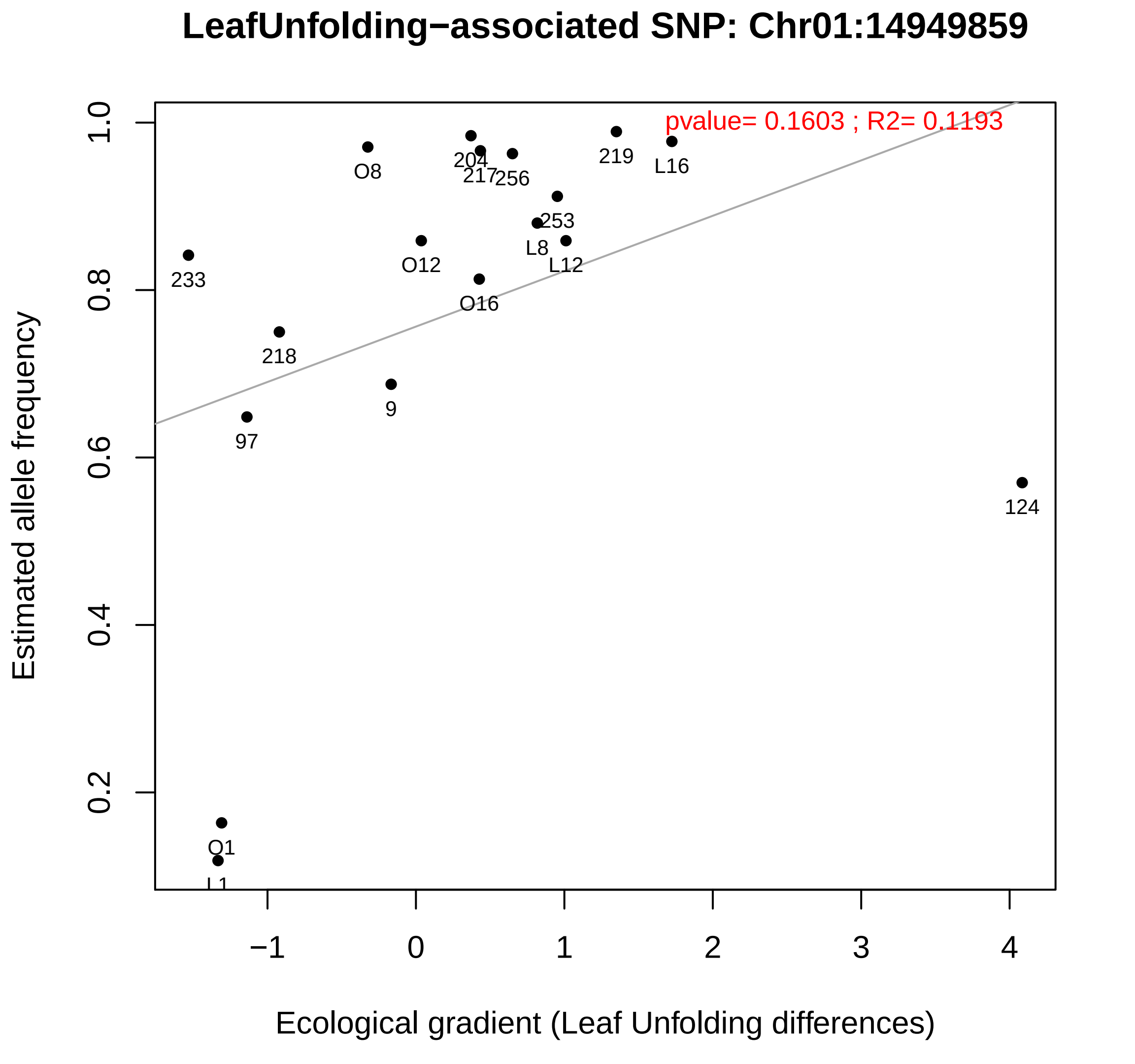

LeafUnfolding−associated SNP: Chr01:14949859
pvalue= 0.1603 ; R2= 0.1193
Estimated allele frequency
Ecological gradient (Leaf Unfolding differences)
1.0
0.8
0.6
0.4
0.2
−1
0
1
2
3
4
233
218
97
O1
O8
O12
9
204
217
256
O16
253
L8
L12
219
L16
124

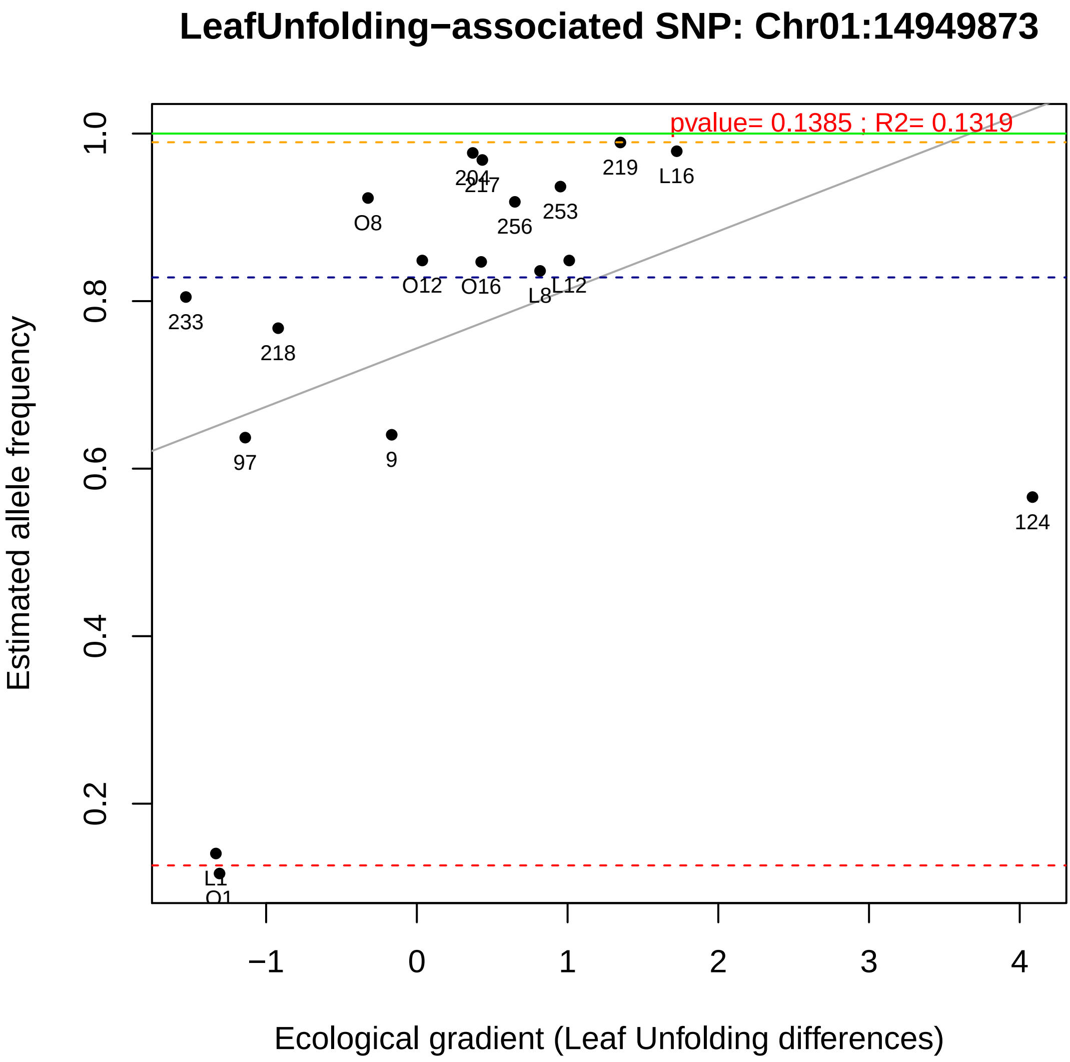

LeafUnfolding−associated SNP: Chr01:14949873
pvalue= 0.1385 ; R2= 0.1319
Estimated allele frequency
Ecological gradient (Leaf Unfolding differences)
0.2
0.4
0.6
0.8
1.0
−1
0
1
2
3
4
219
L16
253
256
O8
O12
O16
L8
L12
233
218
97
9
124
L1
O1

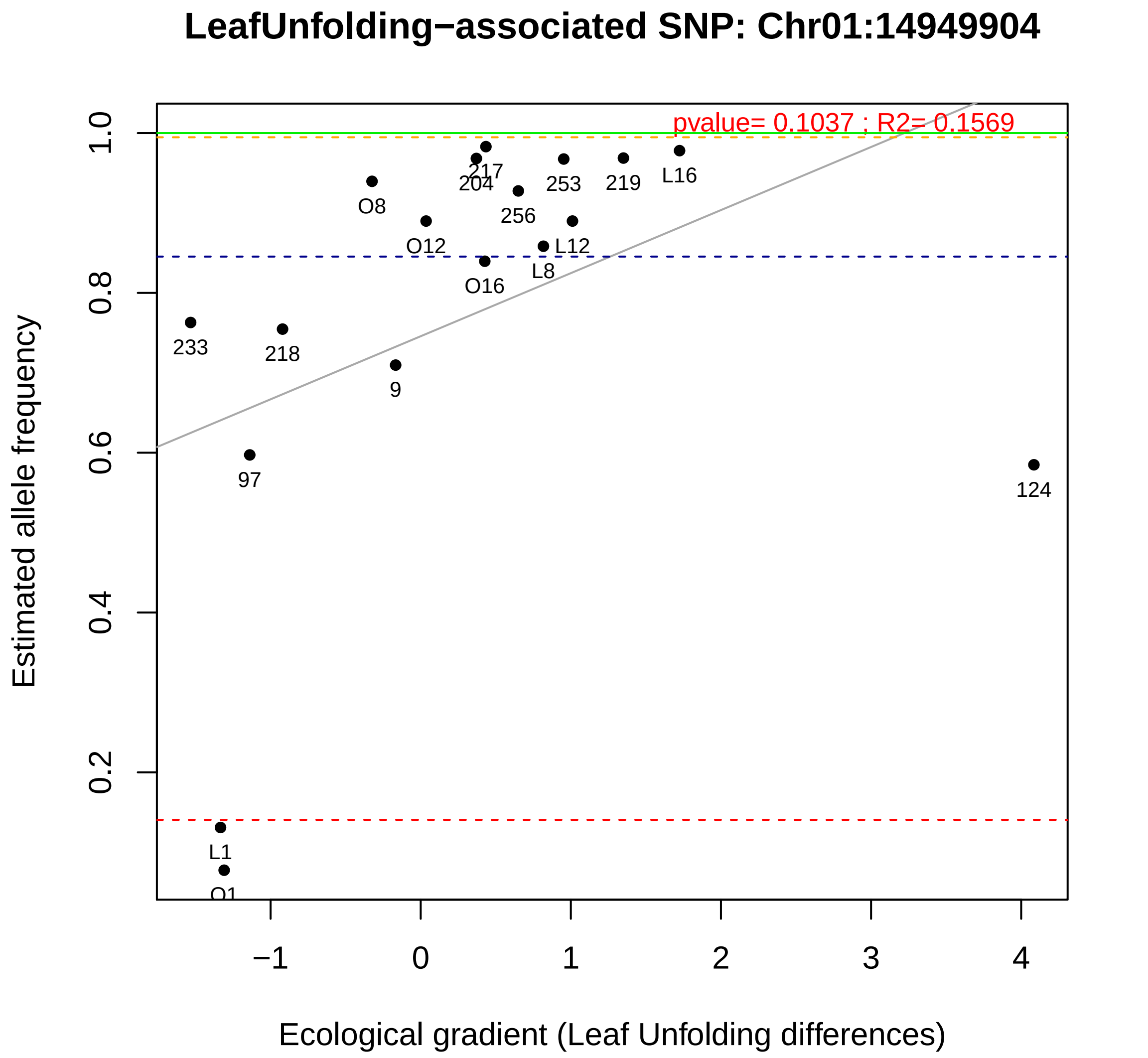

LeafUnfolding−associated SNP: Chr01:14949904
pvalue= 0.1037 ; R2= 0.1569
Estimated allele frequency
Ecological gradient (Leaf Unfolding differences)
0.2
0.4
0.6
0.8
1.0
−1
0
1
2
3
4
217
204
253
219
L16
O8
256
O12
L12
L8
O16
233
218
9
97
124
L1
O1

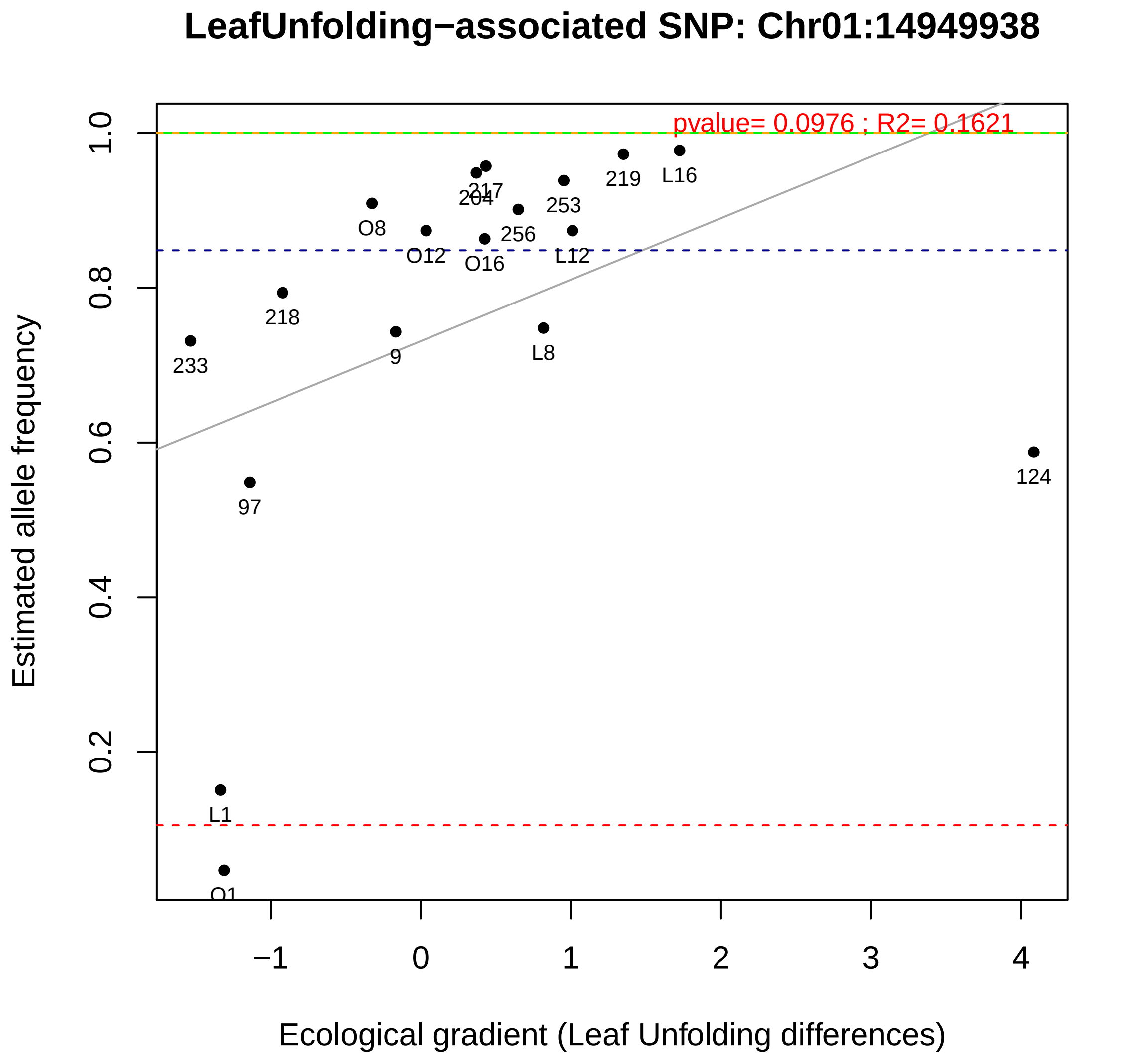
LeafUnfolding−associated SNP: Chr01:14949938
pvalue= 0.0976 ; R2= 0.1621
Estimated allele frequency
Ecological gradient (Leaf Unfolding differences)
1.0
0.8
0.6
0.4
0.2
−1
0
1
2
3
4
219
L16
217
204
253
O8
256
O12
O16
L12
218
233
9
L8
97
124
L1
O1

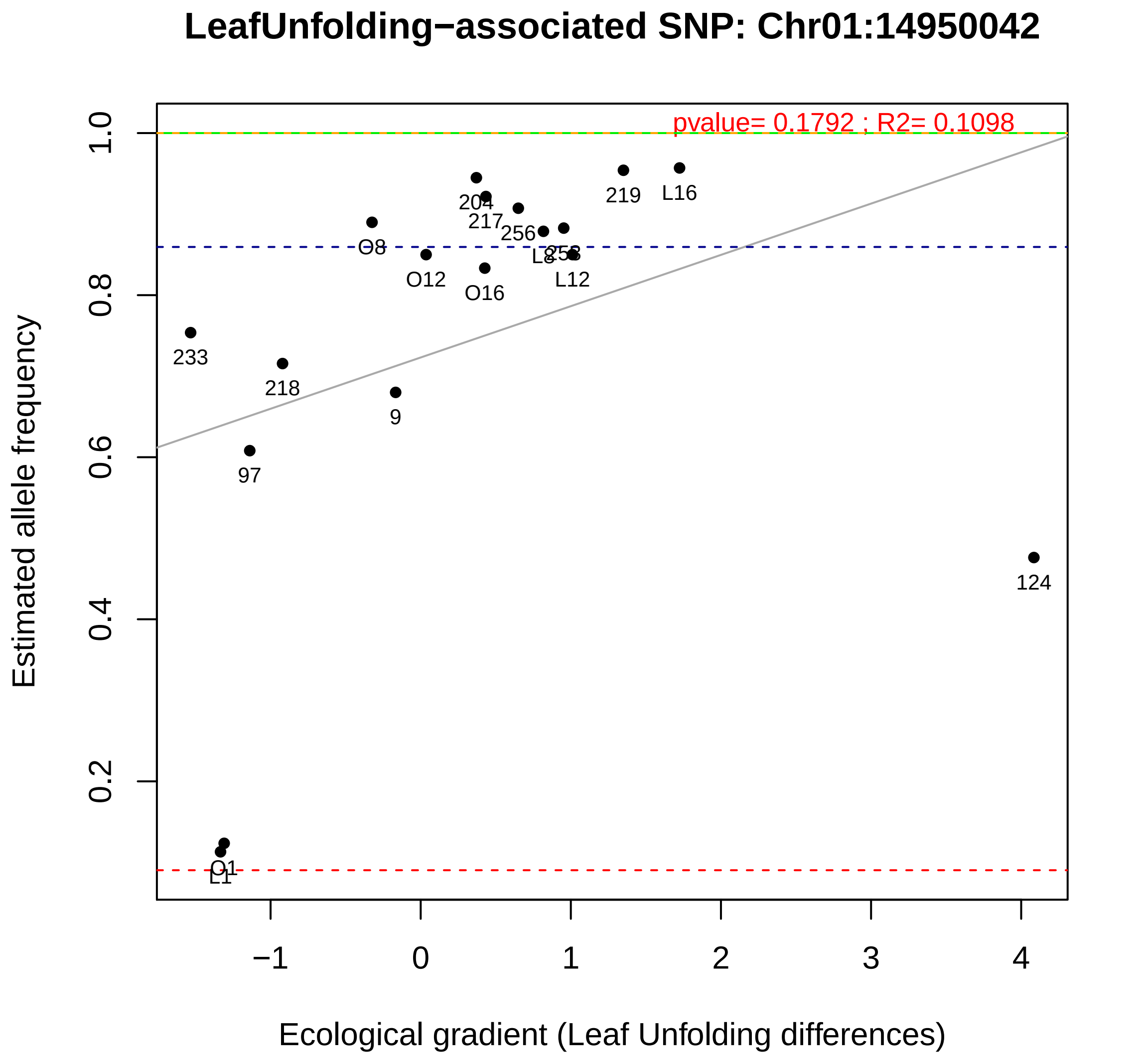
LeafUnfolding−associated SNP: Chr01:14950042
pvalue= 0.1792 ; R2= 0.1098
Estimated allele frequency
Ecological gradient (Leaf Unfolding differences)
1.0
0.8
0.6
0.4
0.2
−1
0
1
2
3
4
204
217
256
219
L16
O8
O12
O16
L8
253
L12
233
218
9
97
124
O1
L1

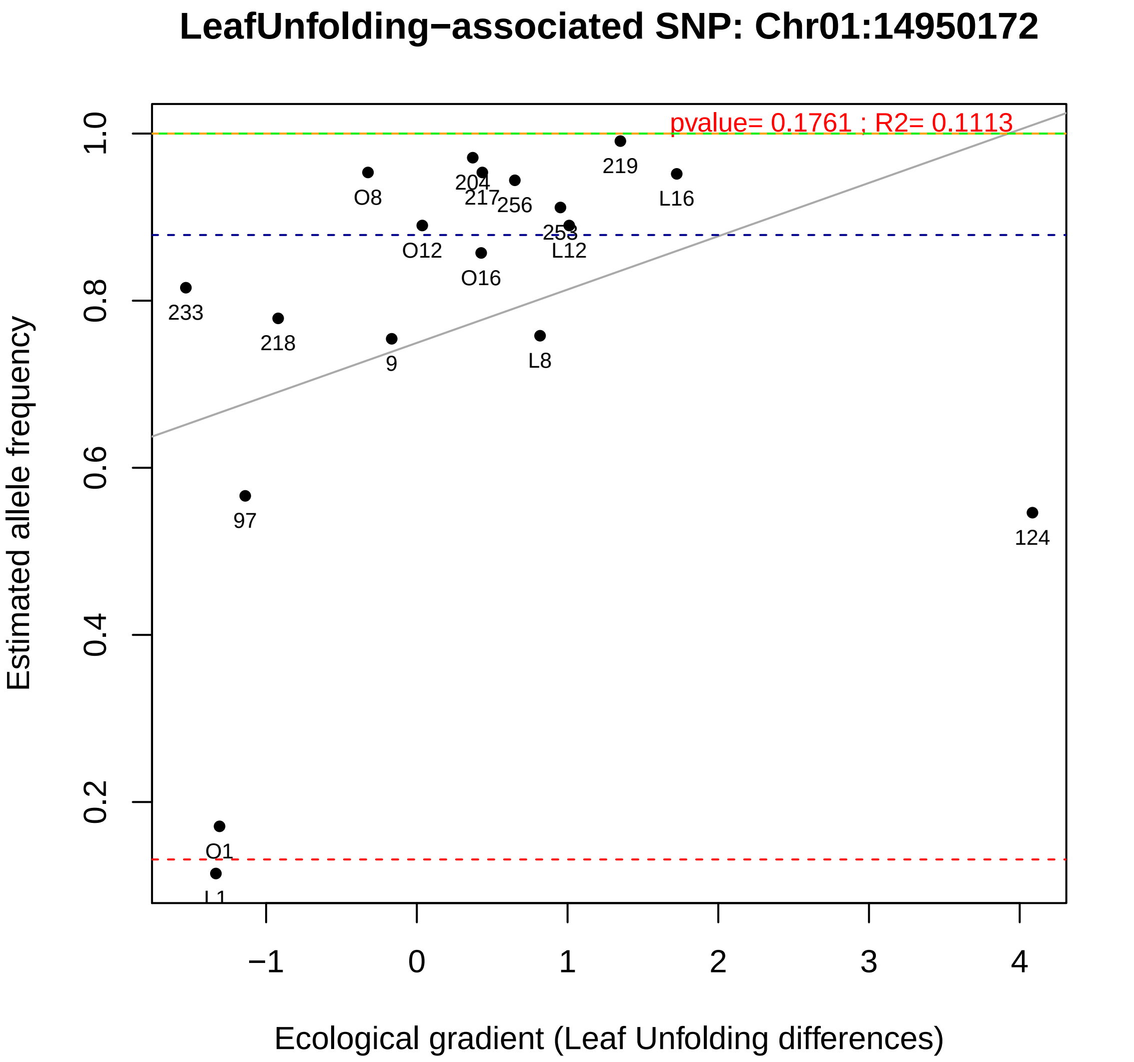

LeafUnfolding−associated SNP: Chr01:14950172
pvalue= 0.1761 ; R2= 0.1113
Estimated allele frequency
Ecological gradient (Leaf Unfolding differences)
1.0
0.8
0.6
0.4
0.2
−1
0
1
2
3
4
219
204
217
256
O8
L16
O12
253
L12
O16
233
218
9
L8
97
124
O1

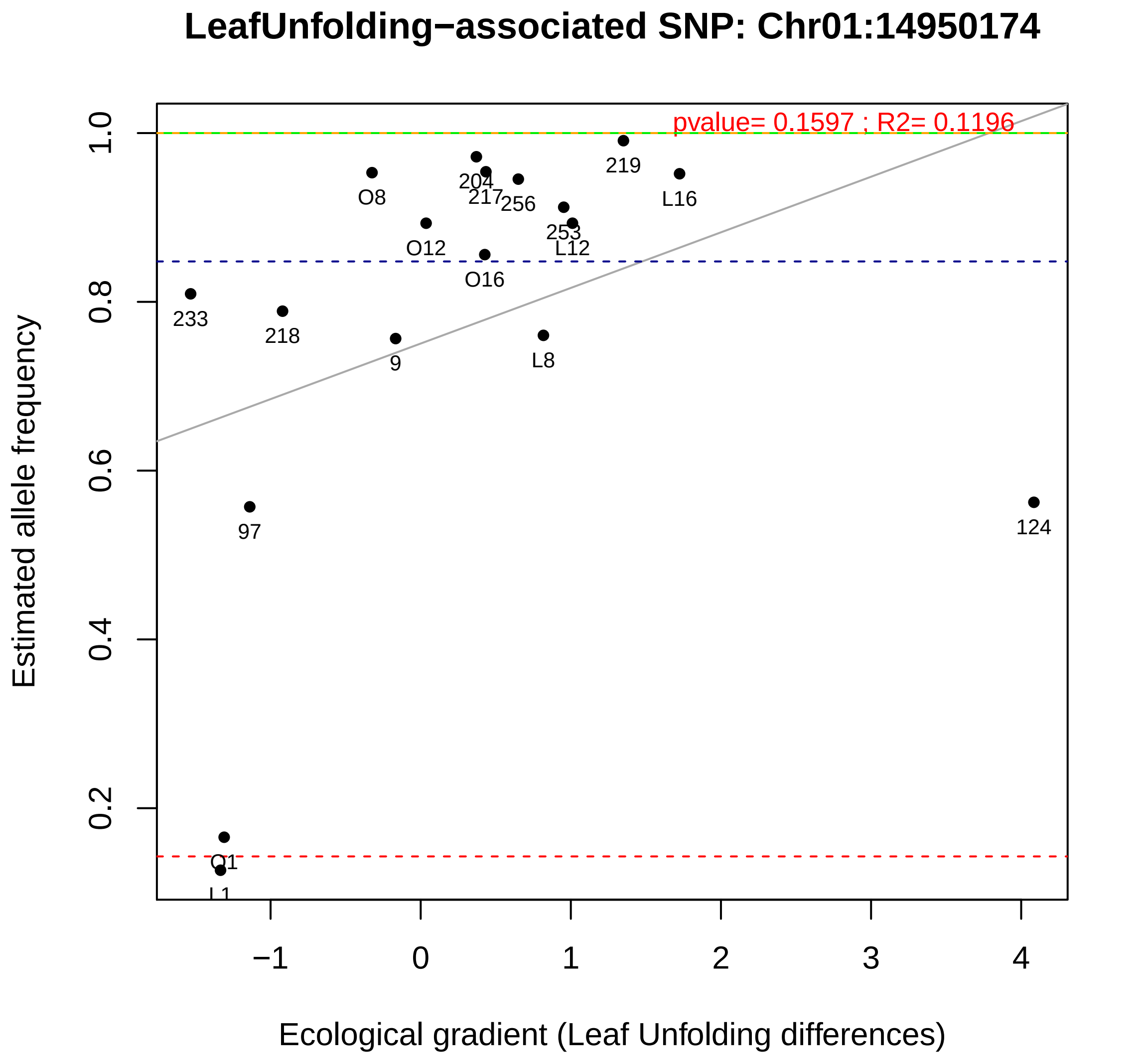

LeafUnfolding−associated SNP: Chr01:14950174
pvalue= 0.1597 ; R2= 0.1196
Estimated allele frequency
Ecological gradient (Leaf Unfolding differences)
1.0
0.8
0.6
0.4
0.2
−1
0
1
2
3
4
219
204
217
256
O8
L16
253
L12
O12
O16
233
218
9
L8
97
124
O1
L1

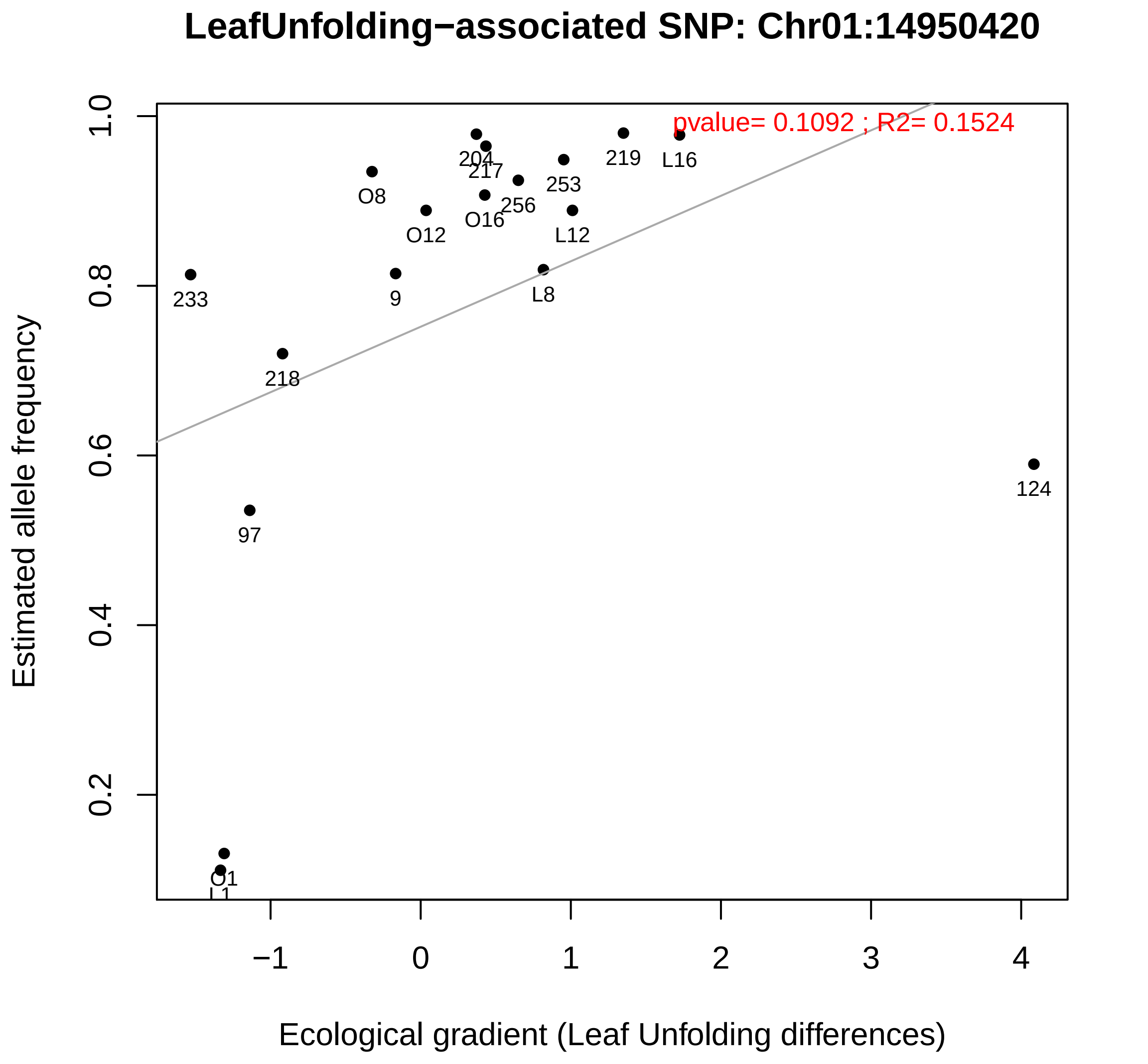

LeafUnfolding−associated SNP: Chr01:14950420
pvalue= 0.1092 ; R2= 0.1524
Estimated allele frequency
0.2
0.4
0.6
0.8
1.0
−1
0
1
2
3
4
Ecological gradient (Leaf Unfolding differences)
204
217
219
L16
O8
253
256
O16
O12
L12
233
9
L8
218
124
97
O1
L1

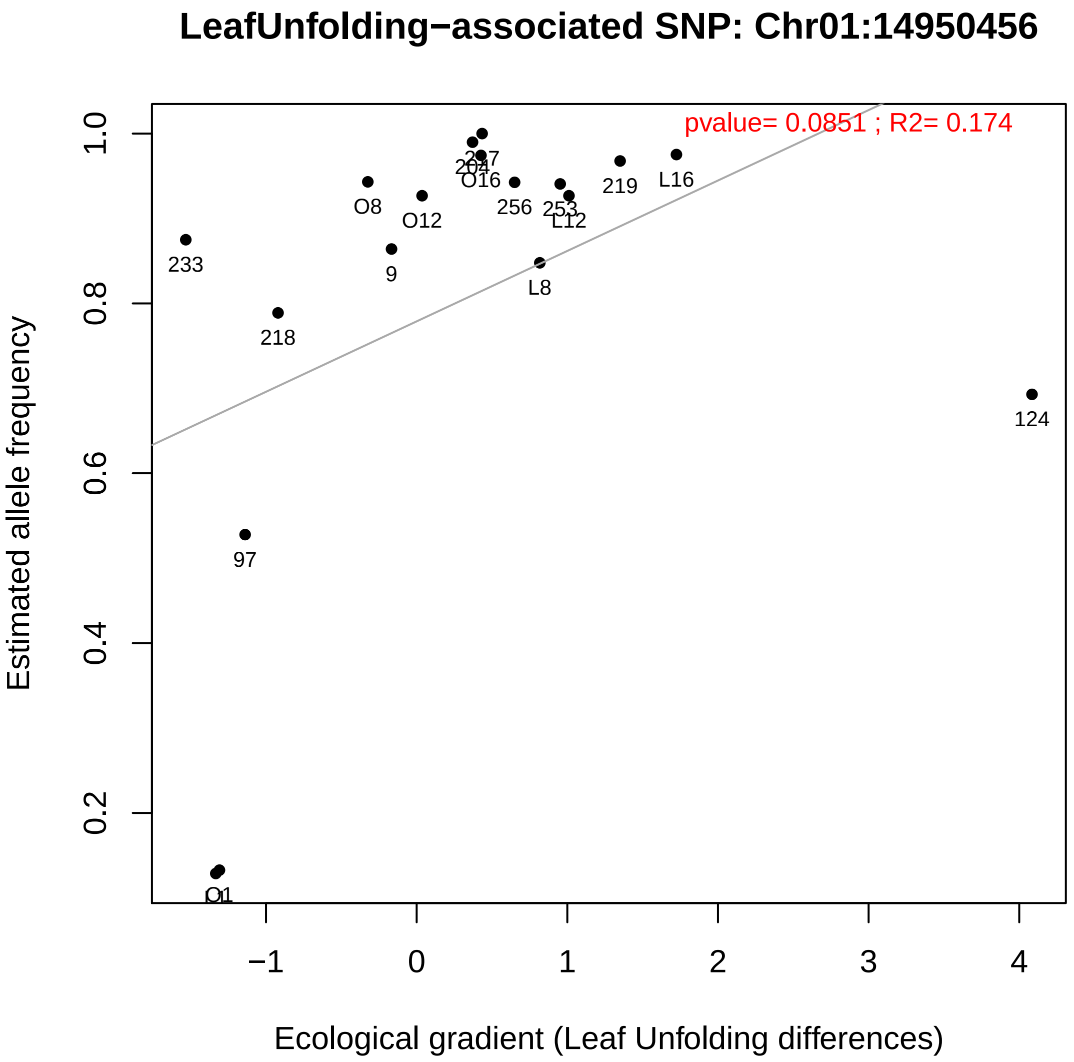
LeafUnfolding−associated SNP: Chr01:14950456
pvalue= 0.0851 ; R2= 0.174
Estimated allele frequency
Ecological gradient (Leaf Unfolding differences)
0.2
0.4
0.6
0.8
1.0
−1
0
1
2
3
4
233
218
97
O8
9
O12
204
O16
256
L8
253
L12
219
L16
124

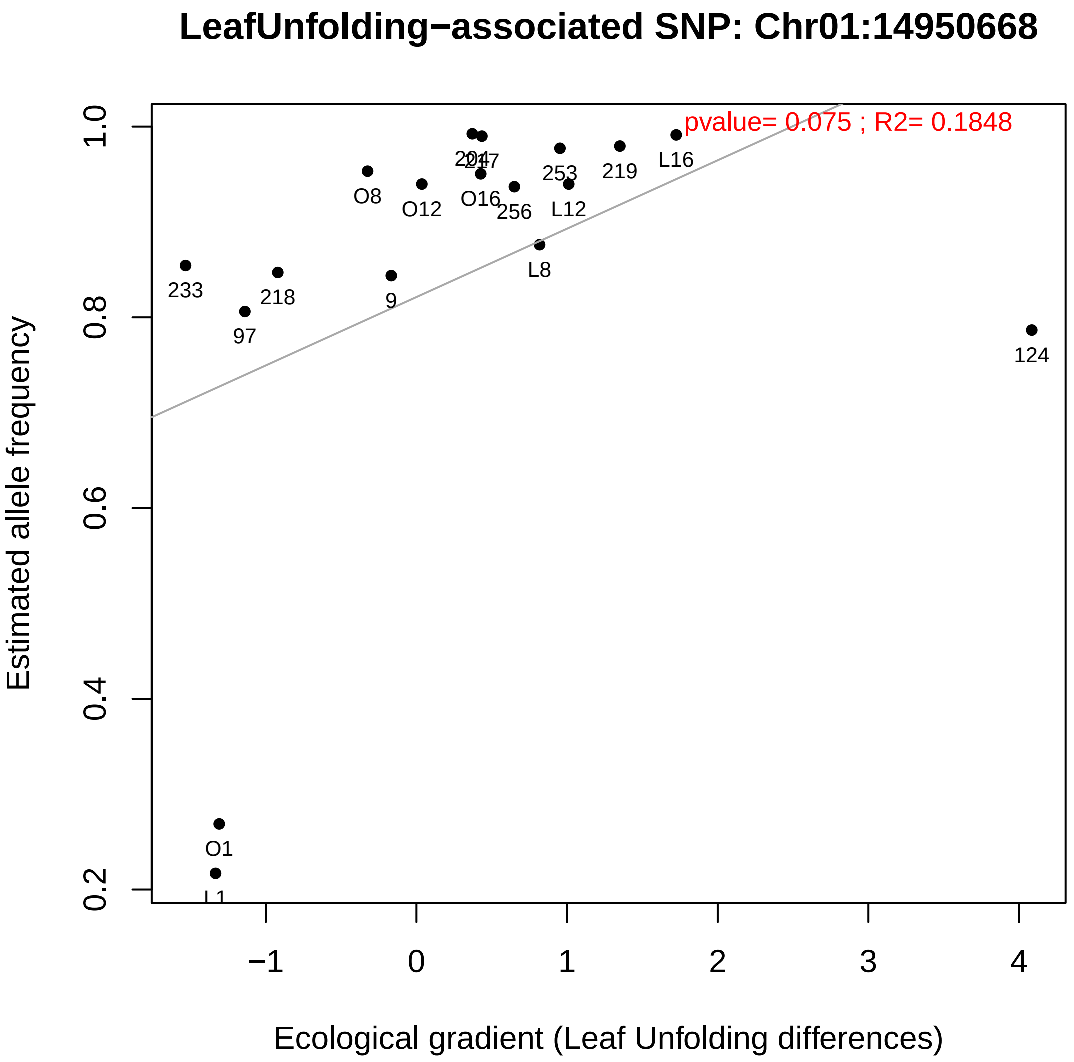

LeafUnfolding−associated SNP: Chr01:14950668
pvalue= 0.075 ; R2= 0.1848
Estimated allele frequency
Ecological gradient (Leaf Unfolding differences)
1.0
0.8
0.6
0.4
0.2
−1
0
1
2
3
4
O8
O12
O16
256
253
L12
219
L16
233
218
97
9
L8
124
O1

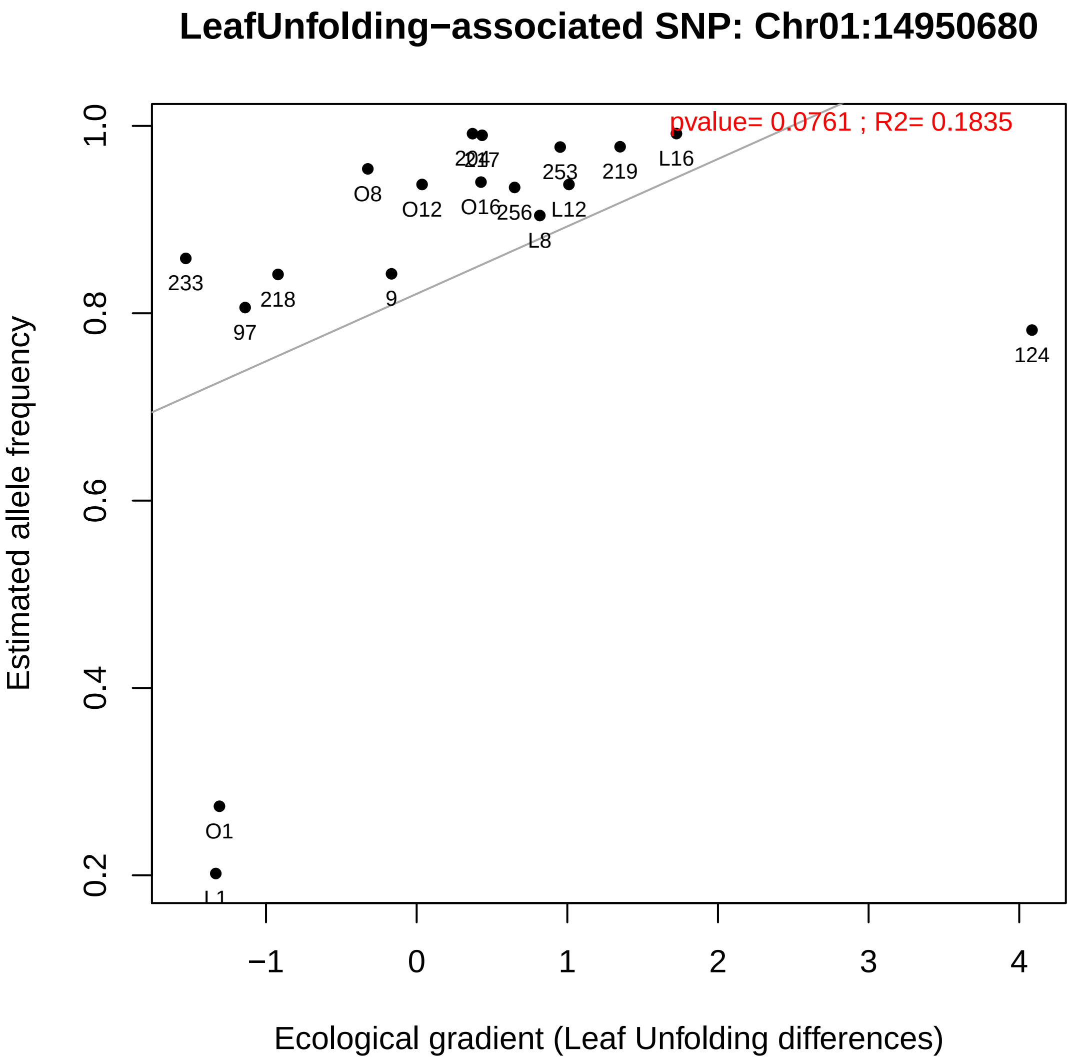

LeafUnfolding−associated SNP: Chr01:14950680
pvalue= 0.0761 ; R2= 0.1835
Estimated allele frequency
Ecological gradient (Leaf Unfolding differences)
0.2
0.4
0.6
0.8
1.0
−1
0
1
2
3
4
233
97
218
O8
9
O12
O16
256
L8
253
L12
219
L16
124
O1

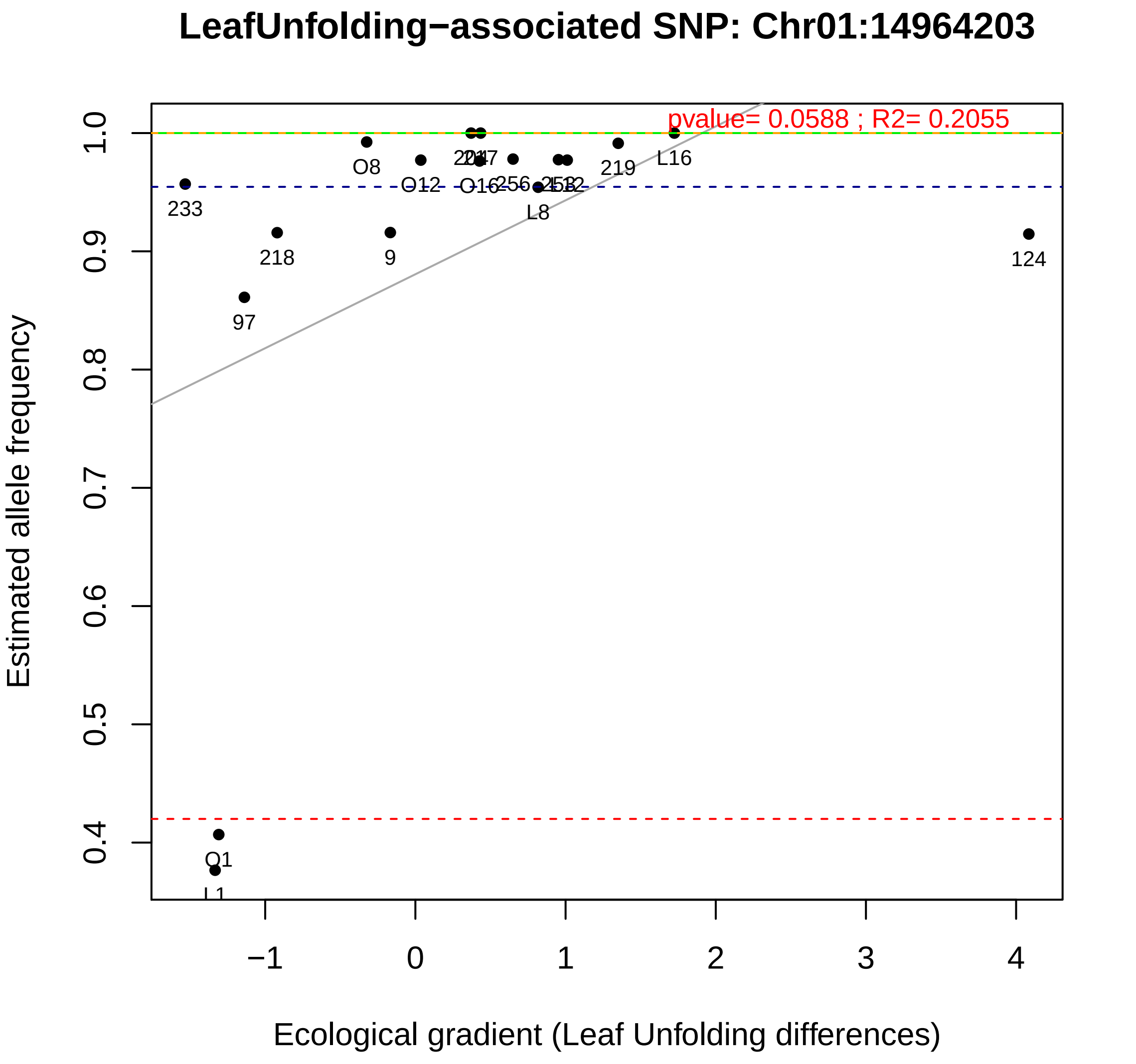
LeafUnfolding−associated SNP: Chr01:14964203
pvalue= 0.0588 ; R2= 0.2055
Estimated allele frequency
Ecological gradient (Leaf Unfolding differences)
0.4
0.5
0.6
0.7
0.8
0.9
1.0
−1
0
1
2
3
4
233
O8
O12
O16
256
L8
219
L16
218
9
97
124
O1

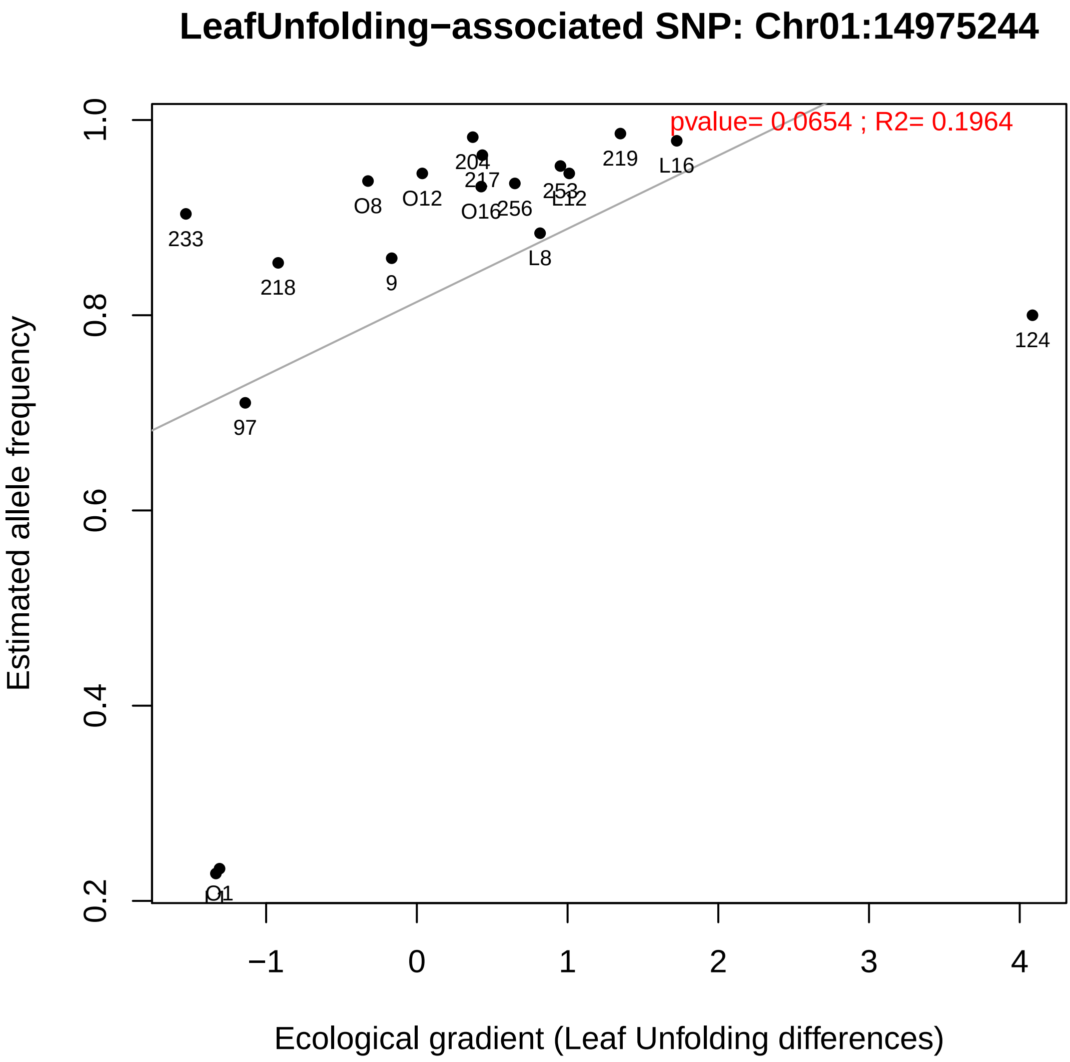

LeafUnfolding−associated SNP: Chr01:14975244
pvalue= 0.0654 ; R2= 0.1964
Estimated allele frequency
Ecological gradient (Leaf Unfolding differences)
0.2
0.4
0.6
0.8
1.0
−1
0
1
2
3
4
233
218
97
O8
9
O12
204
217
O16
256
253
L12
L8
219
L16
124
O1

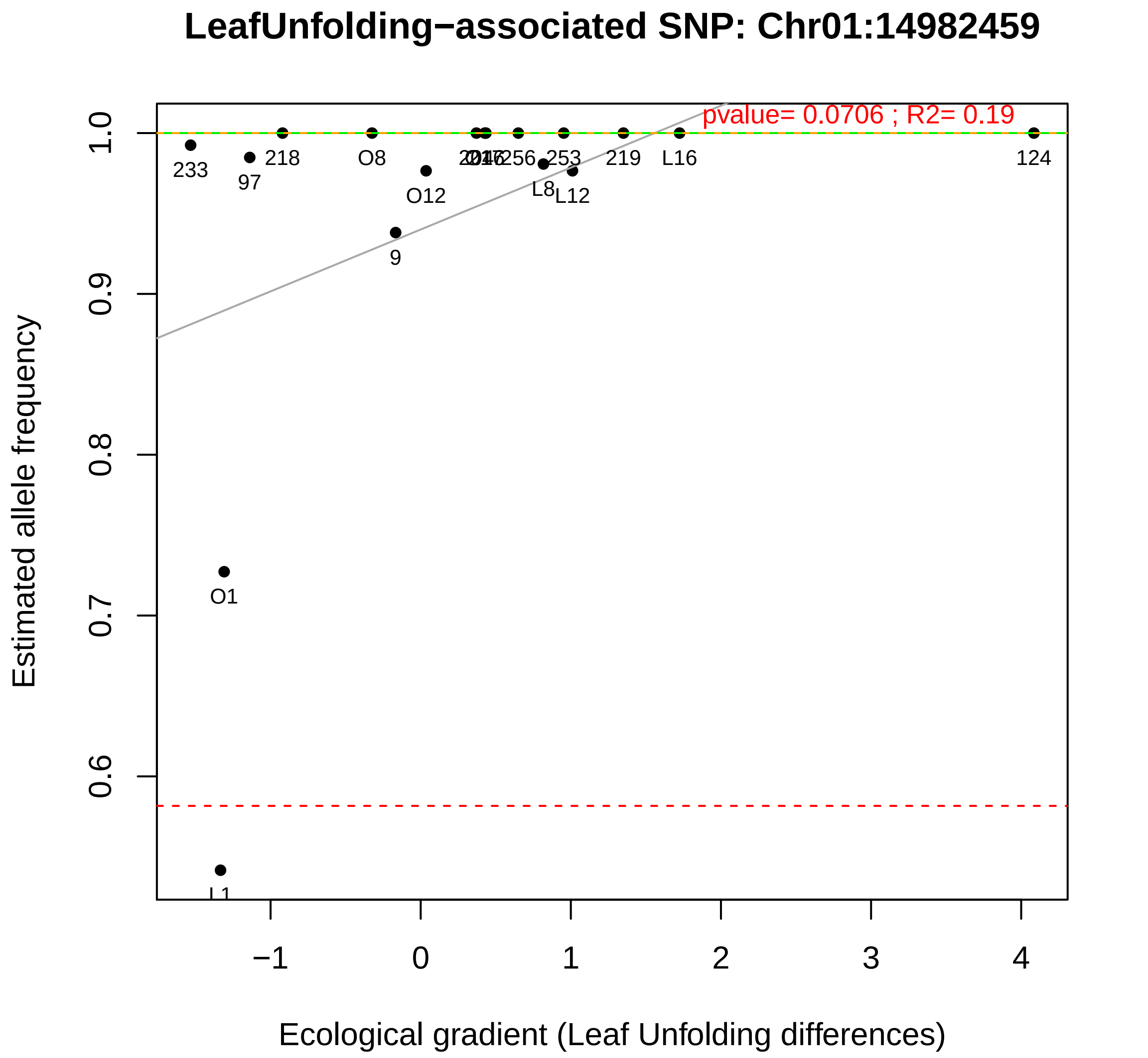
LeafUnfolding−associated SNP: Chr01:14982459
pvalue= 0.0706 ; R2= 0.19
Estimated allele frequency
Ecological gradient (Leaf Unfolding differences)
1.0
0.9
0.8
0.7
0.6
−1
0
1
2
3
4
233
97
218
O8
O12
9
256
253
L8
L12
219
L16
124
O1
L1

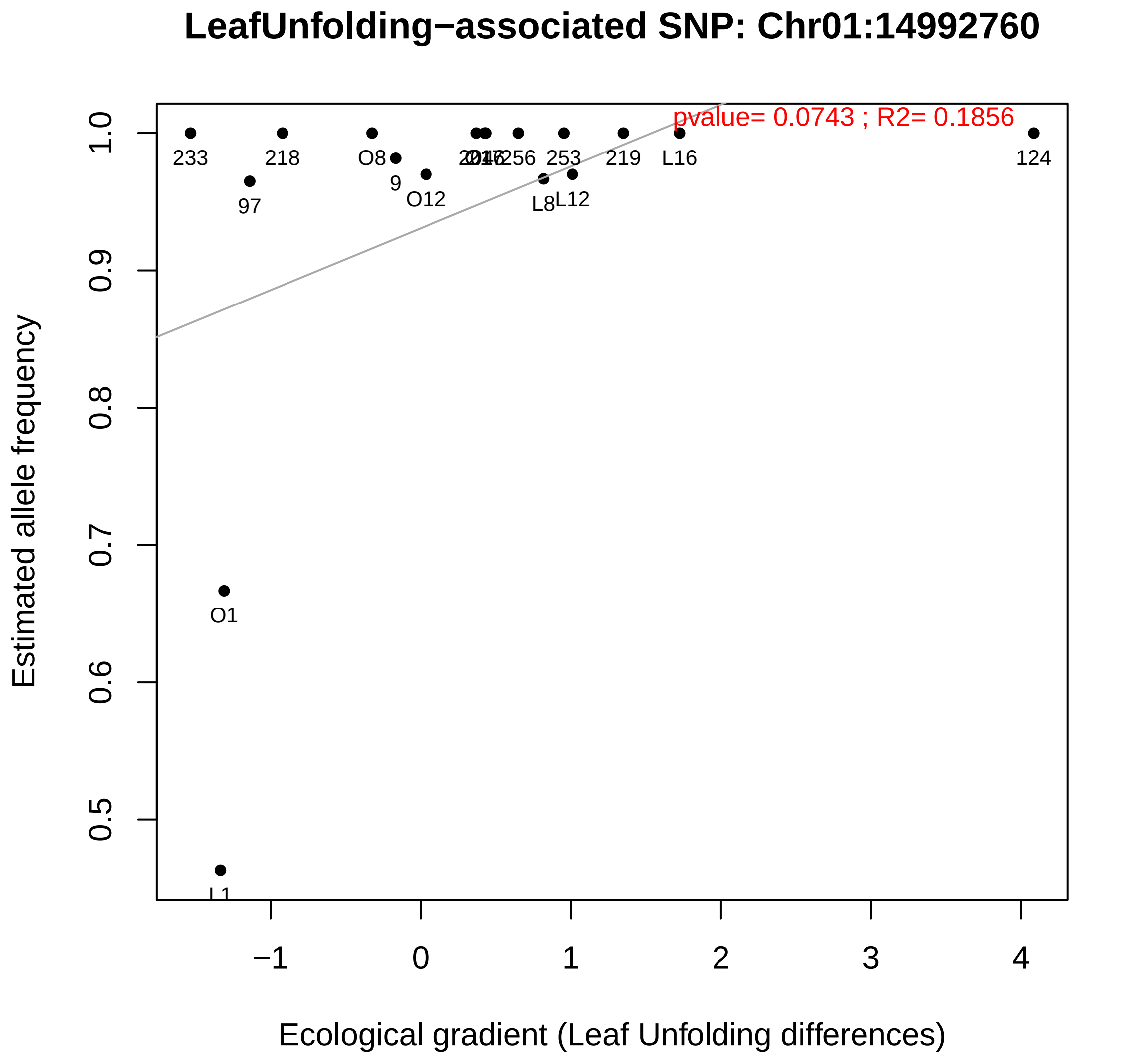

LeafUnfolding−associated SNP: Chr01:14992760
pvalue= 0.0743 ; R2= 0.1856
Estimated allele frequency
Ecological gradient (Leaf Unfolding differences)
233
218
97
O8
9
O12
256
253
219
L16
124
L8
L12
O1
L1

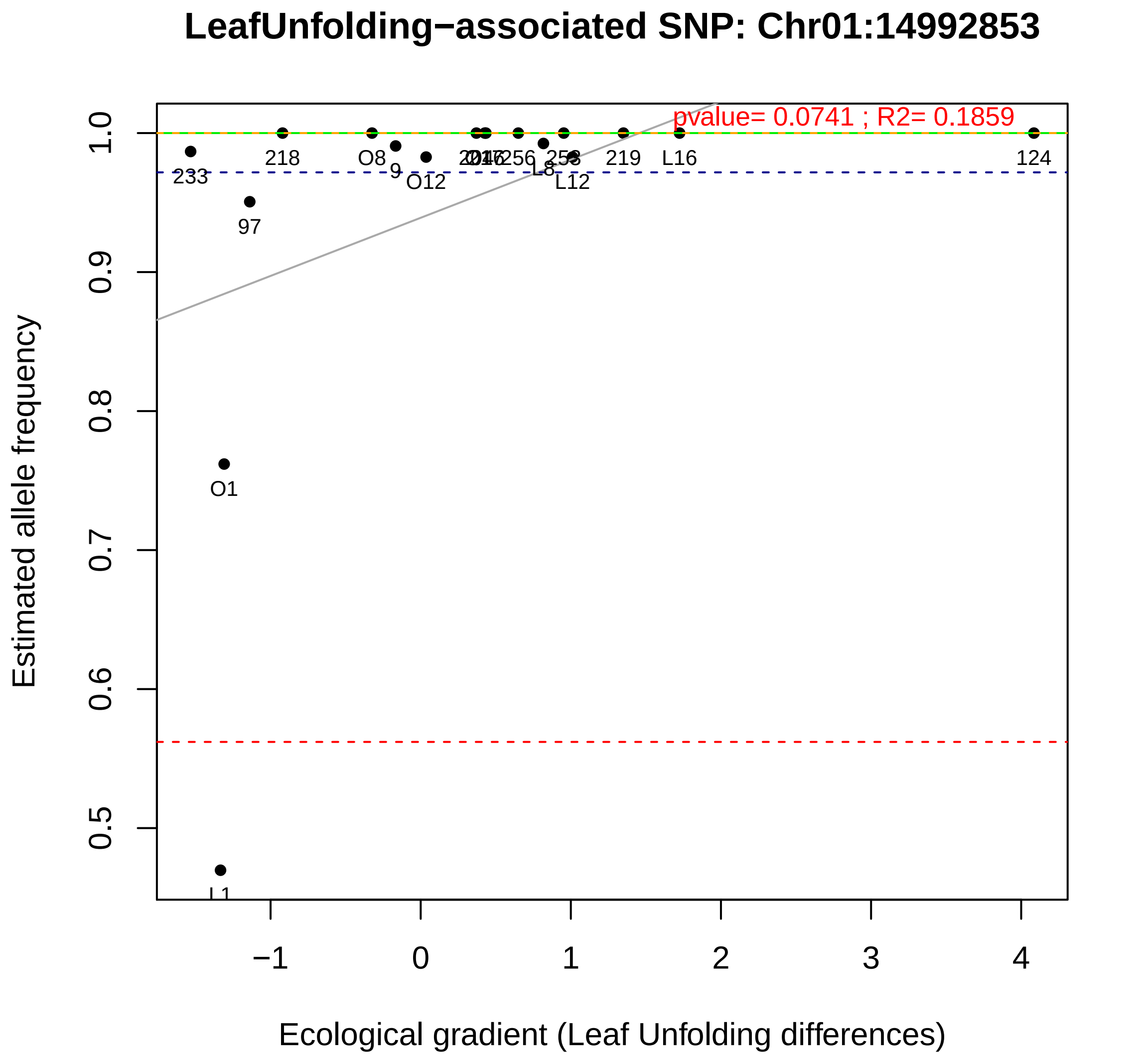

LeafUnfolding−associated SNP: Chr01:14992853
pvalue= 0.0741 ; R2= 0.1859
Estimated allele frequency
Ecological gradient (Leaf Unfolding differences)
1.0
0.9
0.8
0.7
0.6
0.5
−1
0
1
2
3
4
233
218
O8
9
O12
97
O1
L8
L12
219
L16
124

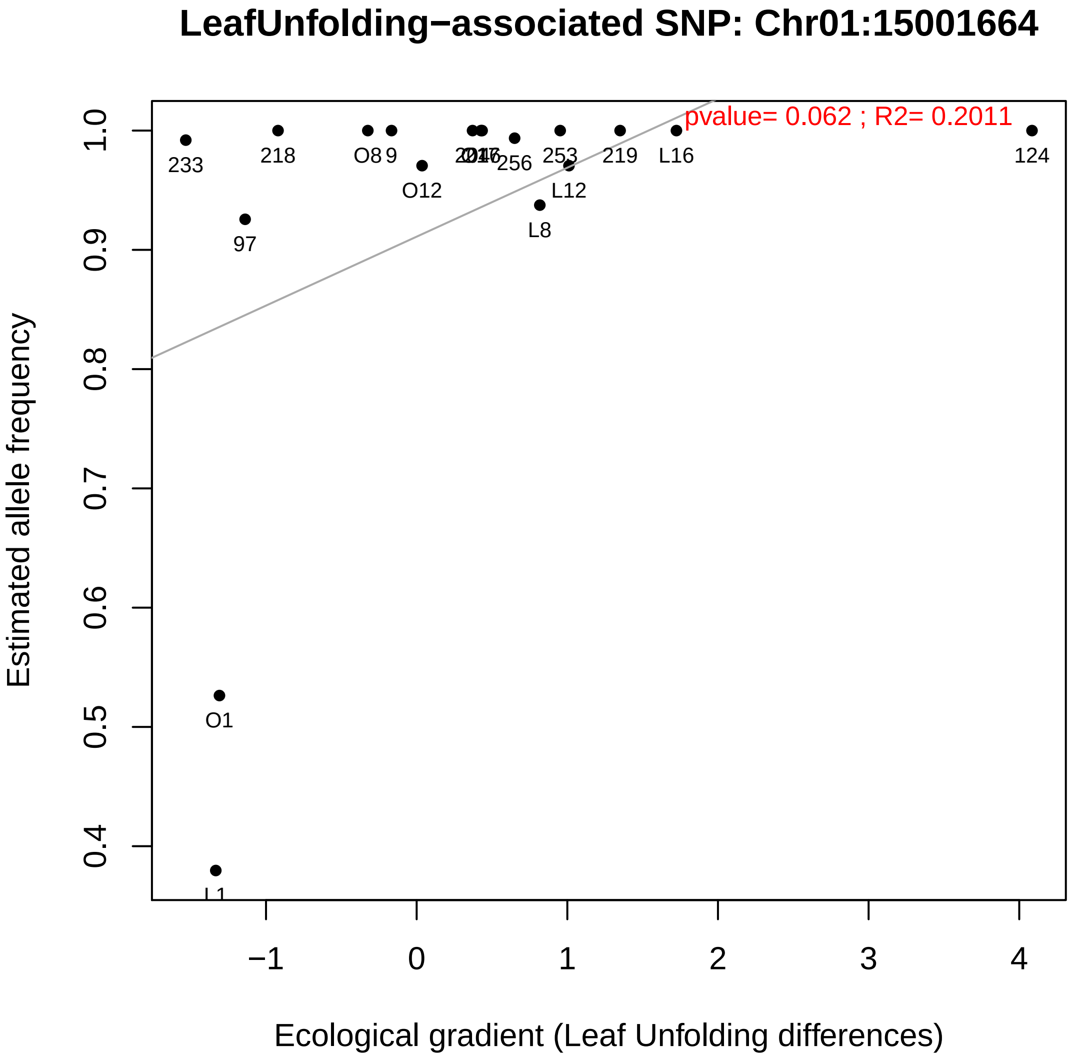
LeafUnfolding−associated SNP: Chr01:15001664
pvalue= 0.062 ; R2= 0.2011
Estimated allele frequency
Ecological gradient (Leaf Unfolding differences)
0.4
0.5
0.6
0.7
0.8
0.9
1.0
−1
0
1
2
3
4
233
218
O8
9
O12
256
253
219
L16
124
L12
L8
97
O1

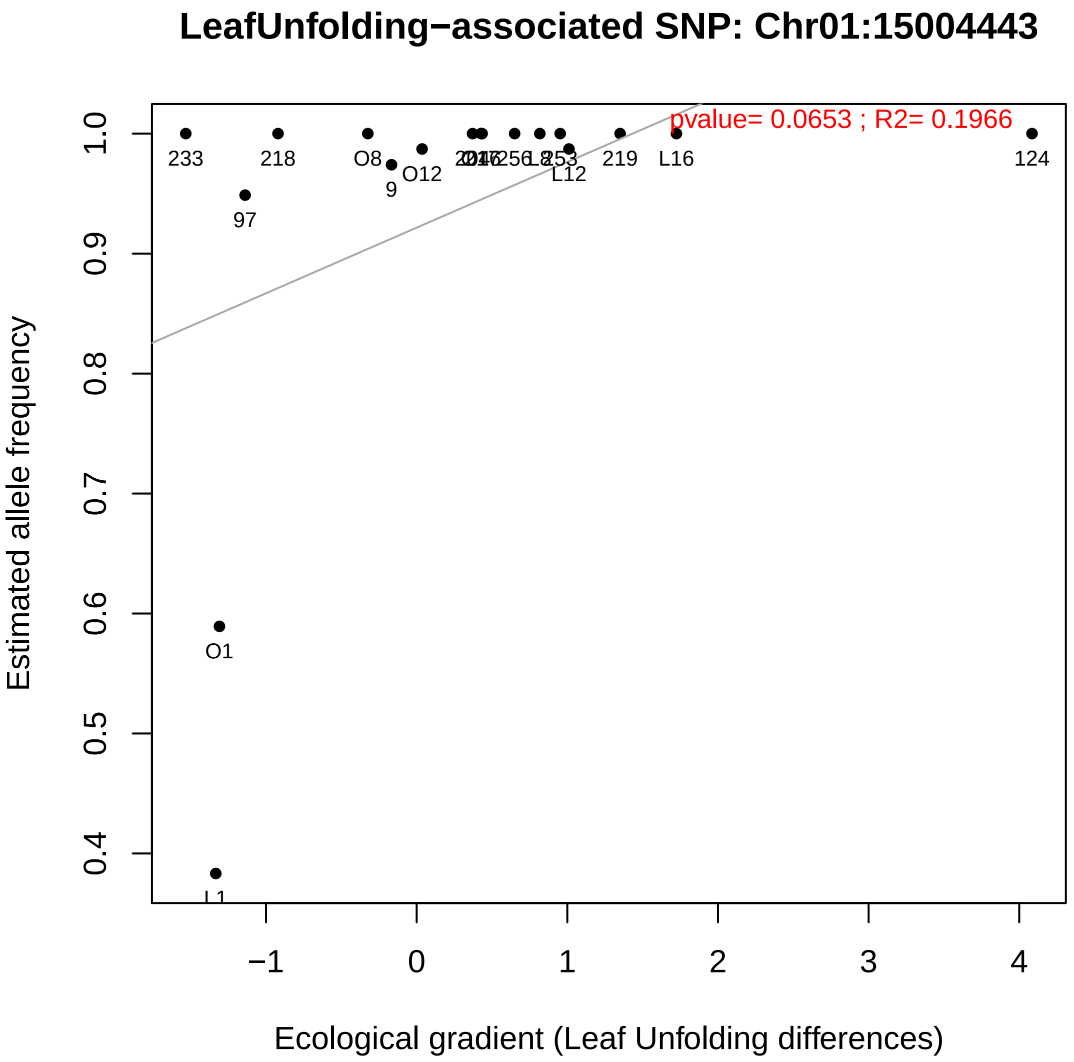
LeafUnfolding−associated SNP: Chr01:15004443
pvalue= 0.0653 ; R2= 0.1966
Estimated allele frequency
Ecological gradient (Leaf Unfolding differences)
0.4
0.5
0.6
0.7
0.8
0.9
1.0
−1
0
1
2
3
4
233
218
O8
9
O12
97
L12
219
L16
124
O1

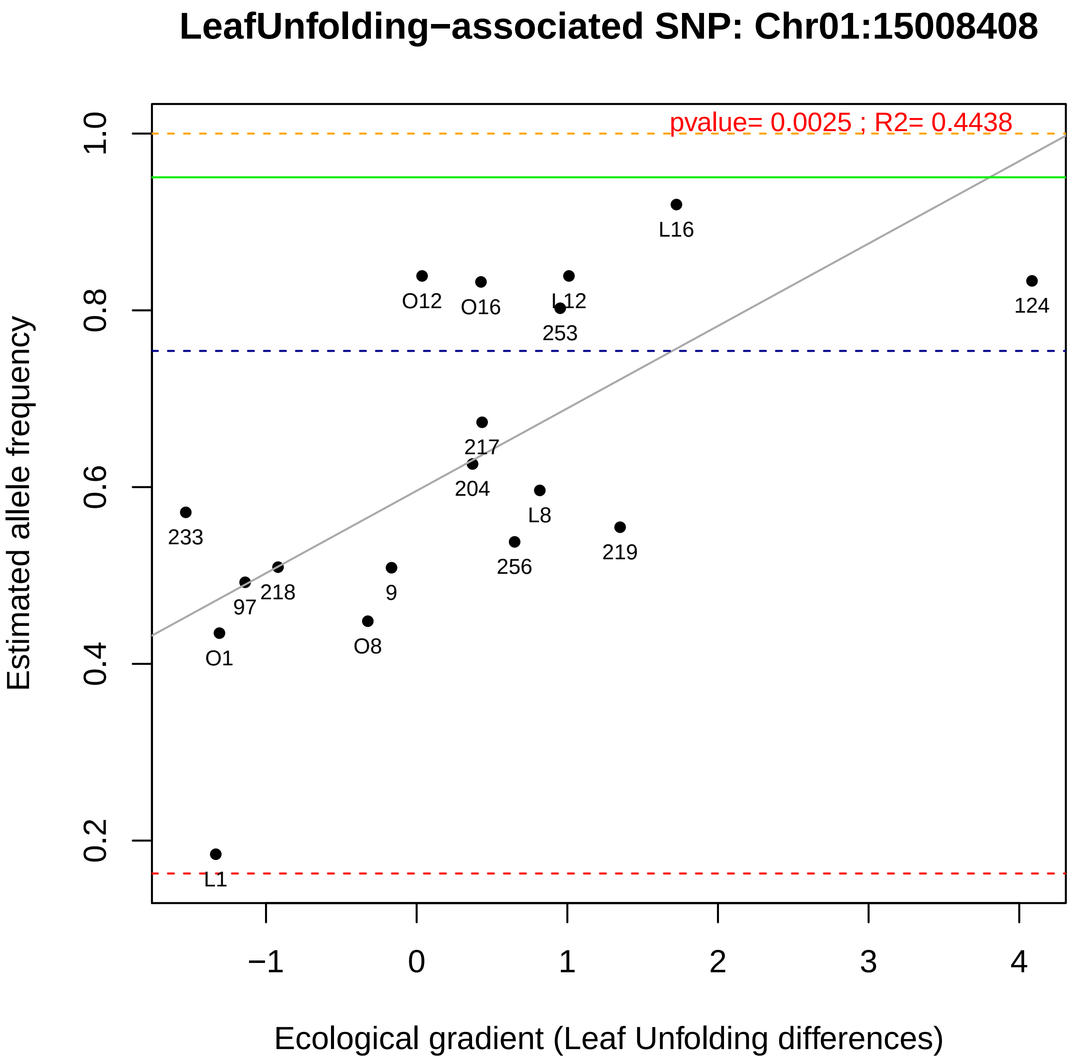
LeafUnfolding−associated SNP: Chr01:15008408
pvalue= 0.0025 ; R2= 0.4438
Estimated allele frequency
Ecological gradient (Leaf Unfolding differences)
L16
O12
O16
L12
253
124
217
204
L8
233
219
256
218
9
97
O1
O8
L1
0.2
0.4
0.6
0.8
1.0
−1
0
1
2
3
4

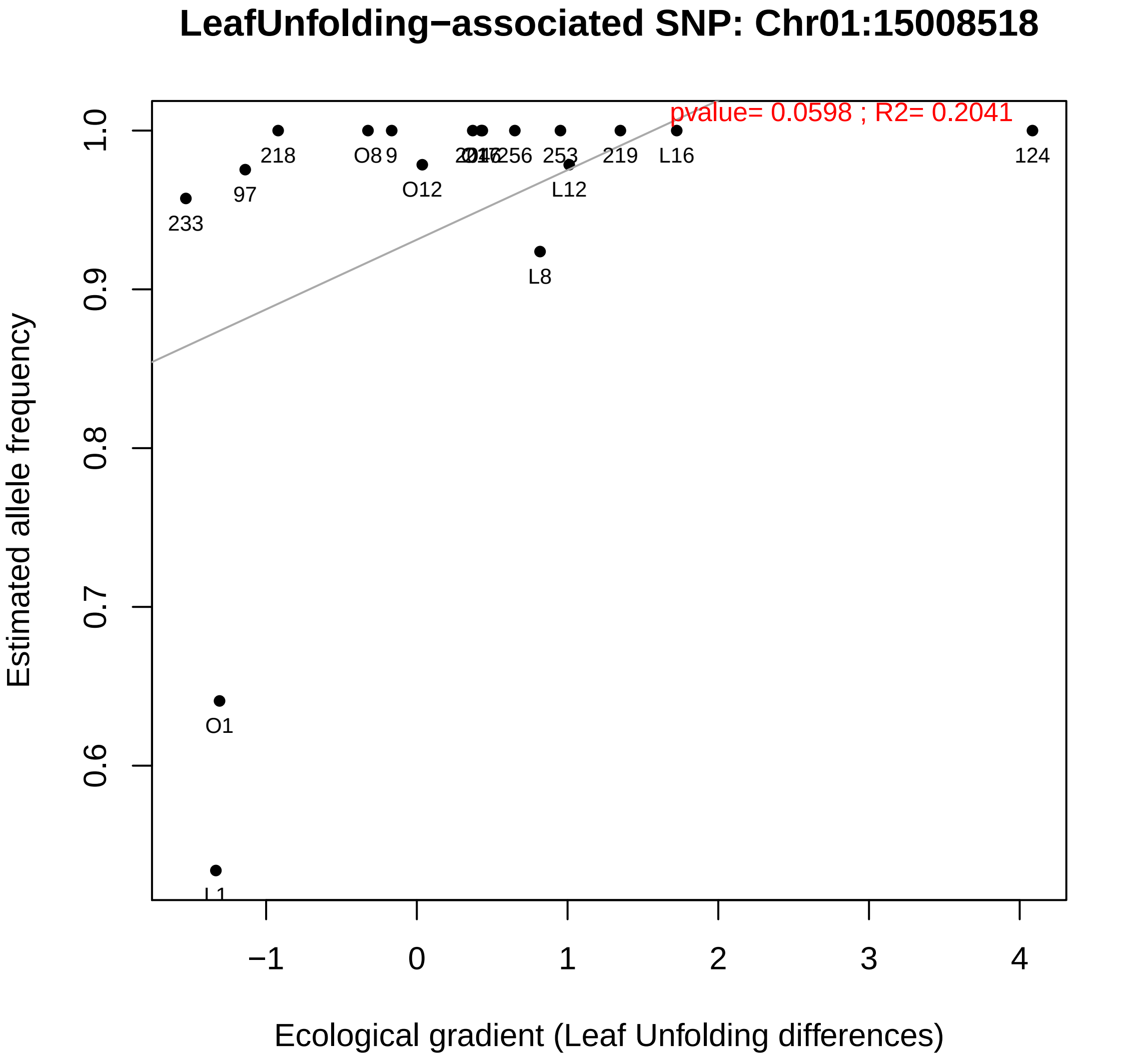

LeafUnfolding−associated SNP: Chr01:15008518
pvalue= 0.0598 ; R2= 0.2041
Estimated allele frequency
Ecological gradient (Leaf Unfolding differences)
0.6
0.7
0.8
0.9
1.0
−1
0
1
2
3
4
218
O8
9
256
253
219
L16
124
97
O12
L12
233
L8
O1

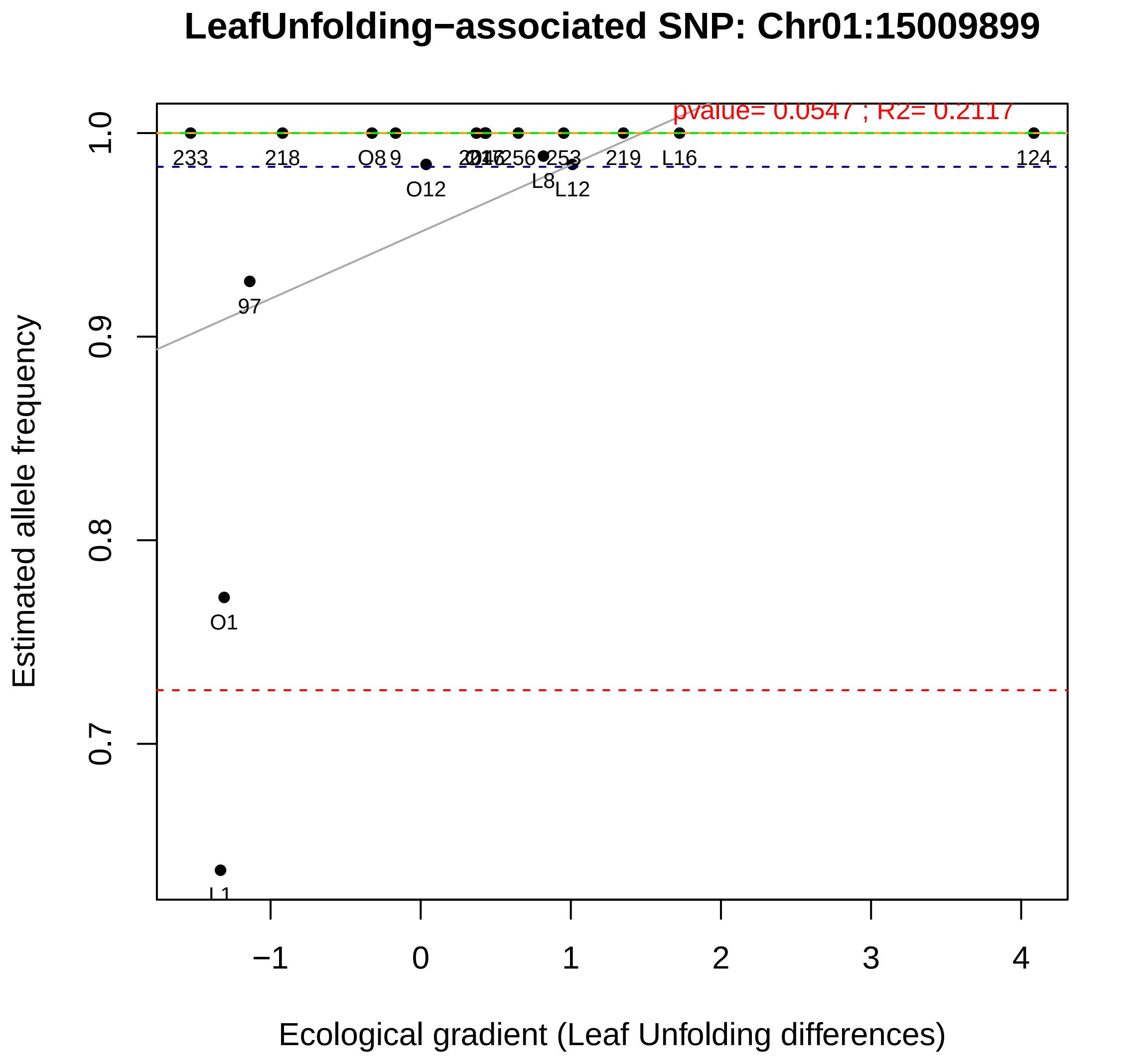

LeafUnfolding−associated SNP: Chr01:15009899
pvalue= 0.0547 ; R2= 0.2117
Estimated allele frequency
1.0
0.9
0.8
0.7
233
218
O8
9
O12
L8
L12
219
L16
124
97
O1
L1
−1
0
1
2
3
4
Ecological gradient (Leaf Unfolding differences)

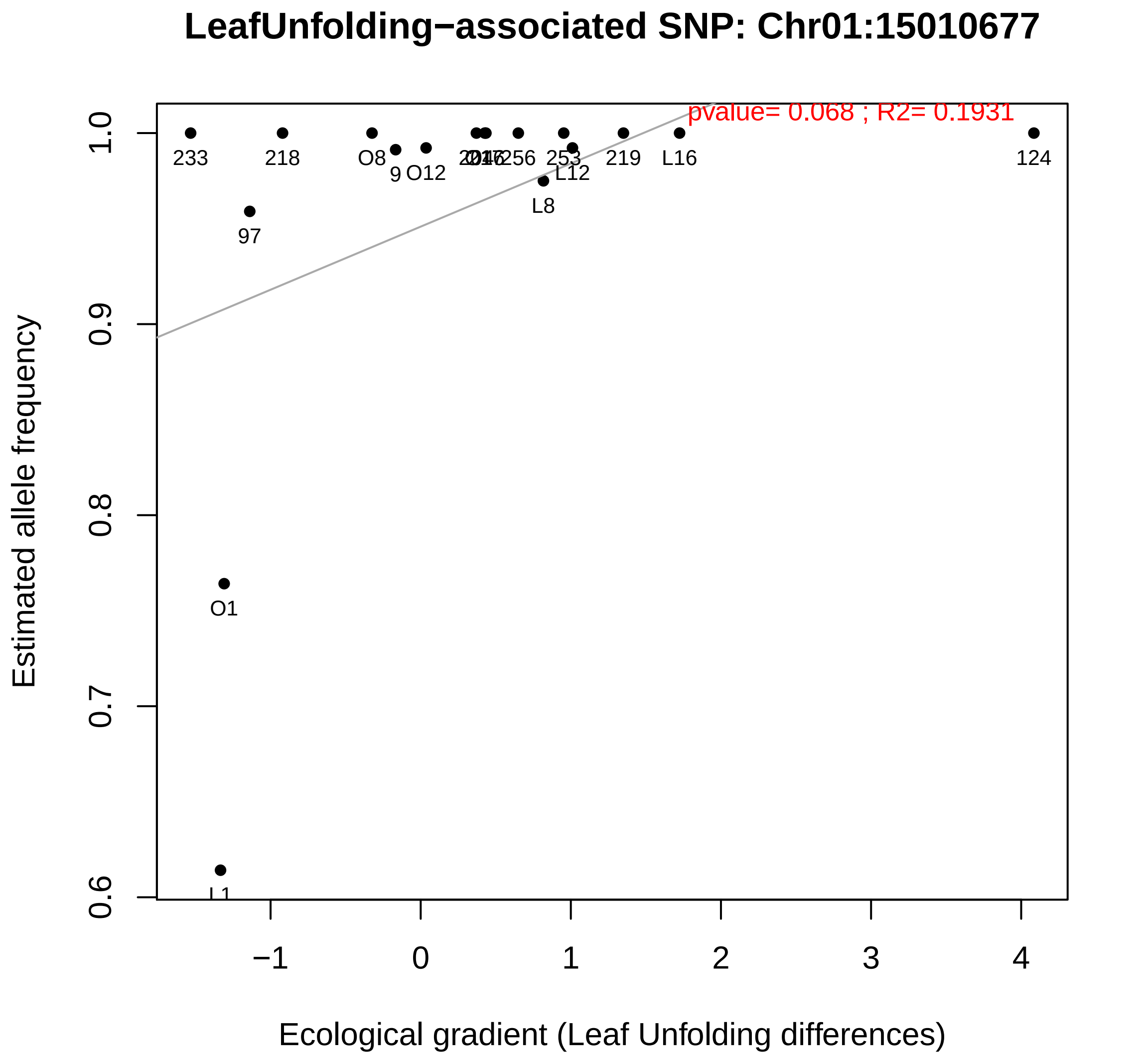

LeafUnfolding−associated SNP: Chr01:15010677
pvalue= 0.068 ; R2= 0.1931
Estimated allele frequency
Ecological gradient (Leaf Unfolding differences)
0.6
0.7
0.8
0.9
1.0
−1
0
1
2
3
4
233
218
O8
9
O12
256
253
L12
219
L16
124
L8
97
O1

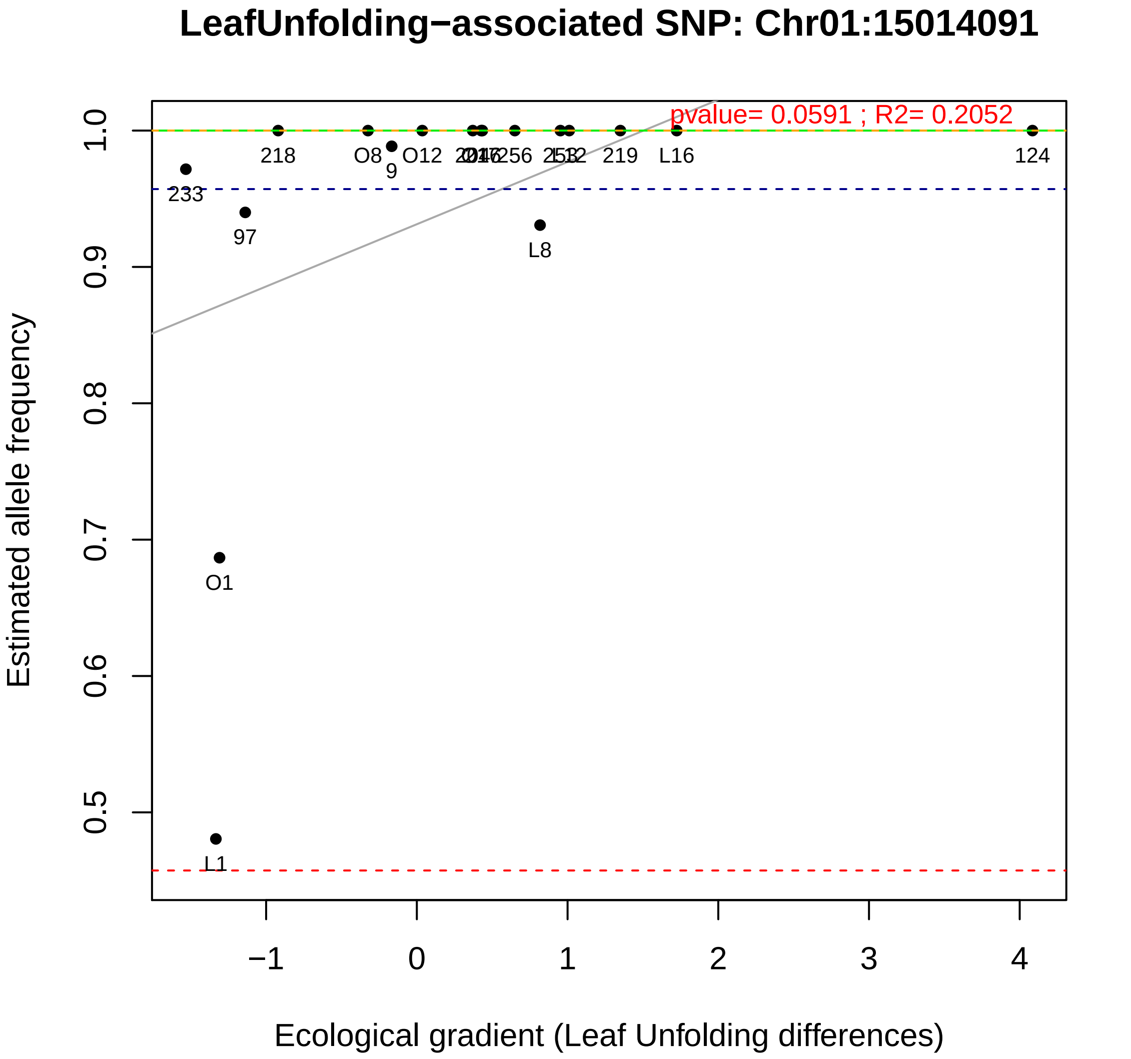

LeafUnfolding−associated SNP: Chr01:15014091
pvalue= 0.0591 ; R2= 0.2052
Estimated allele frequency
Ecological gradient (Leaf Unfolding differences)
1.0
0.9
0.8
0.7
0.6
0.5
−1
0
1
2
3
4
218
O8
O12
9
219
L16
124
233
97
L8
O1
L1

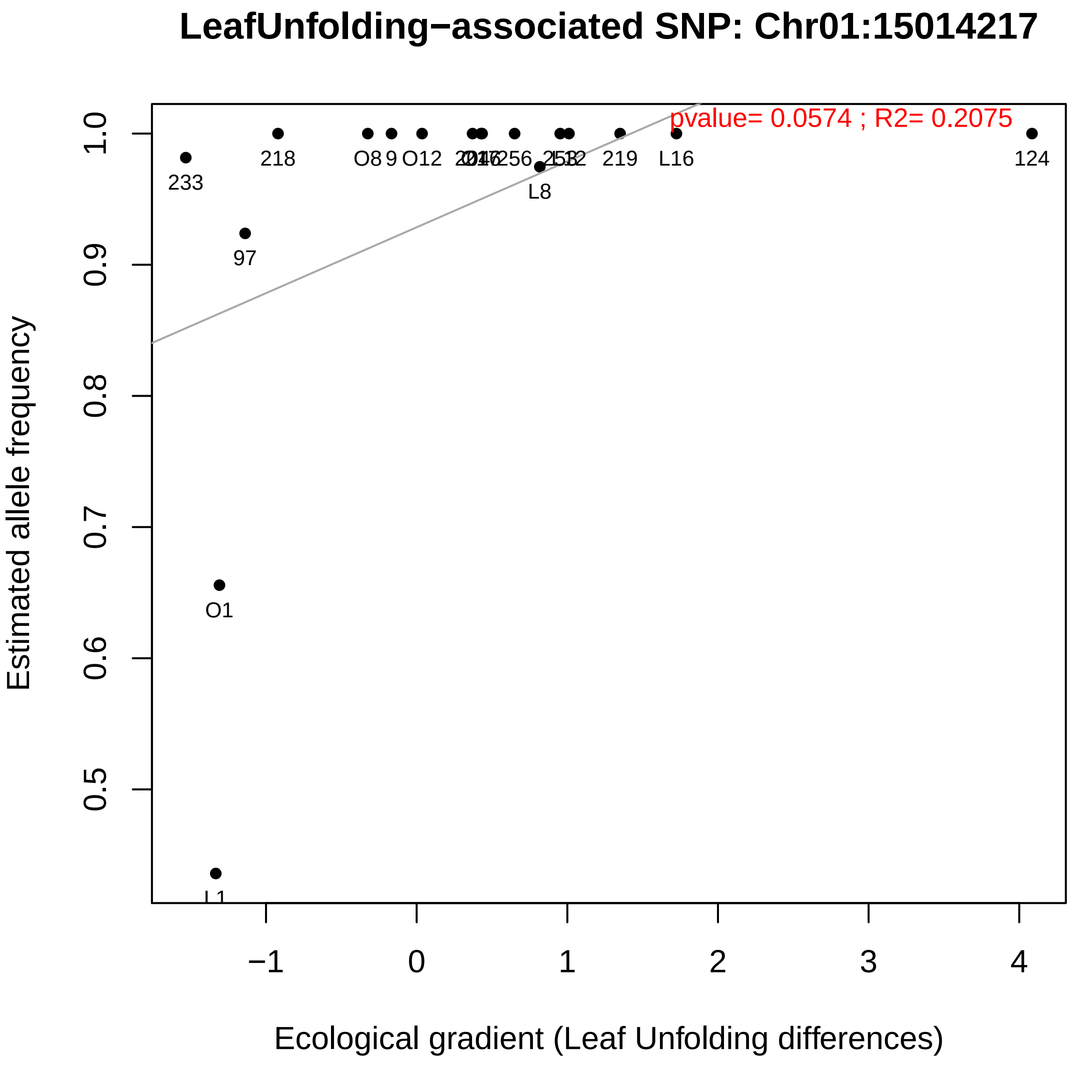

LeafUnfolding−associated SNP: Chr01:15014217
pvalue= 0.0574 ; R2= 0.2075
Estimated allele frequency
Ecological gradient (Leaf Unfolding differences)
1.0
0.9
0.8
0.7
0.6
0.5
−1
0
1
2
3
4
233
218
O8
9
O12
L8
219
L16
124
97
O1

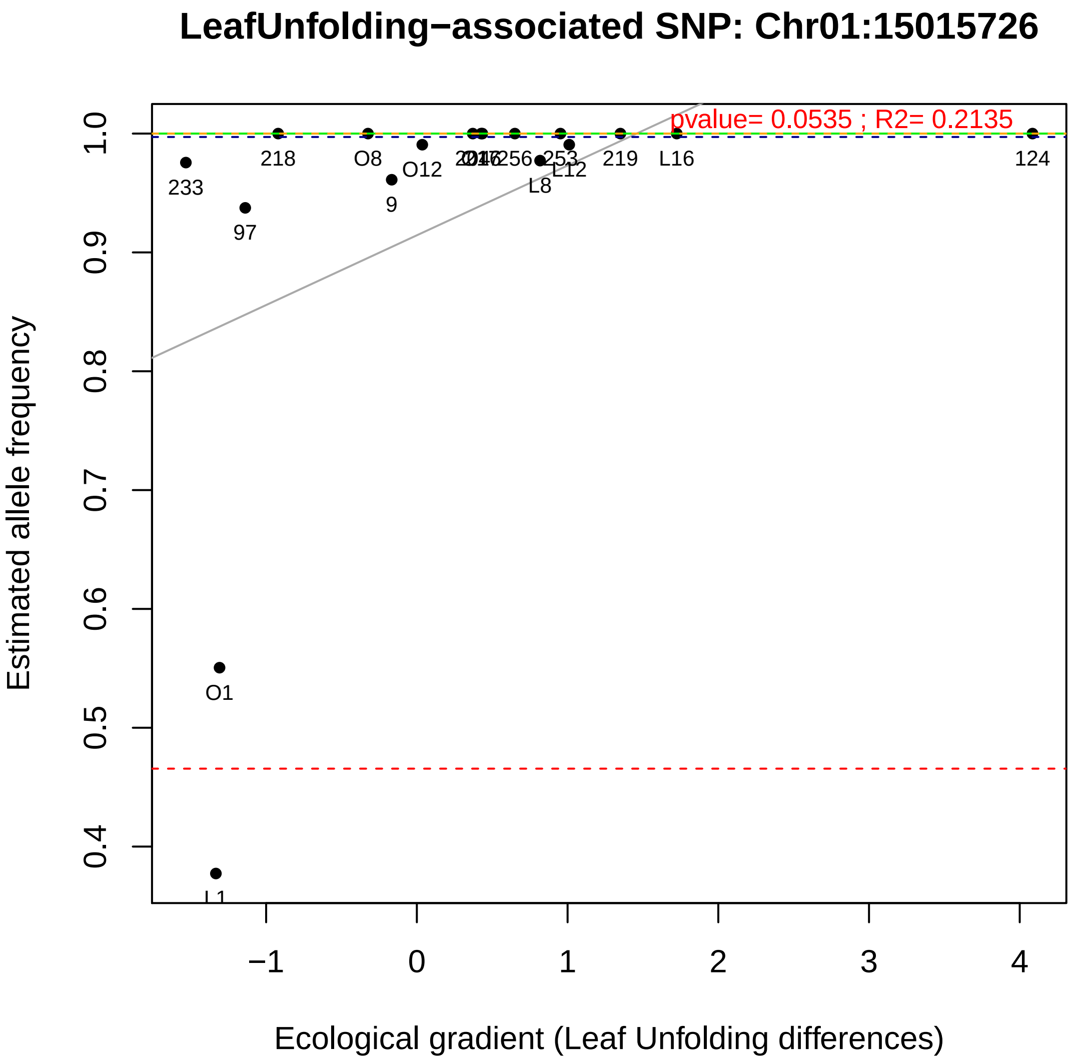
LeafUnfolding−associated SNP: Chr01:15015726
pvalue= 0.0535 ; R2= 0.2135
Estimated allele frequency
Ecological gradient (Leaf Unfolding differences)
1.0
0.9
0.8
0.7
0.6
0.5
0.4
−1
0
1
2
3
4
233
97
218
O8
9
O12
256
L8
253
L12
219
L16
124
O1
L1

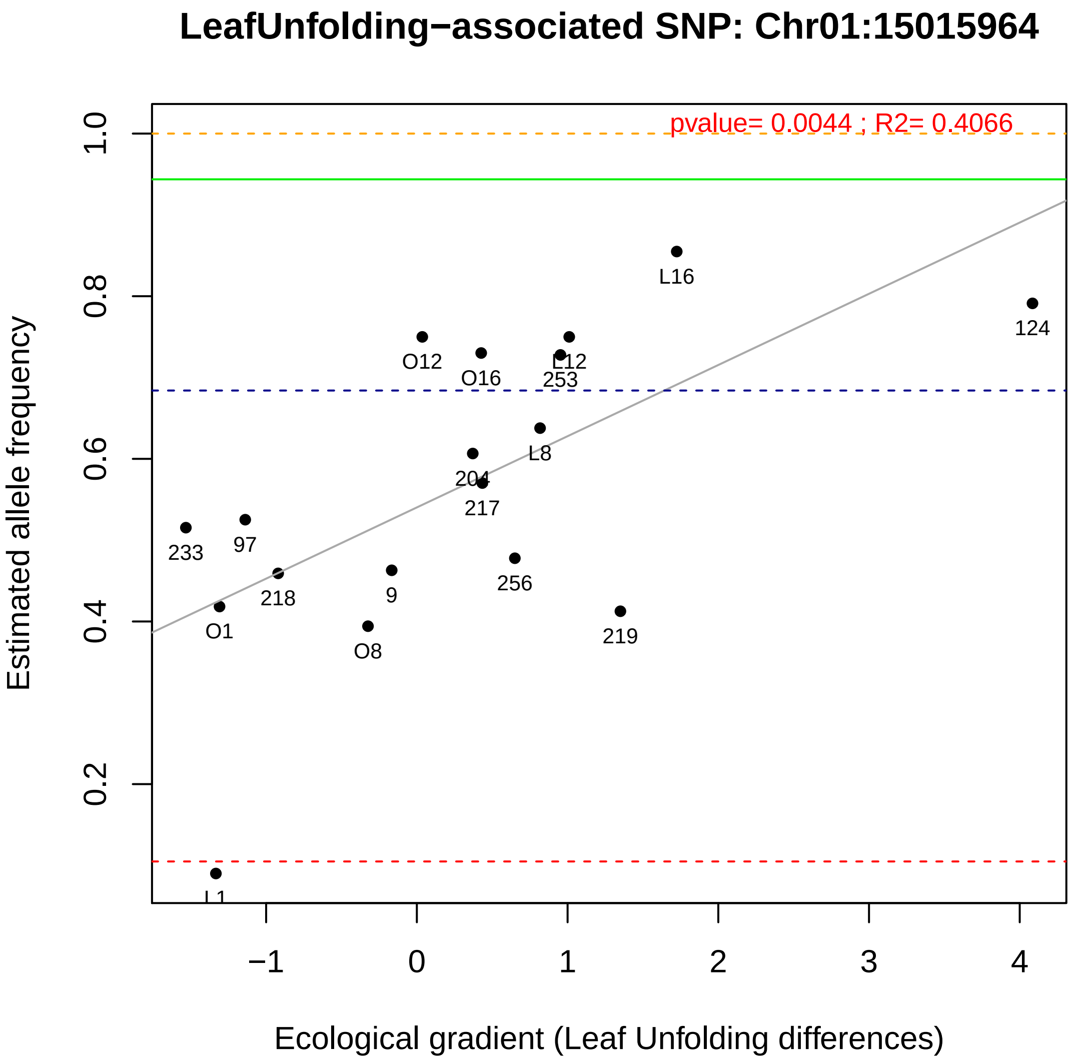
LeafUnfolding−associated SNP: Chr01:15015964
pvalue= 0.0044 ; R2= 0.4066
Estimated allele frequency
Ecological gradient (Leaf Unfolding differences)
0.2
0.4
0.6
0.8
1.0
−1
0
1
2
3
4
L16
124
O12
O16
L12
253
L8
204
217
233
97
218
9
256
O1
O8
219
L1

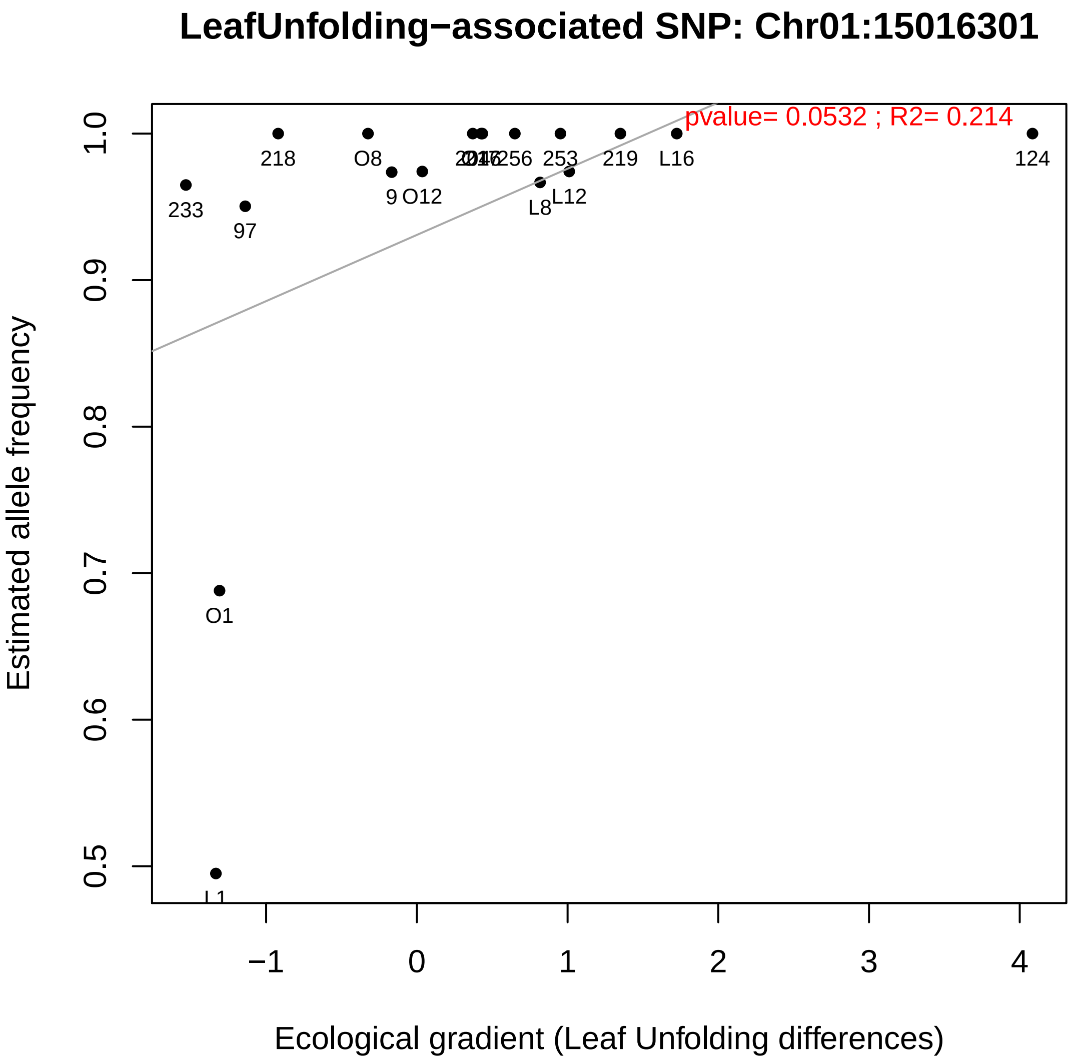
LeafUnfolding−associated SNP: Chr01:15016301
pvalue= 0.0532 ; R2= 0.214
Estimated allele frequency
Ecological gradient (Leaf Unfolding differences)
1.0
0.9
0.8
0.7
0.6
0.5
−1
0
1
2
3
4
218
O8
256
253
219
L16
124
9
O12
L12
L8
233
97
O1
L1

**LeafUnfolding−associated SNP: Chr01:15021395**

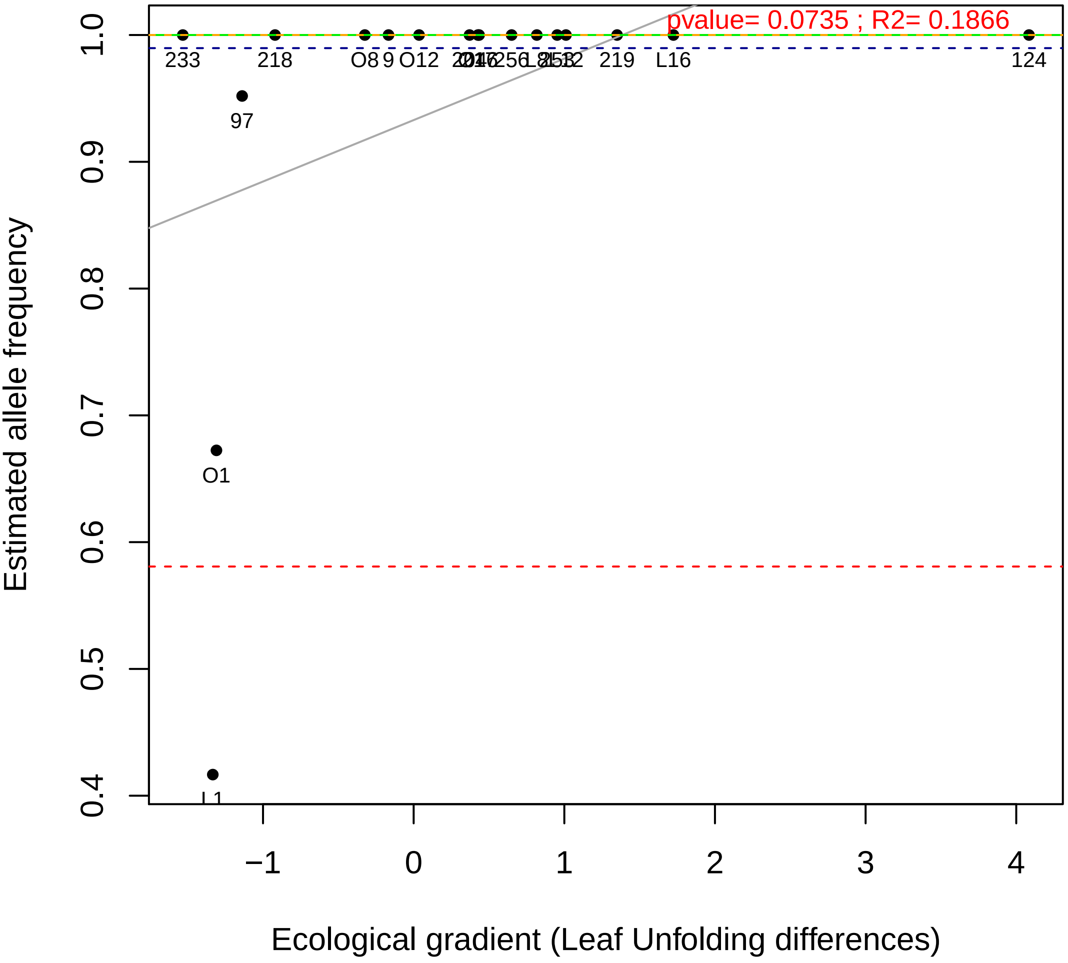

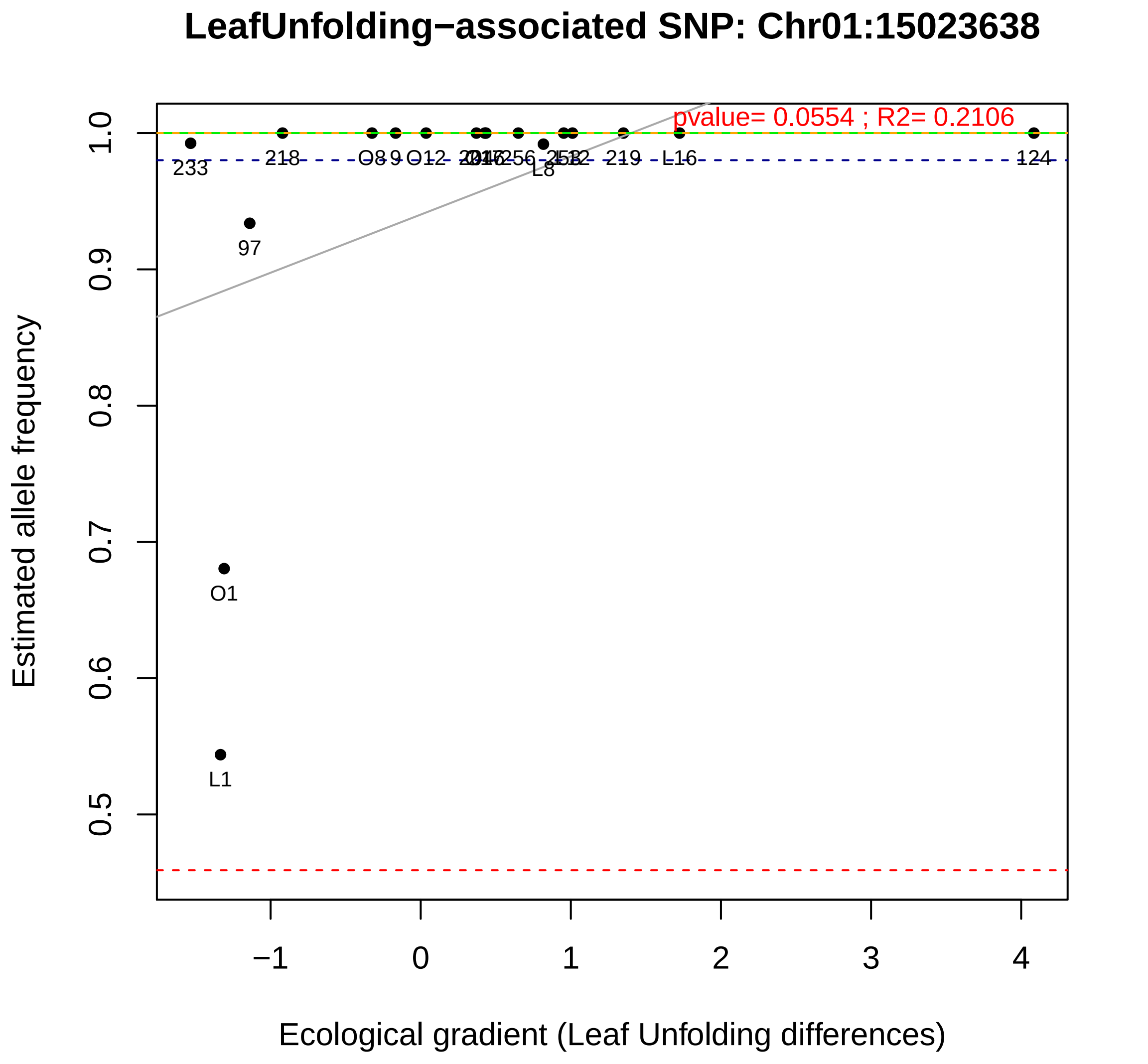

LeafUnfolding−associated SNP: Chr01:15023638
pvalue= 0.0554 ; R2= 0.2106
Estimated allele frequency
Ecological gradient (Leaf Unfolding differences)
0.5
0.6
0.7
0.8
0.9
1.0
−1
0
1
2
3
4
233
218
O8
9
O12
219
L16
124
L8
97
O1
L1

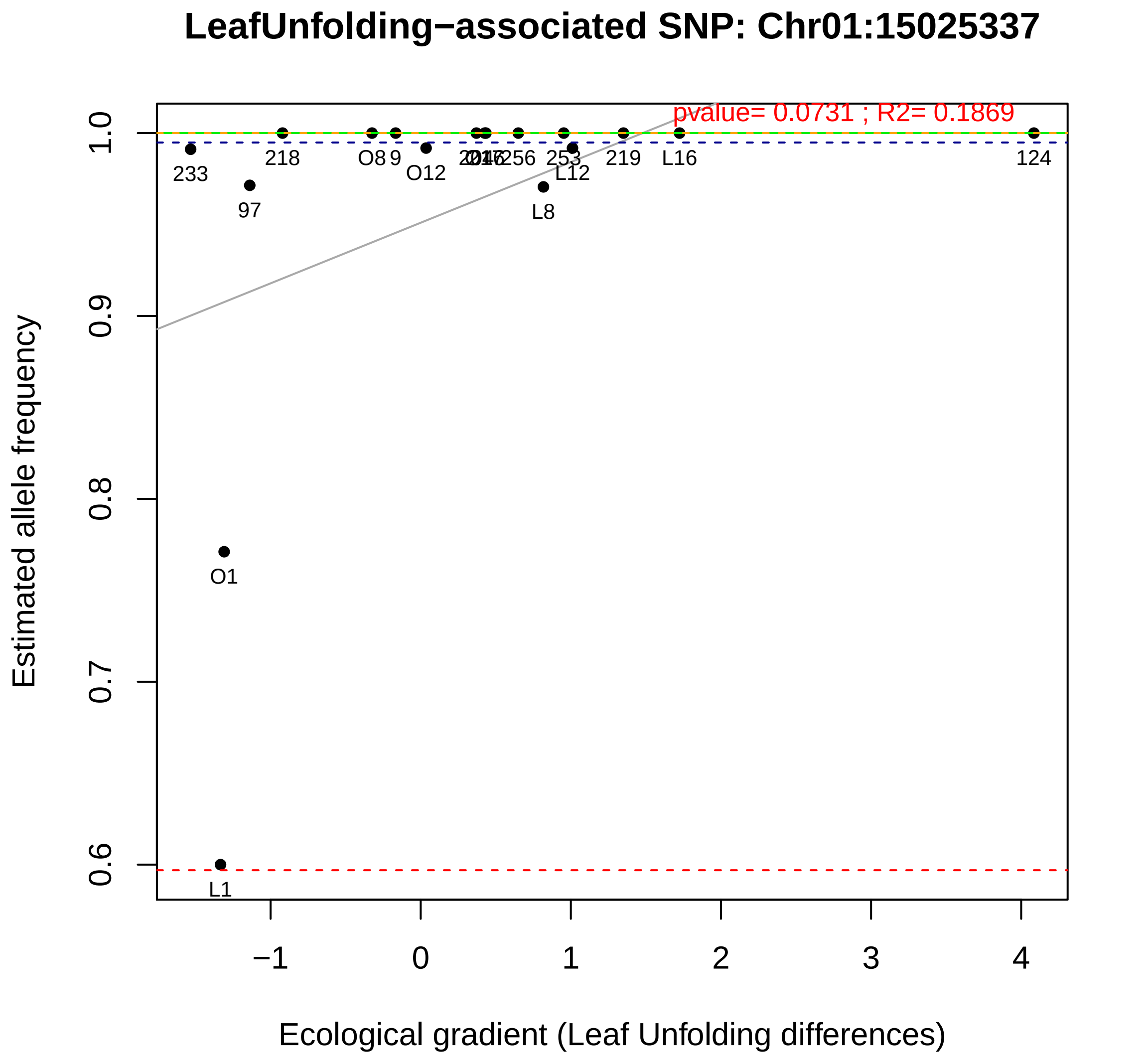

LeafUnfolding−associated SNP: Chr01:15025337
pvalue= 0.0731 ; R2= 0.1869
Estimated allele frequency
Ecological gradient (Leaf Unfolding differences)
1.0
0.9
0.8
0.7
0.6
−1
0
1
2
3
4
233
97
218
O8
9
O12
L8
253
L12
219
L16
124
O1
L1

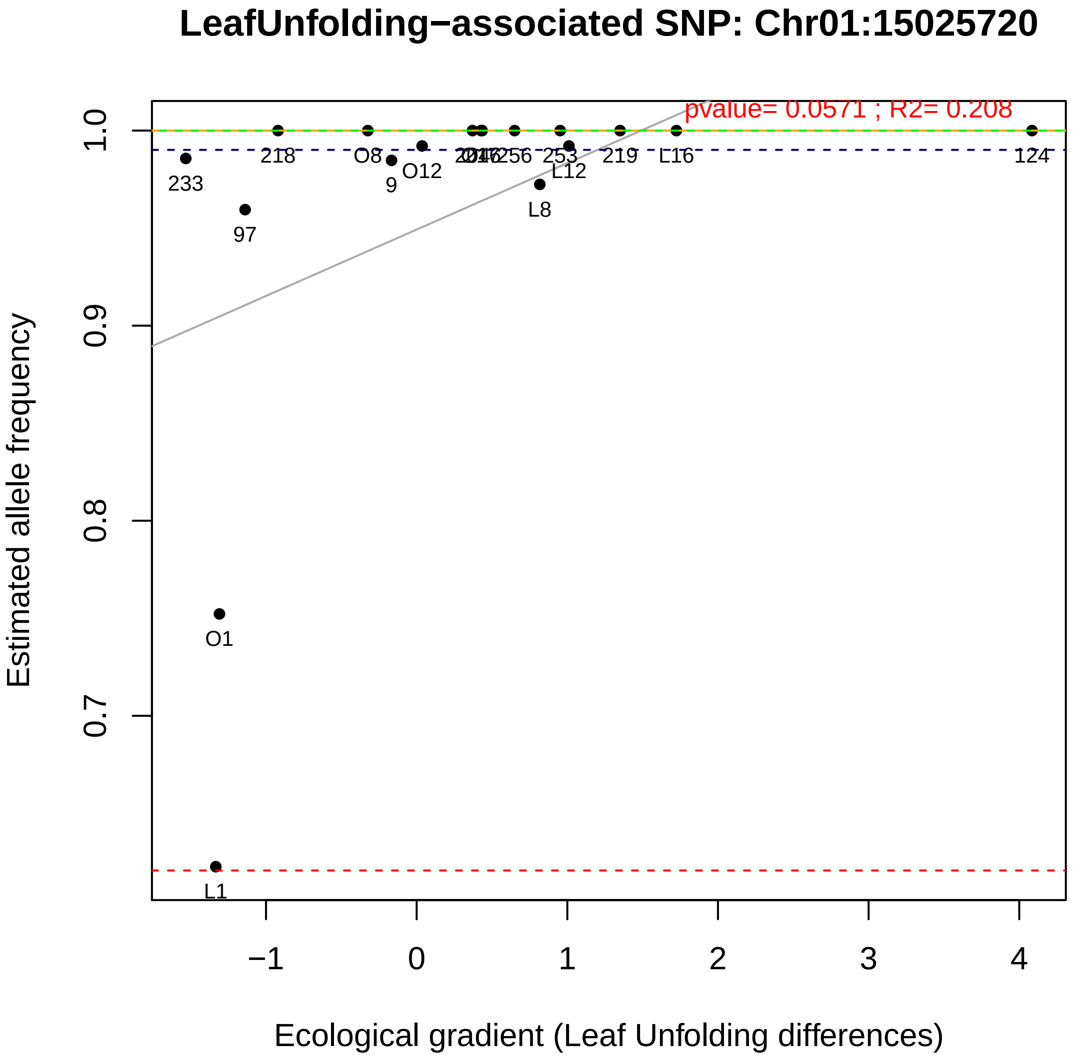
LeafUnfolding−associated SNP: Chr01:15025720
pvalue= 0.0571 ; R2= 0.208
Estimated allele frequency
1.0
0.9
0.8
0.7
233
97
218
O8
9
O12
256
253
L12
L8
219
L16
124
O1
L1
−1
0
1
2
3
4
Ecological gradient (Leaf Unfolding differences)

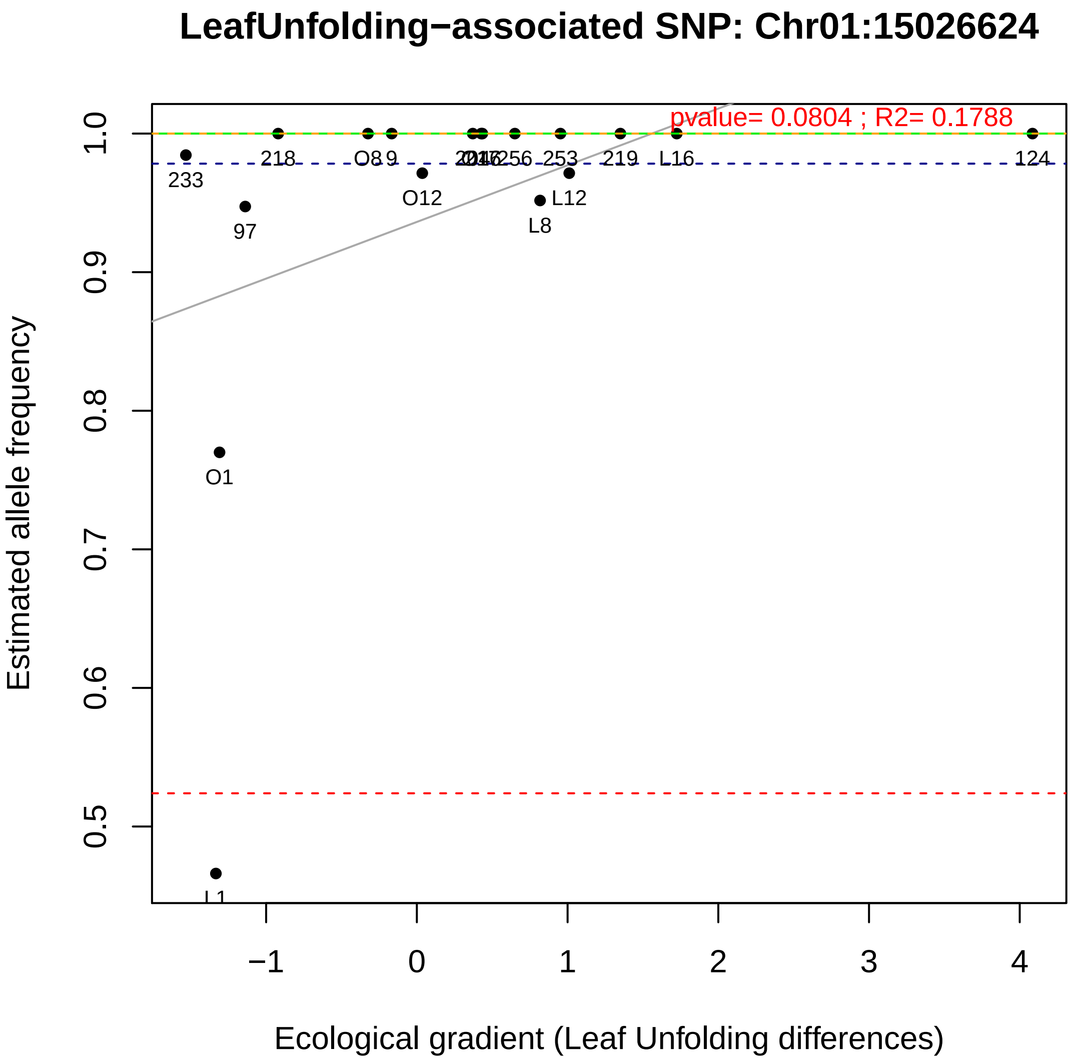
LeafUnfolding−associated SNP: Chr01:15026624
pvalue= 0.0804 ; R2= 0.1788
Estimated allele frequency
Ecological gradient (Leaf Unfolding differences)
1.0
0.9
0.8
0.7
0.6
0.5
−1
0
1
2
3
4
233
218
O8
9
O12
256
253
219
L16
124
97
L12
L8
O1

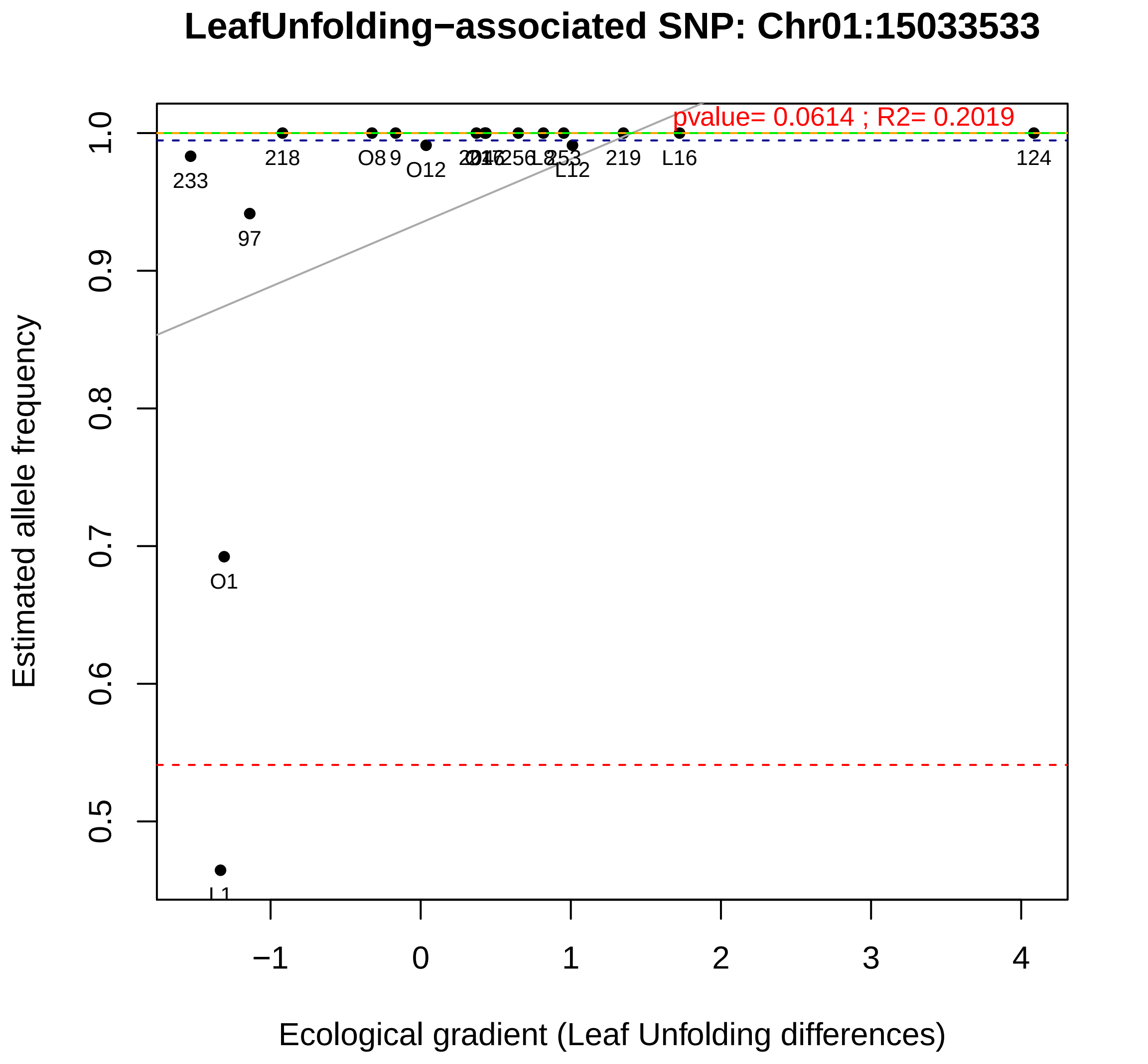
LeafUnfolding−associated SNP: Chr01:15033533
pvalue= 0.0614 ; R2= 0.2019
Estimated allele frequency
Ecological gradient (Leaf Unfolding differences)
1.0
0.9
0.8
0.7
0.6
0.5
−1
0
1
2
3
4
233
218
O8
9
O12
L12
219
L16
124
97
O1
L1

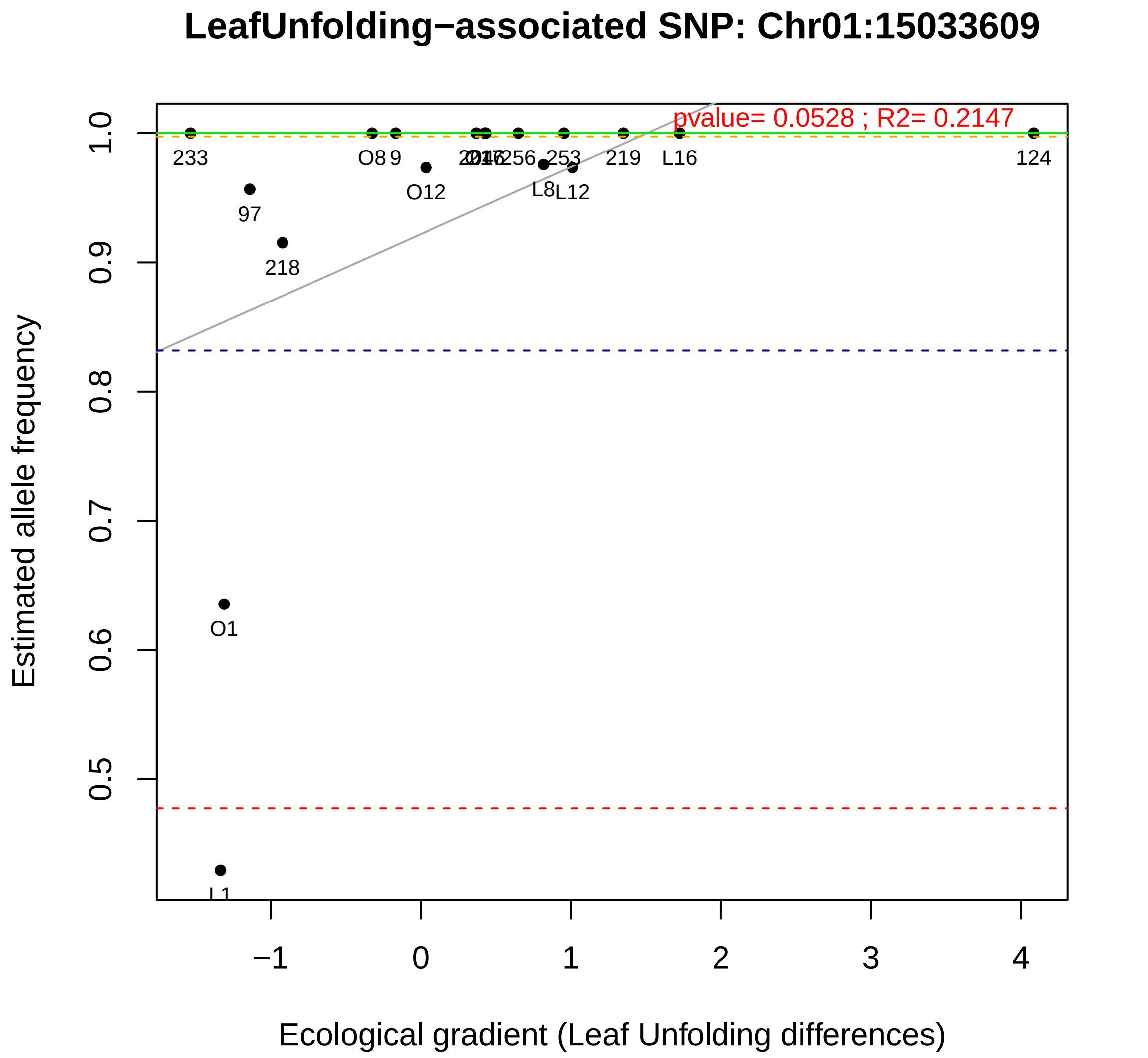
LeafUnfolding−associated SNP: Chr01:15033609
pvalue= 0.0528 ; R2= 0.2147
Estimated allele frequency
Ecological gradient (Leaf Unfolding differences)
233
O8
9
O12
256
253
219
L16
124
97
218
L8
L12
O1
1.0
0.9
0.8
0.7
0.6
0.5
−1
0
1
2
3
4

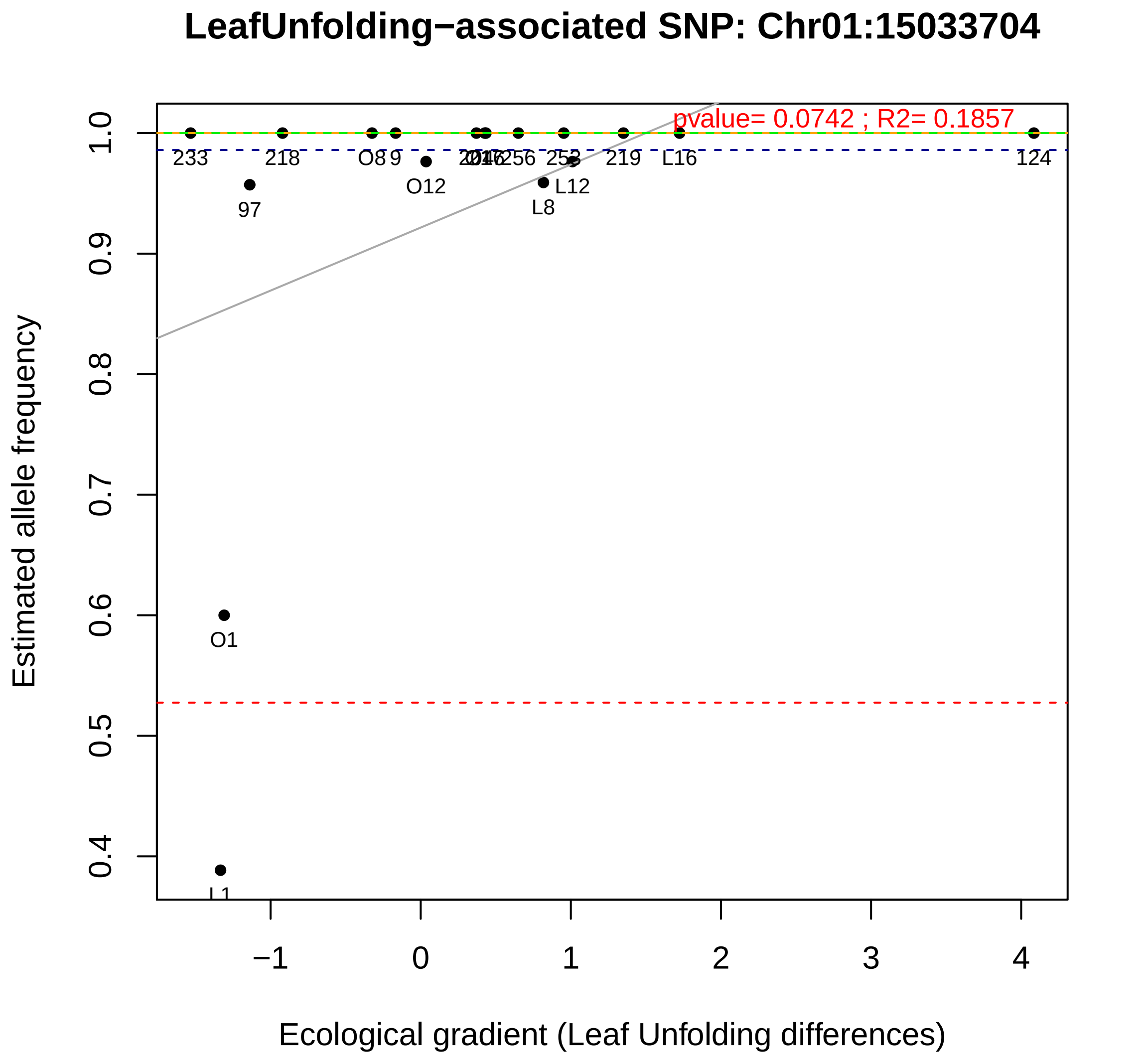
LeafUnfolding−associated SNP: Chr01:15033704
pvalue= 0.0742 ; R2= 0.1857
Estimated allele frequency
Ecological gradient (Leaf Unfolding differences)
233
218
O8
9
O12
97
L8
L12
219
L16
124
O1
256
−1
0
1
2
3
4
0.4
0.5
0.6
0.7
0.8
0.9
1.0

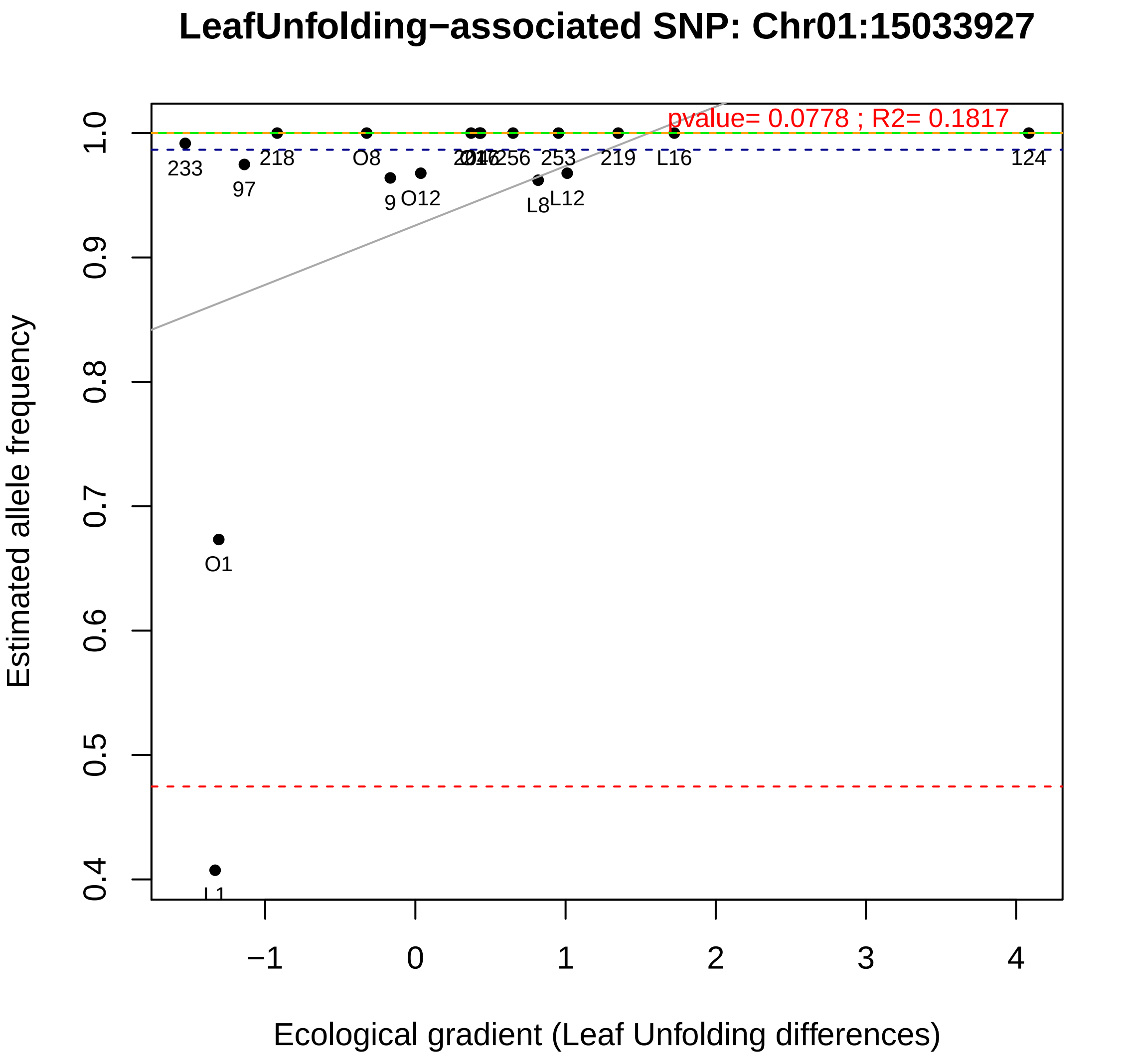
LeafUnfolding−associated SNP: Chr01:15033927
pvalue= 0.0778 ; R2= 0.1817
Estimated allele frequency
Ecological gradient (Leaf Unfolding differences)
0.4
0.5
0.6
0.7
0.8
0.9
1.0
−1
0
1
2
3
4
233
218
97
O8
9
O12
256
253
L8
L12
219
L16
124
O1

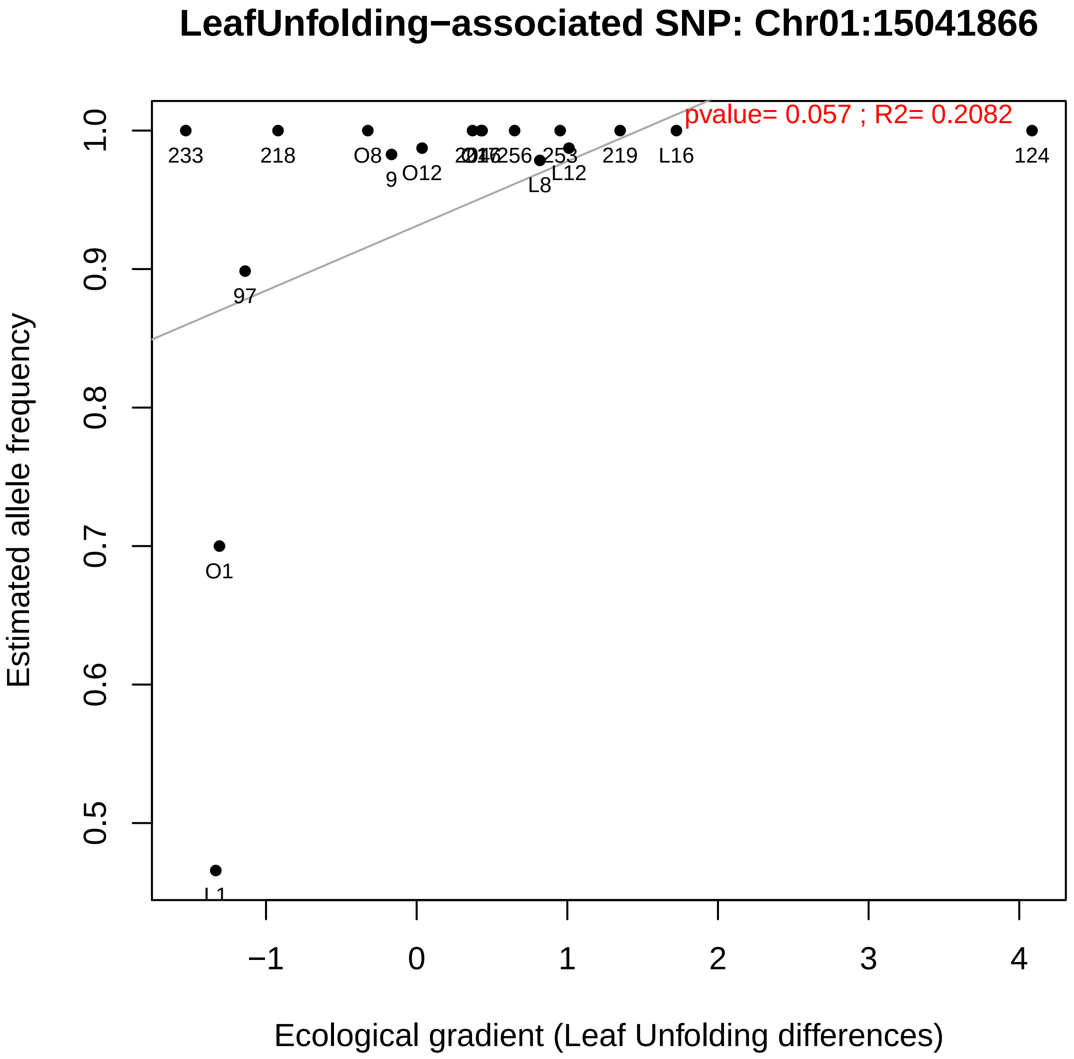
LeafUnfolding−associated SNP: Chr01:15041866
pvalue= 0.057 ; R2= 0.2082
Estimated allele frequency
Ecological gradient (Leaf Unfolding differences)
0.5
0.6
0.7
0.8
0.9
1.0
−1
0
1
2
3
4
233
218
O8
9
O12
256
253
L8
L12
219
L16
124
97
O1
L1

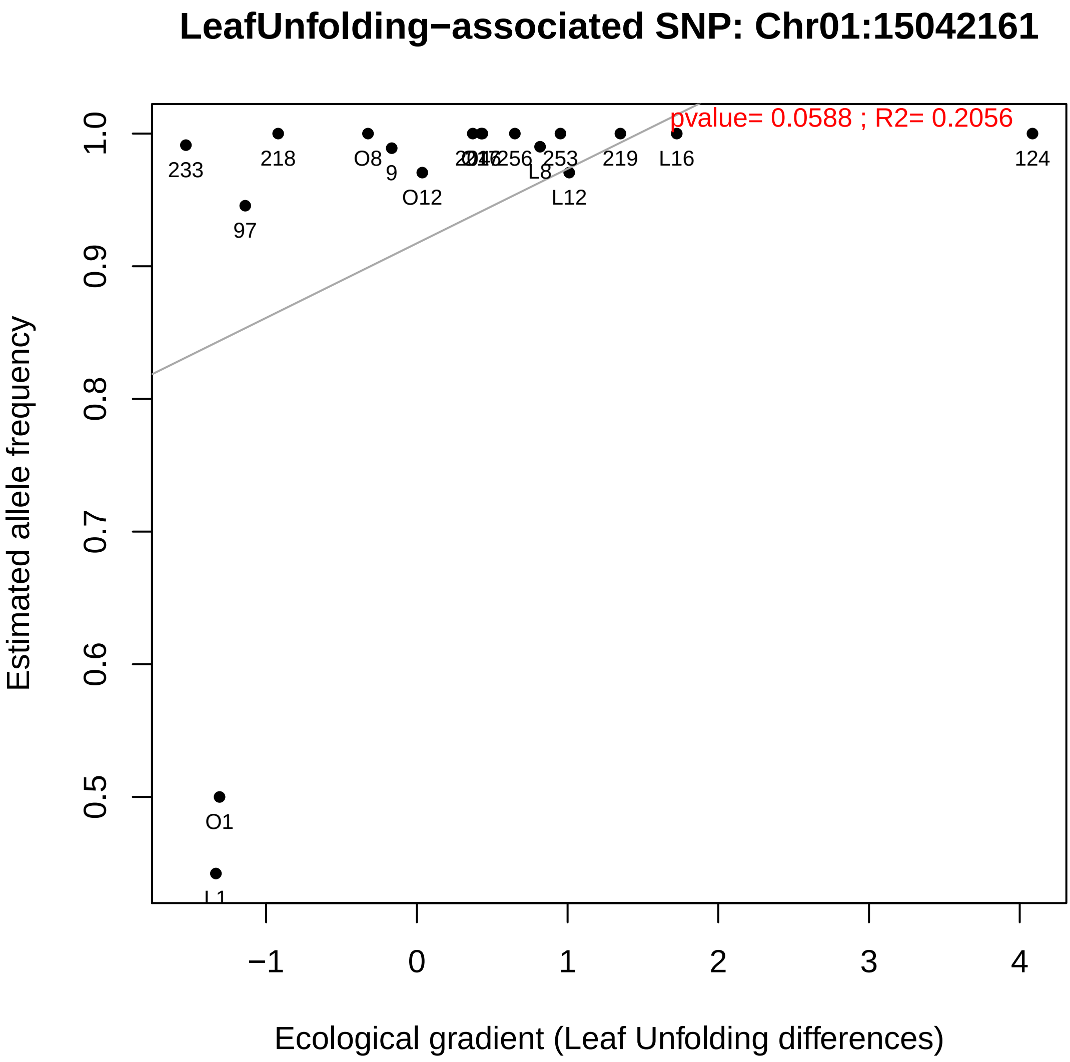
LeafUnfolding−associated SNP: Chr01:15042161
pvalue= 0.0588 ; R2= 0.2056
Estimated allele frequency
Ecological gradient (Leaf Unfolding differences)
1.0
0.9
0.8
0.7
0.6
0.5
−1
0
1
2
3
4
233
218
O8
9
O12
97
256
L8
253
L12
219
L16
124
O1
L1

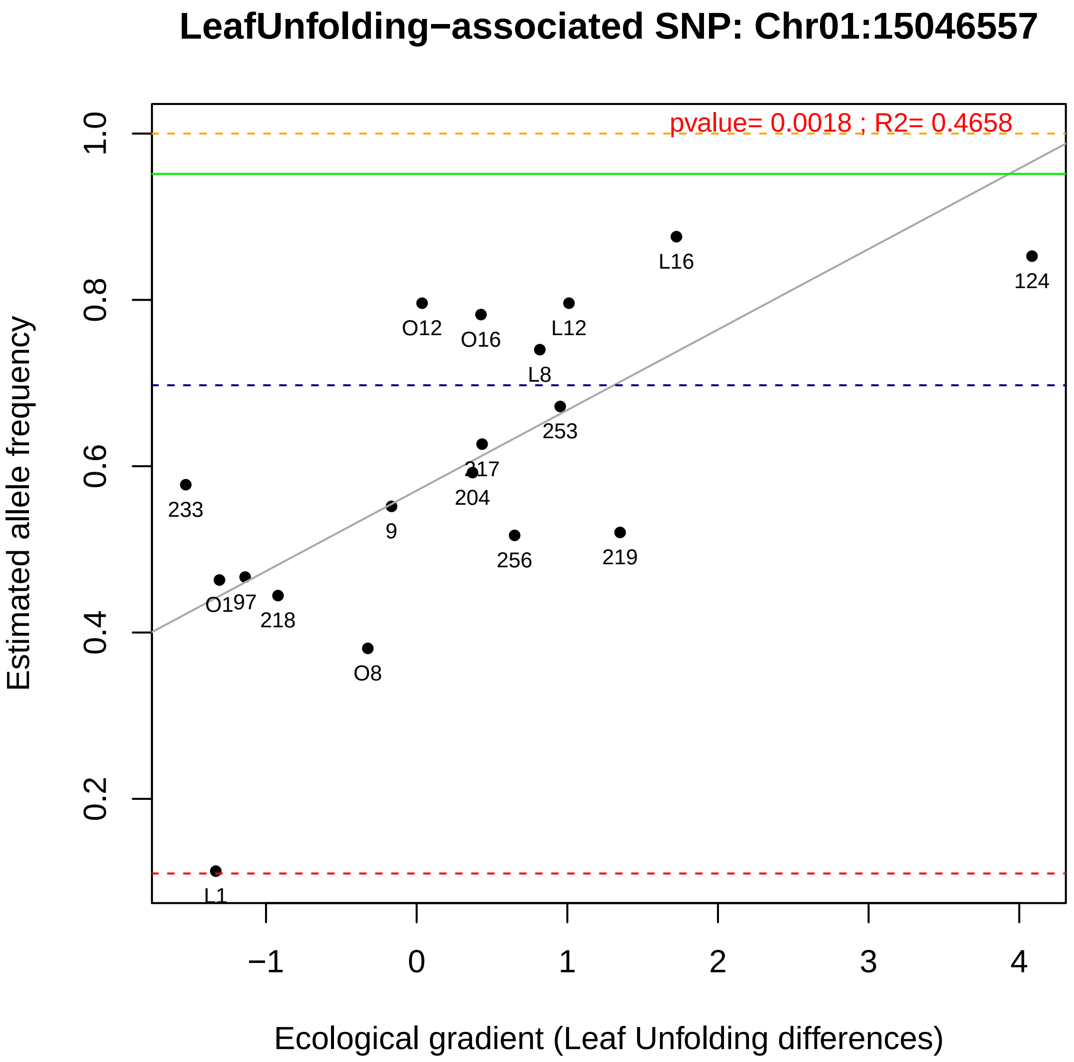

LeafUnfolding−associated SNP: Chr01:15046557
pvalue= 0.0018 ; R2= 0.4658
Estimated allele frequency
Ecological gradient (Leaf Unfolding differences)
1.0
0.8
0.6
0.4
0.2
−1
0
1
2
3
4
L16
124
O12
O16
L12
L8
253
217
204
233
9
256
219
O1
97
218
O8
L1

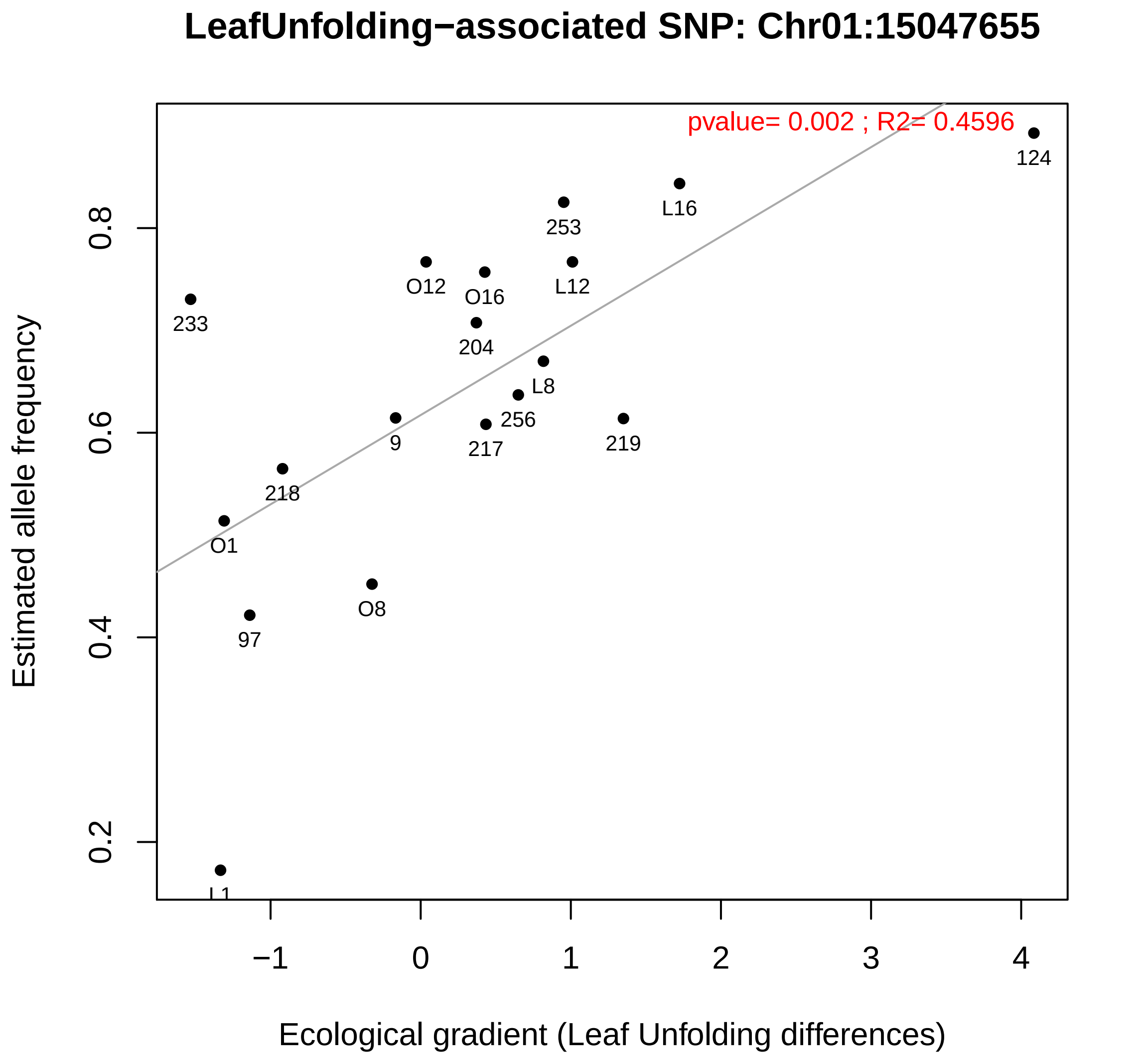

LeafUnfolding−associated SNP: Chr01:15047655
pvalue= 0.002 ; R2= 0.4596
124
L16
253
L12
O12
O16
233
204
L8
256
9
217
219
218
O1
O8
97
L1
Estimated allele frequency
0.2
0.4
0.6
0.8
−1
0
1
2
3
4
Ecological gradient (Leaf Unfolding differences)

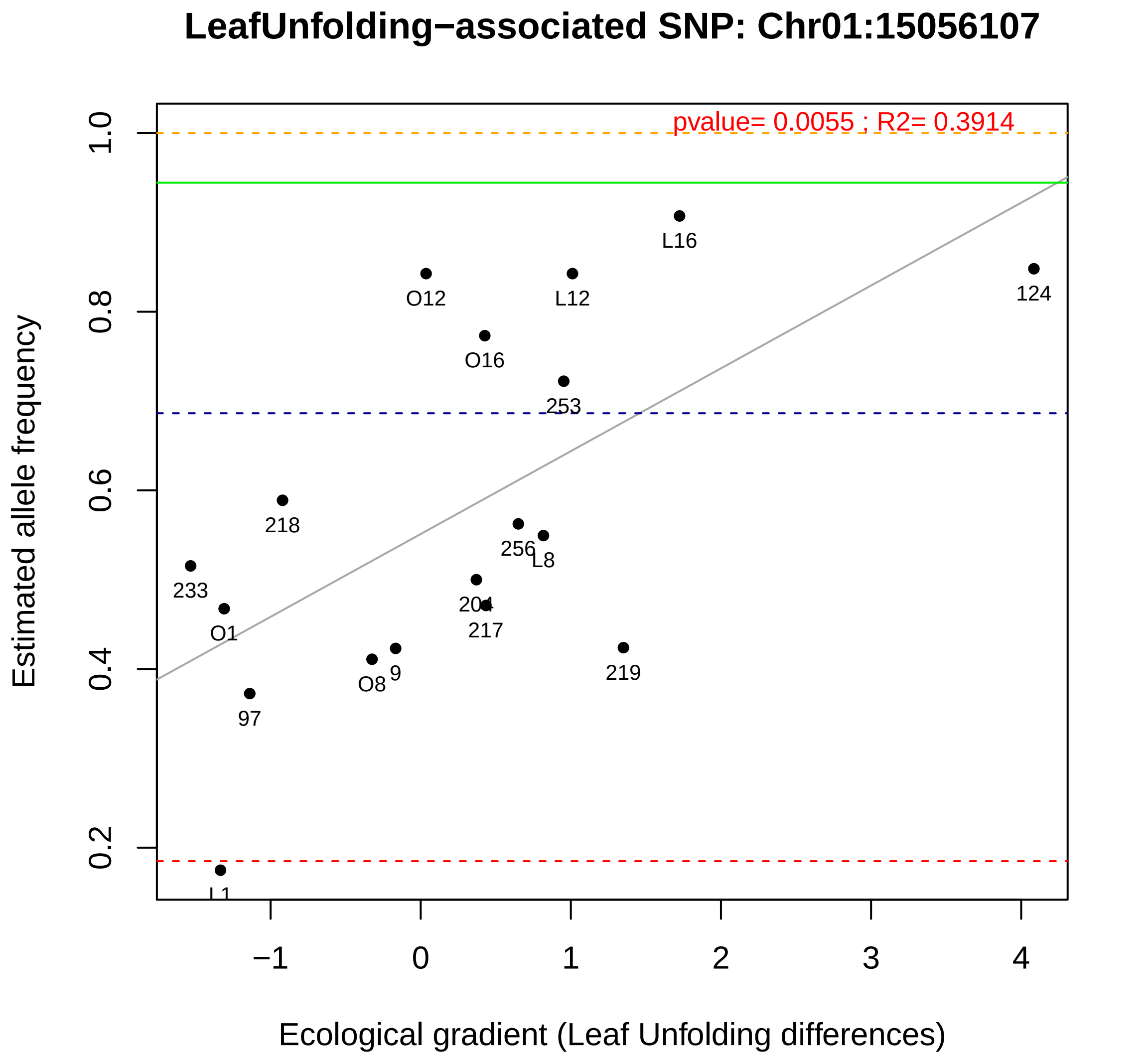

LeafUnfolding−associated SNP: Chr01:15056107
pvalue= 0.0055 ; R2= 0.3914
Estimated allele frequency
0.2
0.4
0.6
0.8
1.0
−1
0
1
2
3
4
Ecological gradient (Leaf Unfolding differences)
L16
O12
L12
124
O16
253
218
256
L8
233
204
217
O1
O8
9
219
97

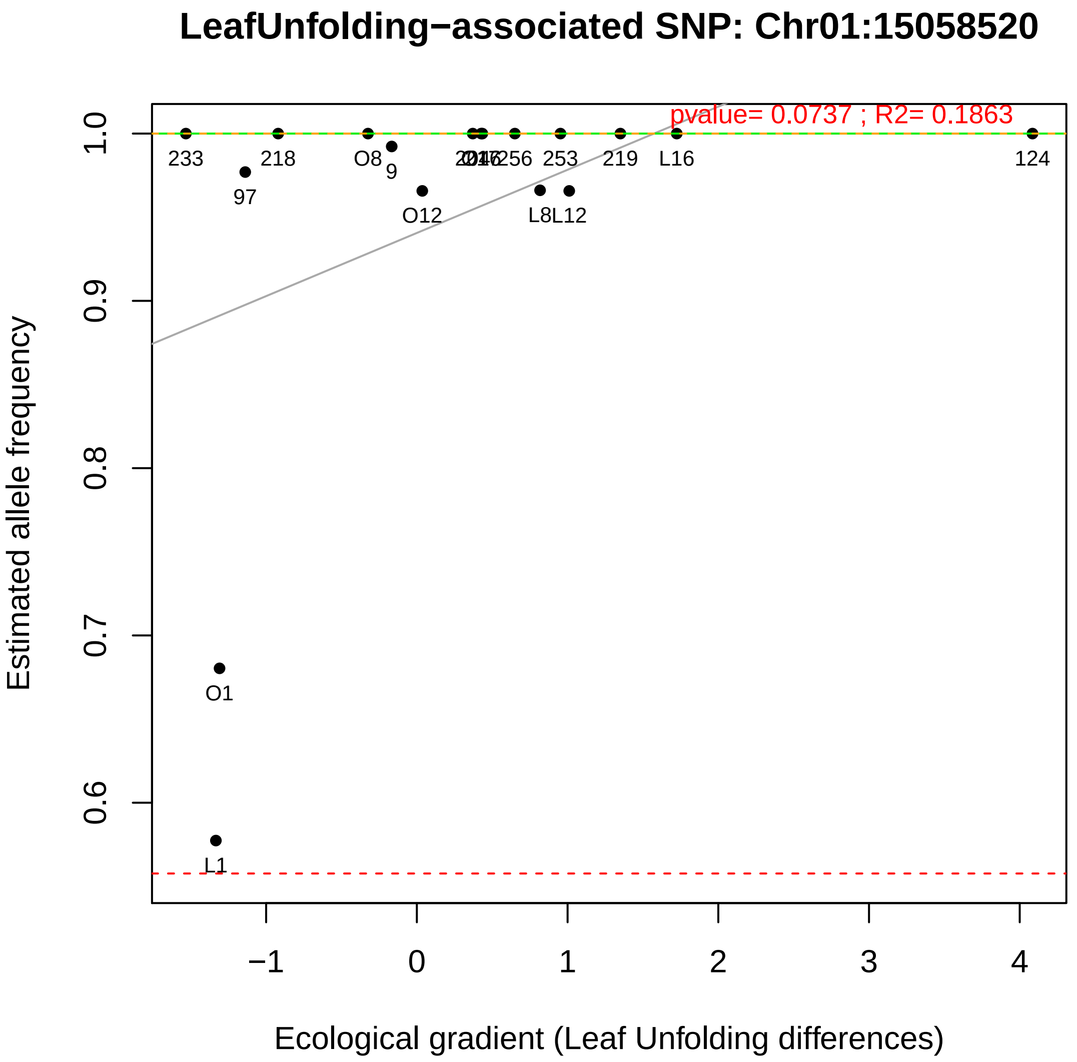
LeafUnfolding−associated SNP: Chr01:15058520
pvalue= 0.0737 ; R2= 0.1863
Estimated allele frequency
Ecological gradient (Leaf Unfolding differences)
1.0
0.9
0.8
0.7
0.6
−1
0
1
2
3
4
233
218
O8
9
256
253
219
L16
124
97
O12
L8
L12
O1
L1

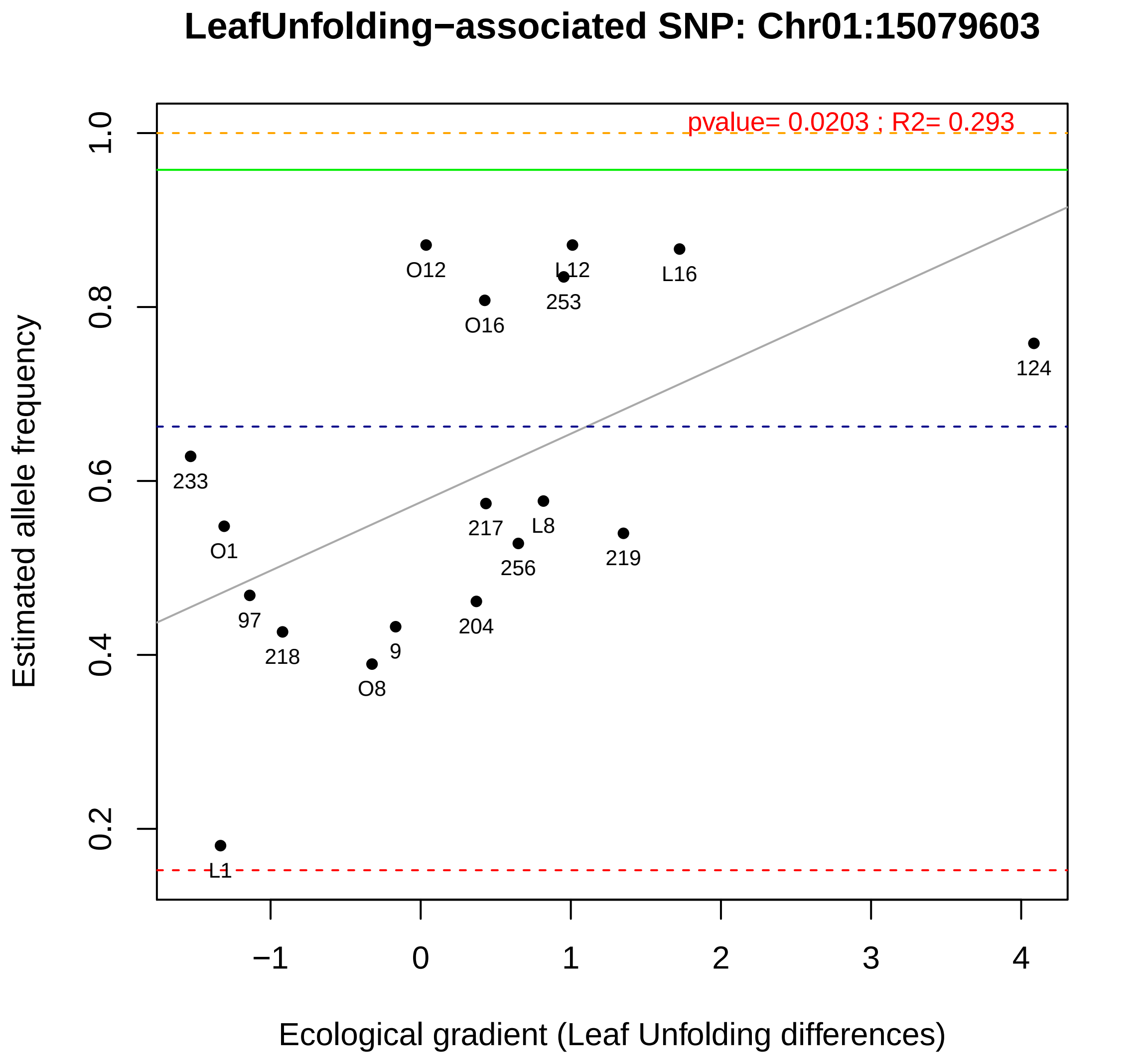

LeafUnfolding−associated SNP: Chr01:15079603
pvalue= 0.0203 ; R2= 0.293
Estimated allele frequency
0.2
0.4
0.6
0.8
1.0
−1
0
1
2
3
4
Ecological gradient (Leaf Unfolding differences)
O12
L12
L16
253
O16
124
233
O1
217
L8
256
219
97
218
9
204
O8
L1

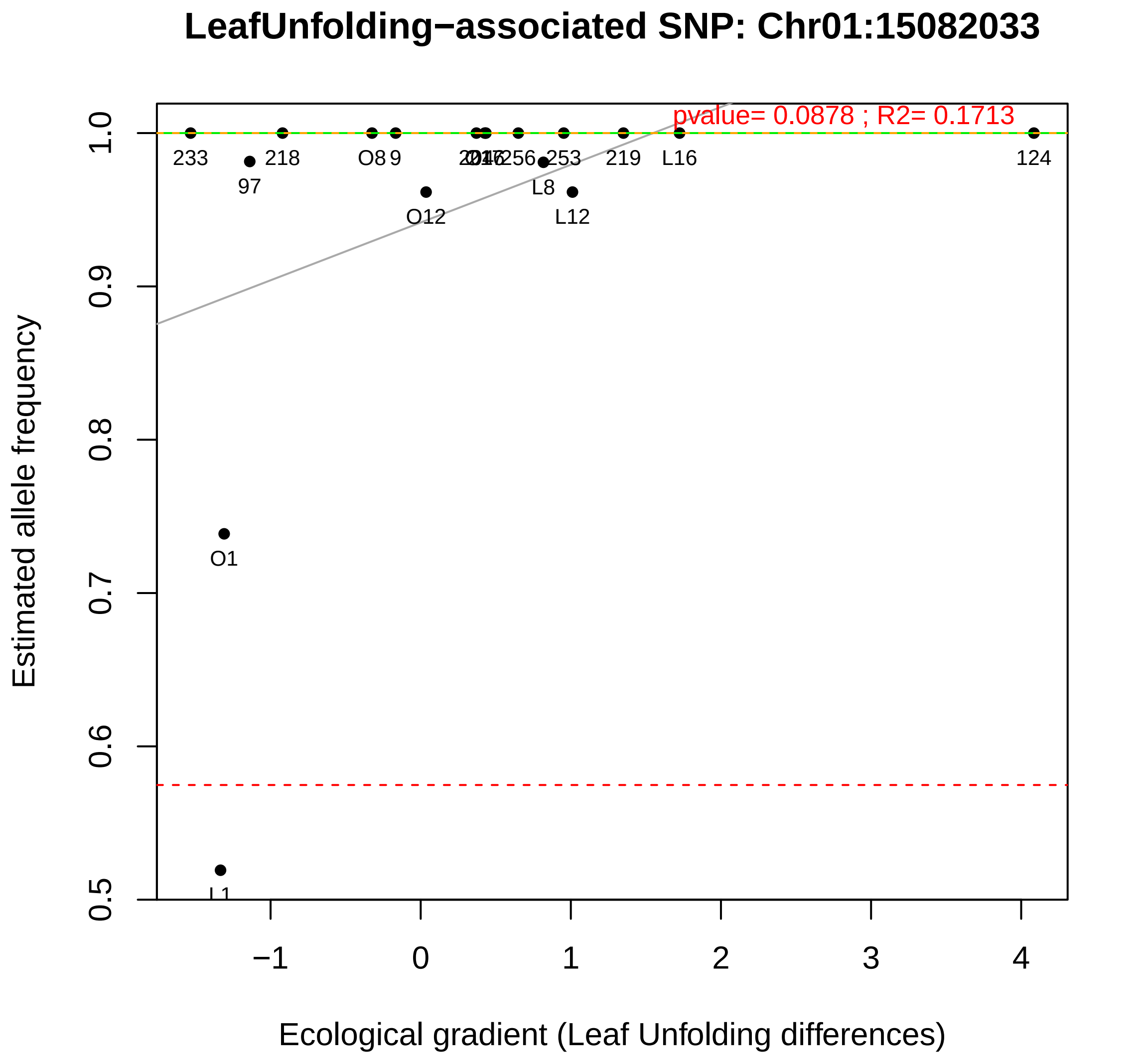
LeafUnfolding−associated SNP: Chr01:15082033
pvalue= 0.0878 ; R2= 0.1713
Estimated allele frequency
Ecological gradient (Leaf Unfolding differences)
233
97
218
O8
9
O12
L8
253
L12
219
L16
124
O1
L1

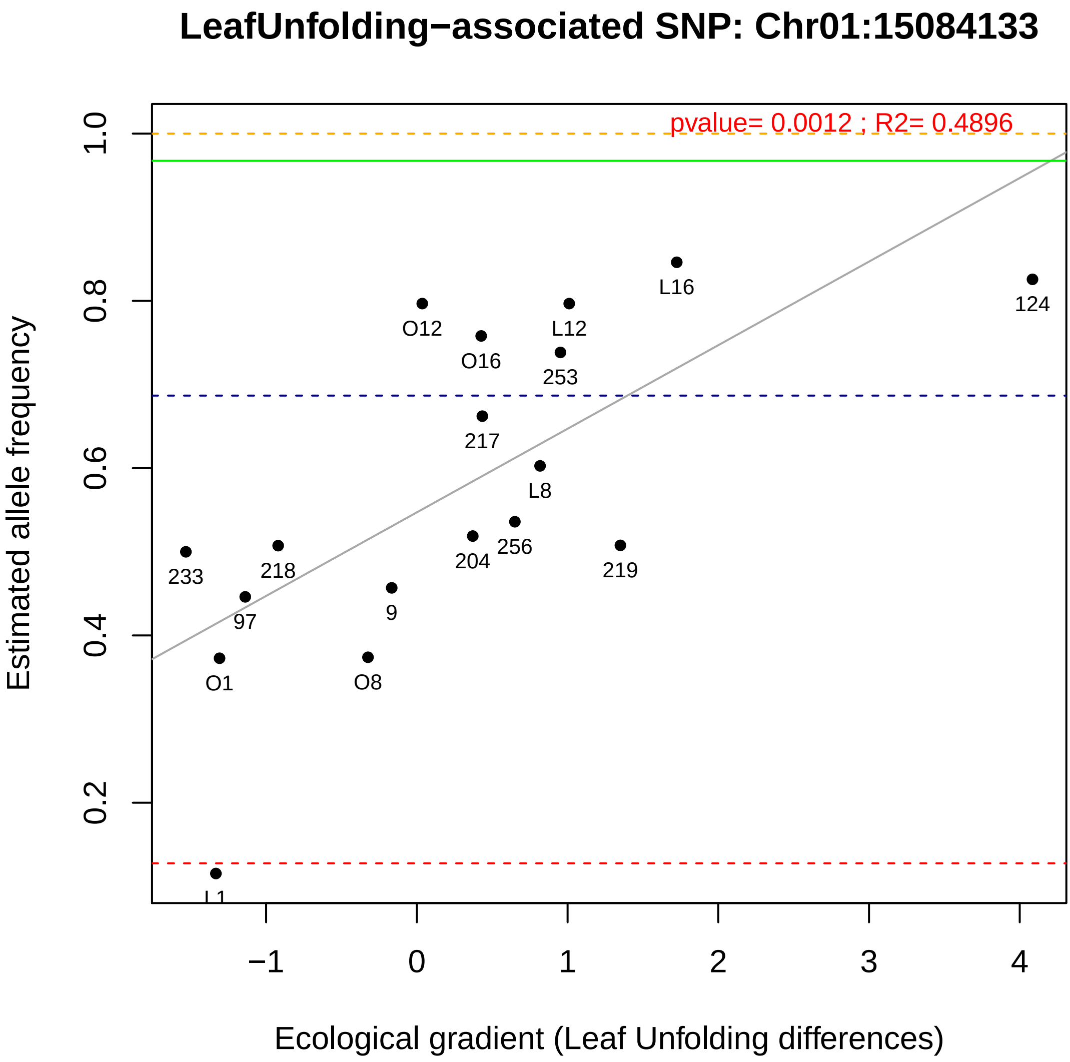

LeafUnfolding−associated SNP: Chr01:15084133
pvalue= 0.0012 ; R2= 0.4896
Estimated allele frequency
0.2
0.4
0.6
0.8
1.0
−1
0
1
2
3
4
Ecological gradient (Leaf Unfolding differences)
233
218
97
O1
O8
9
O12
O16
217
204
256
L8
253
L12
L16
219
124

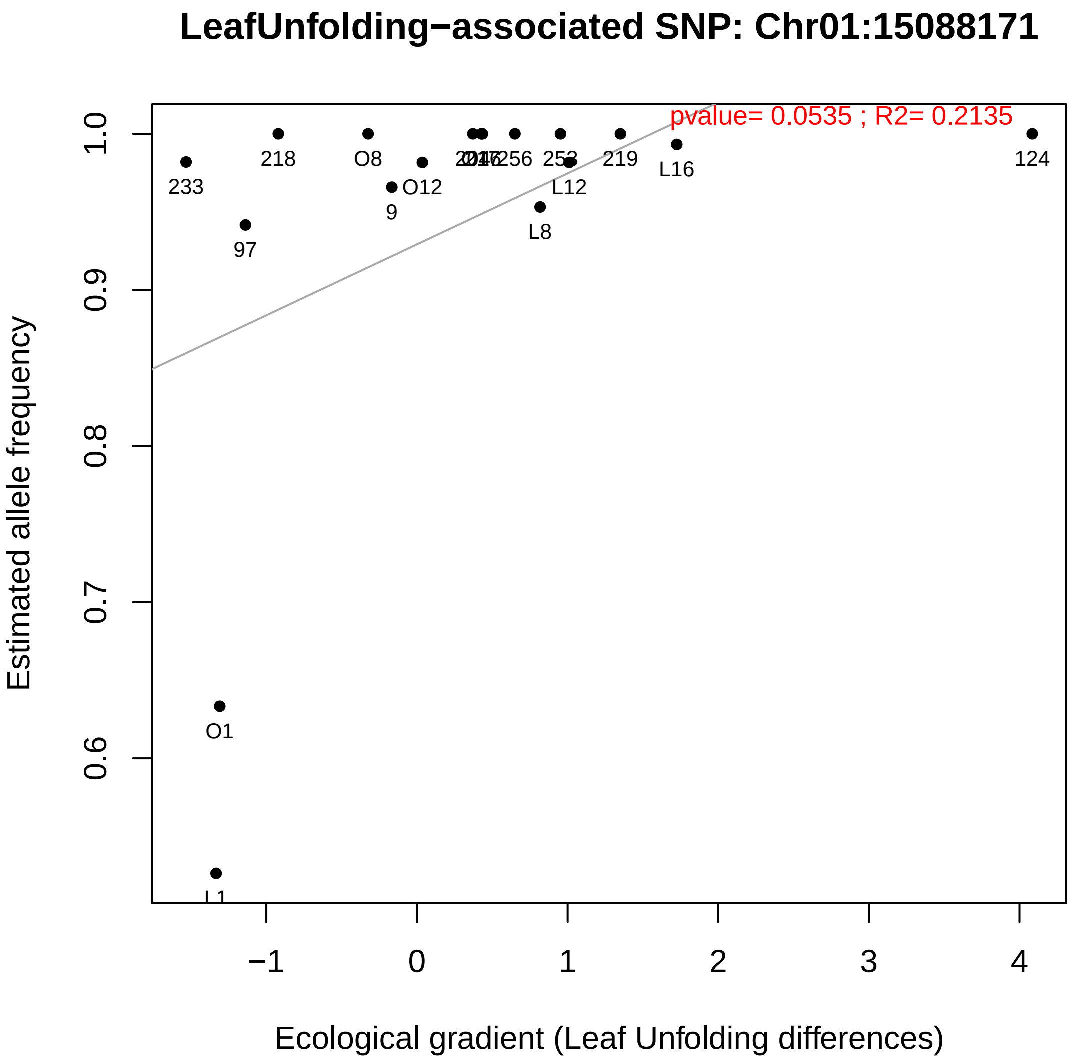
LeafUnfolding−associated SNP: Chr01:15088171
pvalue= 0.0535 ; R2= 0.2135
Estimated allele frequency
Ecological gradient (Leaf Unfolding differences)
1.0
0.9
0.8
0.7
0.6
−1
0
1
2
3
4
233
218
O8
O12
9
97
256
L12
L8
219
L16
124
O1

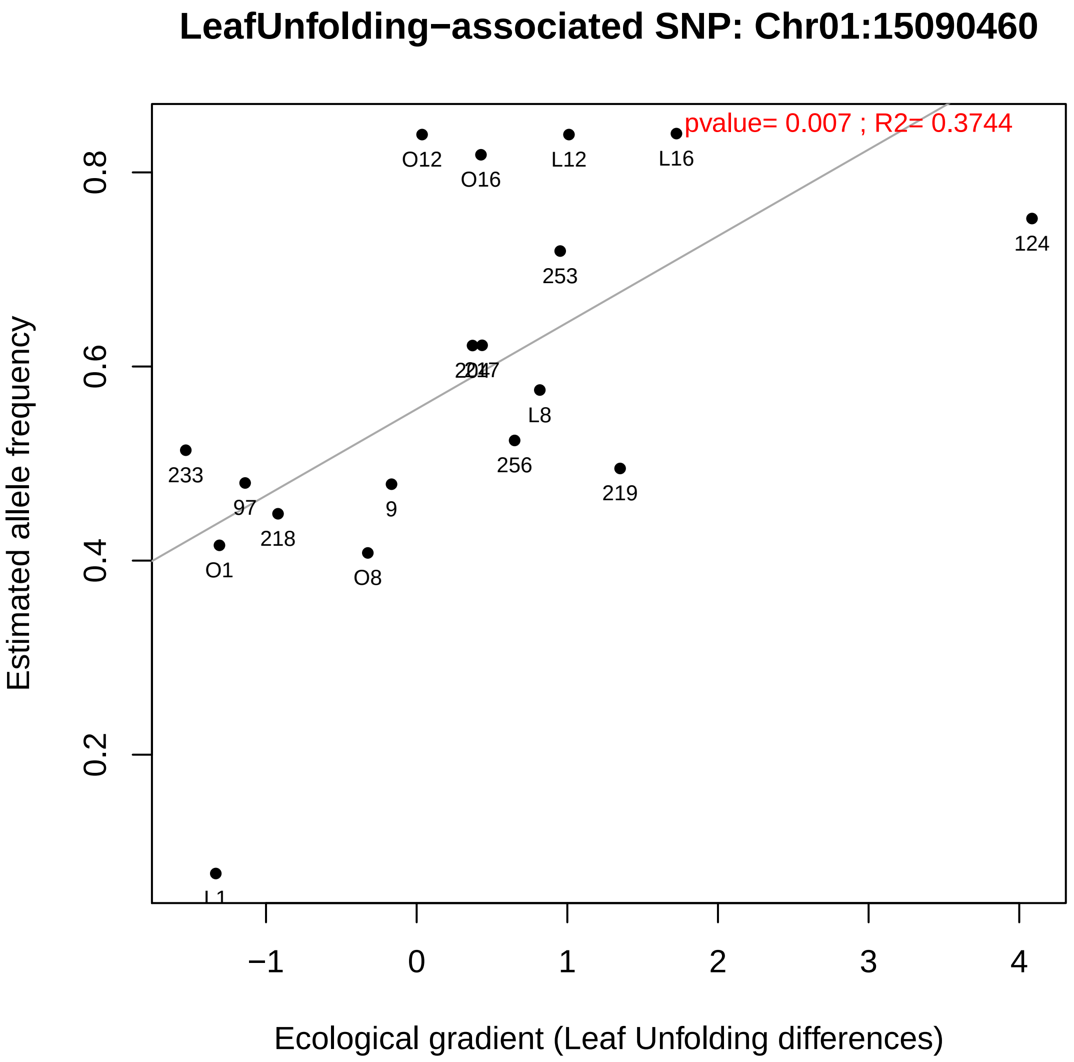

LeafUnfolding−associated SNP: Chr01:15090460
pvalue= 0.007 ; R2= 0.3744
Estimated allele frequency
Ecological gradient (Leaf Unfolding differences)
0.2
0.4
0.6
0.8
−1
0
1
2
3
4
O12
O16
L12
L16
124
253
L8
256
219
233
97
218
9
O1
O8

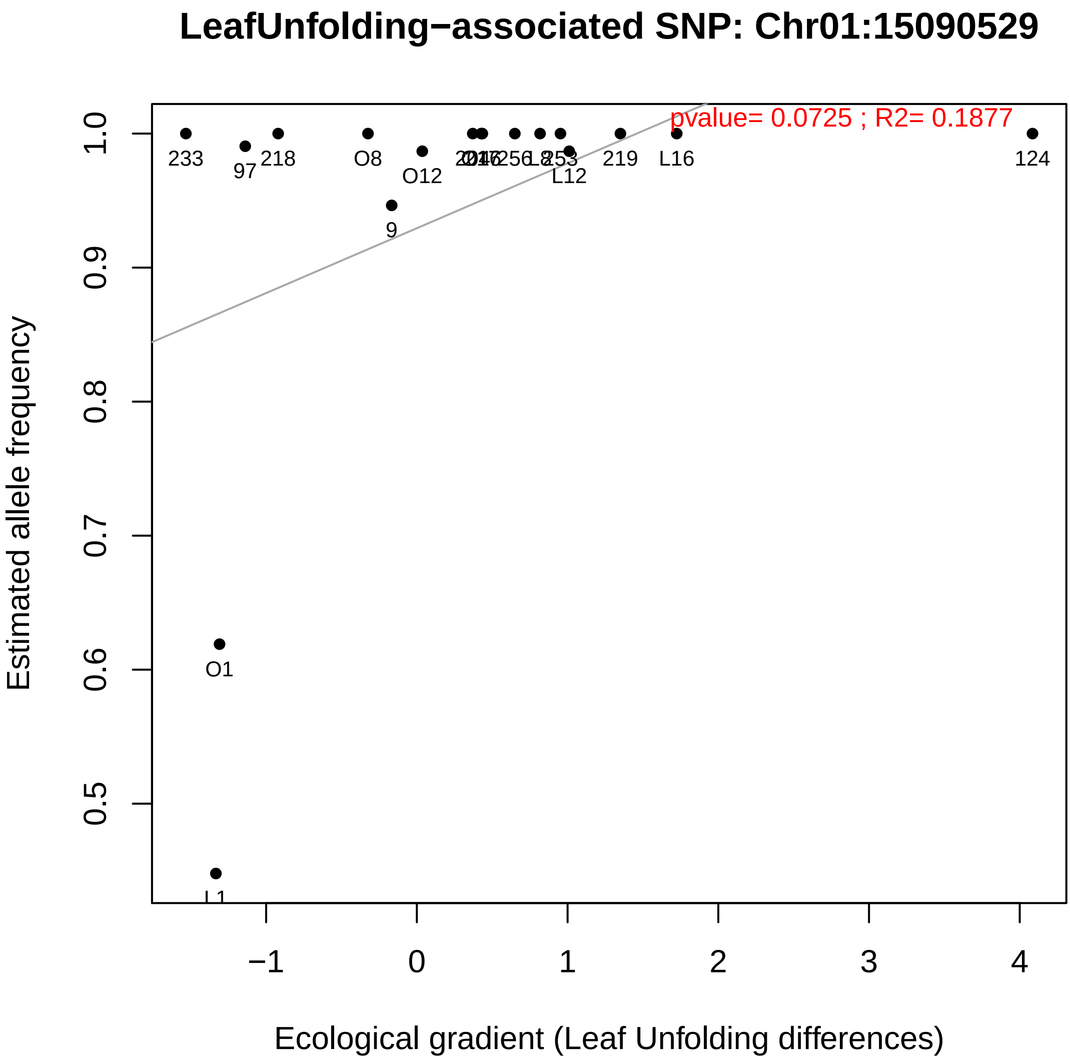
LeafUnfolding−associated SNP: Chr01:15090529
pvalue= 0.0725 ; R2= 0.1877
Estimated allele frequency
0.5
0.6
0.7
0.8
0.9
1.0
−1
0
1
2
3
4
Ecological gradient (Leaf Unfolding differences)
233
97
218
O8
O12
9
219
L16
L12
124
O1
L1

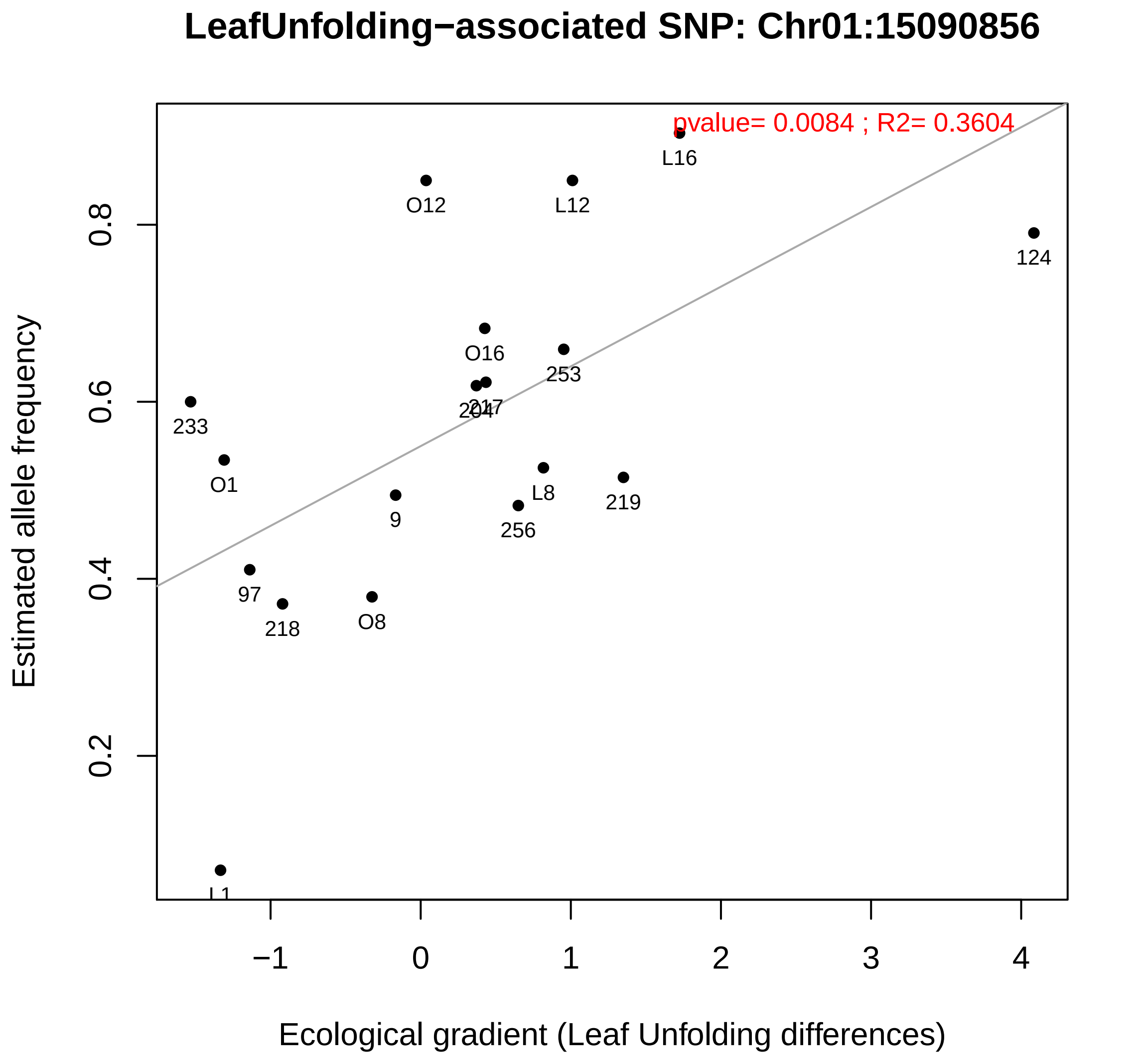
LeafUnfolding−associated SNP: Chr01:15090856
pvalue= 0.0084 ; R2= 0.3604
Estimated allele frequency
Ecological gradient (Leaf Unfolding differences)
0.2
0.4
0.6
0.8
−1
0
1
2
3
4
L16
O12
L12
124
O16
253
217
204
233
O1
L8
219
9
256
97
218
O8

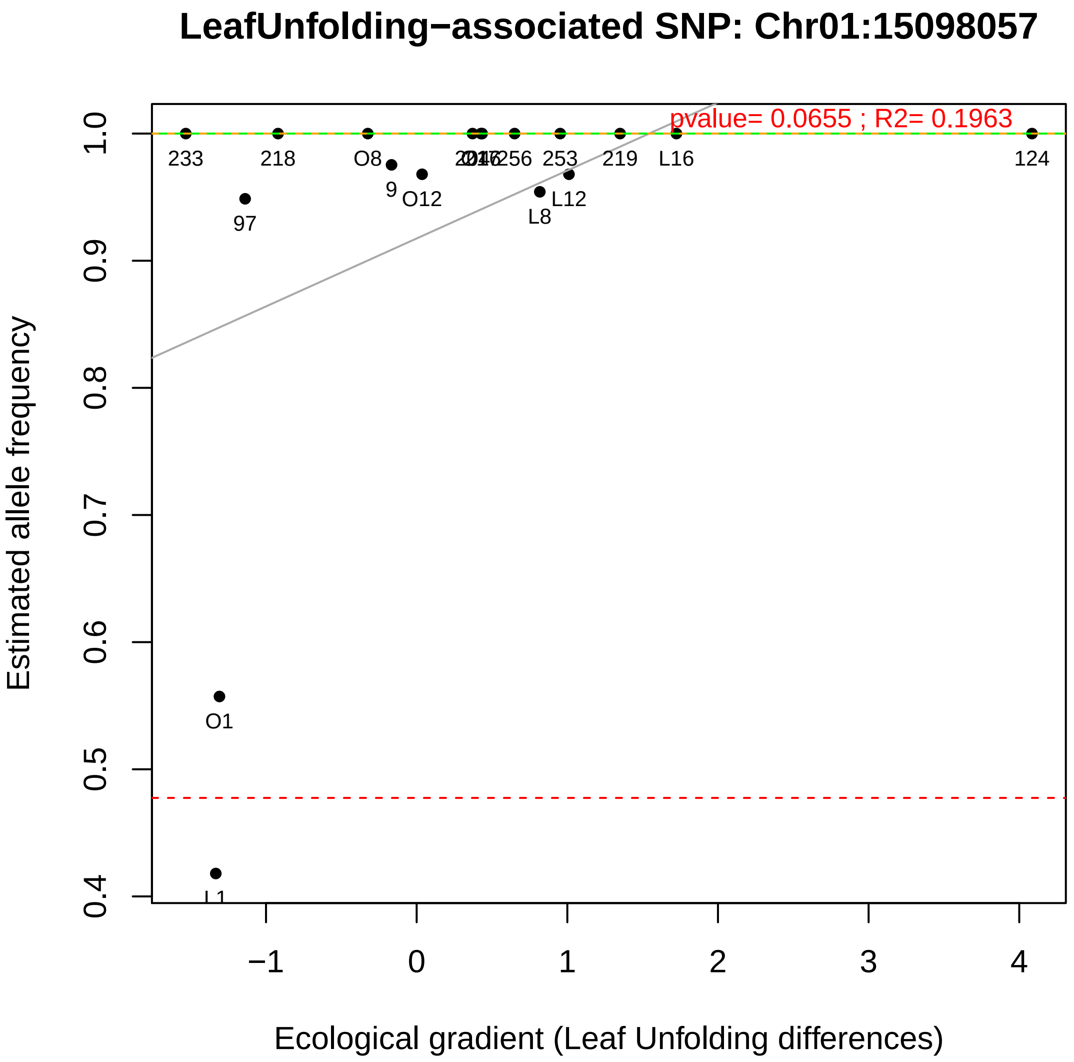

LeafUnfolding−associated SNP: Chr01:15098057
pvalue= 0.0655 ; R2= 0.1963
Estimated allele frequency
Ecological gradient (Leaf Unfolding differences)
233
218
O8
9
O12
256
253
219
L16
124
97
L8
L12
O1
1.0
0.9
0.8
0.7
0.6
0.5
0.4
−1
0
1
2
3
4

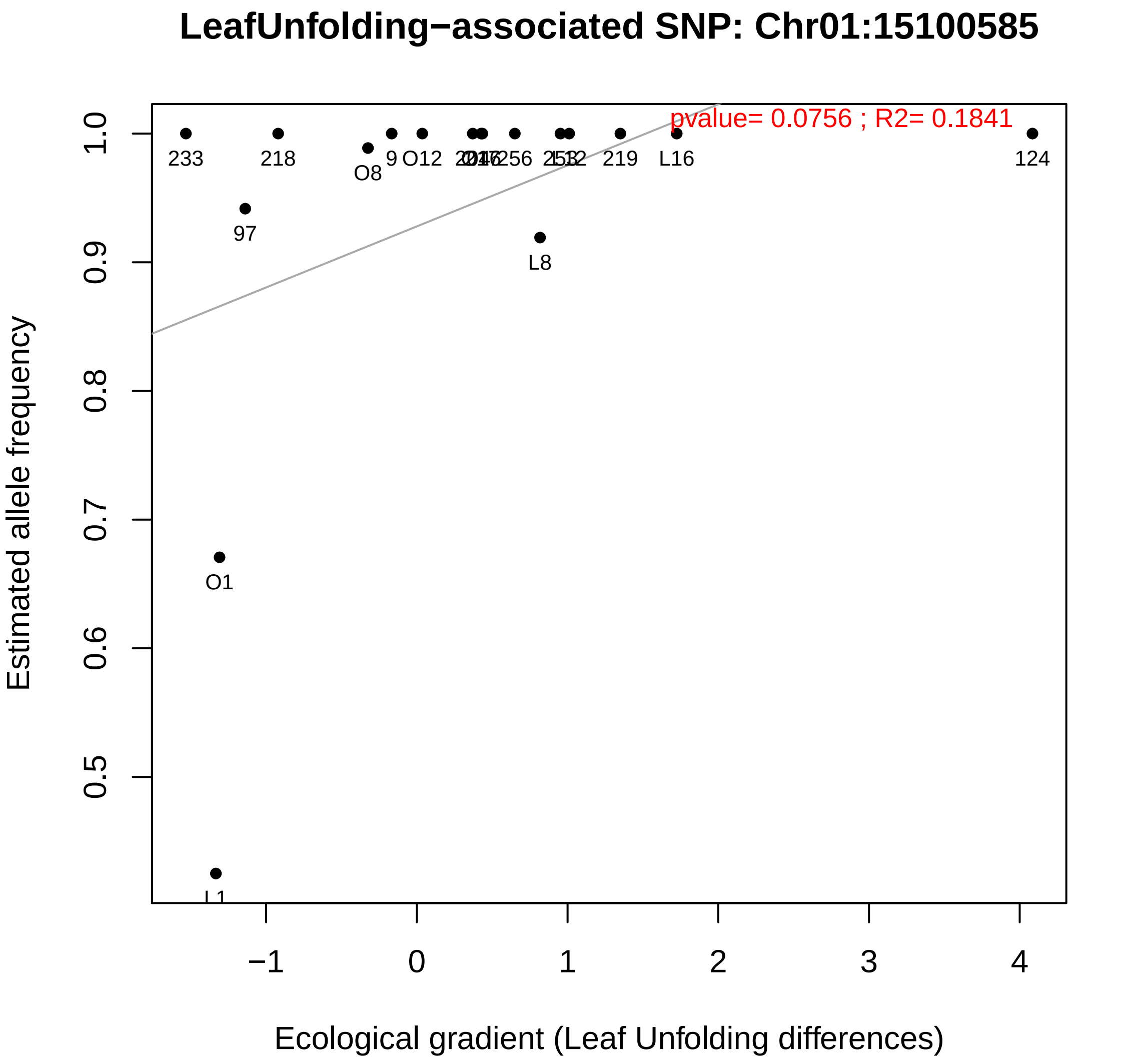

LeafUnfolding−associated SNP: Chr01:15100585
pvalue= 0.0756 ; R2= 0.1841
Estimated allele frequency
Ecological gradient (Leaf Unfolding differences)
1.0
0.9
0.8
0.7
0.6
0.5
−1
0
1
2
3
4
233
218
O8
9
O12
219
L16
124
97
L8
O1
L1

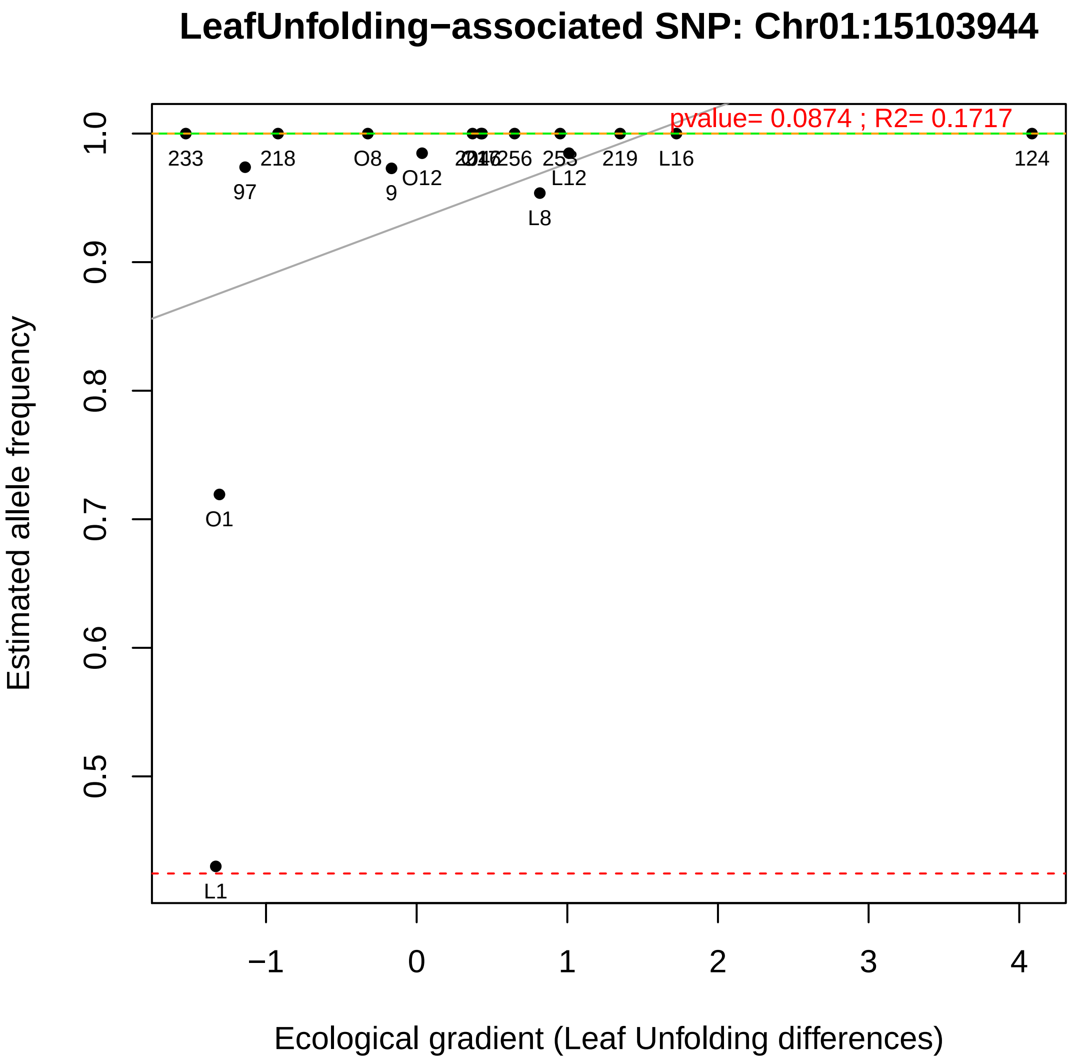
LeafUnfolding−associated SNP: Chr01:15103944
pvalue= 0.0874 ; R2= 0.1717
Estimated allele frequency
Ecological gradient (Leaf Unfolding differences)
1.0
0.9
0.8
0.7
0.6
0.5
−1
0
1
2
3
4
233
218
O8
256
253
219
L16
124
97
9
O12
L12
L8
O1
L1

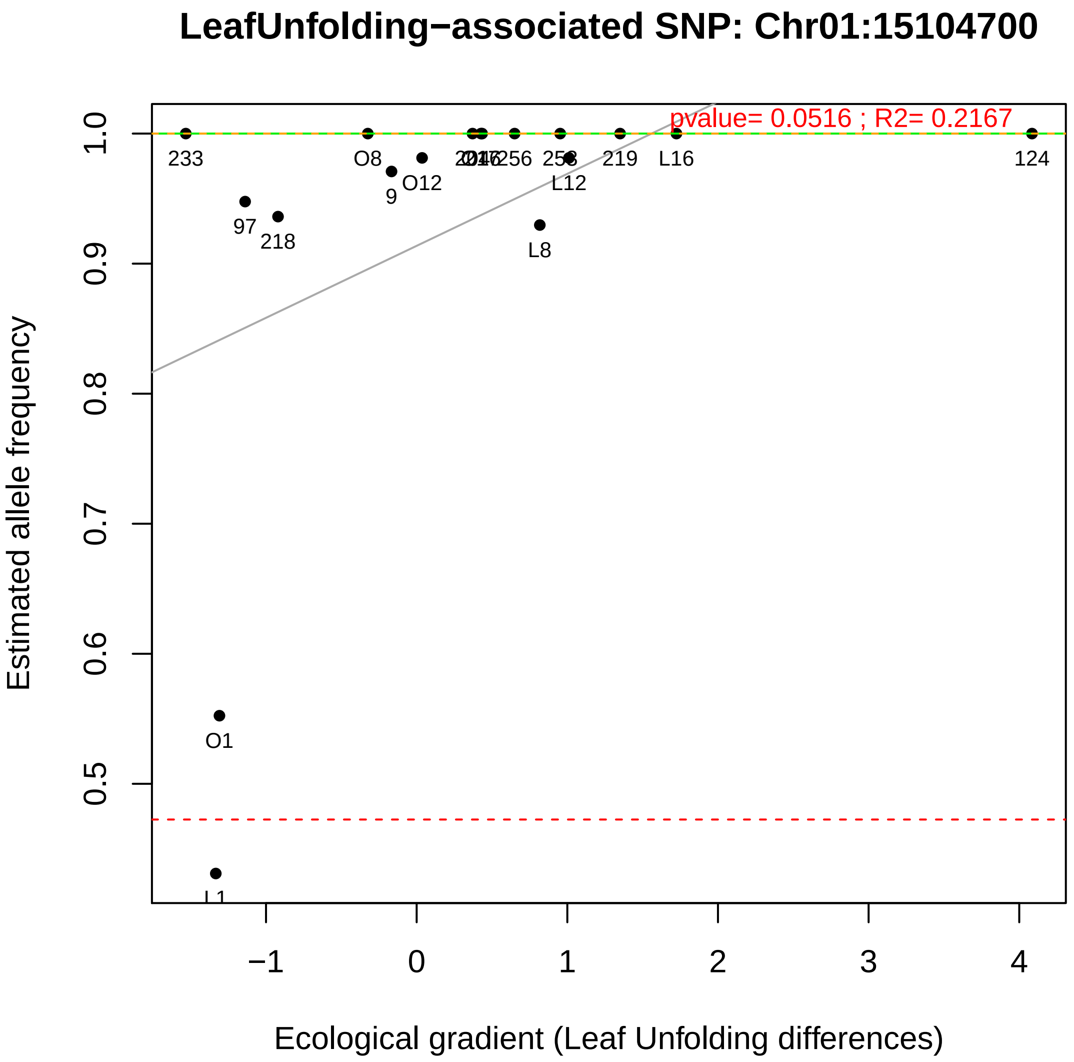

LeafUnfolding−associated SNP: Chr01:15104700
pvalue= 0.0516 ; R2= 0.2167
Estimated allele frequency
Ecological gradient (Leaf Unfolding differences)
1.0
0.9
0.8
0.7
0.6
0.5
−1
0
1
2
3
4
233
O8
O12
9
97
218
L8
L12
219
L16
124
O1

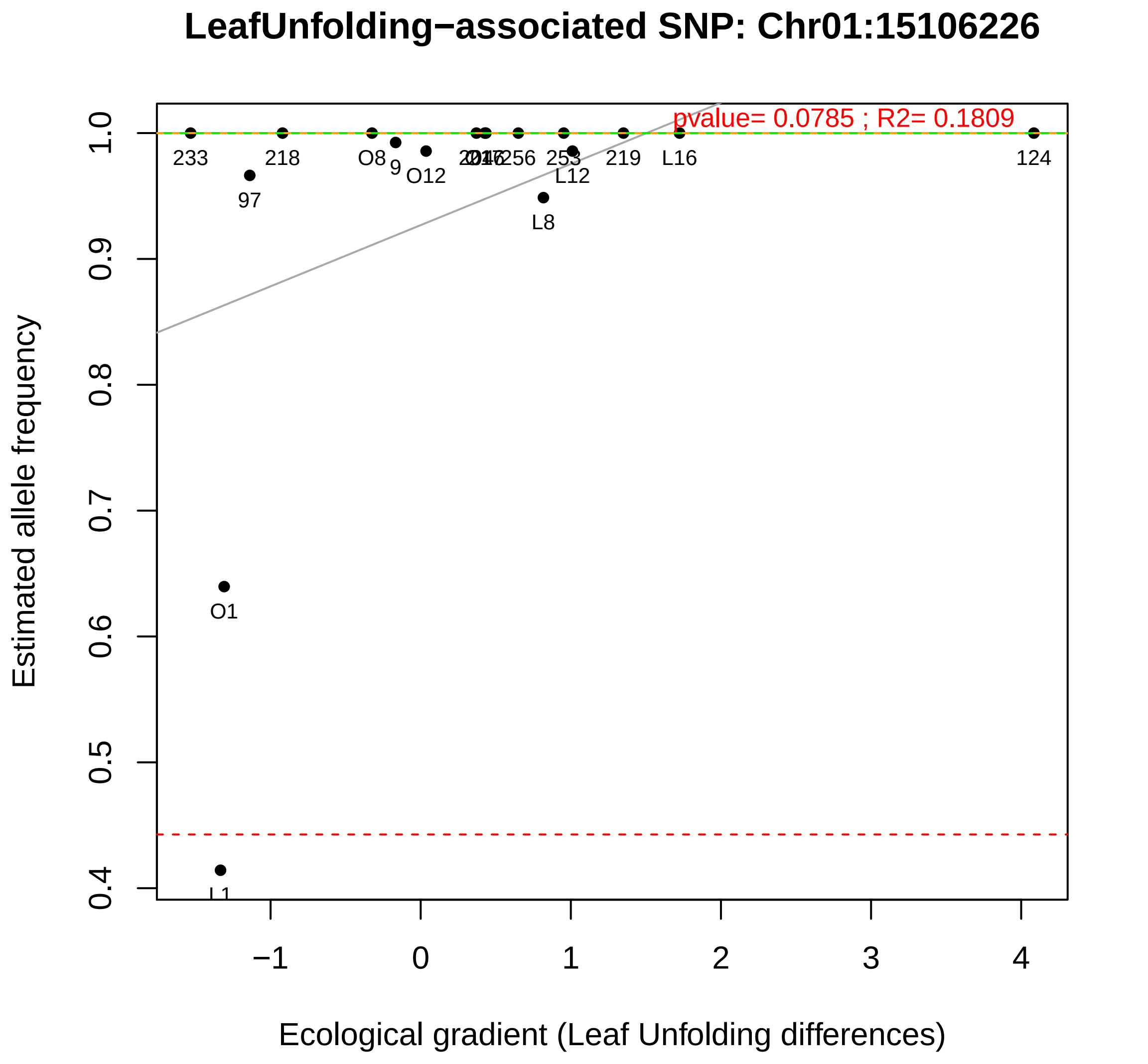

LeafUnfolding−associated SNP: Chr01:15106226
pvalue= 0.0785 ; R2= 0.1809
Estimated allele frequency
Ecological gradient (Leaf Unfolding differences)
1.0
0.9
0.8
0.7
0.6
0.5
0.4
−1
0
1
2
3
4
233
218
O8
9
O12
97
256
253
L12
L8
219
L16
124
O1
L1

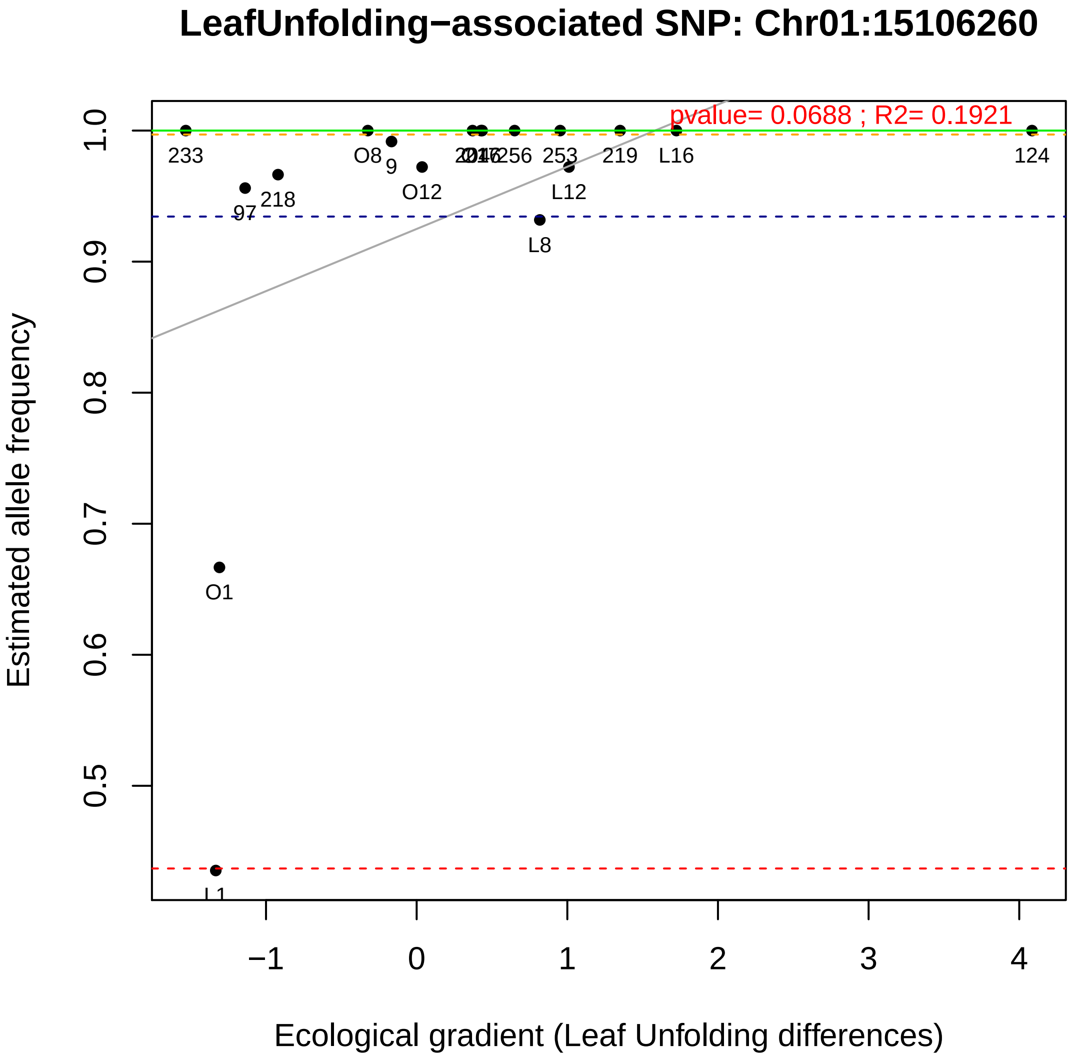
LeafUnfolding−associated SNP: Chr01:15106260
pvalue= 0.0688 ; R2= 0.1921
Estimated allele frequency
Ecological gradient (Leaf Unfolding differences)
1.0
0.9
0.8
0.7
0.6
0.5
−1
0
1
2
3
4
233
O8
9
O12
253
219
L16
124
97
218
L12
L8
O1

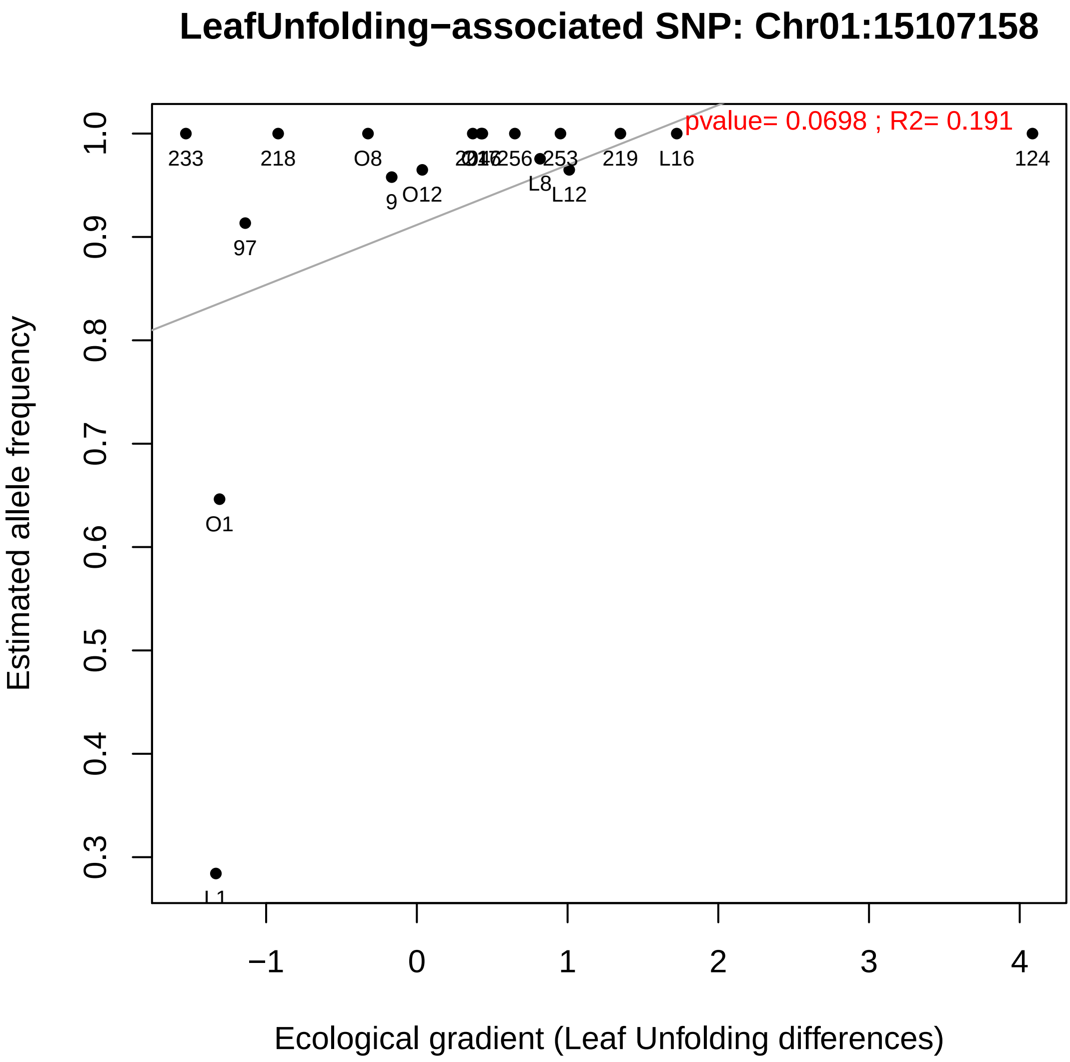
LeafUnfolding−associated SNP: Chr01:15107158
pvalue= 0.0698 ; R2= 0.191
Estimated allele frequency
Ecological gradient (Leaf Unfolding differences)
1.0
0.9
0.8
0.7
0.6
0.5
0.4
0.3
−1
0
1
2
3
4
233
218
O8
9
O12
256
253
L8
L12
219
L16
124
97
O1

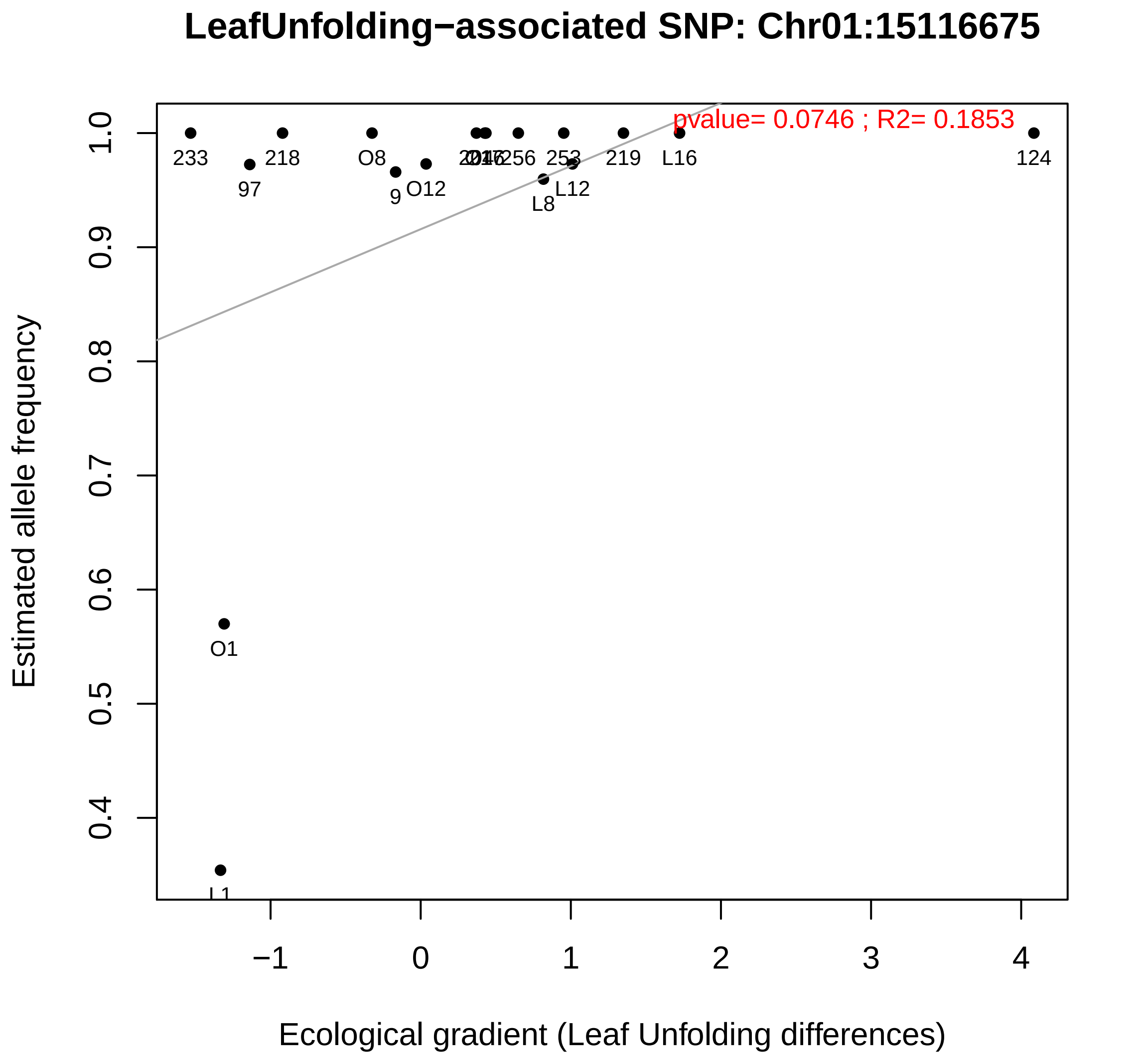

LeafUnfolding−associated SNP: Chr01:15116675
pvalue= 0.0746 ; R2= 0.1853
Estimated allele frequency
Ecological gradient (Leaf Unfolding differences)
1.0
0.9
0.8
0.7
0.6
0.5
0.4
−1
0
1
2
3
4
233
97
218
O8
9
O12
256
253
L8
L12
219
L16
124
O1

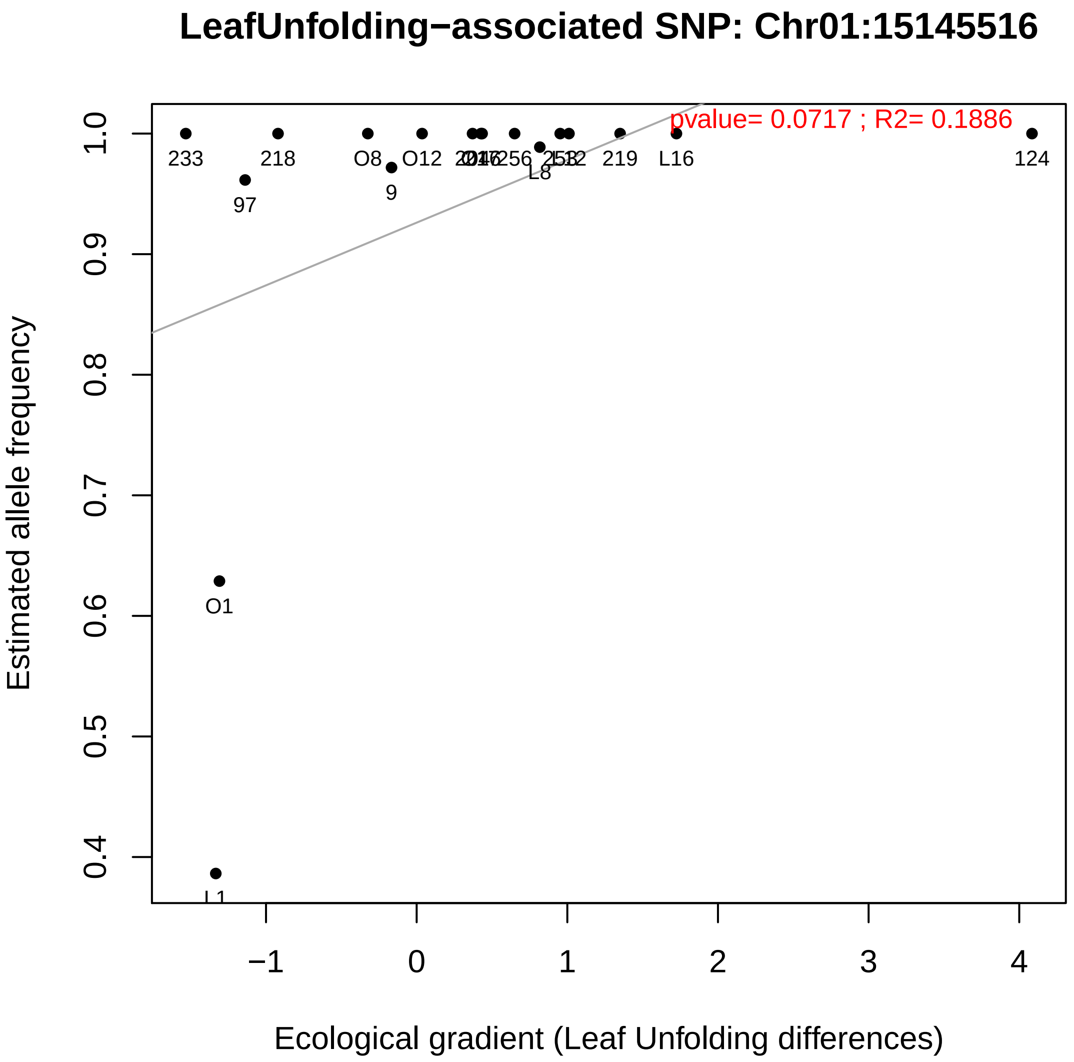

LeafUnfolding−associated SNP: Chr01:15145516
pvalue= 0.0717 ; R2= 0.1886
Estimated allele frequency
Ecological gradient (Leaf Unfolding differences)
1.0
0.9
0.8
0.7
0.6
0.5
0.4
−1
0
1
2
3
4
233
218
O8
O12
219
L16
124
97
9
L8
O1

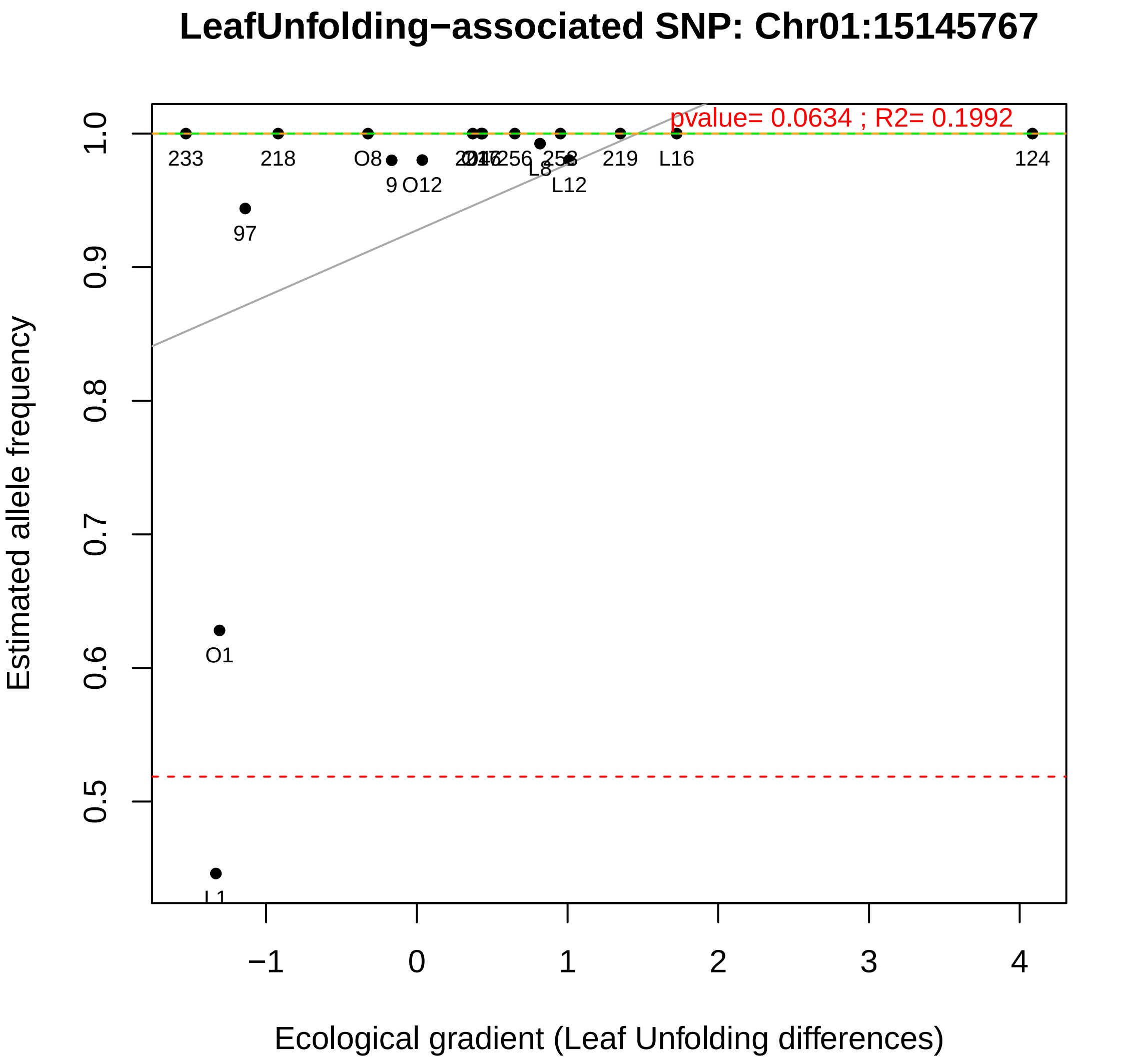

LeafUnfolding−associated SNP: Chr01:15145767
pvalue= 0.0634 ; R2= 0.1992
Estimated allele frequency
Ecological gradient (Leaf Unfolding differences)
1.0
0.9
0.8
0.7
0.6
0.5
−1
0
1
2
3
4
233
218
O8
9
O12
97
219
L16
124
L8
L12
O1
L1

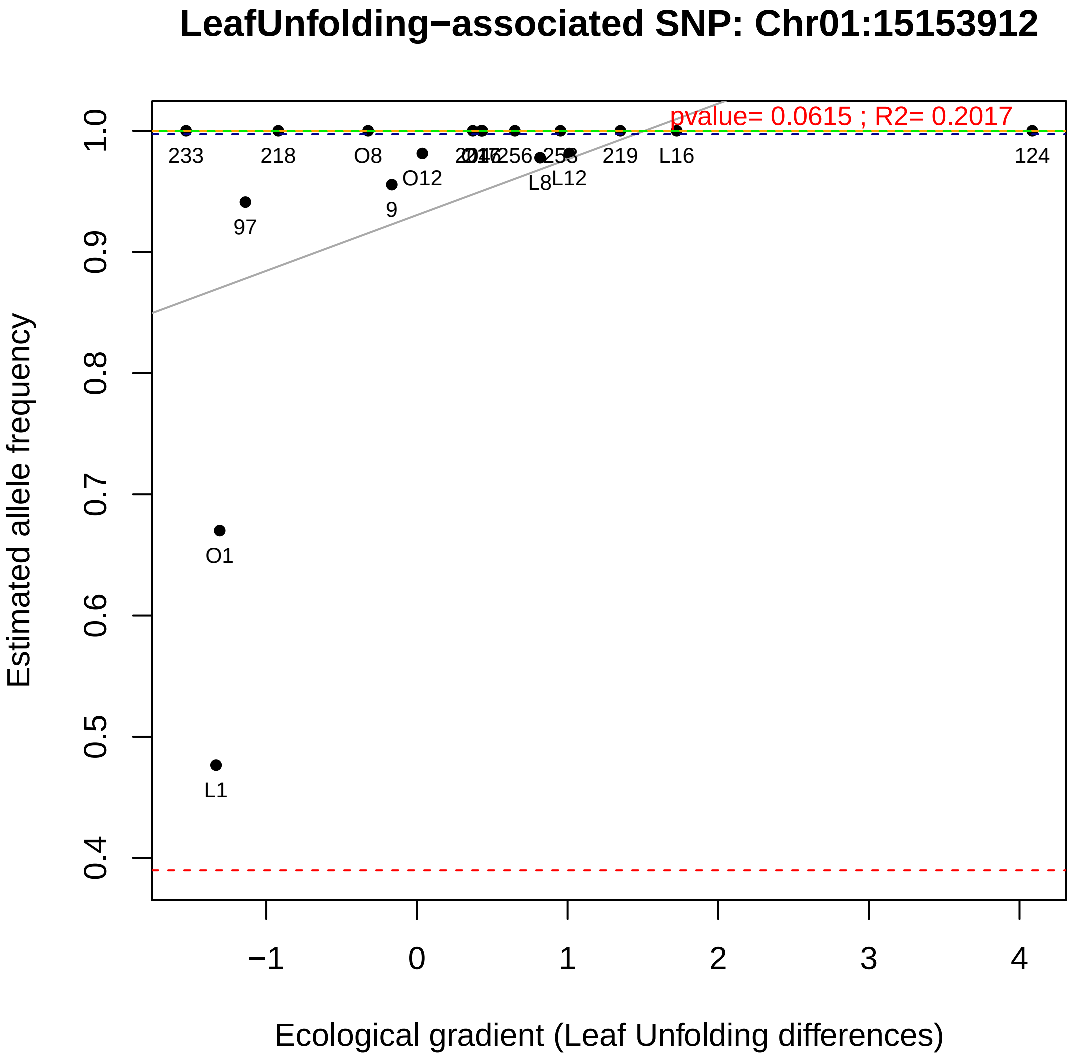
LeafUnfolding−associated SNP: Chr01:15153912
pvalue= 0.0615 ; R2= 0.2017
Estimated allele frequency
Ecological gradient (Leaf Unfolding differences)
1.0
0.9
0.8
0.7
0.6
0.5
0.4
−1
0
1
2
3
4
233
218
O8
256
219
L16
124
O12
L8
L12
97
9
O1
L1

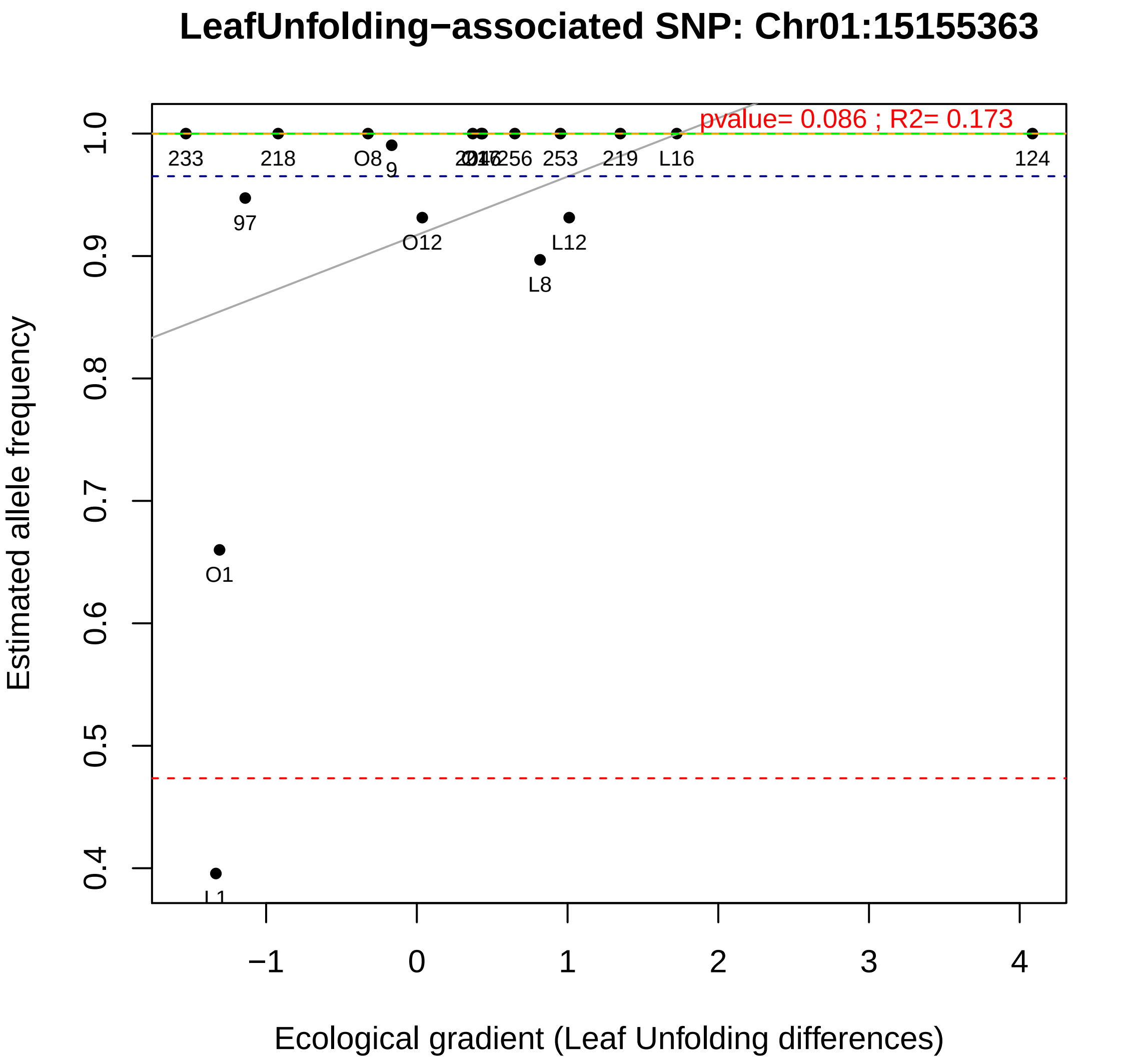

LeafUnfolding−associated SNP: Chr01:15155363
pvalue= 0.086 ; R2= 0.173
Estimated allele frequency
Ecological gradient (Leaf Unfolding differences)
0.4
0.5
0.6
0.7
0.8
0.9
1.0
−1
0
1
2
3
4
233
218
O8
9
253
219
L16
124
97
O12
L12
L8
O1

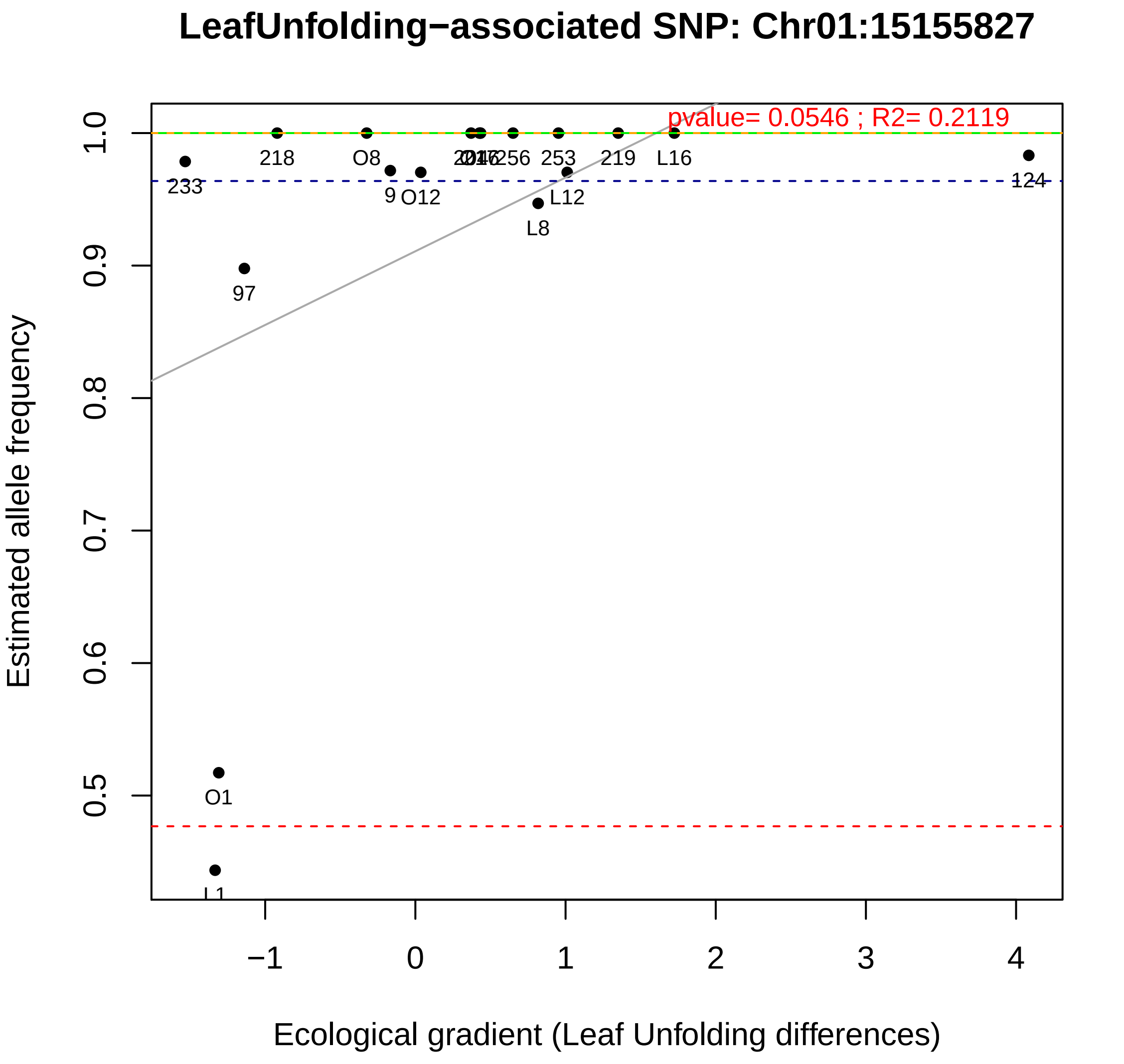

LeafUnfolding−associated SNP: Chr01:15155827
pvalue= 0.0546 ; R2= 0.2119
Estimated allele frequency
Ecological gradient (Leaf Unfolding differences)
1.0
0.9
0.8
0.7
0.6
0.5
−1
0
1
2
3
4
218
O8
256
253
219
L16
233
9
O12
L12
124
L8
97
O1

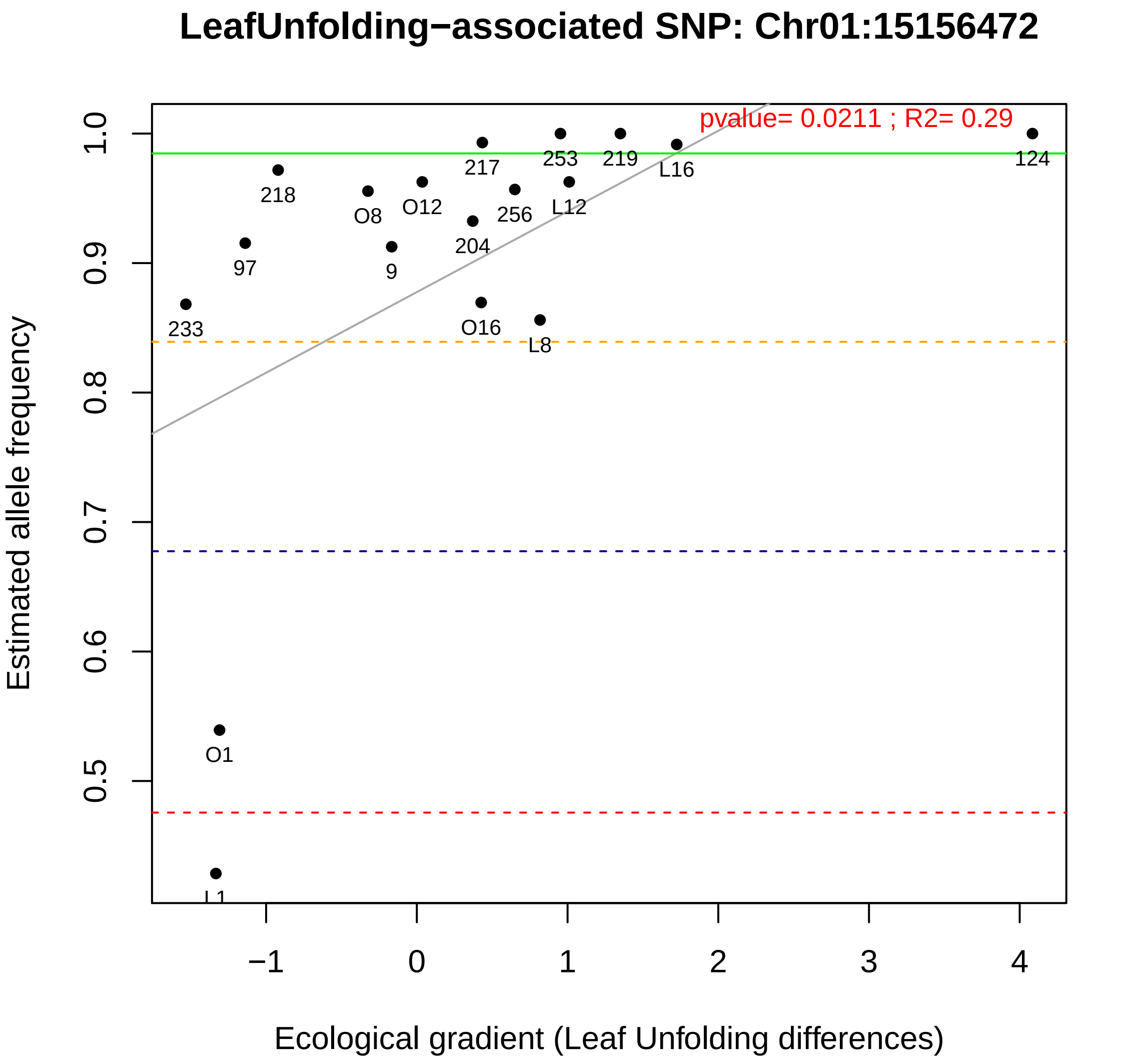
LeafUnfolding−associated SNP: Chr01:15156472
pvalue= 0.0211 ; R2= 0.29
Estimated allele frequency
Ecological gradient (Leaf Unfolding differences)
1.0
0.9
0.8
0.7
0.6
0.5
−1
0
1
2
3
4
233
97
218
O8
9
O12
217
204
O16
256
L8
253
L12
219
L16
124
O1

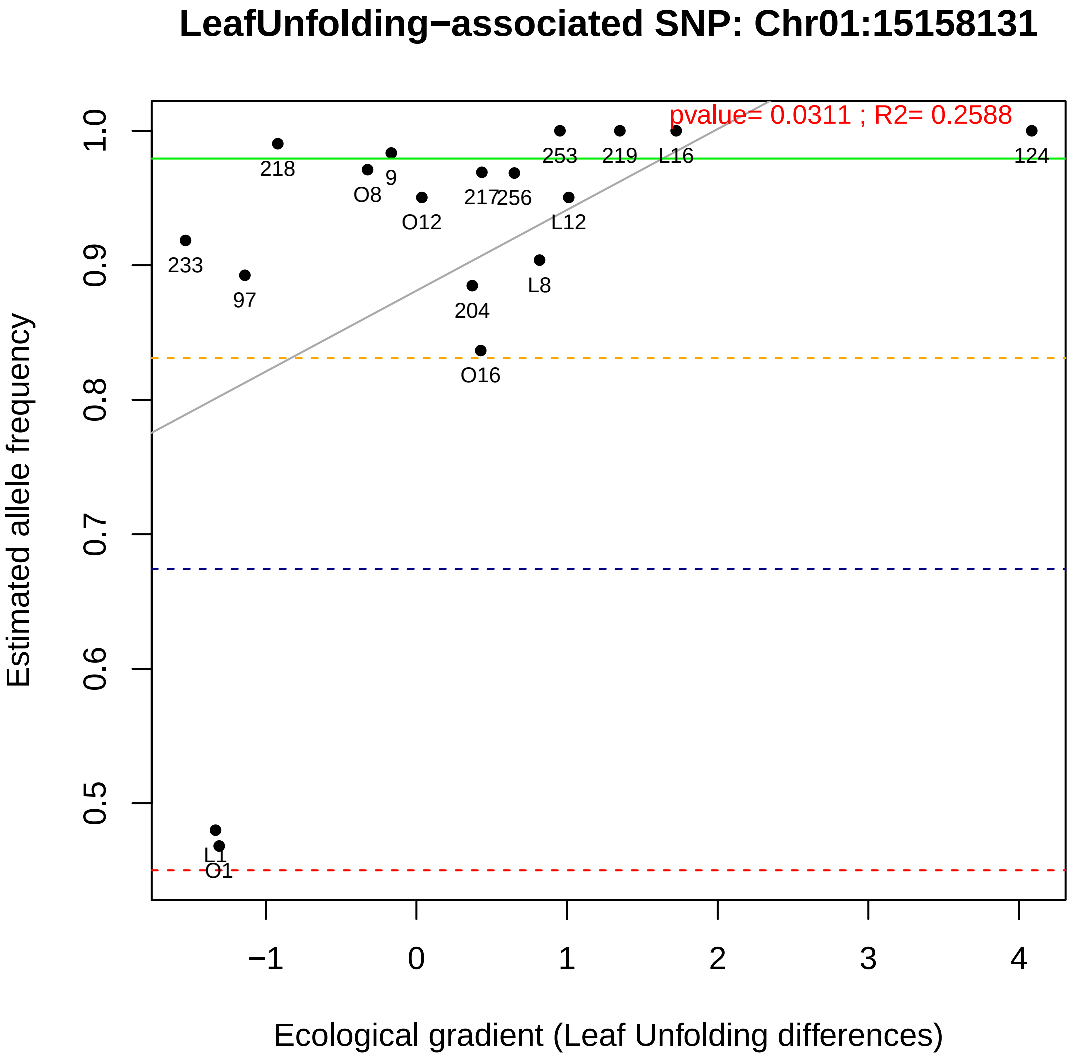
LeafUnfolding−associated SNP: Chr01:15158131
pvalue= 0.0311 ; R2= 0.2588
Estimated allele frequency
Ecological gradient (Leaf Unfolding differences)
1.0
0.9
0.8
0.7
0.6
0.5
−1
0
1
2
3
4
218
O8
9
O12
217
256
253
219
L16
124
L12
233
97
204
L8
O16
L1
O1

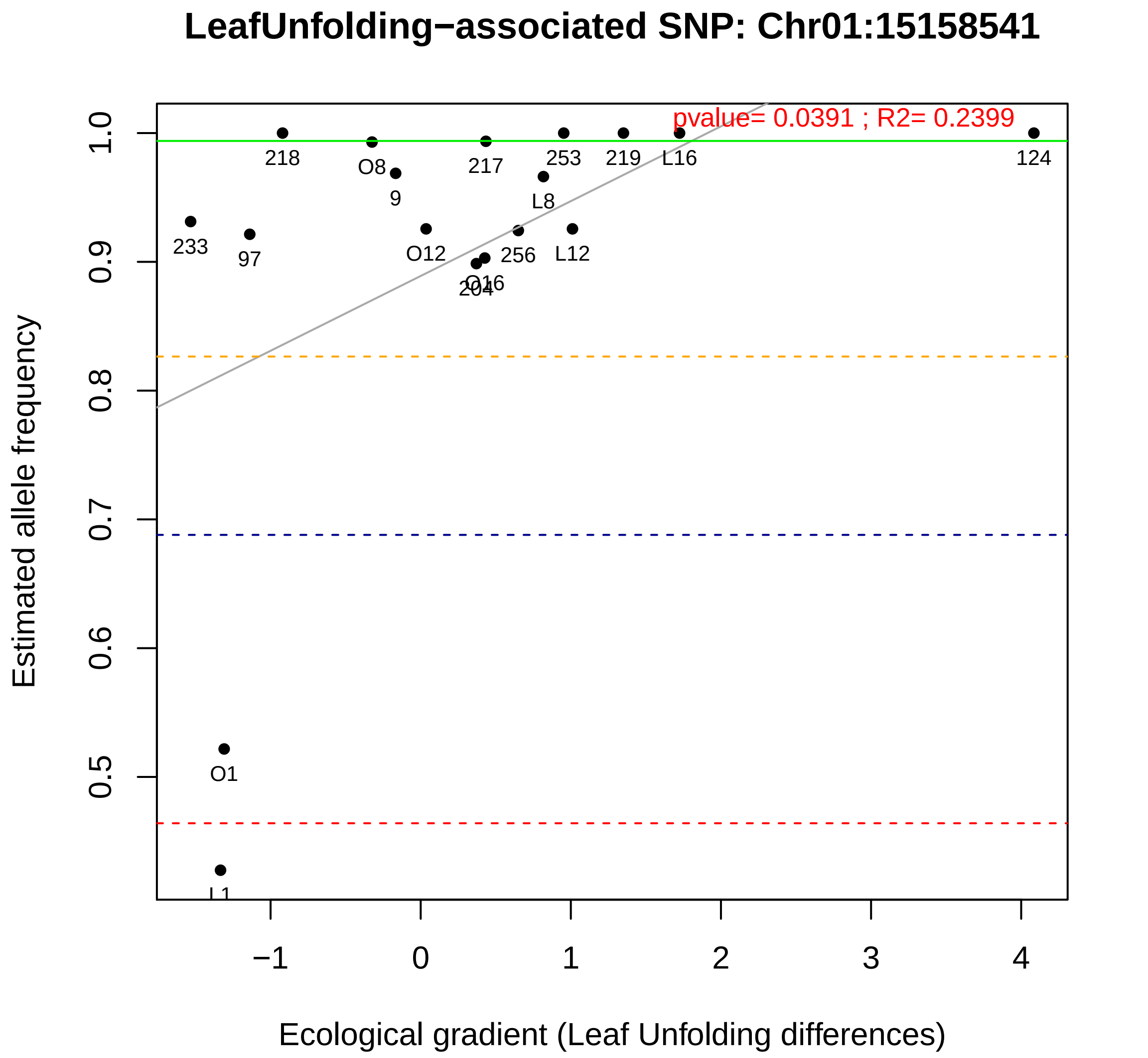
LeafUnfolding−associated SNP: Chr01:15158541
pvalue= 0.0391 ; R2= 0.2399
Estimated allele frequency
0.5
0.6
0.7
0.8
0.9
1.0
−1
0
1
2
3
4
Ecological gradient (Leaf Unfolding differences)
218
O8
9
217
253
219
L16
124
L8
233
97
O12
256
L12
204
O16
O1

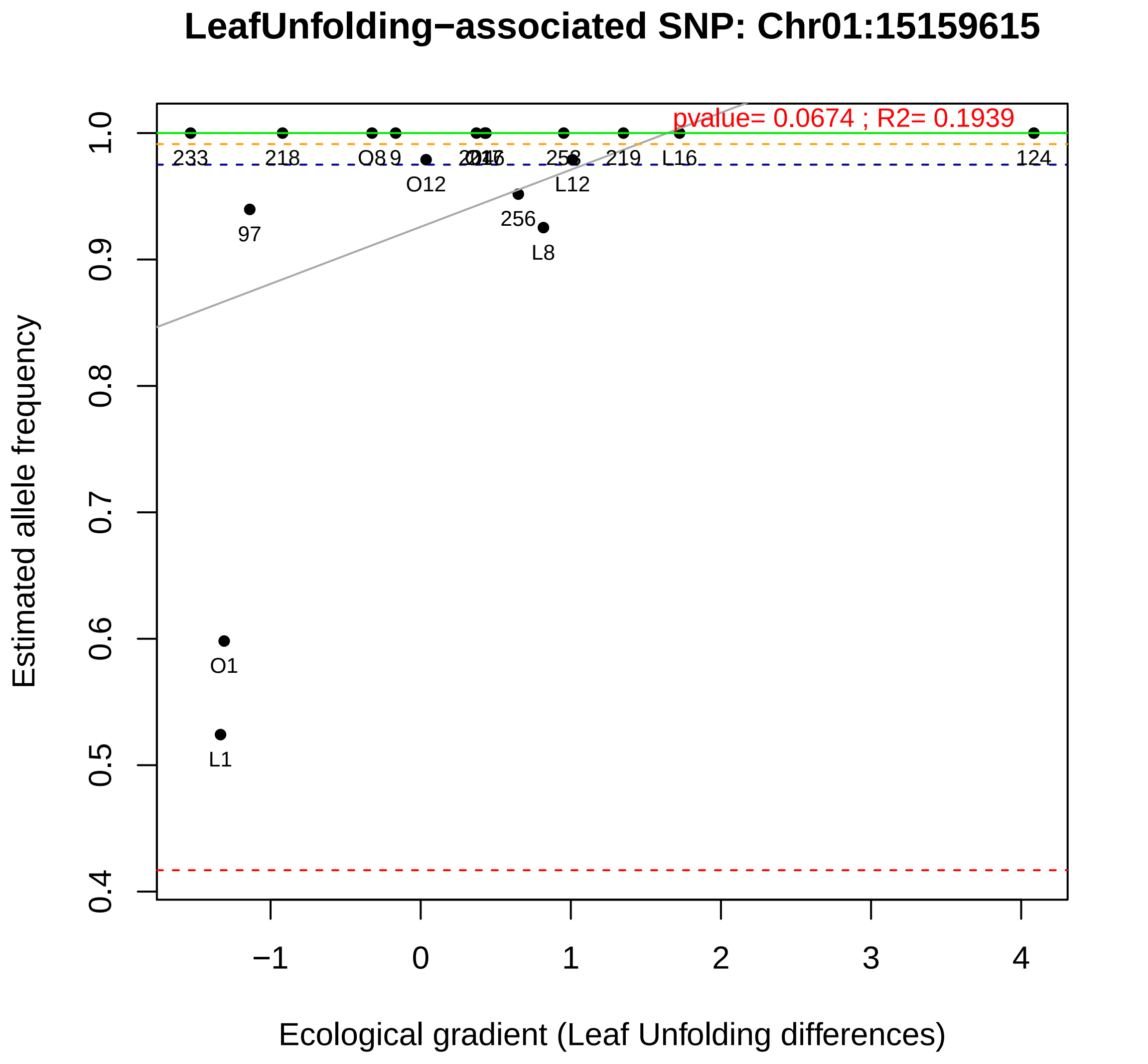

LeafUnfolding−associated SNP: Chr01:15159615
pvalue= 0.0674 ; R2= 0.1939
Estimated allele frequency
Ecological gradient (Leaf Unfolding differences)
1.0
0.9
0.8
0.7
0.6
0.5
0.4
−1
0
1
2
3
4
233
218
O8
9
O12
97
256
L8
L12
219
L16
124
O1
L1

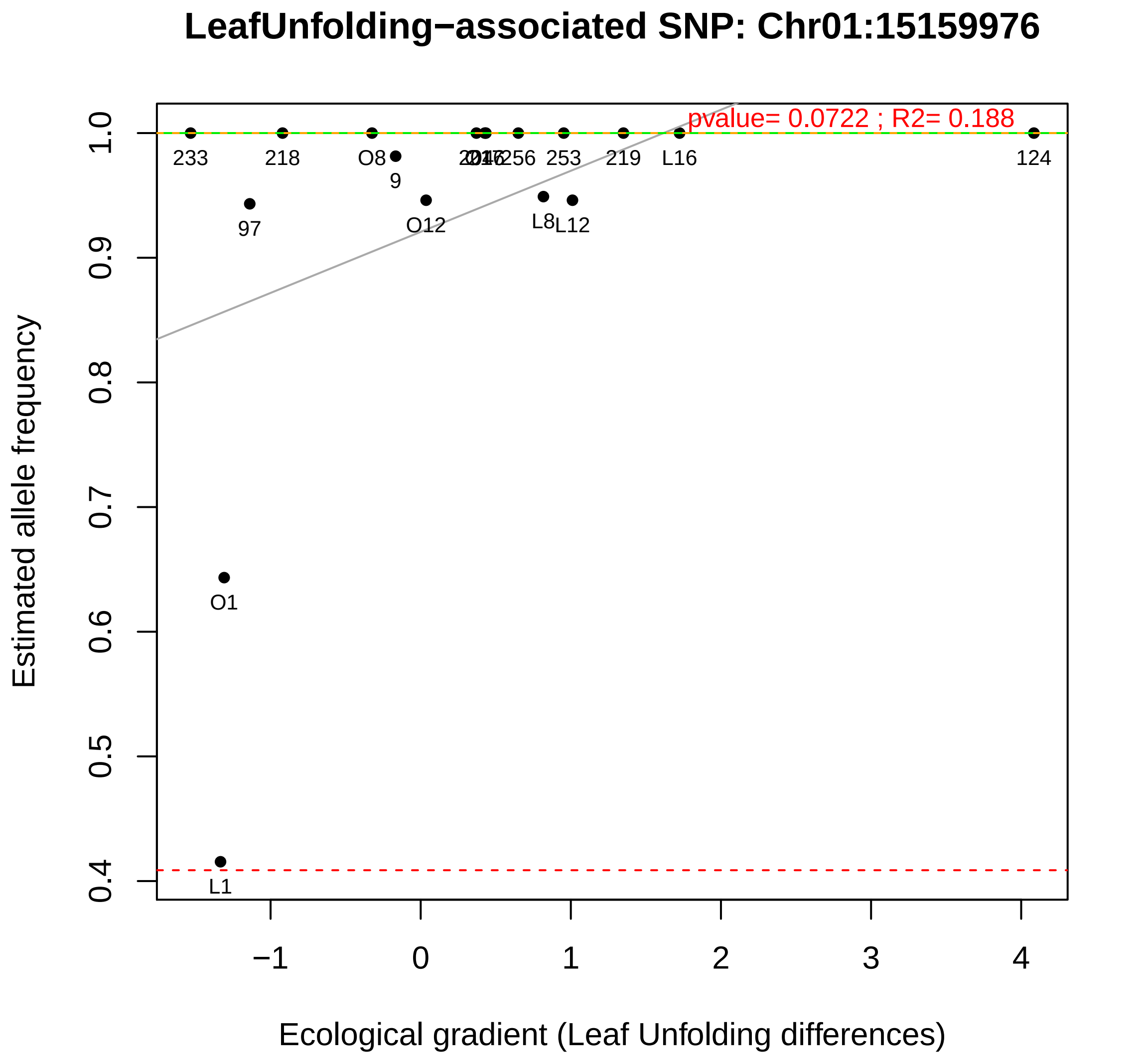

LeafUnfolding−associated SNP: Chr01:15159976
pvalue= 0.0722 ; R2= 0.188
Estimated allele frequency
Ecological gradient (Leaf Unfolding differences)
1.0
0.9
0.8
0.7
0.6
0.5
0.4
−1
0
1
2
3
4
233
218
O8
9
256
253
219
L16
124
97
O12
L8
L12
O1
L1

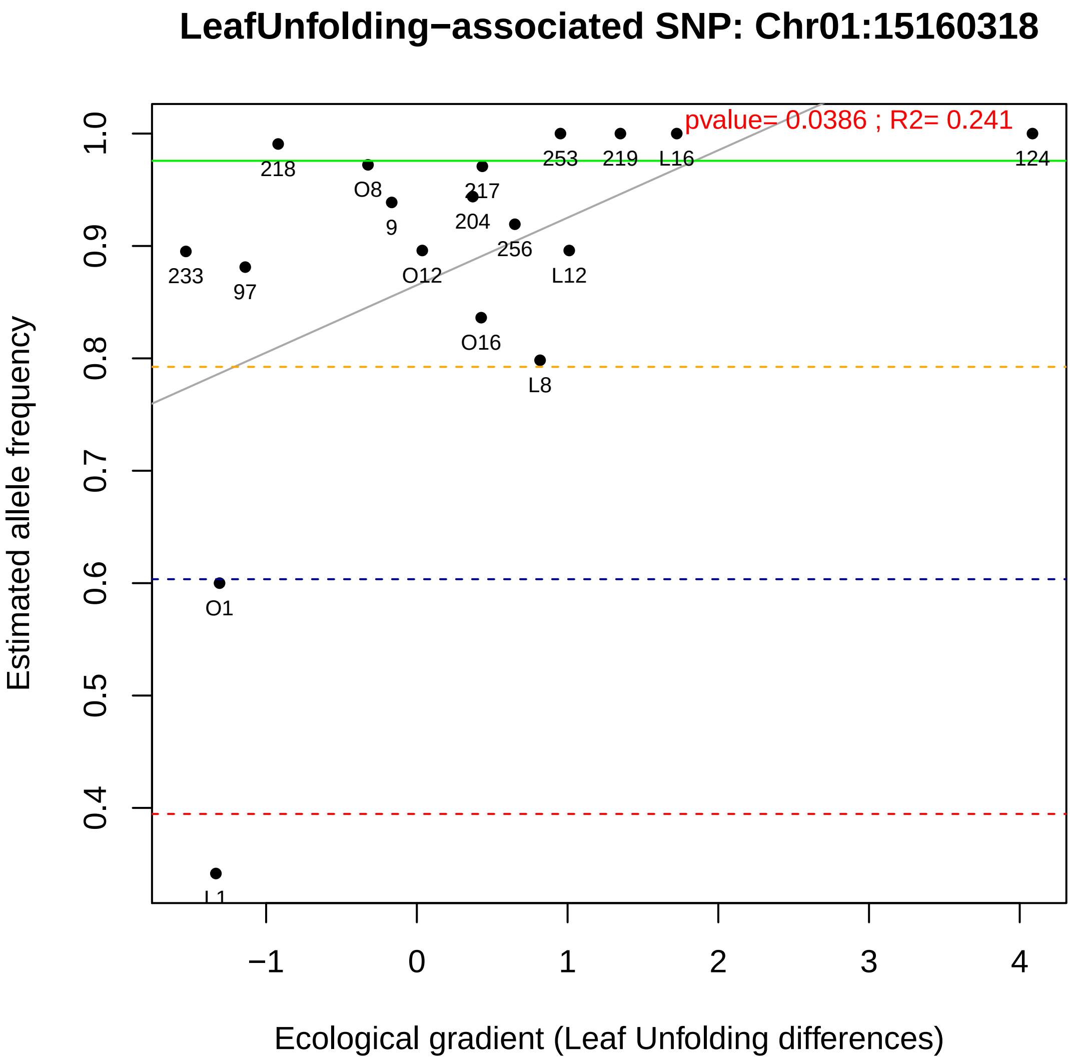
LeafUnfolding−associated SNP: Chr01:15160318
pvalue= 0.0386 ; R2= 0.241
Estimated allele frequency
Ecological gradient (Leaf Unfolding differences)
1.0
0.9
0.8
0.7
0.6
0.5
0.4
−1
0
1
2
3
4
218
O8
9
217
204
253
219
L16
124
256
233
97
O12
L12
O16
L8
O1
L1

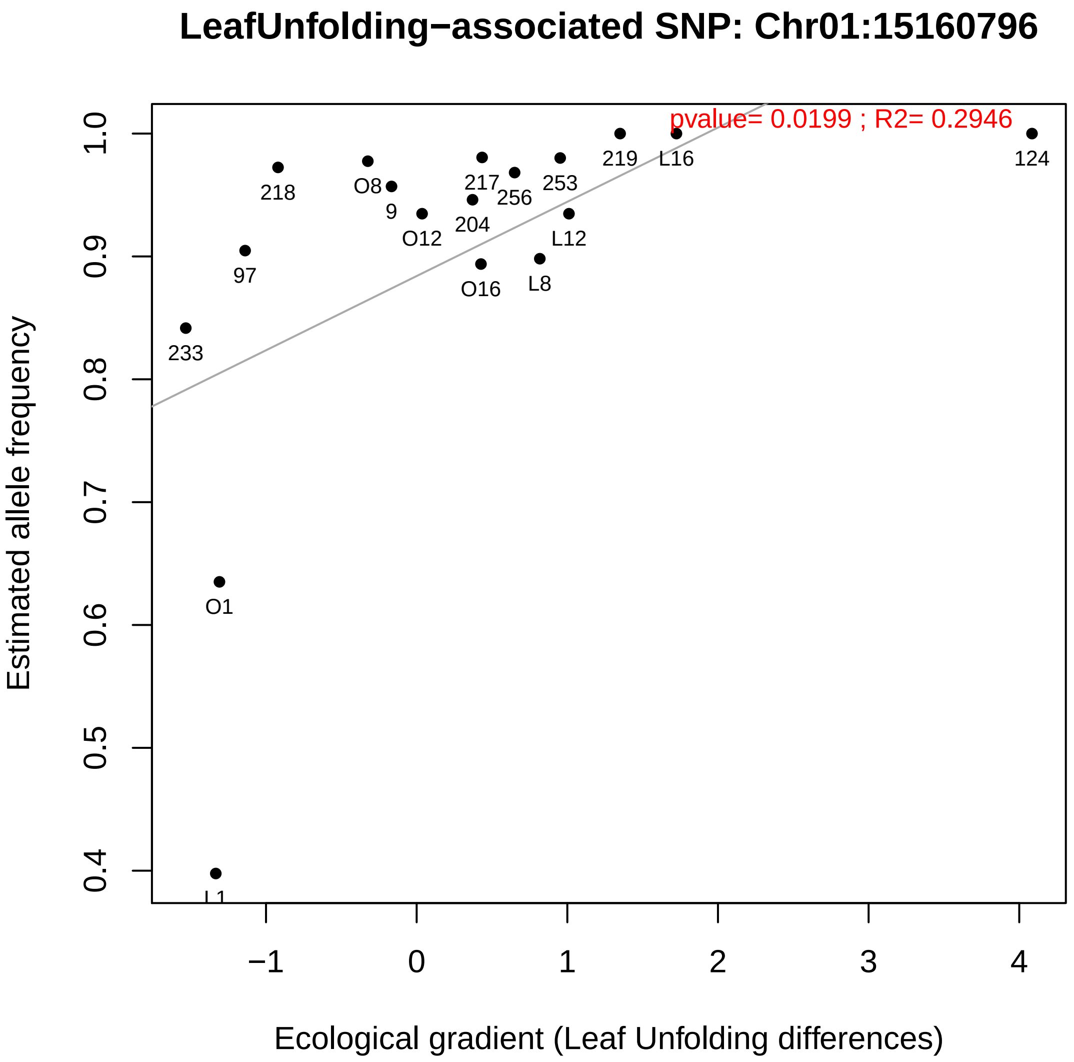

LeafUnfolding−associated SNP: Chr01:15160796
pvalue= 0.0199 ; R2= 0.2946
Estimated allele frequency
Ecological gradient (Leaf Unfolding differences)
0.4
0.5
0.6
0.7
0.8
0.9
1.0
−1
0
1
2
3
4
233
97
218
O8
9
O12
217
204
256
O16
253
L12
L8
219
L16
124
O1
L1

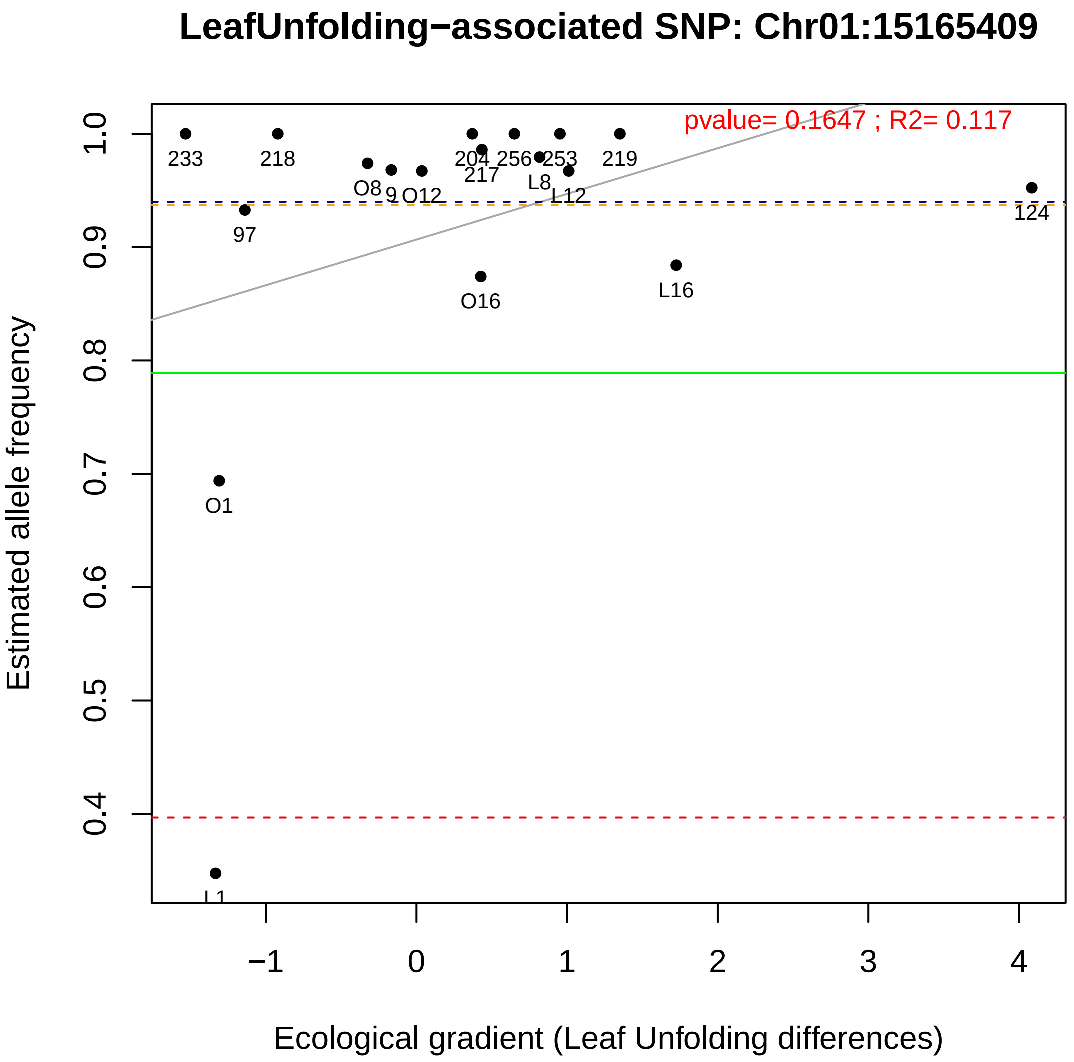
LeafUnfolding−associated SNP: Chr01:15165409
pvalue= 0.1647 ; R2= 0.117
233
218
97
O8
9
O12
204
217
256
L8
253
L12
219
124
O16
L16
O1
L1
1.0
0.9
0.8
0.7
0.6
0.5
0.4
−1
0
1
2
3
4
Estimated allele frequency
Ecological gradient (Leaf Unfolding differences)

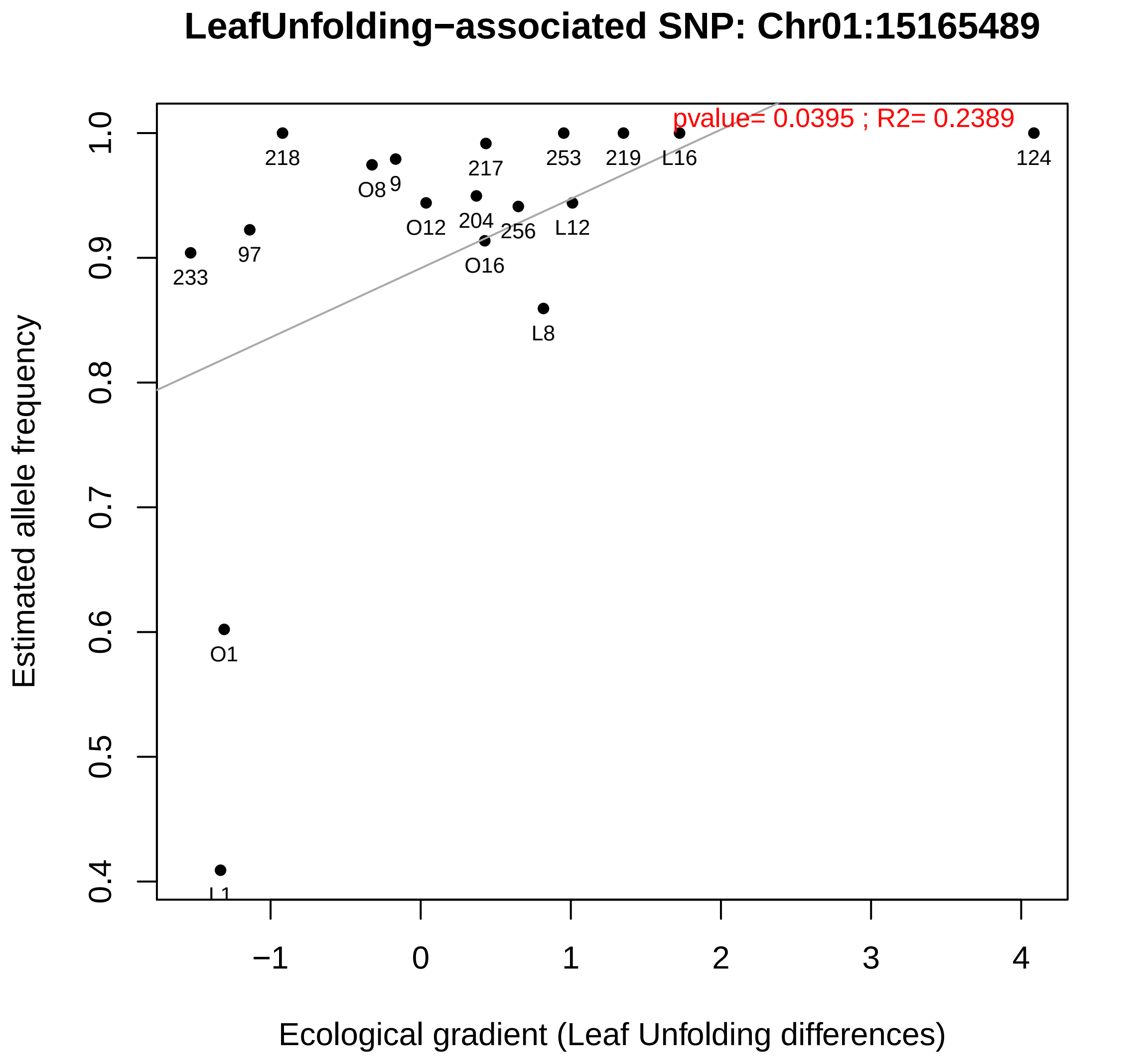
LeafUnfolding−associated SNP: Chr01:15165489
pvalue= 0.0395 ; R2= 0.2389
Estimated allele frequency
Ecological gradient (Leaf Unfolding differences)
0.4
0.5
0.6
0.7
0.8
0.9
1.0
−1
0
1
2
3
4
218
O8
9
217
253
219
L16
124
O12
204
256
L12
97
233
O16
L8
O1
L1

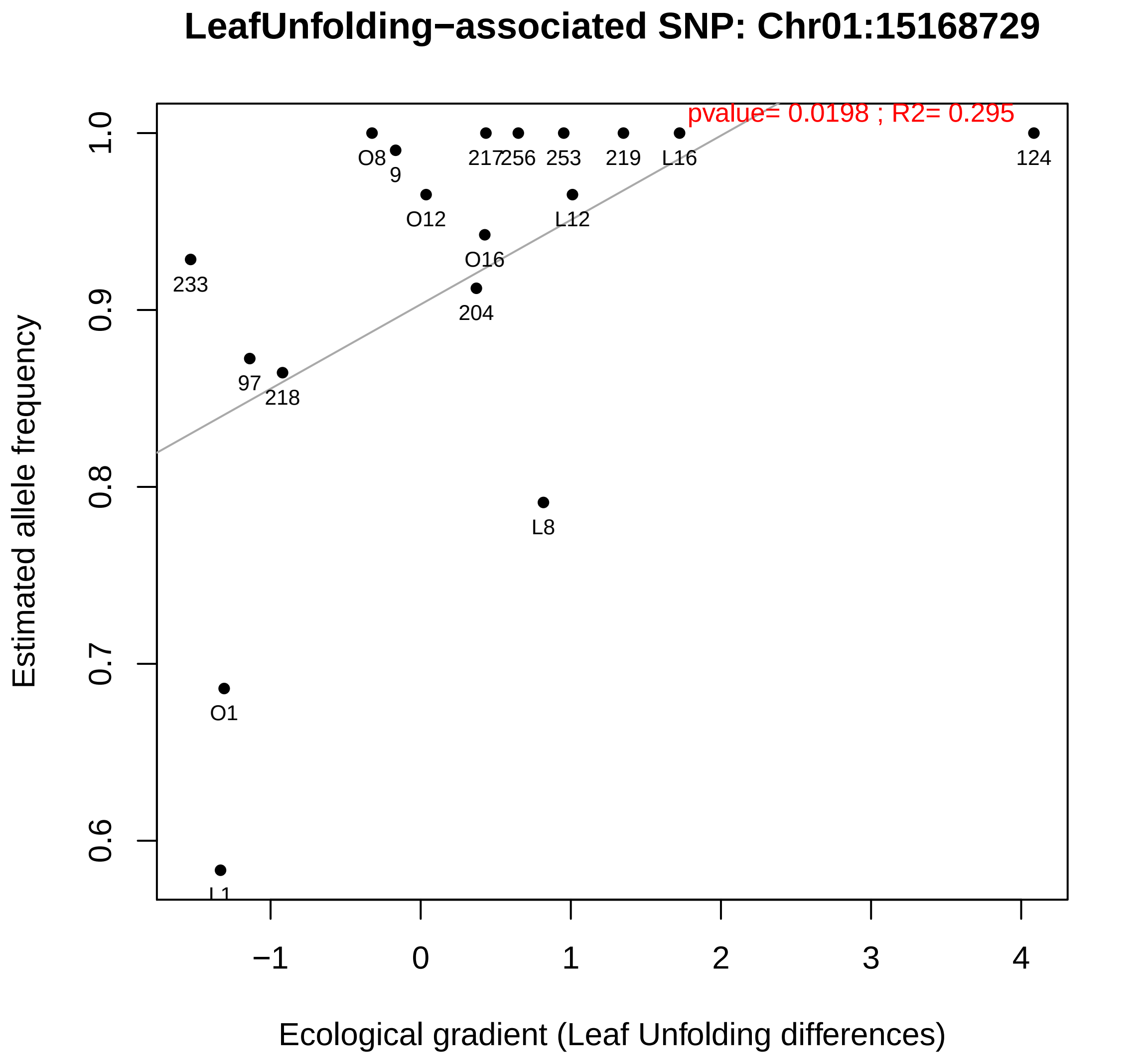

LeafUnfolding−associated SNP: Chr01:15168729
pvalue= 0.0198 ; R2= 0.295
Estimated allele frequency
Ecological gradient (Leaf Unfolding differences)
0.6
0.7
0.8
0.9
1.0
−1
0
1
2
3
4
O8
9
217
256
253
219
L16
124
O12
L12
O16
233
204
97
218
L8
O1

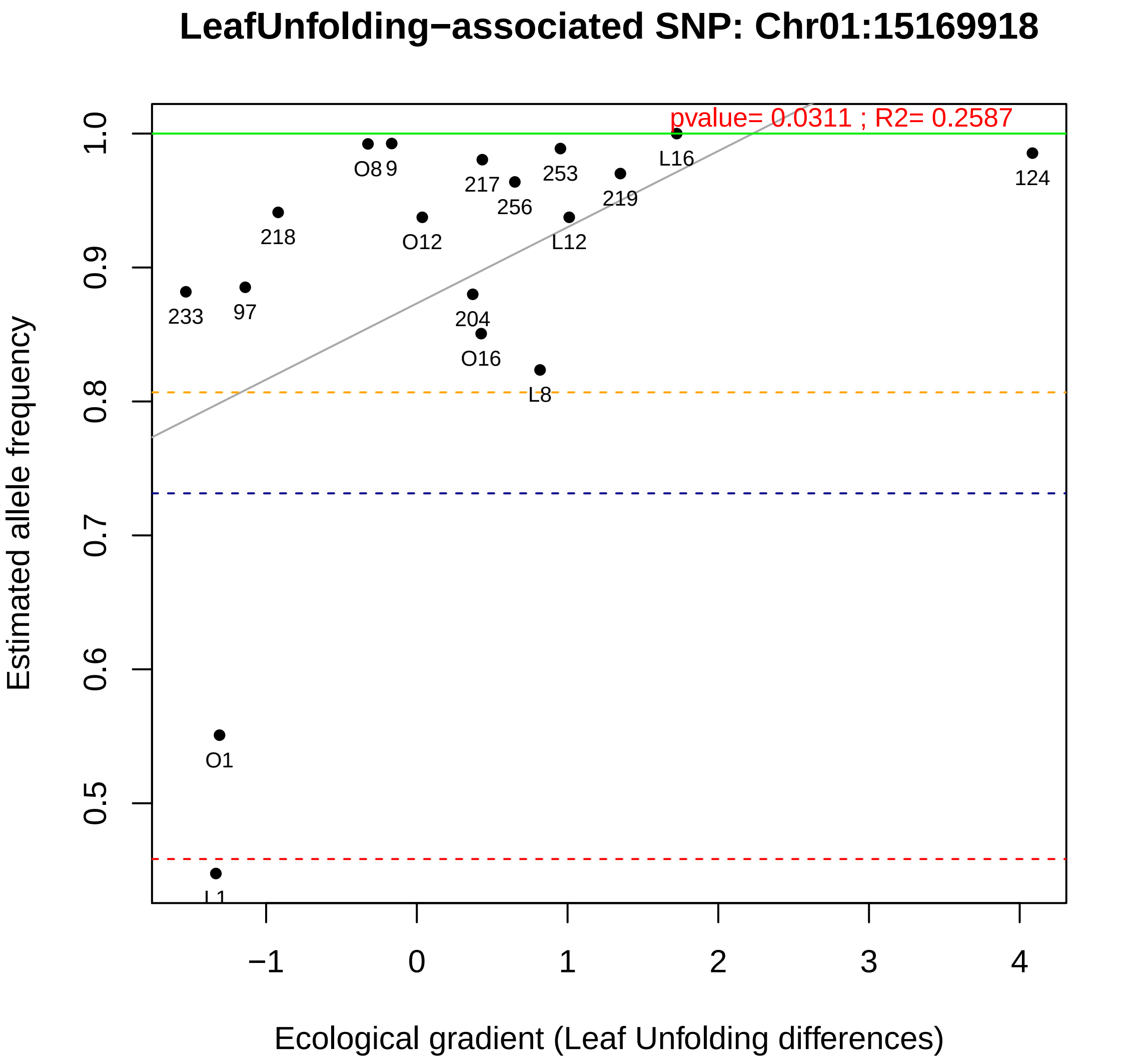
LeafUnfolding−associated SNP: Chr01:15169918
pvalue= 0.0311 ; R2= 0.2587
Estimated allele frequency
Ecological gradient (Leaf Unfolding differences)
1.0
0.9
0.8
0.7
0.6
0.5
−1
0
1
2
3
4
O8
9
217
253
L16
124
256
219
218
O12
L12
233
97
204
O16
L8
O1
L1

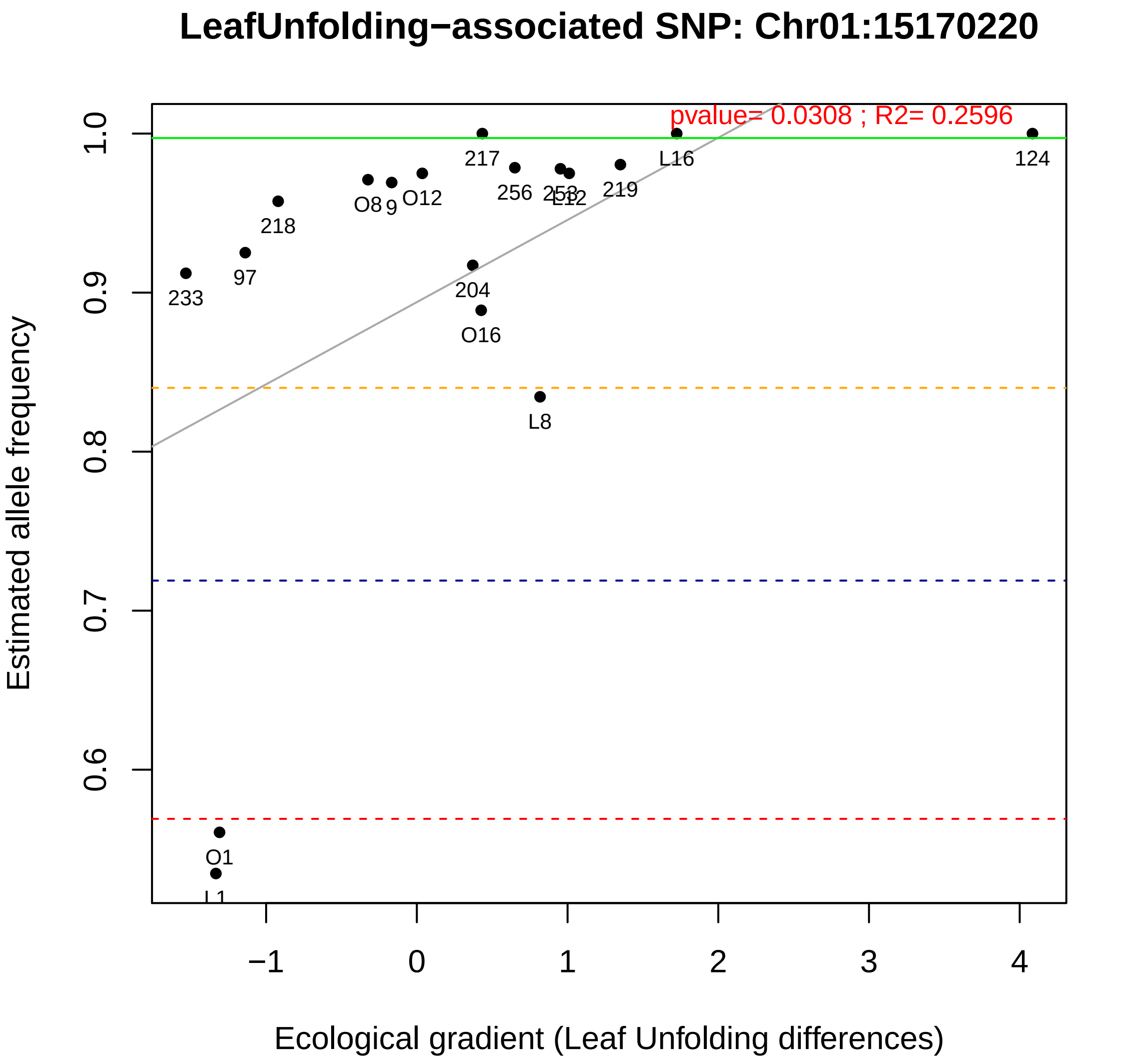
LeafUnfolding−associated SNP: Chr01:15170220
pvalue= 0.0308 ; R2= 0.2596
Estimated allele frequency
Ecological gradient (Leaf Unfolding differences)
1.0
0.9
0.8
0.7
0.6
−1
0
1
2
3
4
217
L16
124
256
253
L12
219
O8
9
O12
218
97
233
204
O16
L8
O1

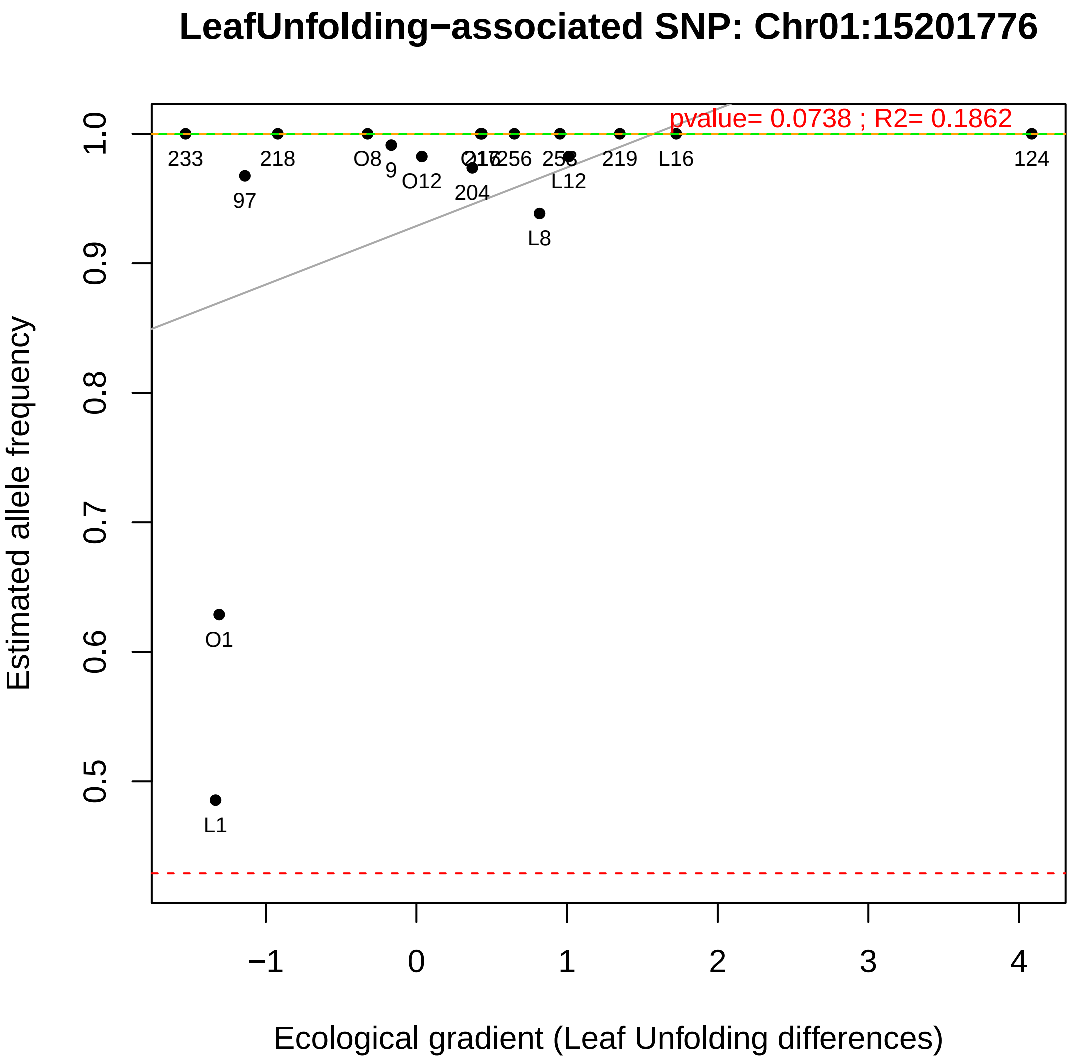

LeafUnfolding−associated SNP: Chr01:15201776
pvalue= 0.0738 ; R2= 0.1862
Estimated allele frequency
0.5
0.6
0.7
0.8
0.9
1.0
−1
0
1
2
3
4
Ecological gradient (Leaf Unfolding differences)
233
218
O8
9
O12
204
256
L12
L8
219
L16
124
97
O1
L1

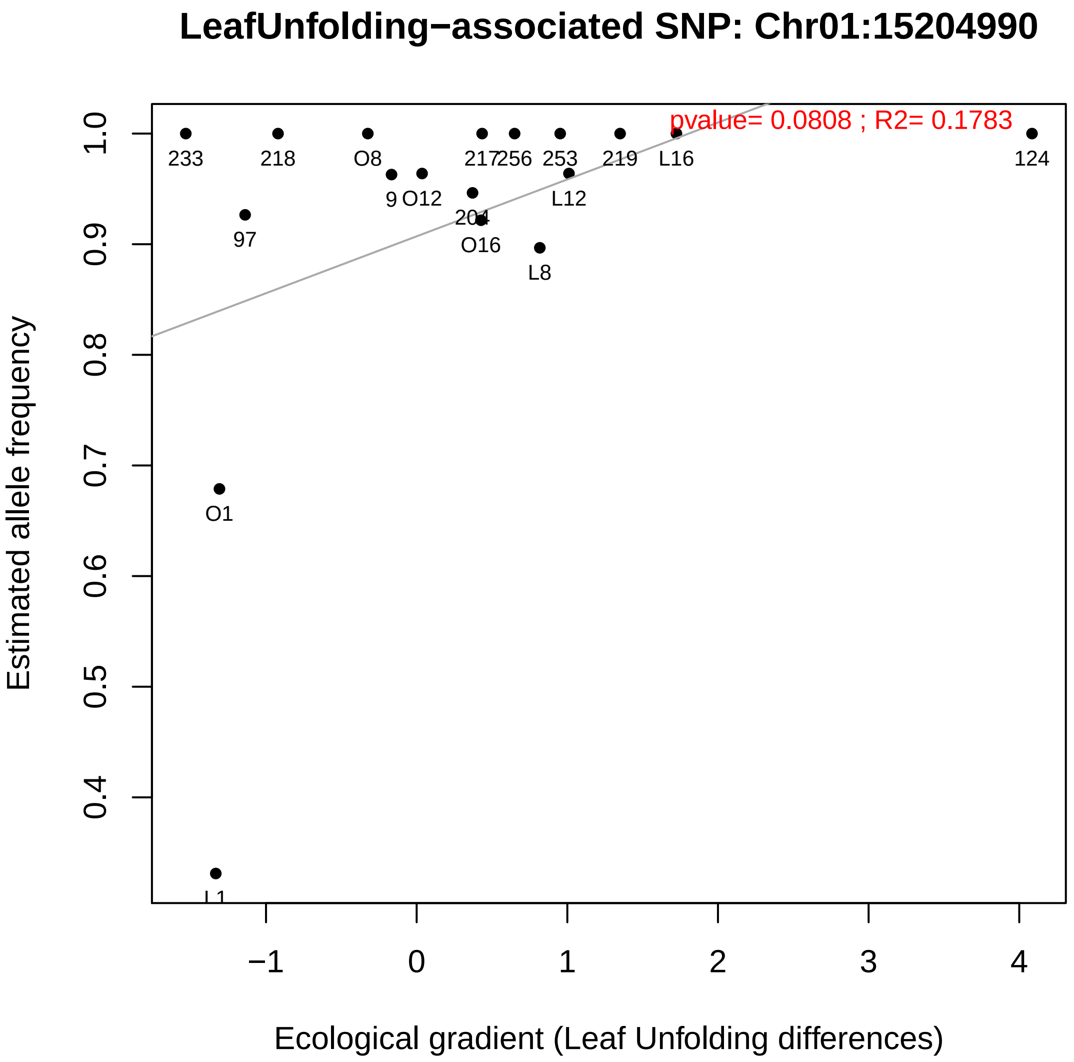
LeafUnfolding−associated SNP: Chr01:15204990
pvalue= 0.0808 ; R2= 0.1783
233
218
O8
9
O12
217
256
253
219
L16
124
204
O16
L12
L8
97
O1
L1
1.0
0.9
0.8
0.7
0.6
0.5
0.4
−1
0
1
2
3
4
Estimated allele frequency
Ecological gradient (Leaf Unfolding differences)

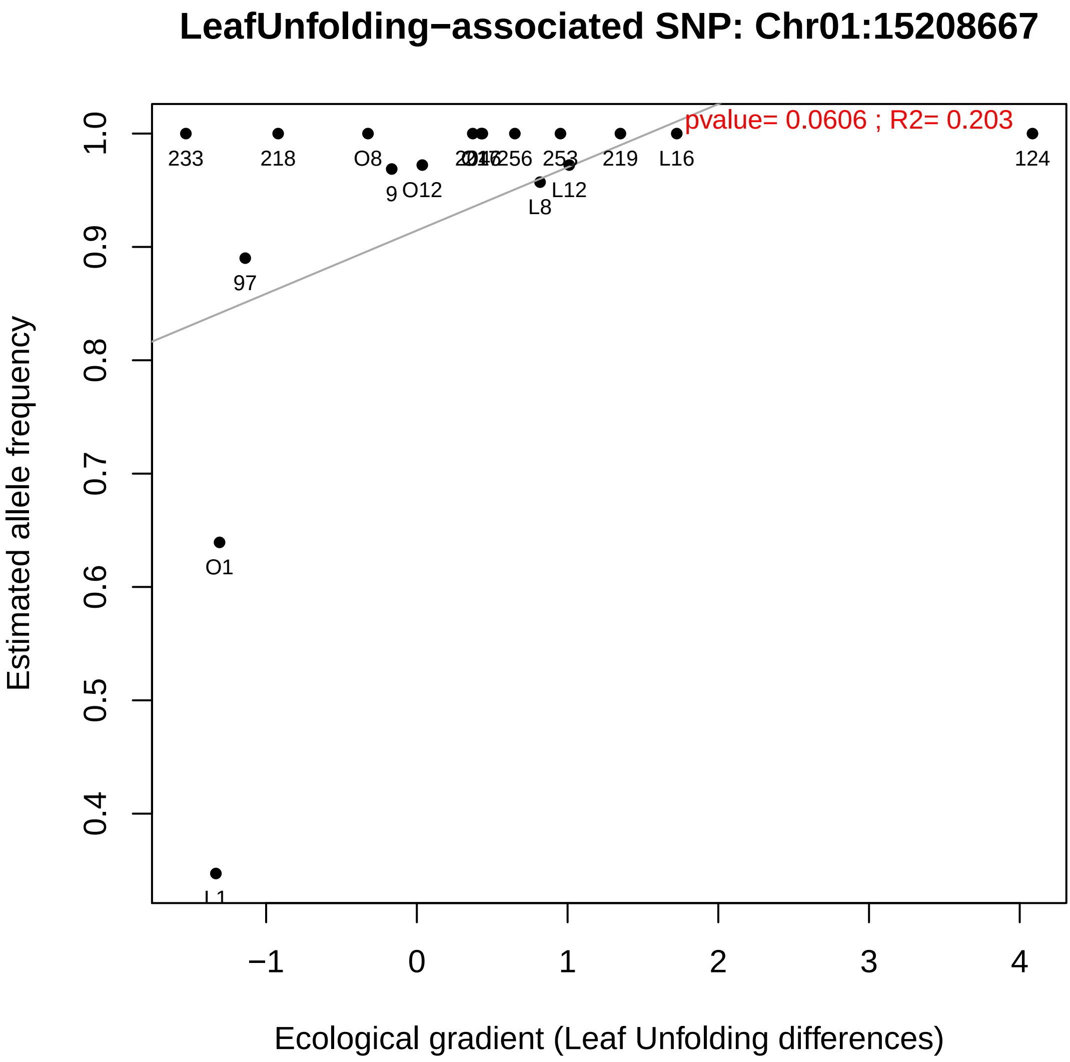
LeafUnfolding−associated SNP: Chr01:15208667
pvalue= 0.0606 ; R2= 0.203
Estimated allele frequency
Ecological gradient (Leaf Unfolding differences)
233
218
O8
9
O12
256
253
L12
L8
219
L16
124
97
O1
L1
1.0
0.9
0.8
0.7
0.6
0.5
0.4
−1
0
1
2
3
4

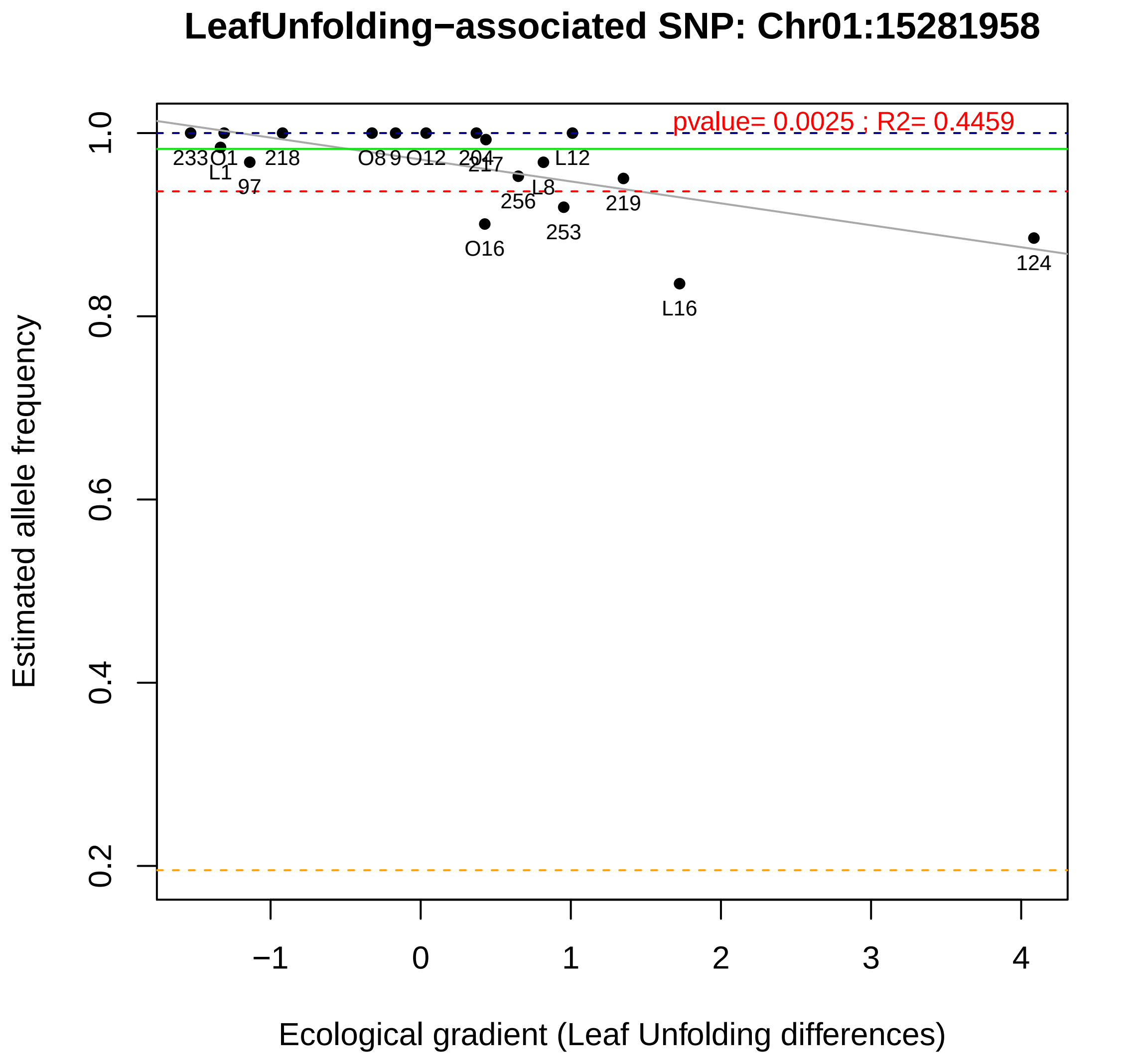

LeafUnfolding−associated SNP: Chr01:15281958
pvalue= 0.0025 ; R2= 0.4459
Estimated allele frequency
1.0
0.8
0.6
0.4
0.2
−1
0
1
2
3
4
Ecological gradient (Leaf Unfolding differences)
233
O1
L1
97
218
O8
9
O12
204
217
256
L8
L12
253
219
O16
L16
124

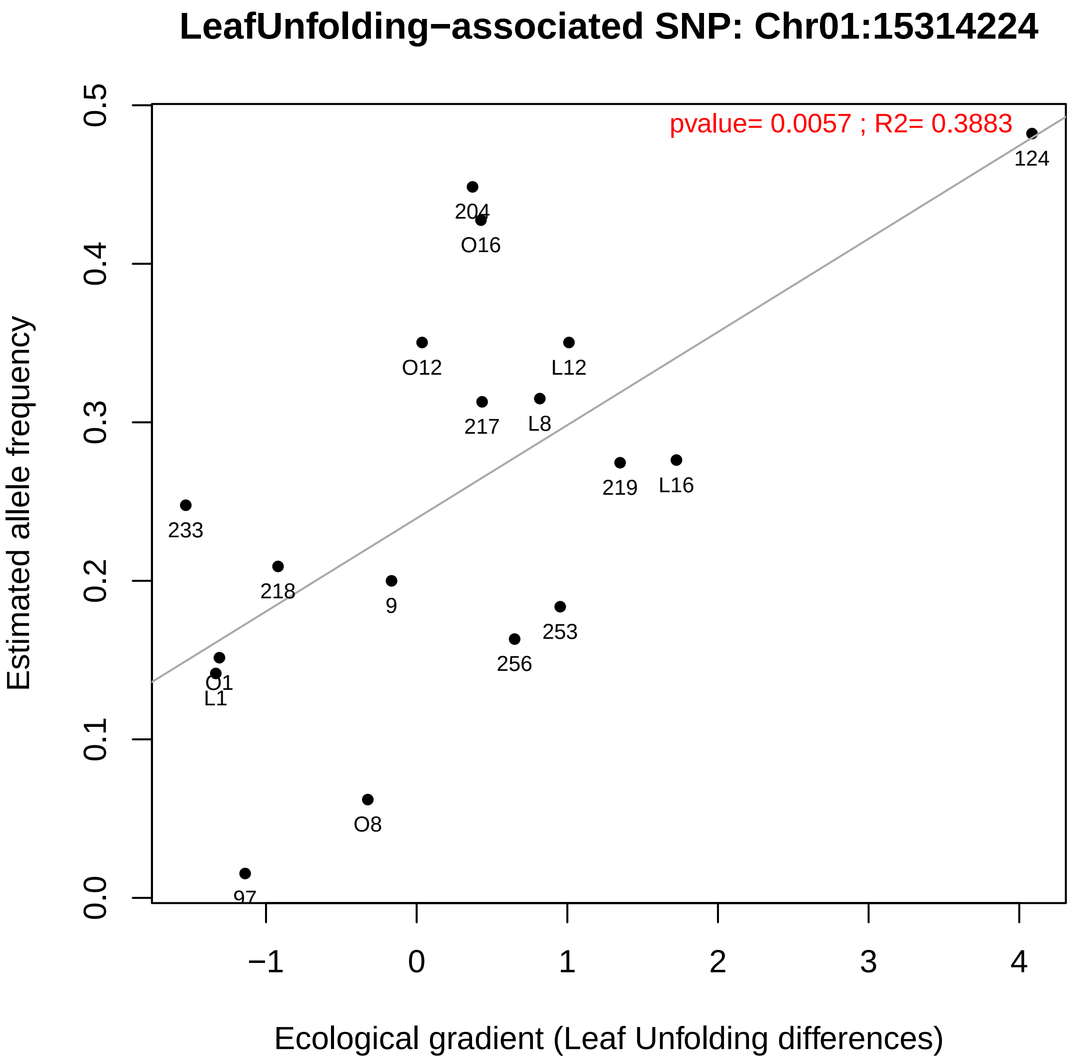

LeafUnfolding−associated SNP: Chr01:15314224
pvalue= 0.0057 ; R2= 0.3883
Estimated allele frequency
0.0
0.1
0.2
0.3
0.4
0.5
Ecological gradient (Leaf Unfolding differences)
−1
0
1
2
3
4
124
204
O16
O12
L12
217
L8
219
L16
233
218
9
253
256
O1
L1
O8
97

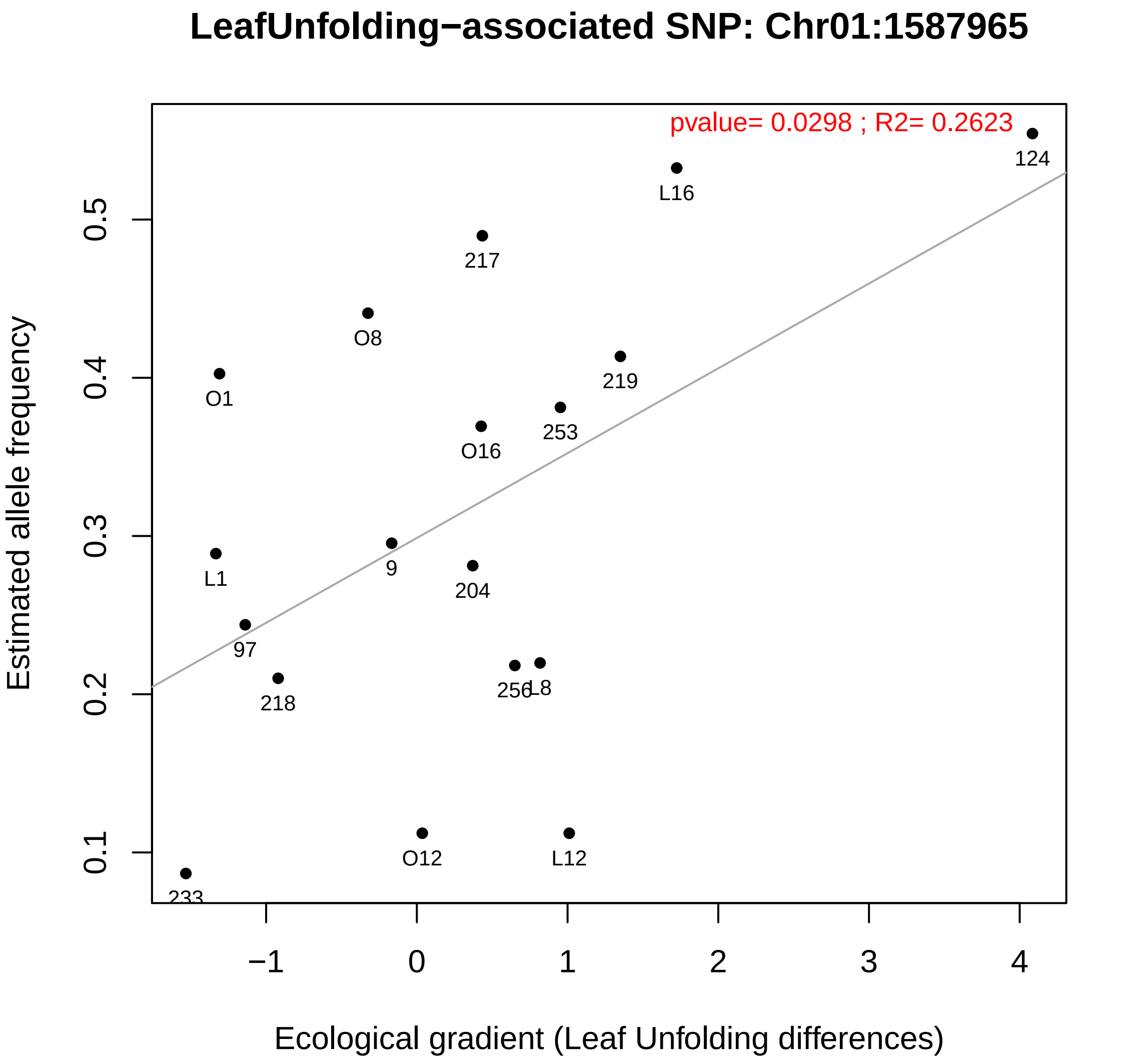

LeafUnfolding−associated SNP: Chr01:1587965
pvalue= 0.0298 ; R2= 0.2623
124
L16
217
O8
219
O1
253
O16
9
L1
204
97
218
256
L8
O12
L12
233
0.1
0.2
0.3
0.4
0.5
−1
0
1
2
3
4
Estimated allele frequency
Ecological gradient (Leaf Unfolding differences)

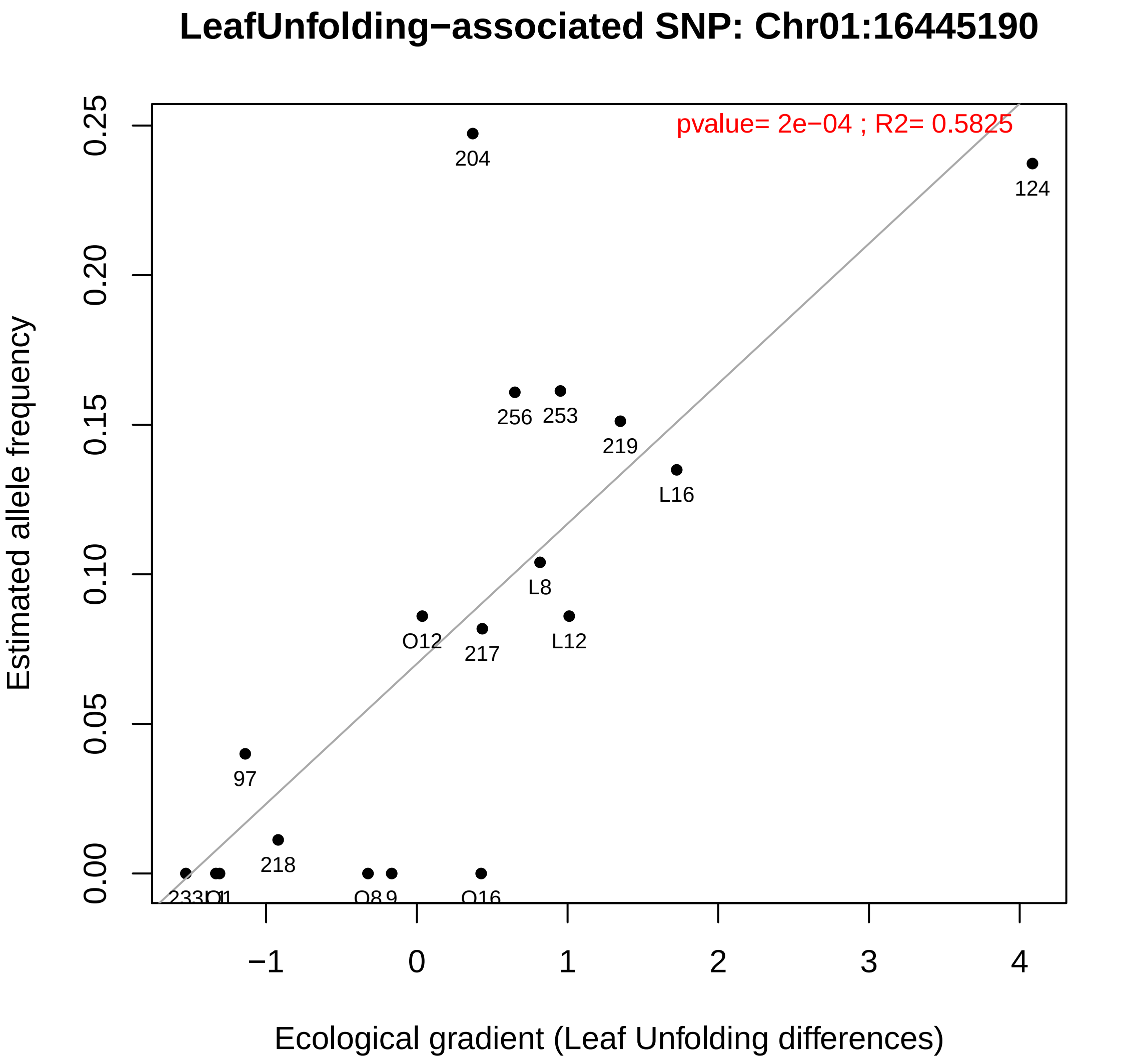
LeafUnfolding−associated SNP: Chr01:16445190
pvalue= 2e−04 ; R2= 0.5825
Estimated allele frequency
0.00
0.05
0.10
0.15
0.20
0.25
Ecological gradient (Leaf Unfolding differences)
−1
0
1
2
3
4
204
124
256
253
219
L16
L8
O12
217
L12
97
218
O8
9
O16

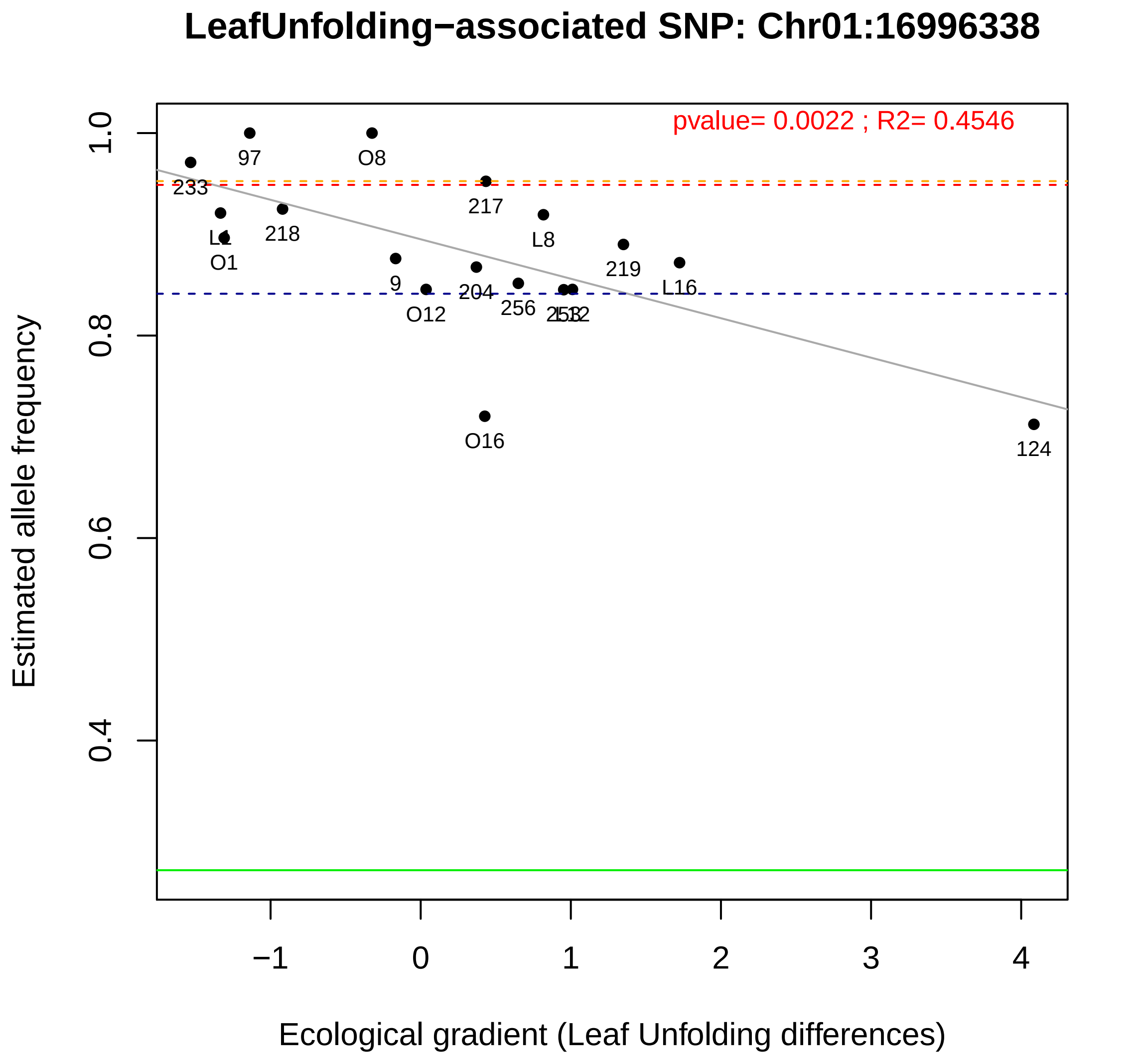

LeafUnfolding−associated SNP: Chr01:16996338
pvalue= 0.0022 ; R2= 0.4546
Estimated allele frequency
Ecological gradient (Leaf Unfolding differences)
1.0
0.8
0.6
0.4
−1
0
1
2
3
4
97
O8
233
217
L1
218
L8
O1
219
9
L16
204
O12
256
O16
124

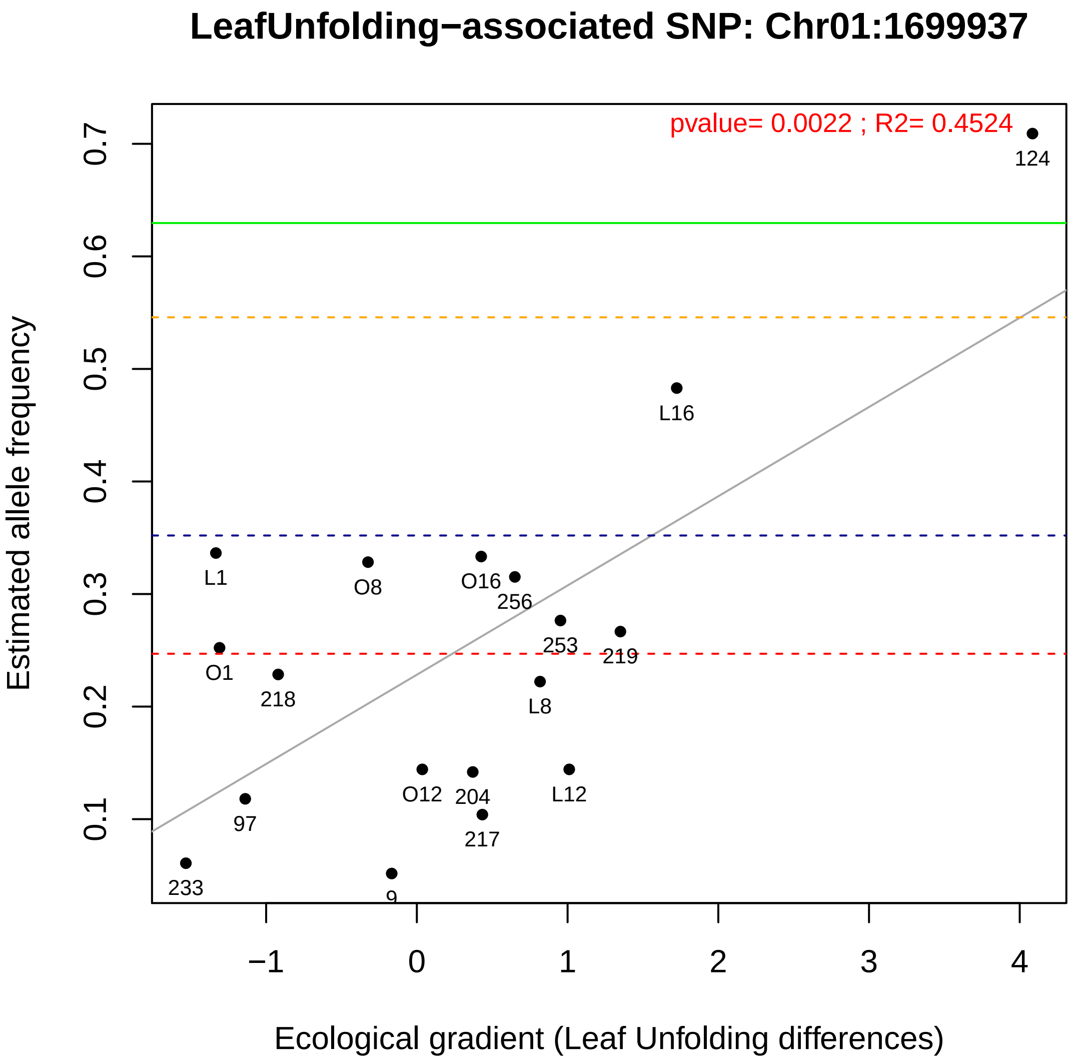

LeafUnfolding−associated SNP: Chr01:1699937
pvalue= 0.0022 ; R2= 0.4524
Estimated allele frequency
Ecological gradient (Leaf Unfolding differences)
0.1
0.2
0.3
0.4
0.5
0.6
0.7
−1
0
1
2
3
4
124
L16
L1
O8
O16
256
253
219
O1
218
L8
O12
204
L12
97
217
233
9

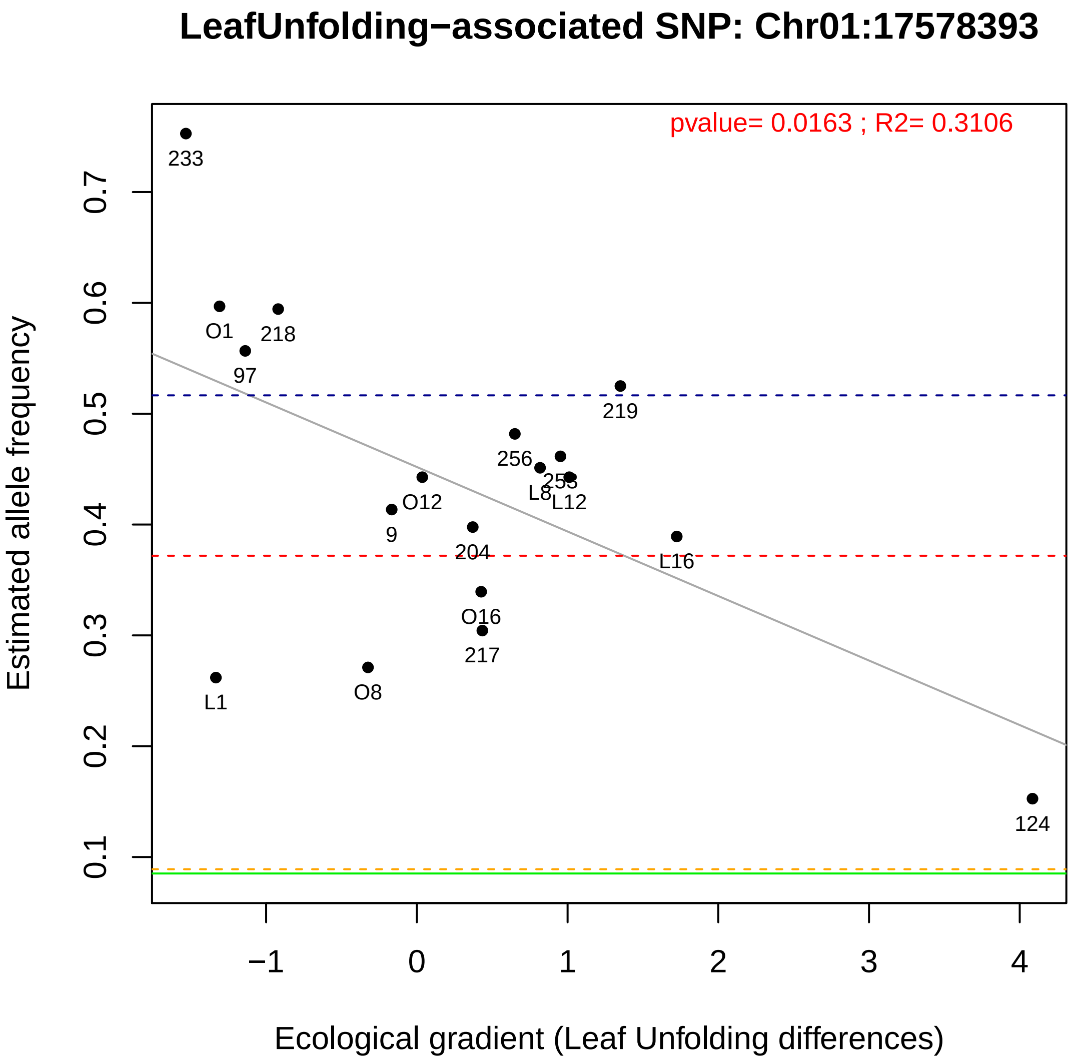

LeafUnfolding−associated SNP: Chr01:17578393
pvalue= 0.0163 ; R2= 0.3106
Estimated allele frequency
0.1
0.2
0.3
0.4
0.5
0.6
0.7
−1
0
1
2
3
4
Ecological gradient (Leaf Unfolding differences)
233
O1
218
97
219
256
253
L8
L12
O12
9
204
L16
O16
217
L1
O8
124

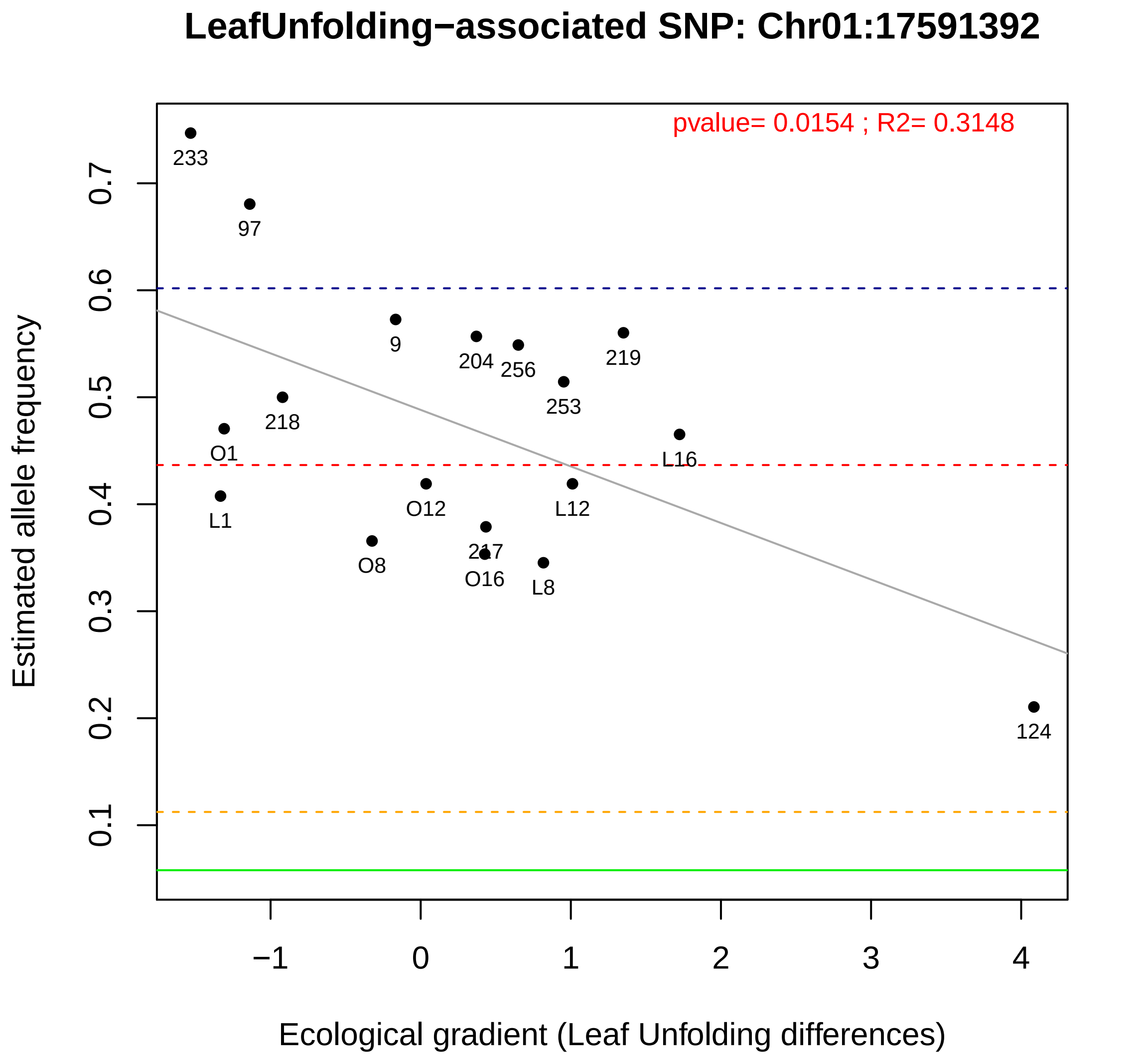
LeafUnfolding−associated SNP: Chr01:17591392
pvalue= 0.0154 ; R2= 0.3148
Estimated allele frequency
Ecological gradient (Leaf Unfolding differences)
0.1
0.2
0.3
0.4
0.5
0.6
0.7
−1
0
1
2
3
4
233
97
9
204
256
219
253
218
O1
L16
L1
O12
L12
O8
217
O16
L8
124

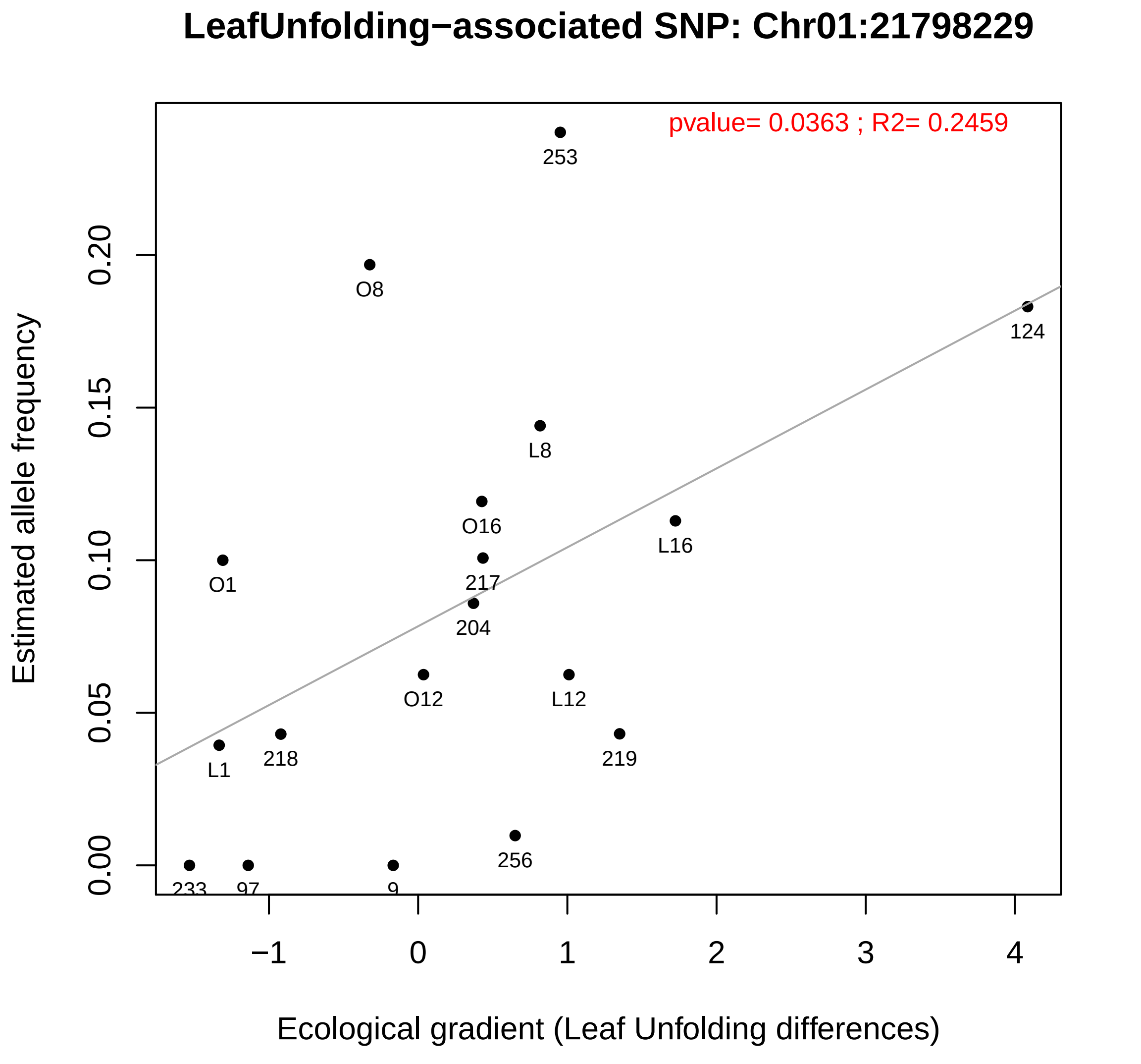
LeafUnfolding−associated SNP: Chr01:21798229
pvalue= 0.0363 ; R2= 0.2459
Estimated allele frequency
Ecological gradient (Leaf Unfolding differences)
0.00
0.05
0.10
0.15
0.20
−1
0
1
2
3
4
253
O8
124
L8
O16
L16
O1
217
204
O12
L12
L1
218
219
256
233
97
9

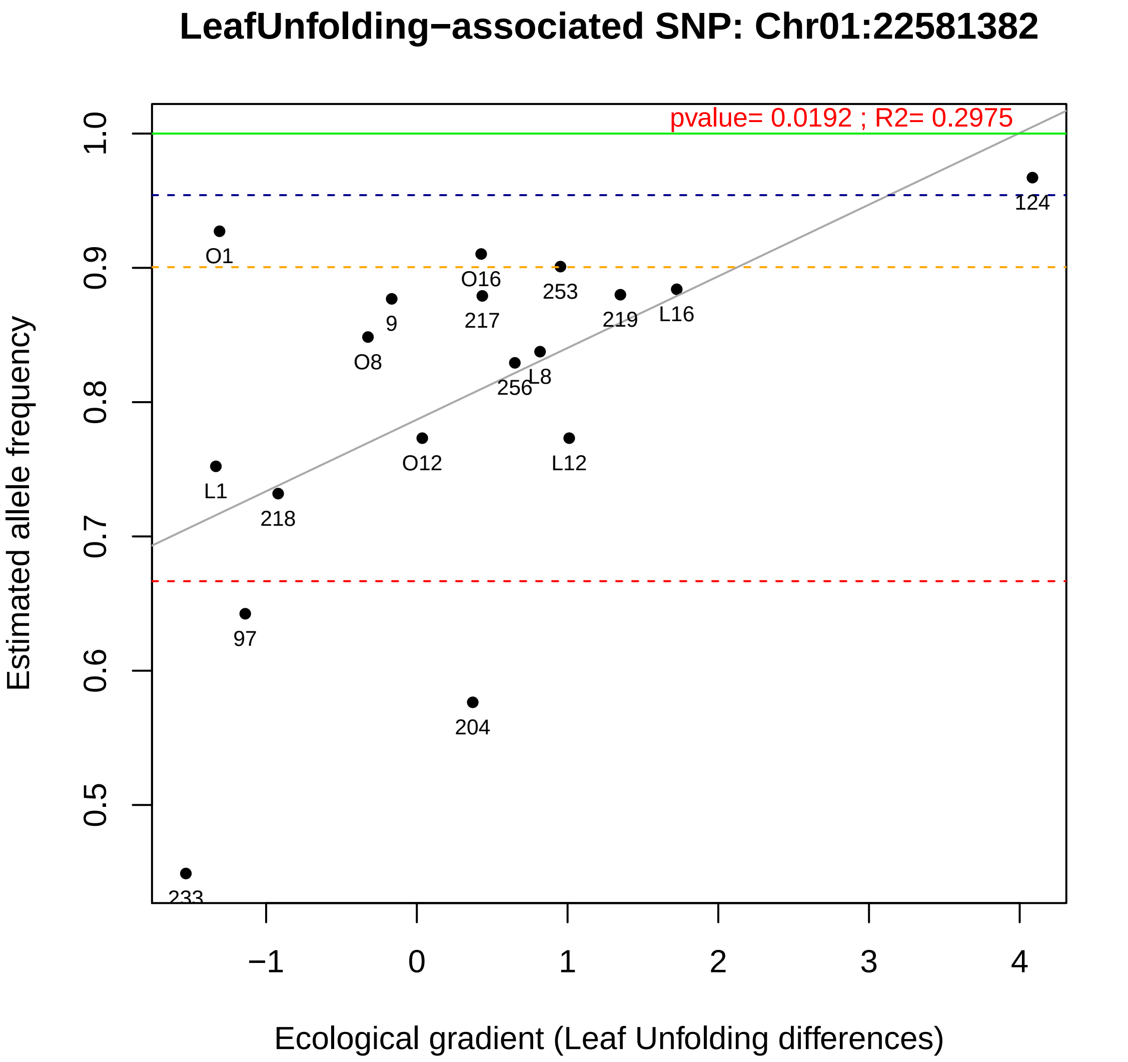

LeafUnfolding−associated SNP: Chr01:22581382
pvalue= 0.0192 ; R2= 0.2975
Estimated allele frequency
0.5
0.6
0.7
0.8
0.9
1.0
Ecological gradient (Leaf Unfolding differences)
−1
0
1
2
3
4
124
O1
O16
253
217
9
219
L16
O8
256
L8
O12
L12
L1
218
97
204
233

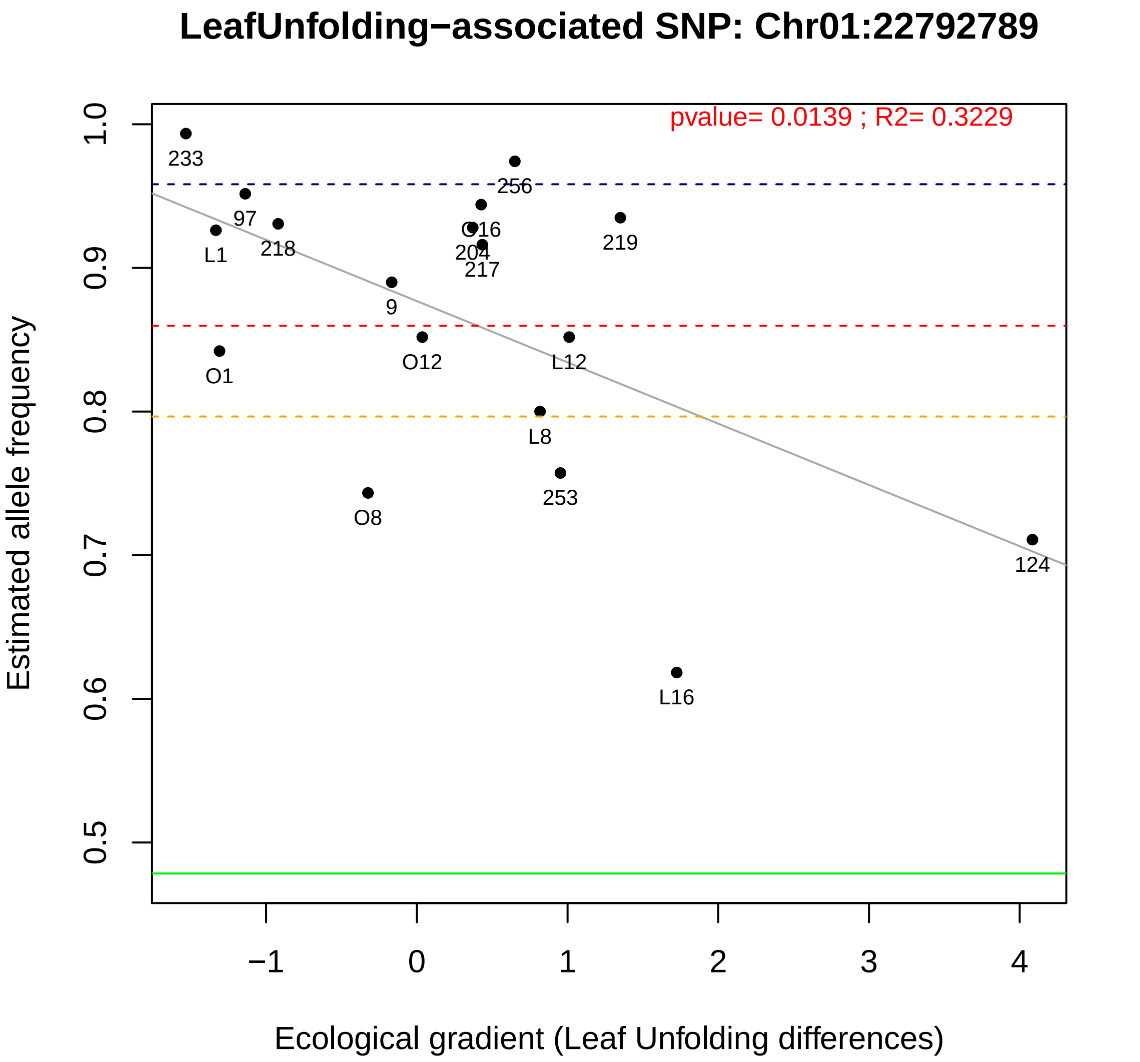
LeafUnfolding−associated SNP: Chr01:22792789
pvalue= 0.0139 ; R2= 0.3229
Estimated allele frequency
Ecological gradient (Leaf Unfolding differences)
1.0
0.9
0.8
0.7
0.6
0.5
−1
0
1
2
3
4
233
97
L1
218
256
O16
204
217
219
9
O12
L12
O1
L8
253
O8
124
L16

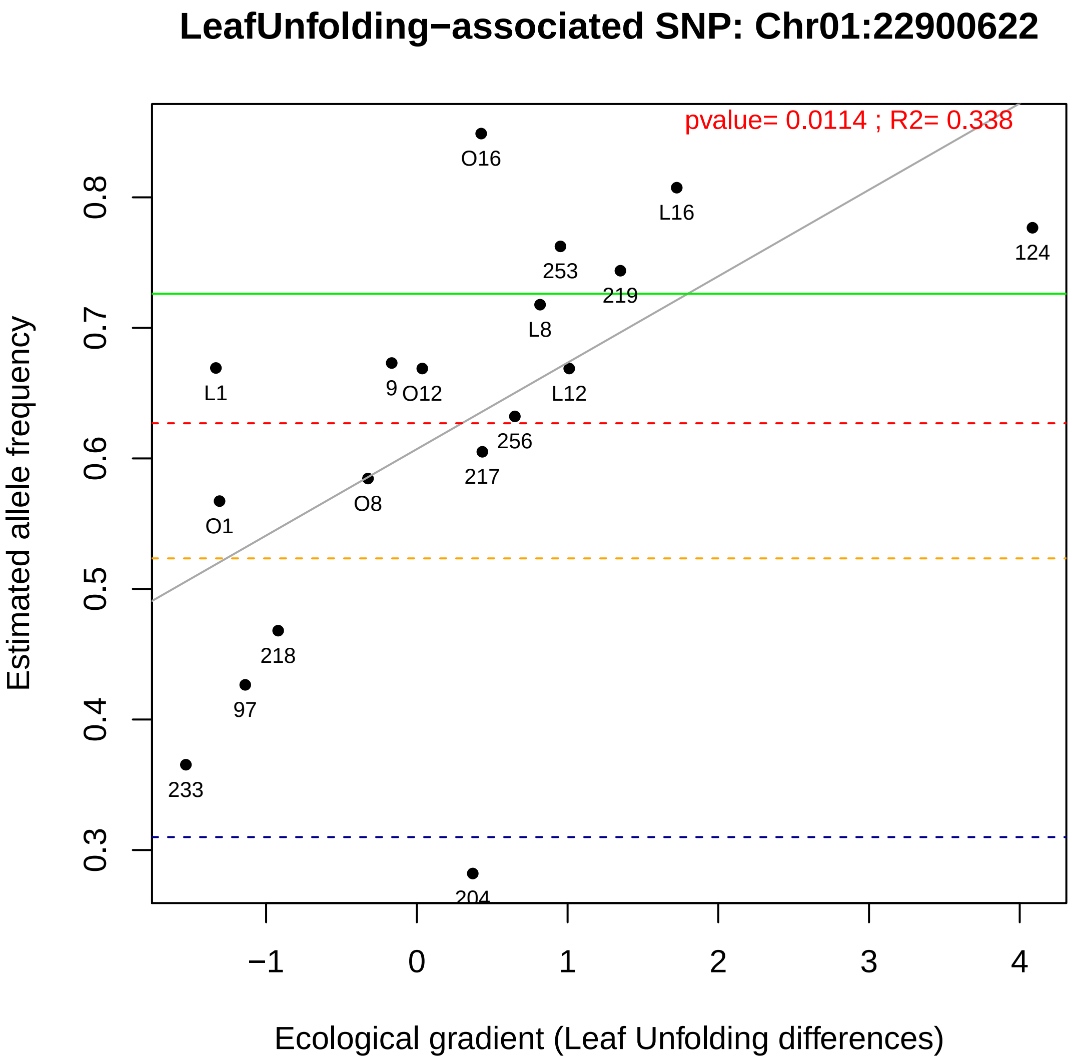
LeafUnfolding−associated SNP: Chr01:22900622
pvalue= 0.0114 ; R2= 0.338
Estimated allele frequency
Ecological gradient (Leaf Unfolding differences)
0.3
0.4
0.5
0.6
0.7
0.8
−1
0
1
2
3
4
O16
L16
124
253
219
L8
L1
9
O12
L12
256
217
O8
O1
218
97
233
204

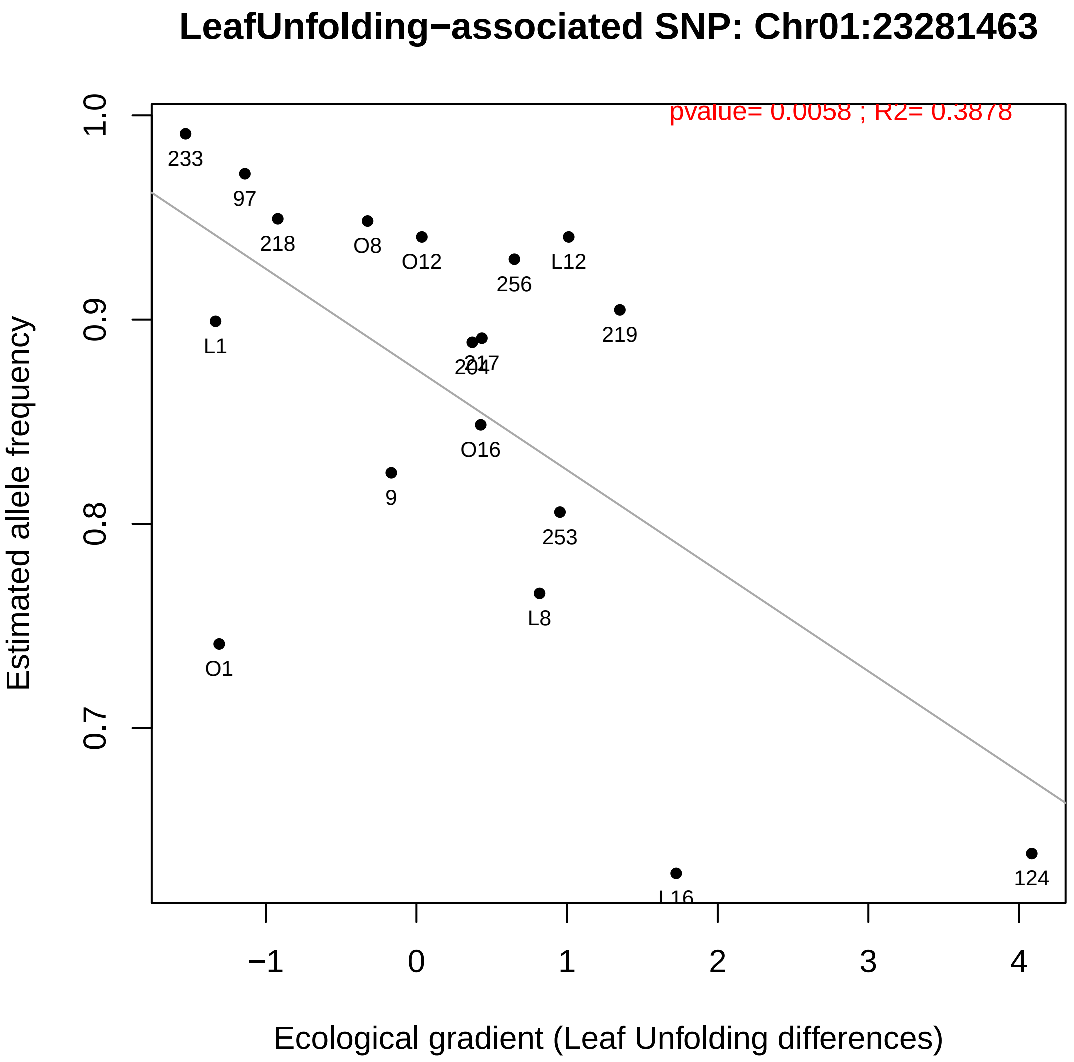

LeafUnfolding−associated SNP: Chr01:23281463
pvalue= 0.0058 ; R2= 0.3878
Estimated allele frequency
Ecological gradient (Leaf Unfolding differences)
1.0
0.9
0.8
0.7
−1
0
1
2
3
4
233
97
218
O8
O12
256
L12
219
L1
217
204
O16
9
253
L8
O1
124
L16

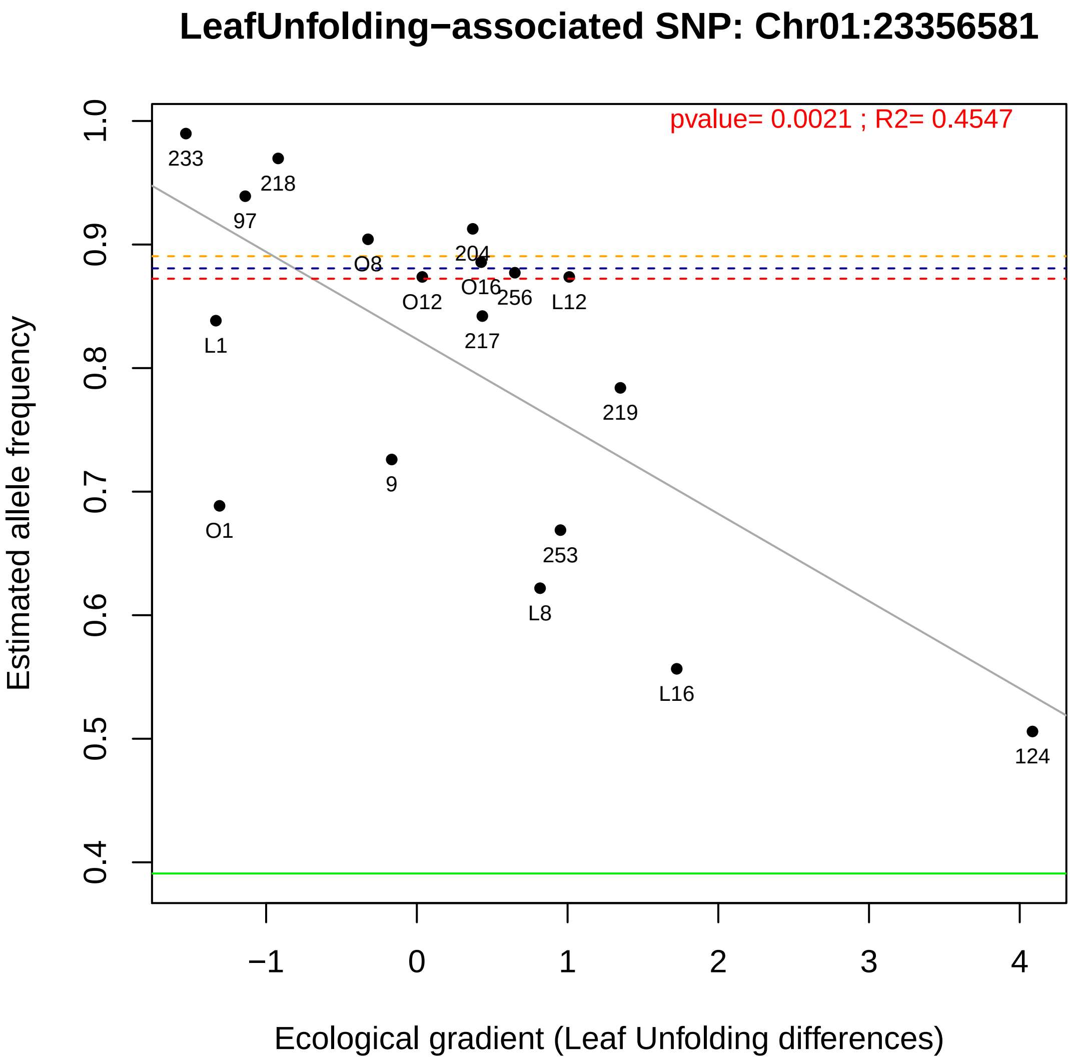
LeafUnfolding−associated SNP: Chr01:23356581
pvalue= 0.0021 ; R2= 0.4547
Estimated allele frequency
Ecological gradient (Leaf Unfolding differences)
1.0
0.9
0.8
0.7
0.6
0.5
0.4
−1
0
1
2
3
4
233
218
97
O8
204
O16
O12
256
L12
217
L1
219
9
O1
253
L8
L16
124

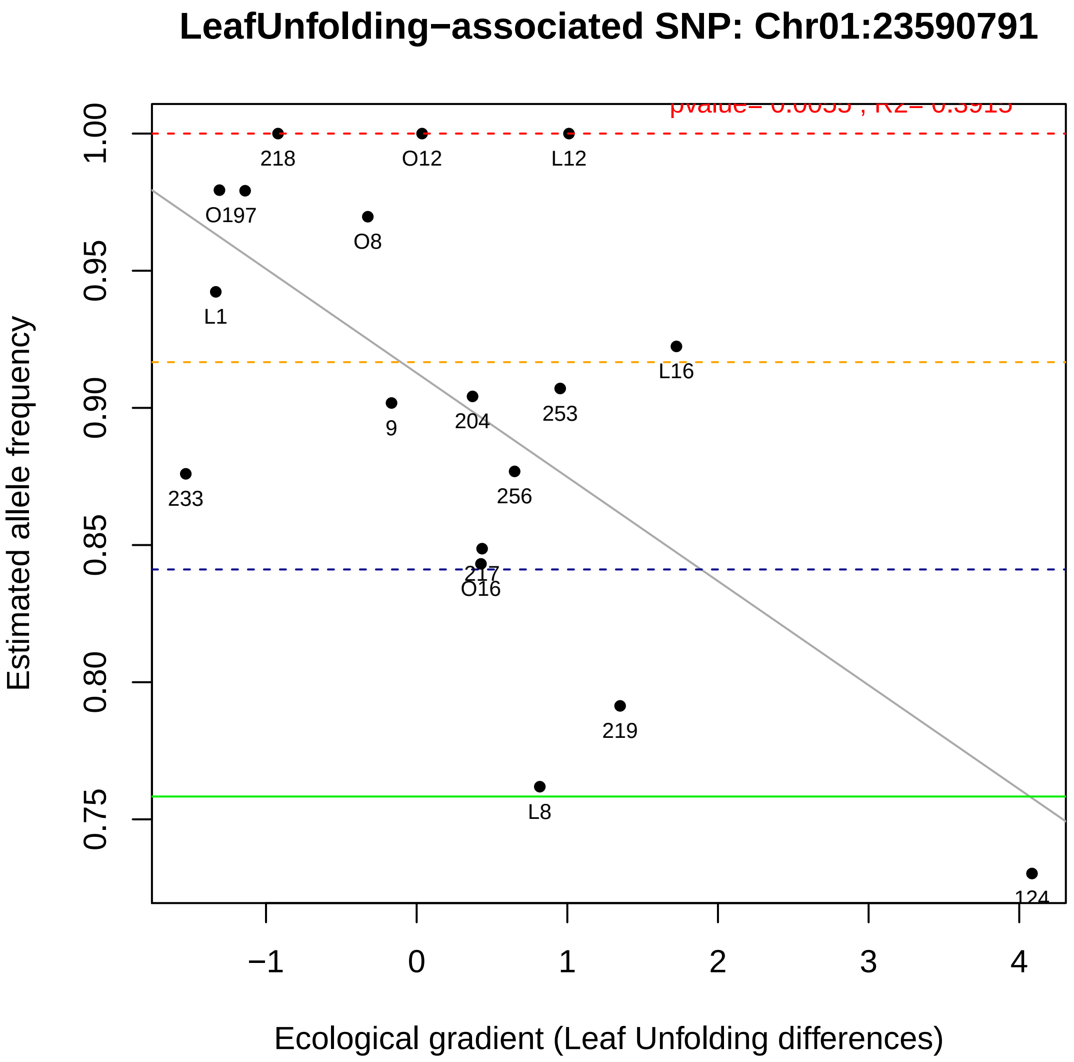
LeafUnfolding−associated SNP: Chr01:23590791
Estimated allele frequency
1.00
0.95
0.90
0.85
0.80
0.75
218
O12
L12
O197
O8
L1
L16
9
204
253
233
256
217
O16
219
L8
124
−1
0
1
2
3
4
Ecological gradient (Leaf Unfolding differences)

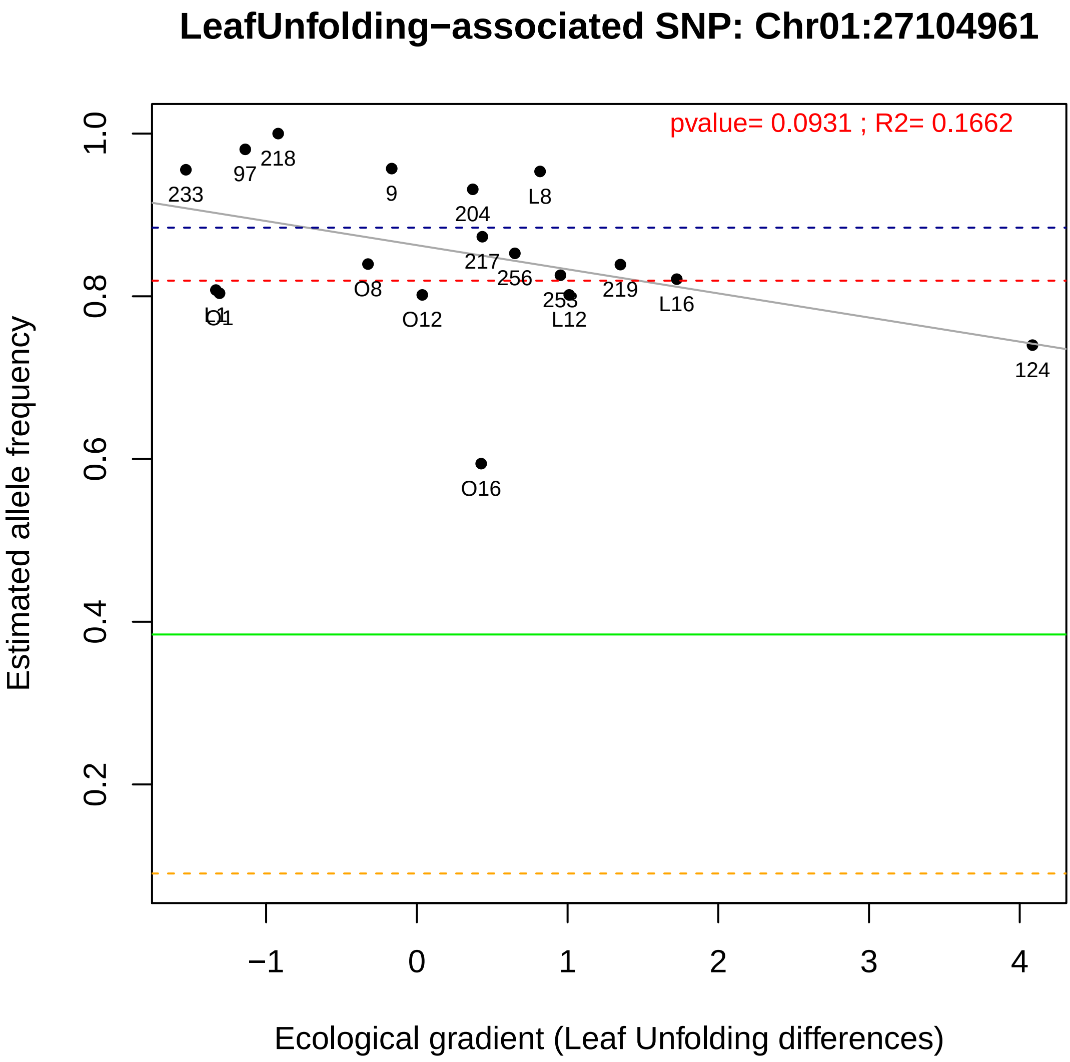

LeafUnfolding−associated SNP: Chr01:27104961
pvalue= 0.0931 ; R2= 0.1662
Estimated allele frequency
Ecological gradient (Leaf Unfolding differences)
0.2
0.4
0.6
0.8
1.0
−1
0
1
2
3
4
233
97
218
9
204
L8
217
256
O8
O12
253
L12
219
L16
124
O16

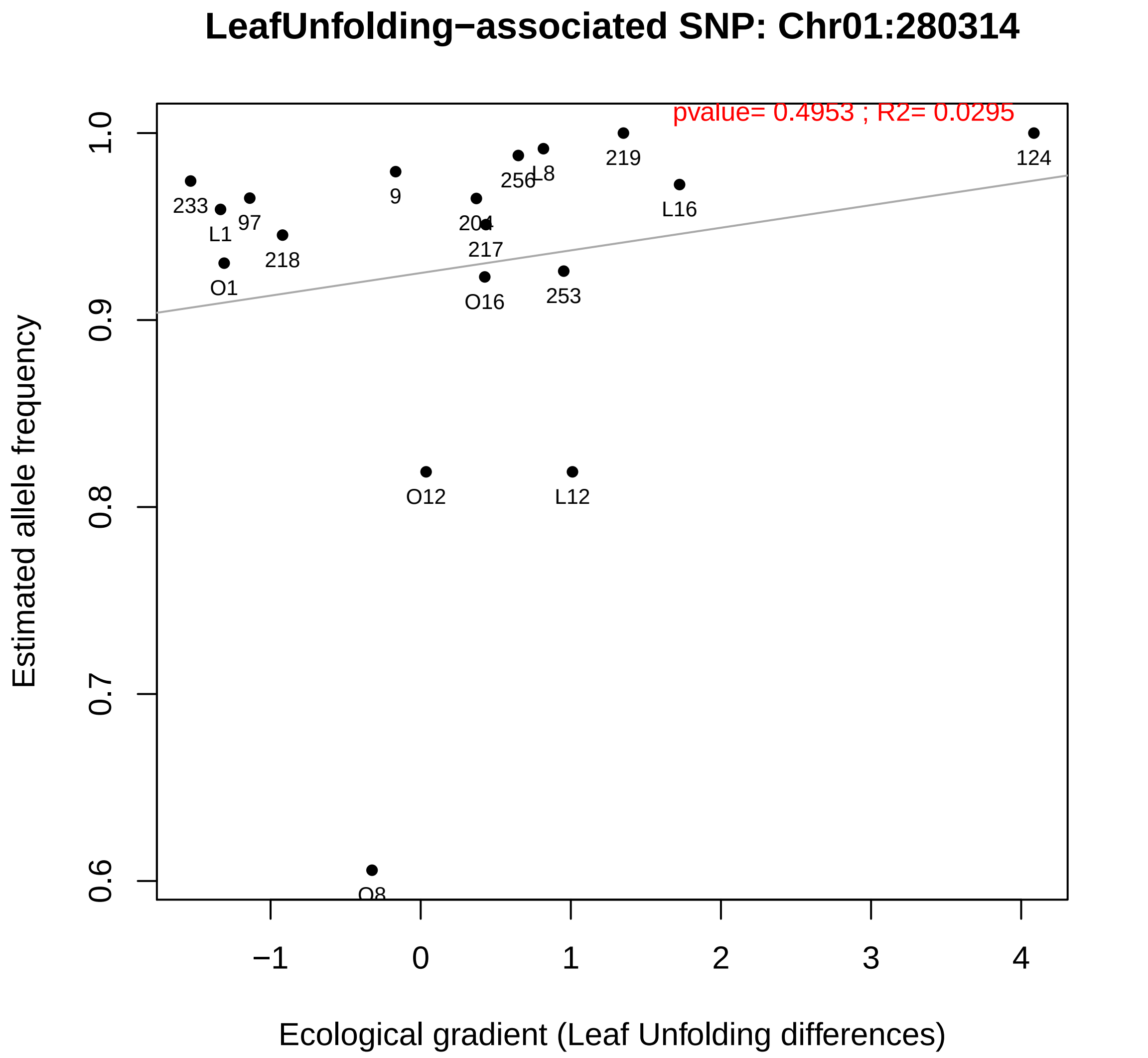

LeafUnfolding−associated SNP: Chr01:280314
pvalue= 0.4953 ; R2= 0.0295
Estimated allele frequency
Ecological gradient (Leaf Unfolding differences)
1.0
0.9
0.8
0.7
0.6
−1
0
1
2
3
4
233
L1
O1
97
218
9
204
217
O16
256
L8
219
253
L16
124
O12
L12
O8

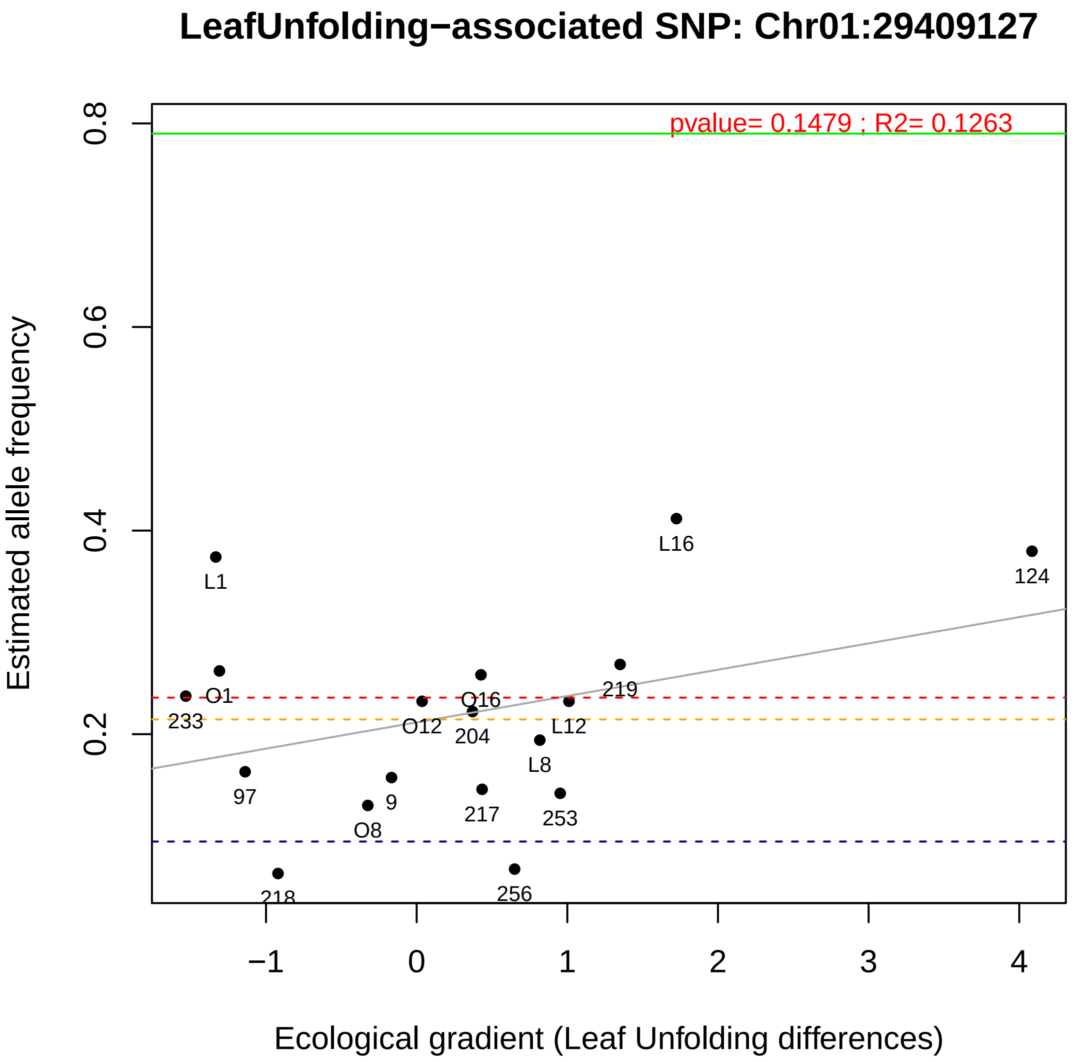

LeafUnfolding−associated SNP: Chr01:29409127
pvalue= 0.1479 ; R2= 0.1263
Estimated allele frequency
0.2
0.4
0.6
0.8
Ecological gradient (Leaf Unfolding differences)
−1
0
1
2
3
4
L16
124
L1
219
O1
233
O16
L12
O12
204
L8
97
9
217
253
O8
218
256

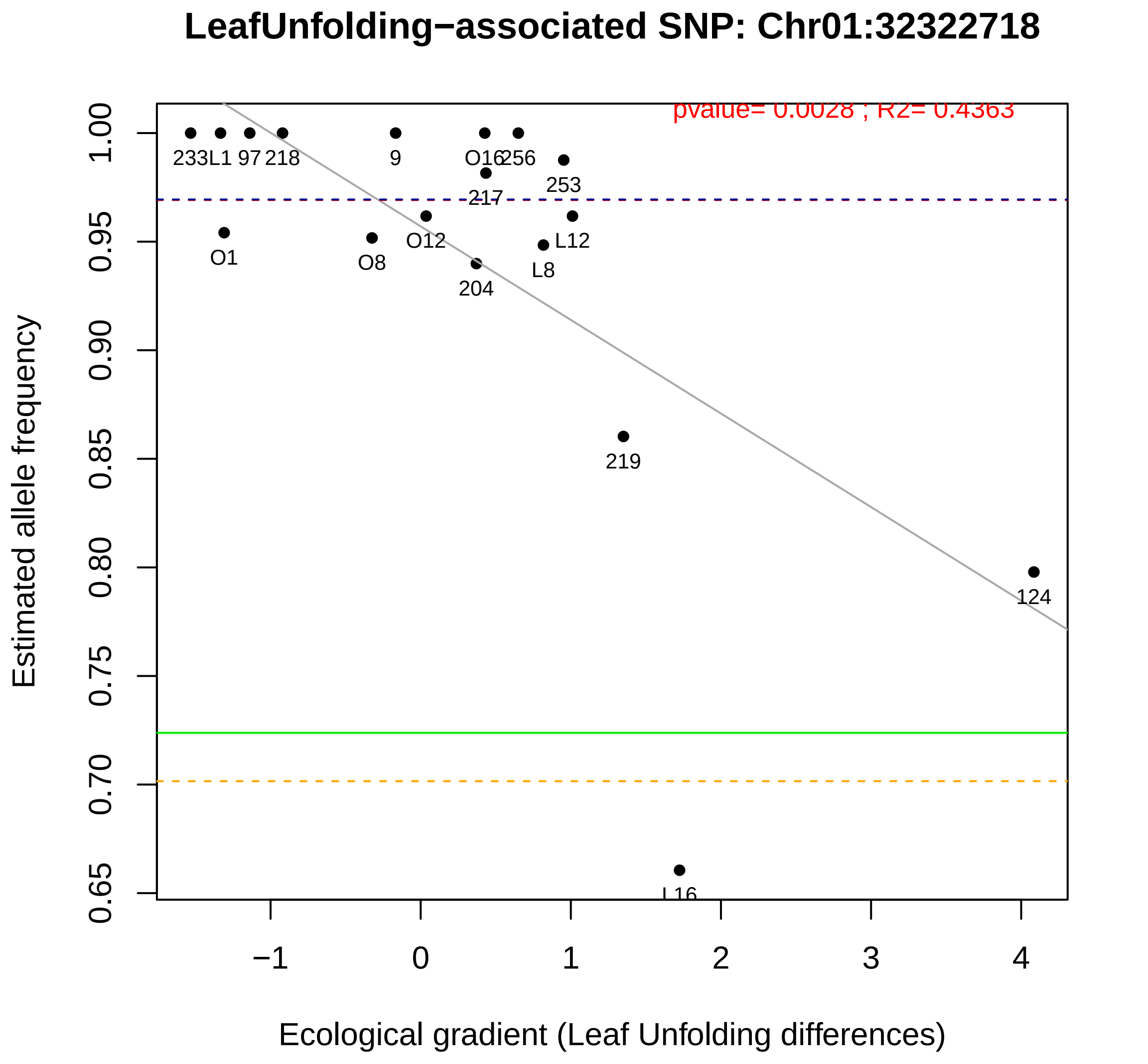

LeafUnfolding−associated SNP: Chr01:32322718
pvalue= 0.0028 ; R2= 0.4363
Estimated allele frequency
Ecological gradient (Leaf Unfolding differences)
1.00
0.95
0.90
0.85
0.80
0.75
0.70
0.65
−1
0
1
2
3
4
233
L1
97
218
9
O16
256
253
217
O1
O8
O12
L12
L8
204
219
124
L16

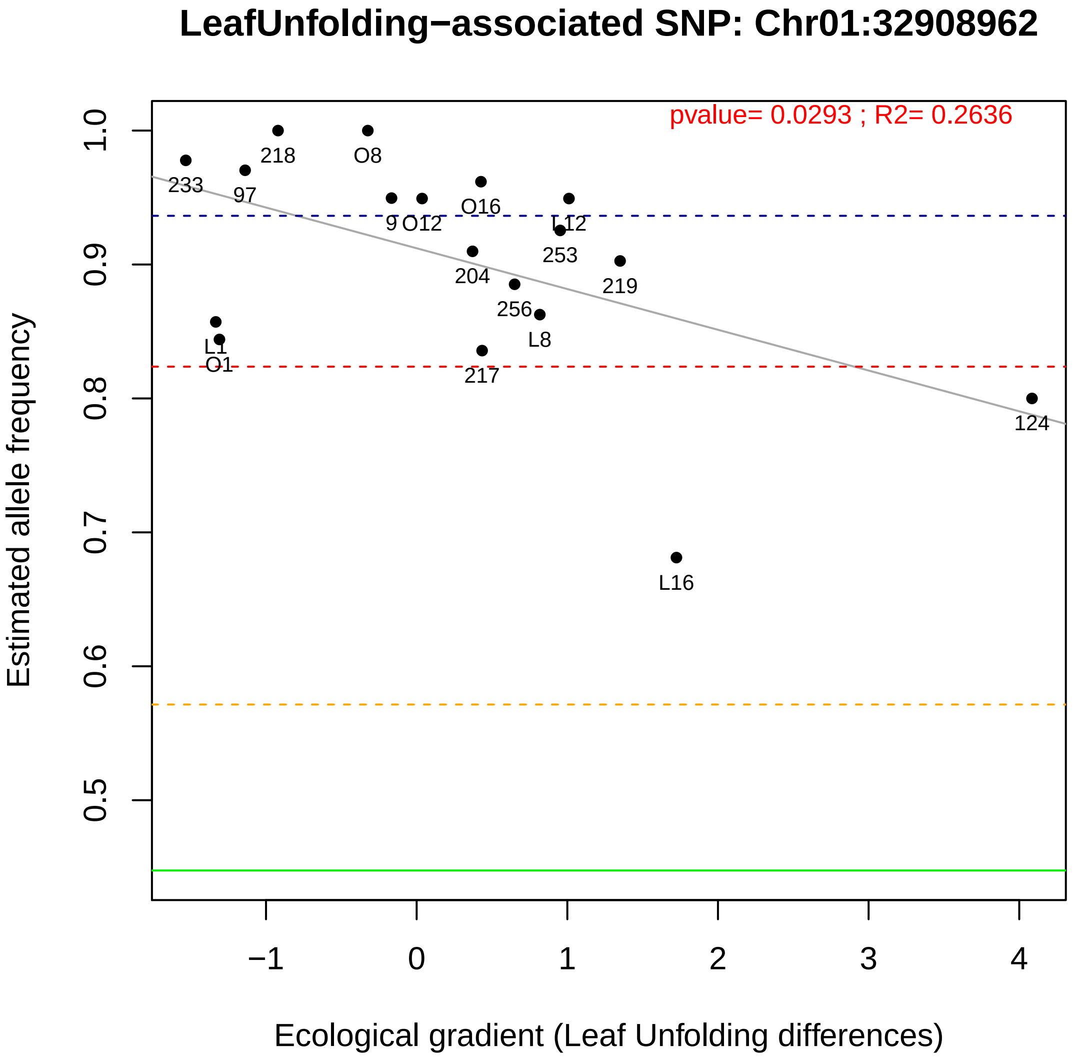
LeafUnfolding−associated SNP: Chr01:32908962
pvalue= 0.0293 ; R2= 0.2636
Estimated allele frequency
Ecological gradient (Leaf Unfolding differences)
233
97
218
O8
9
O12
O16
L12
253
204
219
256
L8
L1
O1
217
124
L16
0.5
0.6
0.7
0.8
0.9
1.0
−1
0
1
2
3
4

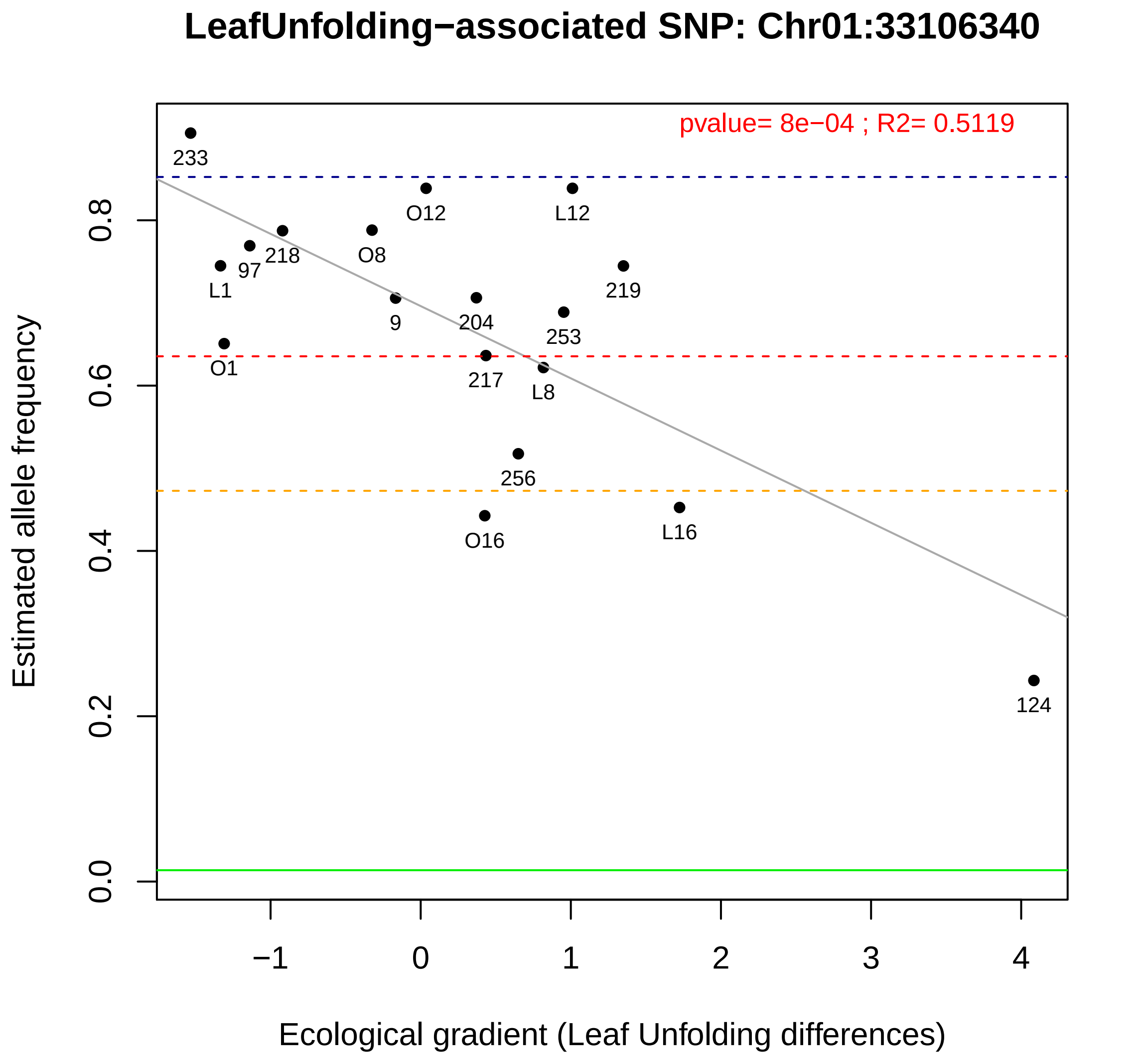
LeafUnfolding−associated SNP: Chr01:33106340
pvalue= 8e−04 ; R2= 0.5119
Estimated allele frequency
Ecological gradient (Leaf Unfolding differences)
0.0
0.2
0.4
0.6
0.8
−1
0
1
2
3
4
233
O12
L12
218
97
L1
O8
9
204
219
253
O1
217
L8
256
O16
L16
124

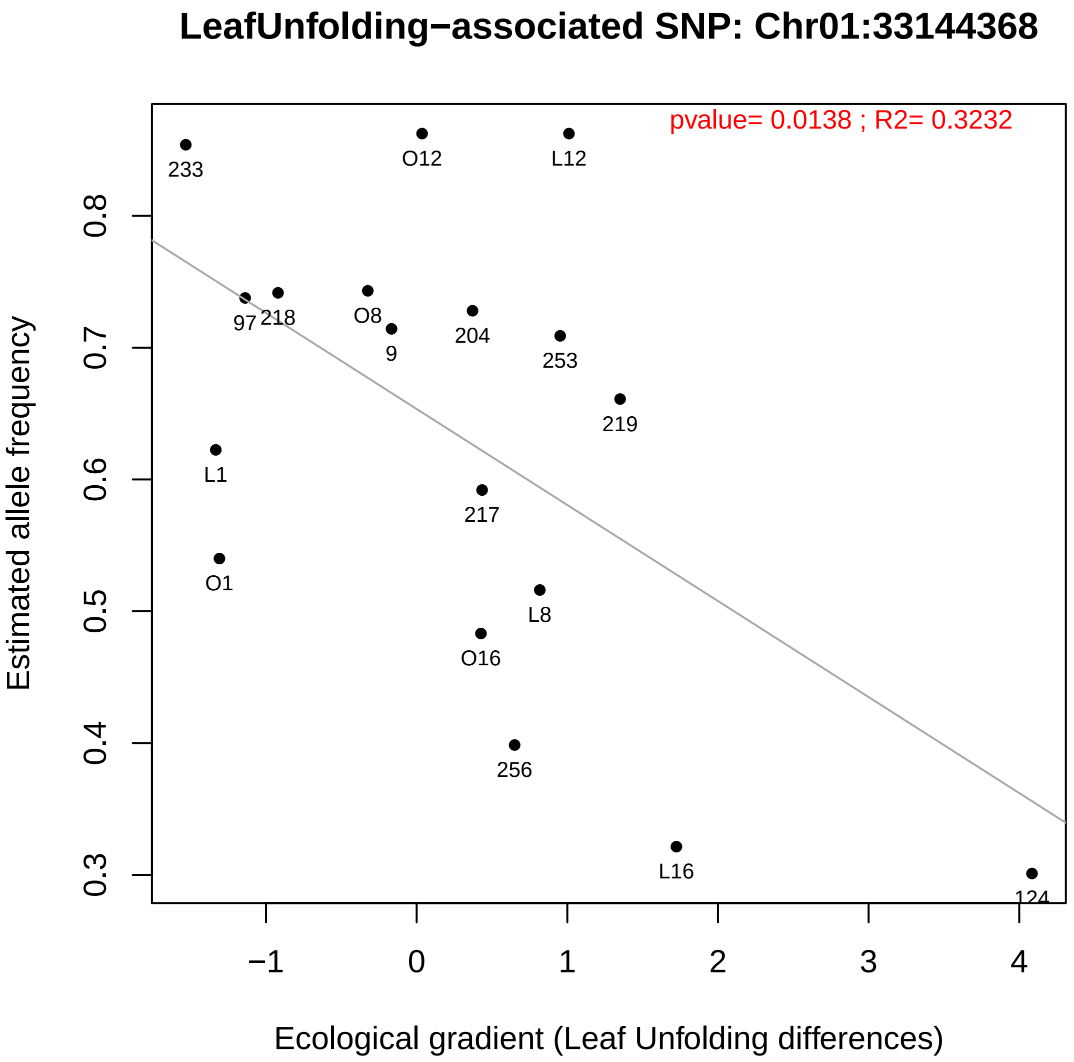
LeafUnfolding−associated SNP: Chr01:33144368
pvalue= 0.0138 ; R2= 0.3232
Estimated allele frequency
0.3
0.4
0.5
0.6
0.7
0.8
−1
0
1
2
3
4
Ecological gradient (Leaf Unfolding differences)
233
O12
L12
97
218
O8
9
204
253
219
L1
217
O1
L8
O16
256
L16
124

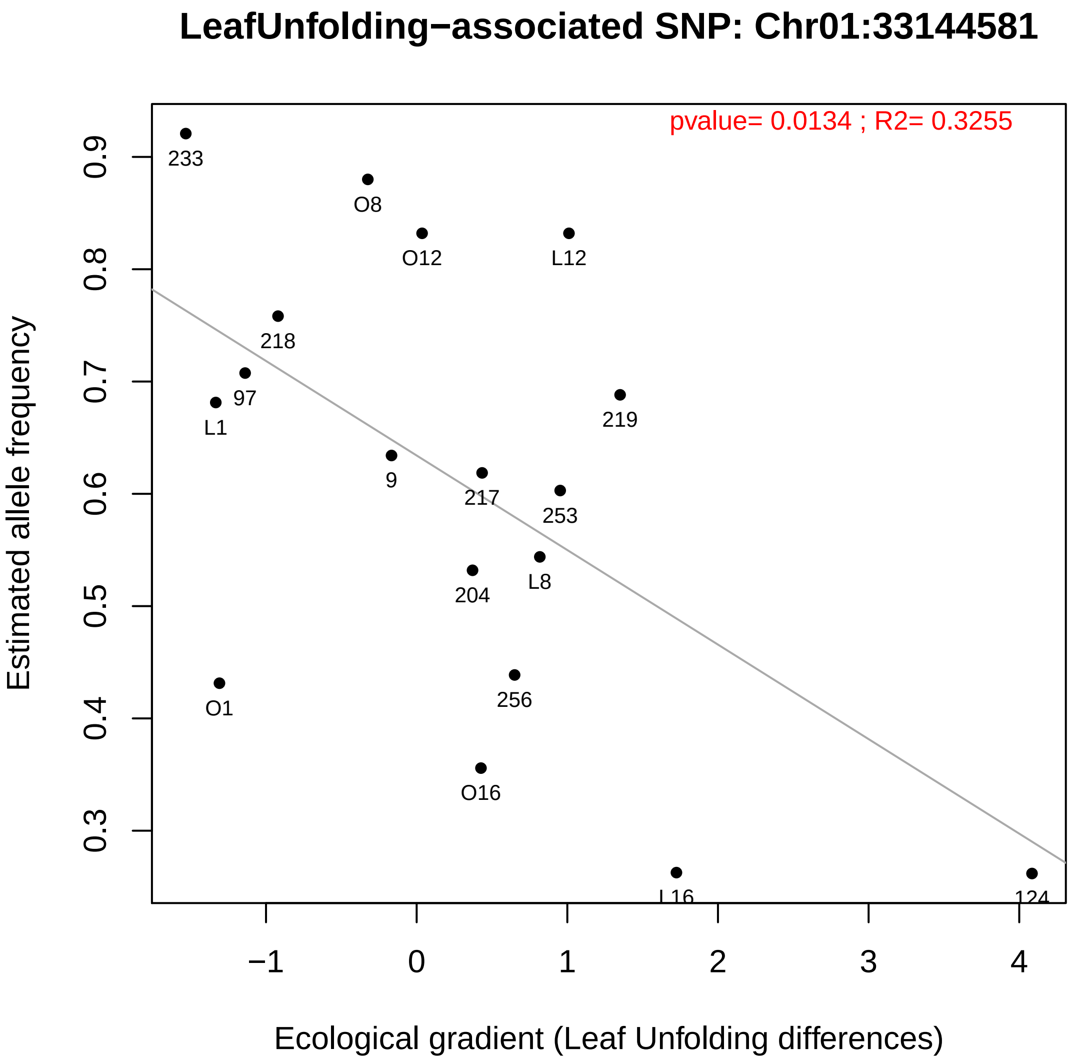
LeafUnfolding−associated SNP: Chr01:33144581
pvalue= 0.0134 ; R2= 0.3255
Estimated allele frequency
0.3
0.4
0.5
0.6
0.7
0.8
0.9
−1
0
1
2
3
4
Ecological gradient (Leaf Unfolding differences)
233
O8
O12
L12
218
97
L1
219
9
217
253
204
L8
256
O1
O16
124

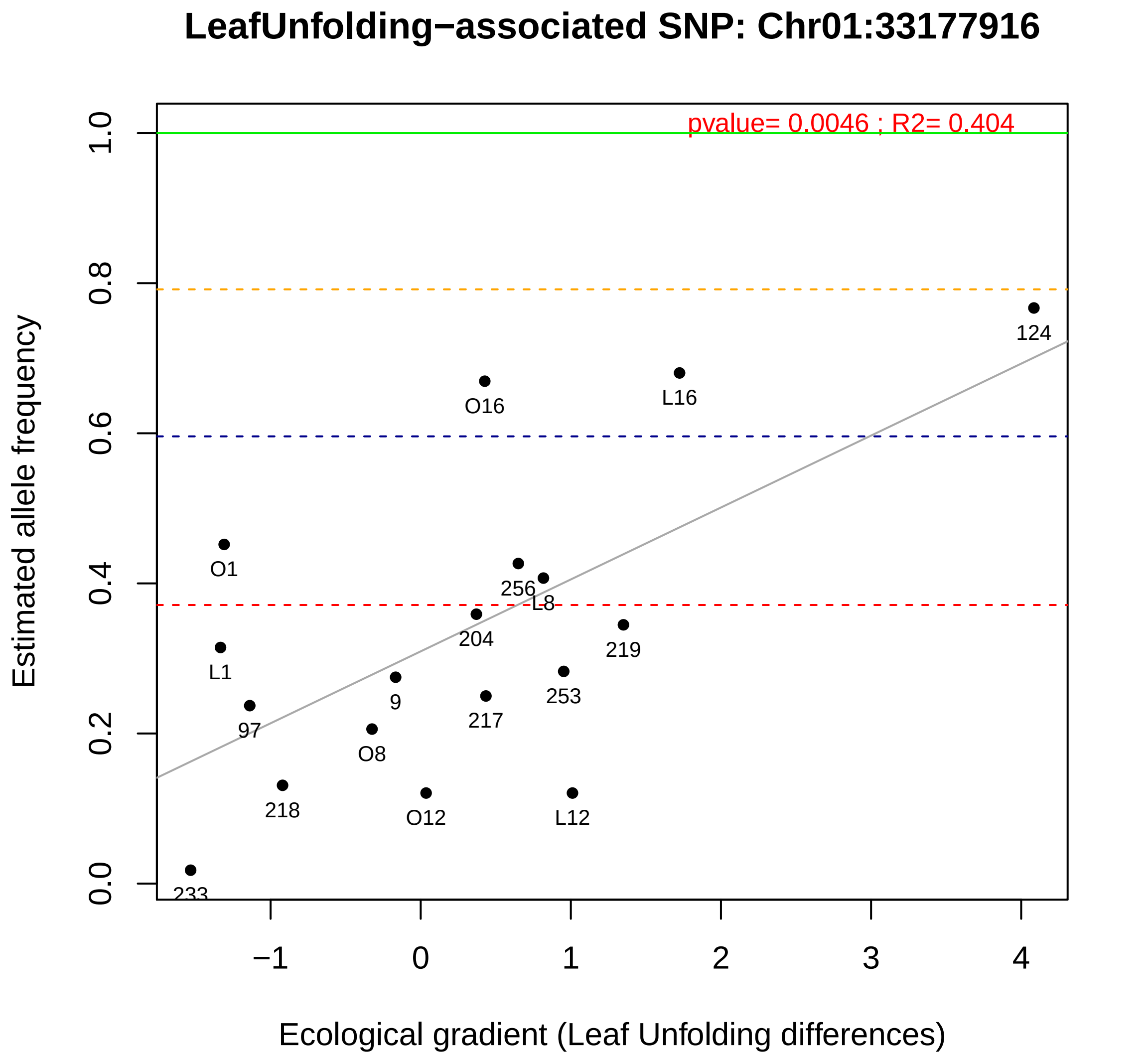

LeafUnfolding−associated SNP: Chr01:33177916
pvalue= 0.0046 ; R2= 0.404
Estimated allele frequency
0.0
0.2
0.4
0.6
0.8
1.0
124
O16
L16
O1
256
L8
204
219
L1
9
253
217
97
O8
218
O12
L12
233
−1
0
1
2
3
4
Ecological gradient (Leaf Unfolding differences)

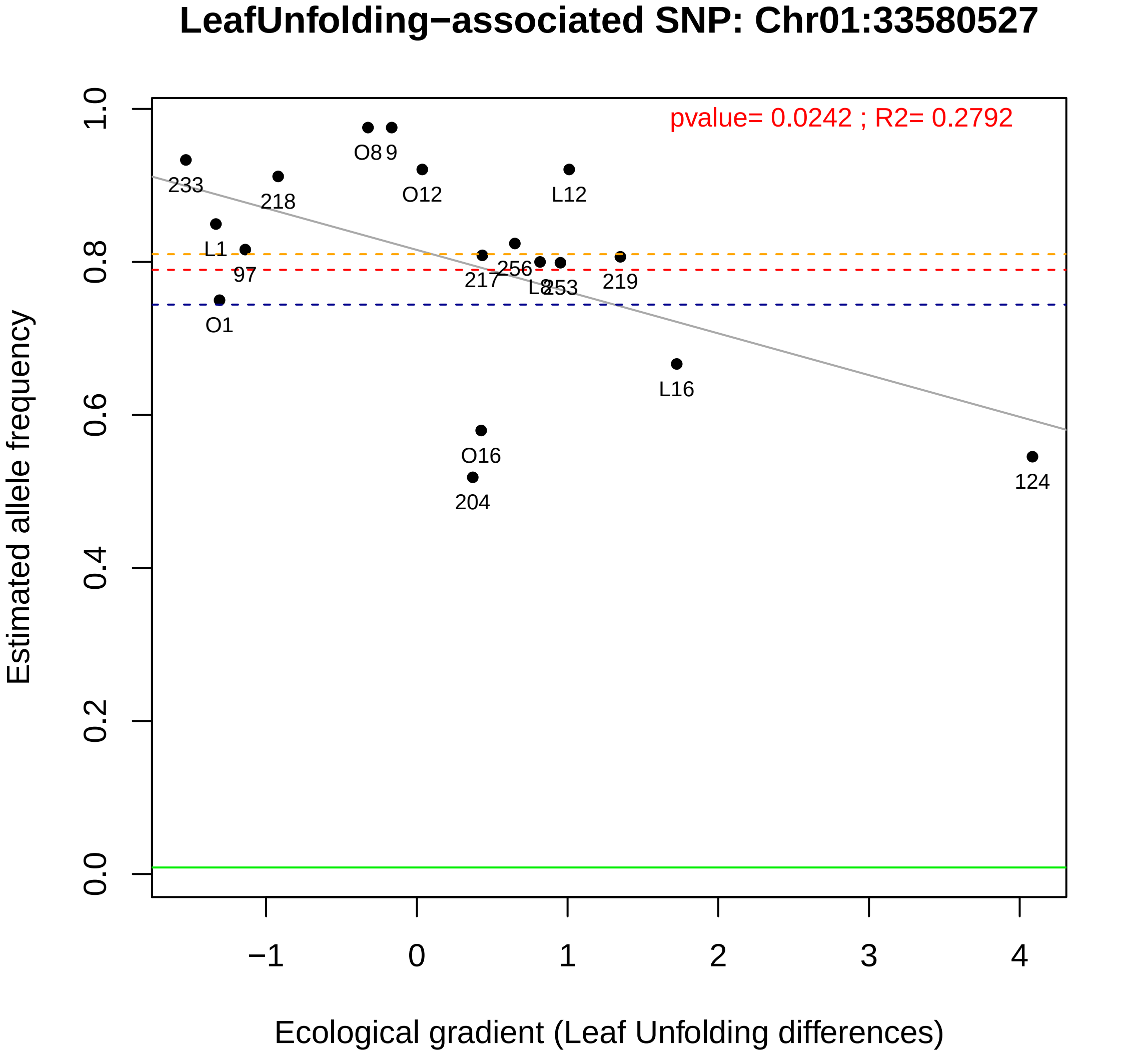
LeafUnfolding−associated SNP: Chr01:33580527
pvalue= 0.0242 ; R2= 0.2792
Estimated allele frequency
Ecological gradient (Leaf Unfolding differences)
0.0
0.2
0.4
0.6
0.8
1.0
−1
0
1
2
3
4
233
218
O8
9
O12
L12
L1
97
217
256
L8
253
219
O1
L16
O16
204
124

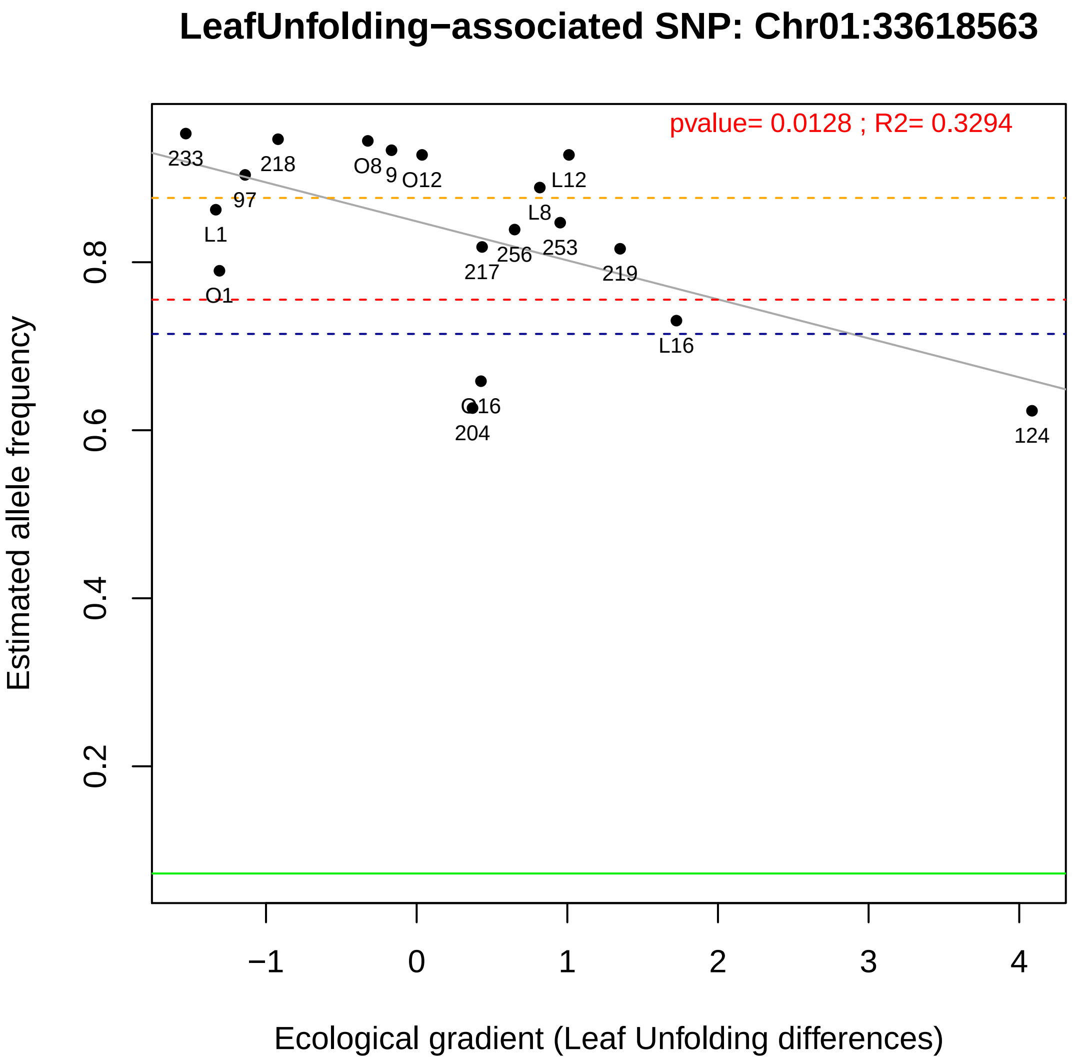
LeafUnfolding−associated SNP: Chr01:33618563
pvalue= 0.0128 ; R2= 0.3294
233
218
97
L1
O1
O8
9
O12
L12
L8
253
256
217
219
L16
O16
204
124
0.2
0.4
0.6
0.8
−1
0
1
2
3
4
Estimated allele frequency
Ecological gradient (Leaf Unfolding differences)

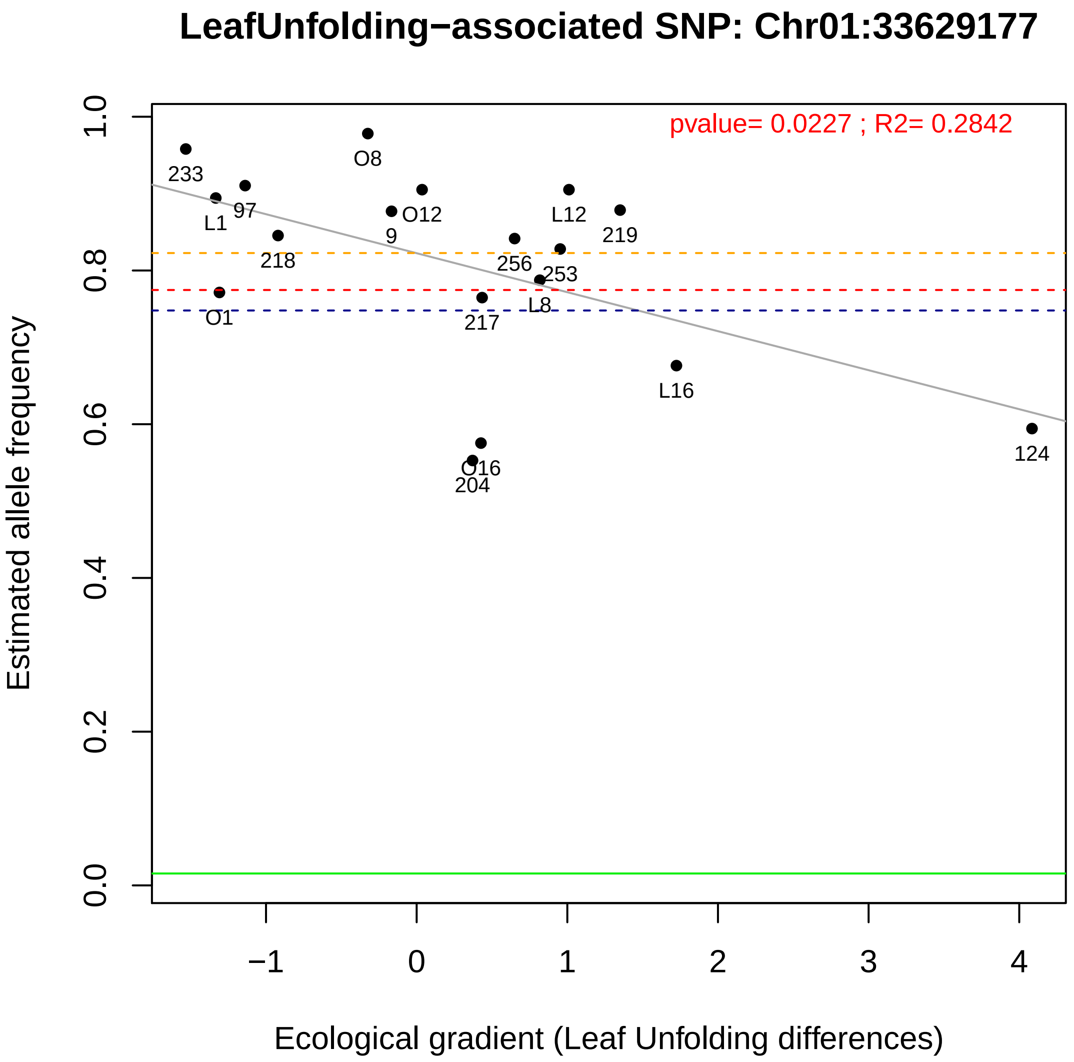
LeafUnfolding−associated SNP: Chr01:33629177
pvalue= 0.0227 ; R2= 0.2842
Estimated allele frequency
Ecological gradient (Leaf Unfolding differences)
0.0
0.2
0.4
0.6
0.8
1.0
−1
0
1
2
3
4
233
O8
97
L1
O12
9
218
L12
219
256
253
L8
O1
217
L16
O16
204
124

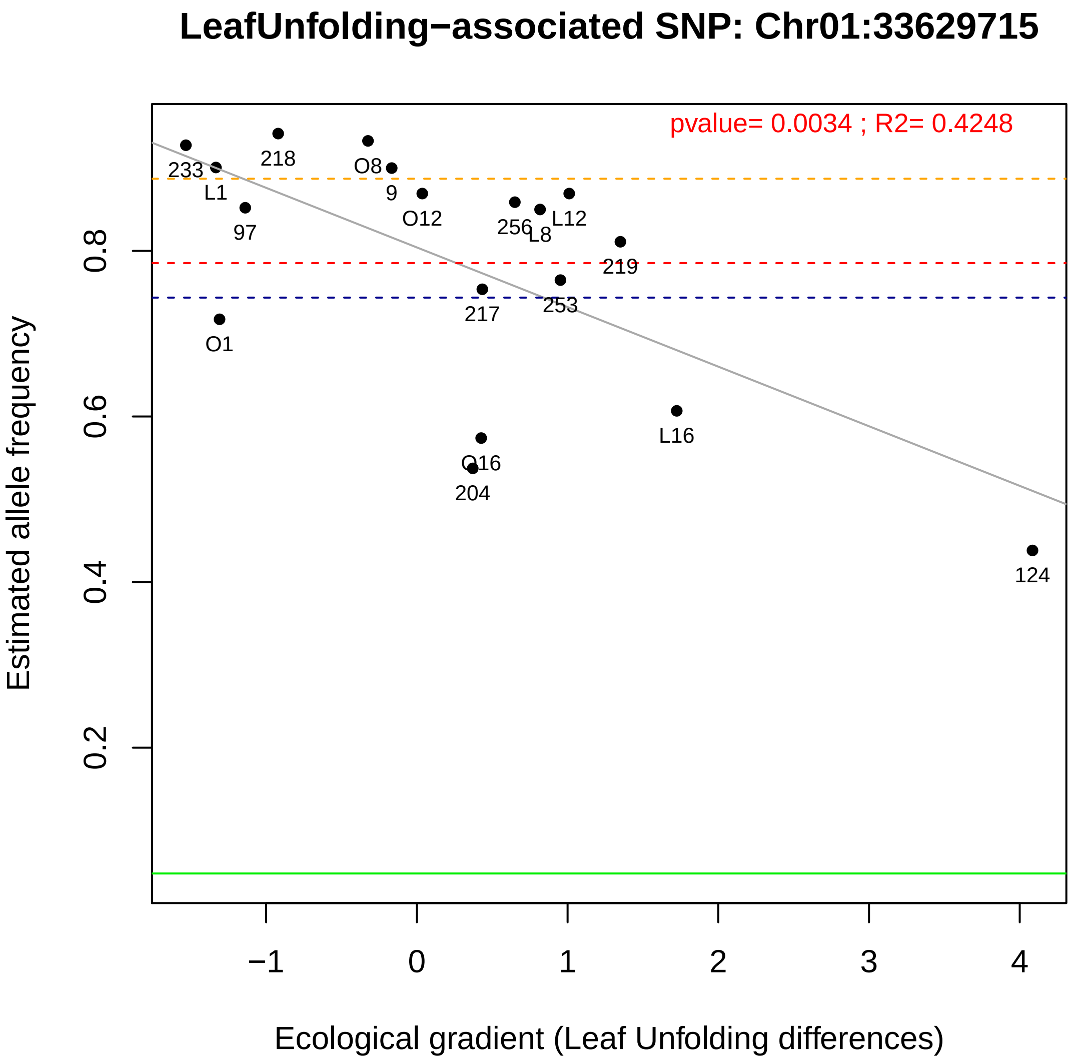
LeafUnfolding−associated SNP: Chr01:33629715
pvalue= 0.0034 ; R2= 0.4248
Estimated allele frequency
0.2
0.4
0.6
0.8
−1
0
1
2
3
4
Ecological gradient (Leaf Unfolding differences)
233
L1
218
97
O8
9
O12
256
L8
L12
219
217
253
O1
O16
204
L16
124

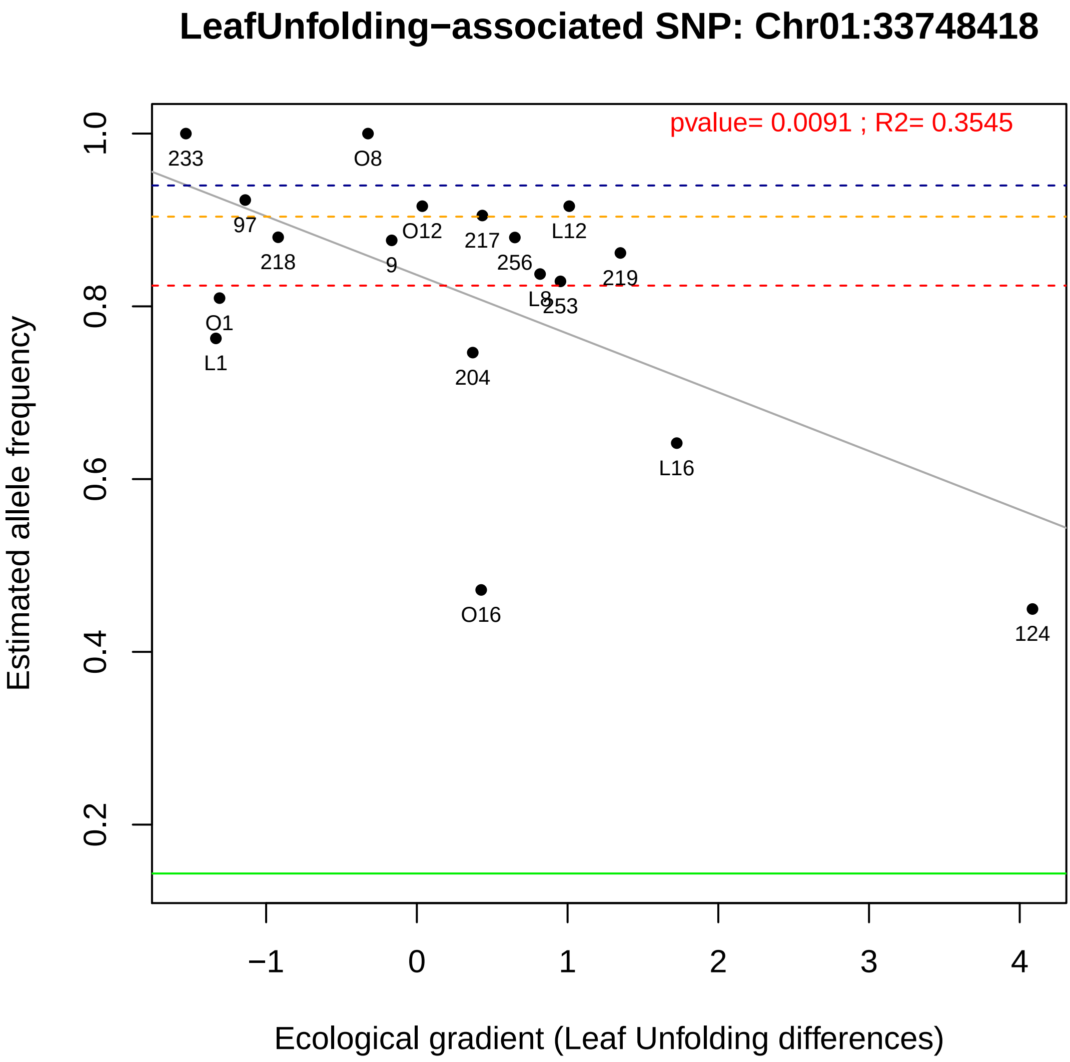

LeafUnfolding−associated SNP: Chr01:33748418
pvalue= 0.0091 ; R2= 0.3545
233
O8
97
218
9
O12
217
256
L12
L8
253
219
O1
L1
204
L16
O16
124
1.0
0.8
0.6
0.4
0.2
−1
0
1
2
3
4
Estimated allele frequency
Ecological gradient (Leaf Unfolding differences)

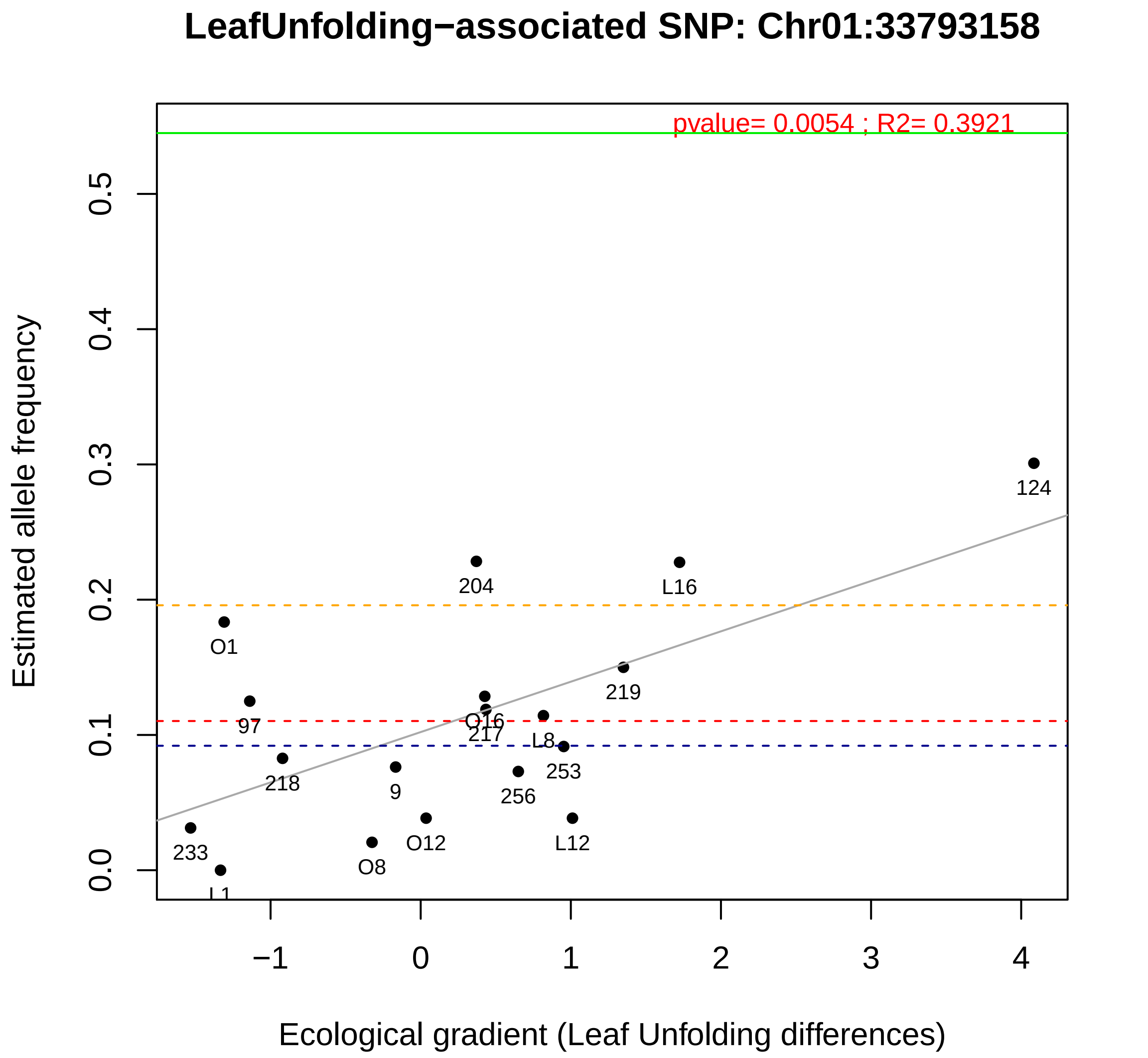
LeafUnfolding−associated SNP: Chr01:33793158
pvalue= 0.0054 ; R2= 0.3921
Estimated allele frequency
Ecological gradient (Leaf Unfolding differences)
0.0
0.1
0.2
0.3
0.4
0.5
−1
0
1
2
3
4
124
204
L16
O1
219
97
O16
217
L8
253
218
9
256
233
O12
L12
O8
L1

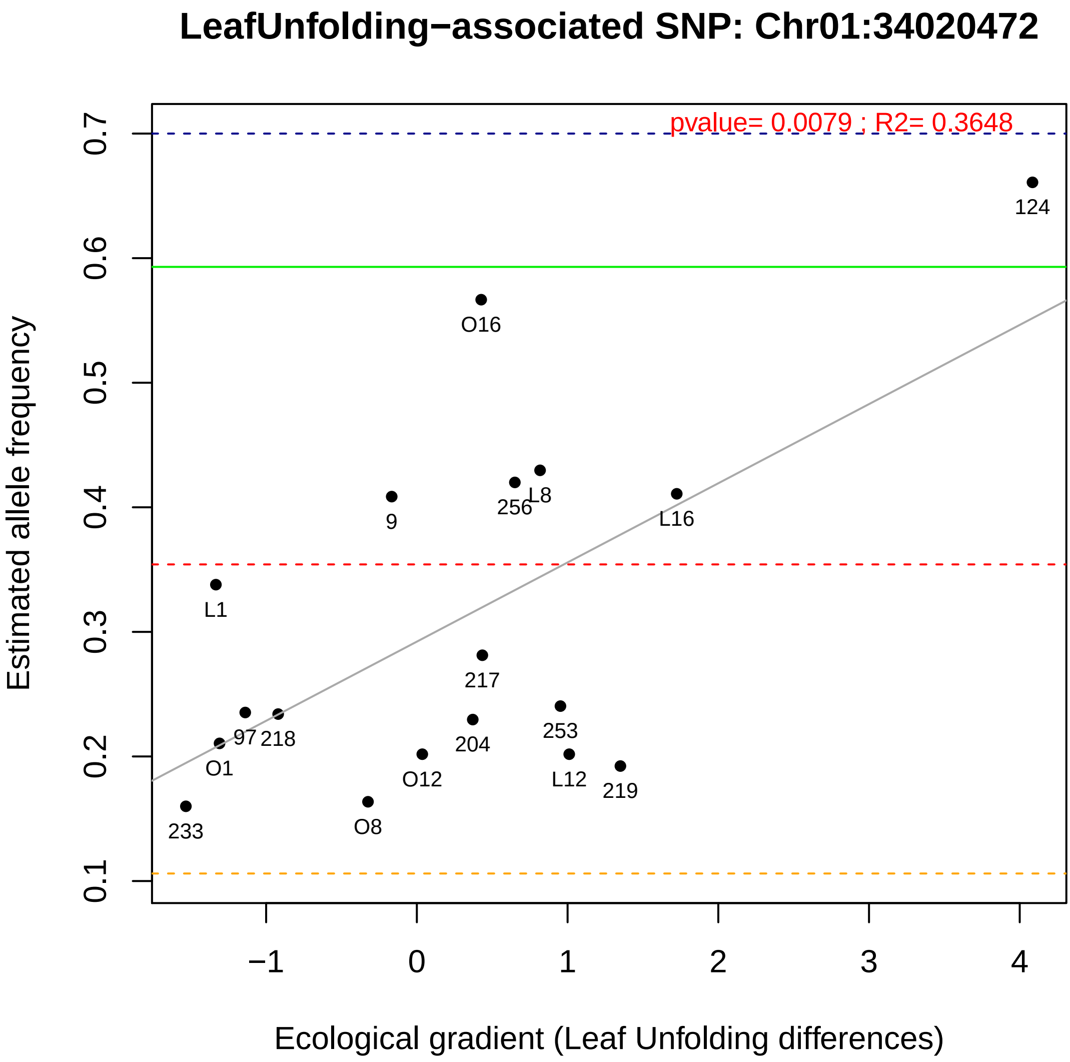
LeafUnfolding−associated SNP: Chr01:34020472
pvalue= 0.0079 ; R2= 0.3648
Estimated allele frequency
Ecological gradient (Leaf Unfolding differences)
0.1
0.2
0.3
0.4
0.5
0.6
0.7
−1
0
1
2
3
4
124
O16
9
256
L8
L16
L1
217
97
218
O1
204
253
O12
L12
219
233
O8

#### LeafUnfolding−associated SNP: Chr01:34142103

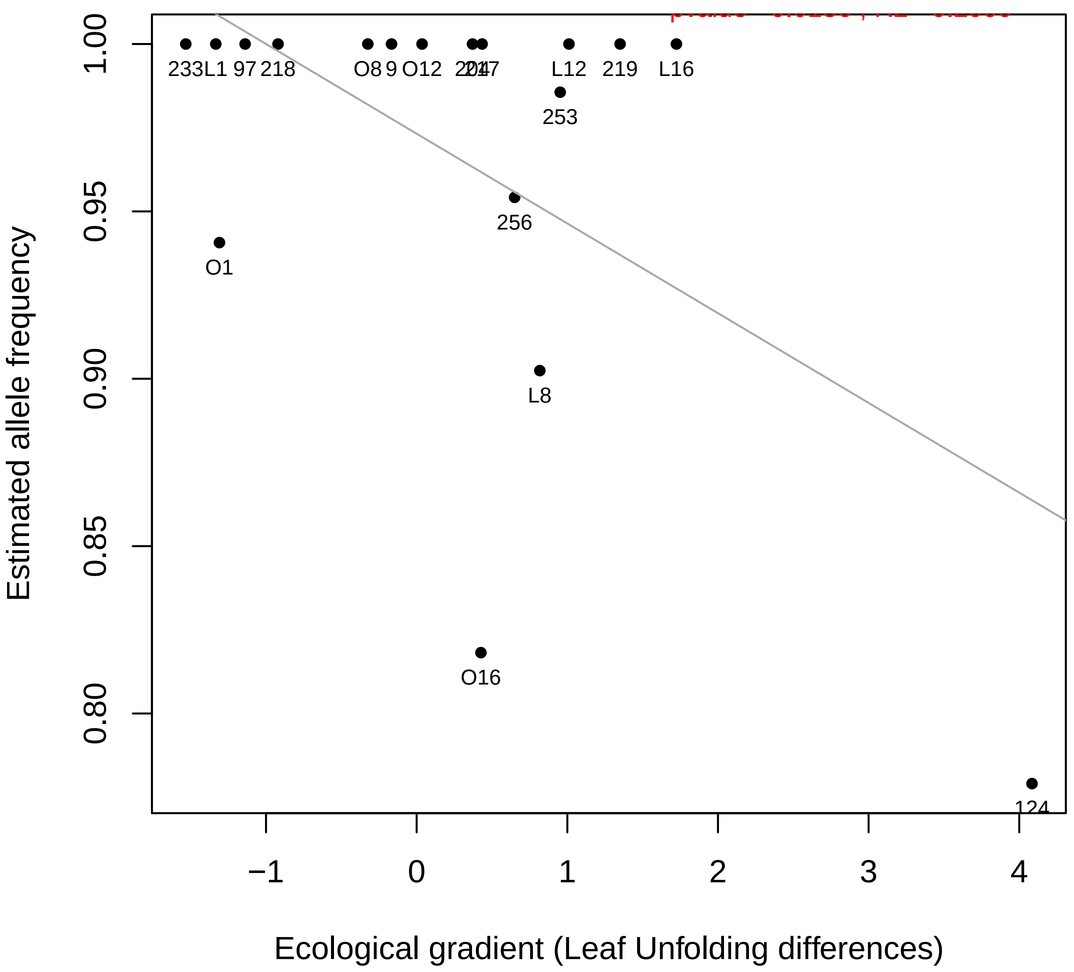

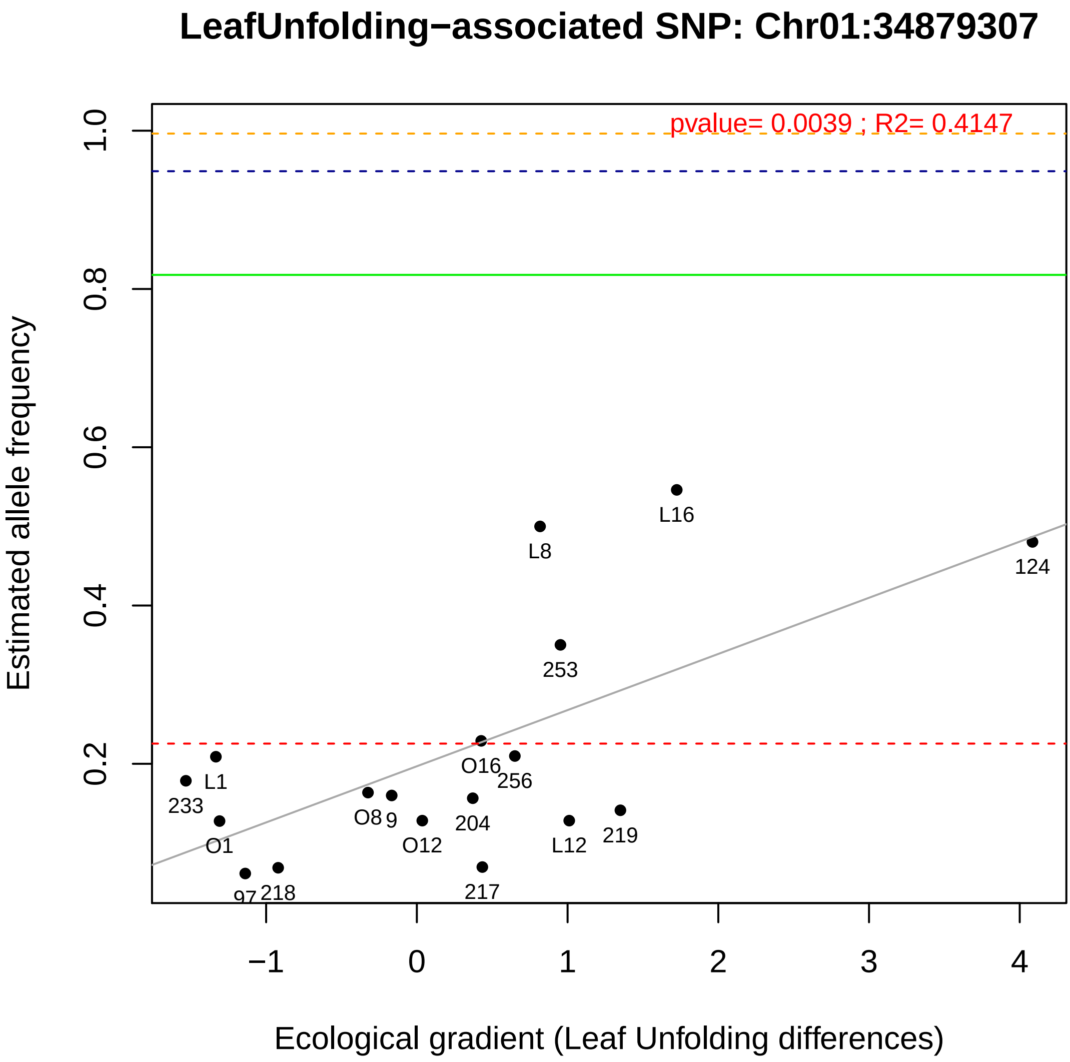
LeafUnfolding−associated SNP: Chr01:34879307
pvalue= 0.0039 ; R2= 0.4147
Estimated allele frequency
1.0
0.8
0.6
0.4
0.2
L16
L8
124
253
O16
256
L1
233
O8
9
204
O1
O12
L12
219
97
218
217
−1
0
1
2
3
4
Ecological gradient (Leaf Unfolding differences)

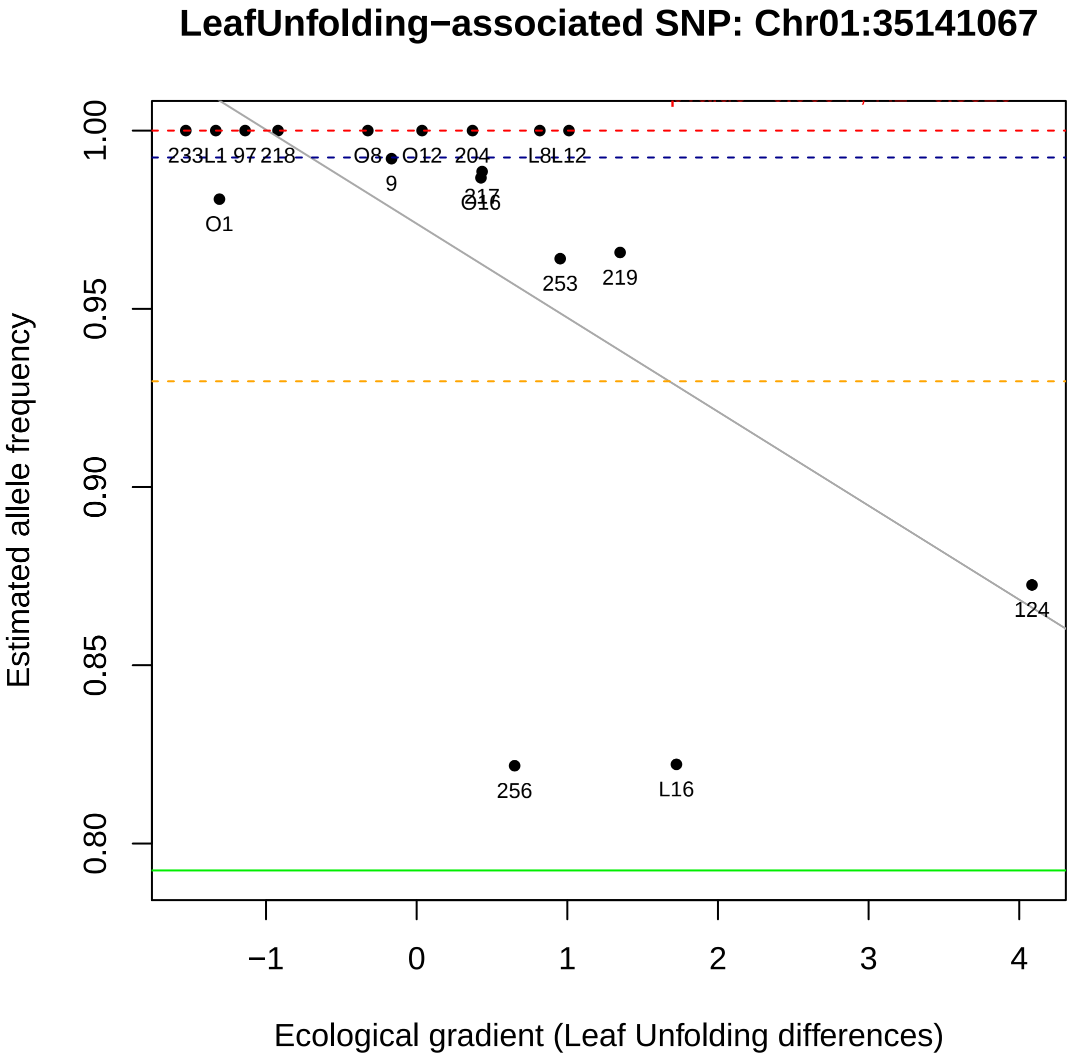
LeafUnfolding−associated SNP: Chr01:35141067
Estimated allele frequency
Ecological gradient (Leaf Unfolding differences)
1.00
0.95
0.90
0.85
0.80
−1
0
1
2
3
4
233
L1
97
218
O8
O12
204
L8
L12
9
217
O16
O1
253
219
124
256
L16

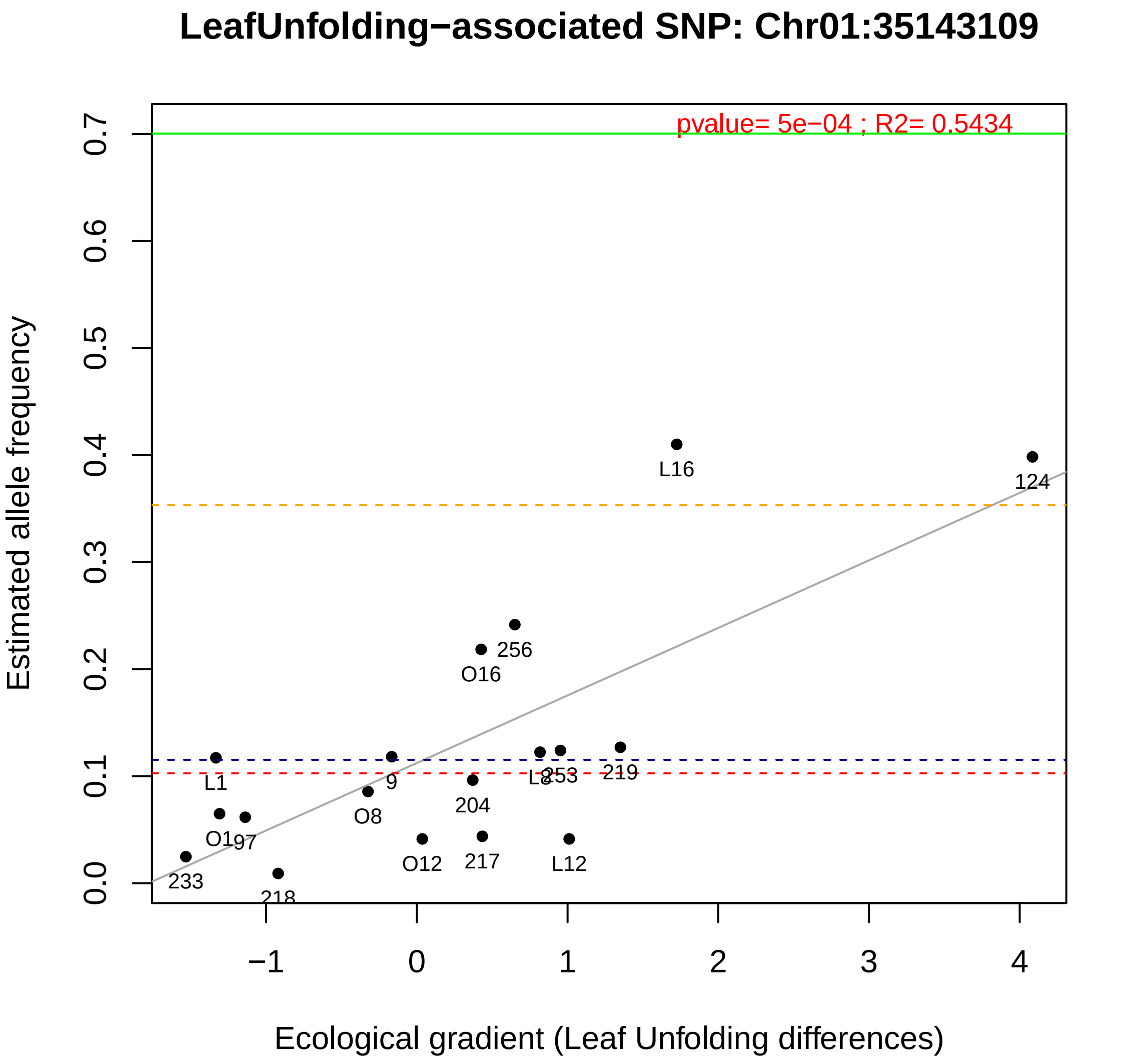
LeafUnfolding−associated SNP: Chr01:35143109
pvalue= 5e−04 ; R2= 0.5434
Estimated allele frequency
Ecological gradient (Leaf Unfolding differences)
0.0
0.1
0.2
0.3
0.4
0.5
0.6
0.7
−1
0
1
2
3
4
L16
124
256
O16
L1
9
204
219
253
O8
O12
217
L12
233
218

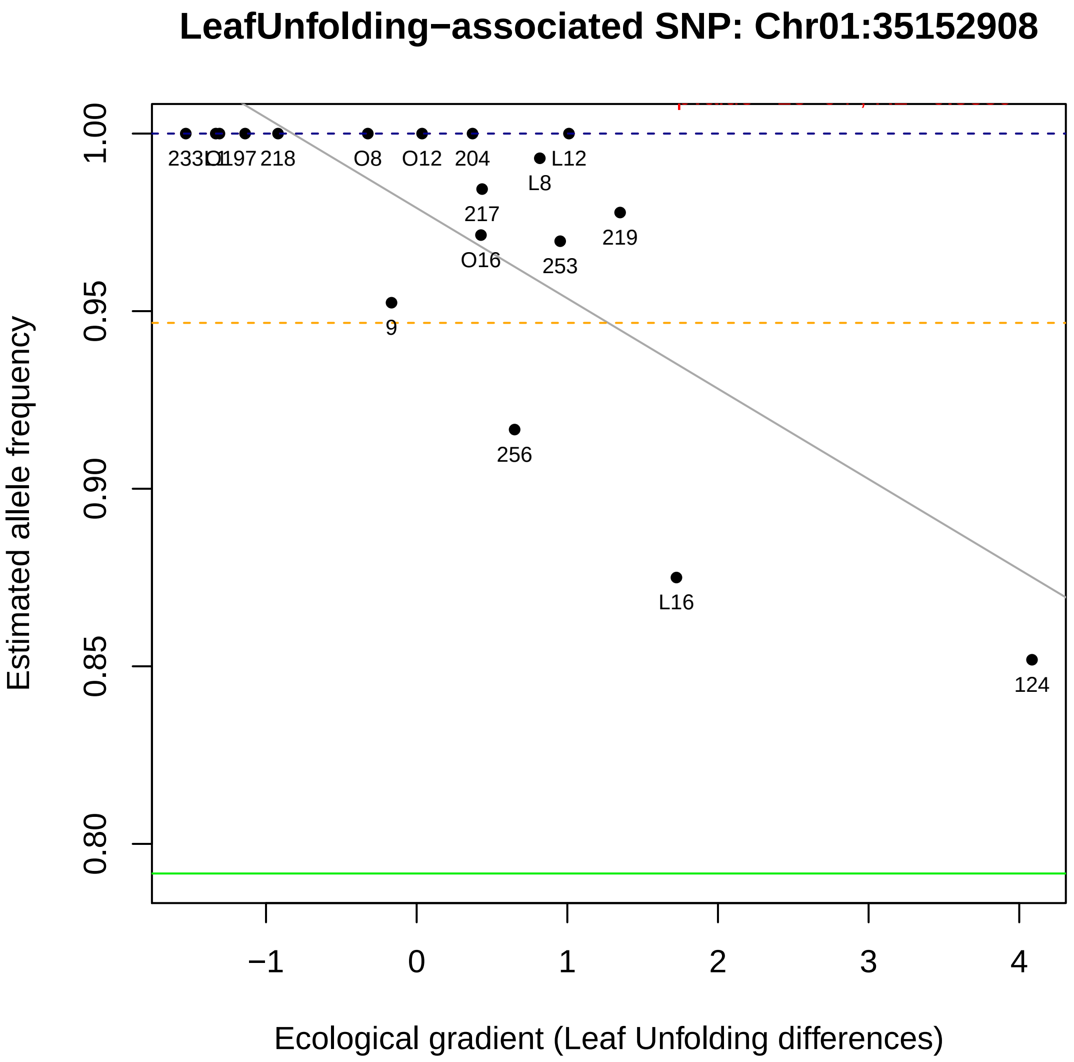

LeafUnfolding−associated SNP: Chr01:35152908
Estimated allele frequency
Ecological gradient (Leaf Unfolding differences)
1.00
0.95
0.90
0.85
0.80
−1
0
1
2
3
4
233
O1
197
218
O8
O12
204
L12
L8
217
219
O16
253
9
256
L16
124

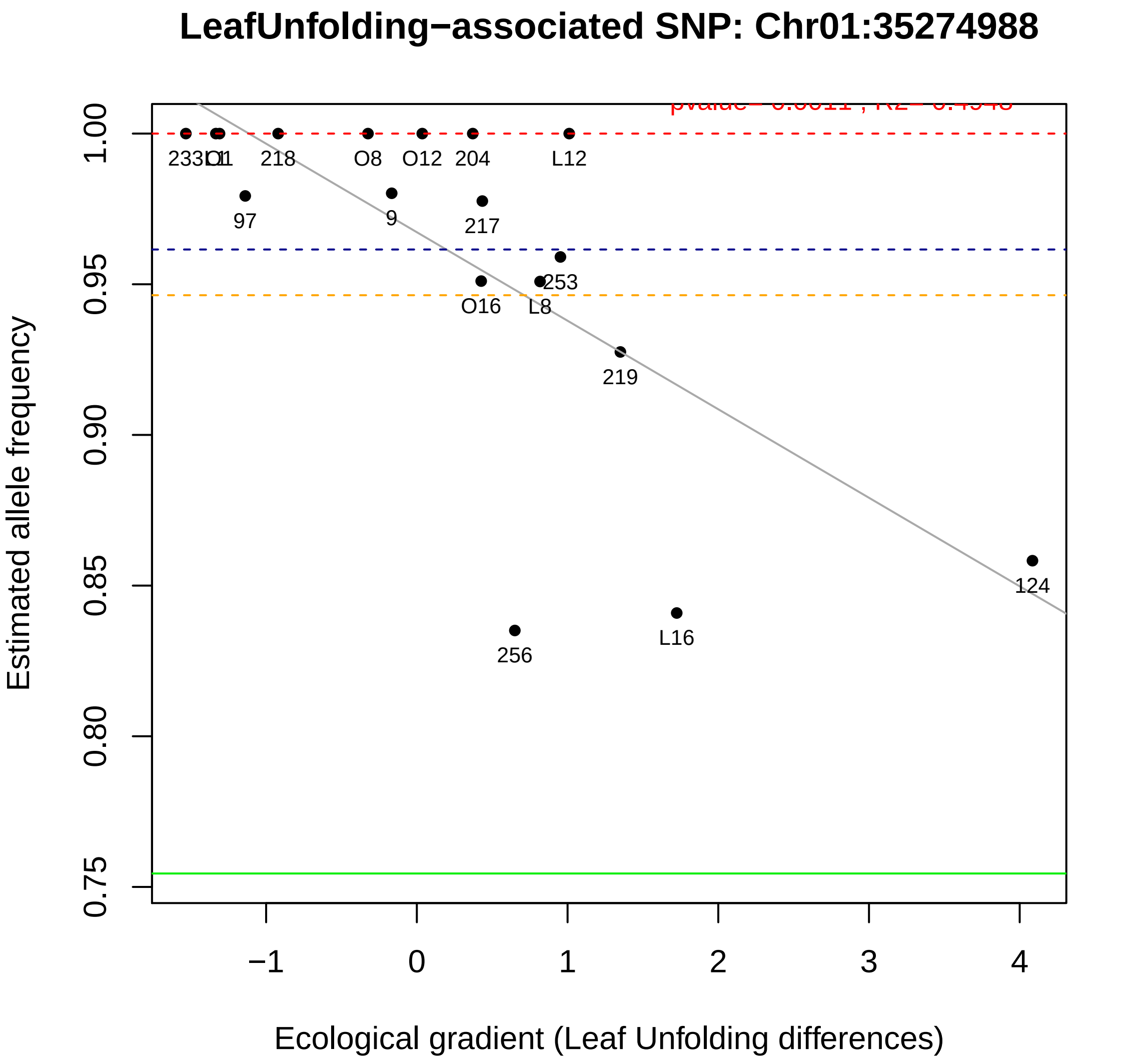
LeafUnfolding−associated SNP: Chr01:35274988
Estimated allele frequency
1.00
0.95
0.90
0.85
0.80
0.75
−1
0
1
2
3
4
Ecological gradient (Leaf Unfolding differences)
233
O1
218
O8
O12
204
L12
97
9
217
253
O16
L8
219
124
256
L16

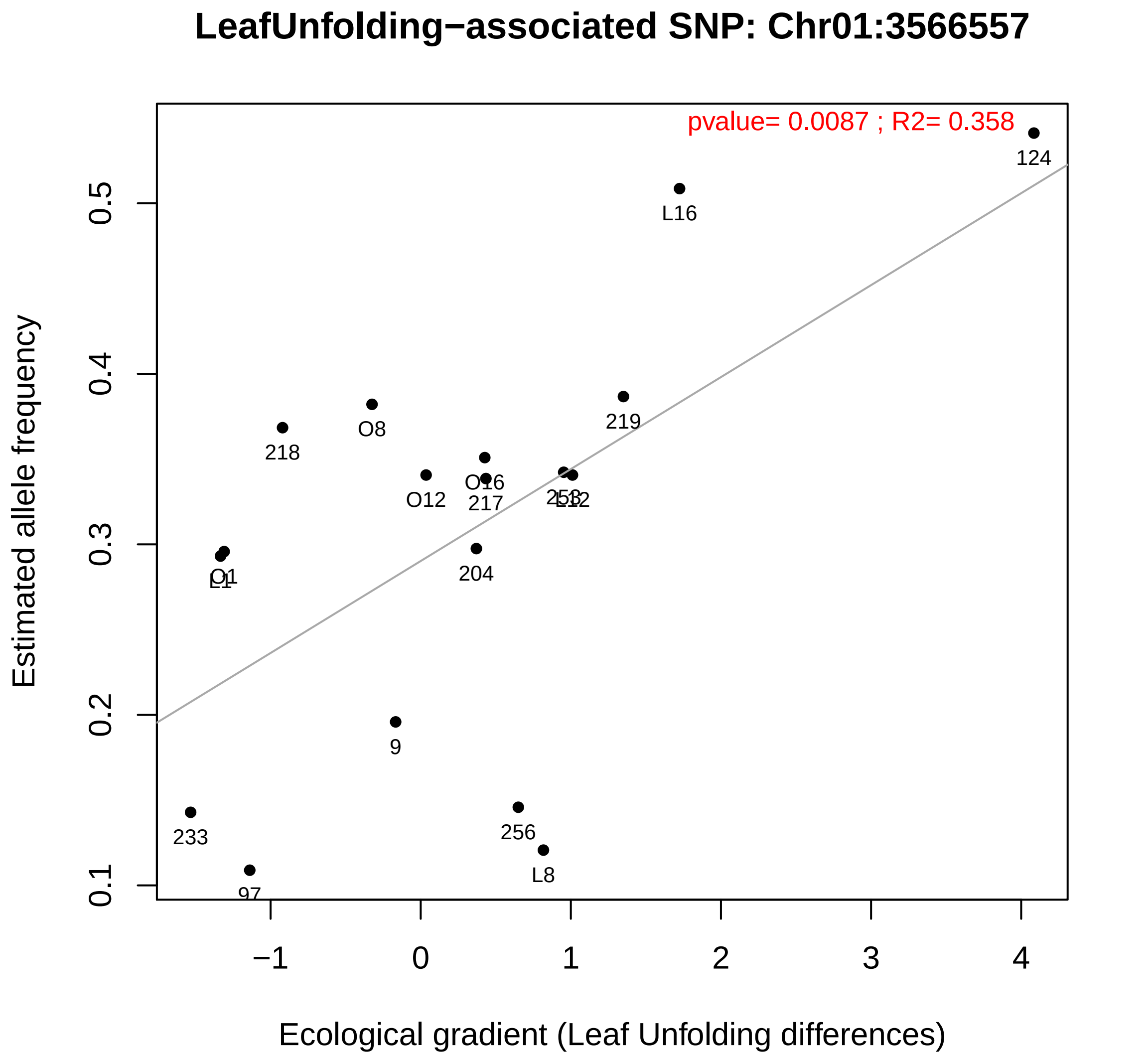
LeafUnfolding−associated SNP: Chr01:3566557
pvalue= 0.0087 ; R2= 0.358
Estimated allele frequency
Ecological gradient (Leaf Unfolding differences)
0.1
0.2
0.3
0.4
0.5
−1
0
1
2
3
4
124
L16
219
O8
218
O12
O16
217
253
L12
O1
L1
204
9
233
97
256
L8

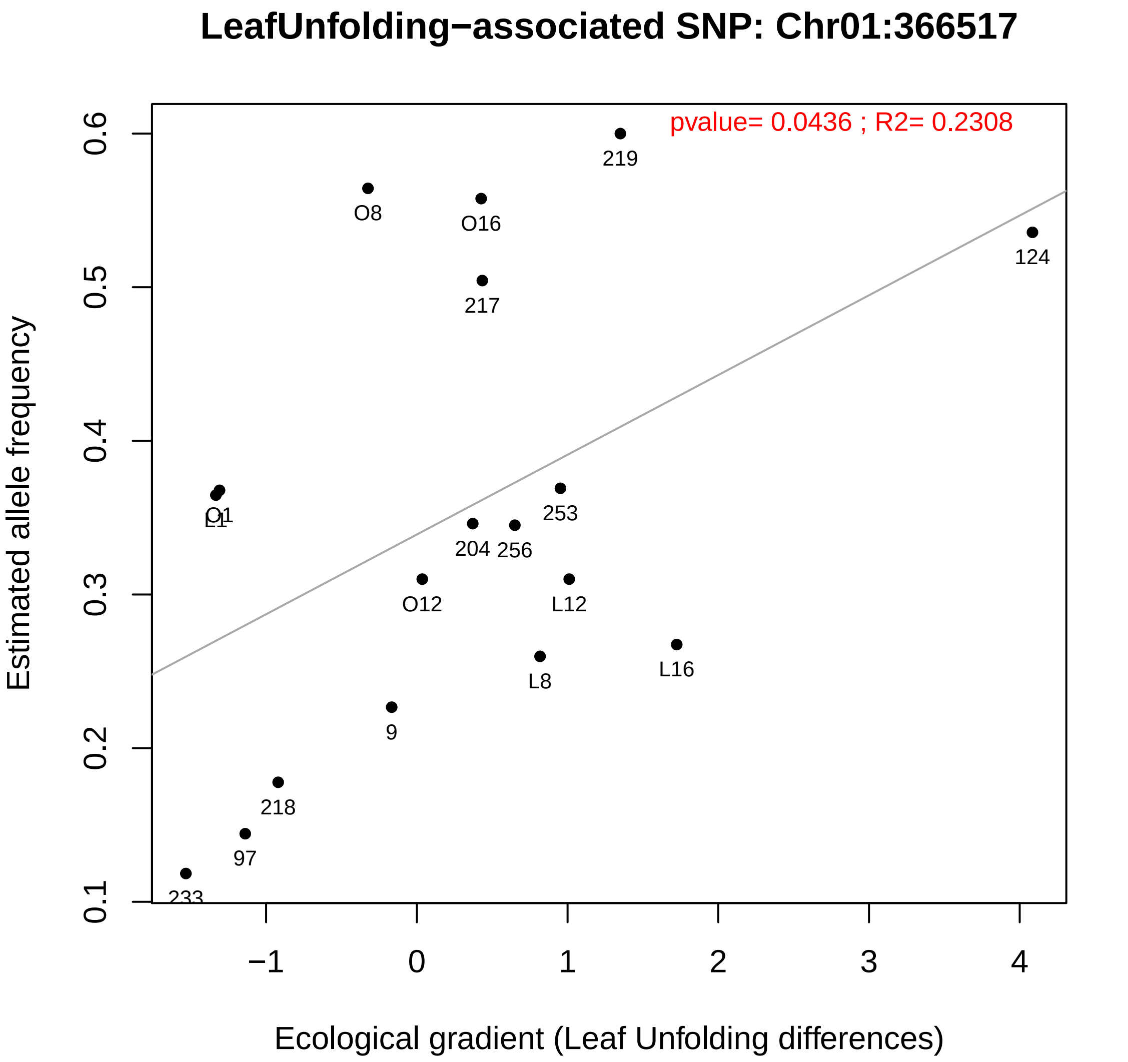

LeafUnfolding−associated SNP: Chr01:366517
pvalue= 0.0436 ; R2= 0.2308
Estimated allele frequency
0.1
0.2
0.3
0.4
0.5
0.6
−1
0
1
2
3
4
Ecological gradient (Leaf Unfolding differences)
219
O8
O16
217
124
O1
L1
253
204
256
O12
L12
L16
L8
9
218
97
233

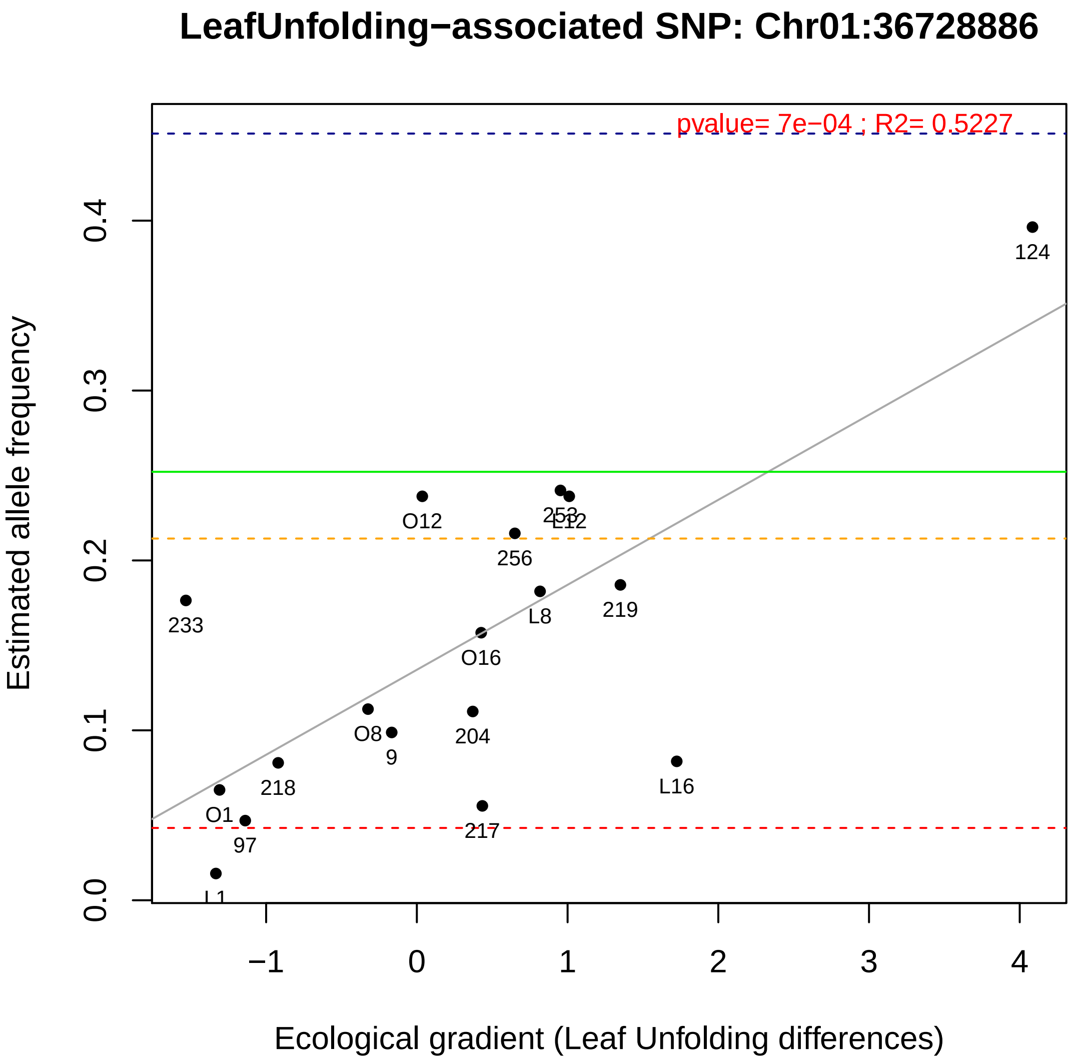

LeafUnfolding−associated SNP: Chr01:36728886
pvalue= 7e−04 ; R2= 0.5227
Estimated allele frequency
0.0
0.1
0.2
0.3
0.4
−1
0
1
2
3
4
Ecological gradient (Leaf Unfolding differences)
124
O12
253
L12
256
233
L8
219
O16
O8
9
204
218
L16
O1
97
217

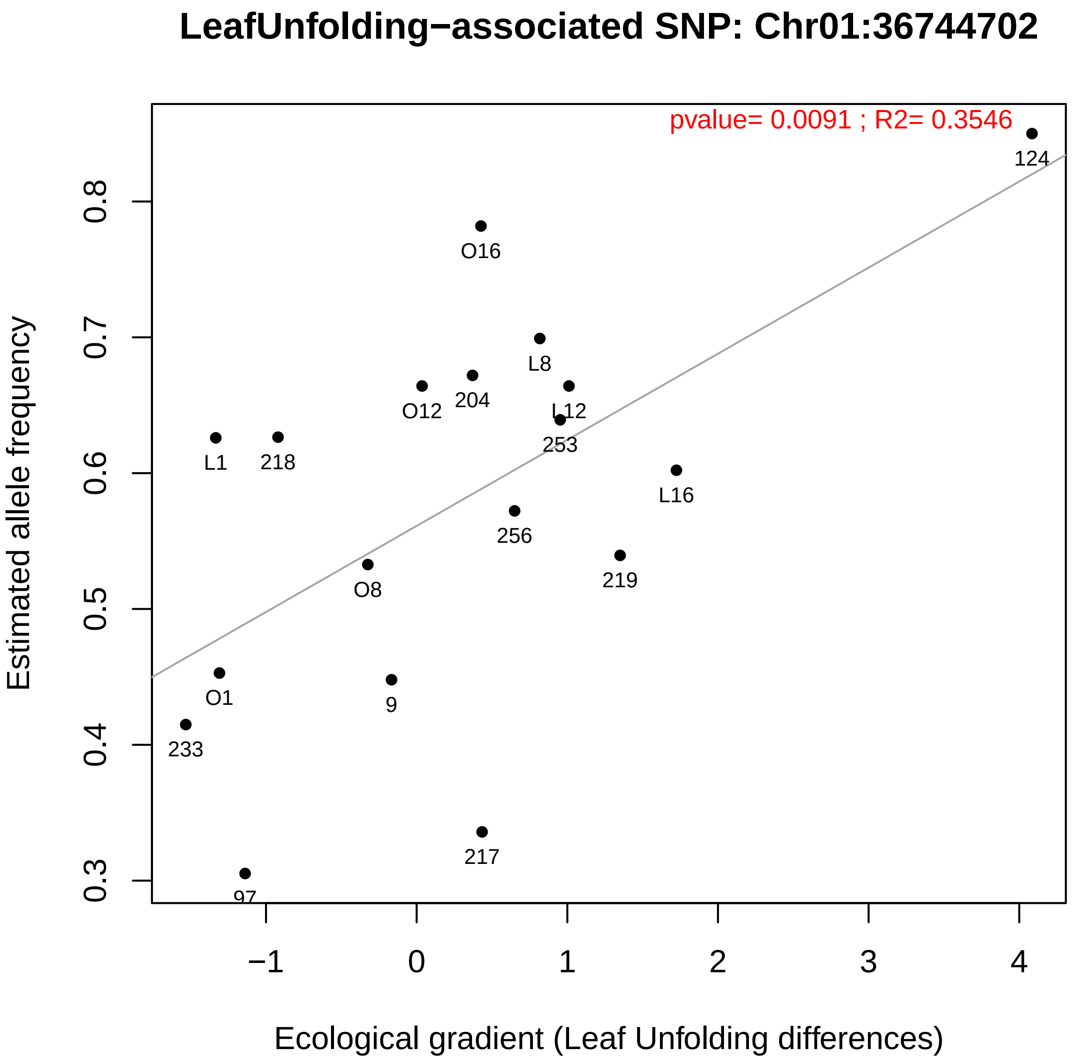
LeafUnfolding−associated SNP: Chr01:36744702
pvalue= 0.0091 ; R2= 0.3546
Estimated allele frequency
Ecological gradient (Leaf Unfolding differences)
0.3
0.4
0.5
0.6
0.7
0.8
−1
0
1
2
3
4
124
O16
L8
L12
204
O12
253
L1
218
L16
256
219
O8
O1
9
233
217
97

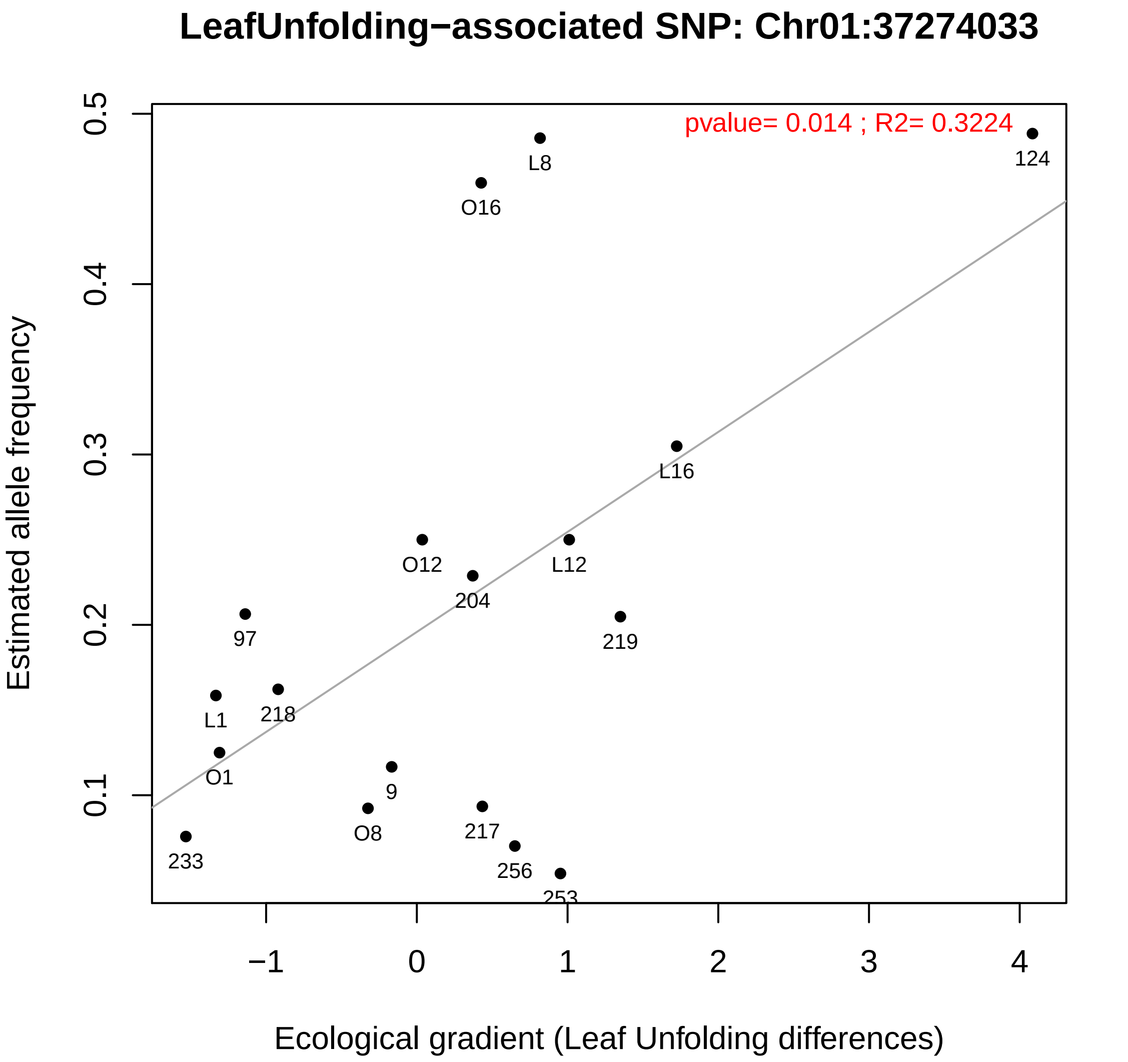

LeafUnfolding−associated SNP: Chr01:37274033
pvalue= 0.014 ; R2= 0.3224
Estimated allele frequency
0.1
0.2
0.3
0.4
0.5
Ecological gradient (Leaf Unfolding differences)
−1
0
1
2
3
4
L8
124
O16
L16
O12
L12
204
97
219
L1
218
O1
9
O8
217
233
256
253

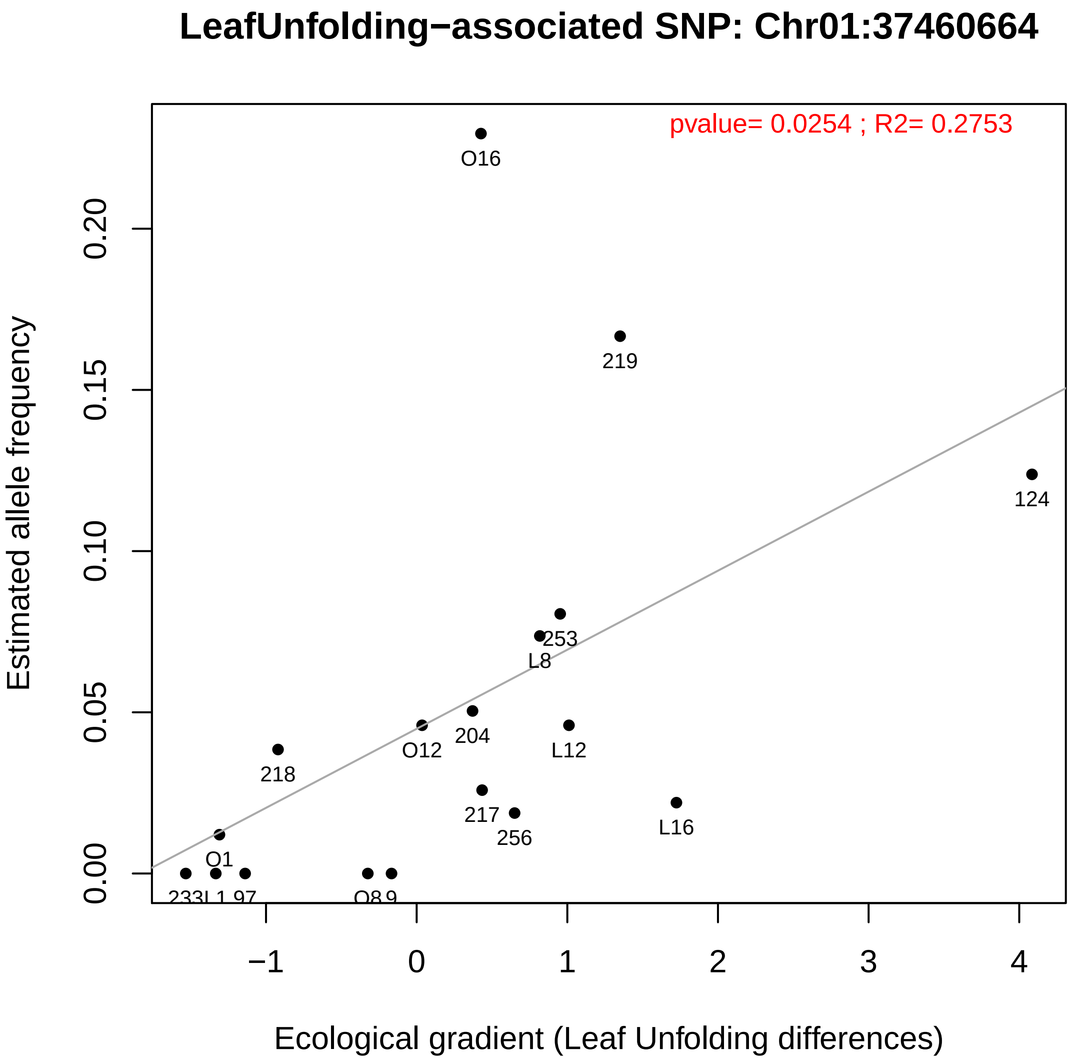

LeafUnfolding−associated SNP: Chr01:37460664
pvalue= 0.0254 ; R2= 0.2753
Estimated allele frequency
Ecological gradient (Leaf Unfolding differences)
0.00
0.05
0.10
0.15
0.20
−1
0
1
2
3
4
O16
219
124
253
L8
204
L12
O12
218
217
256
L16
O1
233
L1
97
O8
9

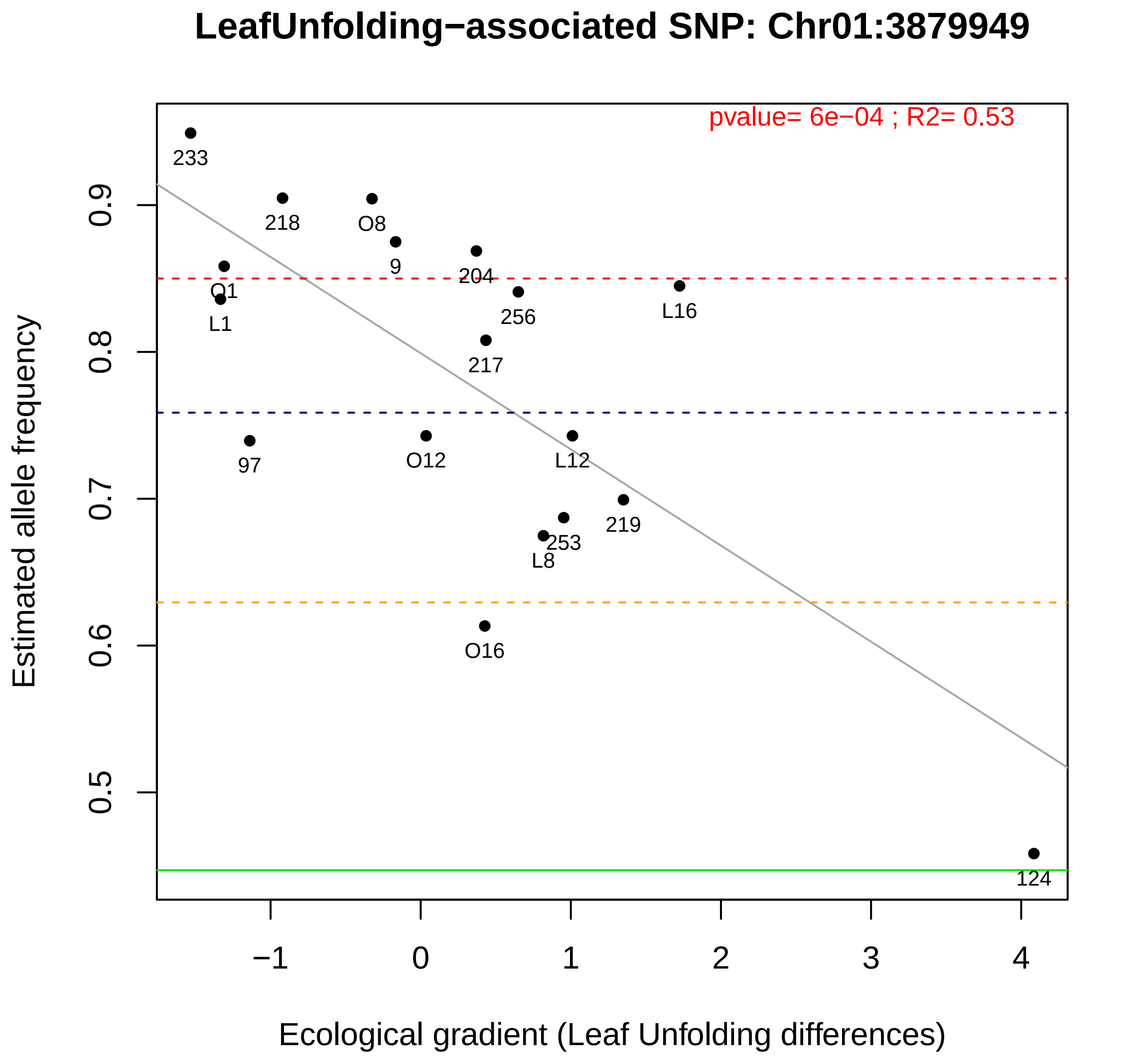
LeafUnfolding−associated SNP: Chr01:3879949
pvalue= 6e−04 ; R2= 0.53
Estimated allele frequency
0.5
0.6
0.7
0.8
0.9
Ecological gradient (Leaf Unfolding differences)
−1
0
1
2
3
4
233
218
O8
9
204
O1
L1
256
L16
217
97
O12
L12
219
253
L8
O16
124

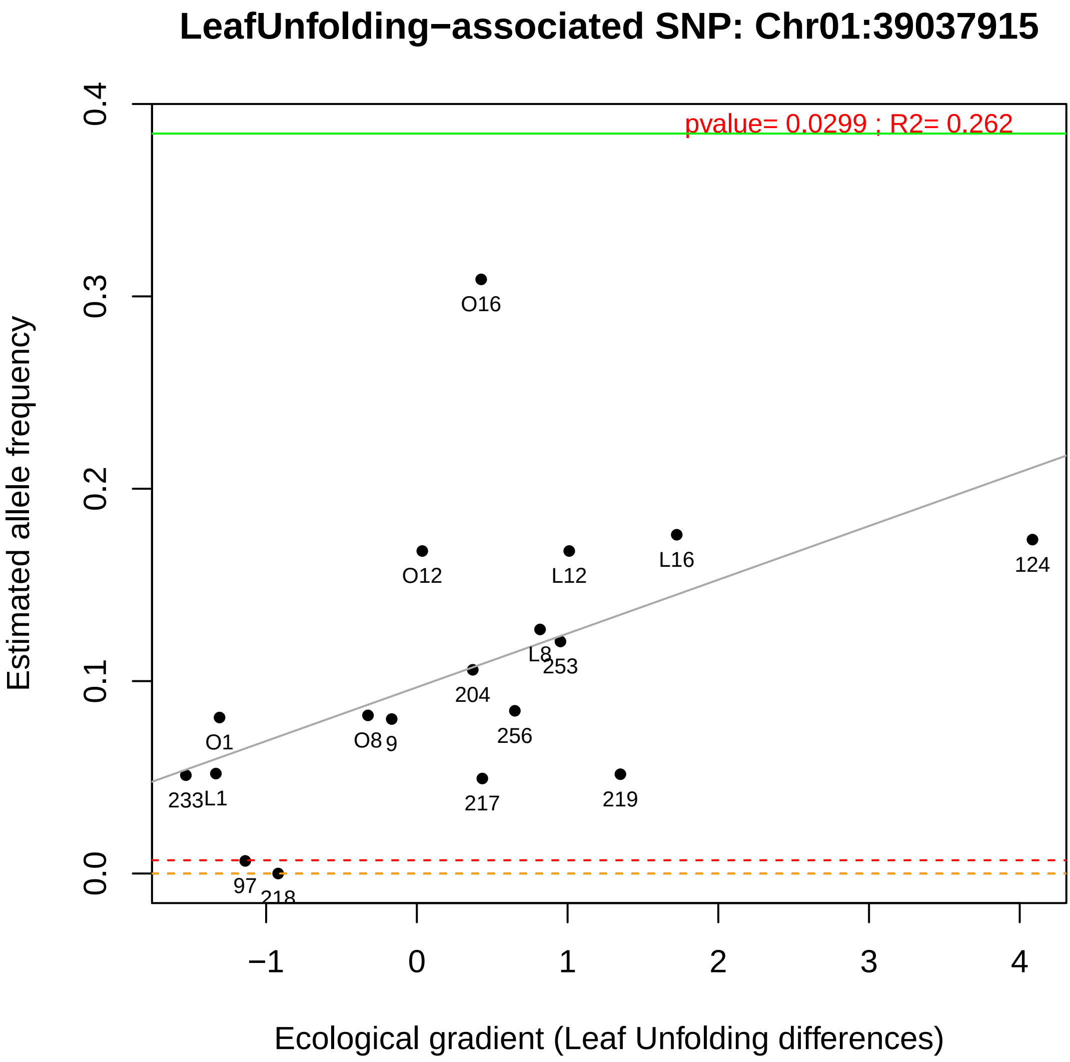

LeafUnfolding−associated SNP: Chr01:39037915
pvalue= 0.0299 ; R2= 0.262
Estimated allele frequency
0.0
0.1
0.2
0.3
0.4
−1
0
1
2
3
4
Ecological gradient (Leaf Unfolding differences)
O16
O12
L12
L16
124
L8
253
204
256
O1
O8
9
233
L1
217
219
97
218

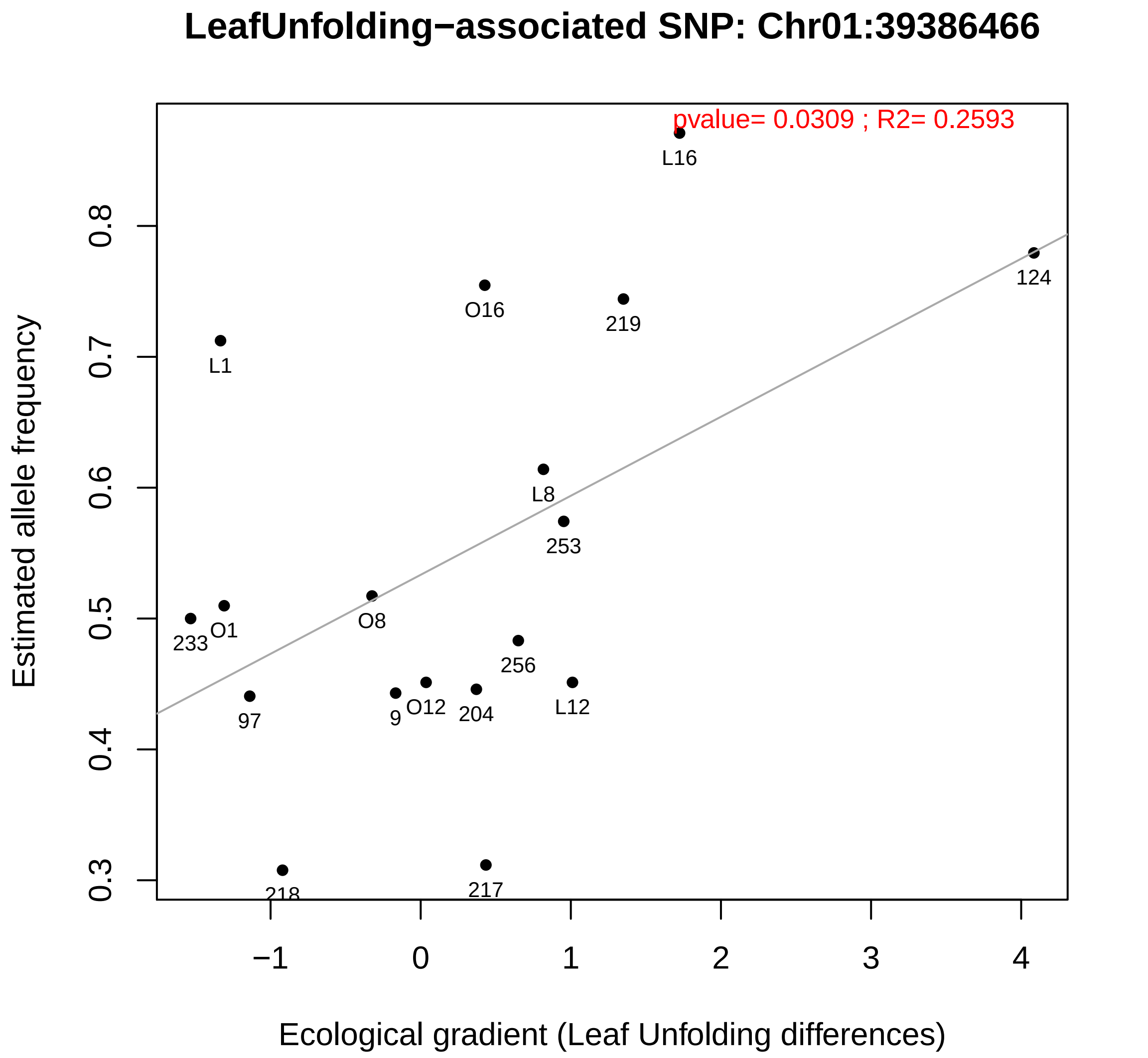

LeafUnfolding−associated SNP: Chr01:39386466
pvalue= 0.0309 ; R2= 0.2593
Estimated allele frequency
Ecological gradient (Leaf Unfolding differences)
0.3
0.4
0.5
0.6
0.7
0.8
−1
0
1
2
3
4
L16
124
O16
219
L1
L8
253
O8
233
O1
256
97
9
O12
204
L12
218
217

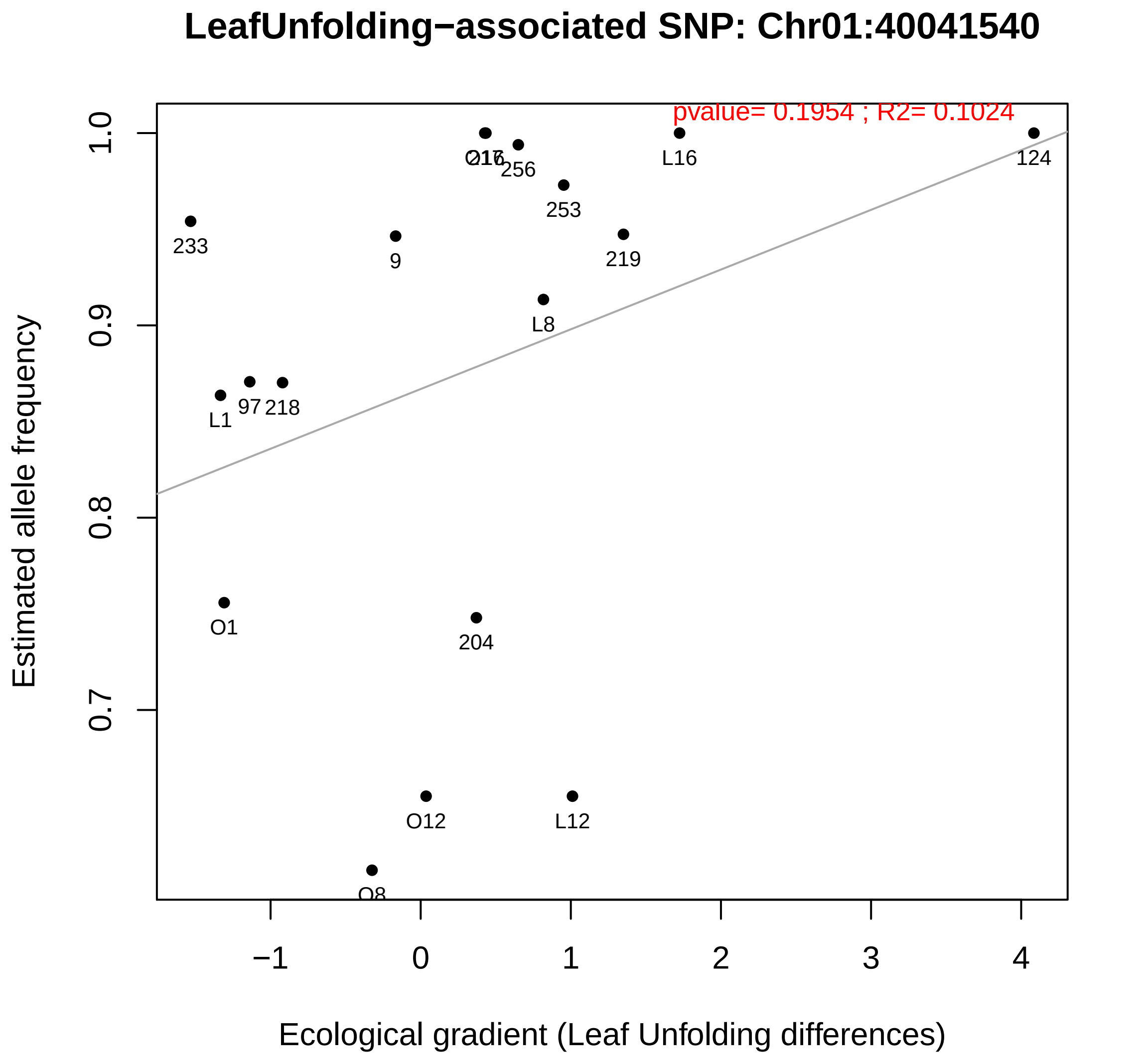
LeafUnfolding−associated SNP: Chr01:40041540
pvalue= 0.1954 ; R2= 0.1024
Estimated allele frequency
Ecological gradient (Leaf Unfolding differences)
1.0
0.9
0.8
0.7
−1
0
1
2
3
4
233
L1
97
218
O1
9
O8
O12
204
L8
256
253
219
L16
L12
124

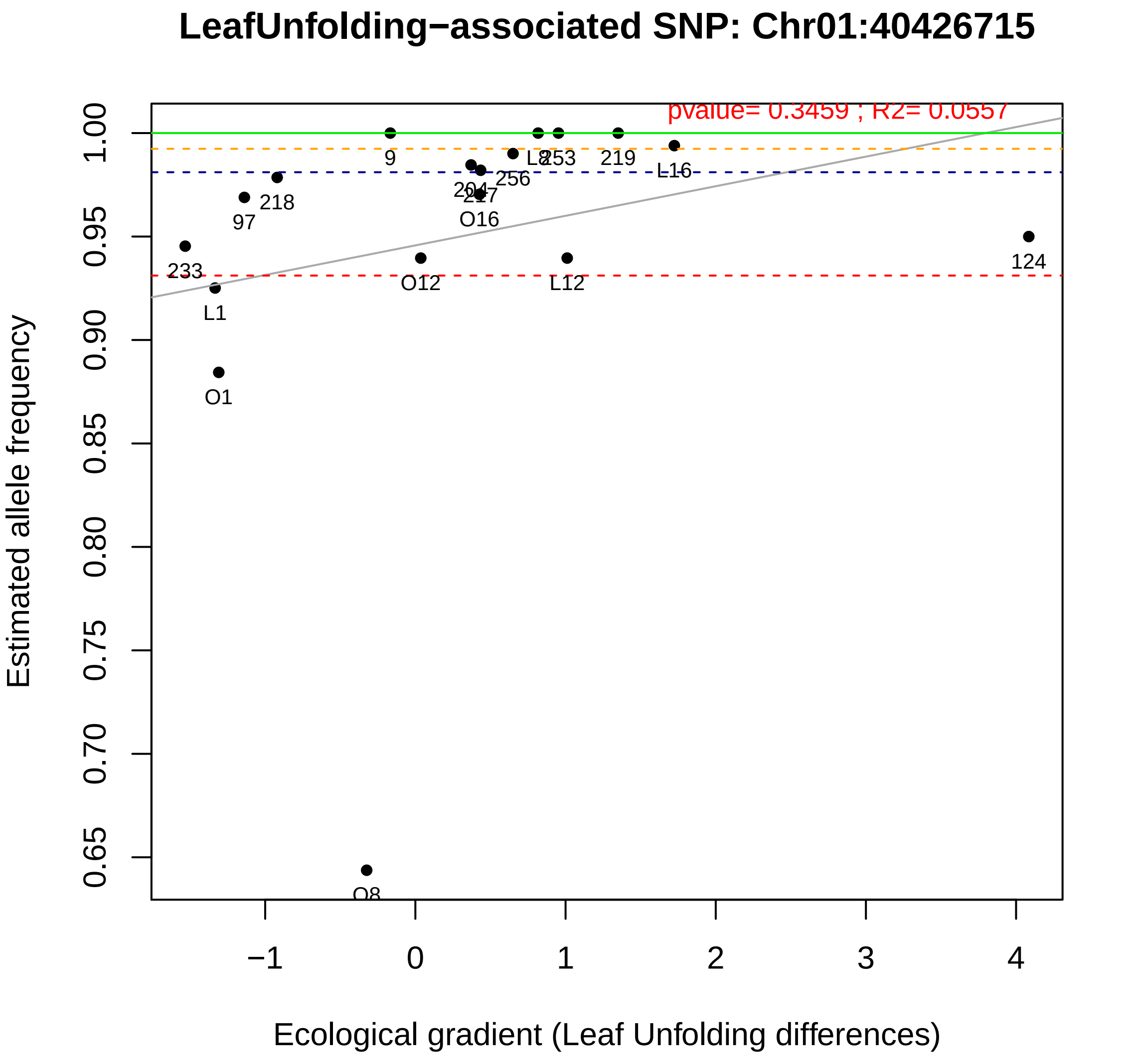

LeafUnfolding−associated SNP: Chr01:40426715
pvalue= 0.3459 ; R2= 0.0557
Estimated allele frequency
Ecological gradient (Leaf Unfolding differences)
1.00
0.95
0.90
0.85
0.80
0.75
0.70
0.65
−1
0
1
2
3
4
9
L8
253
219
L16
256
204
217
O16
218
97
233
L1
O1
O12
L12
124
O8

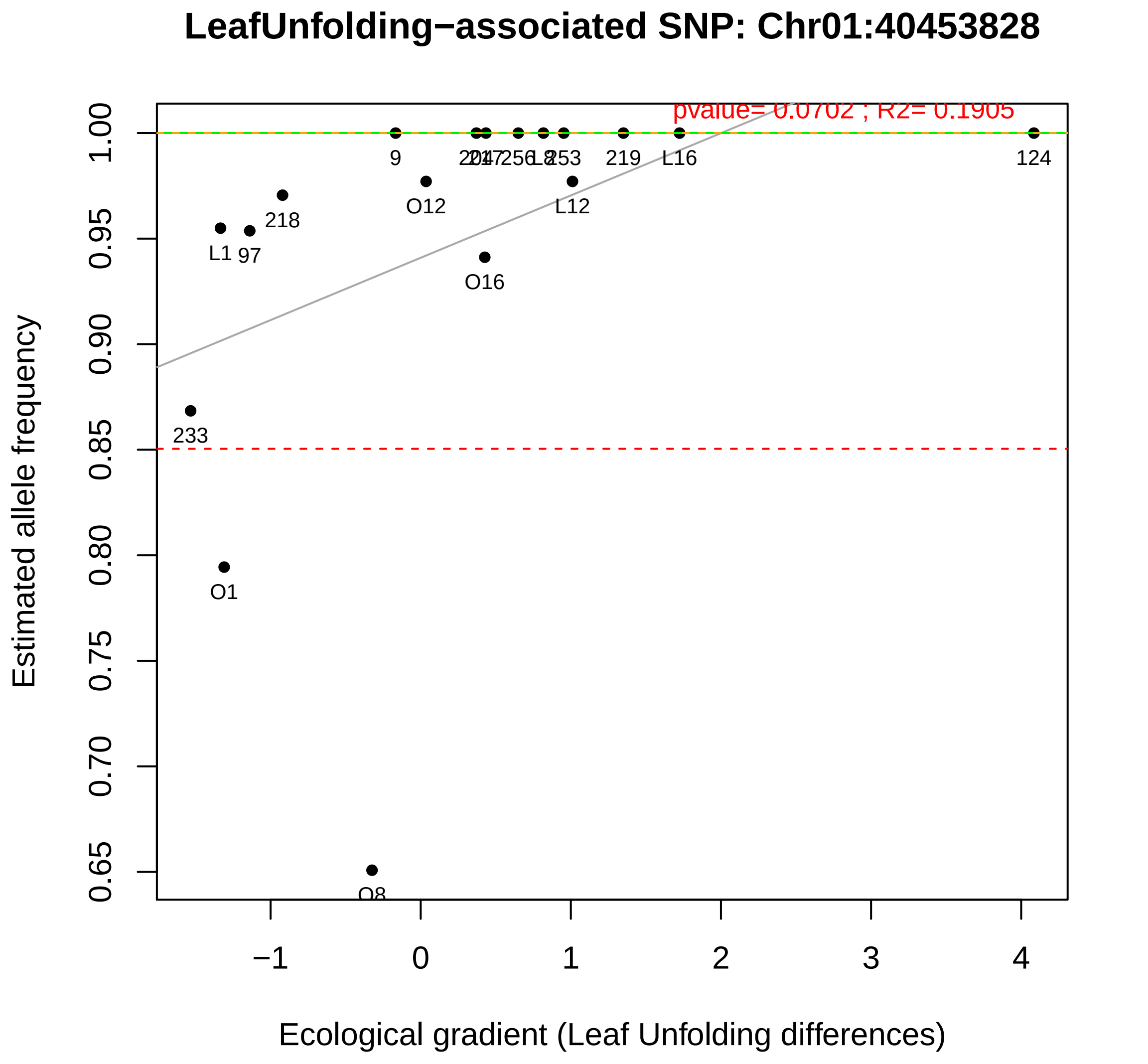

LeafUnfolding−associated SNP: Chr01:40453828
pvalue= 0.0702 ; R2= 0.1905
Estimated allele frequency
Ecological gradient (Leaf Unfolding differences)
1.00
0.95
0.90
0.85
0.80
0.75
0.70
0.65
−1
0
1
2
3
4
9
219
L16
124
O12
L12
218
L1
97
O16
233
O1
O8

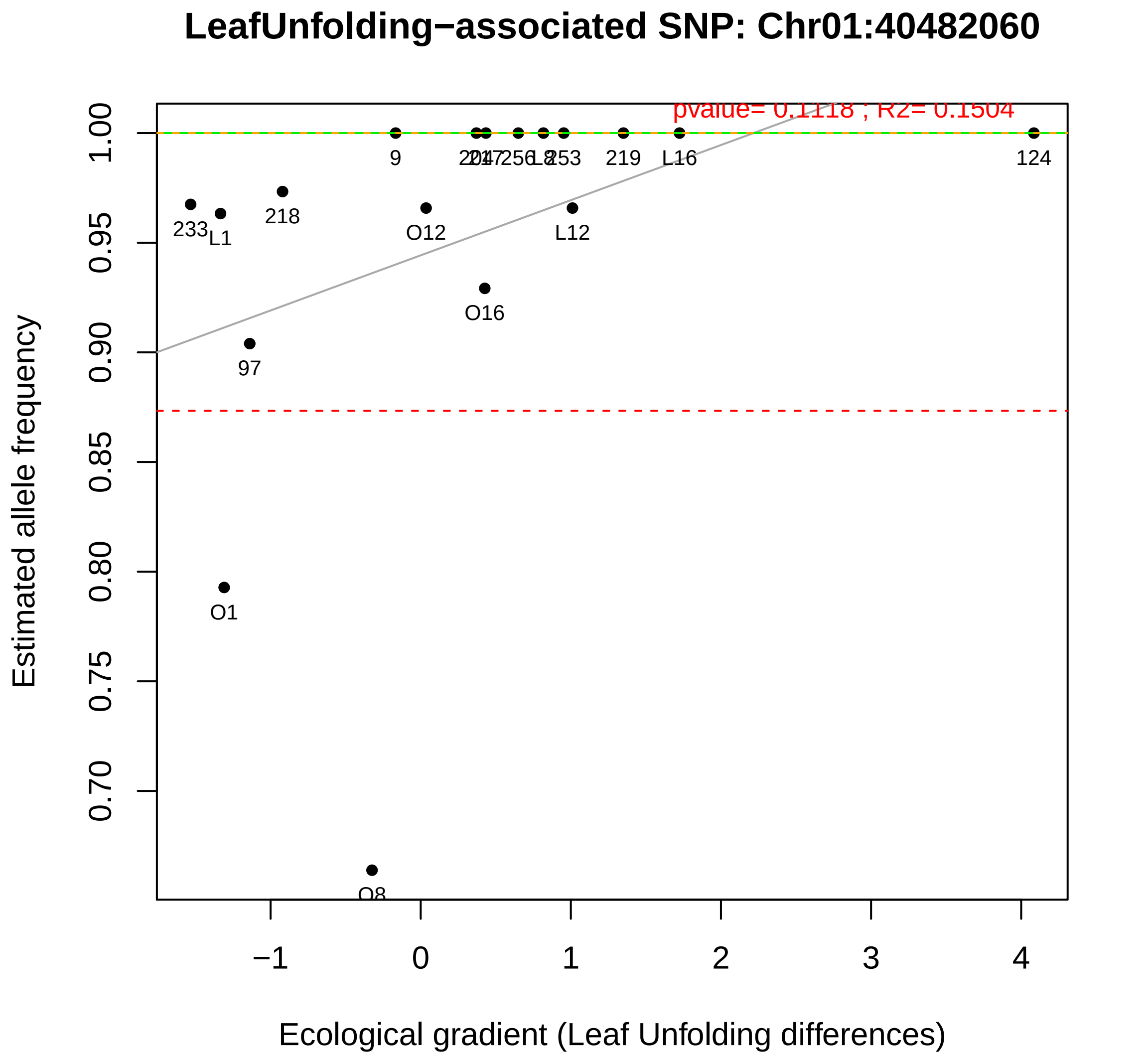

LeafUnfolding−associated SNP: Chr01:40482060
pvalue= 0.1118 ; R2= 0.1504
Estimated allele frequency
1.00
0.95
0.90
0.85
0.80
0.75
0.70
9
219
L16
124
233
L1
218
O12
L12
O16
97
O1
O8
−1
0
1
2
3
4
Ecological gradient (Leaf Unfolding differences)

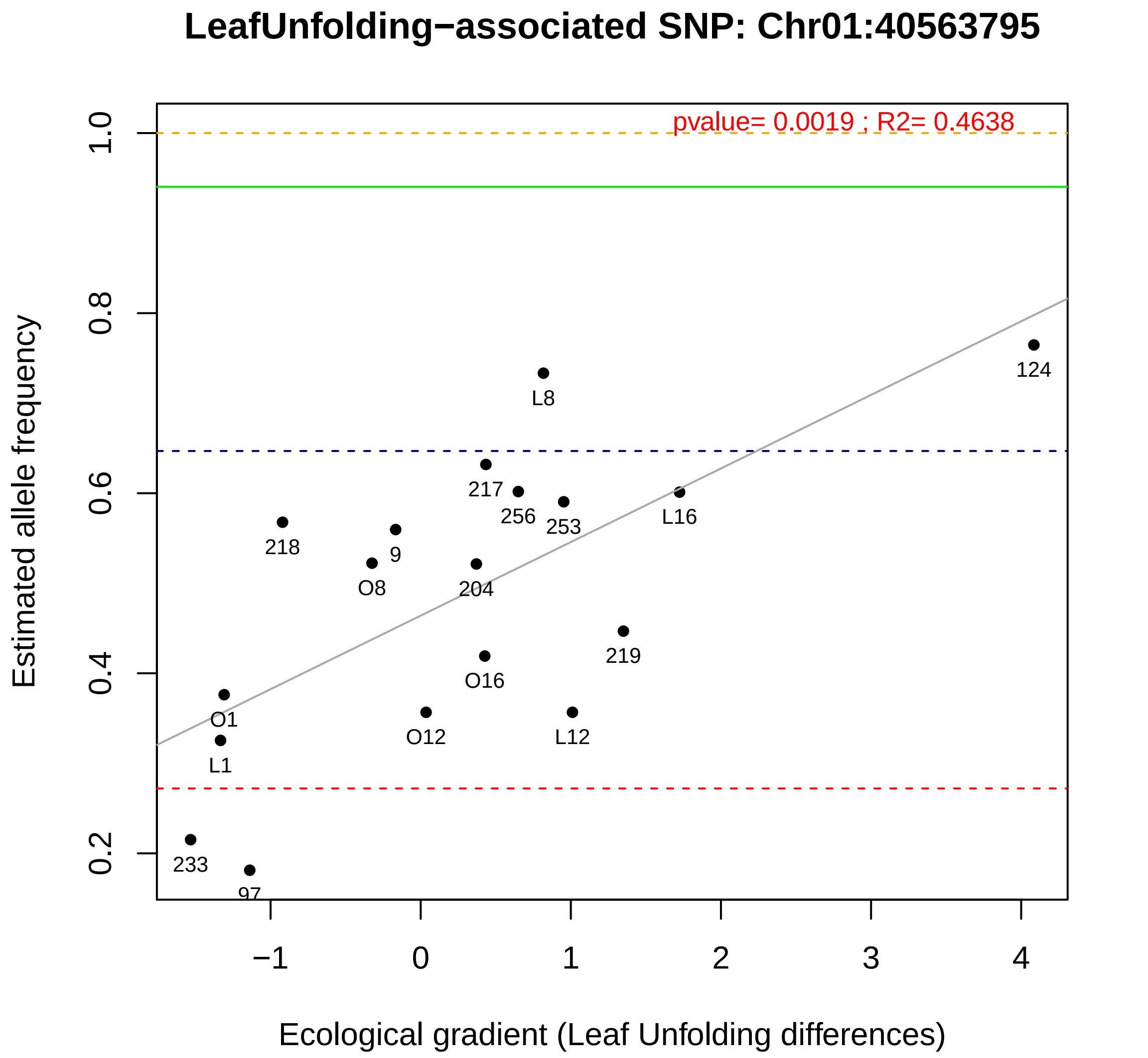

LeafUnfolding−associated SNP: Chr01:40563795
pvalue= 0.0019 ; R2= 0.4638
Estimated allele frequency
Ecological gradient (Leaf Unfolding differences)
0.2
0.4
0.6
0.8
1.0
−1
0
1
2
3
4
124
L8
217
256
253
L16
218
9
O8
204
219
O16
O1
L1
O12
L12
233
97

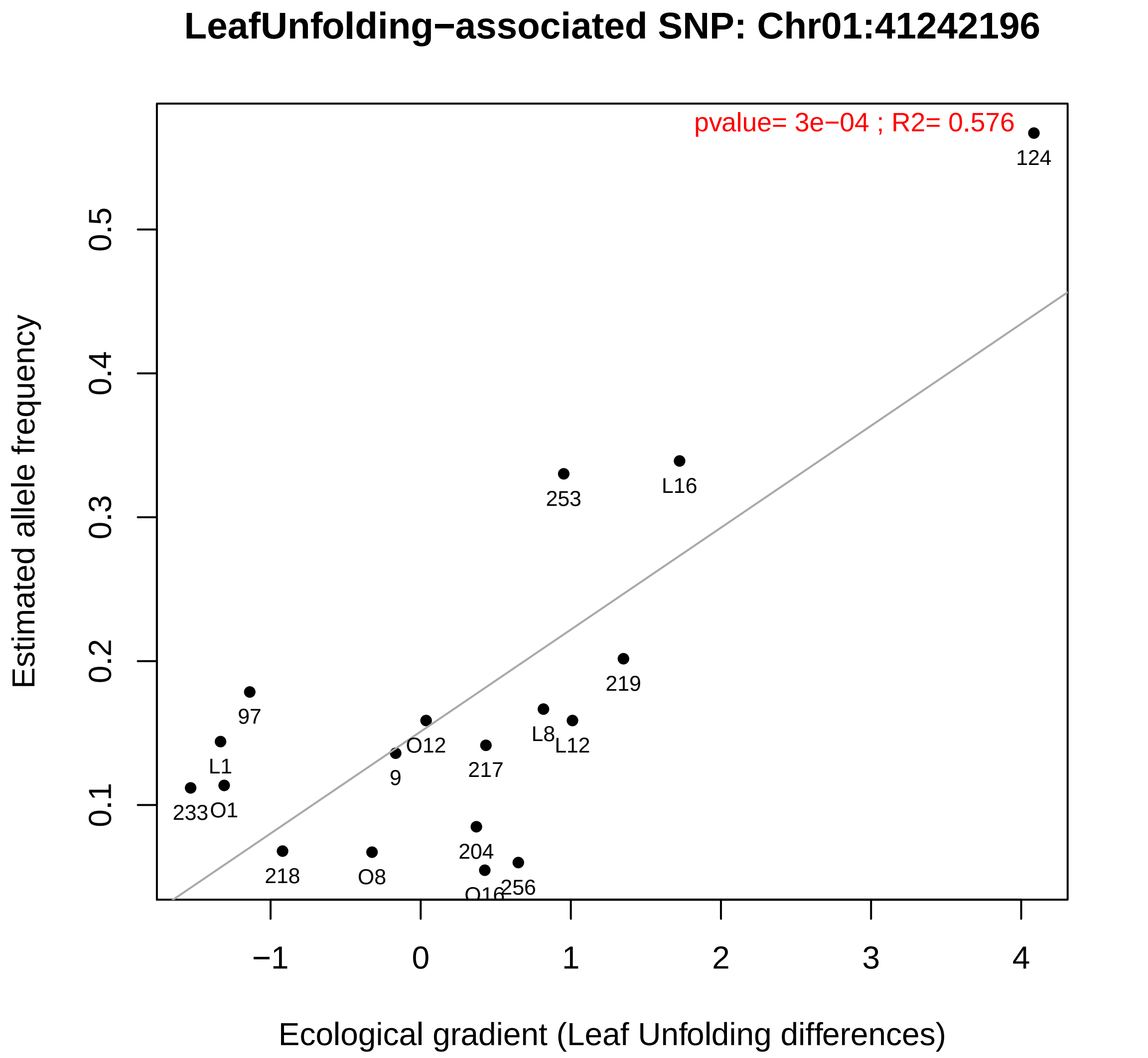
LeafUnfolding−associated SNP: Chr01:41242196
pvalue= 3e−04 ; R2= 0.576
Estimated allele frequency
0.1
0.2
0.3
0.4
0.5
−1
0
1
2
3
4
Ecological gradient (Leaf Unfolding differences)
124
253
L16
219
97
L1
233
O1
O12
9
217
L8
L12
204
218
O8
O16
256

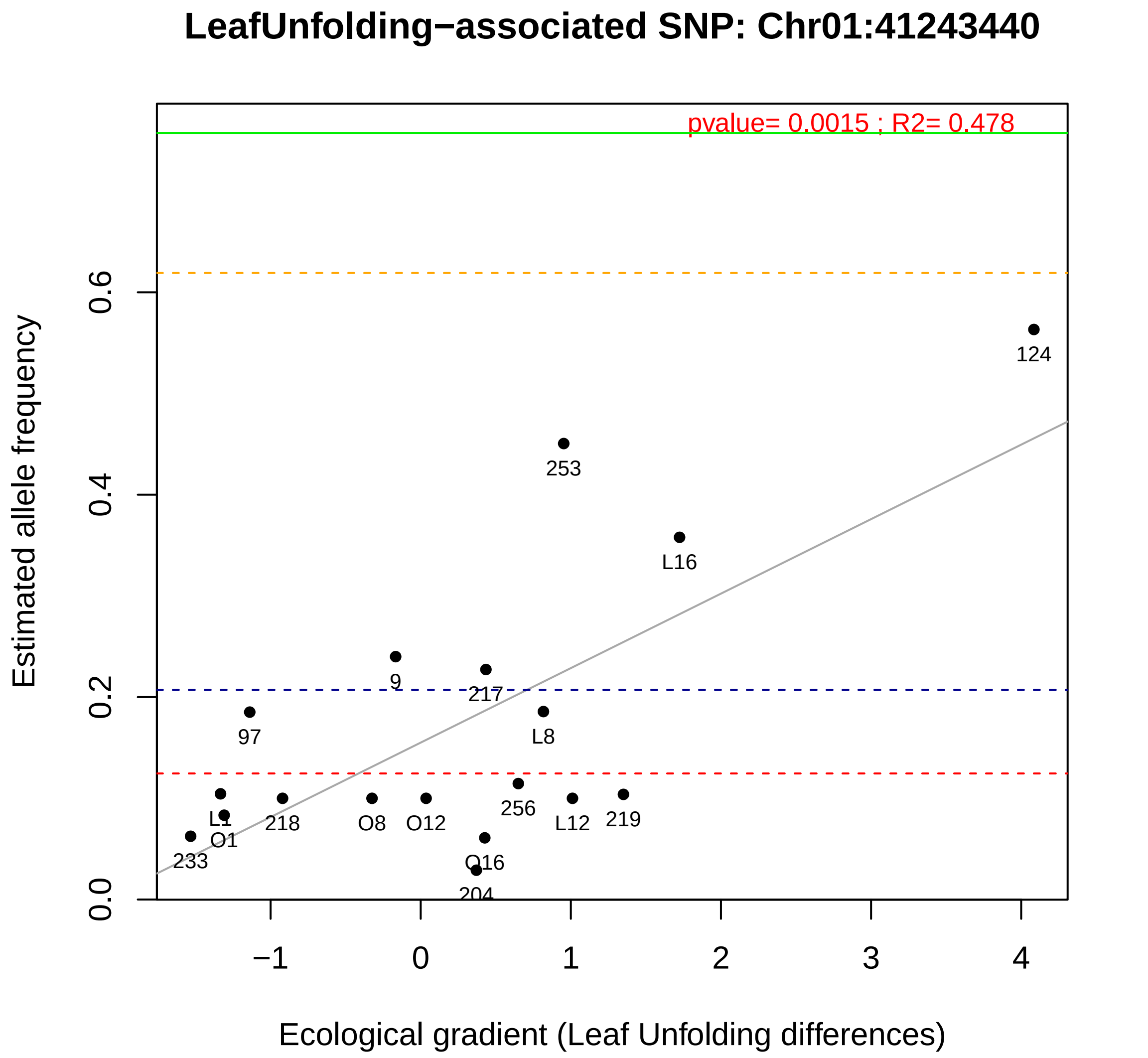

LeafUnfolding−associated SNP: Chr01:41243440
pvalue= 0.0015 ; R2= 0.478
Estimated allele frequency
0.0
0.2
0.4
0.6
−1
0
1
2
3
4
Ecological gradient (Leaf Unfolding differences)
124
253
L16
9
217
97
L8
L1
O1
218
O8
O12
256
L12
219
233
O16
204

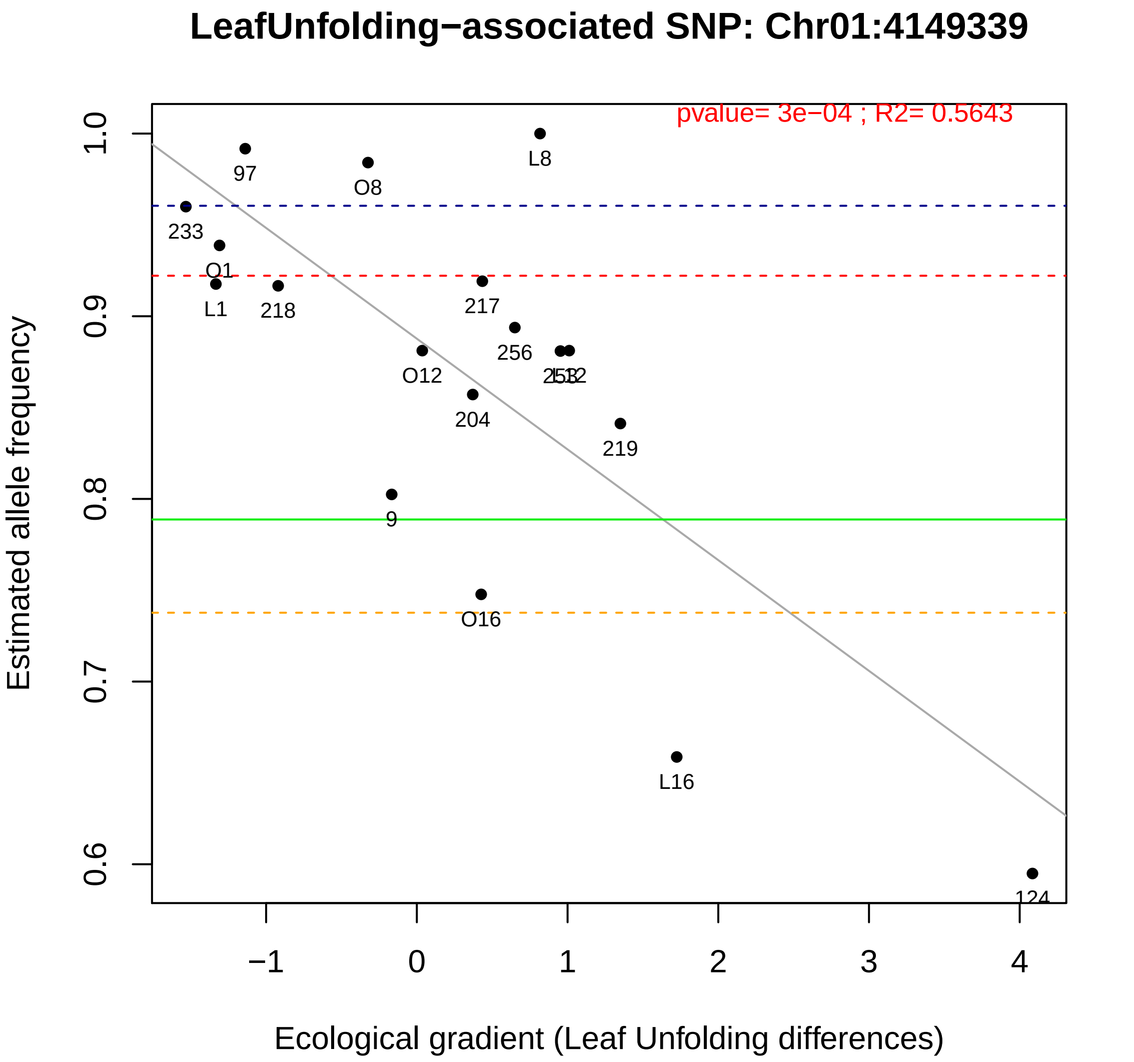
LeafUnfolding−associated SNP: Chr01:4149339
pvalue= 3e−04 ; R2= 0.5643
Estimated allele frequency
Ecological gradient (Leaf Unfolding differences)
1.0
0.9
0.8
0.7
0.6
−1
0
1
2
3
4
97
O8
L8
233
O1
L1
218
217
256
O12
204
219
9
O16
L16
124

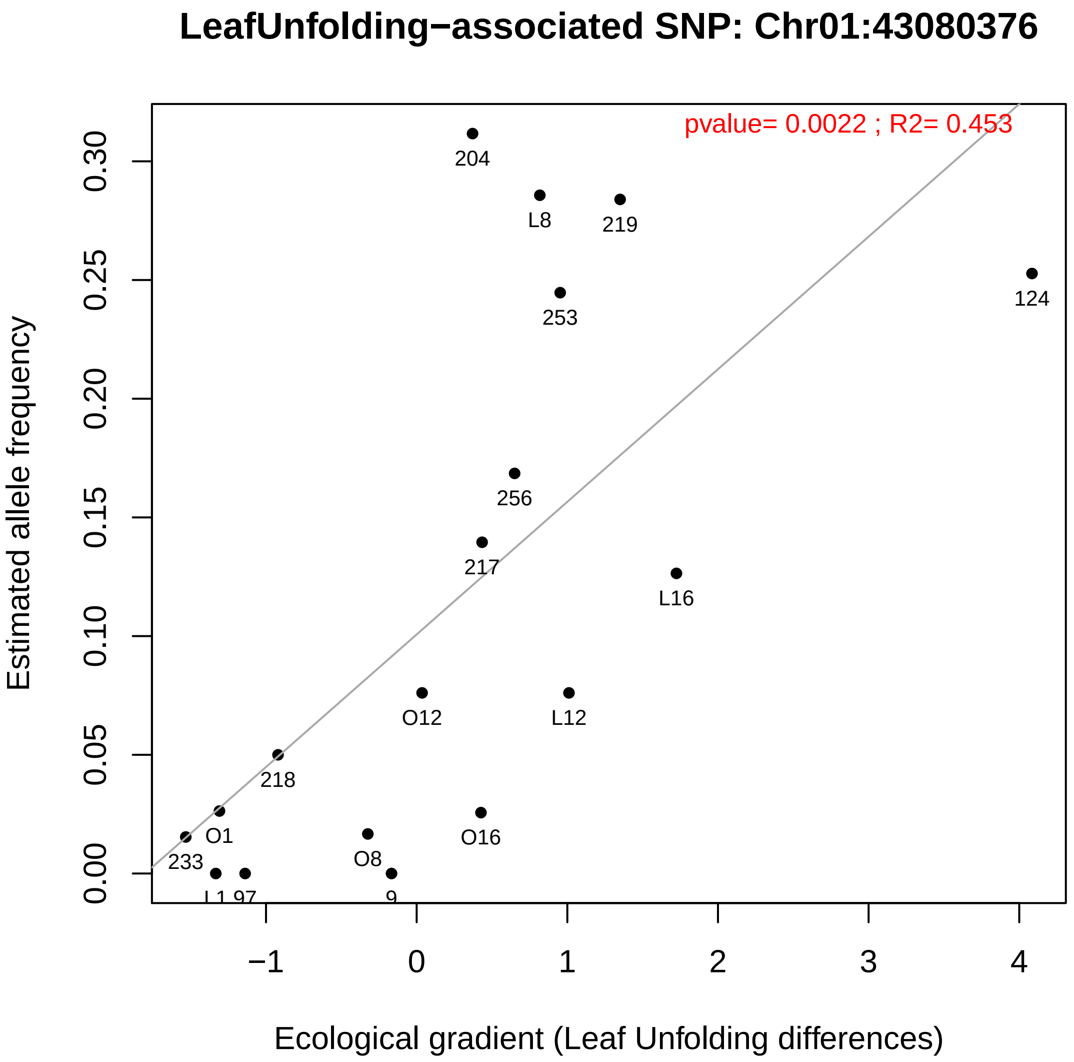
LeafUnfolding−associated SNP: Chr01:43080376
pvalue= 0.0022 ; R2= 0.453
Estimated allele frequency
0.00
0.05
0.10
0.15
0.20
0.25
0.30
Ecological gradient (Leaf Unfolding differences)
−1
0
1
2
3
4
204
L8
219
253
124
256
217
L16
O12
L12
218
O1
233
O8
O16
L1
97
9

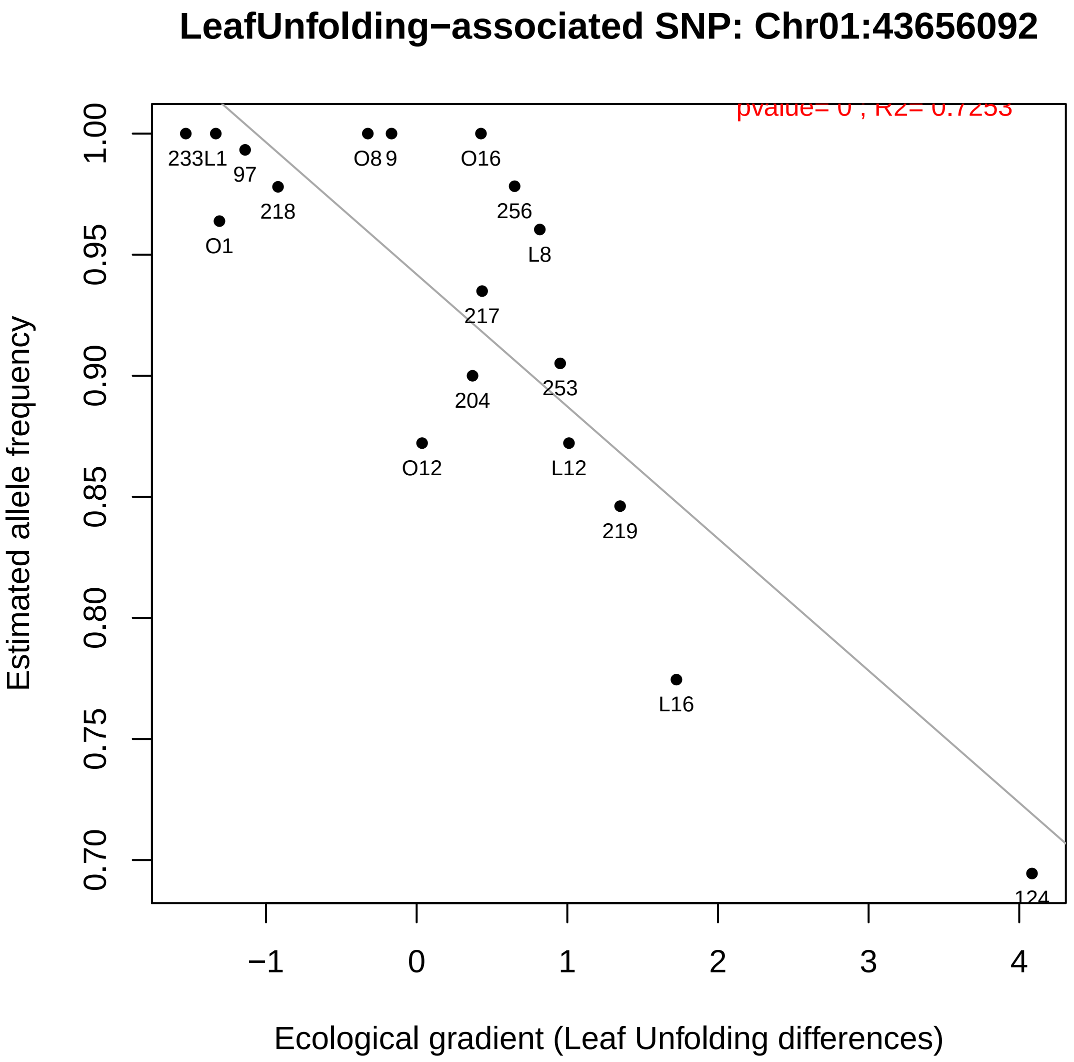

LeafUnfolding−associated SNP: Chr01:43656092
pvalue= 0 ; R2= 0.7253
Estimated allele frequency
1.00
0.95
0.90
0.85
0.80
0.75
0.70
−1
0
1
2
3
4
Ecological gradient (Leaf Unfolding differences)
233
L1
97
218
O1
O8
9
O16
256
L8
217
253
204
O12
L12
219
L16
124

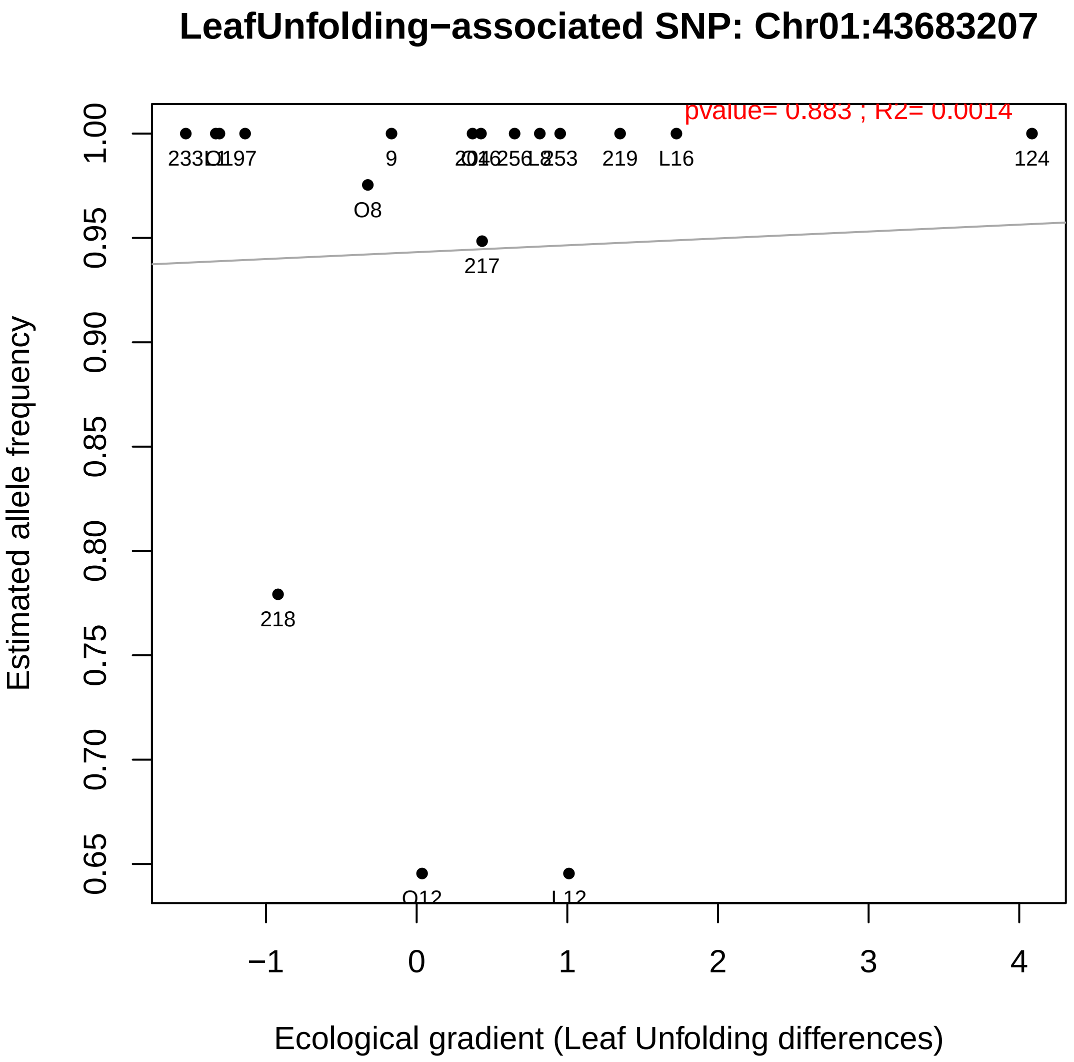

LeafUnfolding−associated SNP: Chr01:43683207
pvalue= 0.883 ; R2= 0.0014
Estimated allele frequency
Ecological gradient (Leaf Unfolding differences)
1.00
0.95
0.90
0.85
0.80
0.75
0.70
0.65
−1
0
1
2
3
4
9
O8
217
218
219
L16
124
O12
L12

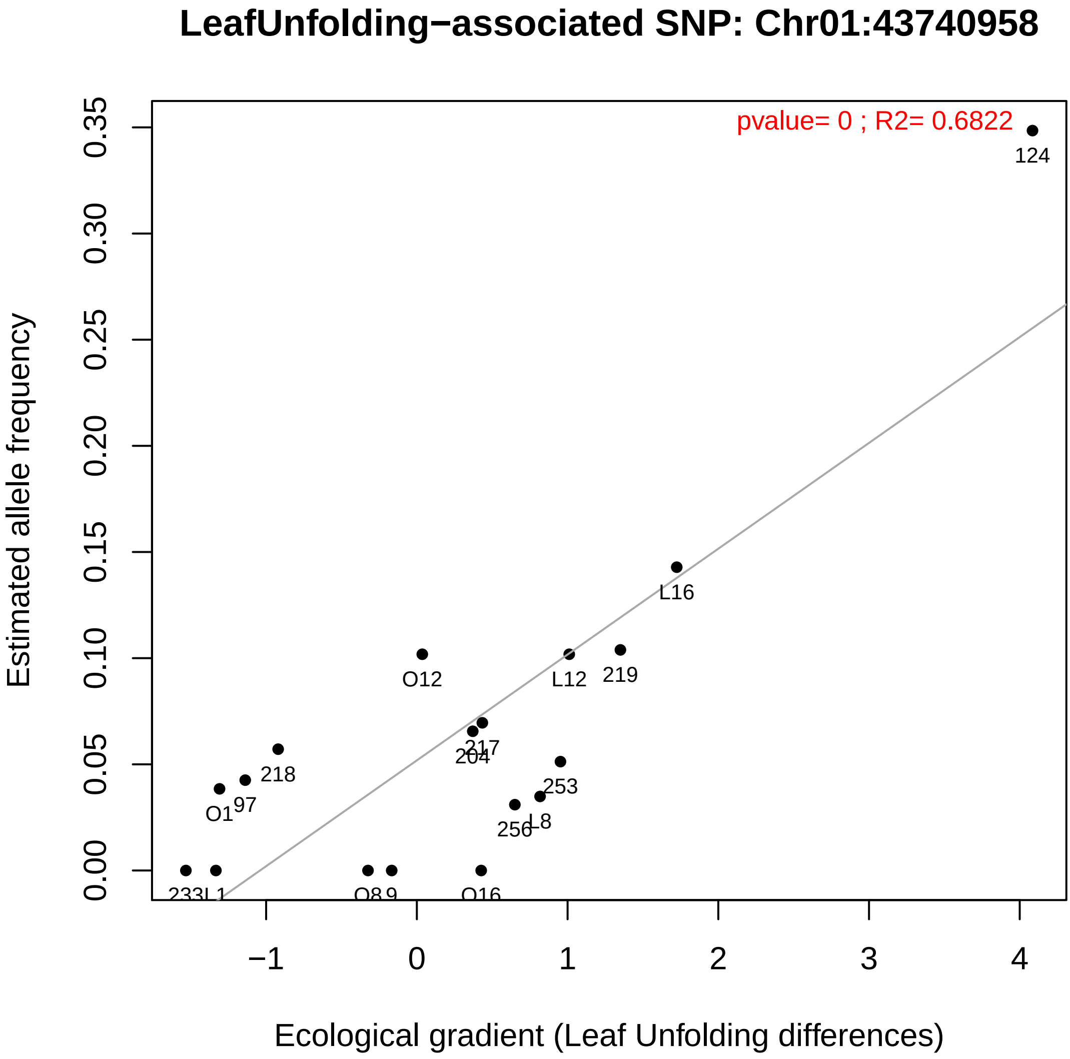
LeafUnfolding−associated SNP: Chr01:43740958
pvalue= 0 ; R2= 0.6822
Estimated allele frequency
Ecological gradient (Leaf Unfolding differences)
0.00
0.05
0.10
0.15
0.20
0.25
0.30
0.35
−1
0
1
2
3
4
124
L16
O12
L12
219
217
204
218
253
O1
97
256
L8
O16
O8
9
233
L1

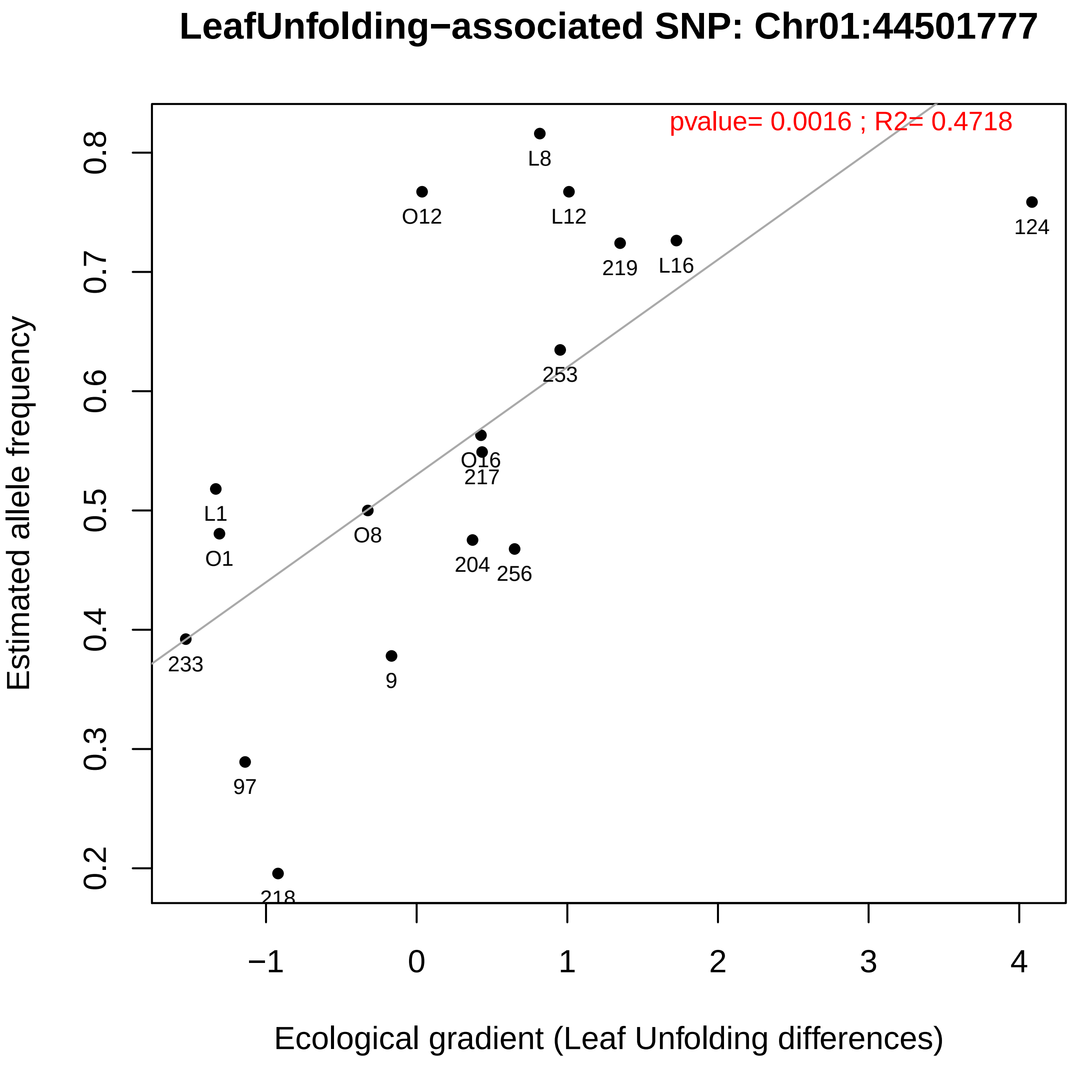
LeafUnfolding−associated SNP: Chr01:44501777
pvalue= 0.0016 ; R2= 0.4718
Estimated allele frequency
Ecological gradient (Leaf Unfolding differences)
L8
O12
L12
124
219
L16
253
O16
217
L1
O8
O1
204
256
233
9
97
218

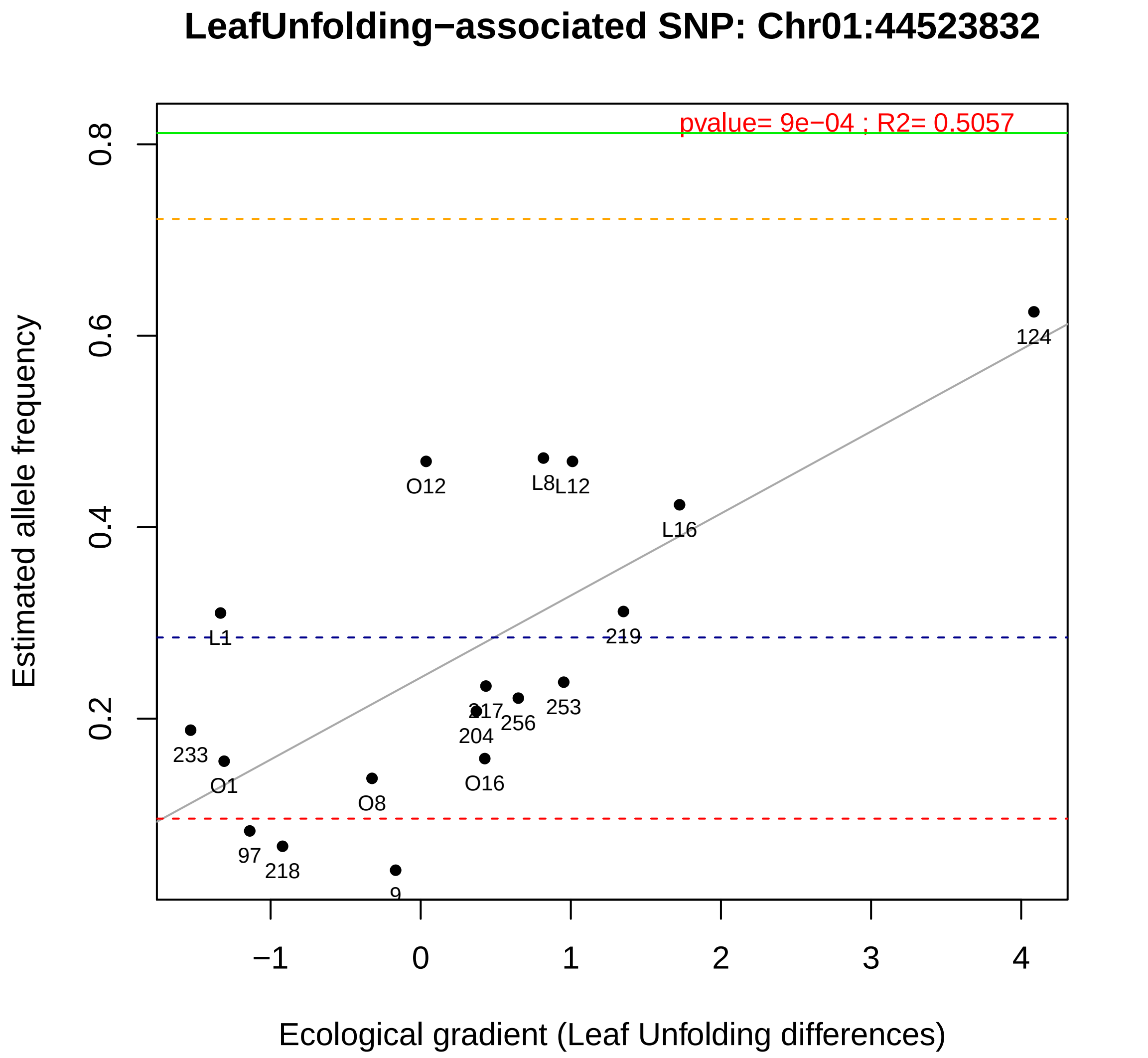

LeafUnfolding−associated SNP: Chr01:44523832
pvalue= 9e−04 ; R2= 0.5057
Estimated allele frequency
0.8
0.6
0.4
0.2
−1
0
1
2
3
4
Ecological gradient (Leaf Unfolding differences)
124
O12
L8
L12
L16
L1
219
217
253
256
204
233
O1
O16
O8
97
218
9

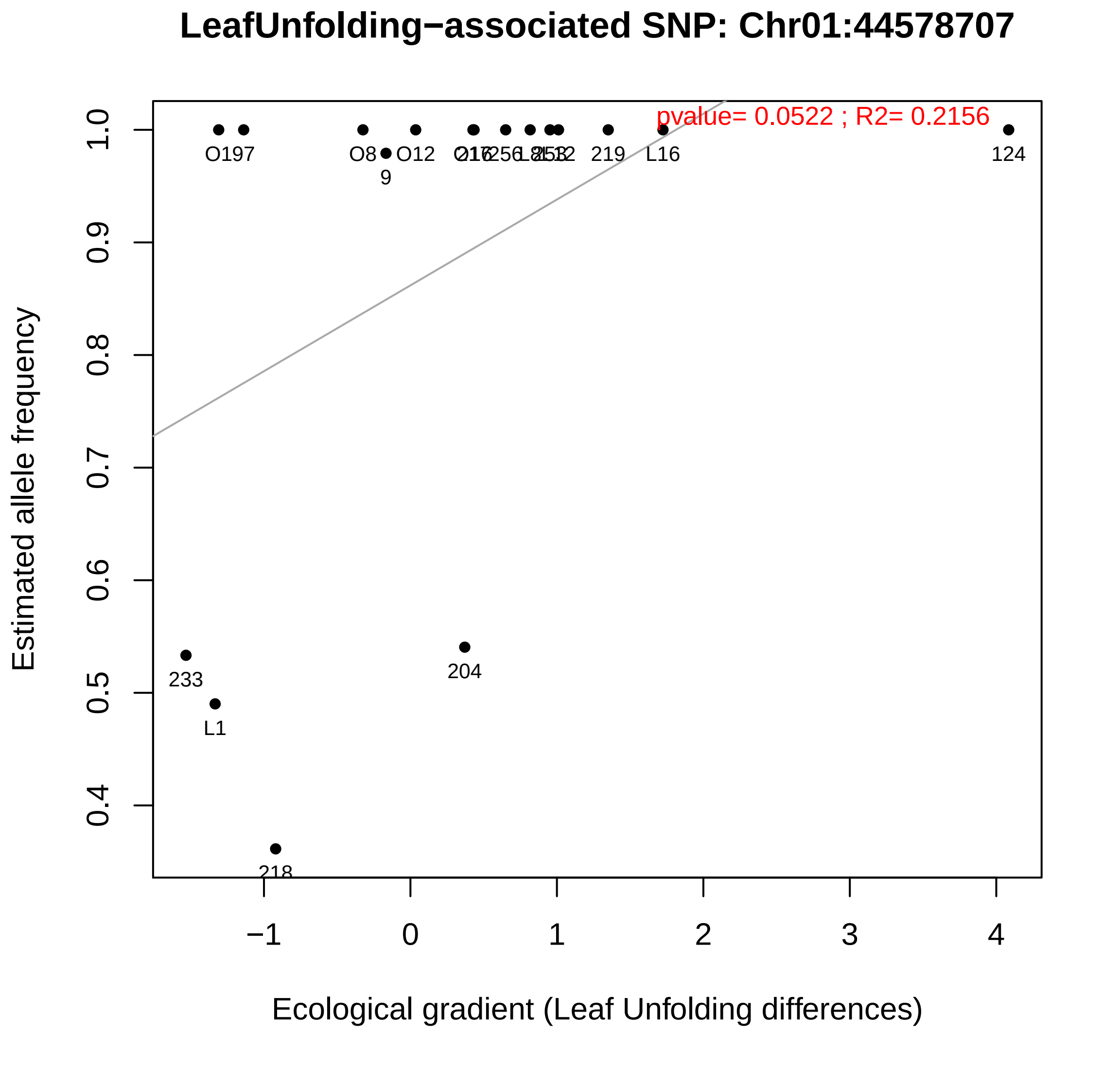
LeafUnfolding−associated SNP: Chr01:44578707
pvalue= 0.0522 ; R2= 0.2156
Estimated allele frequency
Ecological gradient (Leaf Unfolding differences)
1.0
0.9
0.8
0.7
0.6
0.5
0.4
−1
0
1
2
3
4
O197
O8
9
O12
219
L16
124
233
204
L1
218

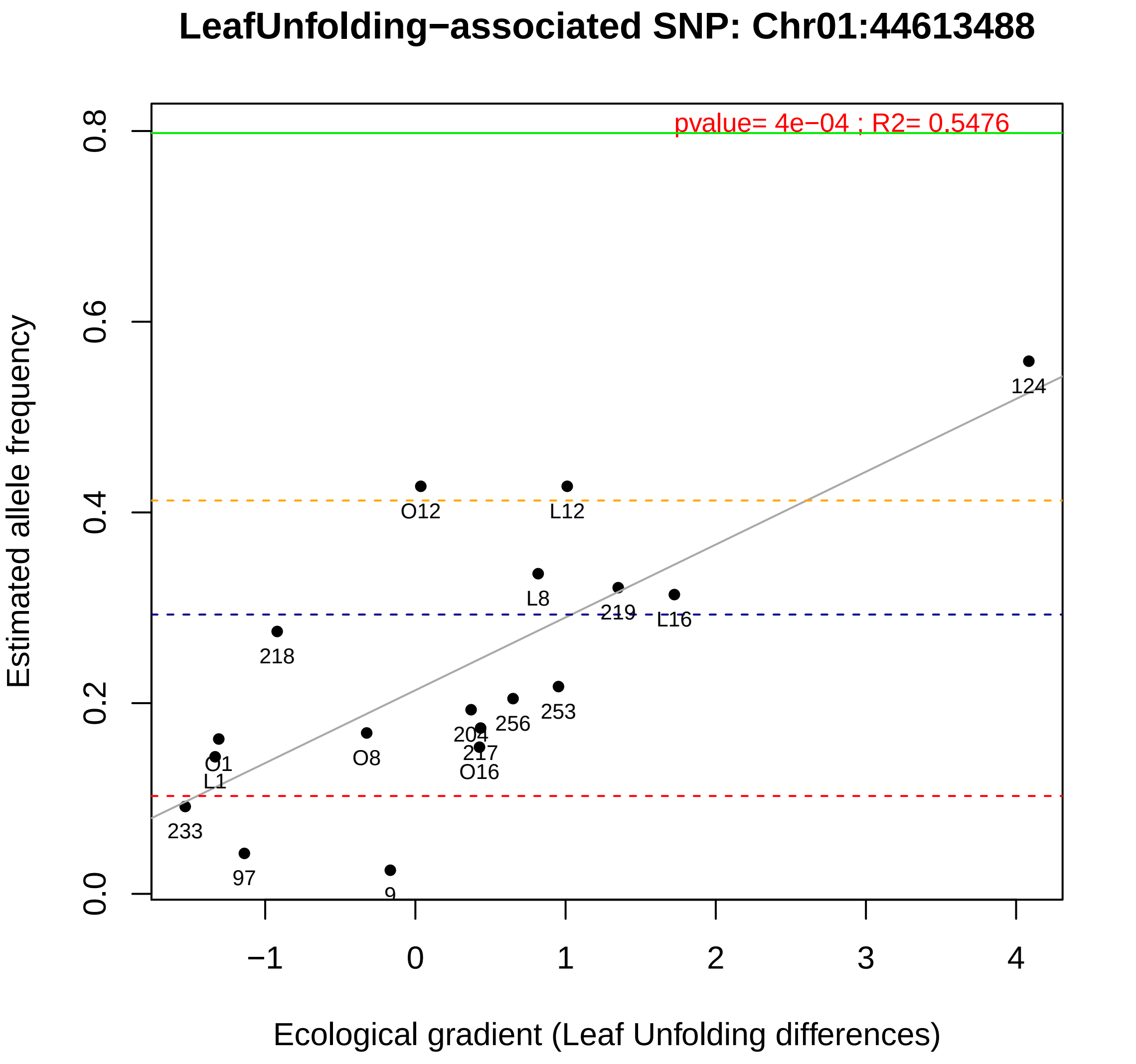
LeafUnfolding−associated SNP: Chr01:44613488
pvalue= 4e−04 ; R2= 0.5476
Estimated allele frequency
0.0
0.2
0.4
0.6
0.8
Ecological gradient (Leaf Unfolding differences)
−1
0
1
2
3
4
124
O12
L12
L8
219
L16
218
253
256
204
217
O16
O8
O1
L1
233
97
9

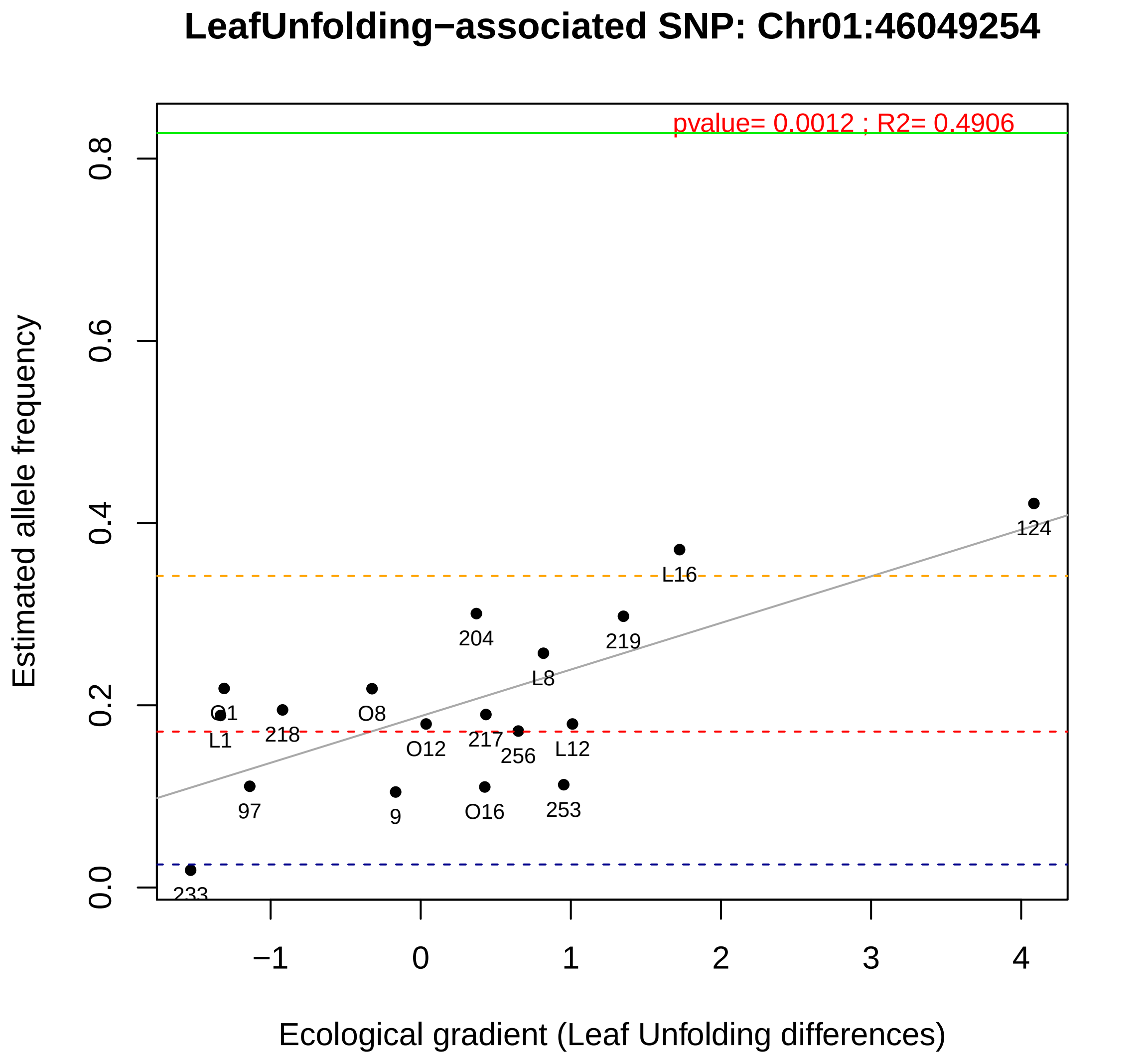

LeafUnfolding−associated SNP: Chr01:46049254
pvalue= 0.0012 ; R2= 0.4906
Estimated allele frequency
0.0
0.2
0.4
0.6
0.8
−1
0
1
2
3
4
Ecological gradient (Leaf Unfolding differences)
124
L16
204
219
L8
O1
L1
218
O8
O12
217
256
L12
97
9
O16
253

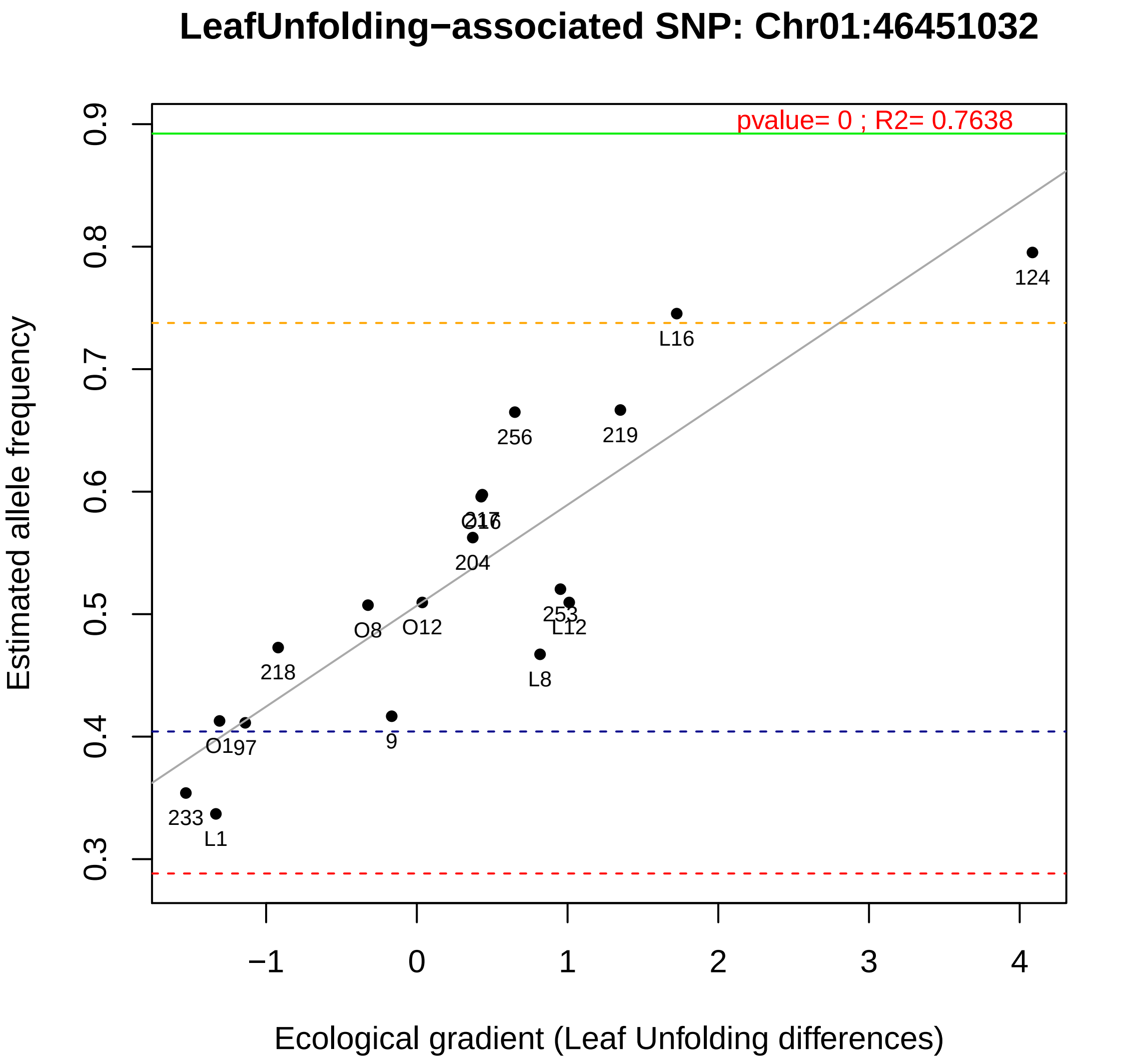

LeafUnfolding−associated SNP: Chr01:46451032
pvalue= 0 ; R2= 0.7638
Estimated allele frequency
Ecological gradient (Leaf Unfolding differences)
0.3
0.4
0.5
0.6
0.7
0.8
0.9
−1
0
1
2
3
4
124
L16
256
219
204
O8
O12
218
L8
253
L12
9
233
L1

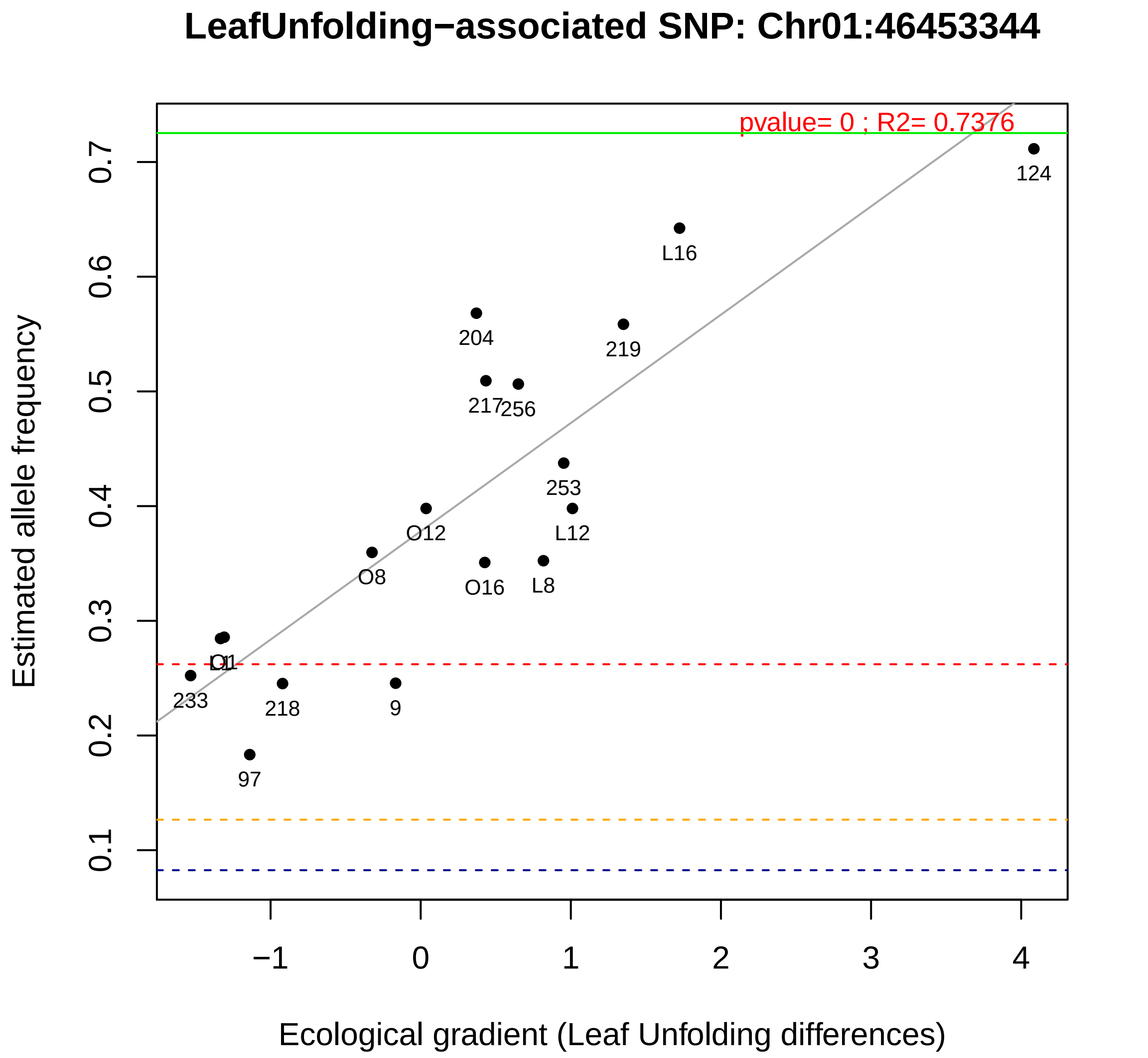

LeafUnfolding−associated SNP: Chr01:46453344
pvalue= 0 ; R2= 0.7376
Estimated allele frequency
Ecological gradient (Leaf Unfolding differences)
0.1
0.2
0.3
0.4
0.5
0.6
0.7
−1
0
1
2
3
4
124
L16
204
219
217
256
253
O12
L12
O8
O16
L8
233
218
9
97

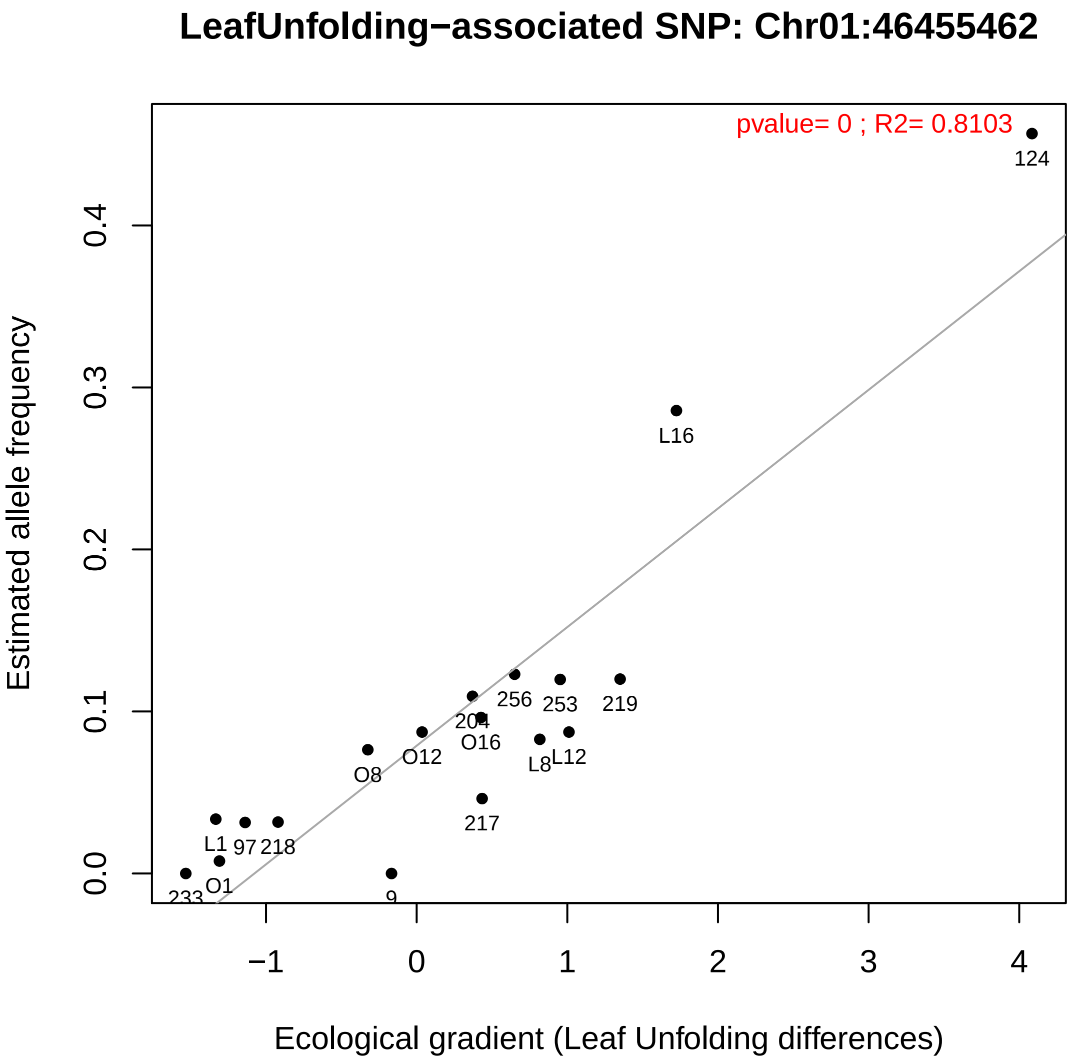

LeafUnfolding−associated SNP: Chr01:46455462
pvalue= 0 ; R2= 0.8103
124
L16
256
253
219
204
O16
O12
L8
L12
O8
217
L1
97
218
O1
233
9
0.0
0.1
0.2
0.3
0.4
−1
0
1
2
3
4
Estimated allele frequency
Ecological gradient (Leaf Unfolding differences)

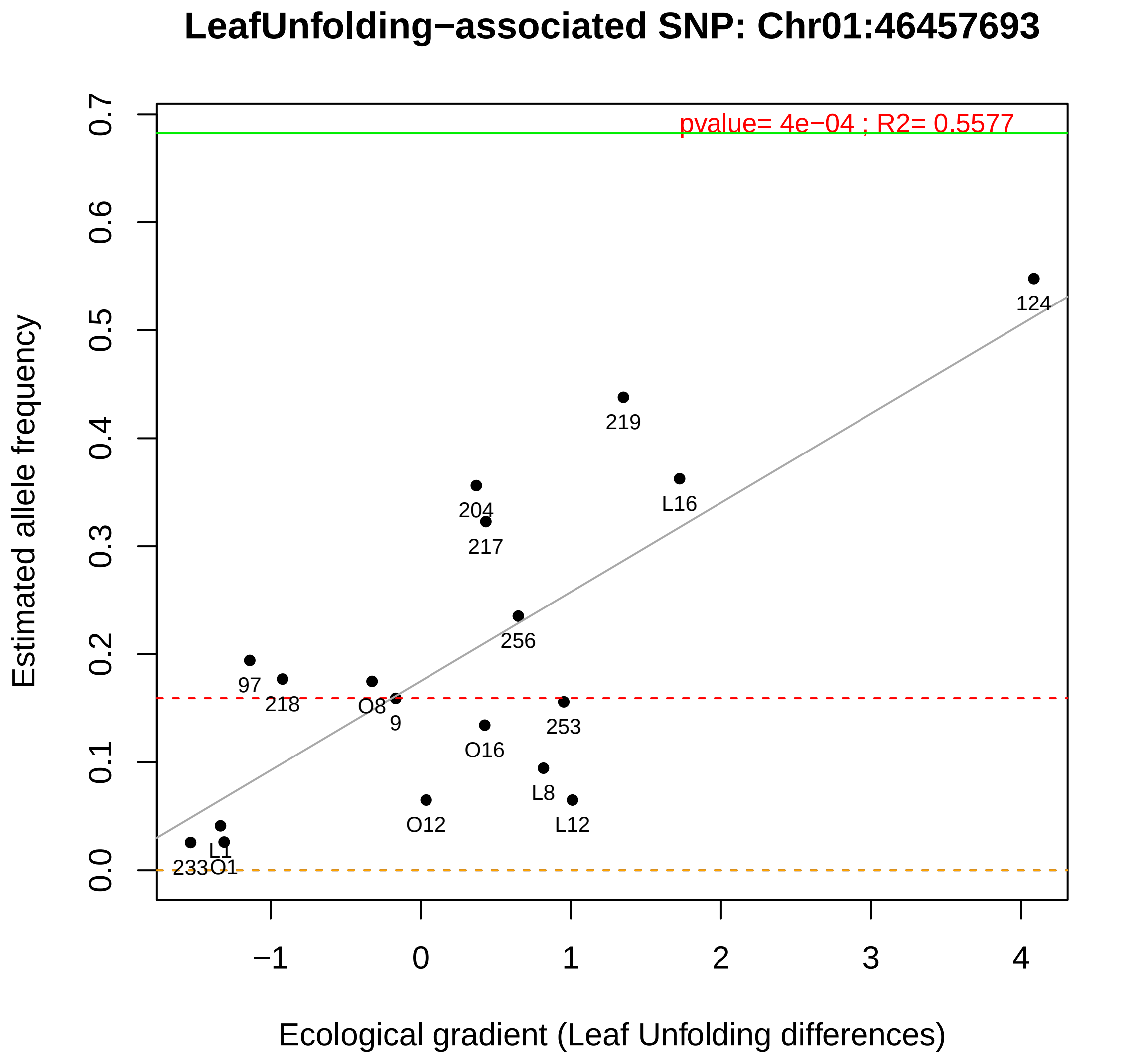

LeafUnfolding−associated SNP: Chr01:46457693
pvalue= 4e−04 ; R2= 0.5577
Estimated allele frequency
Ecological gradient (Leaf Unfolding differences)
0.0
0.1
0.2
0.3
0.4
0.5
0.6
0.7
−1
0
1
2
3
4
124
219
L16
204
217
256
97
218
O8
9
253
O16
L8
L12
O12
L1
233
O1

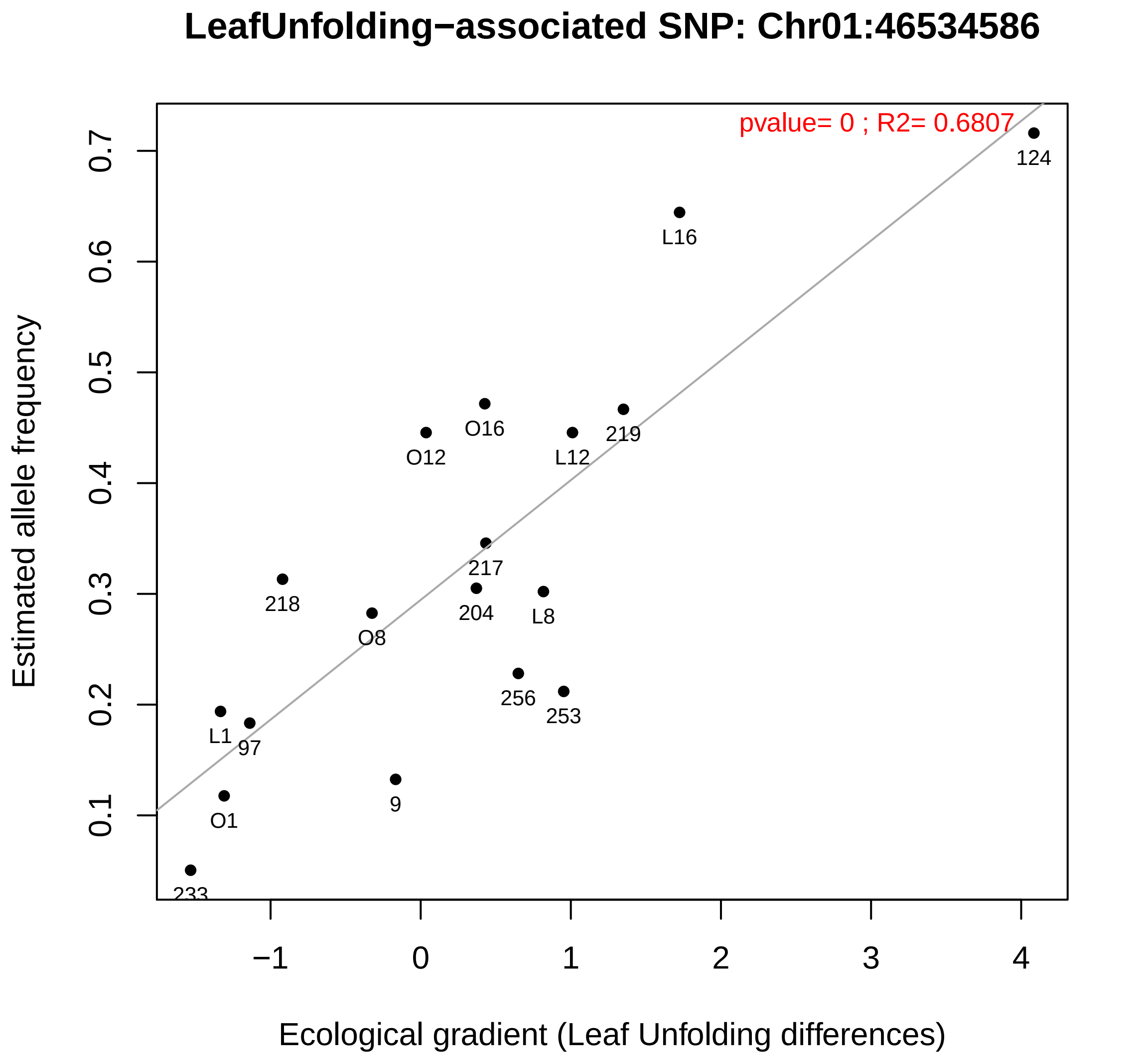
LeafUnfolding−associated SNP: Chr01:46534586
pvalue= 0 ; R2= 0.6807
Estimated allele frequency
Ecological gradient (Leaf Unfolding differences)
0.1
0.2
0.3
0.4
0.5
0.6
0.7
−1
0
1
2
3
4
124
L16
O16
219
O12
L12
217
218
204
L8
O8
256
253
L1
97
O1
9
233

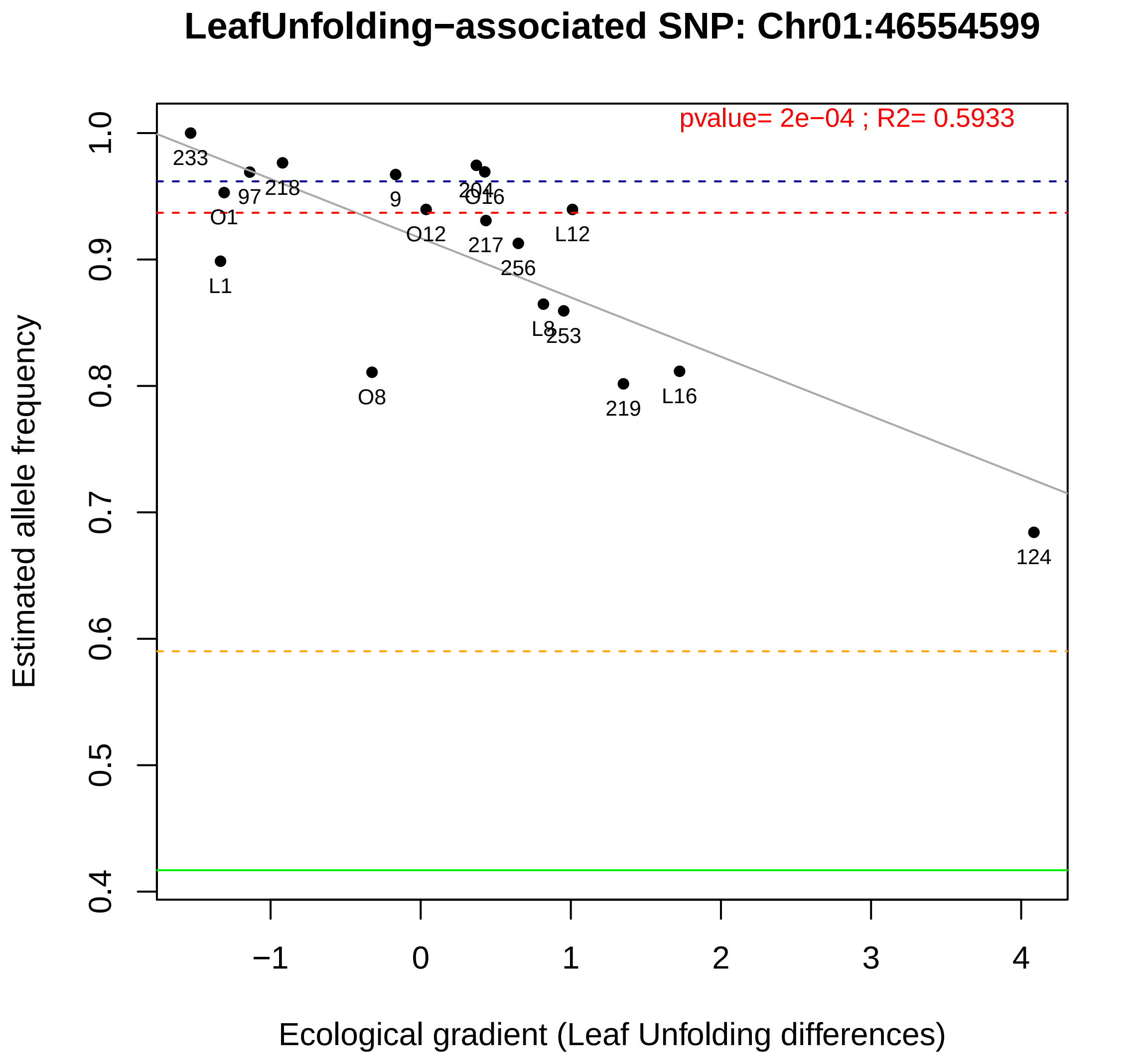

LeafUnfolding−associated SNP: Chr01:46554599
pvalue= 2e−04 ; R2= 0.5933
Estimated allele frequency
Ecological gradient (Leaf Unfolding differences)
0.4
0.5
0.6
0.7
0.8
0.9
1.0
−1
0
1
2
3
4
233
97
218
O1
L1
9
O12
204
O16
217
256
L12
L8
253
O8
219
L16
124

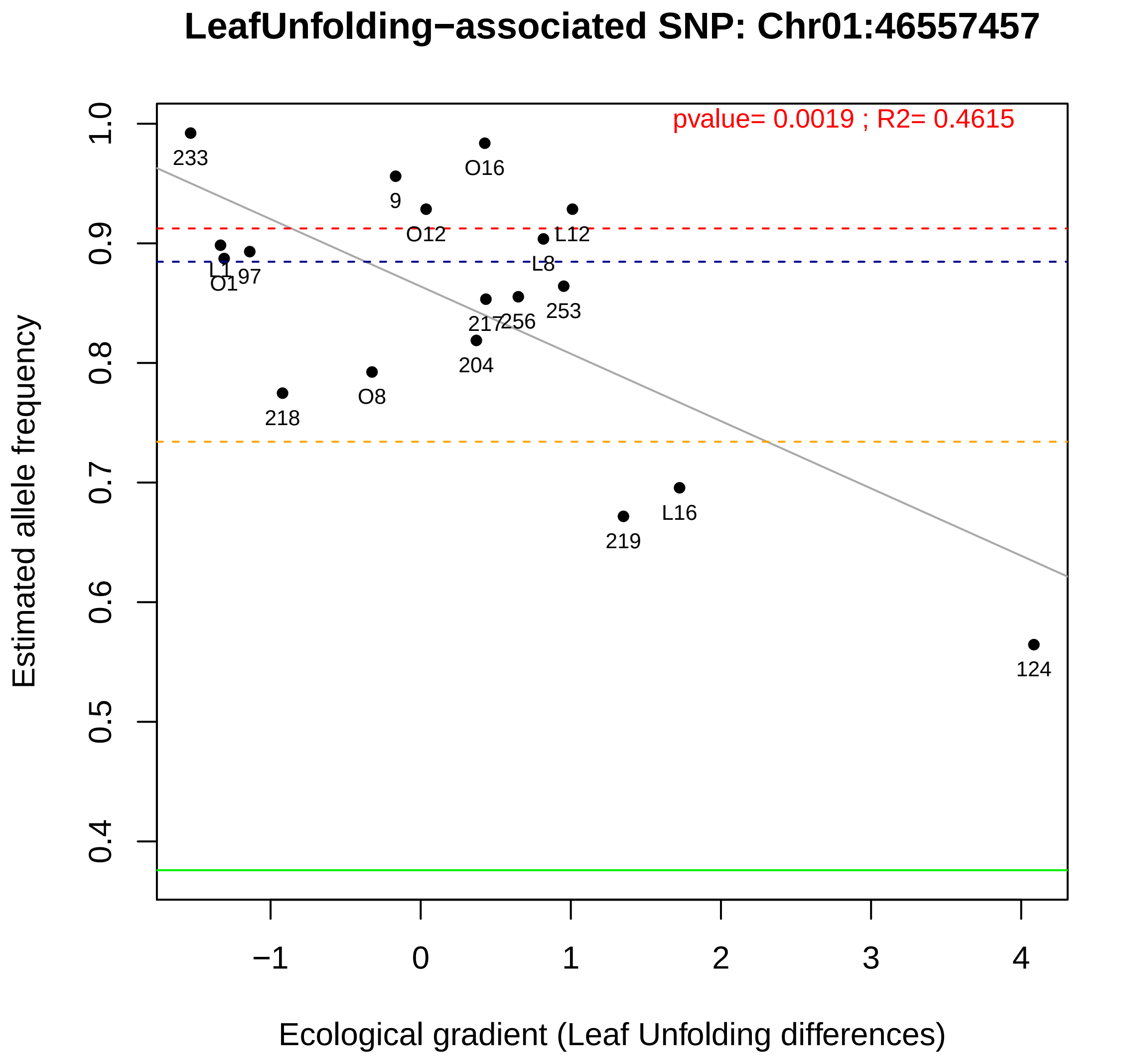

LeafUnfolding−associated SNP: Chr01:46557457
pvalue= 0.0019 ; R2= 0.4615
233
O16
9
O12
L12
L8
L1
O1
97
217
256
253
204
O8
218
L16
219
124
Estimated allele frequency
Ecological gradient (Leaf Unfolding differences)

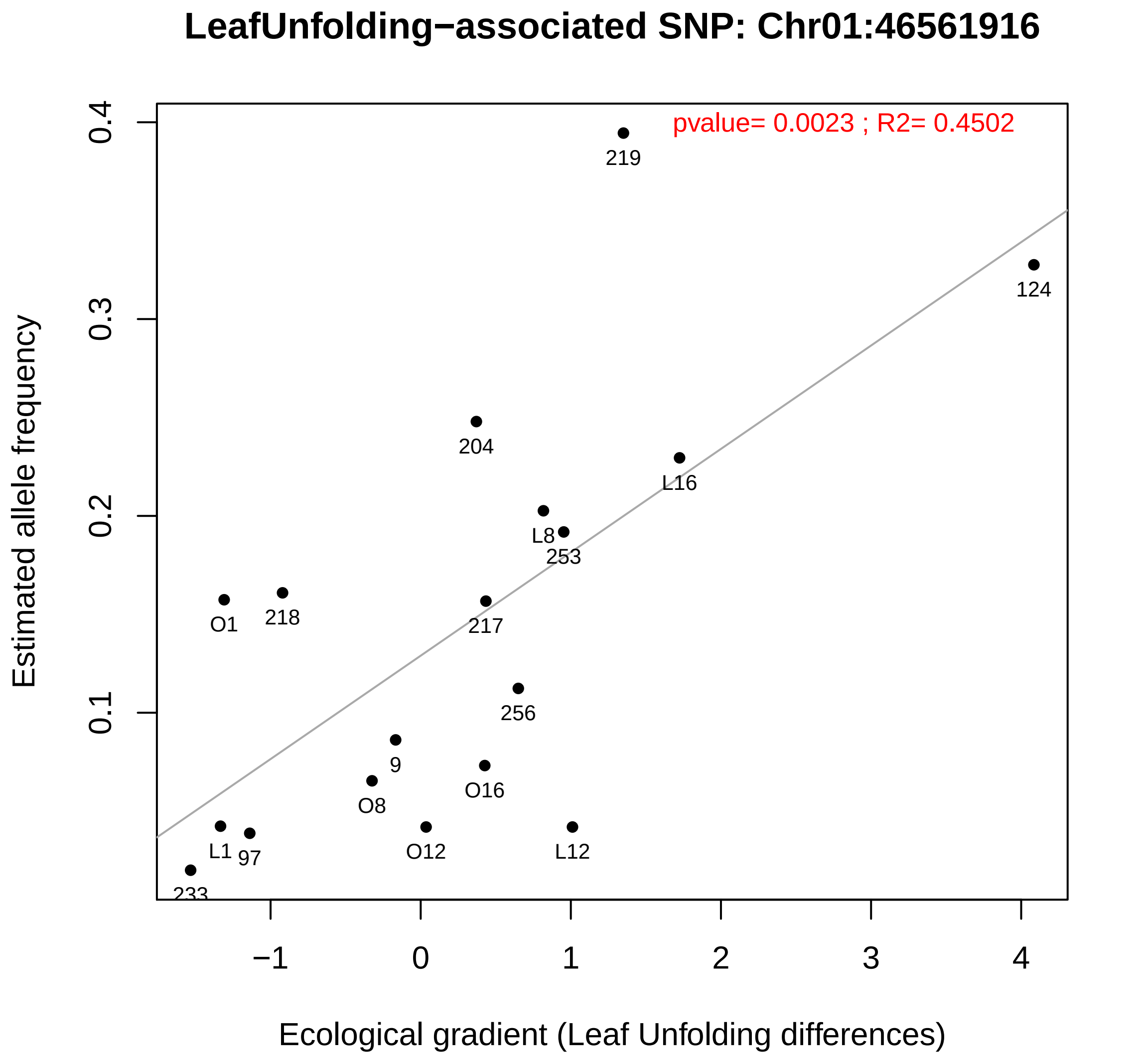
LeafUnfolding−associated SNP: Chr01:46561916
pvalue= 0.0023 ; R2= 0.4502
219
124
204
L16
L8
253
O1
218
217
256
9
O16
O8
L1
97
O12
L12
233
0.1
0.2
0.3
0.4
−1
0
1
2
3
4
Estimated allele frequency
Ecological gradient (Leaf Unfolding differences)

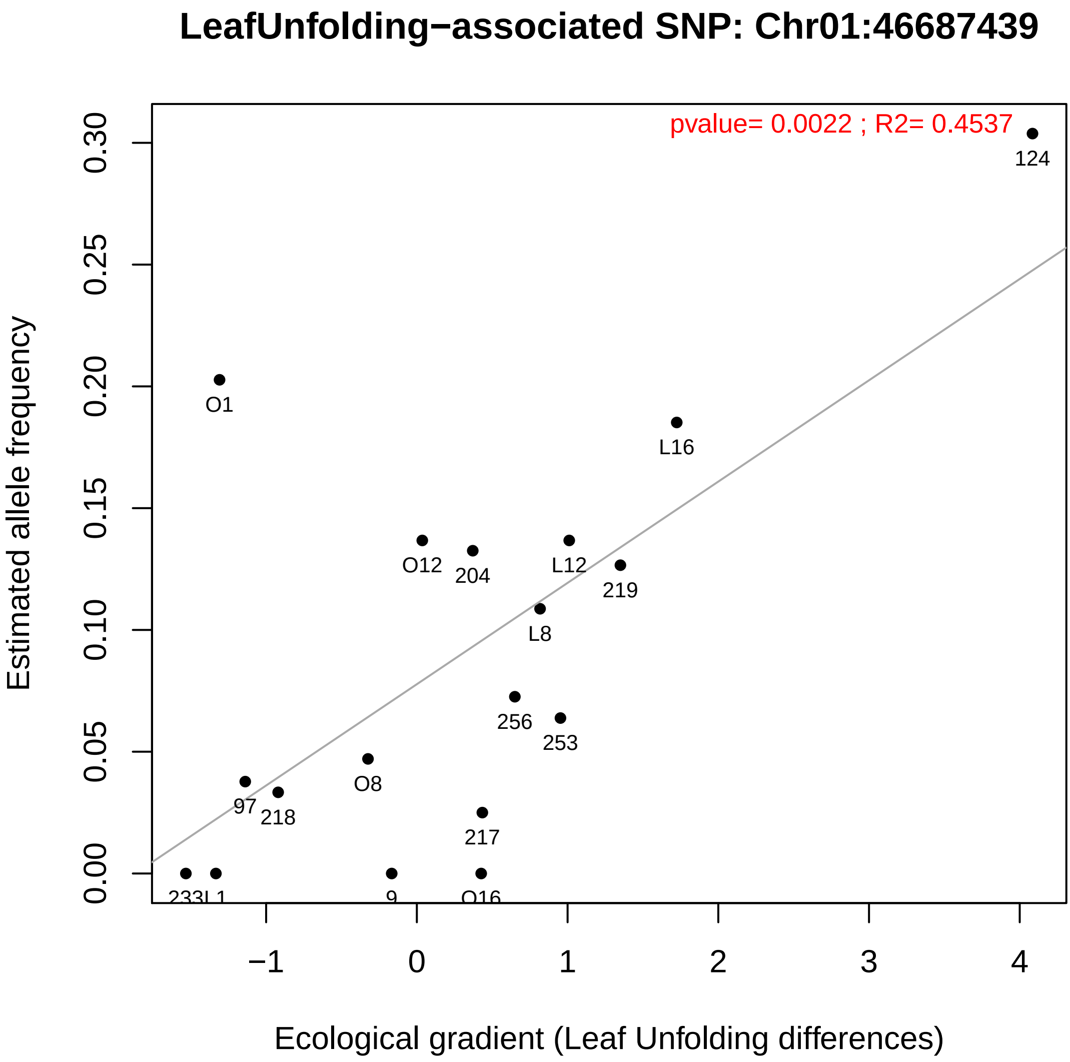

LeafUnfolding−associated SNP: Chr01:46687439
pvalue= 0.0022 ; R2= 0.4537
Estimated allele frequency
0.00
0.05
0.10
0.15
0.20
0.25
0.30
−1
0
1
2
3
4
Ecological gradient (Leaf Unfolding differences)
124
O1
L16
O12
204
L12
219
L8
256
253
O8
97
218
217
233
L1
9
O16

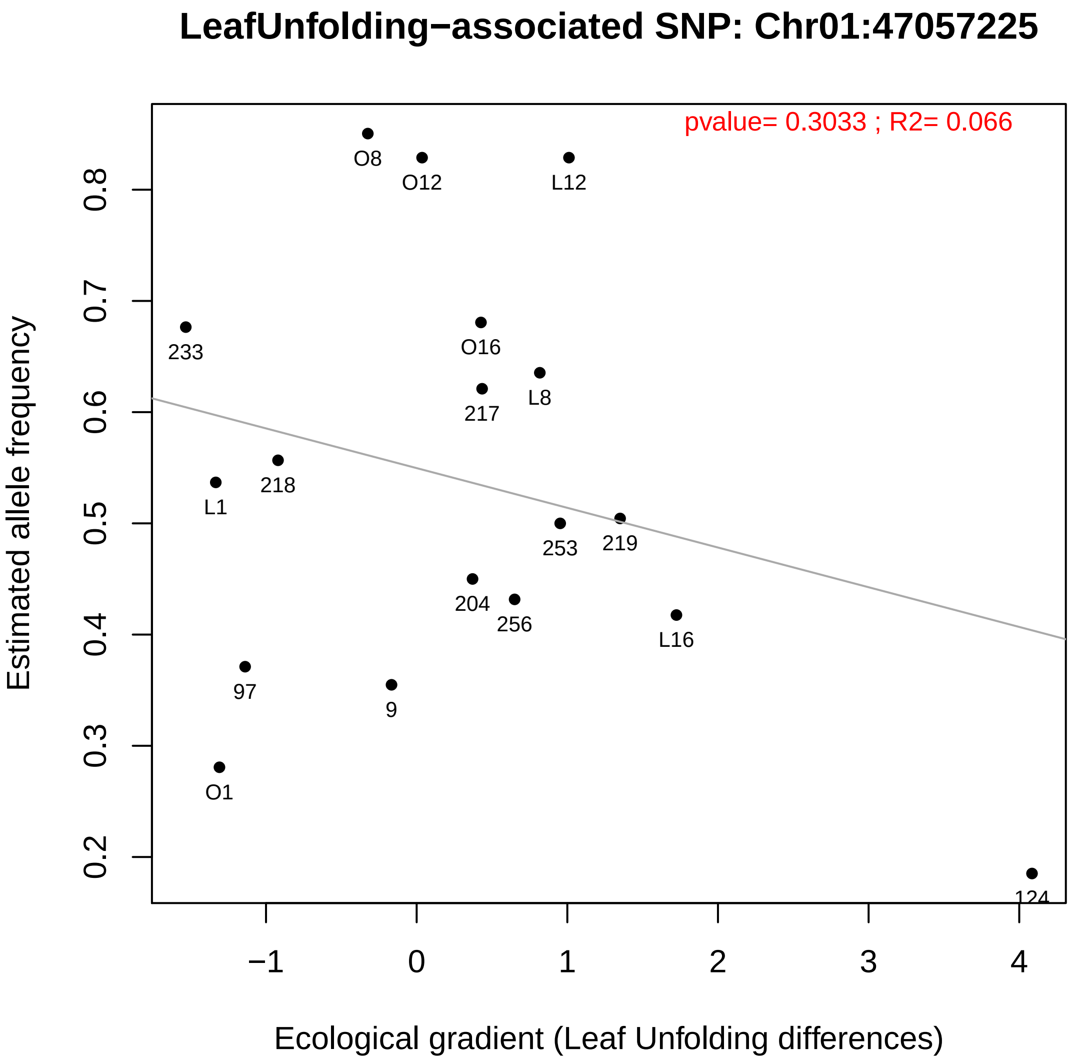
LeafUnfolding−associated SNP: Chr01:47057225
pvalue= 0.3033 ; R2= 0.066
O8
O12
L12
233
O16
L8
217
218
L1
253
219
204
256
L16
97
9
O1
124
0.2
0.3
0.4
0.5
0.6
0.7
0.8
−1
0
1
2
3
4
Estimated allele frequency
Ecological gradient (Leaf Unfolding differences)

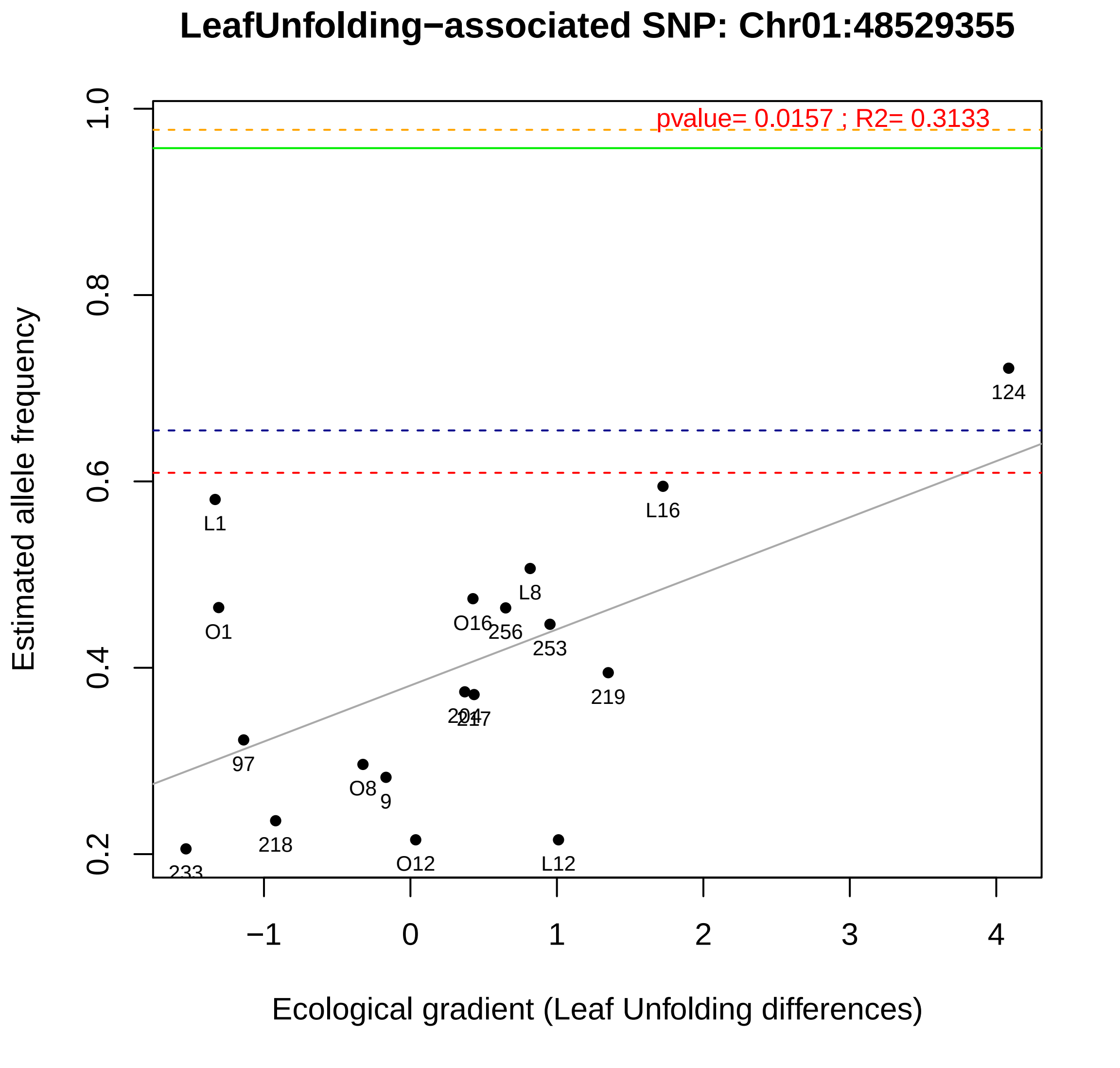

LeafUnfolding−associated SNP: Chr01:48529355
pvalue= 0.0157 ; R2= 0.3133
Estimated allele frequency
Ecological gradient (Leaf Unfolding differences)
1.0
0.8
0.6
0.4
0.2
−1
0
1
2
3
4
124
L16
L1
L8
O16
256
253
O1
219
204
217
97
O8
9
218
O12
L12
233

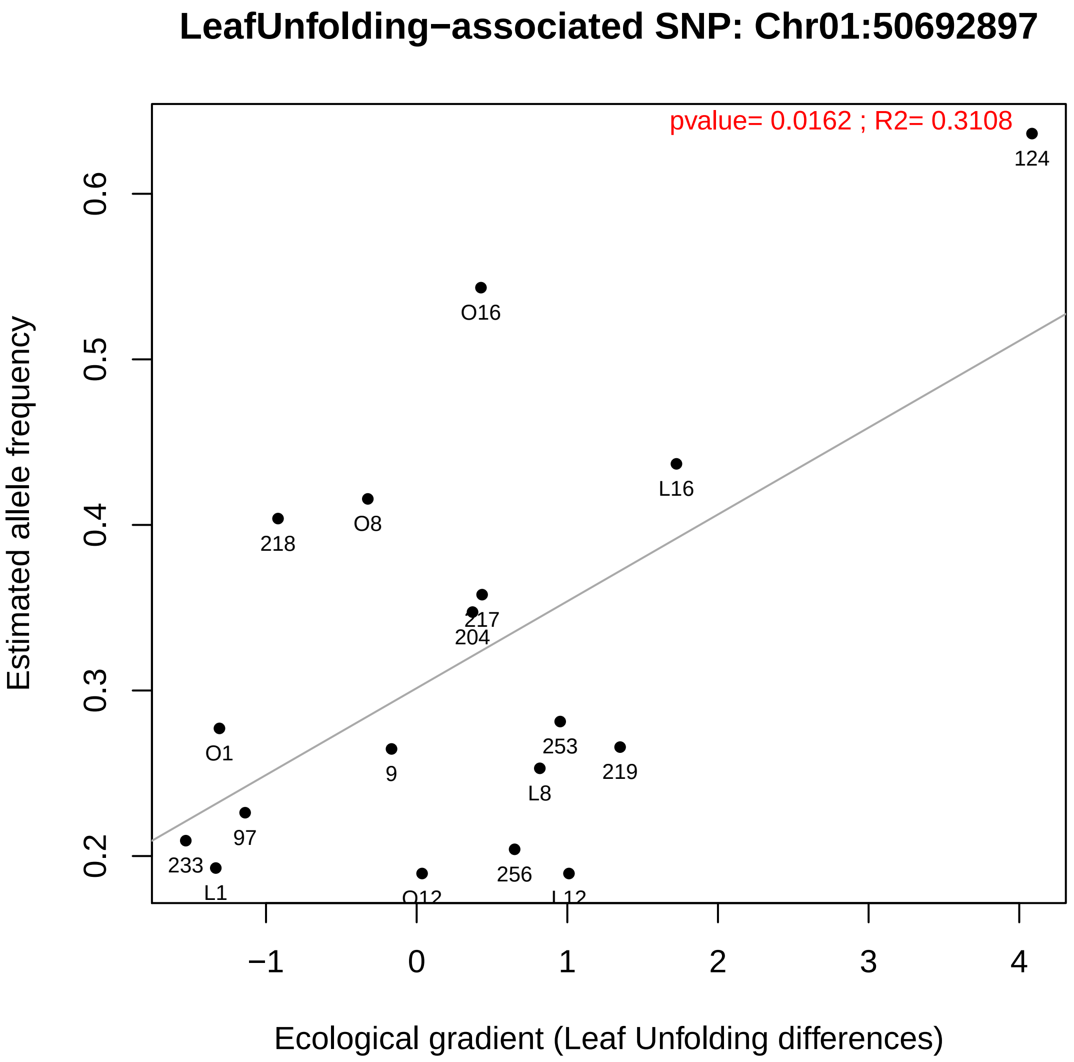

LeafUnfolding−associated SNP: Chr01:50692897
pvalue= 0.0162 ; R2= 0.3108
Estimated allele frequency
Ecological gradient (Leaf Unfolding differences)
0.2
0.3
0.4
0.5
0.6
−1
0
1
2
3
4
124
O16
L16
O8
218
217
204
O1
9
253
219
L8
97
233
L1
256
O12
L12

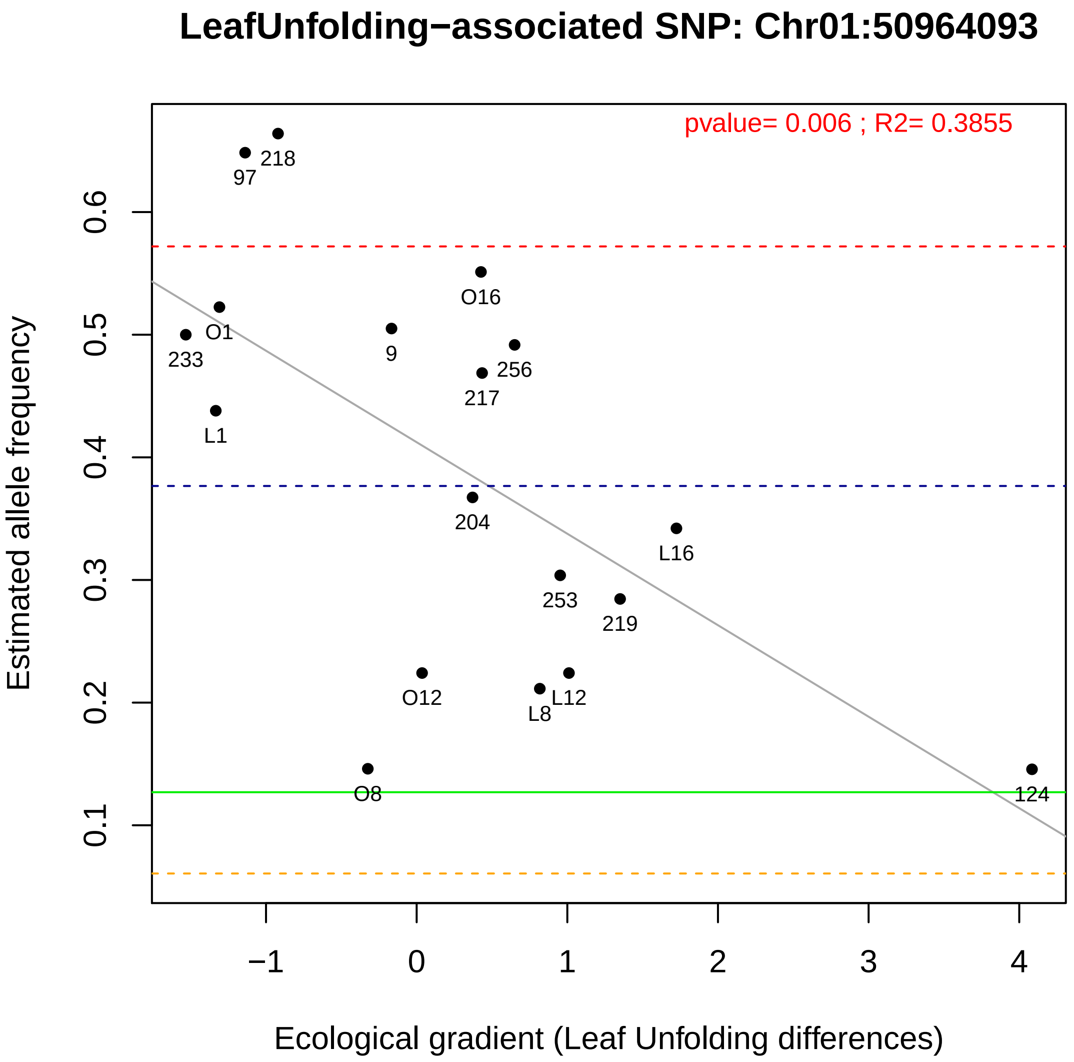
LeafUnfolding−associated SNP: Chr01:50964093
pvalue= 0.006 ; R2= 0.3855
Estimated allele frequency
0.1
0.2
0.3
0.4
0.5
0.6
−1
0
1
2
3
4
Ecological gradient (Leaf Unfolding differences)
218
97
O16
O1
233
9
256
217
L1
204
L16
253
219
O12
L8
L12
O8
124

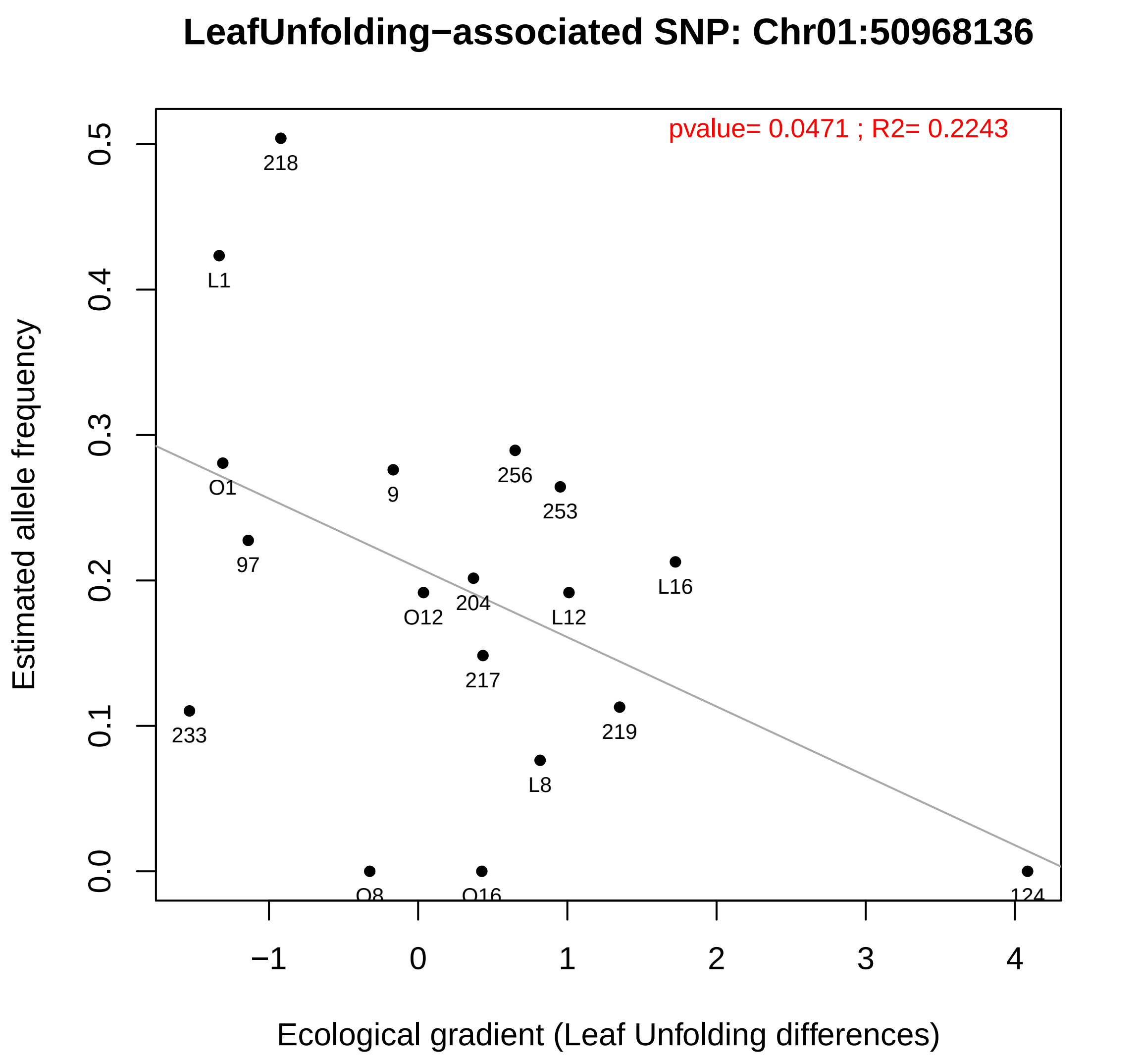
LeafUnfolding−associated SNP: Chr01:50968136
pvalue= 0.0471 ; R2= 0.2243
Estimated allele frequency
0.0
0.1
0.2
0.3
0.4
0.5
218
L1
O1
9
256
253
97
L16
204
O12
L12
217
233
219
L8
O8
O16
124
−1
0
1
2
3
4
Ecological gradient (Leaf Unfolding differences)

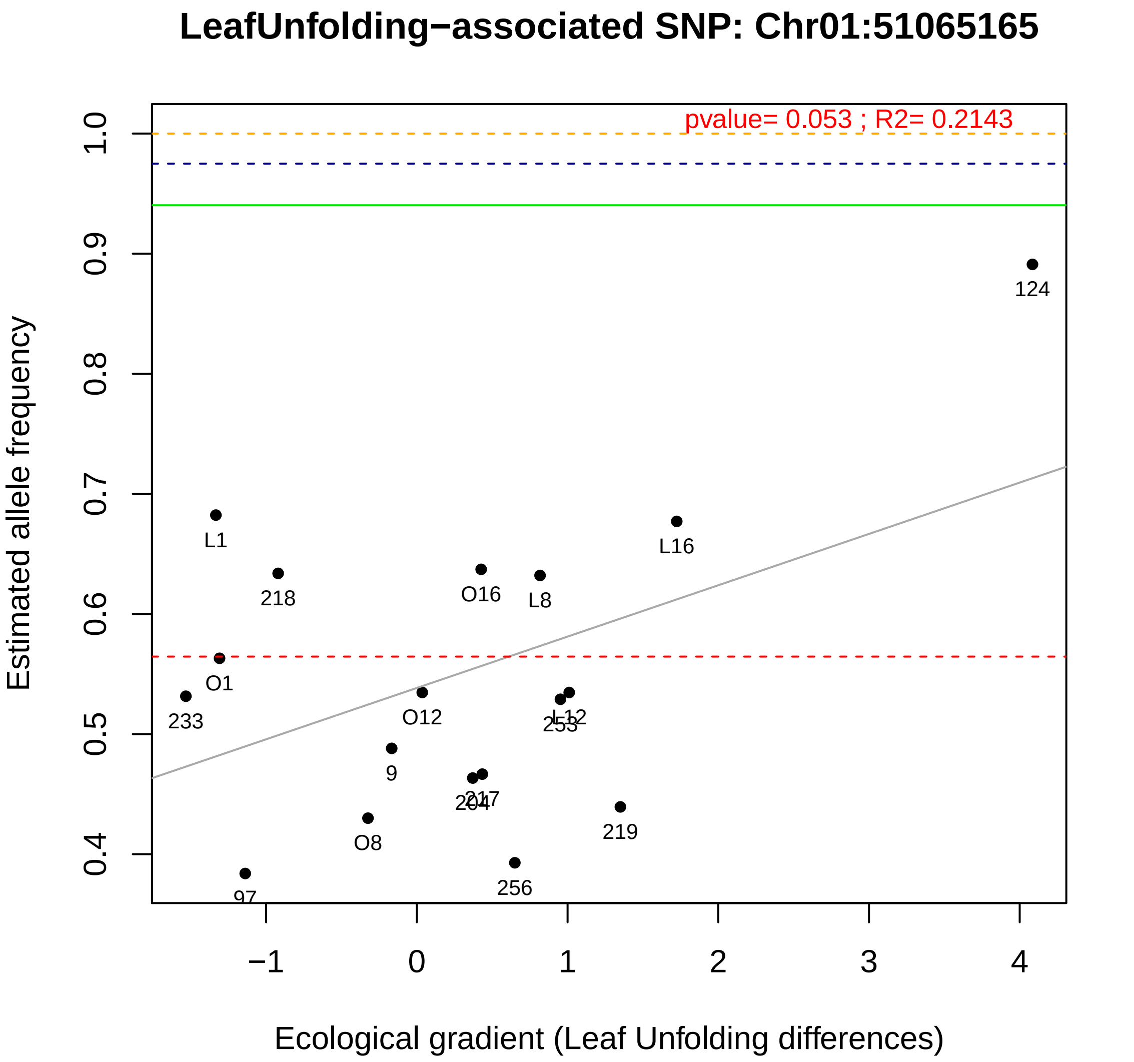
LeafUnfolding−associated SNP: Chr01:51065165
pvalue= 0.053 ; R2= 0.2143
Estimated allele frequency
Ecological gradient (Leaf Unfolding differences)
1.0
0.9
0.8
0.7
0.6
0.5
0.4
−1
0
1
2
3
4
124
L1
L16
218
O16
L8
O1
233
O12
L12
253
9
217
204
O8
219
256
97

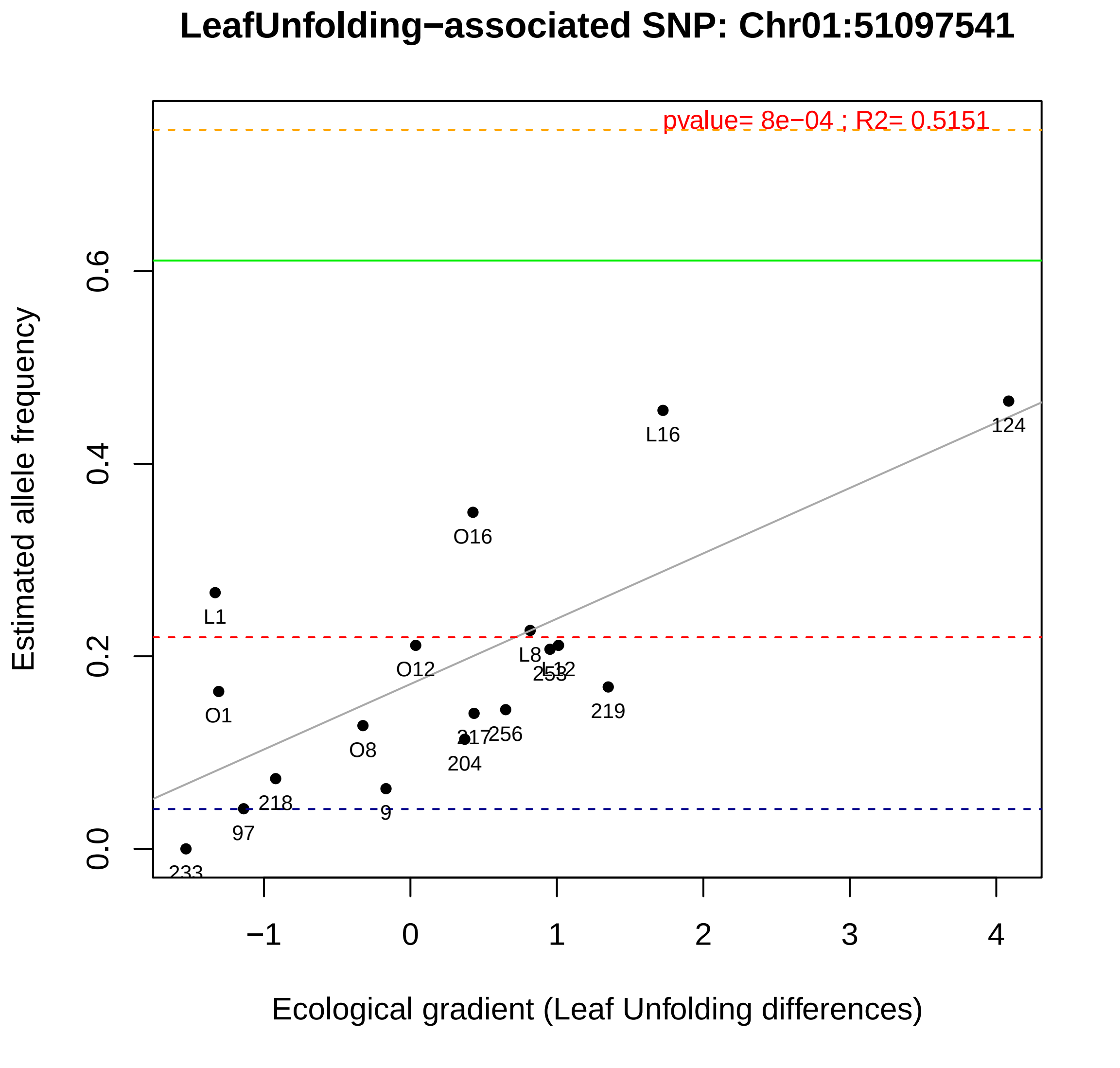

LeafUnfolding−associated SNP: Chr01:51097541
pvalue= 8e−04 ; R2= 0.5151
Estimated allele frequency
0.0
0.2
0.4
0.6
−1
0
1
2
3
4
Ecological gradient (Leaf Unfolding differences)
L16
124
O16
L1
O12
L8
253
L12
219
O1
O8
217
256
204
218
9
97
233

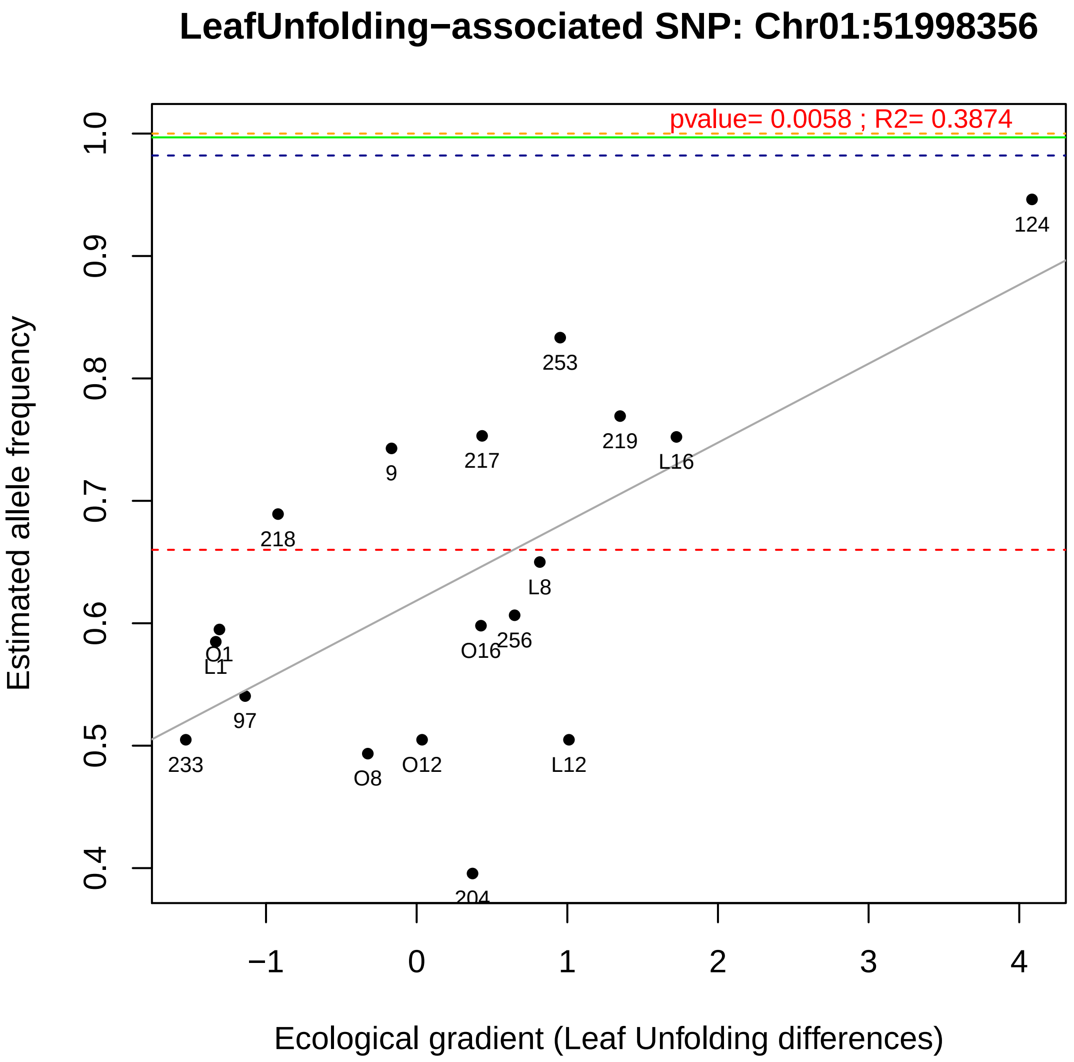

LeafUnfolding−associated SNP: Chr01:51998356
pvalue= 0.0058 ; R2= 0.3874
Estimated allele frequency
1.0
0.9
0.8
0.7
0.6
0.5
0.4
124
253
219
L16
217
9
218
L8
256
O16
O1
L1
97
233
O8
O12
L12
204
−1
0
1
2
3
4
Ecological gradient (Leaf Unfolding differences)

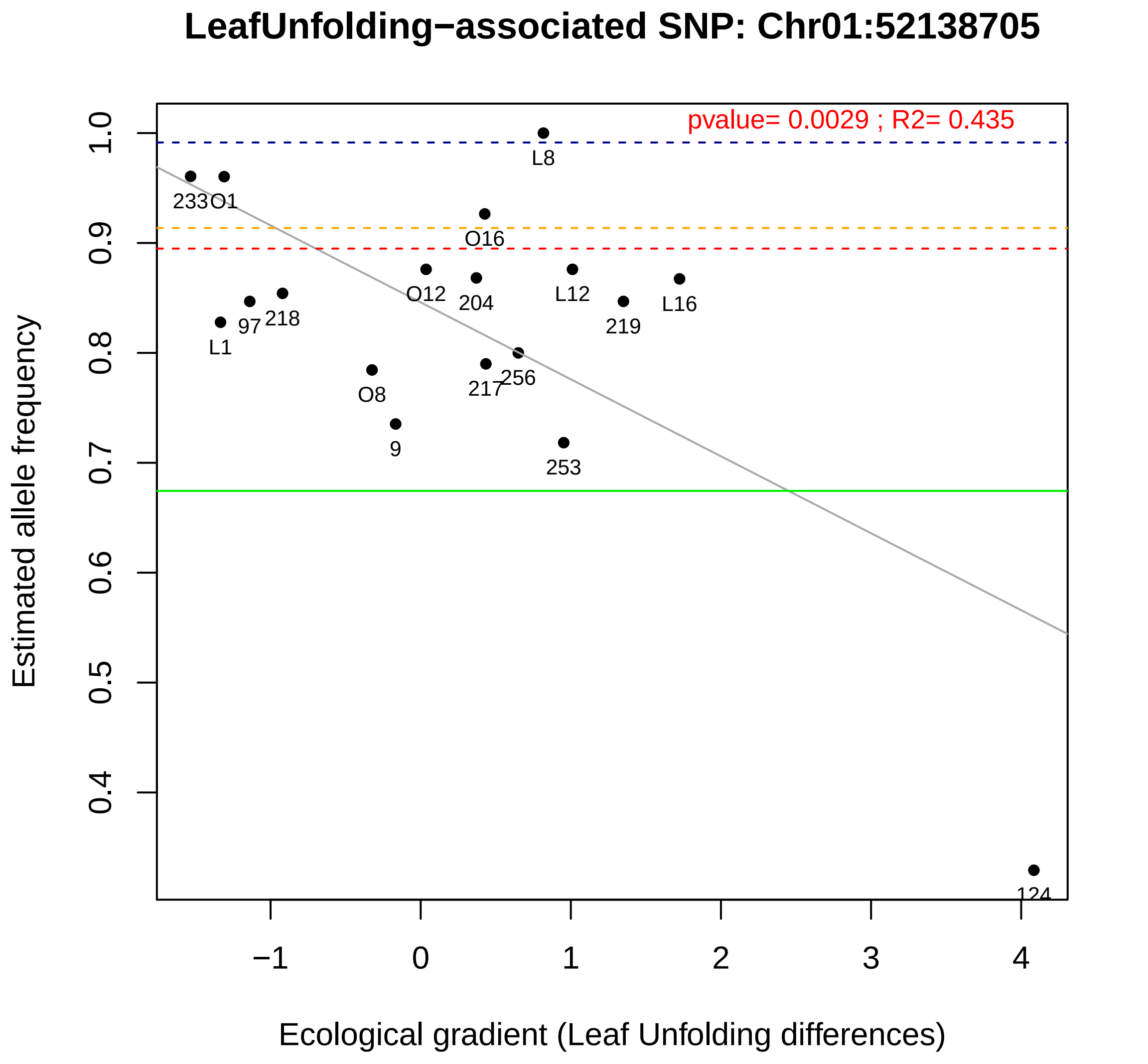
LeafUnfolding−associated SNP: Chr01:52138705
pvalue= 0.0029 ; R2= 0.435
Estimated allele frequency
Ecological gradient (Leaf Unfolding differences)
1.0
0.9
0.8
0.7
0.6
0.5
0.4
−1
0
1
2
3
4
L8
233
O1
O16
O12
204
L12
L16
219
218
97
L1
O8
217
256
9
253
124

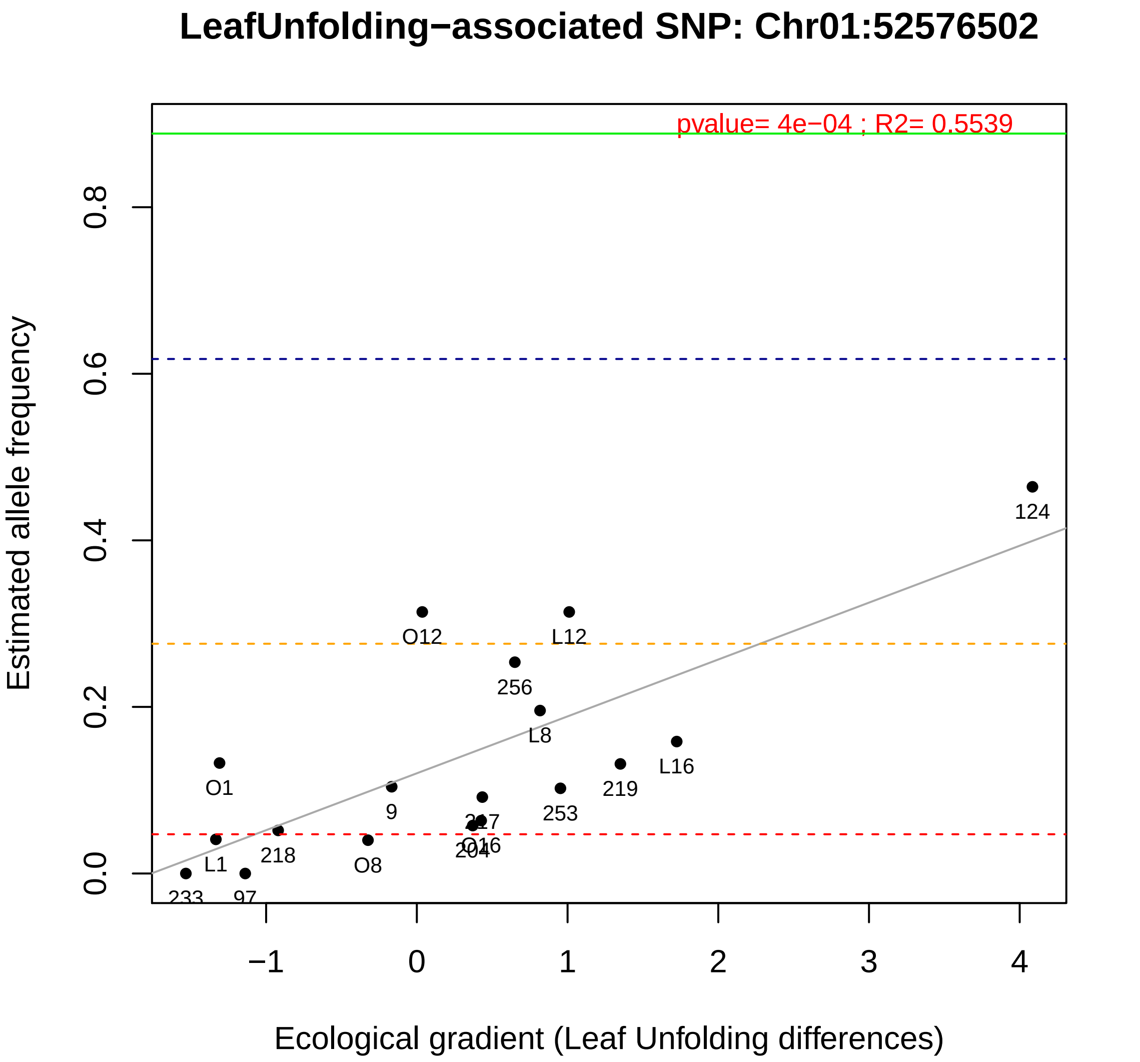
LeafUnfolding−associated SNP: Chr01:52576502
pvalue= 4e−04 ; R2= 0.5539
Estimated allele frequency
0.0
0.2
0.4
0.6
0.8
−1
0
1
2
3
4
Ecological gradient (Leaf Unfolding differences)
124
O12
L12
256
L8
L16
219
O1
9
253
218
L1
O8
233
97

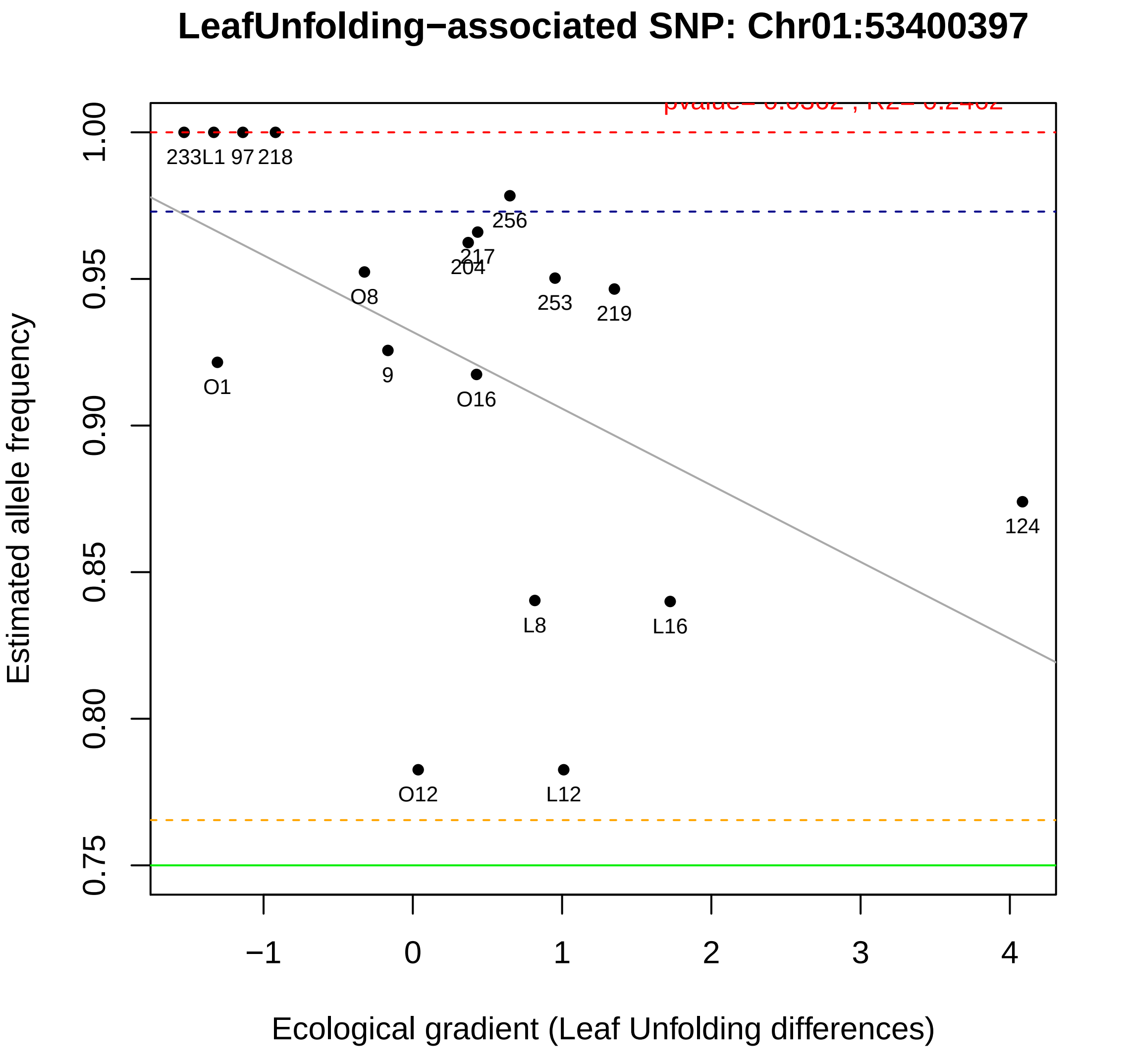
LeafUnfolding−associated SNP: Chr01:53400397
Estimated allele frequency
1.00
0.95
0.90
0.85
0.80
0.75
−1
0
1
2
3
4
Ecological gradient (Leaf Unfolding differences)
233
L1
97
218
256
217
204
O8
253
219
O1
9
O16
124
L8
L16
O12
L12

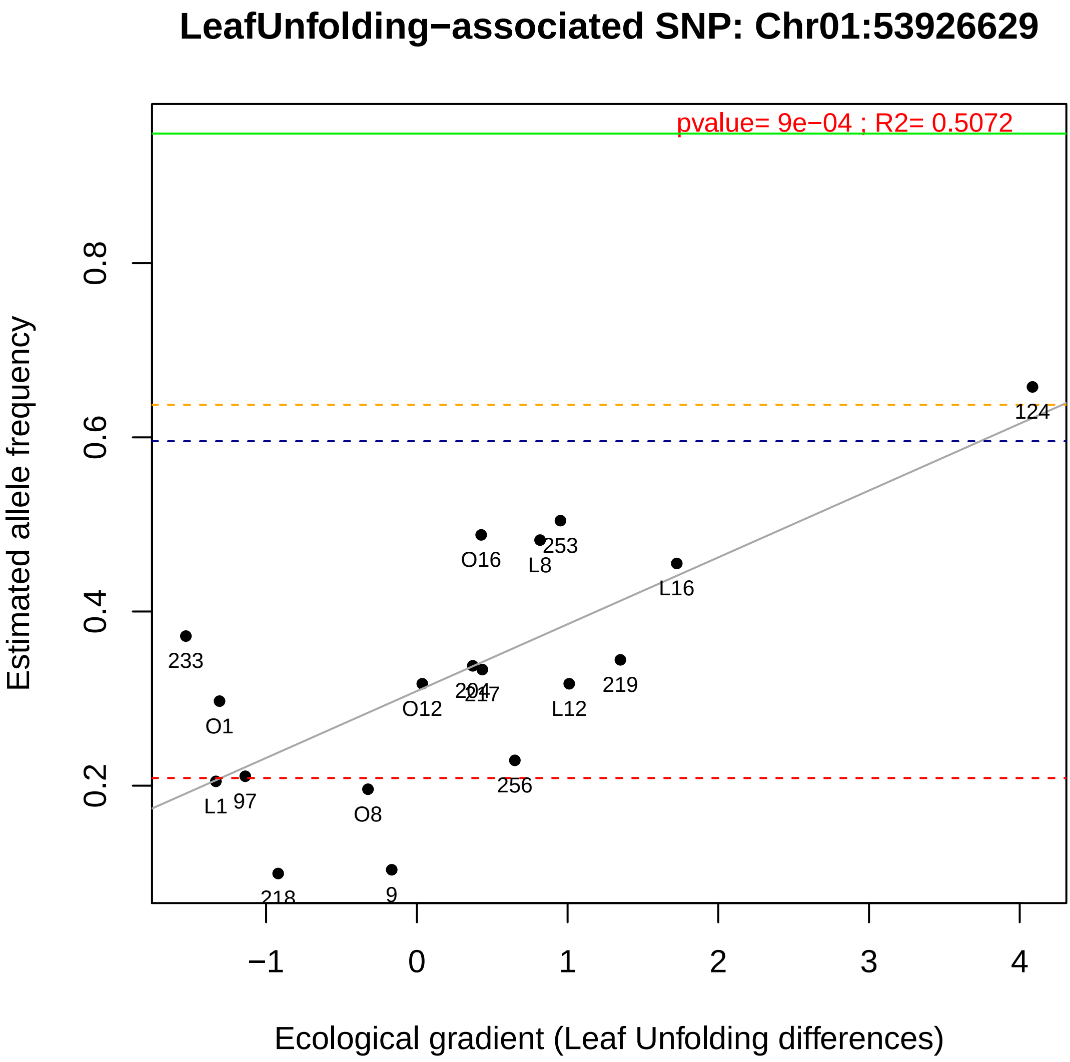

LeafUnfolding−associated SNP: Chr01:53926629
pvalue= 9e−04 ; R2= 0.5072
Estimated allele frequency
0.2
0.4
0.6
0.8
−1
0
1
2
3
4
Ecological gradient (Leaf Unfolding differences)
124
253
O16
L8
L16
233
219
O12
L12
O1
256
L1
97
O8
9

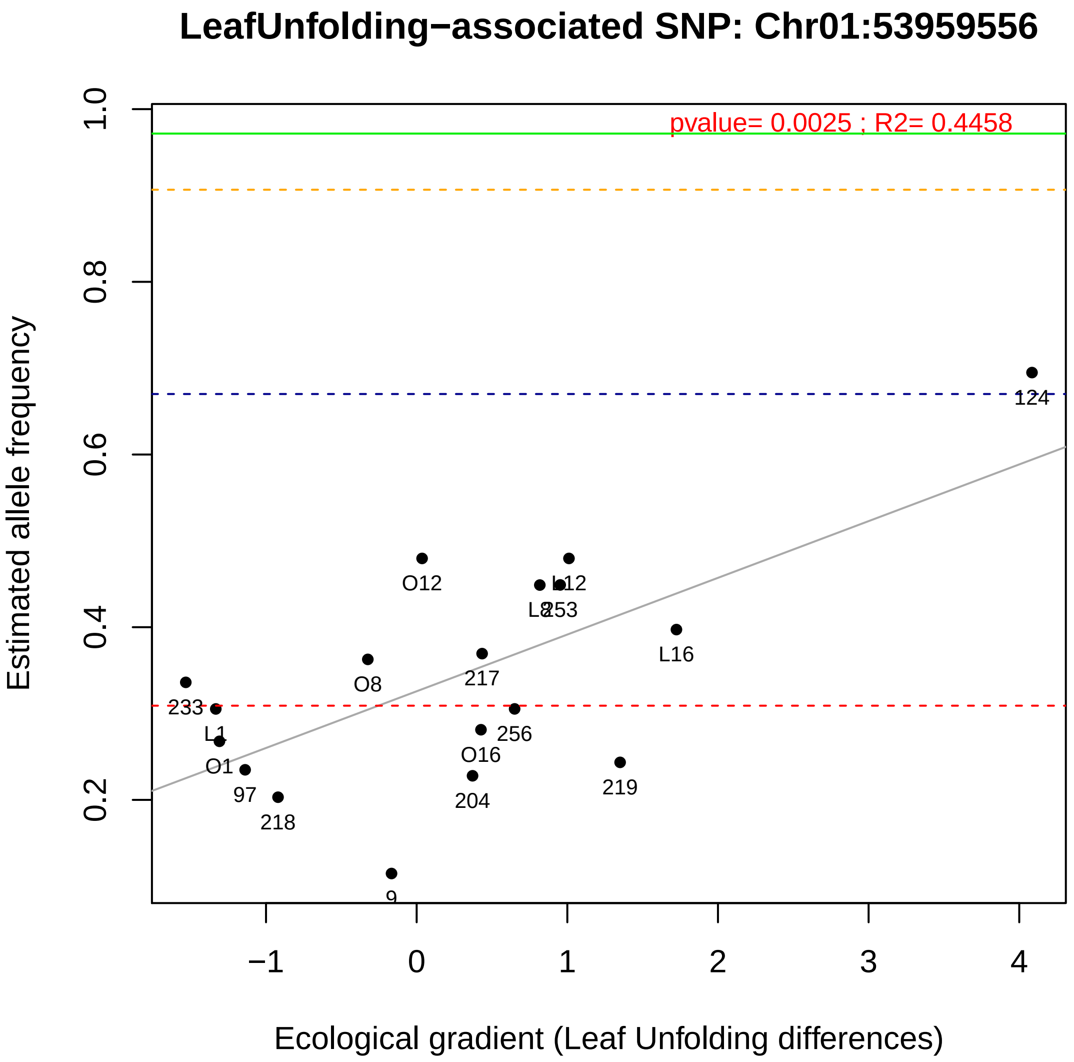
LeafUnfolding−associated SNP: Chr01:53959556
pvalue= 0.0025 ; R2= 0.4458
Estimated allele frequency
Ecological gradient (Leaf Unfolding differences)
0.2
0.4
0.6
0.8
1.0
−1
0
1
2
3
4
124
O12
L12
L8
253
L16
217
O8
233
256
L1
O16
O1
97
218
204
219
9

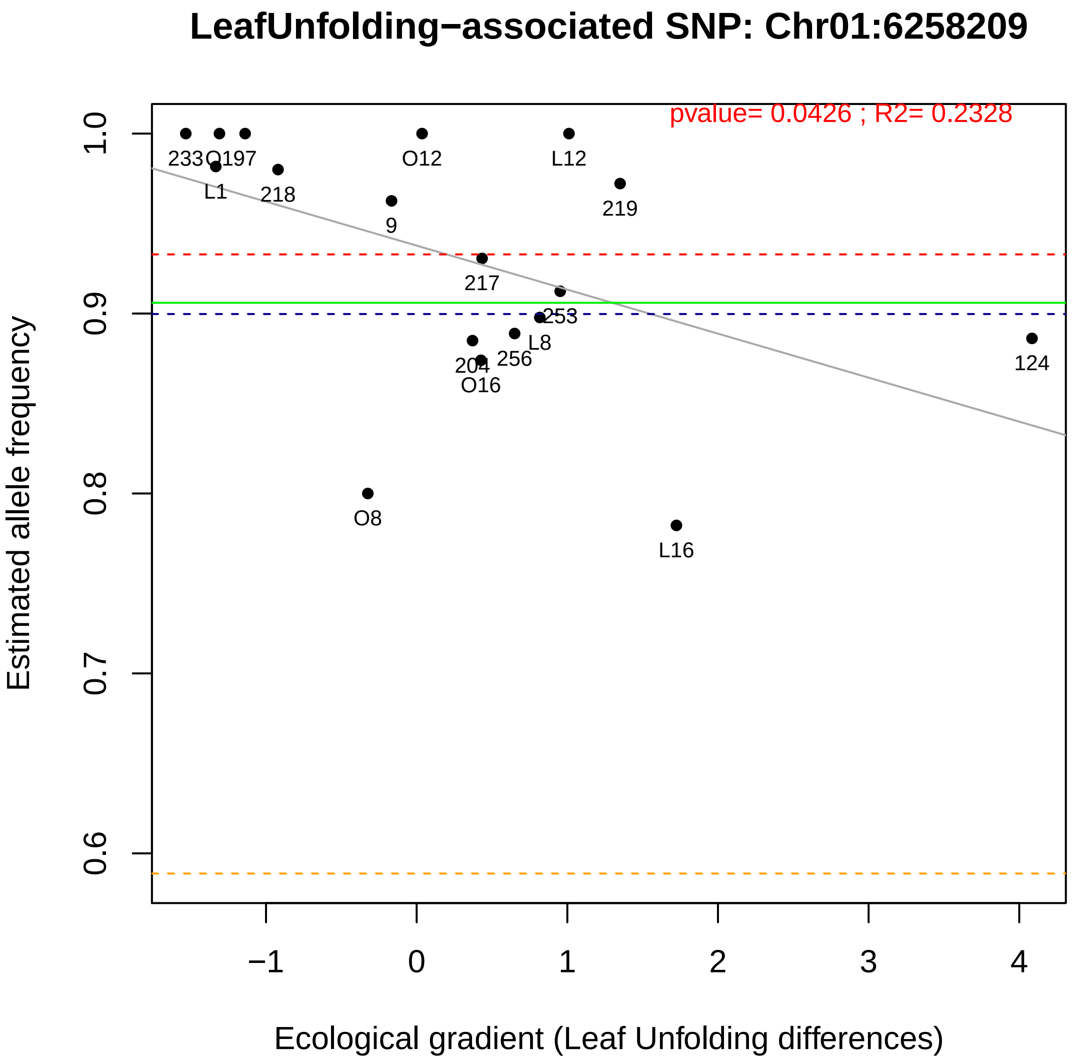

LeafUnfolding−associated SNP: Chr01:6258209
pvalue= 0.0426 ; R2= 0.2328
Estimated allele frequency
Ecological gradient (Leaf Unfolding differences)
1.0
0.9
0.8
0.7
0.6
−1
0
1
2
3
4
233
O197
L1
218
O12
9
L12
219
217
253
L8
256
204
O16
124
O8
L16

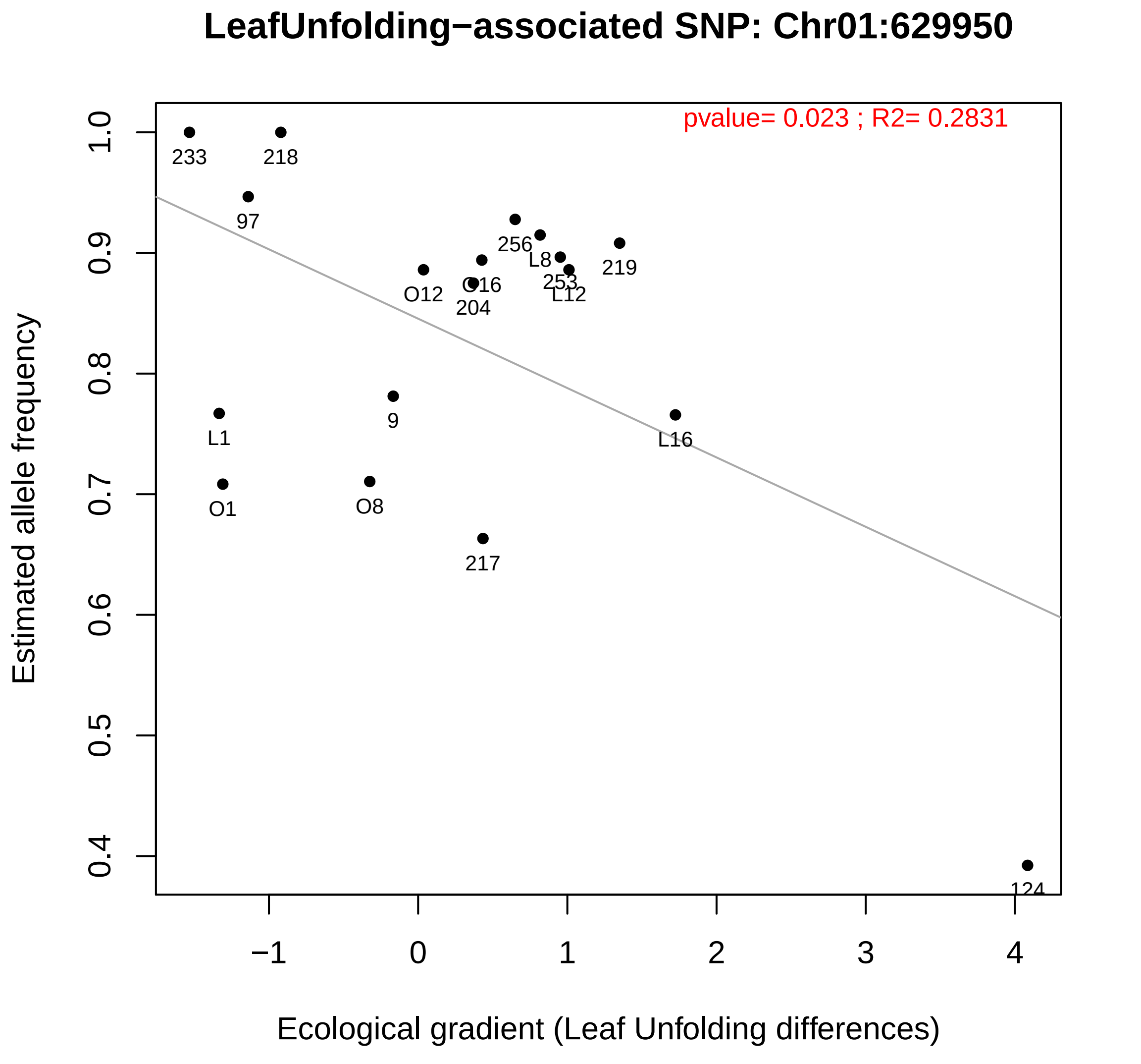

LeafUnfolding−associated SNP: Chr01:629950
pvalue= 0.023 ; R2= 0.2831
Estimated allele frequency
Ecological gradient (Leaf Unfolding differences)
0.4
0.5
0.6
0.7
0.8
0.9
1.0
−1
0
1
2
3
4
233
218
97
256
L8
219
O12
O16
253
204
L12
9
L1
L16
O1
O8
217
124

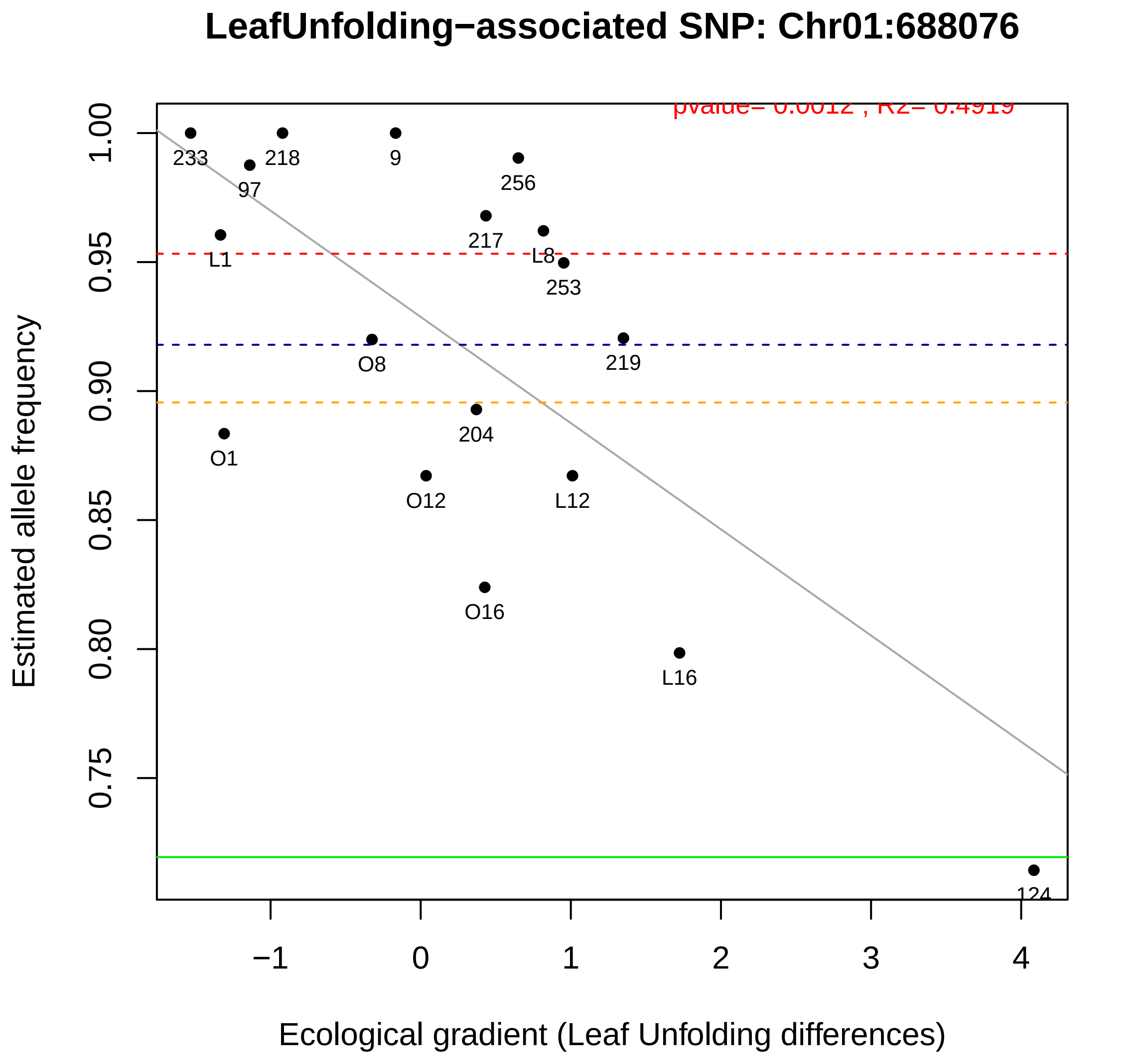

LeafUnfolding−associated SNP: Chr01:688076
pvalue= 0.0012 , R2= 0.4919
Estimated allele frequency
Ecological gradient (Leaf Unfolding differences)
1.00
0.95
0.90
0.85
0.80
0.75
−1
0
1
2
3
4
233
218
9
97
256
217
L1
L8
253
O8
219
204
O1
O12
L12
O16
L16
124

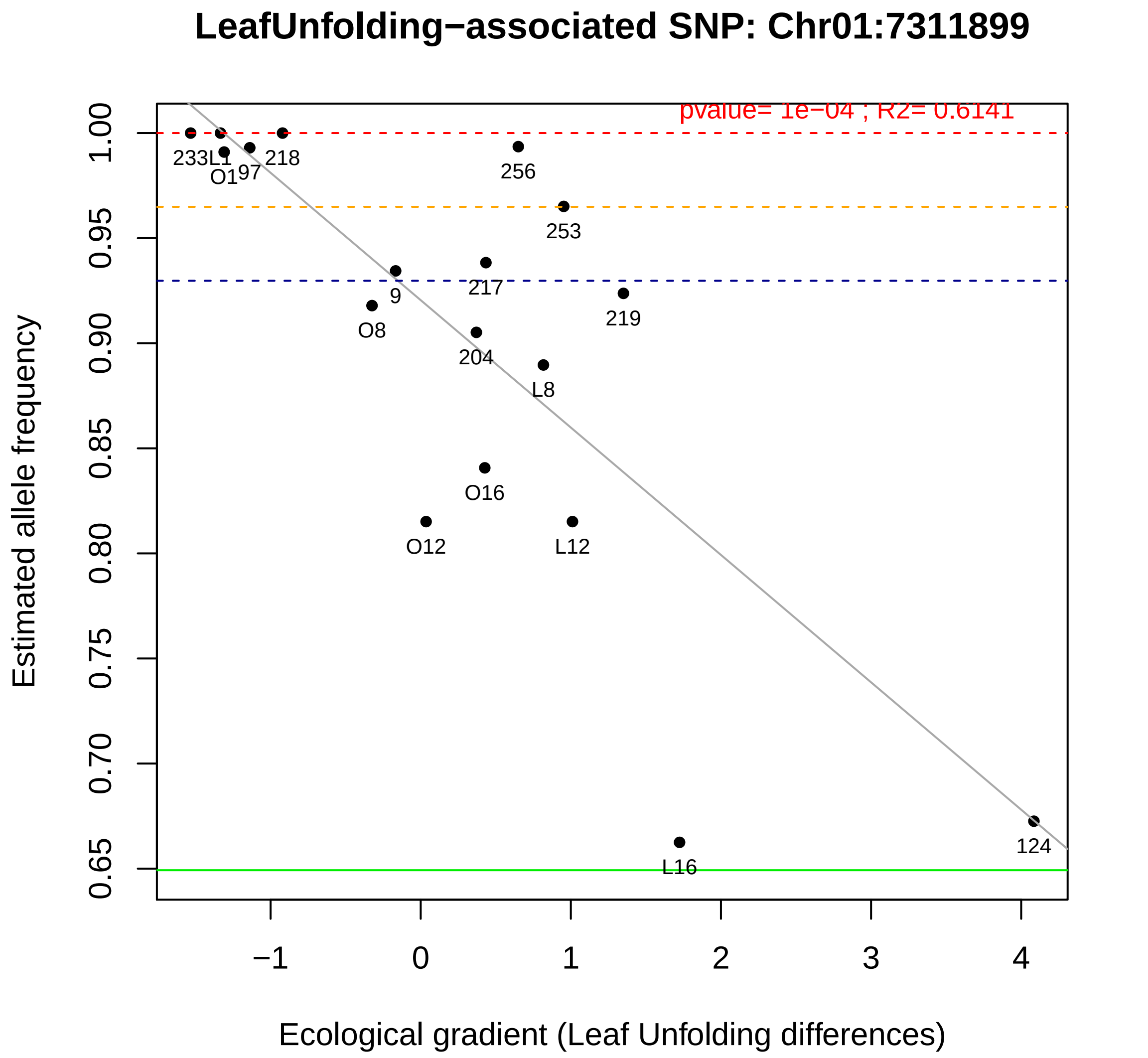

LeafUnfolding−associated SNP: Chr01:7311899
pvalue= 1e−04 ; R2= 0.6141
Estimated allele frequency
Ecological gradient (Leaf Unfolding differences)
0.65
0.70
0.75
0.80
0.85
0.90
0.95
1.00
−1
0
1
2
3
4
233
L1
O1
97
218
256
253
9
217
O8
219
204
L8
O16
O12
L12
L16
124

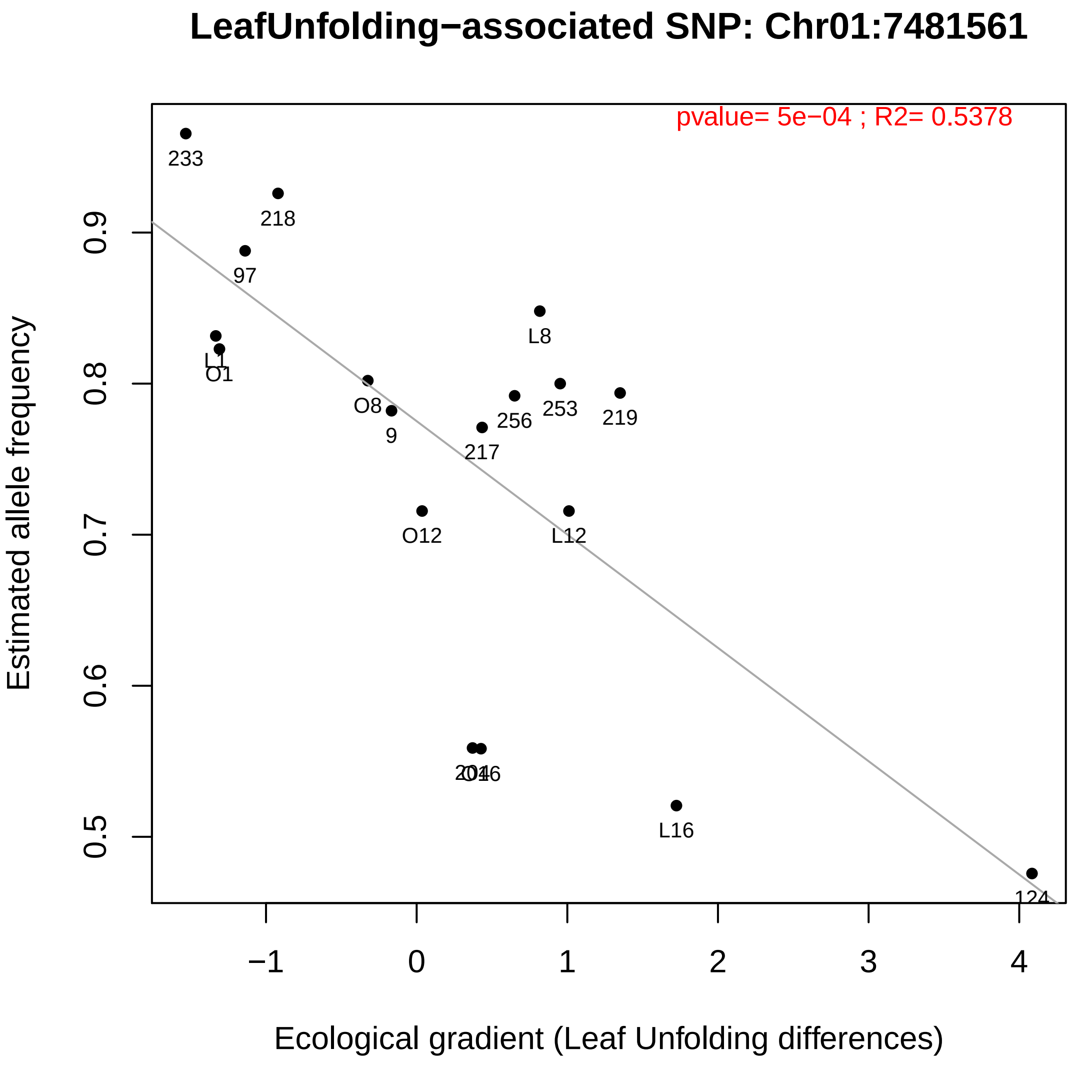

LeafUnfolding−associated SNP: Chr01:7481561
pvalue= 5e−04 ; R2= 0.5378
233
218
97
L1
O1
O8
9
217
256
L8
253
219
O12
L12
L16
124
Estimated allele frequency
0.5
0.6
0.7
0.8
0.9
−1
0
1
2
3
4
Ecological gradient (Leaf Unfolding differences)

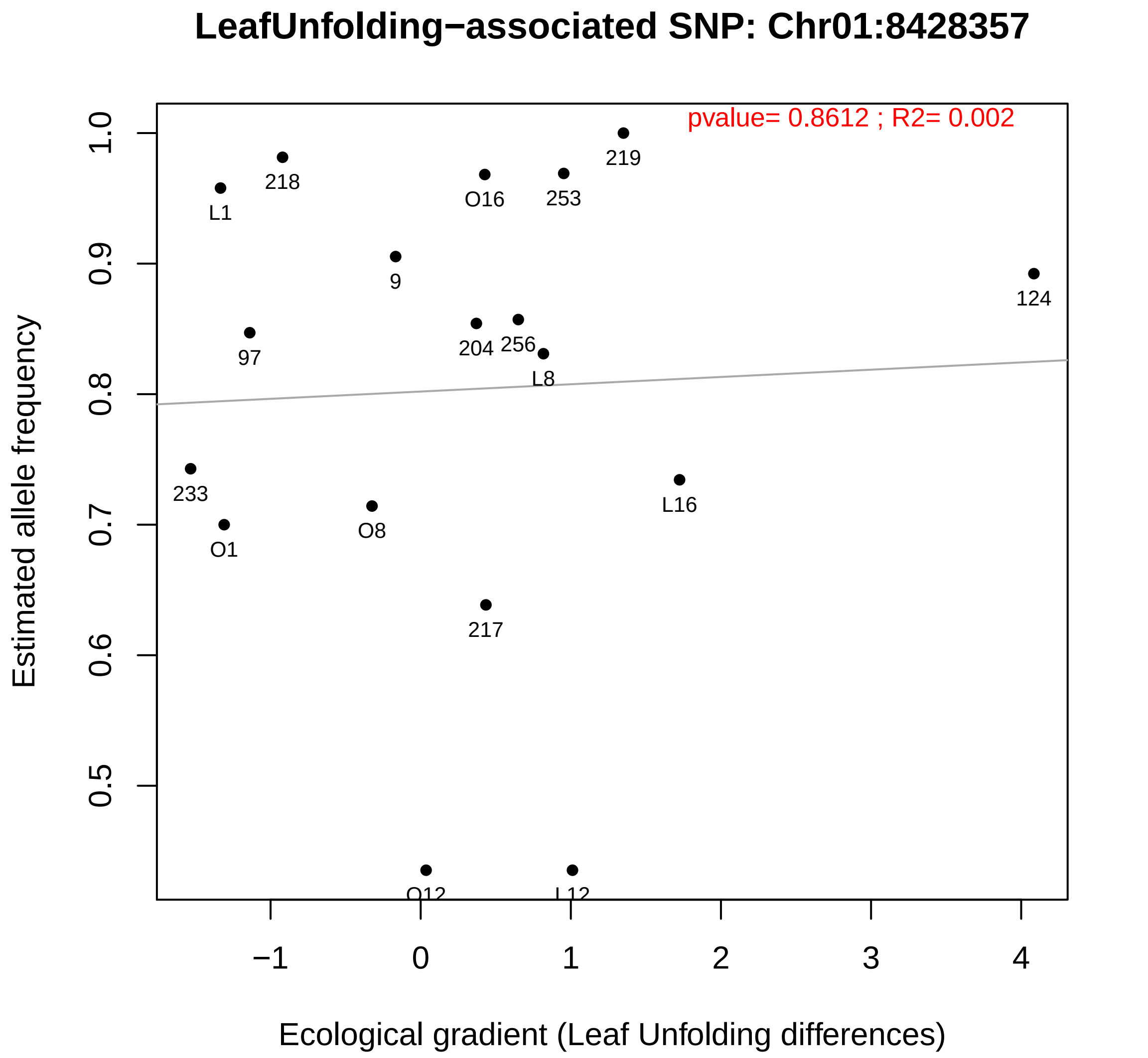
LeafUnfolding−associated SNP: Chr01:8428357
pvalue= 0.8612 ; R2= 0.002
Estimated allele frequency
Ecological gradient (Leaf Unfolding differences)
0.5
0.6
0.7
0.8
0.9
1.0
−1
0
1
2
3
4
219
218
L1
O16
253
9
124
97
204
256
L8
233
O1
O8
L16
217
O12
L12

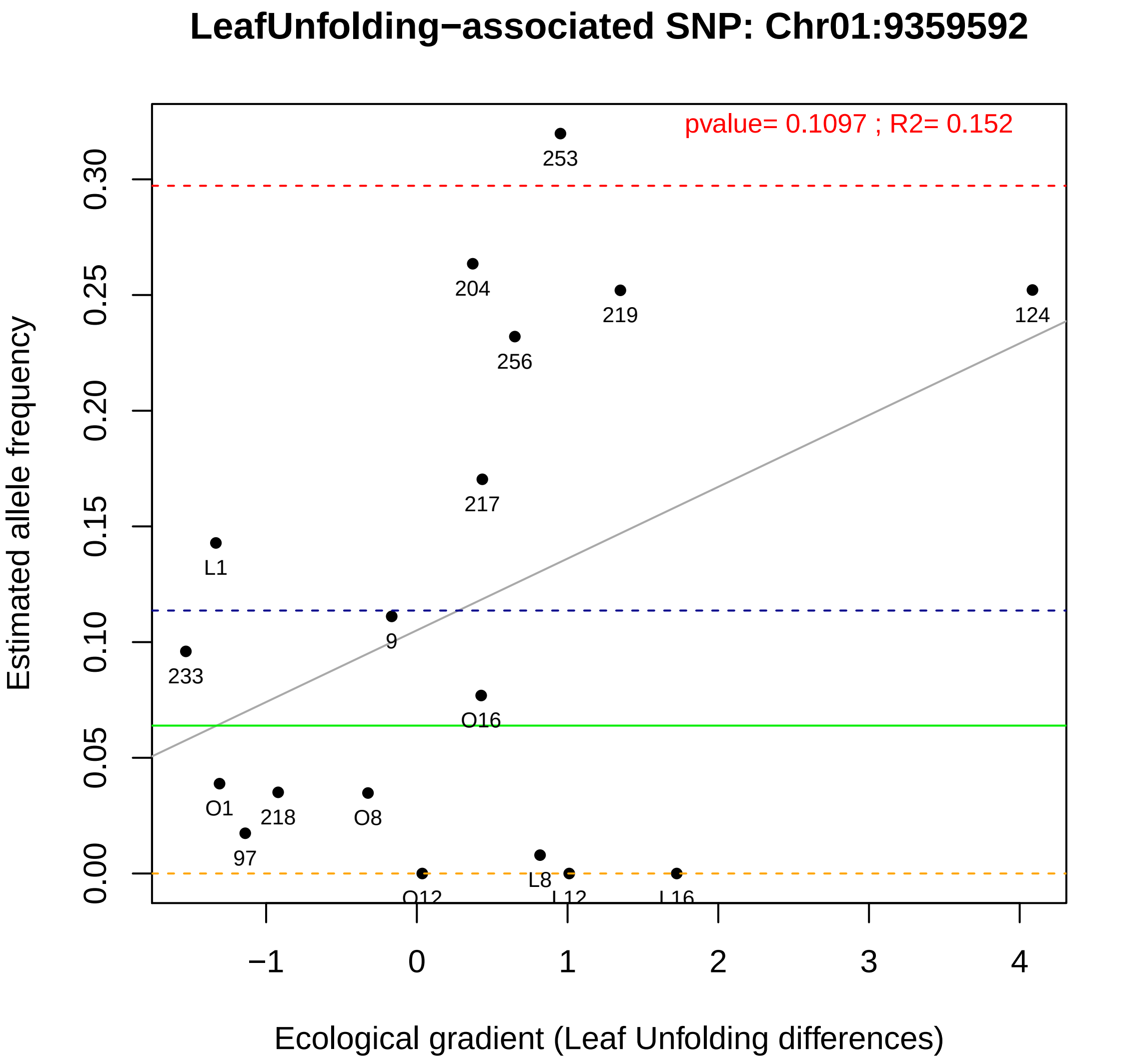

LeafUnfolding−associated SNP: Chr01:9359592
pvalue= 0.1097 ; R2= 0.152
Estimated allele frequency
0.00
0.05
0.10
0.15
0.20
0.25
0.30
−1
0
1
2
3
4
Ecological gradient (Leaf Unfolding differences)
253
204
219
124
256
217
L1
9
233
O16
O1
218
O8
97
L8

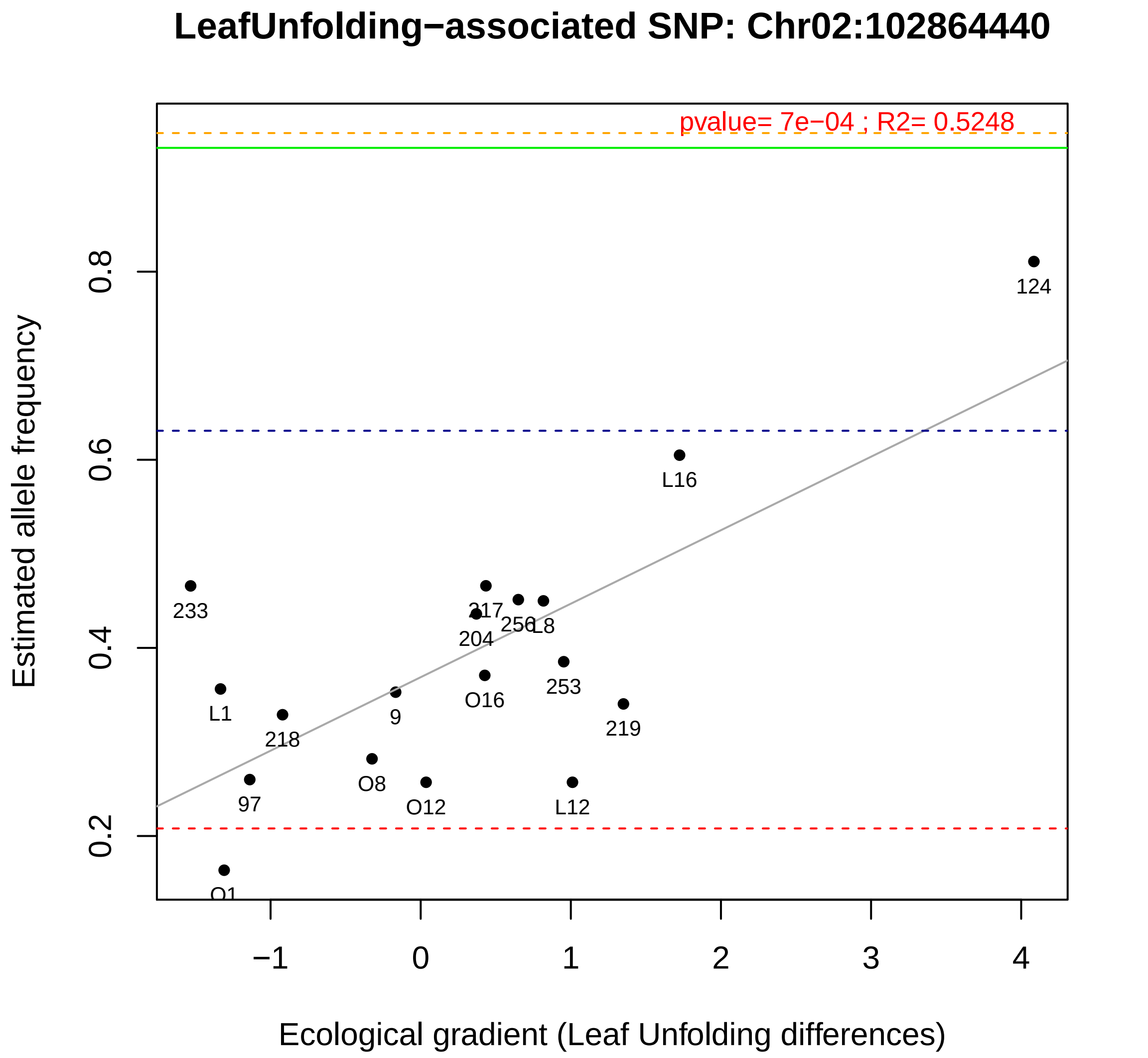
LeafUnfolding−associated SNP: Chr02:102864440
pvalue= 7e−04 ; R2= 0.5248
Estimated allele frequency
Ecological gradient (Leaf Unfolding differences)
0.2
0.4
0.6
0.8
−1
0
1
2
3
4
124
L16
233
217
256
L8
204
253
O16
L1
9
219
218
O8
97
O12
L12
O1

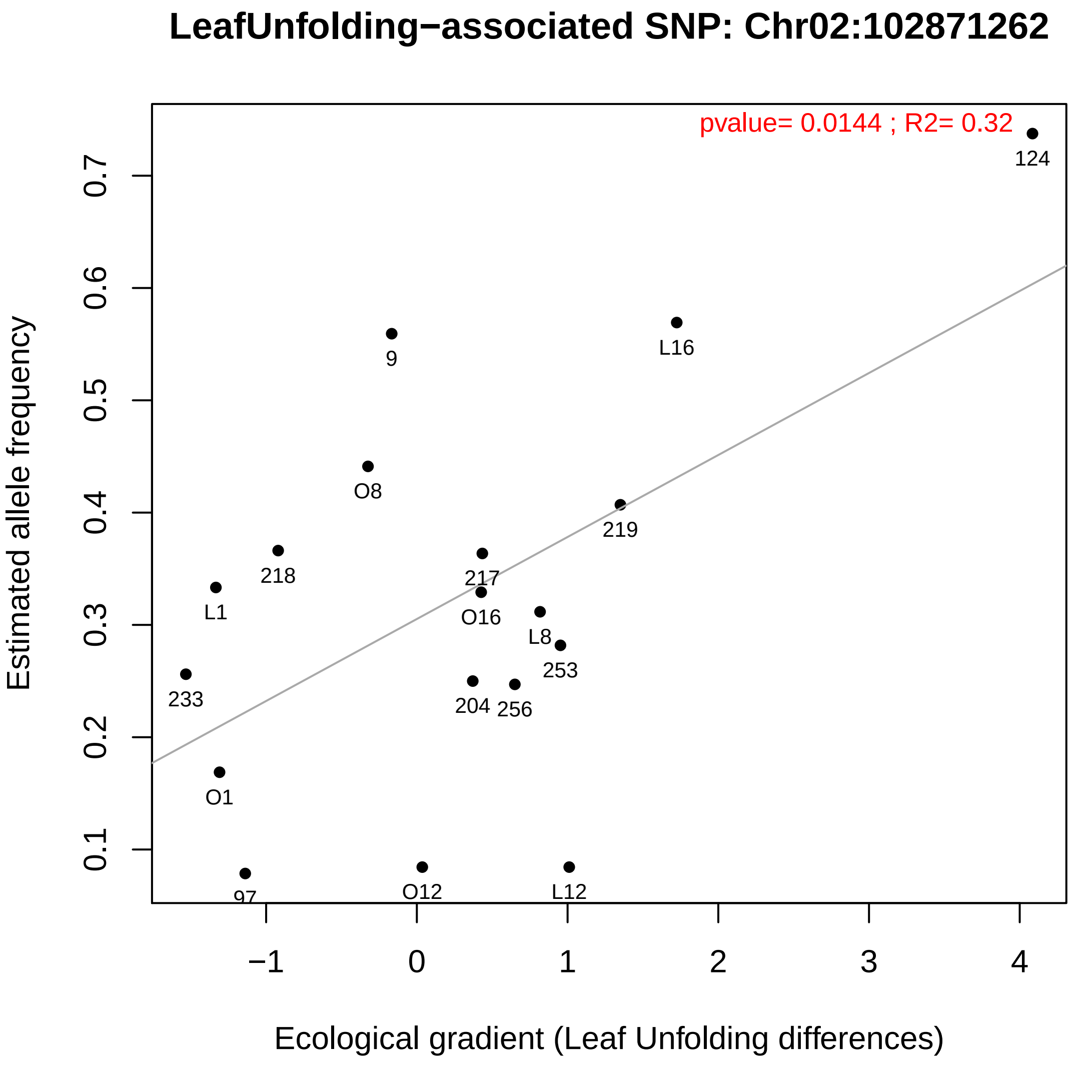
LeafUnfolding−associated SNP: Chr02:102871262
pvalue= 0.0144 ; R2= 0.32
124
9
L16
O8
219
218
217
L1
O16
L8
253
233
204
256
O1
97
O12
L12
0.1
0.2
0.3
0.4
0.5
0.6
0.7
Estimated allele frequency
−1
0
1
2
3
4
Ecological gradient (Leaf Unfolding differences)

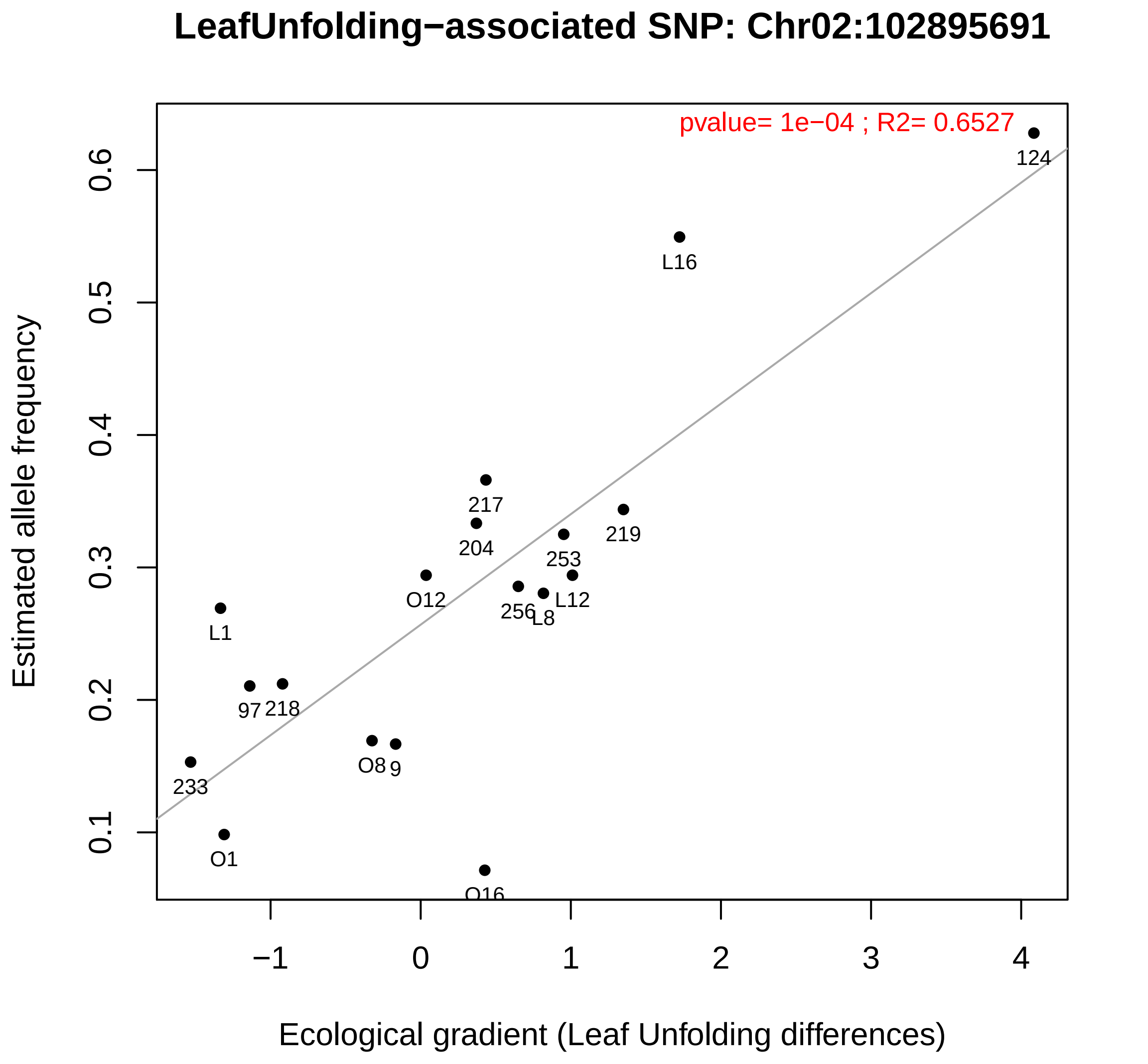
LeafUnfolding−associated SNP: Chr02:102895691
pvalue= 1e−04 ; R2= 0.6527
Estimated allele frequency
Ecological gradient (Leaf Unfolding differences)
0.1
0.2
0.3
0.4
0.5
0.6
−1
0
1
2
3
4
124
L16
217
204
219
253
O12
256
L8
L12
L1
97
218
O8
9
233
O1
O16

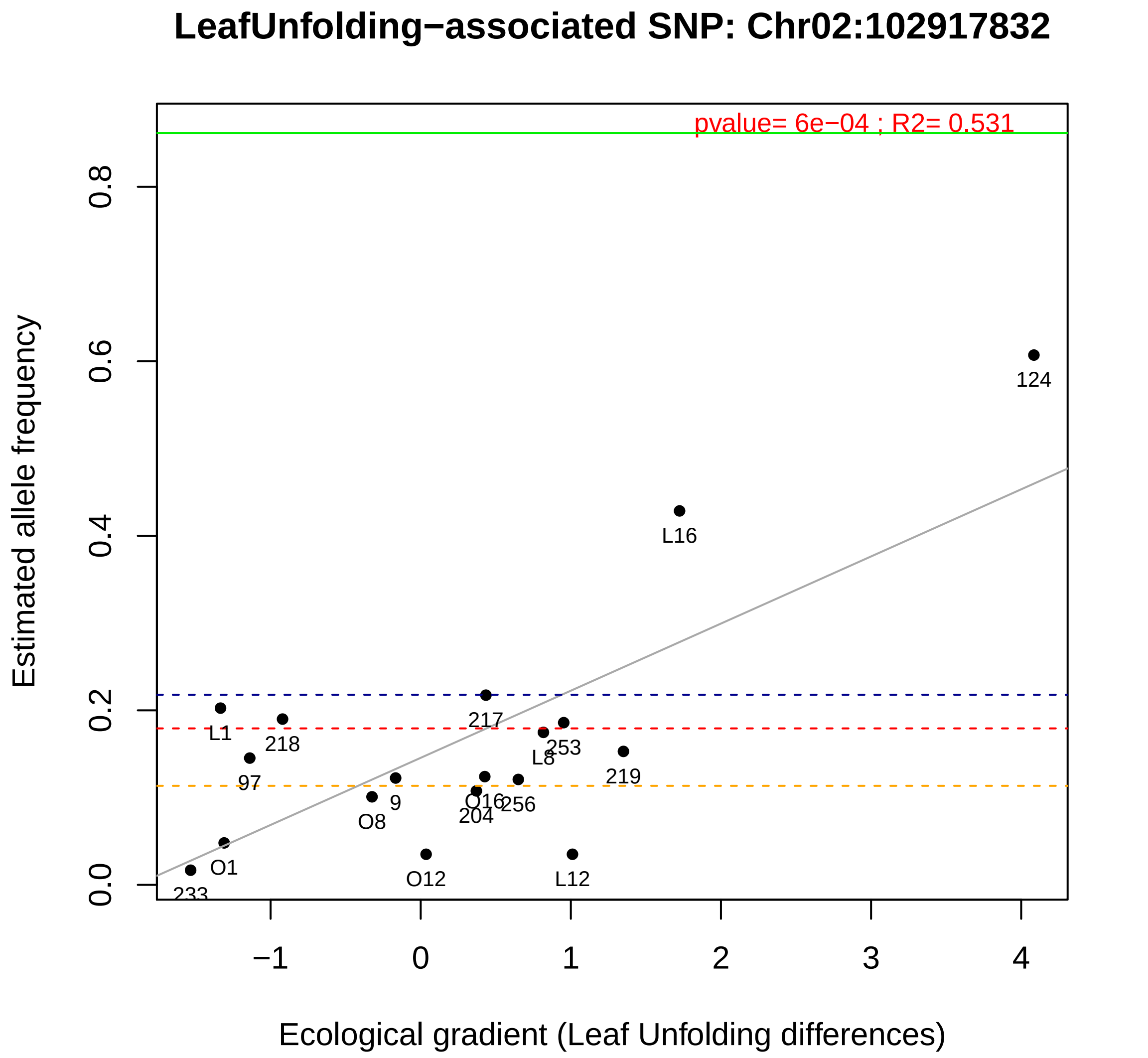

LeafUnfolding−associated SNP: Chr02:102917832
pvalue= 6e−04 ; R2= 0.531
Estimated allele frequency
0.0
0.2
0.4
0.6
0.8
Ecological gradient (Leaf Unfolding differences)
−1
0
1
2
3
4
124
L16
217
L1
218
253
L8
97
219
9
O16
256
204
O8
O1
O12
L12
233

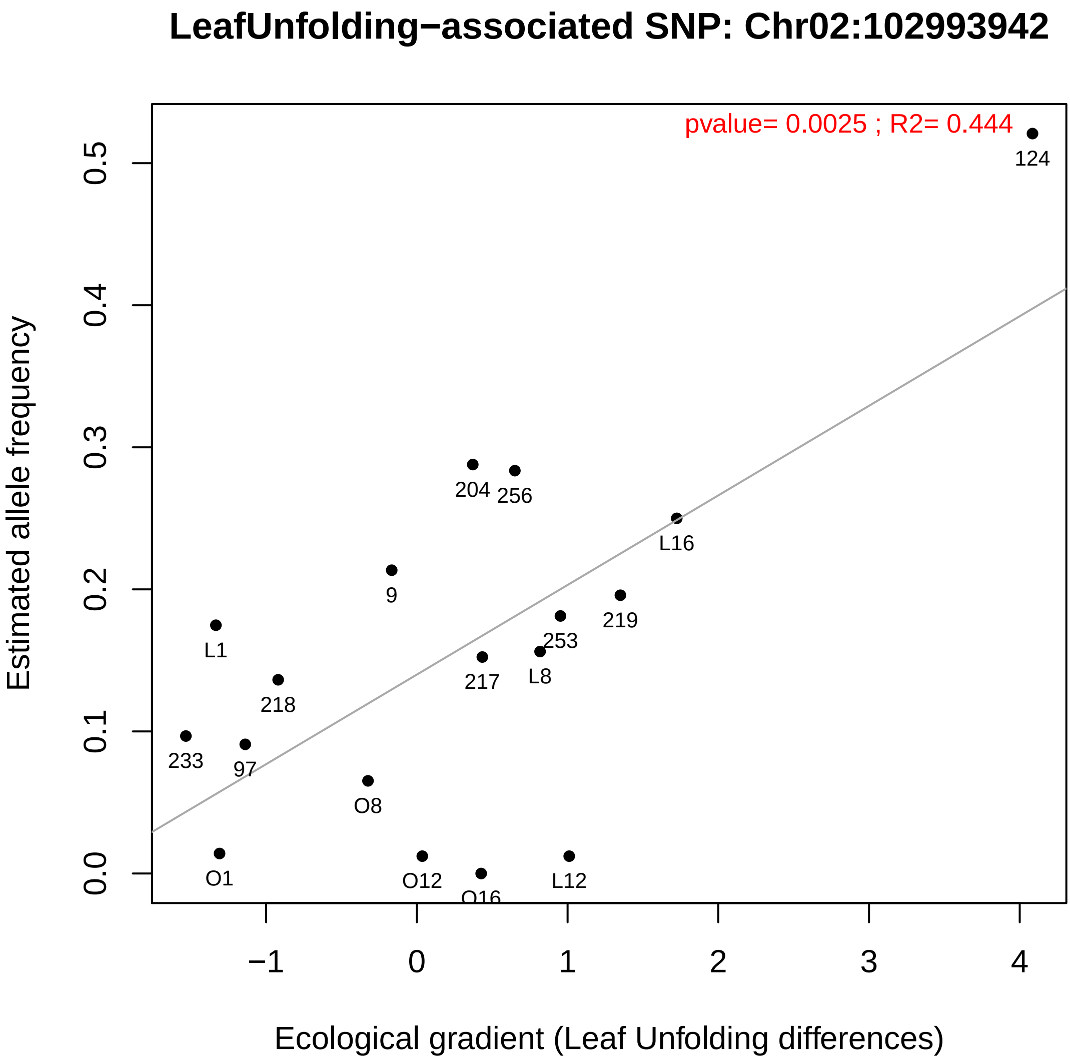
LeafUnfolding−associated SNP: Chr02:102993942
pvalue= 0.0025 ; R2= 0.444
Estimated allele frequency
0.0
0.1
0.2
0.3
0.4
0.5
−1
0
1
2
3
4
Ecological gradient (Leaf Unfolding differences)
124
204
256
L16
9
219
253
L1
217
L8
218
233
97
O8
O1
O12
O16
L12

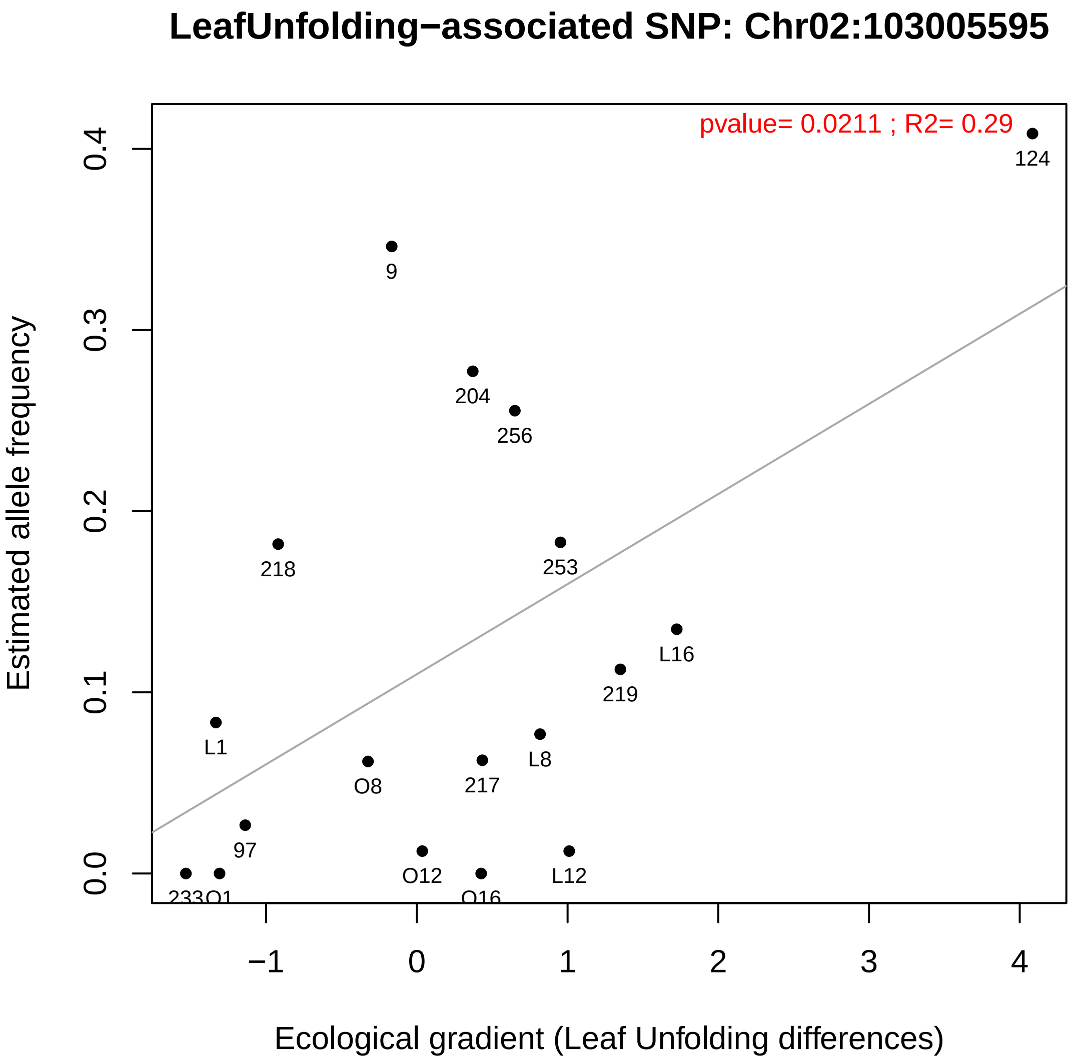

LeafUnfolding−associated SNP: Chr02:103005595
pvalue= 0.0211 ; R2= 0.29
124
9
204
256
218
253
L16
219
L1
L8
O8
217
97
O12
L12
233
O1
O16
Estimated allele frequency
0.0
0.1
0.2
0.3
0.4
−1
0
1
2
3
4
Ecological gradient (Leaf Unfolding differences)

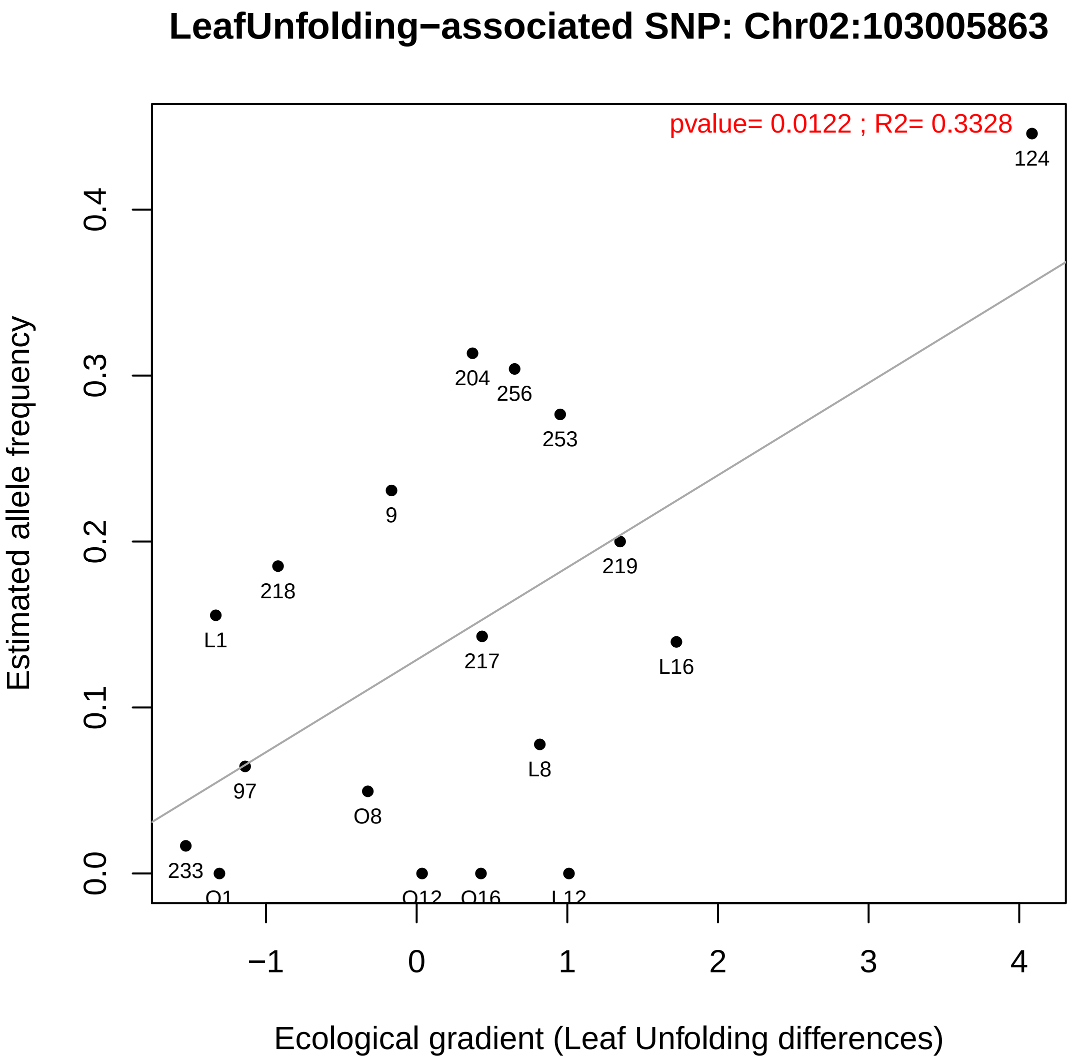
LeafUnfolding−associated SNP: Chr02:103005863
pvalue= 0.0122 ; R2= 0.3328
Estimated allele frequency
Ecological gradient (Leaf Unfolding differences)
0.0
0.1
0.2
0.3
0.4
−1
0
1
2
3
4
124
204
256
253
9
219
218
L1
217
L16
L8
97
O8
233
O1
O12
O16
L12

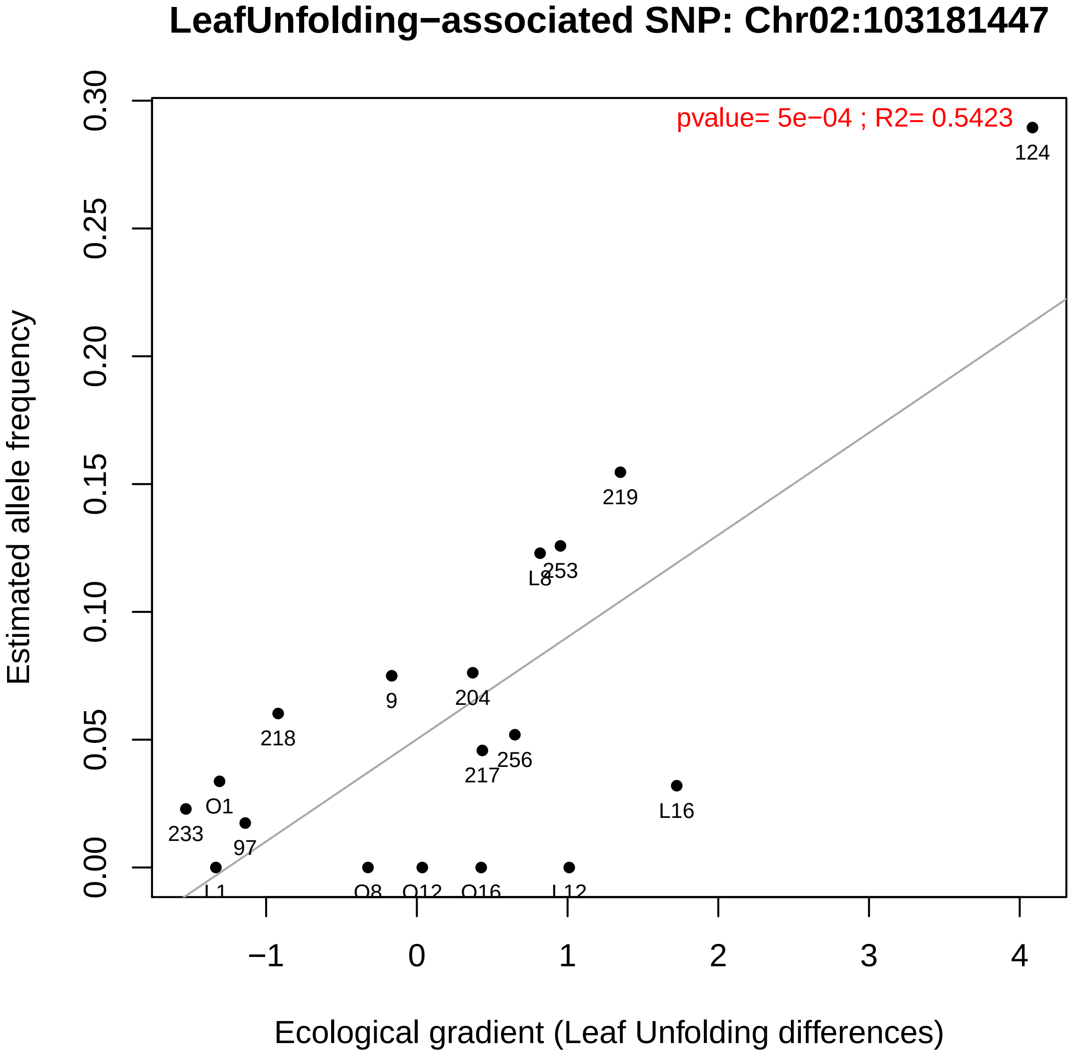

LeafUnfolding−associated SNP: Chr02:103181447
pvalue= 5e−04 ; R2= 0.5423
Estimated allele frequency
0.00
0.05
0.10
0.15
0.20
0.25
0.30
−1
0
1
2
3
4
Ecological gradient (Leaf Unfolding differences)
124
219
L8
253
9
204
218
256
217
L16
O1
233
97
L1
O8
O12
O16
L12

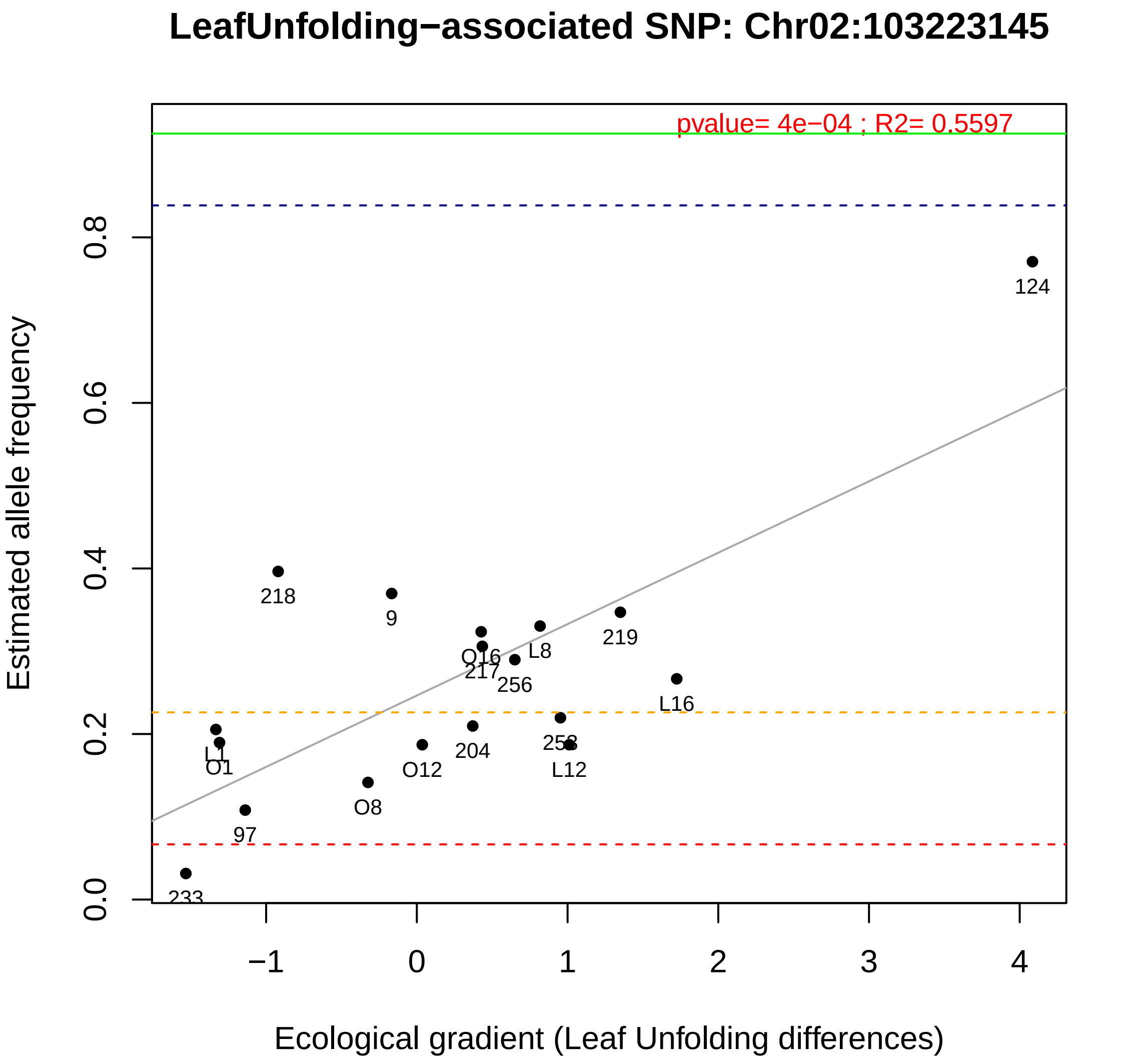

LeafUnfolding−associated SNP: Chr02:103223145
pvalue= 4e−04 ; R2= 0.5597
Estimated allele frequency
0.0
0.2
0.4
0.6
0.8
−1
0
1
2
3
4
Ecological gradient (Leaf Unfolding differences)
124
218
9
O16
217
L8
256
219
L16
L1
O1
O12
204
L12
O8
97
233

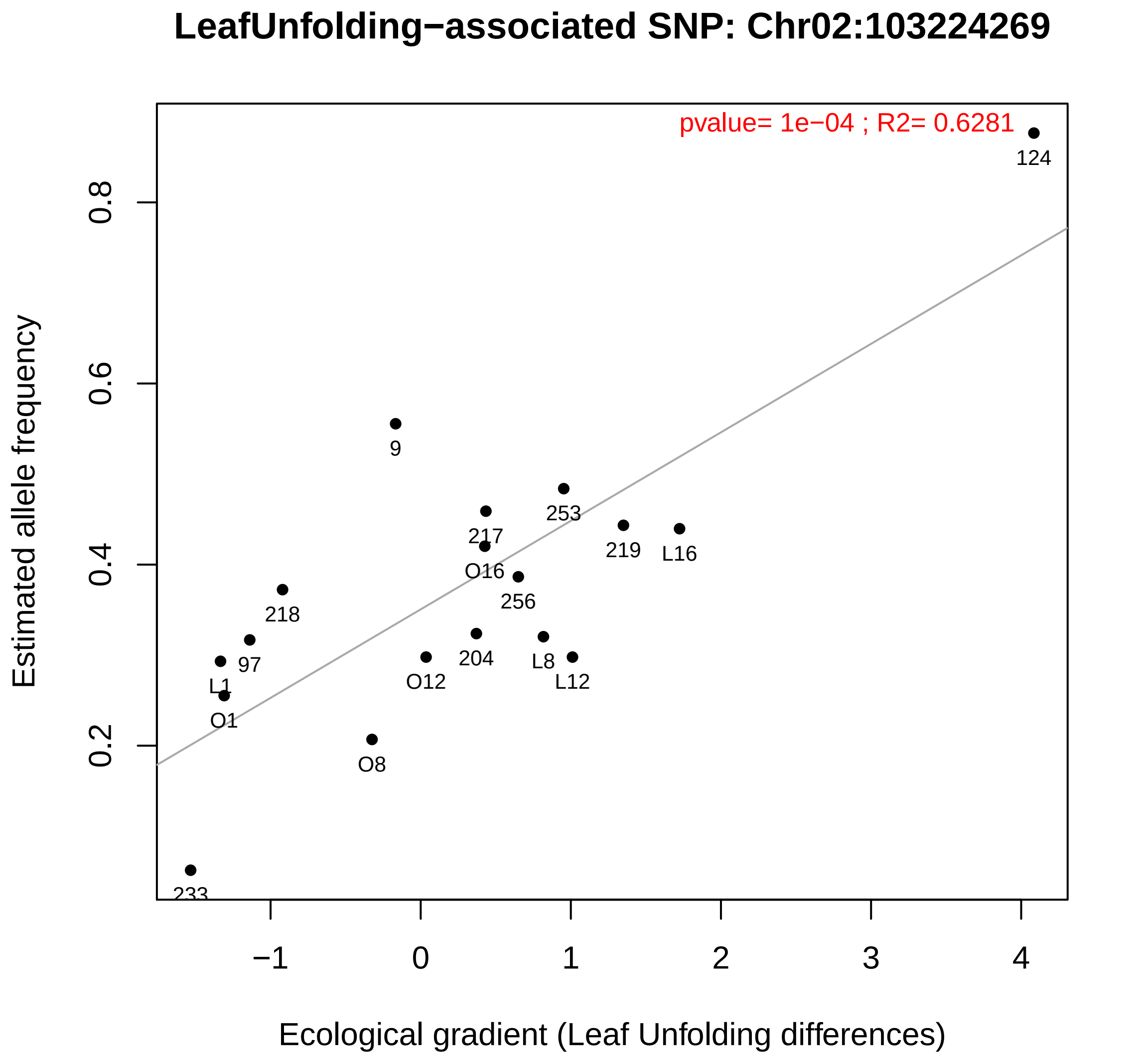

LeafUnfolding−associated SNP: Chr02:103224269
pvalue= 1e−04 ; R2= 0.6281
Estimated allele frequency
0.2
0.4
0.6
0.8
−1
0
1
2
3
4
Ecological gradient (Leaf Unfolding differences)
124
9
253
217
O16
219
L16
256
218
97
L1
O1
204
L8
L12
O12
O8
233

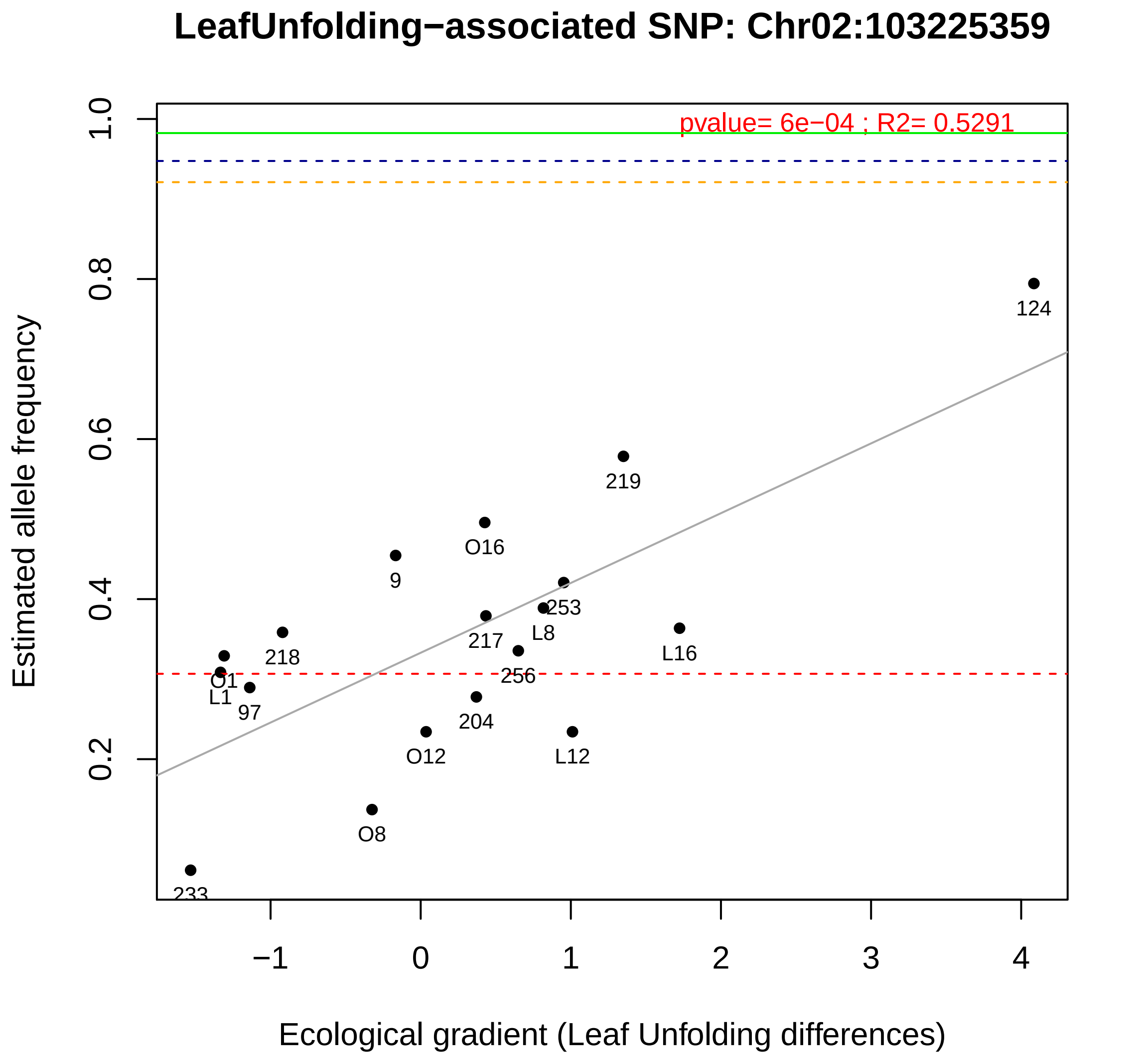
LeafUnfolding−associated SNP: Chr02:103225359
pvalue= 6e−04 ; R2= 0.5291
Estimated allele frequency
Ecological gradient (Leaf Unfolding differences)
0.2
0.4
0.6
0.8
1.0
−1
0
1
2
3
4
124
219
O16
9
253
L8
217
L16
218
256
O1
L1
97
204
O12
L12
O8
233

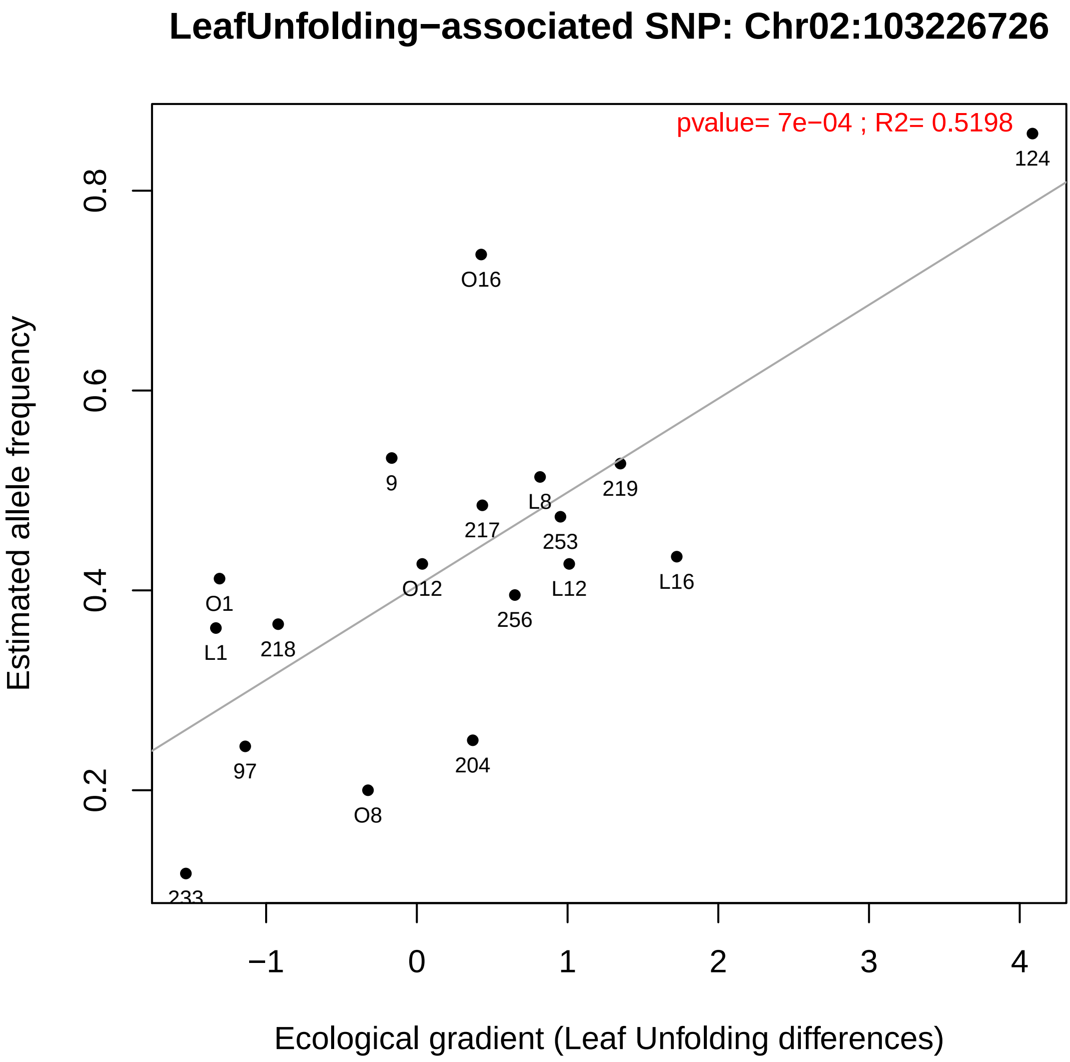

LeafUnfolding−associated SNP: Chr02:103226726
pvalue= 7e−04 ; R2= 0.5198
Estimated allele frequency
0.2
0.4
0.6
0.8
Ecological gradient (Leaf Unfolding differences)
−1
0
1
2
3
4
124
O16
9
L8
219
217
253
O12
L12
L16
O1
256
L1
218
97
204
O8
233

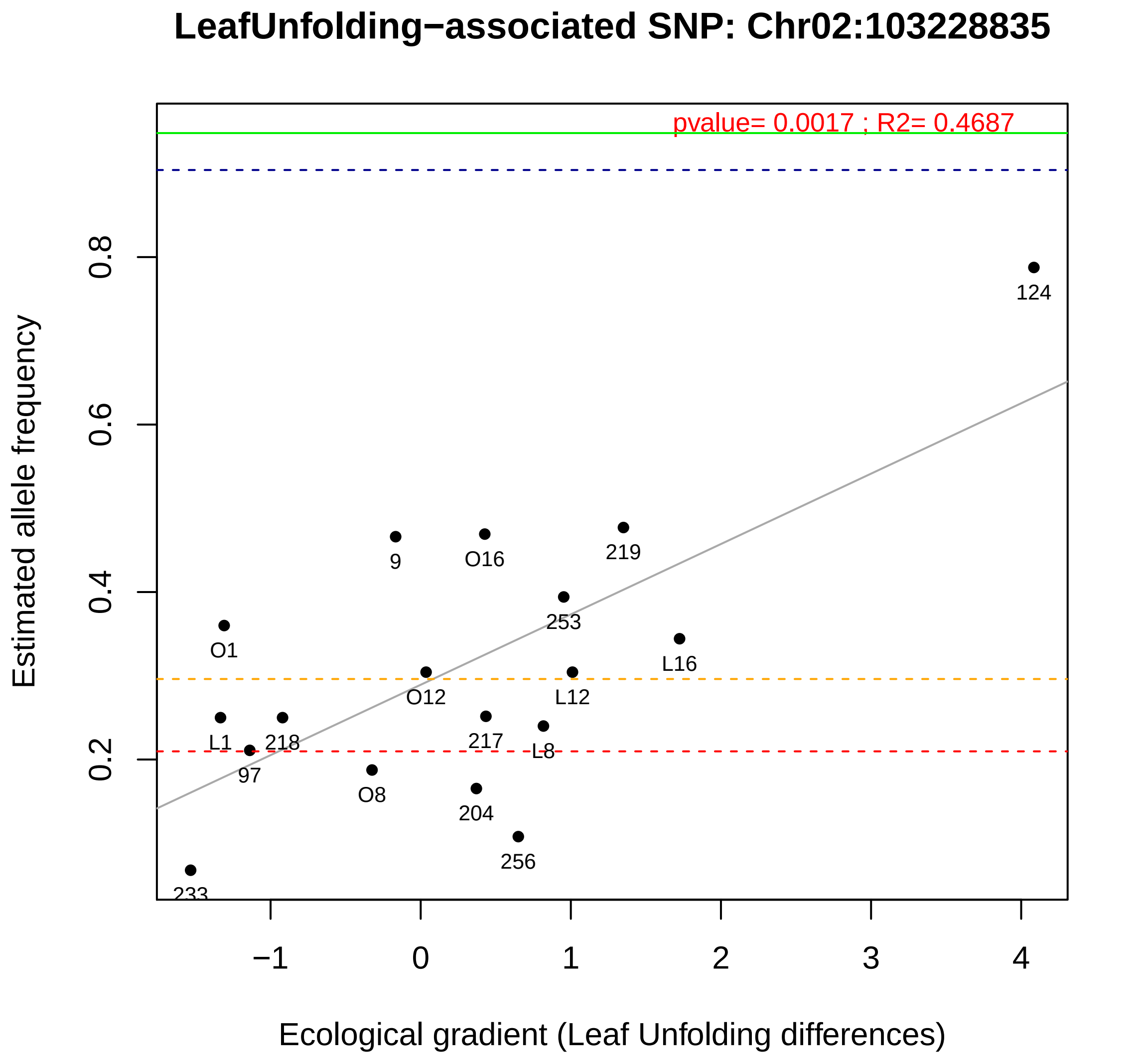

LeafUnfolding−associated SNP: Chr02:103228835
pvalue= 0.0017 ; R2= 0.4687
Estimated allele frequency
0.2
0.4
0.6
0.8
−1
0
1
2
3
4
Ecological gradient (Leaf Unfolding differences)
124
219
O16
9
253
O1
L16
O12
L12
L1
218
97
217
L8
O8
204
256
233

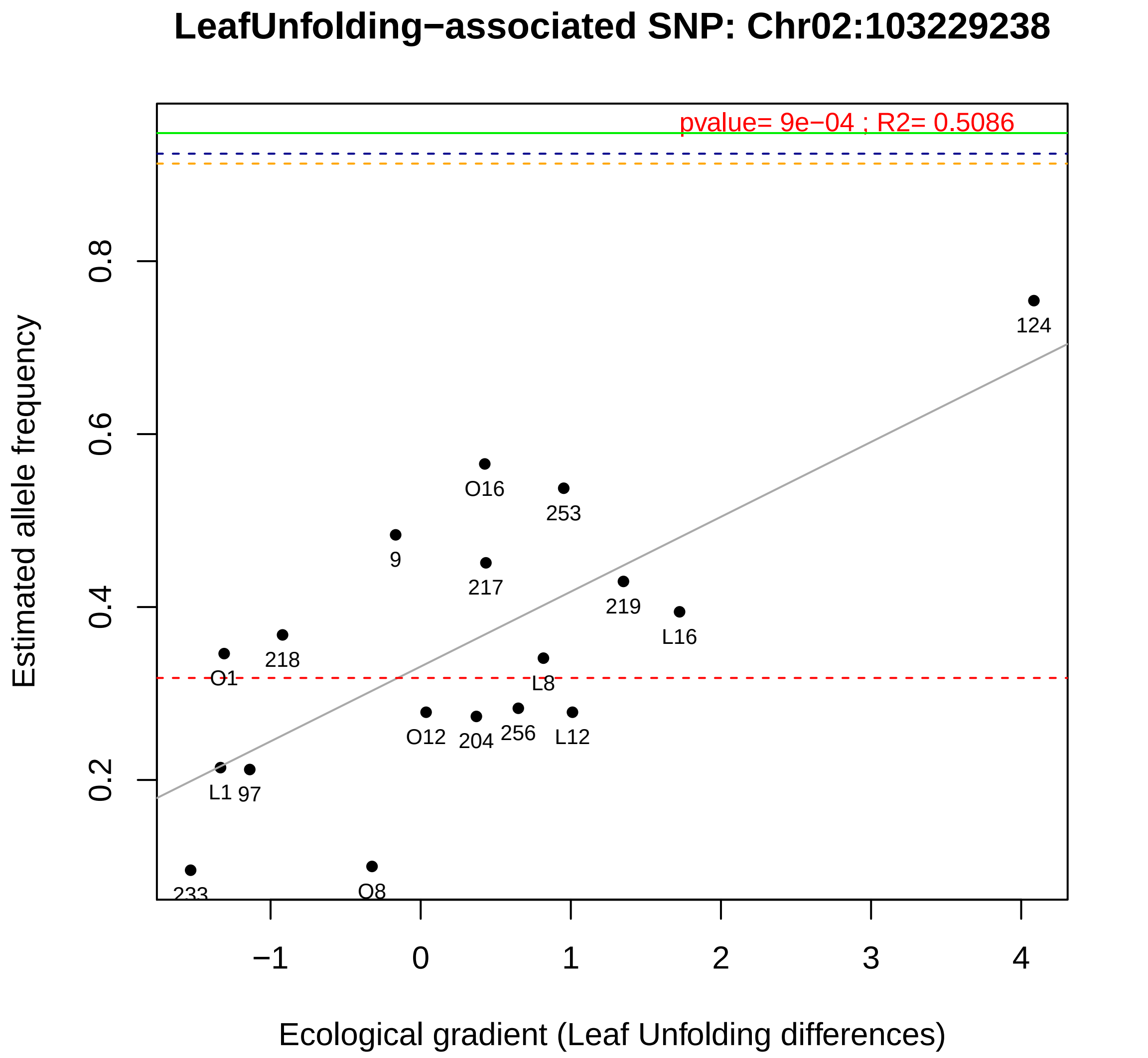

LeafUnfolding−associated SNP: Chr02:103229238
pvalue= 9e−04 ; R2= 0.5086
Estimated allele frequency
0.2
0.4
0.6
0.8
−1
0
1
2
3
4
Ecological gradient (Leaf Unfolding differences)
124
O16
253
9
217
219
L16
218
O1
L8
O12
204
256
L12
L1
97
233
O8

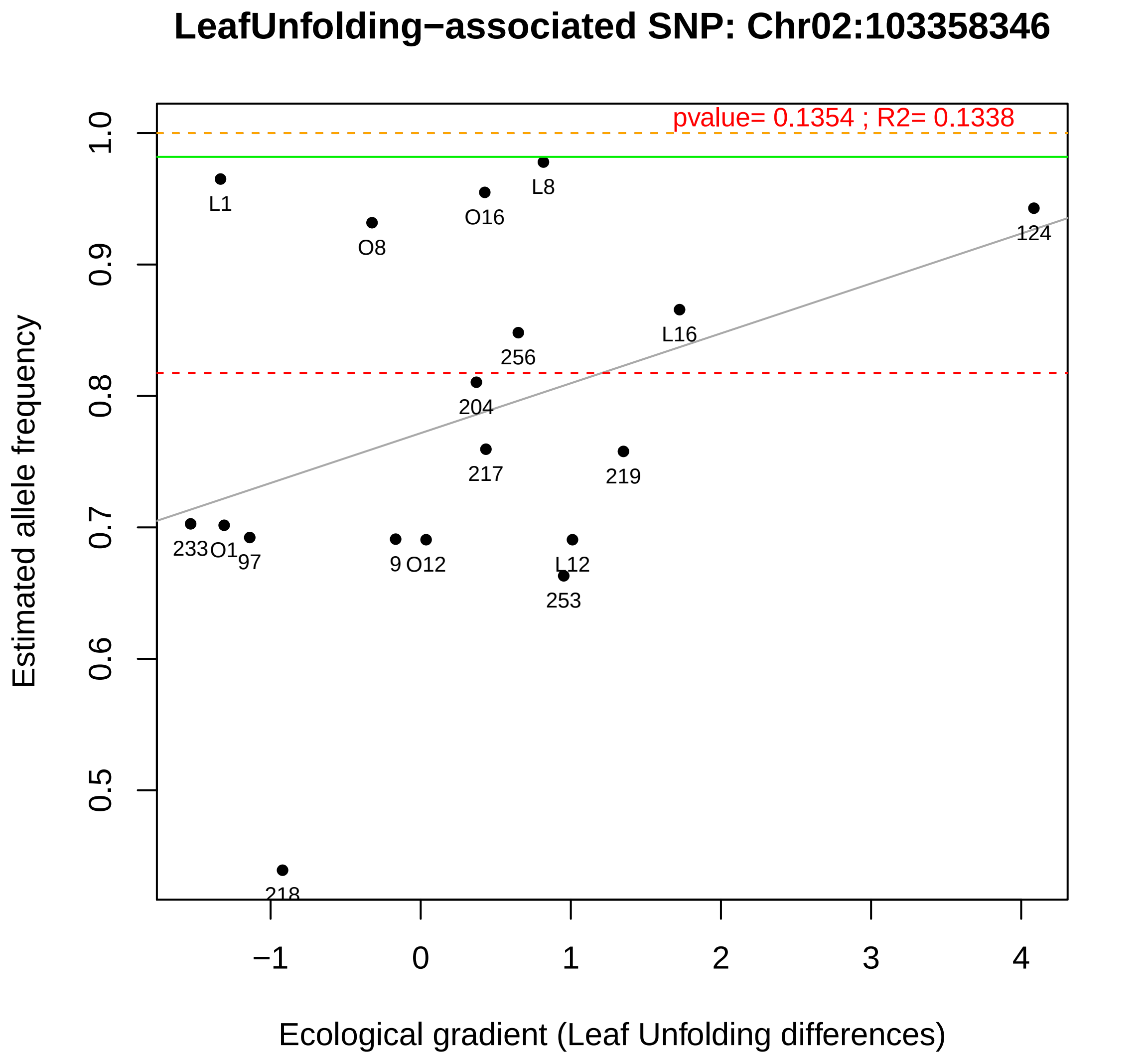

LeafUnfolding−associated SNP: Chr02:103358346
pvalue= 0.1354 ; R2= 0.1338
Estimated allele frequency
0.5
0.6
0.7
0.8
0.9
1.0
L1
L8
O16
O8
124
L16
256
204
217
219
233
O1
97
9
O12
L12
253
218
−1
0
1
2
3
4
Ecological gradient (Leaf Unfolding differences)

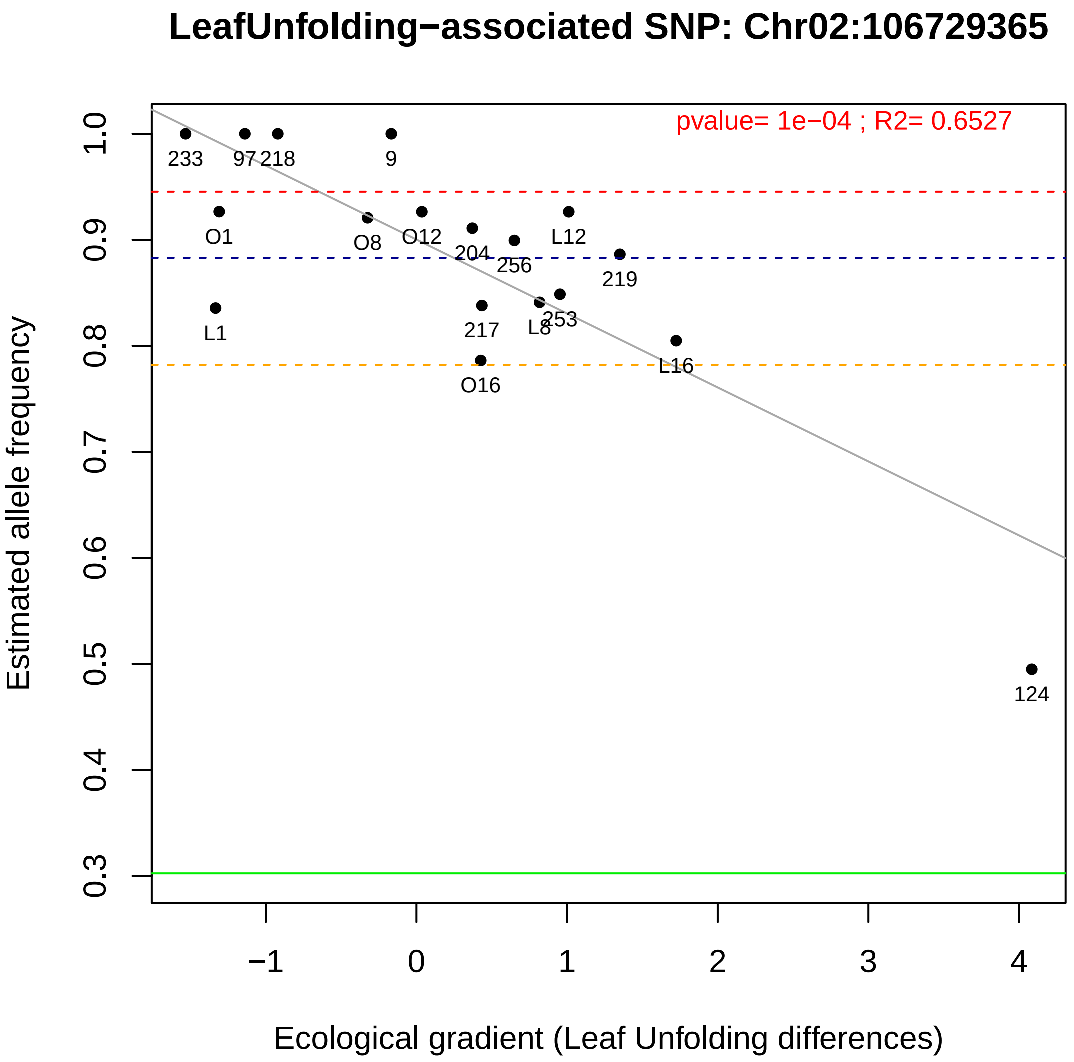
LeafUnfolding−associated SNP: Chr02:106729365
pvalue= 1e−04 ; R2= 0.6527
Estimated allele frequency
Ecological gradient (Leaf Unfolding differences)
1.0
0.9
0.8
0.7
0.6
0.5
0.4
0.3
−1
0
1
2
3
4
233
97
218
9
O1
O8
O12
204
256
L12
219
L1
217
L8
253
L16
O16
124

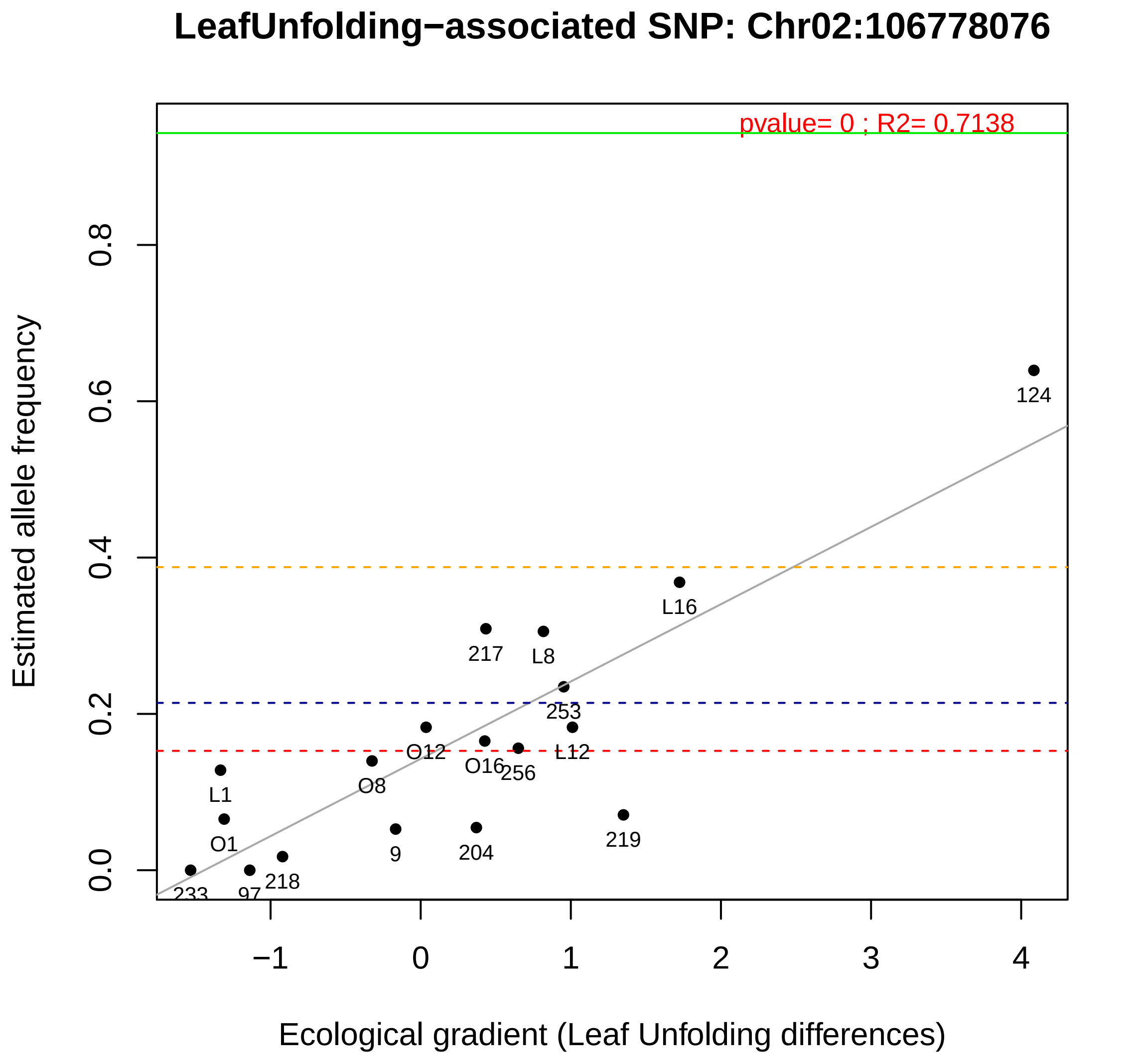
LeafUnfolding−associated SNP: Chr02:106778076
pvalue= 0 ; R2= 0.7138
Estimated allele frequency
0.0
0.2
0.4
0.6
0.8
124
L16
217
L8
253
O12
L12
O16
256
L1
O8
O1
9
204
219
218
233
97
−1
0
1
2
3
4
Ecological gradient (Leaf Unfolding differences)

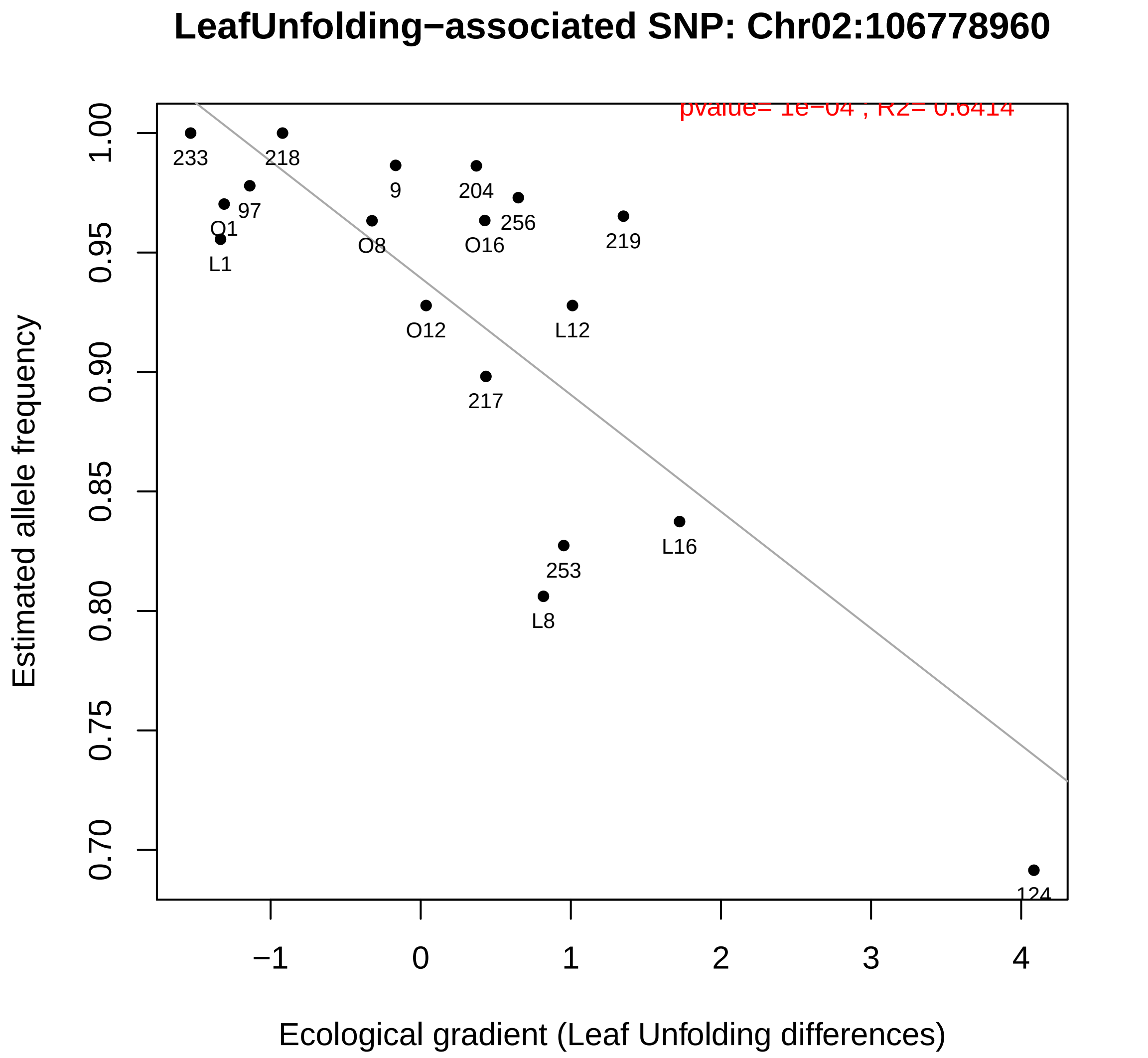
LeafUnfolding−associated SNP: Chr02:106778960
pvalue= 1e−04 ; R2= 0.6414
Estimated allele frequency
Ecological gradient (Leaf Unfolding differences)
1.00
0.95
0.90
0.85
0.80
0.75
0.70
−1
0
1
2
3
4
233
218
97
O1
L1
9
204
256
O8
O16
219
O12
L12
217
L16
253
L8
124

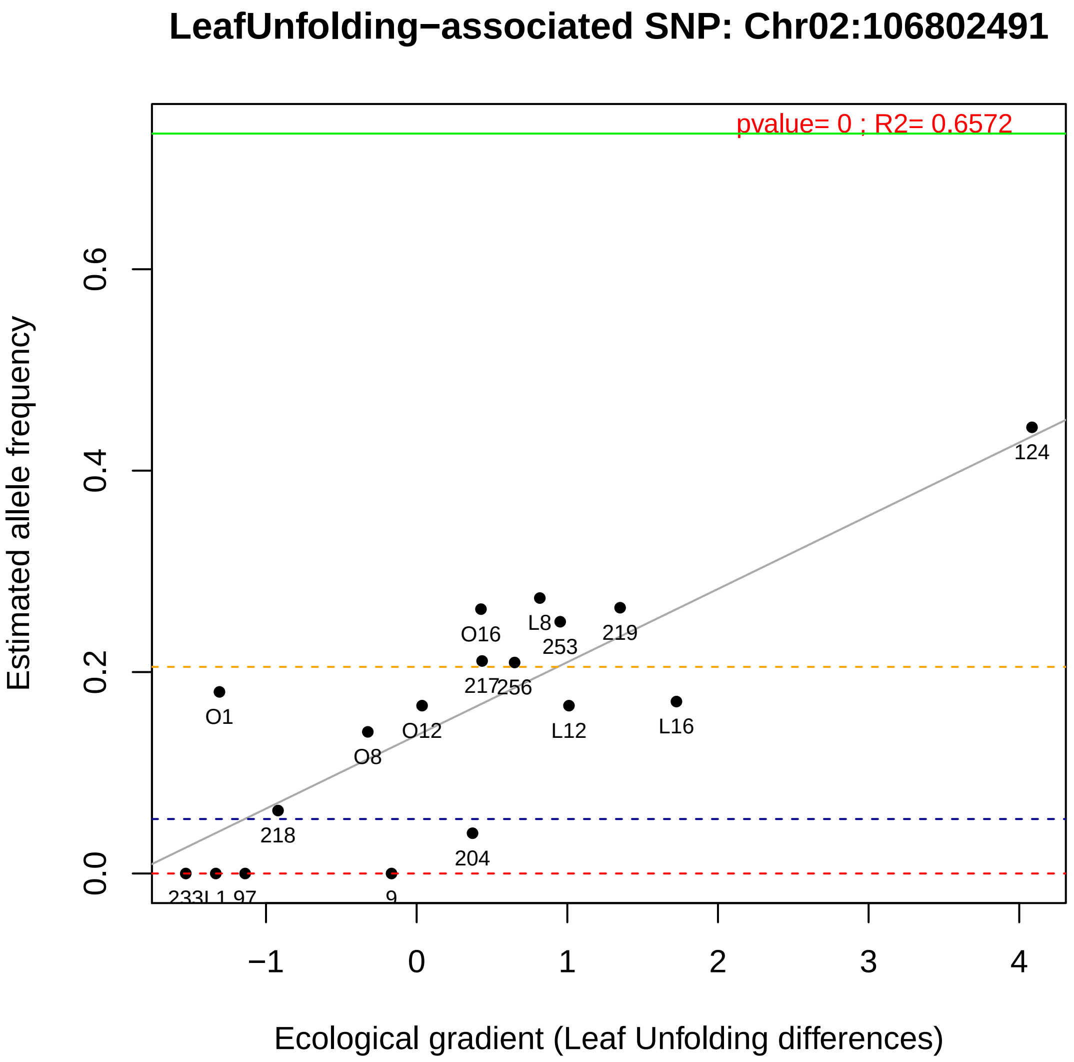

LeafUnfolding−associated SNP: Chr02:106802491
pvalue= 0 ; R2= 0.6572
Estimated allele frequency
0.0
0.2
0.4
0.6
Ecological gradient (Leaf Unfolding differences)
−1
0
1
2
3
4
124
O16
L8
253
219
217
256
O1
O12
L12
L16
O8
218
204
233
L1
97
9

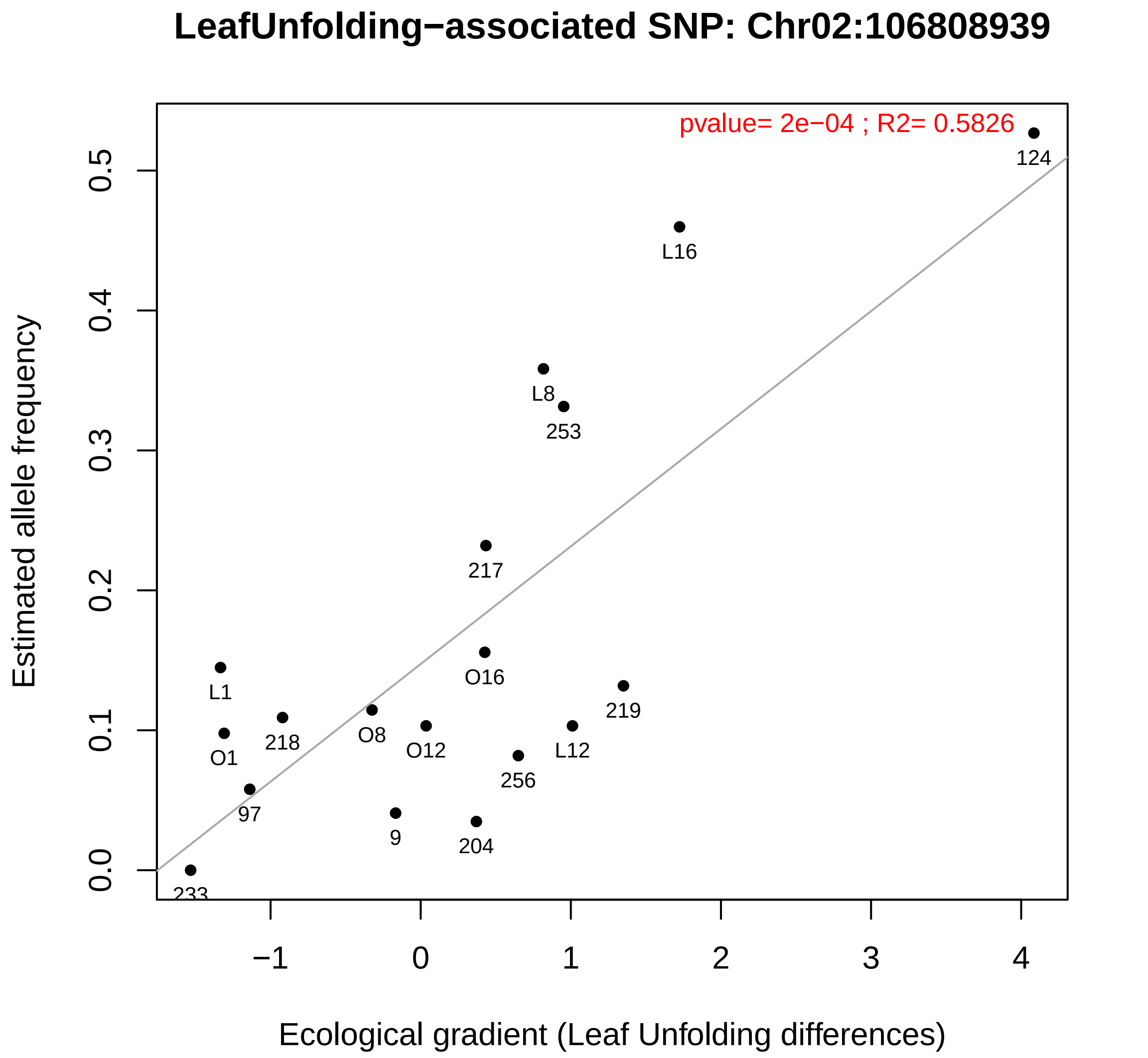
LeafUnfolding−associated SNP: Chr02:106808939
pvalue= 2e−04 ; R2= 0.5826
Estimated allele frequency
Ecological gradient (Leaf Unfolding differences)
0.0
0.1
0.2
0.3
0.4
0.5
−1
0
1
2
3
4
124
L16
L8
253
217
O16
219
L1
218
O8
O12
L12
O1
256
97
9
204
233

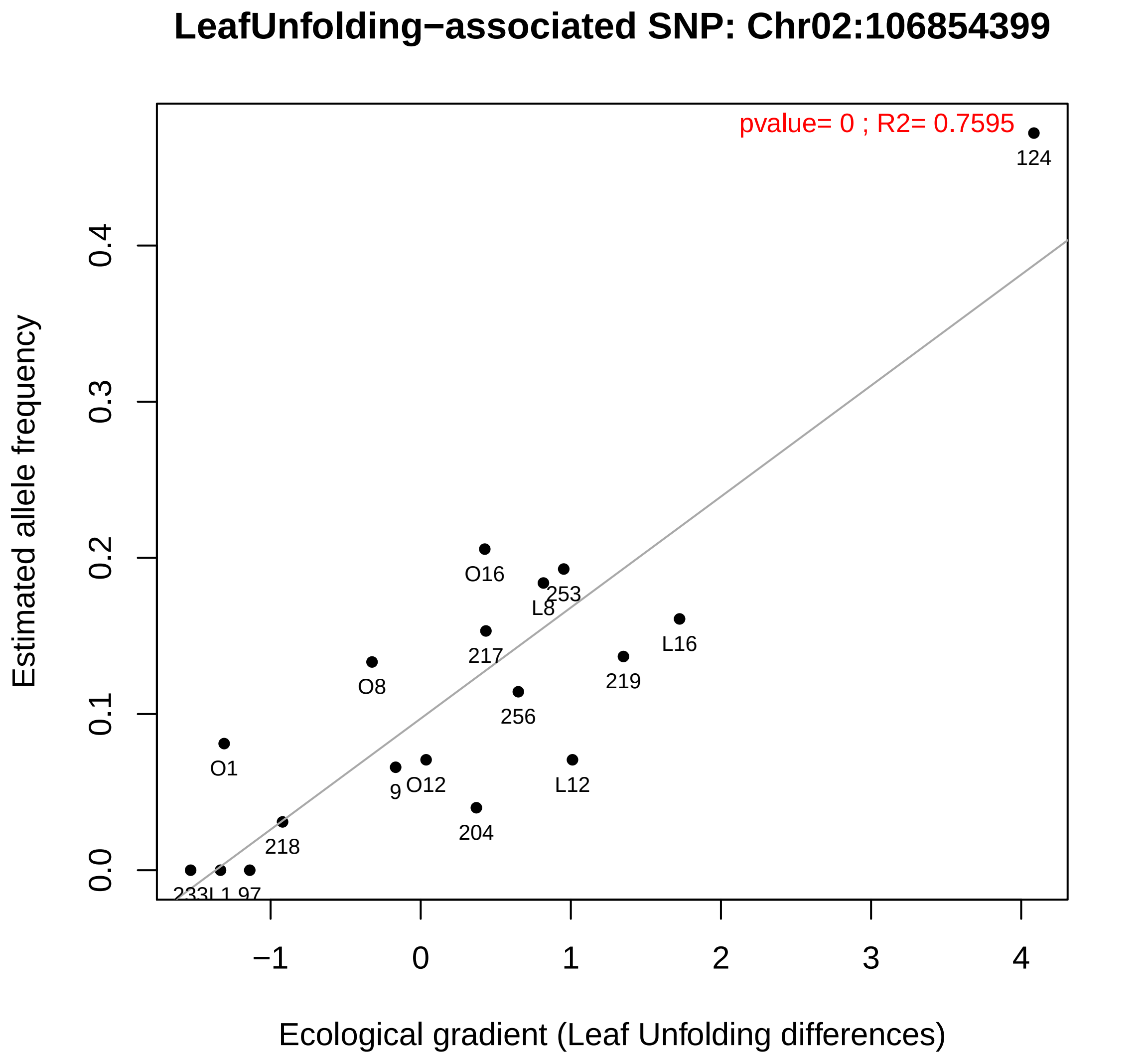

LeafUnfolding−associated SNP: Chr02:106854399
pvalue= 0 ; R2= 0.7595
Estimated allele frequency
Ecological gradient (Leaf Unfolding differences)
0.0
0.1
0.2
0.3
0.4
−1
0
1
2
3
4
124
O16
253
L8
L16
217
219
O8
256
O1
9
O12
L12
204
218
233
L1
97

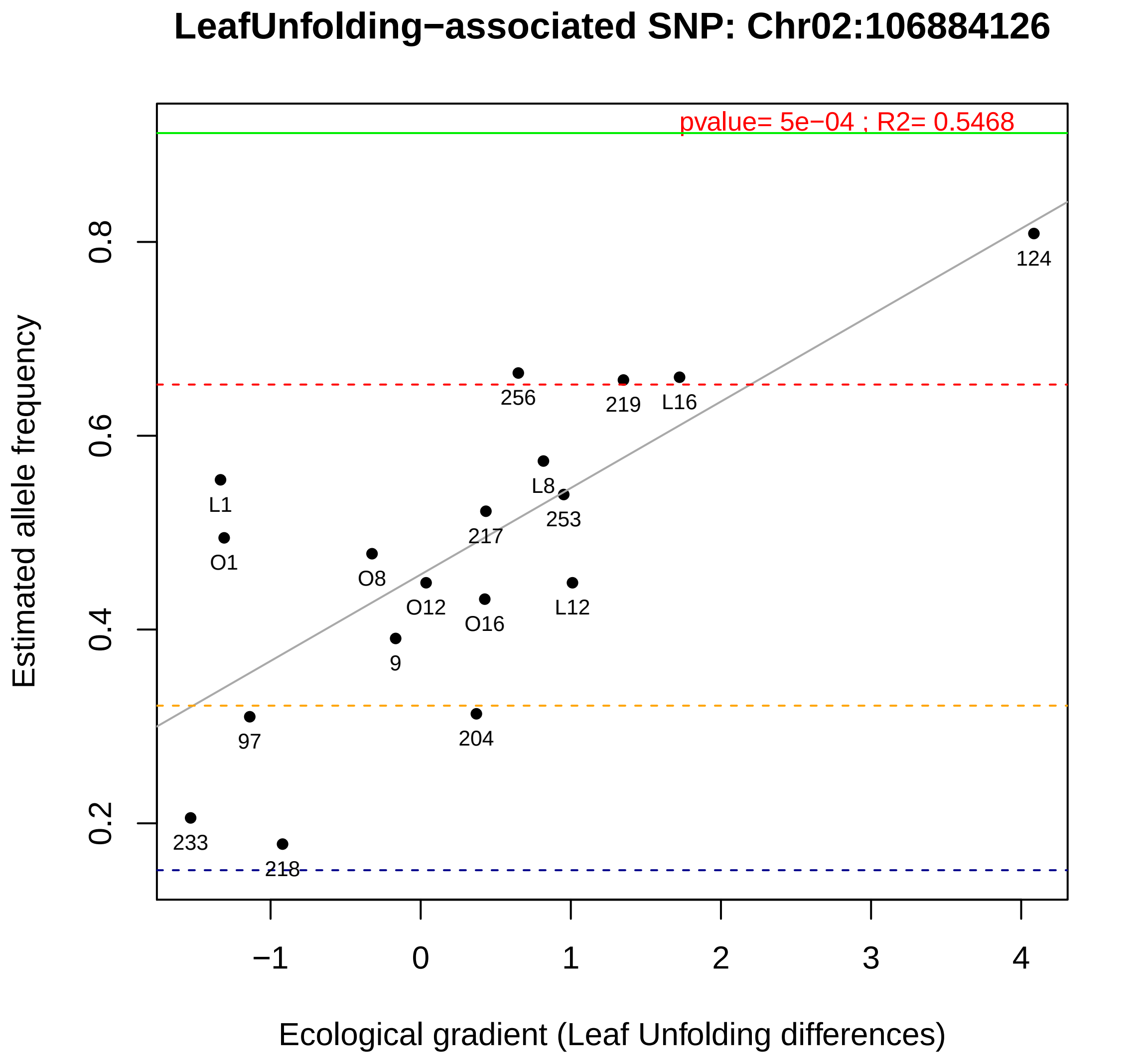

LeafUnfolding−associated SNP: Chr02:106884126
pvalue= 5e−04 ; R2= 0.5468
Estimated allele frequency
0.2
0.4
0.6
0.8
Ecological gradient (Leaf Unfolding differences)
−1
0
1
2
3
4
124
256
219
L16
L8
253
L1
217
O1
O8
O12
O16
L12
9
97
204
233
218

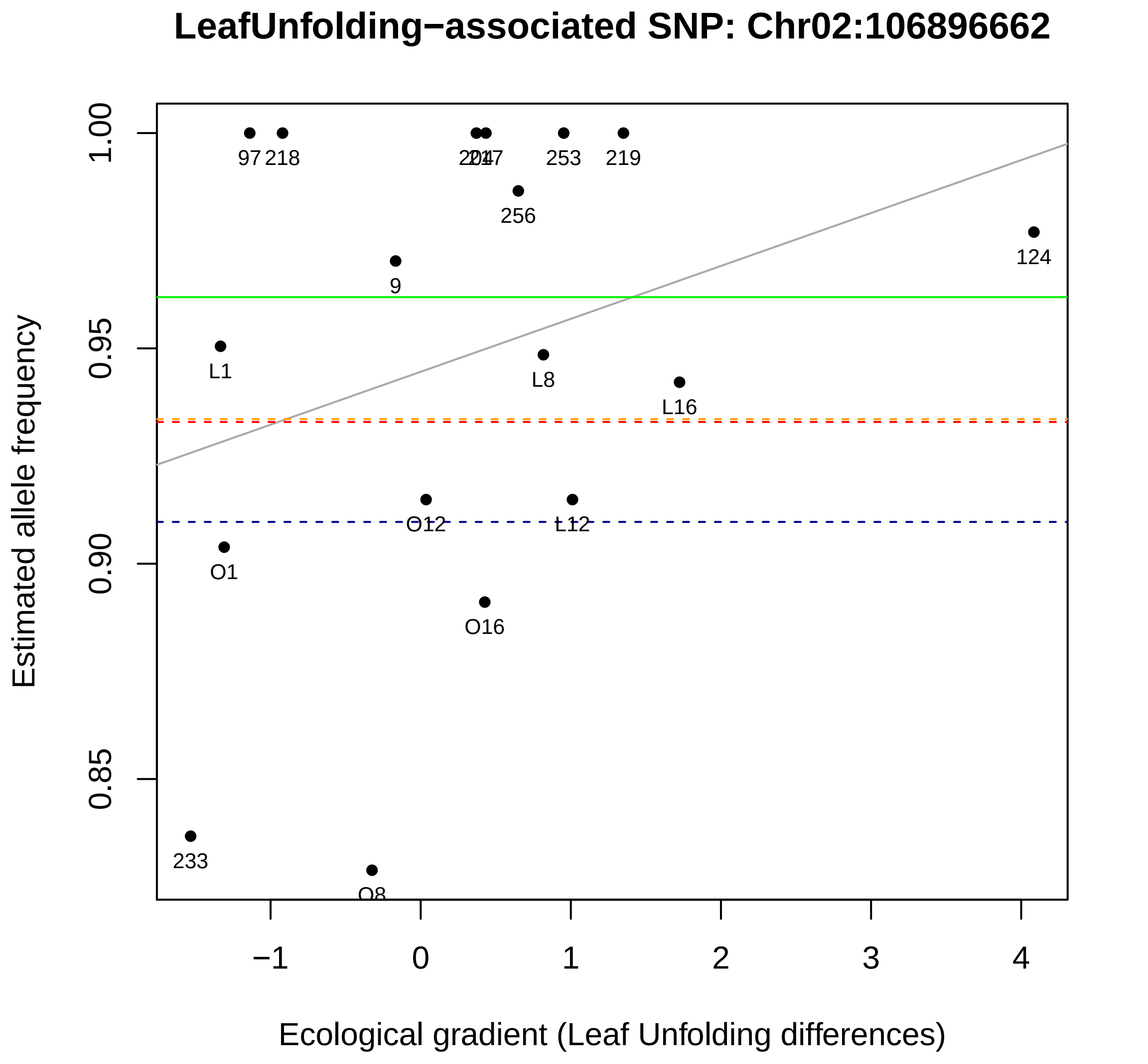
LeafUnfolding−associated SNP: Chr02:106896662
Estimated allele frequency
Ecological gradient (Leaf Unfolding differences)
1.00
0.95
0.90
0.85
−1
0
1
2
3
4
97
218
253
219
256
124
9
L1
L8
L16
O12
L12
O1
O16
233

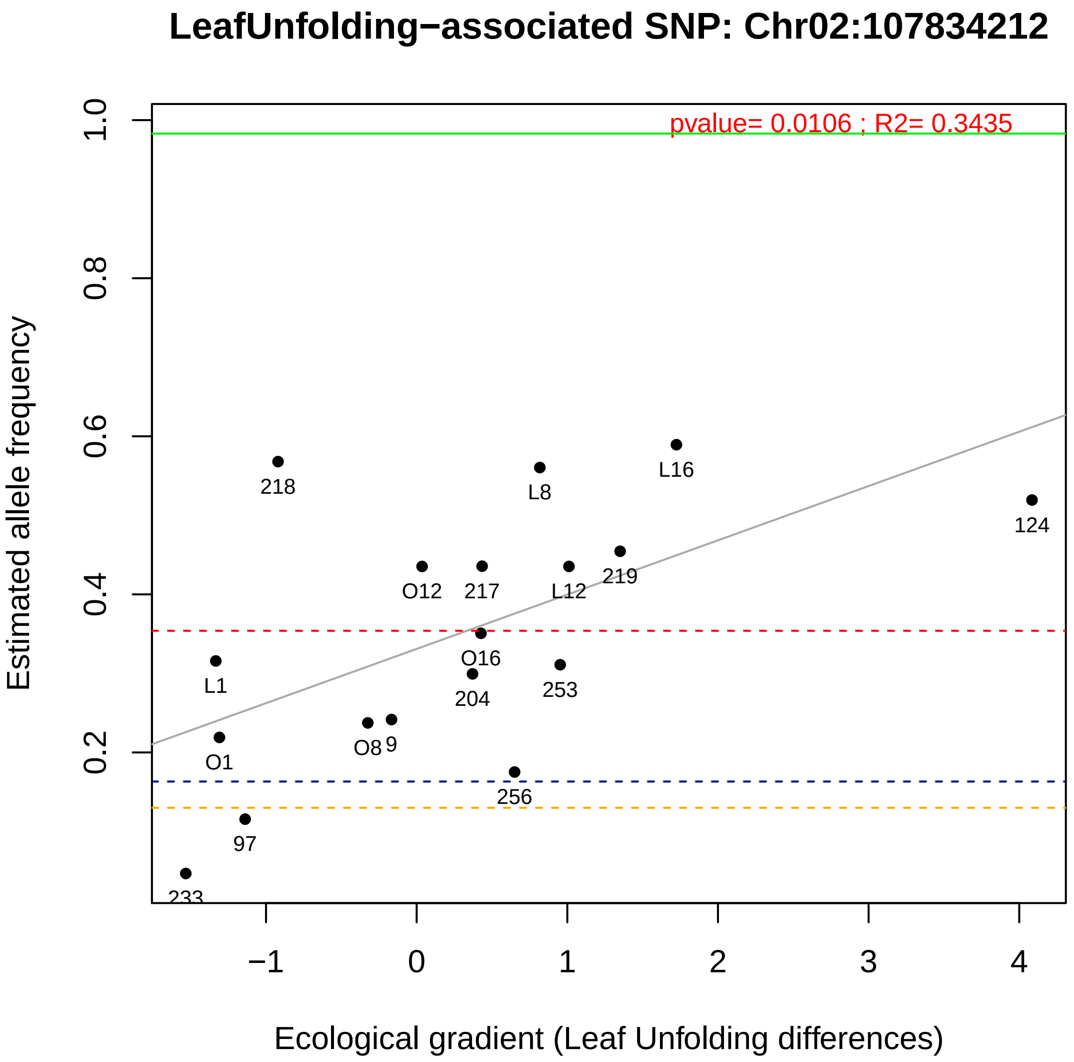
LeafUnfolding−associated SNP: Chr02:107834212
pvalue= 0.0106 ; R2= 0.3435
Estimated allele frequency
Ecological gradient (Leaf Unfolding differences)
1.0
0.8
0.6
0.4
0.2
−1
0
1
2
3
4
218
L8
L16
124
O12
217
L12
219
O16
L1
204
253
O8
9
O1
256
97
233

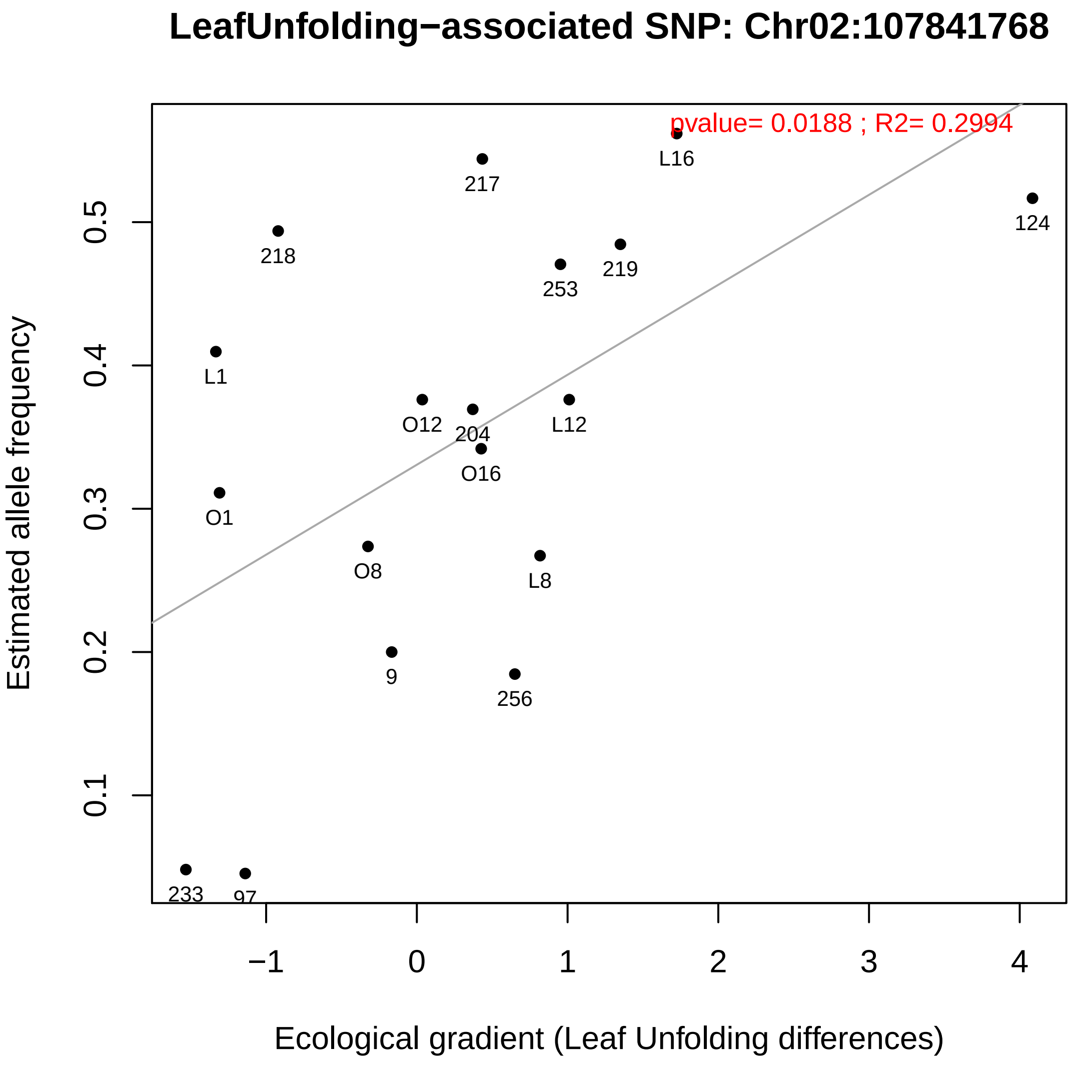

LeafUnfolding−associated SNP: Chr02:107841768
pvalue= 0.0188 ; R2= 0.2994
Estimated allele frequency
0.1
0.2
0.3
0.4
0.5
−1
0
1
2
3
4
Ecological gradient (Leaf Unfolding differences)
L16
217
124
218
219
253
L1
O12
204
L12
O16
O1
O8
L8
9
256
233
97

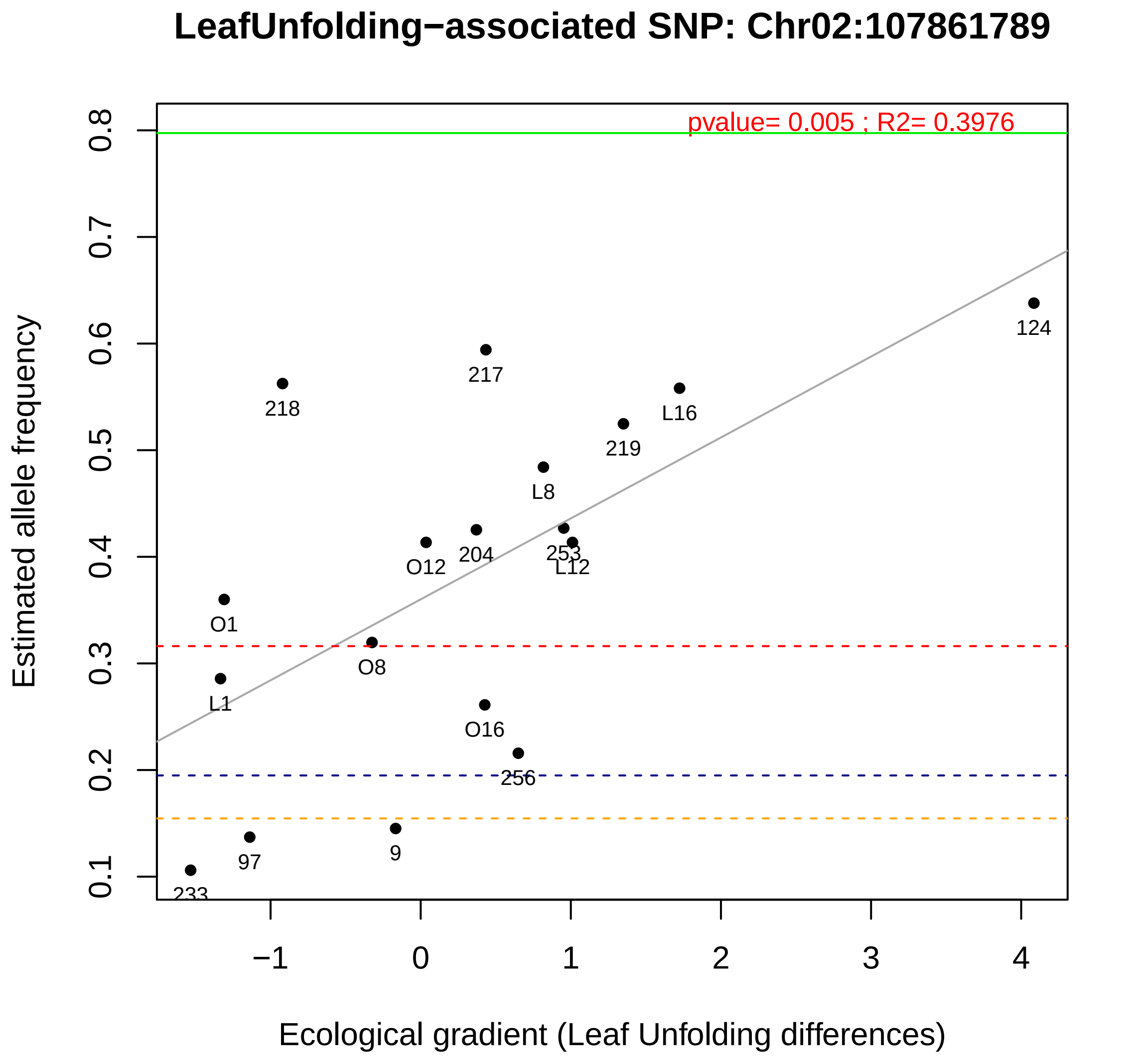
LeafUnfolding−associated SNP: Chr02:107861789
pvalue= 0.005 ; R2= 0.3976
Estimated allele frequency
0.1
0.2
0.3
0.4
0.5
0.6
0.7
0.8
Ecological gradient (Leaf Unfolding differences)
−1
0
1
2
3
4
233
97
9
256
O16
L1
O8
O1
O12
204
253
L12
L8
219
L16
217
218
124

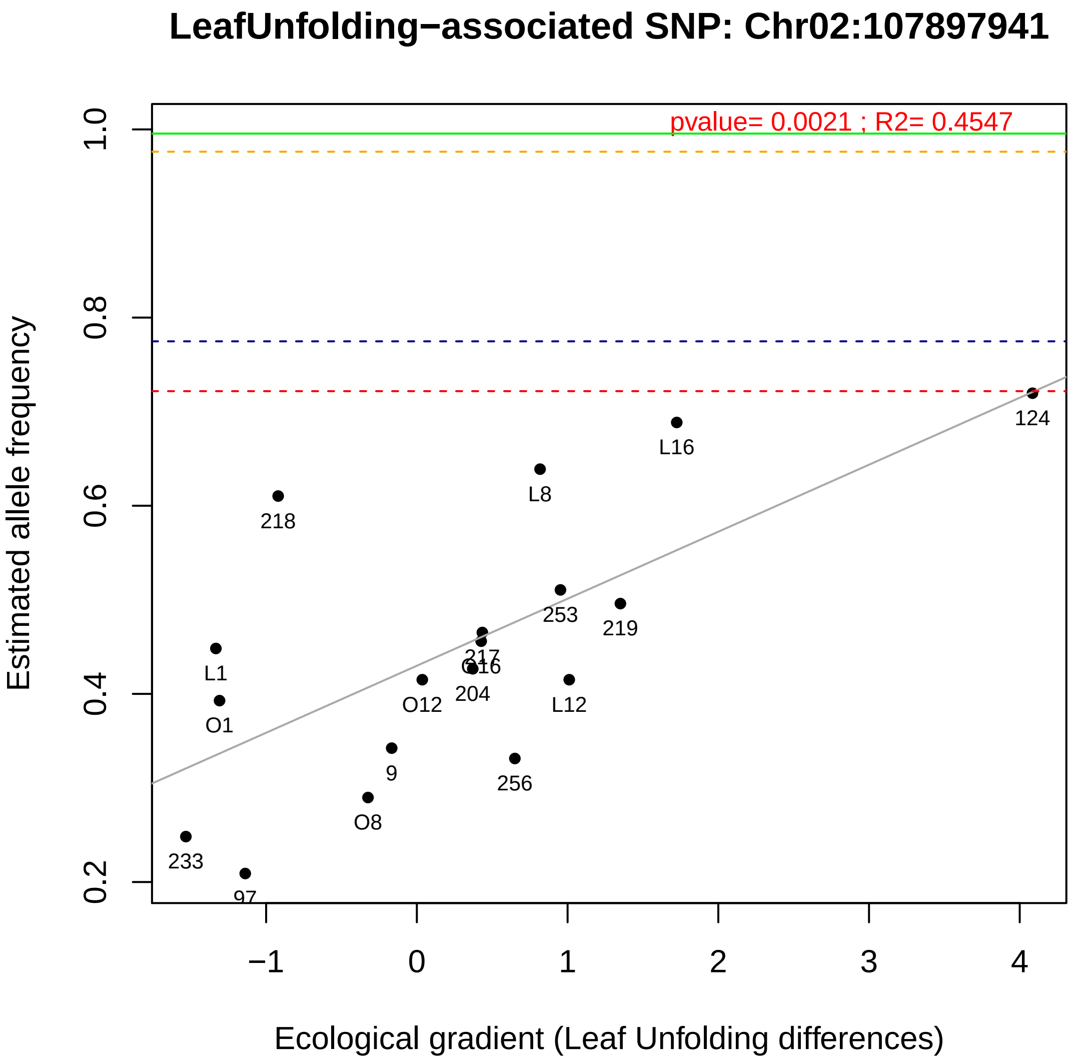
LeafUnfolding−associated SNP: Chr02:107897941
pvalue= 0.0021 ; R2= 0.4547
Estimated allele frequency
Ecological gradient (Leaf Unfolding differences)
0.2
0.4
0.6
0.8
1.0
−1
0
1
2
3
4
124
L16
L8
218
253
219
217
O16
204
L1
O1
O12
L12
9
256
O8
233
97

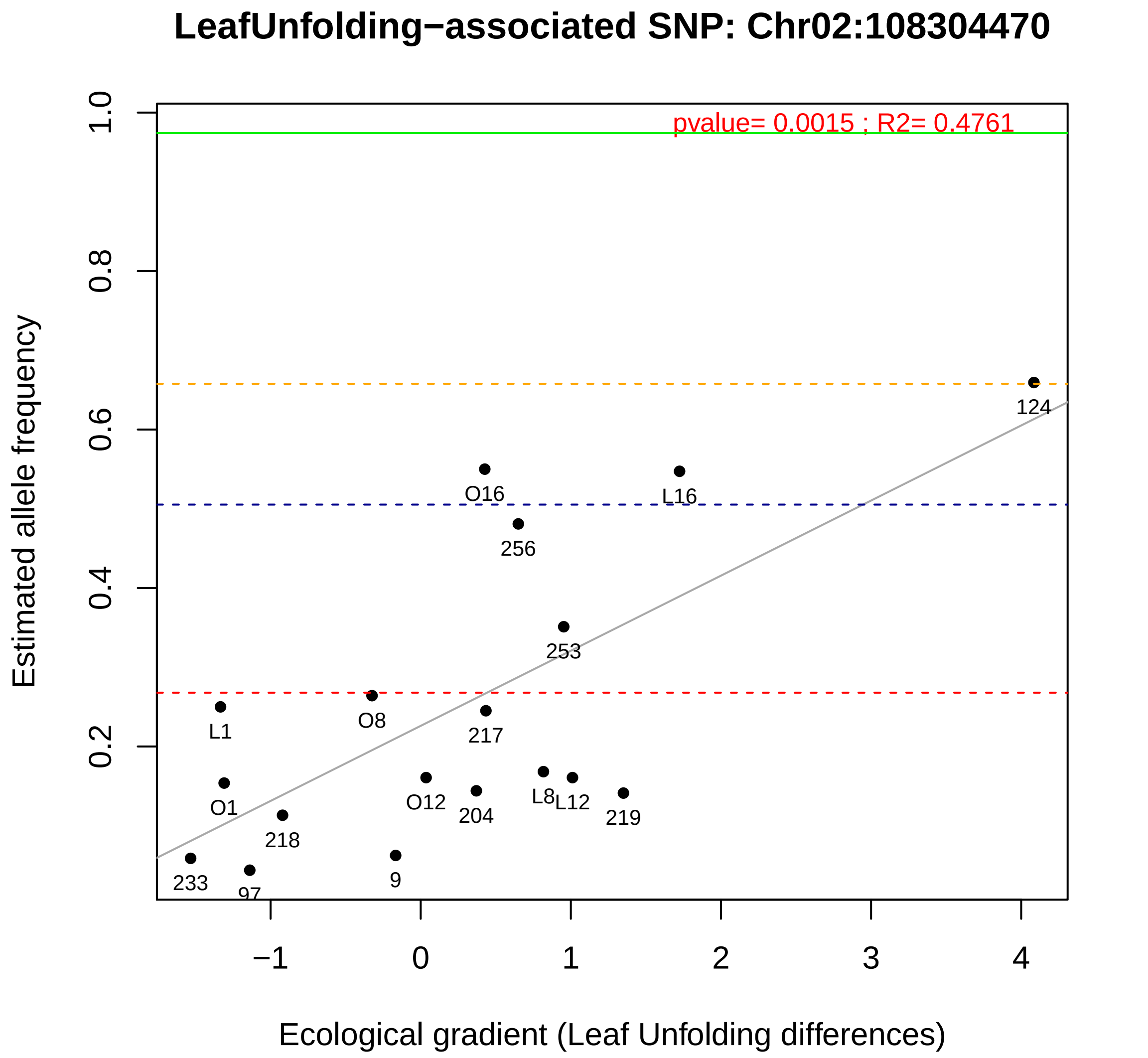
LeafUnfolding−associated SNP: Chr02:108304470
pvalue= 0.0015 ; R2= 0.4761
Estimated allele frequency
1.0
0.8
0.6
0.4
0.2
124
O16
L16
256
253
O8
L1
217
O1
O12
204
L8
L12
219
218
233
97
9
−1
0
1
2
3
4
Ecological gradient (Leaf Unfolding differences)

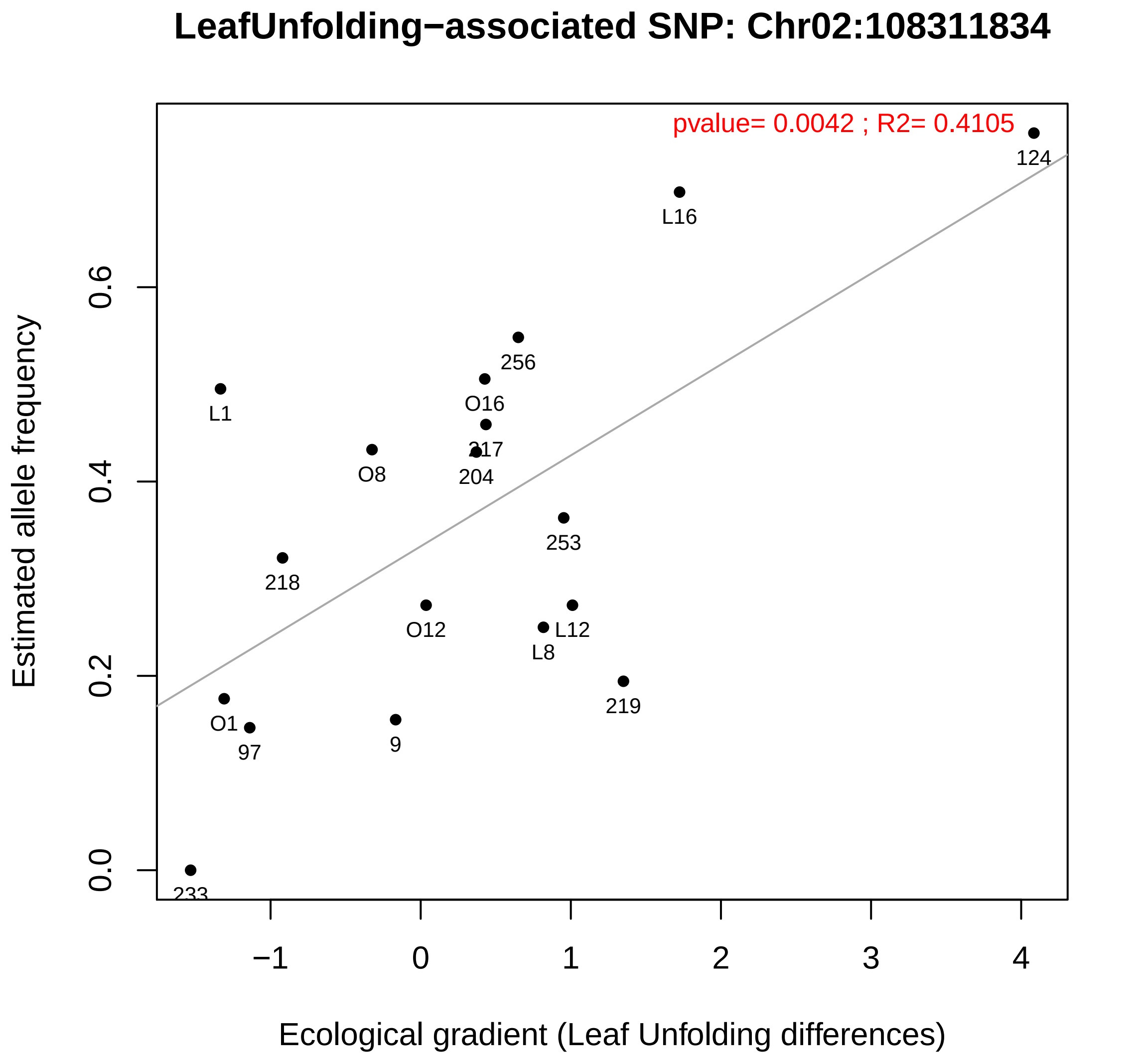

LeafUnfolding−associated SNP: Chr02:108311834
pvalue= 0.0042 ; R2= 0.4105
124
L16
256
L1
O16
217
O8
204
253
218
O12
L12
L8
219
O1
97
9
233
0.0
0.2
0.4
0.6
−1
0
1
2
3
4
Estimated allele frequency
Ecological gradient (Leaf Unfolding differences)

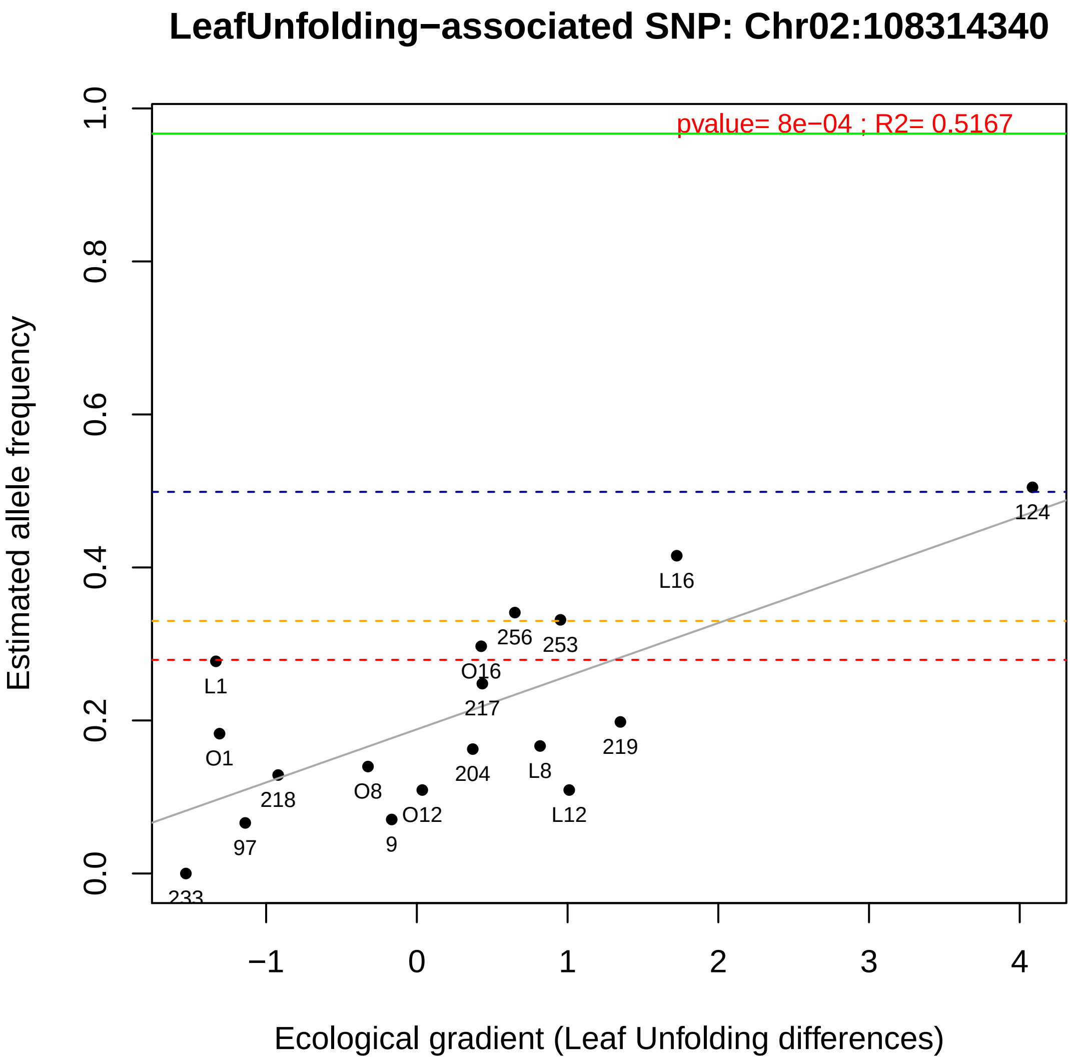
LeafUnfolding−associated SNP: Chr02:108314340
pvalue= 8e−04 ; R2= 0.5167
Estimated allele frequency
0.0
0.2
0.4
0.6
0.8
1.0
−1
0
1
2
3
4
Ecological gradient (Leaf Unfolding differences)
124
L16
256
253
O16
L1
217
O1
219
204
L8
218
O8
O12
L12
97
9
233

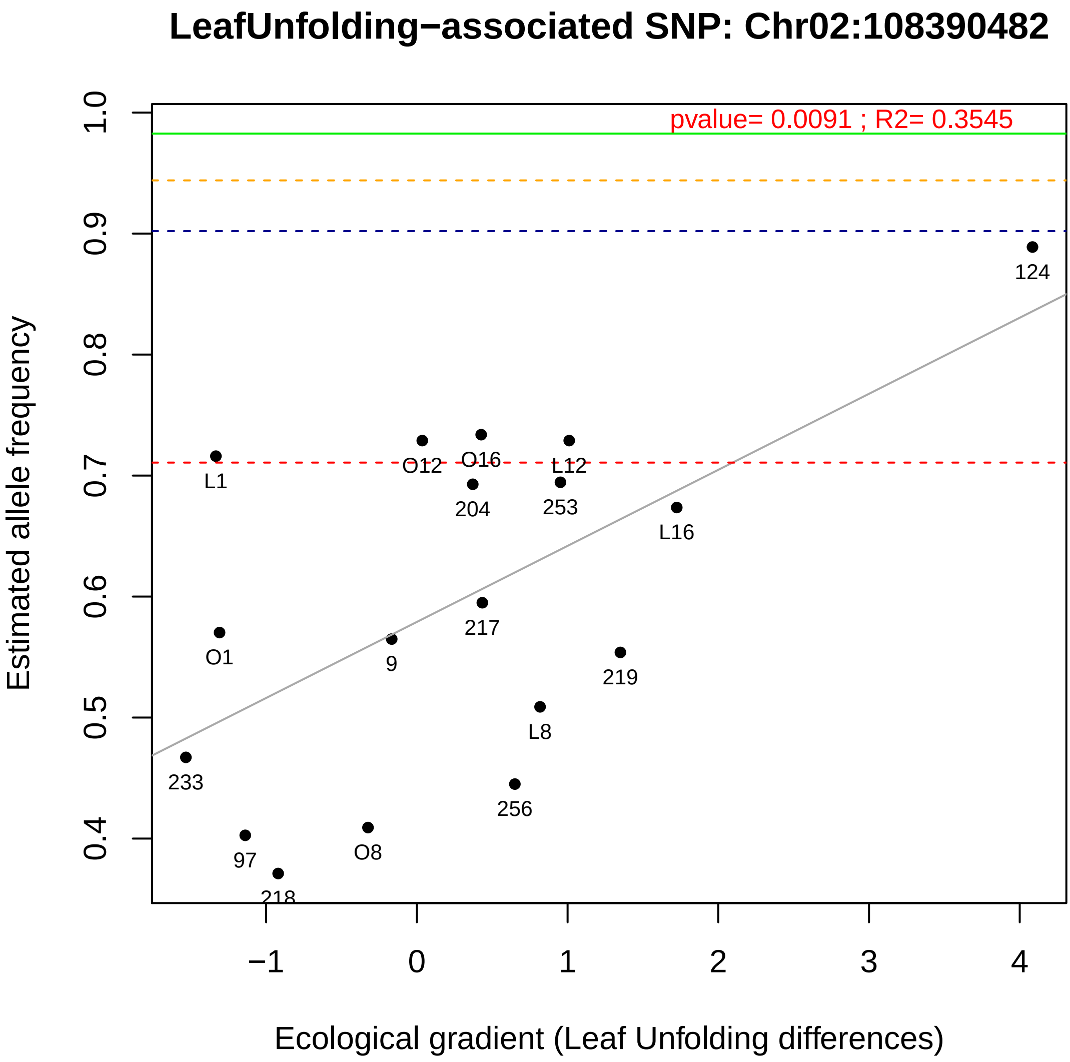

LeafUnfolding−associated SNP: Chr02:108390482
pvalue= 0.0091 ; R2= 0.3545
Estimated allele frequency
Ecological gradient (Leaf Unfolding differences)
0.4
0.5
0.6
0.7
0.8
0.9
1.0
−1
0
1
2
3
4
124
L1
O12
O16
L12
204
253
L16
217
O1
9
219
L8
233
256
97
O8
218

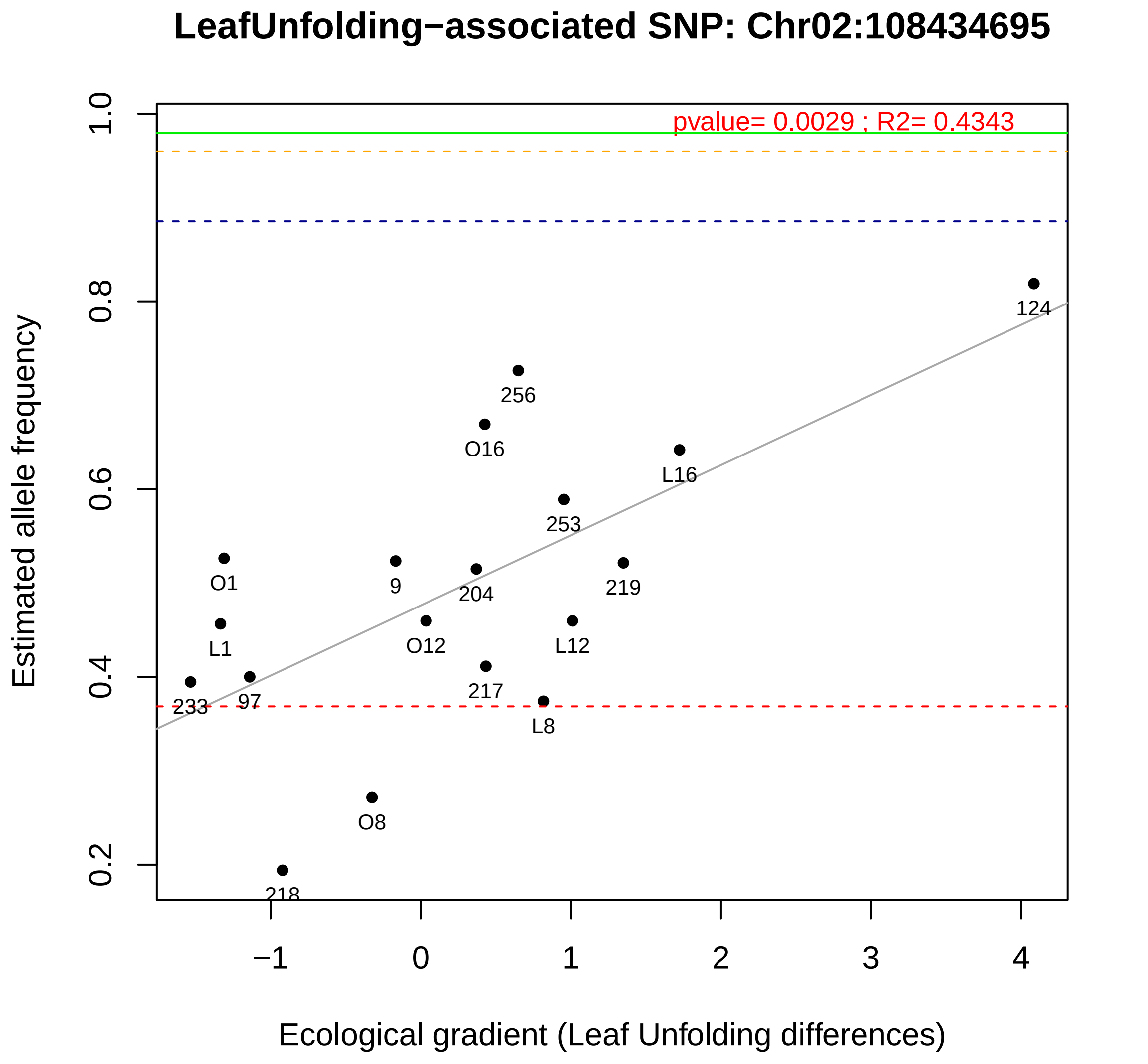
LeafUnfolding−associated SNP: Chr02:108434695
pvalue= 0.0029 ; R2= 0.4343
Estimated allele frequency
0.2
0.4
0.6
0.8
1.0
−1
0
1
2
3
4
Ecological gradient (Leaf Unfolding differences)
124
256
O16
L16
253
O1
9
204
219
L1
O12
L12
217
233
97
L8
O8
218

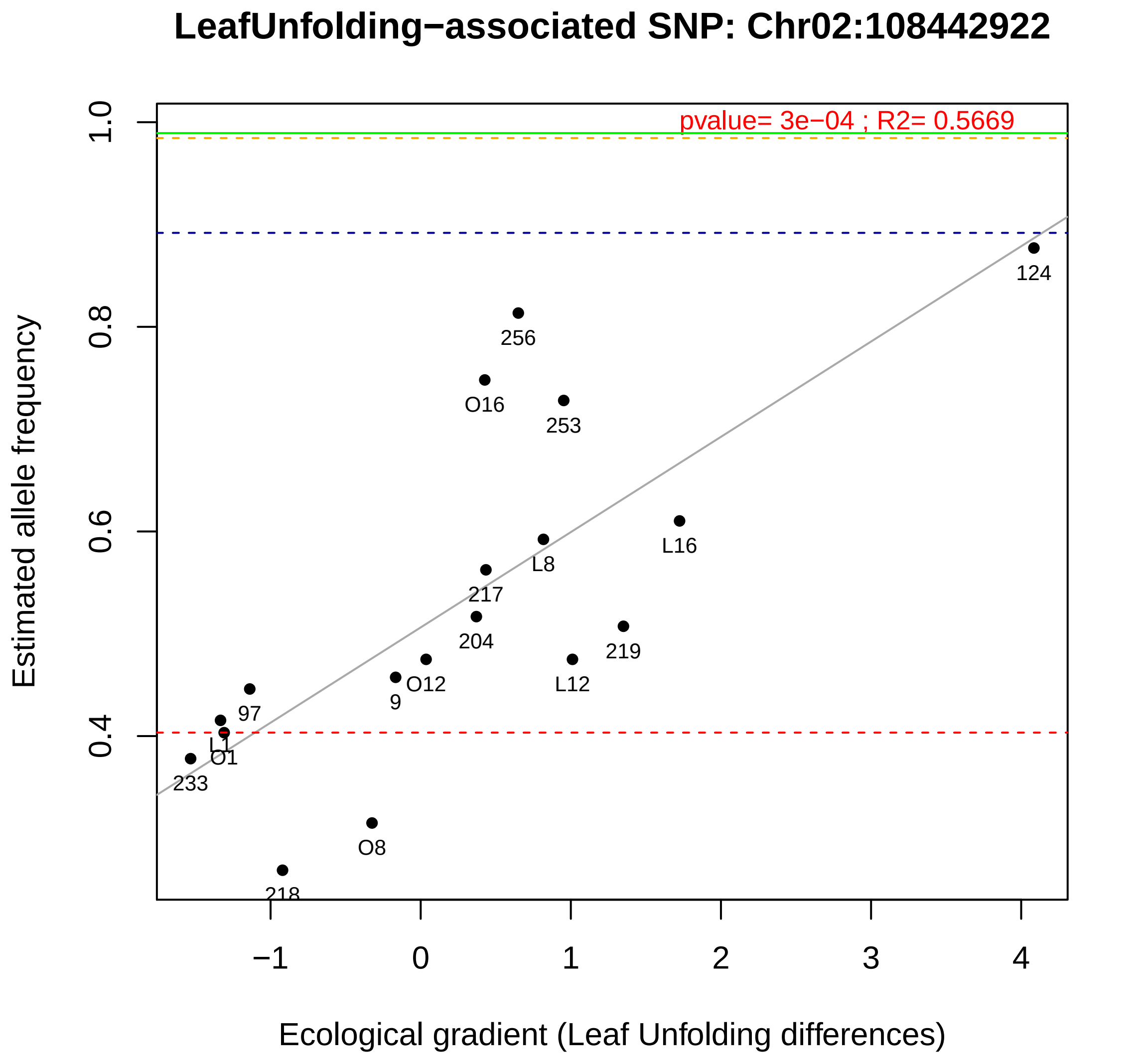

LeafUnfolding−associated SNP: Chr02:108442922
pvalue= 3e−04 ; R2= 0.5669
Estimated allele frequency
0.4
0.6
0.8
1.0
124
256
O16
253
L16
L8
217
204
219
L12
O12
9
97
L1
O1
233
O8
218
−1
0
1
2
3
4
Ecological gradient (Leaf Unfolding differences)

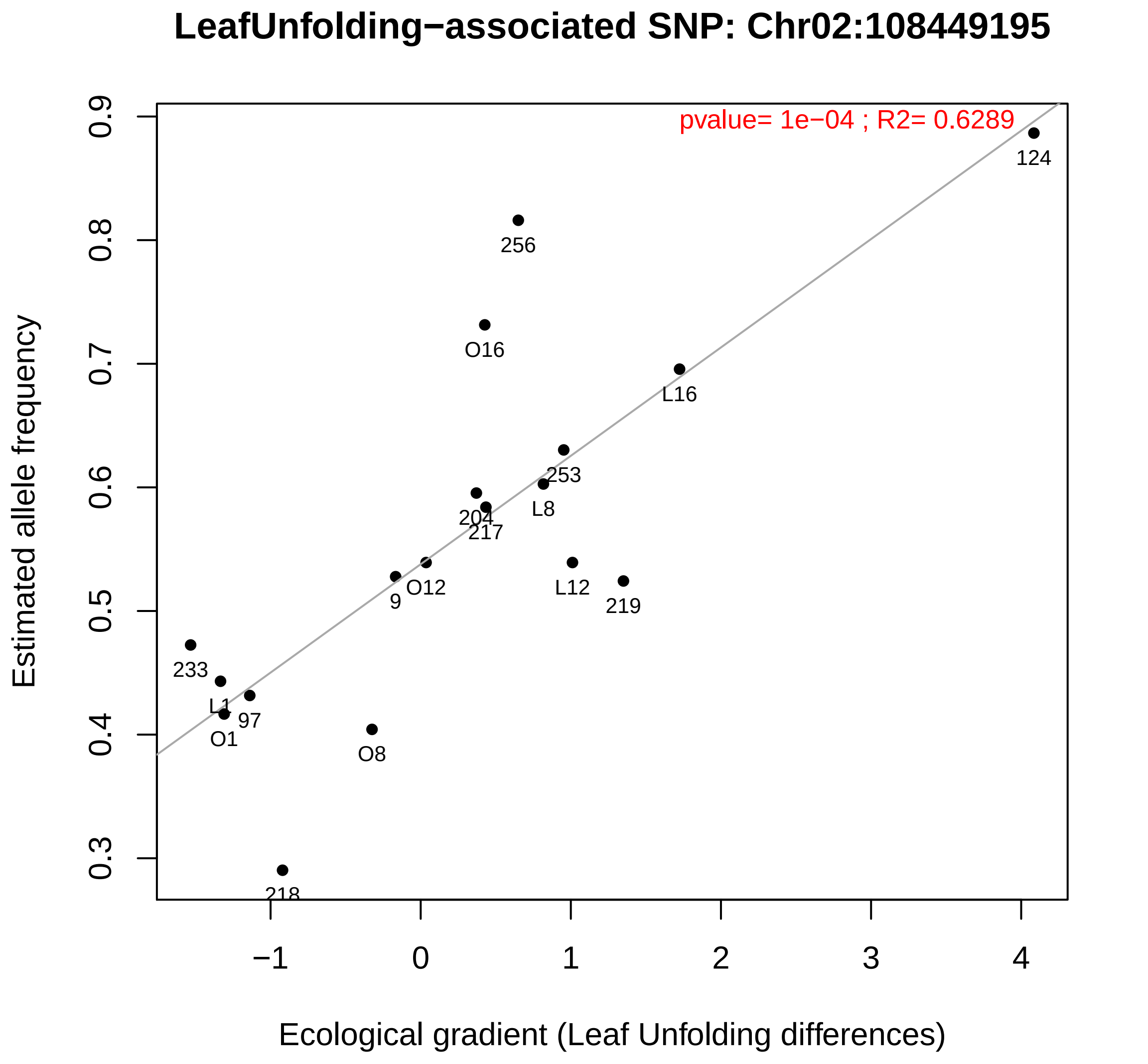
LeafUnfolding−associated SNP: Chr02:108449195
pvalue= 1e−04 ; R2= 0.6289
Estimated allele frequency
Ecological gradient (Leaf Unfolding differences)
0.3
0.4
0.5
0.6
0.7
0.8
0.9
−1
0
1
2
3
4
124
256
O16
L16
253
L8
204
217
O12
9
L12
219
233
L1
97
O1
O8
218

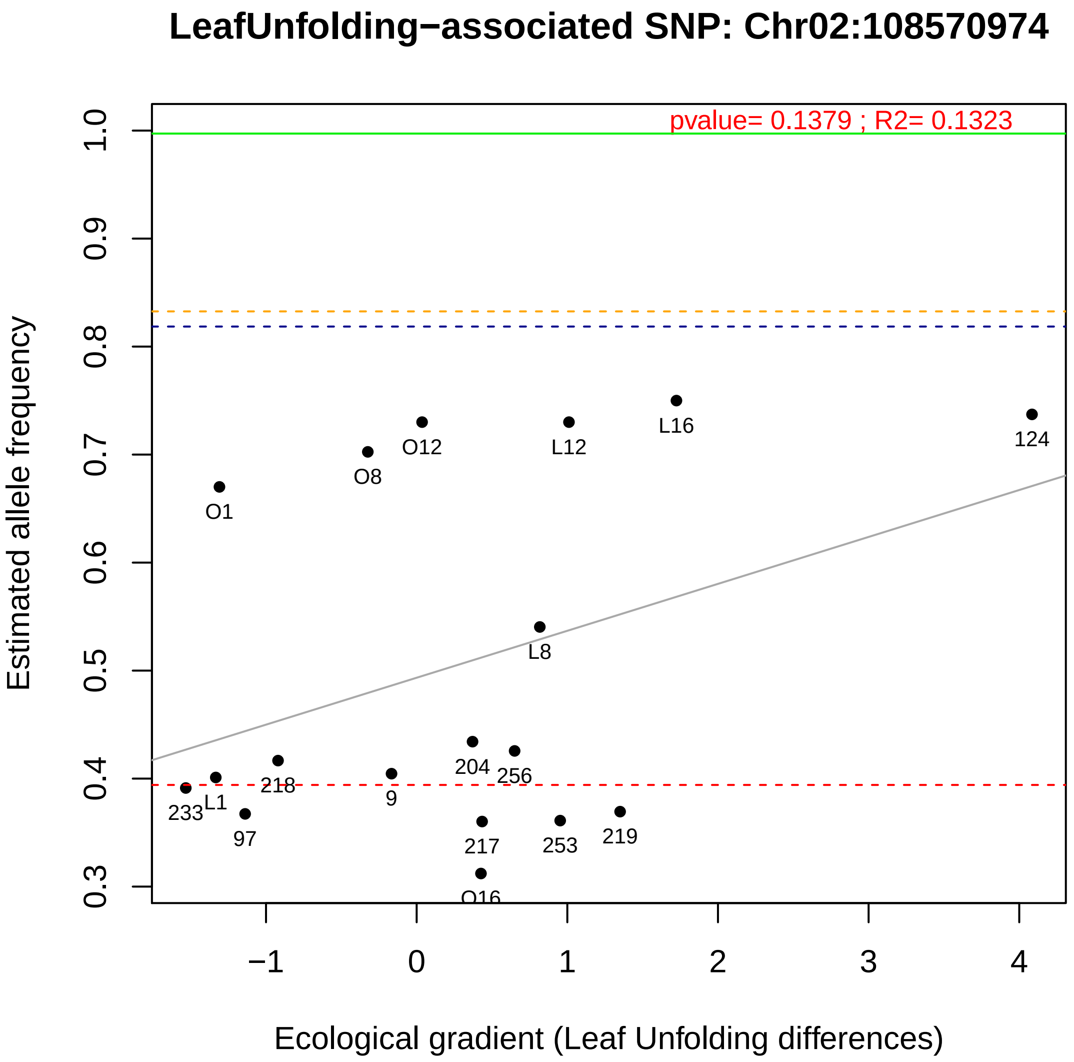
LeafUnfolding−associated SNP: Chr02:108570974
pvalue= 0.1379 ; R2= 0.1323
Estimated allele frequency
0.3
0.4
0.5
0.6
0.7
0.8
0.9
1.0
−1
0
1
2
3
4
Ecological gradient (Leaf Unfolding differences)
O1
O8
O12
L12
L16
124
L8
204
256
218
9
233
L1
97
217
253
219
O16

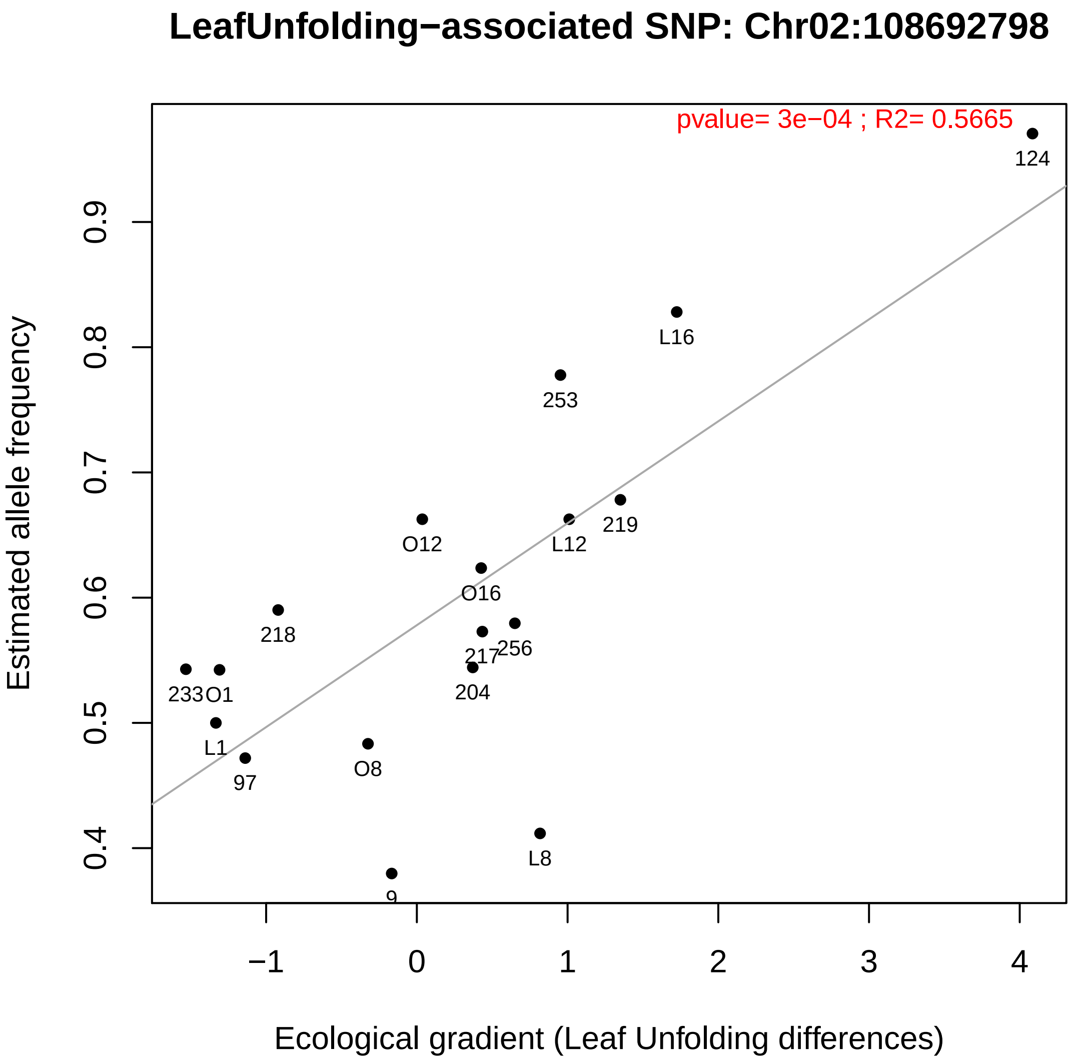

LeafUnfolding−associated SNP: Chr02:108692798
pvalue= 3e−04 ; R2= 0.5665
Estimated allele frequency
Ecological gradient (Leaf Unfolding differences)
0.4
0.5
0.6
0.7
0.8
0.9
−1
0
1
2
3
4
124
L16
253
219
L12
O12
O16
218
217
256
204
233
O1
L1
97
O8
L8
9

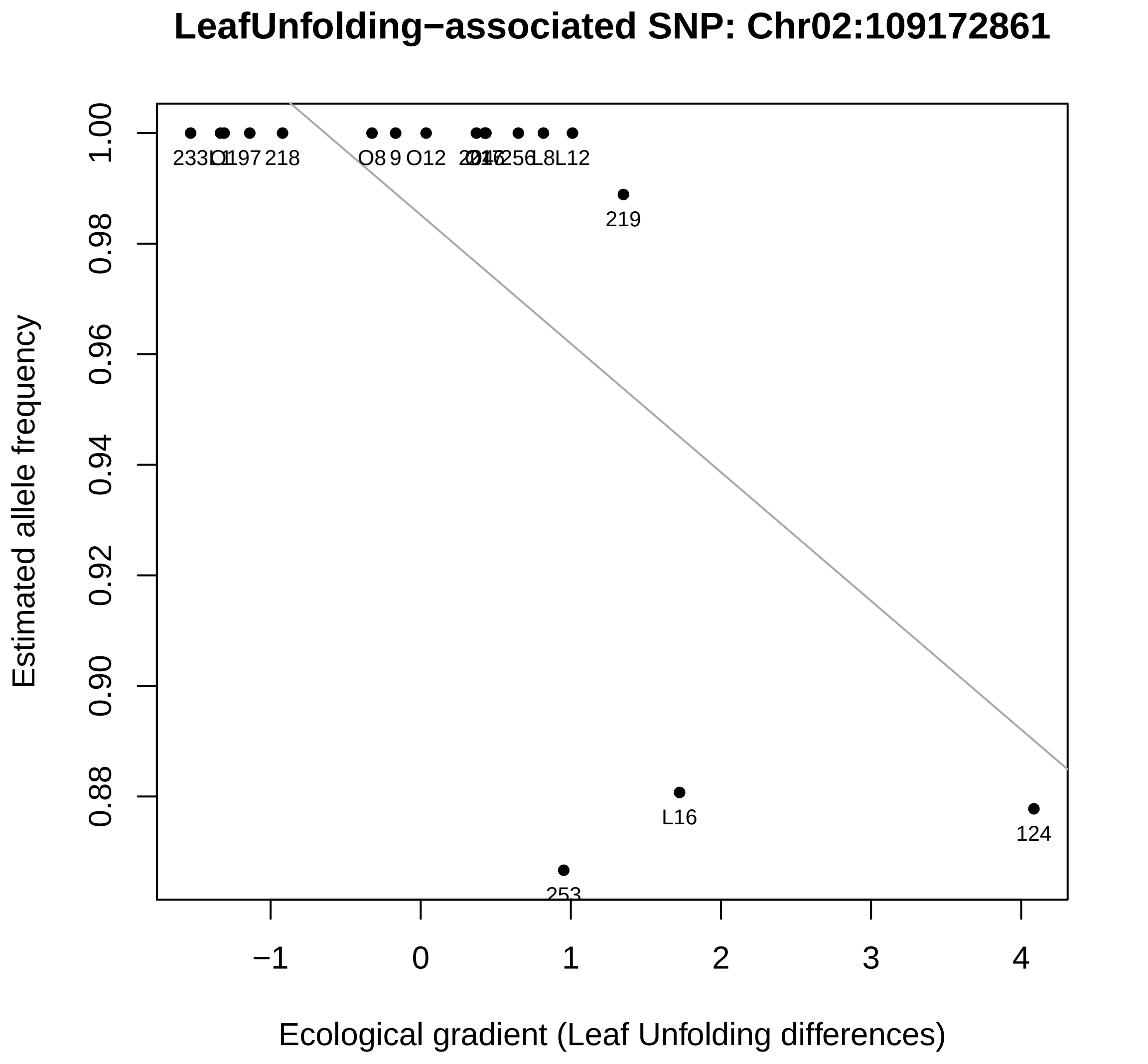
LeafUnfolding−associated SNP: Chr02:109172861
Estimated allele frequency
Ecological gradient (Leaf Unfolding differences)
1.00
0.98
0.96
0.94
0.92
0.90
0.88
−1
0
1
2
3
4
218
O8
9
O12
L8
L12
219
L16
124

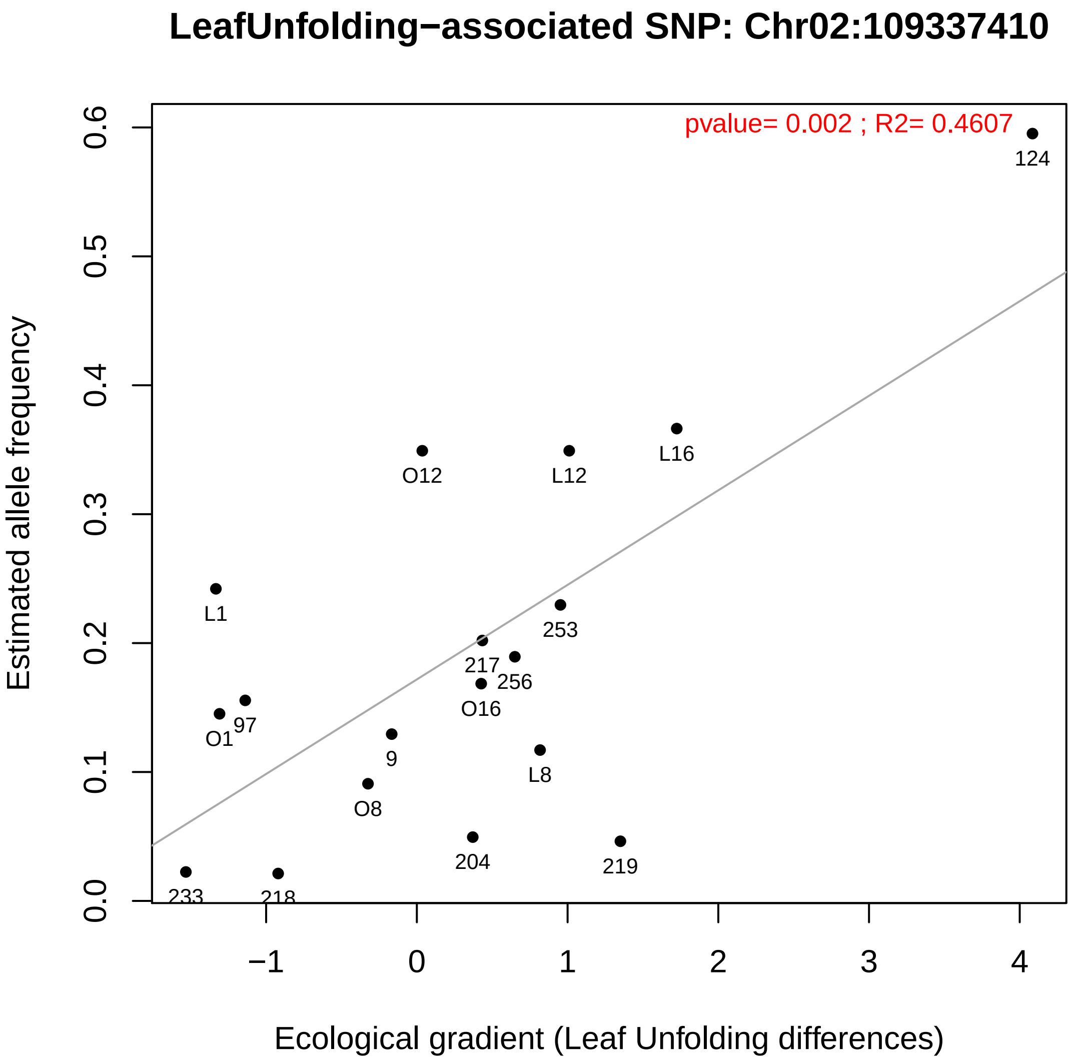

LeafUnfolding−associated SNP: Chr02:109337410
pvalue= 0.002 ; R2= 0.4607
Estimated allele frequency
0.0
0.1
0.2
0.3
0.4
0.5
0.6
Ecological gradient (Leaf Unfolding differences)
−1
0
1
2
3
4
124
L16
O12
L12
L1
253
217
256
O16
97
O1
9
L8
O8
204
219
233
218

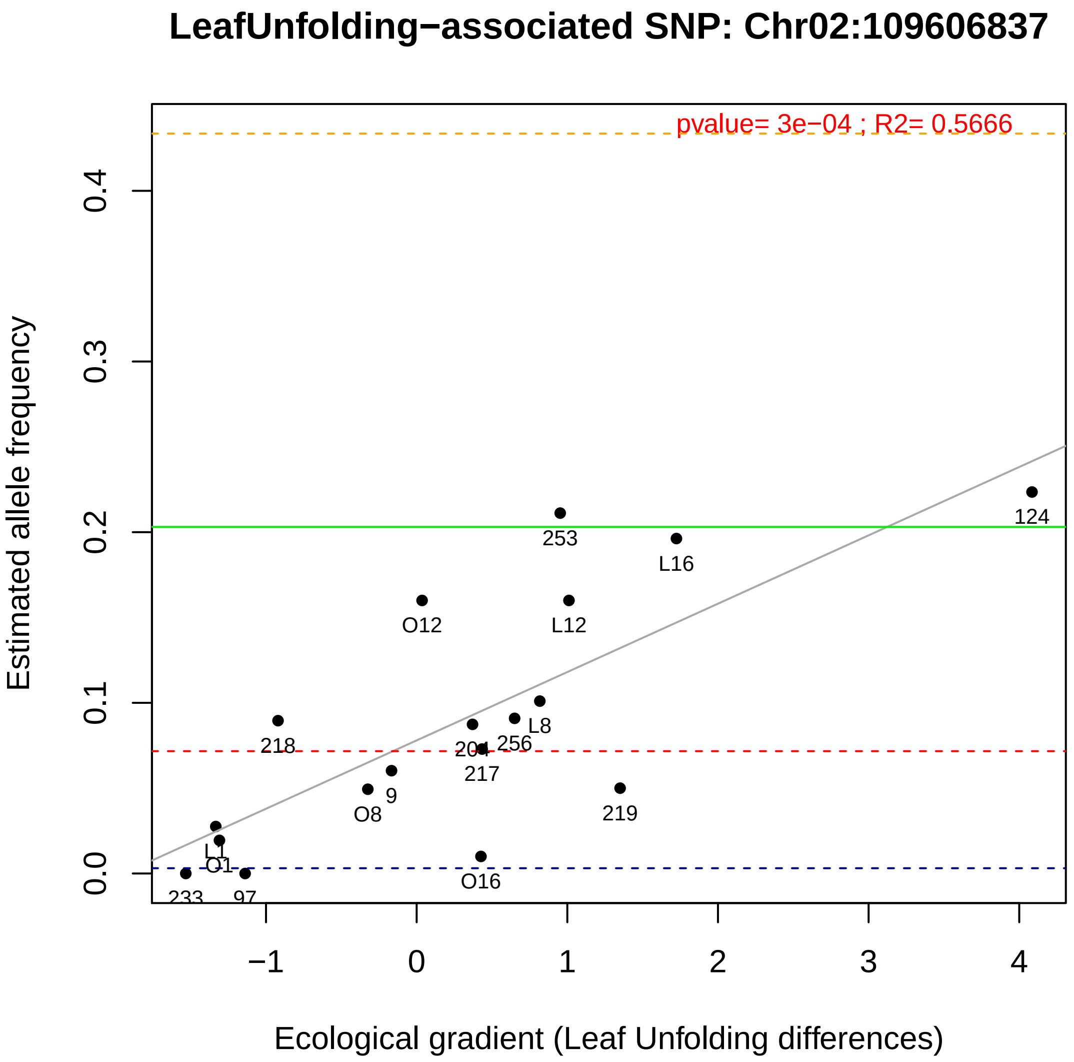

LeafUnfolding−associated SNP: Chr02:109606837
pvalue= 3e−04 ; R2= 0.5666
Estimated allele frequency
0.0
0.1
0.2
0.3
0.4
−1
0
1
2
3
4
Ecological gradient (Leaf Unfolding differences)
124
253
L16
O12
L12
L8
256
204
217
218
9
O8
219
L1
O1
O16
233
97

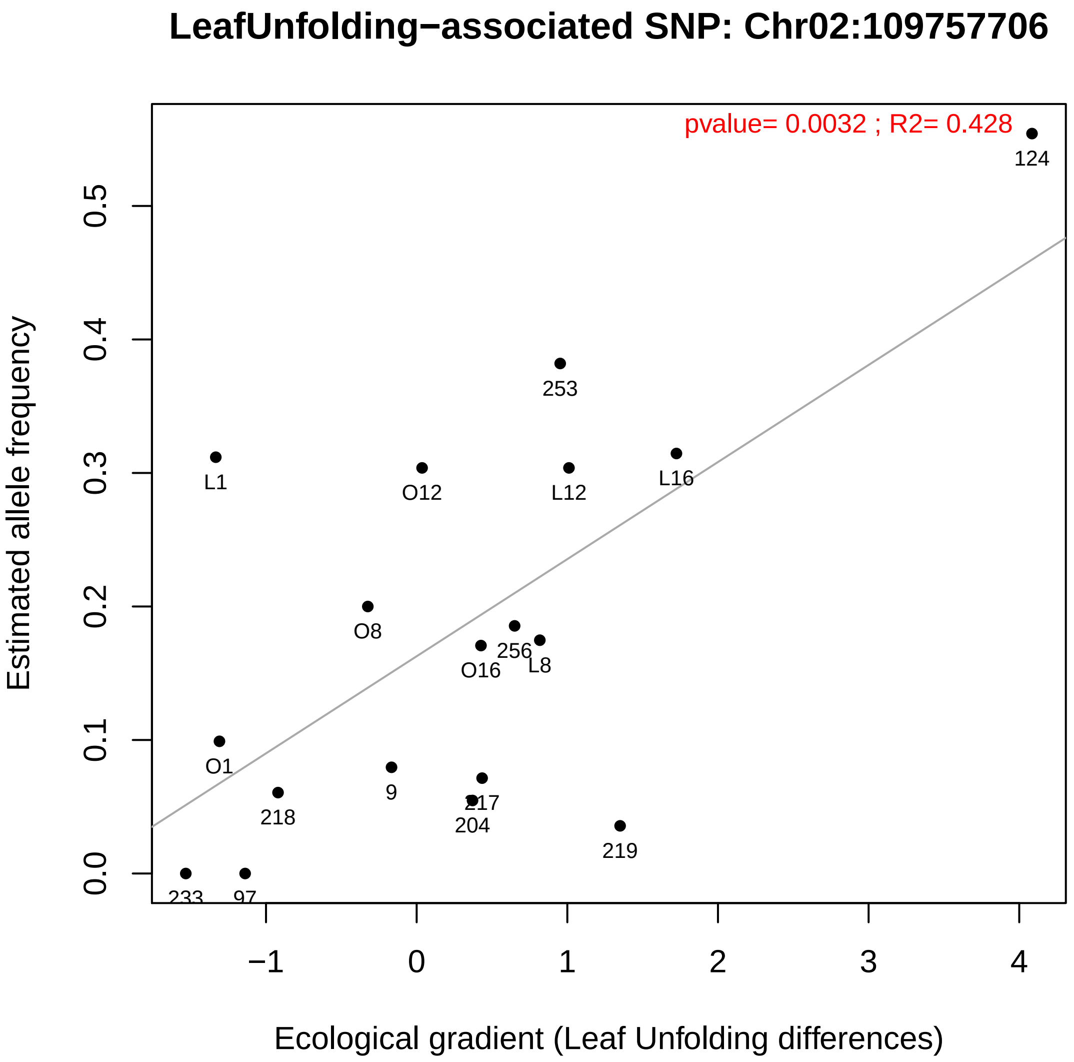

LeafUnfolding−associated SNP: Chr02:109757706
pvalue= 0.0032 ; R2= 0.428
124
253
L1
O12
L12
L16
O8
256
O16
L8
O1
9
217
204
218
219
233
97
0.0
0.1
0.2
0.3
0.4
0.5
−1
0
1
2
3
4
Estimated allele frequency
Ecological gradient (Leaf Unfolding differences)

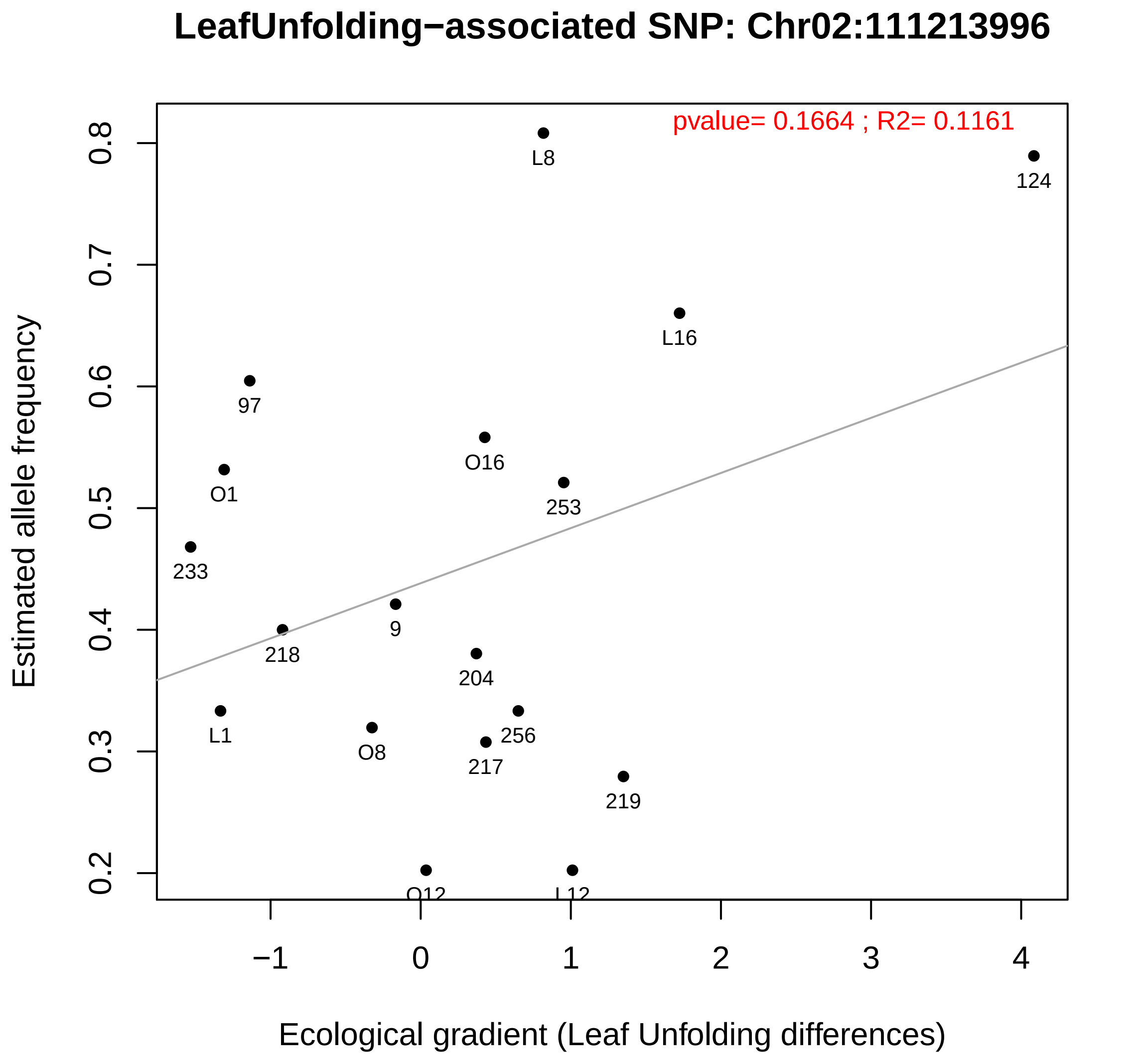

LeafUnfolding−associated SNP: Chr02:111213996
pvalue= 0.1664 ; R2= 0.1161
L8
124
L16
97
O16
O1
253
233
9
218
204
L1
256
O8
217
219
O12
L12
0.2
0.3
0.4
0.5
0.6
0.7
0.8
−1
0
1
2
3
4
Estimated allele frequency
Ecological gradient (Leaf Unfolding differences)

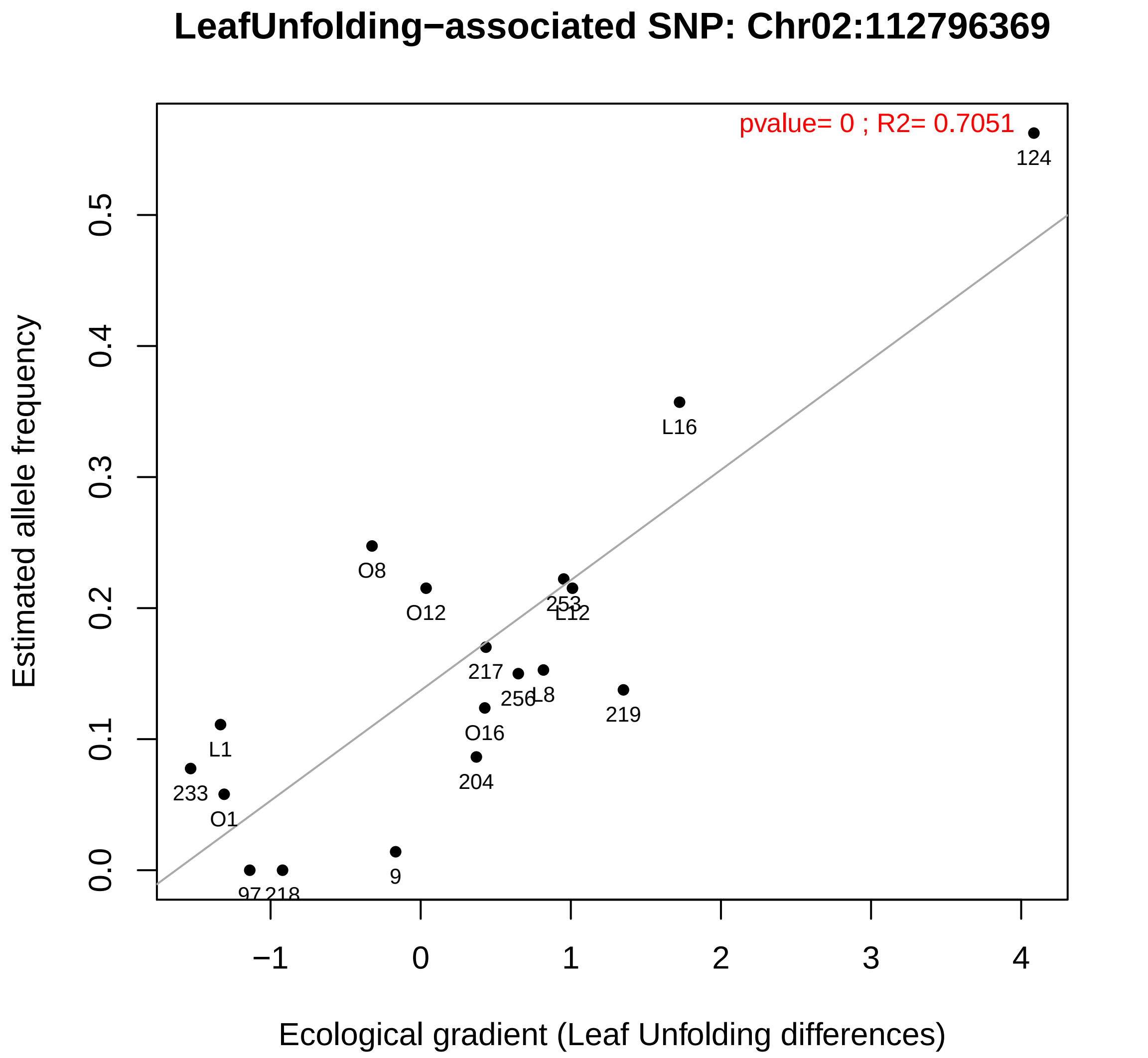

LeafUnfolding−associated SNP: Chr02:112796369
pvalue= 0 ; R2= 0.7051
124
L16
O8
O12
253
L12
217
256
L8
219
O16
L1
204
233
O1
9
97
218
0.0
0.1
0.2
0.3
0.4
0.5
−1
0
1
2
3
4
Estimated allele frequency
Ecological gradient (Leaf Unfolding differences)

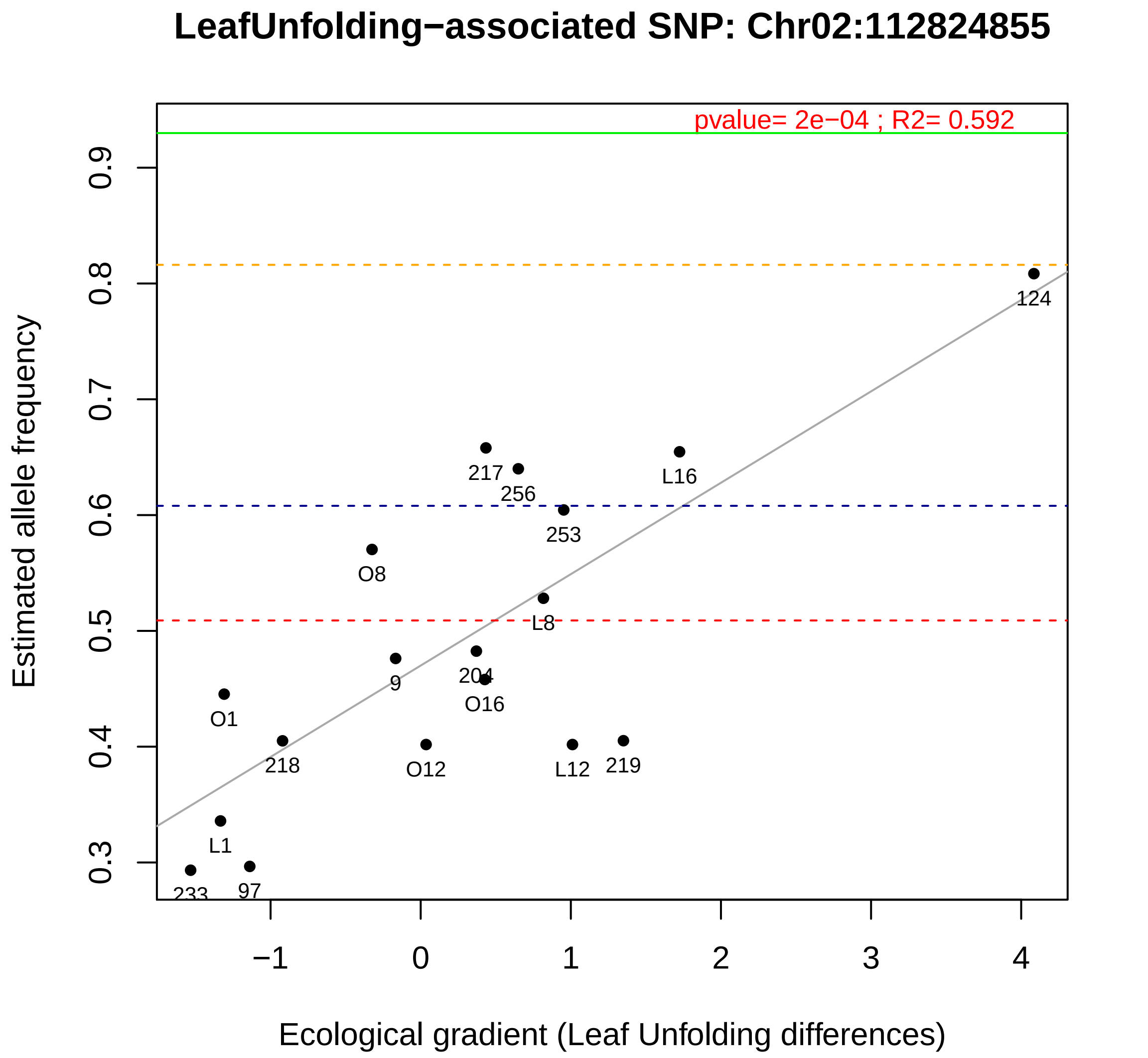
LeafUnfolding−associated SNP: Chr02:112824855
pvalue= 2e−04 ; R2= 0.592
Estimated allele frequency
0.3
0.4
0.5
0.6
0.7
0.8
0.9
−1
0
1
2
3
4
Ecological gradient (Leaf Unfolding differences)
124
217
256
L16
253
O8
L8
9
204
O16
O1
218
O12
L12
219
L1
233
97

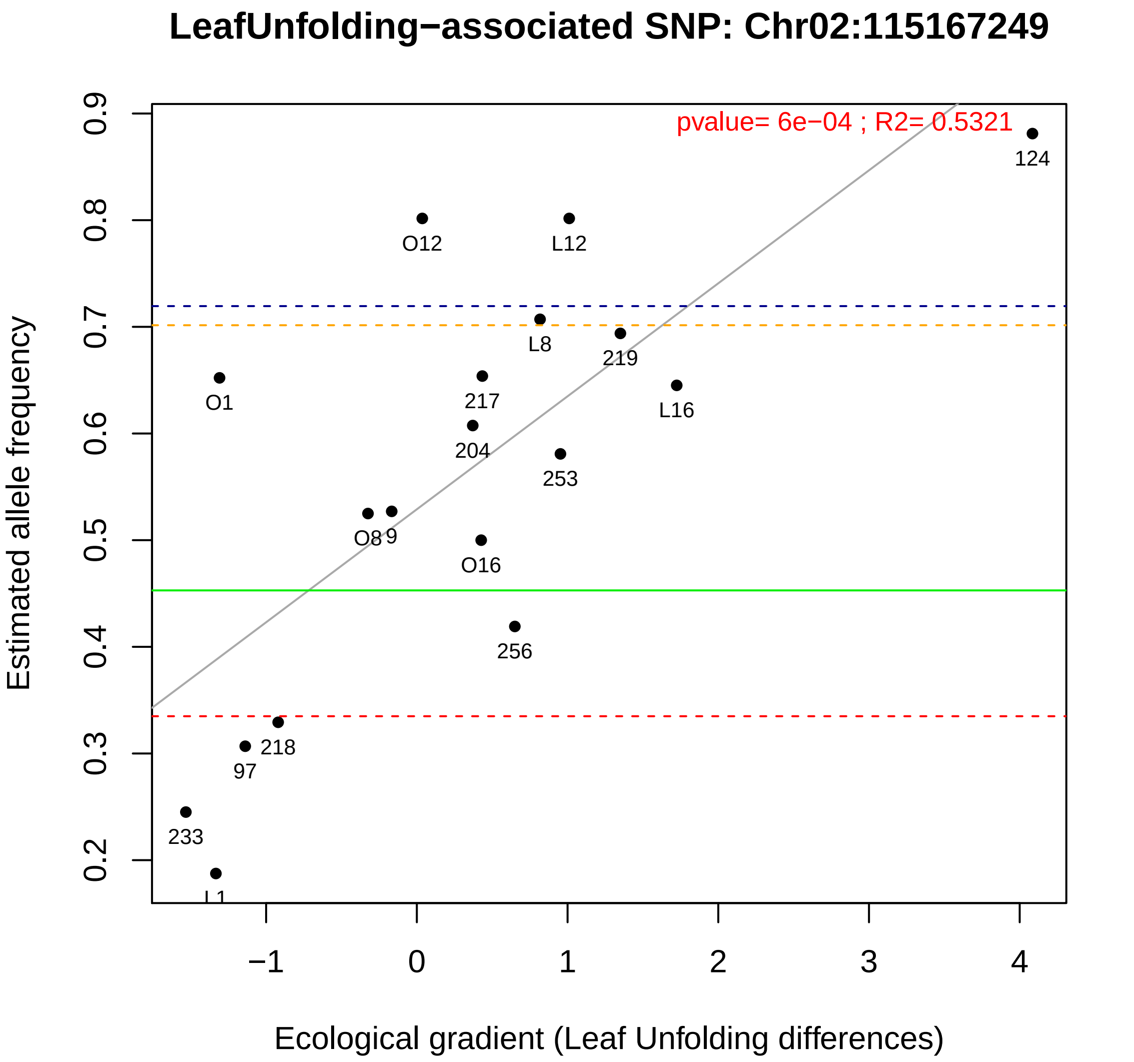

LeafUnfolding−associated SNP: Chr02:115167249
pvalue= 6e−04 ; R2= 0.5321
Estimated allele frequency
Ecological gradient (Leaf Unfolding differences)
0.2
0.3
0.4
0.5
0.6
0.7
0.8
0.9
−1
0
1
2
3
4
124
O12
L12
L8
219
O1
217
L16
204
253
O8
9
O16
256
218
97
233

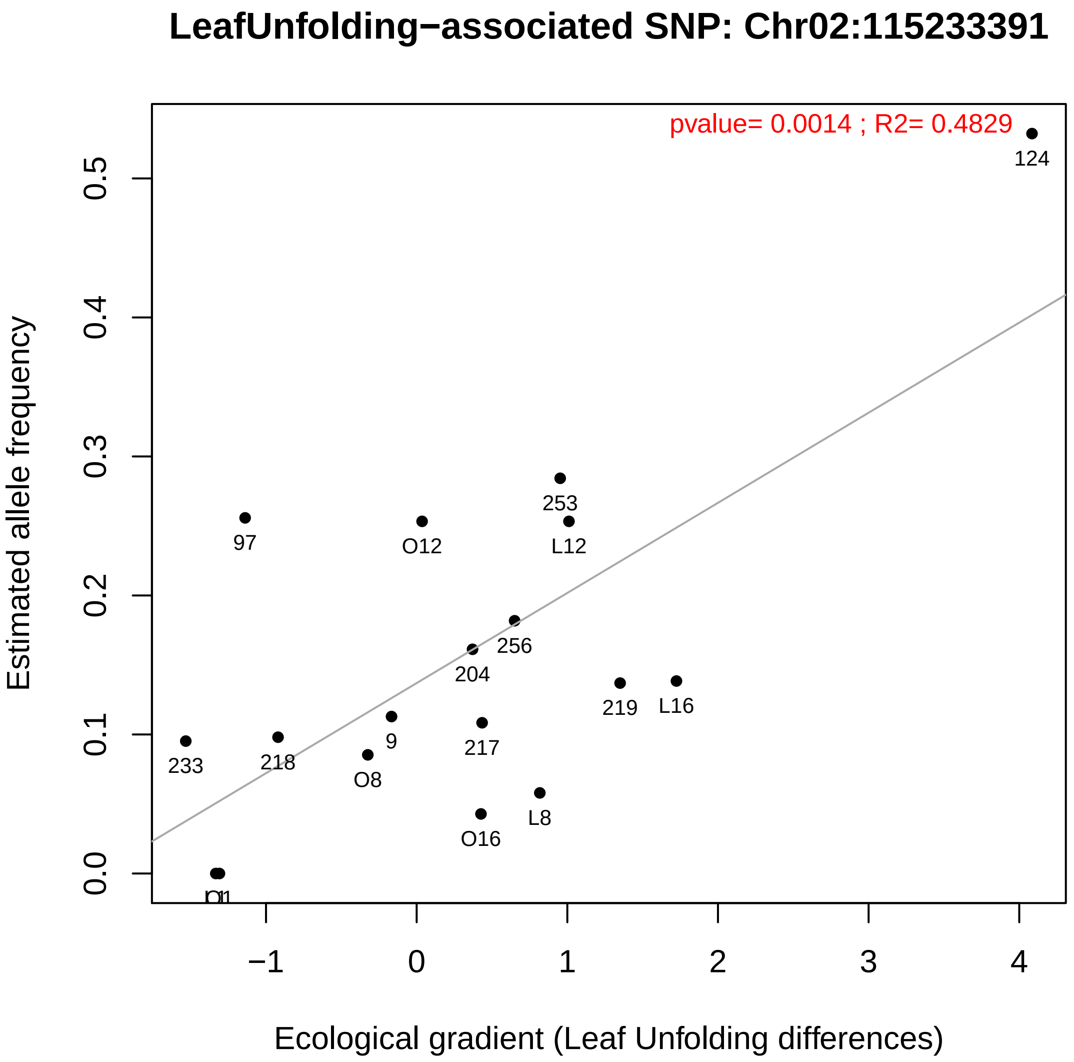

LeafUnfolding−associated SNP: Chr02:115233391
pvalue= 0.0014 ; R2= 0.4829
124
253
97
O12
L12
256
204
219
L16
9
217
233
218
O8
L8
O16
Estimated allele frequency
0.0
0.1
0.2
0.3
0.4
0.5
−1
0
1
2
3
4
Ecological gradient (Leaf Unfolding differences)

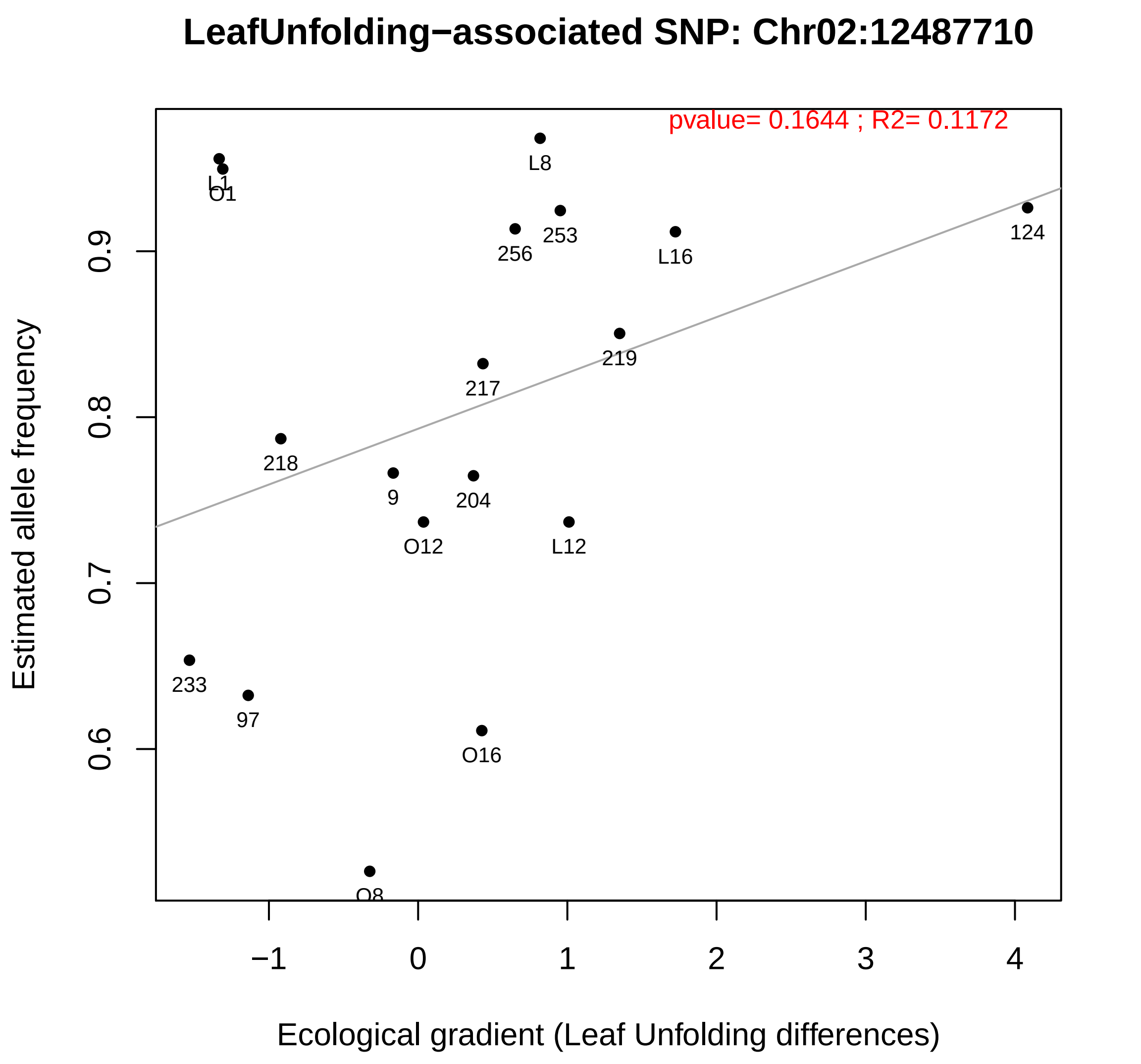

LeafUnfolding−associated SNP: Chr02:12487710
pvalue= 0.1644 ; R2= 0.1172
Estimated allele frequency
Ecological gradient (Leaf Unfolding differences)
0.6
0.7
0.8
0.9
−1
0
1
2
3
4
L8
L1
O1
253
256
L16
124
219
217
218
9
204
O12
L12
233
97
O16
O8

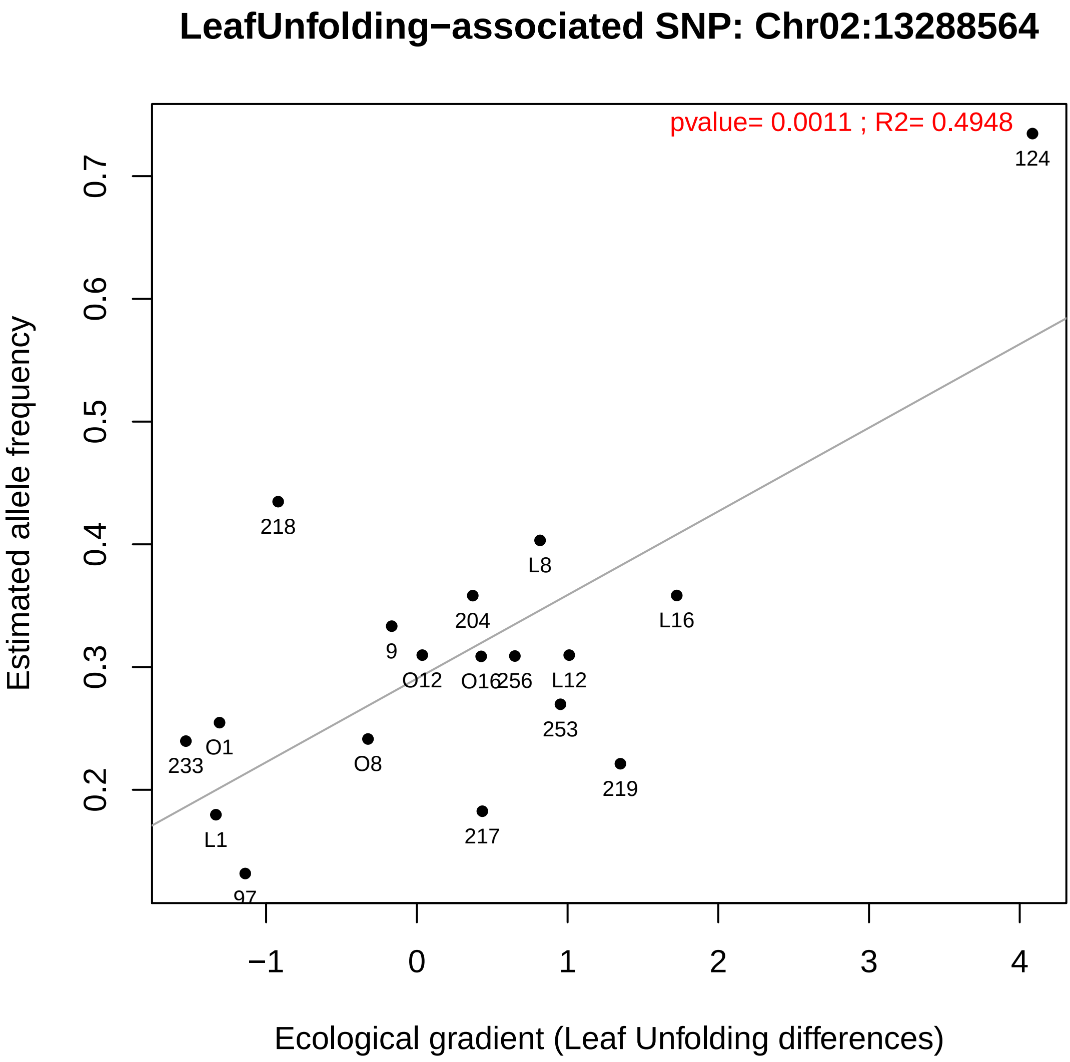
LeafUnfolding−associated SNP: Chr02:13288564
pvalue= 0.0011 ; R2= 0.4948
Estimated allele frequency
Ecological gradient (Leaf Unfolding differences)
0.2
0.3
0.4
0.5
0.6
0.7
−1
0
1
2
3
4
124
218
L8
204
L16
9
O12
O16
256
L12
253
O1
233
O8
219
L1
217
97

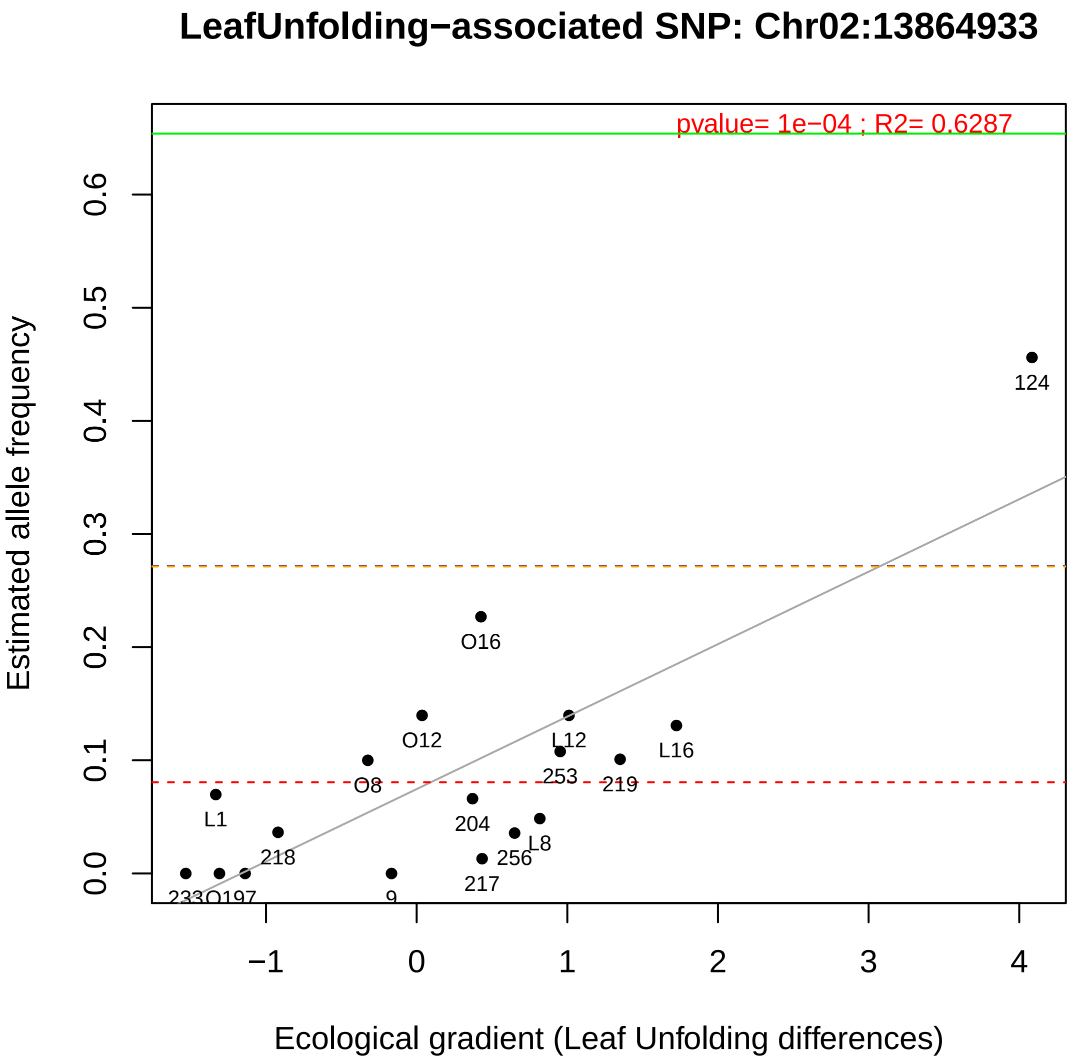

LeafUnfolding−associated SNP: Chr02:13864933
pvalue= 1e−04 ; R2= 0.6287
Estimated allele frequency
0.0
0.1
0.2
0.3
0.4
0.5
0.6
Ecological gradient (Leaf Unfolding differences)
−1
0
1
2
3
4
124
O16
O12
L12
L16
253
219
O8
L1
204
L8
256
218
217
9
233
O197

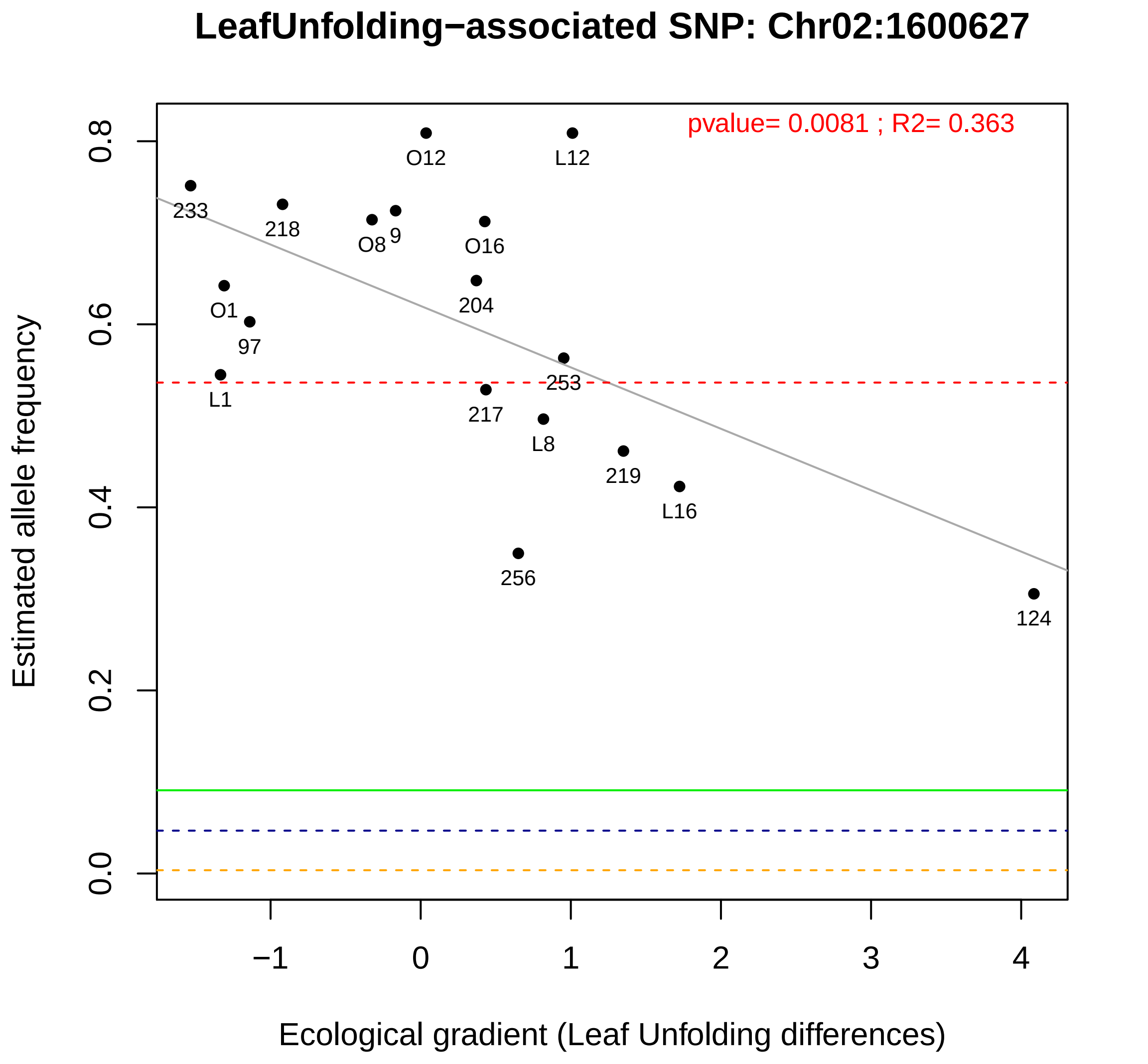
LeafUnfolding−associated SNP: Chr02:1600627
pvalue= 0.0081 ; R2= 0.363
Estimated allele frequency
0.0
0.2
0.4
0.6
0.8
Ecological gradient (Leaf Unfolding differences)
−1
0
1
2
3
4
233
218
O8
9
O12
L12
O16
204
O1
97
L1
253
217
L8
219
L16
256
124

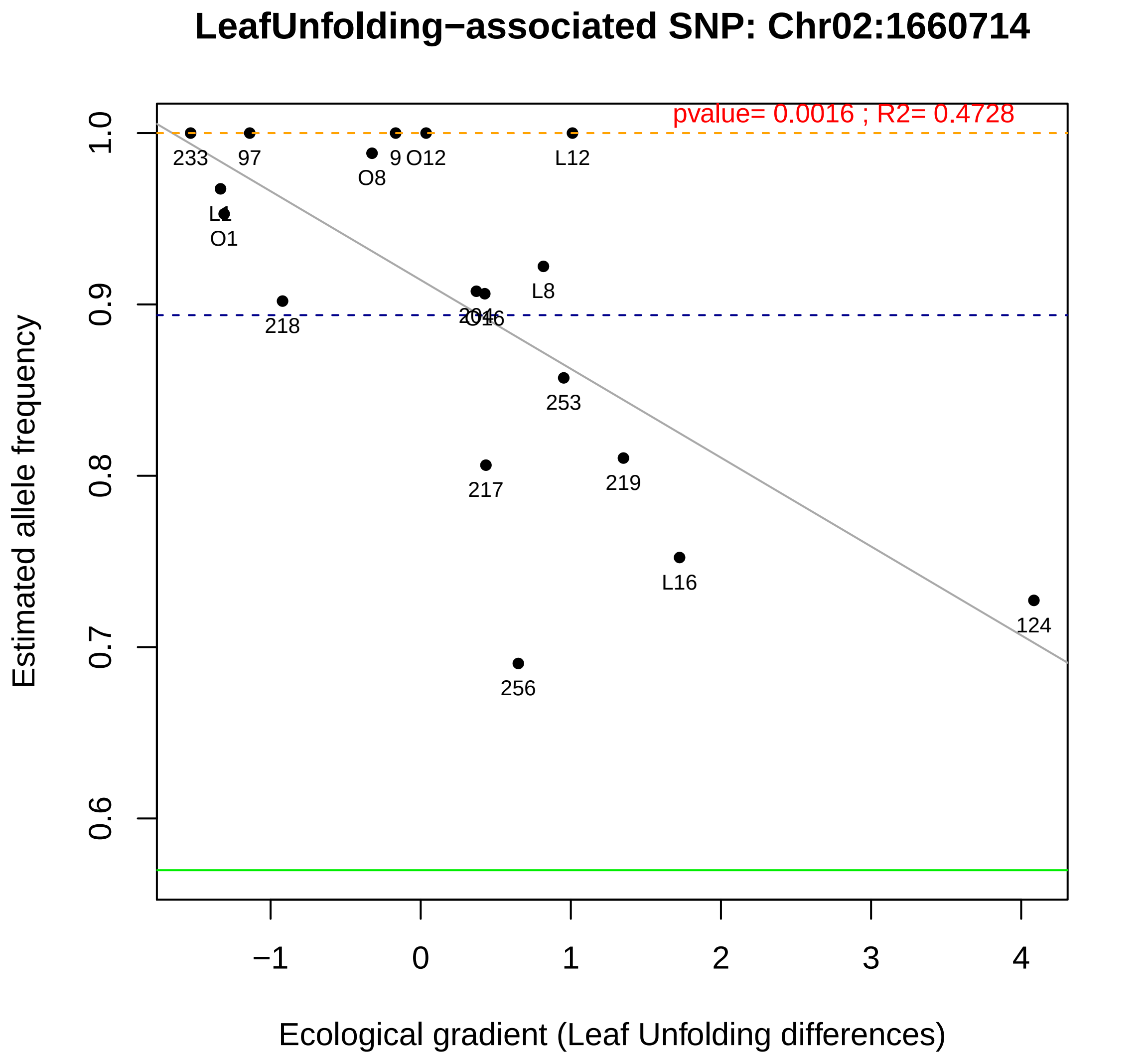

LeafUnfolding−associated SNP: Chr02:1660714
pvalue= 0.0016 ; R2= 0.4728
Estimated allele frequency
Ecological gradient (Leaf Unfolding differences)
1.0
0.9
0.8
0.7
0.6
−1
0
1
2
3
4
233
97
O8
9
O12
L12
L1
O1
218
204
O16
L8
253
217
219
L16
124
256

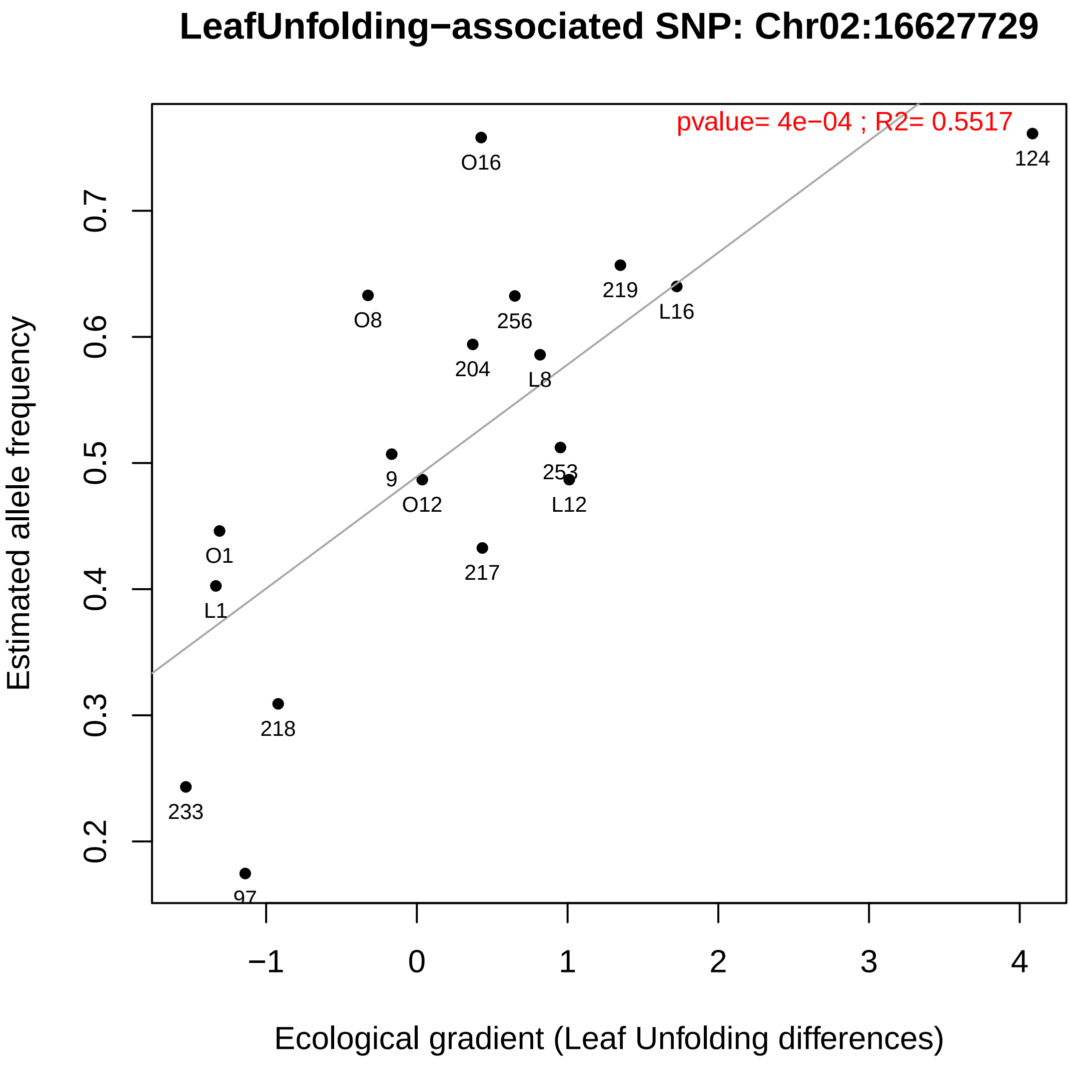

LeafUnfolding−associated SNP: Chr02:16627729
pvalue= 4e−04 ; R2= 0.5517
Estimated allele frequency
0.2
0.3
0.4
0.5
0.6
0.7
−1
0
1
2
3
4
Ecological gradient (Leaf Unfolding differences)
O16
124
219
L16
O8
256
204
L8
9
O12
253
L12
O1
217
L1
218
233
97

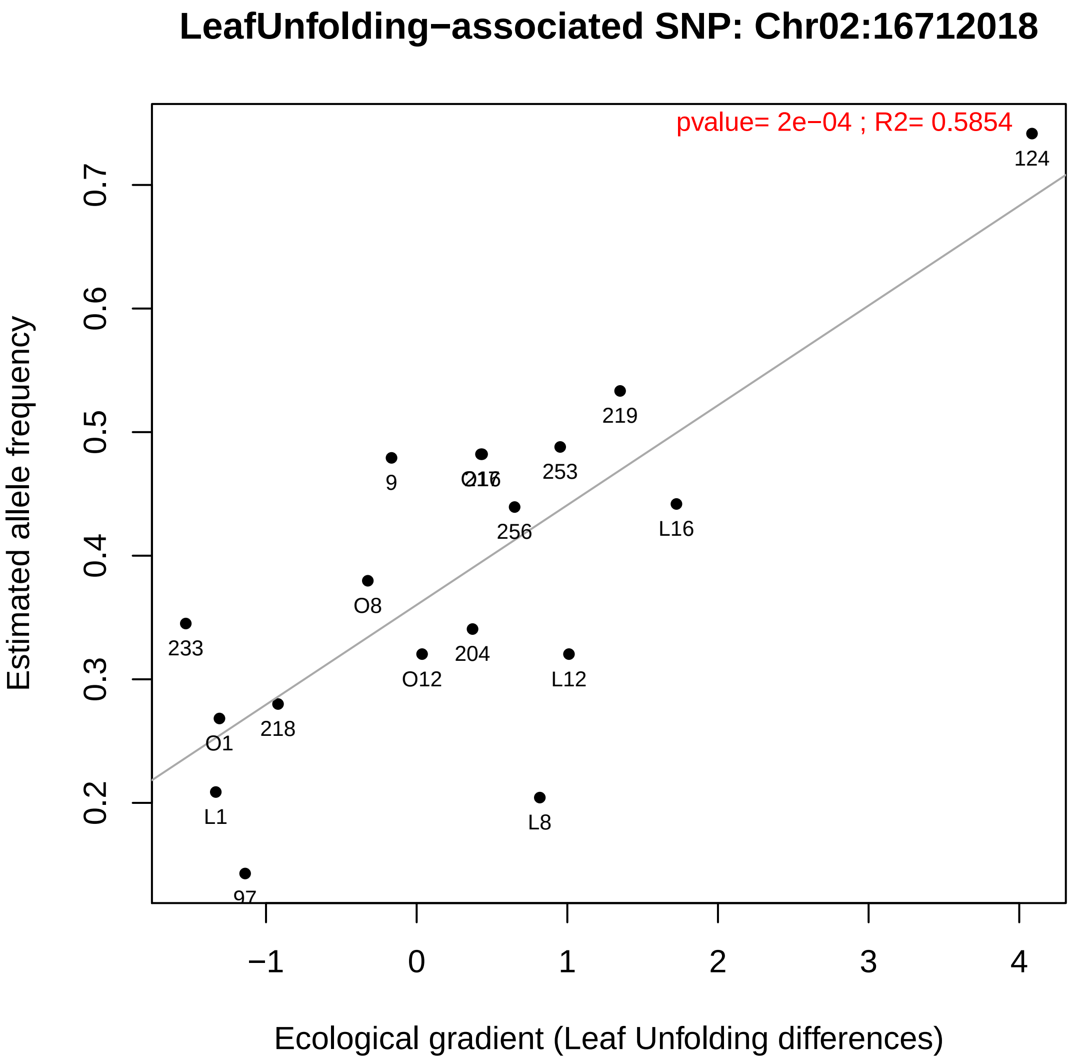

LeafUnfolding−associated SNP: Chr02:16712018
pvalue= 2e−04 ; R2= 0.5854
124
219
253
9
256
L16
O8
233
204
O12
L12
218
O1
L1
L8
97
0.2
0.3
0.4
0.5
0.6
0.7
−1
0
1
2
3
4
Estimated allele frequency
Ecological gradient (Leaf Unfolding differences)

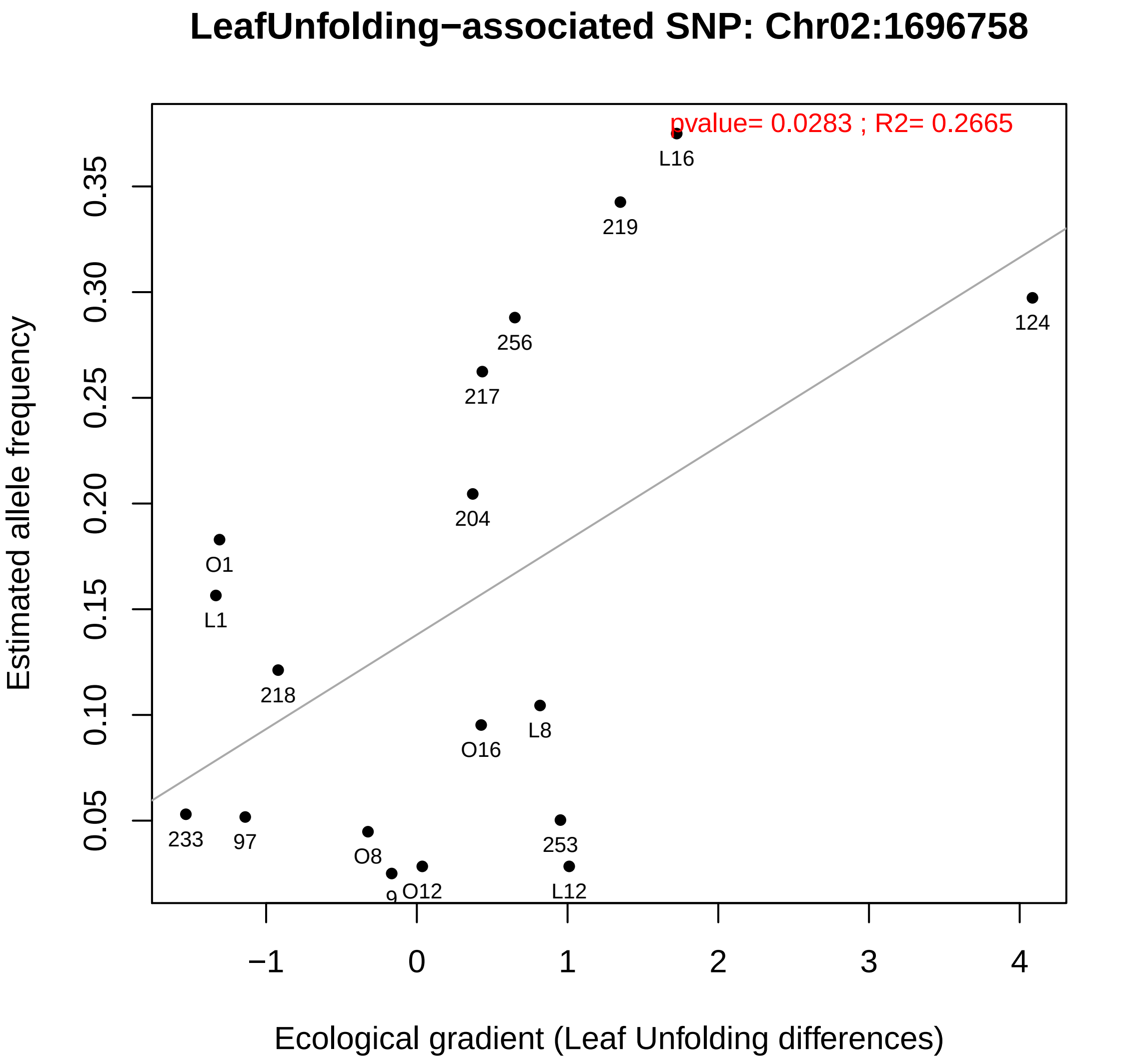

LeafUnfolding−associated SNP: Chr02:1696758
pvalue= 0.0283 ; R2= 0.2665
Estimated allele frequency
0.05
0.10
0.15
0.20
0.25
0.30
0.35
−1
0
1
2
3
4
Ecological gradient (Leaf Unfolding differences)
L16
219
124
256
217
204
O1
L1
218
L8
O16
233
97
O8
9
O12
253
L12

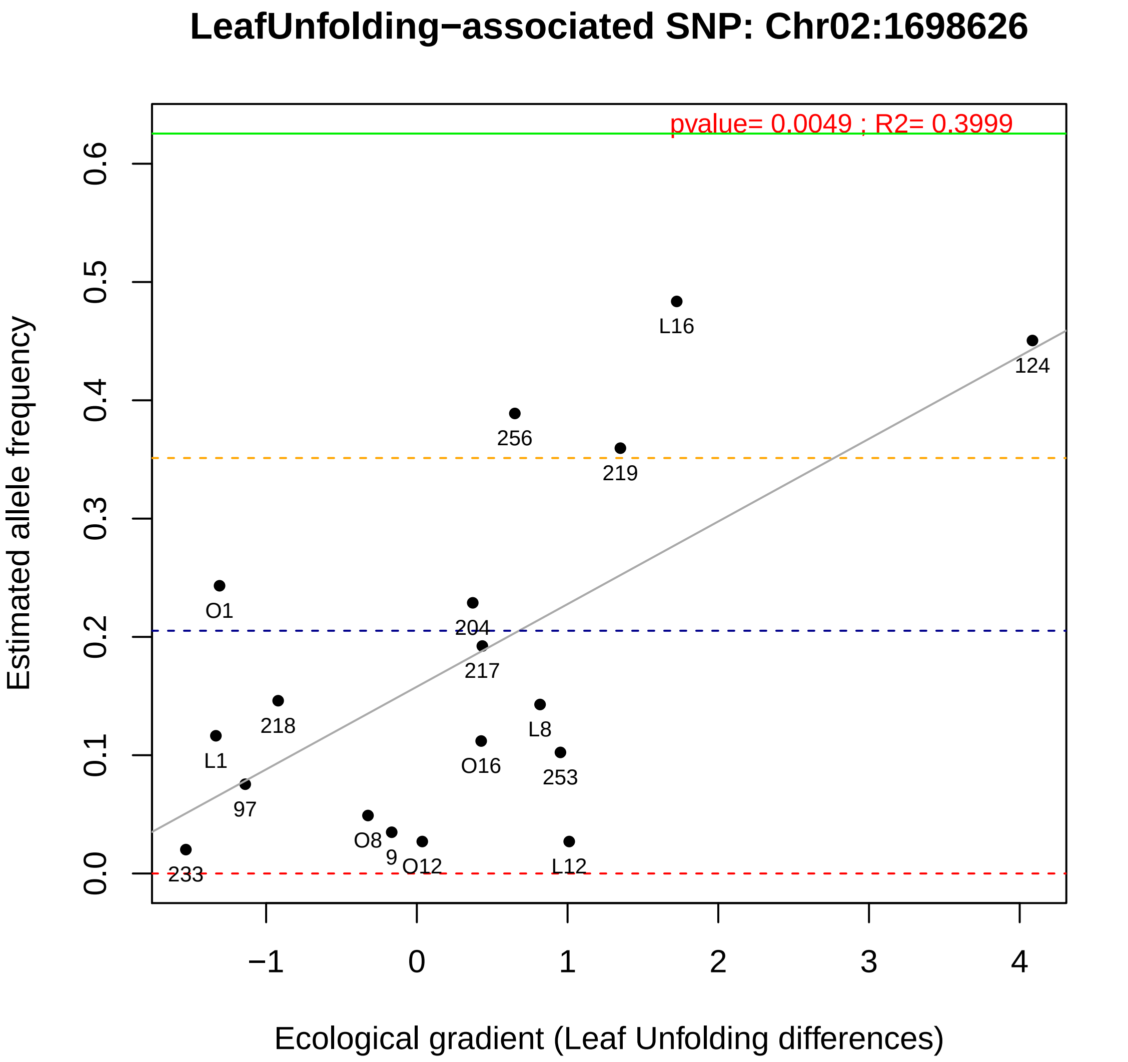

LeafUnfolding−associated SNP: Chr02:1698626
pvalue= 0.0049 ; R2= 0.3999
Estimated allele frequency
0.0
0.1
0.2
0.3
0.4
0.5
0.6
−1
0
1
2
3
4
Ecological gradient (Leaf Unfolding differences)
L16
124
256
219
O1
204
217
218
L8
L1
O16
253
97
O8
9
O12
L12
233

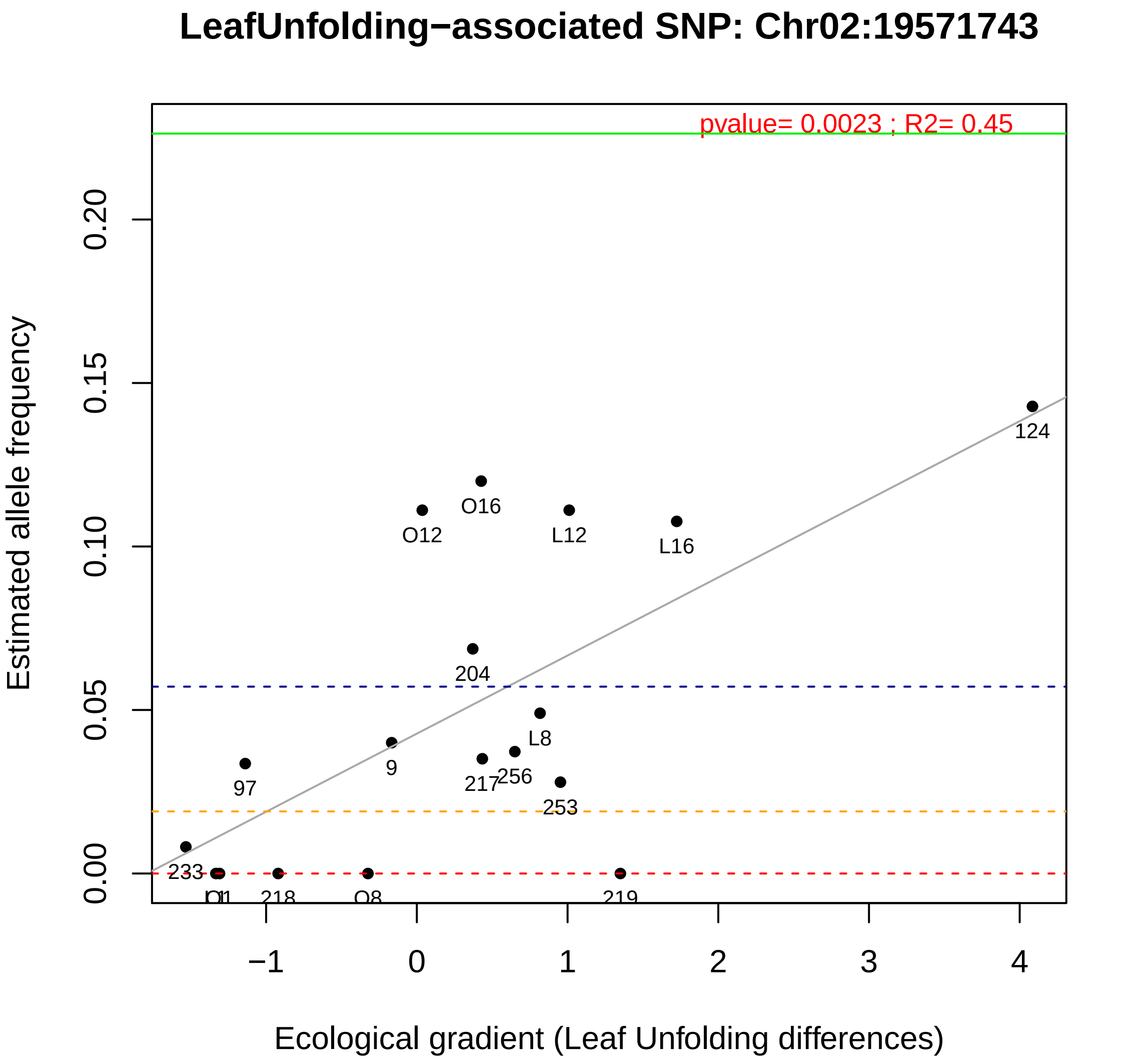

LeafUnfolding−associated SNP: Chr02:19571743
pvalue= 0.0023 ; R2= 0.45
Estimated allele frequency
0.00
0.05
0.10
0.15
0.20
−1
0
1
2
3
4
Ecological gradient (Leaf Unfolding differences)
124
O16
O12
L12
L16
204
L8
9
97
217
256
253
233
218
O8
219

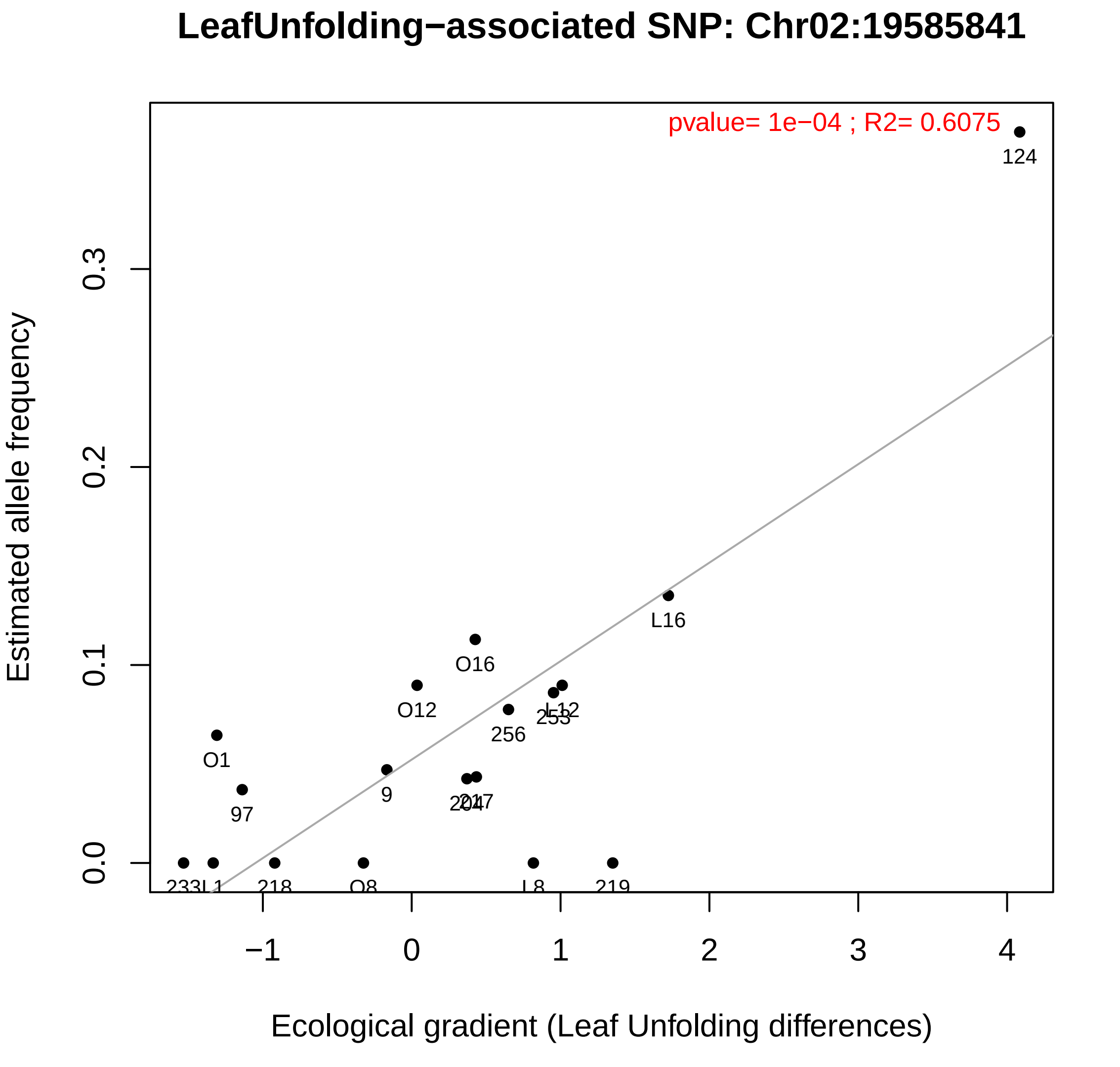

LeafUnfolding−associated SNP: Chr02:19585841
pvalue= 1e−04 ; R2= 0.6075
124
L16
O16
O12
L12
253
256
O1
9
97
204
217
233
L1
218
O8
L8
219
0.0
0.1
0.2
0.3
−1
0
1
2
3
4
Estimated allele frequency
Ecological gradient (Leaf Unfolding differences)

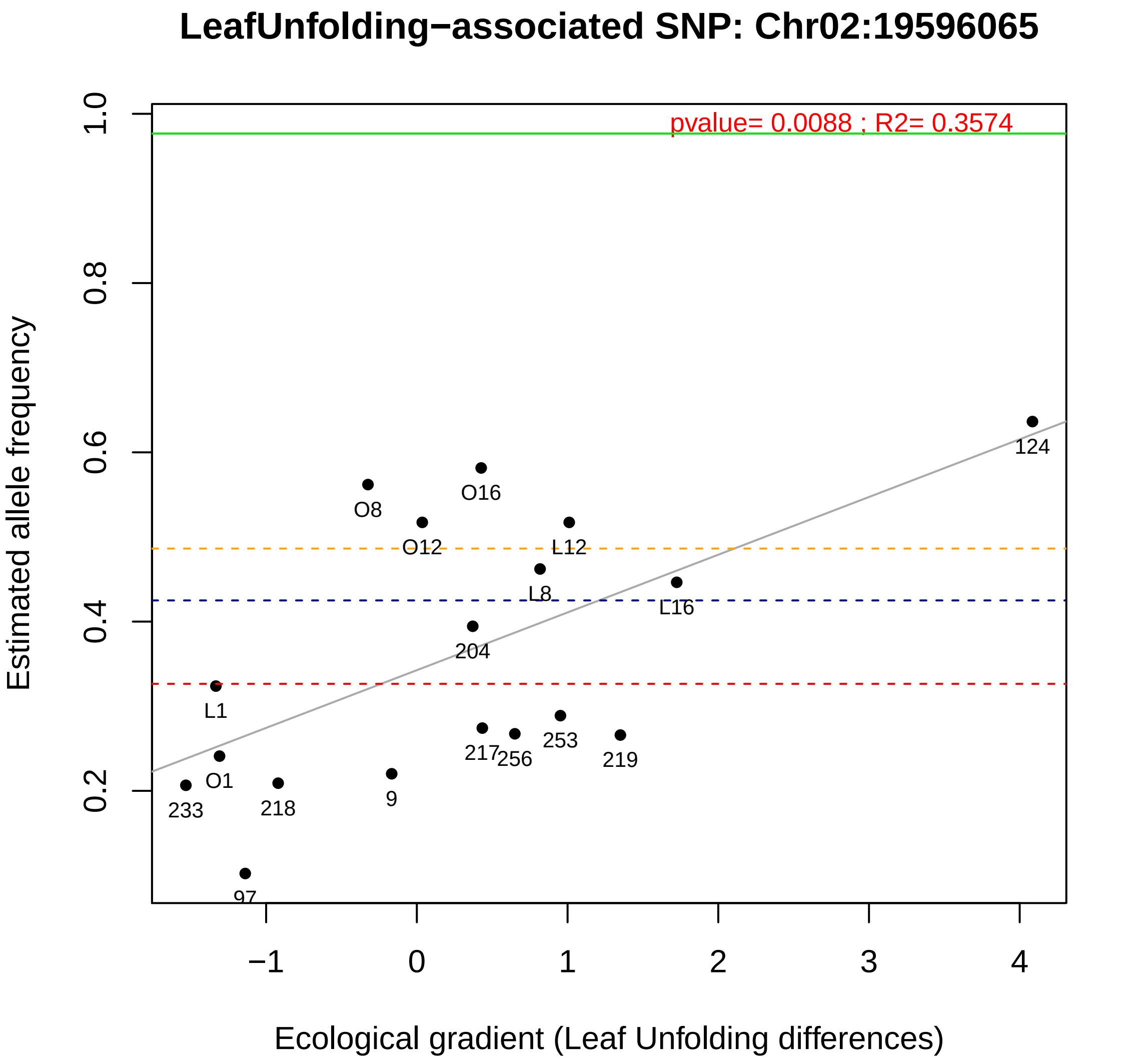
LeafUnfolding−associated SNP: Chr02:19596065
pvalue= 0.0088 ; R2= 0.3574
Estimated allele frequency
Ecological gradient (Leaf Unfolding differences)
0.2
0.4
0.6
0.8
1.0
−1
0
1
2
3
4
124
O16
O8
O12
L12
L8
L16
204
L1
217
256
253
219
O1
233
218
9
97

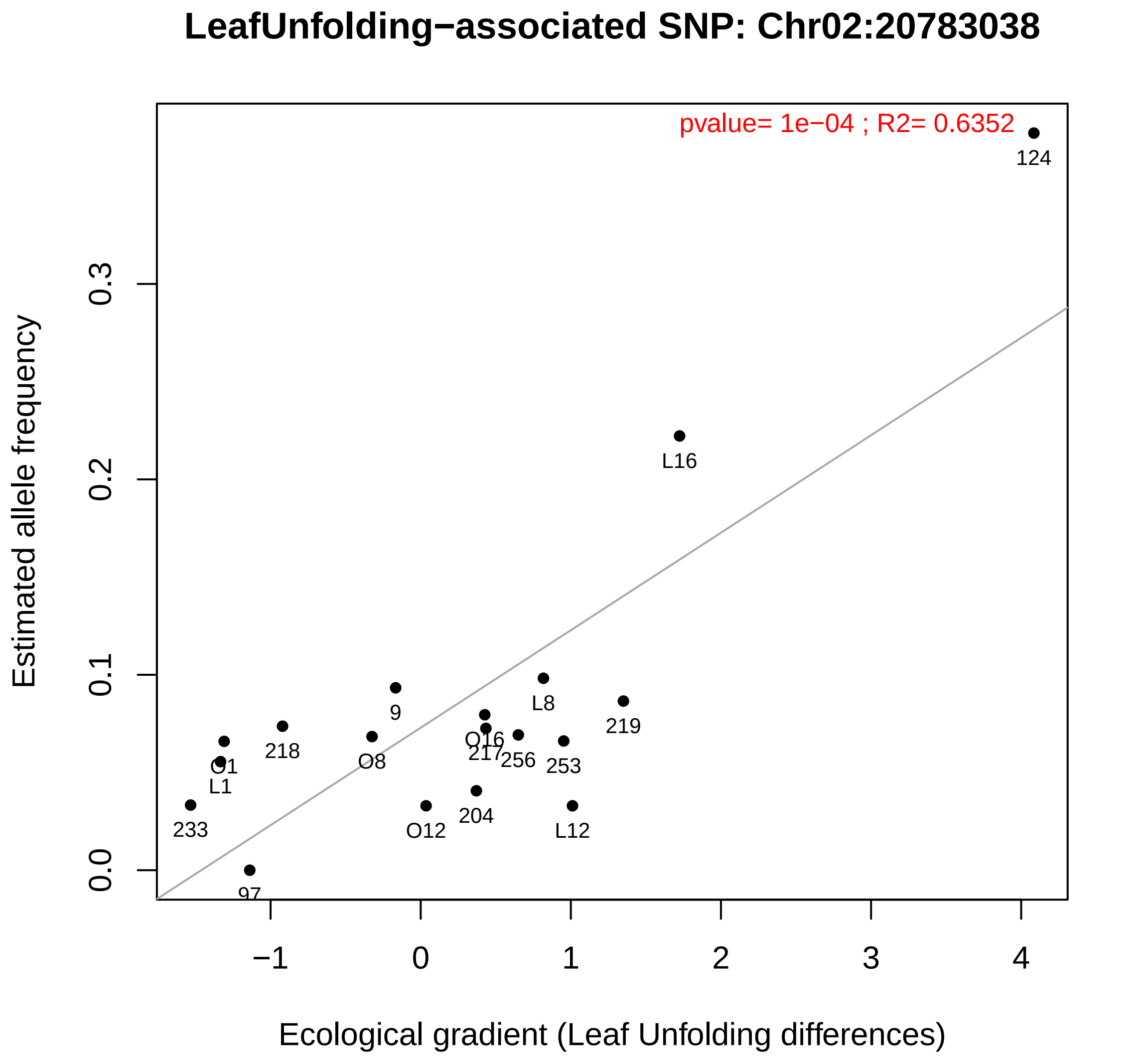

LeafUnfolding−associated SNP: Chr02:20783038
pvalue= 1e−04 ; R2= 0.6352
Estimated allele frequency
0.0
0.1
0.2
0.3
Ecological gradient (Leaf Unfolding differences)
−1
0
1
2
3
4
124
L16
L8
219
9
218
O8
O16
217
256
253
O1
L1
233
97
O12
204
L12

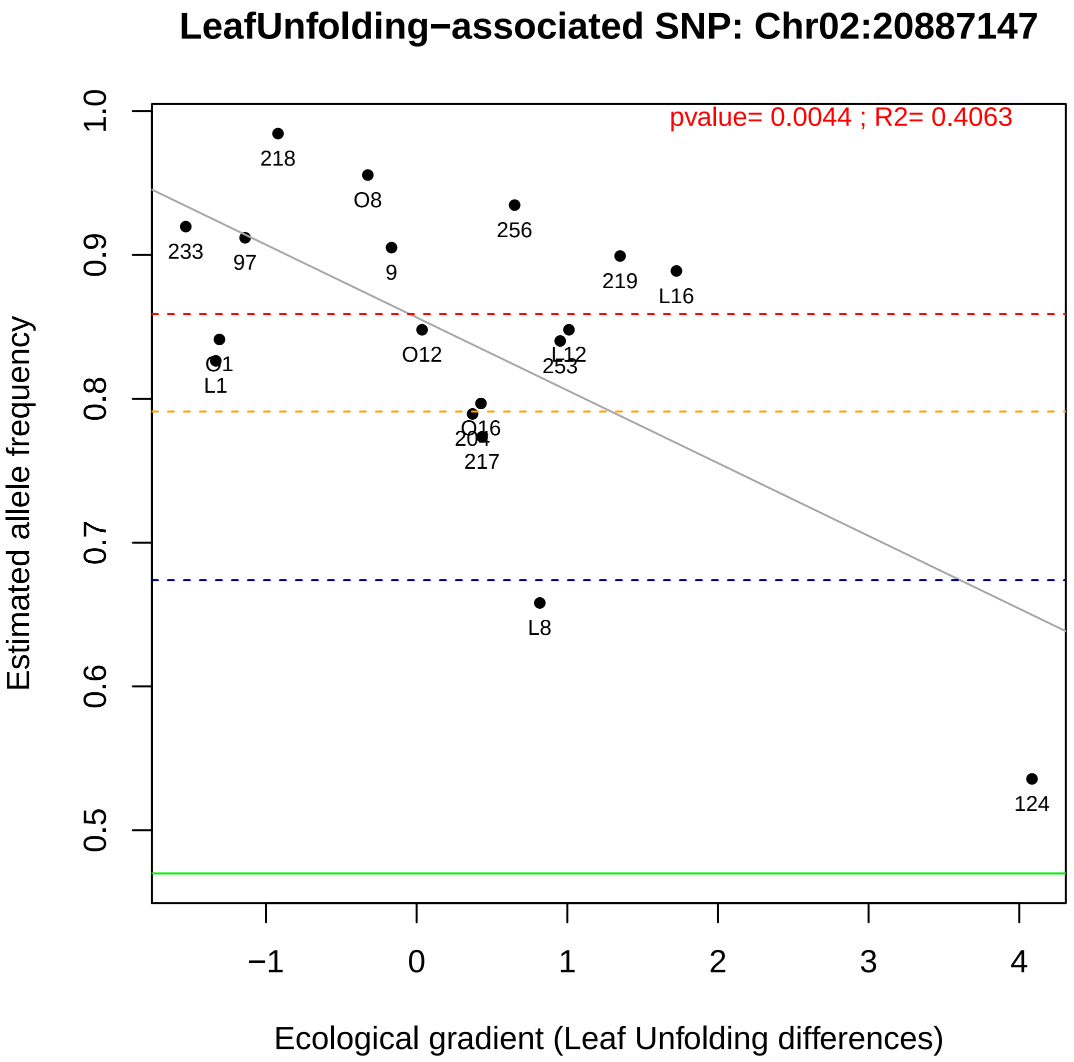

LeafUnfolding−associated SNP: Chr02:20887147
pvalue= 0.0044 ; R2= 0.4063
Estimated allele frequency
Ecological gradient (Leaf Unfolding differences)
0.5
0.6
0.7
0.8
0.9
1.0
−1
0
1
2
3
4
218
O8
256
233
97
9
219
L16
O12
O1
L1
L12
253
O16
204
217
L8
124

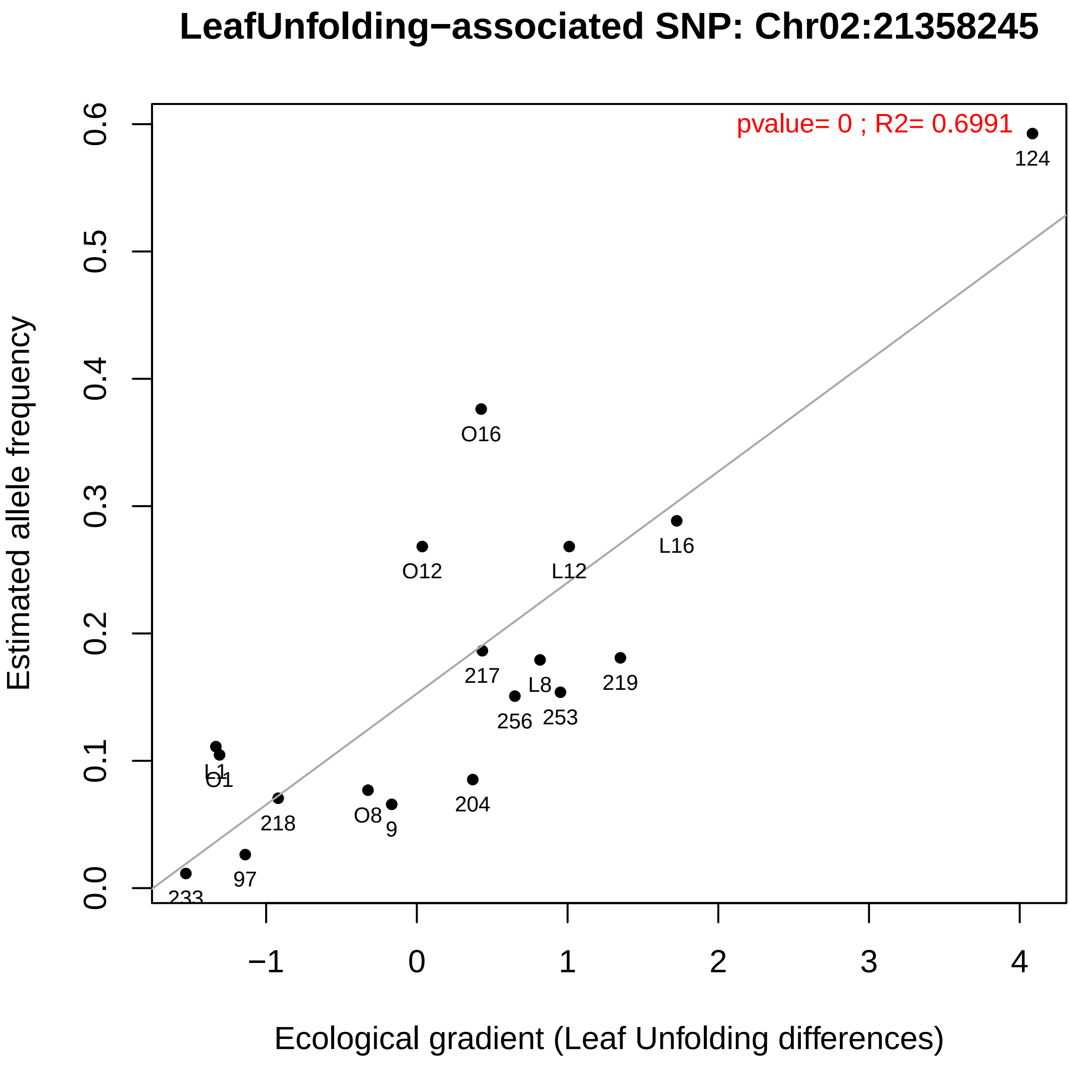

LeafUnfolding−associated SNP: Chr02:21358245
pvalue= 0 ; R2= 0.6991
Estimated allele frequency
Ecological gradient (Leaf Unfolding differences)
0.0
0.1
0.2
0.3
0.4
0.5
0.6
−1
0
1
2
3
4
124
O16
O12
L12
L16
217
L8
219
256
253
L1
O1
218
O8
9
204
97
233

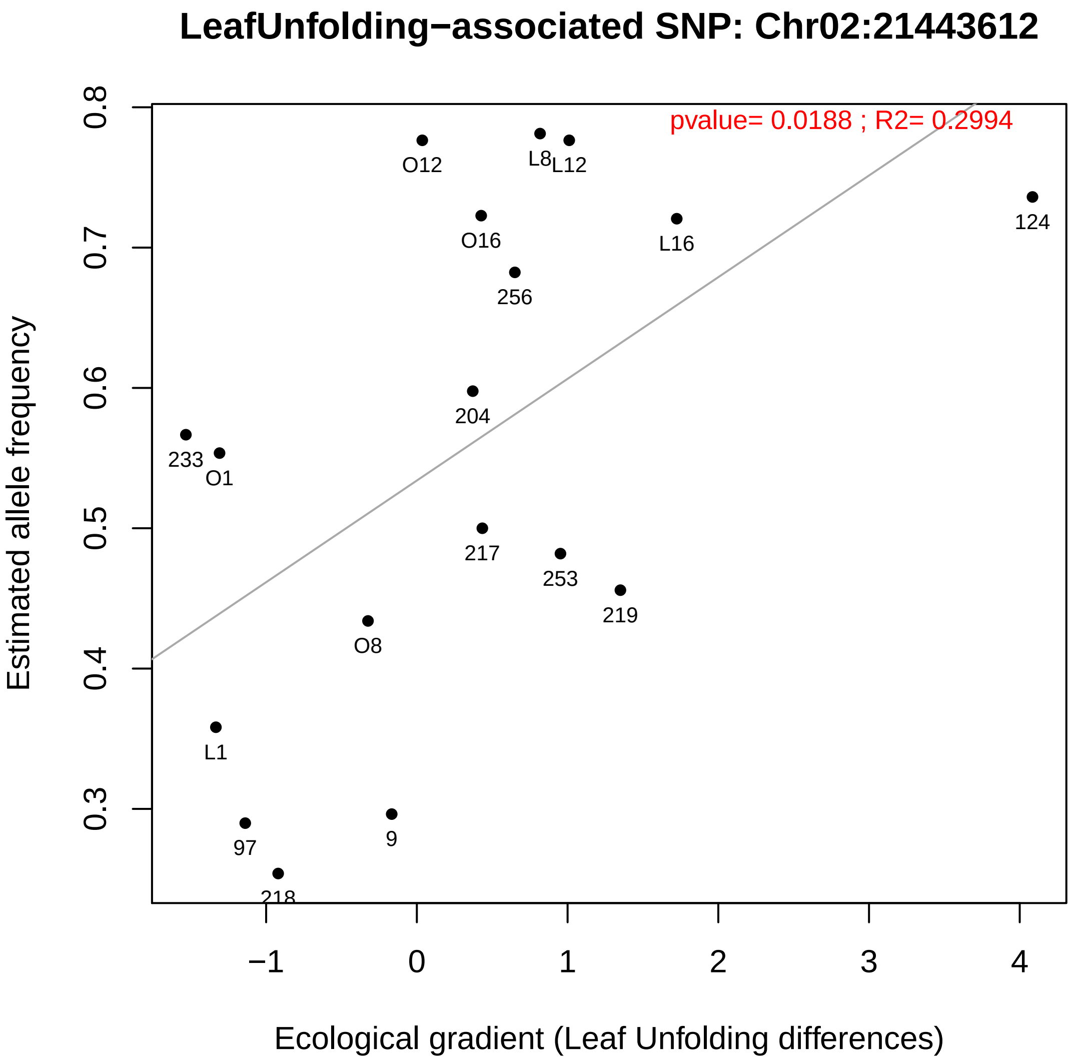
LeafUnfolding−associated SNP: Chr02:21443612
pvalue= 0.0188 ; R2= 0.2994
Estimated allele frequency
Ecological gradient (Leaf Unfolding differences)
0.3
0.4
0.5
0.6
0.7
0.8
−1
0
1
2
3
4
O12
L8
L12
124
O16
L16
256
204
233
O1
217
253
219
O8
L1
97
9
218

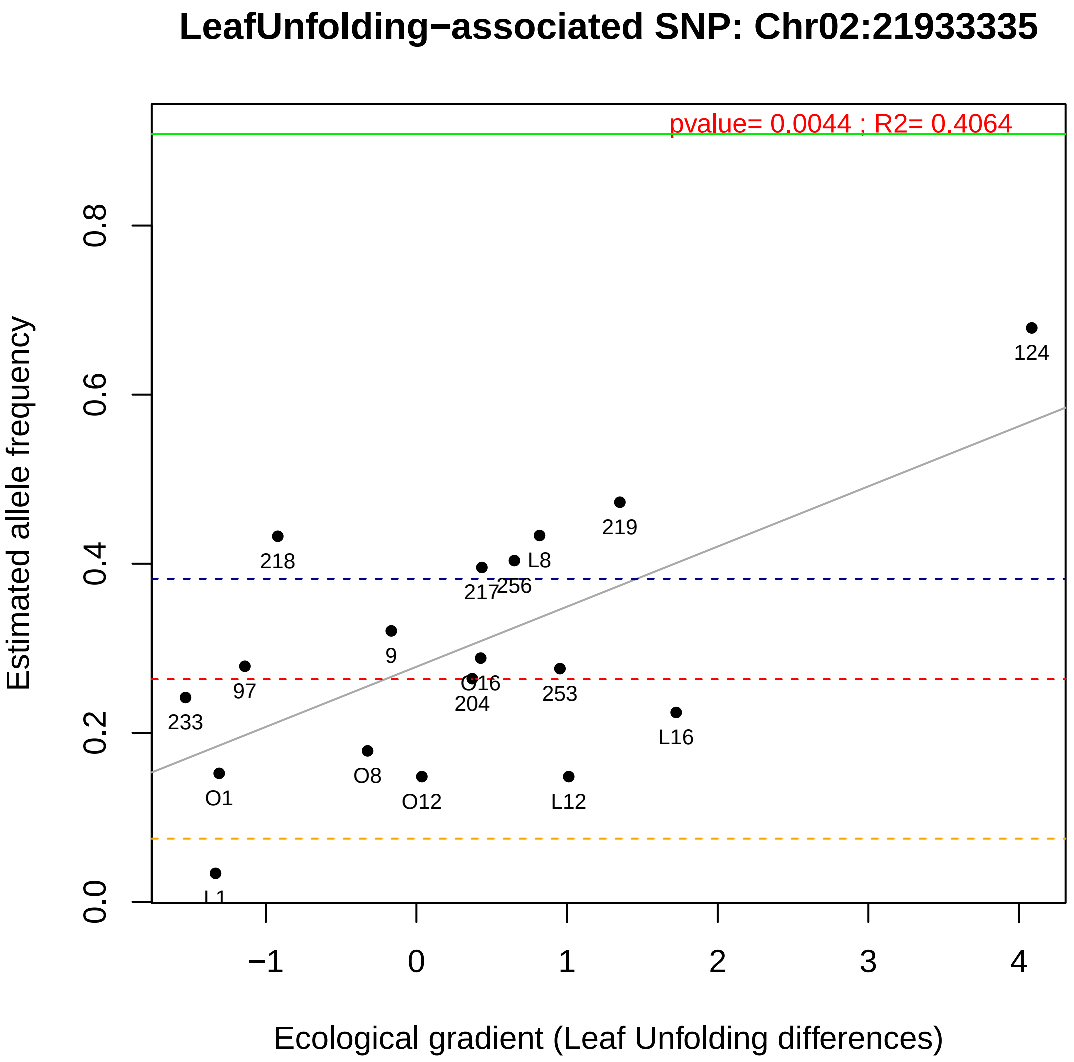

LeafUnfolding−associated SNP: Chr02:21933335
pvalue= 0.0044 ; R2= 0.4064
Estimated allele frequency
0.0
0.2
0.4
0.6
0.8
−1
0
1
2
3
4
Ecological gradient (Leaf Unfolding differences)
124
219
218
L8
256
217
9
97
233
O16
204
253
L16
O8
O1
O12
L12

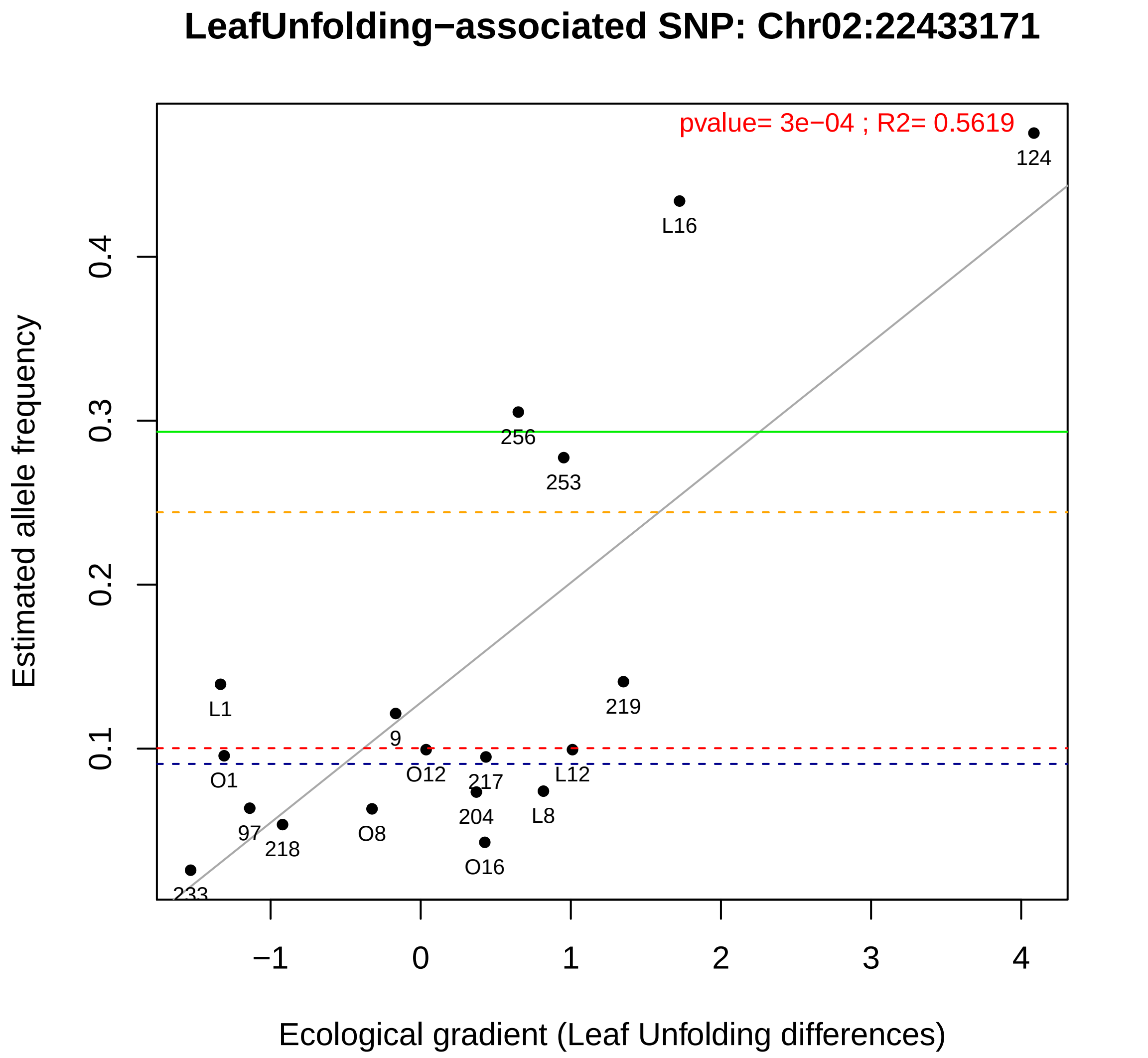

LeafUnfolding−associated SNP: Chr02:22433171
pvalue= 3e−04 ; R2= 0.5619
Estimated allele frequency
0.1
0.2
0.3
0.4
−1
0
1
2
3
4
Ecological gradient (Leaf Unfolding differences)
124
L16
256
253
219
L1
9
O12
217
L12
O1
204
L8
97
218
O8
O16
233

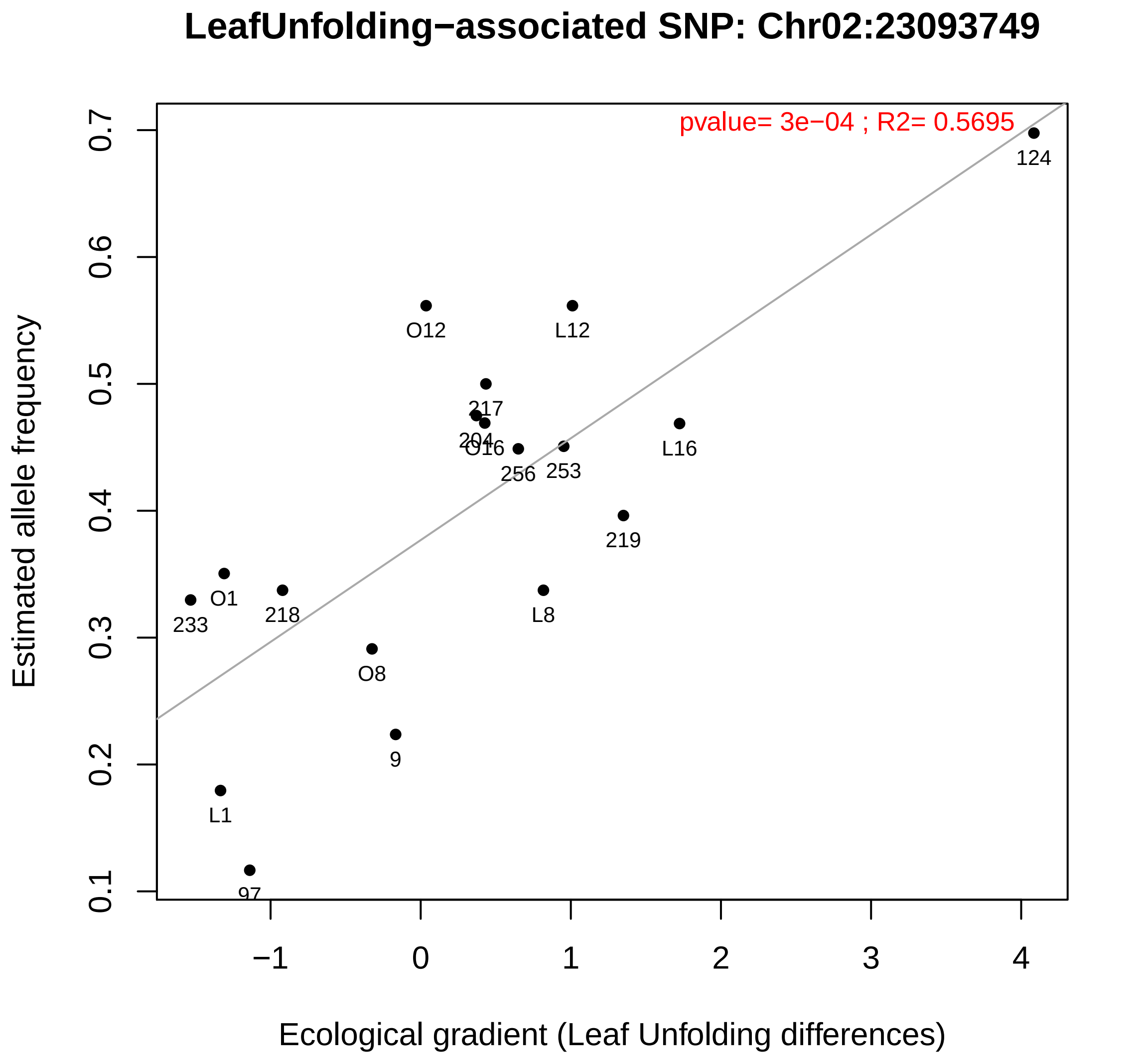
LeafUnfolding−associated SNP: Chr02:23093749
pvalue= 3e−04 ; R2= 0.5695
Estimated allele frequency
Ecological gradient (Leaf Unfolding differences)
0.1
0.2
0.3
0.4
0.5
0.6
0.7
−1
0
1
2
3
4
124
O12
L12
217
204
O16
256
253
L16
219
O1
233
218
L8
O8
9
L1
97

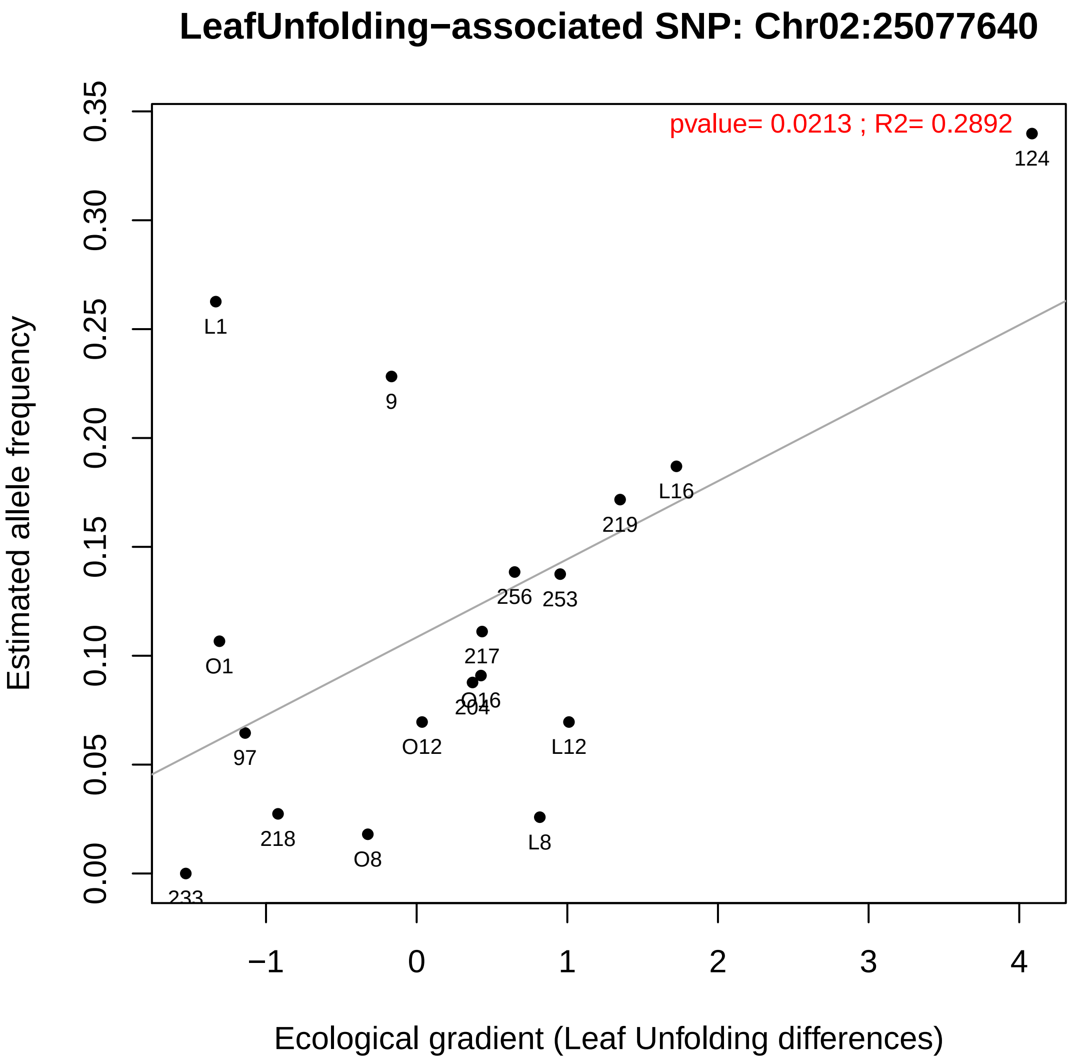
LeafUnfolding−associated SNP: Chr02:25077640
pvalue= 0.0213 ; R2= 0.2892
Estimated allele frequency
Ecological gradient (Leaf Unfolding differences)
0.00
0.05
0.10
0.15
0.20
0.25
0.30
0.35
−1
0
1
2
3
4
124
L1
9
L16
219
256
253
217
O1
O16
204
97
O12
L12
218
O8
L8
233

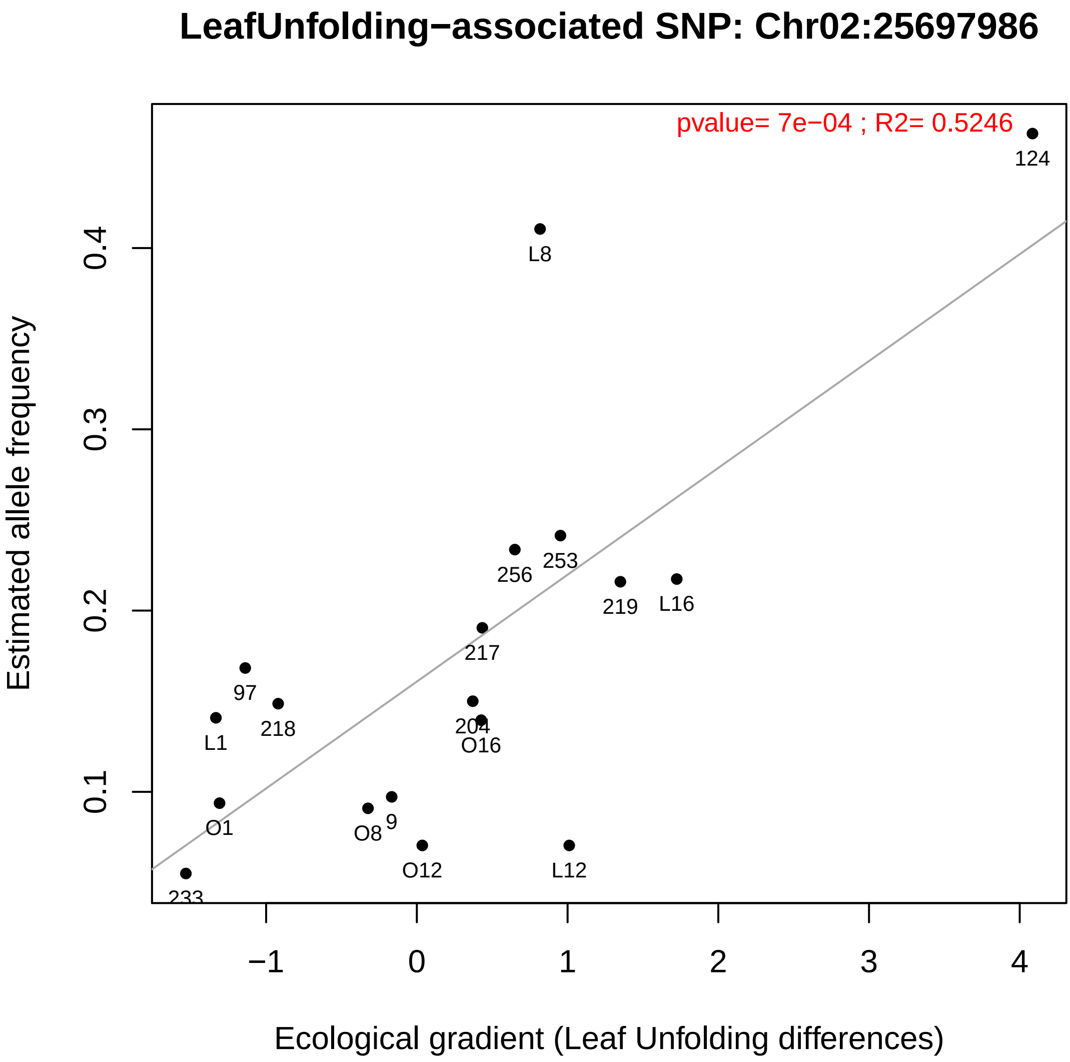

LeafUnfolding−associated SNP: Chr02:25697986
pvalue= 7e−04 ; R2= 0.5246
Estimated allele frequency
0.1
0.2
0.3
0.4
−1
0
1
2
3
4
Ecological gradient (Leaf Unfolding differences)
124
L8
253
256
219
L16
217
97
218
L1
204
O16
O1
O8
9
O12
L12
233

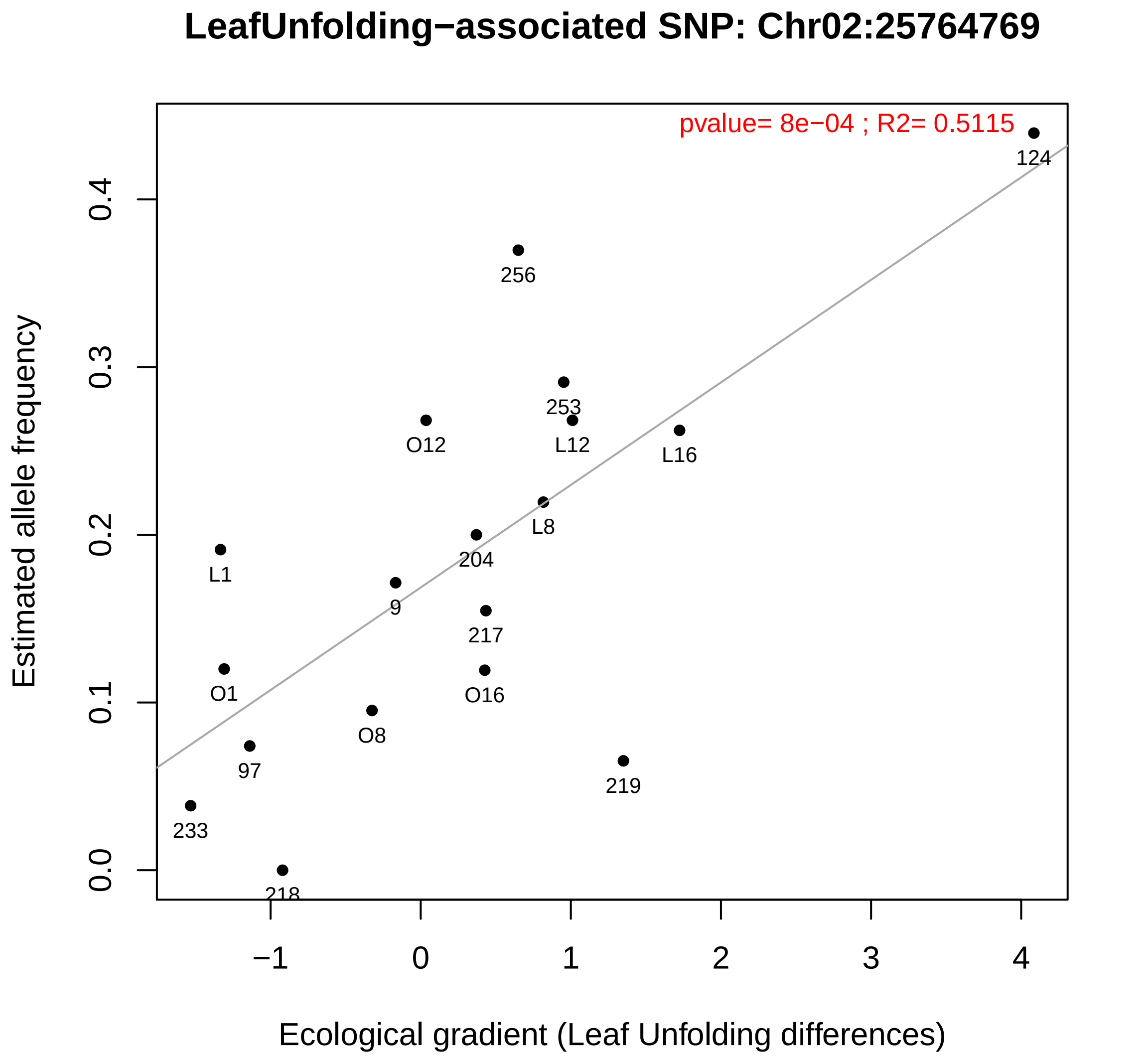

LeafUnfolding−associated SNP: Chr02:25764769
pvalue= 8e−04 ; R2= 0.5115
Estimated allele frequency
0.0
0.1
0.2
0.3
0.4
Ecological gradient (Leaf Unfolding differences)
−1
0
1
2
3
4
124
256
253
O12
L12
L16
L8
204
L1
9
217
O1
O16
O8
97
219
233
218

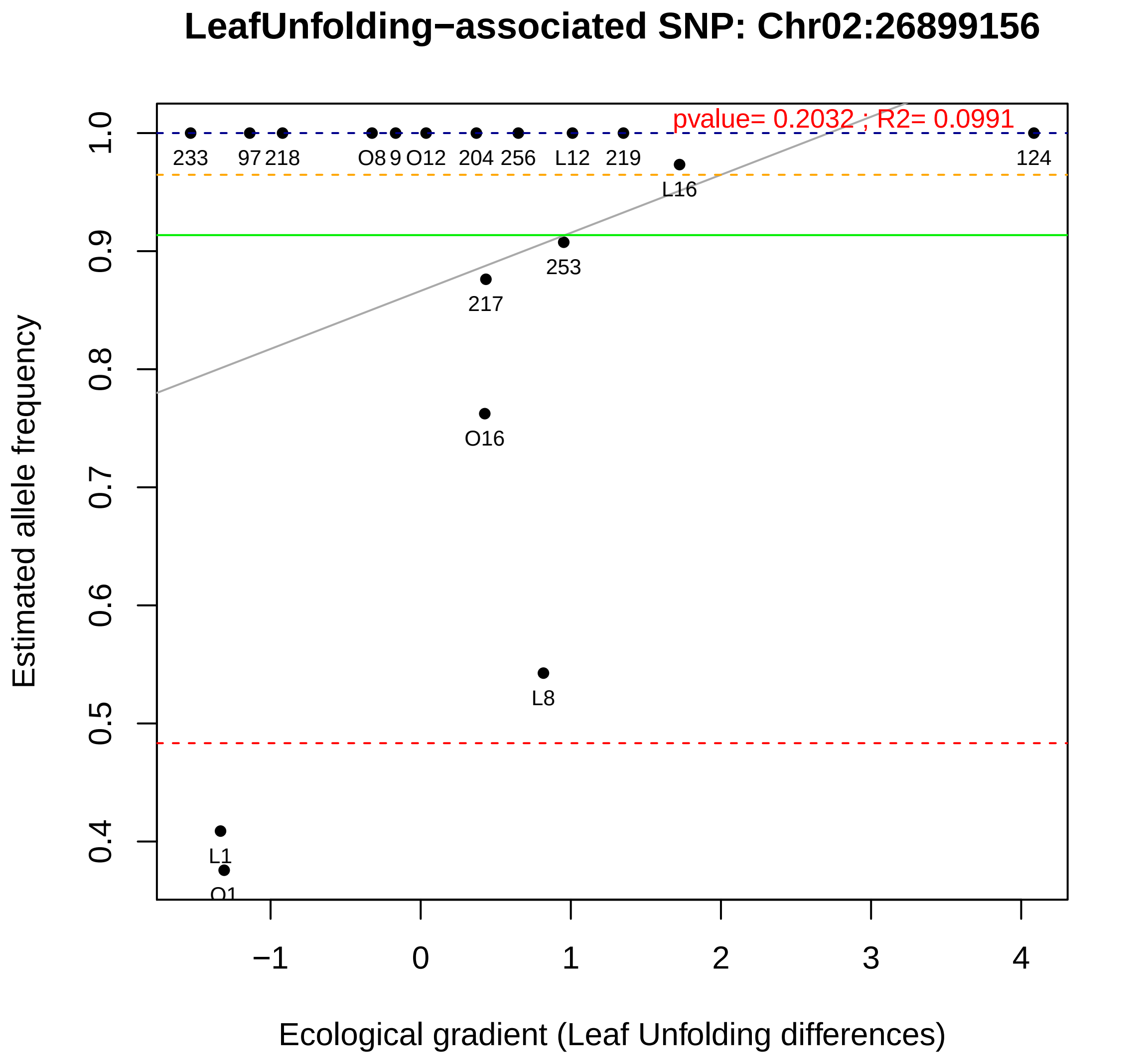
LeafUnfolding−associated SNP: Chr02:26899156
pvalue= 0.2032 ; R2= 0.0991
Estimated allele frequency
Ecological gradient (Leaf Unfolding differences)
1.0
0.9
0.8
0.7
0.6
0.5
0.4
−1
0
1
2
3
4
233
97
218
O8
9
O12
204
256
L12
219
124
L16
253
217
O16
L8
L1
O1

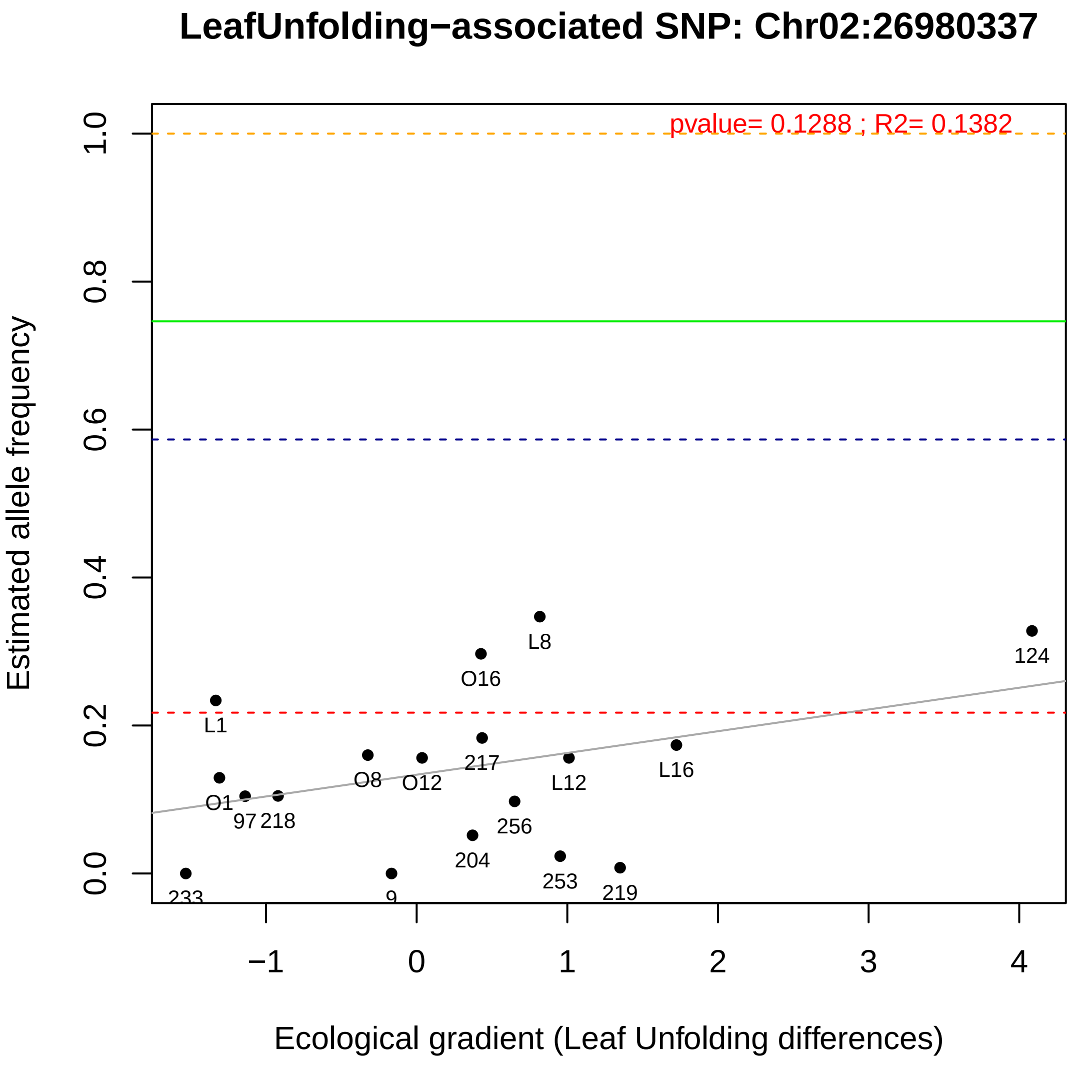

LeafUnfolding−associated SNP: Chr02:26980337
pvalue= 0.1288 ; R2= 0.1382
Estimated allele frequency
Ecological gradient (Leaf Unfolding differences)
0.0
0.2
0.4
0.6
0.8
1.0
−1
0
1
2
3
4
L8
124
O16
L1
217
O8
O12
L12
L16
O1
97
218
256
204
253
219

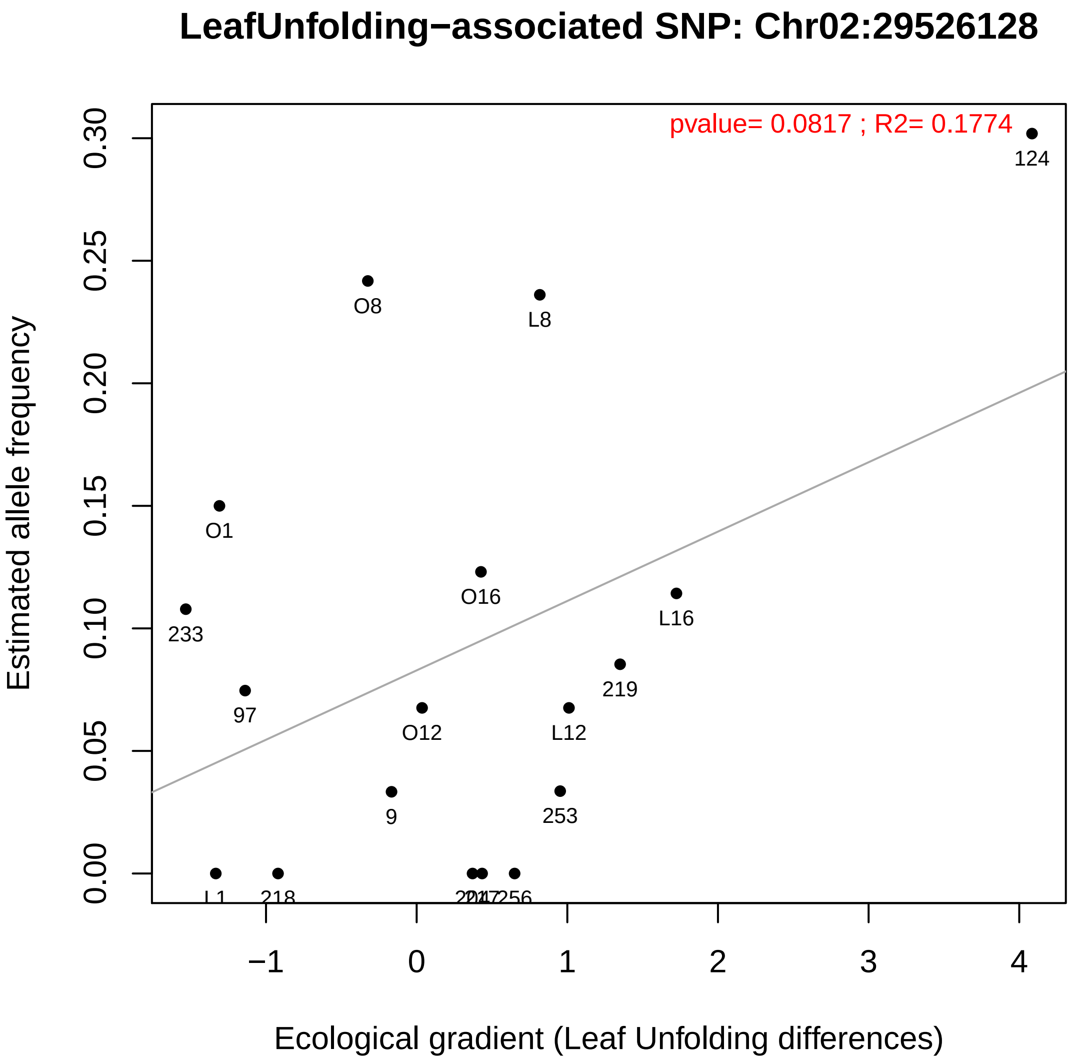
LeafUnfolding−associated SNP: Chr02:29526128
pvalue= 0.0817 ; R2= 0.1774
Estimated allele frequency
Ecological gradient (Leaf Unfolding differences)
0.00
0.05
0.10
0.15
0.20
0.25
0.30
−1
0
1
2
3
4
124
O8
L8
O1
O16
L16
233
219
97
O12
L12
9
253
L1
218
256

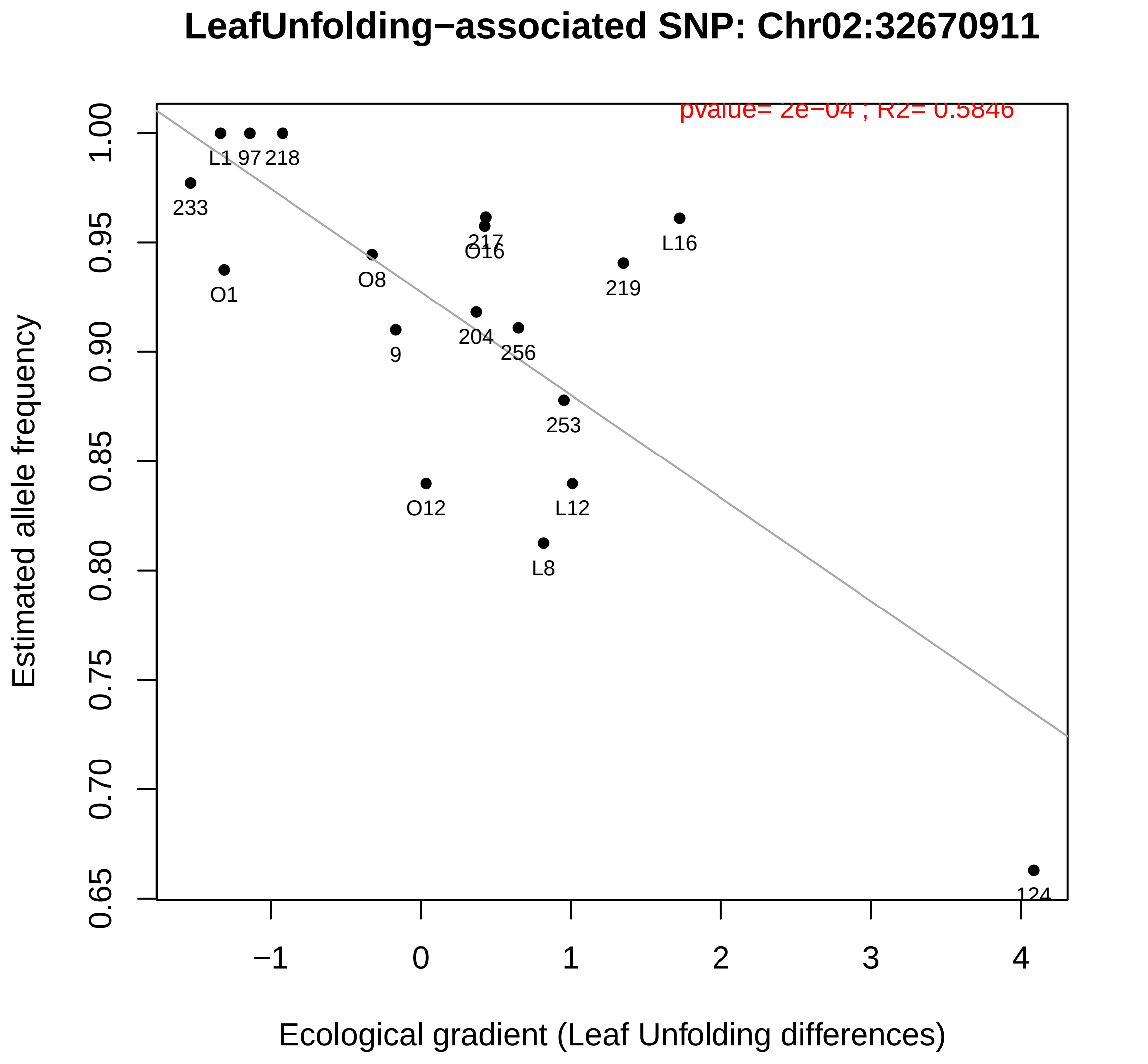

LeafUnfolding−associated SNP: Chr02:32670911
pvalue= 2e−04 ; R2= 0.5846
Estimated allele frequency
Ecological gradient (Leaf Unfolding differences)
0.65
0.70
0.75
0.80
0.85
0.90
0.95
1.00
−1
0
1
2
3
4
L1
97
218
233
O1
O8
9
217
O16
204
256
253
O12
L12
L8
219
L16
124

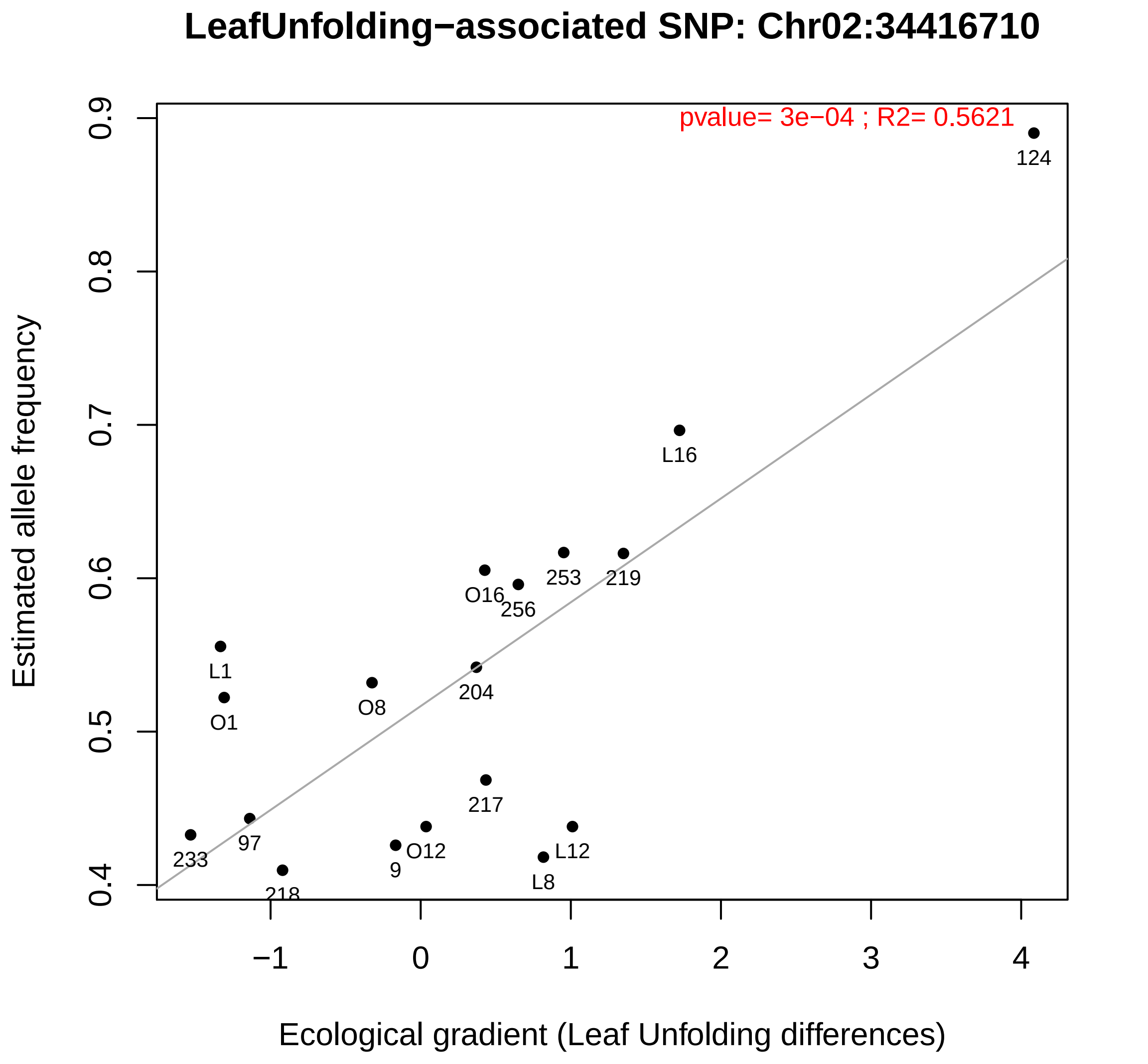
LeafUnfolding−associated SNP: Chr02:34416710
pvalue= 3e−04 ; R2= 0.5621
Estimated allele frequency
Ecological gradient (Leaf Unfolding differences)
0.4
0.5
0.6
0.7
0.8
0.9
−1
0
1
2
3
4
124
L16
253
219
O16
256
L1
O1
O8
204
217
233
97
218
9
O12
L12
L8

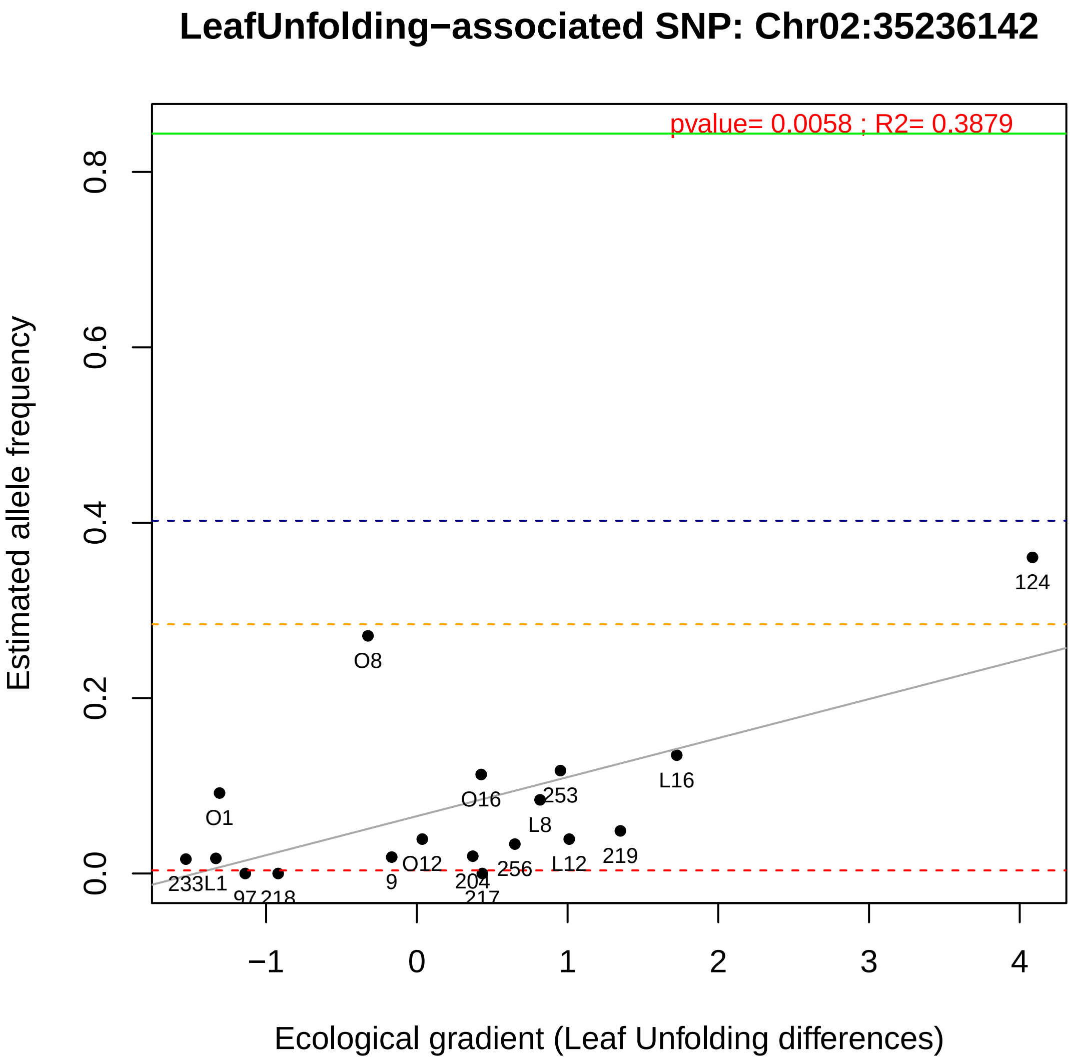
LeafUnfolding−associated SNP: Chr02:35236142
pvalue= 0.0058 ; R2= 0.3879
Estimated allele frequency
0.0
0.2
0.4
0.6
0.8
−1
0
1
2
3
4
Ecological gradient (Leaf Unfolding differences)
124
O8
L16
O16
253
L8
O1
219
L12
O12
256
233
L1
9
204
97
218
217

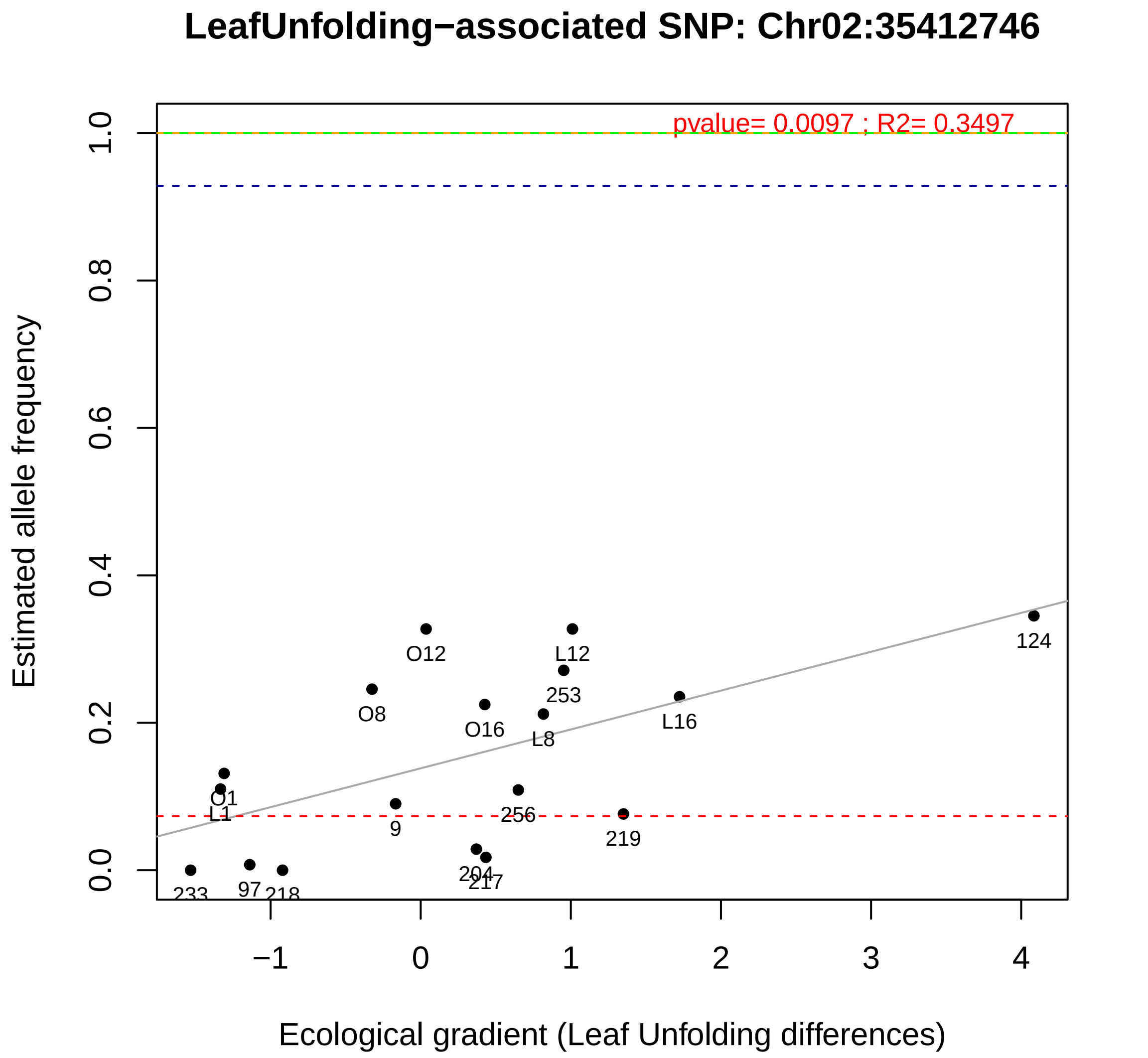

LeafUnfolding−associated SNP: Chr02:35412746
pvalue= 0.0097 ; R2= 0.3497
Estimated allele frequency
0.0
0.2
0.4
0.6
0.8
1.0
−1
0
1
2
3
4
Ecological gradient (Leaf Unfolding differences)
O12
L12
124
253
O8
O16
L8
L16
O1
L1
256
9
219
204
217
233
97
218

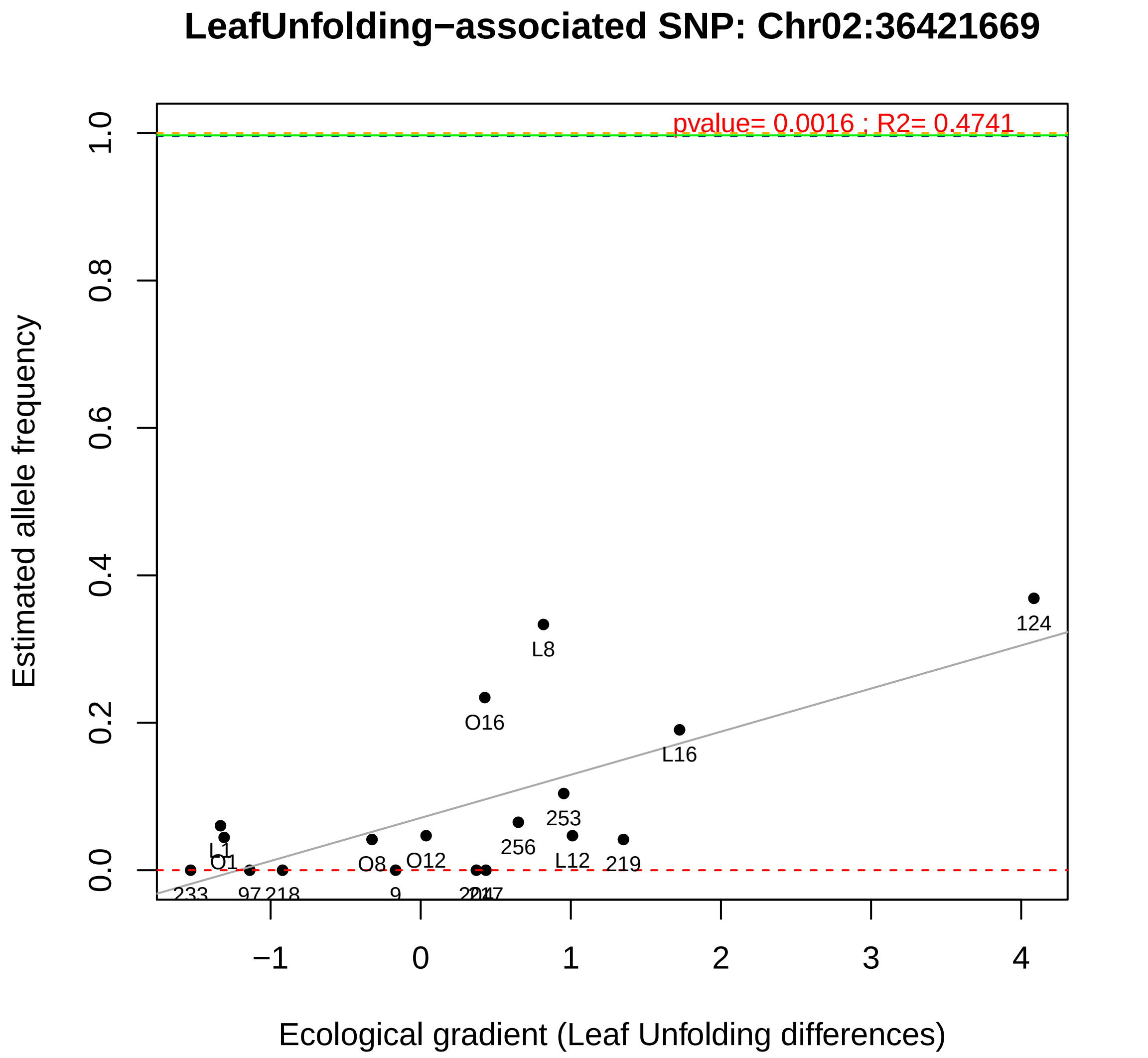

LeafUnfolding−associated SNP: Chr02:36421669
pvalue= 0.0016 ; R2= 0.4741
Estimated allele frequency
Ecological gradient (Leaf Unfolding differences)
0.0
0.2
0.4
0.6
0.8
1.0
−1
0
1
2
3
4
124
L8
O16
L16
253
256
L12
219
O12
O8
9
L1
O1
97
218
233

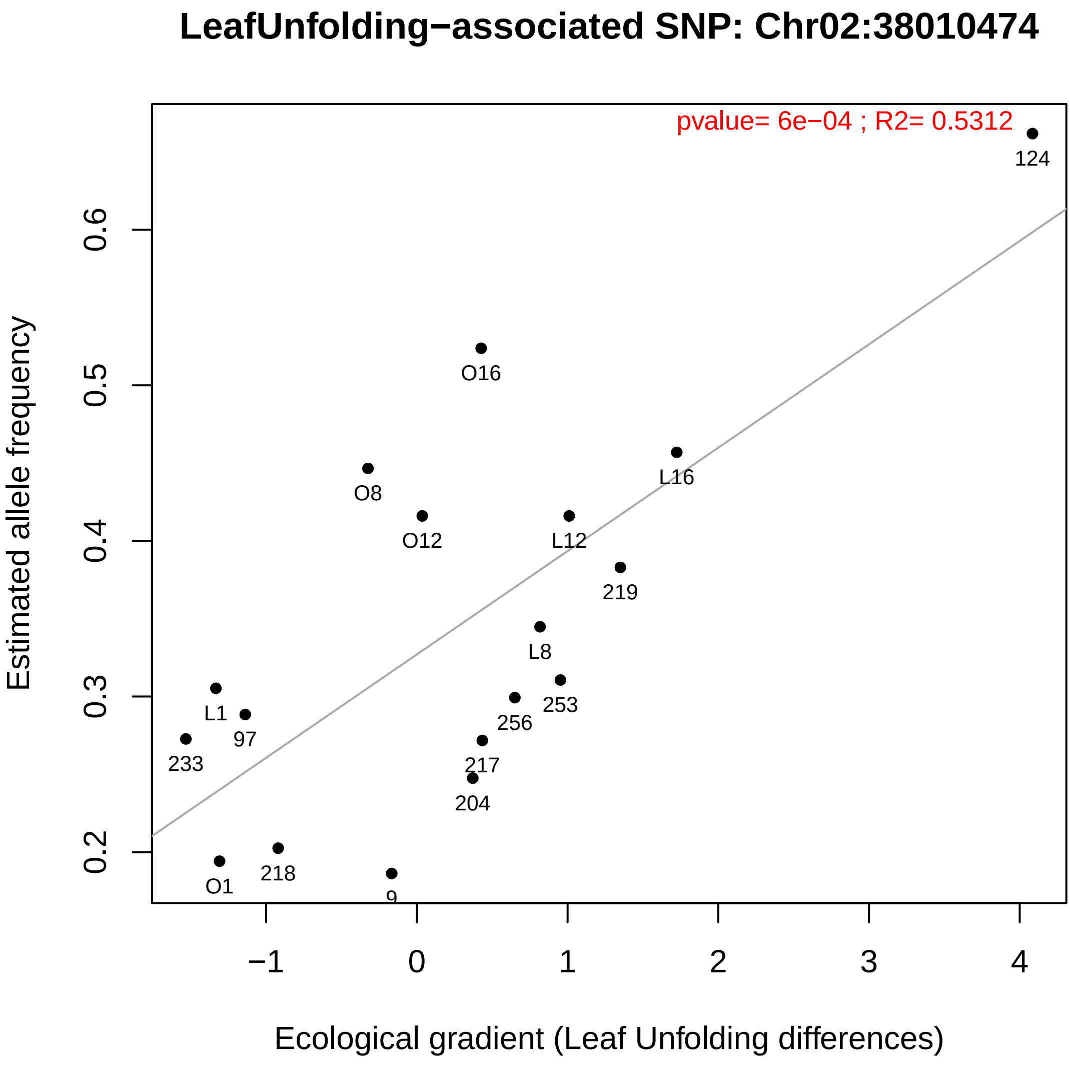
LeafUnfolding−associated SNP: Chr02:38010474
pvalue= 6e−04 ; R2= 0.5312
124
O16
L16
O8
O12
L12
219
L8
253
256
L1
97
233
217
204
218
O1
9
0.2
0.3
0.4
0.5
0.6
−1
0
1
2
3
4
Estimated allele frequency
Ecological gradient (Leaf Unfolding differences)

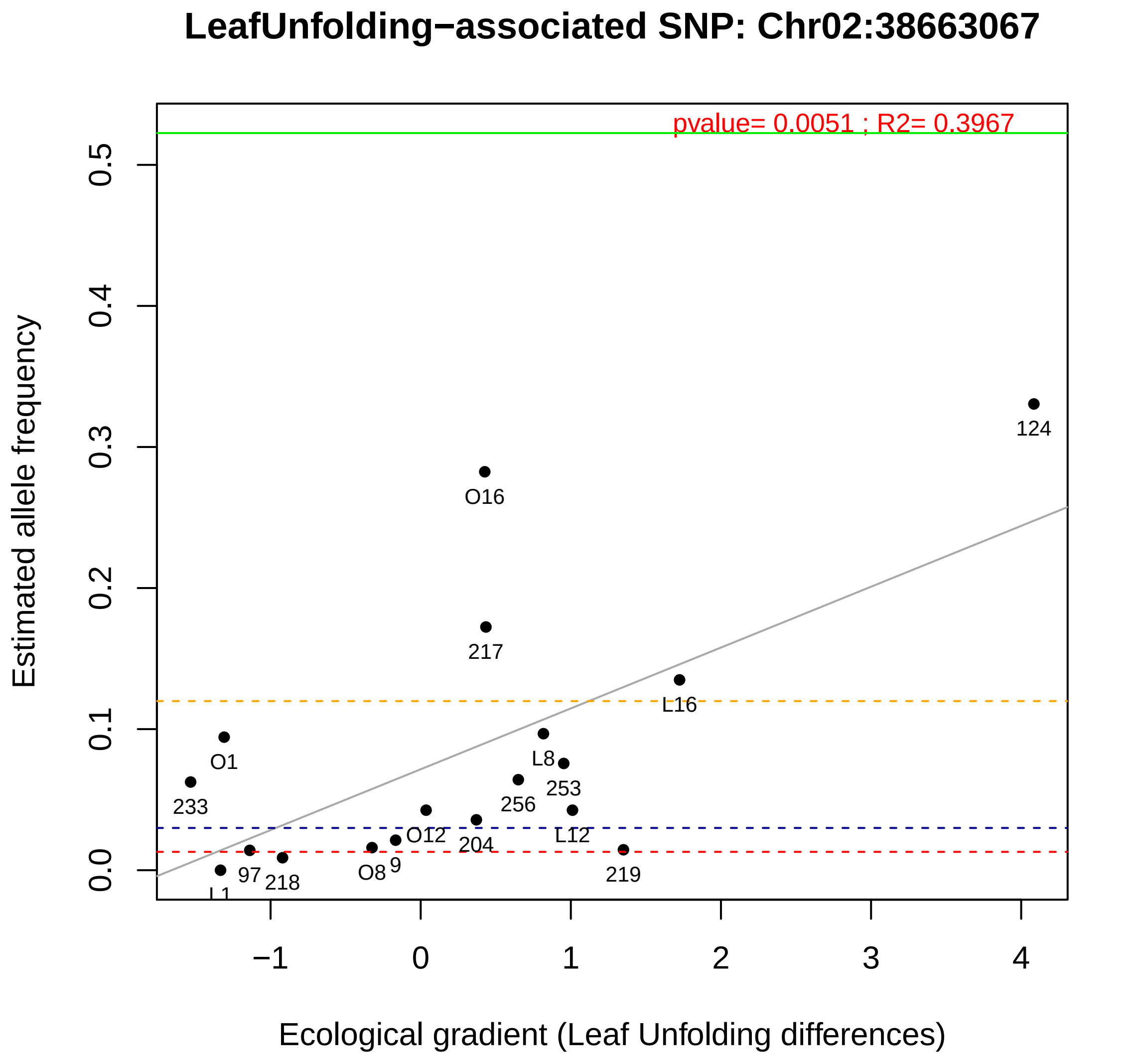

LeafUnfolding−associated SNP: Chr02:38663067
pvalue= 0.0051 ; R2= 0.3967
Estimated allele frequency
0.0
0.1
0.2
0.3
0.4
0.5
Ecological gradient (Leaf Unfolding differences)
−1
0
1
2
3
4
124
O16
217
L16
O1
233
L8
253
256
L12
O12
204
219
97
218
O8
9
L1

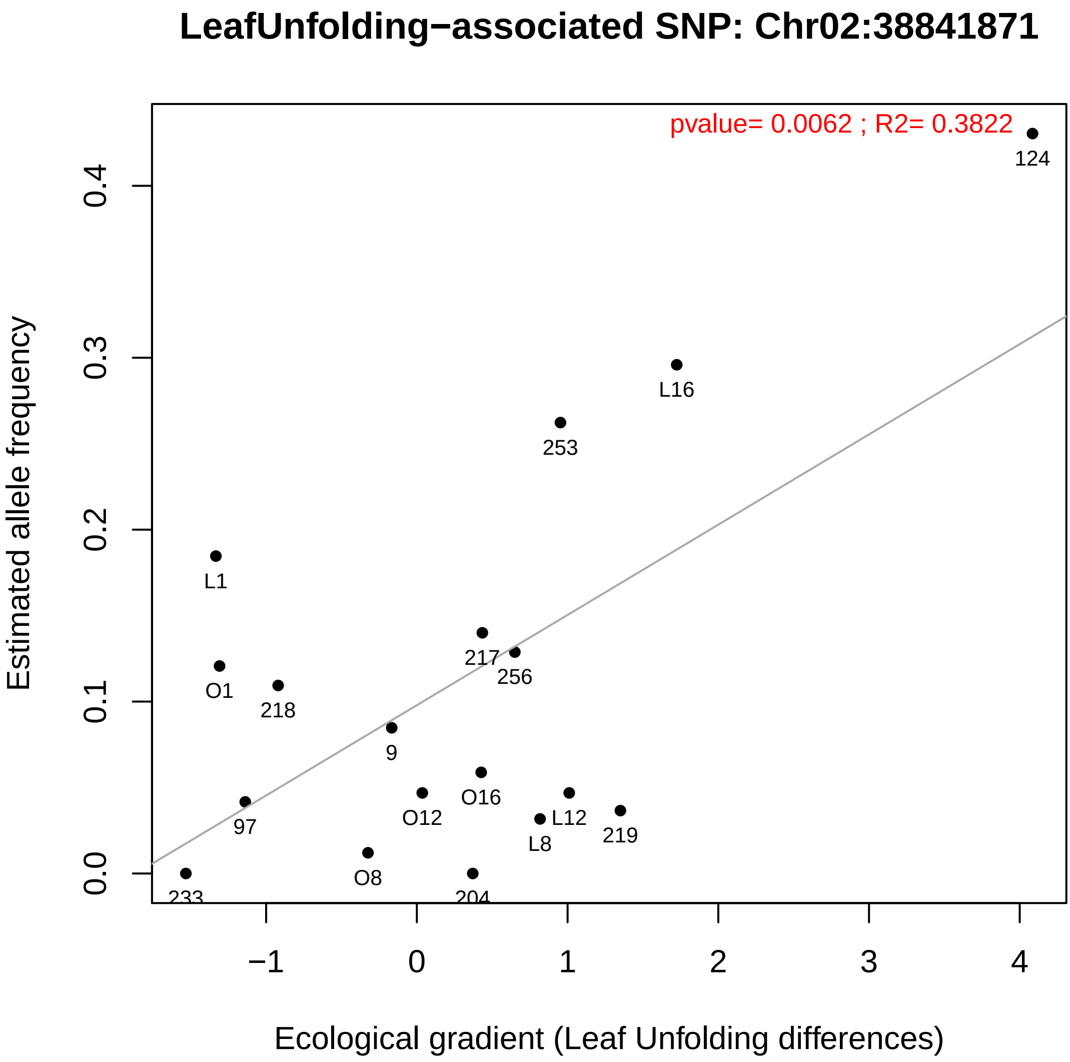
LeafUnfolding−associated SNP: Chr02:38841871
pvalue= 0.0062 ; R2= 0.3822
Estimated allele frequency
Ecological gradient (Leaf Unfolding differences)
0.0
0.1
0.2
0.3
0.4
−1
0
1
2
3
4
124
L16
253
L1
217
256
O1
218
9
O16
O12
L12
219
97
L8
O8
233
204

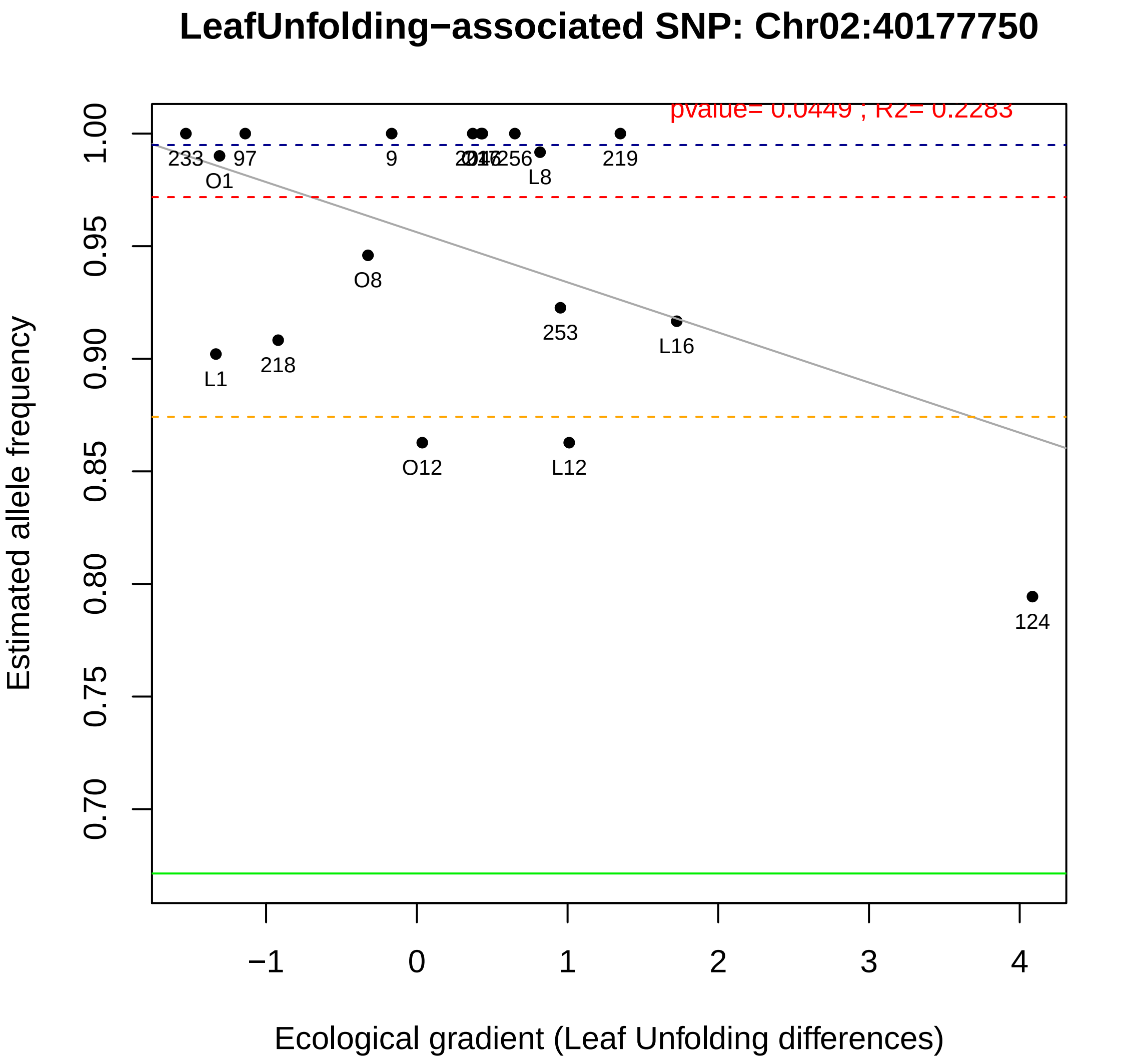

LeafUnfolding−associated SNP: Chr02:40177750
pvalue= 0.0449 ; R2= 0.2283
233
97
O1
9
219
L8
O8
253
L16
L1
218
O12
L12
124
Estimated allele frequency
1.00
0.95
0.90
0.85
0.80
0.75
0.70
−1
0
1
2
3
4
Ecological gradient (Leaf Unfolding differences)

**LeafUnfolding−associated SNP: Chr02:40833124**

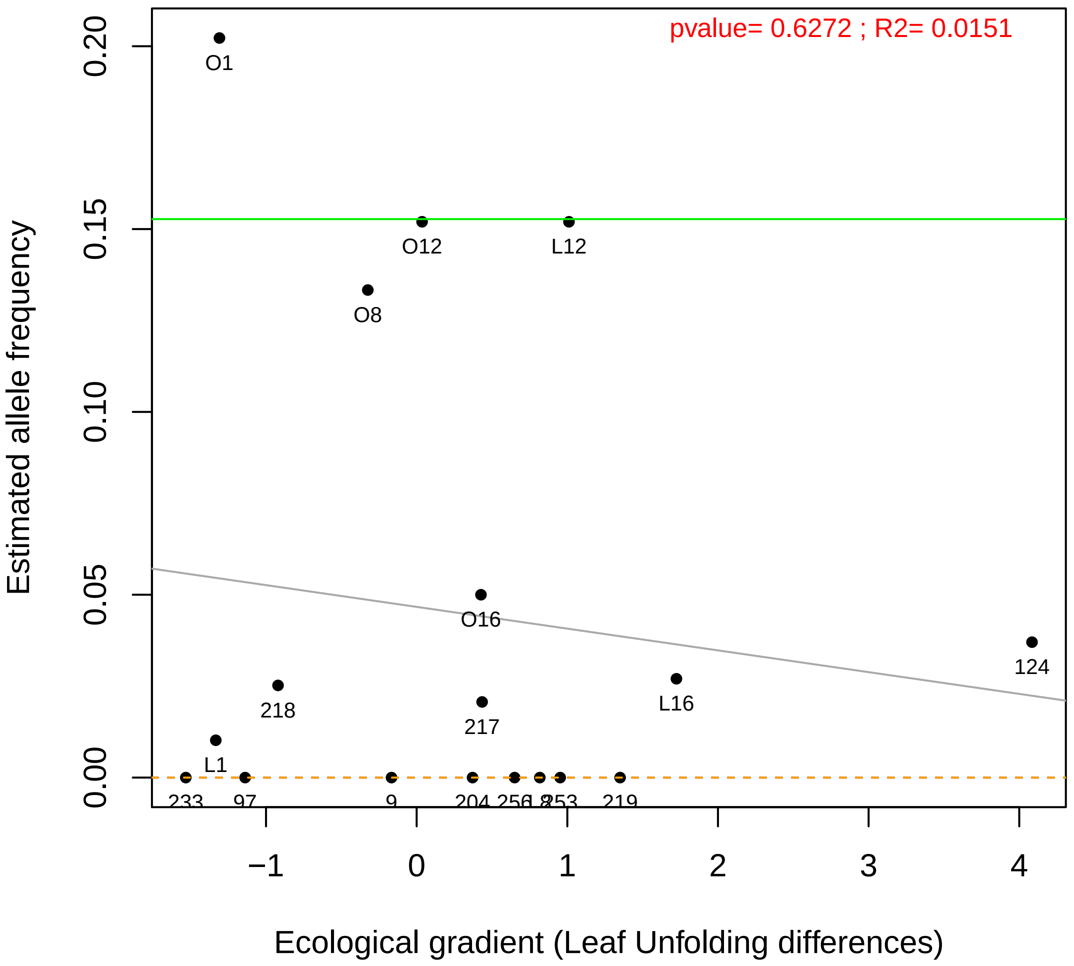

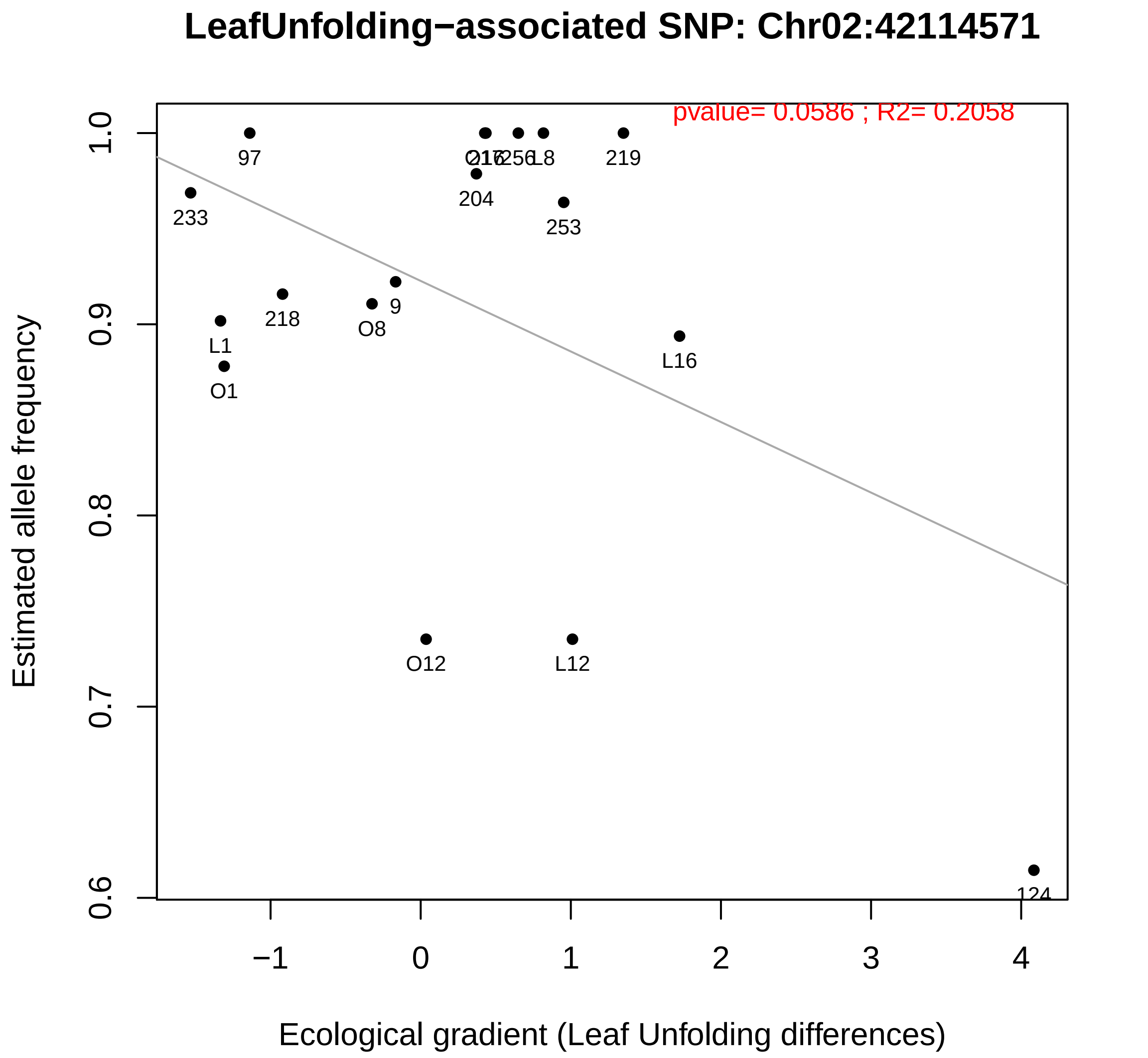
LeafUnfolding−associated SNP: Chr02:42114571
pvalue= 0.0586 ; R2= 0.2058
Estimated allele frequency
Ecological gradient (Leaf Unfolding differences)
1.0
0.9
0.8
0.7
0.6
−1
0
1
2
3
4
97
233
204
253
219
L8
218
L1
O1
O8
9
L16
O12
L12
124

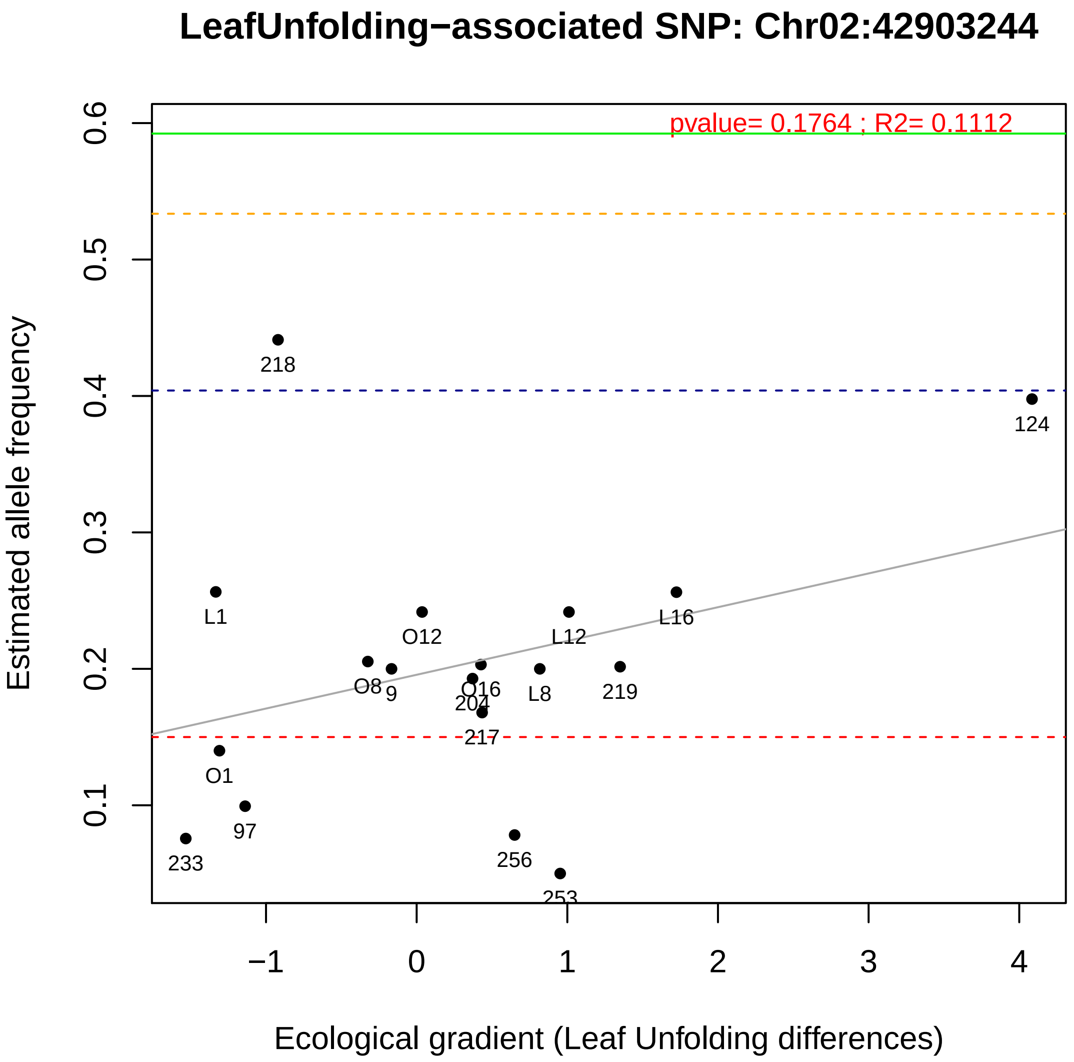

LeafUnfolding−associated SNP: Chr02:42903244
pvalue= 0.1764 ; R2= 0.1112
Estimated allele frequency
0.1
0.2
0.3
0.4
0.5
0.6
218
124
L1
O12
L12
L16
O8
9
O16
204
L8
219
217
O1
97
233
256
253
−1
0
1
2
3
4
Ecological gradient (Leaf Unfolding differences)

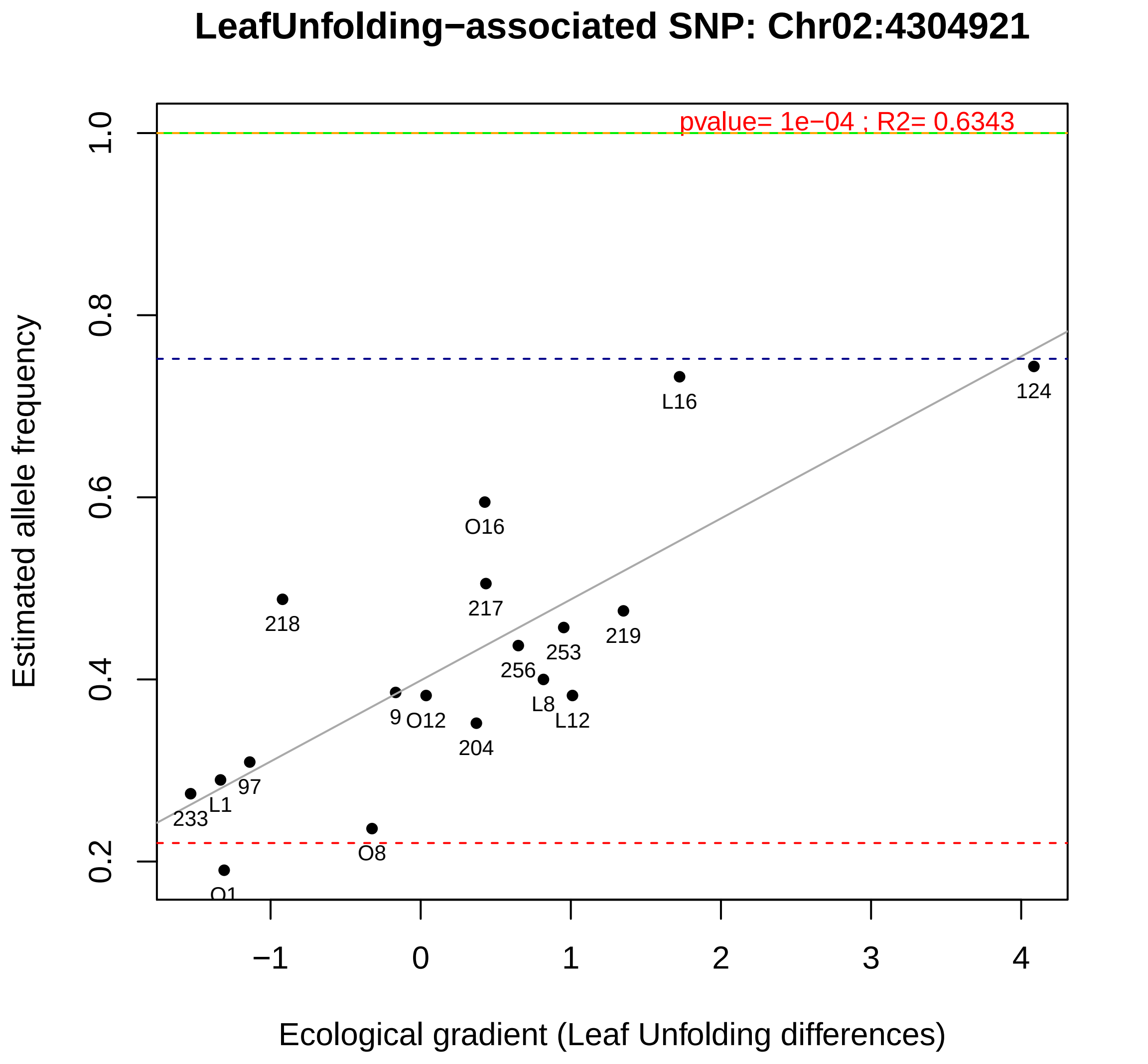

LeafUnfolding−associated SNP: Chr02:4304921
pvalue= 1e−04 ; R2= 0.6343
Estimated allele frequency
Ecological gradient (Leaf Unfolding differences)
0.2
0.4
0.6
0.8
1.0
−1
0
1
2
3
4
L16
124
O16
217
218
219
253
256
L8
L12
9
O12
204
97
L1
233
O8
O1

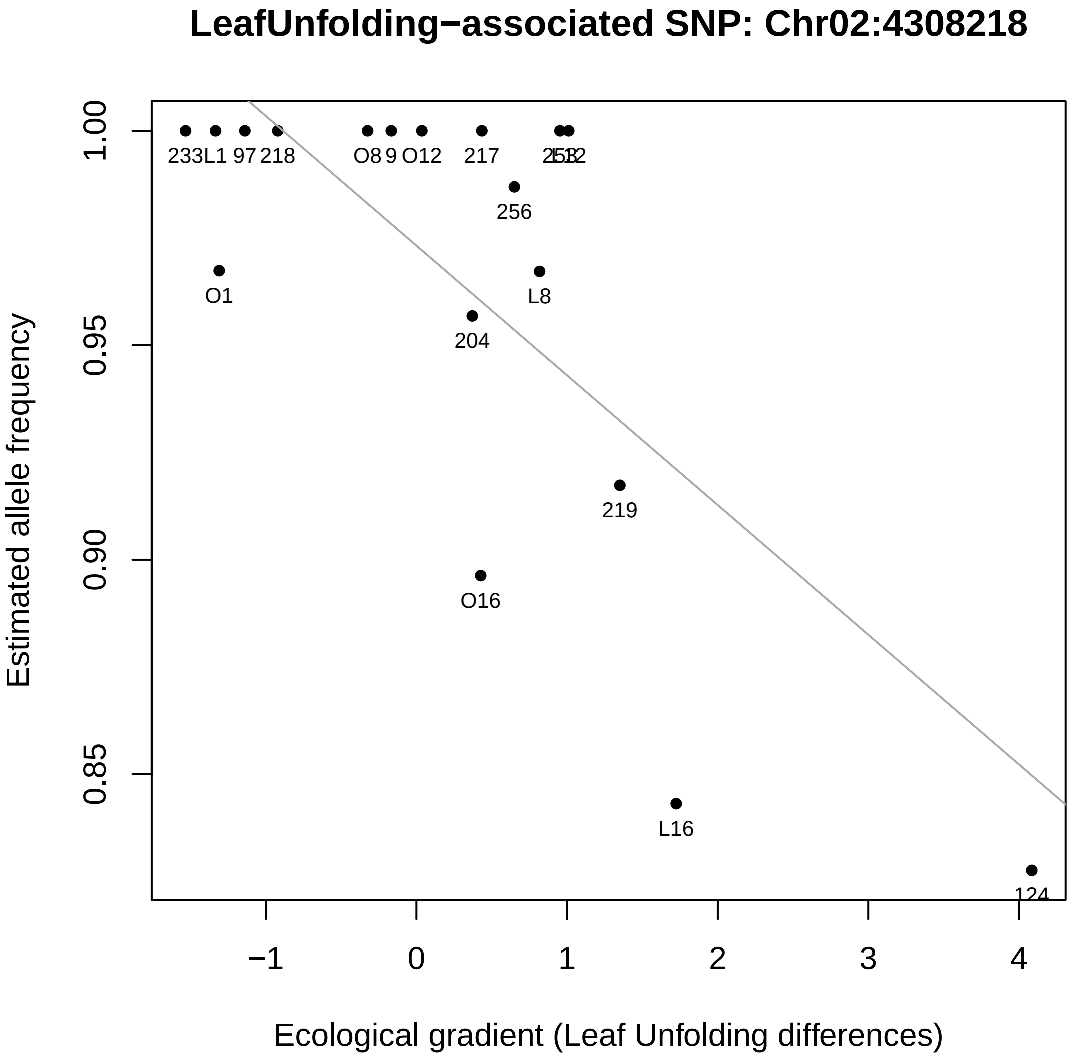
LeafUnfolding−associated SNP: Chr02:4308218
Estimated allele frequency
1.00
0.95
0.90
0.85
−1
0
1
2
3
4
Ecological gradient (Leaf Unfolding differences)
233
L1
97
218
O8
9
O12
217
256
O1
L8
204
219
O16
L16
124

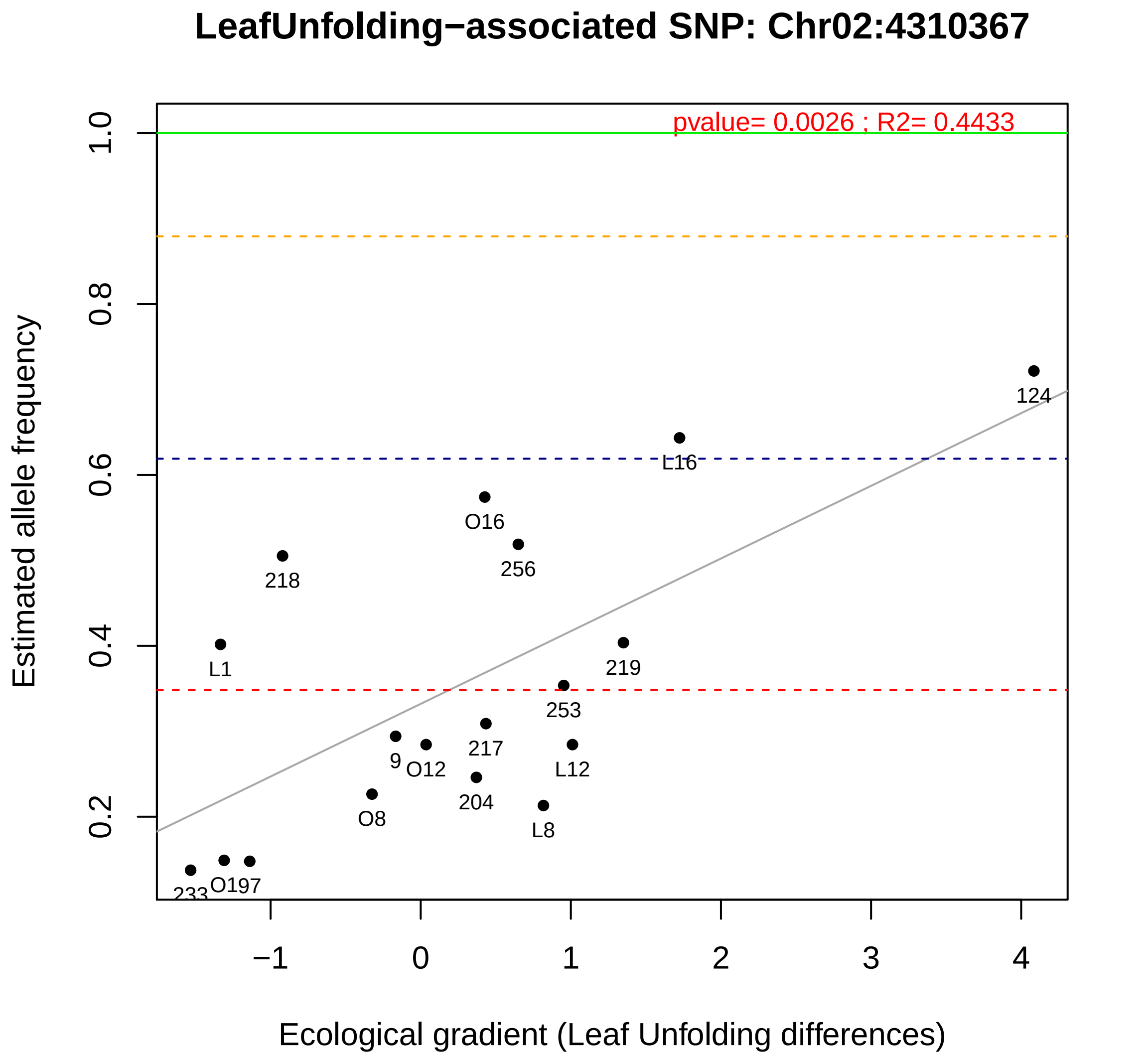
LeafUnfolding−associated SNP: Chr02:4310367
pvalue= 0.0026 ; R2= 0.4433
Estimated allele frequency
1.0
0.8
0.6
0.4
0.2
124
L16
O16
256
218
L1
219
253
217
9
O12
L12
204
O8
L8
233
O197
−1
0
1
2
3
4
Ecological gradient (Leaf Unfolding differences)

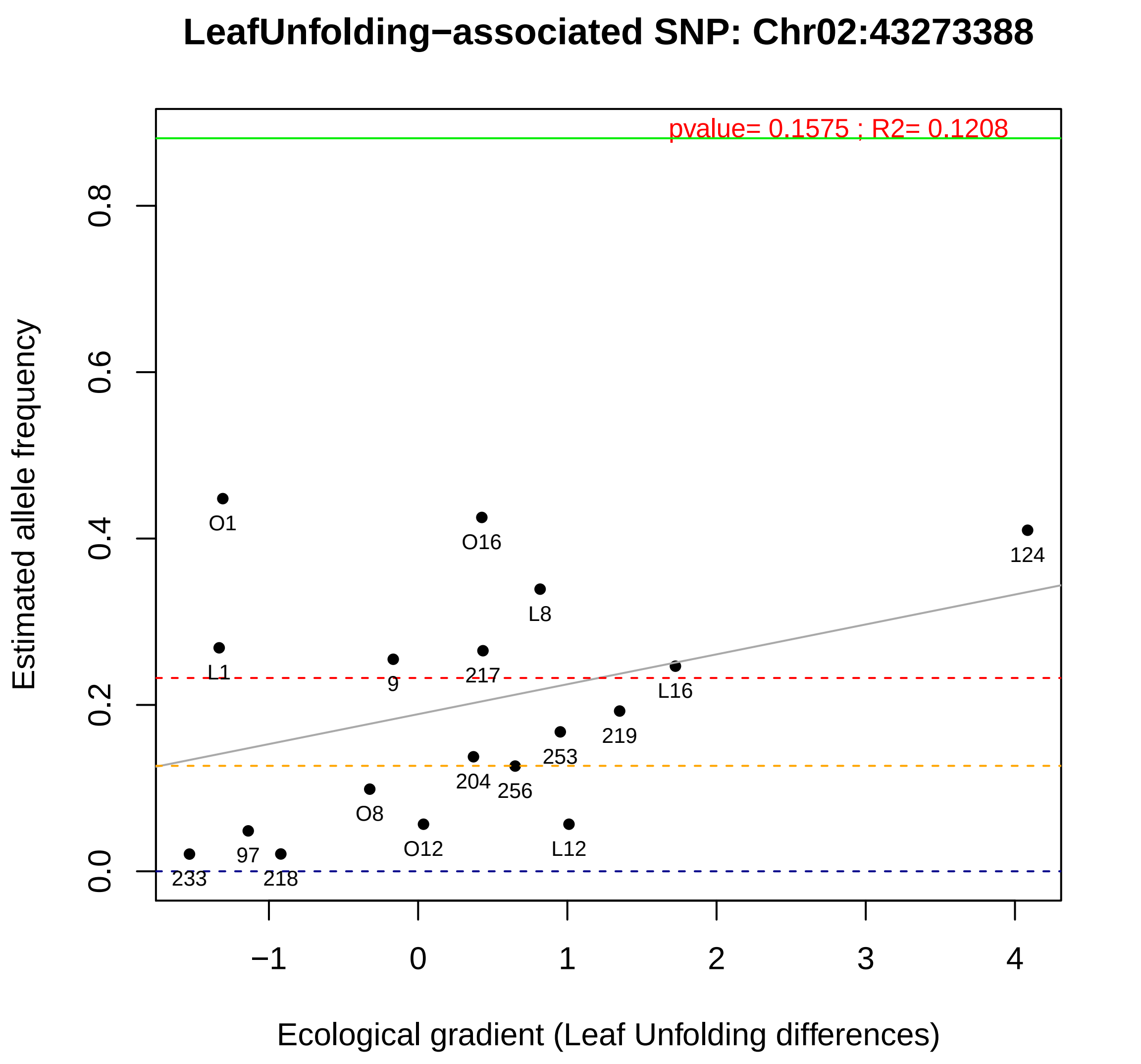

LeafUnfolding−associated SNP: Chr02:43273388
pvalue= 0.1575 ; R2= 0.1208
Estimated allele frequency
0.0
0.2
0.4
0.6
0.8
−1
0
1
2
3
4
Ecological gradient (Leaf Unfolding differences)
O1
O16
124
L8
L1
9
217
L16
219
253
204
256
O8
O12
L12
97
233
218

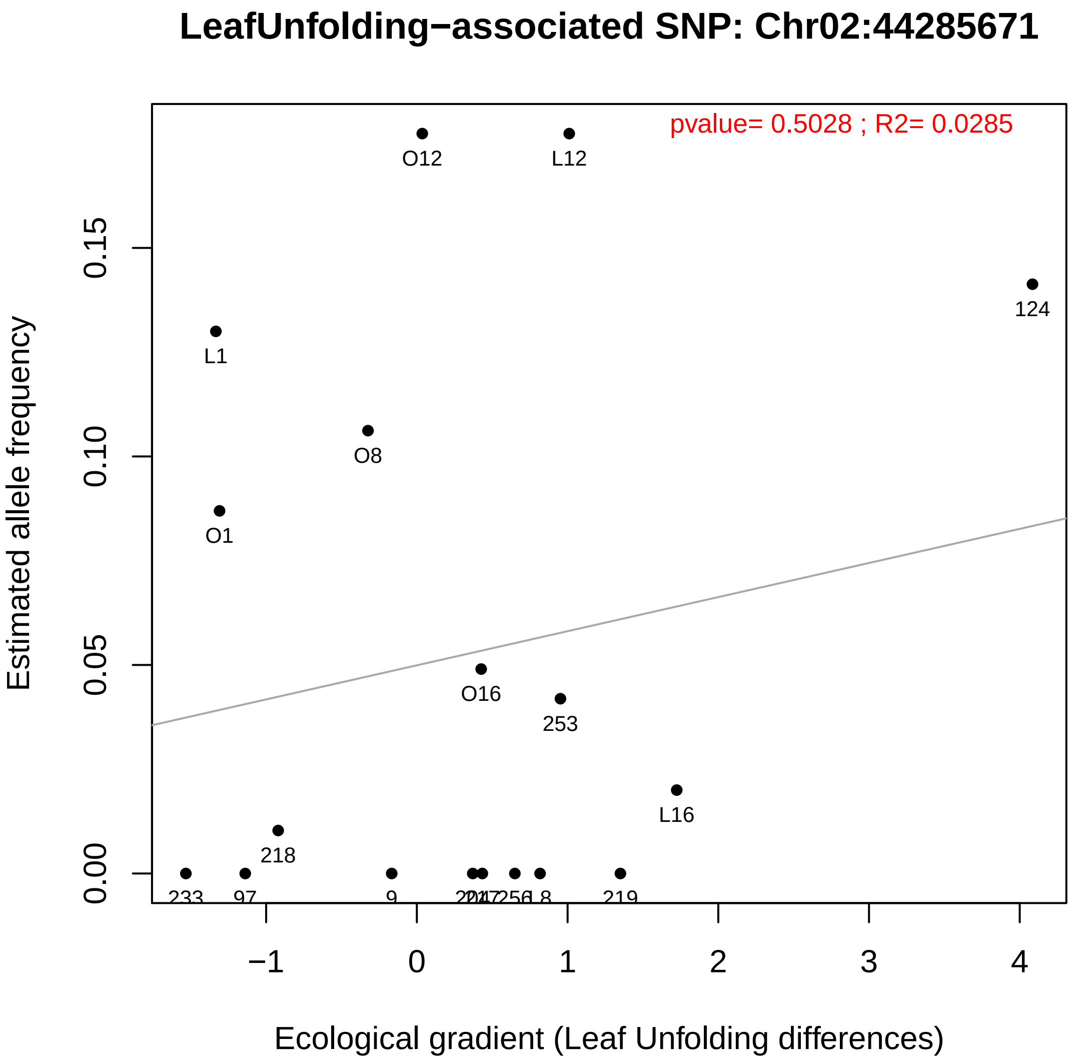
LeafUnfolding−associated SNP: Chr02:44285671
pvalue= 0.5028 ; R2= 0.0285
Estimated allele frequency
0.00
0.05
0.10
0.15
O12
L12
124
L1
O8
O1
O16
253
L16
218
233
97
9
219
−1
0
1
2
3
4
Ecological gradient (Leaf Unfolding differences)

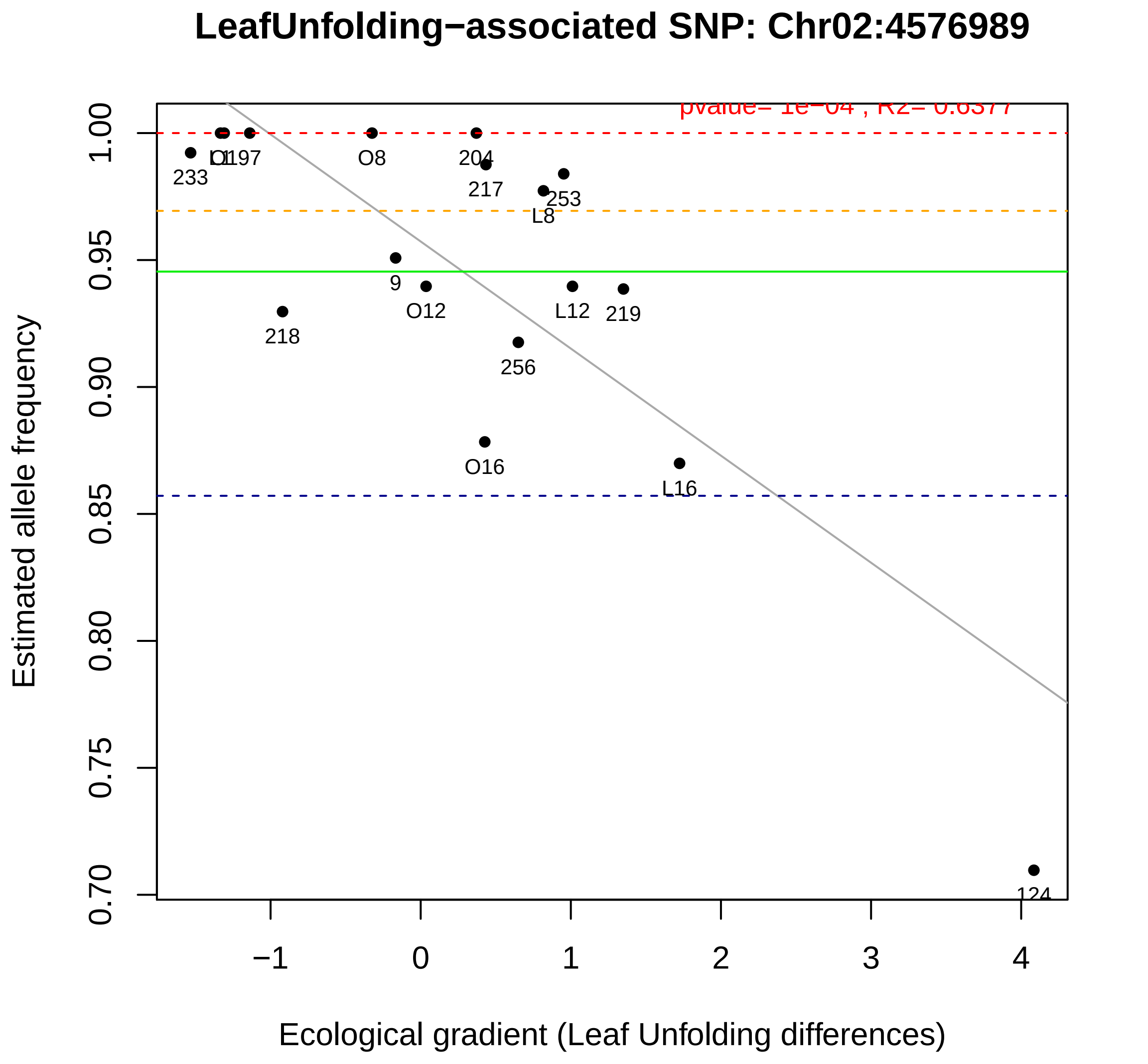

LeafUnfolding−associated SNP: Chr02:4576989
pvalue= 1e−04 , R2= 0.6377
Estimated allele frequency
Ecological gradient (Leaf Unfolding differences)
0.70
0.75
0.80
0.85
0.90
0.95
1.00
−1
0
1
2
3
4
233
O1
197
O8
204
217
253
L8
9
O12
L12
219
218
256
O16
L16
124

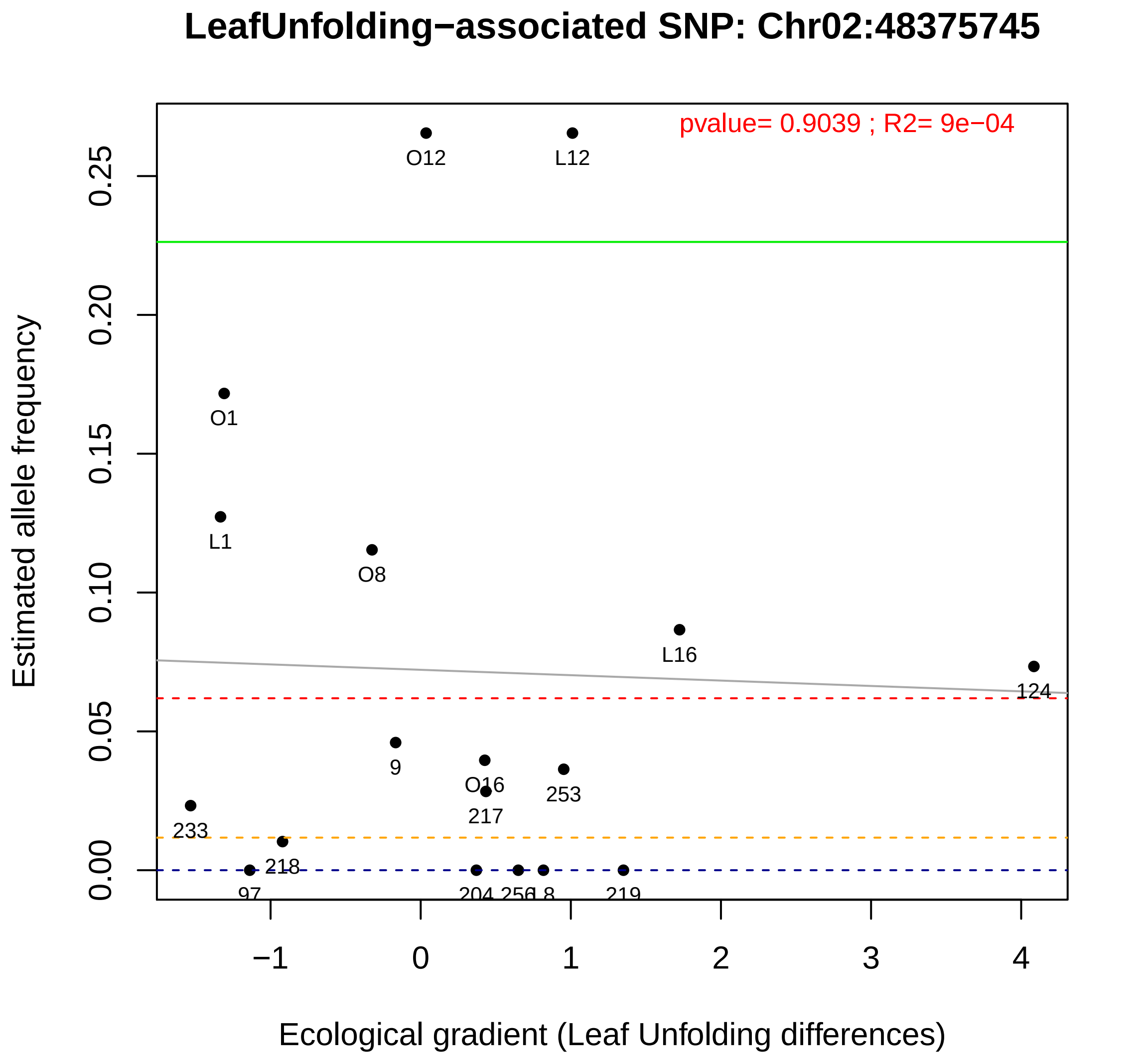

LeafUnfolding−associated SNP: Chr02:48375745
pvalue= 0.9039 ; R2= 9e−04
Estimated allele frequency
Ecological gradient (Leaf Unfolding differences)
0.00
0.05
0.10
0.15
0.20
0.25
−1
0
1
2
3
4
O12
L12
O1
L1
O8
L16
124
9
O16
217
253
233
218
97
204
256
L8
219

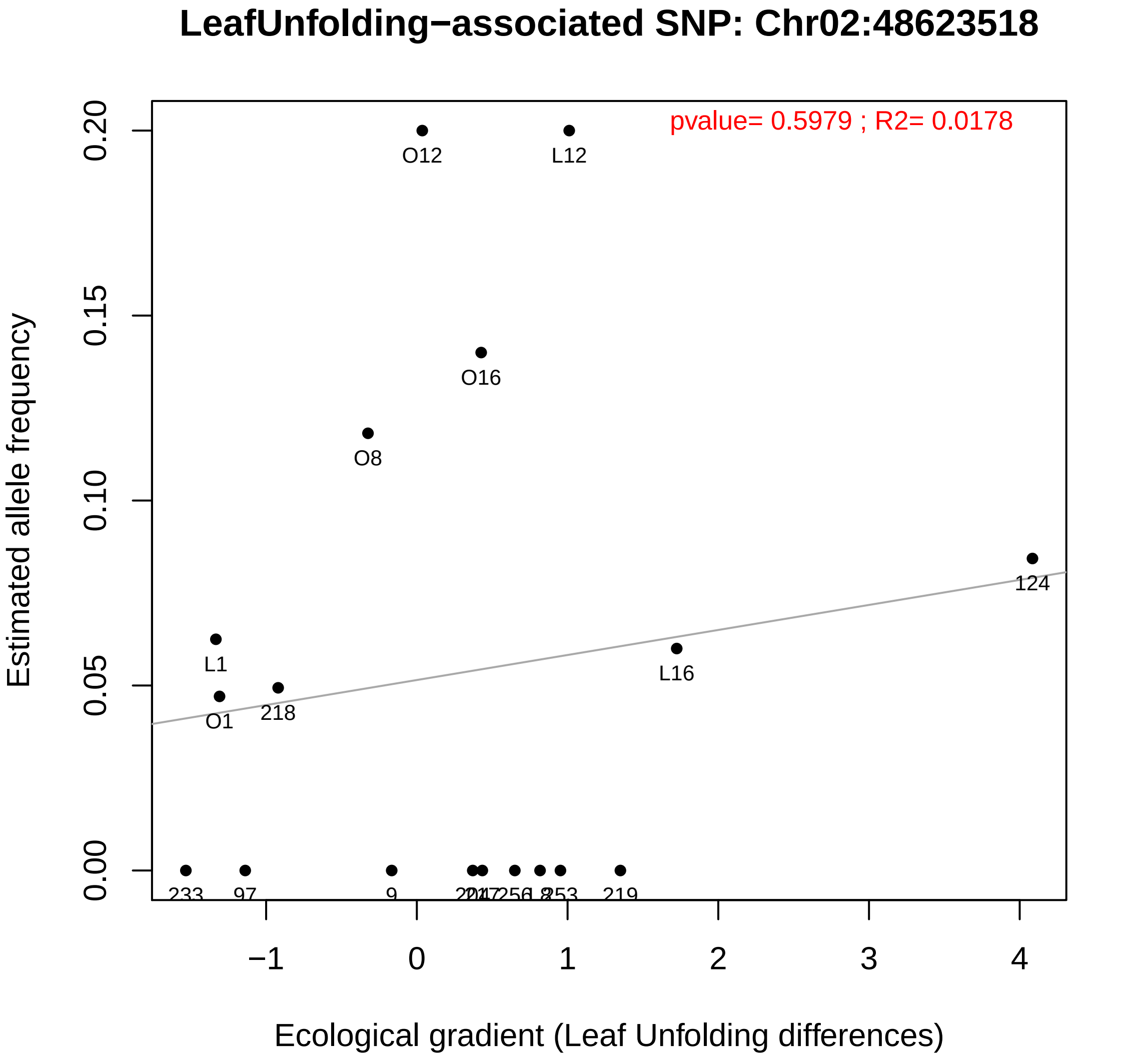
LeafUnfolding−associated SNP: Chr02:48623518
pvalue= 0.5979 ; R2= 0.0178
Estimated allele frequency
0.00
0.05
0.10
0.15
0.20
Ecological gradient (Leaf Unfolding differences)
−1
0
1
2
3
4
O12
L12
O16
O8
124
L1
L16
O1
218
233
97
9
219

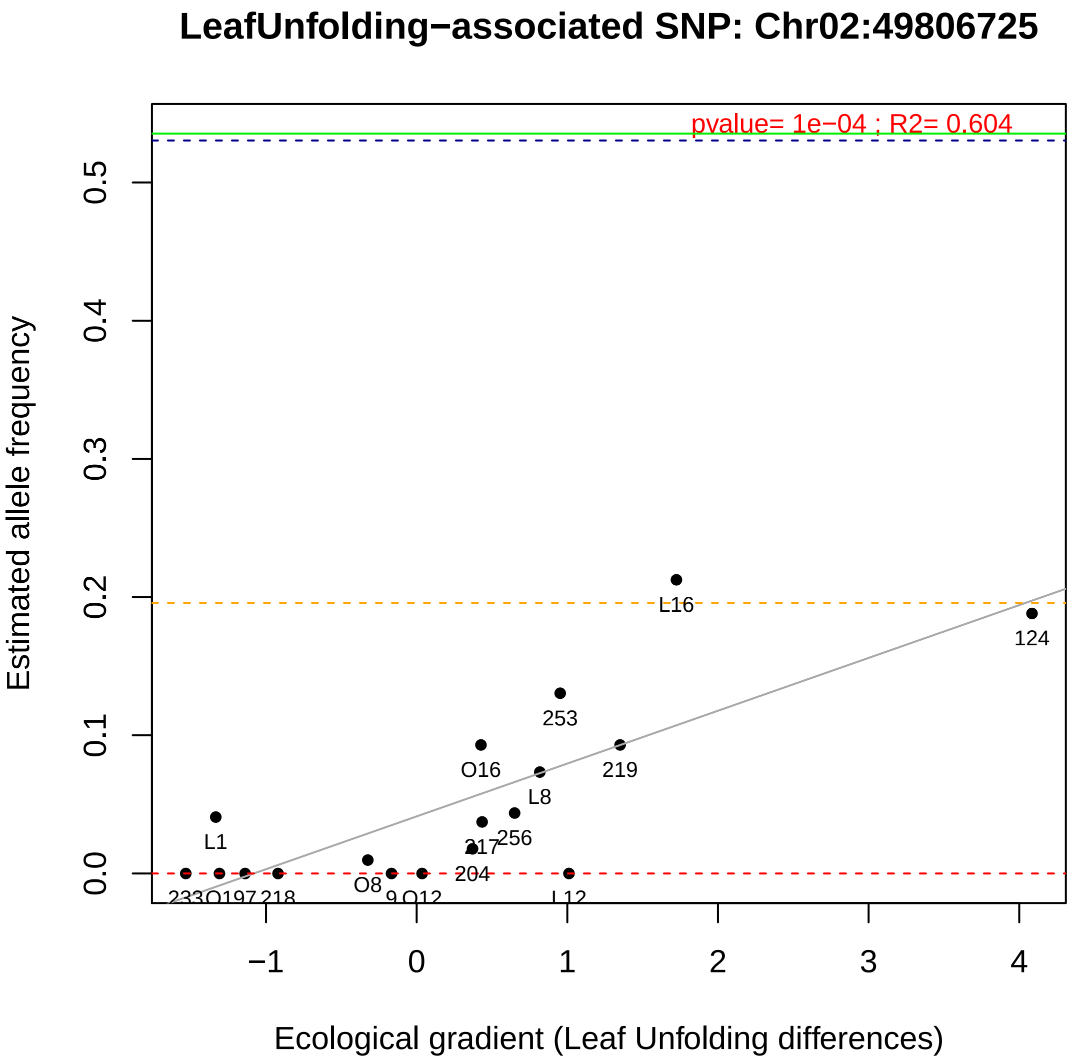
LeafUnfolding−associated SNP: Chr02:49806725
pvalue= 1e−04 ; R2= 0.604
Estimated allele frequency
0.0
0.1
0.2
0.3
0.4
0.5
−1
0
1
2
3
4
Ecological gradient (Leaf Unfolding differences)
L16
124
253
O16
219
L8
256
217
204
L1
O8
233
O197
218
9
O12
L12

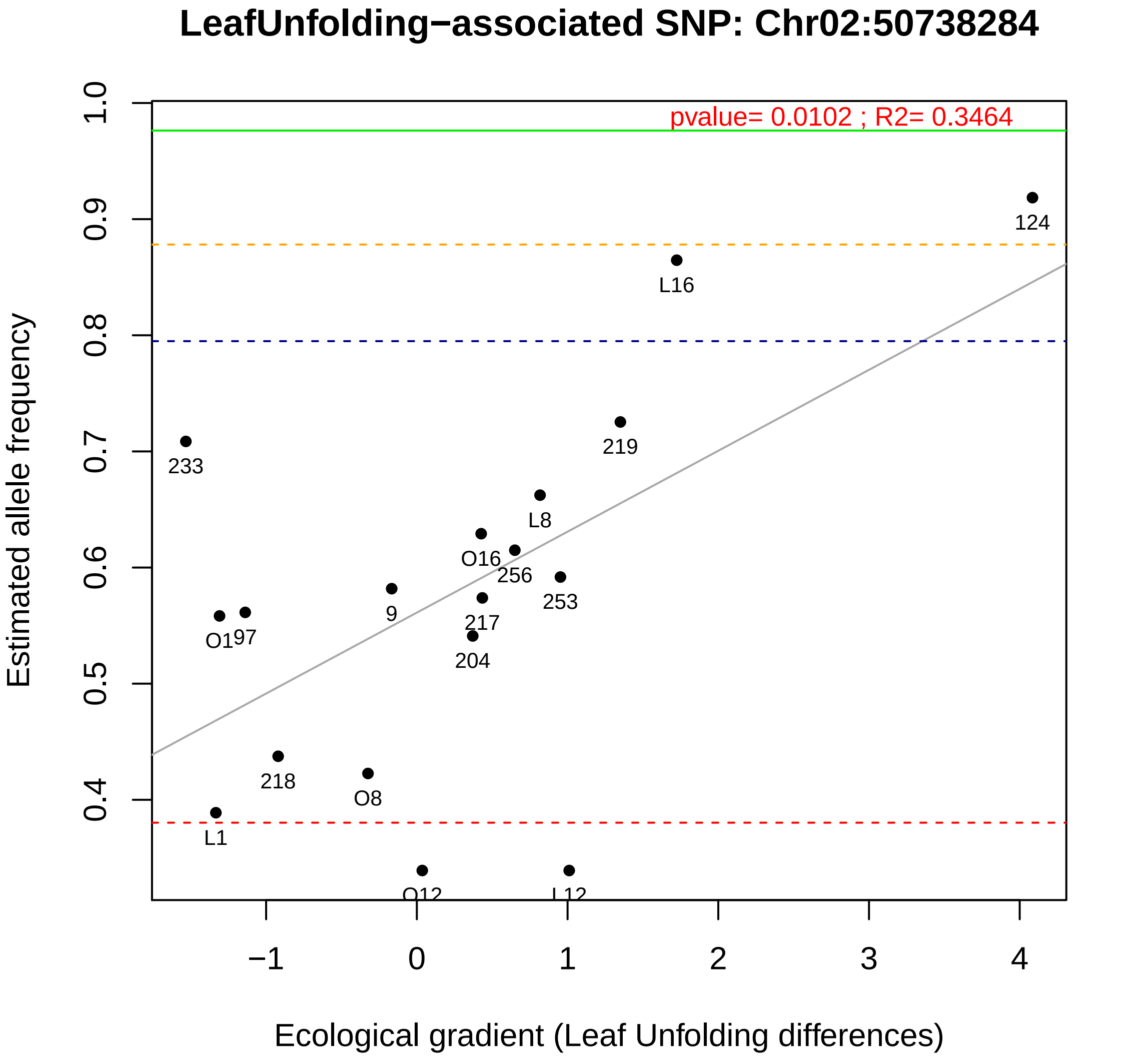

LeafUnfolding−associated SNP: Chr02:50738284
pvalue= 0.0102 ; R2= 0.3464
Estimated allele frequency
Ecological gradient (Leaf Unfolding differences)
1.0
0.9
0.8
0.7
0.6
0.5
0.4
−1
0
1
2
3
4
124
L16
219
233
L8
O16
256
253
9
217
O1
97
204
218
O8
L1

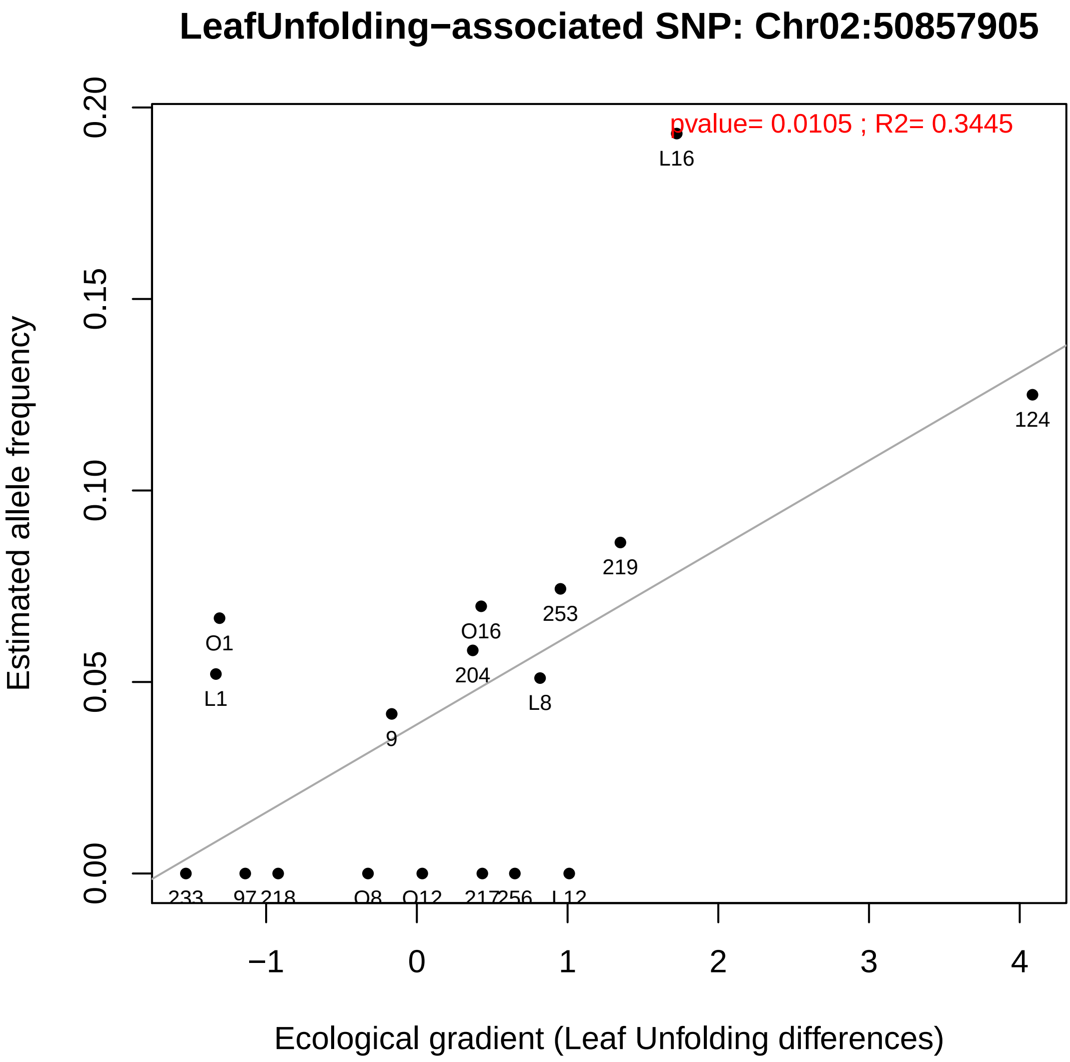
LeafUnfolding−associated SNP: Chr02:50857905
pvalue= 0.0105 ; R2= 0.3445
Estimated allele frequency
0.00
0.05
0.10
0.15
0.20
−1
0
1
2
3
4
Ecological gradient (Leaf Unfolding differences)
L16
124
219
253
O16
O1
204
L1
L8
9
233
97
218
O8
O12
217
256
L12

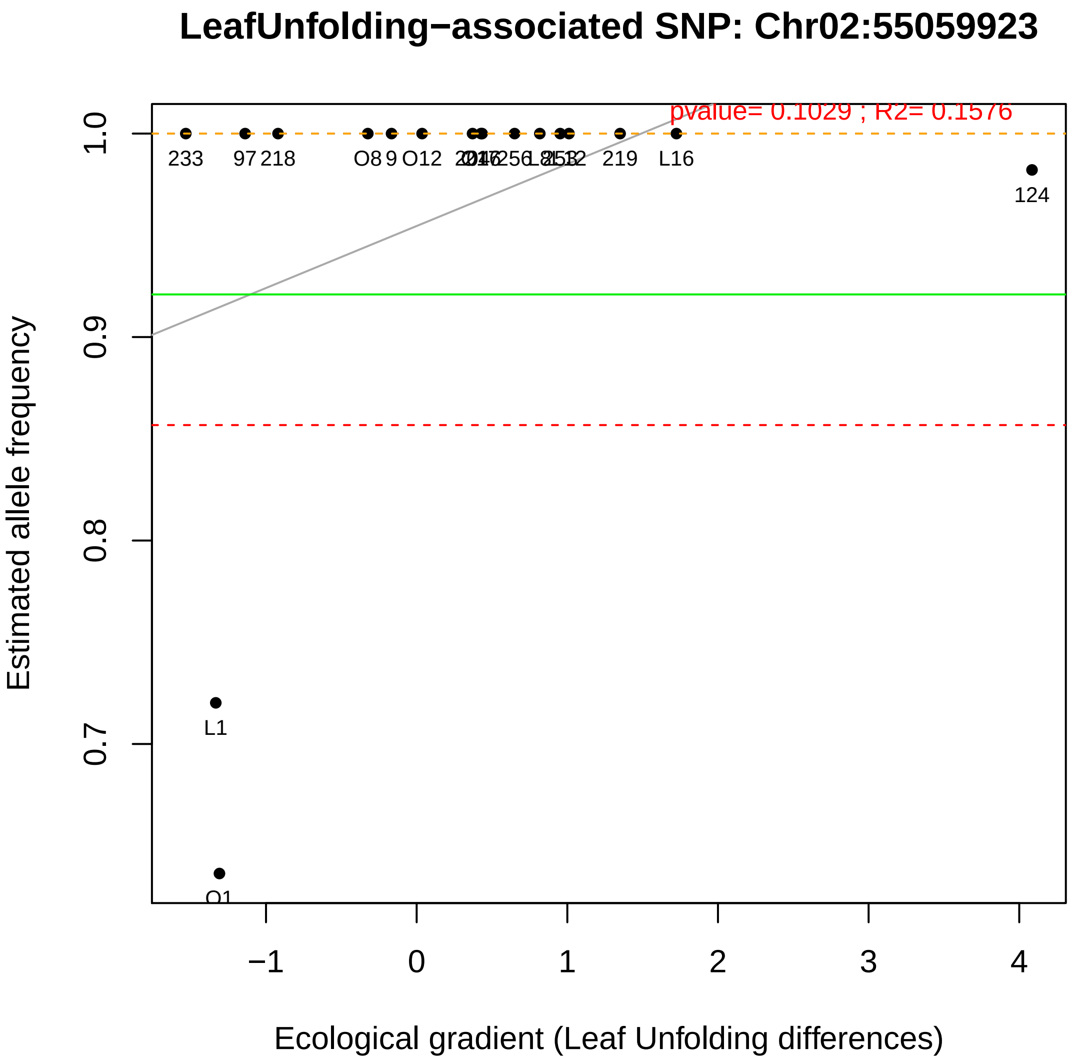
LeafUnfolding−associated SNP: Chr02:55059923
pvalue= 0.1029 ; R2= 0.1576
Estimated allele frequency
1.0
0.9
0.8
0.7
233
97
218
O8
9
O12
219
L16
124
L1
O1
−1
0
1
2
3
4
Ecological gradient (Leaf Unfolding differences)

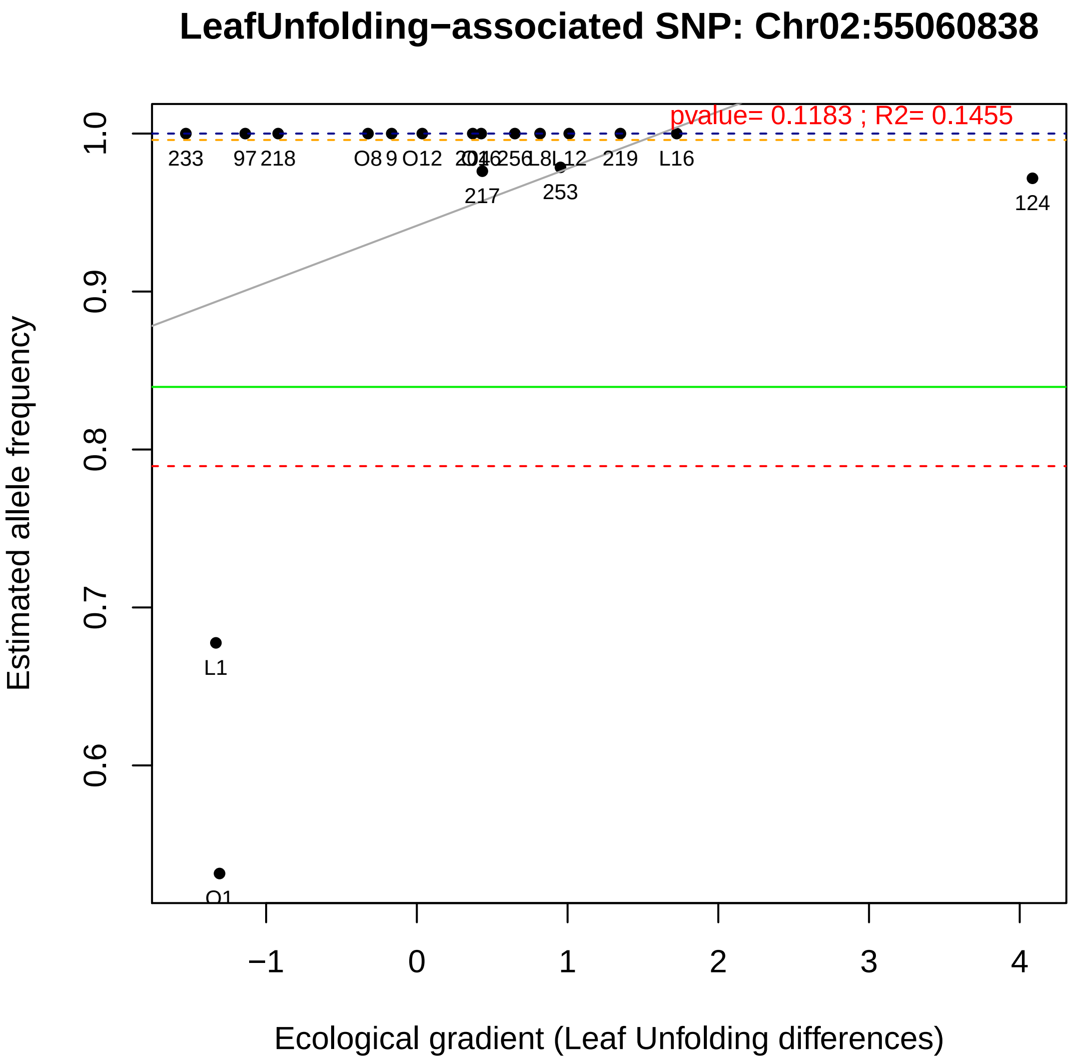

LeafUnfolding−associated SNP: Chr02:55060838
pvalue= 0.1183 ; R2= 0.1455
Estimated allele frequency
Ecological gradient (Leaf Unfolding differences)
233
97
218
O8
9
O12
204
O46
256
L8
L12
219
L16
217
253
124
L1
O1

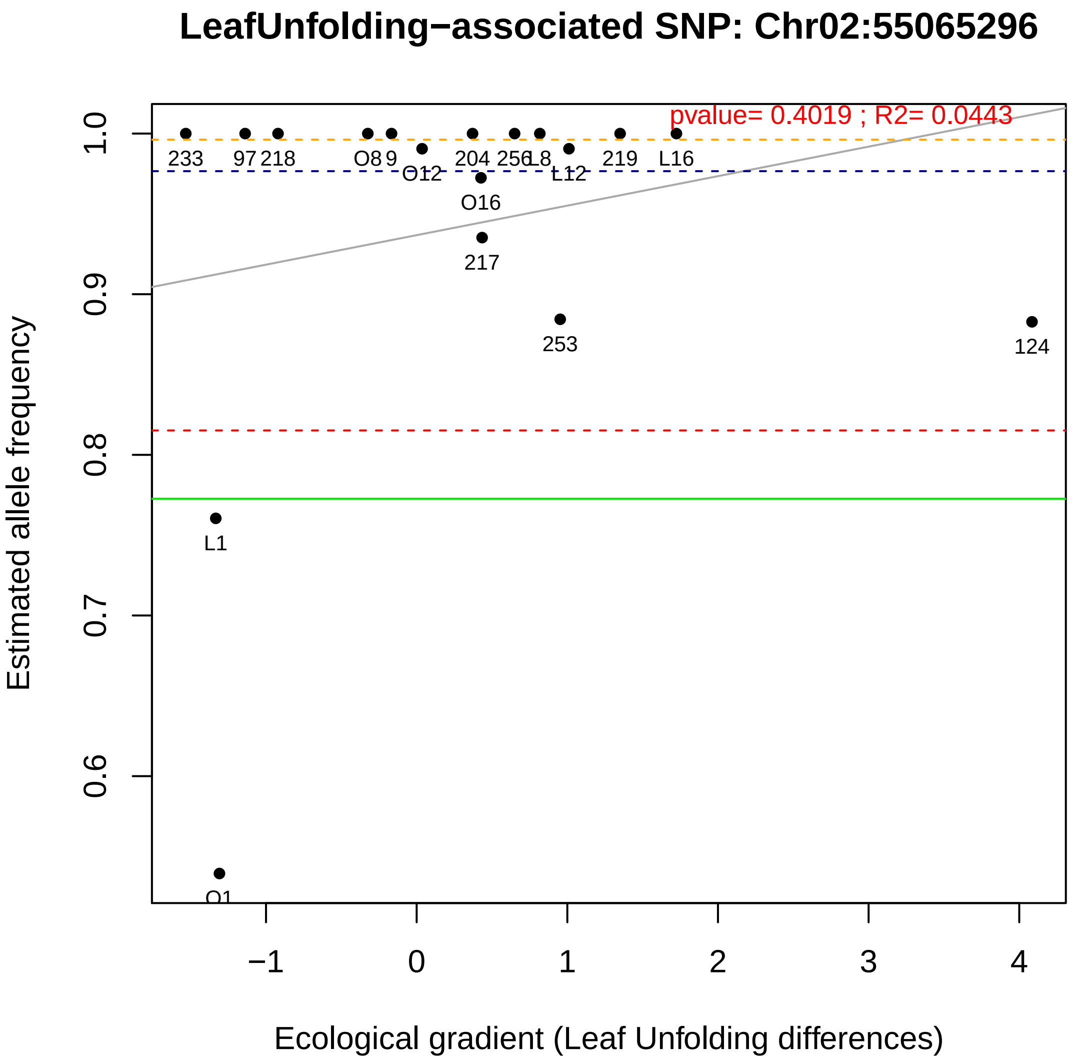

LeafUnfolding−associated SNP: Chr02:55065296
pvalue= 0.4019 ; R2= 0.0443
233
97
218
O8
9
O12
204
256
L8
L12
219
L16
O16
217
253
124
L1
O1
Estimated allele frequency
0.6
0.7
0.8
0.9
1.0
−1
0
1
2
3
4
Ecological gradient (Leaf Unfolding differences)

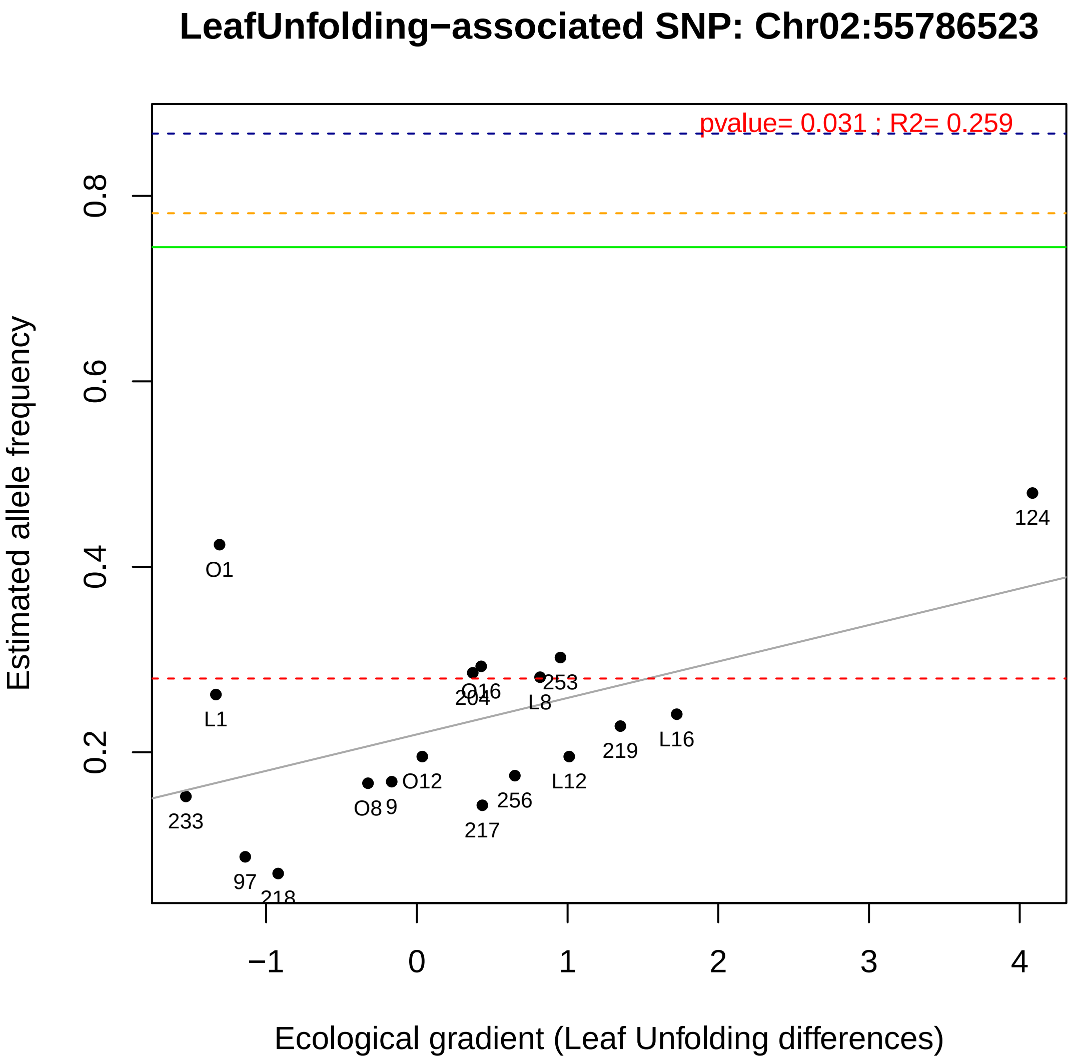

LeafUnfolding−associated SNP: Chr02:55786523
pvalue= 0.031 ; R2= 0.259
Estimated allele frequency
0.2
0.4
0.6
0.8
Ecological gradient (Leaf Unfolding differences)
−1
0
1
2
3
4
124
O1
L1
233
97
218
O8
9
O12
O16
204
217
256
L8
253
L12
219
L16

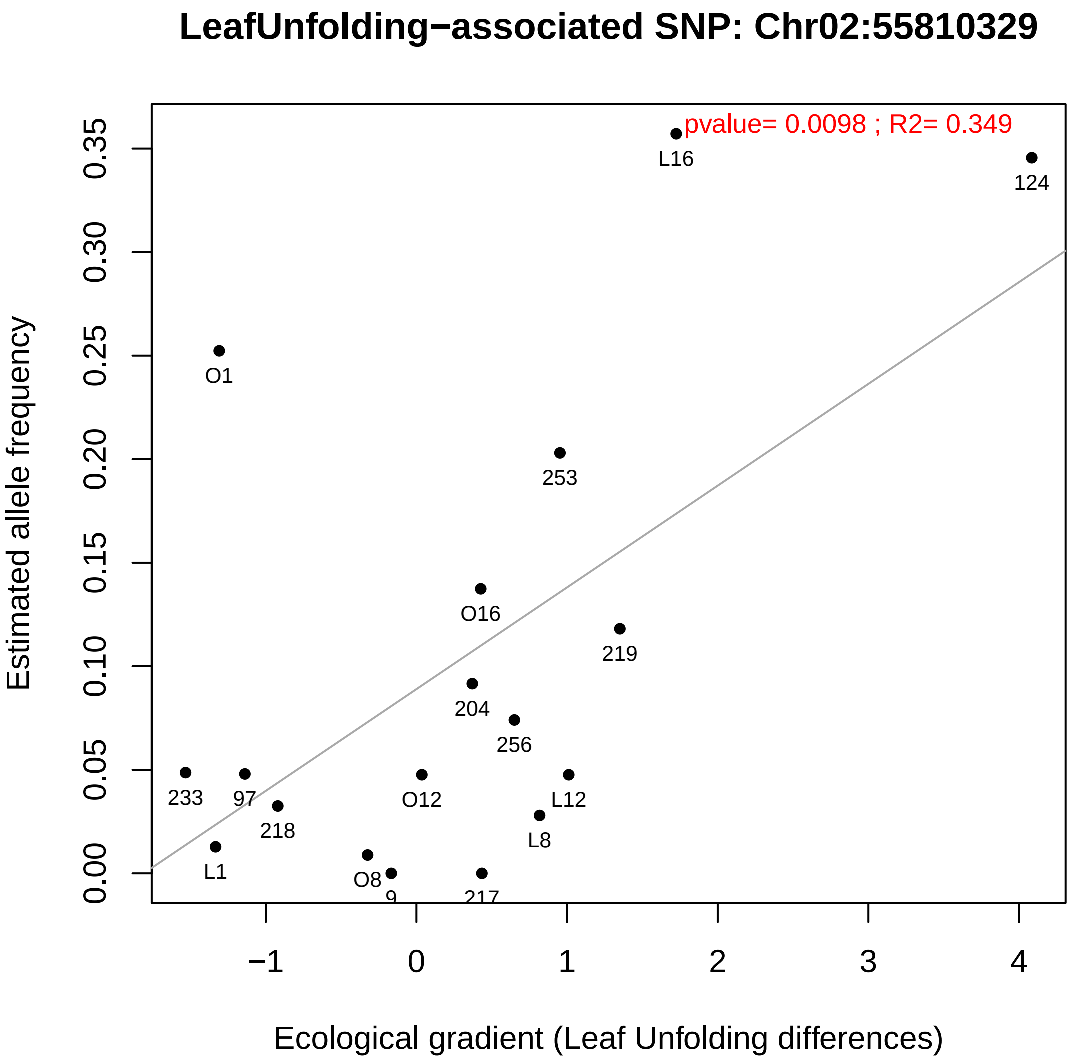
LeafUnfolding−associated SNP: Chr02:55810329
pvalue= 0.0098 ; R2= 0.349
L16
124
O1
253
O16
219
204
256
233
97
218
O12
L12
L8
L1
O8
9
217
0.00
0.05
0.10
0.15
0.20
0.25
0.30
0.35
Estimated allele frequency
−1
0
1
2
3
4
Ecological gradient (Leaf Unfolding differences)

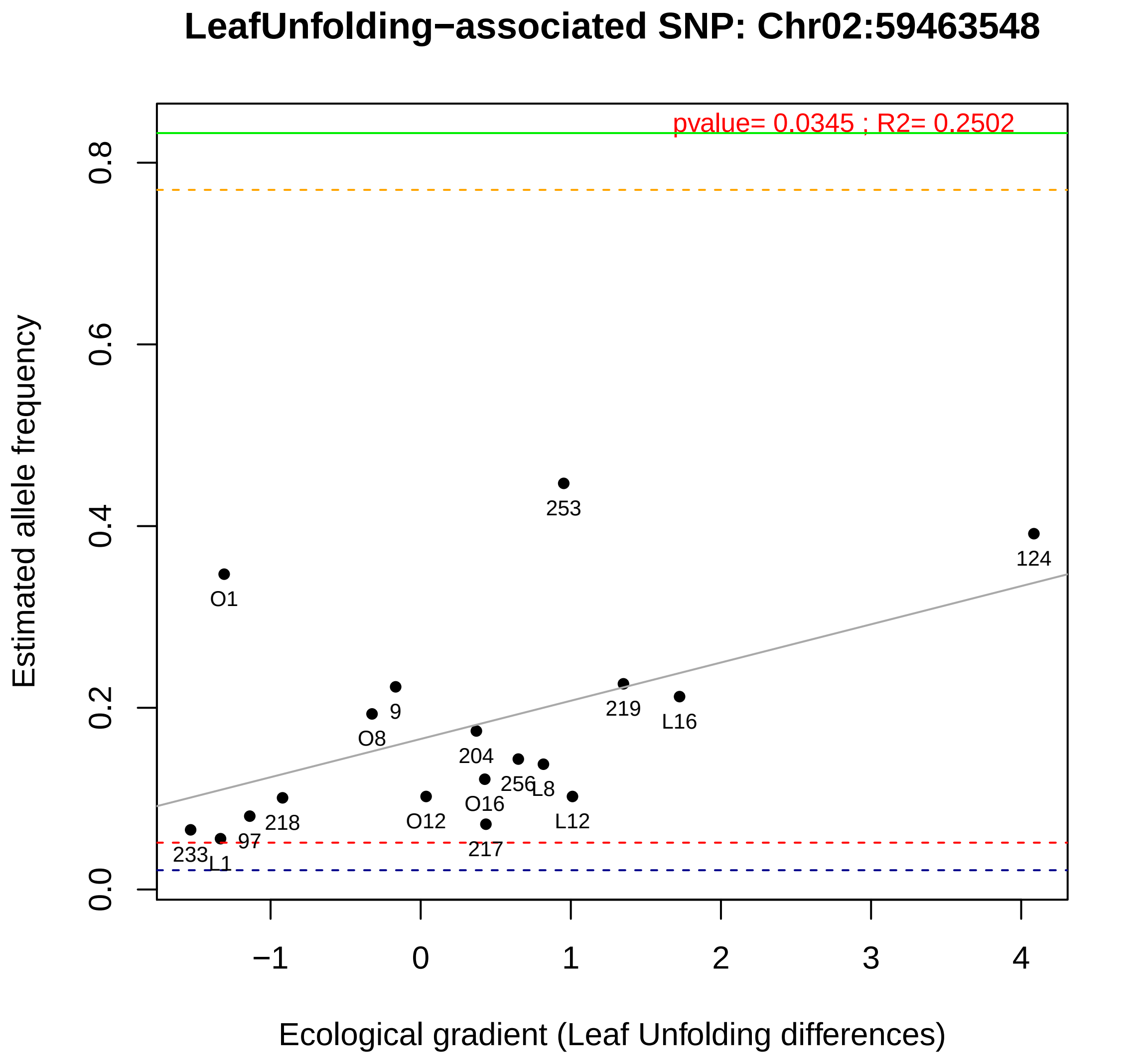

LeafUnfolding−associated SNP: Chr02:59463548
pvalue= 0.0345 ; R2= 0.2502
Estimated allele frequency
0.0
0.2
0.4
0.6
0.8
−1
0
1
2
3
4
Ecological gradient (Leaf Unfolding differences)
253
124
O1
9
O8
219
L16
204
256
L8
O16
L12
O12
218
97
233
L1
217

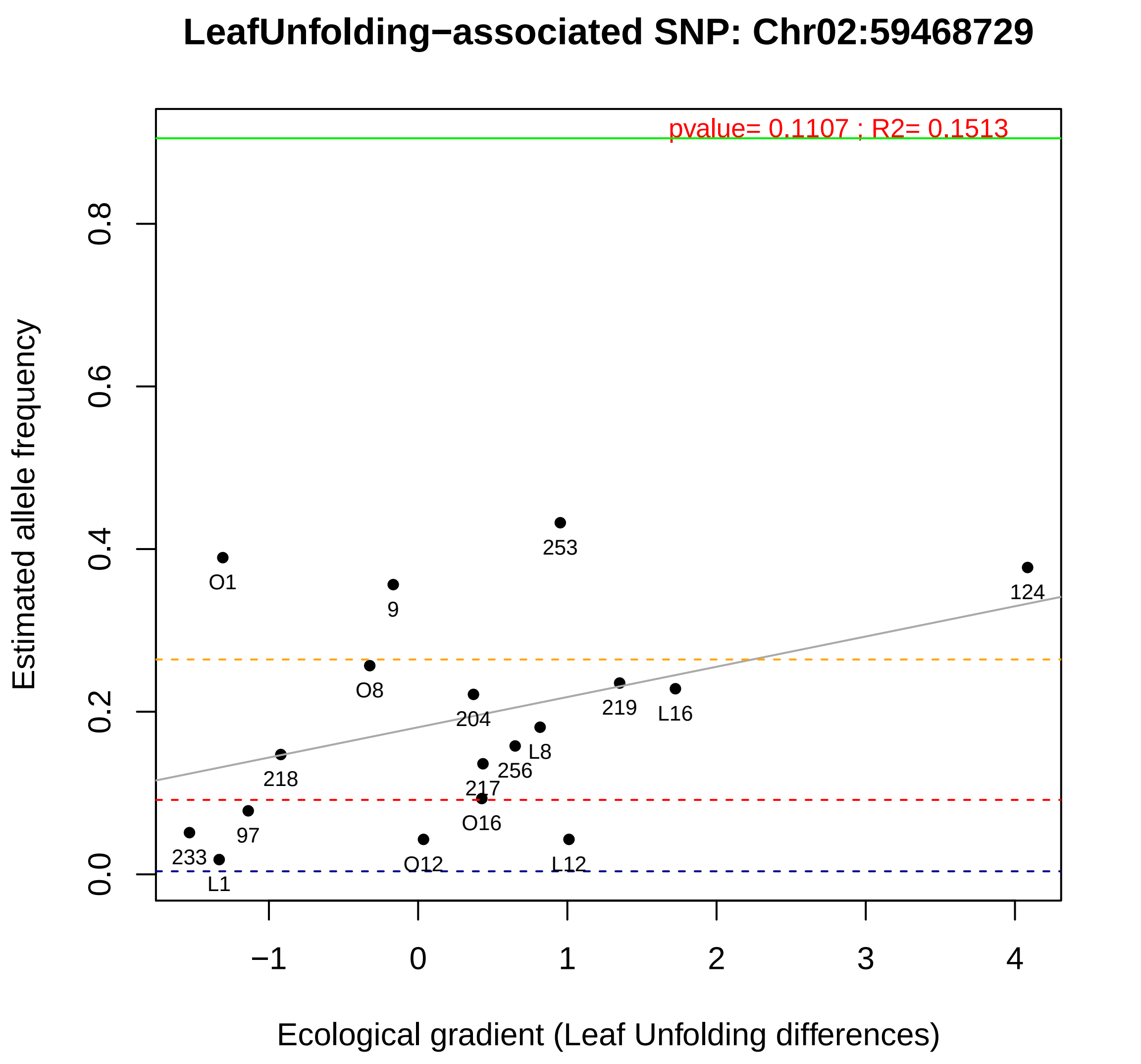

LeafUnfolding−associated SNP: Chr02:59468729
pvalue= 0.1107 ; R2= 0.1513
Estimated allele frequency
0.0
0.2
0.4
0.6
0.8
Ecological gradient (Leaf Unfolding differences)
−1
0
1
2
3
4
O1
9
253
124
O8
204
219
L16
218
L8
256
217
O16
97
233
L1
O12
L12

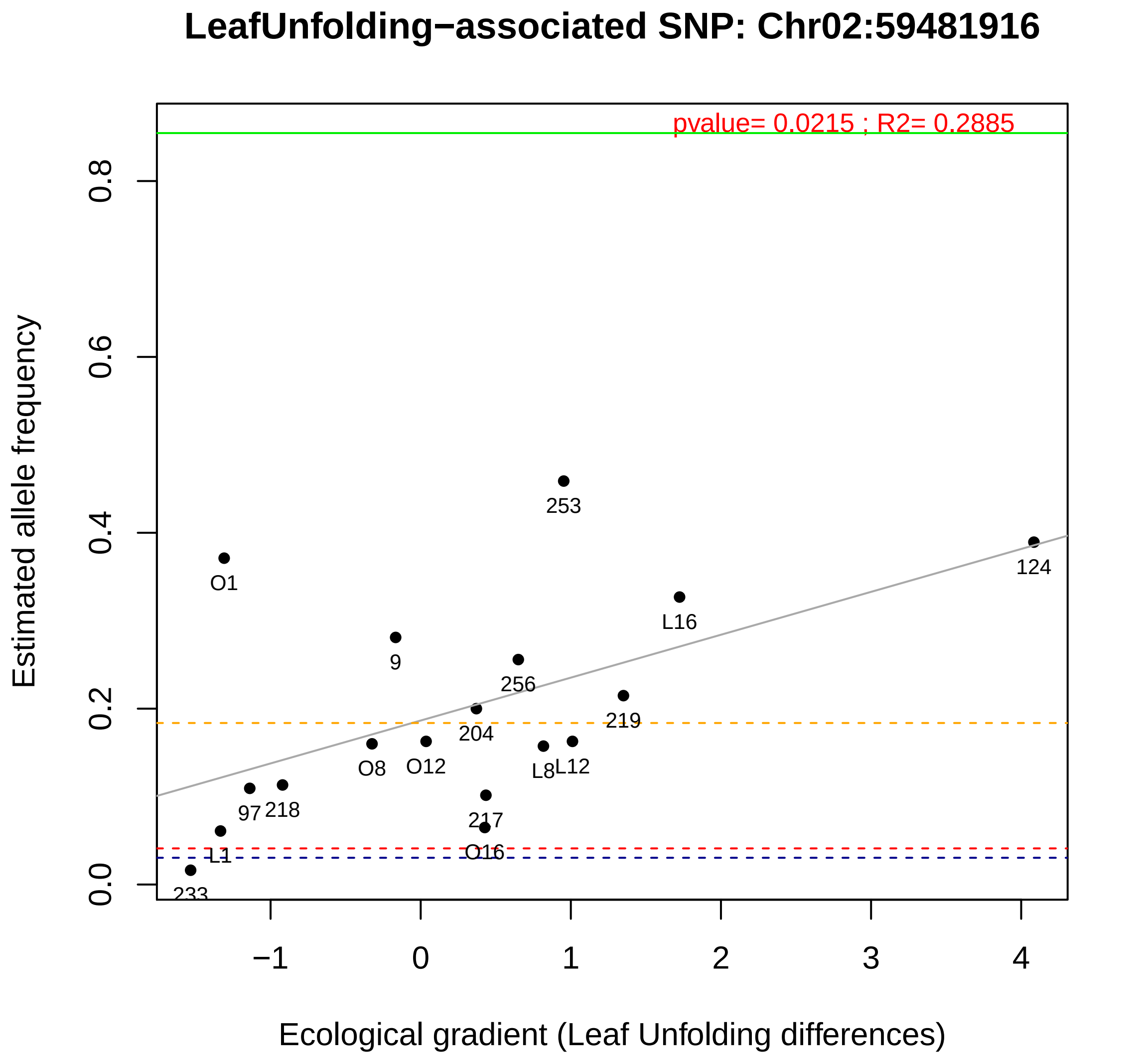
LeafUnfolding−associated SNP: Chr02:59481916
pvalue= 0.0215 ; R2= 0.2885
Estimated allele frequency
Ecological gradient (Leaf Unfolding differences)
0.0
0.2
0.4
0.6
0.8
−1
0
1
2
3
4
253
O1
124
L16
9
256
219
204
O8
O12
L8
L12
97
218
217
O16
L1
233

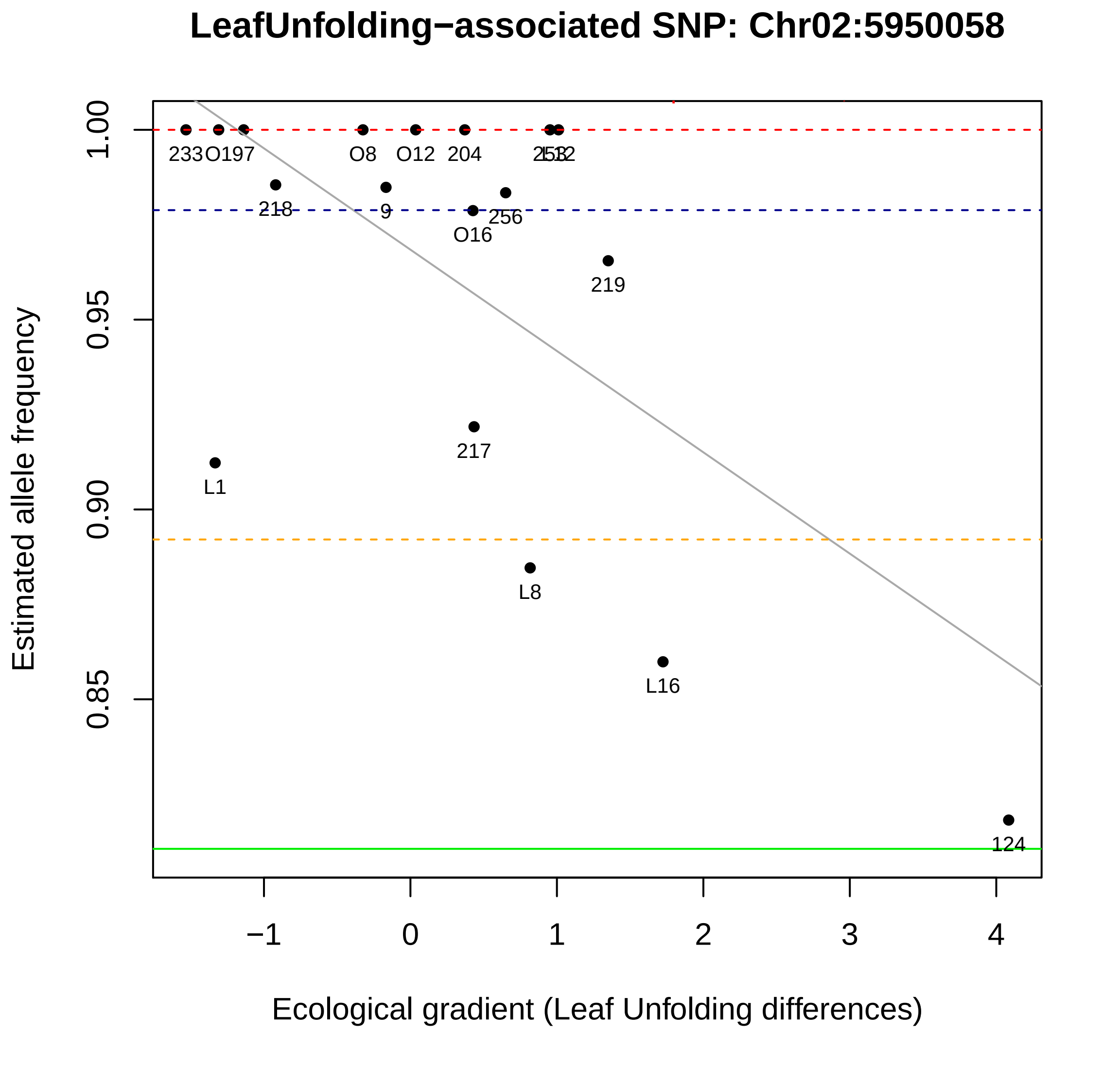

LeafUnfolding−associated SNP: Chr02:5950058
Estimated allele frequency
Ecological gradient (Leaf Unfolding differences)
1.00
0.95
0.90
0.85
−1
0
1
2
3
4
233
O19
7
O8
O12
204
218
9
256
O16
219
217
L1
L8
L16
124

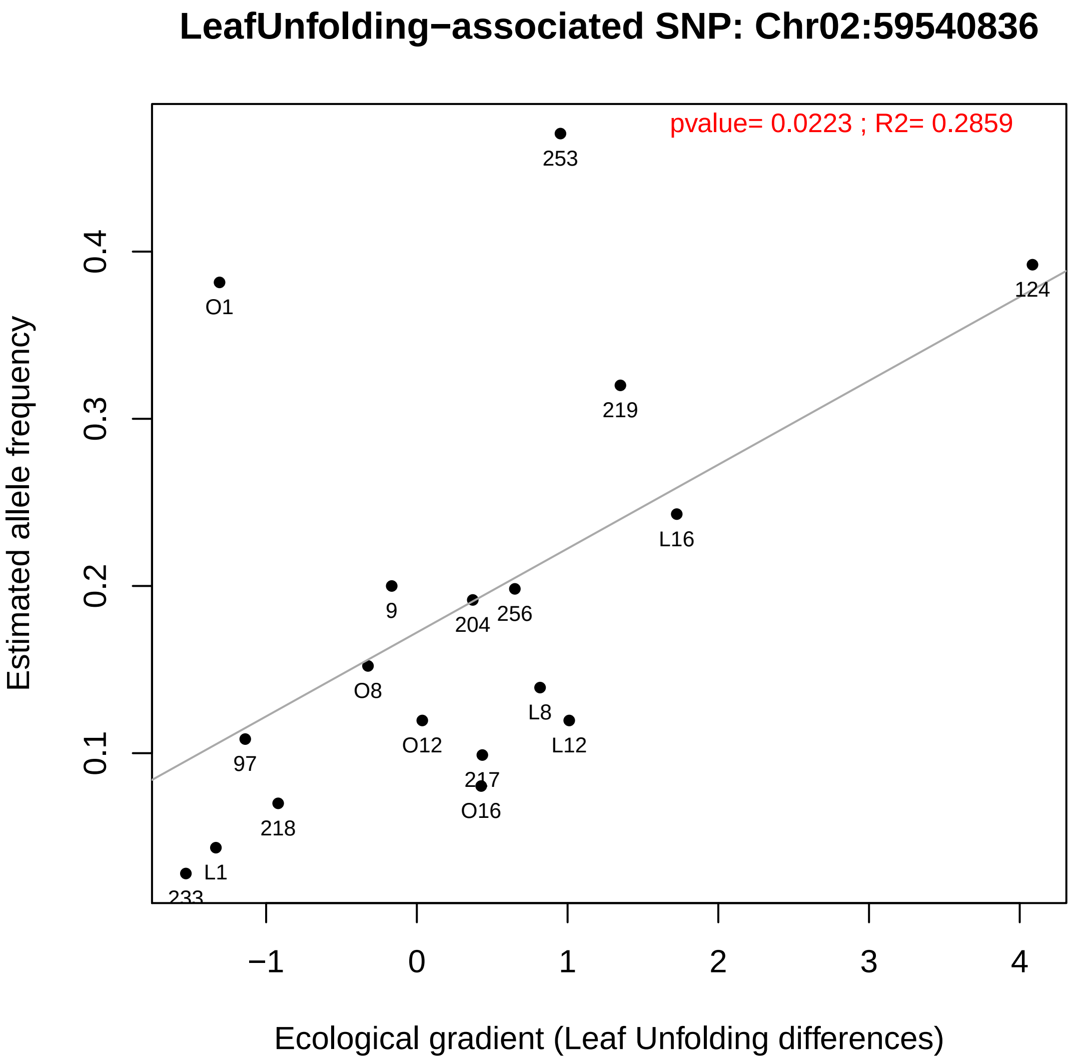
LeafUnfolding−associated SNP: Chr02:59540836
pvalue= 0.0223 ; R2= 0.2859
253
124
O1
219
L16
9
204
256
O8
L8
L12
O12
97
217
O16
218
L1
233
0.1
0.2
0.3
0.4
−1
0
1
2
3
4
Estimated allele frequency
Ecological gradient (Leaf Unfolding differences)

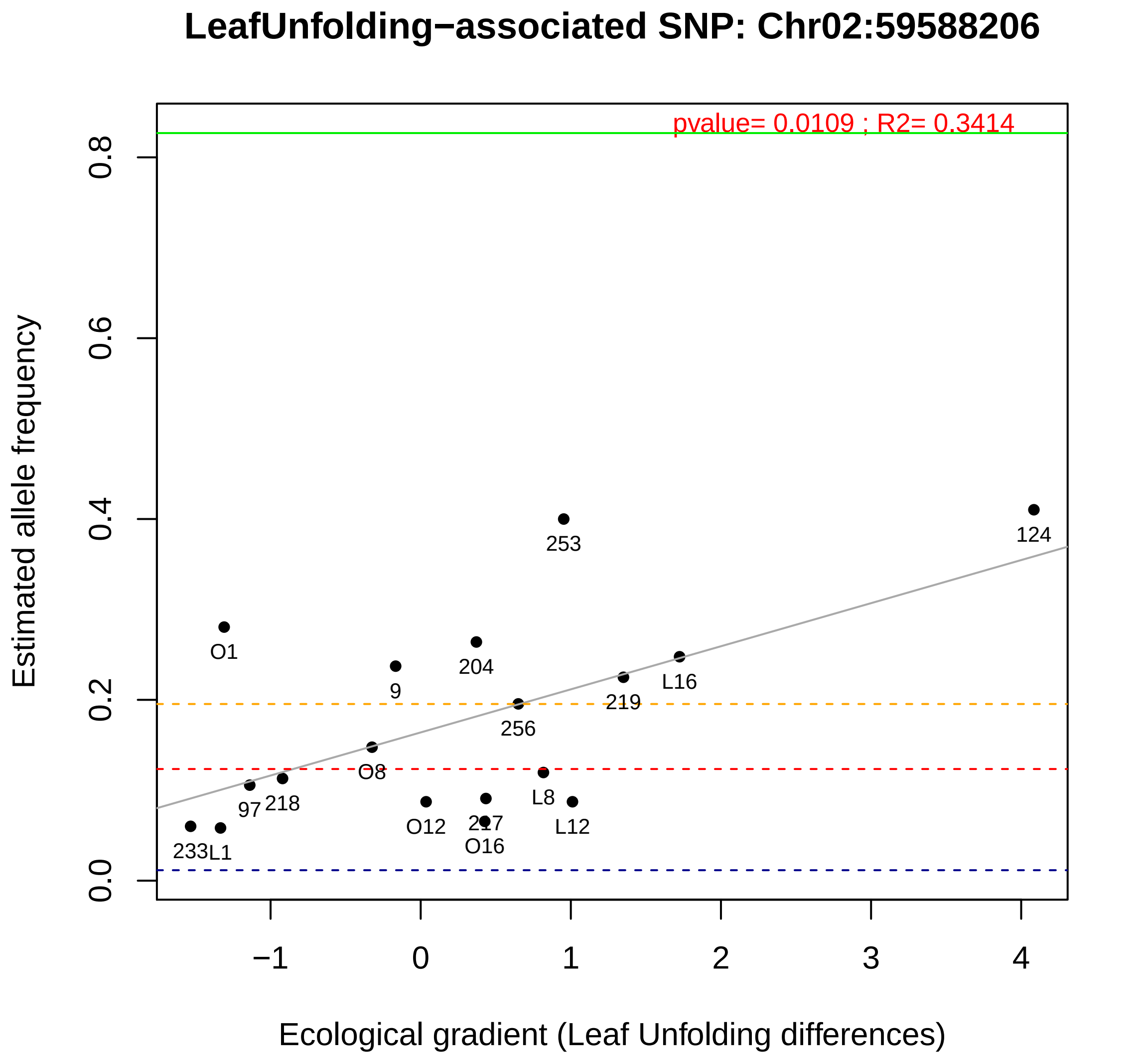
LeafUnfolding−associated SNP: Chr02:59588206
pvalue= 0.0109 ; R2= 0.3414
Estimated allele frequency
0.0
0.2
0.4
0.6
0.8
−1
0
1
2
3
4
Ecological gradient (Leaf Unfolding differences)
O1
9
204
253
124
256
219
L16
O8
97
218
233
L1
O12
217
O16
L8
L12

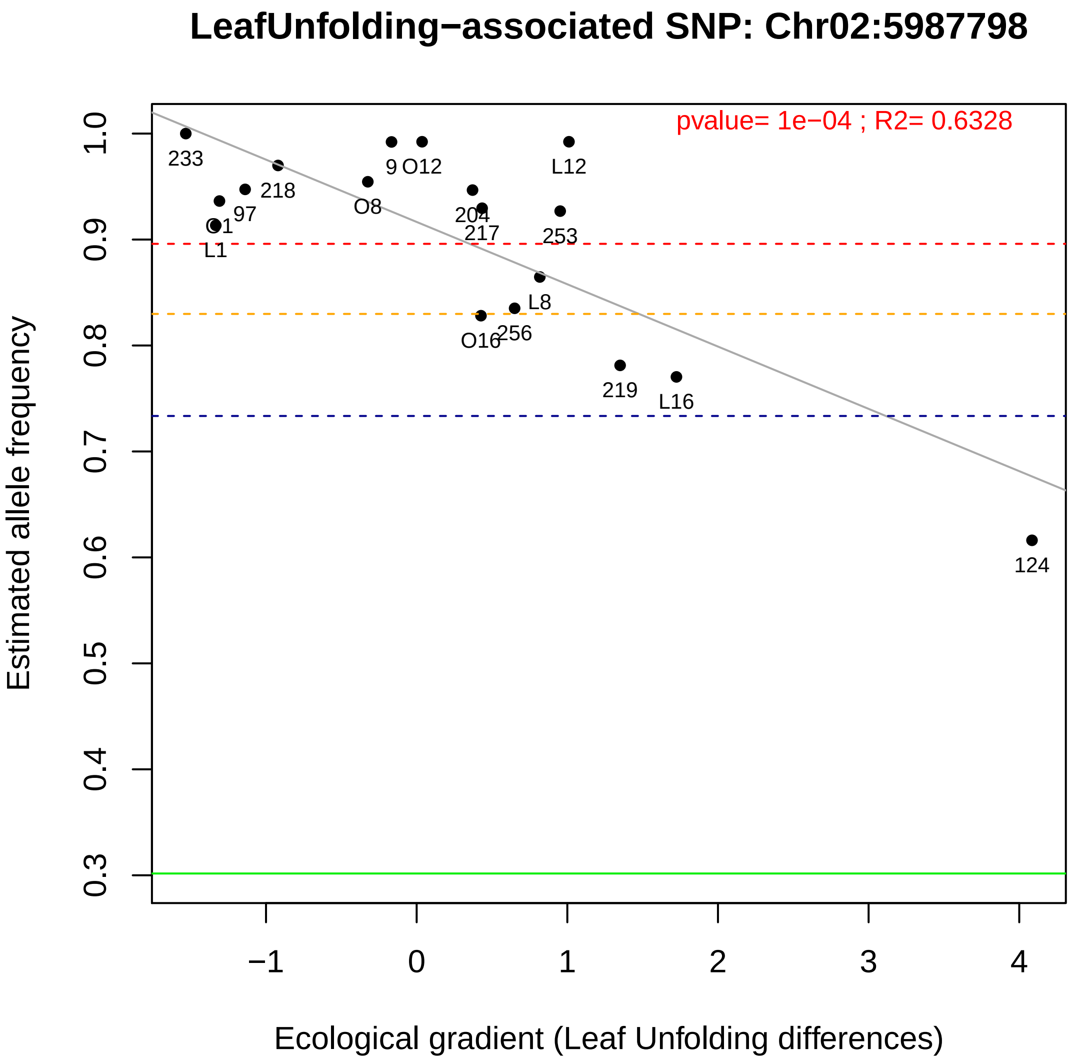

LeafUnfolding−associated SNP: Chr02:5987798
pvalue= 1e−04 ; R2= 0.6328
Estimated allele frequency
Ecological gradient (Leaf Unfolding differences)
1.0
0.9
0.8
0.7
0.6
0.5
0.4
0.3
−1
0
1
2
3
4
233
218
97
O1
L1
O8
9
O12
204
217
L12
253
L8
256
O16
219
L16
124

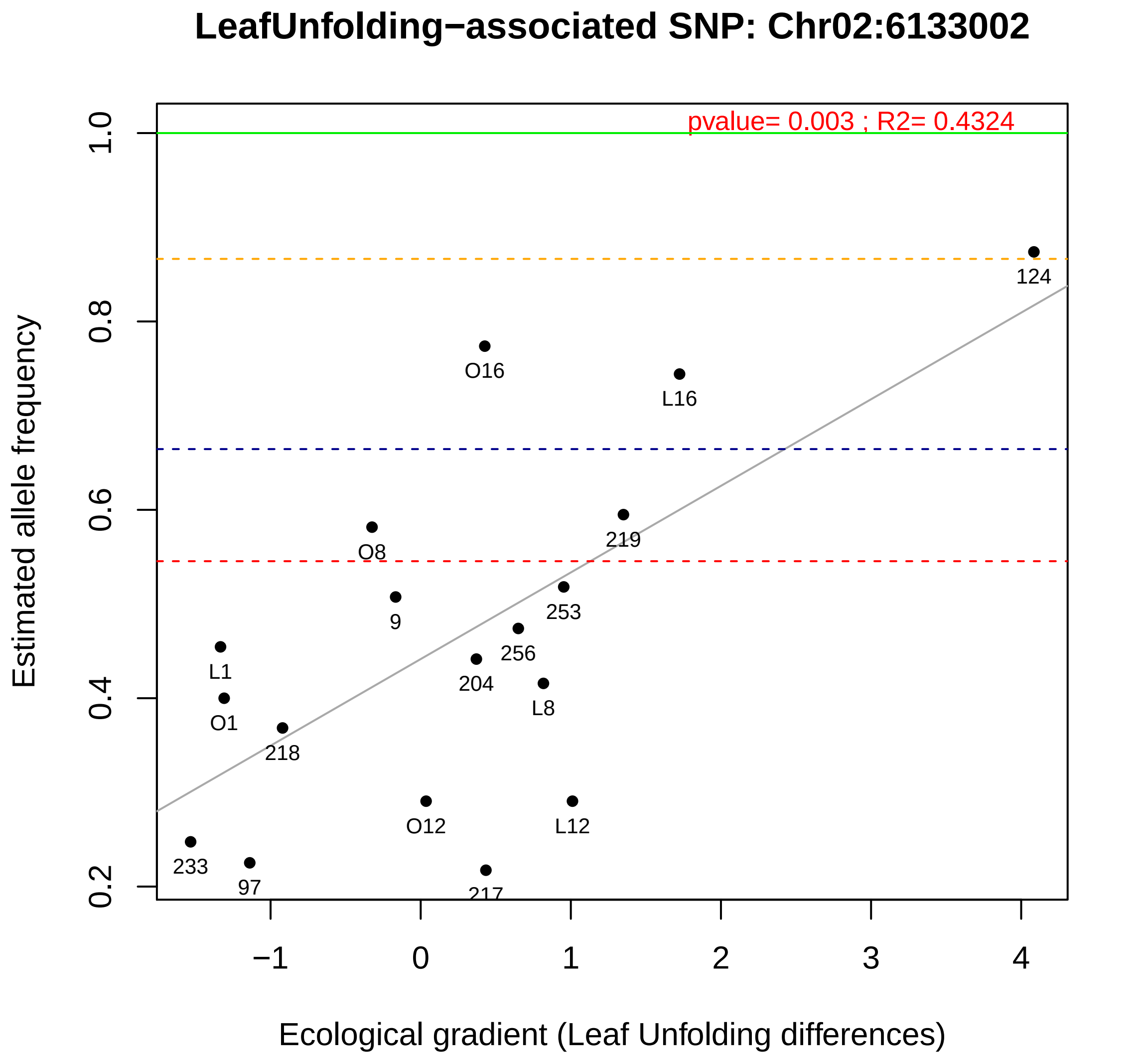
LeafUnfolding−associated SNP: Chr02:6133002
pvalue= 0.003 ; R2= 0.4324
Estimated allele frequency
0.2
0.4
0.6
0.8
1.0
−1
0
1
2
3
4
Ecological gradient (Leaf Unfolding differences)
124
O16
L16
219
O8
9
253
256
L1
204
L8
O1
218
O12
L12
233
97
217

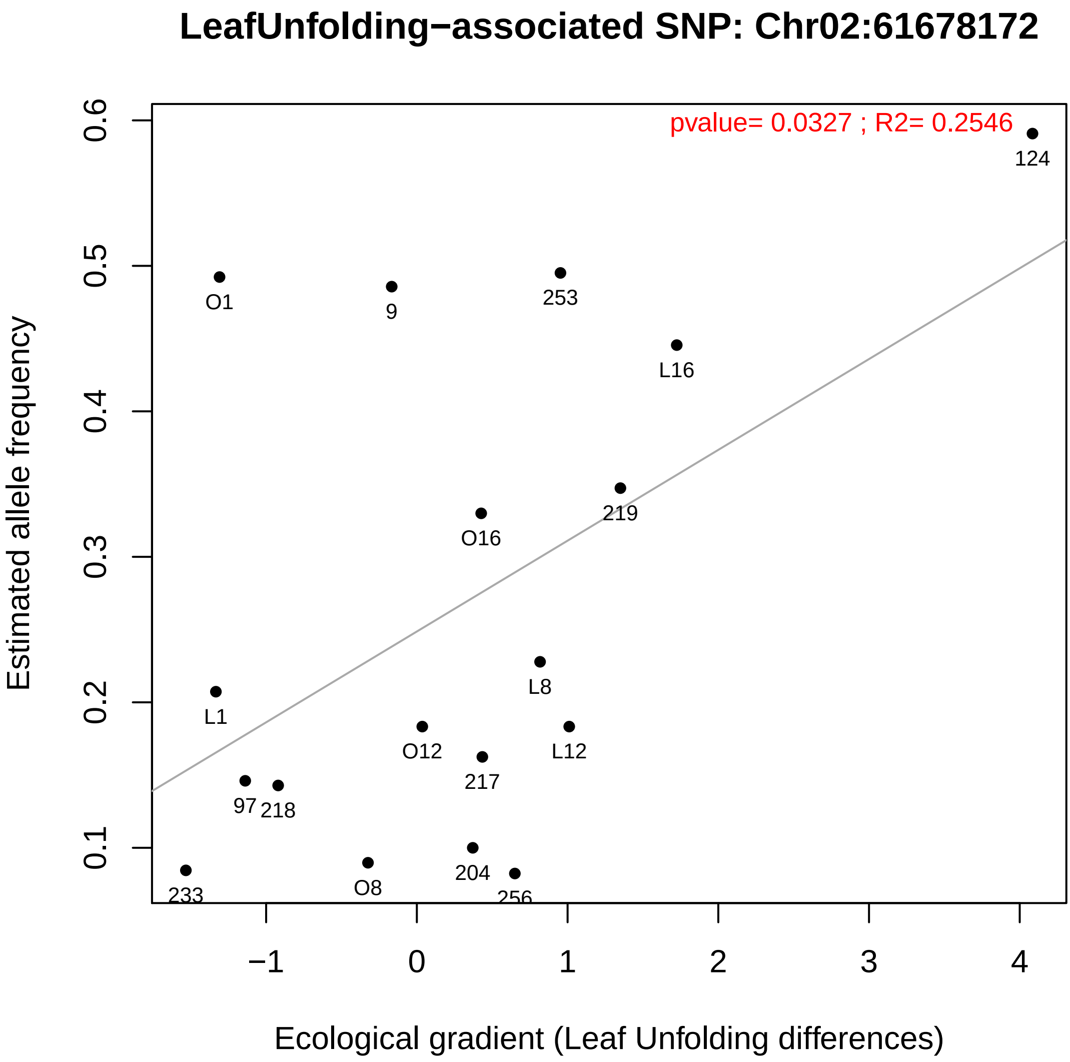
LeafUnfolding−associated SNP: Chr02:61678172
pvalue= 0.0327 ; R2= 0.2546
Estimated allele frequency
Ecological gradient (Leaf Unfolding differences)
0.1
0.2
0.3
0.4
0.5
0.6
−1
0
1
2
3
4
124
O1
9
253
L16
219
O16
L8
L1
O12
L12
217
97
218
233
O8
204
256

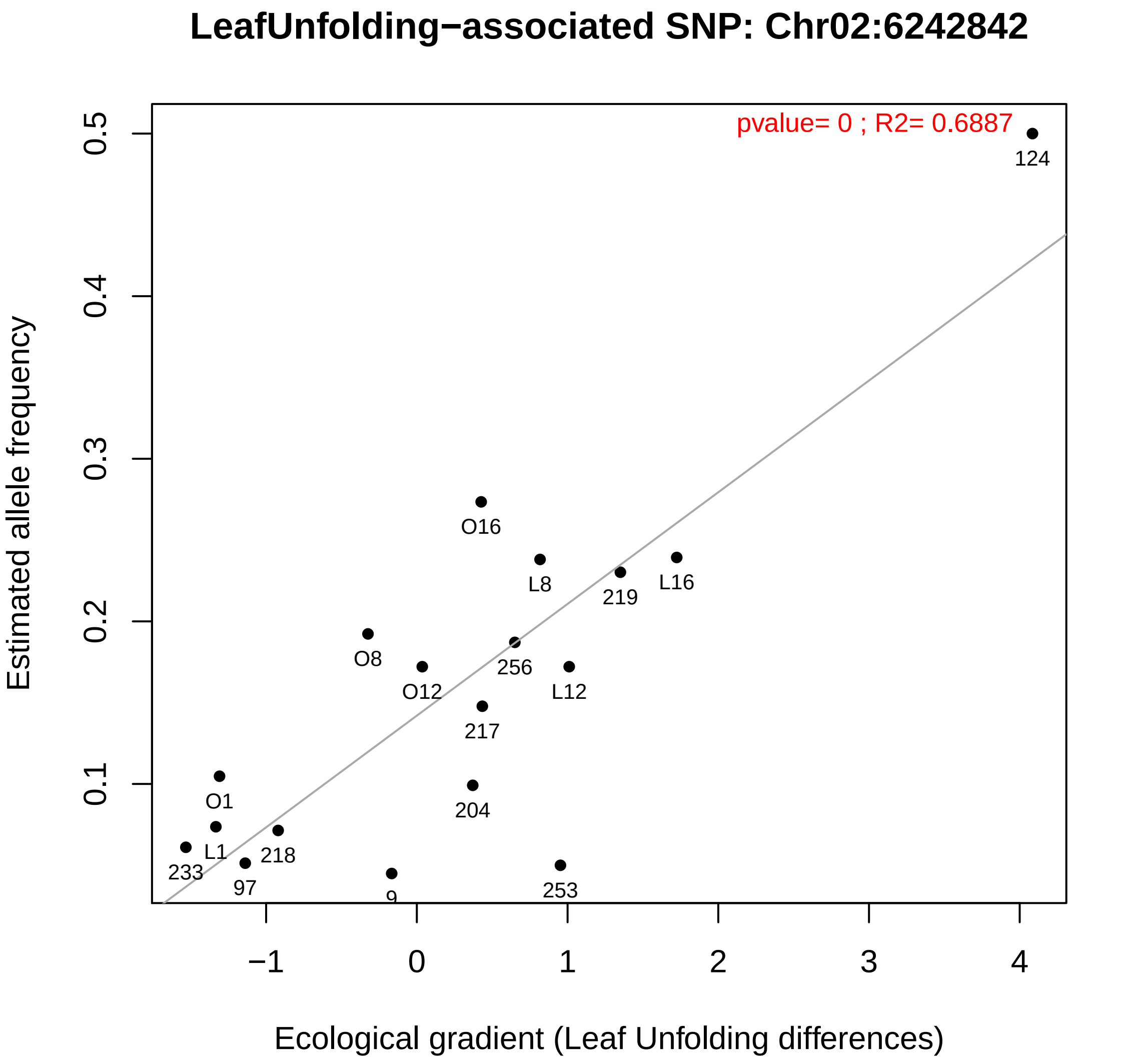
LeafUnfolding−associated SNP: Chr02:6242842
pvalue= 0 ; R2= 0.6887
Estimated allele frequency
0.1
0.2
0.3
0.4
0.5
−1
0
1
2
3
4
Ecological gradient (Leaf Unfolding differences)
124
O16
L8
219
L16
O8
256
O12
L12
217
O1
204
L1
218
233
97
9
253

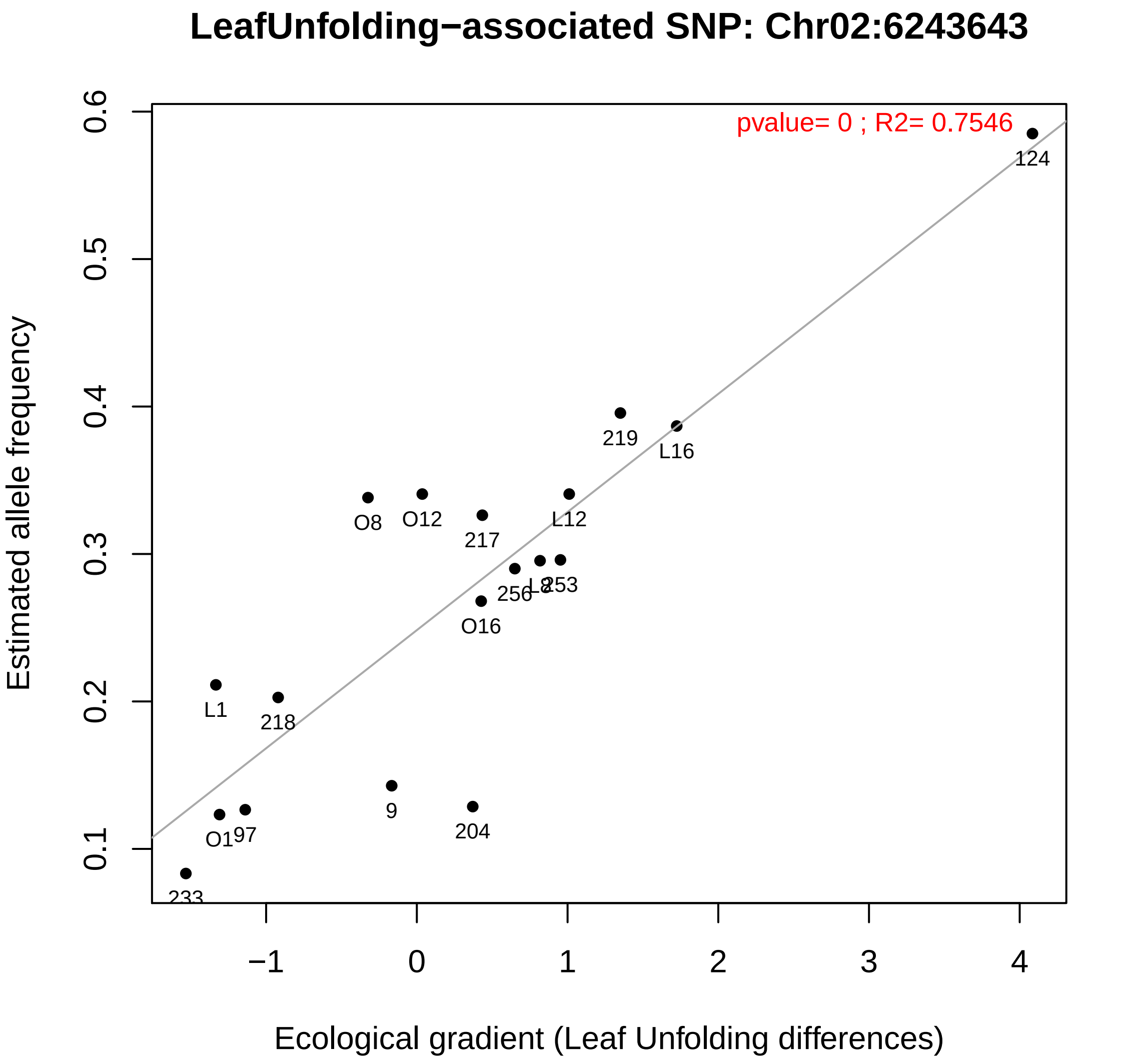
LeafUnfolding−associated SNP: Chr02:6243643
pvalue= 0 ; R2= 0.7546
Estimated allele frequency
Ecological gradient (Leaf Unfolding differences)
0.1
0.2
0.3
0.4
0.5
0.6
−1
0
1
2
3
4
124
219
L16
O8
O12
217
L12
256
L8
253
O16
L1
218
9
204
O1
97
233

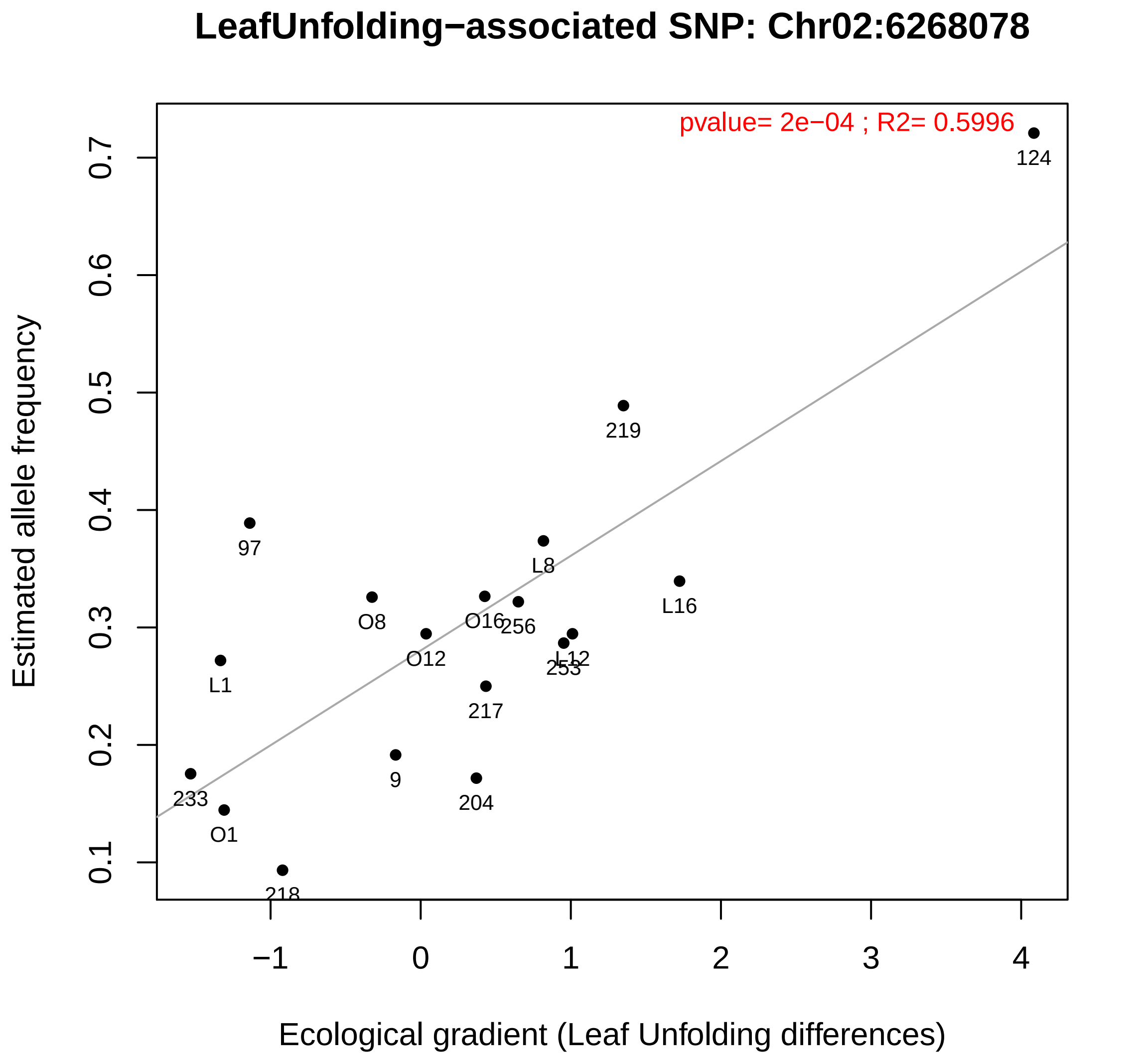
LeafUnfolding−associated SNP: Chr02:6268078
pvalue= 2e−04 ; R2= 0.5996
124
219
97
L8
L16
O8
O16
256
O12
L12
253
L1
217
9
204
233
O1
218
0.1
0.2
0.3
0.4
0.5
0.6
0.7
−1
0
1
2
3
4
Estimated allele frequency
Ecological gradient (Leaf Unfolding differences)

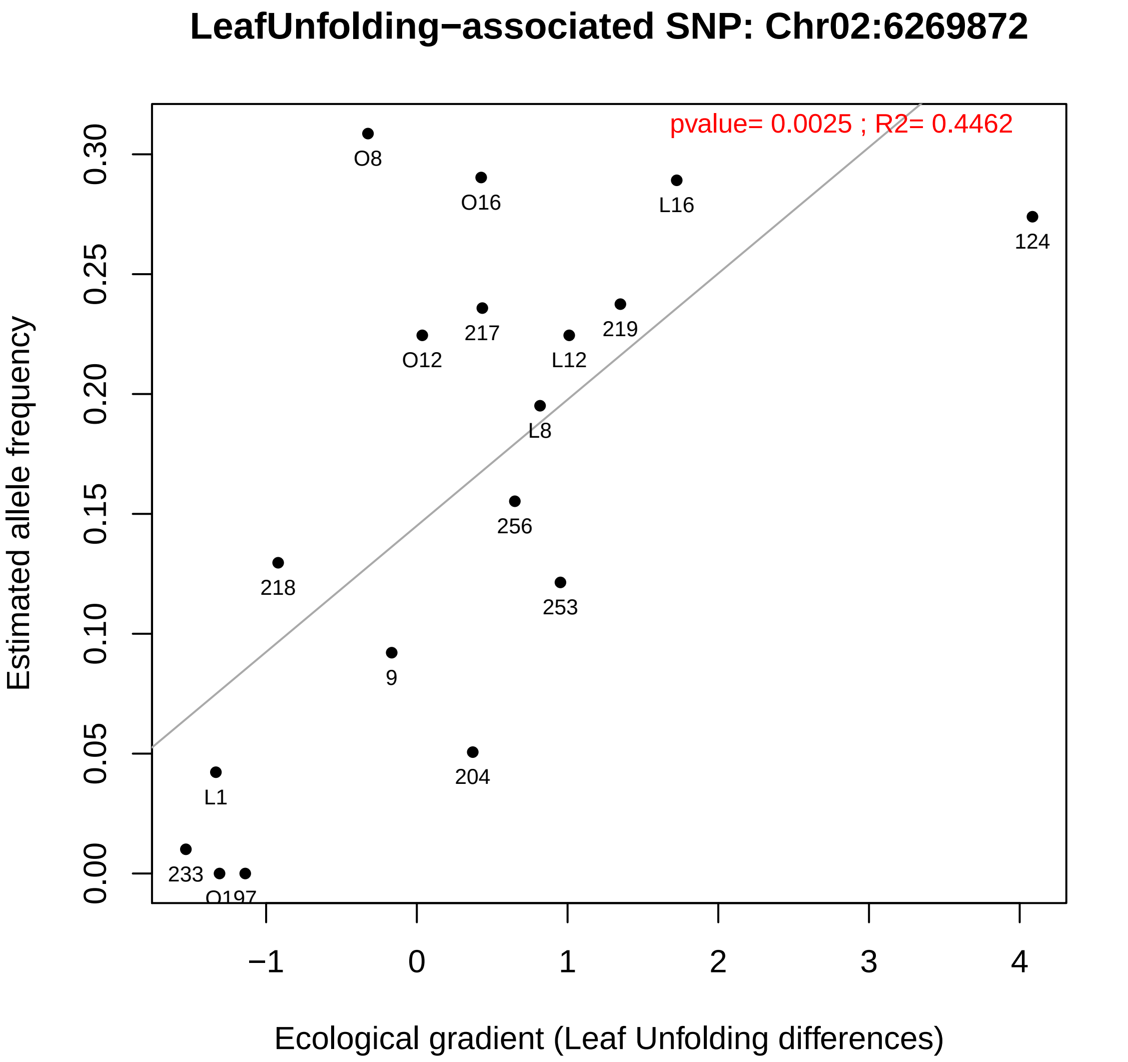
LeafUnfolding−associated SNP: Chr02:6269872
pvalue= 0.0025 ; R2= 0.4462
Estimated allele frequency
0.00
0.05
0.10
0.15
0.20
0.25
0.30
−1
0
1
2
3
4
Ecological gradient (Leaf Unfolding differences)
O8
O16
L16
124
217
219
O12
L12
L8
256
218
253
9
204
L1
233
O197

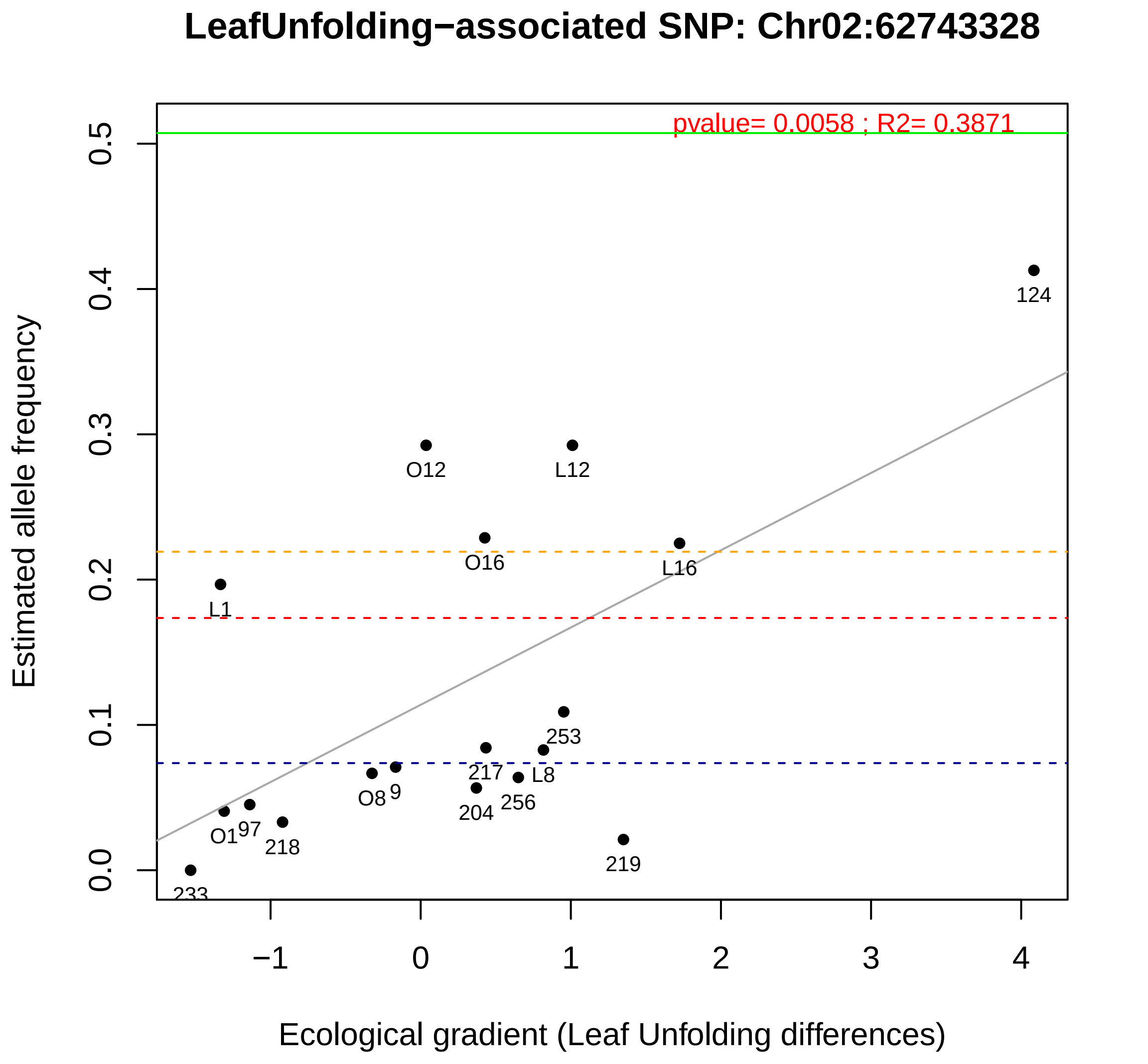

LeafUnfolding−associated SNP: Chr02:62743328
pvalue= 0.0058 ; R2= 0.3871
Estimated allele frequency
Ecological gradient (Leaf Unfolding differences)
0.0
0.1
0.2
0.3
0.4
0.5
−1
0
1
2
3
4
124
O12
L12
O16
L16
L1
253
217
L8
256
204
O8
9
O1
97
218
219
233

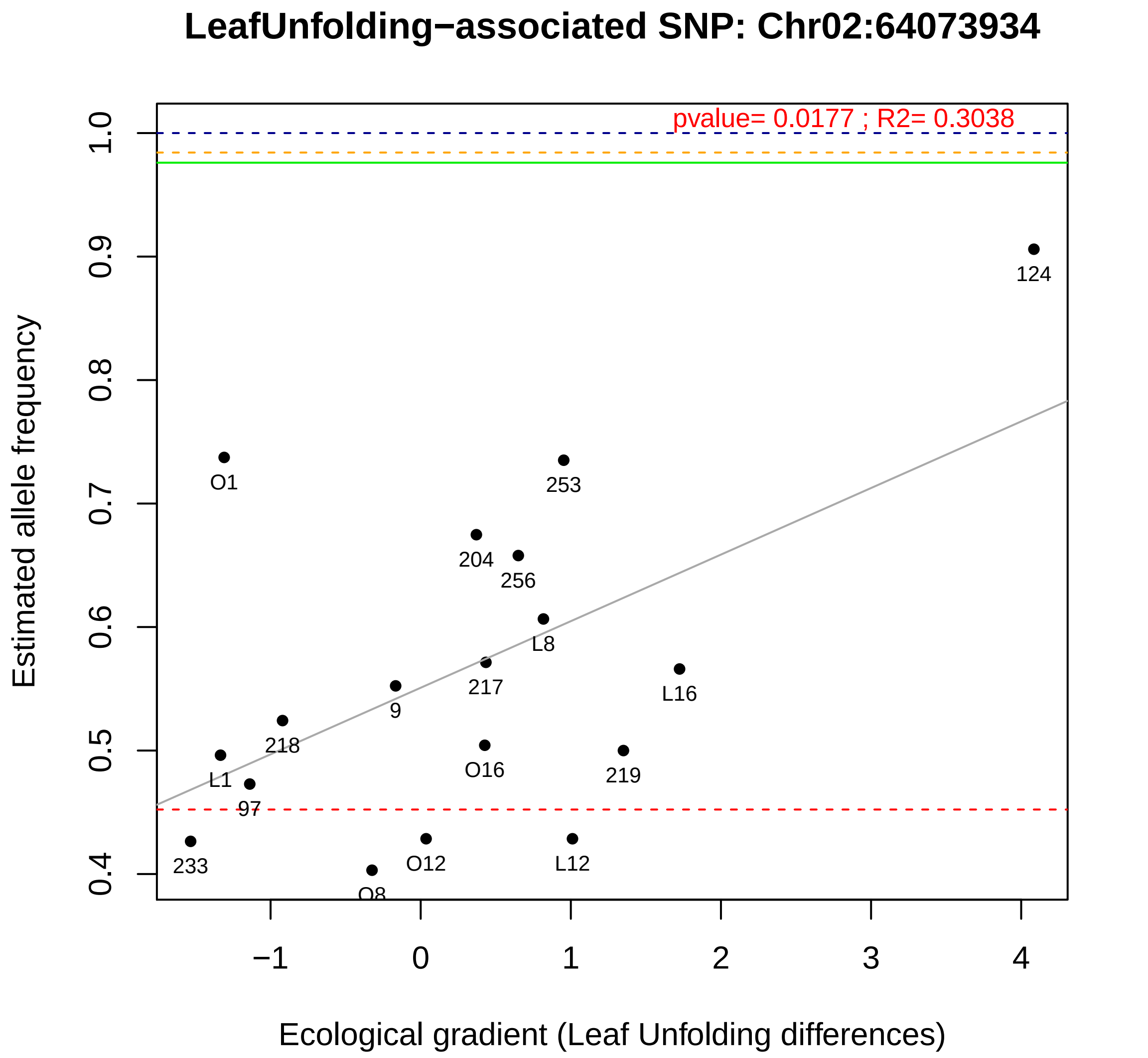

LeafUnfolding−associated SNP: Chr02:64073934
pvalue= 0.0177 ; R2= 0.3038
Estimated allele frequency
0.4
0.5
0.6
0.7
0.8
0.9
1.0
Ecological gradient (Leaf Unfolding differences)
−1
0
1
2
3
4
124
O1
253
204
256
L8
217
L16
9
218
O16
219
L1
97
233
O12
L12
O8

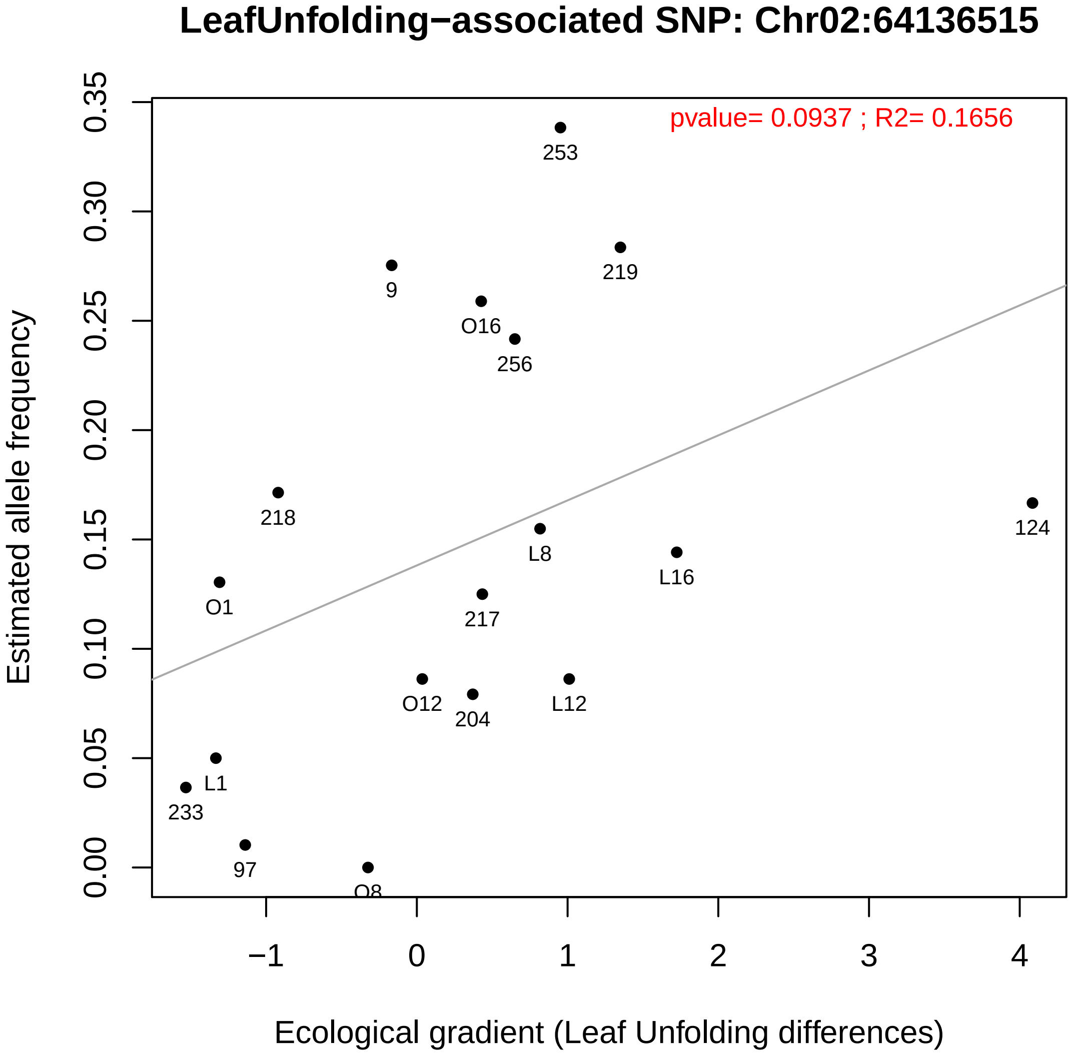

LeafUnfolding−associated SNP: Chr02:64136515
pvalue= 0.0937 ; R2= 0.1656
Estimated allele frequency
Ecological gradient (Leaf Unfolding differences)
0.00
0.05
0.10
0.15
0.20
0.25
0.30
0.35
−1
0
1
2
3
4
253
219
9
O16
256
218
124
L8
L16
O1
217
O12
204
L12
L1
233
97
O8

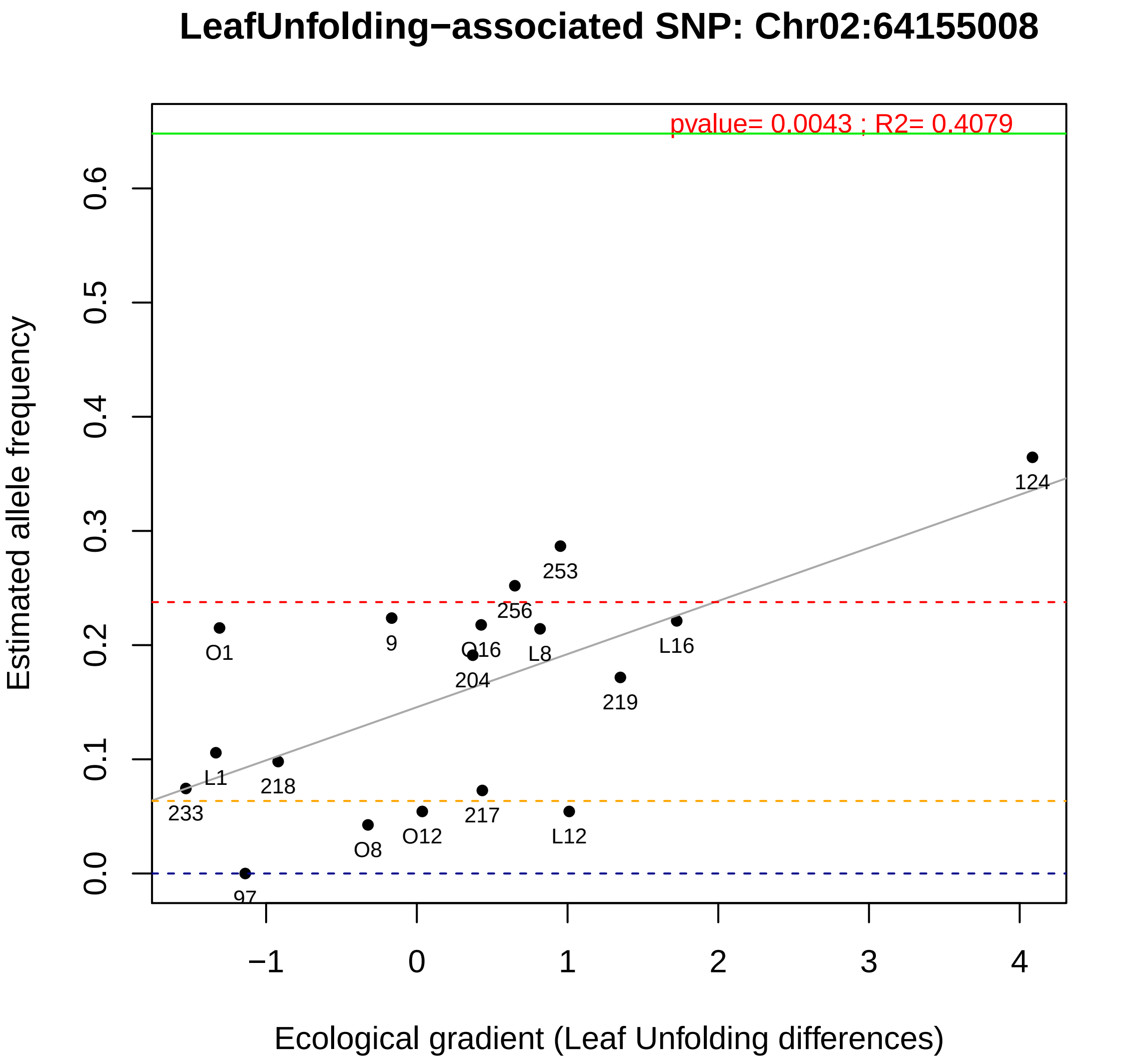

LeafUnfolding−associated SNP: Chr02:64155008
pvalue= 0.0043 ; R2= 0.4079
Estimated allele frequency
0.0
0.1
0.2
0.3
0.4
0.5
0.6
−1
0
1
2
3
4
Ecological gradient (Leaf Unfolding differences)
124
253
256
9
O16
L8
L16
O1
204
219
L1
218
233
217
O12
L12
O8
97

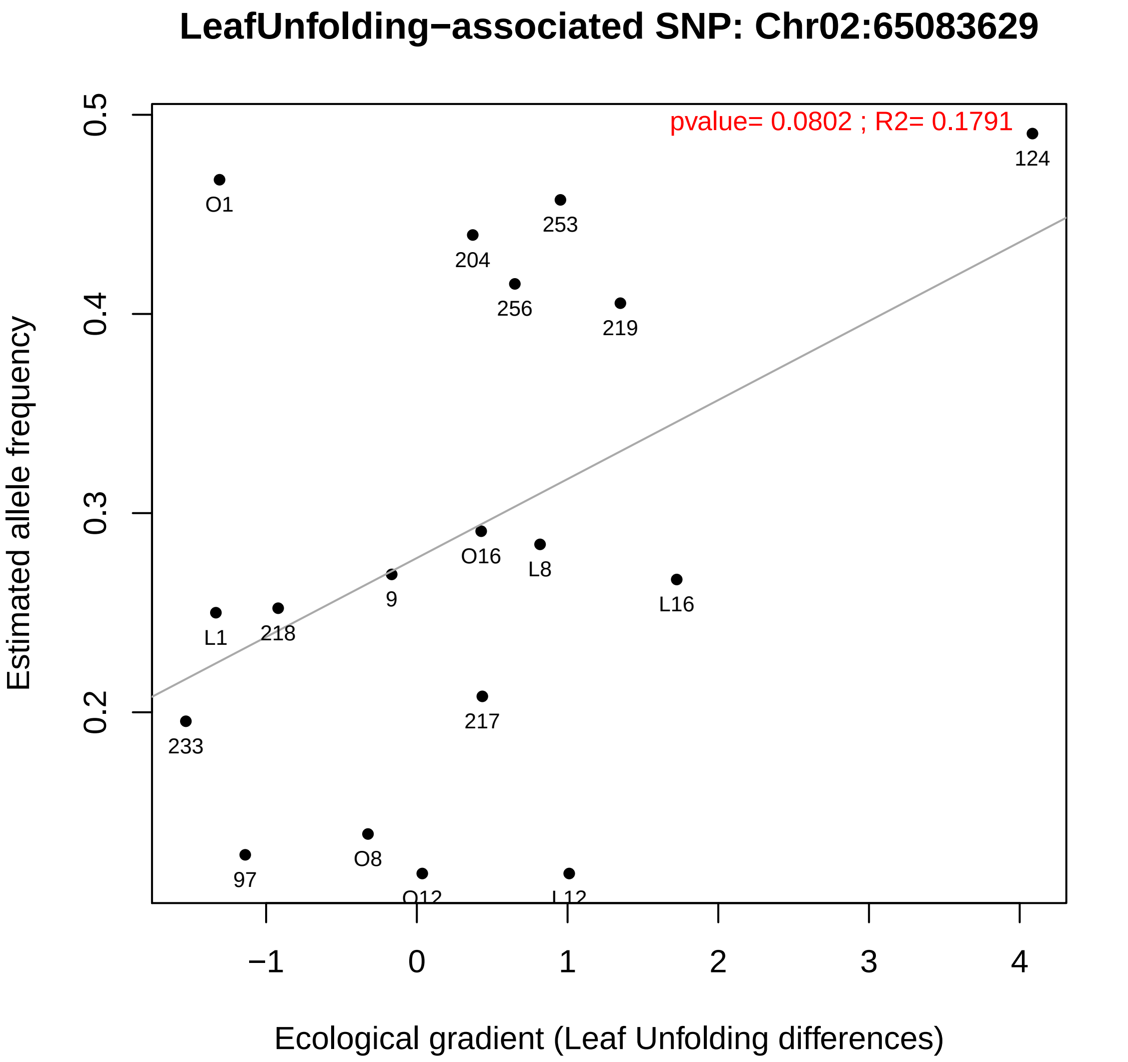
LeafUnfolding−associated SNP: Chr02:65083629
pvalue= 0.0802 ; R2= 0.1791
Estimated allele frequency
Ecological gradient (Leaf Unfolding differences)
0.2
0.3
0.4
0.5
−1
0
1
2
3
4
124
O1
253
204
256
219
O16
L8
9
L16
L1
218
217
233
O8
97
O12
L12

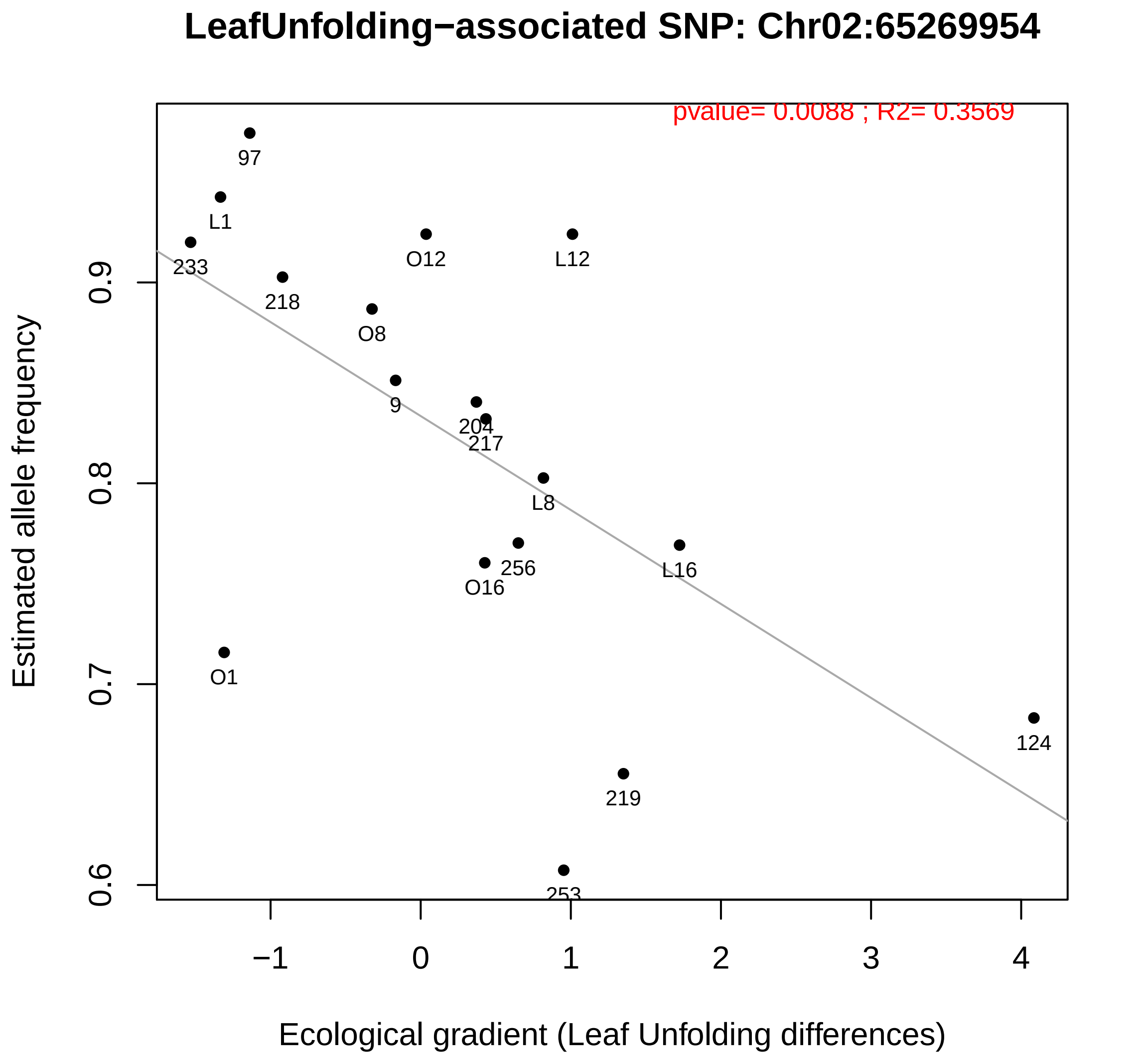

LeafUnfolding−associated SNP: Chr02:65269954
pvalue= 0.0088 ; R2= 0.3569
Estimated allele frequency
Ecological gradient (Leaf Unfolding differences)
0.6
0.7
0.8
0.9
−1
0
1
2
3
4
97
L1
233
218
O12
L12
O8
9
204
217
L8
256
O16
L16
O1
124
219
253

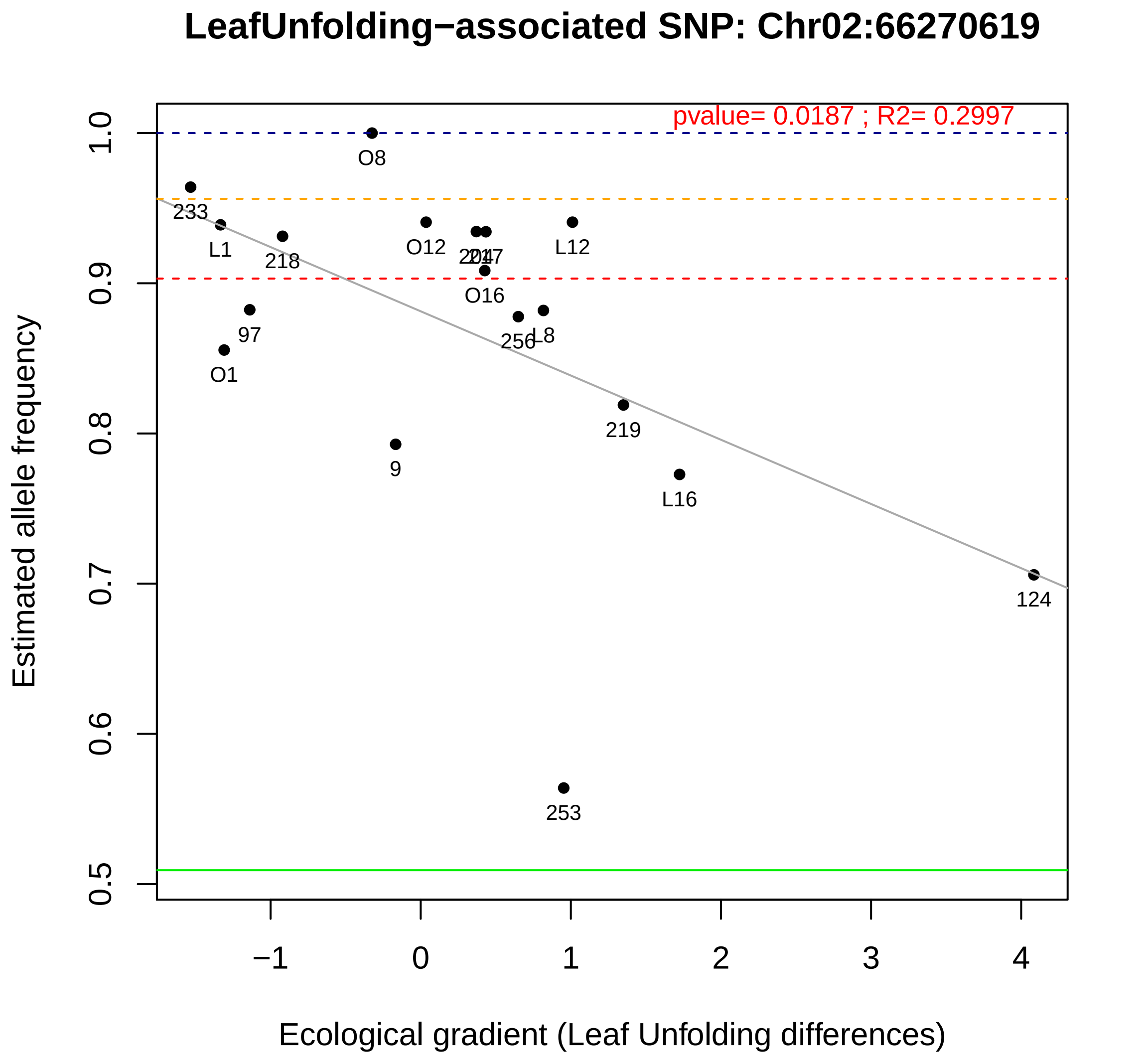

LeafUnfolding−associated SNP: Chr02:66270619
pvalue= 0.0187 ; R2= 0.2997
Estimated allele frequency
Ecological gradient (Leaf Unfolding differences)
1.0
0.9
0.8
0.7
0.6
0.5
−1
0
1
2
3
4
O8
233
L1
218
O12
L12
O16
97
O1
256
L8
219
9
L16
124
253

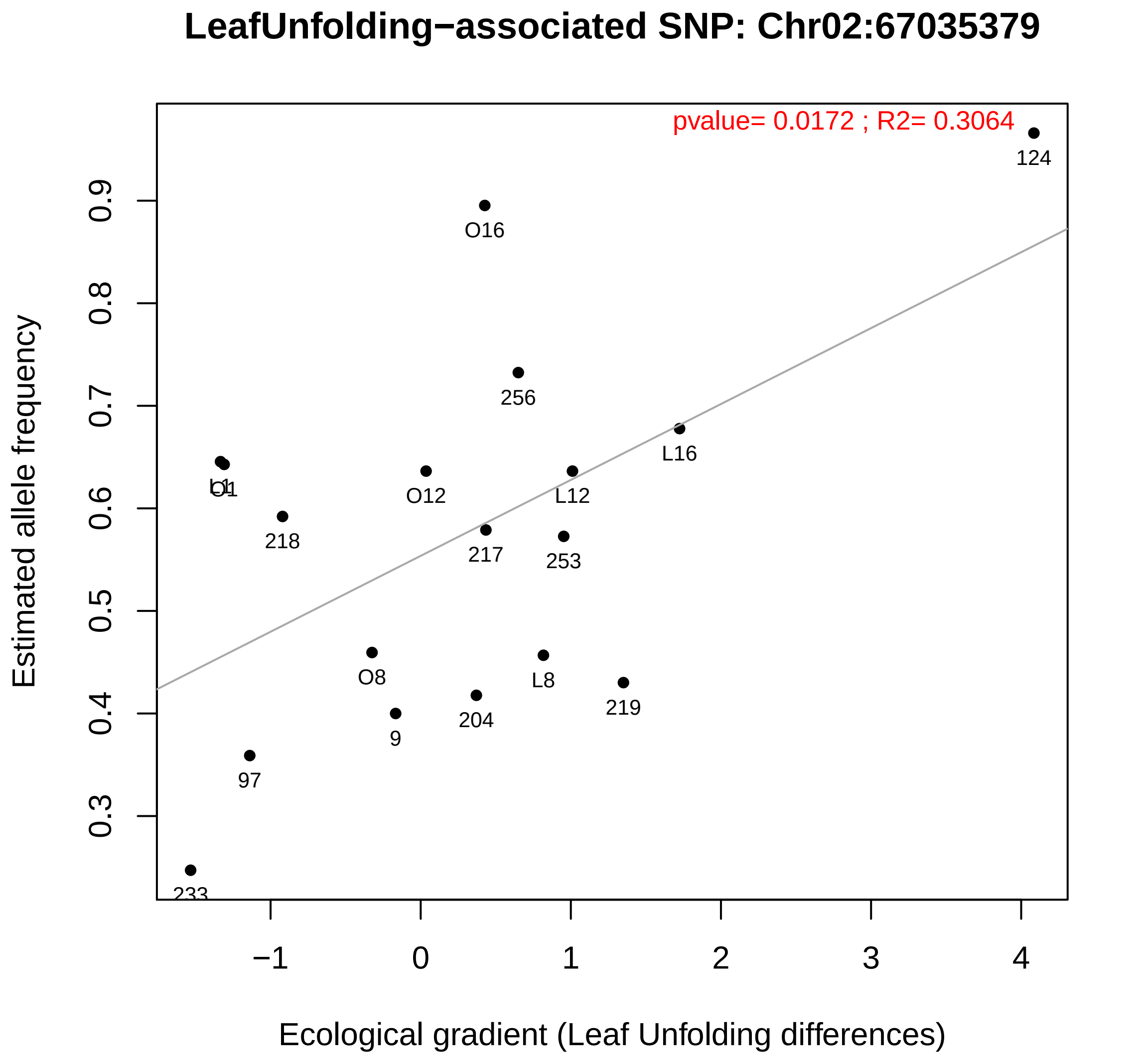
LeafUnfolding−associated SNP: Chr02:67035379
pvalue= 0.0172 ; R2= 0.3064
124
O16
256
L16
O12
L12
O1
218
217
253
O8
L8
219
204
9
97
233
0.3
0.4
0.5
0.6
0.7
0.8
0.9
−1
0
1
2
3
4
Estimated allele frequency
Ecological gradient (Leaf Unfolding differences)

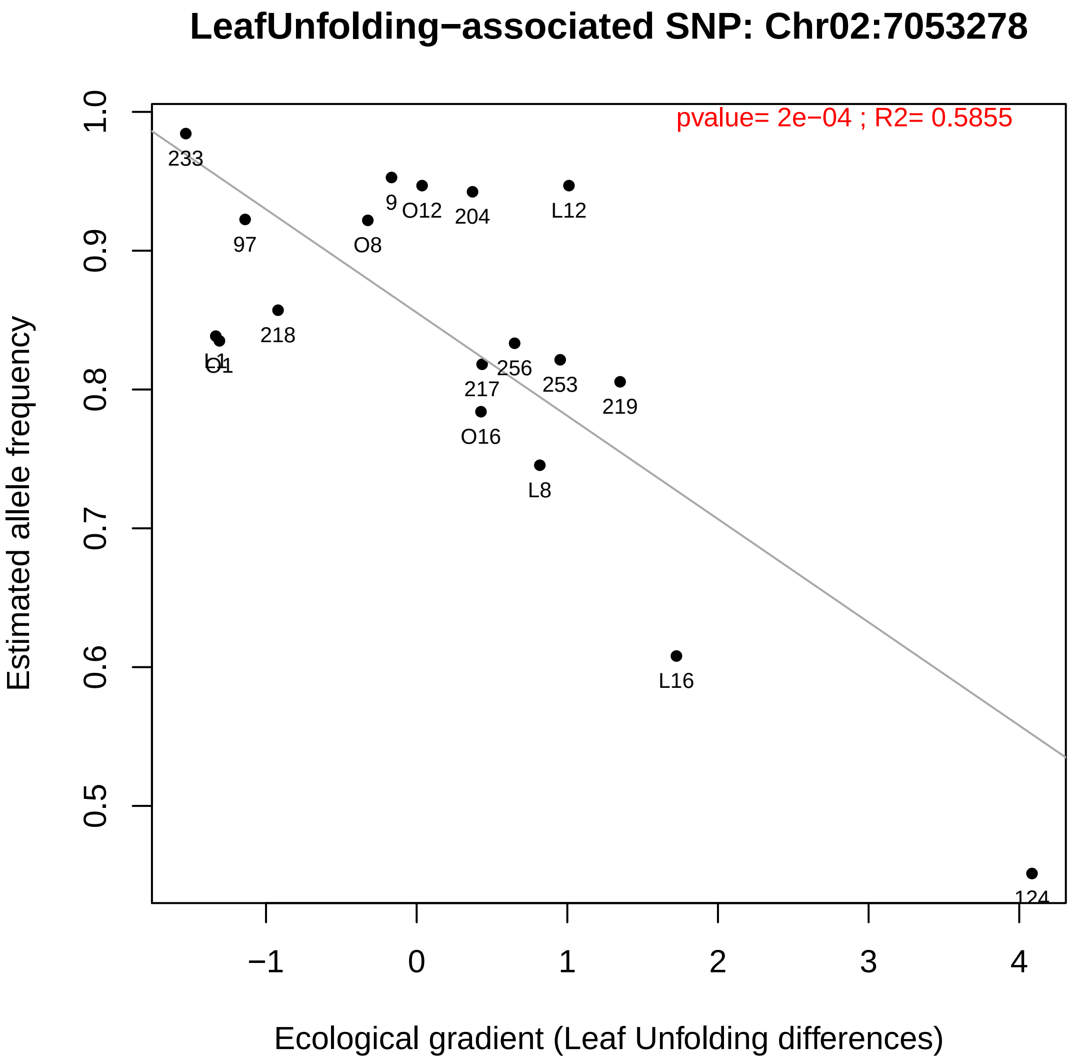
LeafUnfolding−associated SNP: Chr02:7053278
pvalue= 2e−04 ; R2= 0.5855
Estimated allele frequency
Ecological gradient (Leaf Unfolding differences)
233
97
218
L1
O1
9
O12
204
L12
O8
256
217
253
219
O16
L8
L16
124
1.0
0.9
0.8
0.7
0.6
0.5
−1
0
1
2
3
4

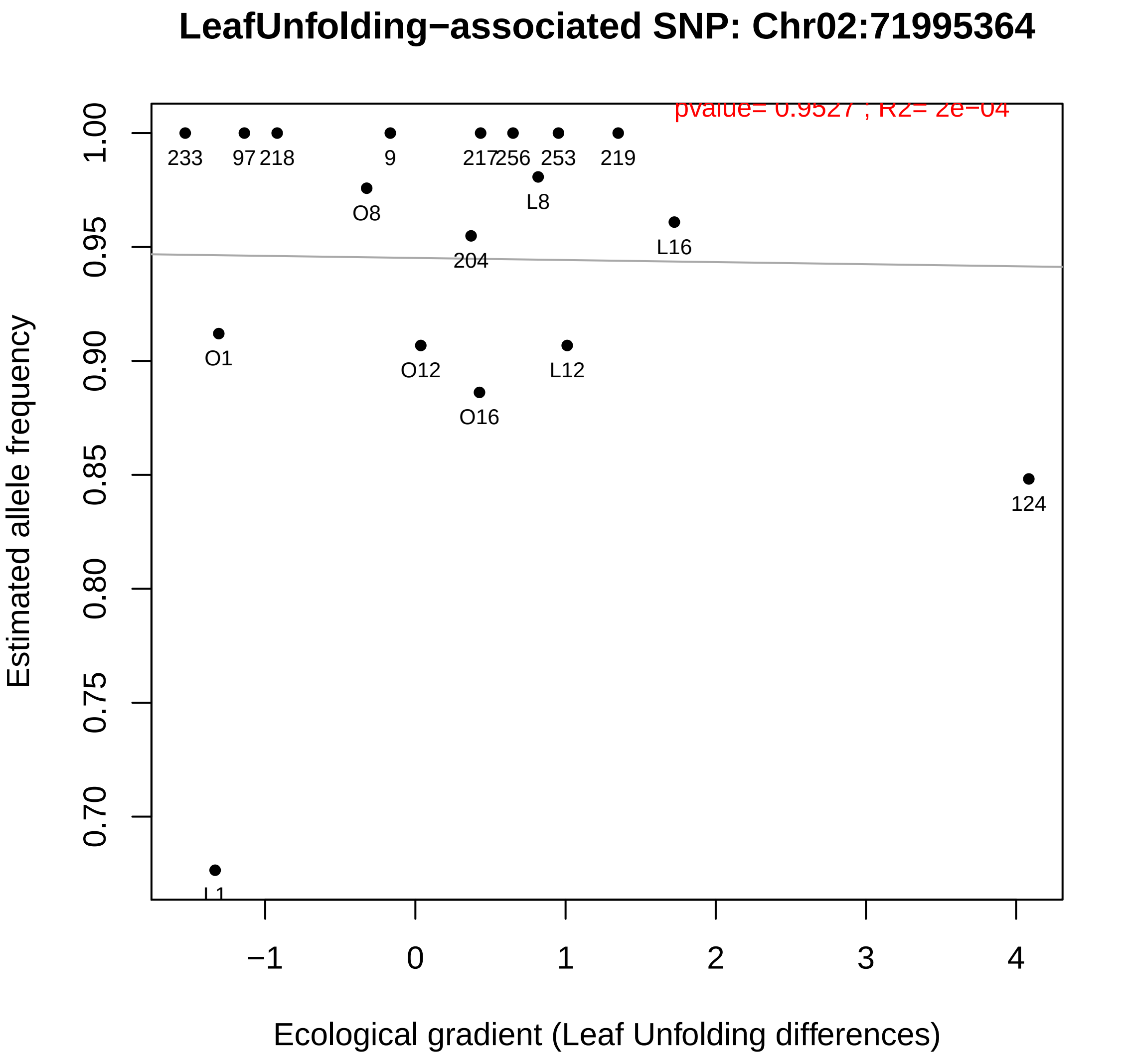
LeafUnfolding−associated SNP: Chr02:71995364
pvalue= 0.9527 ; R2= 2e−04
Estimated allele frequency
Ecological gradient (Leaf Unfolding differences)
0.70
0.75
0.80
0.85
0.90
0.95
1.00
−1
0
1
2
3
4
233
97
218
9
217
256
253
219
O8
L8
204
L16
O1
O12
L12
O16
124
L1

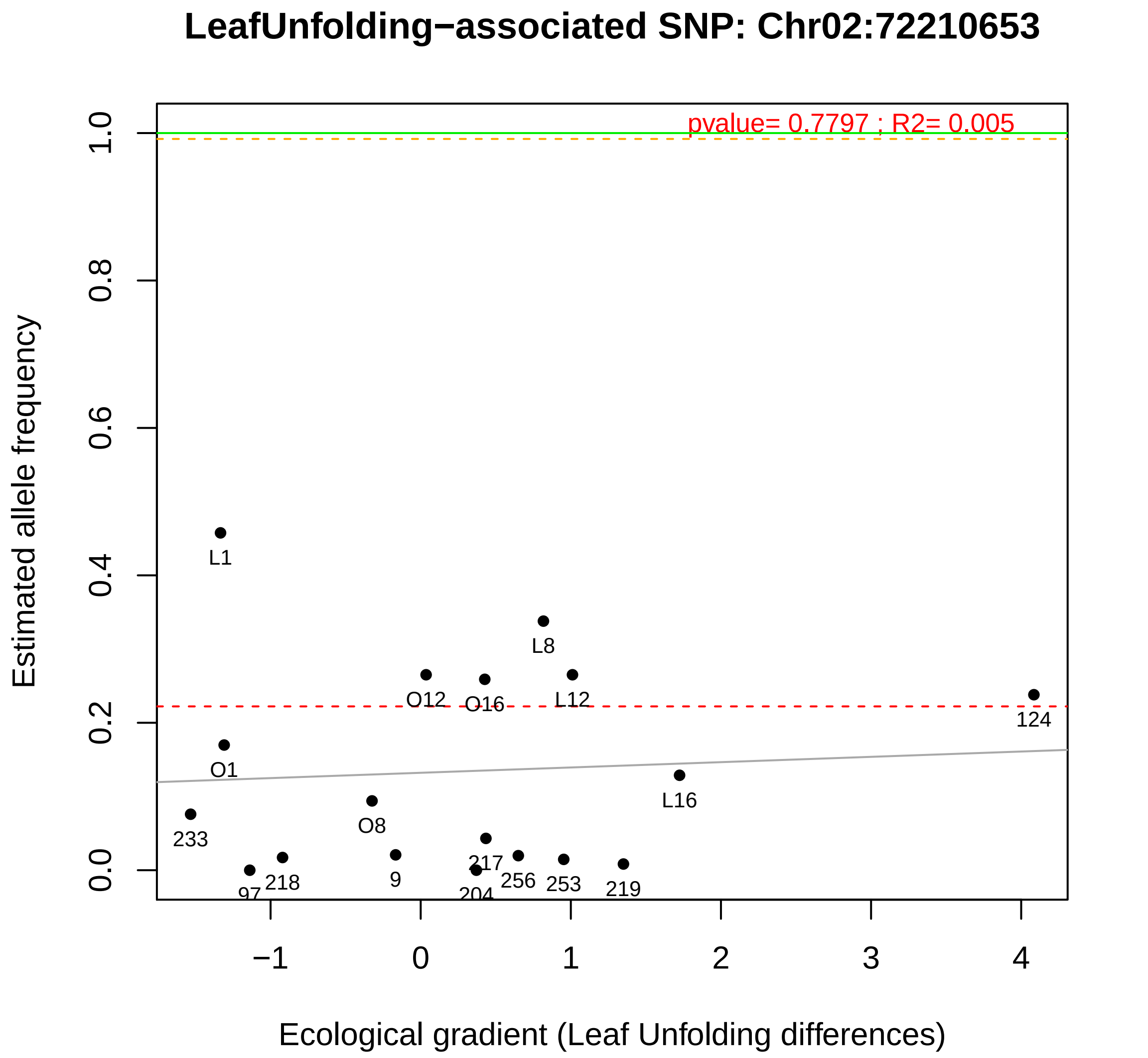
LeafUnfolding−associated SNP: Chr02:72210653
pvalue= 0.7797 ; R2= 0.005
Estimated allele frequency
Ecological gradient (Leaf Unfolding differences)
0.0
0.2
0.4
0.6
0.8
1.0
−1
0
1
2
3
4
L1
L8
O12
O16
L12
124
O1
L16
233
O8
217
256
253
219
218
9
97
204

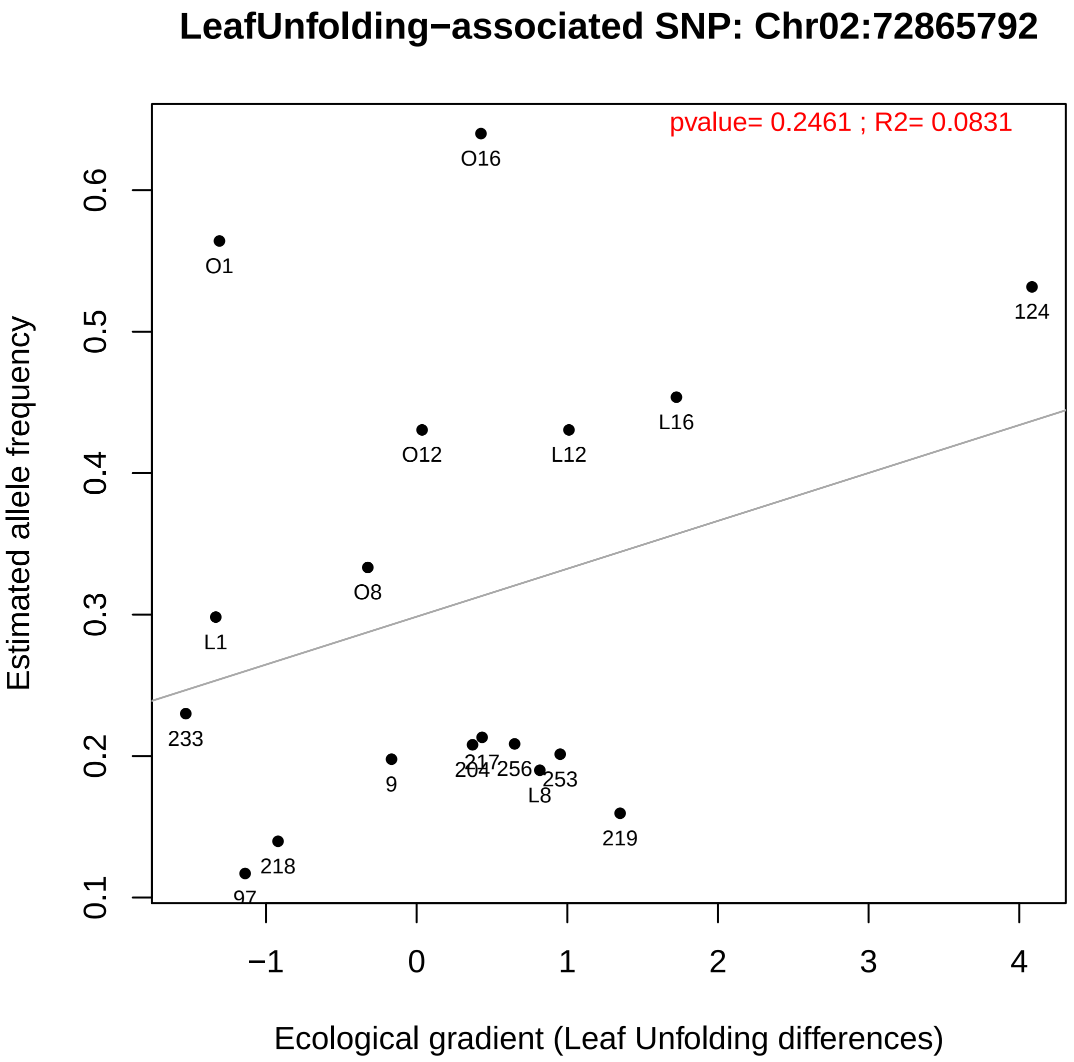
LeafUnfolding−associated SNP: Chr02:72865792
pvalue= 0.2461 ; R2= 0.0831
O16
O1
124
L16
O12
L12
O8
L1
233
217
204
256
253
L8
9
219
218
97
0.1
0.2
0.3
0.4
0.5
0.6
−1
0
1
2
3
4
Estimated allele frequency
Ecological gradient (Leaf Unfolding differences)

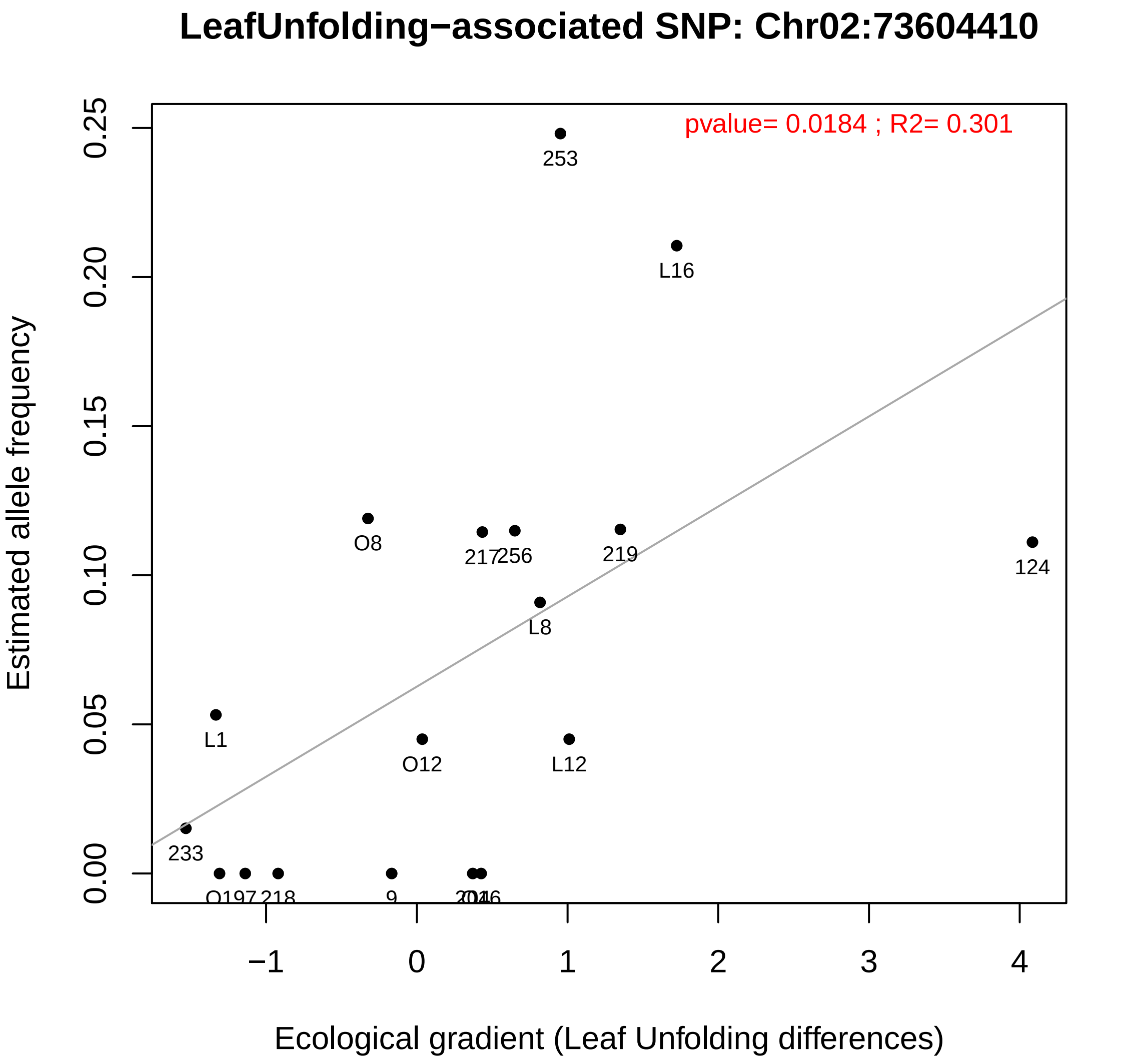

LeafUnfolding−associated SNP: Chr02:73604410
pvalue= 0.0184 ; R2= 0.301
253
L16
O8
217
256
219
124
L8
L1
O12
L12
233
218
9
Estimated allele frequency
0.00
0.05
0.10
0.15
0.20
0.25
−1
0
1
2
3
4
Ecological gradient (Leaf Unfolding differences)

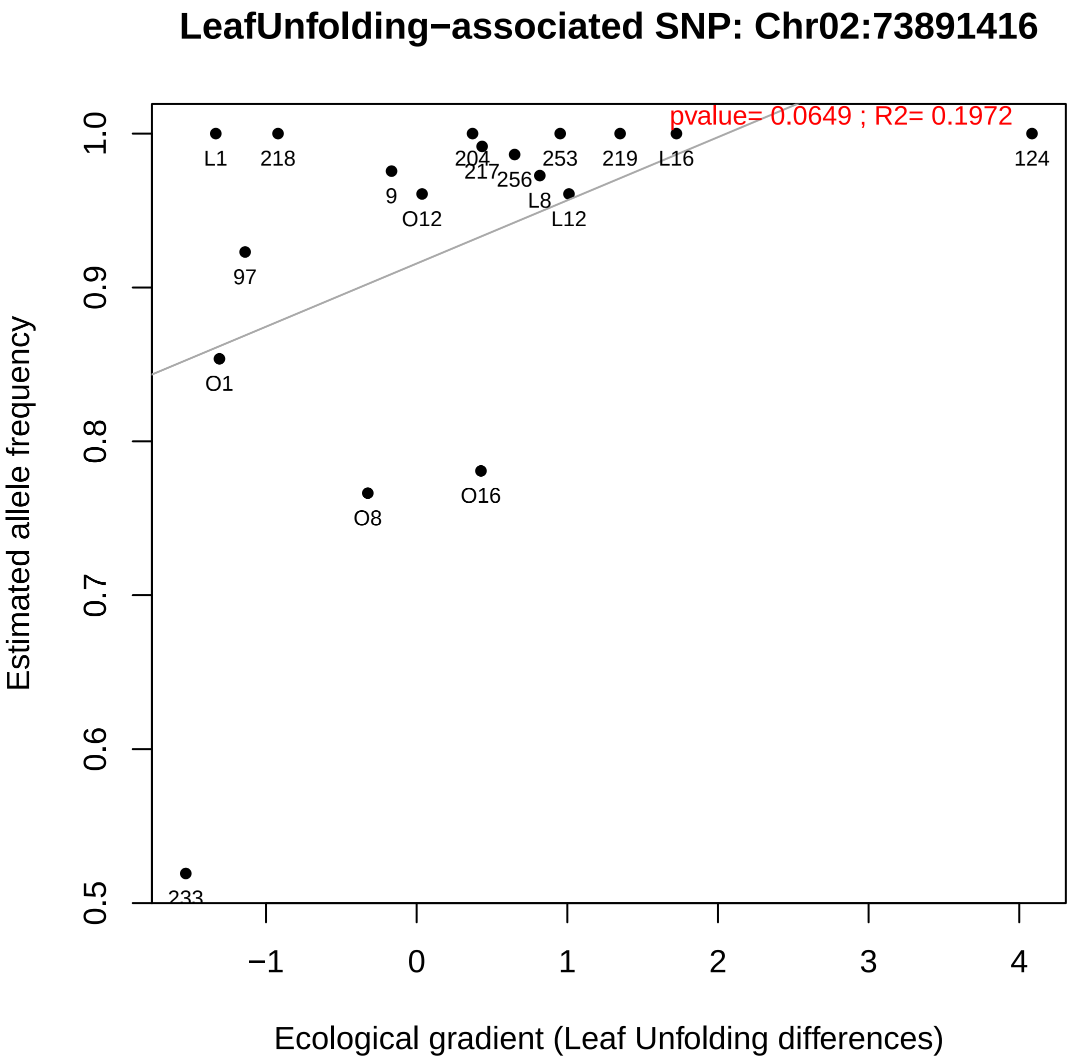
LeafUnfolding−associated SNP: Chr02:73891416
pvalue= 0.0649 ; R2= 0.1972
Estimated allele frequency
0.5
0.6
0.7
0.8
0.9
1.0
−1
0
1
2
3
4
Ecological gradient (Leaf Unfolding differences)
L1
218
204
217
256
253
219
L16
124
9
O12
L8
L12
97
O1
O8
O16
233

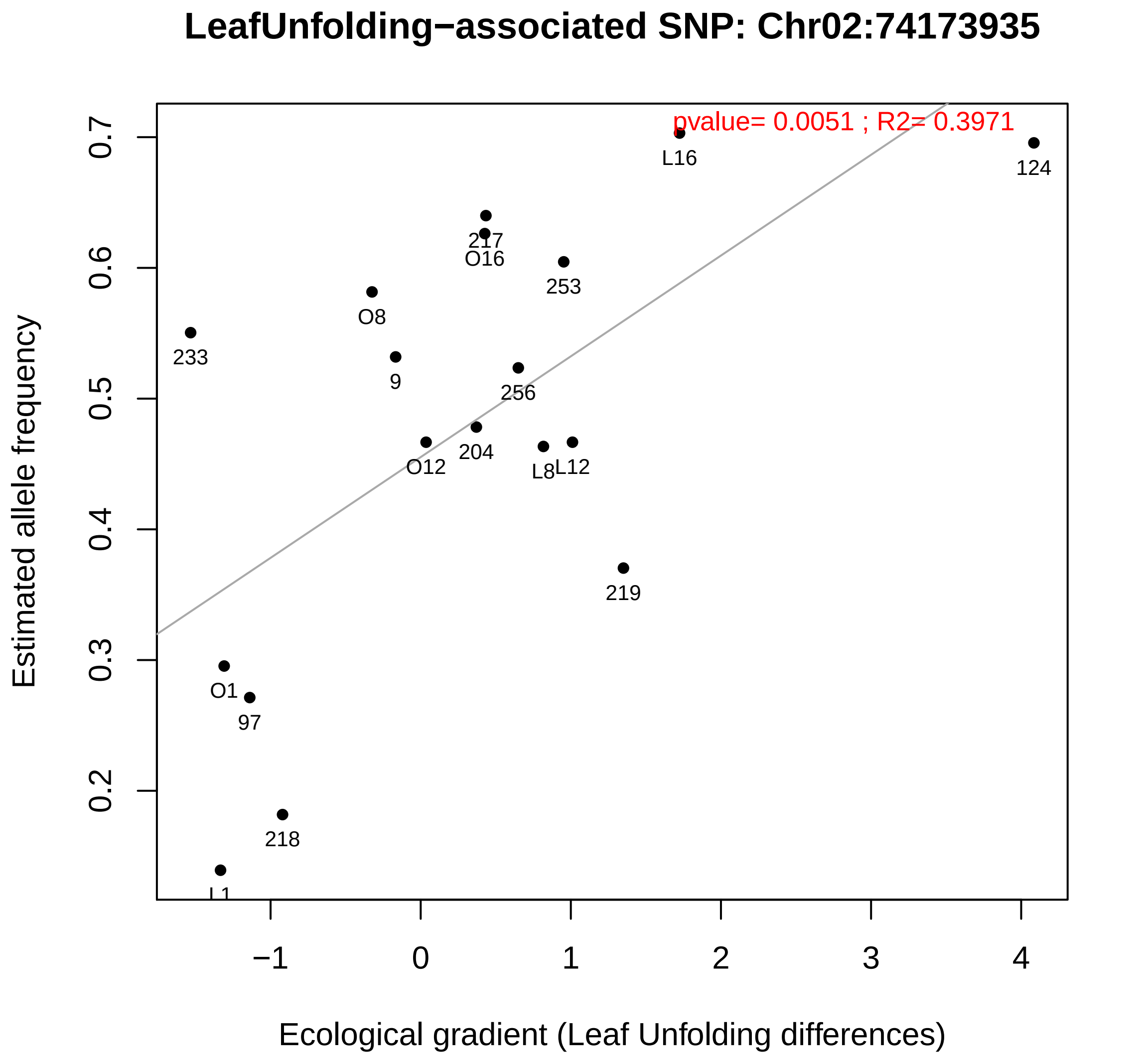

LeafUnfolding−associated SNP: Chr02:74173935
pvalue= 0.0051 ; R2= 0.3971
Estimated allele frequency
Ecological gradient (Leaf Unfolding differences)
0.2
0.3
0.4
0.5
0.6
0.7
−1
0
1
2
3
4
L16
124
217
O16
253
O8
233
9
256
204
O12
L8
L12
219
O1
97
218
L1

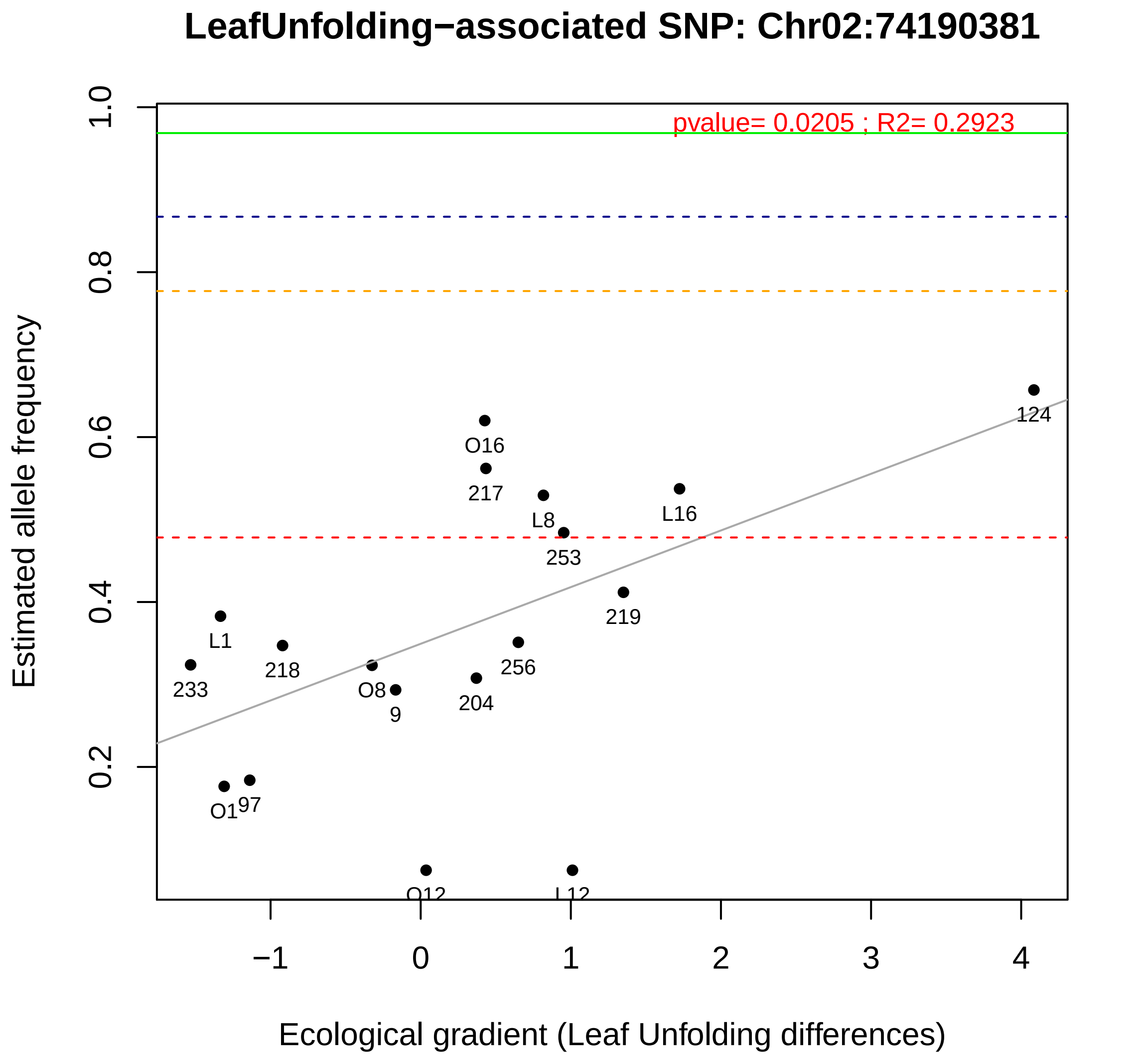

LeafUnfolding−associated SNP: Chr02:74190381
pvalue= 0.0205 ; R2= 0.2923
Estimated allele frequency
Ecological gradient (Leaf Unfolding differences)
1.0
0.8
0.6
0.4
0.2
−1
0
1
2
3
4
124
O16
217
L8
253
L16
219
L1
218
233
O8
9
204
256
O1
97
O12
L12

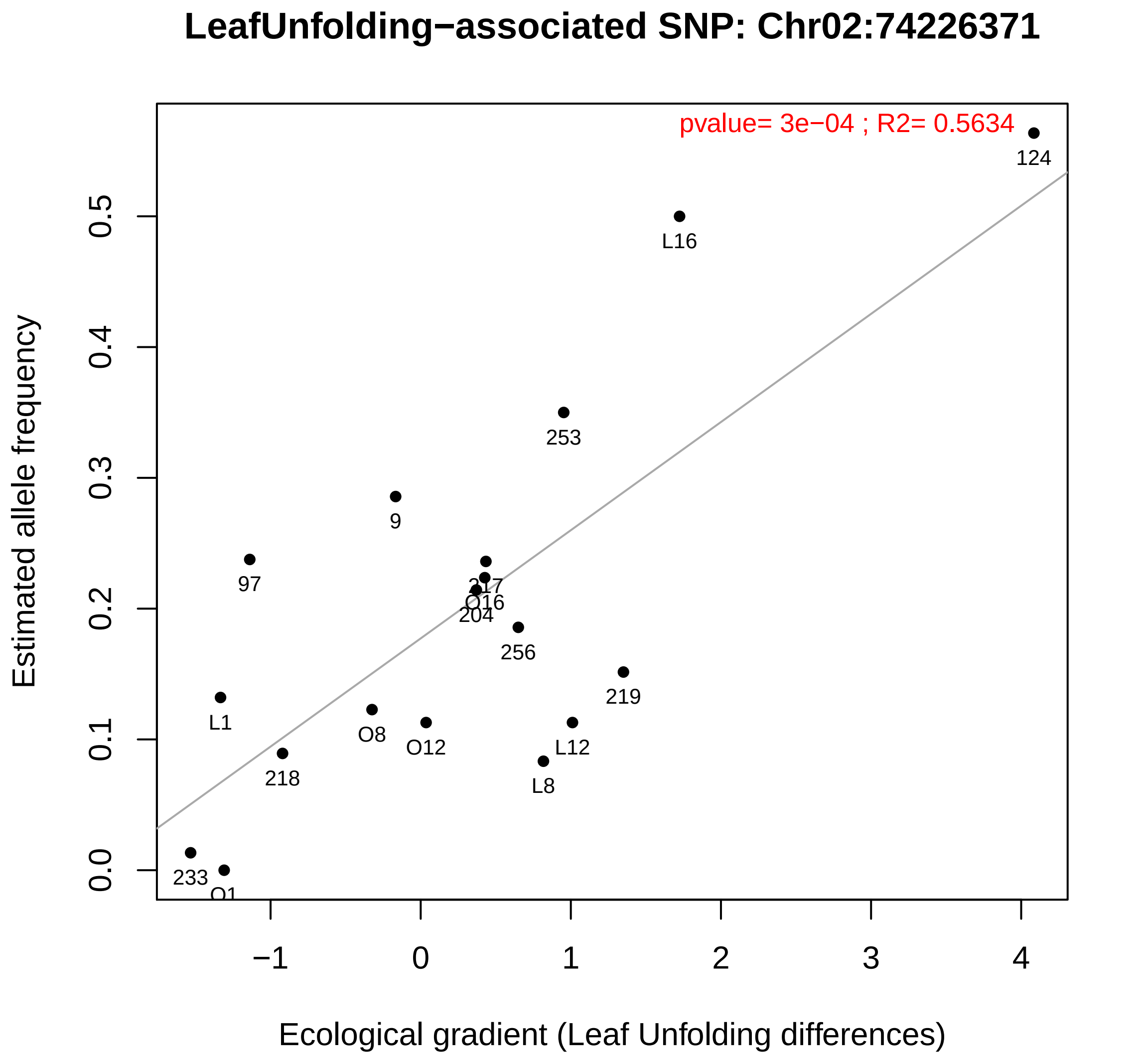

LeafUnfolding−associated SNP: Chr02:74226371
pvalue= 3e−04 ; R2= 0.5634
124
L16
253
9
97
217
O16
204
256
219
L1
O8
O12
L12
218
L8
233
O1
0.0
0.1
0.2
0.3
0.4
0.5
−1
0
1
2
3
4
Estimated allele frequency
Ecological gradient (Leaf Unfolding differences)

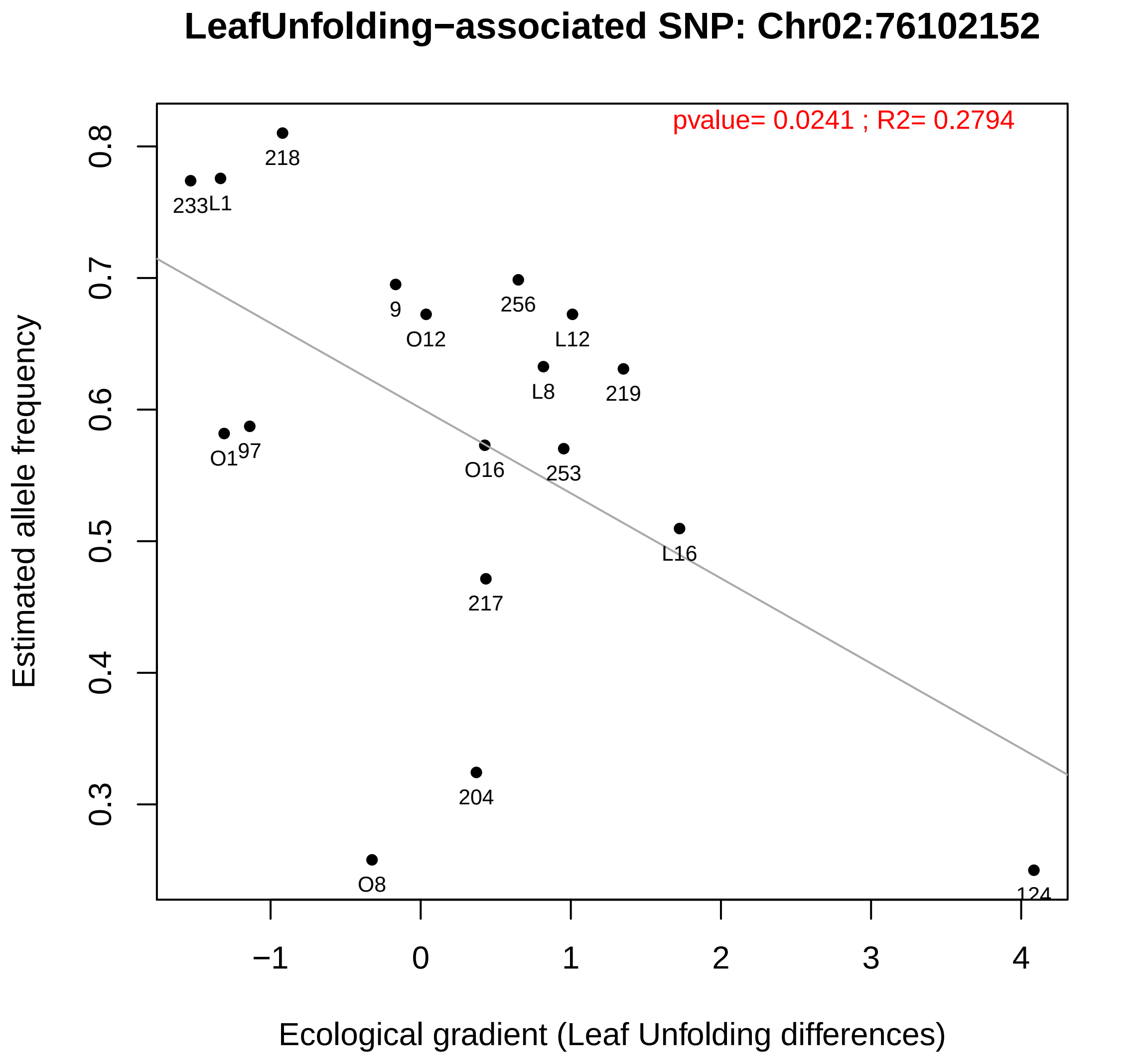

LeafUnfolding−associated SNP: Chr02:76102152
pvalue= 0.0241 ; R2= 0.2794
Estimated allele frequency
0.3
0.4
0.5
0.6
0.7
0.8
−1
0
1
2
3
4
Ecological gradient (Leaf Unfolding differences)
218
233
L1
9
O12
256
L12
L8
219
O1
97
O16
253
L16
217
204
O8
124

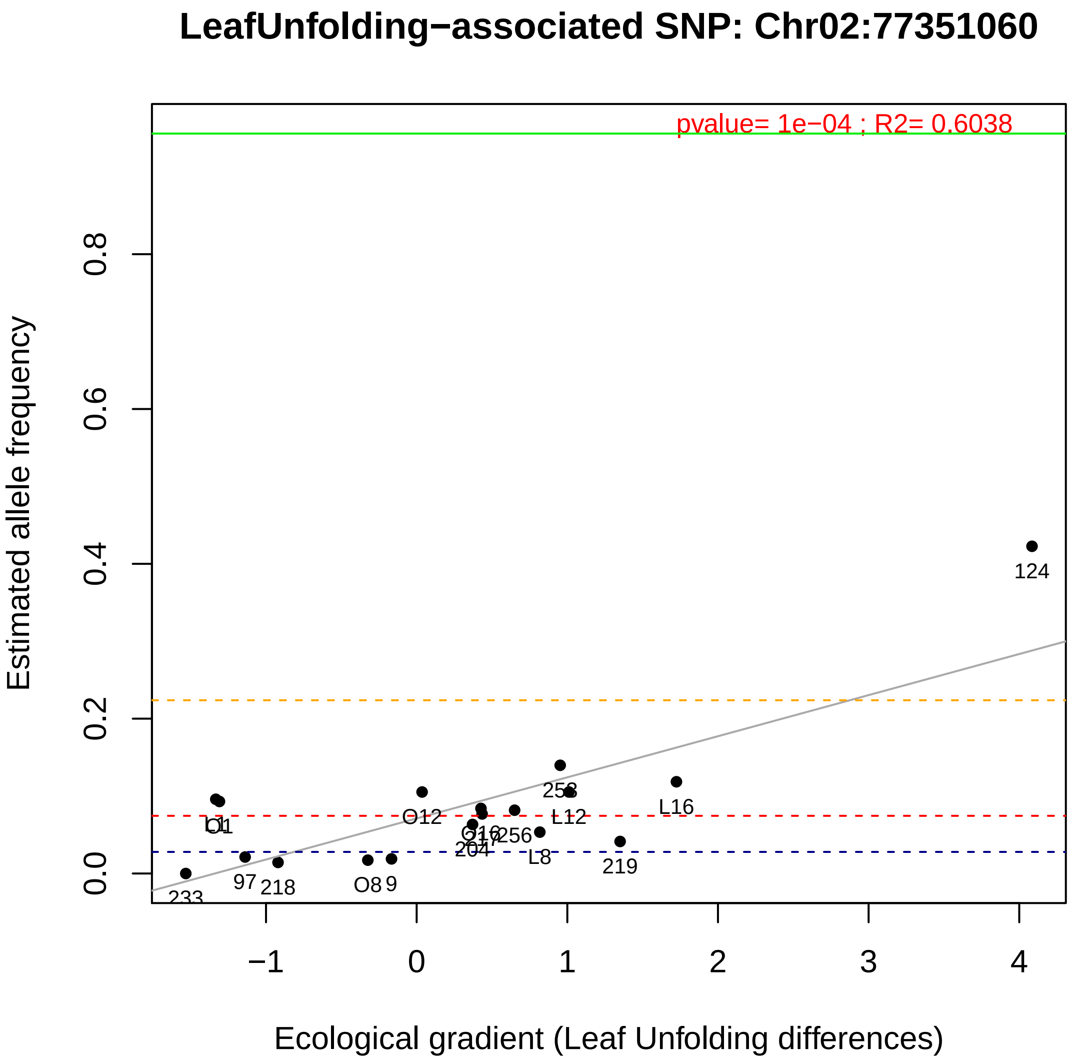
LeafUnfolding−associated SNP: Chr02:77351060
pvalue= 1e−04 ; R2= 0.6038
Estimated allele frequency
0.0
0.2
0.4
0.6
0.8
Ecological gradient (Leaf Unfolding differences)
−1
0
1
2
3
4
124
L16
O12
L12
256
L8
219
97
218
O8
9
233
204

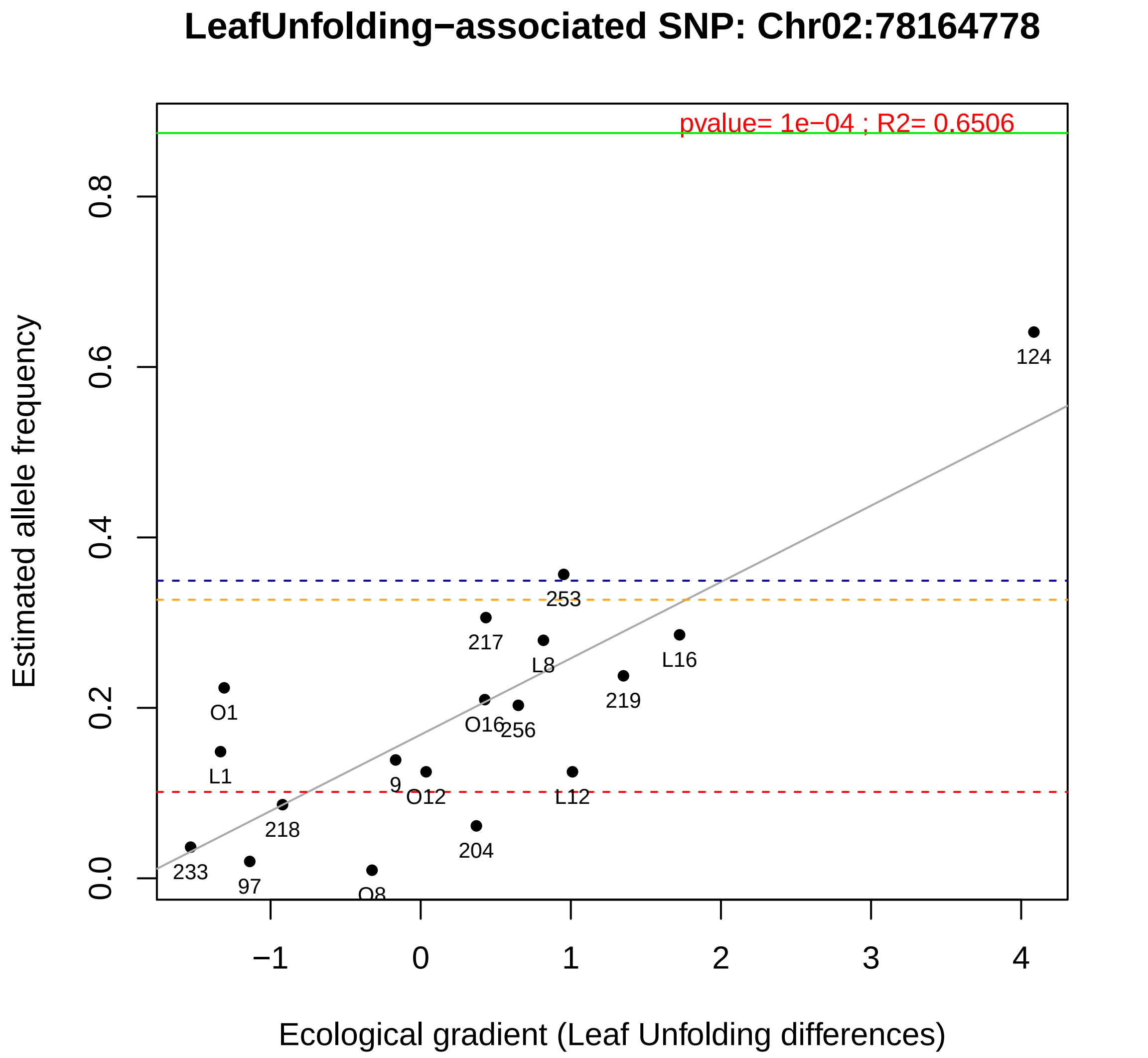
LeafUnfolding−associated SNP: Chr02:78164778
pvalue= 1e−04 ; R2= 0.6506
Estimated allele frequency
0.0
0.2
0.4
0.6
0.8
−1
0
1
2
3
4
Ecological gradient (Leaf Unfolding differences)
124
253
217
L8
L16
219
O1
O16
256
L1
9
O12
L12
218
204
233
97
O8

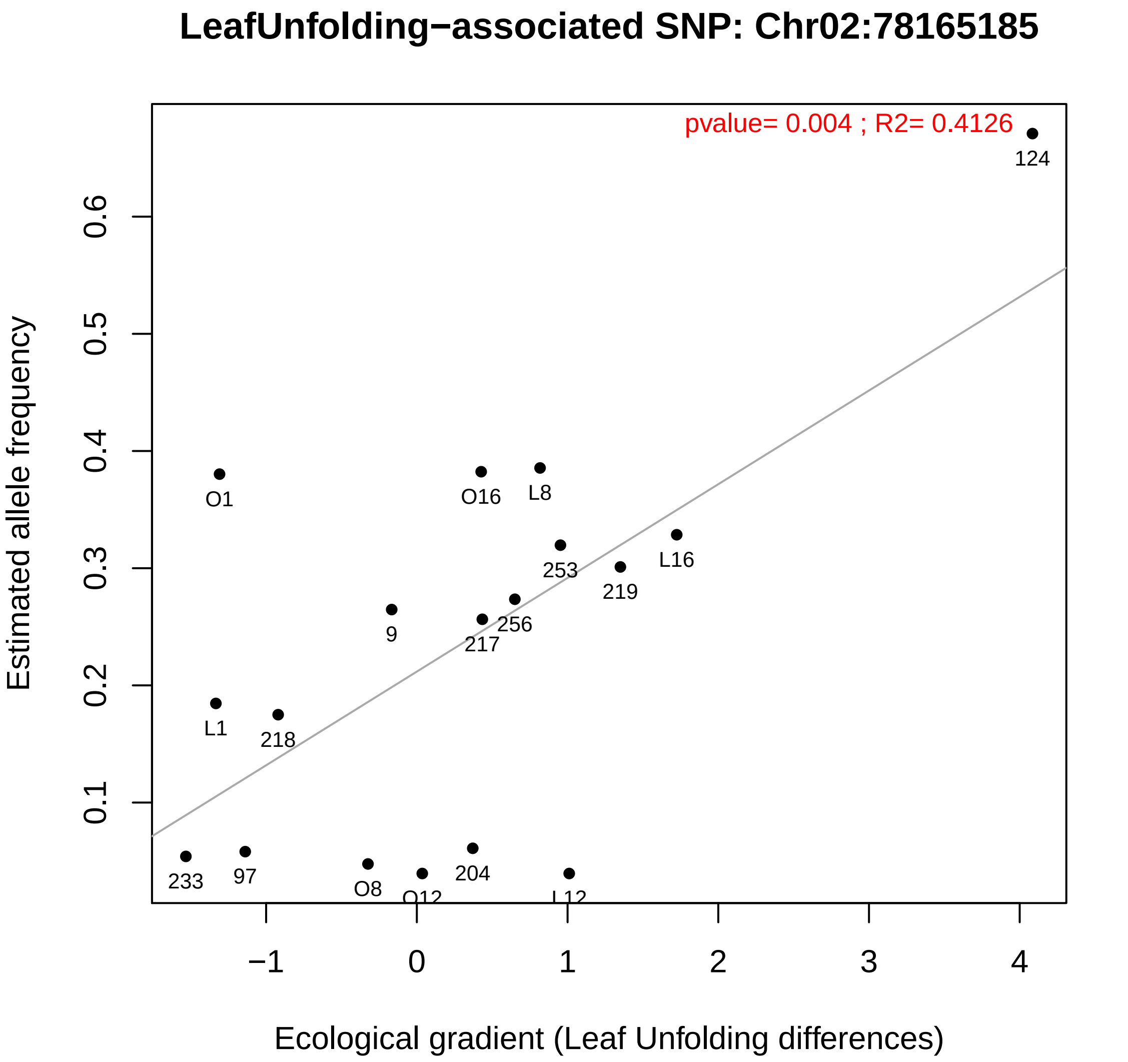

LeafUnfolding−associated SNP: Chr02:78165185
pvalue= 0.004 ; R2= 0.4126
Estimated allele frequency
Ecological gradient (Leaf Unfolding differences)
0.1
0.2
0.3
0.4
0.5
0.6
−1
0
1
2
3
4
124
O1
O16
L8
253
L16
219
256
9
217
L1
218
233
97
O8
O12
204
L12

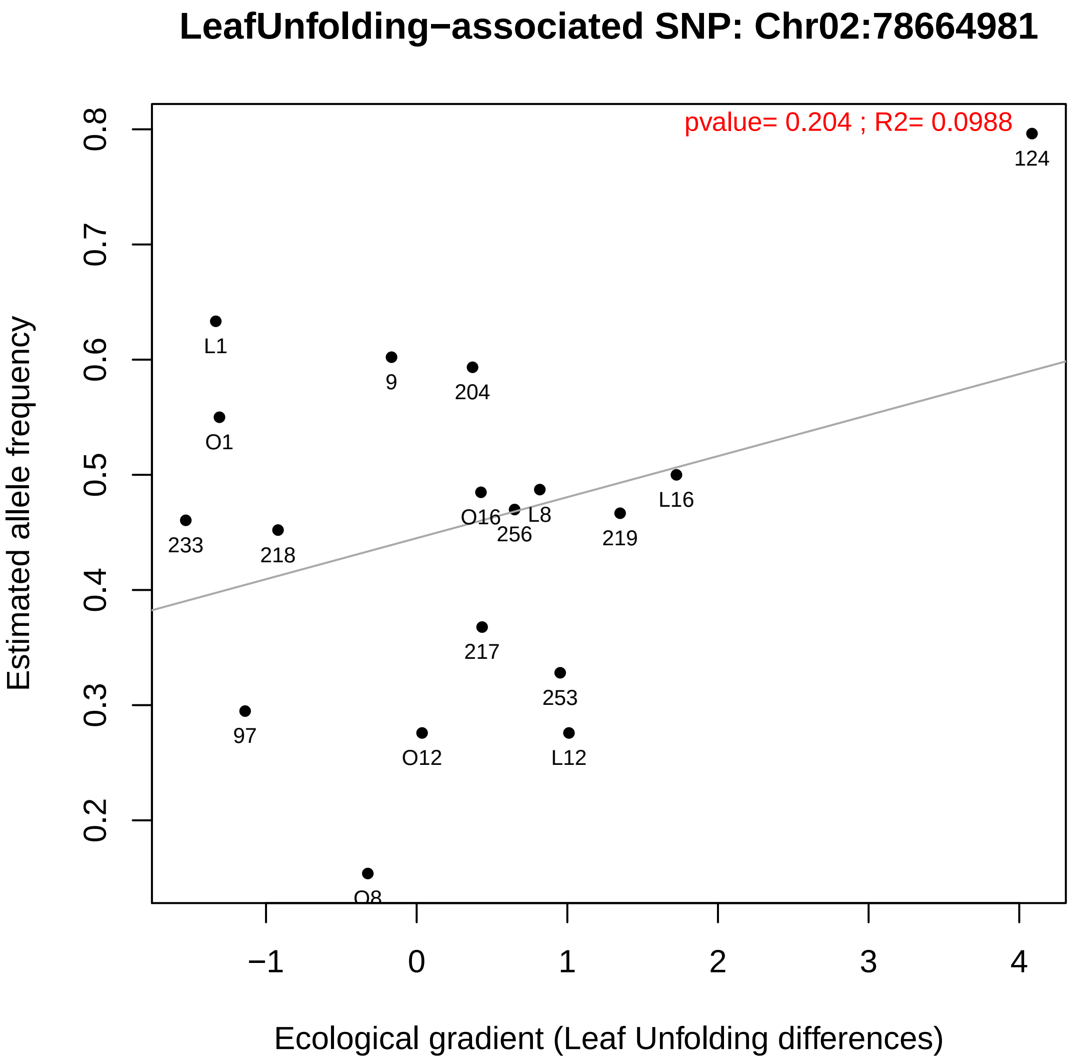

LeafUnfolding−associated SNP: Chr02:78664981
pvalue= 0.204 ; R2= 0.0988
Estimated allele frequency
Ecological gradient (Leaf Unfolding differences)
0.2
0.3
0.4
0.5
0.6
0.7
0.8
−1
0
1
2
3
4
124
L1
9
204
O1
L16
O16
L8
256
219
233
218
217
253
97
O12
L12
O8

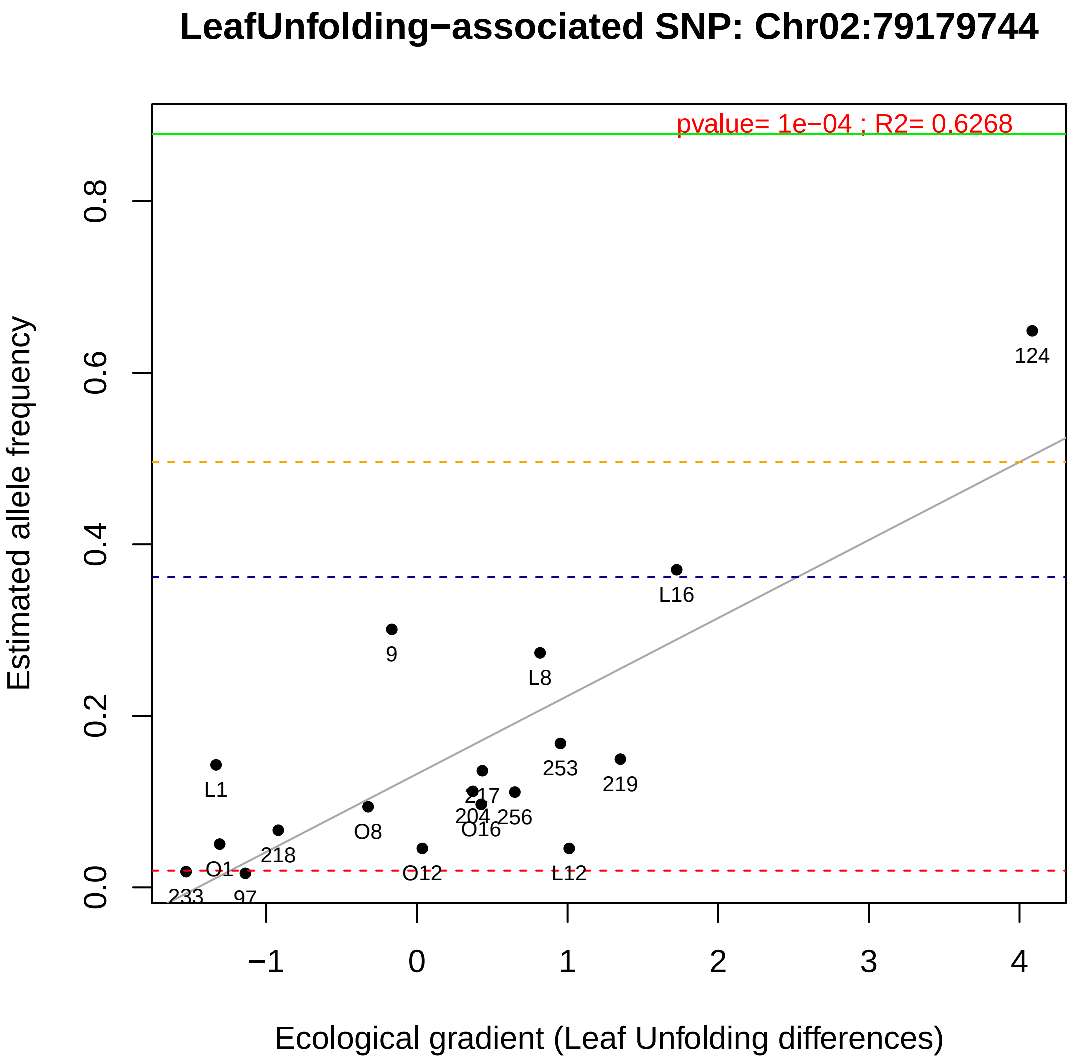

LeafUnfolding−associated SNP: Chr02:79179744
pvalue= 1e−04 ; R2= 0.6268
Estimated allele frequency
Ecological gradient (Leaf Unfolding differences)
0.0
0.2
0.4
0.6
0.8
−1
0
1
2
3
4
124
L16
9
L8
253
219
L1
217
204
256
O16
O8
218
O12
L12
O1
233
97

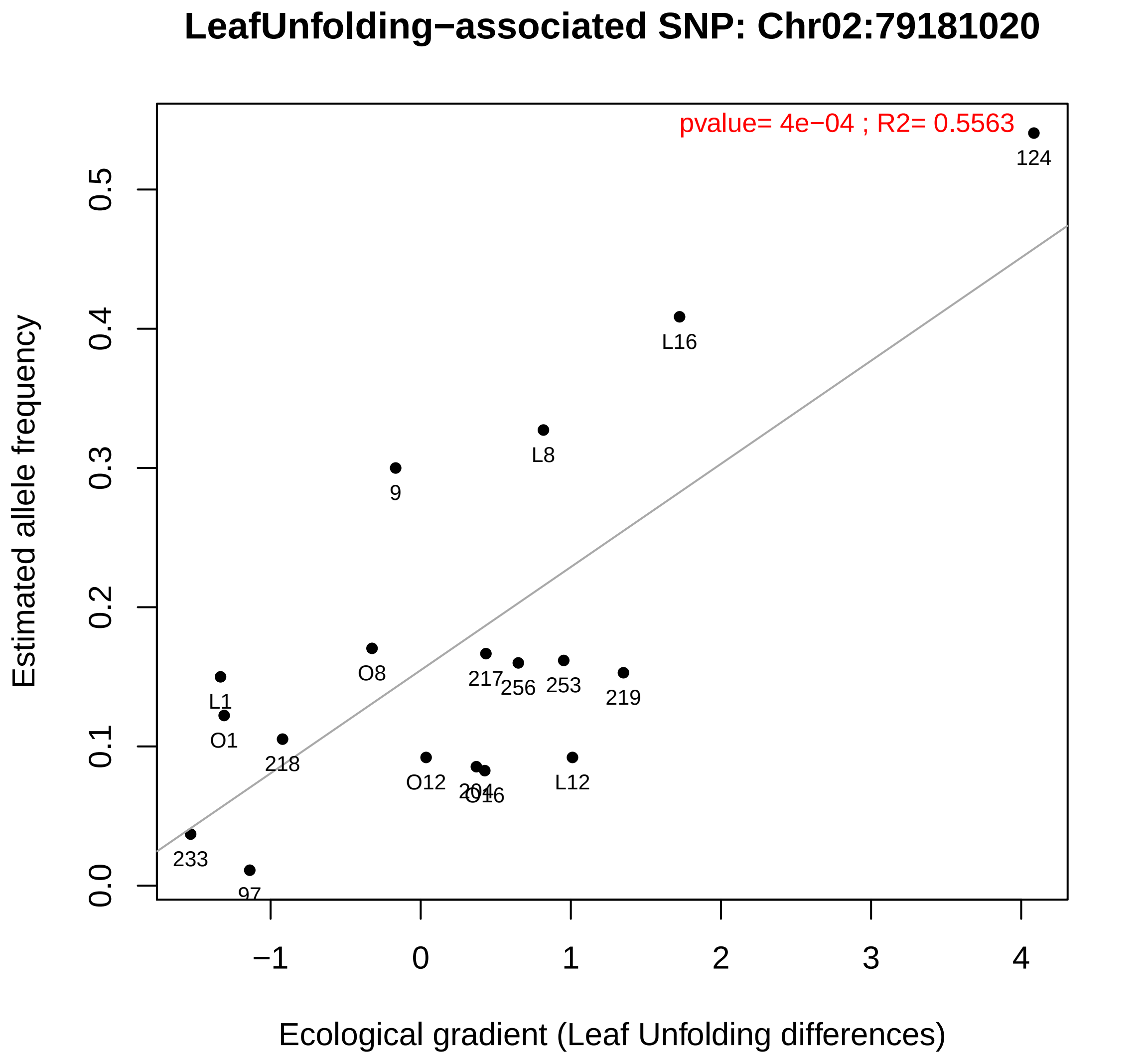
LeafUnfolding−associated SNP: Chr02:79181020
pvalue= 4e−04 ; R2= 0.5563
Estimated allele frequency
Ecological gradient (Leaf Unfolding differences)
0.0
0.1
0.2
0.3
0.4
0.5
−1
0
1
2
3
4
124
L16
L8
9
O8
217
256
253
219
L1
O1
218
O12
204
O16
L12
233
97

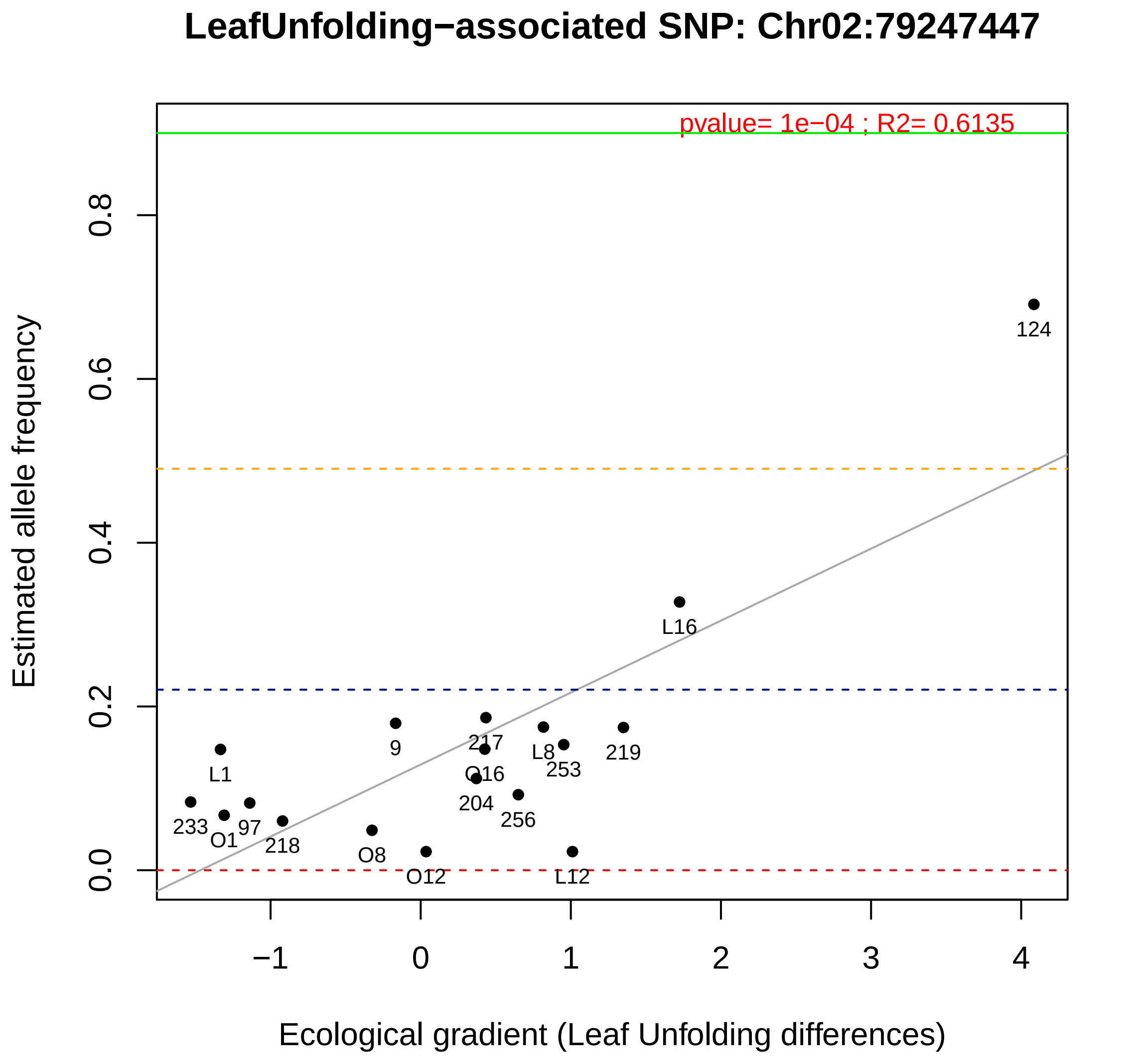
LeafUnfolding−associated SNP: Chr02:79247447
pvalue= 1e−04 ; R2= 0.6135
Estimated allele frequency
0.0
0.2
0.4
0.6
0.8
−1
0
1
2
3
4
Ecological gradient (Leaf Unfolding differences)
124
L16
9
217
L8
253
219
L1
O16
204
256
233
O1
97
218
O8
O12
L12

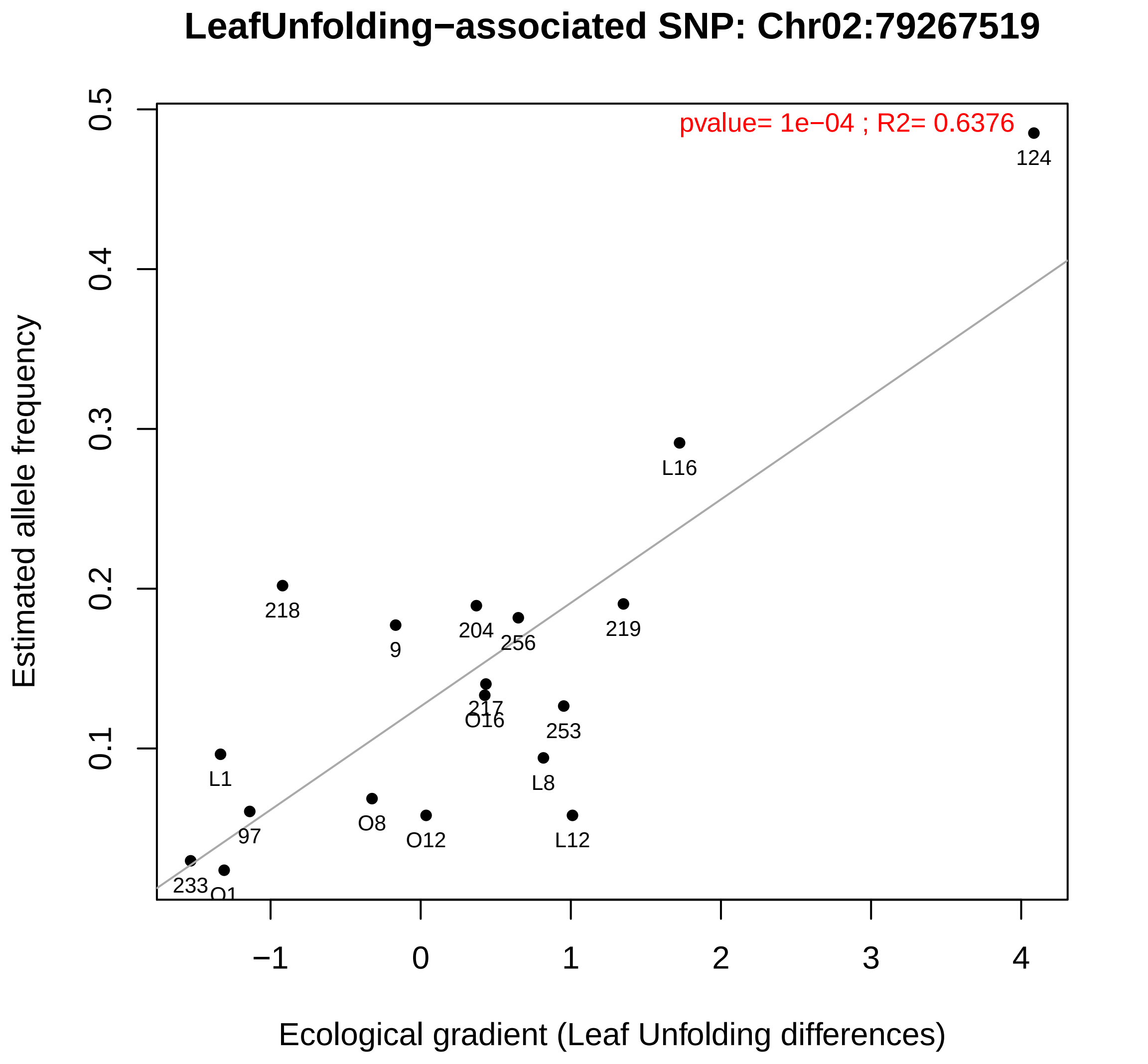

LeafUnfolding−associated SNP: Chr02:79267519
pvalue= 1e−04 ; R2= 0.6376
Estimated allele frequency
Ecological gradient (Leaf Unfolding differences)
0.1
0.2
0.3
0.4
0.5
−1
0
1
2
3
4
124
L16
218
9
204
256
219
217
O16
253
L1
L8
97
O8
O12
L12
233
O1

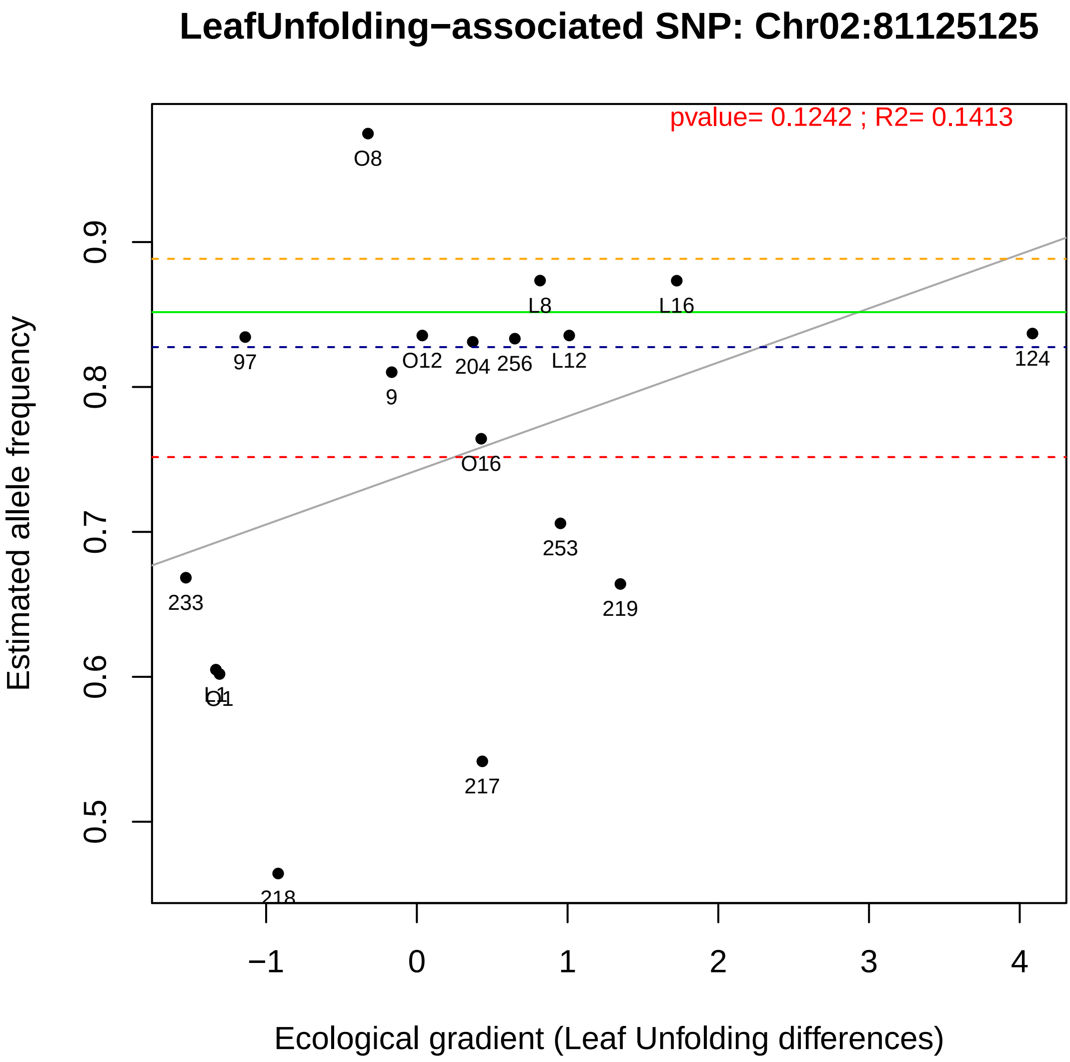
LeafUnfolding−associated SNP: Chr02:81125125
pvalue= 0.1242 ; R2= 0.1413
Estimated allele frequency
Ecological gradient (Leaf Unfolding differences)
0.5
0.6
0.7
0.8
0.9
−1
0
1
2
3
4
O8
L8
L16
97
O12
204
256
L12
124
9
O16
253
233
219
O1
217
218

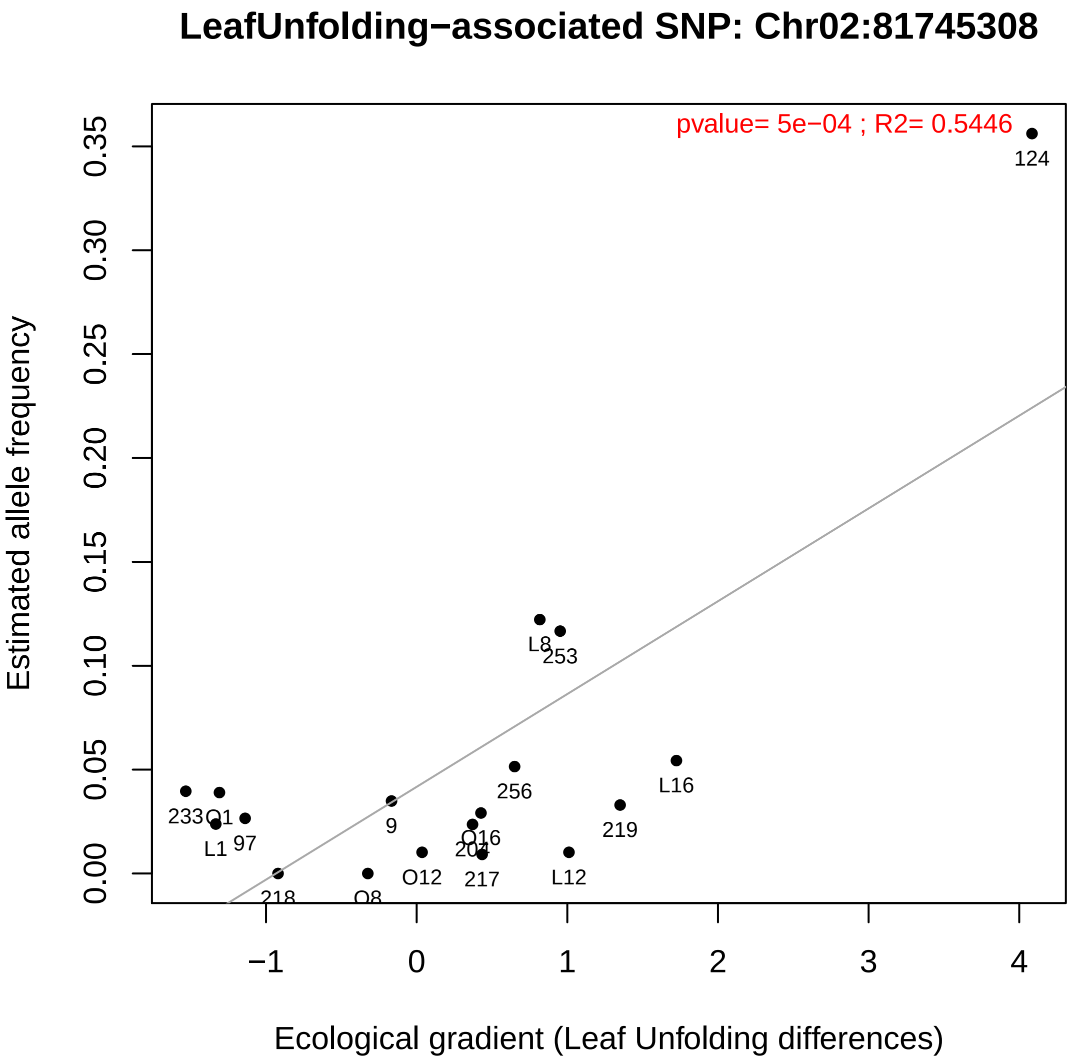
LeafUnfolding−associated SNP: Chr02:81745308
pvalue= 5e−04 ; R2= 0.5446
Estimated allele frequency
Ecological gradient (Leaf Unfolding differences)
0.00
0.05
0.10
0.15
0.20
0.25
0.30
0.35
−1
0
1
2
3
4
124
L8
253
256
L16
233
O1
L1
97
9
219
O16
204
217
O12
L12
218
O8

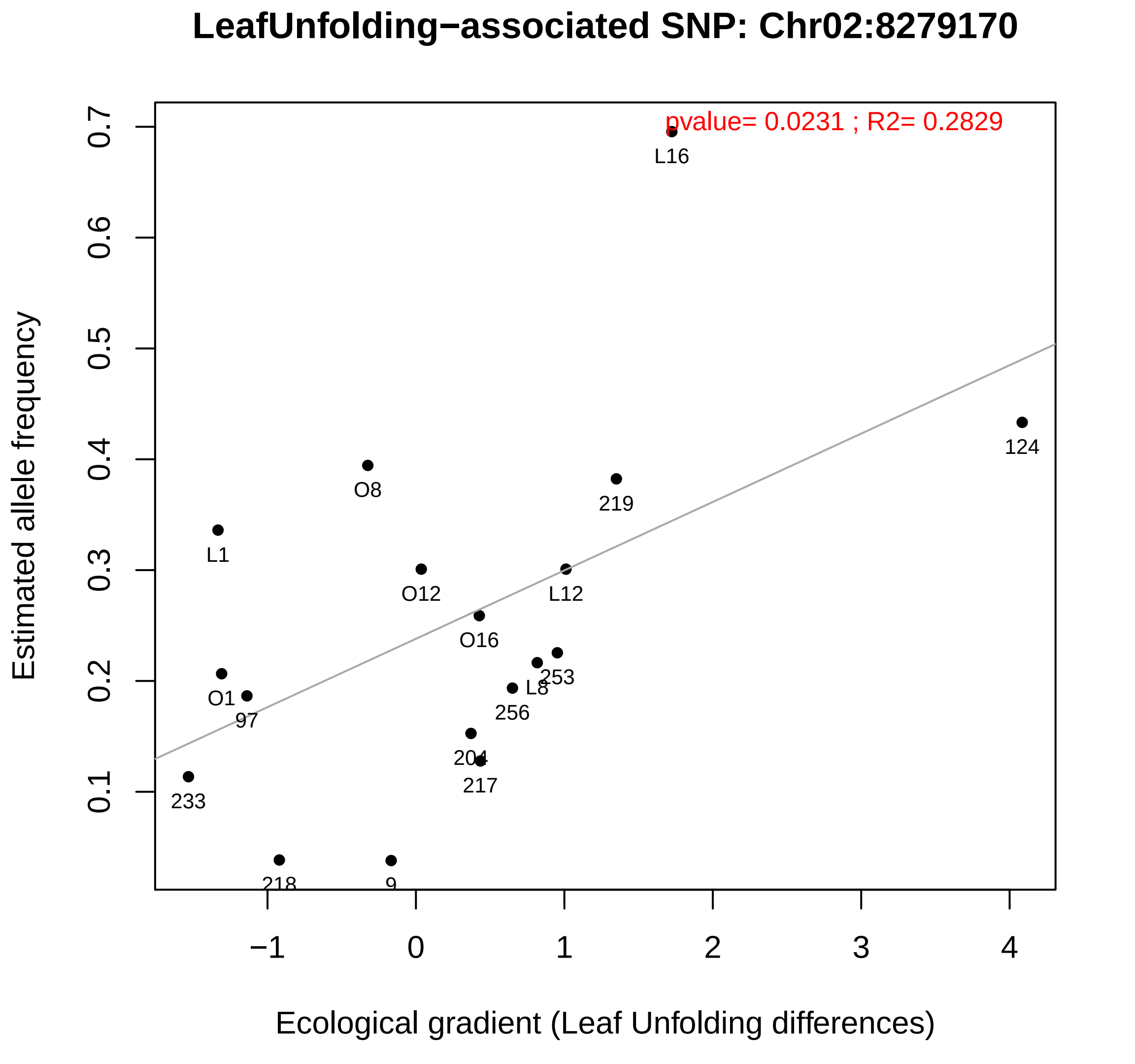
LeafUnfolding−associated SNP: Chr02:8279170
pvalue= 0.0231 ; R2= 0.2829
Estimated allele frequency
Ecological gradient (Leaf Unfolding differences)
0.1
0.2
0.3
0.4
0.5
0.6
0.7
−1
0
1
2
3
4
L16
124
O8
219
L1
O12
L12
O16
253
L8
O1
97
256
204
217
233
218
9

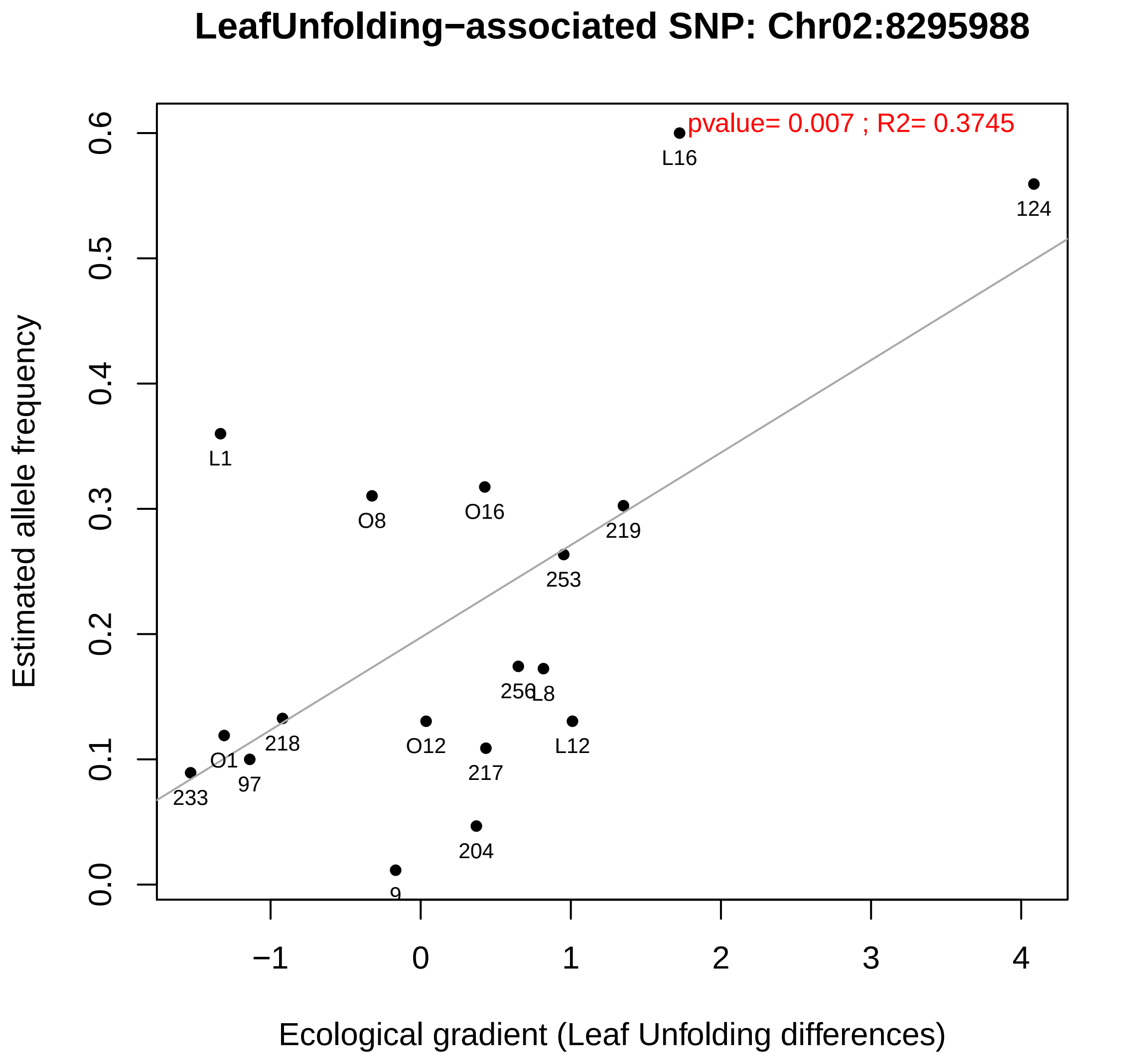

LeafUnfolding−associated SNP: Chr02:8295988
pvalue= 0.007 ; R2= 0.3745
L16
124
L1
O8
O16
219
253
256
L8
218
O12
L12
O1
97
217
233
204
9
0.0
0.1
0.2
0.3
0.4
0.5
0.6
−1
0
1
2
3
4
Estimated allele frequency
Ecological gradient (Leaf Unfolding differences)

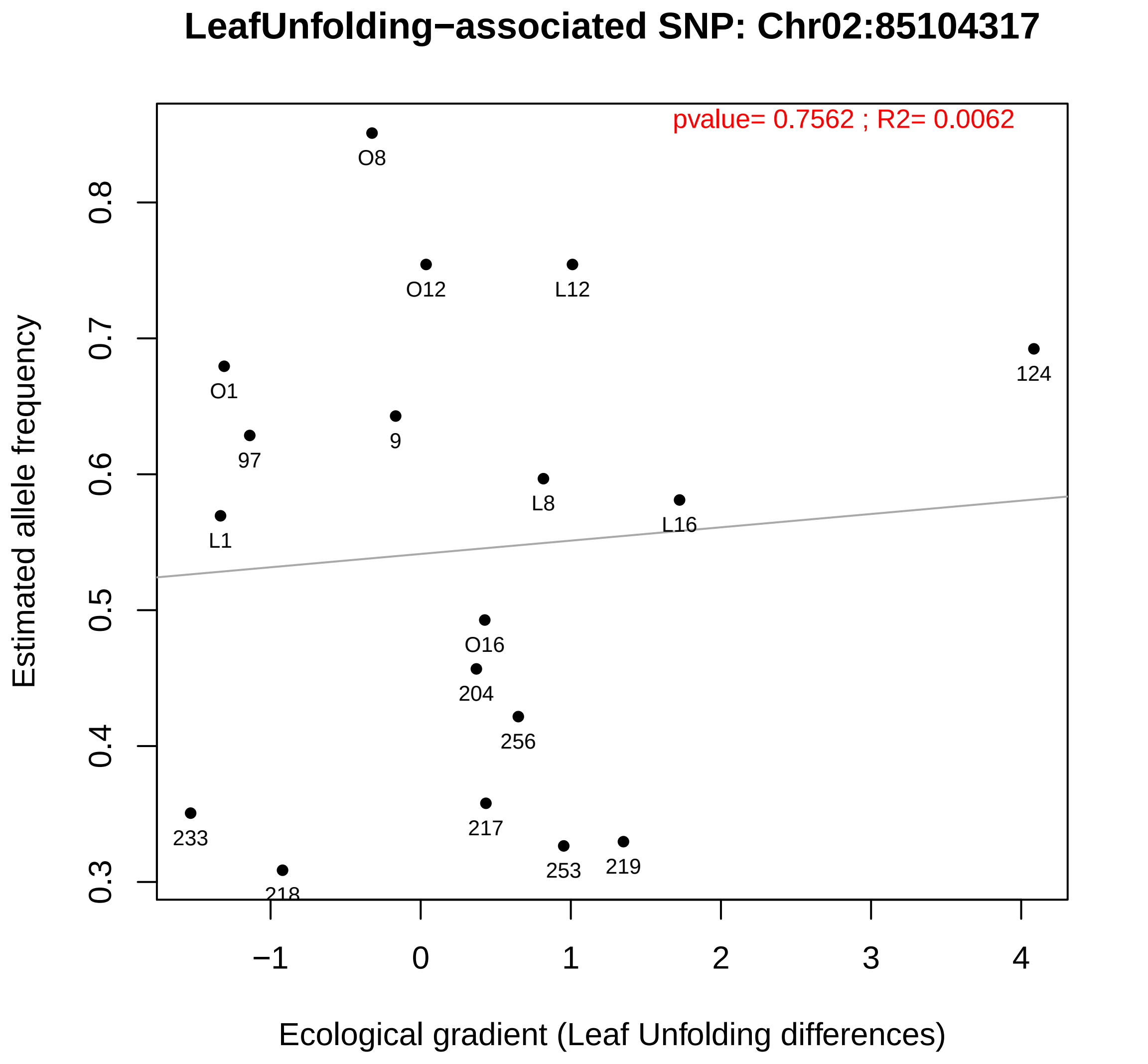
LeafUnfolding−associated SNP: Chr02:85104317
pvalue= 0.7562 ; R2= 0.0062
Estimated allele frequency
Ecological gradient (Leaf Unfolding differences)
0.3
0.4
0.5
0.6
0.7
0.8
−1
0
1
2
3
4
O8
O12
L12
124
O1
9
97
L8
L16
L1
O16
204
256
233
217
253
219
218

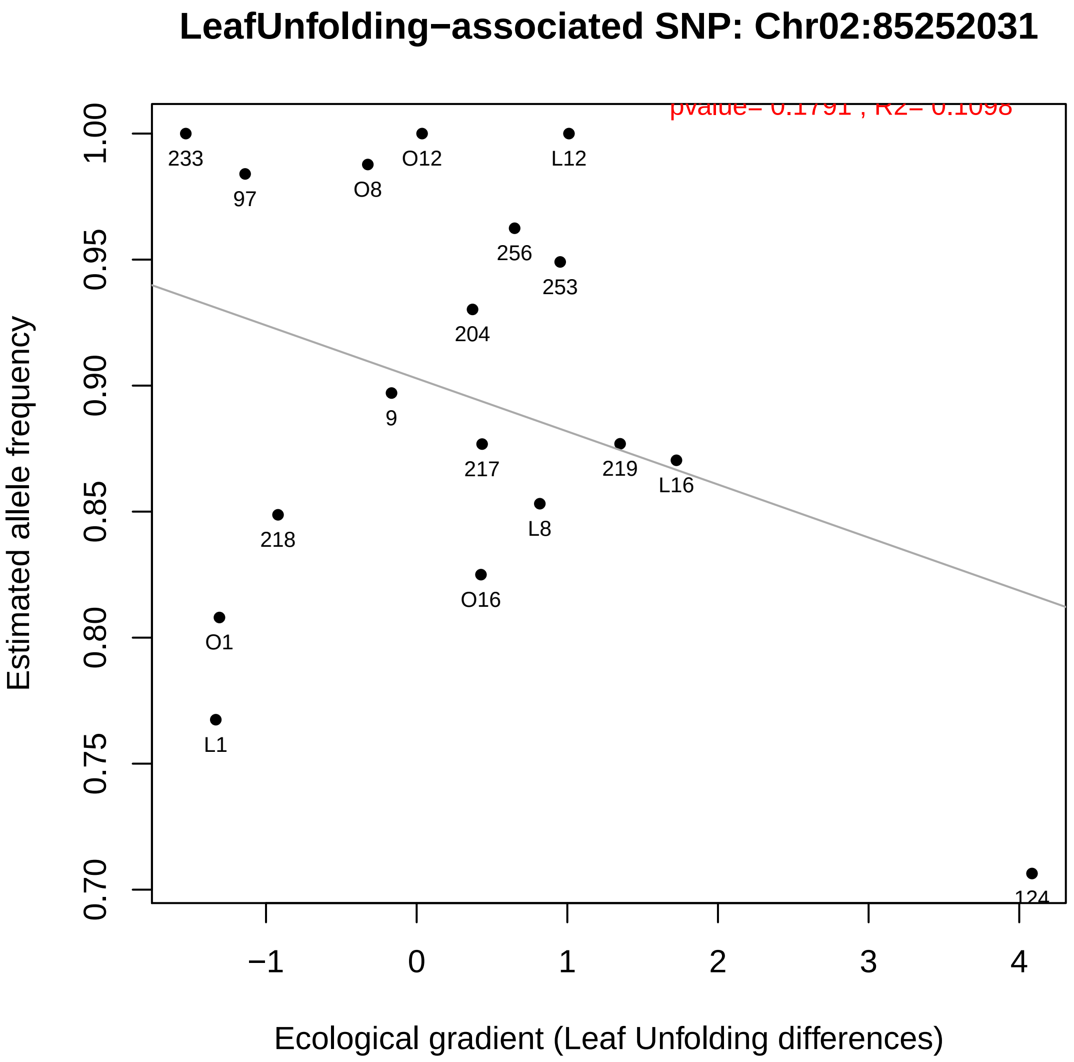

LeafUnfolding−associated SNP: Chr02:85252031
pvalue= 0.1791 , R2= 0.1098
Estimated allele frequency
Ecological gradient (Leaf Unfolding differences)
0.70
0.75
0.80
0.85
0.90
0.95
1.00
−1
0
1
2
3
4
233
97
O8
O12
L12
256
253
204
9
217
219
L16
L8
218
O16
O1
L1
124

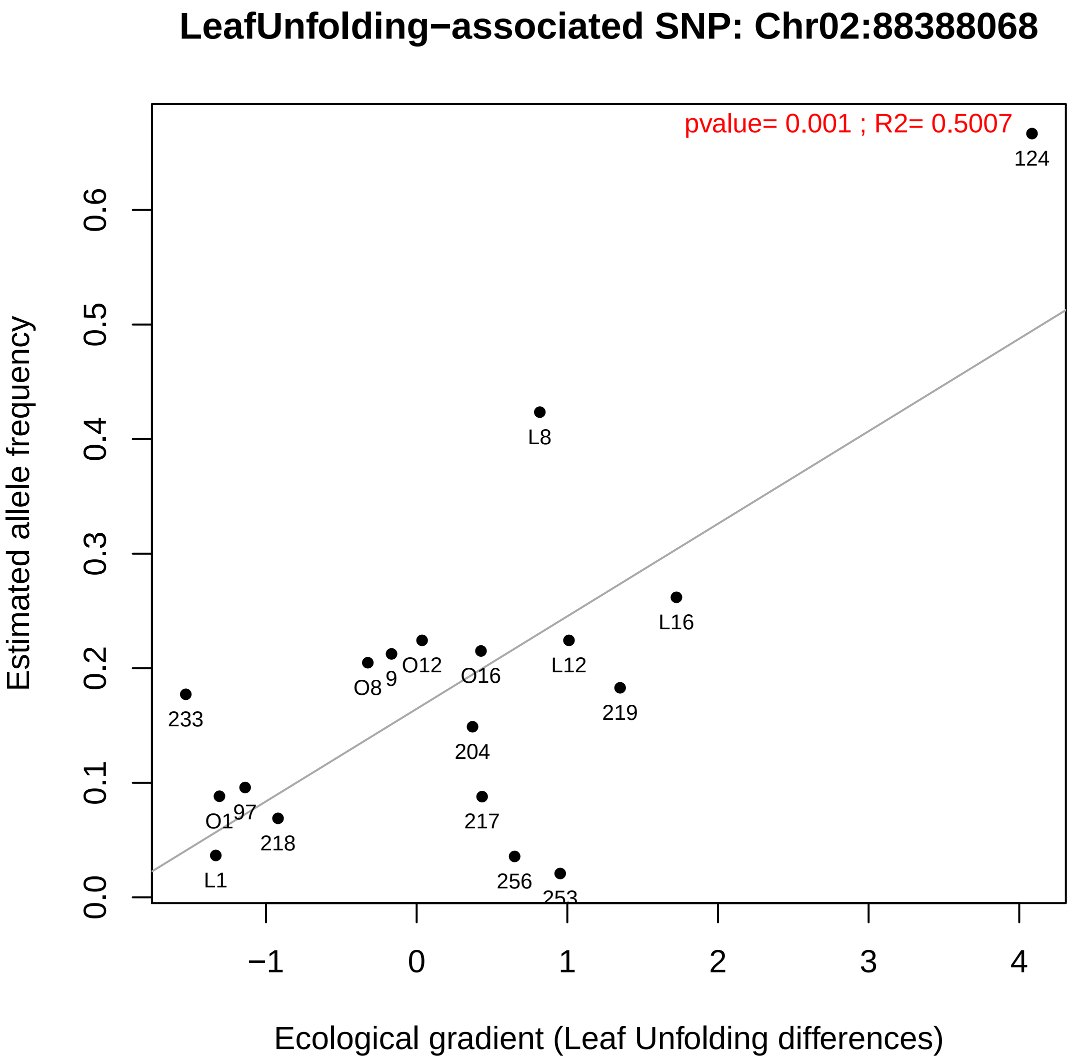
LeafUnfolding−associated SNP: Chr02:88388068
pvalue= 0.001 ; R2= 0.5007
Estimated allele frequency
Ecological gradient (Leaf Unfolding differences)
0.0
0.1
0.2
0.3
0.4
0.5
0.6
−1
0
1
2
3
4
124
L8
L16
L12
219
O12
O16
9
O8
233
204
217
97
O1
218
L1
256

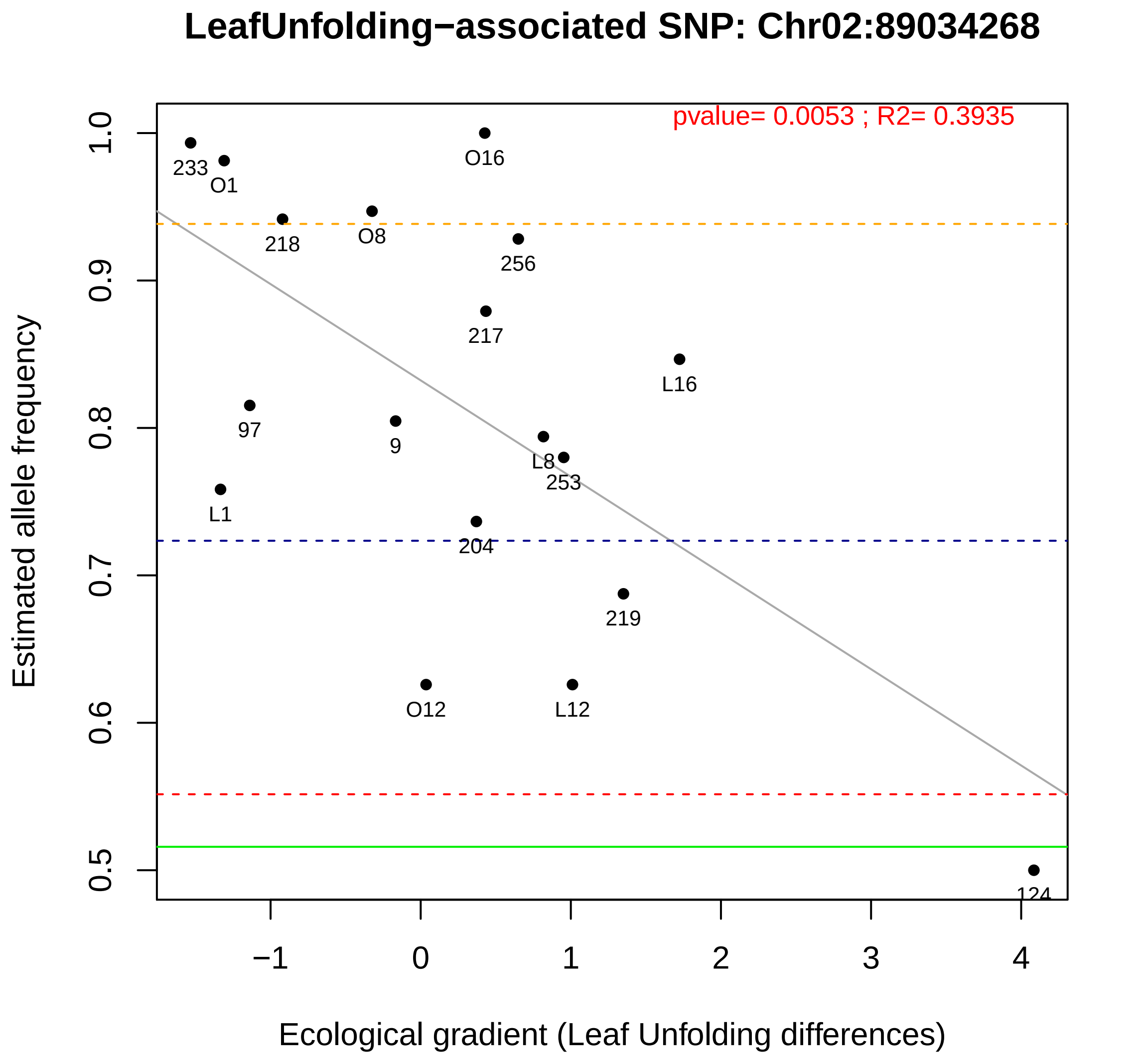

LeafUnfolding−associated SNP: Chr02:89034268
pvalue= 0.0053 ; R2= 0.3935
Estimated allele frequency
Ecological gradient (Leaf Unfolding differences)
1.0
0.9
0.8
0.7
0.6
0.5
−1
0
1
2
3
4
233
O1
O16
218
O8
256
217
L16
97
9
L8
253
L1
204
219
O12
L12
124

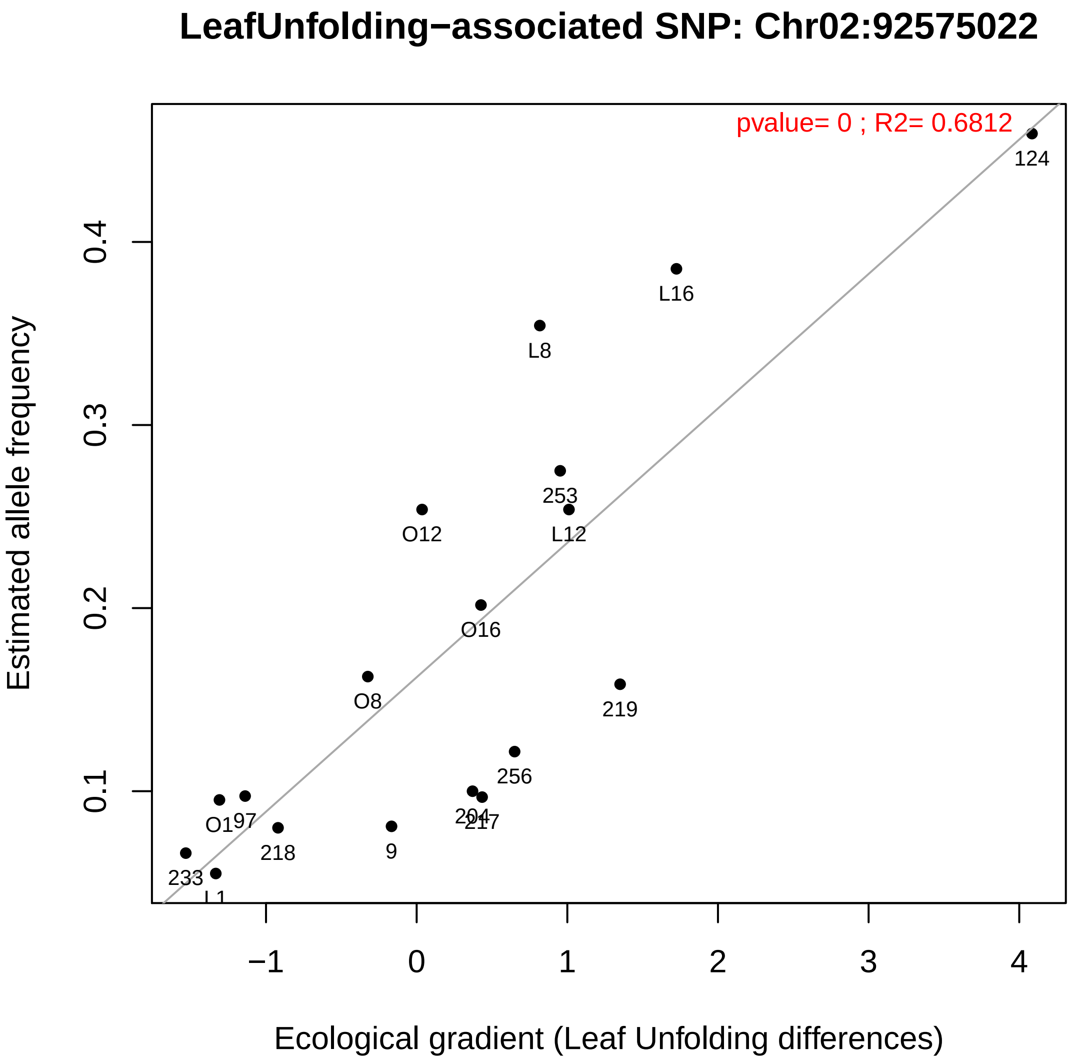

LeafUnfolding−associated SNP: Chr02:92575022
pvalue= 0 ; R2= 0.6812
Estimated allele frequency
Ecological gradient (Leaf Unfolding differences)
0.1
0.2
0.3
0.4
−1
0
1
2
3
4
124
L16
L8
253
L12
O12
O16
O8
219
256
204
217
O1
97
218
9
233
L1

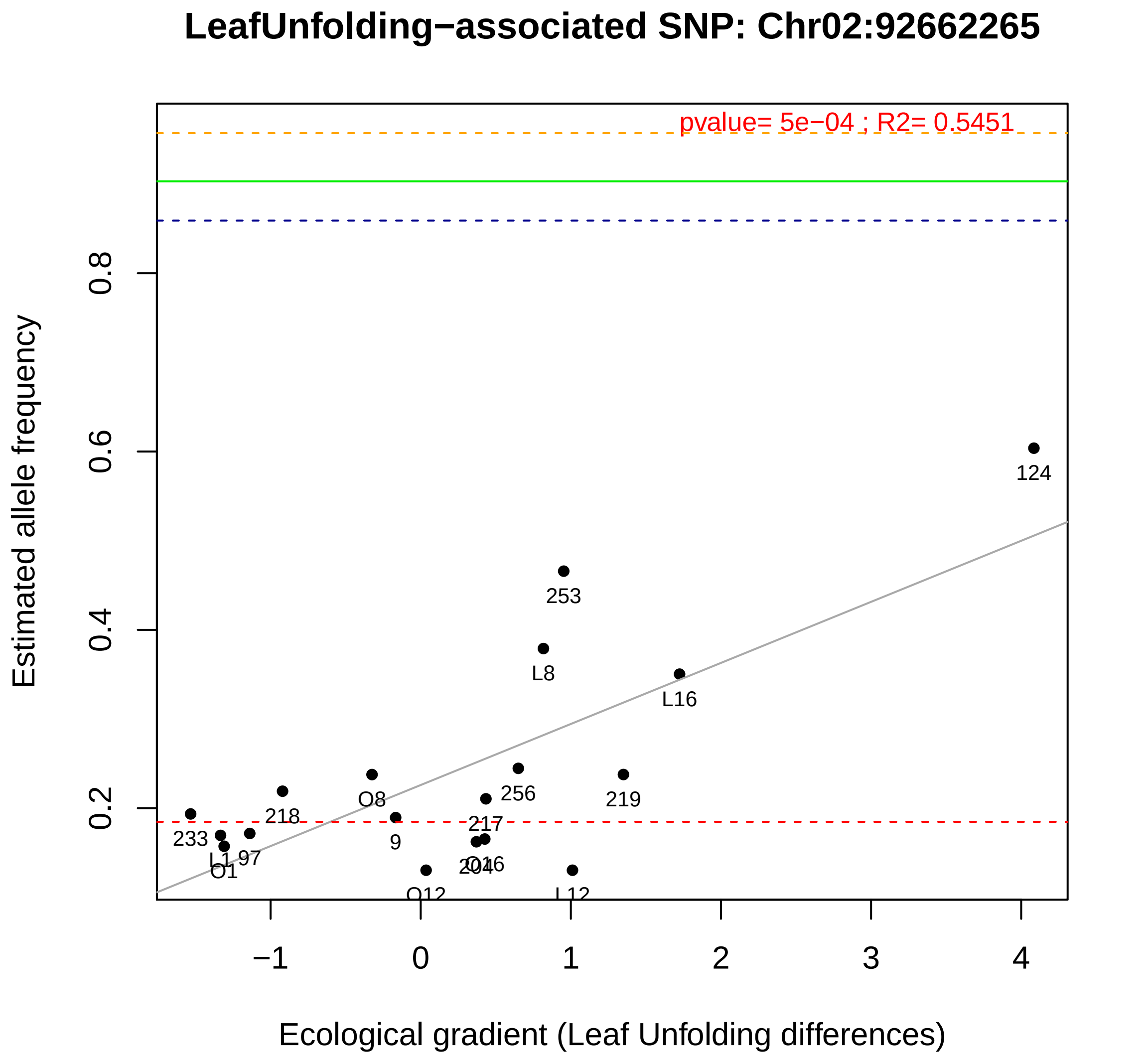

LeafUnfolding−associated SNP: Chr02:92662265
pvalue= 5e−04 ; R2= 0.5451
Estimated allele frequency
0.2
0.4
0.6
0.8
Ecological gradient (Leaf Unfolding differences)
−1
0
1
2
3
4
124
253
L8
L16
256
219
O8
218
217
233
9
O16
204
L1
97
O1
O12
L12

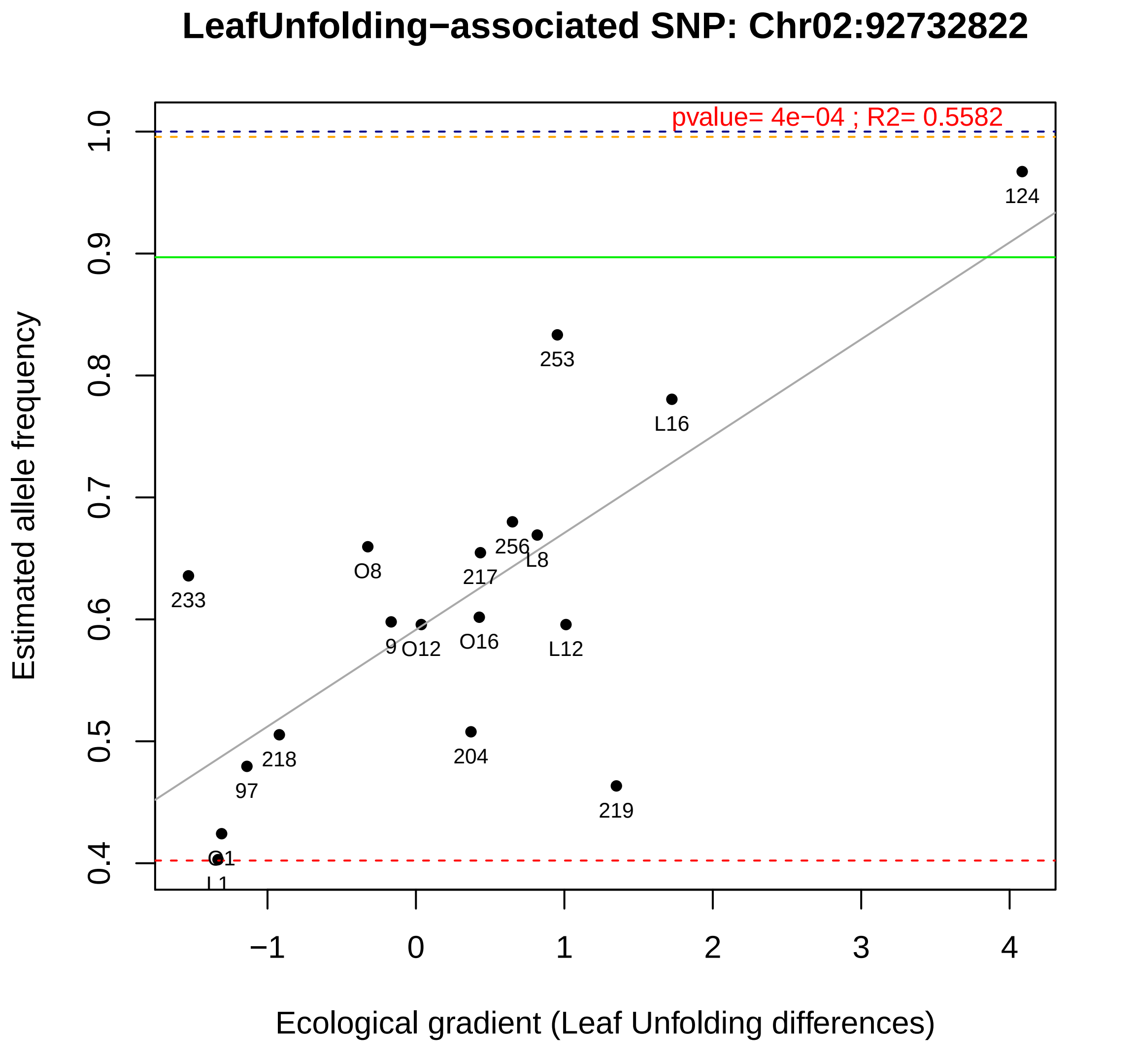
LeafUnfolding−associated SNP: Chr02:92732822
pvalue= 4e−04 ; R2= 0.5582
Estimated allele frequency
0.4
0.5
0.6
0.7
0.8
0.9
1.0
−1
0
1
2
3
4
Ecological gradient (Leaf Unfolding differences)
124
253
L16
256
L8
O8
217
233
9
O12
O16
L12
218
204
97
219
O1
L1

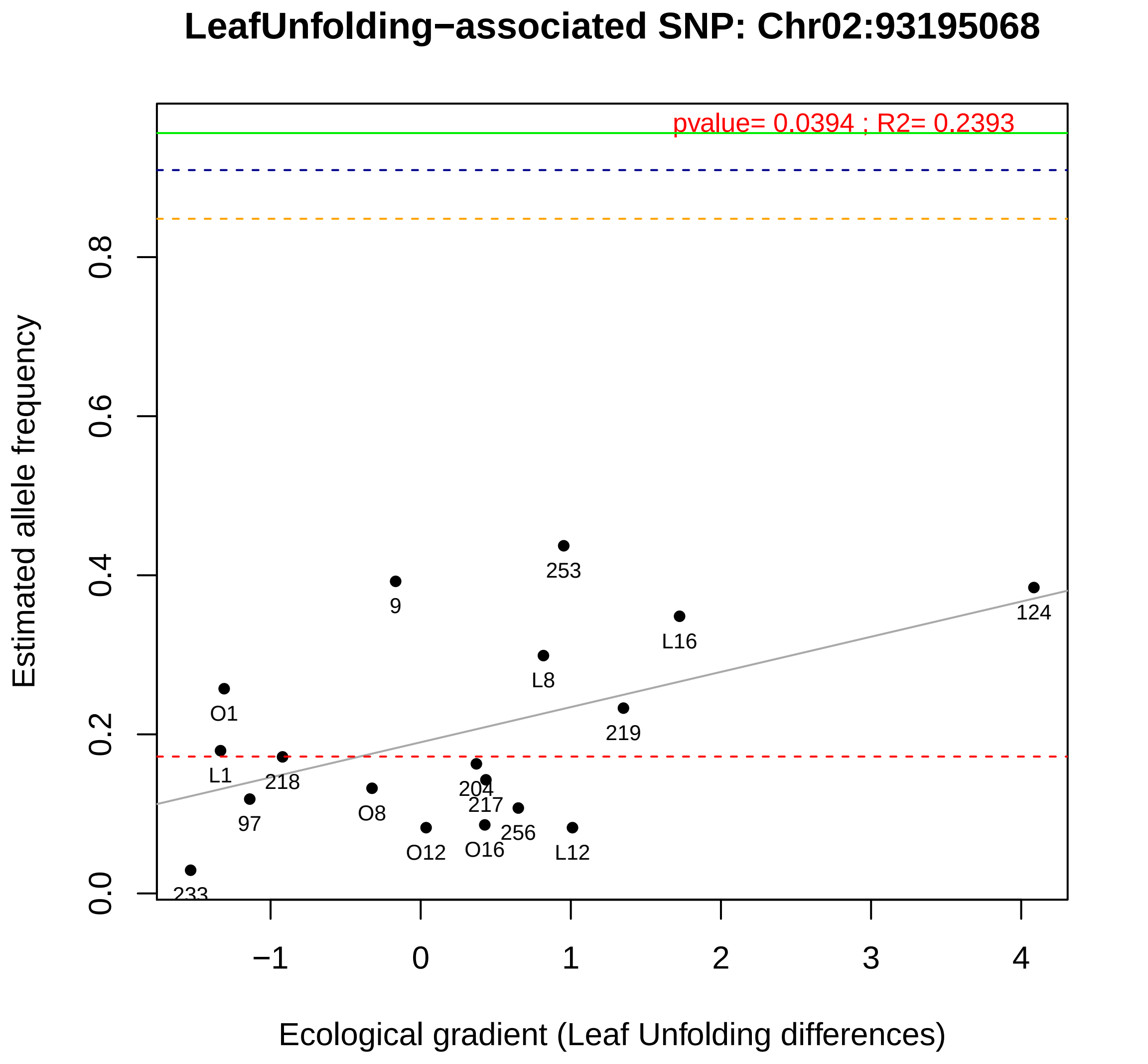
LeafUnfolding−associated SNP: Chr02:93195068
pvalue= 0.0394 ; R2= 0.2393
Estimated allele frequency
0.0
0.2
0.4
0.6
0.8
−1
0
1
2
3
4
Ecological gradient (Leaf Unfolding differences)
253
9
124
L16
L8
O1
219
L1
218
204
217
97
O8
256
O12
O16
L12
233

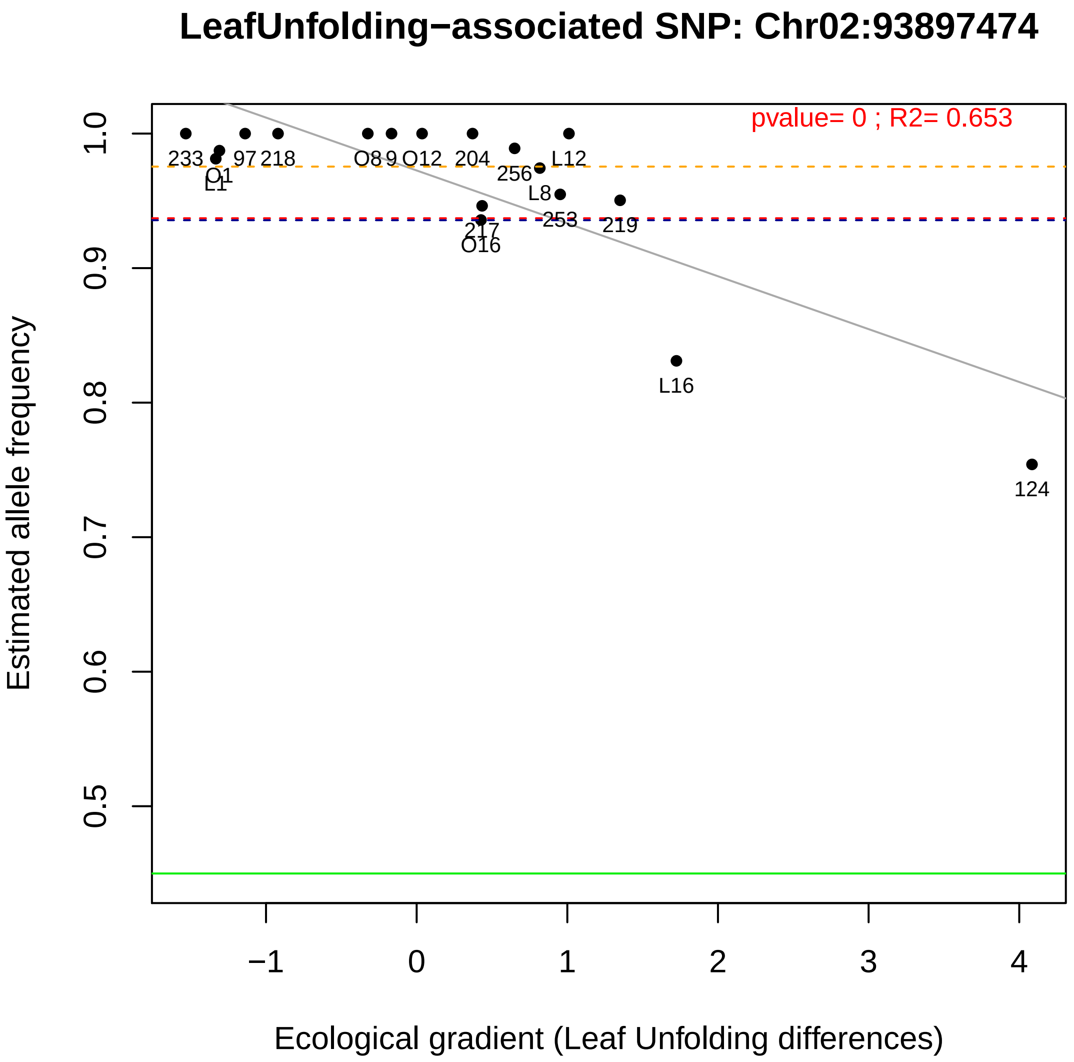
LeafUnfolding−associated SNP: Chr02:93897474
pvalue= 0 ; R2= 0.653
Estimated allele frequency
Ecological gradient (Leaf Unfolding differences)
1.0
0.9
0.8
0.7
0.6
0.5
−1
0
1
2
3
4
233
O1
L1
97
218
O8
9
O12
204
256
L12
L8
253
217
O16
219
L16
124

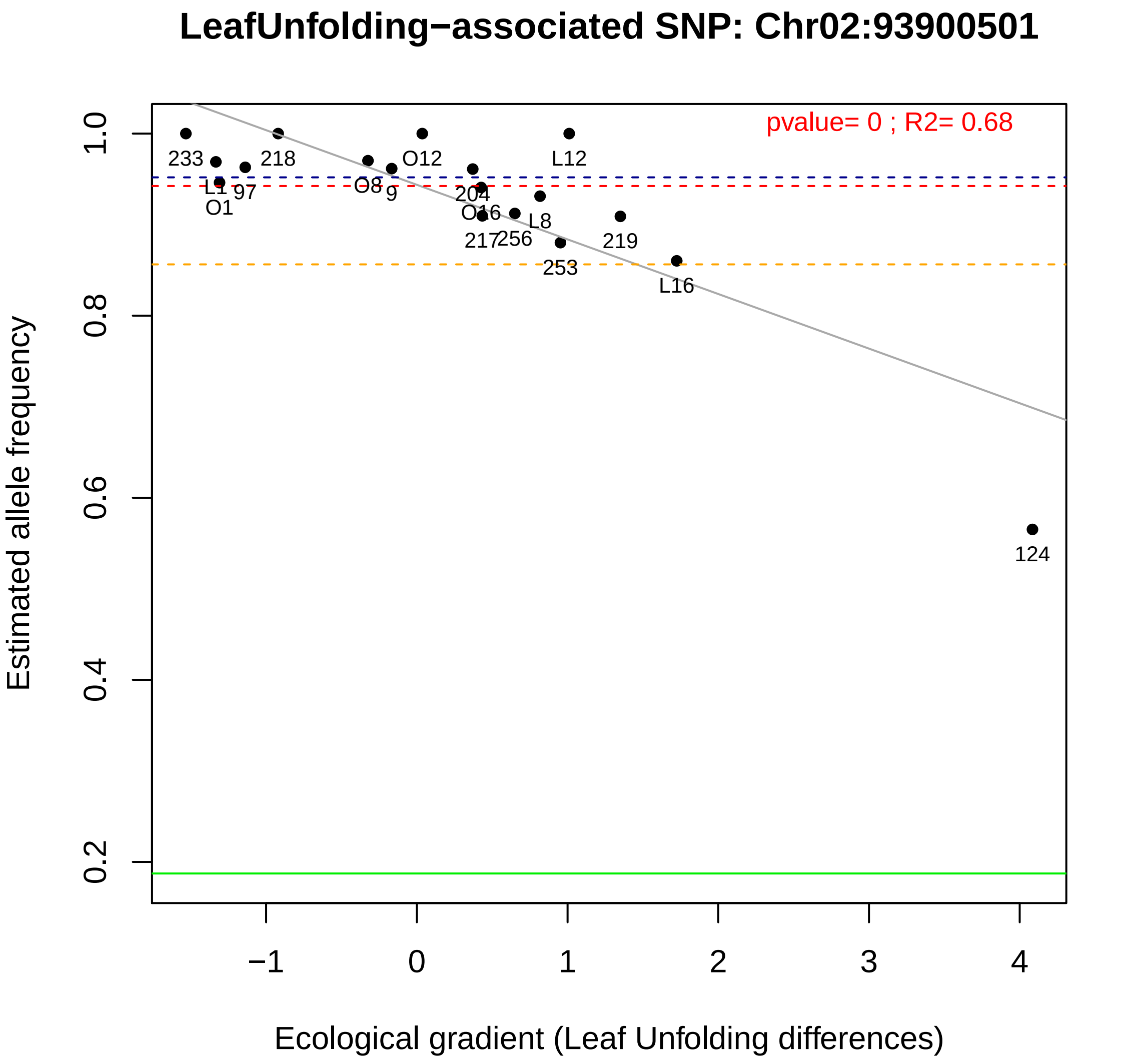

LeafUnfolding−associated SNP: Chr02:93900501
pvalue= 0 ; R2= 0.68
Estimated allele frequency
Ecological gradient (Leaf Unfolding differences)
1.0
0.8
0.6
0.4
0.2
−1
0
1
2
3
4
233
218
O12
L12
L1
97
O1
O8
9
204
O16
L8
217
256
219
253
L16
124

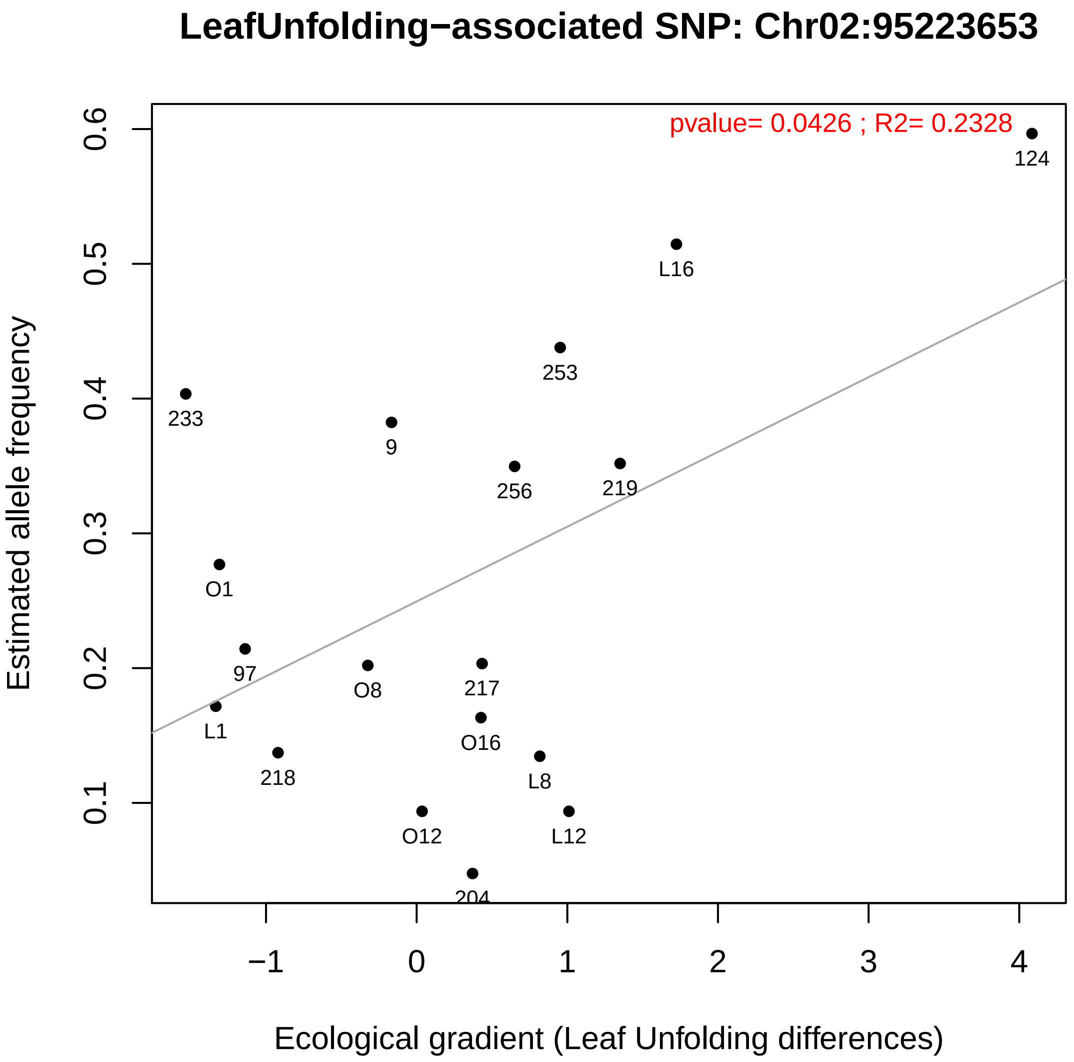
LeafUnfolding−associated SNP: Chr02:95223653
pvalue= 0.0426 ; R2= 0.2328
Estimated allele frequency
Ecological gradient (Leaf Unfolding differences)
0.1
0.2
0.3
0.4
0.5
0.6
−1
0
1
2
3
4
124
L16
253
233
9
256
219
O1
97
O8
217
L1
O16
218
L8
O12
L12
204

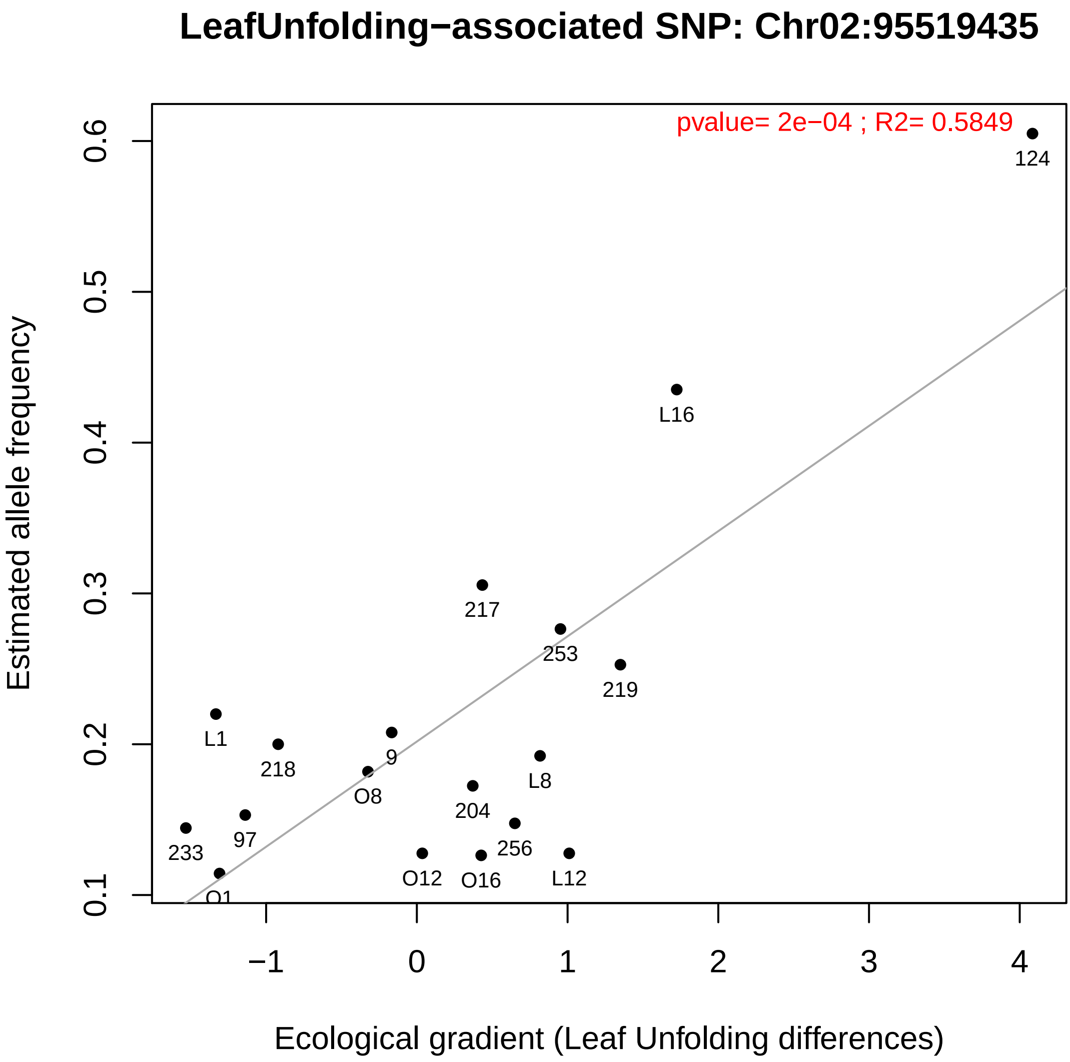
LeafUnfolding−associated SNP: Chr02:95519435
pvalue= 2e−04 ; R2= 0.5849
Estimated allele frequency
Ecological gradient (Leaf Unfolding differences)
0.1
0.2
0.3
0.4
0.5
0.6
−1
0
1
2
3
4
124
L16
217
253
219
L1
218
9
O8
L8
204
97
233
256
O12
O16
L12
O1

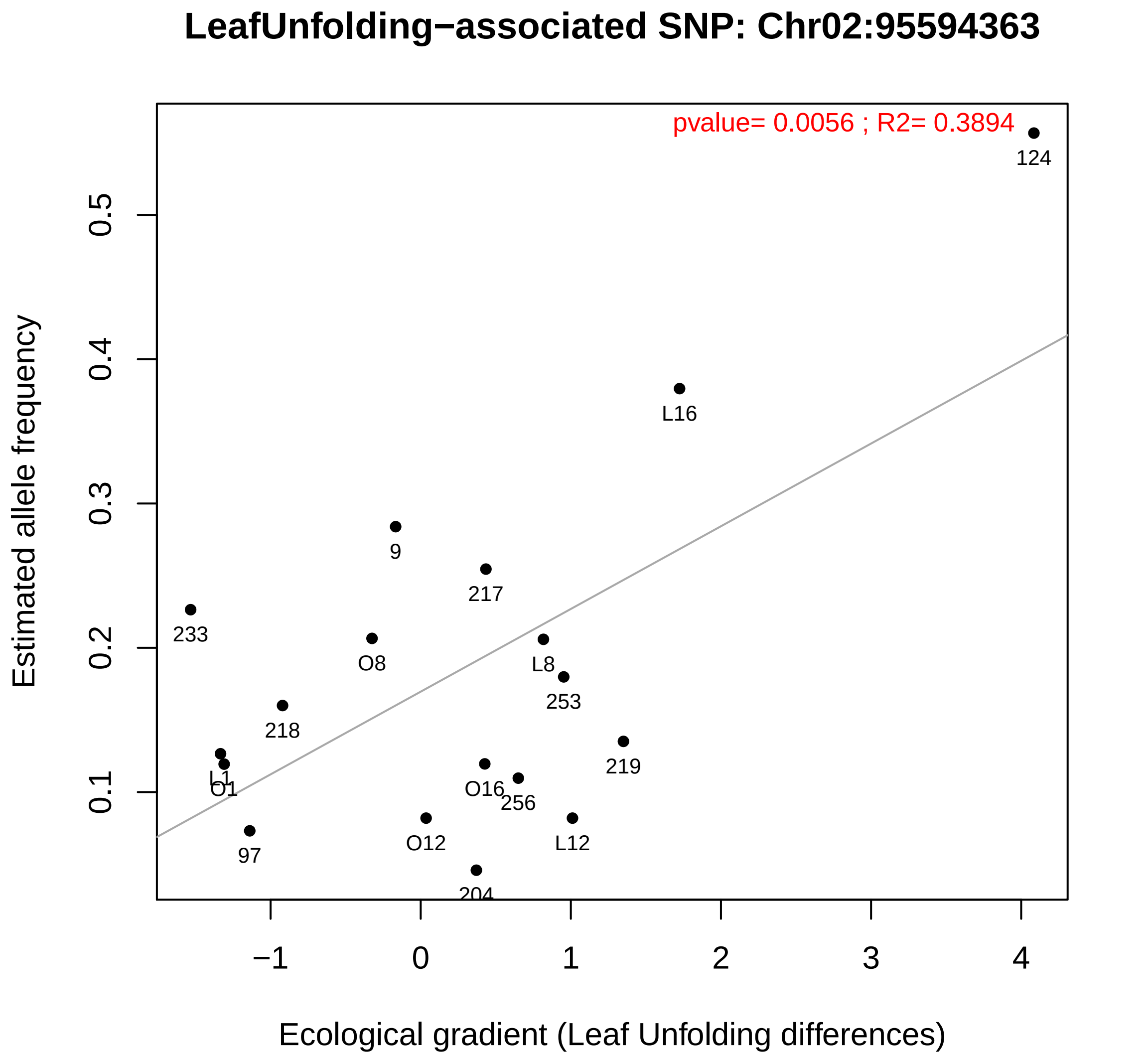

LeafUnfolding−associated SNP: Chr02:95594363
pvalue= 0.0056 ; R2= 0.3894
Estimated allele frequency
0.1
0.2
0.3
0.4
0.5
Ecological gradient (Leaf Unfolding differences)
−1
0
1
2
3
4
124
L16
9
217
233
O8
L8
253
218
219
L1
O1
O16
256
97
O12
L12
204

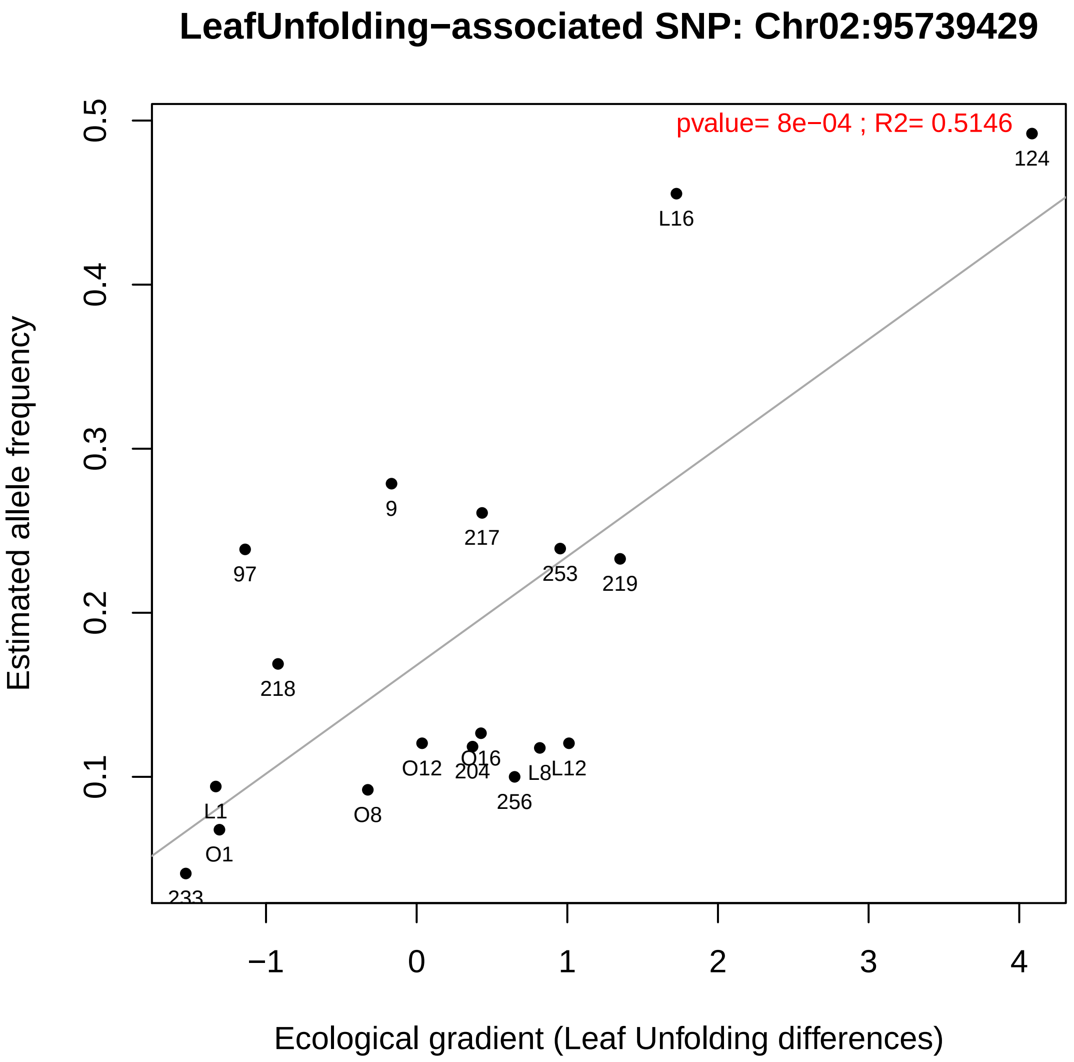
LeafUnfolding−associated SNP: Chr02:95739429
pvalue= 8e−04 ; R2= 0.5146
Estimated allele frequency
0.1
0.2
0.3
0.4
0.5
−1
0
1
2
3
4
Ecological gradient (Leaf Unfolding differences)
124
L16
9
217
97
253
219
218
O12
O16
204
L8
L12
256
L1
O8
O1
233

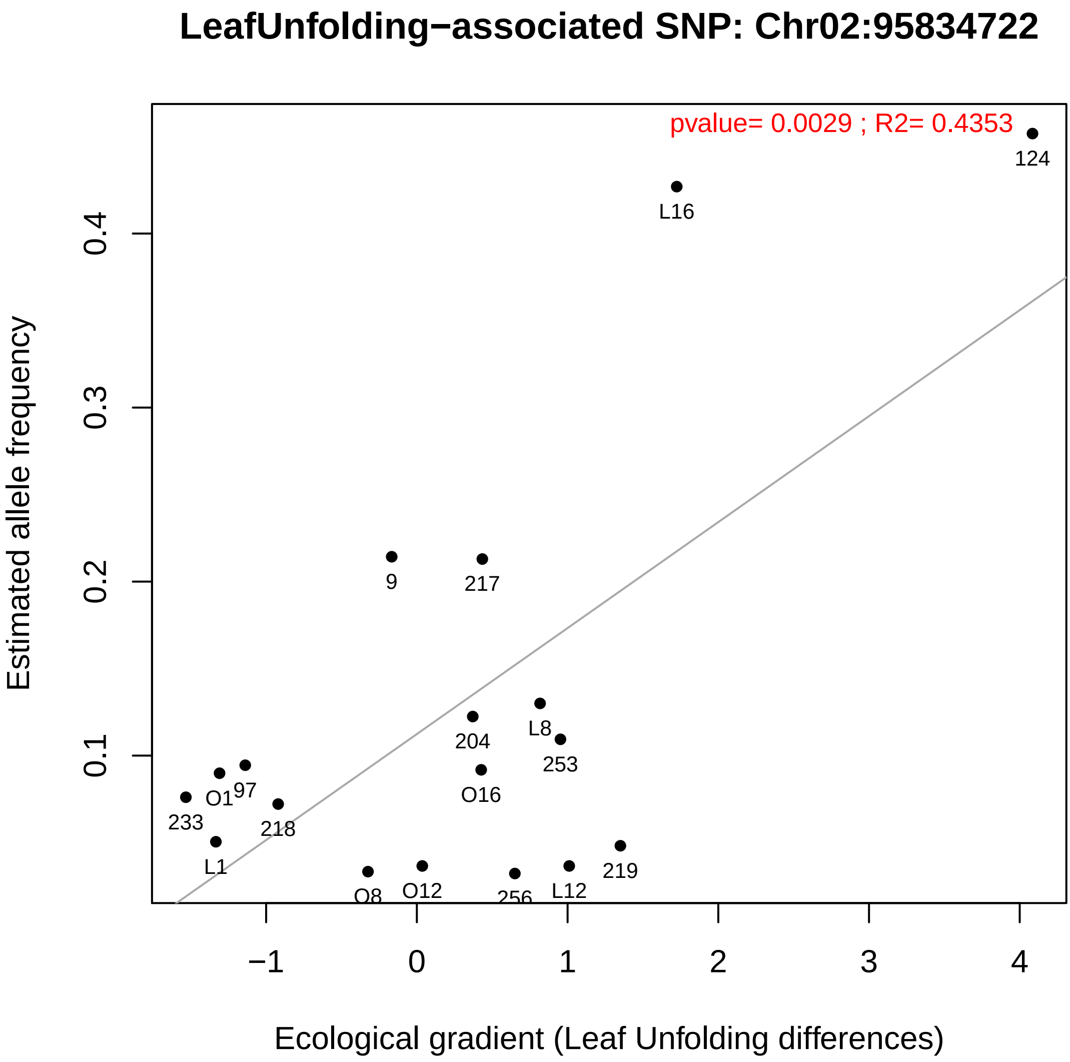

LeafUnfolding−associated SNP: Chr02:95834722
pvalue= 0.0029 ; R2= 0.4353
124
L16
9
217
204
L8
253
O16
O1
97
233
218
L1
O8
O12
256
L12
219
0.1
0.2
0.3
0.4
−1
0
1
2
3
4
Estimated allele frequency
Ecological gradient (Leaf Unfolding differences)

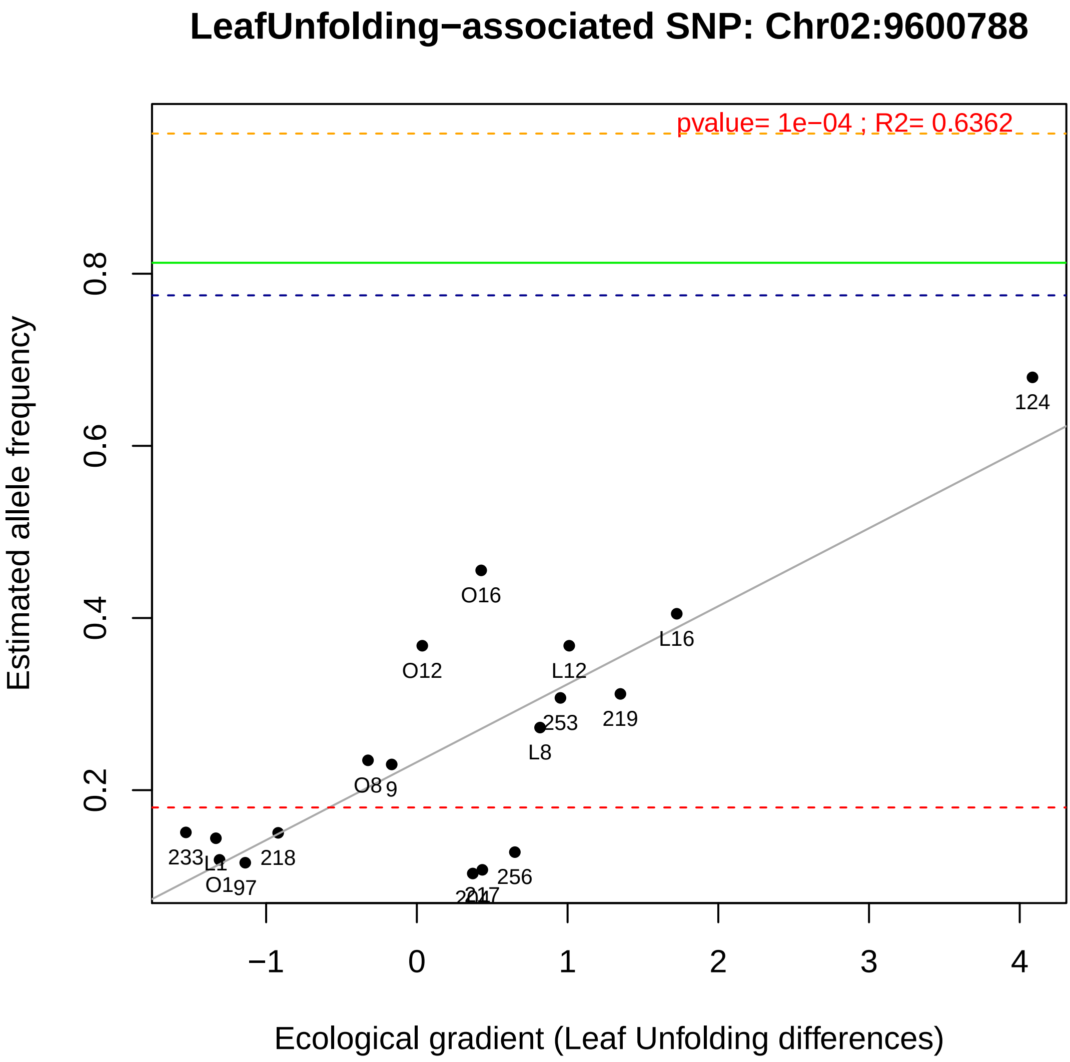

LeafUnfolding−associated SNP: Chr02:9600788
pvalue= 1e−04 ; R2= 0.6362
Estimated allele frequency
0.2
0.4
0.6
0.8
−1
0
1
2
3
4
Ecological gradient (Leaf Unfolding differences)
124
O16
L16
O12
L12
253
219
L8
O8
9
233
L1
O197
218
256
217

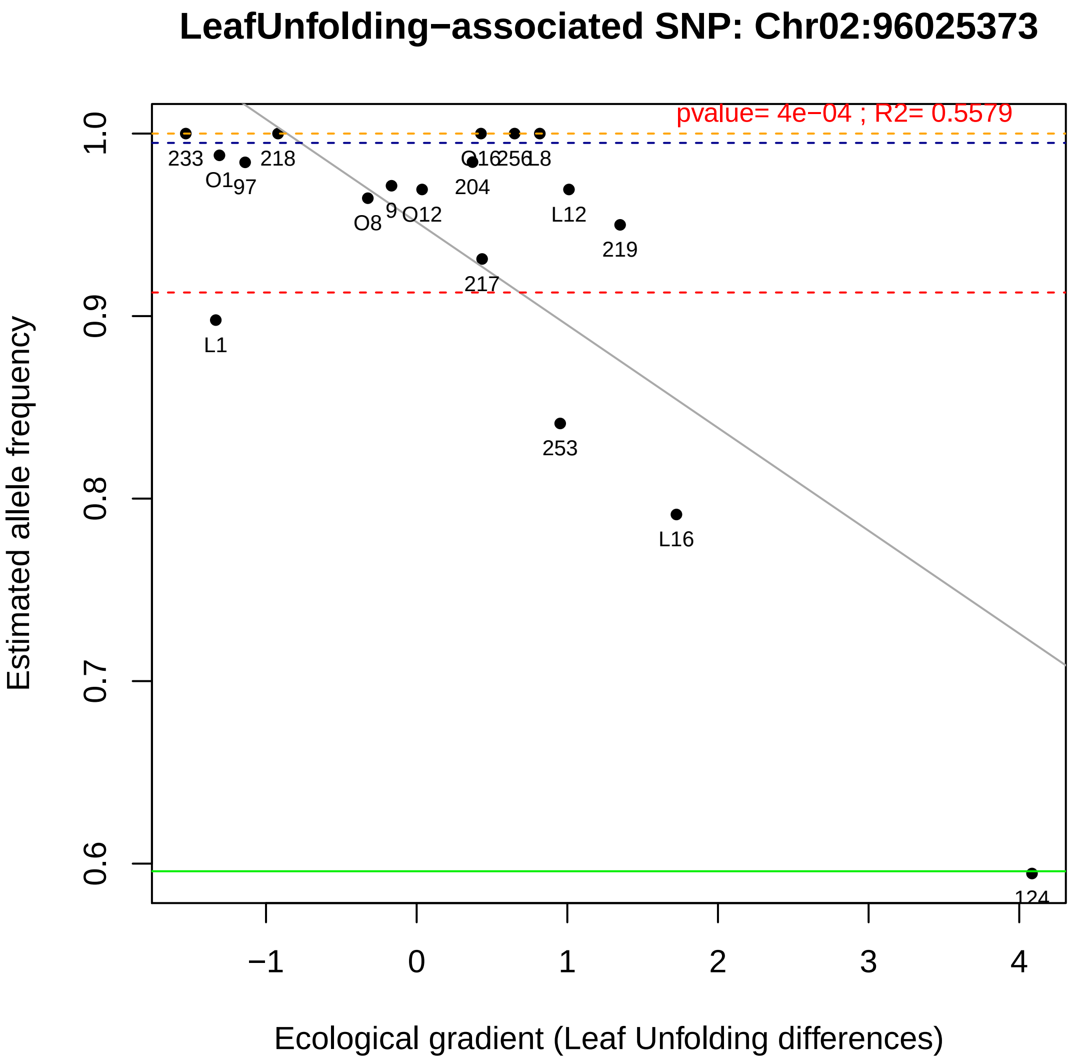
LeafUnfolding−associated SNP: Chr02:96025373
pvalue= 4e−04 ; R2= 0.5579
Estimated allele frequency
0.6
0.7
0.8
0.9
1.0
−1
0
1
2
3
4
Ecological gradient (Leaf Unfolding differences)
233
O1
97
218
O8
9
O12
204
O16
256
L8
L12
219
217
L1
253
L16
124

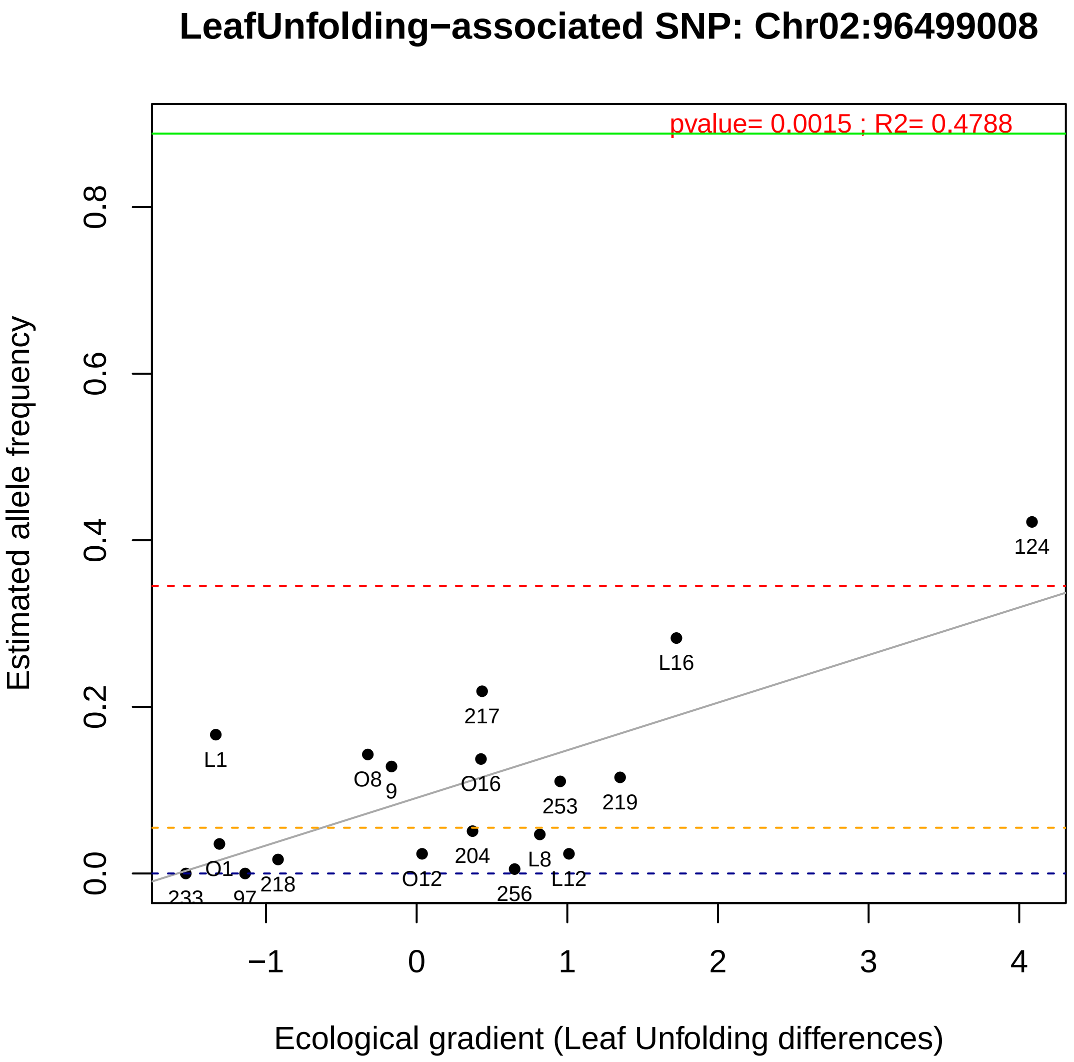

LeafUnfolding−associated SNP: Chr02:96499008
pvalue= 0.0015 ; R2= 0.4788
Estimated allele frequency
0.0
0.2
0.4
0.6
0.8
−1
0
1
2
3
4
Ecological gradient (Leaf Unfolding differences)
124
L16
217
L1
O8
9
O16
253
219
204
L8
L12
O12
256
O1
218
233
97

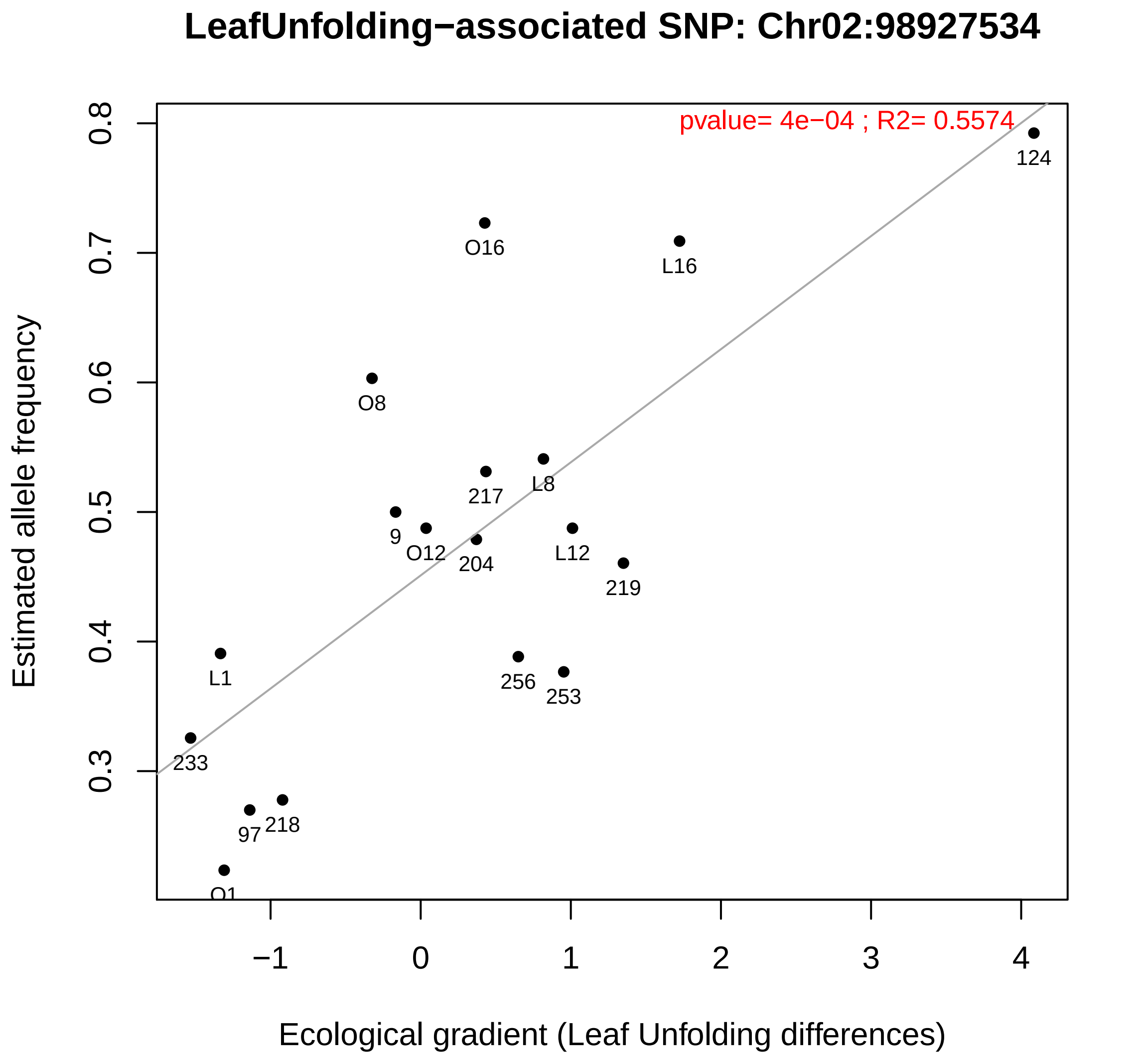
LeafUnfolding−associated SNP: Chr02:98927534
pvalue= 4e−04 ; R2= 0.5574
Estimated allele frequency
Ecological gradient (Leaf Unfolding differences)
0.3
0.4
0.5
0.6
0.7
0.8
−1
0
1
2
3
4
124
O16
L16
O8
217
L8
9
O12
204
L12
219
L1
256
253
233
97
218
O1

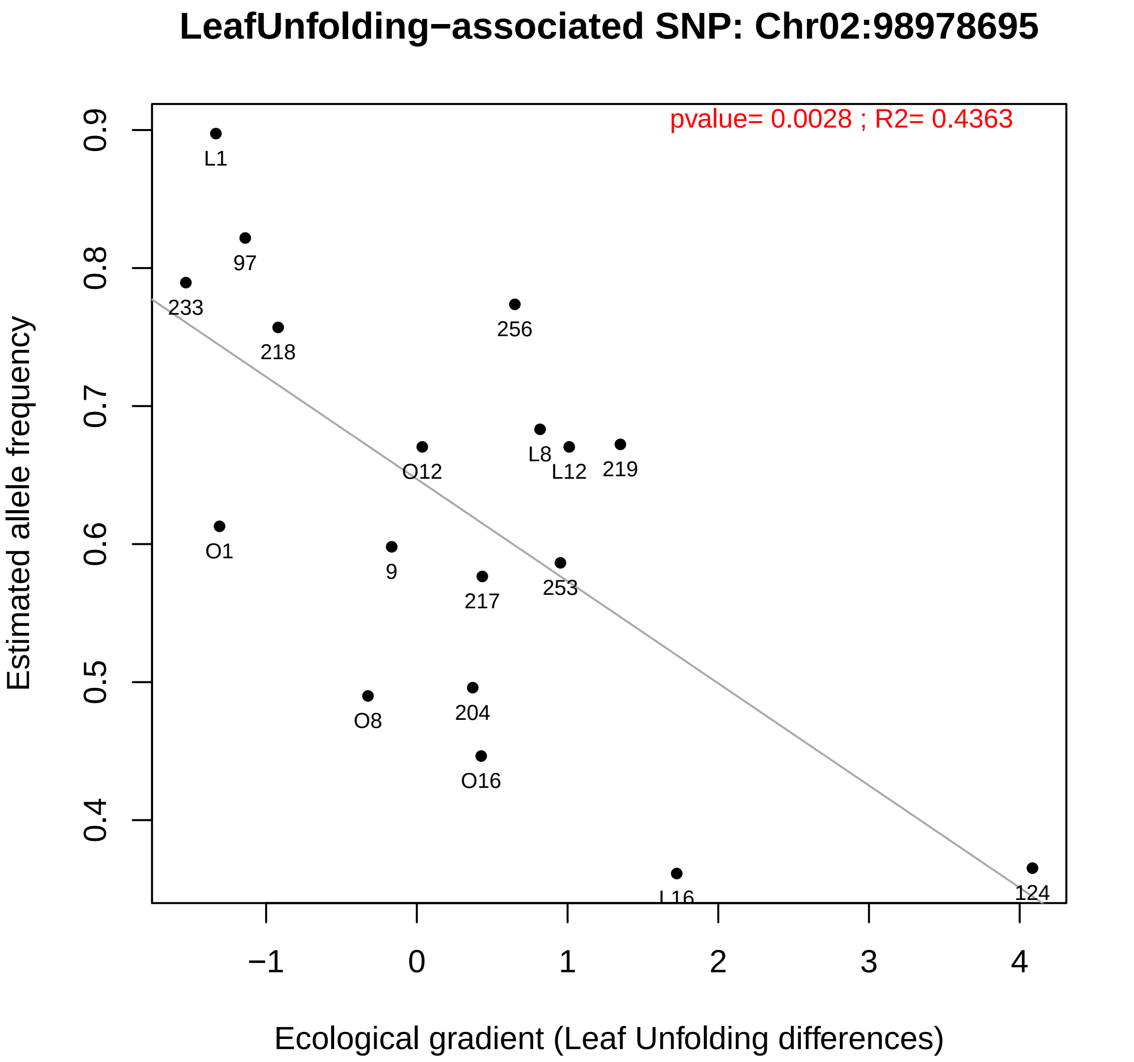
LeafUnfolding−associated SNP: Chr02:98978695
pvalue= 0.0028 ; R2= 0.4363
Estimated allele frequency
0.9
0.8
0.7
0.6
0.5
0.4
L1
97
233
218
256
O12
L8
L12
219
O1
9
217
253
O8
204
O16
124
−1
0
1
2
3
4
Ecological gradient (Leaf Unfolding differences)

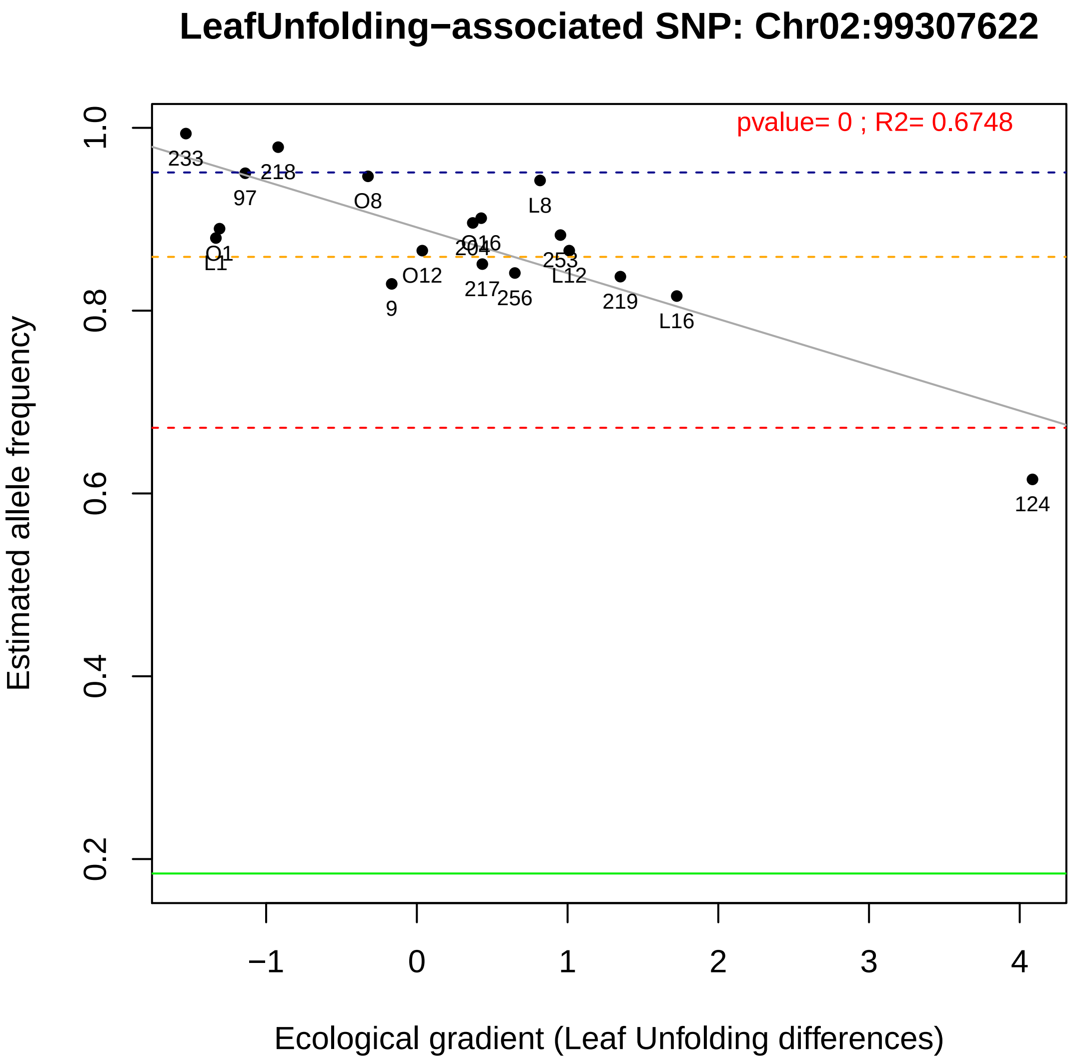
LeafUnfolding−associated SNP: Chr02:99307622
pvalue= 0 ; R2= 0.6748
Estimated allele frequency
Ecological gradient (Leaf Unfolding differences)
0.2
0.4
0.6
0.8
1.0
−1
0
1
2
3
4
233
218
97
O8
L8
O1
L1
O16
204
O12
9
217
256
253
L12
219
L16
124

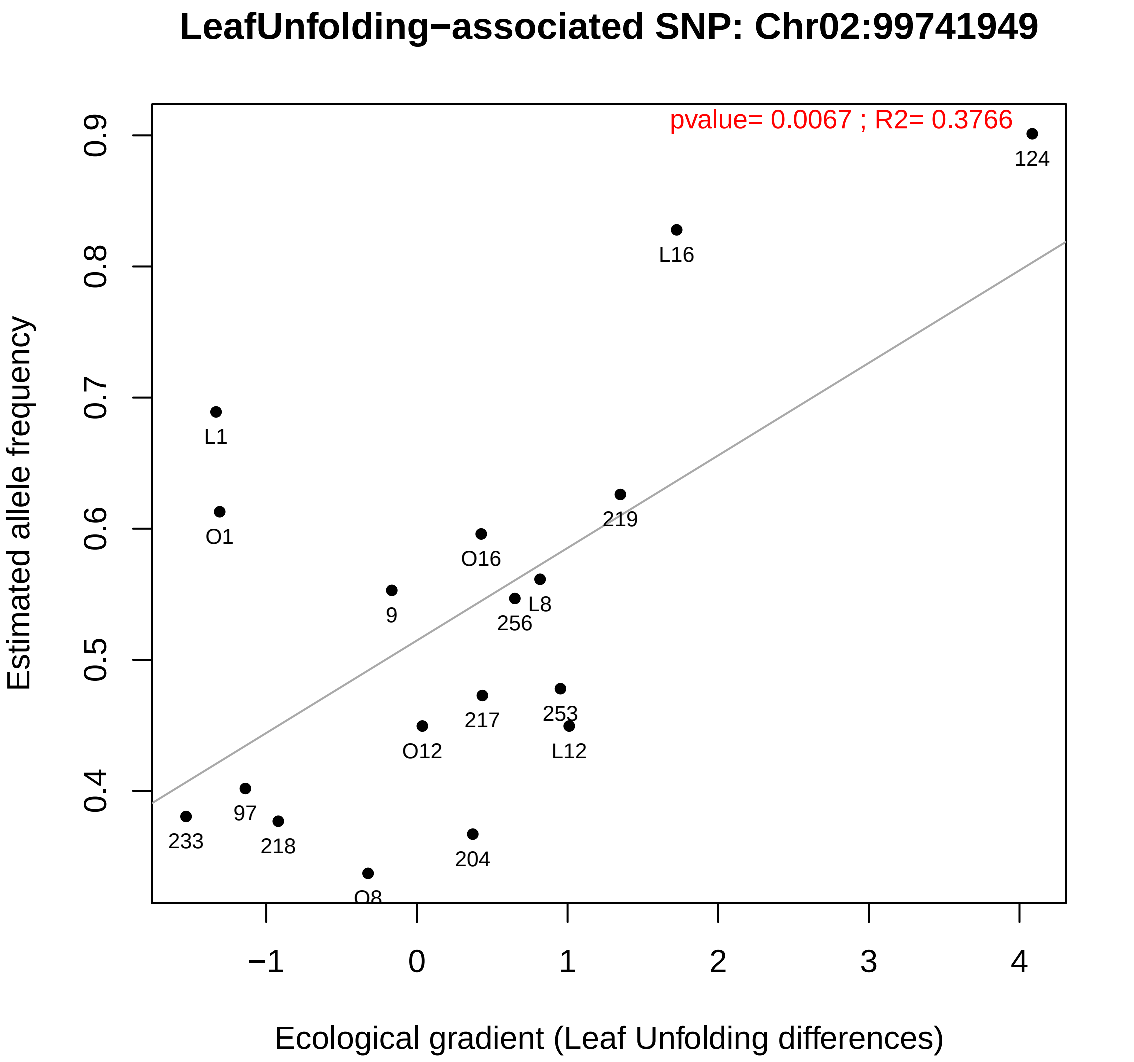

LeafUnfolding−associated SNP: Chr02:99741949
pvalue= 0.0067 ; R2= 0.3766
Estimated allele frequency
Ecological gradient (Leaf Unfolding differences)
0.4
0.5
0.6
0.7
0.8
0.9
−1
0
1
2
3
4
124
L16
L1
O1
219
O16
L8
256
9
217
253
L12
O12
97
233
218
204
O8

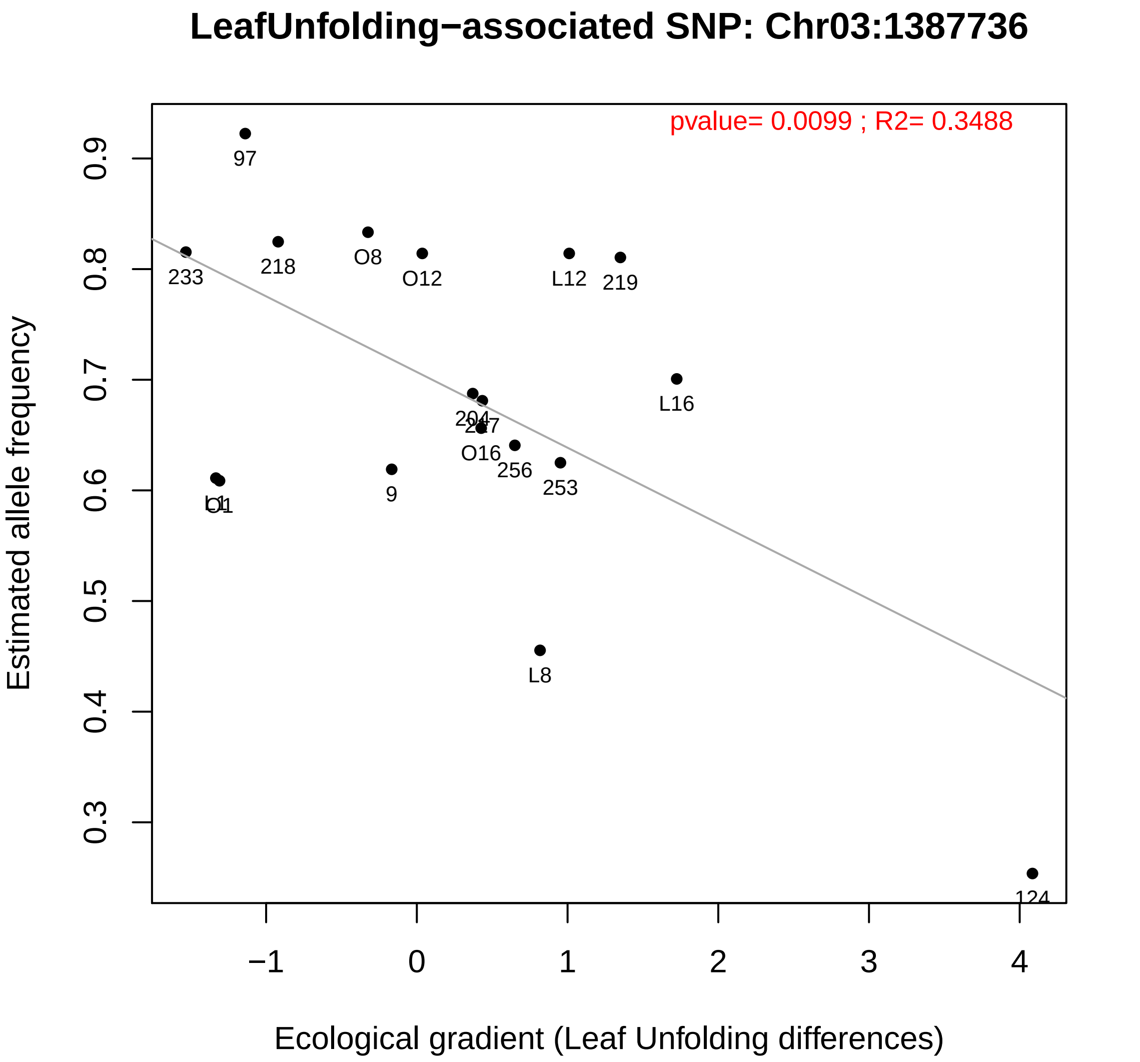
LeafUnfolding−associated SNP: Chr03:1387736
pvalue= 0.0099 ; R2= 0.3488
Estimated allele frequency
Ecological gradient (Leaf Unfolding differences)
0.3
0.4
0.5
0.6
0.7
0.8
0.9
−1
0
1
2
3
4
97
233
218
O8
O12
L12
219
L16
204
247
O16
256
253
9
L1
O1
L8
124

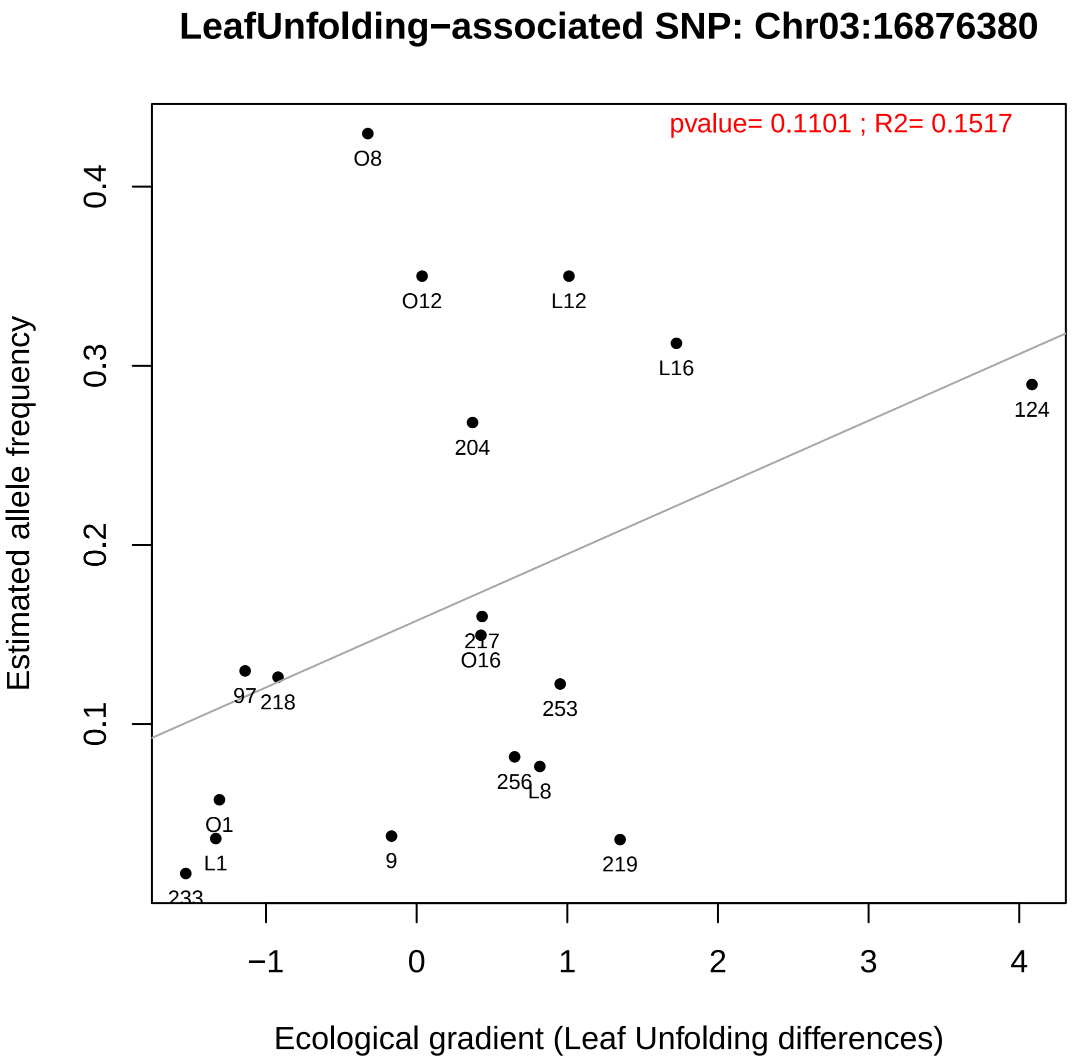

LeafUnfolding−associated SNP: Chr03:16876380
pvalue= 0.1101 ; R2= 0.1517
Estimated allele frequency
0.1
0.2
0.3
0.4
Ecological gradient (Leaf Unfolding differences)
−1
0
1
2
3
4
O8
O12
L12
L16
124
204
217
O16
97
218
253
256
L8
O1
L1
9
219
233

**LeafUnfolding−associated SNP: Chr03:1748374**

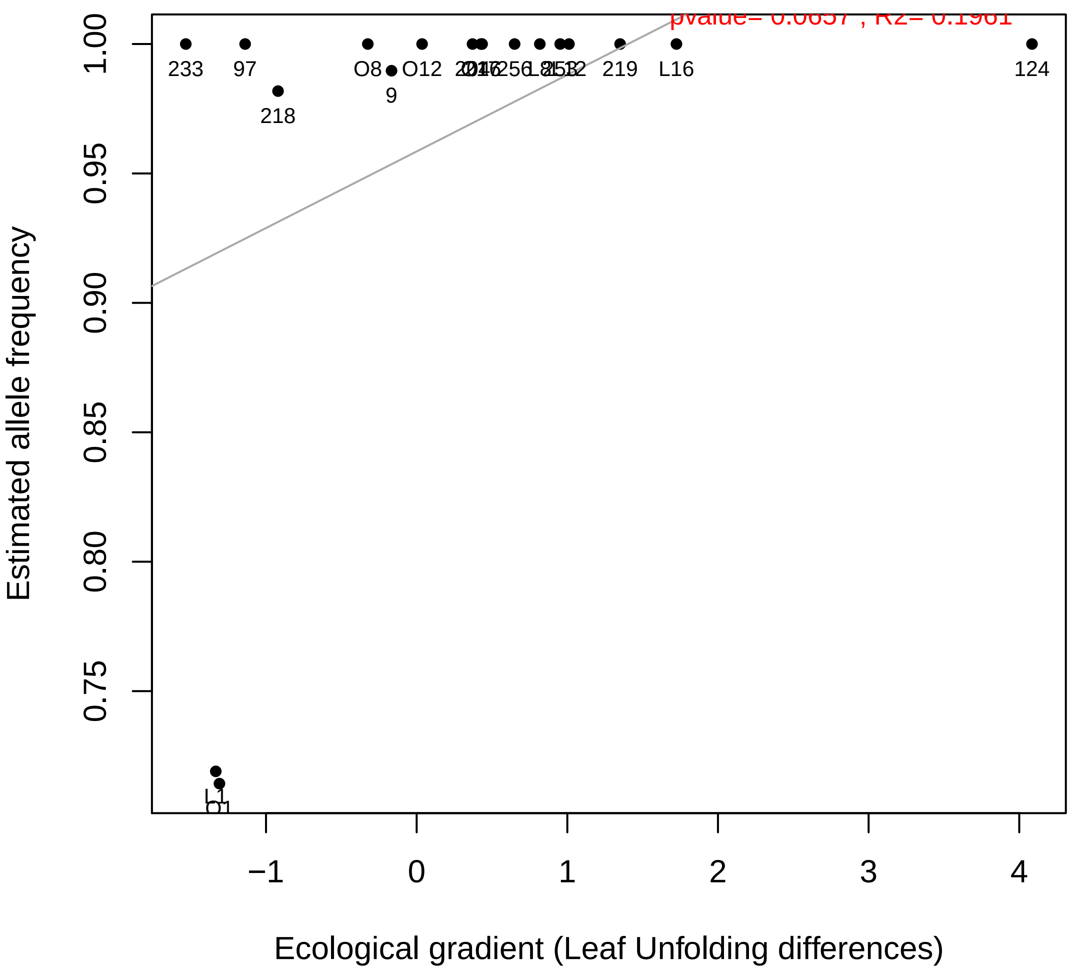

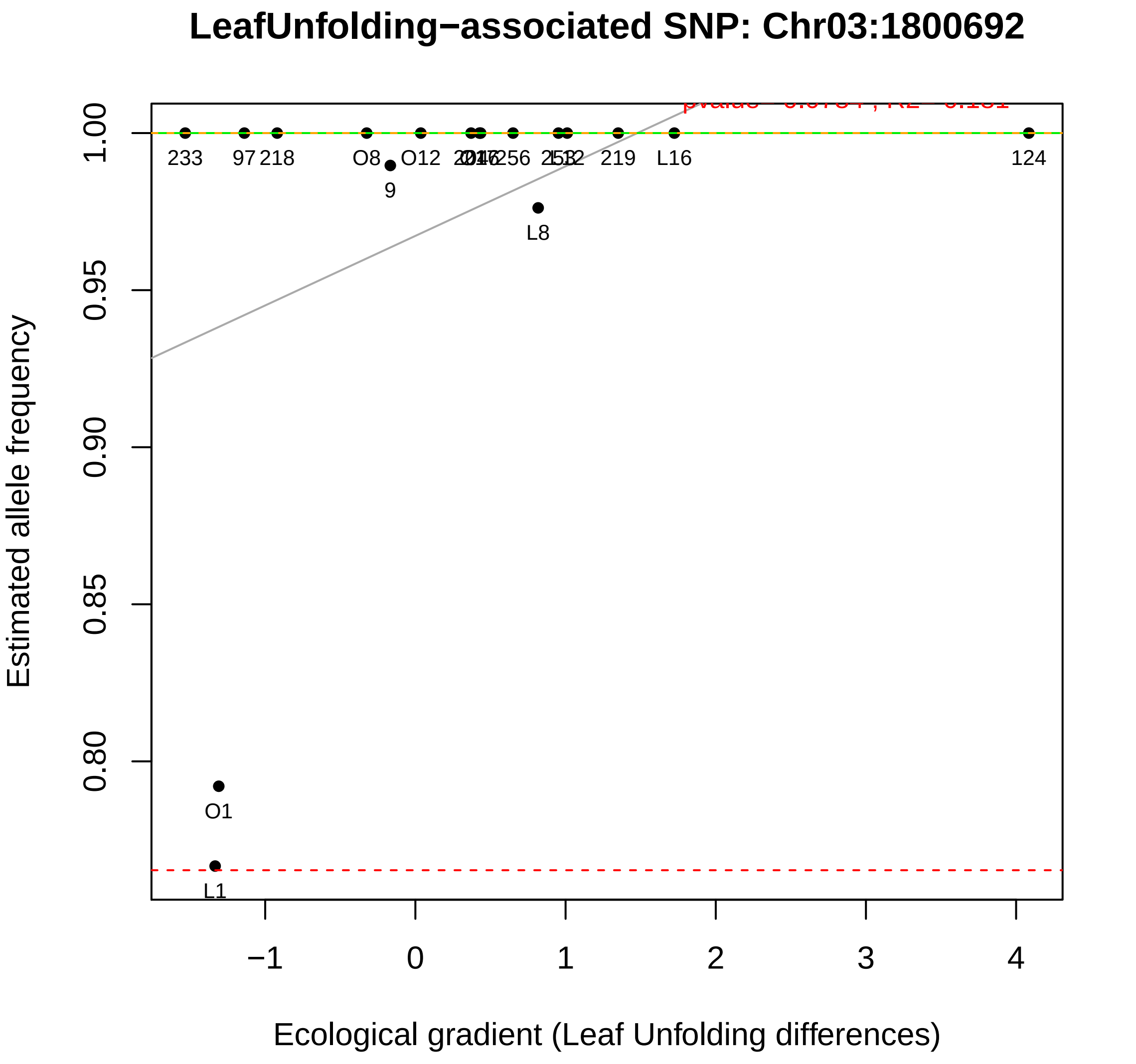
LeafUnfolding−associated SNP: Chr03:1800692
Estimated allele frequency
1.00
0.95
0.90
0.85
0.80
233
97
218
O8
O12
256
219
L16
124
9
L8
O1
L1
−1
0
1
2
3
4
Ecological gradient (Leaf Unfolding differences)

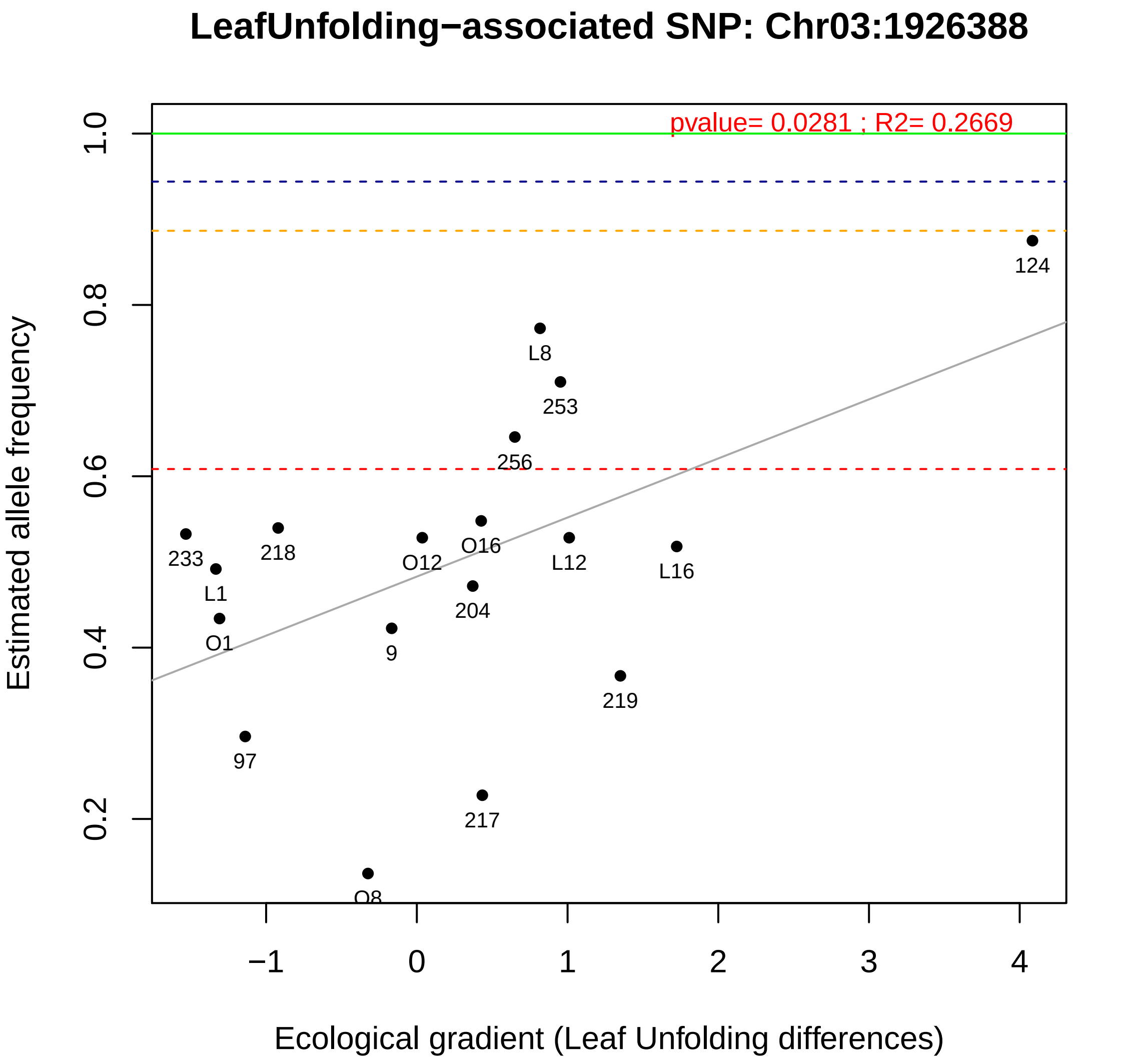
LeafUnfolding−associated SNP: Chr03:1926388
pvalue= 0.0281 ; R2= 0.2669
Estimated allele frequency
1.0
0.8
0.6
0.4
0.2
124
L8
253
256
233
218
O12
O16
L12
L16
L1
204
O1
9
219
97
217
O8
−1
0
1
2
3
4
Ecological gradient (Leaf Unfolding differences)

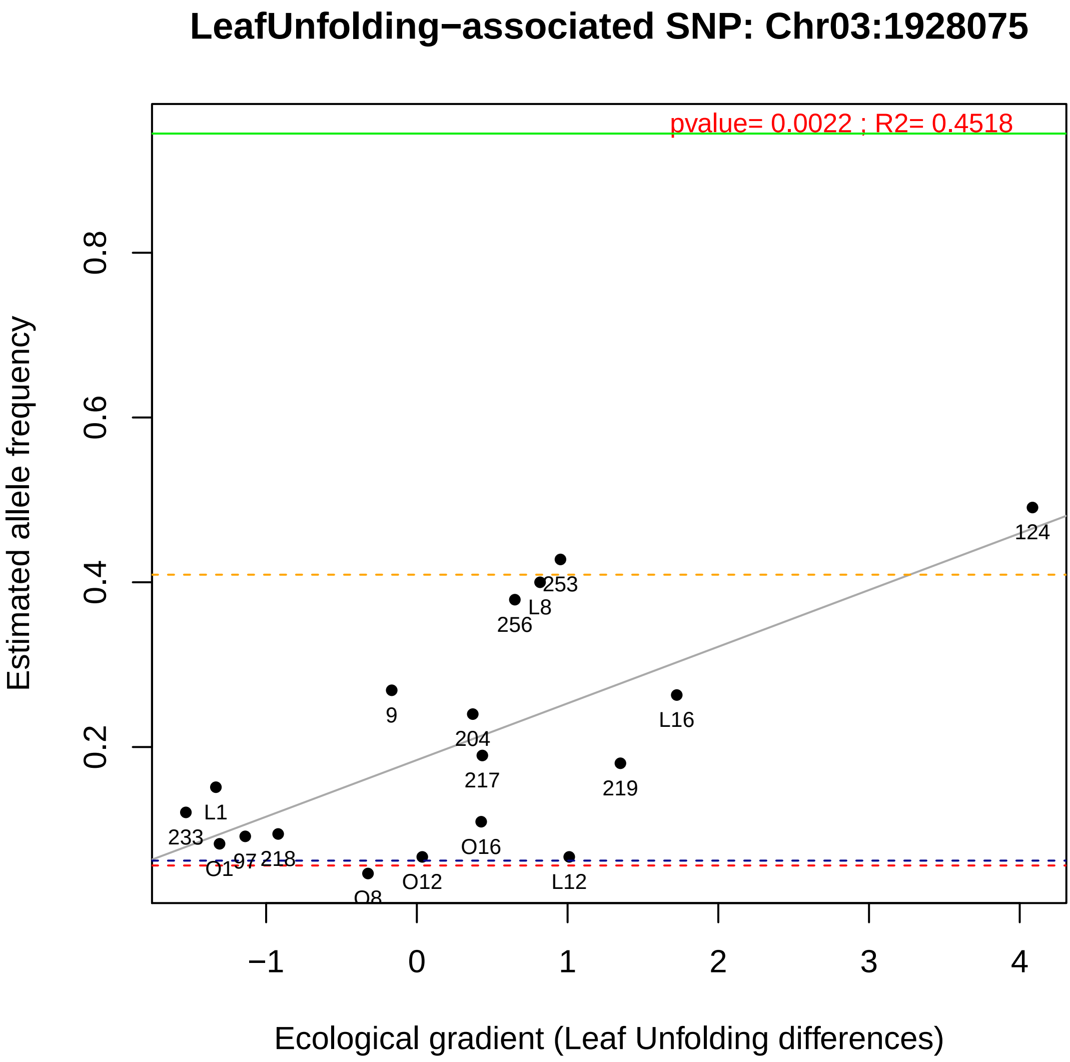
LeafUnfolding−associated SNP: Chr03:1928075
pvalue= 0.0022 ; R2= 0.4518
Estimated allele frequency
0.2
0.4
0.6
0.8
−1
0
1
2
3
4
Ecological gradient (Leaf Unfolding differences)
124
253
L8
256
9
204
L16
217
219
L1
233
O16
O1
97
218
O12
L12
O8

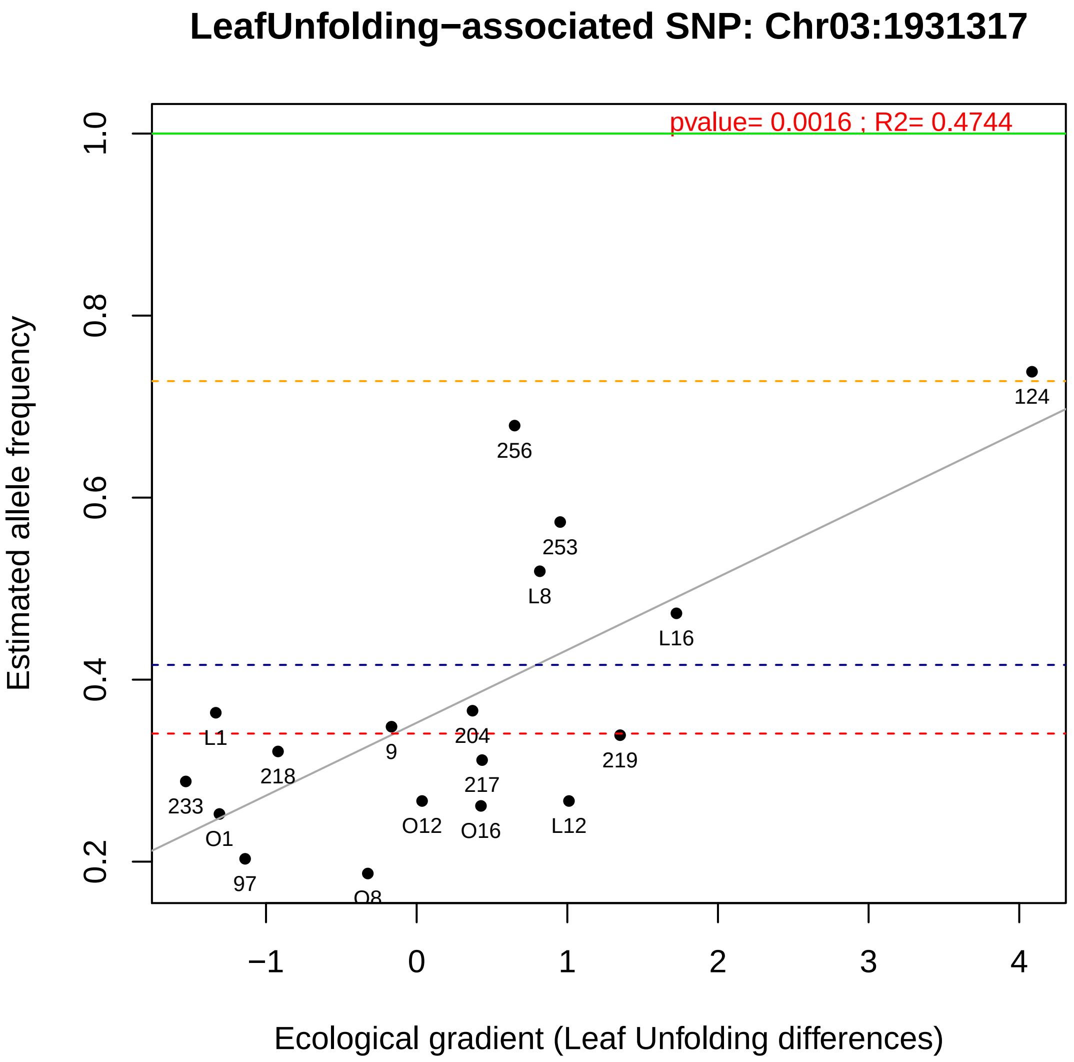

LeafUnfolding−associated SNP: Chr03:1931317
pvalue= 0.0016 ; R2= 0.4744
Estimated allele frequency
0.2
0.4
0.6
0.8
1.0
Ecological gradient (Leaf Unfolding differences)
−1
0
1
2
3
4
124
256
253
L8
L16
L1
218
9
204
219
217
233
O1
O12
O16
L12
97
O8

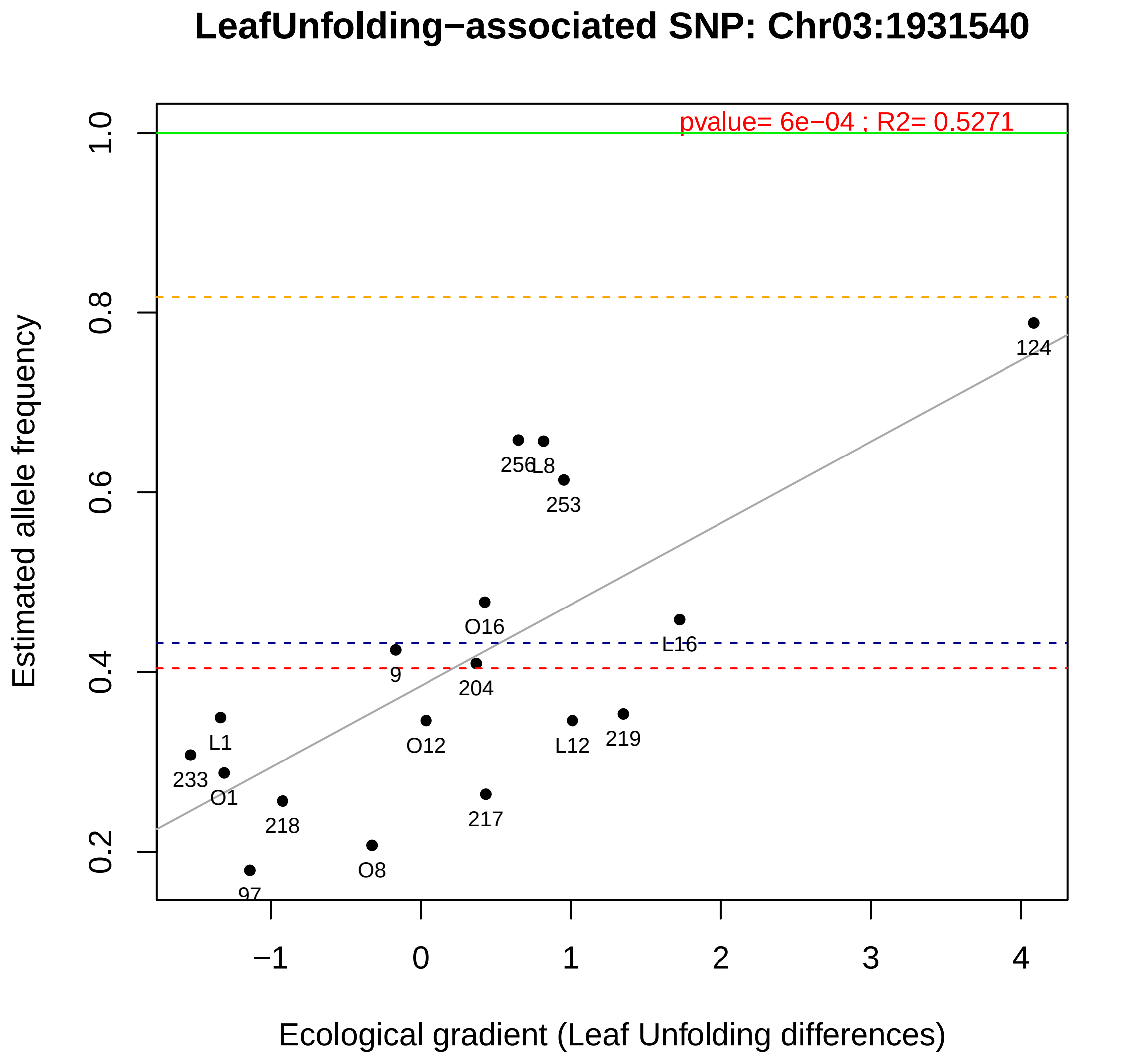
LeafUnfolding−associated SNP: Chr03:1931540
pvalue= 6e−04 ; R2= 0.5271
Estimated allele frequency
0.2
0.4
0.6
0.8
1.0
−1
0
1
2
3
4
Ecological gradient (Leaf Unfolding differences)
124
256
L8
253
O16
L16
9
204
L1
O12
L12
219
233
O1
218
217
97
O8

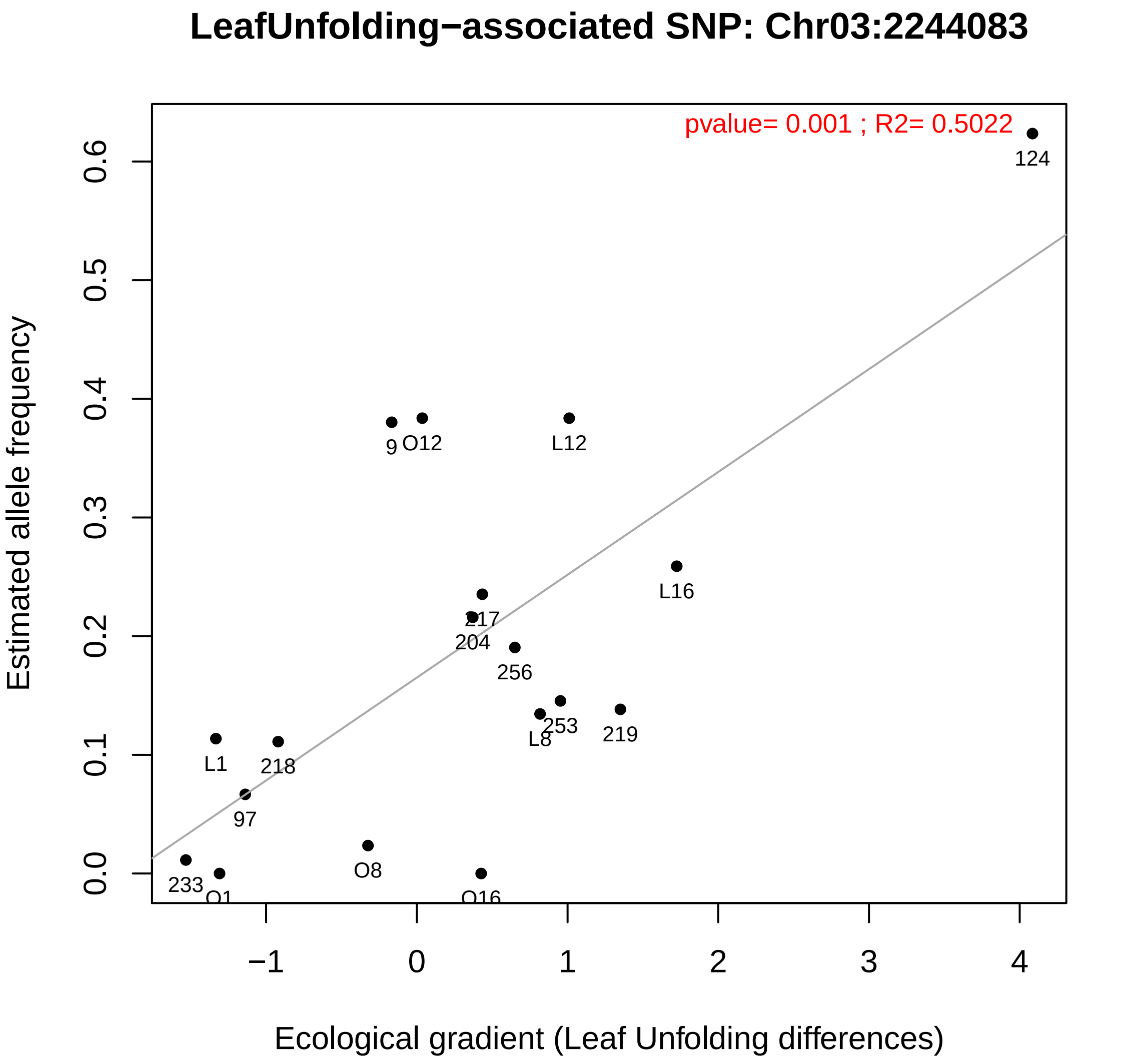
LeafUnfolding−associated SNP: Chr03:2244083
pvalue= 0.001 ; R2= 0.5022
Estimated allele frequency
Ecological gradient (Leaf Unfolding differences)
0.0
0.1
0.2
0.3
0.4
0.5
0.6
−1
0
1
2
3
4
124
9
O12
L12
L16
217
204
256
253
L8
219
L1
218
97
O8
233
O1
O16

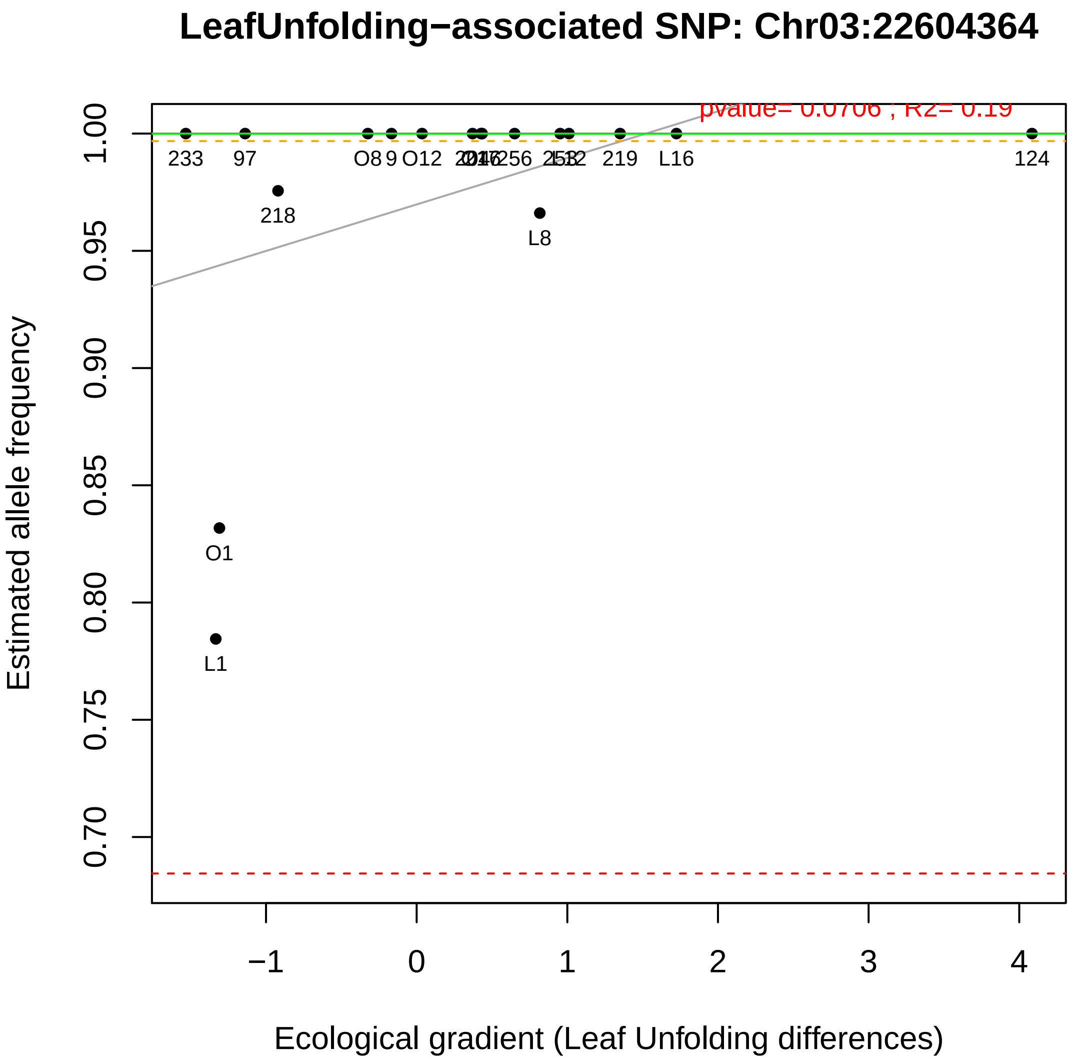
LeafUnfolding−associated SNP: Chr03:22604364
pvalue= 0.0706 ; R2= 0.19
Estimated allele frequency
Ecological gradient (Leaf Unfolding differences)
1.00
0.95
0.90
0.85
0.80
0.75
0.70
−1
0
1
2
3
4
233
97
218
O8
9
O12
L16
219
124
L8
O1
L1

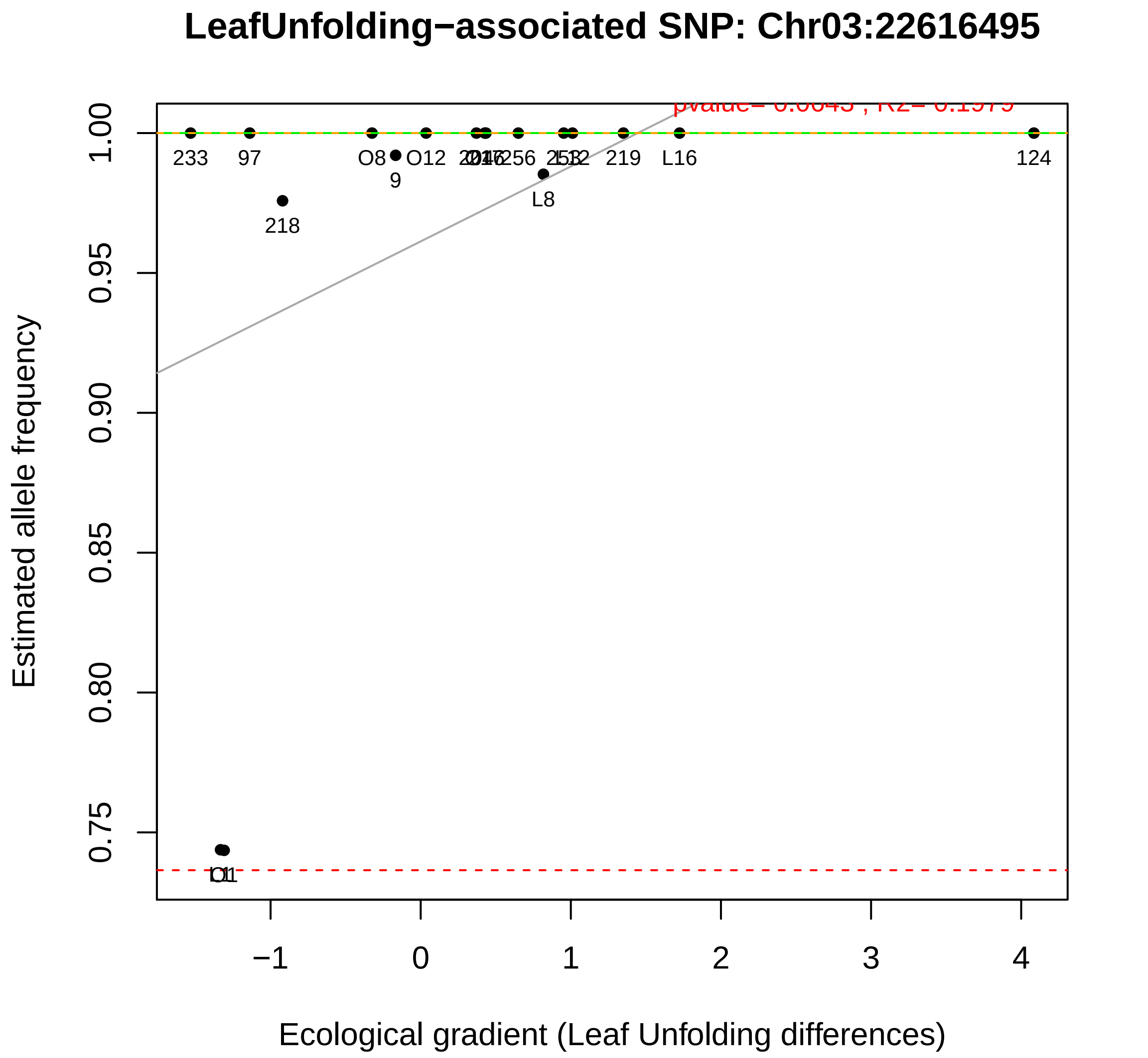

LeafUnfolding−associated SNP: Chr03:22616495
Estimated allele frequency
Ecological gradient (Leaf Unfolding differences)
1.00
0.95
0.90
0.85
0.80
0.75
−1
0
1
2
3
4
233
97
218
O8
9
O12
L8
219
L16
124

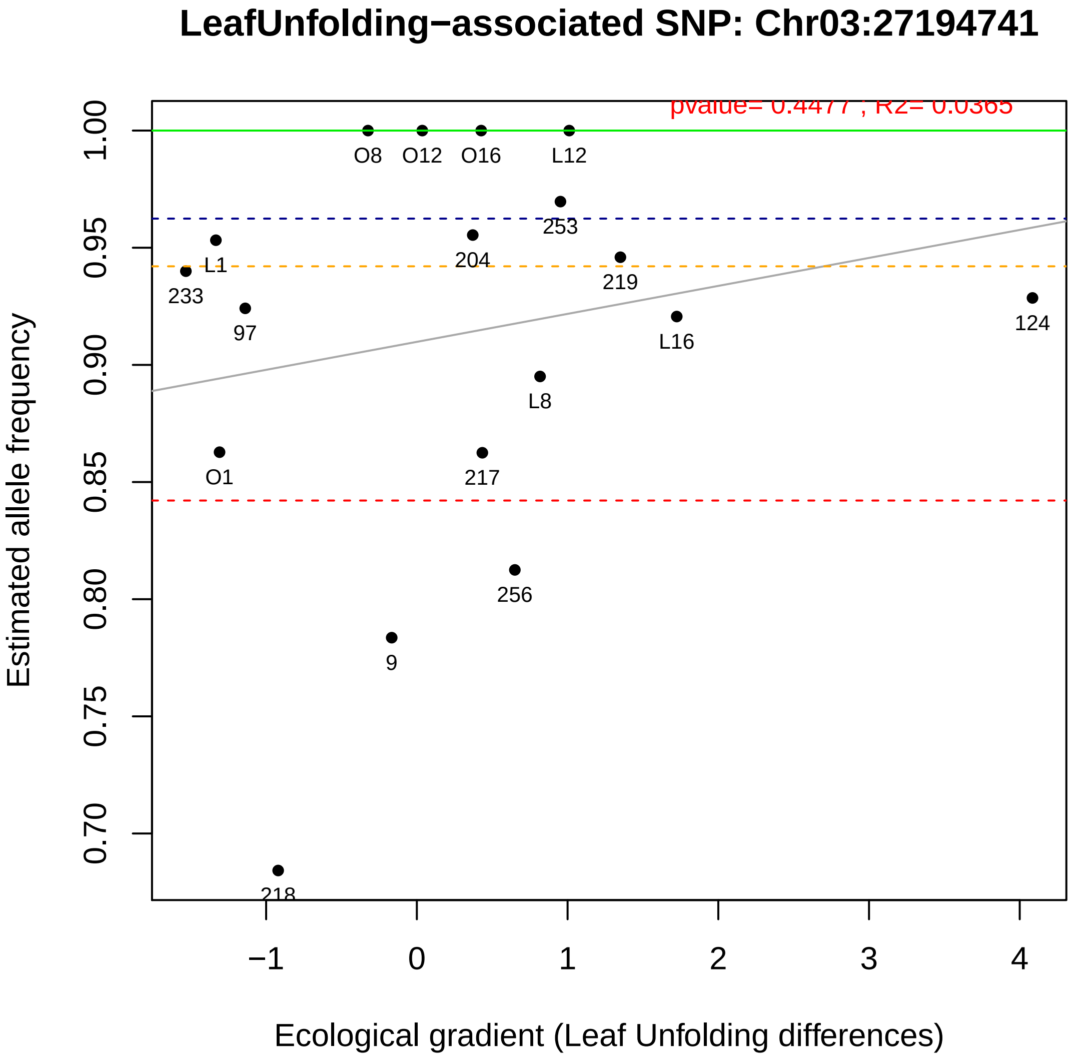

LeafUnfolding−associated SNP: Chr03:27194741
pvalue= 0.4477 ; R2= 0.0365
Estimated allele frequency
1.00
0.95
0.90
0.85
0.80
0.75
0.70
O8
O12
O16
L12
253
204
L1
233
219
97
L16
124
L8
O1
217
256
9
218
−1
0
1
2
3
4
Ecological gradient (Leaf Unfolding differences)

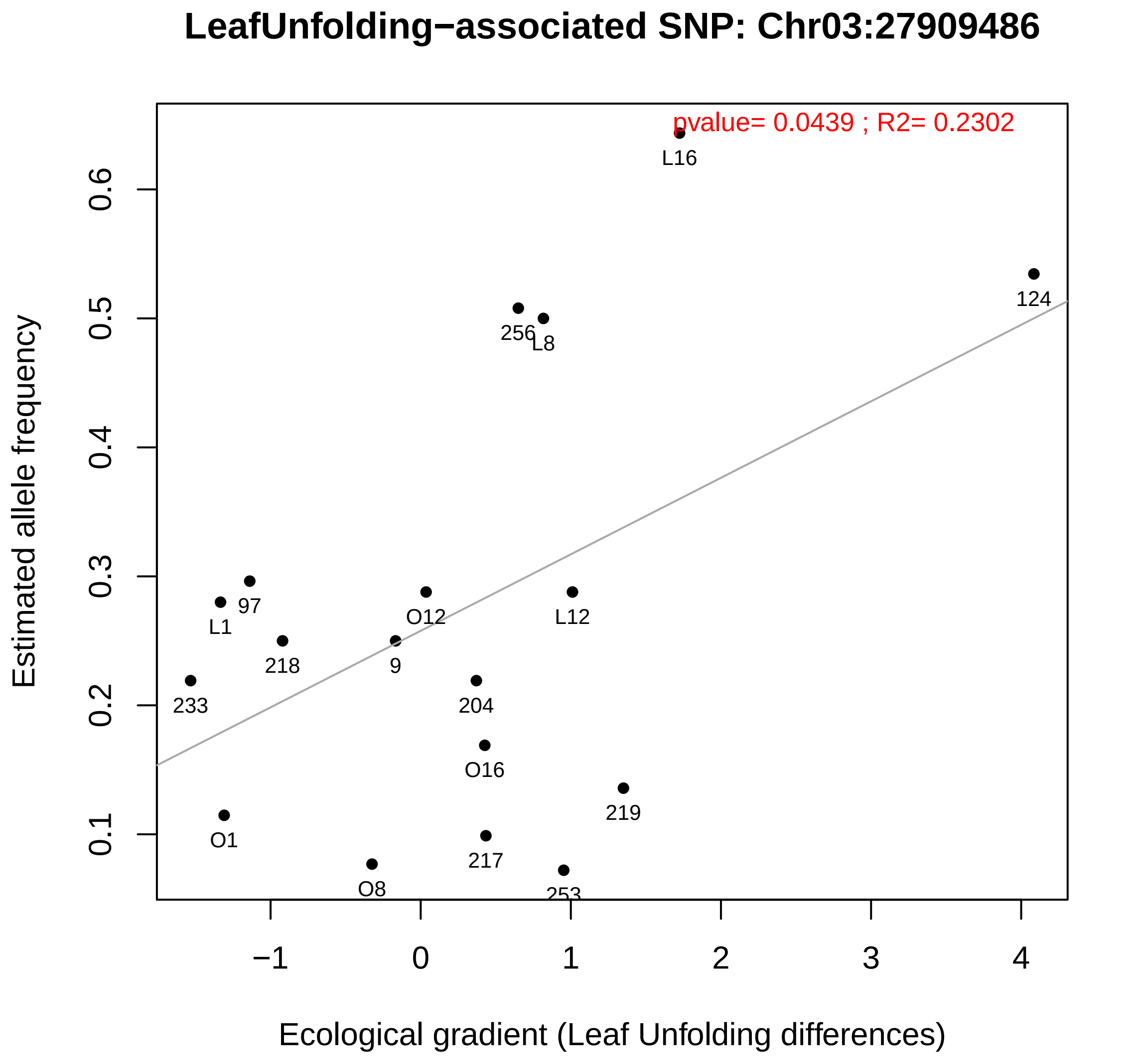
LeafUnfolding−associated SNP: Chr03:27909486
pvalue= 0.0439 ; R2= 0.2302
Estimated allele frequency
Ecological gradient (Leaf Unfolding differences)
0.1
0.2
0.3
0.4
0.5
0.6
−1
0
1
2
3
4
L16
124
256
L8
97
L1
O12
L12
218
9
233
204
O16
219
O1
217
O8
253

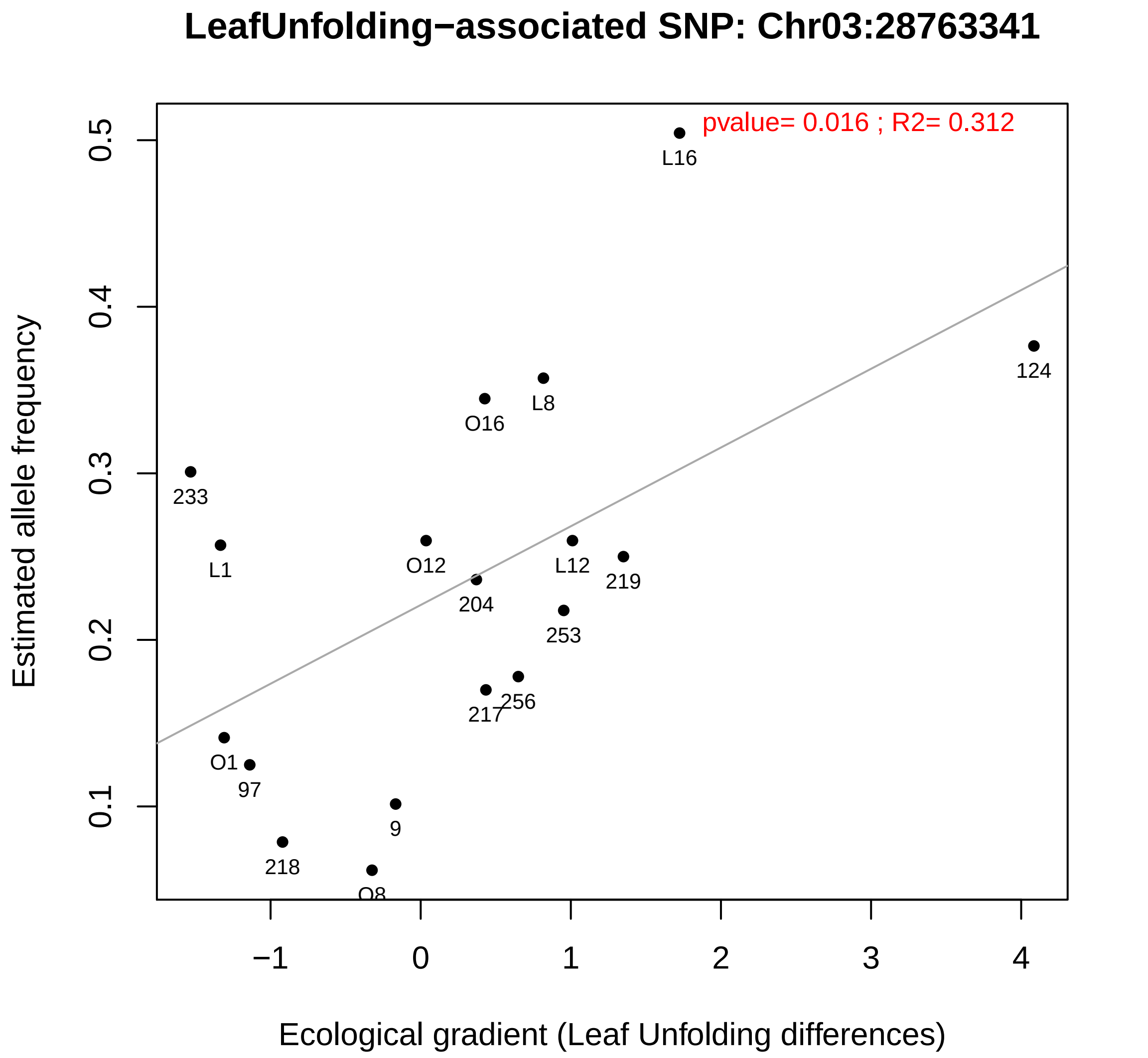
LeafUnfolding−associated SNP: Chr03:28763341
pvalue= 0.016 ; R2= 0.312
Estimated allele frequency
Ecological gradient (Leaf Unfolding differences)
0.1
0.2
0.3
0.4
0.5
−1
0
1
2
3
4
L16
124
L8
O16
233
L1
O12
204
L12
219
253
256
217
O1
97
9
218
O8

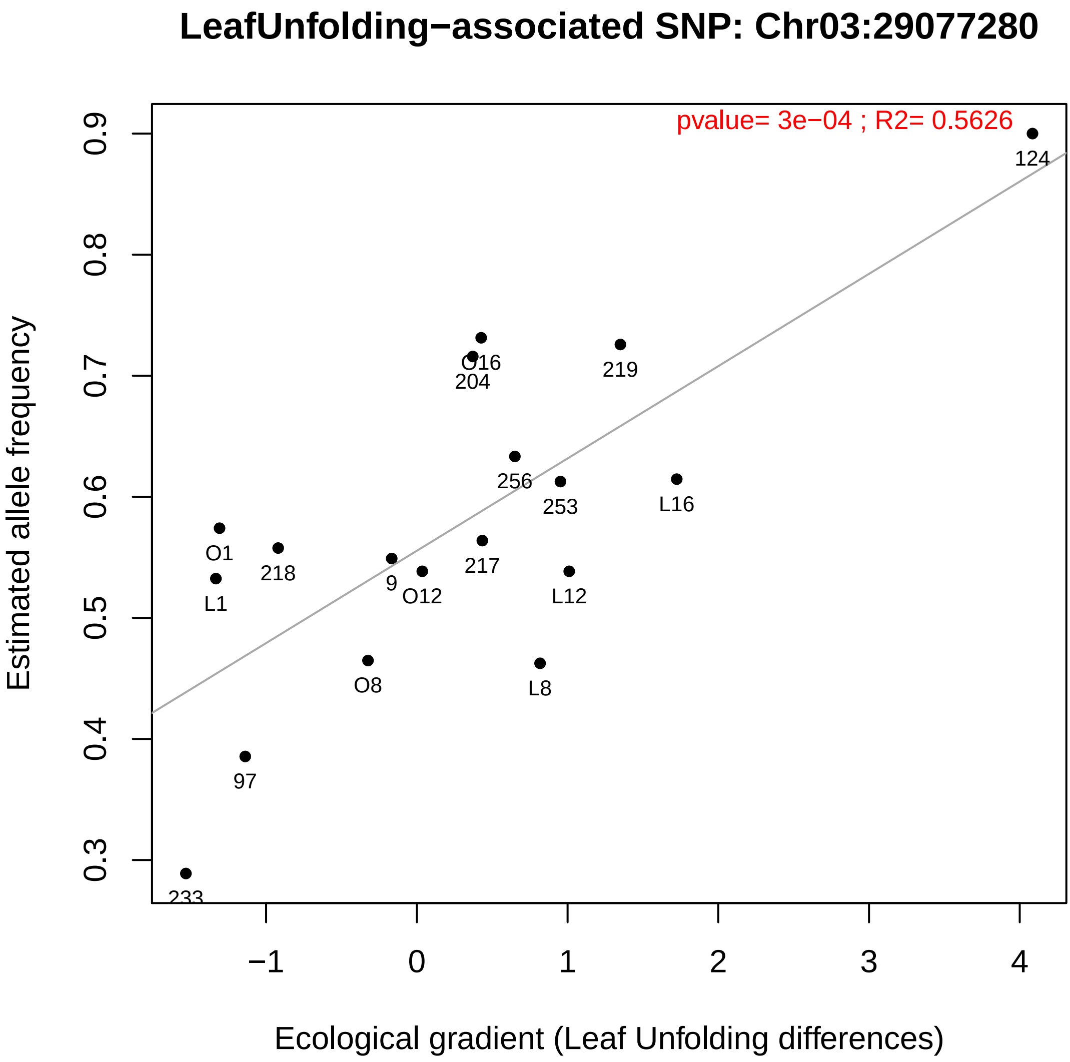
LeafUnfolding−associated SNP: Chr03:29077280
pvalue= 3e−04 ; R2= 0.5626
Estimated allele frequency
Ecological gradient (Leaf Unfolding differences)
0.3
0.4
0.5
0.6
0.7
0.8
0.9
−1
0
1
2
3
4
124
O16
204
219
256
253
L16
O1
218
9
O12
217
L12
L1
O8
L8
97
233

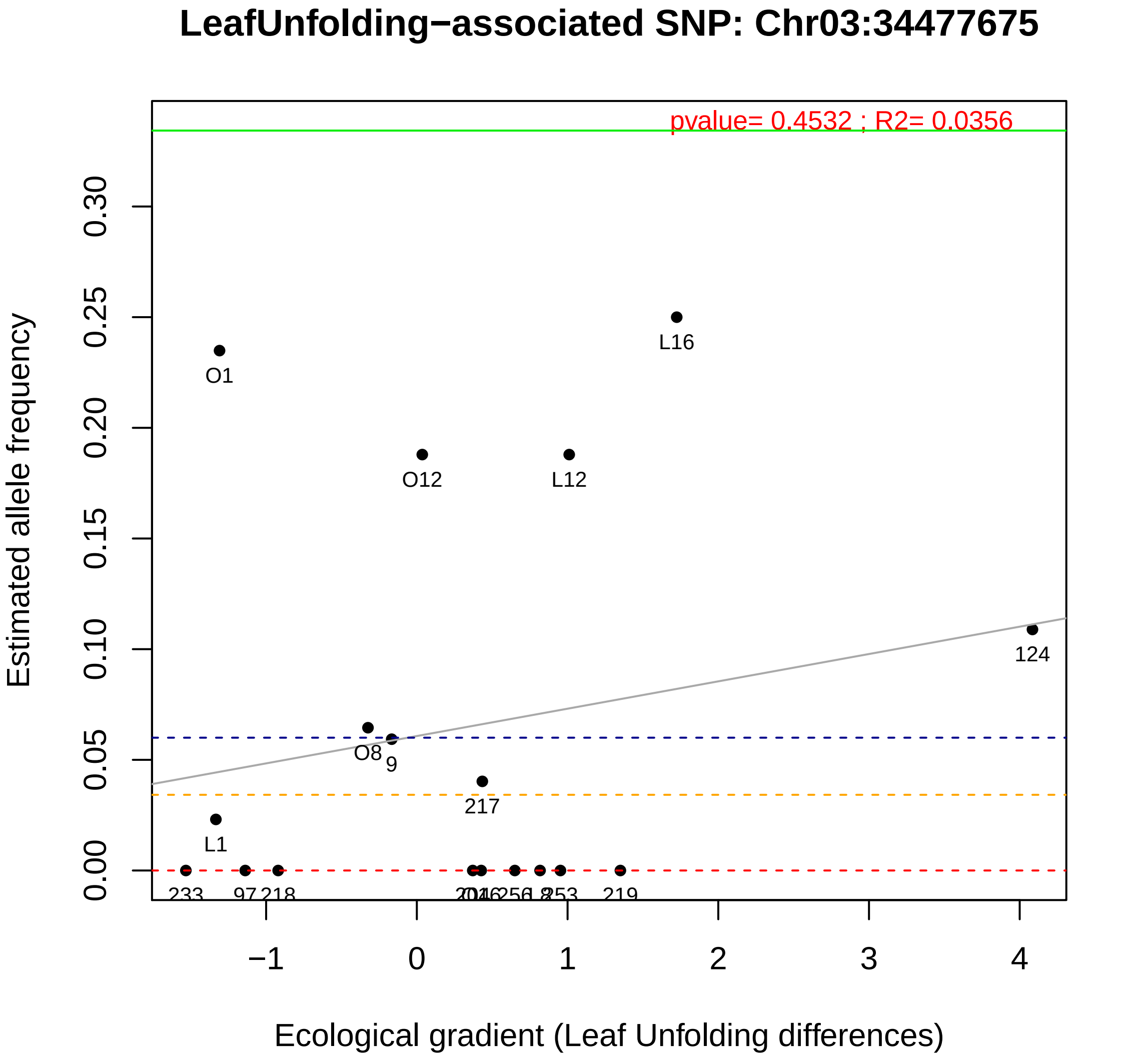
LeafUnfolding−associated SNP: Chr03:34477675
pvalue= 0.4532 ; R2= 0.0356
Estimated allele frequency
0.00
0.05
0.10
0.15
0.20
0.25
0.30
−1
0
1
2
3
4
Ecological gradient (Leaf Unfolding differences)
O1
L16
O12
L12
124
O8
9
217
L1
233
97
218
219

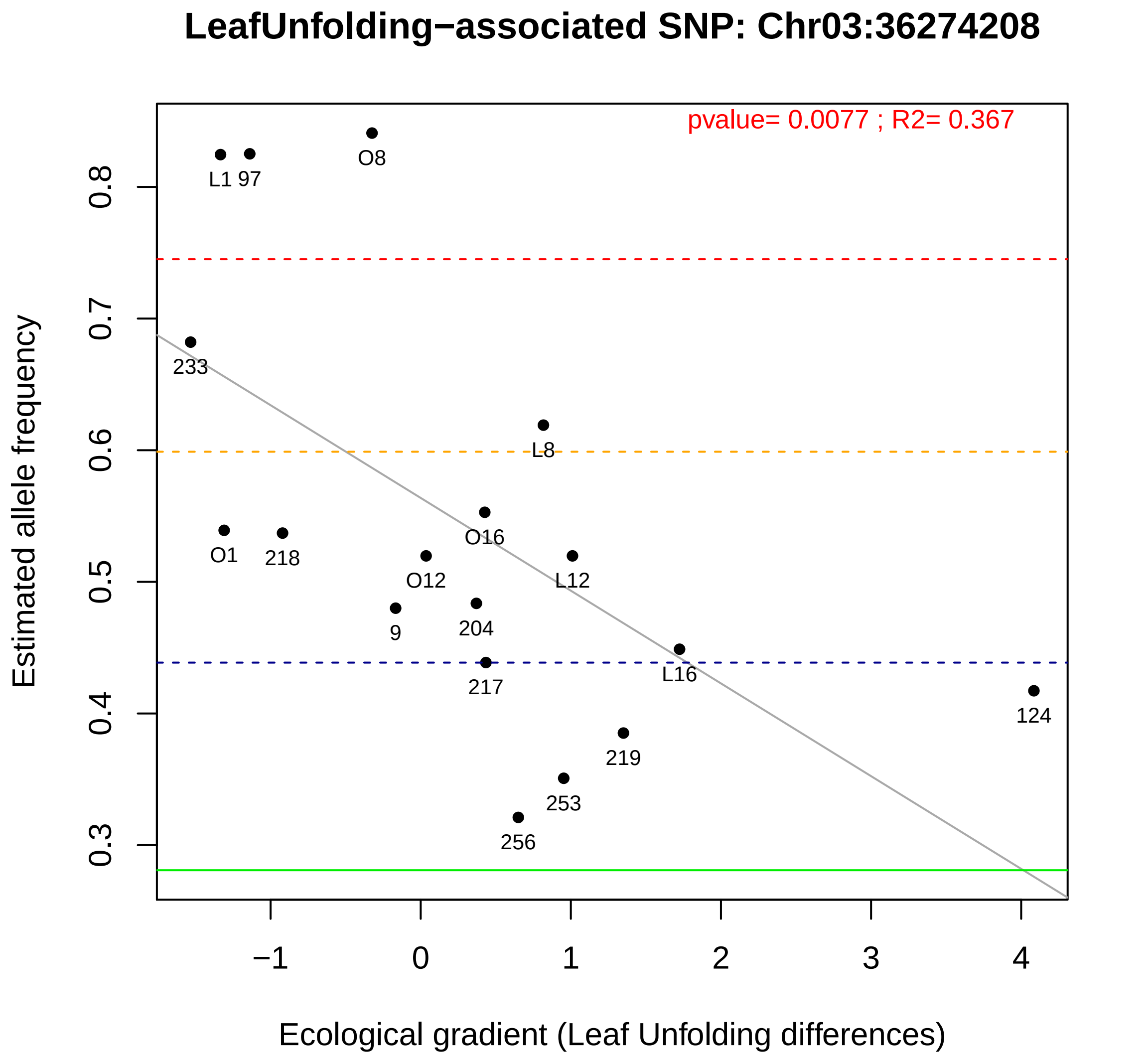

LeafUnfolding−associated SNP: Chr03:36274208
pvalue= 0.0077 ; R2= 0.367
Estimated allele frequency
Ecological gradient (Leaf Unfolding differences)
0.3
0.4
0.5
0.6
0.7
0.8
−1
0
1
2
3
4
L1
97
O8
233
L8
O16
O1
218
O12
L12
9
204
217
L16
124
219
253
256

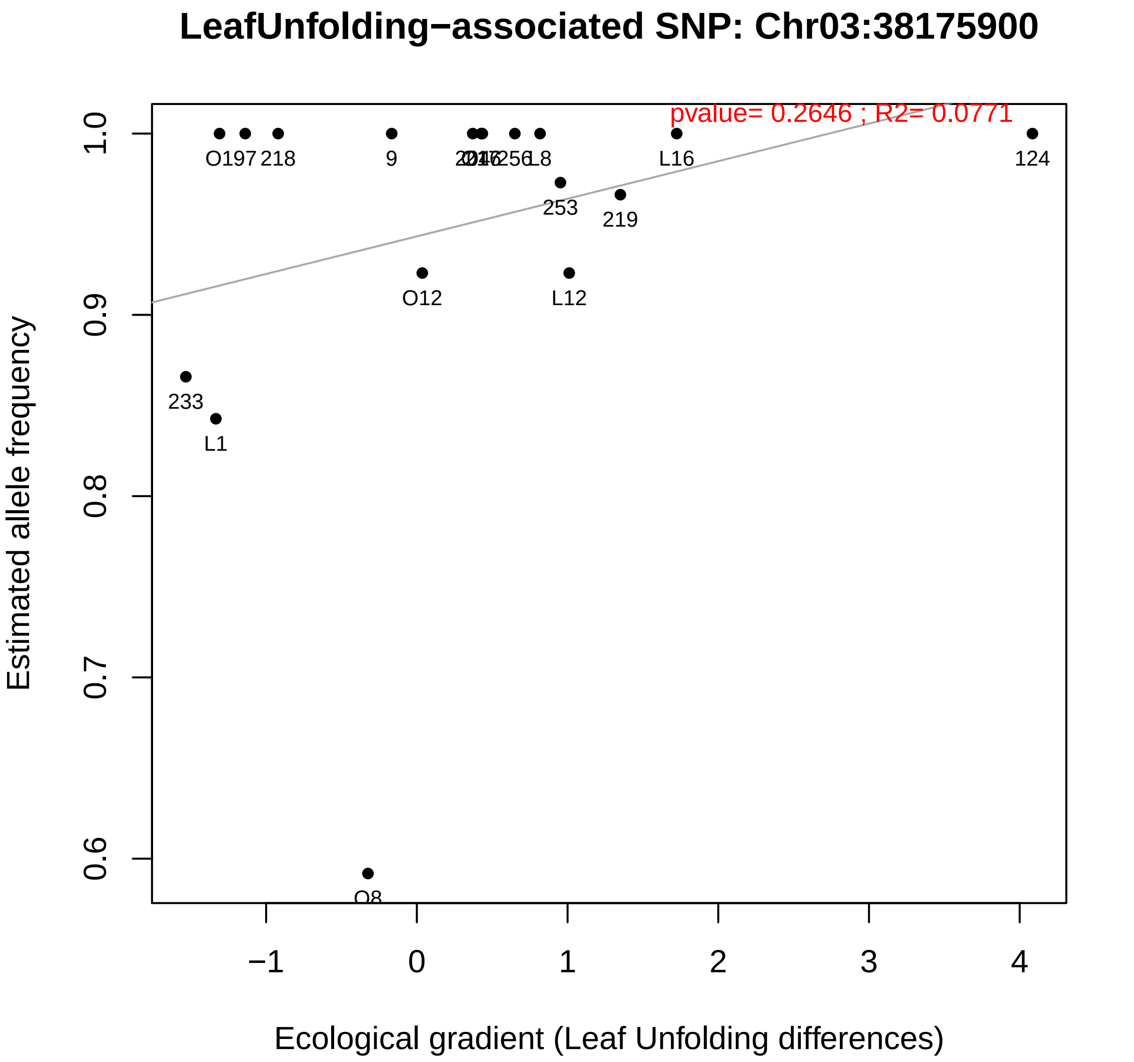
LeafUnfolding−associated SNP: Chr03:38175900
pvalue= 0.2646 ; R2= 0.0771
Estimated allele frequency
Ecological gradient (Leaf Unfolding differences)
0.6
0.7
0.8
0.9
1.0
−1
0
1
2
3
4
O197
218
9
L8
L16
124
253
219
O12
L12
233
L1

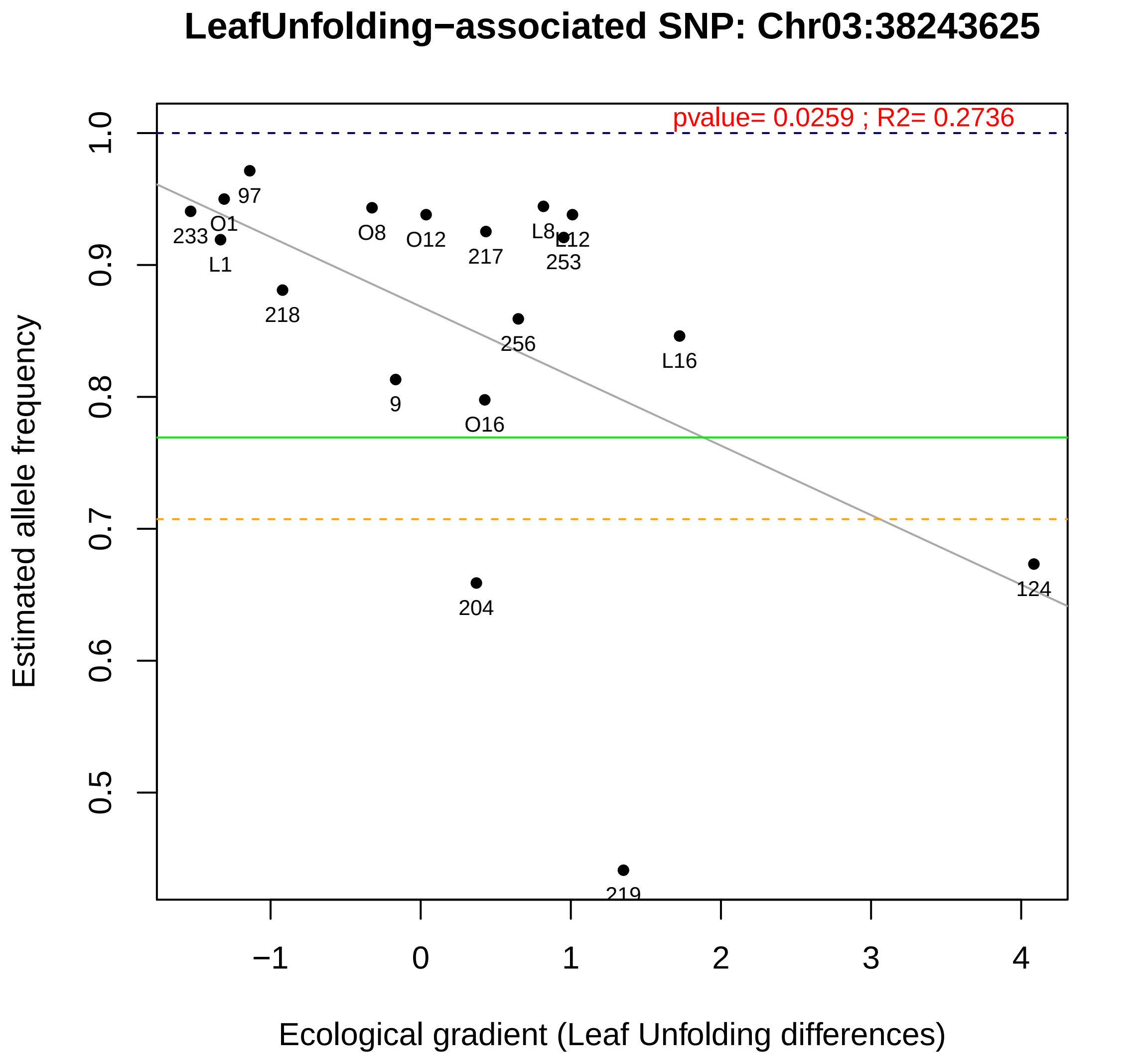

LeafUnfolding−associated SNP: Chr03:38243625
pvalue= 0.0259 ; R2= 0.2736
Estimated allele frequency
Ecological gradient (Leaf Unfolding differences)
1.0
0.9
0.8
0.7
0.6
0.5
−1
0
1
2
3
4
97
233
O1
L1
218
O8
O12
217
L8
L12
253
256
L16
9
O16
204
124
219

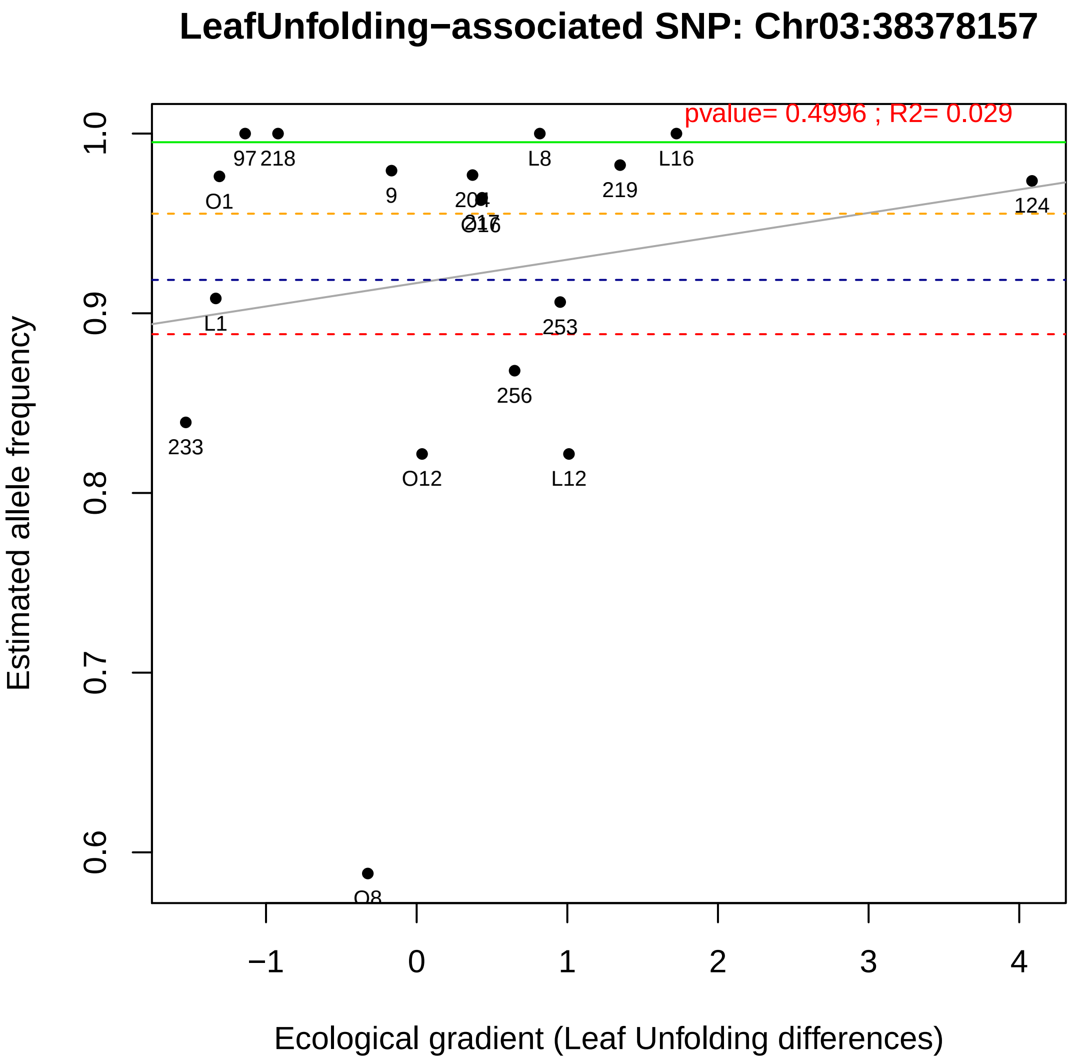

LeafUnfolding−associated SNP: Chr03:38378157
pvalue= 0.4996 ; R2= 0.029
Estimated allele frequency
0.6
0.7
0.8
0.9
1.0
Ecological gradient (Leaf Unfolding differences)
−1
0
1
2
3
4
97
218
L8
L16
O1
9
204
217
O16
219
124
L1
253
256
233
O12
L12
O8

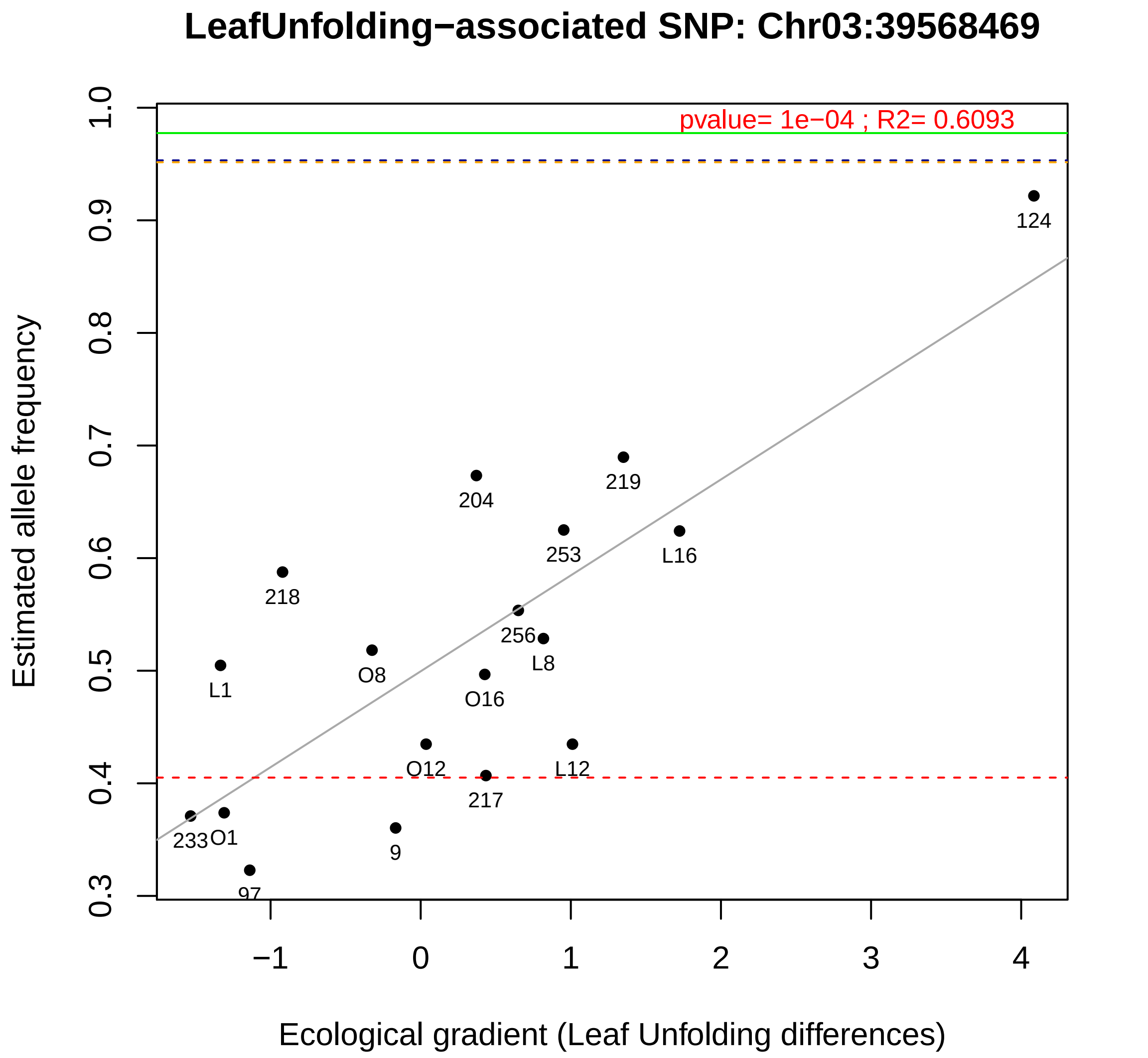

LeafUnfolding−associated SNP: Chr03:39568469
pvalue= 1e−04 ; R2= 0.6093
Estimated allele frequency
Ecological gradient (Leaf Unfolding differences)
0.3
0.4
0.5
0.6
0.7
0.8
0.9
1.0
−1
0
1
2
3
4
124
219
204
253
L16
218
256
L8
O8
L1
O16
O12
L12
217
233
O1
9
97

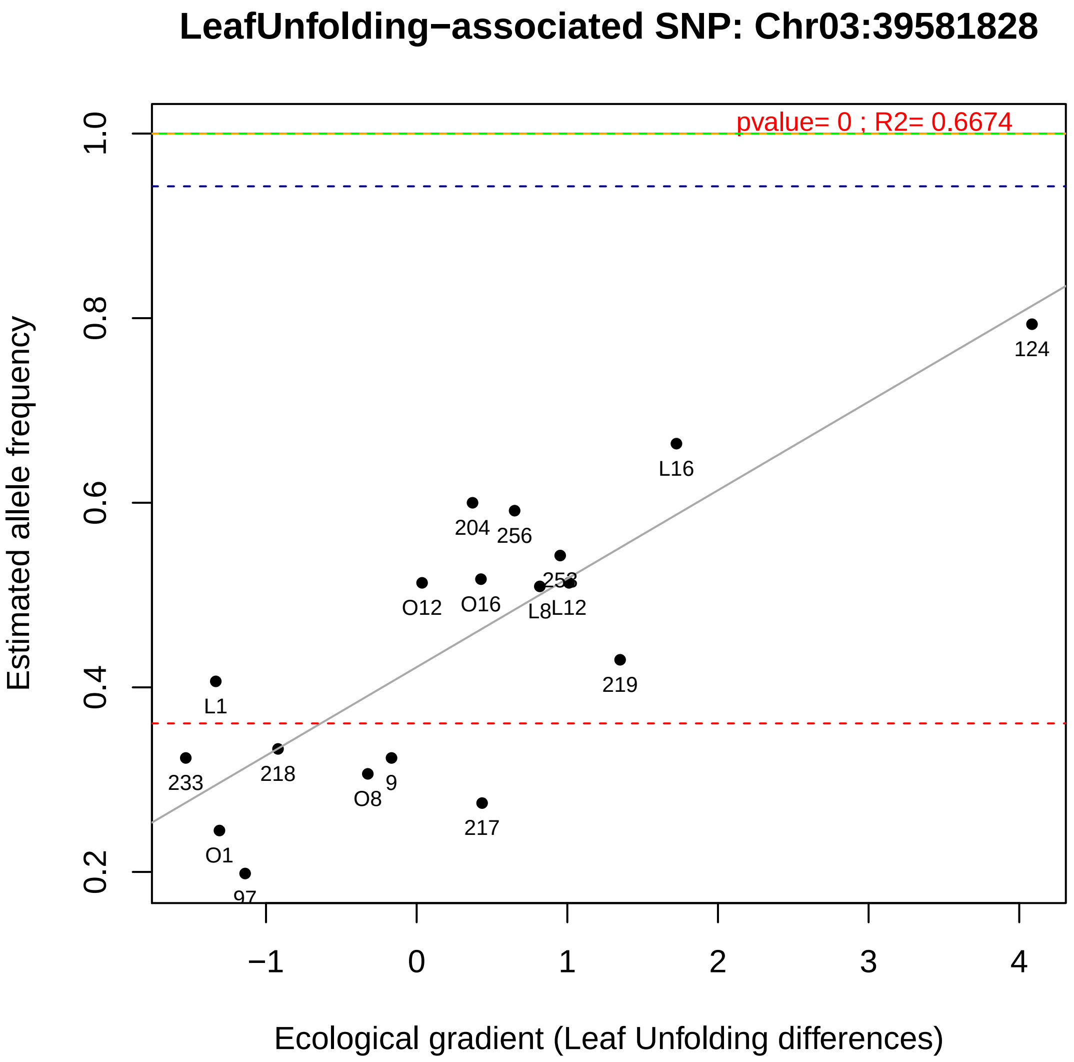
LeafUnfolding−associated SNP: Chr03:39581828
pvalue= 0 ; R2= 0.6674
Estimated allele frequency
Ecological gradient (Leaf Unfolding differences)
0.2
0.4
0.6
0.8
1.0
−1
0
1
2
3
4
124
L16
204
256
253
O12
O16
L8
L12
219
L1
233
218
O8
9
217
O1
97

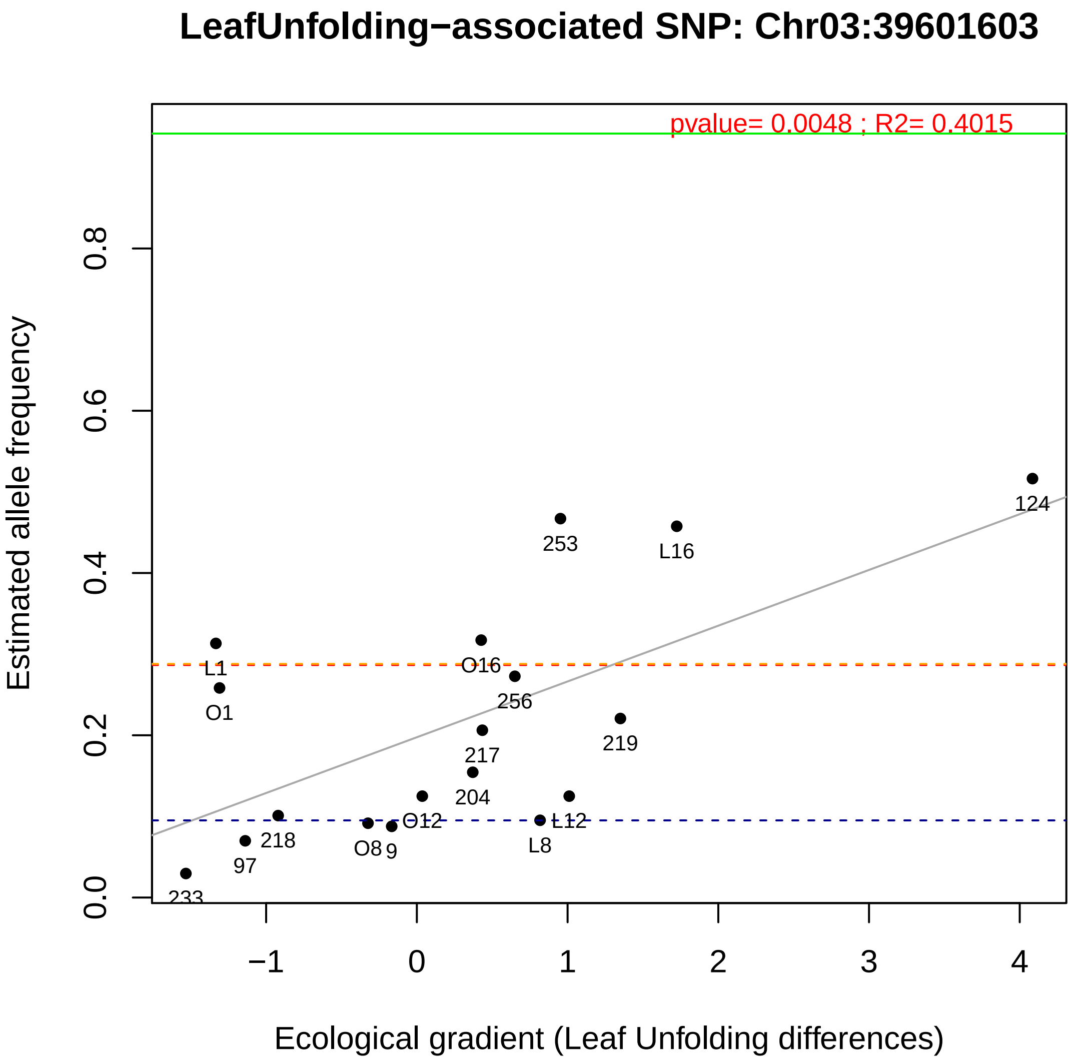

LeafUnfolding−associated SNP: Chr03:39601603
pvalue= 0.0048 ; R2= 0.4015
Estimated allele frequency
0.0
0.2
0.4
0.6
0.8
−1
0
1
2
3
4
Ecological gradient (Leaf Unfolding differences)
124
253
L16
L1
O16
256
O1
219
217
204
O12
L12
218
O8
9
L8
97
233

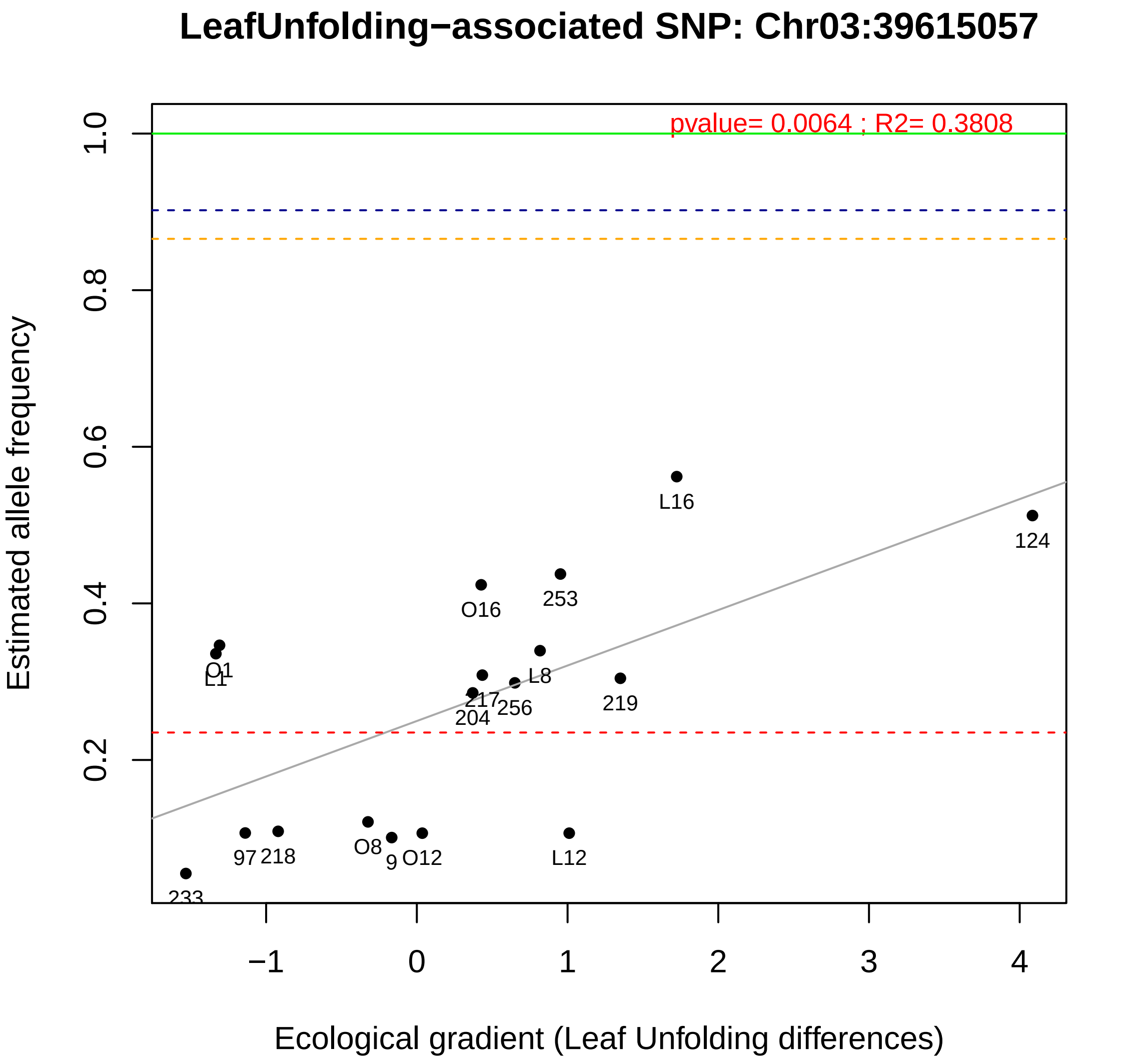

LeafUnfolding−associated SNP: Chr03:39615057
pvalue= 0.0064 ; R2= 0.3808
Estimated allele frequency
0.2
0.4
0.6
0.8
1.0
Ecological gradient (Leaf Unfolding differences)
−1
0
1
2
3
4
L16
124
O16
253
O1
L1
L8
217
256
204
219
97
218
O8
9
O12
L12
233

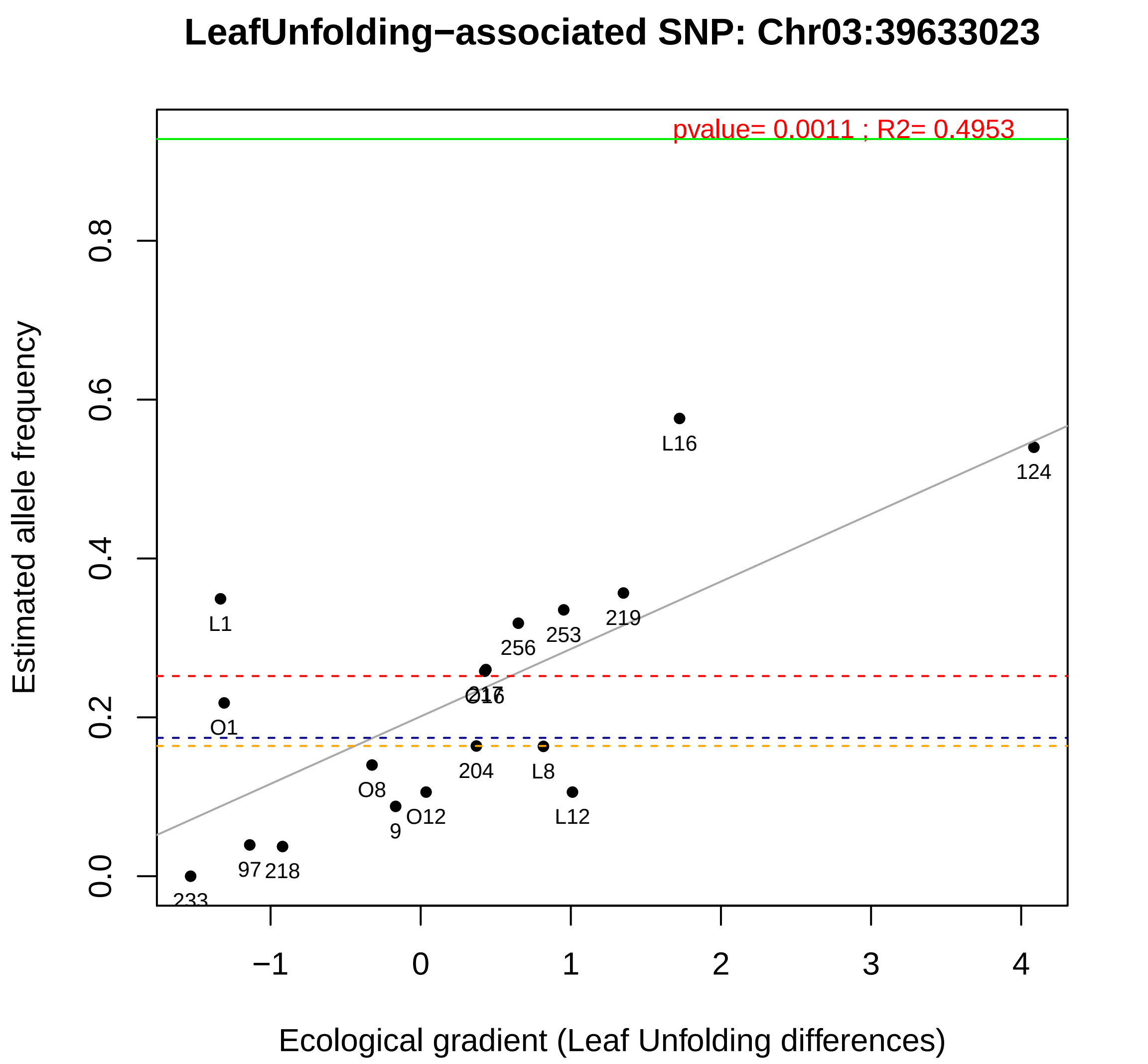
LeafUnfolding−associated SNP: Chr03:39633023
pvalue= 0.0011 ; R2= 0.4953
Estimated allele frequency
0.0
0.2
0.4
0.6
0.8
Ecological gradient (Leaf Unfolding differences)
−1
0
1
2
3
4
L16
124
L1
219
253
256
O1
204
L8
O8
O12
L12
9
97
218

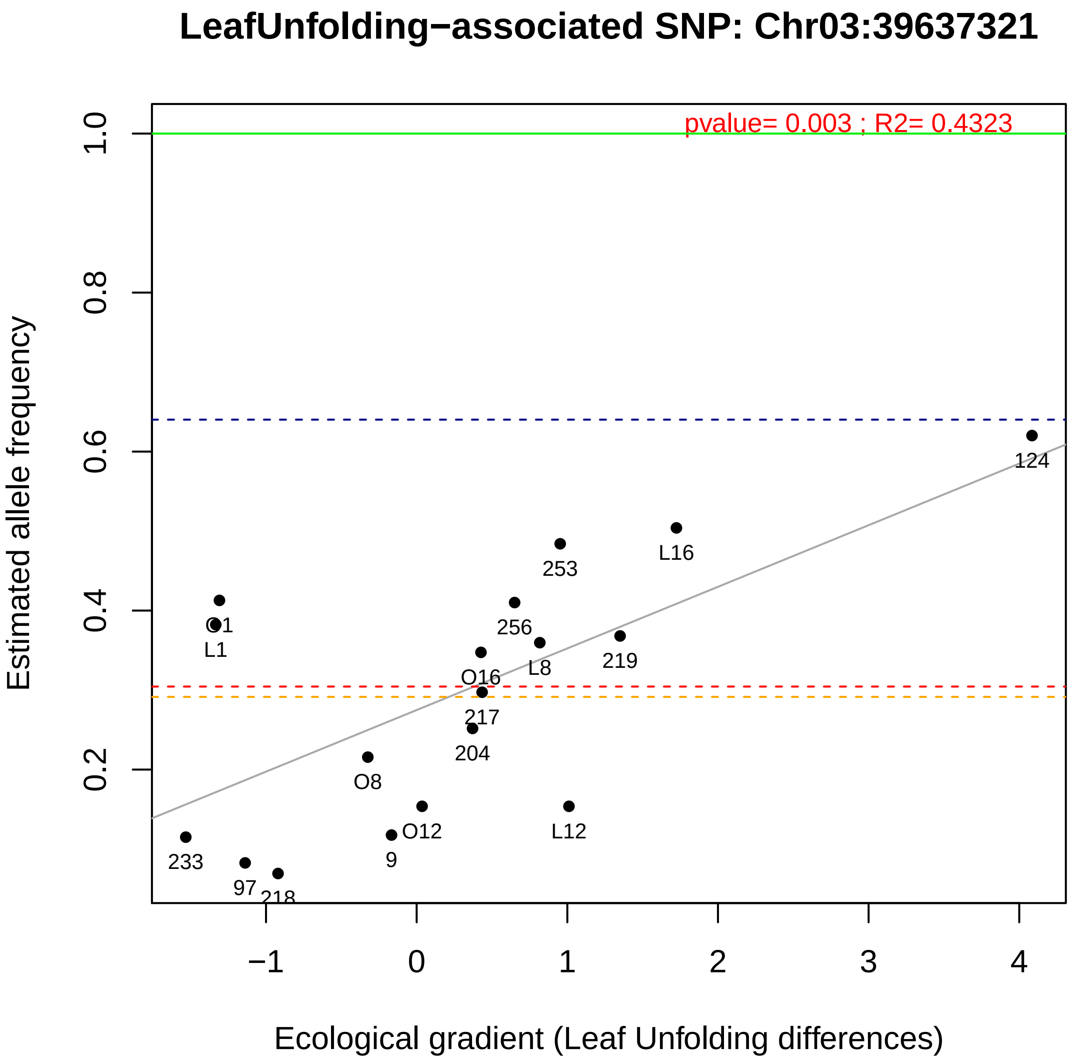
LeafUnfolding−associated SNP: Chr03:39637321
pvalue= 0.003 ; R2= 0.4323
Estimated allele frequency
Ecological gradient (Leaf Unfolding differences)
1.0
0.8
0.6
0.4
0.2
−1
0
1
2
3
4
124
L16
253
256
O1
L1
L8
219
O16
217
204
O8
O12
L12
233
9
97
218

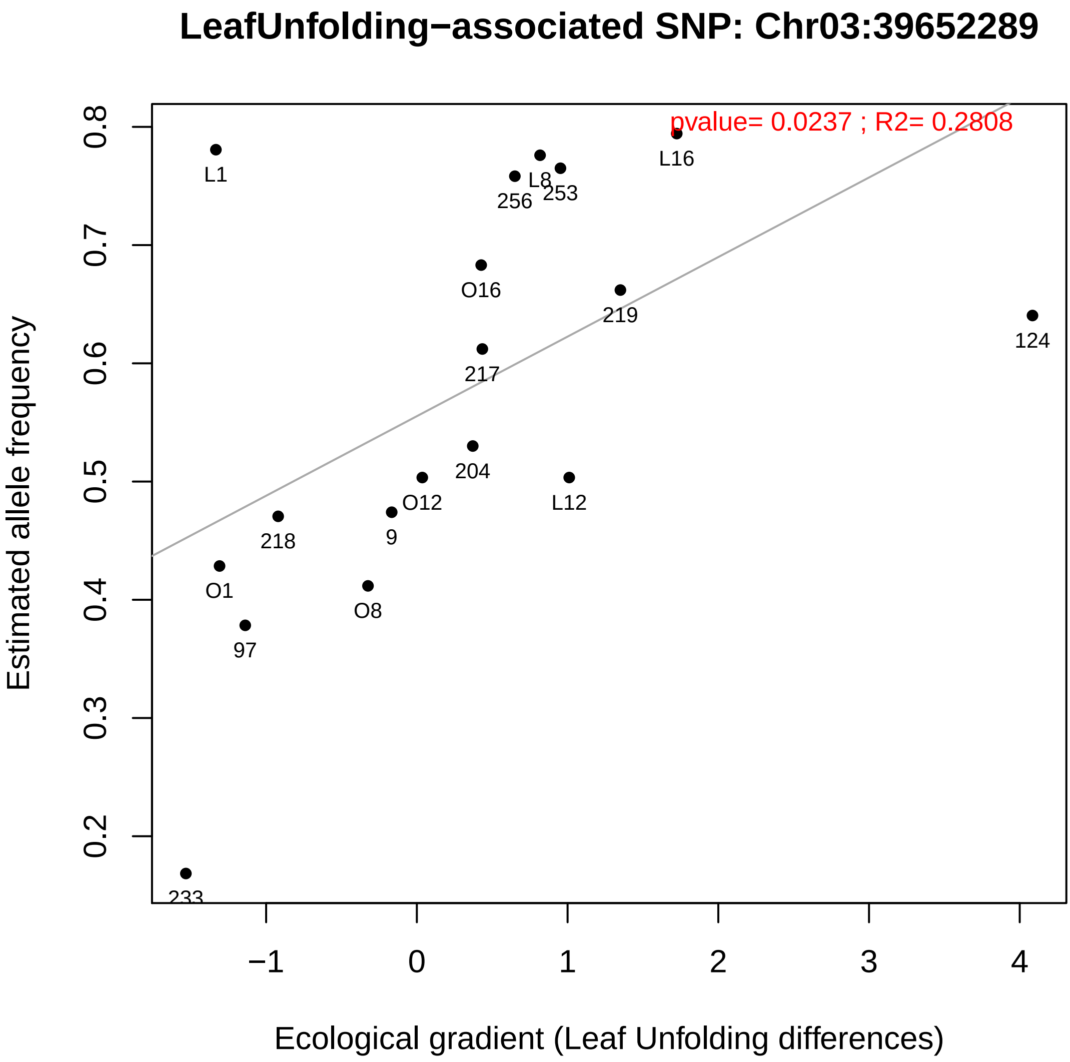

LeafUnfolding−associated SNP: Chr03:39652289
pvalue= 0.0237 ; R2= 0.2808
Estimated allele frequency
0.2
0.3
0.4
0.5
0.6
0.7
0.8
Ecological gradient (Leaf Unfolding differences)
−1
0
1
2
3
4
L1
L8
256
253
L16
O16
219
124
217
204
L12
O12
218
9
O1
O8
97
233

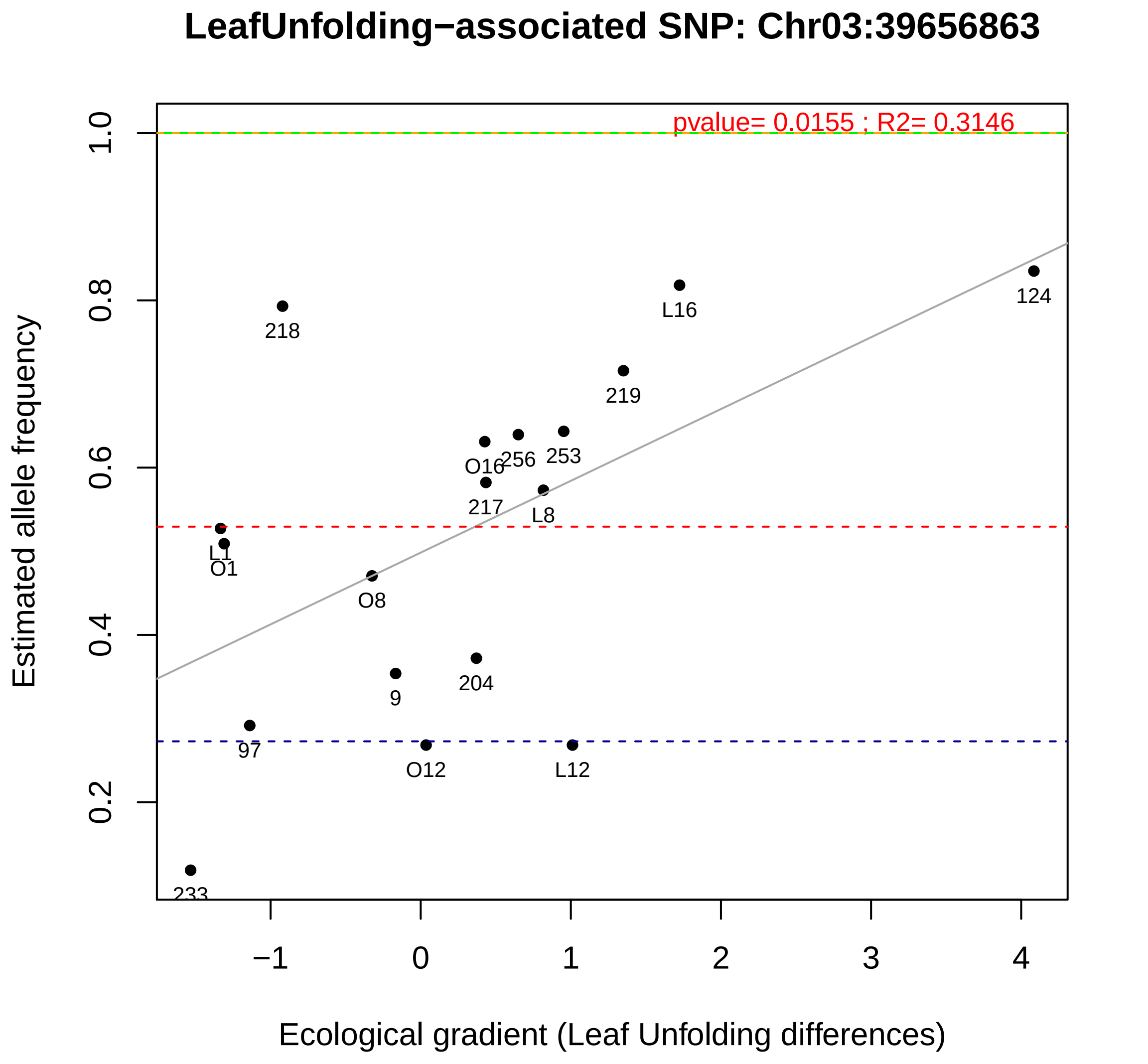

LeafUnfolding−associated SNP: Chr03:39656863
pvalue= 0.0155 ; R2= 0.3146
Estimated allele frequency
Ecological gradient (Leaf Unfolding differences)
1.0
0.8
0.6
0.4
0.2
−1
0
1
2
3
4
218
L16
124
219
O16
256
253
217
L8
L1
O1
O8
9
204
97
O12
L12
233

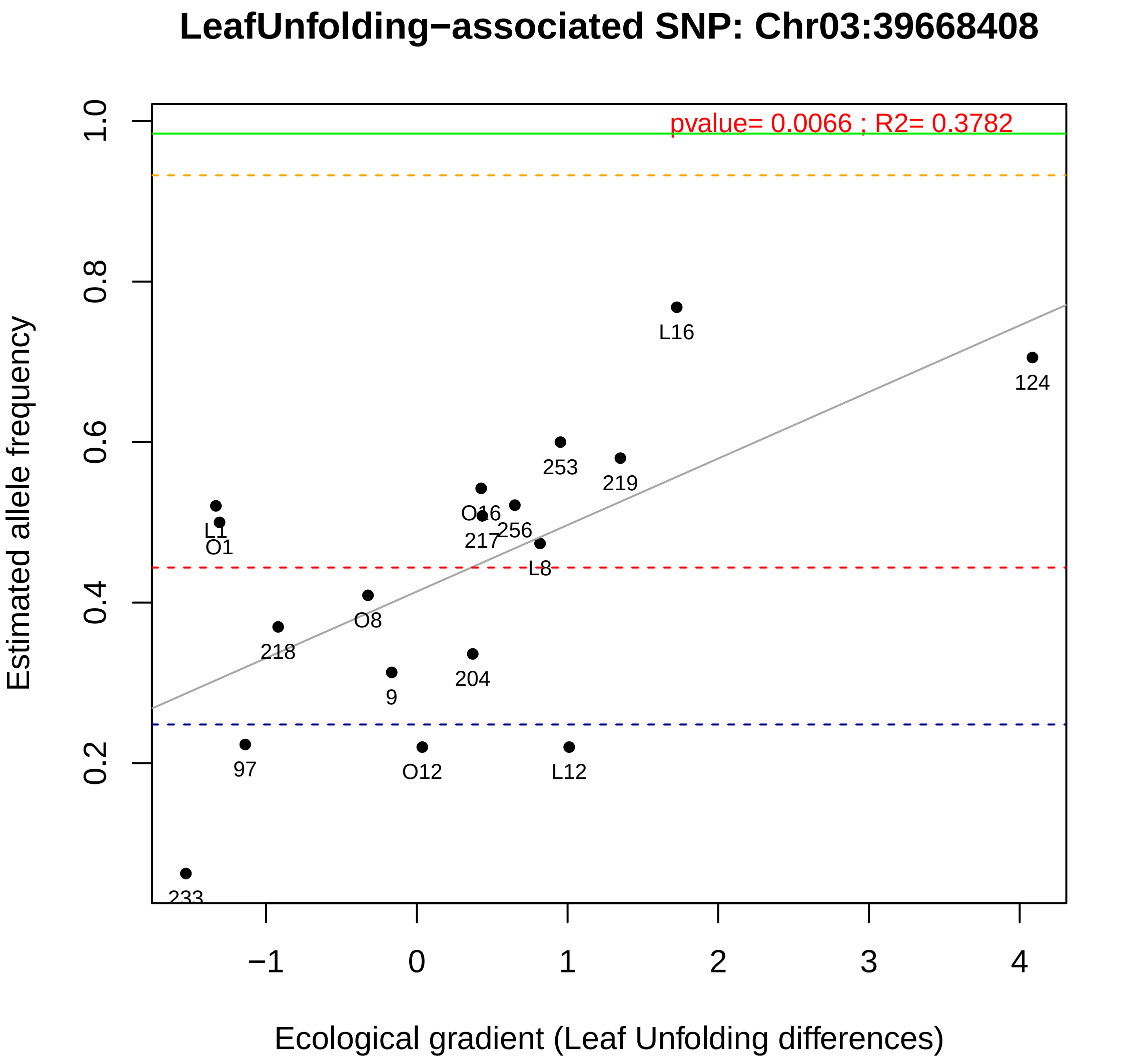
LeafUnfolding−associated SNP: Chr03:39668408
pvalue= 0.0066 ; R2= 0.3782
Estimated allele frequency
Ecological gradient (Leaf Unfolding differences)
0.2
0.4
0.6
0.8
1.0
−1
0
1
2
3
4
L16
124
253
219
O16
256
217
L1
O1
L8
O8
218
204
9
97
O12
L12
233

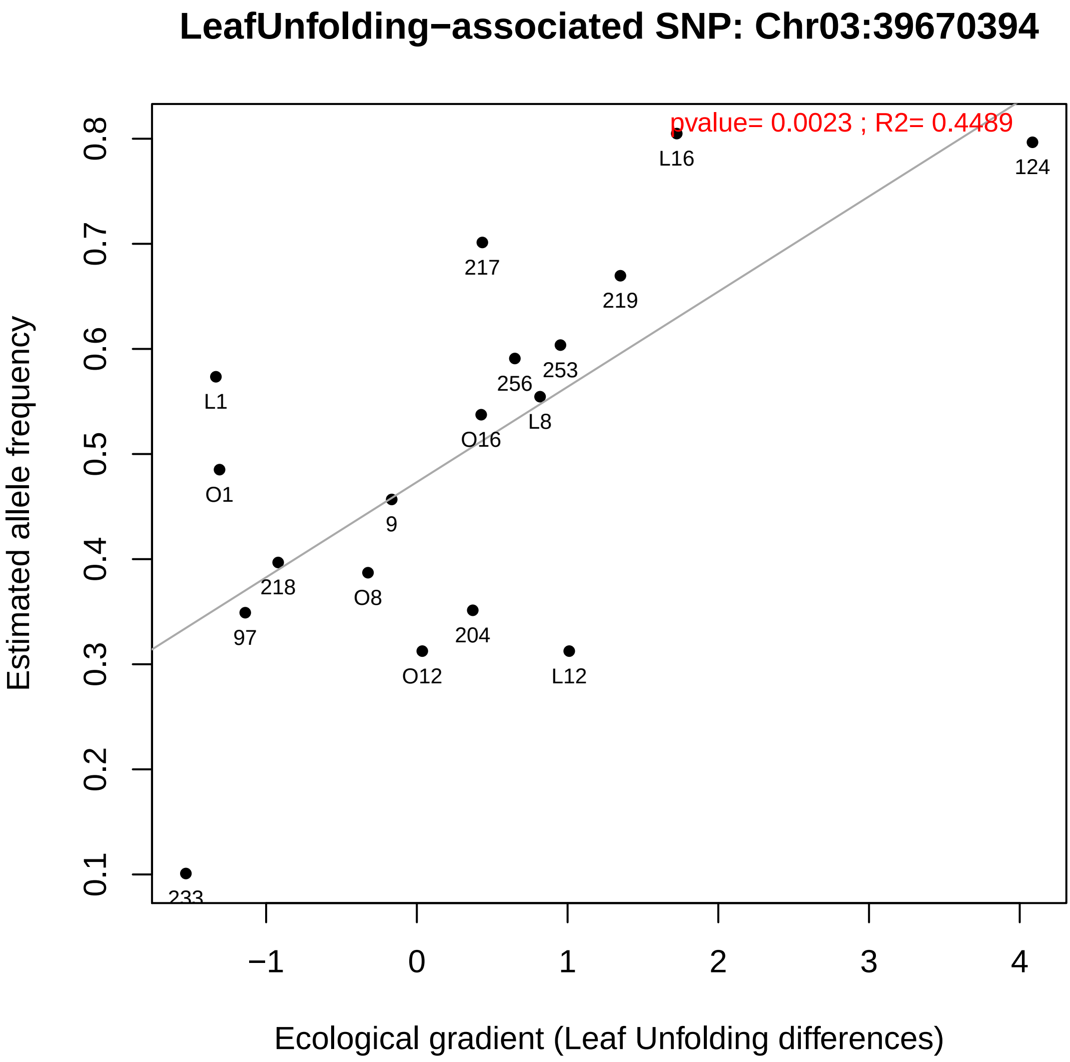

LeafUnfolding−associated SNP: Chr03:39670394
pvalue= 0.0023 ; R2= 0.4489
Estimated allele frequency
Ecological gradient (Leaf Unfolding differences)
L16
124
217
219
253
256
L8
L1
O16
O1
9
218
O8
97
204
O12
L12
233
0.1
0.2
0.3
0.4
0.5
0.6
0.7
0.8
−1
0
1
2
3
4

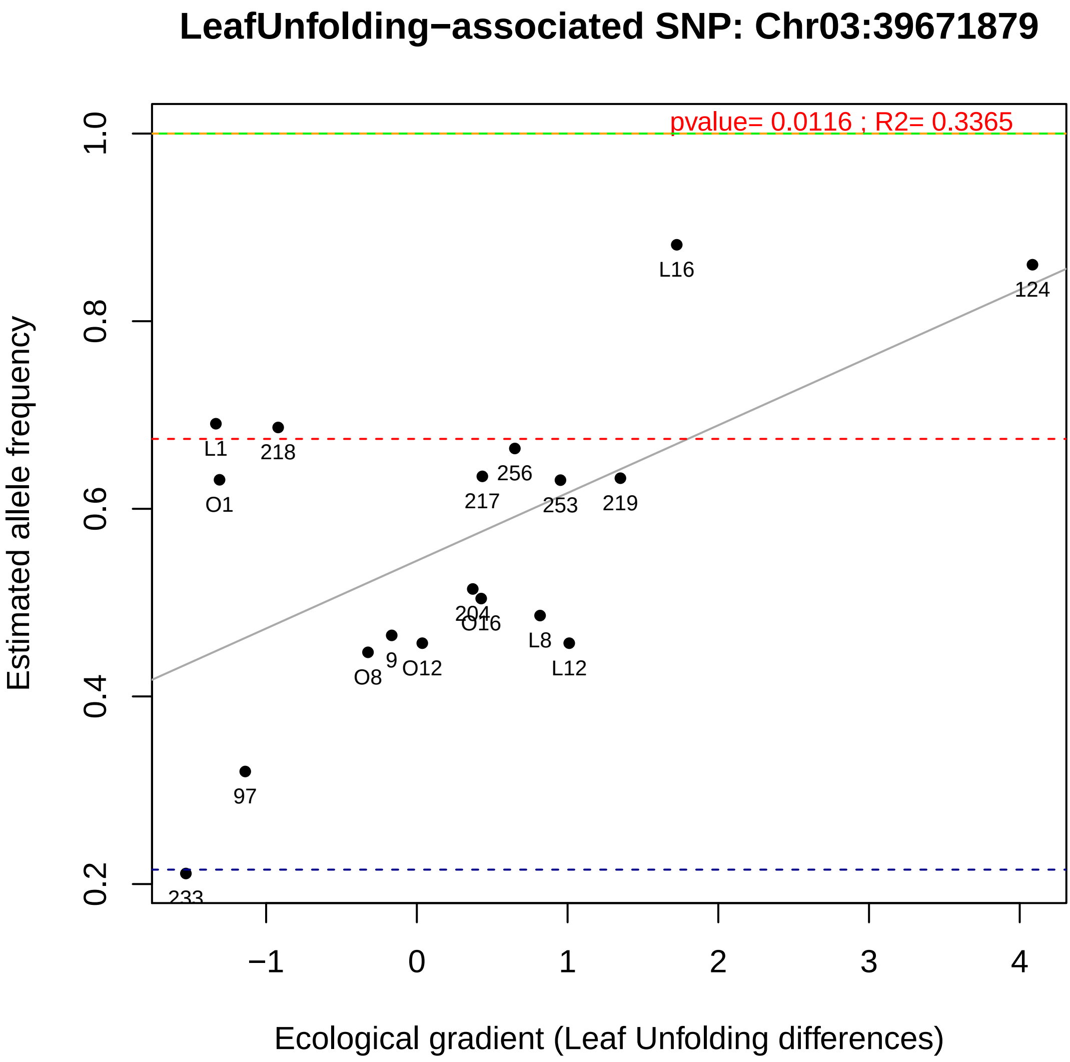

LeafUnfolding−associated SNP: Chr03:39671879
pvalue= 0.0116 ; R2= 0.3365
Estimated allele frequency
Ecological gradient (Leaf Unfolding differences)
0.2
0.4
0.6
0.8
1.0
−1
0
1
2
3
4
L16
124
L1
218
O1
256
217
253
219
204
O16
L8
L12
9
O8
O12
97
233

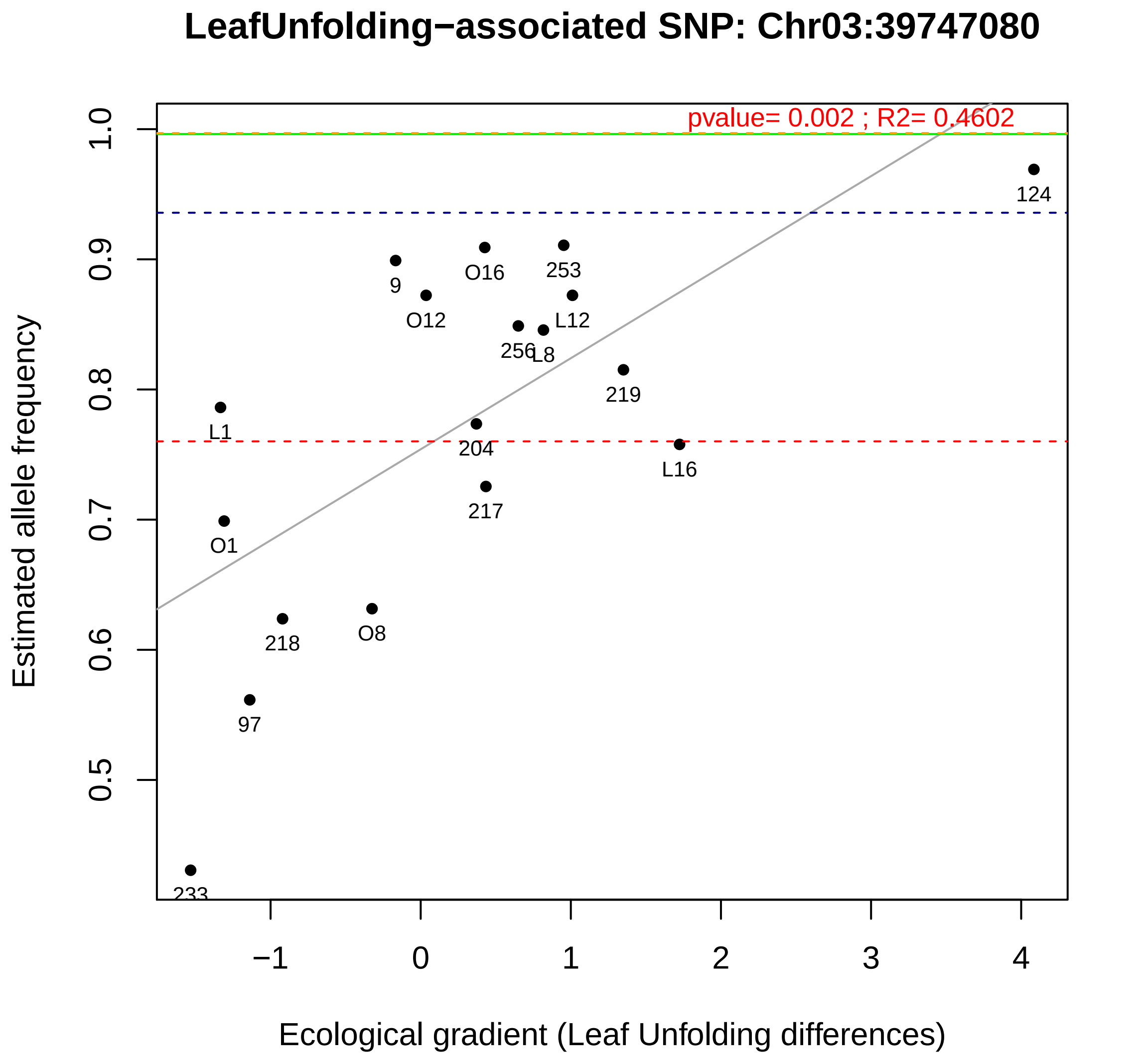
LeafUnfolding−associated SNP: Chr03:39747080
pvalue= 0.002 ; R2= 0.4602
Estimated allele frequency
Ecological gradient (Leaf Unfolding differences)
1.0
0.9
0.8
0.7
0.6
0.5
−1
0
1
2
3
4
124
9
O16
253
O12
L12
256
L8
219
L1
204
L16
217
O1
218
O8
97
233

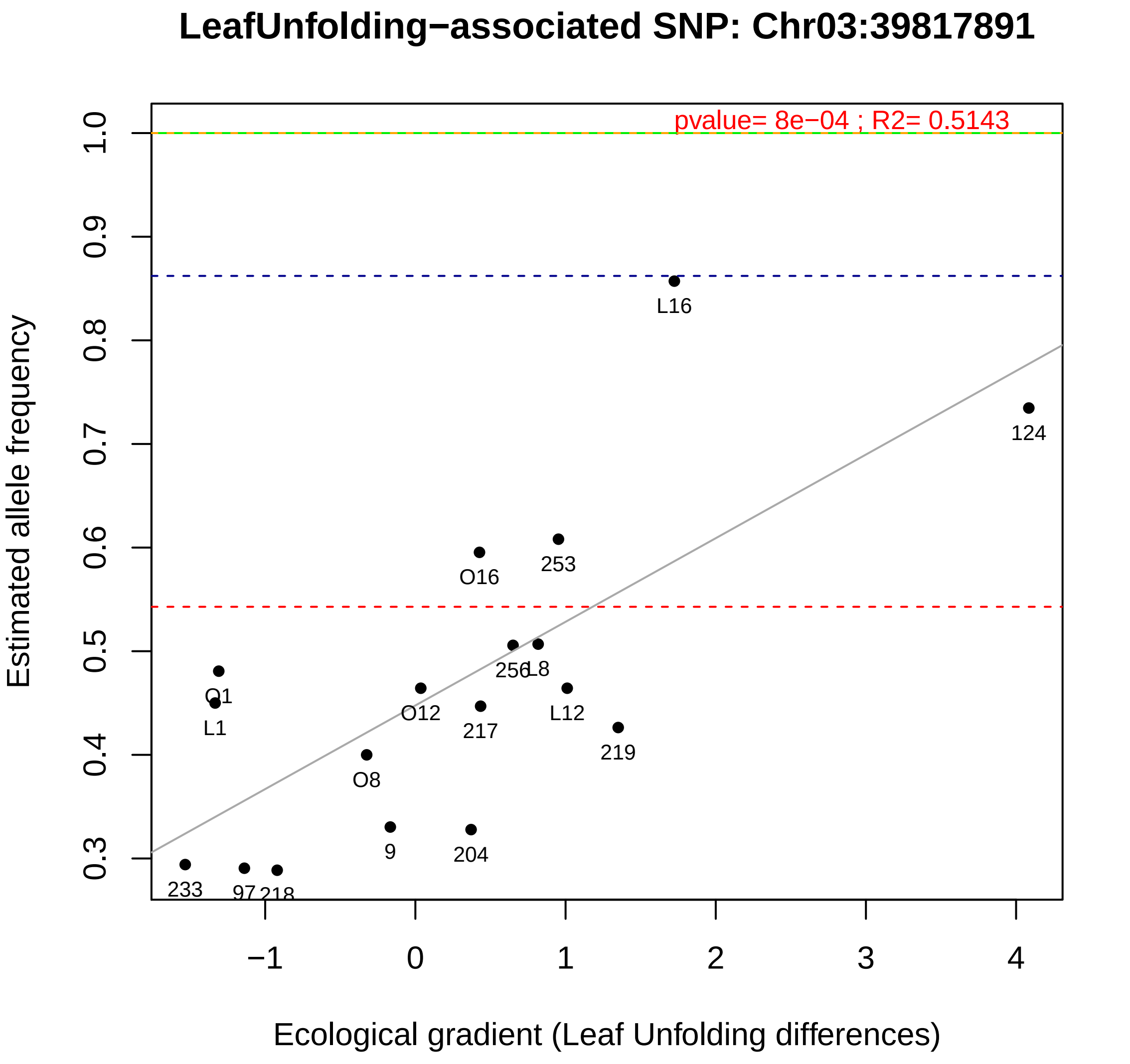

LeafUnfolding−associated SNP: Chr03:39817891
pvalue= 8e−04 ; R2= 0.5143
Estimated allele frequency
Ecological gradient (Leaf Unfolding differences)
1.0
0.9
0.8
0.7
0.6
0.5
0.4
0.3
−1
0
1
2
3
4
L16
124
253
O16
256
L8
O1
L1
O12
217
L12
219
O8
9
204
233
97
218

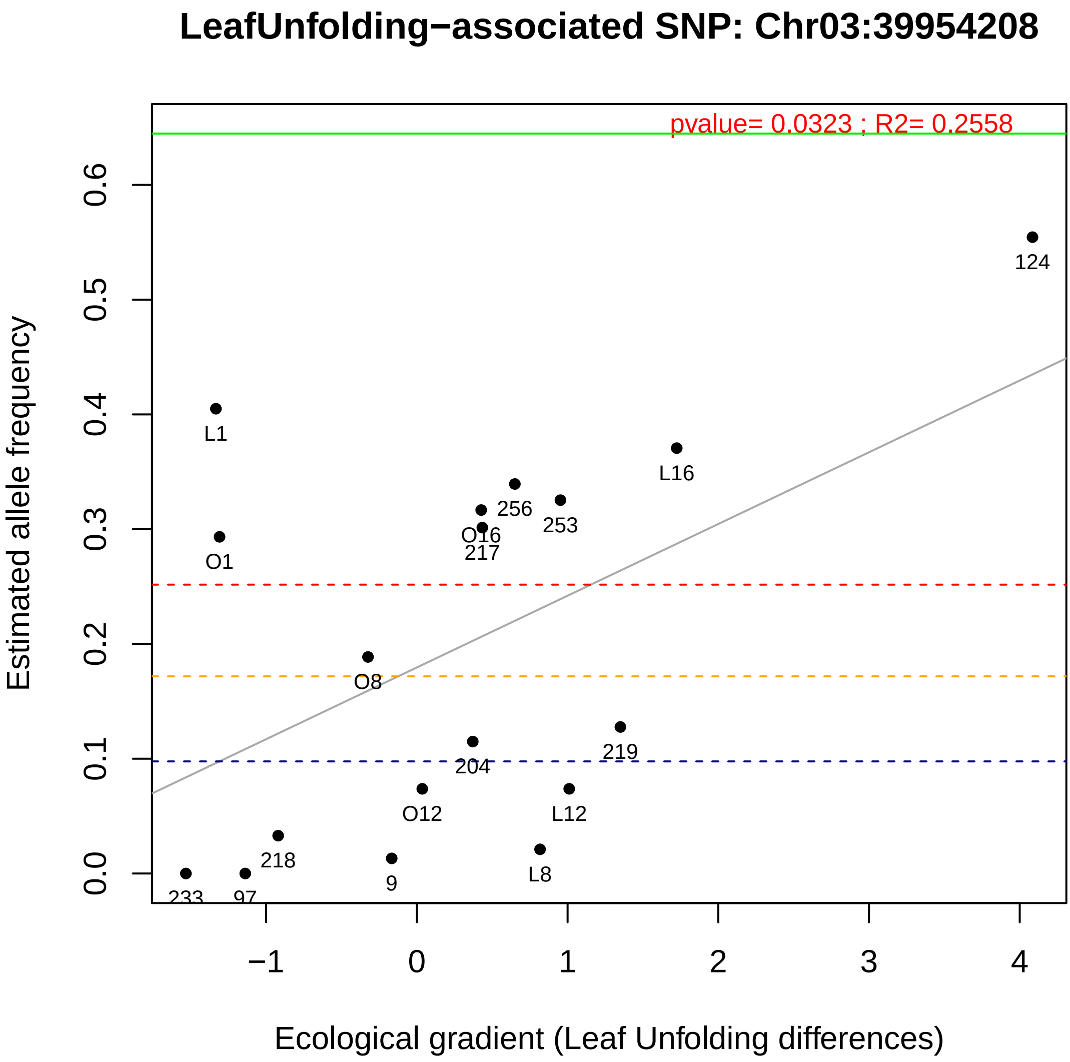

LeafUnfolding−associated SNP: Chr03:39954208
pvalue= 0.0323 ; R2= 0.2558
Estimated allele frequency
0.0
0.1
0.2
0.3
0.4
0.5
0.6
−1
0
1
2
3
4
Ecological gradient (Leaf Unfolding differences)
124
L1
L16
256
253
O16
217
O1
O8
204
219
O12
L12
218
9
L8
233
97

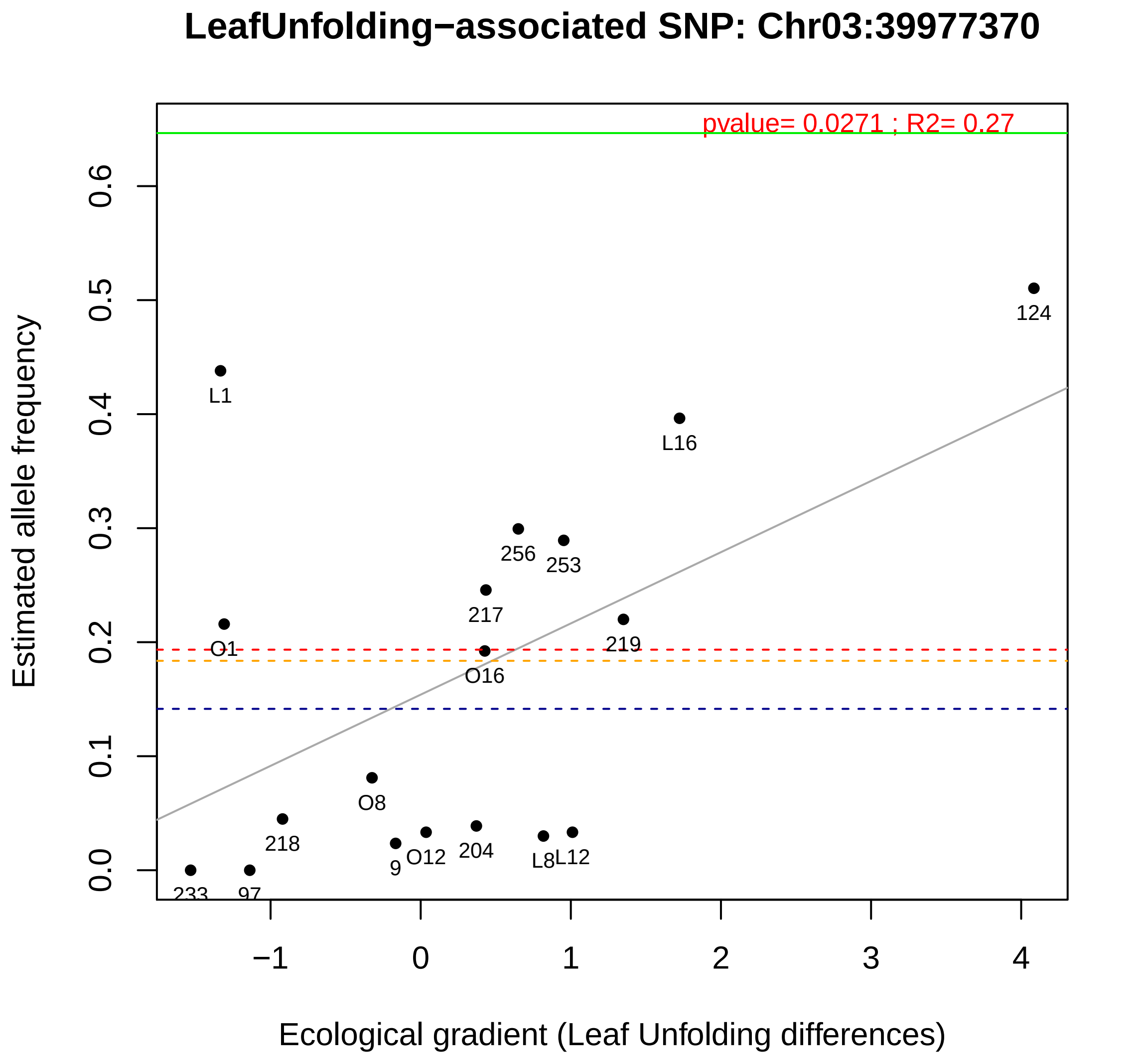

LeafUnfolding−associated SNP: Chr03:39977370
pvalue= 0.0271 ; R2= 0.27
Estimated allele frequency
0.0
0.1
0.2
0.3
0.4
0.5
0.6
−1
0
1
2
3
4
Ecological gradient (Leaf Unfolding differences)
124
L1
L16
256
253
217
219
O1
O16
O8
218
9
O12
204
L8
L12
233
97

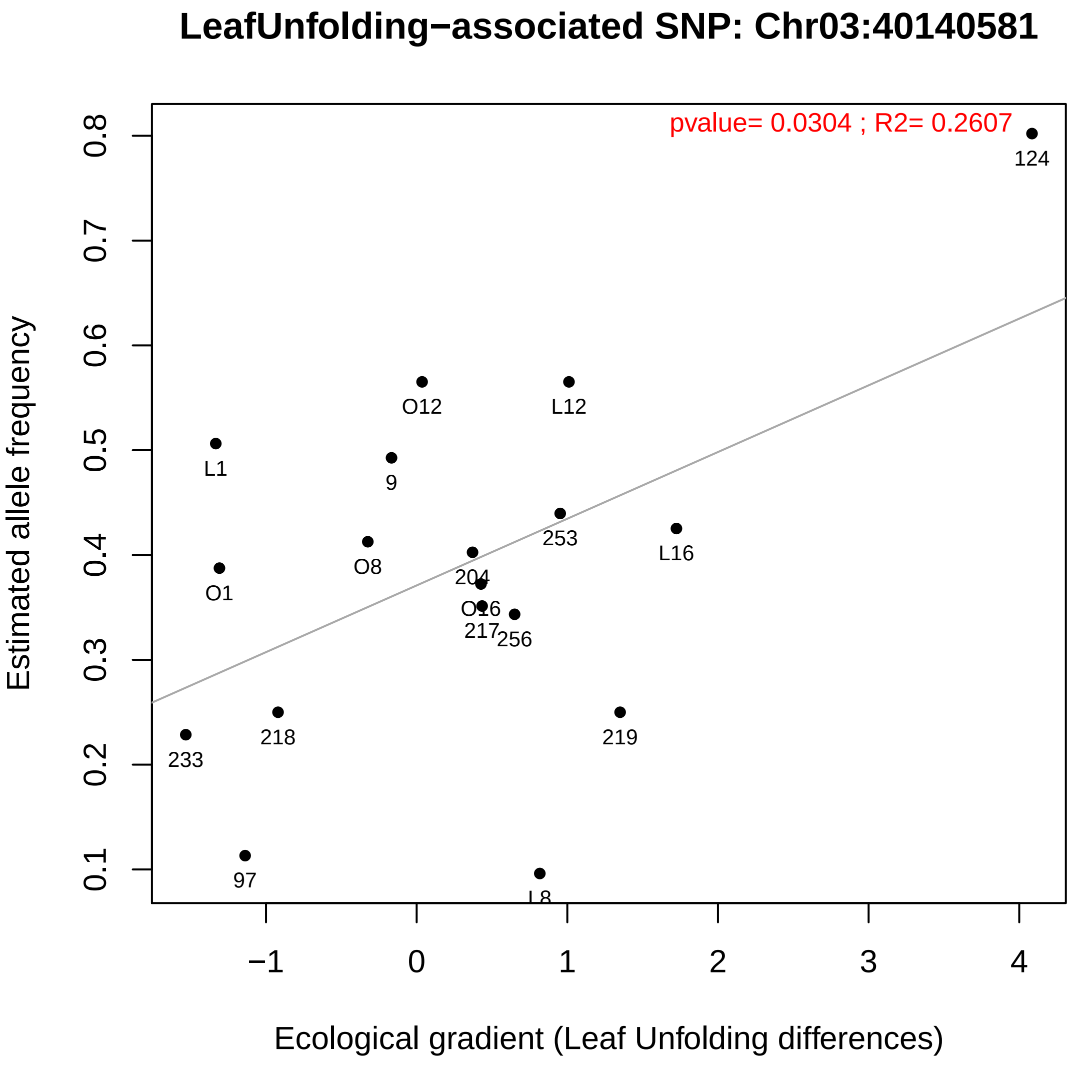

LeafUnfolding−associated SNP: Chr03:40140581
pvalue= 0.0304 ; R2= 0.2607
Estimated allele frequency
Ecological gradient (Leaf Unfolding differences)
0.1
0.2
0.3
0.4
0.5
0.6
0.7
0.8
−1
0
1
2
3
4
124
O12
L12
L1
9
253
L16
O8
204
O1
O16
217
256
218
219
233
97
L8

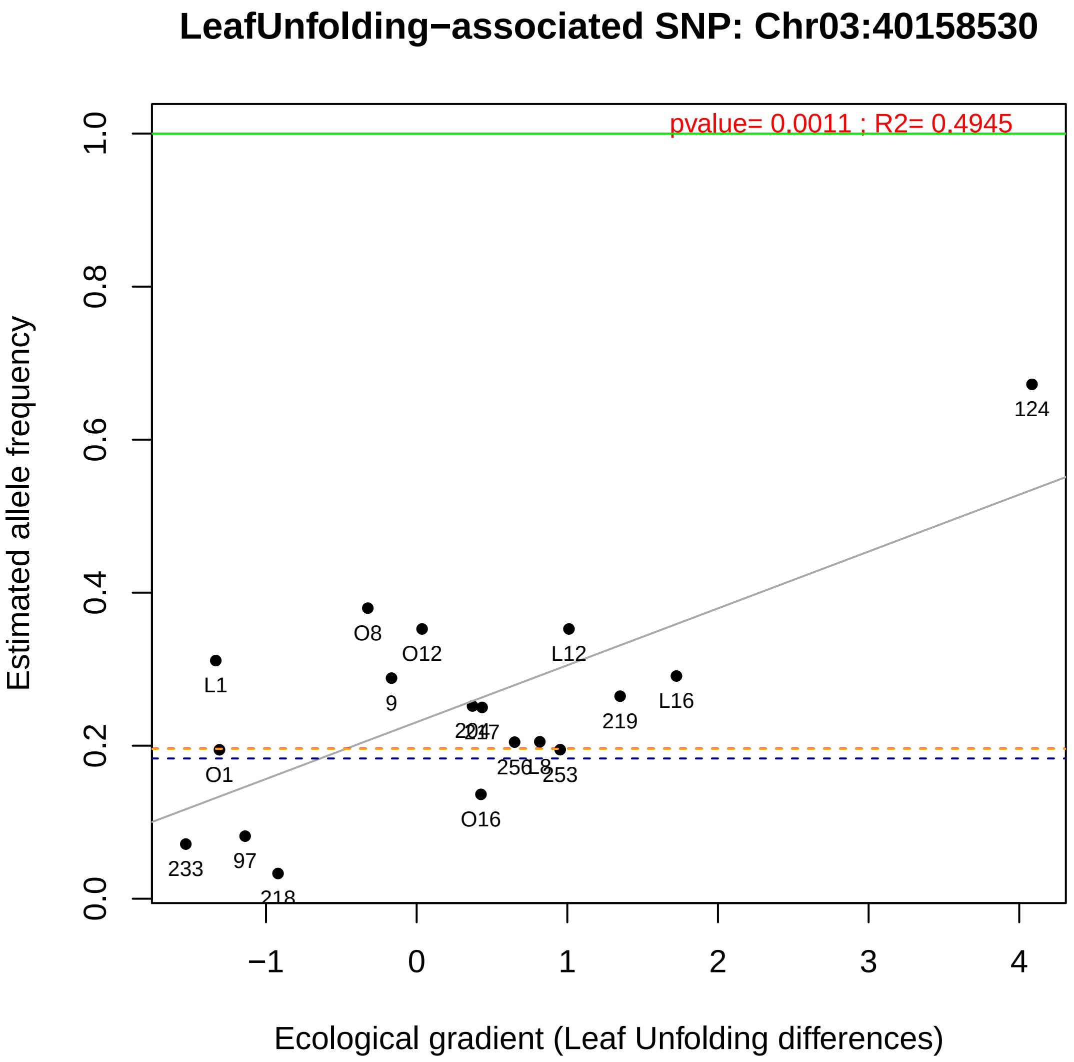
LeafUnfolding−associated SNP: Chr03:40158530
pvalue= 0.0011 ; R2= 0.4945
Estimated allele frequency
0.0
0.2
0.4
0.6
0.8
1.0
−1
0
1
2
3
4
Ecological gradient (Leaf Unfolding differences)
124
O8
O12
L12
L1
9
L16
219
O1
256
L8
253
O16
233
97
218

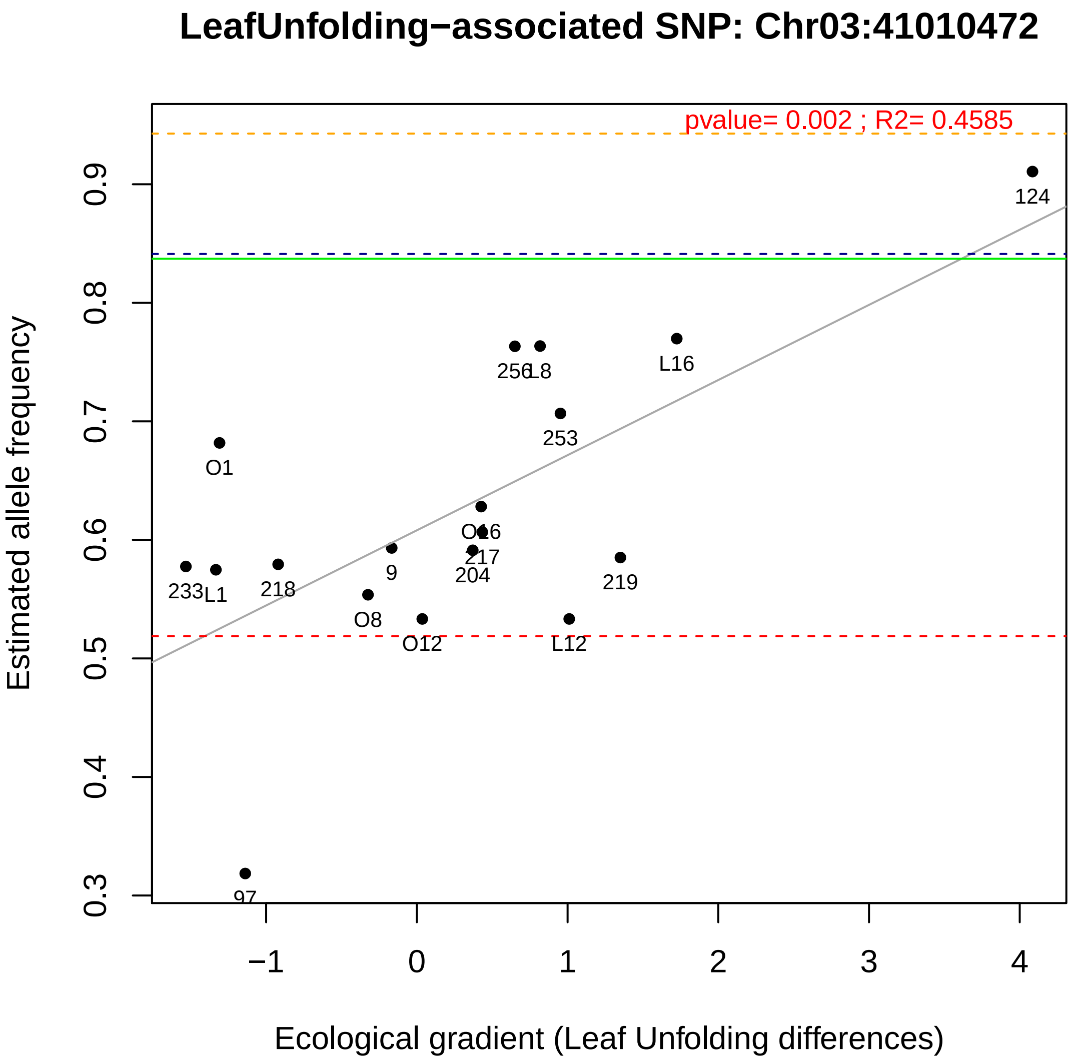
LeafUnfolding−associated SNP: Chr03:41010472
pvalue= 0.002 ; R2= 0.4585
Estimated allele frequency
0.3
0.4
0.5
0.6
0.7
0.8
0.9
124
256
L8
L16
253
O1
O16
217
204
9
233
L1
218
219
O8
O12
L12
97
−1
0
1
2
3
4
Ecological gradient (Leaf Unfolding differences)

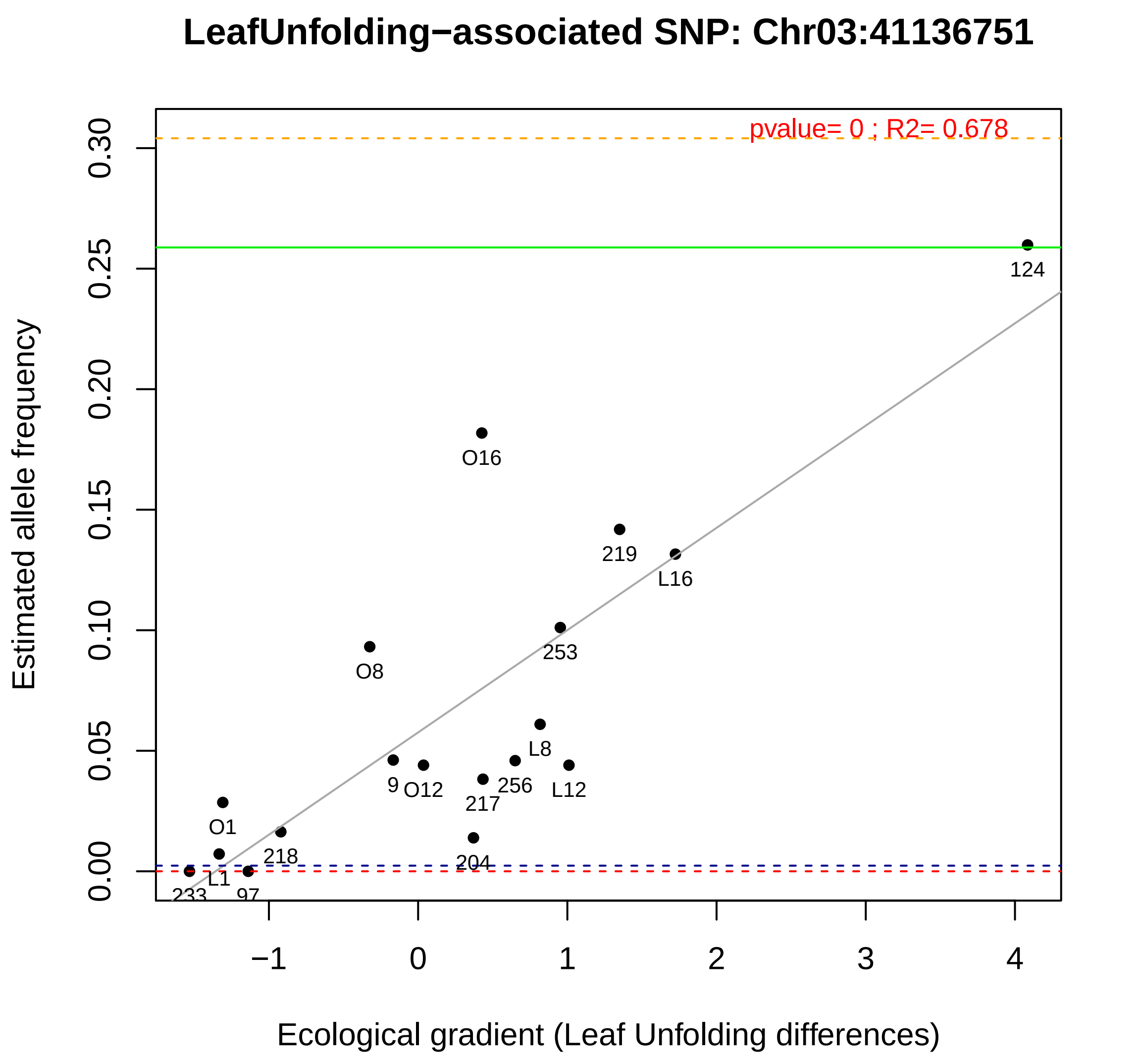

LeafUnfolding−associated SNP: Chr03:41136751
pvalue= 0 ; R2= 0.678
Estimated allele frequency
Ecological gradient (Leaf Unfolding differences)
0.00
0.05
0.10
0.15
0.20
0.25
0.30
−1
0
1
2
3
4
124
O16
219
L16
253
O8
L8
9
O12
256
L12
217
O1
218
204
L1
233
97

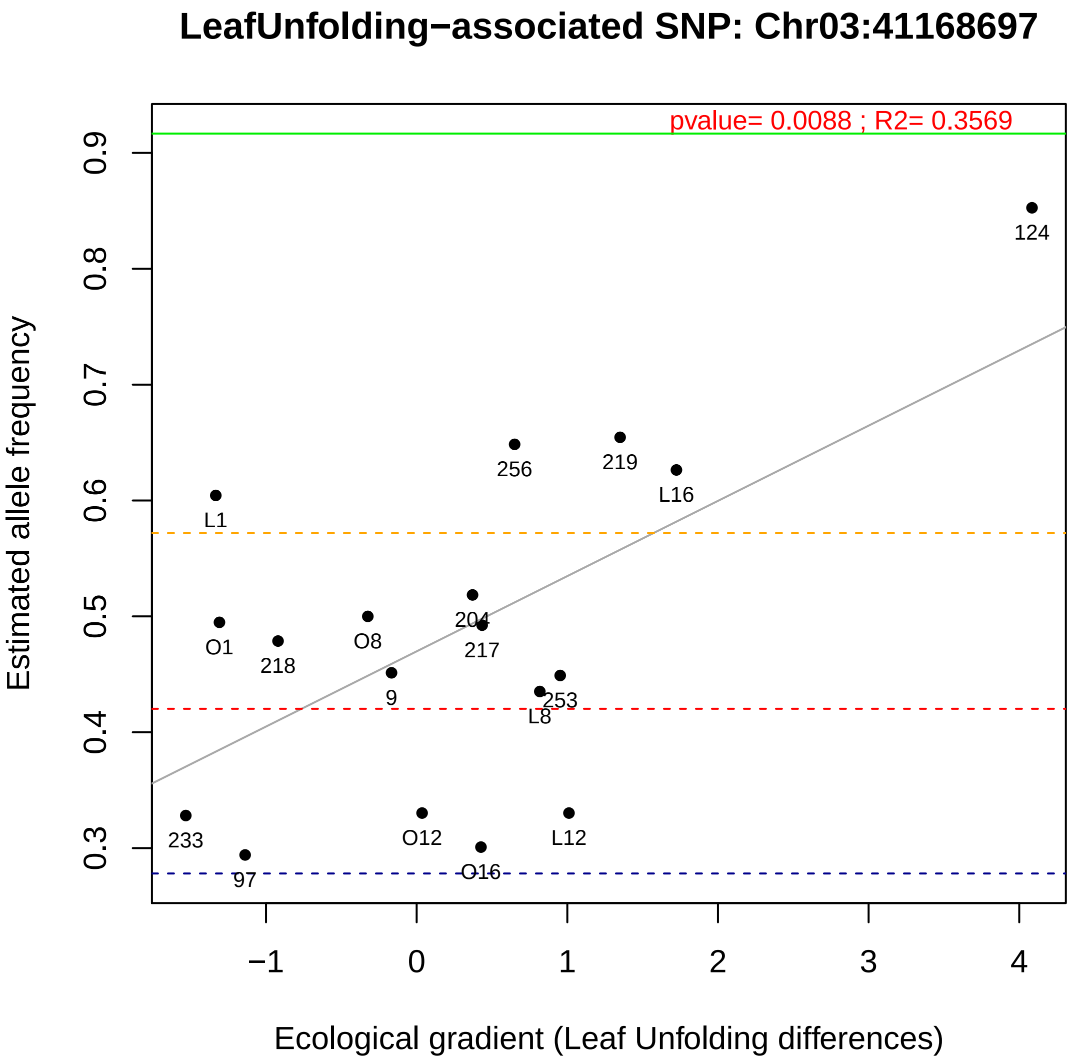
LeafUnfolding−associated SNP: Chr03:41168697
pvalue= 0.0088 ; R2= 0.3569
Estimated allele frequency
0.3
0.4
0.5
0.6
0.7
0.8
0.9
−1
0
1
2
3
4
Ecological gradient (Leaf Unfolding differences)
124
256
219
L16
L1
204
217
O1
218
O8
9
253
L8
233
O12
L12
97
O16

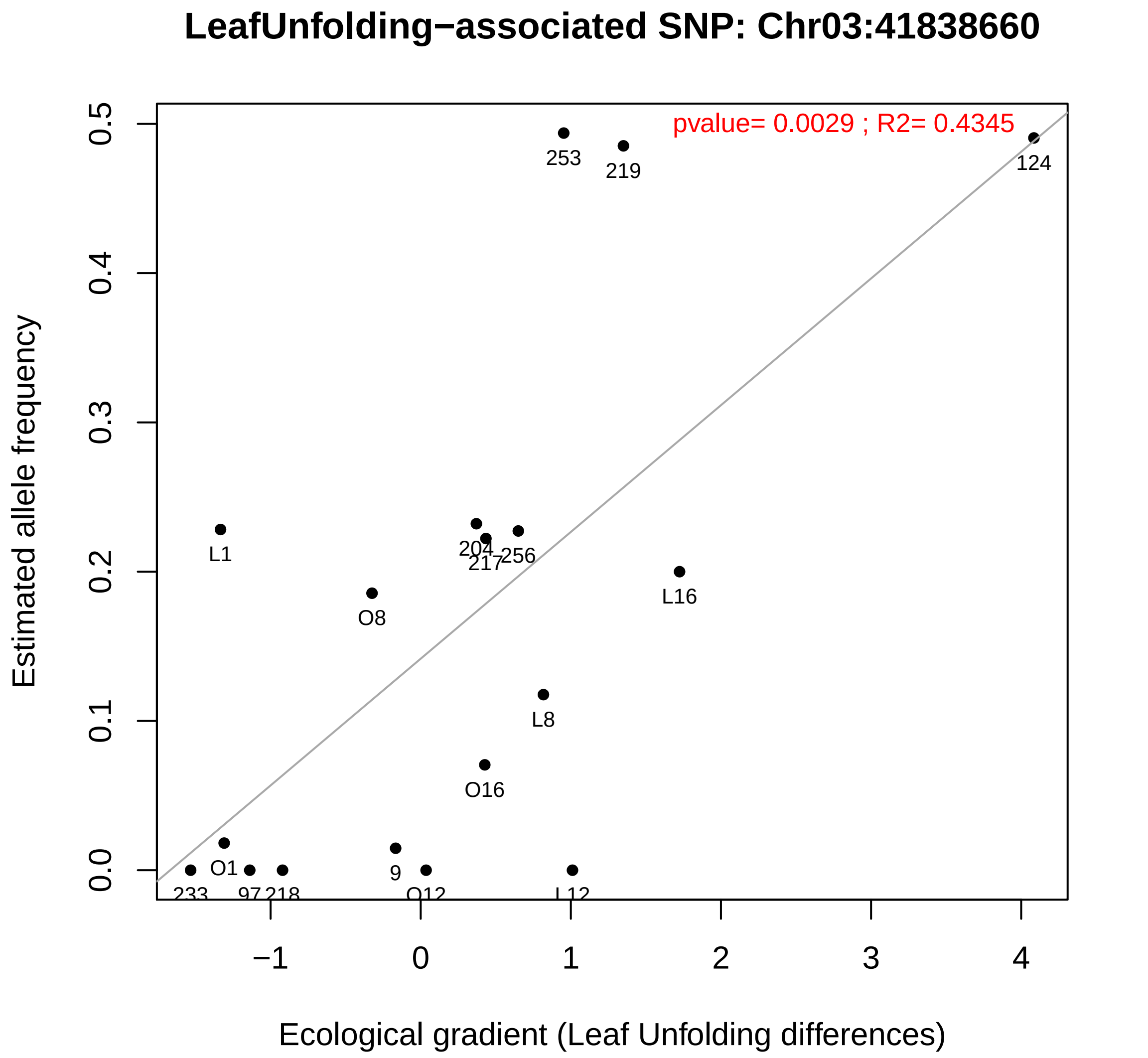
LeafUnfolding−associated SNP: Chr03:41838660
pvalue= 0.0029 ; R2= 0.4345
Estimated allele frequency
0.0
0.1
0.2
0.3
0.4
0.5
−1
0
1
2
3
4
Ecological gradient (Leaf Unfolding differences)
253
219
124
L1
204
217
256
L16
O8
L8
O16
O1
9
97
218
233
O12
L12

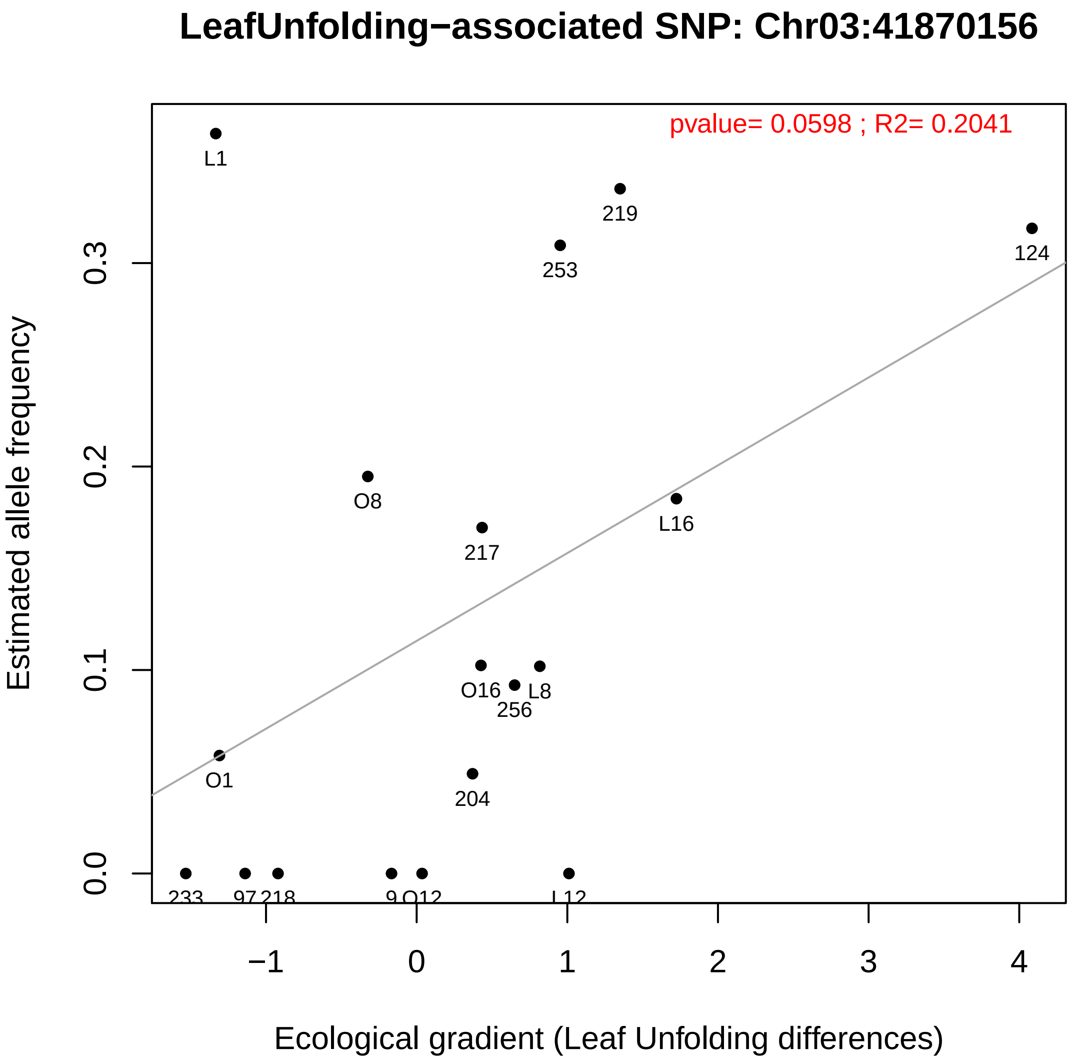
LeafUnfolding−associated SNP: Chr03:41870156
pvalue= 0.0598 ; R2= 0.2041
Estimated allele frequency
Ecological gradient (Leaf Unfolding differences)
0.0
0.1
0.2
0.3
−1
0
1
2
3
4
L1
219
124
253
O8
217
L16
O16
L8
256
O1
204
233
97
218
9
O12
L12

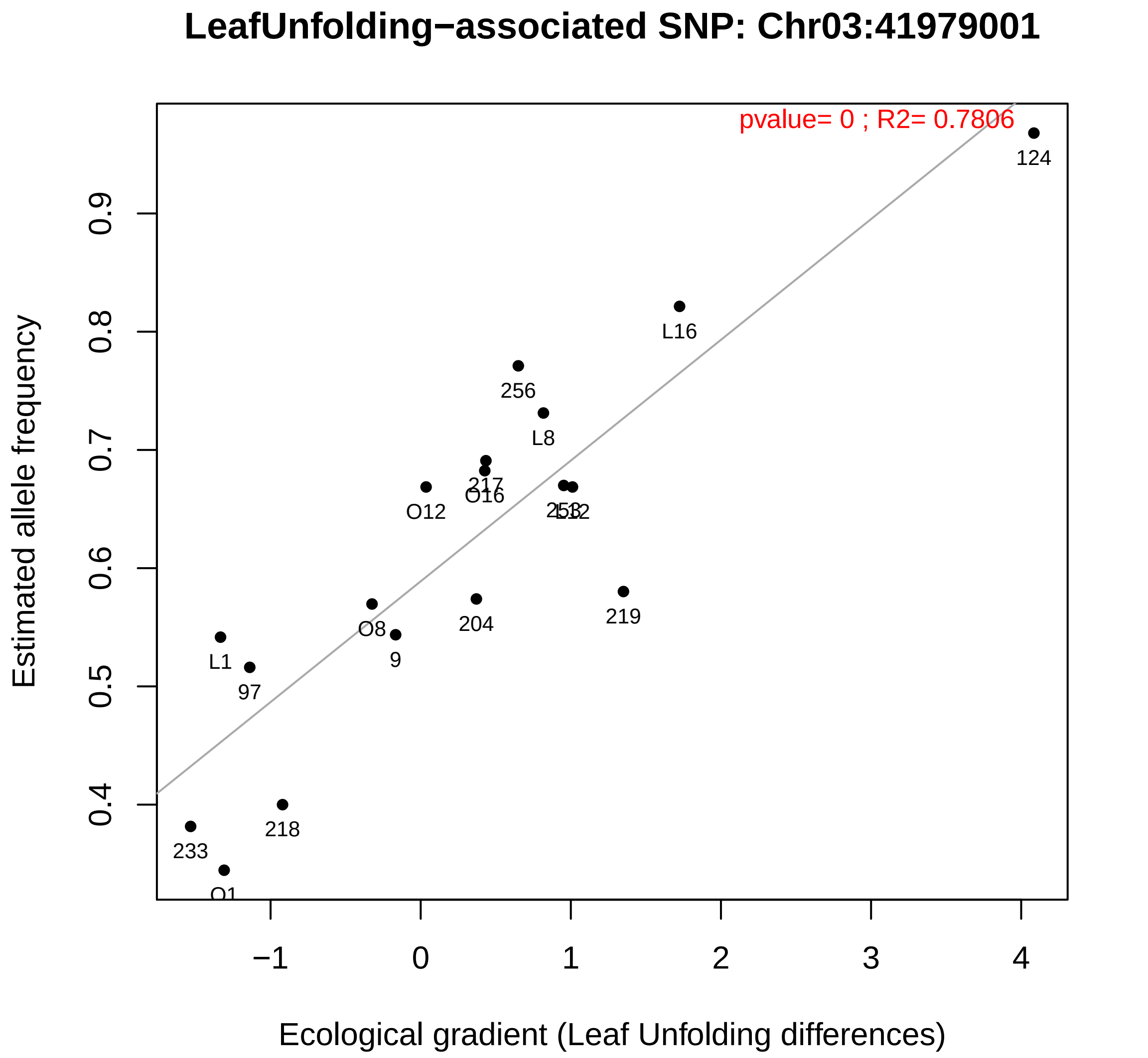
LeafUnfolding−associated SNP: Chr03:41979001
pvalue= 0 ; R2= 0.7806
Estimated allele frequency
Ecological gradient (Leaf Unfolding differences)
0.4
0.5
0.6
0.7
0.8
0.9
−1
0
1
2
3
4
124
L16
256
L8
217
O16
O12
251
L12
O8
9
204
219
L1
97
218
233
O1

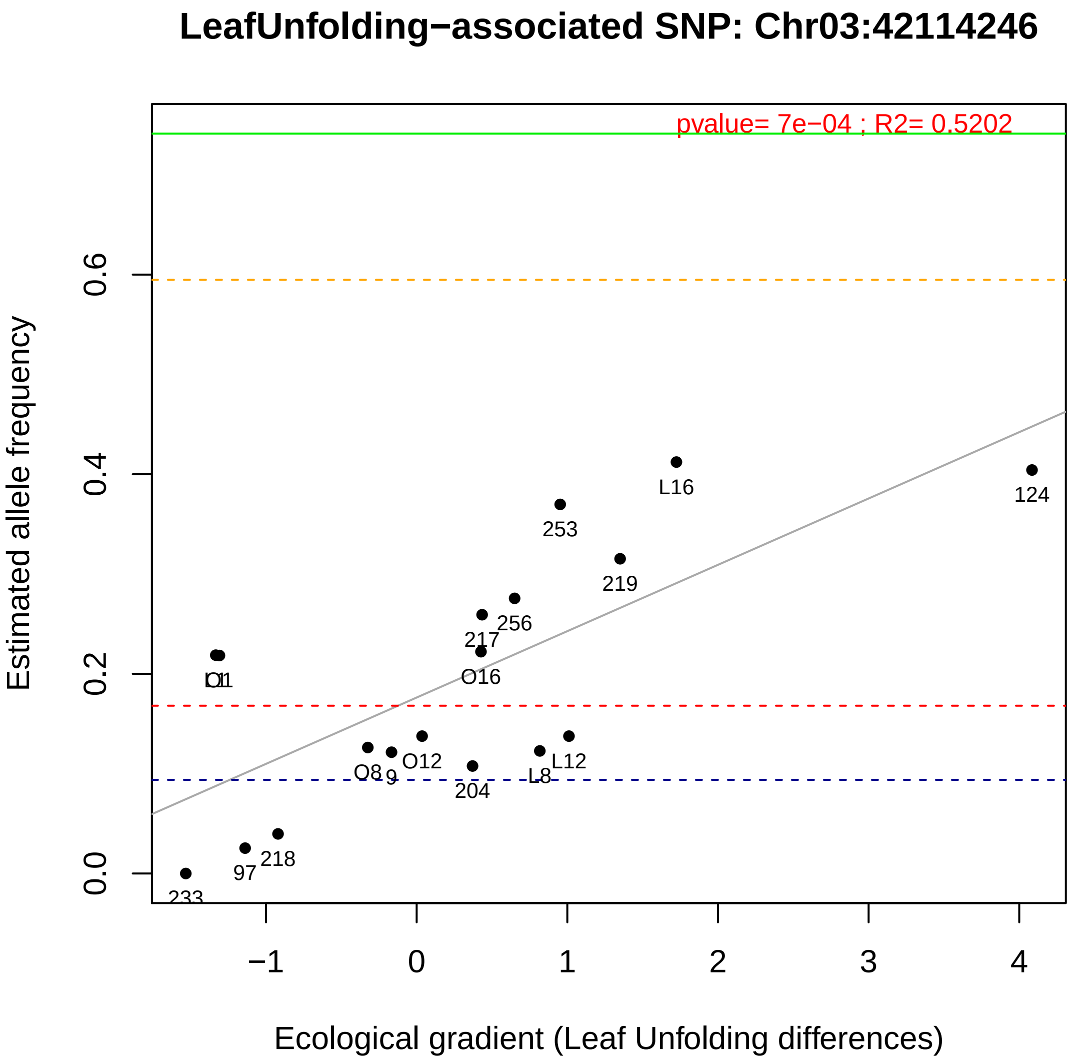
LeafUnfolding−associated SNP: Chr03:42114246
pvalue= 7e−04 ; R2= 0.5202
Estimated allele frequency
0.0
0.2
0.4
0.6
Ecological gradient (Leaf Unfolding differences)
−1
0
1
2
3
4
233
97
218
O8
9
O12
204
L8
L12
O16
217
256
253
219
L16
124

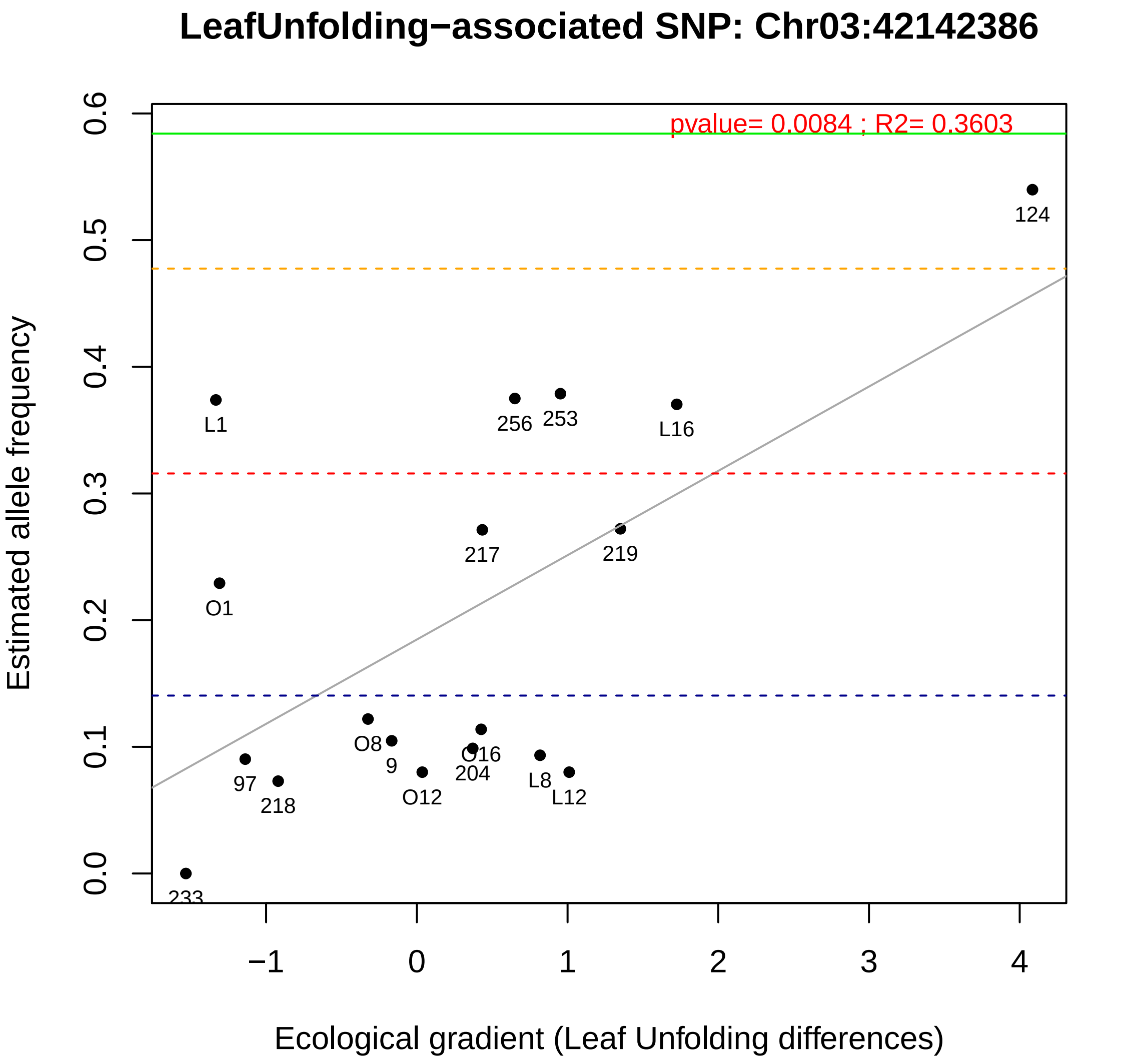

LeafUnfolding−associated SNP: Chr03:42142386
pvalue= 0.0084 ; R2= 0.3603
Estimated allele frequency
Ecological gradient (Leaf Unfolding differences)
0.0
0.1
0.2
0.3
0.4
0.5
0.6
−1
0
1
2
3
4
124
L1
256
253
L16
217
219
O1
O8
9
O16
204
L8
L12
97
218
O12
233

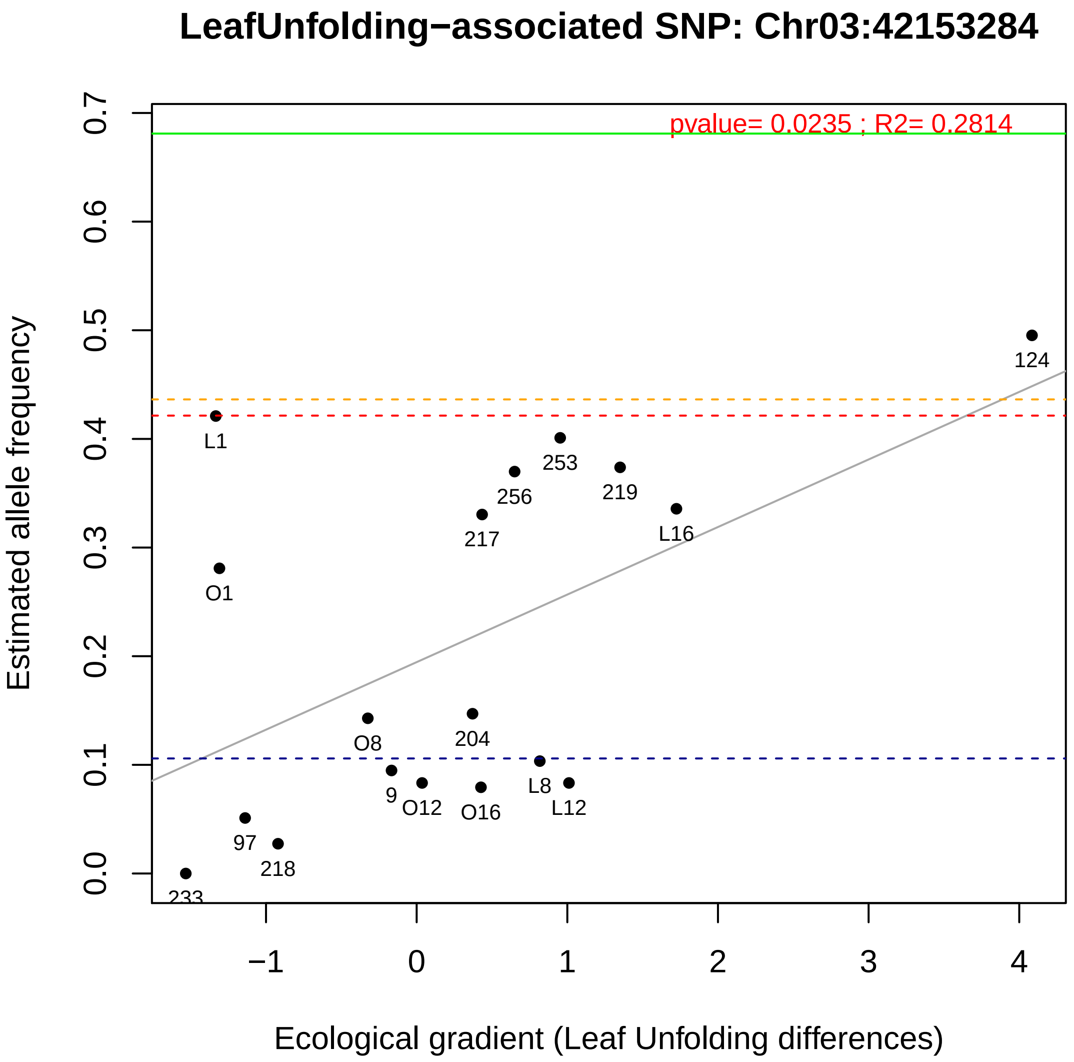
LeafUnfolding−associated SNP: Chr03:42153284
pvalue= 0.0235 ; R2= 0.2814
Estimated allele frequency
0.0
0.1
0.2
0.3
0.4
0.5
0.6
0.7
−1
0
1
2
3
4
Ecological gradient (Leaf Unfolding differences)
124
L1
253
256
219
217
L16
O1
O8
204
9
O12
O16
L8
L12
97
218
233

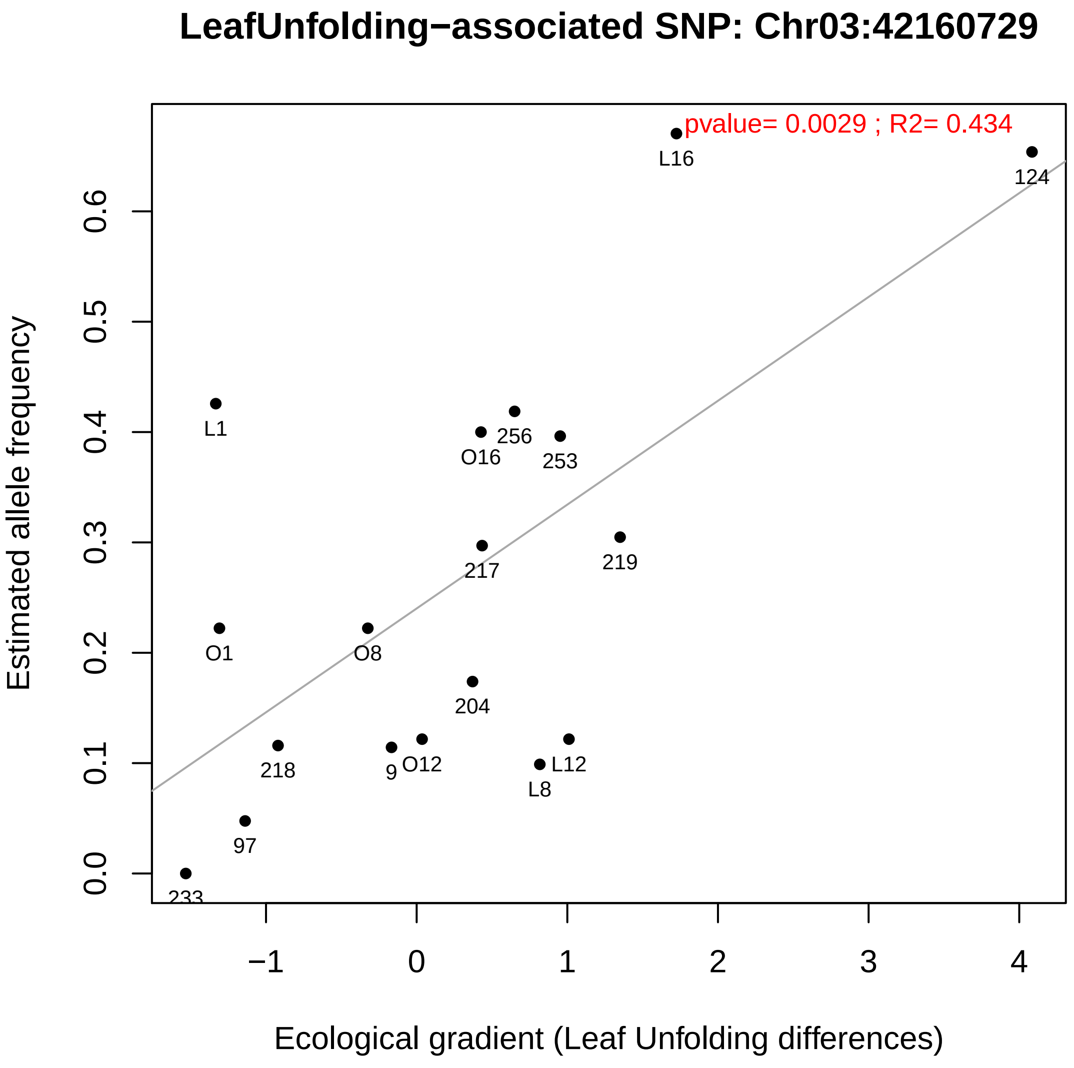

LeafUnfolding−associated SNP: Chr03:42160729
pvalue= 0.0029 ; R2= 0.434
L16
124
L1
256
O16
253
217
219
O1
O8
204
218
9
O12
L12
L8
97
233
Estimated allele frequency
0.0
0.1
0.2
0.3
0.4
0.5
0.6
−1
0
1
2
3
4
Ecological gradient (Leaf Unfolding differences)

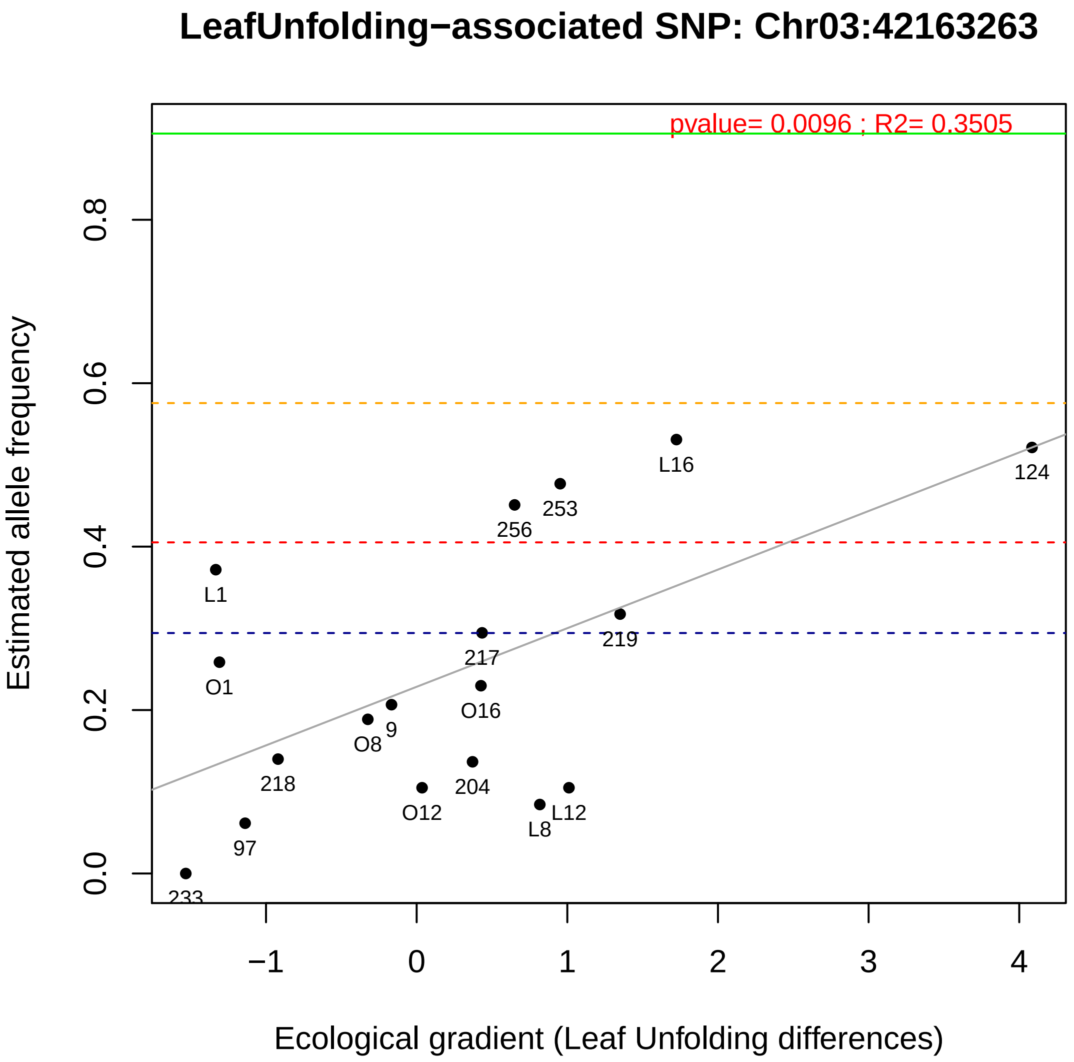
LeafUnfolding−associated SNP: Chr03:42163263
pvalue= 0.0096 ; R2= 0.3505
Estimated allele frequency
0.0
0.2
0.4
0.6
0.8
−1
0
1
2
3
4
Ecological gradient (Leaf Unfolding differences)
L16
124
253
256
L1
219
217
O1
O16
9
O8
218
204
O12
L12
L8
97
233

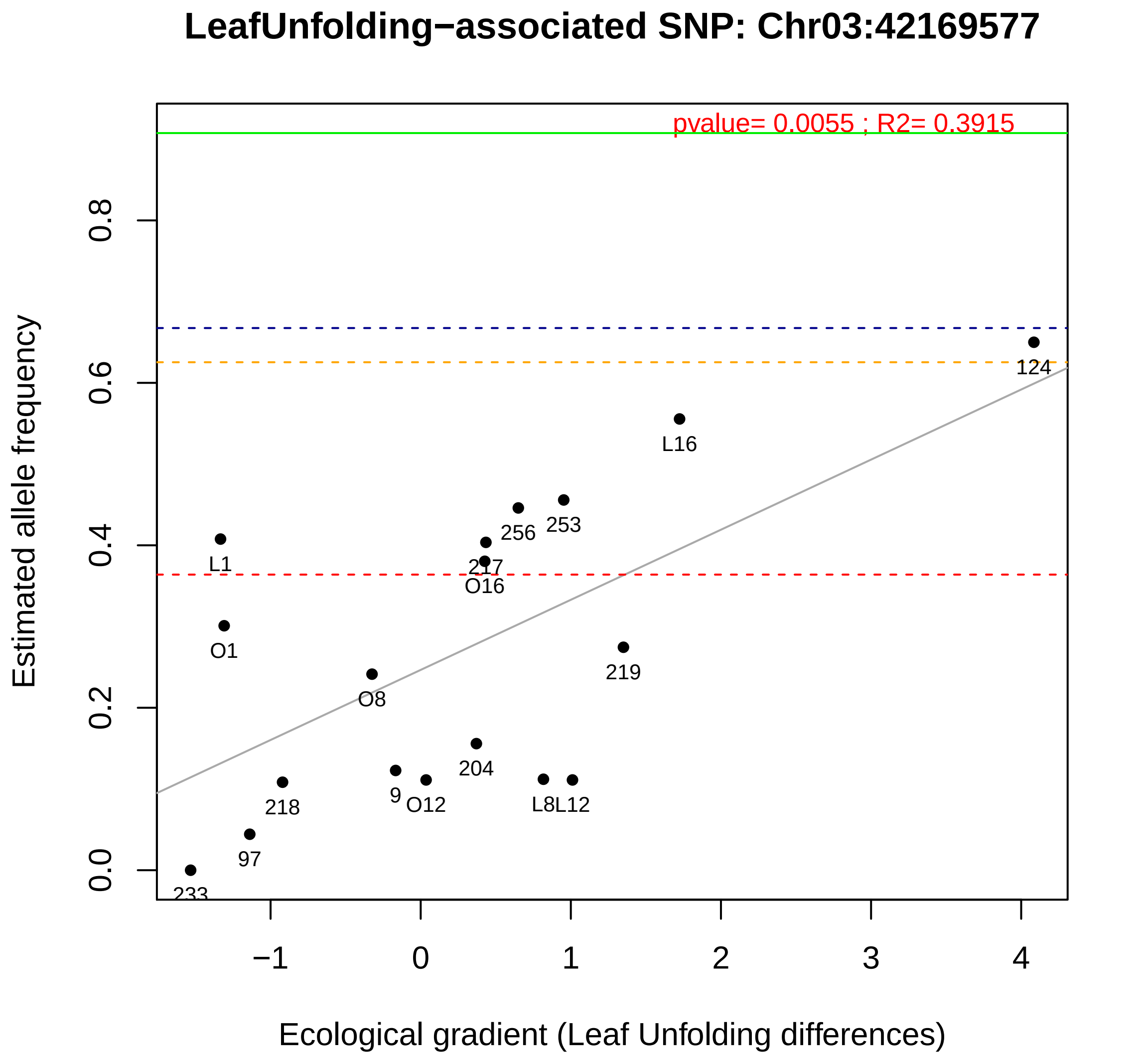

LeafUnfolding−associated SNP: Chr03:42169577
pvalue= 0.0055 ; R2= 0.3915
Estimated allele frequency
0.0
0.2
0.4
0.6
0.8
−1
0
1
2
3
4
Ecological gradient (Leaf Unfolding differences)
124
L16
256
253
L1
217
O16
O1
219
O8
204
218
9
O12
L8
L12
97
233

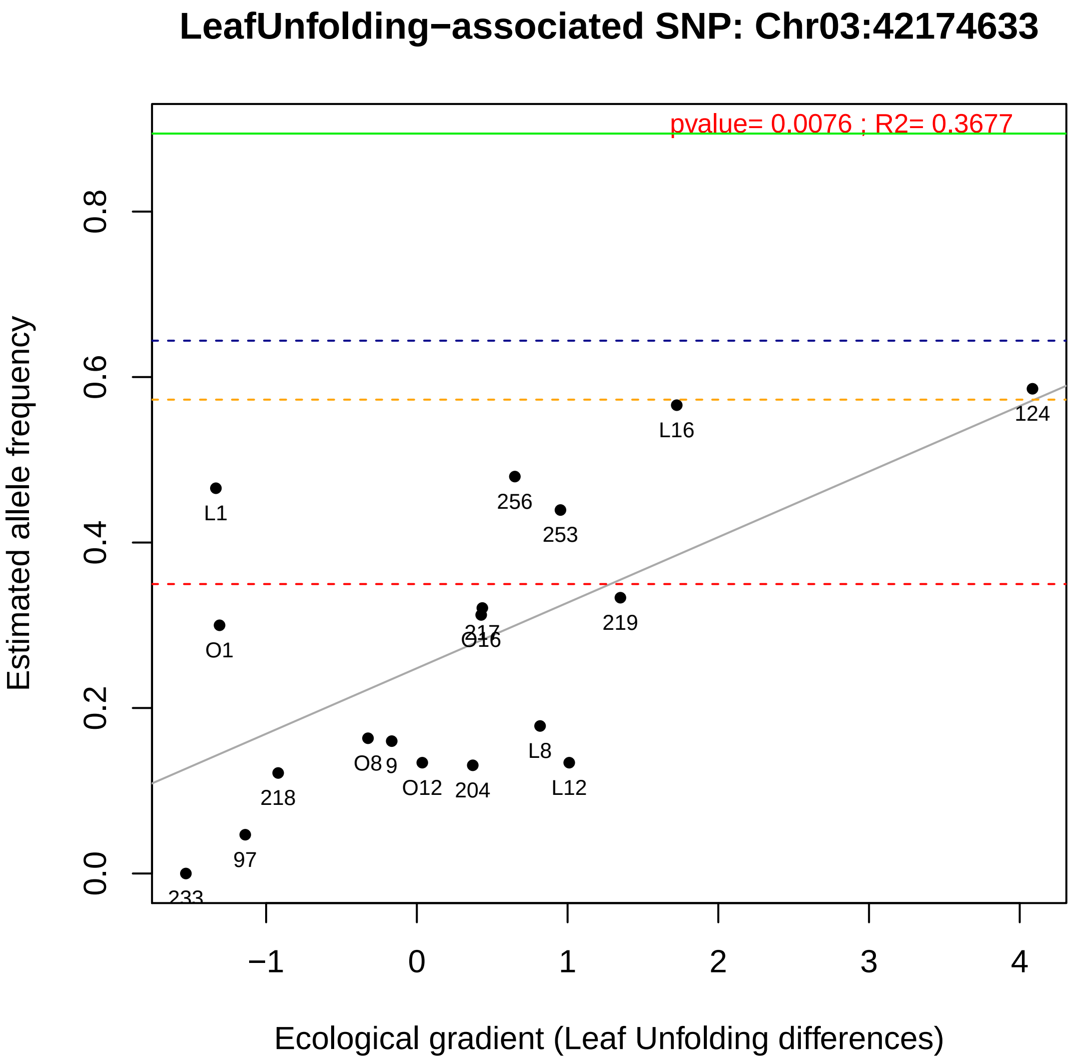
LeafUnfolding−associated SNP: Chr03:42174633
pvalue= 0.0076 ; R2= 0.3677
Estimated allele frequency
0.0
0.2
0.4
0.6
0.8
−1
0
1
2
3
4
Ecological gradient (Leaf Unfolding differences)
124
L16
L1
256
253
219
217
O16
O1
L8
O8
9
O12
204
L12
218
97
233

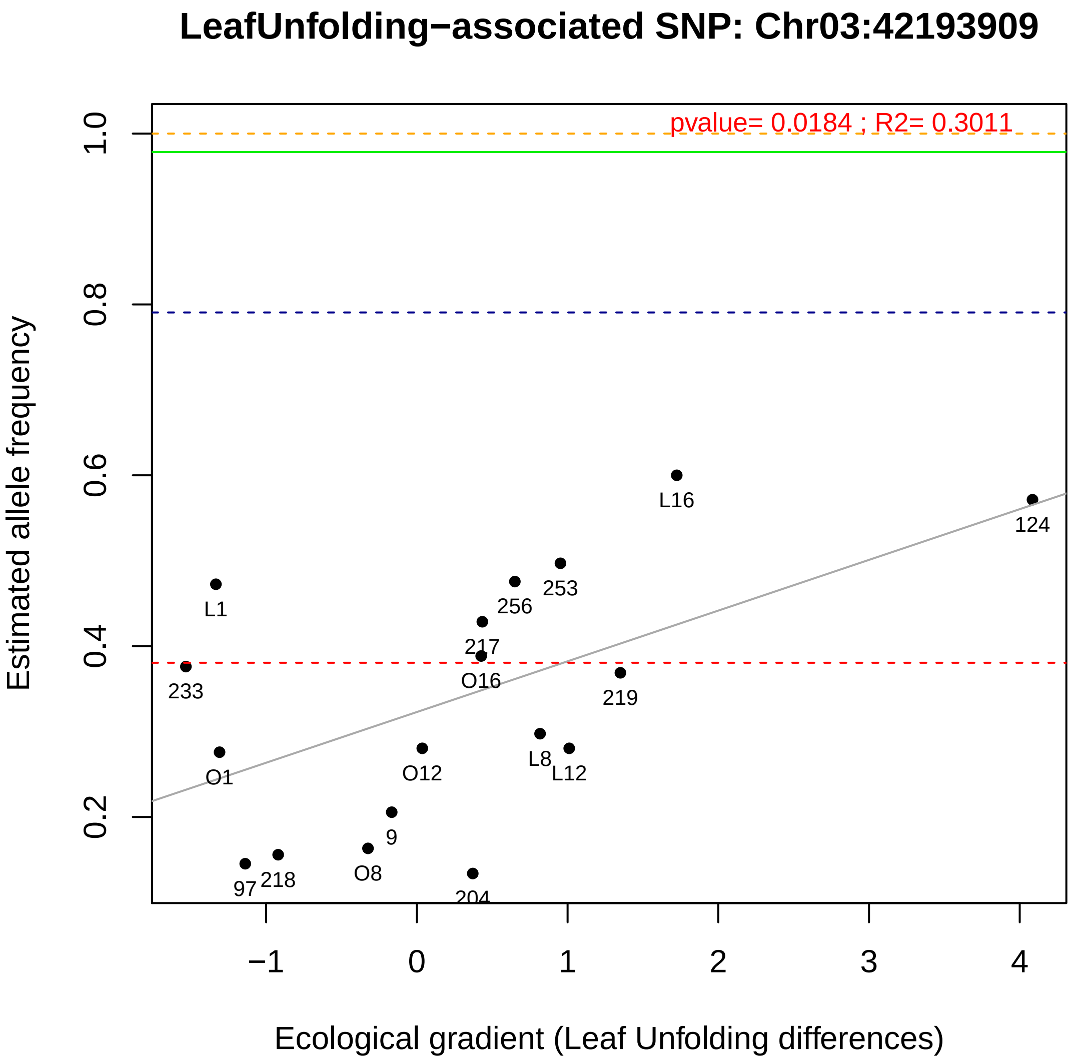

LeafUnfolding−associated SNP: Chr03:42193909
pvalue= 0.0184 ; R2= 0.3011
Estimated allele frequency
0.2
0.4
0.6
0.8
1.0
Ecological gradient (Leaf Unfolding differences)
−1
0
1
2
3
4
L16
124
253
256
L1
217
O16
233
219
O1
O12
L8
L12
9
O8
97
218
204

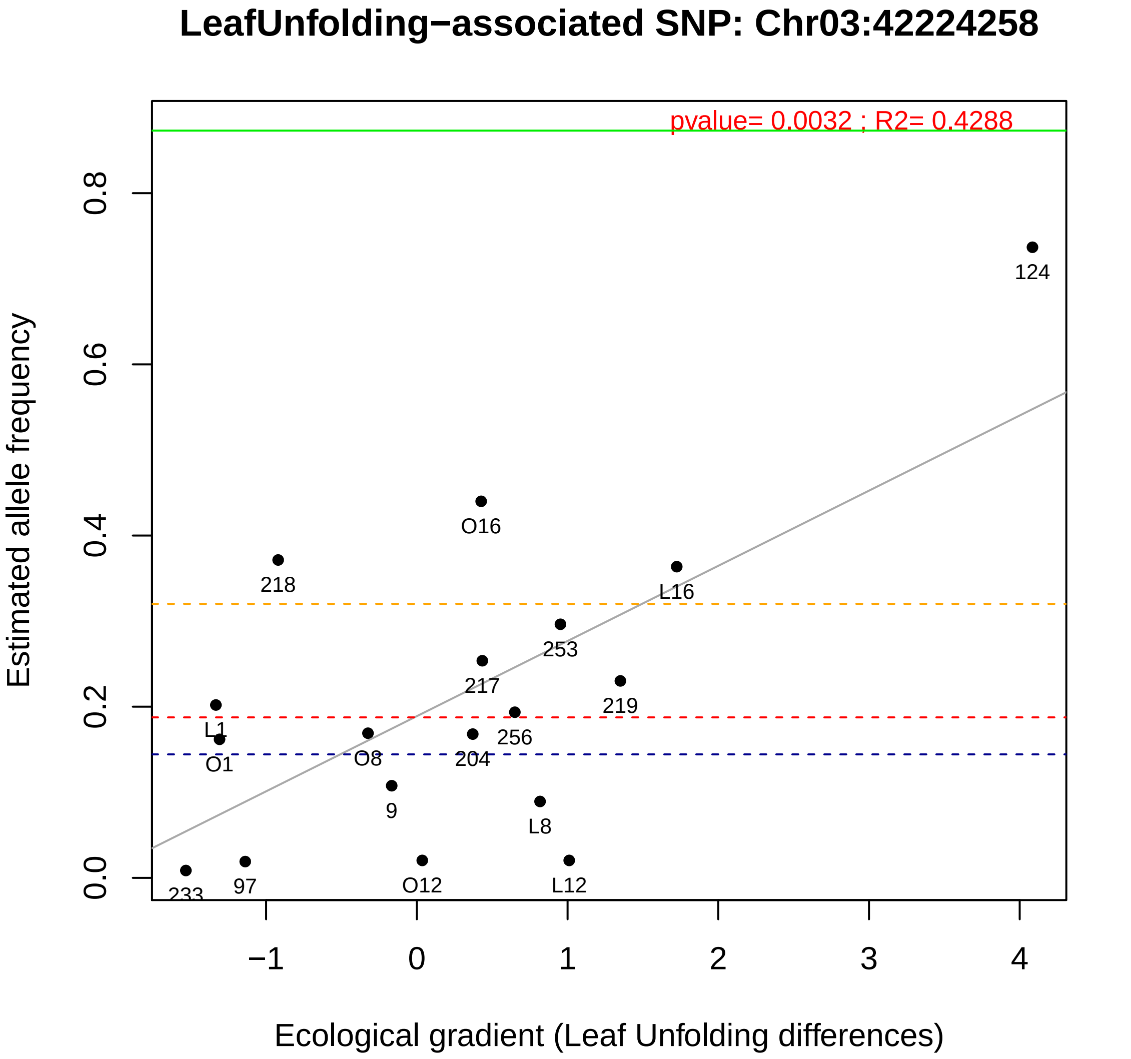
LeafUnfolding−associated SNP: Chr03:42224258
pvalue= 0.0032 ; R2= 0.4288
Estimated allele frequency
0.0
0.2
0.4
0.6
0.8
−1
0
1
2
3
4
Ecological gradient (Leaf Unfolding differences)
124
O16
218
L16
253
217
219
256
L1
O1
O8
204
9
L8
233
97
O12
L12

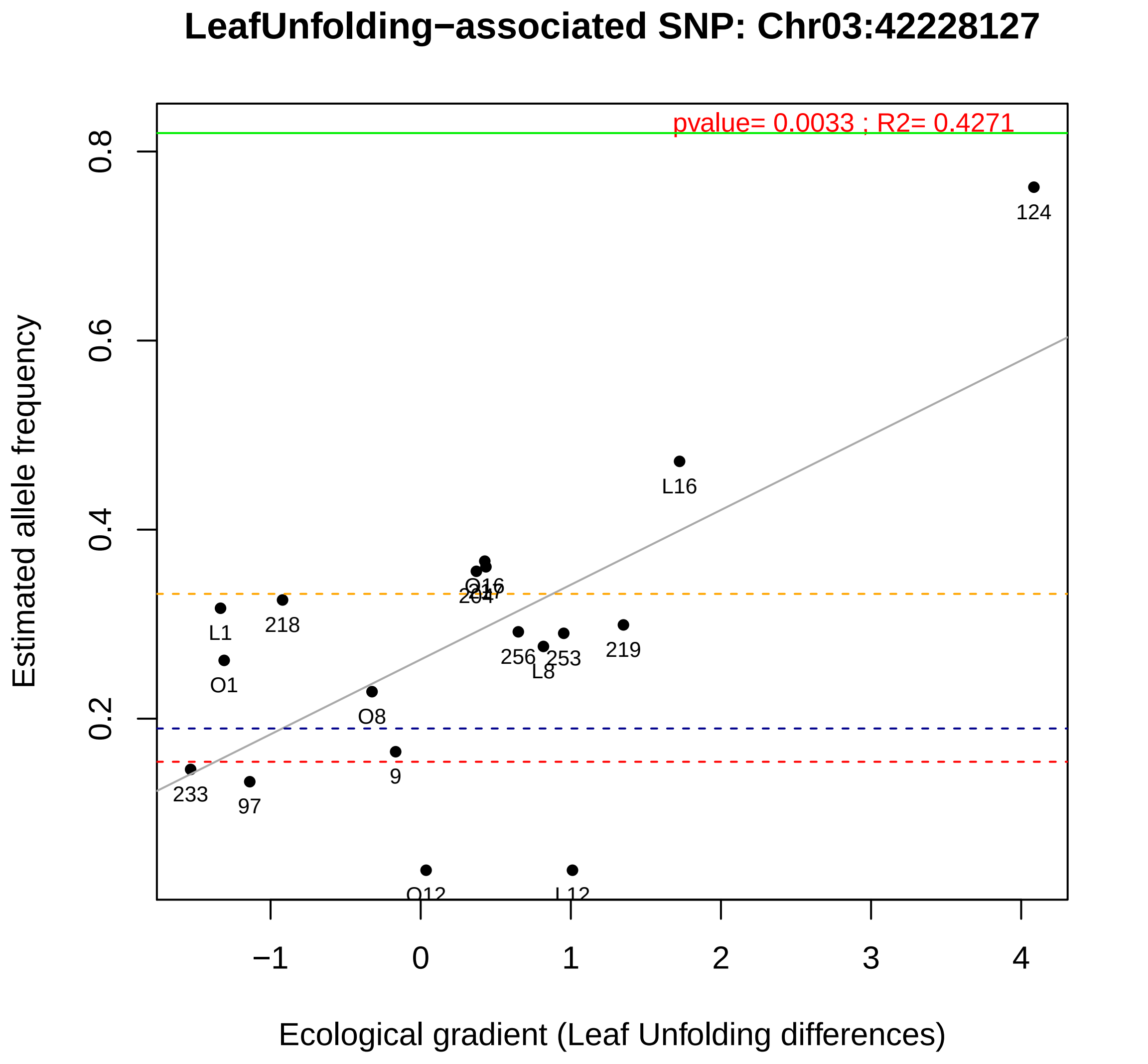
LeafUnfolding−associated SNP: Chr03:42228127
pvalue= 0.0033 ; R2= 0.4271
Estimated allele frequency
0.2
0.4
0.6
0.8
Ecological gradient (Leaf Unfolding differences)
−1
0
1
2
3
4
124
L16
218
L1
O1
O8
9
233
97
256
L8
253
219
O12
L12

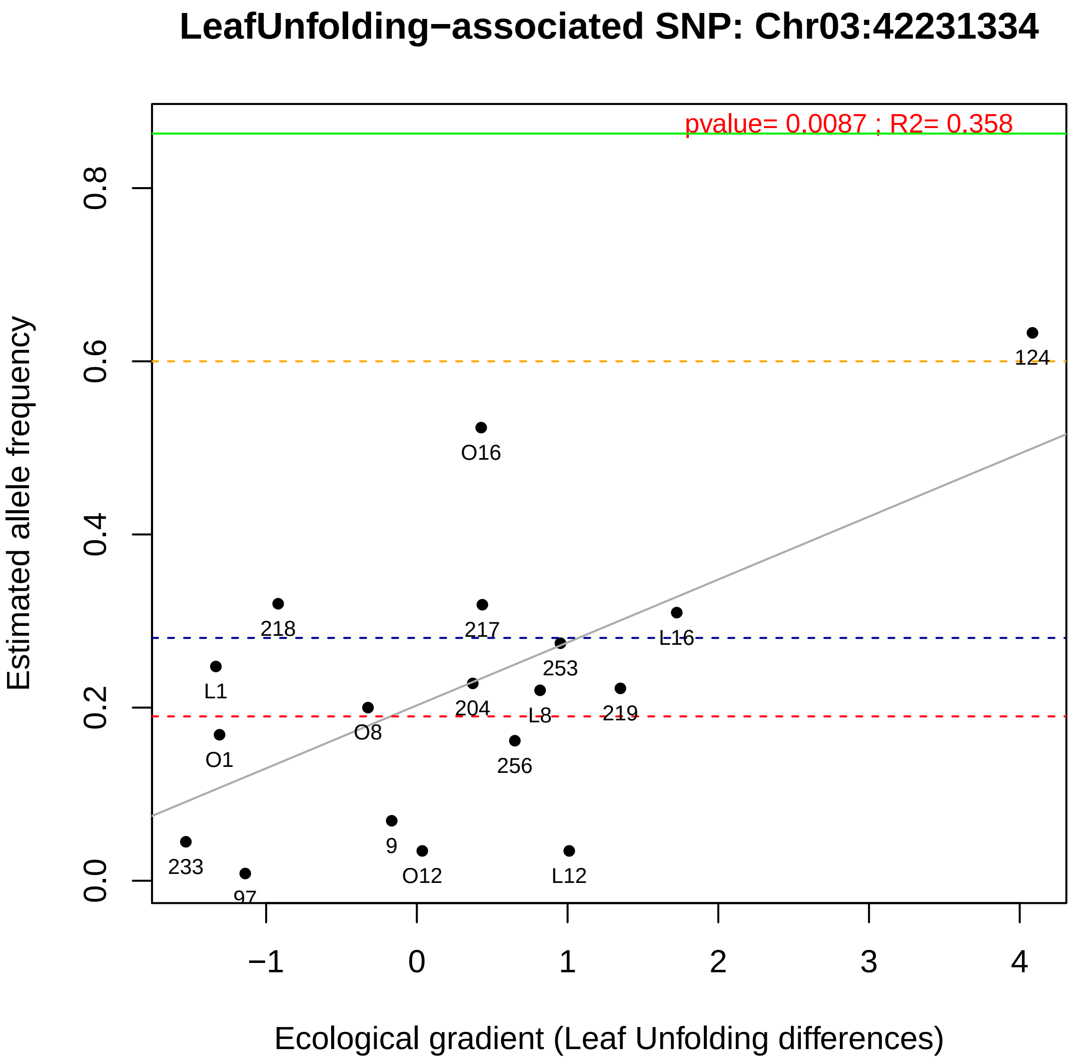
LeafUnfolding−associated SNP: Chr03:42231334
pvalue= 0.0087 ; R2= 0.358
Estimated allele frequency
0.0
0.2
0.4
0.6
0.8
−1
0
1
2
3
4
Ecological gradient (Leaf Unfolding differences)
124
O16
218
217
L16
253
L1
204
L8
219
O8
O1
256
9
233
O12
L12
97

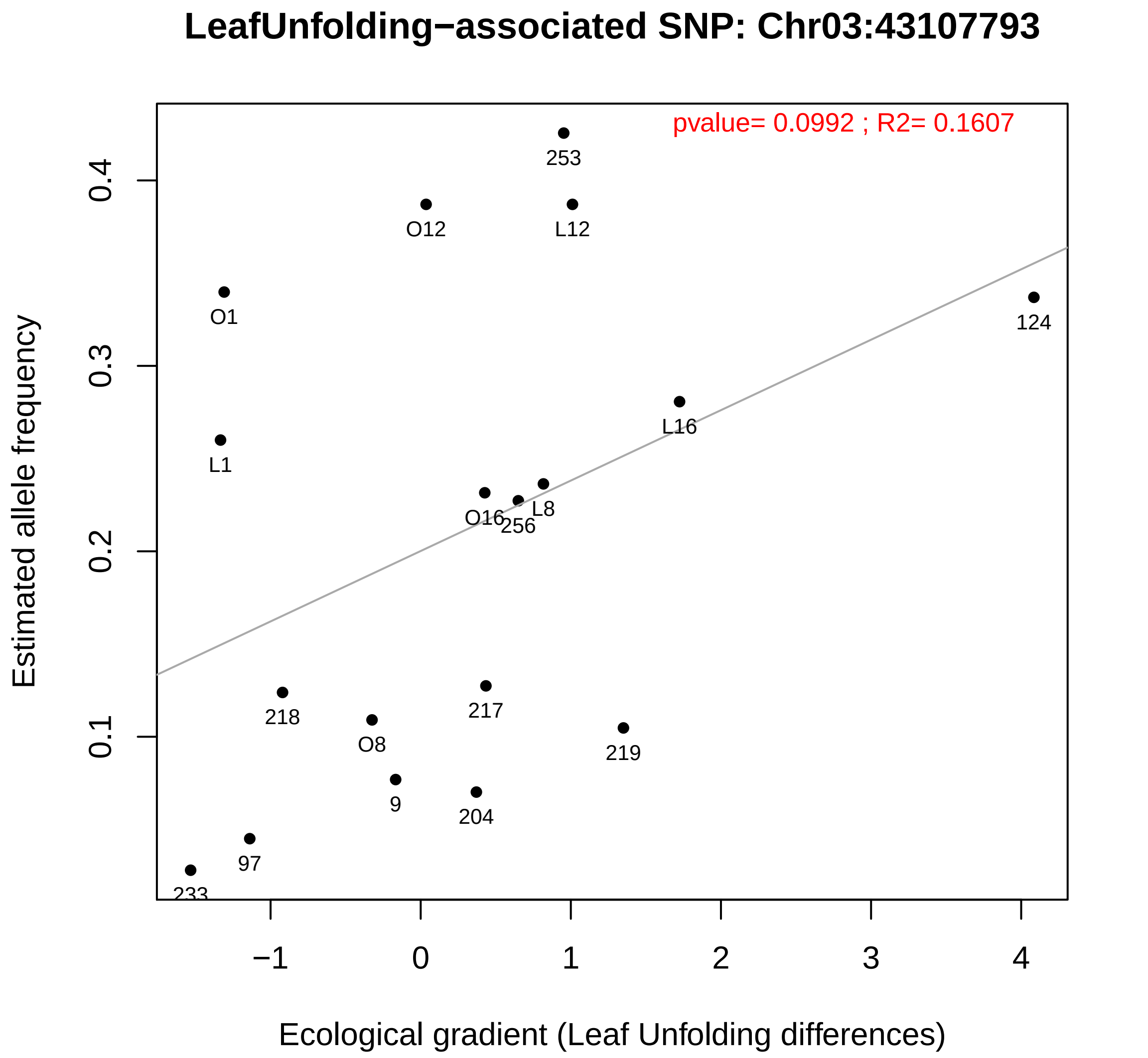
LeafUnfolding−associated SNP: Chr03:43107793
pvalue= 0.0992 ; R2= 0.1607
Estimated allele frequency
Ecological gradient (Leaf Unfolding differences)
0.1
0.2
0.3
0.4
−1
0
1
2
3
4
253
O12
L12
O1
124
L16
L1
O16
256
L8
218
217
O8
219
9
204
97
233

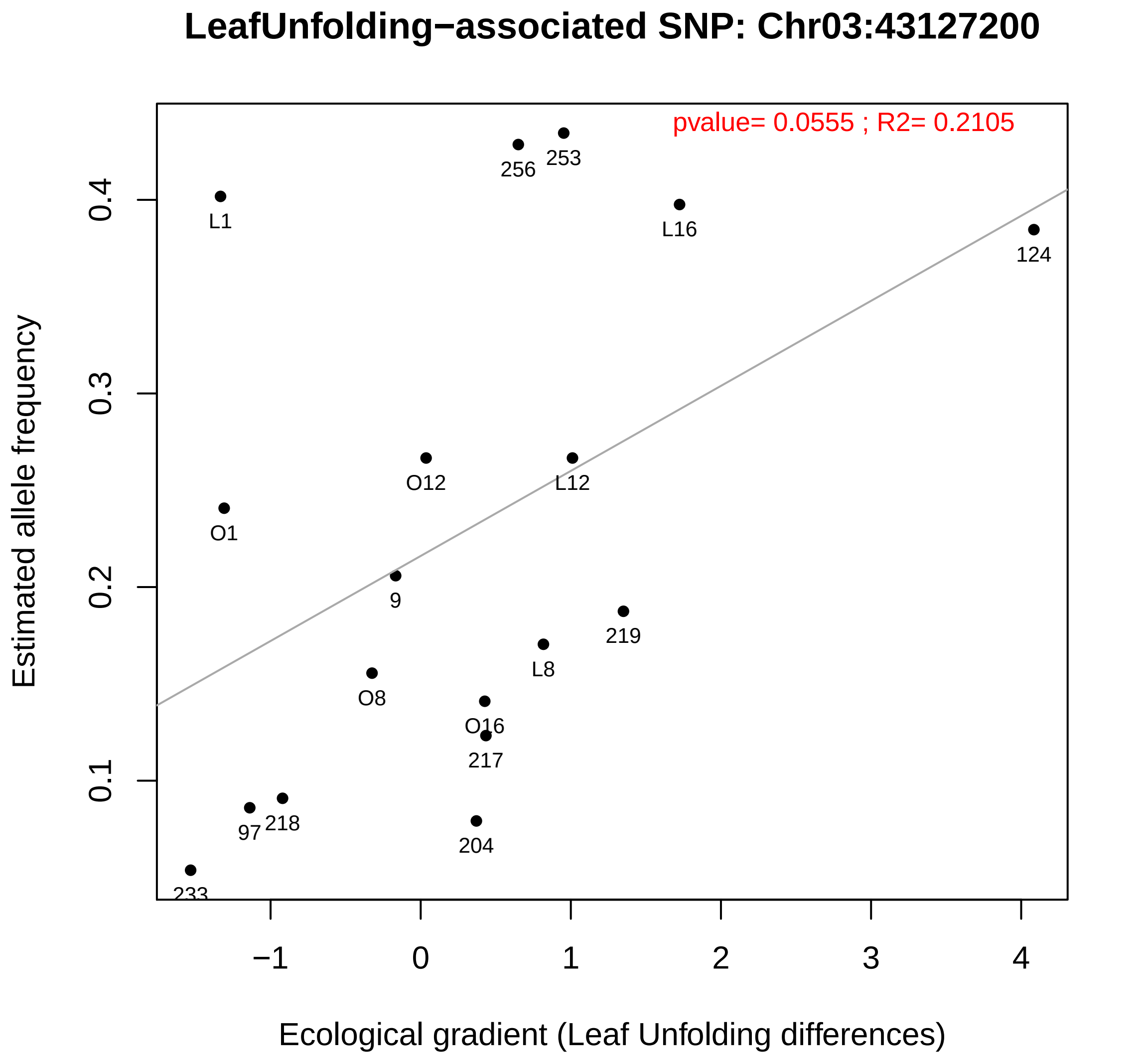
LeafUnfolding−associated SNP: Chr03:43127200
pvalue= 0.0555 ; R2= 0.2105
Estimated allele frequency
Ecological gradient (Leaf Unfolding differences)
0.1
0.2
0.3
0.4
−1
0
1
2
3
4
256
253
L1
L16
124
O12
L12
O1
9
219
L8
O8
O16
217
97
218
204
233

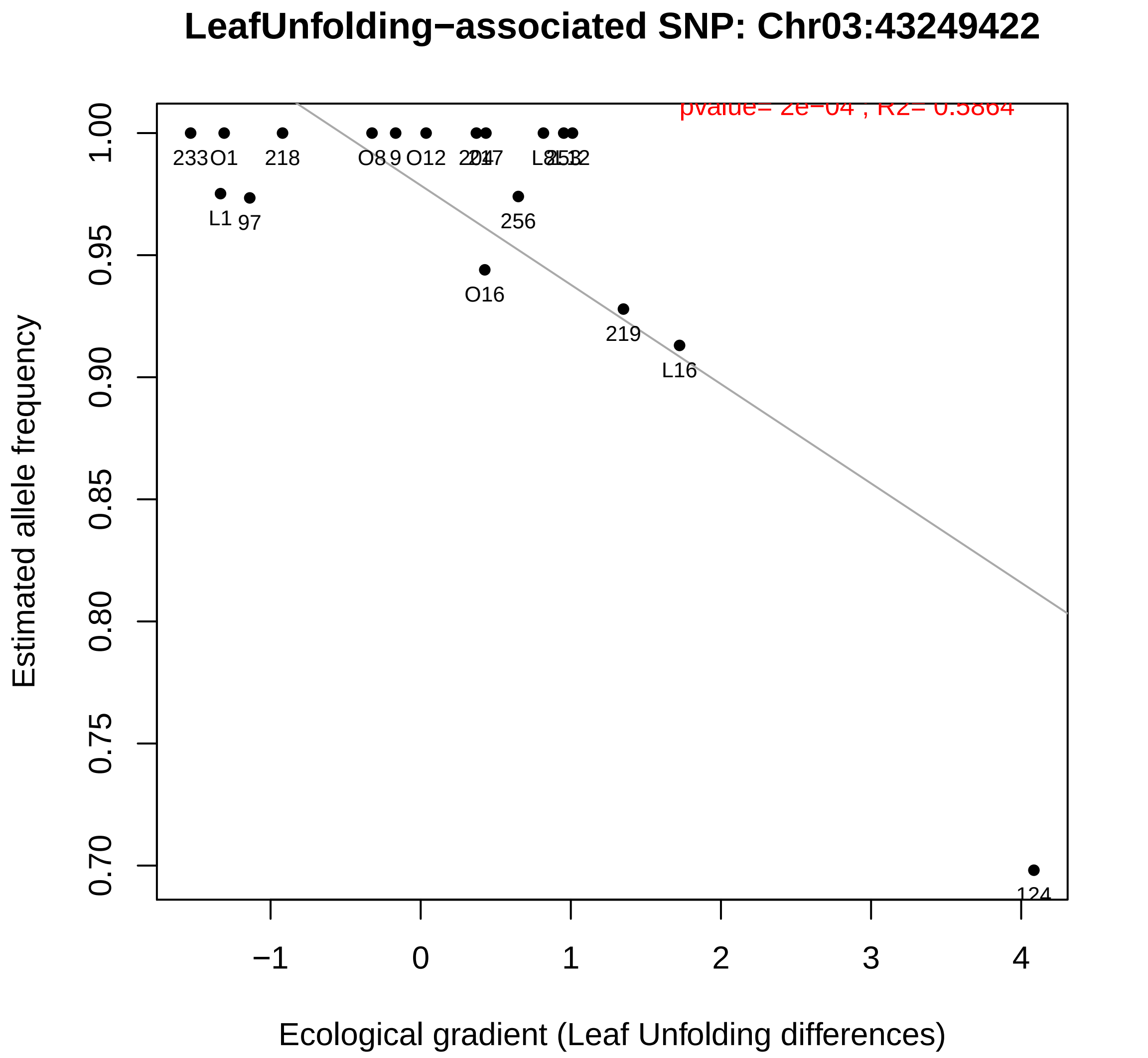

LeafUnfolding−associated SNP: Chr03:43249422
pvalue= 2e−04 , R2= 0.5864
233
O1
218
O8
9
O12
L1
97
256
O16
219
L16
124
Estimated allele frequency
Ecological gradient (Leaf Unfolding differences)
1.00
0.95
0.90
0.85
0.80
0.75
0.70
−1
0
1
2
3
4

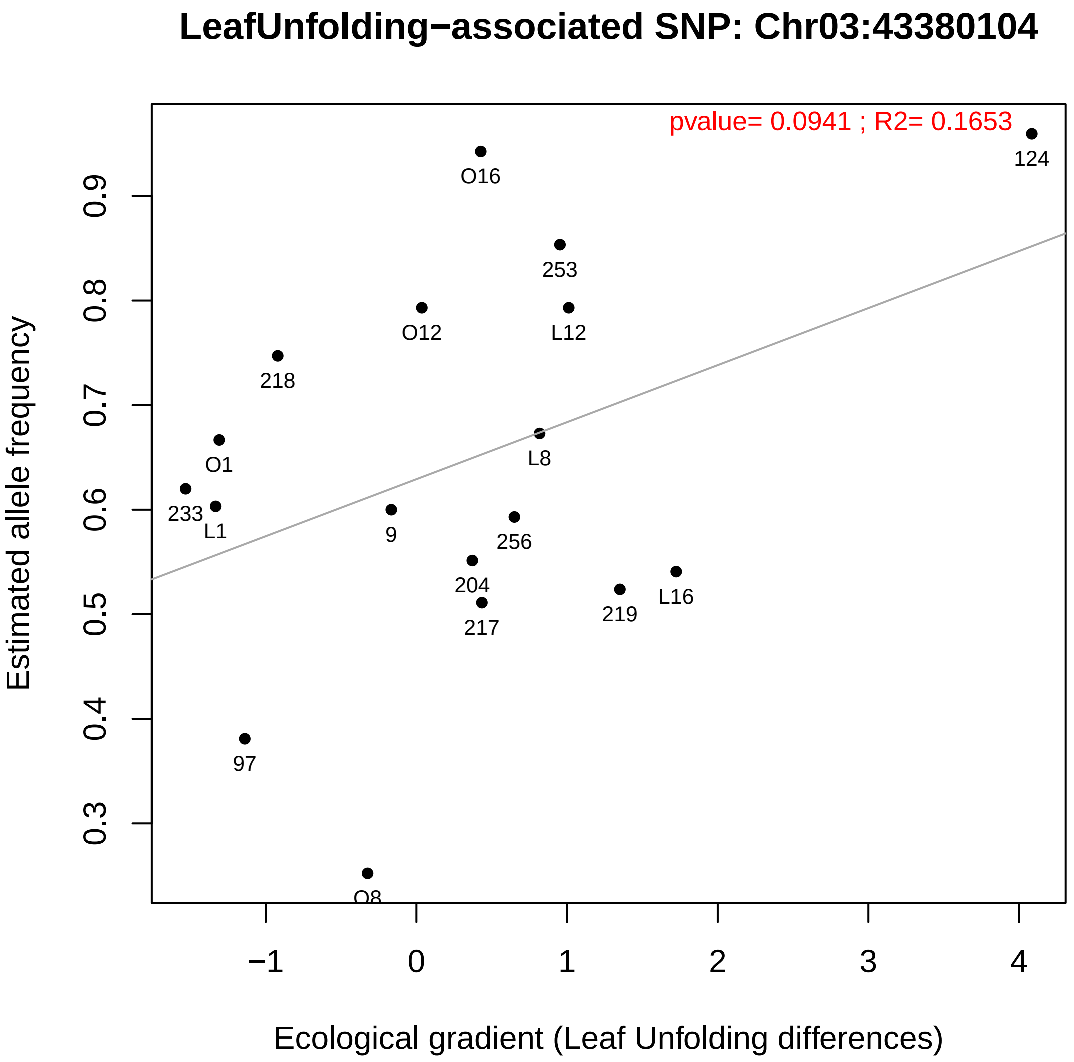

LeafUnfolding−associated SNP: Chr03:43380104
pvalue= 0.0941 ; R2= 0.1653
Estimated allele frequency
Ecological gradient (Leaf Unfolding differences)
0.3
0.4
0.5
0.6
0.7
0.8
0.9
−1
0
1
2
3
4
124
O16
253
O12
L12
218
O1
L8
233
L1
9
256
204
217
219
L16
97
O8

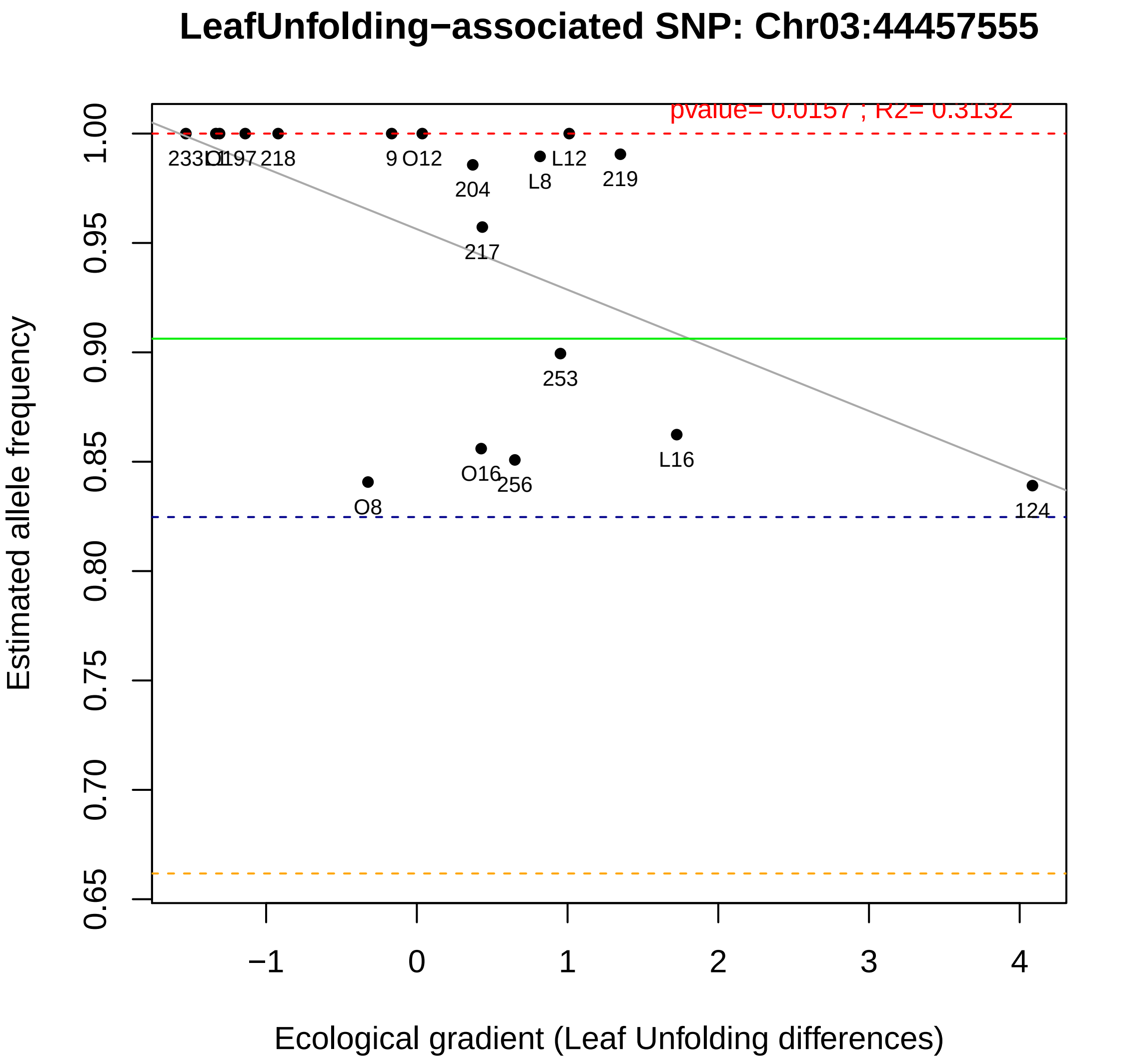
LeafUnfolding−associated SNP: Chr03:44457555
pvalue= 0.0157 ; R2= 0.3132
Estimated allele frequency
1.00
0.95
0.90
0.85
0.80
0.75
0.70
0.65
233
O1
197
218
9
O12
204
L12
L8
219
217
253
O16
256
L16
O8
124
−1
0
1
2
3
4
Ecological gradient (Leaf Unfolding differences)

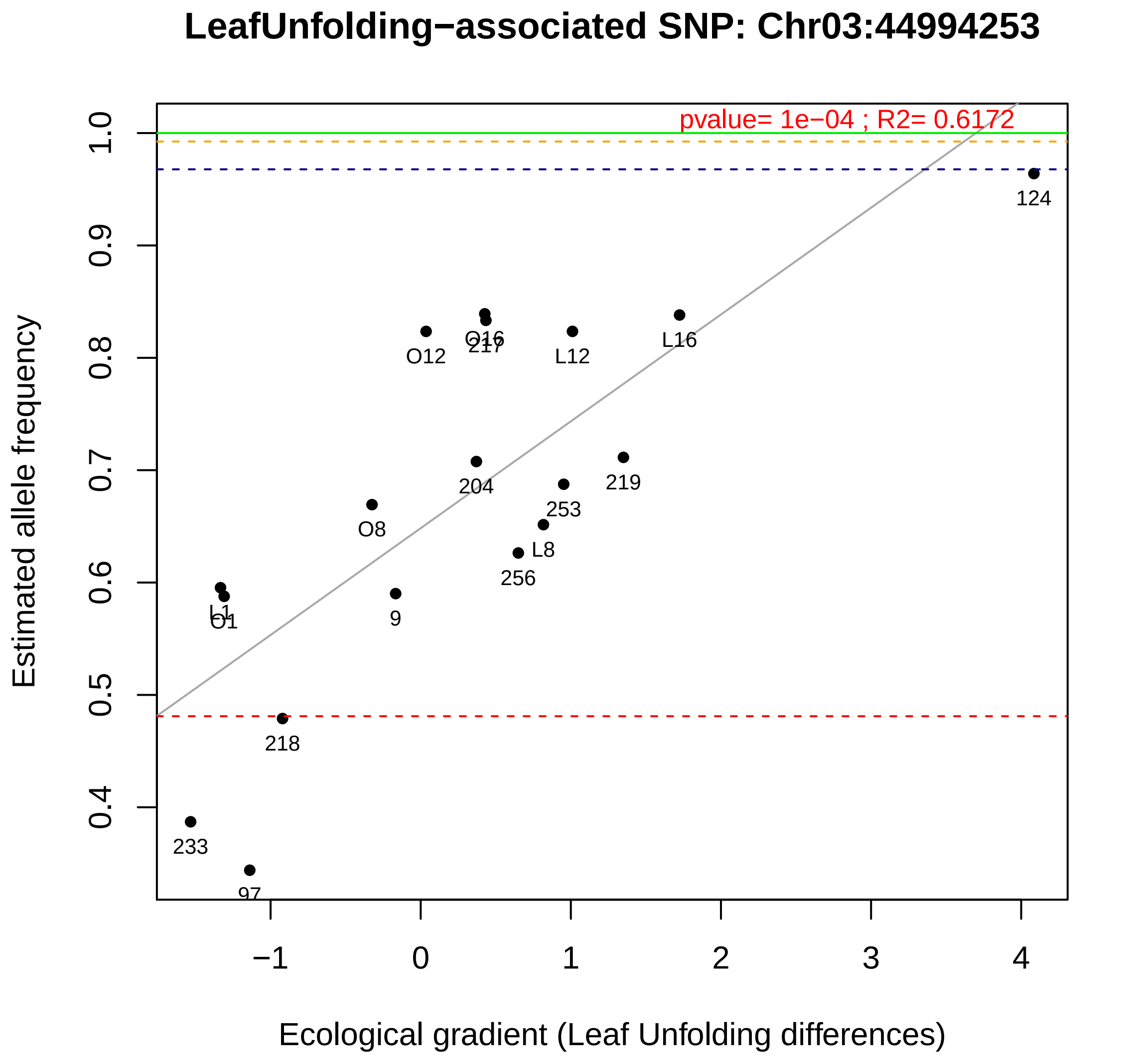
LeafUnfolding−associated SNP: Chr03:44994253
pvalue= 1e−04 ; R2= 0.6172
Estimated allele frequency
Ecological gradient (Leaf Unfolding differences)
0.4
0.5
0.6
0.7
0.8
0.9
1.0
−1
0
1
2
3
4
124
O16
217
O12
L12
L16
204
219
253
O8
L8
256
L1
O1
9
218
233
97

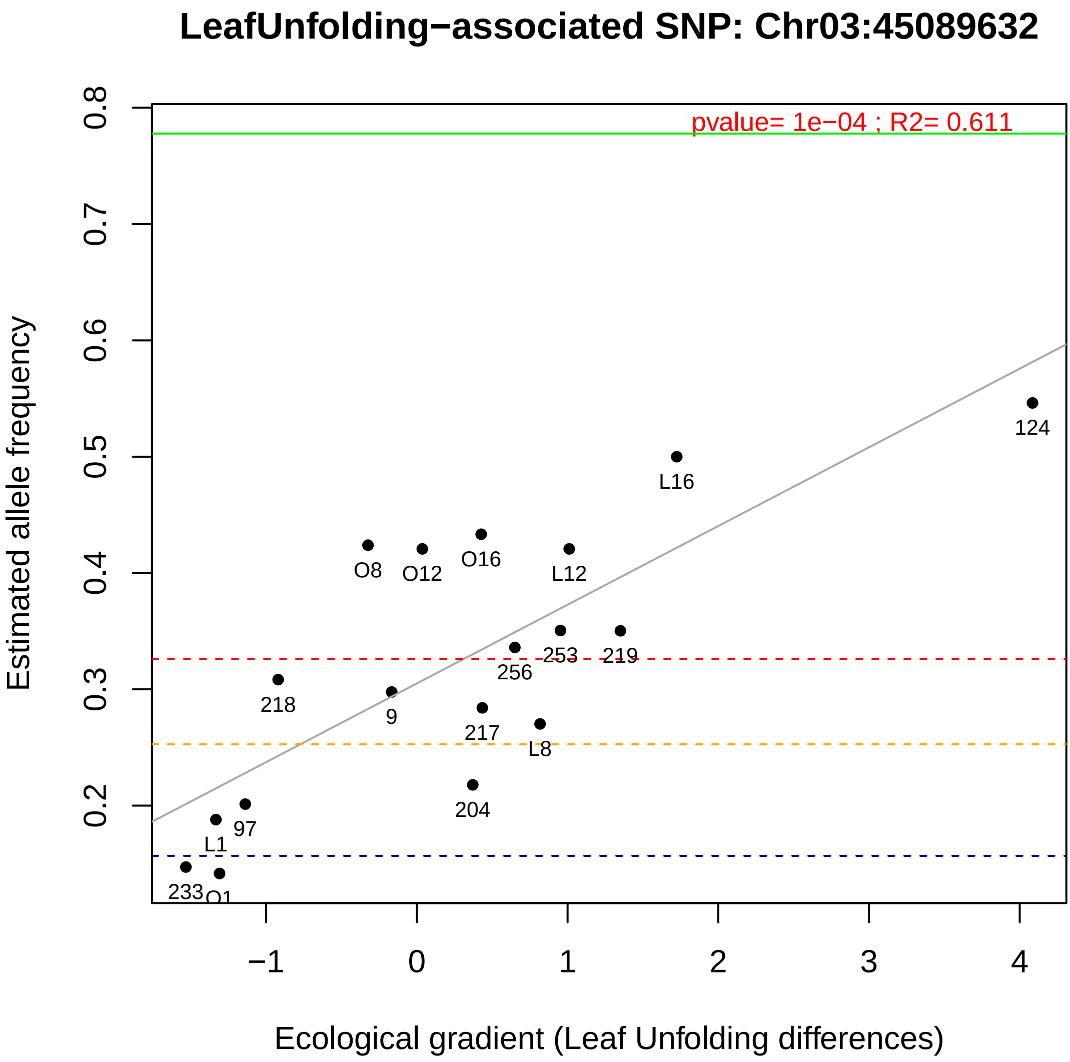

LeafUnfolding−associated SNP: Chr03:45089632
pvalue= 1e−04 ; R2= 0.611
Estimated allele frequency
Ecological gradient (Leaf Unfolding differences)
0.8
0.7
0.6
0.5
0.4
0.3
0.2
−1
0
1
2
3
4
124
L16
O8
O12
O16
L12
253
219
256
218
9
217
L8
204
97
L1
233
O1

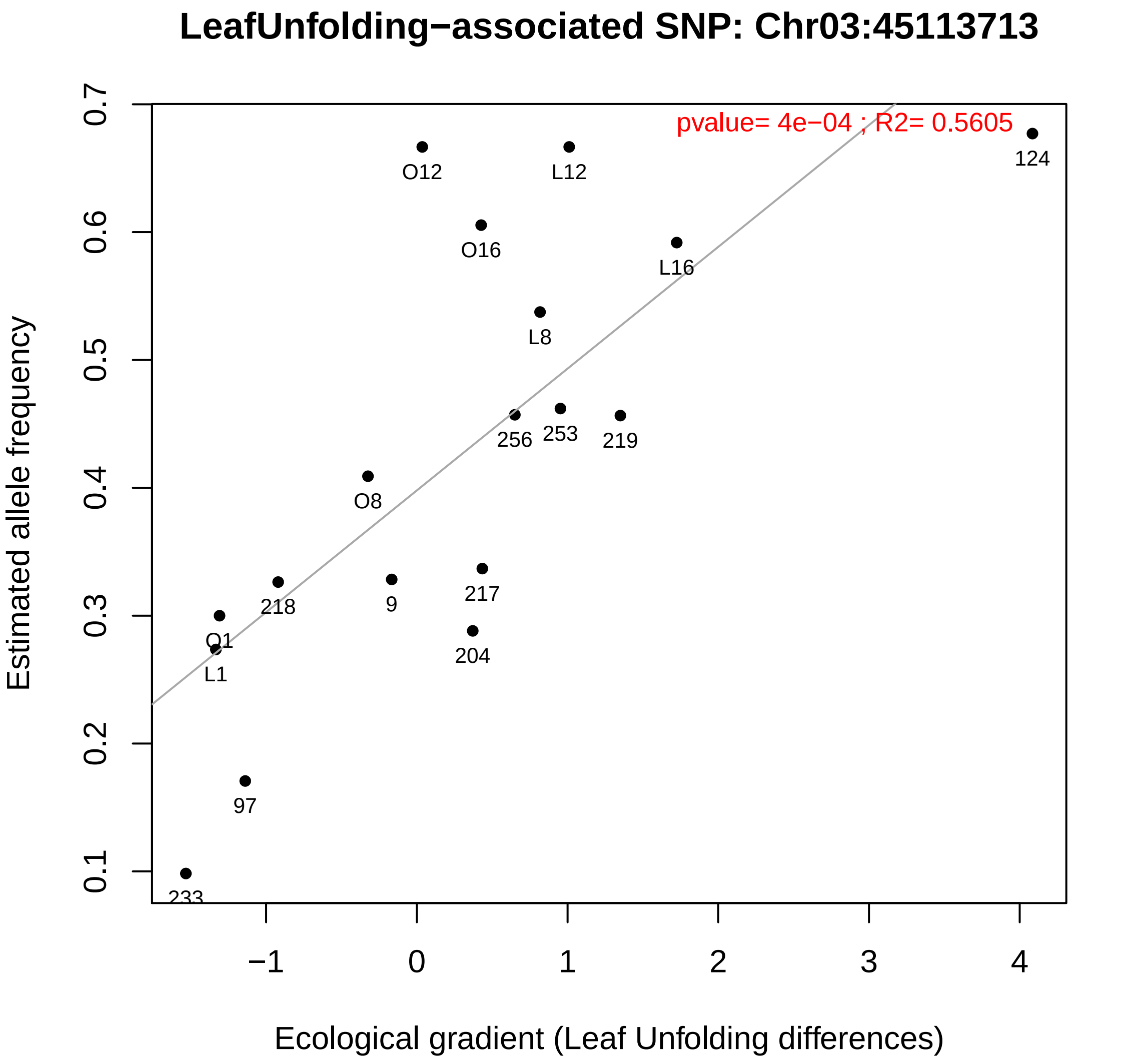

LeafUnfolding−associated SNP: Chr03:45113713
pvalue= 4e−04 ; R2= 0.5605
Estimated allele frequency
Ecological gradient (Leaf Unfolding differences)
0.1
0.2
0.3
0.4
0.5
0.6
0.7
−1
0
1
2
3
4
O12
L12
124
O16
L16
L8
256
253
219
O8
218
9
217
O1
L1
204
97
233

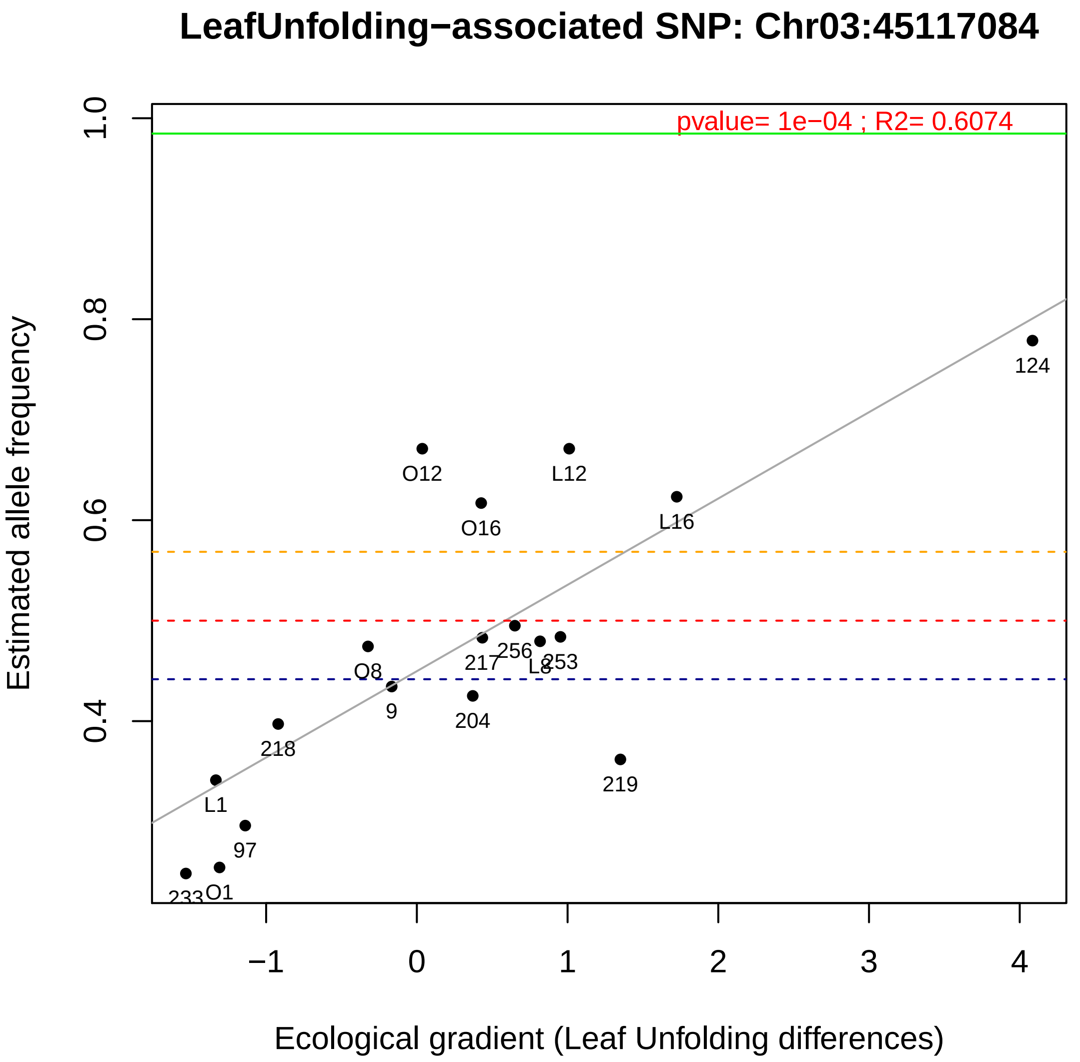
LeafUnfolding−associated SNP: Chr03:45117084
pvalue= 1e−04 ; R2= 0.6074
Estimated allele frequency
Ecological gradient (Leaf Unfolding differences)
0.4
0.6
0.8
1.0
−1
0
1
2
3
4
124
O12
L12
O16
L16
256
217
L8
253
O8
9
204
218
219
L1
97
233
O1

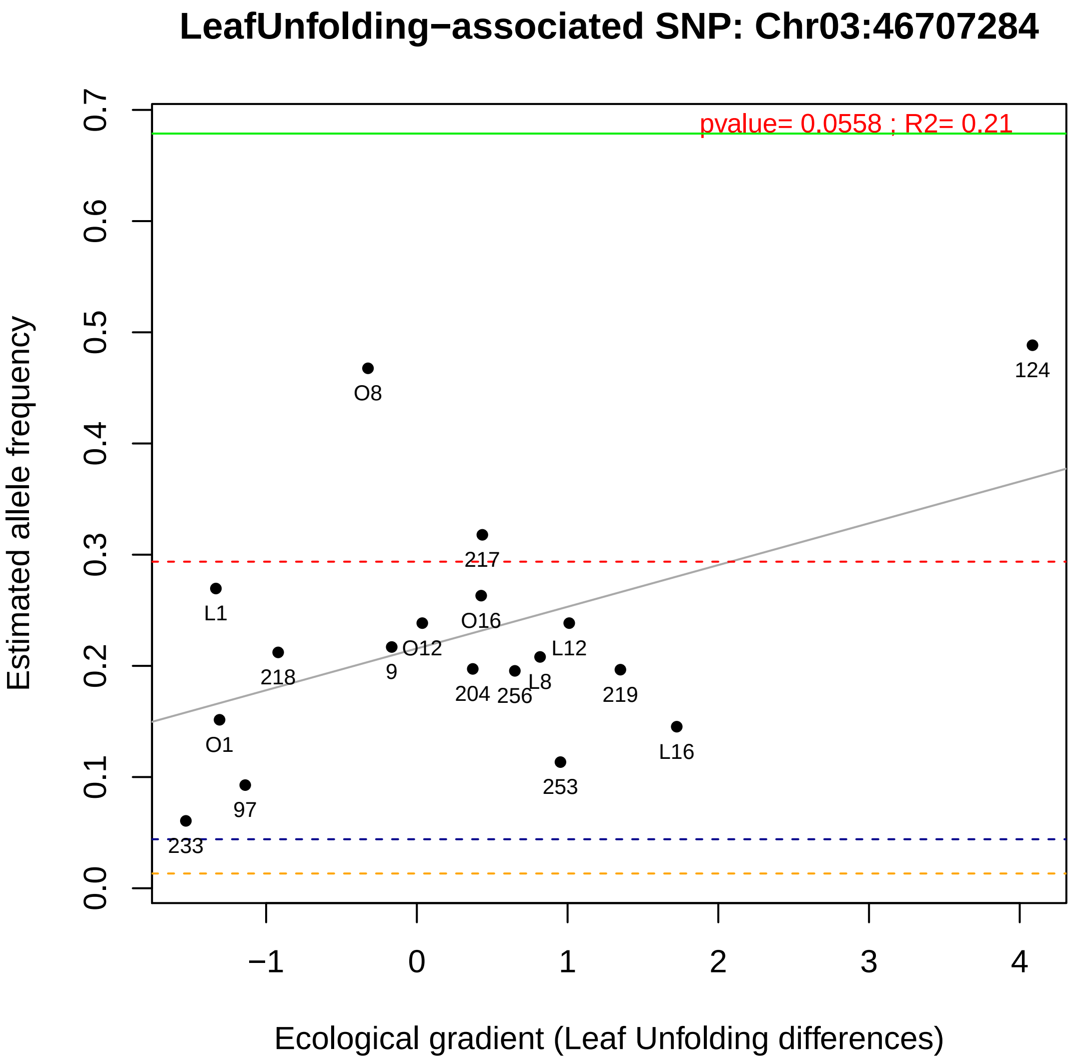
LeafUnfolding−associated SNP: Chr03:46707284
pvalue= 0.0558 ; R2= 0.21
Estimated allele frequency
0.0
0.1
0.2
0.3
0.4
0.5
0.6
0.7
−1
0
1
2
3
4
Ecological gradient (Leaf Unfolding differences)
O8
124
217
L1
O16
O12
9
218
L12
204
256
L8
219
O1
L16
253
97
233

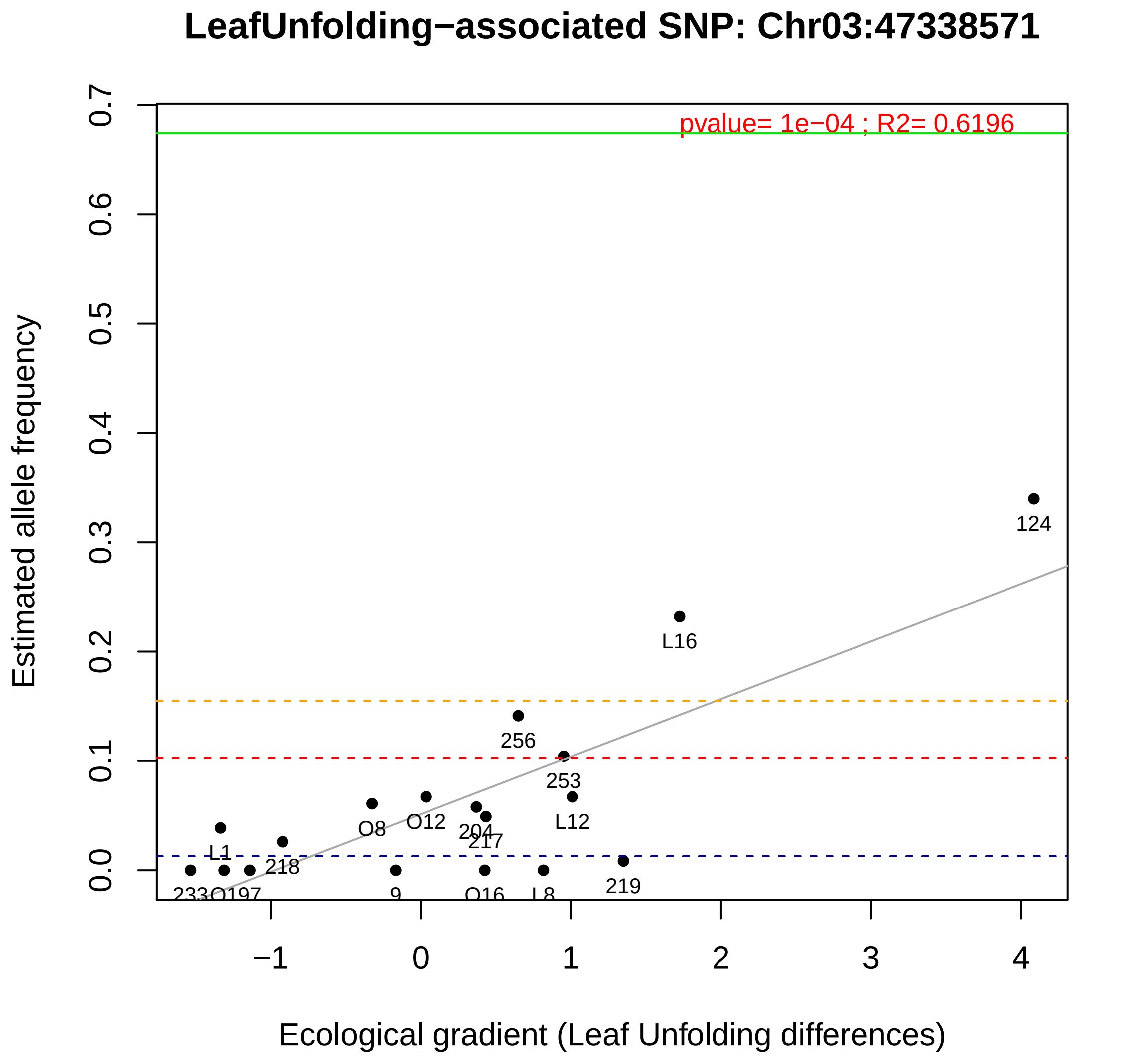
LeafUnfolding−associated SNP: Chr03:47338571
pvalue= 1e−04 ; R2= 0.6196
Estimated allele frequency
Ecological gradient (Leaf Unfolding differences)
0.0
0.1
0.2
0.3
0.4
0.5
0.6
0.7
−1
0
1
2
3
4
124
L16
256
253
L12
O12
O8
204
217
L1
218
219
9
O16
L8

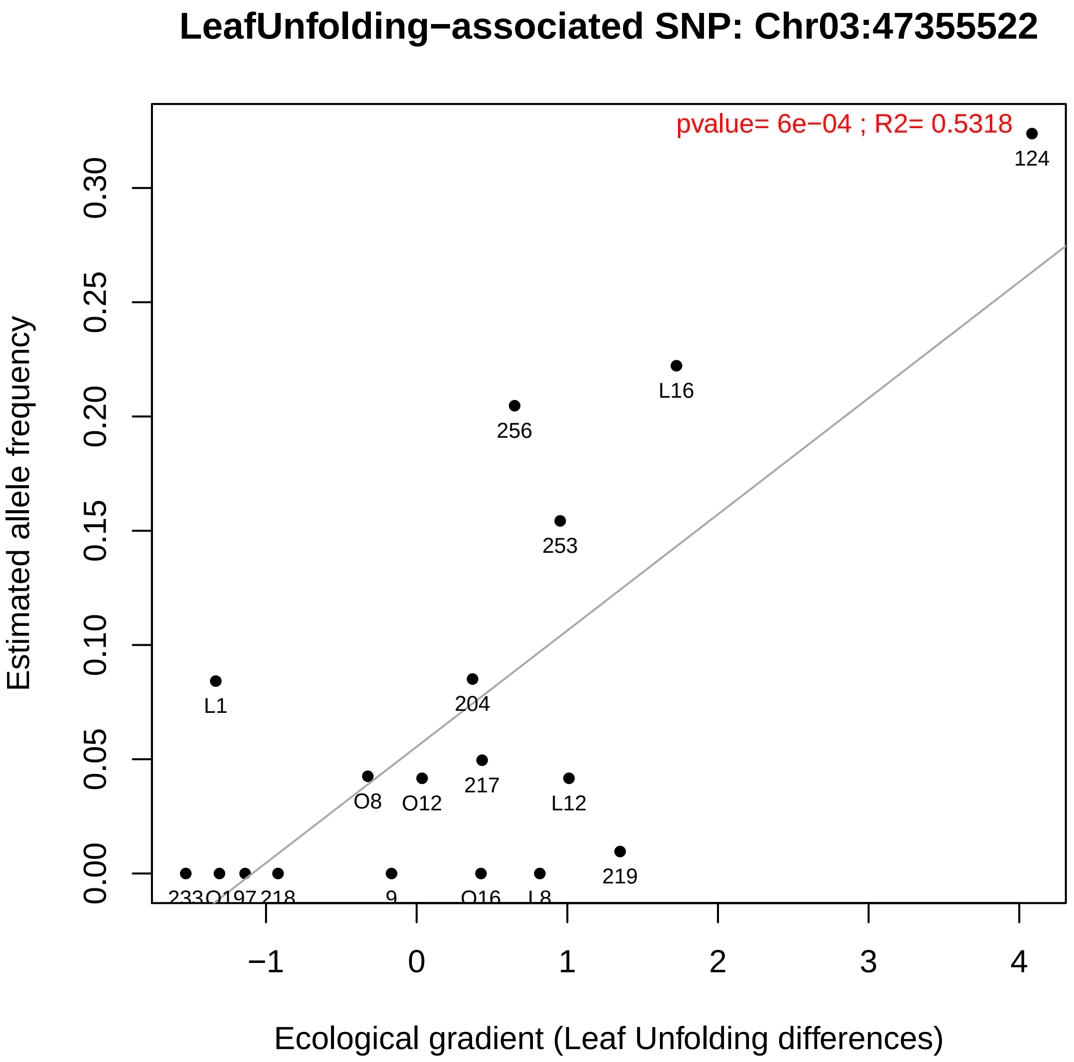
LeafUnfolding−associated SNP: Chr03:47355522
pvalue= 6e−04 ; R2= 0.5318
Estimated allele frequency
0.00
0.05
0.10
0.15
0.20
0.25
0.30
Ecological gradient (Leaf Unfolding differences)
−1
0
1
2
3
4
124
L16
256
253
L1
204
217
O8
O12
L12
219
233
O197
218
9
O16
L8

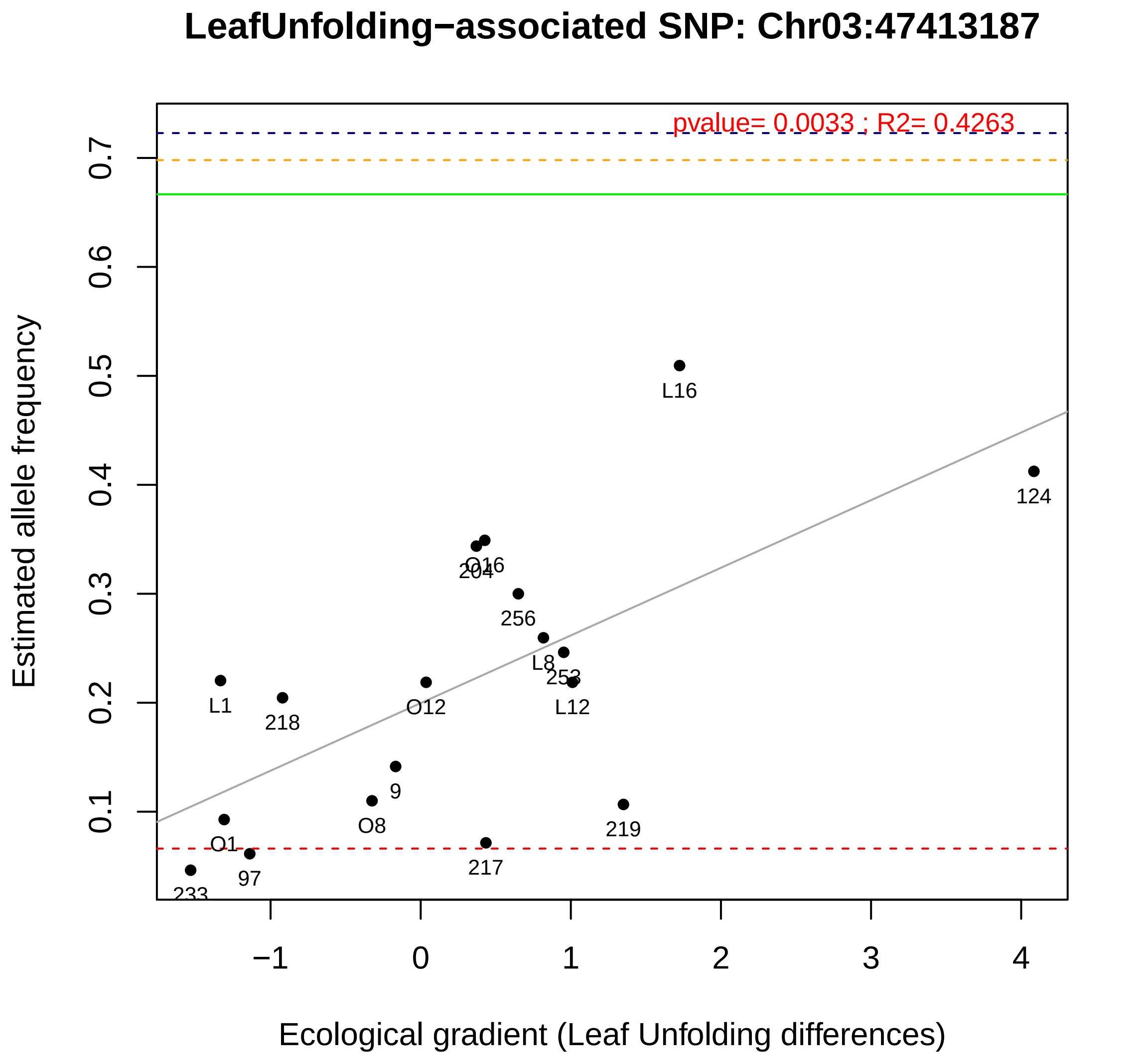

LeafUnfolding−associated SNP: Chr03:47413187
pvalue= 0.0033 ; R2= 0.4263
Estimated allele frequency
0.1
0.2
0.3
0.4
0.5
0.6
0.7
−1
0
1
2
3
4
Ecological gradient (Leaf Unfolding differences)
L16
124
O16
204
256
L8
253
L12
L1
218
O12
9
O8
O1
97
217
219
233

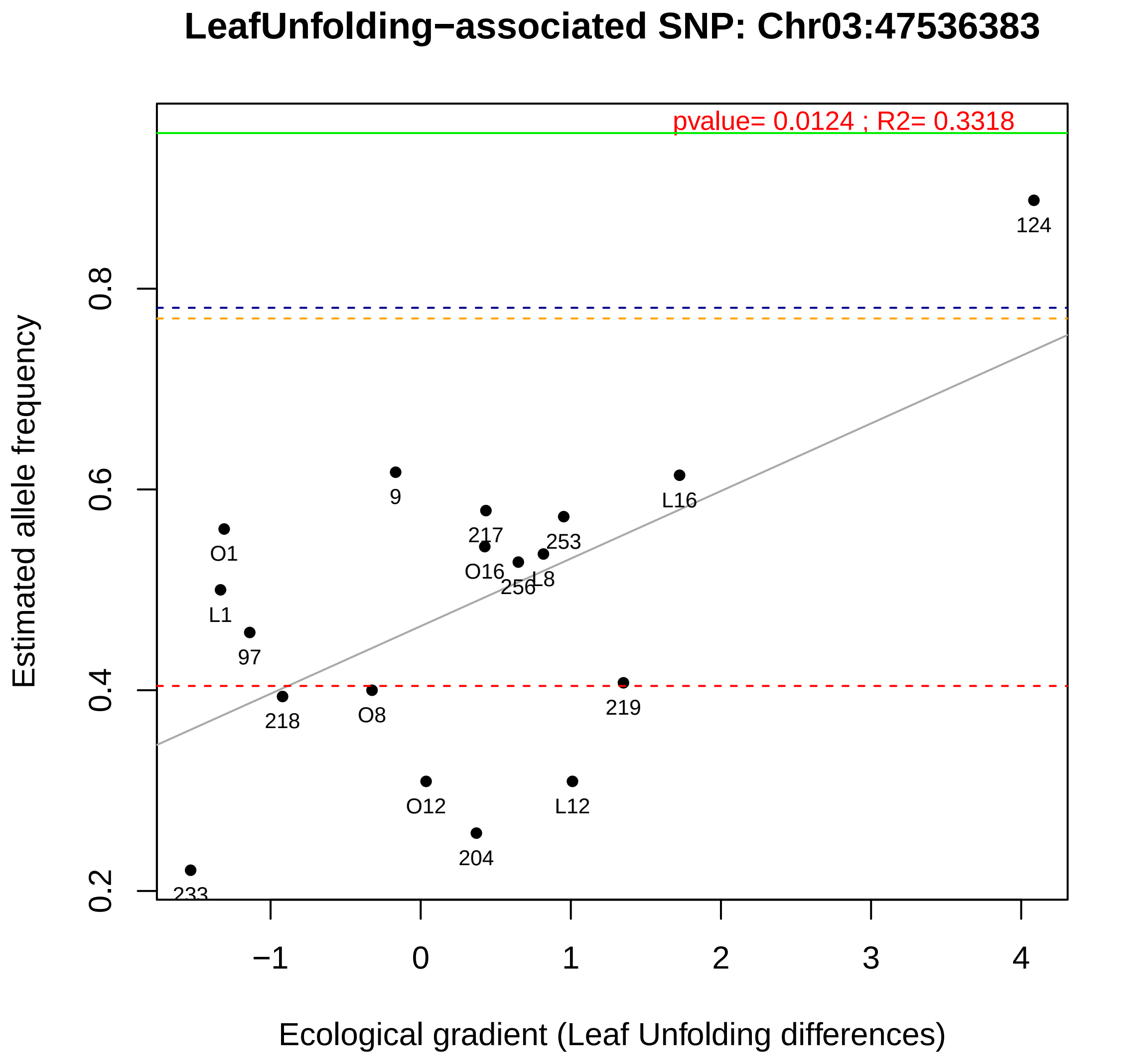
LeafUnfolding−associated SNP: Chr03:47536383
pvalue= 0.0124 ; R2= 0.3318
Estimated allele frequency
0.2
0.4
0.6
0.8
−1
0
1
2
3
4
Ecological gradient (Leaf Unfolding differences)
124
L16
9
217
253
O1
O16
256
L8
L1
97
218
O8
219
O12
L12
204
233

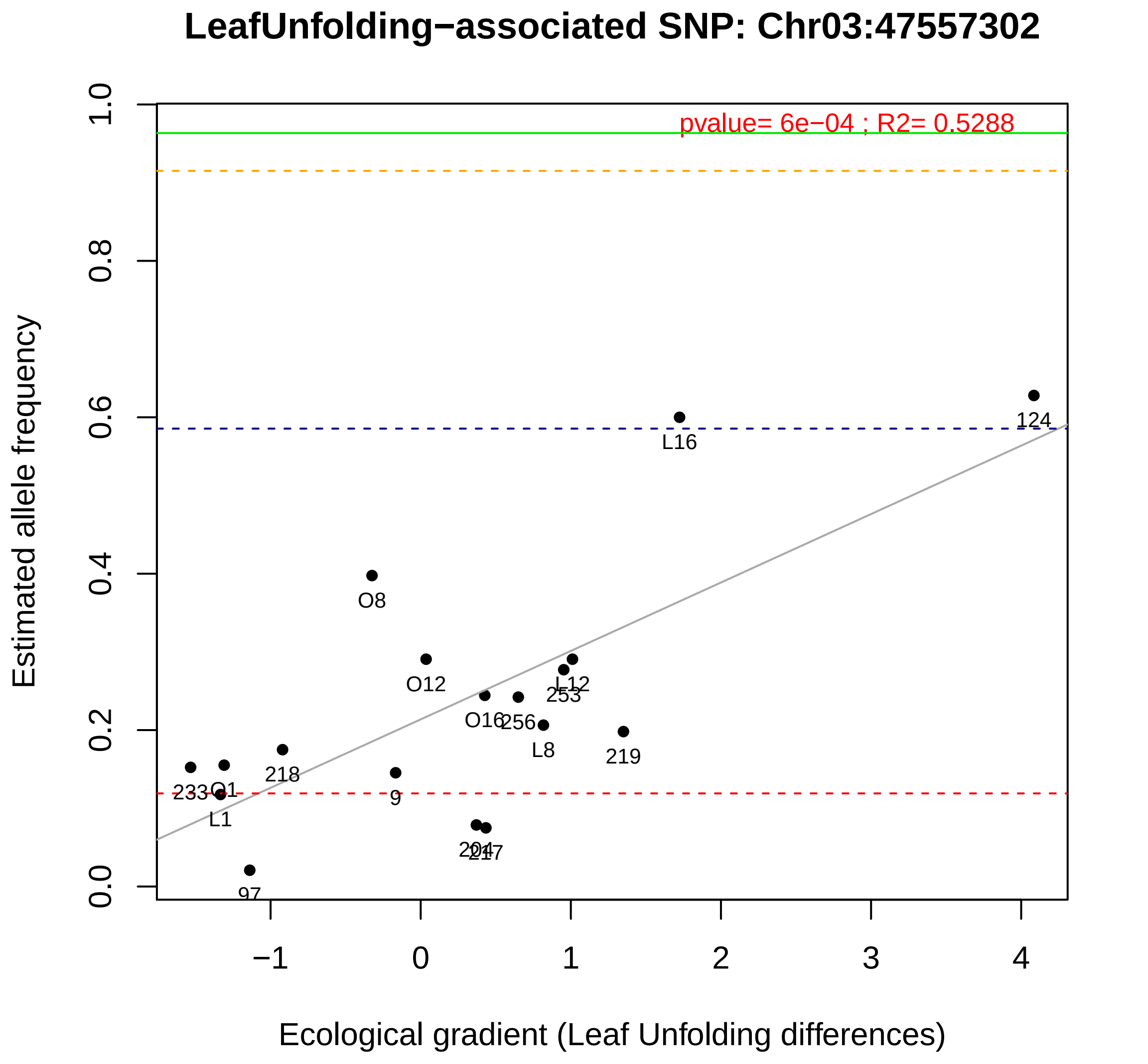

LeafUnfolding−associated SNP: Chr03:47557302
pvalue= 6e−04 ; R2= 0.5288
Estimated allele frequency
Ecological gradient (Leaf Unfolding differences)
0.0
0.2
0.4
0.6
0.8
1.0
−1
0
1
2
3
4
124
L16
O8
O12
L12
253
O16
256
L8
219
218
233
O1
L1
9
204
217
97

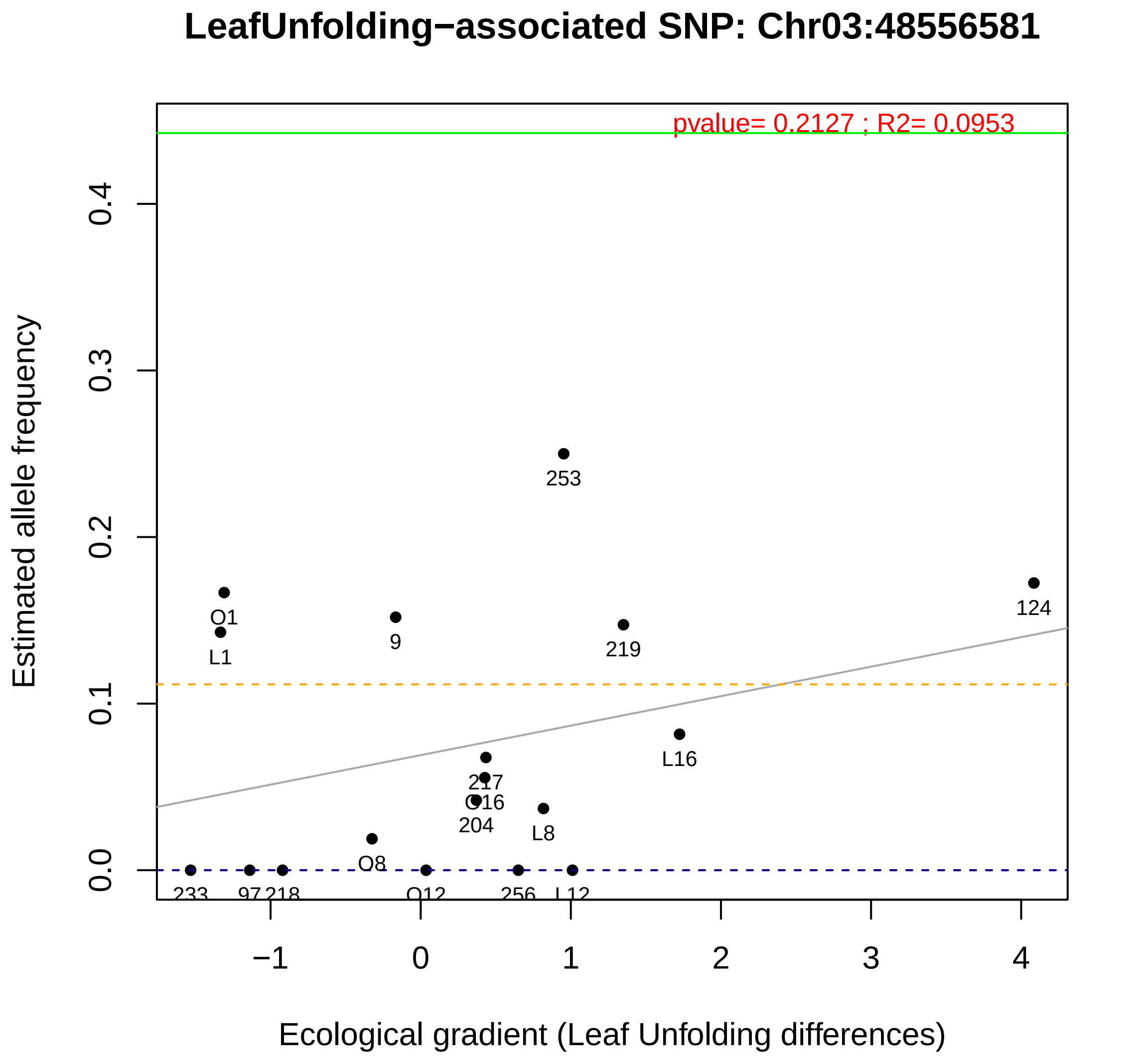

LeafUnfolding−associated SNP: Chr03:48556581
pvalue= 0.2127 ; R2= 0.0953
Estimated allele frequency
0.0
0.1
0.2
0.3
0.4
−1
0
1
2
3
4
Ecological gradient (Leaf Unfolding differences)
253
O1
L1
9
219
124
L16
217
O16
204
L8
O8
233
97
218
O12
256
L12

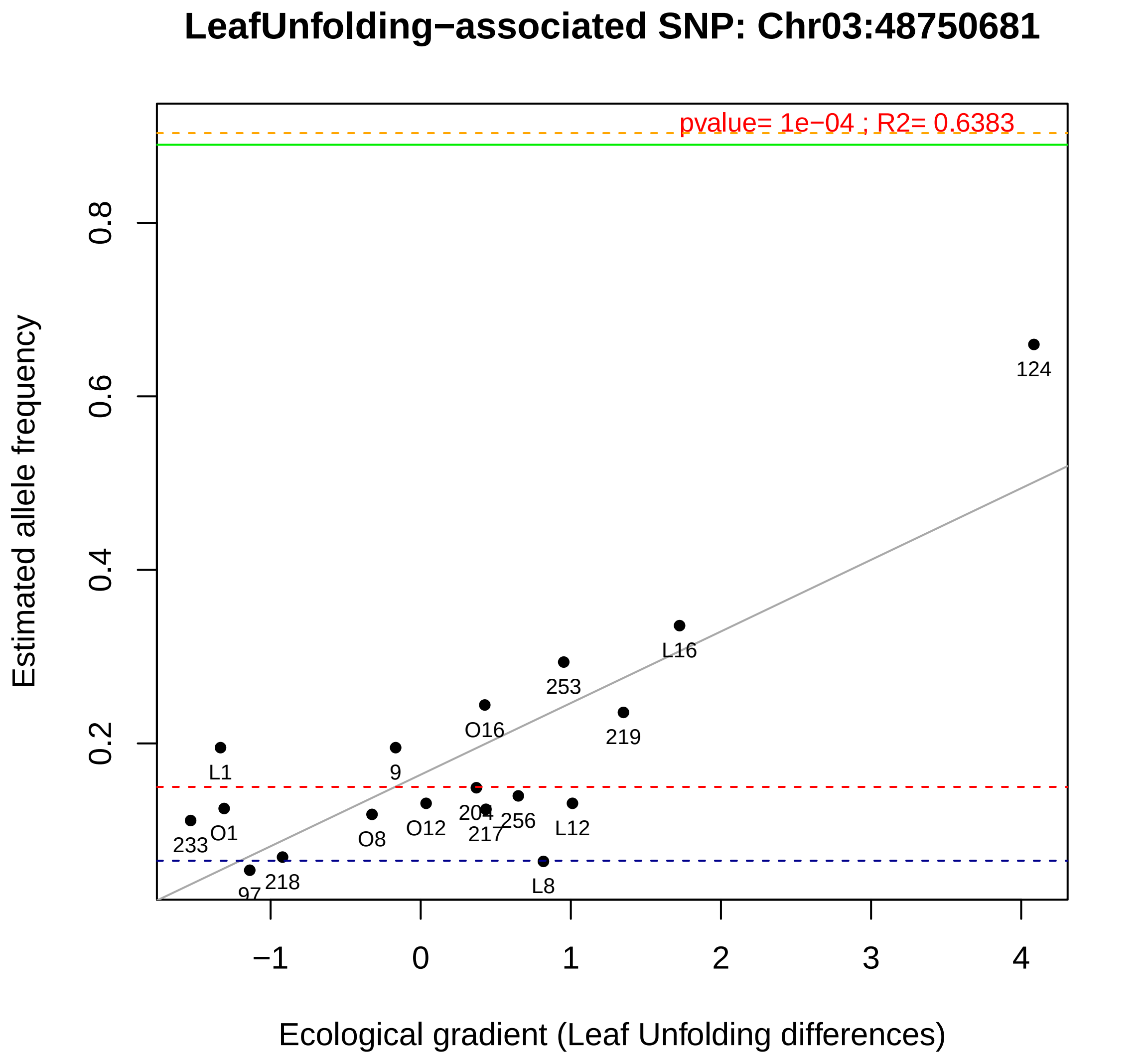

LeafUnfolding−associated SNP: Chr03:48750681
pvalue= 1e−04 ; R2= 0.6383
Estimated allele frequency
0.2
0.4
0.6
0.8
−1
0
1
2
3
4
Ecological gradient (Leaf Unfolding differences)
124
L16
253
219
O16
L1
9
204
217
256
L12
233
O1
O8
O12
218
97
L8

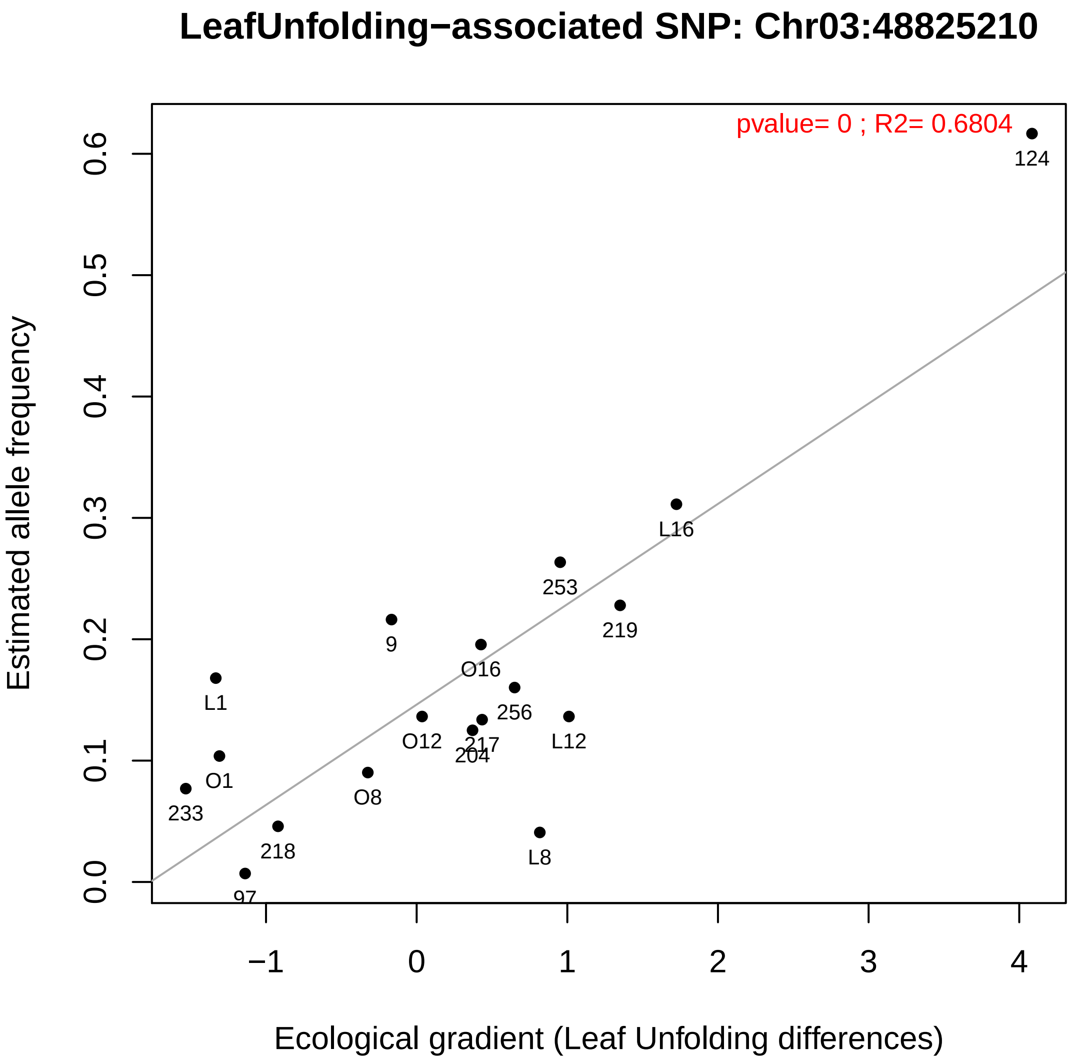

LeafUnfolding−associated SNP: Chr03:48825210
pvalue= 0 ; R2= 0.6804
124
L16
253
219
9
O16
256
L1
O12
217
204
L12
O1
O8
233
218
L8
97
0.0
0.1
0.2
0.3
0.4
0.5
0.6
−1
0
1
2
3
4
Estimated allele frequency
Ecological gradient (Leaf Unfolding differences)

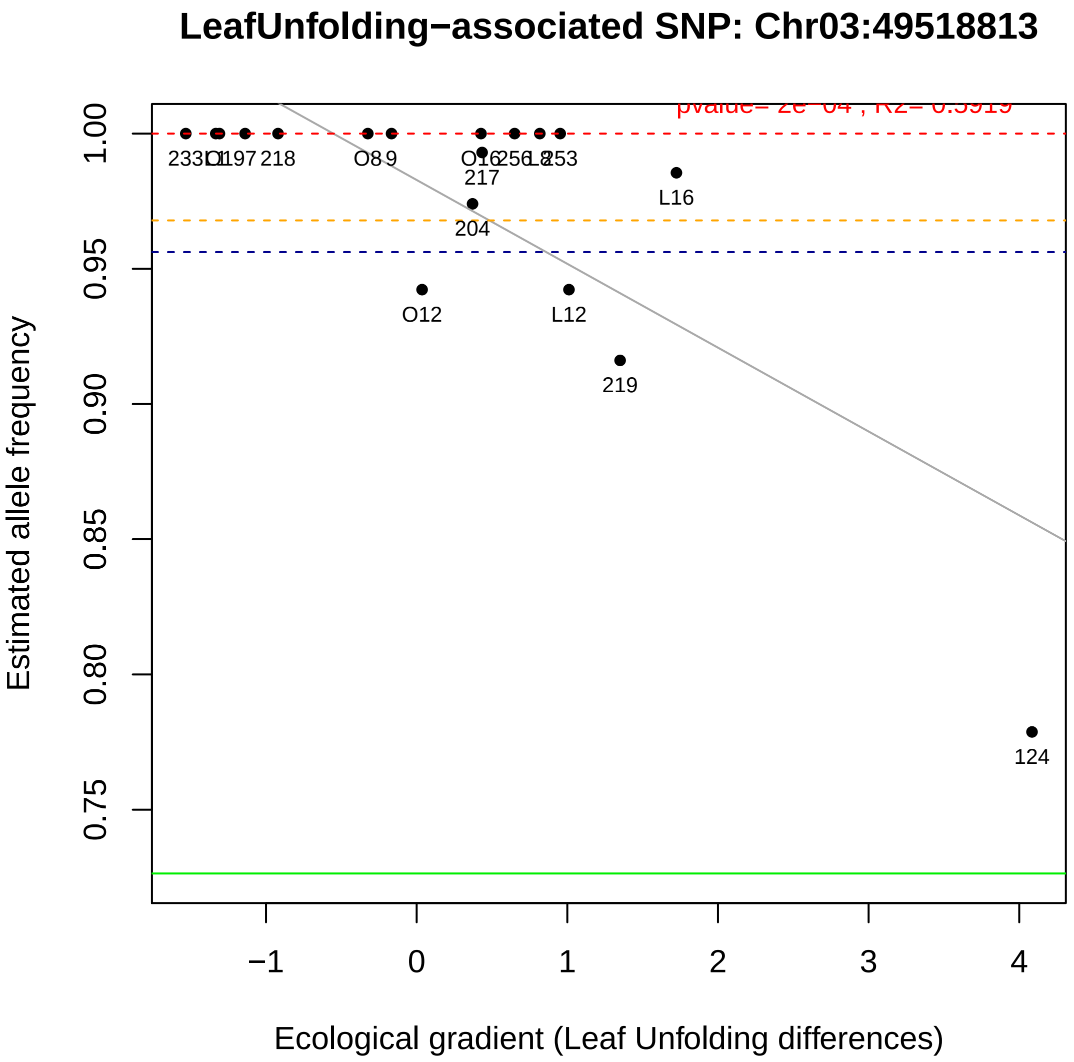

LeafUnfolding−associated SNP: Chr03:49518813
pvalue= 2e−04 , R2= 0.5919
Estimated allele frequency
1.00
0.95
0.90
0.85
0.80
0.75
−1
0
1
2
3
4
Ecological gradient (Leaf Unfolding differences)
233
O1
197
218
O8
9
O16
256
L8
253
217
L16
204
O12
L12
219
124

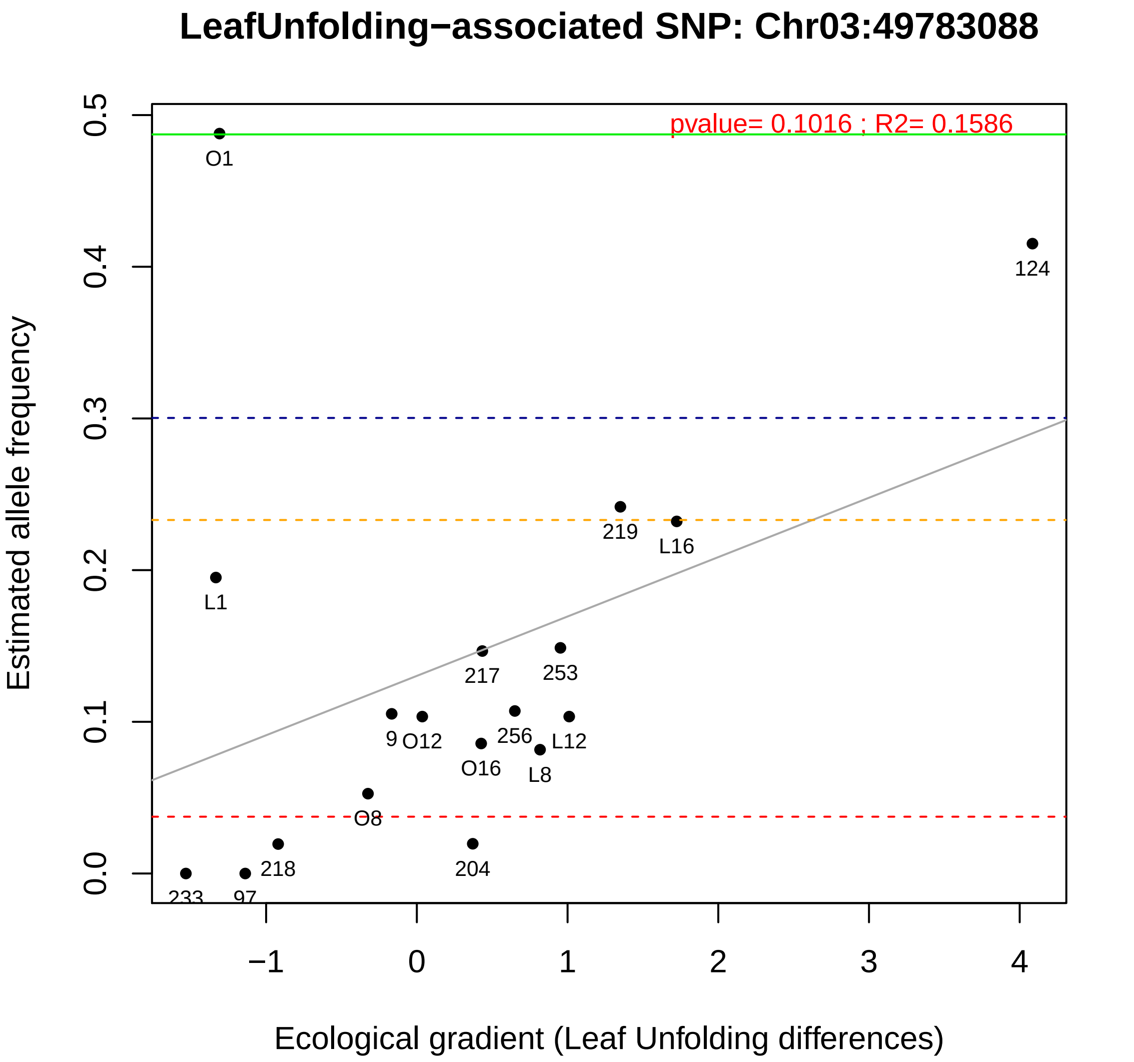
LeafUnfolding−associated SNP: Chr03:49783088
pvalue= 0.1016 ; R2= 0.1586
Estimated allele frequency
0.0
0.1
0.2
0.3
0.4
0.5
Ecological gradient (Leaf Unfolding differences)
−1
0
1
2
3
4
O1
124
219
L16
L1
217
253
9
O12
256
L12
O16
L8
O8
218
204
233
97

**LeafUnfolding−associated SNP: Chr03:50311687**

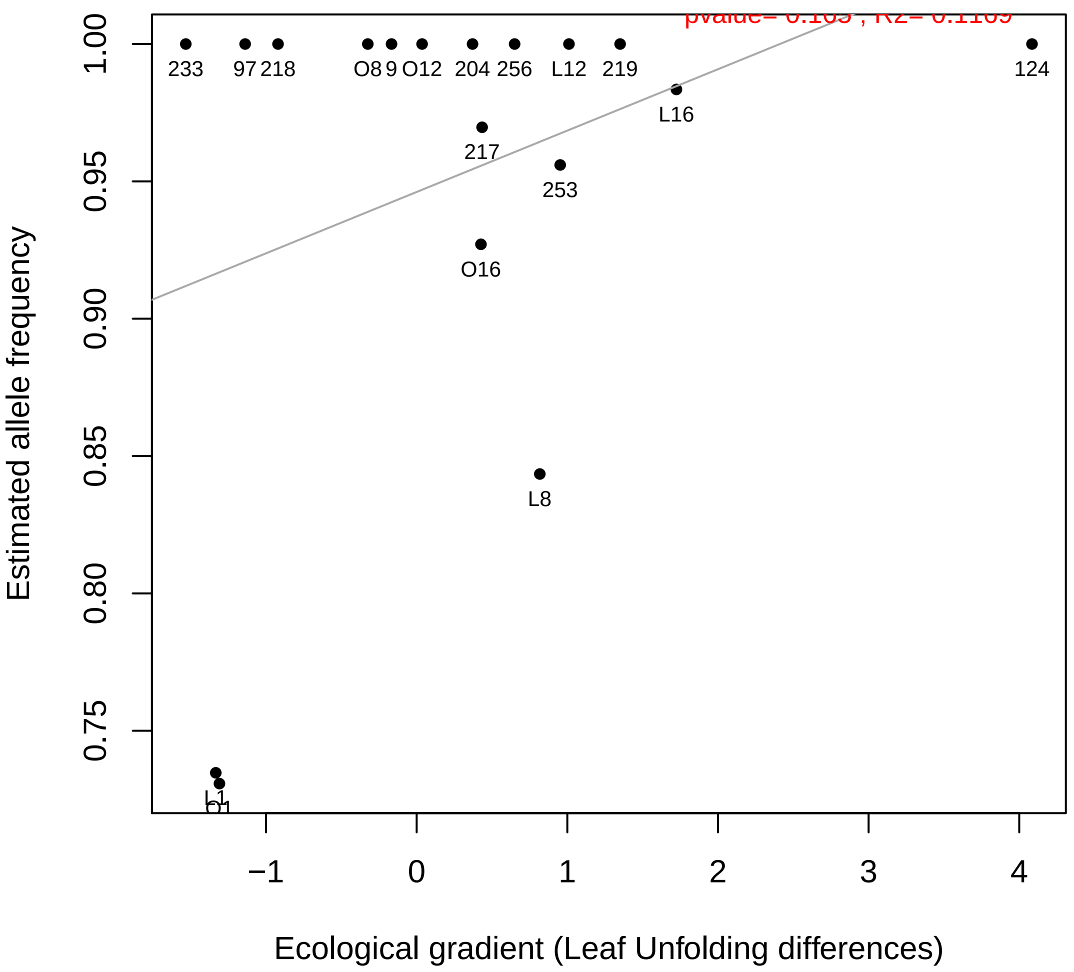

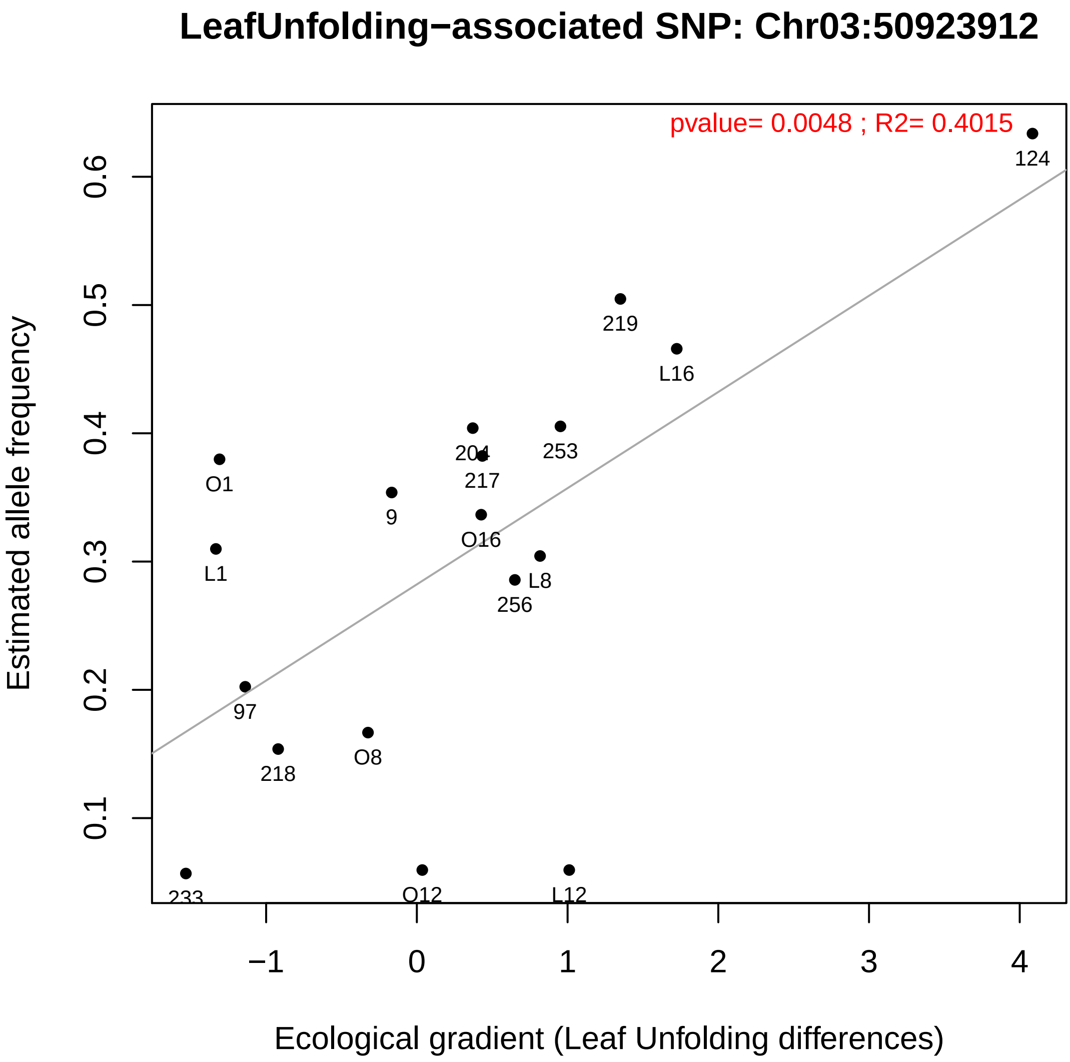
LeafUnfolding−associated SNP: Chr03:50923912
pvalue= 0.0048 ; R2= 0.4015
Estimated allele frequency
Ecological gradient (Leaf Unfolding differences)
0.1
0.2
0.3
0.4
0.5
0.6
−1
0
1
2
3
4
124
219
L16
204
253
217
O1
9
O16
L1
L8
256
97
218
O8
233
O12
L12

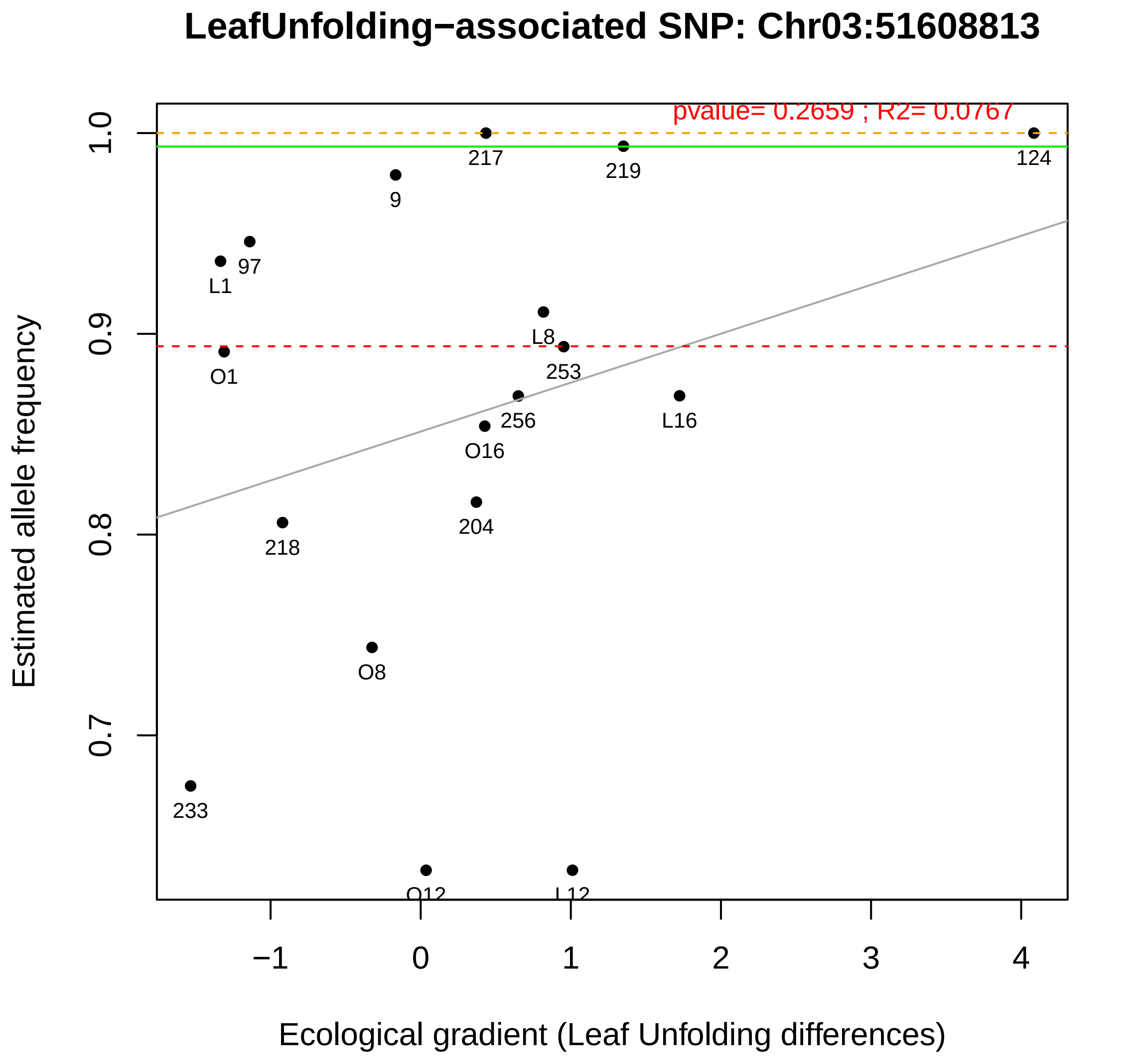

LeafUnfolding−associated SNP: Chr03:51608813
pvalue= 0.2659 ; R2= 0.0767
Estimated allele frequency
1.0
0.9
0.8
0.7
−1
0
1
2
3
4
217
219
124
9
97
L1
L8
253
O1
256
L16
O16
204
218
O8
233
O12
L12
Ecological gradient (Leaf Unfolding differences)

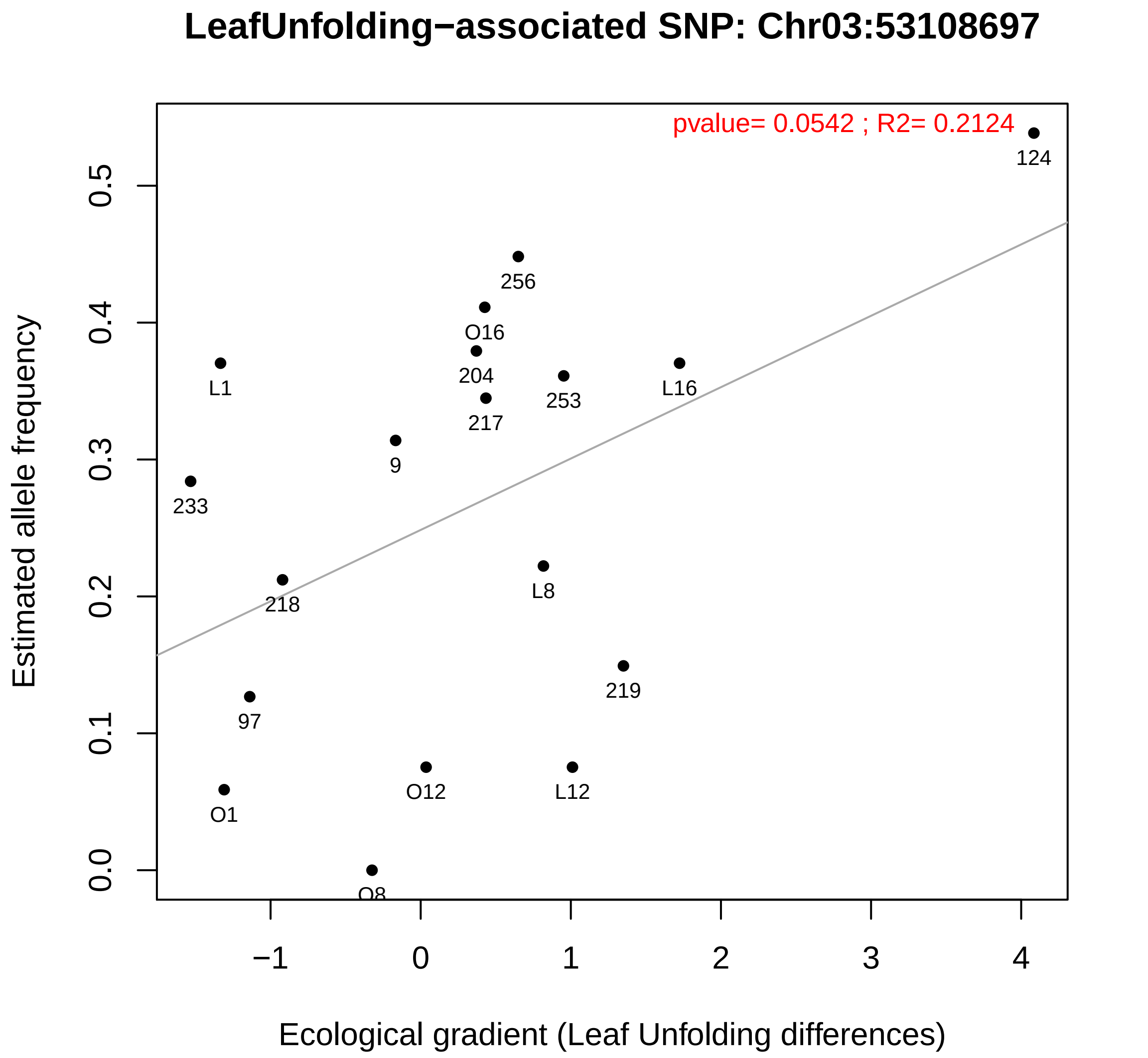
LeafUnfolding−associated SNP: Chr03:53108697
pvalue= 0.0542 ; R2= 0.2124
Estimated allele frequency
0.0
0.1
0.2
0.3
0.4
0.5
124
256
O16
204
L1
253
L16
217
9
233
218
L8
219
97
O12
L12
O1
O8
−1
0
1
2
3
4
Ecological gradient (Leaf Unfolding differences)

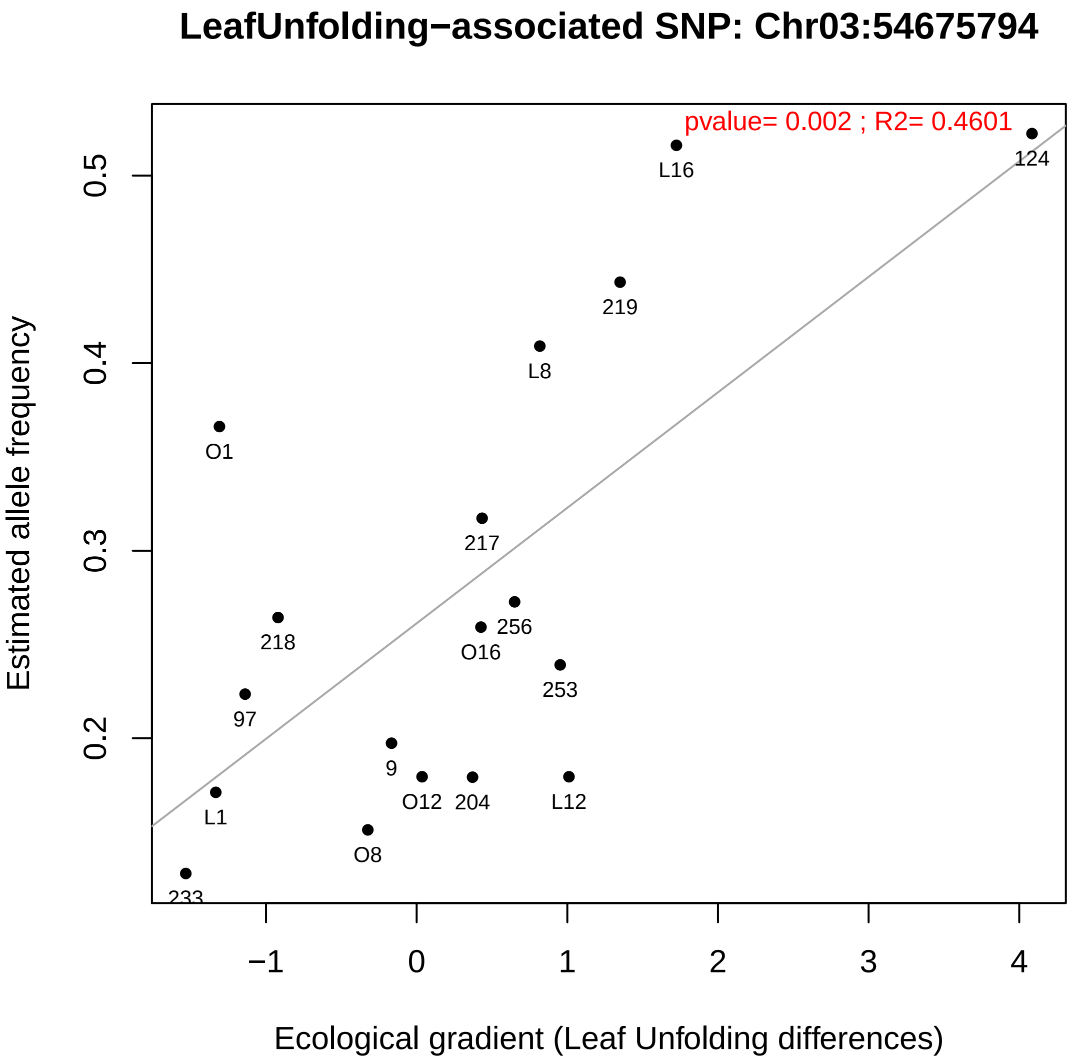

LeafUnfolding−associated SNP: Chr03:54675794
pvalue= 0.002 ; R2= 0.4601
Estimated allele frequency
0.2
0.3
0.4
0.5
−1
0
1
2
3
4
Ecological gradient (Leaf Unfolding differences)
L16
124
219
L8
O1
217
256
O16
218
253
97
9
O12
204
L12
L1
O8
233

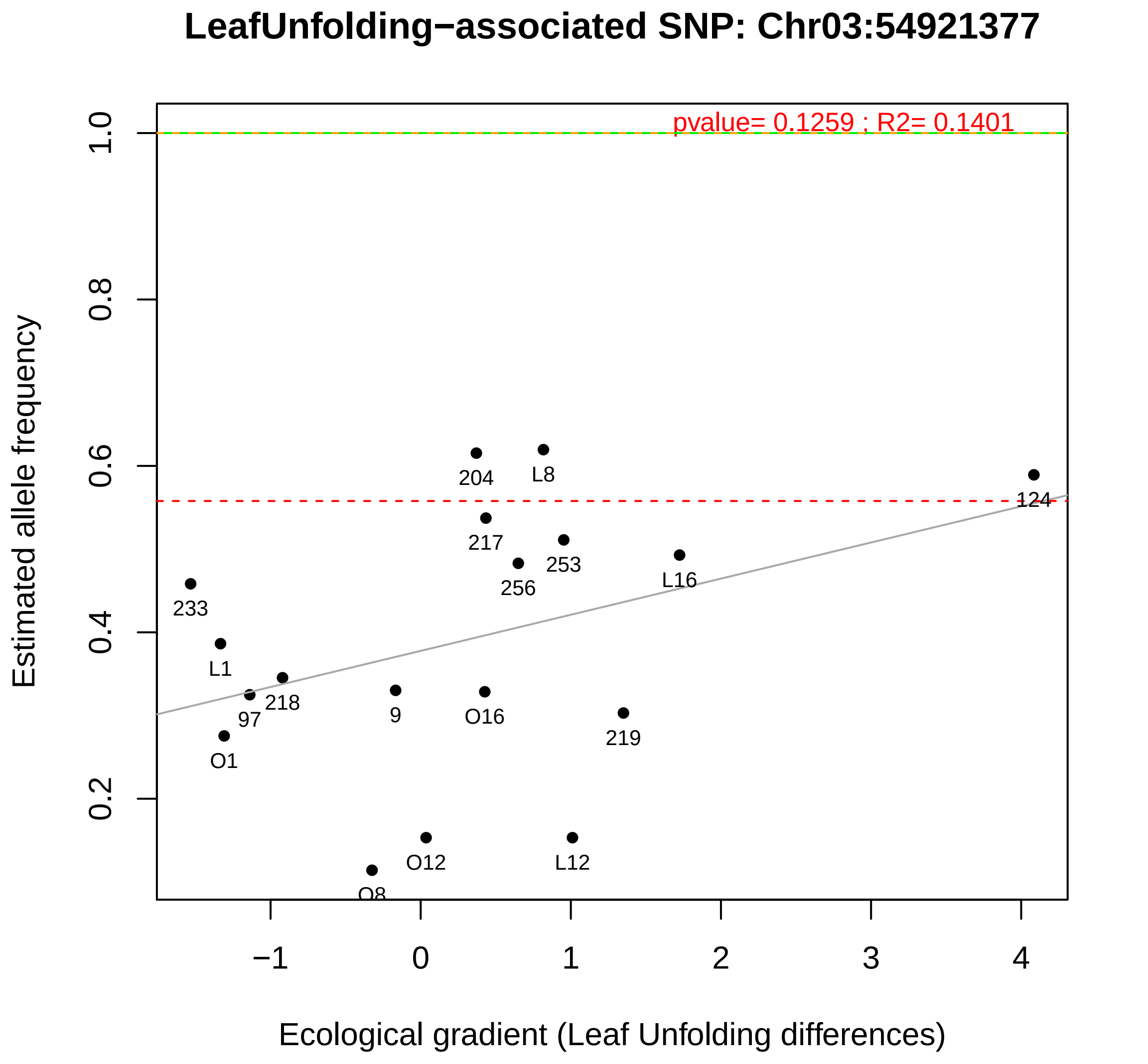
LeafUnfolding−associated SNP: Chr03:54921377
pvalue= 0.1259 ; R2= 0.1401
Estimated allele frequency
Ecological gradient (Leaf Unfolding differences)
0.2
0.4
0.6
0.8
1.0
−1
0
1
2
3
4
204
L8
124
217
253
256
L16
233
L1
218
97
9
O16
219
O1
O12
L12
O8

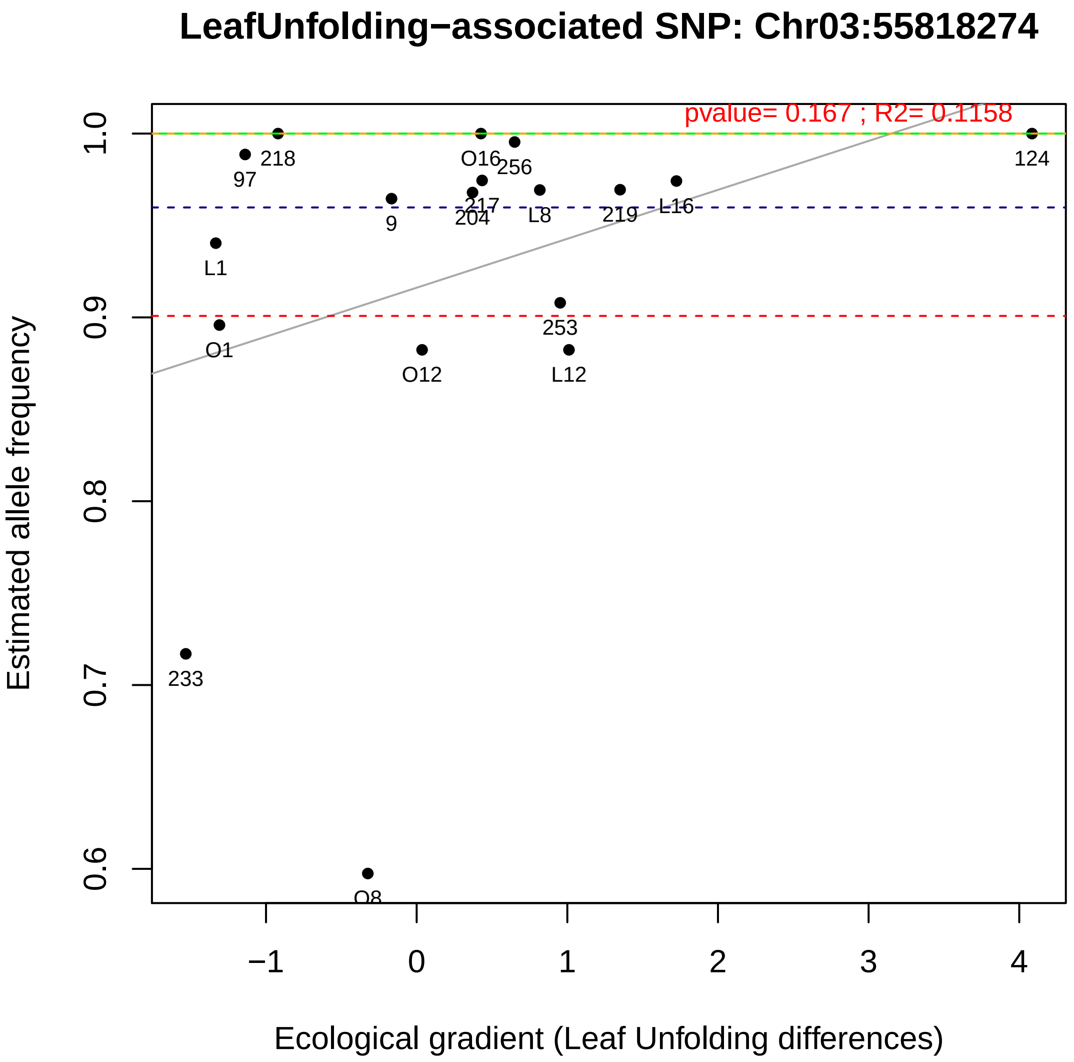

LeafUnfolding−associated SNP: Chr03:55818274
pvalue= 0.167 ; R2= 0.1158
Estimated allele frequency
Ecological gradient (Leaf Unfolding differences)
1.0
0.9
0.8
0.7
0.6
−1
0
1
2
3
4
218
97
O16
256
124
9
217
204
L8
219
L16
L1
253
O1
O12
L12
233
O8

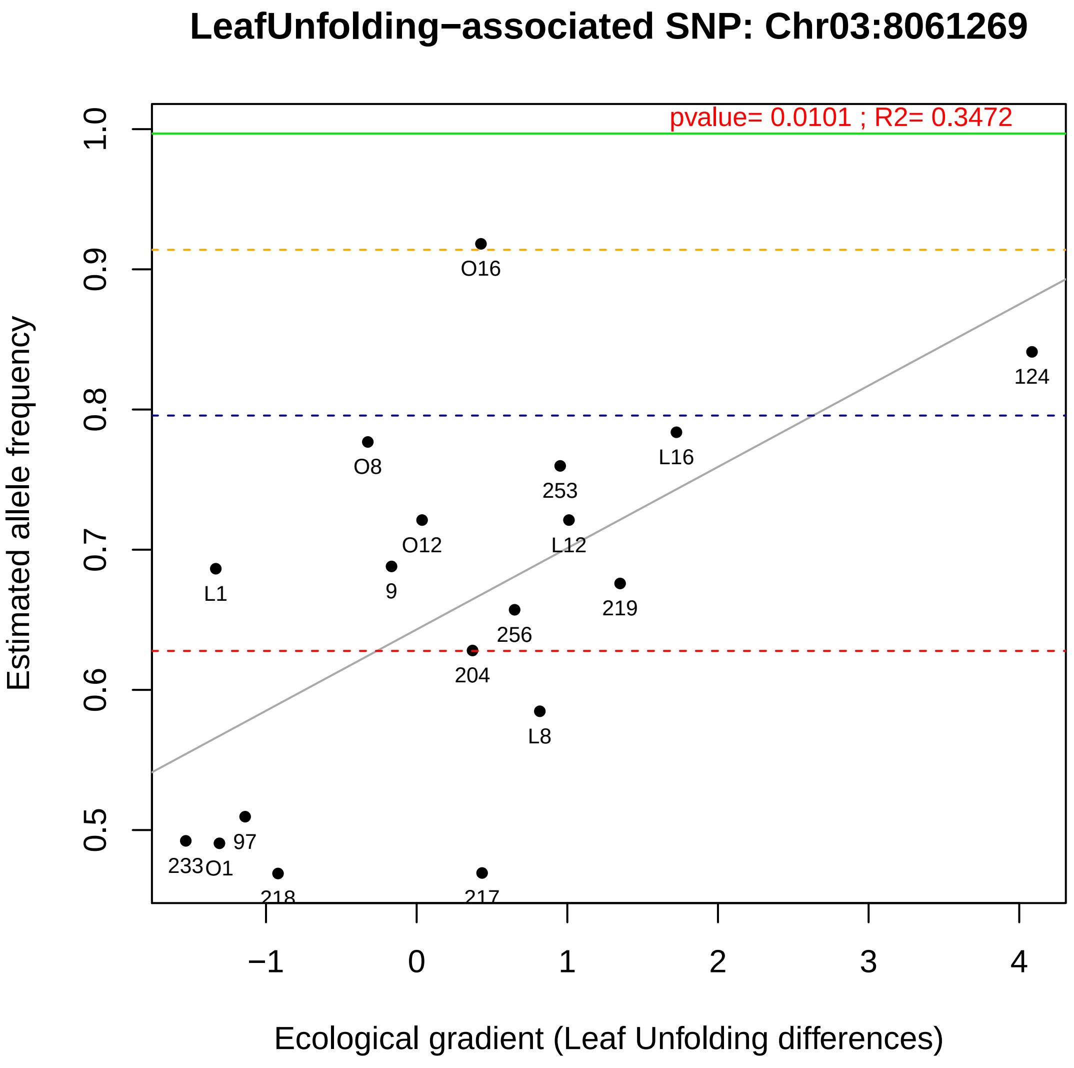

LeafUnfolding−associated SNP: Chr03:8061269
pvalue= 0.0101 ; R2= 0.3472
Estimated allele frequency
Ecological gradient (Leaf Unfolding differences)
0.5
0.6
0.7
0.8
0.9
1.0
−1
0
1
2
3
4
O16
124
L16
O8
253
O12
L12
L1
9
219
256
204
L8
97
233
O1
218
217

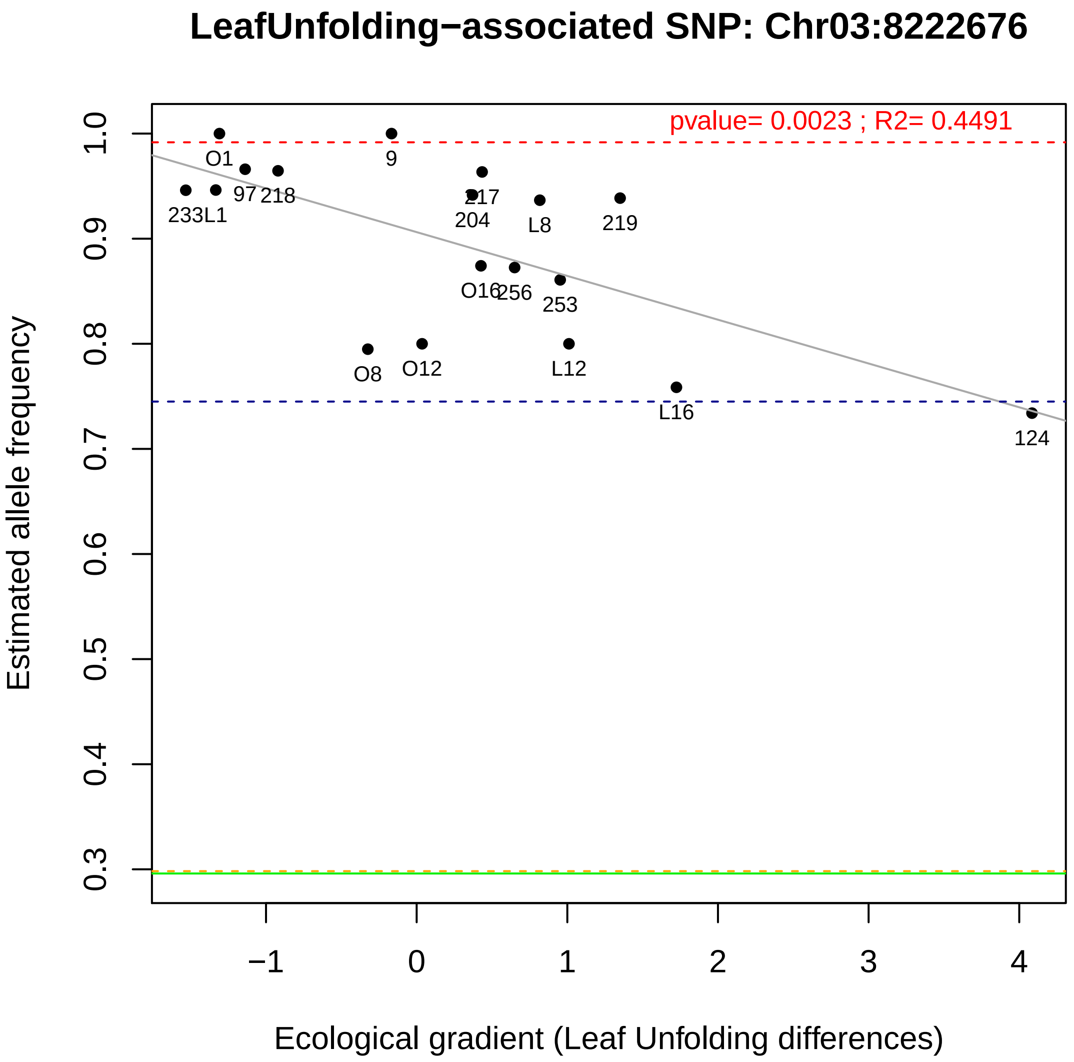

LeafUnfolding−associated SNP: Chr03:8222676
pvalue= 0.0023 ; R2= 0.4491
Estimated allele frequency
Ecological gradient (Leaf Unfolding differences)
1.0
0.9
0.8
0.7
0.6
0.5
0.4
0.3
−1
0
1
2
3
4
O1
9
97
218
217
204
233
L1
L8
219
O16
256
253
O8
O12
L12
L16
124

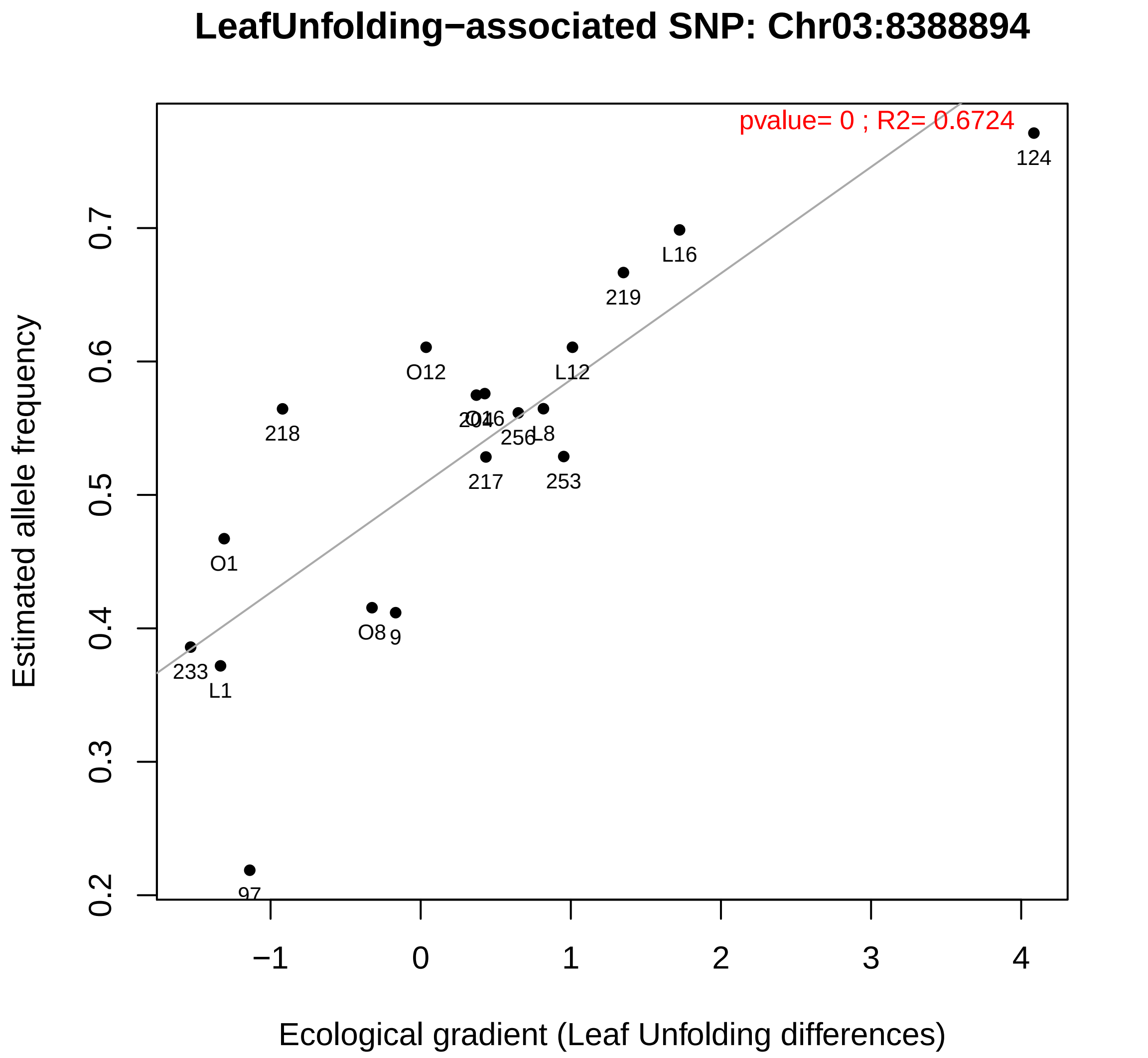

LeafUnfolding−associated SNP: Chr03:8388894
pvalue= 0 ; R2= 0.6724
124
L16
219
O12
L12
218
204
O16
256
L8
217
253
O1
O8
9
233
L1
97
Estimated allele frequency
0.2
0.3
0.4
0.5
0.6
0.7
−1
0
1
2
3
4
Ecological gradient (Leaf Unfolding differences)

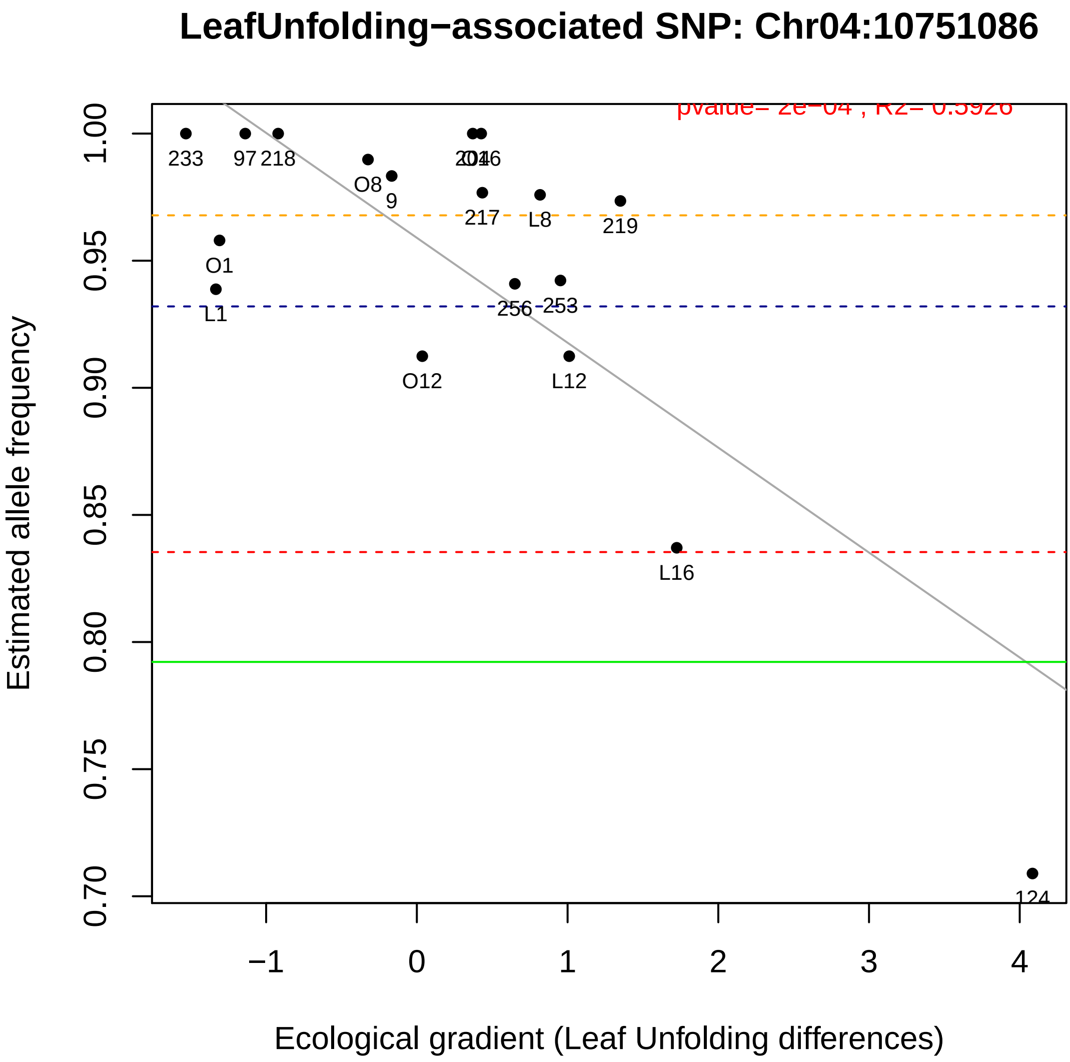
LeafUnfolding−associated SNP: Chr04:10751086
pvalue= 2e−04 , R2= 0.5926
Estimated allele frequency
Ecological gradient (Leaf Unfolding differences)
0.70
0.75
0.80
0.85
0.90
0.95
1.00
−1
0
1
2
3
4
233
97
218
O8
9
217
L8
219
O1
L1
256
253
O12
L12
L16
124

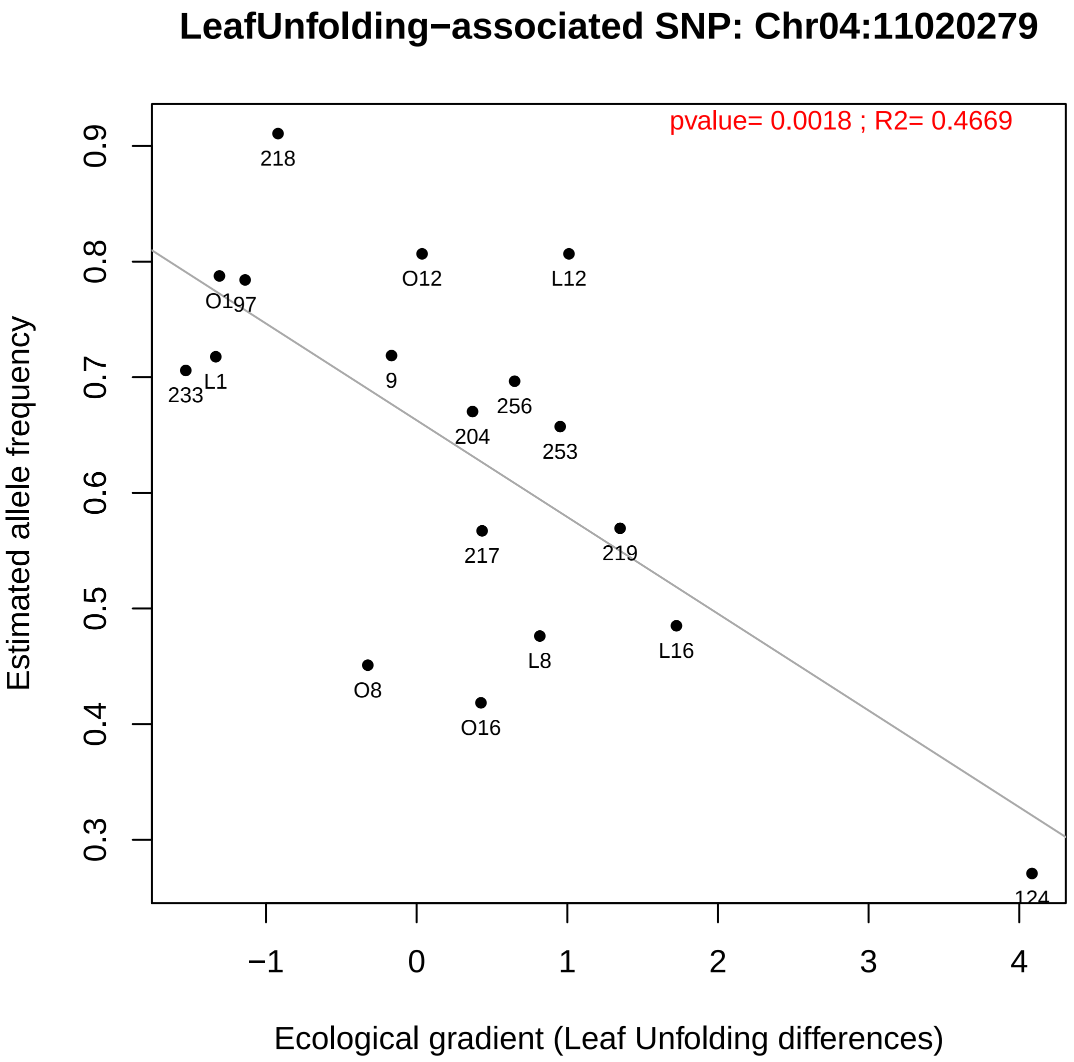

LeafUnfolding−associated SNP: Chr04:11020279
pvalue= 0.0018 ; R2= 0.4669
218
O12
L12
O1
97
233
L1
9
256
204
253
217
219
L8
L16
O8
O16
124
0.3
0.4
0.5
0.6
0.7
0.8
0.9
−1
0
1
2
3
4
Estimated allele frequency
Ecological gradient (Leaf Unfolding differences)

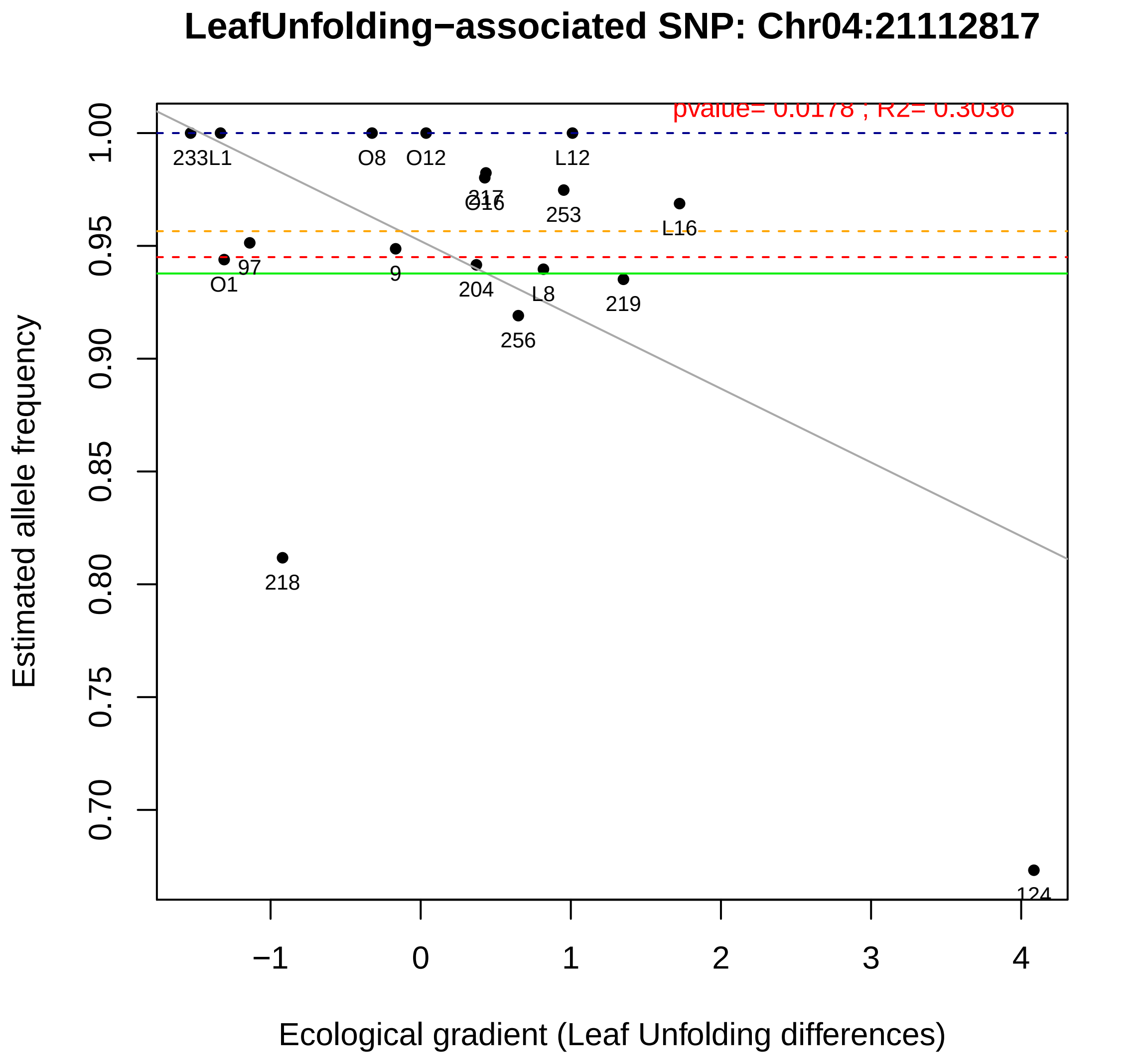

LeafUnfolding−associated SNP: Chr04:21112817
pvalue= 0.0178 ; R2= 0.3036
Estimated allele frequency
Ecological gradient (Leaf Unfolding differences)
1.00
0.95
0.90
0.85
0.80
0.75
0.70
−1
0
1
2
3
4
233
L1
O8
O12
L12
217
O16
253
L16
97
O1
9
204
L8
219
256
218
124

#### LeafUnfolding−associated SNP: Chr04:24393593

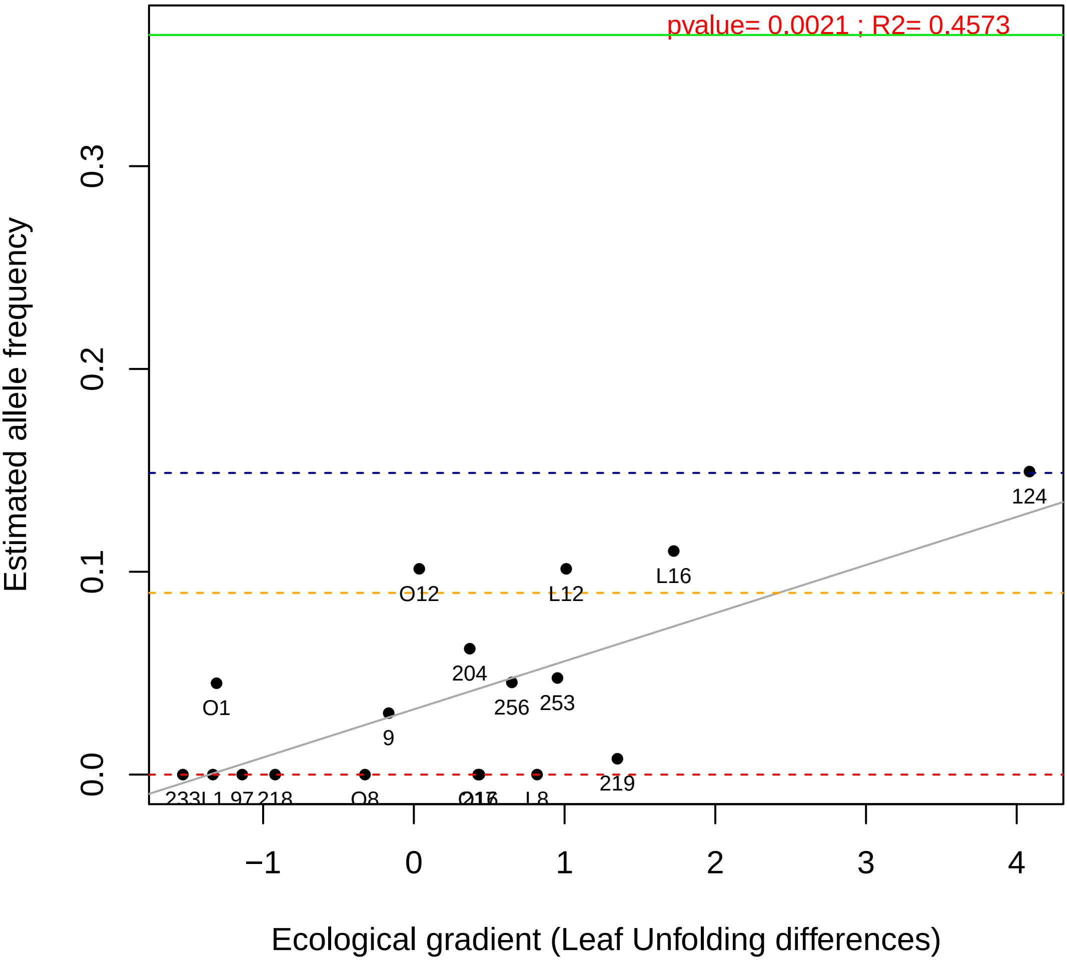

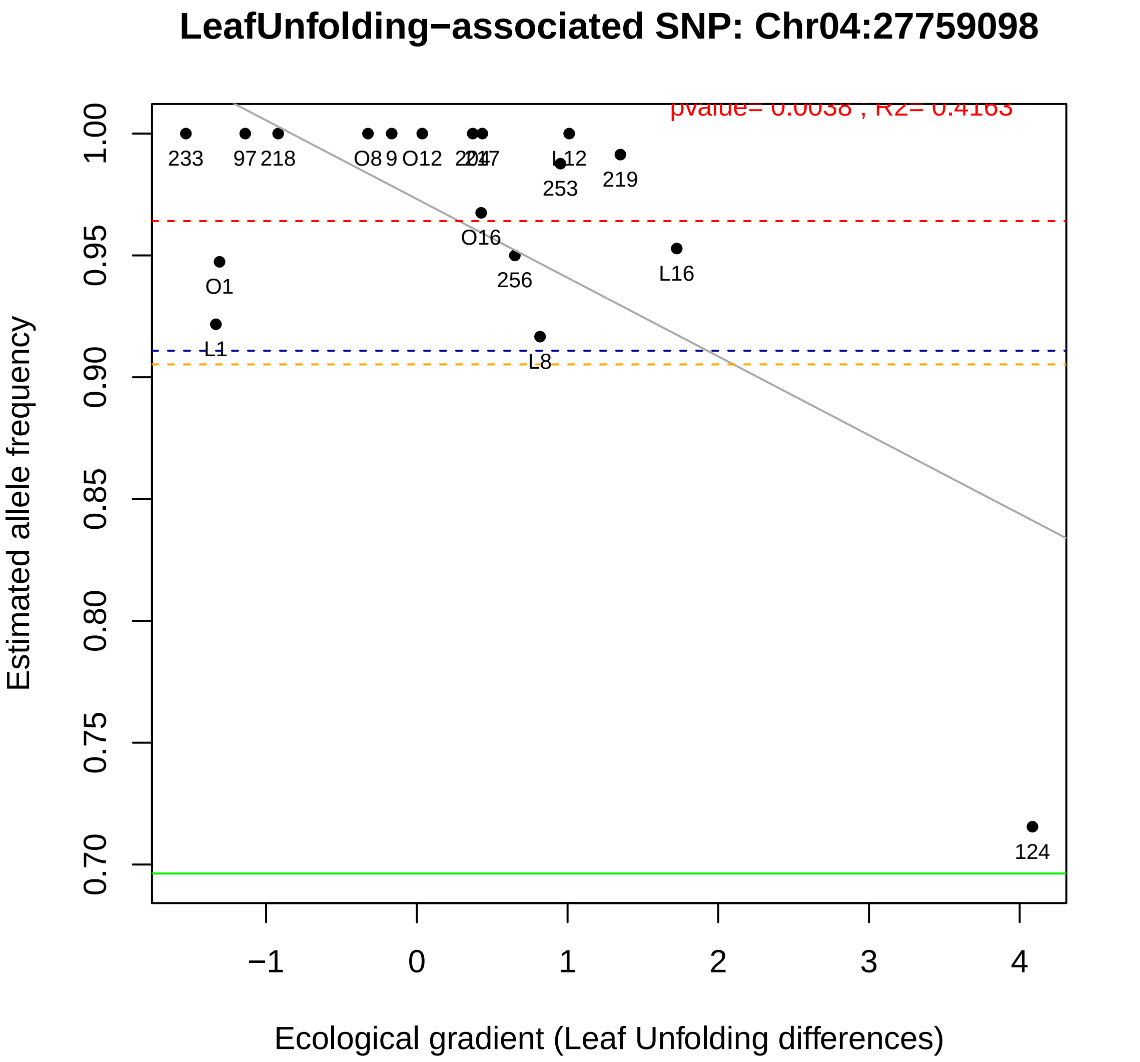

LeafUnfolding−associated SNP: Chr04:27759098
pvalue= 0.0038 ; R2= 0.4163
Estimated allele frequency
Ecological gradient (Leaf Unfolding differences)
0.70
0.75
0.80
0.85
0.90
0.95
1.00
−1
0
1
2
3
4
233
97
218
O8
9
O12
L12
253
219
O16
256
L16
O1
L1
L8
124

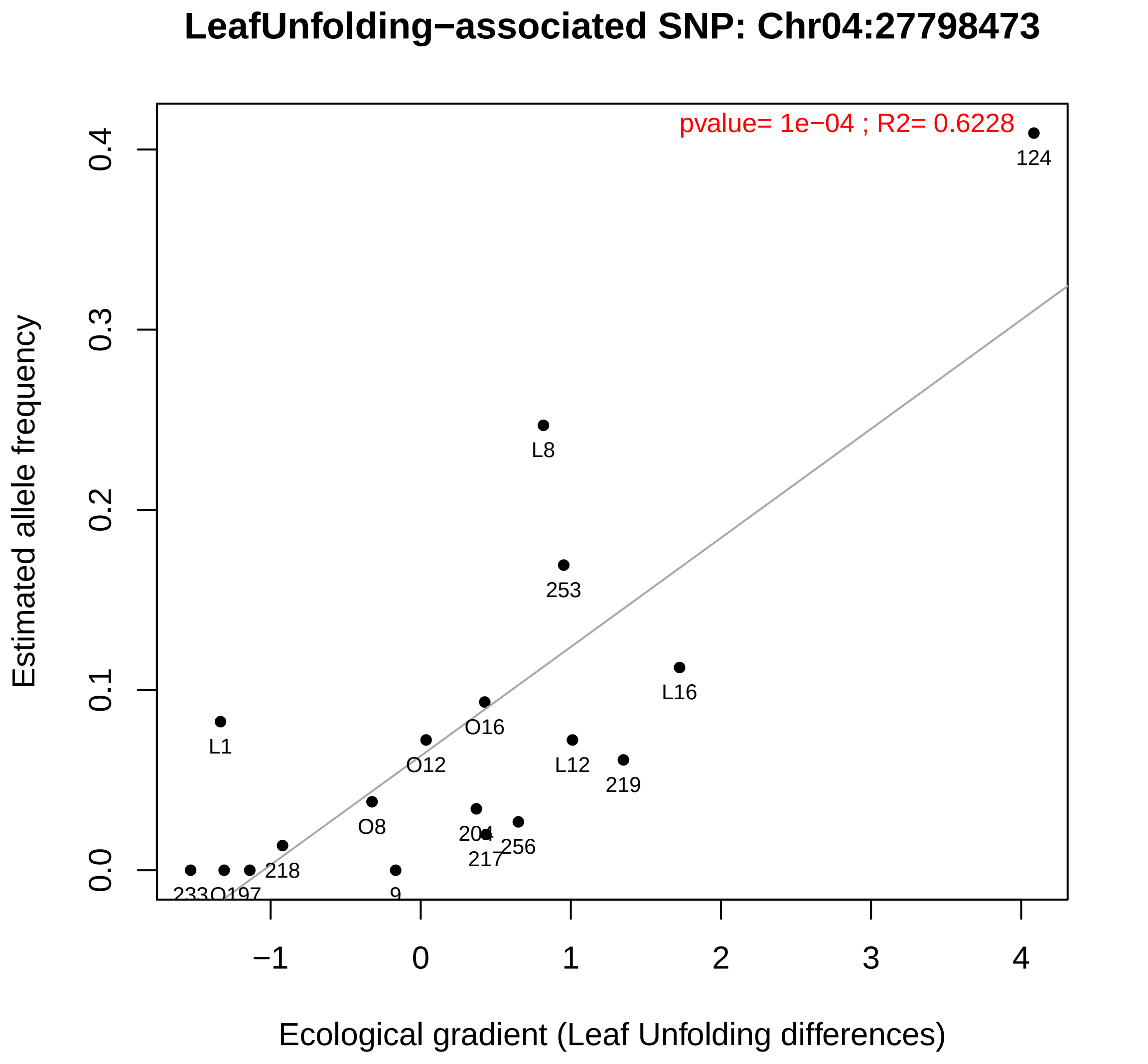
LeafUnfolding−associated SNP: Chr04:27798473
pvalue= 1e−04 ; R2= 0.6228
124
L8
253
L16
O16
L1
O12
L12
219
O8
204
217
256
218
233
O197
9
0.0
0.1
0.2
0.3
0.4
−1
0
1
2
3
4
Estimated allele frequency
Ecological gradient (Leaf Unfolding differences)

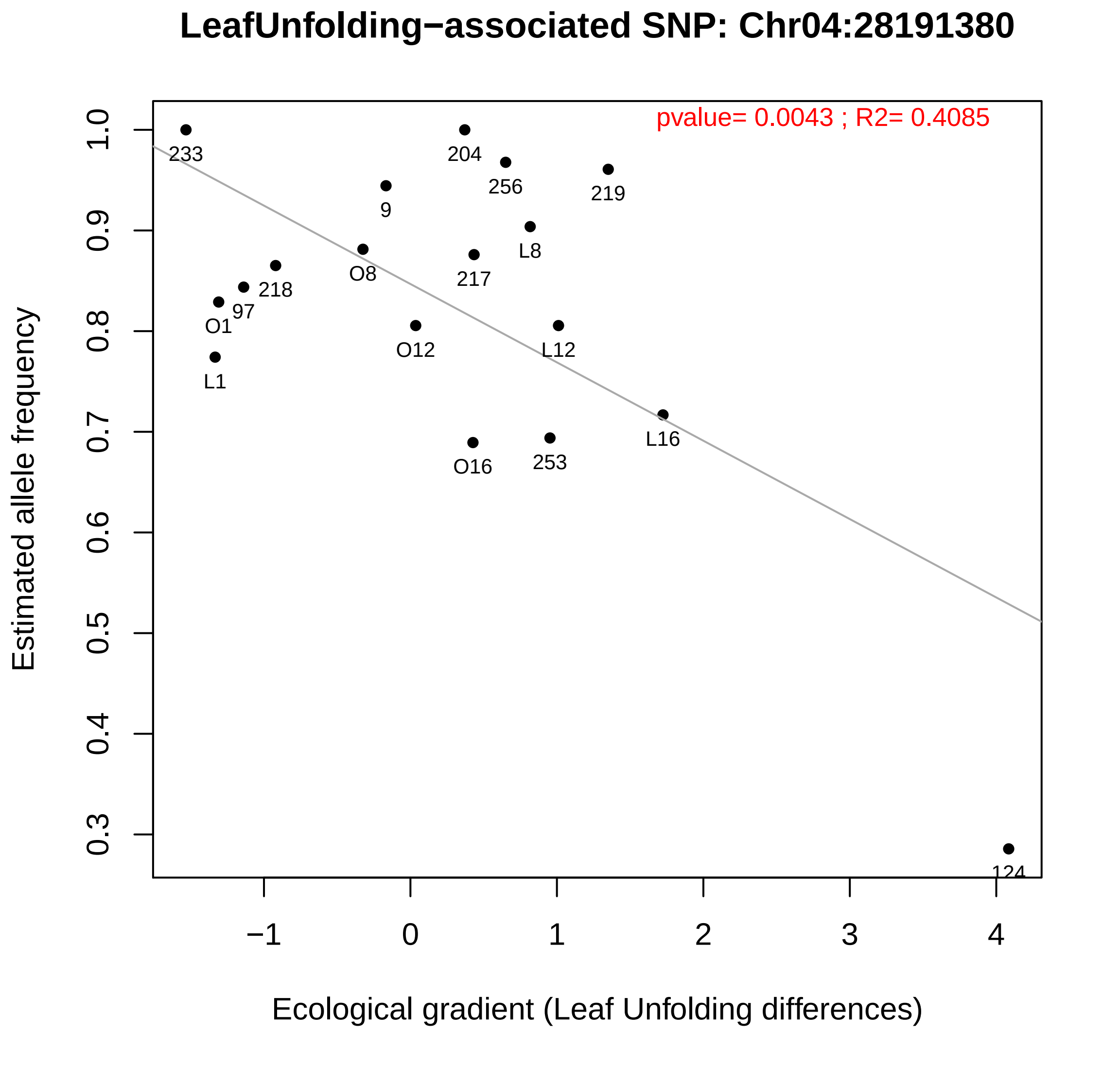

LeafUnfolding−associated SNP: Chr04:28191380
pvalue= 0.0043 ; R2= 0.4085
Estimated allele frequency
Ecological gradient (Leaf Unfolding differences)
0.3
0.4
0.5
0.6
0.7
0.8
0.9
1.0
−1
0
1
2
3
4
233
204
256
219
9
L8
O8
217
218
97
O1
O12
L12
L1
L16
O16
253
124

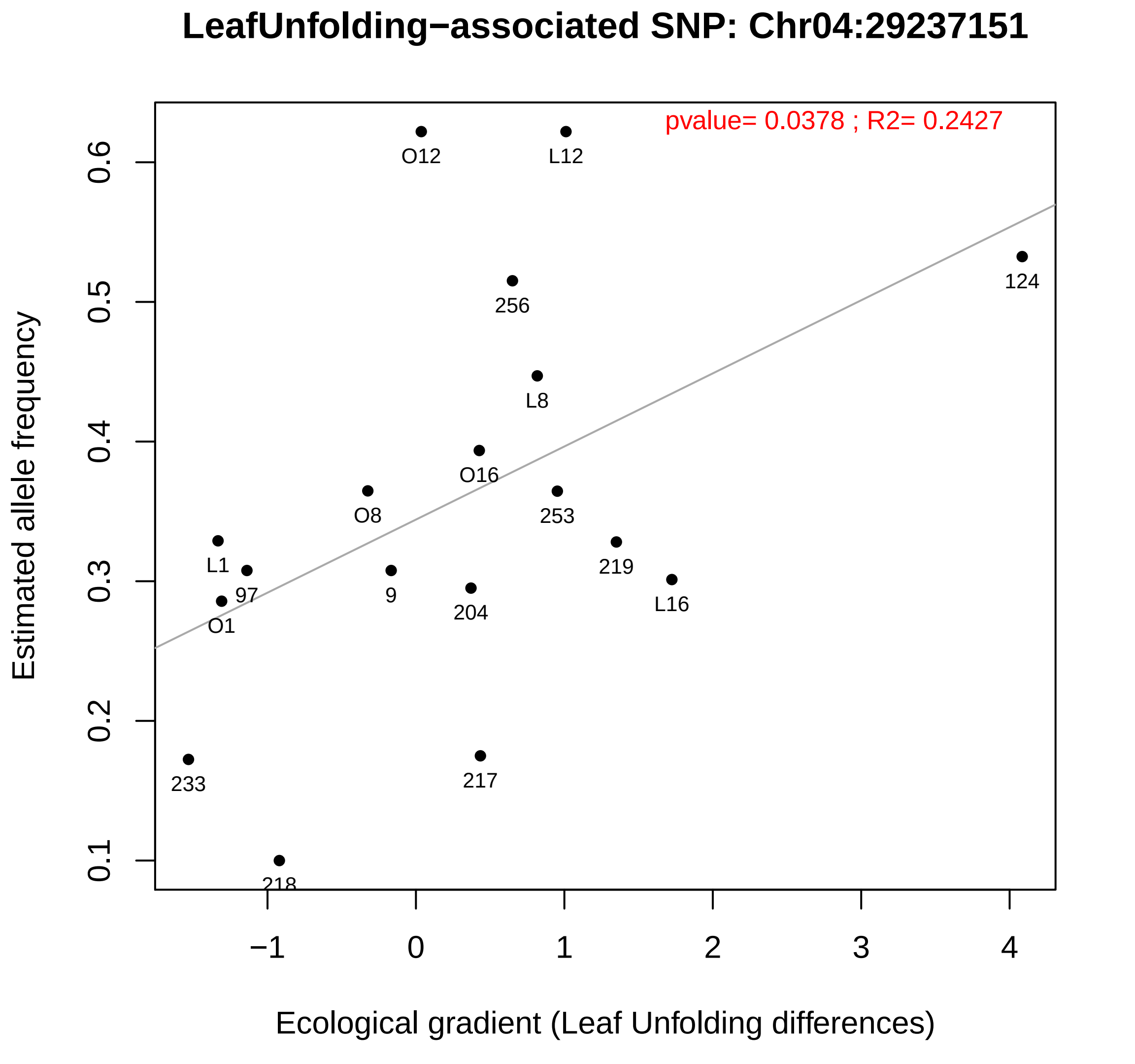
LeafUnfolding−associated SNP: Chr04:29237151
pvalue= 0.0378 ; R2= 0.2427
Estimated allele frequency
Ecological gradient (Leaf Unfolding differences)
0.1
0.2
0.3
0.4
0.5
0.6
−1
0
1
2
3
4
O12
L12
124
256
L8
O16
O8
253
L1
97
9
219
L16
O1
204
233
217
218

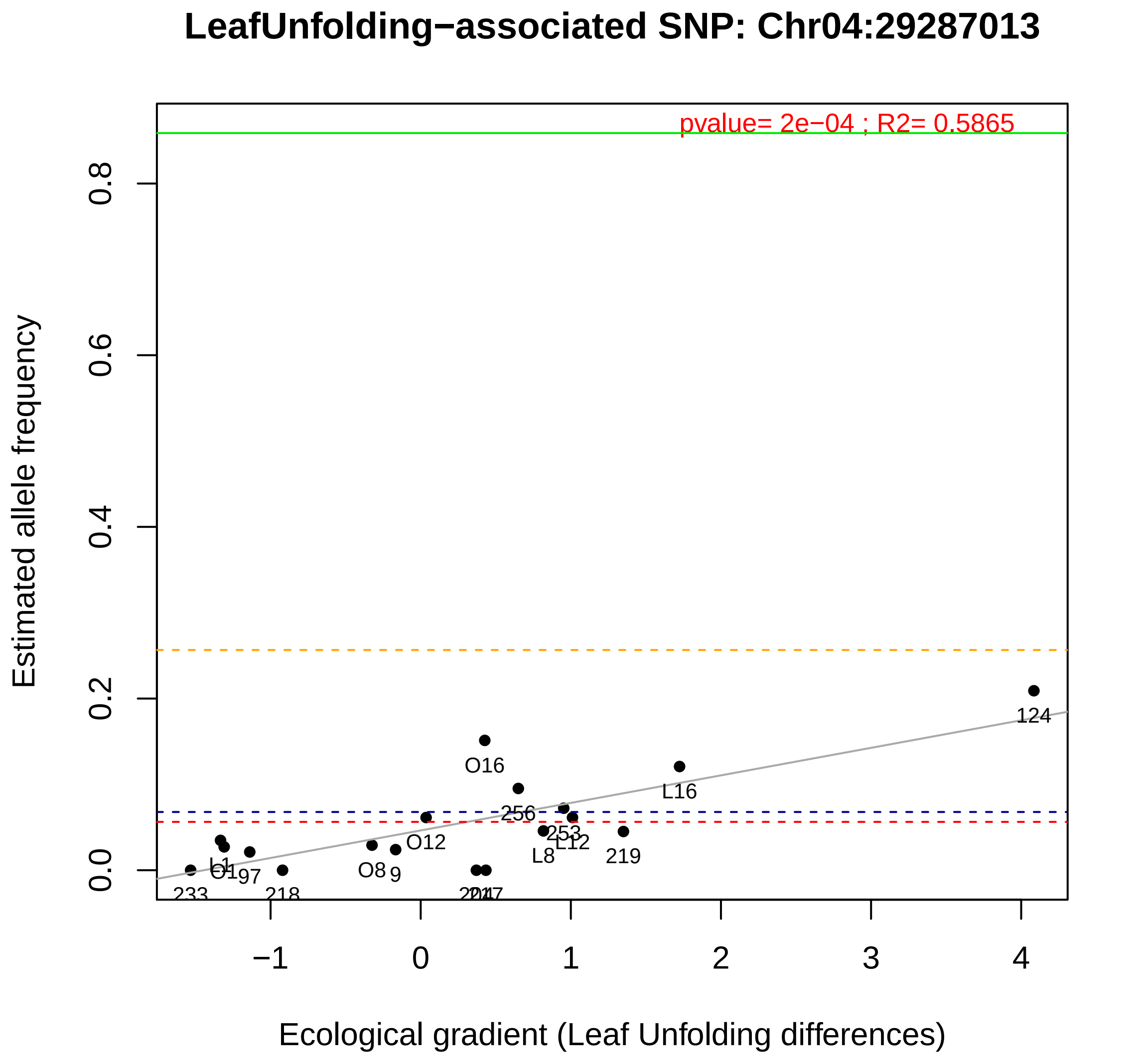

LeafUnfolding−associated SNP: Chr04:29287013
pvalue= 2e−04 ; R2= 0.5865
Estimated allele frequency
Ecological gradient (Leaf Unfolding differences)
0.0
0.2
0.4
0.6
0.8
−1
0
1
2
3
4
124
O16
L16
256
253
L12
L8
219
O12
O8
9
L1
O1
97
218
233

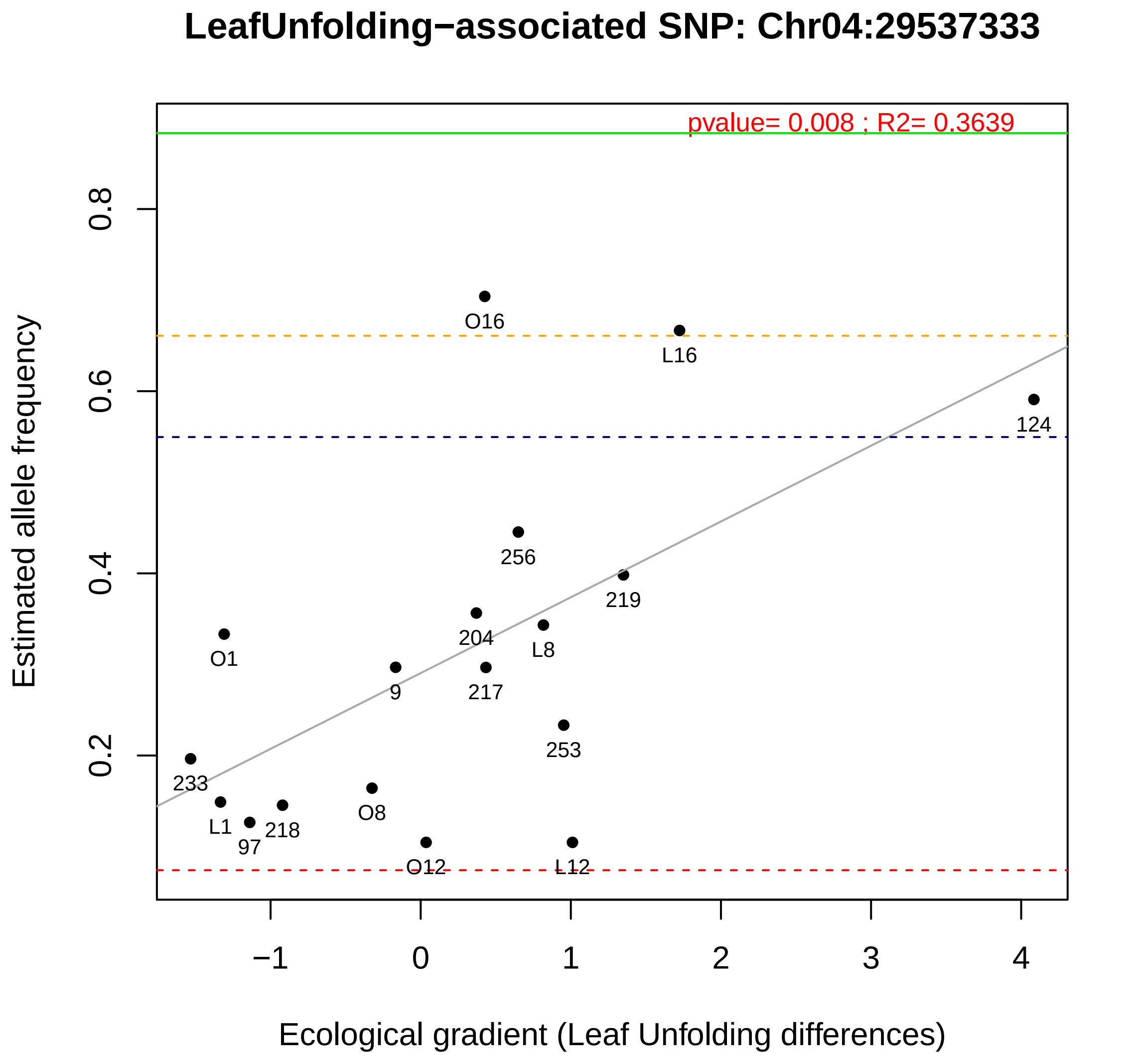

LeafUnfolding−associated SNP: Chr04:29537333
pvalue= 0.008 ; R2= 0.3639
Estimated allele frequency
Ecological gradient (Leaf Unfolding differences)
0.2
0.4
0.6
0.8
−1
0
1
2
3
4
O16
L16
124
256
219
204
L8
O1
9
217
253
233
L1
97
218
O8
O12
L12

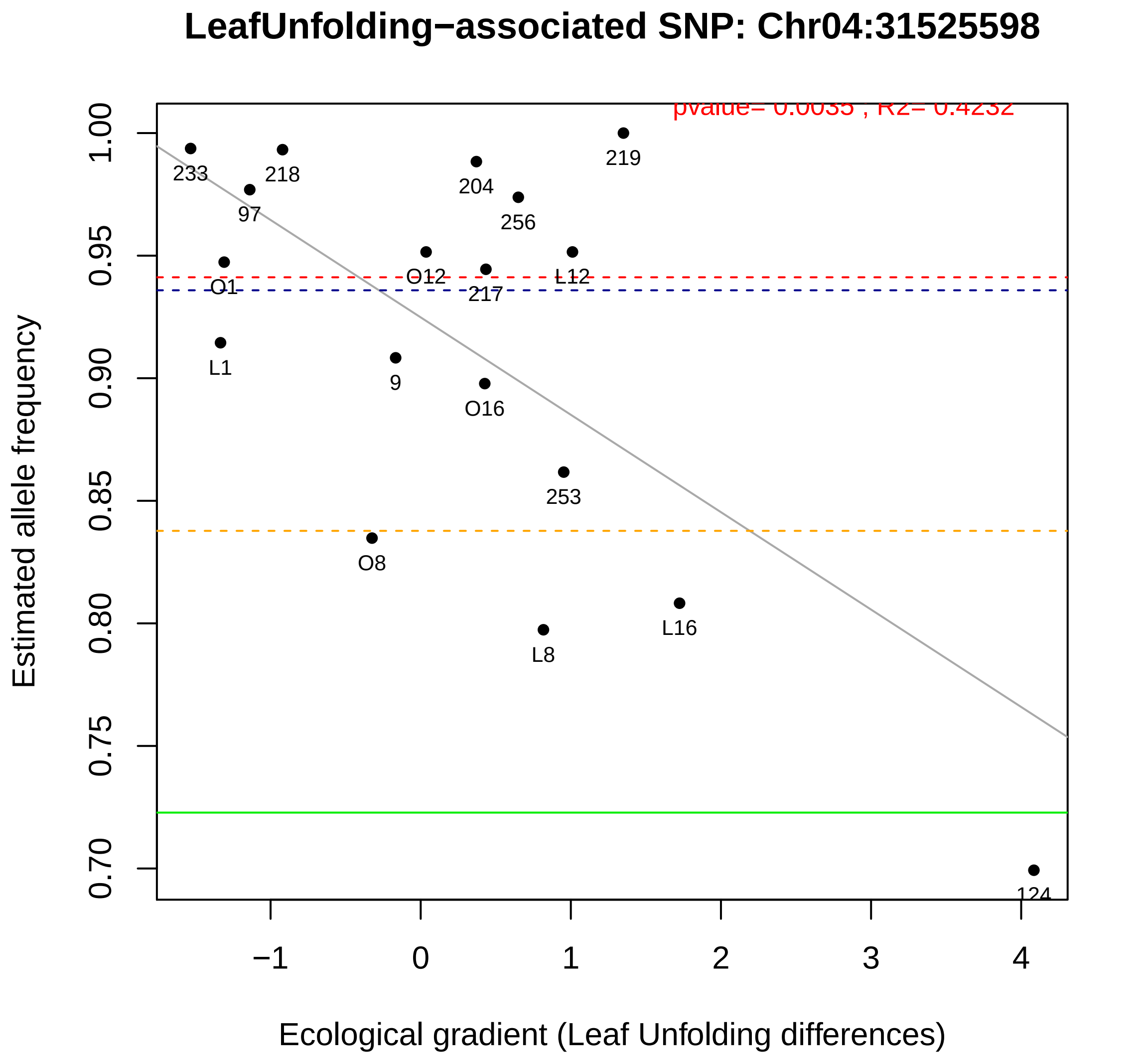

LeafUnfolding−associated SNP: Chr04:31525598
pvalue= 0.0035 , R2= 0.4232
Estimated allele frequency
Ecological gradient (Leaf Unfolding differences)
0.70
0.75
0.80
0.85
0.90
0.95
1.00
−1
0
1
2
3
4
233
218
97
204
256
219
O1
O12
217
L12
L1
9
O16
253
O8
L8
L16
124

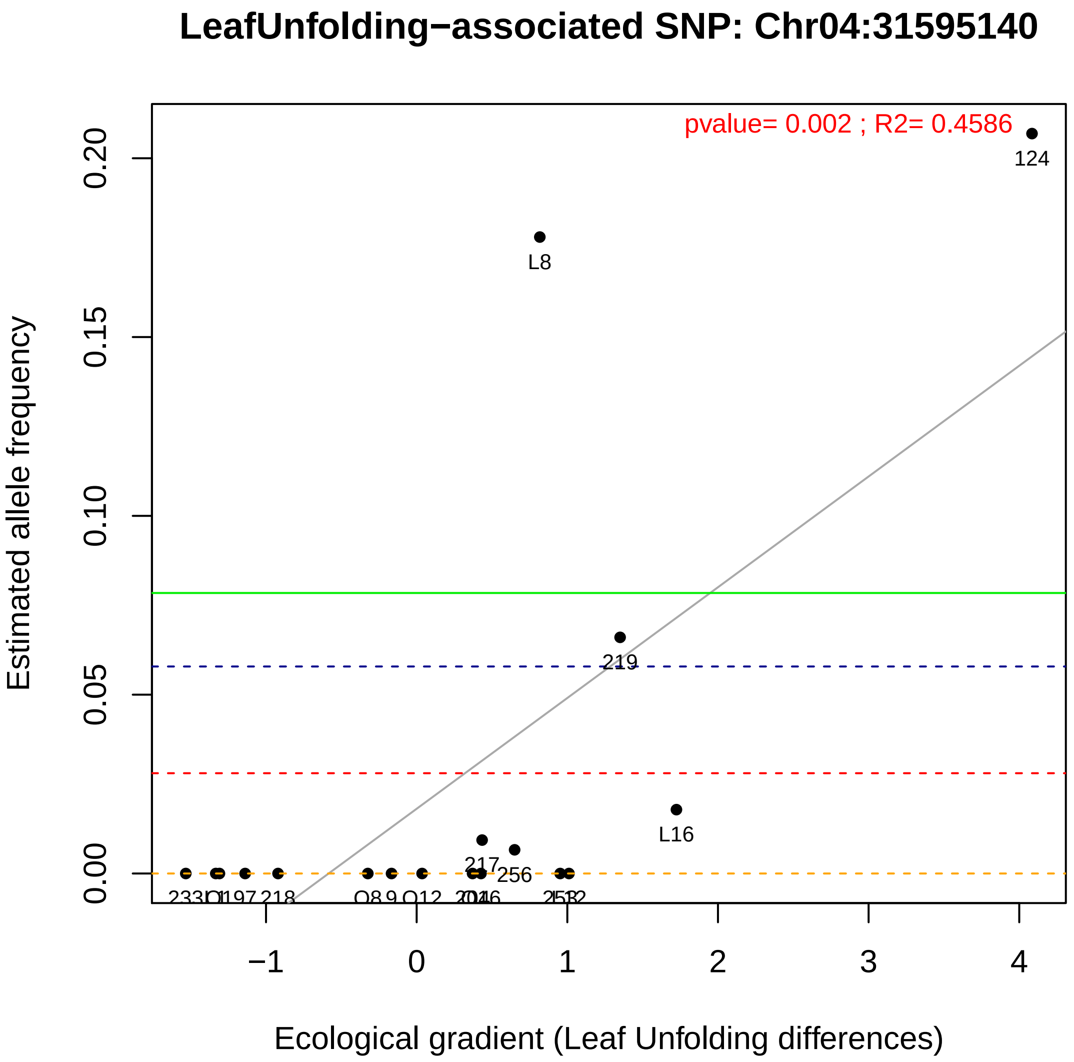

LeafUnfolding−associated SNP: Chr04:31595140
pvalue= 0.002 ; R2= 0.4586
Estimated allele frequency
0.00
0.05
0.10
0.15
0.20
Ecological gradient (Leaf Unfolding differences)
−1
0
1
2
3
4
124
L8
219
L16
217
256
218
O8
9
O12

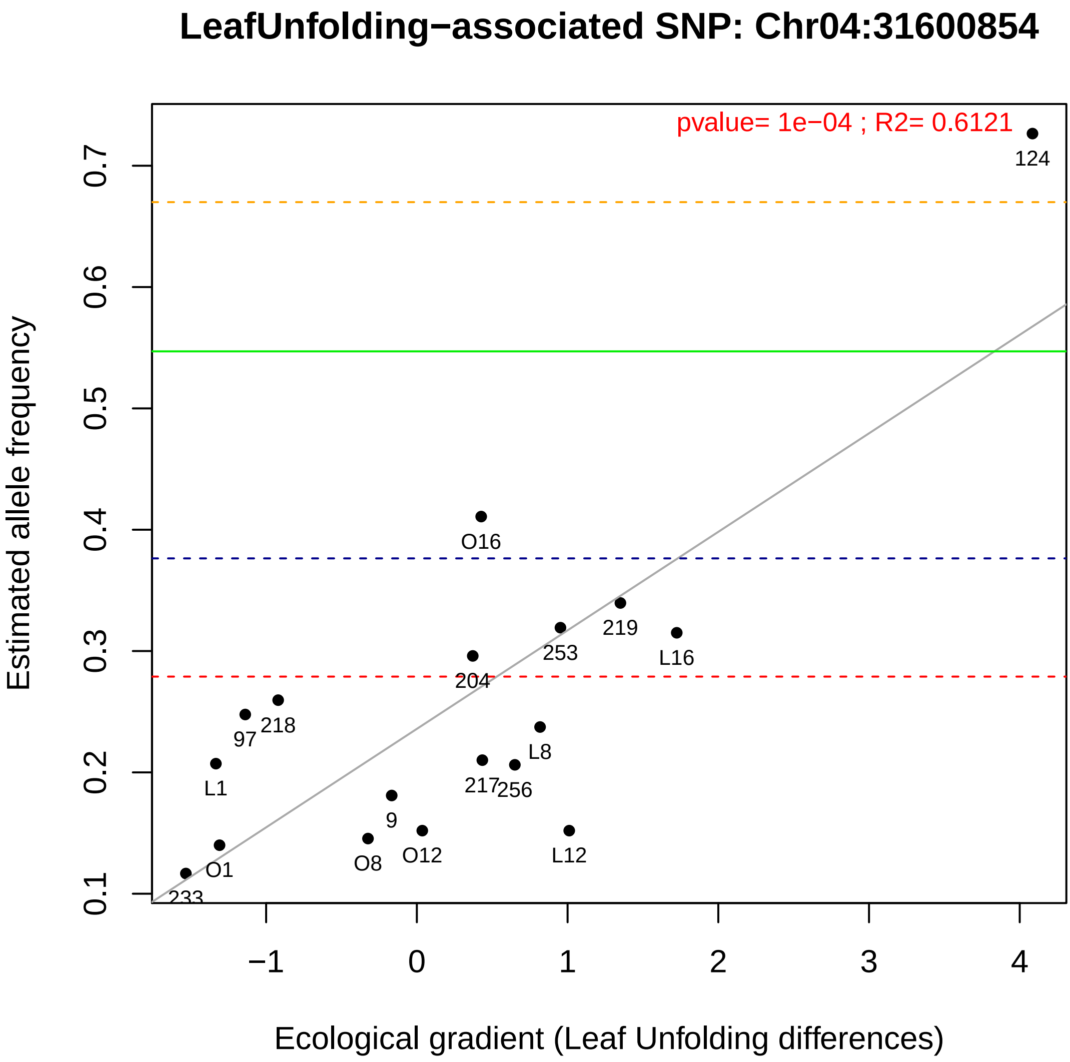

LeafUnfolding−associated SNP: Chr04:31600854
pvalue= 1e−04 ; R2= 0.6121
Estimated allele frequency
0.1
0.2
0.3
0.4
0.5
0.6
0.7
−1
0
1
2
3
4
Ecological gradient (Leaf Unfolding differences)
124
O16
219
253
L16
204
218
97
L1
L8
217
256
9
O12
O8
L12
O1
233

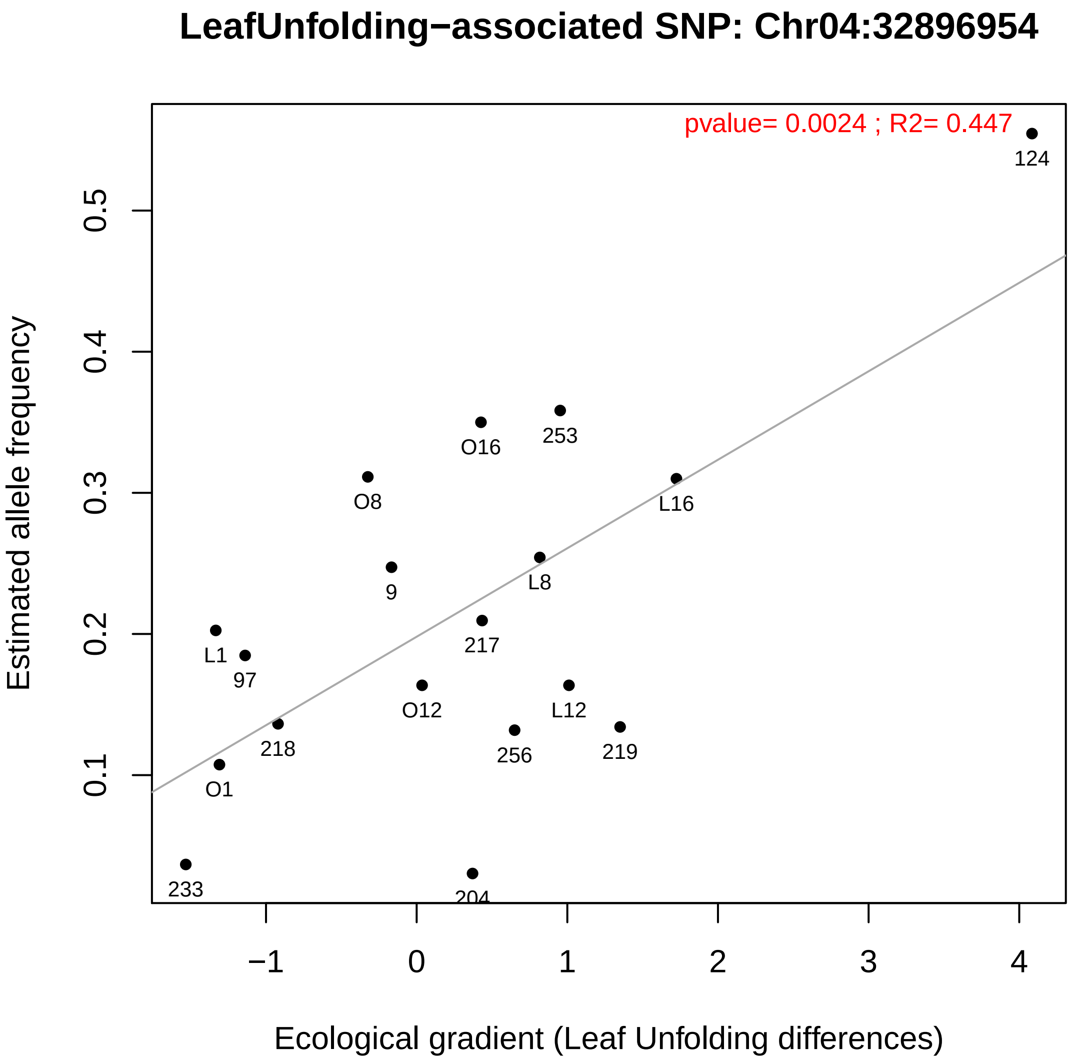
LeafUnfolding−associated SNP: Chr04:32896954
pvalue= 0.0024 ; R2= 0.447
124
253
O16
O8
L16
9
L8
217
L1
97
O12
L12
218
256
219
O1
233
204
0.1
0.2
0.3
0.4
0.5
−1
0
1
2
3
4
Estimated allele frequency
Ecological gradient (Leaf Unfolding differences)

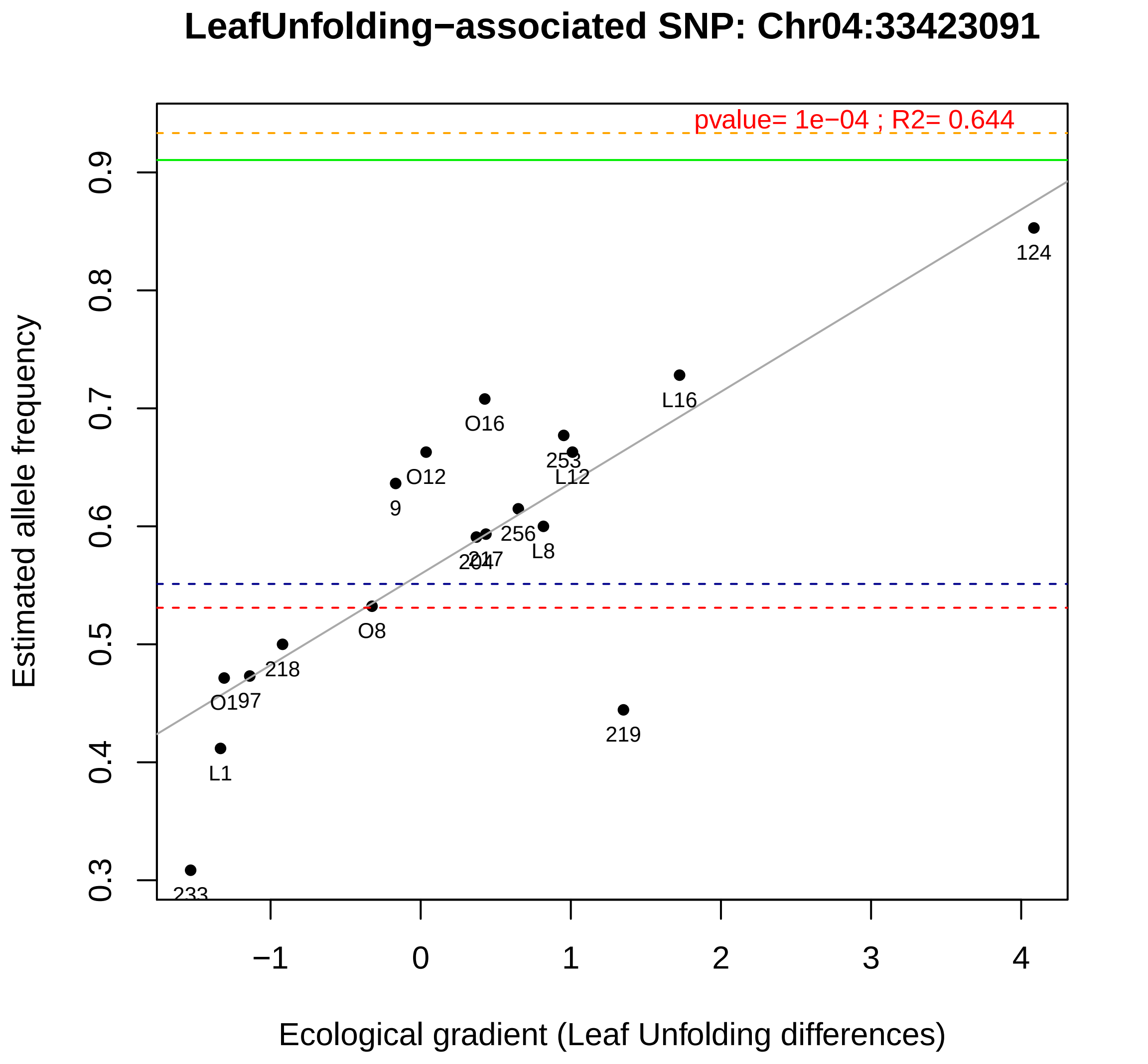

LeafUnfolding−associated SNP: Chr04:33423091
pvalue= 1e−04 ; R2= 0.644
Estimated allele frequency
0.3
0.4
0.5
0.6
0.7
0.8
0.9
−1
0
1
2
3
4
Ecological gradient (Leaf Unfolding differences)
124
L16
O16
253
L12
O12
9
256
L8
217
204
O8
218
O1
97
219
L1
233

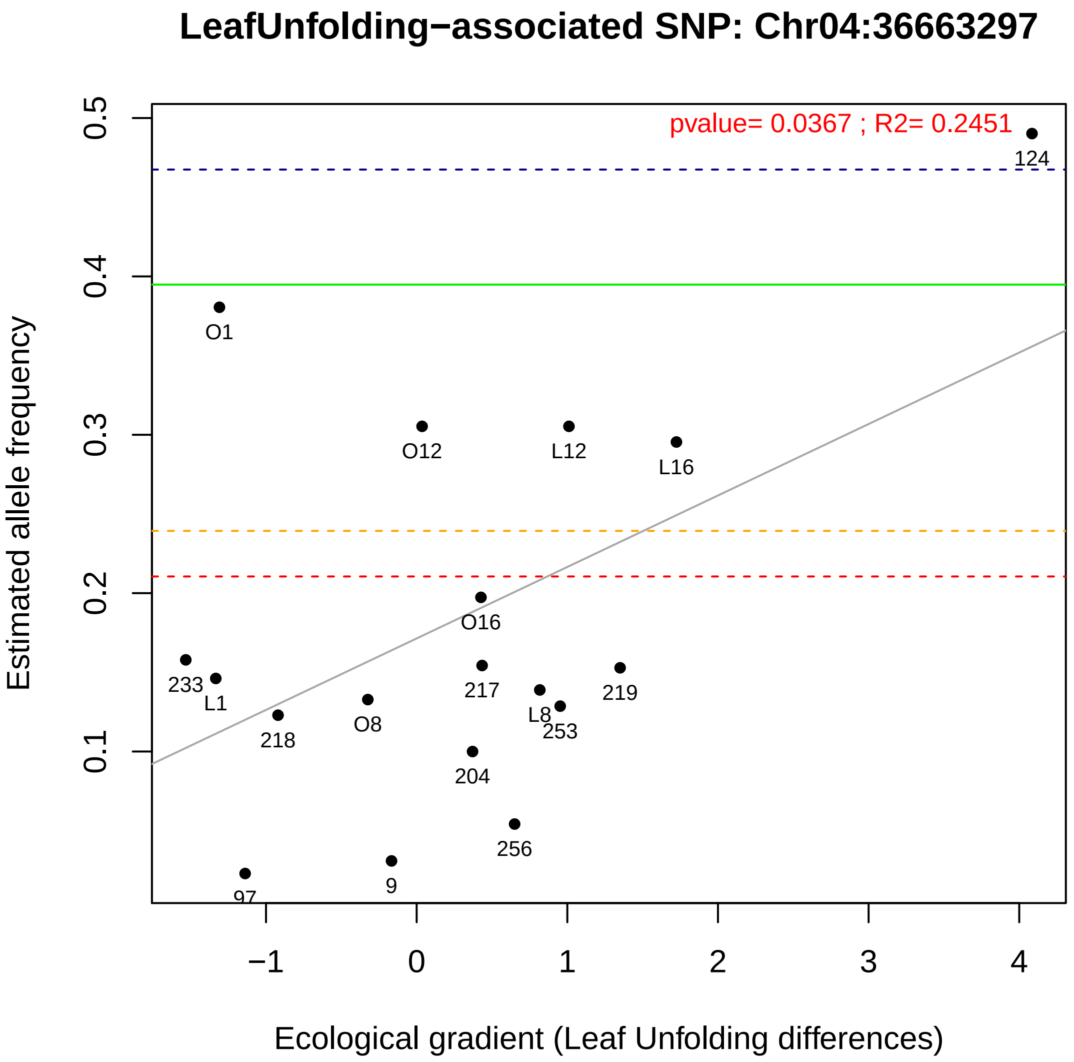
LeafUnfolding−associated SNP: Chr04:36663297
pvalue= 0.0367 ; R2= 0.2451
Estimated allele frequency
Ecological gradient (Leaf Unfolding differences)
0.1
0.2
0.3
0.4
0.5
−1
0
1
2
3
4
124
O1
O12
L12
L16
O16
233
L1
217
219
L8
253
218
O8
204
256
9
97

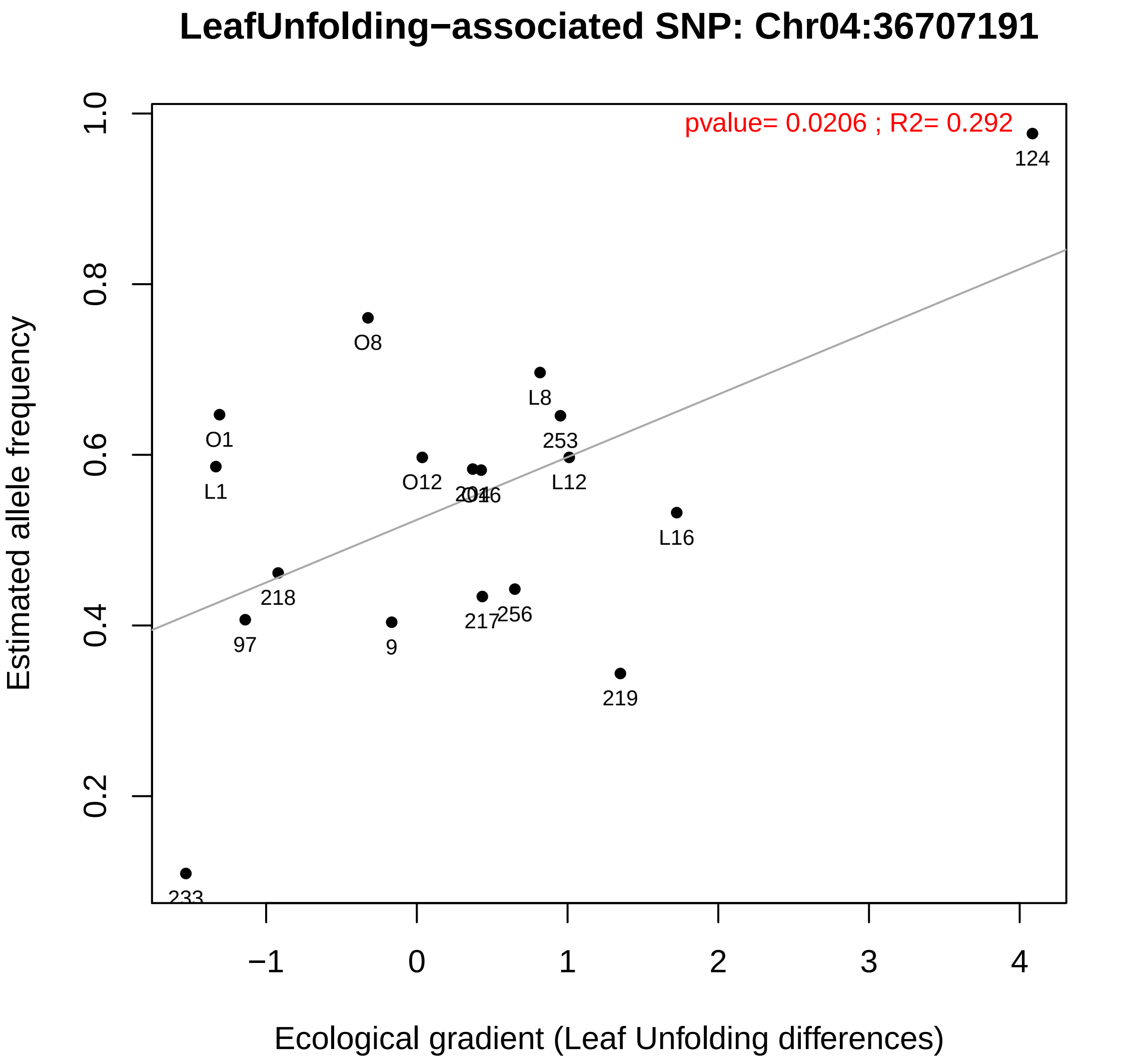

LeafUnfolding−associated SNP: Chr04:36707191
pvalue= 0.0206 ; R2= 0.292
Estimated allele frequency
Ecological gradient (Leaf Unfolding differences)
1.0
0.8
0.6
0.4
0.2
−1
0
1
2
3
4
124
O8
L8
253
O1
L1
O12
L12
L16
218
97
9
217
256
219
233

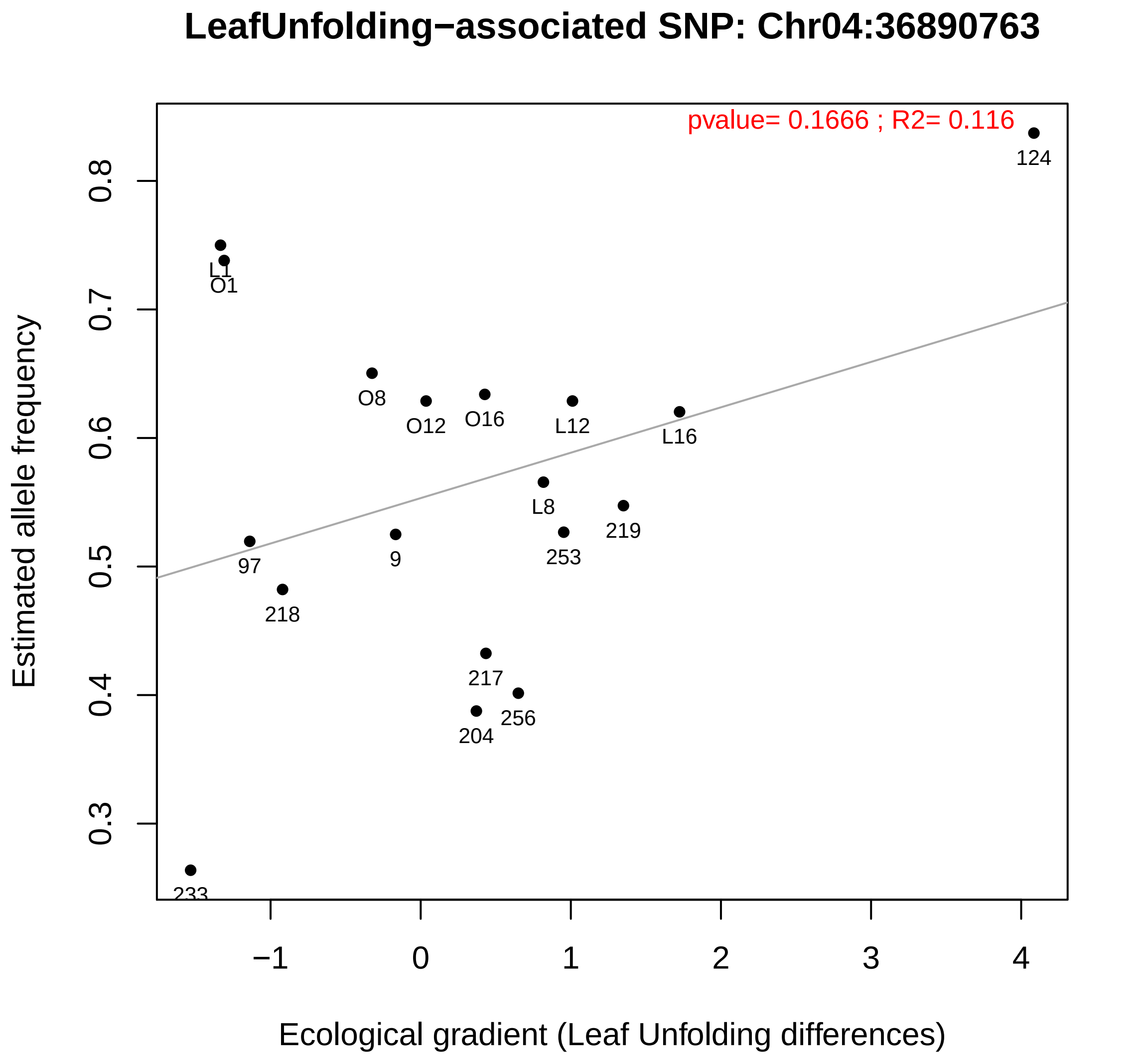
LeafUnfolding−associated SNP: Chr04:36890763
pvalue= 0.1666 ; R2= 0.116
124
L1
O1
O8
O12
O16
L12
L16
L8
219
97
9
253
218
217
256
204
233
0.3
0.4
0.5
0.6
0.7
0.8
−1
0
1
2
3
4
Estimated allele frequency
Ecological gradient (Leaf Unfolding differences)

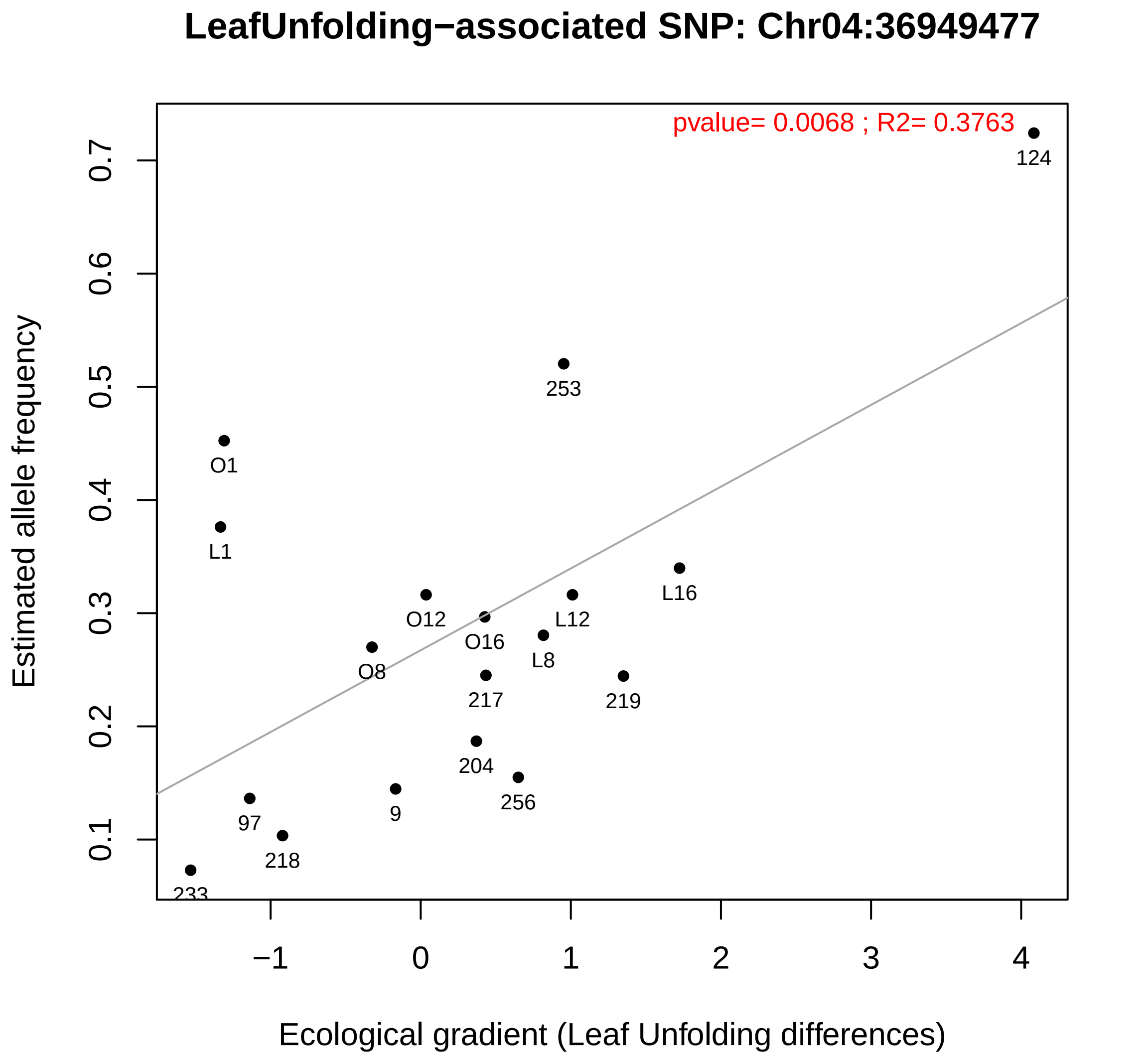

LeafUnfolding−associated SNP: Chr04:36949477
pvalue= 0.0068 ; R2= 0.3763
Estimated allele frequency
Ecological gradient (Leaf Unfolding differences)
0.1
0.2
0.3
0.4
0.5
0.6
0.7
−1
0
1
2
3
4
124
253
O1
L1
L16
O12
L12
O16
L8
O8
217
219
204
256
9
97
218
233

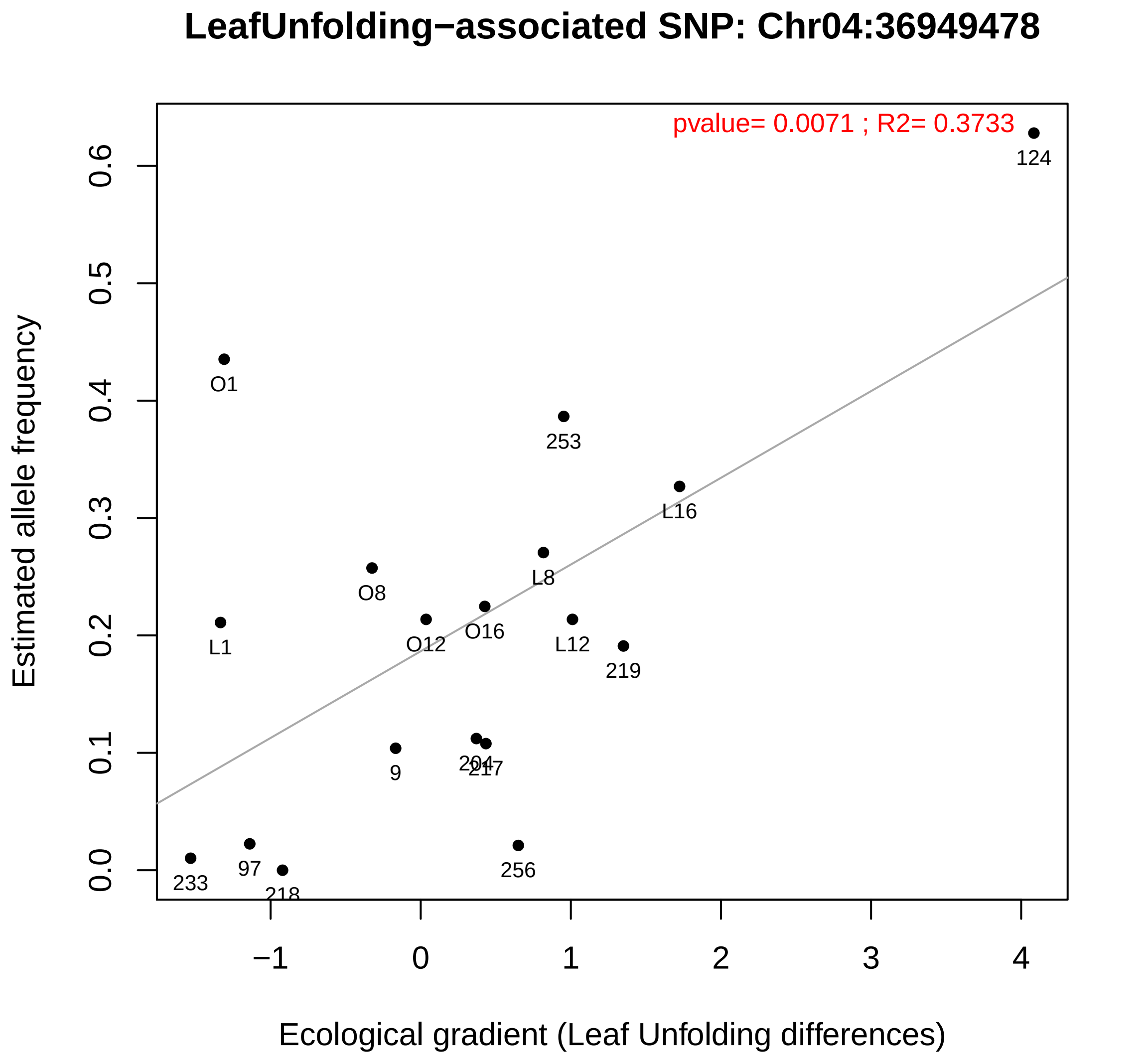

LeafUnfolding−associated SNP: Chr04:36949478
pvalue= 0.0071 ; R2= 0.3733
Estimated allele frequency
Ecological gradient (Leaf Unfolding differences)
0.0
0.1
0.2
0.3
0.4
0.5
0.6
−1
0
1
2
3
4
124
O1
253
L16
L8
O8
O16
L1
O12
L12
219
9
204
217
256
233
97
218

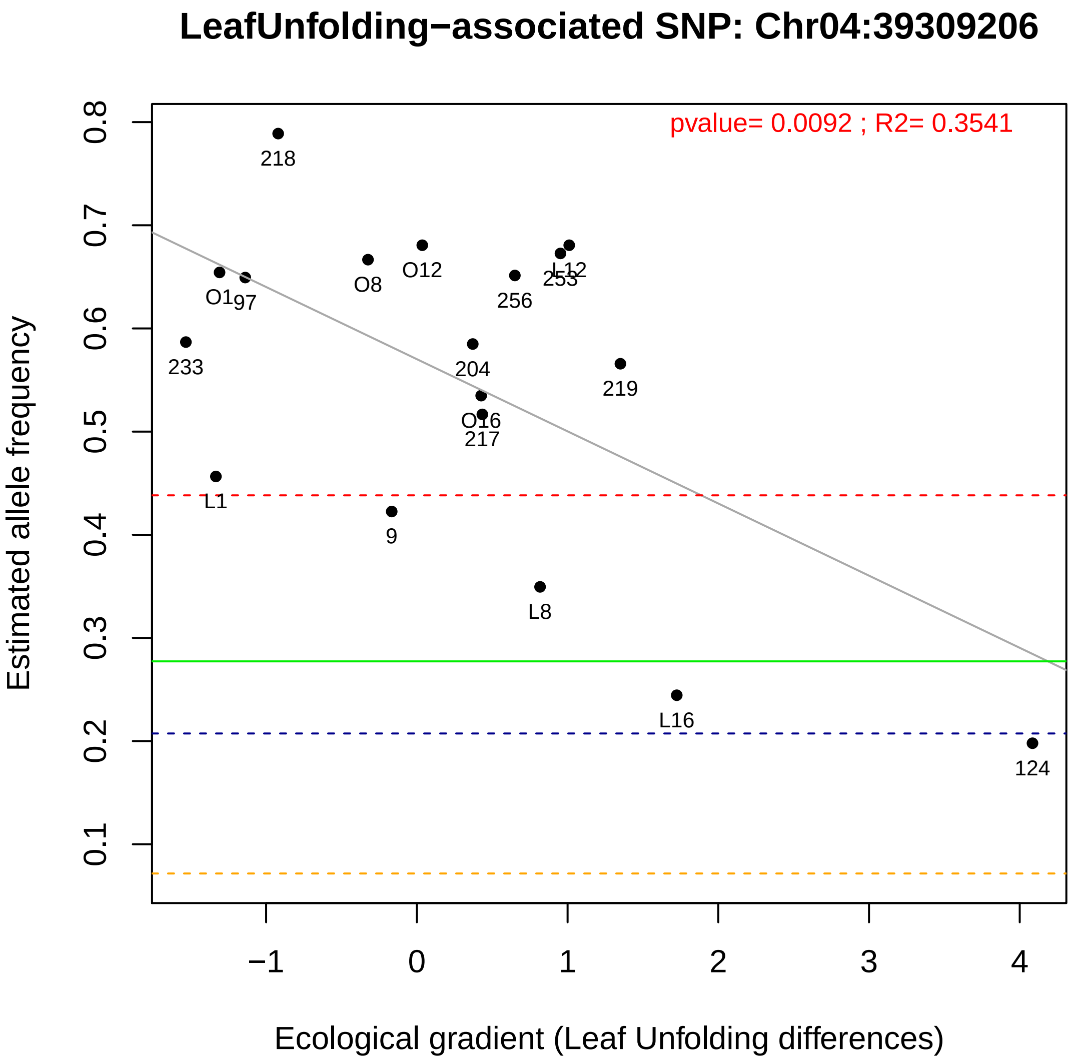

LeafUnfolding−associated SNP: Chr04:39309206
pvalue= 0.0092 ; R2= 0.3541
Estimated allele frequency
Ecological gradient (Leaf Unfolding differences)
0.1
0.2
0.3
0.4
0.5
0.6
0.7
0.8
−1
0
1
2
3
4
218
O12
O8
L12
253
256
O1
97
233
204
219
O16
217
L1
9
L8
L16
124

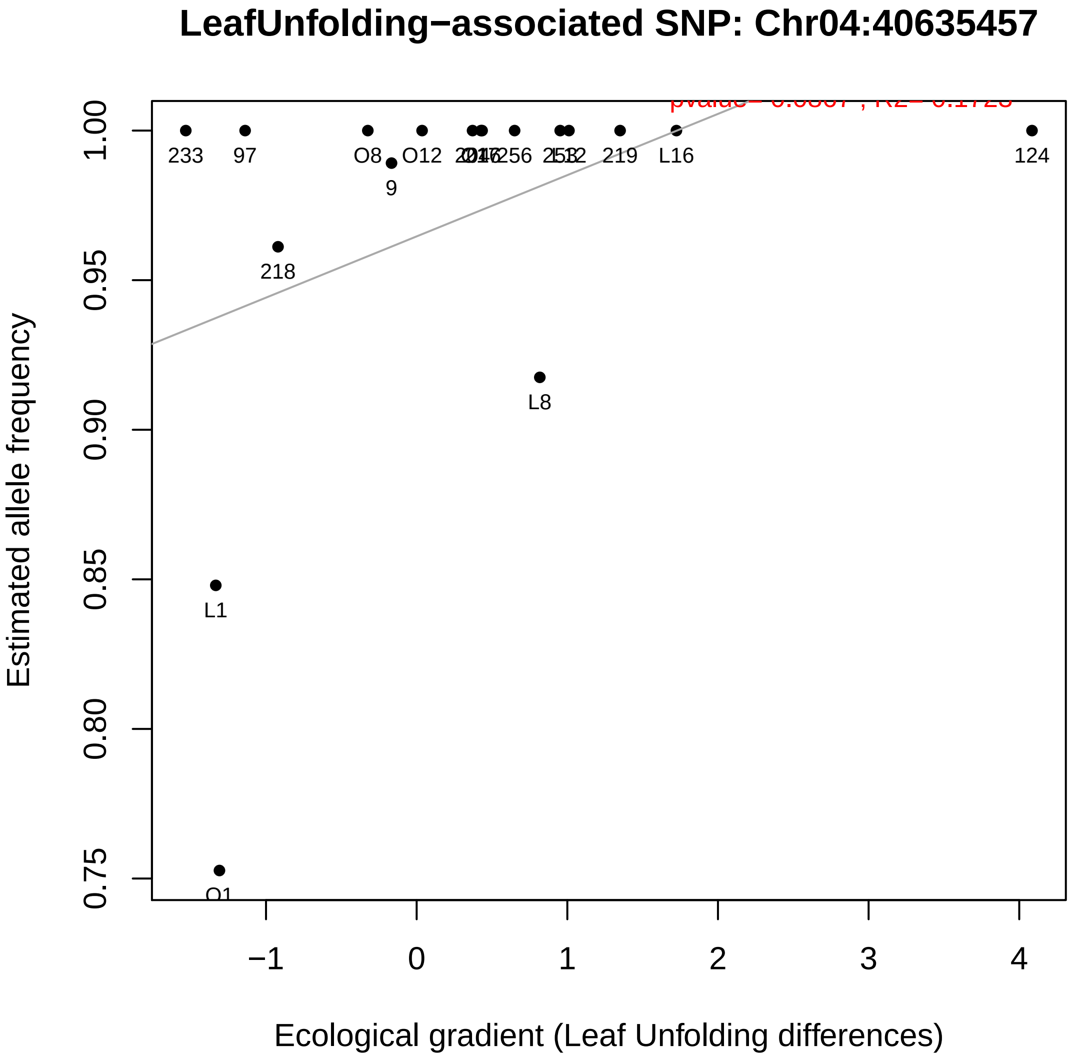
LeafUnfolding−associated SNP: Chr04:40635457
Estimated allele frequency
Ecological gradient (Leaf Unfolding differences)
1.00
0.95
0.90
0.85
0.80
0.75
−1
0
1
2
3
4
233
97
O8
9
O12
219
L16
124
218
L8
L1

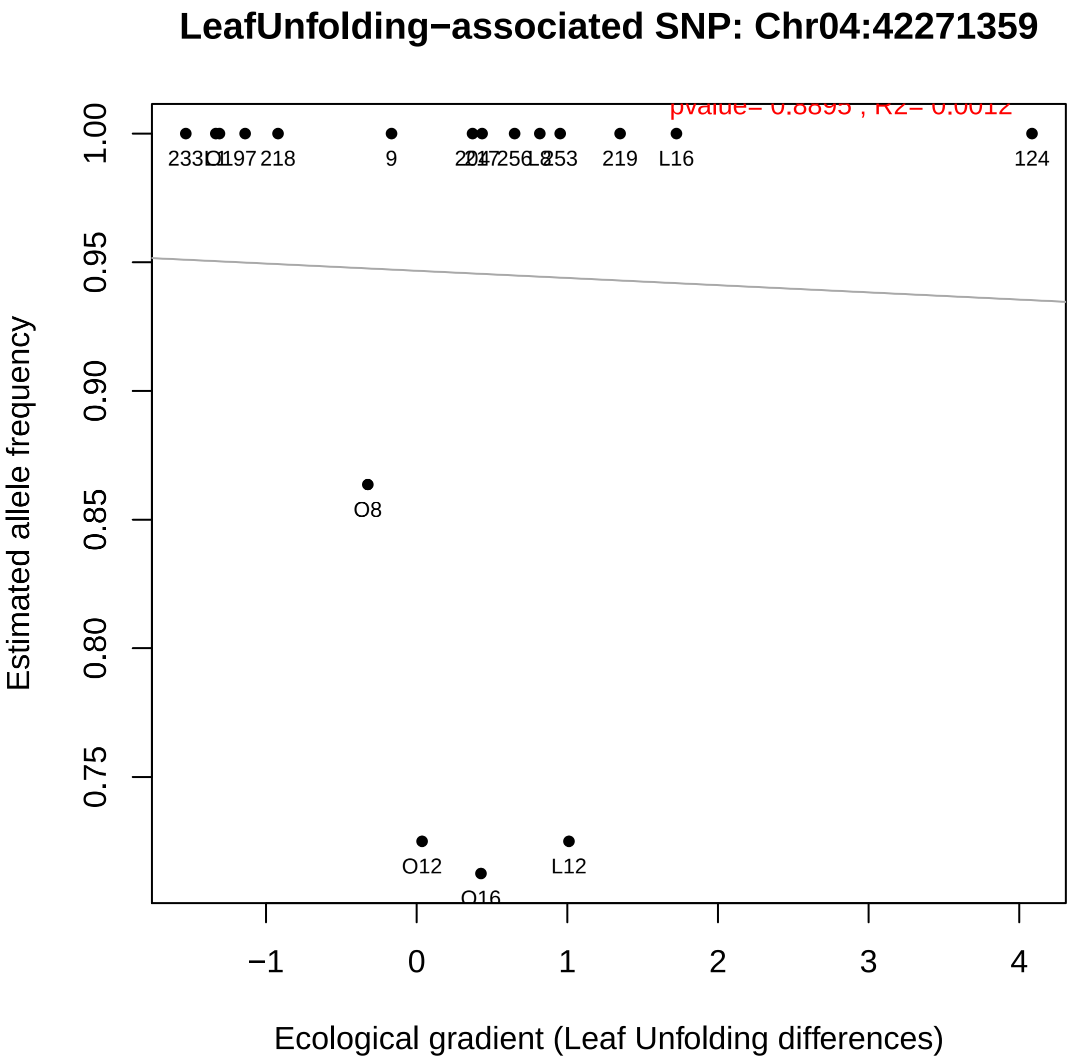

LeafUnfolding−associated SNP: Chr04:42271359
pvalue= 0.8895 , R2= 0.0012
Estimated allele frequency
0.75
0.80
0.85
0.90
0.95
1.00
Ecological gradient (Leaf Unfolding differences)
−1
0
1
2
3
4
218
9
219
L16
124
O8
O12
O16
L12

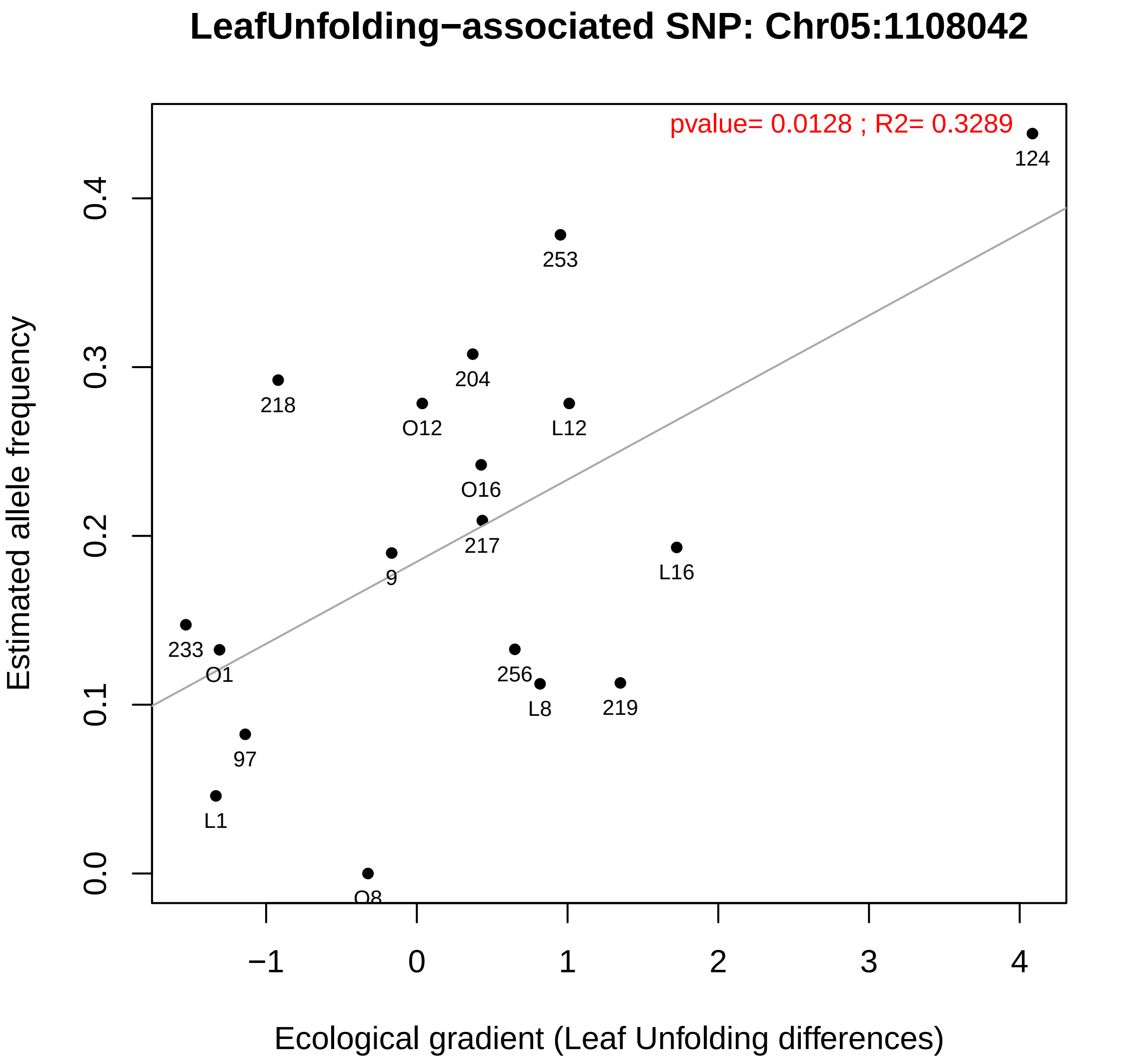

LeafUnfolding−associated SNP: Chr05:1108042
pvalue= 0.0128 ; R2= 0.3289
124
253
204
218
O12
L12
O16
217
9
L16
233
O1
256
L8
219
97
L1
O8
0.0
0.1
0.2
0.3
0.4
−1
0
1
2
3
4
Estimated allele frequency
Ecological gradient (Leaf Unfolding differences)

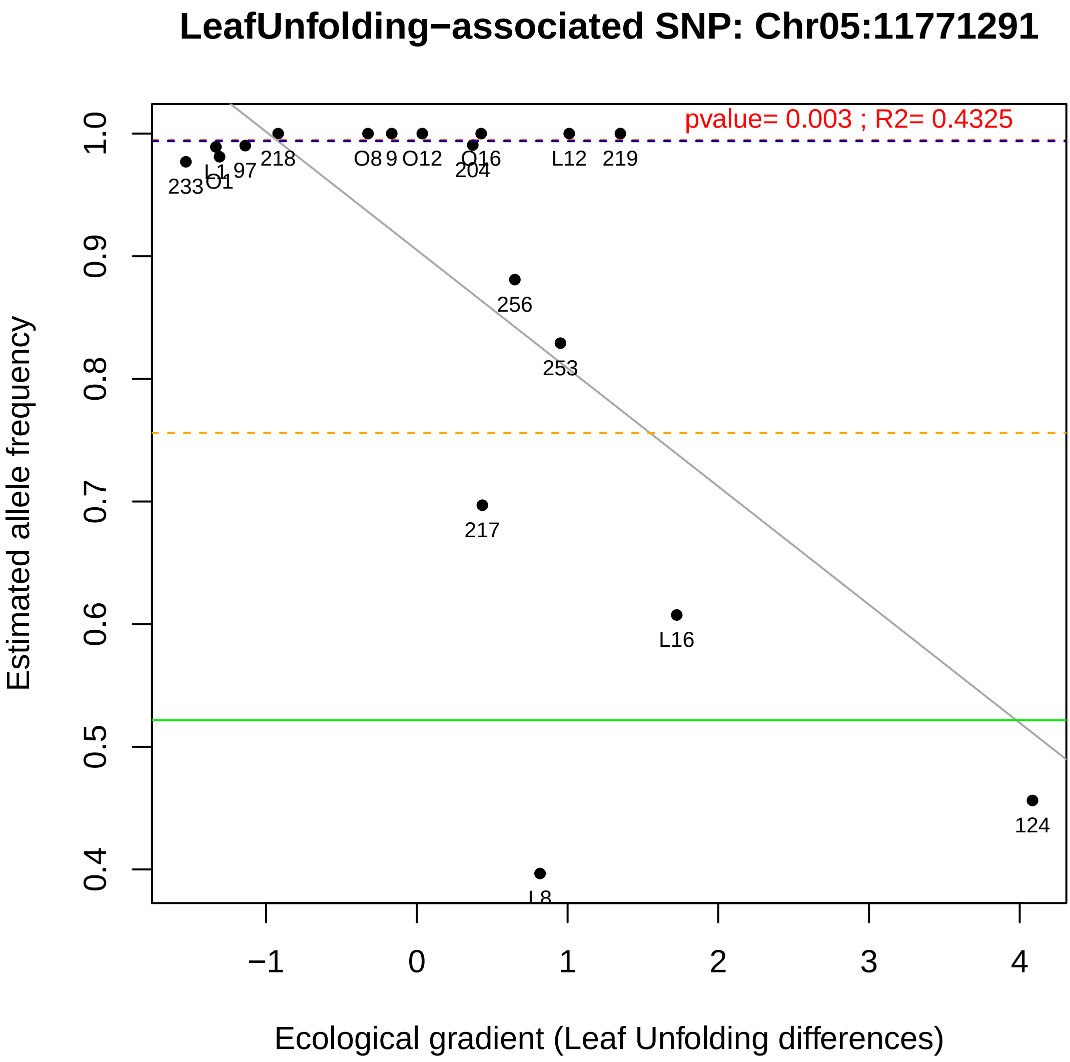
LeafUnfolding−associated SNP: Chr05:11771291
pvalue= 0.003 ; R2= 0.4325
Estimated allele frequency
Ecological gradient (Leaf Unfolding differences)
1.0
0.9
0.8
0.7
0.6
0.5
0.4
−1
0
1
2
3
4
233
L1
O1
97
218
O8
9
O12
O16
204
L12
219
256
253
217
L16
124
L8

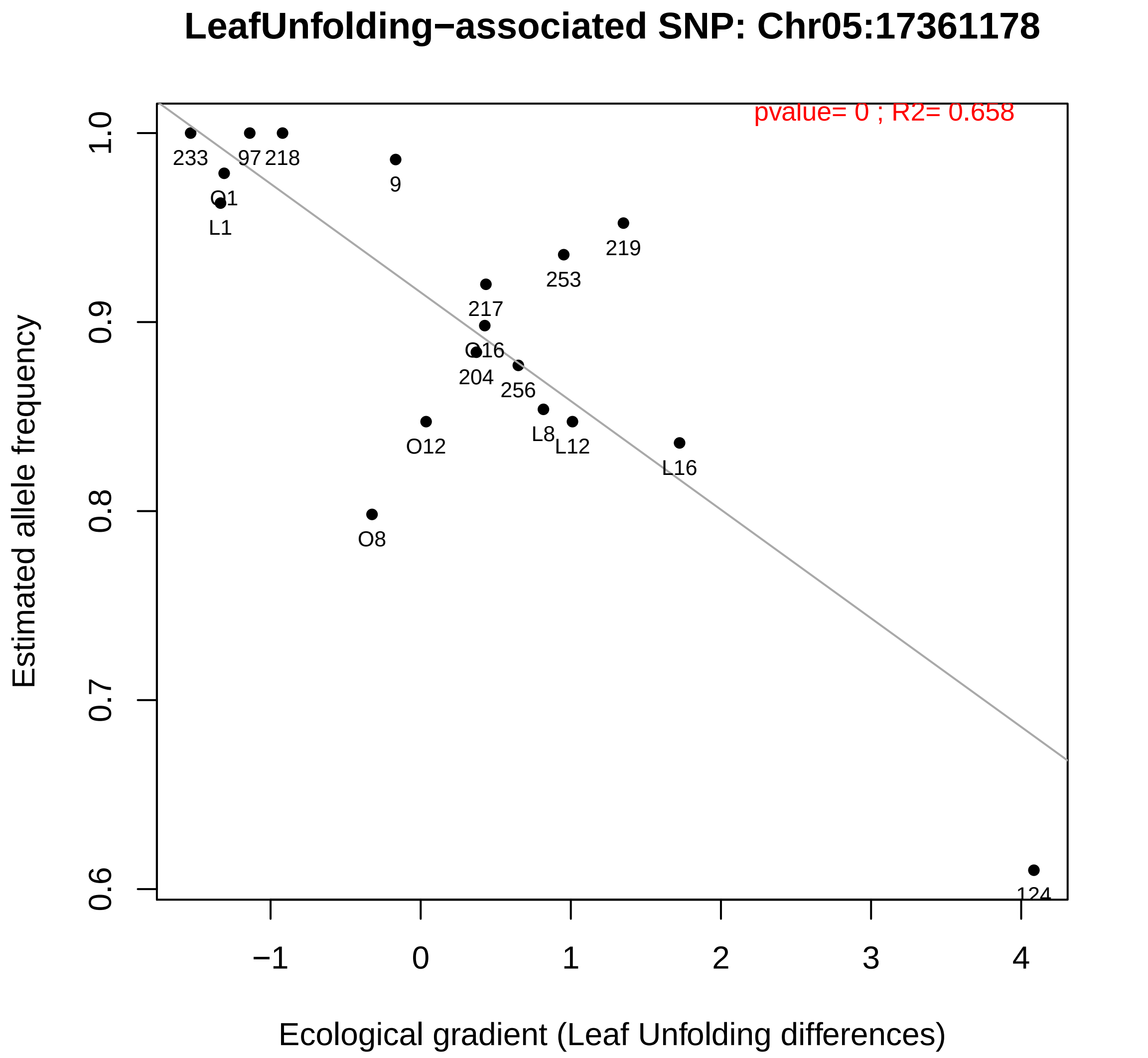

LeafUnfolding−associated SNP: Chr05:17361178
pvalue= 0 ; R2= 0.658
Estimated allele frequency
Ecological gradient (Leaf Unfolding differences)
1.0
0.9
0.8
0.7
0.6
−1
0
1
2
3
4
233
97
218
O1
L1
9
219
253
217
O16
204
256
L8
L12
O12
L16
O8
124

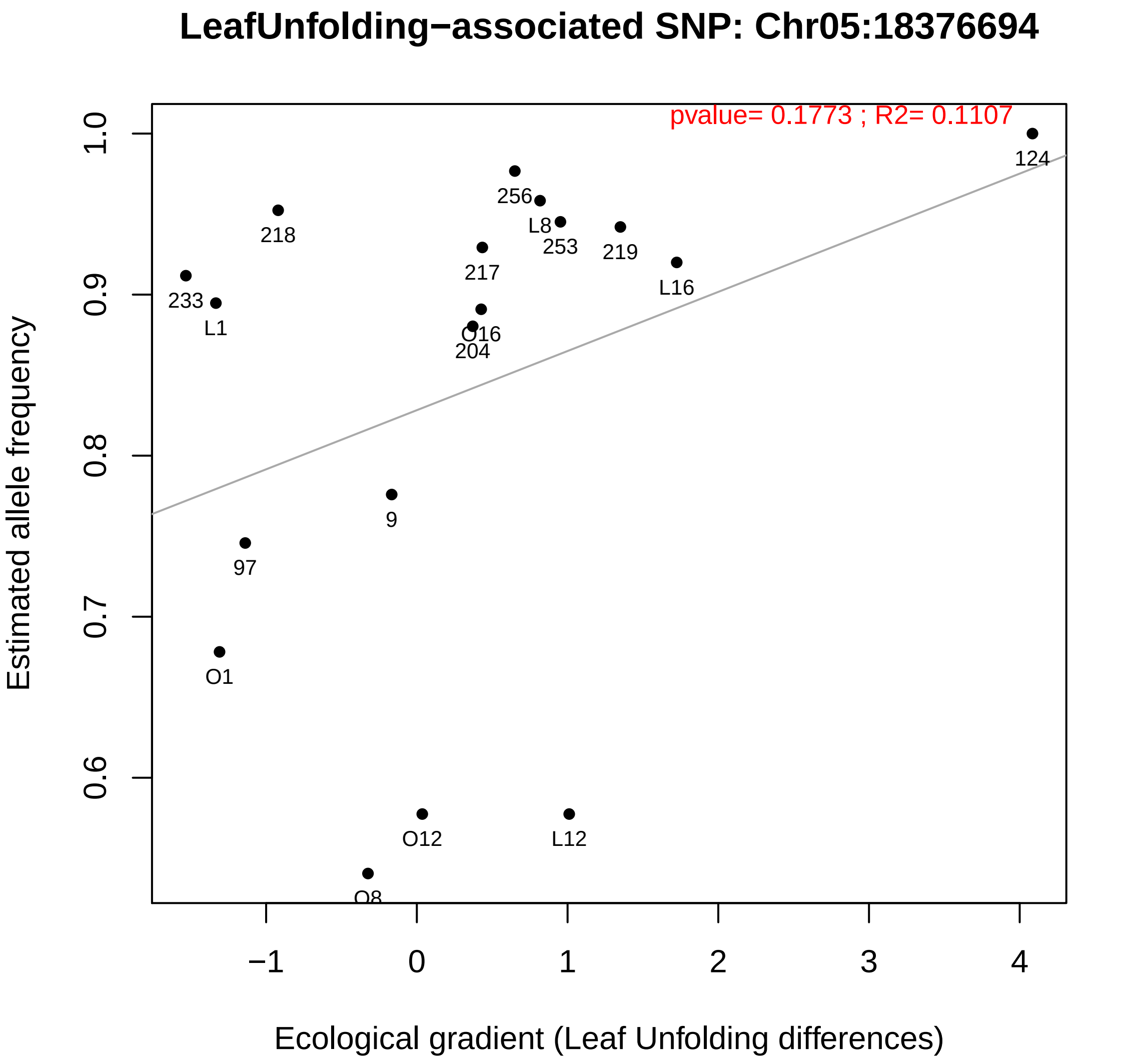

LeafUnfolding−associated SNP: Chr05:18376694
pvalue= 0.1773 ; R2= 0.1107
Estimated allele frequency
Ecological gradient (Leaf Unfolding differences)
0.6
0.7
0.8
0.9
1.0
−1
0
1
2
3
4
124
256
L8
253
219
218
217
L16
233
L1
O16
204
9
97
O1
O12
L12
O8

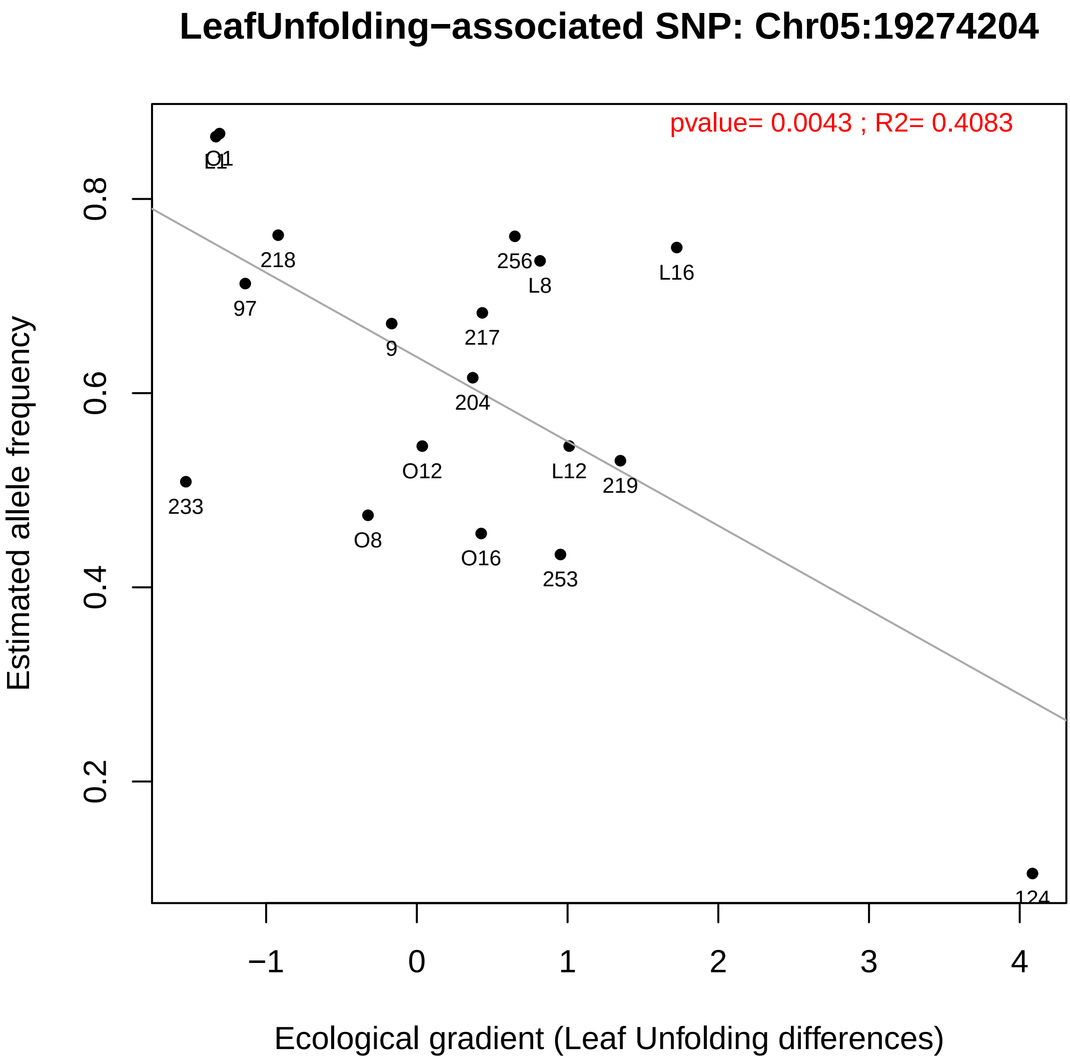

LeafUnfolding−associated SNP: Chr05:19274204
pvalue= 0.0043 ; R2= 0.4083
Estimated allele frequency
0.2
0.4
0.6
0.8
−1
0
1
2
3
4
Ecological gradient (Leaf Unfolding differences)
O1
L1
218
97
9
217
256
L8
L16
204
O12
L12
219
233
O8
O16
253
124

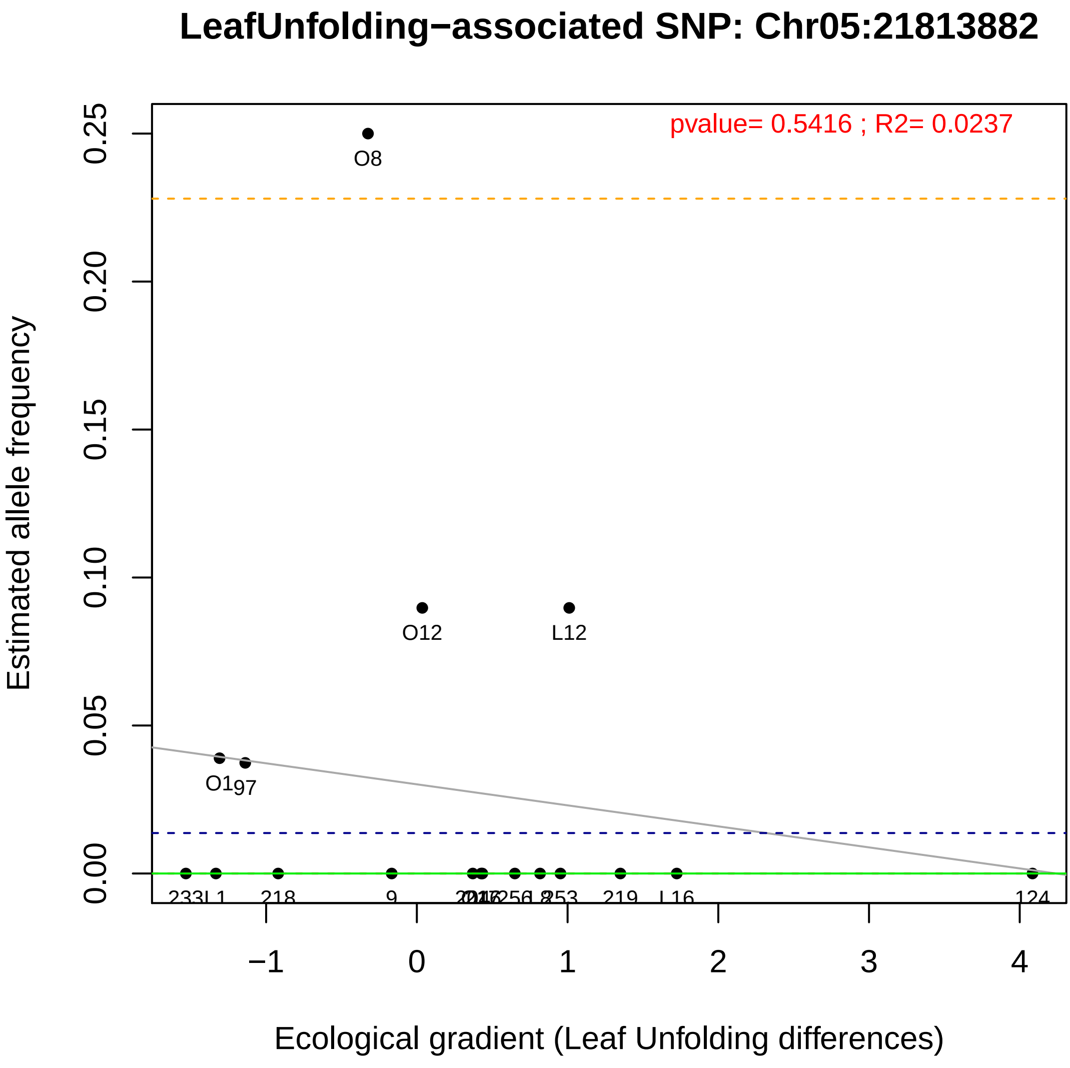

LeafUnfolding−associated SNP: Chr05:21813882
pvalue= 0.5416 ; R2= 0.0237
Estimated allele frequency
0.00
0.05
0.10
0.15
0.20
0.25
O8
O12
L12
O1
97
233
L1
218
9
124
219
L16
−1
0
1
2
3
4
Ecological gradient (Leaf Unfolding differences)

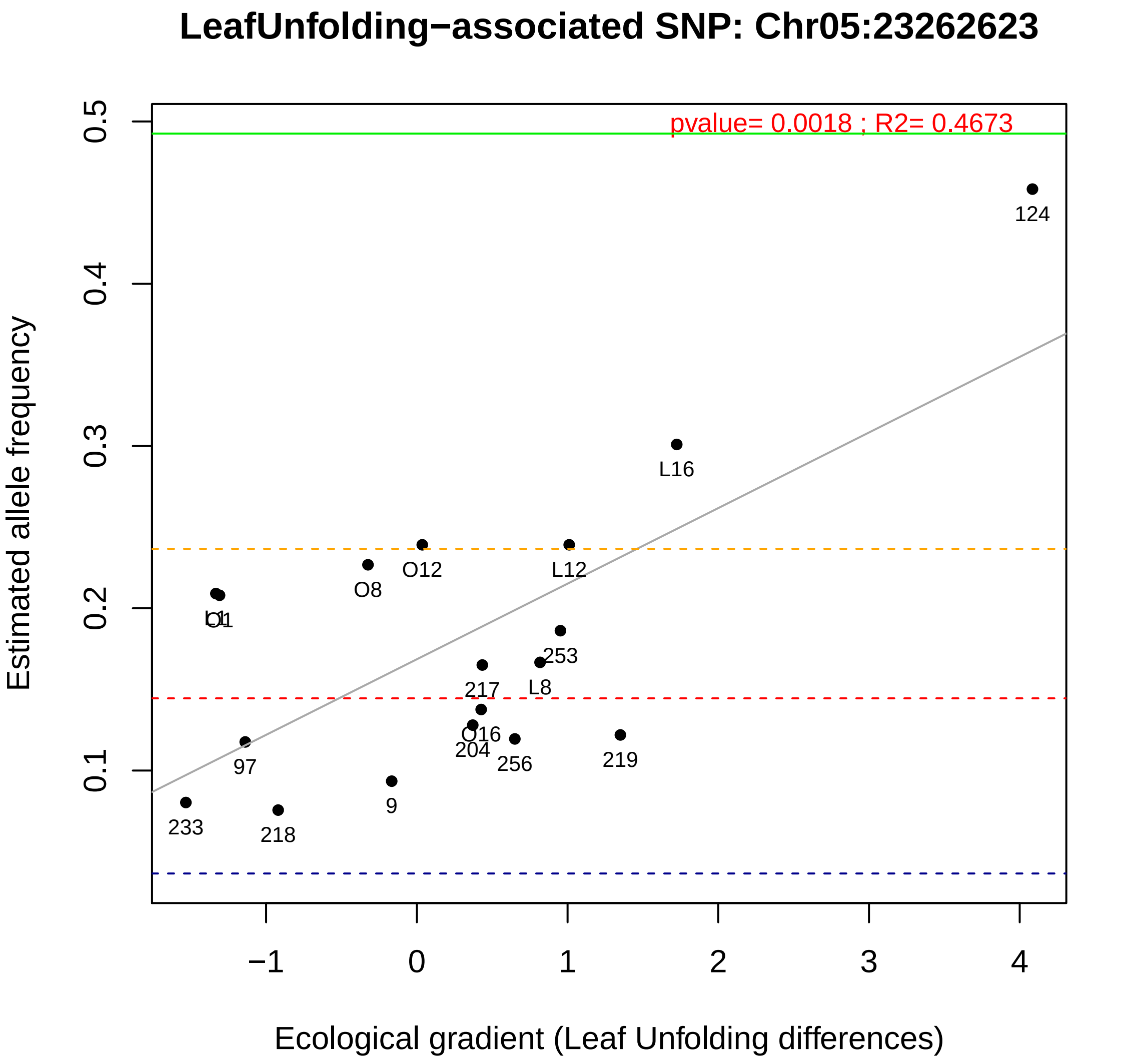
LeafUnfolding−associated SNP: Chr05:23262623
pvalue= 0.0018 ; R2= 0.4673
Estimated allele frequency
0.1
0.2
0.3
0.4
0.5
−1
0
1
2
3
4
Ecological gradient (Leaf Unfolding differences)
124
L16
O12
L12
O8
253
217
L8
O16
204
256
219
97
9
233
218

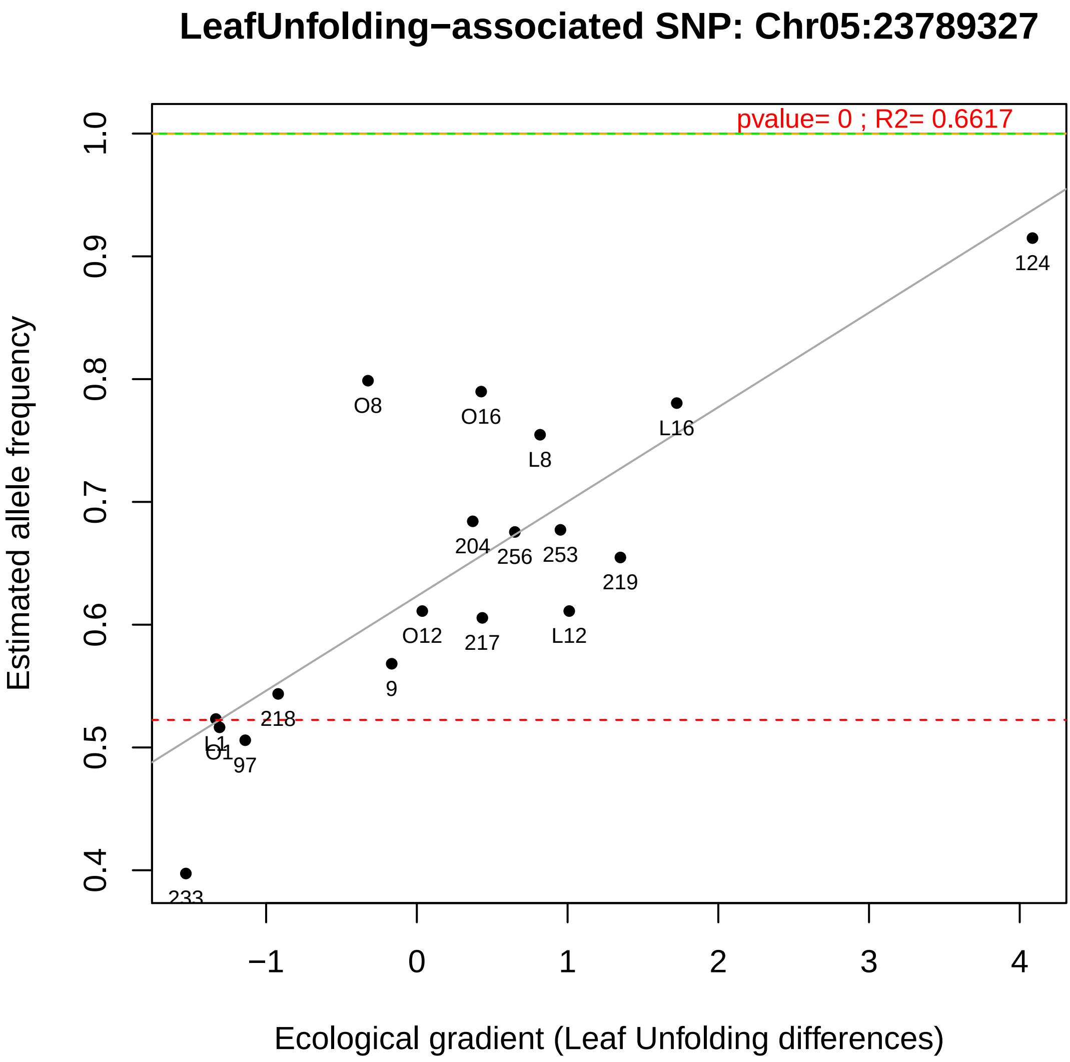

LeafUnfolding−associated SNP: Chr05:23789327
pvalue= 0 ; R2= 0.6617
Estimated allele frequency
Ecological gradient (Leaf Unfolding differences)
0.4
0.5
0.6
0.7
0.8
0.9
1.0
−1
0
1
2
3
4
124
O8
O16
L16
L8
204
256
253
219
O12
217
L12
9
218
L1
O1
97
233

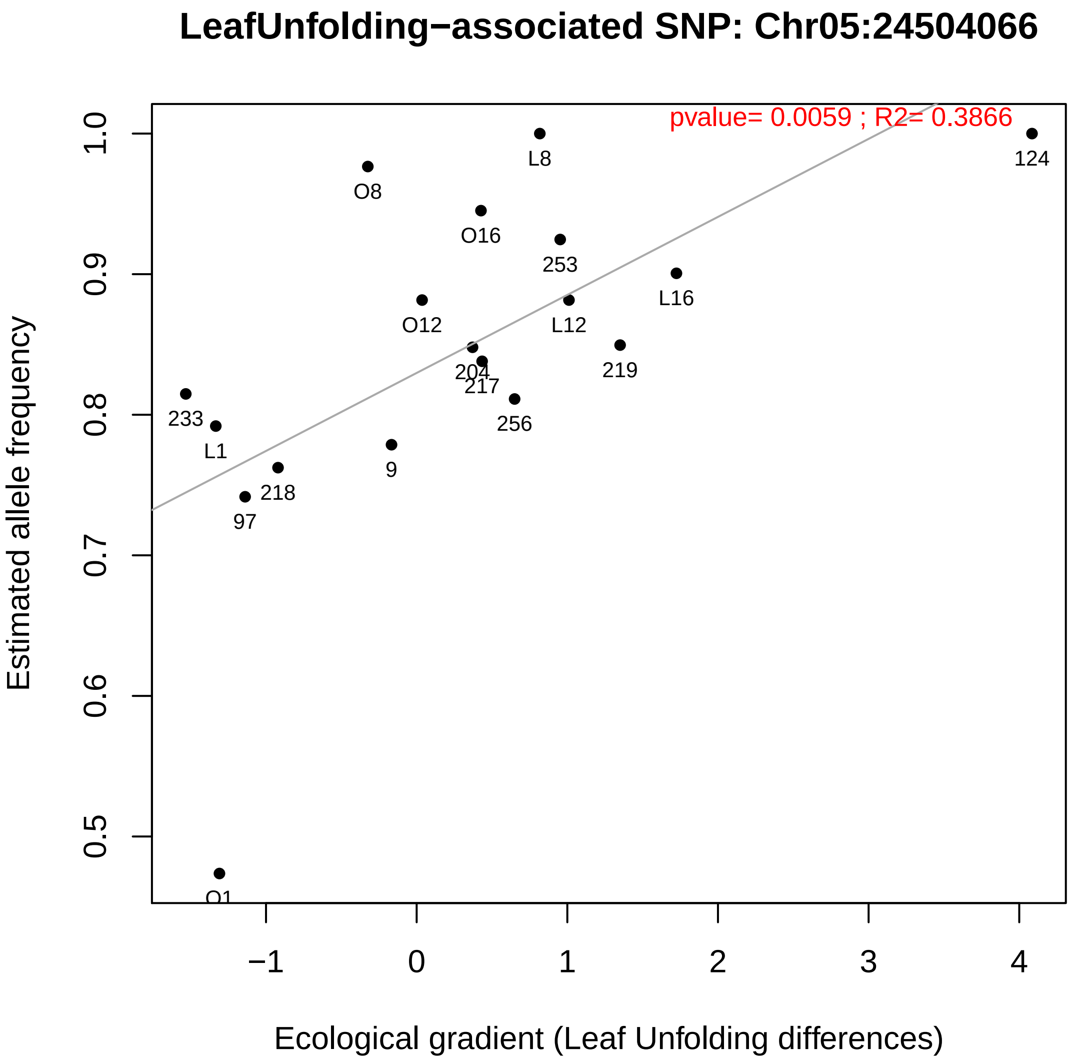

LeafUnfolding−associated SNP: Chr05:24504066
pvalue= 0.0059 ; R2= 0.3866
Estimated allele frequency
Ecological gradient (Leaf Unfolding differences)
0.5
0.6
0.7
0.8
0.9
1.0
−1
0
1
2
3
4
L8
124
O8
O16
253
L16
O12
L12
219
204
217
256
233
L1
218
9
97
O1

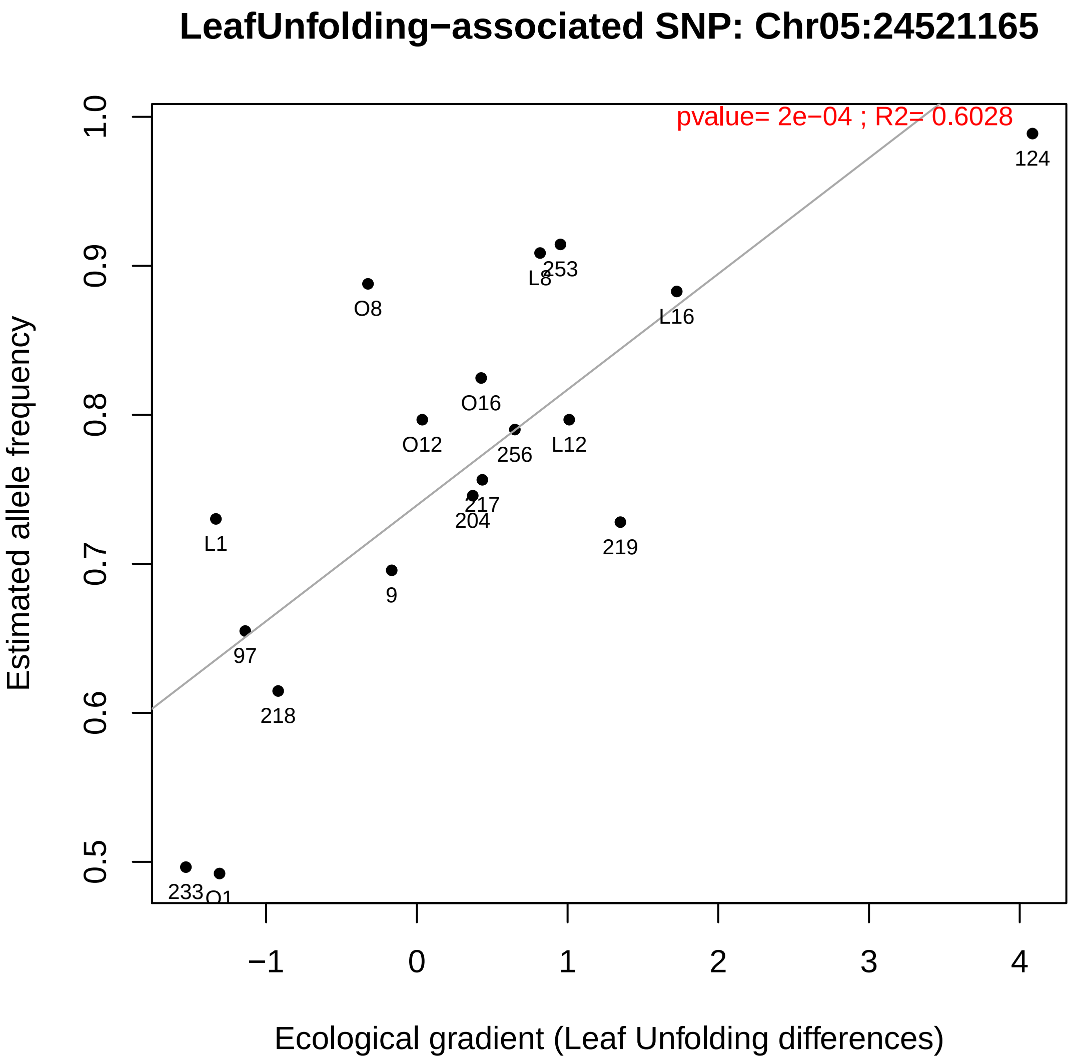
LeafUnfolding−associated SNP: Chr05:24521165
pvalue= 2e−04 ; R2= 0.6028
Estimated allele frequency
Ecological gradient (Leaf Unfolding differences)
1.0
0.9
0.8
0.7
0.6
0.5
−1
0
1
2
3
4
124
253
L8
O8
L16
O16
O12
256
L12
217
204
L1
219
9
97
218
233
O1

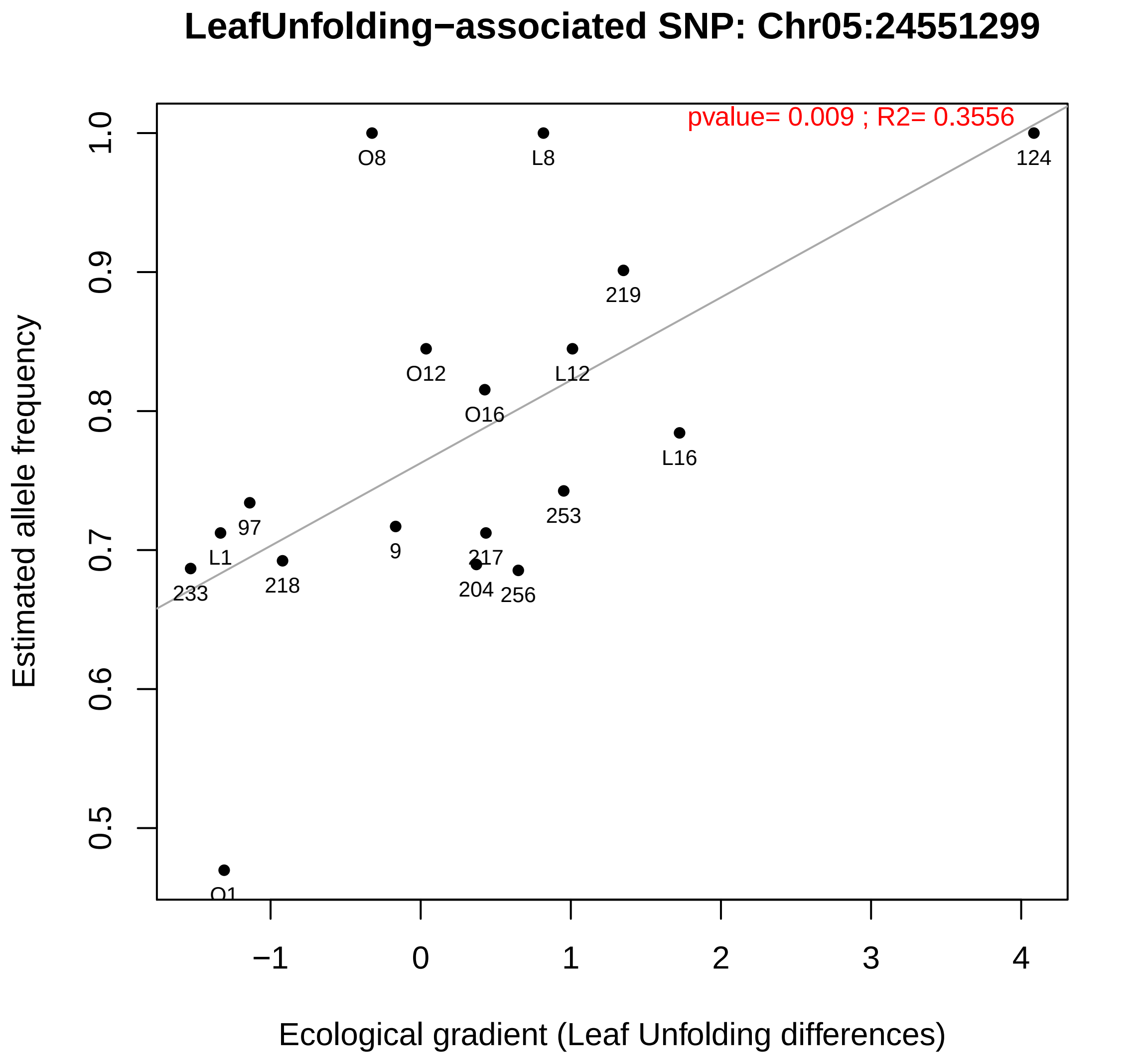

LeafUnfolding−associated SNP: Chr05:24551299
pvalue= 0.009 ; R2= 0.3556
Estimated allele frequency
Ecological gradient (Leaf Unfolding differences)
1.0
0.9
0.8
0.7
0.6
0.5
−1
0
1
2
3
4
O8
L8
124
219
O12
L12
O16
L16
253
97
L1
9
217
233
218
204
256
O1

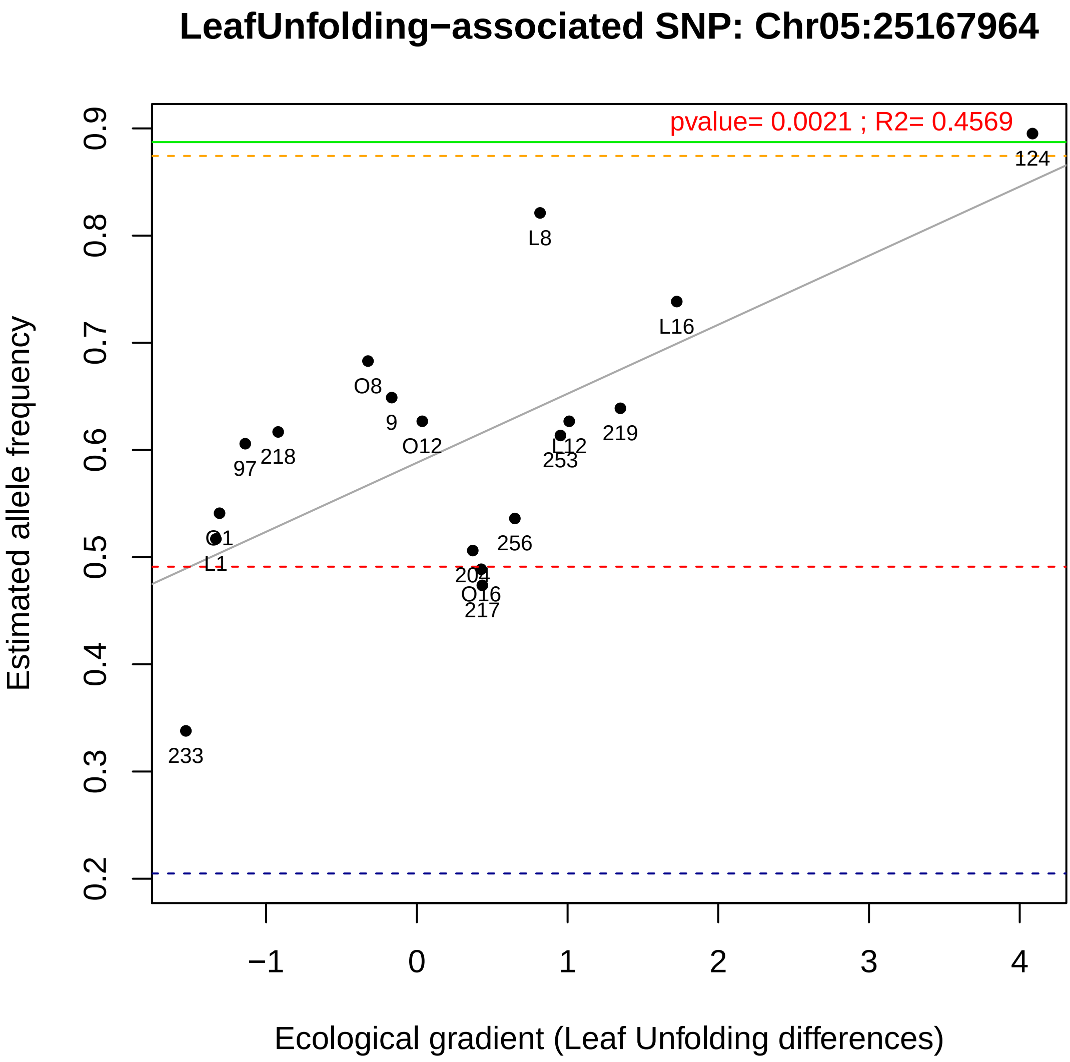

LeafUnfolding−associated SNP: Chr05:25167964
pvalue= 0.0021 ; R2= 0.4569
Estimated allele frequency
Ecological gradient (Leaf Unfolding differences)
0.2
0.3
0.4
0.5
0.6
0.7
0.8
0.9
−1
0
1
2
3
4
124
L8
L16
O8
9
O12
219
L12
253
97
218
O1
L1
256
204
O16
217
233

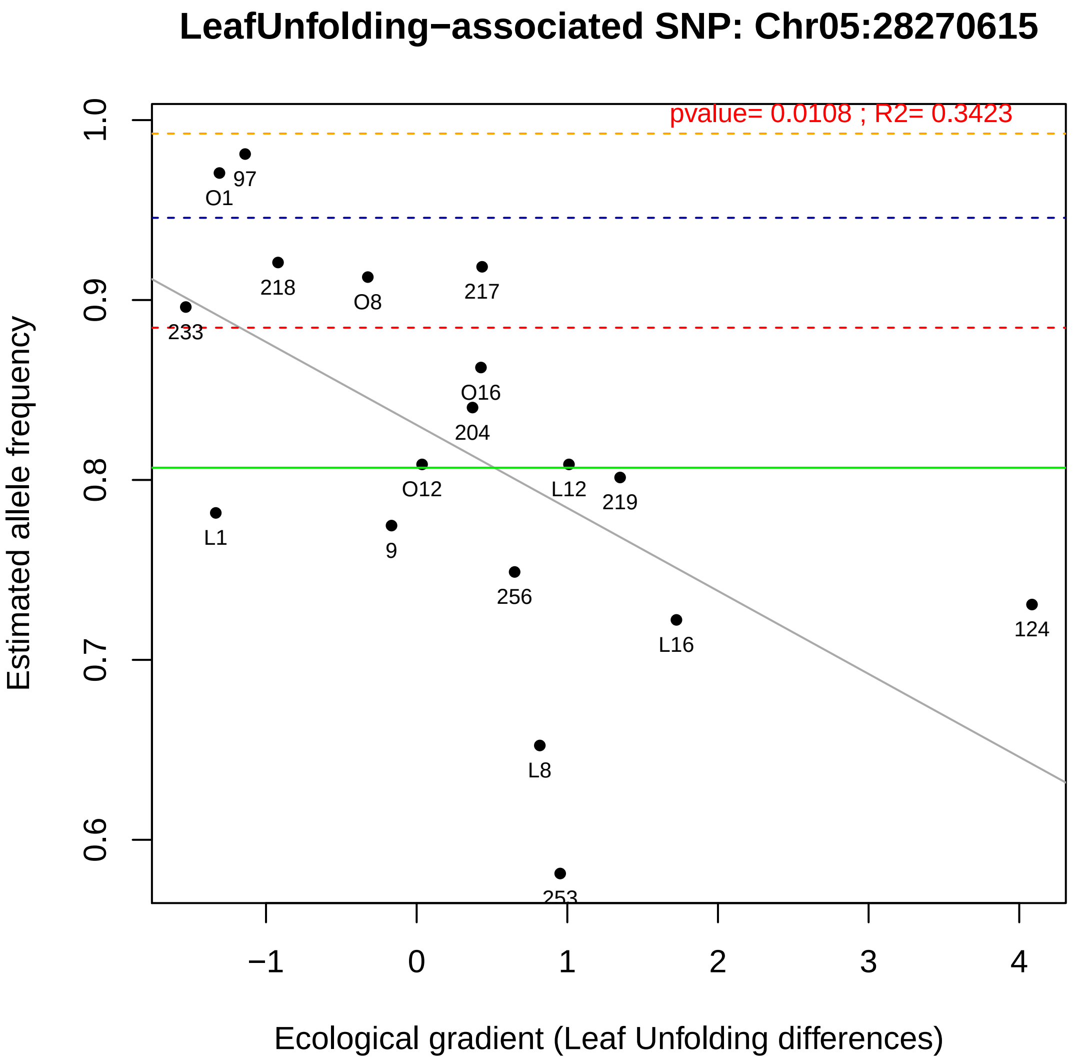
LeafUnfolding−associated SNP: Chr05:28270615
pvalue= 0.0108 ; R2= 0.3423
Estimated allele frequency
Ecological gradient (Leaf Unfolding differences)
0.6
0.7
0.8
0.9
1.0
−1
0
1
2
3
4
97
O1
218
O8
217
233
O16
204
O12
L12
219
L1
9
256
L16
124
L8
253

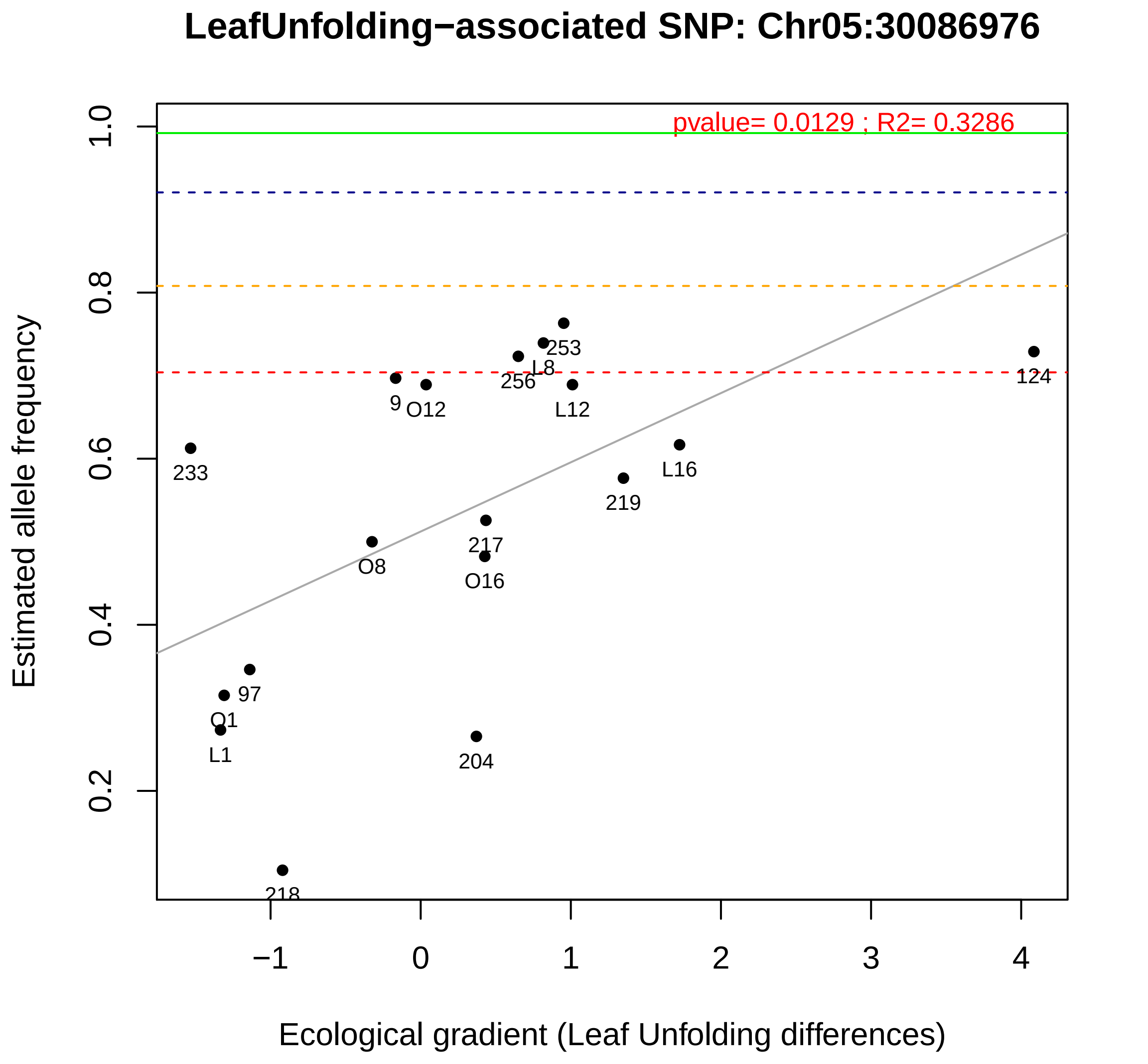

LeafUnfolding−associated SNP: Chr05:30086976
pvalue= 0.0129 ; R2= 0.3286
Estimated allele frequency
Ecological gradient (Leaf Unfolding differences)
0.2
0.4
0.6
0.8
1.0
−1
0
1
2
3
4
253
L8
256
L12
9
O12
124
233
L16
219
217
O8
O16
97
O1
L1
204
218

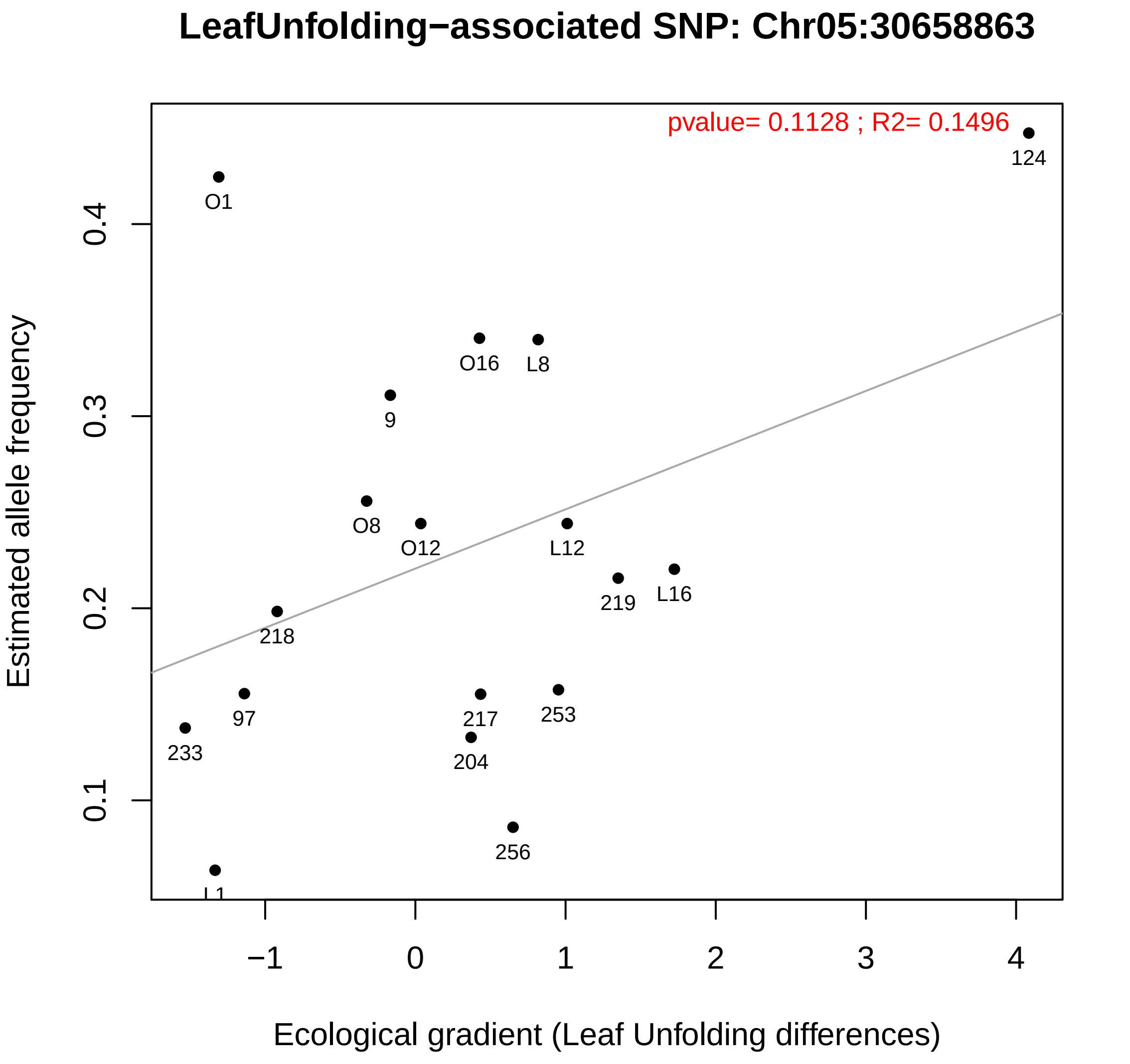
LeafUnfolding−associated SNP: Chr05:30658863
pvalue= 0.1128 ; R2= 0.1496
124
O1
O16
L8
9
O8
O12
L12
219
L16
218
97
233
217
253
204
256
L1
0.1
0.2
0.3
0.4
−1
0
1
2
3
4
Estimated allele frequency
Ecological gradient (Leaf Unfolding differences)

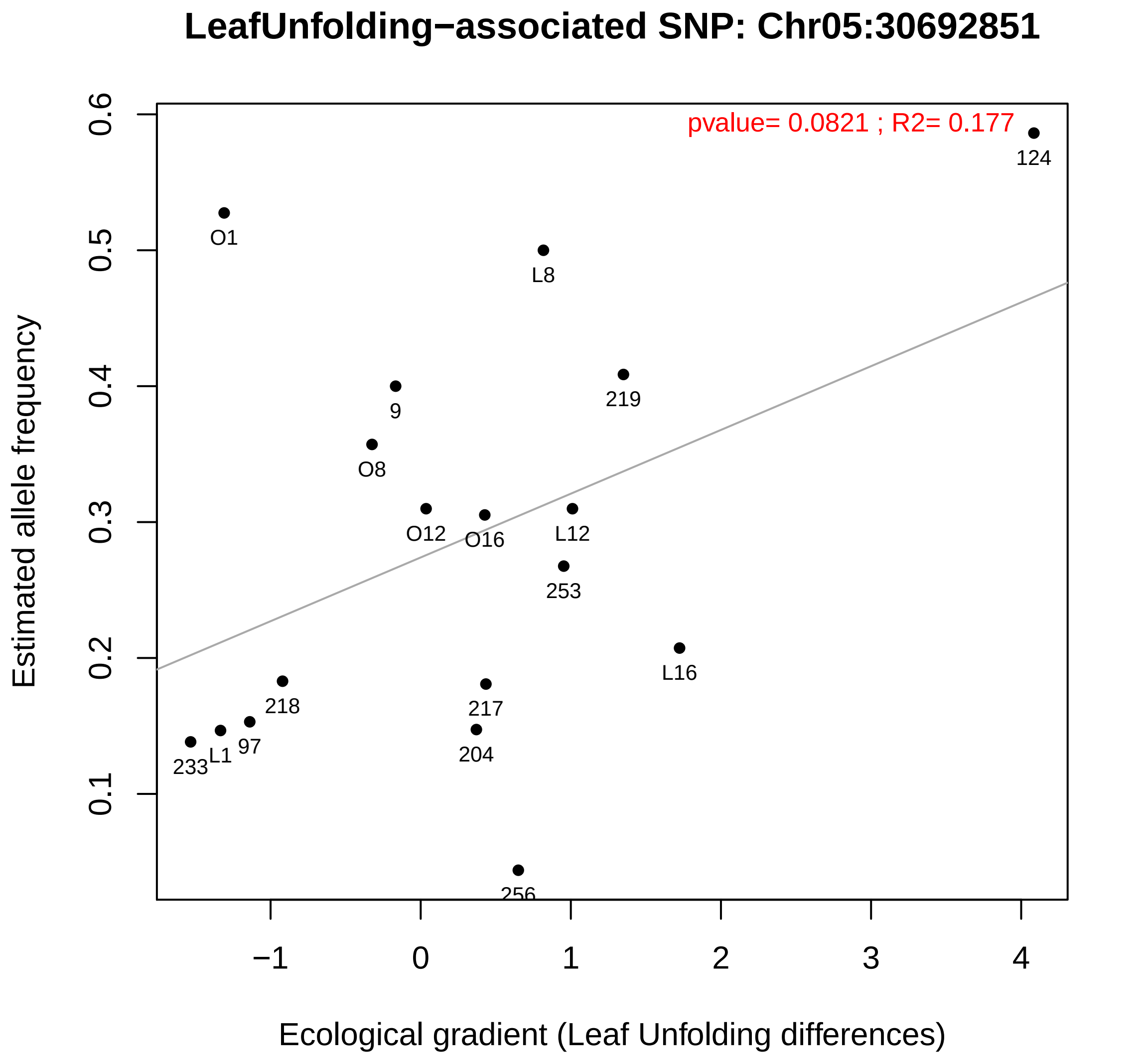
LeafUnfolding−associated SNP: Chr05:30692851
pvalue= 0.0821 ; R2= 0.177
Estimated allele frequency
Ecological gradient (Leaf Unfolding differences)
0.1
0.2
0.3
0.4
0.5
0.6
−1
0
1
2
3
4
124
O1
L8
219
9
O8
O12
O16
L12
253
L16
218
217
233
L1
97
204
256

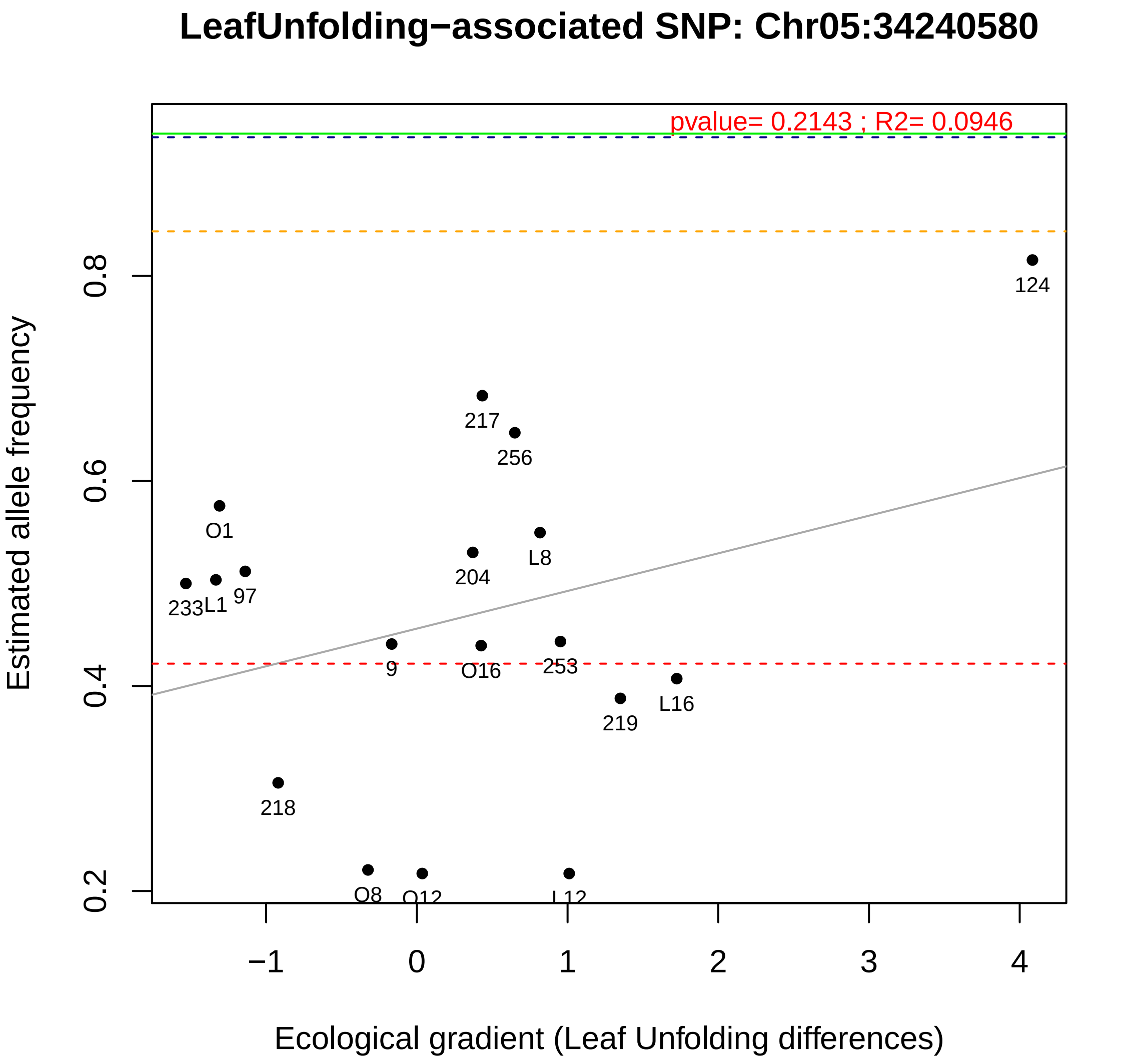

LeafUnfolding−associated SNP: Chr05:34240580
pvalue= 0.2143 ; R2= 0.0946
Estimated allele frequency
0.2
0.4
0.6
0.8
Ecological gradient (Leaf Unfolding differences)
−1
0
1
2
3
4
124
217
256
O1
L8
204
233
L1
97
9
O16
253
L16
219
218
O8
O12
L12

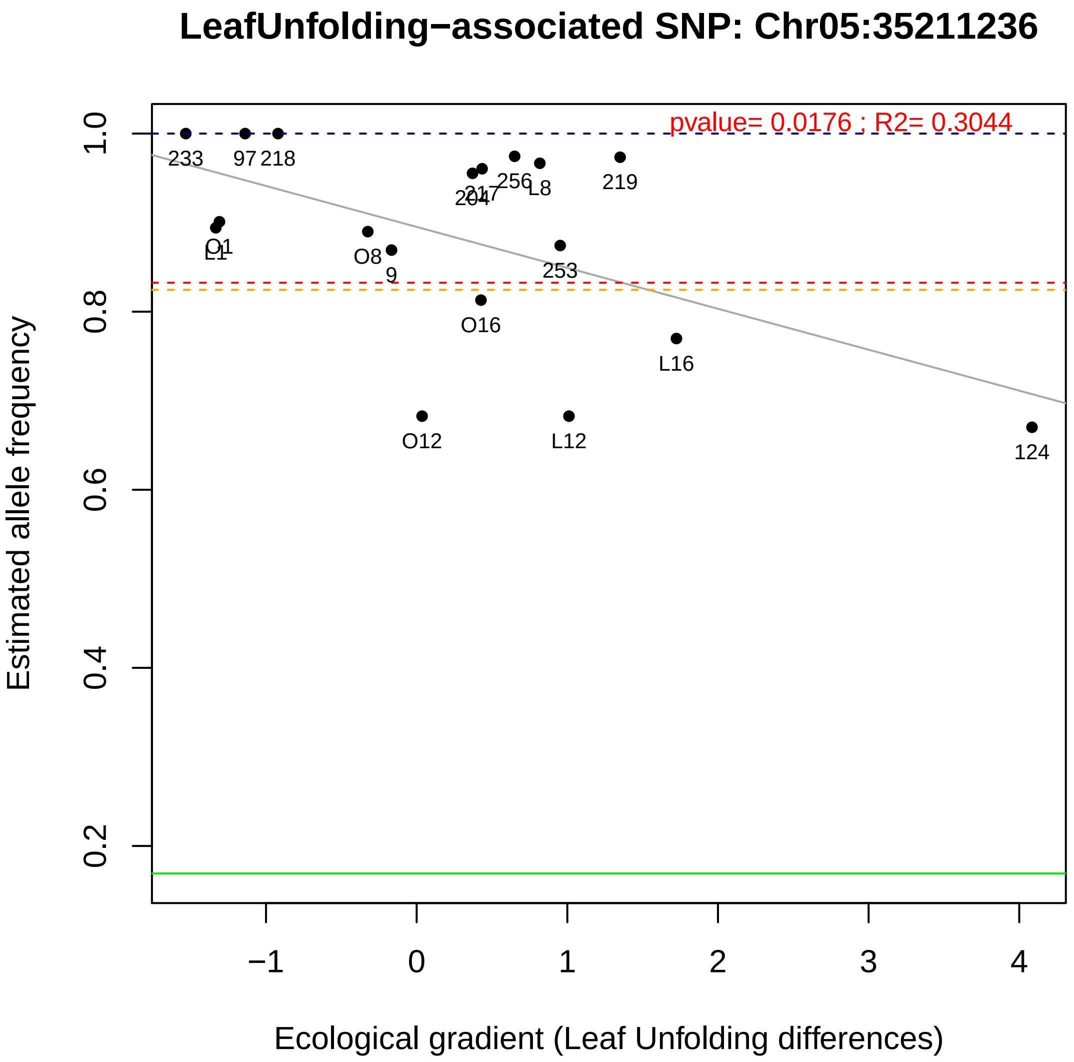

LeafUnfolding−associated SNP: Chr05:35211236
pvalue= 0.0176 ; R2= 0.3044
Estimated allele frequency
Ecological gradient (Leaf Unfolding differences)
0.2
0.4
0.6
0.8
1.0
−1
0
1
2
3
4
233
97
218
256
L8
219
217
204
O1
L1
O8
9
253
O16
L16
O12
L12
124

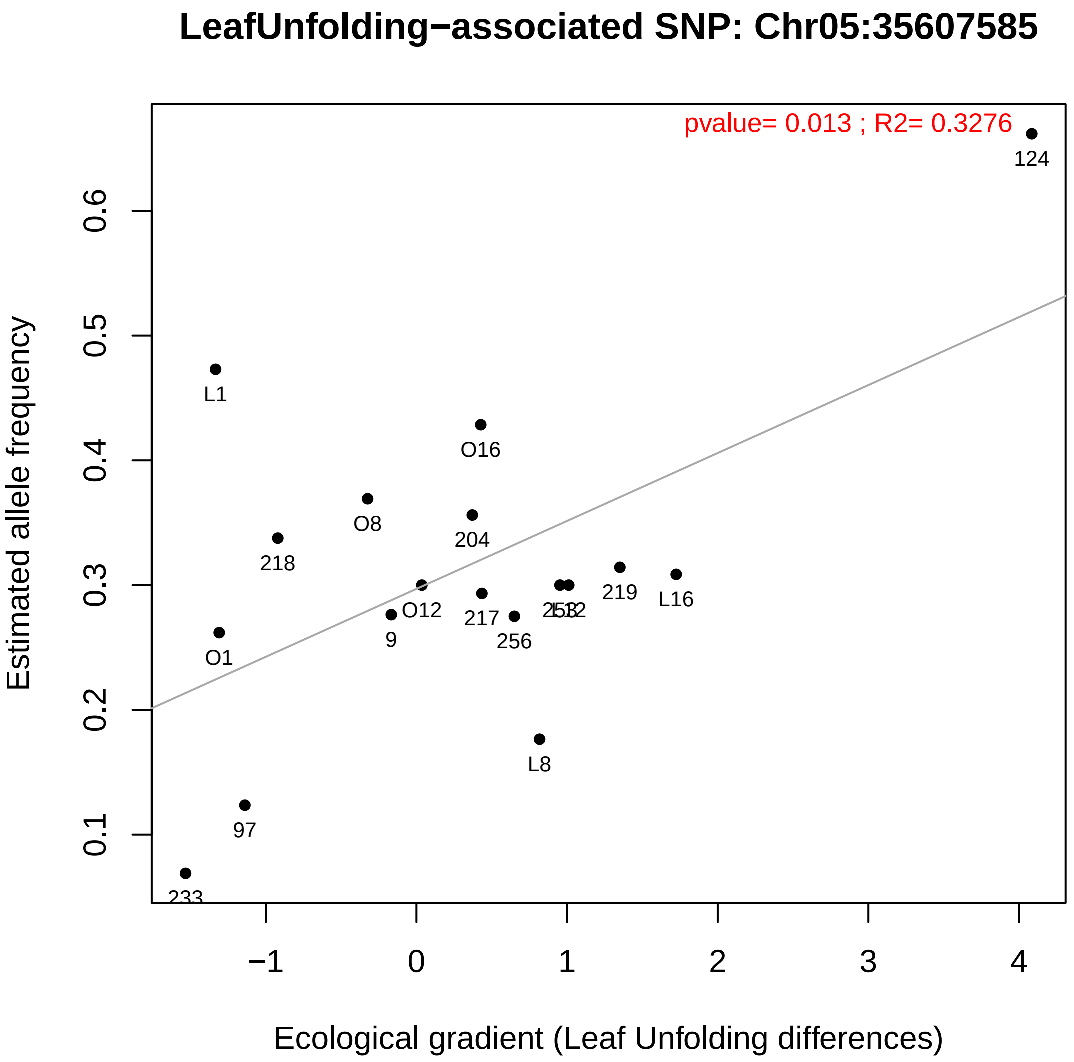
LeafUnfolding−associated SNP: Chr05:35607585
pvalue= 0.013 ; R2= 0.3276
Estimated allele frequency
0.1
0.2
0.3
0.4
0.5
0.6
Ecological gradient (Leaf Unfolding differences)
−1
0
1
2
3
4
124
L1
O16
O8
204
218
219
L16
O12
217
9
256
O1
L8
97
233

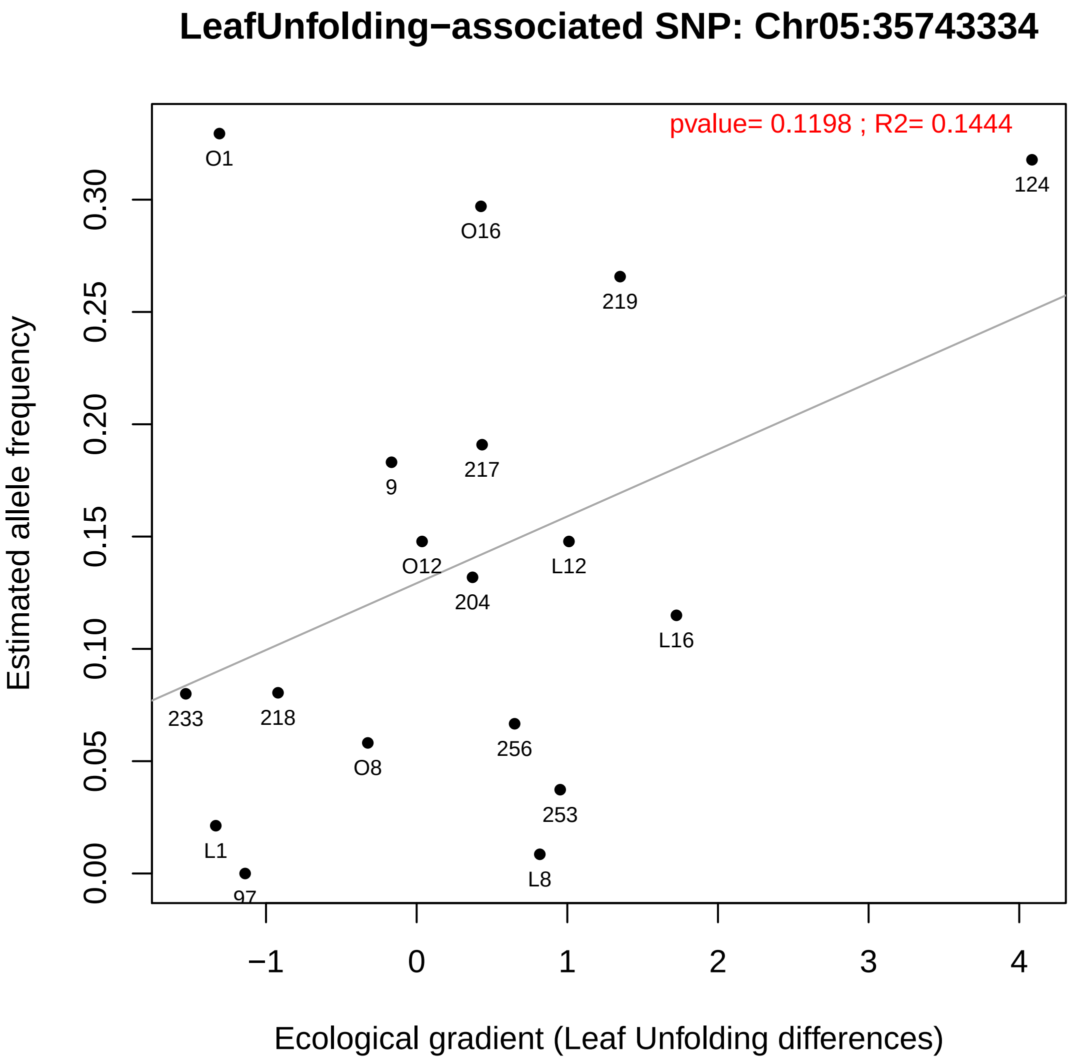
LeafUnfolding−associated SNP: Chr05:35743334
pvalue= 0.1198 ; R2= 0.1444
O1
124
O16
219
217
9
O12
L12
204
L16
233
218
O8
256
253
L1
97
L8
0.00
0.05
0.10
0.15
0.20
0.25
0.30
−1
0
1
2
3
4
Estimated allele frequency
Ecological gradient (Leaf Unfolding differences)

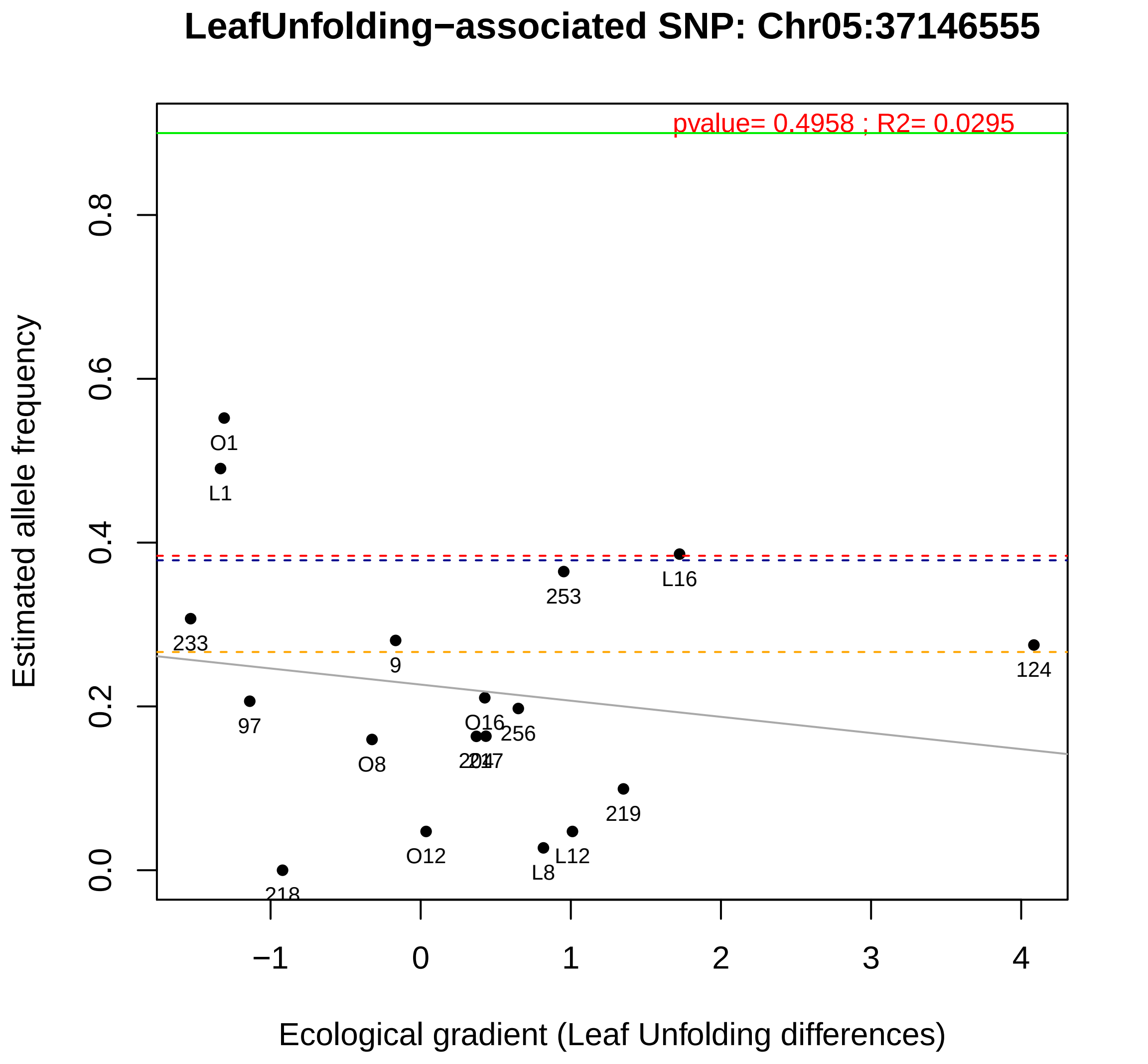

LeafUnfolding−associated SNP: Chr05:37146555
pvalue= 0.4958 ; R2= 0.0295
Estimated allele frequency
0.0
0.2
0.4
0.6
0.8
−1
0
1
2
3
4
Ecological gradient (Leaf Unfolding differences)
O1
L1
L16
253
233
9
124
97
O16
256
O8
219
O12
L12
L8
218

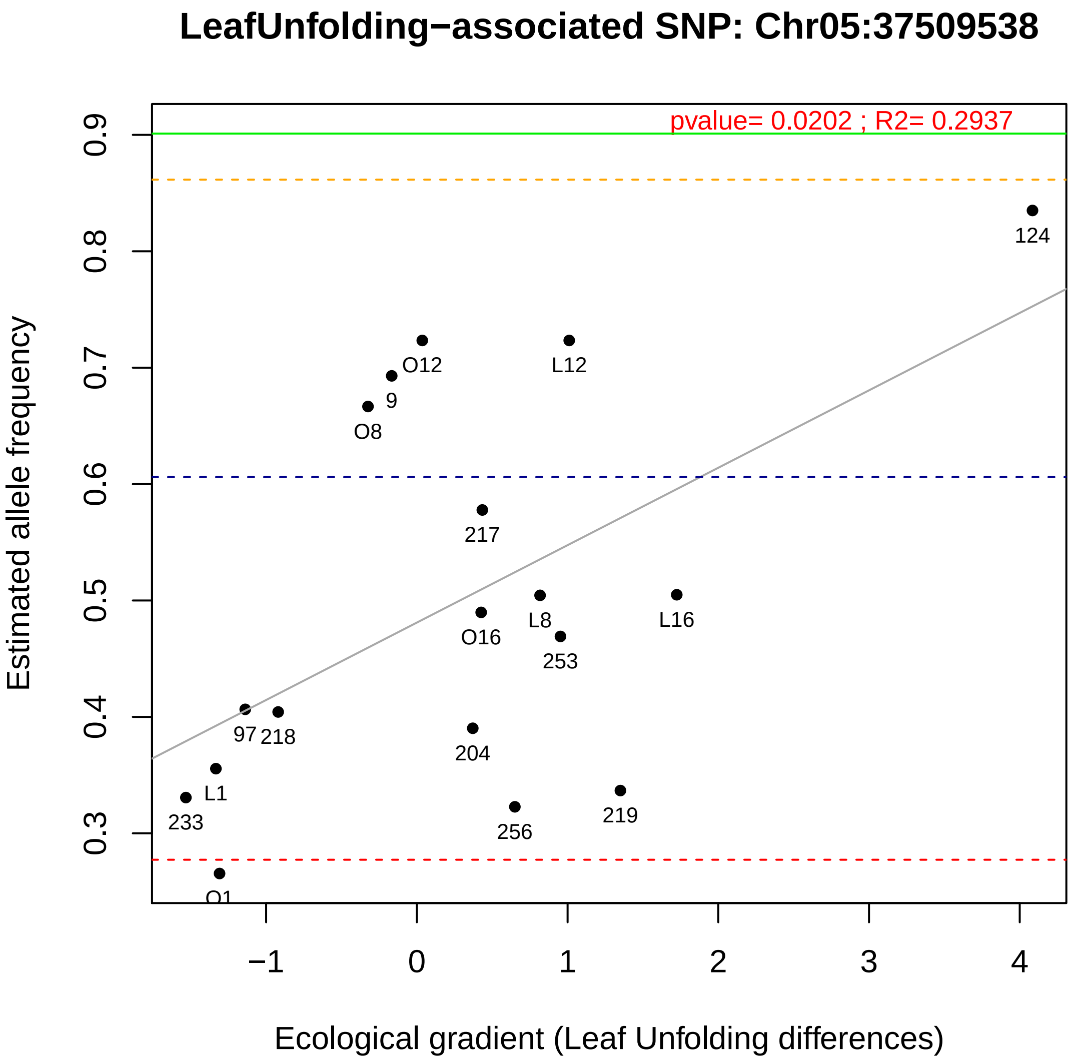

LeafUnfolding−associated SNP: Chr05:37509538
pvalue= 0.0202 ; R2= 0.2937
Estimated allele frequency
0.3
0.4
0.5
0.6
0.7
0.8
0.9
−1
0
1
2
3
4
Ecological gradient (Leaf Unfolding differences)
124
O12
L12
9
O8
217
L8
L16
O16
253
97
218
204
L1
233
256
219
O1

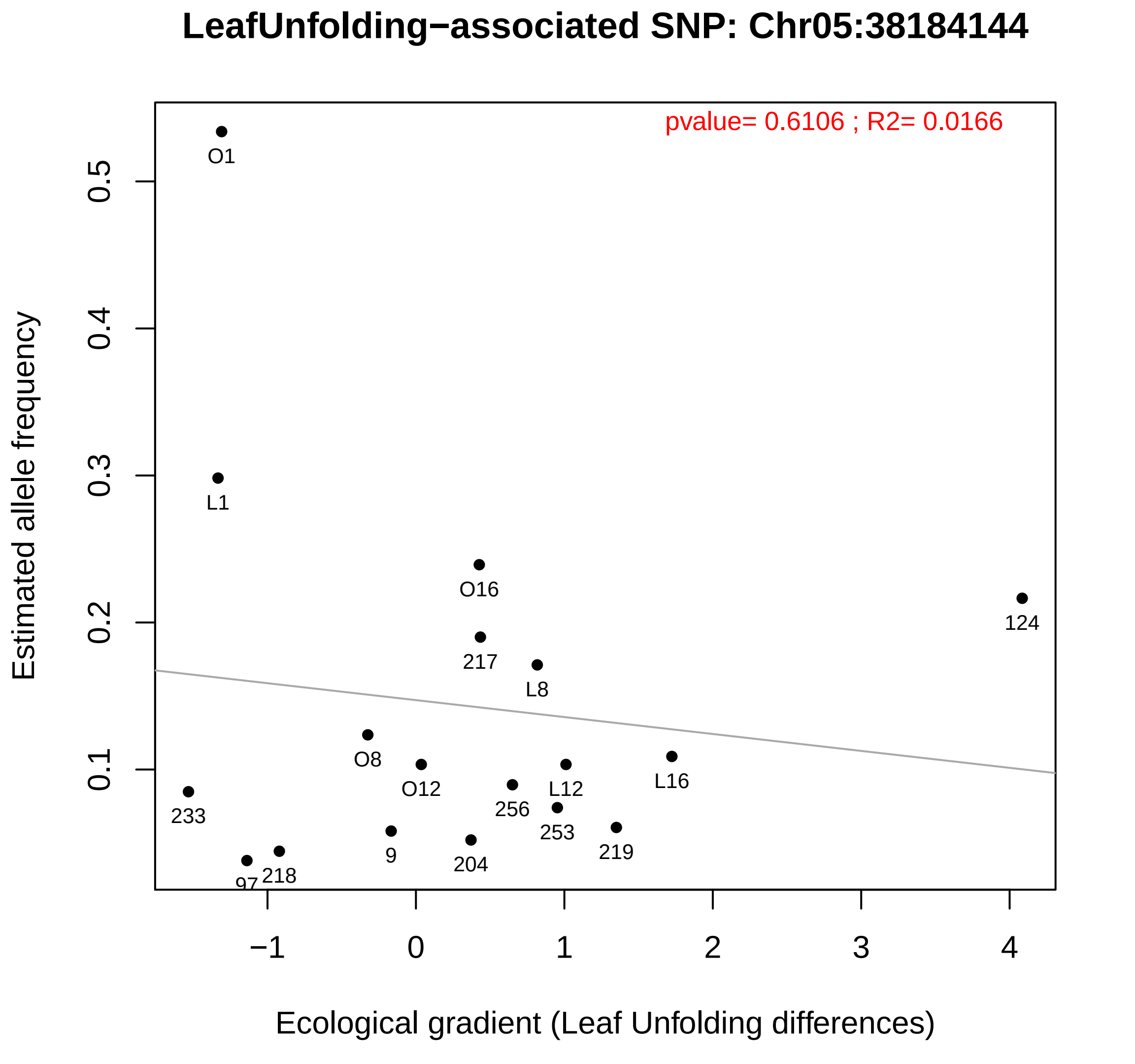

LeafUnfolding−associated SNP: Chr05:38184144
pvalue= 0.6106 ; R2= 0.0166
Estimated allele frequency
Ecological gradient (Leaf Unfolding differences)
0.1
0.2
0.3
0.4
0.5
−1
0
1
2
3
4
O1
L1
O16
124
217
L8
O8
O12
L12
L16
233
256
253
219
9
204
97
218

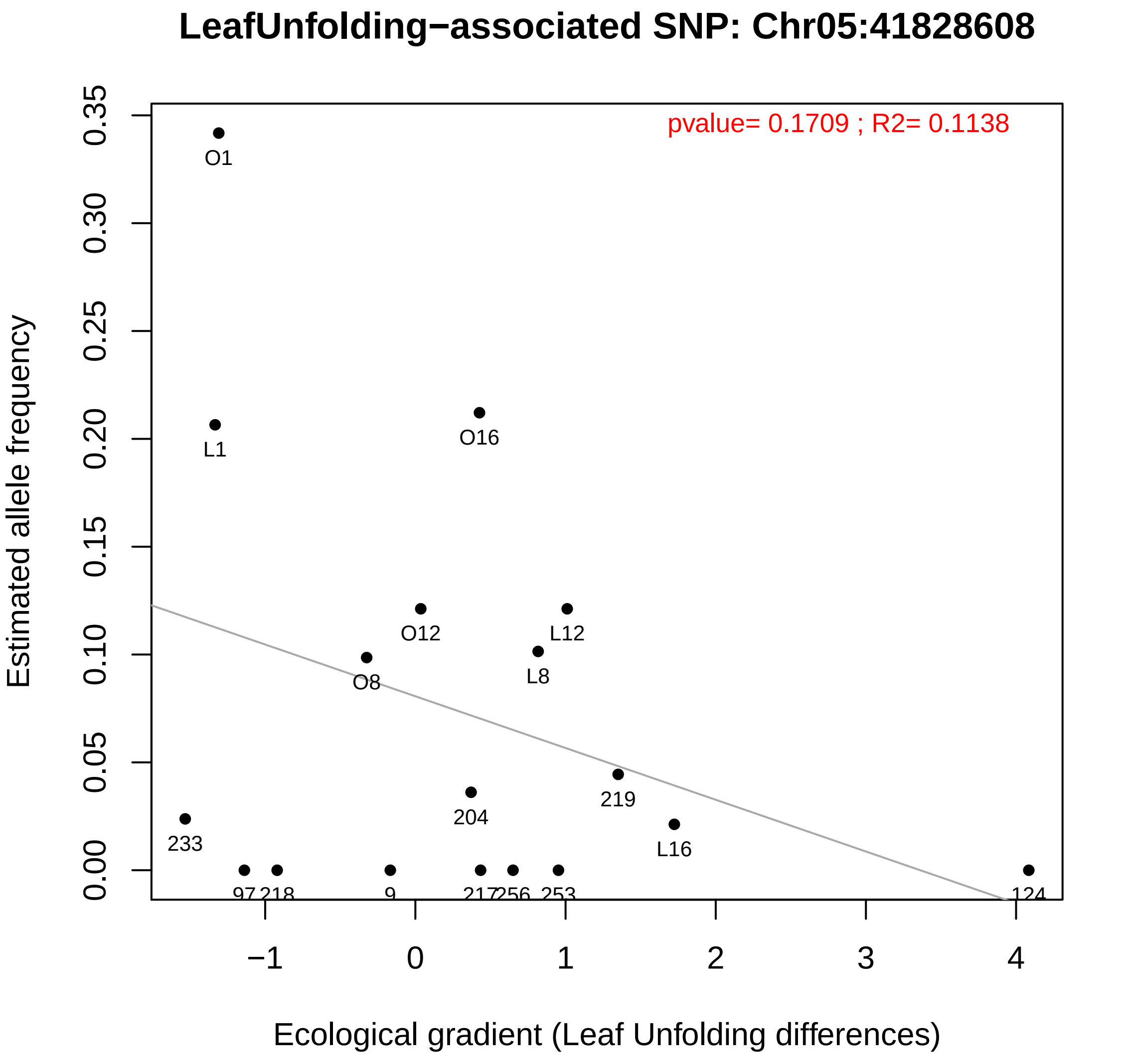

LeafUnfolding−associated SNP: Chr05:41828608
pvalue= 0.1709 ; R2= 0.1138
Estimated allele frequency
0.00
0.05
0.10
0.15
0.20
0.25
0.30
0.35
−1
0
1
2
3
4
Ecological gradient (Leaf Unfolding differences)
O1
L1
O16
O12
L12
O8
L8
219
204
233
L16
97
218
9
217
256
253
124

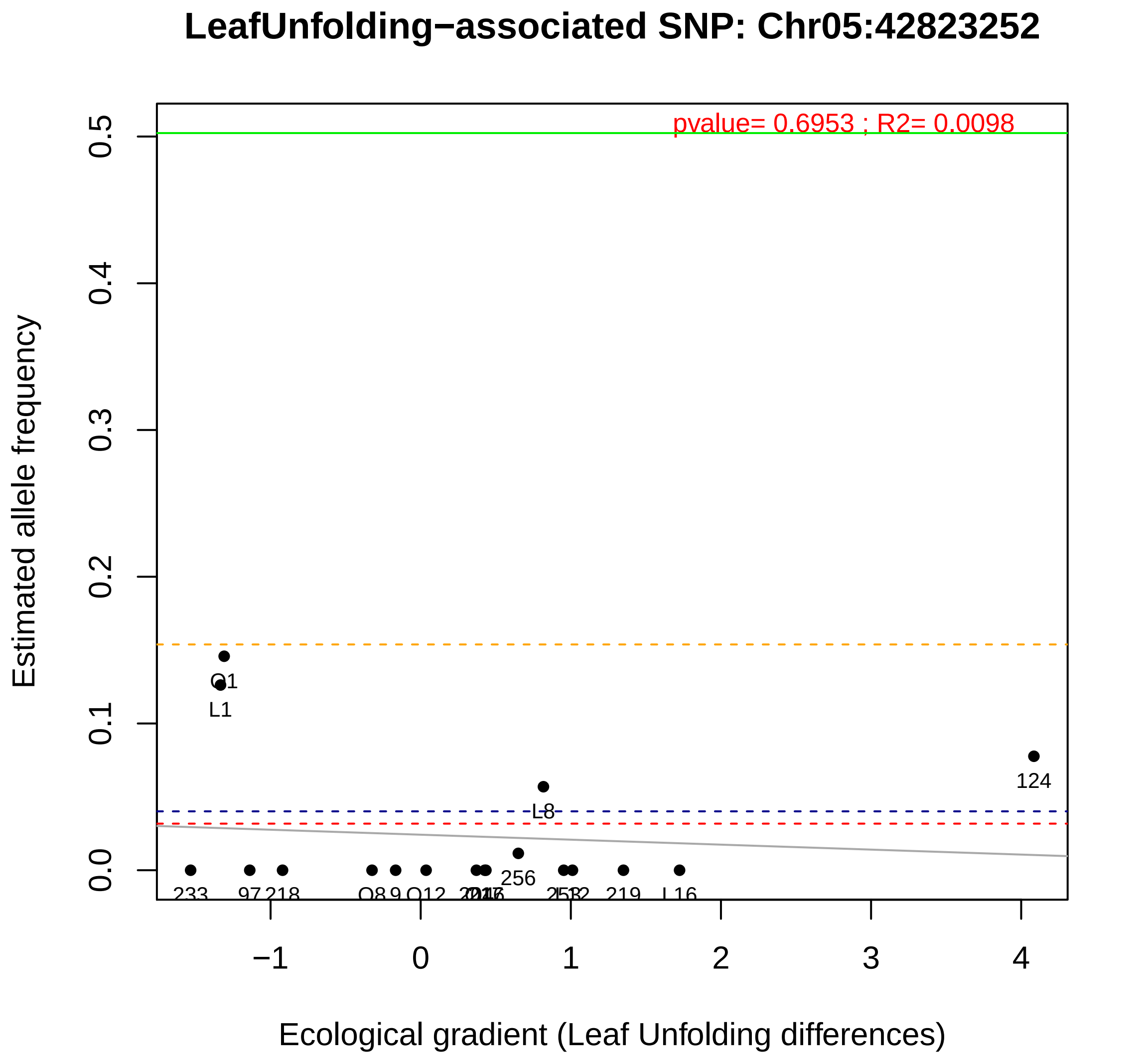

LeafUnfolding−associated SNP: Chr05:42823252
pvalue= 0.6953 ; R2= 0.0098
Estimated allele frequency
0.0
0.1
0.2
0.3
0.4
0.5
Q1
L1
124
L8
256
233
97
218
Q8
9
Q12
219
L16
−1
0
1
2
3
4
Ecological gradient (Leaf Unfolding differences)

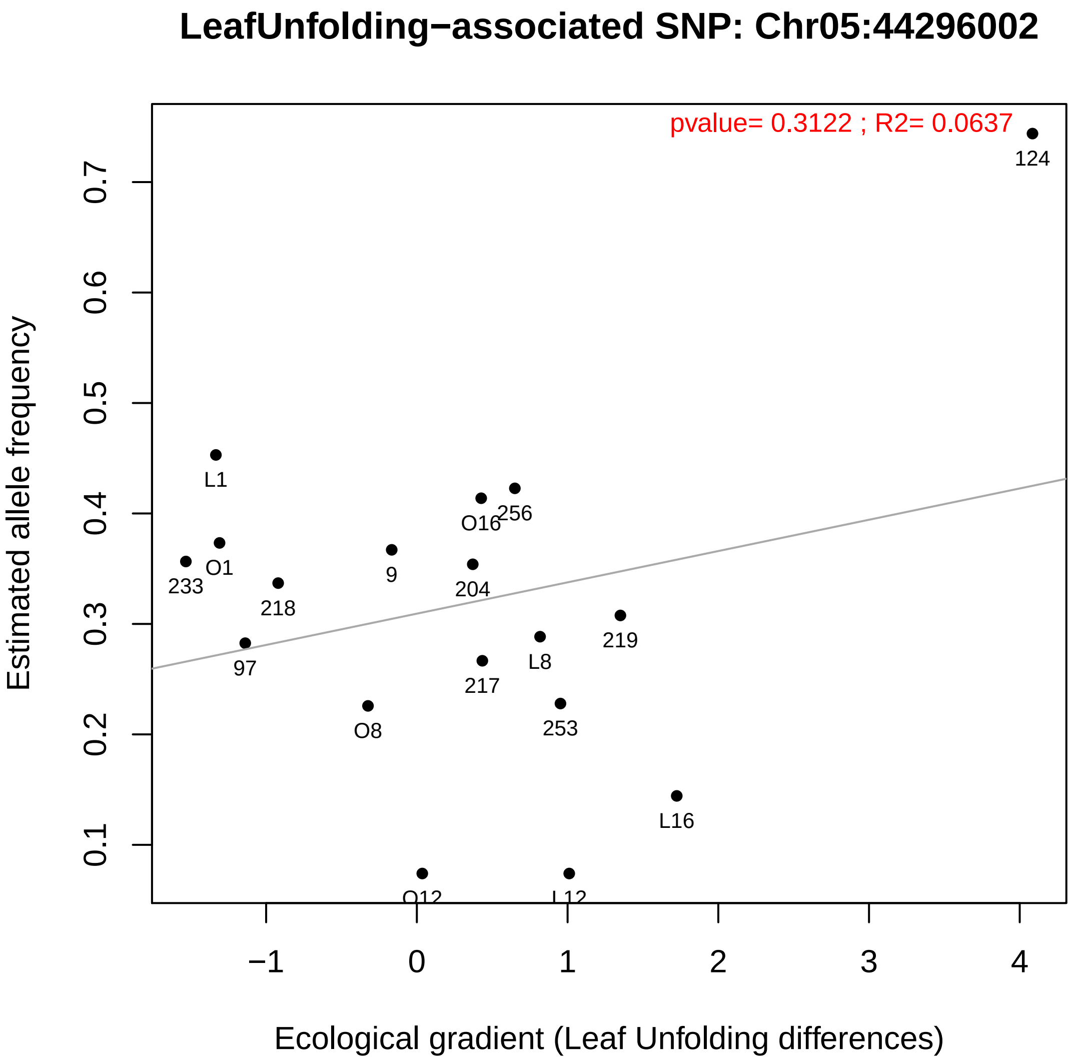
LeafUnfolding−associated SNP: Chr05:44296002
pvalue= 0.3122 ; R2= 0.0637
Estimated allele frequency
Ecological gradient (Leaf Unfolding differences)
0.1
0.2
0.3
0.4
0.5
0.6
0.7
−1
0
1
2
3
4
124
L1
O16
256
O1
233
9
204
218
97
219
L8
217
O8
253
L16

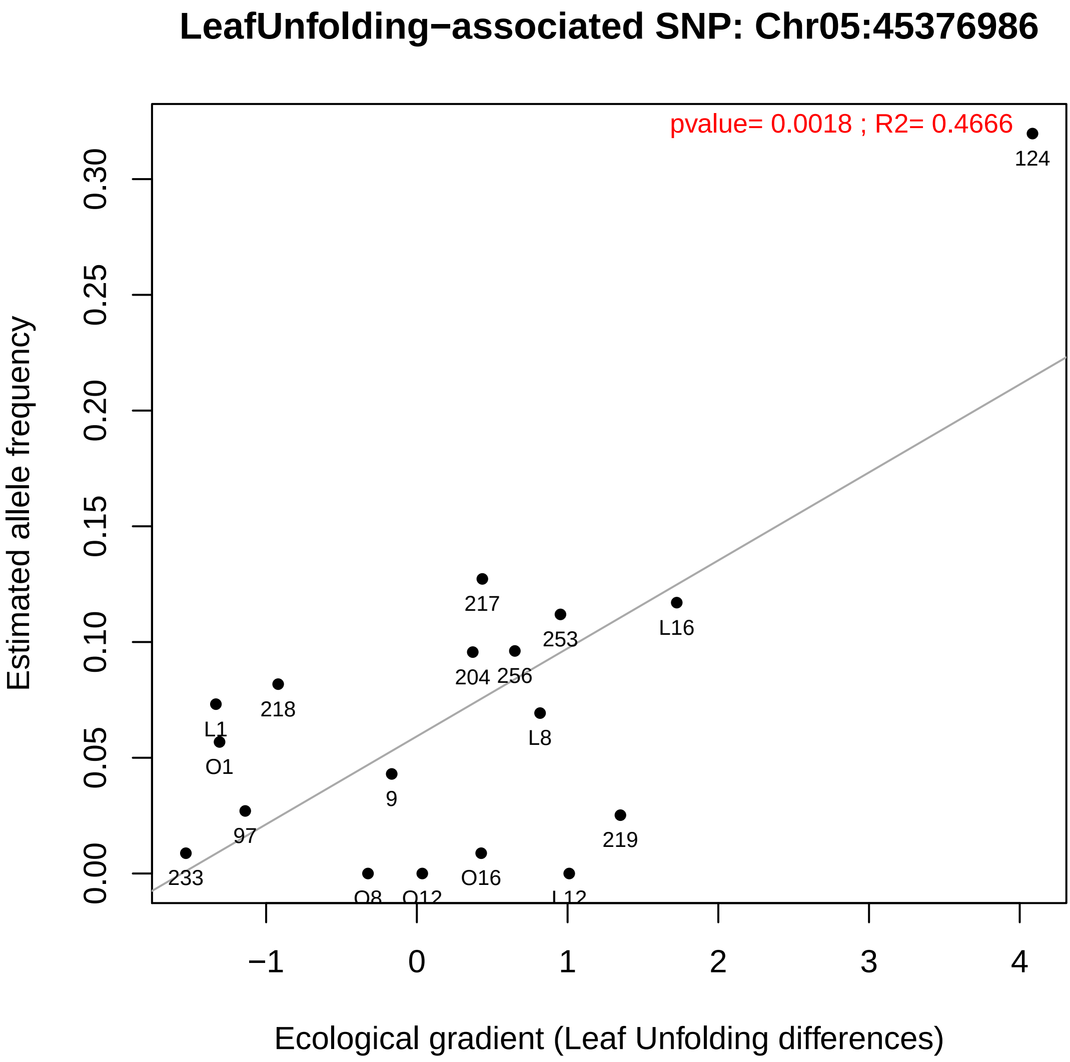

LeafUnfolding−associated SNP: Chr05:45376986
pvalue= 0.0018 ; R2= 0.4666
Estimated allele frequency
0.00
0.05
0.10
0.15
0.20
0.25
0.30
−1
0
1
2
3
4
Ecological gradient (Leaf Unfolding differences)
124
217
253
L16
204
256
218
L1
O1
L8
9
97
219
233
O8
O12
O16
L12

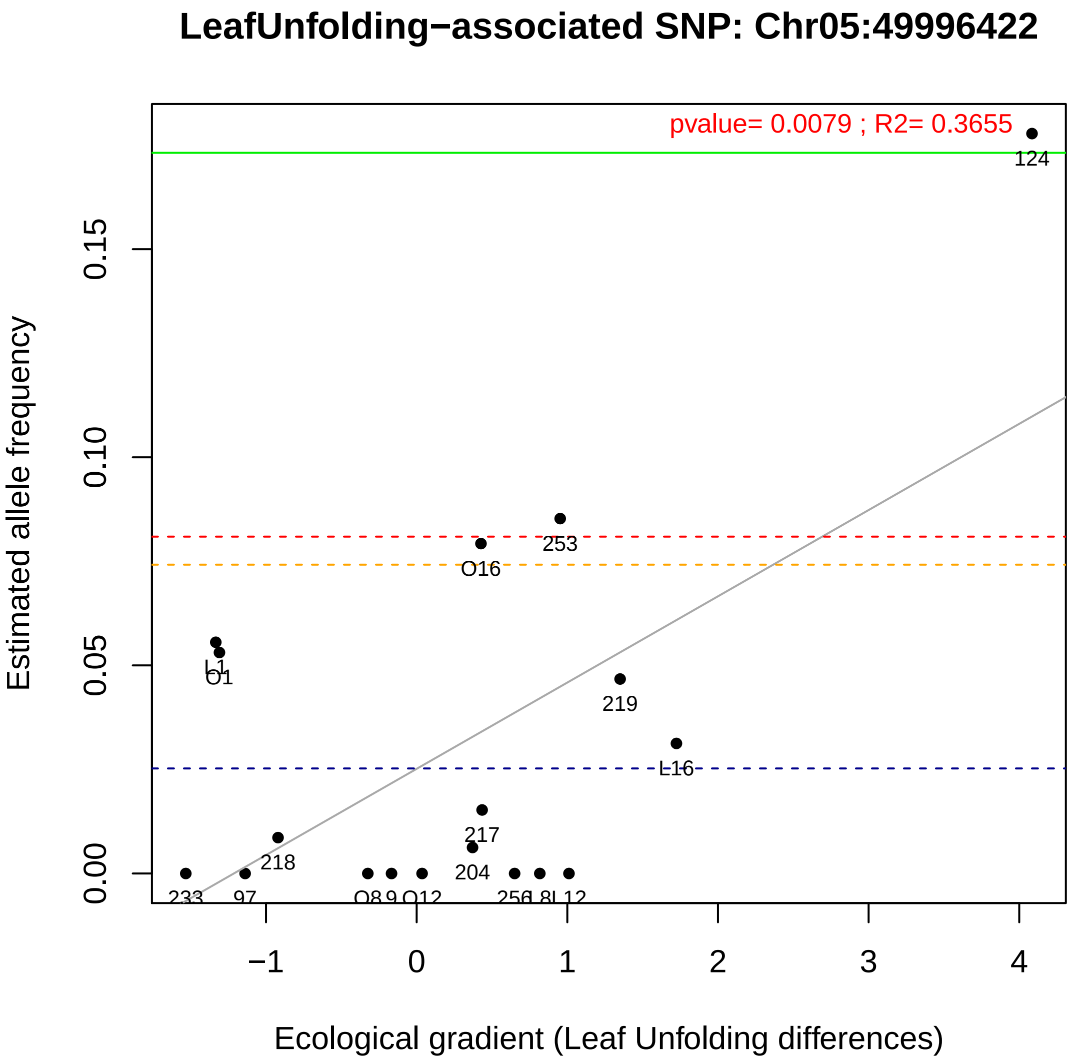

LeafUnfolding−associated SNP: Chr05:49996422
pvalue= 0.0079 ; R2= 0.3655
Estimated allele frequency
0.00
0.05
0.10
0.15
Ecological gradient (Leaf Unfolding differences)
−1
0
1
2
3
4
124
253
O16
L1
O1
219
L16
217
218
204
233
97
O8
9
O12
256
L8
L12

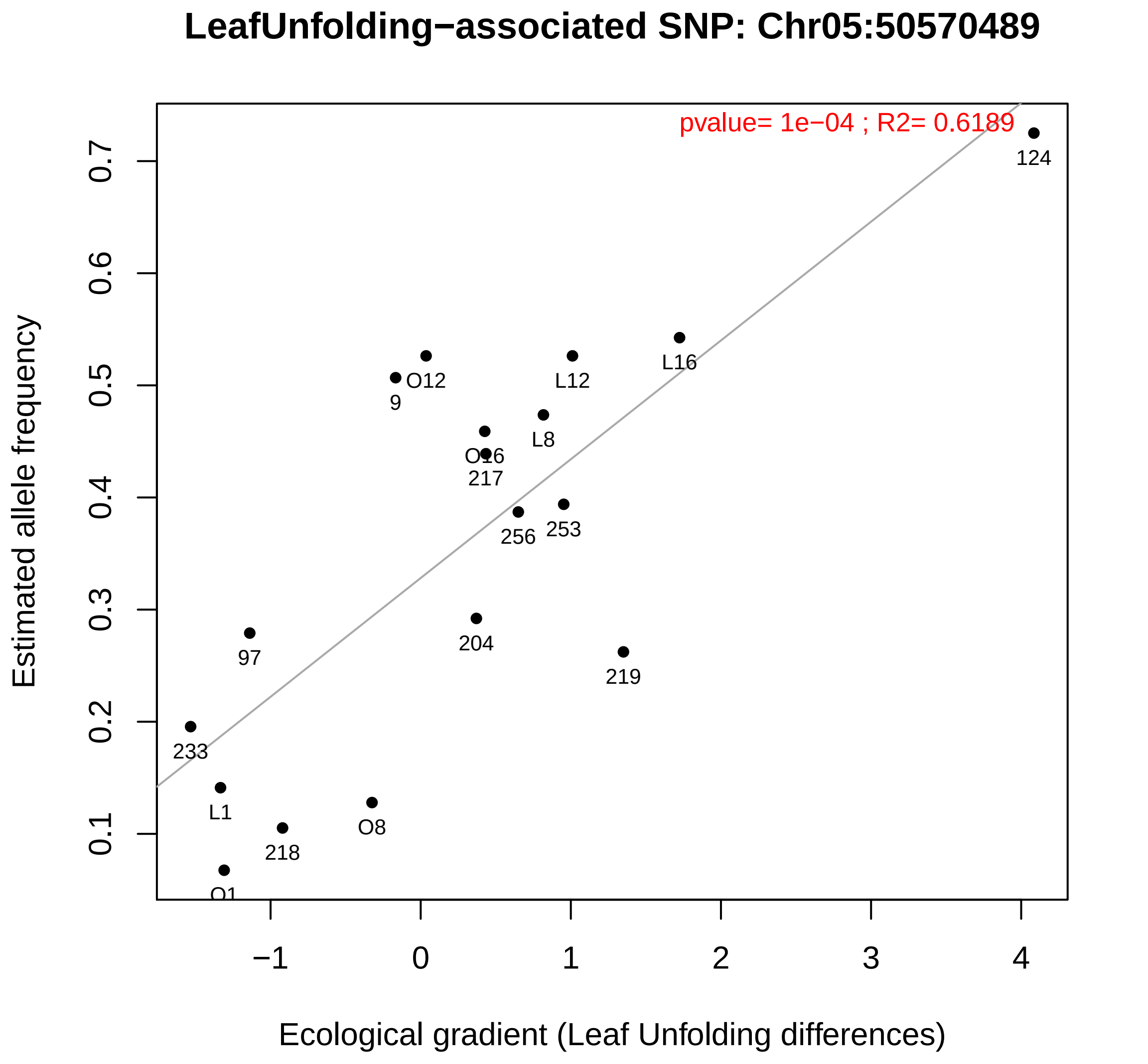

LeafUnfolding−associated SNP: Chr05:50570489
pvalue= 1e−04 ; R2= 0.6189
Estimated allele frequency
0.1
0.2
0.3
0.4
0.5
0.6
0.7
−1
0
1
2
3
4
Ecological gradient (Leaf Unfolding differences)
124
L16
L12
O12
9
L8
O16
217
256
253
204
219
97
233
L1
O8
218
O1

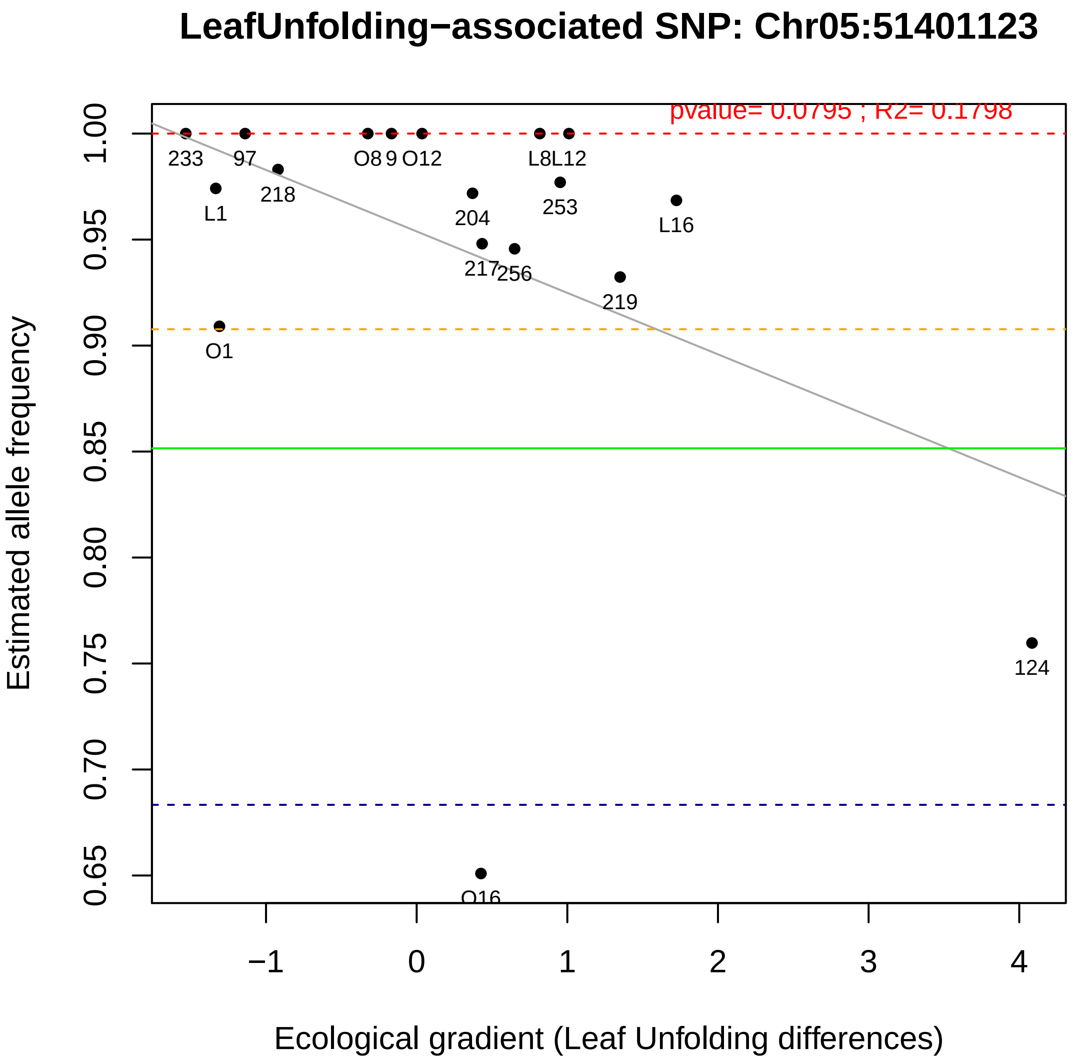

LeafUnfolding−associated SNP: Chr05:51401123
pvalue= 0.0795 ; R2= 0.1798
Estimated allele frequency
Ecological gradient (Leaf Unfolding differences)
1.00
0.95
0.90
0.85
0.80
0.75
0.70
0.65
−1
0
1
2
3
4
233
97
218
L1
O1
O8
9
O12
204
217
256
L8
L12
253
219
L16
124
O16

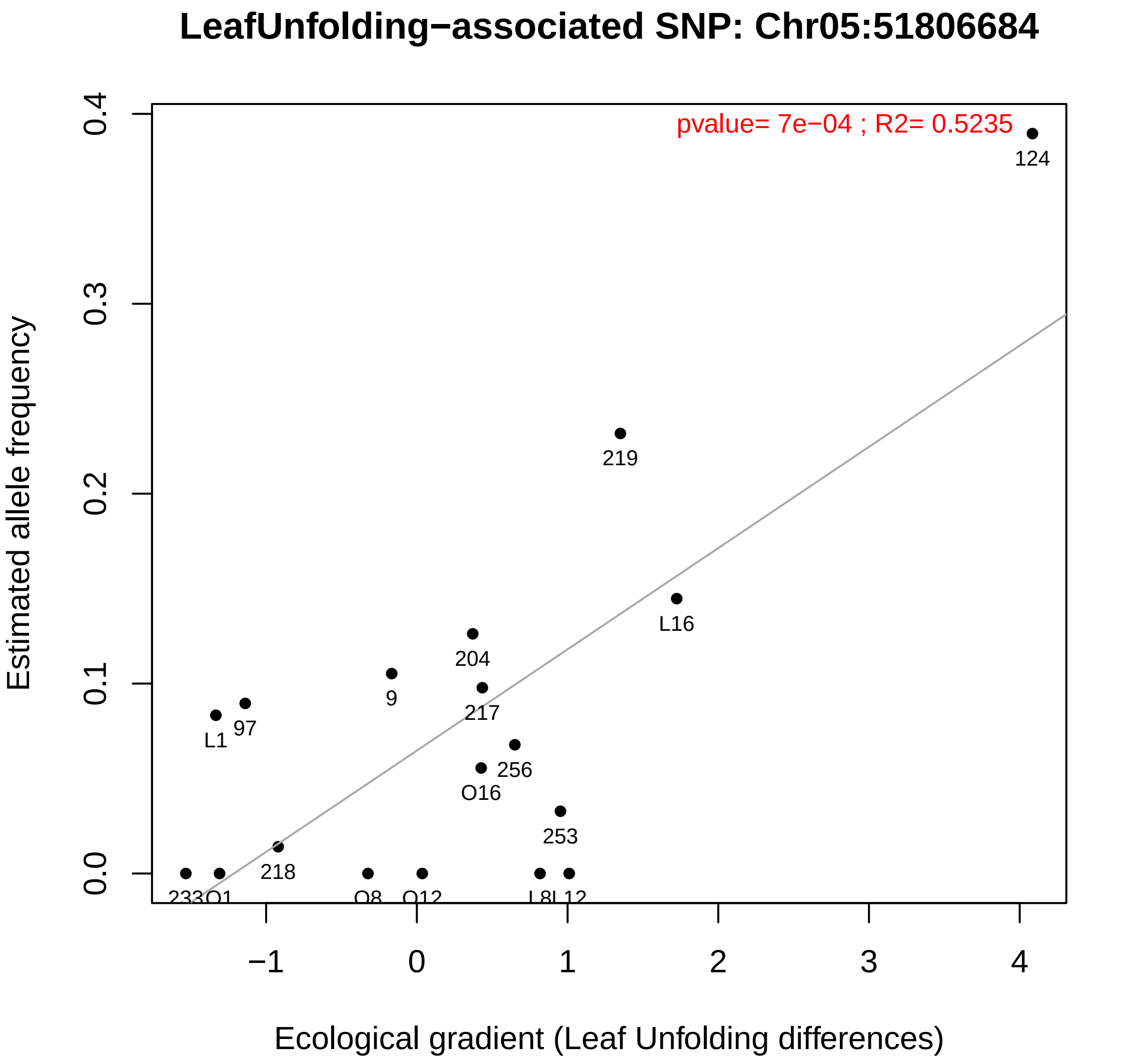

LeafUnfolding−associated SNP: Chr05:51806684
pvalue= 7e−04 ; R2= 0.5235
Estimated allele frequency
Ecological gradient (Leaf Unfolding differences)
0.0
0.1
0.2
0.3
0.4
−1
0
1
2
3
4
124
219
L16
204
9
217
97
L1
256
O16
253
218
O8
O12

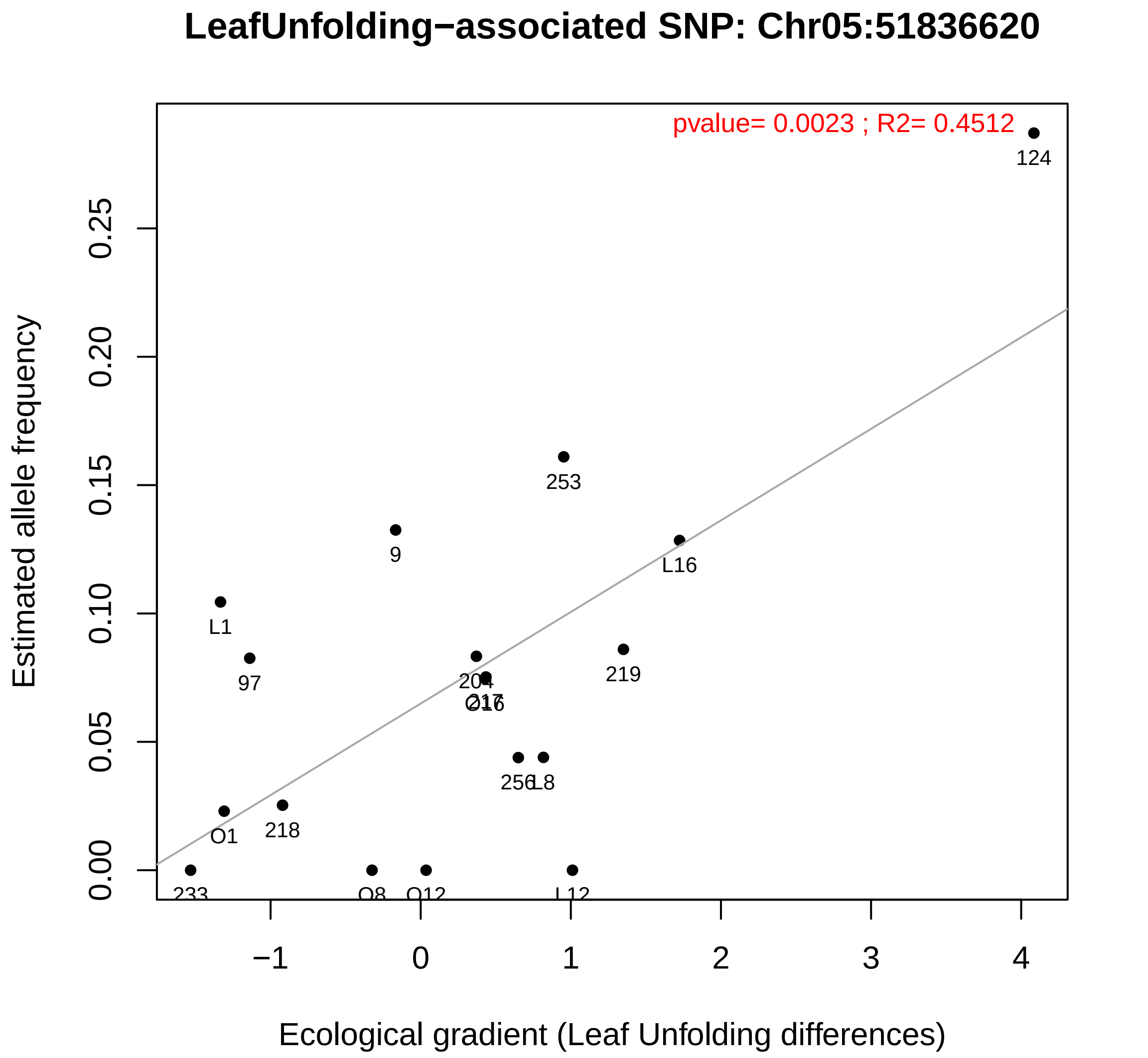
LeafUnfolding−associated SNP: Chr05:51836620
pvalue= 0.0023 ; R2= 0.4512
Estimated allele frequency
Ecological gradient (Leaf Unfolding differences)
0.00
0.05
0.10
0.15
0.20
0.25
−1
0
1
2
3
4
124
253
9
L16
L1
97
204
219
256
L8
218
O1
233
O8
O12
L12

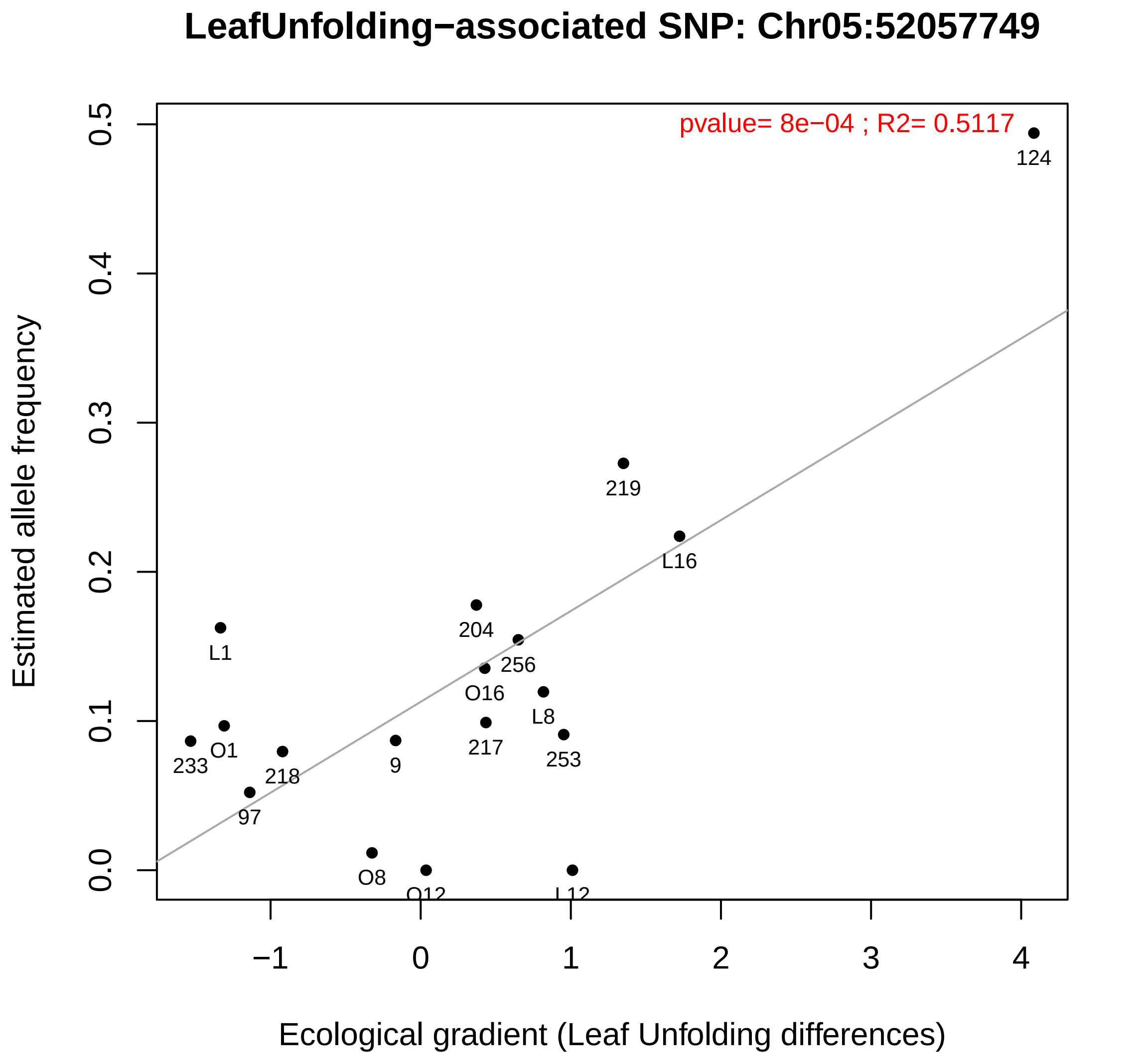

LeafUnfolding−associated SNP: Chr05:52057749
pvalue= 8e−04 ; R2= 0.5117
Estimated allele frequency
Ecological gradient (Leaf Unfolding differences)
0.0
0.1
0.2
0.3
0.4
0.5
−1
0
1
2
3
4
124
219
L16
204
L1
256
O16
L8
217
253
O1
233
218
9
97
O8

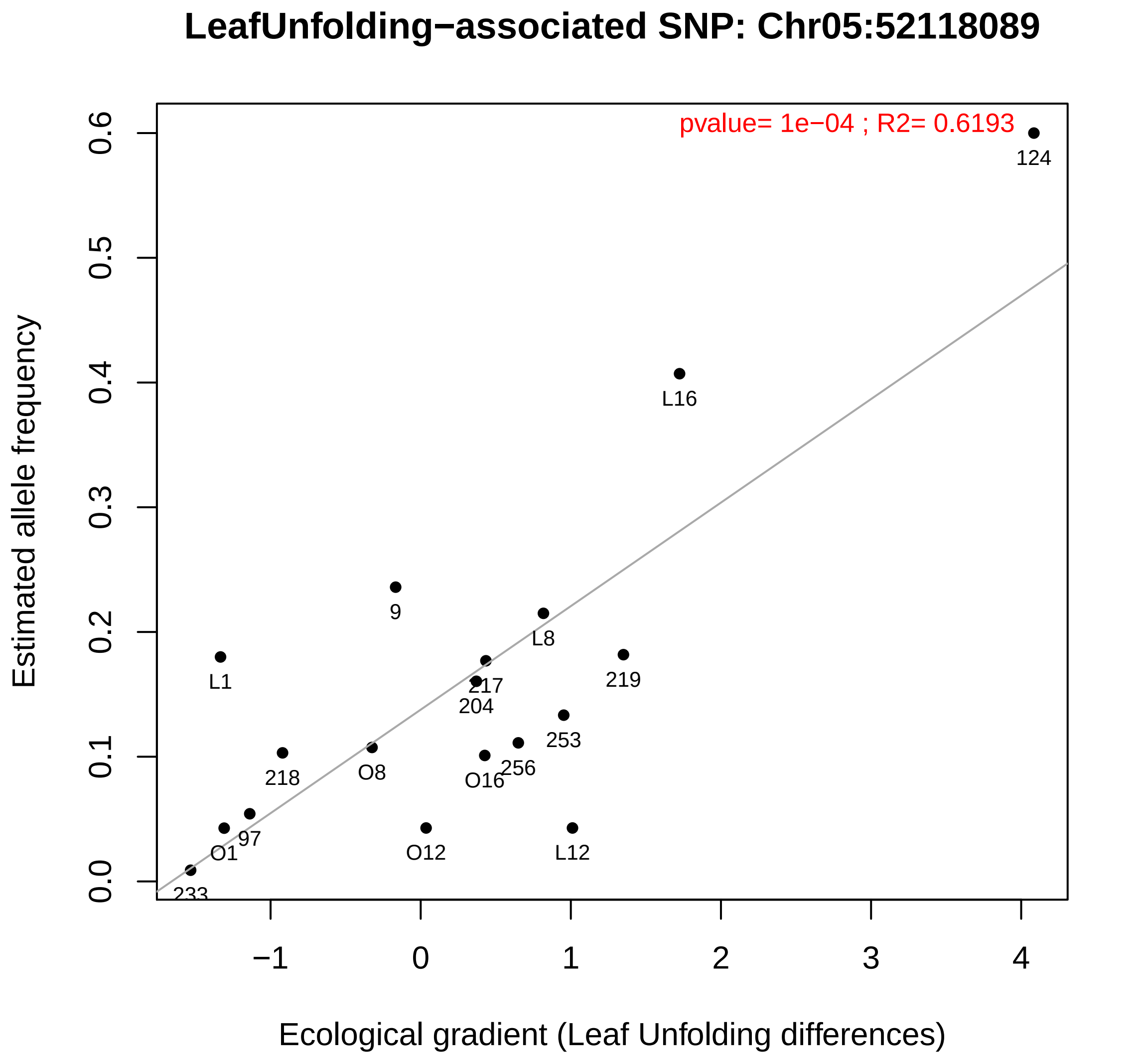

LeafUnfolding−associated SNP: Chr05:52118089
pvalue= 1e−04 ; R2= 0.6193
Estimated allele frequency
Ecological gradient (Leaf Unfolding differences)
0.0
0.1
0.2
0.3
0.4
0.5
0.6
−1
0
1
2
3
4
124
L16
9
L8
L1
217
204
219
253
218
O8
256
O16
97
O1
O12
L12
233

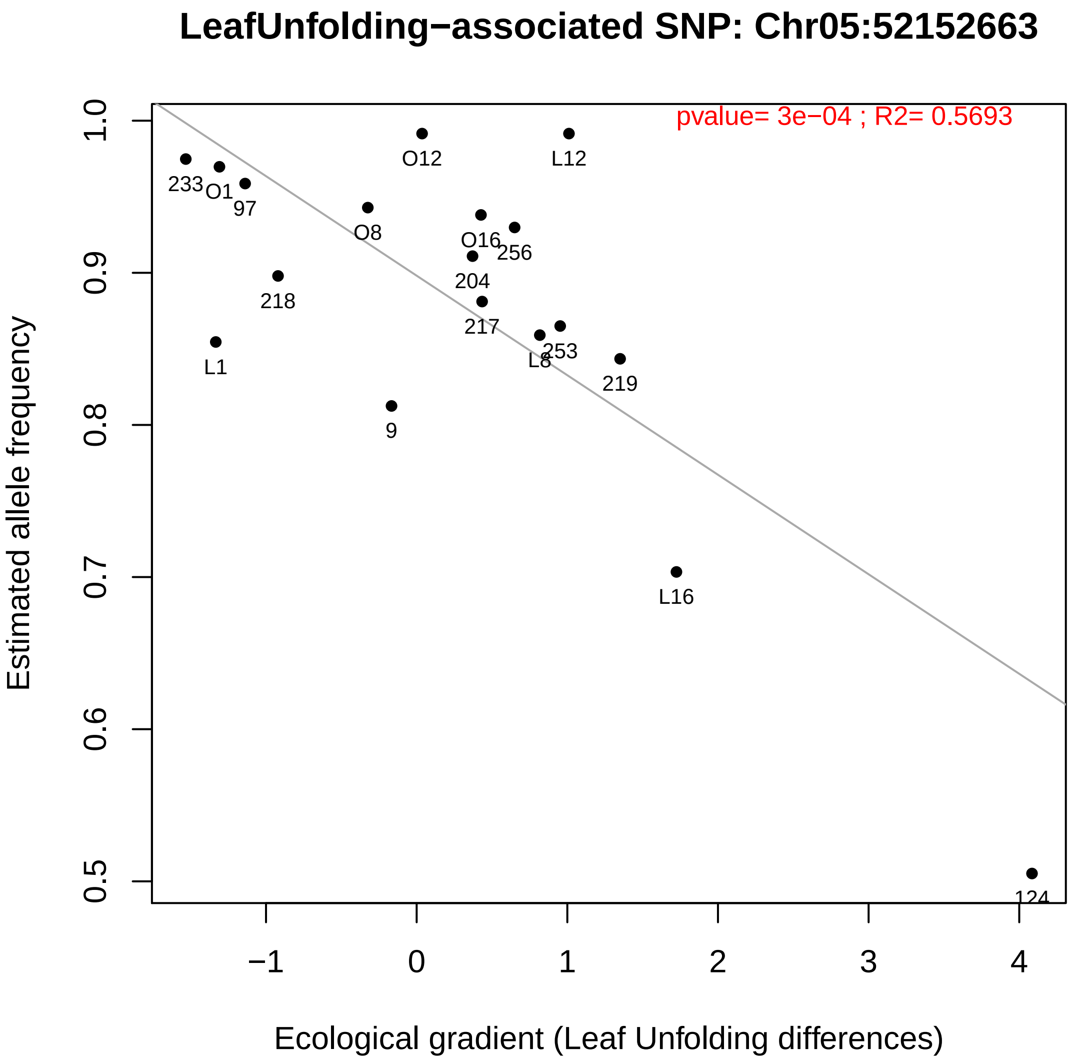
LeafUnfolding−associated SNP: Chr05:52152663
pvalue= 3e−04 ; R2= 0.5693
Estimated allele frequency
Ecological gradient (Leaf Unfolding differences)
0.5
0.6
0.7
0.8
0.9
1.0
−1
0
1
2
3
4
233
O1
97
O8
O12
L12
O16
256
204
218
217
253
L8
L1
219
9
L16
124

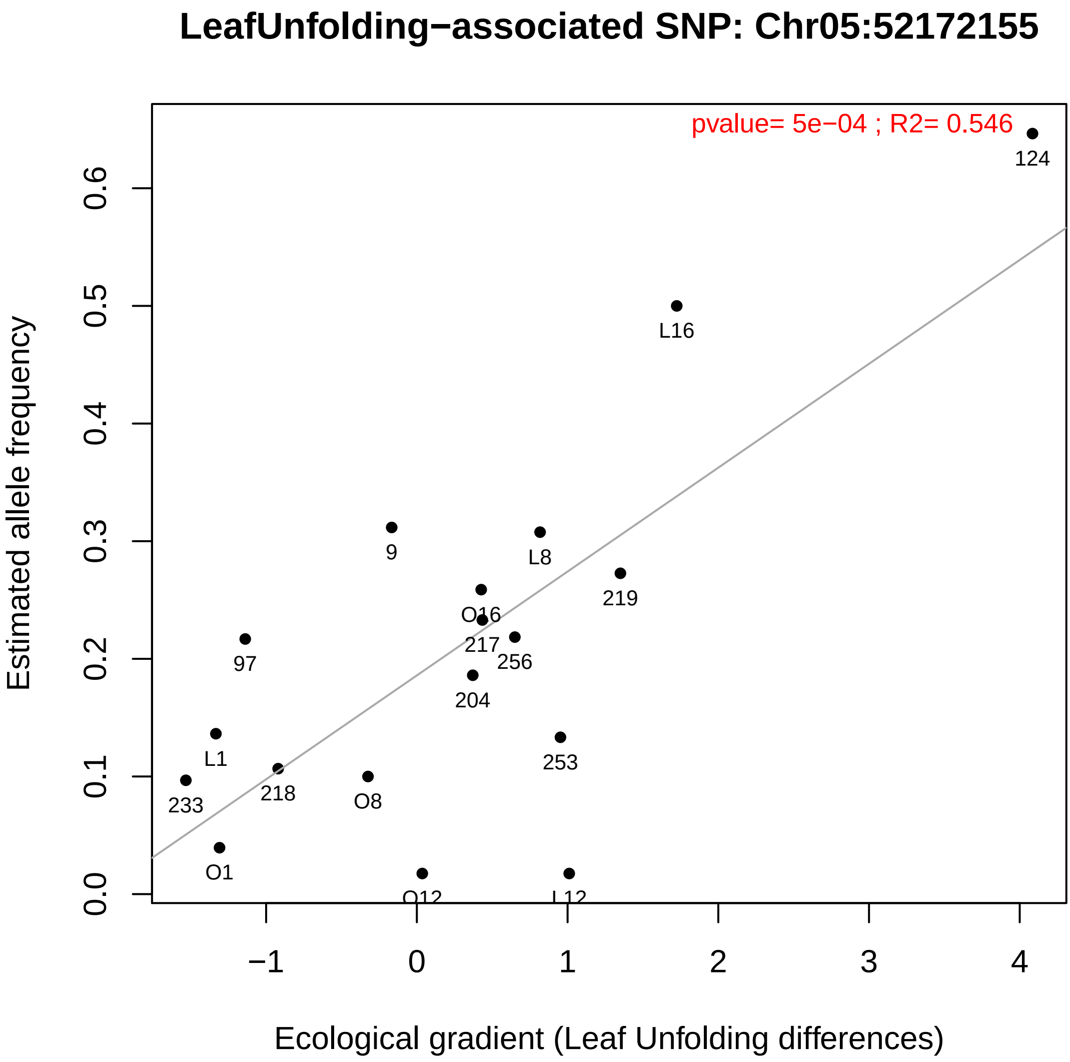
LeafUnfolding−associated SNP: Chr05:52172155
pvalue= 5e−04 ; R2= 0.546
Estimated allele frequency
Ecological gradient (Leaf Unfolding differences)
0.0
0.1
0.2
0.3
0.4
0.5
0.6
−1
0
1
2
3
4
124
L16
9
L8
219
O16
217
256
97
204
253
L1
218
233
O8
O1
O12
L12

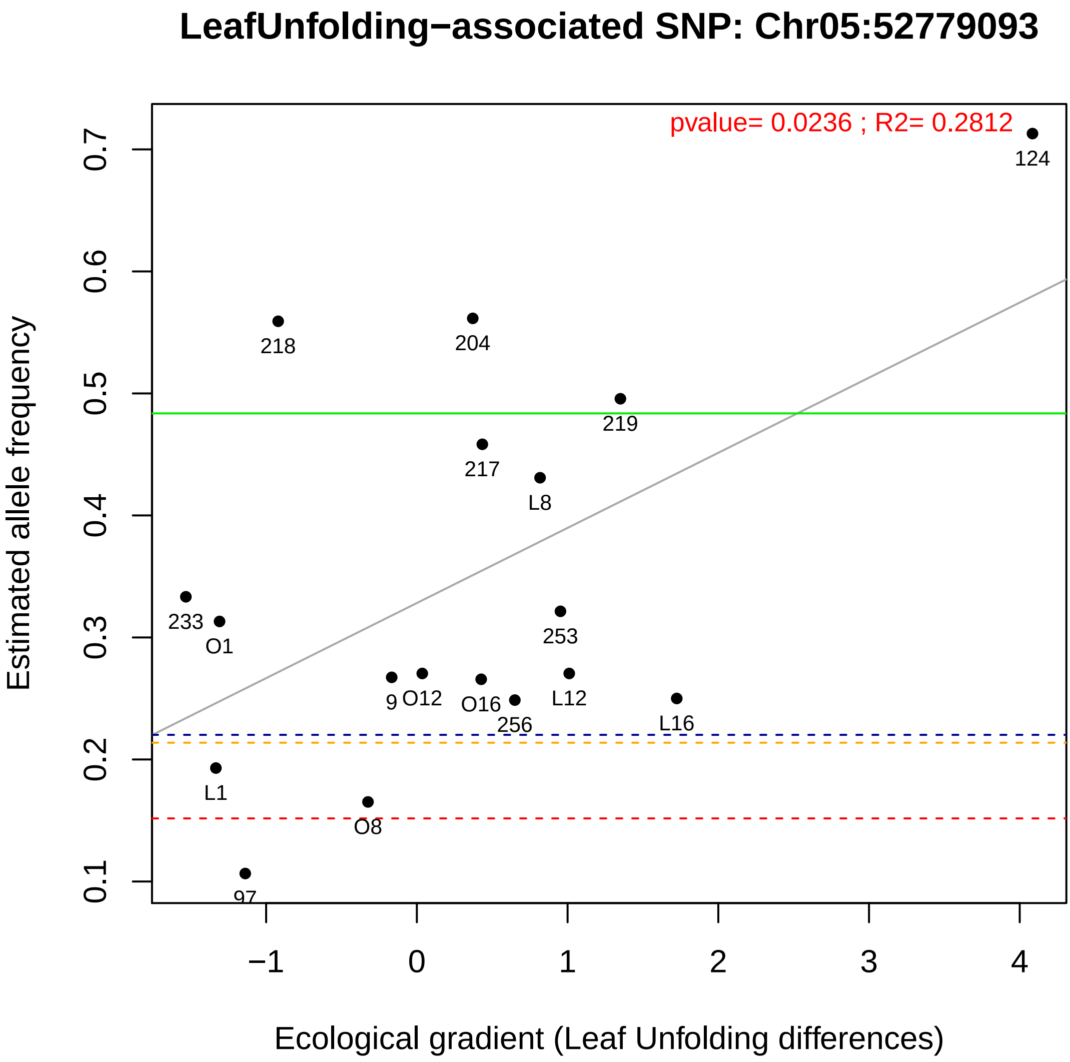
LeafUnfolding−associated SNP: Chr05:52779093
pvalue= 0.0236 ; R2= 0.2812
Estimated allele frequency
Ecological gradient (Leaf Unfolding differences)
0.1
0.2
0.3
0.4
0.5
0.6
0.7
−1
0
1
2
3
4
124
218
204
219
217
L8
233
O1
253
9
O12
O16
256
L12
L16
L1
O8
97

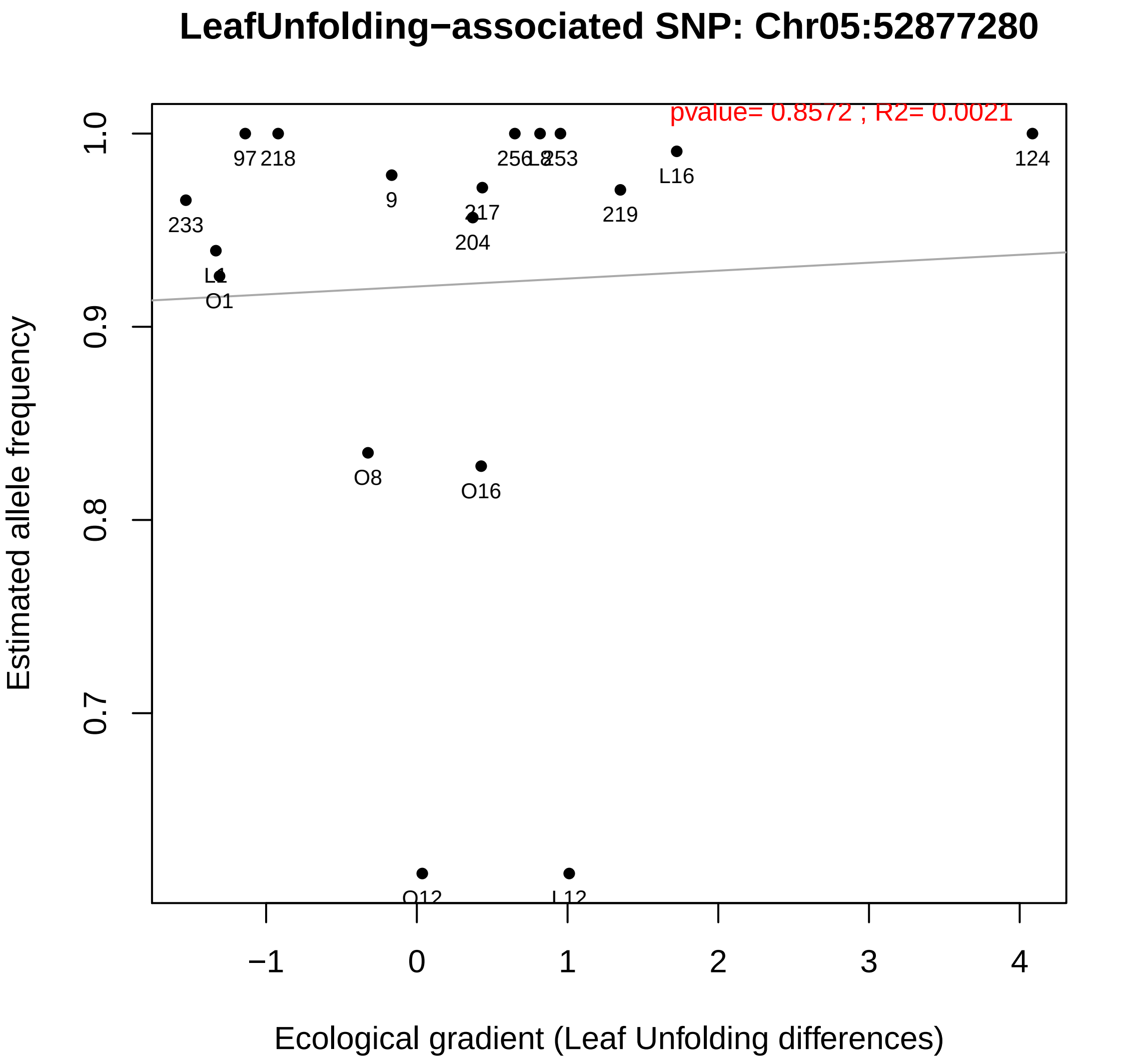
LeafUnfolding−associated SNP: Chr05:52877280
pvalue= 0.8572 ; R2= 0.0021
Estimated allele frequency
Ecological gradient (Leaf Unfolding differences)
1.0
0.9
0.8
0.7
−1
0
1
2
3
4
97
218
233
9
217
204
219
L16
124
O1
O8
O16
O12
L12

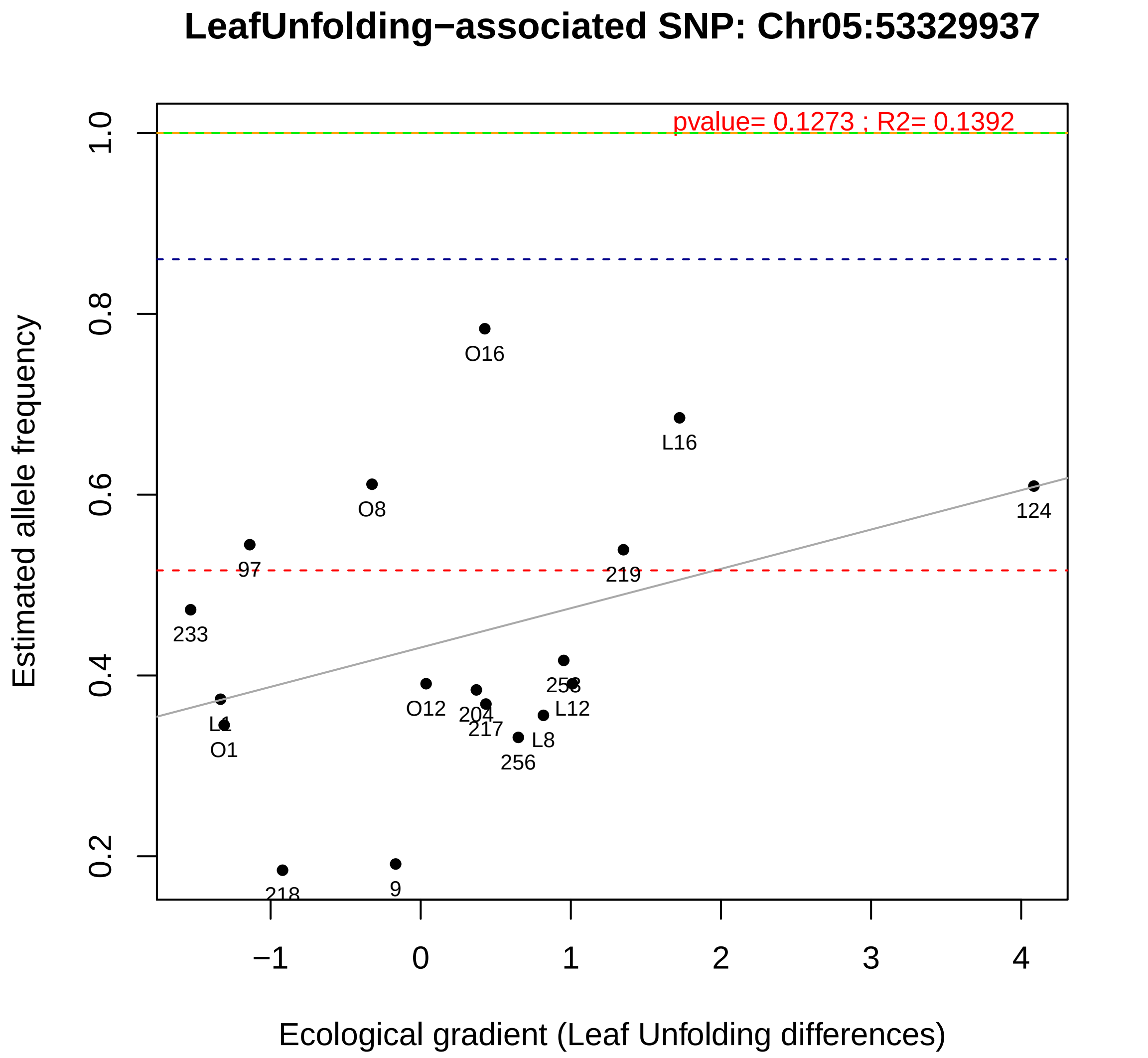

LeafUnfolding−associated SNP: Chr05:53329937
pvalue= 0.1273 ; R2= 0.1392
Estimated allele frequency
0.2
0.4
0.6
0.8
1.0
Ecological gradient (Leaf Unfolding differences)
−1
0
1
2
3
4
O16
L16
O8
124
97
219
233
O12
204
217
L12
L8
256
L1
O1
218
9

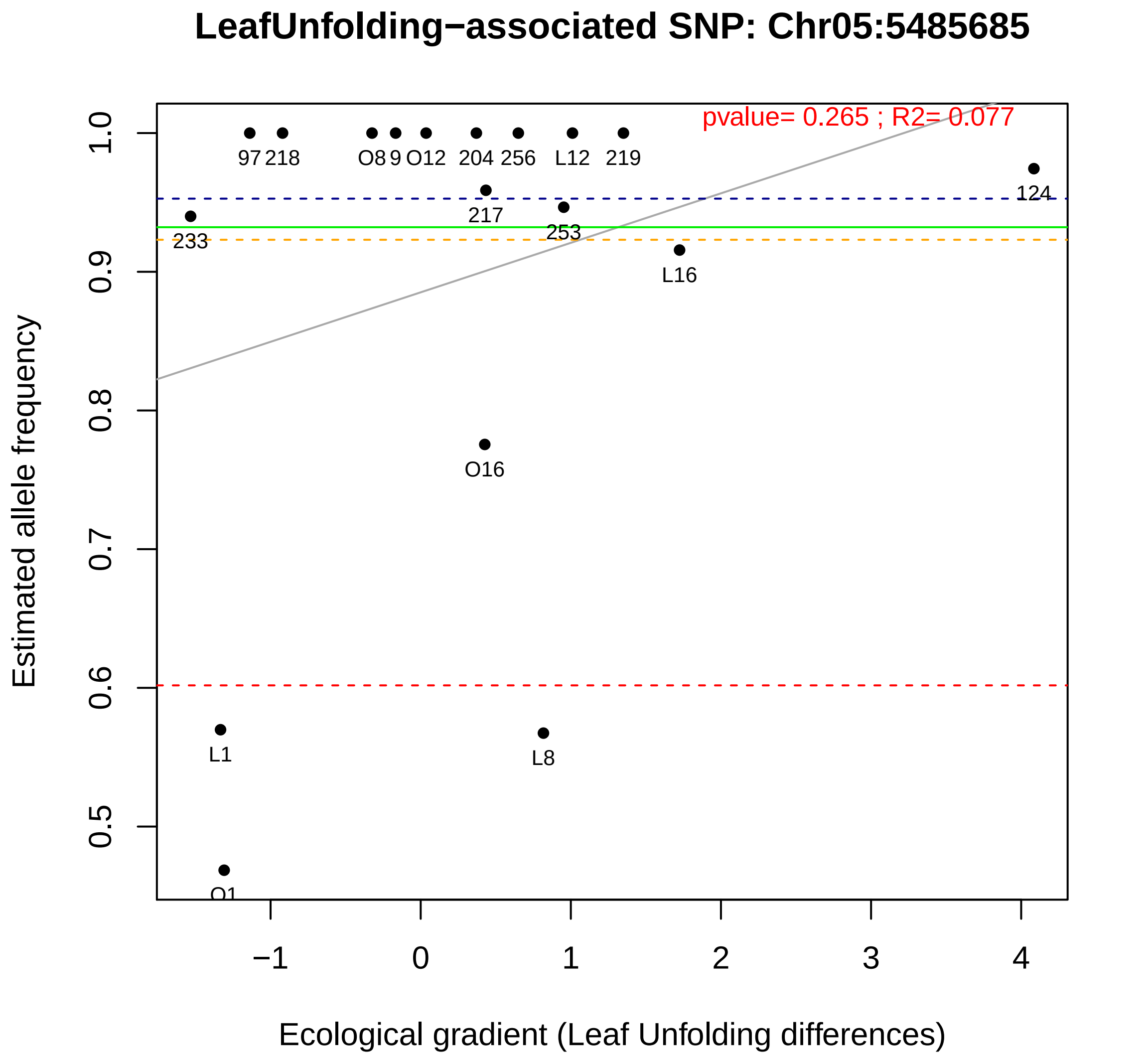
LeafUnfolding−associated SNP: Chr05:5485685
pvalue= 0.265 ; R2= 0.077
Estimated allele frequency
Ecological gradient (Leaf Unfolding differences)
1.0
0.9
0.8
0.7
0.6
0.5
−1
0
1
2
3
4
97
218
O8
9
O12
204
256
L12
219
124
217
233
253
L16
O16
L1
L8
O1

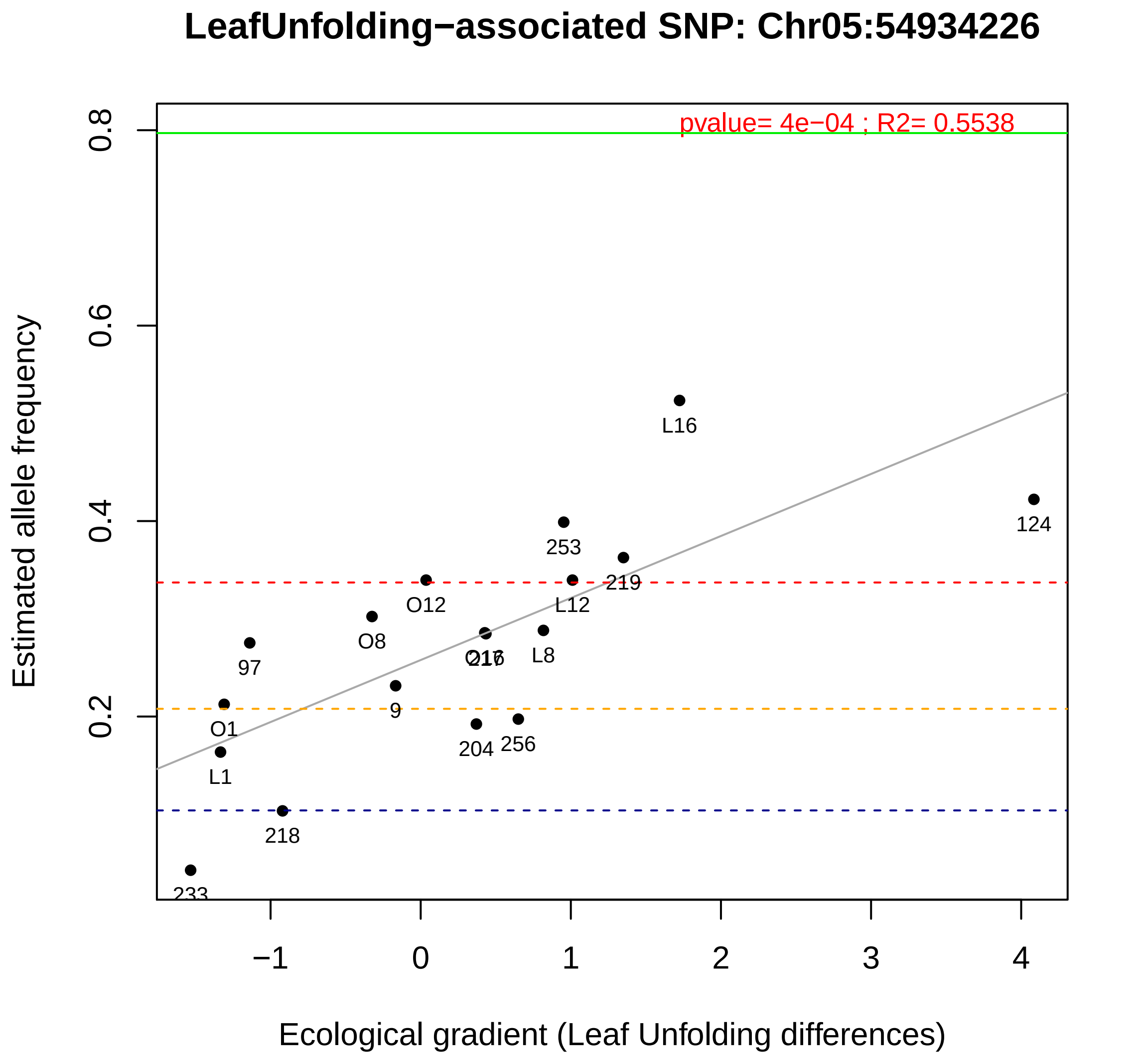

LeafUnfolding−associated SNP: Chr05:54934226
pvalue= 4e−04 ; R2= 0.5538
Estimated allele frequency
0.8
0.6
0.4
0.2
Ecological gradient (Leaf Unfolding differences)
−1
0
1
2
3
4
L16
124
253
219
L12
O12
O8
L8
97
9
O1
204
256
L1
218
233

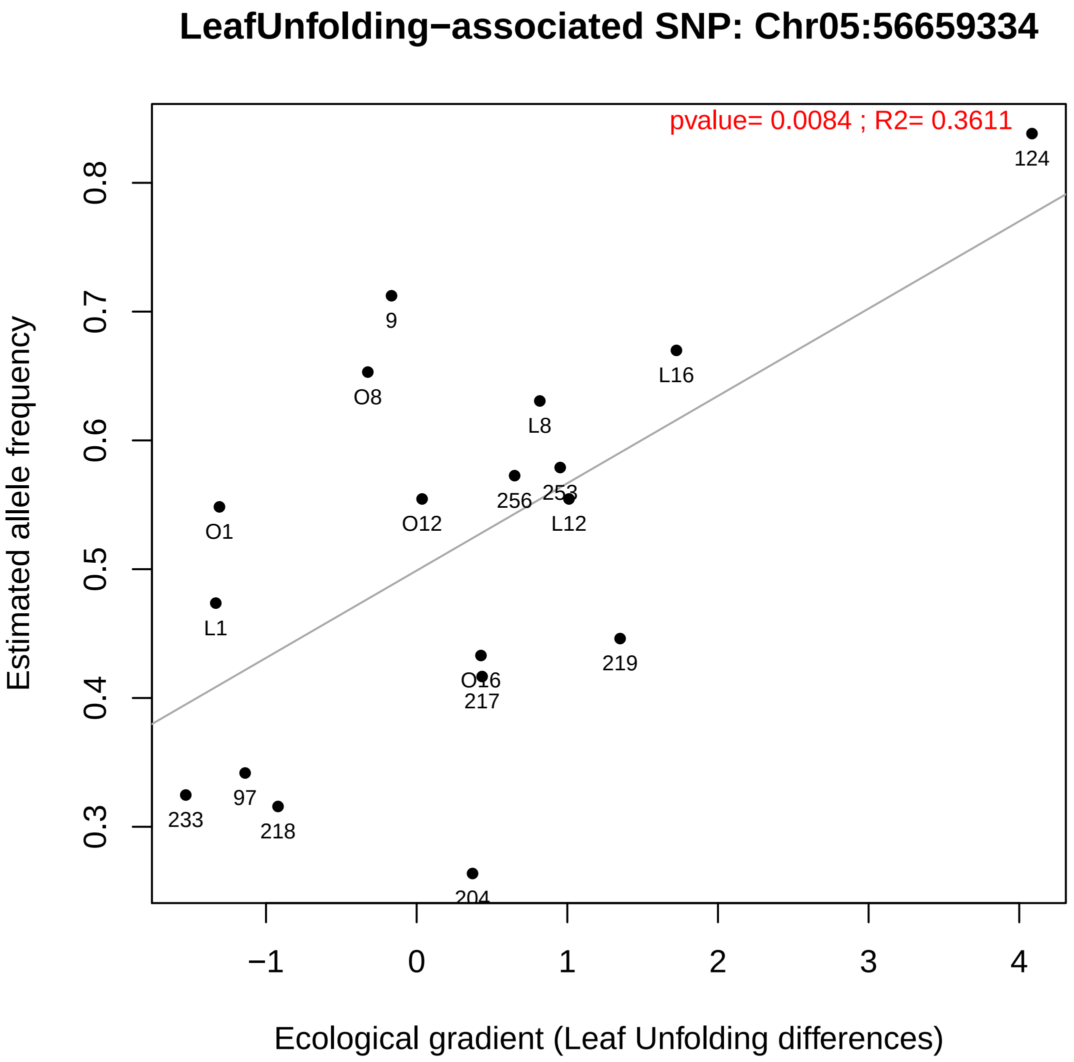

LeafUnfolding−associated SNP: Chr05:56659334
pvalue= 0.0084 ; R2= 0.3611
124
9
L16
O8
L8
256
253
L12
O12
O1
L1
219
O16
217
97
233
218
204
0.3
0.4
0.5
0.6
0.7
0.8
−1
0
1
2
3
4
Estimated allele frequency
Ecological gradient (Leaf Unfolding differences)

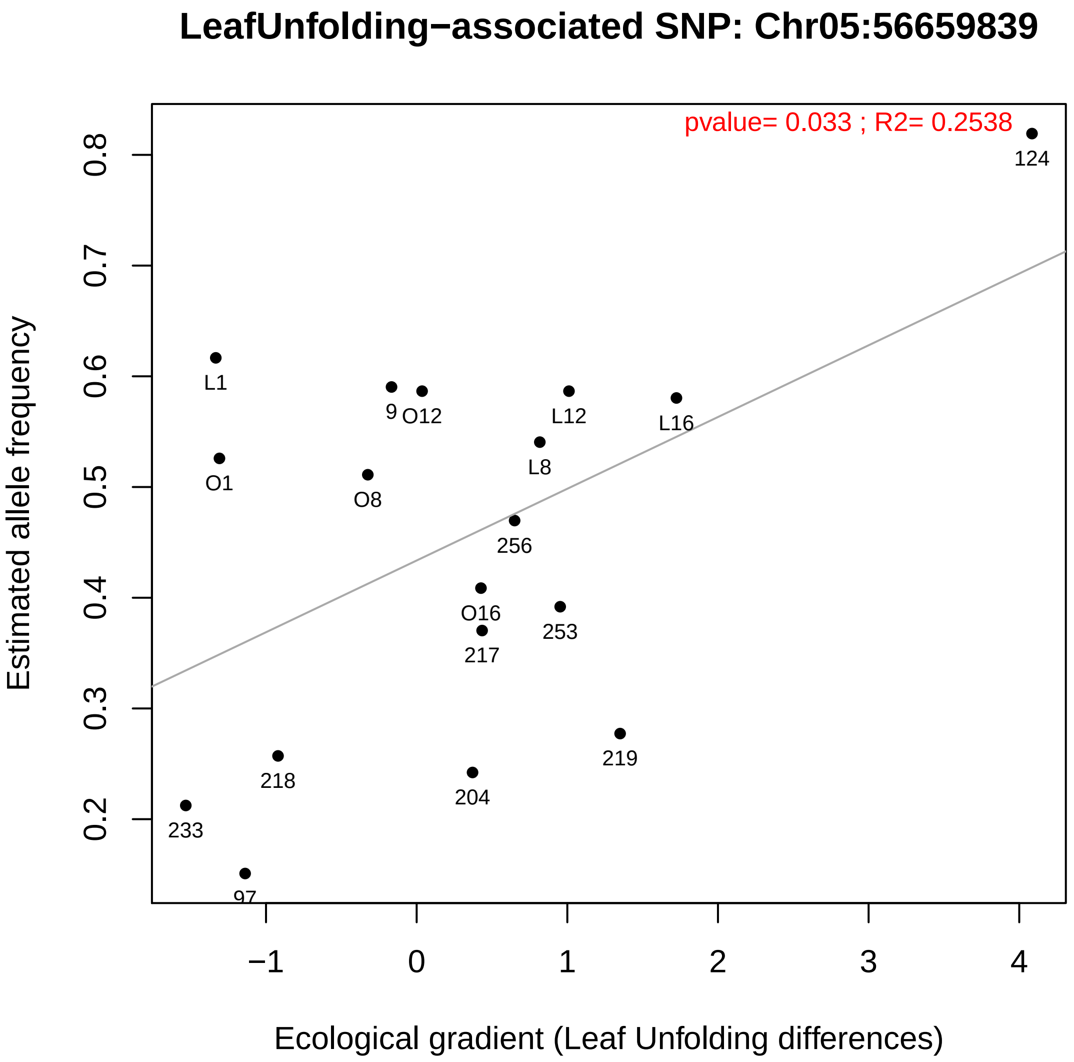

LeafUnfolding−associated SNP: Chr05:56659839
pvalue= 0.033 ; R2= 0.2538
Estimated allele frequency
Ecological gradient (Leaf Unfolding differences)
0.2
0.3
0.4
0.5
0.6
0.7
0.8
−1
0
1
2
3
4
124
L1
9
O12
L12
L16
L8
O1
O8
256
O16
253
217
219
218
204
233
97

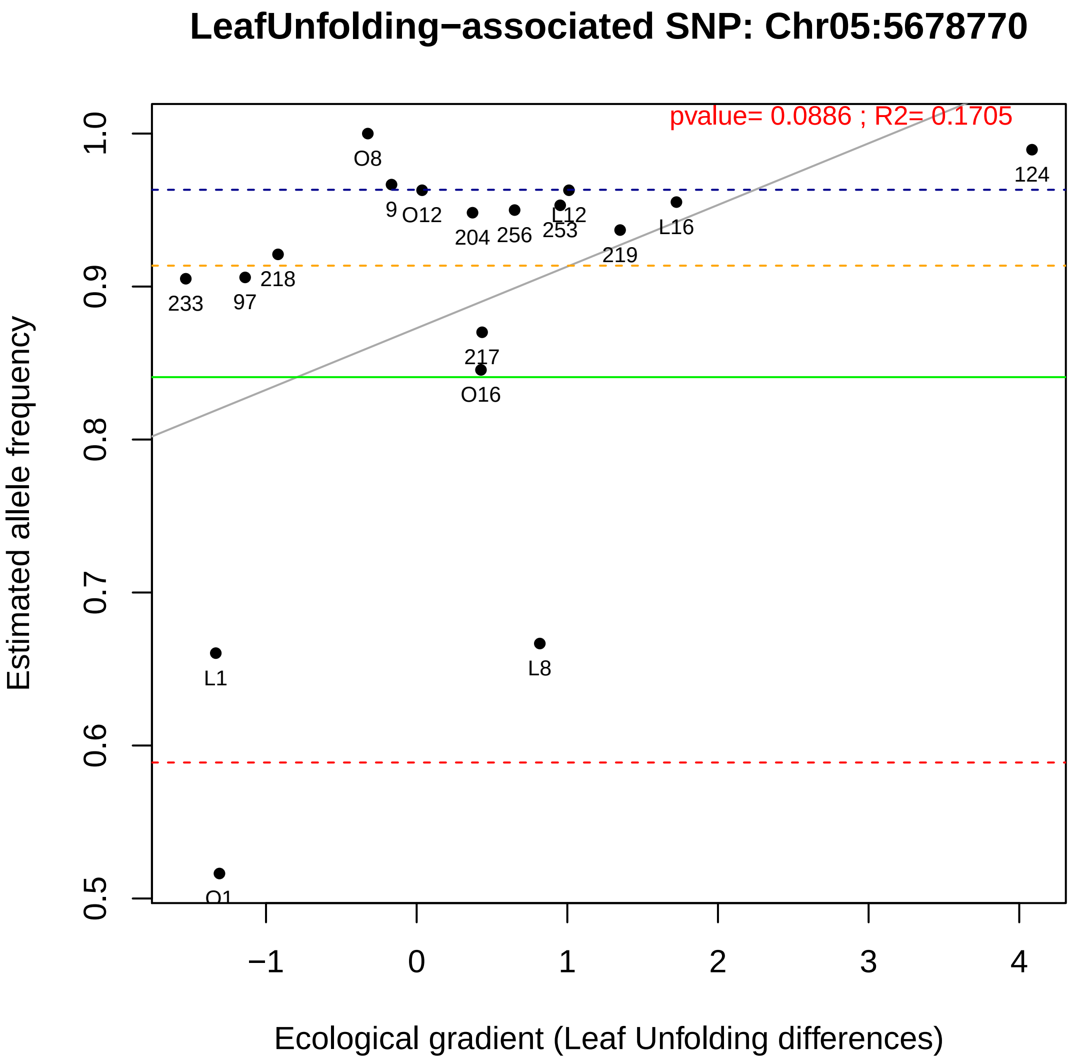

LeafUnfolding−associated SNP: Chr05:5678770
pvalue= 0.0886 ; R2= 0.1705
Estimated allele frequency
Ecological gradient (Leaf Unfolding differences)
1.0
0.9
0.8
0.7
0.6
0.5
−1
0
1
2
3
4
O8
9
O12
204
256
253
L12
219
L16
124
233
97
218
217
O16
L1
L8
O1

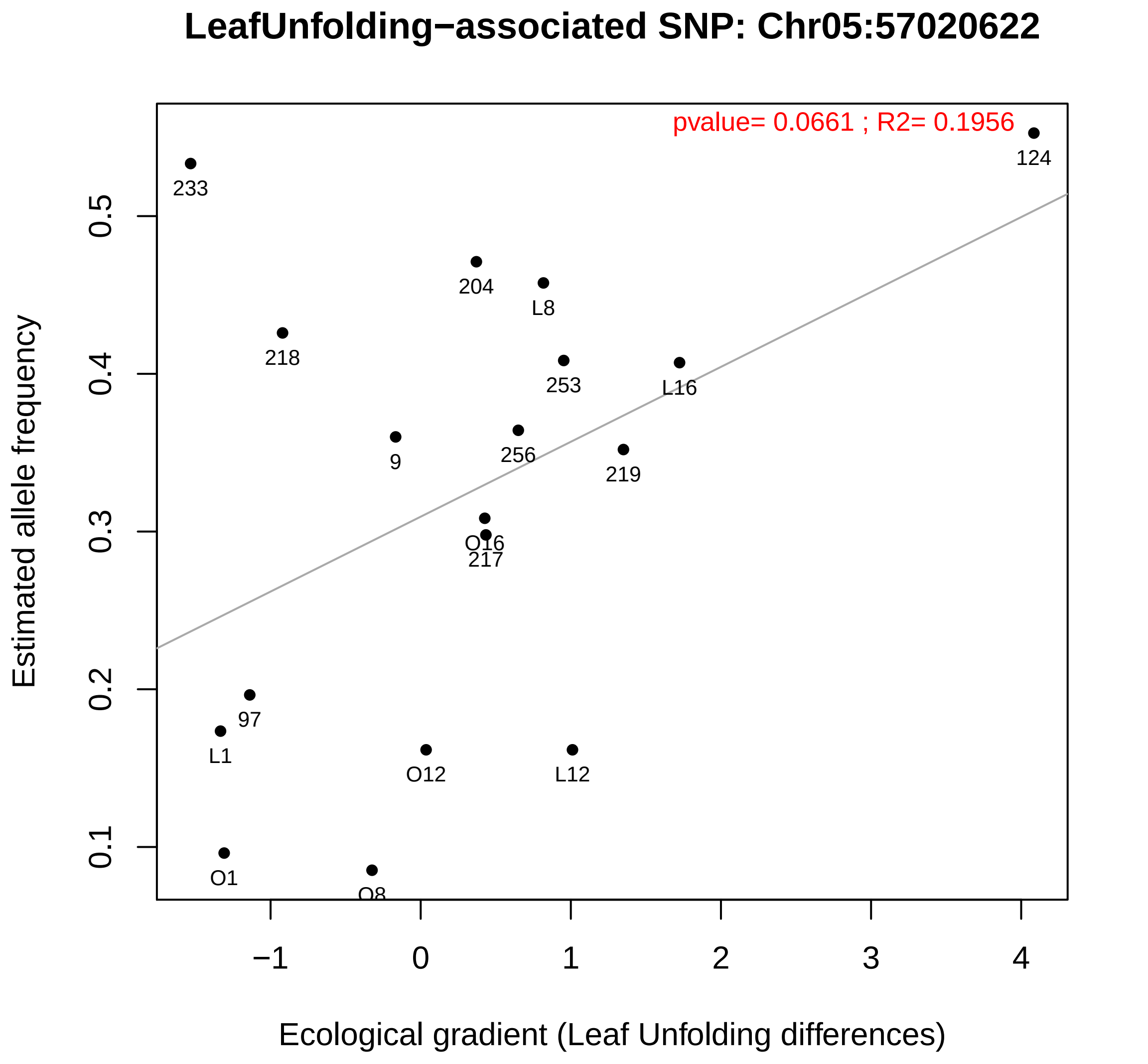

LeafUnfolding−associated SNP: Chr05:57020622
pvalue= 0.0661 ; R2= 0.1956
Estimated allele frequency
Ecological gradient (Leaf Unfolding differences)
0.1
0.2
0.3
0.4
0.5
−1
0
1
2
3
4
233
124
204
L8
218
253
L16
9
256
219
O16
217
97
L1
O12
L12
O1
O8

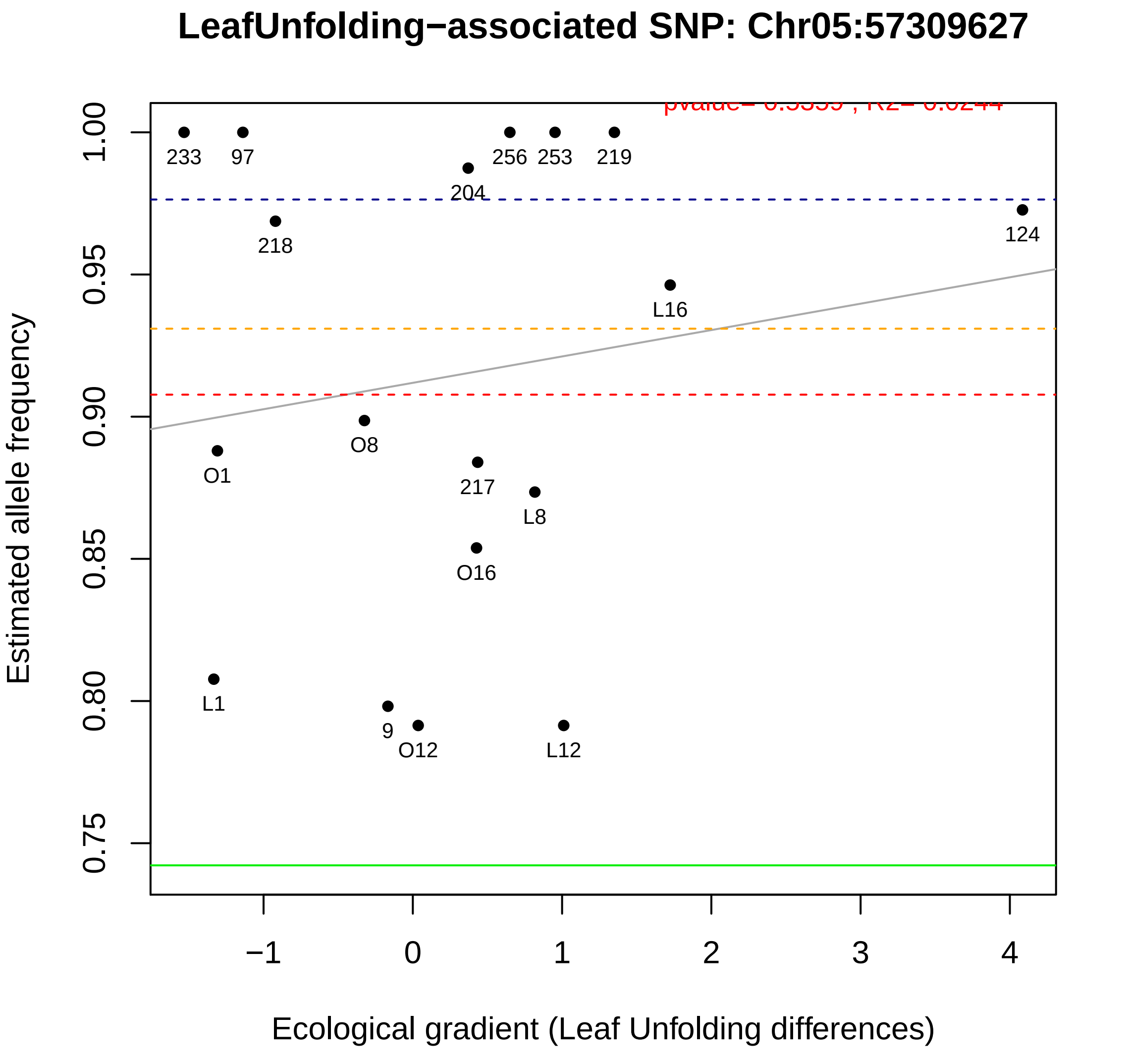
LeafUnfolding−associated SNP: Chr05:57309627
Estimated allele frequency
0.75
0.80
0.85
0.90
0.95
1.00
−1
0
1
2
3
4
Ecological gradient (Leaf Unfolding differences)
233
97
204
256
253
219
218
124
L16
O8
O1
217
L8
O16
L1
9
O12
L12

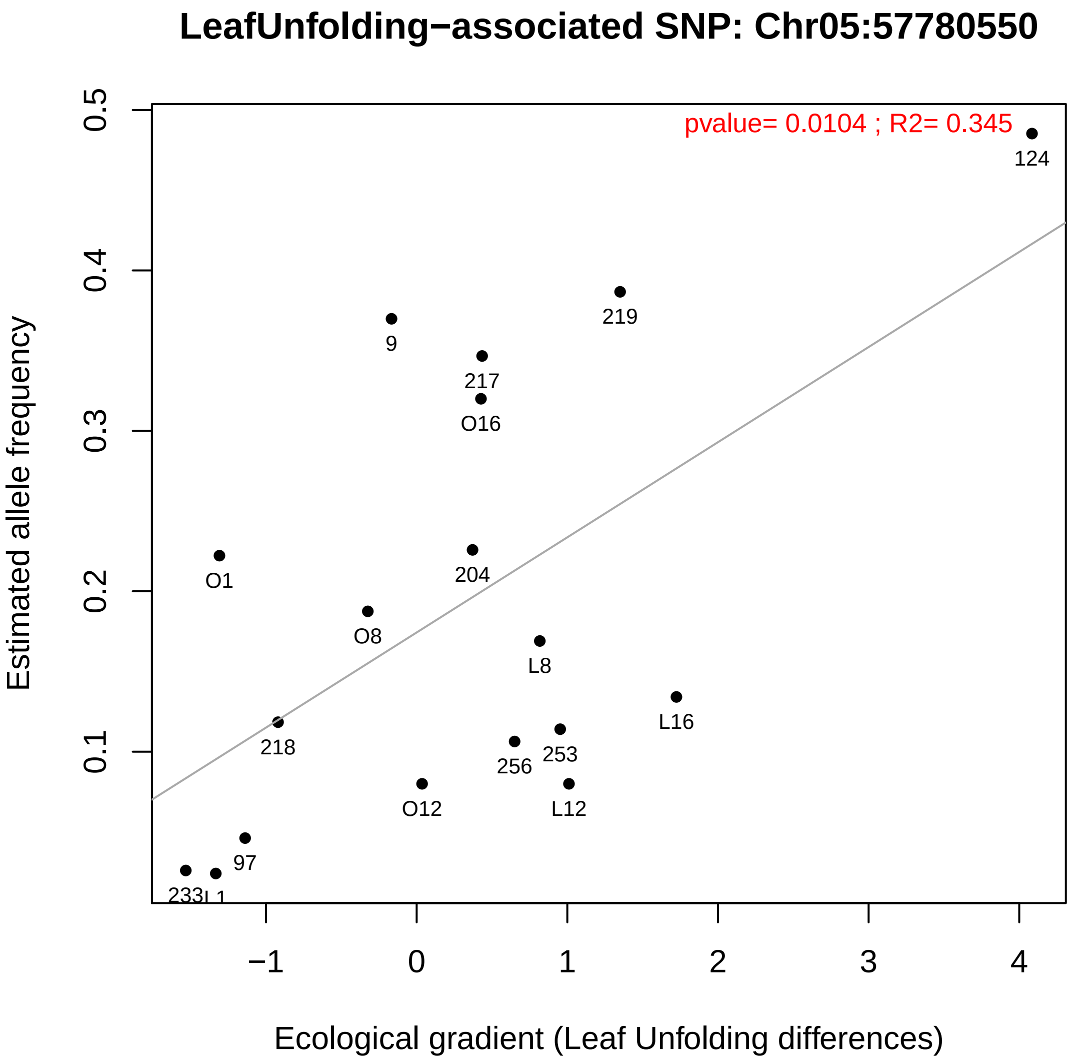

LeafUnfolding−associated SNP: Chr05:57780550
pvalue= 0.0104 ; R2= 0.345
124
219
9
217
O16
204
O1
O8
L8
L16
218
256
253
O12
L12
97
233
L1
0.1
0.2
0.3
0.4
0.5
−1
0
1
2
3
4
Estimated allele frequency
Ecological gradient (Leaf Unfolding differences)

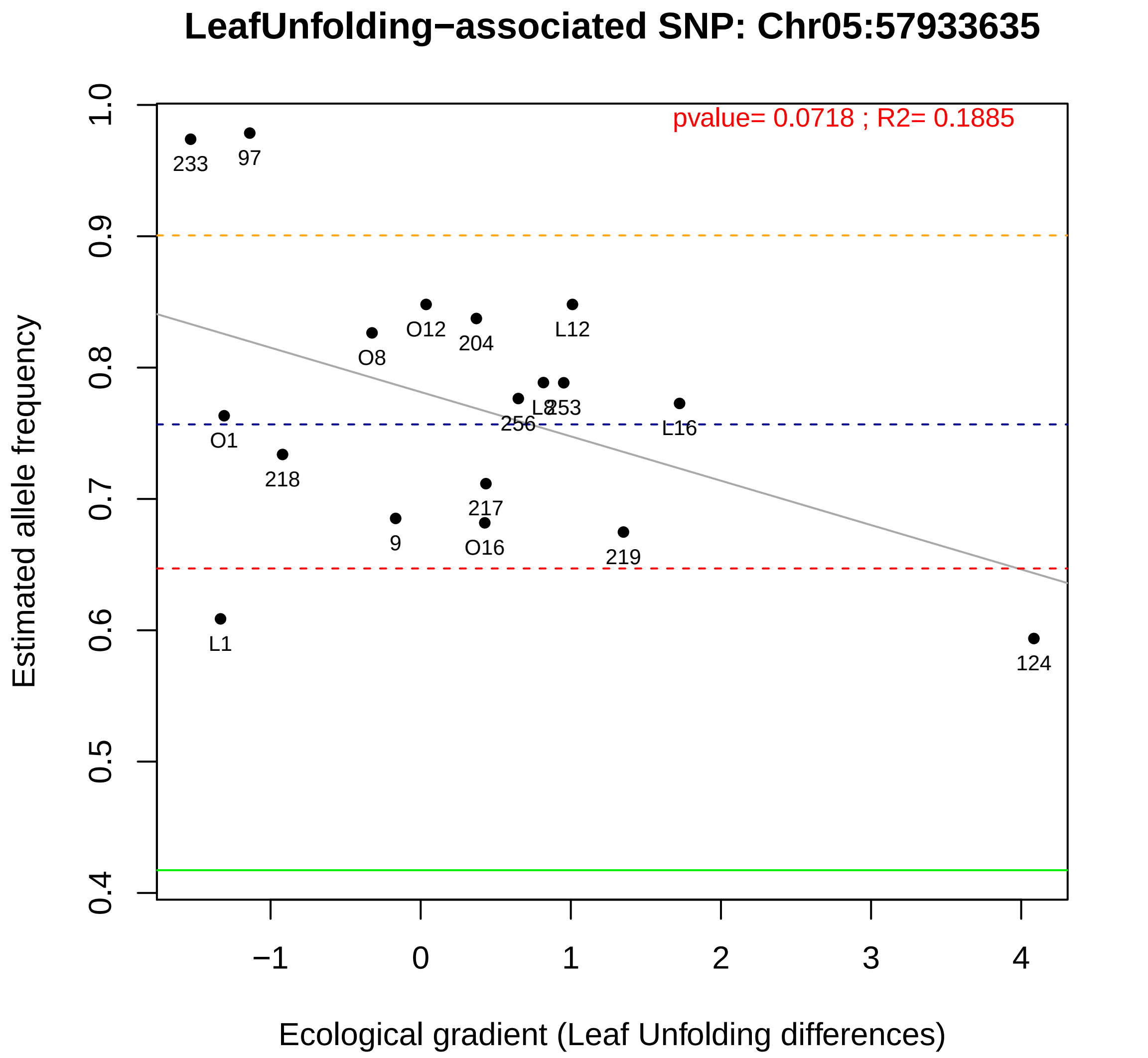

LeafUnfolding−associated SNP: Chr05:57933635
pvalue= 0.0718 ; R2= 0.1885
Estimated allele frequency
Ecological gradient (Leaf Unfolding differences)
0.4
0.5
0.6
0.7
0.8
0.9
1.0
−1
0
1
2
3
4
233
97
O12
204
L12
O8
L8
253
256
L16
O1
218
217
9
O16
219
L1
124

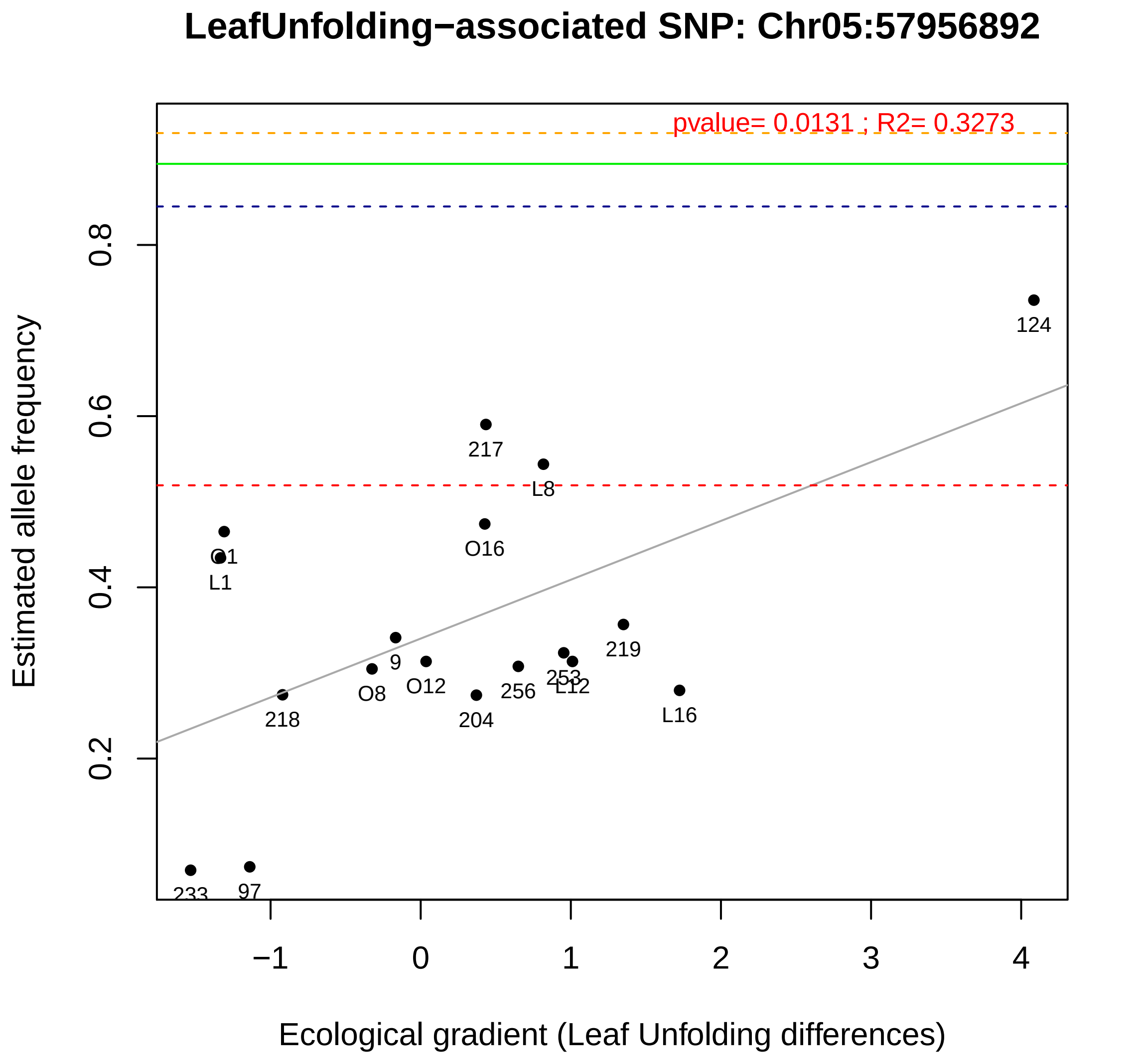

LeafUnfolding−associated SNP: Chr05:57956892
pvalue= 0.0131 ; R2= 0.3273
Estimated allele frequency
0.2
0.4
0.6
0.8
−1
0
1
2
3
4
Ecological gradient (Leaf Unfolding differences)
124
217
L8
O16
O1
L1
9
O12
O8
218
204
256
253
L12
219
L16
233
97

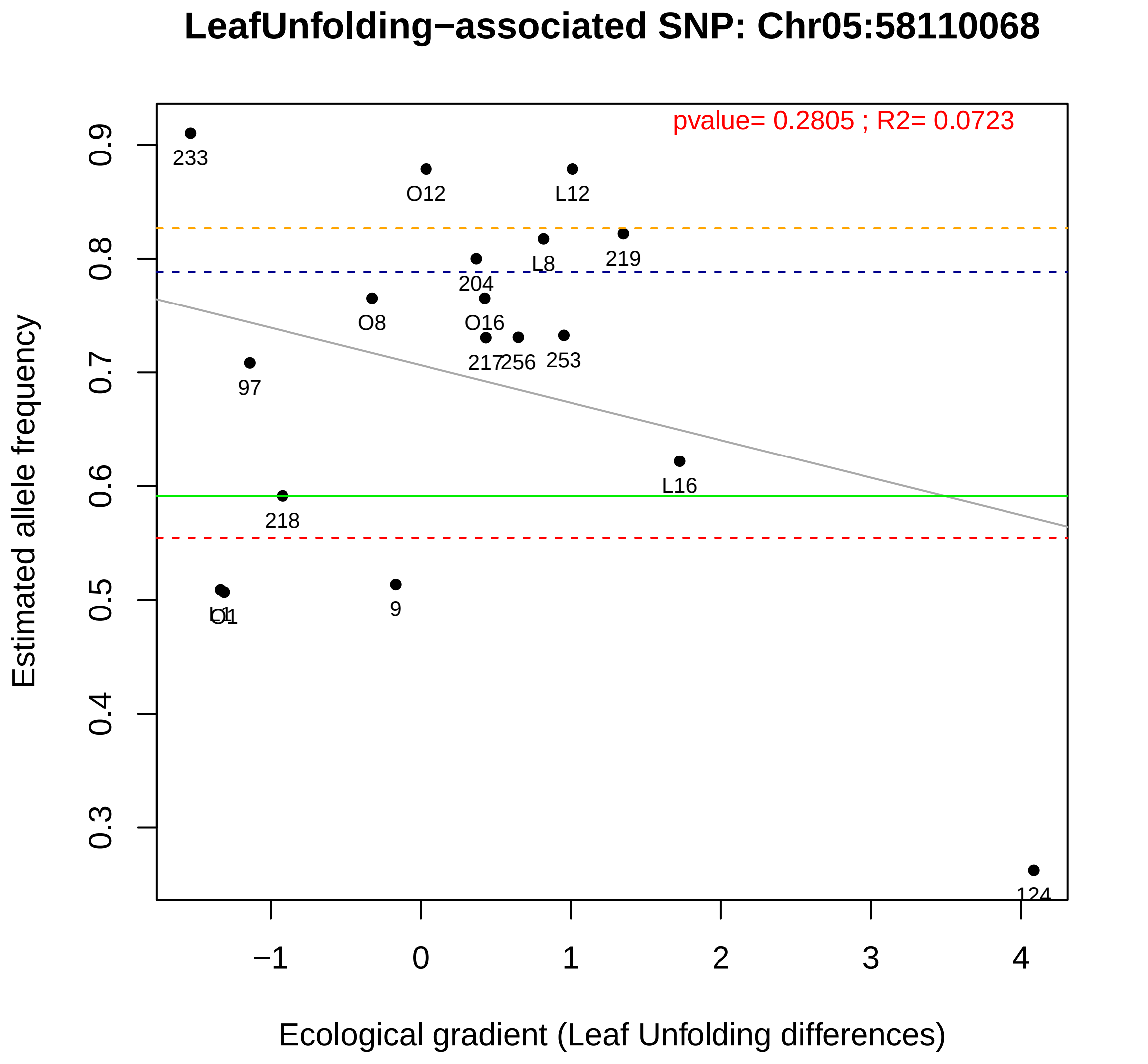
LeafUnfolding−associated SNP: Chr05:58110068
pvalue= 0.2805 ; R2= 0.0723
Estimated allele frequency
Ecological gradient (Leaf Unfolding differences)
233
O12
L12
219
L8
204
O8
O16
217
256
253
97
L16
218
9
124
0.3
0.4
0.5
0.6
0.7
0.8
0.9
−1
0
1
2
3
4

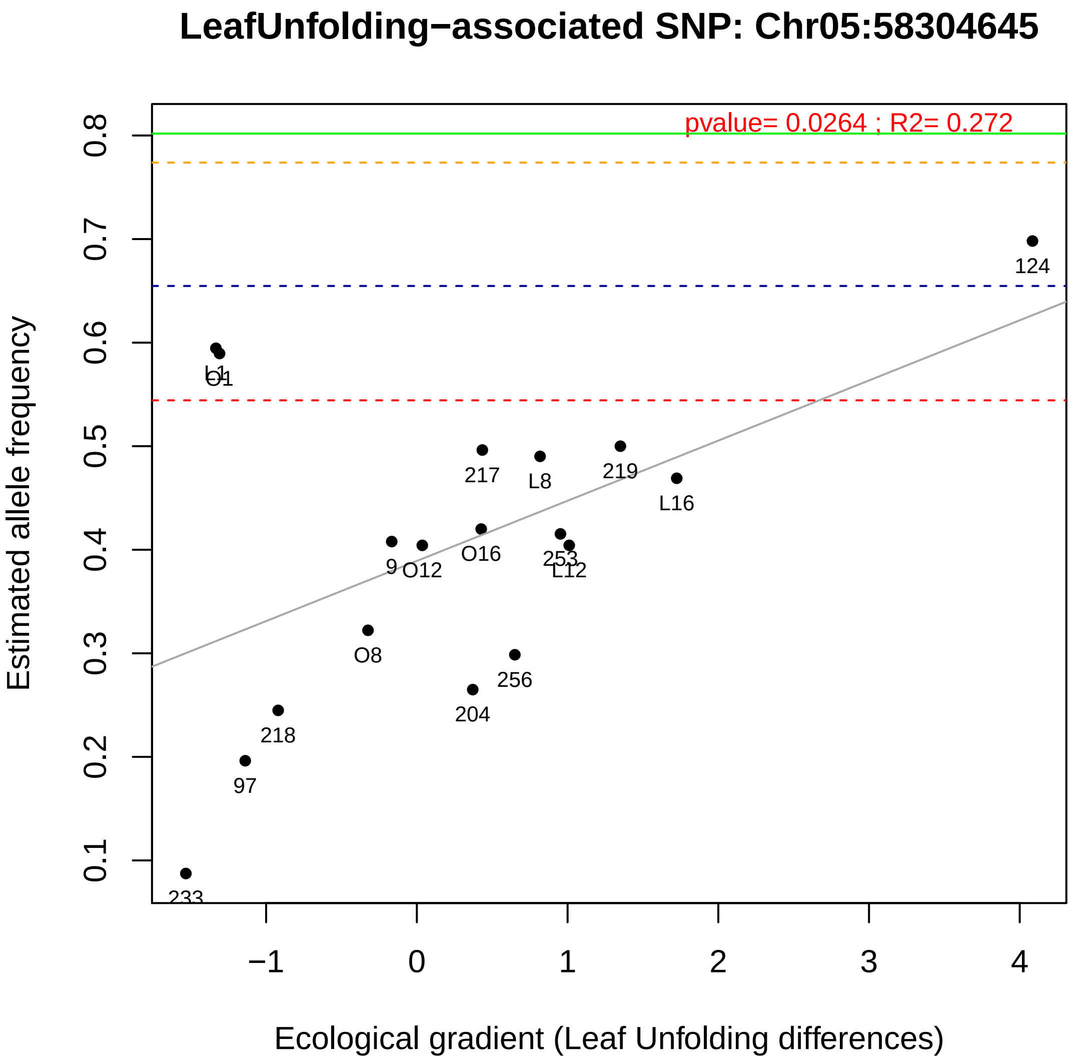

LeafUnfolding−associated SNP: Chr05:58304645
pvalue= 0.0264 ; R2= 0.272
Estimated allele frequency
0.1
0.2
0.3
0.4
0.5
0.6
0.7
0.8
−1
0
1
2
3
4
Ecological gradient (Leaf Unfolding differences)
124
L1
O1
217
L8
219
L16
9
O12
O16
253
L12
O8
256
204
218
97
233

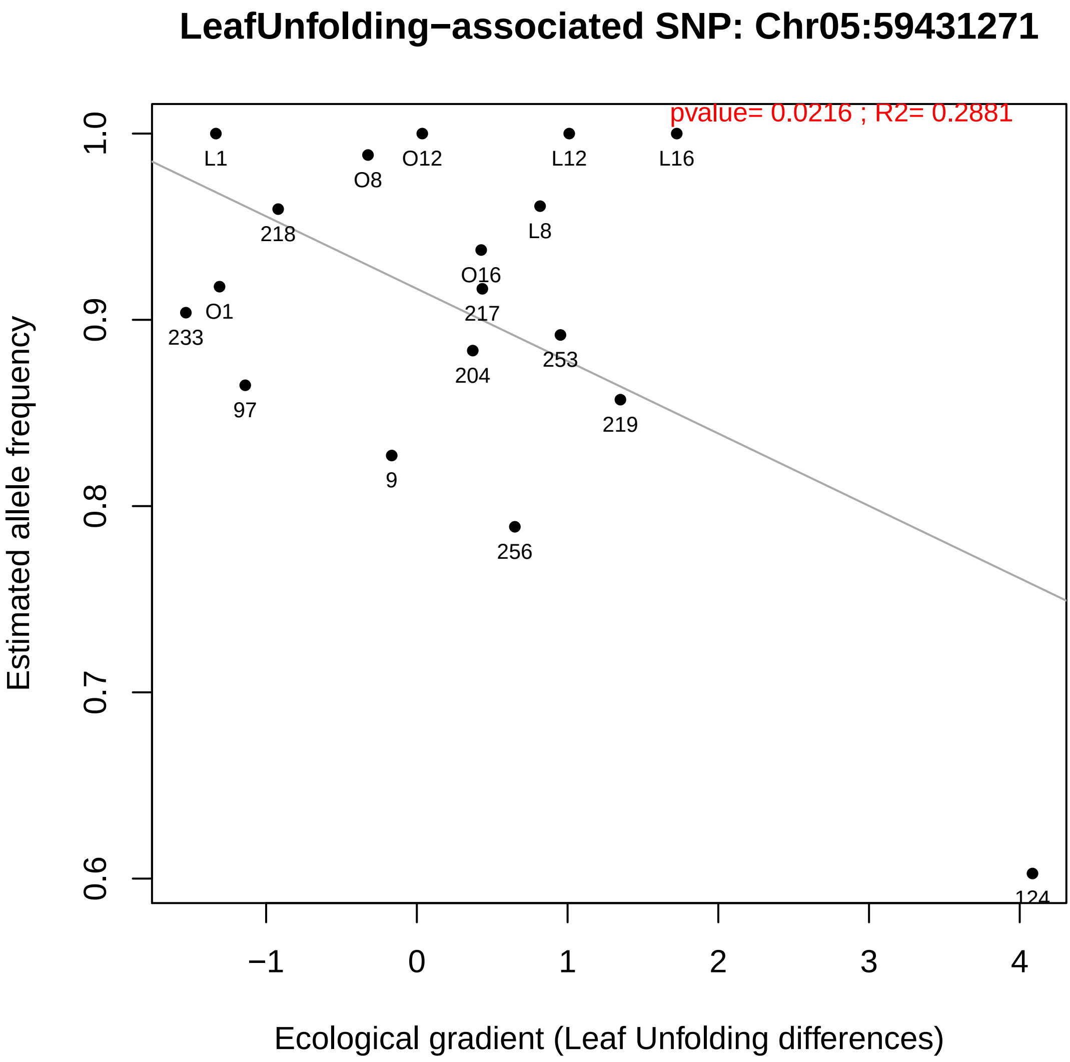
LeafUnfolding−associated SNP: Chr05:59431271
pvalue= 0.0216 ; R2= 0.2881
Estimated allele frequency
Ecological gradient (Leaf Unfolding differences)
0.6
0.7
0.8
0.9
1.0
−1
0
1
2
3
4
L1
O8
O12
L12
L16
218
L8
O16
217
O1
233
253
204
97
219
9
256
124

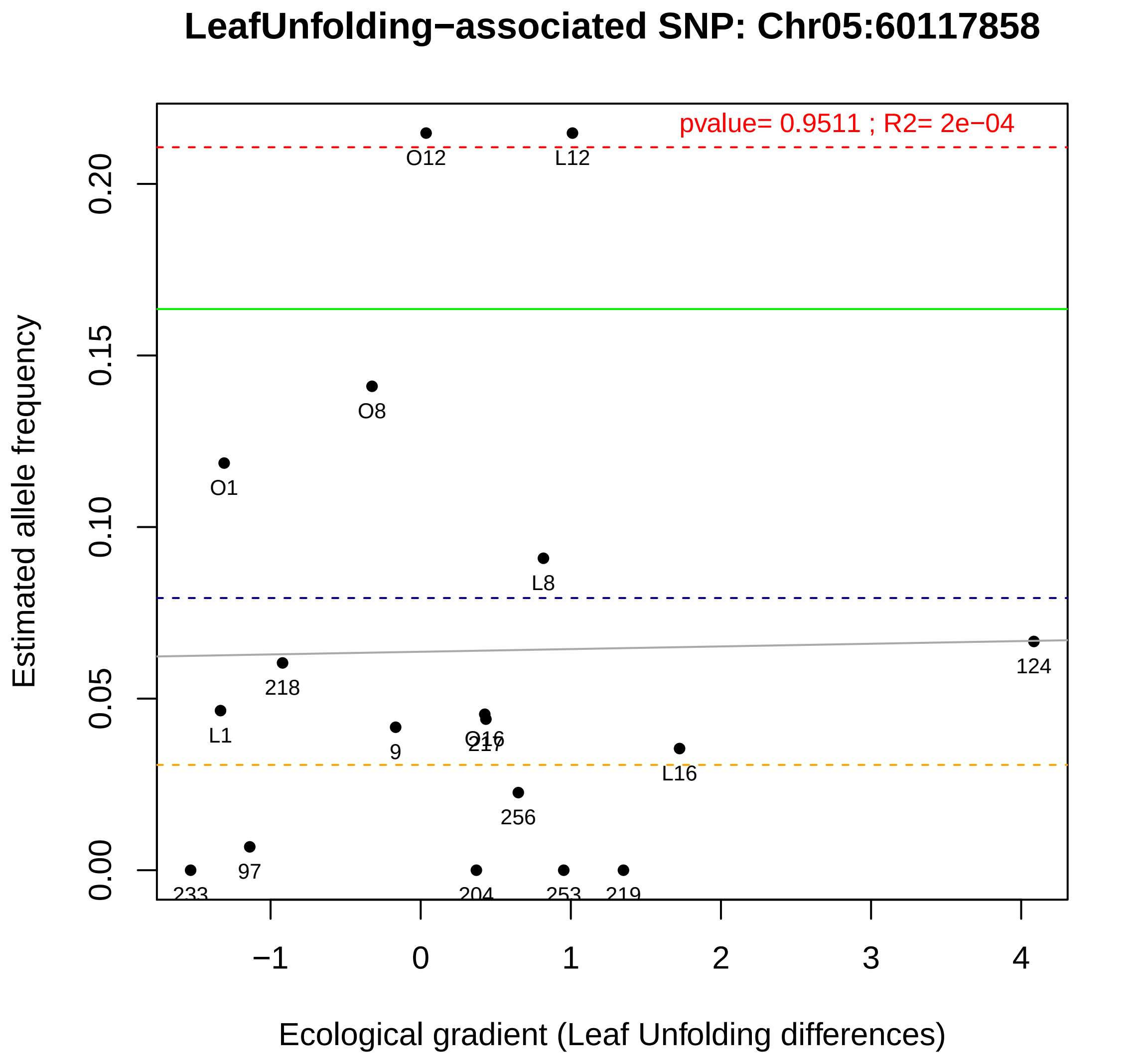
LeafUnfolding−associated SNP: Chr05:60117858
pvalue= 0.9511 ; R2= 2e−04
Estimated allele frequency
0.00
0.05
0.10
0.15
0.20
O12
L12
O8
O1
L8
124
218
L1
9
O16
217
L16
256
97
233
204
253
219
−1
0
1
2
3
4
Ecological gradient (Leaf Unfolding differences)

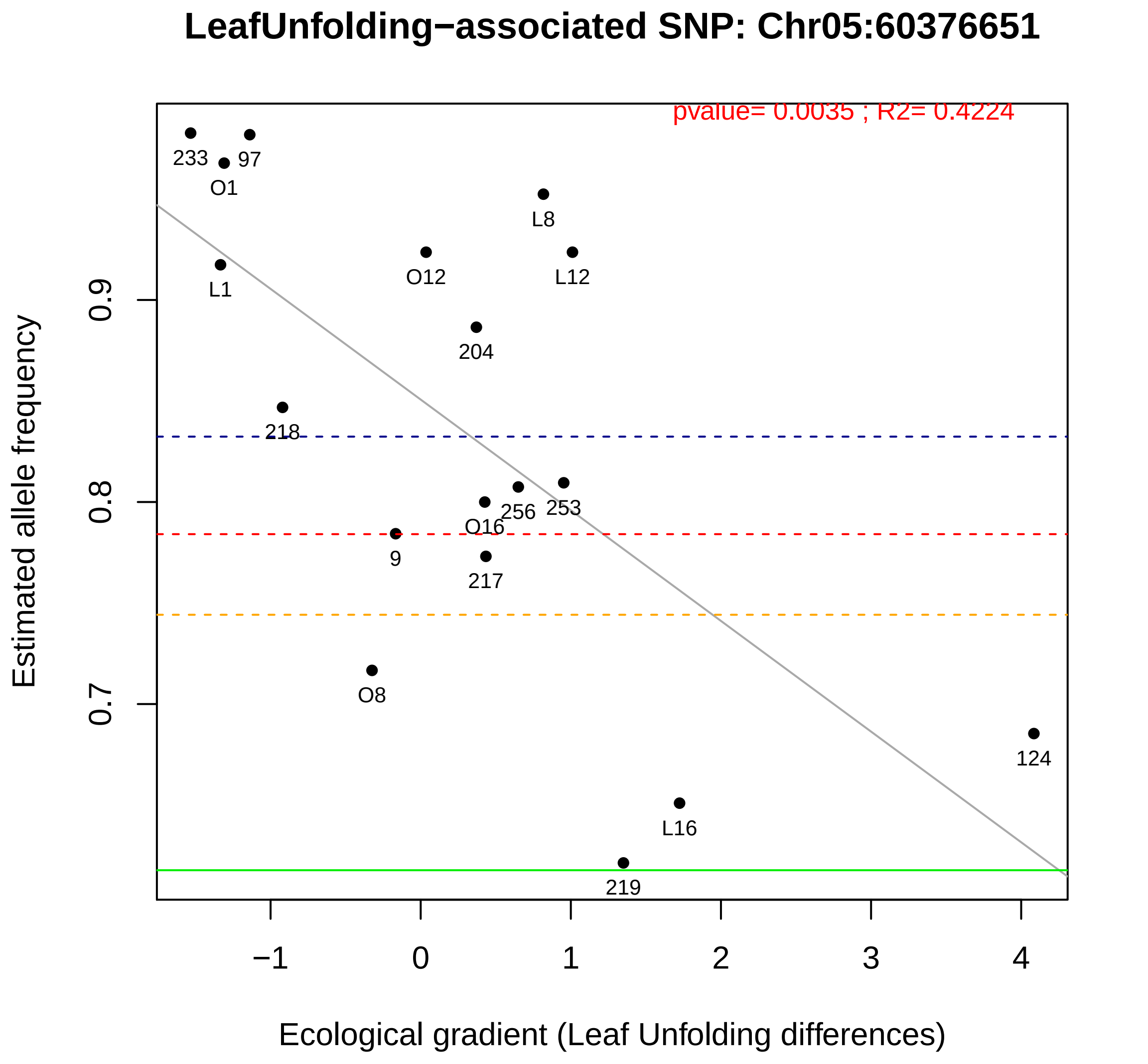
LeafUnfolding−associated SNP: Chr05:60376651
pvalue= 0.0035 ; R2= 0.4224
Estimated allele frequency
0.7
0.8
0.9
−1
0
1
2
3
4
Ecological gradient (Leaf Unfolding differences)
233
97
O1
L8
O12
L12
L1
204
218
256
253
O16
9
217
O8
124
L16
219

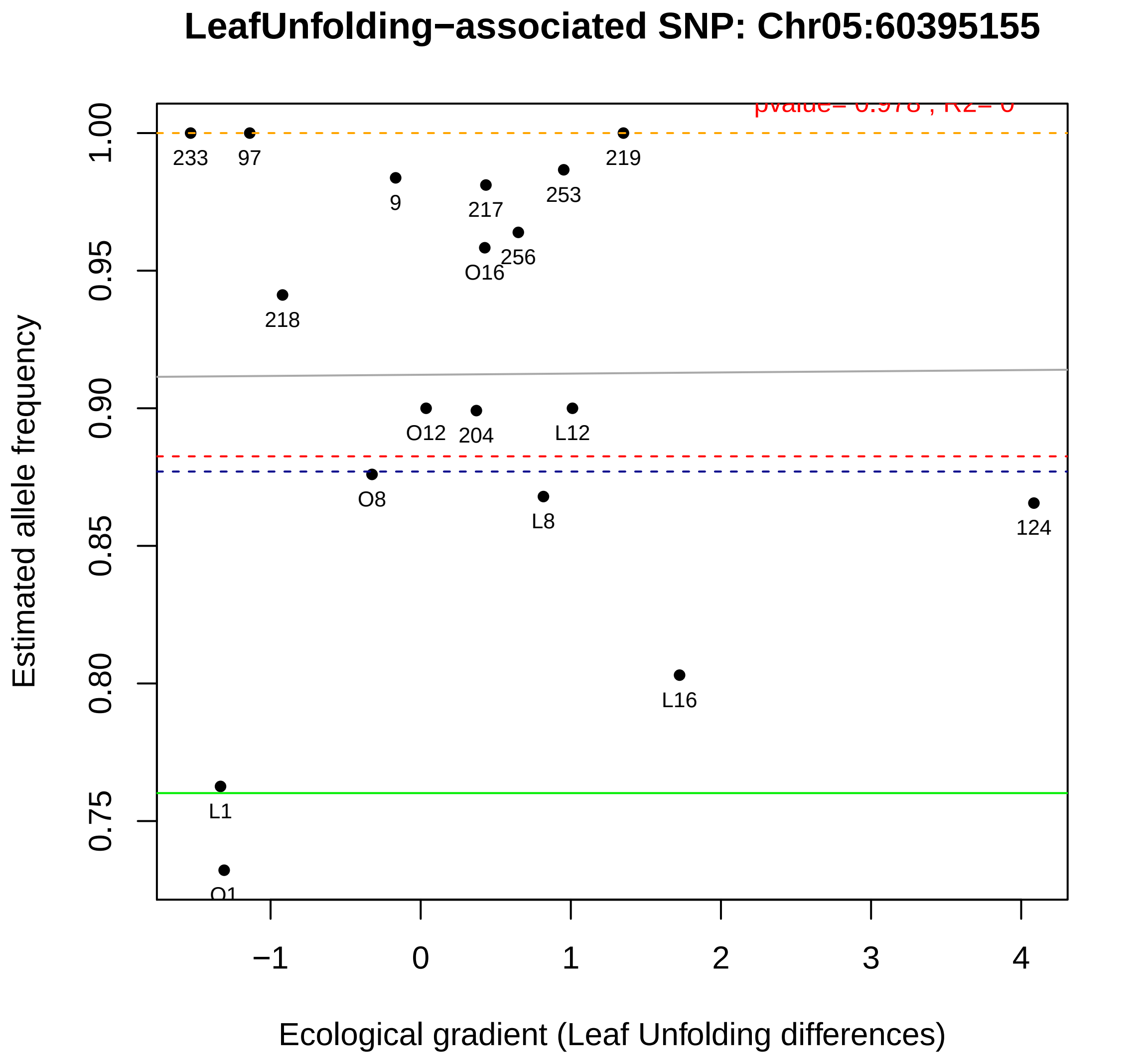

LeafUnfolding−associated SNP: Chr05:60395155
pvalue= 0.978 ; R2= 0
233
97
219
9
217
253
256
O16
218
O12
204
L12
O8
L8
124
L16
L1
O1
Estimated allele frequency
0.75
0.80
0.85
0.90
0.95
1.00
−1
0
1
2
3
4
Ecological gradient (Leaf Unfolding differences)

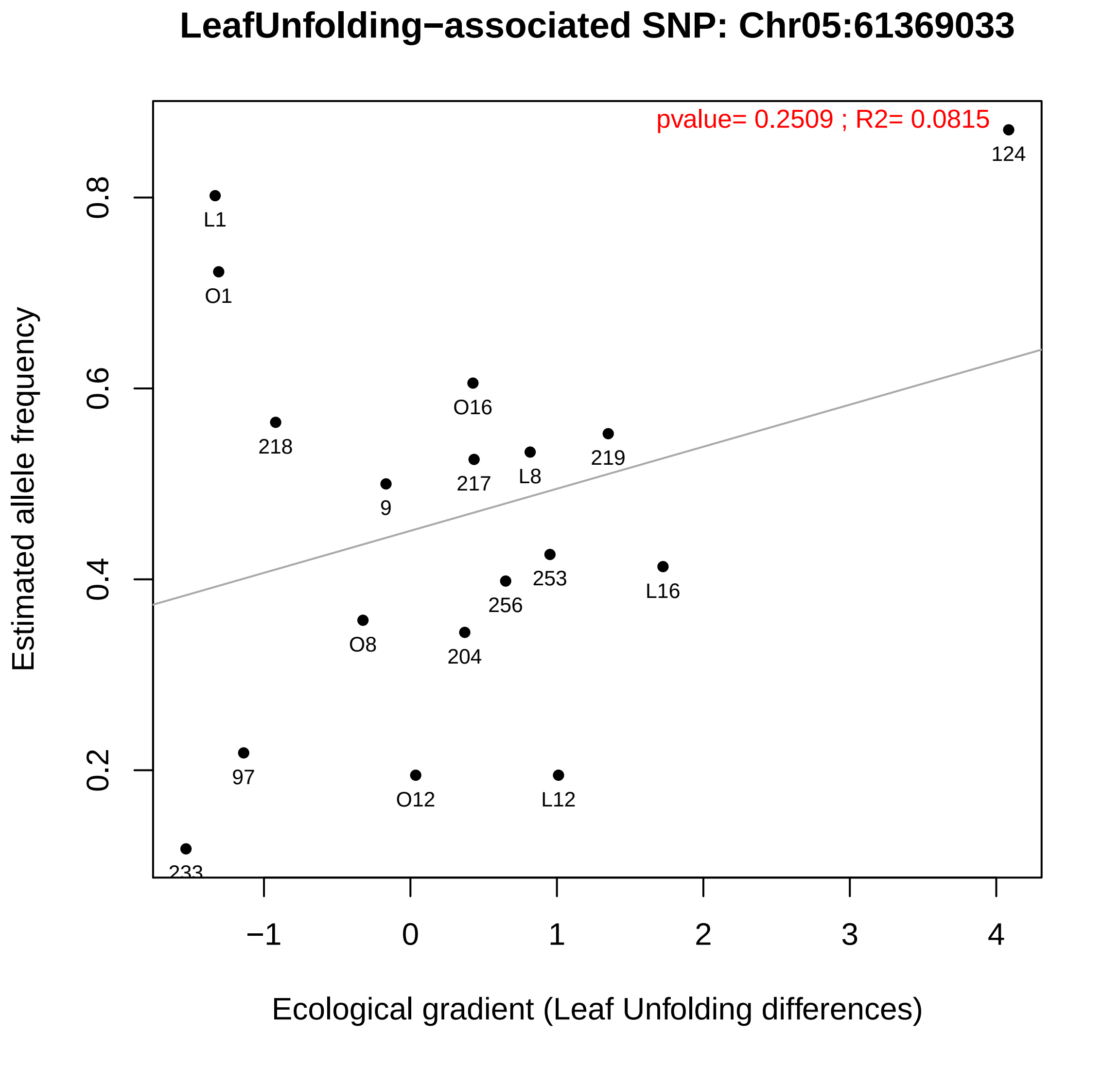

LeafUnfolding−associated SNP: Chr05:61369033
pvalue= 0.2509 ; R2= 0.0815
Estimated allele frequency
Ecological gradient (Leaf Unfolding differences)
0.2
0.4
0.6
0.8
−1
0
1
2
3
4
124
L1
O1
O16
218
219
217
L8
9
253
256
L16
O8
204
97
O12
L12
233

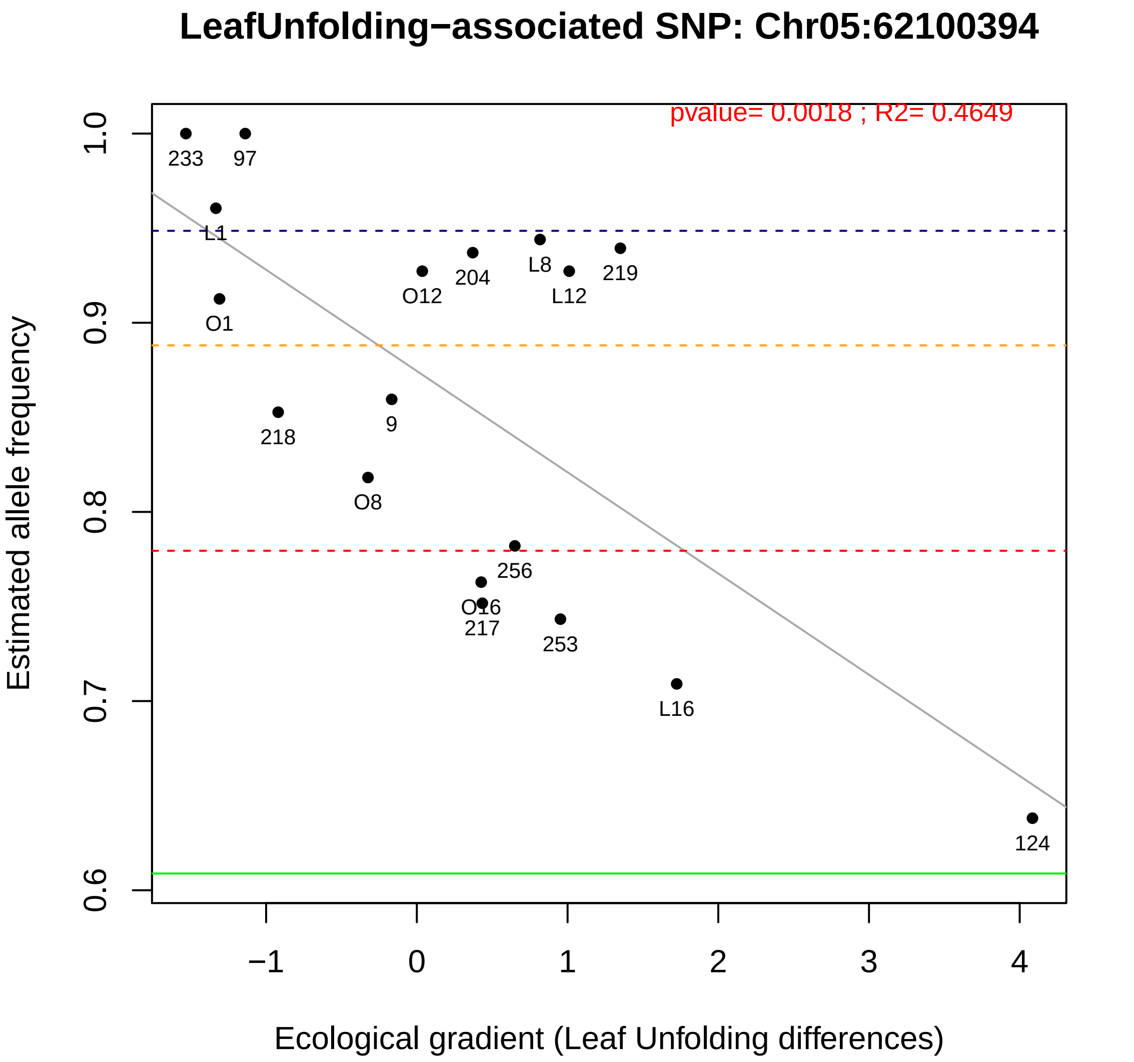

LeafUnfolding−associated SNP: Chr05:62100394
pvalue= 0.0018 ; R2= 0.4649
233
97
L1
O1
O12
204
L8
L12
219
218
9
O8
256
O16
217
253
L16
124
Estimated allele frequency
0.6
0.7
0.8
0.9
1.0
−1
0
1
2
3
4
Ecological gradient (Leaf Unfolding differences)

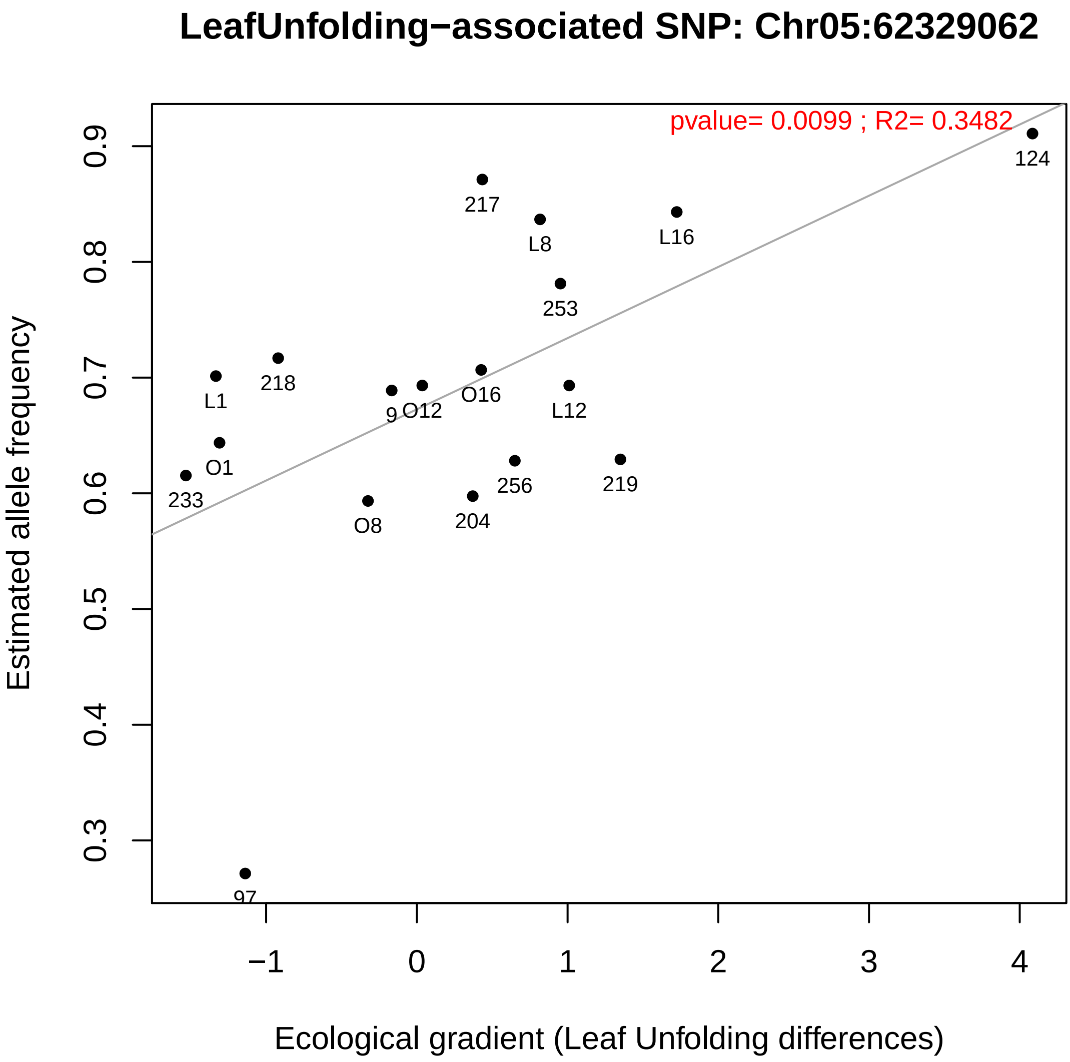
LeafUnfolding−associated SNP: Chr05:62329062
pvalue= 0.0099 ; R2= 0.3482
Estimated allele frequency
Ecological gradient (Leaf Unfolding differences)
0.3
0.4
0.5
0.6
0.7
0.8
0.9
−1
0
1
2
3
4
124
217
L8
L16
253
218
L1
9
O12
O16
L12
O1
233
256
219
O8
204
97

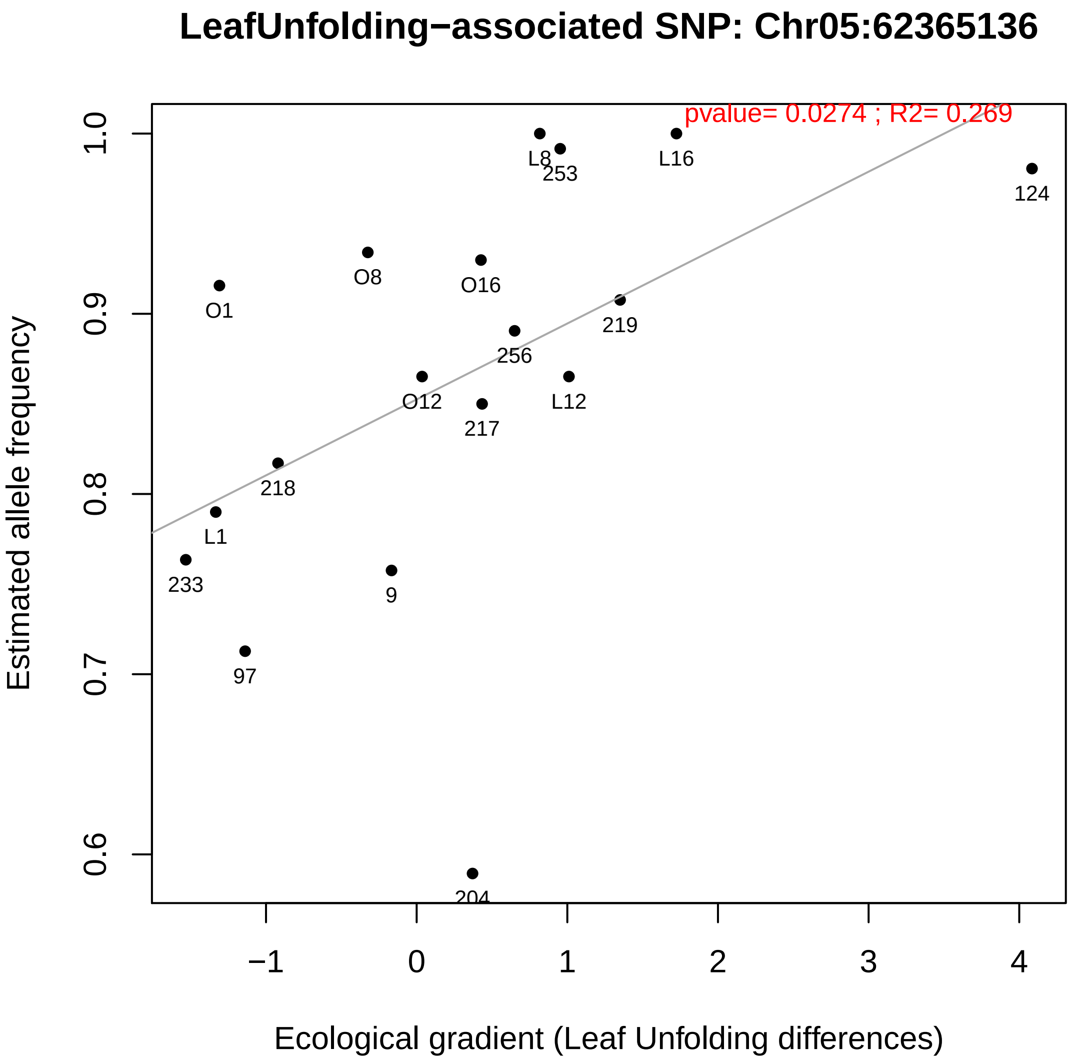
LeafUnfolding−associated SNP: Chr05:62365136
pvalue= 0.0274 ; R2= 0.269
Estimated allele frequency
Ecological gradient (Leaf Unfolding differences)
1.0
0.9
0.8
0.7
0.6
−1
0
1
2
3
4
L8
253
L16
124
O8
O16
O1
219
256
O12
L12
217
218
L1
233
9
97
204

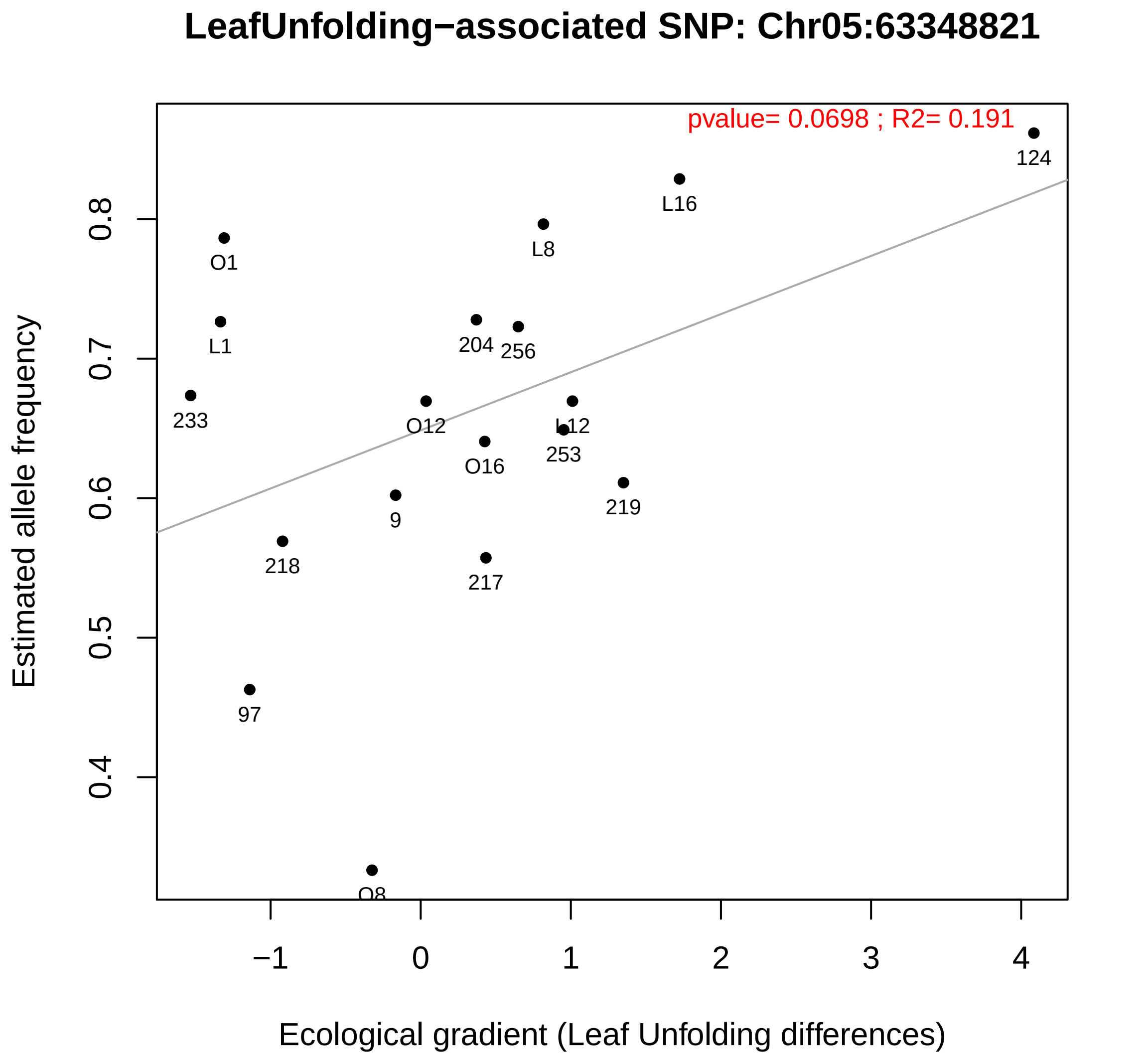
LeafUnfolding−associated SNP: Chr05:63348821
pvalue= 0.0698 ; R2= 0.191
124
L16
L8
O1
L1
204
256
233
O12
L12
253
O16
219
9
218
217
97
O8
0.4
0.5
0.6
0.7
0.8
−1
0
1
2
3
4
Estimated allele frequency
Ecological gradient (Leaf Unfolding differences)

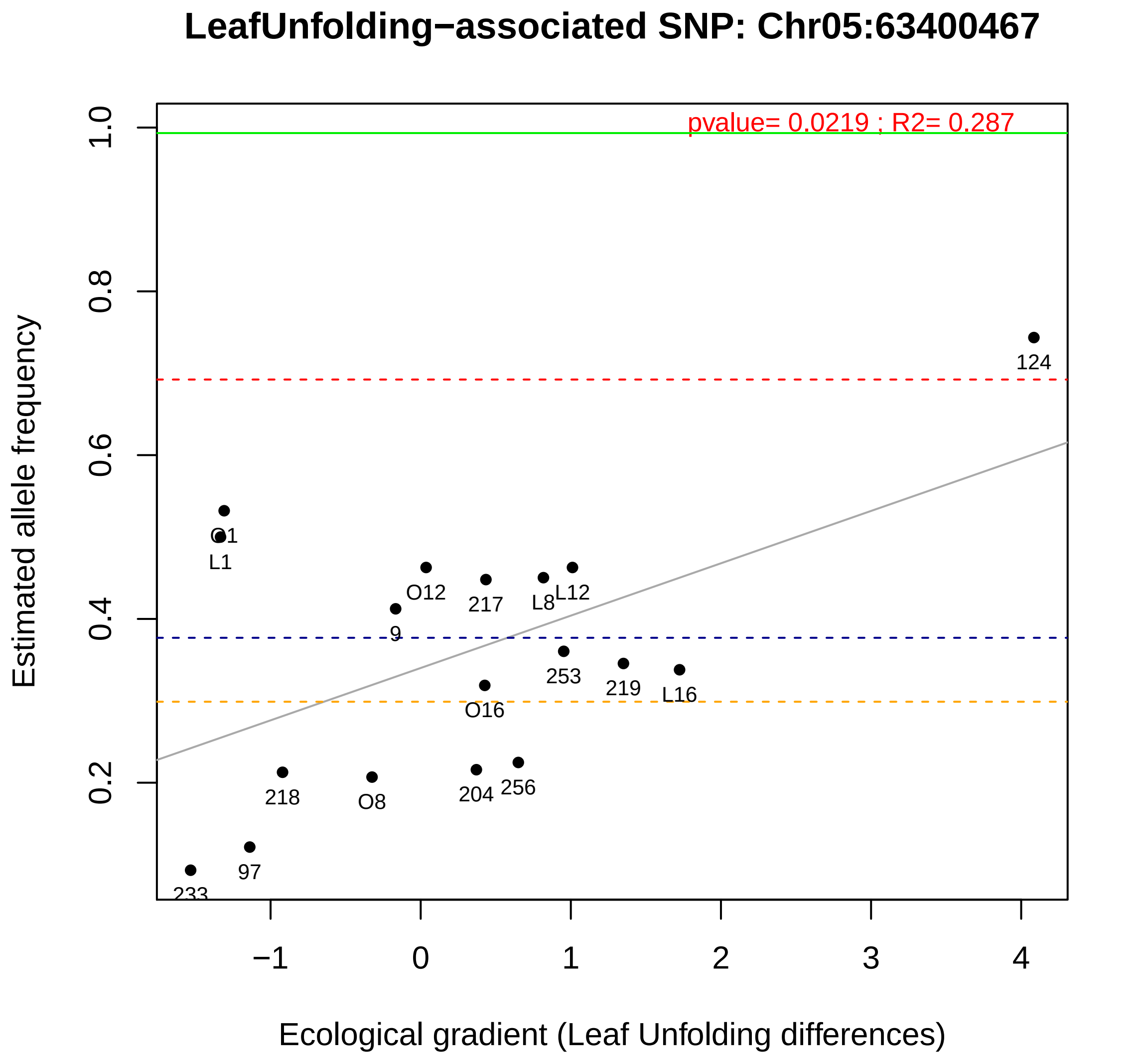
LeafUnfolding−associated SNP: Chr05:63400467
pvalue= 0.0219 ; R2= 0.287
Estimated allele frequency
0.2
0.4
0.6
0.8
1.0
Ecological gradient (Leaf Unfolding differences)
−1
0
1
2
3
4
124
O1
L1
O12
217
L8
L12
9
253
219
L16
O16
218
O8
204
256
97
233

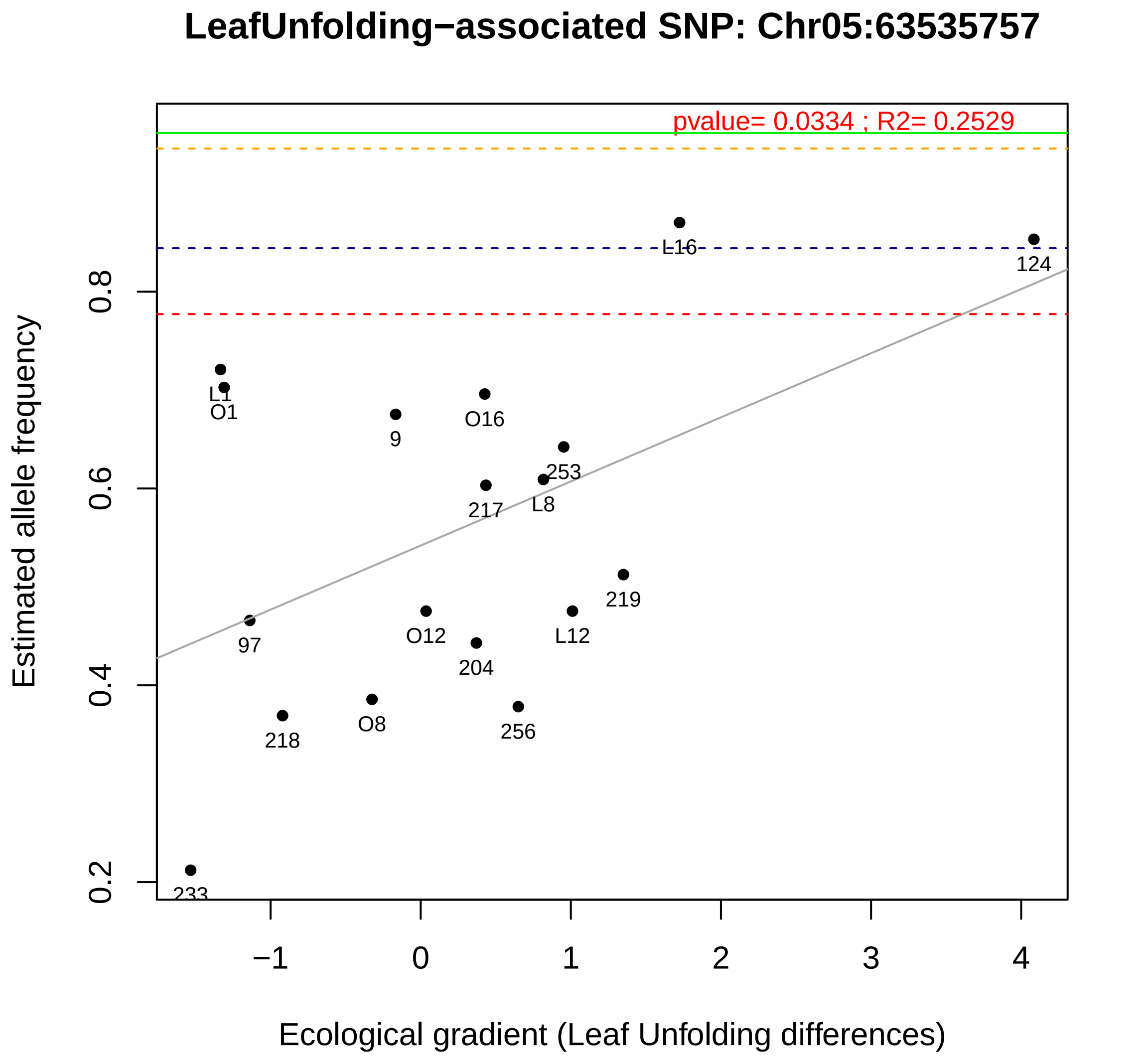
LeafUnfolding−associated SNP: Chr05:63535757
pvalue= 0.0334 ; R2= 0.2529
Estimated allele frequency
0.2
0.4
0.6
0.8
−1
0
1
2
3
4
Ecological gradient (Leaf Unfolding differences)
L16
124
L1
O1
9
O16
253
L8
217
219
97
O12
L12
204
218
O8
256
233

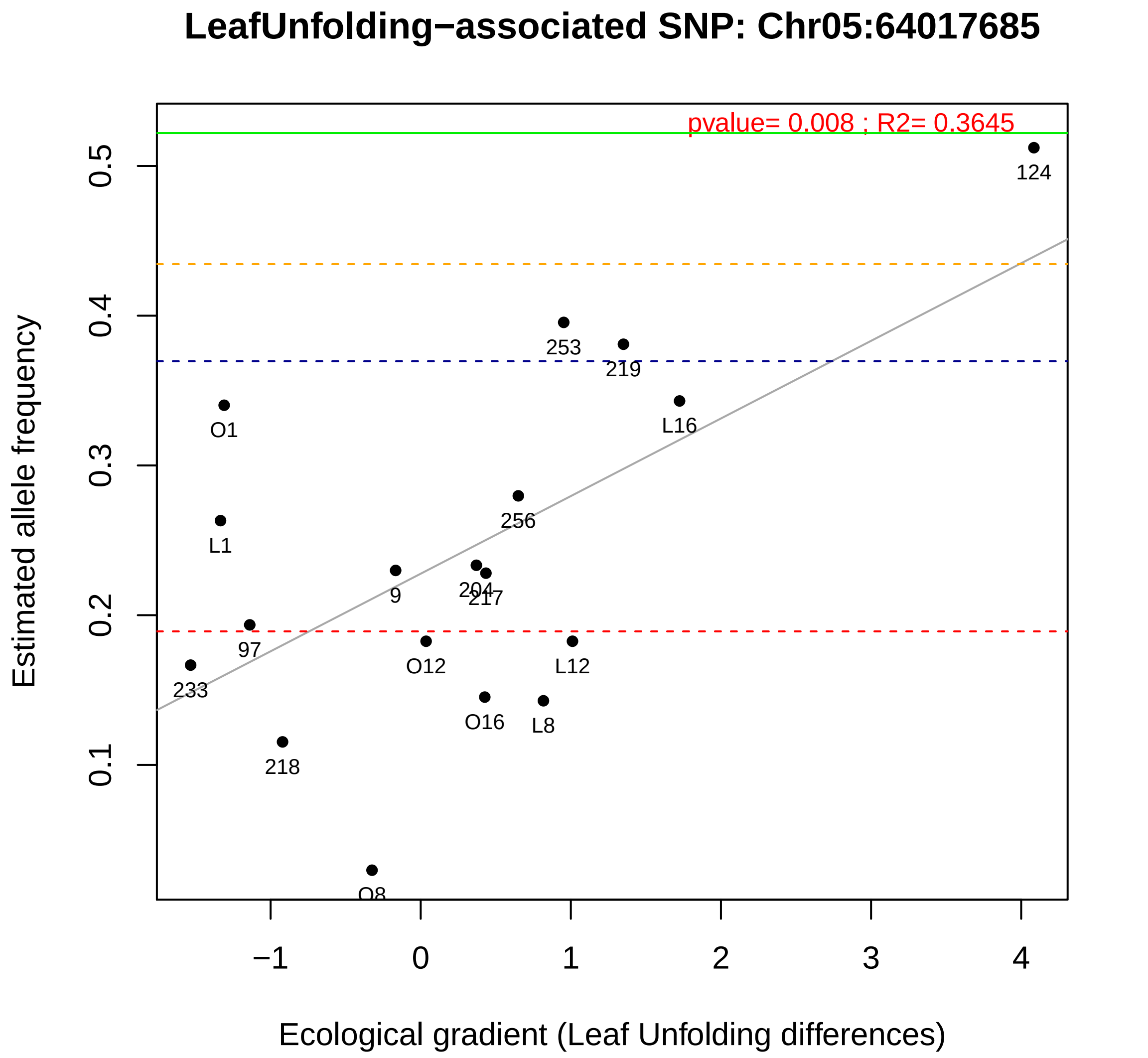

LeafUnfolding−associated SNP: Chr05:64017685
pvalue= 0.008 ; R2= 0.3645
Estimated allele frequency
0.1
0.2
0.3
0.4
0.5
−1
0
1
2
3
4
Ecological gradient (Leaf Unfolding differences)
124
253
219
L16
O1
256
L1
9
204
217
97
O12
L12
233
O16
L8
218
O8

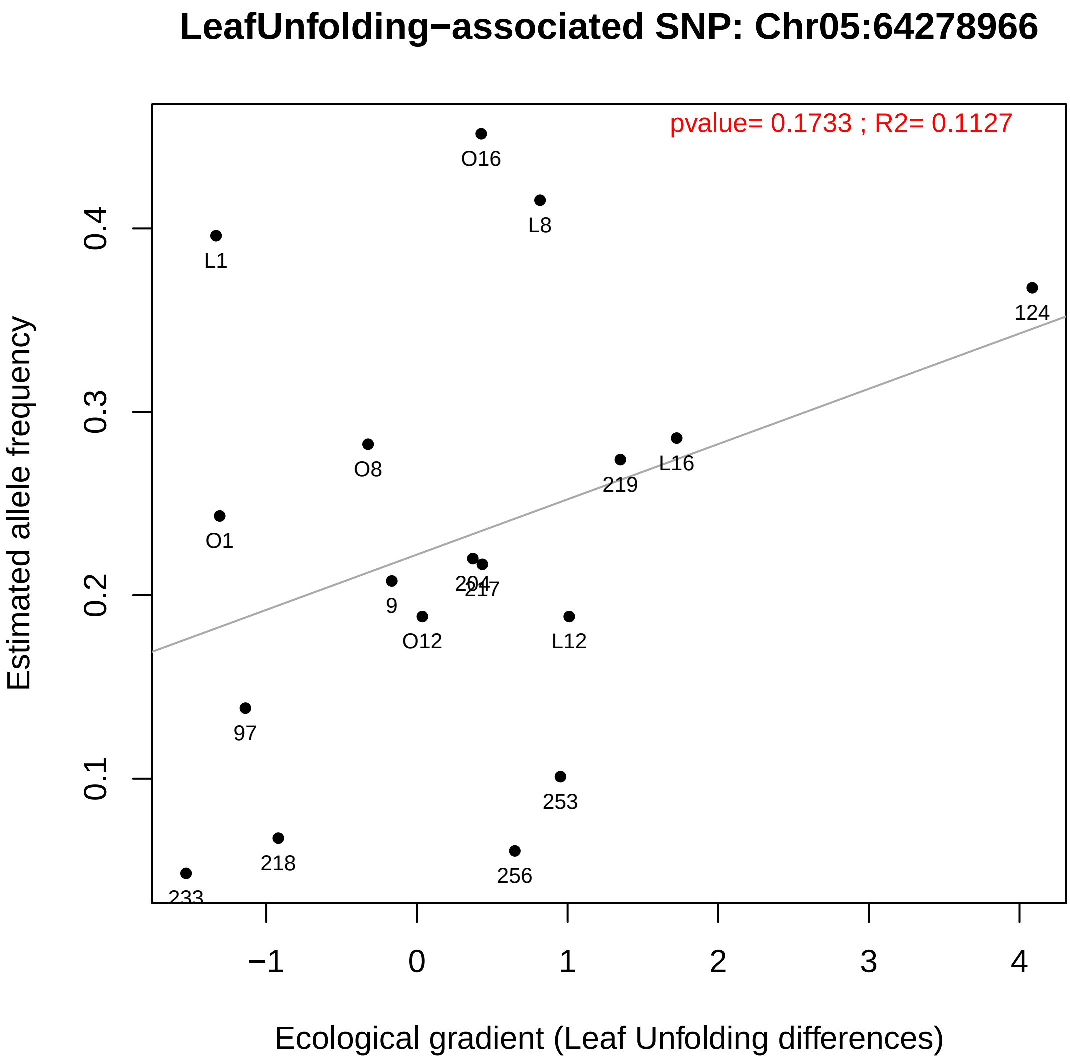

LeafUnfolding−associated SNP: Chr05:64278966
pvalue= 0.1733 ; R2= 0.1127
O16
L8
L1
124
O8
219
L16
O1
204
217
9
O12
L12
97
253
218
256
233
0.1
0.2
0.3
0.4
−1
0
1
2
3
4
Estimated allele frequency
Ecological gradient (Leaf Unfolding differences)

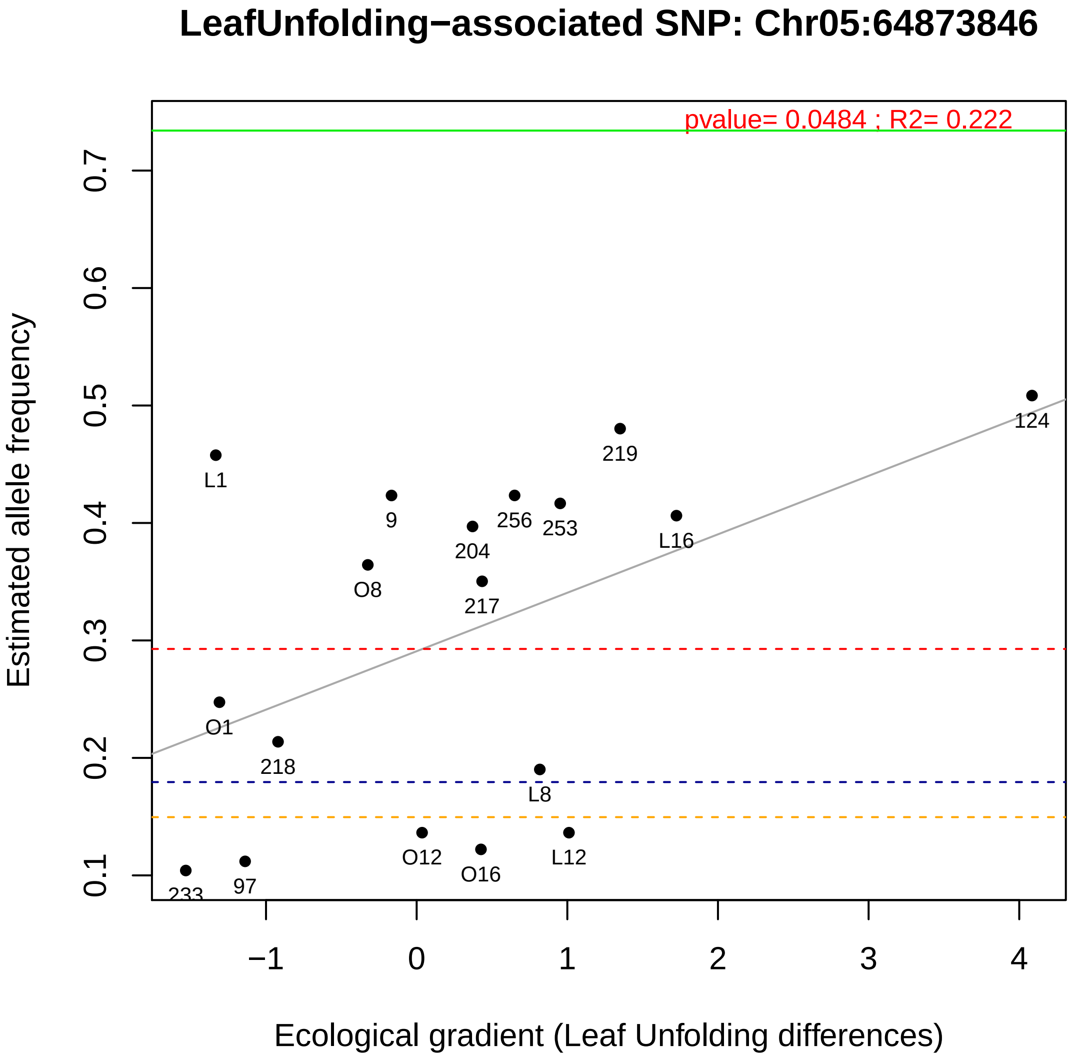

LeafUnfolding−associated SNP: Chr05:64873846
pvalue= 0.0484 ; R2= 0.222
Estimated allele frequency
0.1
0.2
0.3
0.4
0.5
0.6
0.7
−1
0
1
2
3
4
Ecological gradient (Leaf Unfolding differences)
L1
9
256
253
219
124
204
L16
O8
217
O1
218
L8
O12
O16
L12
233
97

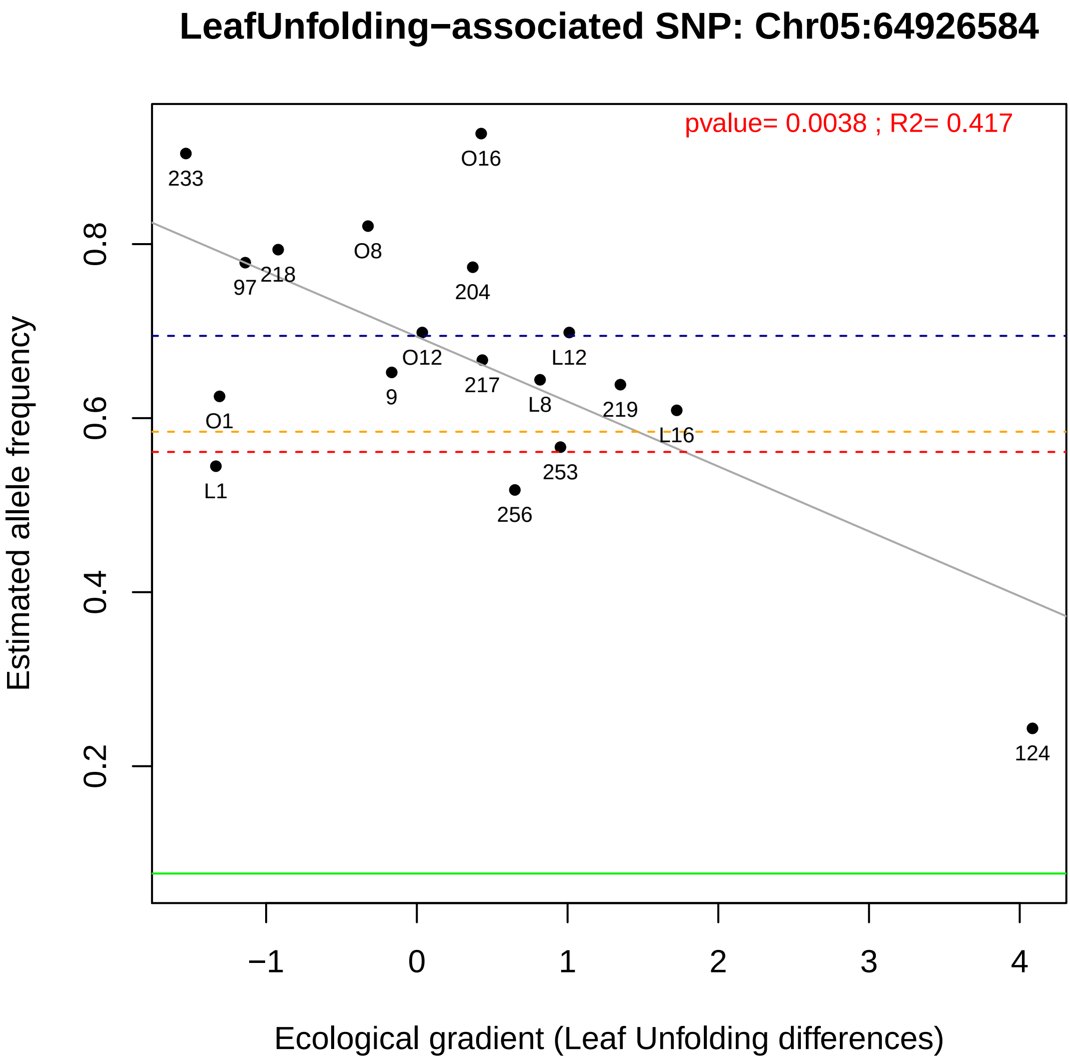
LeafUnfolding−associated SNP: Chr05:64926584
pvalue= 0.0038 ; R2= 0.417
Estimated allele frequency
0.2
0.4
0.6
0.8
−1
0
1
2
3
4
Ecological gradient (Leaf Unfolding differences)
233
O16
97
218
O8
204
O12
L12
217
9
O1
L8
219
L16
253
L1
256
124

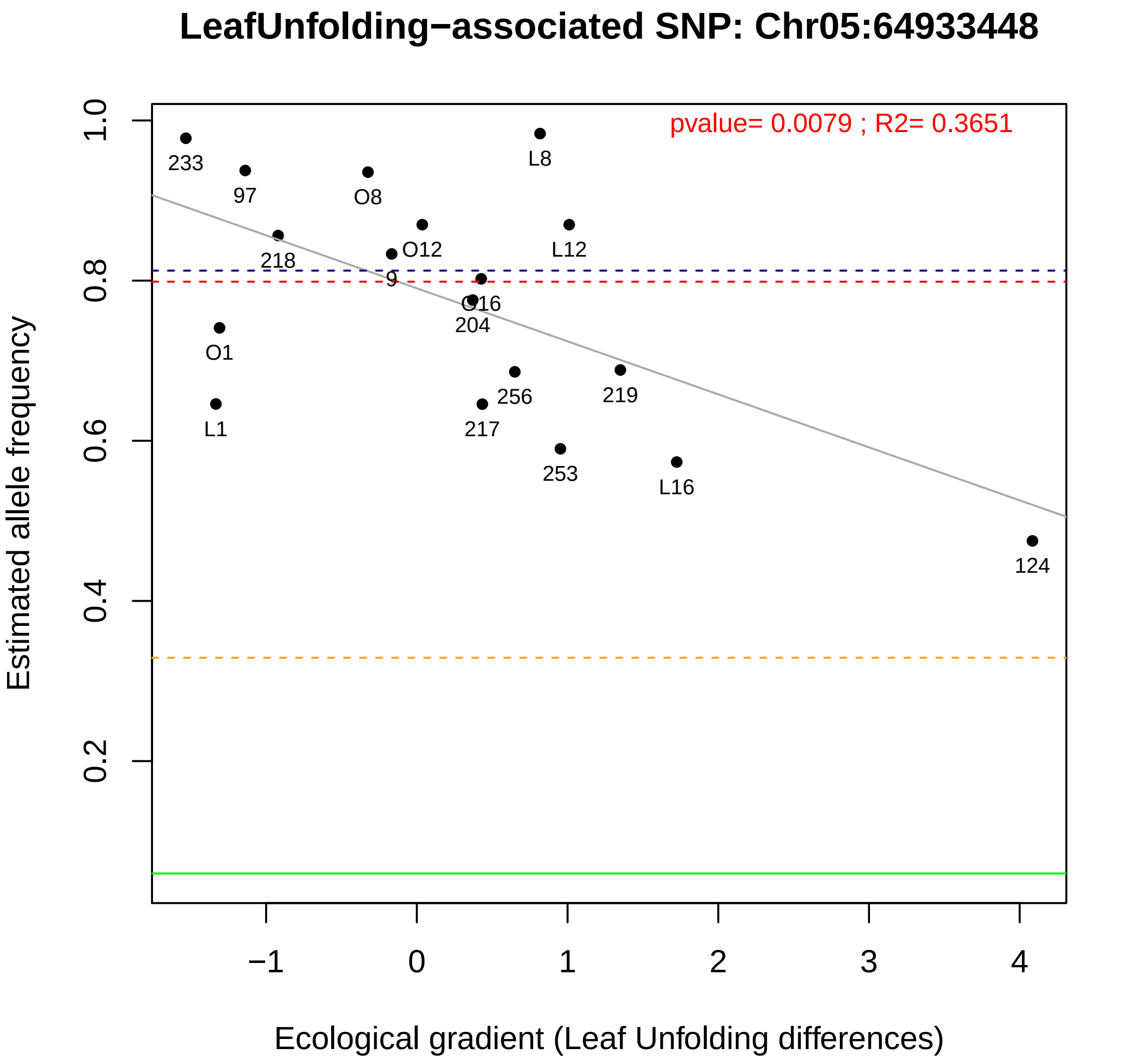
LeafUnfolding−associated SNP: Chr05:64933448
pvalue= 0.0079 ; R2= 0.3651
233
97
O8
L8
218
O12
L12
9
O16
204
O1
L1
256
217
219
253
L16
124
1.0
0.8
0.6
0.4
0.2
−1
0
1
2
3
4
Estimated allele frequency
Ecological gradient (Leaf Unfolding differences)

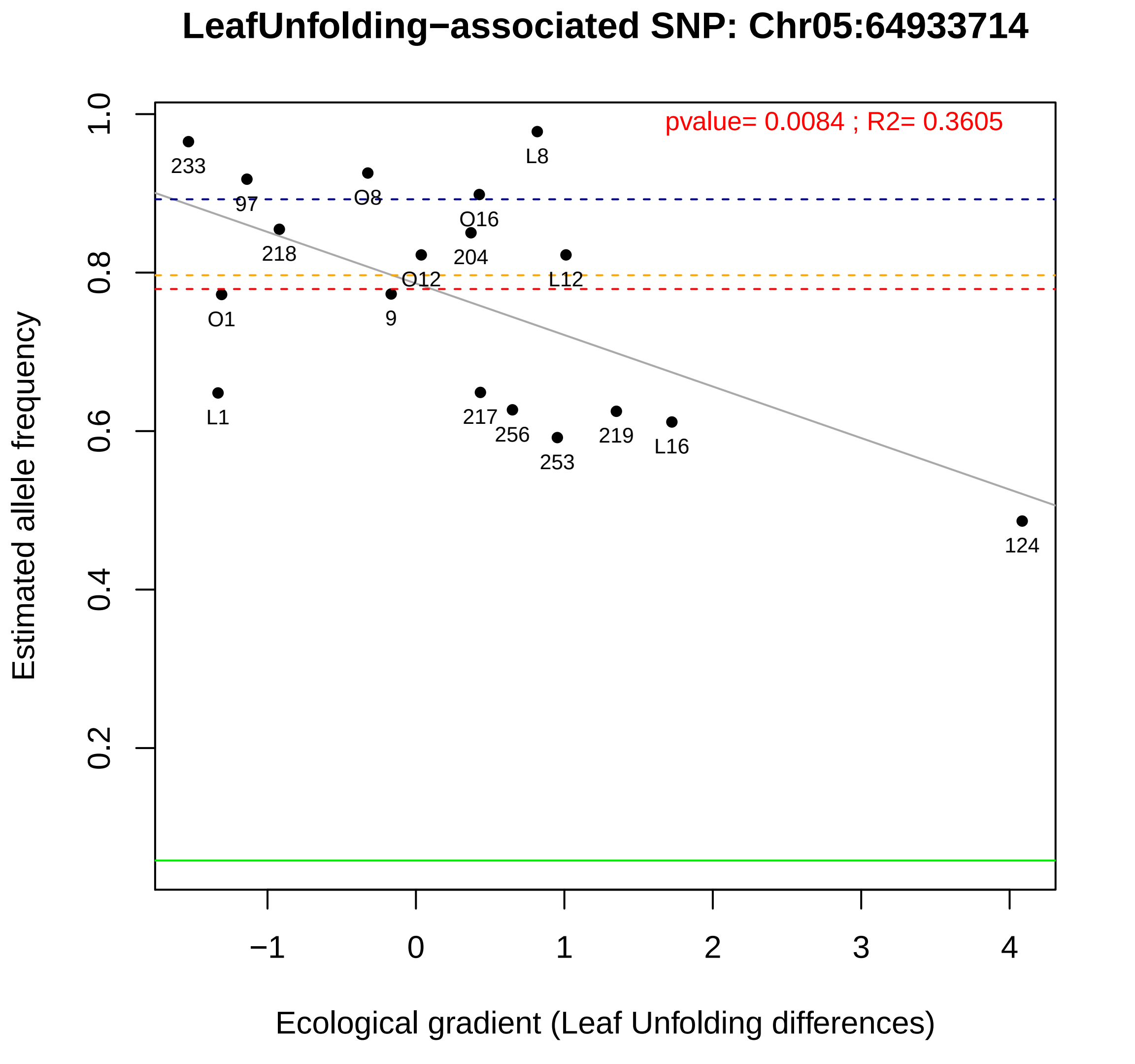

LeafUnfolding−associated SNP: Chr05:64933714
pvalue= 0.0084 ; R2= 0.3605
Estimated allele frequency
Ecological gradient (Leaf Unfolding differences)
0.2
0.4
0.6
0.8
1.0
−1
0
1
2
3
4
233
97
O8
L8
O16
218
204
O12
L12
O1
9
L1
217
256
253
219
L16
124

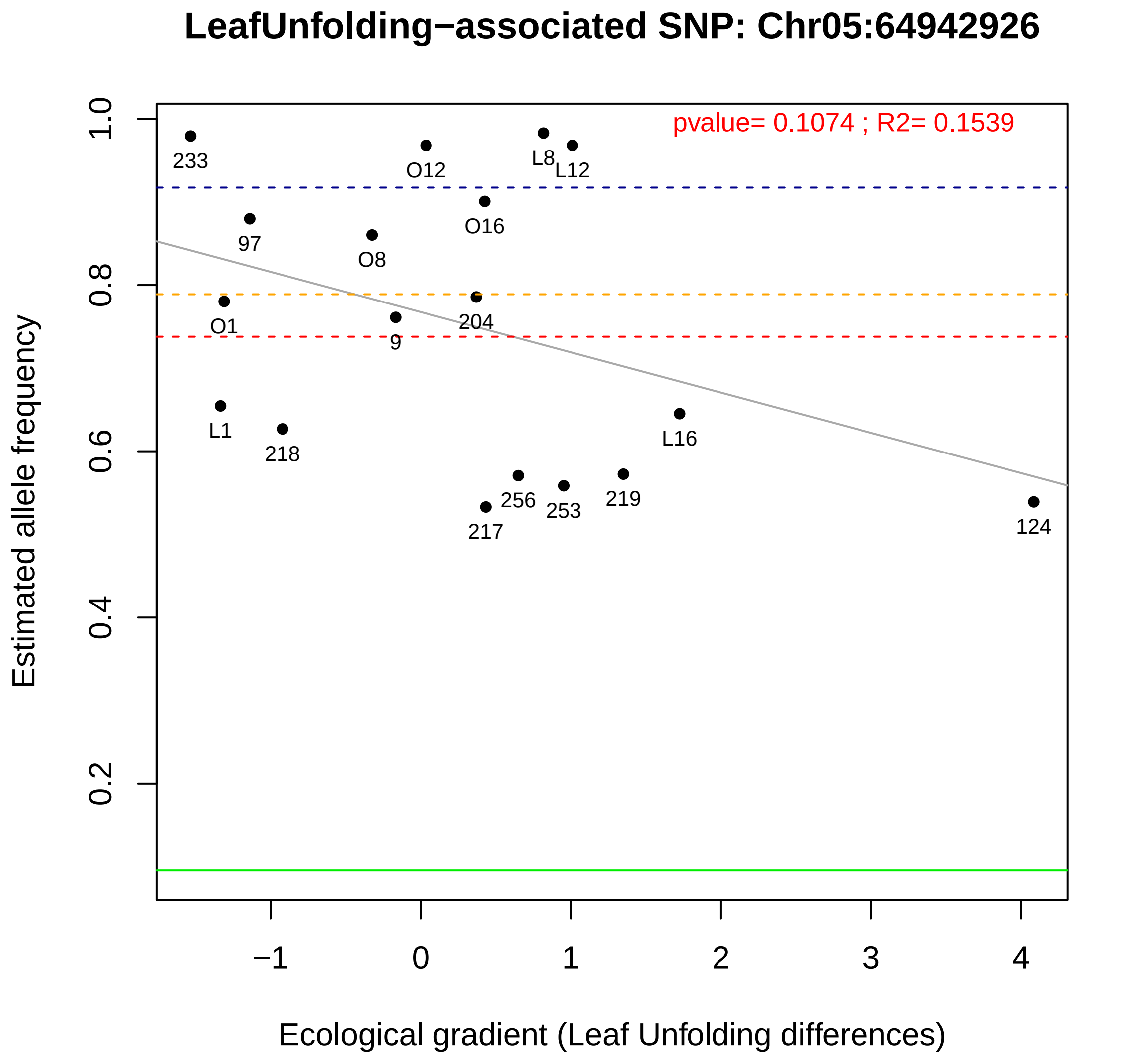

LeafUnfolding−associated SNP: Chr05:64942926
pvalue= 0.1074 ; R2= 0.1539
Estimated allele frequency
Ecological gradient (Leaf Unfolding differences)
0.2
0.4
0.6
0.8
1.0
−1
0
1
2
3
4
233
O12
L8
L12
97
O8
O16
O1
9
204
L1
218
L16
256
253
219
217
124

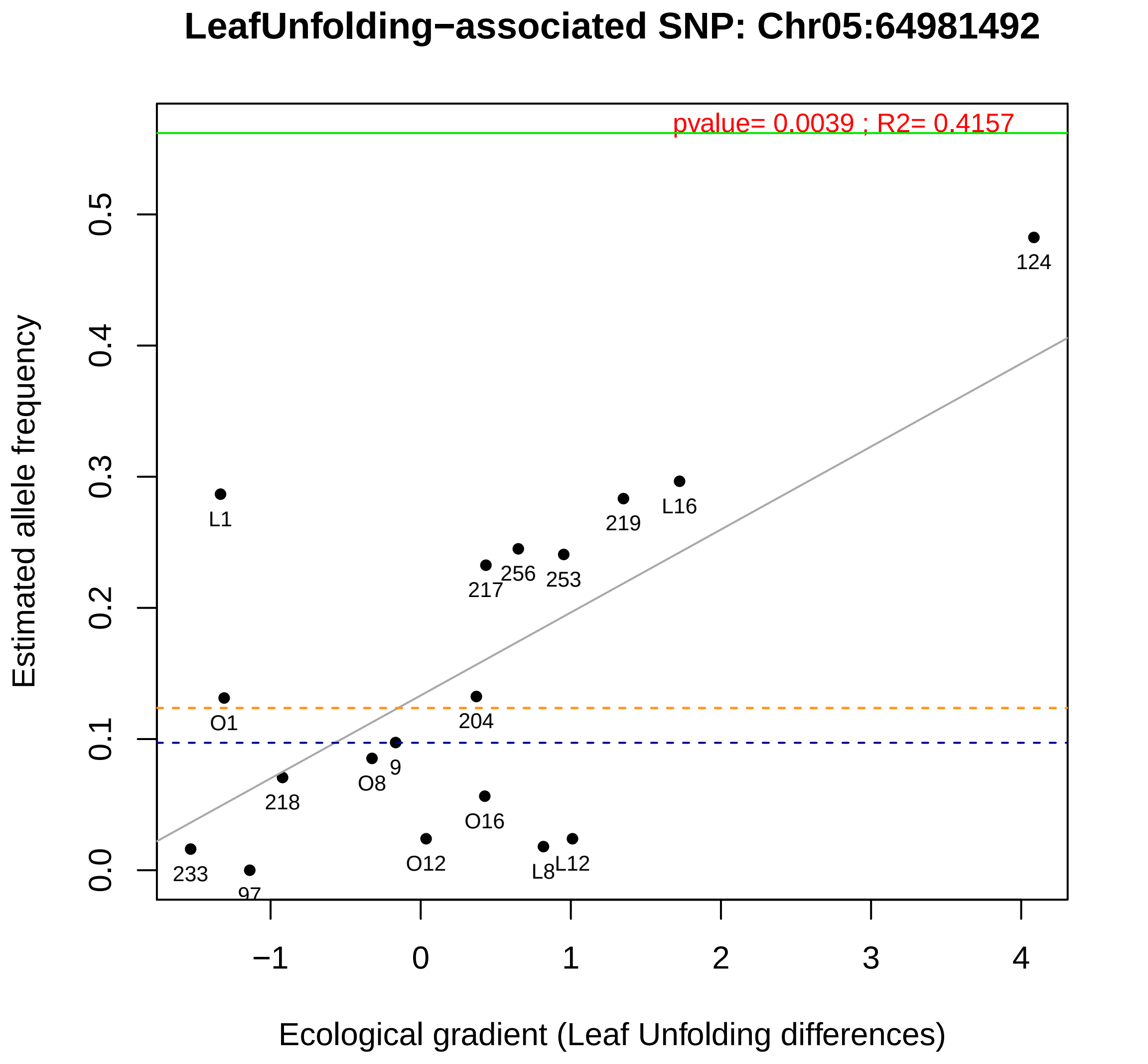

LeafUnfolding−associated SNP: Chr05:64981492
pvalue= 0.0039 ; R2= 0.4157
Estimated allele frequency
0.0
0.1
0.2
0.3
0.4
0.5
−1
0
1
2
3
4
Ecological gradient (Leaf Unfolding differences)
124
L1
219
L16
256
217
253
204
O1
9
O8
218
O16
O12
L8
L12
233
97

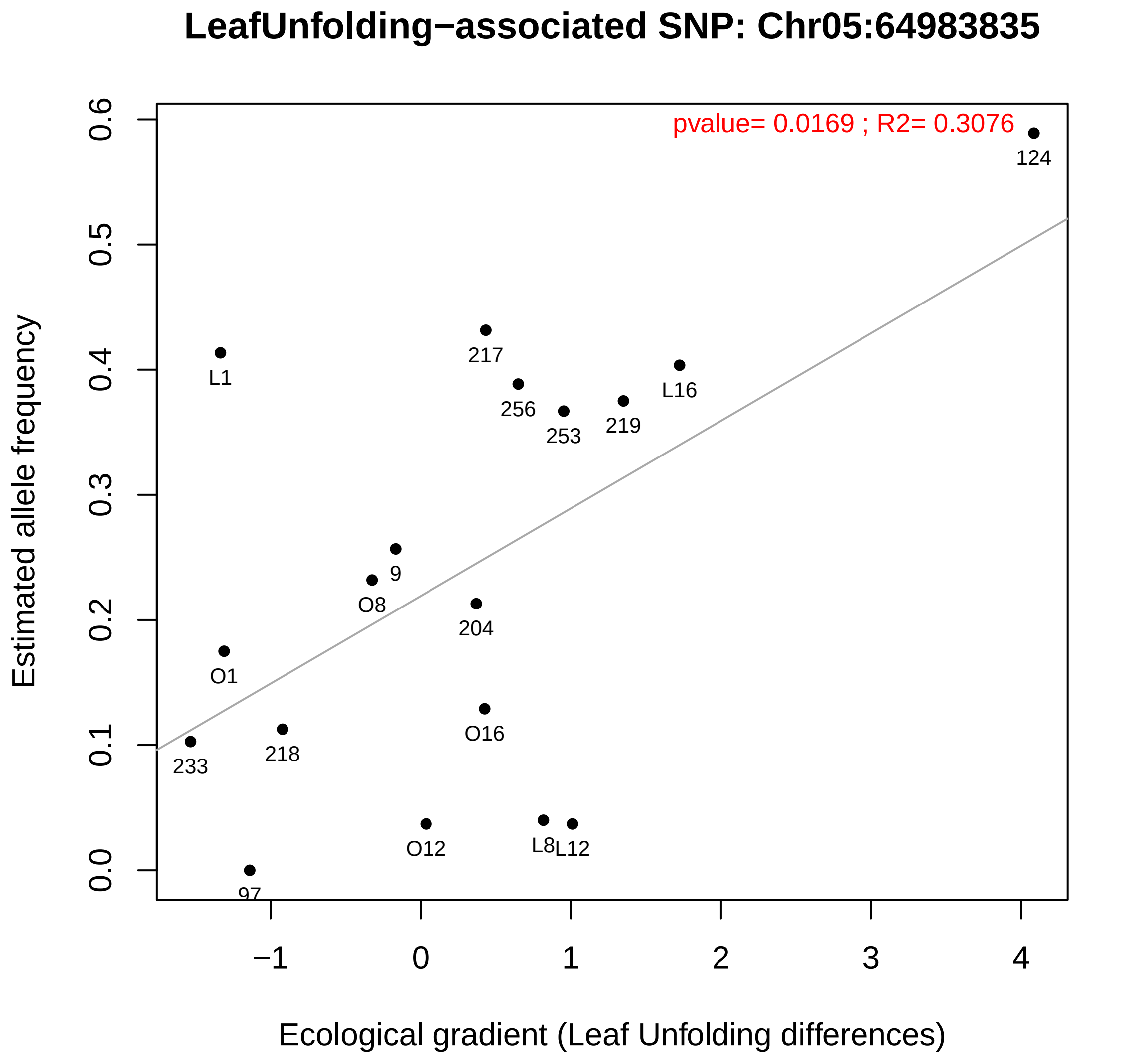
LeafUnfolding−associated SNP: Chr05:64983835
pvalue= 0.0169 ; R2= 0.3076
Estimated allele frequency
Ecological gradient (Leaf Unfolding differences)
0.0
0.1
0.2
0.3
0.4
0.5
0.6
−1
0
1
2
3
4
124
217
L1
256
L16
253
219
9
O8
204
O1
O16
233
218
O12
L8
L12
97

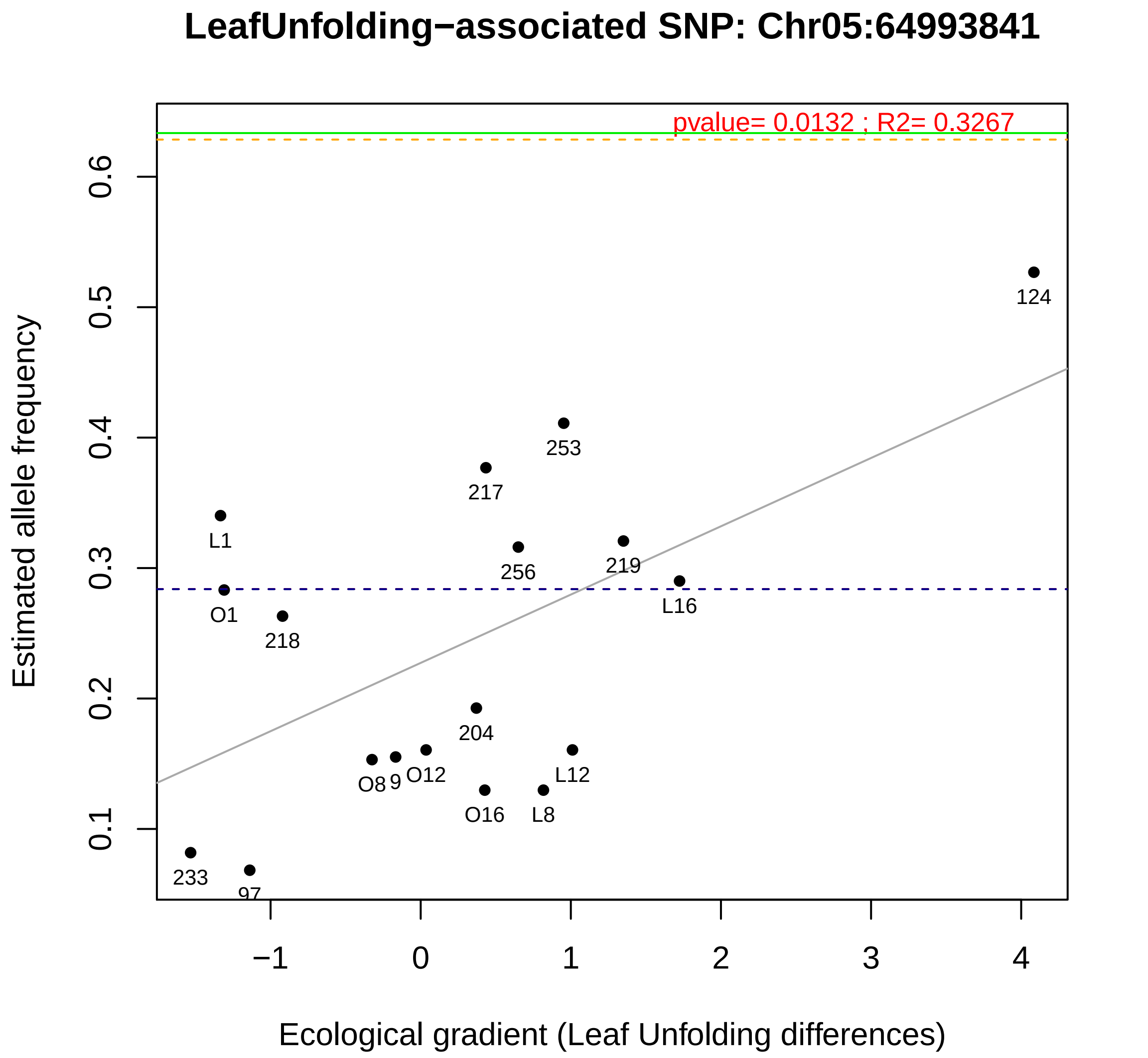

LeafUnfolding−associated SNP: Chr05:64993841
pvalue= 0.0132 ; R2= 0.3267
Estimated allele frequency
0.1
0.2
0.3
0.4
0.5
0.6
Ecological gradient (Leaf Unfolding differences)
−1
0
1
2
3
4
124
253
217
L1
256
219
L16
O1
218
204
O8
9
O12
L12
O16
L8
233
97

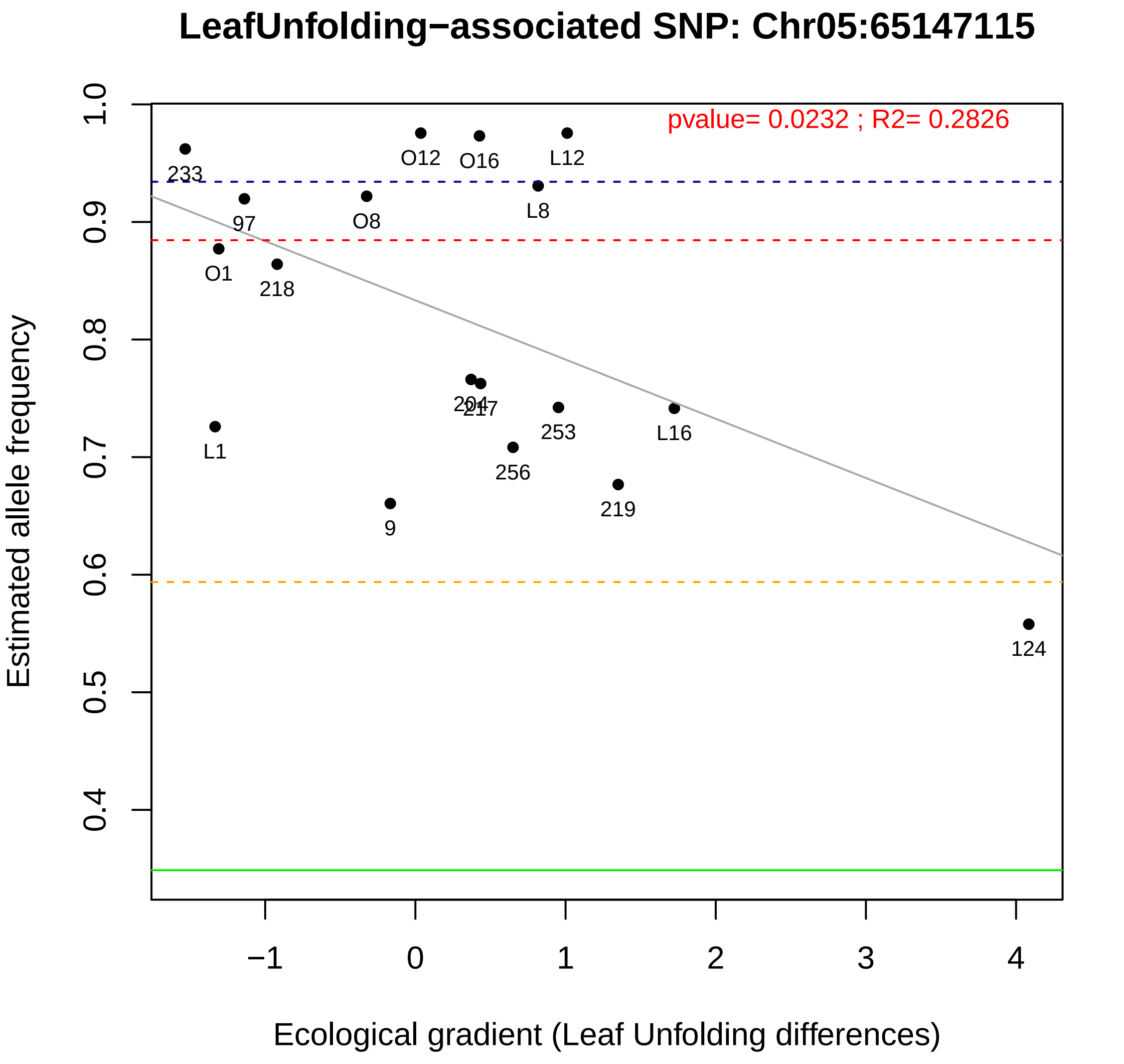
LeafUnfolding−associated SNP: Chr05:65147115
pvalue= 0.0232 ; R2= 0.2826
Estimated allele frequency
Ecological gradient (Leaf Unfolding differences)
1.0
0.9
0.8
0.7
0.6
0.5
0.4
−1
0
1
2
3
4
233
O12
O16
L12
97
O8
L8
O1
218
204
217
253
L16
L1
256
219
9
124

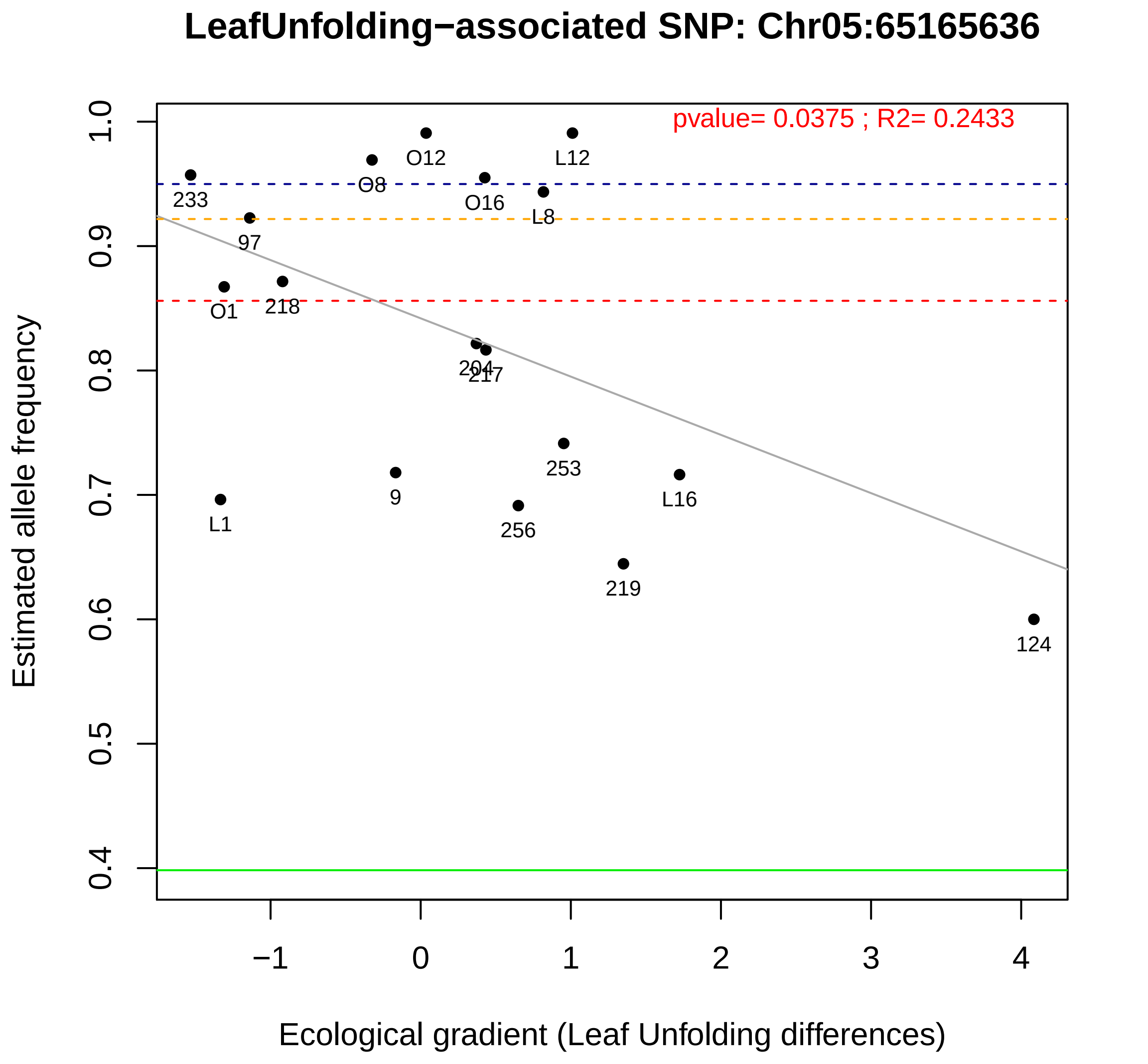

LeafUnfolding−associated SNP: Chr05:65165636
pvalue= 0.0375 ; R2= 0.2433
Estimated allele frequency
Ecological gradient (Leaf Unfolding differences)
0.4
0.5
0.6
0.7
0.8
0.9
1.0
−1
0
1
2
3
4
233
97
O1
218
L1
O8
O12
9
O16
204
217
256
253
L12
L8
219
L16
124

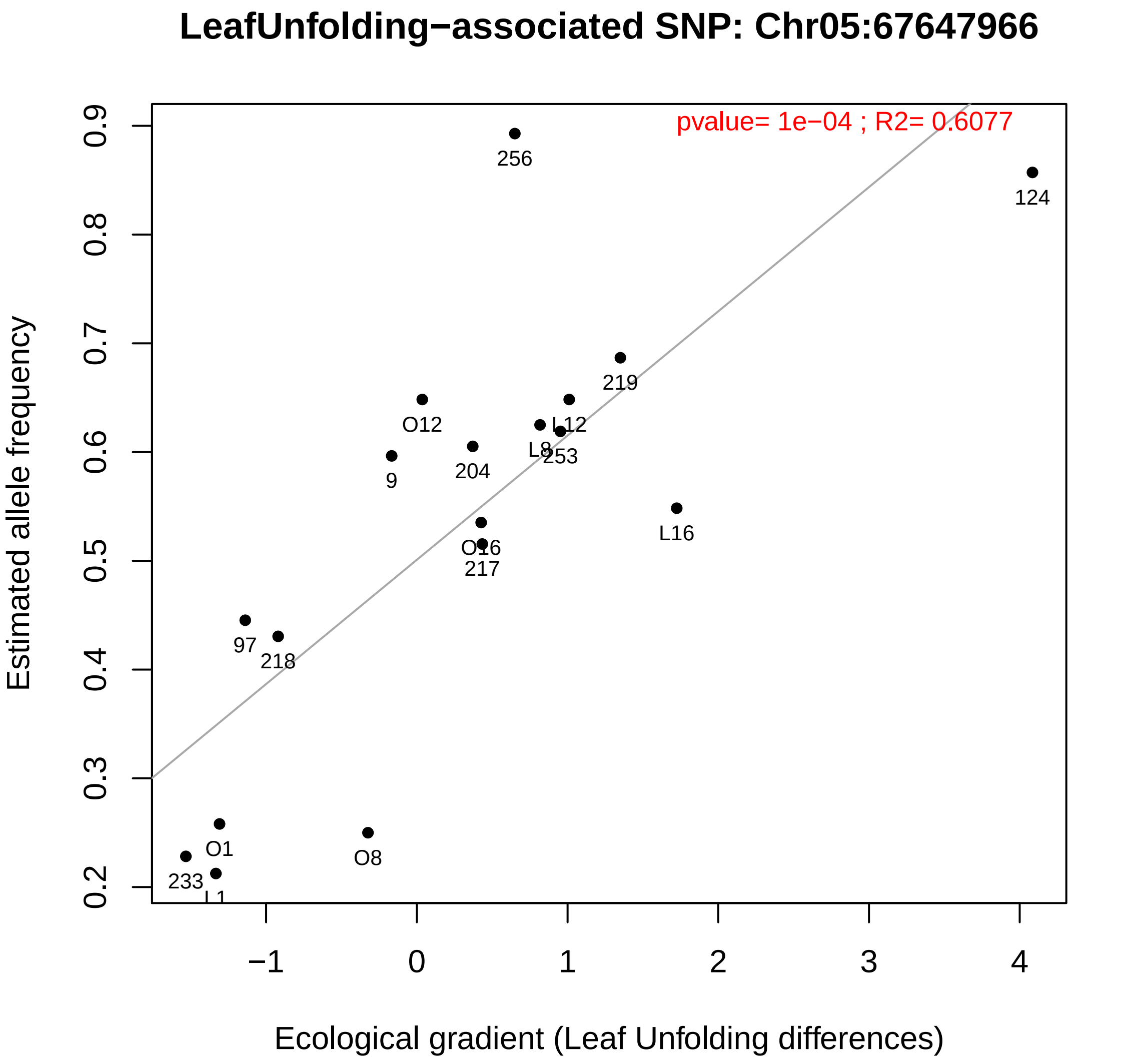
LeafUnfolding−associated SNP: Chr05:67647966
pvalue= 1e−04 ; R2= 0.6077
256
124
219
O12
L12
L8
253
204
9
L16
O16
217
97
218
O1
233
L1
O8
0.2
0.3
0.4
0.5
0.6
0.7
0.8
0.9
−1
0
1
2
3
4
Estimated allele frequency
Ecological gradient (Leaf Unfolding differences)

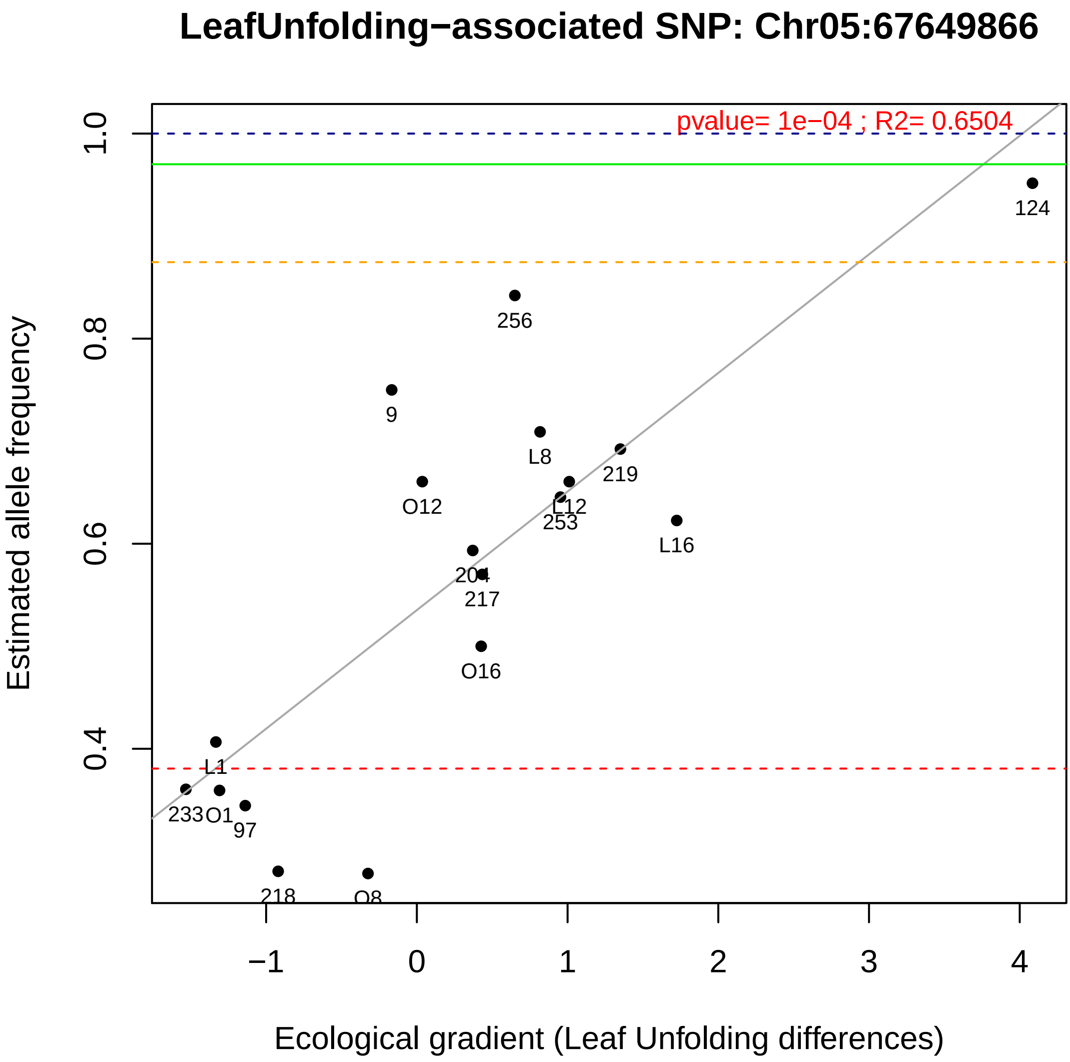

LeafUnfolding−associated SNP: Chr05:67649866
pvalue= 1e−04 ; R2= 0.6504
Estimated allele frequency
0.4
0.6
0.8
1.0
−1
0
1
2
3
4
Ecological gradient (Leaf Unfolding differences)
124
256
9
L8
219
O12
L12
253
L16
204
217
O16
L1
233
O1
97
218
O8

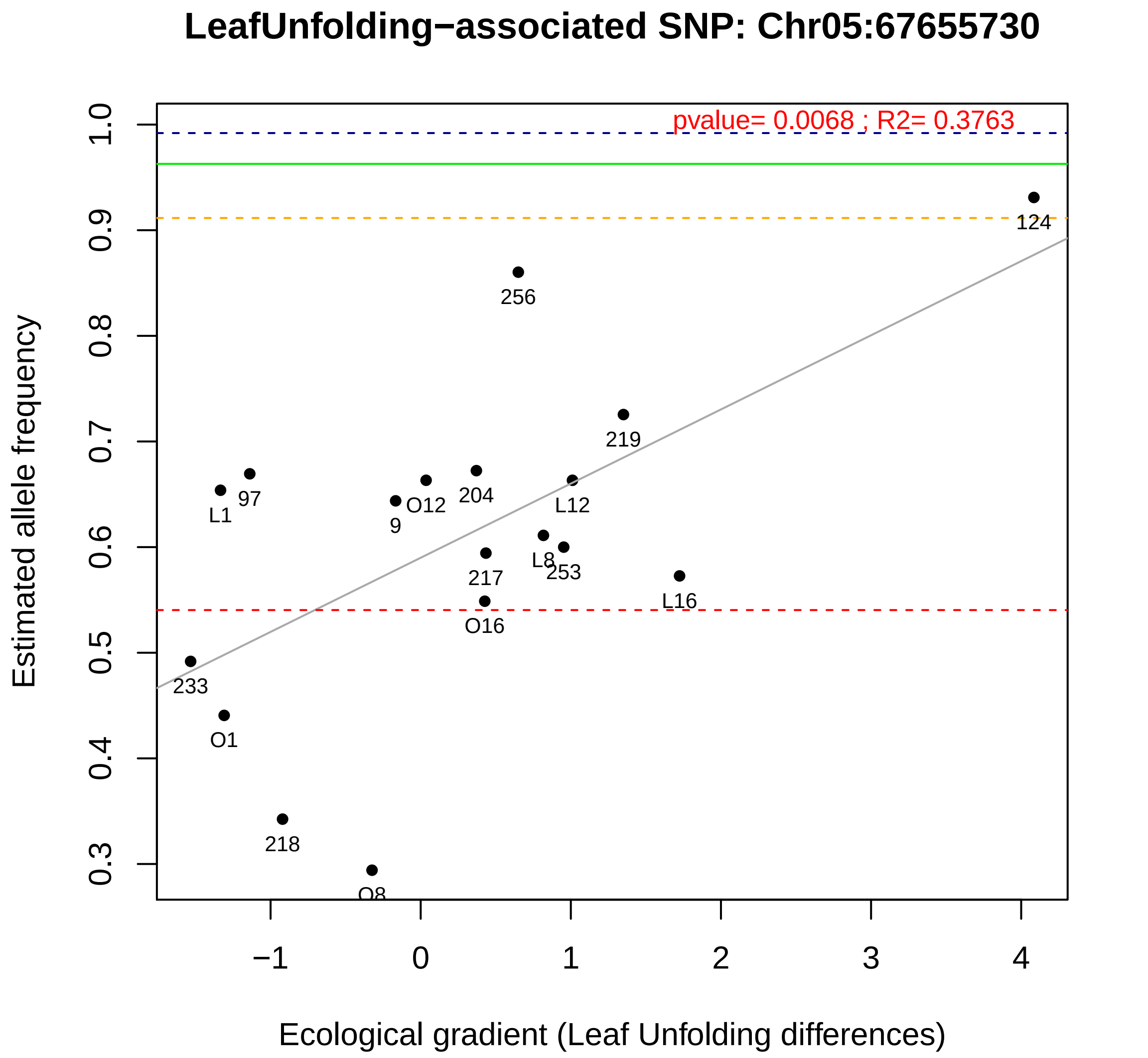

LeafUnfolding−associated SNP: Chr05:67655730
pvalue= 0.0068 ; R2= 0.3763
Estimated allele frequency
Ecological gradient (Leaf Unfolding differences)
0.3
0.4
0.5
0.6
0.7
0.8
0.9
1.0
−1
0
1
2
3
4
124
256
219
204
O12
9
97
L1
L12
L8
253
217
L16
O16
233
O1
218
O8

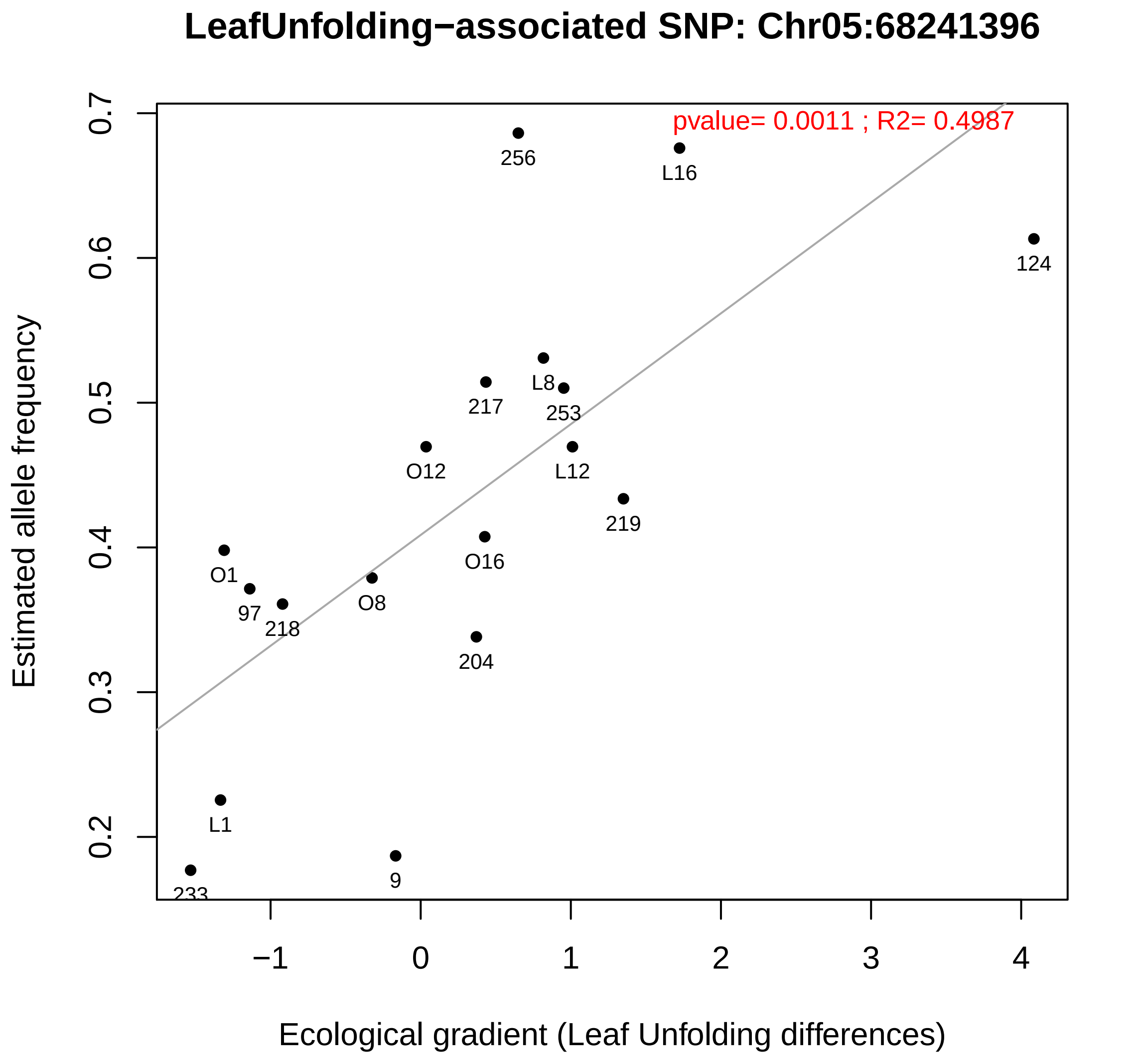
LeafUnfolding−associated SNP: Chr05:68241396
pvalue= 0.0011 ; R2= 0.4987
Estimated allele frequency
Ecological gradient (Leaf Unfolding differences)
0.2
0.3
0.4
0.5
0.6
0.7
−1
0
1
2
3
4
256
L16
124
L8
217
253
O12
L12
219
O16
O1
97
218
O8
204
L1
233
9

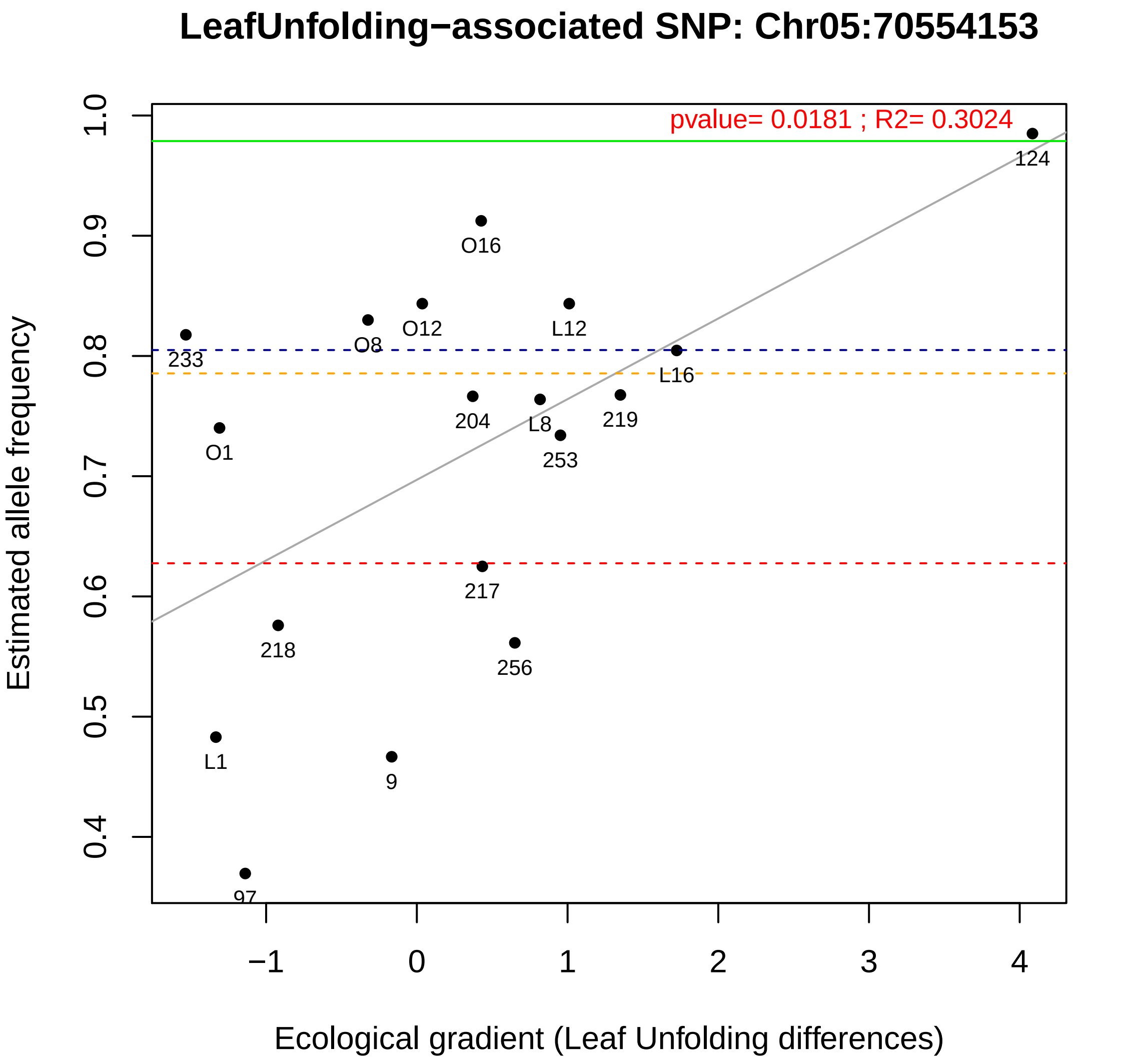
LeafUnfolding−associated SNP: Chr05:70554153
pvalue= 0.0181 ; R2= 0.3024
Estimated allele frequency
Ecological gradient (Leaf Unfolding differences)
0.4
0.5
0.6
0.7
0.8
0.9
1.0
−1
0
1
2
3
4
124
O16
O12
O8
233
L12
L16
204
L8
219
253
O1
217
218
256
L1
9
97

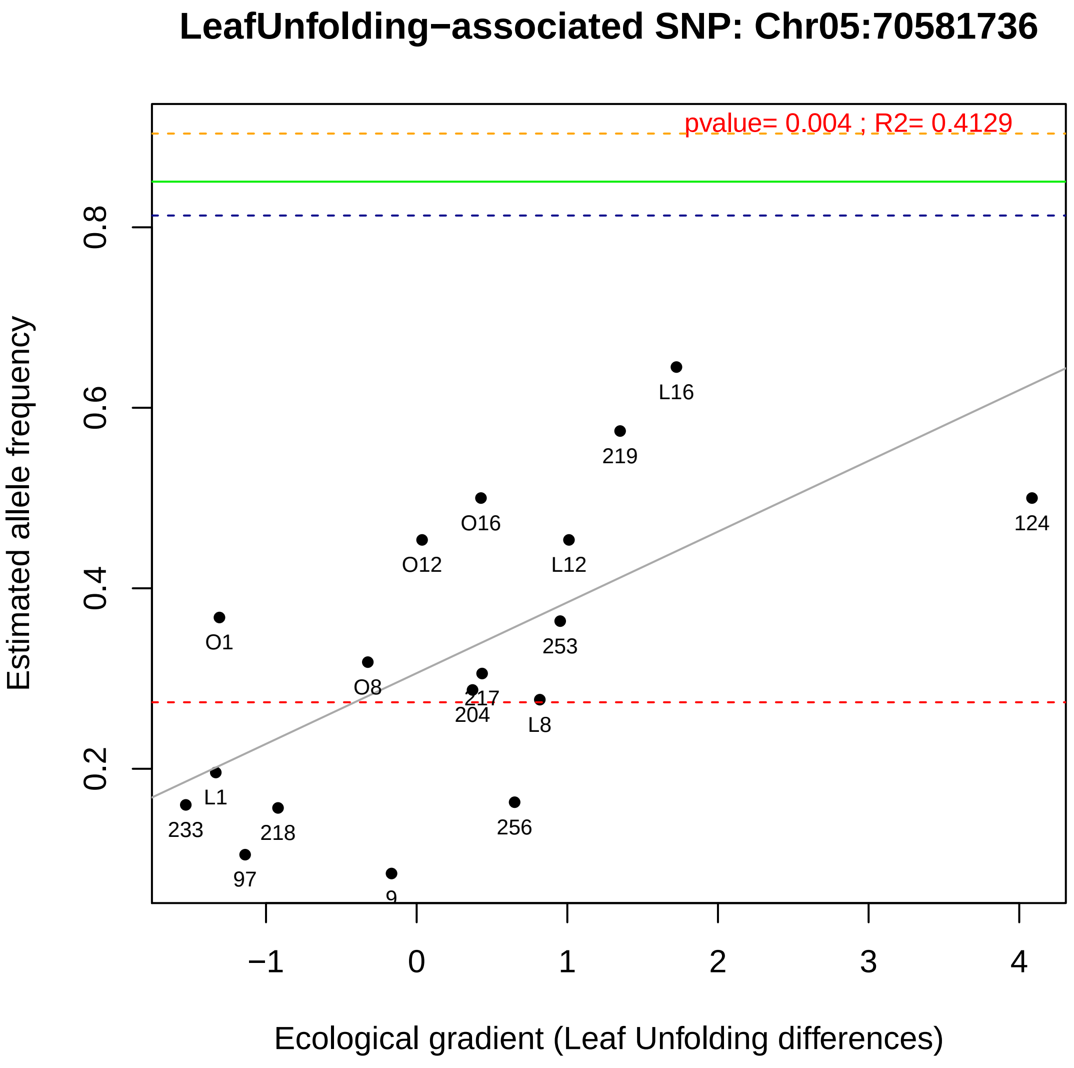

LeafUnfolding−associated SNP: Chr05:70581736
pvalue= 0.004 ; R2= 0.4129
Estimated allele frequency
0.2
0.4
0.6
0.8
−1
0
1
2
3
4
Ecological gradient (Leaf Unfolding differences)
L16
219
O16
124
O12
L12
O1
253
O8
217
204
L8
L1
233
218
256
97
9

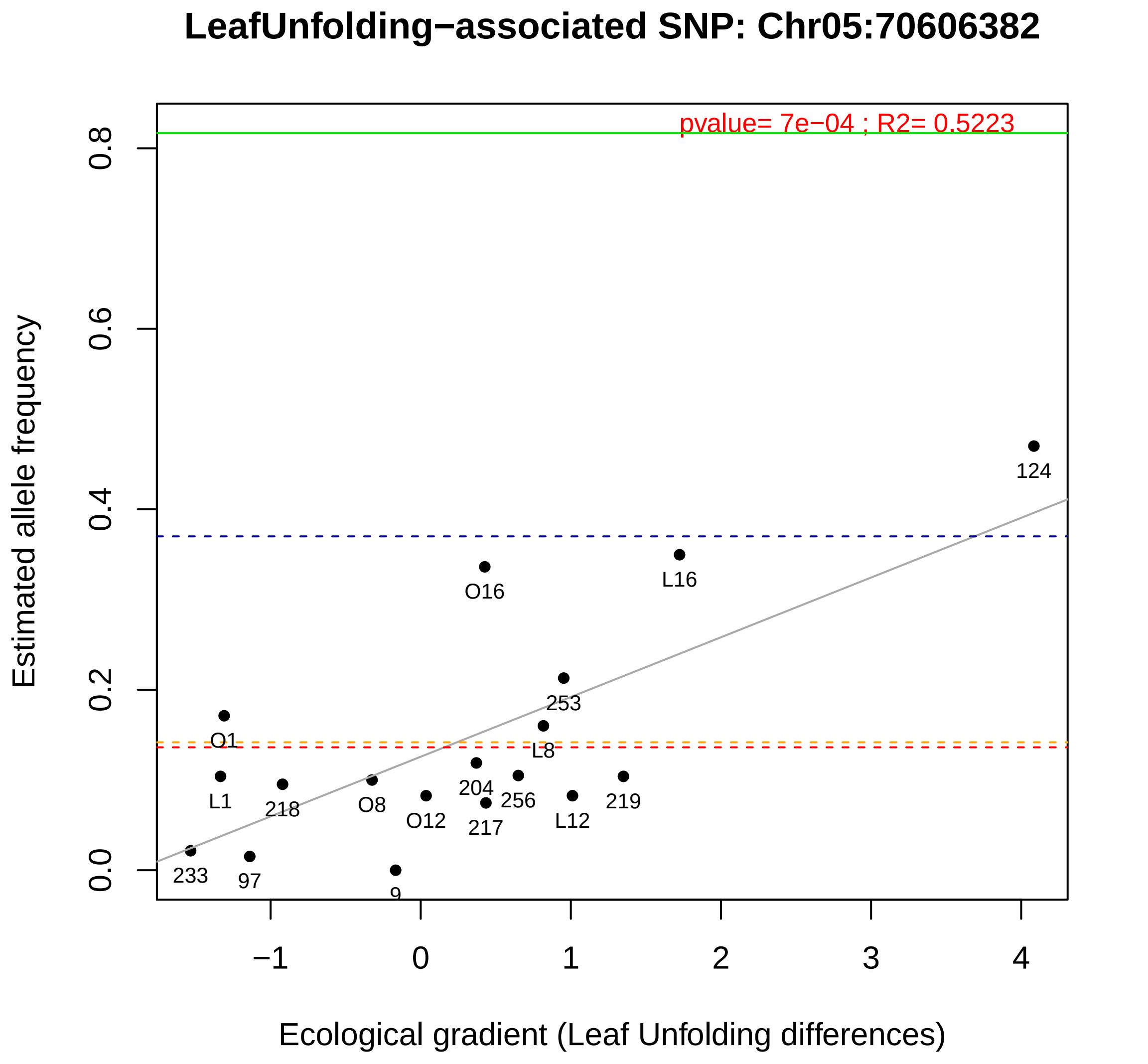
LeafUnfolding−associated SNP: Chr05:70606382
pvalue= 7e−04 ; R2= 0.5223
Estimated allele frequency
Ecological gradient (Leaf Unfolding differences)
0.0
0.2
0.4
0.6
0.8
−1
0
1
2
3
4
124
O16
L16
253
O1
L8
L1
218
O8
204
256
O12
217
L12
219
233
97
9

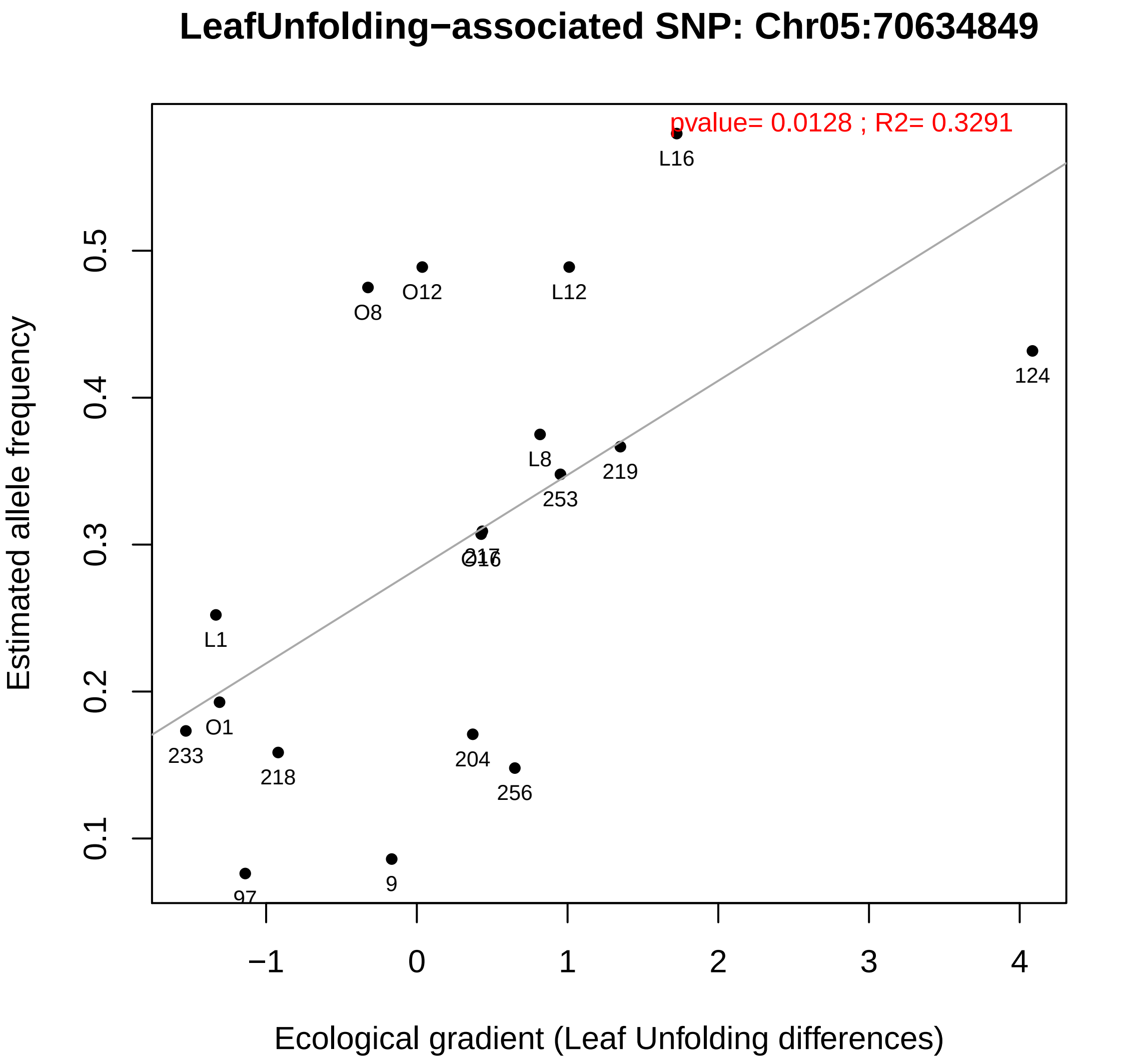
LeafUnfolding−associated SNP: Chr05:70634849
pvalue= 0.0128 ; R2= 0.3291
Estimated allele frequency
0.1
0.2
0.3
0.4
0.5
−1
0
1
2
3
4
Ecological gradient (Leaf Unfolding differences)
L16
O12
O8
L12
124
L8
219
253
217
O16
L1
O1
233
218
204
256
9
97

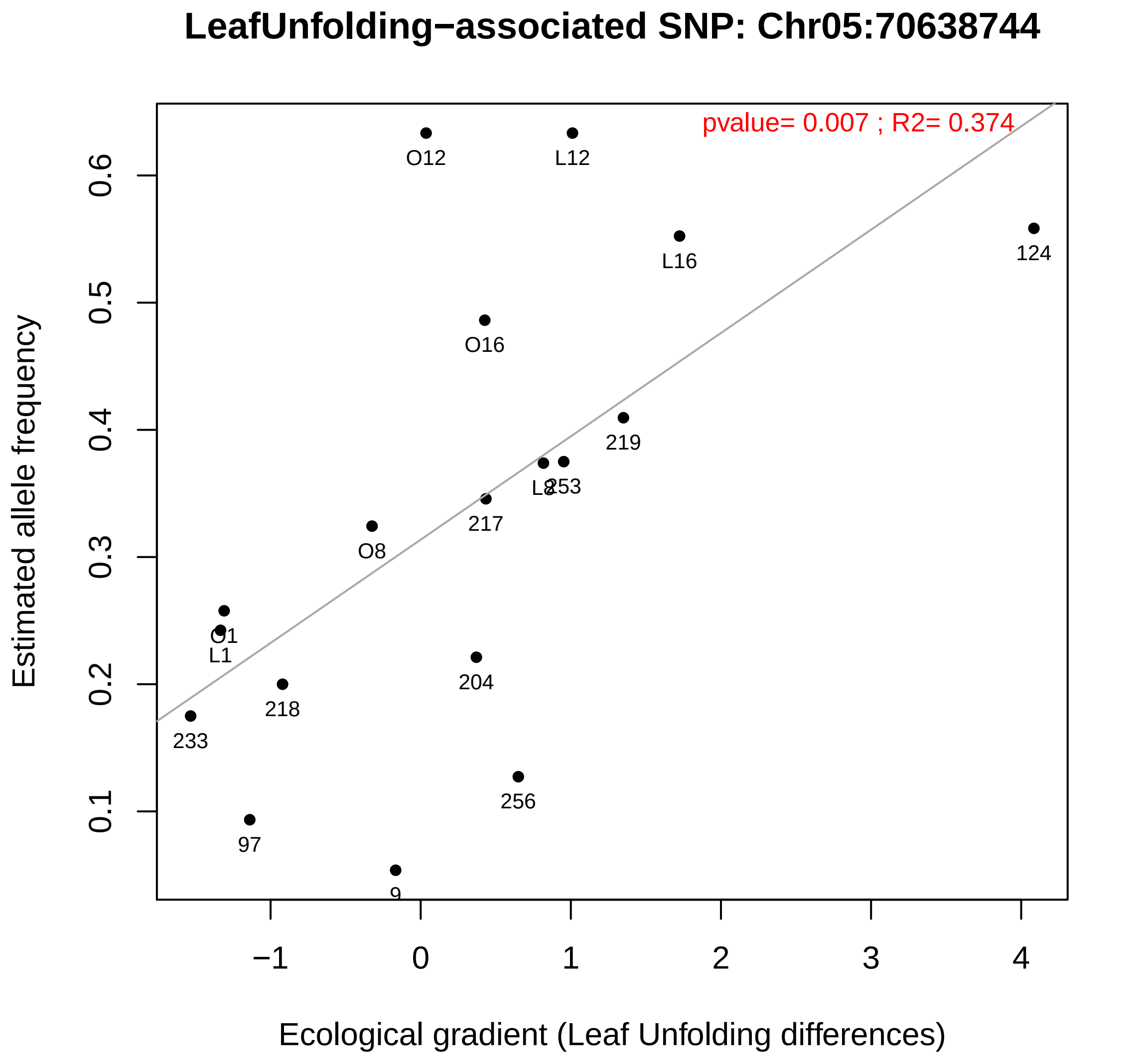
LeafUnfolding−associated SNP: Chr05:70638744
pvalue= 0.007 ; R2= 0.374
Estimated allele frequency
Ecological gradient (Leaf Unfolding differences)
0.1
0.2
0.3
0.4
0.5
0.6
−1
0
1
2
3
4
O12
L12
L16
124
O16
219
L8
253
217
O8
O1
L1
218
204
233
256
97
9

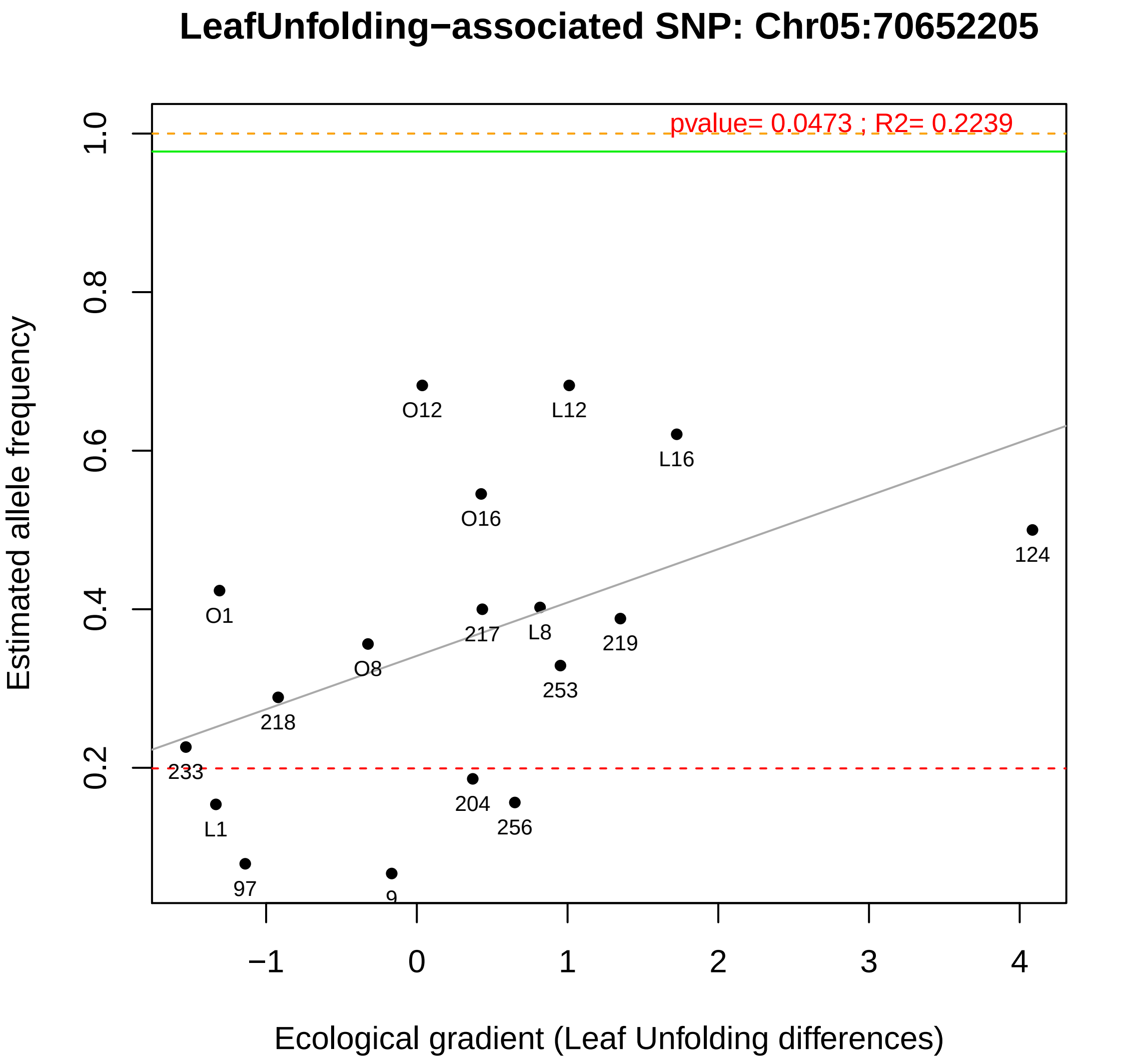

LeafUnfolding−associated SNP: Chr05:70652205
pvalue= 0.0473 ; R2= 0.2239
Estimated allele frequency
0.2
0.4
0.6
0.8
1.0
−1
0
1
2
3
4
Ecological gradient (Leaf Unfolding differences)
O12
L12
L16
O16
124
O1
217
L8
219
O8
253
218
233
204
256
L1
97
9

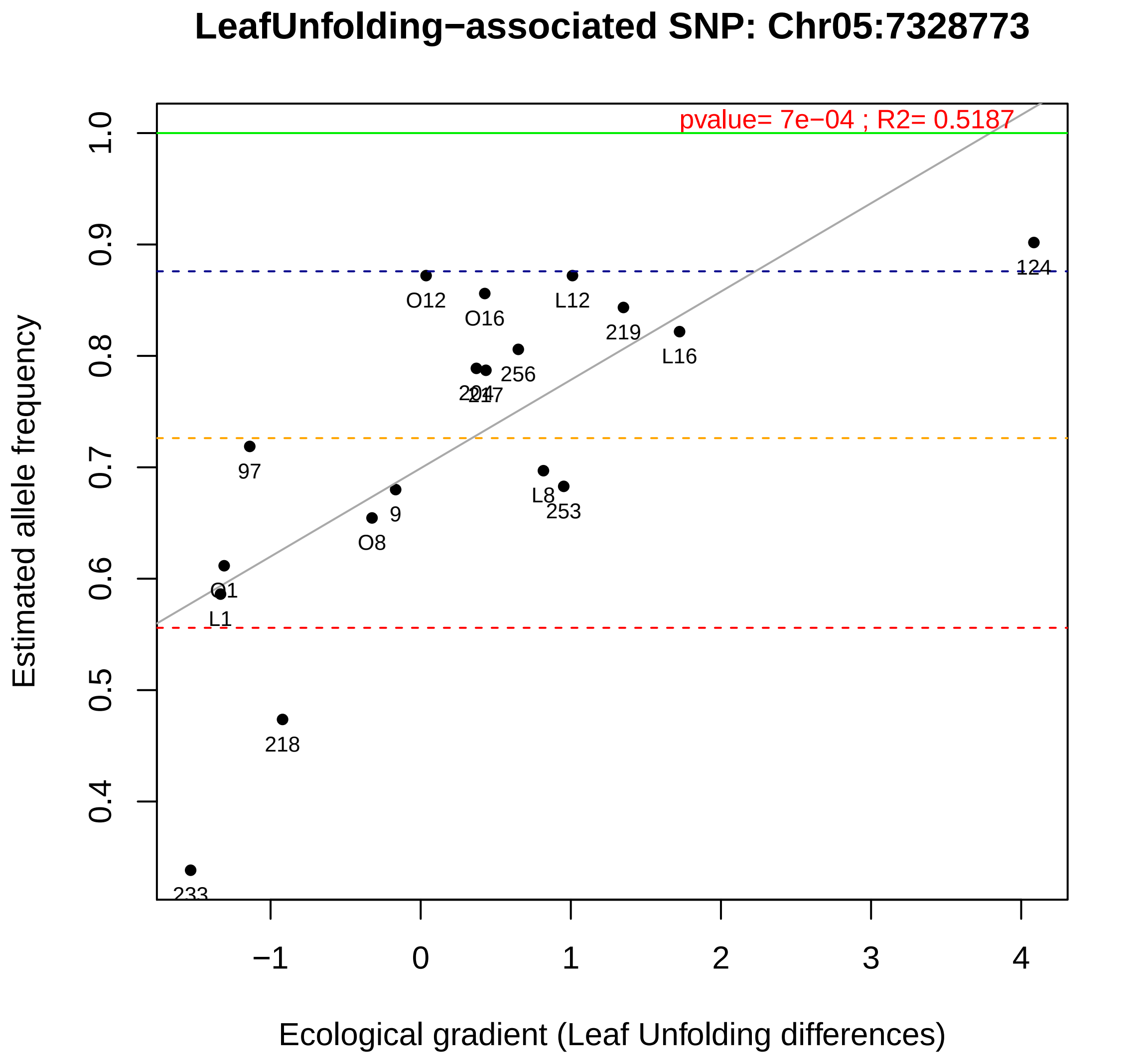
LeafUnfolding−associated SNP: Chr05:7328773
pvalue= 7e−04 ; R2= 0.5187
Estimated allele frequency
Ecological gradient (Leaf Unfolding differences)
0.4
0.5
0.6
0.7
0.8
0.9
1.0
−1
0
1
2
3
4
124
O12
L12
O16
219
L16
256
97
9
O8
O1
L1
L8
253
218
233

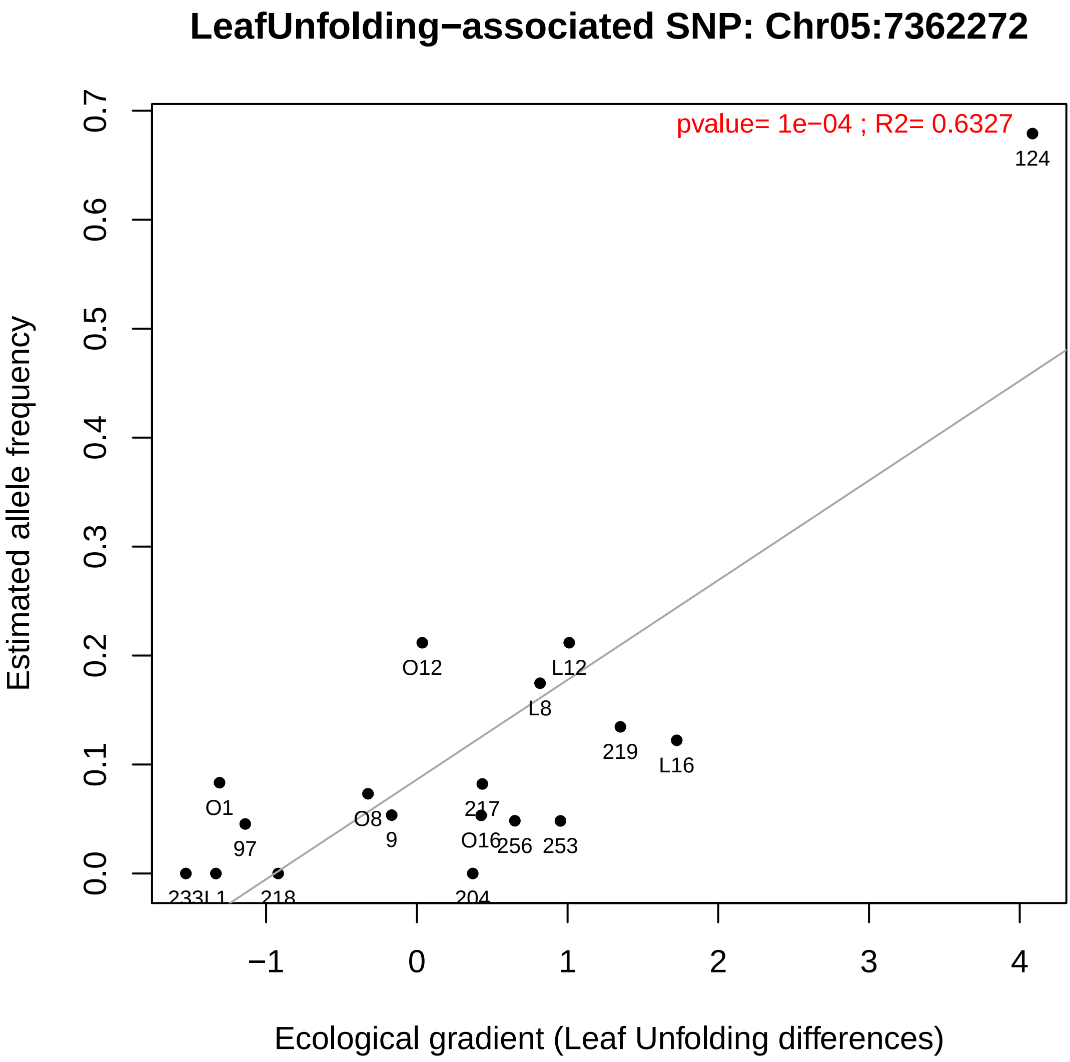

LeafUnfolding−associated SNP: Chr05:7362272
pvalue= 1e−04 ; R2= 0.6327
Estimated allele frequency
Ecological gradient (Leaf Unfolding differences)
0.0
0.1
0.2
0.3
0.4
0.5
0.6
0.7
−1
0
1
2
3
4
124
O12
L12
L8
219
L16
O1
97
O8
9
217
O16
256
253
233
L1
218
204

#### LeafUnfolding−associated SNP: Chr05:7395743

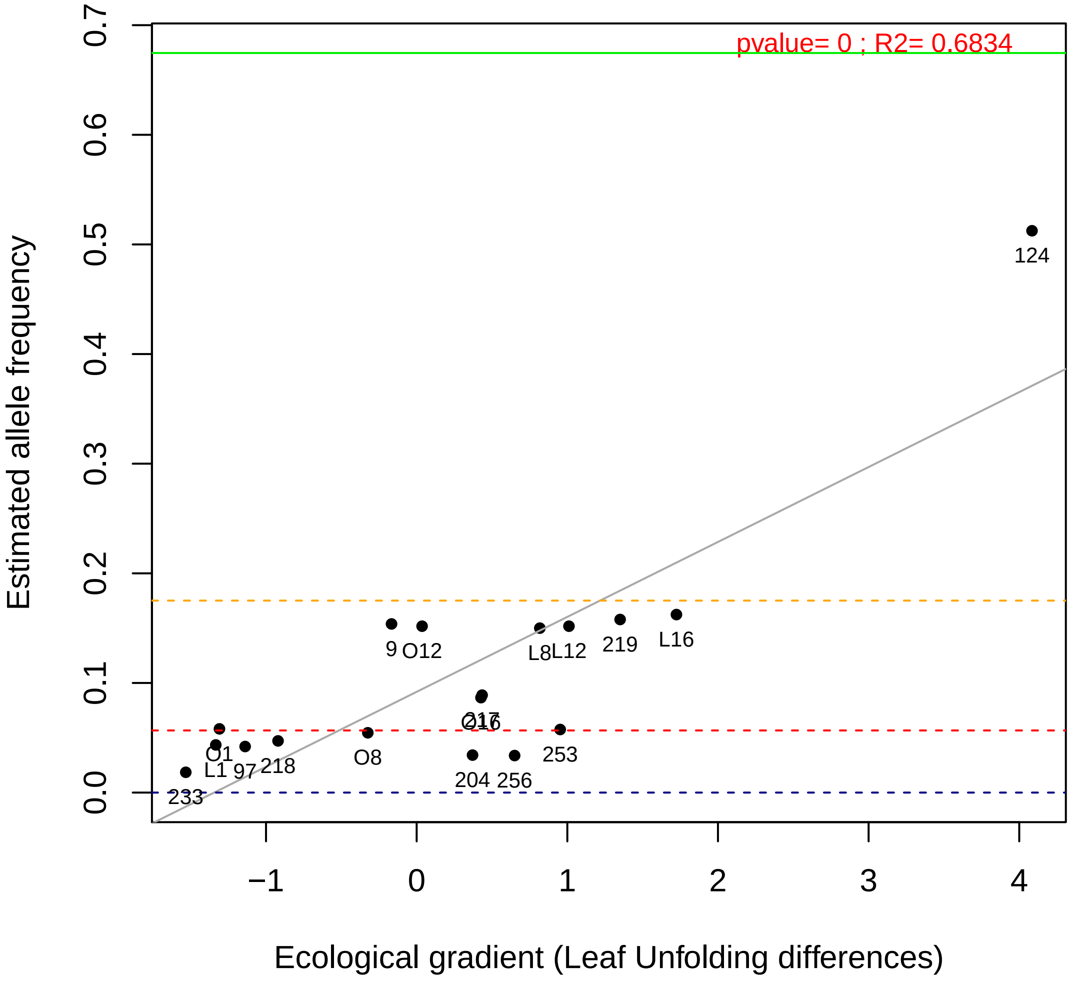

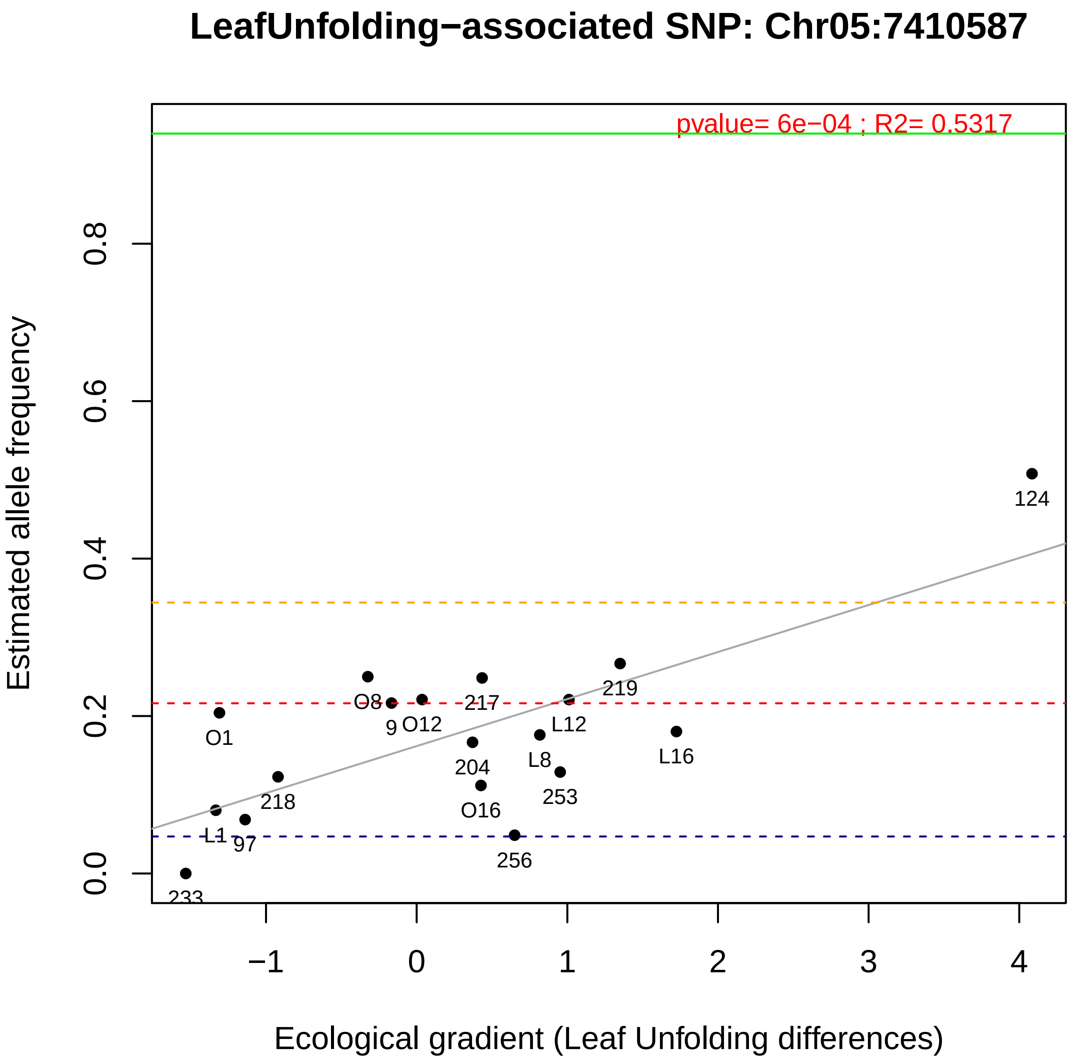
LeafUnfolding−associated SNP: Chr05:7410587
pvalue= 6e−04 ; R2= 0.5317
Estimated allele frequency
0.0
0.2
0.4
0.6
0.8
−1
0
1
2
3
4
Ecological gradient (Leaf Unfolding differences)
124
219
O8
217
O12
9
L12
O1
L16
204
L8
253
218
O16
L1
97
256
233

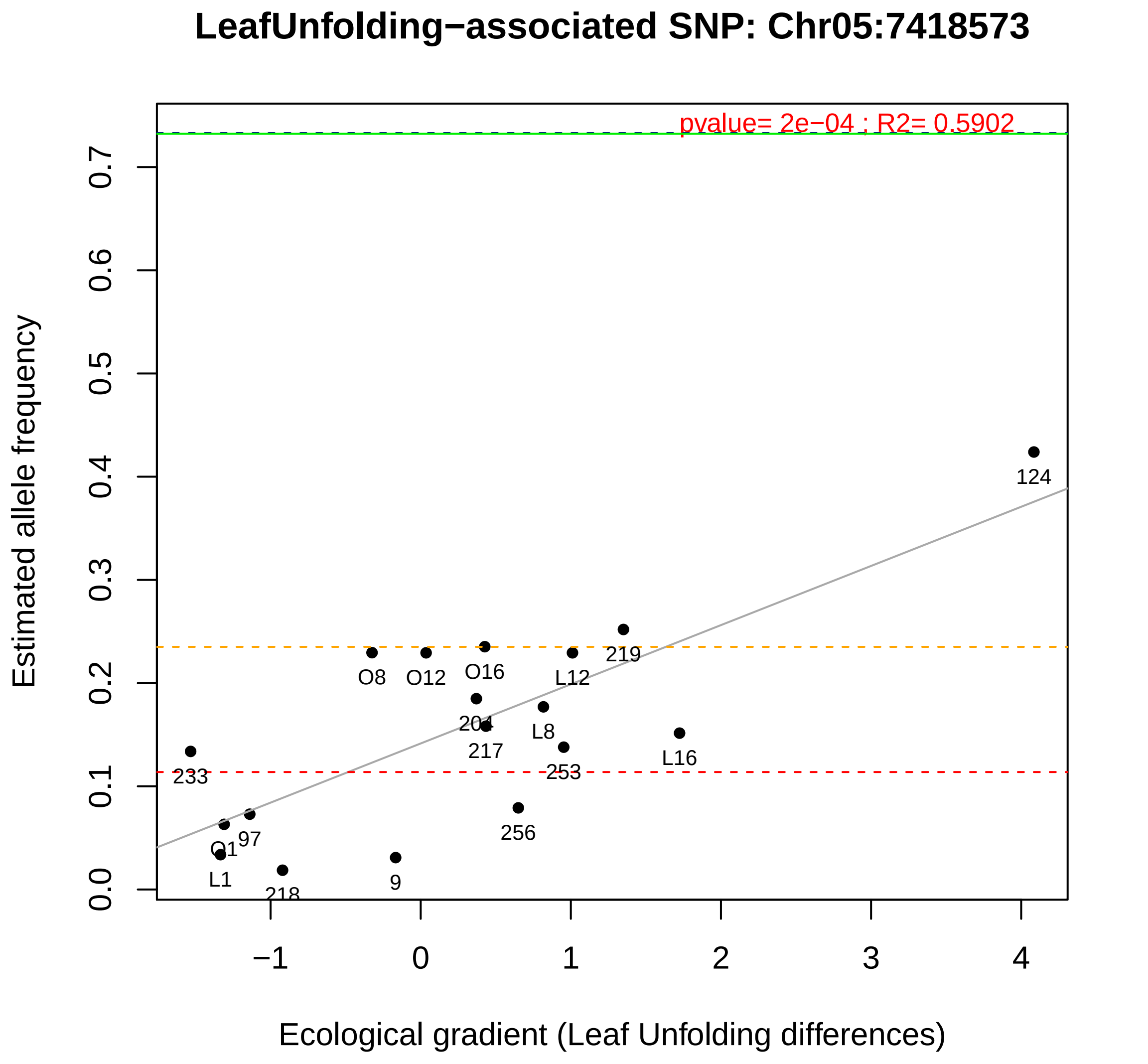

LeafUnfolding−associated SNP: Chr05:7418573
pvalue= 2e−04 ; R2= 0.5902
Estimated allele frequency
Ecological gradient (Leaf Unfolding differences)
0.0
0.1
0.2
0.3
0.4
0.5
0.6
0.7
−1
0
1
2
3
4
124
219
O8
O12
O16
L12
204
217
L8
253
L16
233
256
O1
97
L1
218
9

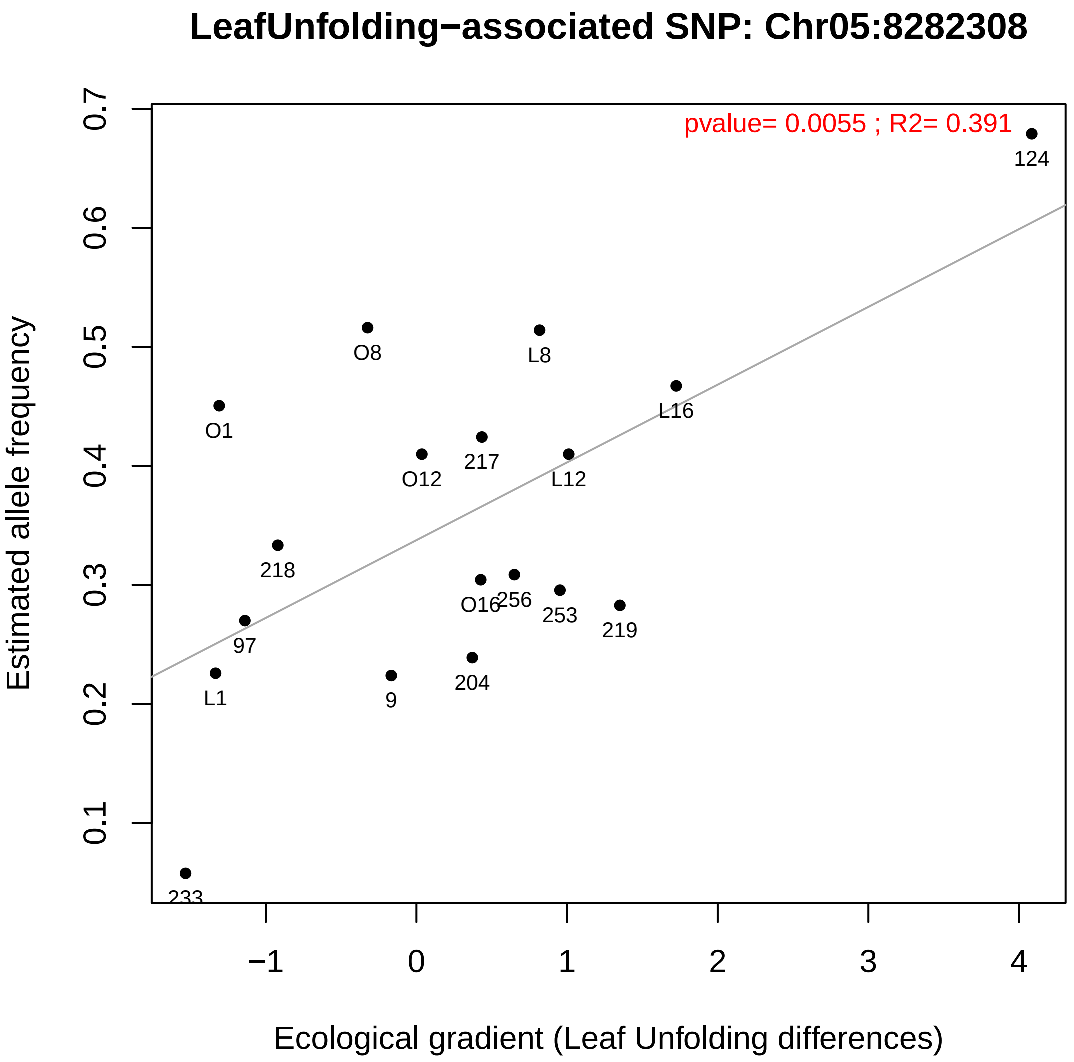

LeafUnfolding−associated SNP: Chr05:8282308
pvalue= 0.0055 ; R2= 0.391
Estimated allele frequency
Ecological gradient (Leaf Unfolding differences)
0.1
0.2
0.3
0.4
0.5
0.6
0.7
−1
0
1
2
3
4
124
O8
L8
L16
O1
217
O12
L12
218
O16
256
253
219
97
L1
9
204
233

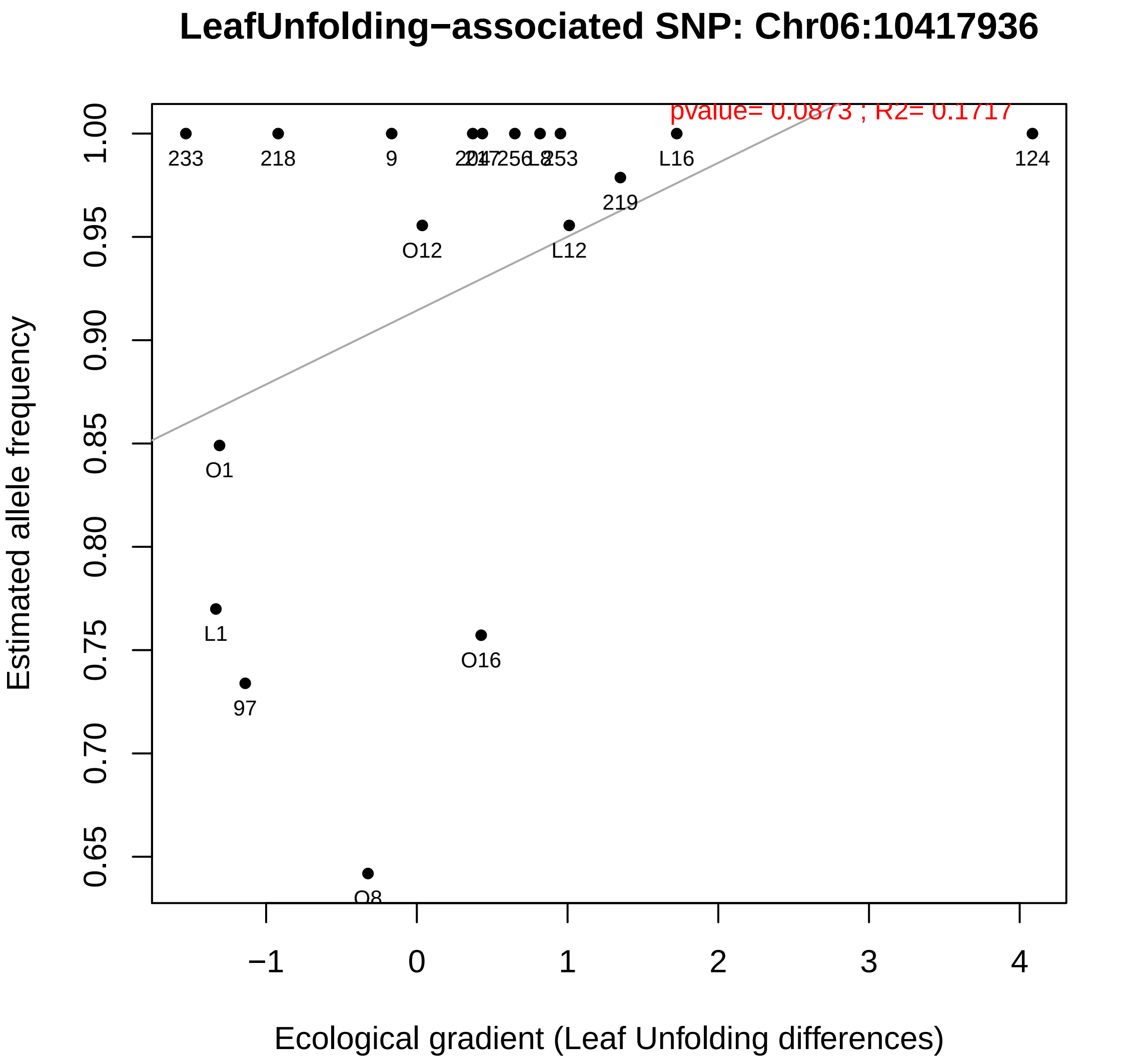

LeafUnfolding−associated SNP: Chr06:10417936
pvalue= 0.0873 ; R2= 0.1717
Estimated allele frequency
1.00
0.95
0.90
0.85
0.80
0.75
0.70
0.65
233
218
9
L16
124
219
O12
L12
O1
L1
O16
97
O8
−1
0
1
2
3
4
Ecological gradient (Leaf Unfolding differences)

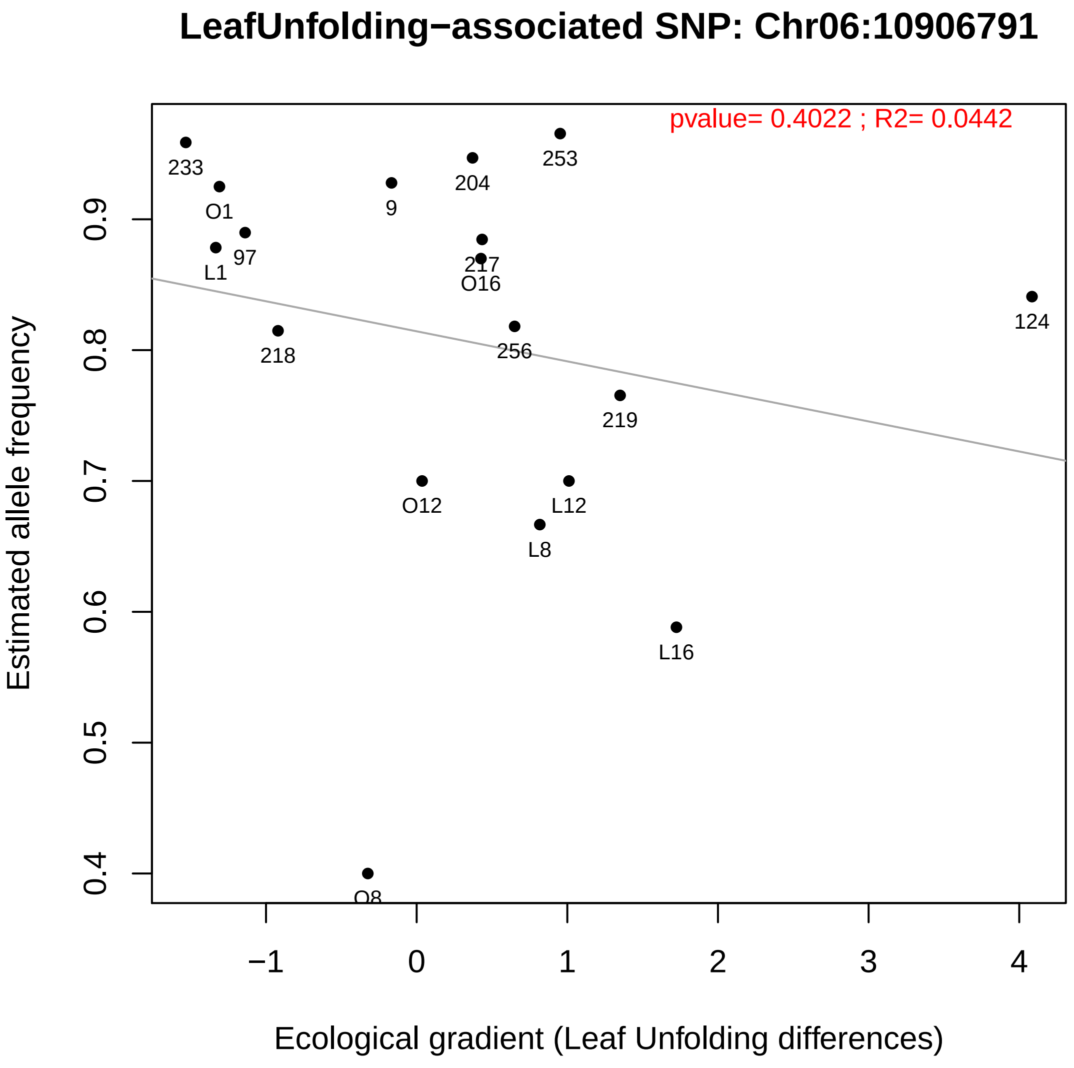
LeafUnfolding−associated SNP: Chr06:10906791
pvalue= 0.4022 ; R2= 0.0442
Estimated allele frequency
0.4
0.5
0.6
0.7
0.8
0.9
−1
0
1
2
3
4
Ecological gradient (Leaf Unfolding differences)
233
O1
97
L1
218
9
204
253
217
O16
256
124
219
O12
L12
L8
L16
O8

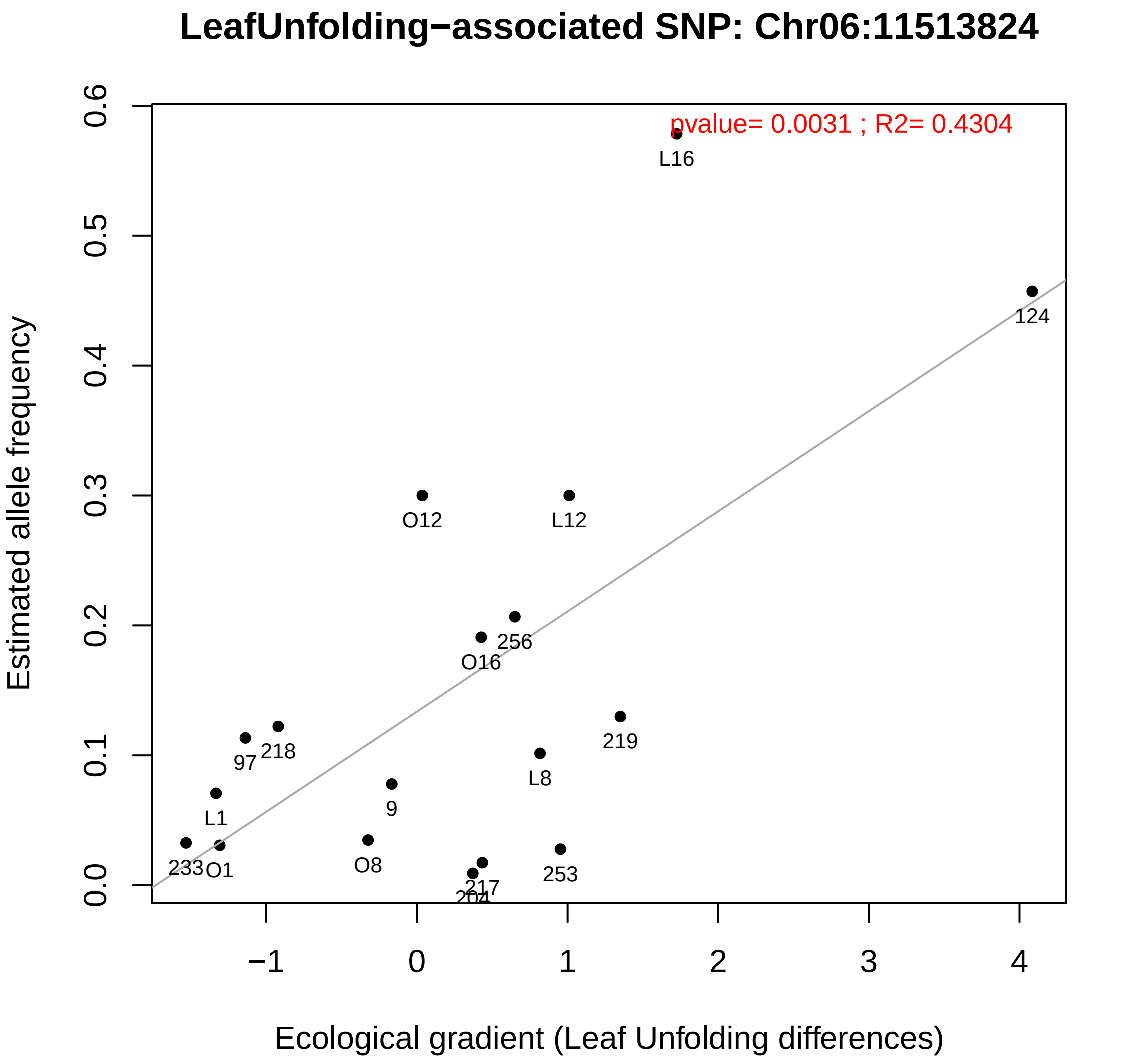
LeafUnfolding−associated SNP: Chr06:11513824
pvalue= 0.0031 ; R2= 0.4304
Estimated allele frequency
0.0
0.1
0.2
0.3
0.4
0.5
0.6
−1
0
1
2
3
4
Ecological gradient (Leaf Unfolding differences)
L16
124
O12
L12
256
O16
219
97
218
L8
L1
9
233
O1
O8
253
217
204

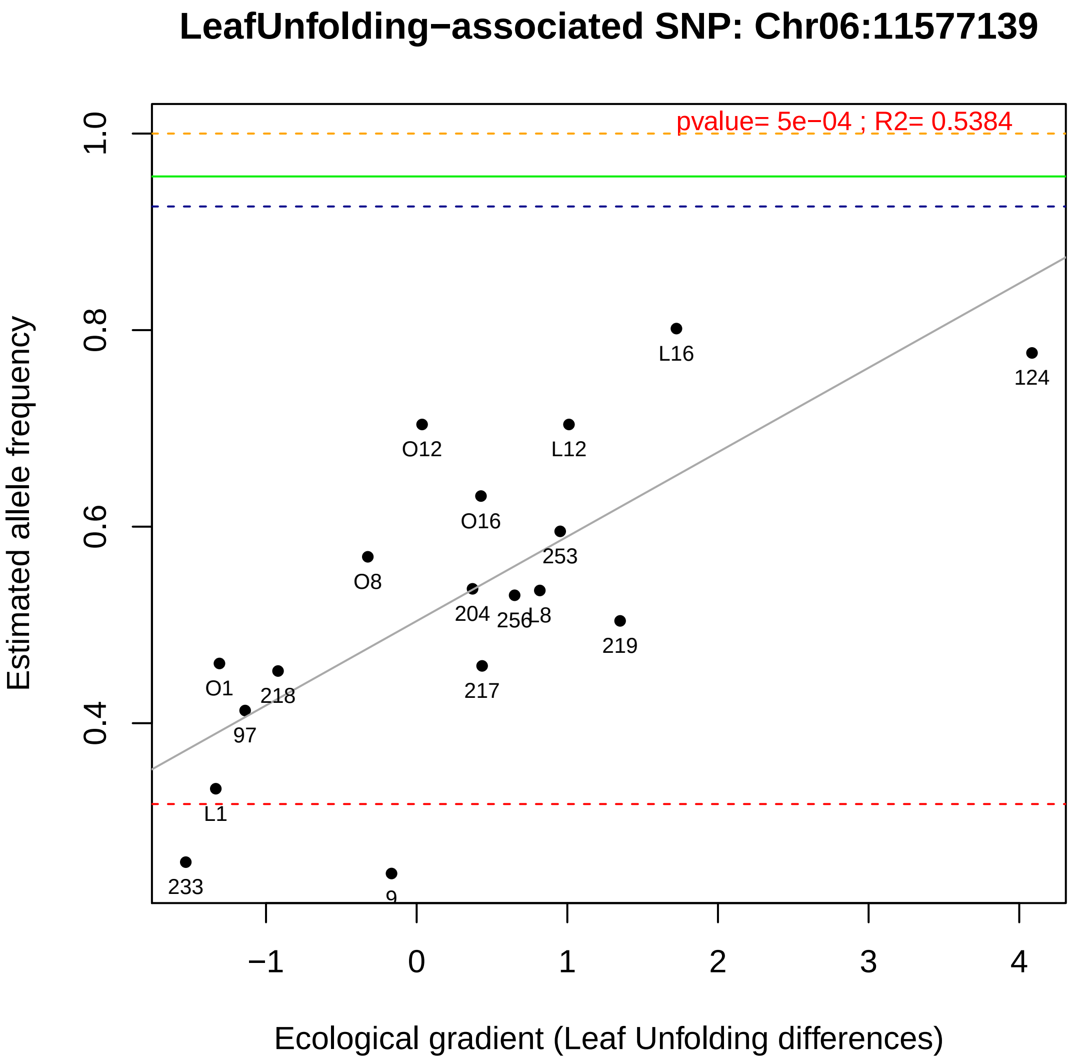

LeafUnfolding−associated SNP: Chr06:11577139
pvalue= 5e−04 ; R2= 0.5384
Estimated allele frequency
Ecological gradient (Leaf Unfolding differences)
1.0
0.8
0.6
0.4
−1
0
1
2
3
4
L16
124
O12
L12
O16
253
O8
204
256
L8
219
O1
218
217
97
L1
233
9

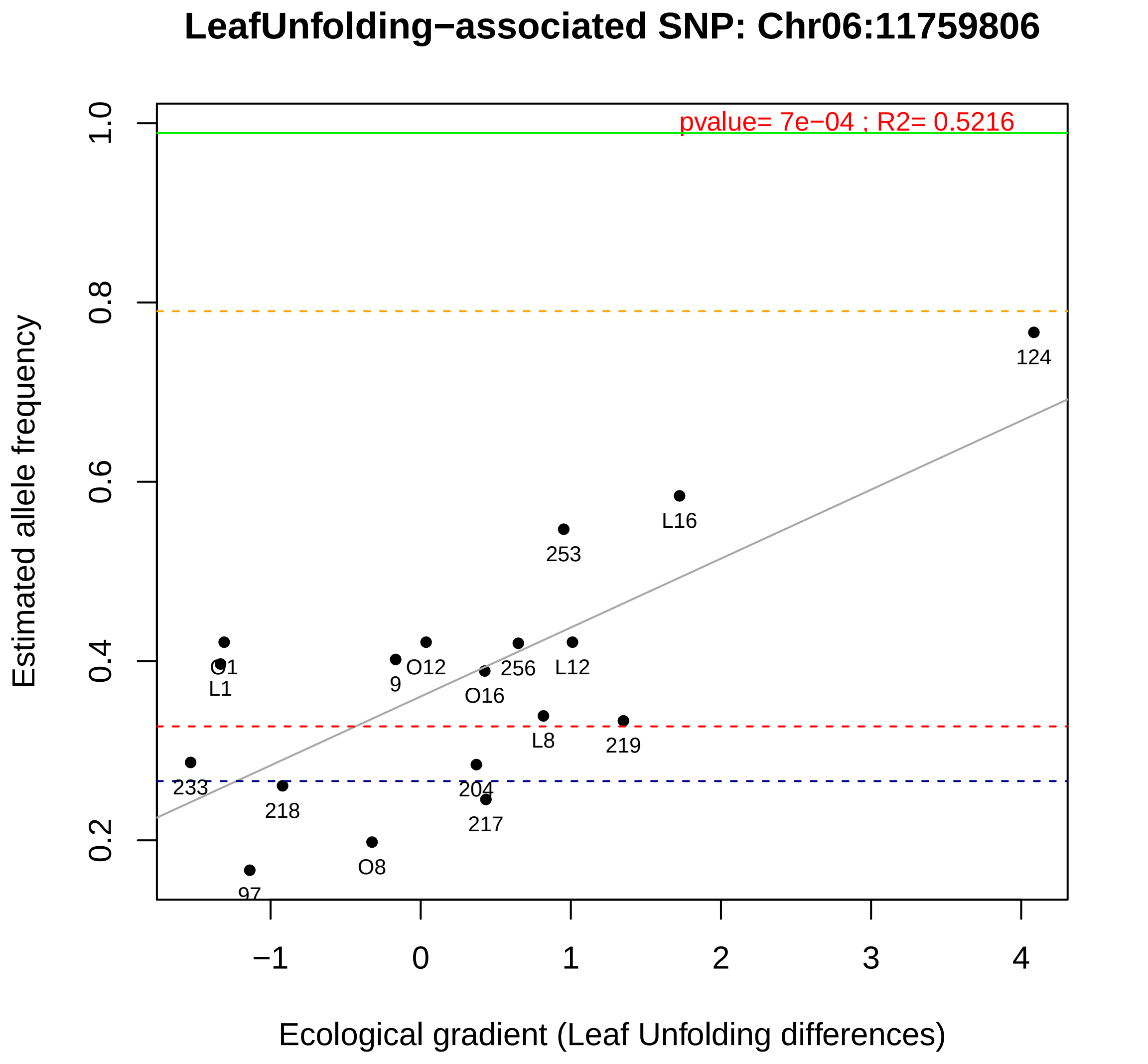

LeafUnfolding−associated SNP: Chr06:11759806
pvalue= 7e−04 ; R2= 0.5216
Estimated allele frequency
0.2
0.4
0.6
0.8
1.0
−1
0
1
2
3
4
Ecological gradient (Leaf Unfolding differences)
124
L16
253
O1
L1
9
O12
256
L12
O16
L8
219
233
218
204
217
O8
97

**LeafUnfolding−associated SNP: Chr06:11811775**

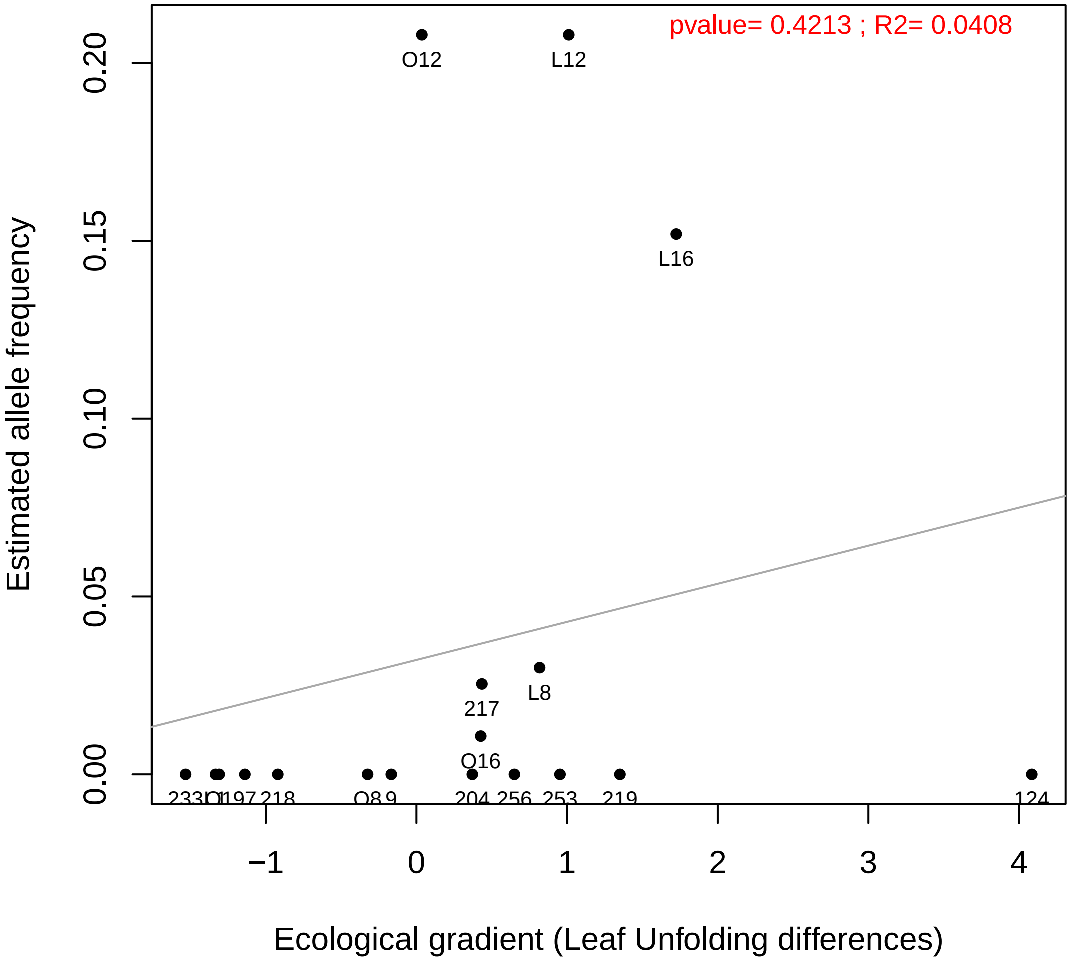

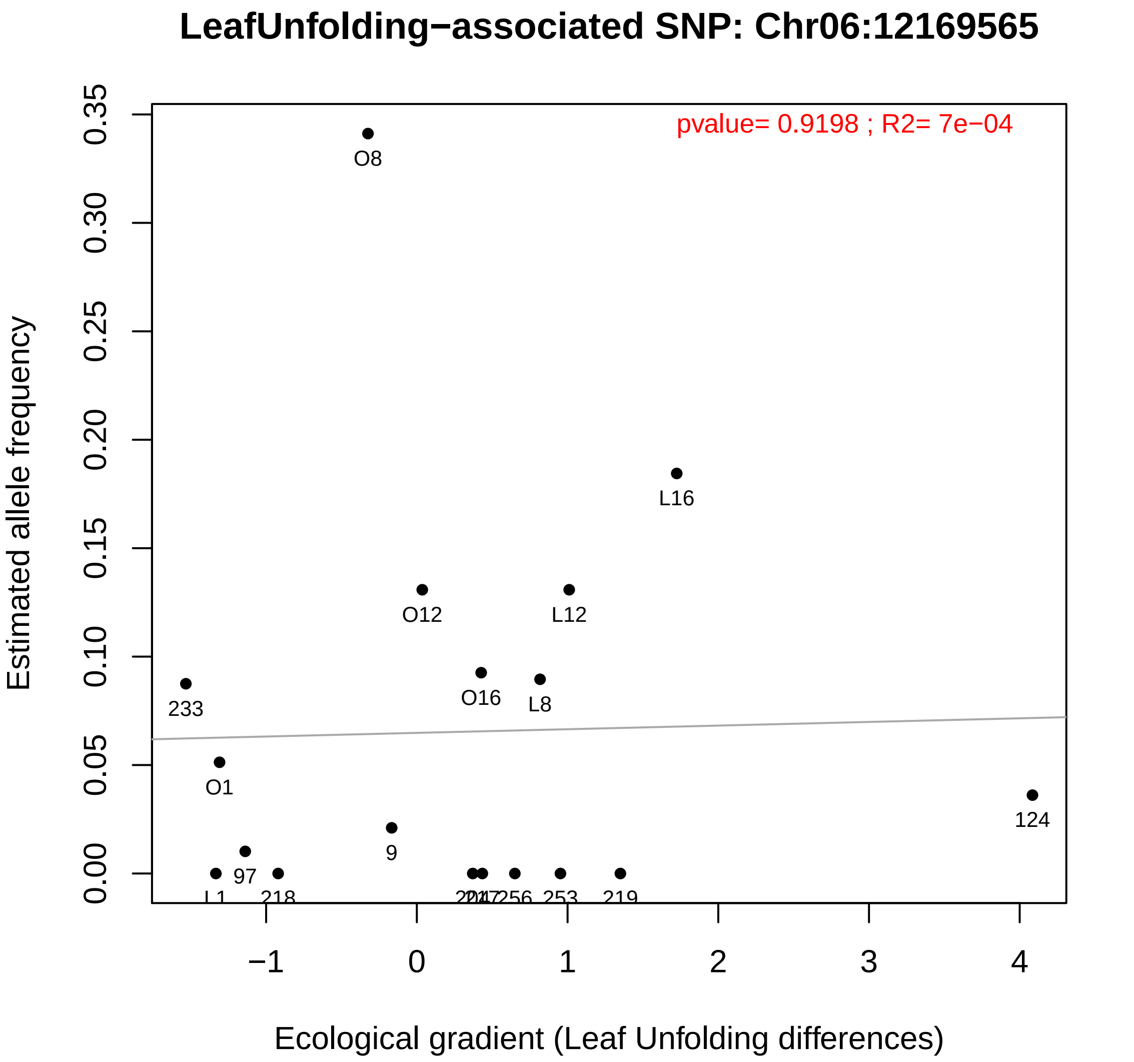
LeafUnfolding−associated SNP: Chr06:12169565
pvalue= 0.9198 ; R2= 7e−04
Estimated allele frequency
Ecological gradient (Leaf Unfolding differences)
0.00
0.05
0.10
0.15
0.20
0.25
0.30
0.35
−1
0
1
2
3
4
O8
L16
O12
L12
233
O16
L8
O1
124
9
97
L1
218
256
253
219

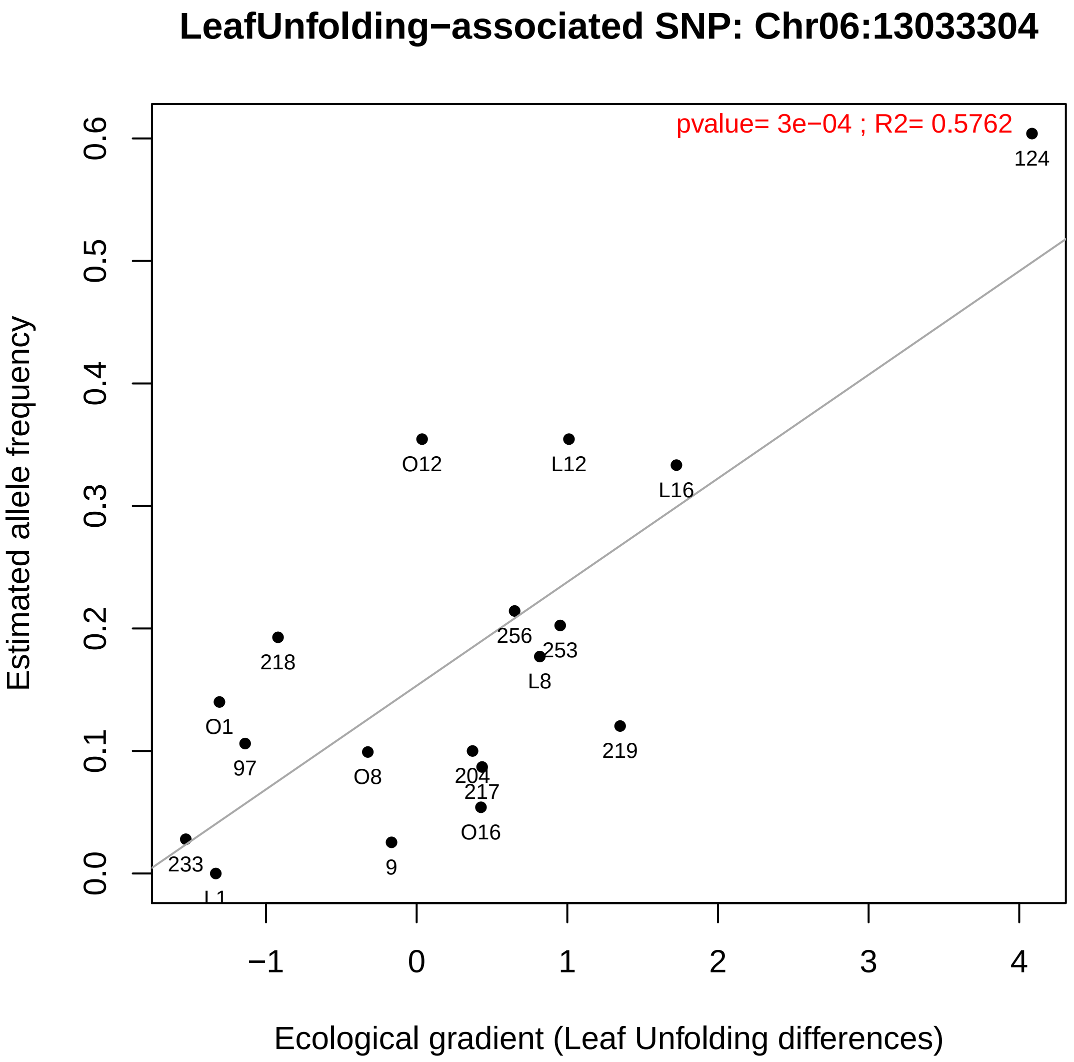
LeafUnfolding−associated SNP: Chr06:13033304
pvalue= 3e−04 ; R2= 0.5762
124
O12
L12
L16
256
253
L8
218
O1
97
O8
204
217
219
O16
233
9
0.0
0.1
0.2
0.3
0.4
0.5
0.6
−1
0
1
2
3
4
Estimated allele frequency
Ecological gradient (Leaf Unfolding differences)

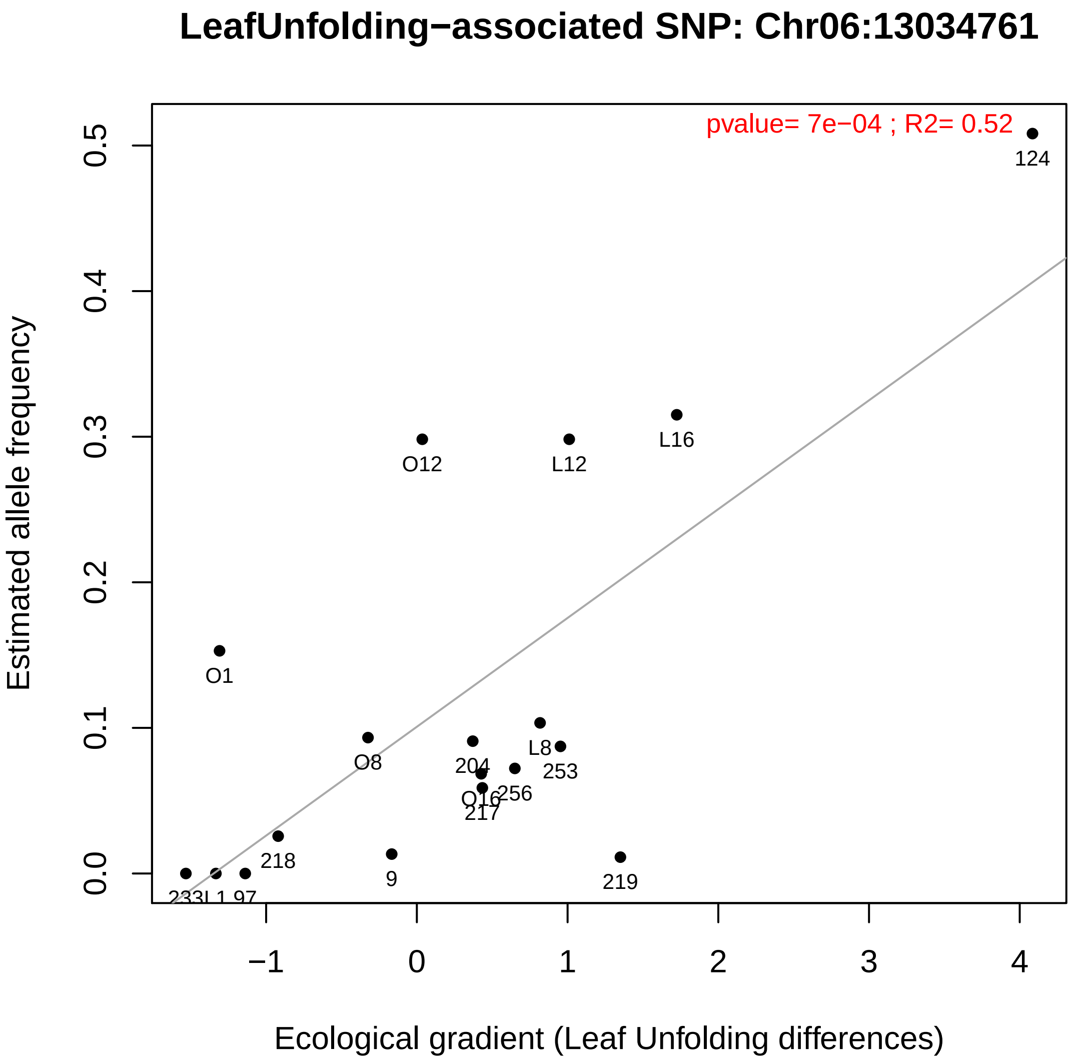
LeafUnfolding−associated SNP: Chr06:13034761
pvalue= 7e−04 ; R2= 0.52
124
O12
L12
L16
O1
O8
204
L8
253
256
O16
217
218
9
219
233
L1
97
0.0
0.1
0.2
0.3
0.4
0.5
−1
0
1
2
3
4
Estimated allele frequency
Ecological gradient (Leaf Unfolding differences)

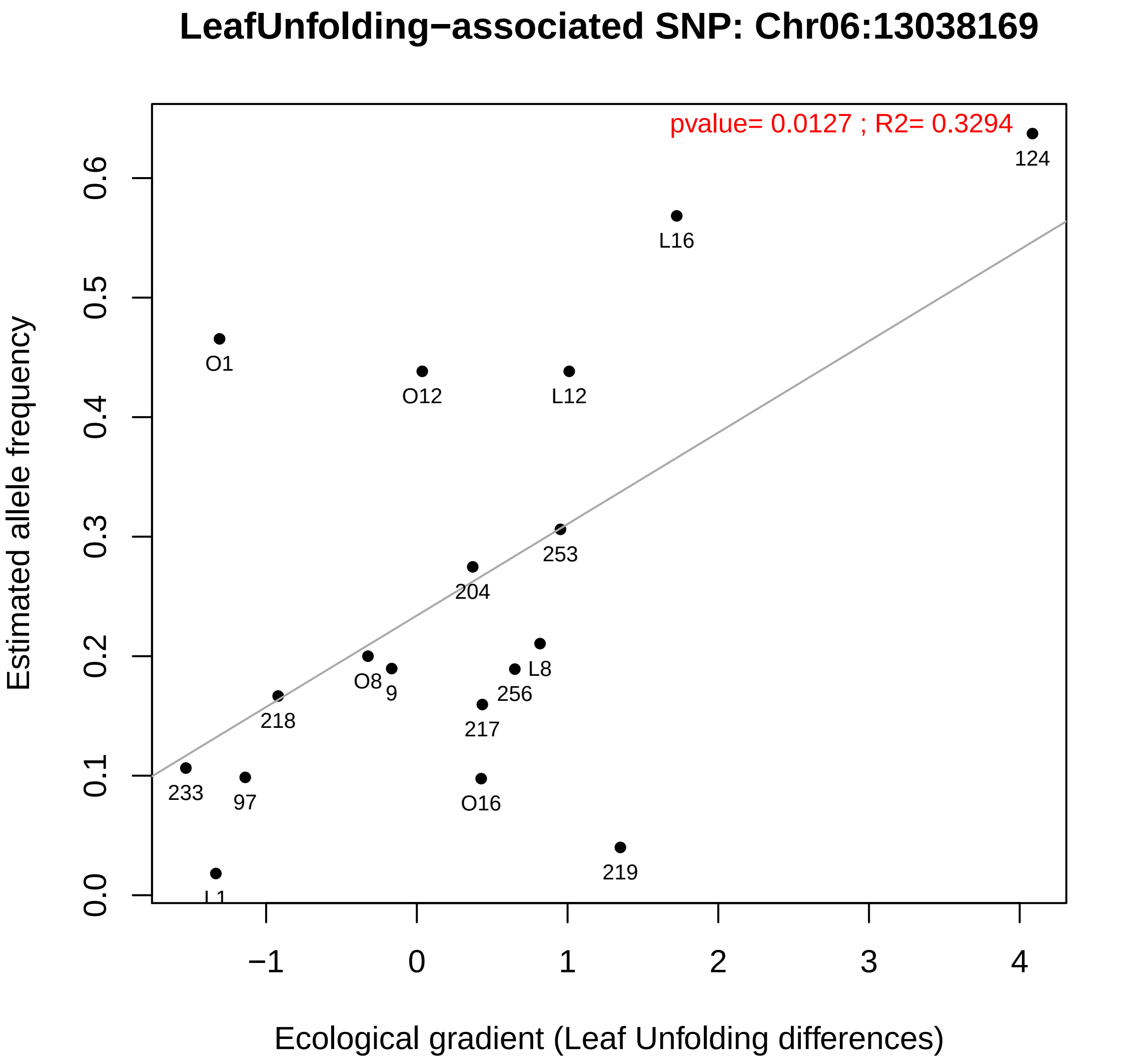

LeafUnfolding−associated SNP: Chr06:13038169
pvalue= 0.0127 ; R2= 0.3294
Estimated allele frequency
Ecological gradient (Leaf Unfolding differences)
0.0
0.1
0.2
0.3
0.4
0.5
0.6
−1
0
1
2
3
4
124
L16
O1
O12
L12
253
204
O8
9
L8
256
218
217
233
97
O16
219

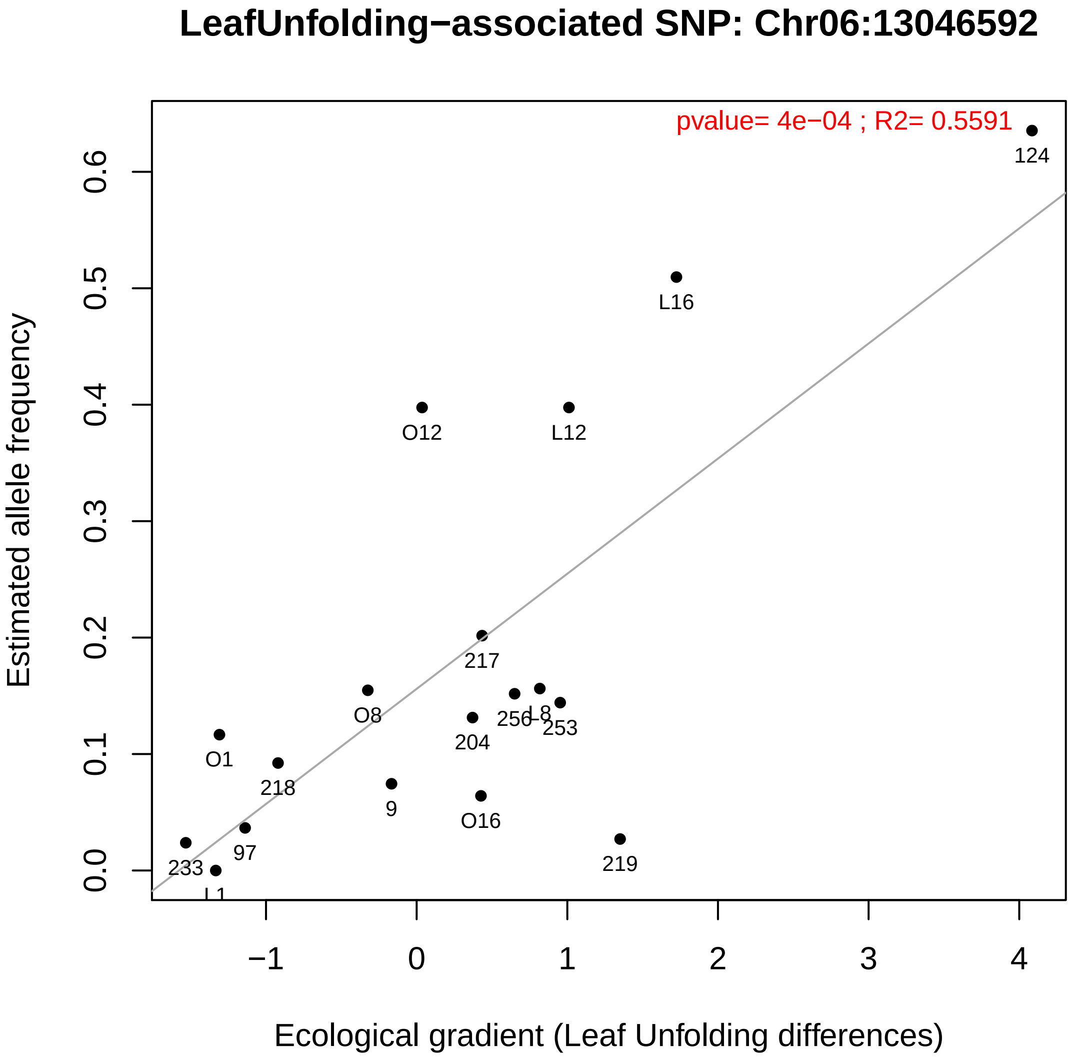
LeafUnfolding−associated SNP: Chr06:13046592
pvalue= 4e−04 ; R2= 0.5591
Estimated allele frequency
0.0
0.1
0.2
0.3
0.4
0.5
0.6
Ecological gradient (Leaf Unfolding differences)
−1
0
1
2
3
4
124
L16
O12
L12
217
O8
256
L8
253
204
O1
218
9
O16
97
233
219

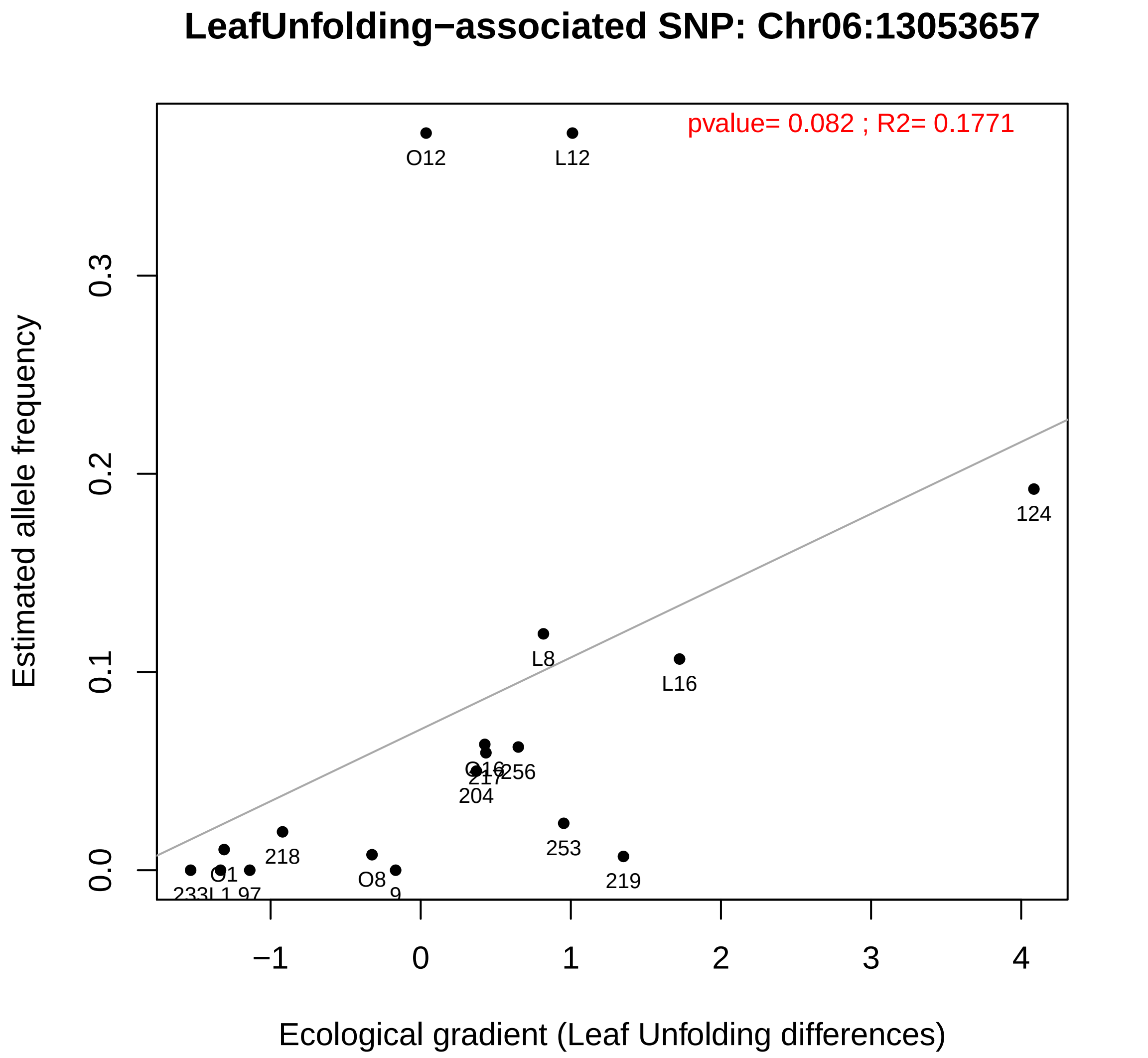
LeafUnfolding−associated SNP: Chr06:13053657
pvalue= 0.082 ; R2= 0.1771
O12
L12
124
L8
L16
O16
217
256
204
253
219
218
O1
O8
233
L1
97
9
Estimated allele frequency
0.0
0.1
0.2
0.3
−1
0
1
2
3
4
Ecological gradient (Leaf Unfolding differences)

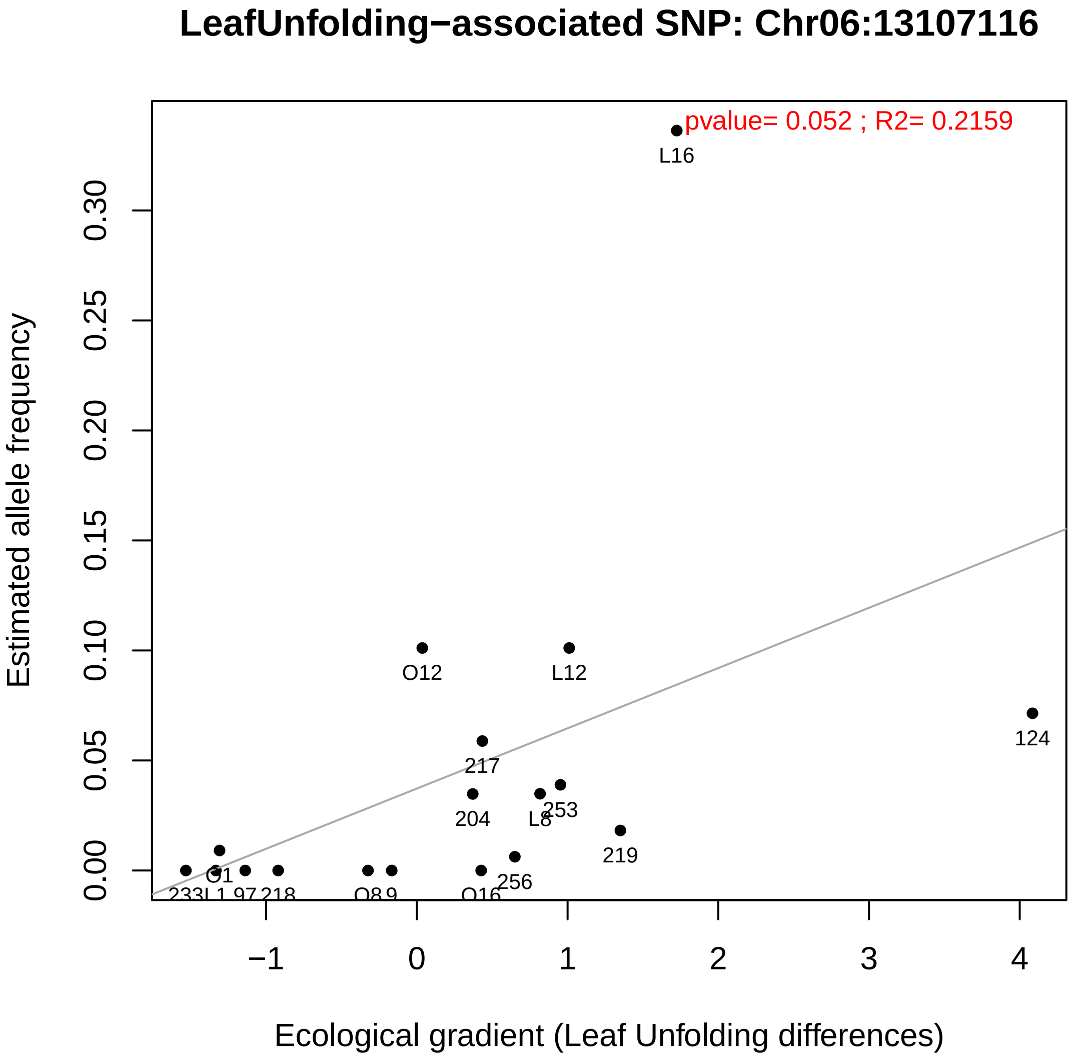
LeafUnfolding−associated SNP: Chr06:13107116
pvalue= 0.052 ; R2= 0.2159
L16
O12
L12
124
217
204
L8
253
219
256
O1
233
L1
97
218
O8
9
O16
Estimated allele frequency
0.00
0.05
0.10
0.15
0.20
0.25
0.30
−1
0
1
2
3
4
Ecological gradient (Leaf Unfolding differences)

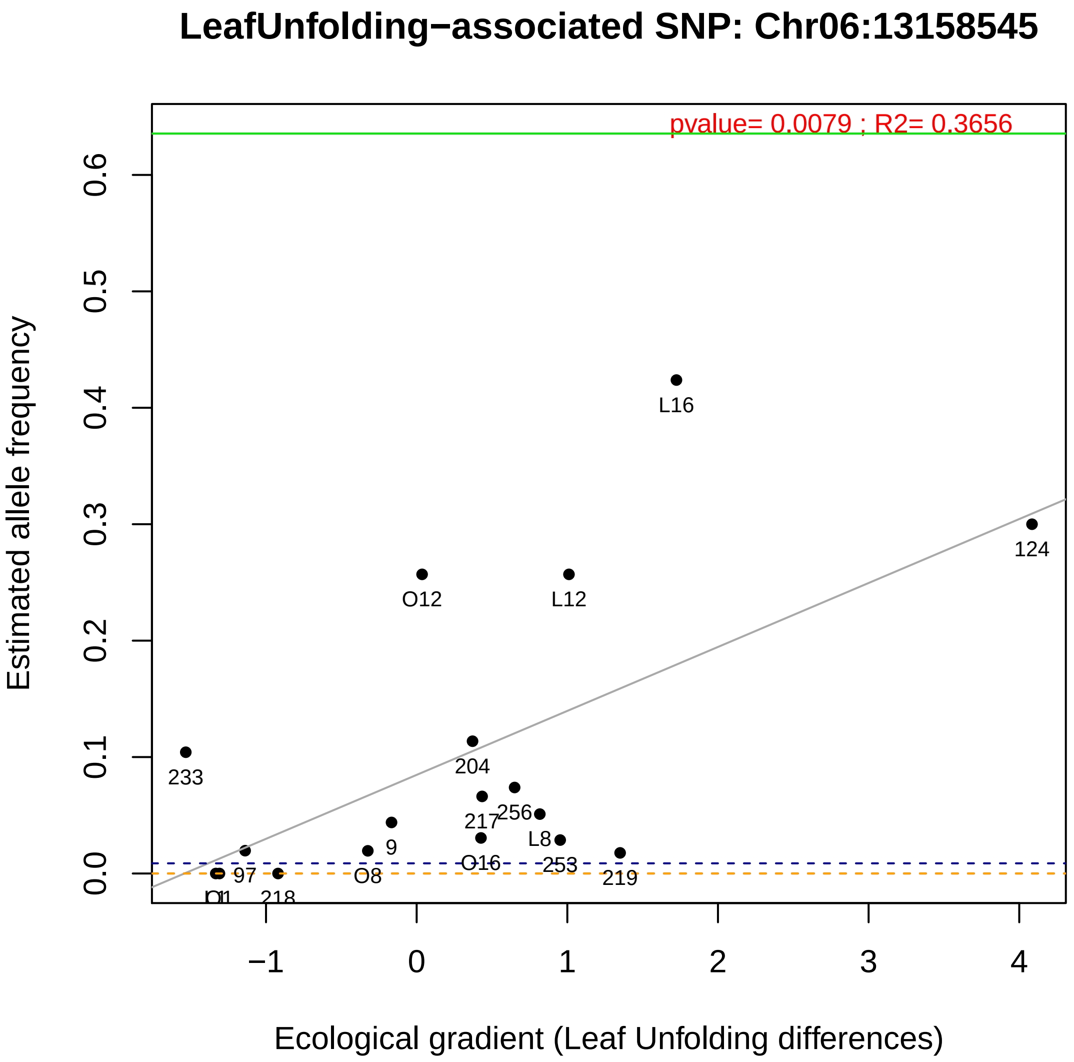

LeafUnfolding−associated SNP: Chr06:13158545
pvalue= 0.0079 ; R2= 0.3656
Estimated allele frequency
Ecological gradient (Leaf Unfolding differences)
0.0
0.1
0.2
0.3
0.4
0.5
0.6
−1
0
1
2
3
4
L16
124
O12
L12
233
204
217
256
L8
9
O16
253
97
O8
219
218

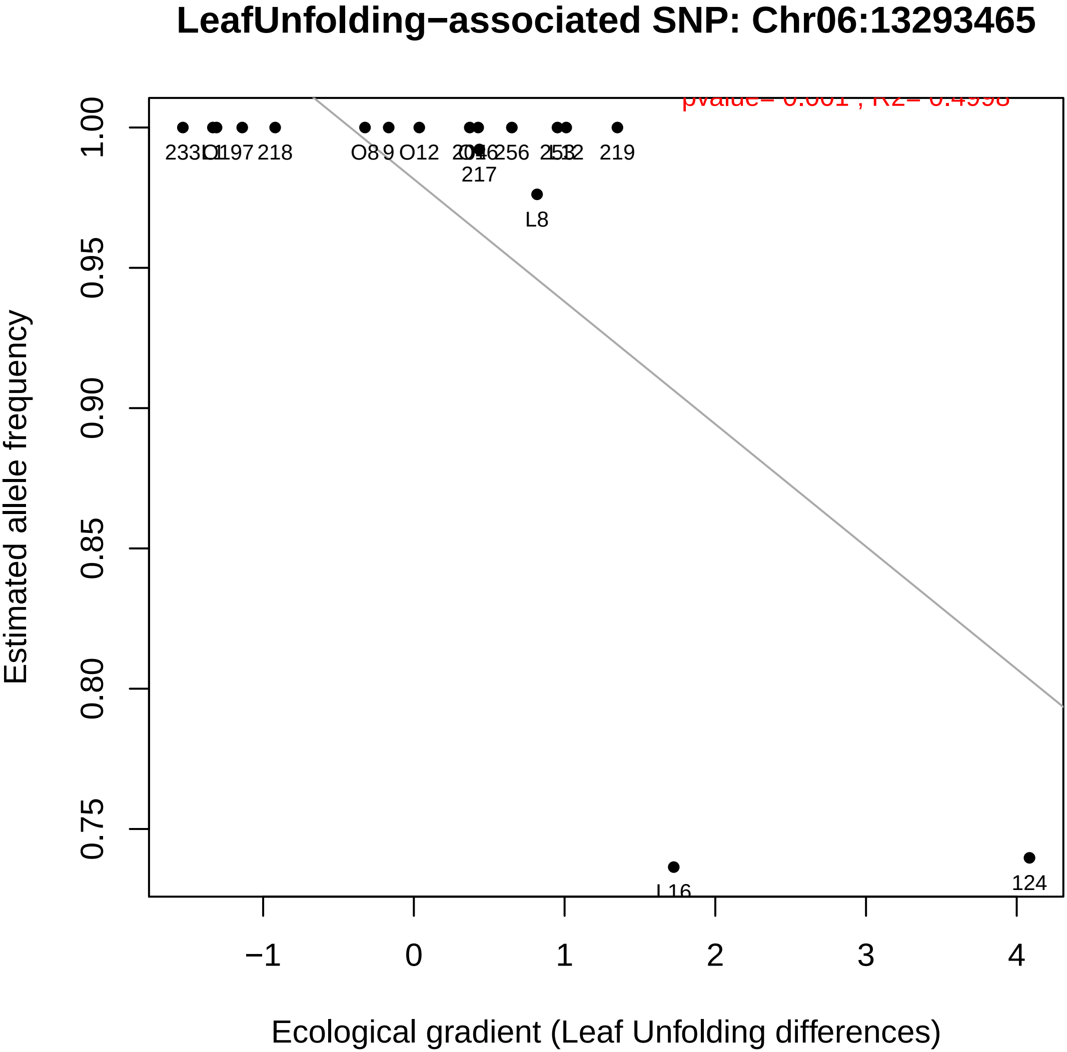
LeafUnfolding−associated SNP: Chr06:13293465
pvalue= 0.001 ; R2= 0.4998
Estimated allele frequency
0.75
0.80
0.85
0.90
0.95
1.00
−1
0
1
2
3
4
Ecological gradient (Leaf Unfolding differences)
218
O8
9
O12
256
219
217
L8
124

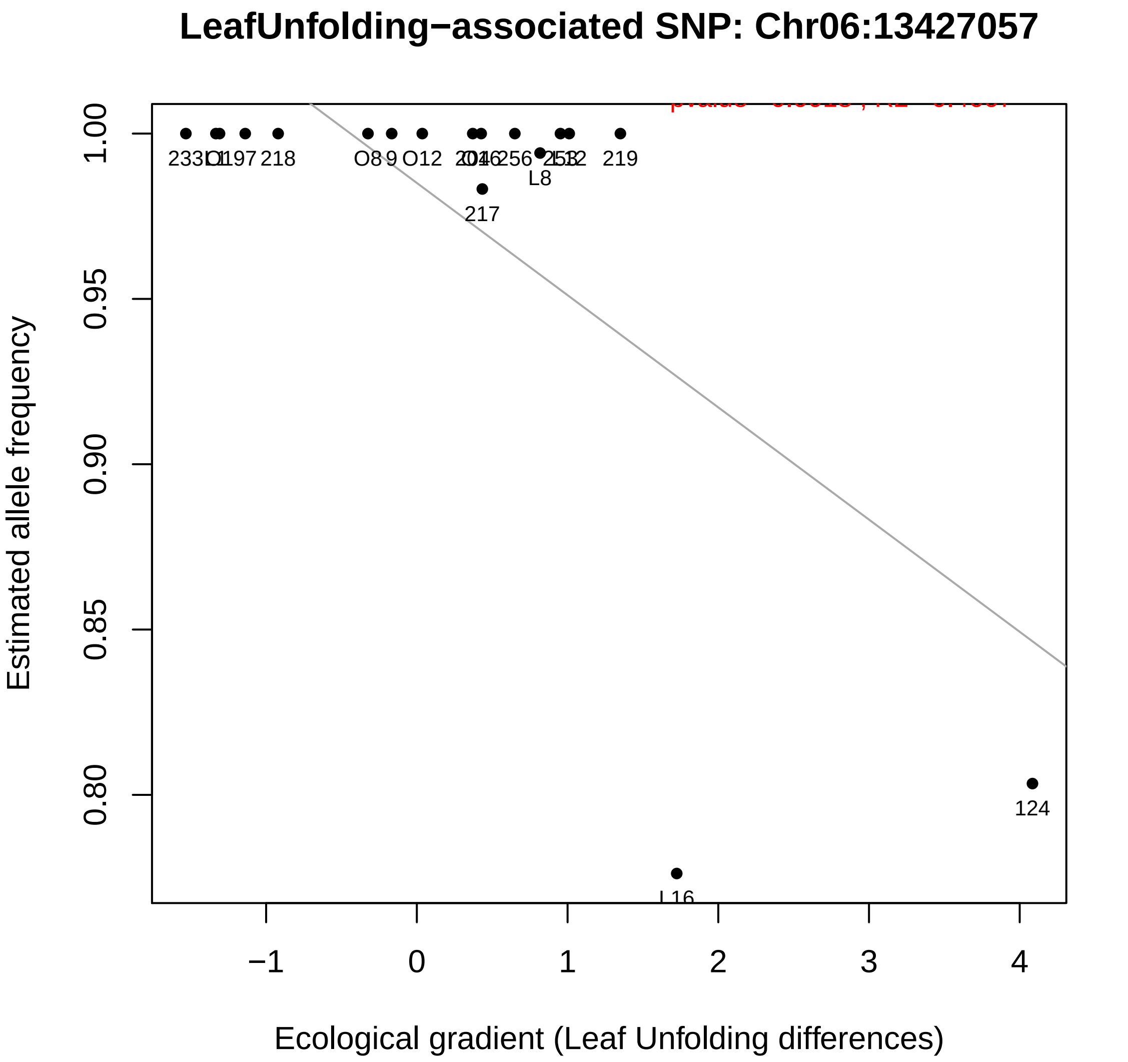
LeafUnfolding−associated SNP: Chr06:13427057
Estimated allele frequency
Ecological gradient (Leaf Unfolding differences)
1.00
0.95
0.90
0.85
0.80
−1
0
1
2
3
4
218
O8
9
O12
256
219
L8
217
124

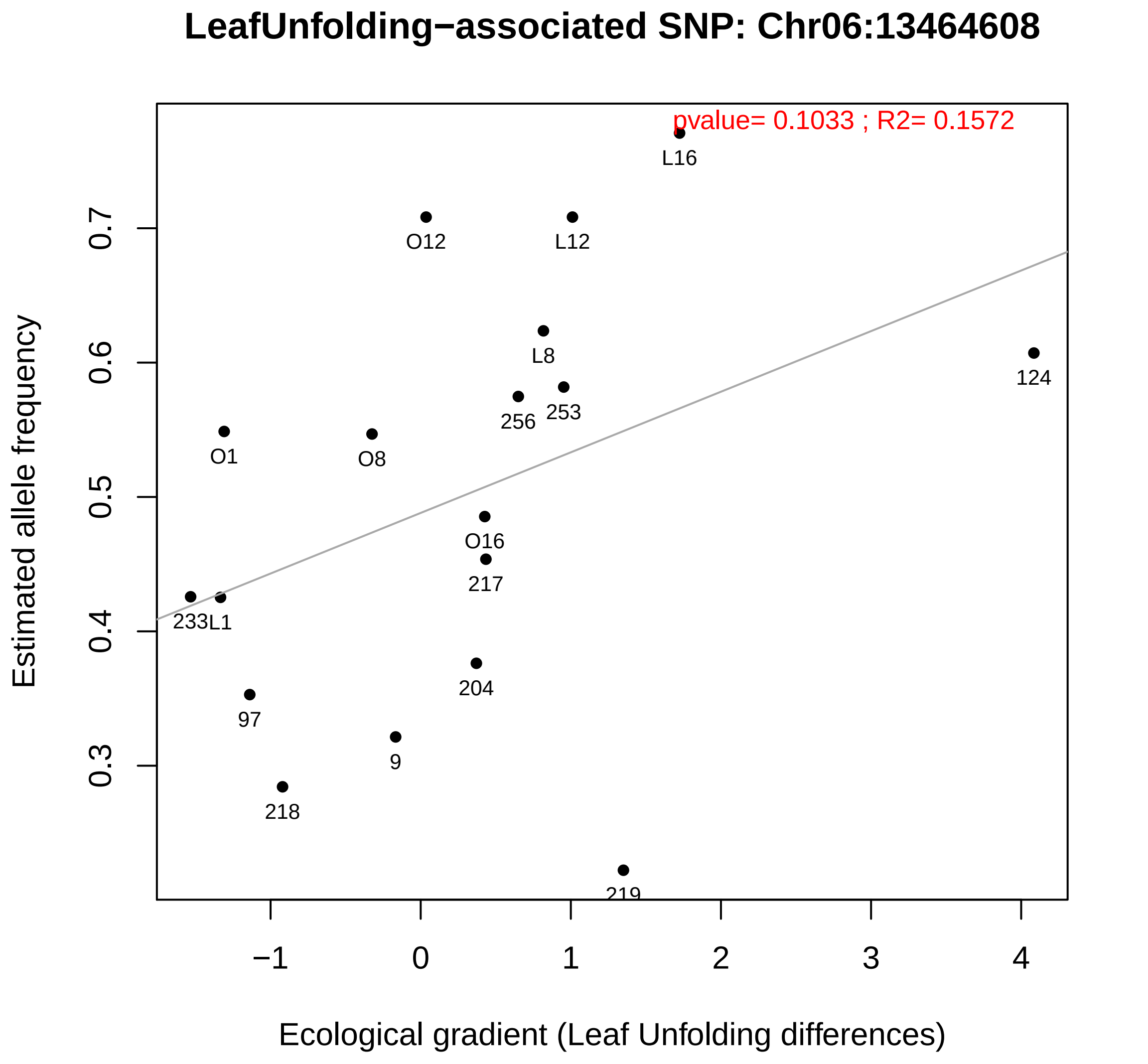

LeafUnfolding−associated SNP: Chr06:13464608
pvalue= 0.1033 ; R2= 0.1572
L16
O12
L12
L8
124
256
253
O1
O8
O16
217
233
L1
204
97
9
218
219
0.3
0.4
0.5
0.6
0.7
−1
0
1
2
3
4
Estimated allele frequency
Ecological gradient (Leaf Unfolding differences)

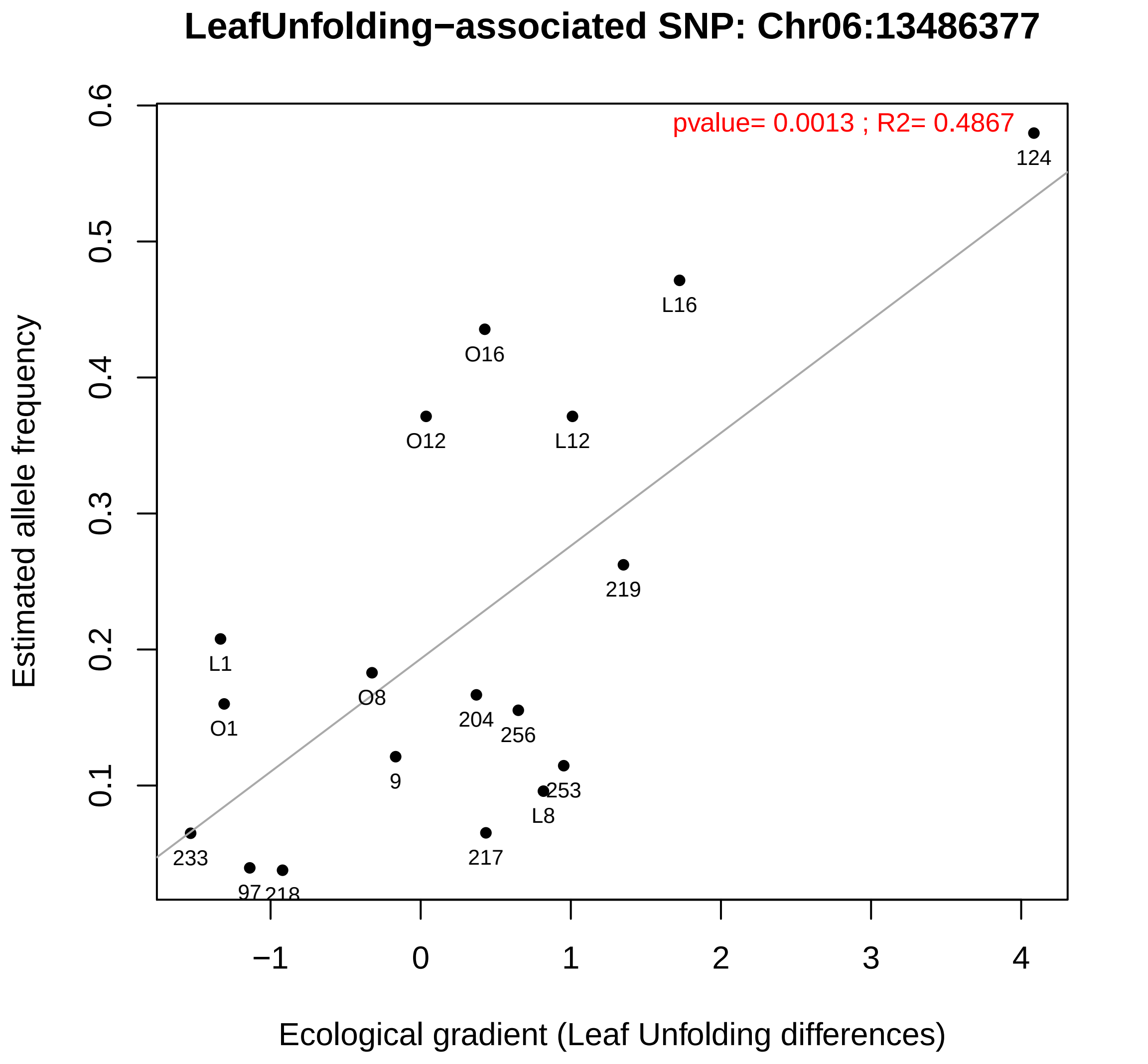
LeafUnfolding−associated SNP: Chr06:13486377
pvalue= 0.0013 ; R2= 0.4867
Estimated allele frequency
Ecological gradient (Leaf Unfolding differences)
0.1
0.2
0.3
0.4
0.5
0.6
−1
0
1
2
3
4
124
L16
O16
O12
L12
219
L1
O8
O1
204
256
9
253
L8
233
217
97
218

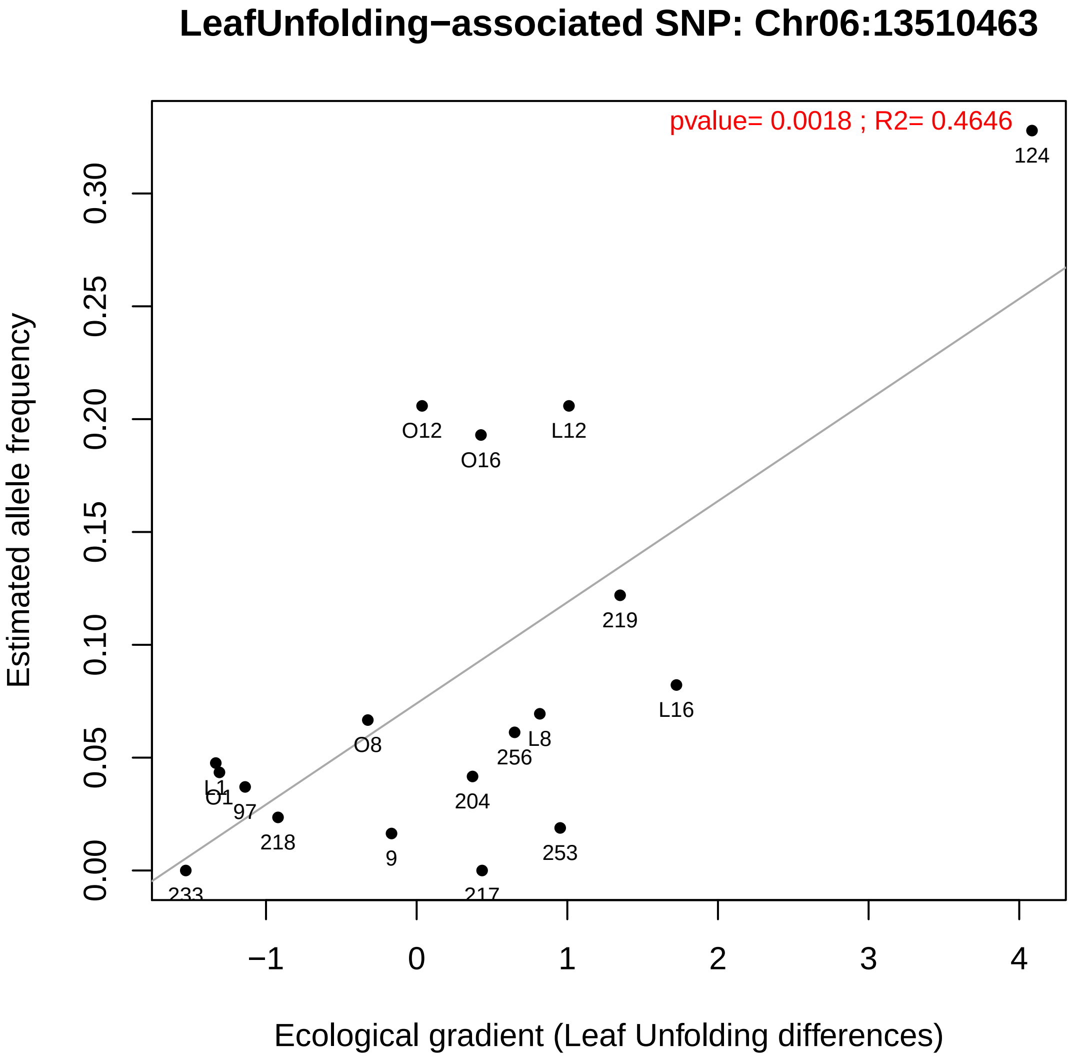
LeafUnfolding−associated SNP: Chr06:13510463
pvalue= 0.0018 ; R2= 0.4646
Estimated allele frequency
0.00
0.05
0.10
0.15
0.20
0.25
0.30
−1
0
1
2
3
4
Ecological gradient (Leaf Unfolding differences)
124
O12
O16
L12
219
L16
L8
256
O8
204
L1
O1
97
218
9
253
233
217

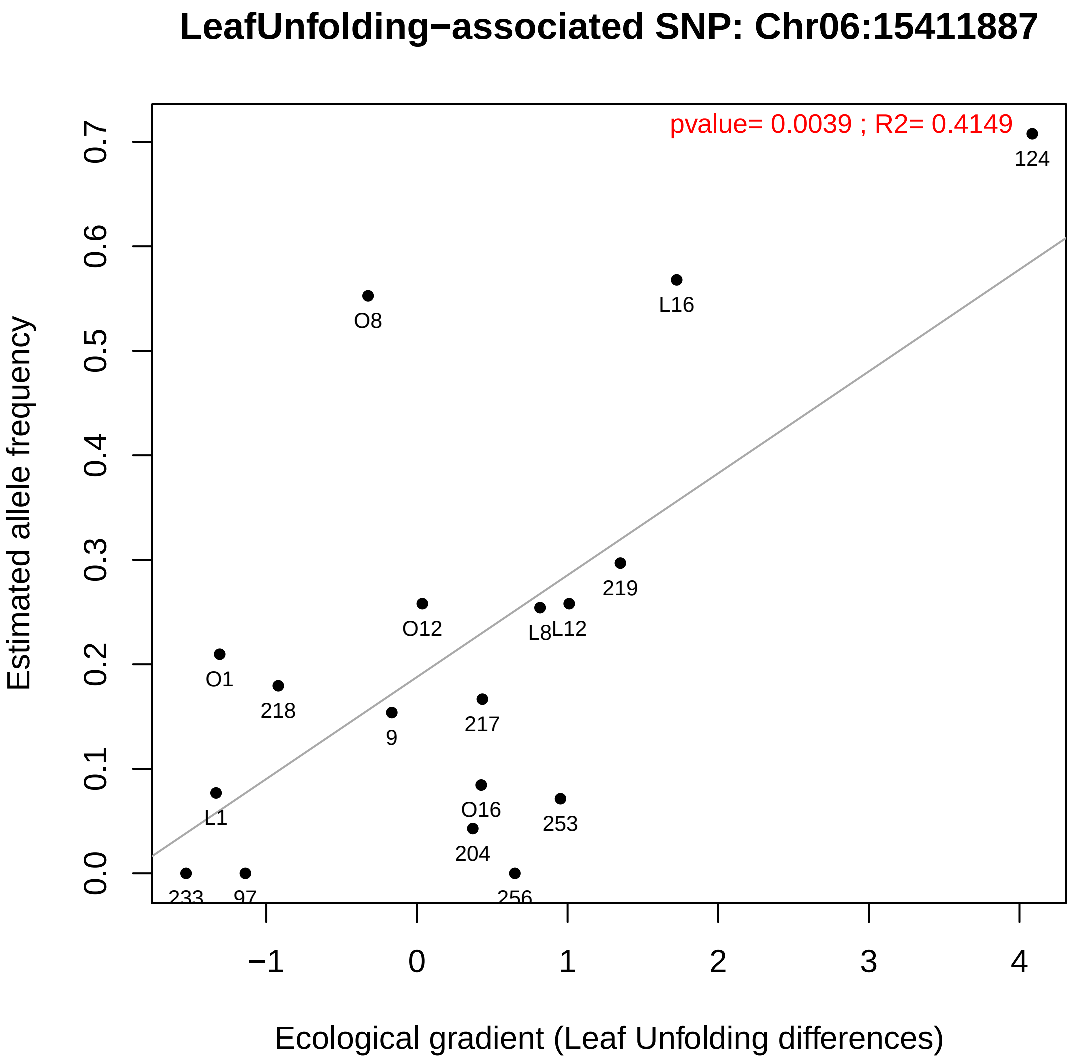

LeafUnfolding−associated SNP: Chr06:15411887
pvalue= 0.0039 ; R2= 0.4149
Estimated allele frequency
Ecological gradient (Leaf Unfolding differences)
0.0
0.1
0.2
0.3
0.4
0.5
0.6
0.7
−1
0
1
2
3
4
124
L16
O8
219
O12
L8
L12
O1
218
9
217
L1
O16
253
204
233
97
256

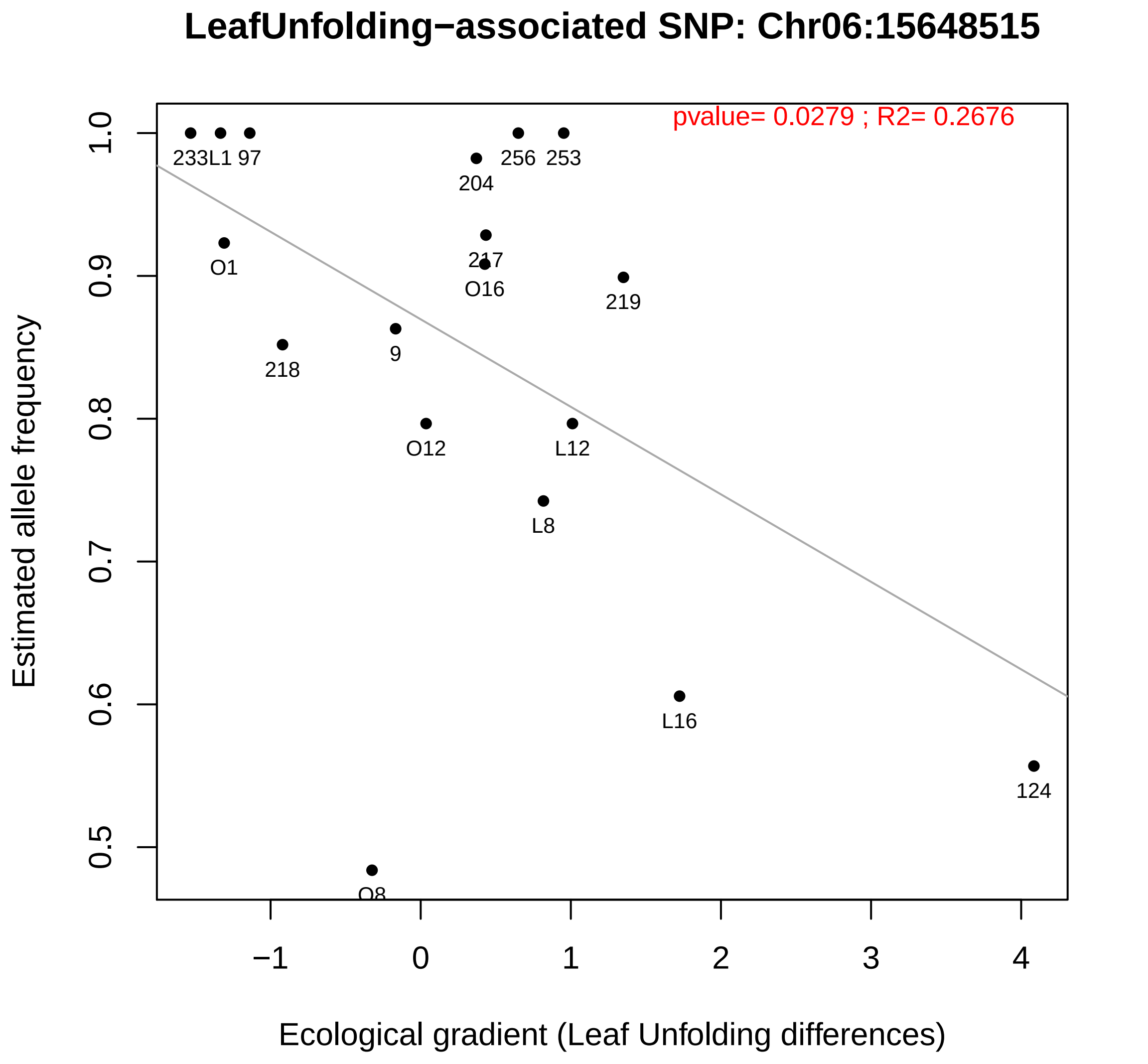
LeafUnfolding−associated SNP: Chr06:15648515
pvalue= 0.0279 ; R2= 0.2676
Estimated allele frequency
Ecological gradient (Leaf Unfolding differences)
0.5
0.6
0.7
0.8
0.9
1.0
−1
0
1
2
3
4
233
L1
97
256
253
204
O1
217
O16
219
218
9
O12
L12
L8
L16
124
O8

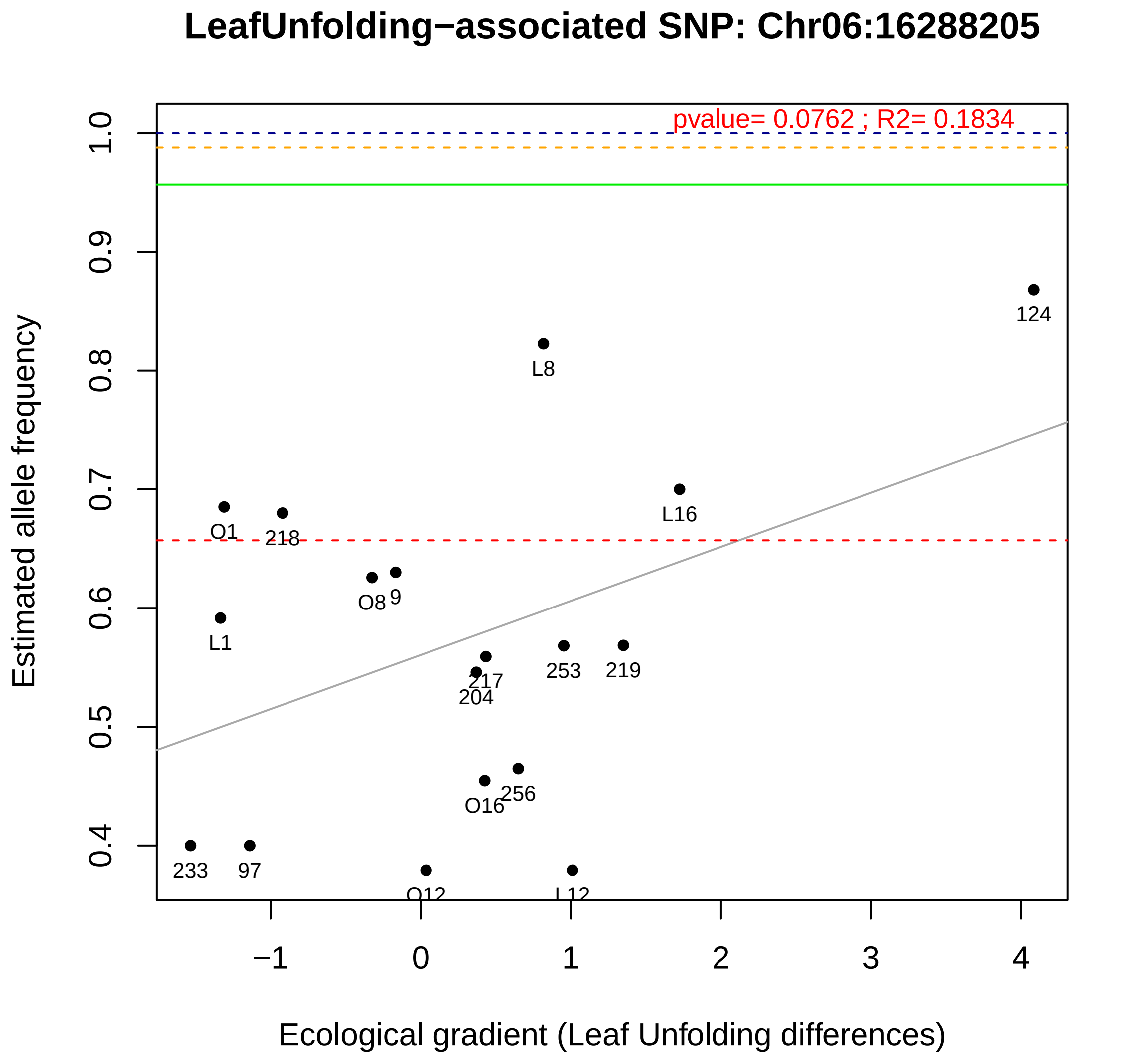

LeafUnfolding−associated SNP: Chr06:16288205
pvalue= 0.0762 ; R2= 0.1834
Estimated allele frequency
Ecological gradient (Leaf Unfolding differences)
0.4
0.5
0.6
0.7
0.8
0.9
1.0
−1
0
1
2
3
4
124
L8
L16
O1
218
O8
9
L1
217
204
253
219
256
O16
233
97

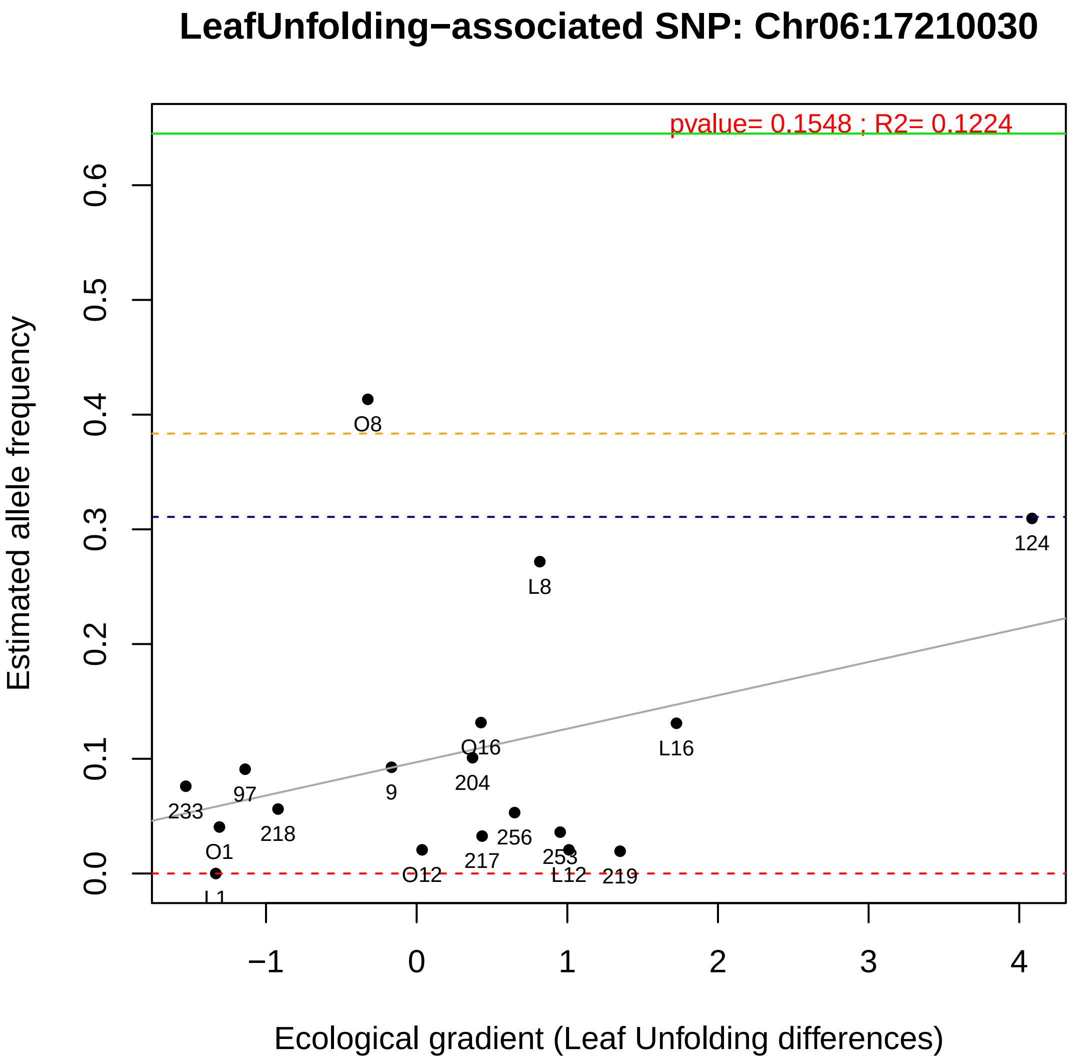

LeafUnfolding−associated SNP: Chr06:17210030
pvalue= 0.1548 ; R2= 0.1224
Estimated allele frequency
0.0
0.1
0.2
0.3
0.4
0.5
0.6
−1
0
1
2
3
4
Ecological gradient (Leaf Unfolding differences)
O8
124
L8
L16
O16
204
9
97
233
218
O1
L1
O12
217
256
253
L12
219

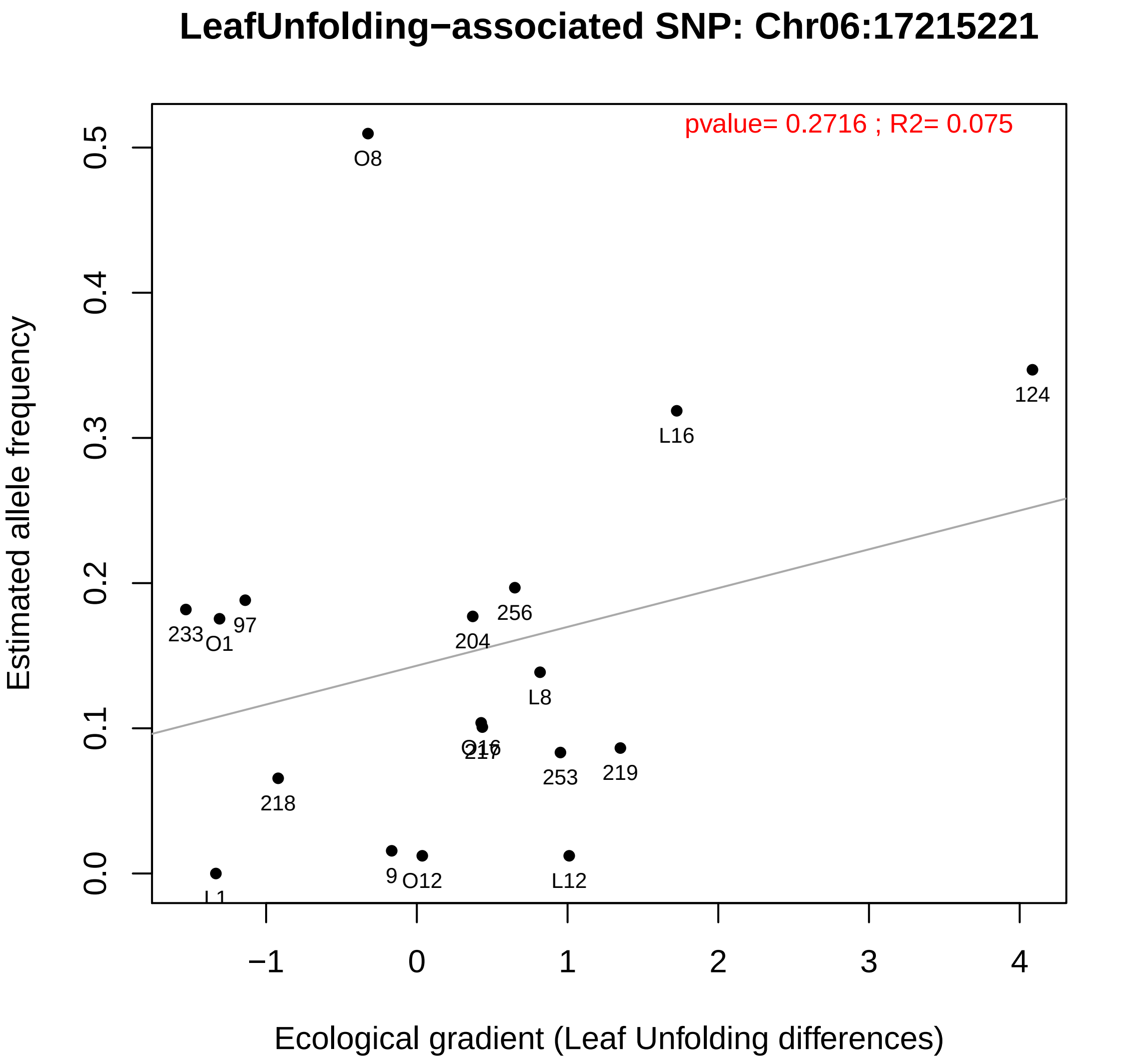

LeafUnfolding−associated SNP: Chr06:17215221
pvalue= 0.2716 ; R2= 0.075
Estimated allele frequency
Ecological gradient (Leaf Unfolding differences)
0.0
0.1
0.2
0.3
0.4
0.5
−1
0
1
2
3
4
O8
124
L16
256
233
O1
97
204
L8
O16
217
253
219
218
9
O12
L12
L1

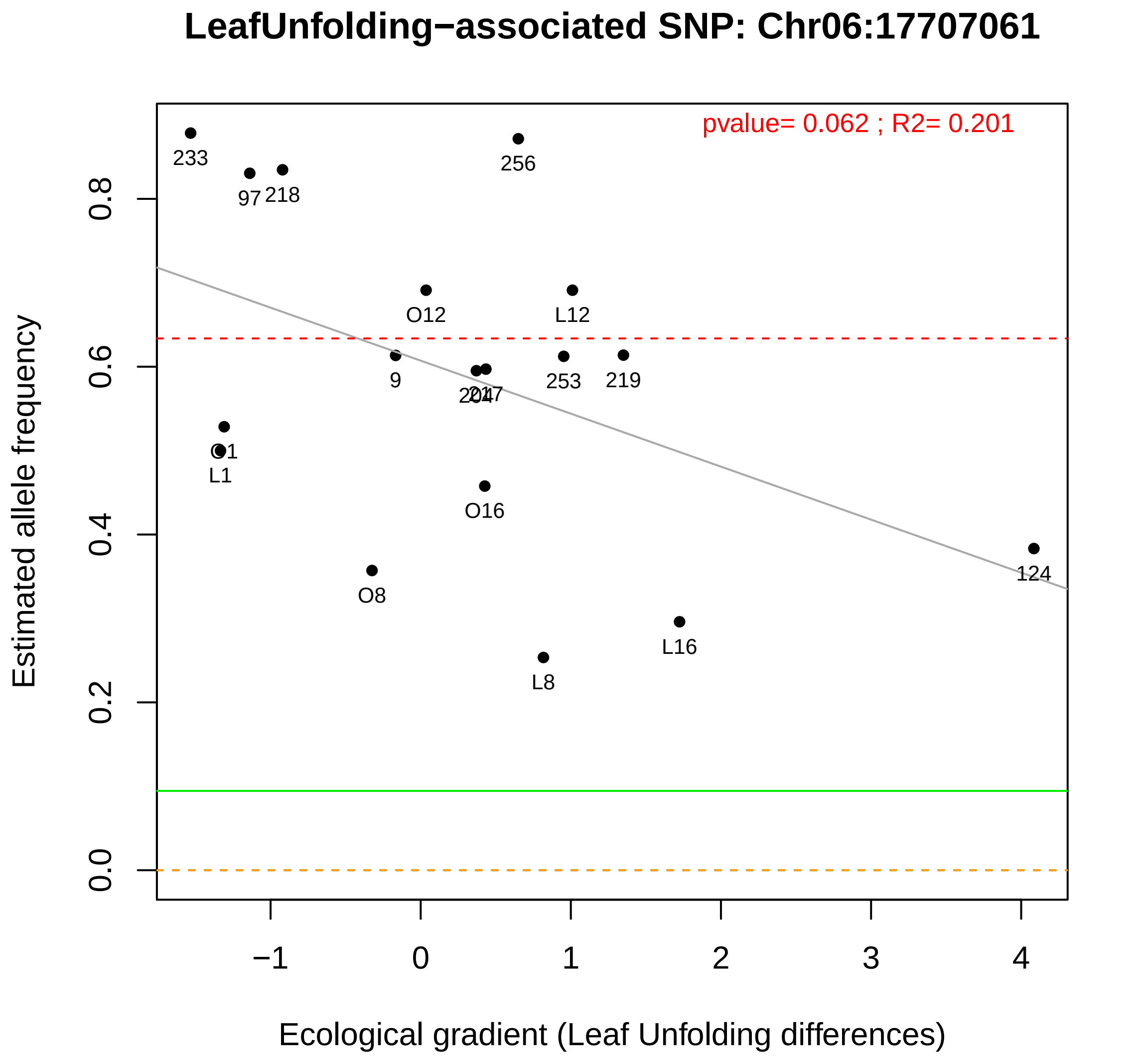
LeafUnfolding−associated SNP: Chr06:17707061
pvalue= 0.062 ; R2= 0.201
Estimated allele frequency
Ecological gradient (Leaf Unfolding differences)
0.0
0.2
0.4
0.6
0.8
−1
0
1
2
3
4
233
97
218
256
O12
L12
9
204
217
253
219
O1
L1
O16
O8
124
L16
L8

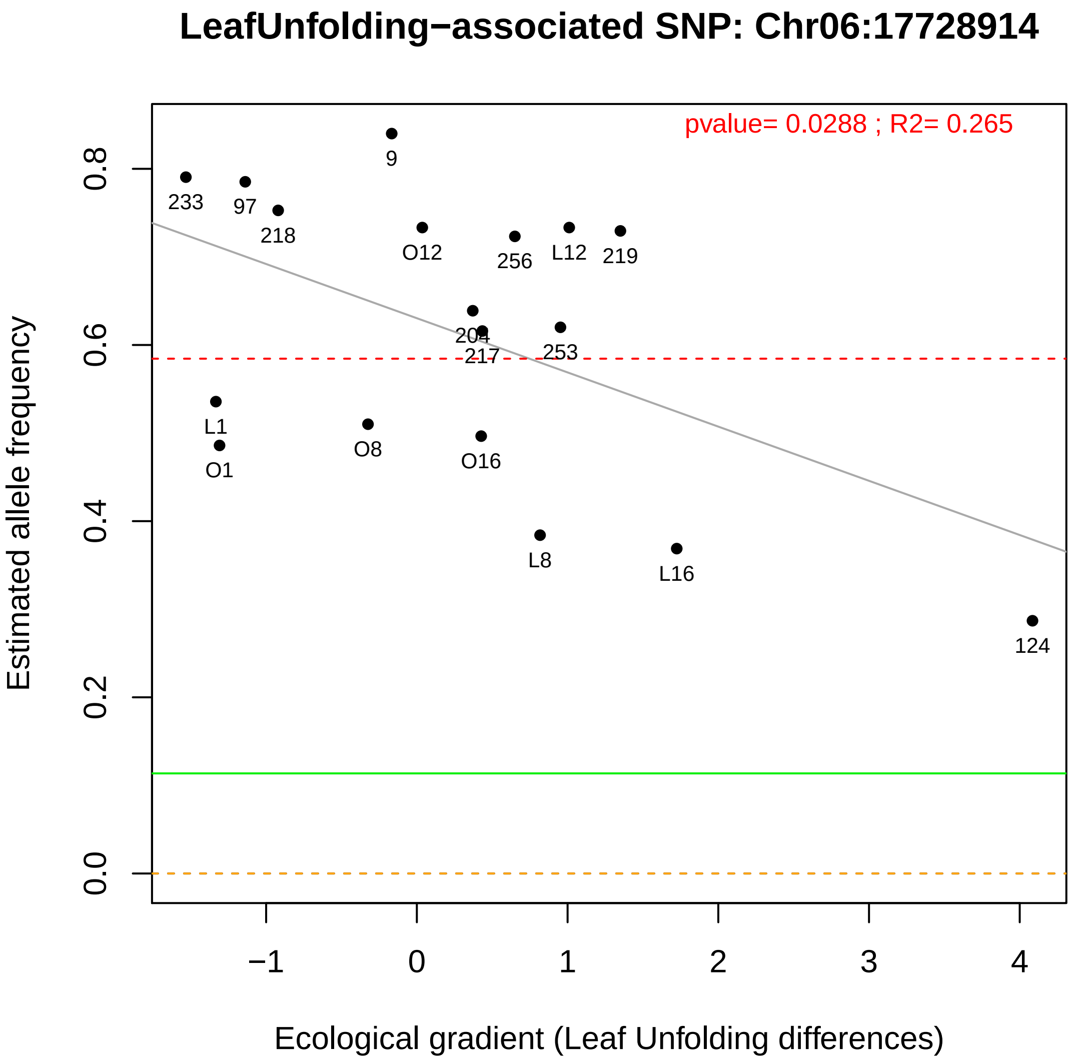
LeafUnfolding−associated SNP: Chr06:17728914
pvalue= 0.0288 ; R2= 0.265
Estimated allele frequency
0.0
0.2
0.4
0.6
0.8
−1
0
1
2
3
4
Ecological gradient (Leaf Unfolding differences)
9
233
97
218
O12
256
L12
219
204
217
253
L1
O1
O8
O16
L8
L16
124

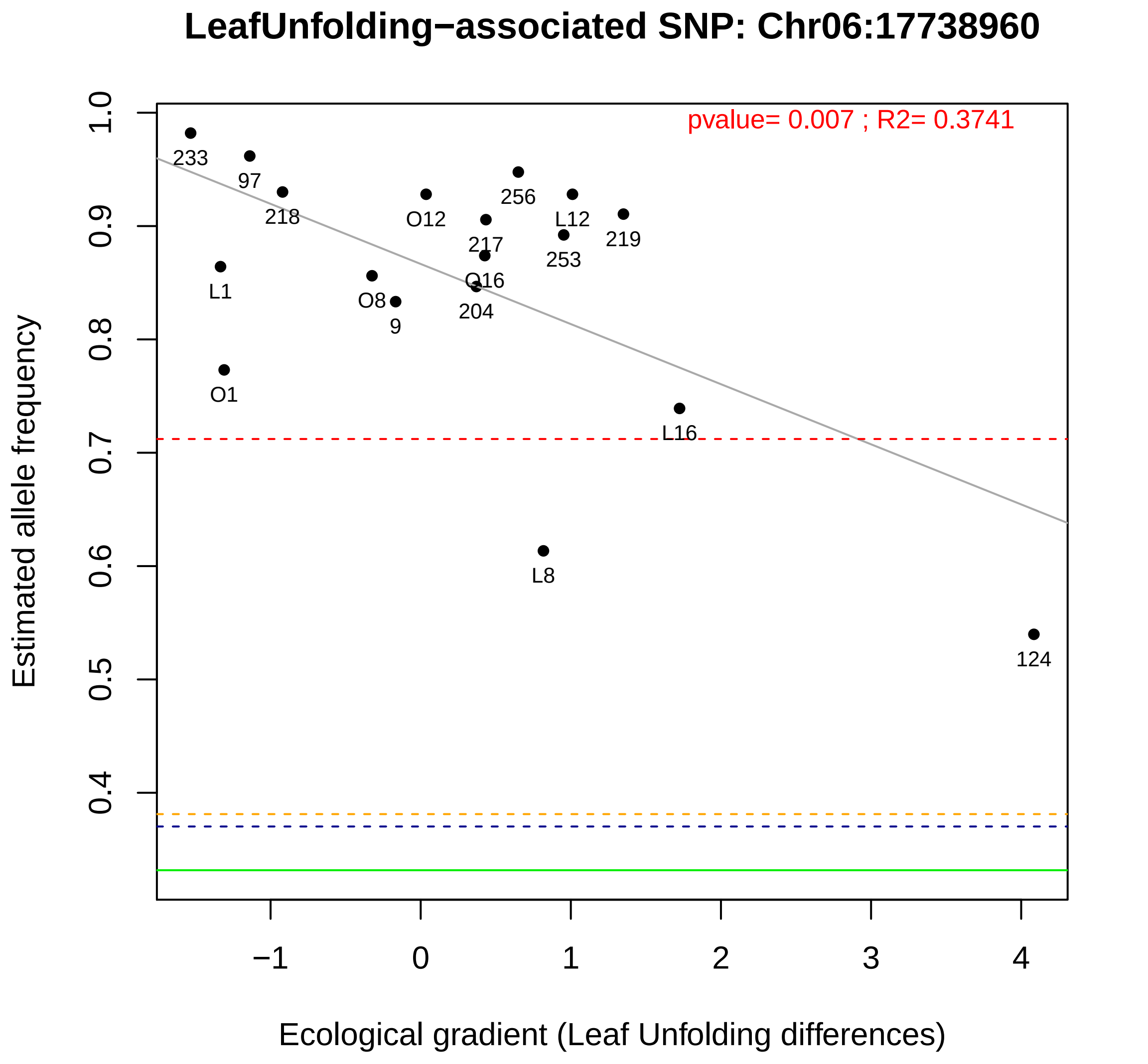
LeafUnfolding−associated SNP: Chr06:17738960
pvalue= 0.007 ; R2= 0.3741
Estimated allele frequency
Ecological gradient (Leaf Unfolding differences)
233
97
218
L1
O1
O8
9
O12
217
O16
204
256
L12
253
219
L16
L8
124

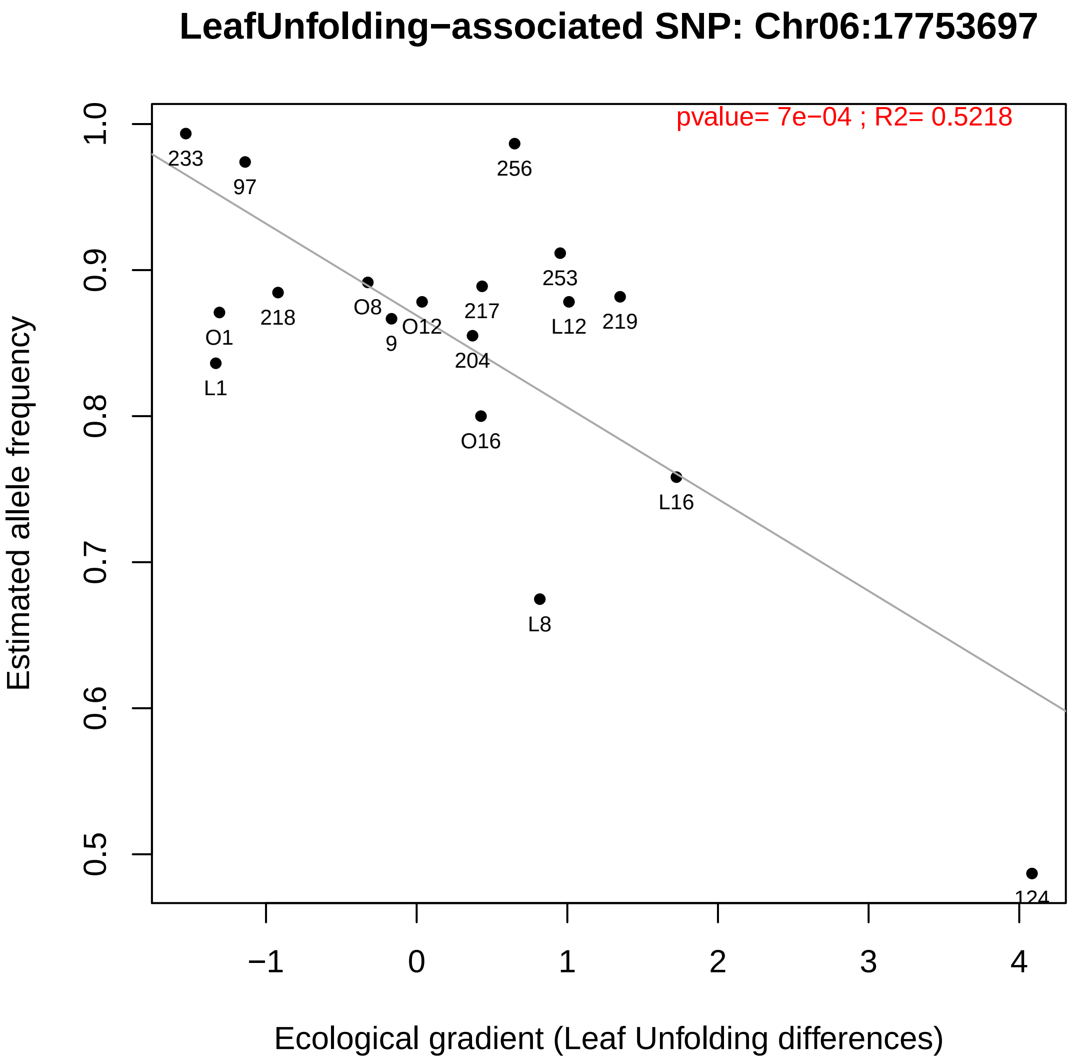
LeafUnfolding−associated SNP: Chr06:17753697
pvalue= 7e−04 ; R2= 0.5218
Estimated allele frequency
Ecological gradient (Leaf Unfolding differences)
233
97
256
253
217
L12
219
218
O1
L1
O8
9
O12
204
O16
L16
L8
124
1.0
0.9
0.8
0.7
0.6
0.5
−1
0
1
2
3
4

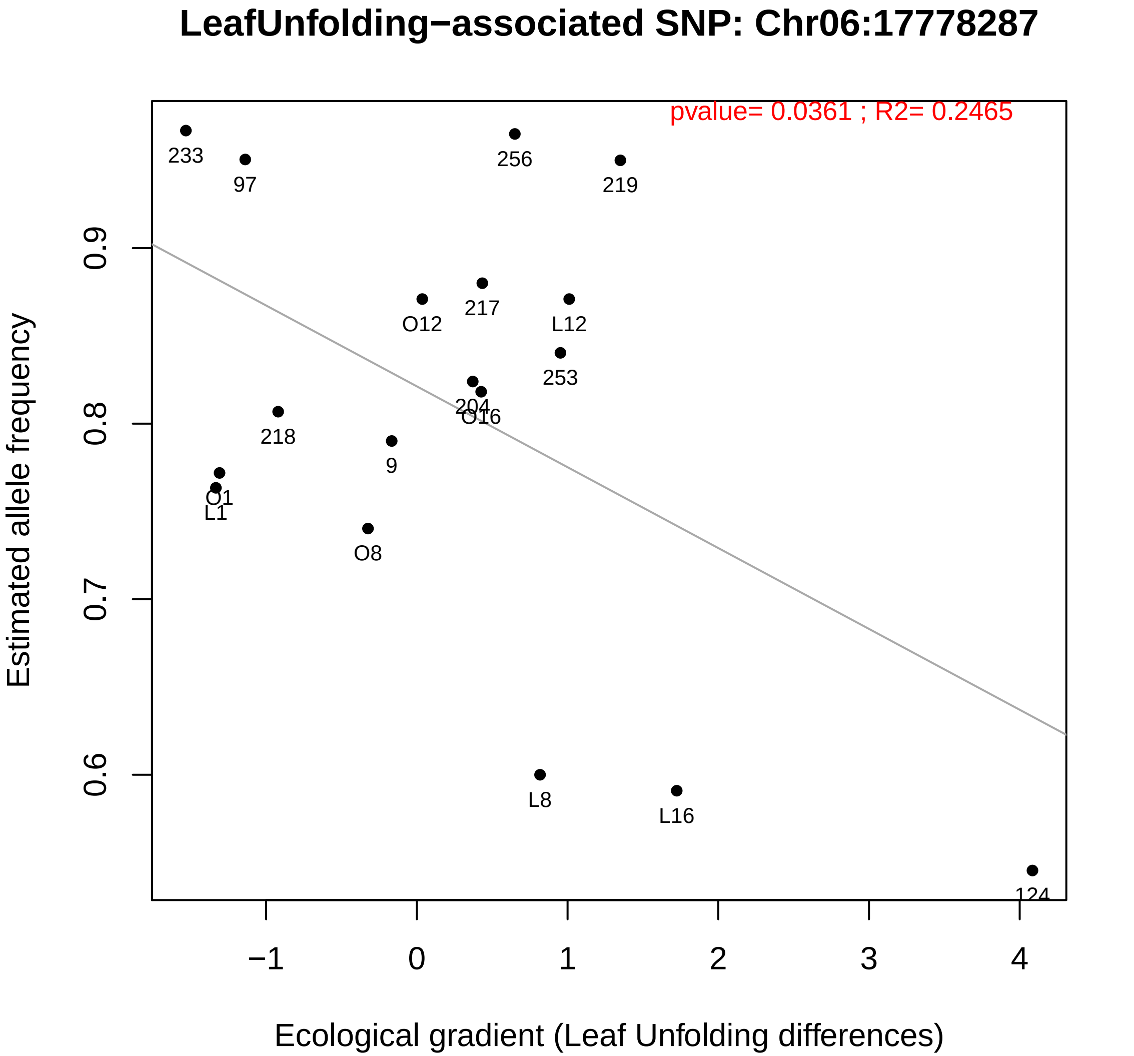
LeafUnfolding−associated SNP: Chr06:17778287
pvalue= 0.0361 ; R2= 0.2465
Estimated allele frequency
0.6
0.7
0.8
0.9
−1
0
1
2
3
4
Ecological gradient (Leaf Unfolding differences)
233
97
256
219
217
O12
L12
253
204
O16
218
9
O1
L1
O8
L8
L16
124

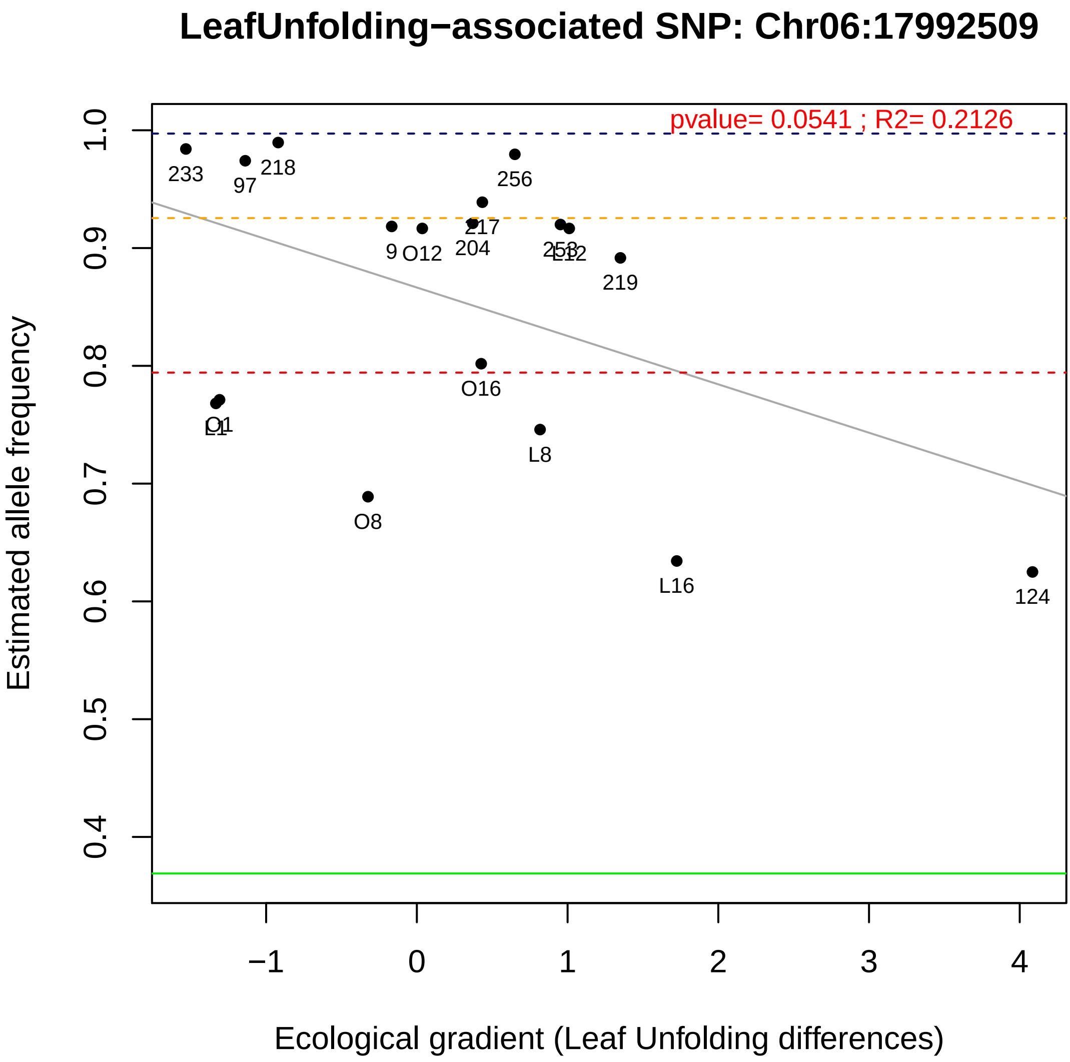

LeafUnfolding−associated SNP: Chr06:17992509
pvalue= 0.0541 ; R2= 0.2126
Estimated allele frequency
Ecological gradient (Leaf Unfolding differences)
0.4
0.5
0.6
0.7
0.8
0.9
1.0
−1
0
1
2
3
4
233
97
218
256
9
O12
217
204
219
O16
L8
O8
L16
124

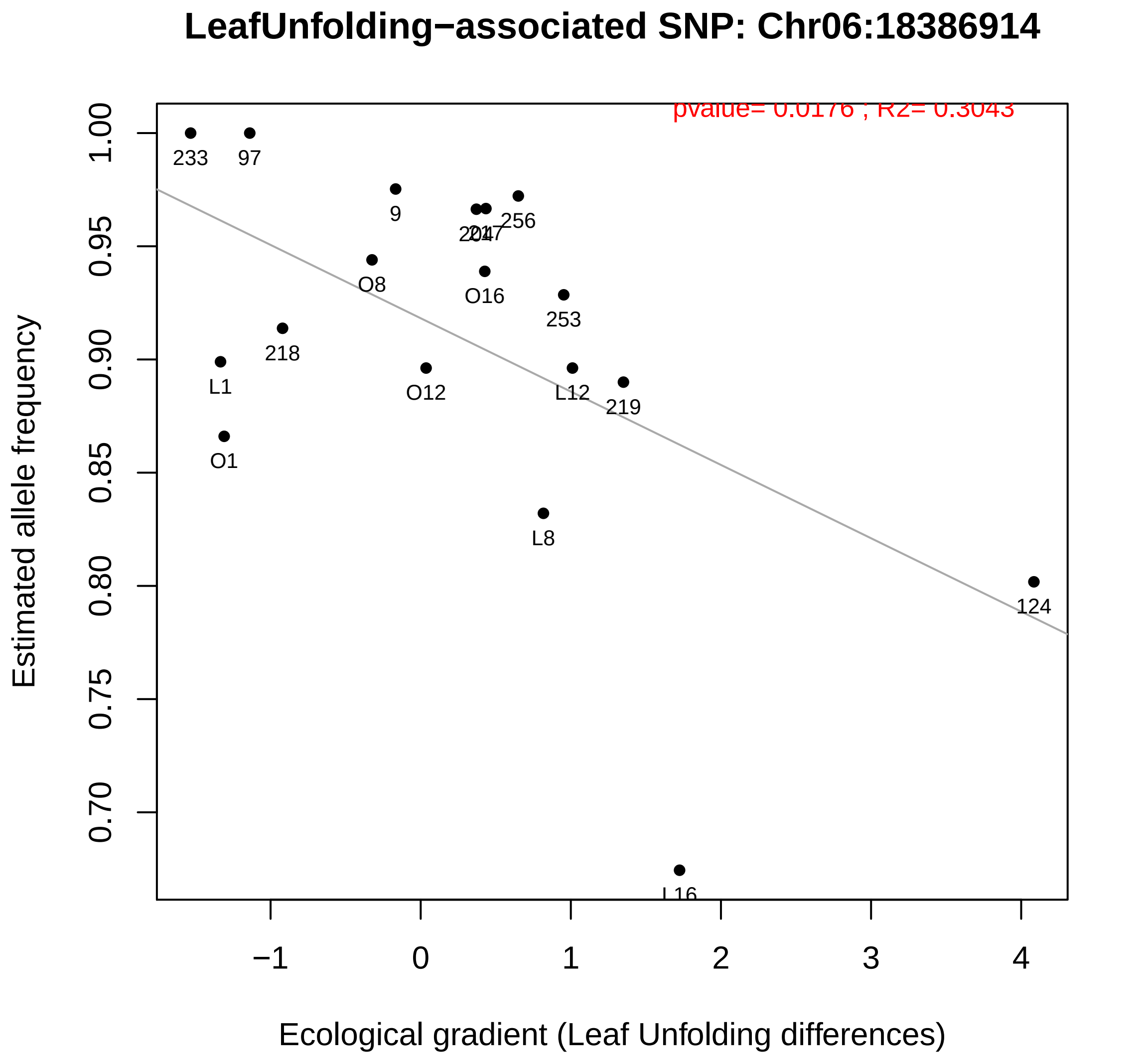

LeafUnfolding−associated SNP: Chr06:18386914
pvalue= 0.0176 ; R2= 0.3043
Estimated allele frequency
Ecological gradient (Leaf Unfolding differences)
1.00
0.95
0.90
0.85
0.80
0.75
0.70
−1
0
1
2
3
4
233
97
9
256
O8
O16
253
218
L1
O12
L12
219
O1
L8
124

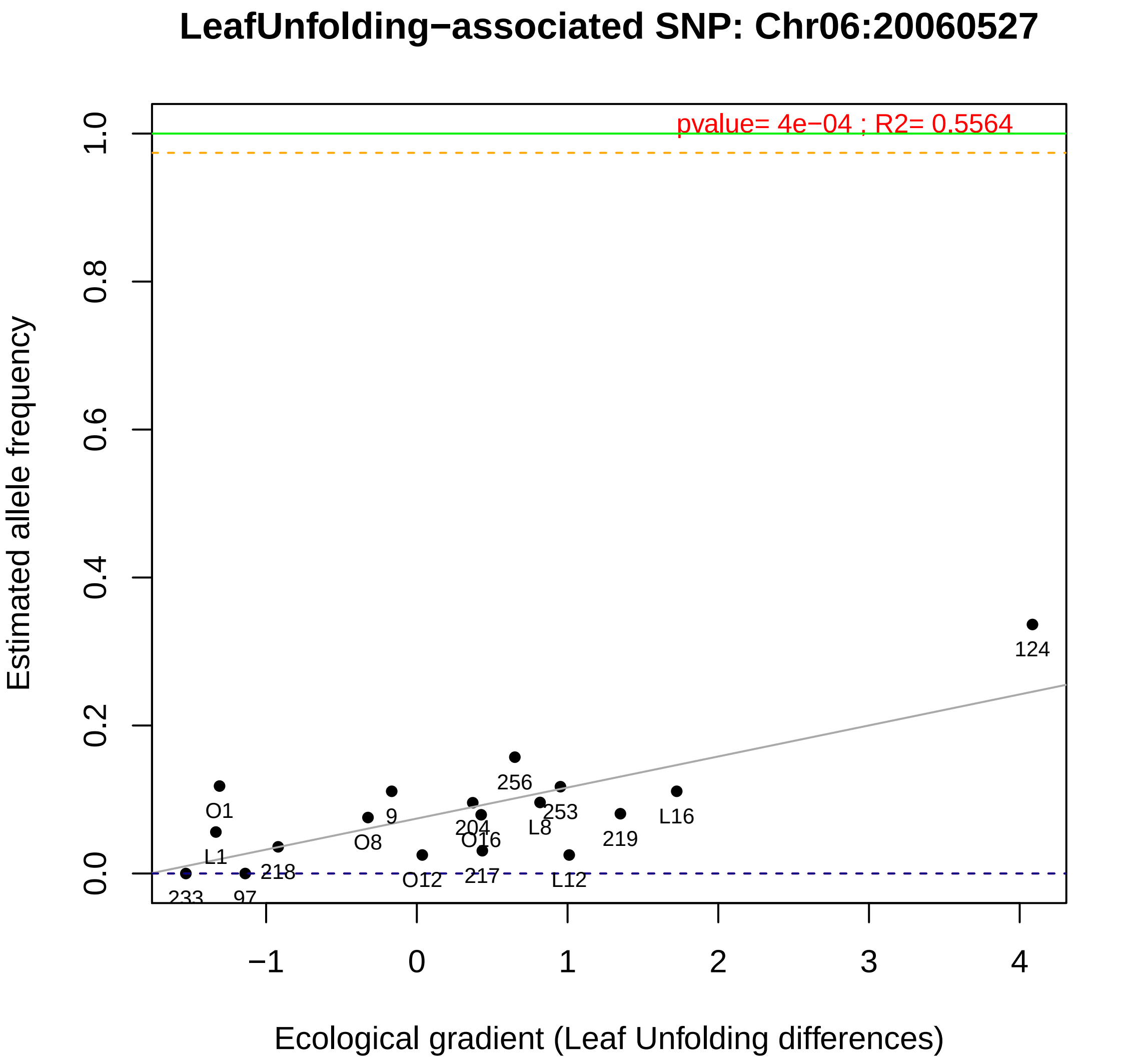

LeafUnfolding−associated SNP: Chr06:20060527
pvalue= 4e−04 ; R2= 0.5564
Estimated allele frequency
Ecological gradient (Leaf Unfolding differences)
0.0
0.2
0.4
0.6
0.8
1.0
−1
0
1
2
3
4
124
256
O1
9
253
L16
L1
O8
204
O16
L8
219
218
O12
217
L12
233
97

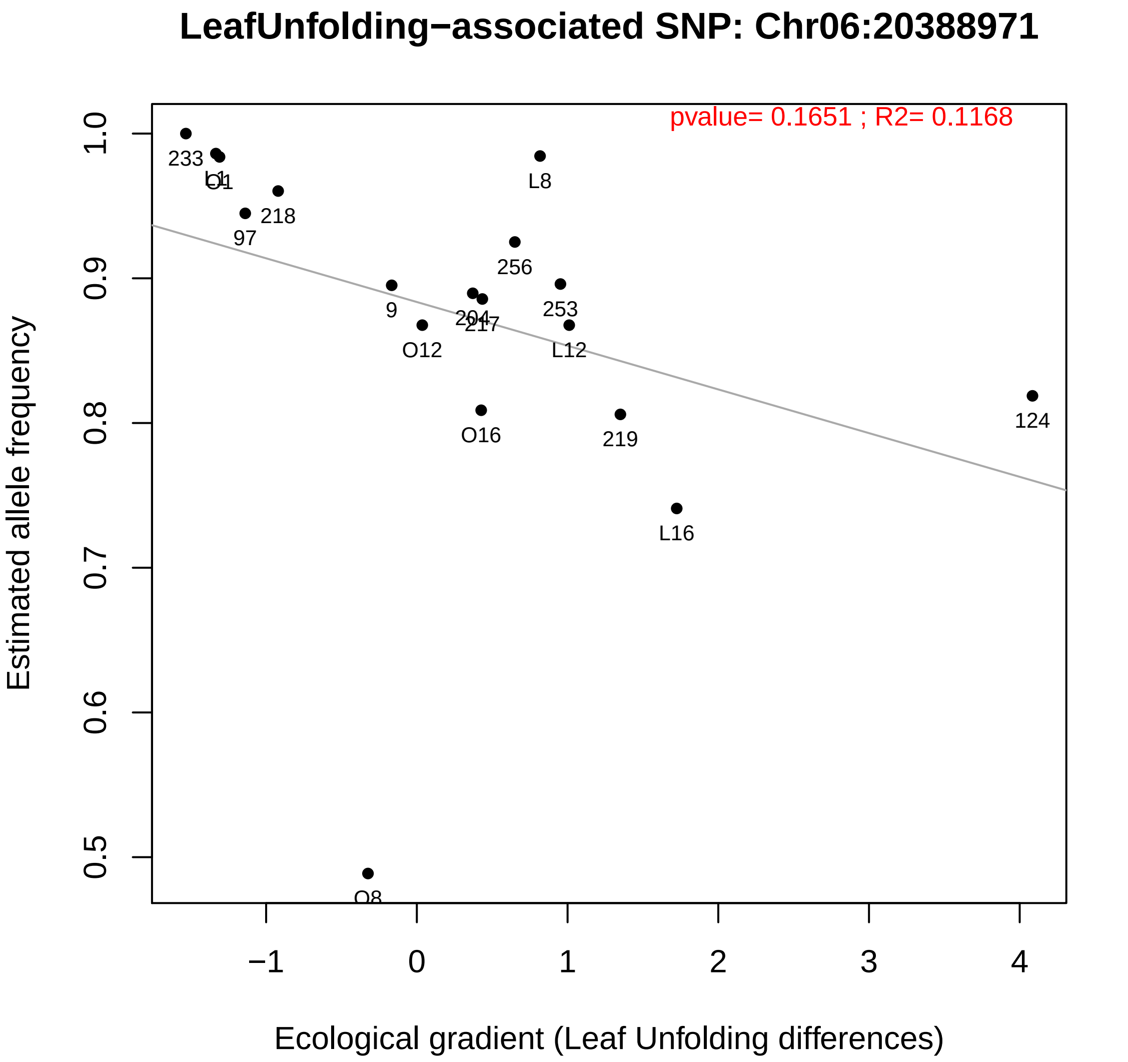
LeafUnfolding−associated SNP: Chr06:20388971
pvalue= 0.1651 ; R2= 0.1168
Estimated allele frequency
Ecological gradient (Leaf Unfolding differences)
1.0
0.9
0.8
0.7
0.6
0.5
−1
0
1
2
3
4
233
L1
O1
218
97
L8
256
9
204
217
253
O12
L12
124
O16
219
L16
O8

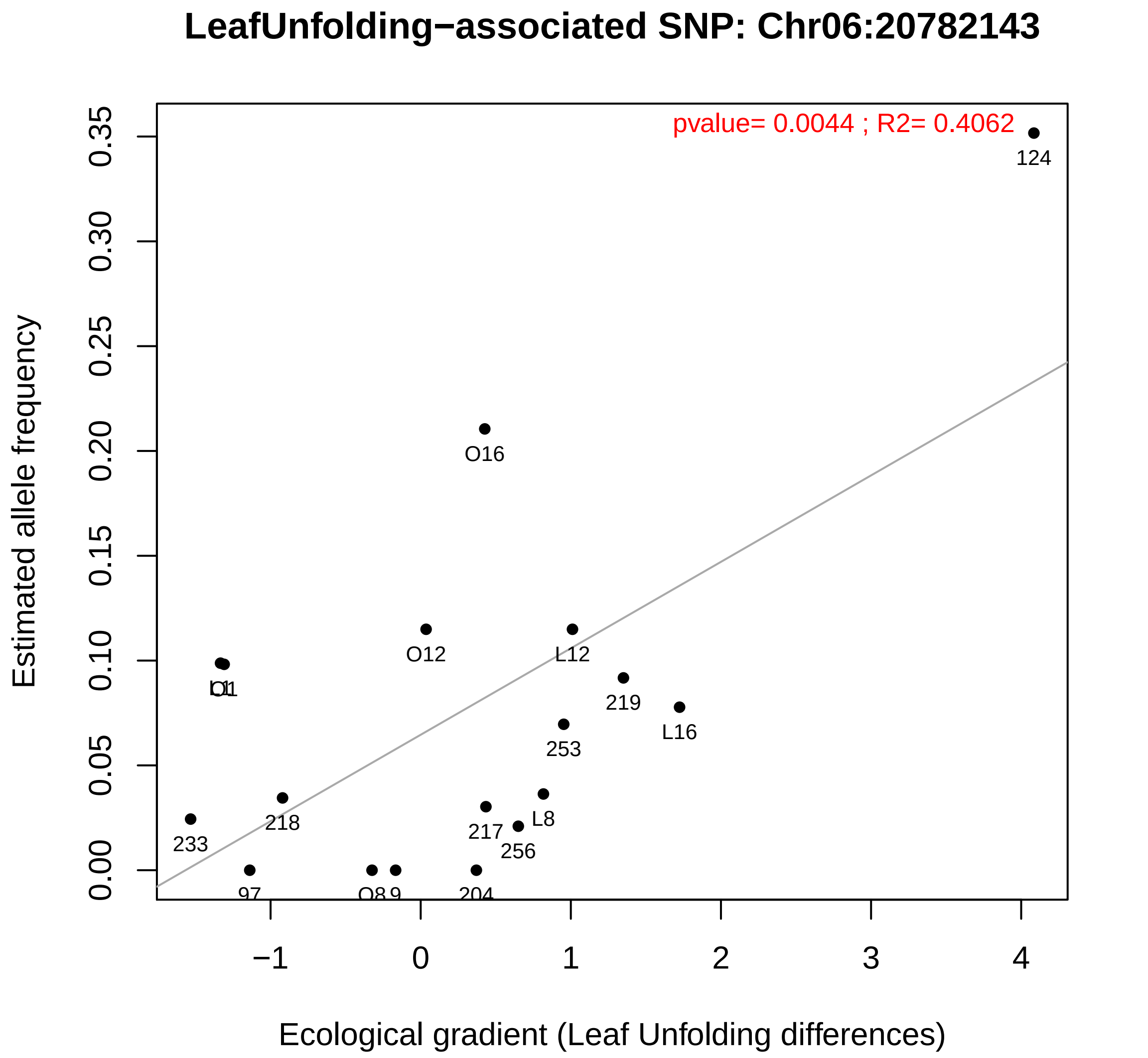

LeafUnfolding−associated SNP: Chr06:20782143
pvalue= 0.0044 ; R2= 0.4062
Estimated allele frequency
Ecological gradient (Leaf Unfolding differences)
0.00
0.05
0.10
0.15
0.20
0.25
0.30
0.35
−1
0
1
2
3
4
124
O16
O12
L12
219
L16
253
218
233
217
L8
256
97
204

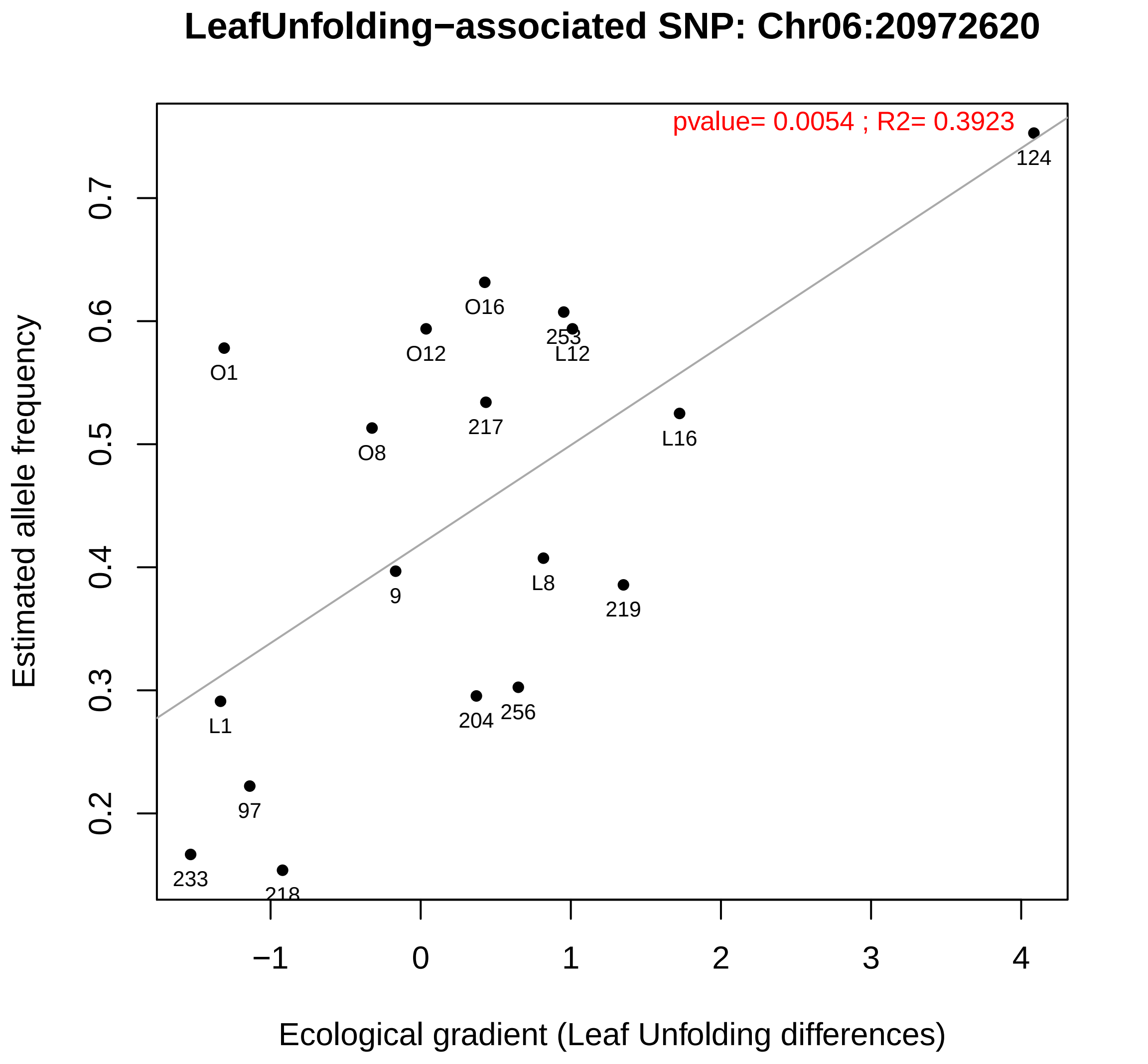

LeafUnfolding−associated SNP: Chr06:20972620
pvalue= 0.0054 ; R2= 0.3923
124
O16
253
L12
O12
O1
217
L16
O8
9
L8
219
L1
204
256
97
233
218
0.7
0.6
0.5
0.4
0.3
0.2
−1
0
1
2
3
4
Estimated allele frequency
Ecological gradient (Leaf Unfolding differences)

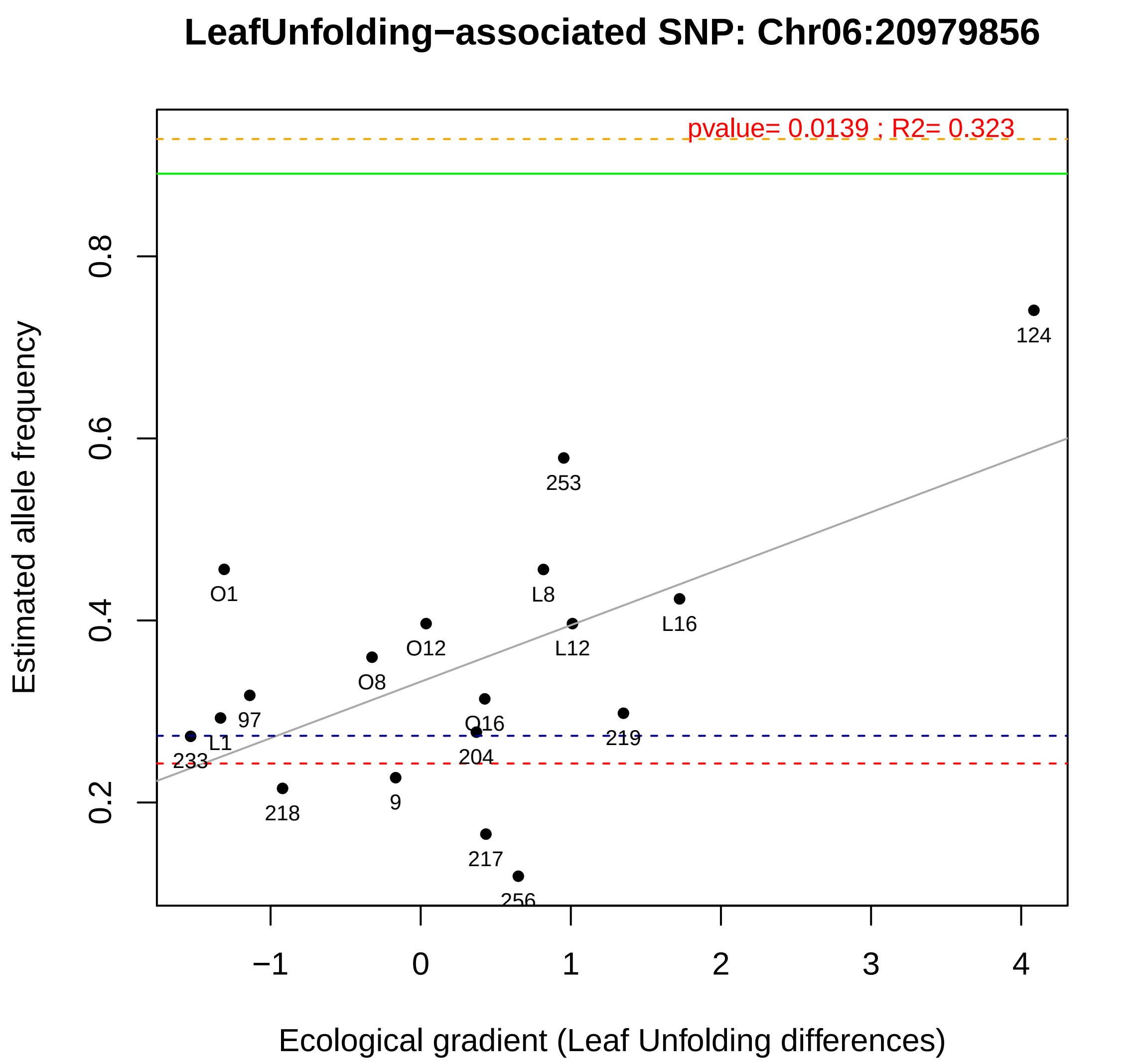

LeafUnfolding−associated SNP: Chr06:20979856
pvalue= 0.0139 ; R2= 0.323
Estimated allele frequency
0.2
0.4
0.6
0.8
−1
0
1
2
3
4
Ecological gradient (Leaf Unfolding differences)
124
253
O1
L8
L16
O12
L12
O8
97
L1
233
O16
204
219
218
9
217
256

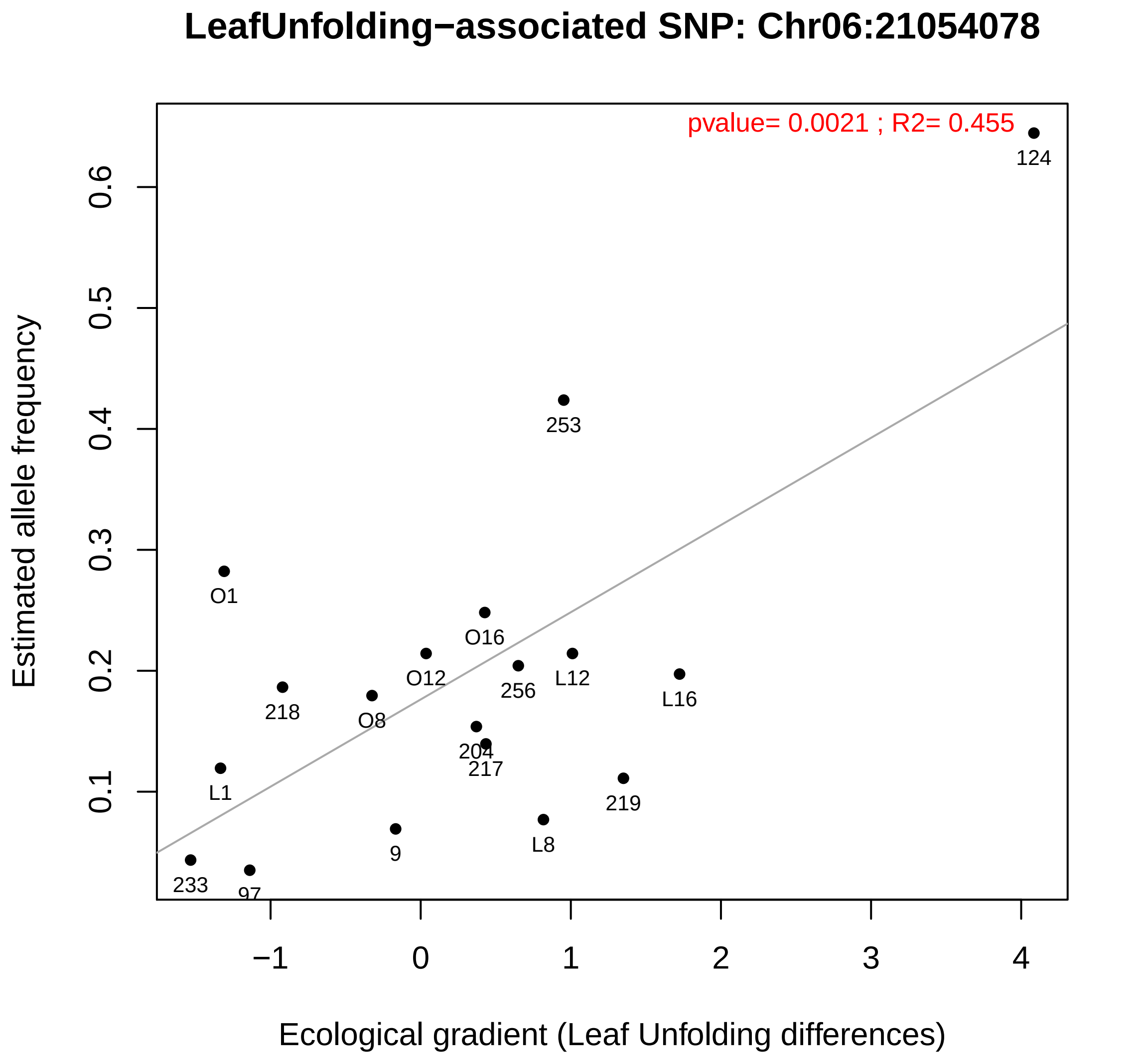

LeafUnfolding−associated SNP: Chr06:21054078
pvalue= 0.0021 ; R2= 0.455
124
253
O1
O16
O12
L12
256
L16
218
O8
204
217
L1
219
9
L8
233
97
0.1
0.2
0.3
0.4
0.5
0.6
−1
0
1
2
3
4
Estimated allele frequency
Ecological gradient (Leaf Unfolding differences)

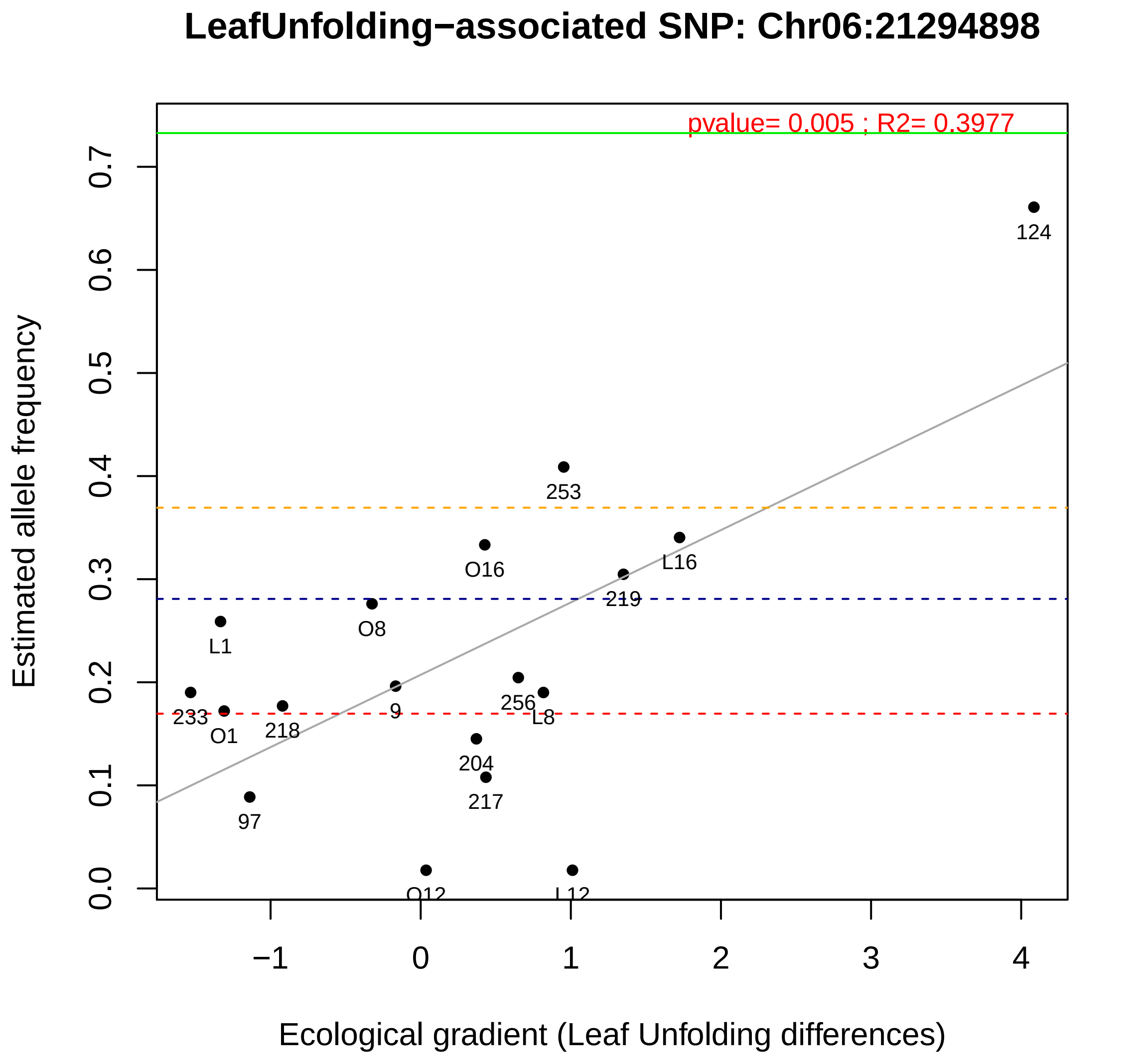

LeafUnfolding−associated SNP: Chr06:21294898
pvalue= 0.005 ; R2= 0.3977
Estimated allele frequency
Ecological gradient (Leaf Unfolding differences)
0.0
0.1
0.2
0.3
0.4
0.5
0.6
0.7
−1
0
1
2
3
4
124
253
O16
L16
219
O8
L1
233
O1
218
9
256
L8
204
217
97
O12
L12

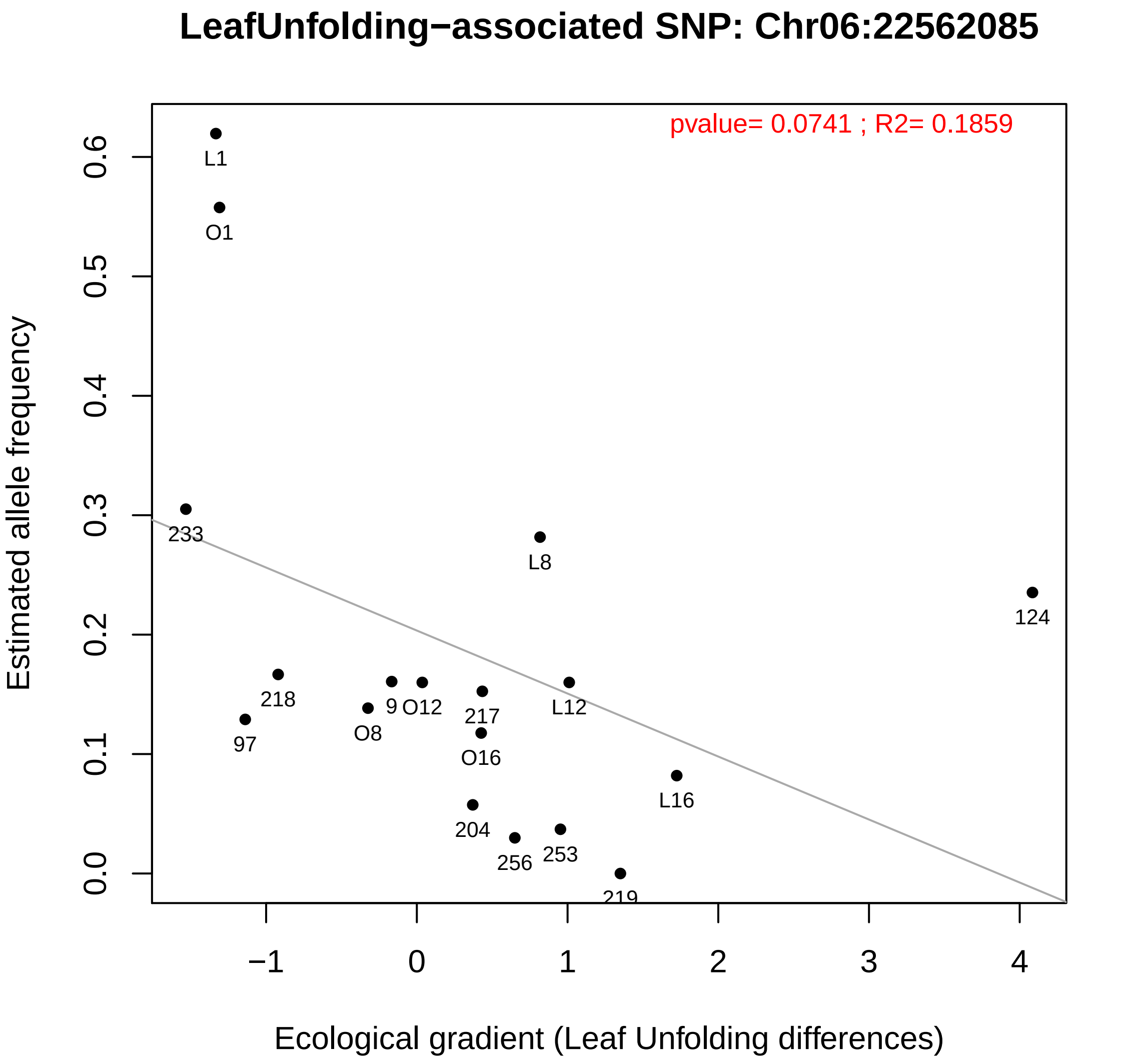

LeafUnfolding−associated SNP: Chr06:22562085
pvalue= 0.0741 ; R2= 0.1859
Estimated allele frequency
0.0
0.1
0.2
0.3
0.4
0.5
0.6
−1
0
1
2
3
4
Ecological gradient (Leaf Unfolding differences)
L1
O1
233
L8
124
218
97
O8
9
O12
217
O16
L12
L16
204
256
253
219

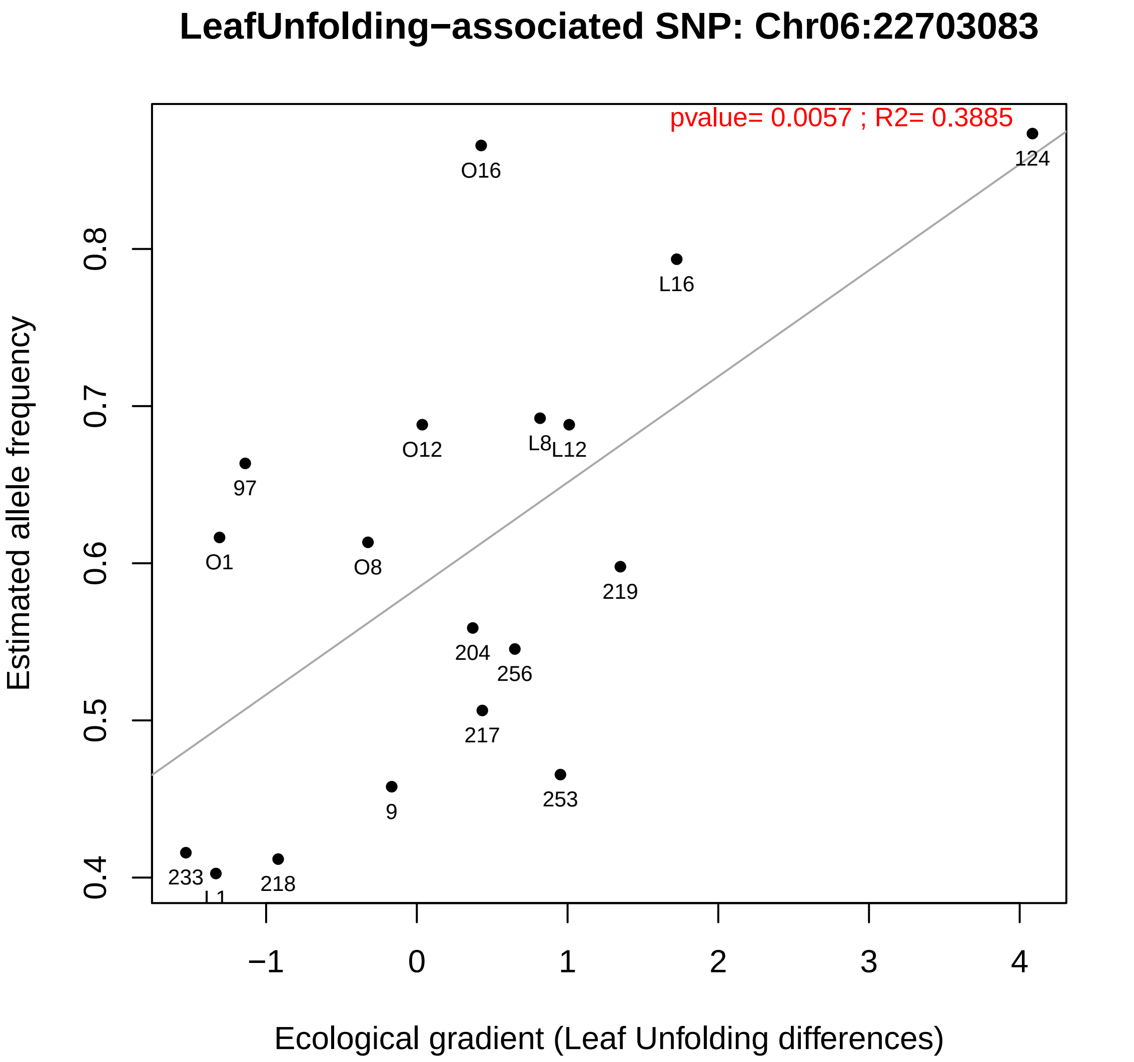

LeafUnfolding−associated SNP: Chr06:22703083
pvalue= 0.0057 ; R2= 0.3885
Estimated allele frequency
Ecological gradient (Leaf Unfolding differences)
0.4
0.5
0.6
0.7
0.8
−1
0
1
2
3
4
O16
124
L16
O12
L8
L12
97
O1
O8
219
204
256
217
253
9
233
218

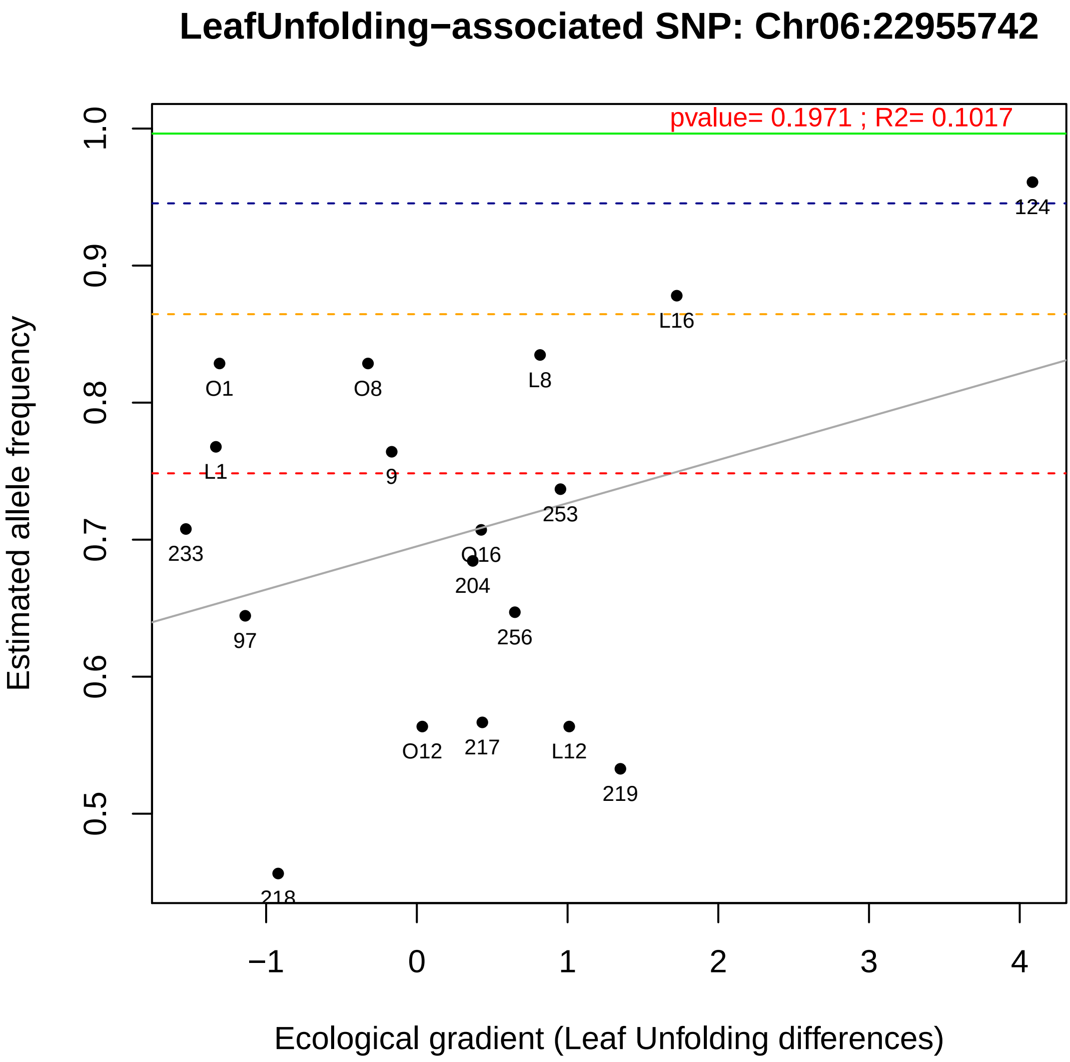

LeafUnfolding−associated SNP: Chr06:22955742
pvalue= 0.1971 ; R2= 0.1017
Estimated allele frequency
Ecological gradient (Leaf Unfolding differences)
1.0
0.9
0.8
0.7
0.6
0.5
−1
0
1
2
3
4
124
L16
O1
O8
L8
L1
9
253
233
O16
204
97
256
O12
217
L12
219
218

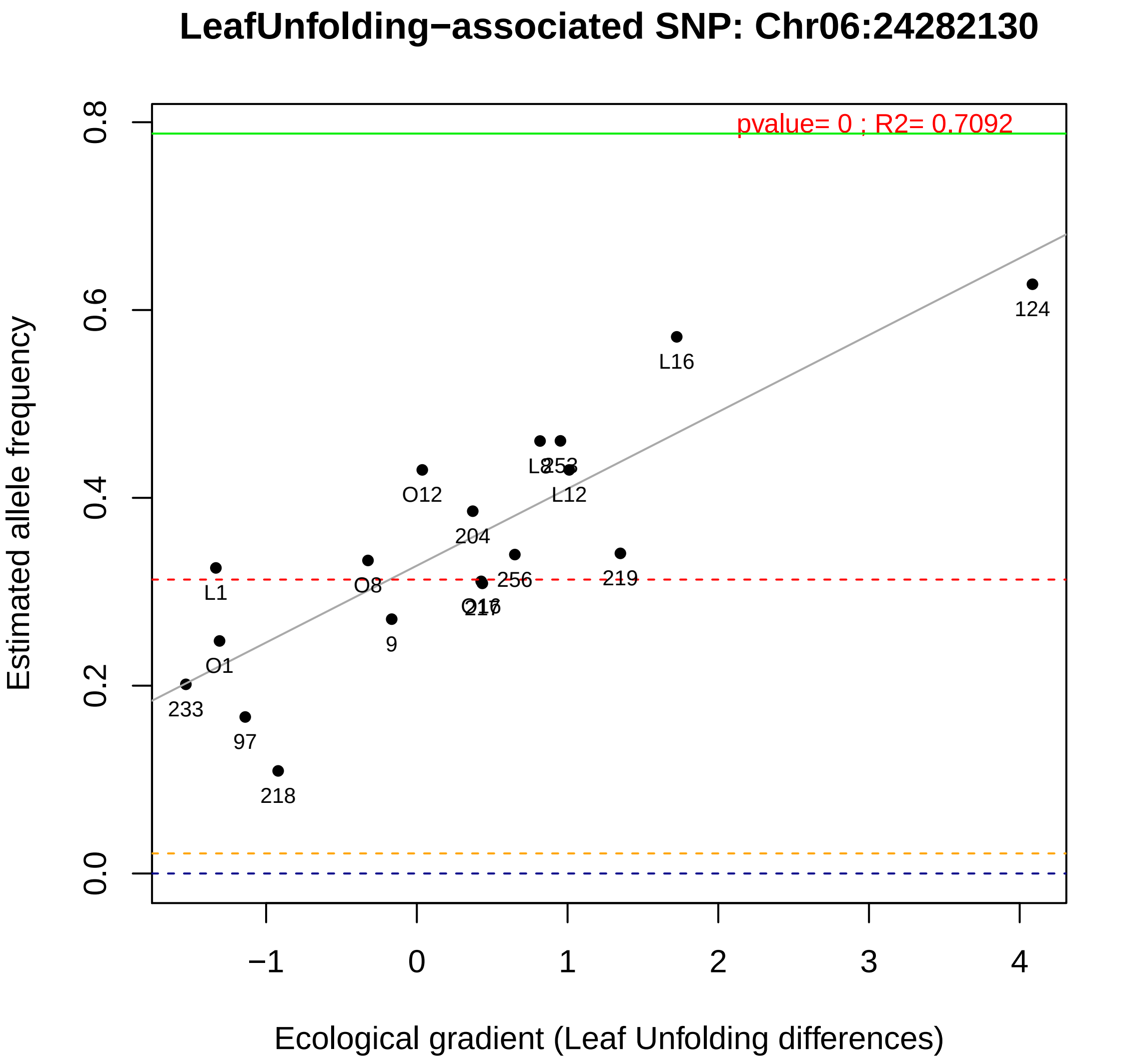
LeafUnfolding−associated SNP: Chr06:24282130
pvalue= 0 ; R2= 0.7092
Estimated allele frequency
Ecological gradient (Leaf Unfolding differences)
0.0
0.2
0.4
0.6
0.8
−1
0
1
2
3
4
124
L16
L8
253
L12
O12
204
256
219
L1
O8
O16
217
9
O1
233
97
218

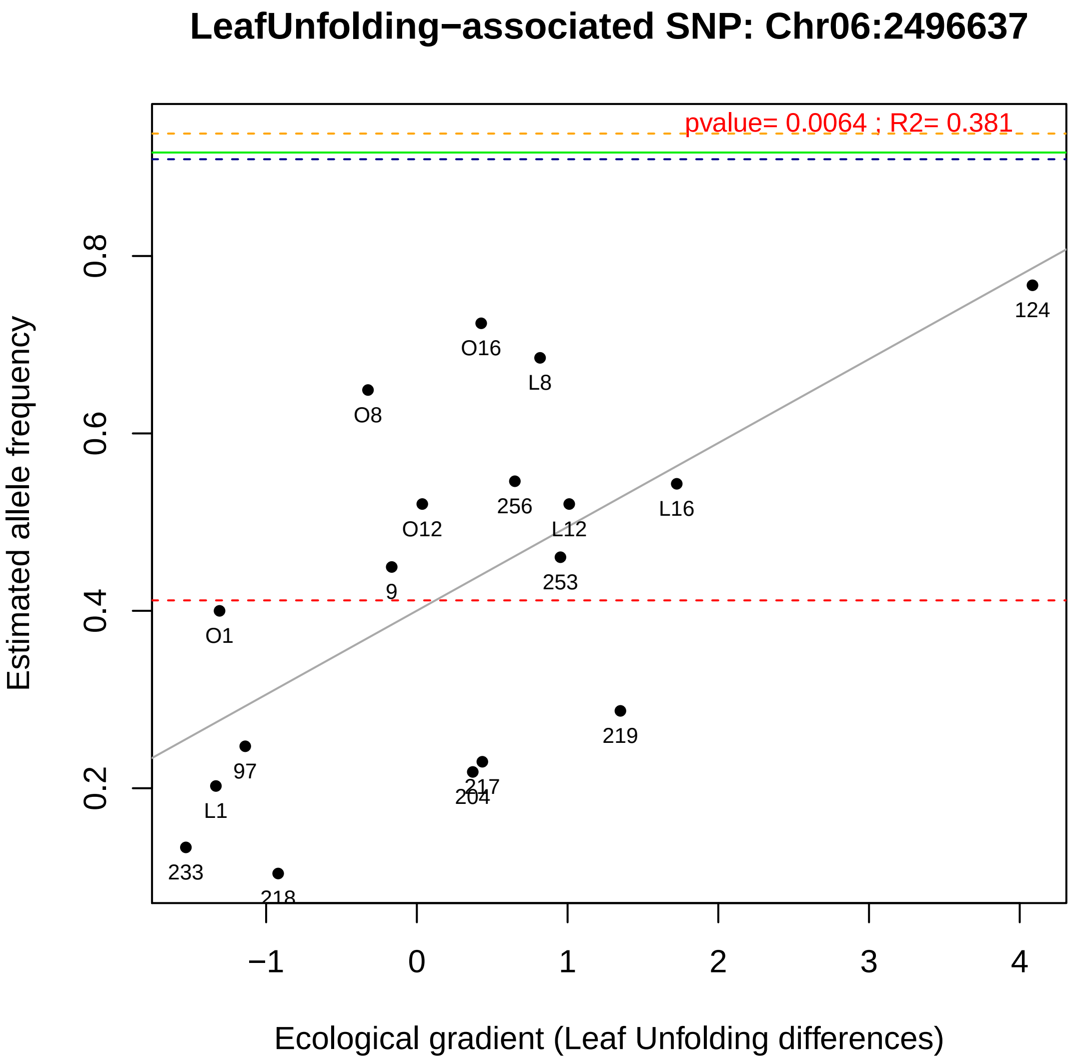

LeafUnfolding−associated SNP: Chr06:2496637
pvalue= 0.0064 ; R2= 0.381
Estimated allele frequency
0.2
0.4
0.6
0.8
−1
0
1
2
3
4
Ecological gradient (Leaf Unfolding differences)
124
O16
L8
O8
256
L16
O12
L12
253
9
O1
219
97
217
204
L1
233
218

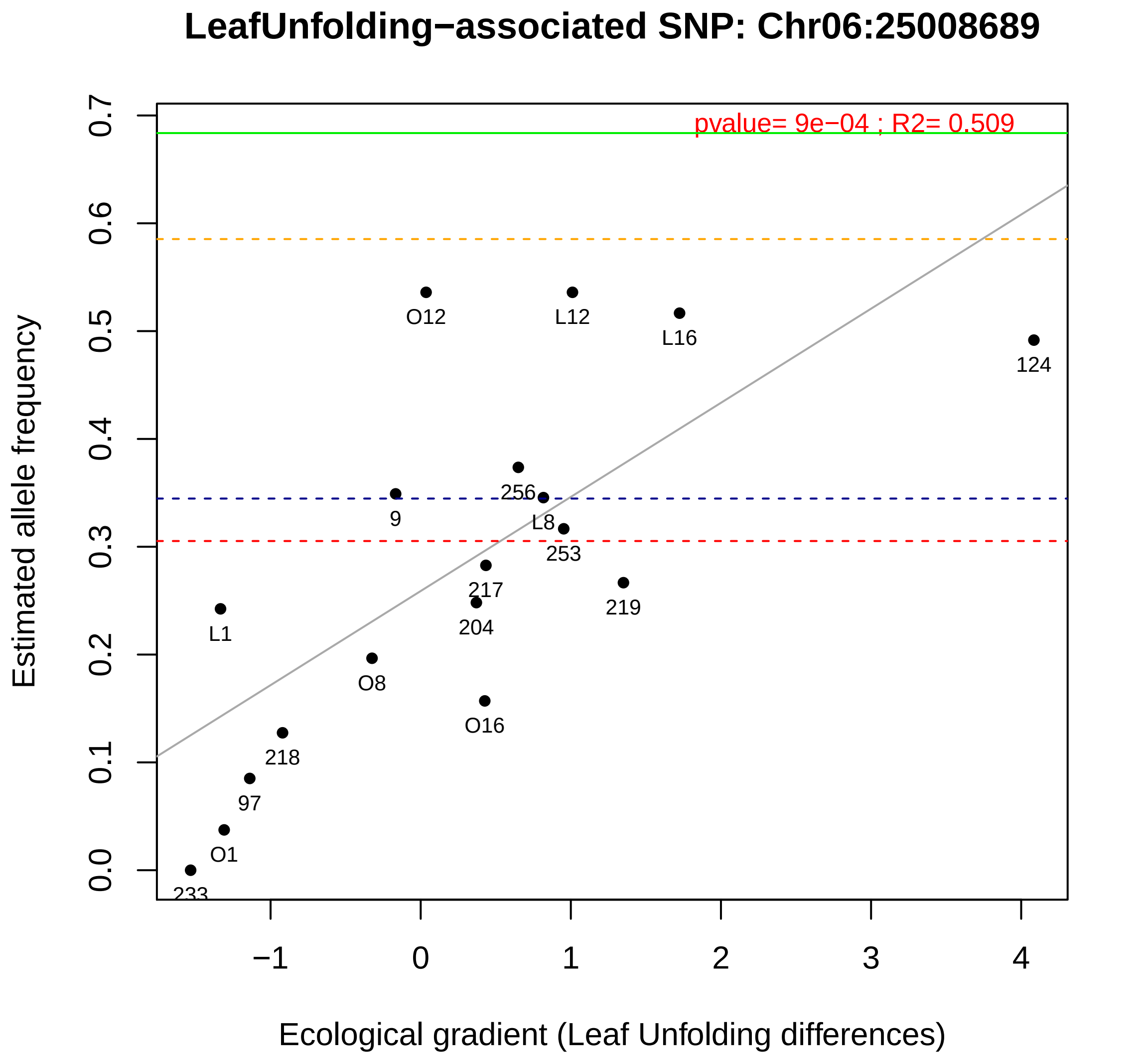
LeafUnfolding−associated SNP: Chr06:25008689
pvalue= 9e−04 ; R2= 0.509
Estimated allele frequency
Ecological gradient (Leaf Unfolding differences)
0.0
0.1
0.2
0.3
0.4
0.5
0.6
0.7
−1
0
1
2
3
4
O12
L12
L16
124
256
9
L8
253
217
219
L1
204
O8
O16
218
97
O1
233

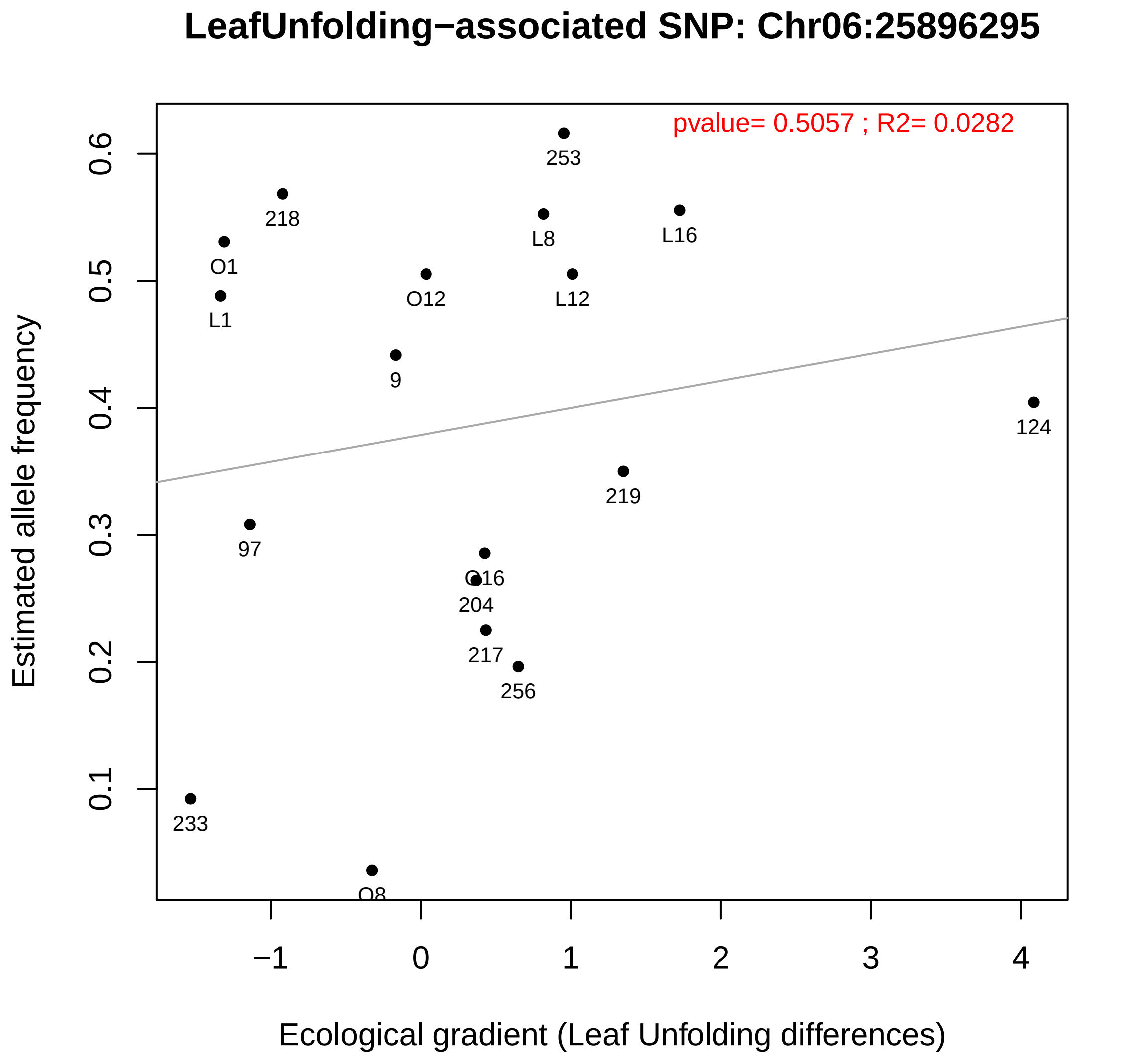
LeafUnfolding−associated SNP: Chr06:25896295
pvalue= 0.5057 ; R2= 0.0282
Estimated allele frequency
Ecological gradient (Leaf Unfolding differences)
0.1
0.2
0.3
0.4
0.5
0.6
−1
0
1
2
3
4
253
218
L8
L16
O1
O12
L12
L1
9
124
219
97
O16
204
217
256
233
O8

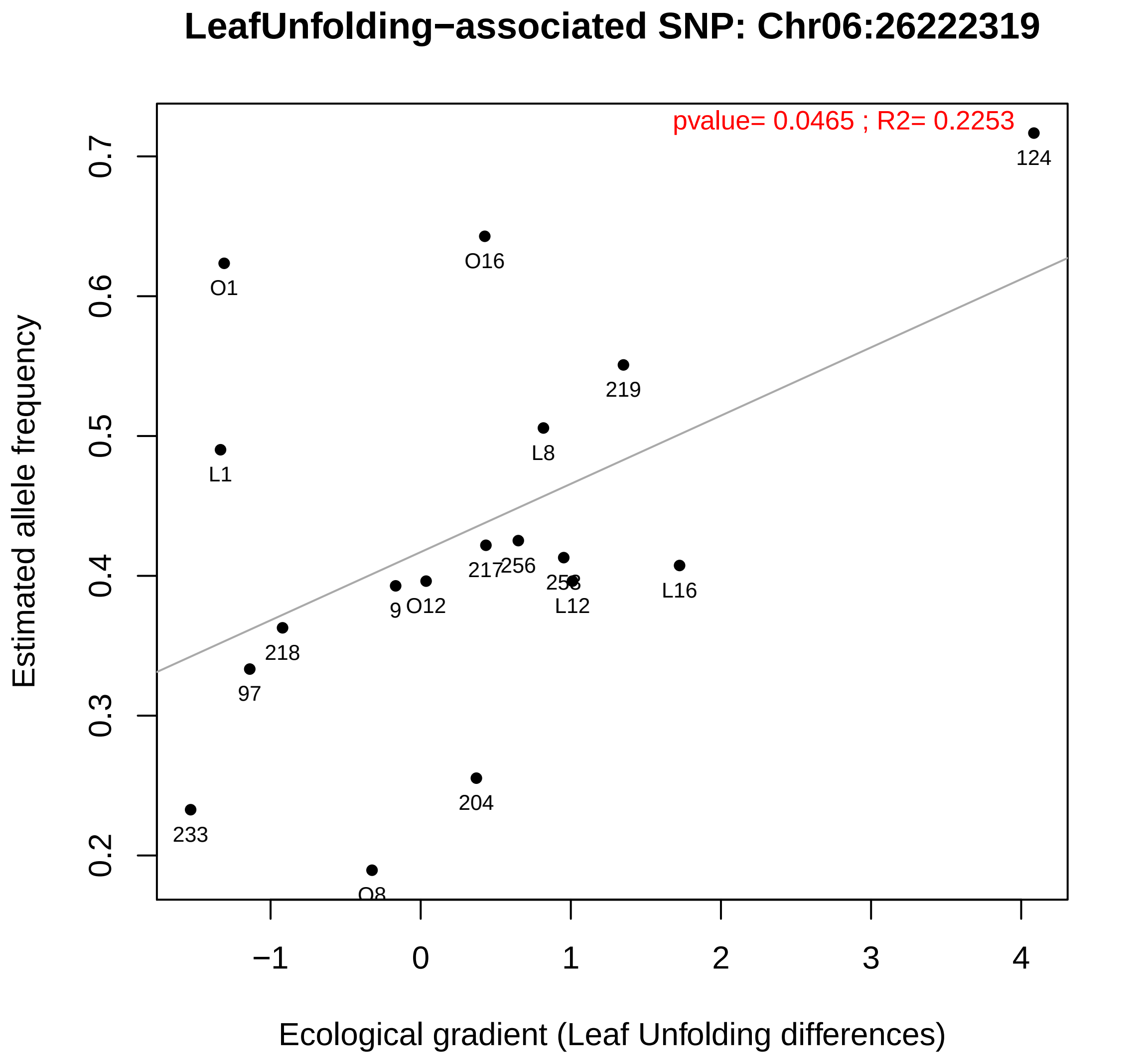
LeafUnfolding−associated SNP: Chr06:26222319
pvalue= 0.0465 ; R2= 0.2253
Estimated allele frequency
Ecological gradient (Leaf Unfolding differences)
0.2
0.3
0.4
0.5
0.6
0.7
−1
0
1
2
3
4
124
O16
O1
219
L8
L1
217
256
L12
L16
9
O12
218
97
204
233
O8

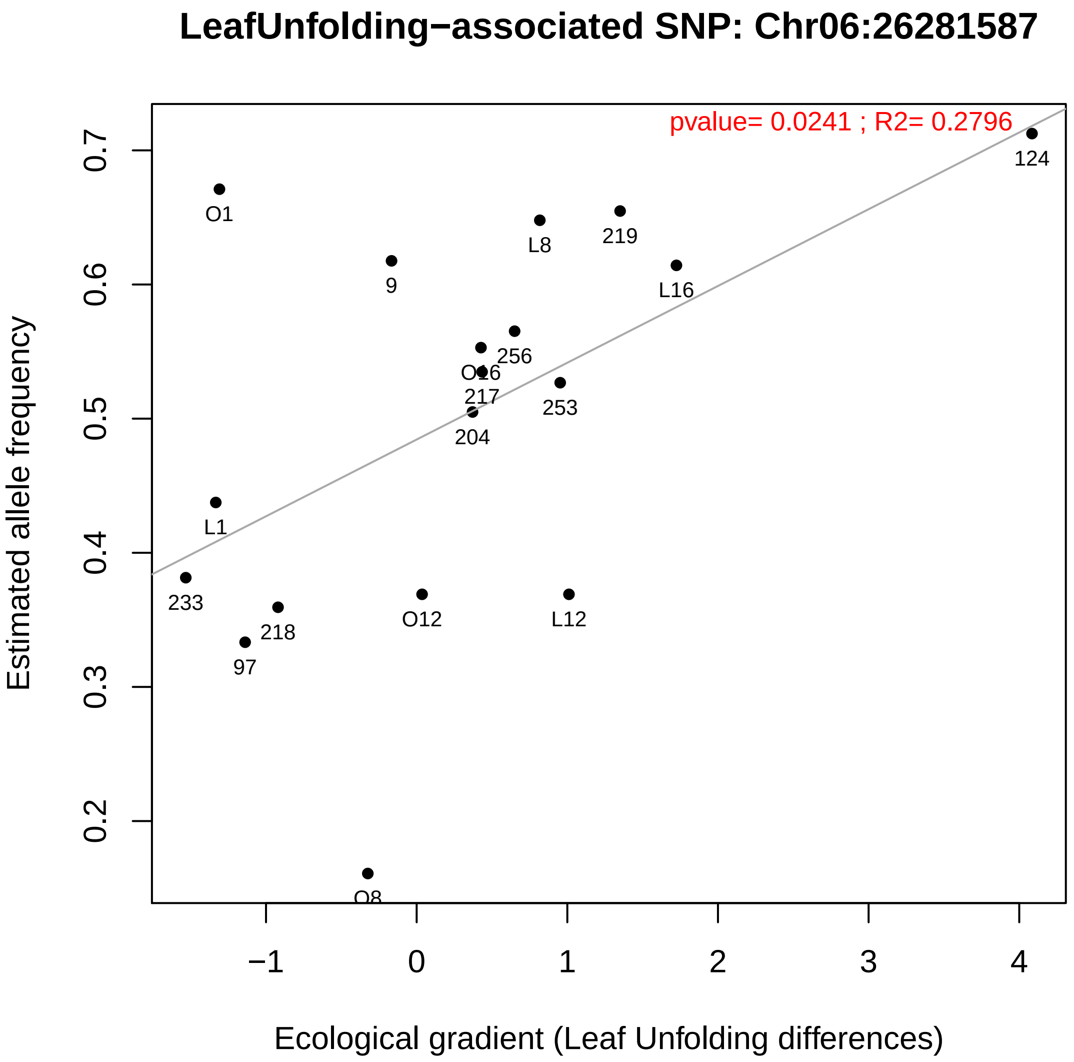
LeafUnfolding−associated SNP: Chr06:26281587
pvalue= 0.0241 ; R2= 0.2796
124
O1
L8
219
9
L16
256
O16
217
253
204
L1
233
218
O12
L12
97
O8
Estimated allele frequency
0.2
0.3
0.4
0.5
0.6
0.7
−1
0
1
2
3
4
Ecological gradient (Leaf Unfolding differences)

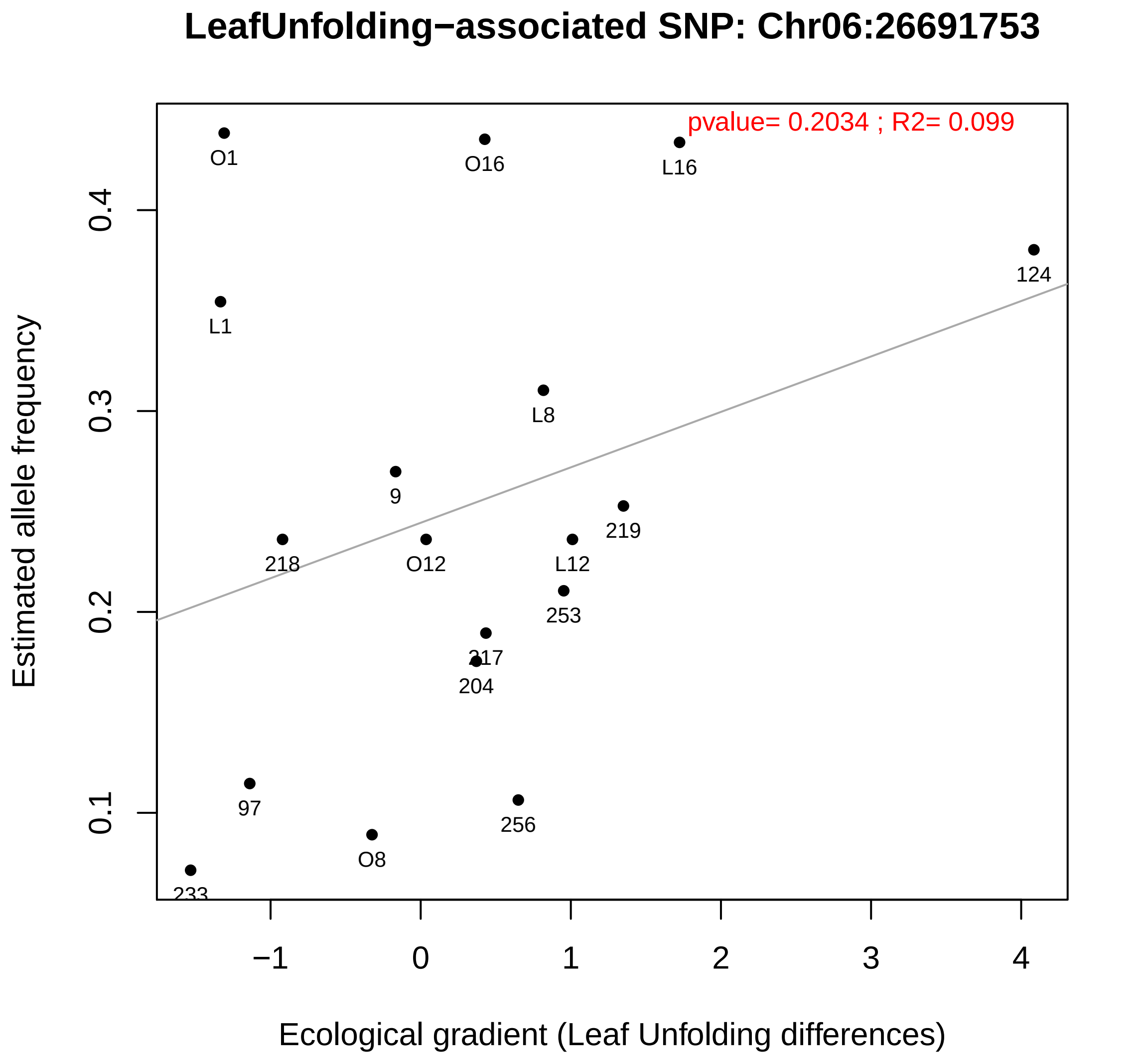

LeafUnfolding−associated SNP: Chr06:26691753
pvalue= 0.2034 ; R2= 0.099
Estimated allele frequency
Ecological gradient (Leaf Unfolding differences)
0.1
0.2
0.3
0.4
−1
0
1
2
3
4
O1
O16
L16
124
L1
L8
9
218
O12
L12
219
253
217
204
97
256
O8
233

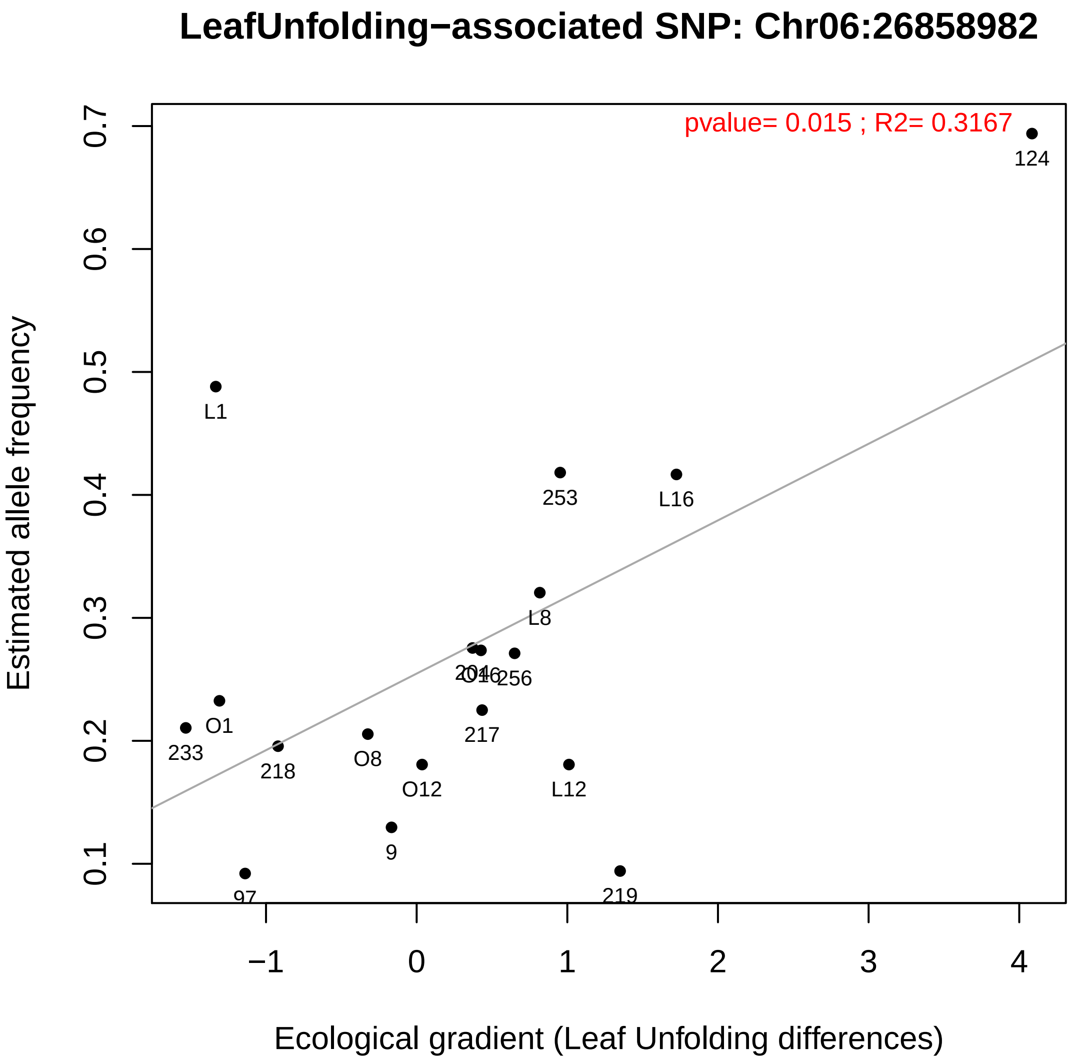
LeafUnfolding−associated SNP: Chr06:26858982
pvalue= 0.015 ; R2= 0.3167
Estimated allele frequency
Ecological gradient (Leaf Unfolding differences)
0.1
0.2
0.3
0.4
0.5
0.6
0.7
−1
0
1
2
3
4
124
L1
253
L16
L8
204
O16
256
217
O1
233
218
O8
O12
L12
9
97
219

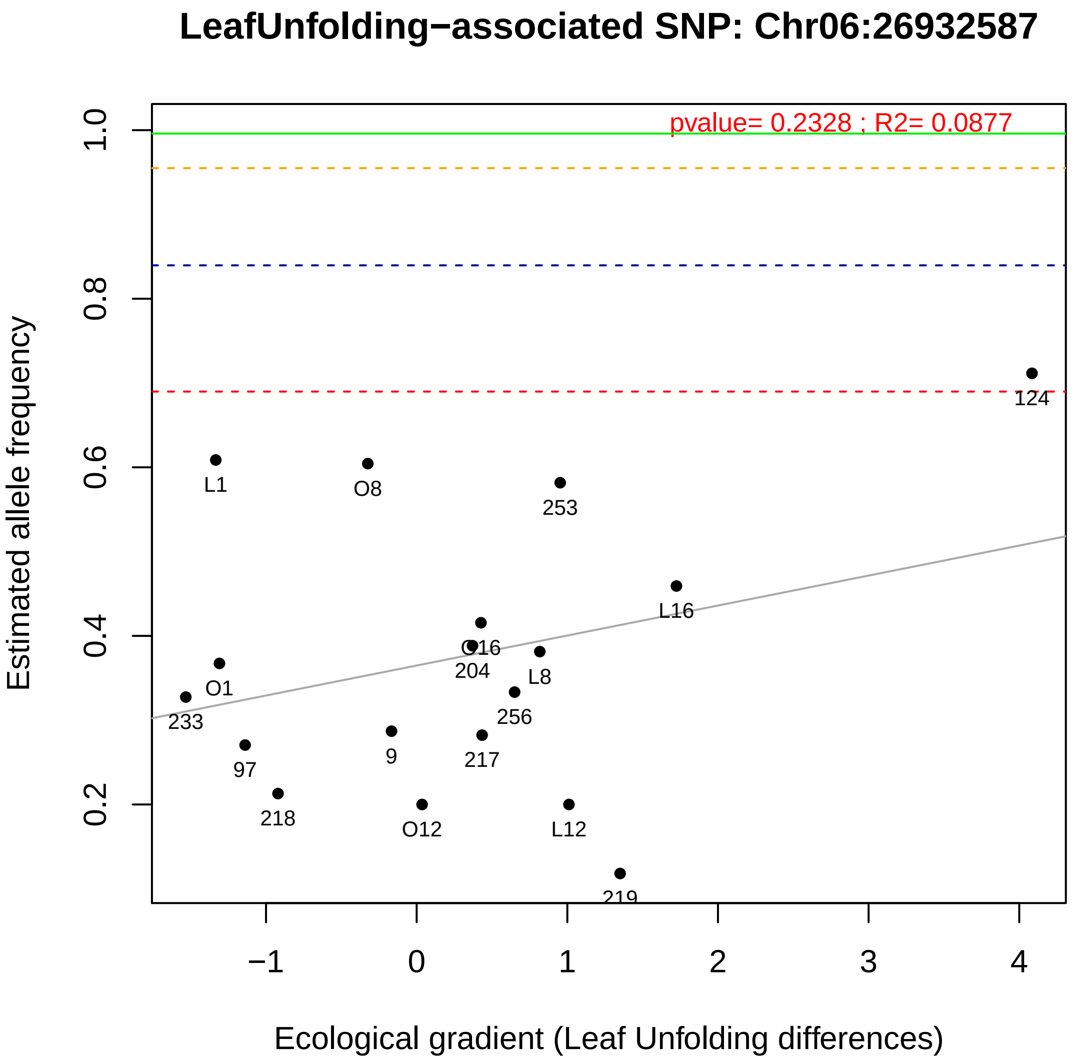

LeafUnfolding−associated SNP: Chr06:26932587
pvalue= 0.2328 ; R2= 0.0877
Estimated allele frequency
1.0
0.8
0.6
0.4
0.2
124
L1
O8
253
L16
O16
204
L8
O1
233
256
97
9
217
218
O12
L12
219
−1
0
1
2
3
4
Ecological gradient (Leaf Unfolding differences)

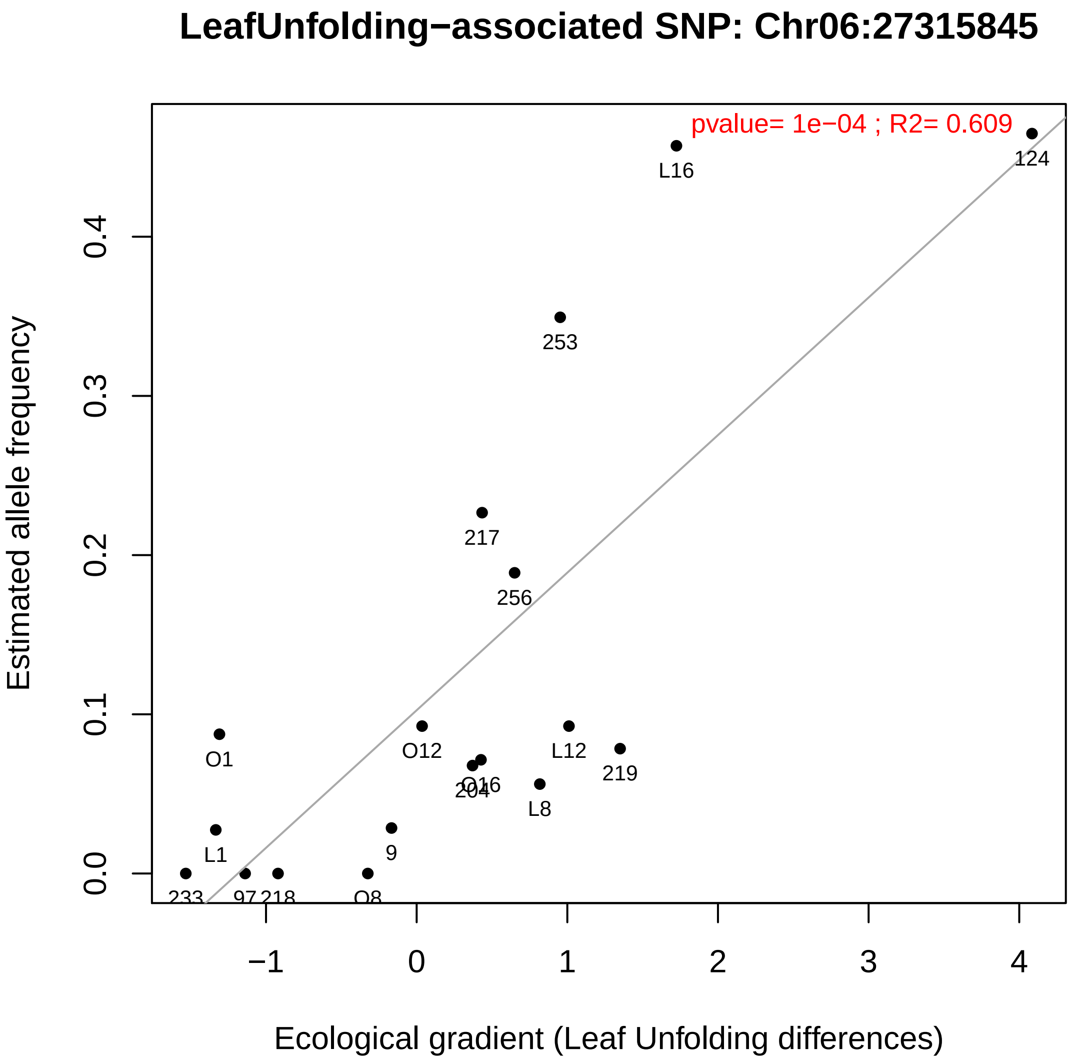

LeafUnfolding−associated SNP: Chr06:27315845
pvalue= 1e−04 ; R2= 0.609
L16
124
253
217
256
O1
O12
L12
219
O16
204
L8
L1
9
233
97
218
O8
0.0
0.1
0.2
0.3
0.4
−1
0
1
2
3
4
Estimated allele frequency
Ecological gradient (Leaf Unfolding differences)

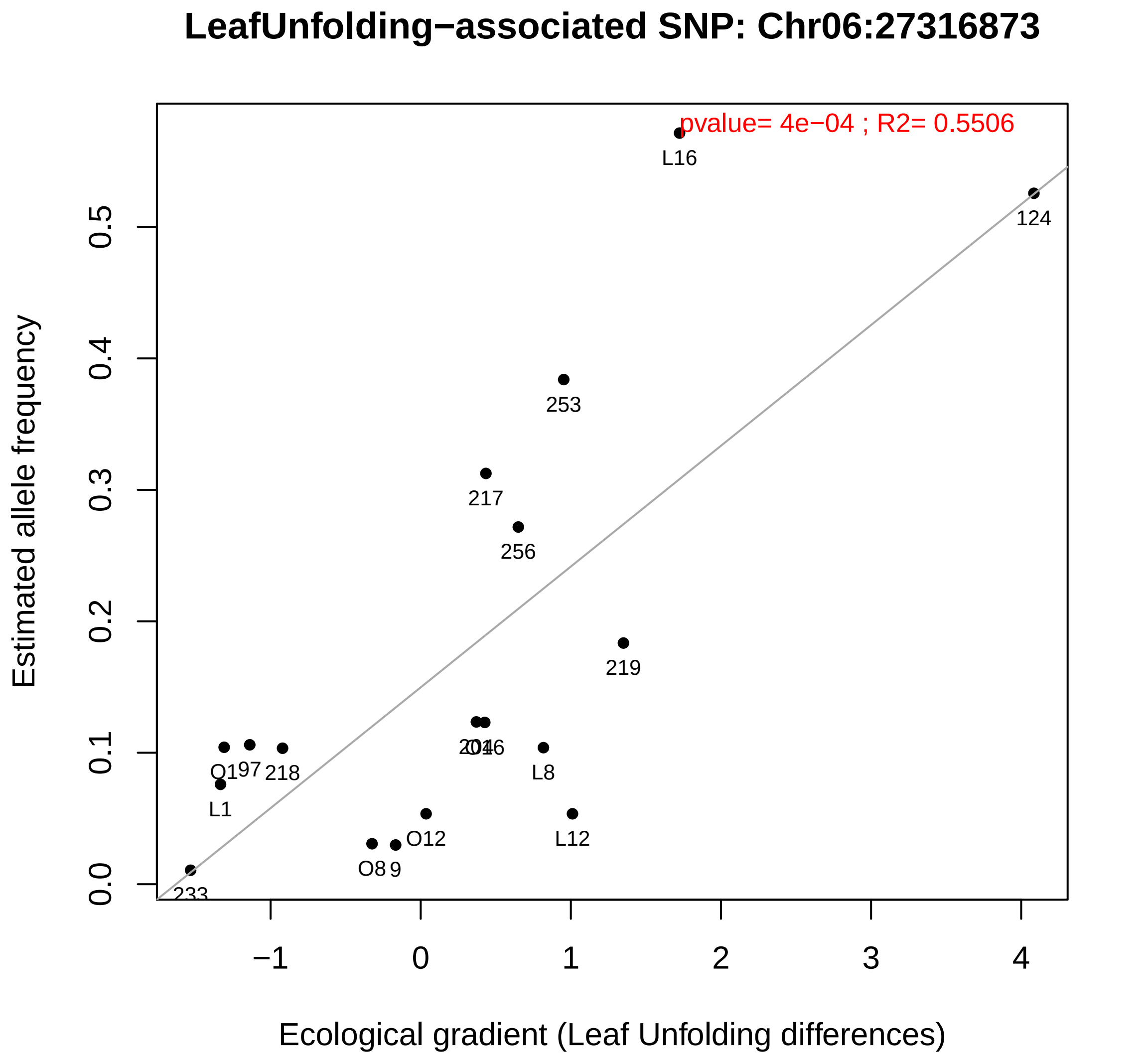

LeafUnfolding−associated SNP: Chr06:27316873
pvalue= 4e−04 ; R2= 0.5506
Estimated allele frequency
Ecological gradient (Leaf Unfolding differences)
0.0
0.1
0.2
0.3
0.4
0.5
−1
0
1
2
3
4
L16
124
253
217
256
219
L8
L12
O12
O8
9
218
L1
233

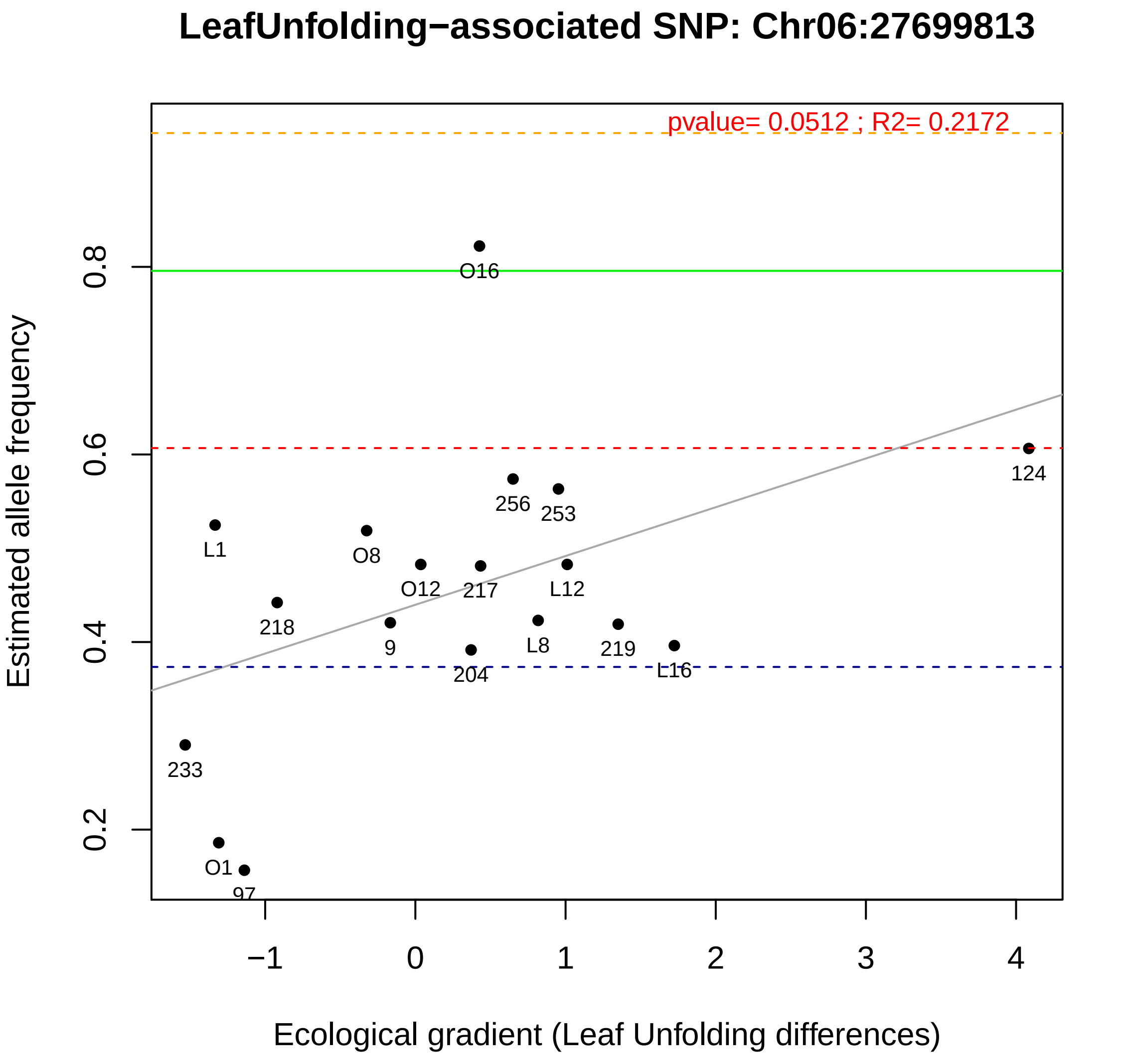

LeafUnfolding−associated SNP: Chr06:27699813
pvalue= 0.0512 ; R2= 0.2172
Estimated allele frequency
0.2
0.4
0.6
0.8
−1
0
1
2
3
4
Ecological gradient (Leaf Unfolding differences)
O16
124
256
253
L1
O8
O12
217
L12
218
9
L8
219
L16
204
233
O1
97

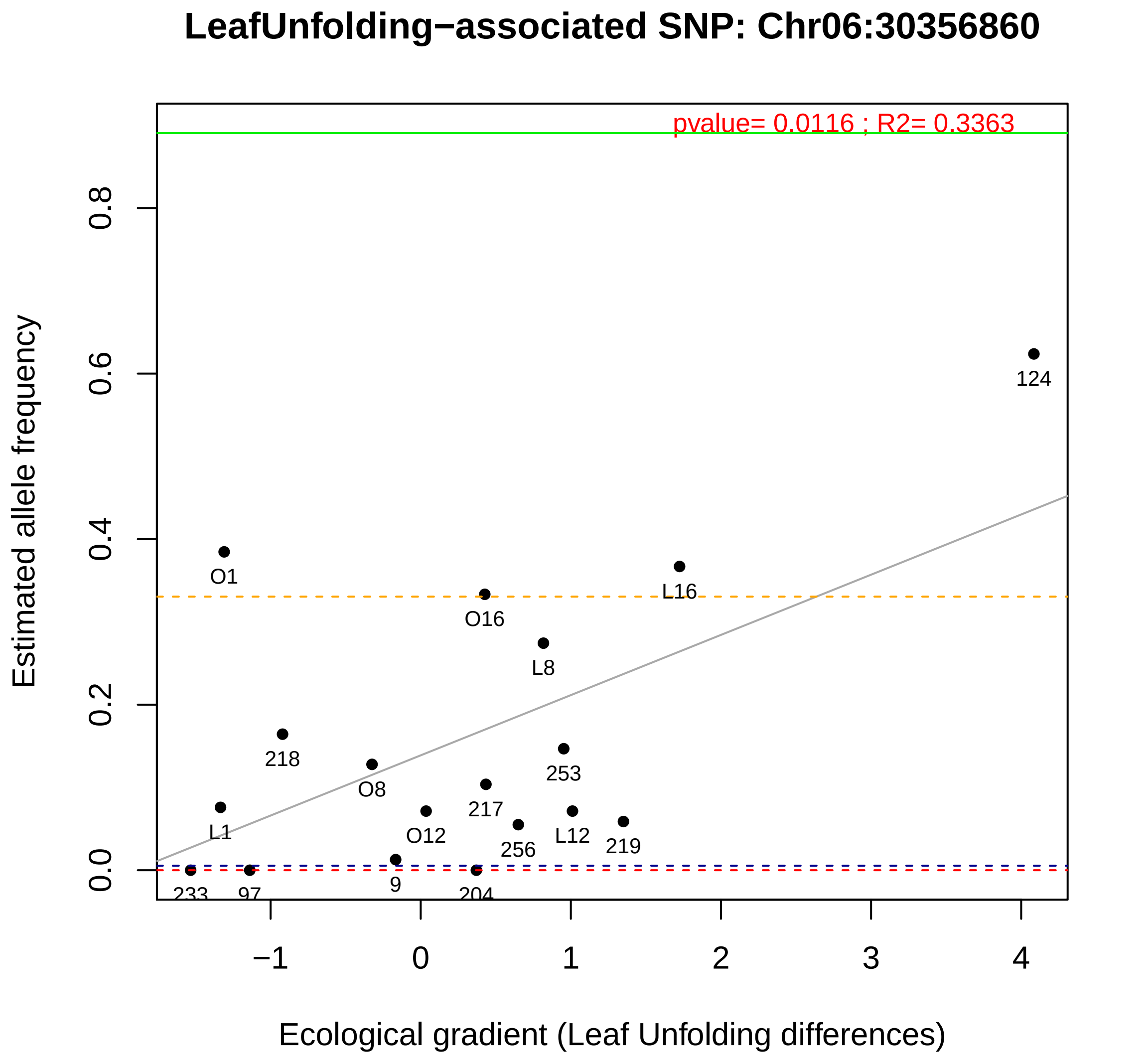

LeafUnfolding−associated SNP: Chr06:30356860
pvalue= 0.0116 ; R2= 0.3363
Estimated allele frequency
0.0
0.2
0.4
0.6
0.8
−1
0
1
2
3
4
Ecological gradient (Leaf Unfolding differences)
124
O1
L16
O16
L8
218
253
O8
217
L1
O12
256
L12
219
9

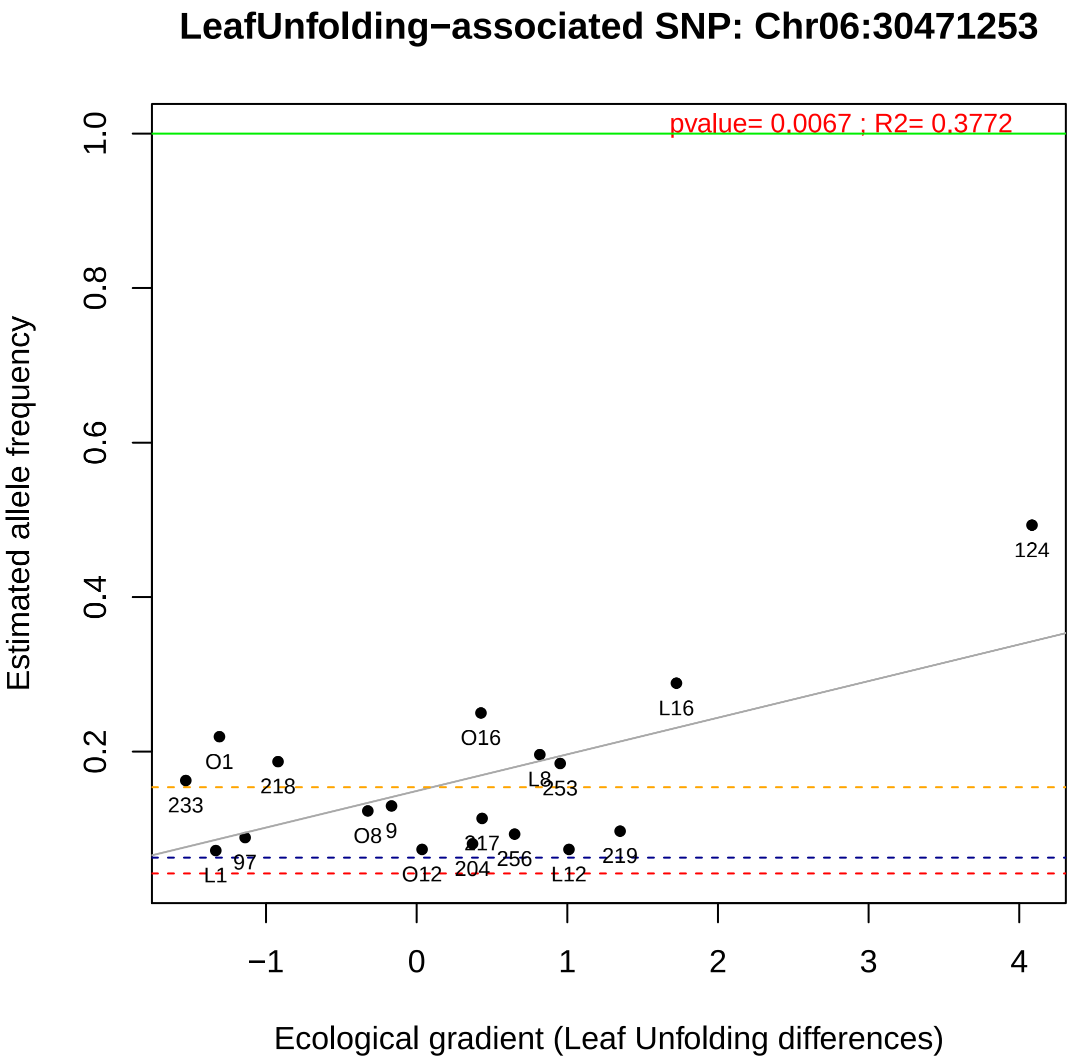
LeafUnfolding−associated SNP: Chr06:30471253
pvalue= 0.0067 ; R2= 0.3772
Estimated allele frequency
Ecological gradient (Leaf Unfolding differences)
0.2
0.4
0.6
0.8
1.0
−1
0
1
2
3
4
124
L16
O16
O1
218
233
L8
253
O8
9
217
204
256
219
L12
O12
97
L1

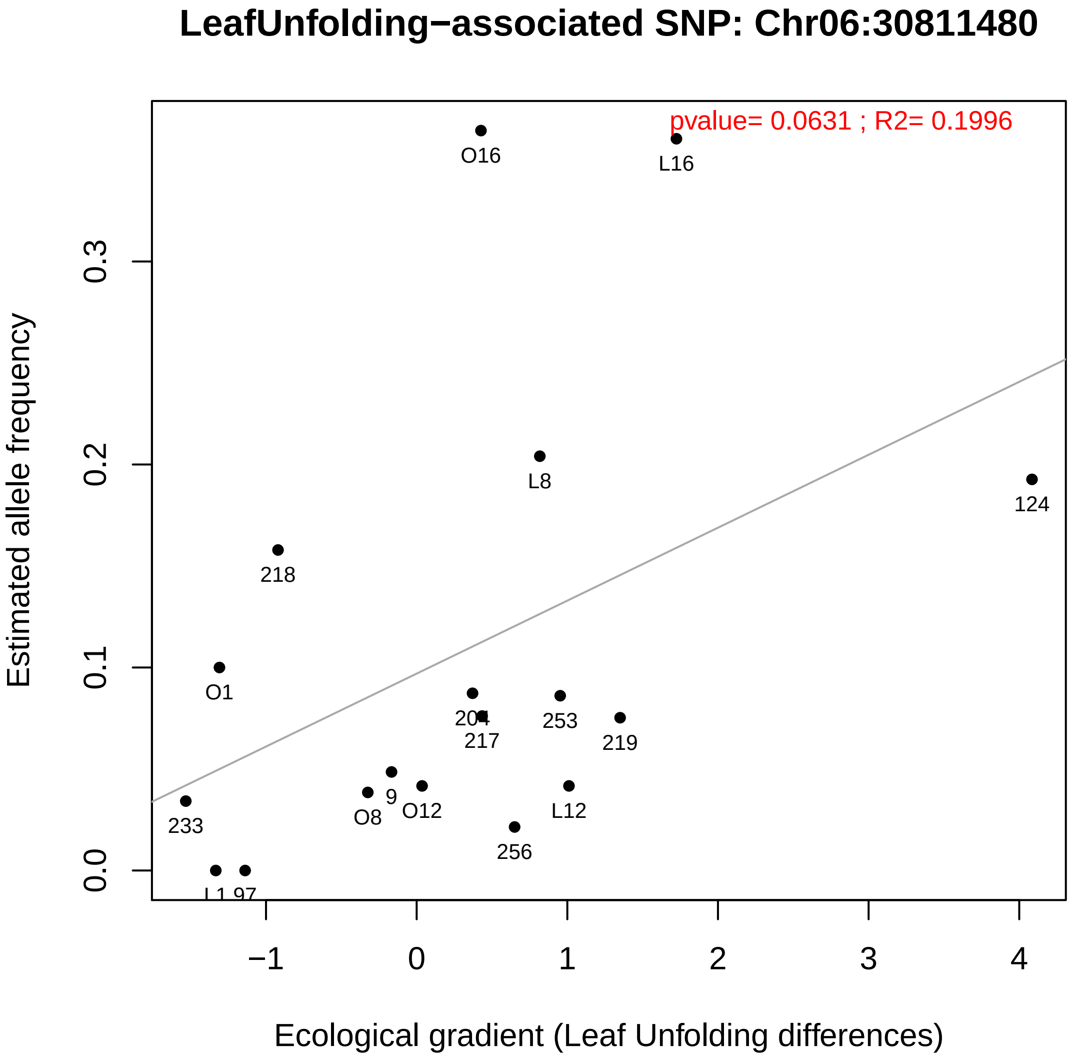
LeafUnfolding−associated SNP: Chr06:30811480
pvalue= 0.0631 ; R2= 0.1996
O16
L16
L8
124
218
O1
204
217
253
219
9
O12
O8
233
L12
256
Estimated allele frequency
0.0
0.1
0.2
0.3
−1
0
1
2
3
4
Ecological gradient (Leaf Unfolding differences)

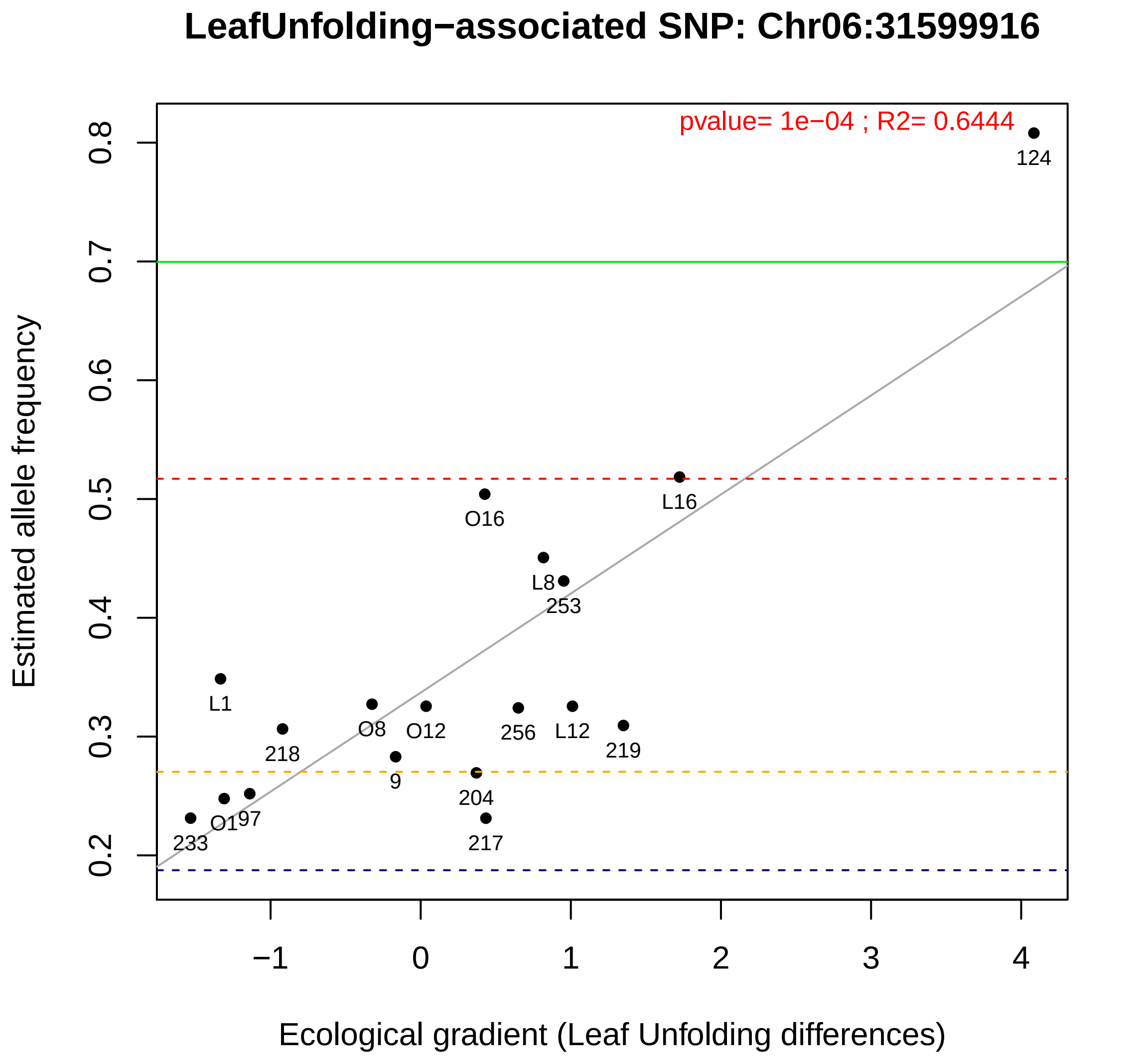

LeafUnfolding−associated SNP: Chr06:31599916
pvalue= 1e−04 ; R2= 0.6444
Estimated allele frequency
Ecological gradient (Leaf Unfolding differences)
0.2
0.3
0.4
0.5
0.6
0.7
0.8
−1
0
1
2
3
4
124
L16
O16
L8
253
L1
218
O8
O12
256
L12
219
9
204
233
O1
97
217

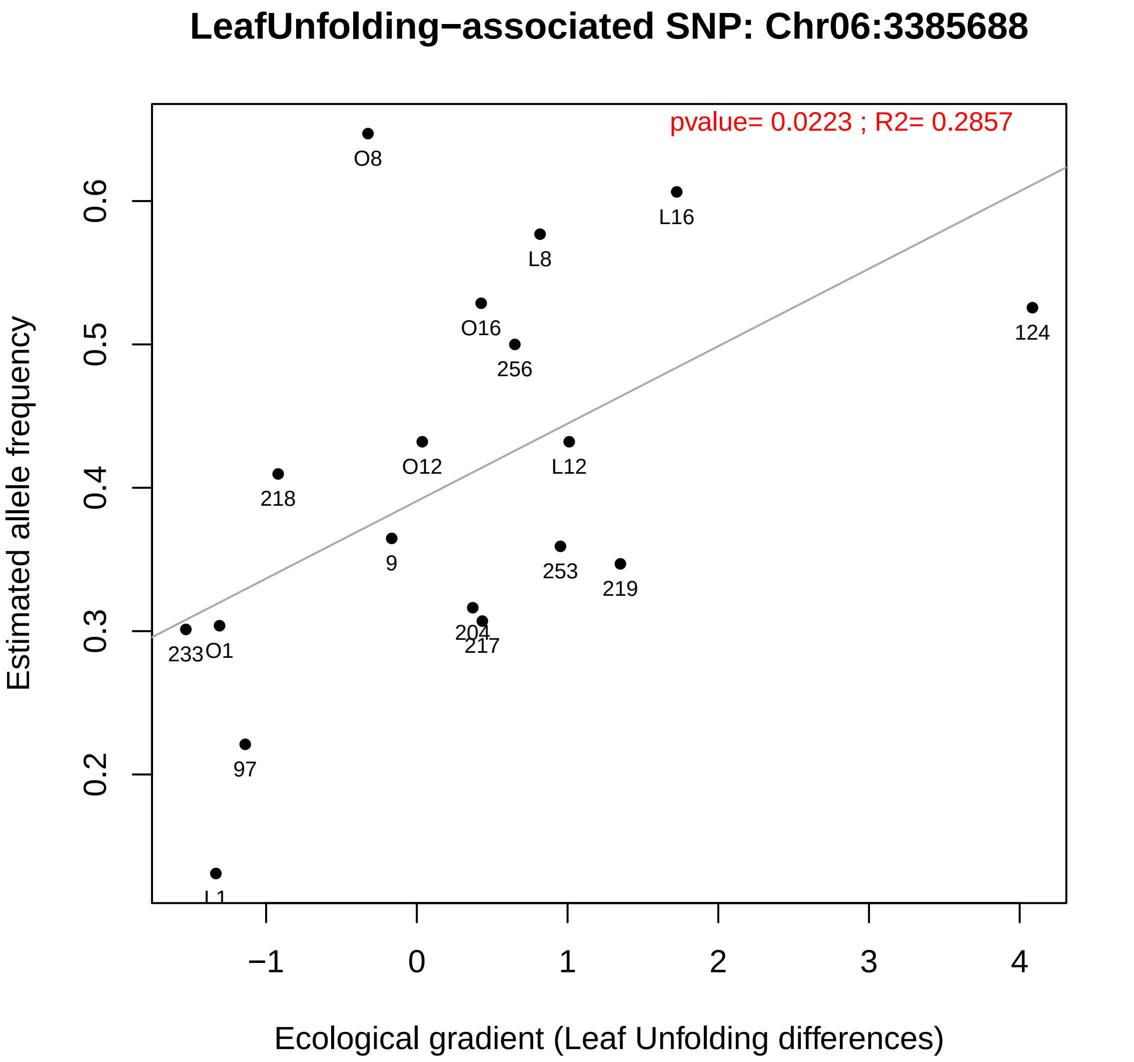
LeafUnfolding−associated SNP: Chr06:3385688
pvalue= 0.0223 ; R2= 0.2857
Estimated allele frequency
0.2
0.3
0.4
0.5
0.6
−1
0
1
2
3
4
Ecological gradient (Leaf Unfolding differences)
O8
L16
L8
O16
124
256
O12
L12
218
9
253
219
204
217
233
O1
97
L1

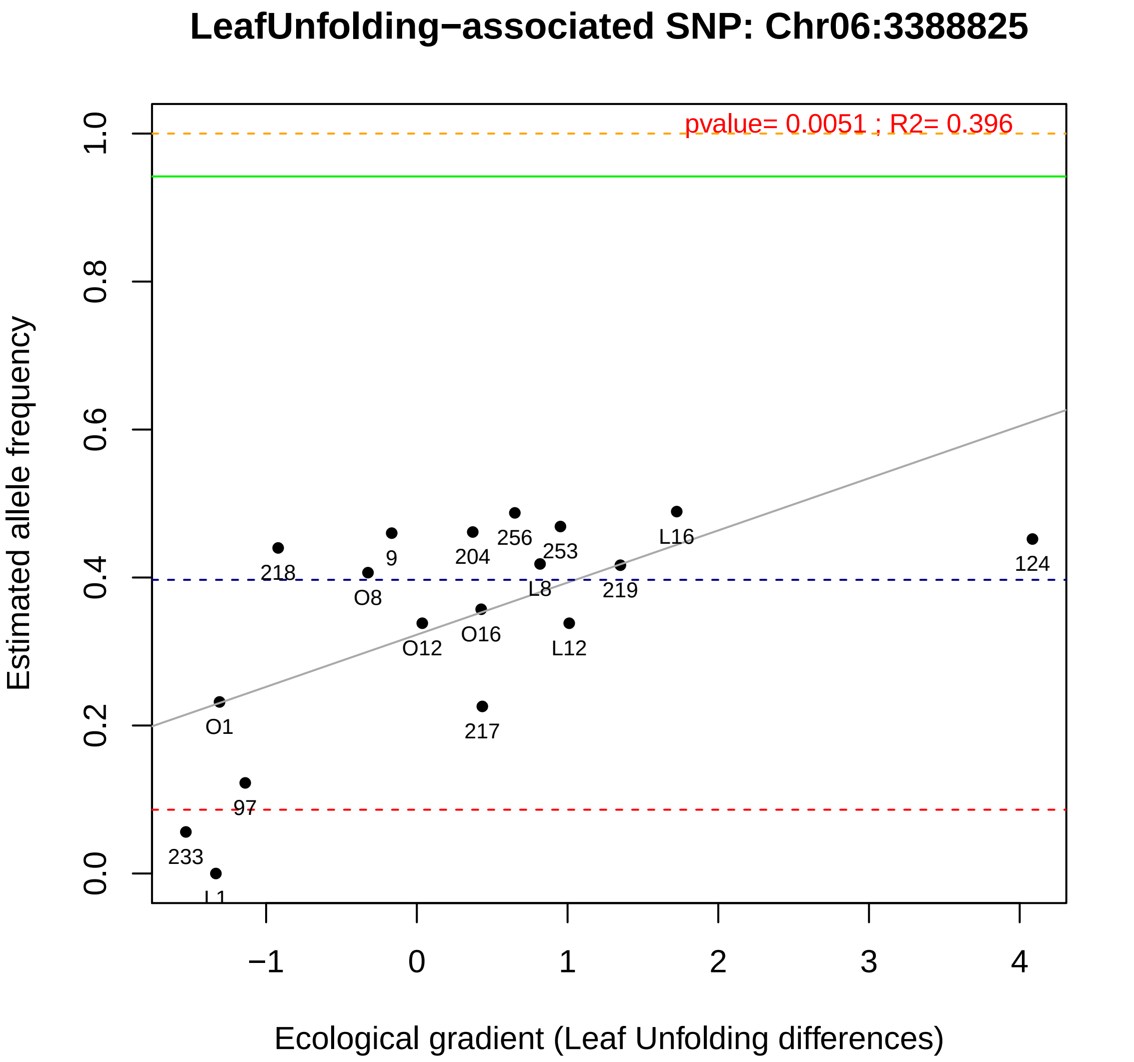

LeafUnfolding−associated SNP: Chr06:3388825
pvalue= 0.0051 ; R2= 0.396
Estimated allele frequency
Ecological gradient (Leaf Unfolding differences)
0.0
0.2
0.4
0.6
0.8
1.0
−1
0
1
2
3
4
218
9
O8
204
256
253
L8
219
L16
124
O12
O16
L12
O1
217
97
233
L1

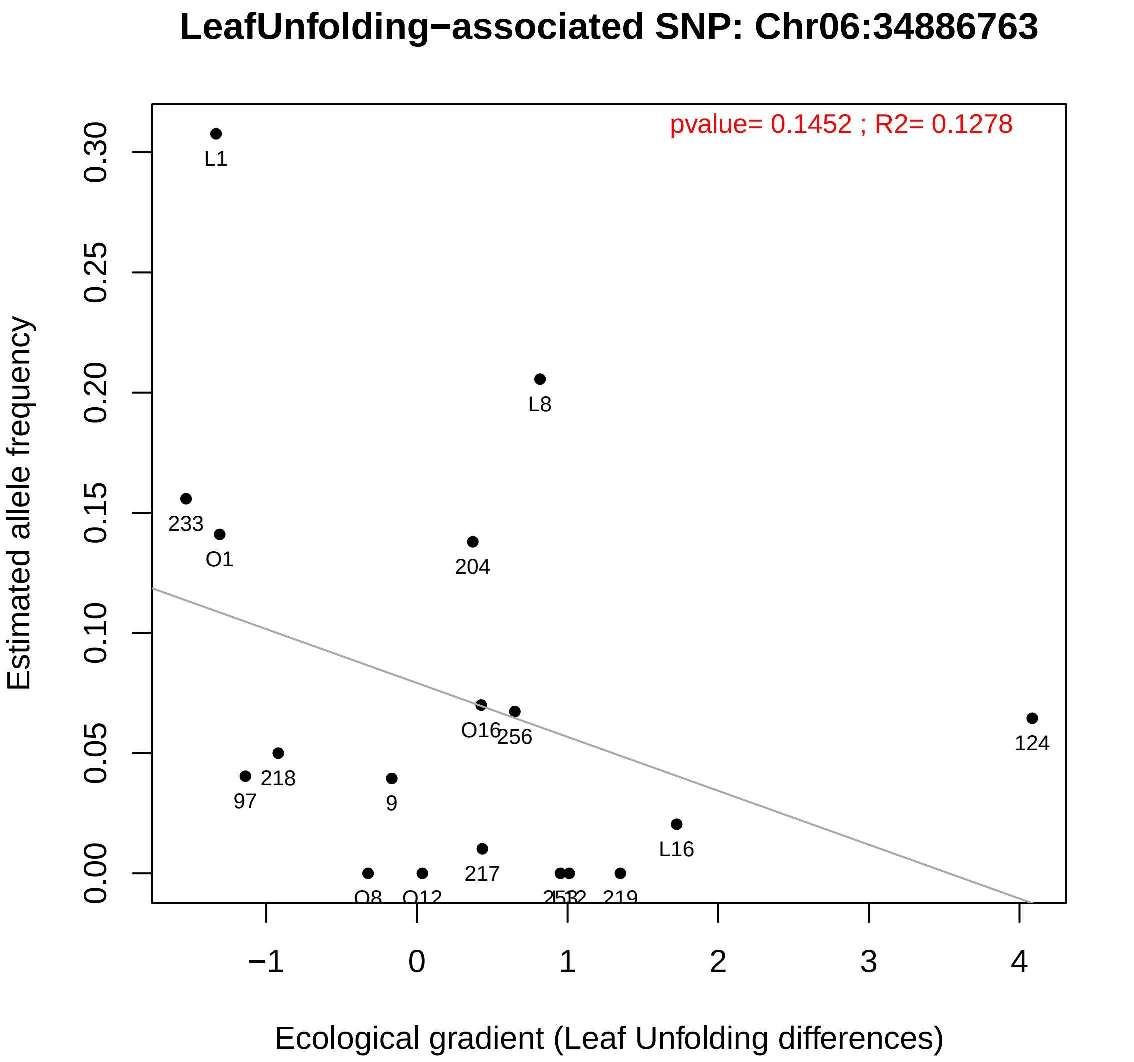

LeafUnfolding−associated SNP: Chr06:34886763
pvalue= 0.1452 ; R2= 0.1278
Estimated allele frequency
0.00
0.05
0.10
0.15
0.20
0.25
0.30
Ecological gradient (Leaf Unfolding differences)
−1
0
1
2
3
4
L1
L8
233
O1
204
O16
256
124
218
97
9
217
L16
O8
O12
219

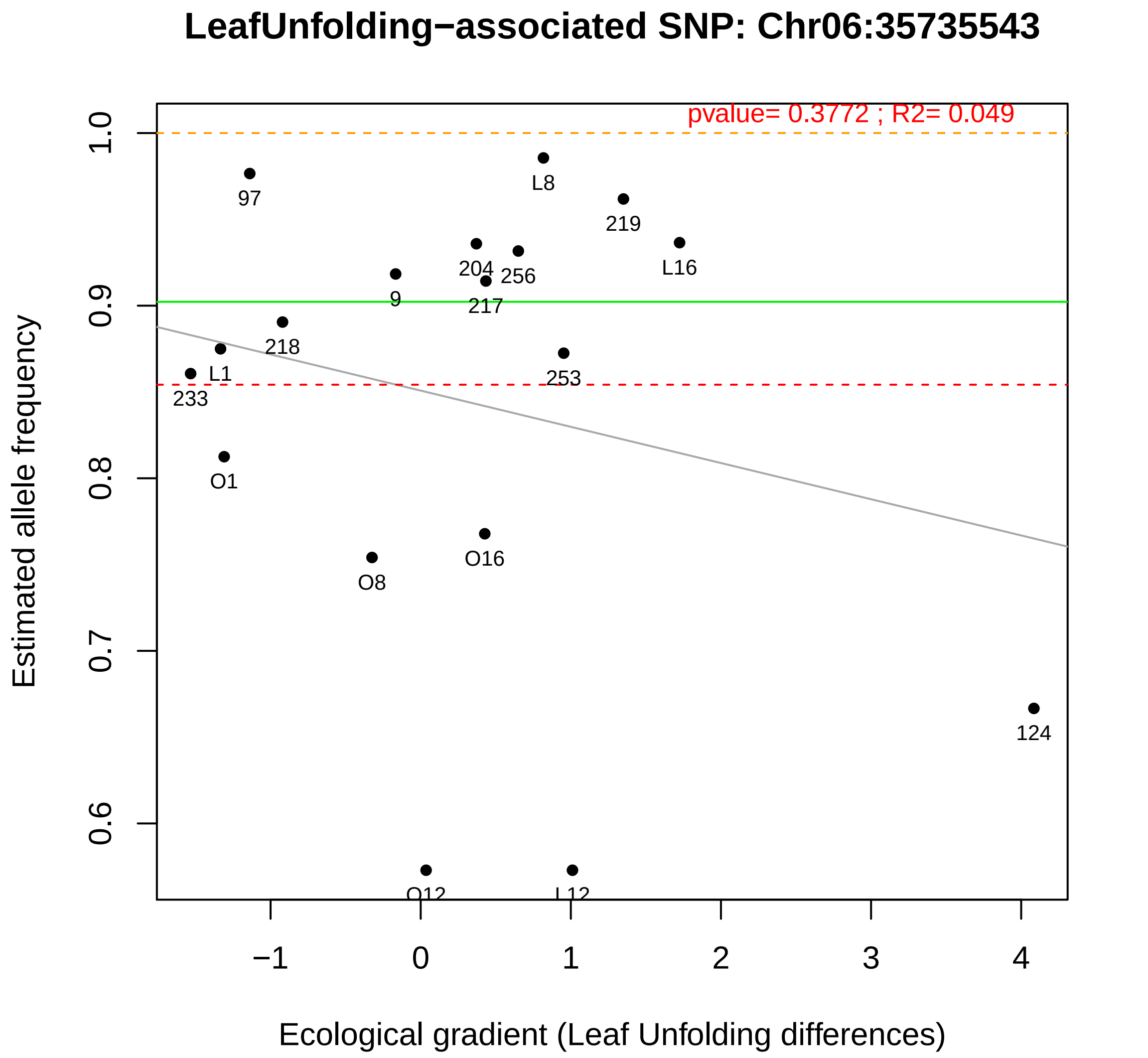
LeafUnfolding−associated SNP: Chr06:35735543
pvalue= 0.3772 ; R2= 0.049
Estimated allele frequency
Ecological gradient (Leaf Unfolding differences)
0.6
0.7
0.8
0.9
1.0
−1
0
1
2
3
4
97
L8
219
L16
204
256
9
217
218
L1
233
253
O1
O16
O8
124
O12
L12

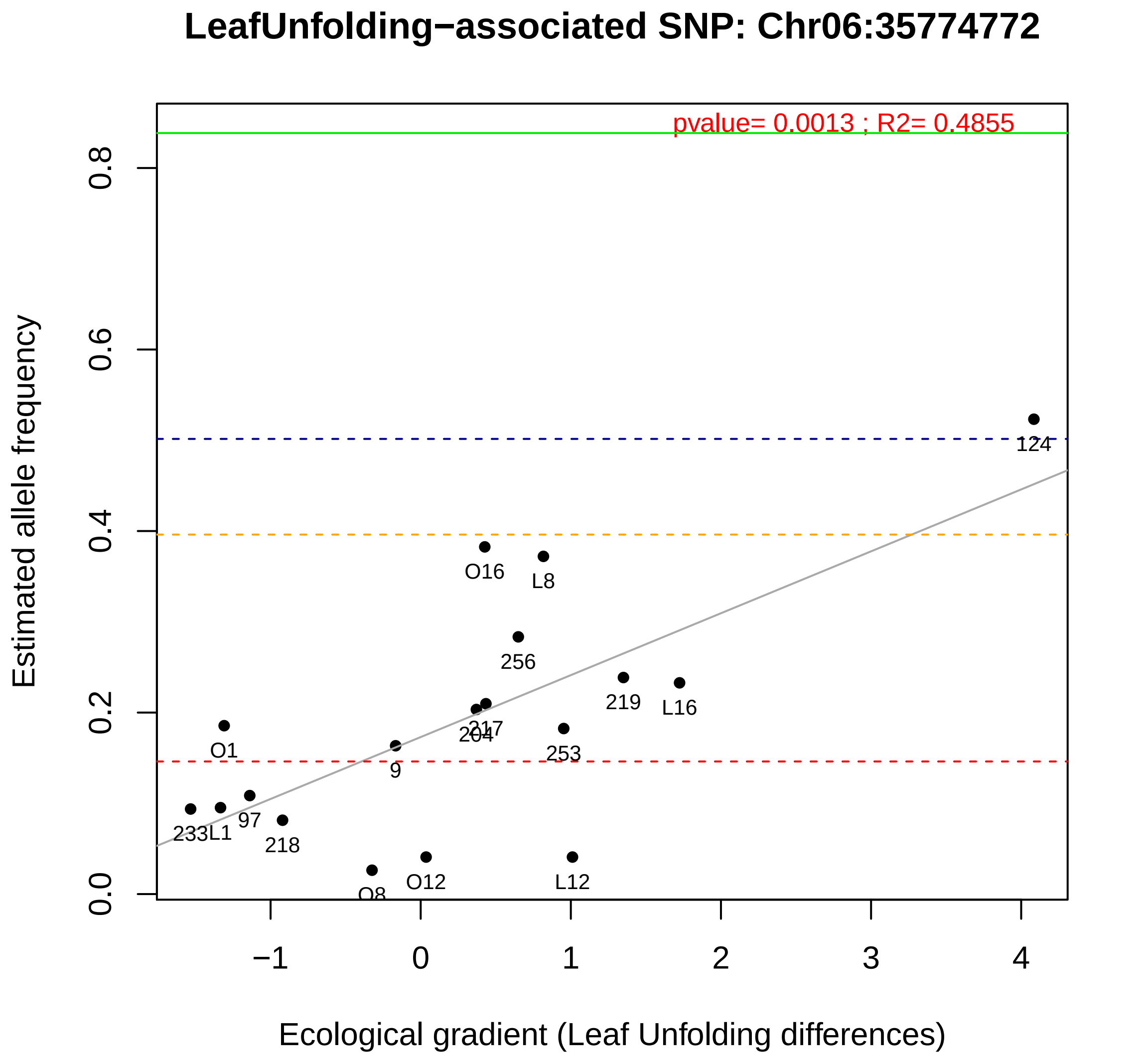

LeafUnfolding−associated SNP: Chr06:35774772
pvalue= 0.0013 ; R2= 0.4855
Estimated allele frequency
Ecological gradient (Leaf Unfolding differences)
0.0
0.2
0.4
0.6
0.8
−1
0
1
2
3
4
124
O16
L8
256
219
L16
O1
217
204
253
9
97
233
L1
218
O12
L12
O8

**LeafUnfolding−associated SNP: Chr06:36084089**

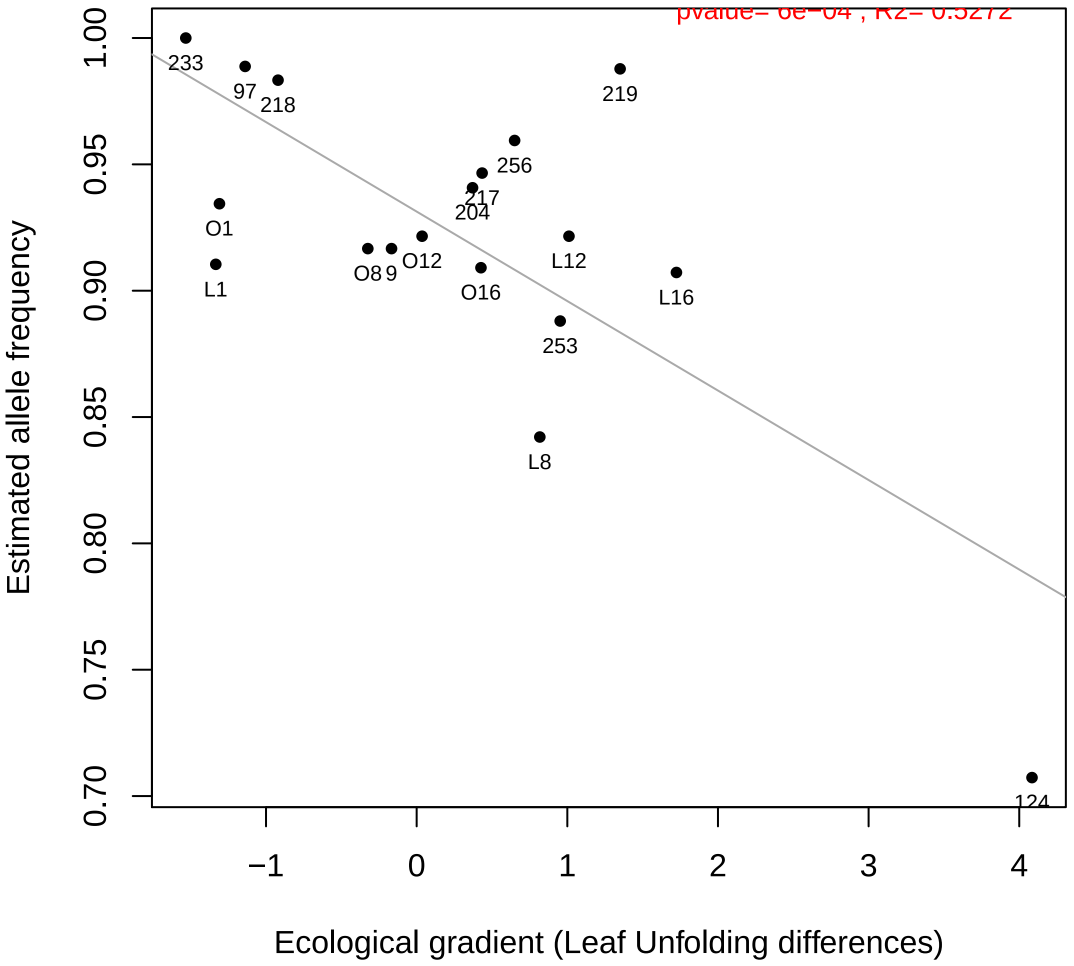

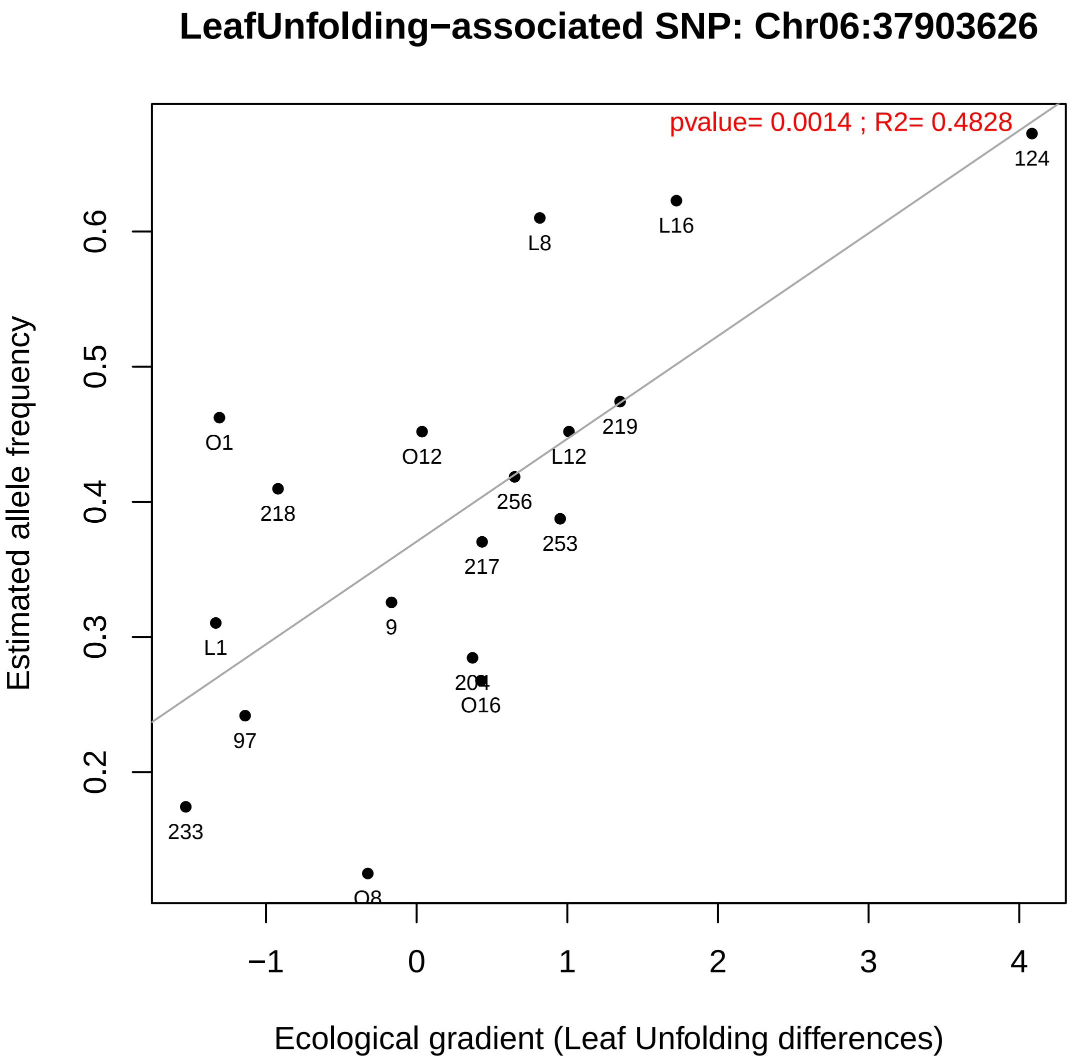
LeafUnfolding−associated SNP: Chr06:37903626
pvalue= 0.0014 ; R2= 0.4828
Estimated allele frequency
0.2
0.3
0.4
0.5
0.6
−1
0
1
2
3
4
Ecological gradient (Leaf Unfolding differences)
124
L16
L8
O1
O12
219
L12
256
218
253
217
9
L1
204
O16
97
233
O8

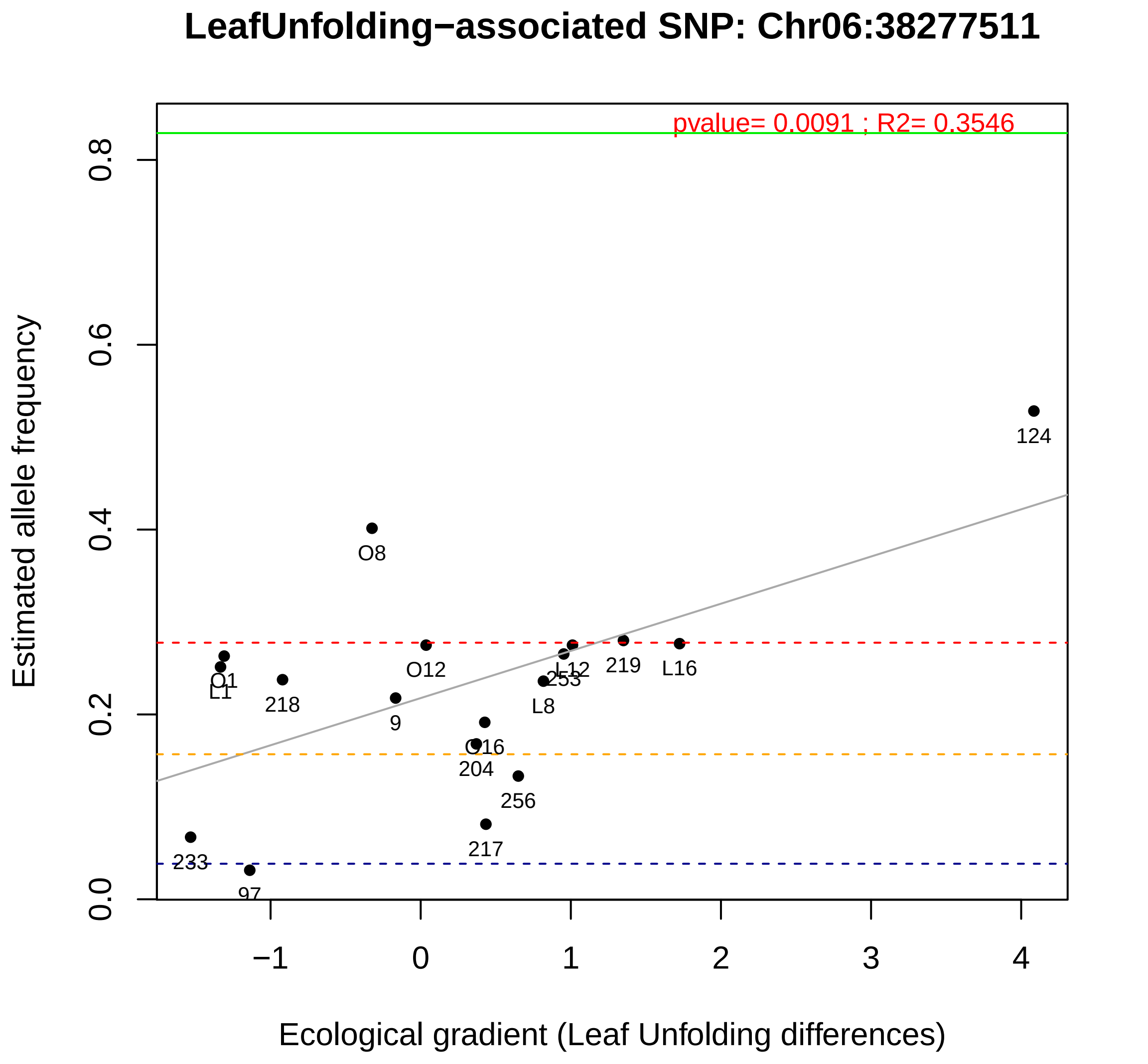
LeafUnfolding−associated SNP: Chr06:38277511
pvalue= 0.0091 ; R2= 0.3546
Estimated allele frequency
0.0
0.2
0.4
0.6
0.8
−1
0
1
2
3
4
Ecological gradient (Leaf Unfolding differences)
124
O8
O12
O1
L1
218
9
L12
253
L8
219
L16
O16
204
256
217
233
97

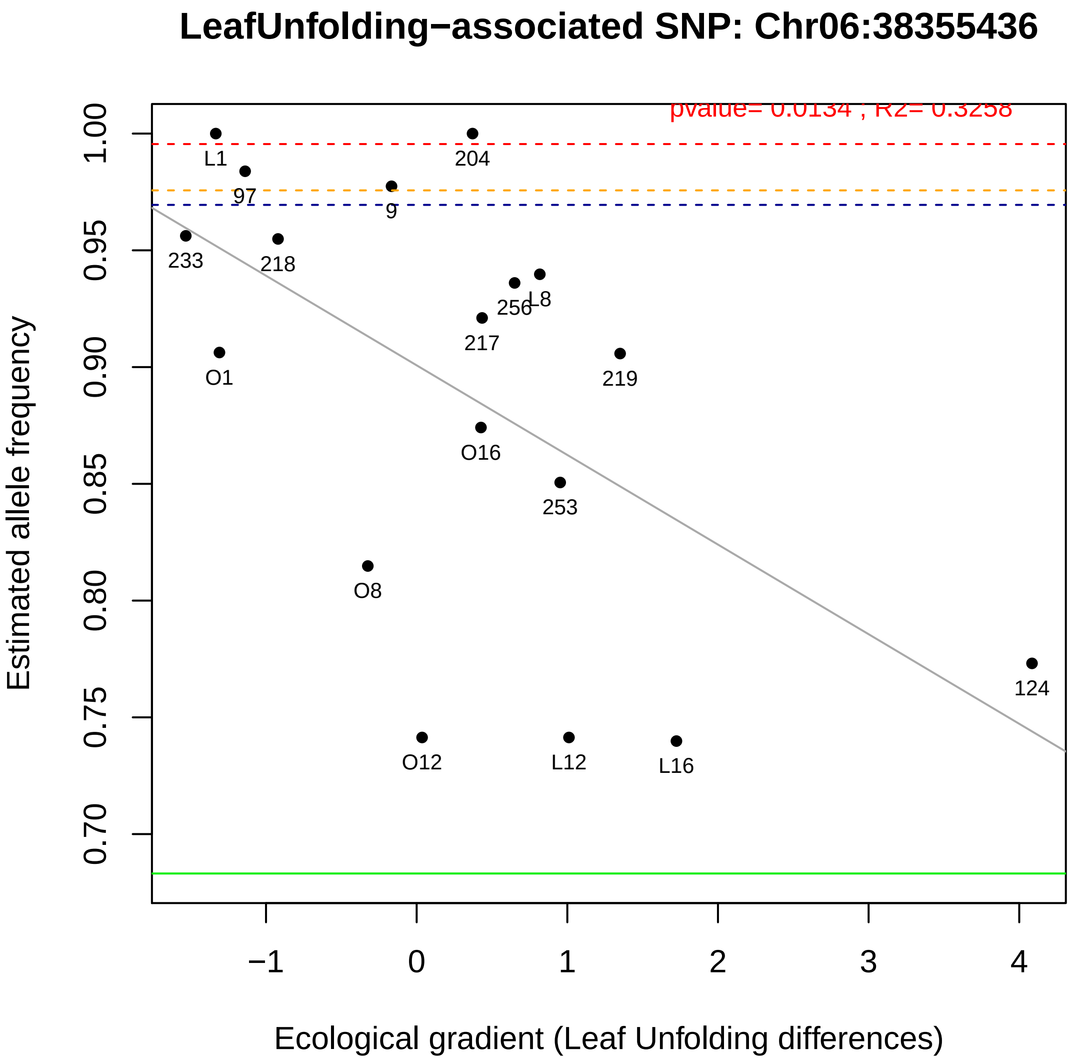

LeafUnfolding−associated SNP: Chr06:38355436
pvalue= 0.0134 ; R2= 0.3258
Estimated allele frequency
Ecological gradient (Leaf Unfolding differences)
1.00
0.95
0.90
0.85
0.80
0.75
0.70
−1
0
1
2
3
4
L1
204
97
9
233
218
256
L8
217
O1
219
O16
253
O8
124
O12
L12
L16

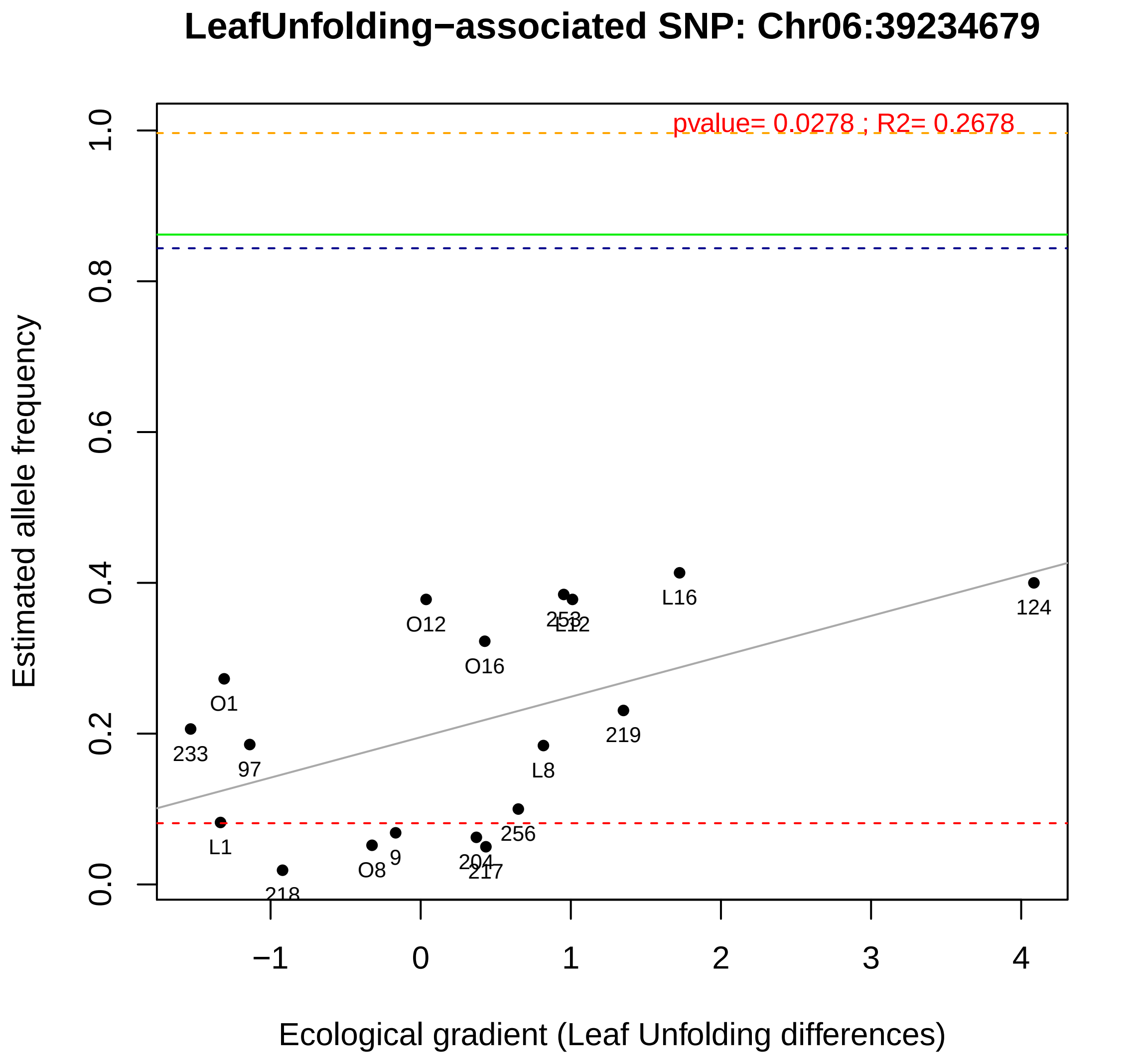

LeafUnfolding−associated SNP: Chr06:39234679
pvalue= 0.0278 ; R2= 0.2678
Estimated allele frequency
Ecological gradient (Leaf Unfolding differences)
0.0
0.2
0.4
0.6
0.8
1.0
−1
0
1
2
3
4
O12
O16
L12
L16
124
O1
233
97
L8
219
L1
218
O8
9
204
217
256

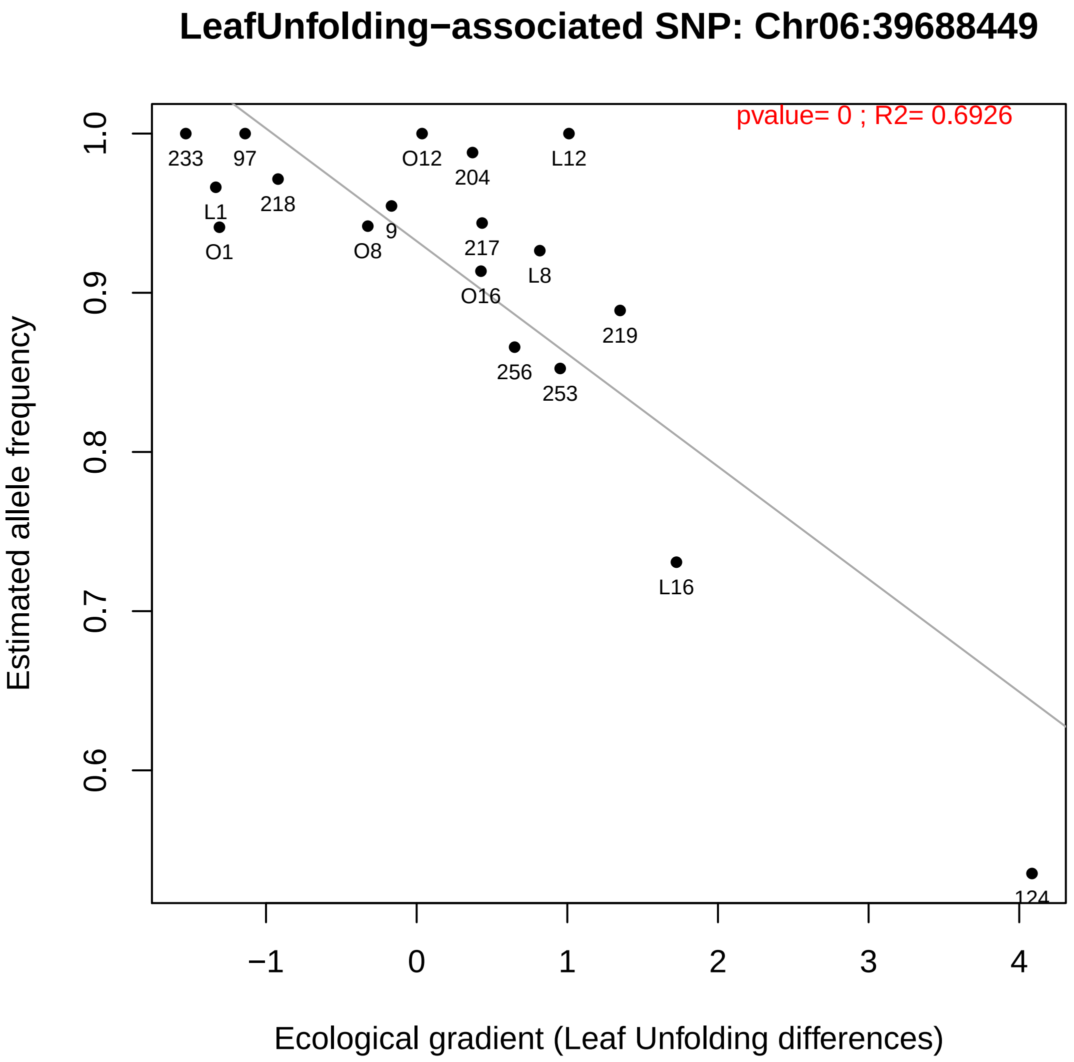
LeafUnfolding−associated SNP: Chr06:39688449
pvalue= 0 ; R2= 0.6926
Estimated allele frequency
Ecological gradient (Leaf Unfolding differences)
1.0
0.9
0.8
0.7
0.6
−1
0
1
2
3
4
233
97
O12
204
L12
L1
218
O1
9
O8
217
L8
O16
219
256
253
L16
124

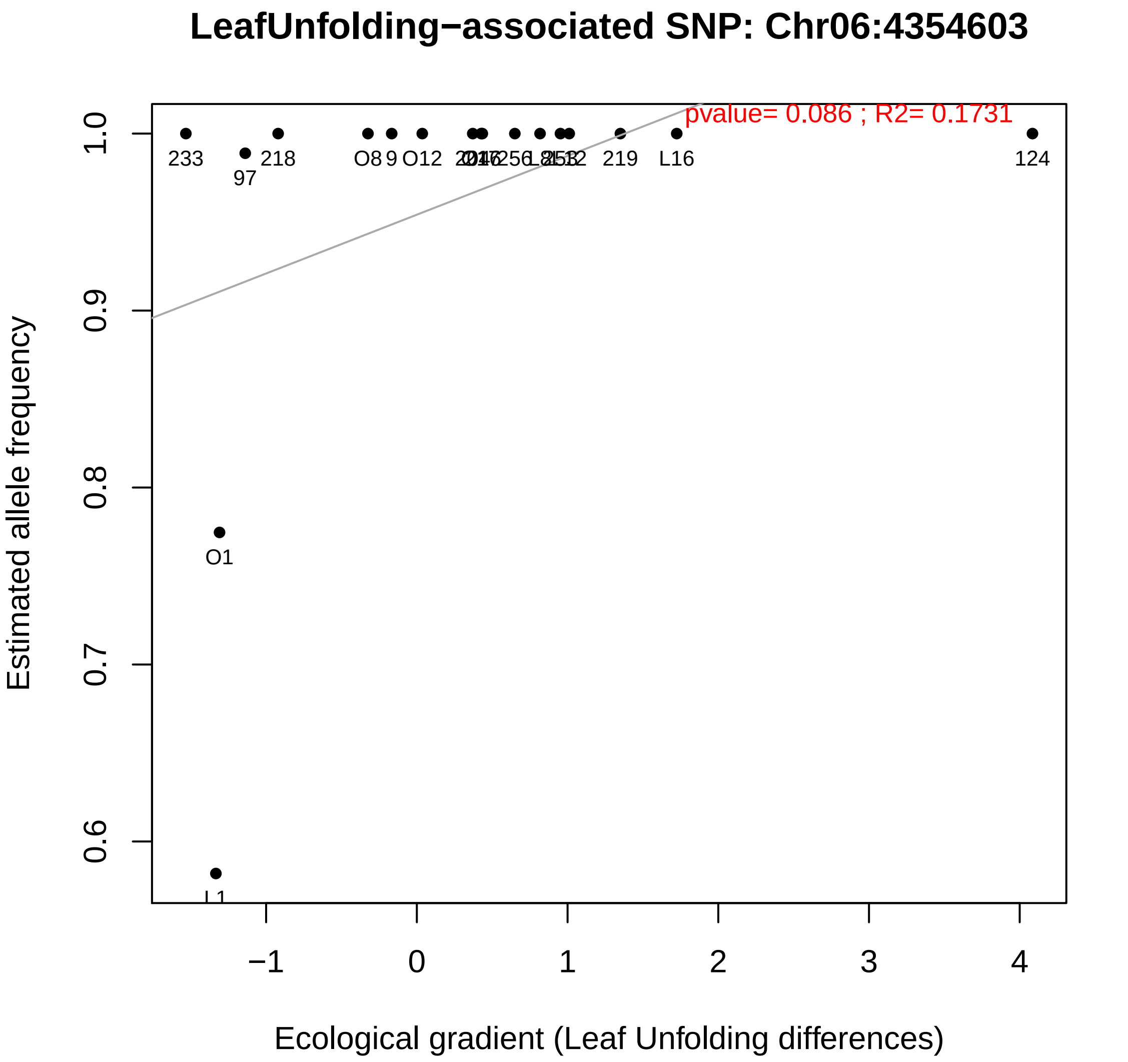
LeafUnfolding−associated SNP: Chr06:4354603
pvalue= 0.086 ; R2= 0.1731
Estimated allele frequency
Ecological gradient (Leaf Unfolding differences)
0.6
0.7
0.8
0.9
1.0
−1
0
1
2
3
4
233
97
218
O8
9
O12
219
L16
124
O1
L1

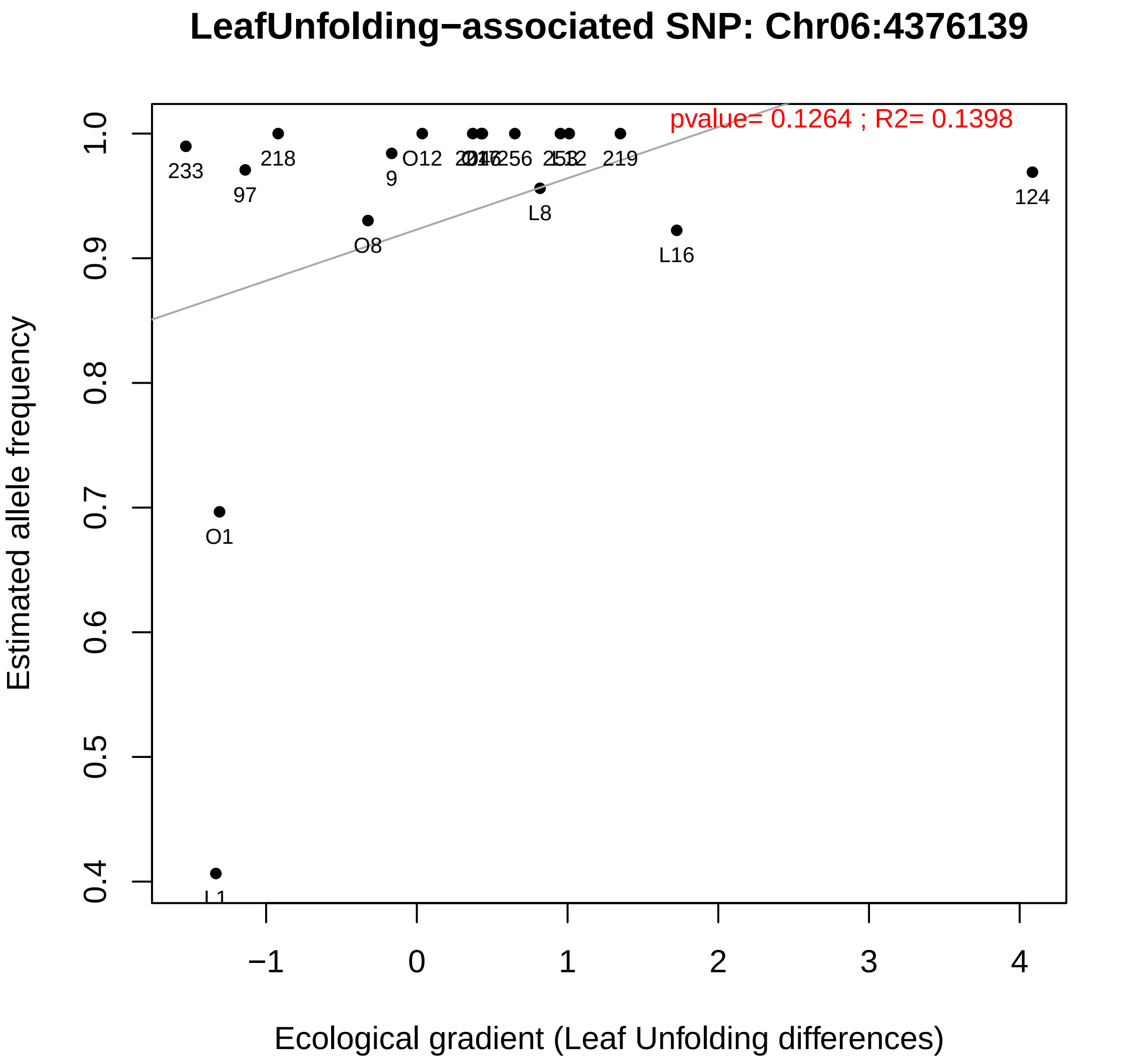
LeafUnfolding−associated SNP: Chr06:4376139
pvalue= 0.1264 ; R2= 0.1398
Estimated allele frequency
Ecological gradient (Leaf Unfolding differences)
0.4
0.5
0.6
0.7
0.8
0.9
1.0
−1
0
1
2
3
4
233
218
97
9
O12
256
219
L8
O8
L16
124
O1

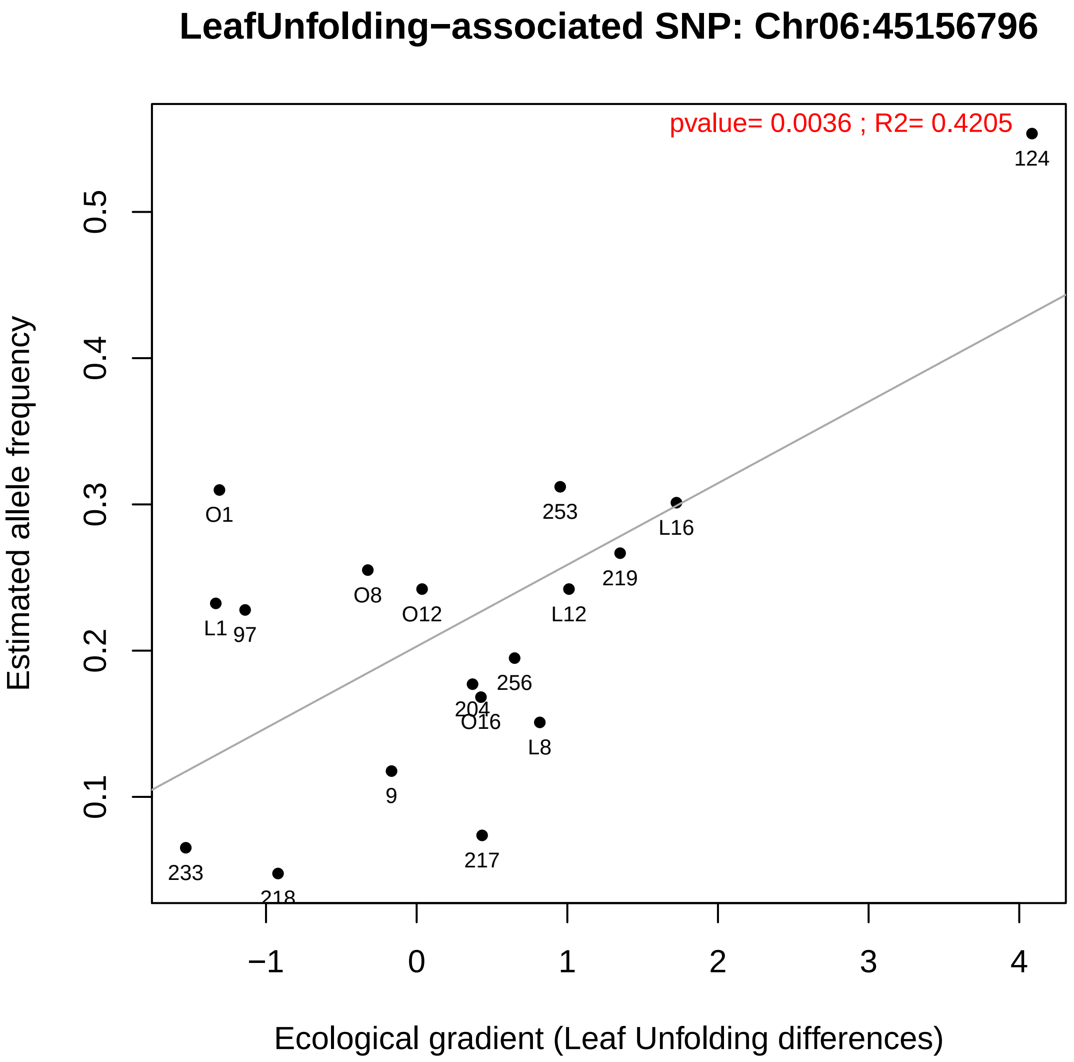

LeafUnfolding−associated SNP: Chr06:45156796
pvalue= 0.0036 ; R2= 0.4205
124
O1
253
L16
219
O8
O12
L12
L1
97
256
204
O16
L8
9
217
233
218
0.1
0.2
0.3
0.4
0.5
−1
0
1
2
3
4
Estimated allele frequency
Ecological gradient (Leaf Unfolding differences)

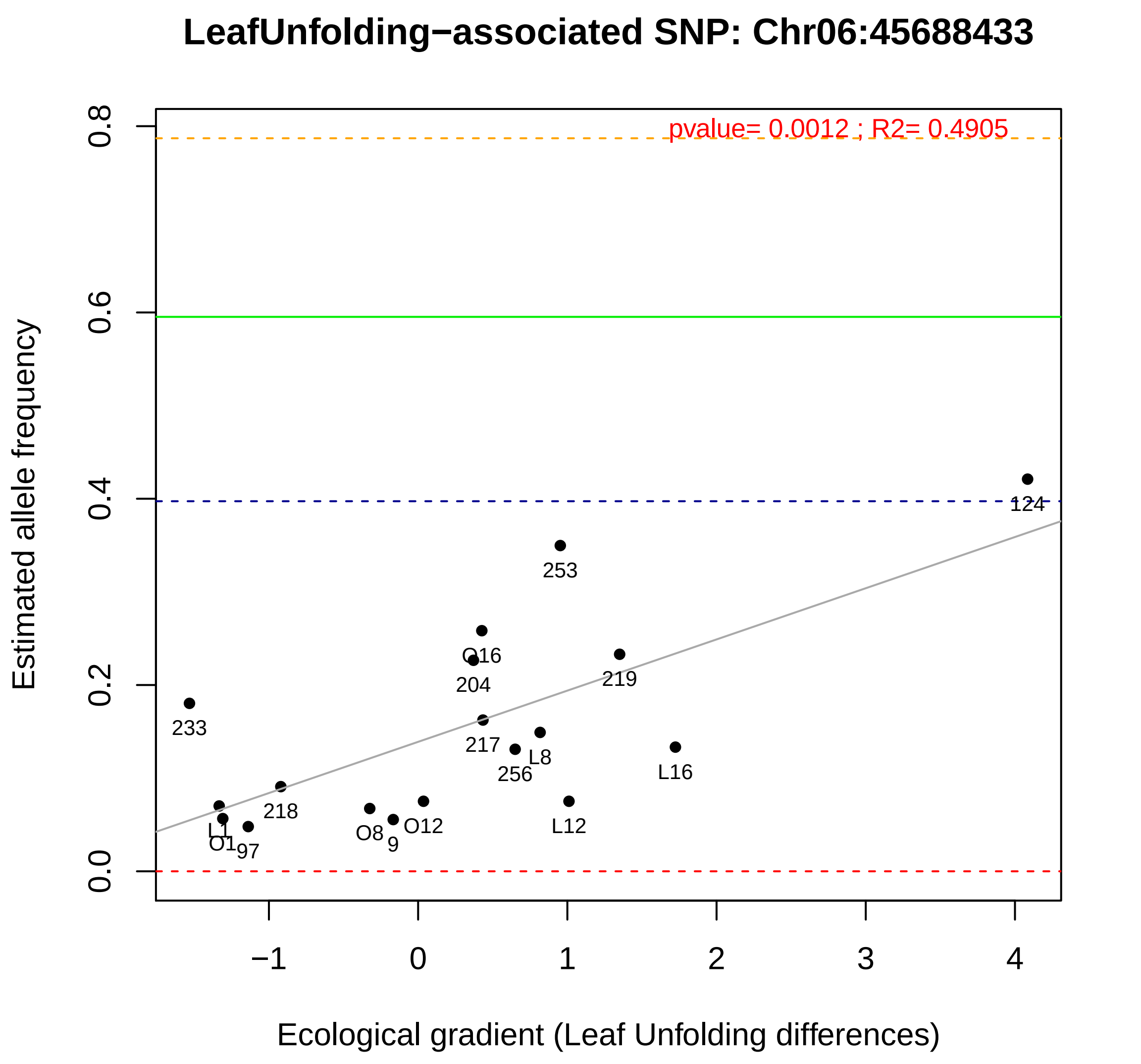

LeafUnfolding−associated SNP: Chr06:45688433
pvalue= 0.0012 ; R2= 0.4905
Estimated allele frequency
0.0
0.2
0.4
0.6
0.8
Ecological gradient (Leaf Unfolding differences)
−1
0
1
2
3
4
124
253
O16
204
219
233
217
L8
256
L16
218
L1
O1
97
O8
9
O12
L12

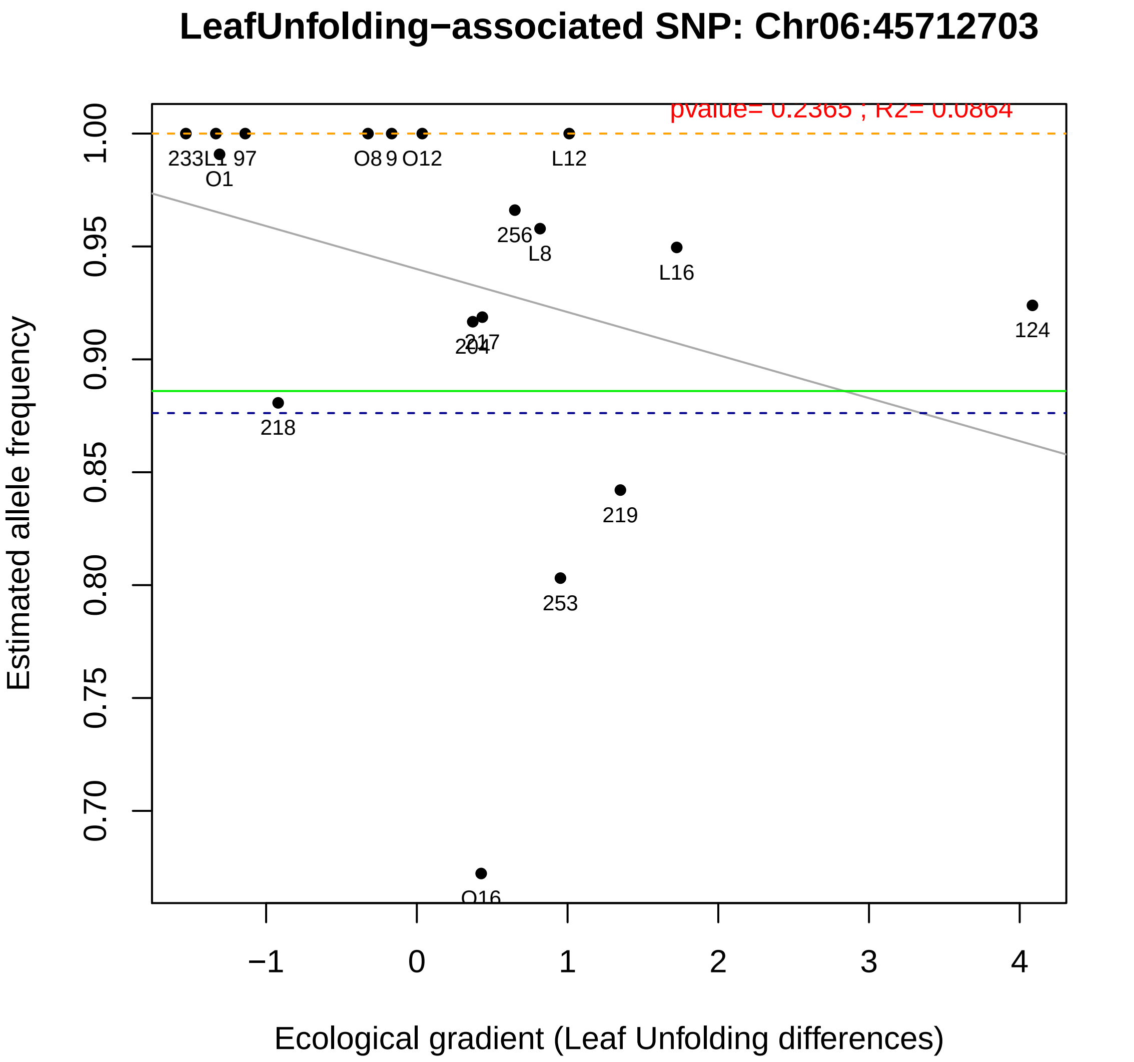

LeafUnfolding−associated SNP: Chr06:45712703
pvalue= 0.2365 ; R2= 0.0864
Estimated allele frequency
Ecological gradient (Leaf Unfolding differences)
0.70
0.75
0.80
0.85
0.90
0.95
1.00
−1
0
1
2
3
4
233
L1
97
O1
O8
9
O12
L12
256
L8
L16
217
204
124
218
219
253
O16

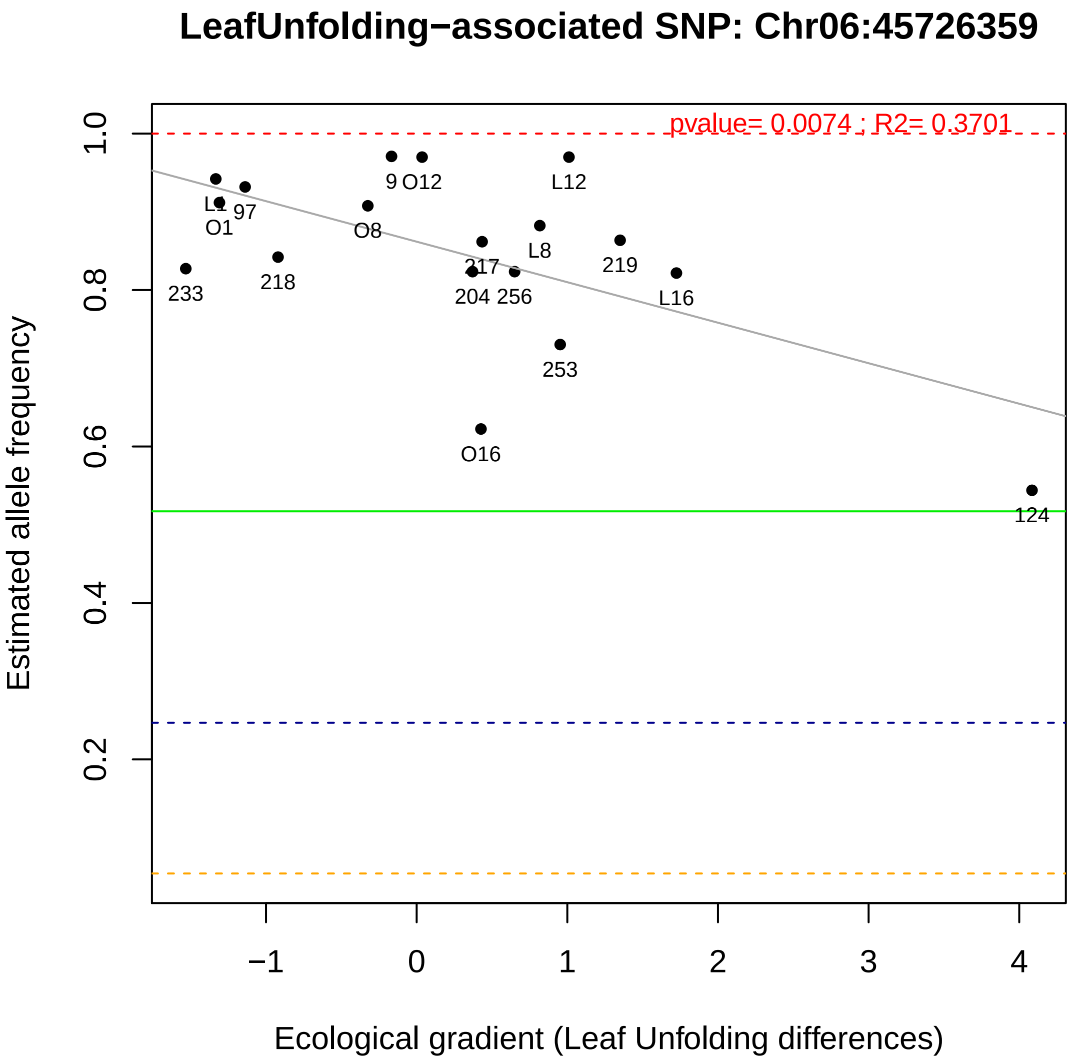
LeafUnfolding−associated SNP: Chr06:45726359
pvalue= 0.0074 ; R2= 0.3701
Estimated allele frequency
1.0
0.8
0.6
0.4
0.2
−1
0
1
2
3
4
Ecological gradient (Leaf Unfolding differences)
9
O12
L12
L1
97
O1
O8
218
233
217
L8
219
204
256
L16
253
O16
124

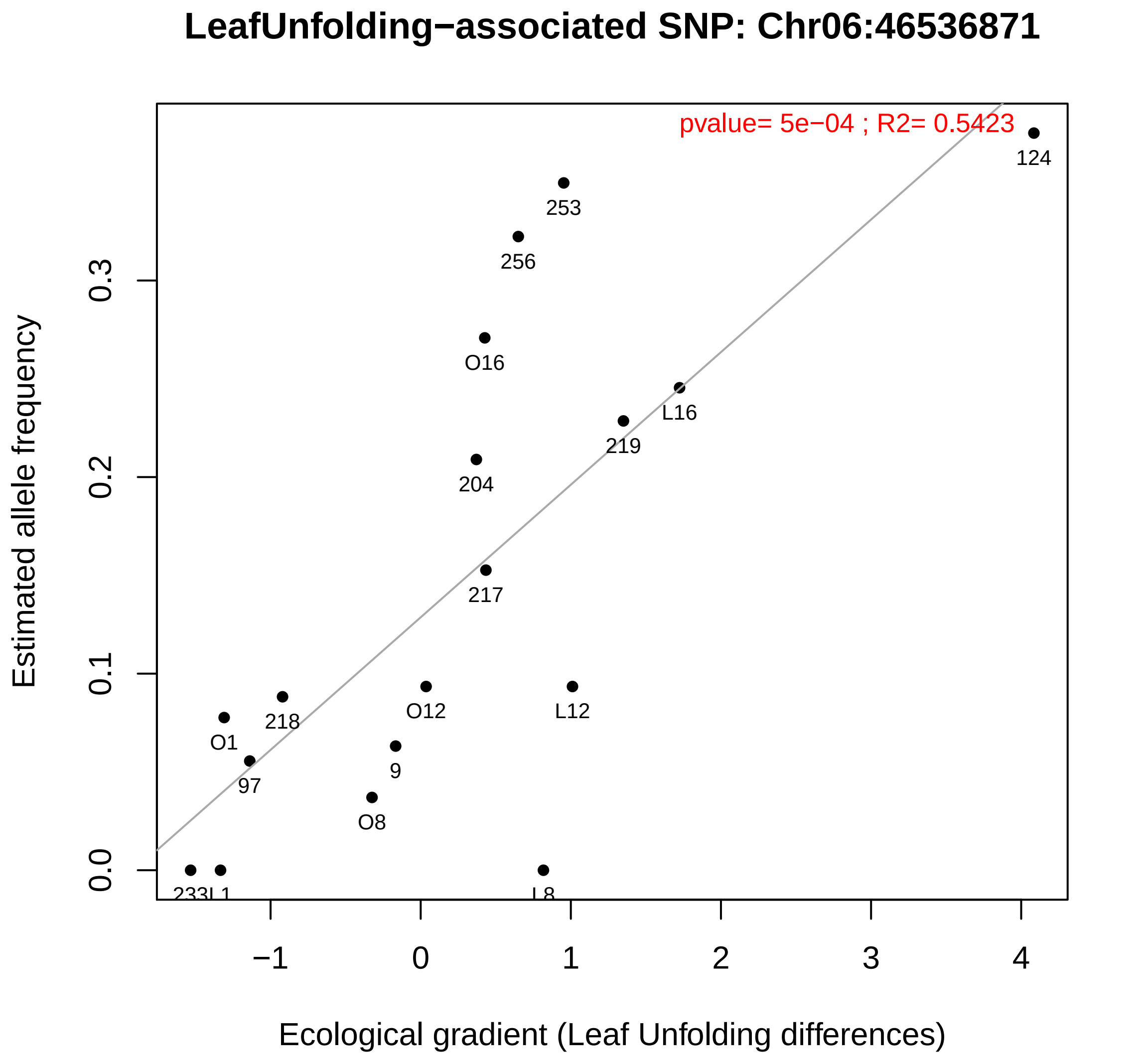

LeafUnfolding−associated SNP: Chr06:46536871
pvalue= 5e−04 ; R2= 0.5423
Estimated allele frequency
Ecological gradient (Leaf Unfolding differences)
0.0
0.1
0.2
0.3
−1
0
1
2
3
4
124
253
256
O16
L16
219
204
217
218
O12
L12
O1
97
9
O8
233
L1
L8

### LeafUnfolding−associated SNP: Chr06:47017727

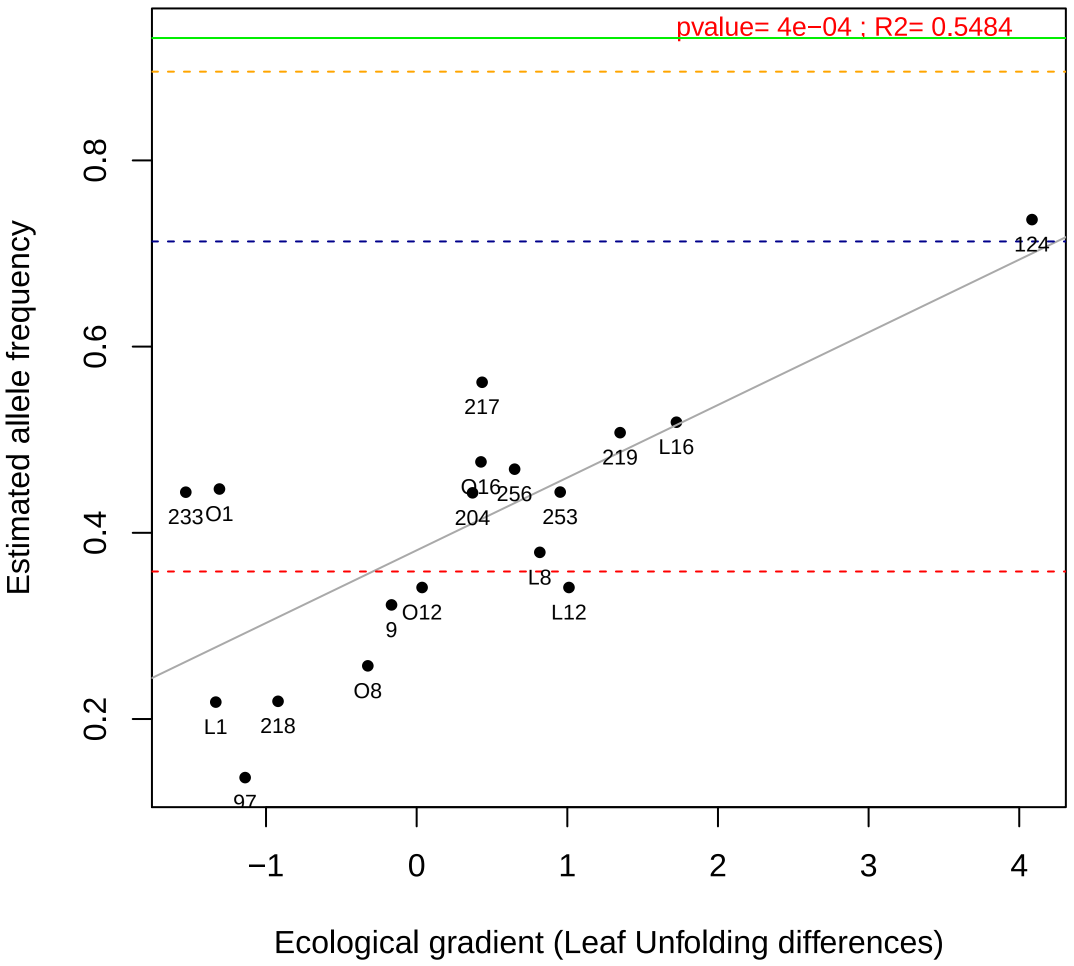

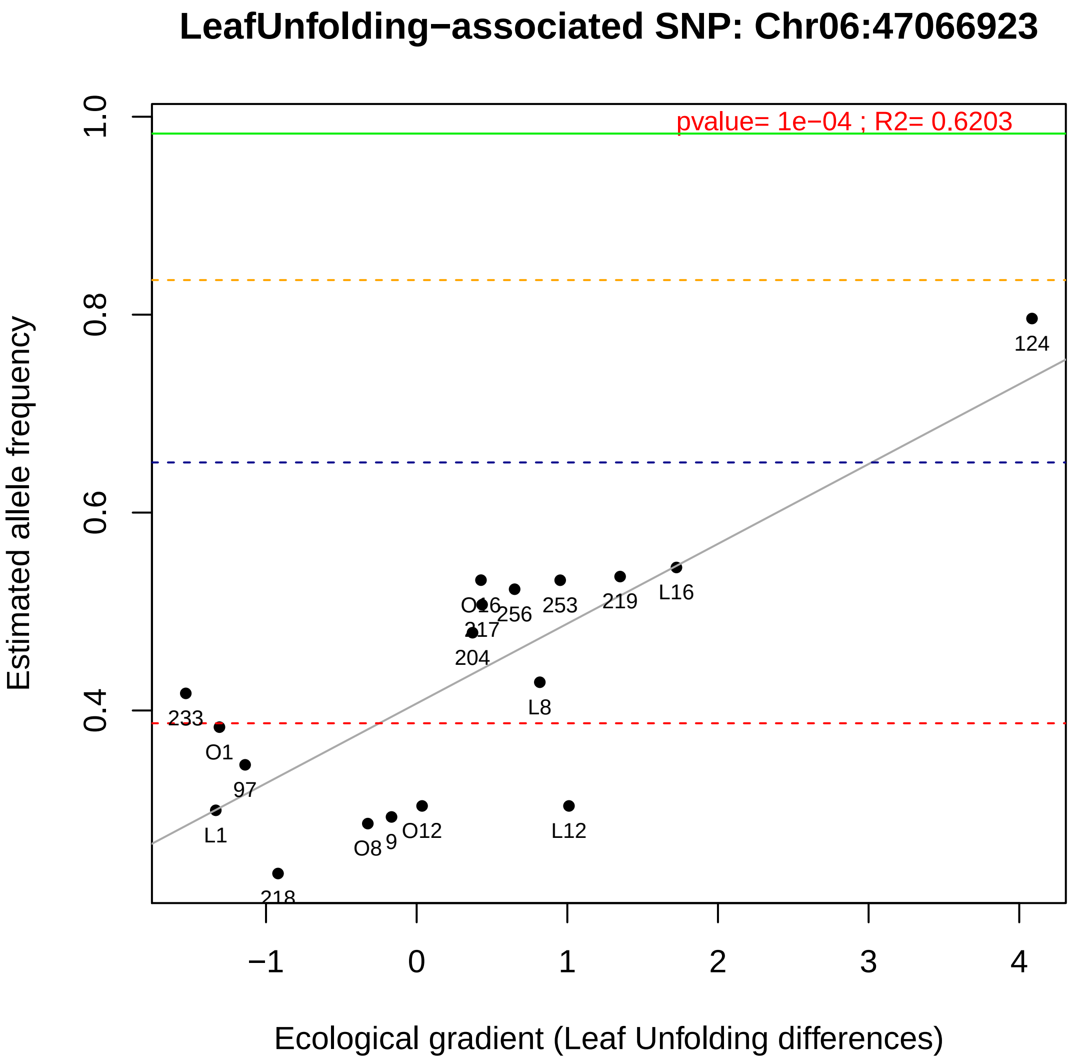

LeafUnfolding−associated SNP: Chr06:47066923
pvalue= 1e−04 ; R2= 0.6203
Estimated allele frequency
0.4
0.6
0.8
1.0
124
L16
219
253
256
O16
217
204
L8
233
O1
97
L1
O8
9
O12
L12
218
−1
0
1
2
3
4
Ecological gradient (Leaf Unfolding differences)

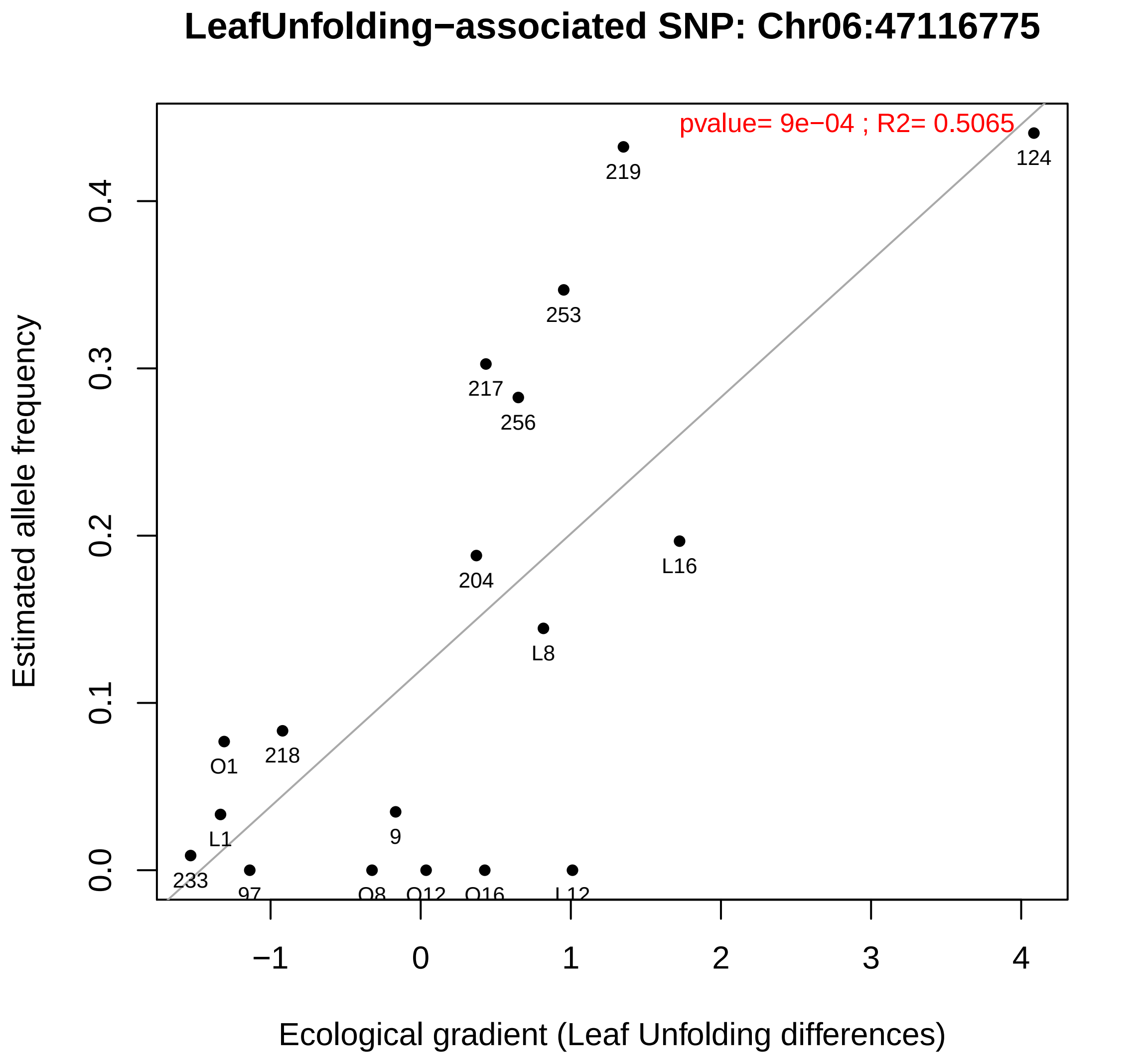

LeafUnfolding−associated SNP: Chr06:47116775
pvalue= 9e−04 ; R2= 0.5065
Estimated allele frequency
0.0
0.1
0.2
0.3
0.4
−1
0
1
2
3
4
Ecological gradient (Leaf Unfolding differences)
219
124
253
217
256
204
L16
L8
O1
218
L1
9
233
97
O8
O12
O16
L12

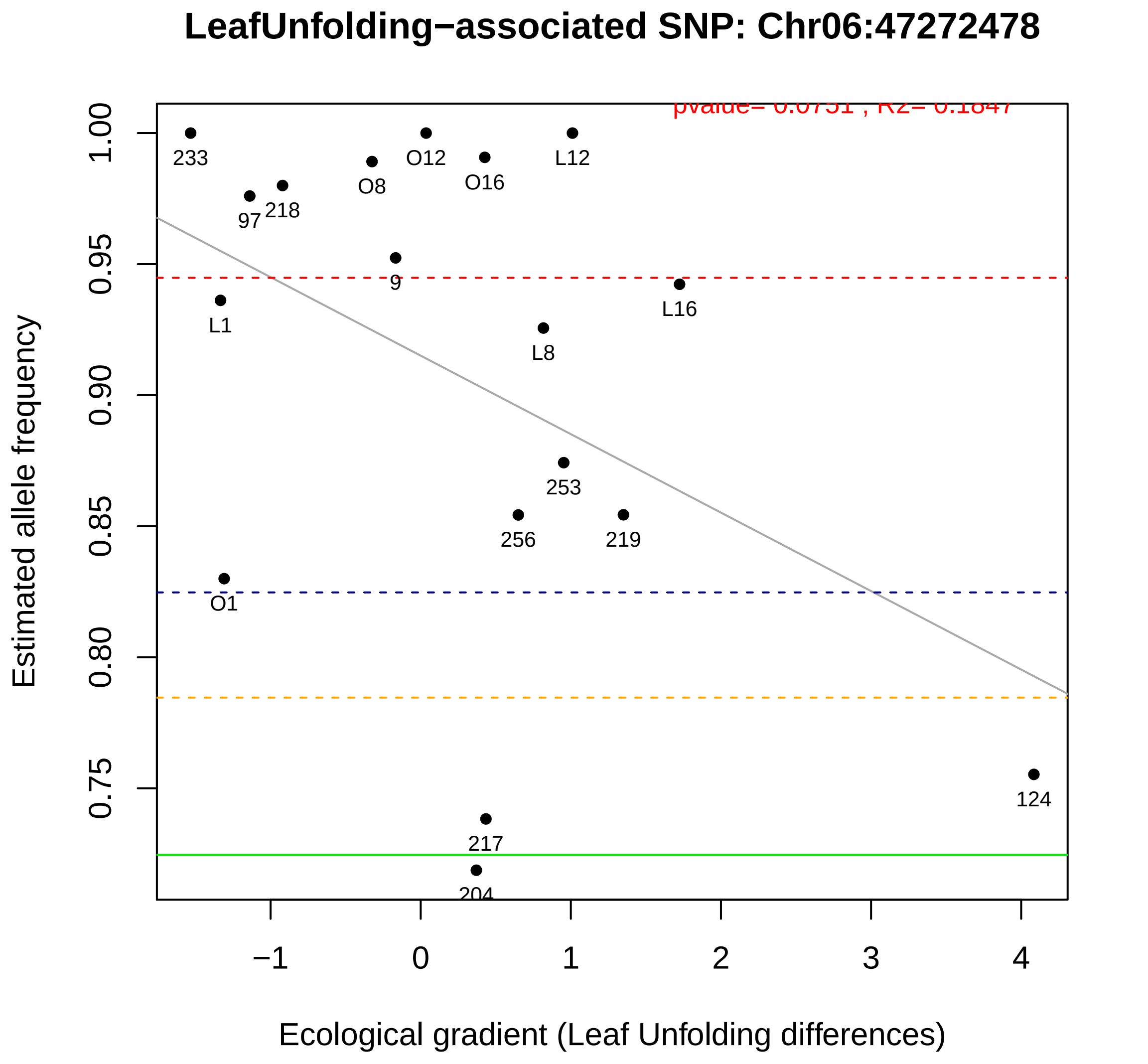

LeafUnfolding−associated SNP: Chr06:47272478
pvalue= 0.0731 , R2= 0.1847
Estimated allele frequency
Ecological gradient (Leaf Unfolding differences)
1.00
0.95
0.90
0.85
0.80
0.75
−1
0
1
2
3
4
233
O12
L12
O8
O16
97
218
9
L16
L1
L8
253
256
219
O1
124
217
204

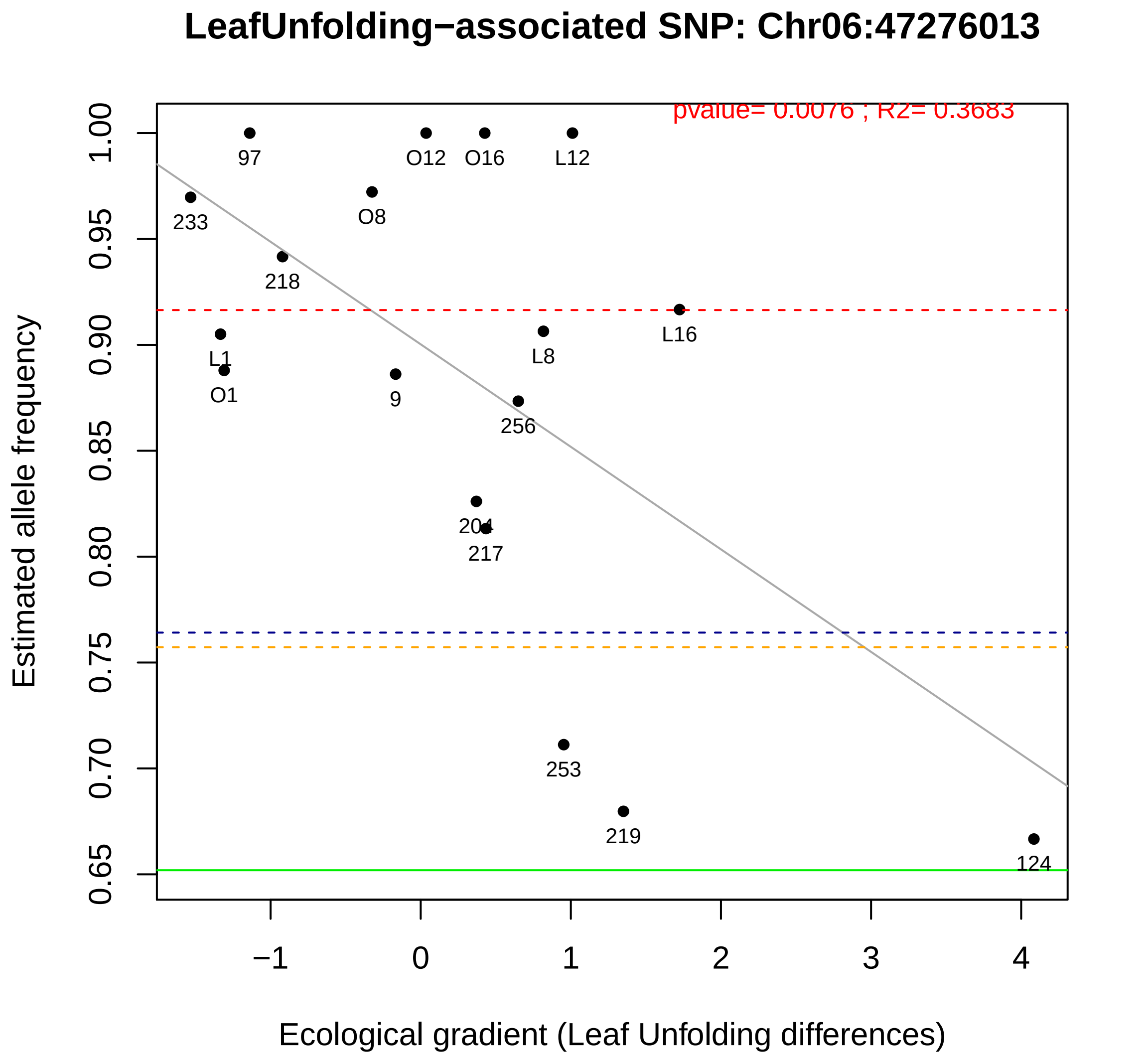

LeafUnfolding−associated SNP: Chr06:47276013
pvalue= 0.0076 ; R2= 0.3683
Estimated allele frequency
Ecological gradient (Leaf Unfolding differences)
1.00
0.95
0.90
0.85
0.80
0.75
0.70
0.65
−1
0
1
2
3
4
97
O12
O16
L12
233
O8
218
L16
L1
L8
O1
9
256
204
217
253
219
124

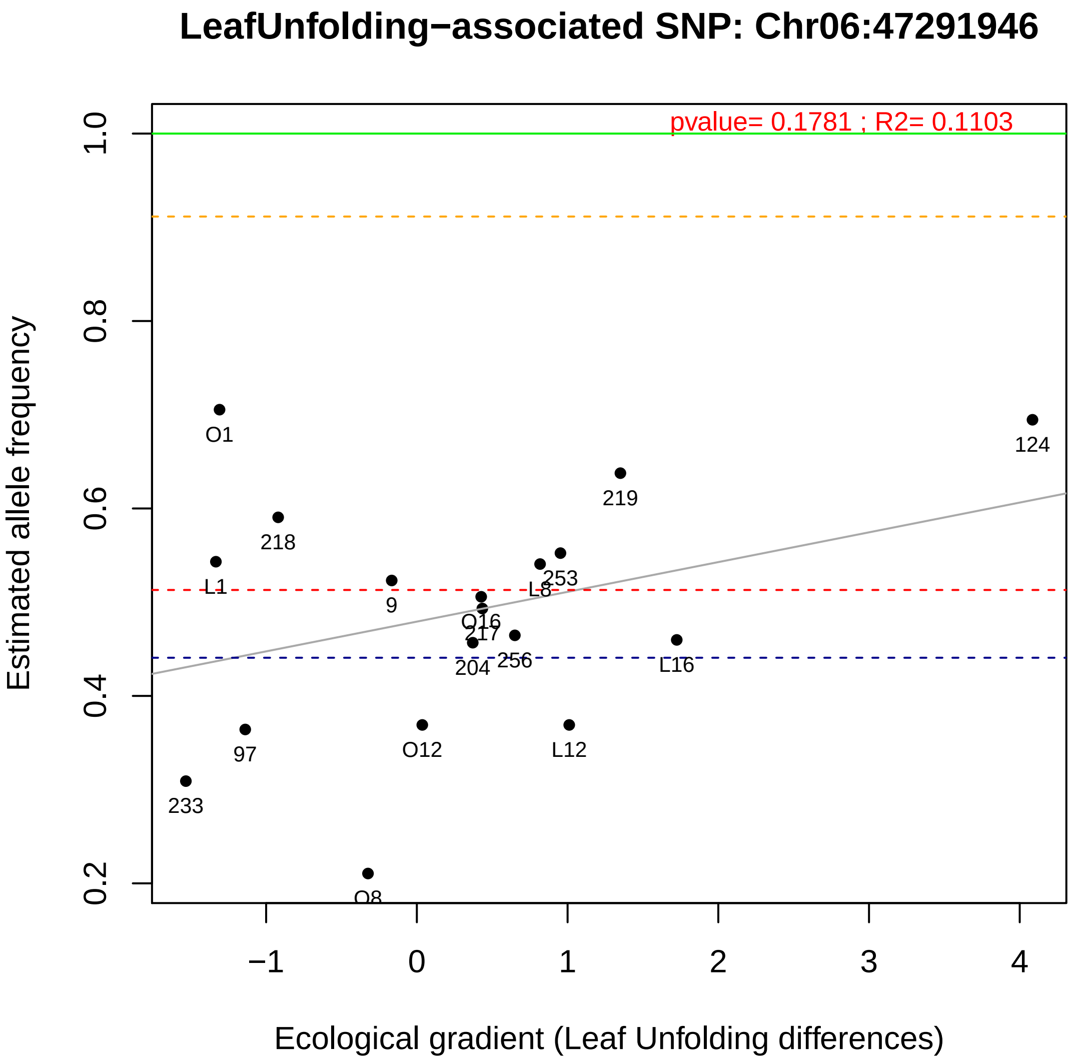
LeafUnfolding−associated SNP: Chr06:47291946
pvalue= 0.1781 ; R2= 0.1103
Estimated allele frequency
0.2
0.4
0.6
0.8
1.0
−1
0
1
2
3
4
Ecological gradient (Leaf Unfolding differences)
O1
124
219
218
L1
253
L8
9
O16
217
204
256
L16
97
O12
L12
233
O8

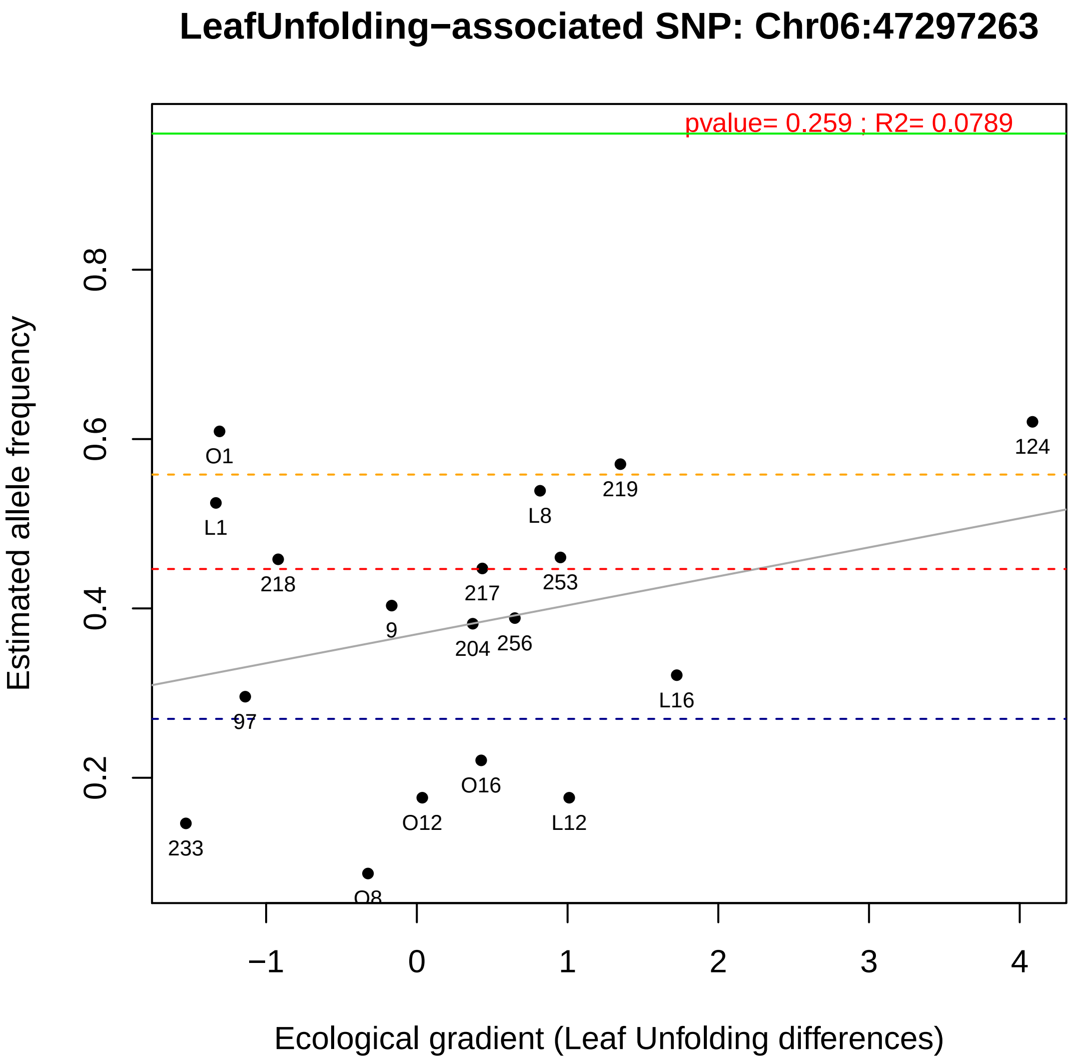

LeafUnfolding−associated SNP: Chr06:47297263
pvalue= 0.259 ; R2= 0.0789
Estimated allele frequency
0.2
0.4
0.6
0.8
−1
0
1
2
3
4
Ecological gradient (Leaf Unfolding differences)
O1
L1
218
9
217
204
256
253
L8
219
124
L16
97
233
O8
O12
O16
L12

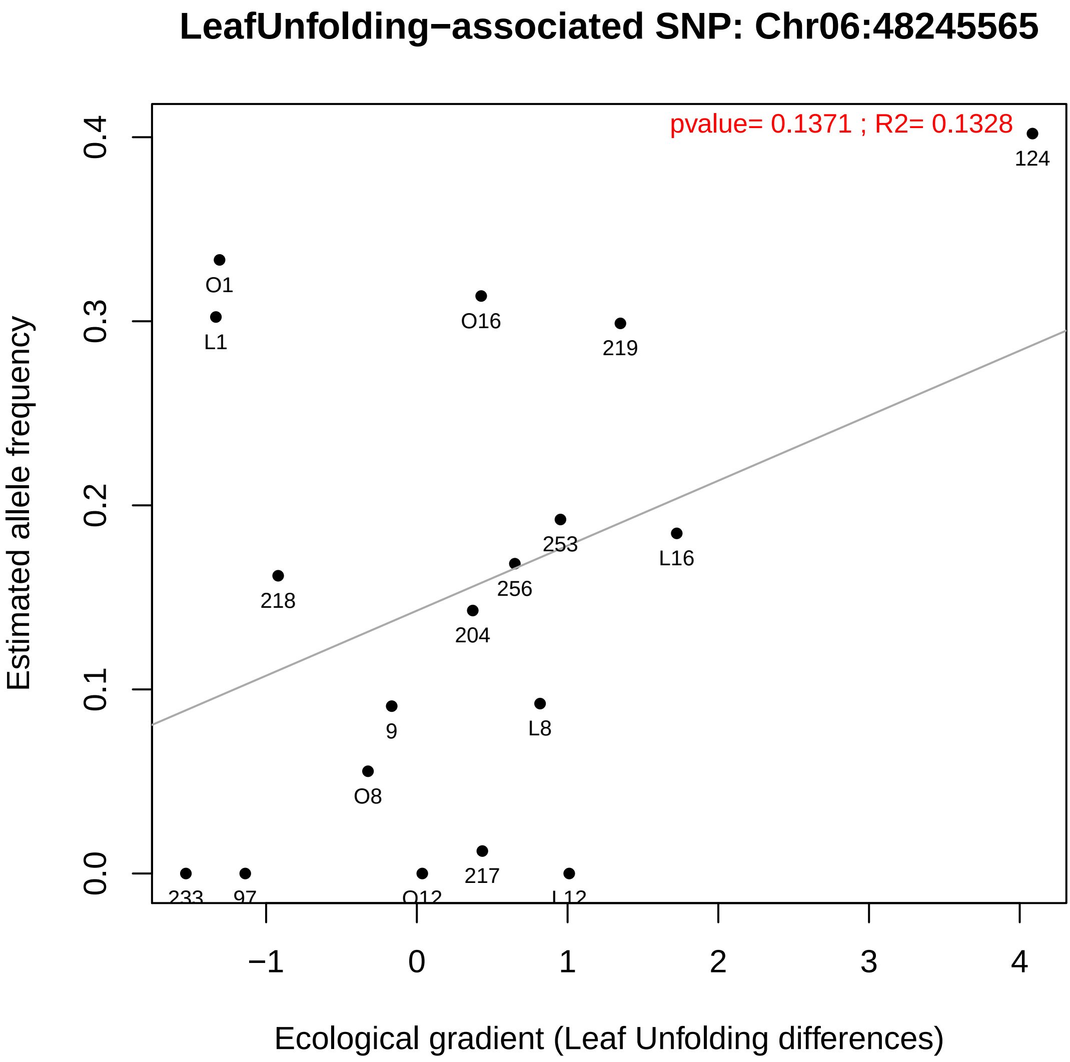
LeafUnfolding−associated SNP: Chr06:48245565
pvalue= 0.1371 ; R2= 0.1328
Estimated allele frequency
Ecological gradient (Leaf Unfolding differences)
0.0
0.1
0.2
0.3
0.4
−1
0
1
2
3
4
124
O1
L1
O16
219
253
L16
256
218
204
9
L8
O8
217
233
97
O12
L12

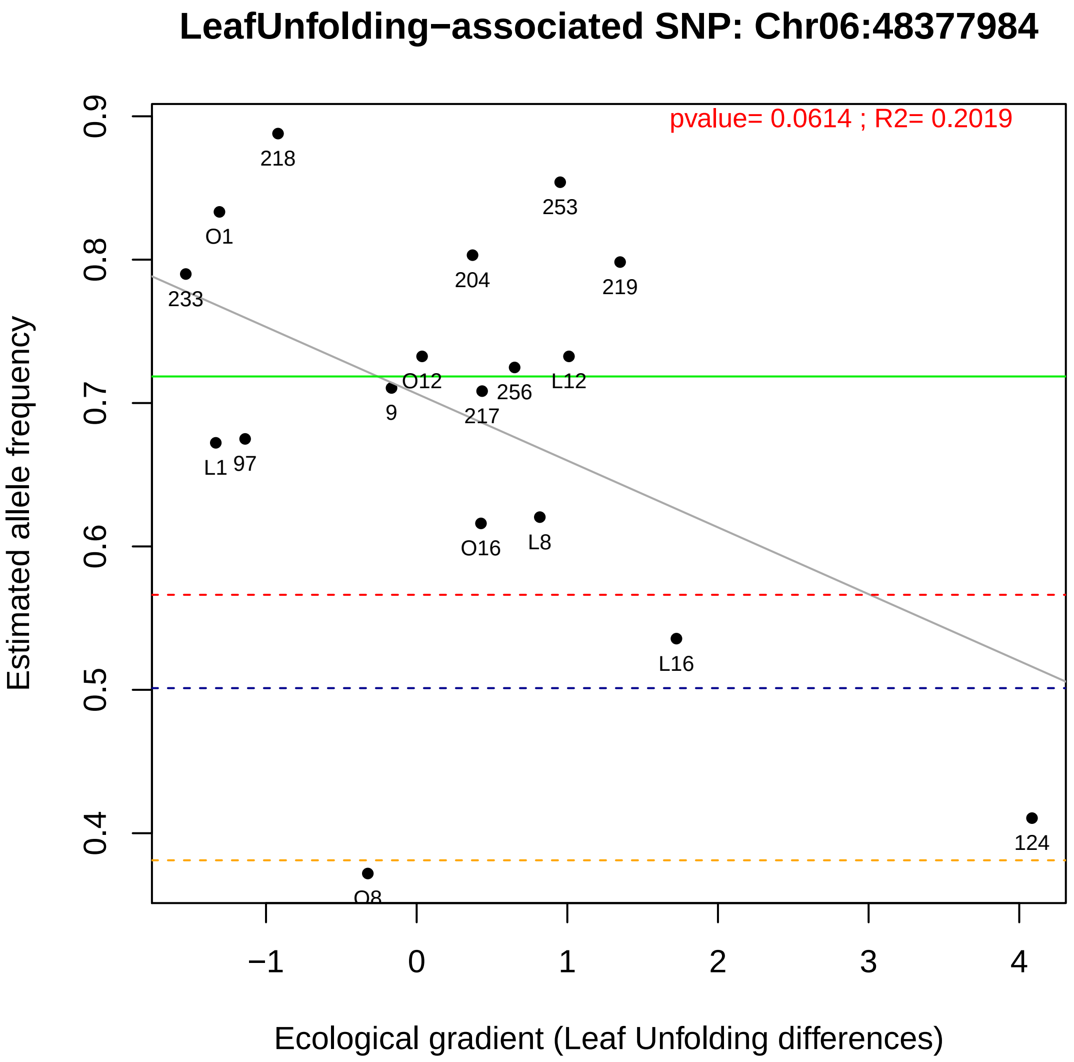

LeafUnfolding−associated SNP: Chr06:48377984
pvalue= 0.0614 ; R2= 0.2019
Estimated allele frequency
0.4
0.5
0.6
0.7
0.8
0.9
−1
0
1
2
3
4
Ecological gradient (Leaf Unfolding differences)
218
O1
233
253
204
219
O12
9
217
256
L12
L1
97
O16
L8
L16
124
O8

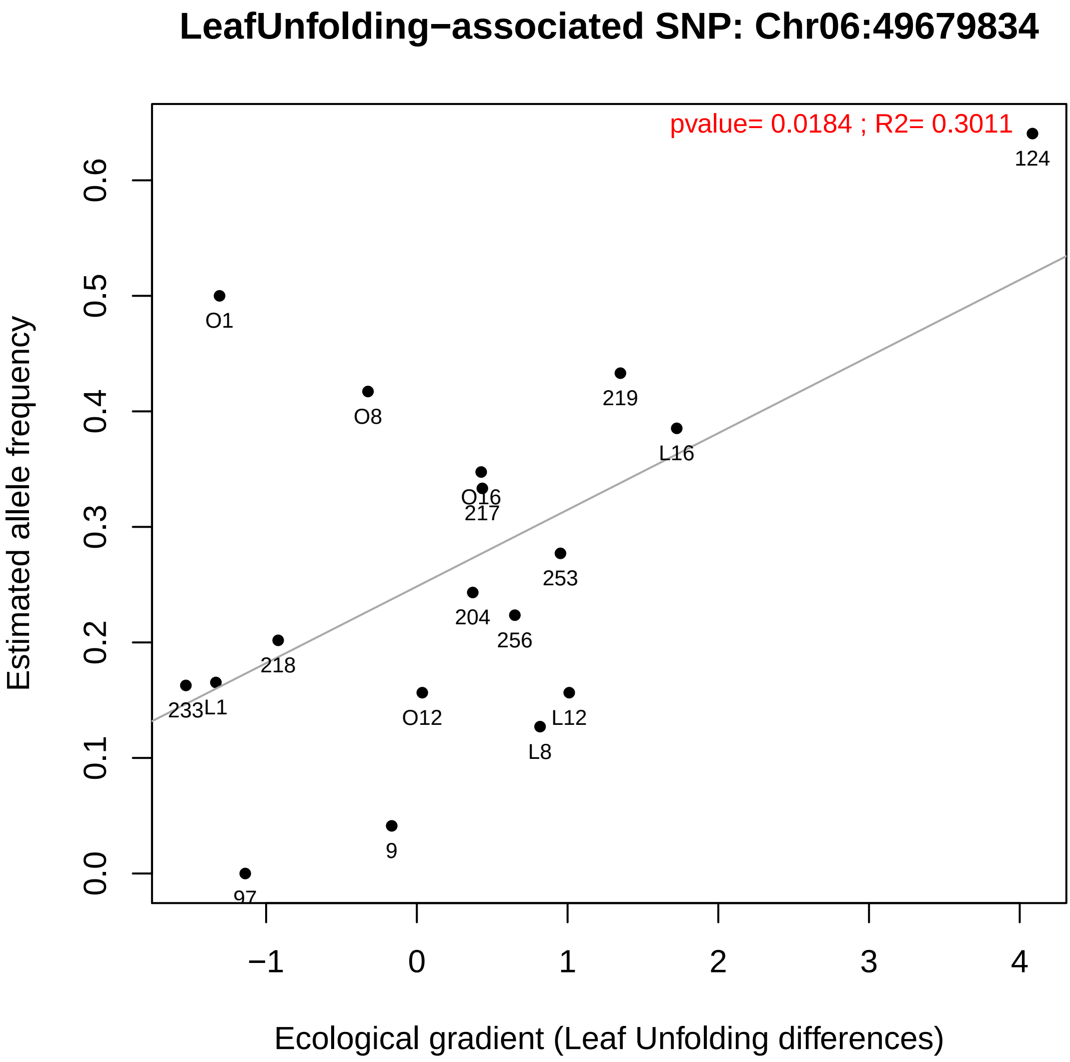
LeafUnfolding−associated SNP: Chr06:49679834
pvalue= 0.0184 ; R2= 0.3011
Estimated allele frequency
Ecological gradient (Leaf Unfolding differences)
0.0
0.1
0.2
0.3
0.4
0.5
0.6
−1
0
1
2
3
4
124
O1
O8
219
L16
O16
217
253
204
256
218
233
L1
O12
L12
L8
9
97

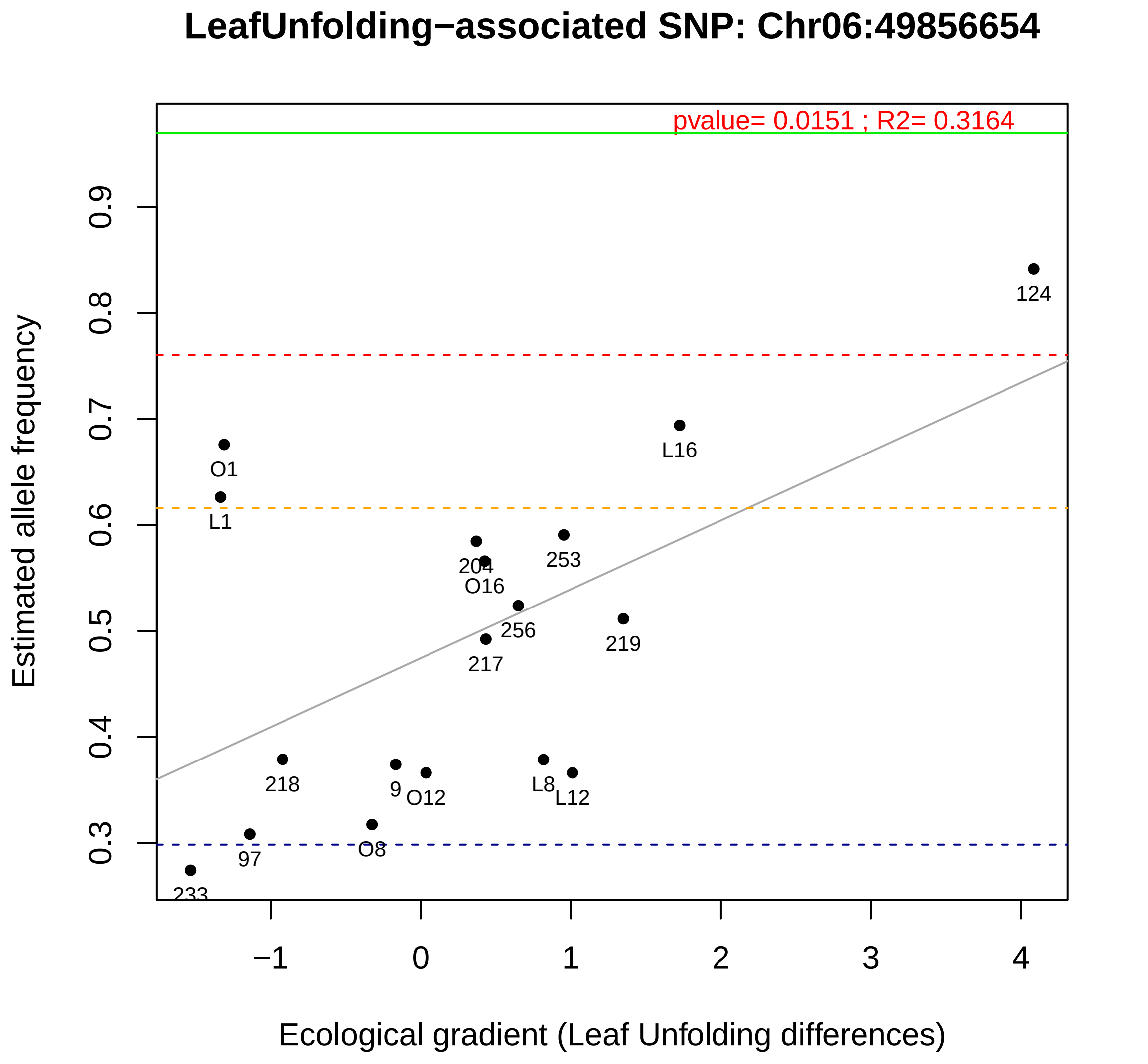
LeafUnfolding−associated SNP: Chr06:49856654
pvalue= 0.0151 ; R2= 0.3164
Estimated allele frequency
0.3
0.4
0.5
0.6
0.7
0.8
0.9
−1
0
1
2
3
4
Ecological gradient (Leaf Unfolding differences)
124
L16
O1
L1
204
O16
253
256
219
217
218
9
O12
L8
L12
97
O8
233

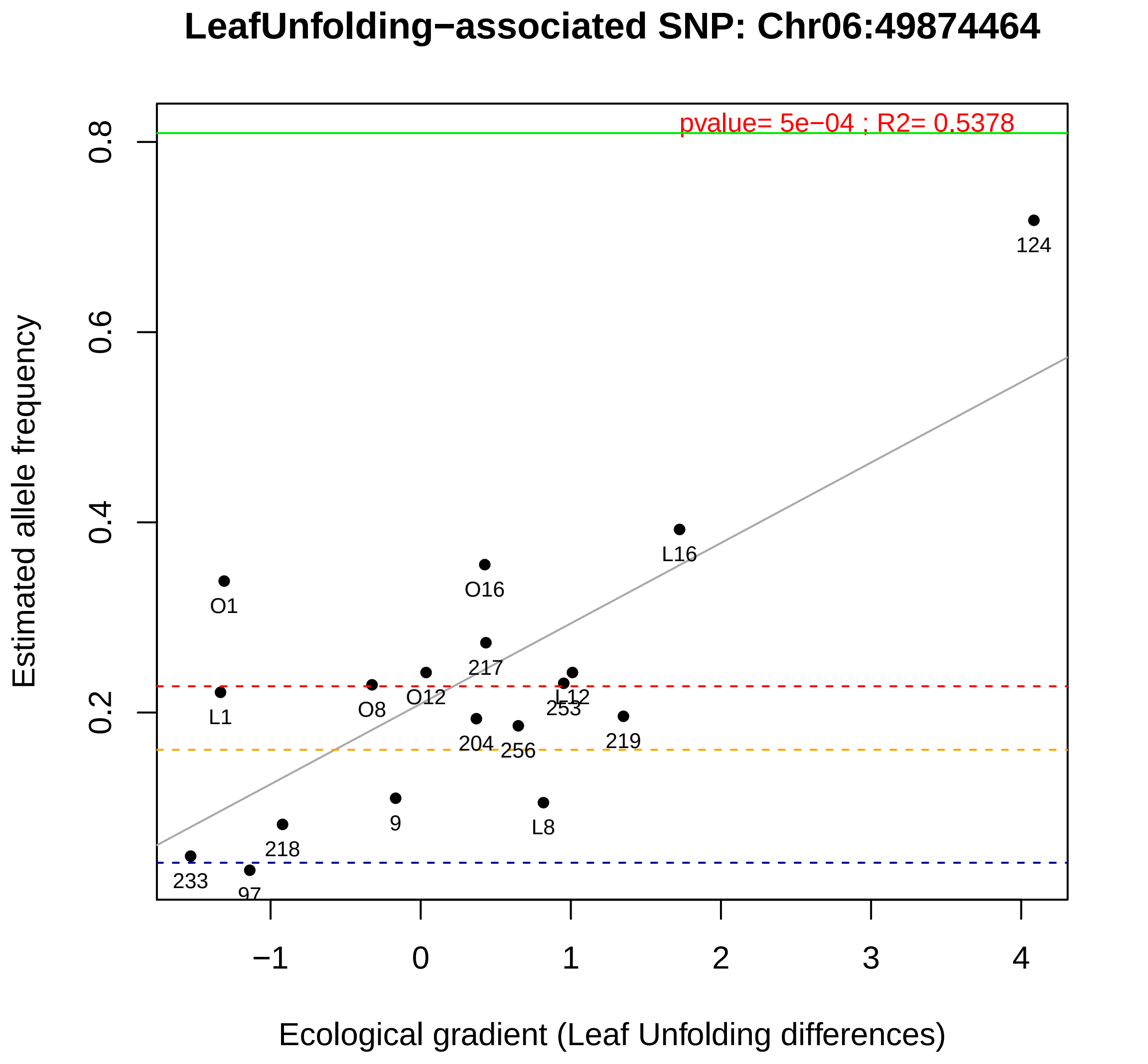

LeafUnfolding−associated SNP: Chr06:49874464
pvalue= 5e−04 ; R2= 0.5378
Estimated allele frequency
Ecological gradient (Leaf Unfolding differences)
0.2
0.4
0.6
0.8
−1
0
1
2
3
4
124
L16
O16
O1
217
L1
O8
O12
L12
253
204
256
219
9
L8
218
233
97

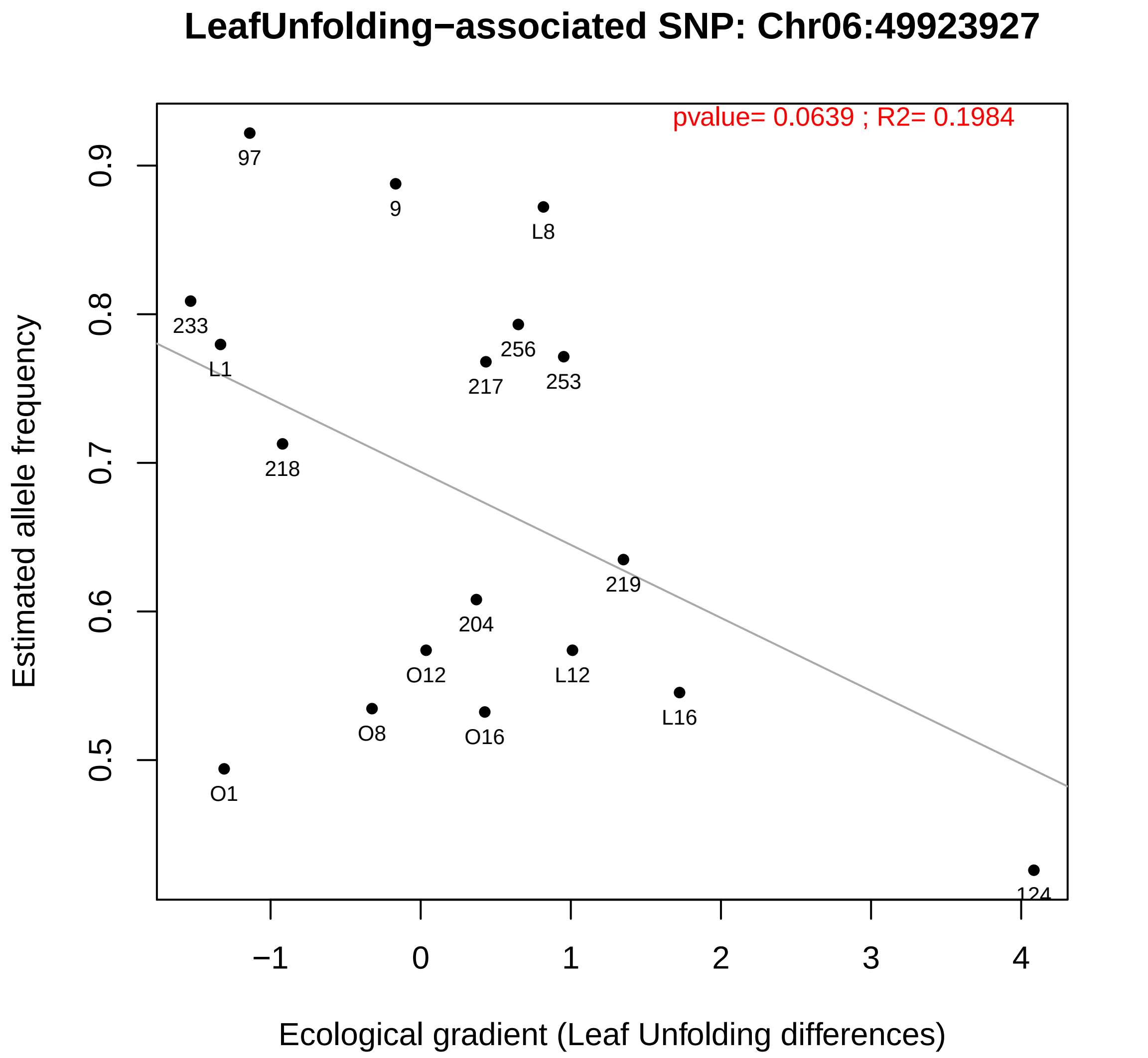
LeafUnfolding−associated SNP: Chr06:49923927
pvalue= 0.0639 ; R2= 0.1984
Estimated allele frequency
Ecological gradient (Leaf Unfolding differences)
0.5
0.6
0.7
0.8
0.9
−1
0
1
2
3
4
97
9
L8
233
L1
256
217
253
218
219
204
O12
L12
L16
O8
O16
O1
124

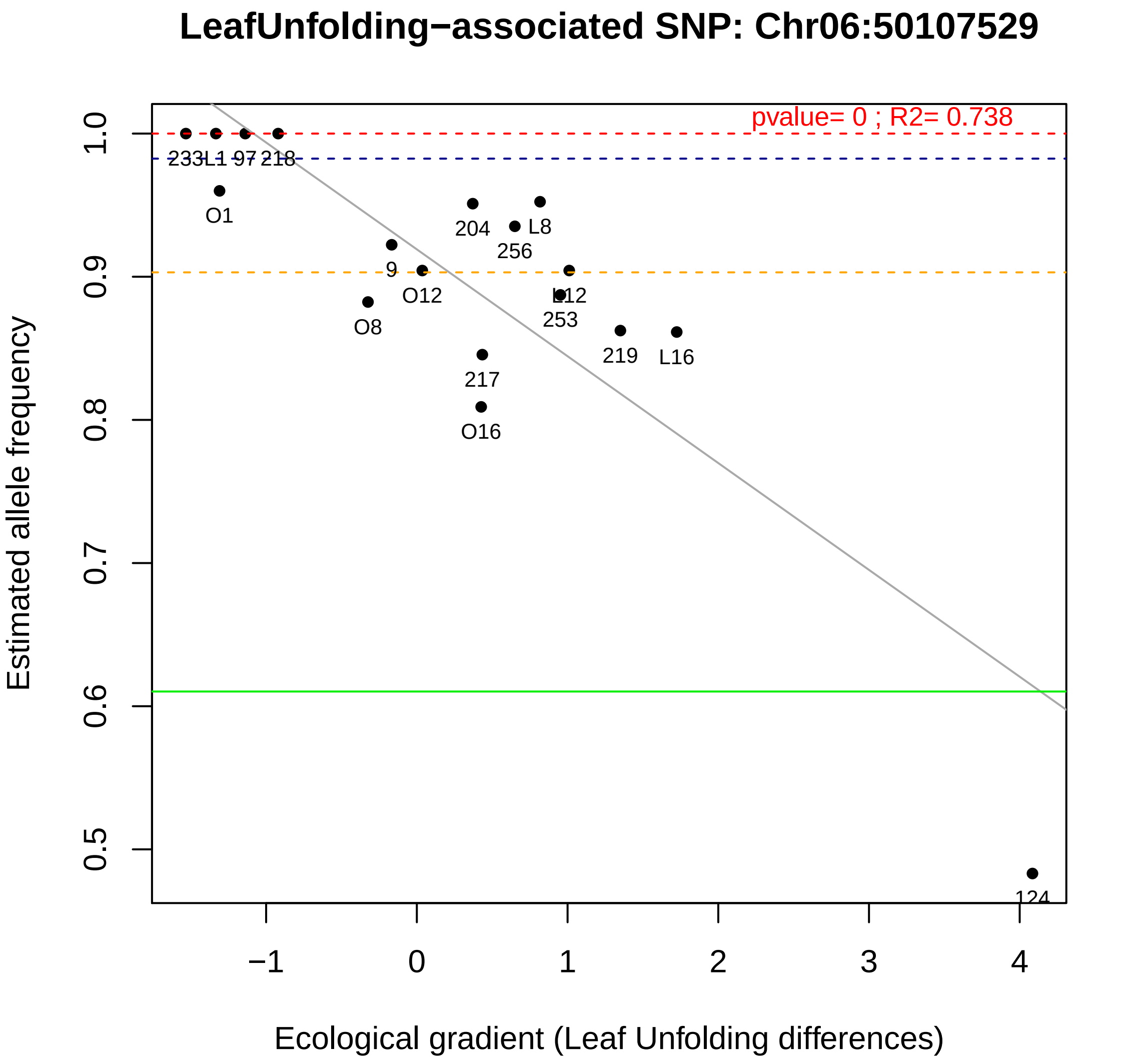
LeafUnfolding−associated SNP: Chr06:50107529
pvalue= 0 ; R2= 0.738
Estimated allele frequency
Ecological gradient (Leaf Unfolding differences)
1.0
0.9
0.8
0.7
0.6
0.5
−1
0
1
2
3
4
233
L1
97
218
O1
204
L8
256
9
O12
L12
253
O8
219
L16
217
O16
124

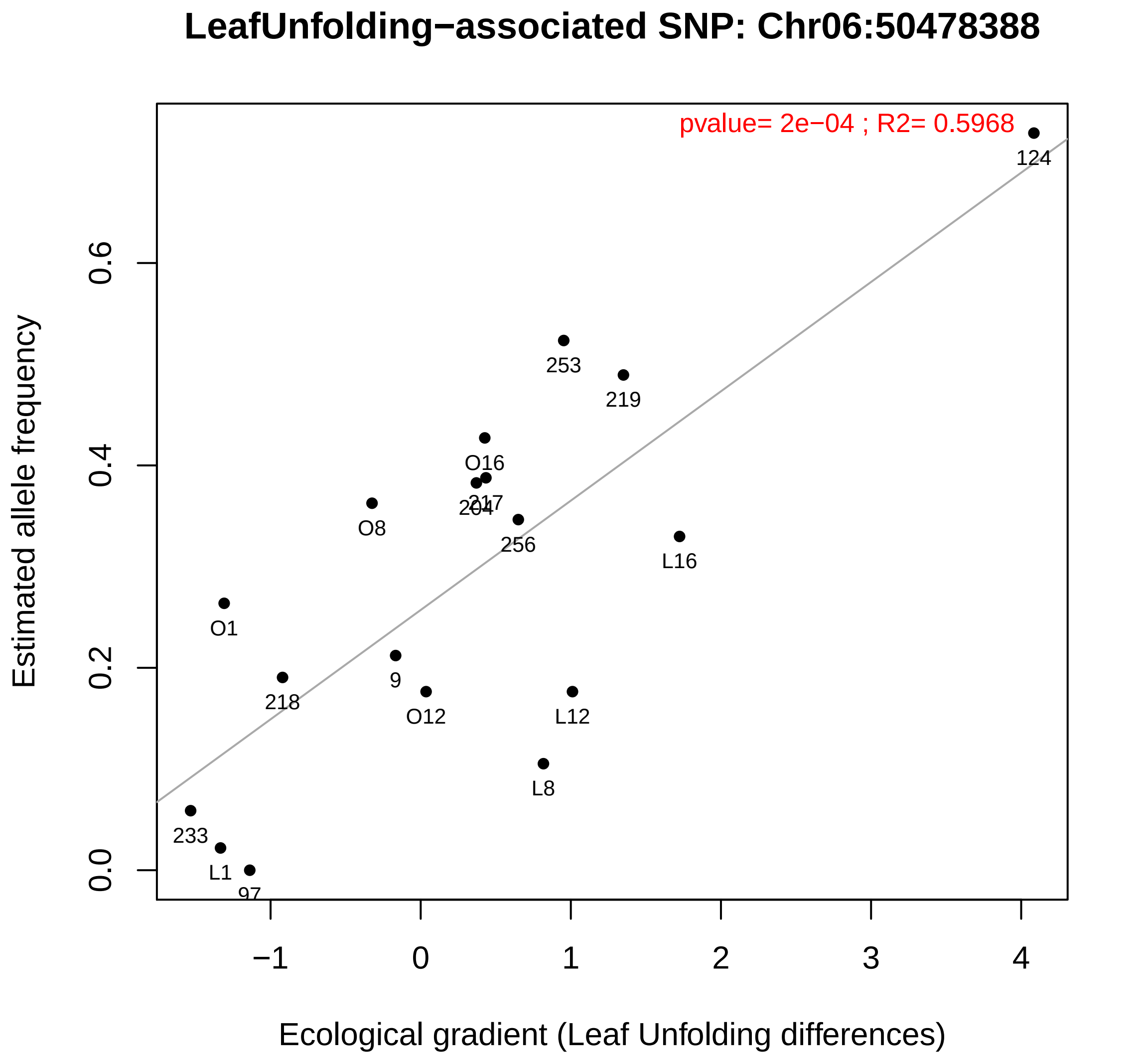

LeafUnfolding−associated SNP: Chr06:50478388
pvalue= 2e−04 ; R2= 0.5968
Estimated allele frequency
0.0
0.2
0.4
0.6
Ecological gradient (Leaf Unfolding differences)
−1
0
1
2
3
4
124
253
219
O16
217
204
O8
256
L16
O1
9
218
O12
L12
L8
233
L1
97

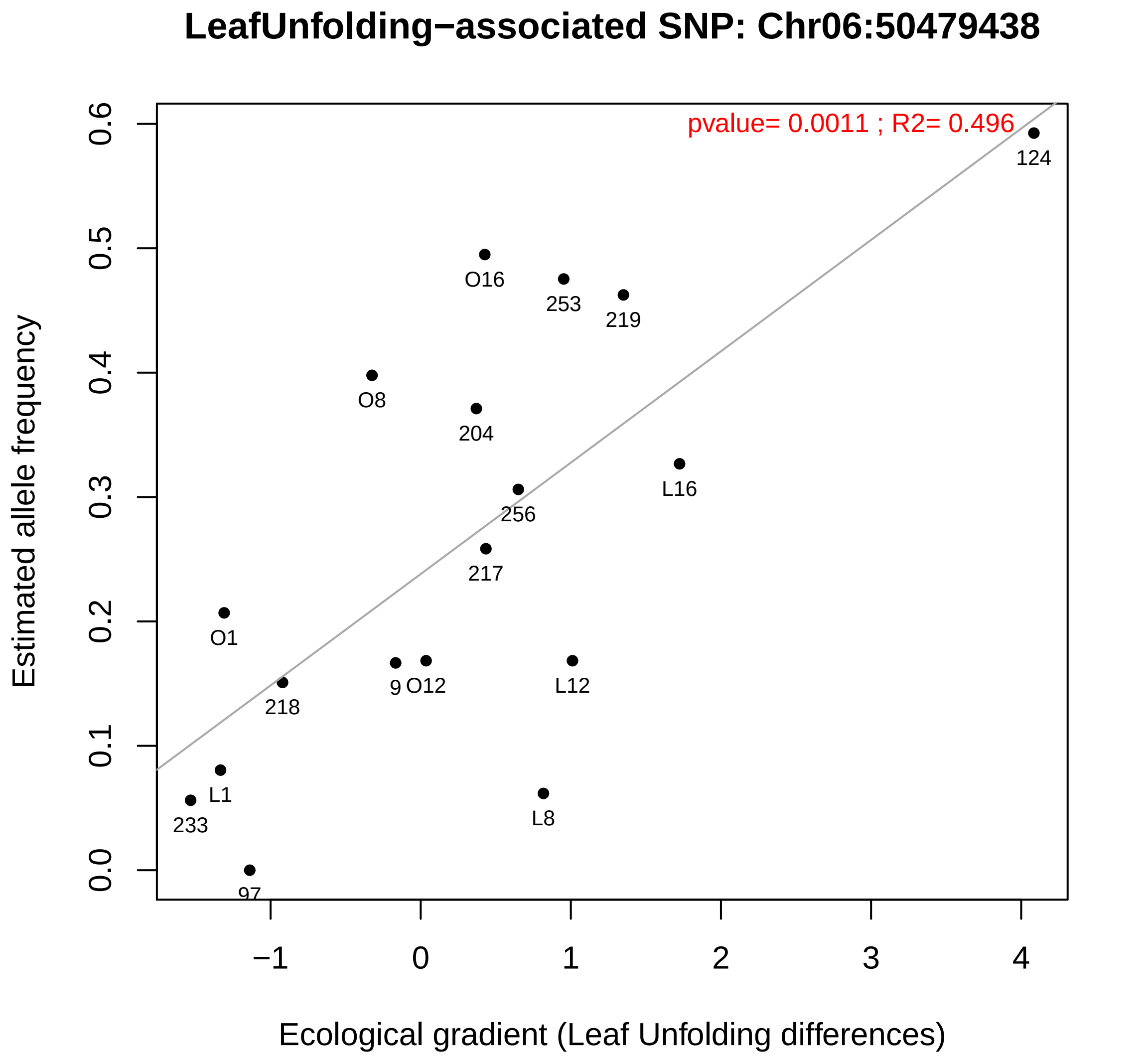

LeafUnfolding−associated SNP: Chr06:50479438
pvalue= 0.0011 ; R2= 0.496
Estimated allele frequency
Ecological gradient (Leaf Unfolding differences)
0.0
0.1
0.2
0.3
0.4
0.5
0.6
−1
0
1
2
3
4
124
O16
253
219
O8
204
L16
256
217
O1
218
9
O12
L12
L1
233
L8
97

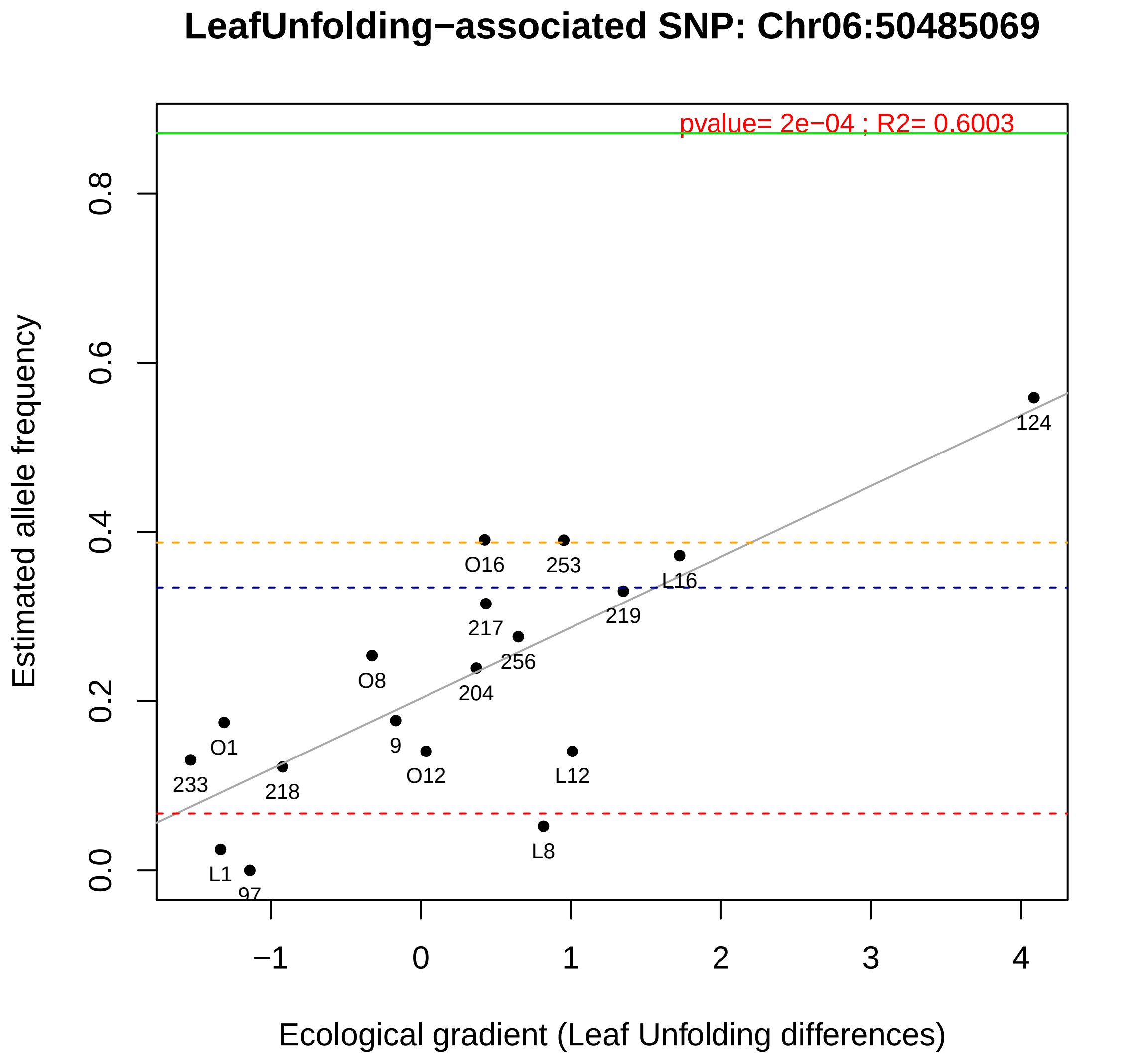
LeafUnfolding−associated SNP: Chr06:50485069
pvalue= 2e−04 ; R2= 0.6003
Estimated allele frequency
0.0
0.2
0.4
0.6
0.8
Ecological gradient (Leaf Unfolding differences)
−1
0
1
2
3
4
124
O16
253
L16
219
217
256
O8
204
O1
9
233
218
O12
L12
L8
L1
97

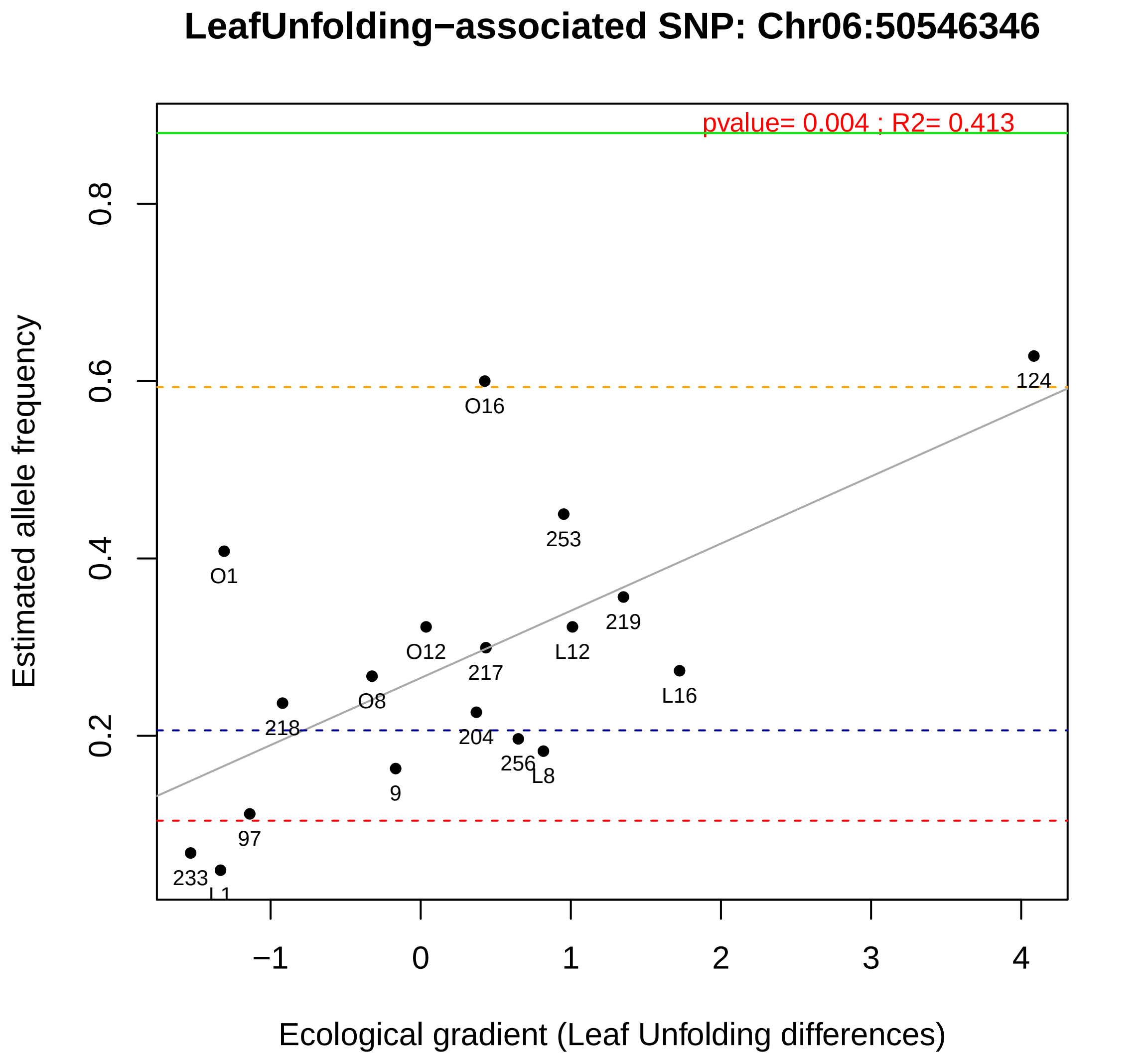

LeafUnfolding−associated SNP: Chr06:50546346
pvalue= 0.004 ; R2= 0.413
Estimated allele frequency
0.2
0.4
0.6
0.8
Ecological gradient (Leaf Unfolding differences)
−1
0
1
2
3
4
124
O16
253
O1
219
O12
L12
217
L16
O8
218
204
256
L8
9
97
233
L1

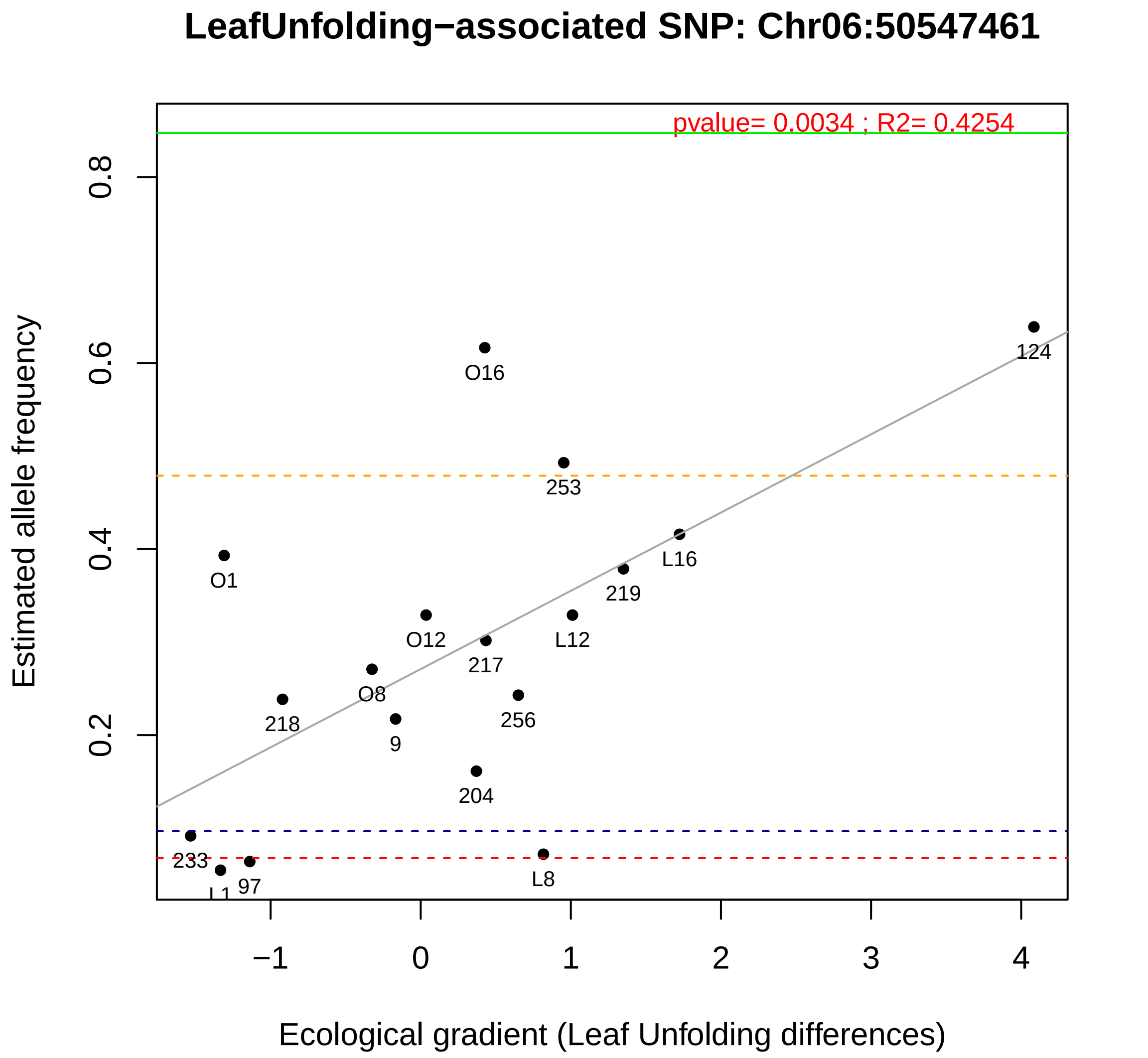

LeafUnfolding−associated SNP: Chr06:50547461
pvalue= 0.0034 ; R2= 0.4254
Estimated allele frequency
0.2
0.4
0.6
0.8
Ecological gradient (Leaf Unfolding differences)
−1
0
1
2
3
4
124
O16
253
L16
219
O1
O12
L12
217
O8
256
218
9
204
233
97
L8

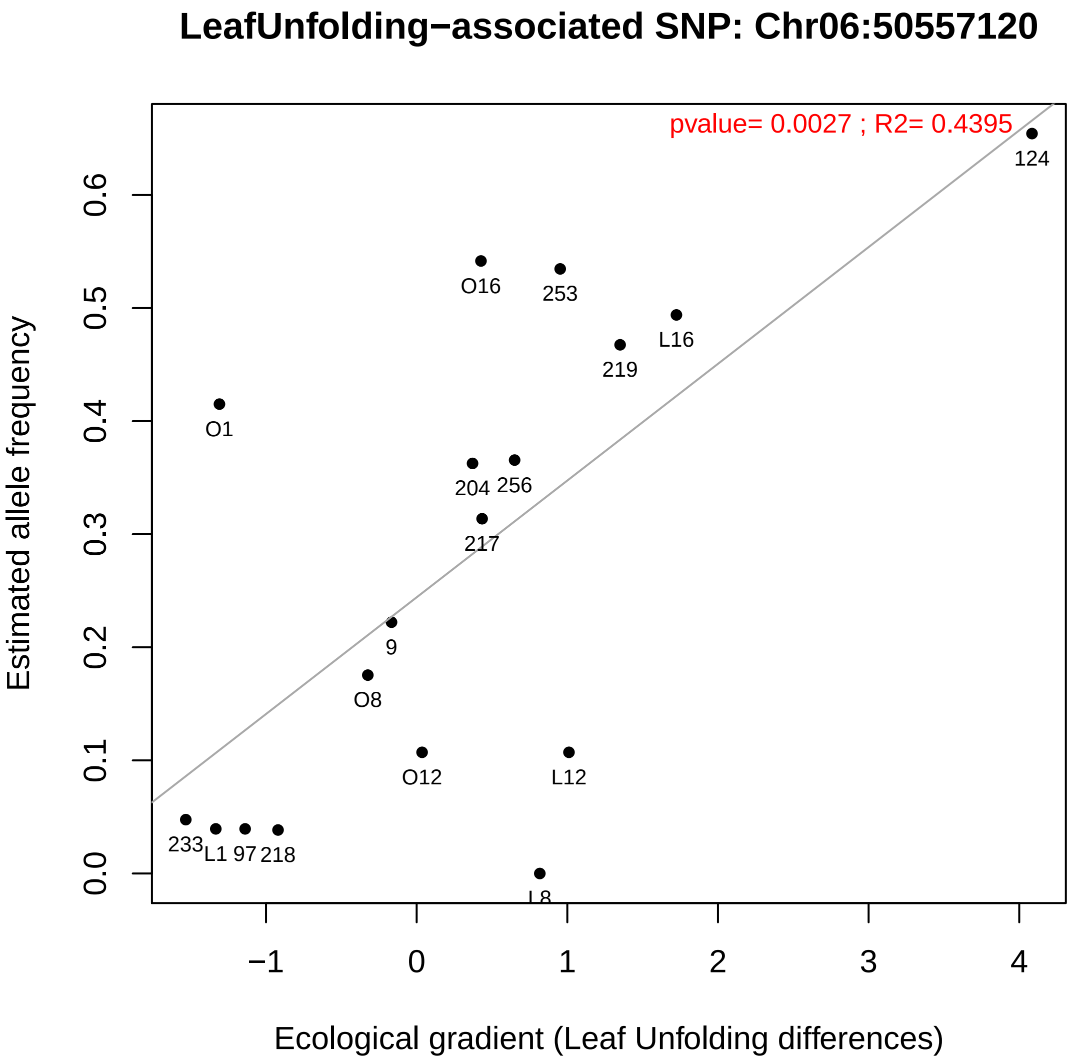

LeafUnfolding−associated SNP: Chr06:50557120
pvalue= 0.0027 ; R2= 0.4395
Estimated allele frequency
Ecological gradient (Leaf Unfolding differences)
0.0
0.1
0.2
0.3
0.4
0.5
0.6
−1
0
1
2
3
4
124
O16
253
L16
219
O1
204
256
217
9
O8
O12
L12
233
L1
97
218

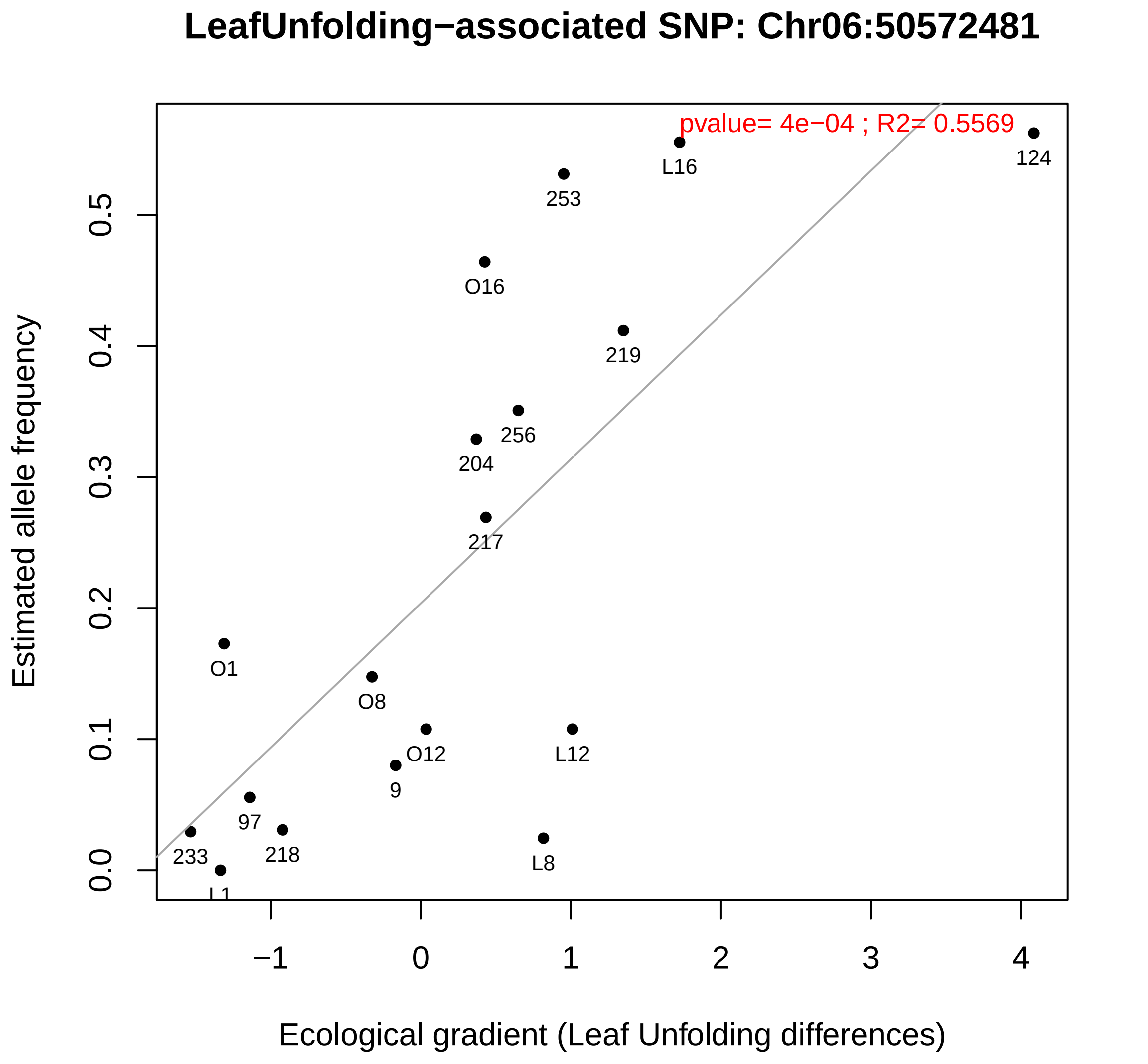
LeafUnfolding−associated SNP: Chr06:50572481
pvalue= 4e−04 ; R2= 0.5569
Estimated allele frequency
Ecological gradient (Leaf Unfolding differences)
0.0
0.1
0.2
0.3
0.4
0.5
−1
0
1
2
3
4
124
L16
253
O16
219
256
204
217
O1
O8
O12
L12
9
97
218
233
L8
L1

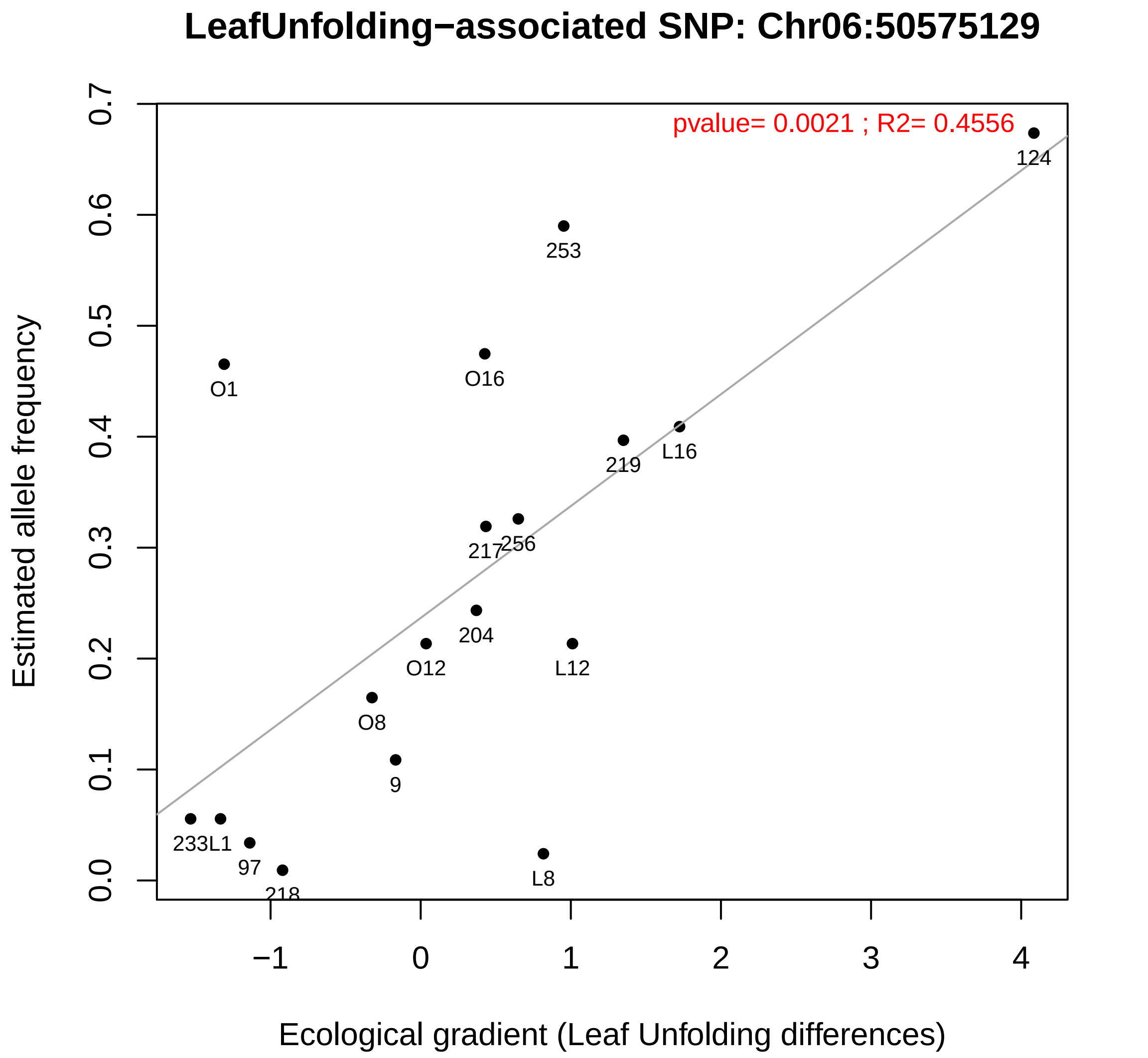

LeafUnfolding−associated SNP: Chr06:50575129
pvalue= 0.0021 ; R2= 0.4556
124
253
O16
O1
219
L16
256
217
204
O12
L12
O8
9
233
L1
97
218
L8
0.0
0.1
0.2
0.3
0.4
0.5
0.6
0.7
−1
0
1
2
3
4
Estimated allele frequency
Ecological gradient (Leaf Unfolding differences)

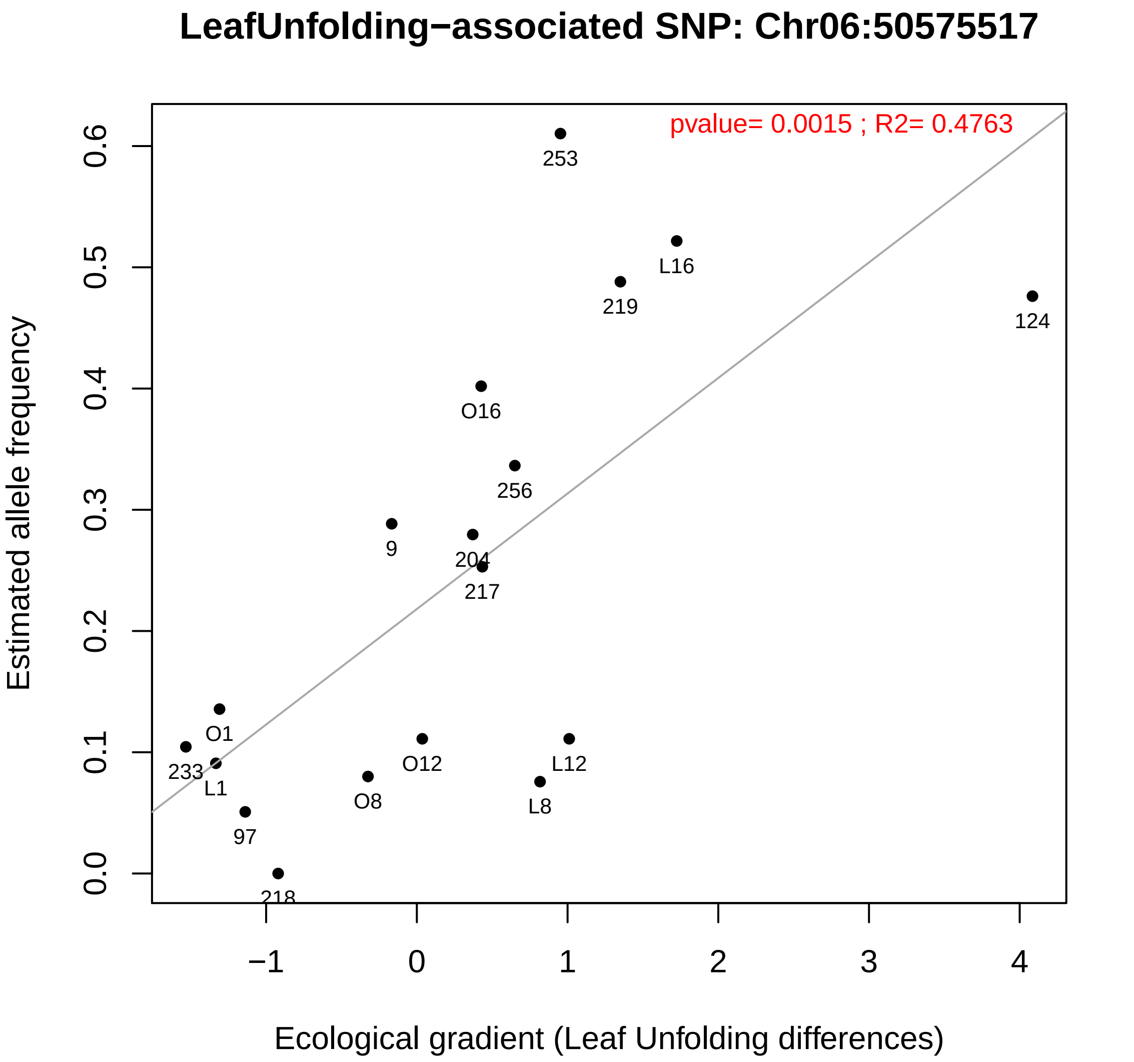
LeafUnfolding−associated SNP: Chr06:50575517
pvalue= 0.0015 ; R2= 0.4763
Estimated allele frequency
0.0
0.1
0.2
0.3
0.4
0.5
0.6
253
L16
219
124
O16
256
9
204
217
O1
233
L1
O12
L12
O8
L8
97
218
−1
0
1
2
3
4
Ecological gradient (Leaf Unfolding differences)

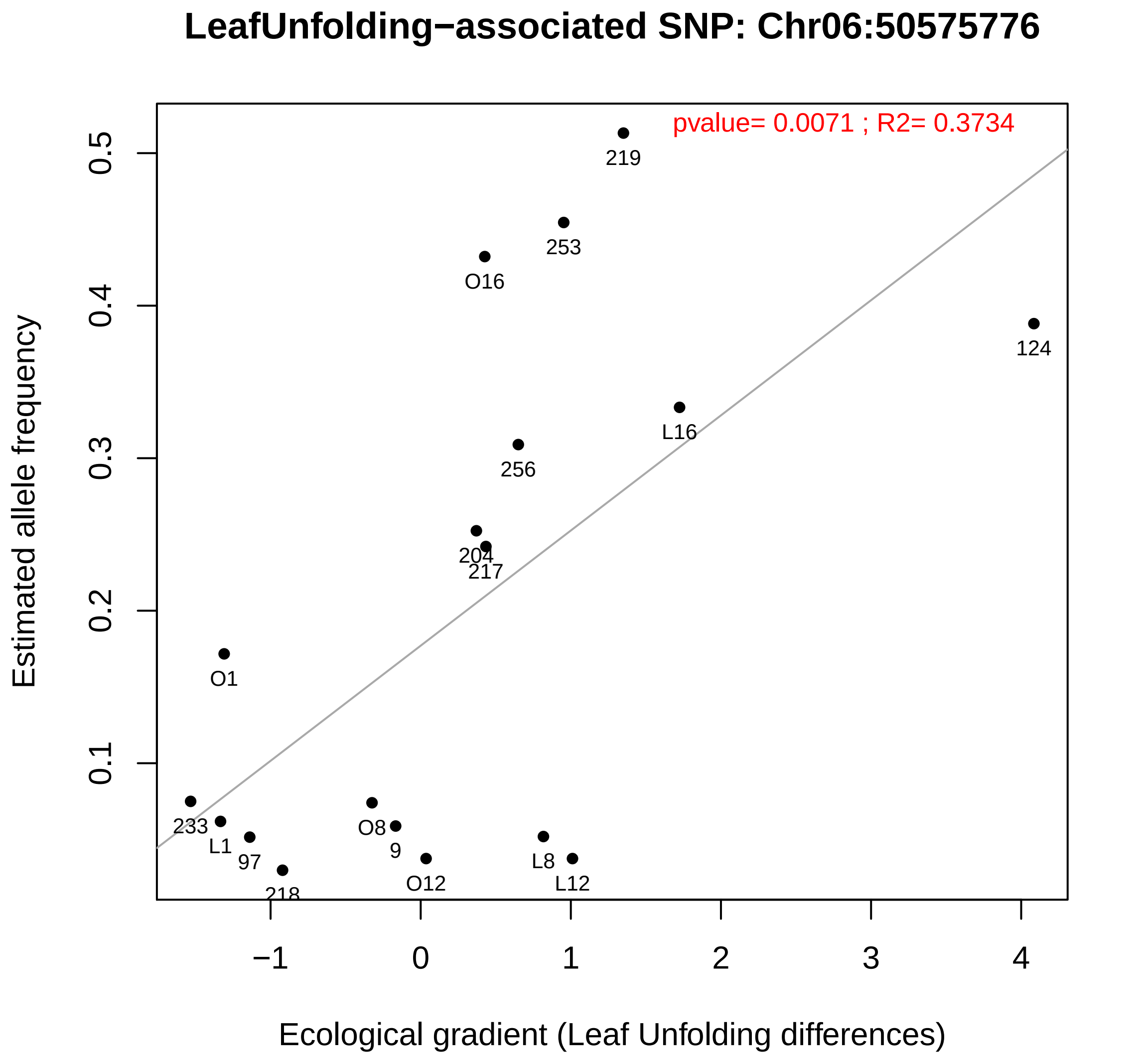

LeafUnfolding−associated SNP: Chr06:50575776
pvalue= 0.0071 ; R2= 0.3734
Estimated allele frequency
Ecological gradient (Leaf Unfolding differences)
0.1
0.2
0.3
0.4
0.5
−1
0
1
2
3
4
219
253
O16
124
L16
256
204
217
O1
233
L1
97
218
O8
9
O12
L8
L12

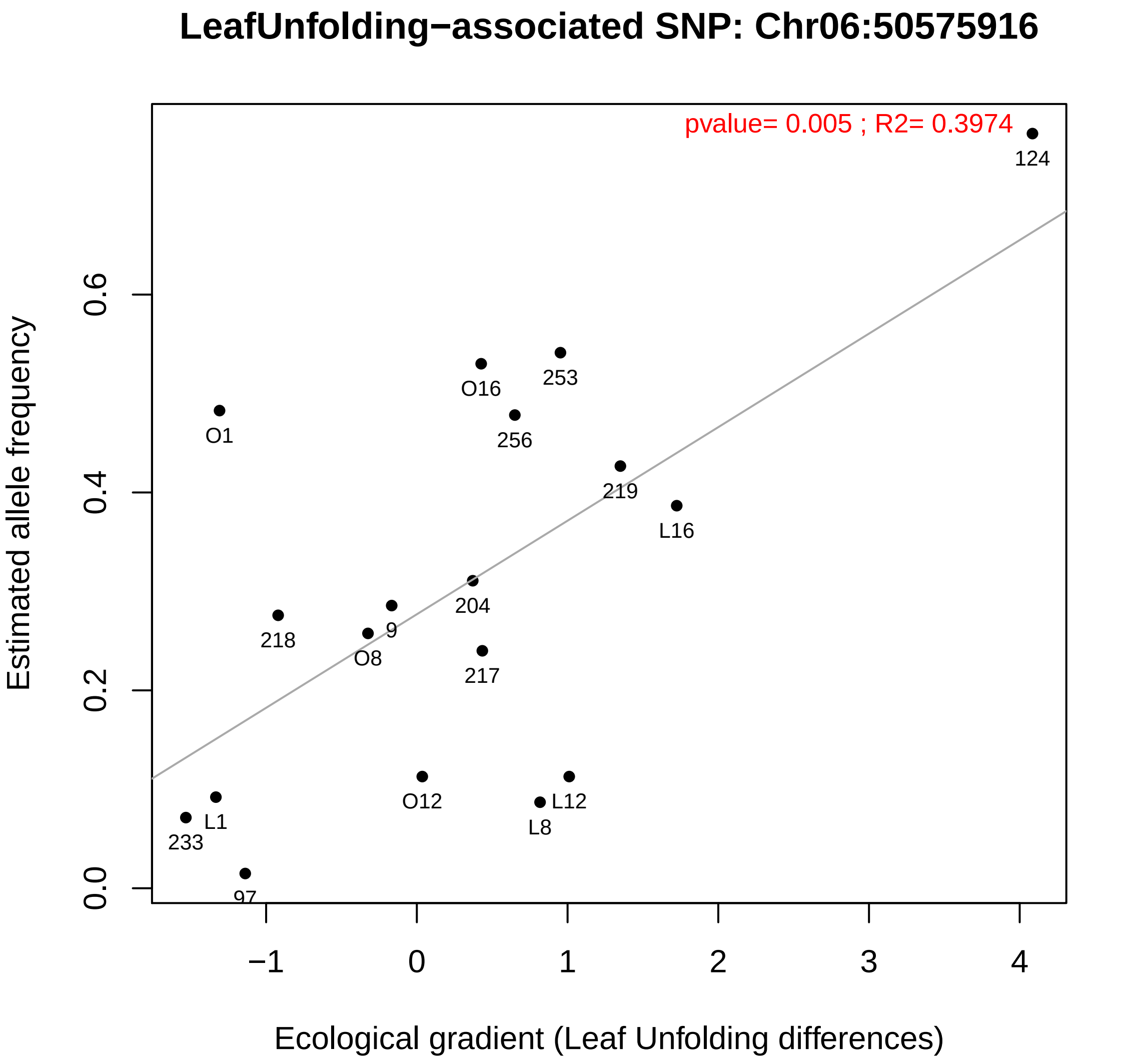
LeafUnfolding−associated SNP: Chr06:50575916
pvalue= 0.005 ; R2= 0.3974
124
253
O16
O1
256
219
L16
204
9
218
O8
217
O12
L12
L1
L8
233
97
0.0
0.2
0.4
0.6
−1
0
1
2
3
4
Estimated allele frequency
Ecological gradient (Leaf Unfolding differences)

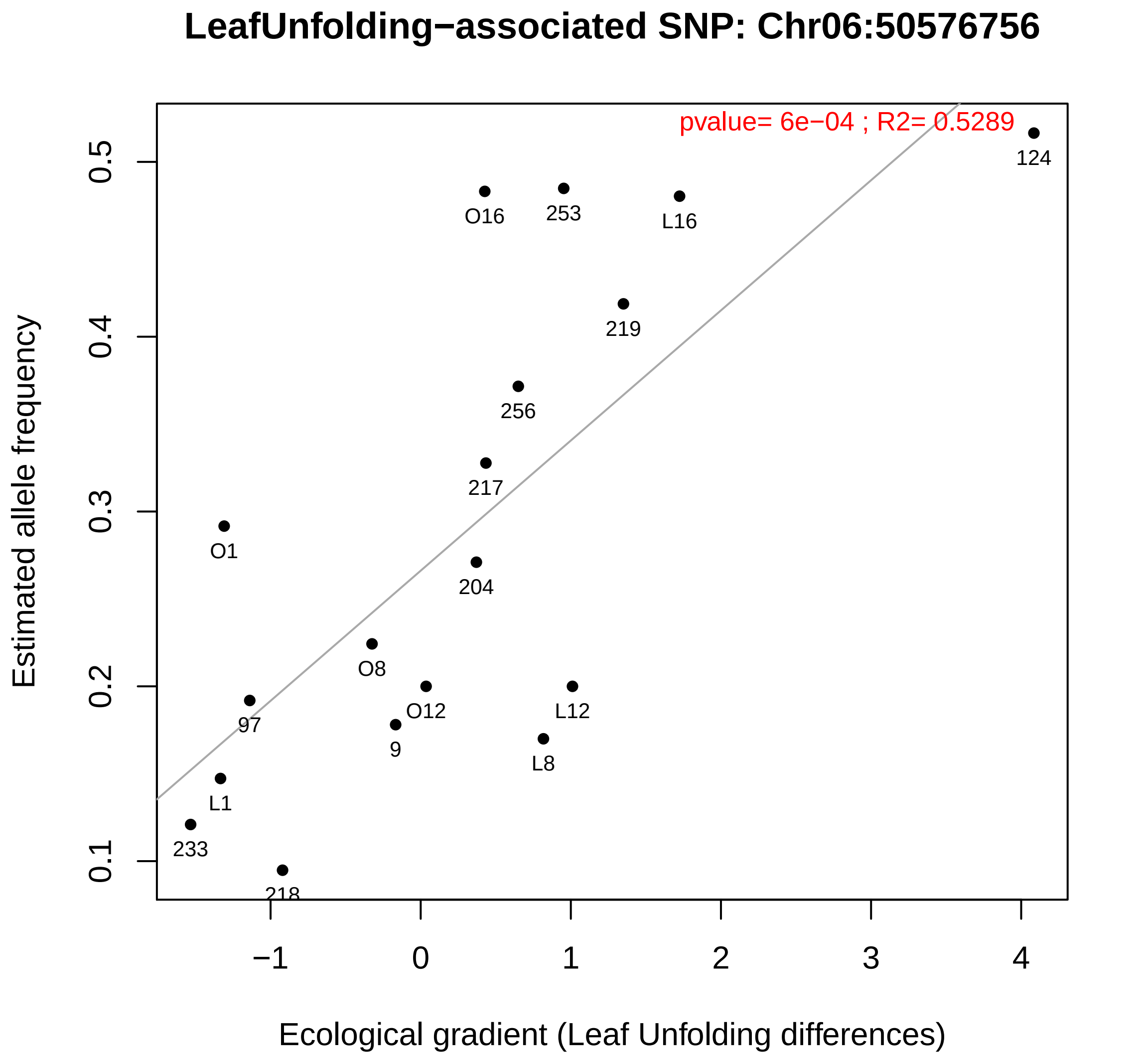

LeafUnfolding−associated SNP: Chr06:50576756
pvalue= 6e−04 ; R2= 0.5289
124
O16
253
L16
219
256
217
O1
204
O8
O12
L12
97
9
L8
L1
233
218
0.1
0.2
0.3
0.4
0.5
−1
0
1
2
3
4
Estimated allele frequency
Ecological gradient (Leaf Unfolding differences)

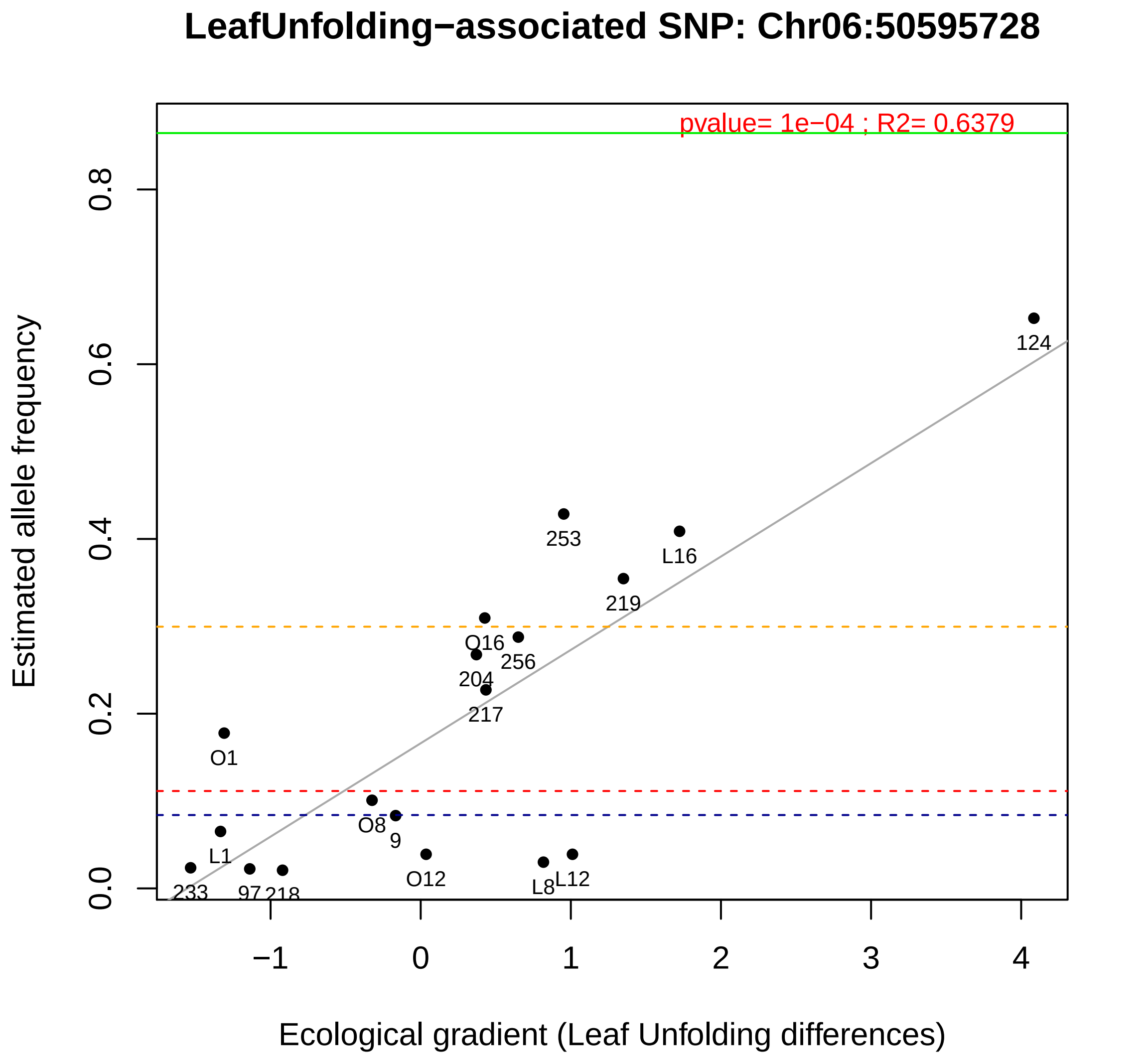

LeafUnfolding−associated SNP: Chr06:50595728
pvalue= 1e−04 ; R2= 0.6379
Estimated allele frequency
0.0
0.2
0.4
0.6
0.8
−1
0
1
2
3
4
Ecological gradient (Leaf Unfolding differences)
124
253
L16
219
O16
256
204
217
O1
O8
9
L1
O12
L8
L12
233
97
218

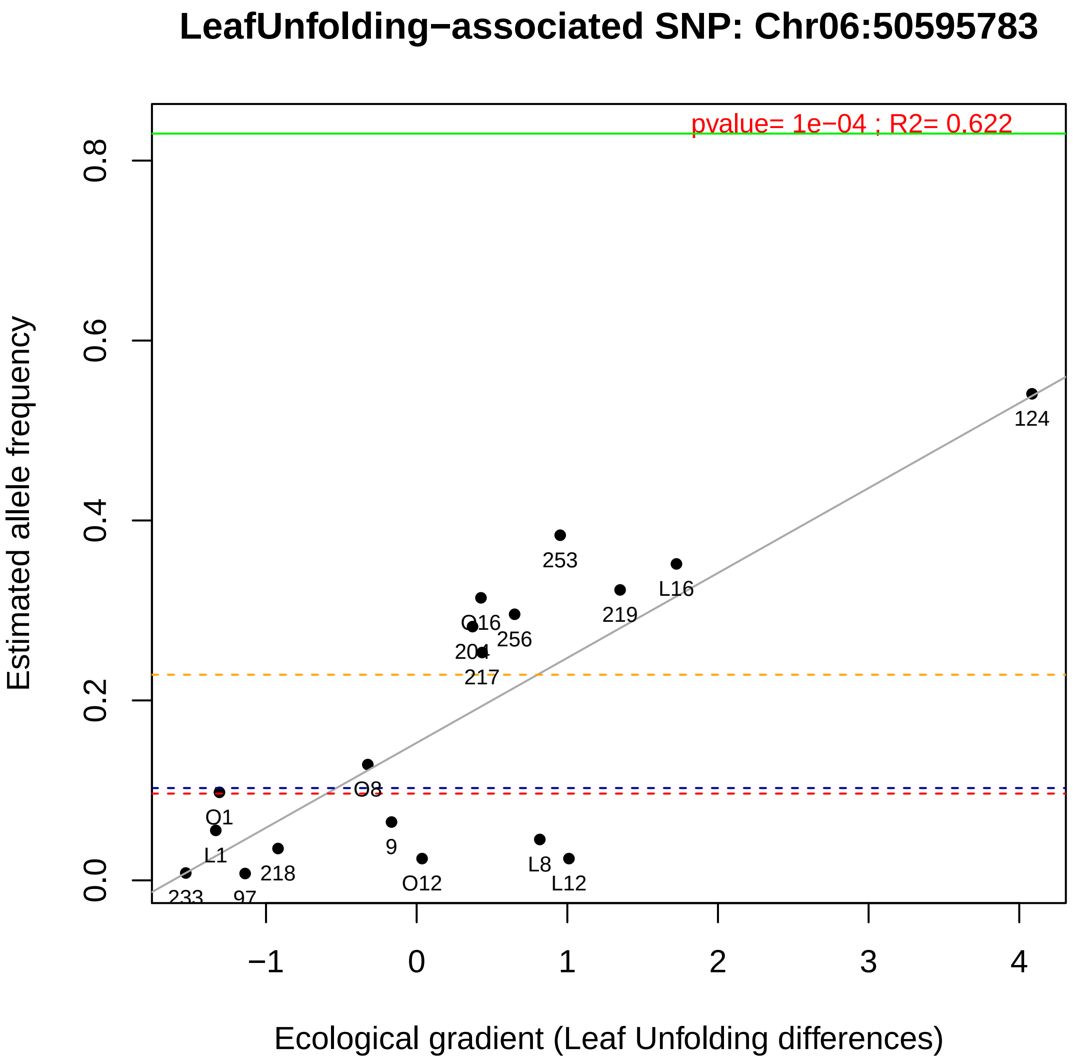

LeafUnfolding−associated SNP: Chr06:50595783
pvalue= 1e−04 ; R2= 0.622
Estimated allele frequency
0.0
0.2
0.4
0.6
0.8
−1
0
1
2
3
4
Ecological gradient (Leaf Unfolding differences)
124
253
L16
219
O16
256
204
217
O8
O1
L1
9
218
O12
L8
L12
233
97

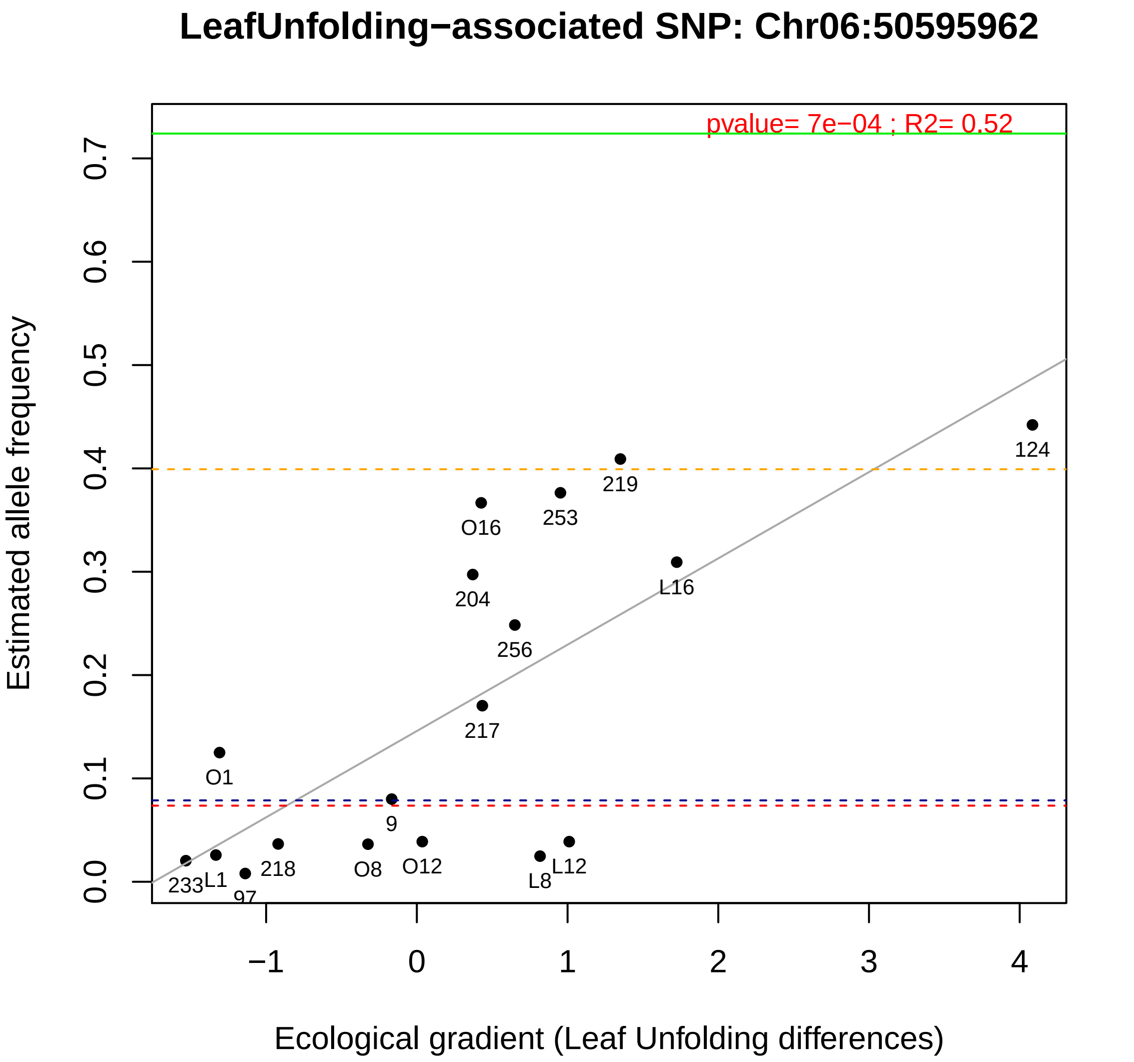

LeafUnfolding−associated SNP: Chr06:50595962
pvalue= 7e−04 ; R2= 0.52
Estimated allele frequency
0.0
0.1
0.2
0.3
0.4
0.5
0.6
0.7
−1
0
1
2
3
4
Ecological gradient (Leaf Unfolding differences)
124
219
253
O16
L16
204
256
217
O1
9
233
L1
97
218
O8
O12
L8
L12

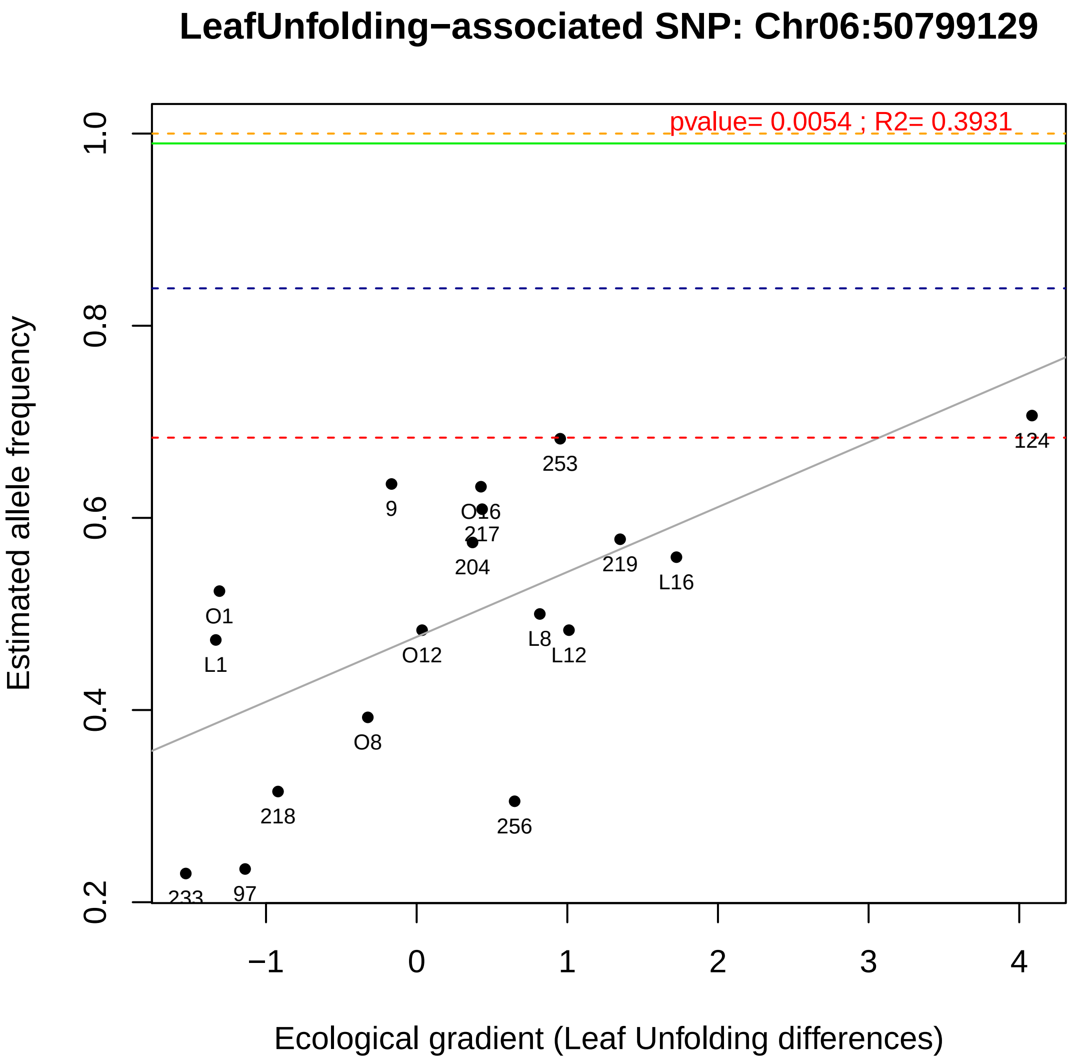

LeafUnfolding−associated SNP: Chr06:50799129
pvalue= 0.0054 ; R2= 0.3931
Estimated allele frequency
Ecological gradient (Leaf Unfolding differences)
0.2
0.4
0.6
0.8
1.0
−1
0
1
2
3
4
233
97
218
O8
L1
O1
9
O12
204
217
O16
256
L8
L12
253
219
L16
124

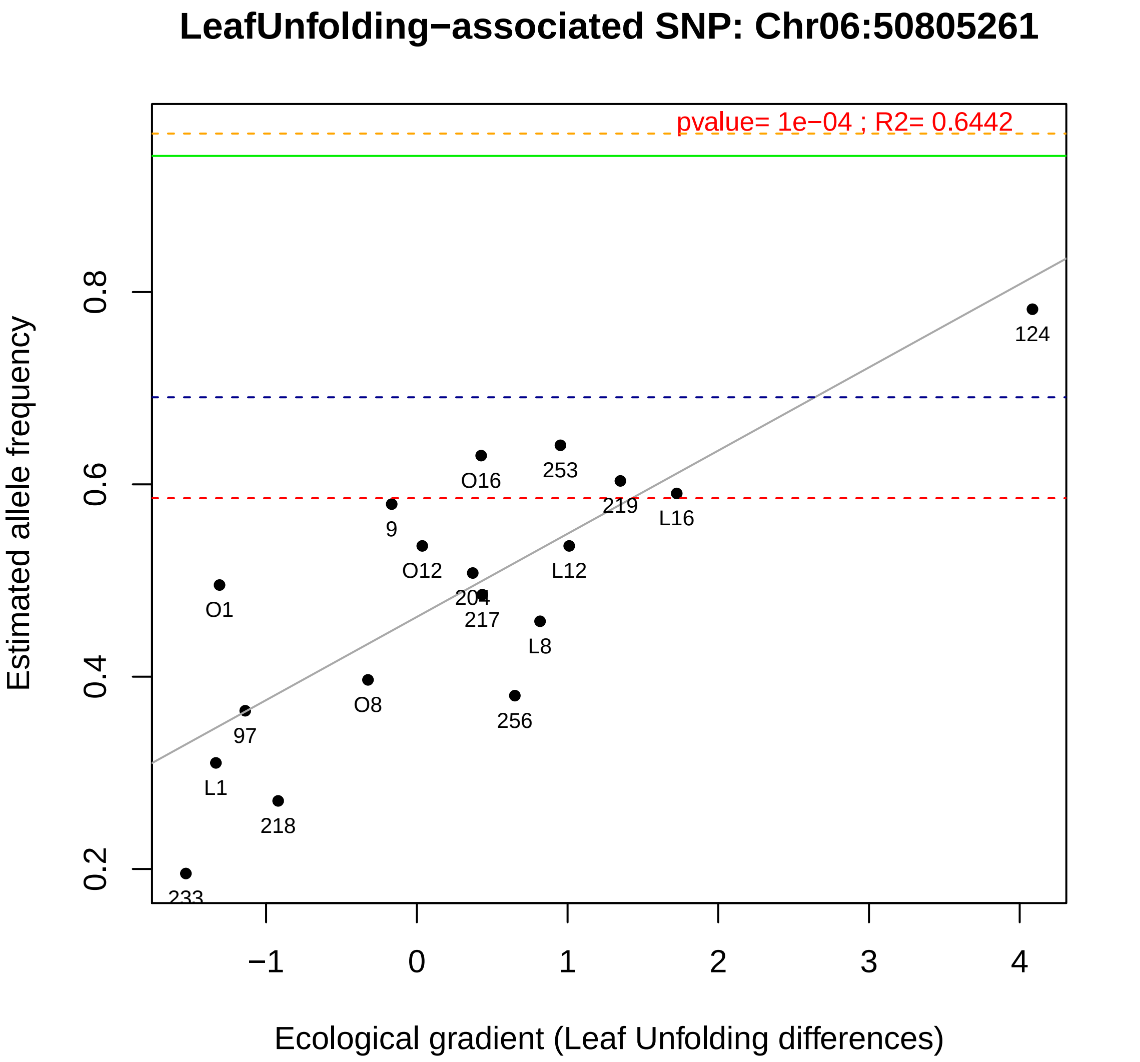
LeafUnfolding−associated SNP: Chr06:50805261
pvalue= 1e−04 ; R2= 0.6442
Estimated allele frequency
0.2
0.4
0.6
0.8
Ecological gradient (Leaf Unfolding differences)
−1
0
1
2
3
4
124
253
O16
219
L16
9
O12
L12
O1
204
217
L8
O8
256
97
L1
218
233

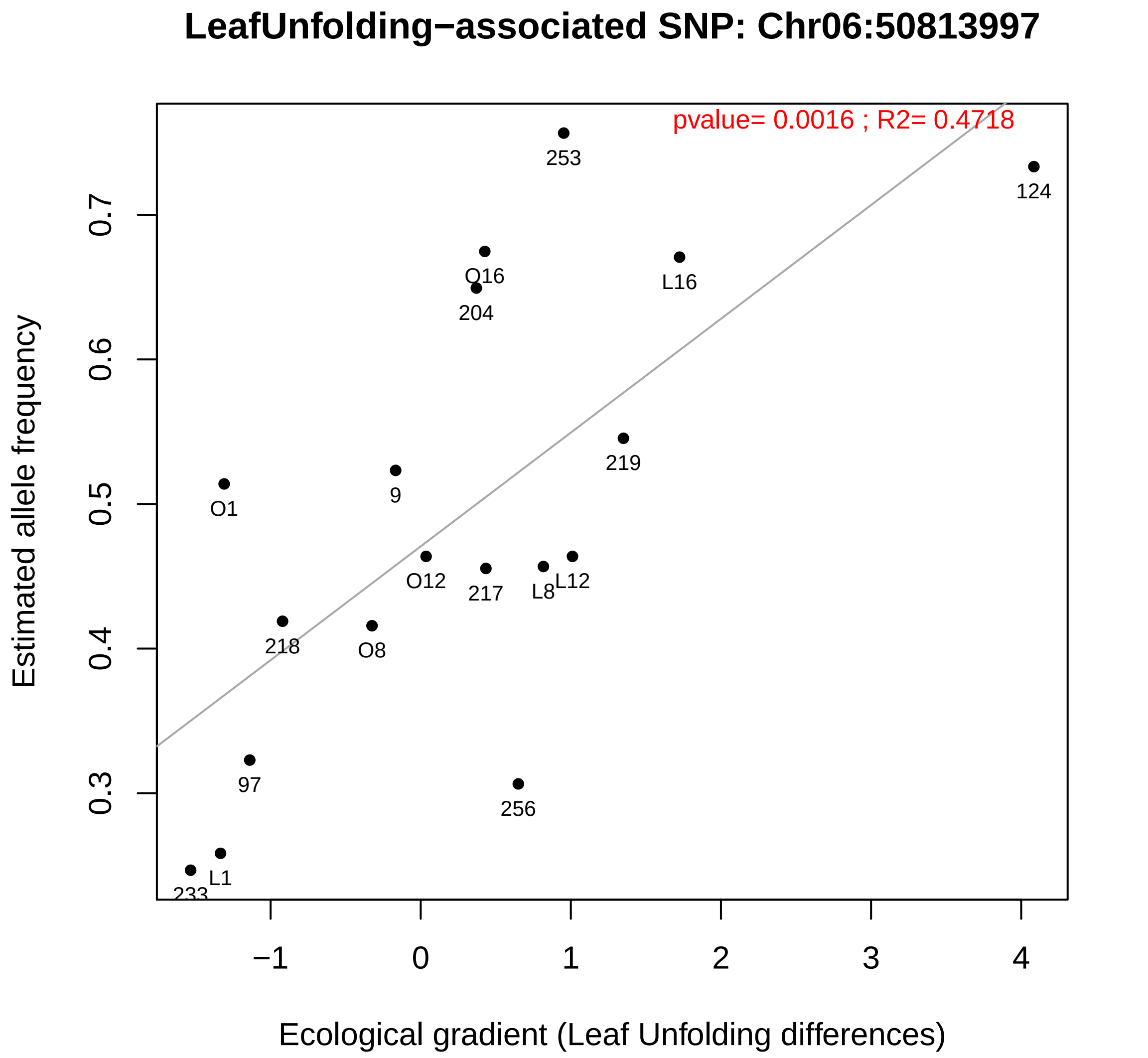
LeafUnfolding−associated SNP: Chr06:50813997
pvalue= 0.0016 ; R2= 0.4718
Estimated allele frequency
Ecological gradient (Leaf Unfolding differences)
0.3
0.4
0.5
0.6
0.7
−1
0
1
2
3
4
253
124
O16
204
L16
219
O1
9
O12
217
L8
L12
218
O8
97
256
L1
233

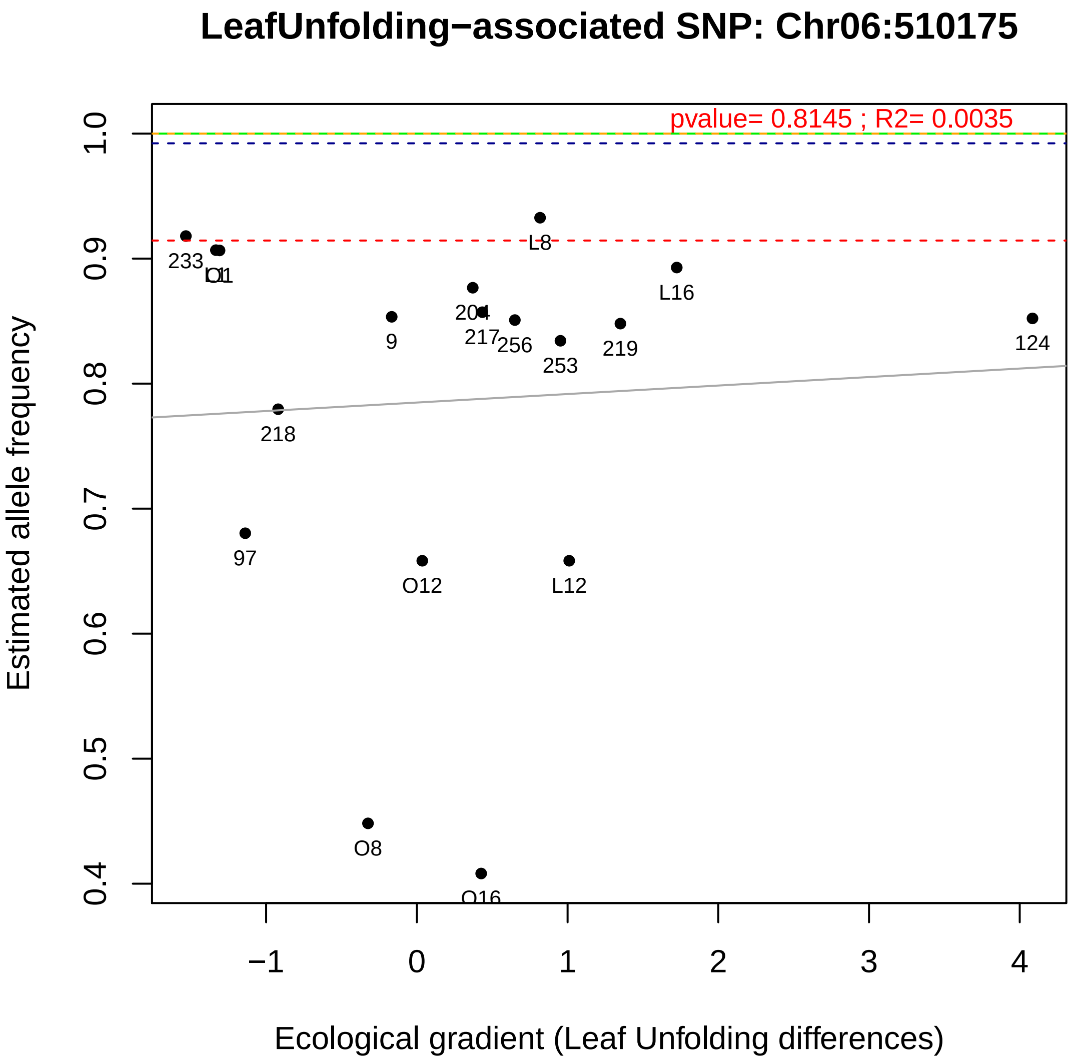

LeafUnfolding−associated SNP: Chr06:510175
pvalue= 0.8145 ; R2= 0.0035
Estimated allele frequency
Ecological gradient (Leaf Unfolding differences)
1.0
0.9
0.8
0.7
0.6
0.5
0.4
−1
0
1
2
3
4
233
L1
O1
L8
L16
204
217
9
256
253
219
124
218
97
O12
L12
O8
O16

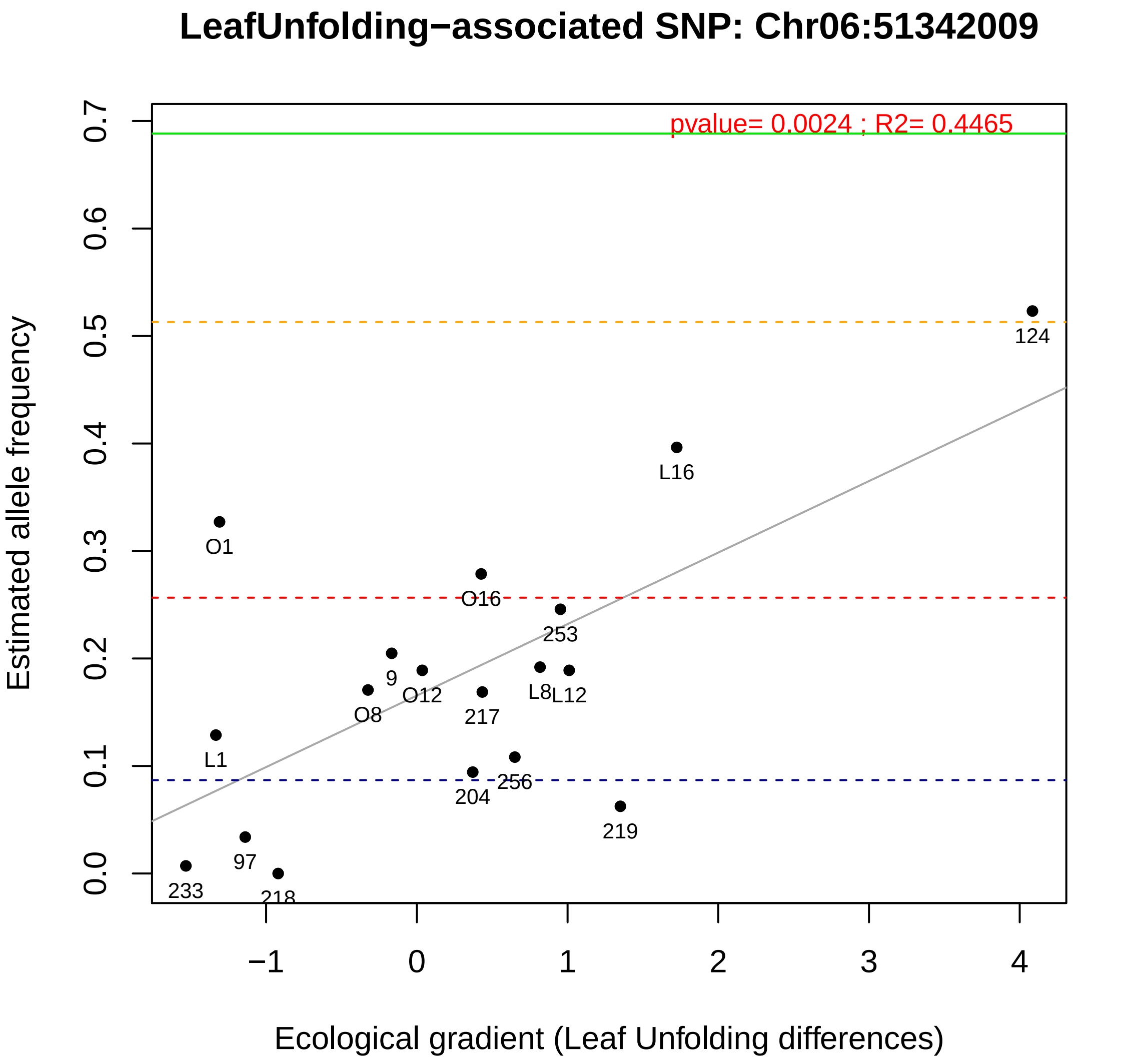

LeafUnfolding−associated SNP: Chr06:51342009
pvalue= 0.0024 ; R2= 0.4465
Estimated allele frequency
Ecological gradient (Leaf Unfolding differences)
0.0
0.1
0.2
0.3
0.4
0.5
0.6
0.7
−1
0
1
2
3
4
124
L16
O1
O16
253
9
O12
O8
217
L8
L12
L1
256
204
219
97
233
218

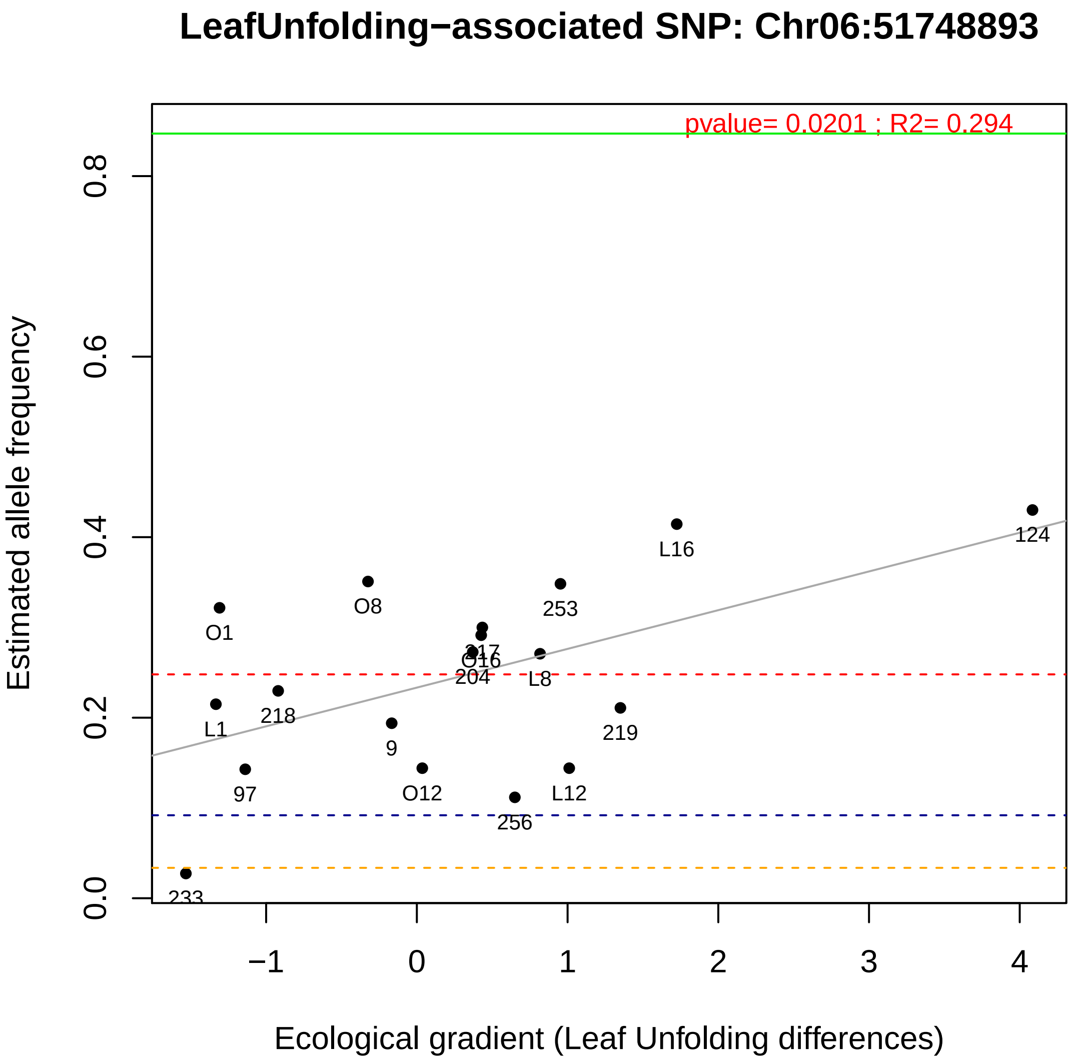

LeafUnfolding−associated SNP: Chr06:51748893
pvalue= 0.0201 ; R2= 0.294
Estimated allele frequency
0.0
0.2
0.4
0.6
0.8
Ecological gradient (Leaf Unfolding differences)
−1
0
1
2
3
4
124
L16
253
O8
O1
217
O16
204
L8
218
L1
219
9
97
O12
L12
256
233

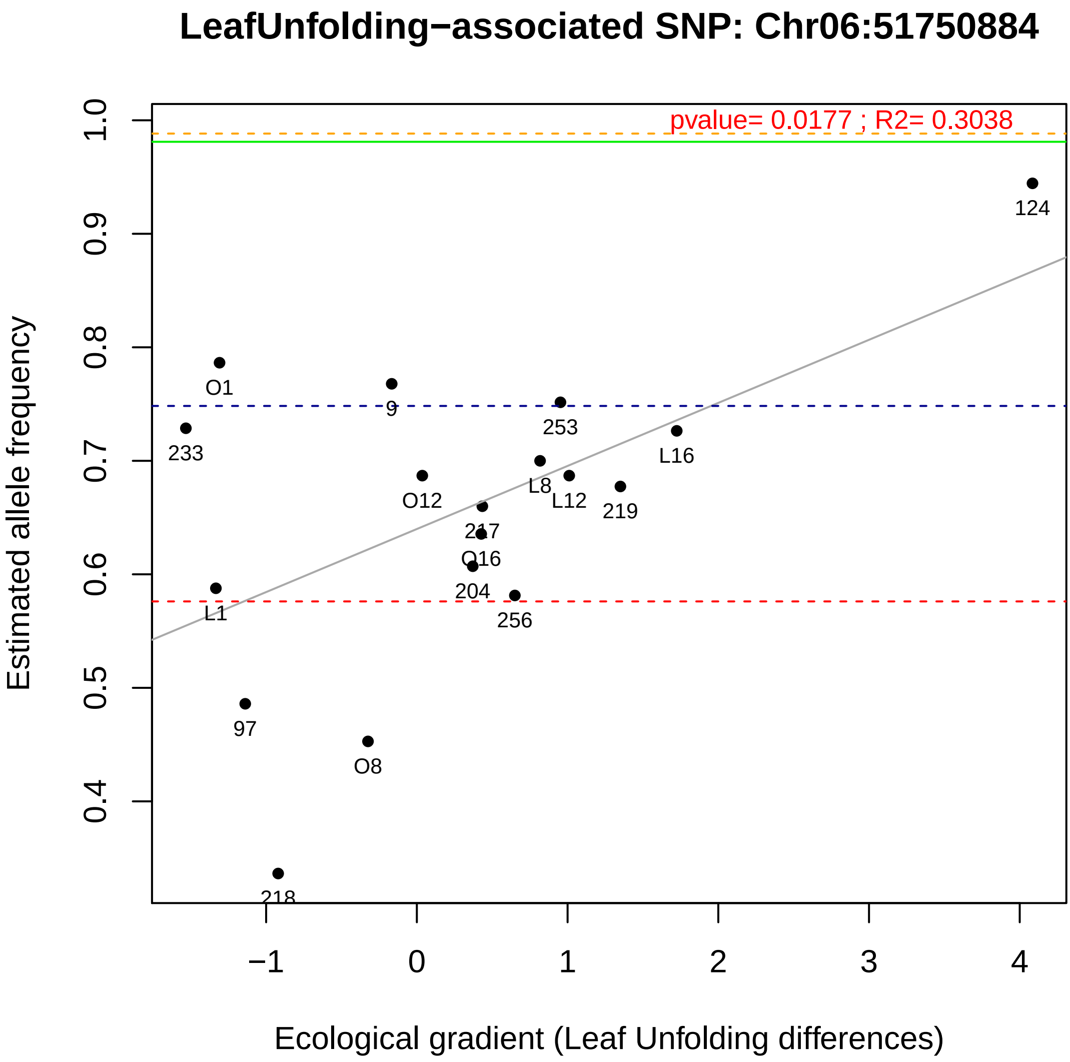

LeafUnfolding−associated SNP: Chr06:51750884
pvalue= 0.0177 ; R2= 0.3038
Estimated allele frequency
Ecological gradient (Leaf Unfolding differences)
1.0
0.9
0.8
0.7
0.6
0.5
0.4
−1
0
1
2
3
4
124
O1
9
253
233
L16
O12
L8
L12
219
217
O16
204
256
L1
97
O8
218

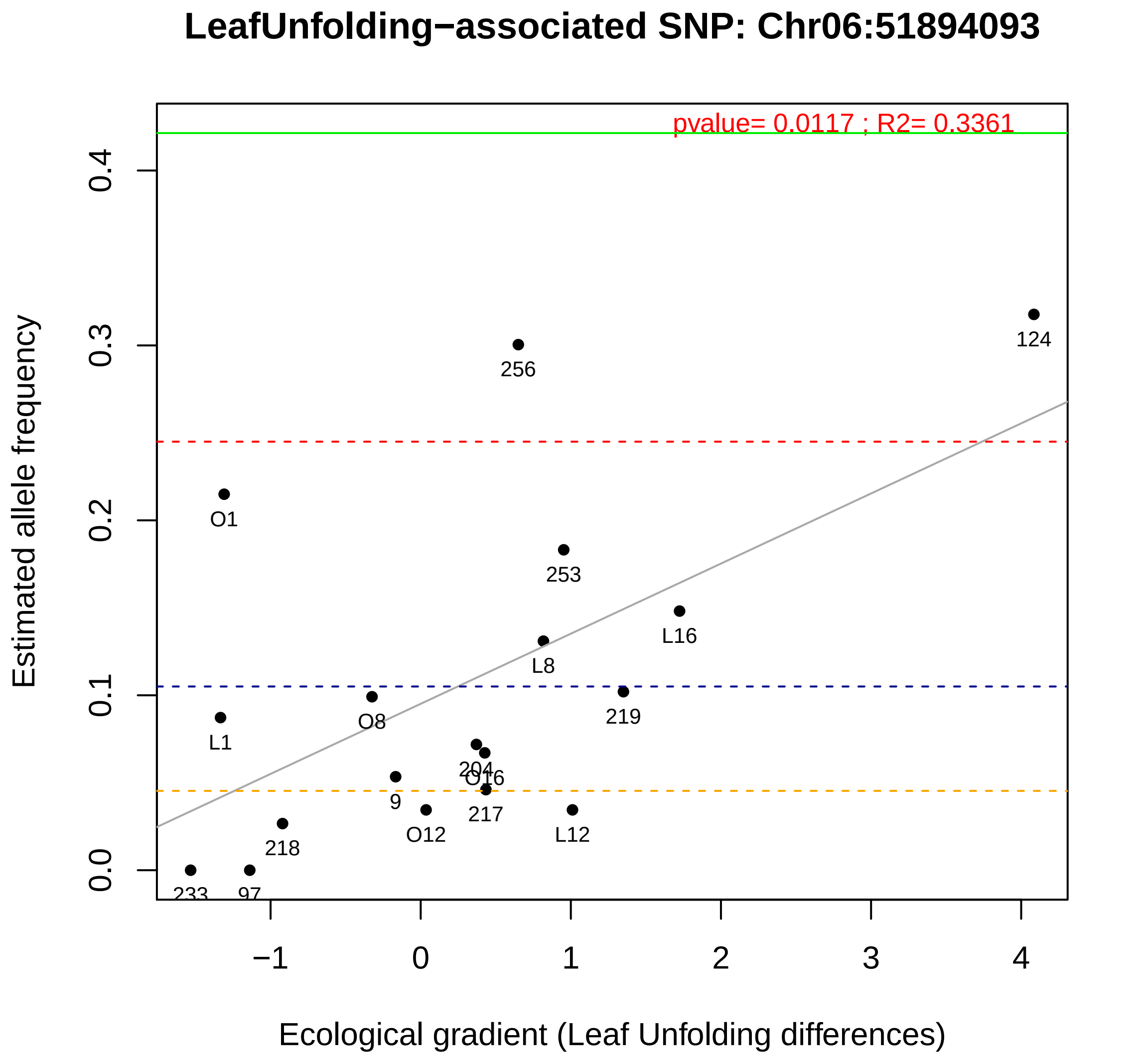
LeafUnfolding−associated SNP: Chr06:51894093
pvalue= 0.0117 ; R2= 0.3361
Estimated allele frequency
0.0
0.1
0.2
0.3
0.4
−1
0
1
2
3
4
Ecological gradient (Leaf Unfolding differences)
124
256
O1
253
L16
L8
219
O8
L1
204
O16
9
217
O12
L12
218
233
97

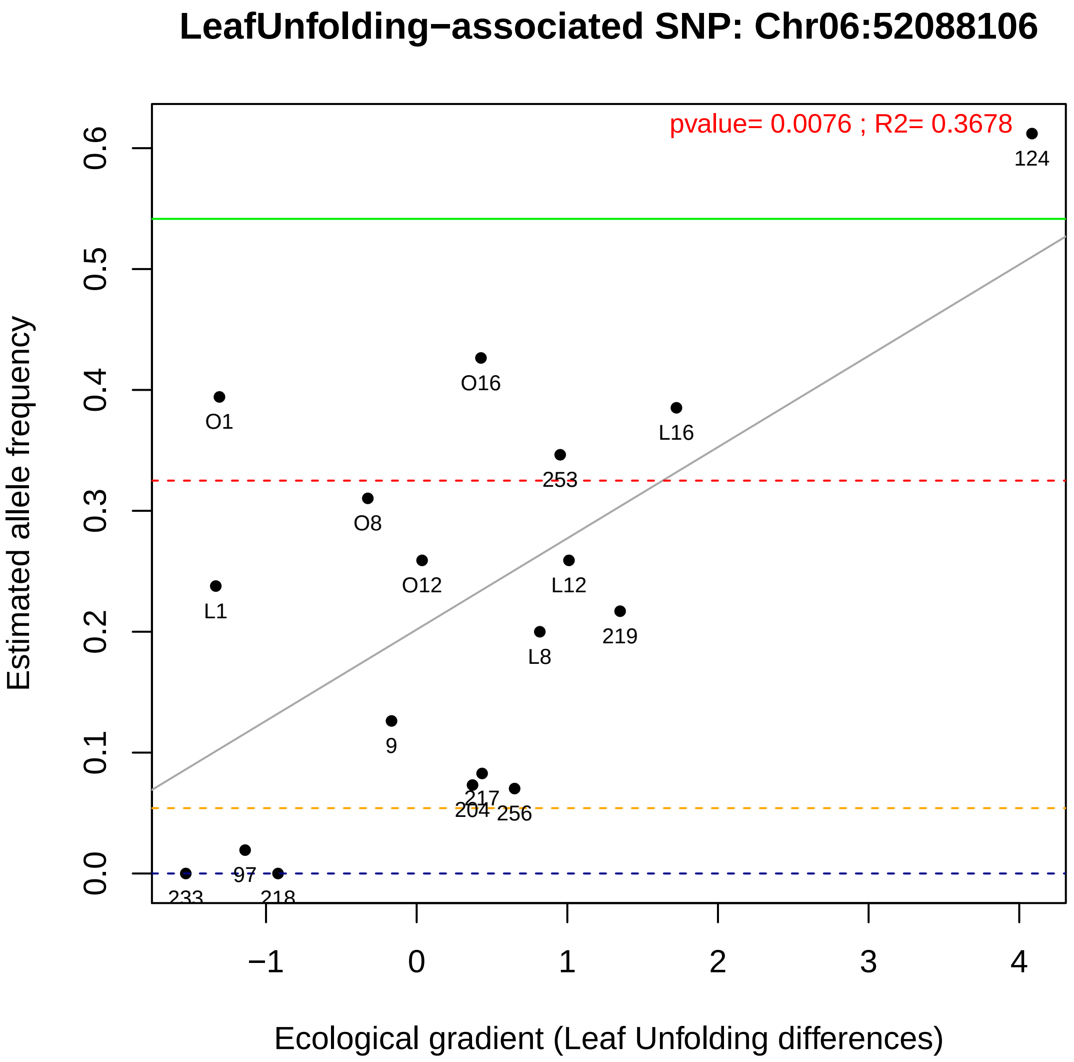
LeafUnfolding−associated SNP: Chr06:52088106
pvalue= 0.0076 ; R2= 0.3678
Estimated allele frequency
0.0
0.1
0.2
0.3
0.4
0.5
0.6
−1
0
1
2
3
4
Ecological gradient (Leaf Unfolding differences)
124
O16
O1
L16
253
O8
O12
L12
L1
219
L8
9
217
204
256
97
233
218

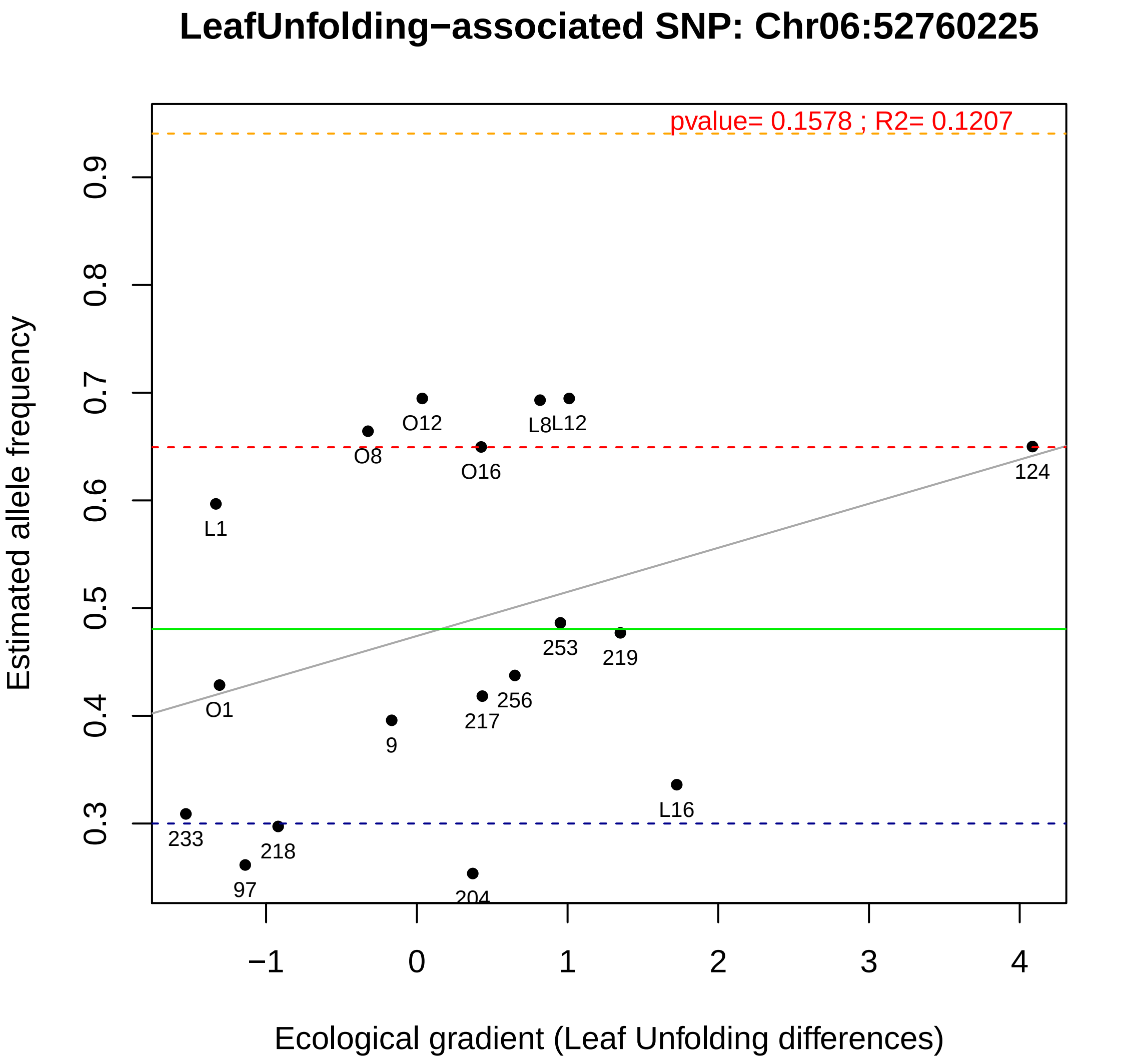
LeafUnfolding−associated SNP: Chr06:52760225
pvalue= 0.1578 ; R2= 0.1207
Estimated allele frequency
0.3
0.4
0.5
0.6
0.7
0.8
0.9
−1
0
1
2
3
4
Ecological gradient (Leaf Unfolding differences)
O12
L8
L12
O8
O16
124
L1
253
219
256
O1
217
9
L16
233
218
97
204

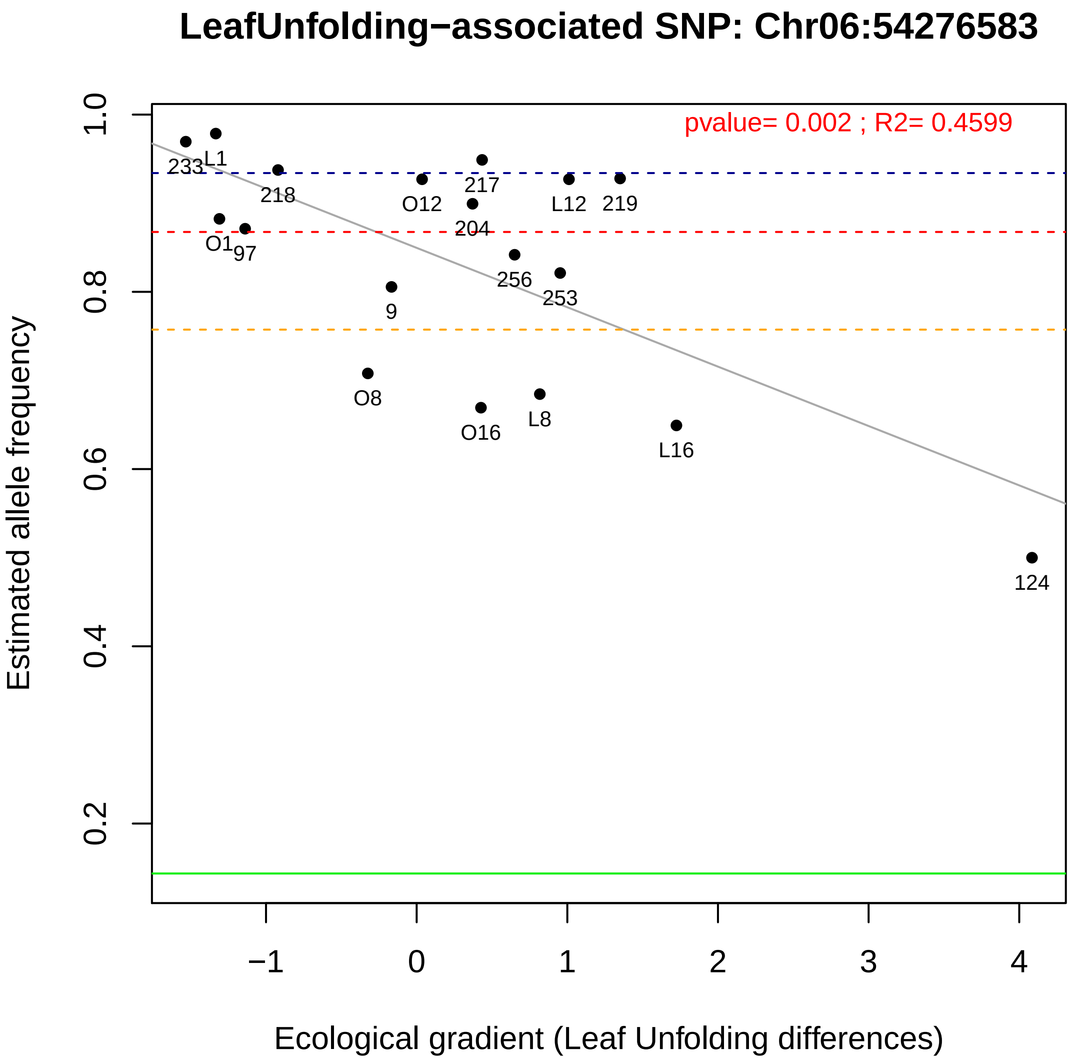
LeafUnfolding−associated SNP: Chr06:54276583
pvalue= 0.002 ; R2= 0.4599
Estimated allele frequency
Ecological gradient (Leaf Unfolding differences)
0.2
0.4
0.6
0.8
1.0
−1
0
1
2
3
4
233
L1
218
O12
217
L12
219
204
O1
97
256
253
9
O8
O16
L8
L16
124

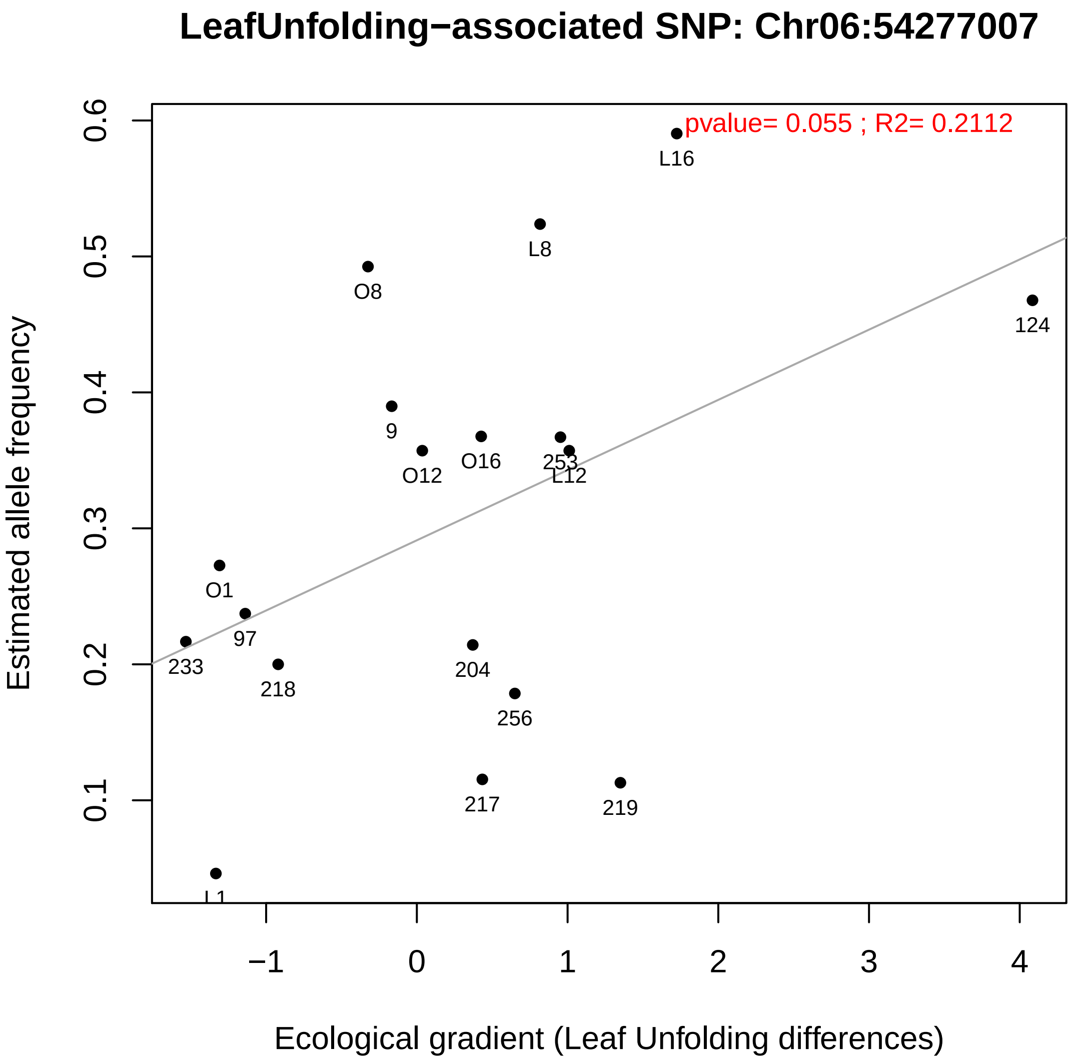
LeafUnfolding−associated SNP: Chr06:54277007
pvalue= 0.055 ; R2= 0.2112
L16
L8
O8
124
9
O12
O16
253
L12
O1
97
233
218
204
256
217
219
L1
Estimated allele frequency
0.1
0.2
0.3
0.4
0.5
0.6
−1
0
1
2
3
4
Ecological gradient (Leaf Unfolding differences)

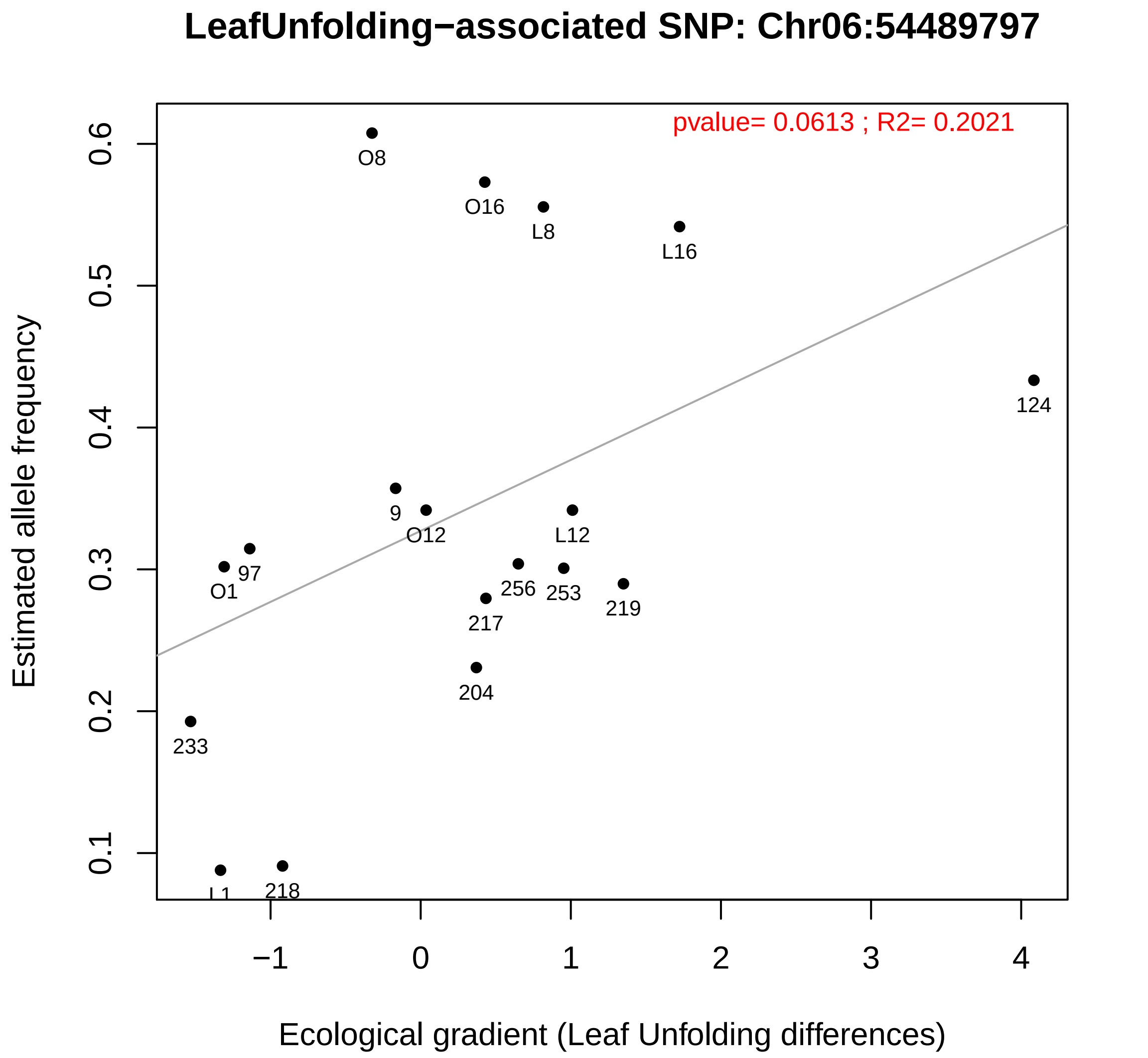
LeafUnfolding−associated SNP: Chr06:54489797
pvalue= 0.0613 ; R2= 0.2021
Estimated allele frequency
Ecological gradient (Leaf Unfolding differences)
0.1
0.2
0.3
0.4
0.5
0.6
−1
0
1
2
3
4
O8
O16
L8
L16
124
9
O12
L12
97
O1
256
253
219
217
204
233
218

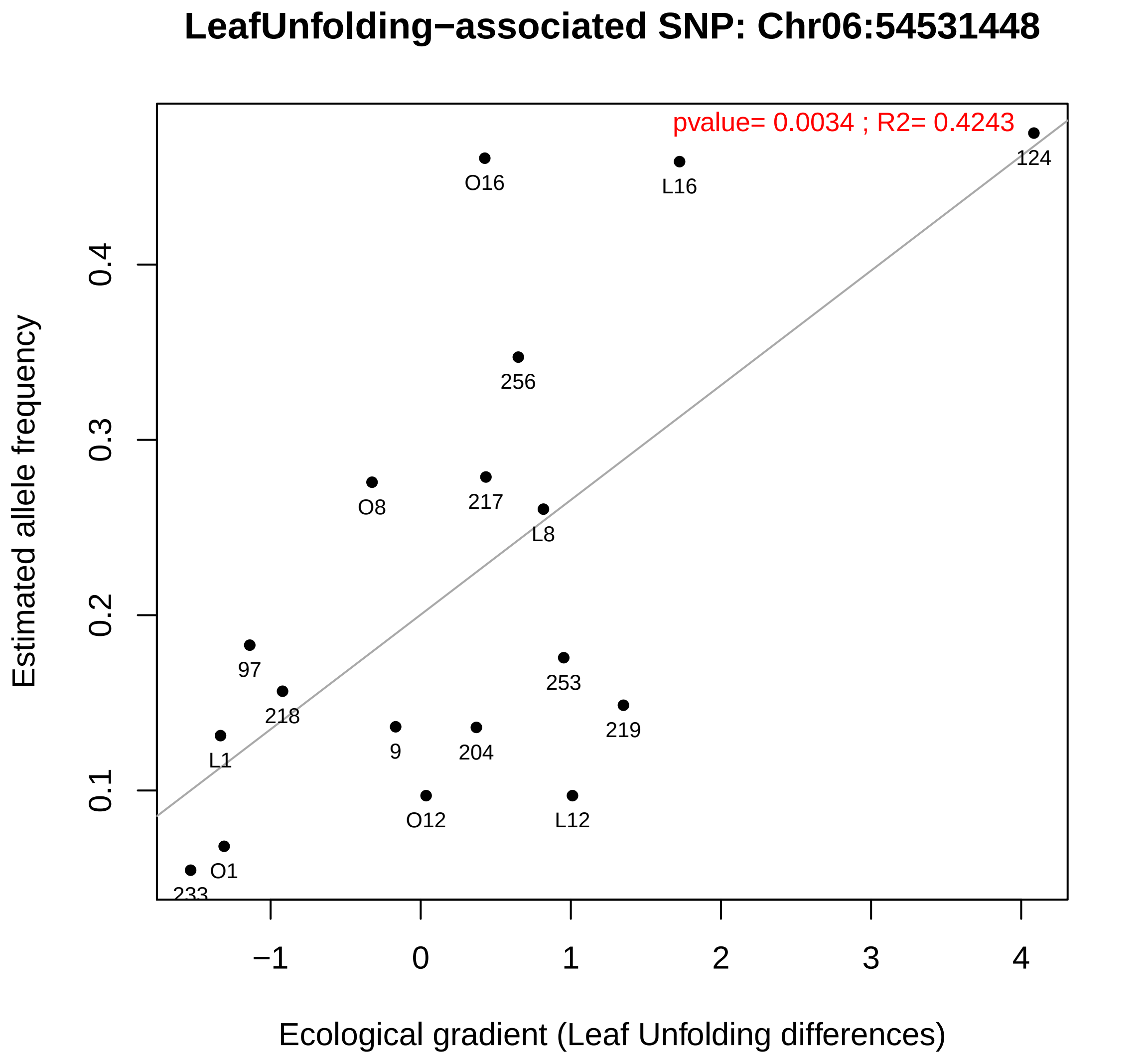
LeafUnfolding−associated SNP: Chr06:54531448
pvalue= 0.0034 ; R2= 0.4243
Estimated allele frequency
Ecological gradient (Leaf Unfolding differences)
0.1
0.2
0.3
0.4
−1
0
1
2
3
4
124
O16
L16
256
O8
217
L8
97
218
253
219
L1
9
204
O12
L12
O1
233

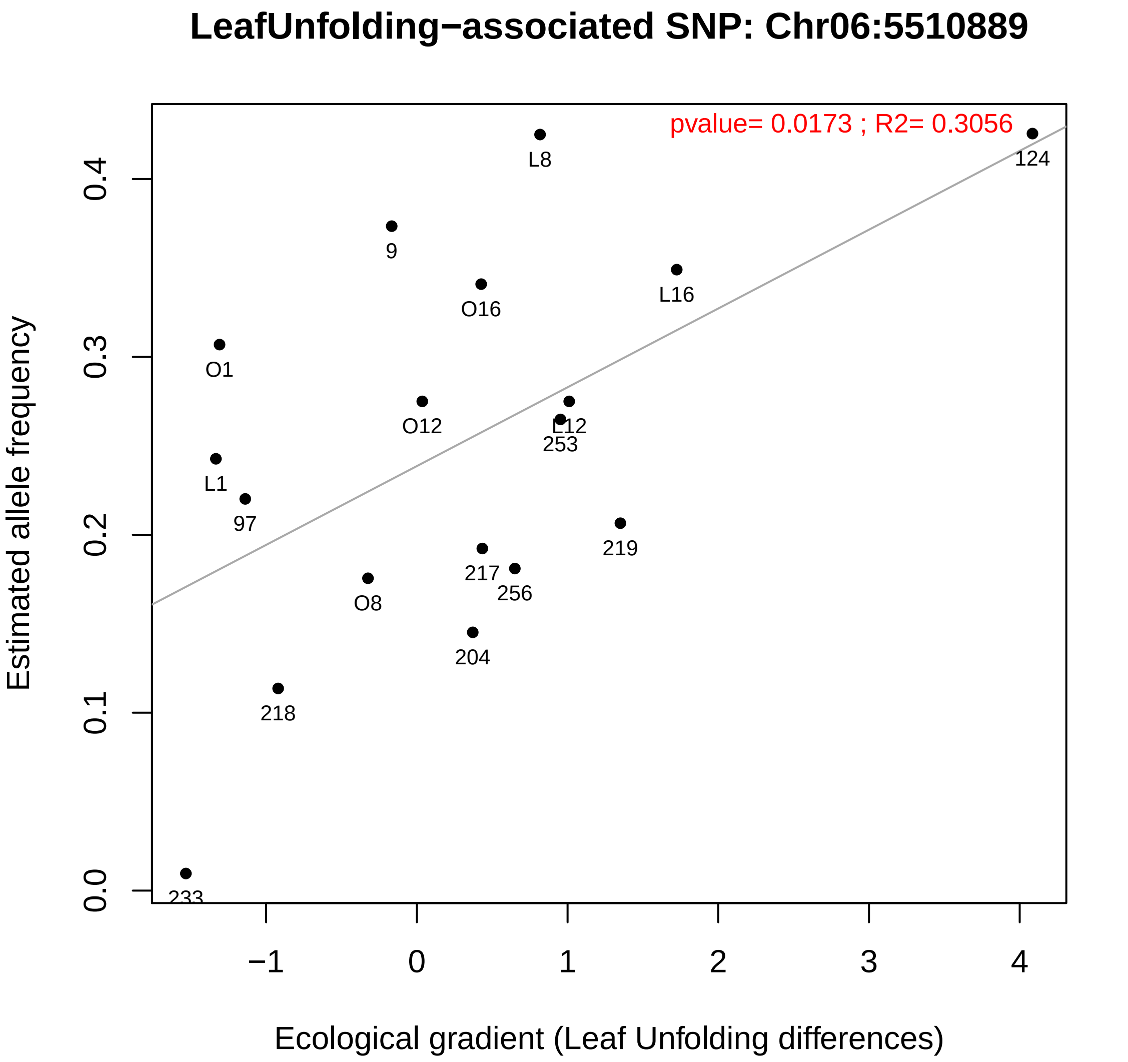

LeafUnfolding−associated SNP: Chr06:5510889
pvalue= 0.0173 ; R2= 0.3056
L8
124
9
O16
L16
O1
O12
L12
253
L1
97
219
217
256
O8
204
218
233
0.0
0.1
0.2
0.3
0.4
−1
0
1
2
3
4
Estimated allele frequency
Ecological gradient (Leaf Unfolding differences)

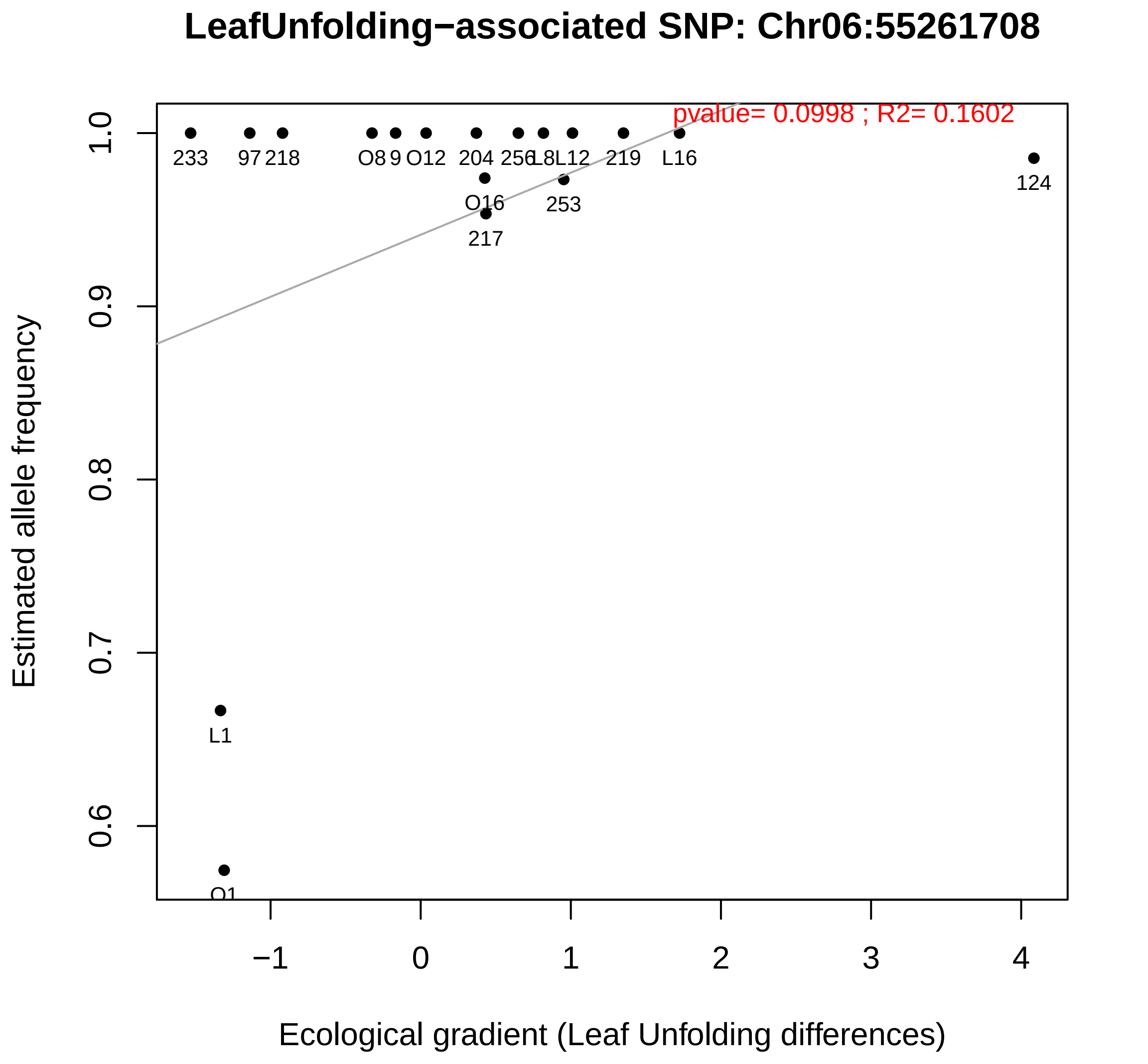
LeafUnfolding−associated SNP: Chr06:55261708
pvalue= 0.0998 ; R2= 0.1602
Estimated allele frequency
Ecological gradient (Leaf Unfolding differences)
1.0
0.9
0.8
0.7
0.6
−1
0
1
2
3
4
233
97
218
O8
9
O12
204
256
L8
L12
219
L16
124
O16
253
217
L1
O1

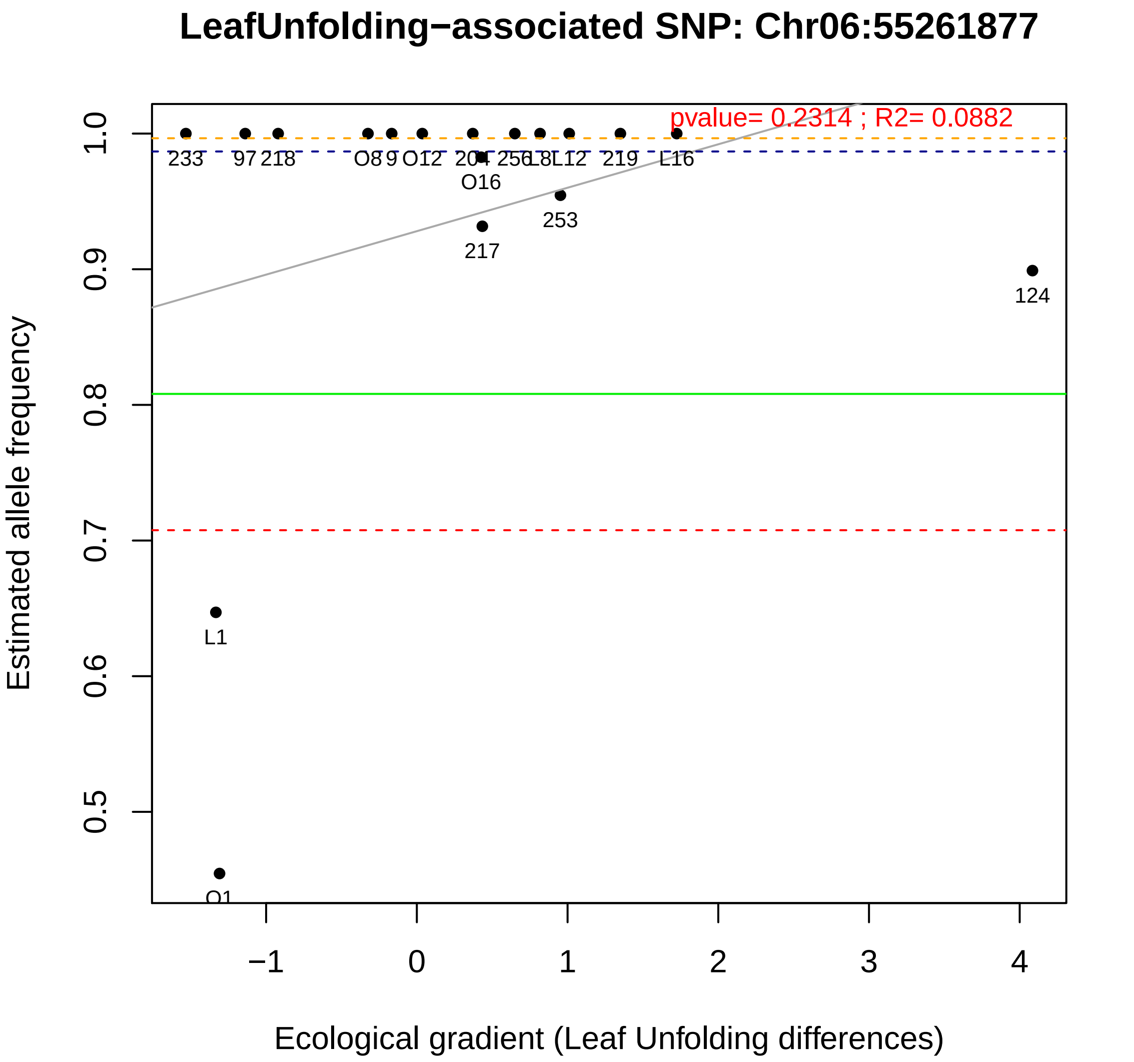

LeafUnfolding−associated SNP: Chr06:55261877
pvalue= 0.2314 ; R2= 0.0882
Estimated allele frequency
Ecological gradient (Leaf Unfolding differences)
0.5
0.6
0.7
0.8
0.9
1.0
−1
0
1
2
3
4
233
97
218
O8
9
O12
204
256
L8
L12
219
L16
O16
253
217
124
L1
O1

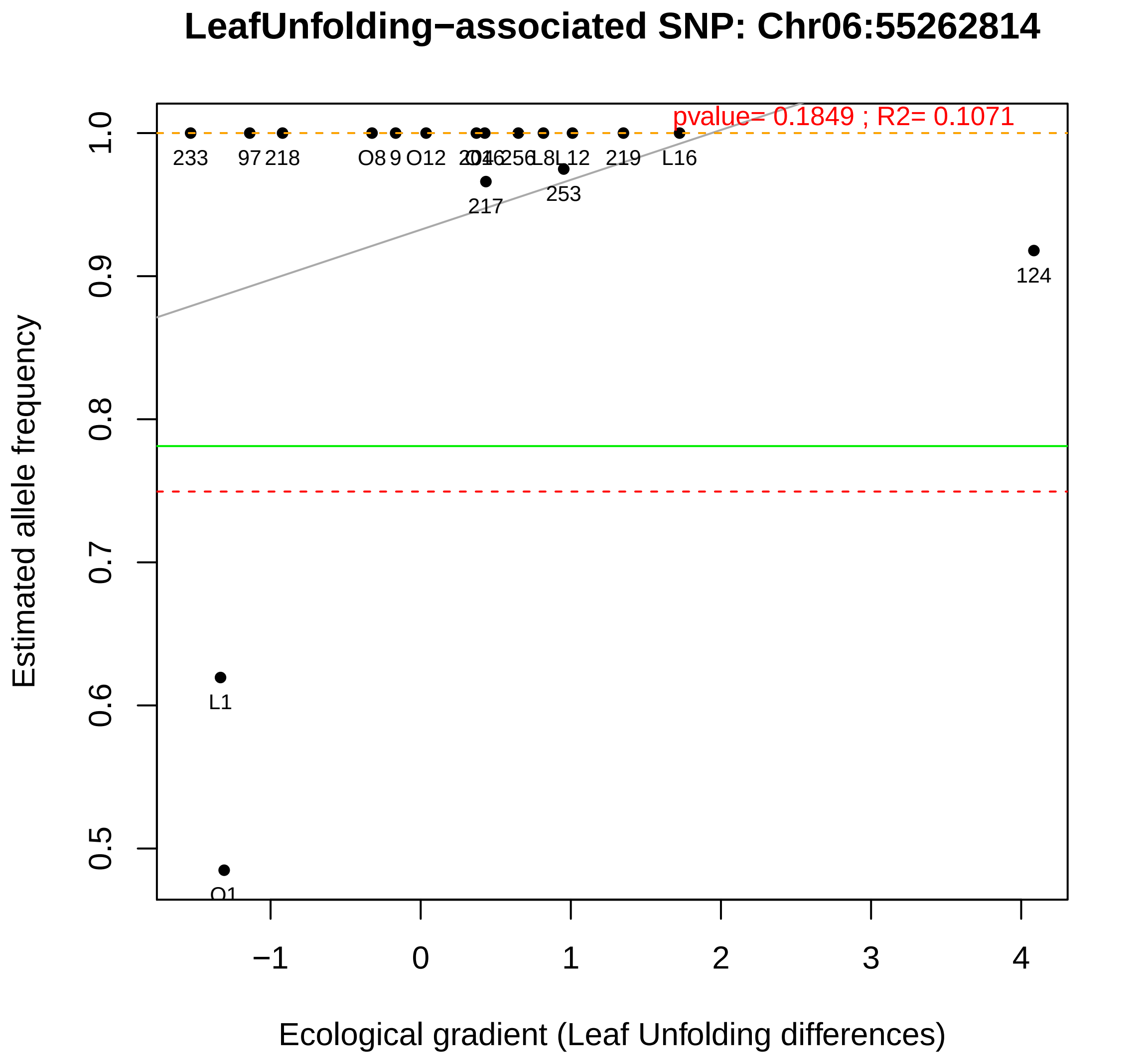

LeafUnfolding−associated SNP: Chr06:55262814
pvalue= 0.1849 ; R2= 0.1071
Estimated allele frequency
Ecological gradient (Leaf Unfolding differences)
0.5
0.6
0.7
0.8
0.9
1.0
−1
0
1
2
3
4
233
97
218
O8
9
O12
L8
L12
219
L16
217
253
124
L1

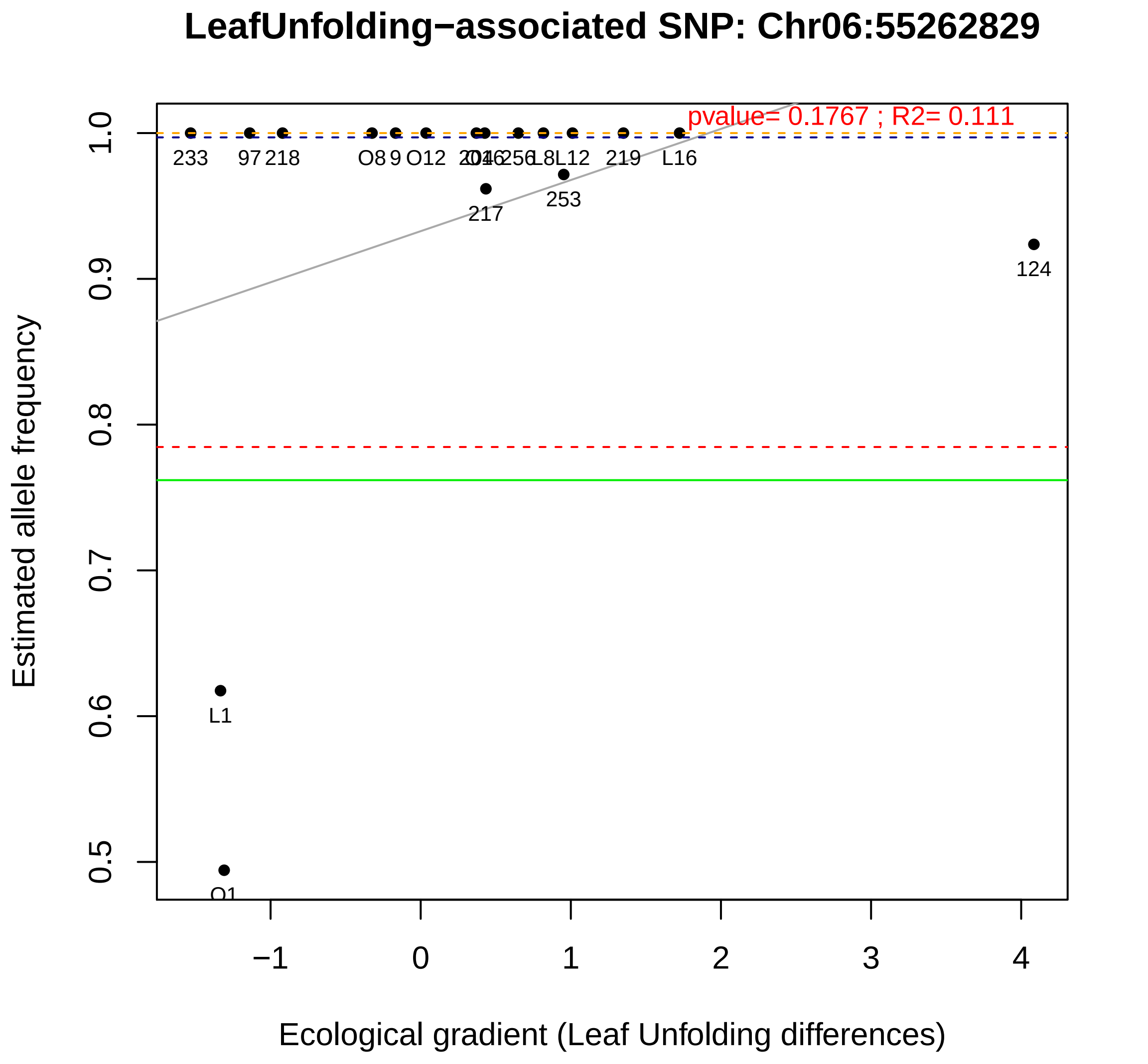

LeafUnfolding−associated SNP: Chr06:55262829
pvalue= 0.1767 ; R2= 0.111
Estimated allele frequency
Ecological gradient (Leaf Unfolding differences)
1.0
0.9
0.8
0.7
0.6
0.5
−1
0
1
2
3
4
233
97
218
O8
9
O12
219
L16
217
253
124
L1
O1

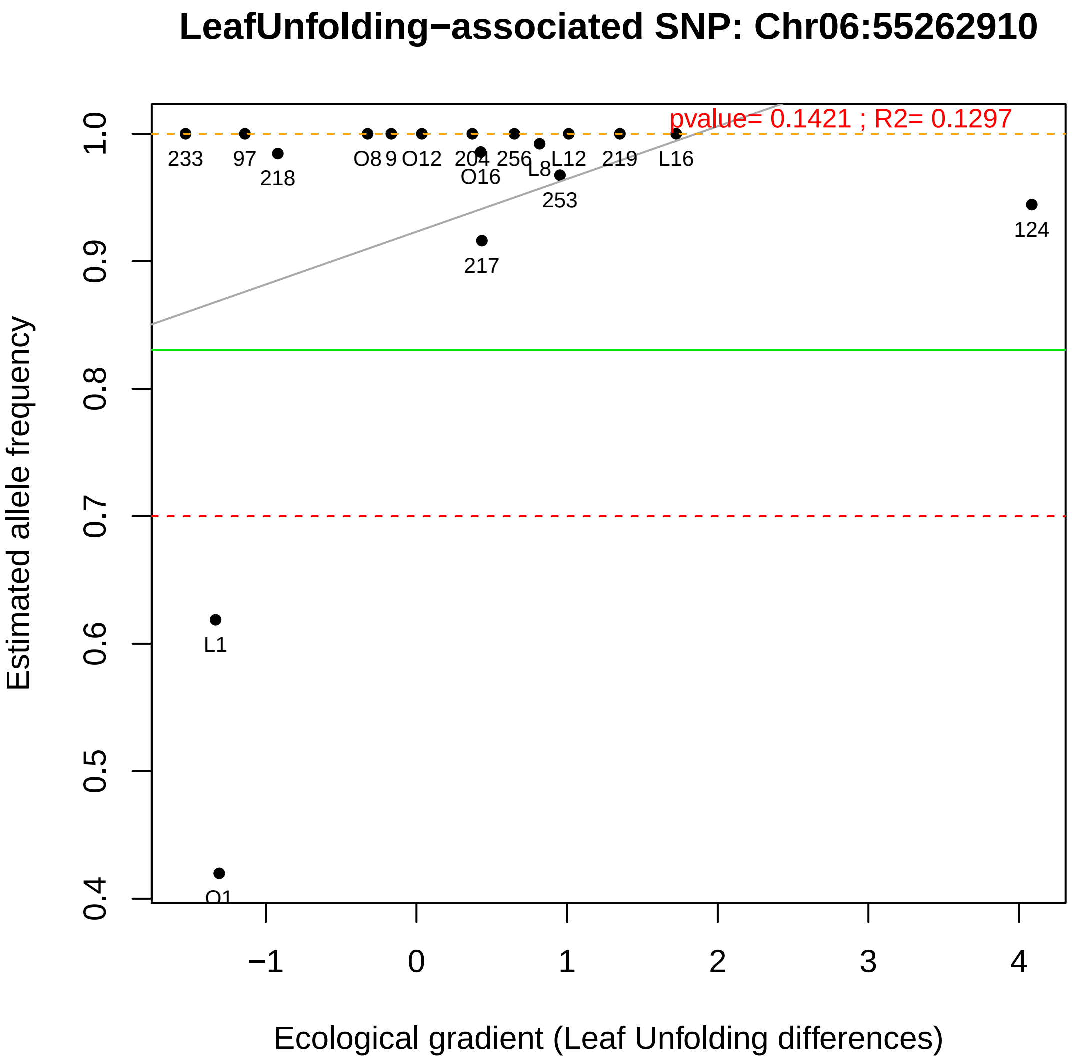

LeafUnfolding−associated SNP: Chr06:55262910
pvalue= 0.1421 ; R2= 0.1297
Estimated allele frequency
Ecological gradient (Leaf Unfolding differences)
1.0
0.9
0.8
0.7
0.6
0.5
0.4
−1
0
1
2
3
4
233
97
218
O8
9
O12
204
O16
256
L8
L12
253
219
L16
217
124
L1
O1

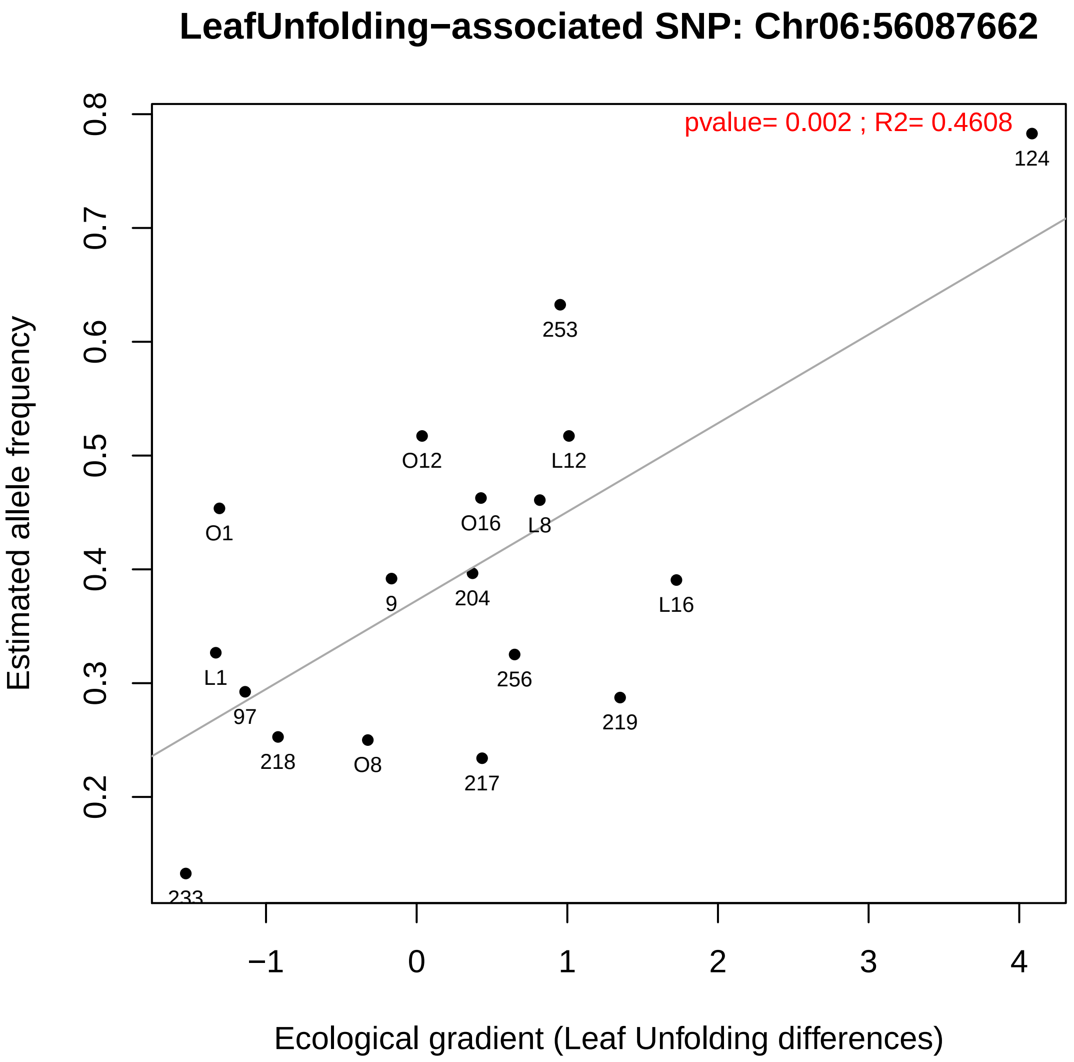

LeafUnfolding−associated SNP: Chr06:56087662
pvalue= 0.002 ; R2= 0.4608
124
253
O12
L12
O1
O16
L8
9
204
L16
L1
256
97
219
218
O8
217
233
0.2
0.3
0.4
0.5
0.6
0.7
0.8
−1
0
1
2
3
4
Estimated allele frequency
Ecological gradient (Leaf Unfolding differences)

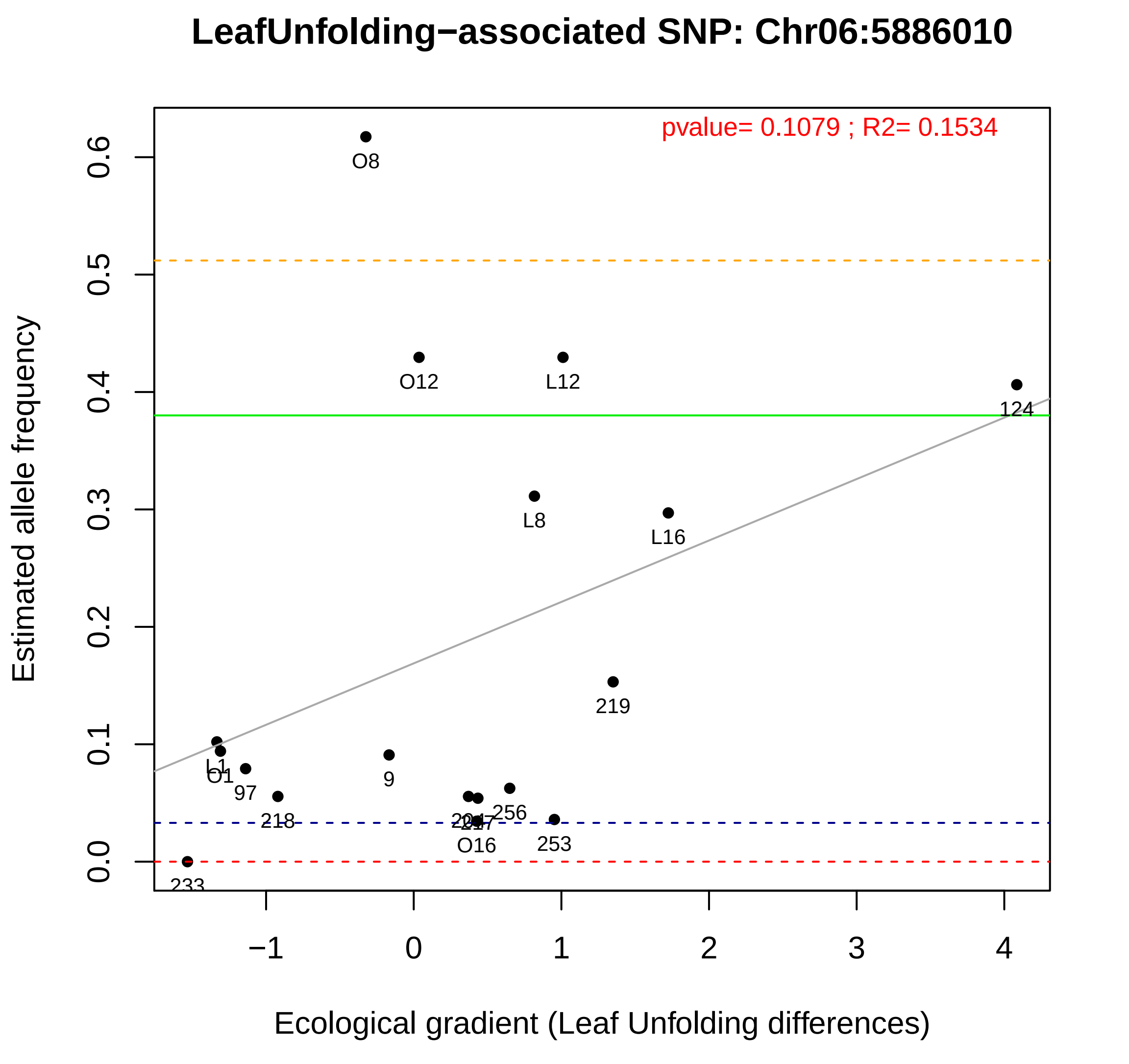
LeafUnfolding−associated SNP: Chr06:5886010
pvalue= 0.1079 ; R2= 0.1534
Estimated allele frequency
Ecological gradient (Leaf Unfolding differences)
0.0
0.1
0.2
0.3
0.4
0.5
0.6
−1
0
1
2
3
4
O8
O12
L12
124
L8
L16
219
L1
O1
97
218
9
256
253
O16
233

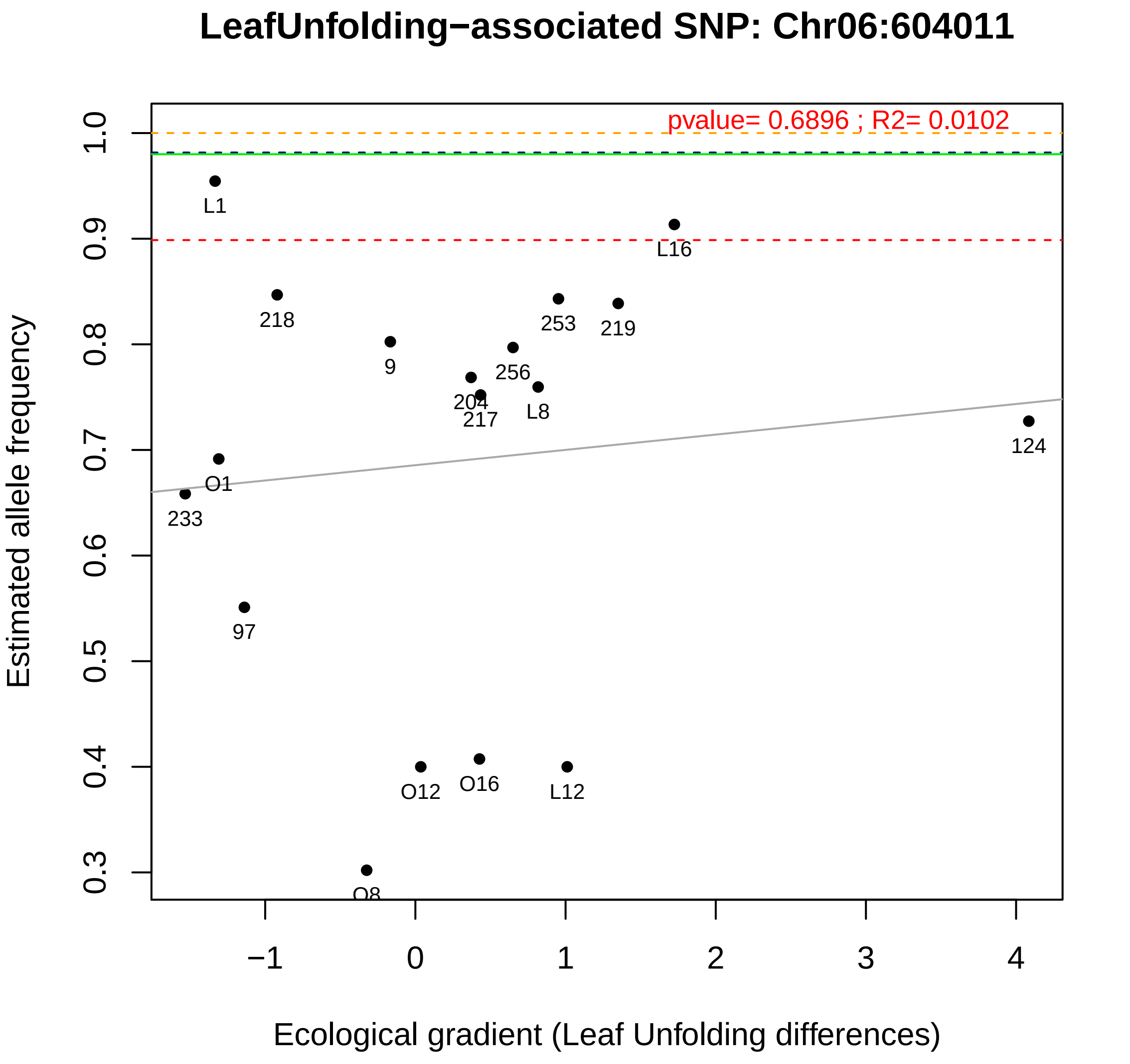
LeafUnfolding−associated SNP: Chr06:604011
pvalue= 0.6896 ; R2= 0.0102
Estimated allele frequency
Ecological gradient (Leaf Unfolding differences)
1.0
0.9
0.8
0.7
0.6
0.5
0.4
0.3
−1
0
1
2
3
4
L1
L16
218
253
219
9
256
204
217
L8
124
O1
233
97
O12
O16
L12
O8

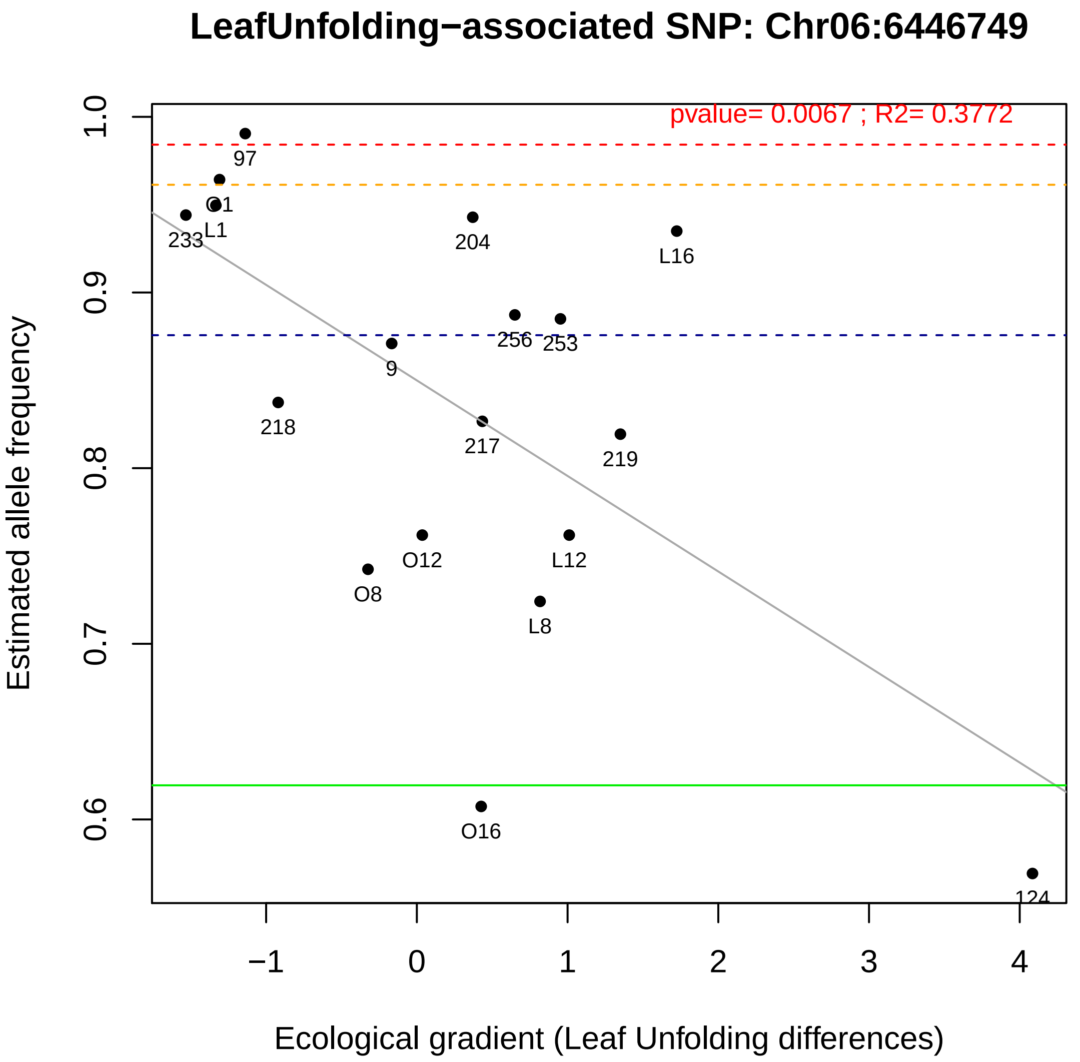

LeafUnfolding−associated SNP: Chr06:6446749
pvalue= 0.0067 ; R2= 0.3772
Estimated allele frequency
Ecological gradient (Leaf Unfolding differences)
1.0
0.9
0.8
0.7
0.6
−1
0
1
2
3
4
97
O1
L1
233
204
L16
256
253
9
218
217
219
O12
L12
O8
L8
O16
124

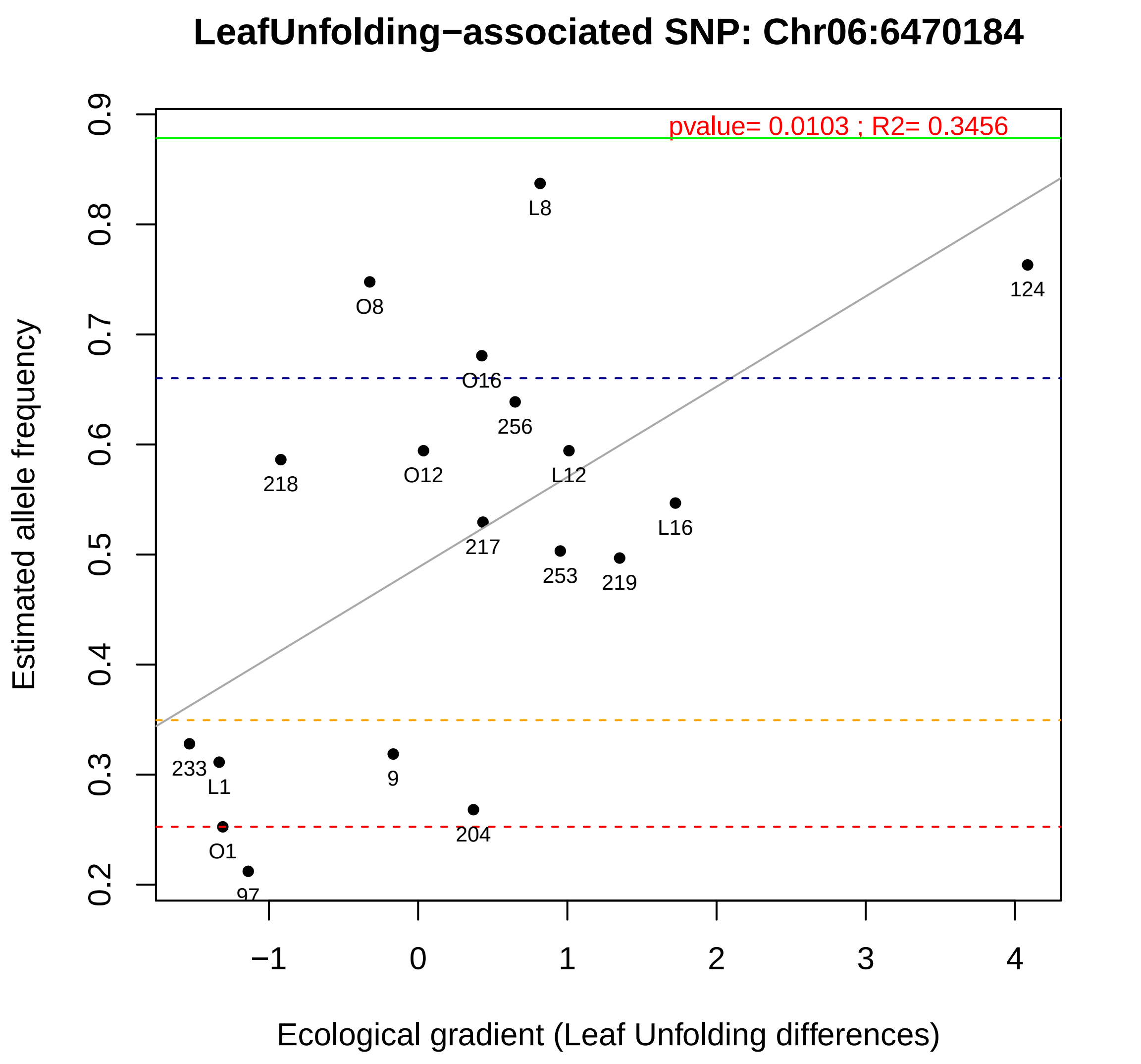
LeafUnfolding−associated SNP: Chr06:6470184
pvalue= 0.0103 ; R2= 0.3456
Estimated allele frequency
Ecological gradient (Leaf Unfolding differences)
0.2
0.3
0.4
0.5
0.6
0.7
0.8
0.9
−1
0
1
2
3
4
L8
124
O8
O16
256
218
O12
L12
L16
217
253
219
233
L1
9
204
O1
97

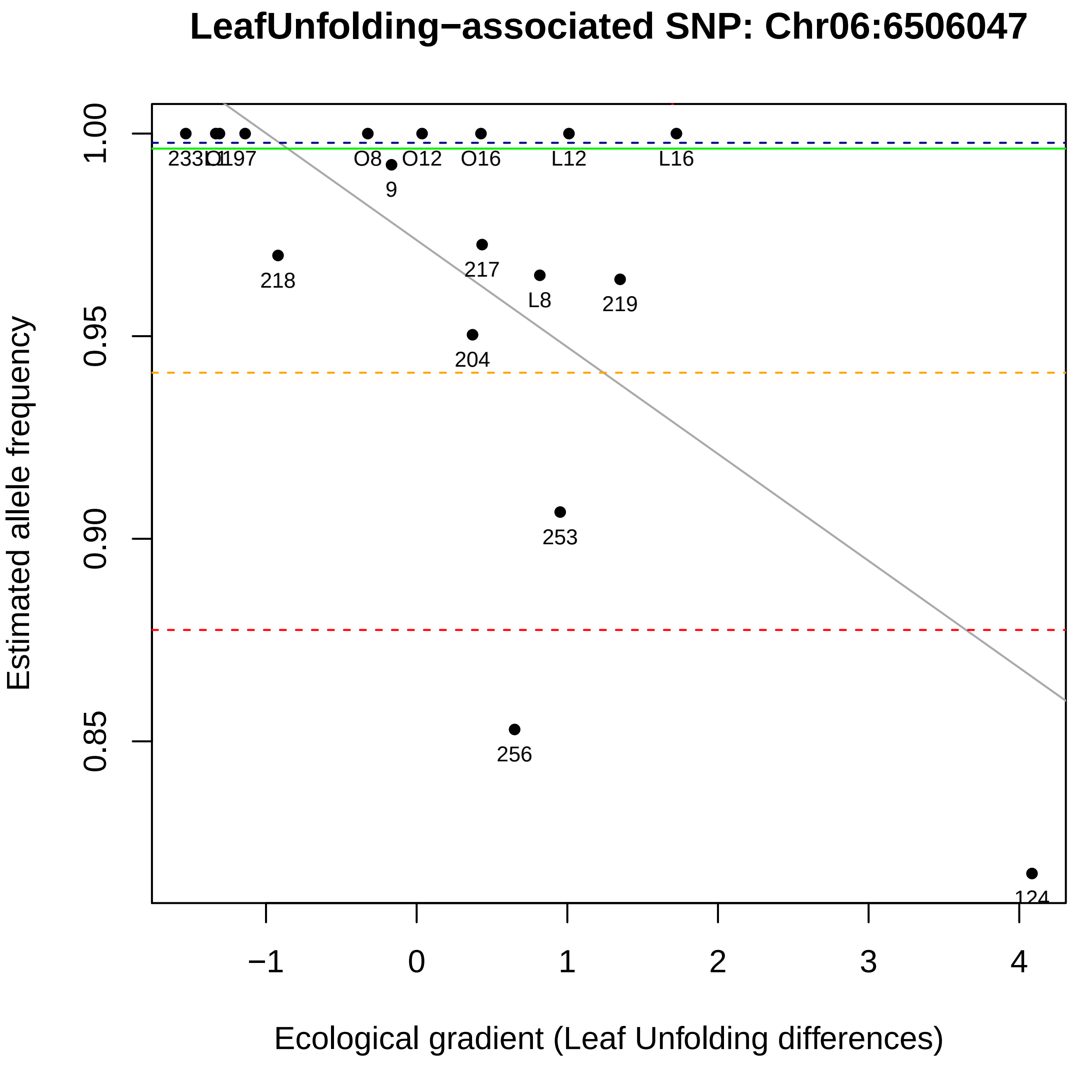

LeafUnfolding−associated SNP: Chr06:6506047
Estimated allele frequency
Ecological gradient (Leaf Unfolding differences)
1.00
0.95
0.90
0.85
−1
0
1
2
3
4
233
O1
197
O8
9
O12
O16
L12
L16
218
217
L8
219
204
253
256
124

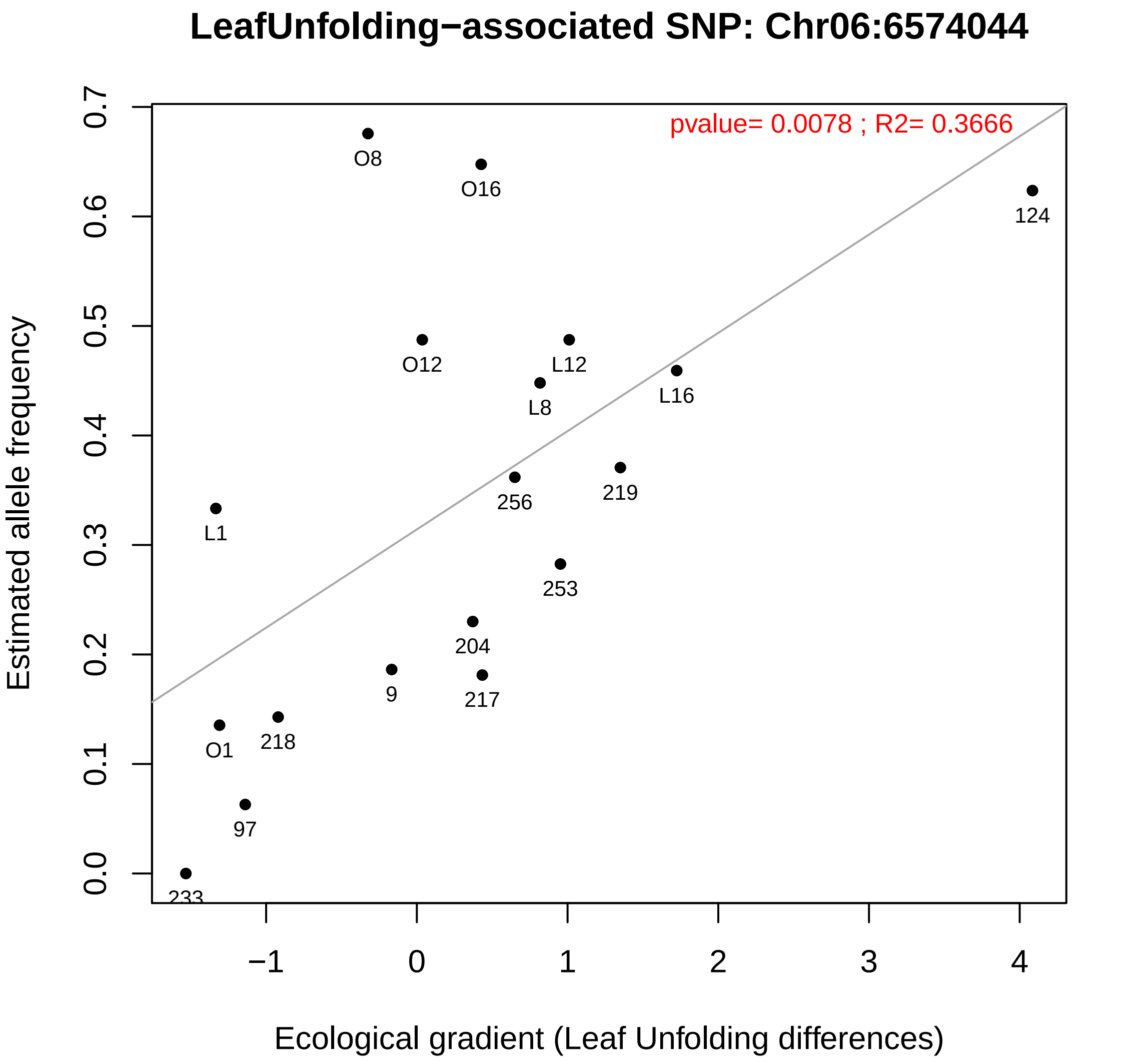

LeafUnfolding−associated SNP: Chr06:6574044
pvalue= 0.0078 ; R2= 0.3666
Estimated allele frequency
Ecological gradient (Leaf Unfolding differences)
0.0
0.1
0.2
0.3
0.4
0.5
0.6
0.7
−1
0
1
2
3
4
O8
O16
124
O12
L12
L8
L16
256
219
L1
253
204
9
217
O1
218
97
233

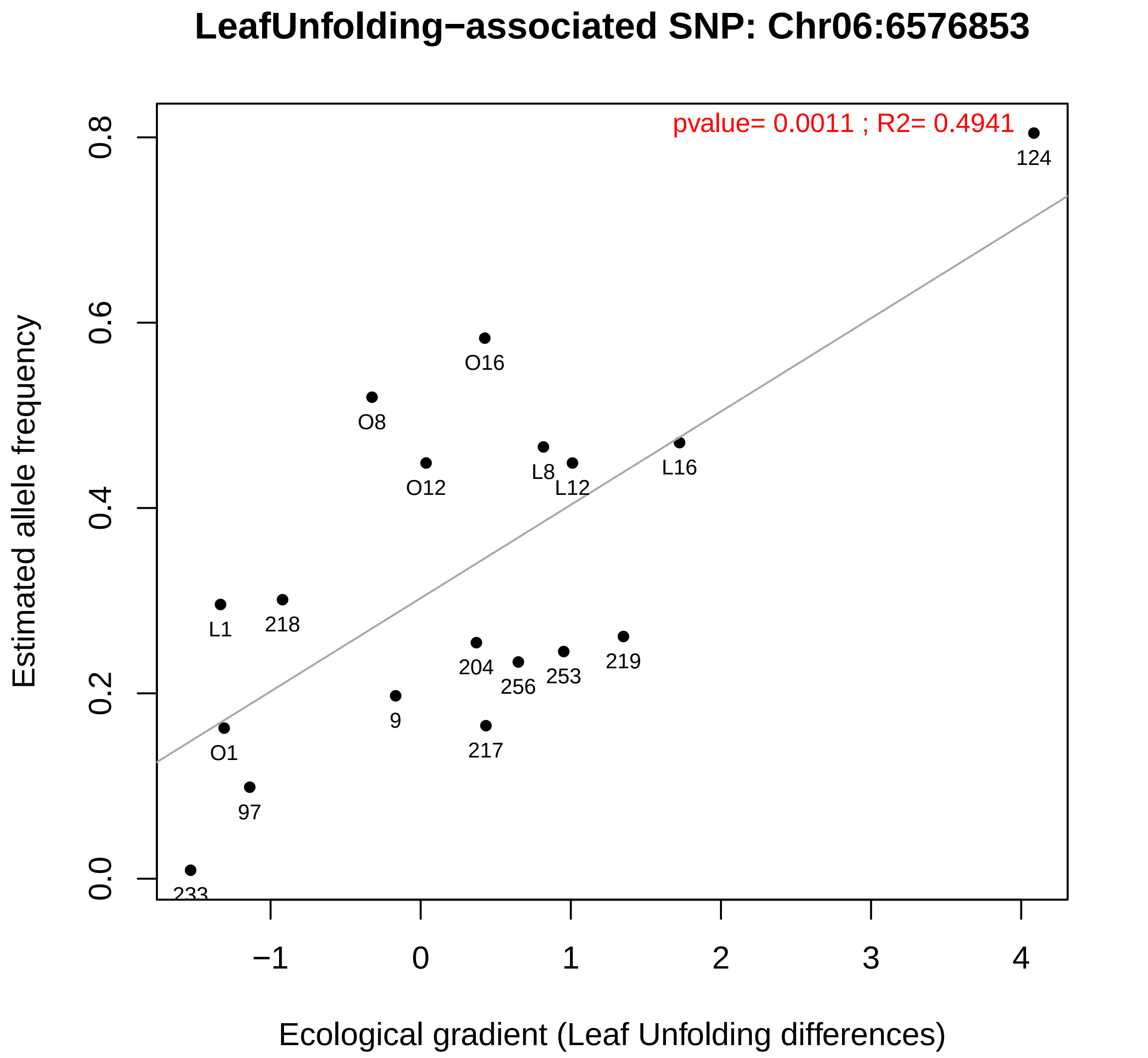
LeafUnfolding−associated SNP: Chr06:6576853
pvalue= 0.0011 ; R2= 0.4941
Estimated allele frequency
0.0
0.2
0.4
0.6
0.8
Ecological gradient (Leaf Unfolding differences)
−1
0
1
2
3
4
124
O16
O8
O12
L8
L12
L16
L1
218
204
256
253
219
9
217
O1
97
233

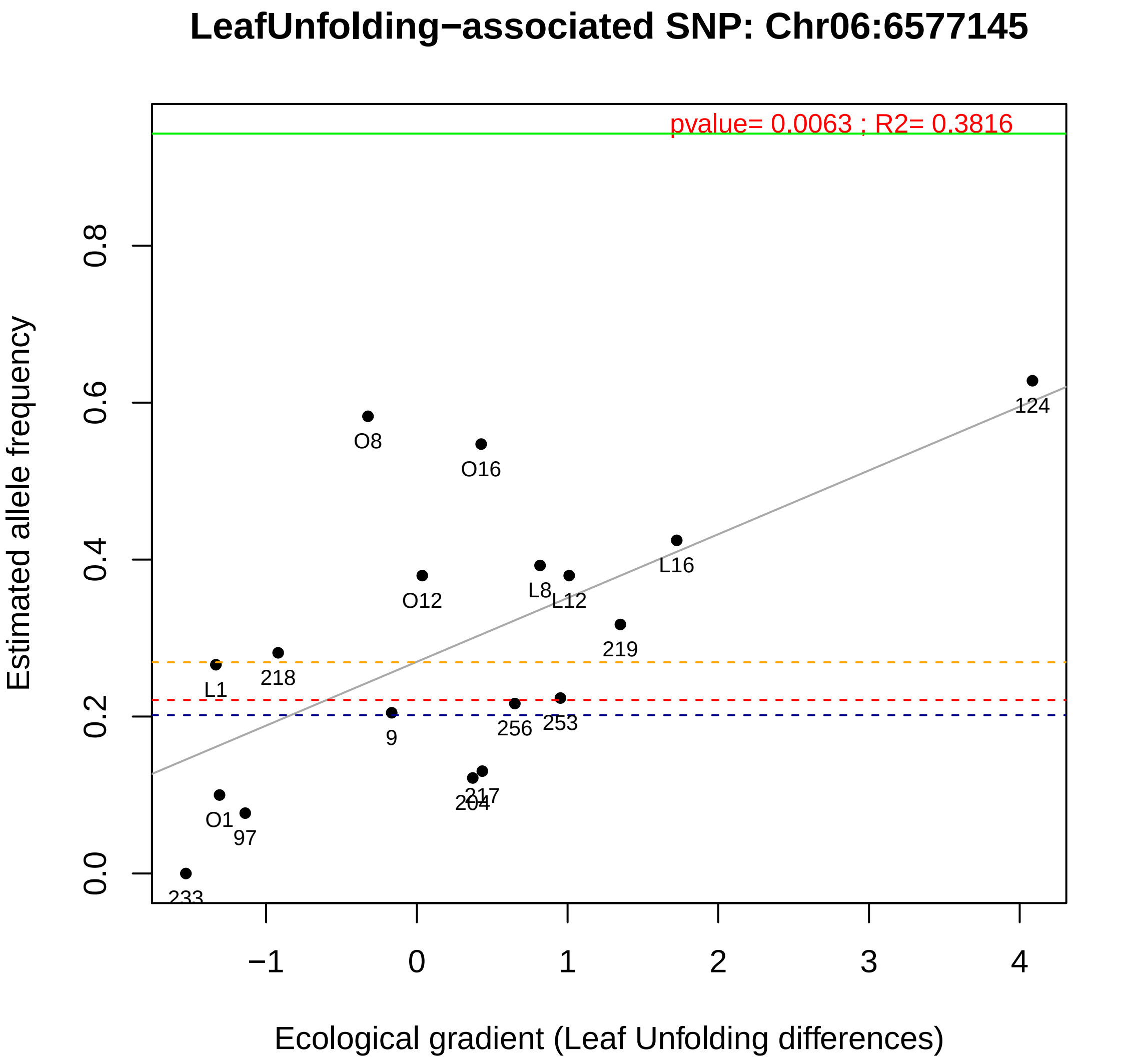
LeafUnfolding−associated SNP: Chr06:6577145
pvalue= 0.0063 ; R2= 0.3816
Estimated allele frequency
0.0
0.2
0.4
0.6
0.8
−1
0
1
2
3
4
Ecological gradient (Leaf Unfolding differences)
124
O8
O16
L16
O12
L8
L12
219
218
L1
9
256
253
217
204
O1
97
233

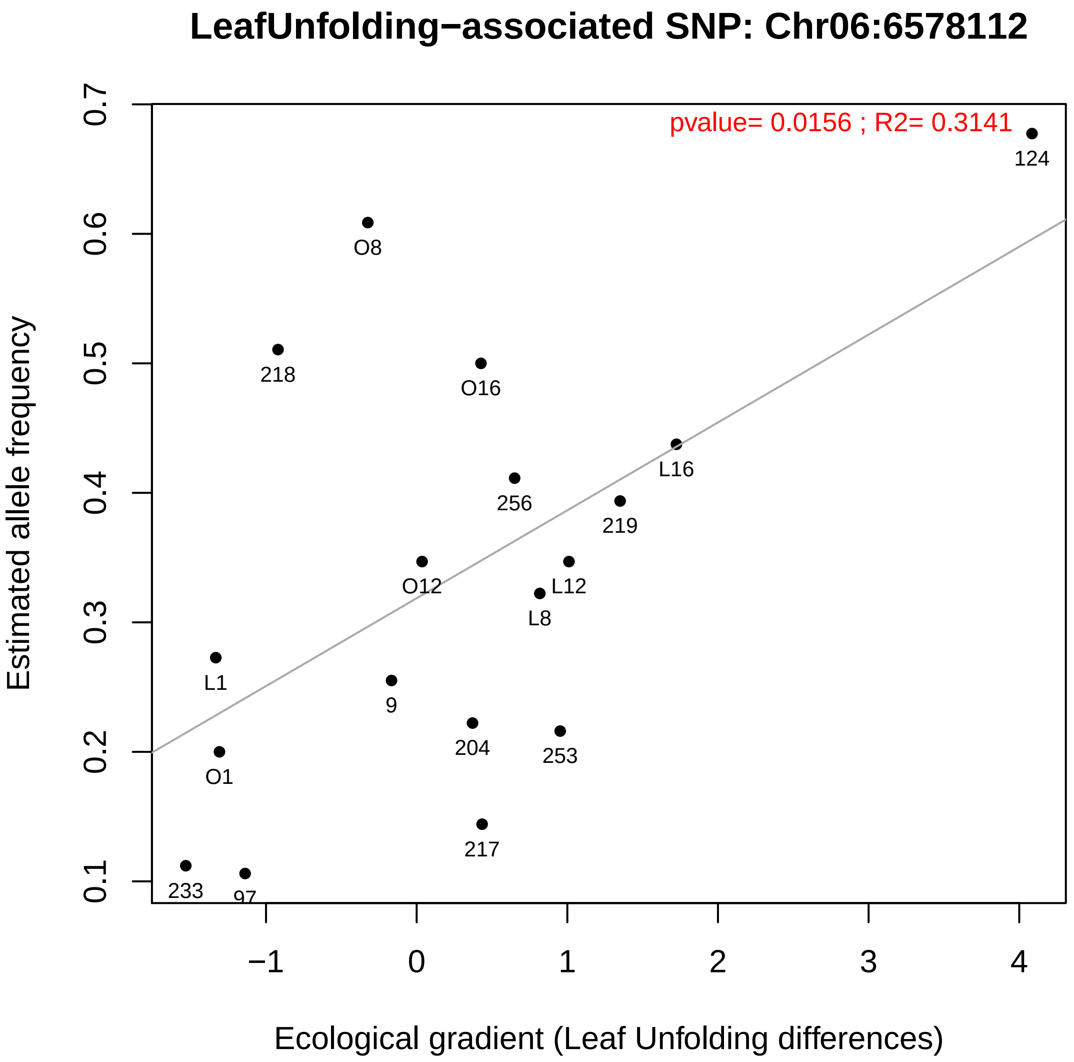
LeafUnfolding−associated SNP: Chr06:6578112
pvalue= 0.0156 ; R2= 0.3141
Estimated allele frequency
Ecological gradient (Leaf Unfolding differences)
0.1
0.2
0.3
0.4
0.5
0.6
0.7
−1
0
1
2
3
4
124
O8
218
O16
L16
256
219
O12
L12
L8
L1
9
204
253
O1
217
233
97

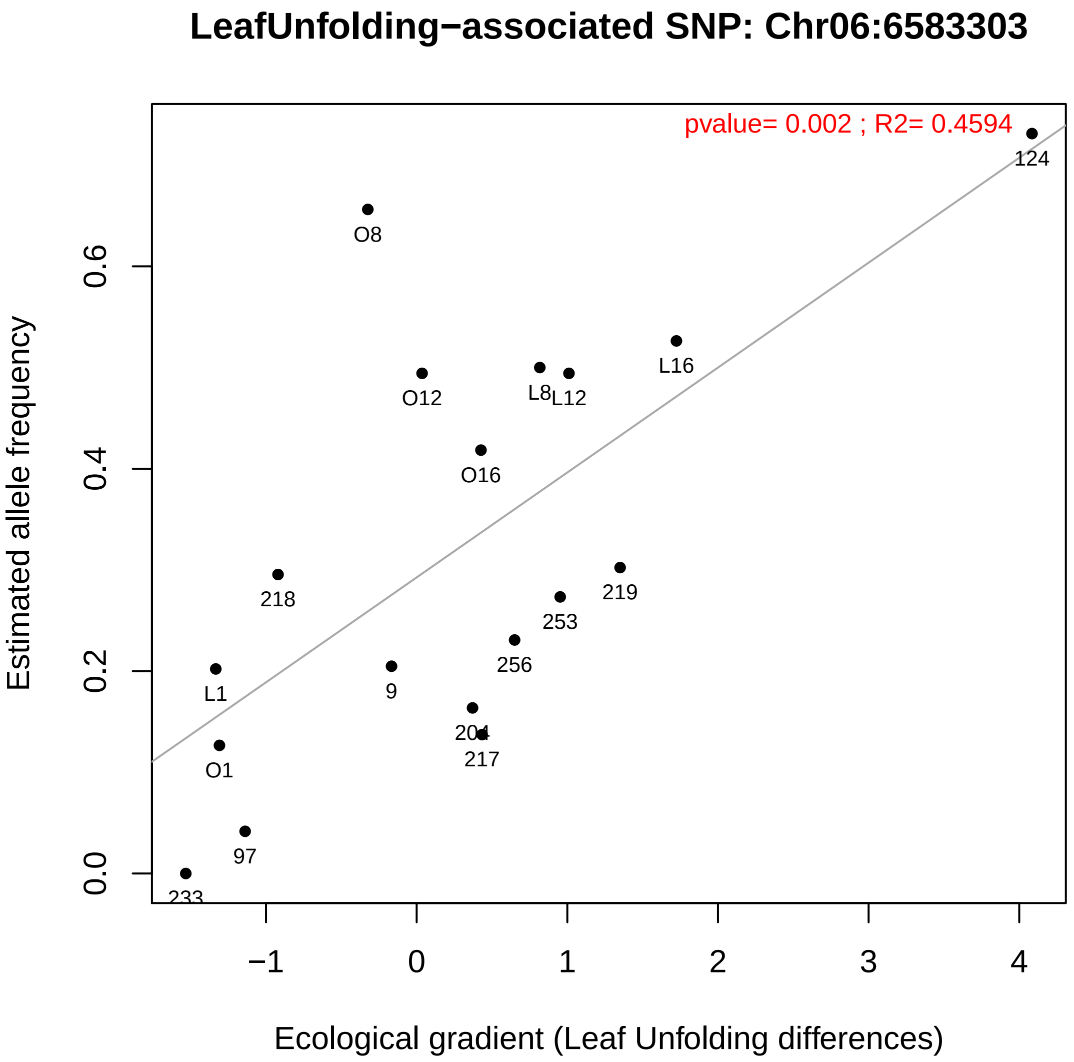
LeafUnfolding−associated SNP: Chr06:6583303
pvalue= 0.002 ; R2= 0.4594
Estimated allele frequency
Ecological gradient (Leaf Unfolding differences)
0.0
0.2
0.4
0.6
−1
0
1
2
3
4
124
O8
L16
O12
L8
L12
O16
218
219
253
256
L1
9
204
217
O1
97
233

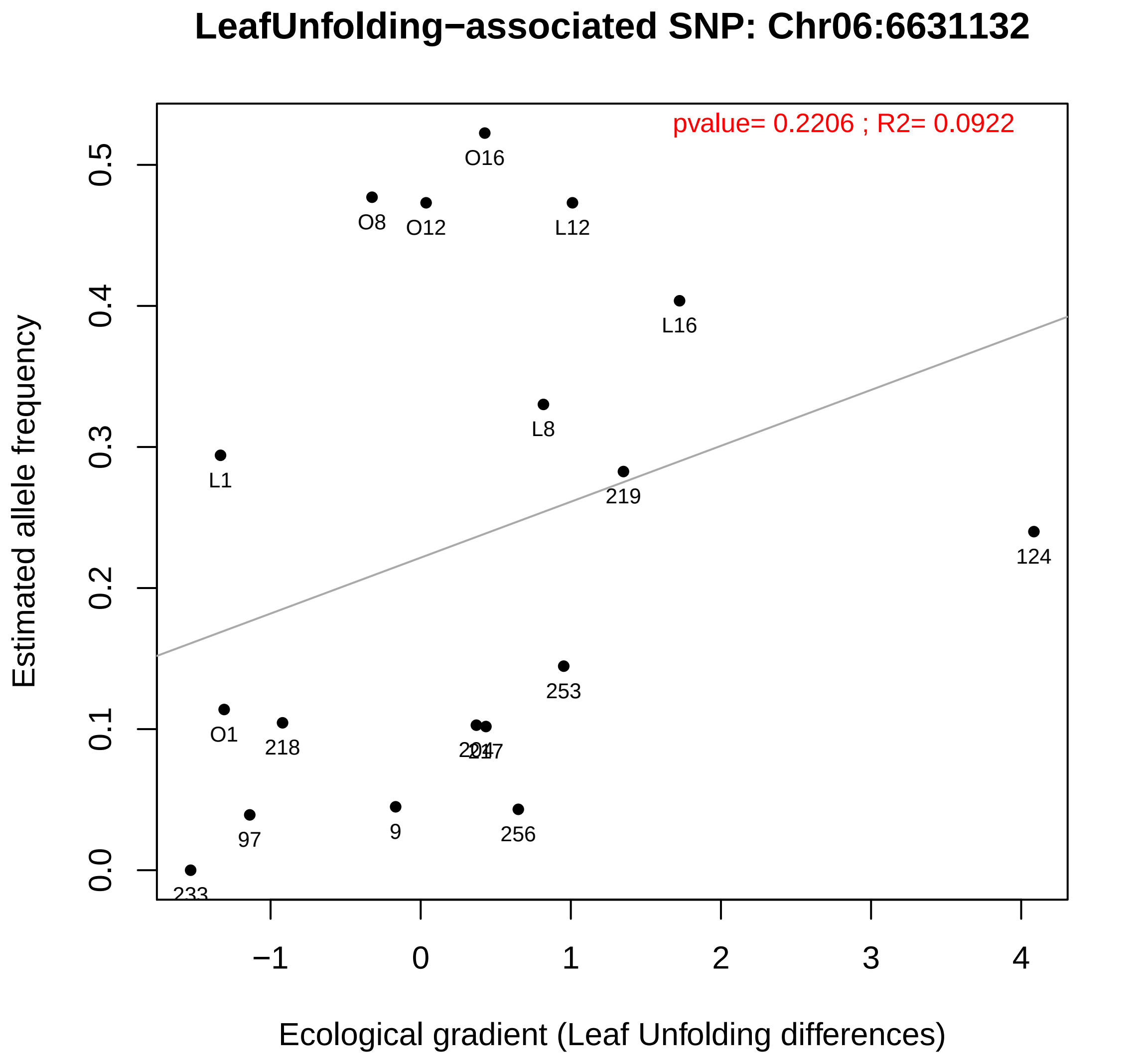
LeafUnfolding−associated SNP: Chr06:6631132
pvalue= 0.2206 ; R2= 0.0922
Estimated allele frequency
0.0
0.1
0.2
0.3
0.4
0.5
Ecological gradient (Leaf Unfolding differences)
−1
0
1
2
3
4
O16
O8
O12
L12
L16
L8
L1
219
124
253
O1
218
97
9
256
233

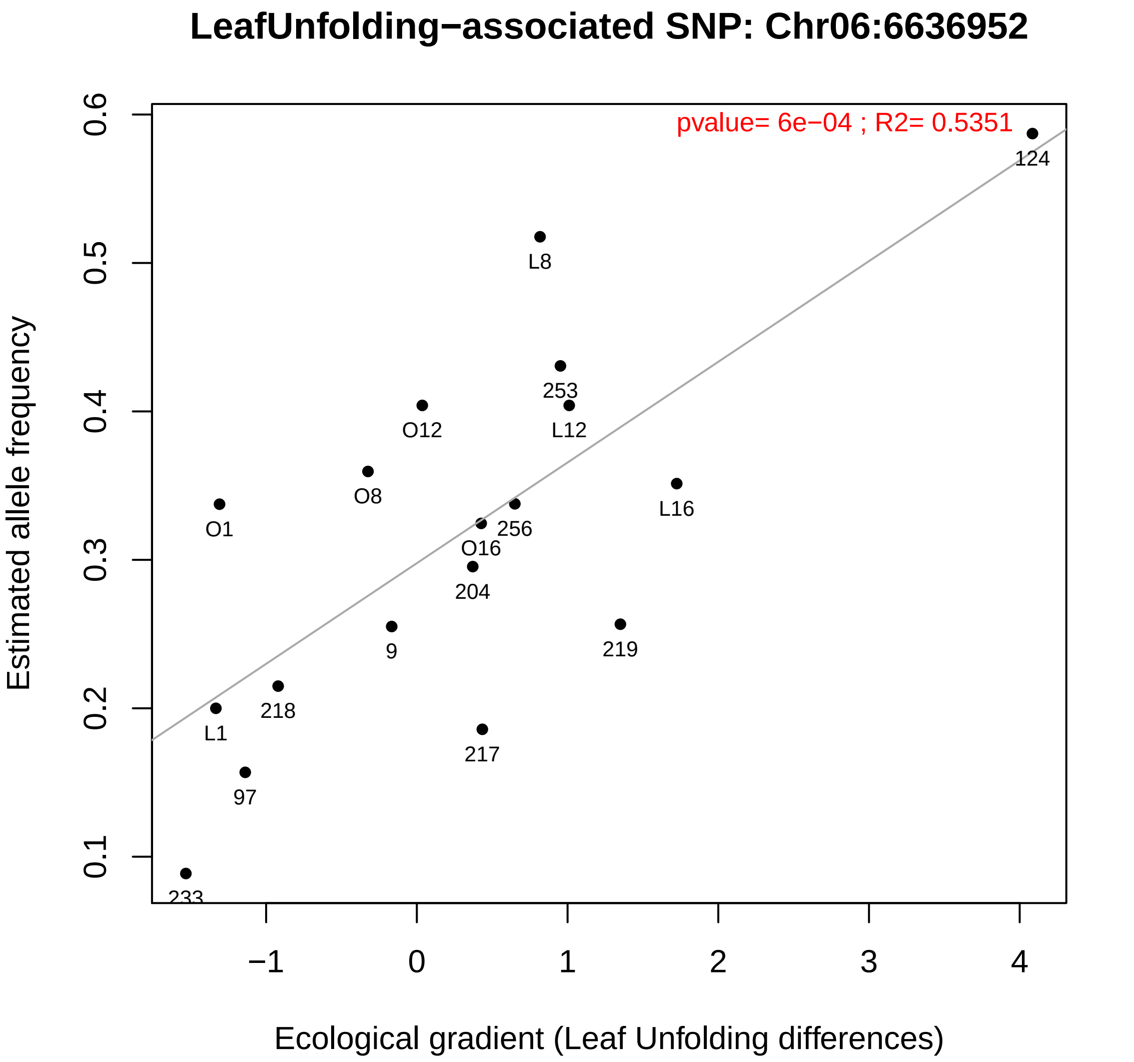
LeafUnfolding−associated SNP: Chr06:6636952
pvalue= 6e−04 ; R2= 0.5351
Estimated allele frequency
Ecological gradient (Leaf Unfolding differences)
0.1
0.2
0.3
0.4
0.5
0.6
−1
0
1
2
3
4
124
L8
253
L12
O12
O8
L16
O1
256
O16
204
9
219
218
L1
217
97
233

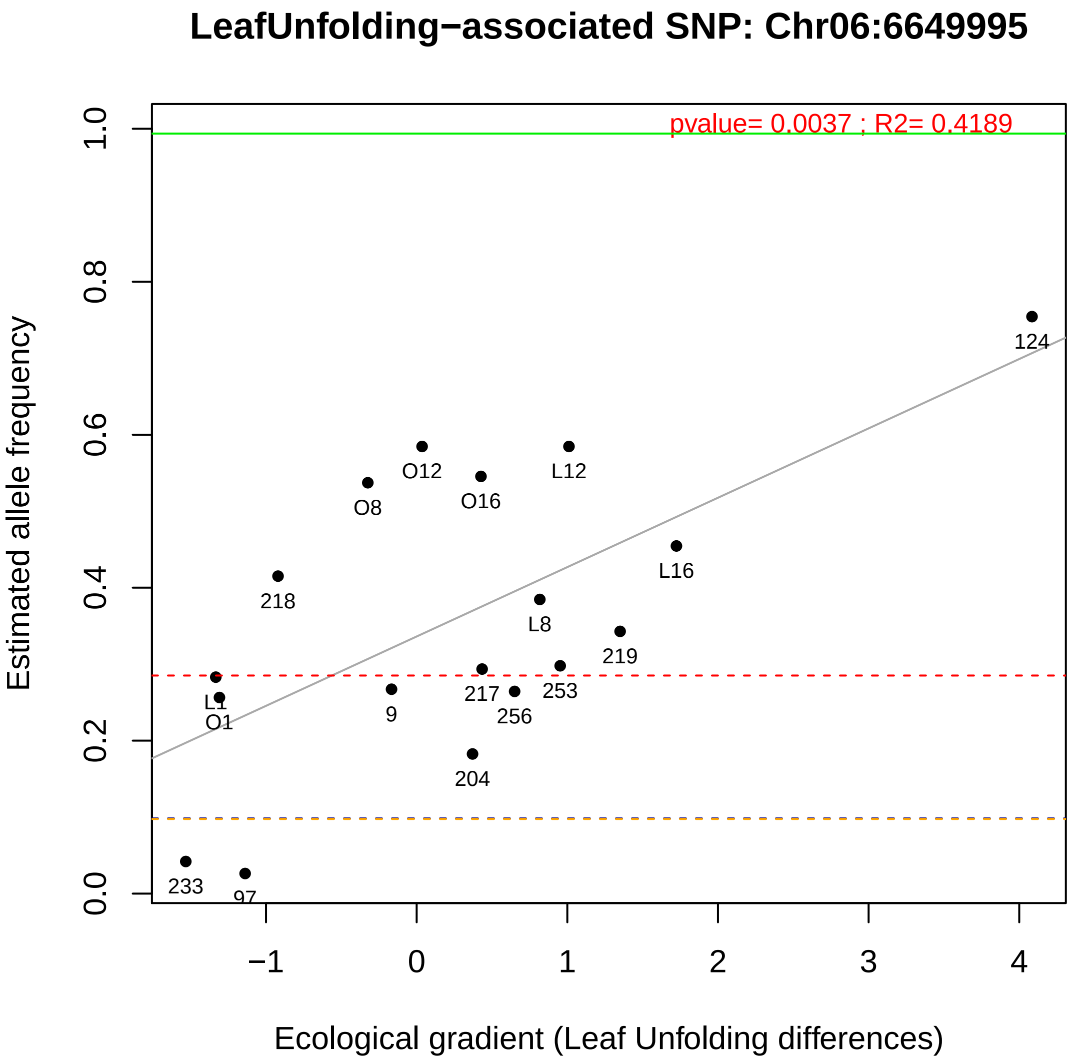

LeafUnfolding−associated SNP: Chr06:6649995
pvalue= 0.0037 ; R2= 0.4189
Estimated allele frequency
0.0
0.2
0.4
0.6
0.8
1.0
−1
0
1
2
3
4
Ecological gradient (Leaf Unfolding differences)
124
O12
L12
O8
O16
L16
218
L8
219
L1
O1
9
217
256
253
204
233
97

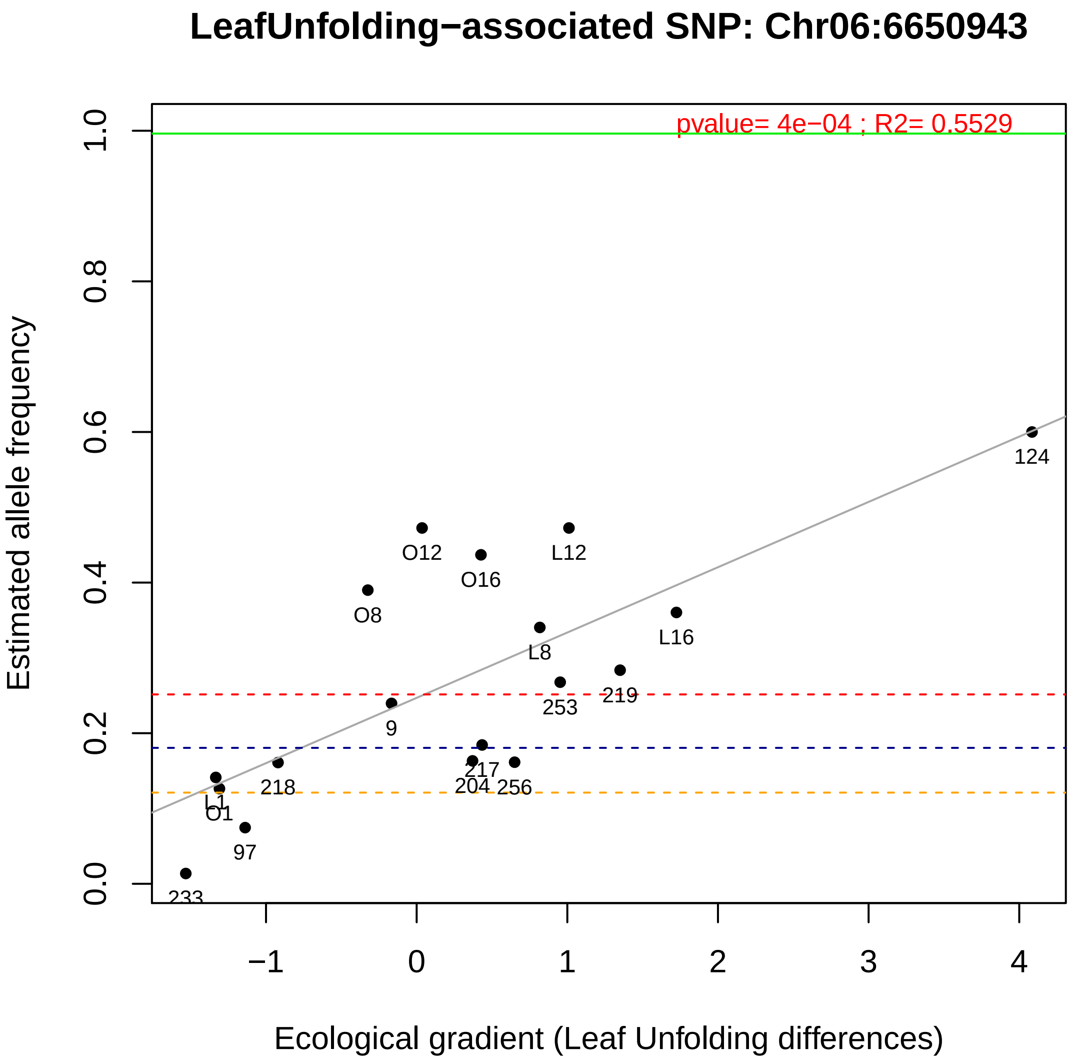
LeafUnfolding−associated SNP: Chr06:6650943
pvalue= 4e−04 ; R2= 0.5529
Estimated allele frequency
Ecological gradient (Leaf Unfolding differences)
0.0
0.2
0.4
0.6
0.8
1.0
−1
0
1
2
3
4
124
O12
L12
O16
O8
L16
L8
219
253
9
217
204
256
218
L1
O1
97
233

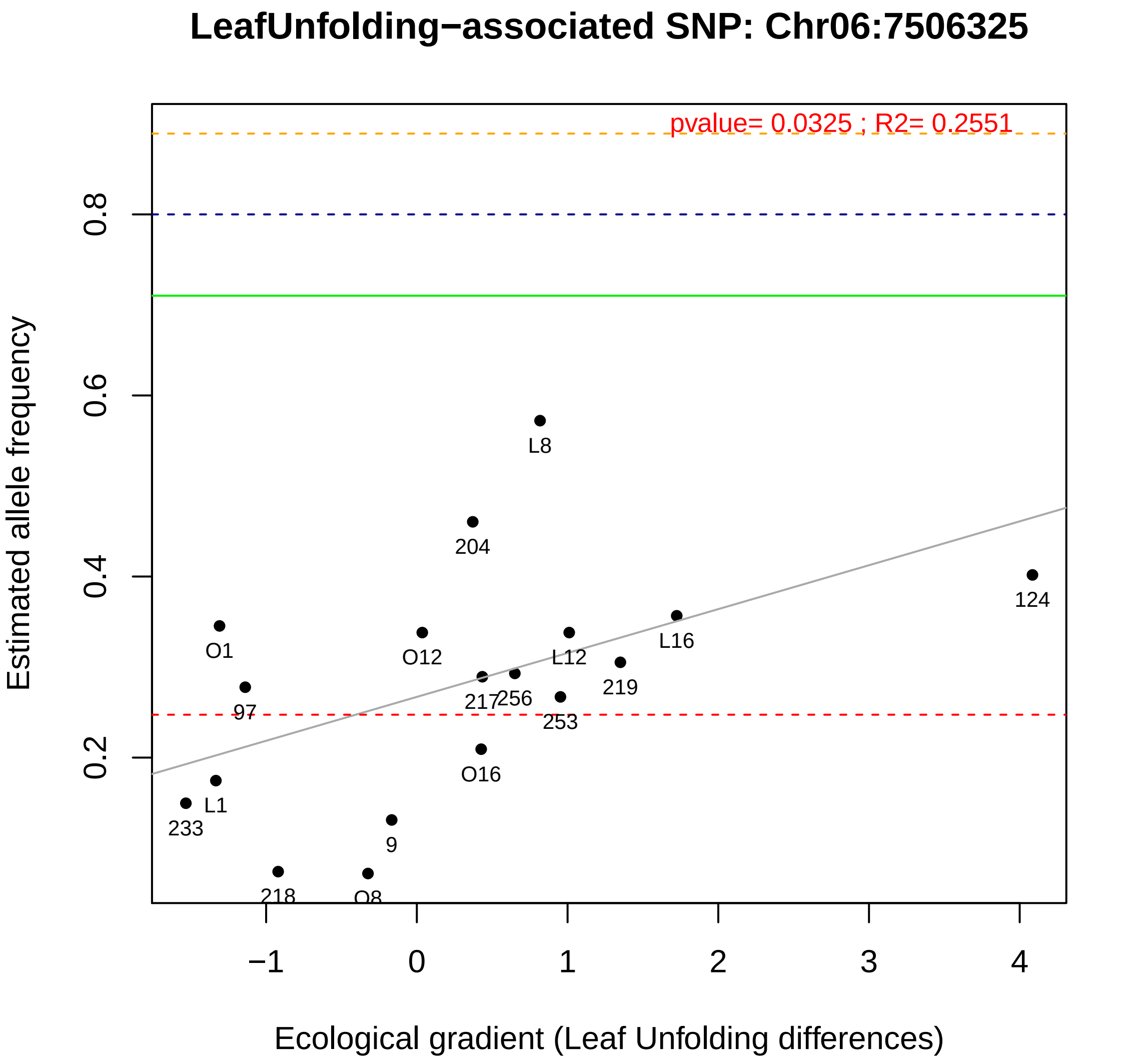

LeafUnfolding−associated SNP: Chr06:7506325
pvalue= 0.0325 ; R2= 0.2551
Estimated allele frequency
0.2
0.4
0.6
0.8
−1
0
1
2
3
4
Ecological gradient (Leaf Unfolding differences)
L8
204
124
L16
O1
O12
L12
219
97
217
256
253
O16
L1
233
9
218
O8

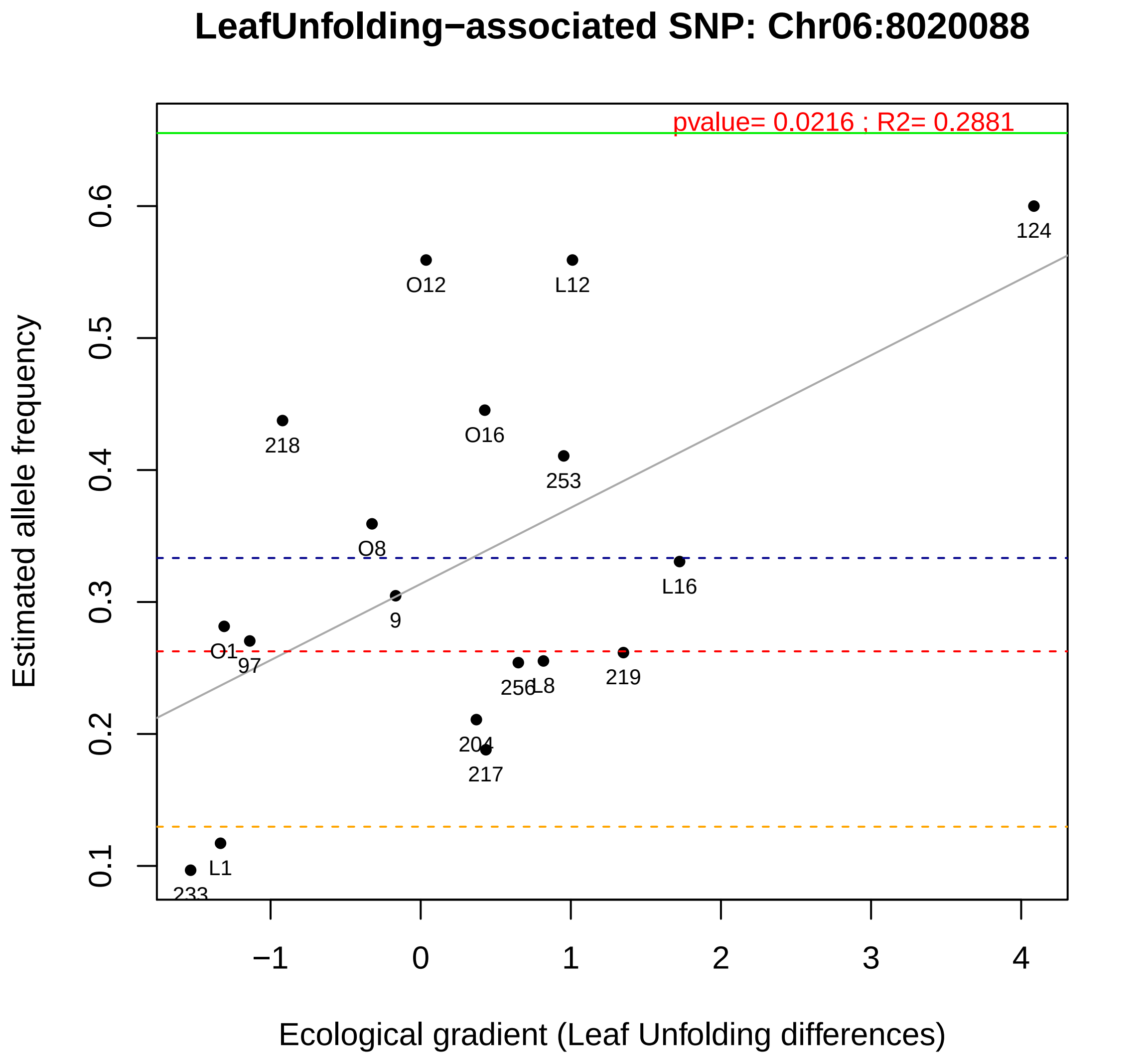

LeafUnfolding−associated SNP: Chr06:8020088
pvalue= 0.0216 ; R2= 0.2881
Estimated allele frequency
0.1
0.2
0.3
0.4
0.5
0.6
−1
0
1
2
3
4
Ecological gradient (Leaf Unfolding differences)
124
O12
L12
218
O16
253
O8
L16
9
O1
97
256
L8
219
204
217
L1
233

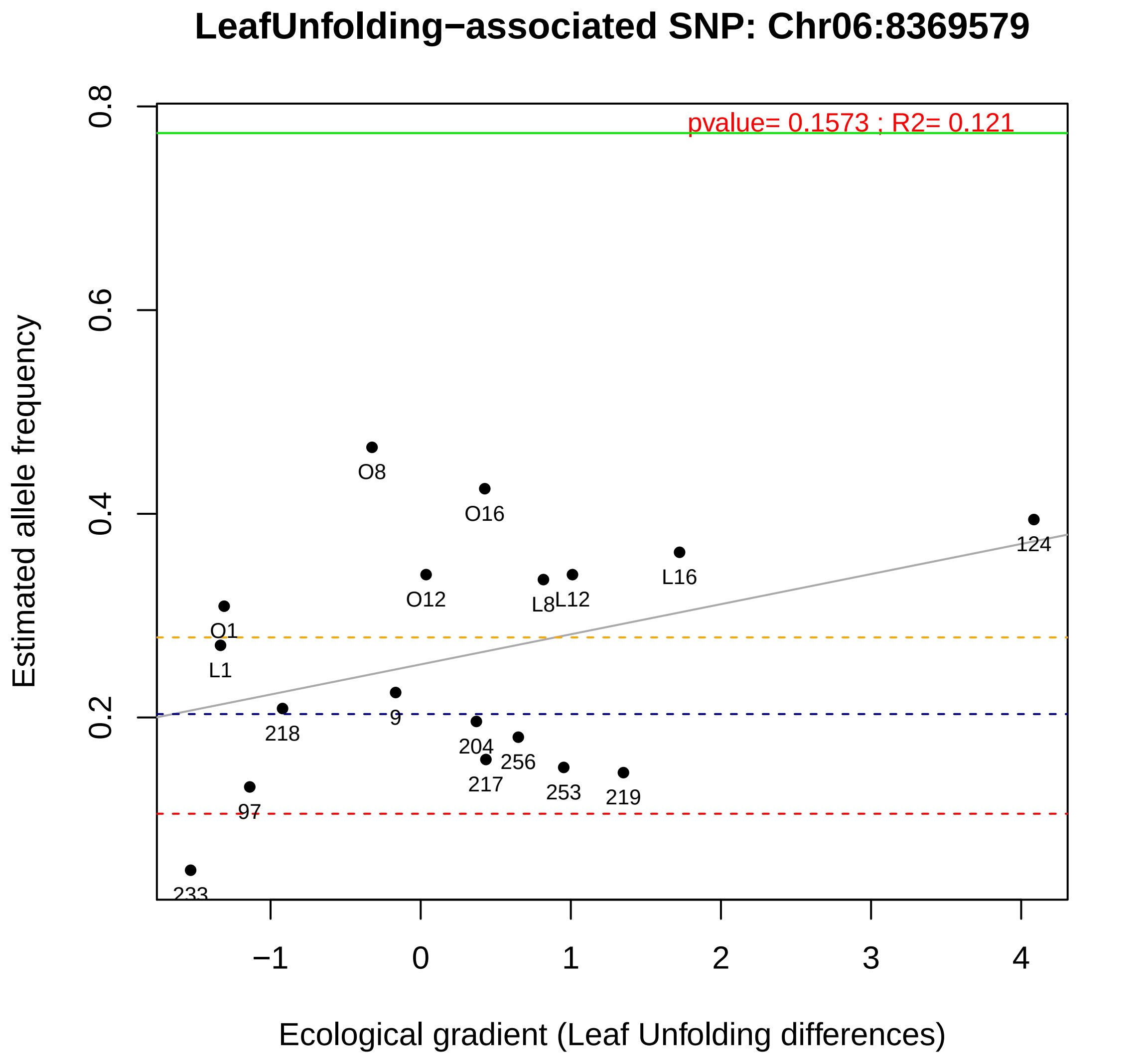

LeafUnfolding−associated SNP: Chr06:8369579
pvalue= 0.1573 ; R2= 0.121
Estimated allele frequency
0.2
0.4
0.6
0.8
−1
0
1
2
3
4
Ecological gradient (Leaf Unfolding differences)
O8
O16
124
L16
O12
L8
L12
O1
L1
218
9
204
256
217
253
219
97
233

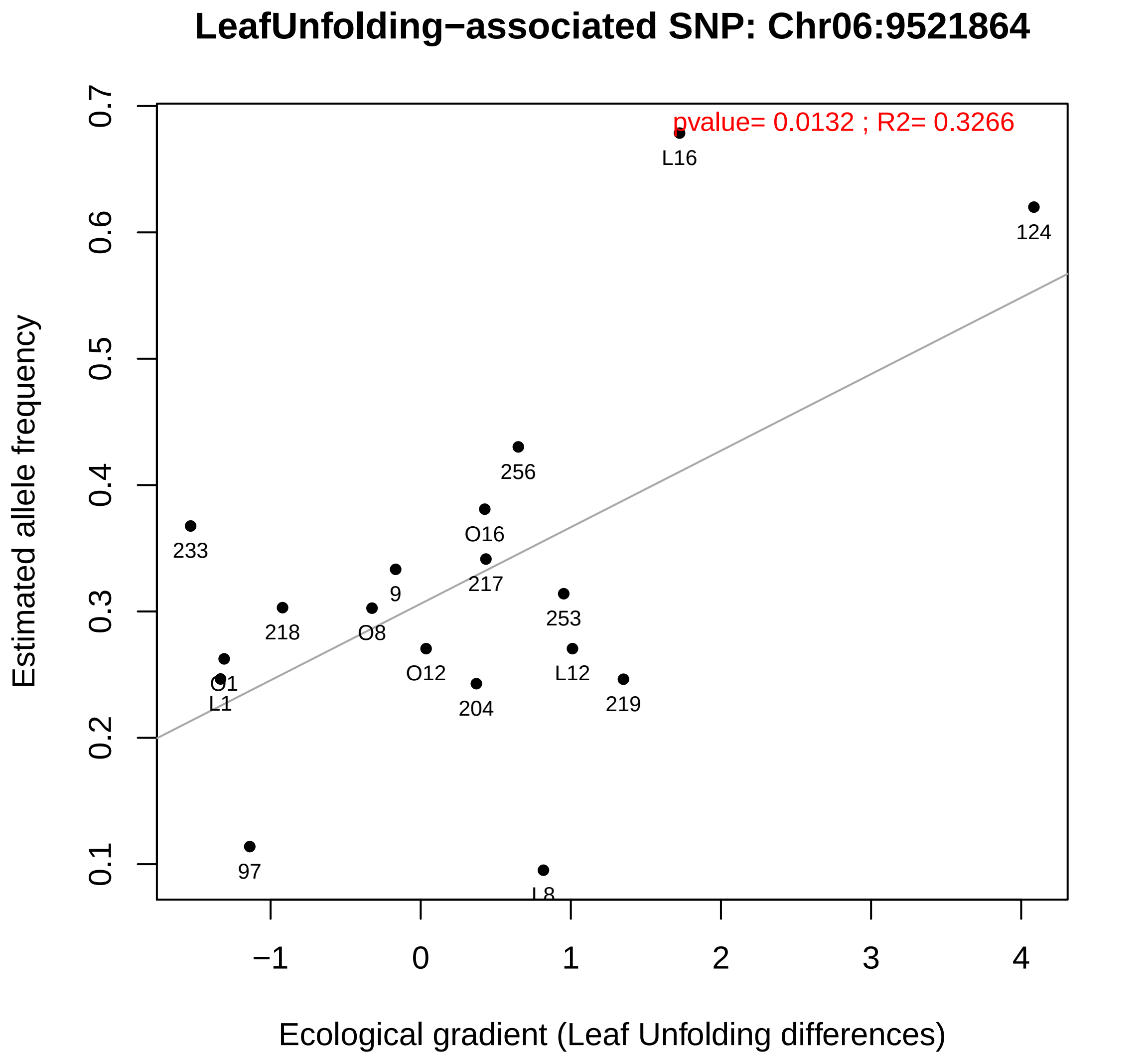

LeafUnfolding−associated SNP: Chr06:9521864
pvalue= 0.0132 ; R2= 0.3266
Estimated allele frequency
0.1
0.2
0.3
0.4
0.5
0.6
0.7
Ecological gradient (Leaf Unfolding differences)
−1
0
1
2
3
4
L16
124
256
O16
233
217
9
253
218
O8
O12
L12
O1
L1
204
219
97
L8

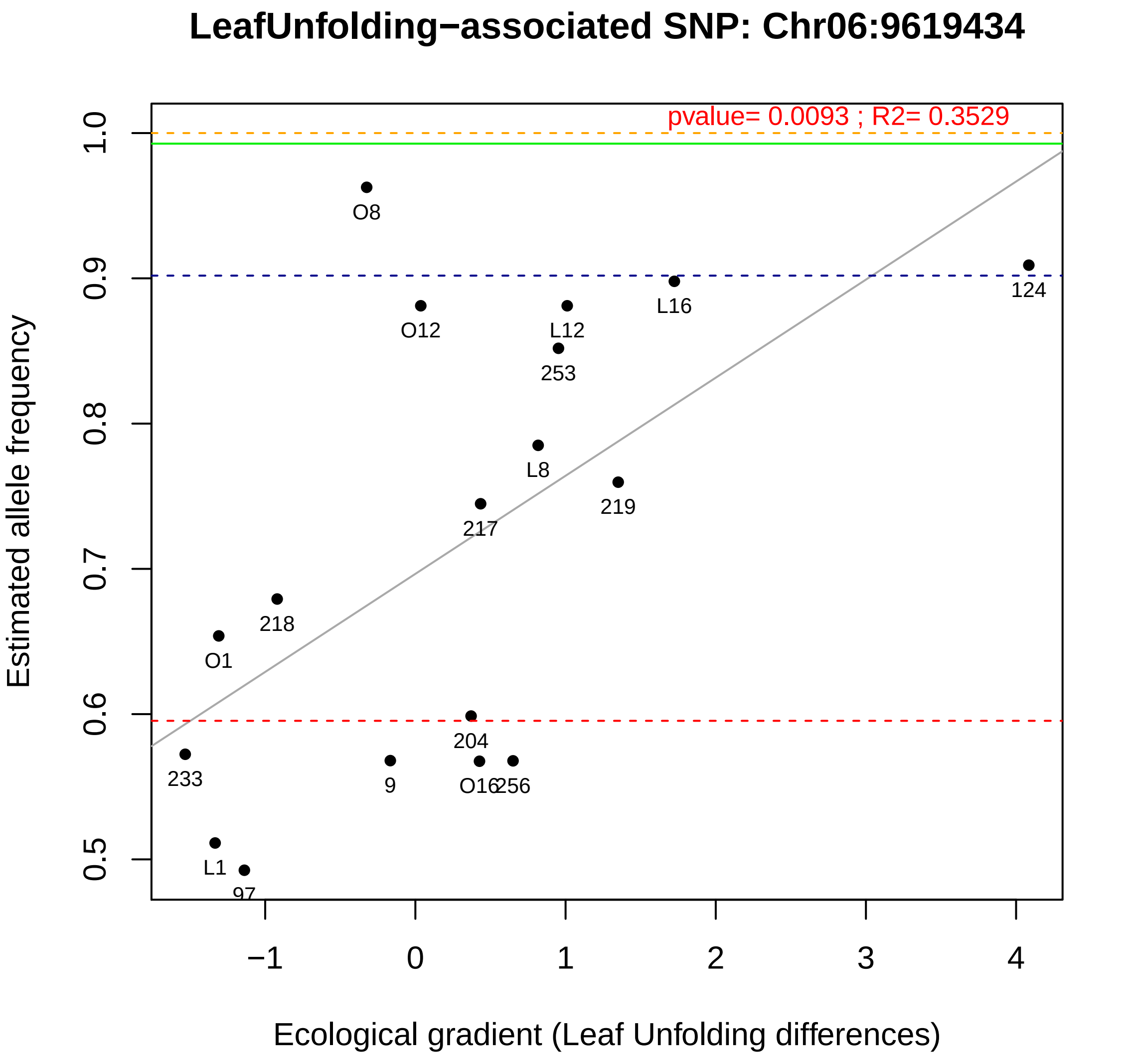

LeafUnfolding−associated SNP: Chr06:9619434
pvalue= 0.0093 ; R2= 0.3529
Estimated allele frequency
Ecological gradient (Leaf Unfolding differences)
0.5
0.6
0.7
0.8
0.9
1.0
−1
0
1
2
3
4
O8
O12
L12
253
L16
124
L8
219
217
218
O1
204
233
9
O16
256
L1
97

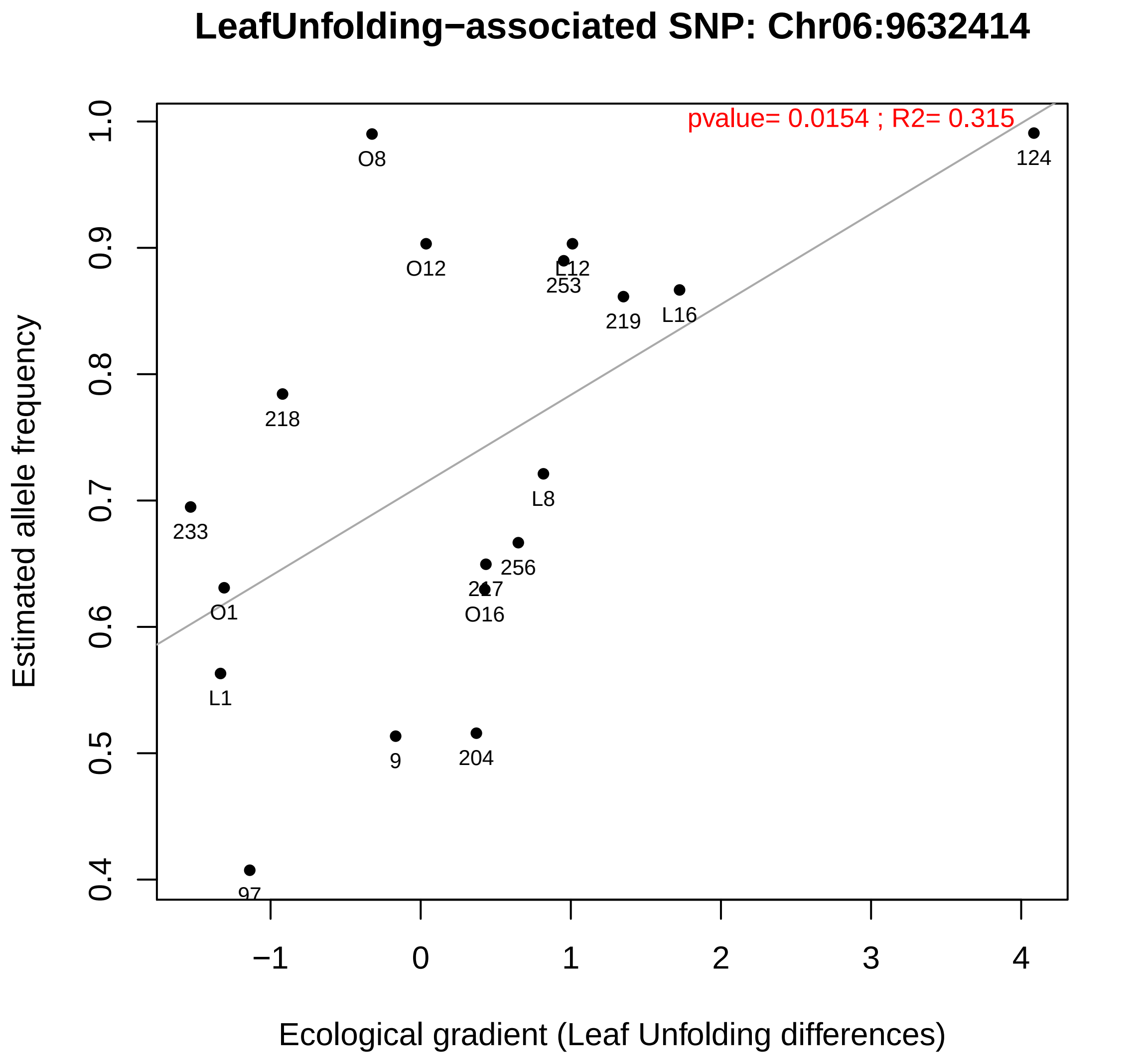

LeafUnfolding−associated SNP: Chr06:9632414
pvalue= 0.0154 ; R2= 0.315
O8
124
O12
L12
253
219
L16
218
L8
233
256
217
O1
O16
L1
9
204
97
Estimated allele frequency
1.0
0.9
0.8
0.7
0.6
0.5
0.4
−1
0
1
2
3
4
Ecological gradient (Leaf Unfolding differences)

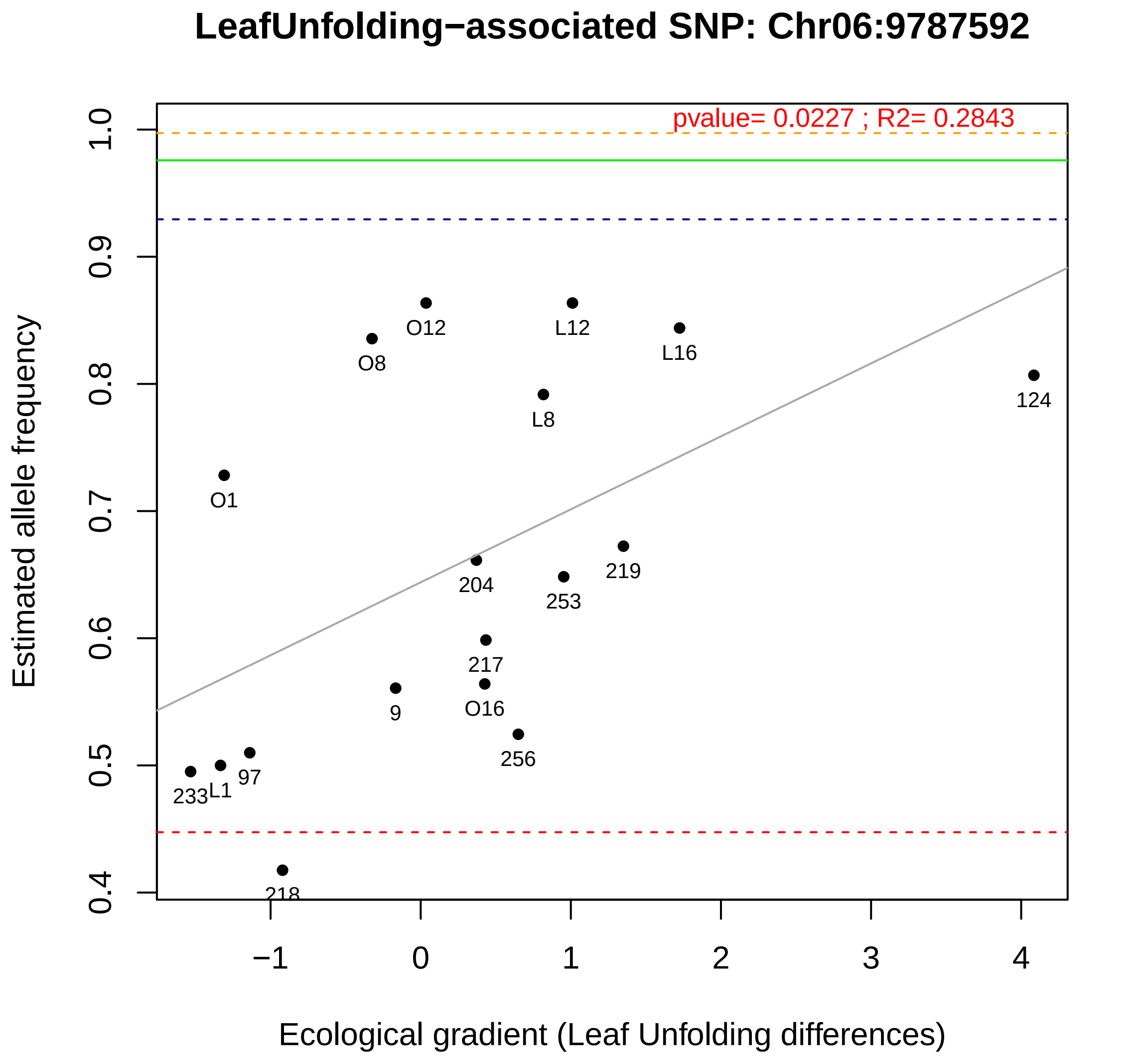

LeafUnfolding−associated SNP: Chr06:9787592
pvalue= 0.0227 ; R2= 0.2843
Estimated allele frequency
Ecological gradient (Leaf Unfolding differences)
1.0
0.9
0.8
0.7
0.6
0.5
0.4
−1
0
1
2
3
4
O12
L12
L16
O8
124
L8
O1
219
204
253
217
9
O16
256
233
L1
97
218

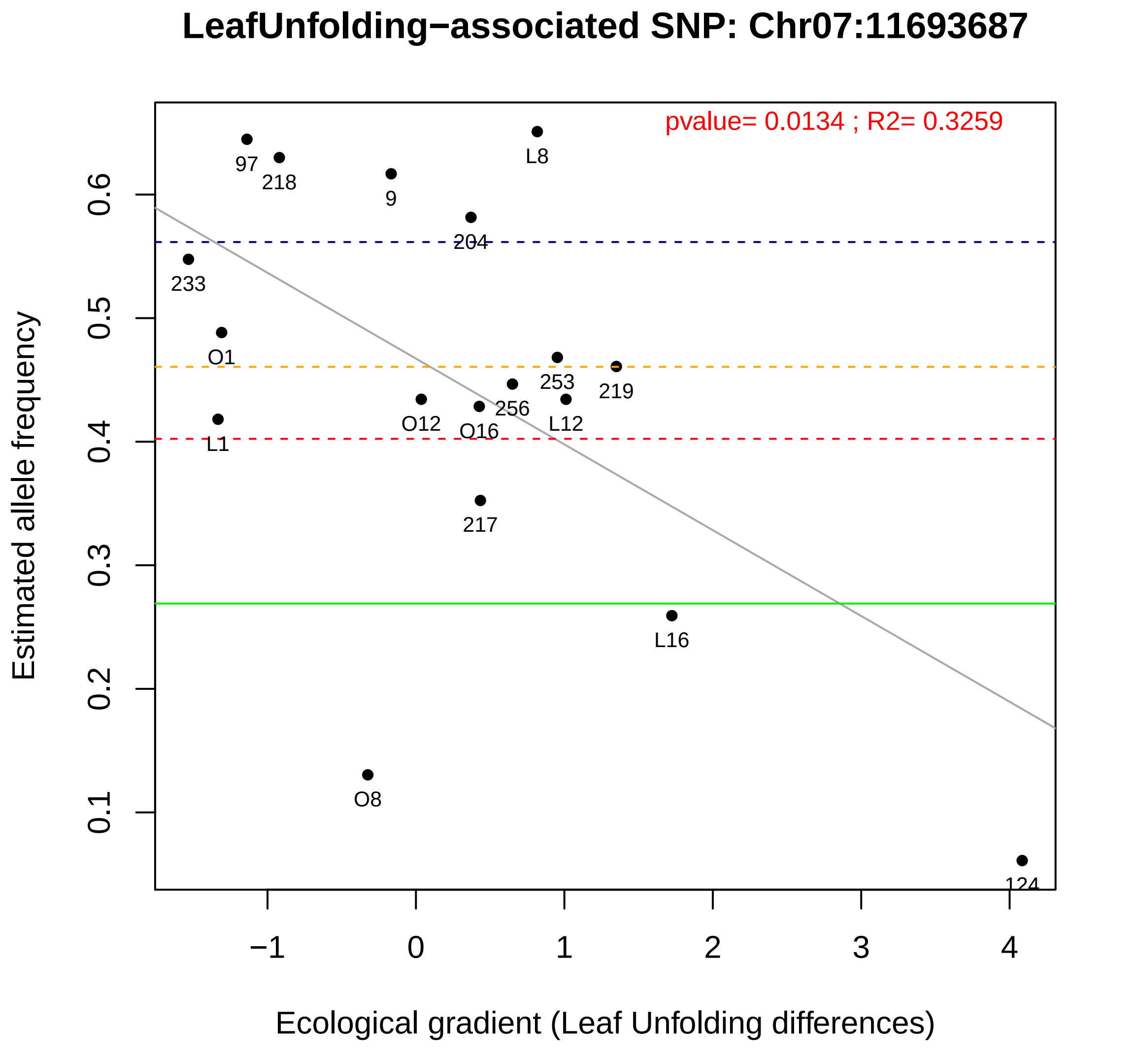
LeafUnfolding−associated SNP: Chr07:11693687
pvalue= 0.0134 ; R2= 0.3259
Estimated allele frequency
Ecological gradient (Leaf Unfolding differences)
0.1
0.2
0.3
0.4
0.5
0.6
−1
0
1
2
3
4
97
218
9
L8
204
233
O1
253
219
256
O12
O16
L12
L1
217
L16
O8
124

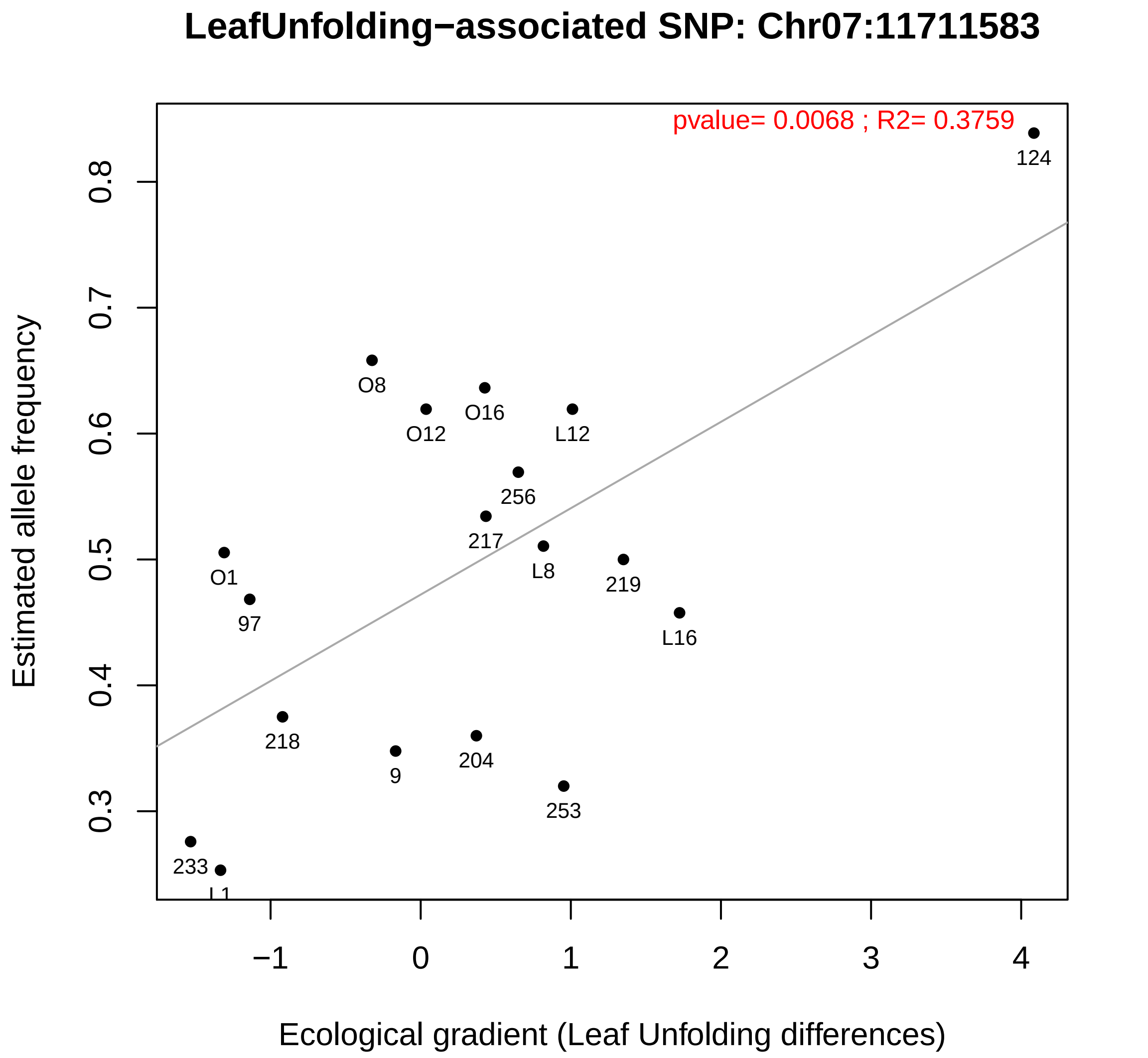
LeafUnfolding−associated SNP: Chr07:11711583
pvalue= 0.0068 ; R2= 0.3759
Estimated allele frequency
Ecological gradient (Leaf Unfolding differences)
0.3
0.4
0.5
0.6
0.7
0.8
−1
0
1
2
3
4
124
O8
O12
O16
L12
256
217
L8
219
L16
O1
97
218
9
204
253
233

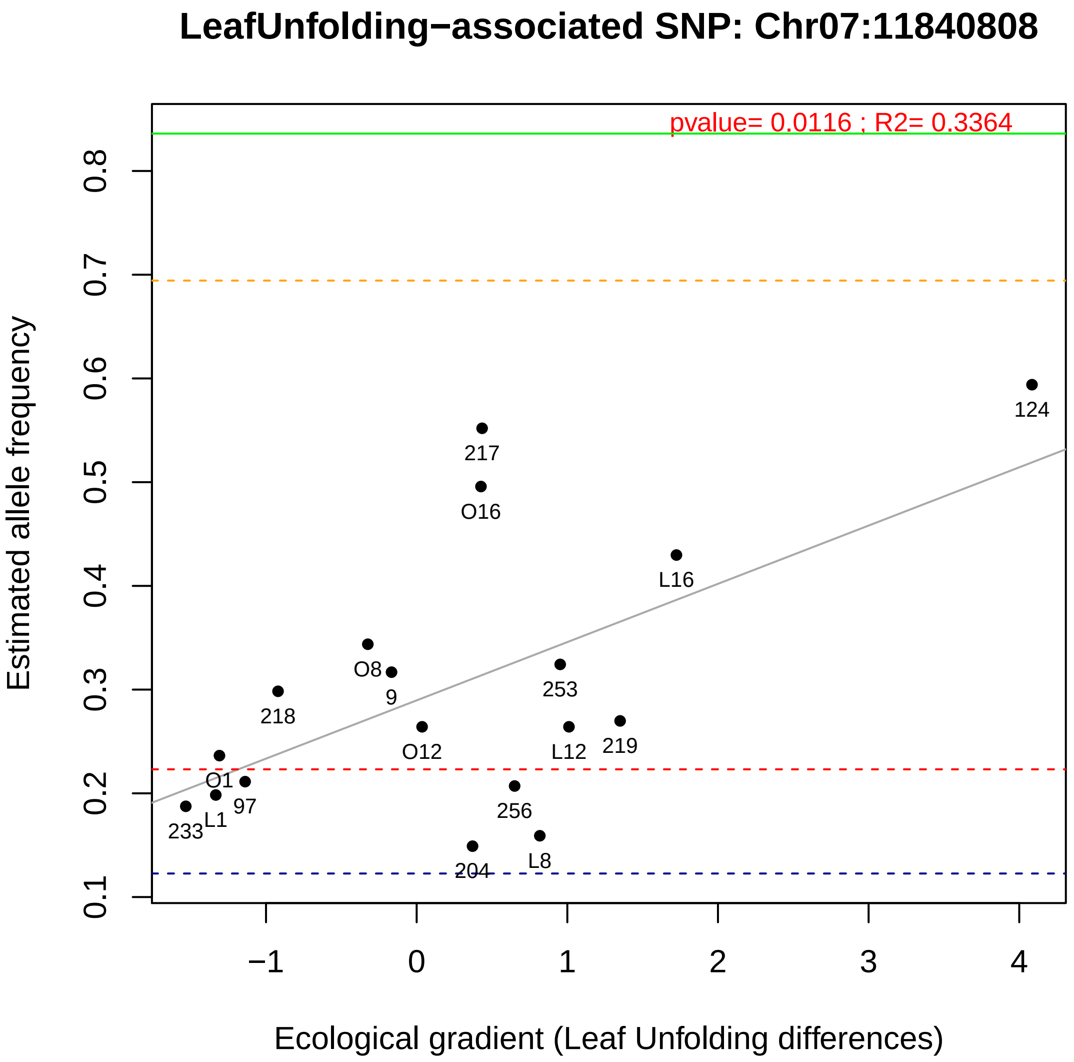

LeafUnfolding−associated SNP: Chr07:11840808
pvalue= 0.0116 ; R2= 0.3364
Estimated allele frequency
0.1
0.2
0.3
0.4
0.5
0.6
0.7
0.8
−1
0
1
2
3
4
Ecological gradient (Leaf Unfolding differences)
124
217
O16
L16
O8
9
253
218
O12
L12
219
O1
L1
97
233
256
204
L8

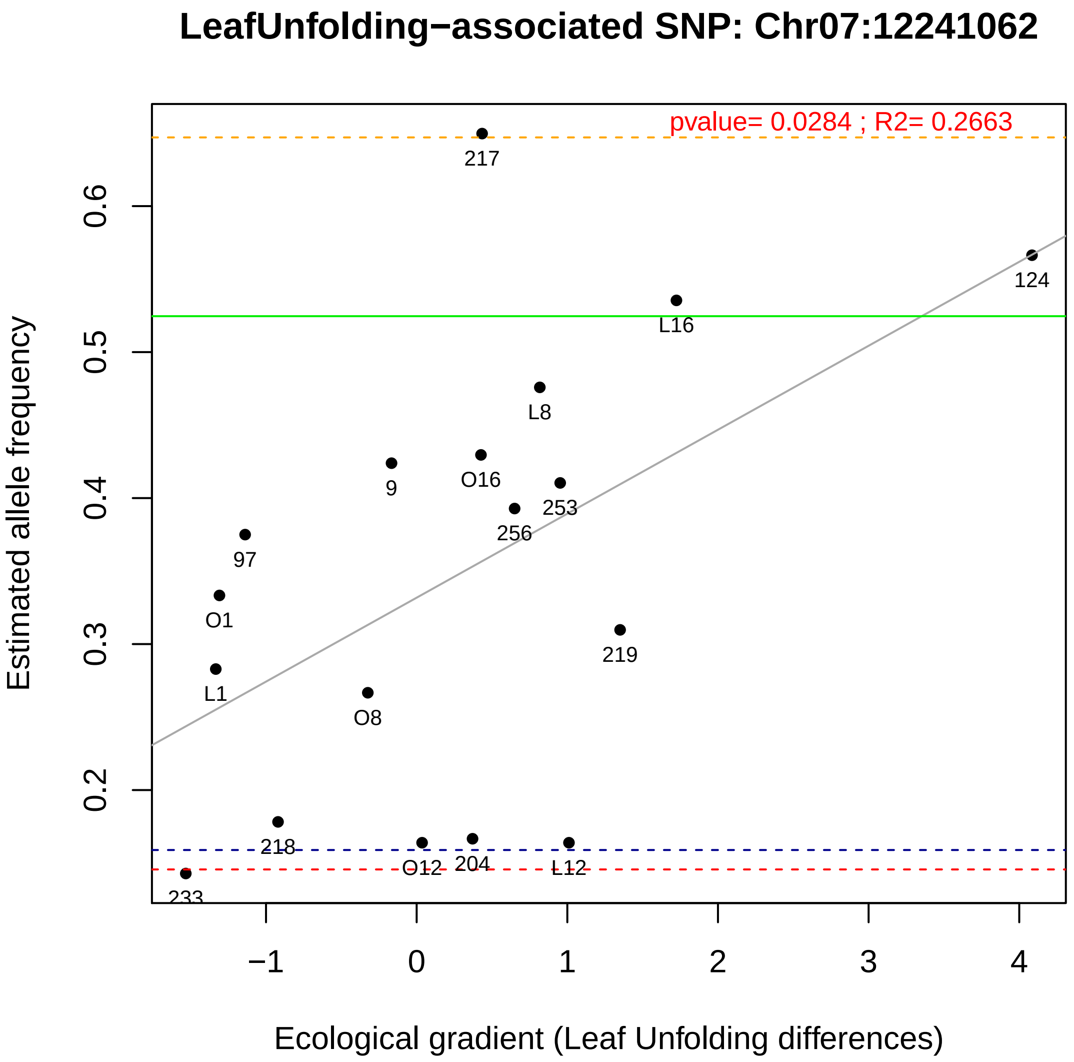

LeafUnfolding−associated SNP: Chr07:12241062
pvalue= 0.0284 ; R2= 0.2663
217
124
L16
L8
9
O16
253
256
97
O1
219
L1
O8
218
O12
204
L12
233
Estimated allele frequency
0.2
0.3
0.4
0.5
0.6
−1
0
1
2
3
4
Ecological gradient (Leaf Unfolding differences)

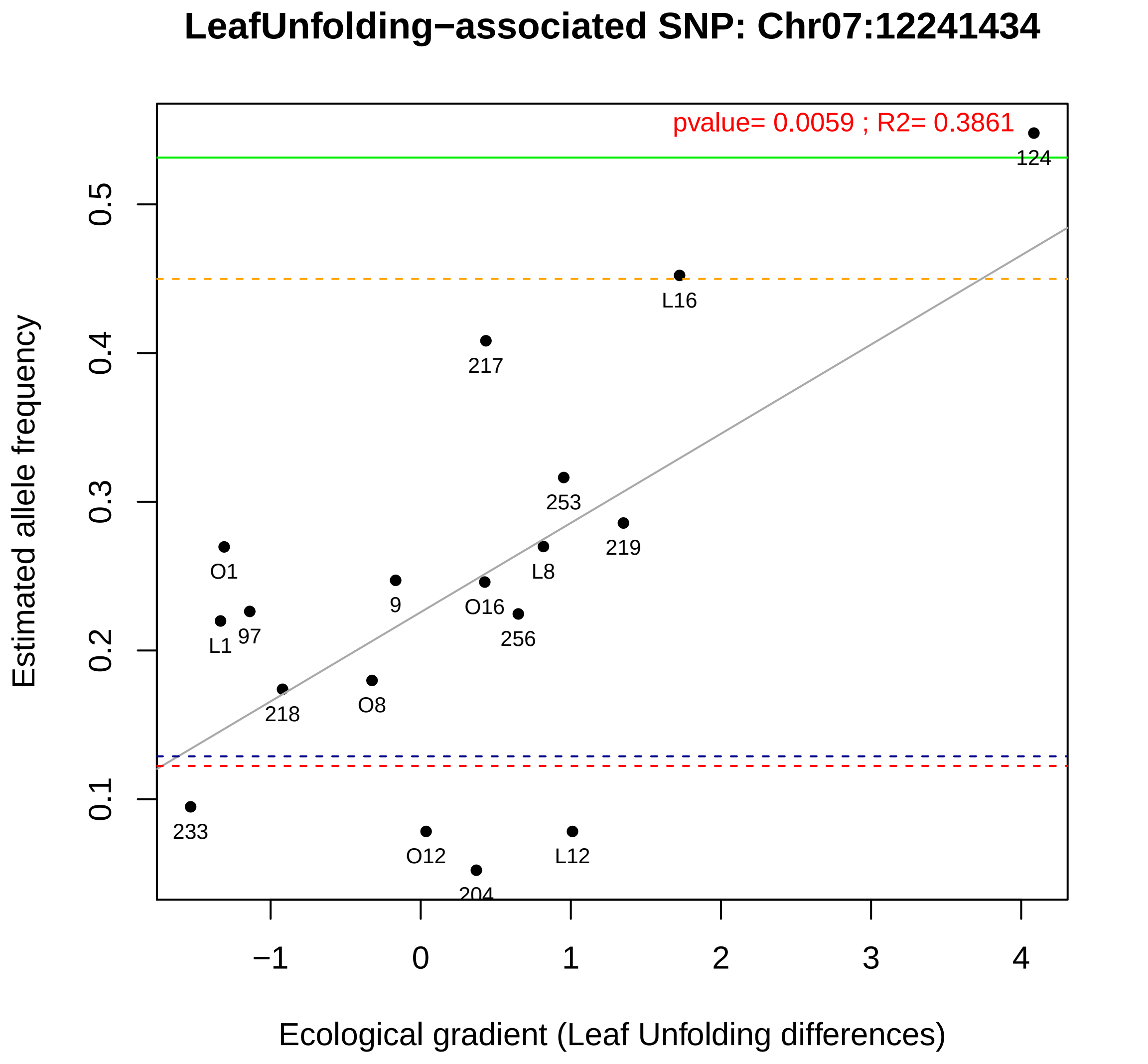
LeafUnfolding−associated SNP: Chr07:12241434
pvalue= 0.0059 ; R2= 0.3861
Estimated allele frequency
0.1
0.2
0.3
0.4
0.5
−1
0
1
2
3
4
Ecological gradient (Leaf Unfolding differences)
124
L16
217
253
219
O1
L8
9
O16
256
L1
97
218
O8
233
O12
L12
204

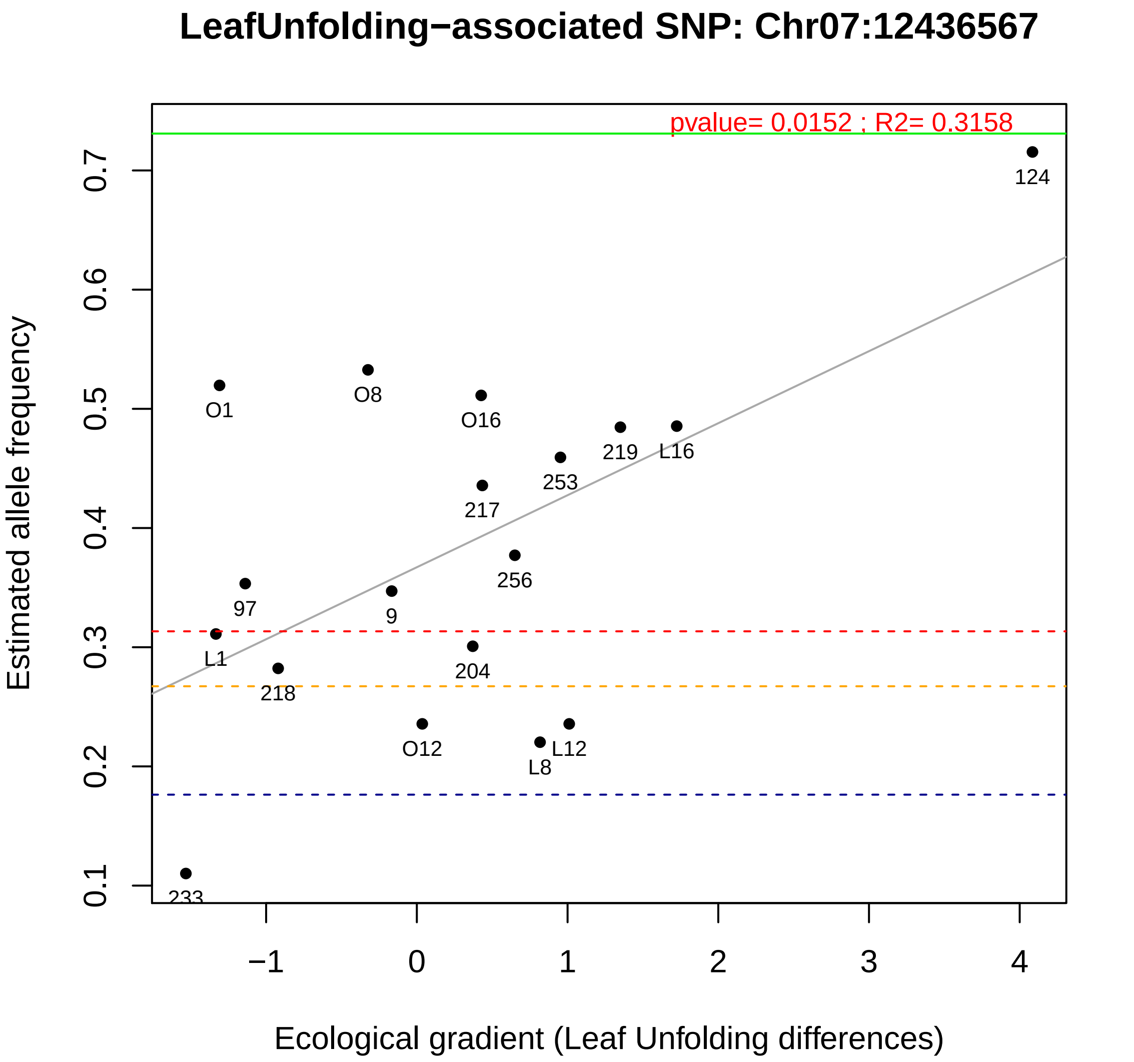
LeafUnfolding−associated SNP: Chr07:12436567
pvalue= 0.0152 ; R2= 0.3158
Estimated allele frequency
0.1
0.2
0.3
0.4
0.5
0.6
0.7
Ecological gradient (Leaf Unfolding differences)
−1
0
1
2
3
4
124
O1
O8
O16
219
L16
253
217
256
97
9
L1
218
204
O12
L8
L12
233

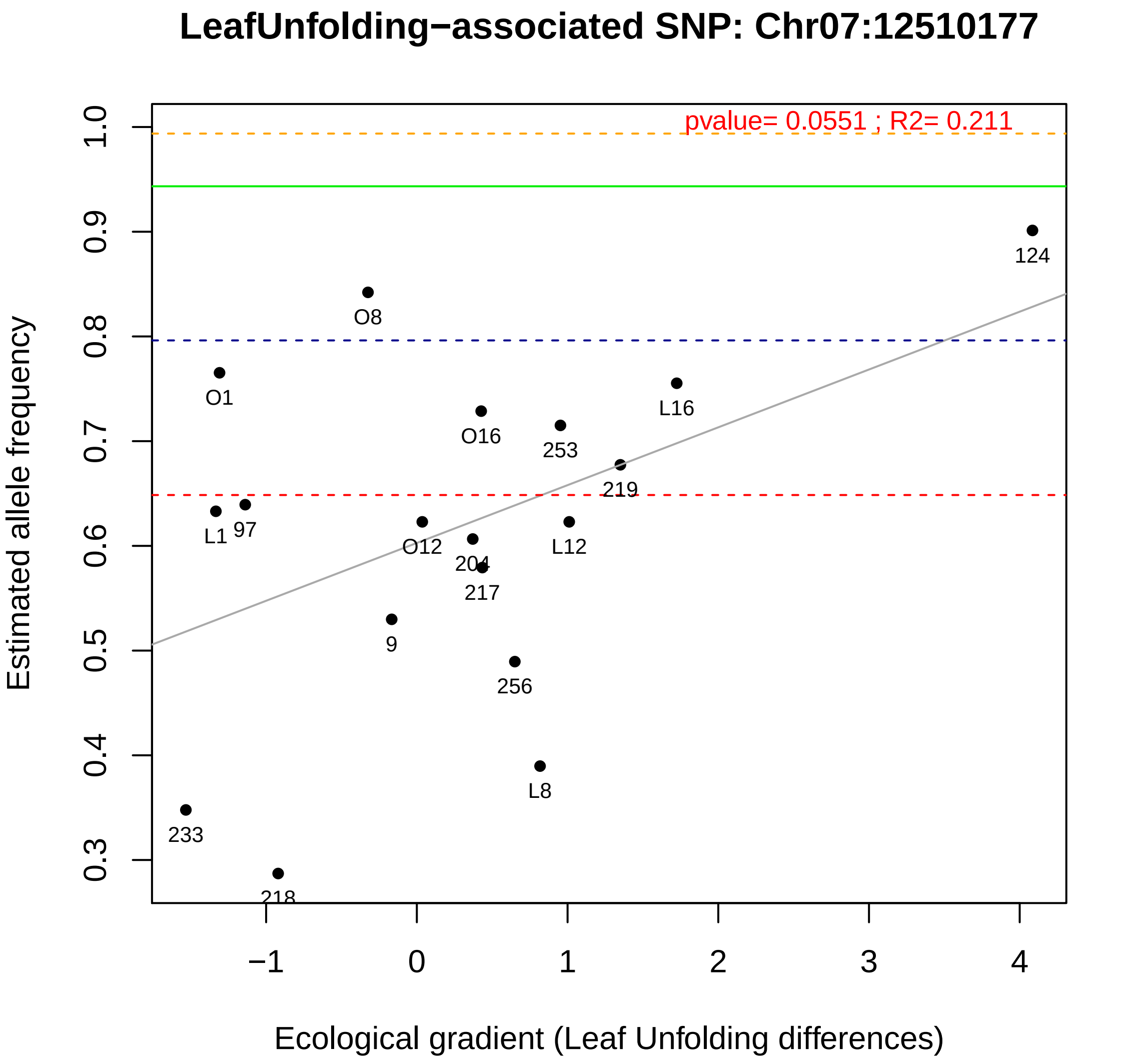

LeafUnfolding−associated SNP: Chr07:12510177
pvalue= 0.0551 ; R2= 0.211
Estimated allele frequency
Ecological gradient (Leaf Unfolding differences)
0.3
0.4
0.5
0.6
0.7
0.8
0.9
1.0
−1
0
1
2
3
4
124
O8
O1
L16
O16
253
219
L1
97
O12
L12
204
217
9
256
L8
233
218

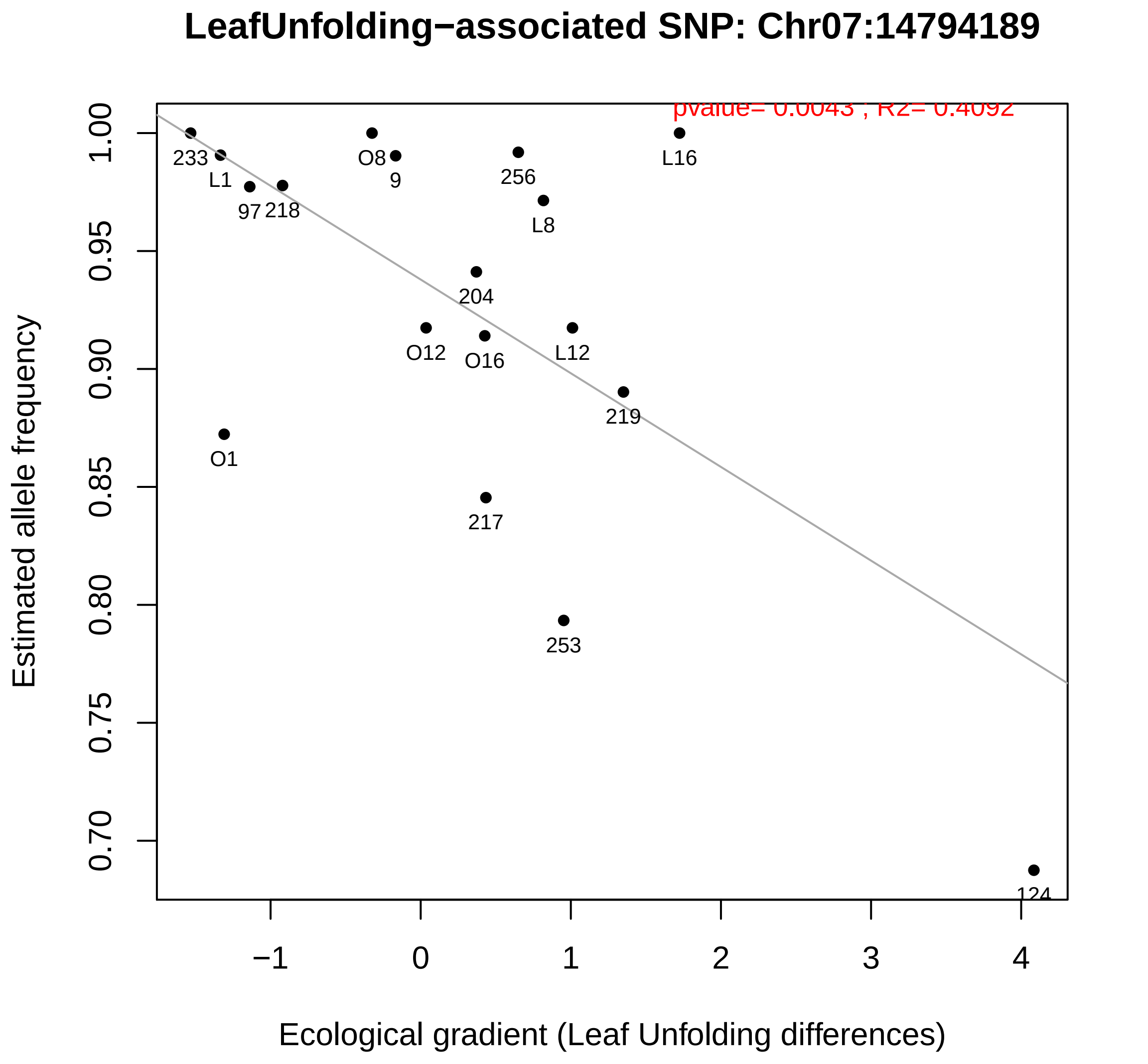
LeafUnfolding−associated SNP: Chr07:14794189
pvalue= 0.0043 ; R2= 0.4092
Estimated allele frequency
Ecological gradient (Leaf Unfolding differences)
1.00
0.95
0.90
0.85
0.80
0.75
0.70
−1
0
1
2
3
4
233
L1
97
218
O8
9
256
L16
L8
204
O12
O16
L12
219
O1
217
253
124

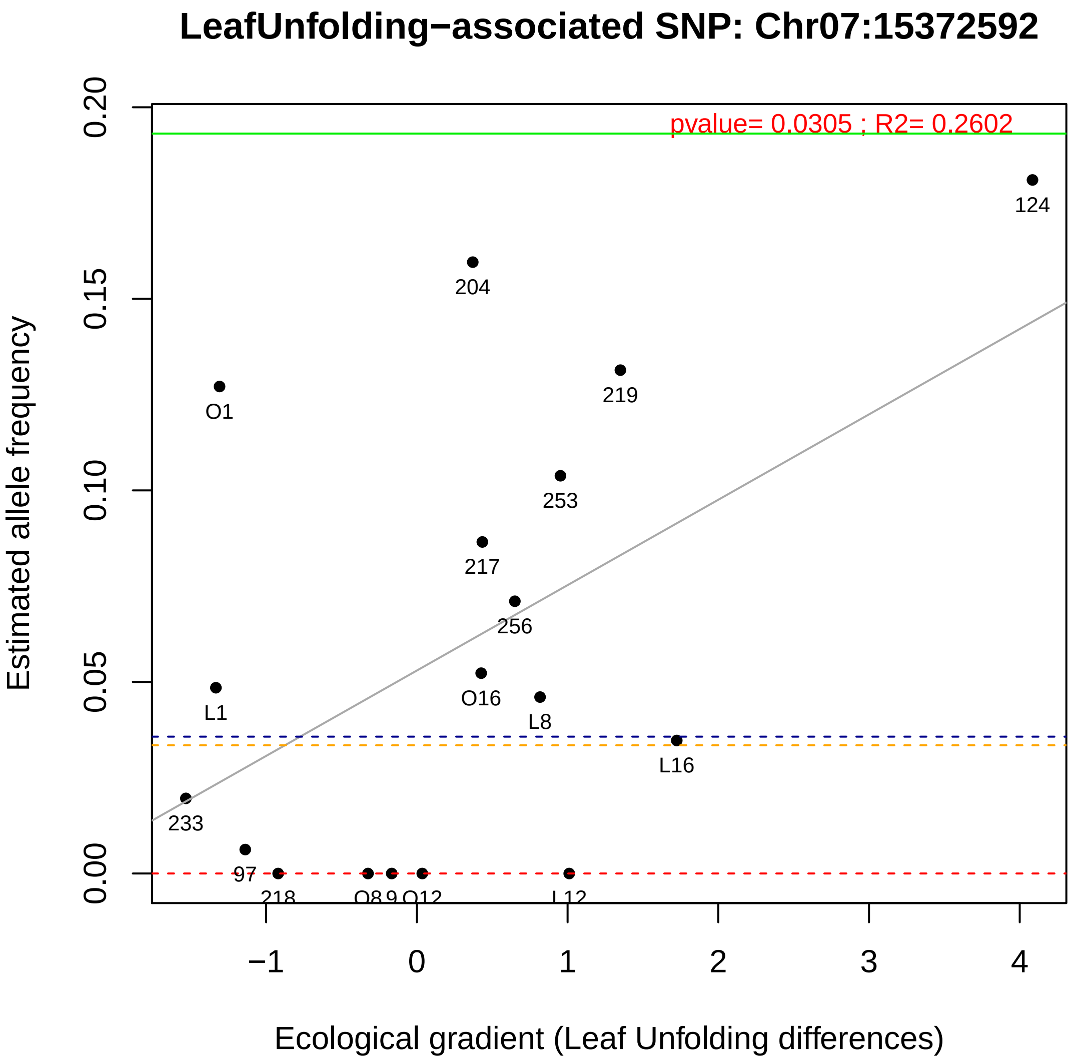

LeafUnfolding−associated SNP: Chr07:15372592
pvalue= 0.0305 ; R2= 0.2602
Estimated allele frequency
0.00
0.05
0.10
0.15
0.20
Ecological gradient (Leaf Unfolding differences)
−1
0
1
2
3
4
124
204
219
O1
253
217
256
O16
L8
L1
L16
233
97
218
O8
9
O12
L12

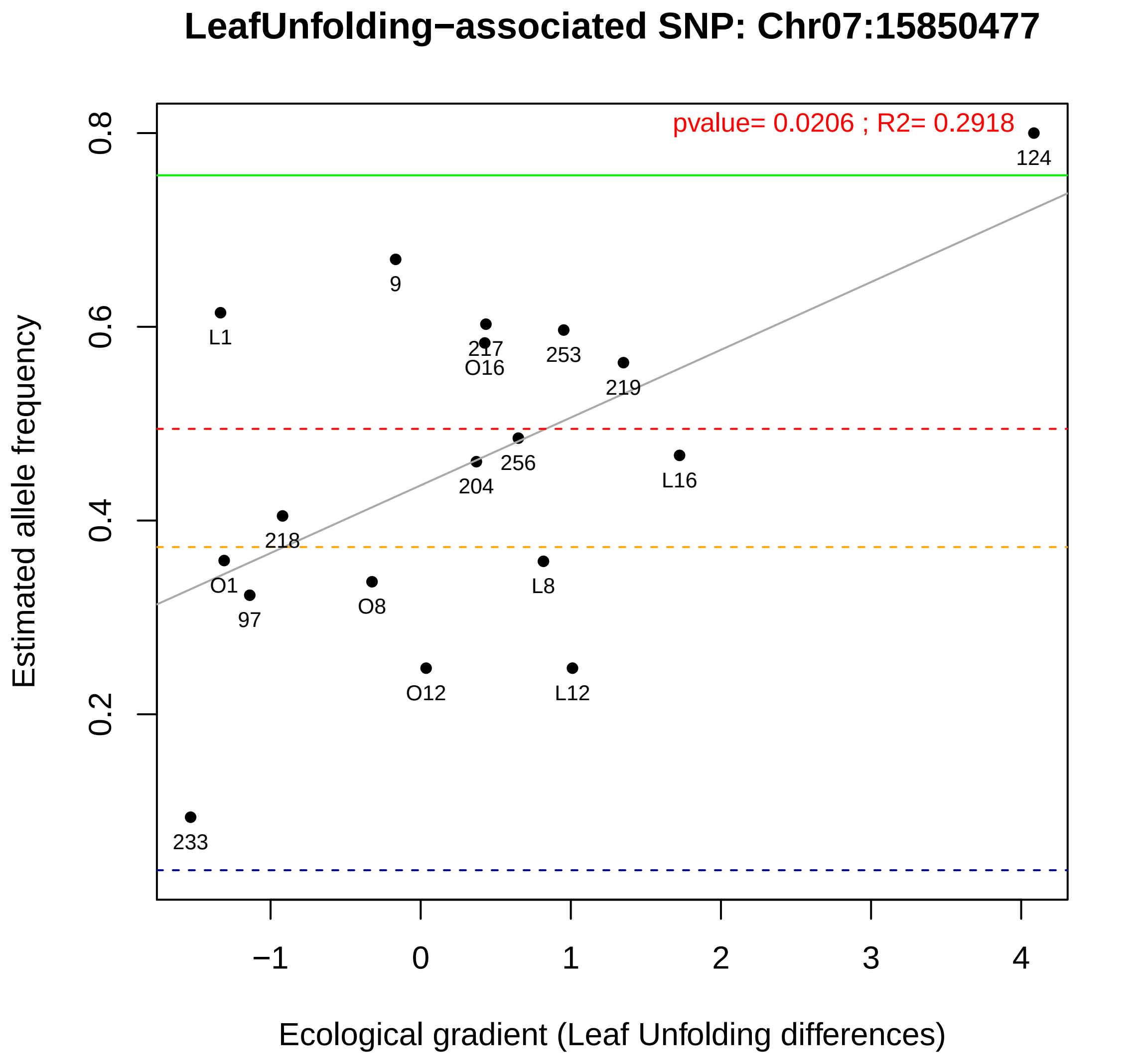

LeafUnfolding−associated SNP: Chr07:15850477
pvalue= 0.0206 ; R2= 0.2918
Estimated allele frequency
0.2
0.4
0.6
0.8
124
9
L1
217
O16
253
219
256
204
L16
218
O1
97
O8
L8
O12
L12
233
−1
0
1
2
3
4
Ecological gradient (Leaf Unfolding differences)

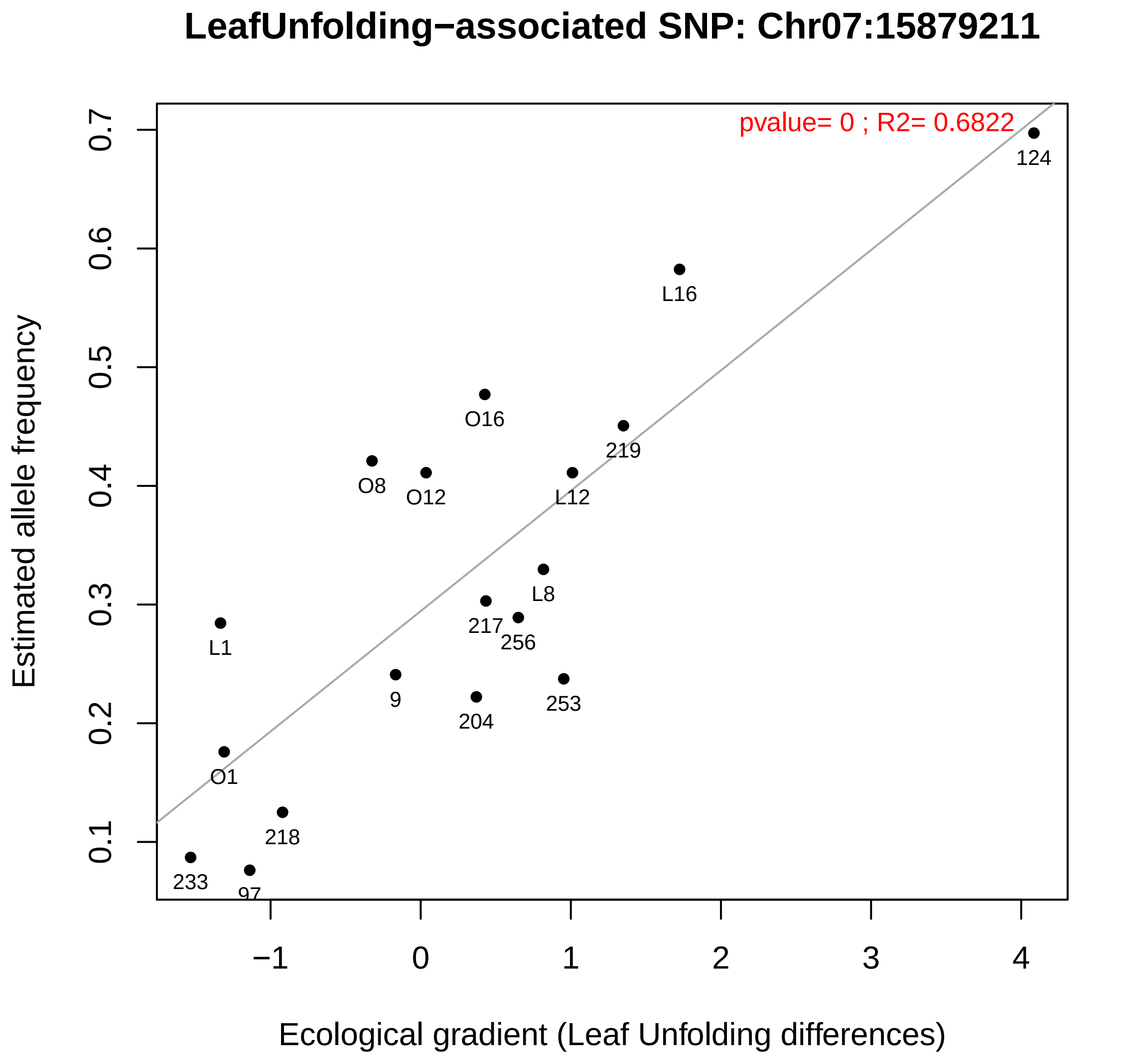

LeafUnfolding−associated SNP: Chr07:15879211
pvalue= 0 ; R2= 0.6822
Estimated allele frequency
Ecological gradient (Leaf Unfolding differences)
0.1
0.2
0.3
0.4
0.5
0.6
0.7
−1
0
1
2
3
4
124
L16
O16
219
O8
O12
L12
L8
217
256
L1
9
204
253
O1
218
233
97

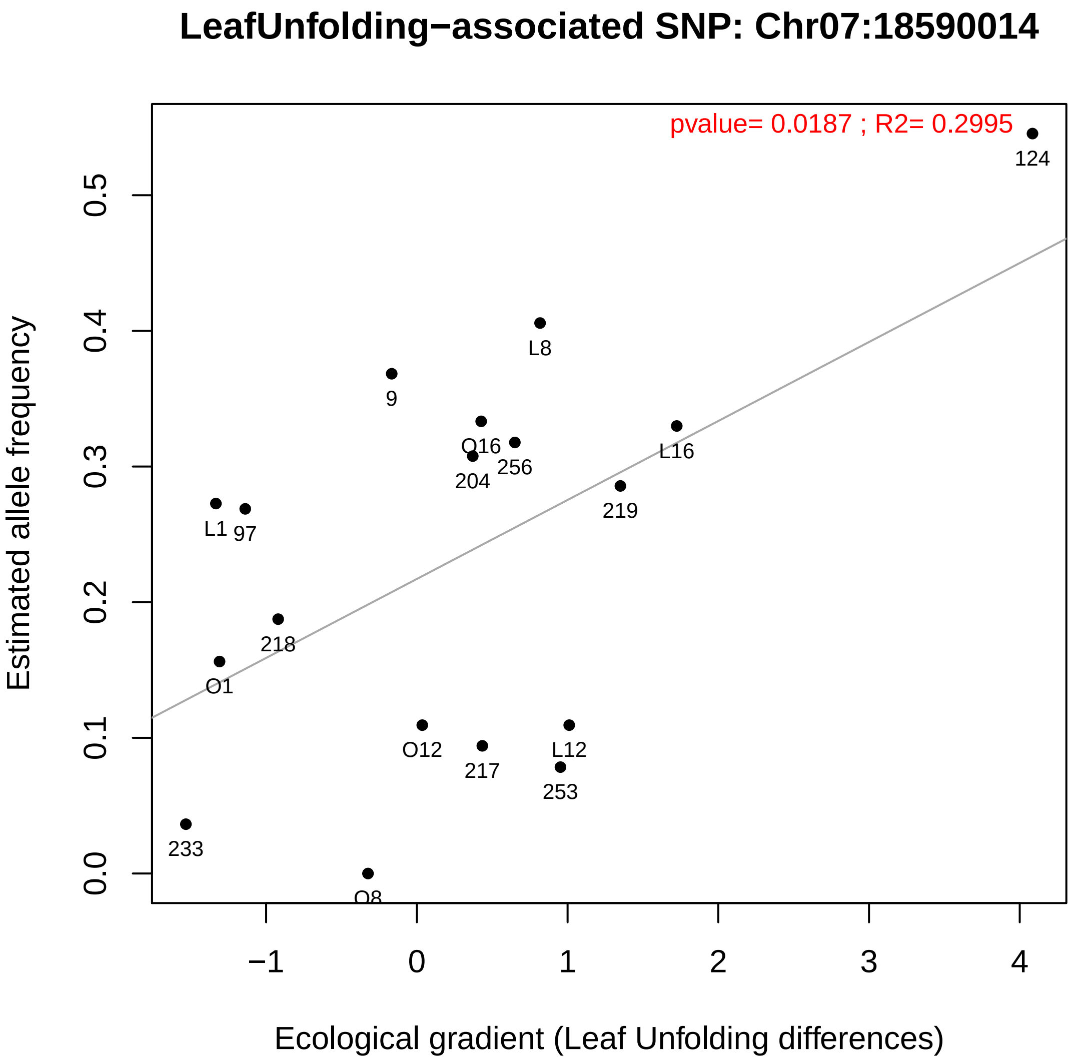
LeafUnfolding−associated SNP: Chr07:18590014
pvalue= 0.0187 ; R2= 0.2995
Estimated allele frequency
Ecological gradient (Leaf Unfolding differences)
0.0
0.1
0.2
0.3
0.4
0.5
−1
0
1
2
3
4
124
L8
9
O16
256
204
L16
219
L1
97
218
O1
O12
217
L12
253
233
O8

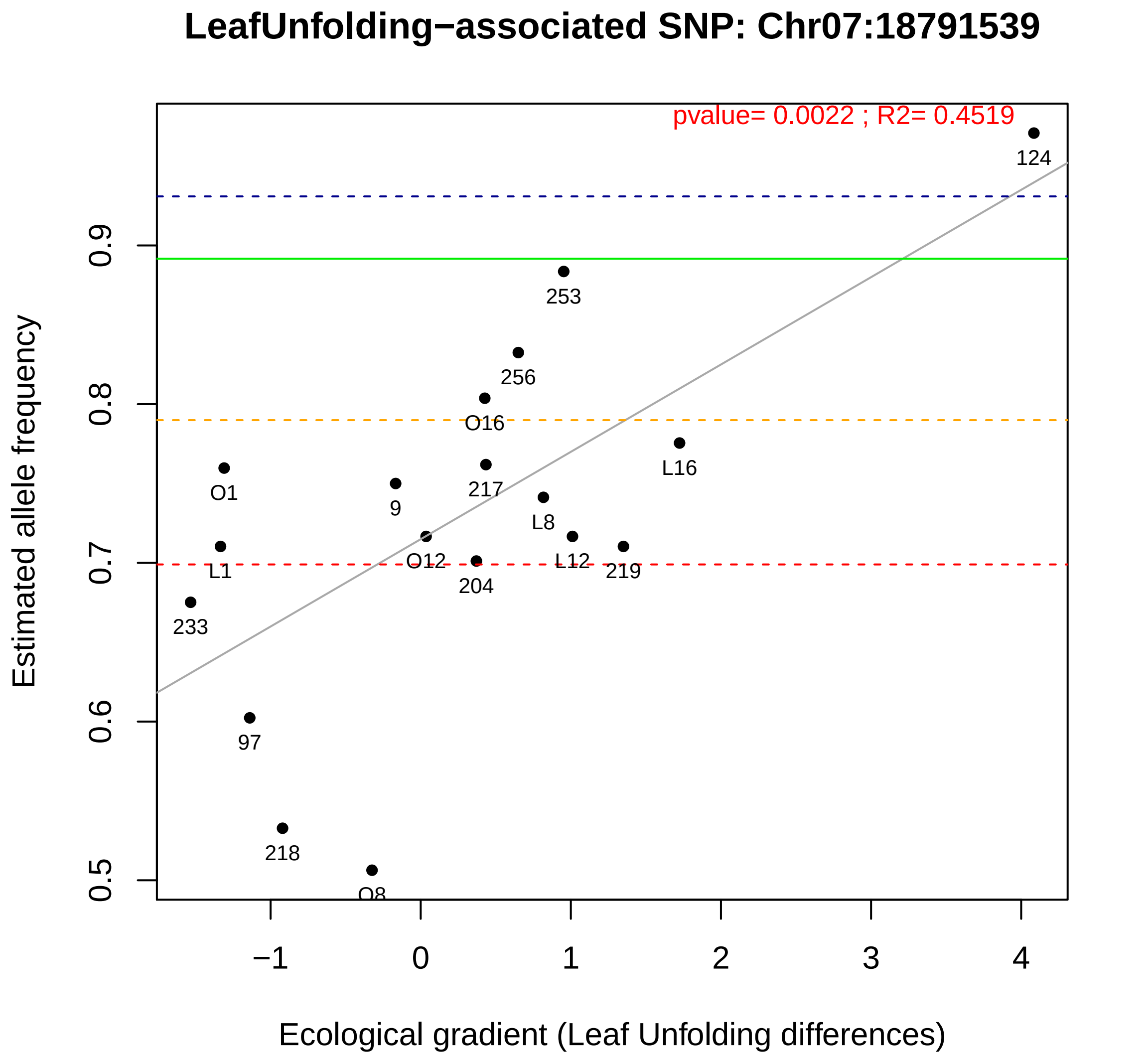

LeafUnfolding−associated SNP: Chr07:18791539
pvalue= 0.0022 ; R2= 0.4519
Estimated allele frequency
Ecological gradient (Leaf Unfolding differences)
0.5
0.6
0.7
0.8
0.9
−1
0
1
2
3
4
124
253
256
O16
L16
O1
9
217
L8
O12
L12
219
L1
204
233
97
218
O8

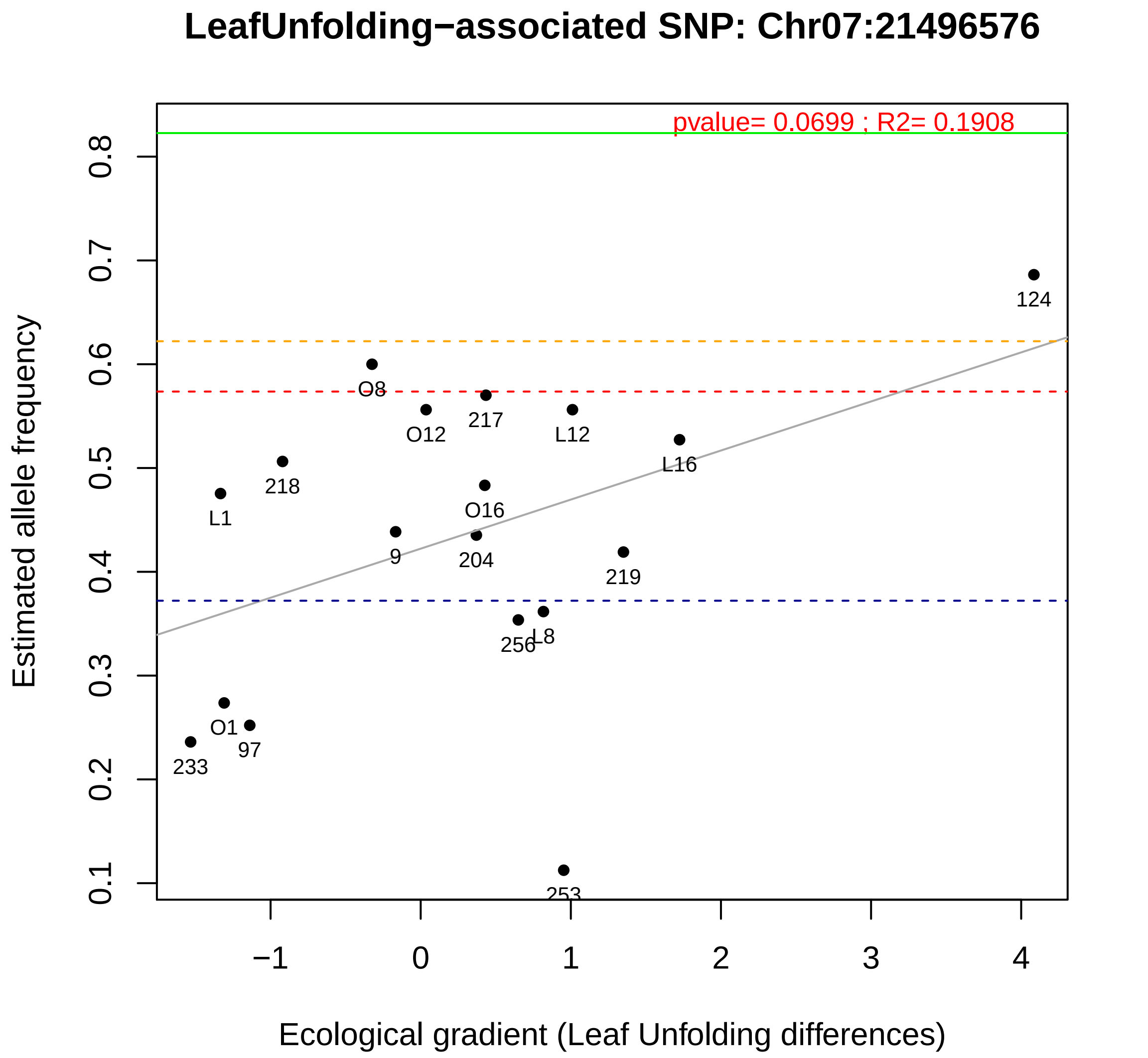
LeafUnfolding−associated SNP: Chr07:21496576
pvalue= 0.0699 ; R2= 0.1908
Estimated allele frequency
0.1
0.2
0.3
0.4
0.5
0.6
0.7
0.8
−1
0
1
2
3
4
Ecological gradient (Leaf Unfolding differences)
124
O8
217
O12
L12
L16
218
L1
O16
9
204
219
256
L8
O1
97
233
253

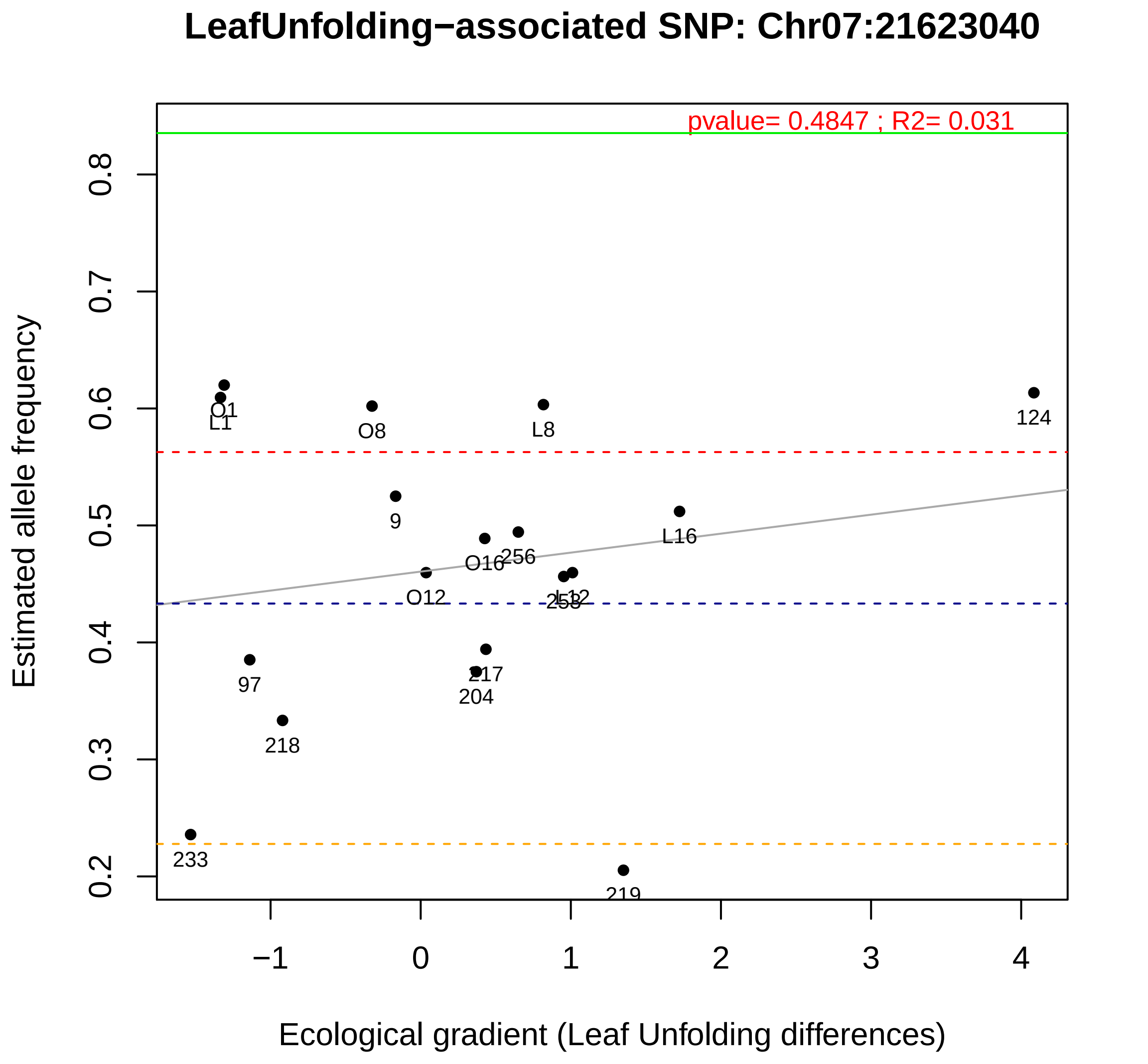
LeafUnfolding−associated SNP: Chr07:21623040
pvalue= 0.4847 ; R2= 0.031
Estimated allele frequency
0.2
0.3
0.4
0.5
0.6
0.7
0.8
Ecological gradient (Leaf Unfolding differences)
−1
0
1
2
3
4
O1
L1
O8
L8
124
9
L16
O16
256
O12
L12
253
97
217
204
218
233
219

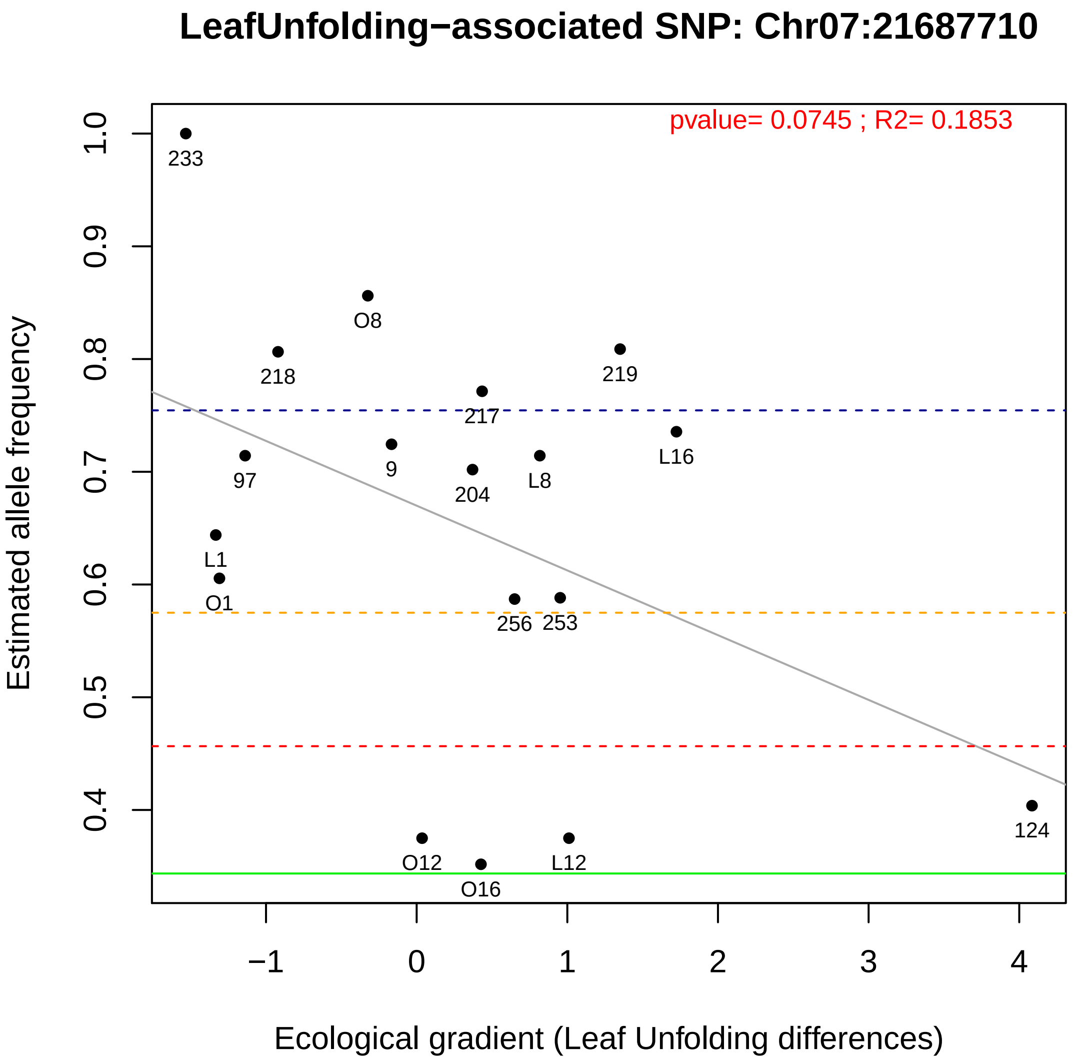

LeafUnfolding−associated SNP: Chr07:21687710
pvalue= 0.0745 ; R2= 0.1853
Estimated allele frequency
1.0
0.9
0.8
0.7
0.6
0.5
0.4
233
O8
218
219
217
L16
9
97
204
L8
L1
O1
256
253
124
O12
L12
O16
−1
0
1
2
3
4
Ecological gradient (Leaf Unfolding differences)

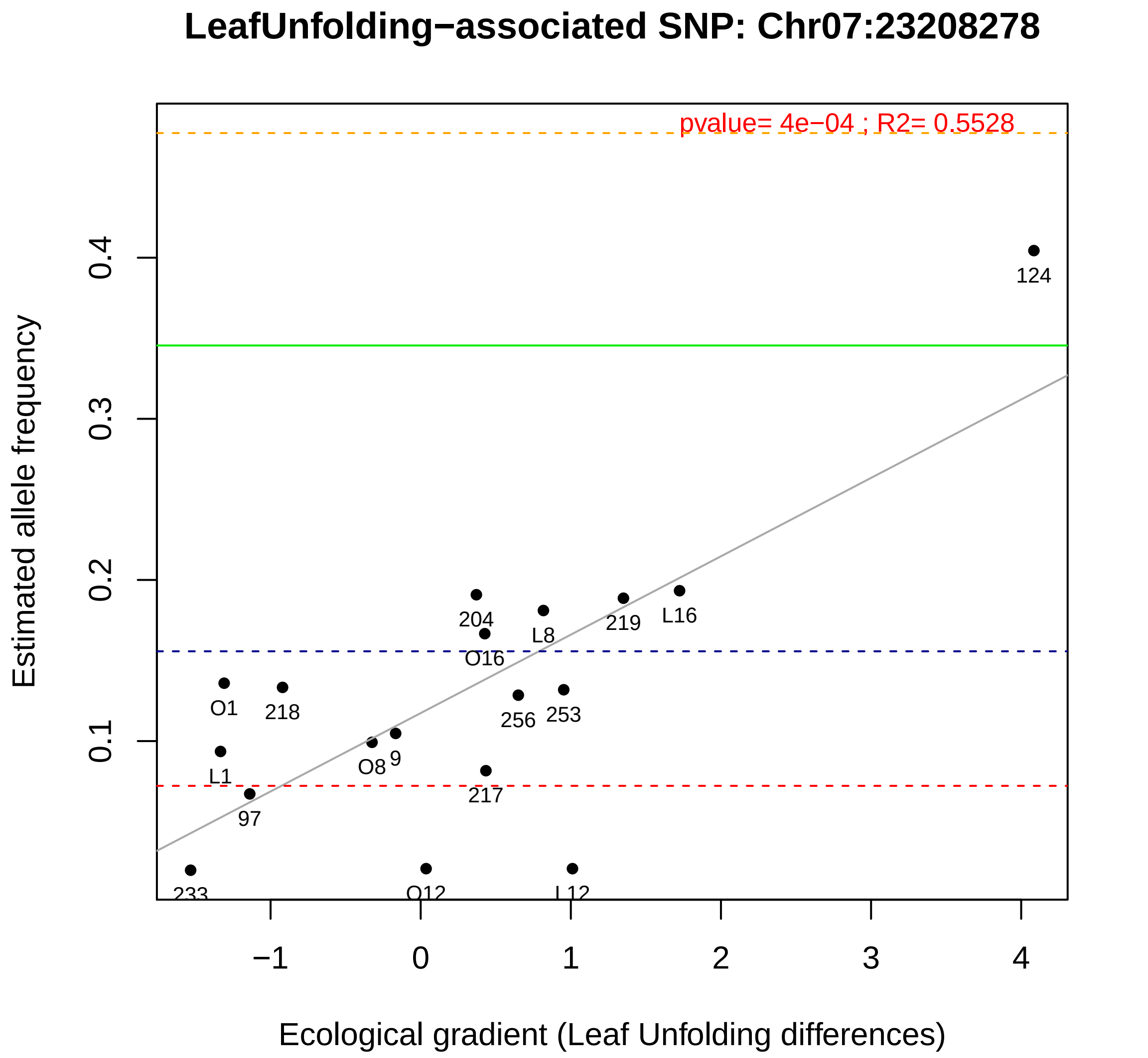

LeafUnfolding−associated SNP: Chr07:23208278
pvalue= 4e−04 ; R2= 0.5528
Estimated allele frequency
0.1
0.2
0.3
0.4
124
204
L8
219
L16
O16
O1
218
256
253
O8
9
L1
217
97
233
O12
L12
−1
0
1
2
3
4
Ecological gradient (Leaf Unfolding differences)

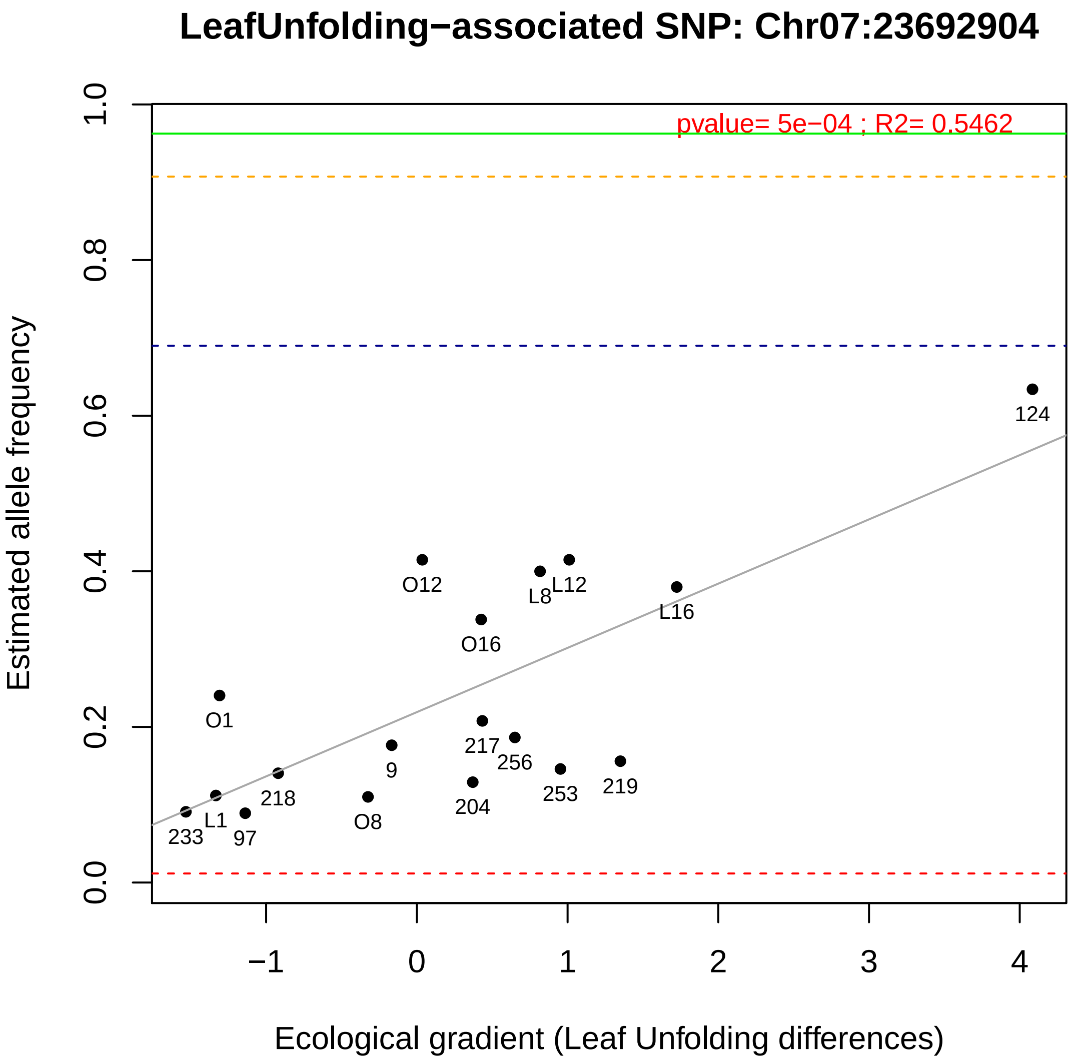
LeafUnfolding−associated SNP: Chr07:23692904
pvalue= 5e−04 ; R2= 0.5462
Estimated allele frequency
0.0
0.2
0.4
0.6
0.8
1.0
−1
0
1
2
3
4
Ecological gradient (Leaf Unfolding differences)
124
O12
L8
L12
L16
O16
O1
217
256
9
218
253
219
204
O8
233
L1
97

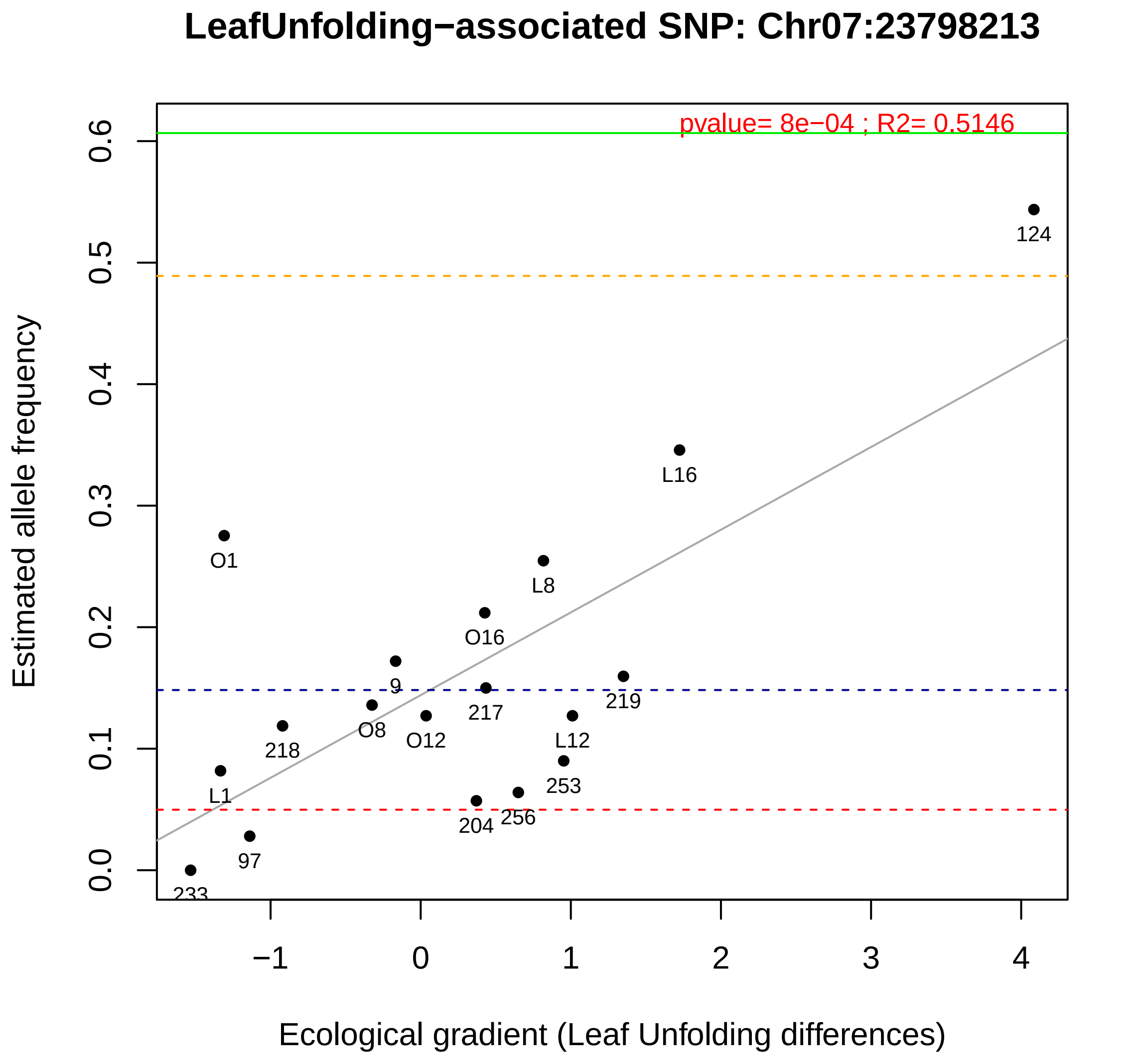

LeafUnfolding−associated SNP: Chr07:23798213
pvalue= 8e−04 ; R2= 0.5146
Estimated allele frequency
0.0
0.1
0.2
0.3
0.4
0.5
0.6
Ecological gradient (Leaf Unfolding differences)
−1
0
1
2
3
4
124
L16
O1
L8
O16
9
217
219
O8
O12
L12
218
253
L1
204
256
97
233

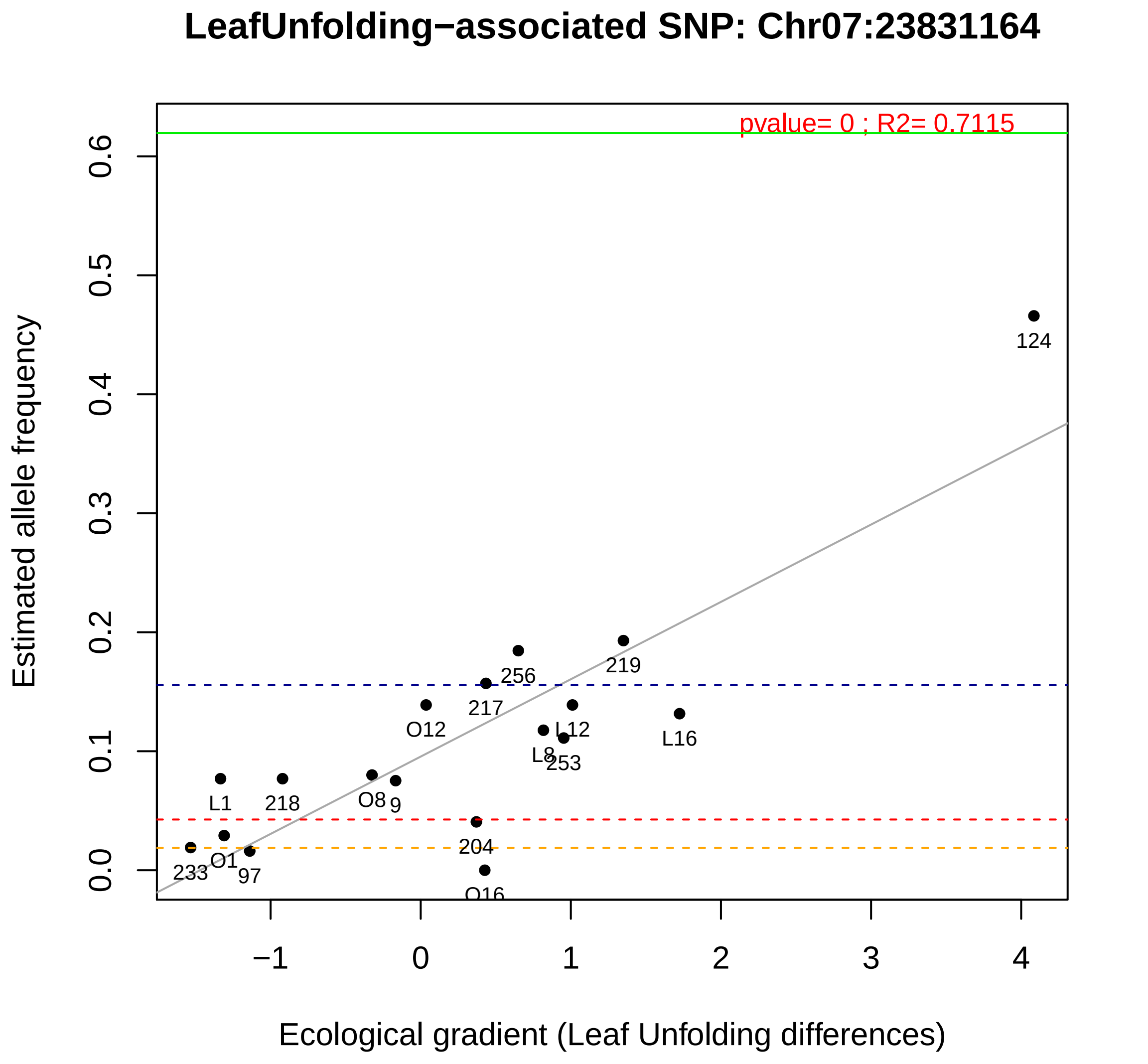

LeafUnfolding−associated SNP: Chr07:23831164
pvalue= 0 ; R2= 0.7115
Estimated allele frequency
0.0
0.1
0.2
0.3
0.4
0.5
0.6
−1
0
1
2
3
4
Ecological gradient (Leaf Unfolding differences)
124
219
256
217
O12
L12
L16
L8
253
L1
218
O8
9
204
O16
233
O1
97

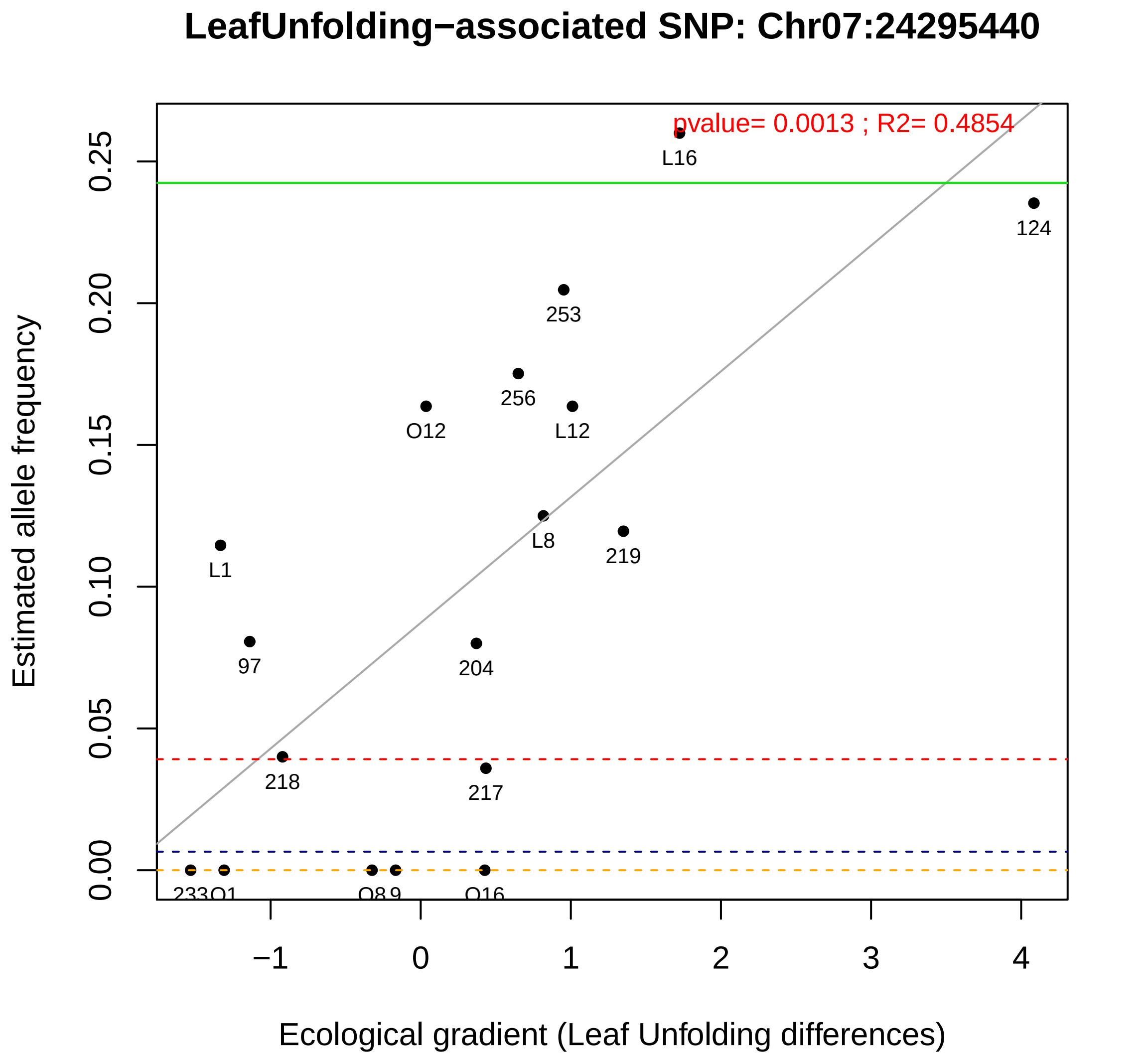
LeafUnfolding−associated SNP: Chr07:24295440
pvalue= 0.0013 ; R2= 0.4854
Estimated allele frequency
0.00
0.05
0.10
0.15
0.20
0.25
−1
0
1
2
3
4
Ecological gradient (Leaf Unfolding differences)
L16
124
253
256
O12
L12
L8
219
L1
97
204
218
217
233
O1
O8
9
O16

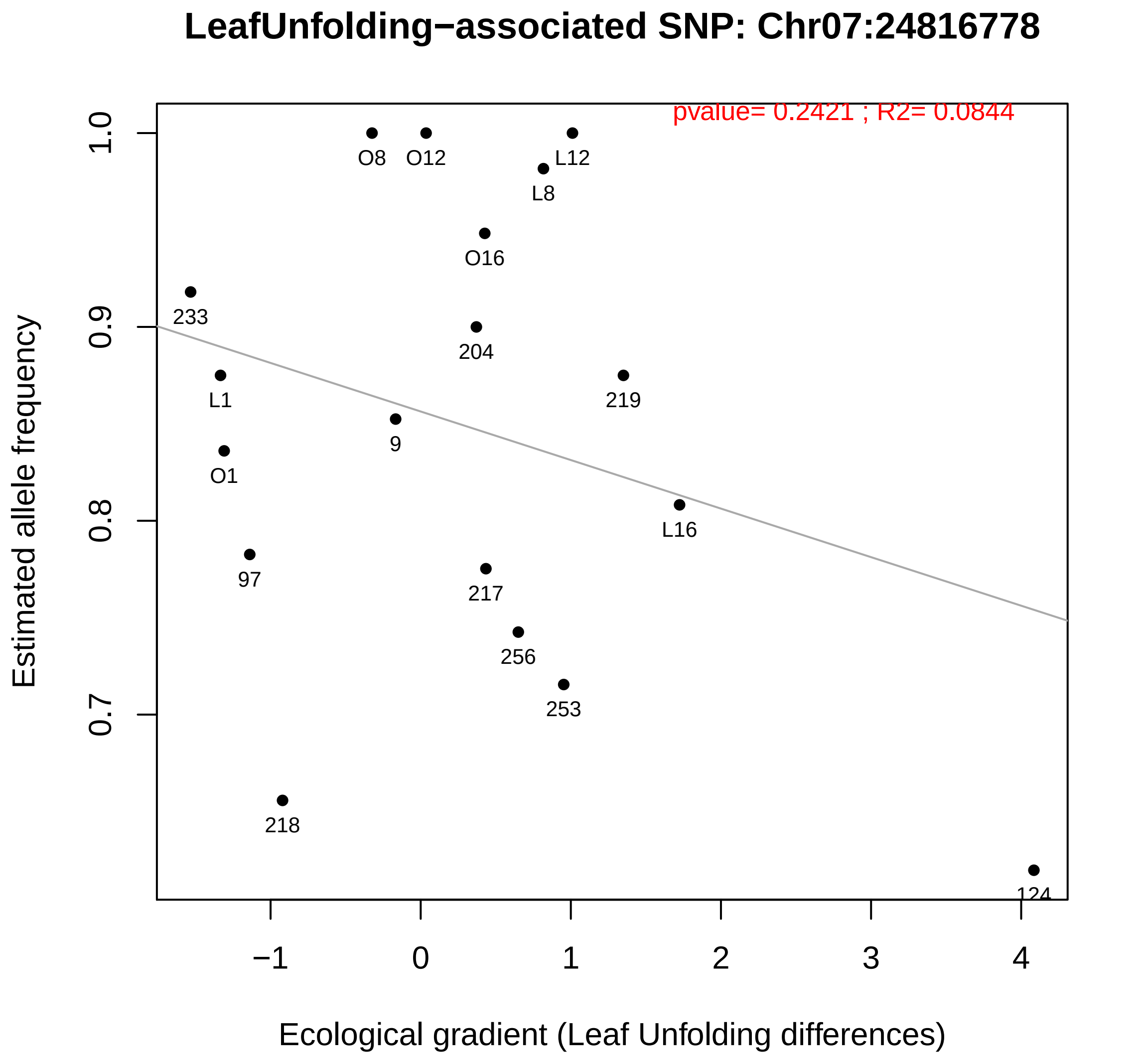
LeafUnfolding−associated SNP: Chr07:24816778
pvalue= 0.2421 ; R2= 0.0844
Estimated allele frequency
1.0
0.9
0.8
0.7
−1
0
1
2
3
4
Ecological gradient (Leaf Unfolding differences)
O8
O12
L12
L8
O16
233
204
L1
219
9
O1
L16
97
217
256
253
218
124

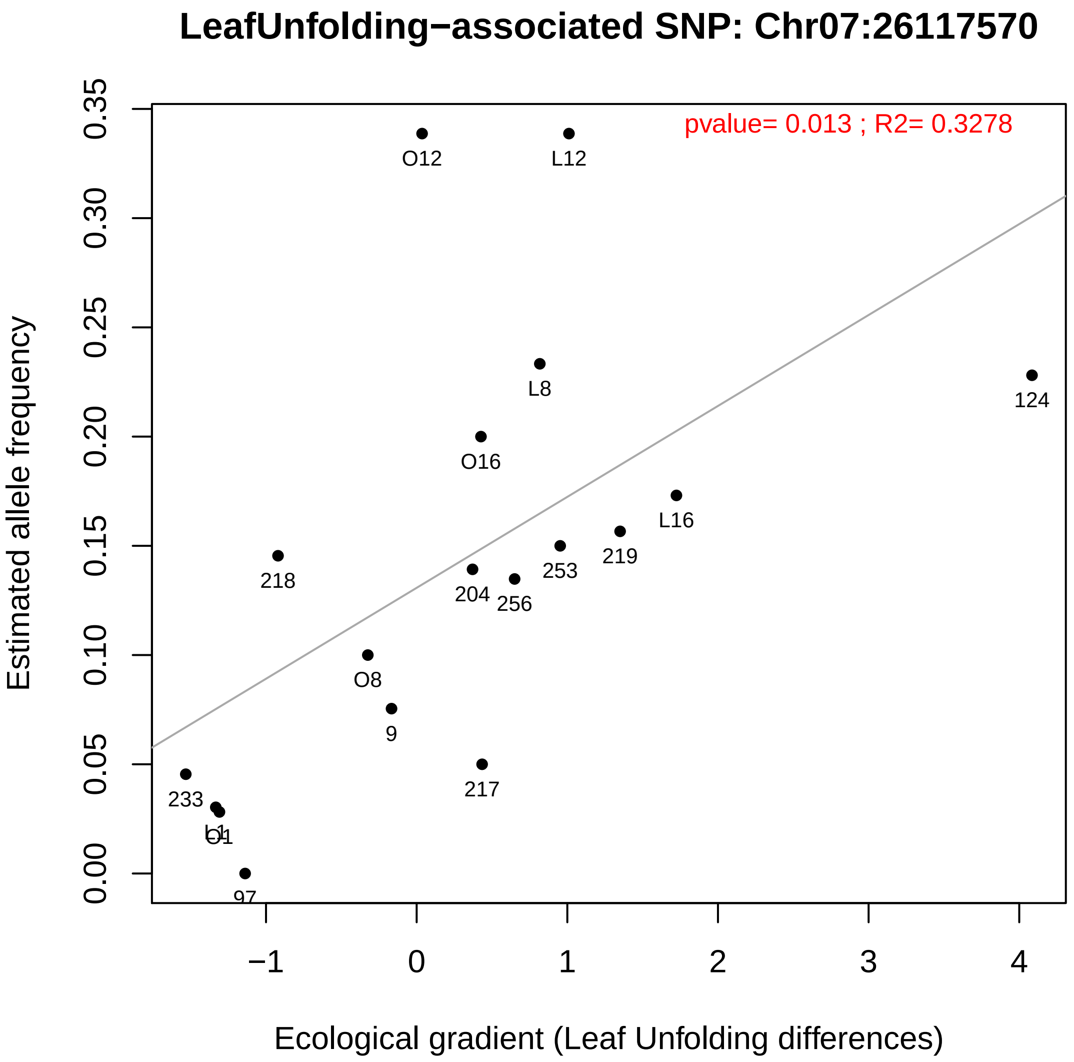

LeafUnfolding−associated SNP: Chr07:26117570
pvalue= 0.013 ; R2= 0.3278
Estimated allele frequency
Ecological gradient (Leaf Unfolding differences)
0.00
0.05
0.10
0.15
0.20
0.25
0.30
0.35
−1
0
1
2
3
4
O12
L12
L8
124
O16
L16
219
253
218
204
256
O8
9
217
233
L1
O1
97

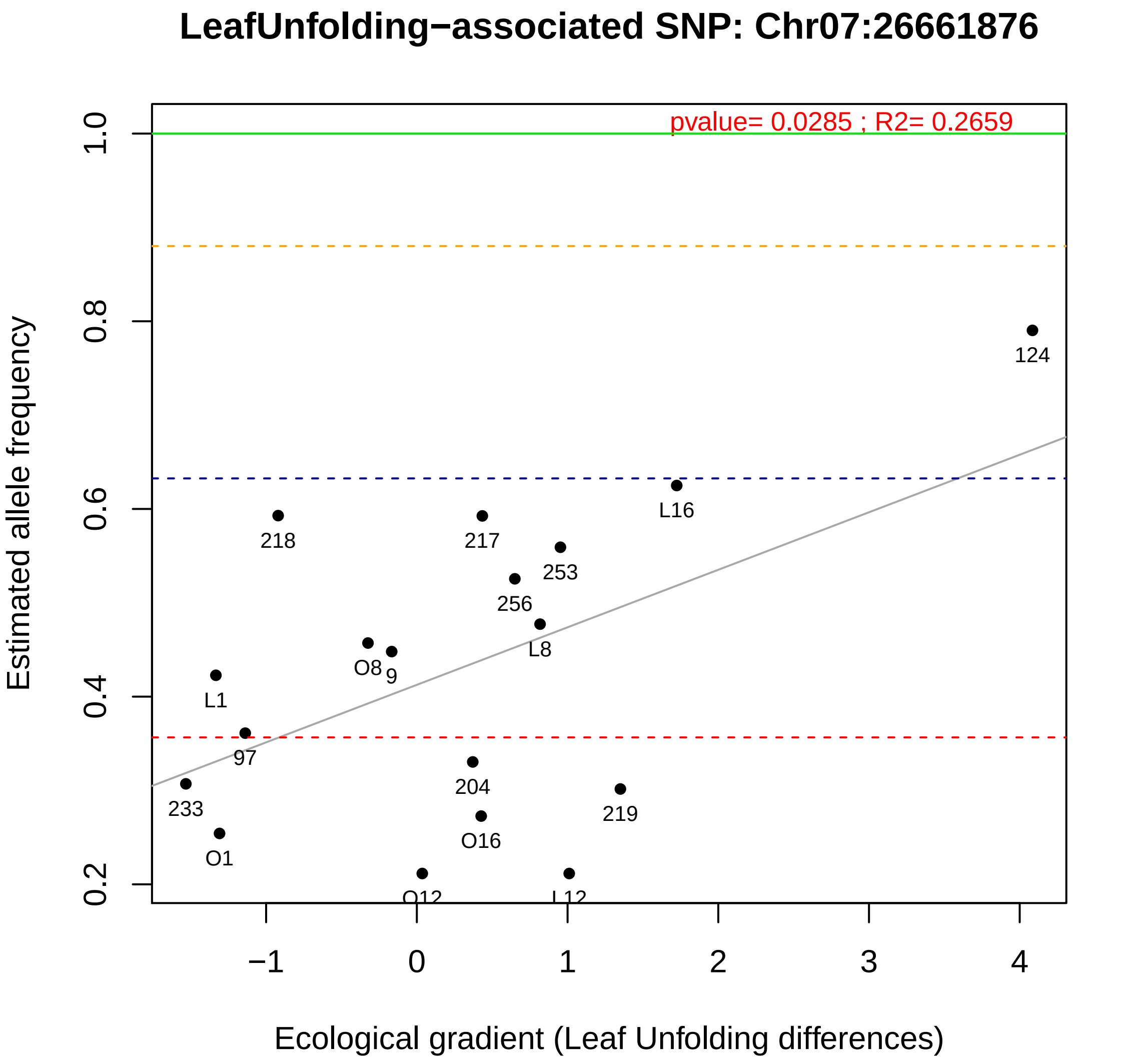
LeafUnfolding−associated SNP: Chr07:26661876
pvalue= 0.0285 ; R2= 0.2659
Estimated allele frequency
Ecological gradient (Leaf Unfolding differences)
0.2
0.4
0.6
0.8
1.0
−1
0
1
2
3
4
124
L16
218
217
253
256
L8
O8
9
L1
97
204
233
219
O16
O1
O12
L12

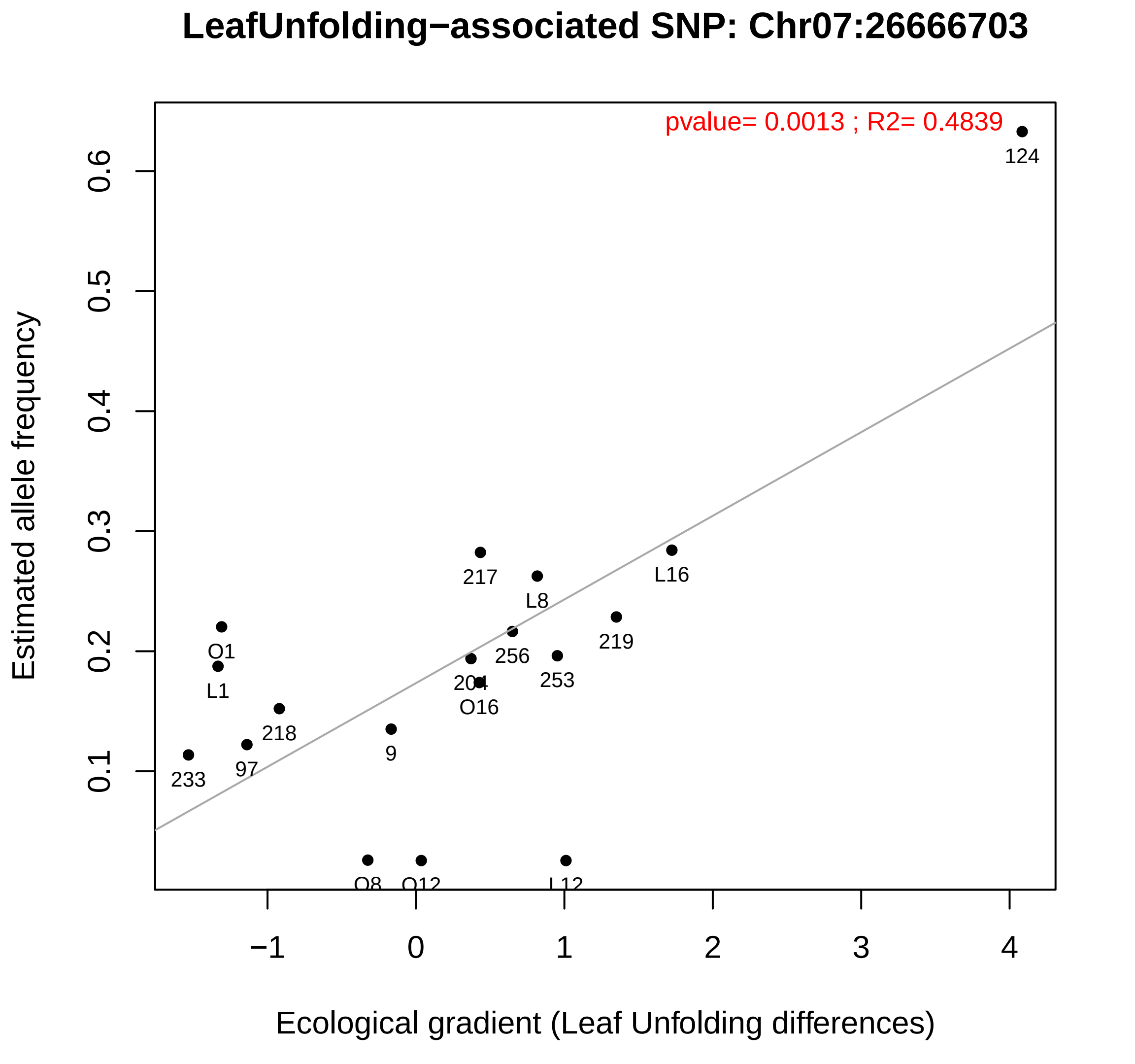

LeafUnfolding−associated SNP: Chr07:26666703
pvalue= 0.0013 ; R2= 0.4839
Estimated allele frequency
0.1
0.2
0.3
0.4
0.5
0.6
Ecological gradient (Leaf Unfolding differences)
−1
0
1
2
3
4
124
217
L8
L16
219
256
253
204
O16
O1
L1
218
97
233
9
O8
O12
L12

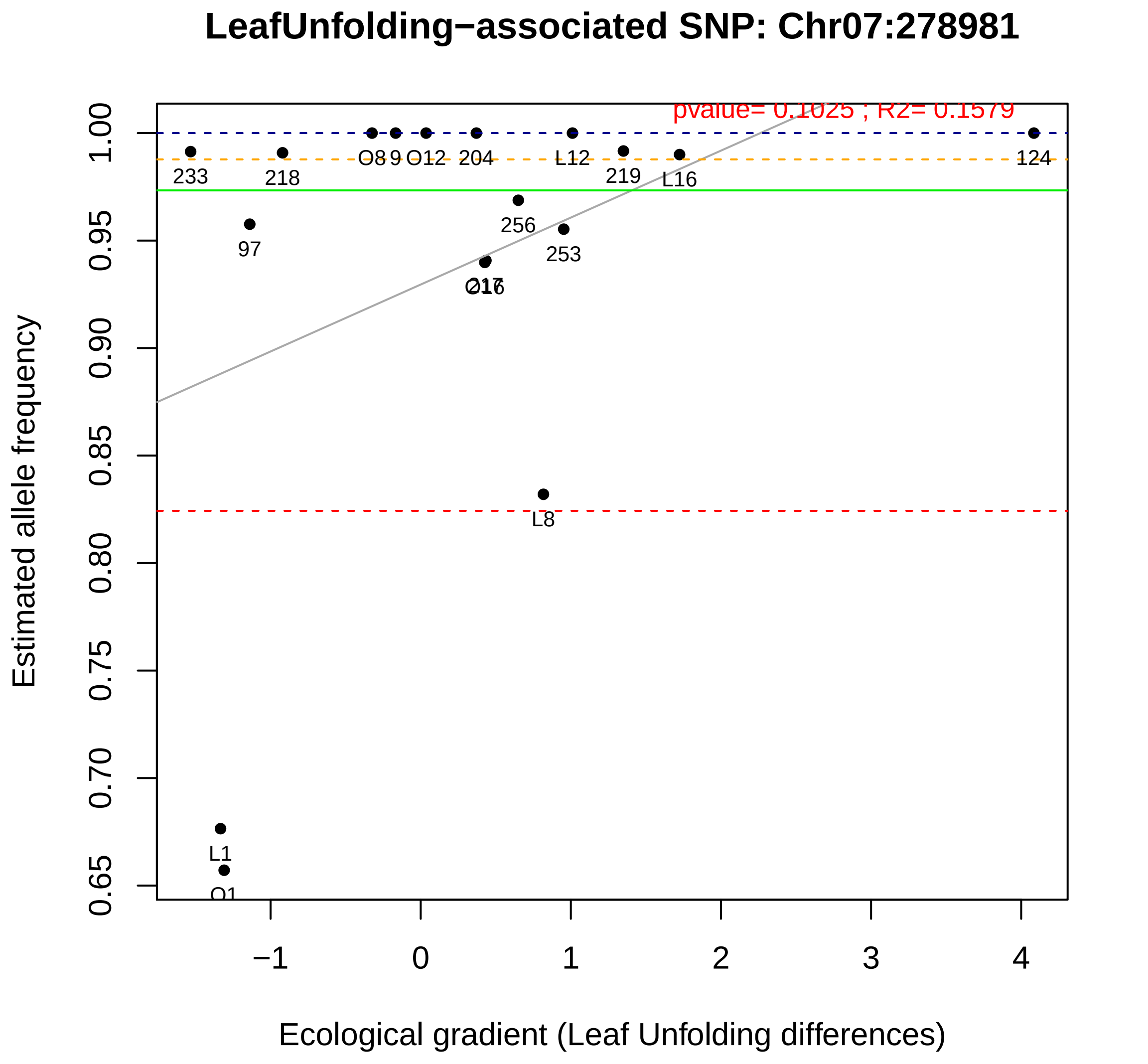

LeafUnfolding−associated SNP: Chr07:278981
pvalue= 0.1025 ; R2= 0.1579
Estimated allele frequency
Ecological gradient (Leaf Unfolding differences)
1.00
0.95
0.90
0.85
0.80
0.75
0.70
0.65
−1
0
1
2
3
4
233
218
97
O8
9
O12
204
L12
219
L16
124
256
253
L8
L1
O1

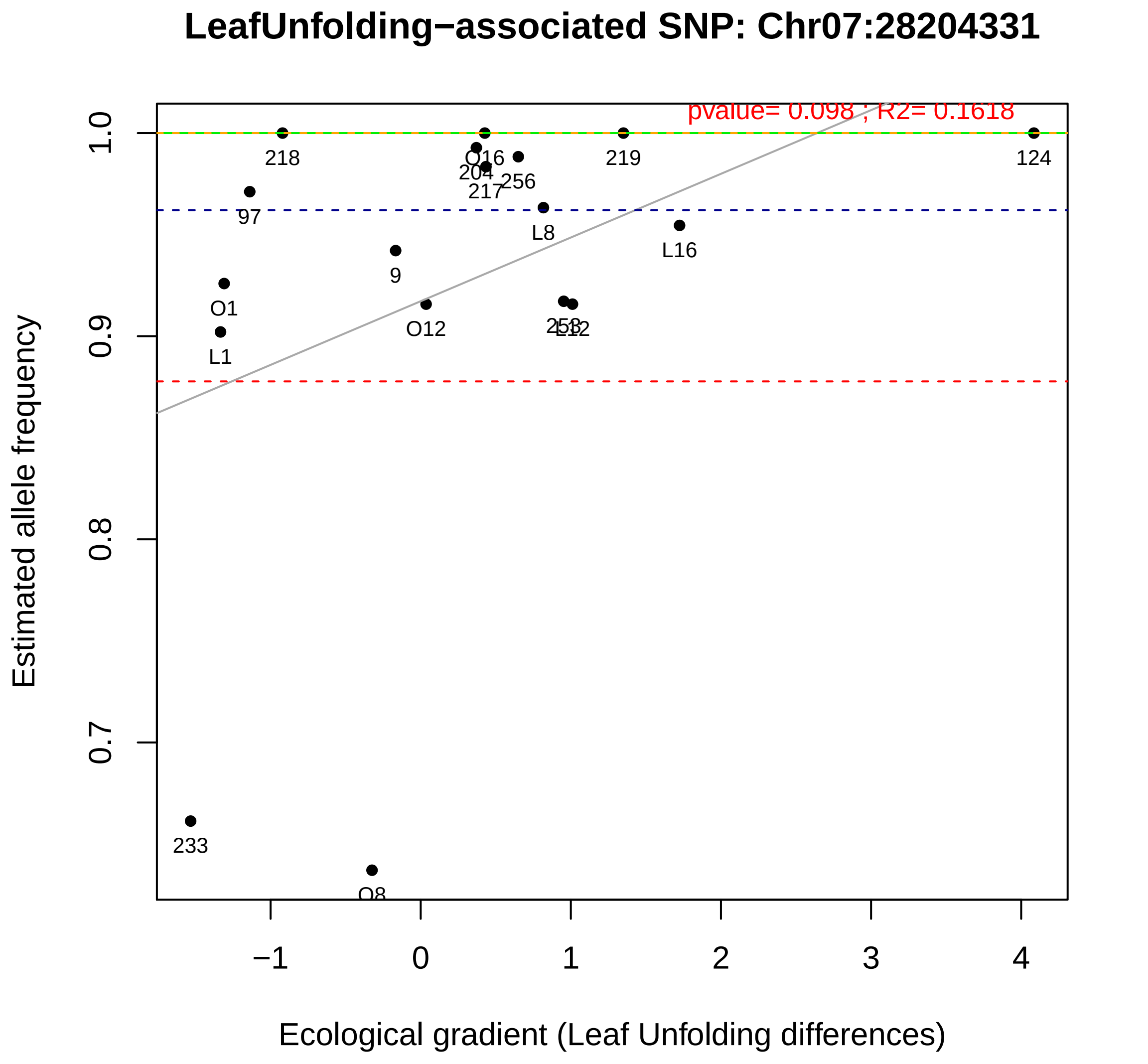

LeafUnfolding−associated SNP: Chr07:28204331
pvalue= 0.098 ; R2= 0.1618
Estimated allele frequency
Ecological gradient (Leaf Unfolding differences)
1.0
0.9
0.8
0.7
−1
0
1
2
3
4
218
O16
204
217
256
219
124
97
L8
L16
9
O1
O12
253
L12
L1
233
O8

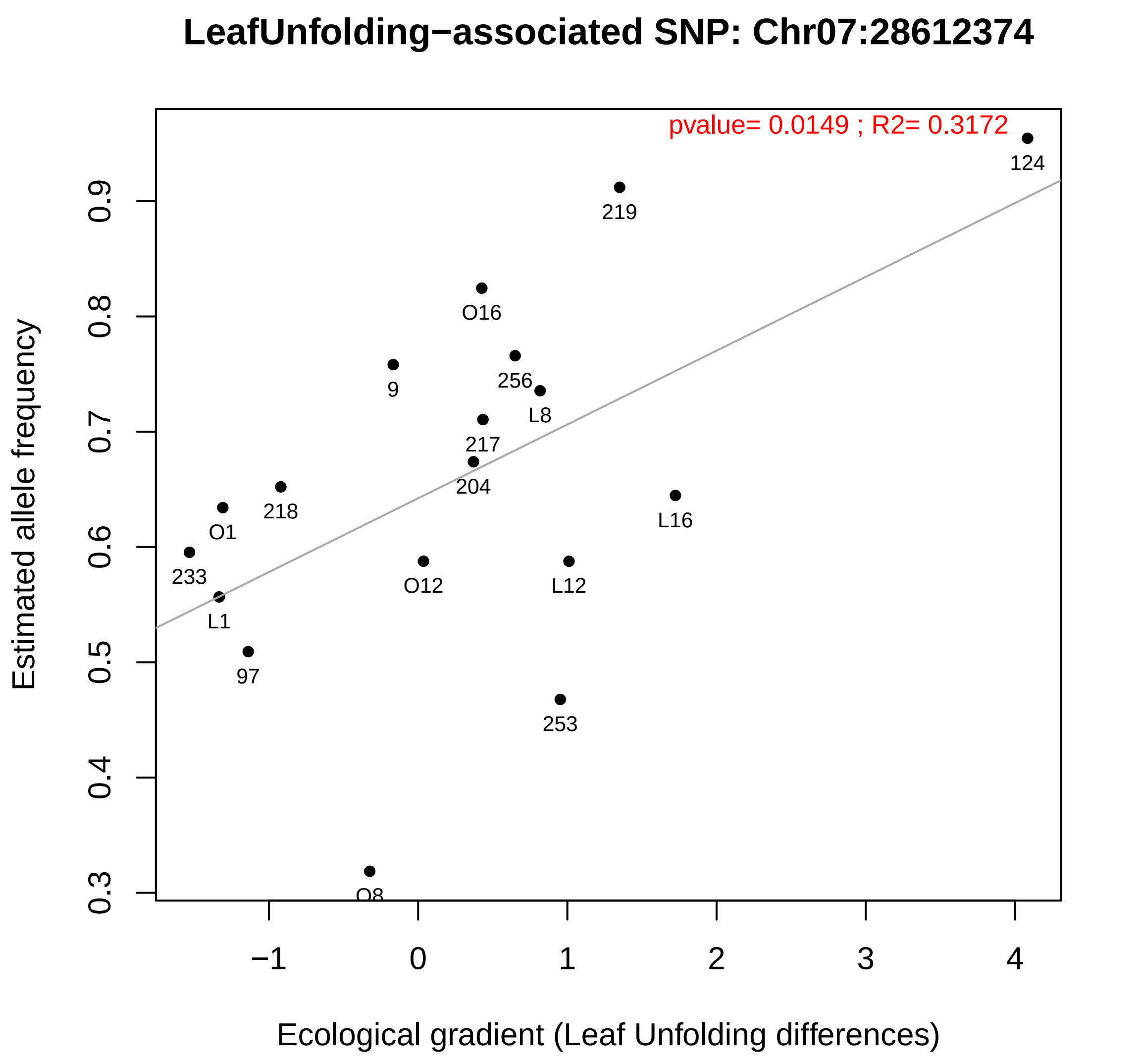
LeafUnfolding−associated SNP: Chr07:28612374
pvalue= 0.0149 ; R2= 0.3172
Estimated allele frequency
Ecological gradient (Leaf Unfolding differences)
0.3
0.4
0.5
0.6
0.7
0.8
0.9
−1
0
1
2
3
4
124
219
O16
256
9
L8
217
204
218
O1
L16
233
O12
L12
L1
97
253
O8

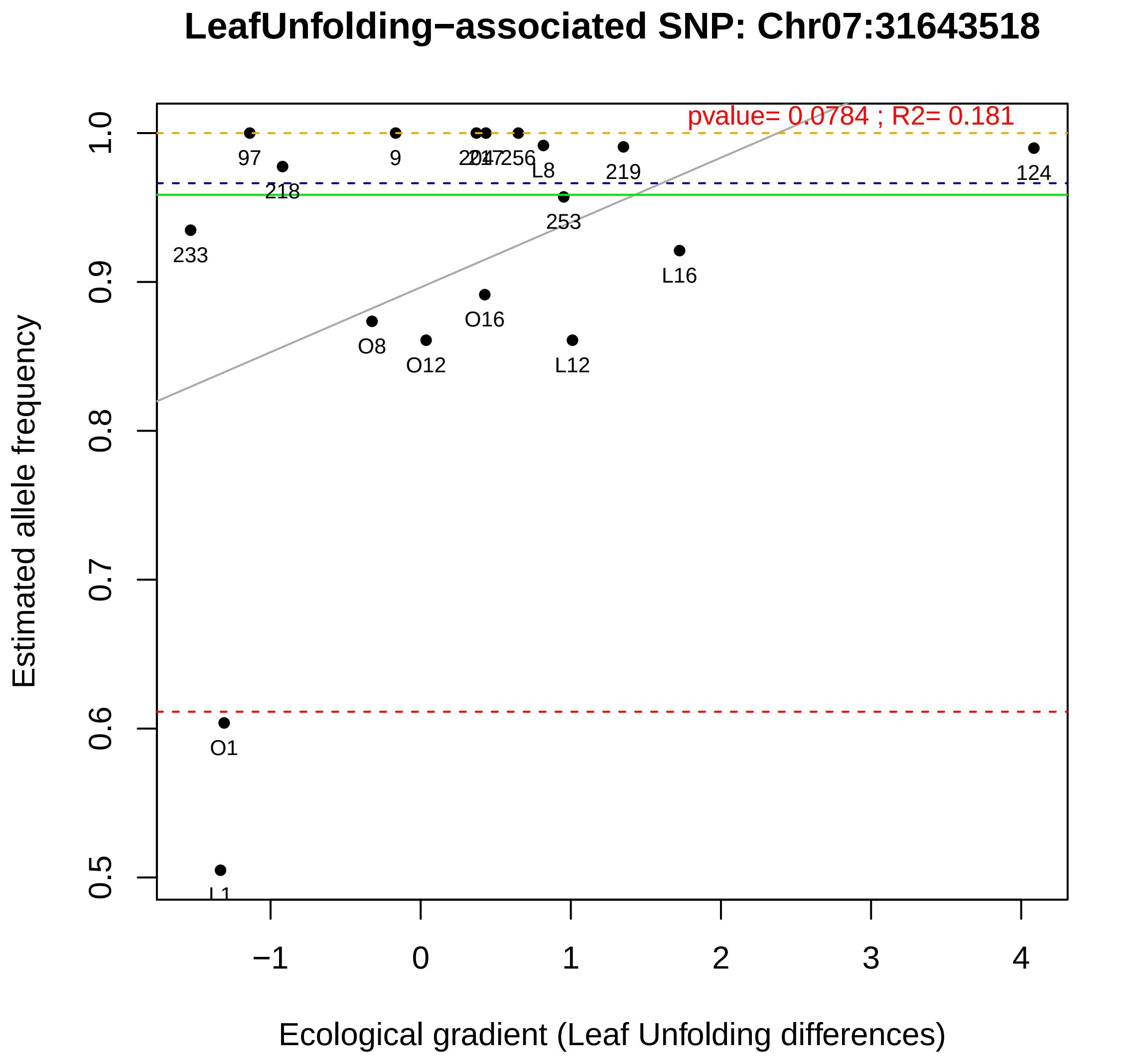

LeafUnfolding−associated SNP: Chr07:31643518
pvalue= 0.0784 ; R2= 0.181
Estimated allele frequency
Ecological gradient (Leaf Unfolding differences)
1.0
0.9
0.8
0.7
0.6
0.5
−1
0
1
2
3
4
97
9
256
L8
219
124
218
253
233
L16
O16
O8
O12
L12
O1
L1

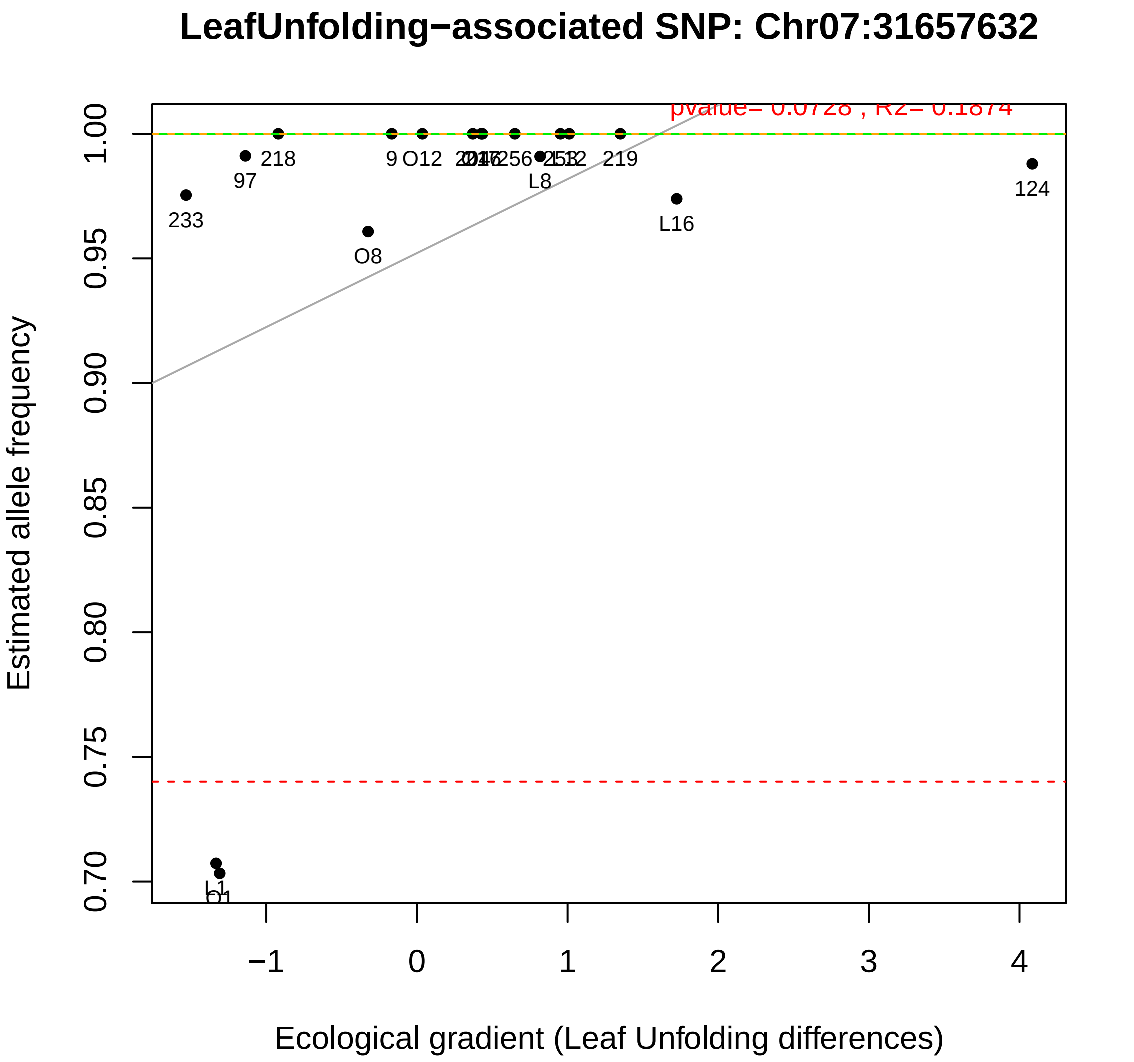
LeafUnfolding−associated SNP: Chr07:31657632
pvalue= 0.0728 , R2= 0.1874
Estimated allele frequency
Ecological gradient (Leaf Unfolding differences)
1.00
0.95
0.90
0.85
0.80
0.75
0.70
−1
0
1
2
3
4
218
97
233
9
O12
O8
219
L8
L16
124
L1
O1

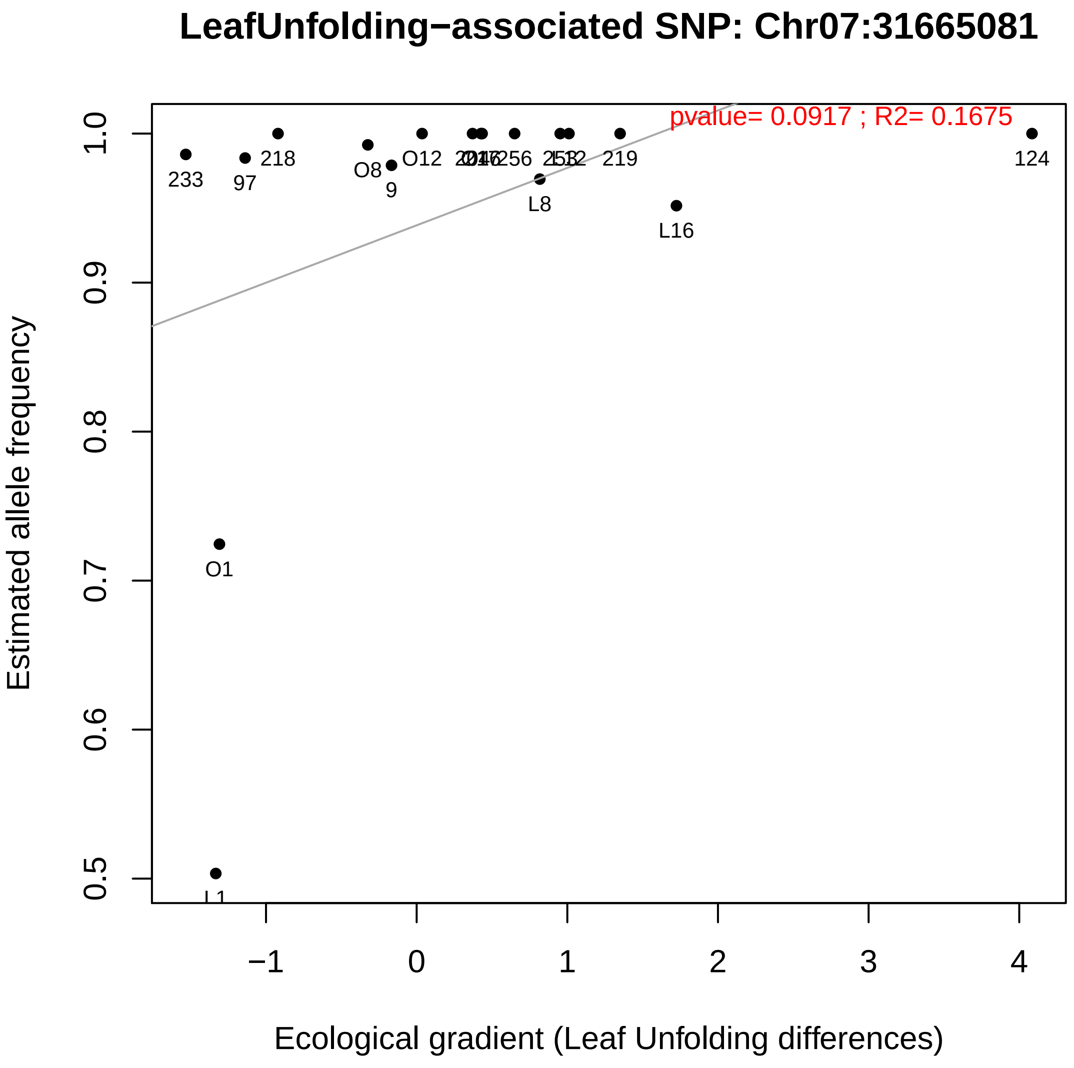

LeafUnfolding−associated SNP: Chr07:31665081
pvalue= 0.0917 ; R2= 0.1675
Estimated allele frequency
Ecological gradient (Leaf Unfolding differences)
0.5
0.6
0.7
0.8
0.9
1.0
−1
0
1
2
3
4
233
97
218
O8
9
O12
256
L8
219
L16
124
O1

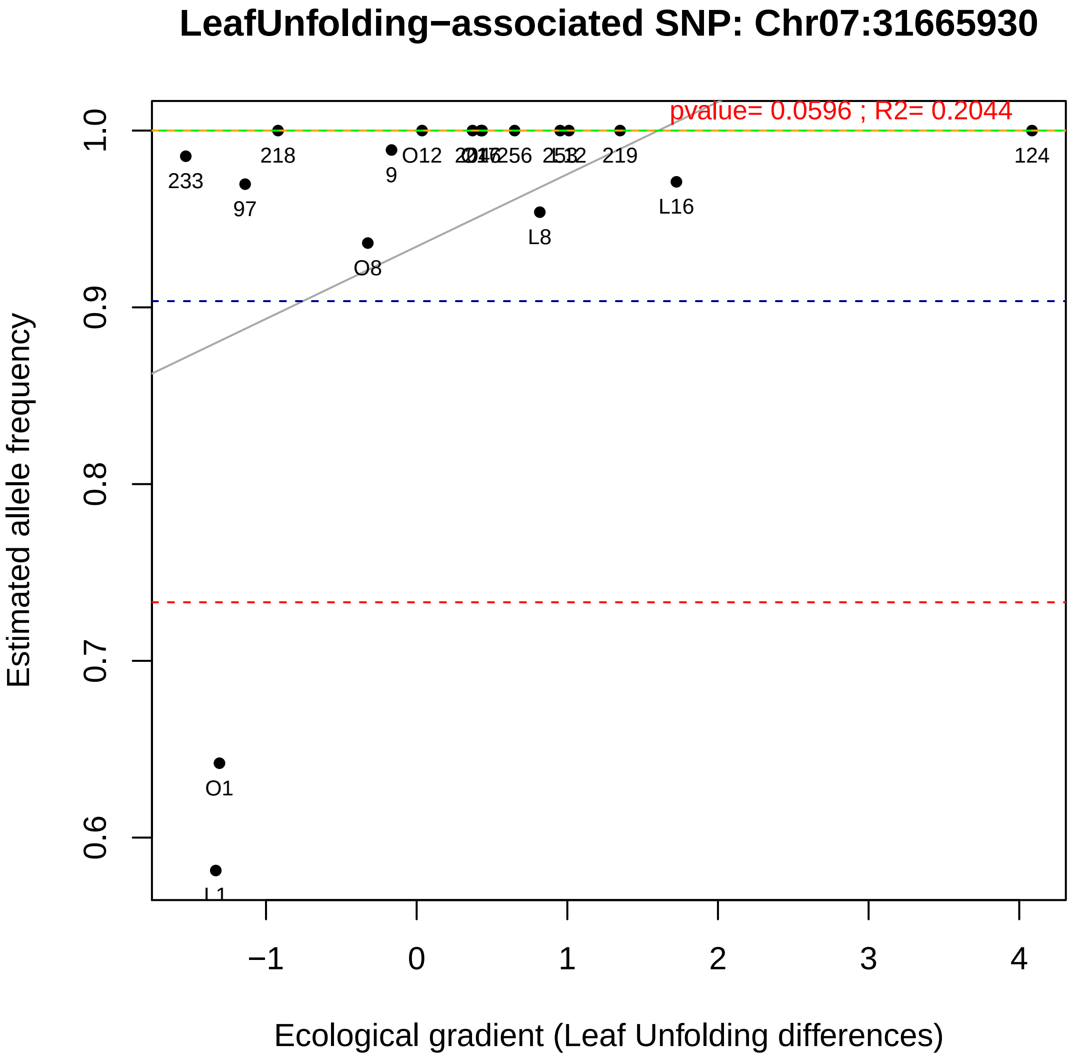
LeafUnfolding−associated SNP: Chr07:31665930
pvalue= 0.0596 ; R2= 0.2044
Estimated allele frequency
Ecological gradient (Leaf Unfolding differences)
1.0
0.9
0.8
0.7
0.6
−1
0
1
2
3
4
233
218
97
9
O12
256
219
124
O8
L8
L16
O1

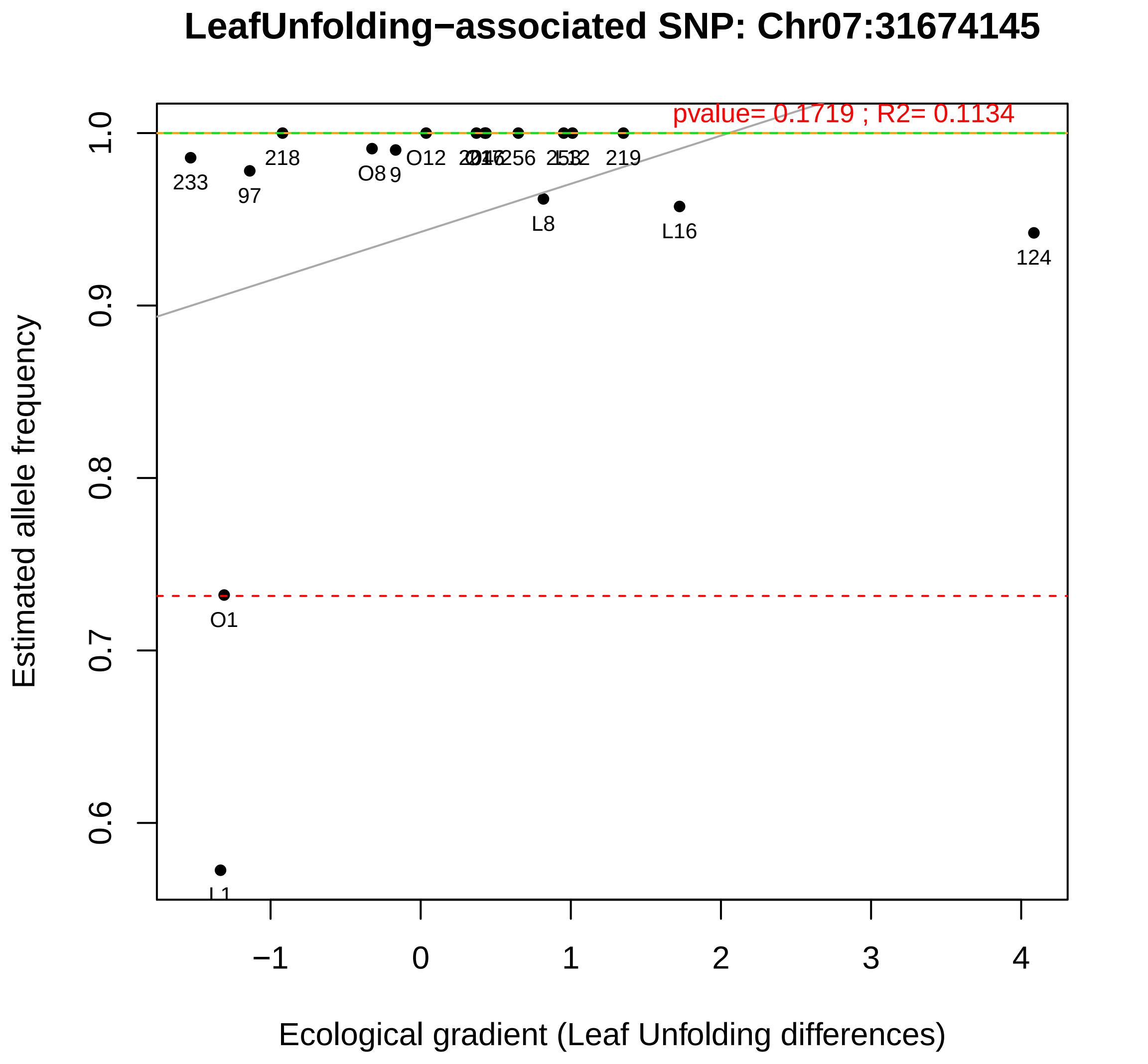

LeafUnfolding−associated SNP: Chr07:31674145
pvalue= 0.1719 ; R2= 0.1134
Estimated allele frequency
Ecological gradient (Leaf Unfolding differences)
1.0
0.9
0.8
0.7
0.6
−1
0
1
2
3
4
233
97
218
O8
9
O12
L8
219
L16
124
O1

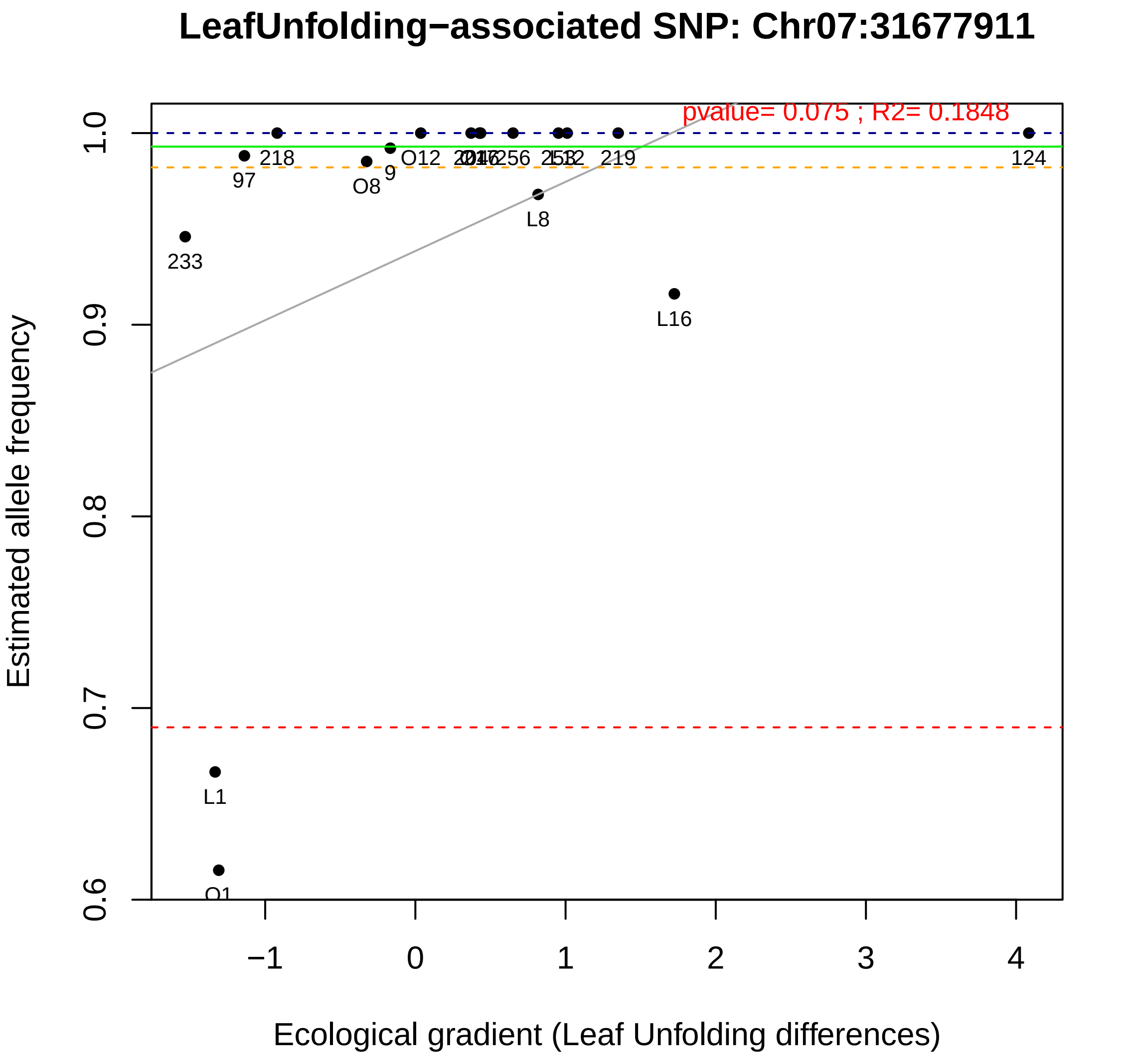
LeafUnfolding−associated SNP: Chr07:31677911
pvalue= 0.075 ; R2= 0.1848
Estimated allele frequency
Ecological gradient (Leaf Unfolding differences)
1.0
0.9
0.8
0.7
0.6
−1
0
1
2
3
4
218
97
O8
9
O12
233
L8
219
124
L16
L1
O1

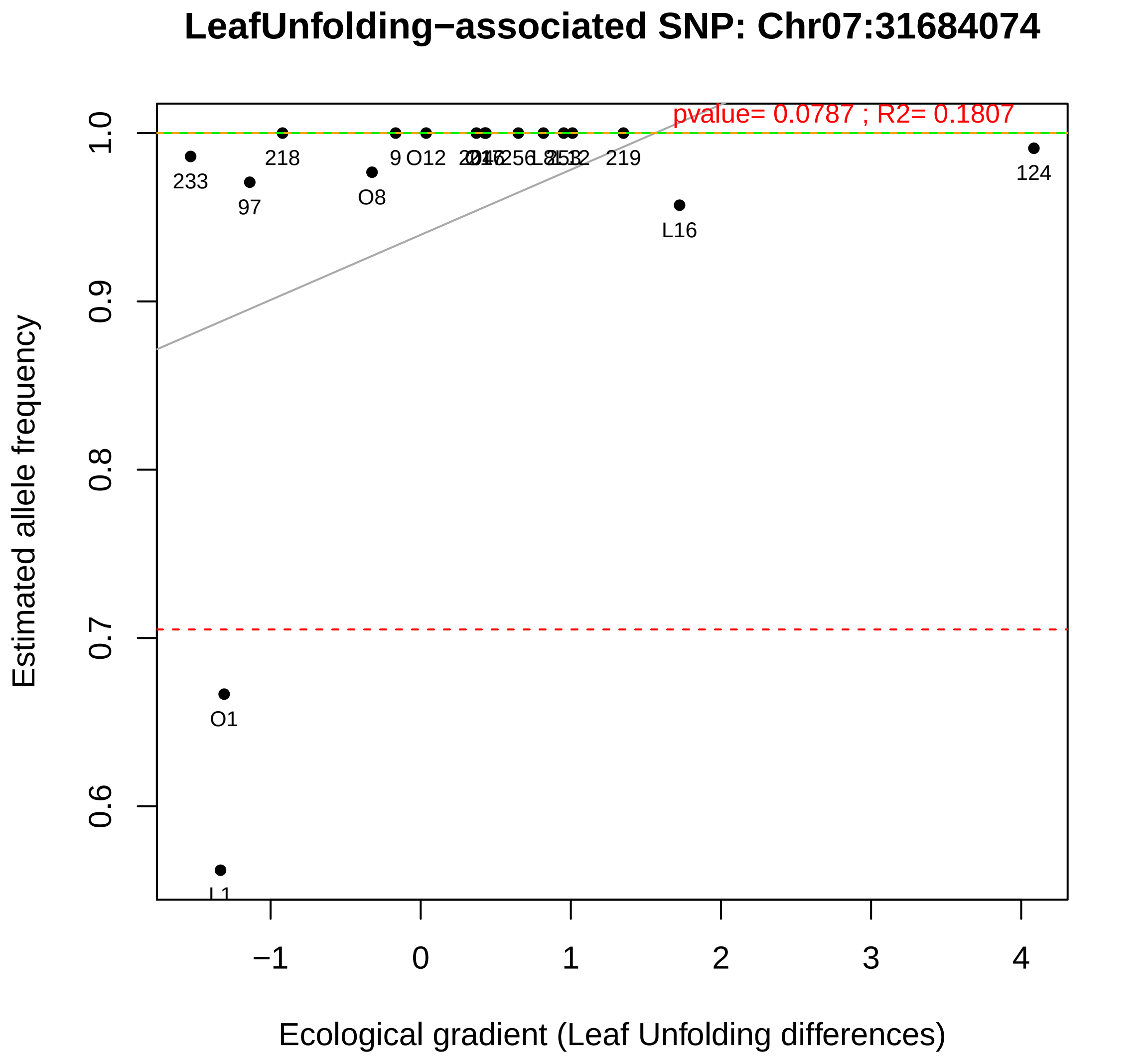

LeafUnfolding−associated SNP: Chr07:31684074
pvalue= 0.0787 ; R2= 0.1807
Estimated allele frequency
1.0
0.9
0.8
0.7
0.6
233
218
97
O8
9
O12
219
124
L16
O1
L1
−1
0
1
2
3
4
Ecological gradient (Leaf Unfolding differences)

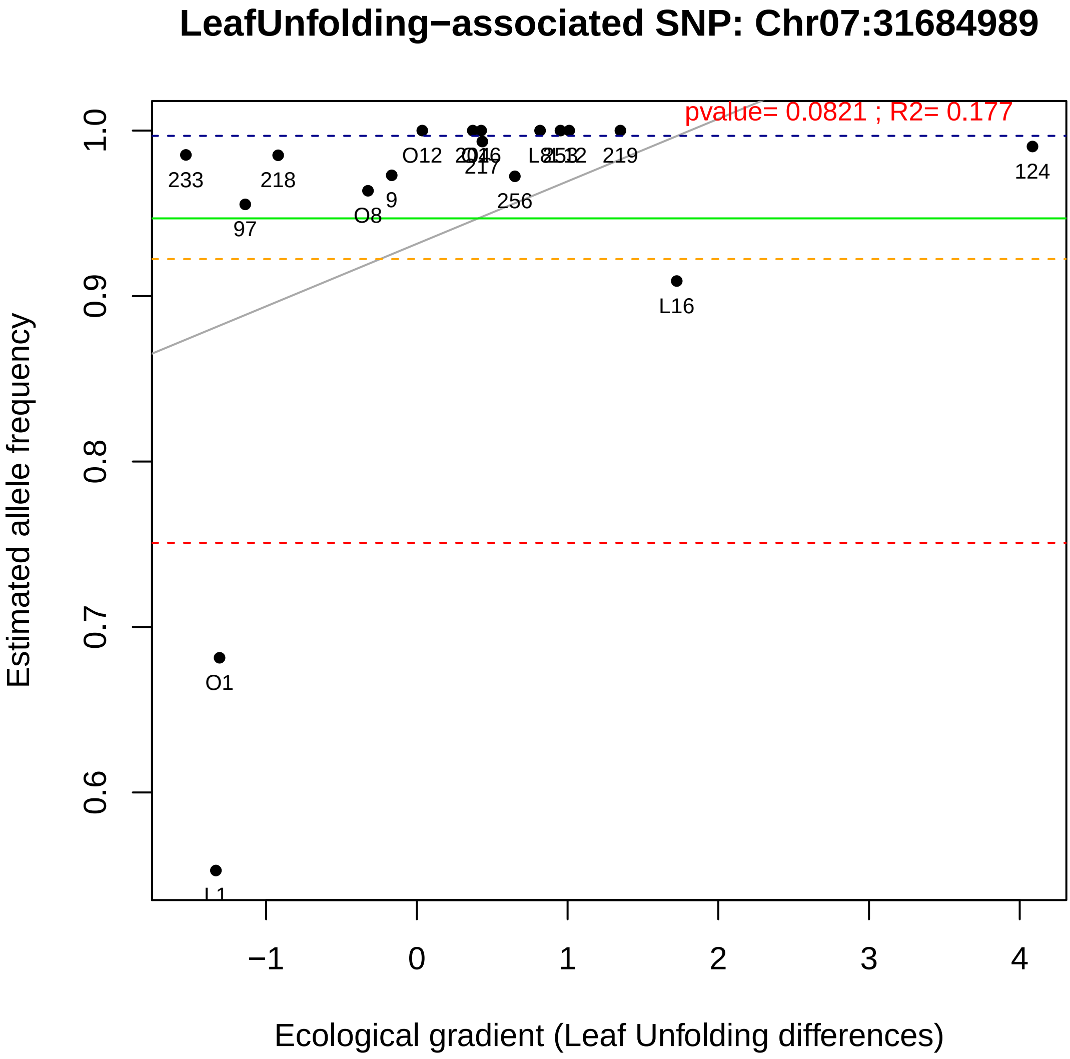

LeafUnfolding−associated SNP: Chr07:31684989
pvalue= 0.0821 ; R2= 0.177
Estimated allele frequency
Ecological gradient (Leaf Unfolding differences)
1.0
0.9
0.8
0.7
0.6
−1
0
1
2
3
4
233
218
97
O8
9
O12
217
256
219
124
L16
O1

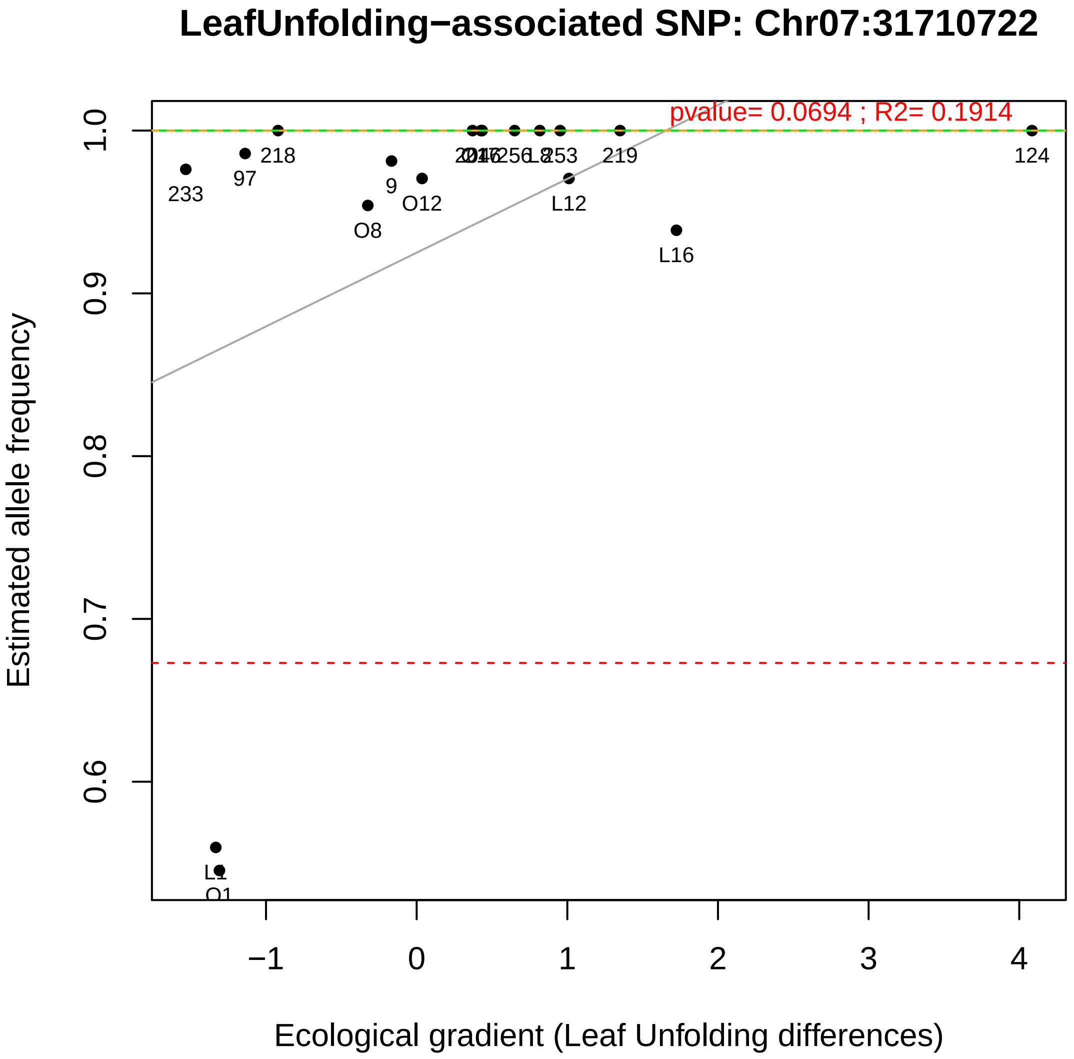

LeafUnfolding−associated SNP: Chr07:31710722
pvalue= 0.0694 ; R2= 0.1914
Estimated allele frequency
1.0
0.9
0.8
0.7
0.6
218
97
233
9
O12
O8
219
124
L12
L16
L1
O1
−1
0
1
2
3
4
Ecological gradient (Leaf Unfolding differences)

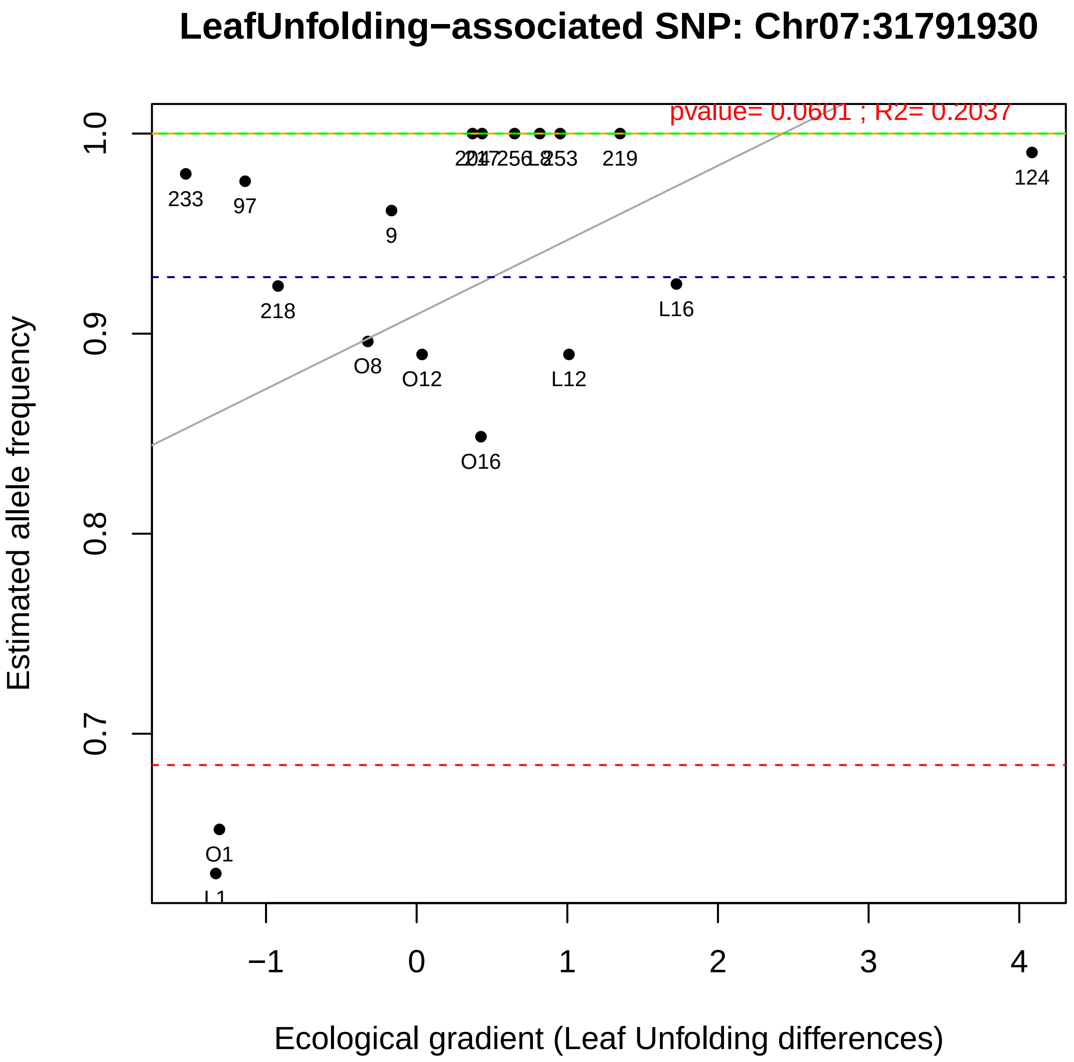
LeafUnfolding−associated SNP: Chr07:31791930
pvalue= 0.0601 ; R2= 0.2037
Estimated allele frequency
Ecological gradient (Leaf Unfolding differences)
1.0
0.9
0.8
0.7
−1
0
1
2
3
4
233
97
218
9
O8
O12
O16
L12
219
L16
124
O1
L1
253
256

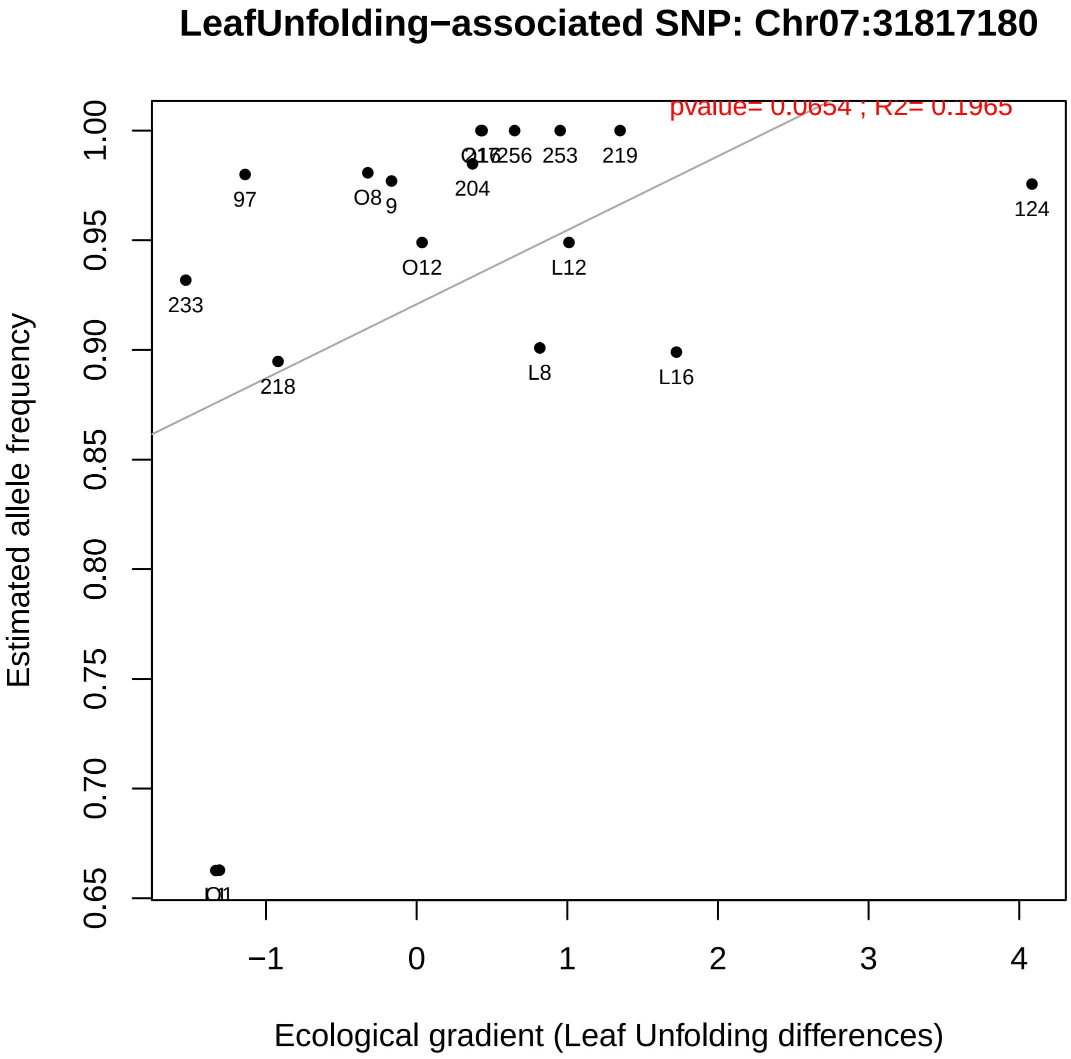
LeafUnfolding−associated SNP: Chr07:31817180
pvalue= 0.0654 ; R2= 0.1965
Estimated allele frequency
Ecological gradient (Leaf Unfolding differences)
0.65
0.70
0.75
0.80
0.85
0.90
0.95
1.00
−1
0
1
2
3
4
256
253
219
204
97
O8
9
124
O12
L12
233
218
L8
L16

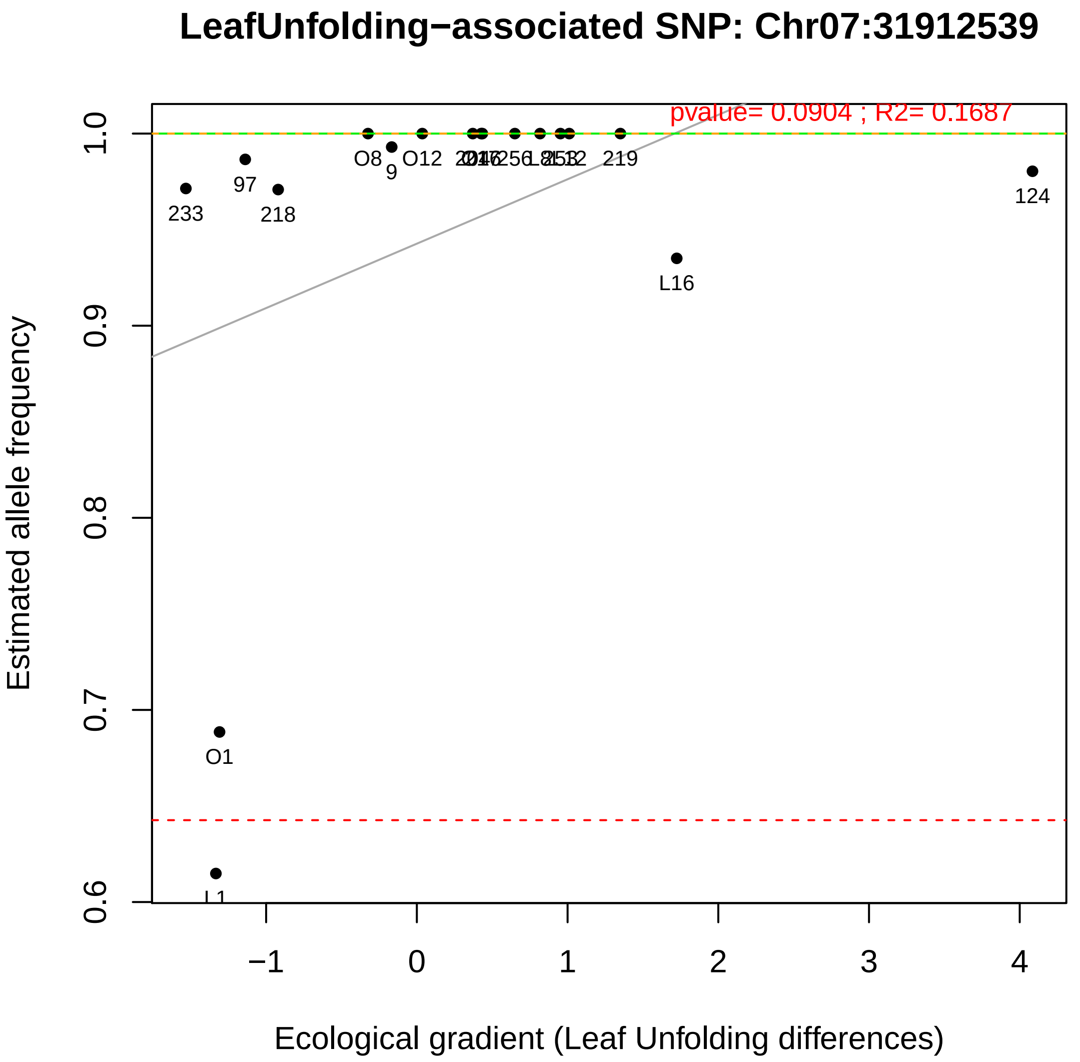
LeafUnfolding−associated SNP: Chr07:31912539
pvalue= 0.0904 ; R2= 0.1687
Estimated allele frequency
Ecological gradient (Leaf Unfolding differences)
1.0
0.9
0.8
0.7
0.6
−1
0
1
2
3
4
O8
9
O12
219
97
233
218
124
L16
O1

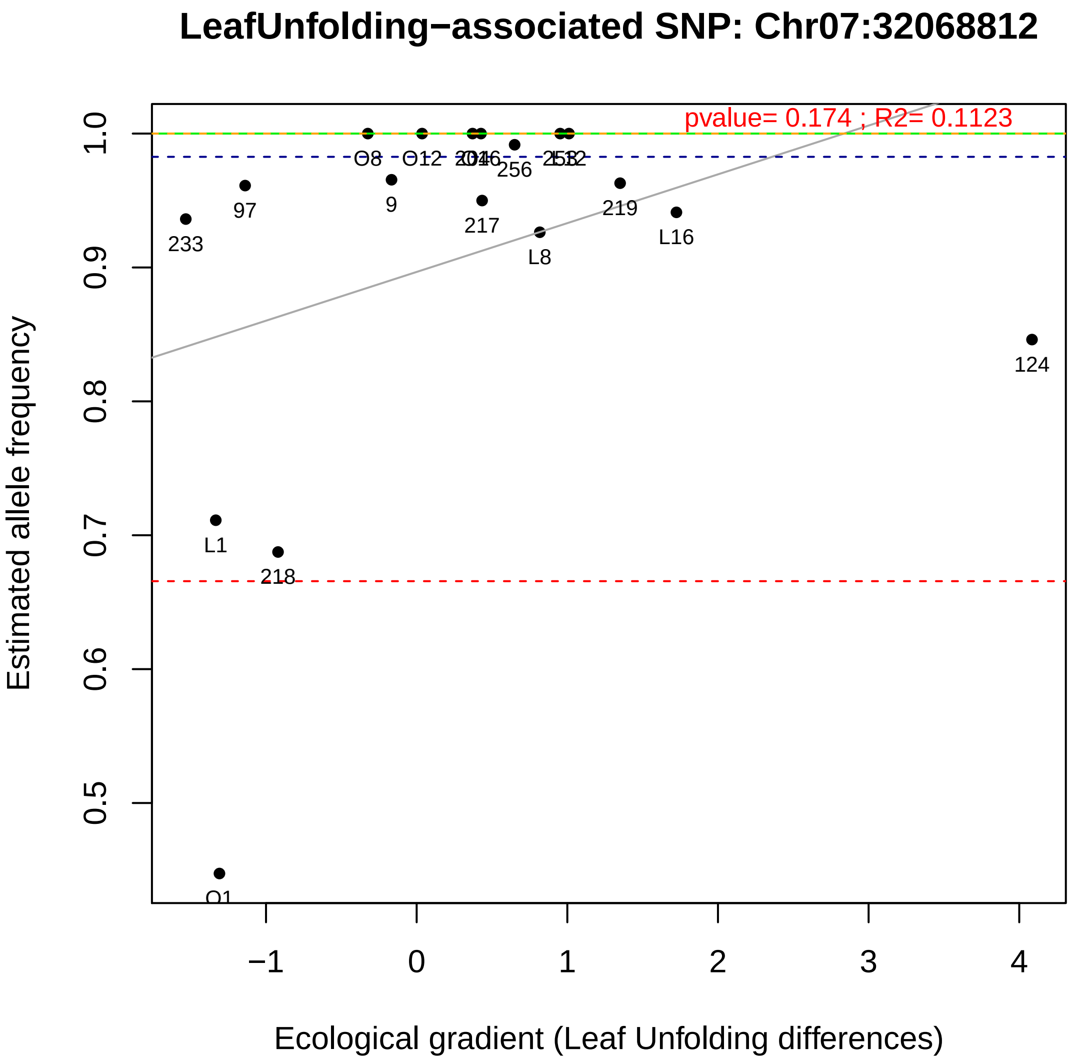
LeafUnfolding−associated SNP: Chr07:32068812
pvalue= 0.174 ; R2= 0.1123
Estimated allele frequency
1.0
0.9
0.8
0.7
0.6
0.5
−1
0
1
2
3
4
Ecological gradient (Leaf Unfolding differences)
O8
O12
256
9
97
233
217
L8
219
L16
124
L1
218
O1

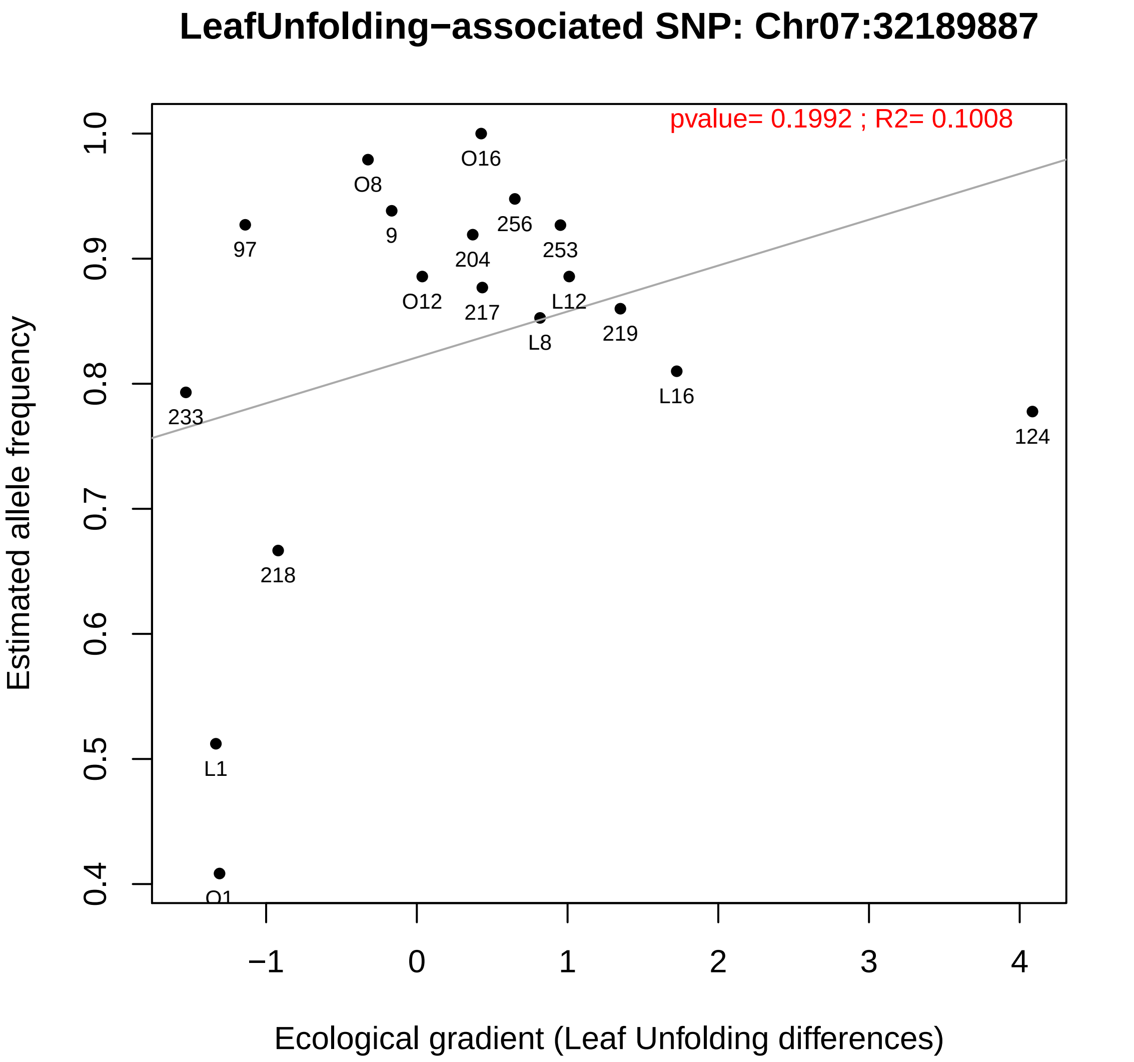

LeafUnfolding−associated SNP: Chr07:32189887
pvalue= 0.1992 ; R2= 0.1008
Estimated allele frequency
Ecological gradient (Leaf Unfolding differences)
1.0
0.9
0.8
0.7
0.6
0.5
0.4
−1
0
1
2
3
4
O16
O8
256
9
97
204
253
O12
217
L12
L8
219
L16
233
124
218
L1
O1

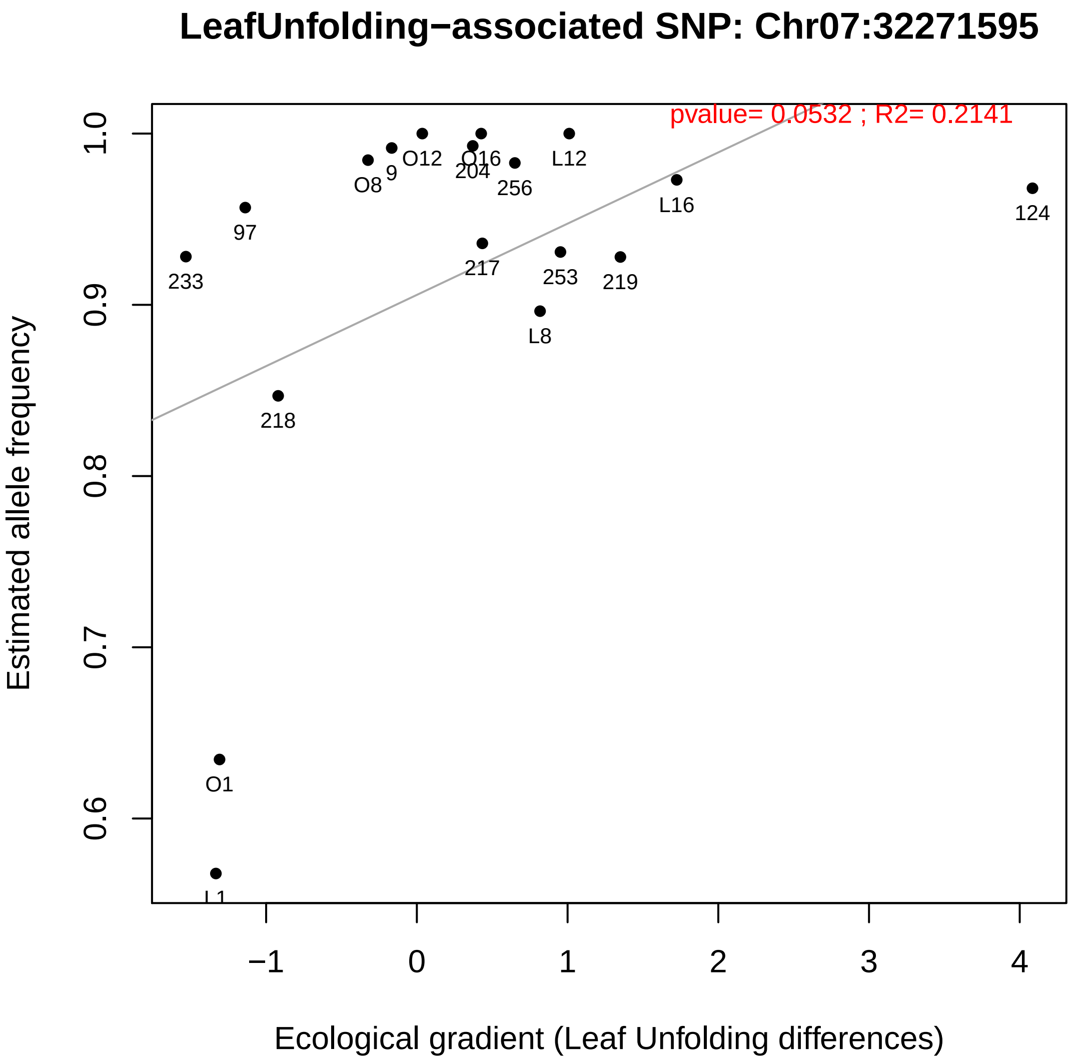
LeafUnfolding−associated SNP: Chr07:32271595
pvalue= 0.0532 ; R2= 0.2141
Estimated allele frequency
Ecological gradient (Leaf Unfolding differences)
0.6
0.7
0.8
0.9
1.0
−1
0
1
2
3
4
O12
O16
L12
204
9
O8
256
L16
124
97
233
217
253
219
L8
218
O1
L1

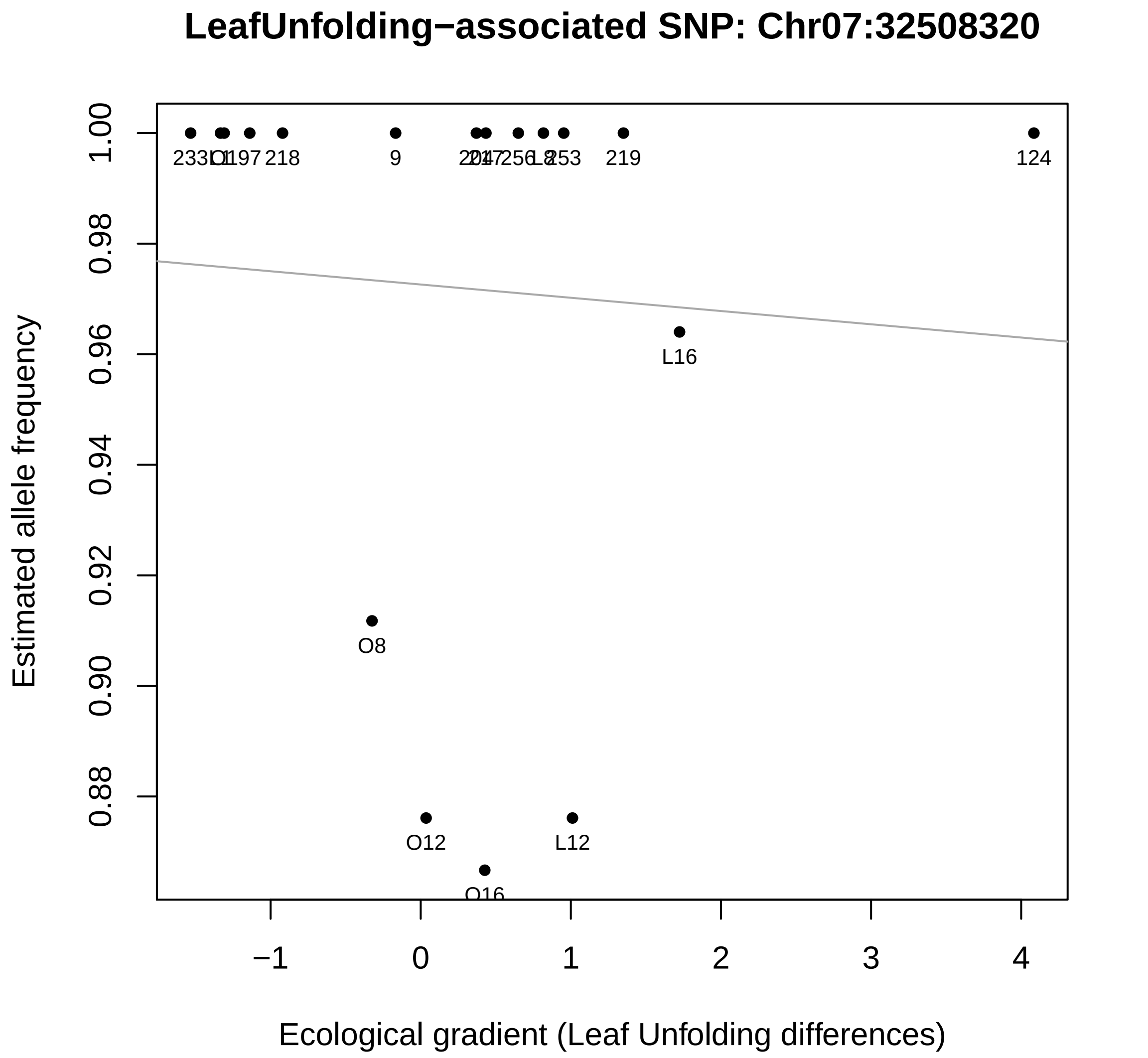
LeafUnfolding−associated SNP: Chr07:32508320
Estimated allele frequency
Ecological gradient (Leaf Unfolding differences)
1.00
0.98
0.96
0.94
0.92
0.90
0.88
−1
0
1
2
3
4
218
9
219
124
L16
O8
O12
L12
O16

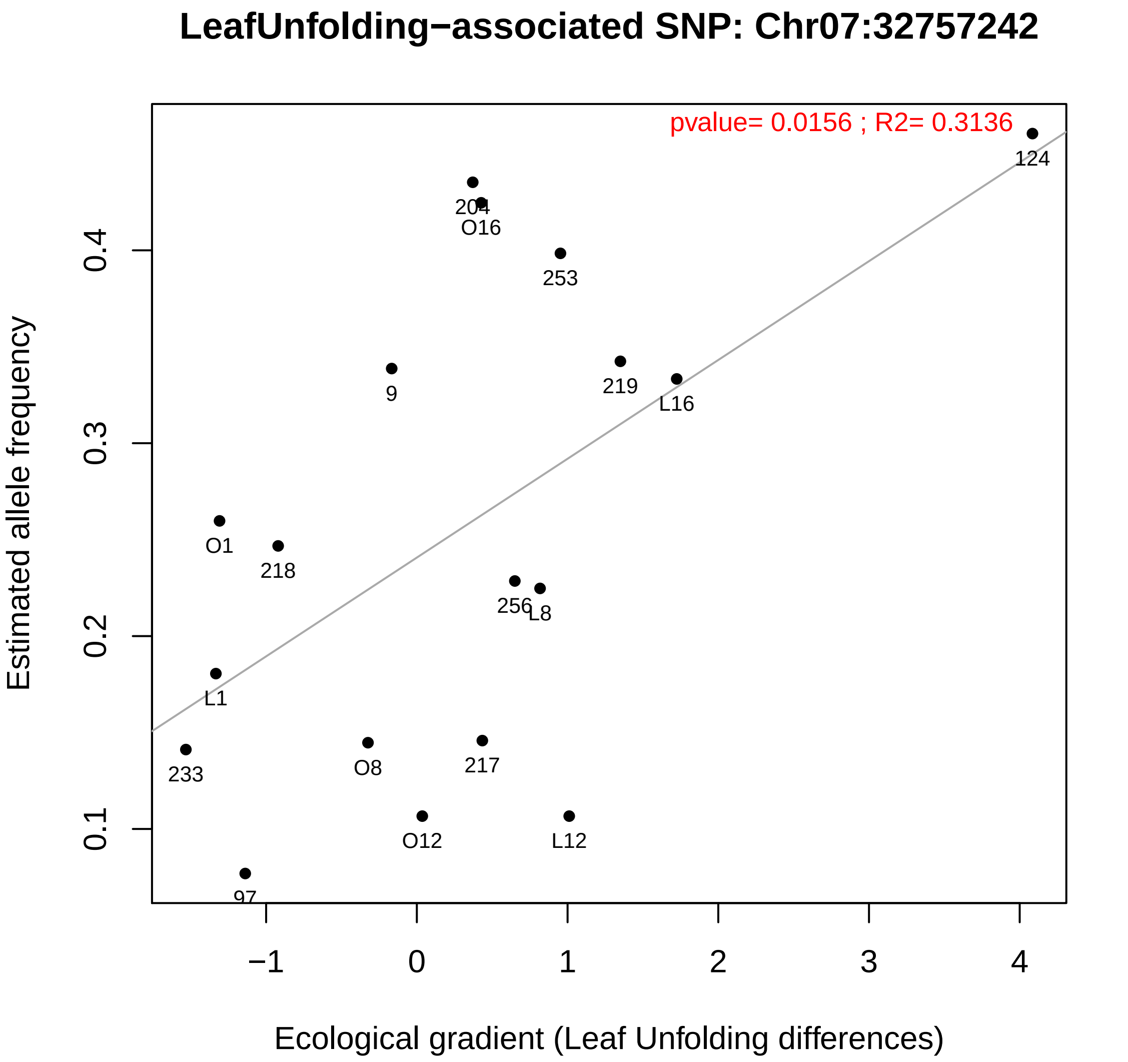

LeafUnfolding−associated SNP: Chr07:32757242
pvalue= 0.0156 ; R2= 0.3136
Estimated allele frequency
0.1
0.2
0.3
0.4
−1
0
1
2
3
4
Ecological gradient (Leaf Unfolding differences)
124
204
O16
253
9
219
L16
O1
218
256
L8
L1
233
O8
217
O12
L12
97

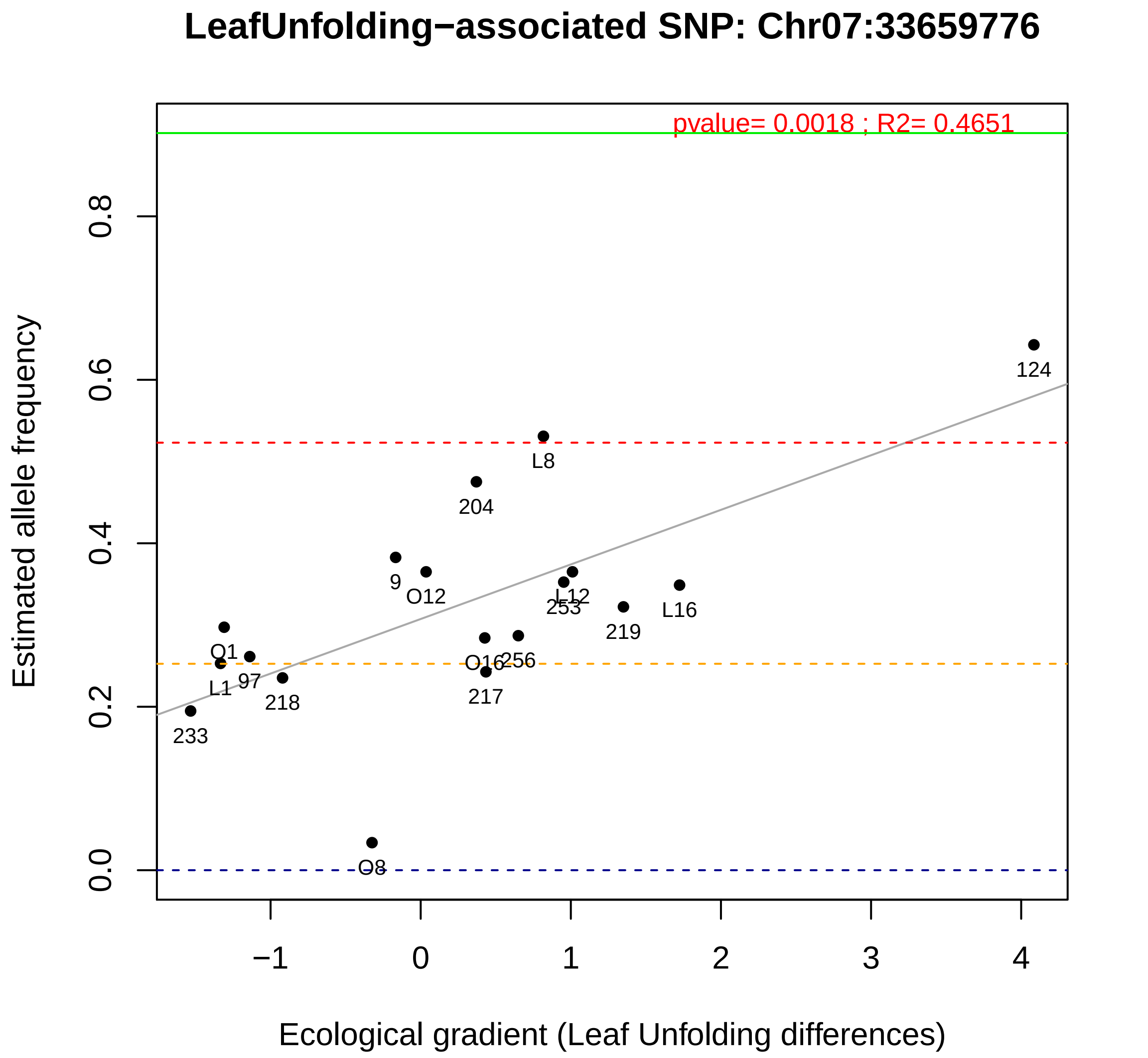
LeafUnfolding−associated SNP: Chr07:33659776
pvalue= 0.0018 ; R2= 0.4651
Estimated allele frequency
0.0
0.2
0.4
0.6
0.8
Ecological gradient (Leaf Unfolding differences)
−1
0
1
2
3
4
124
L8
204
9
O12
L12
253
L16
219
O1
L1
97
218
O16
256
217
233
O8

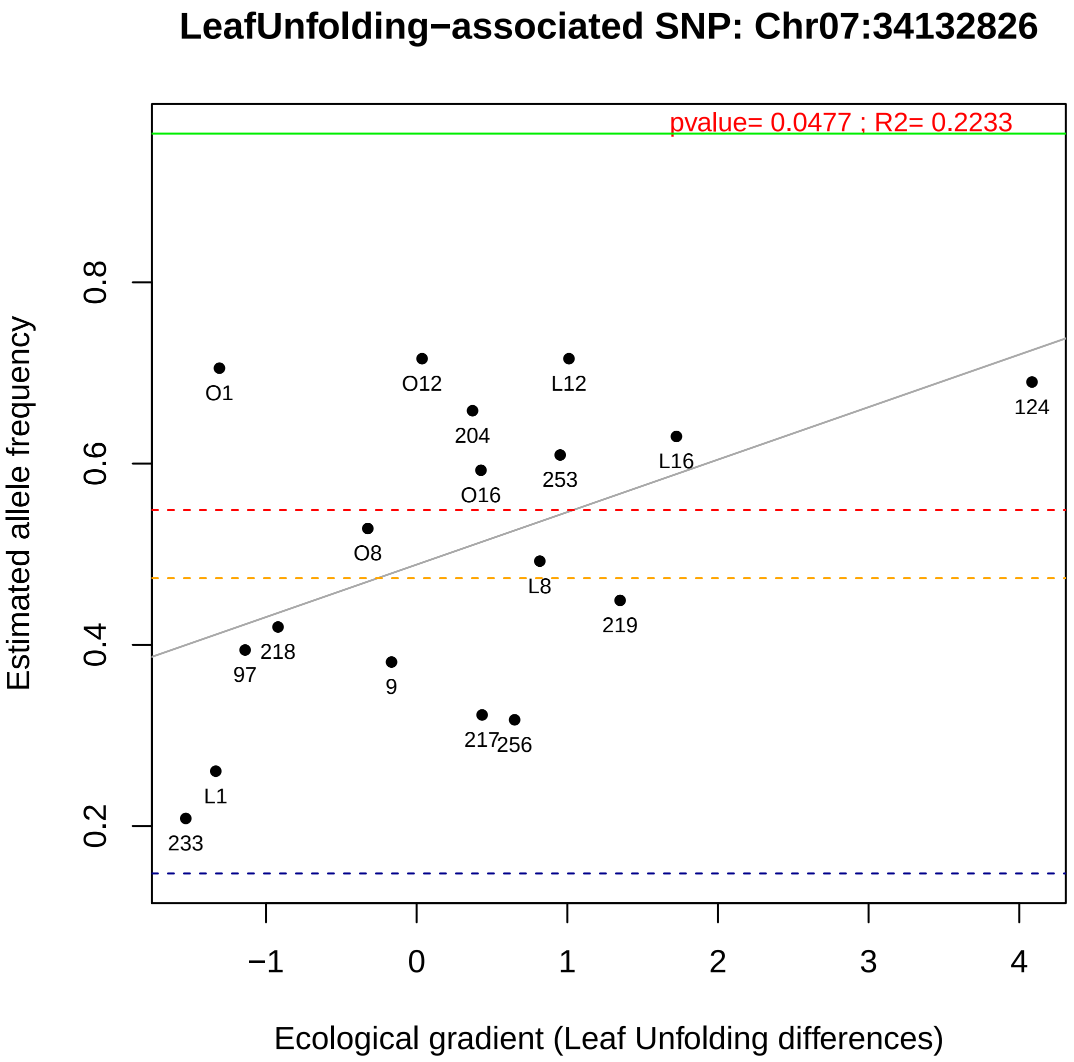

LeafUnfolding−associated SNP: Chr07:34132826
pvalue= 0.0477 ; R2= 0.2233
Estimated allele frequency
0.2
0.4
0.6
0.8
−1
0
1
2
3
4
Ecological gradient (Leaf Unfolding differences)
O1
O12
L12
124
204
L16
253
O16
O8
L8
219
218
97
9
217
256
L1
233

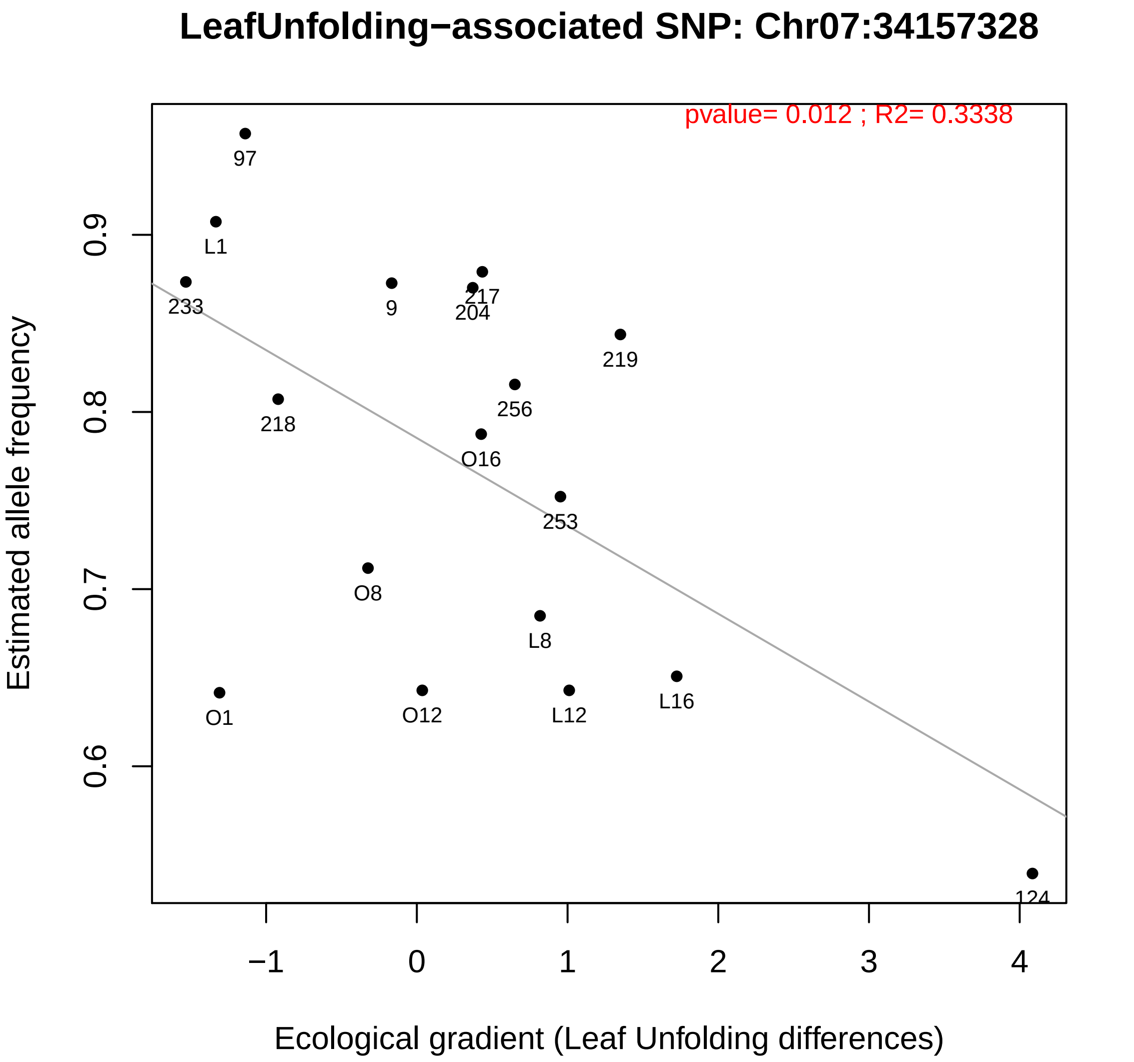
LeafUnfolding−associated SNP: Chr07:34157328
pvalue= 0.012 ; R2= 0.3338
Estimated allele frequency
Ecological gradient (Leaf Unfolding differences)
0.6
0.7
0.8
0.9
−1
0
1
2
3
4
97
L1
233
9
217
204
219
256
218
O16
253
O8
L8
O1
O12
L12
L16
124

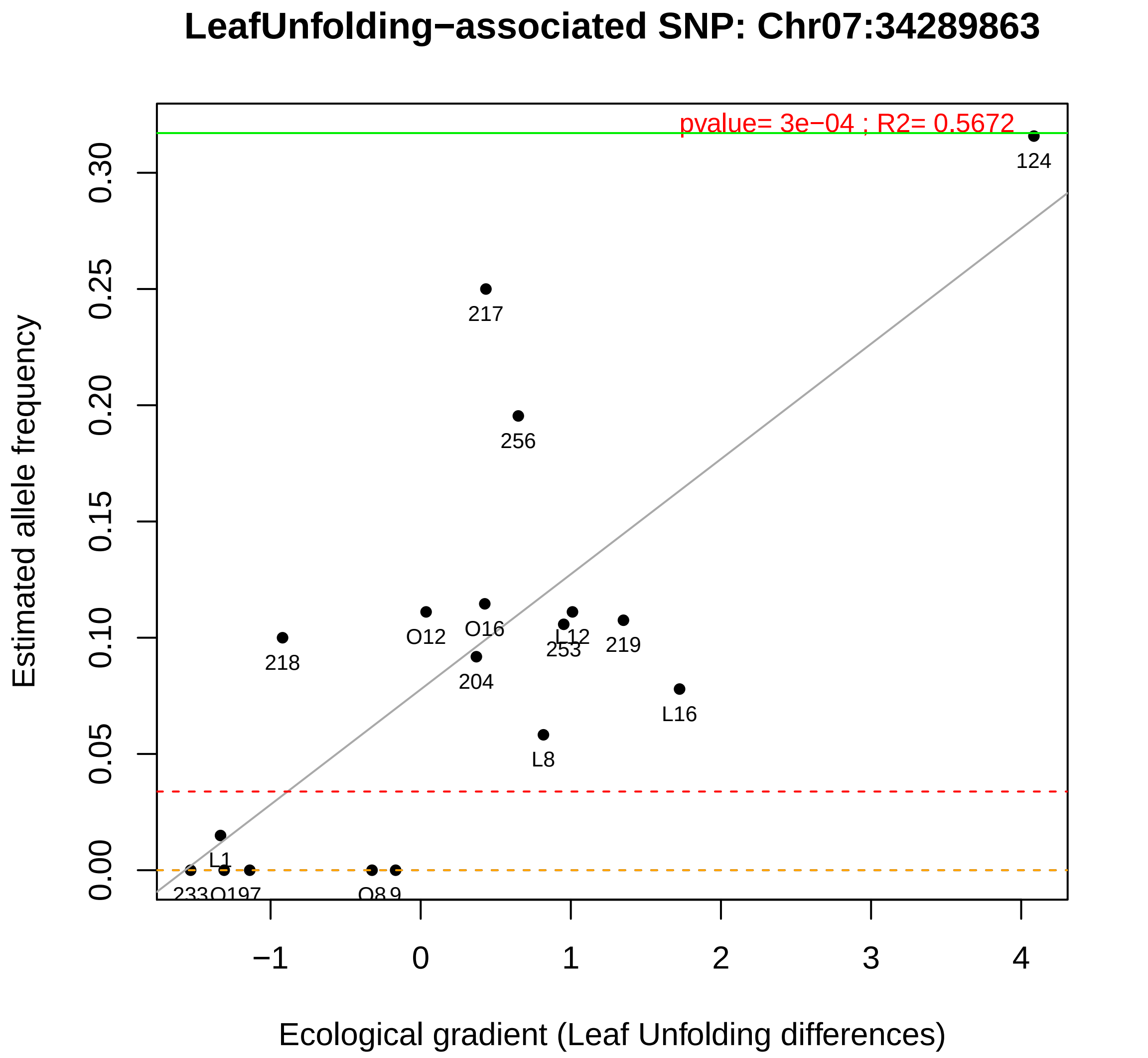
LeafUnfolding−associated SNP: Chr07:34289863
pvalue= 3e−04 ; R2= 0.5672
Estimated allele frequency
0.00
0.05
0.10
0.15
0.20
0.25
0.30
−1
0
1
2
3
4
Ecological gradient (Leaf Unfolding differences)
124
217
256
218
O12
O16
204
L12
253
219
L16
L8
L1
233
O197
O8
9

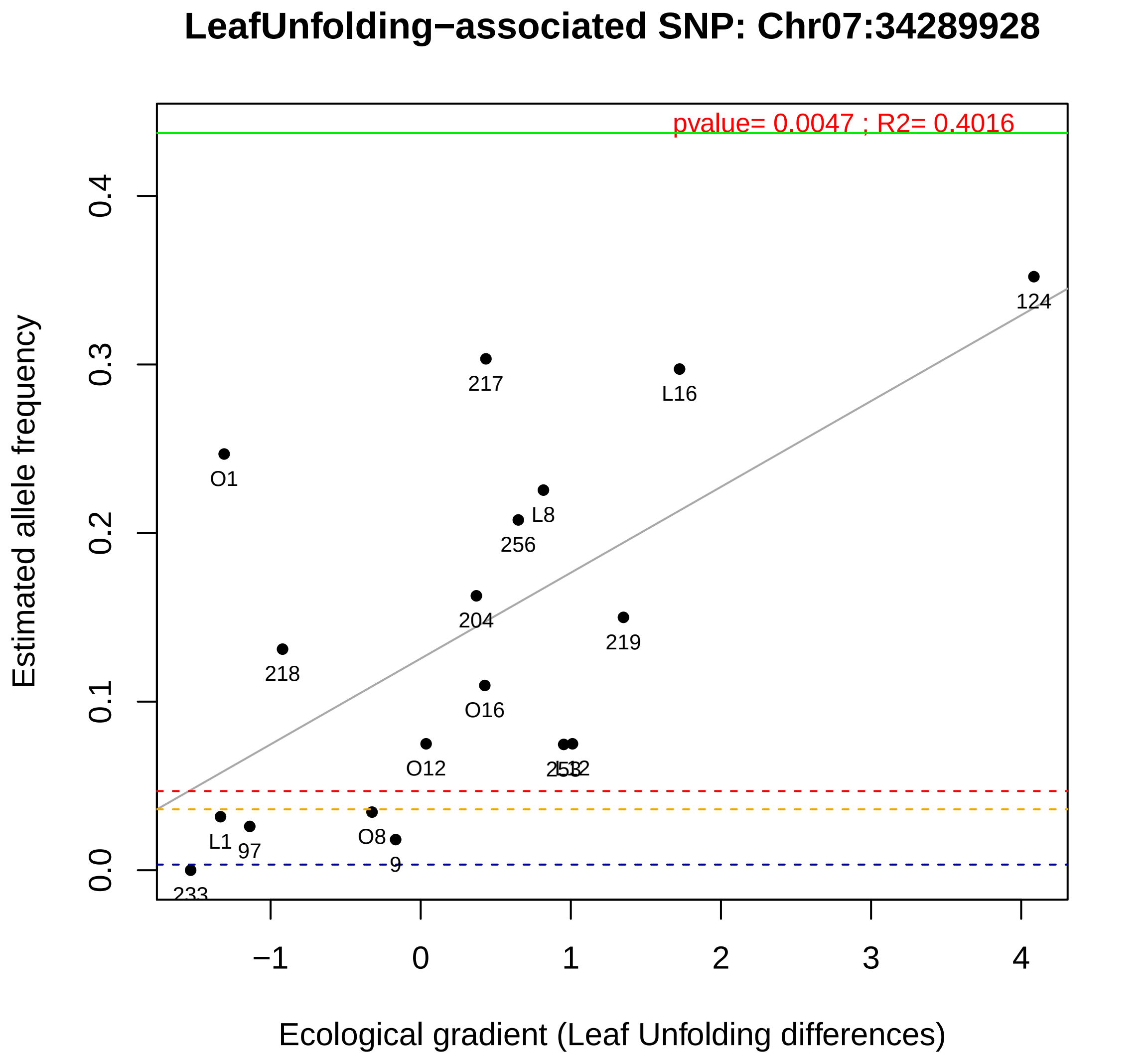
LeafUnfolding−associated SNP: Chr07:34289928
pvalue= 0.0047 ; R2= 0.4016
Estimated allele frequency
0.0
0.1
0.2
0.3
0.4
Ecological gradient (Leaf Unfolding differences)
−1
0
1
2
3
4
124
217
L16
O1
L8
256
204
219
218
O16
O12
L1
97
O8
9
233

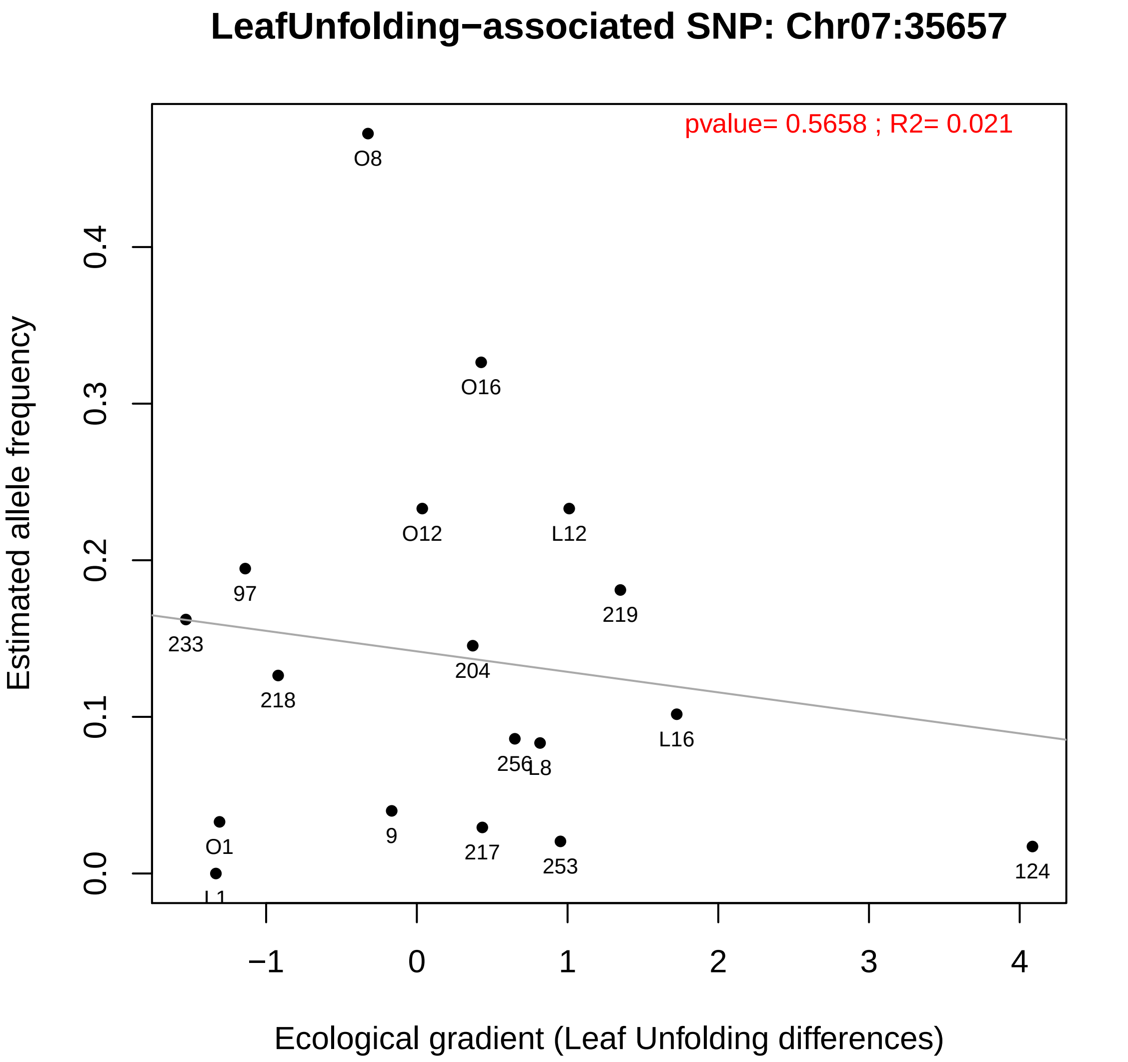
LeafUnfolding−associated SNP: Chr07:35657
pvalue= 0.5658 ; R2= 0.021
Estimated allele frequency
0.0
0.1
0.2
0.3
0.4
−1
0
1
2
3
4
Ecological gradient (Leaf Unfolding differences)
O8
O16
O12
L12
97
219
233
204
218
L16
256
L8
O1
9
217
253
124

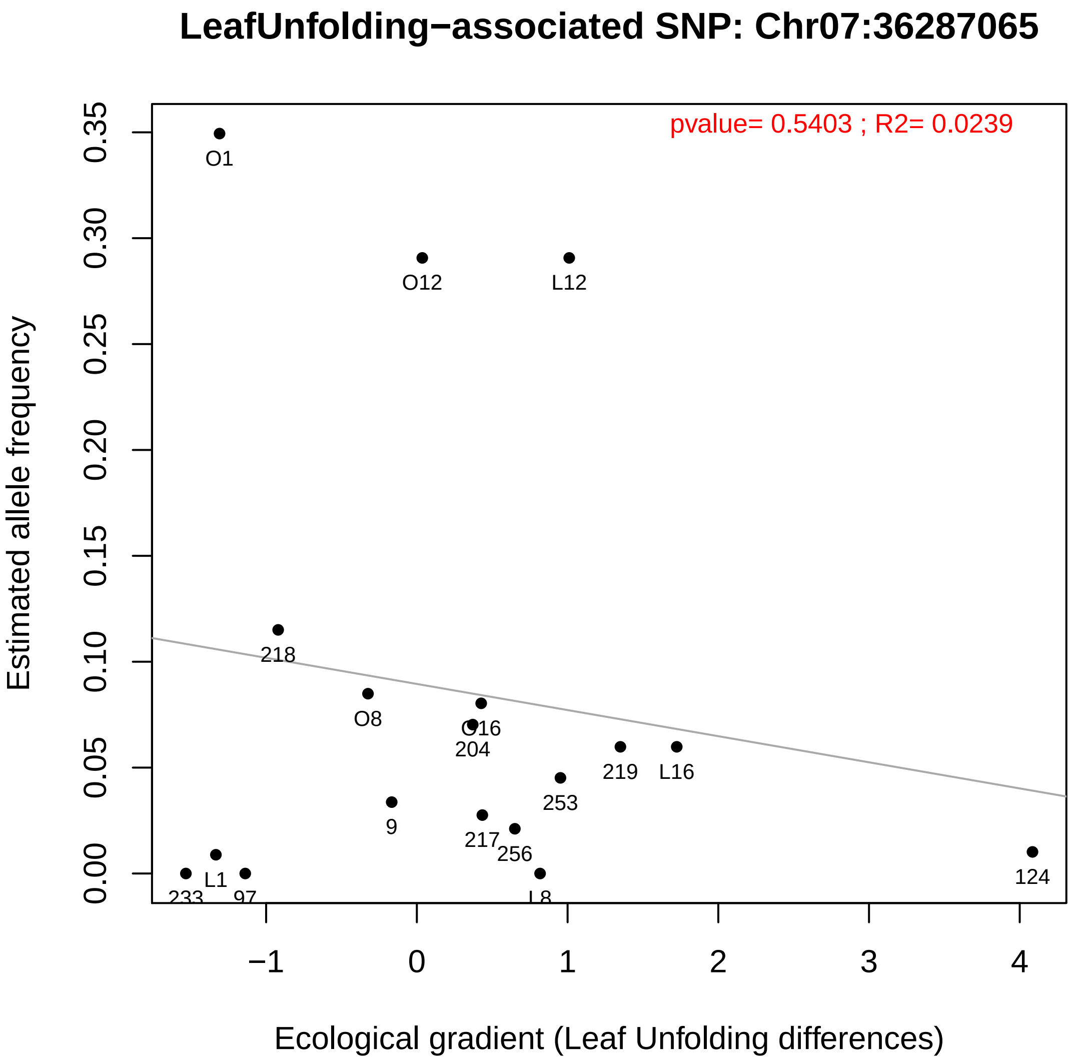
LeafUnfolding−associated SNP: Chr07:36287065
pvalue= 0.5403 ; R2= 0.0239
Estimated allele frequency
Ecological gradient (Leaf Unfolding differences)
0.00
0.05
0.10
0.15
0.20
0.25
0.30
0.35
−1
0
1
2
3
4
O1
O12
L12
218
O8
O16
204
219
L16
253
9
217
256
L1
124

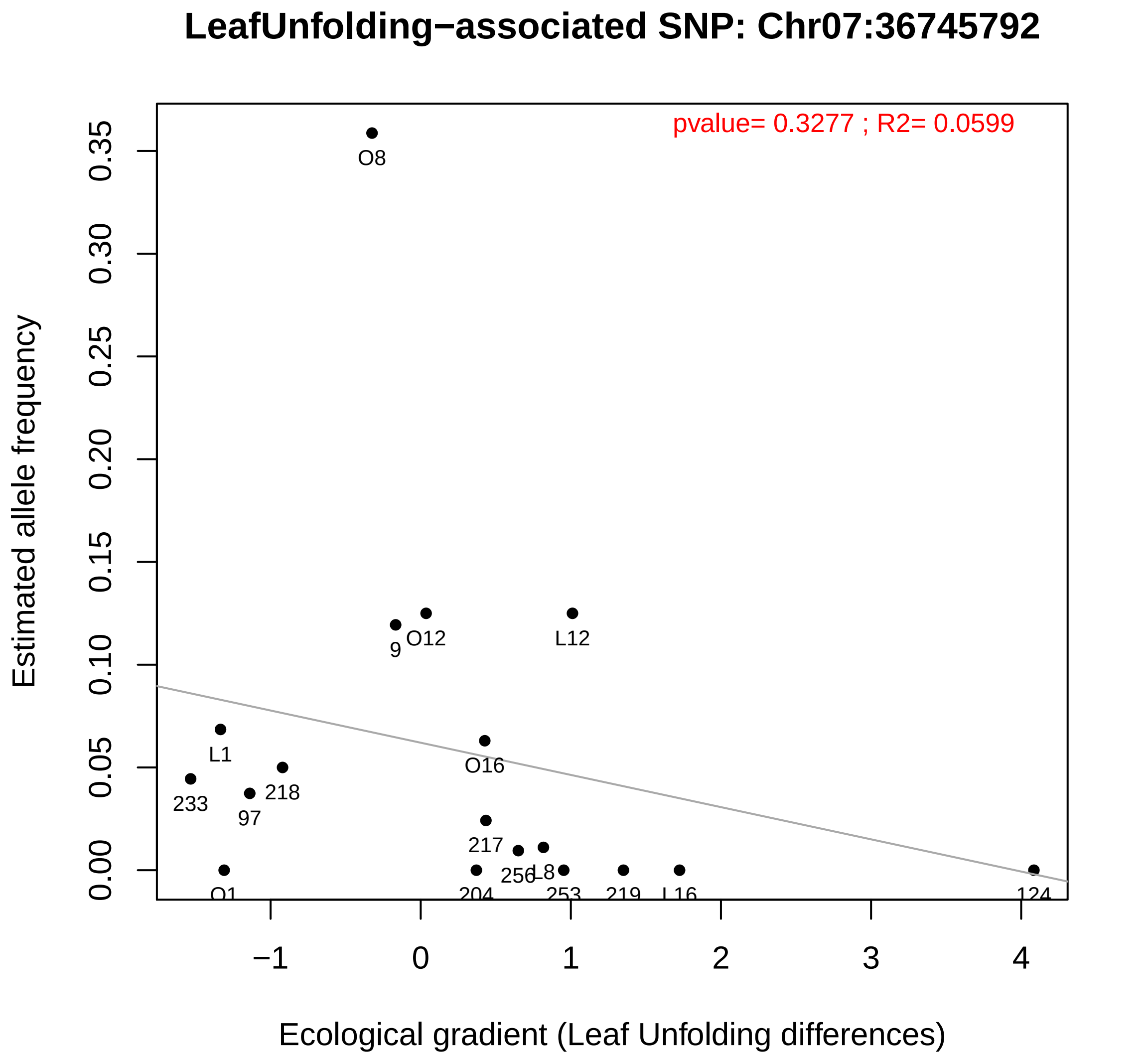
LeafUnfolding−associated SNP: Chr07:36745792
pvalue= 0.3277 ; R2= 0.0599
Estimated allele frequency
0.00
0.05
0.10
0.15
0.20
0.25
0.30
0.35
Ecological gradient (Leaf Unfolding differences)
−1
0
1
2
3
4
O8
9
O12
L12
L1
O16
233
218
97
217
256
L8
O1
204
253
219
L16
124

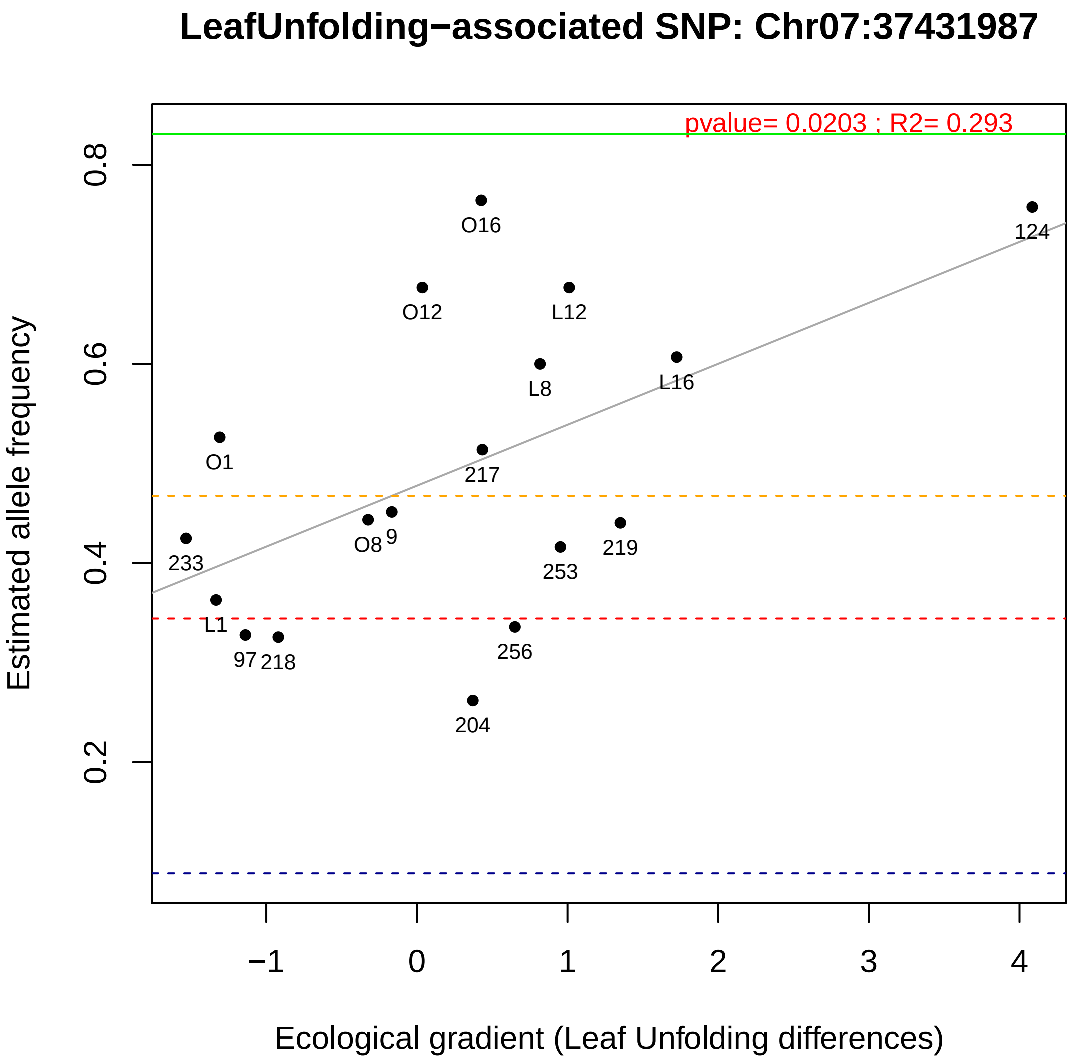

LeafUnfolding−associated SNP: Chr07:37431987
pvalue= 0.0203 ; R2= 0.293
Estimated allele frequency
0.2
0.4
0.6
0.8
−1
0
1
2
3
4
Ecological gradient (Leaf Unfolding differences)
O16
124
O12
L12
L8
L16
O1
217
O8
9
233
219
253
L1
97
218
256
204

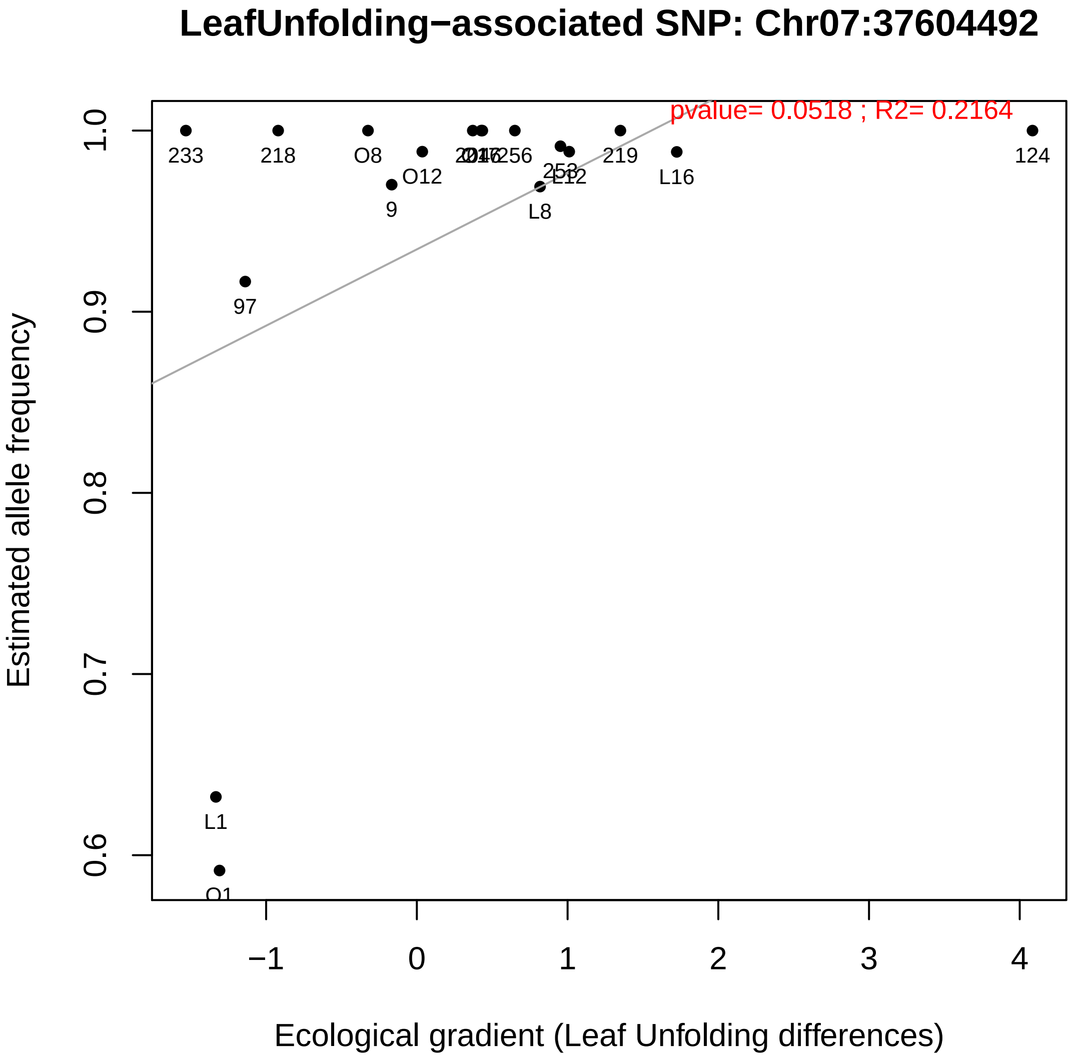
LeafUnfolding−associated SNP: Chr07:37604492
pvalue= 0.0518 ; R2= 0.2164
Estimated allele frequency
Ecological gradient (Leaf Unfolding differences)
0.6
0.7
0.8
0.9
1.0
−1
0
1
2
3
4
233
218
O8
O12
9
256
253
L12
L8
219
L16
124
97
L1
O1

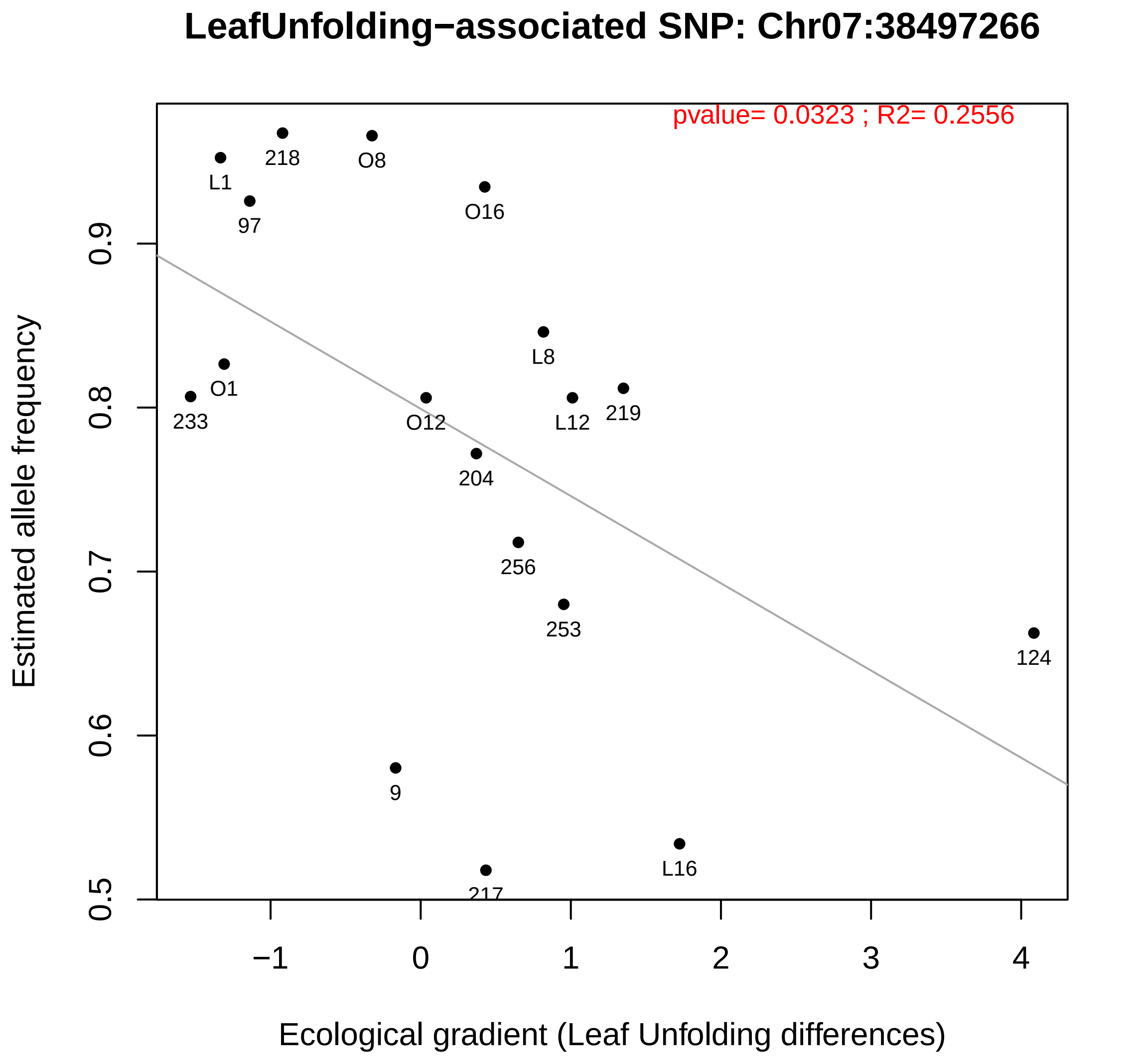
LeafUnfolding−associated SNP: Chr07:38497266
pvalue= 0.0323 ; R2= 0.2556
Estimated allele frequency
Ecological gradient (Leaf Unfolding differences)
0.5
0.6
0.7
0.8
0.9
−1
0
1
2
3
4
218
O8
L1
97
O16
L8
O1
233
O12
L12
219
204
256
253
124
9
L16
217

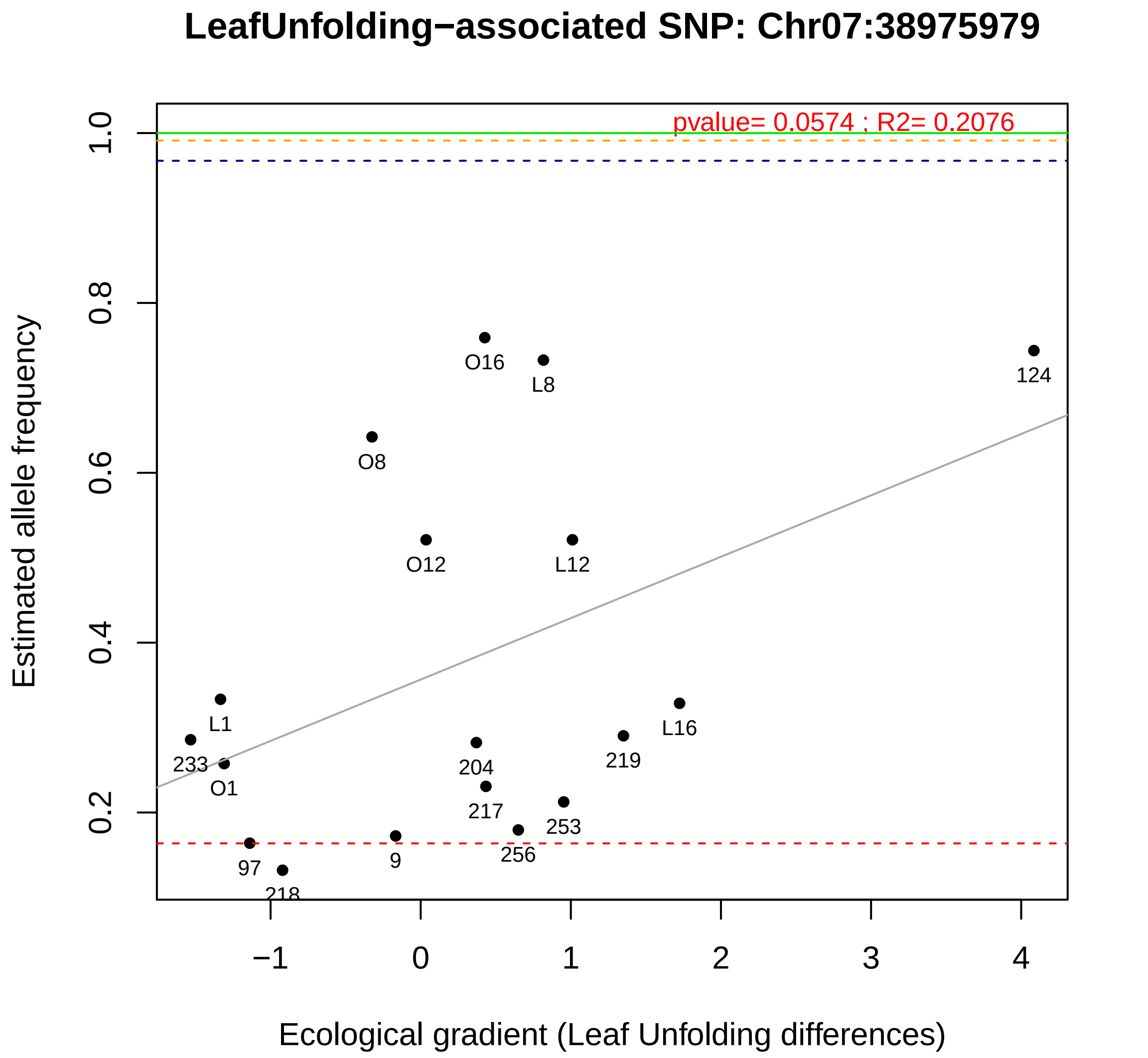

LeafUnfolding−associated SNP: Chr07:38975979
pvalue= 0.0574 ; R2= 0.2076
Estimated allele frequency
1.0
0.8
0.6
0.4
0.2
O16
L8
124
O8
O12
L12
L1
233
O1
204
217
219
L16
253
256
97
9
218
−1
0
1
2
3
4
Ecological gradient (Leaf Unfolding differences)

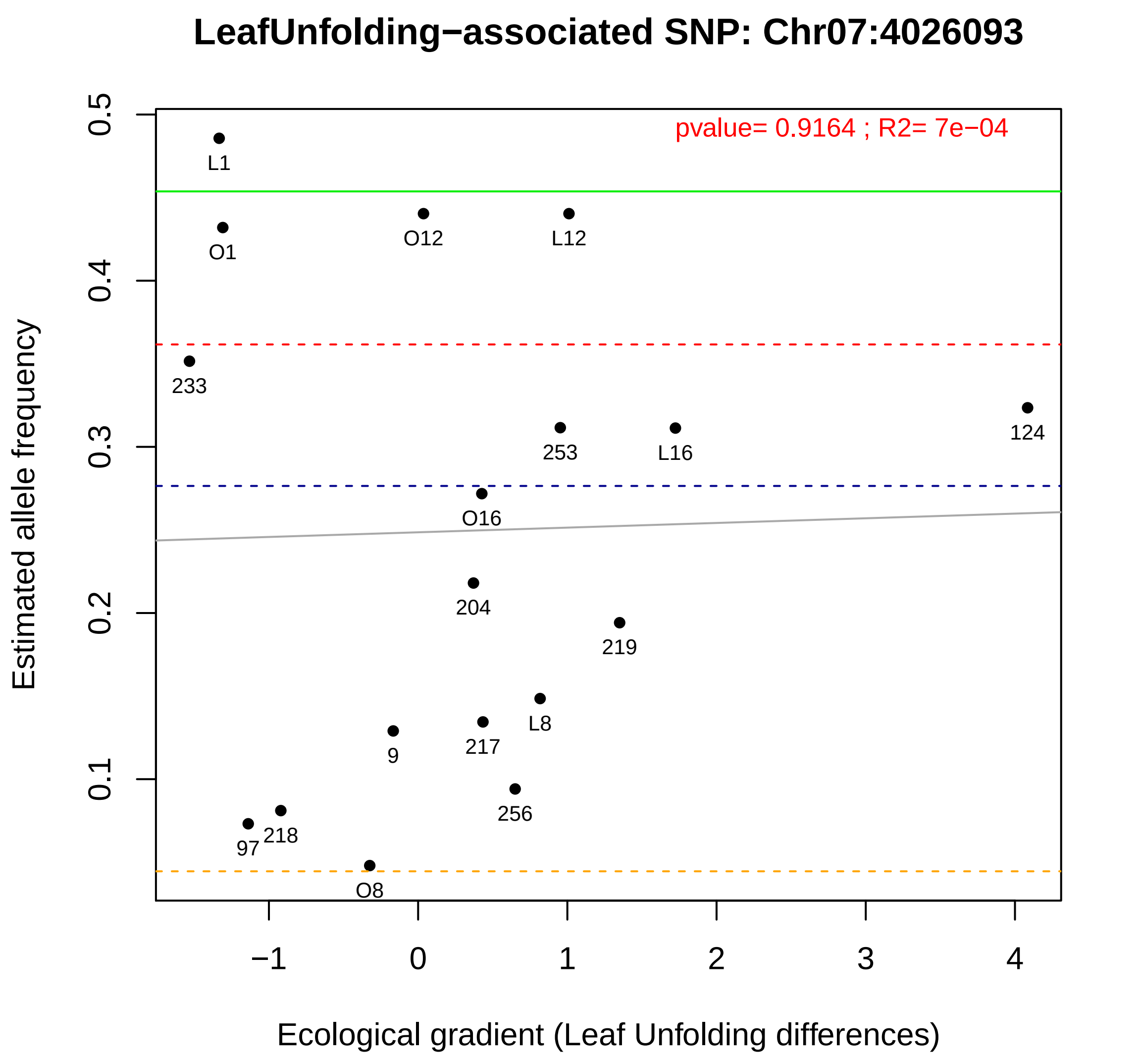
LeafUnfolding−associated SNP: Chr07:4026093
pvalue= 0.9164 ; R2= 7e−04
Estimated allele frequency
0.1
0.2
0.3
0.4
0.5
Ecological gradient (Leaf Unfolding differences)
−1
0
1
2
3
4
L1
O1
O12
L12
233
253
L16
124
O16
204
219
L8
217
9
256
97
218
O8

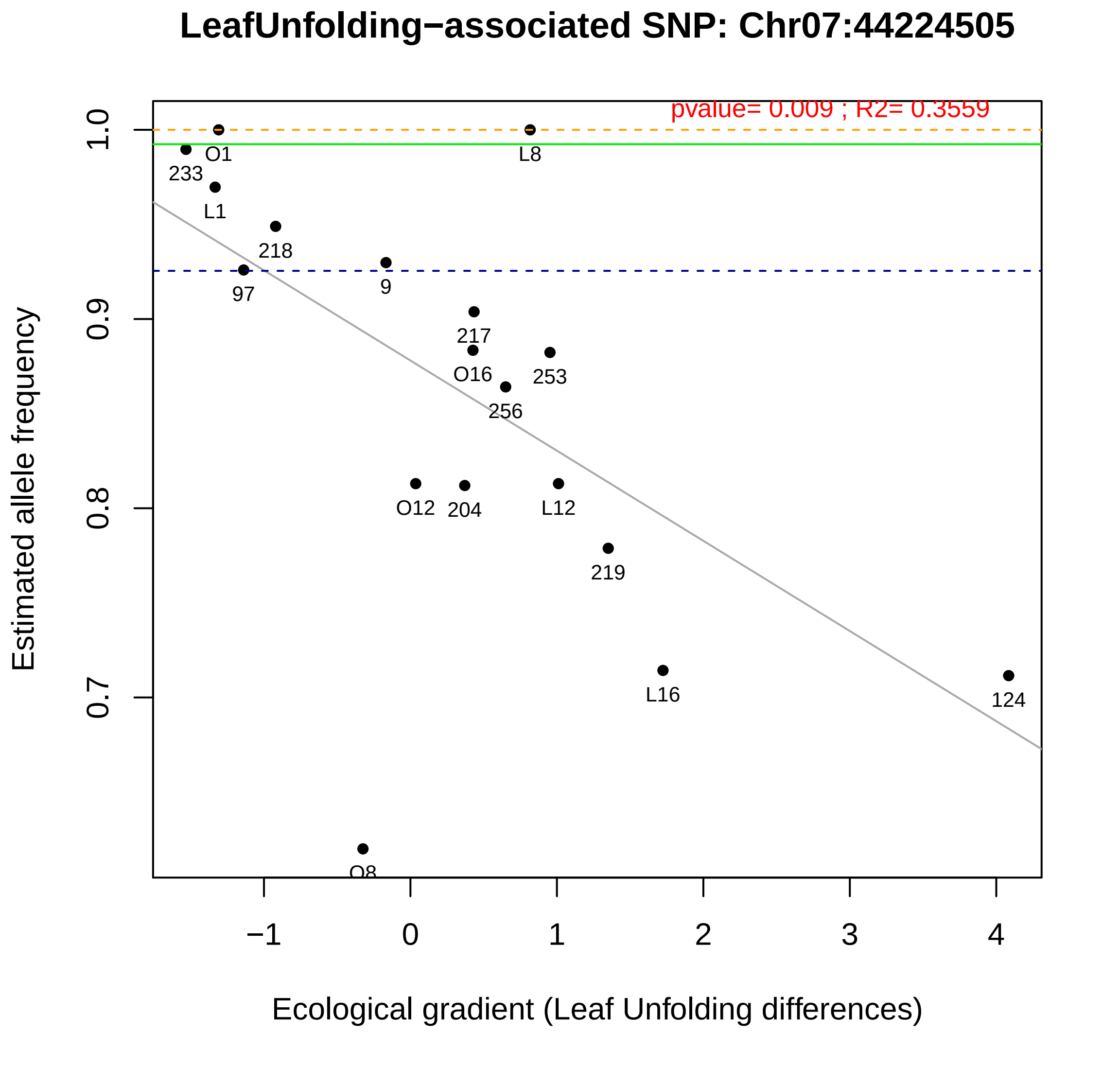

LeafUnfolding−associated SNP: Chr07:44224505
pvalue= 0.009 ; R2= 0.3559
Estimated allele frequency
1.0
0.9
0.8
0.7
O1
L8
233
L1
218
97
9
217
O16
253
256
O12
204
L12
219
L16
124
O8
−1
0
1
2
3
4
Ecological gradient (Leaf Unfolding differences)

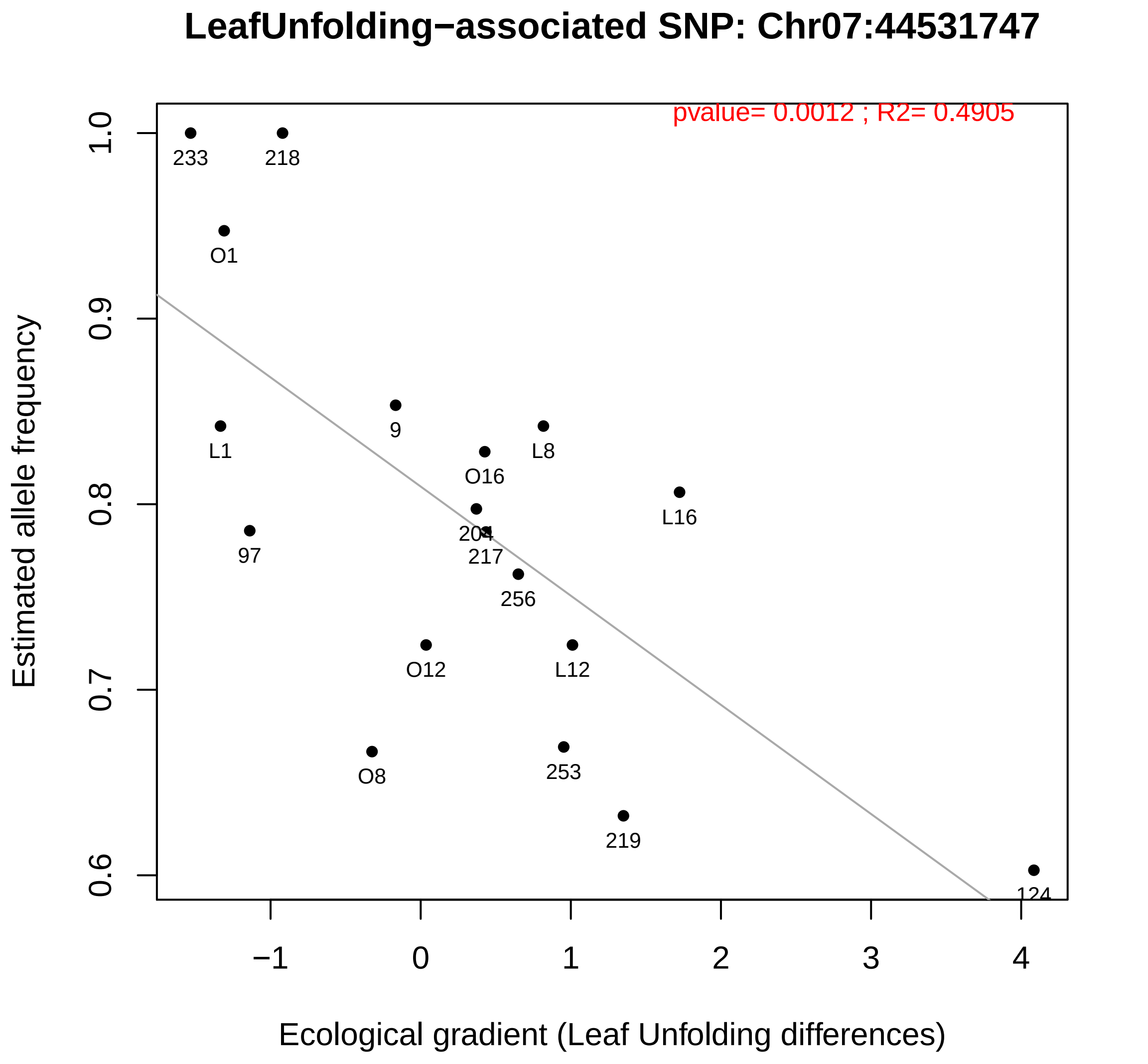
LeafUnfolding−associated SNP: Chr07:44531747
pvalue= 0.0012 ; R2= 0.4905
Estimated allele frequency
Ecological gradient (Leaf Unfolding differences)
1.0
0.9
0.8
0.7
0.6
−1
0
1
2
3
4
233
218
O1
L1
9
L8
O16
L16
204
217
97
256
O12
L12
O8
253
219
124

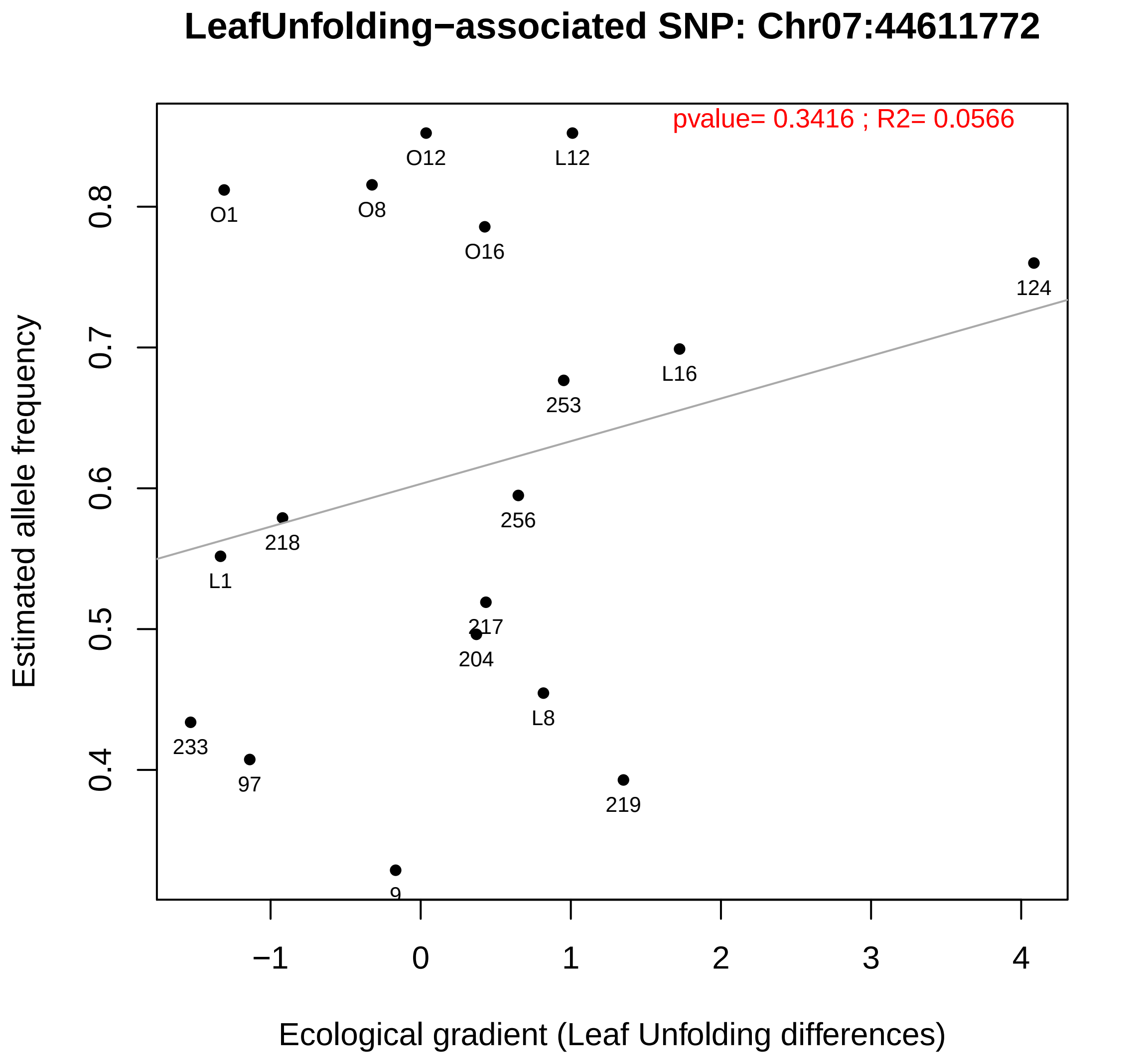
LeafUnfolding−associated SNP: Chr07:44611772
pvalue= 0.3416 ; R2= 0.0566
O12
L12
O1
O8
O16
124
L16
253
256
218
L1
217
204
L8
233
97
219
9
0.4
0.5
0.6
0.7
0.8
−1
0
1
2
3
4
Estimated allele frequency
Ecological gradient (Leaf Unfolding differences)

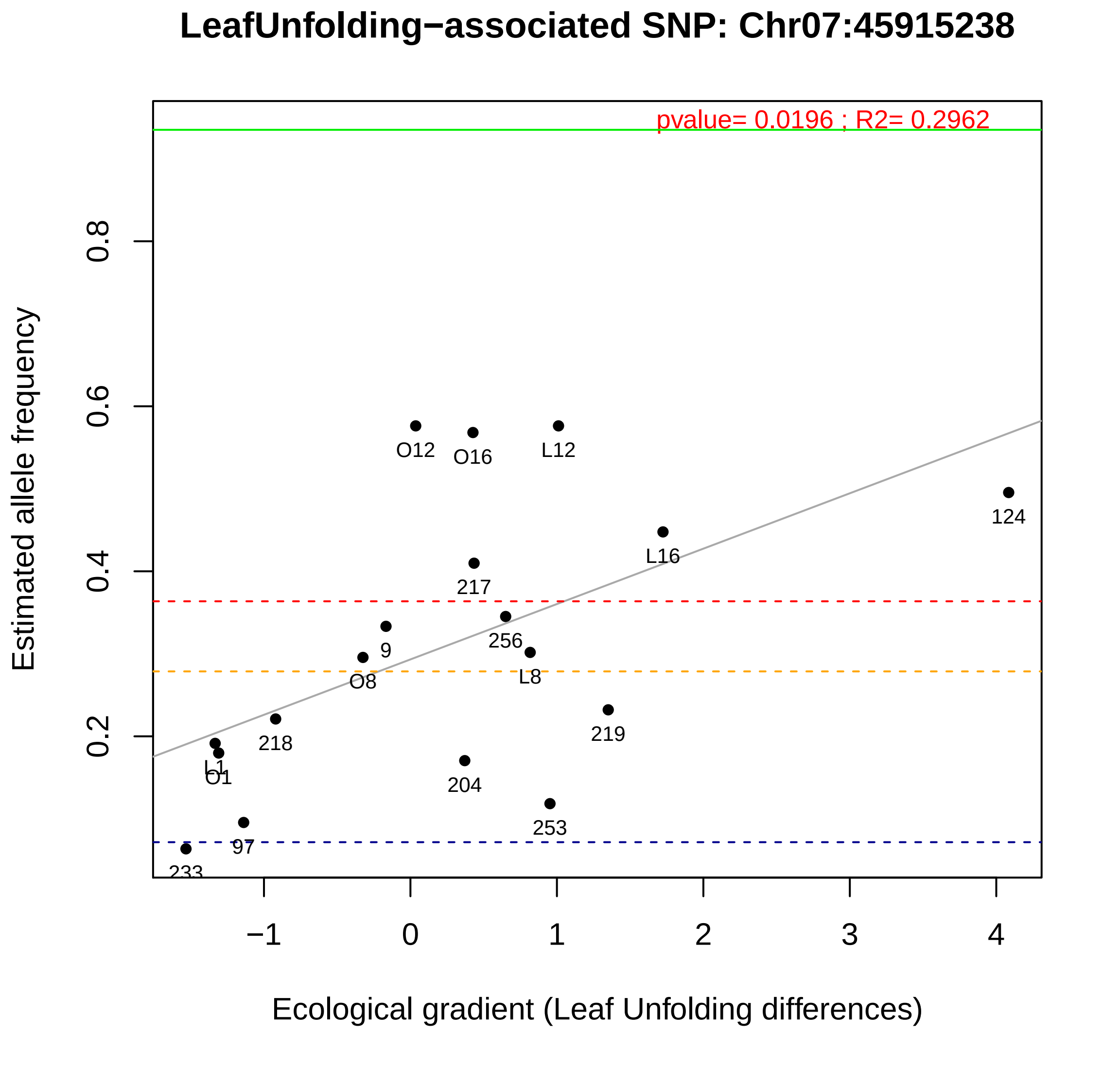
LeafUnfolding−associated SNP: Chr07:45915238
pvalue= 0.0196 ; R2= 0.2962
Estimated allele frequency
0.2
0.4
0.6
0.8
−1
0
1
2
3
4
Ecological gradient (Leaf Unfolding differences)
O12
O16
L12
124
L16
217
256
9
O8
L8
219
218
L1
O1
204
253
97
233

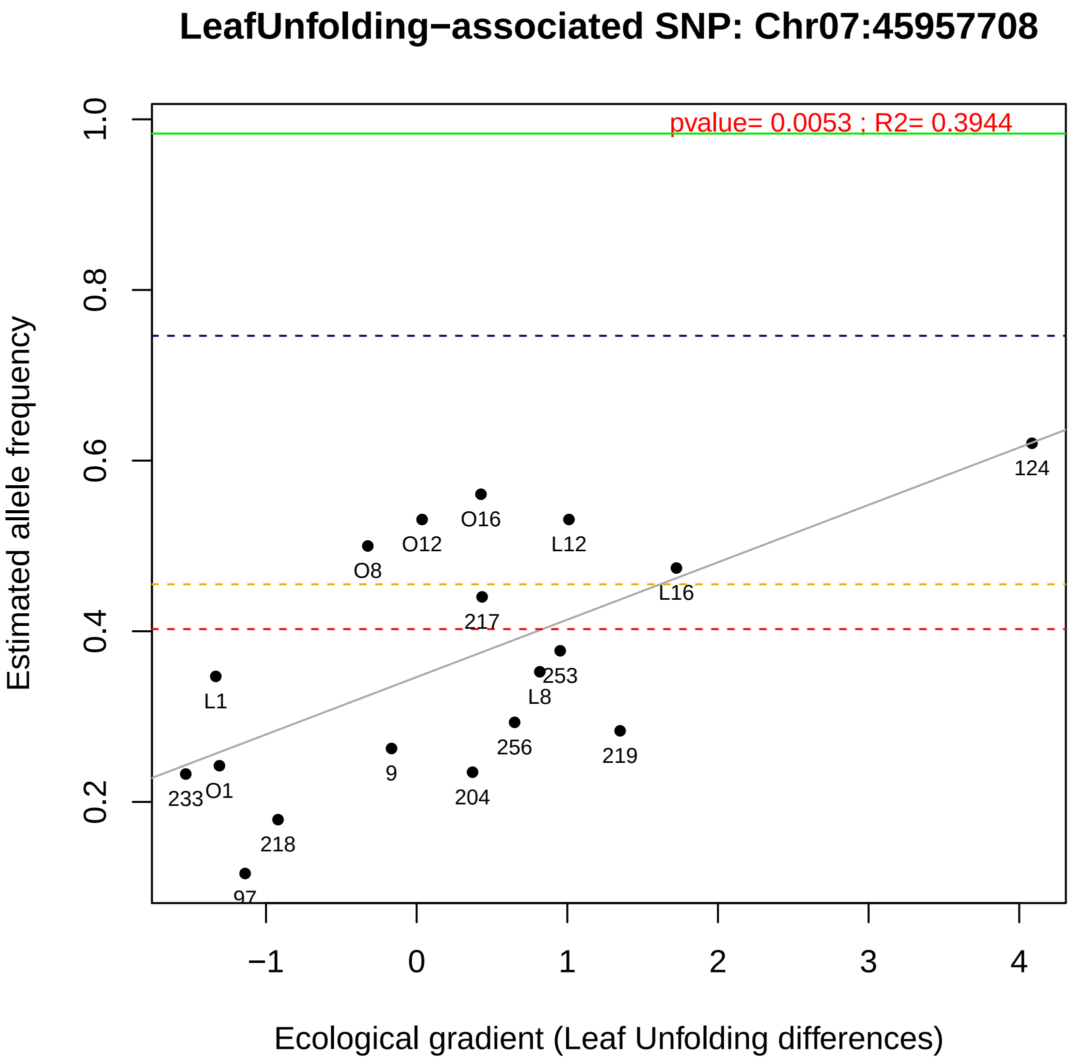

LeafUnfolding−associated SNP: Chr07:45957708
pvalue= 0.0053 ; R2= 0.3944
Estimated allele frequency
1.0
0.8
0.6
0.4
0.2
124
O16
O12
L12
O8
L16
217
253
L1
L8
256
219
9
O1
233
204
218
97
−1
0
1
2
3
4
Ecological gradient (Leaf Unfolding differences)

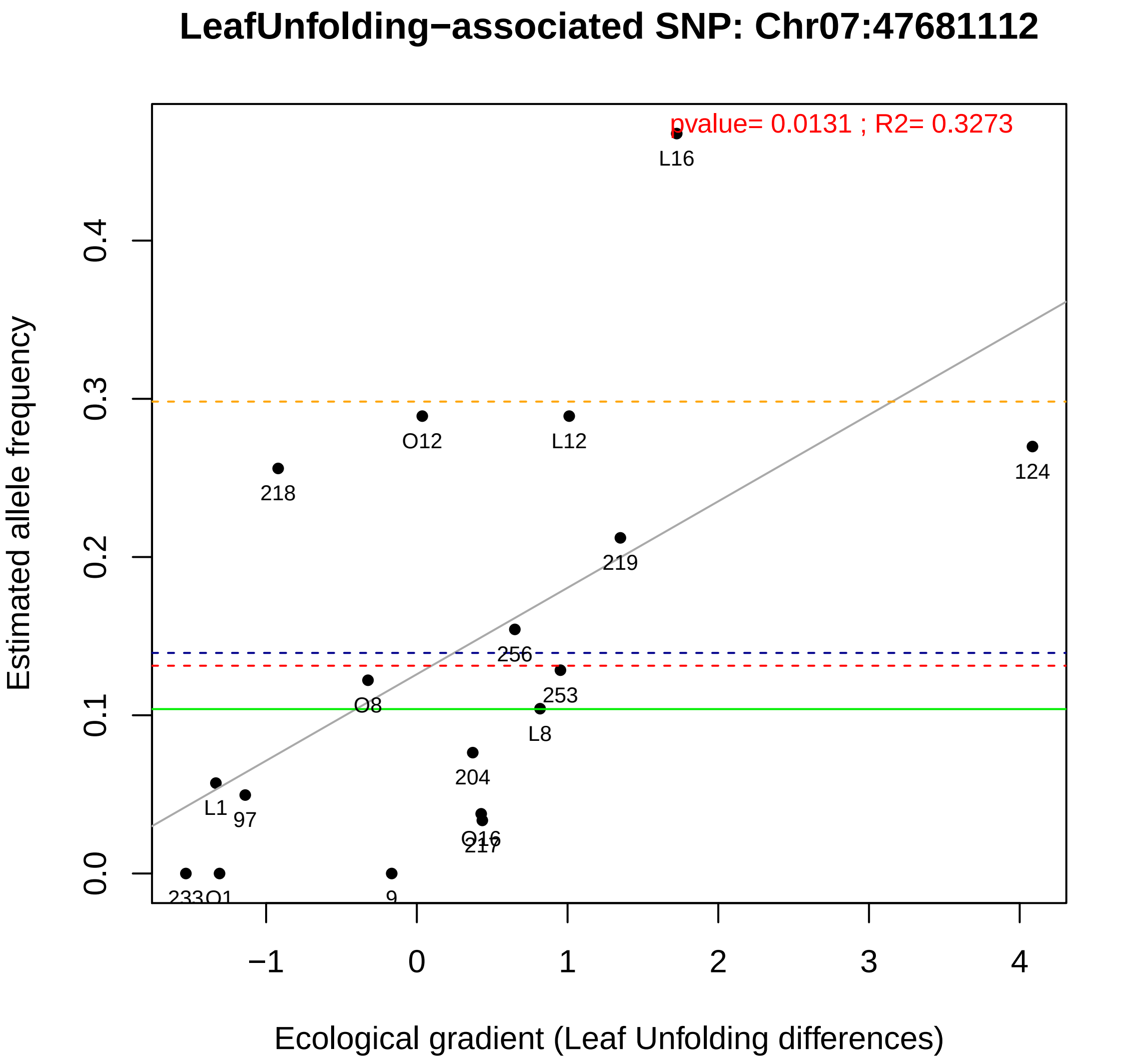

LeafUnfolding−associated SNP: Chr07:47681112
pvalue= 0.0131 ; R2= 0.3273
Estimated allele frequency
Ecological gradient (Leaf Unfolding differences)
0.0
0.1
0.2
0.3
0.4
−1
0
1
2
3
4
L16
O12
L12
124
218
219
256
253
O8
L8
204
L1
97
O16
217
233
O1
9

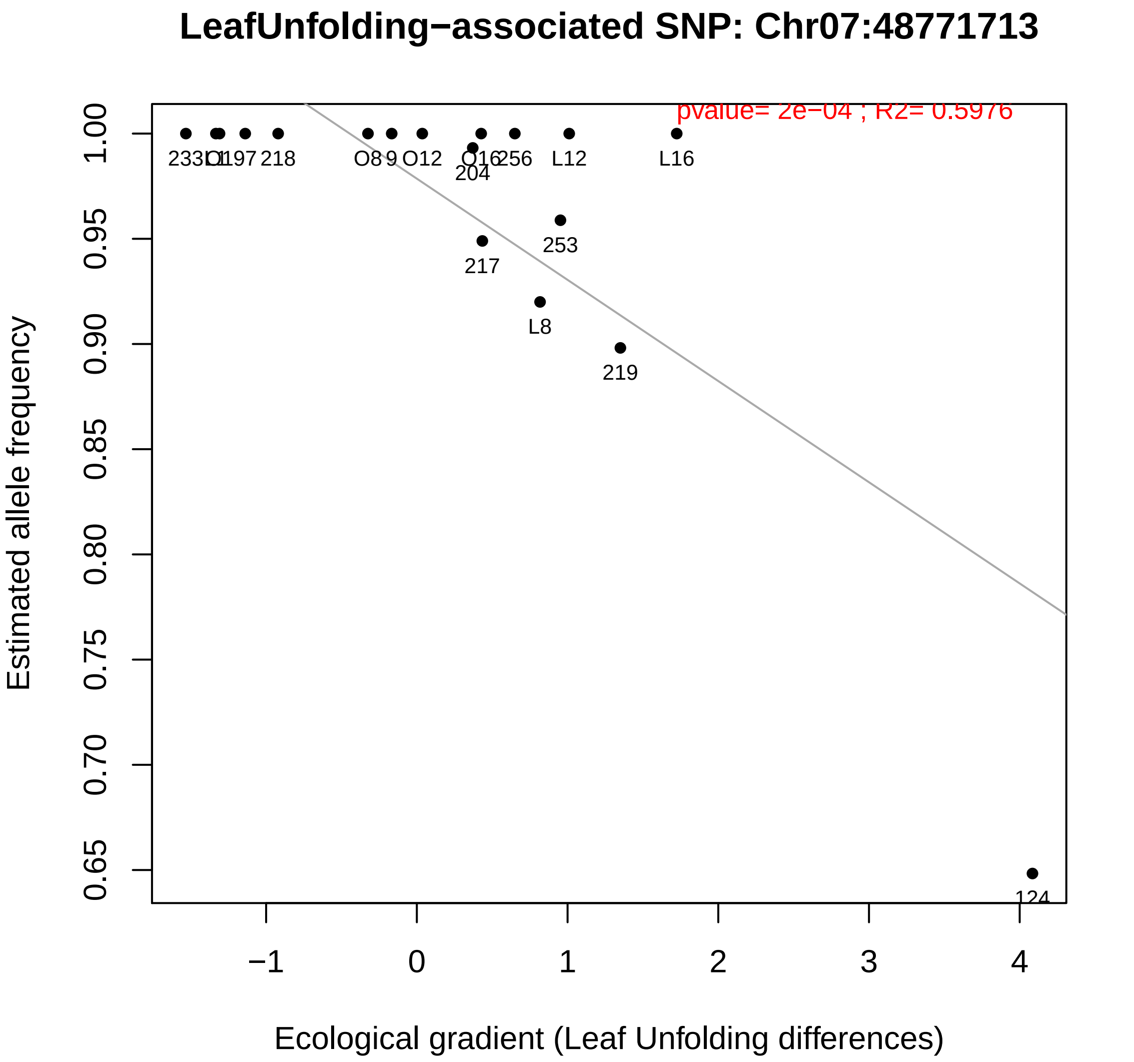

LeafUnfolding−associated SNP: Chr07:48771713
pvalue= 2e−04 ; R2= 0.5976
Estimated allele frequency
Ecological gradient (Leaf Unfolding differences)
1.00
0.95
0.90
0.85
0.80
0.75
0.70
0.65
−1
0
1
2
3
4
233
O1
197
218
O8
9
O12
O16
204
256
L12
L16
253
217
L8
219
124

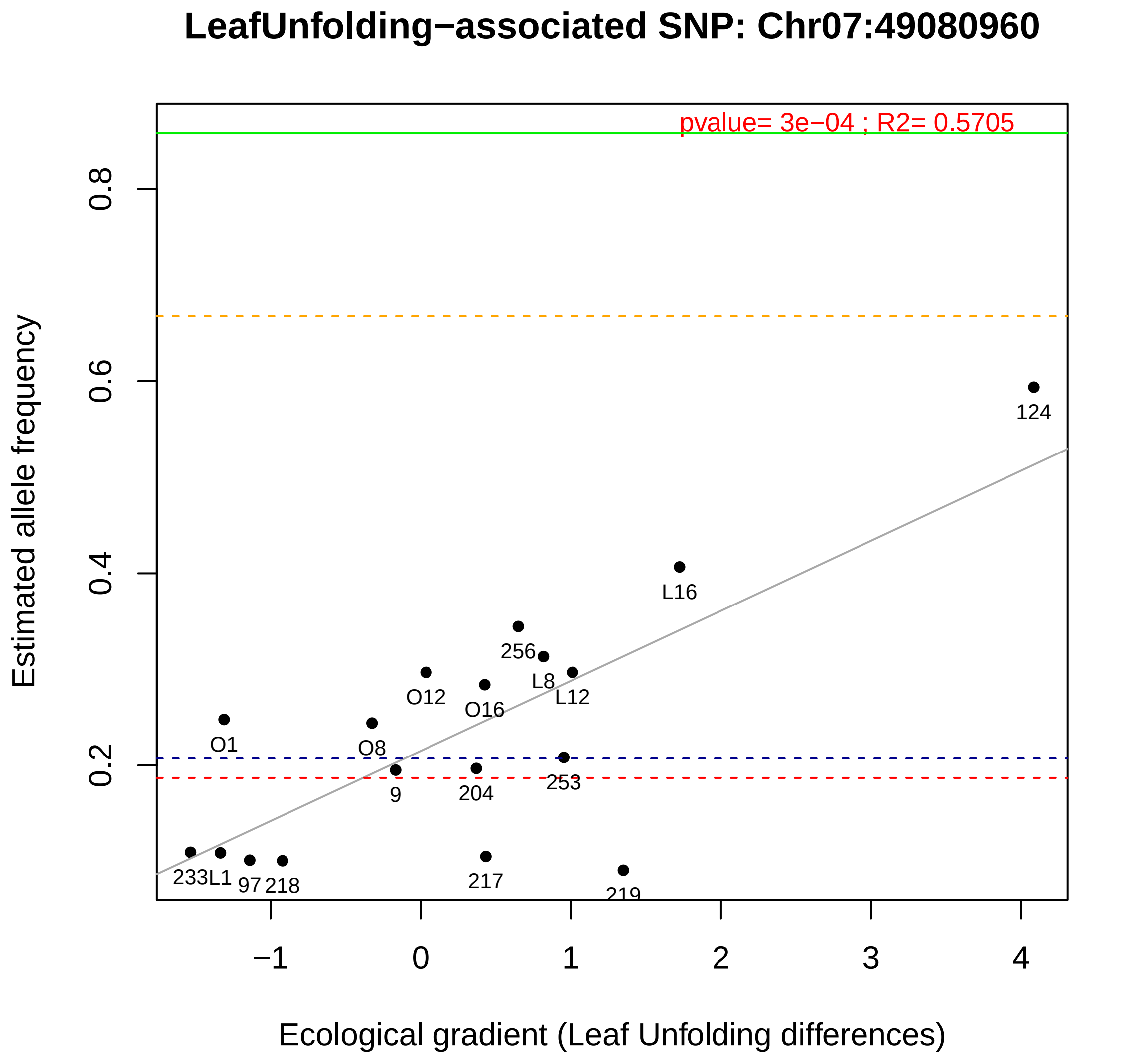

LeafUnfolding−associated SNP: Chr07:49080960
pvalue= 3e−04 ; R2= 0.5705
Estimated allele frequency
0.2
0.4
0.6
0.8
−1
0
1
2
3
4
Ecological gradient (Leaf Unfolding differences)
124
L16
256
L8
L12
O12
O16
O1
O8
9
204
253
233
L1
97
218
217
219

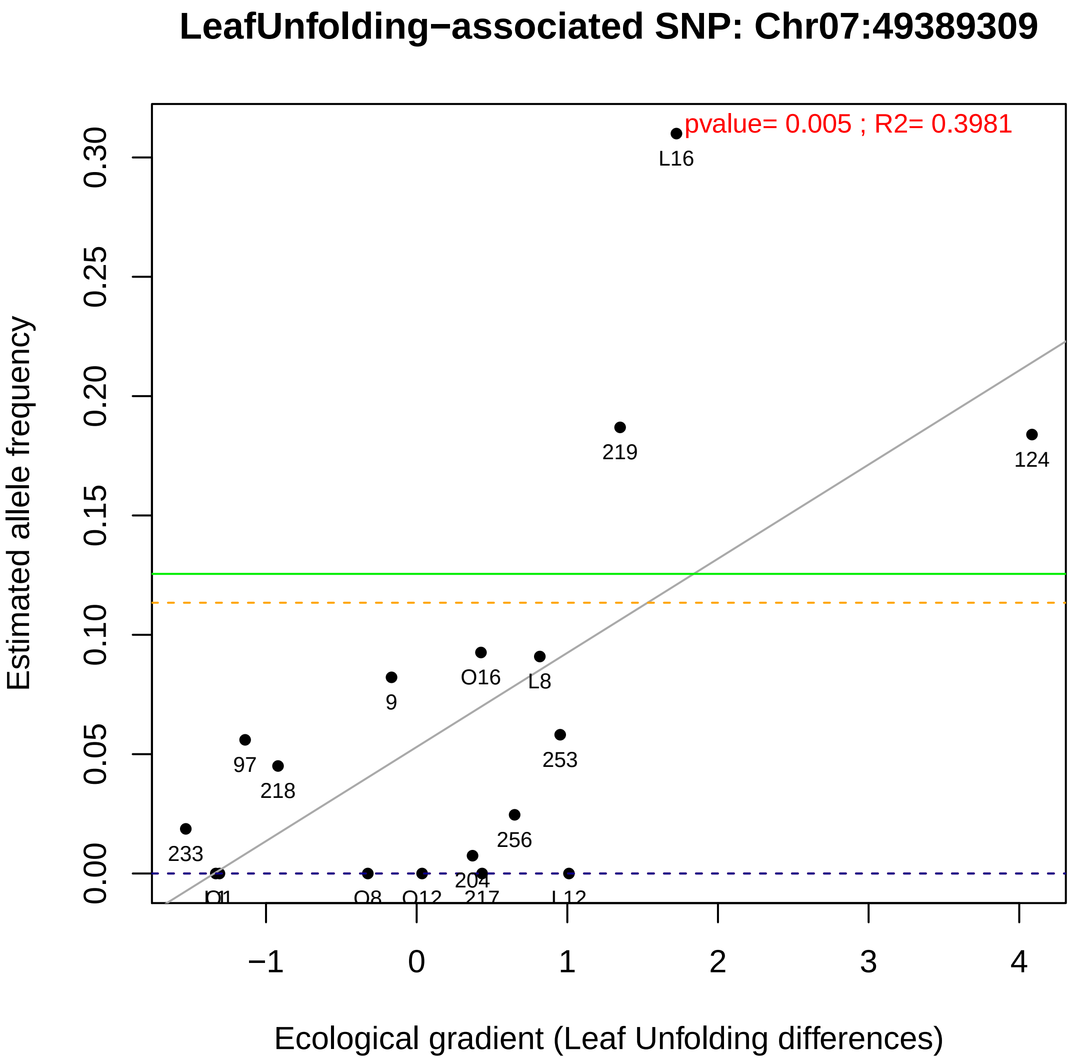

LeafUnfolding−associated SNP: Chr07:49389309
pvalue= 0.005 ; R2= 0.3981
L16
219
124
O16
L8
9
253
97
218
256
233
204
O8
O12
217
L12
0.00
0.05
0.10
0.15
0.20
0.25
0.30
−1
0
1
2
3
4
Estimated allele frequency
Ecological gradient (Leaf Unfolding differences)

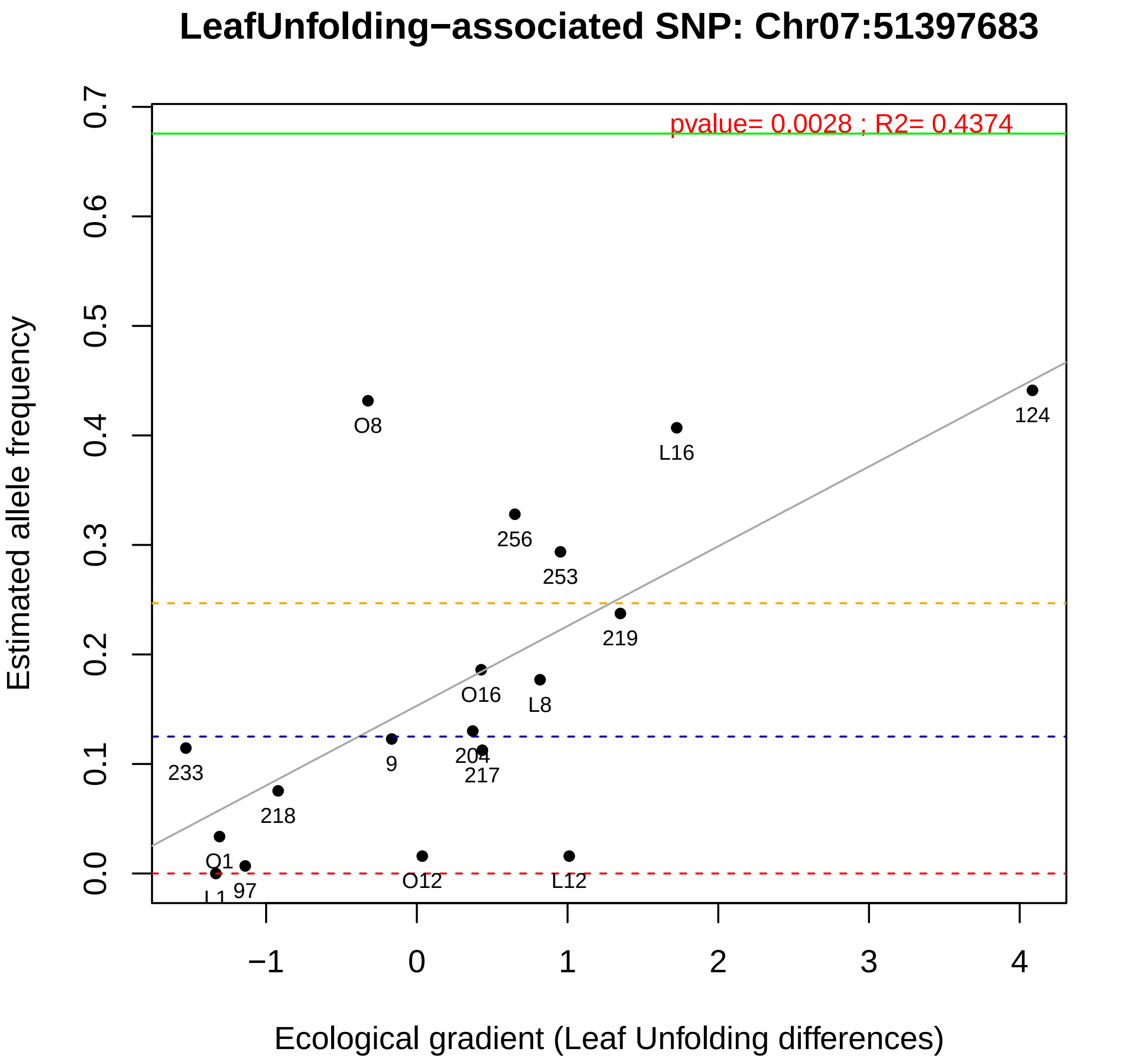
LeafUnfolding−associated SNP: Chr07:51397683
pvalue= 0.0028 ; R2= 0.4374
Estimated allele frequency
0.0
0.1
0.2
0.3
0.4
0.5
0.6
0.7
Ecological gradient (Leaf Unfolding differences)
−1
0
1
2
3
4
O8
124
L16
256
253
219
O16
L8
233
9
204
217
218
O1
97
L1
O12
L12

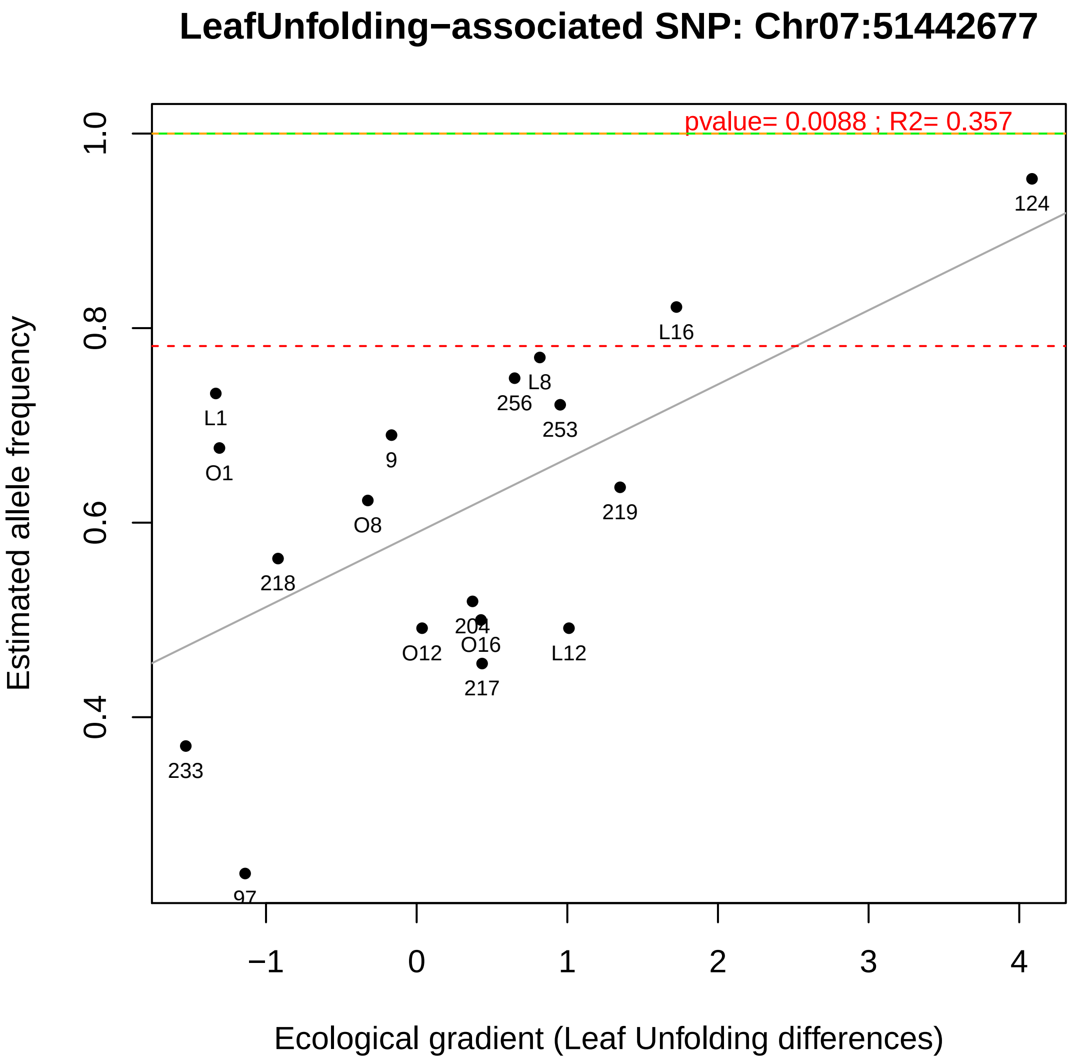

LeafUnfolding−associated SNP: Chr07:51442677
pvalue= 0.0088 ; R2= 0.357
124
L16
L8
256
253
L1
O1
9
219
O8
218
204
O16
O12
L12
217
233
97
1.0
0.8
0.6
0.4
−1
0
1
2
3
4
Estimated allele frequency
Ecological gradient (Leaf Unfolding differences)

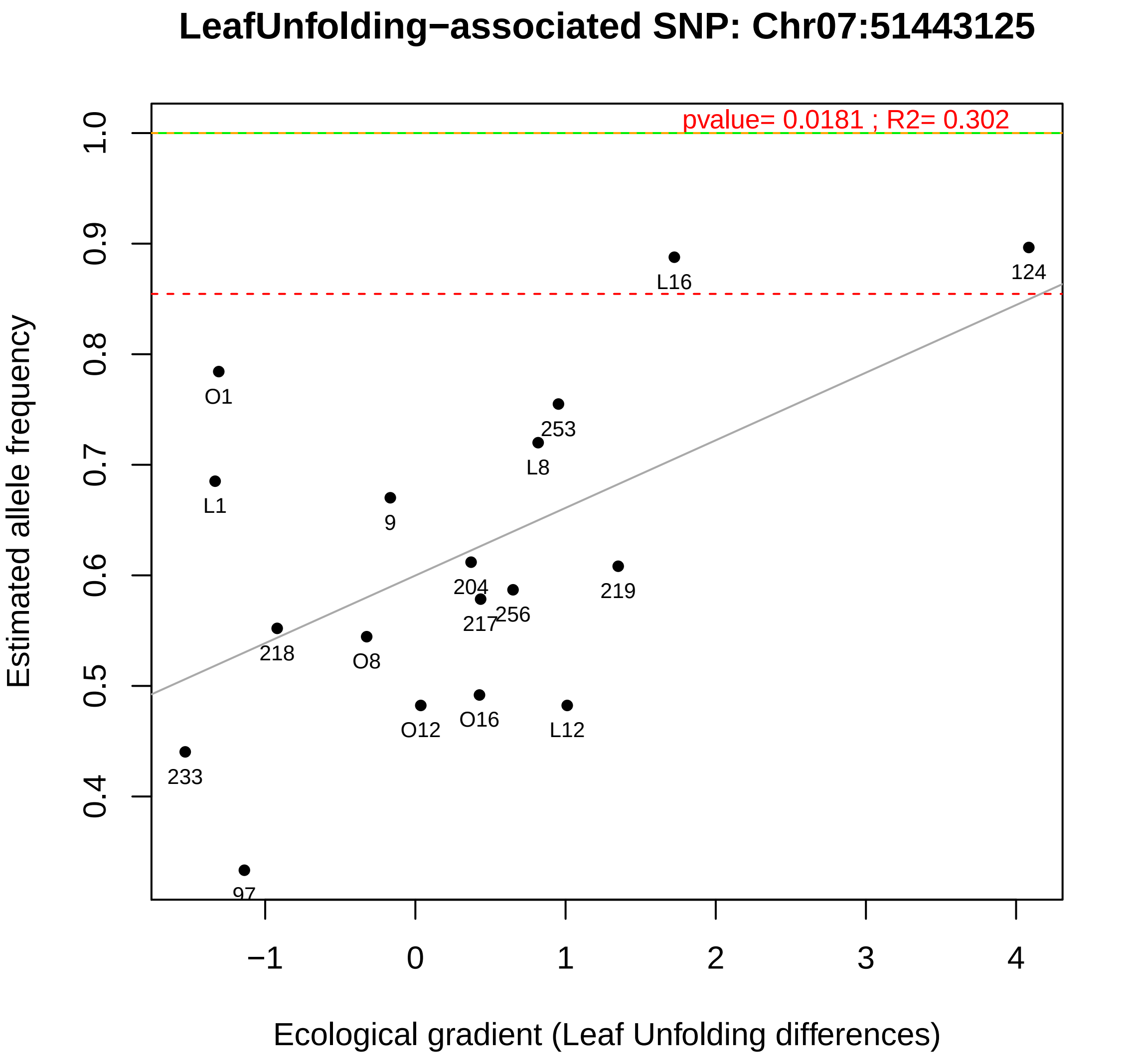
LeafUnfolding−associated SNP: Chr07:51443125
pvalue= 0.0181 ; R2= 0.302
Estimated allele frequency
Ecological gradient (Leaf Unfolding differences)
1.0
0.9
0.8
0.7
0.6
0.5
0.4
−1
0
1
2
3
4
L16
124
O1
253
L8
L1
9
204
219
217
256
218
O8
O12
O16
L12
233
97

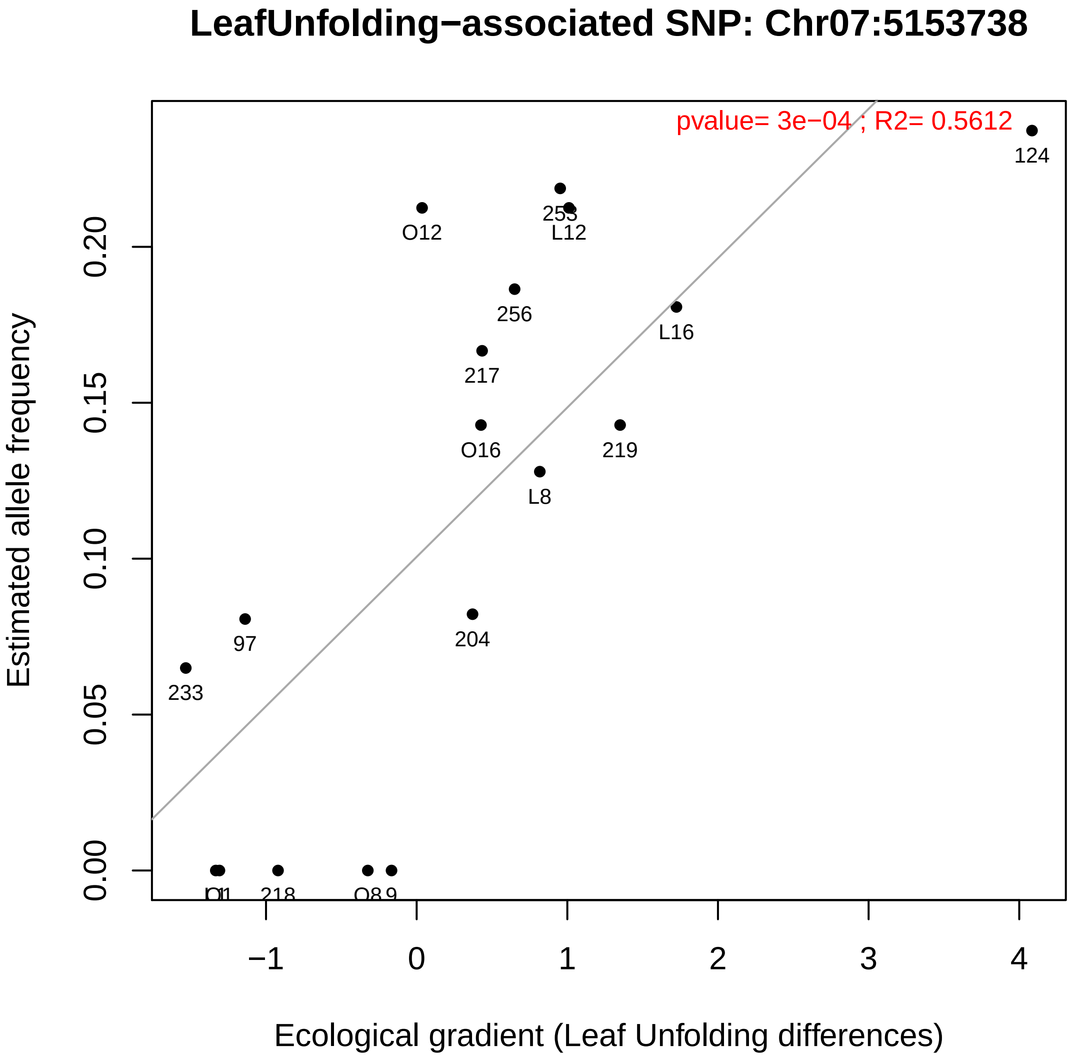

LeafUnfolding−associated SNP: Chr07:5153738
pvalue= 3e−04 ; R2= 0.5612
Estimated allele frequency
Ecological gradient (Leaf Unfolding differences)
0.00
0.05
0.10
0.15
0.20
−1
0
1
2
3
4
124
O12
253
L12
256
L16
217
O16
219
L8
97
204
233
218

**LeafUnfolding−associated SNP: Chr07:51576806**

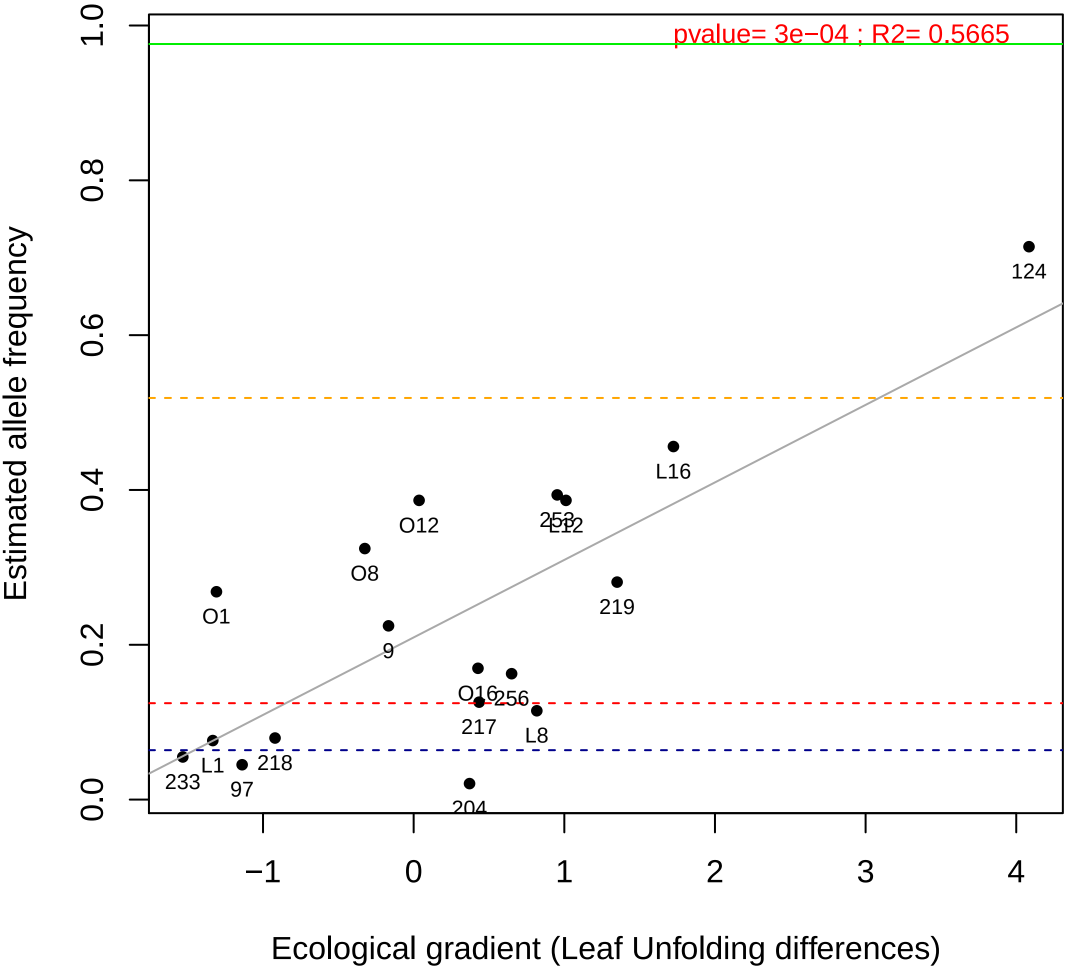

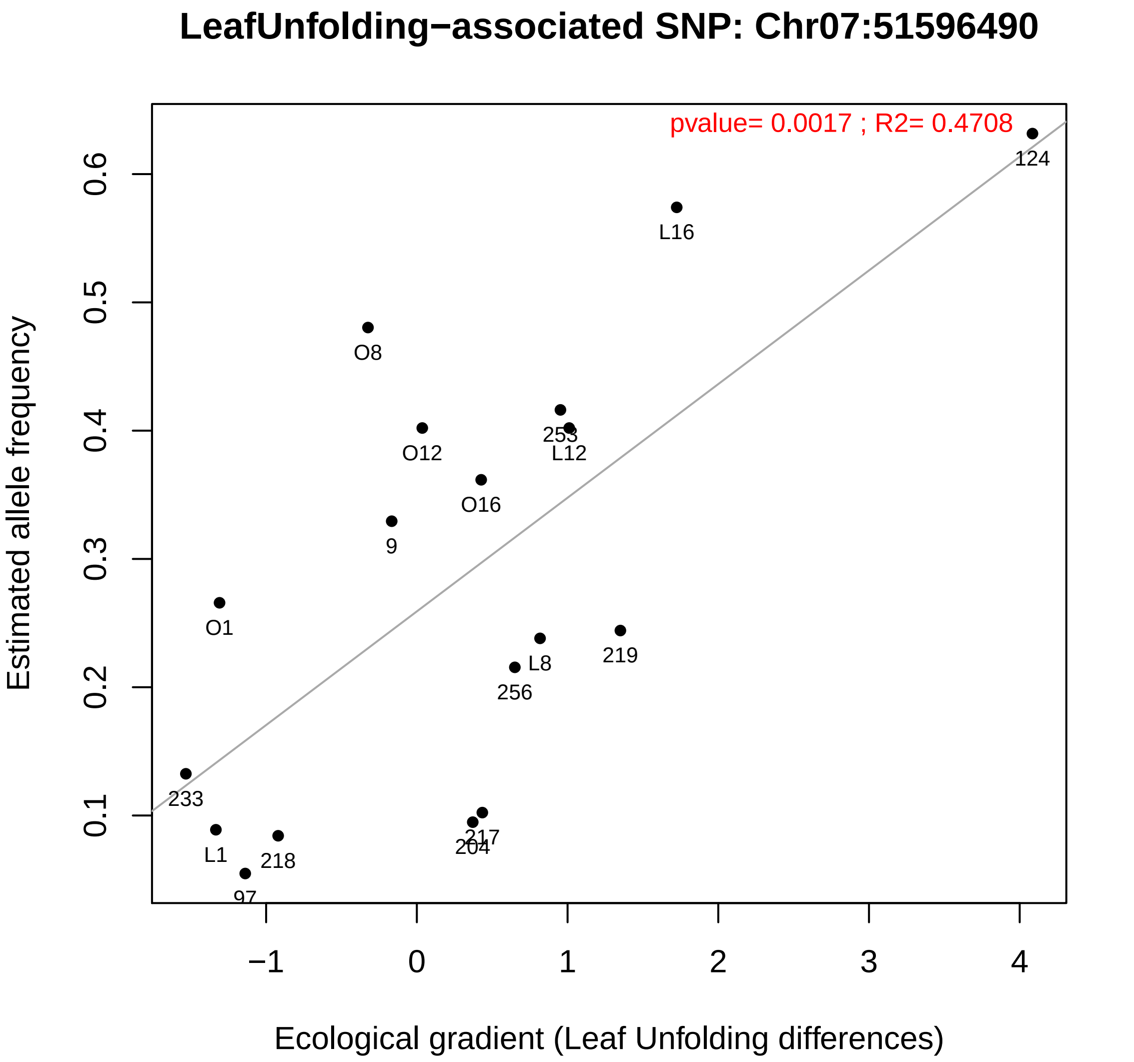
LeafUnfolding−associated SNP: Chr07:51596490
pvalue= 0.0017 ; R2= 0.4708
124
L16
O8
O12
253
L12
O16
9
O1
L8
219
256
233
L1
218
97
217
204
0.1
0.2
0.3
0.4
0.5
0.6
−1
0
1
2
3
4
Estimated allele frequency
Ecological gradient (Leaf Unfolding differences)

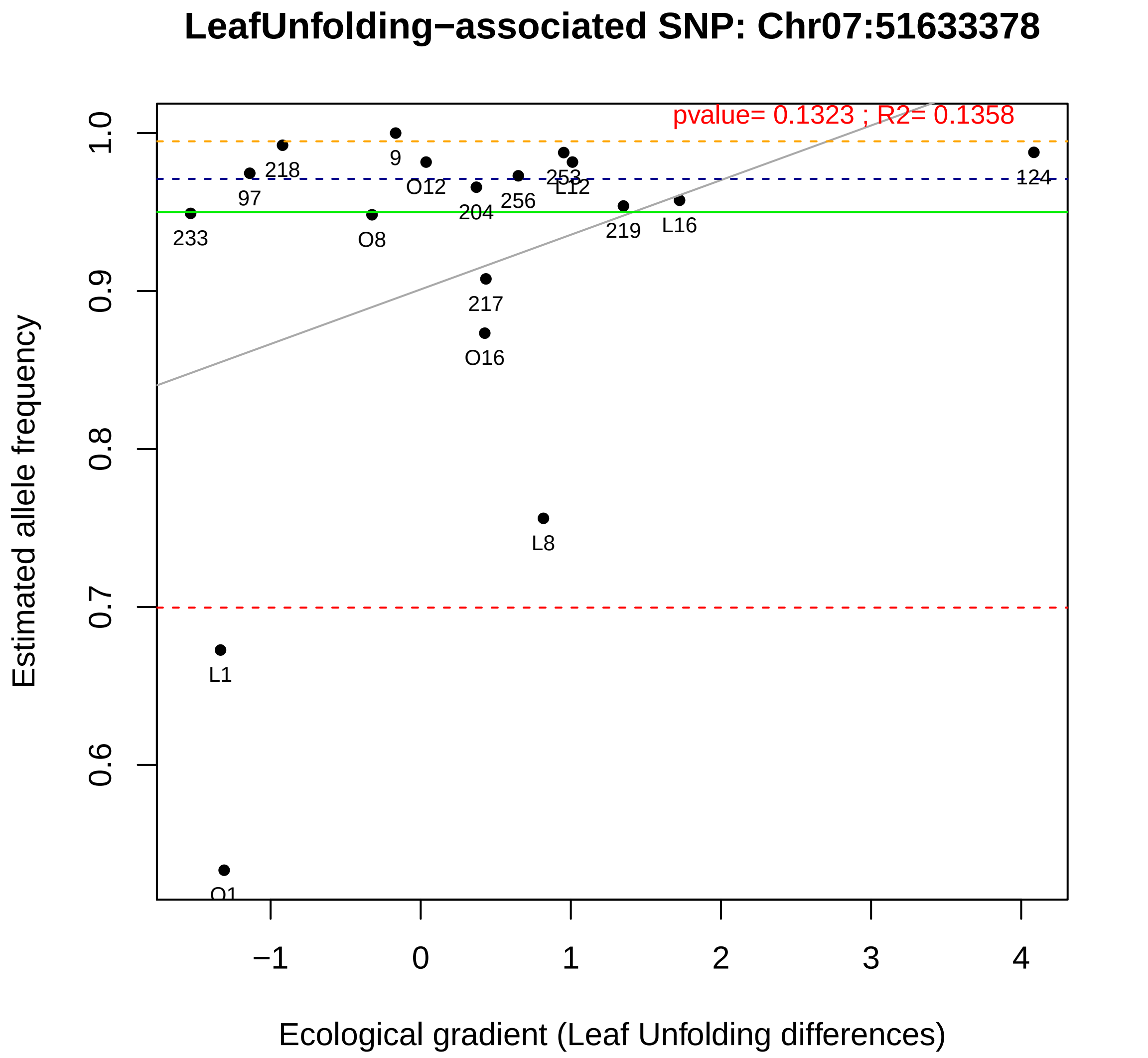

LeafUnfolding−associated SNP: Chr07:51633378
pvalue= 0.1323 ; R2= 0.1358
Estimated allele frequency
Ecological gradient (Leaf Unfolding differences)
0.6
0.7
0.8
0.9
1.0
−1
0
1
2
3
4
218
97
9
O12
204
256
253
L12
124
233
O8
219
L16
217
O16
L8
L1
O1

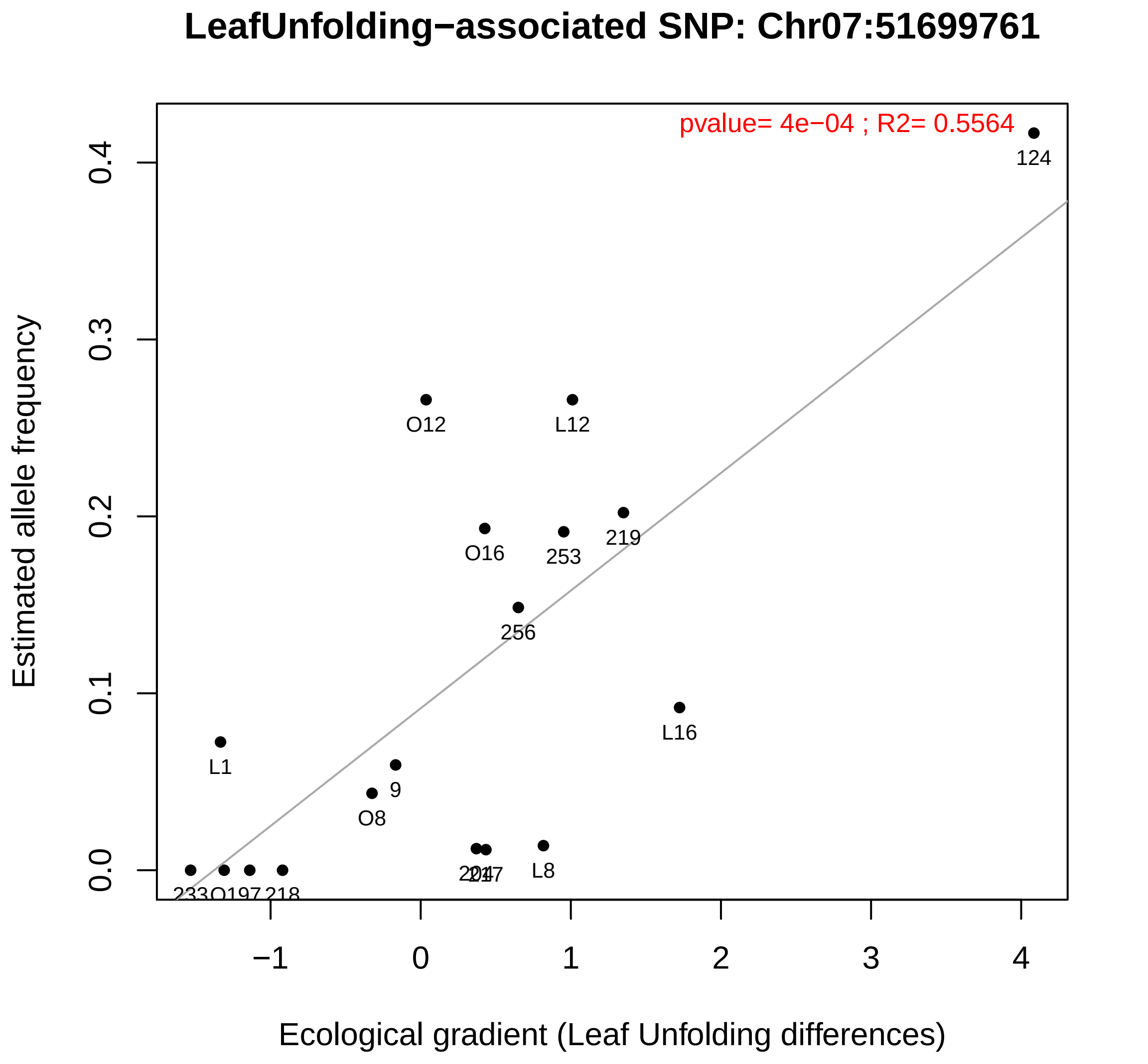
LeafUnfolding−associated SNP: Chr07:51699761
pvalue= 4e−04 ; R2= 0.5564
Estimated allele frequency
Ecological gradient (Leaf Unfolding differences)
0.0
0.1
0.2
0.3
0.4
−1
0
1
2
3
4
124
O12
L12
O16
253
219
256
L16
L1
9
O8
L8
218

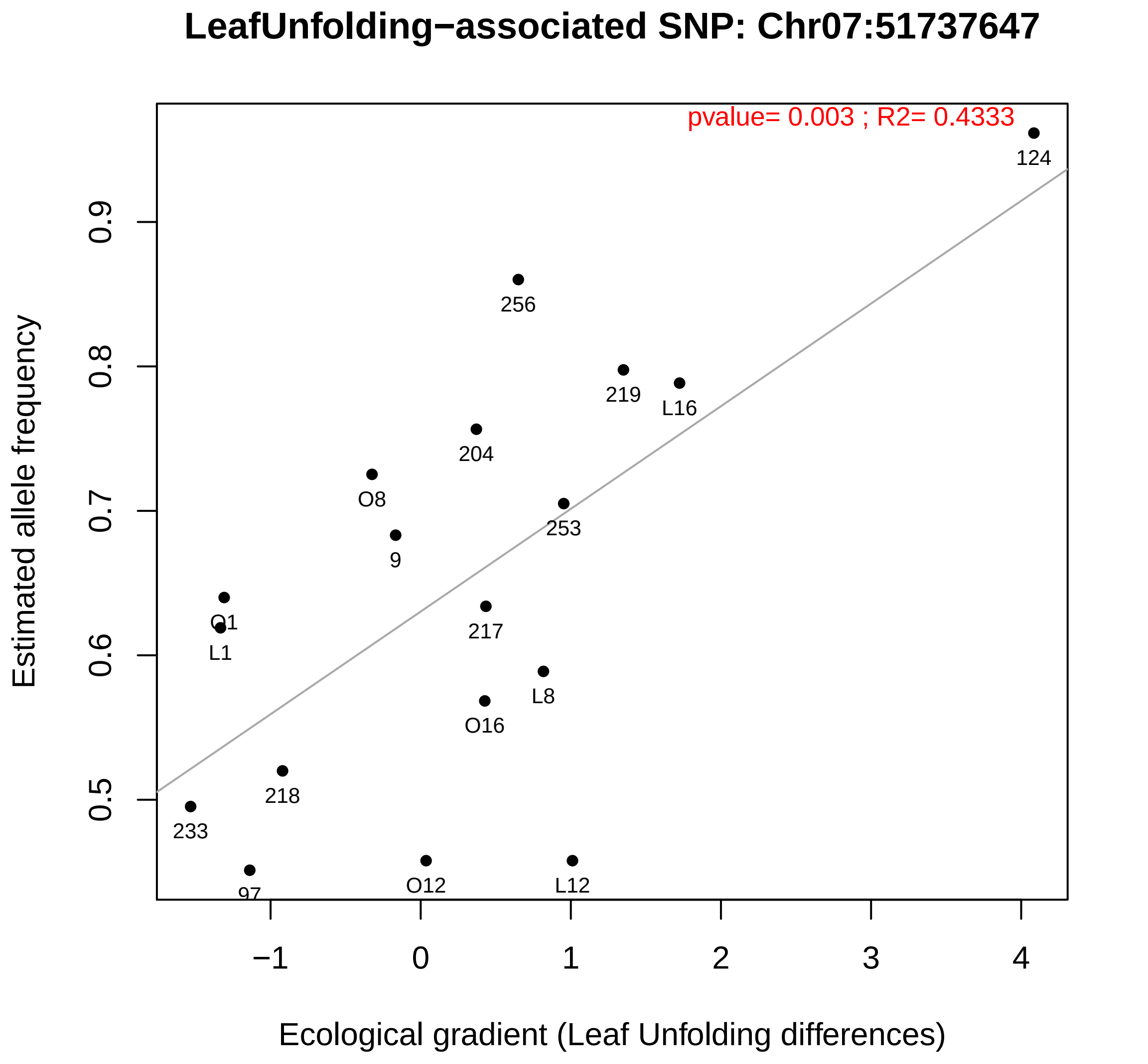

LeafUnfolding−associated SNP: Chr07:51737647
pvalue= 0.003 ; R2= 0.4333
124
256
219
L16
204
O8
253
9
O1
L1
217
L8
O16
218
233
97
O12
L12
0.5
0.6
0.7
0.8
0.9
−1
0
1
2
3
4
Estimated allele frequency
Ecological gradient (Leaf Unfolding differences)

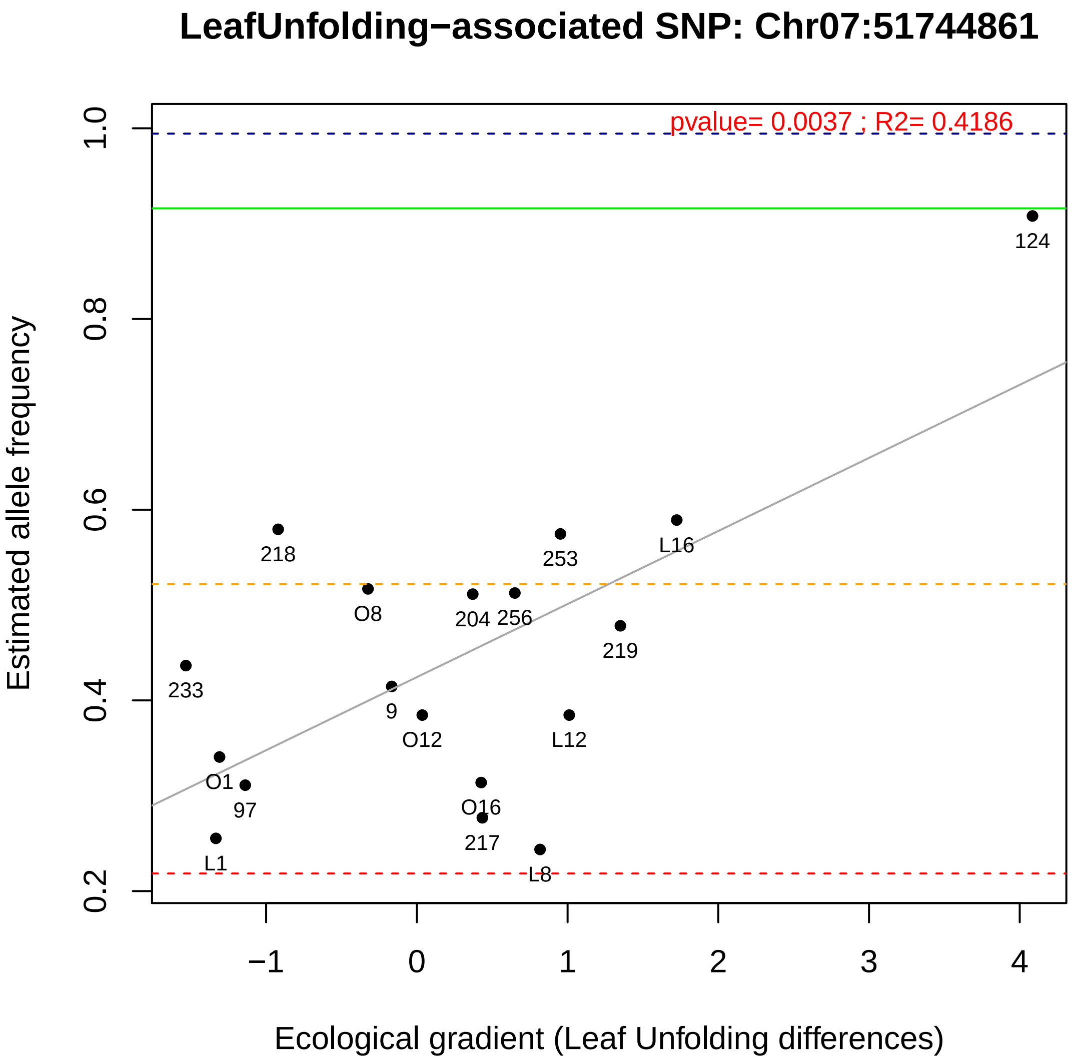
LeafUnfolding−associated SNP: Chr07:51744861
pvalue= 0.0037 ; R2= 0.4186
Estimated allele frequency
0.2
0.4
0.6
0.8
1.0
Ecological gradient (Leaf Unfolding differences)
−1
0
1
2
3
4
124
218
253
L16
O8
204
256
219
233
9
O12
L12
O1
97
O16
217
L1
L8

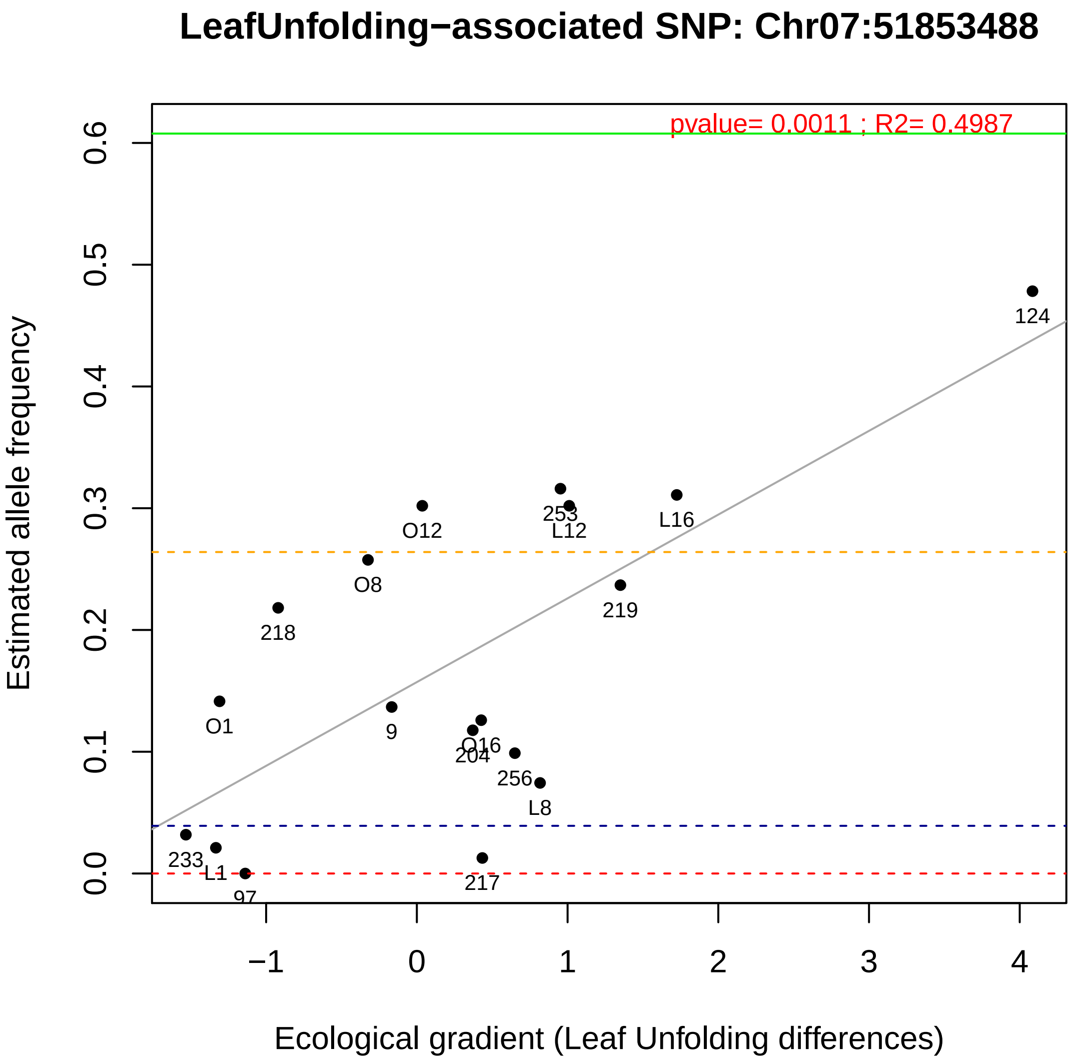

LeafUnfolding−associated SNP: Chr07:51853488
pvalue= 0.0011 ; R2= 0.4987
Estimated allele frequency
0.0
0.1
0.2
0.3
0.4
0.5
0.6
−1
0
1
2
3
4
Ecological gradient (Leaf Unfolding differences)
124
253
L12
L16
O12
O8
219
218
O1
9
O16
204
256
L8
233
L1
97
217

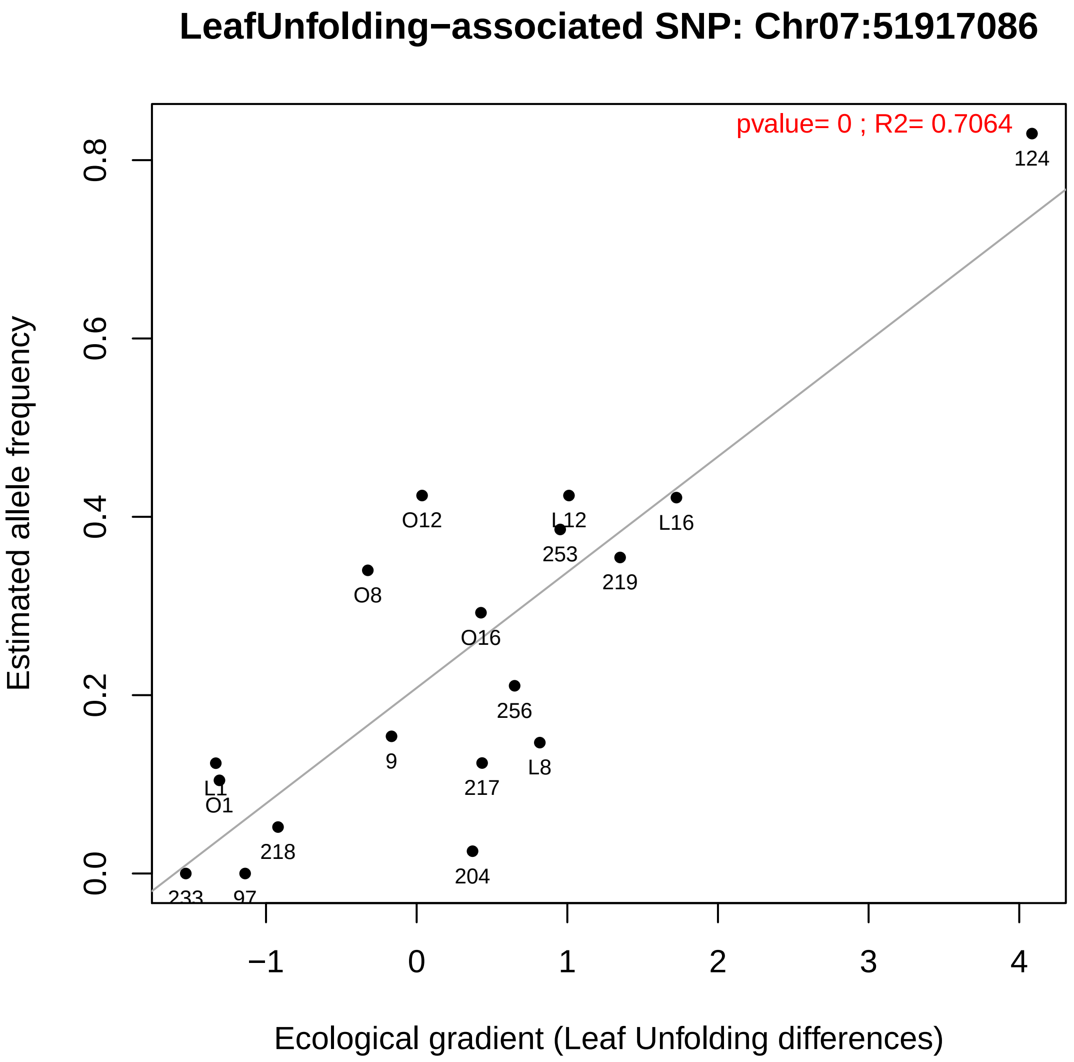

LeafUnfolding−associated SNP: Chr07:51917086
pvalue= 0 ; R2= 0.7064
Estimated allele frequency
Ecological gradient (Leaf Unfolding differences)
0.0
0.2
0.4
0.6
0.8
−1
0
1
2
3
4
124
O12
L12
253
L16
219
O8
O16
256
9
217
L8
L1
O1
218
204
233
97

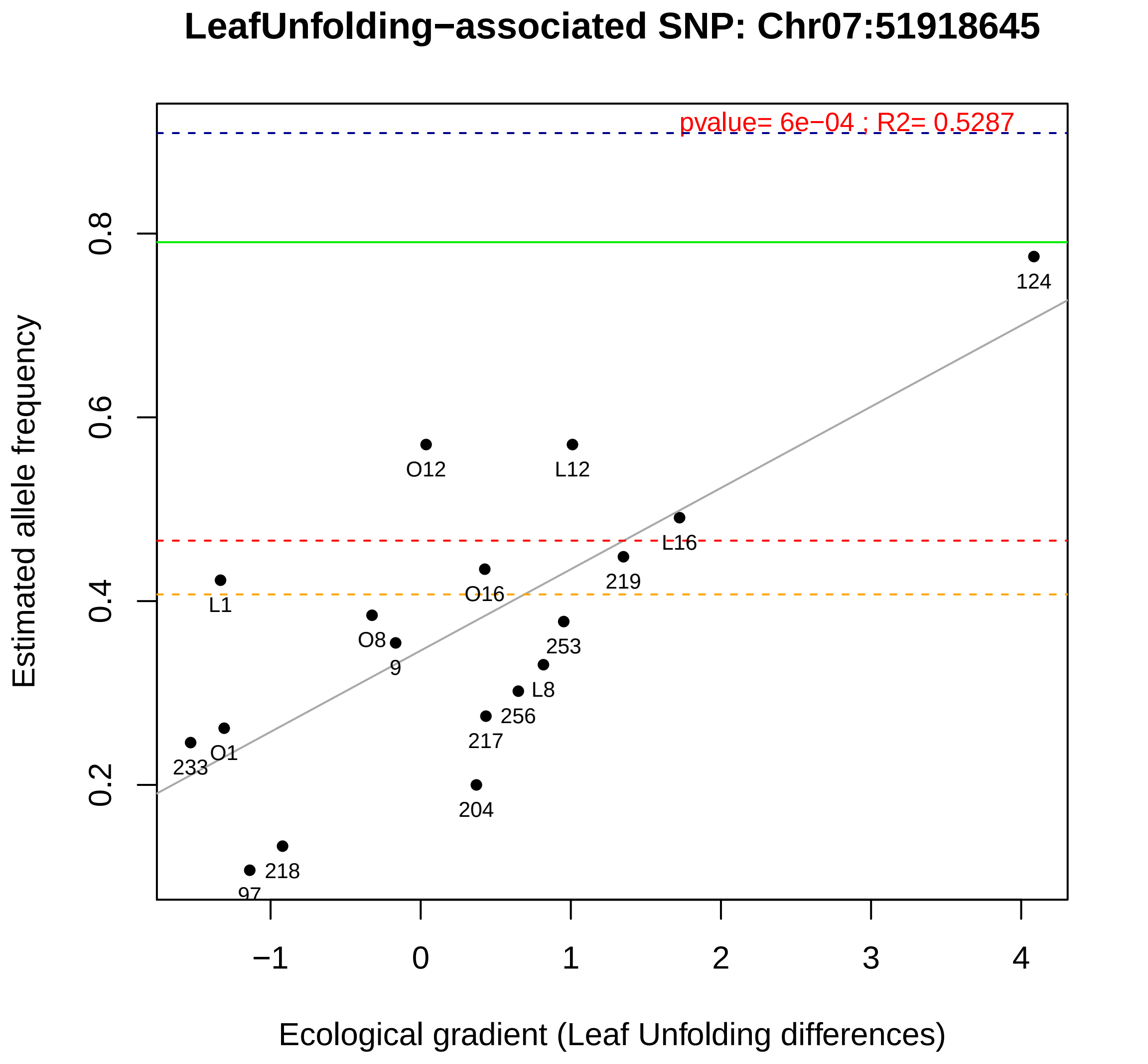
LeafUnfolding−associated SNP: Chr07:51918645
pvalue= 6e−04 ; R2= 0.5287
Estimated allele frequency
0.2
0.4
0.6
0.8
−1
0
1
2
3
4
Ecological gradient (Leaf Unfolding differences)
124
O12
L12
L16
219
O16
L1
253
O8
9
L8
256
217
233
O1
204
218
97

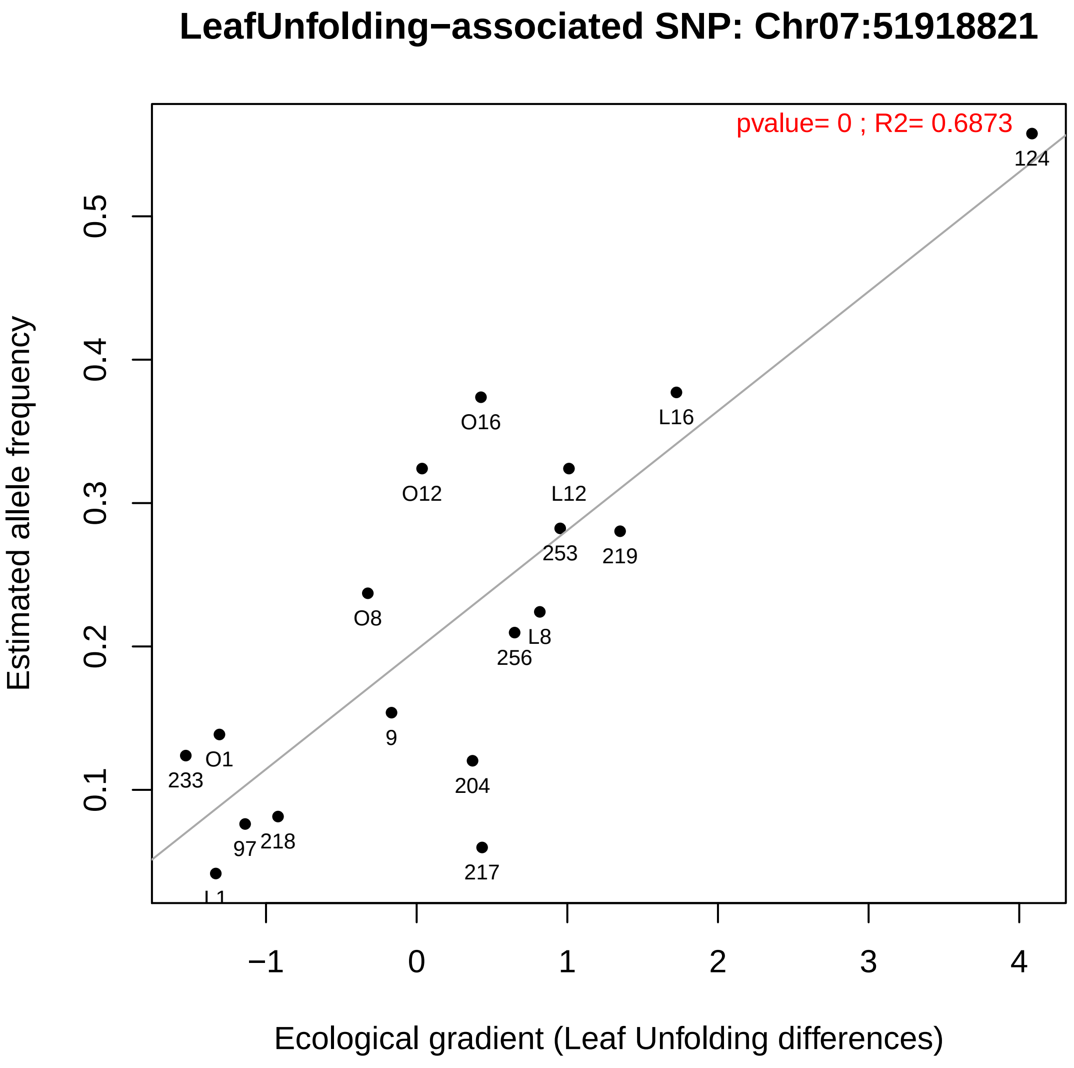

LeafUnfolding−associated SNP: Chr07:51918821
pvalue= 0 ; R2= 0.6873
124
O16
L16
O12
L12
253
219
O8
L8
256
9
O1
233
204
97
218
217
L1
0.1
0.2
0.3
0.4
0.5
−1
0
1
2
3
4
Estimated allele frequency
Ecological gradient (Leaf Unfolding differences)

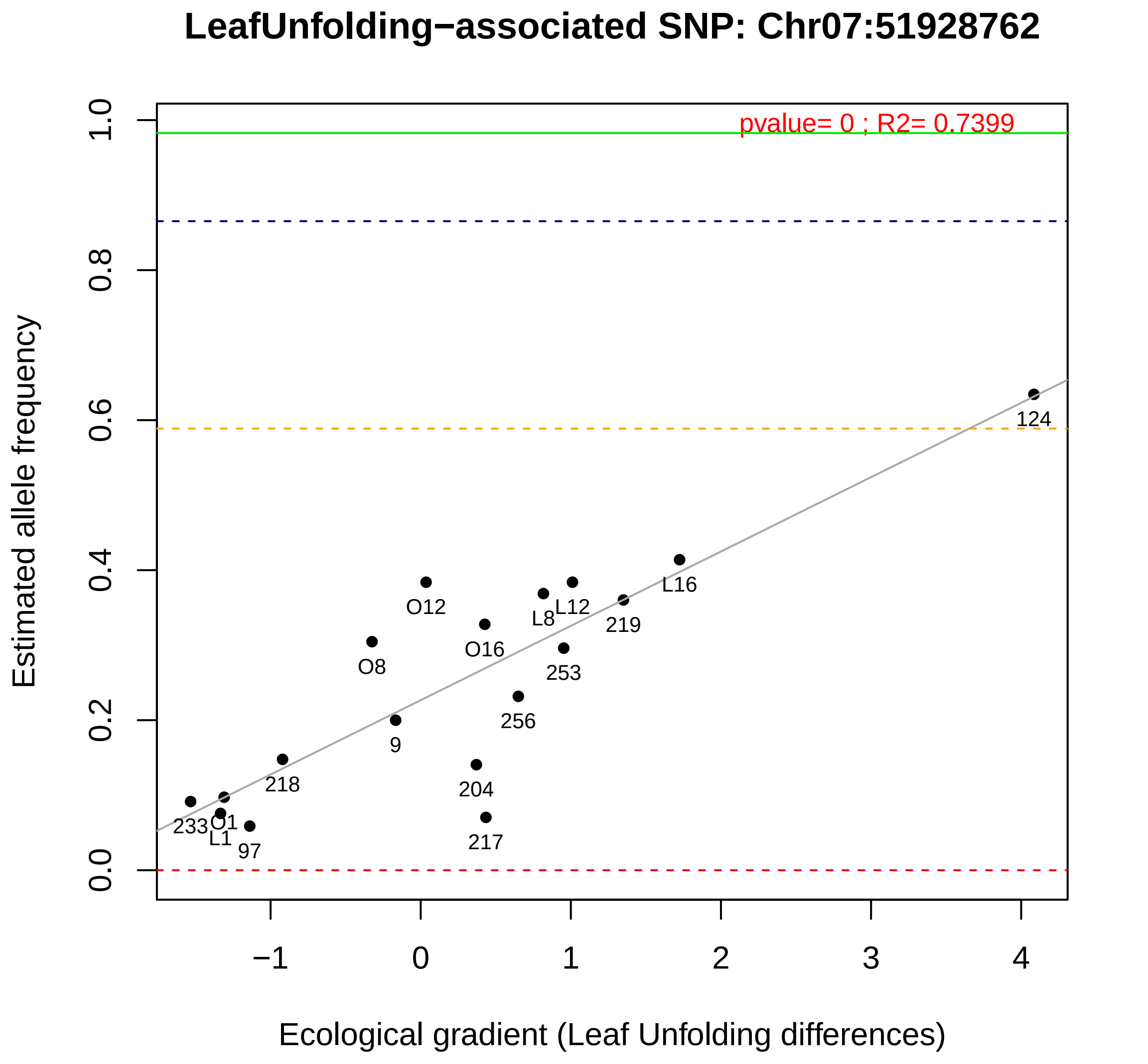
LeafUnfolding−associated SNP: Chr07:51928762
pvalue= 0 ; R2= 0.7399
Estimated allele frequency
0.0
0.2
0.4
0.6
0.8
1.0
−1
0
1
2
3
4
Ecological gradient (Leaf Unfolding differences)
124
L16
O12
L8
L12
219
O16
253
O8
256
9
218
204
233
O1
L1
97
217

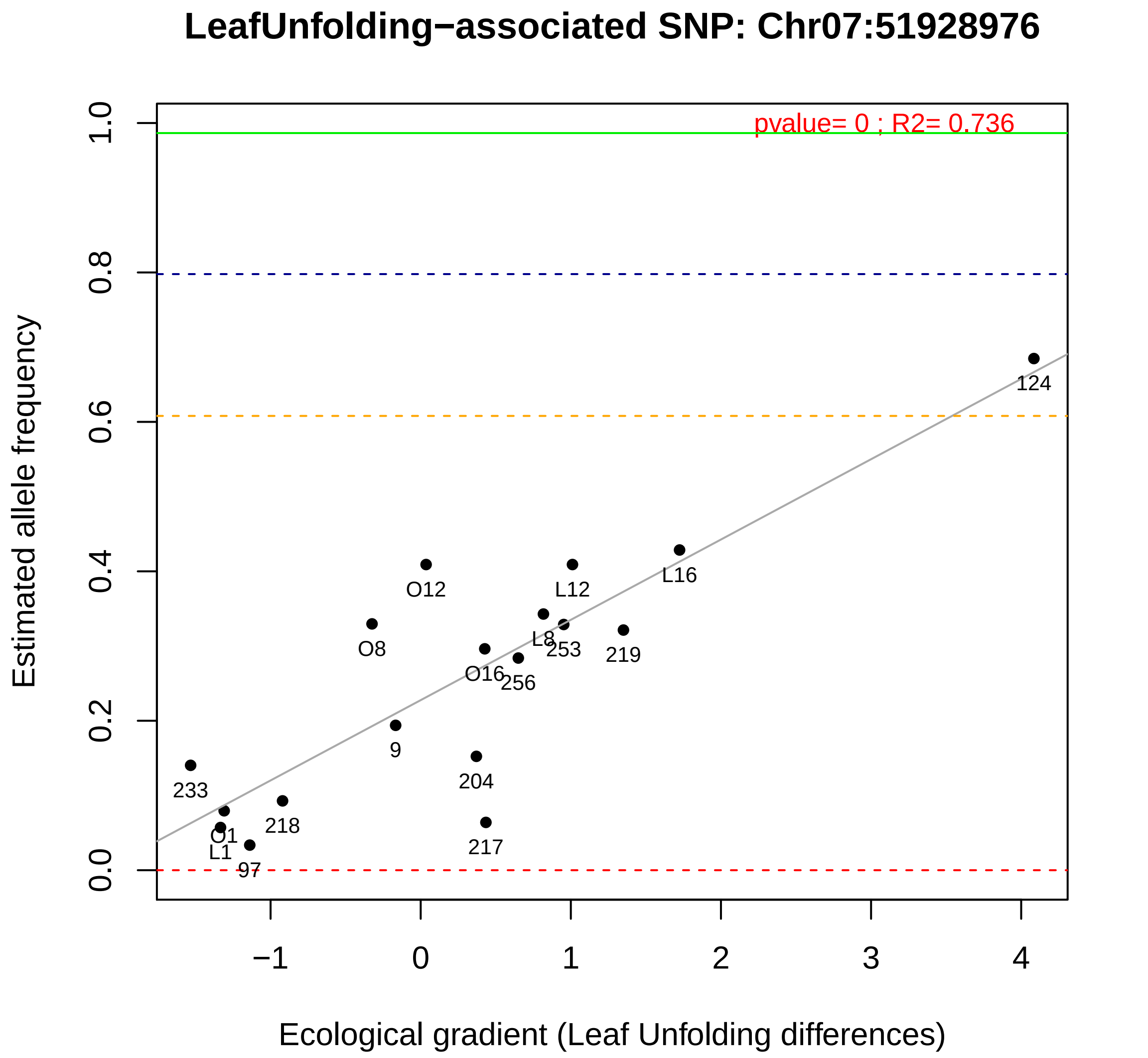

LeafUnfolding−associated SNP: Chr07:51928976
pvalue= 0 ; R2= 0.736
Estimated allele frequency
0.0
0.2
0.4
0.6
0.8
1.0
−1
0
1
2
3
4
Ecological gradient (Leaf Unfolding differences)
124
L16
O12
L12
O8
L8
253
219
O16
256
9
204
233
218
O1
L1
97
217

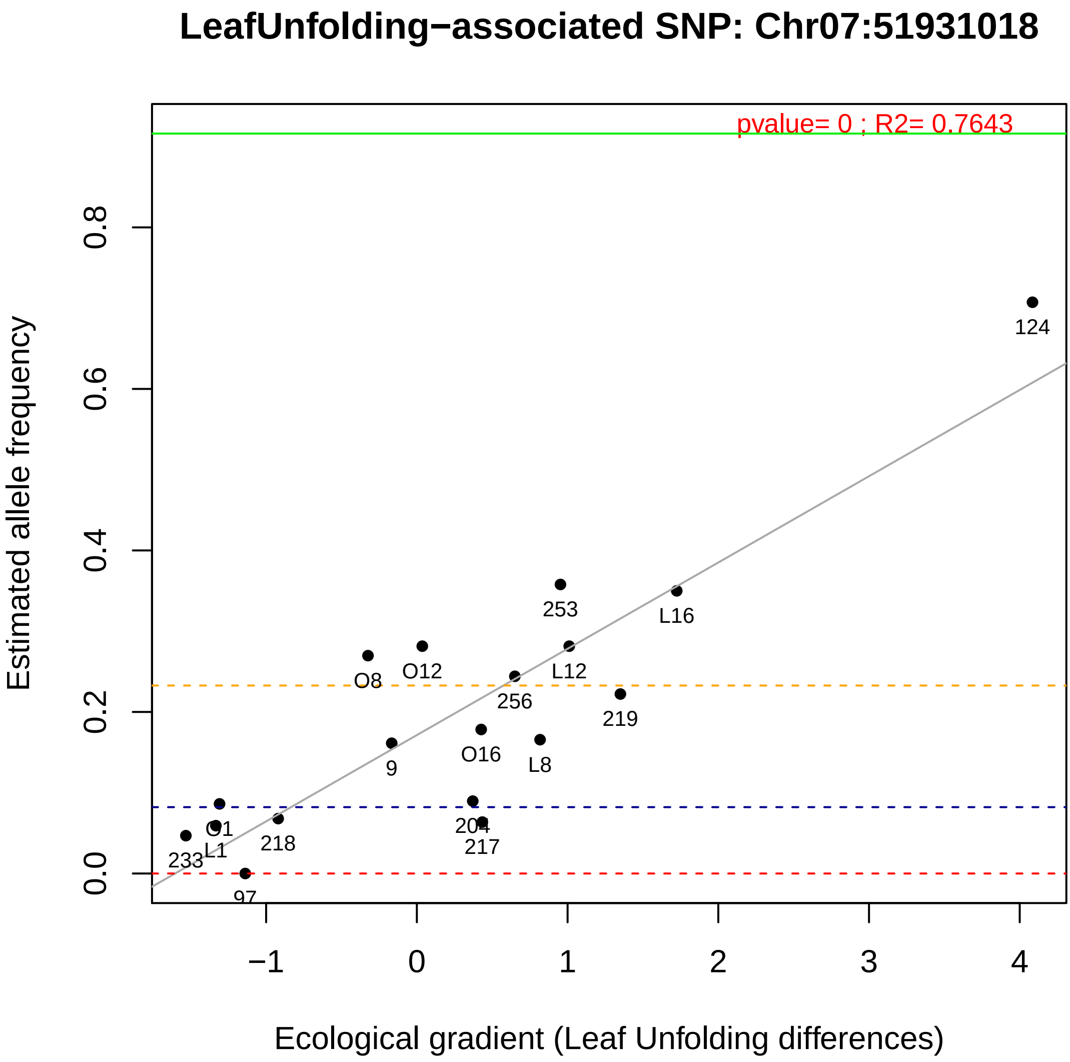

LeafUnfolding−associated SNP: Chr07:51931018
pvalue= 0 ; R2= 0.7643
Estimated allele frequency
0.0
0.2
0.4
0.6
0.8
−1
0
1
2
3
4
Ecological gradient (Leaf Unfolding differences)
124
253
L16
O8
O12
L12
256
219
9
O16
L8
O1
L1
218
204
217
233
97

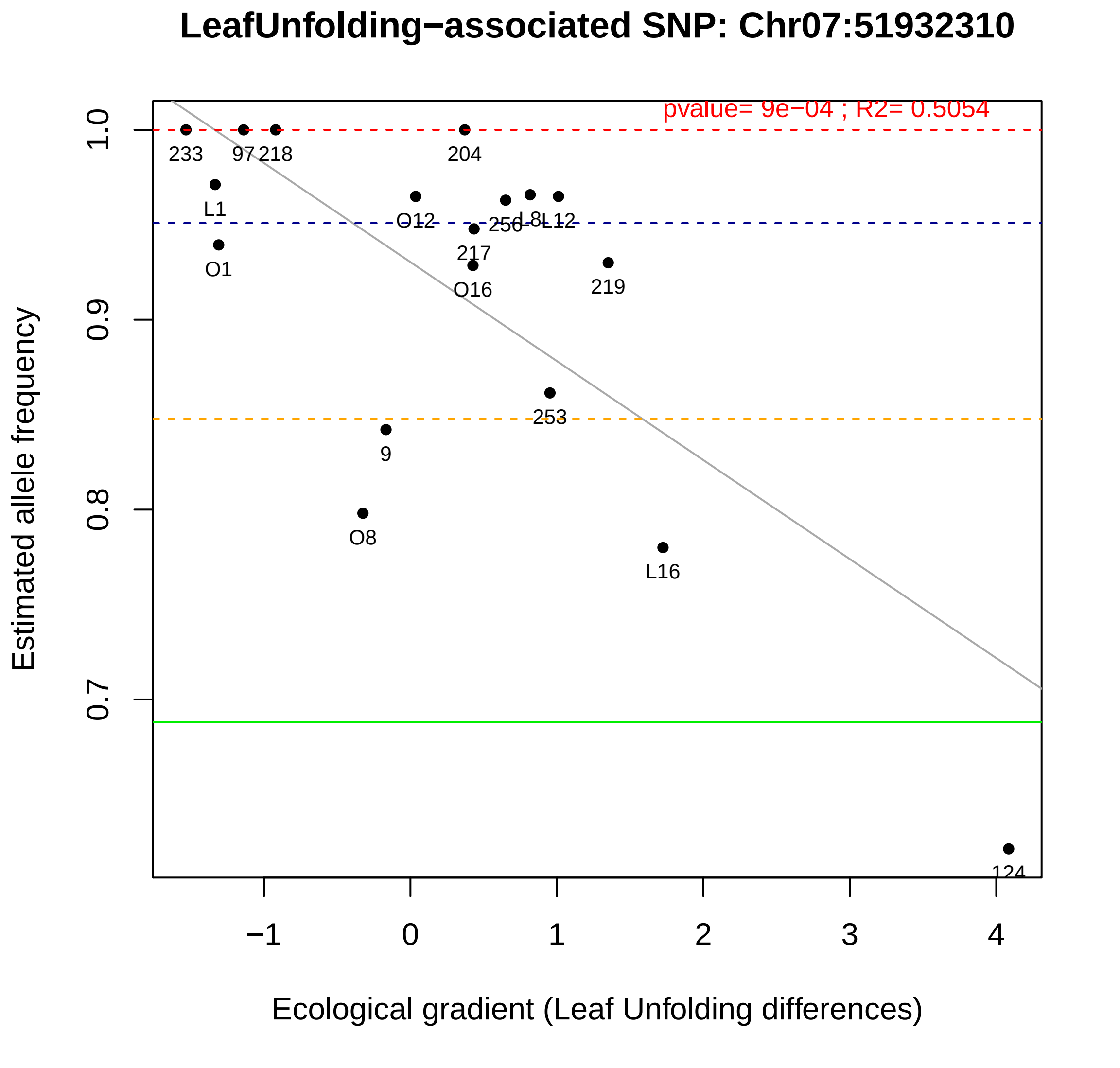

LeafUnfolding−associated SNP: Chr07:51932310
pvalue= 9e−04 ; R2= 0.5054
Estimated allele frequency
Ecological gradient (Leaf Unfolding differences)
1.0
0.9
0.8
0.7
−1
0
1
2
3
4
233
97
218
204
L1
O12
256
L8
L12
217
O1
O16
219
253
9
O8
L16
124

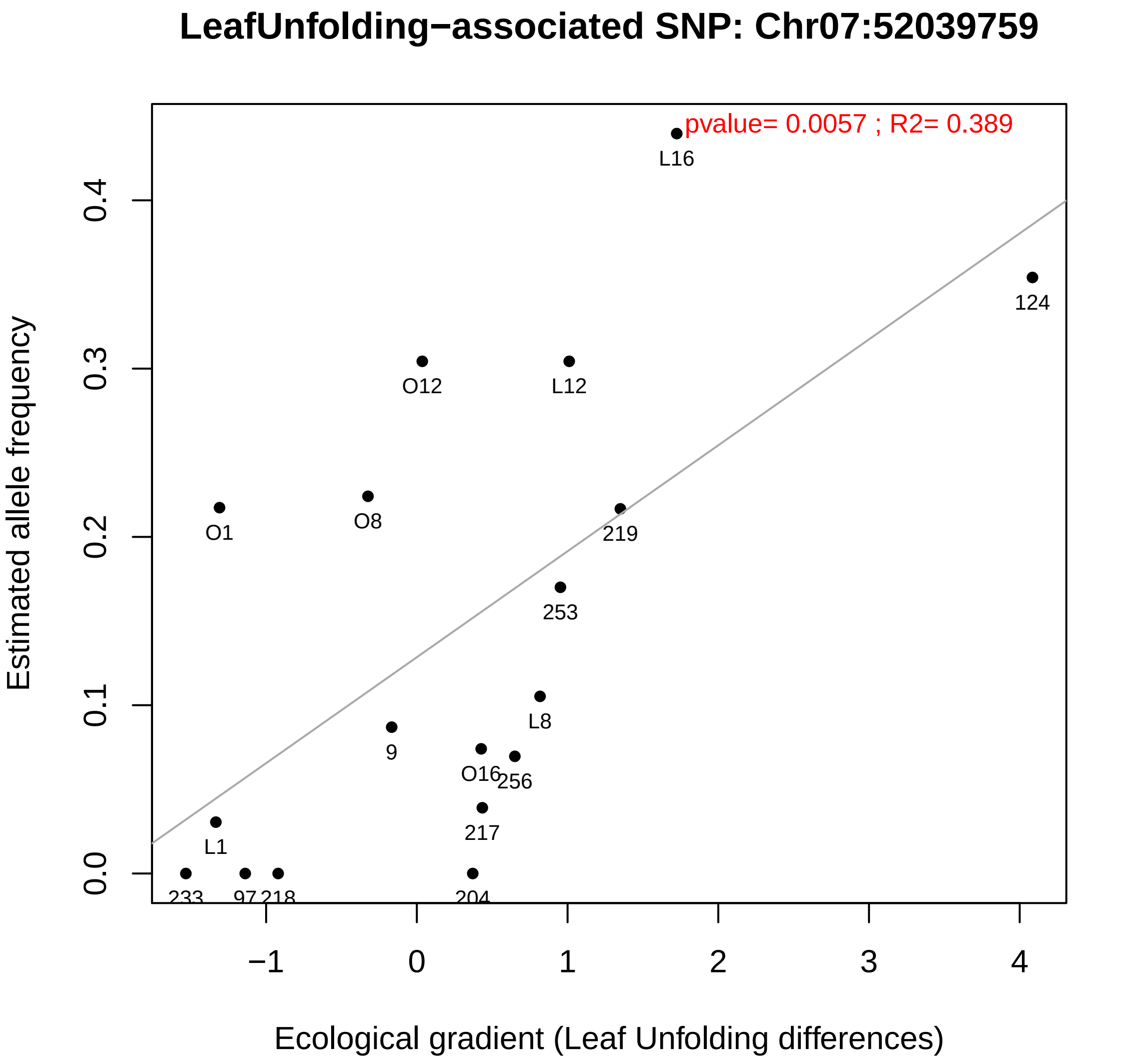
LeafUnfolding−associated SNP: Chr07:52039759
pvalue= 0.0057 ; R2= 0.389
L16
124
O12
L12
O1
O8
219
253
L8
9
O16
256
217
L1
233
97
218
204
0.0
0.1
0.2
0.3
0.4
−1
0
1
2
3
4
Estimated allele frequency
Ecological gradient (Leaf Unfolding differences)

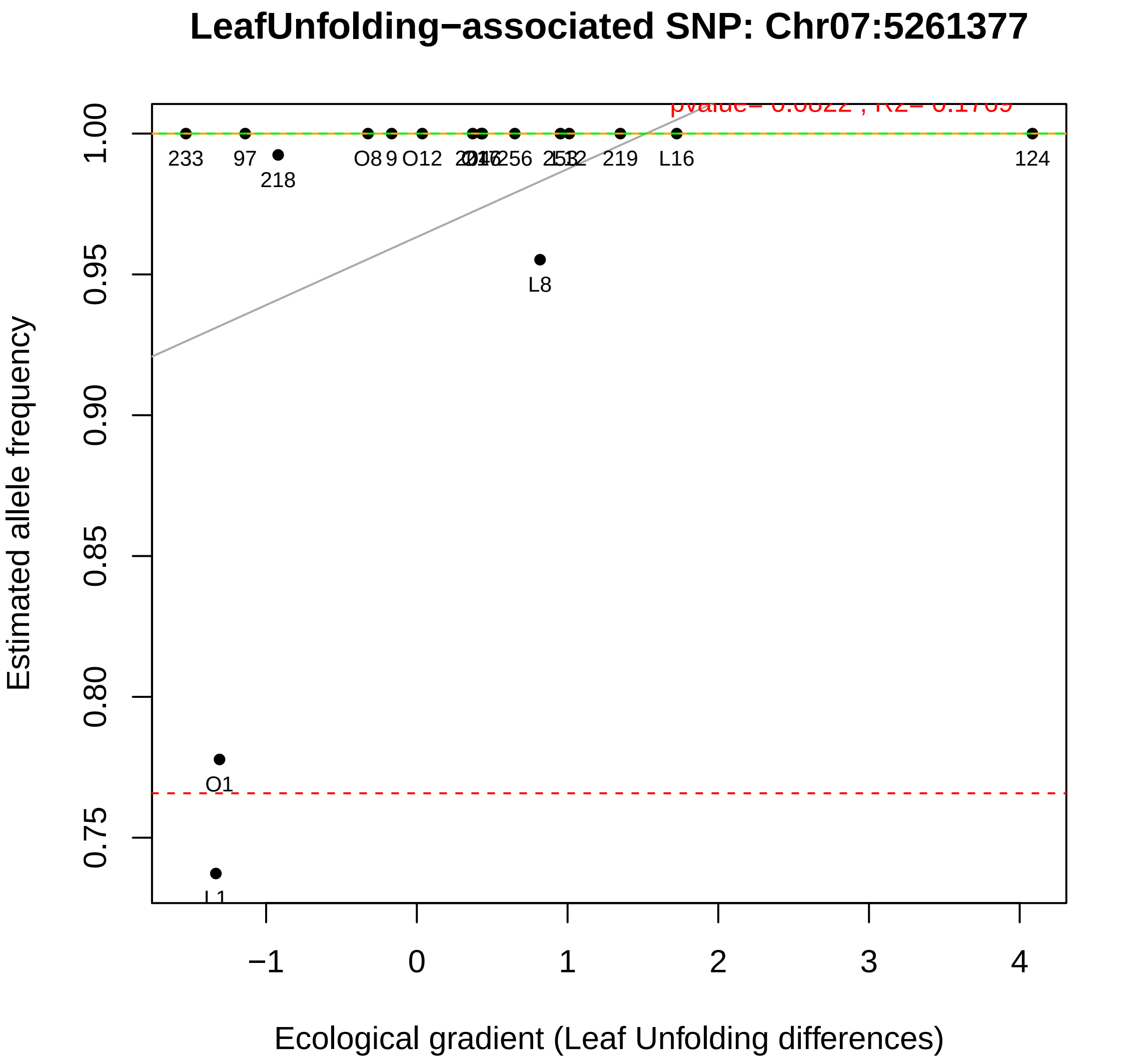

LeafUnfolding−associated SNP: Chr07:5261377
pvalue= 0.0822 ; R2= 0.1769
Estimated allele frequency
Ecological gradient (Leaf Unfolding differences)
1.00
0.95
0.90
0.85
0.80
0.75
−1
0
1
2
3
4
233
97
218
O8
9
O12
219
L16
124
L8
O1
L1

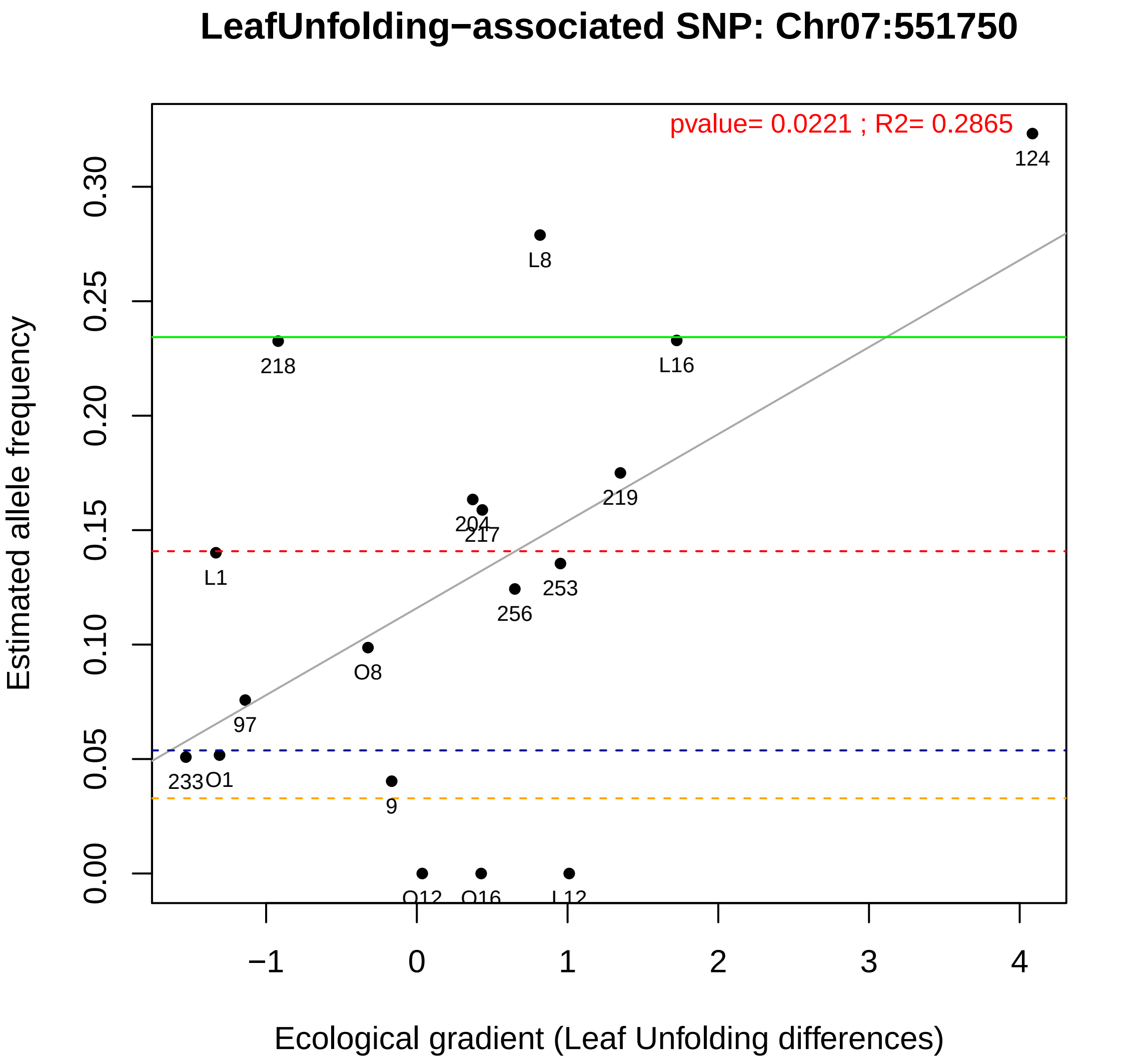

LeafUnfolding−associated SNP: Chr07:551750
pvalue= 0.0221 ; R2= 0.2865
Estimated allele frequency
Ecological gradient (Leaf Unfolding differences)
0.00
0.05
0.10
0.15
0.20
0.25
0.30
−1
0
1
2
3
4
124
L8
218
L16
219
204
217
L1
253
256
O8
97
233
O1
9
O12
O16
L12

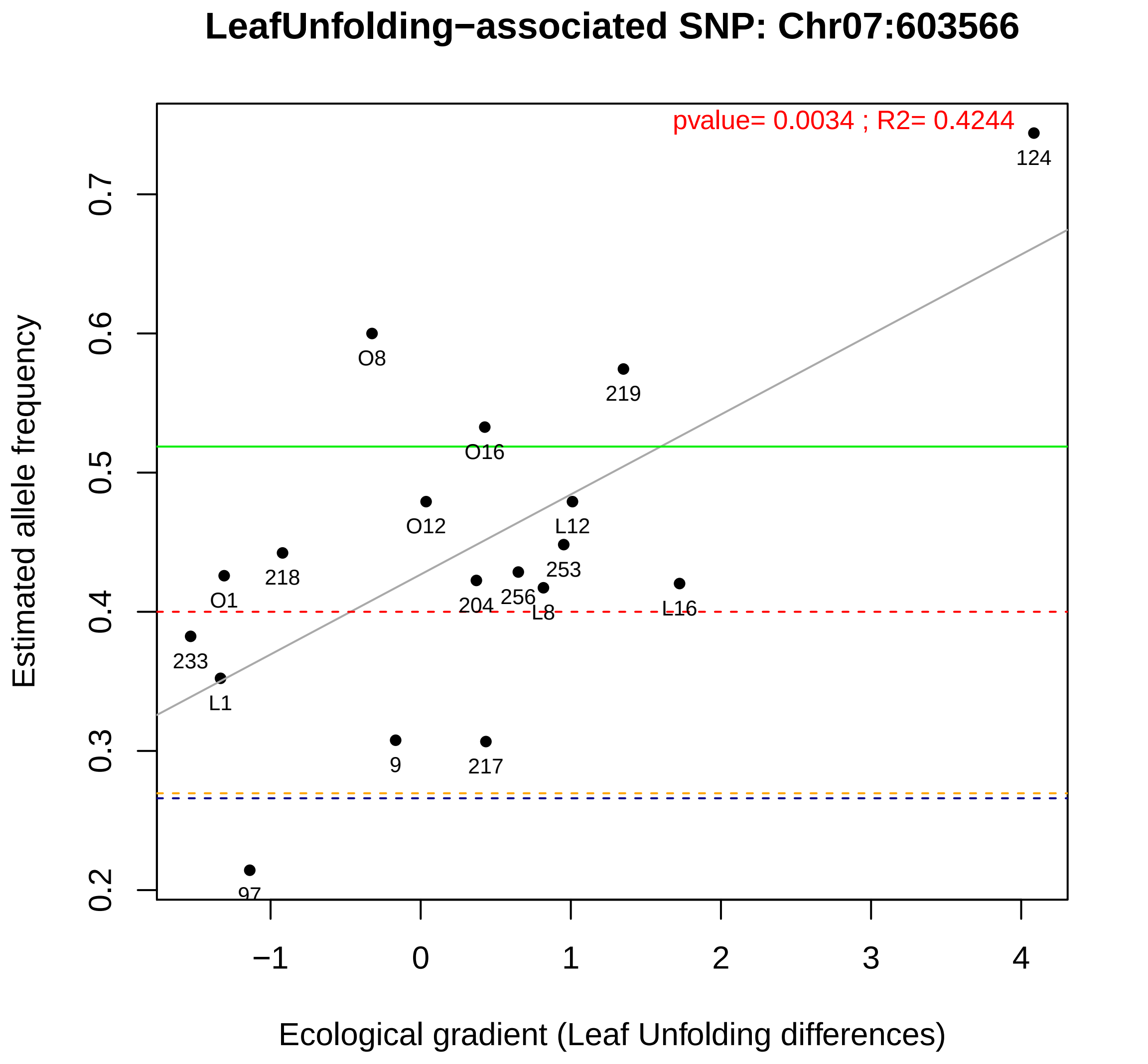

LeafUnfolding−associated SNP: Chr07:603566
pvalue= 0.0034 ; R2= 0.4244
Estimated allele frequency
Ecological gradient (Leaf Unfolding differences)
0.2
0.3
0.4
0.5
0.6
0.7
−1
0
1
2
3
4
124
O8
219
O16
O12
L12
253
218
O1
204
256
L8
L16
233
L1
9
217
97

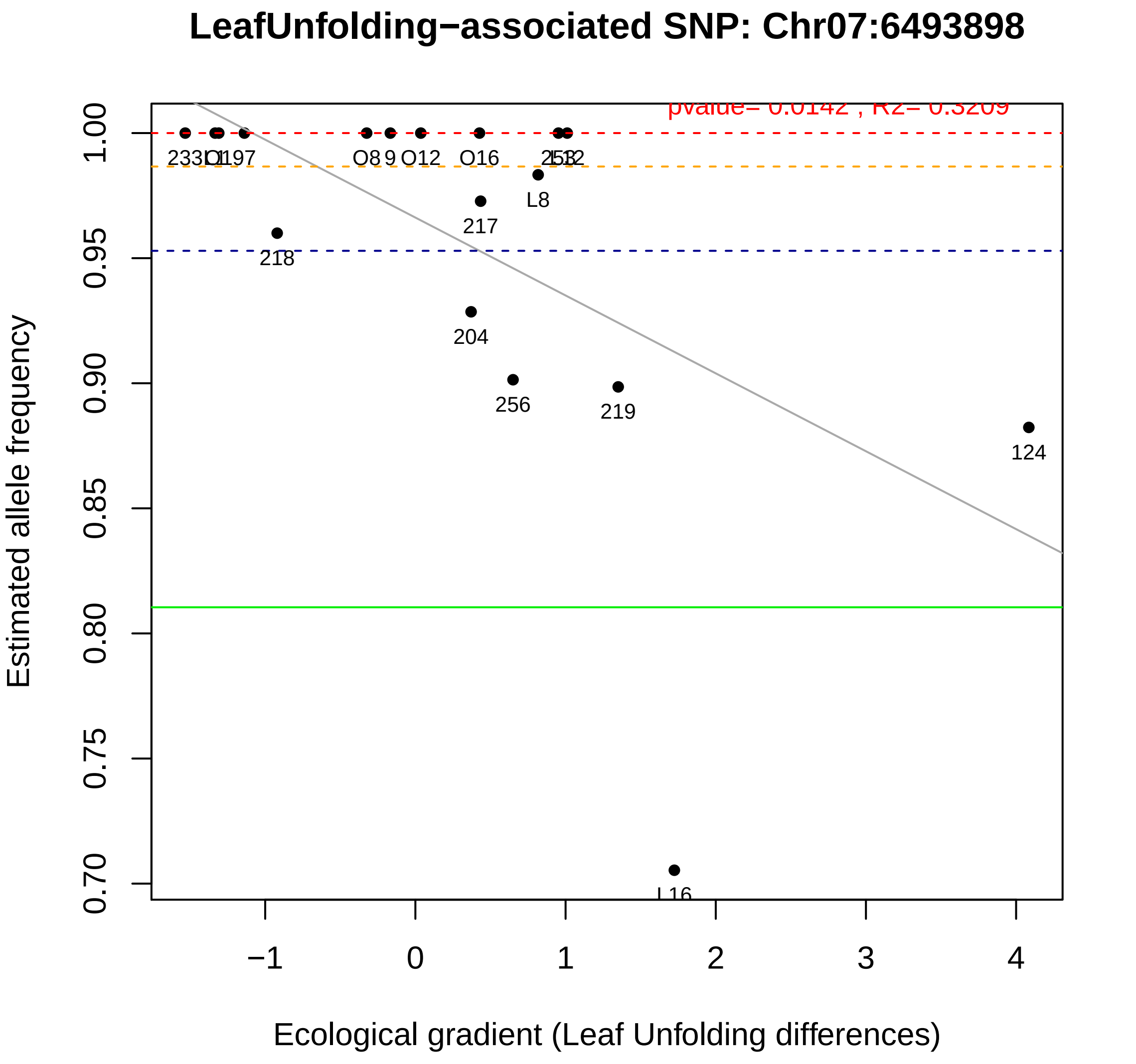

LeafUnfolding−associated SNP: Chr07:6493898
pvalue= 0.0142 , R2= 0.3209
Estimated allele frequency
Ecological gradient (Leaf Unfolding differences)
1.00
0.95
0.90
0.85
0.80
0.75
0.70
−1
0
1
2
3
4
O8
9
O12
O16
L8
217
218
204
256
219
124

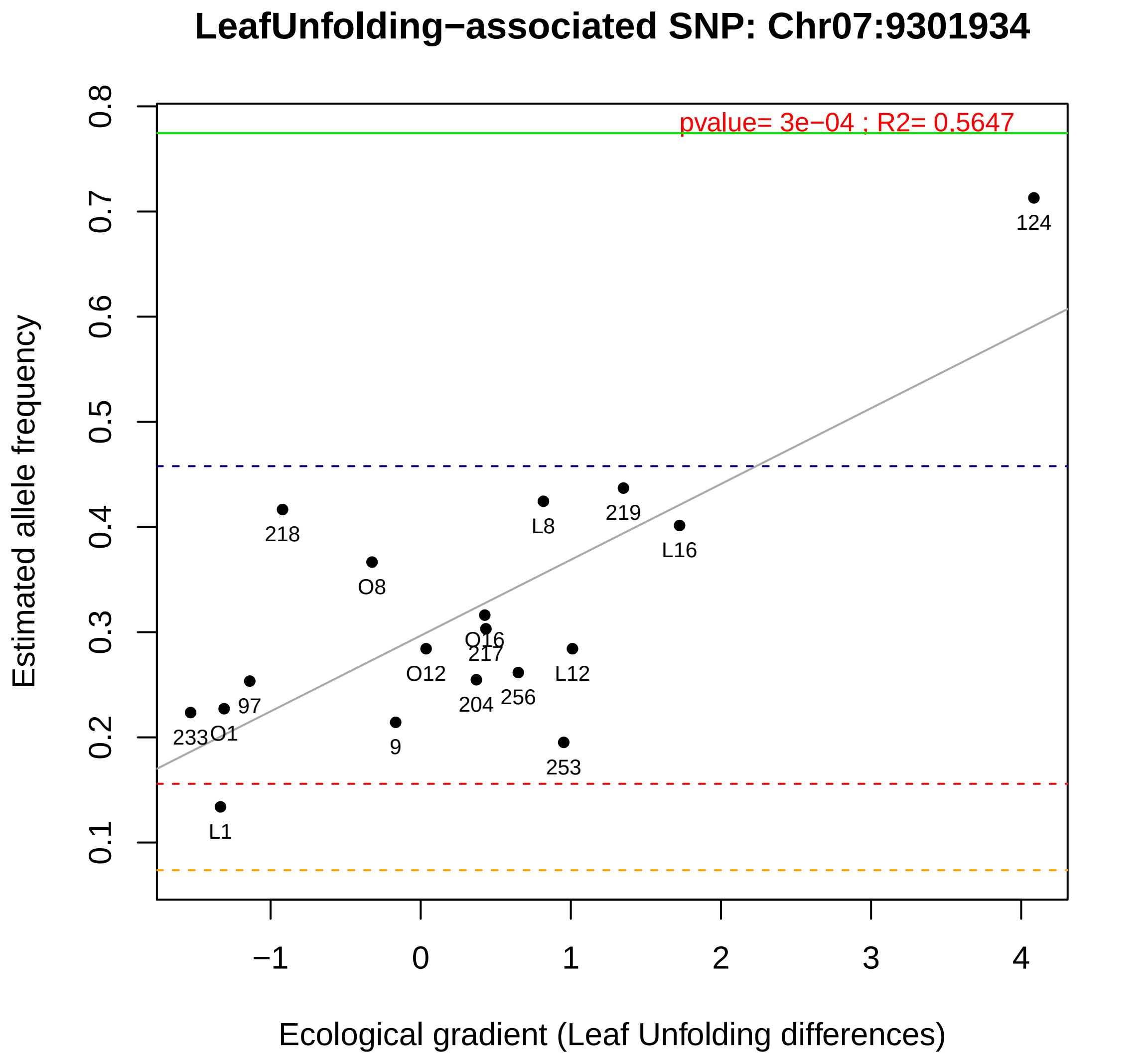
LeafUnfolding−associated SNP: Chr07:9301934
pvalue= 3e−04 ; R2= 0.5647
Estimated allele frequency
Ecological gradient (Leaf Unfolding differences)
0.1
0.2
0.3
0.4
0.5
0.6
0.7
0.8
−1
0
1
2
3
4
124
218
L8
219
L16
O8
O16
217
O12
L12
204
256
97
233
O1
9
253
L1

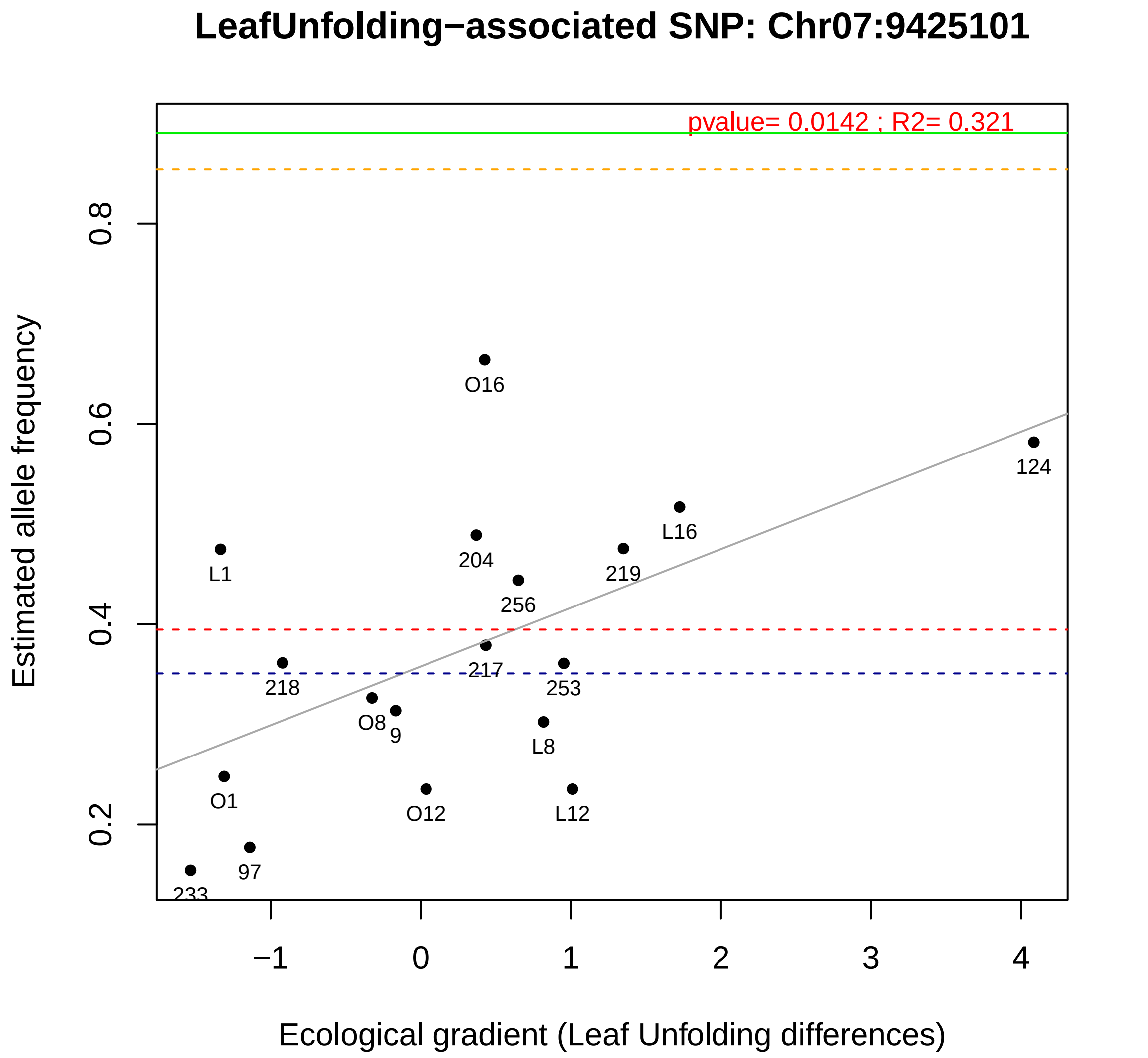
LeafUnfolding−associated SNP: Chr07:9425101
pvalue= 0.0142 ; R2= 0.321
Estimated allele frequency
0.2
0.4
0.6
0.8
−1
0
1
2
3
4
Ecological gradient (Leaf Unfolding differences)
O16
124
L16
204
L1
219
256
217
218
253
O8
9
L8
O1
O12
L12
97
233

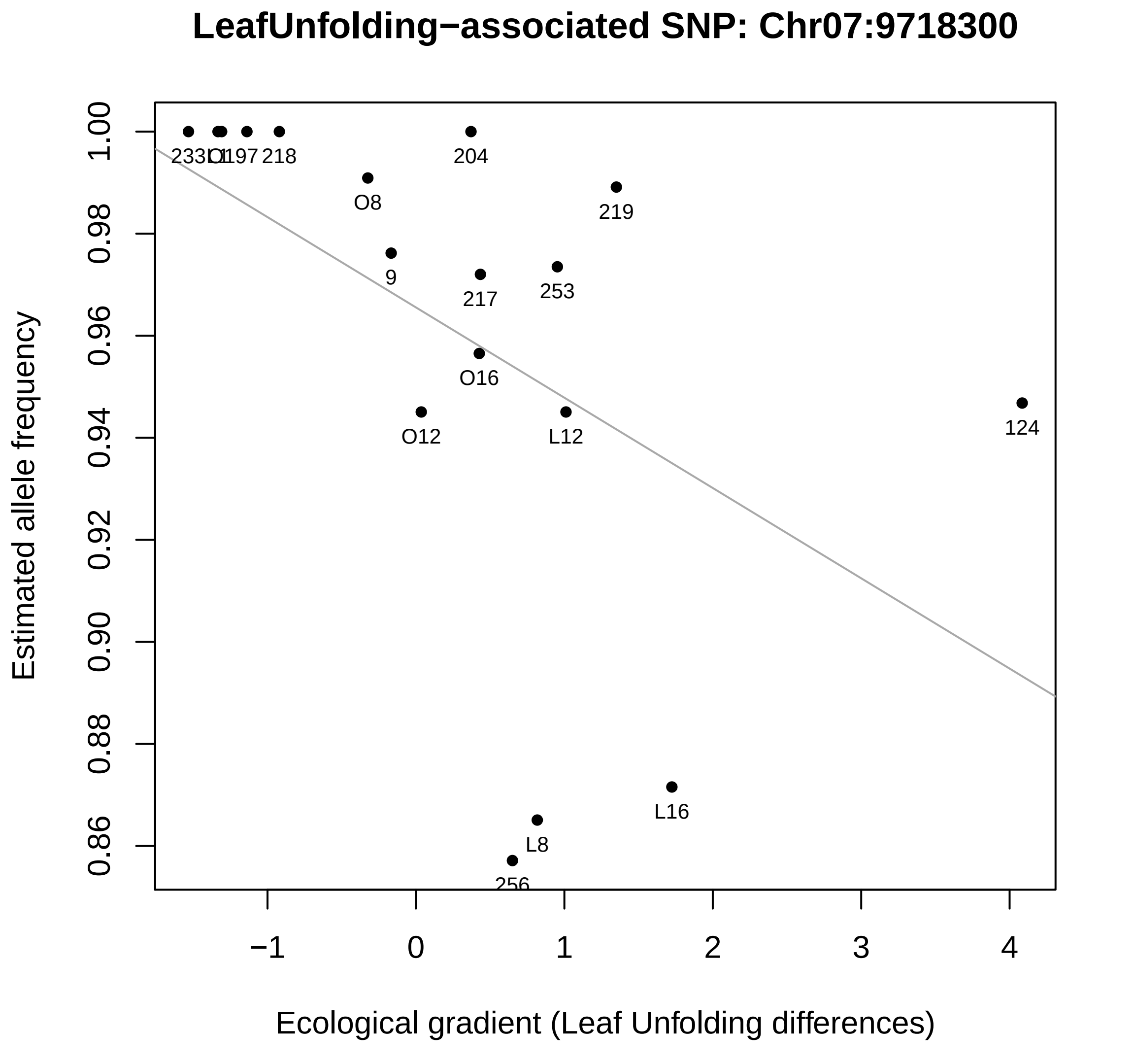

LeafUnfolding−associated SNP: Chr07:9718300
Estimated allele frequency
Ecological gradient (Leaf Unfolding differences)
1.00
0.98
0.96
0.94
0.92
0.90
0.88
0.86
−1
0
1
2
3
4
233
O1
197
218
204
O8
219
9
217
253
O16
O12
L12
124
L16
L8
256

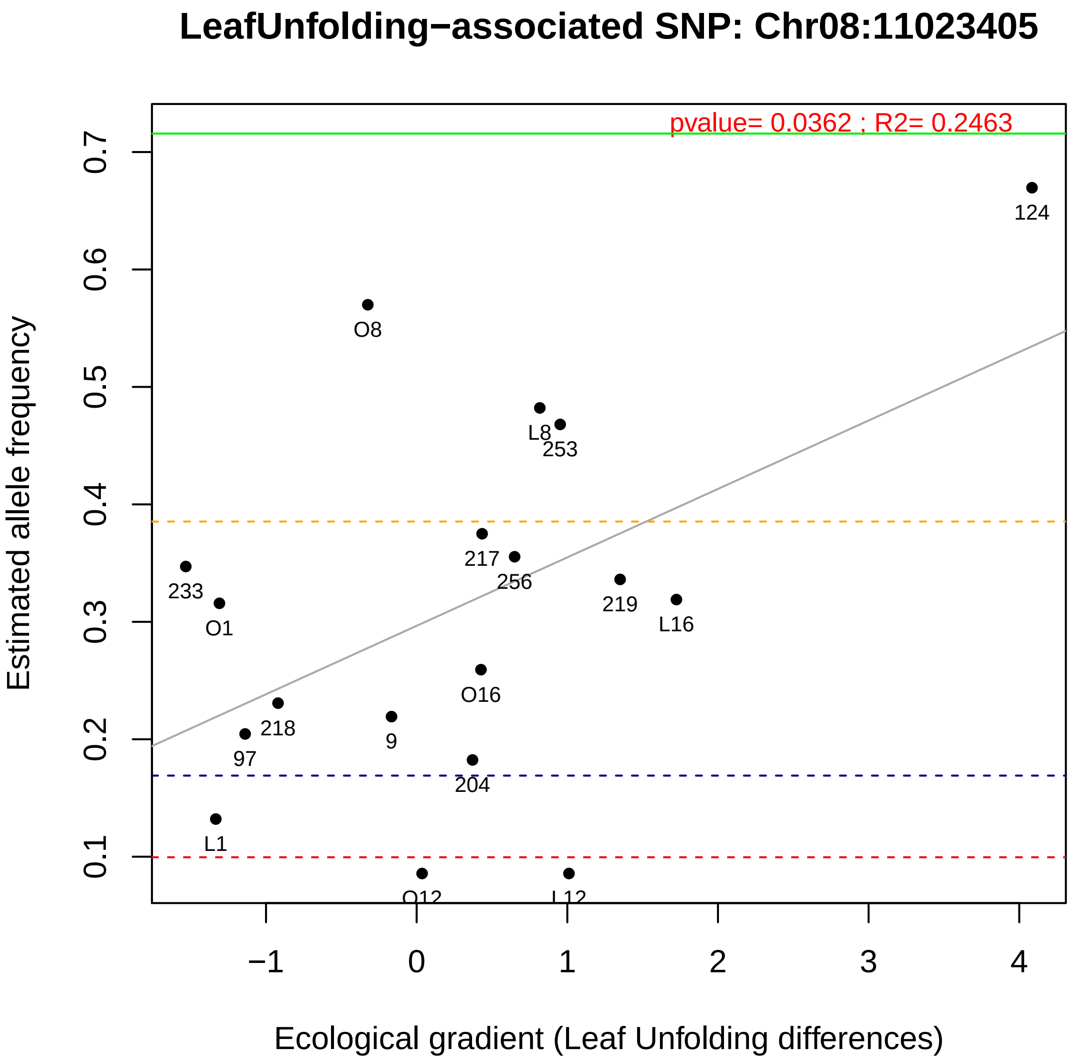

LeafUnfolding−associated SNP: Chr08:11023405
pvalue= 0.0362 ; R2= 0.2463
Estimated allele frequency
0.1
0.2
0.3
0.4
0.5
0.6
0.7
−1
0
1
2
3
4
Ecological gradient (Leaf Unfolding differences)
124
O8
L8
253
217
256
233
O1
219
L16
O16
218
97
9
204
L1
O12
L12

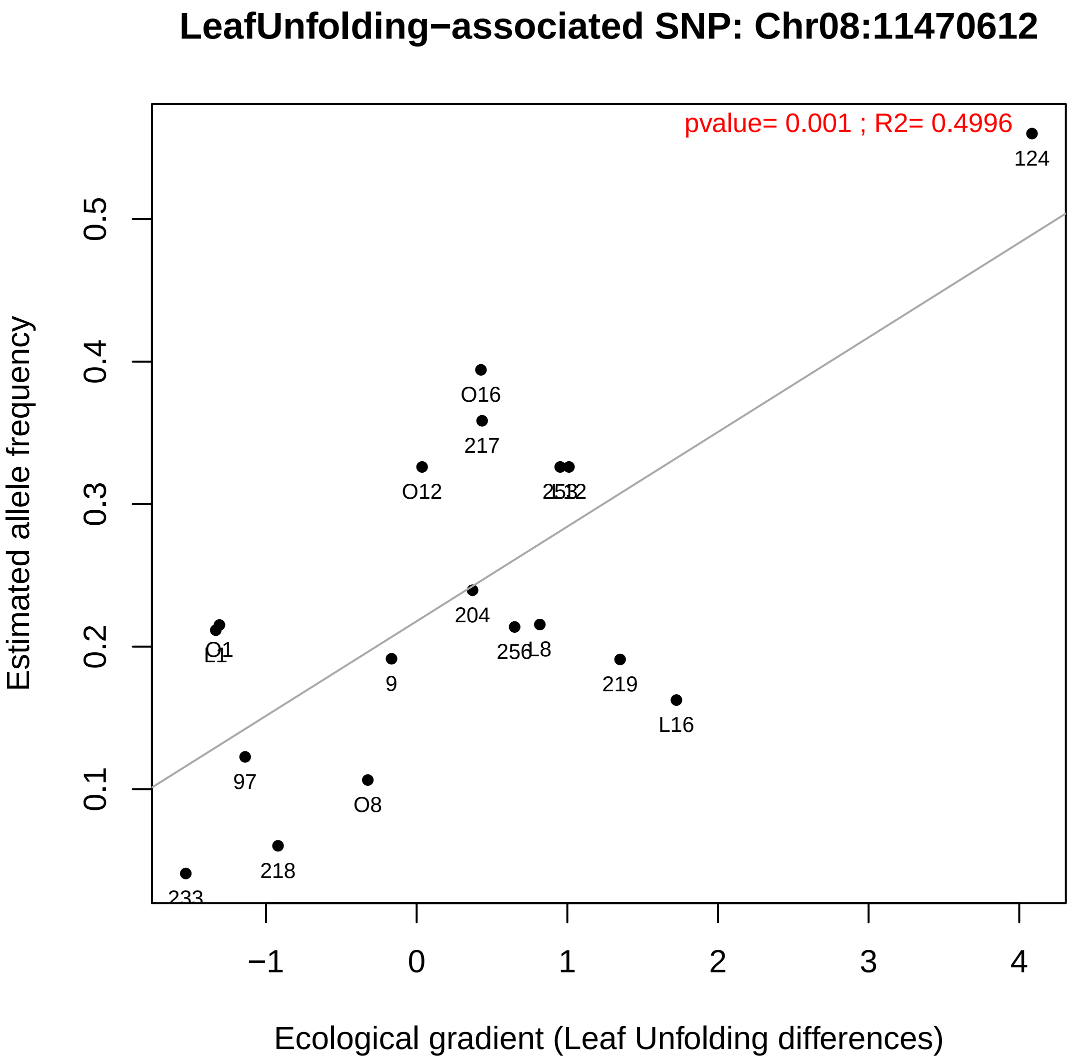

LeafUnfolding−associated SNP: Chr08:11470612
pvalue= 0.001 ; R2= 0.4996
Estimated allele frequency
Ecological gradient (Leaf Unfolding differences)
0.1
0.2
0.3
0.4
0.5
−1
0
1
2
3
4
124
O16
217
O12
204
256
L8
219
L16
9
97
O8
218
233
O1
L1

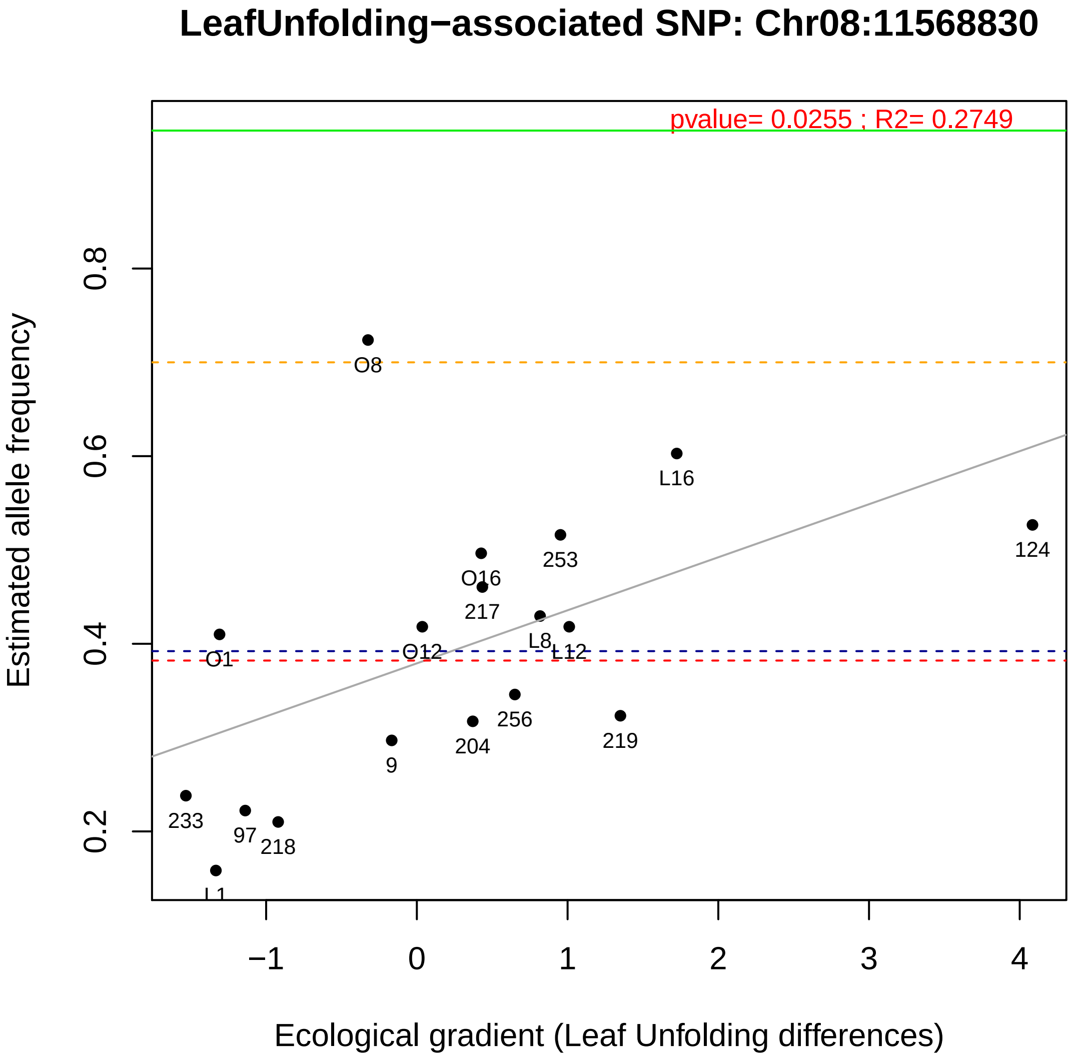
LeafUnfolding−associated SNP: Chr08:11568830
pvalue= 0.0255 ; R2= 0.2749
Estimated allele frequency
0.2
0.4
0.6
0.8
−1
0
1
2
3
4
Ecological gradient (Leaf Unfolding differences)
O8
L16
124
253
O16
217
L8
L12
O12
O1
256
219
204
9
233
97
218

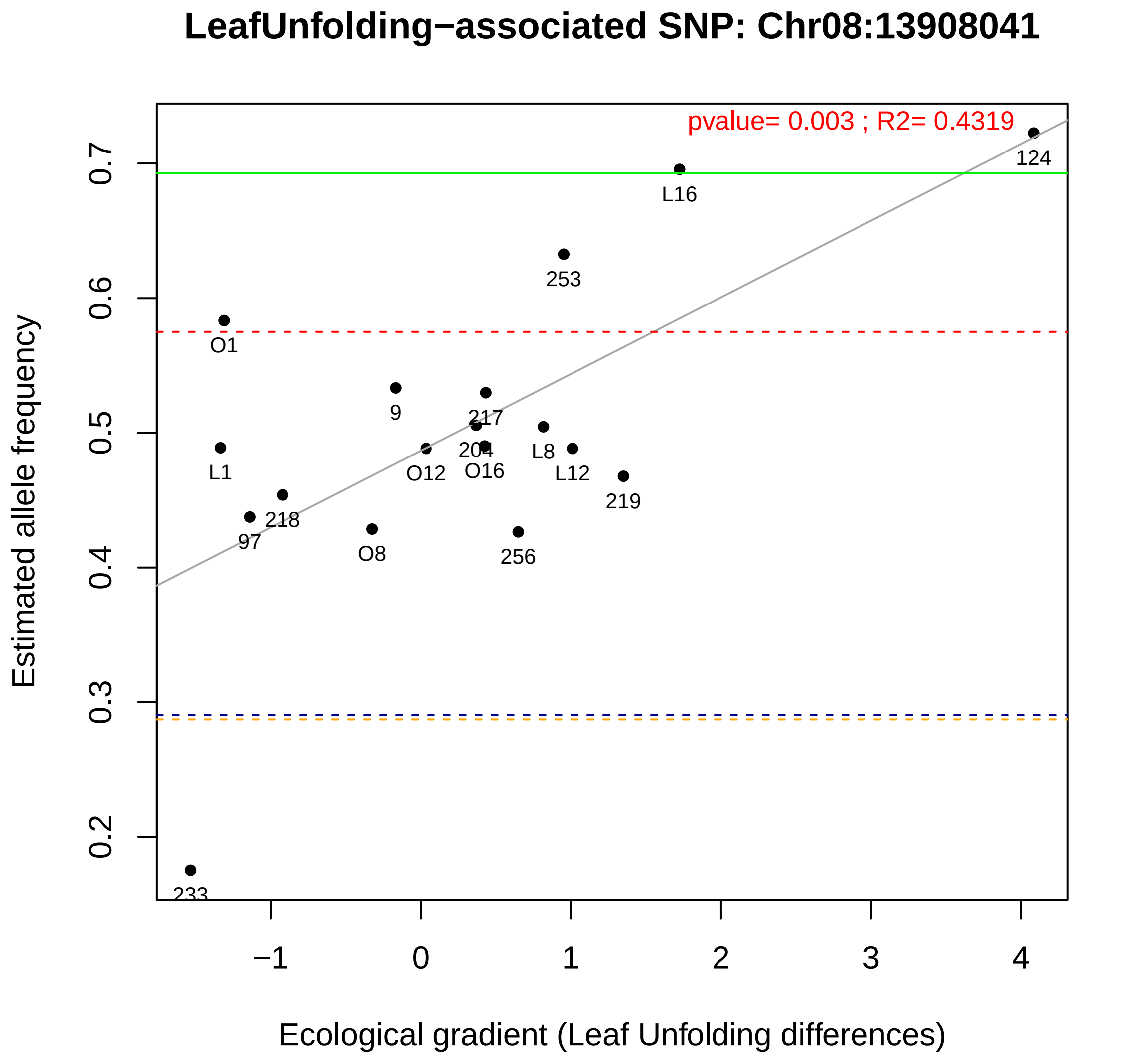
LeafUnfolding−associated SNP: Chr08:13908041
pvalue= 0.003 ; R2= 0.4319
Estimated allele frequency
Ecological gradient (Leaf Unfolding differences)
0.2
0.3
0.4
0.5
0.6
0.7
−1
0
1
2
3
4
124
L16
253
O1
9
217
204
O16
L8
L1
O12
L12
219
218
97
O8
256
233

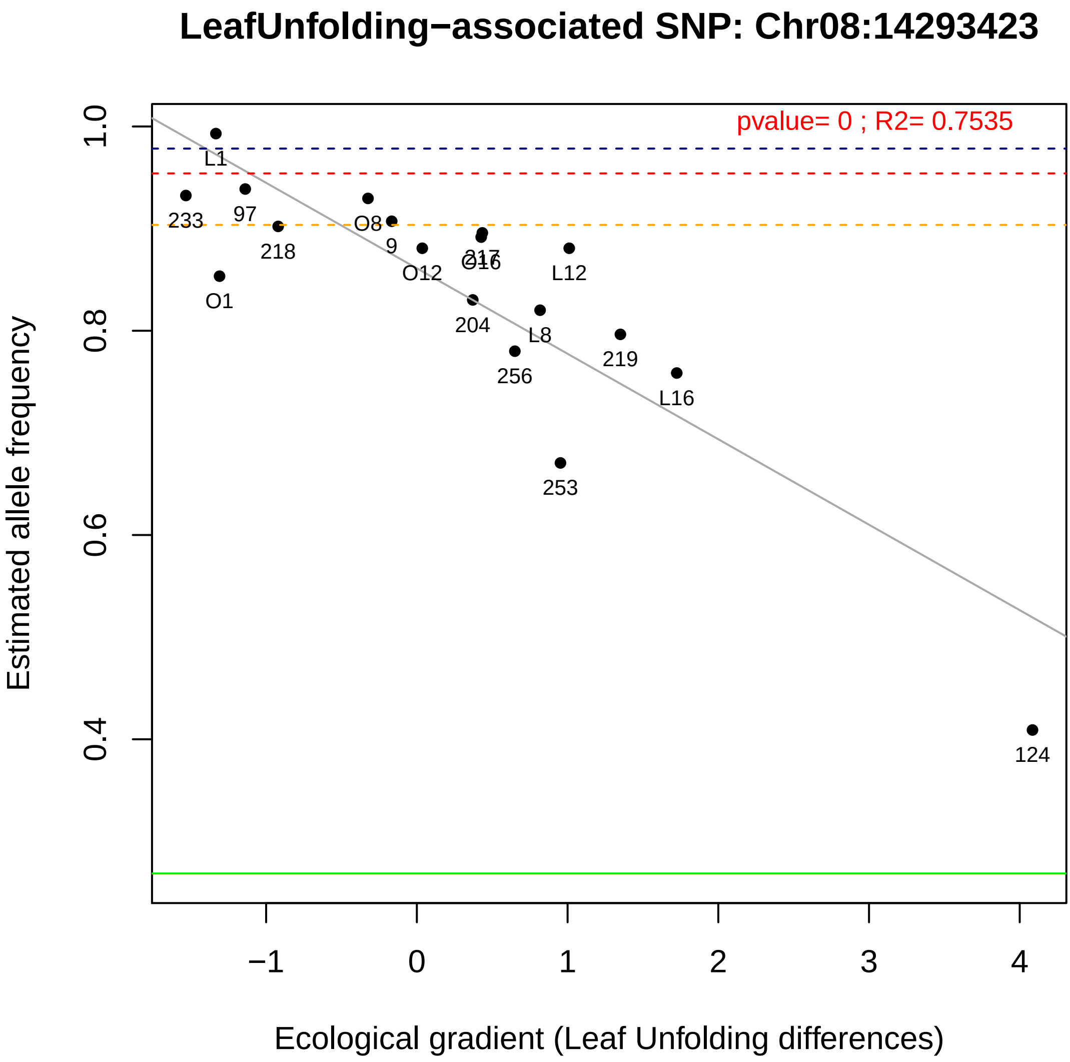

LeafUnfolding−associated SNP: Chr08:14293423
pvalue= 0 ; R2= 0.7535
Estimated allele frequency
Ecological gradient (Leaf Unfolding differences)
0.4
0.6
0.8
1.0
−1
0
1
2
3
4
L1
233
97
218
O1
O8
9
O12
217
O16
204
L12
L8
256
219
L16
253
124

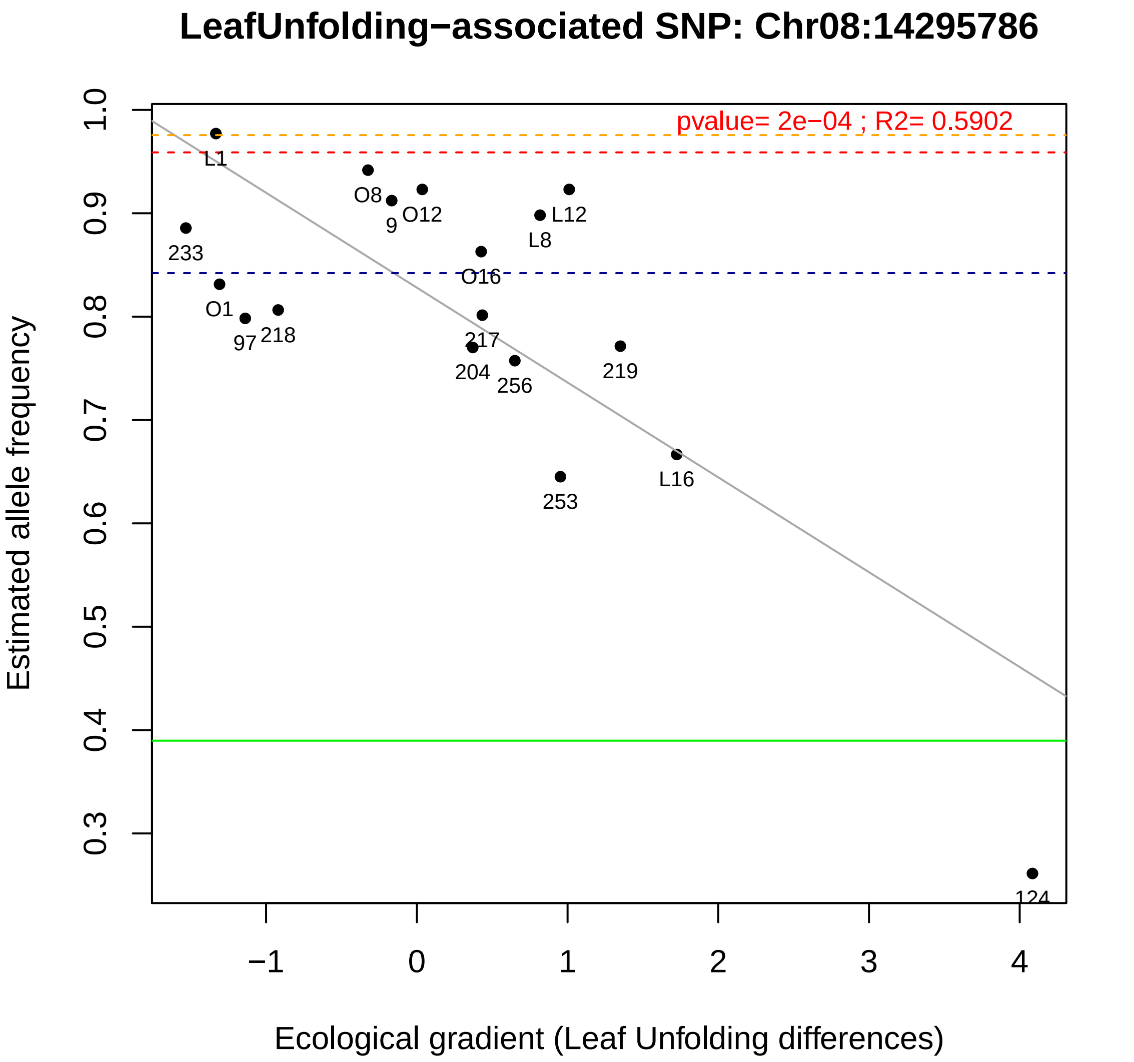
LeafUnfolding−associated SNP: Chr08:14295786
pvalue= 2e−04 ; R2= 0.5902
Estimated allele frequency
Ecological gradient (Leaf Unfolding differences)
0.3
0.4
0.5
0.6
0.7
0.8
0.9
1.0
−1
0
1
2
3
4
L1
O8
9
O12
L12
L8
233
O16
O1
97
218
217
204
256
219
L16
253
124

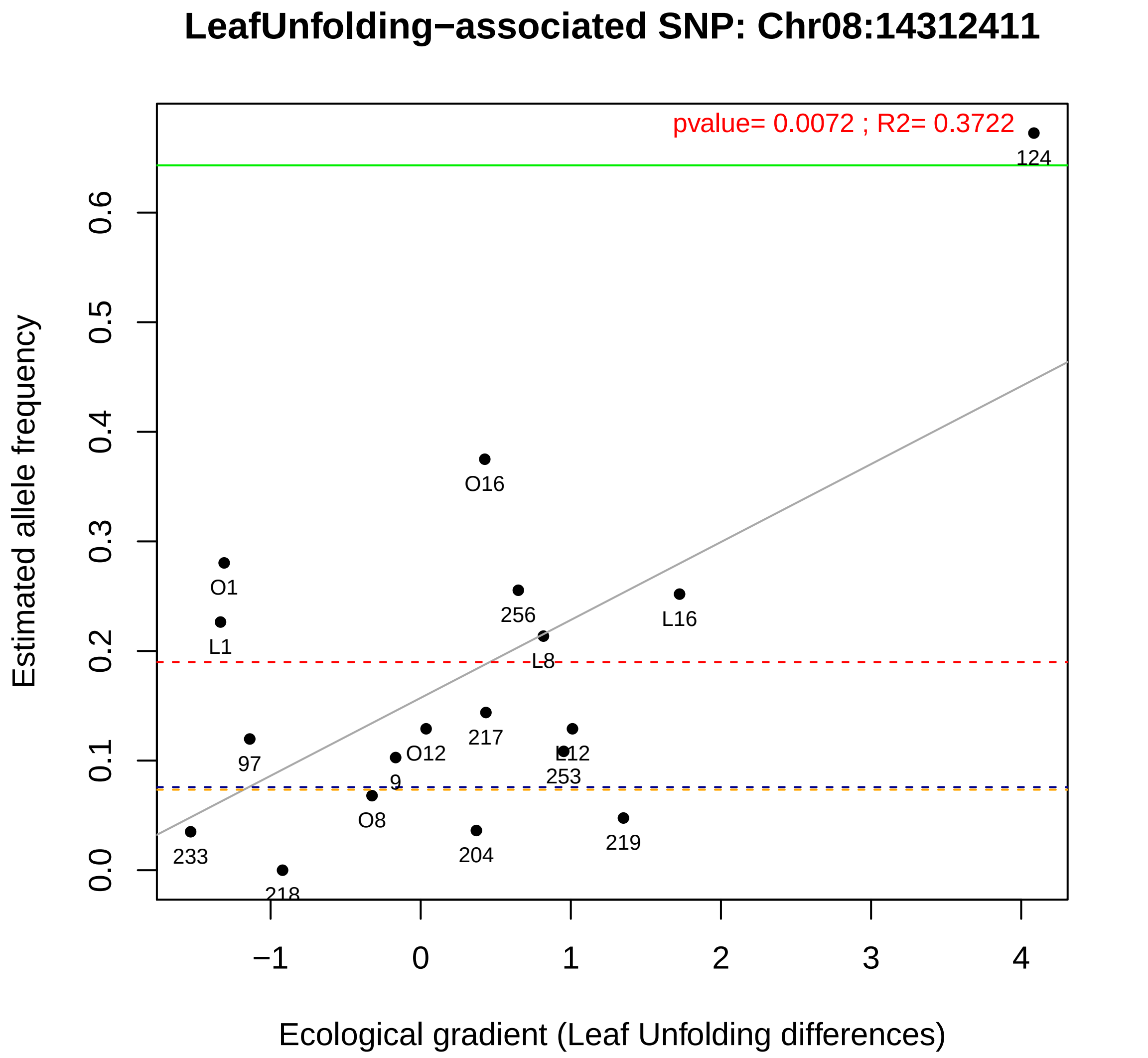
LeafUnfolding−associated SNP: Chr08:14312411
pvalue= 0.0072 ; R2= 0.3722
Estimated allele frequency
Ecological gradient (Leaf Unfolding differences)
0.0
0.1
0.2
0.3
0.4
0.5
0.6
−1
0
1
2
3
4
124
O16
O1
L1
256
L8
L16
97
O12
9
217
L12
253
O8
233
218
204
219

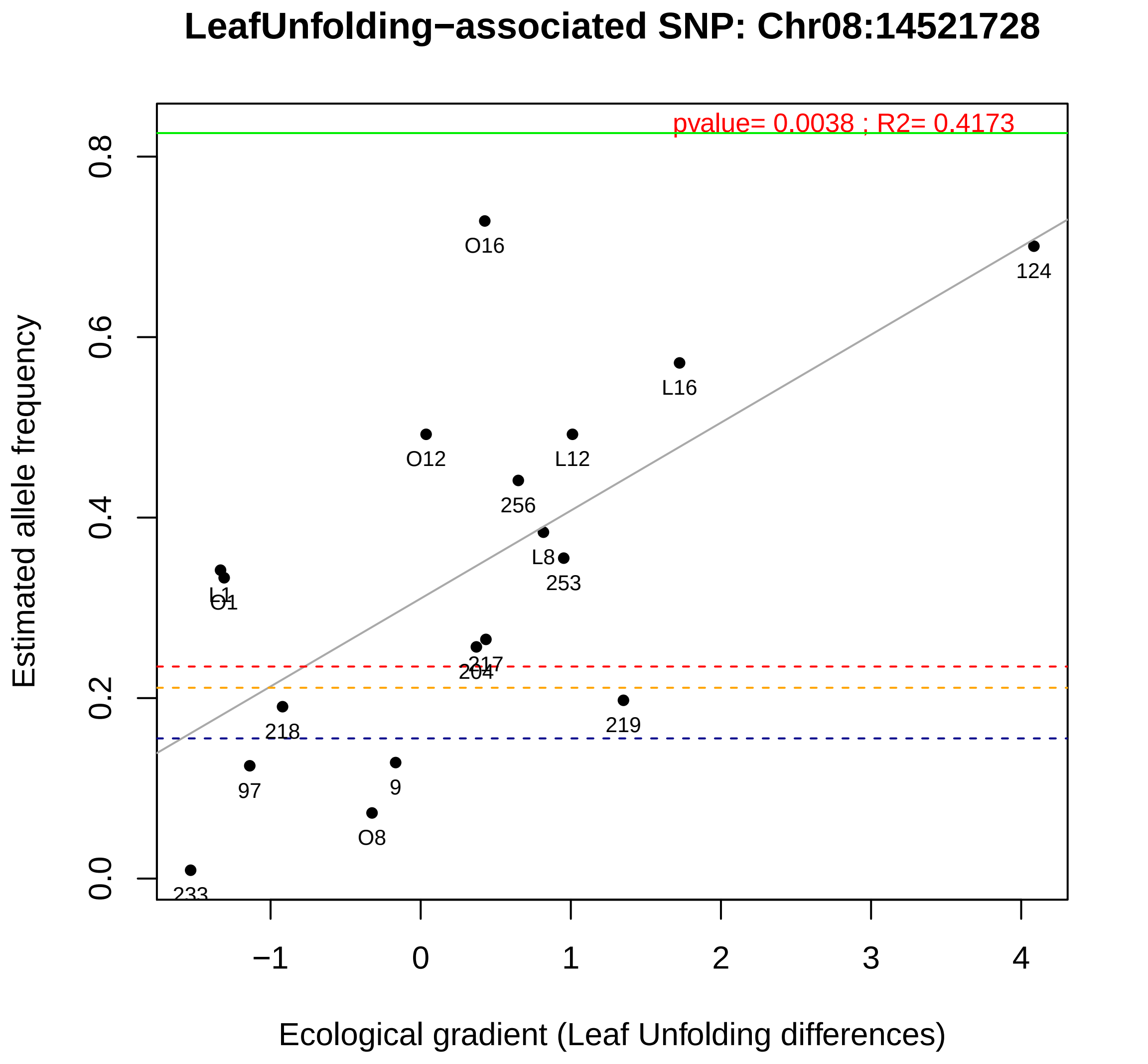
LeafUnfolding−associated SNP: Chr08:14521728
pvalue= 0.0038 ; R2= 0.4173
Estimated allele frequency
Ecological gradient (Leaf Unfolding differences)
0.0
0.2
0.4
0.6
0.8
−1
0
1
2
3
4
O16
124
L16
O12
L12
256
L8
253
L1
O1
217
204
219
218
97
9
O8
233

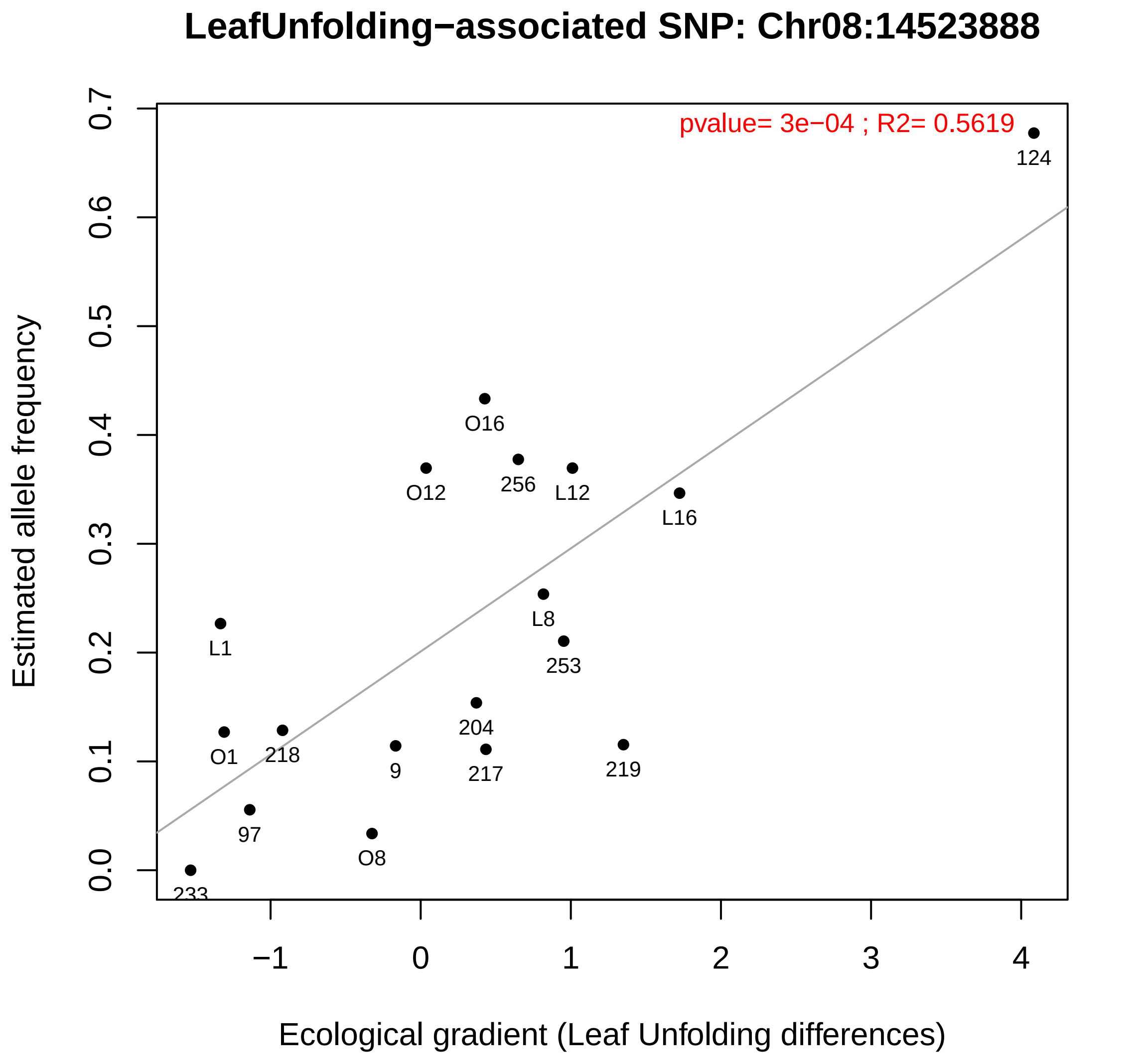

LeafUnfolding−associated SNP: Chr08:14523888
pvalue= 3e−04 ; R2= 0.5619
Estimated allele frequency
Ecological gradient (Leaf Unfolding differences)
0.0
0.1
0.2
0.3
0.4
0.5
0.6
0.7
−1
0
1
2
3
4
124
O16
O12
256
L12
L16
L8
L1
253
204
O1
218
9
217
219
97
O8
233

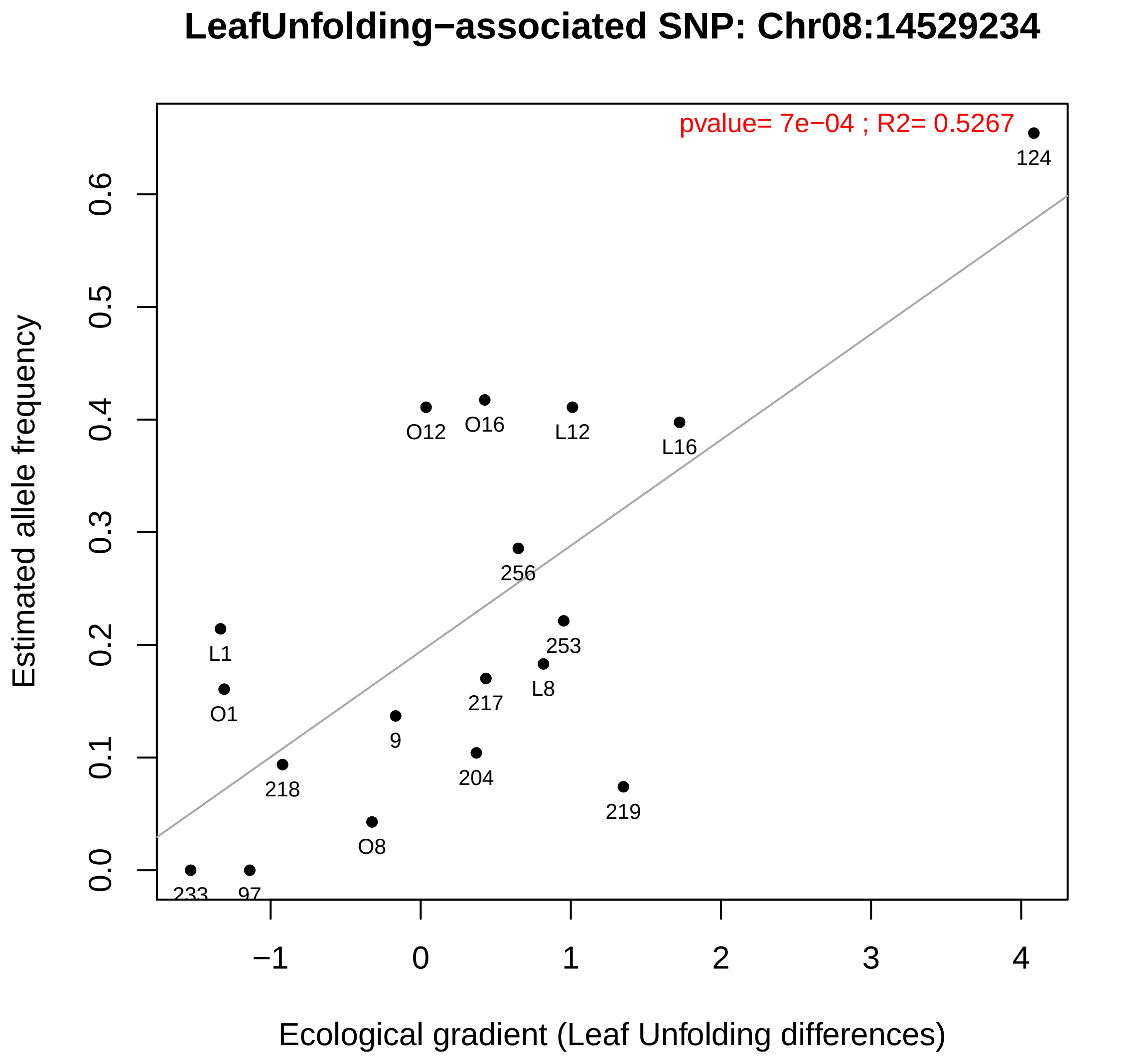
LeafUnfolding−associated SNP: Chr08:14529234
pvalue= 7e−04 ; R2= 0.5267
Estimated allele frequency
Ecological gradient (Leaf Unfolding differences)
0.0
0.1
0.2
0.3
0.4
0.5
0.6
−1
0
1
2
3
4
124
O12
O16
L12
L16
256
253
L1
L8
217
O1
9
204
218
219
O8
233
97

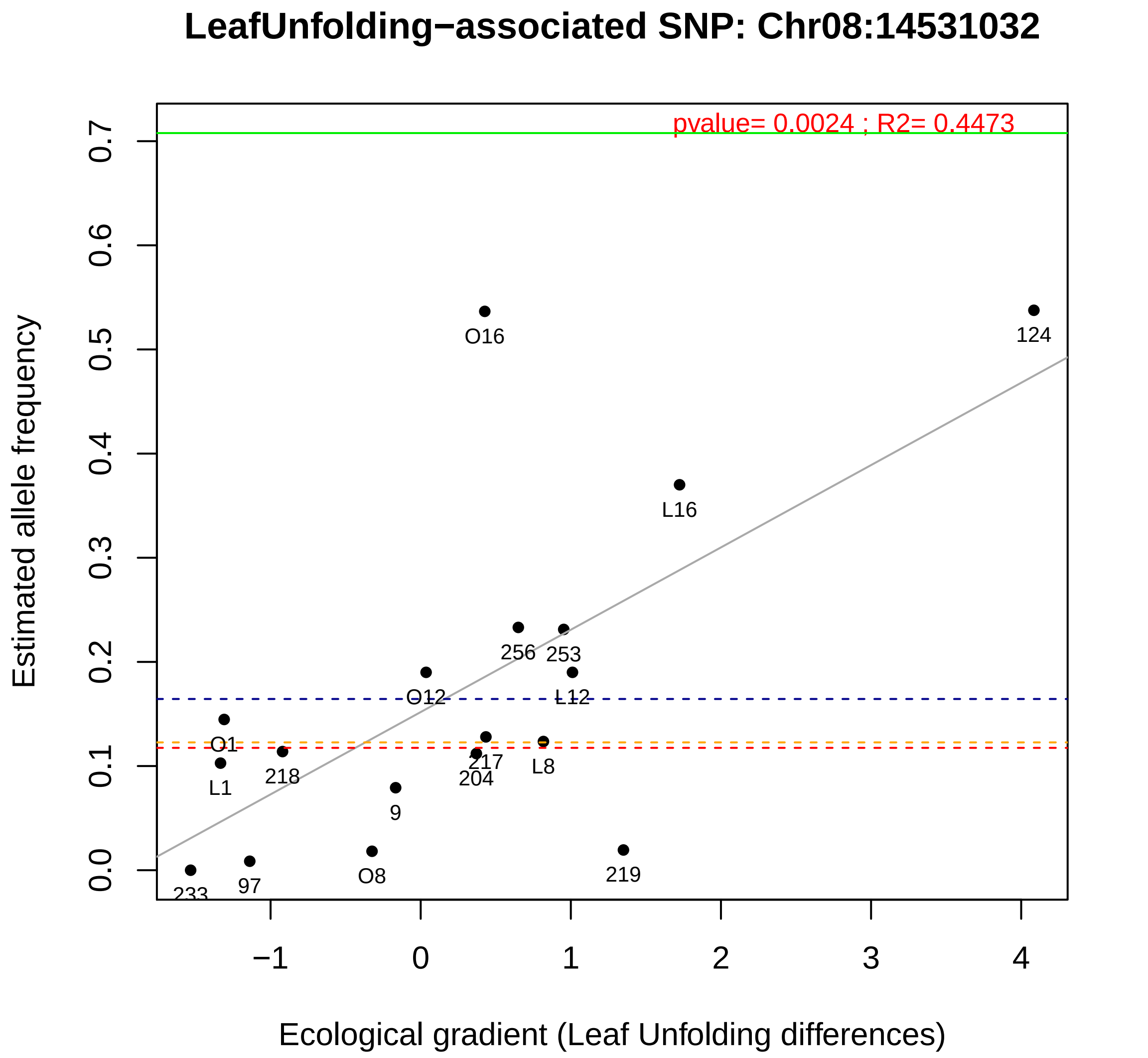
LeafUnfolding−associated SNP: Chr08:14531032
pvalue= 0.0024 ; R2= 0.4473
Estimated allele frequency
0.0
0.1
0.2
0.3
0.4
0.5
0.6
0.7
Ecological gradient (Leaf Unfolding differences)
−1
0
1
2
3
4
O16
124
L16
256
253
O12
L12
O1
L1
218
217
204
L8
9
233
97
O8
219

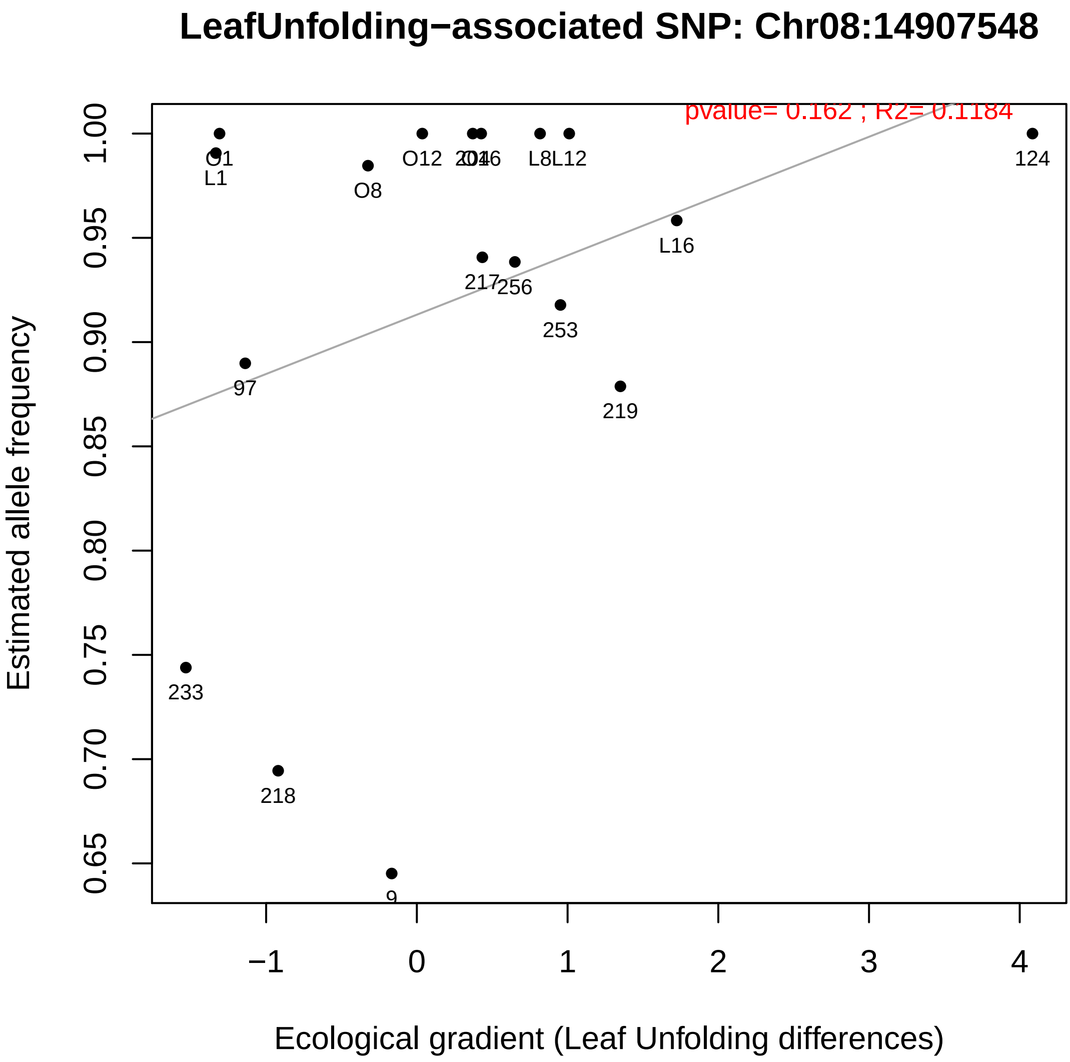

LeafUnfolding−associated SNP: Chr08:14907548
pvalue= 0.162 ; R2= 0.1184
Estimated allele frequency
Ecological gradient (Leaf Unfolding differences)
1.00
0.95
0.90
0.85
0.80
0.75
0.70
0.65
−1
0
1
2
3
4
O1
L1
O8
O12
O16
L8
L12
124
L16
217
256
253
97
219
233
218
9

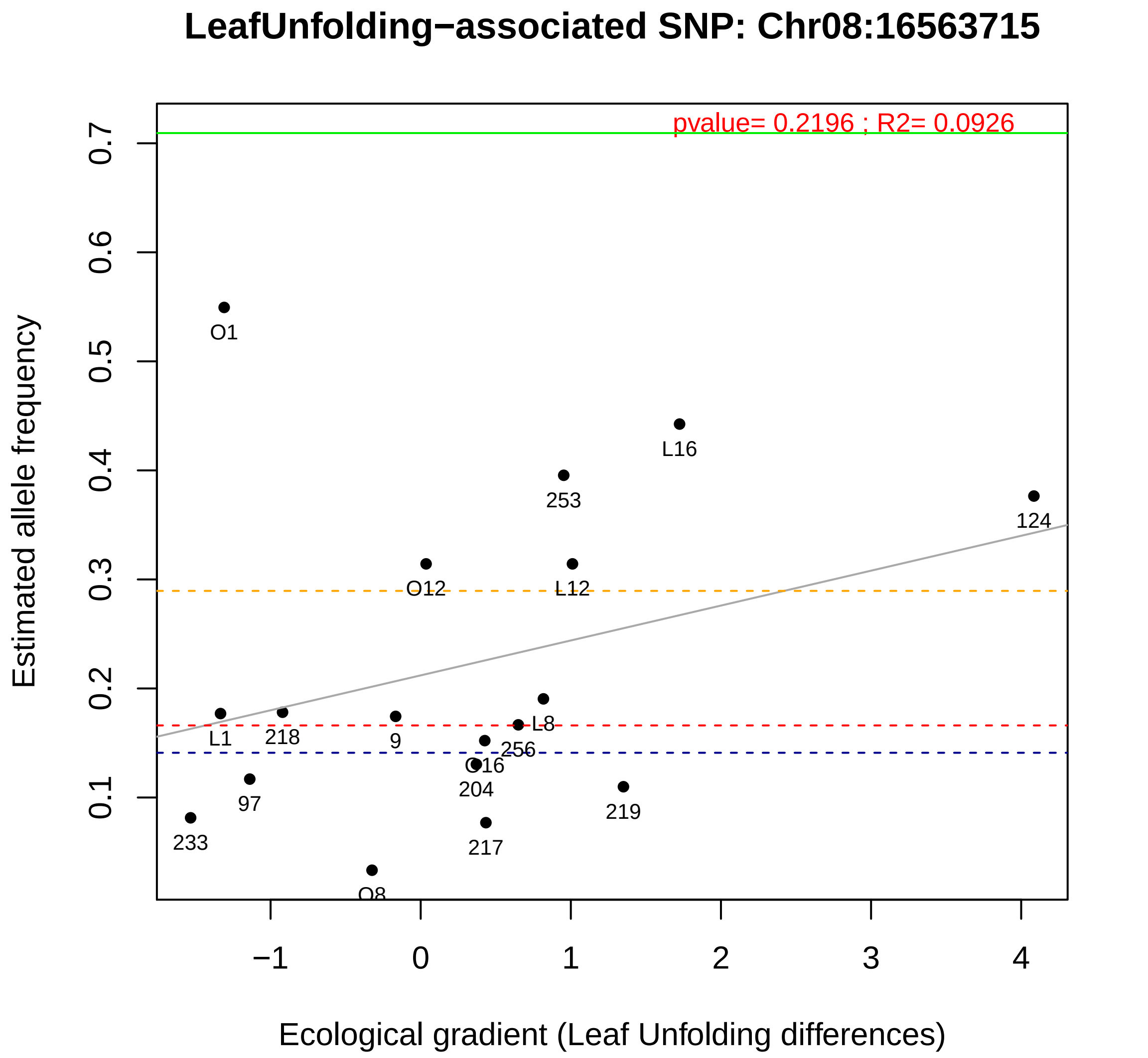
LeafUnfolding−associated SNP: Chr08:16563715
pvalue= 0.2196 ; R2= 0.0926
Estimated allele frequency
0.1
0.2
0.3
0.4
0.5
0.6
0.7
O1
L16
253
124
O12
L12
L1
218
9
L8
256
O16
204
97
219
233
217
O8
−1
0
1
2
3
4
Ecological gradient (Leaf Unfolding differences)

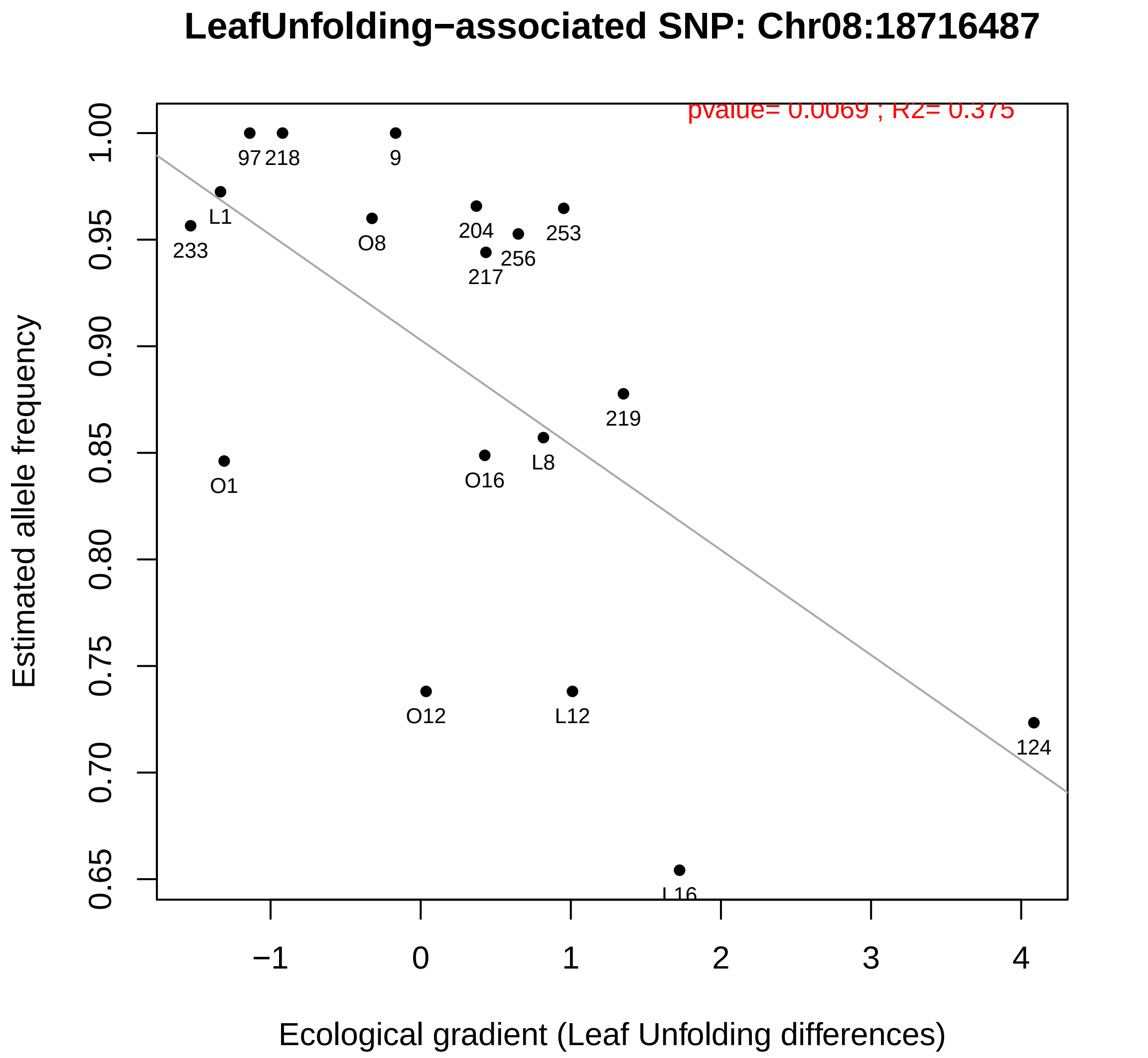

LeafUnfolding−associated SNP: Chr08:18716487
pvalue= 0.0069 ; R2= 0.375
Estimated allele frequency
Ecological gradient (Leaf Unfolding differences)
0.65
0.70
0.75
0.80
0.85
0.90
0.95
1.00
−1
0
1
2
3
4
97
218
9
L1
233
O8
204
253
256
217
219
L8
O1
O16
O12
L12
124

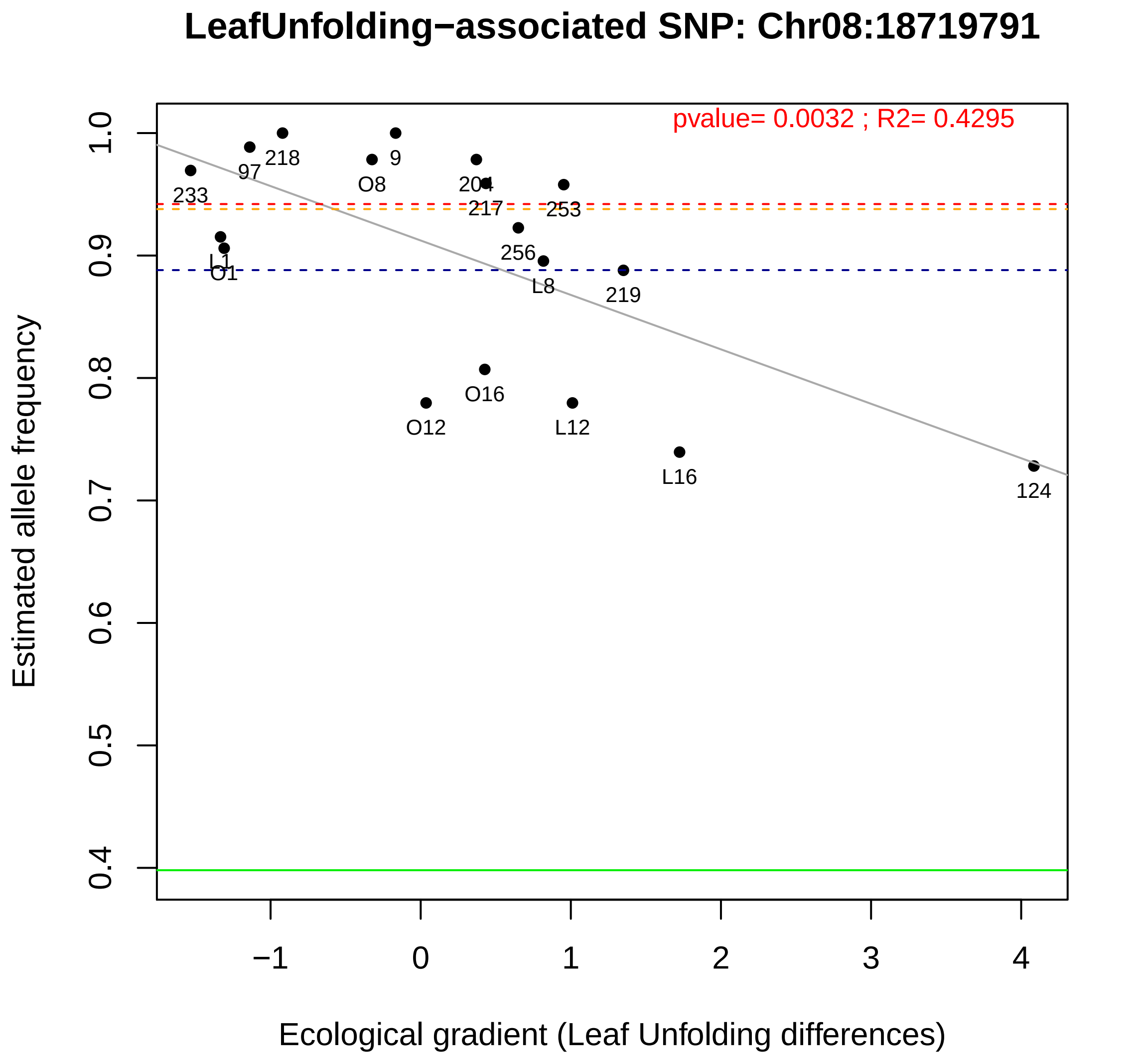
LeafUnfolding−associated SNP: Chr08:18719791
pvalue= 0.0032 ; R2= 0.4295
Estimated allele frequency
Ecological gradient (Leaf Unfolding differences)
1.0
0.9
0.8
0.7
0.6
0.5
0.4
−1
0
1
2
3
4
233
97
218
O8
9
204
217
253
256
L1
O1
L8
219
O16
O12
L12
L16
124

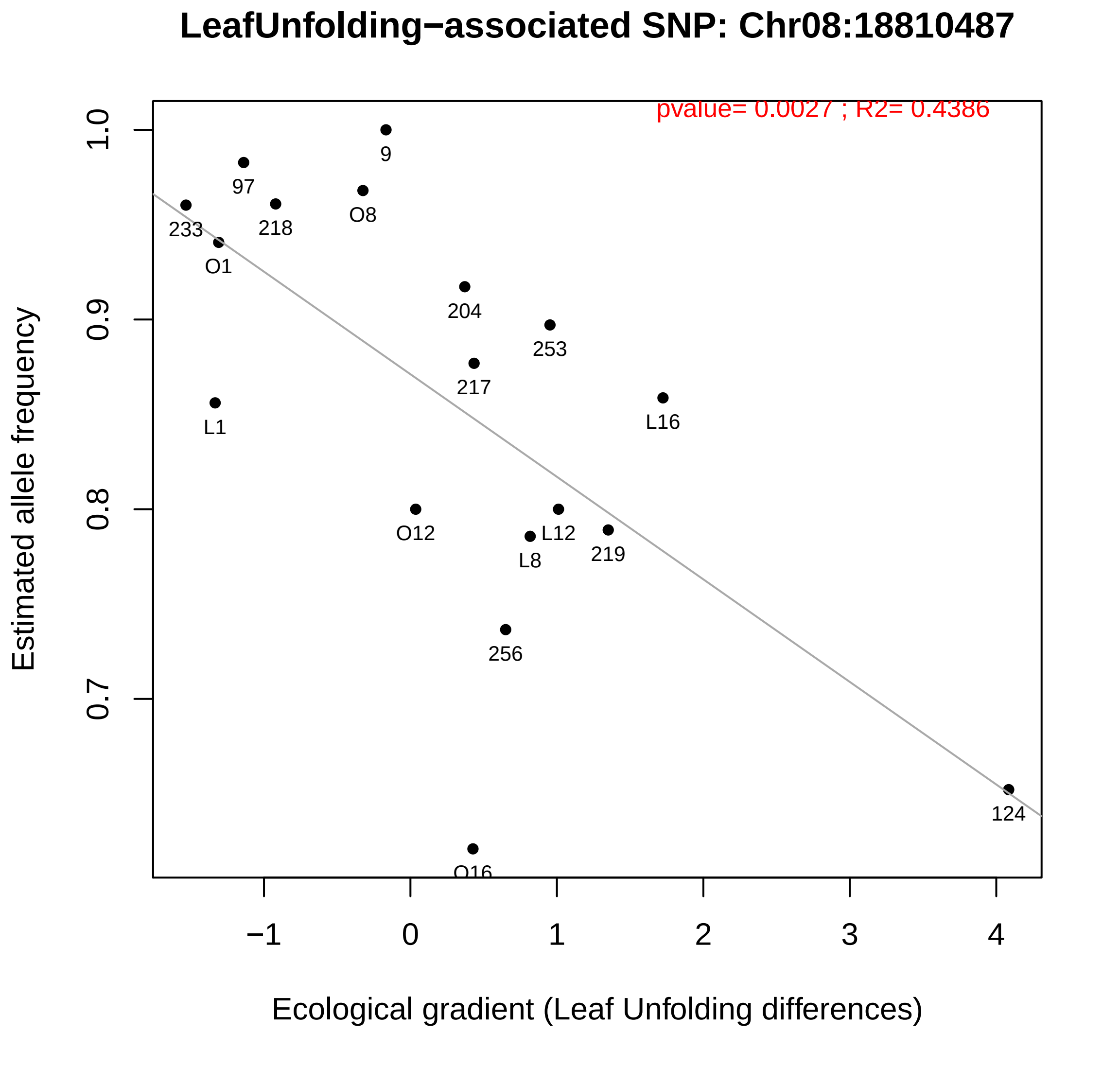

LeafUnfolding−associated SNP: Chr08:18810487
pvalue= 0.0027 ; R2= 0.4386
Estimated allele frequency
Ecological gradient (Leaf Unfolding differences)
1.0
0.9
0.8
0.7
−1
0
1
2
3
4
9
97
O8
233
218
O1
204
253
217
L1
L16
O12
L12
L8
219
256
124
O16

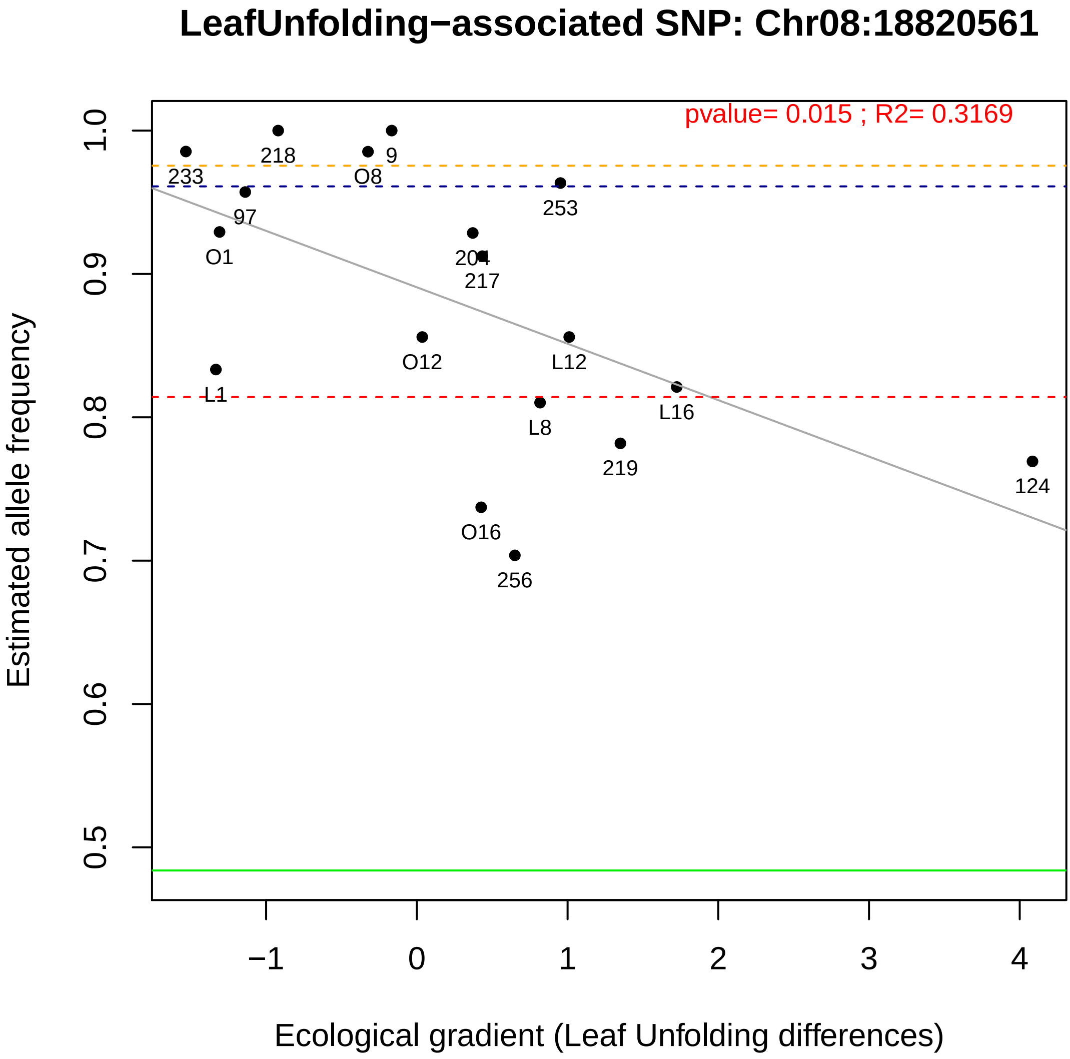
LeafUnfolding−associated SNP: Chr08:18820561
pvalue= 0.015 ; R2= 0.3169
Estimated allele frequency
Ecological gradient (Leaf Unfolding differences)
0.5
0.6
0.7
0.8
0.9
1.0
−1
0
1
2
3
4
233
218
O8
9
97
253
O1
204
217
O12
L12
L1
L16
L8
219
124
O16
256

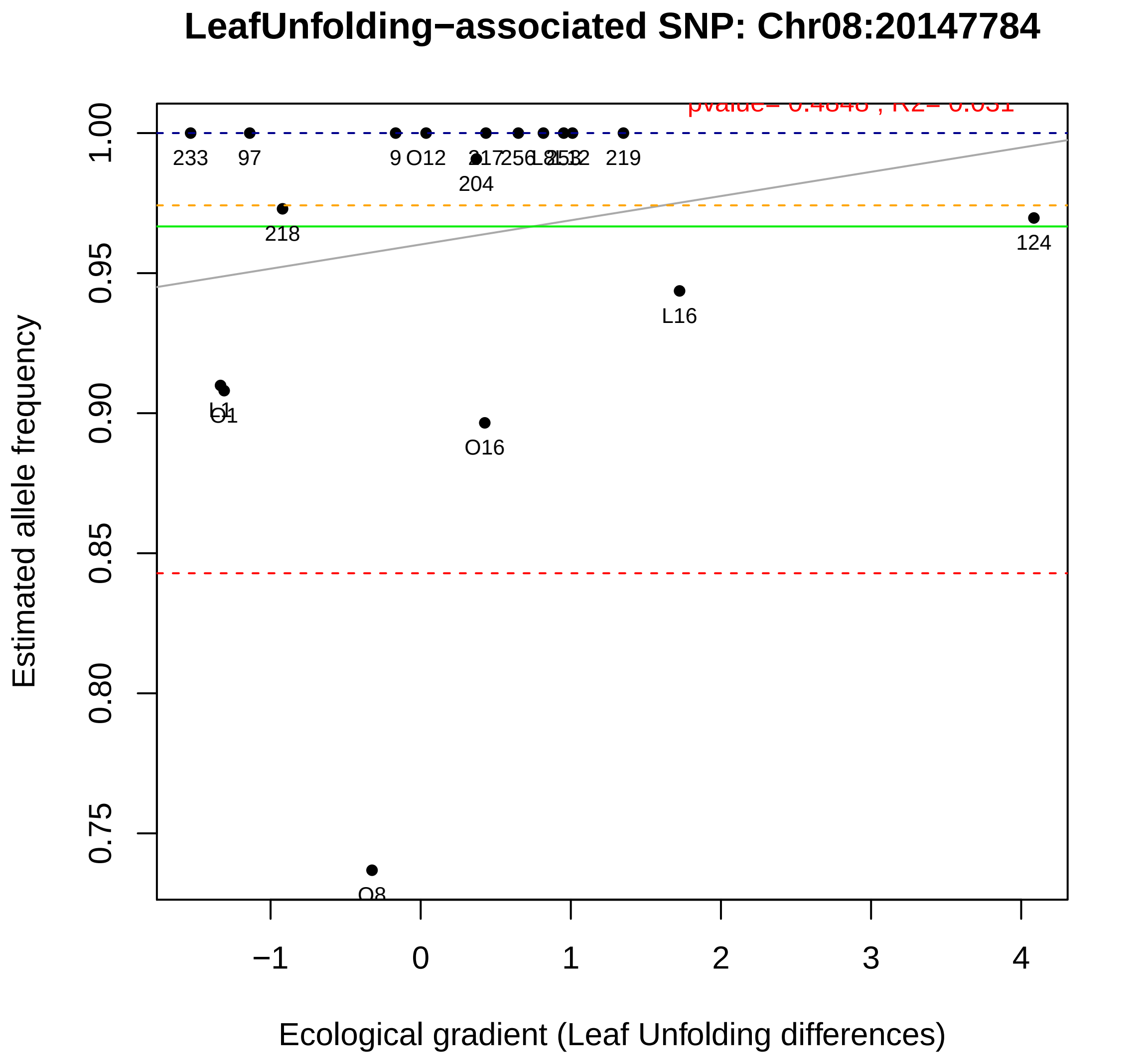

LeafUnfolding−associated SNP: Chr08:20147784
pvalue= 0.4848 ; R2= 0.031
Estimated allele frequency
Ecological gradient (Leaf Unfolding differences)
233
97
9
O12
217
204
256
219
218
124
L16
L1
O1
O16
O8

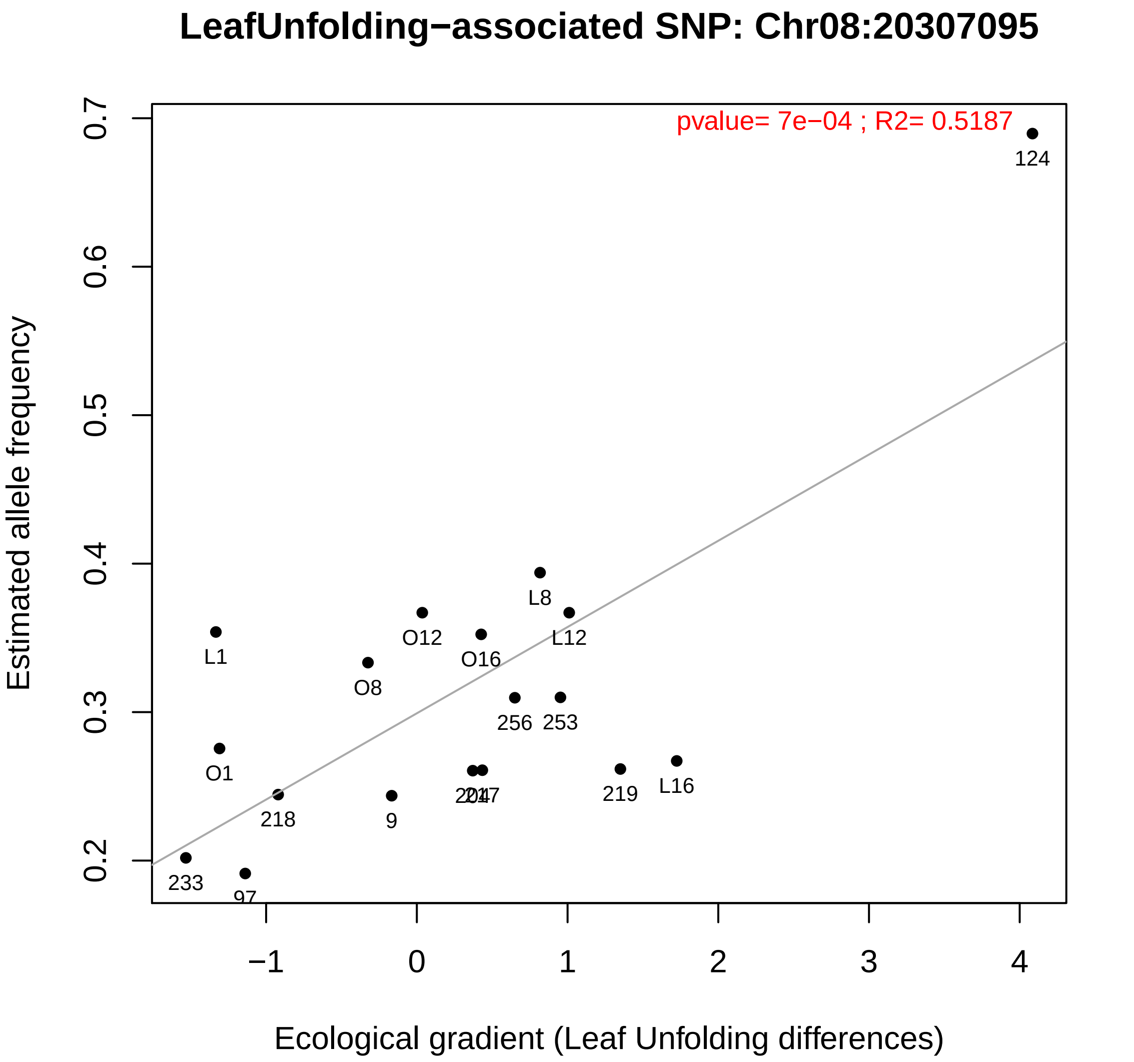

LeafUnfolding−associated SNP: Chr08:20307095
pvalue= 7e−04 ; R2= 0.5187
Estimated allele frequency
Ecological gradient (Leaf Unfolding differences)
0.2
0.3
0.4
0.5
0.6
0.7
−1
0
1
2
3
4
124
L8
L12
O12
O16
L1
O8
256
253
O1
218
9
219
L16
233

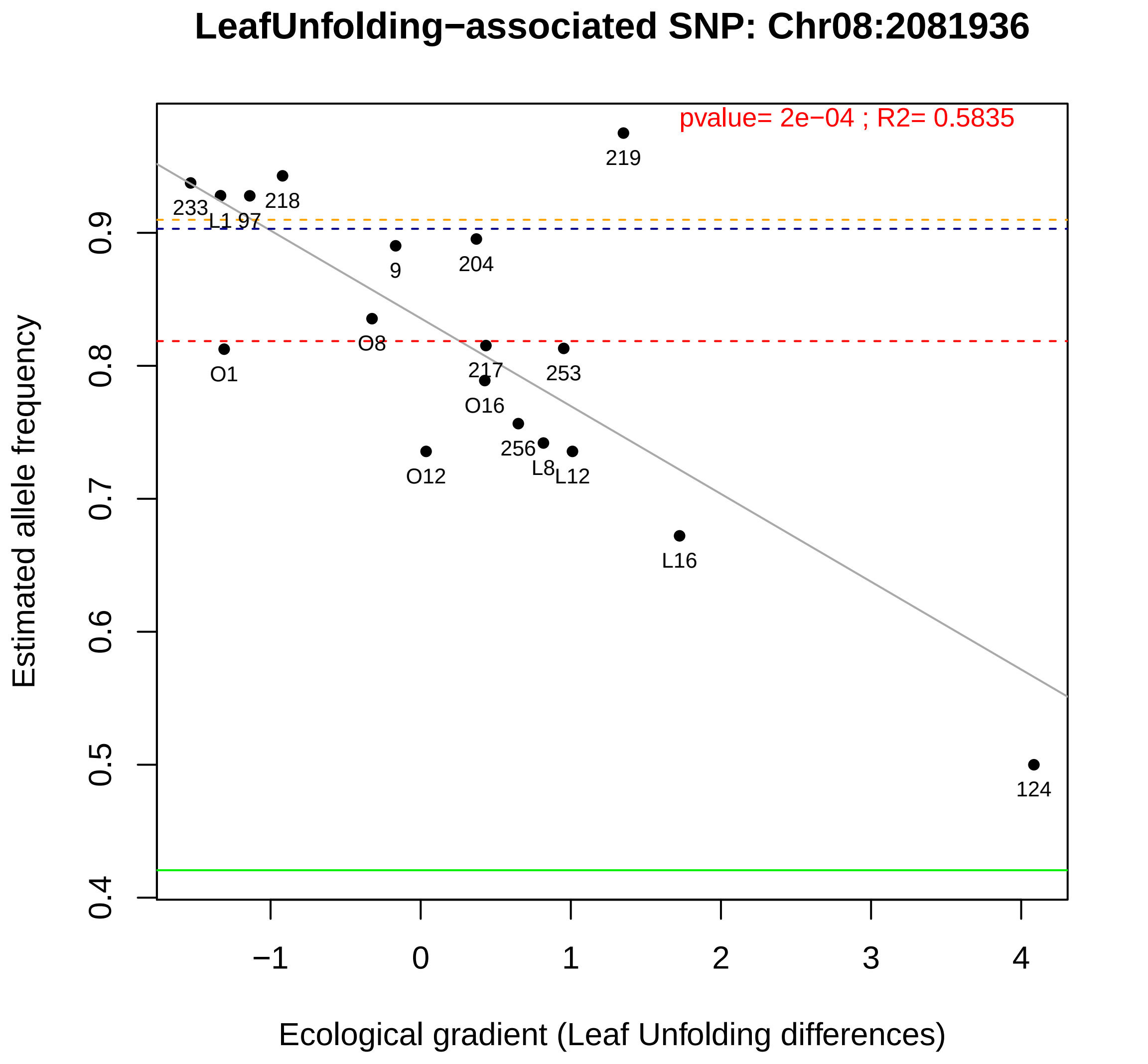

LeafUnfolding−associated SNP: Chr08:2081936
pvalue= 2e−04 ; R2= 0.5835
219
233
L1
97
218
9
204
O8
O1
217
253
O16
256
L8
L12
O12
L16
124
0.4
0.5
0.6
0.7
0.8
0.9
−1
0
1
2
3
4
Estimated allele frequency
Ecological gradient (Leaf Unfolding differences)

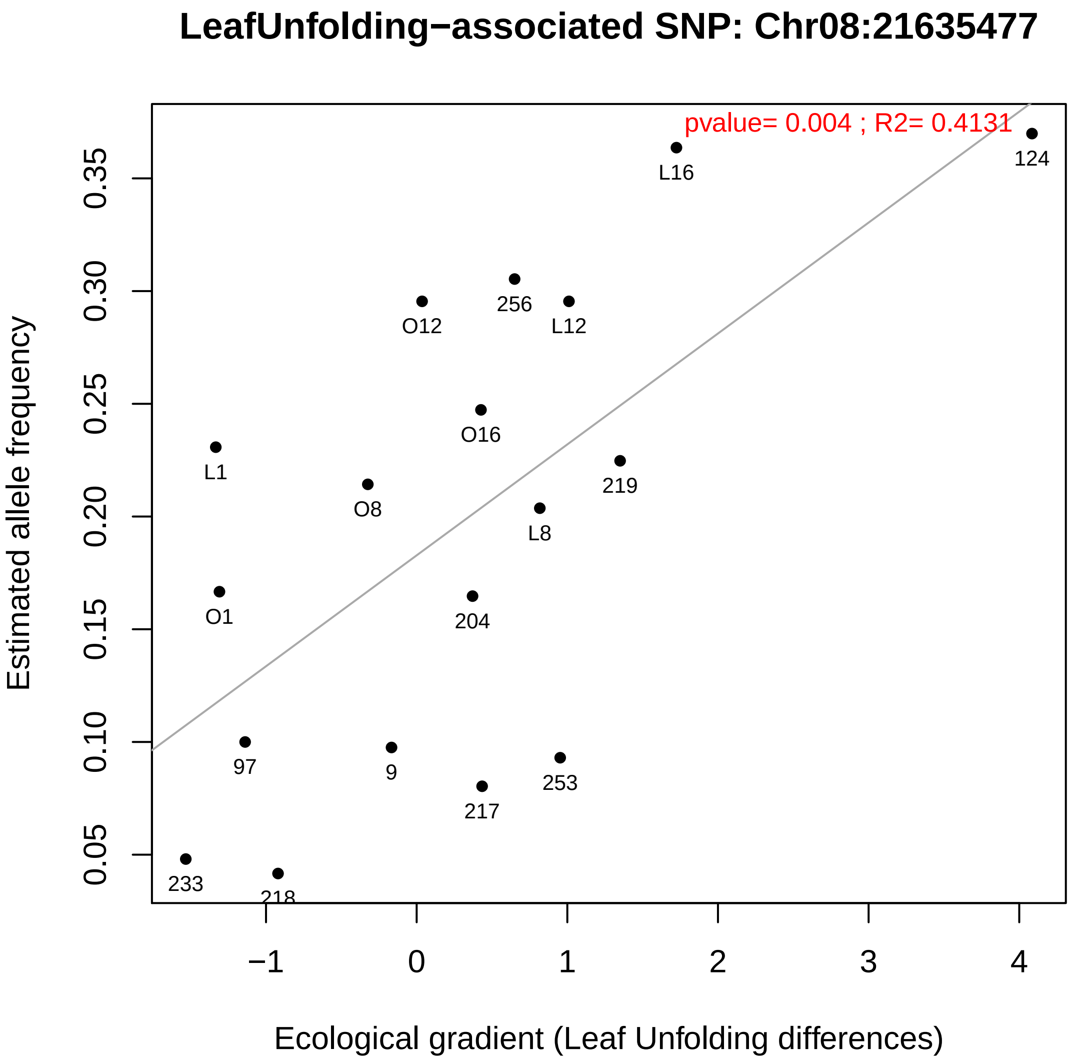
LeafUnfolding−associated SNP: Chr08:21635477
pvalue= 0.004 ; R2= 0.4131
Estimated allele frequency
Ecological gradient (Leaf Unfolding differences)
0.05
0.10
0.15
0.20
0.25
0.30
0.35
−1
0
1
2
3
4
L16
124
256
O12
L12
O16
L1
219
O8
L8
O1
204
97
9
253
217
233
218

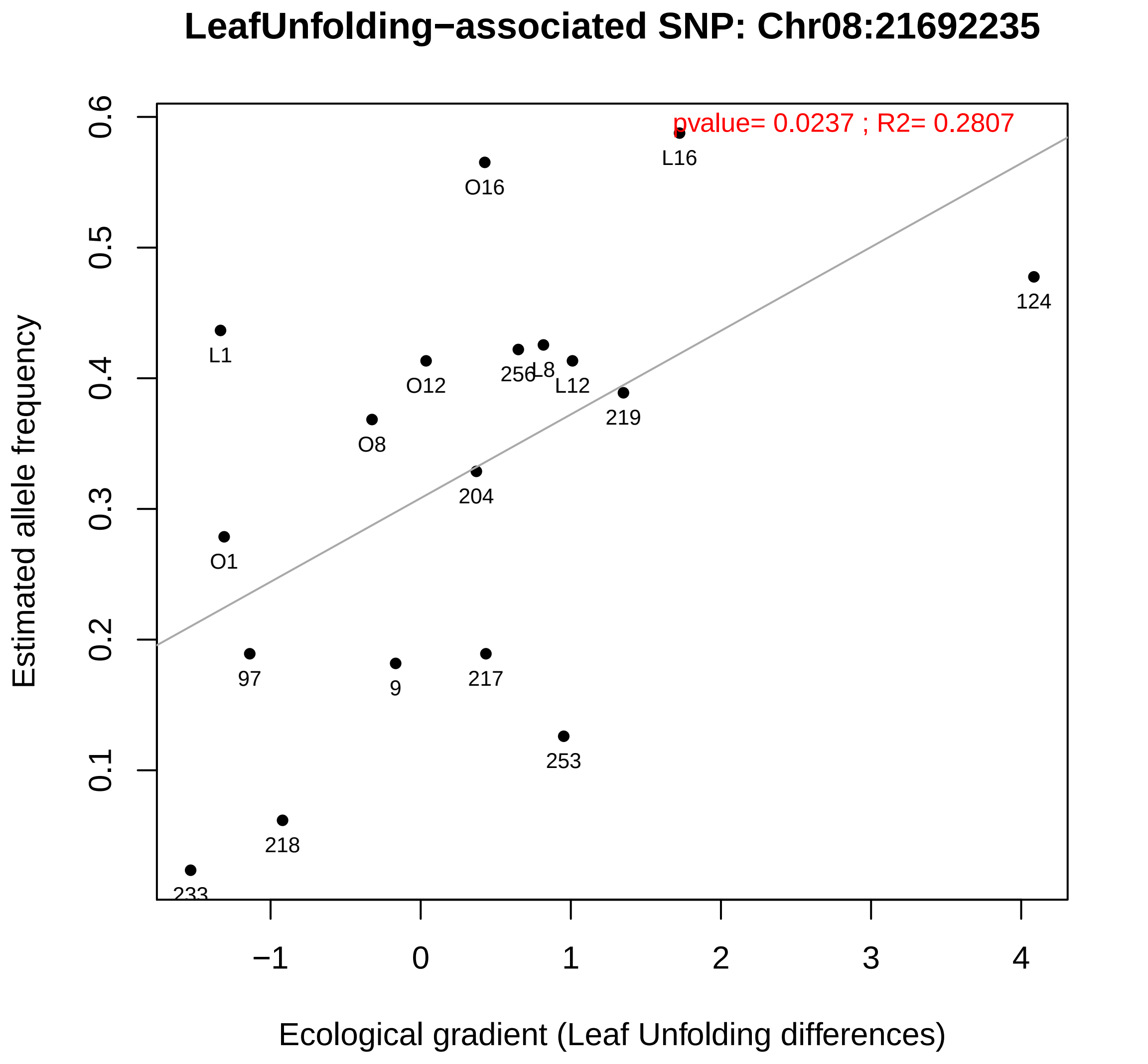

LeafUnfolding−associated SNP: Chr08:21692235
pvalue= 0.0237 ; R2= 0.2807
Estimated allele frequency
Ecological gradient (Leaf Unfolding differences)
0.1
0.2
0.3
0.4
0.5
0.6
−1
0
1
2
3
4
L16
O16
124
L1
O12
256
L8
L12
219
O8
204
O1
97
9
217
253
218
233

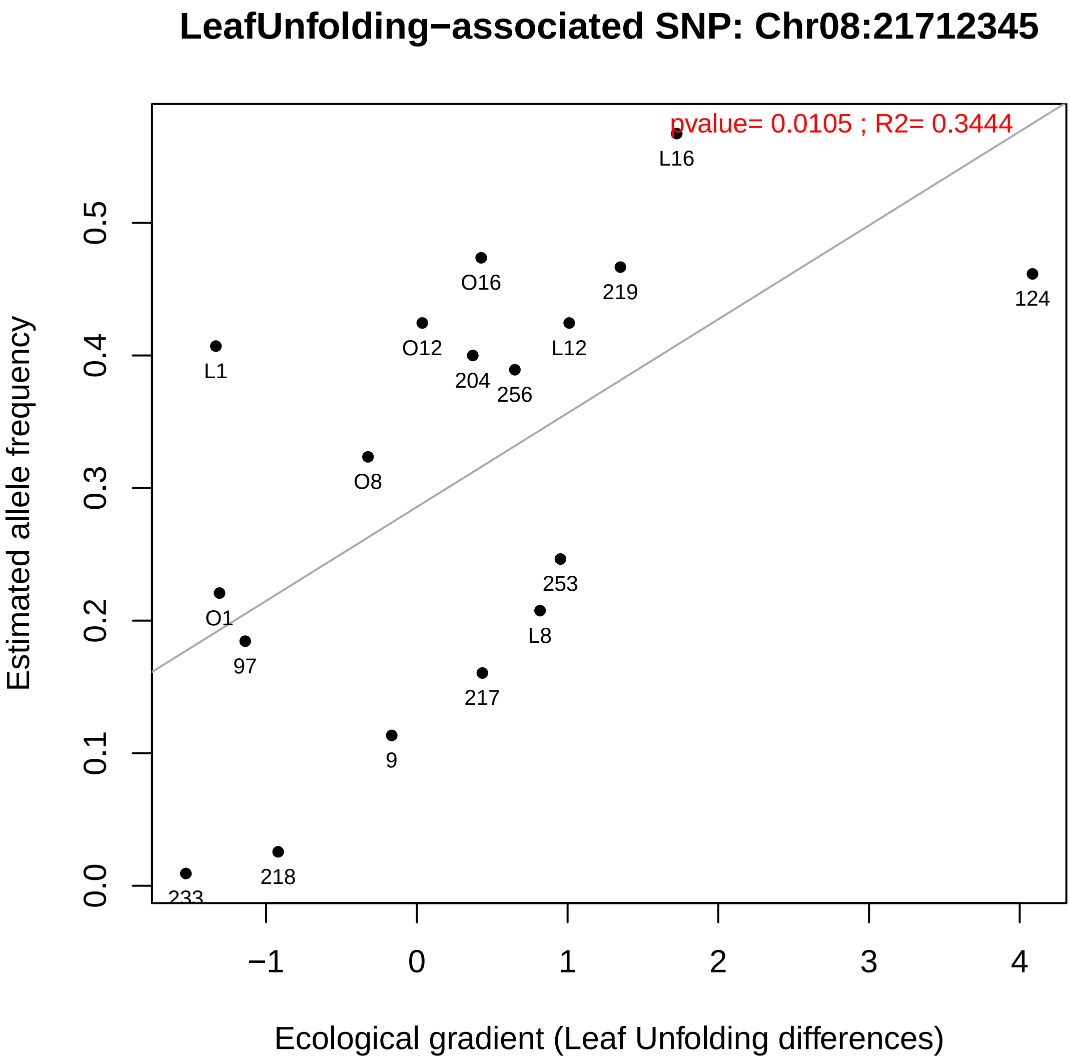
LeafUnfolding−associated SNP: Chr08:21712345
pvalue= 0.0105 ; R2= 0.3444
Estimated allele frequency
Ecological gradient (Leaf Unfolding differences)
0.0
0.1
0.2
0.3
0.4
0.5
−1
0
1
2
3
4
L16
O16
219
124
O12
L12
L1
204
256
O8
253
O1
L8
97
217
9
218
233

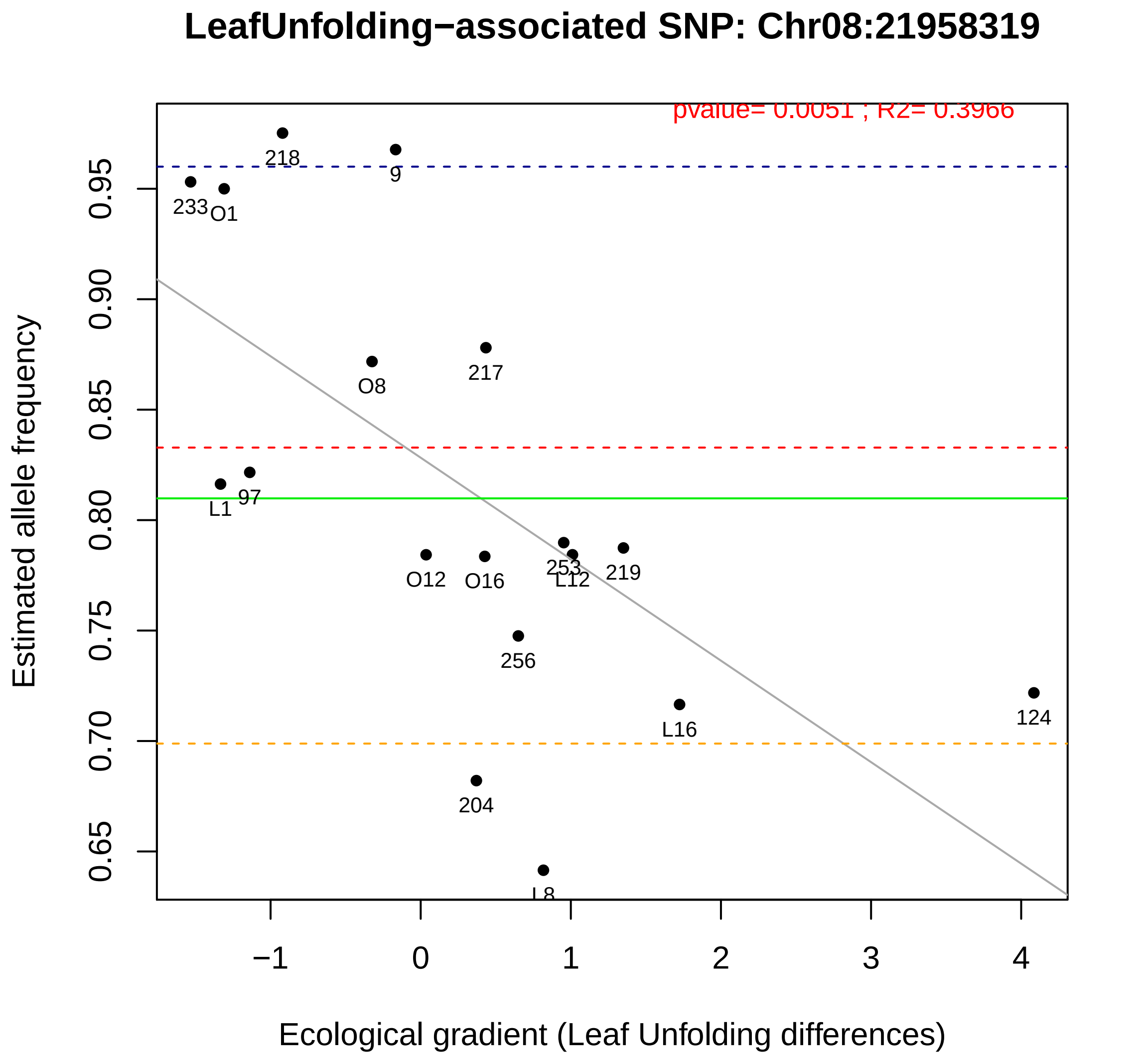
LeafUnfolding−associated SNP: Chr08:21958319
pvalue= 0.0051 ; R2= 0.3966
Estimated allele frequency
0.65
0.70
0.75
0.80
0.85
0.90
0.95
Ecological gradient (Leaf Unfolding differences)
−1
0
1
2
3
4
218
9
233
O1
O8
217
L1
97
O12
O16
253
L12
219
256
L16
124
204
L8

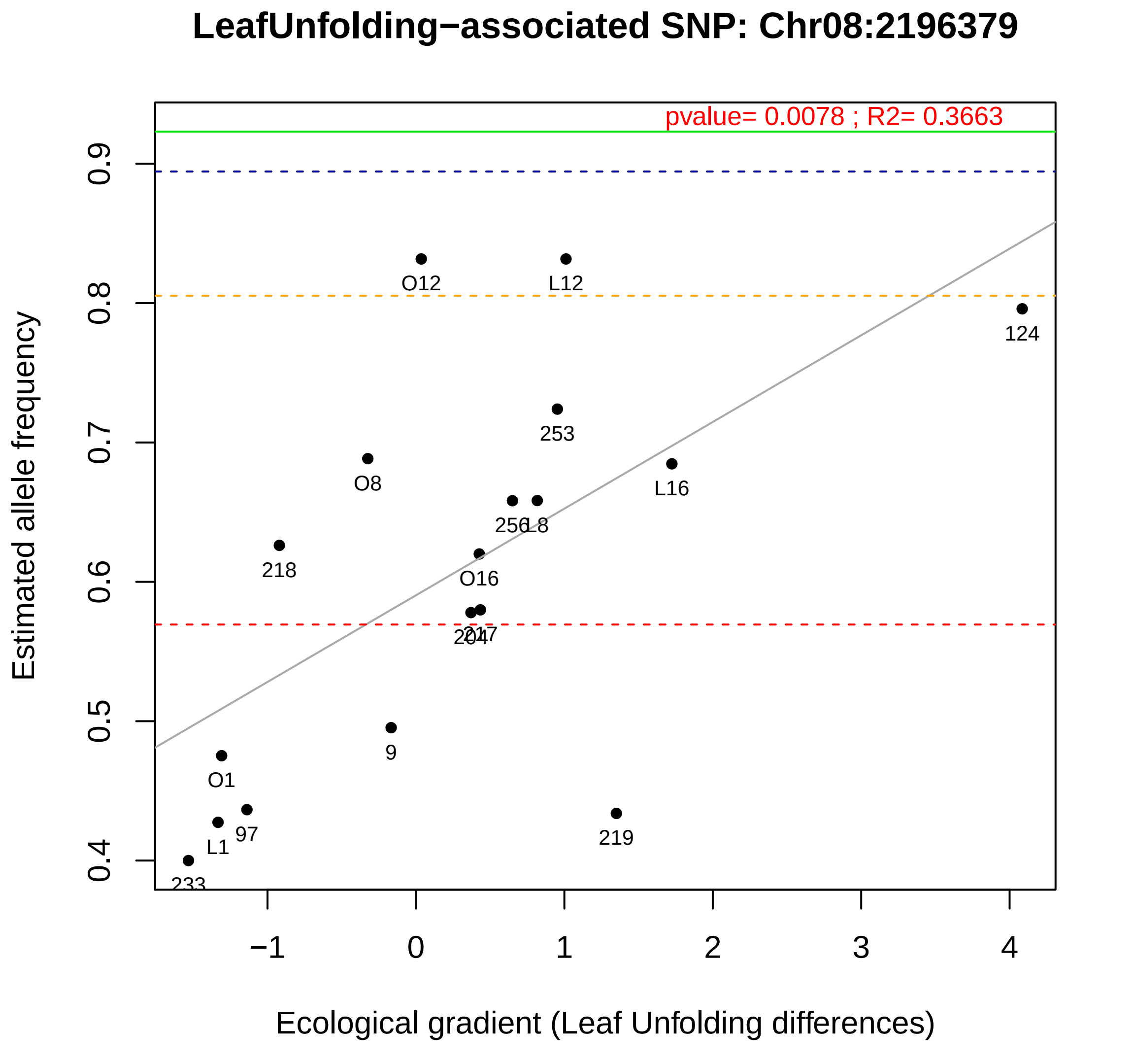
LeafUnfolding−associated SNP: Chr08:2196379
pvalue= 0.0078 ; R2= 0.3663
Estimated allele frequency
Ecological gradient (Leaf Unfolding differences)
0.4
0.5
0.6
0.7
0.8
0.9
−1
0
1
2
3
4
O12
L12
124
253
O8
L16
256
L8
218
O16
204
217
9
O1
97
L1
219
233

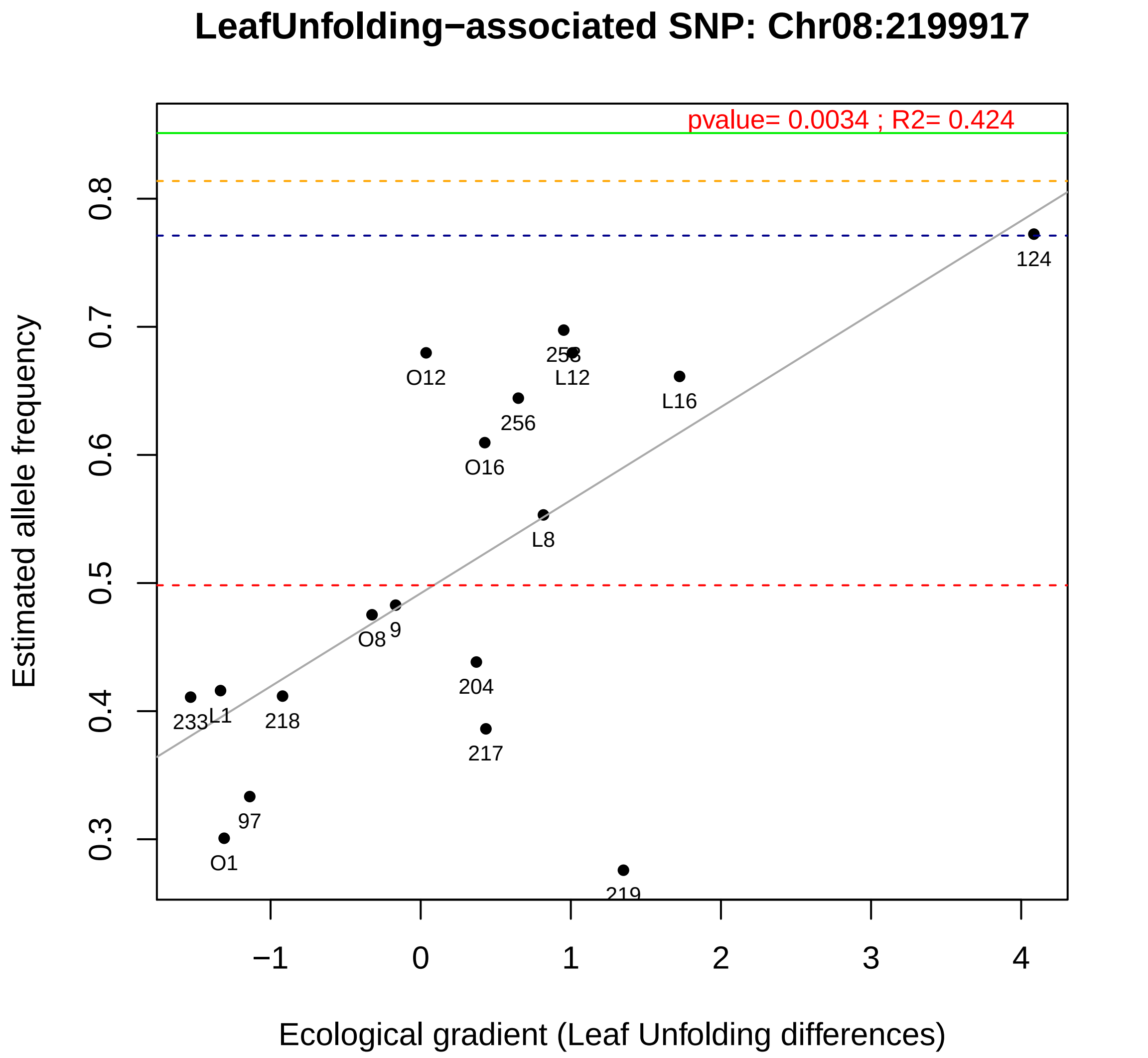

LeafUnfolding−associated SNP: Chr08:2199917
pvalue= 0.0034 ; R2= 0.424
Estimated allele frequency
0.3
0.4
0.5
0.6
0.7
0.8
Ecological gradient (Leaf Unfolding differences)
−1
0
1
2
3
4
124
O12
L12
L16
256
O16
L8
O8
9
204
233
L1
218
217
97
O1
219

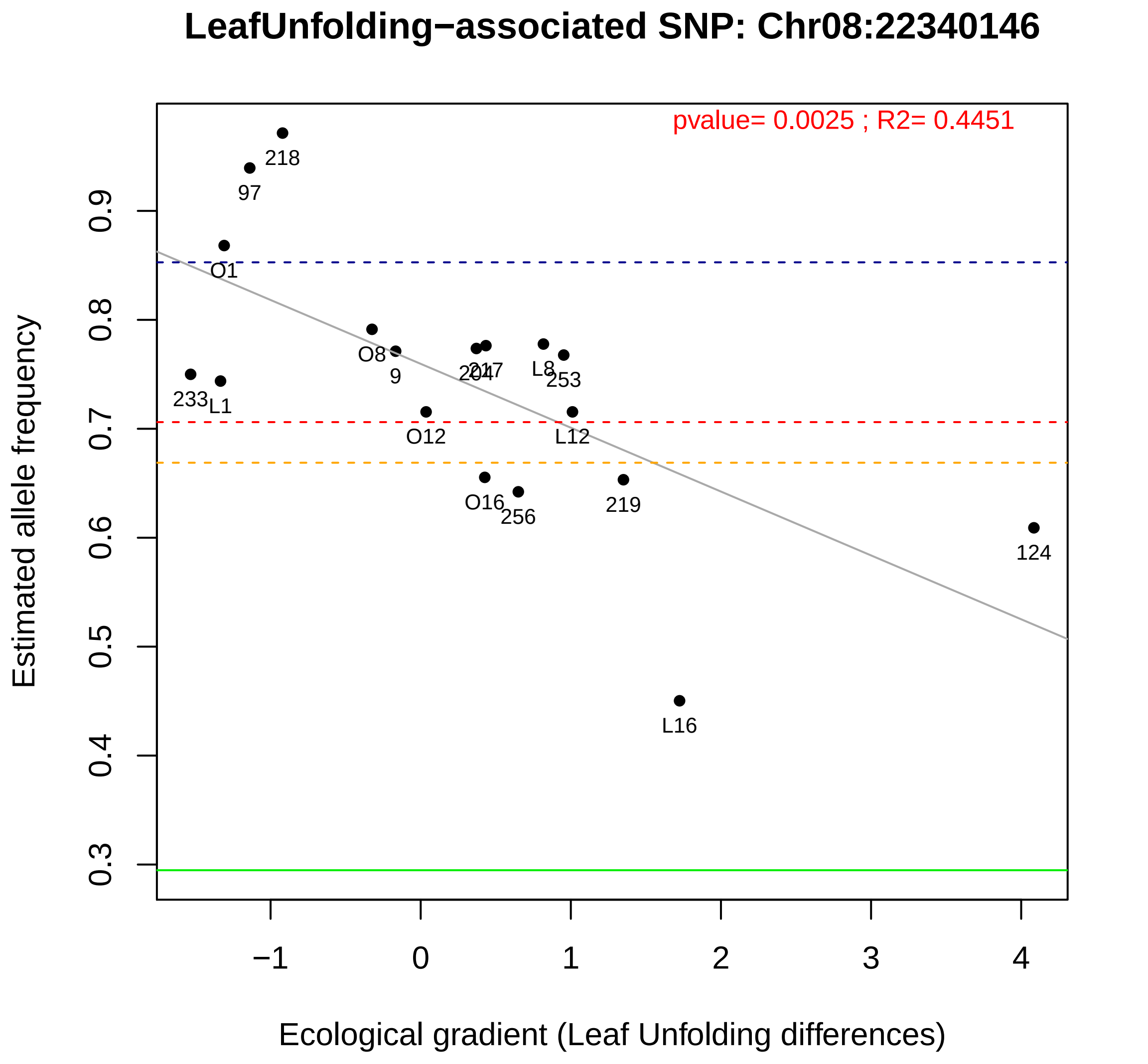
LeafUnfolding−associated SNP: Chr08:22340146
pvalue= 0.0025 ; R2= 0.4451
Estimated allele frequency
Ecological gradient (Leaf Unfolding differences)
0.3
0.4
0.5
0.6
0.7
0.8
0.9
−1
0
1
2
3
4
218
97
O1
O8
9
204
217
L8
253
233
L1
O12
L12
O16
256
219
124
L16

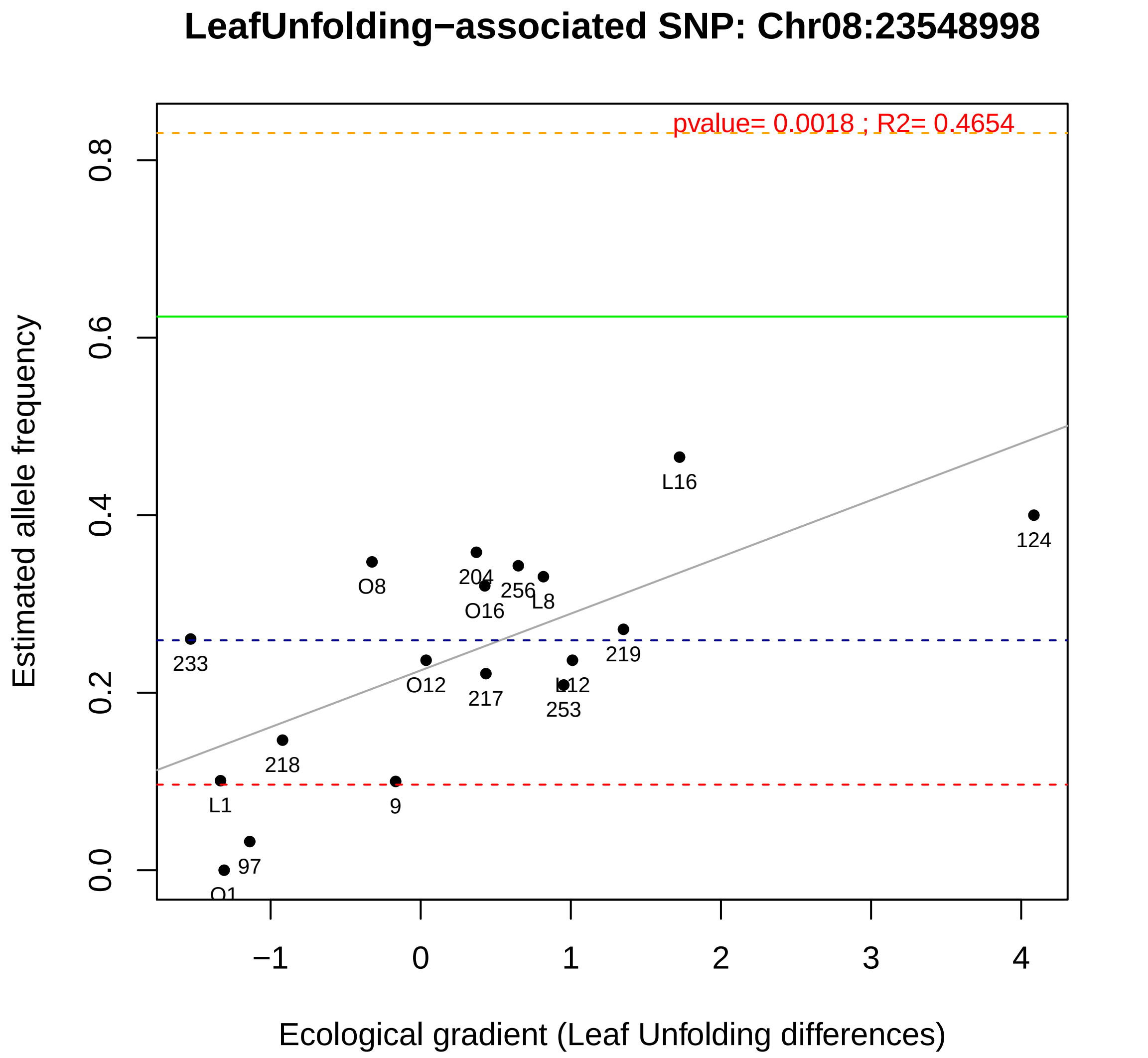

LeafUnfolding−associated SNP: Chr08:23548998
pvalue= 0.0018 ; R2= 0.4654
Estimated allele frequency
0.0
0.2
0.4
0.6
0.8
−1
0
1
2
3
4
Ecological gradient (Leaf Unfolding differences)
L16
124
O8
204
256
O16
L8
233
219
O12
217
L12
253
218
L1
9
97

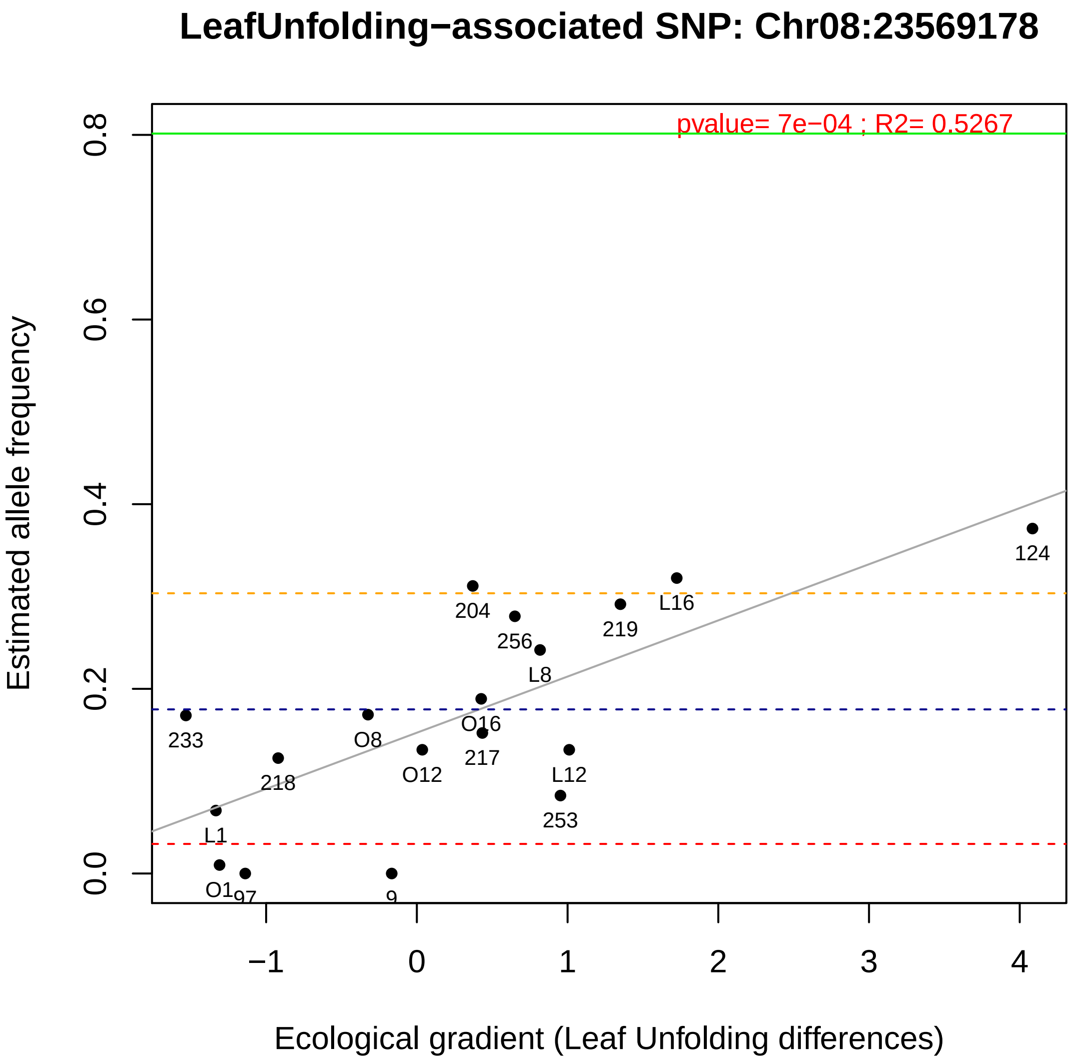
LeafUnfolding−associated SNP: Chr08:23569178
pvalue= 7e−04 ; R2= 0.5267
Estimated allele frequency
0.0
0.2
0.4
0.6
0.8
−1
0
1
2
3
4
Ecological gradient (Leaf Unfolding differences)
124
L16
204
219
256
L8
O16
233
O8
217
L12
O12
218
253
L1
O1
97
9

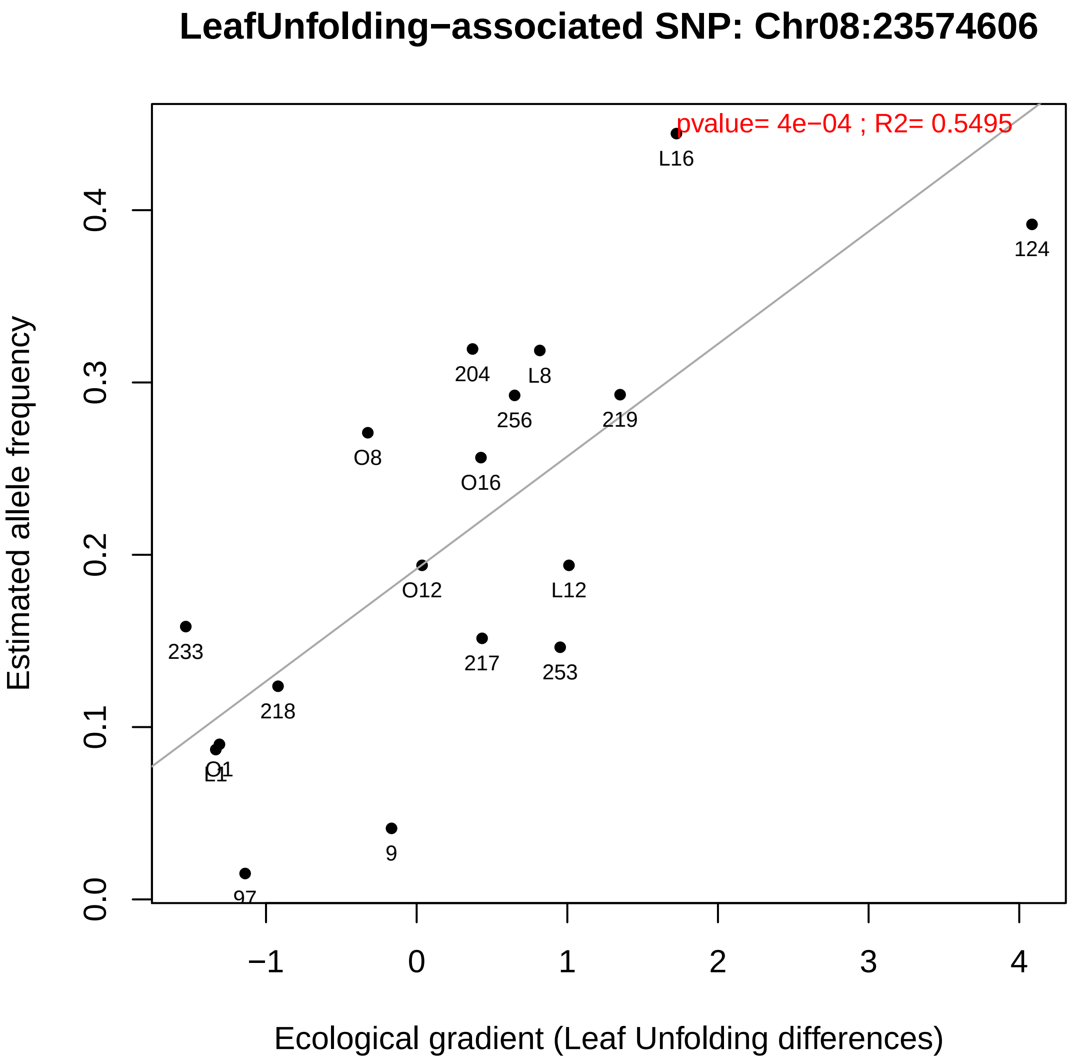

LeafUnfolding−associated SNP: Chr08:23574606
pvalue= 4e−04 ; R2= 0.5495
Estimated allele frequency
Ecological gradient (Leaf Unfolding differences)
0.0
0.1
0.2
0.3
0.4
−1
0
1
2
3
4
L16
124
204
L8
256
219
O8
O16
O12
L12
233
217
253
218
O1
L1
9
97

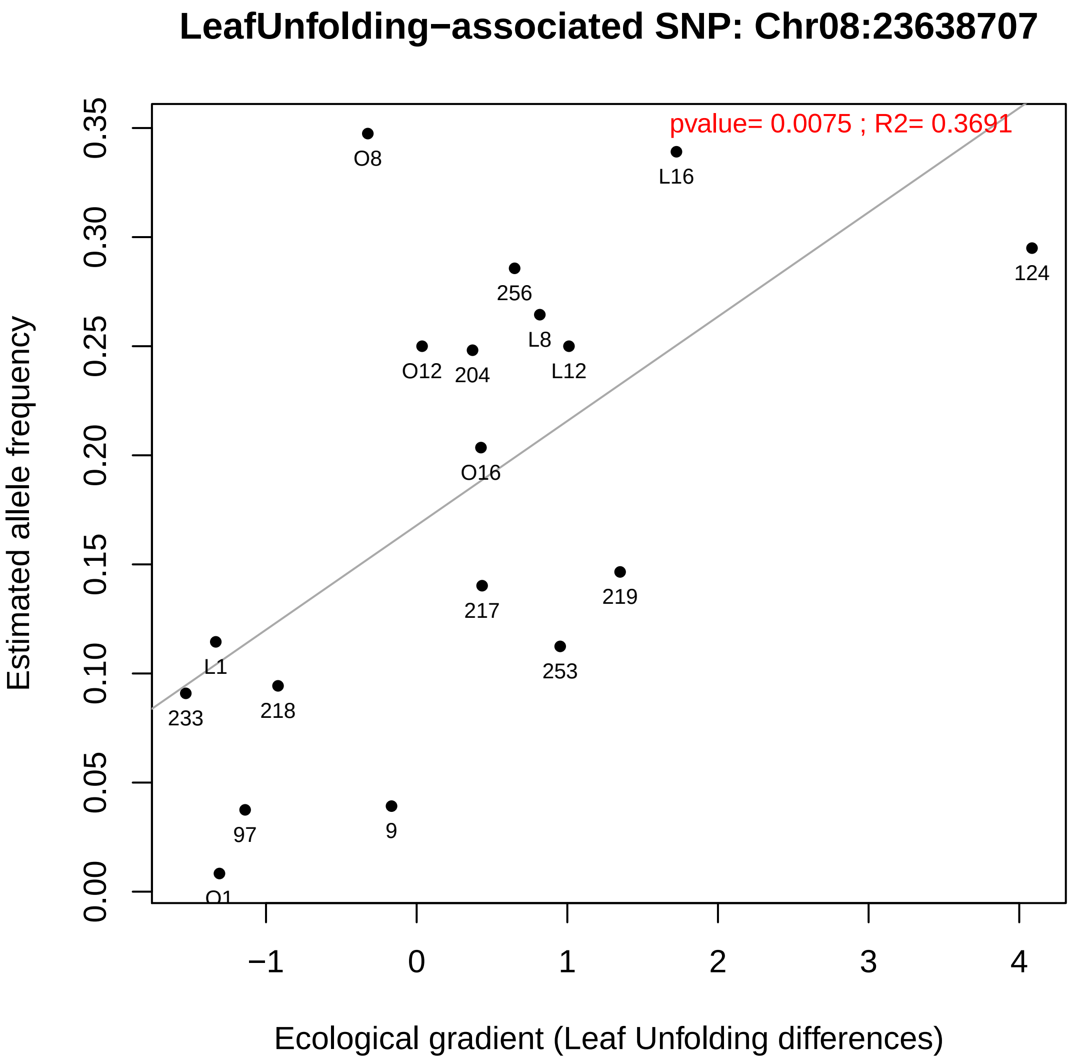
LeafUnfolding−associated SNP: Chr08:23638707
pvalue= 0.0075 ; R2= 0.3691
O8
L16
124
256
L8
O12
204
L12
O16
217
219
L1
253
233
218
97
9
O1
Estimated allele frequency
0.00
0.05
0.10
0.15
0.20
0.25
0.30
0.35
−1
0
1
2
3
4
Ecological gradient (Leaf Unfolding differences)

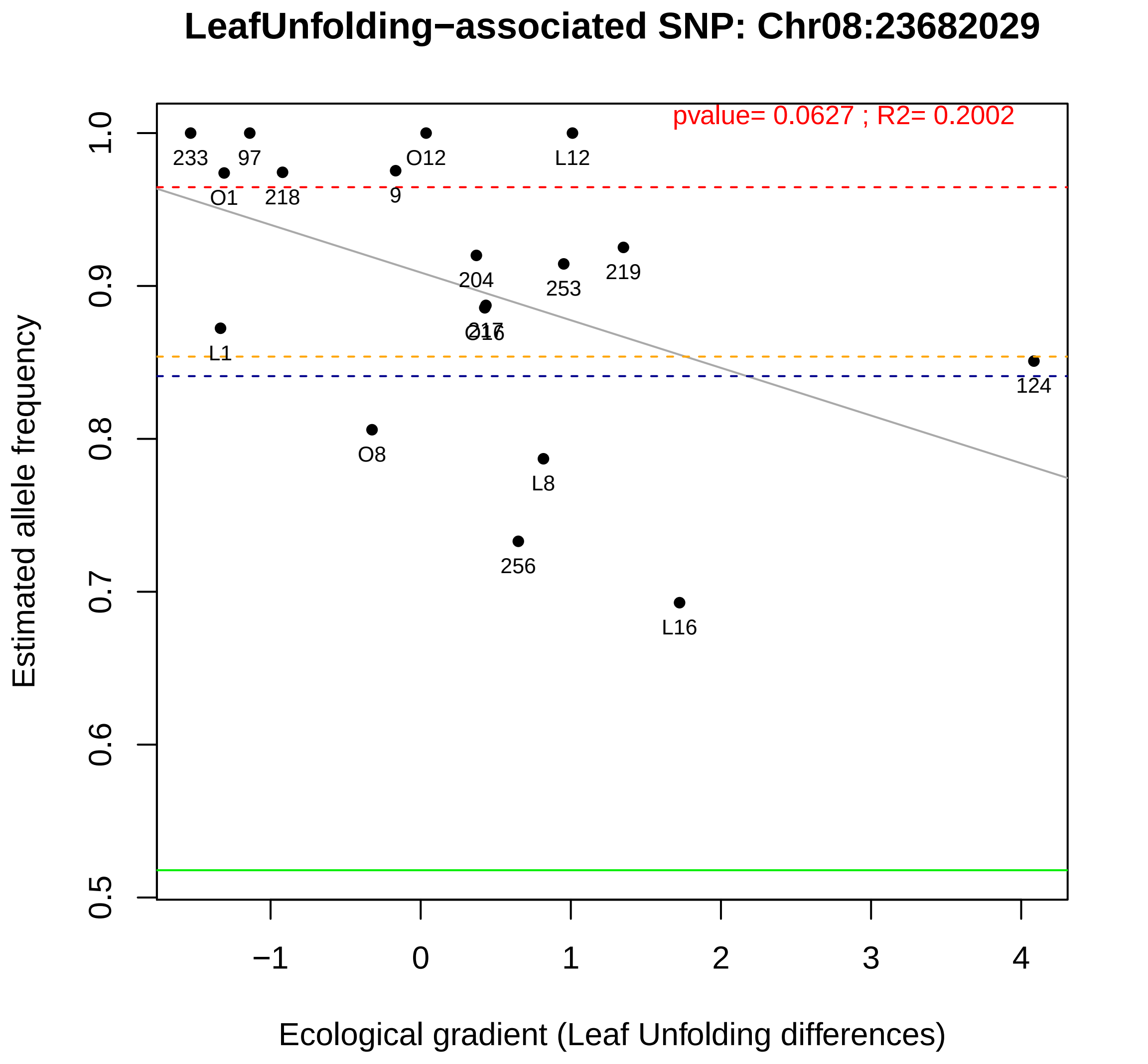

LeafUnfolding−associated SNP: Chr08:23682029
pvalue= 0.0627 ; R2= 0.2002
Estimated allele frequency
Ecological gradient (Leaf Unfolding differences)
1.0
0.9
0.8
0.7
0.6
0.5
−1
0
1
2
3
4
233
97
O12
L12
O1
218
9
204
253
219
L1
124
O8
L8
256
L16

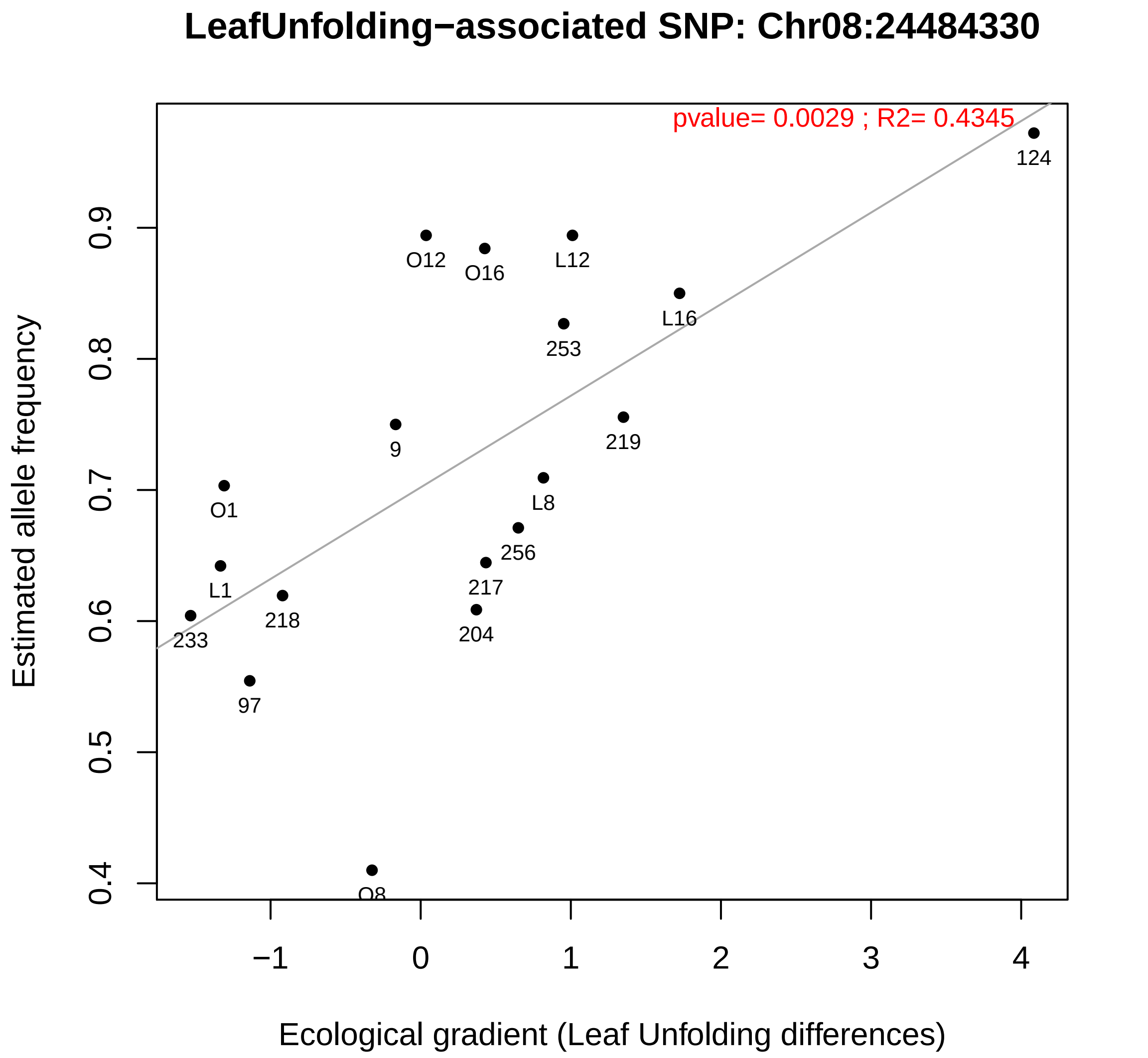
LeafUnfolding−associated SNP: Chr08:24484330
pvalue= 0.0029 ; R2= 0.4345
124
O12
O16
L12
L16
253
9
219
O1
L8
256
217
L1
218
233
204
97
O8
0.4
0.5
0.6
0.7
0.8
0.9
−1
0
1
2
3
4
Estimated allele frequency
Ecological gradient (Leaf Unfolding differences)

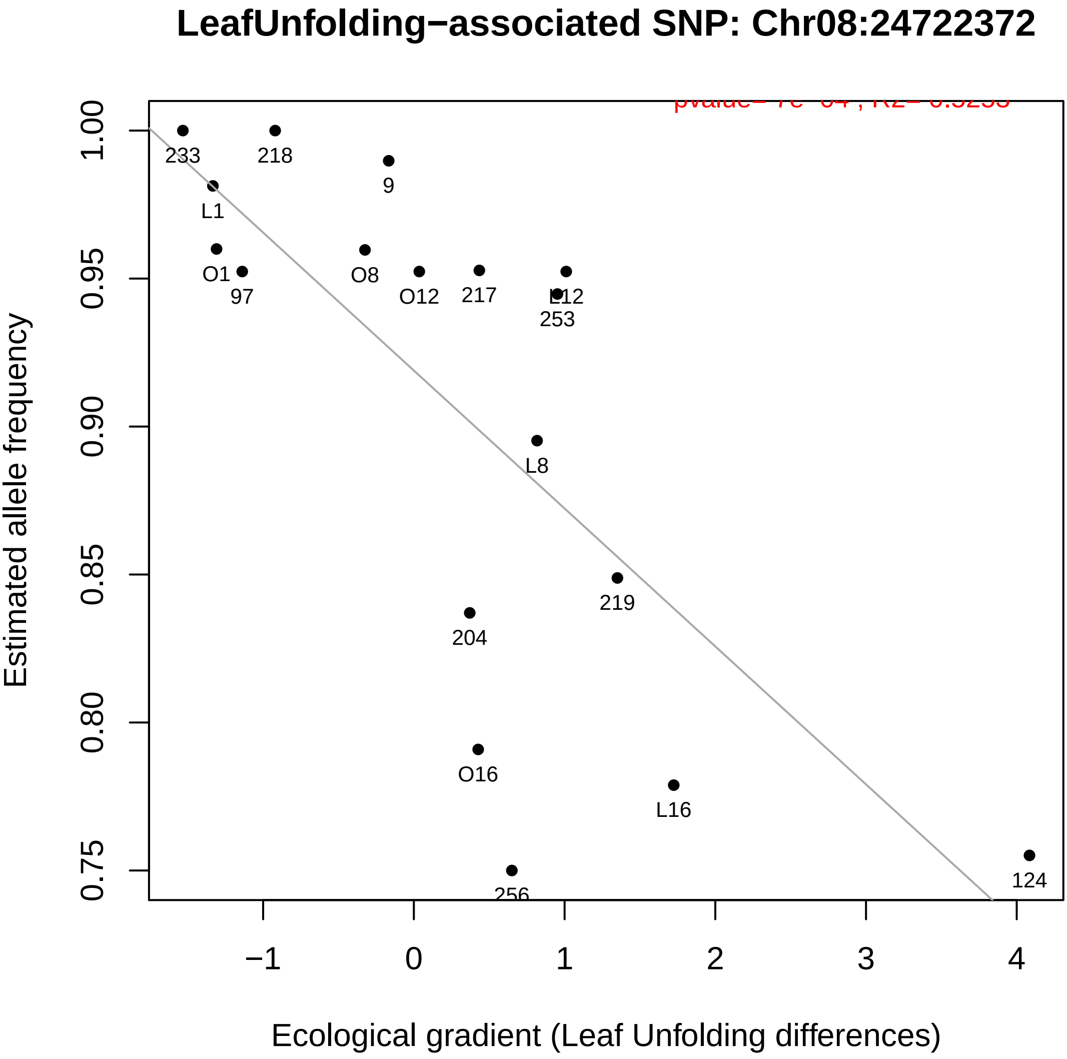
LeafUnfolding−associated SNP: Chr08:24722372
Estimated allele frequency
1.00
0.95
0.90
0.85
0.80
0.75
233
218
9
L1
O1
97
O8
O12
217
L12
253
L8
219
204
O16
L16
256
124
−1
0
1
2
3
4
Ecological gradient (Leaf Unfolding differences)

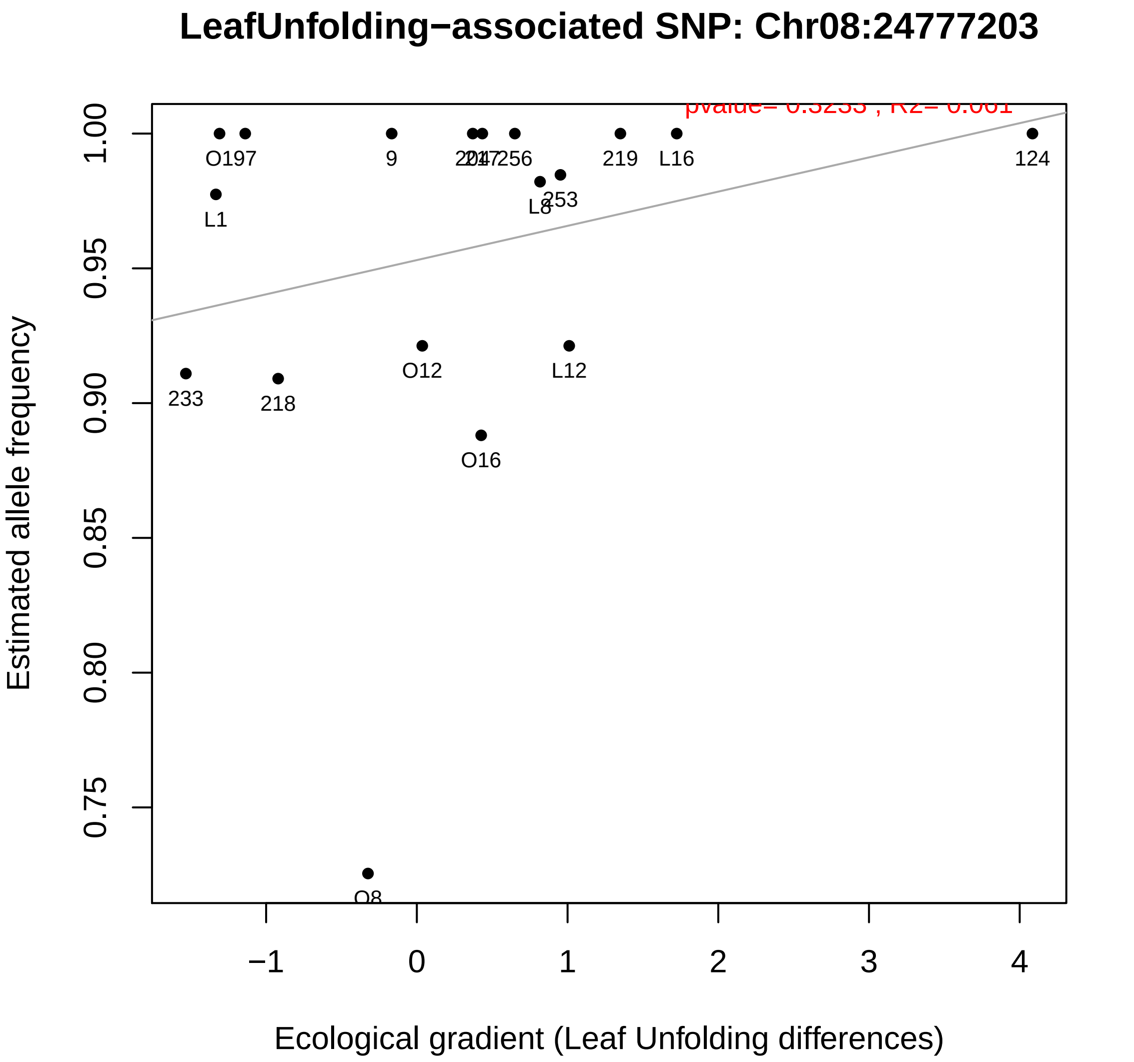

LeafUnfolding−associated SNP: Chr08:24777203
pvalue= 0.3233 ; R2= 0.061
Estimated allele frequency
Ecological gradient (Leaf Unfolding differences)
1.00
0.95
0.90
0.85
0.80
0.75
−1
0
1
2
3
4
O197
9
256
219
L16
124
L1
L8
253
233
218
O12
L12
O16
O8

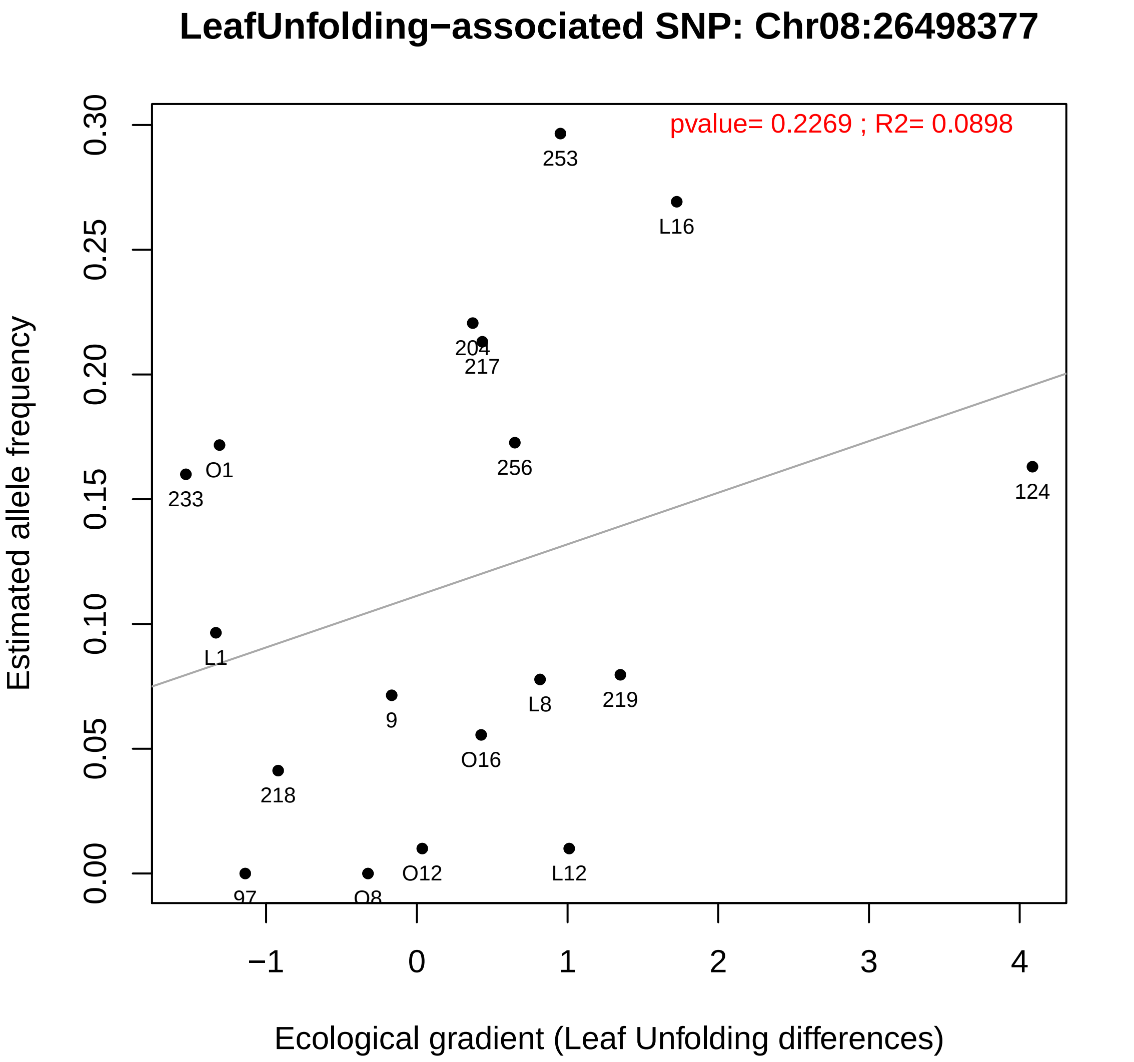
LeafUnfolding−associated SNP: Chr08:26498377
pvalue= 0.2269 ; R2= 0.0898
Estimated allele frequency
0.00
0.05
0.10
0.15
0.20
0.25
0.30
Ecological gradient (Leaf Unfolding differences)
−1
0
1
2
3
4
253
L16
204
217
256
O1
233
124
L1
9
L8
219
O16
218
O12
L12
97
O8

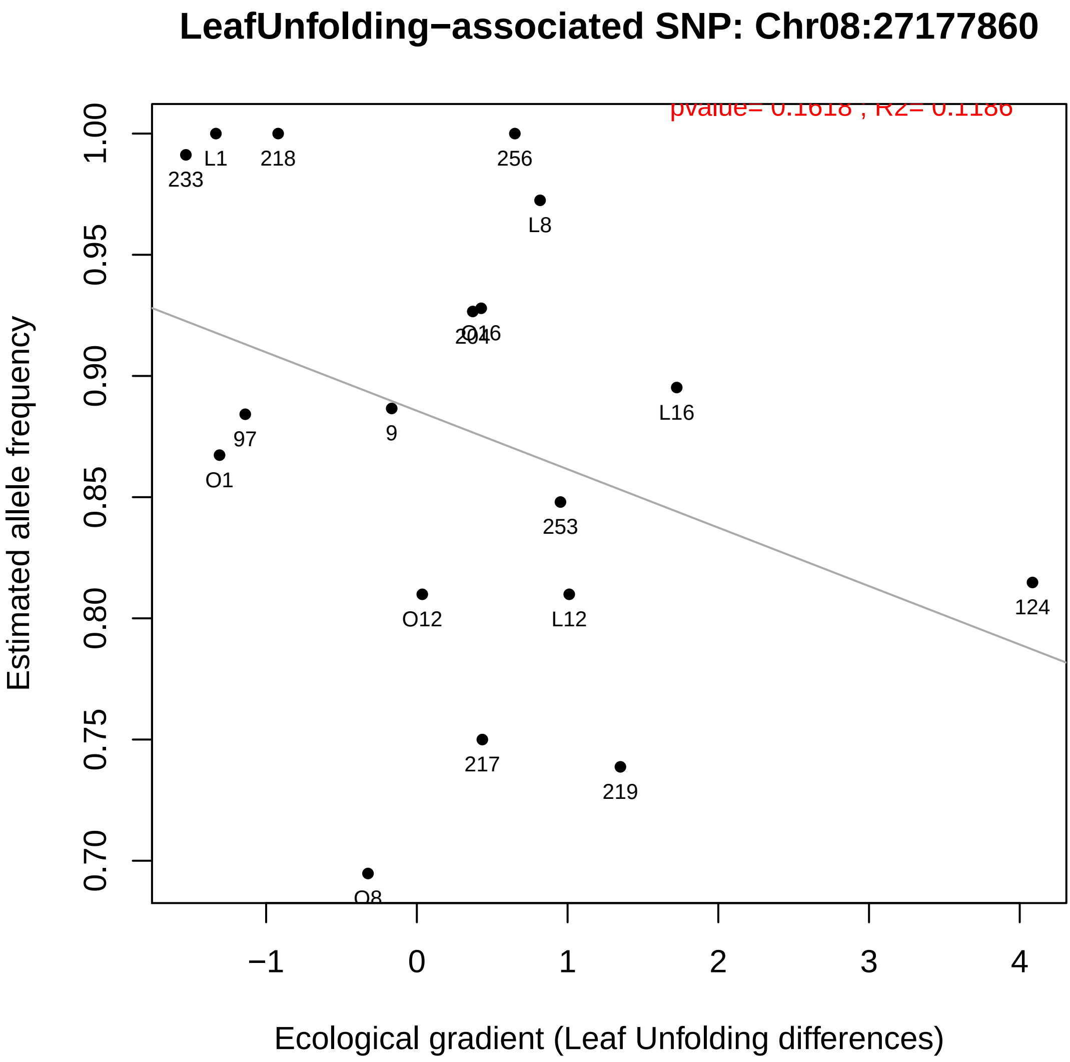

LeafUnfolding−associated SNP: Chr08:27177860
pvalue= 0.1618 ; R2= 0.1186
Estimated allele frequency
Ecological gradient (Leaf Unfolding differences)
1.00
0.95
0.90
0.85
0.80
0.75
0.70
−1
0
1
2
3
4
233
L1
218
256
L8
204
O16
L16
97
9
O1
253
124
O12
L12
217
219
O8

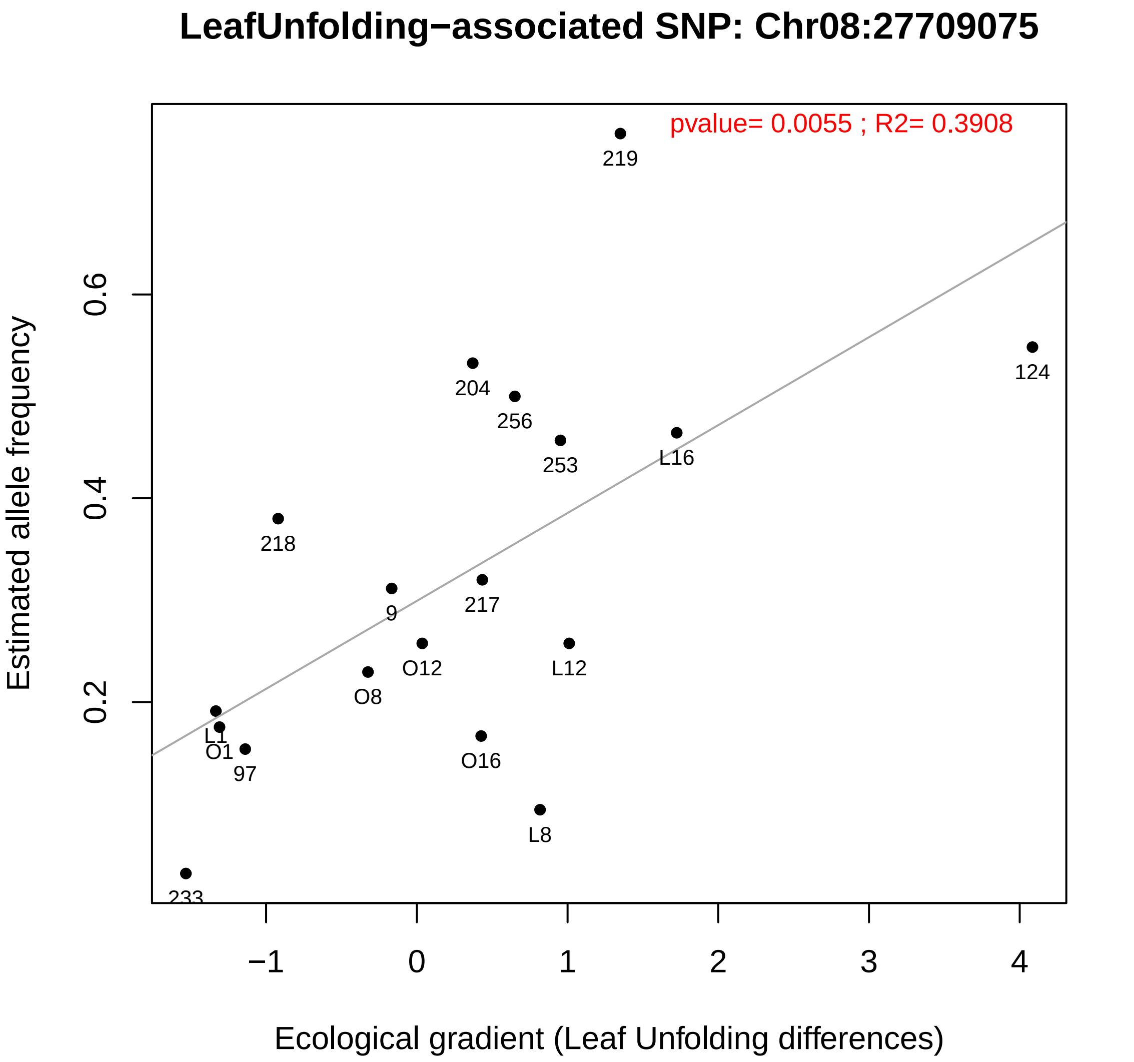
LeafUnfolding−associated SNP: Chr08:27709075
pvalue= 0.0055 ; R2= 0.3908
219
204
256
253
L16
124
218
9
217
O12
L12
O8
L1
O1
97
O16
L8
233
0.2
0.4
0.6
−1
0
1
2
3
4
Estimated allele frequency
Ecological gradient (Leaf Unfolding differences)

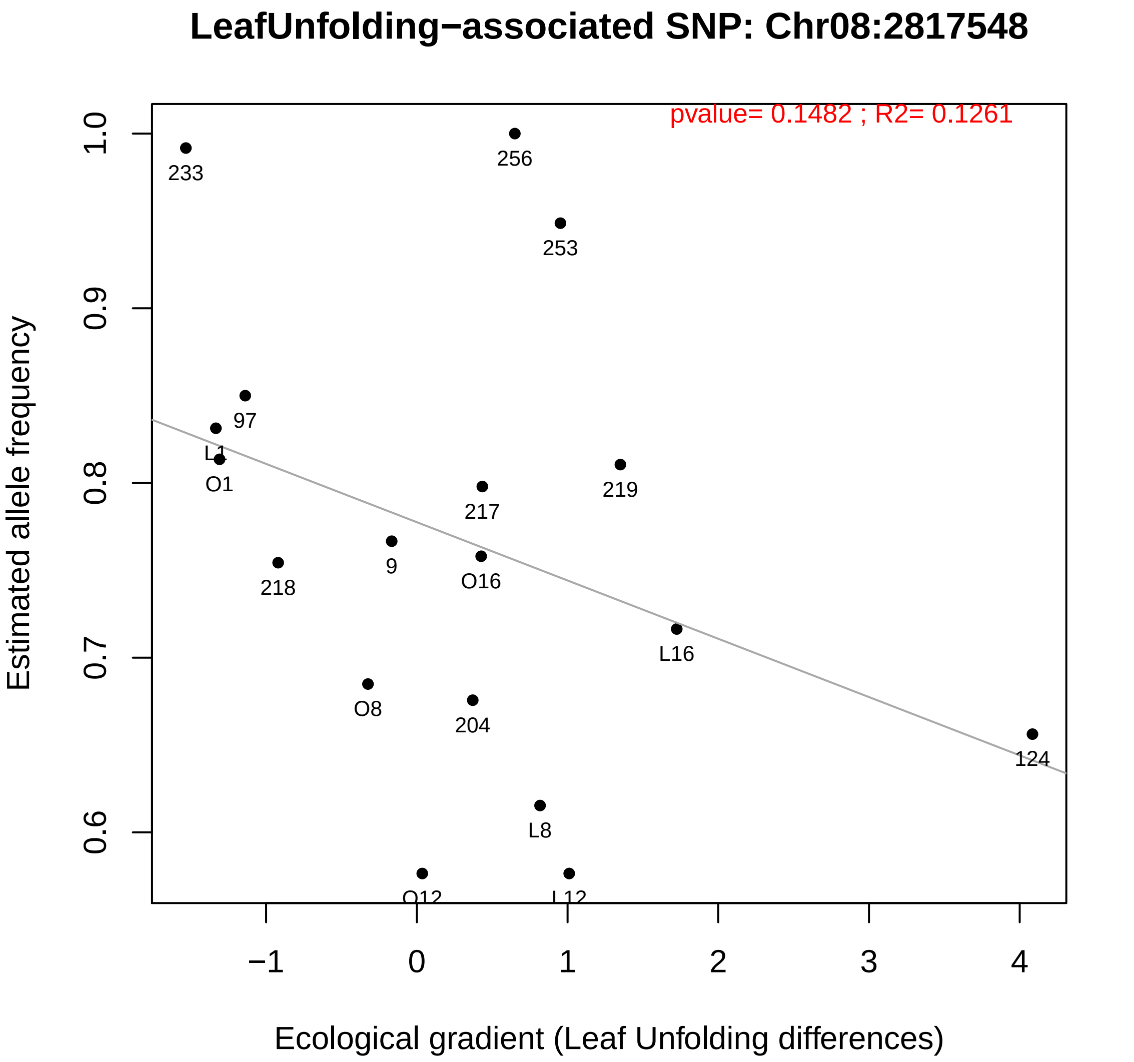

LeafUnfolding−associated SNP: Chr08:2817548
pvalue= 0.1482 ; R2= 0.1261
Estimated allele frequency
0.6
0.7
0.8
0.9
1.0
Ecological gradient (Leaf Unfolding differences)
−1
0
1
2
3
4
233
256
253
97
L1
O1
219
217
9
O16
218
L16
O8
204
124
L8
O12
L12

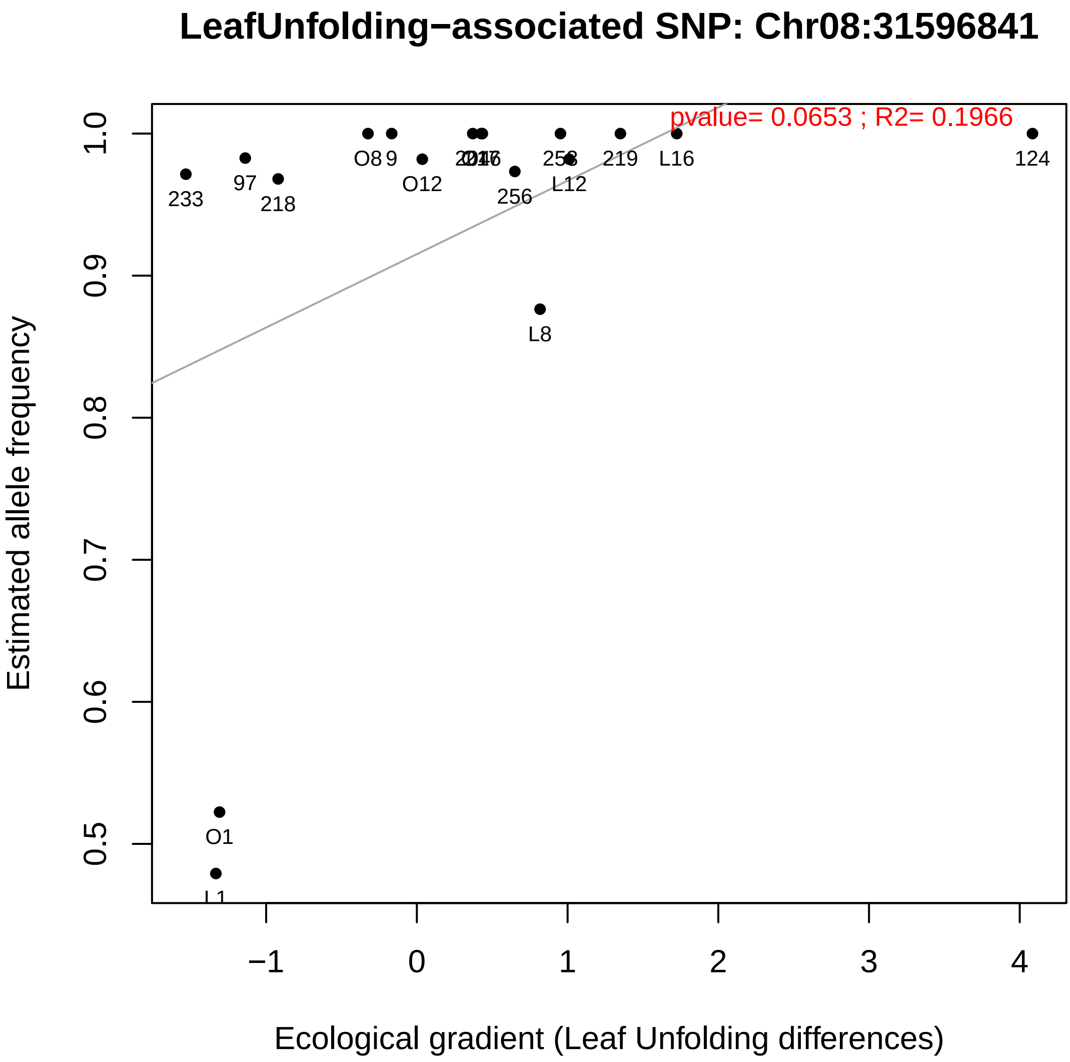

LeafUnfolding−associated SNP: Chr08:31596841
pvalue= 0.0653 ; R2= 0.1966
Estimated allele frequency
Ecological gradient (Leaf Unfolding differences)
0.5
0.6
0.7
0.8
0.9
1.0
−1
0
1
2
3
4
233
97
218
O8
9
O12
256
L8
219
L16
124
L12
O1

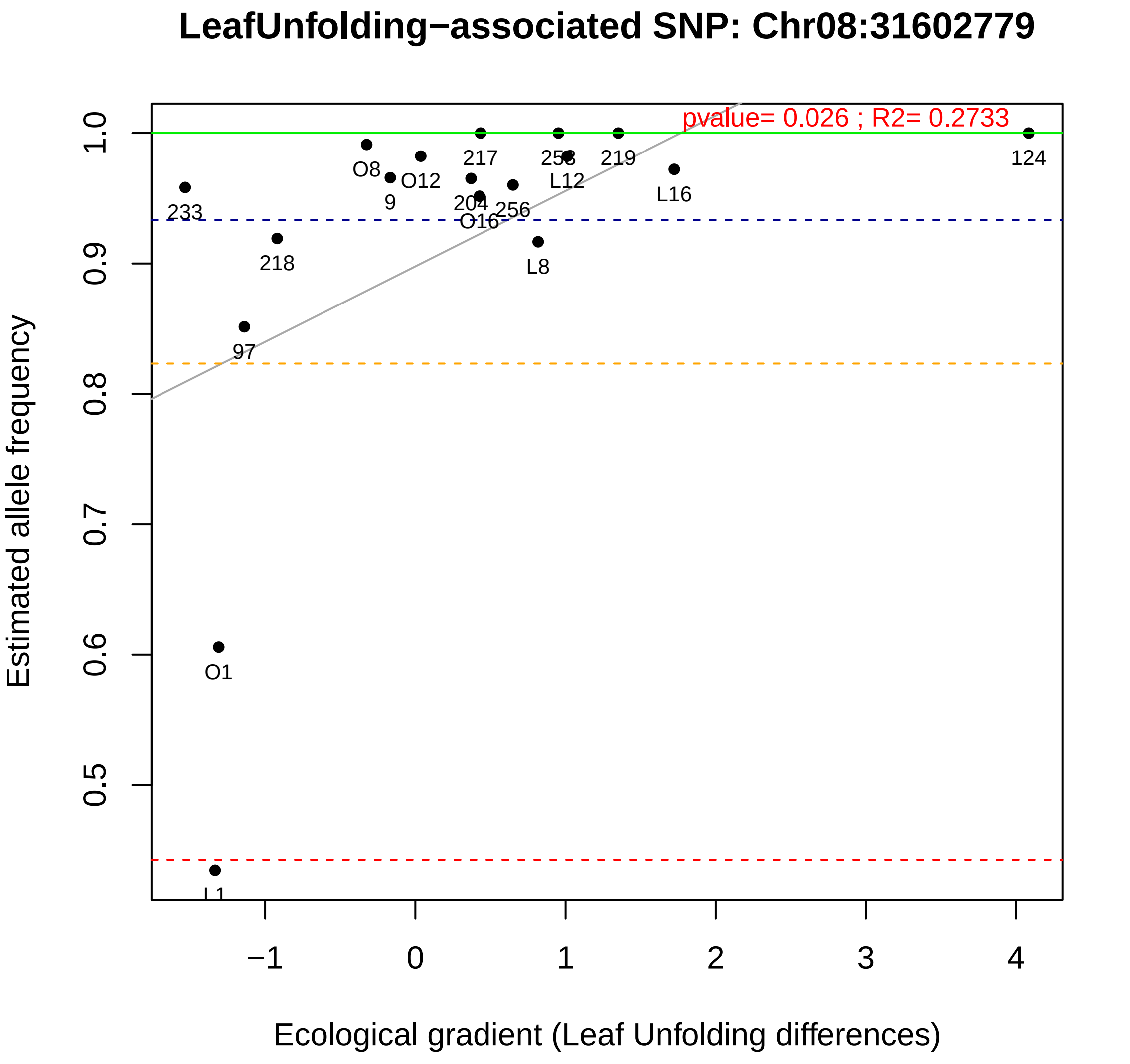
LeafUnfolding−associated SNP: Chr08:31602779
pvalue= 0.026 ; R2= 0.2733
Estimated allele frequency
Ecological gradient (Leaf Unfolding differences)
1.0
0.9
0.8
0.7
0.6
0.5
−1
0
1
2
3
4
233
218
97
O1
L1
O8
9
O12
217
204
O16
256
L8
L12
219
L16
124

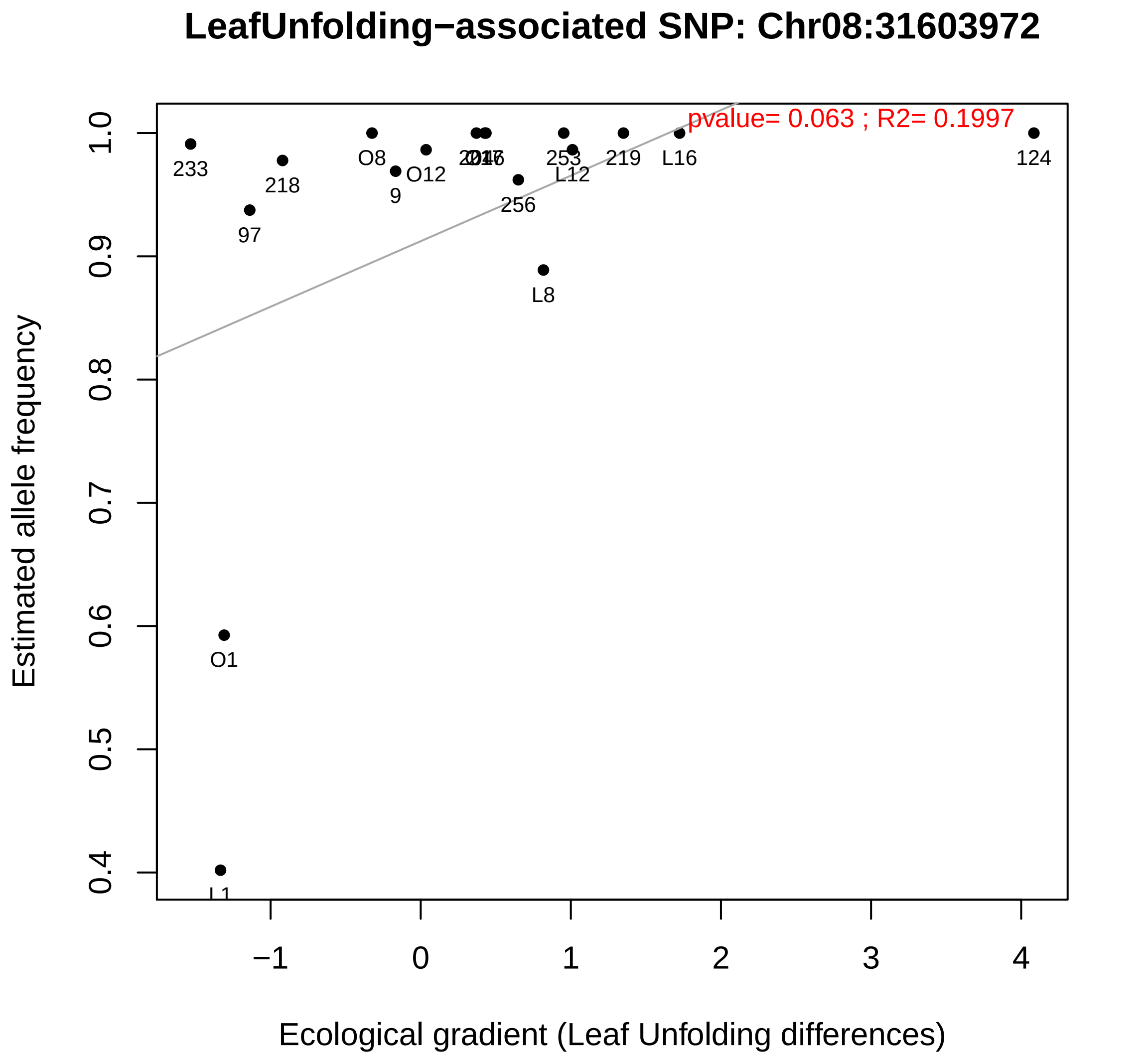

LeafUnfolding−associated SNP: Chr08:31603972
pvalue= 0.063 ; R2= 0.1997
Estimated allele frequency
Ecological gradient (Leaf Unfolding differences)
1.0
0.9
0.8
0.7
0.6
0.5
0.4
−1
0
1
2
3
4
233
218
97
O8
9
O12
256
253
L12
219
L16
124
L8
O1

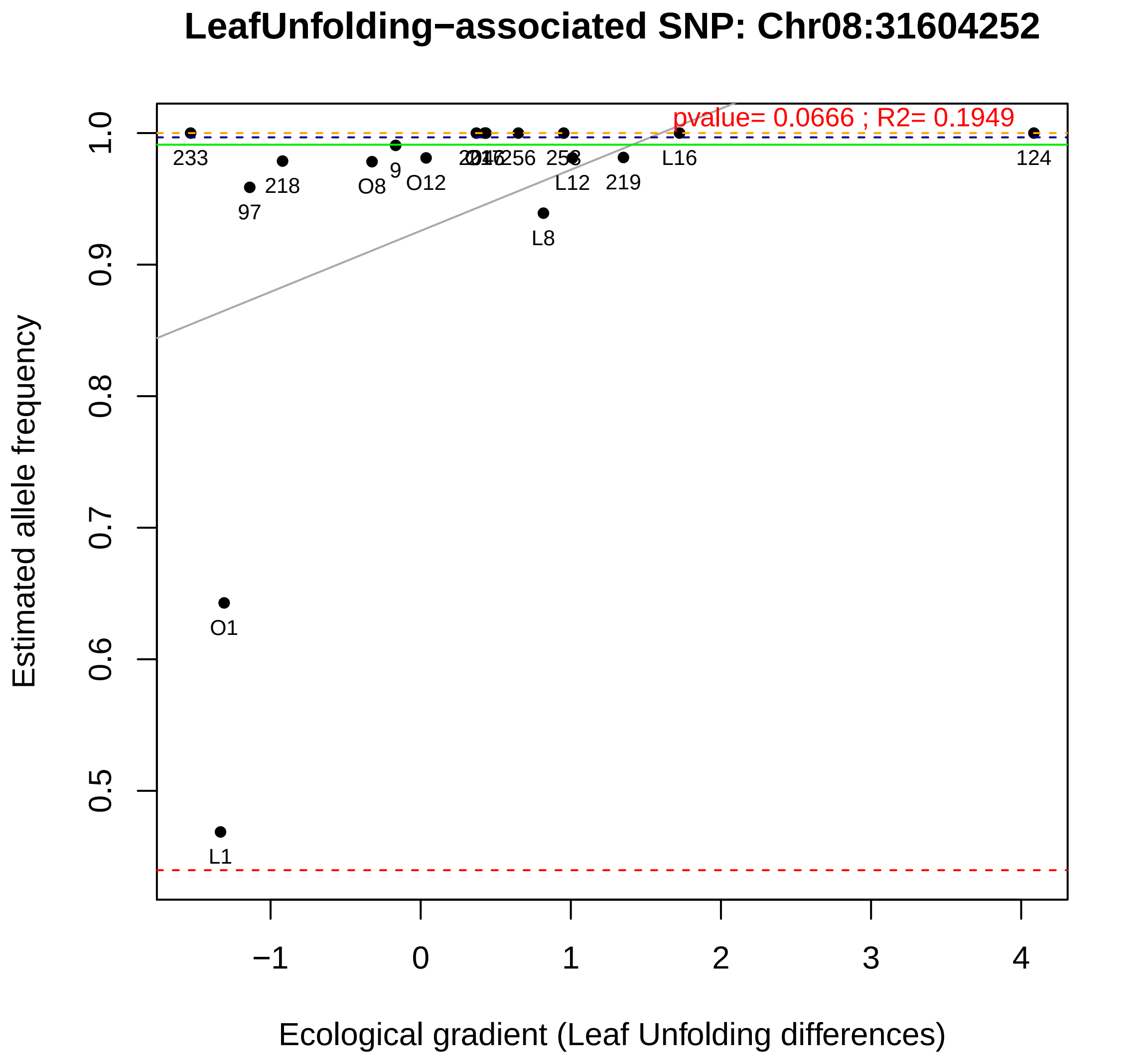

LeafUnfolding−associated SNP: Chr08:31604252
pvalue= 0.0666 ; R2= 0.1949
Estimated allele frequency
Ecological gradient (Leaf Unfolding differences)
1.0
0.9
0.8
0.7
0.6
0.5
−1
0
1
2
3
4
233
218
97
O8
9
O12
256
L12
219
L16
124
L8
O1
L1

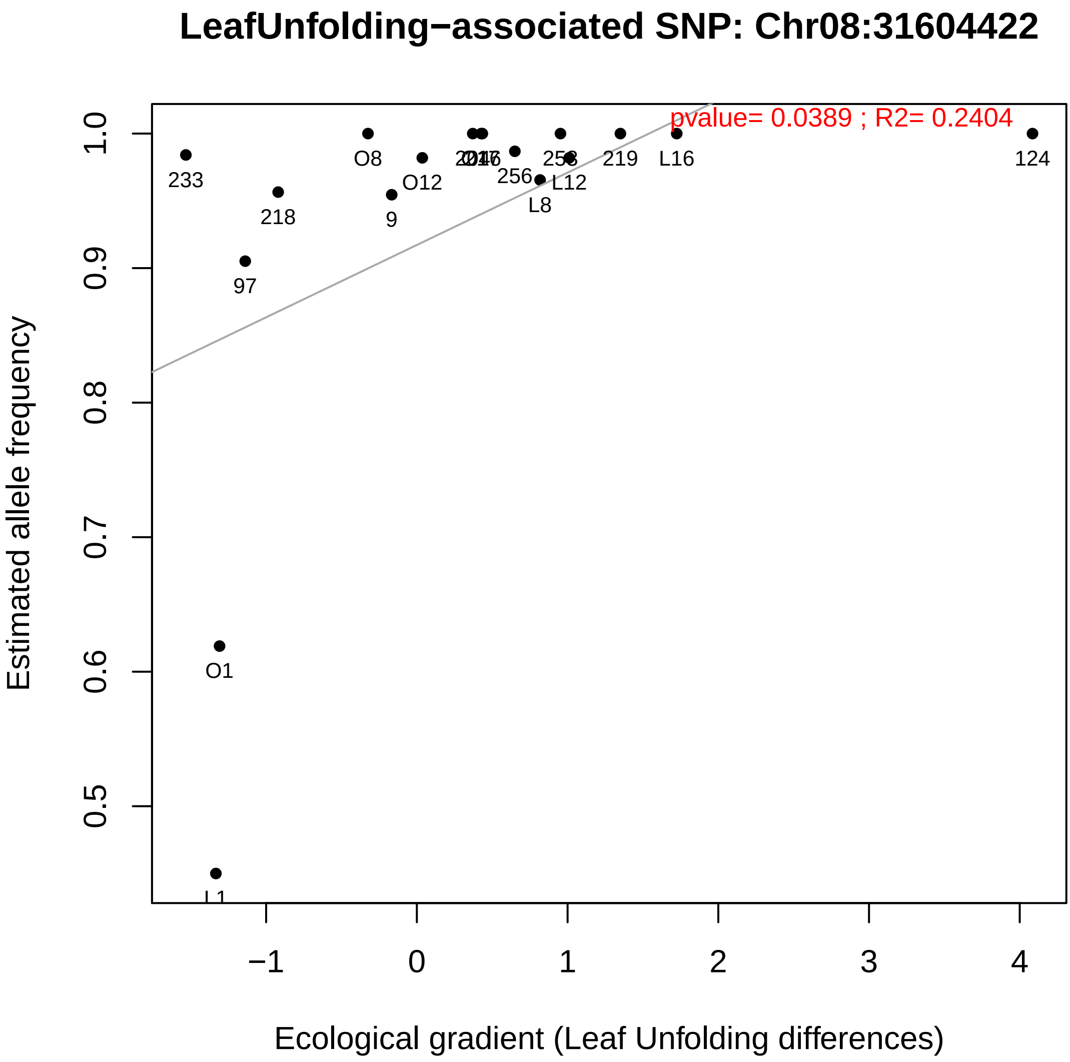

LeafUnfolding−associated SNP: Chr08:31604422
pvalue= 0.0389 ; R2= 0.2404
Estimated allele frequency
Ecological gradient (Leaf Unfolding differences)
1.0
0.9
0.8
0.7
0.6
0.5
−1
0
1
2
3
4
233
218
97
O8
O12
9
256
L8
L12
219
L16
124
O1

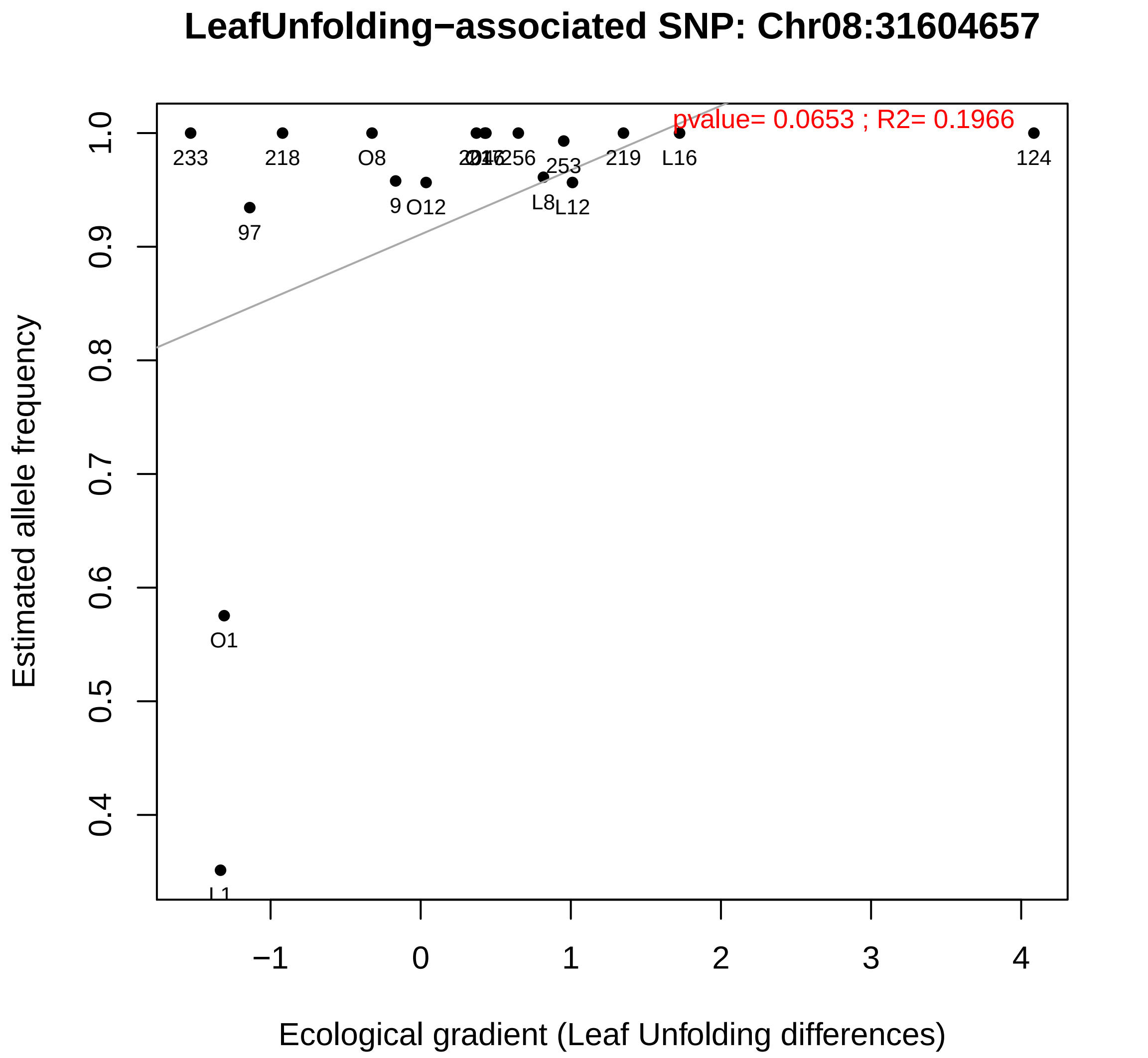

LeafUnfolding−associated SNP: Chr08:31604657
pvalue= 0.0653 ; R2= 0.1966
Estimated allele frequency
Ecological gradient (Leaf Unfolding differences)
233
218
O8
256
253
219
L16
124
97
9
O12
L8
L12
O1
L1
1.0
0.9
0.8
0.7
0.6
0.5
0.4
−1
0
1
2
3
4

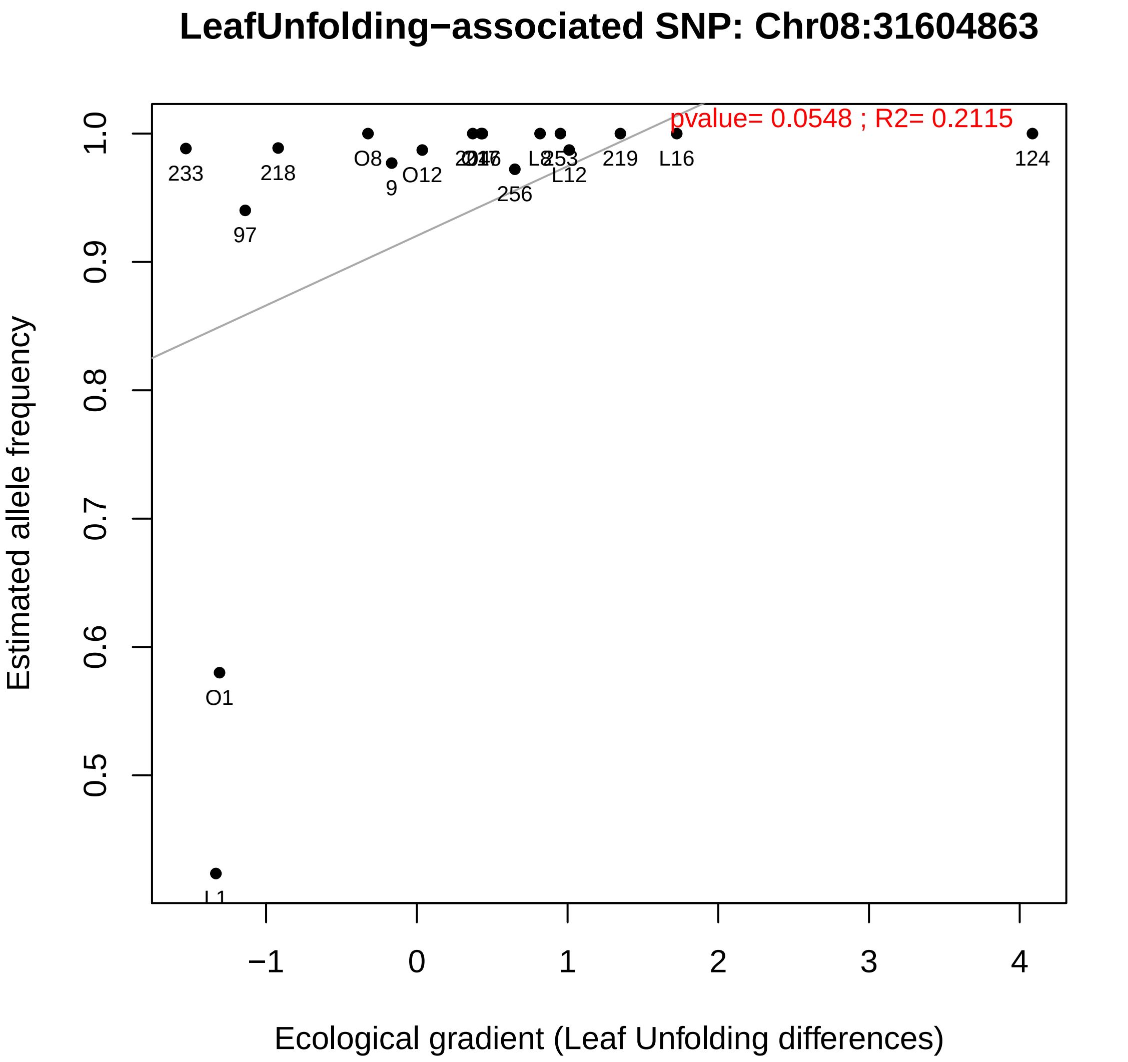

LeafUnfolding−associated SNP: Chr08:31604863
pvalue= 0.0548 ; R2= 0.2115
Estimated allele frequency
Ecological gradient (Leaf Unfolding differences)
1.0
0.9
0.8
0.7
0.6
0.5
−1
0
1
2
3
4
233
218
97
O8
9
O12
256
L12
219
L16
124
O1
L1

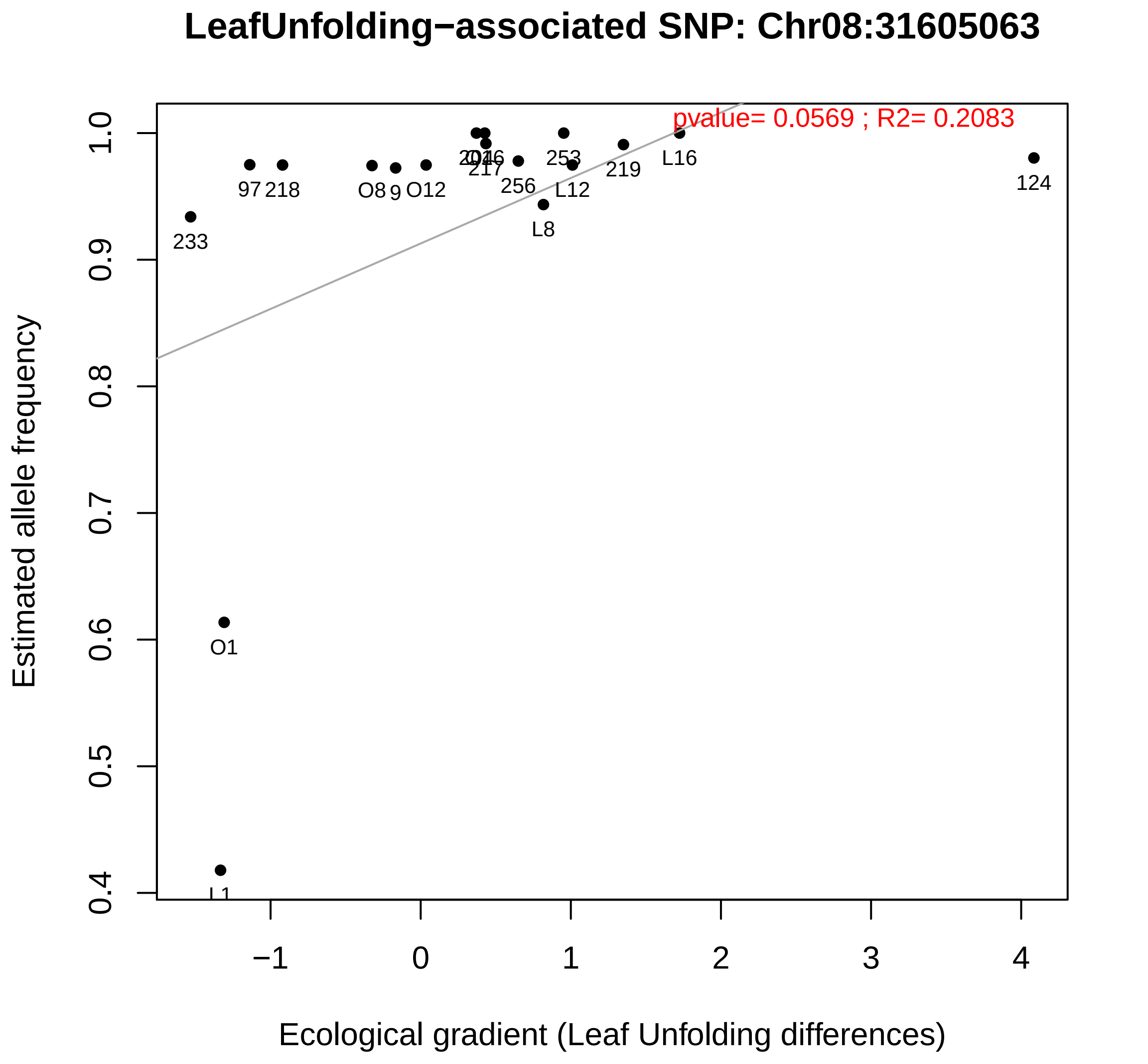

LeafUnfolding−associated SNP: Chr08:31605063
pvalue= 0.0569 ; R2= 0.2083
Estimated allele frequency
Ecological gradient (Leaf Unfolding differences)
1.0
0.9
0.8
0.7
0.6
0.5
0.4
−1
0
1
2
3
4
233
97
218
O8
9
O12
O16
217
256
253
L12
L8
219
L16
124
O1
L1

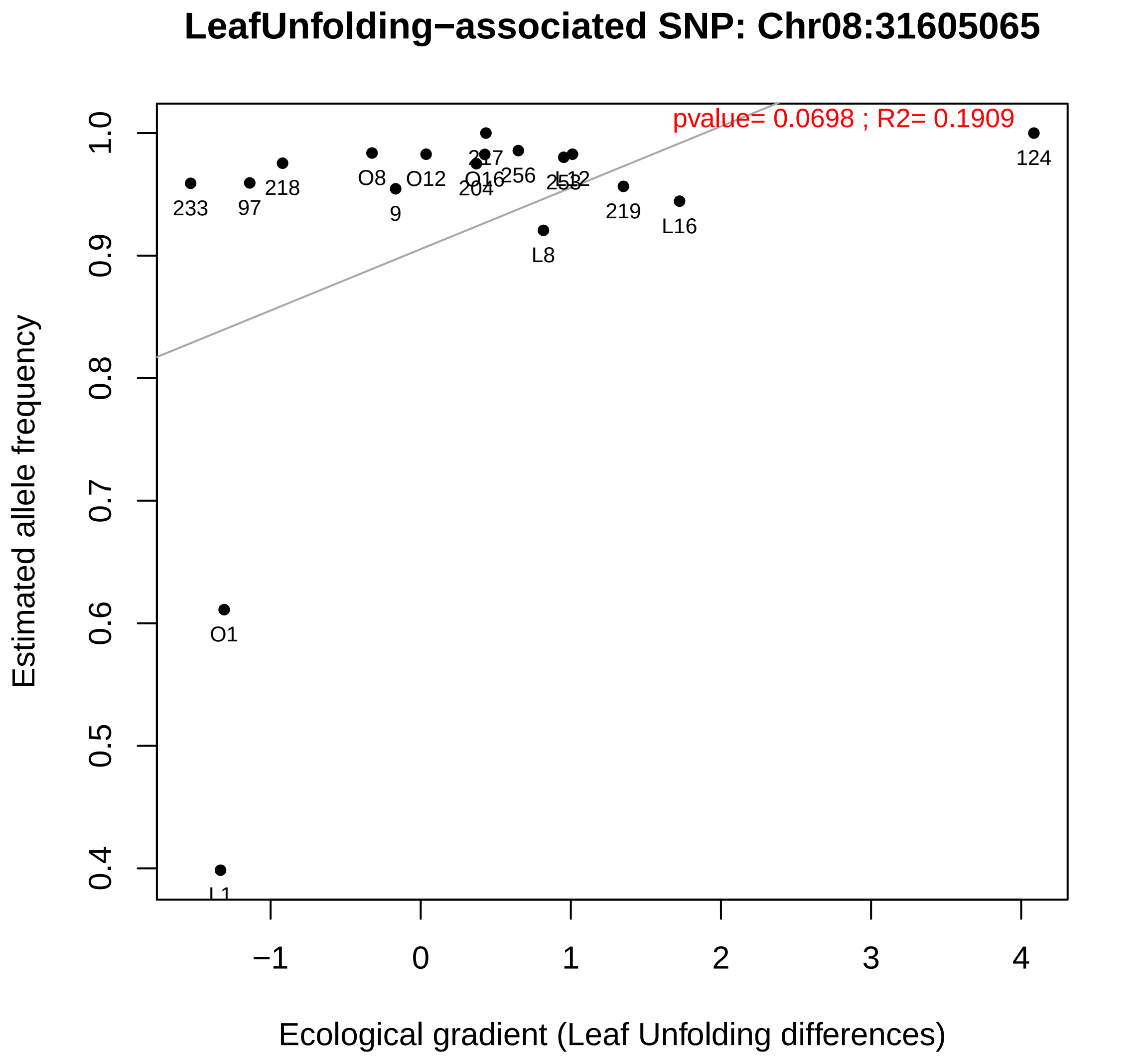

LeafUnfolding−associated SNP: Chr08:31605065
pvalue= 0.0698 ; R2= 0.1909
Estimated allele frequency
Ecological gradient (Leaf Unfolding differences)
0.4
0.5
0.6
0.7
0.8
0.9
1.0
−1
0
1
2
3
4
233
97
218
O8
9
O12
217
O16
204
256
253
L12
L8
219
L16
124
O1

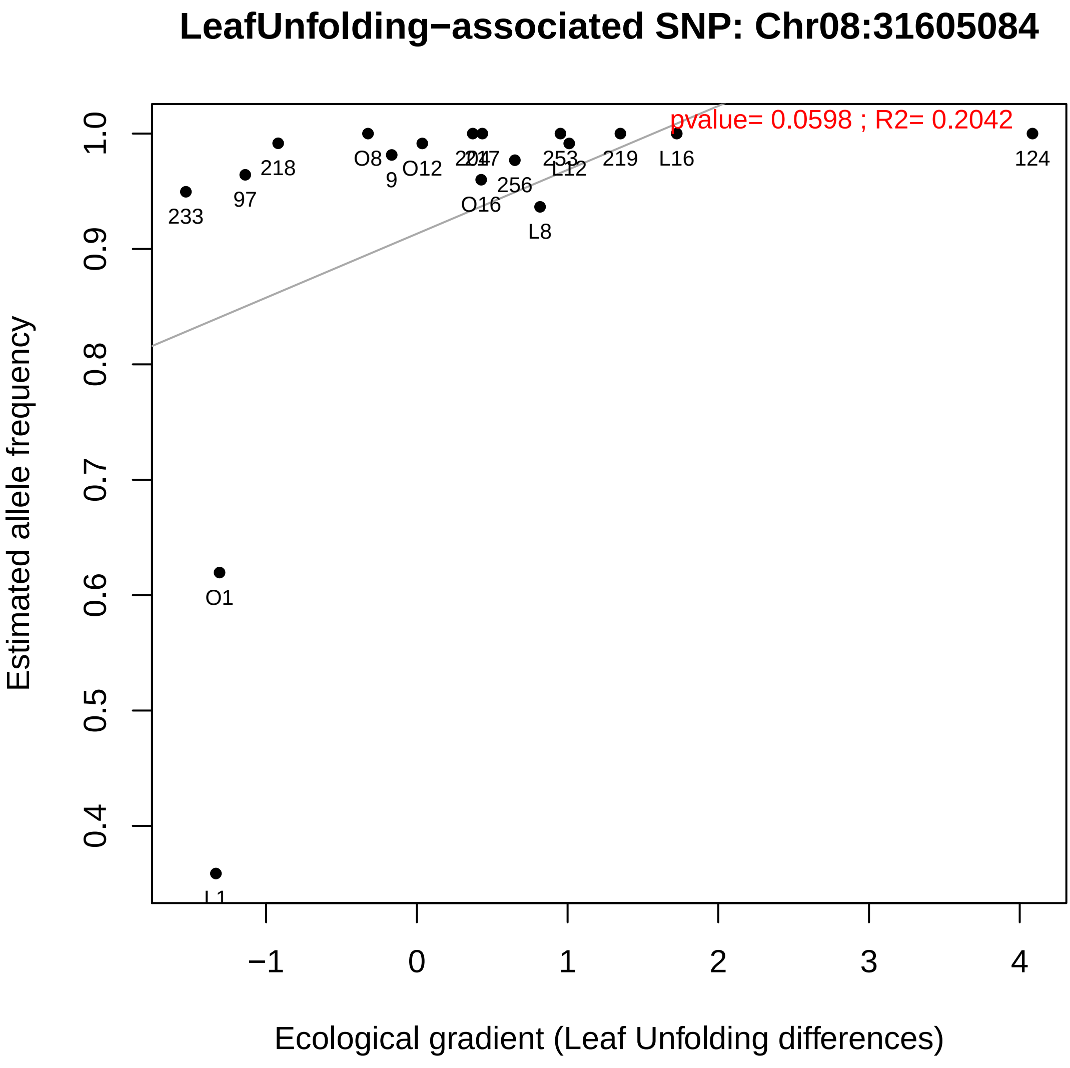

LeafUnfolding−associated SNP: Chr08:31605084
pvalue= 0.0598 ; R2= 0.2042
Estimated allele frequency
Ecological gradient (Leaf Unfolding differences)
0.4
0.5
0.6
0.7
0.8
0.9
1.0
−1
0
1
2
3
4
233
97
218
O8
9
O12
O16
256
L8
253
L12
219
L16
124
O1

**LeafUnfolding−associated SNP: Chr08:31605134**

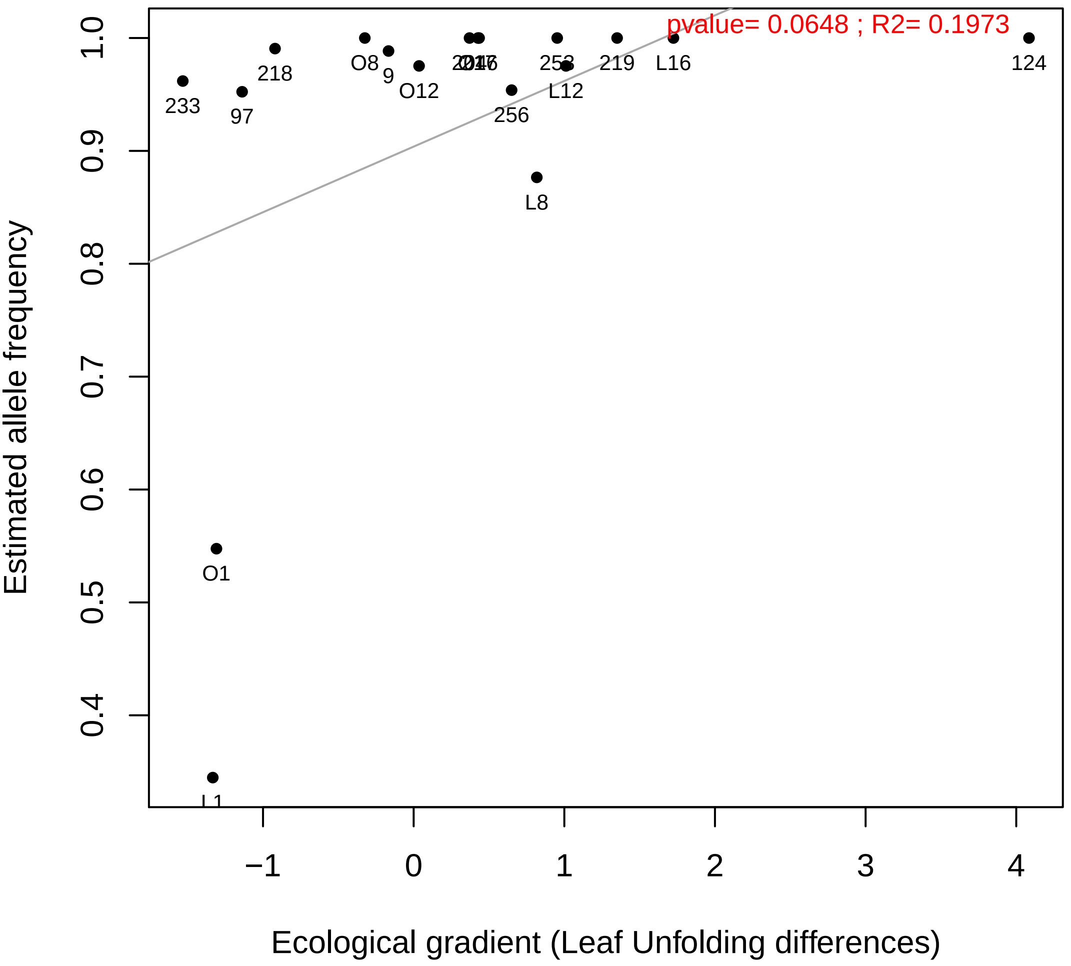

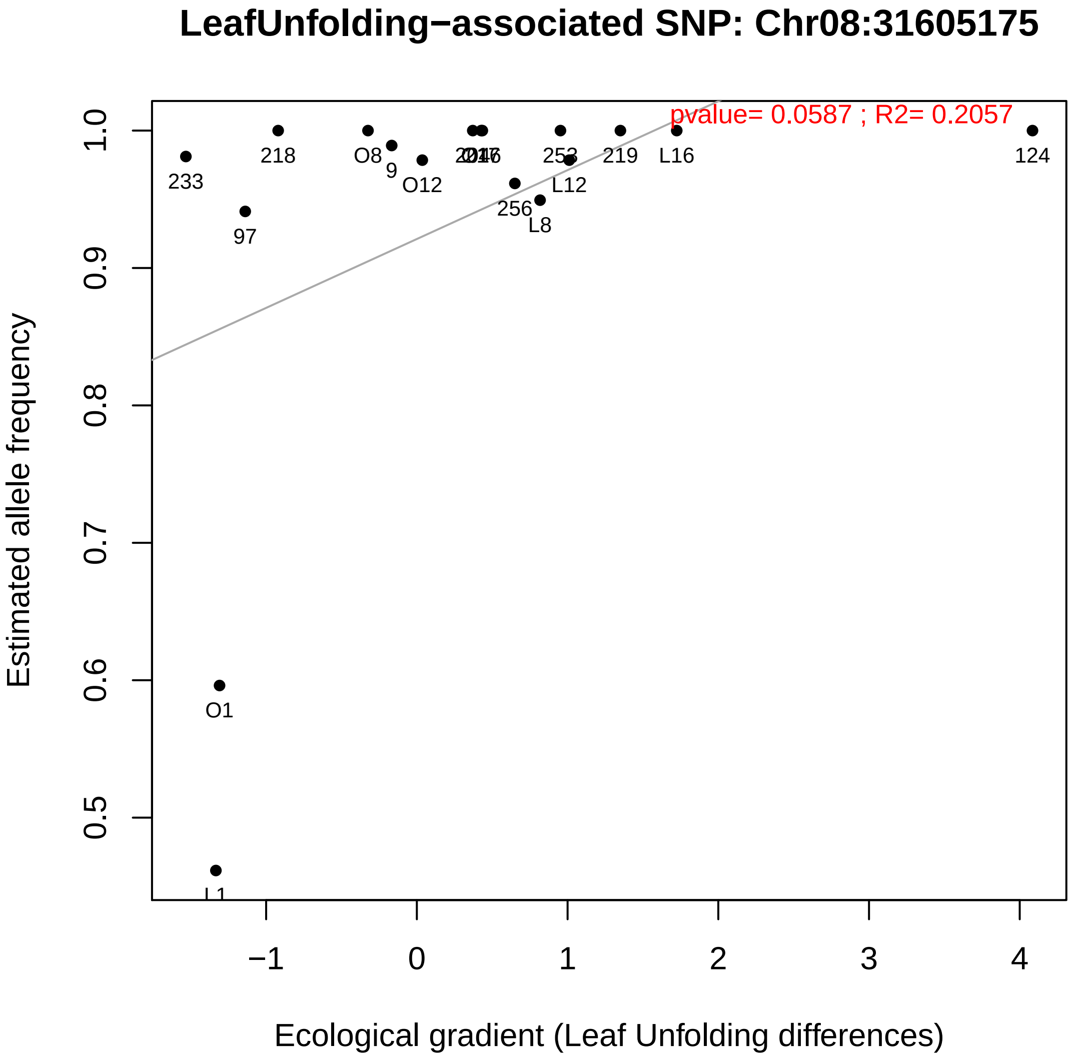

LeafUnfolding−associated SNP: Chr08:31605175
pvalue= 0.0587 ; R2= 0.2057
Estimated allele frequency
Ecological gradient (Leaf Unfolding differences)
0.5
0.6
0.7
0.8
0.9
1.0
−1
0
1
2
3
4
233
218
O8
9
O12
256
L8
253
L12
219
L16
124
97
O1

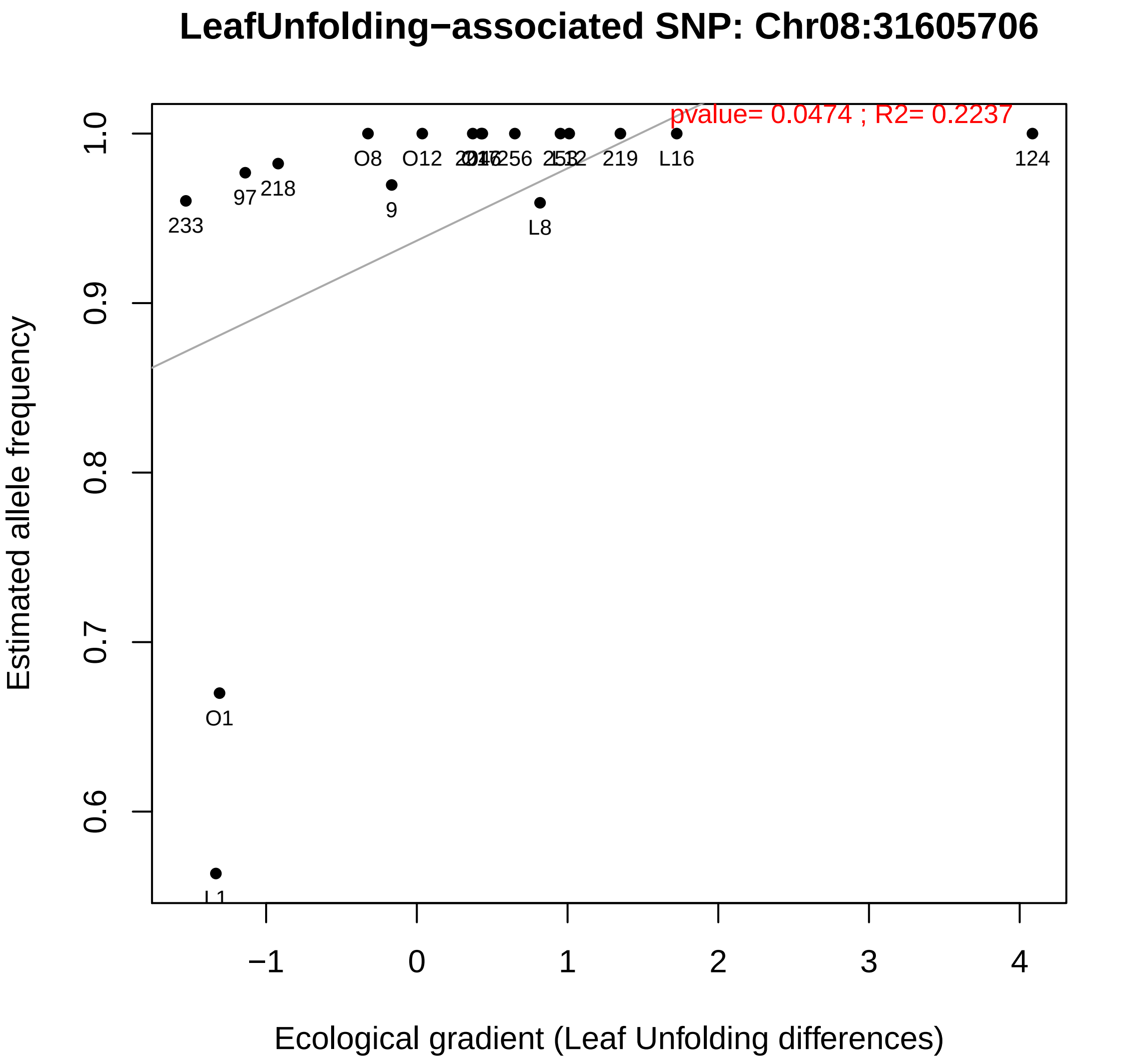
LeafUnfolding−associated SNP: Chr08:31605706
pvalue= 0.0474 ; R2= 0.2237
Estimated allele frequency
1.0
0.9
0.8
0.7
0.6
−1
0
1
2
3
4
Ecological gradient (Leaf Unfolding differences)
O8
O12
256
219
L16
124
233
97
218
9
L8
O1

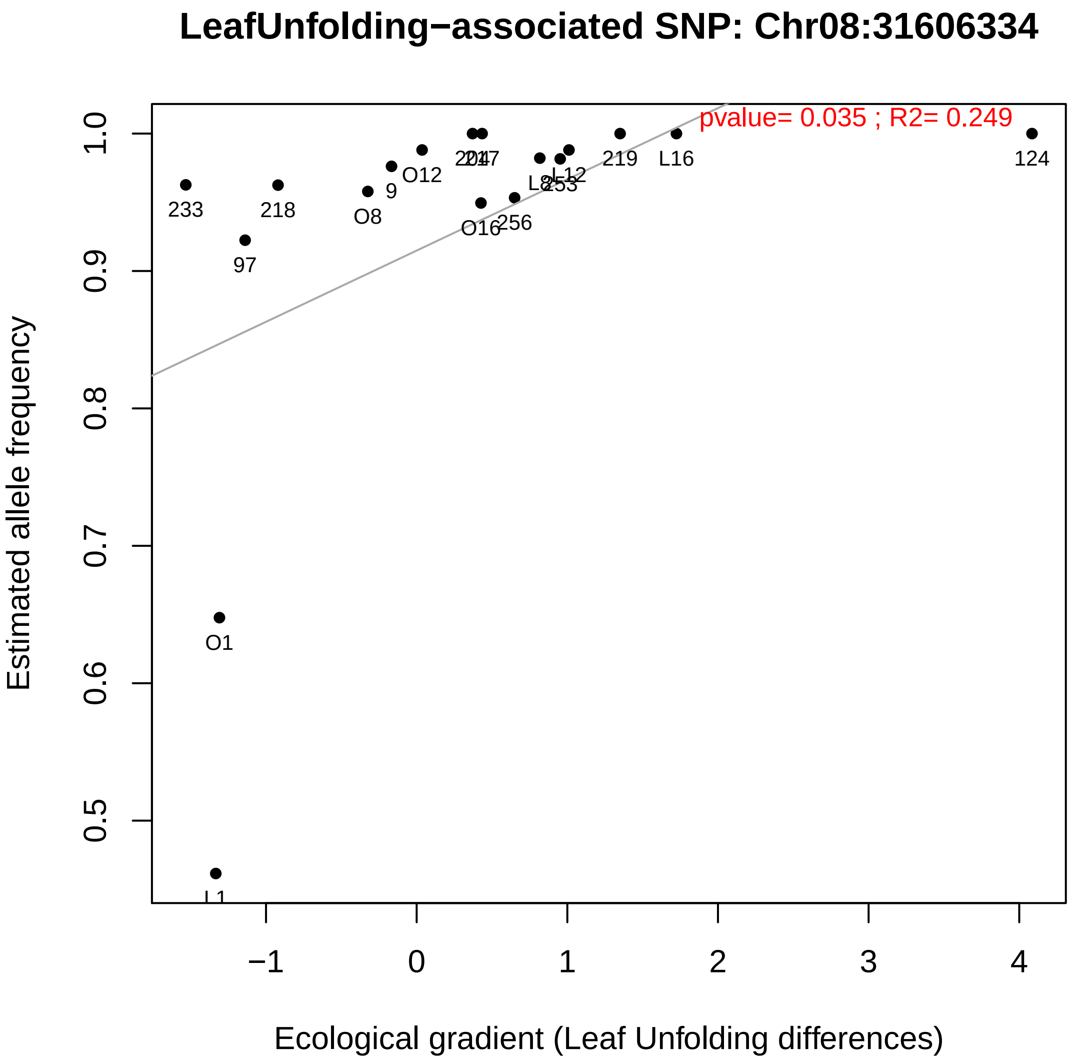

LeafUnfolding−associated SNP: Chr08:31606334
pvalue= 0.035 ; R2= 0.249
Estimated allele frequency
Ecological gradient (Leaf Unfolding differences)
1.0
0.9
0.8
0.7
0.6
0.5
−1
0
1
2
3
4
233
218
97
O8
9
O12
204
217
O16
256
L8
253
L12
219
L16
124
O1
L1

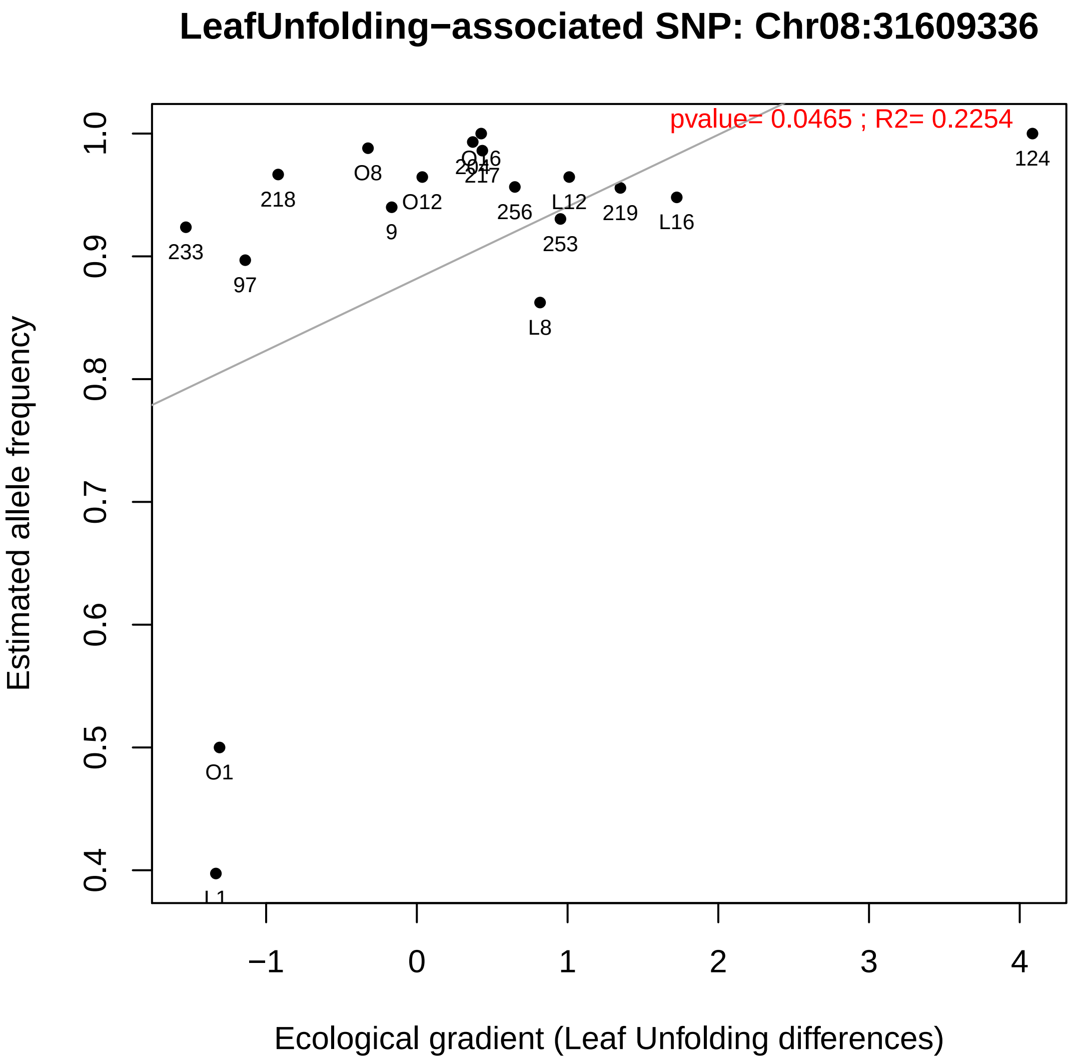
LeafUnfolding−associated SNP: Chr08:31609336
pvalue= 0.0465 ; R2= 0.2254
Estimated allele frequency
Ecological gradient (Leaf Unfolding differences)
1.0
0.9
0.8
0.7
0.6
0.5
0.4
−1
0
1
2
3
4
124
O8
218
233
97
O12
9
256
L12
253
219
L16
L8
O1
O16
217

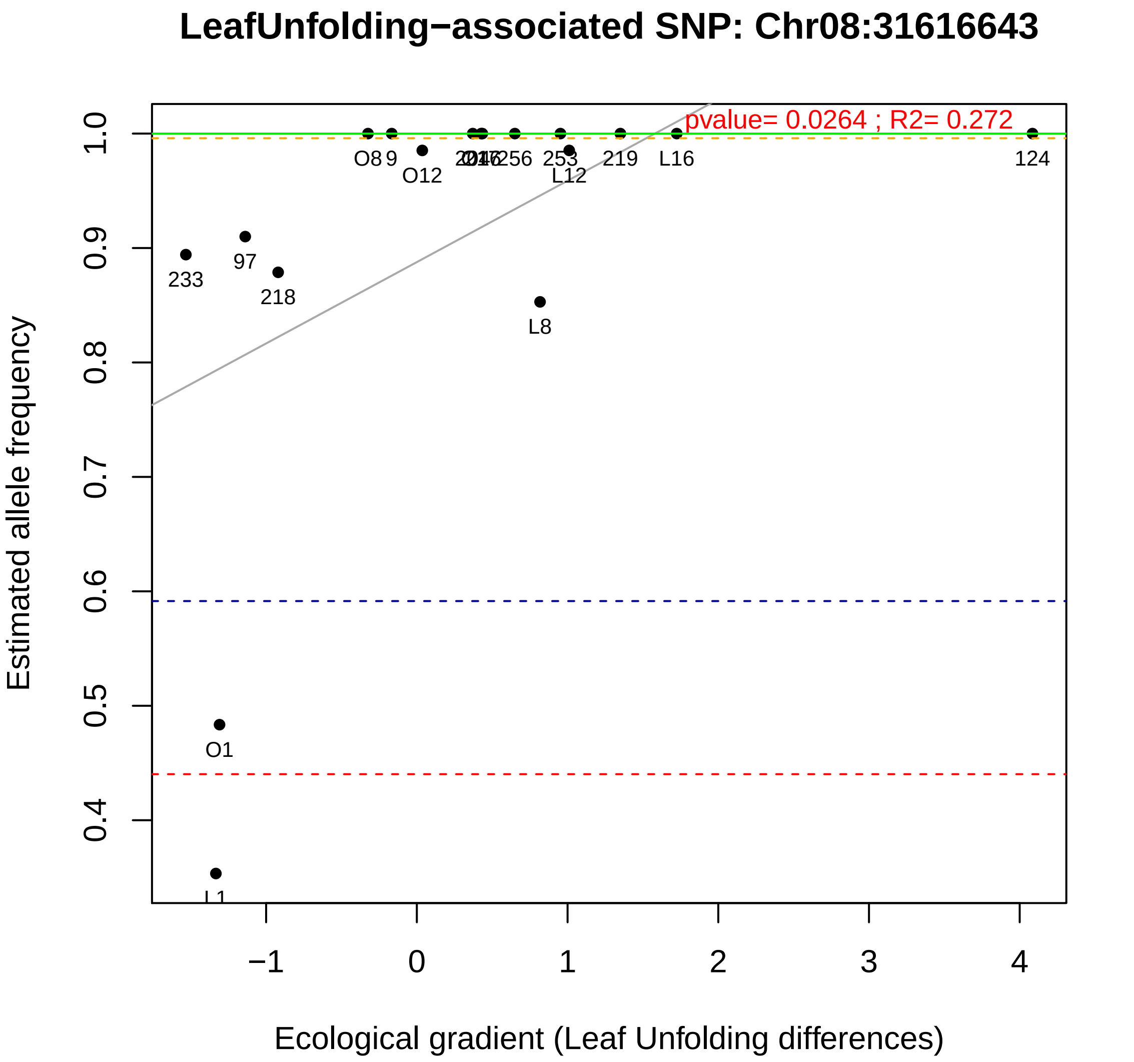
LeafUnfolding−associated SNP: Chr08:31616643
pvalue= 0.0264 ; R2= 0.272
Estimated allele frequency
Ecological gradient (Leaf Unfolding differences)
0.4
0.5
0.6
0.7
0.8
0.9
1.0
−1
0
1
2
3
4
O8
9
O12
256
253
L12
219
L16
124
233
97
218
L8
O1

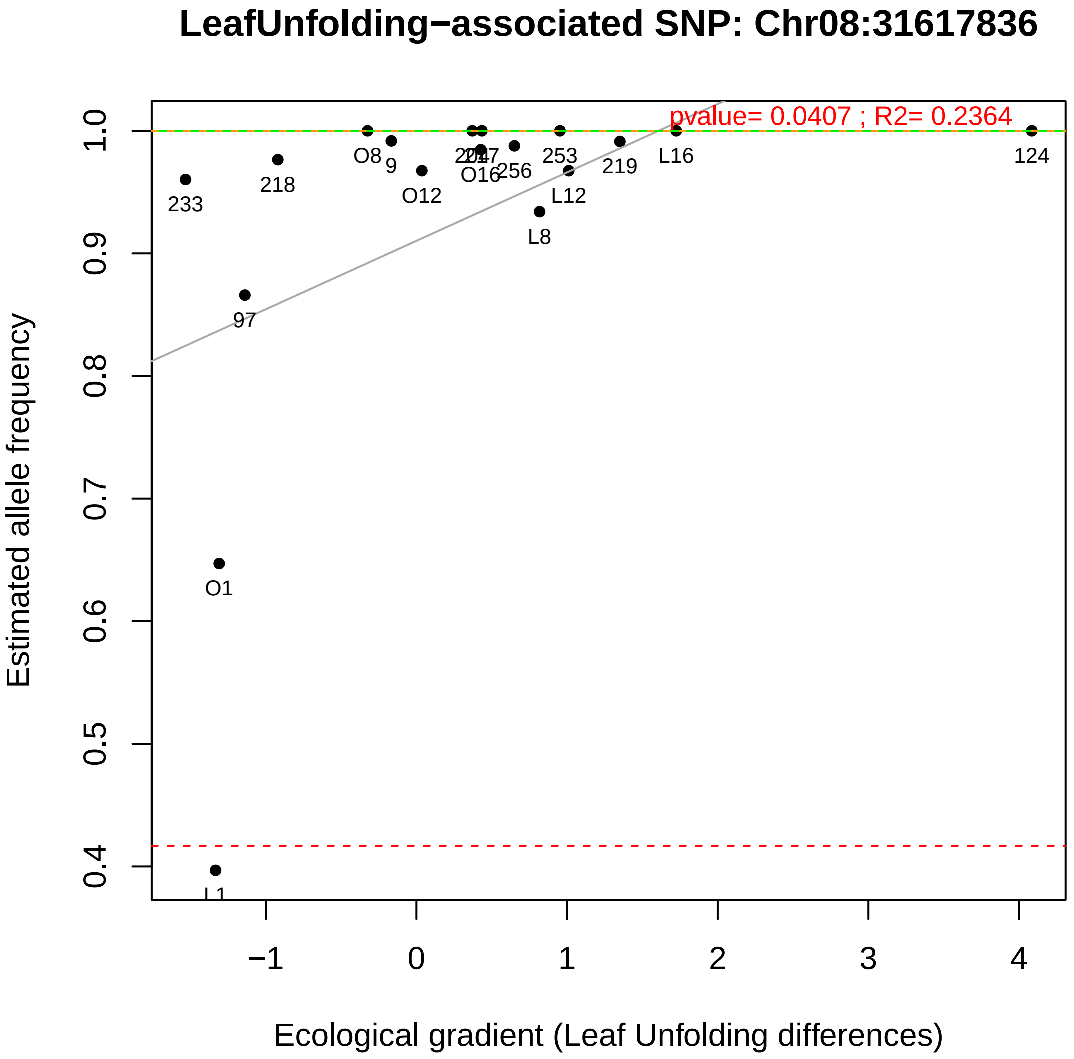
LeafUnfolding−associated SNP: Chr08:31617836
pvalue= 0.0407 ; R2= 0.2364
Estimated allele frequency
Ecological gradient (Leaf Unfolding differences)
0.4
0.5
0.6
0.7
0.8
0.9
1.0
−1
0
1
2
3
4
233
218
97
O1
O8
9
O12
O16
256
253
L12
L8
219
L16
124

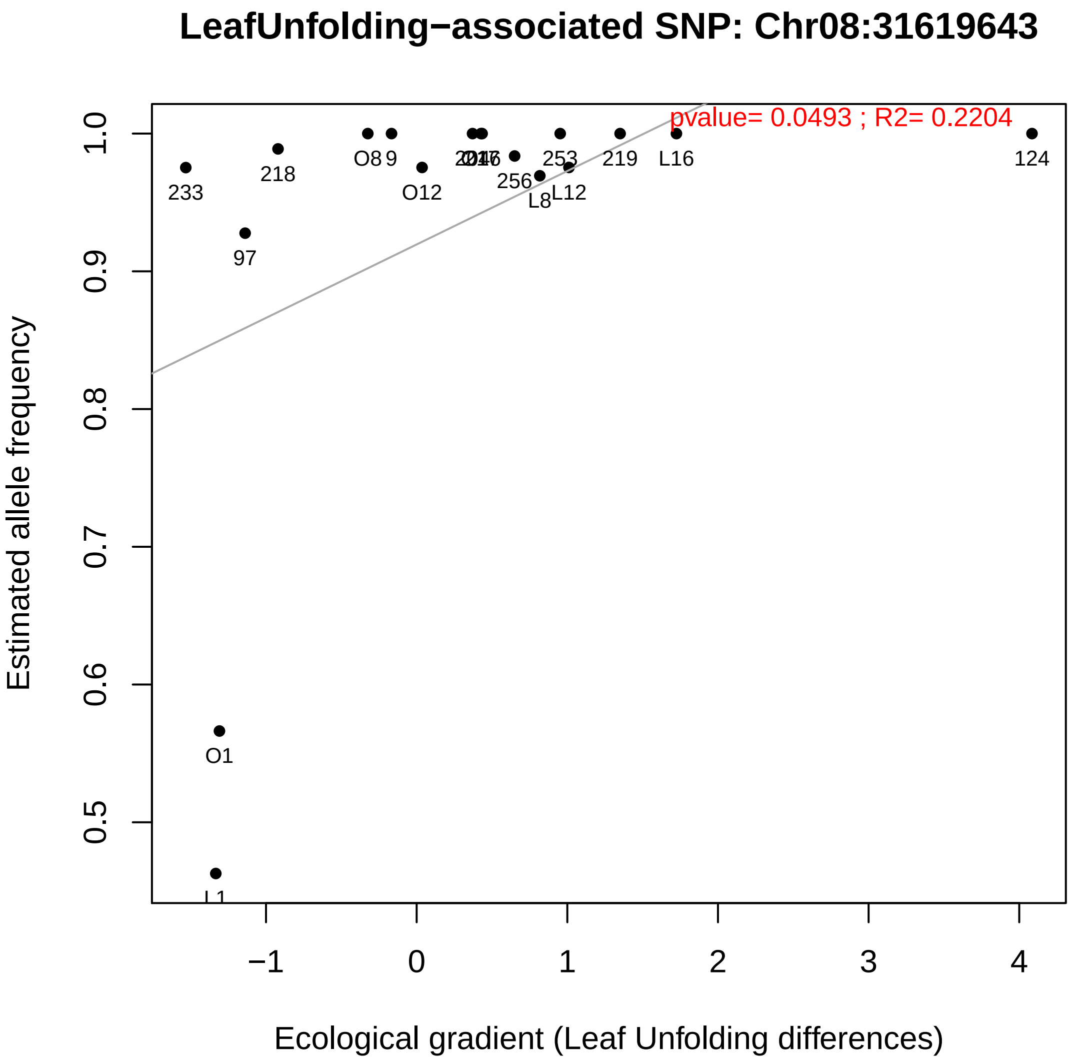
LeafUnfolding−associated SNP: Chr08:31619643
pvalue= 0.0493 ; R2= 0.2204
Estimated allele frequency
Ecological gradient (Leaf Unfolding differences)
1.0
0.9
0.8
0.7
0.6
0.5
−1
0
1
2
3
4
233
97
218
O8
9
O12
256
L8
253
L12
219
L16
124
O1
L1

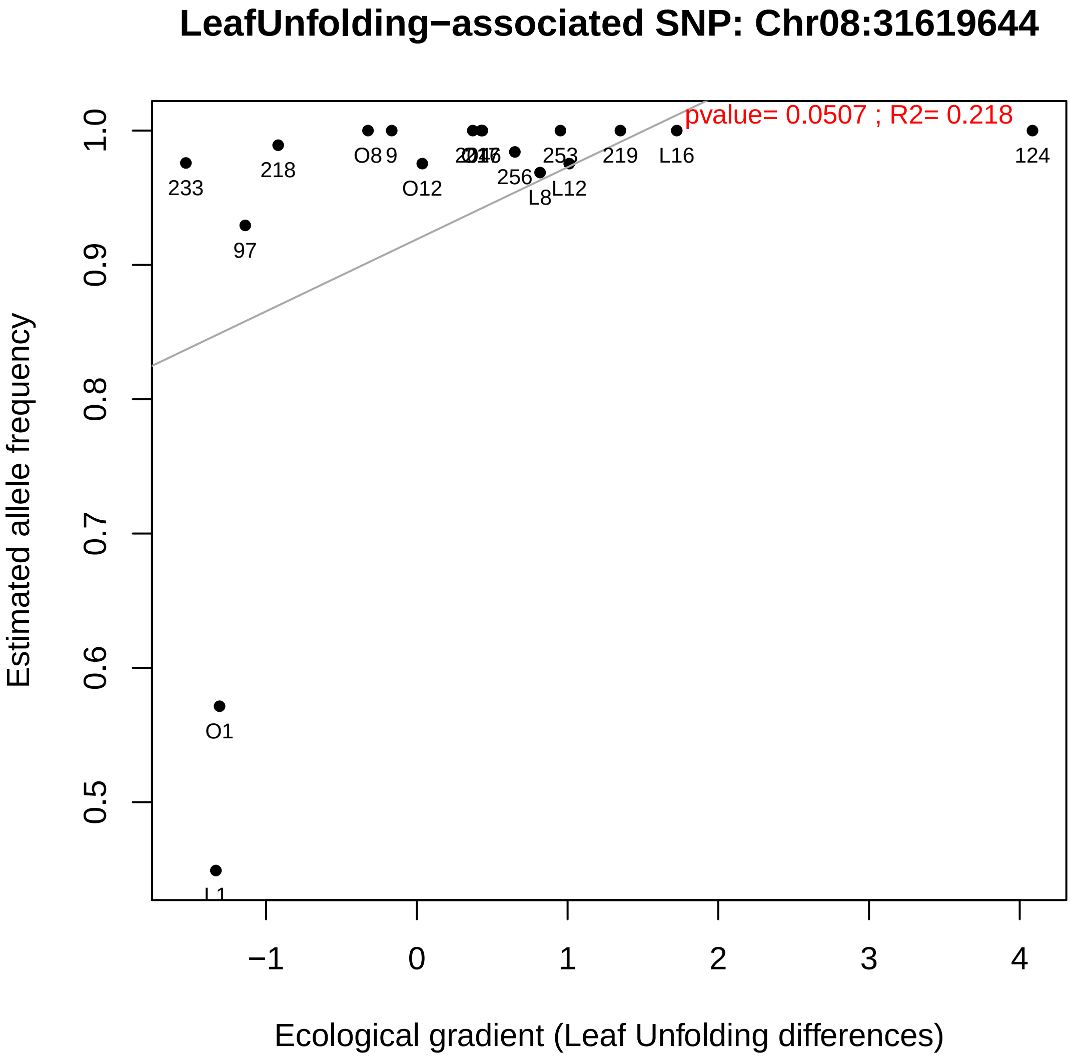

LeafUnfolding−associated SNP: Chr08:31619644
pvalue= 0.0507 ; R2= 0.218
Estimated allele frequency
Ecological gradient (Leaf Unfolding differences)
0.5
0.6
0.7
0.8
0.9
1.0
−1
0
1
2
3
4
233
97
218
O8
9
O12
256
L8
L12
253
219
L16
124
O1

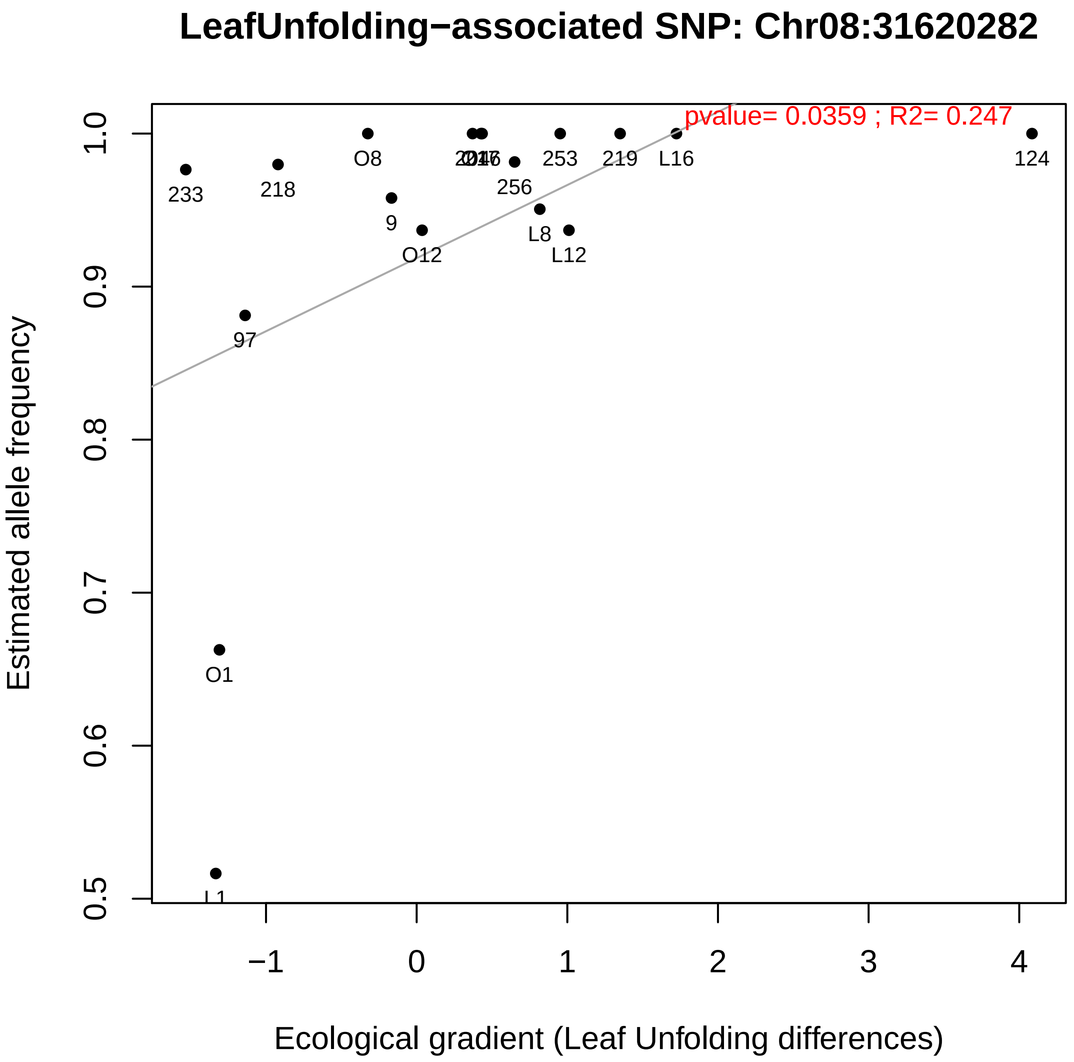

LeafUnfolding−associated SNP: Chr08:31620282
pvalue= 0.0359 ; R2= 0.247
Estimated allele frequency
Ecological gradient (Leaf Unfolding differences)
1.0
0.9
0.8
0.7
0.6
0.5
−1
0
1
2
3
4
233
218
O8
9
O12
97
O1
256
L8
L12
253
219
L16
124

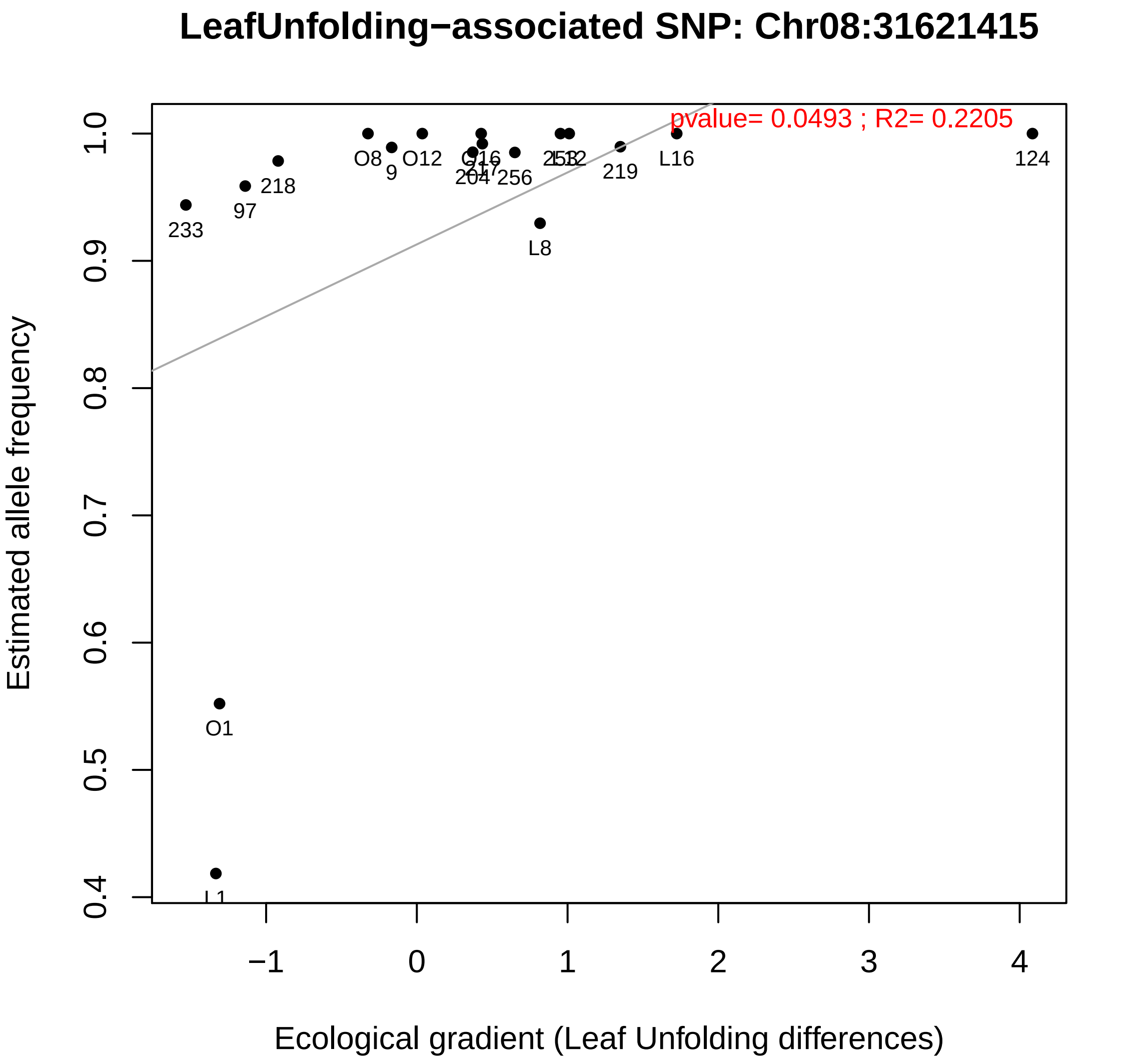

LeafUnfolding−associated SNP: Chr08:31621415
pvalue= 0.0493 ; R2= 0.2205
Estimated allele frequency
Ecological gradient (Leaf Unfolding differences)
1.0
0.9
0.8
0.7
0.6
0.5
0.4
−1
0
1
2
3
4
233
97
218
O8
9
O12
O16
217
204
256
L8
251
L12
219
L16
124
O1
L1

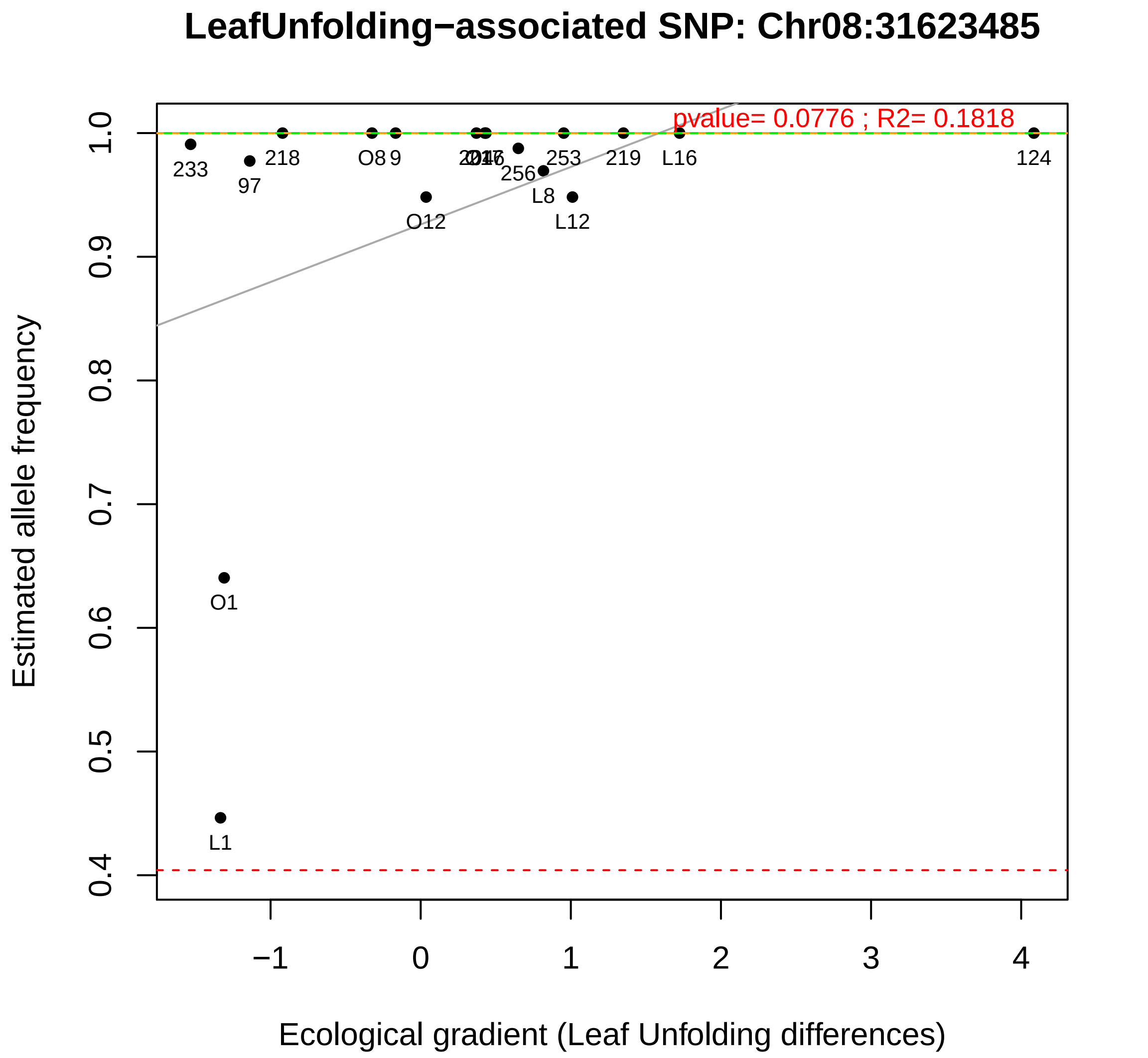
LeafUnfolding−associated SNP: Chr08:31623485
pvalue= 0.0776 ; R2= 0.1818
Estimated allele frequency
Ecological gradient (Leaf Unfolding differences)
1.0
0.9
0.8
0.7
0.6
0.5
0.4
−1
0
1
2
3
4
233
97
218
O8
9
O12
256
253
L8
L12
219
L16
124
O1
L1

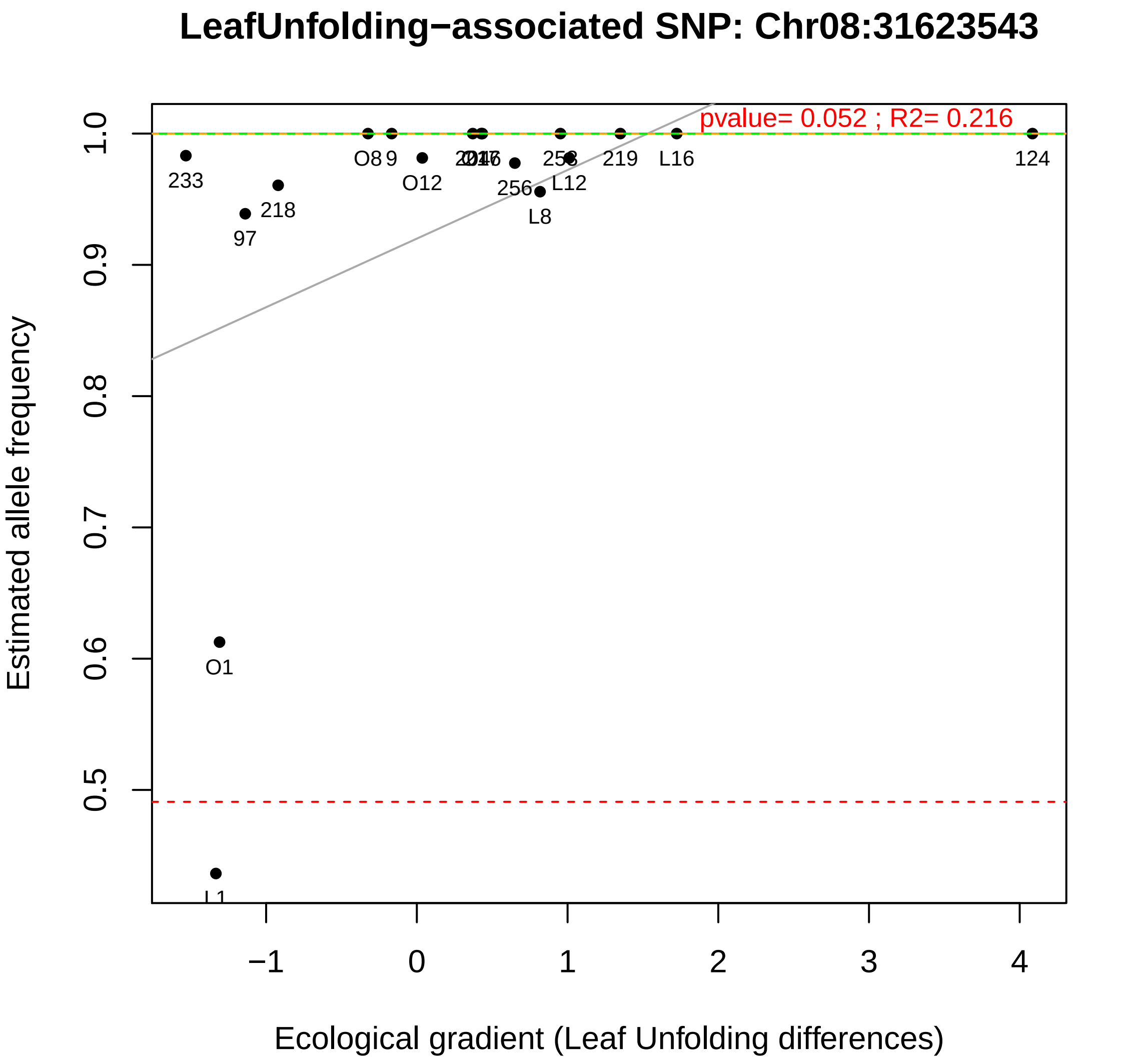

LeafUnfolding−associated SNP: Chr08:31623543
pvalue= 0.052 ; R2= 0.216
Estimated allele frequency
Ecological gradient (Leaf Unfolding differences)
1.0
0.9
0.8
0.7
0.6
0.5
−1
0
1
2
3
4
233
97
218
O8
9
O12
256
L12
L8
219
L16
124
O1

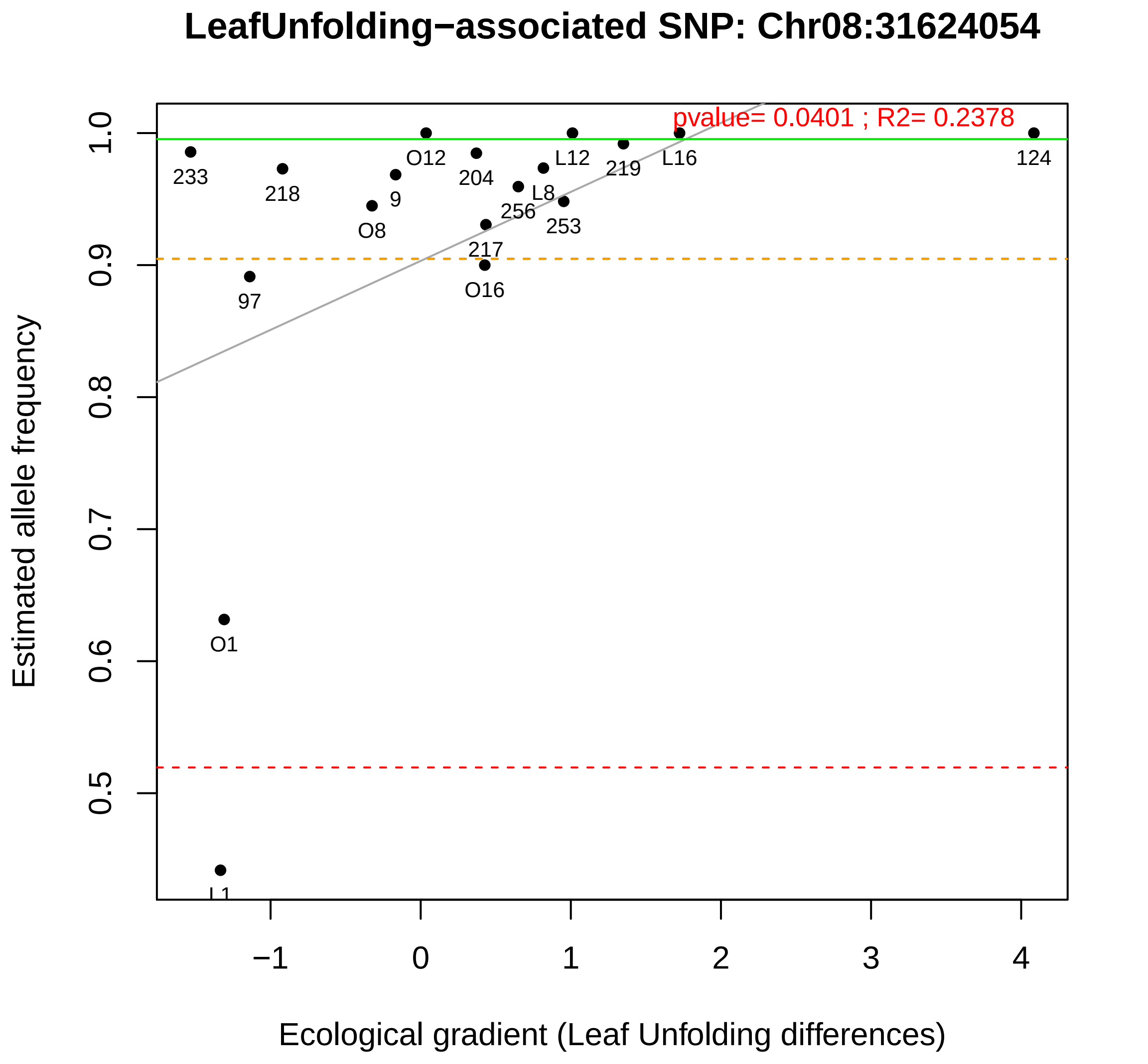
LeafUnfolding−associated SNP: Chr08:31624054
pvalue= 0.0401 ; R2= 0.2378
Estimated allele frequency
Ecological gradient (Leaf Unfolding differences)
0.5
0.6
0.7
0.8
0.9
1.0
−1
0
1
2
3
4
233
218
97
O1
O8
9
O12
204
217
O16
256
L8
253
L12
219
L16
124

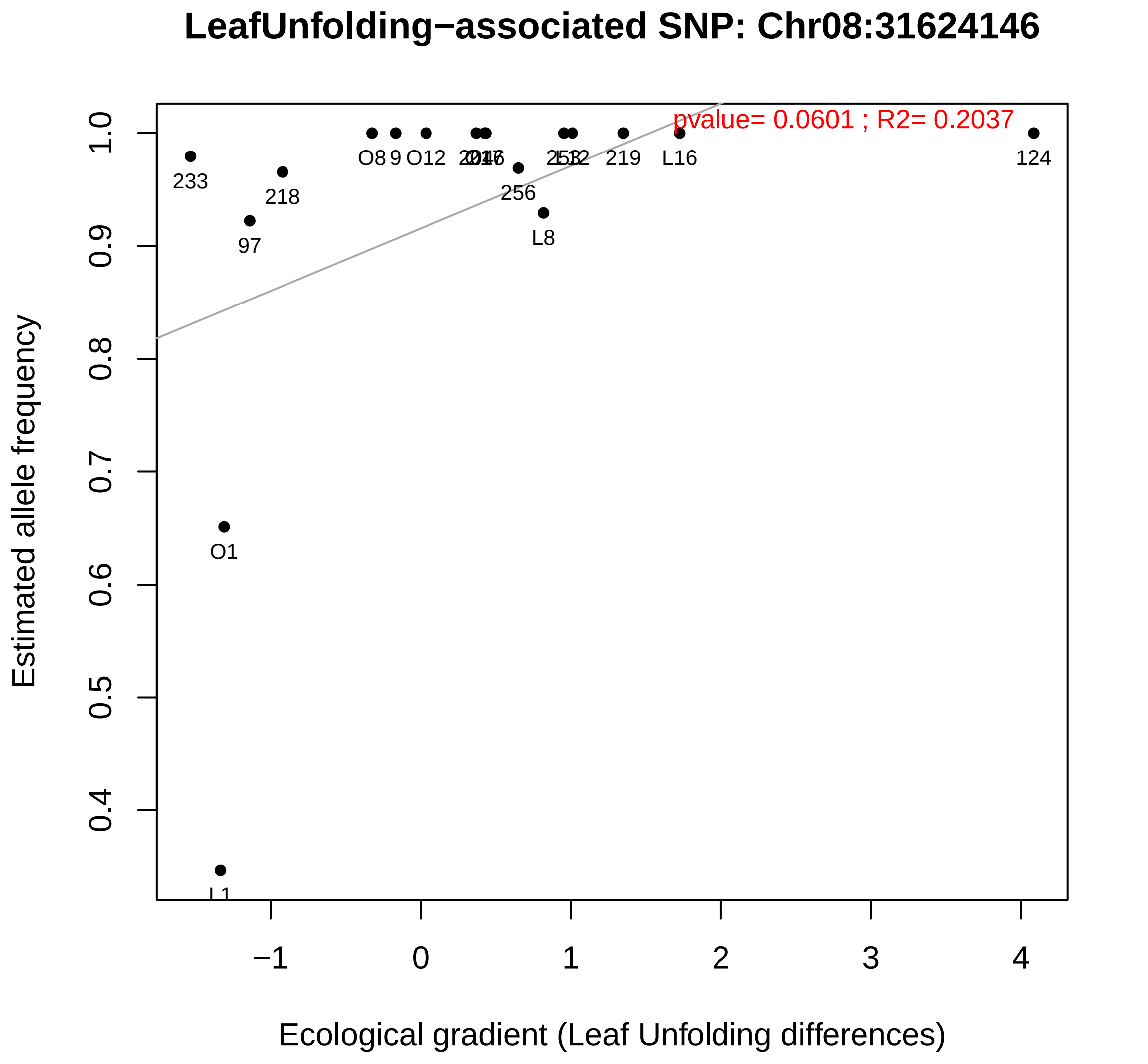
LeafUnfolding−associated SNP: Chr08:31624146
pvalue= 0.0601 ; R2= 0.2037
Estimated allele frequency
Ecological gradient (Leaf Unfolding differences)
1.0
0.9
0.8
0.7
0.6
0.5
0.4
−1
0
1
2
3
4
233
218
97
O8
9
O12
256
L8
219
L16
124
O1

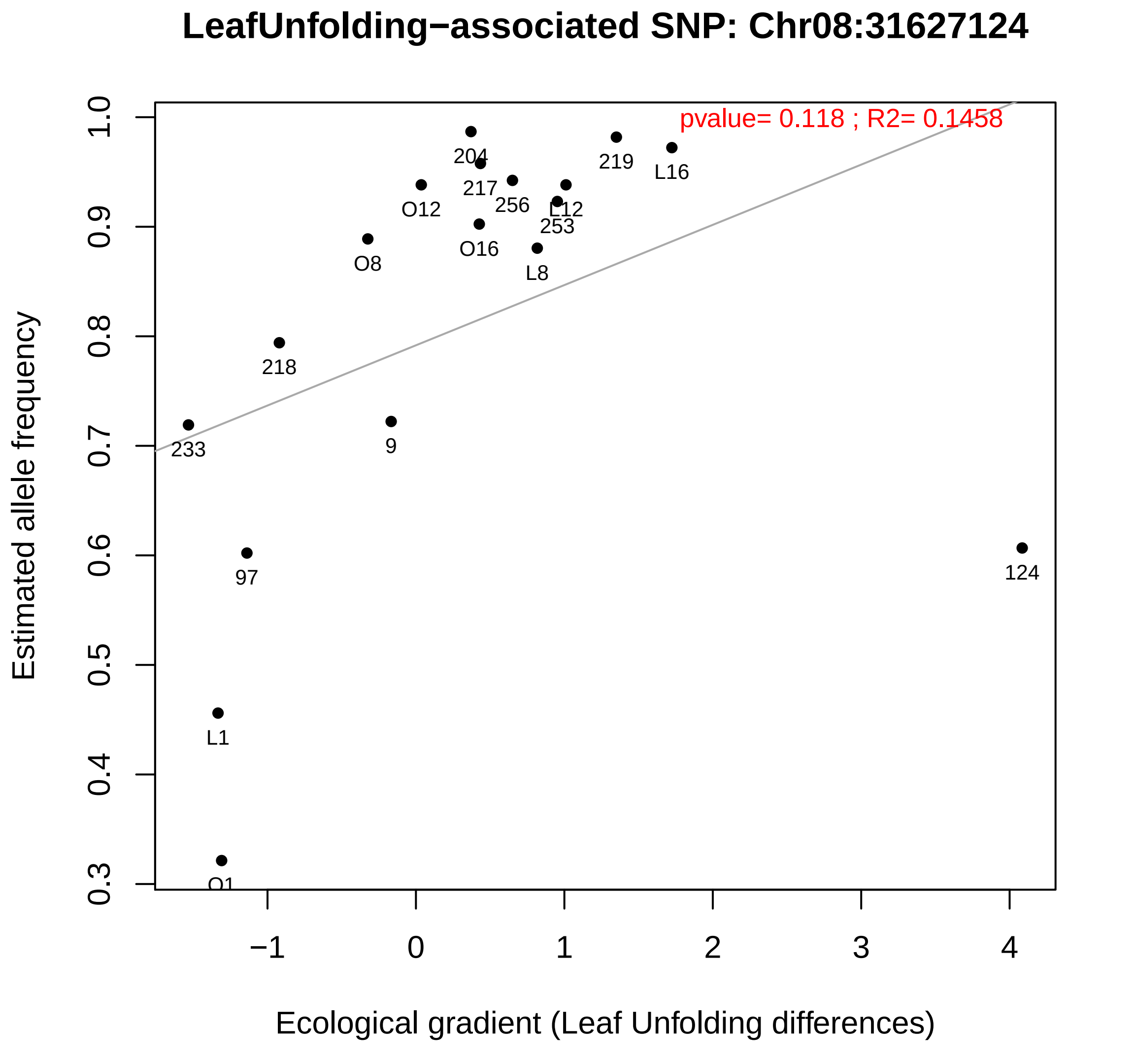

LeafUnfolding−associated SNP: Chr08:31627124
pvalue= 0.118 ; R2= 0.1458
Estimated allele frequency
Ecological gradient (Leaf Unfolding differences)
0.3
0.4
0.5
0.6
0.7
0.8
0.9
1.0
−1
0
1
2
3
4
204
219
L16
217
O12
256
L12
253
O16
O8
L8
218
233
9
97
124
L1
O1

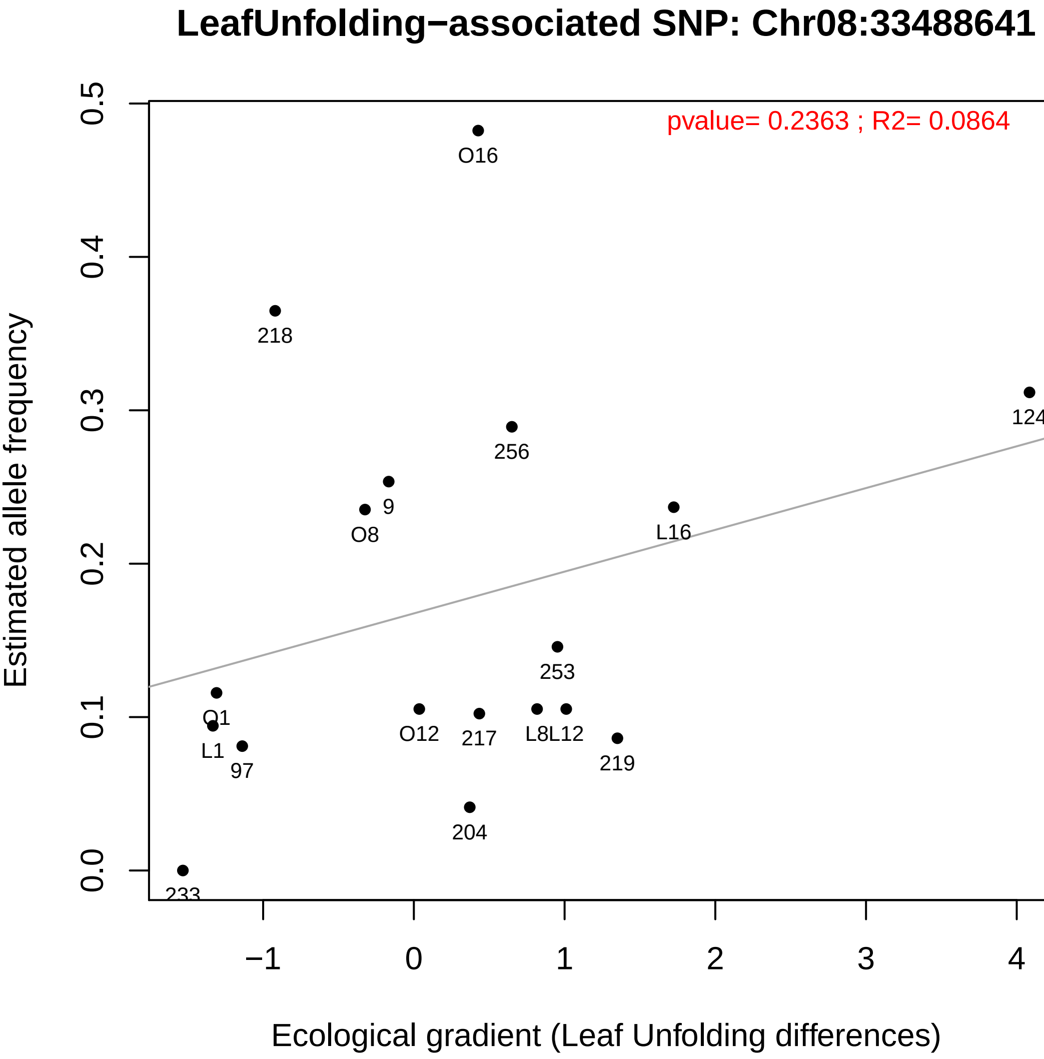
LeafUnfolding−associated SNP: Chr08:33488641
pvalue= 0.2363 ; R2= 0.0864
Estimated allele frequency
Ecological gradient (Leaf Unfolding differences)
0.0
0.1
0.2
0.3
0.4
0.5
−1
0
1
2
3
4
O16
218
124
256
9
O8
L16
253
O1
L1
97
O12
217
L8
L12
219
204
233

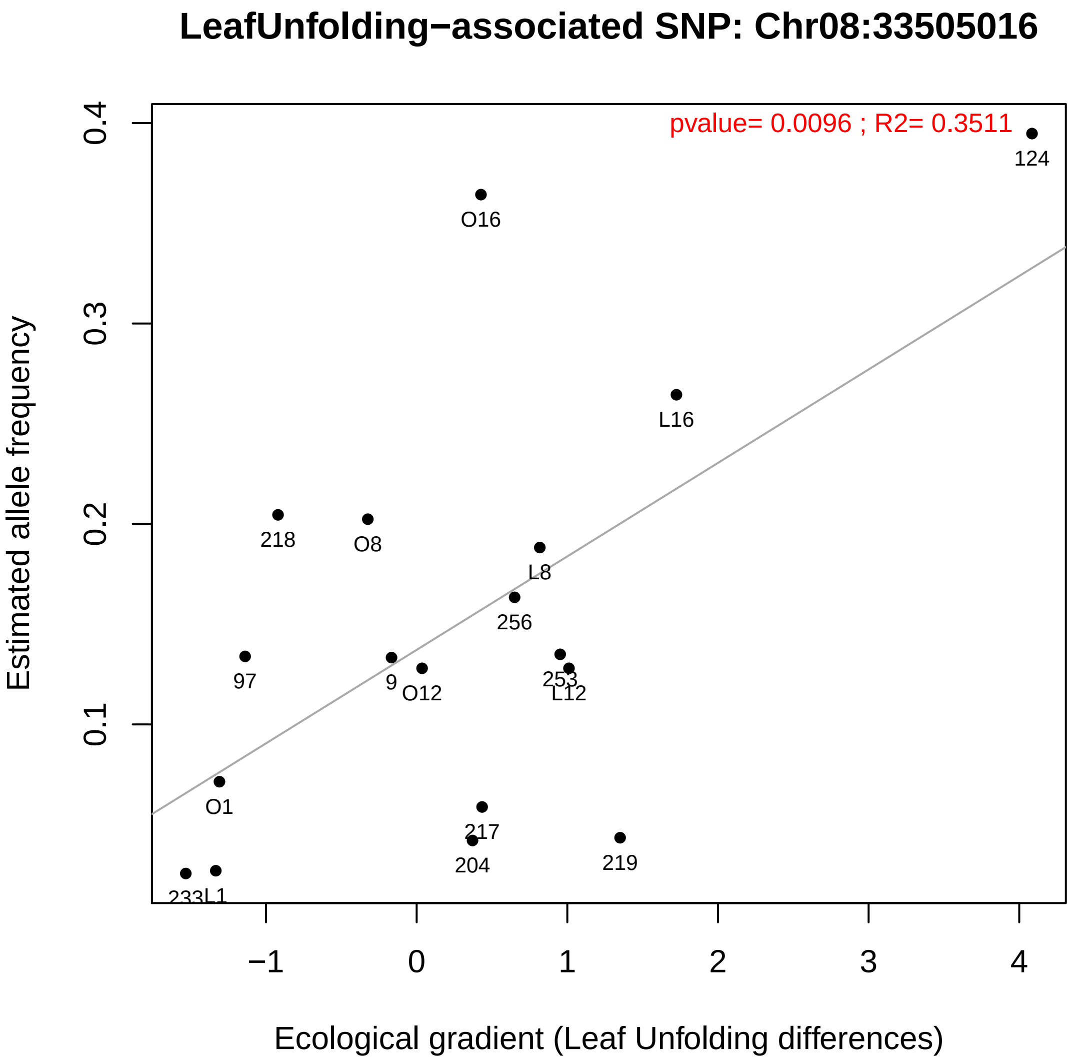

LeafUnfolding−associated SNP: Chr08:33505016
pvalue= 0.0096 ; R2= 0.3511
Estimated allele frequency
Ecological gradient (Leaf Unfolding differences)
0.1
0.2
0.3
0.4
−1
0
1
2
3
4
124
O16
L16
218
O8
L8
256
97
9
O12
253
L12
O1
217
204
219
233
L1

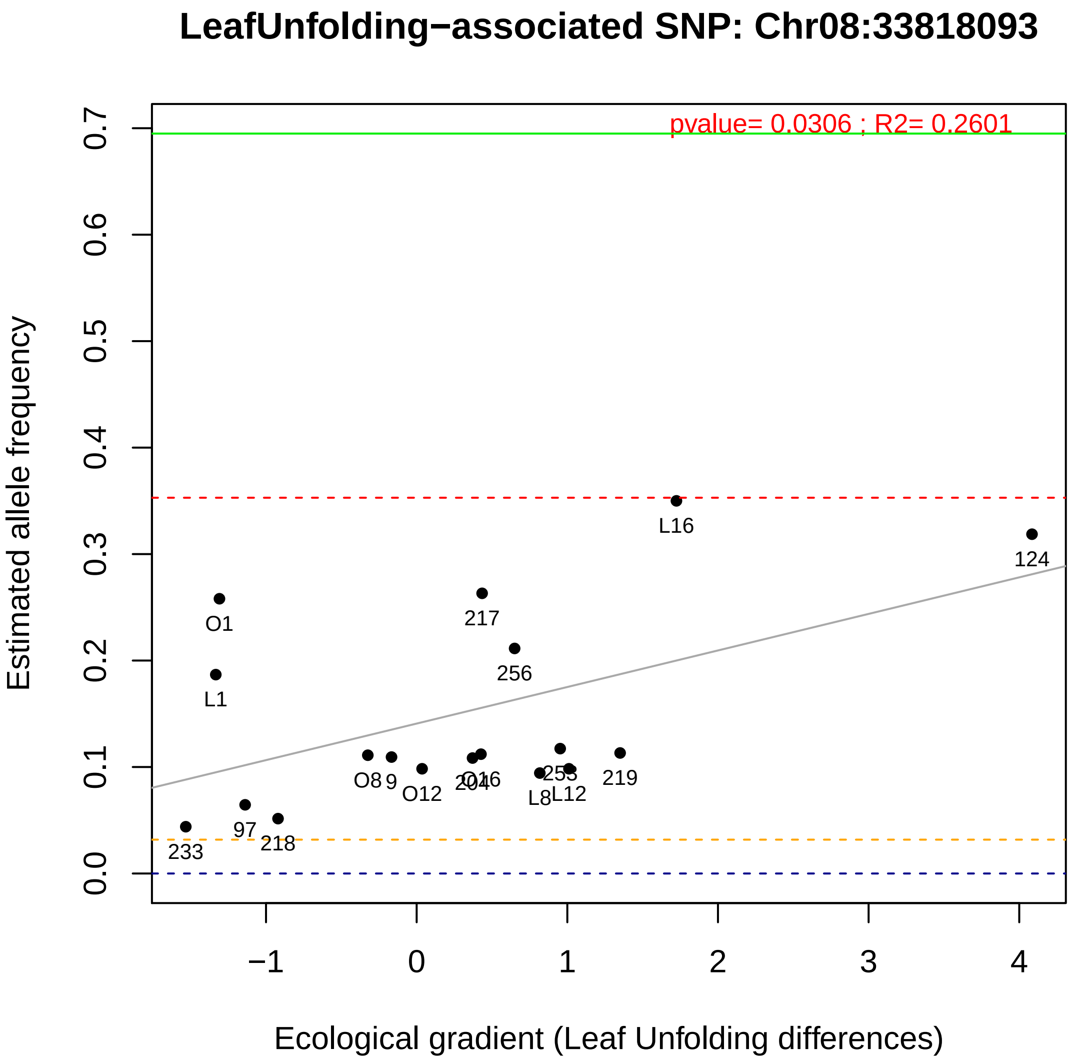
LeafUnfolding−associated SNP: Chr08:33818093
pvalue= 0.0306 ; R2= 0.2601
Estimated allele frequency
0.0
0.1
0.2
0.3
0.4
0.5
0.6
0.7
Ecological gradient (Leaf Unfolding differences)
−1
0
1
2
3
4
L16
124
O1
217
256
L1
O8
9
O12
204
O16
253
L8
L12
219
97
218
233

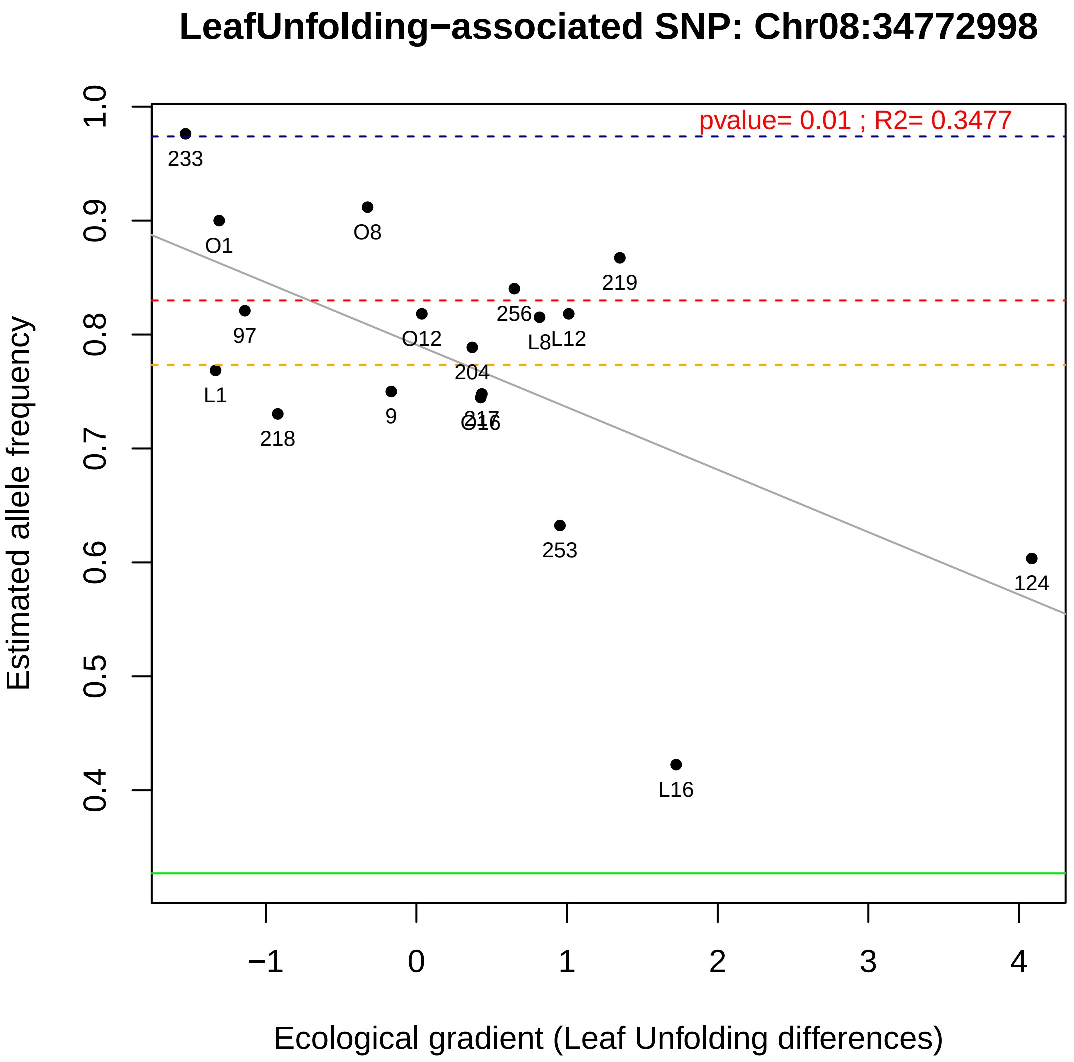

LeafUnfolding−associated SNP: Chr08:34772998
pvalue= 0.01 ; R2= 0.3477
Estimated allele frequency
Ecological gradient (Leaf Unfolding differences)
0.4
0.5
0.6
0.7
0.8
0.9
1.0
−1
0
1
2
3
4
233
O1
O8
219
256
97
O12
L8
L12
204
L1
9
218
217
O16
253
124
L16

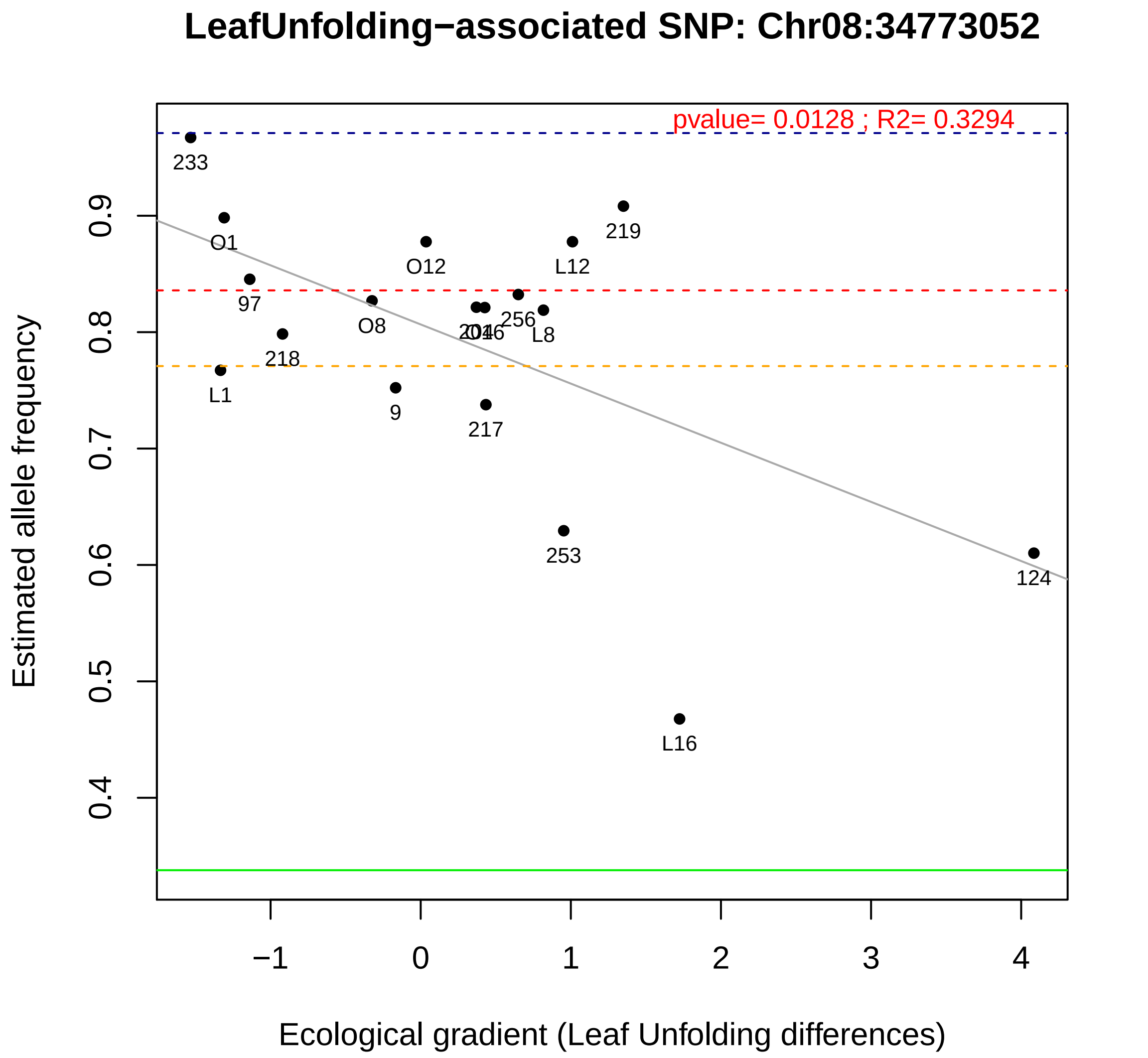

LeafUnfolding−associated SNP: Chr08:34773052
pvalue= 0.0128 ; R2= 0.3294
233
O1
97
218
L1
O8
O12
9
217
204
O16
256
L8
L12
219
253
L16
124
0.4
0.5
0.6
0.7
0.8
0.9
−1
0
1
2
3
4
Estimated allele frequency
Ecological gradient (Leaf Unfolding differences)

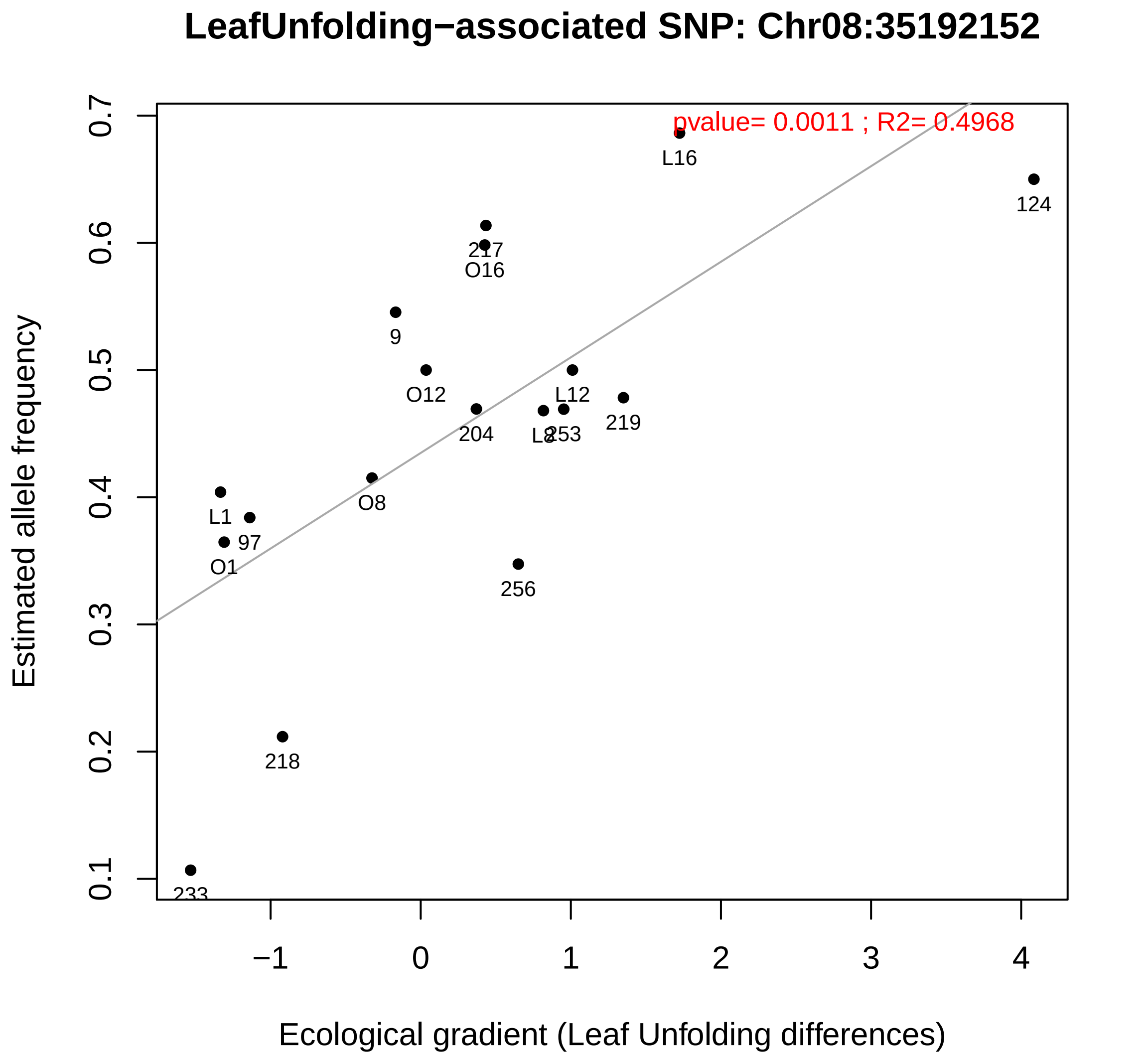
LeafUnfolding−associated SNP: Chr08:35192152
pvalue= 0.0011 ; R2= 0.4968
Estimated allele frequency
Ecological gradient (Leaf Unfolding differences)
0.1
0.2
0.3
0.4
0.5
0.6
0.7
−1
0
1
2
3
4
L16
124
217
O16
9
O12
L12
204
L8
253
219
O8
L1
97
O1
256
218
233

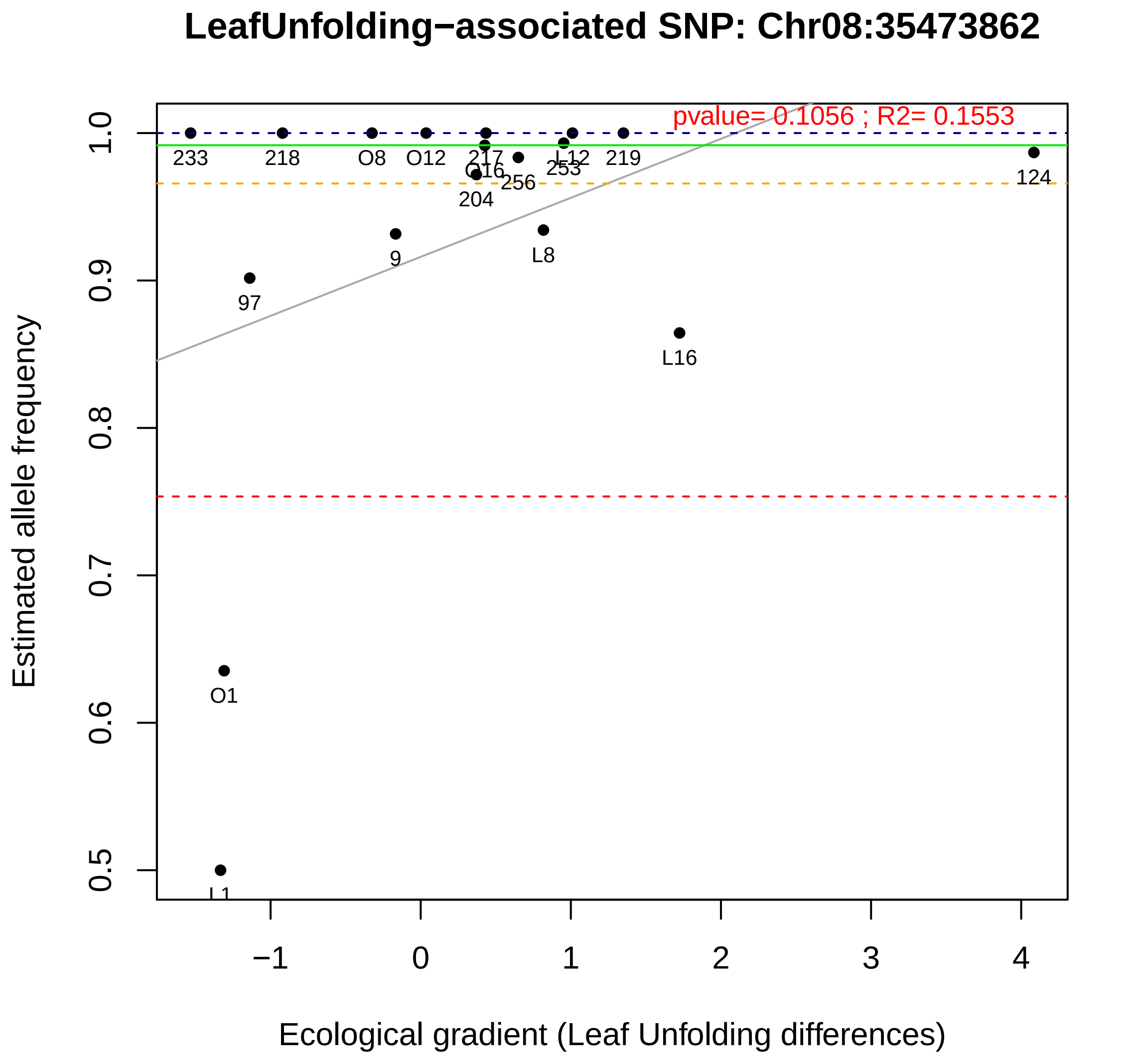
LeafUnfolding−associated SNP: Chr08:35473862
pvalue= 0.1056 ; R2= 0.1553
Estimated allele frequency
Ecological gradient (Leaf Unfolding differences)
0.5
0.6
0.7
0.8
0.9
1.0
−1
0
1
2
3
4
233
218
O8
O12
217
O16
204
256
L12
253
219
124
9
L8
97
L16
O1

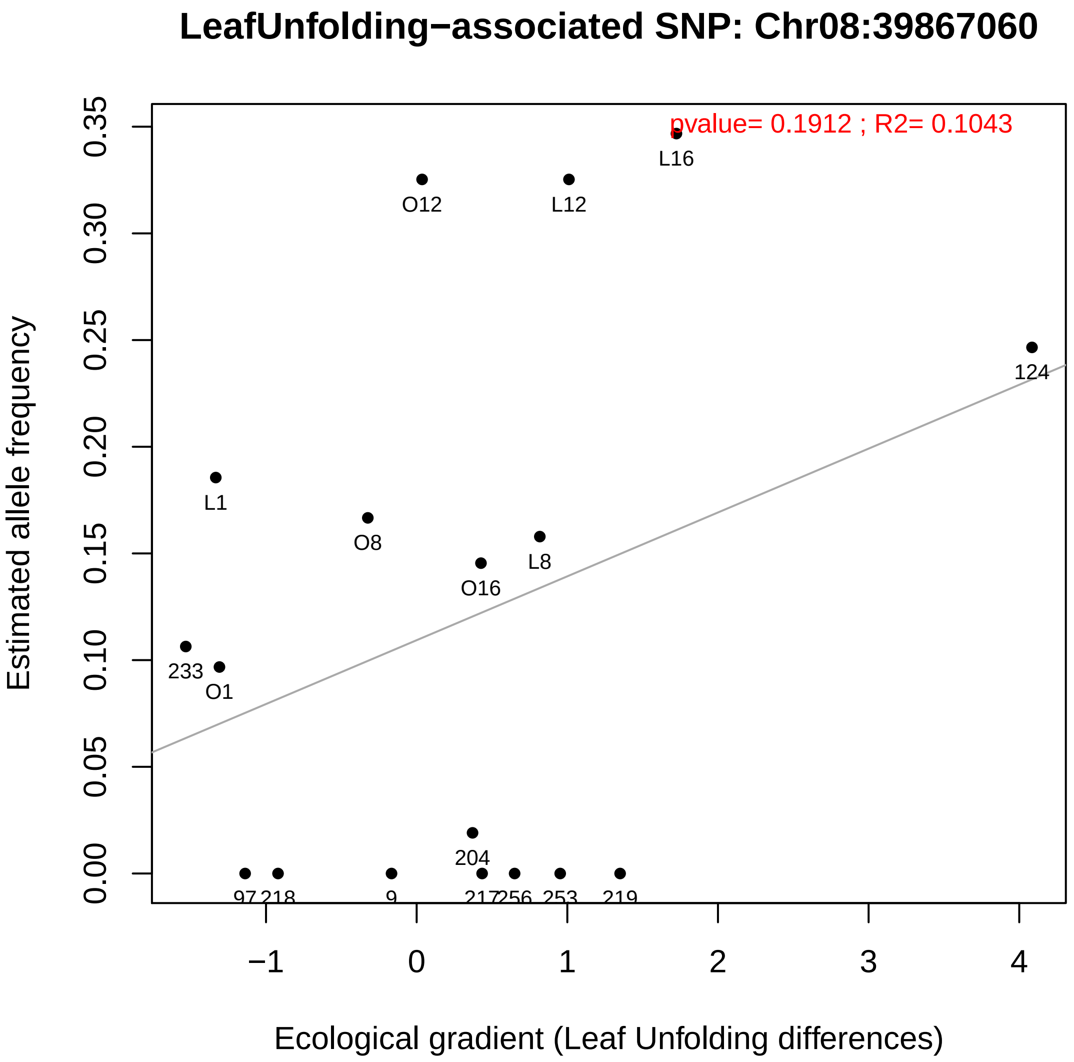

LeafUnfolding−associated SNP: Chr08:39867060
pvalue= 0.1912 ; R2= 0.1043
Estimated allele frequency
Ecological gradient (Leaf Unfolding differences)
0.00
0.05
0.10
0.15
0.20
0.25
0.30
0.35
−1
0
1
2
3
4
L16
O12
L12
124
L1
O8
L8
O16
233
O1
204
97
218
9
217
256
253
219

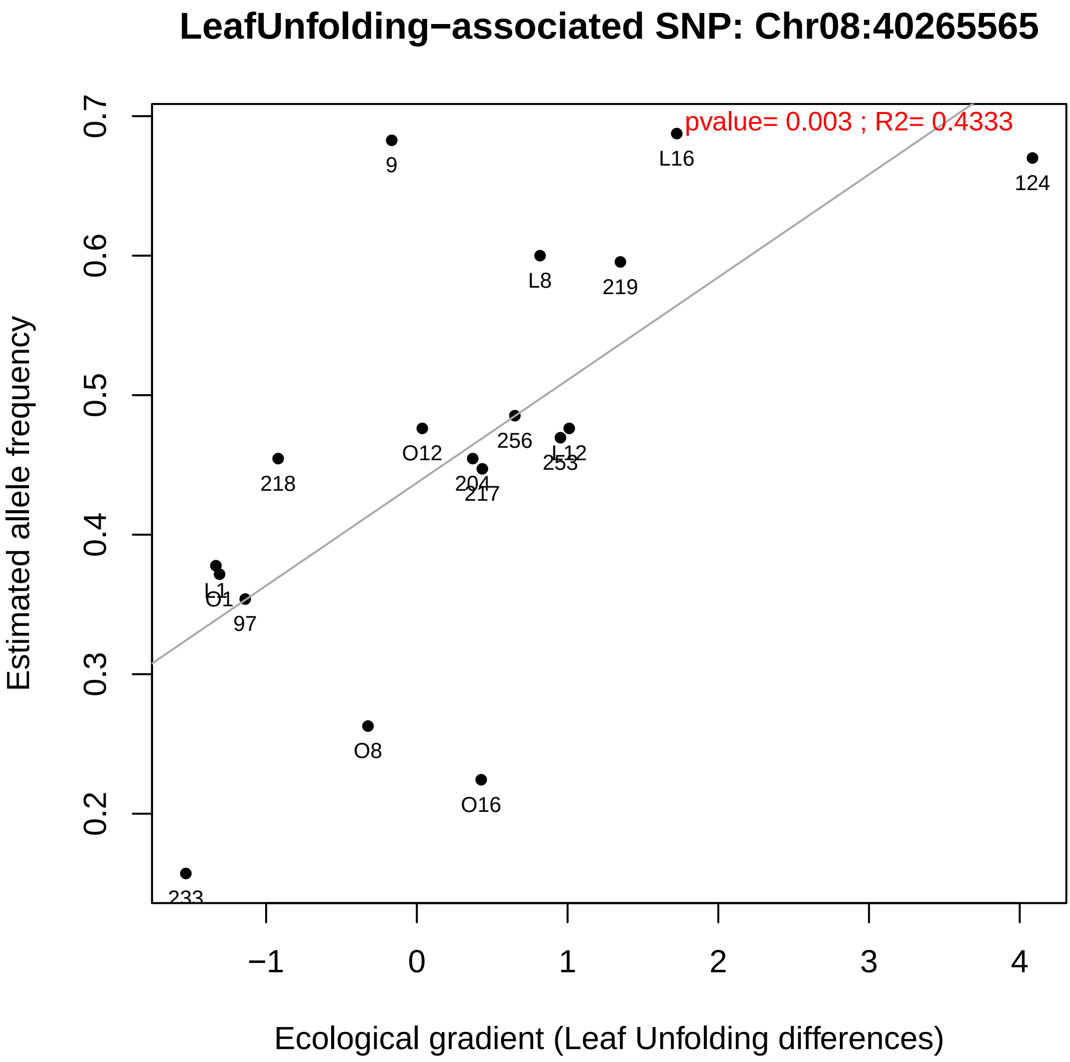
LeafUnfolding−associated SNP: Chr08:40265565
pvalue= 0.003 ; R2= 0.4333
Estimated allele frequency
Ecological gradient (Leaf Unfolding differences)
0.2
0.3
0.4
0.5
0.6
0.7
−1
0
1
2
3
4
9
L16
124
L8
219
256
O12
L12
253
204
217
218
L1
O1
97
O8
O16
233

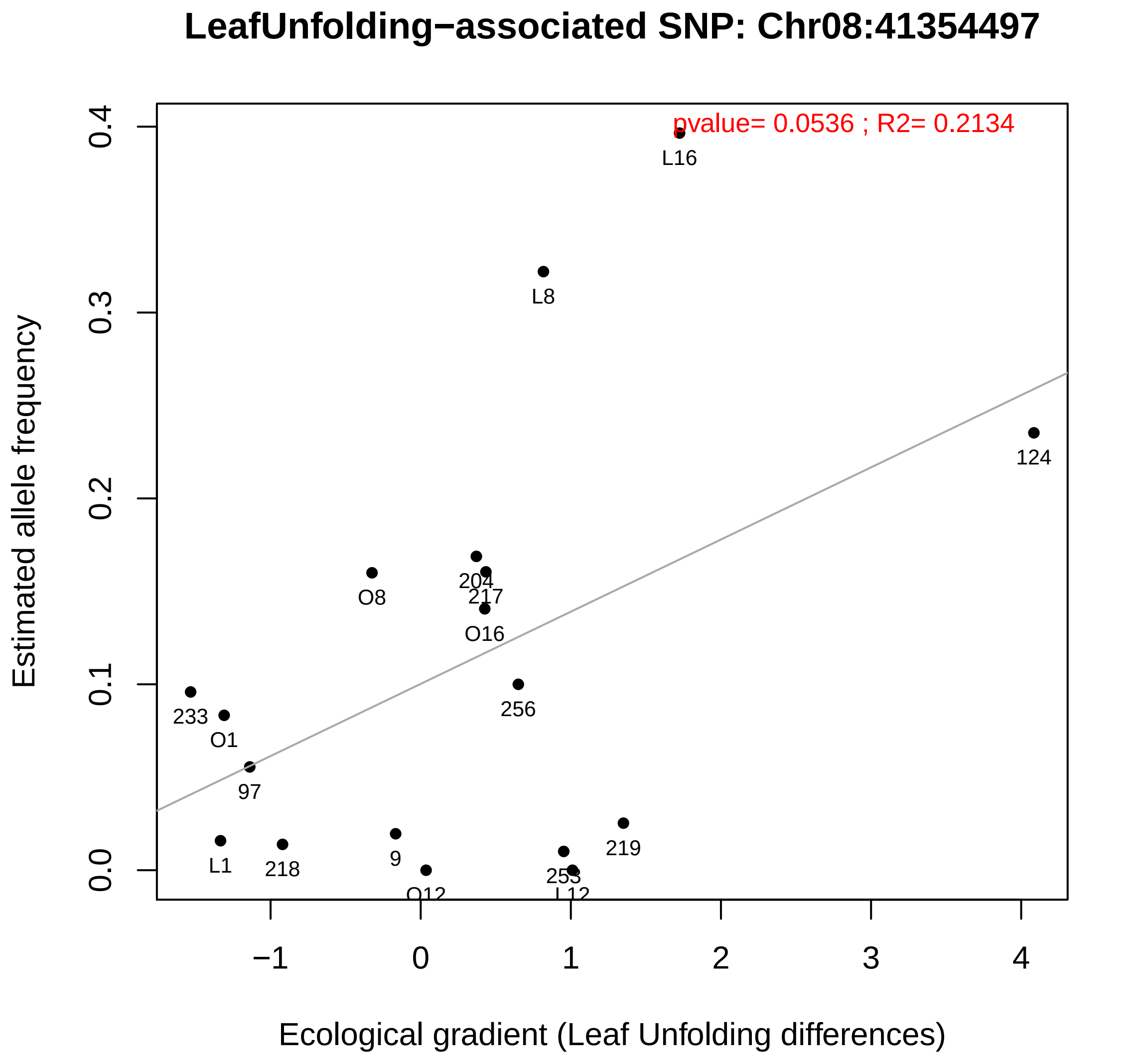

LeafUnfolding−associated SNP: Chr08:41354497
pvalue= 0.0536 ; R2= 0.2134
Estimated allele frequency
Ecological gradient (Leaf Unfolding differences)
0.0
0.1
0.2
0.3
0.4
−1
0
1
2
3
4
L16
L8
124
O8
204
217
O16
256
233
O1
97
L1
218
9
O12
253
L12
219

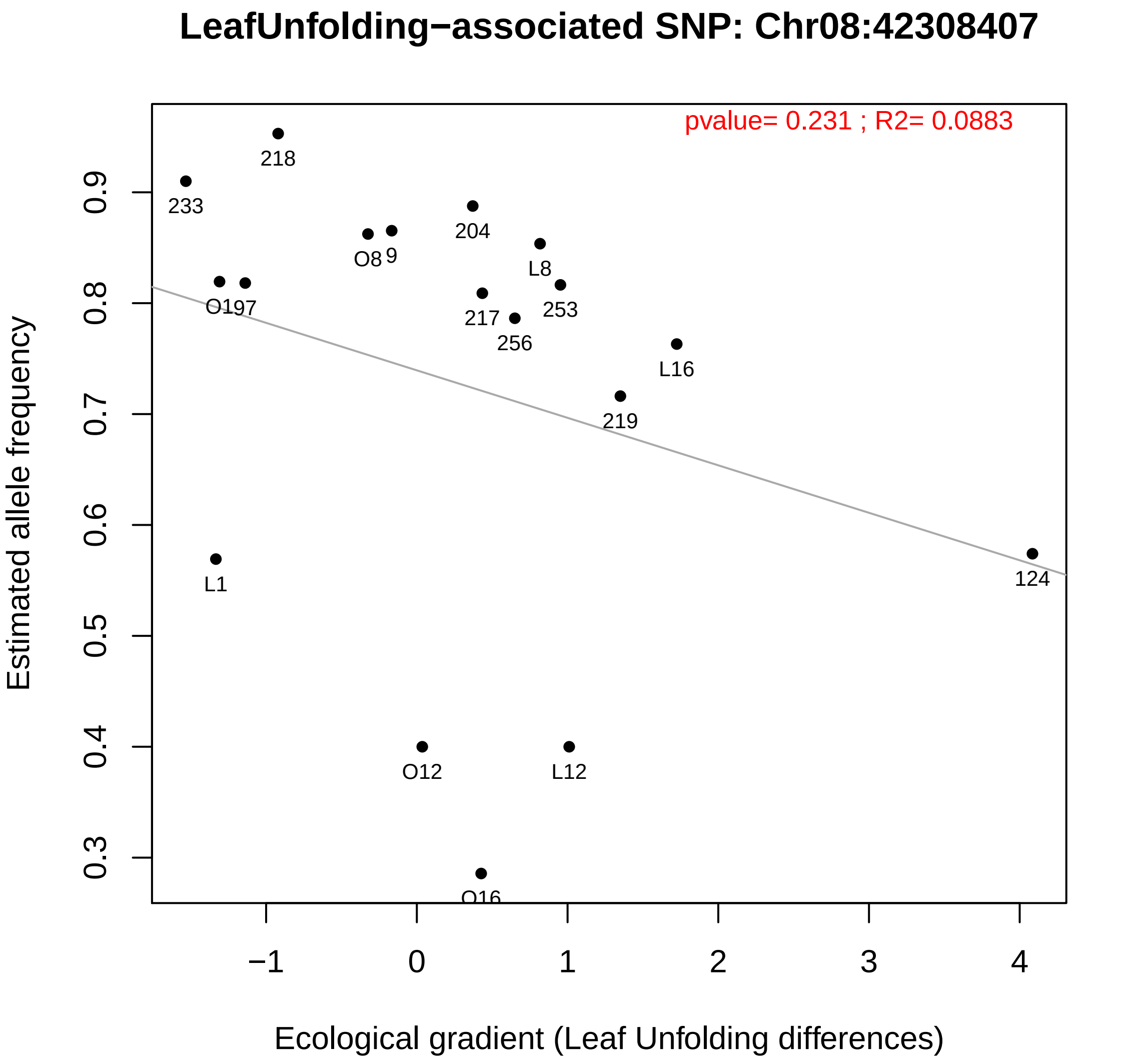
LeafUnfolding−associated SNP: Chr08:42308407
pvalue= 0.231 ; R2= 0.0883
218
233
204
O8
9
L8
O197
217
253
256
L16
219
L1
124
O12
L12
O16
0.3
0.4
0.5
0.6
0.7
0.8
0.9
−1
0
1
2
3
4
Estimated allele frequency
Ecological gradient (Leaf Unfolding differences)

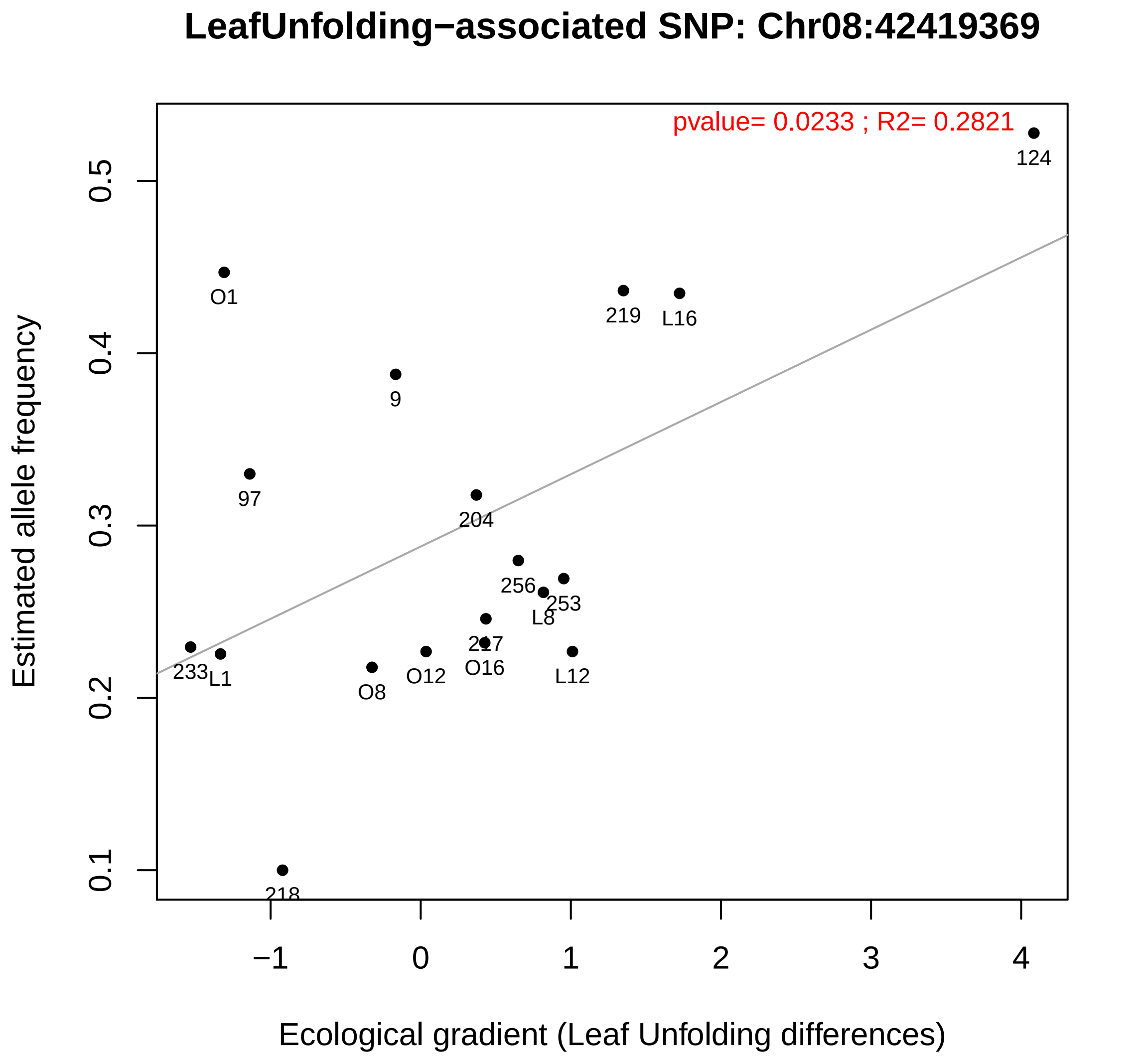

LeafUnfolding−associated SNP: Chr08:42419369
pvalue= 0.0233 ; R2= 0.2821
Estimated allele frequency
0.1
0.2
0.3
0.4
0.5
−1
0
1
2
3
4
Ecological gradient (Leaf Unfolding differences)
124
O1
219
L16
9
97
204
256
253
L8
217
O16
L12
O12
O8
233
L1
218

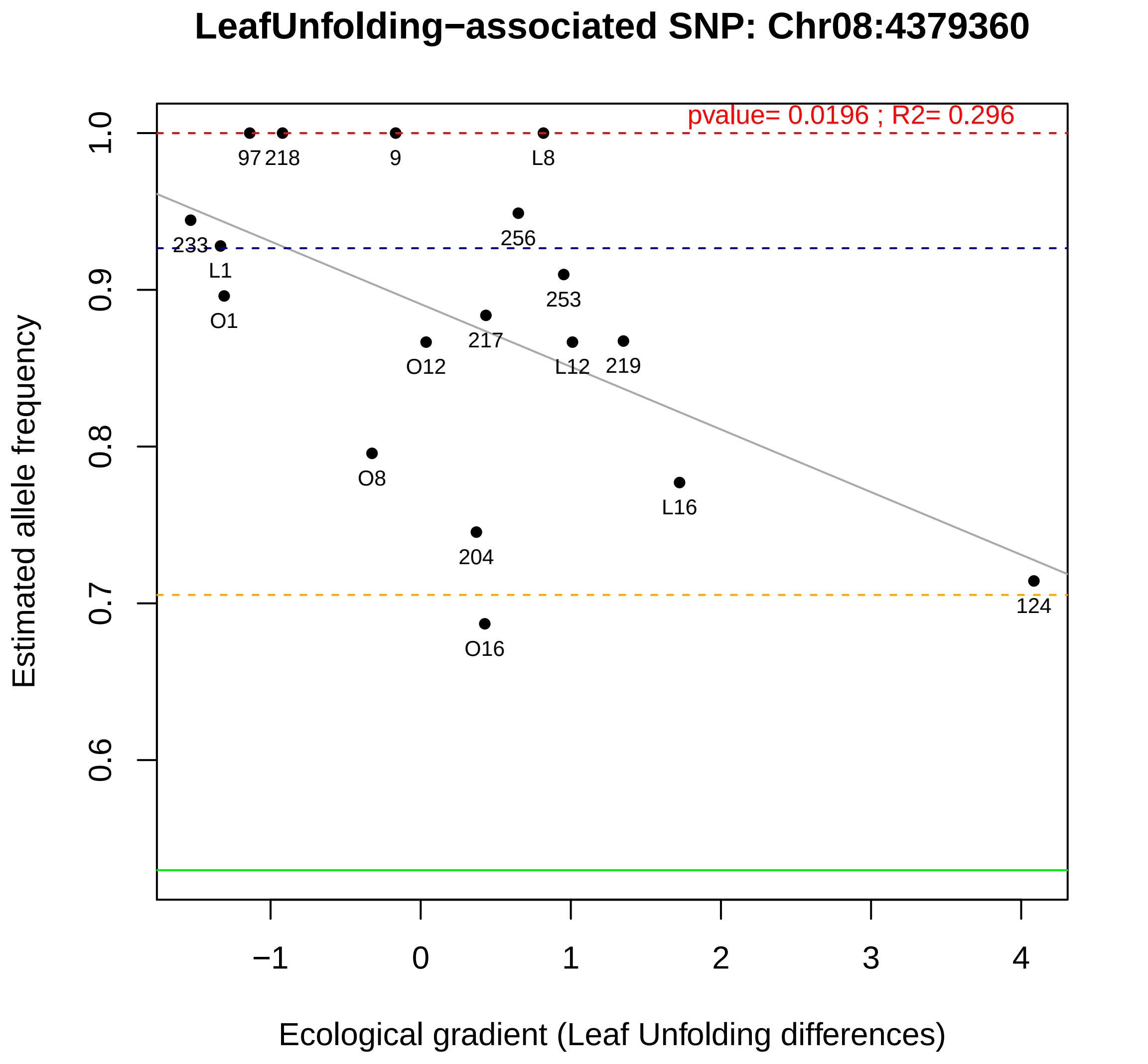

LeafUnfolding−associated SNP: Chr08:4379360
pvalue= 0.0196 ; R2= 0.296
Estimated allele frequency
Ecological gradient (Leaf Unfolding differences)
1.0
0.9
0.8
0.7
0.6
−1
0
1
2
3
4
97
218
9
L8
233
L1
256
O1
253
217
O12
L12
219
O8
L16
204
124
O16

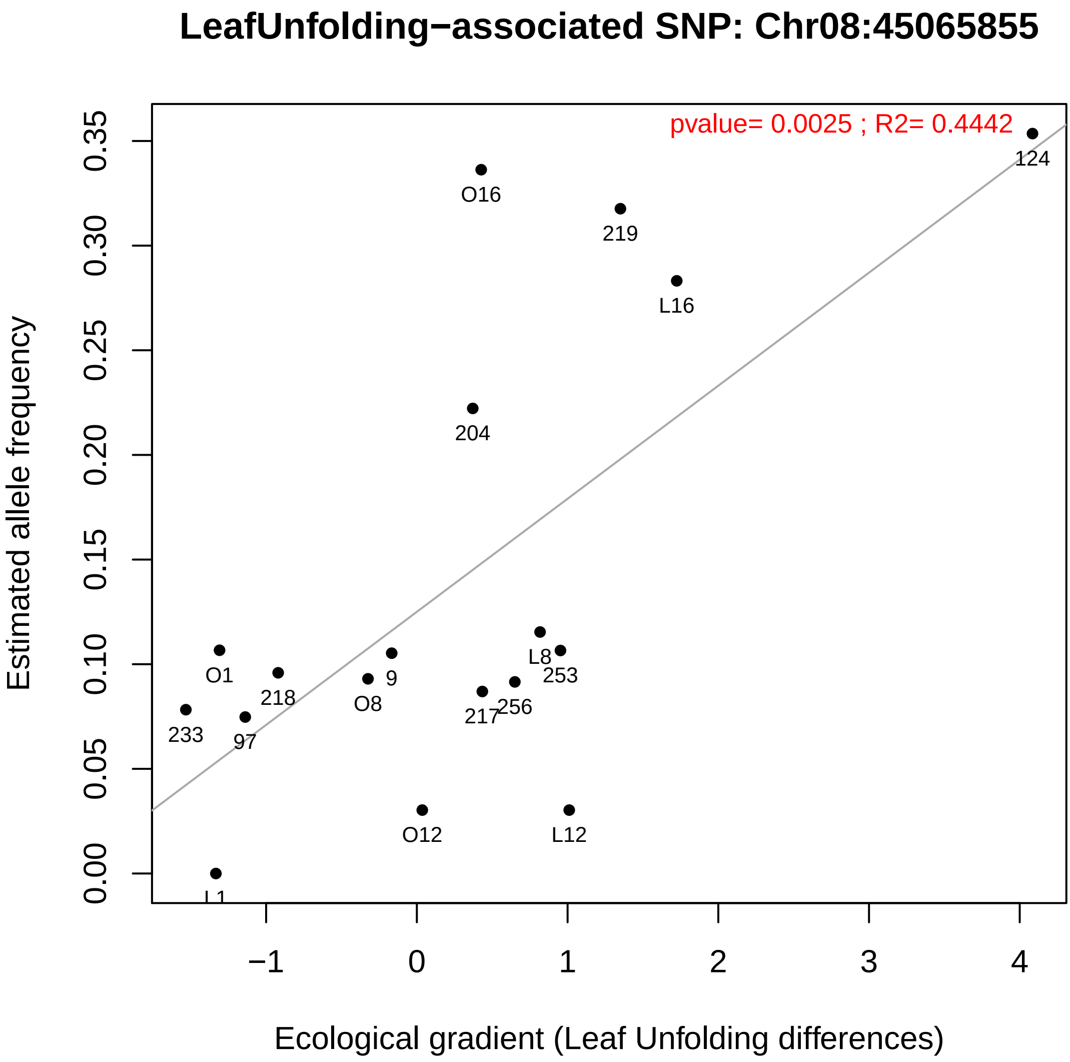

LeafUnfolding−associated SNP: Chr08:45065855
pvalue= 0.0025 ; R2= 0.4442
Estimated allele frequency
Ecological gradient (Leaf Unfolding differences)
0.00
0.05
0.10
0.15
0.20
0.25
0.30
0.35
−1
0
1
2
3
4
124
O16
219
L16
204
L8
253
O1
9
218
O8
256
217
233
97
O12
L12
L1

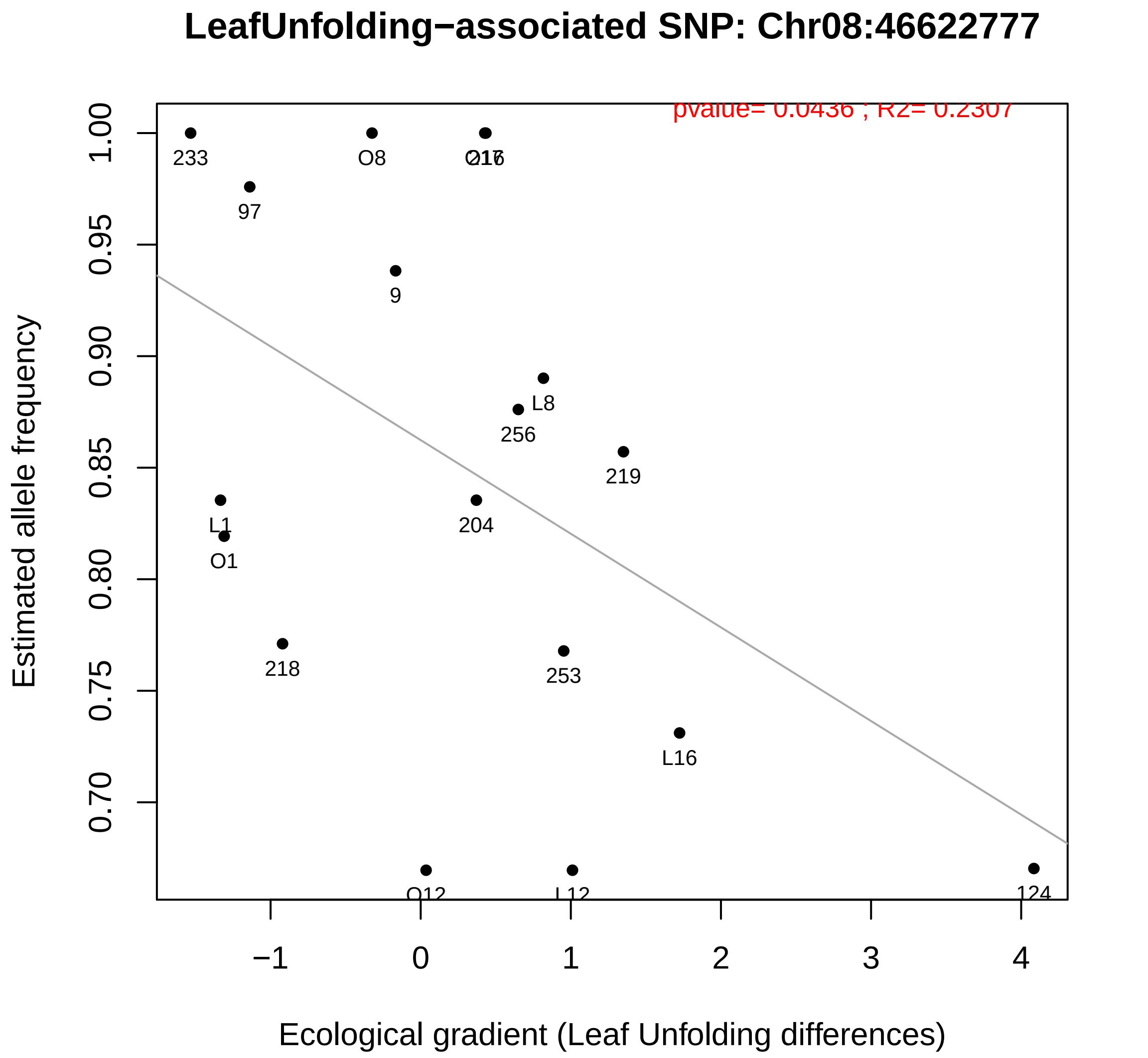

LeafUnfolding−associated SNP: Chr08:46622777
pvalue= 0.0436 ; R2= 0.2307
Estimated allele frequency
1.00
0.95
0.90
0.85
0.80
0.75
0.70
−1
0
1
2
3
4
Ecological gradient (Leaf Unfolding differences)
233
O8
97
9
L8
256
219
L1
204
O1
218
253
L16
O12
L12
124

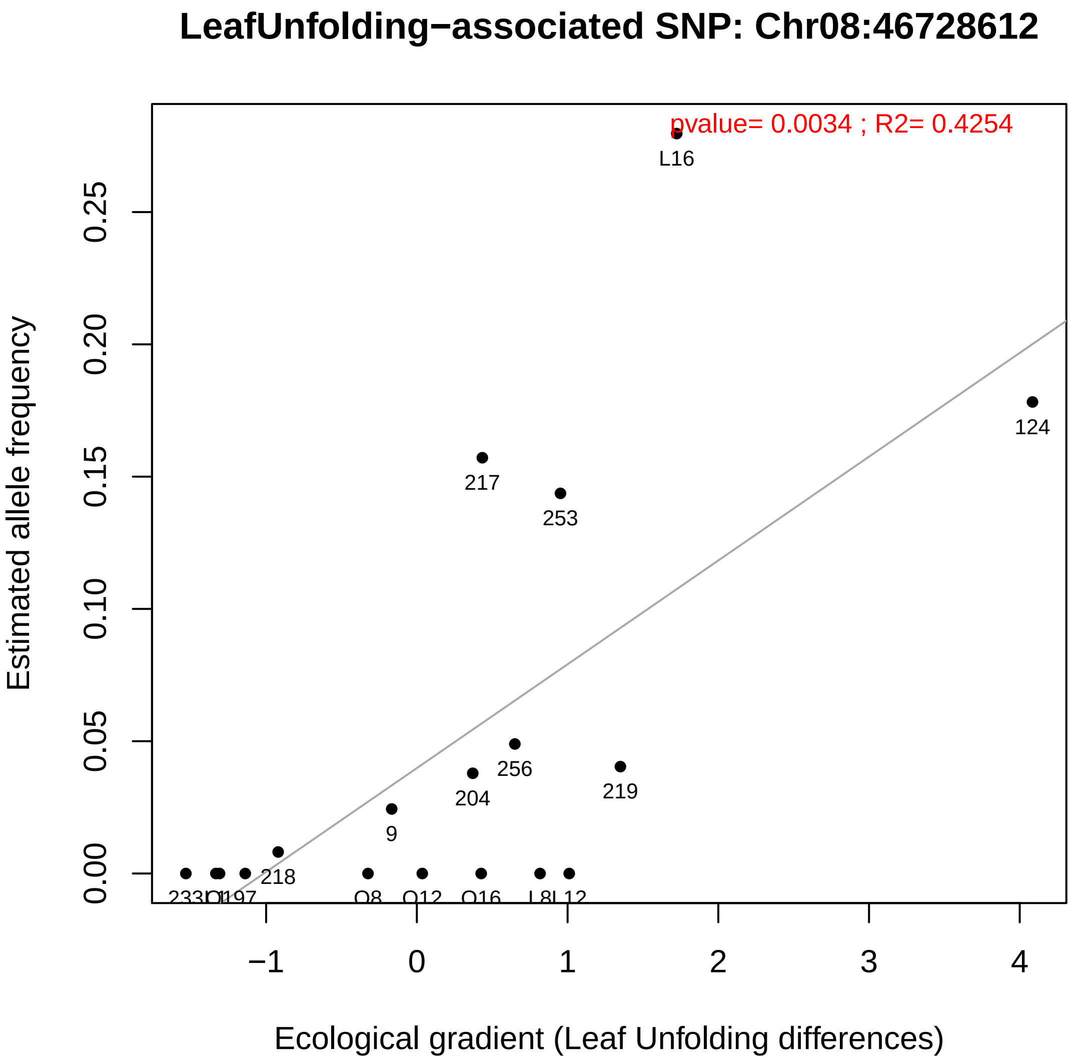
LeafUnfolding−associated SNP: Chr08:46728612
pvalue= 0.0034 ; R2= 0.4254
Estimated allele frequency
0.00
0.05
0.10
0.15
0.20
0.25
−1
0
1
2
3
4
Ecological gradient (Leaf Unfolding differences)
L16
124
217
253
256
204
219
9
218
O8
O12
O16

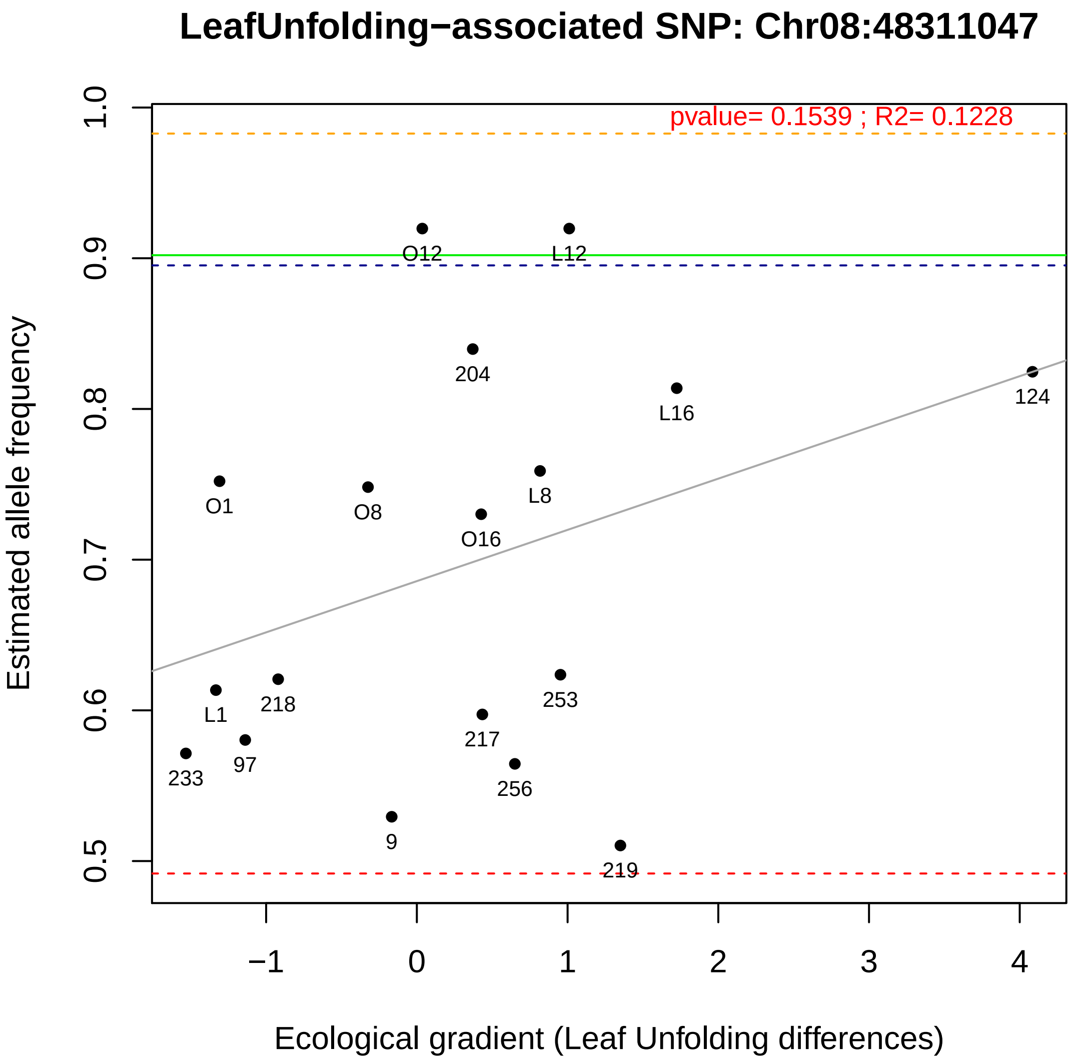

LeafUnfolding−associated SNP: Chr08:48311047
pvalue= 0.1539 ; R2= 0.1228
Estimated allele frequency
Ecological gradient (Leaf Unfolding differences)
0.5
0.6
0.7
0.8
0.9
1.0
−1
0
1
2
3
4
O12
L12
204
L16
124
O1
O8
O16
L8
L1
218
253
217
233
97
256
9
219

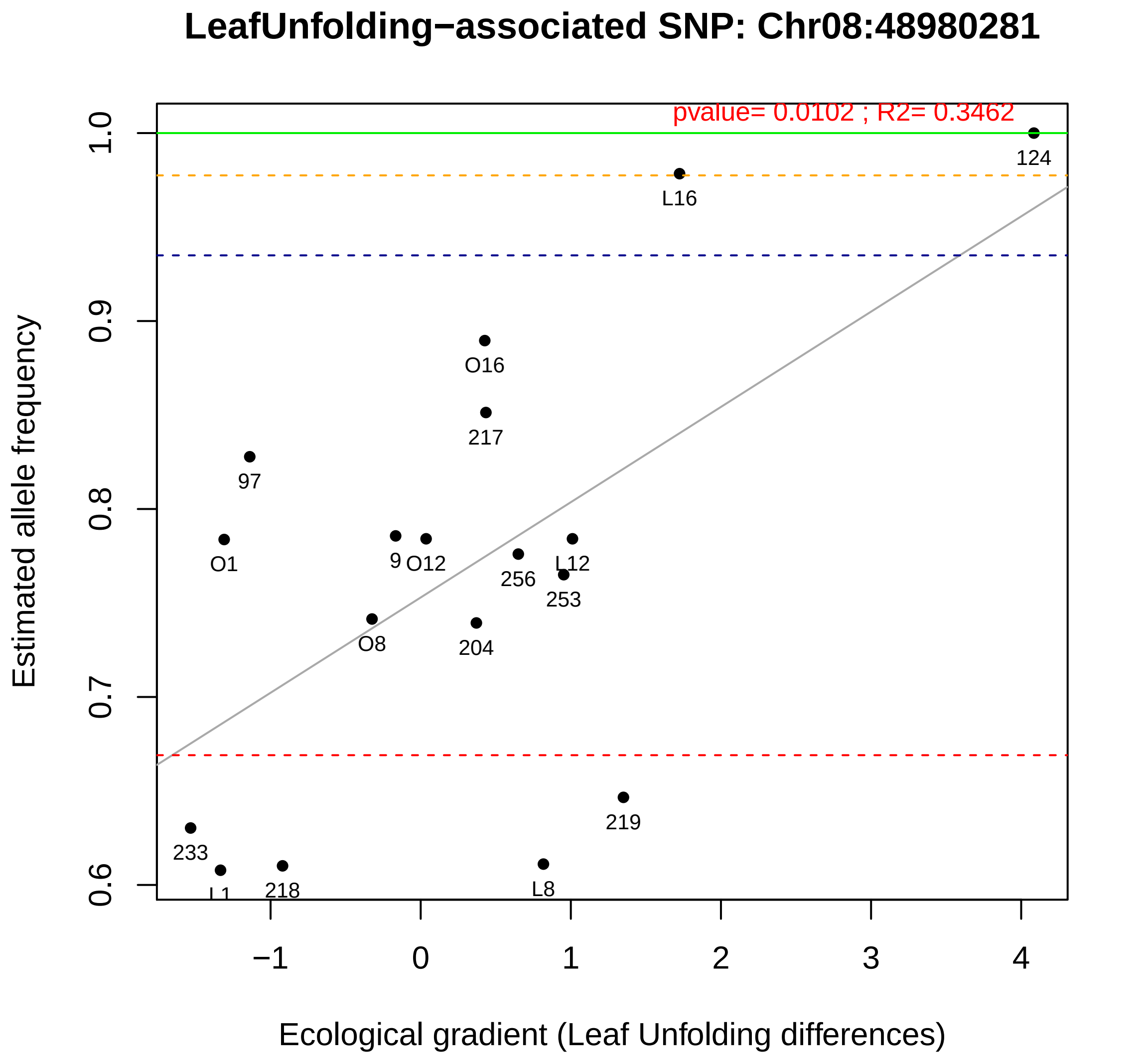
LeafUnfolding−associated SNP: Chr08:48980281
pvalue= 0.0102 ; R2= 0.3462
Estimated allele frequency
Ecological gradient (Leaf Unfolding differences)
0.6
0.7
0.8
0.9
1.0
−1
0
1
2
3
4
124
L16
O16
217
97
O1
9
O12
L12
256
253
O8
204
219
233
218
L8

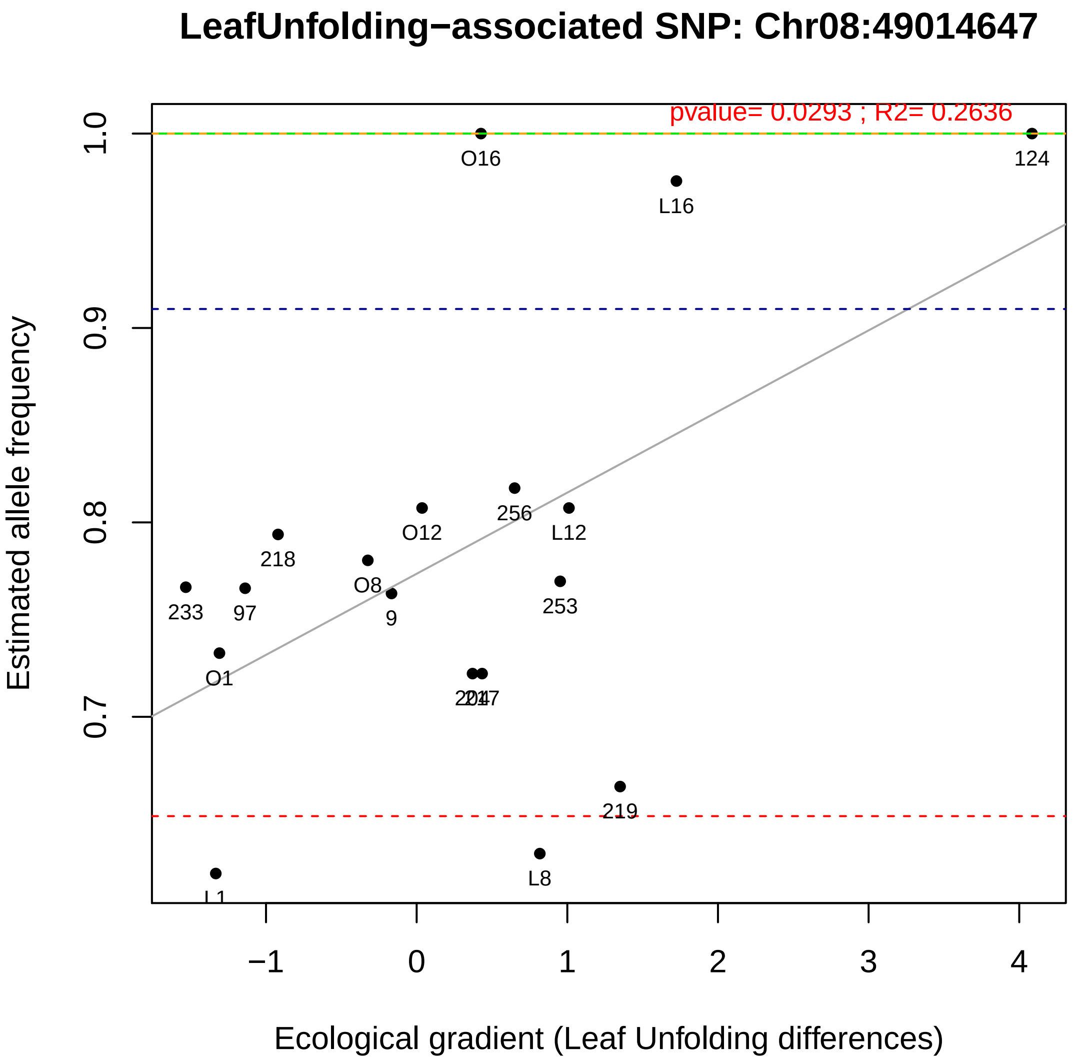

LeafUnfolding−associated SNP: Chr08:49014647
pvalue= 0.0293 ; R2= 0.2636
Estimated allele frequency
Ecological gradient (Leaf Unfolding differences)
1.0
0.9
0.8
0.7
−1
0
1
2
3
4
O16
124
L16
256
O12
L12
218
O8
233
97
9
253
O1
219
L8

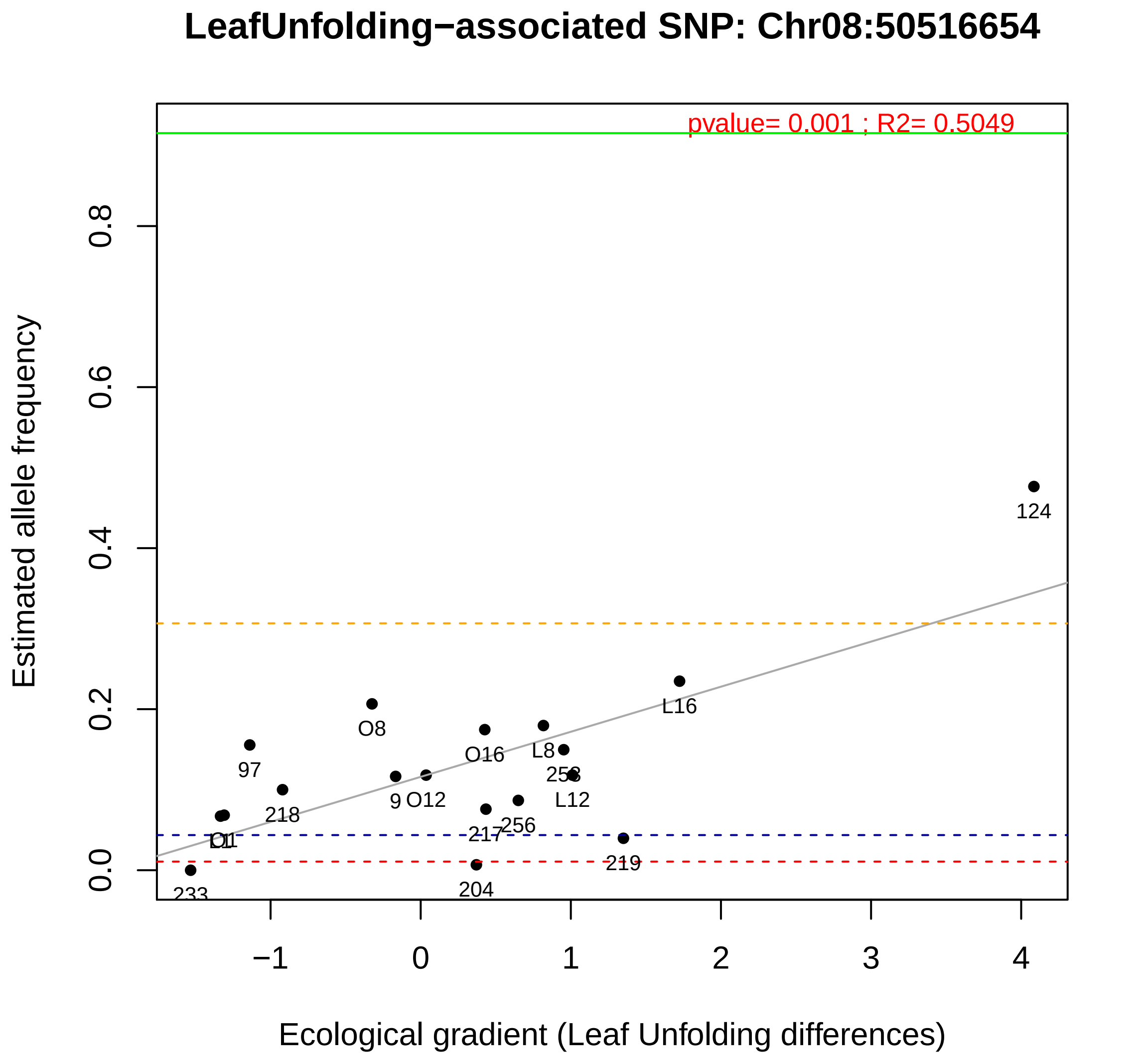
LeafUnfolding−associated SNP: Chr08:50516654
pvalue= 0.001 ; R2= 0.5049
Estimated allele frequency
0.0
0.2
0.4
0.6
0.8
−1
0
1
2
3
4
Ecological gradient (Leaf Unfolding differences)
124
L16
O8
O16
L8
97
9
O12
L12
218
217
256
219
204
233

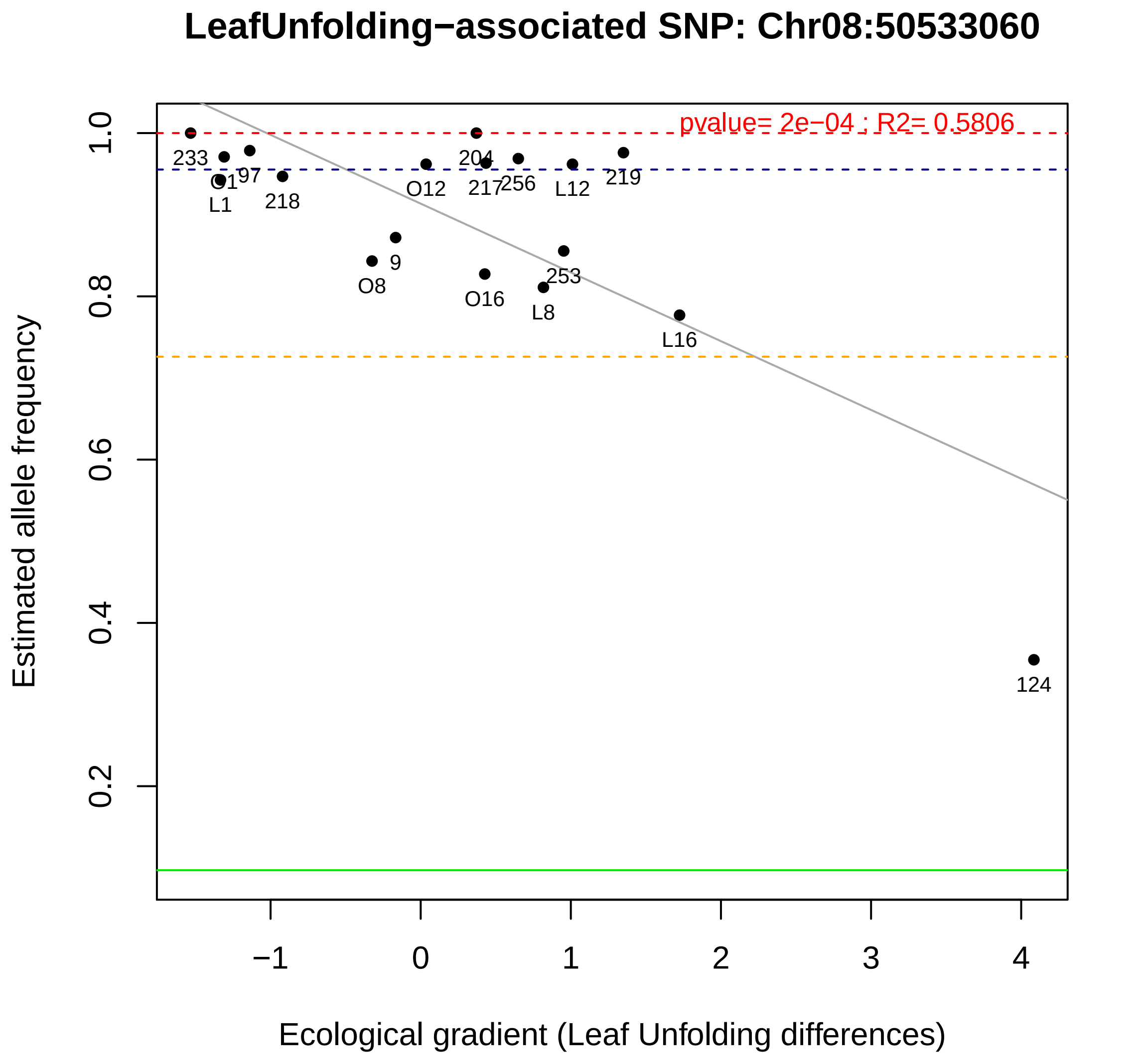

LeafUnfolding−associated SNP: Chr08:50533060
pvalue= 2e−04 ; R2= 0.5806
Estimated allele frequency
Ecological gradient (Leaf Unfolding differences)
0.2
0.4
0.6
0.8
1.0
−1
0
1
2
3
4
233
O1
L1
97
218
O12
204
217
256
L12
219
9
O8
O16
253
L8
L16
124

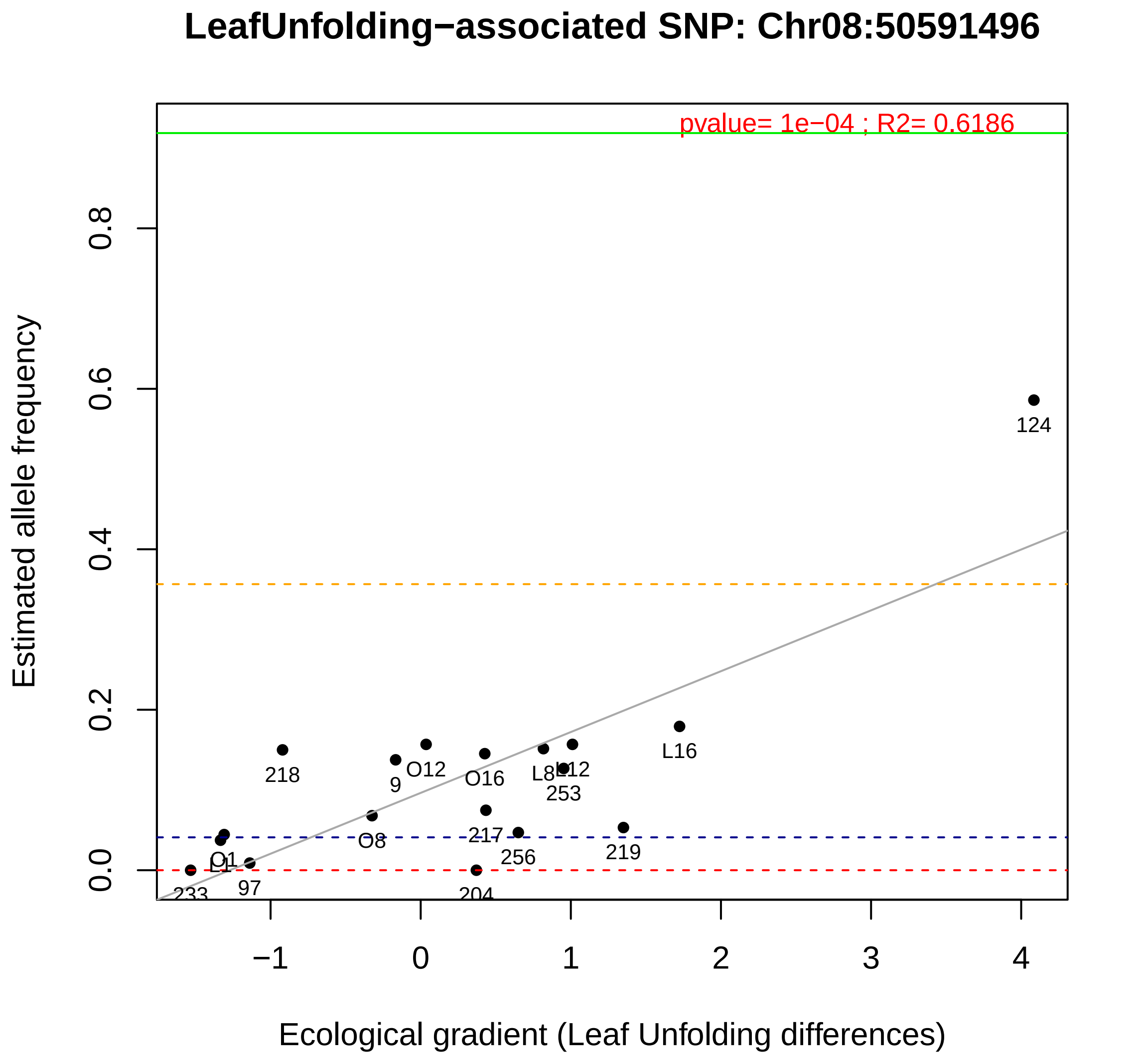

LeafUnfolding−associated SNP: Chr08:50591496
pvalue= 1e−04 ; R2= 0.6186
Estimated allele frequency
0.0
0.2
0.4
0.6
0.8
−1
0
1
2
3
4
Ecological gradient (Leaf Unfolding differences)
124
L16
218
O12
9
O16
L8
L12
253
O8
217
256
219
O1
L1
97
233
204

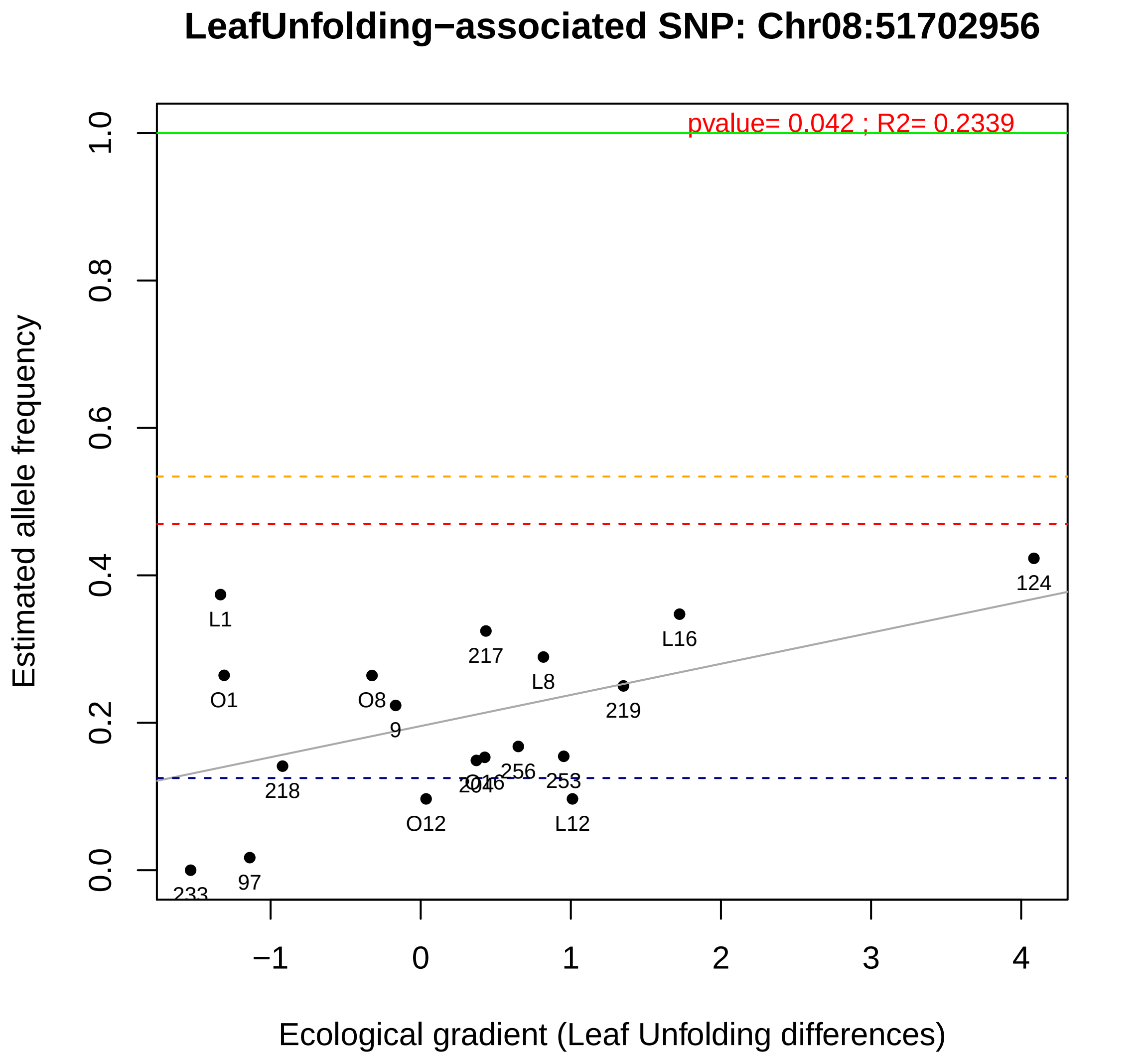

LeafUnfolding−associated SNP: Chr08:51702956
pvalue= 0.042 ; R2= 0.2339
Estimated allele frequency
0.0
0.2
0.4
0.6
0.8
1.0
−1
0
1
2
3
4
Ecological gradient (Leaf Unfolding differences)
L1
O1
O8
9
218
233
97
O12
217
L8
256
253
L12
219
L16
124

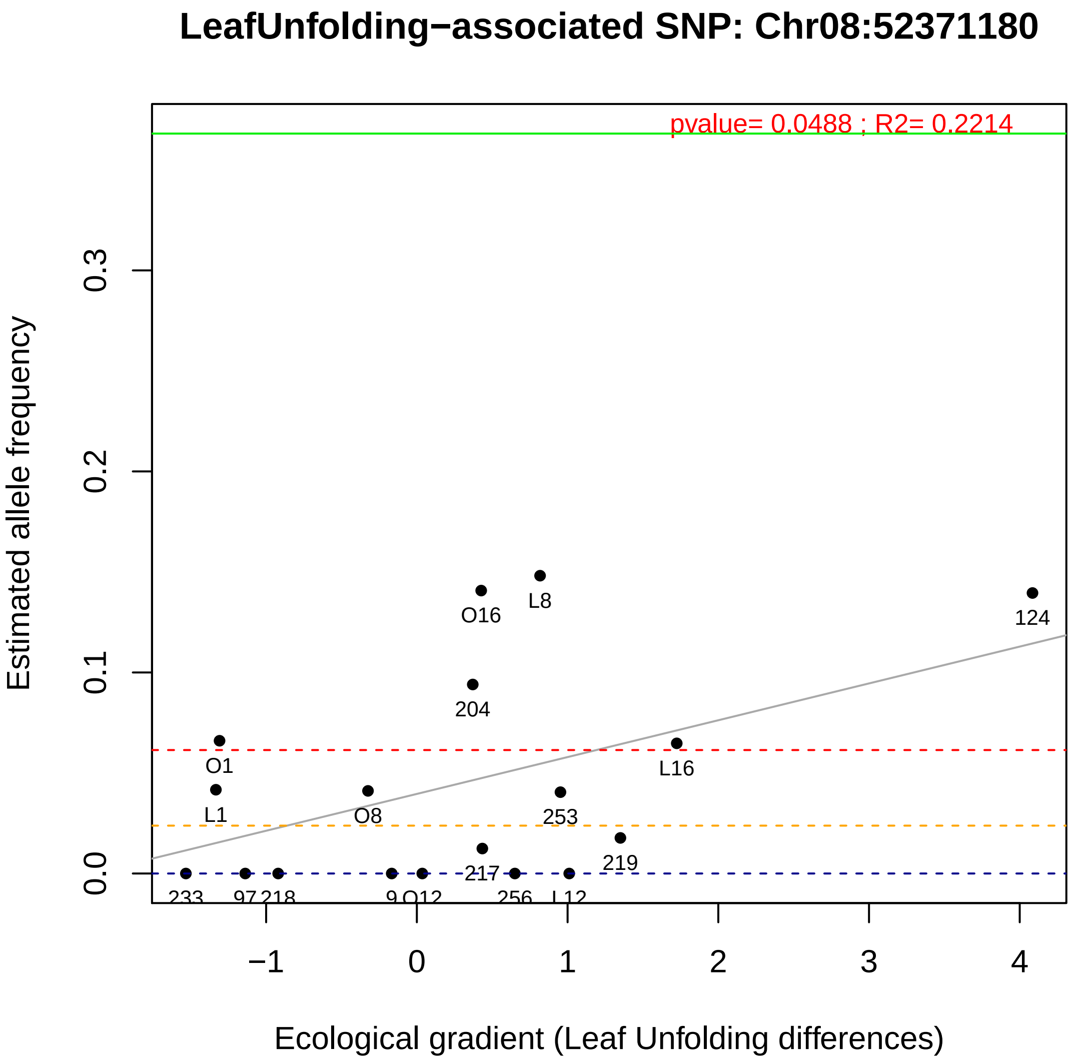

LeafUnfolding−associated SNP: Chr08:52371180
pvalue= 0.0488 ; R2= 0.2214
Estimated allele frequency
0.0
0.1
0.2
0.3
−1
0
1
2
3
4
Ecological gradient (Leaf Unfolding differences)
O16
L8
124
204
O1
L16
L1
O8
253
219
217
233
97
218
9
O12
256
L12

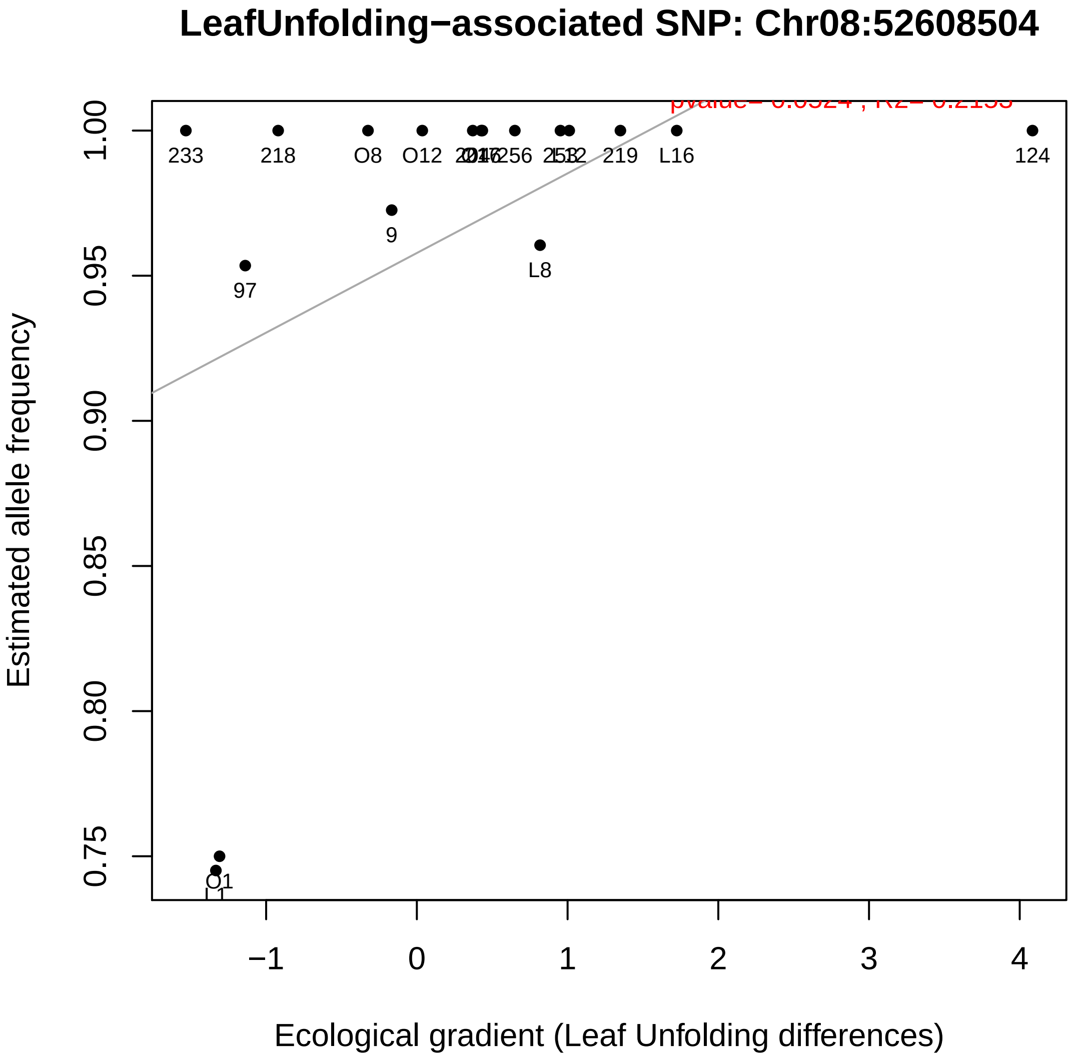
LeafUnfolding−associated SNP: Chr08:52608504
Estimated allele frequency
1.00
0.95
0.90
0.85
0.80
0.75
233
218
O8
O12
219
L16
124
9
97
L8
O1
−1
0
1
2
3
4
Ecological gradient (Leaf Unfolding differences)

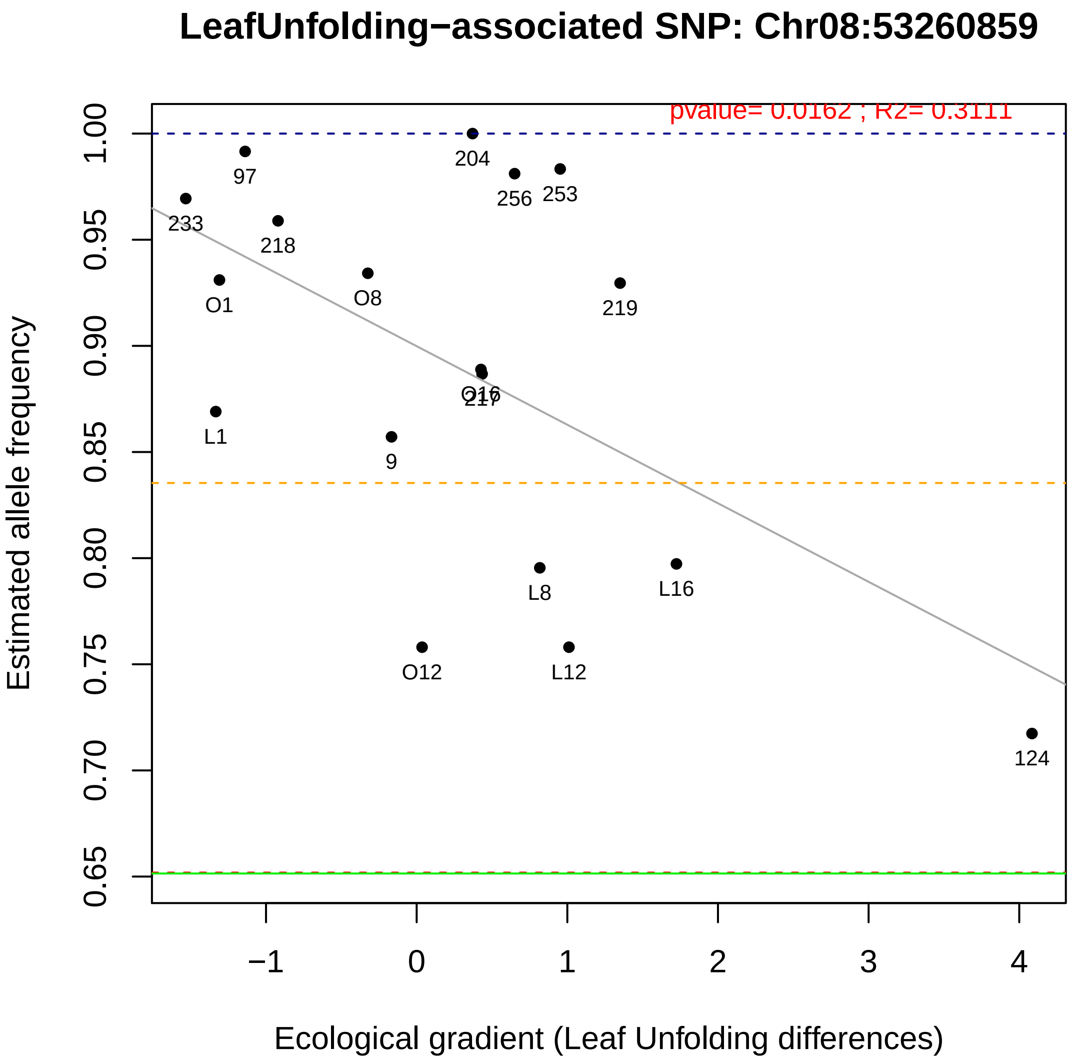

LeafUnfolding−associated SNP: Chr08:53260859
pvalue= 0.0162 ; R2= 0.3111
Estimated allele frequency
Ecological gradient (Leaf Unfolding differences)
1.00
0.95
0.90
0.85
0.80
0.75
0.70
0.65
−1
0
1
2
3
4
233
97
218
O1
L1
O8
9
204
256
253
219
O16
217
L8
L16
O12
L12
124

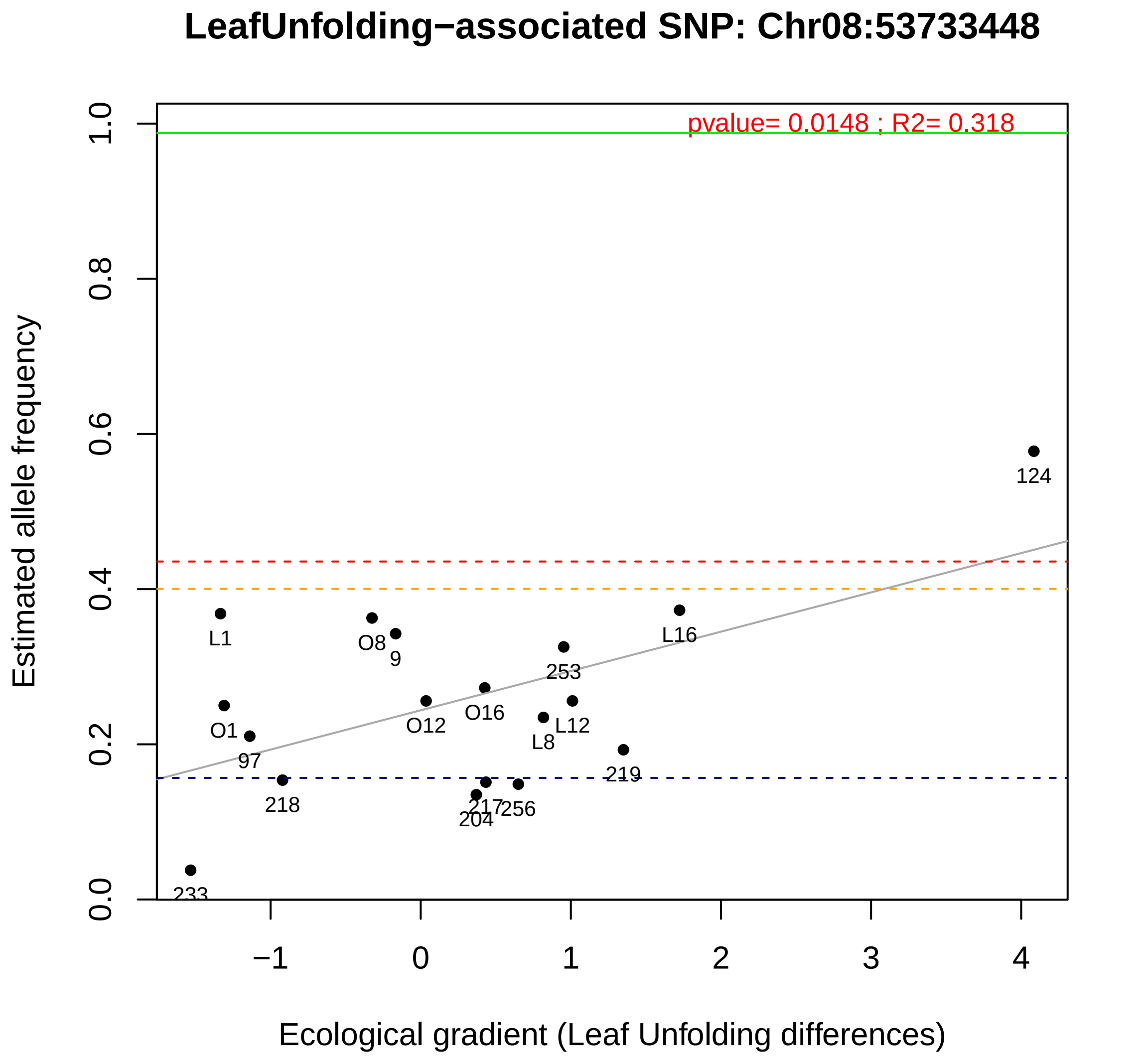
LeafUnfolding−associated SNP: Chr08:53733448
pvalue= 0.0148 ; R2= 0.318
Estimated allele frequency
0.0
0.2
0.4
0.6
0.8
1.0
−1
0
1
2
3
4
Ecological gradient (Leaf Unfolding differences)
L1
O1
97
218
233
O8
9
O12
O16
217
204
256
L8
L12
253
219
L16
124

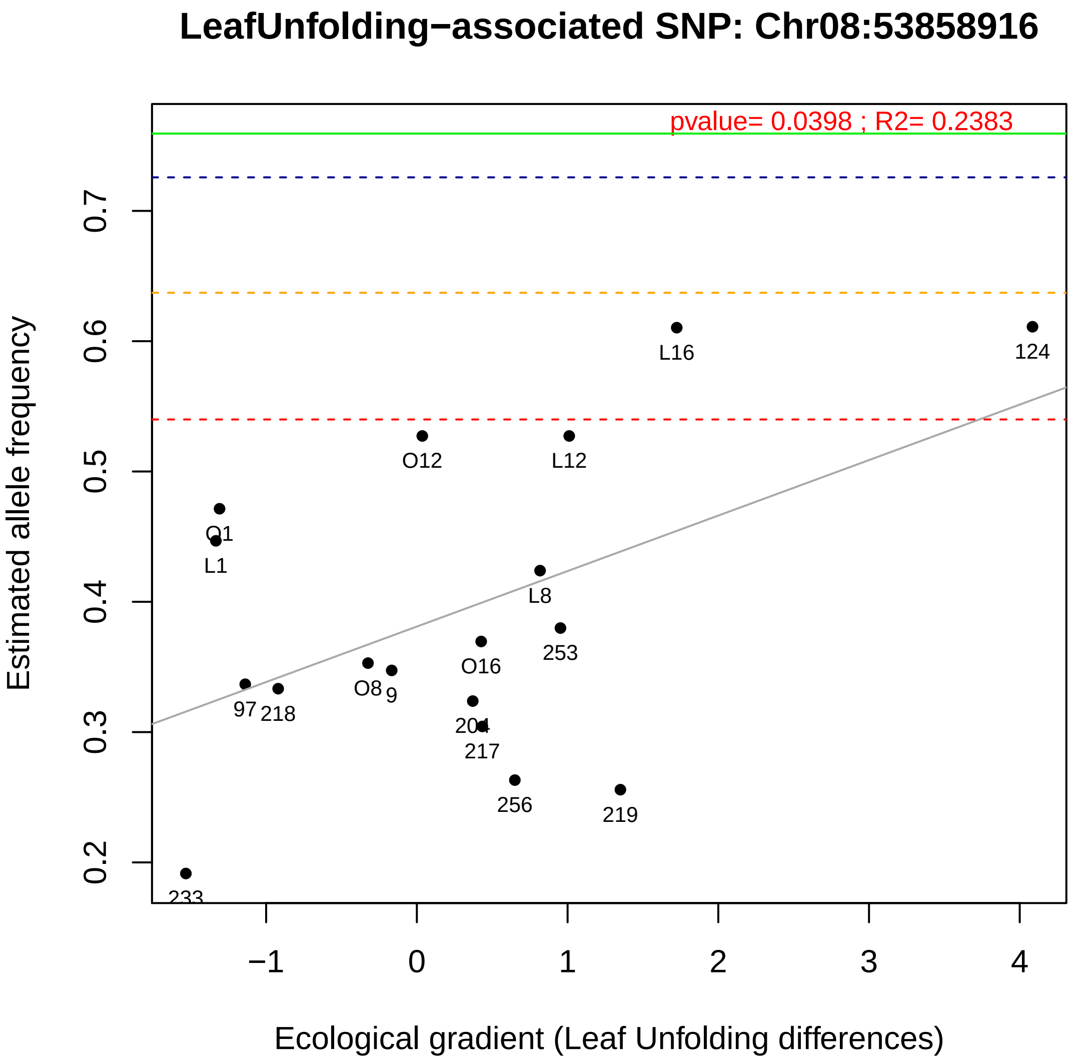

LeafUnfolding−associated SNP: Chr08:53858916
pvalue= 0.0398 ; R2= 0.2383
Estimated allele frequency
0.2
0.3
0.4
0.5
0.6
0.7
Ecological gradient (Leaf Unfolding differences)
−1
0
1
2
3
4
L16
124
O12
L12
O1
L1
L8
253
O16
O8
9
97
218
204
217
256
219
233

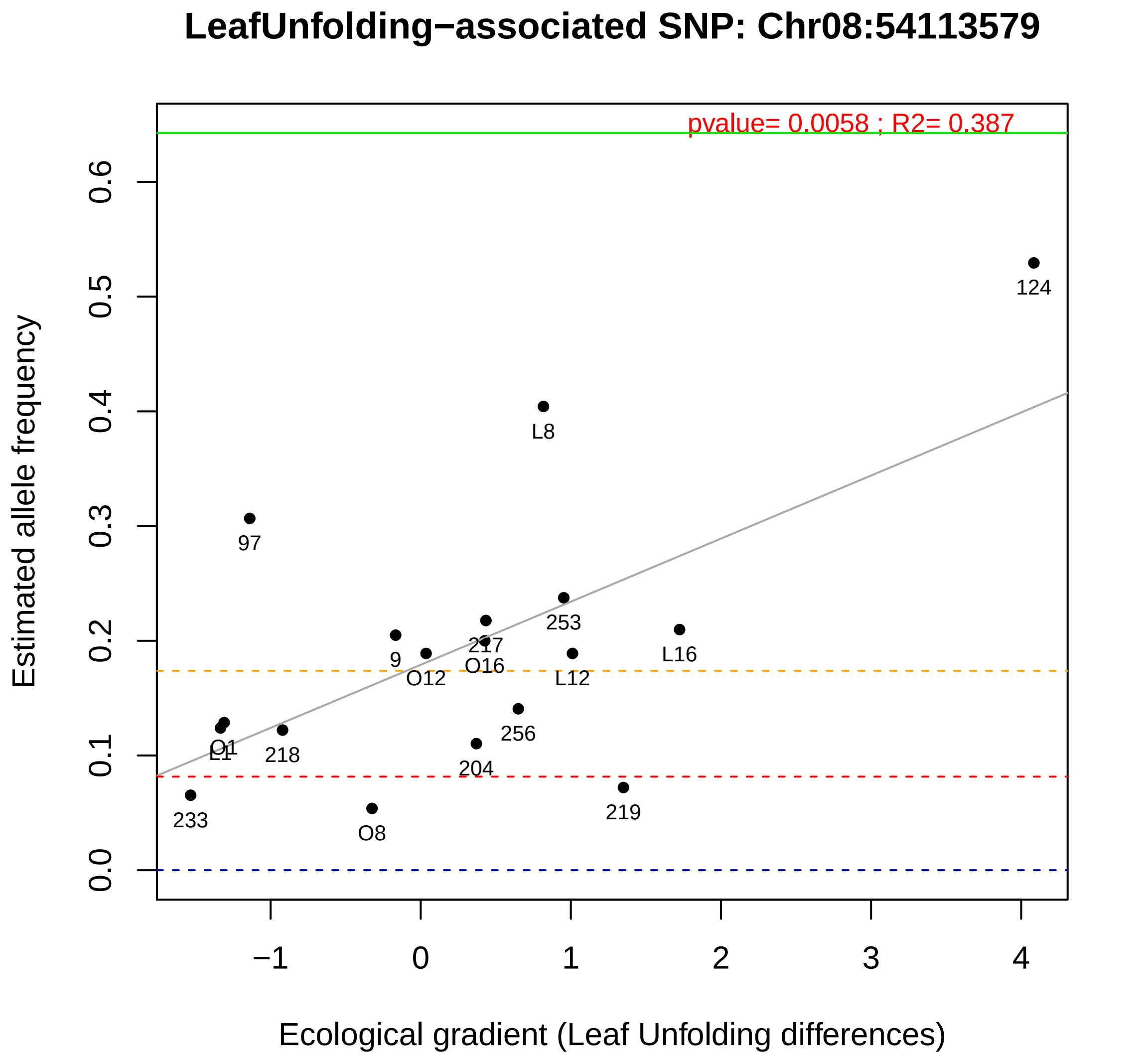
LeafUnfolding−associated SNP: Chr08:54113579
pvalue= 0.0058 ; R2= 0.387
Estimated allele frequency
0.0
0.1
0.2
0.3
0.4
0.5
0.6
−1
0
1
2
3
4
Ecological gradient (Leaf Unfolding differences)
124
L8
97
253
217
O16
9
O12
L12
L16
256
O1
L1
218
204
233
O8
219

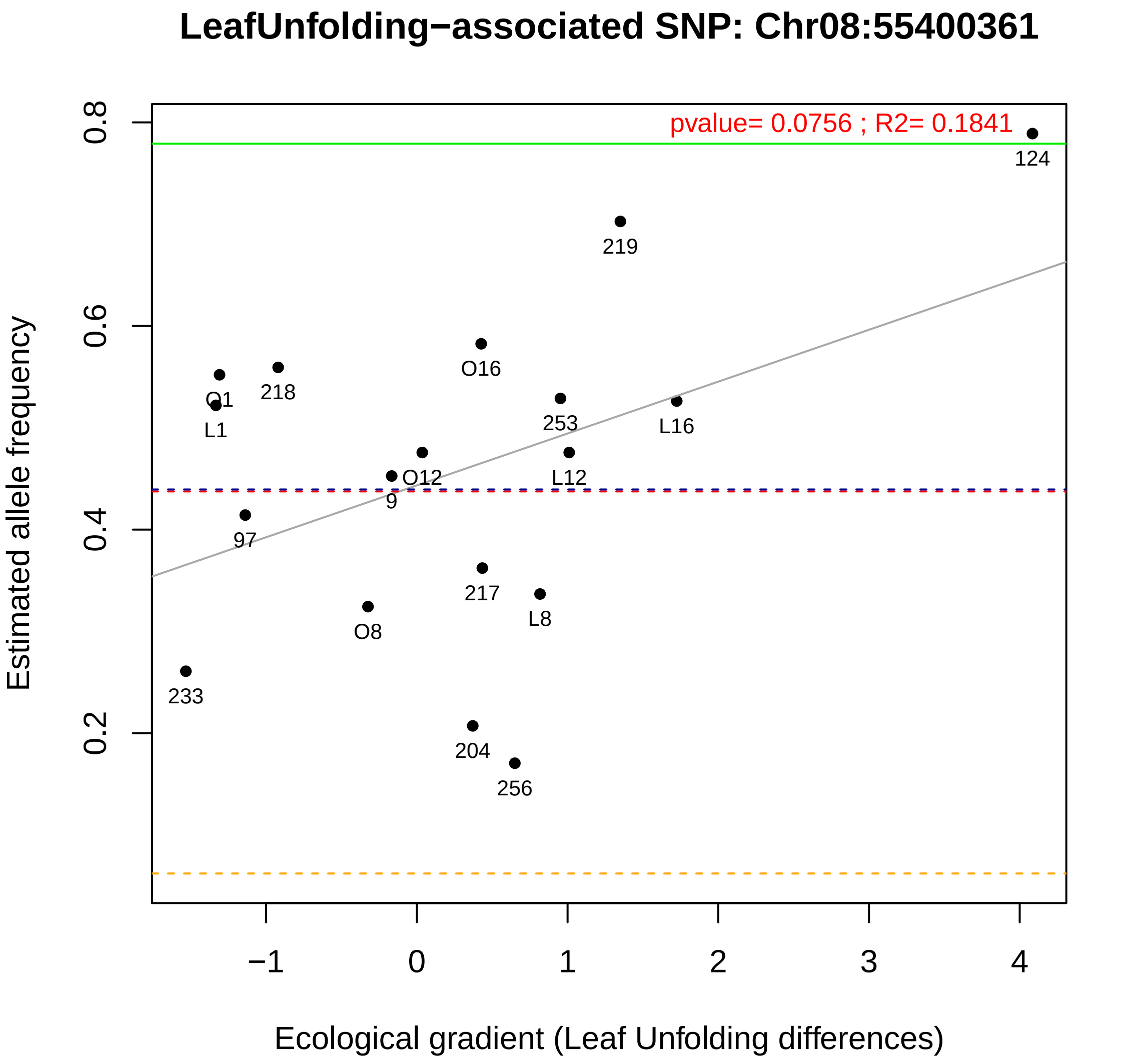

LeafUnfolding−associated SNP: Chr08:55400361
pvalue= 0.0756 ; R2= 0.1841
Estimated allele frequency
0.2
0.4
0.6
0.8
−1
0
1
2
3
4
Ecological gradient (Leaf Unfolding differences)
124
219
O16
218
O1
L1
253
L16
O12
9
L12
97
217
L8
O8
233
204
256

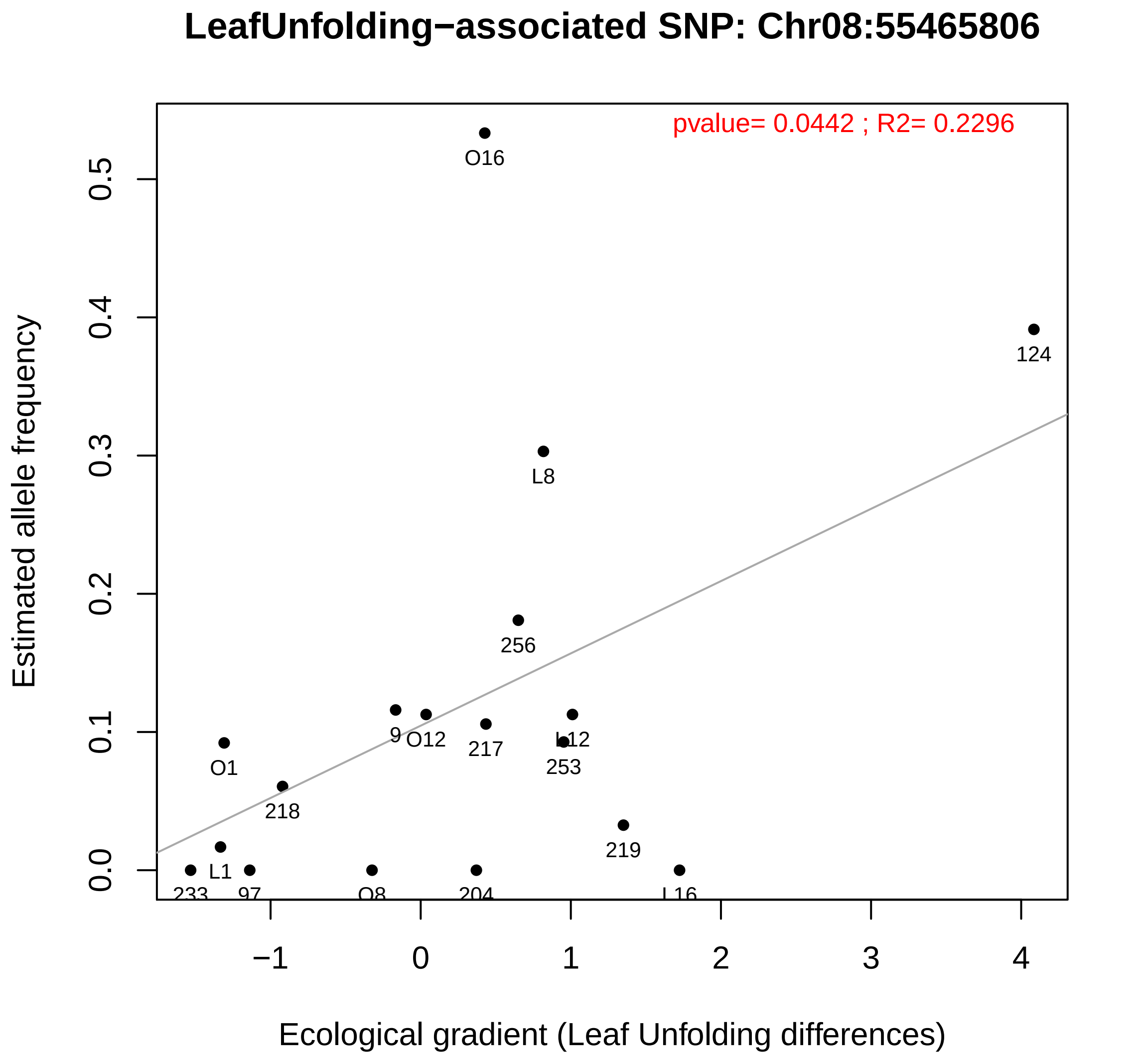

LeafUnfolding−associated SNP: Chr08:55465806
pvalue= 0.0442 ; R2= 0.2296
Estimated allele frequency
Ecological gradient (Leaf Unfolding differences)
0.0
0.1
0.2
0.3
0.4
0.5
−1
0
1
2
3
4
O16
124
L8
256
9
O12
217
L12
253
O1
218
219
L1
233
97
O8
204
L16

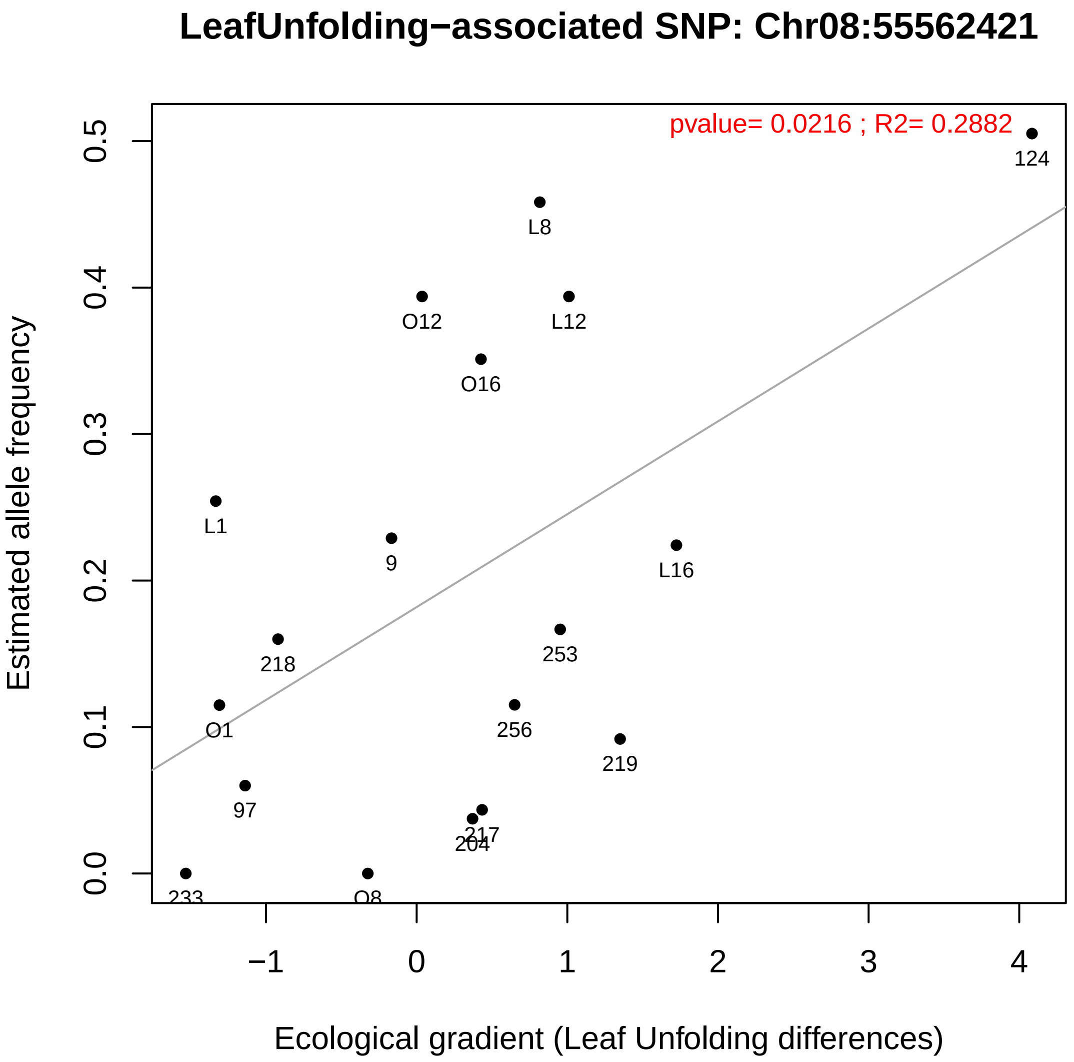
LeafUnfolding−associated SNP: Chr08:55562421
pvalue= 0.0216 ; R2= 0.2882
Estimated allele frequency
Ecological gradient (Leaf Unfolding differences)
0.0
0.1
0.2
0.3
0.4
0.5
−1
0
1
2
3
4
124
L8
O12
L12
O16
L1
9
L16
218
253
O1
256
219
97
217
204
233
O8

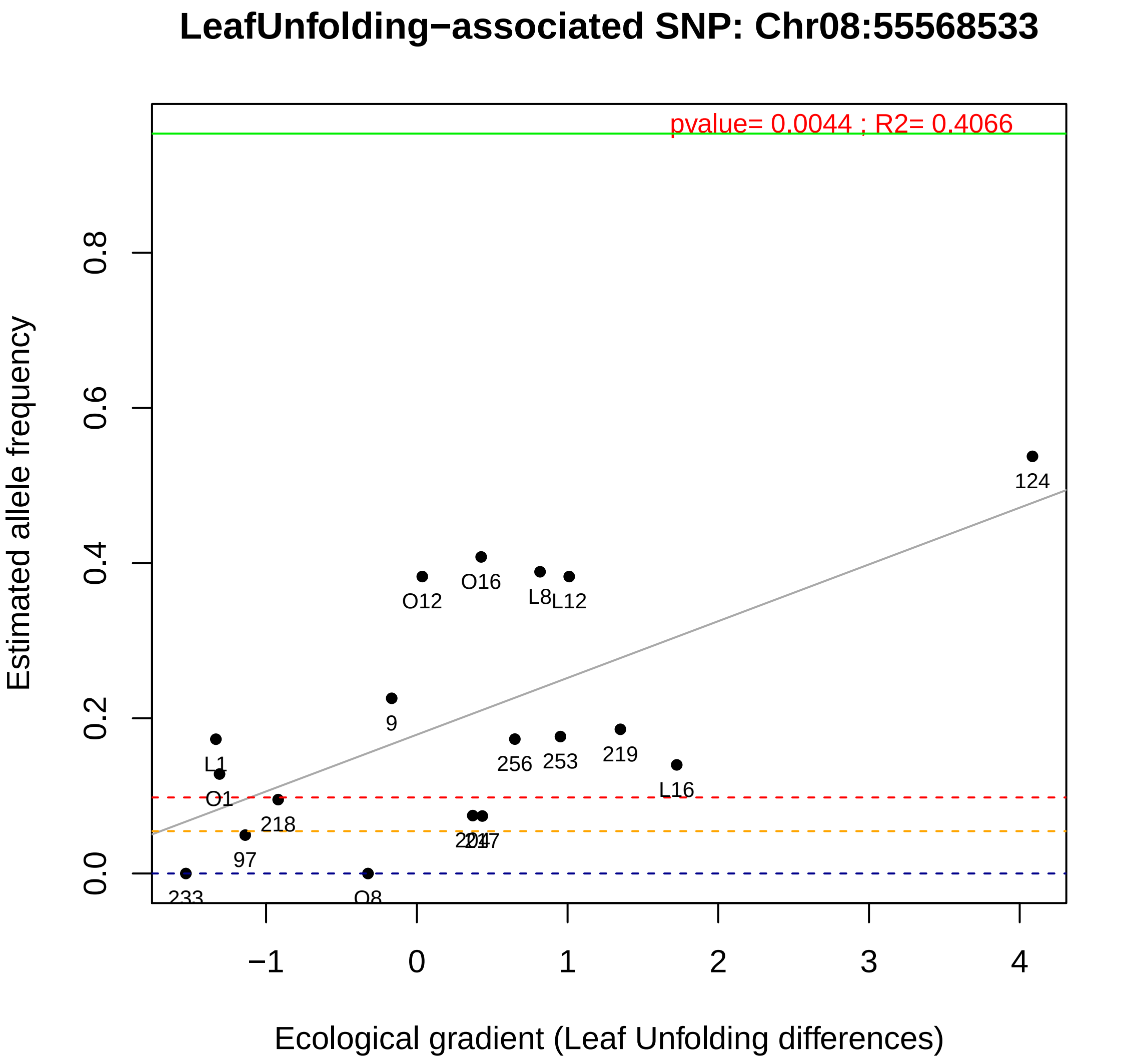

LeafUnfolding−associated SNP: Chr08:55568533
pvalue= 0.0044 ; R2= 0.4066
Estimated allele frequency
0.0
0.2
0.4
0.6
0.8
−1
0
1
2
3
4
Ecological gradient (Leaf Unfolding differences)
124
O16
O12
L8
L12
9
L1
O1
218
97
256
253
219
L16

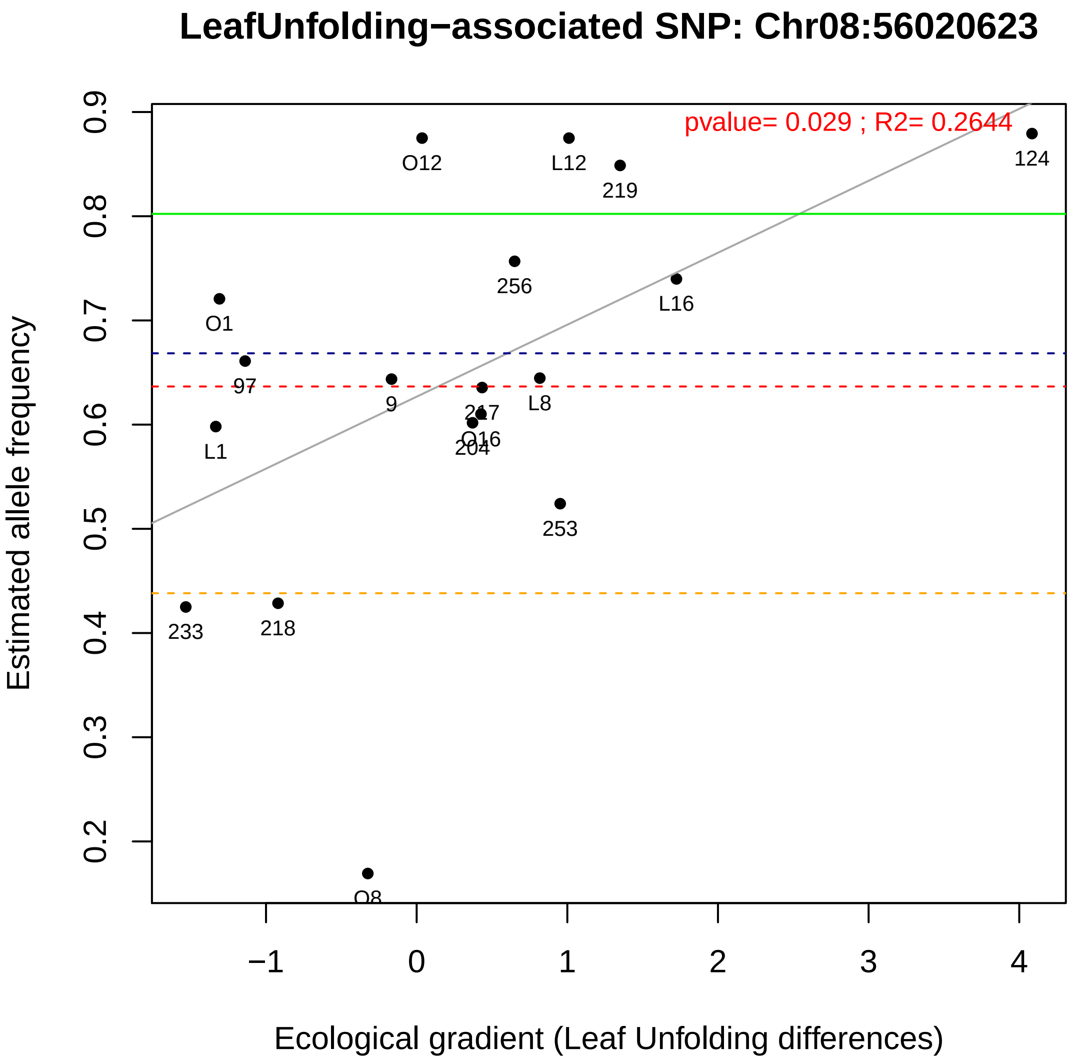

LeafUnfolding−associated SNP: Chr08:56020623
pvalue= 0.029 ; R2= 0.2644
Estimated allele frequency
Ecological gradient (Leaf Unfolding differences)
0.2
0.3
0.4
0.5
0.6
0.7
0.8
0.9
−1
0
1
2
3
4
O12
L12
219
124
256
L16
O1
97
9
217
L8
O16
204
L1
253
233
218
O8

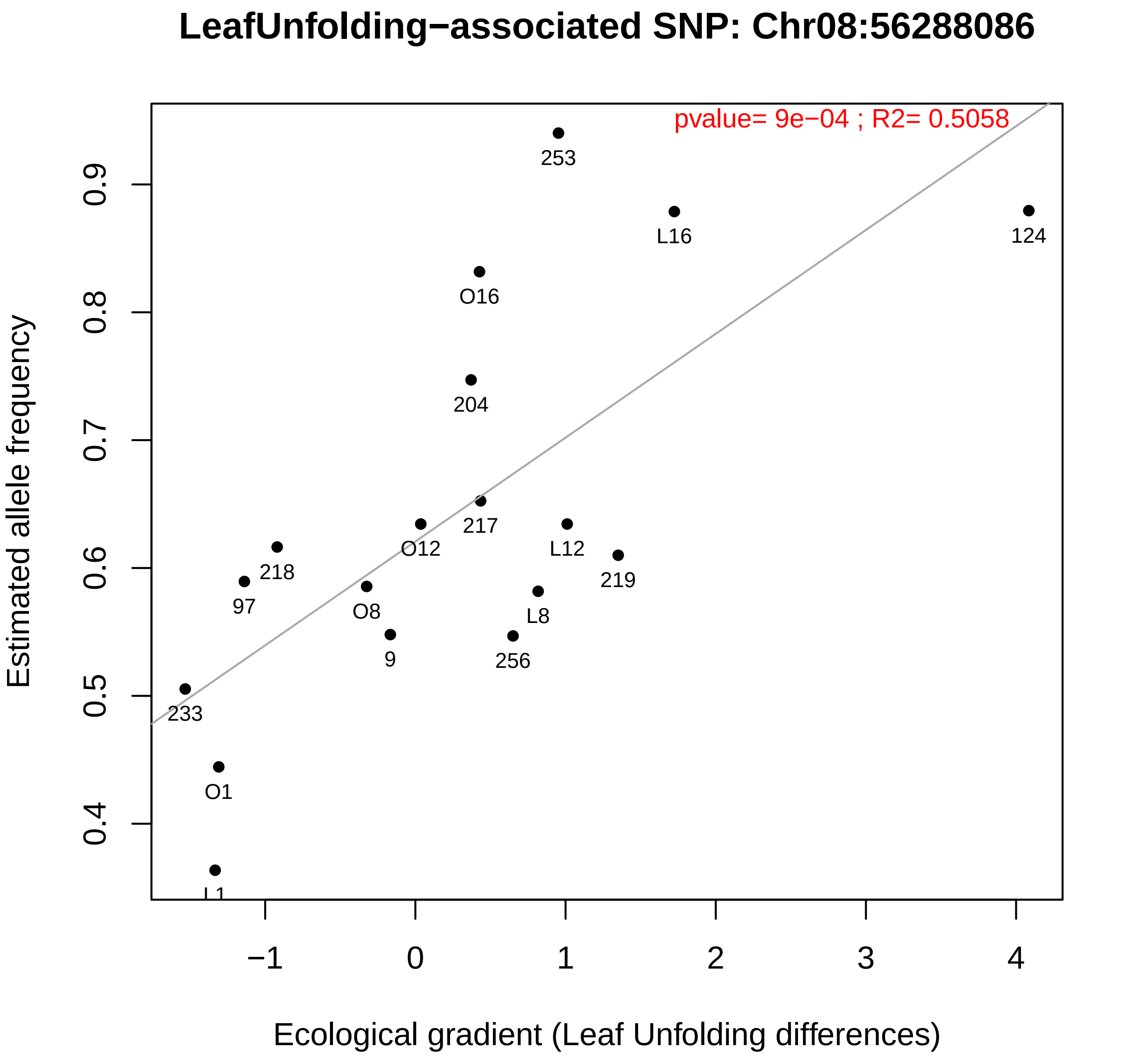
LeafUnfolding−associated SNP: Chr08:56288086
pvalue= 9e−04 ; R2= 0.5058
253
L16
124
O16
204
217
O12
L12
218
219
97
O8
L8
9
256
233
O1
L1
0.4
0.5
0.6
0.7
0.8
0.9
−1
0
1
2
3
4
Estimated allele frequency
Ecological gradient (Leaf Unfolding differences)

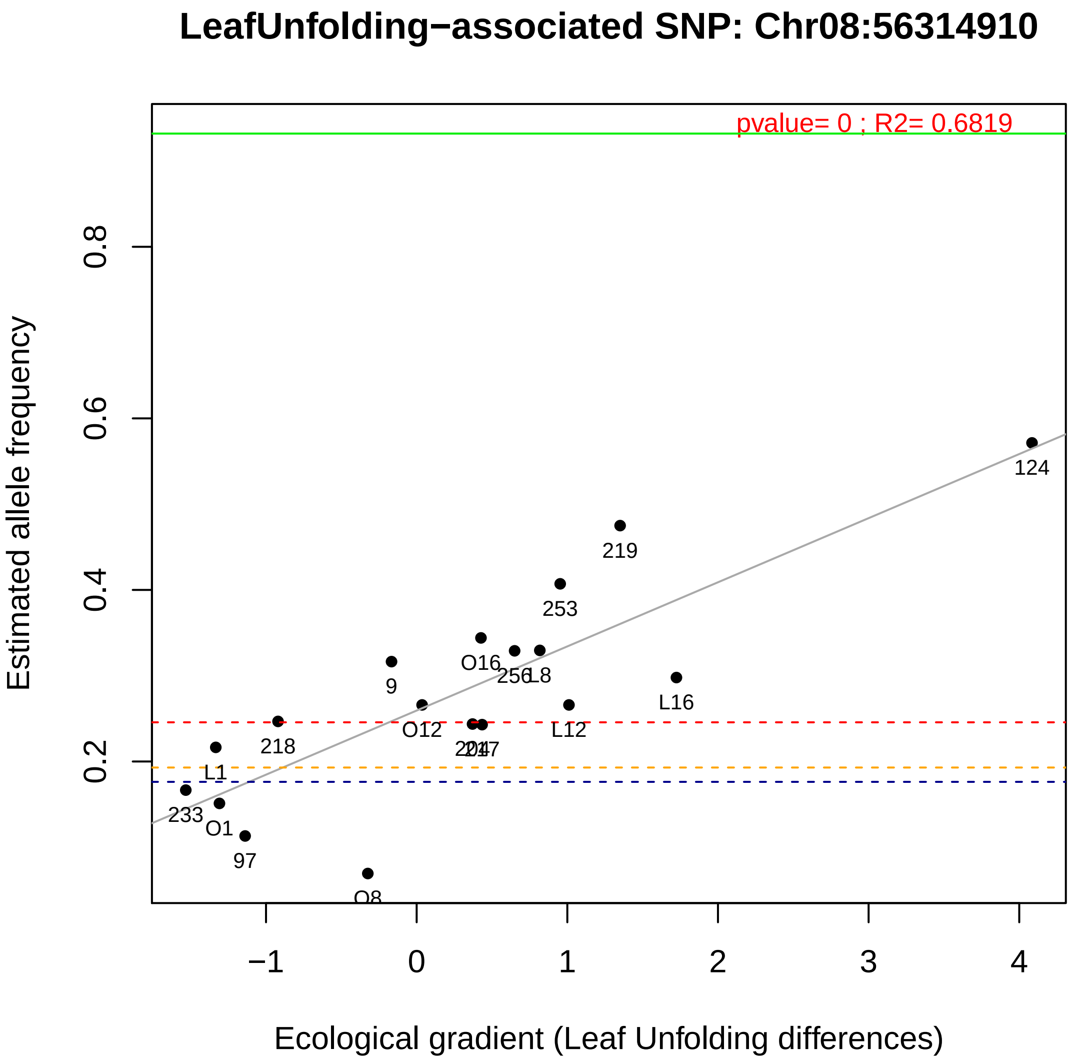
LeafUnfolding−associated SNP: Chr08:56314910
pvalue= 0 ; R2= 0.6819
Estimated allele frequency
0.2
0.4
0.6
0.8
Ecological gradient (Leaf Unfolding differences)
−1
0
1
2
3
4
124
219
253
O16
256
L8
9
L16
O12
L12
218
L1
233
O1
97
O8

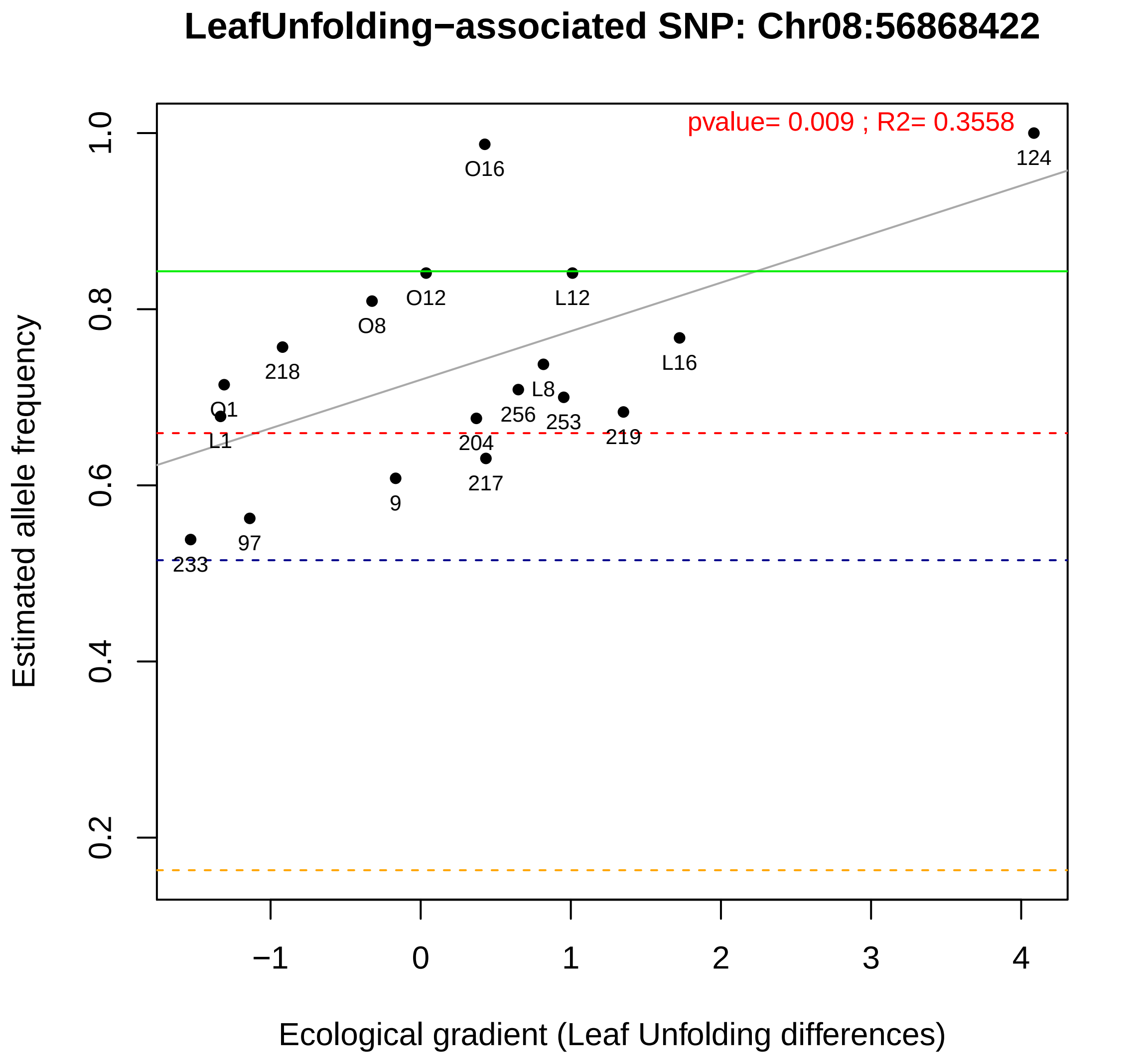

LeafUnfolding−associated SNP: Chr08:56868422
pvalue= 0.009 ; R2= 0.3558
Estimated allele frequency
Ecological gradient (Leaf Unfolding differences)
0.2
0.4
0.6
0.8
1.0
−1
0
1
2
3
4
124
O16
O12
L12
O8
218
L16
O1
L8
256
253
L1
204
219
217
9
97
233

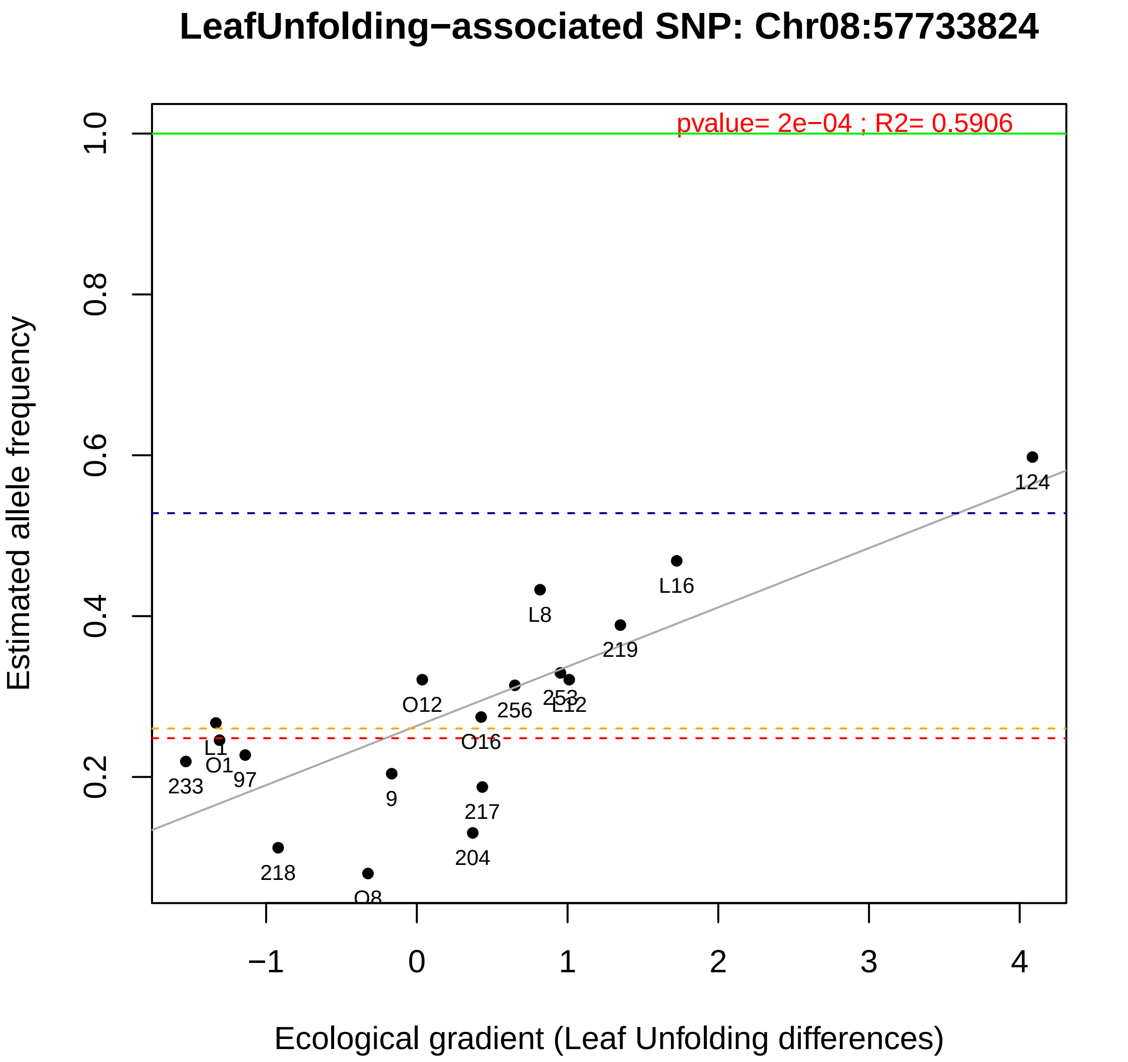

LeafUnfolding−associated SNP: Chr08:57733824
pvalue= 2e−04 ; R2= 0.5906
Estimated allele frequency
Ecological gradient (Leaf Unfolding differences)
1.0
0.8
0.6
0.4
0.2
−1
0
1
2
3
4
124
L16
L8
219
253
L12
256
O12
O16
L1
O1
97
233
9
217
204
218
O8

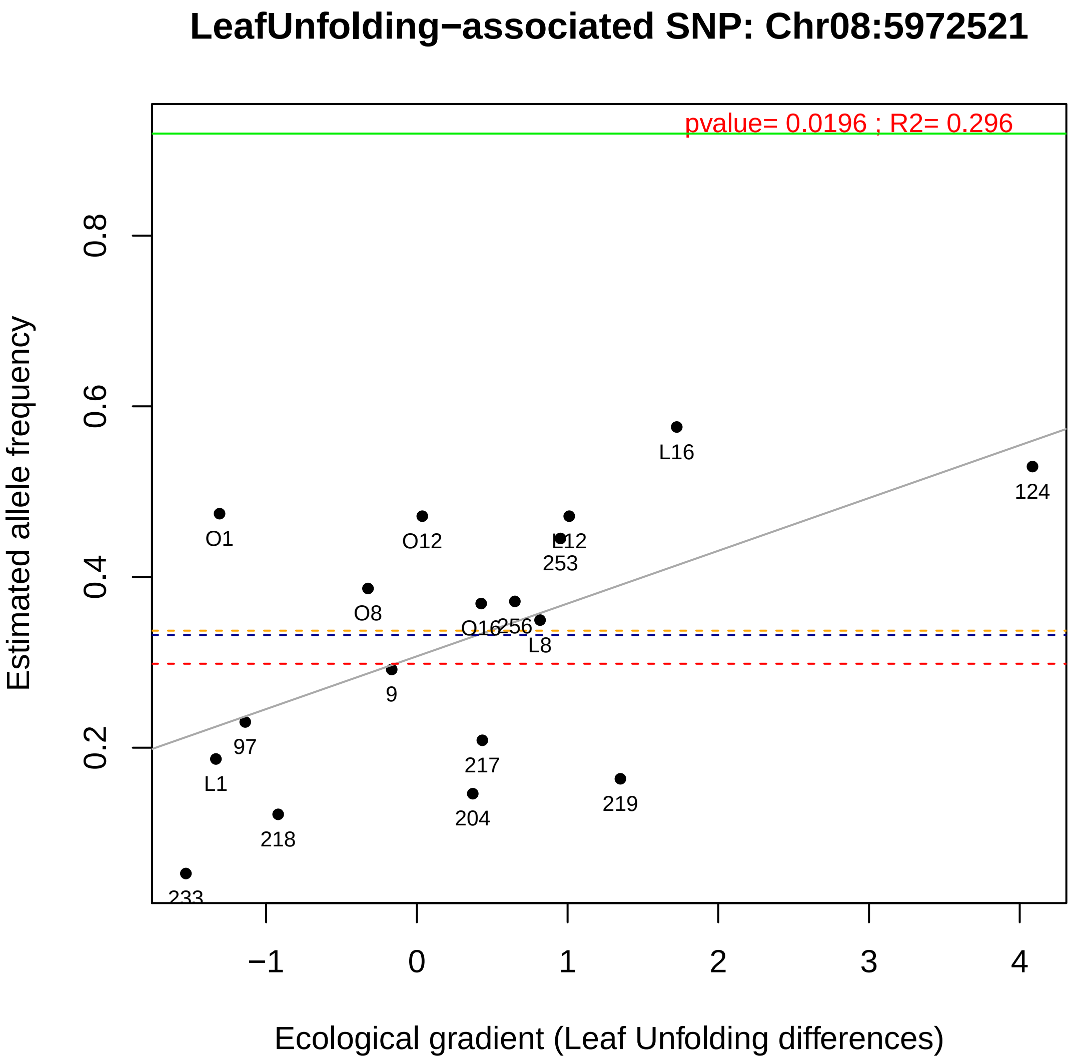
LeafUnfolding−associated SNP: Chr08:5972521
pvalue= 0.0196 ; R2= 0.296
Estimated allele frequency
0.2
0.4
0.6
0.8
−1
0
1
2
3
4
Ecological gradient (Leaf Unfolding differences)
L16
124
O1
O12
L12
253
O8
O16
256
L8
9
97
L1
217
219
204
218
233

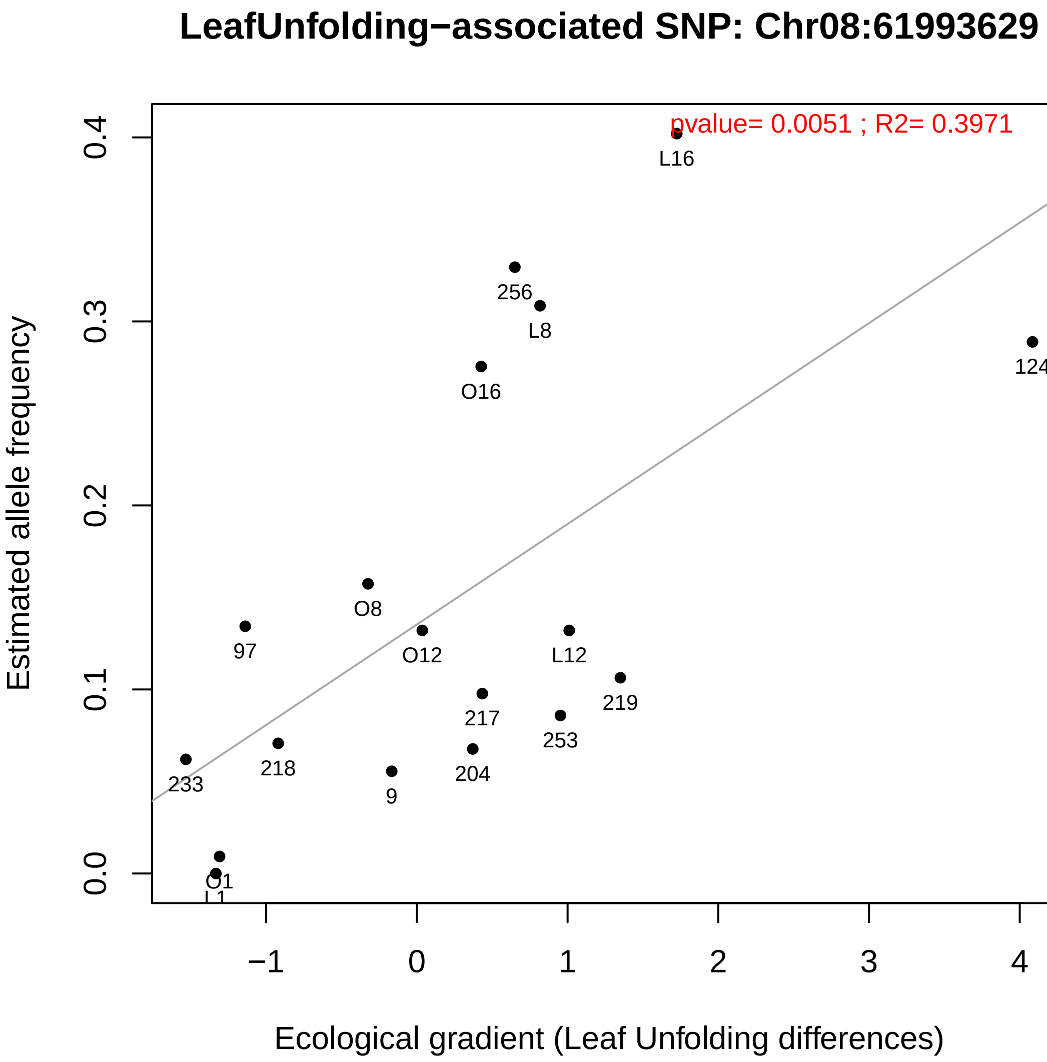
LeafUnfolding−associated SNP: Chr08:61993629
pvalue= 0.0051 ; R2= 0.3971
Estimated allele frequency
0.0
0.1
0.2
0.3
0.4
−1
0
1
2
3
4
Ecological gradient (Leaf Unfolding differences)
L16
256
L8
124
O16
O8
97
O12
L12
219
217
253
233
218
204
9
O1
L1

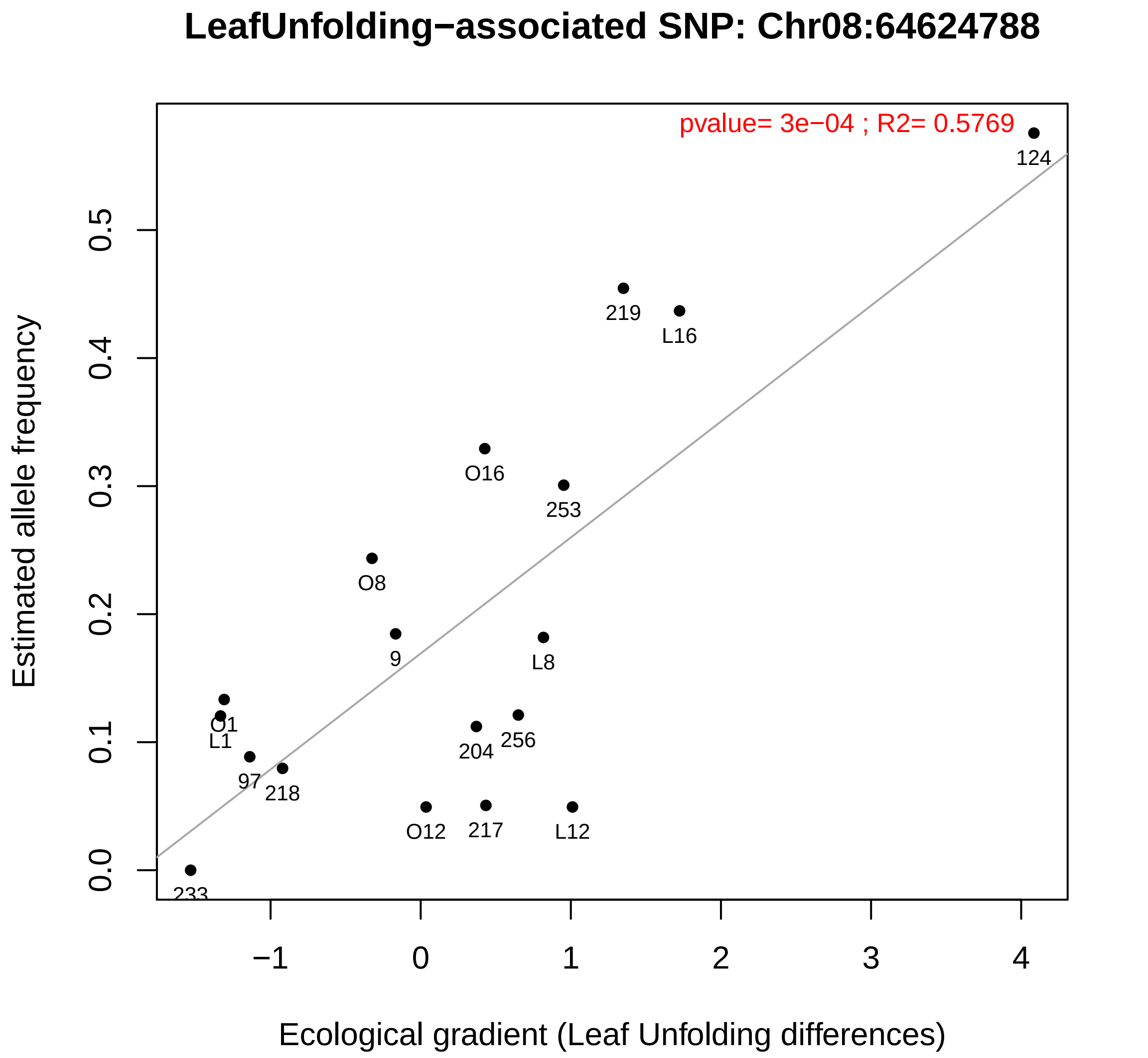

LeafUnfolding−associated SNP: Chr08:64624788
pvalue= 3e−04 ; R2= 0.5769
Estimated allele frequency
Ecological gradient (Leaf Unfolding differences)
0.0
0.1
0.2
0.3
0.4
0.5
−1
0
1
2
3
4
124
219
L16
O16
253
O8
9
L8
O1
L1
97
218
204
256
O12
217
L12
233

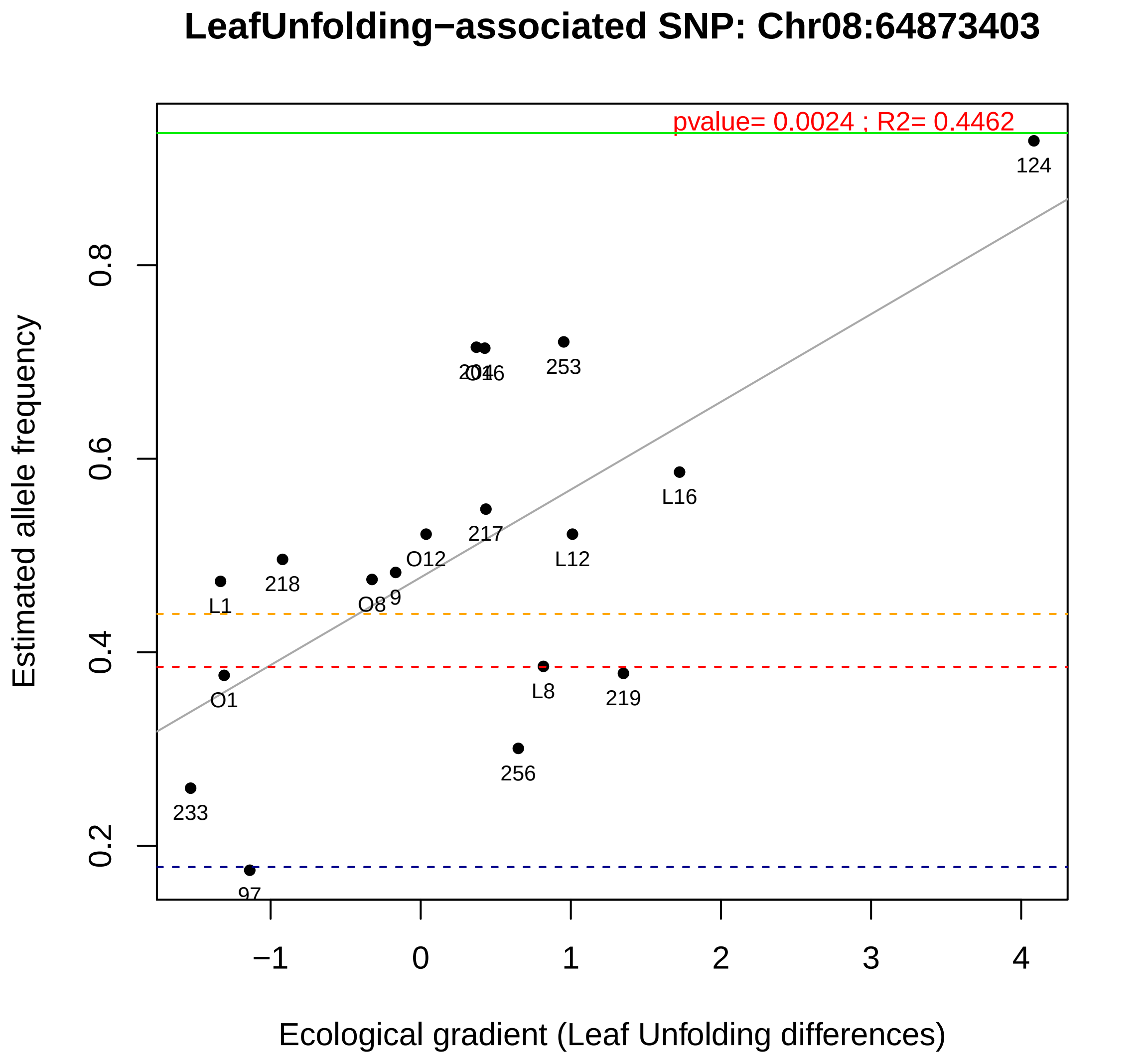
LeafUnfolding−associated SNP: Chr08:64873403
pvalue= 0.0024 ; R2= 0.4462
124
204
O16
253
L16
217
L12
O12
218
L1
O8
9
O1
L8
219
256
233
97
0.2
0.4
0.6
0.8
−1
0
1
2
3
4
Estimated allele frequency
Ecological gradient (Leaf Unfolding differences)

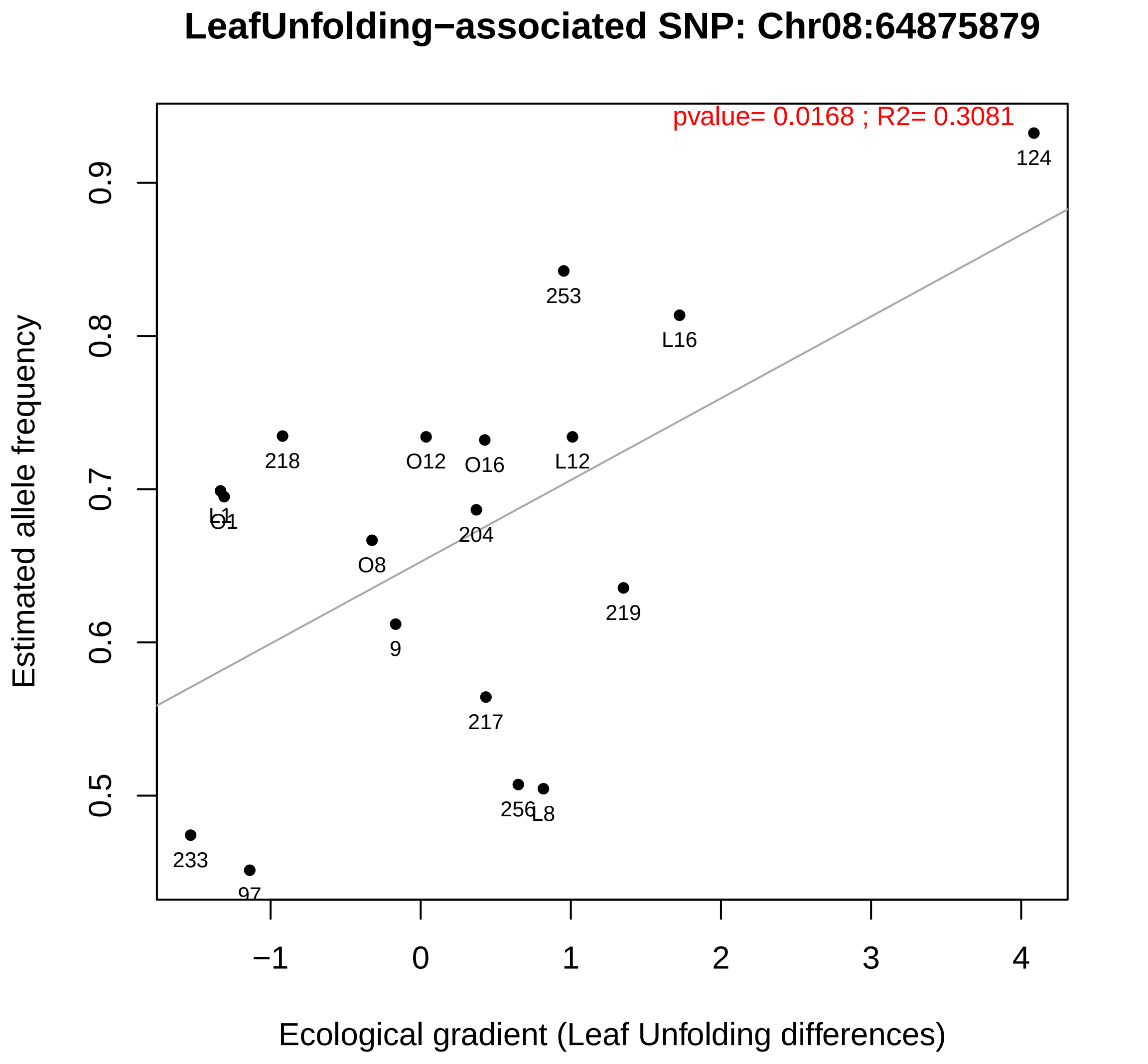
LeafUnfolding−associated SNP: Chr08:64875879
pvalue= 0.0168 ; R2= 0.3081
Estimated allele frequency
0.5
0.6
0.7
0.8
0.9
−1
0
1
2
3
4
Ecological gradient (Leaf Unfolding differences)
124
253
L16
218
O12
O16
L12
L1
O1
204
O8
219
9
217
256
L8
233
97

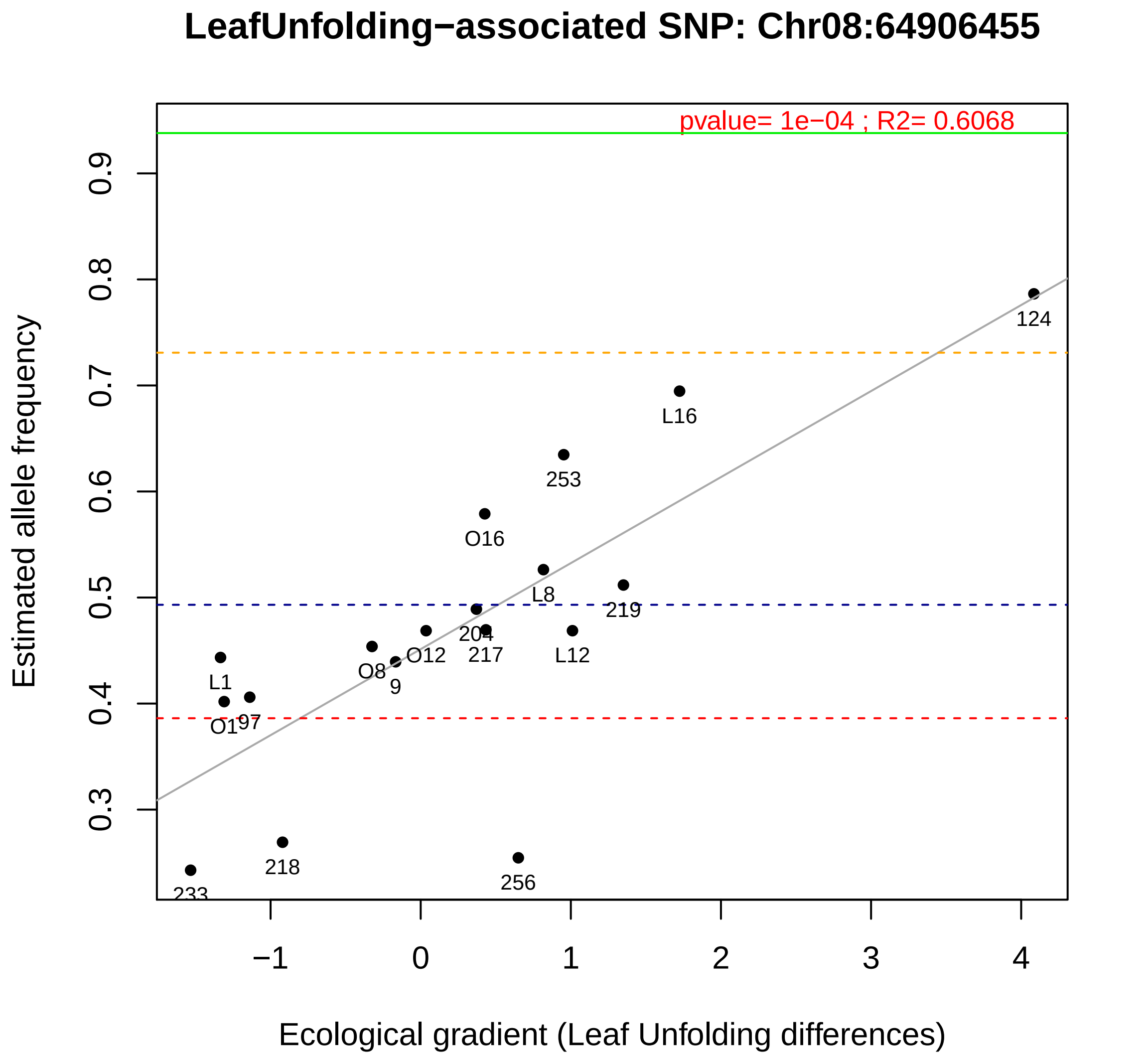
LeafUnfolding−associated SNP: Chr08:64906455
pvalue= 1e−04 ; R2= 0.6068
Estimated allele frequency
0.3
0.4
0.5
0.6
0.7
0.8
0.9
−1
0
1
2
3
4
Ecological gradient (Leaf Unfolding differences)
124
L16
253
O16
L8
219
204
217
L12
O12
O8
9
L1
O1
97
218
233
256

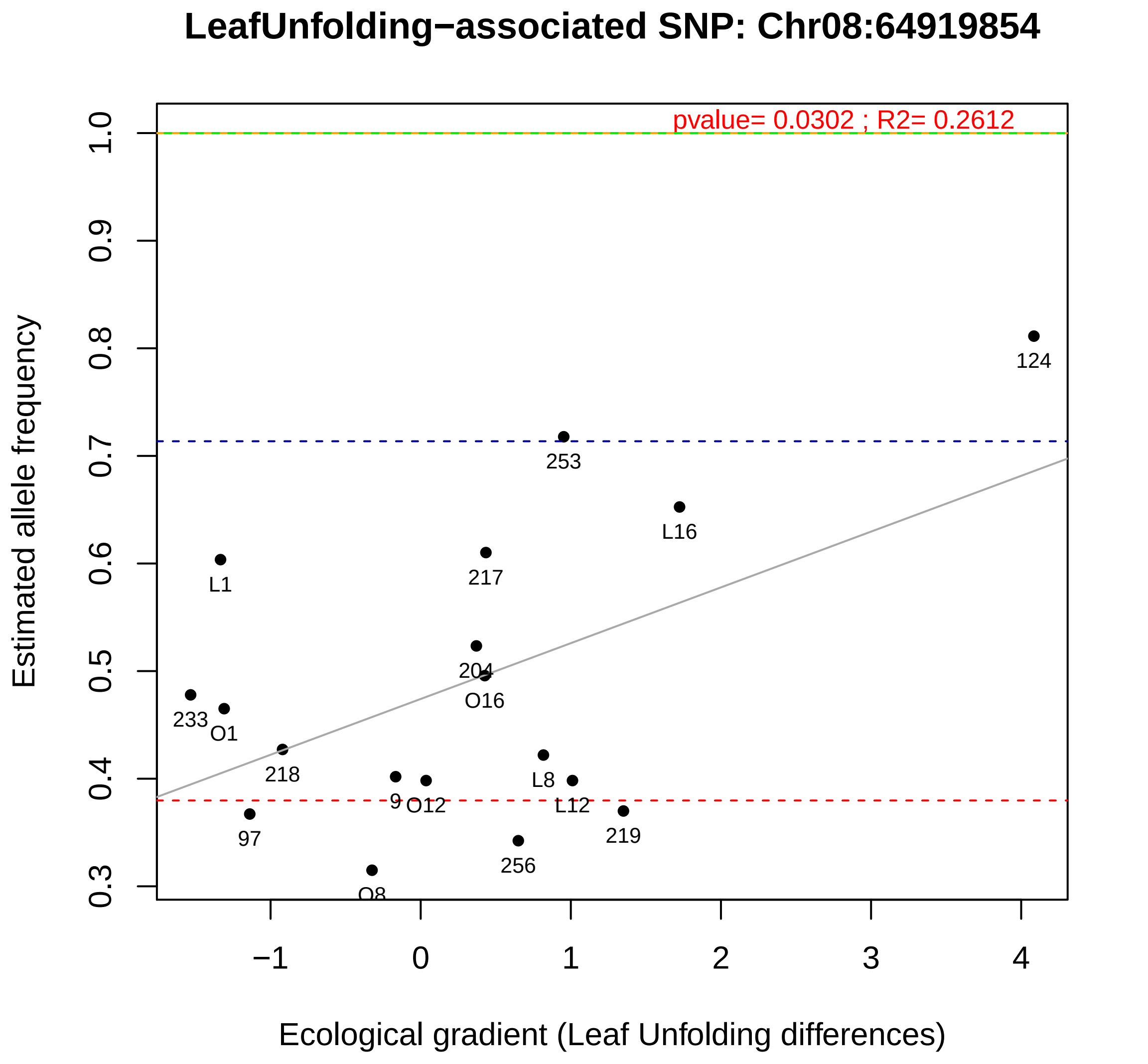

LeafUnfolding−associated SNP: Chr08:64919854
pvalue= 0.0302 ; R2= 0.2612
Estimated allele frequency
Ecological gradient (Leaf Unfolding differences)
0.3
0.4
0.5
0.6
0.7
0.8
0.9
1.0
−1
0
1
2
3
4
124
253
L16
L1
217
204
O16
233
O1
218
L8
9
O12
L12
97
219
256
O8

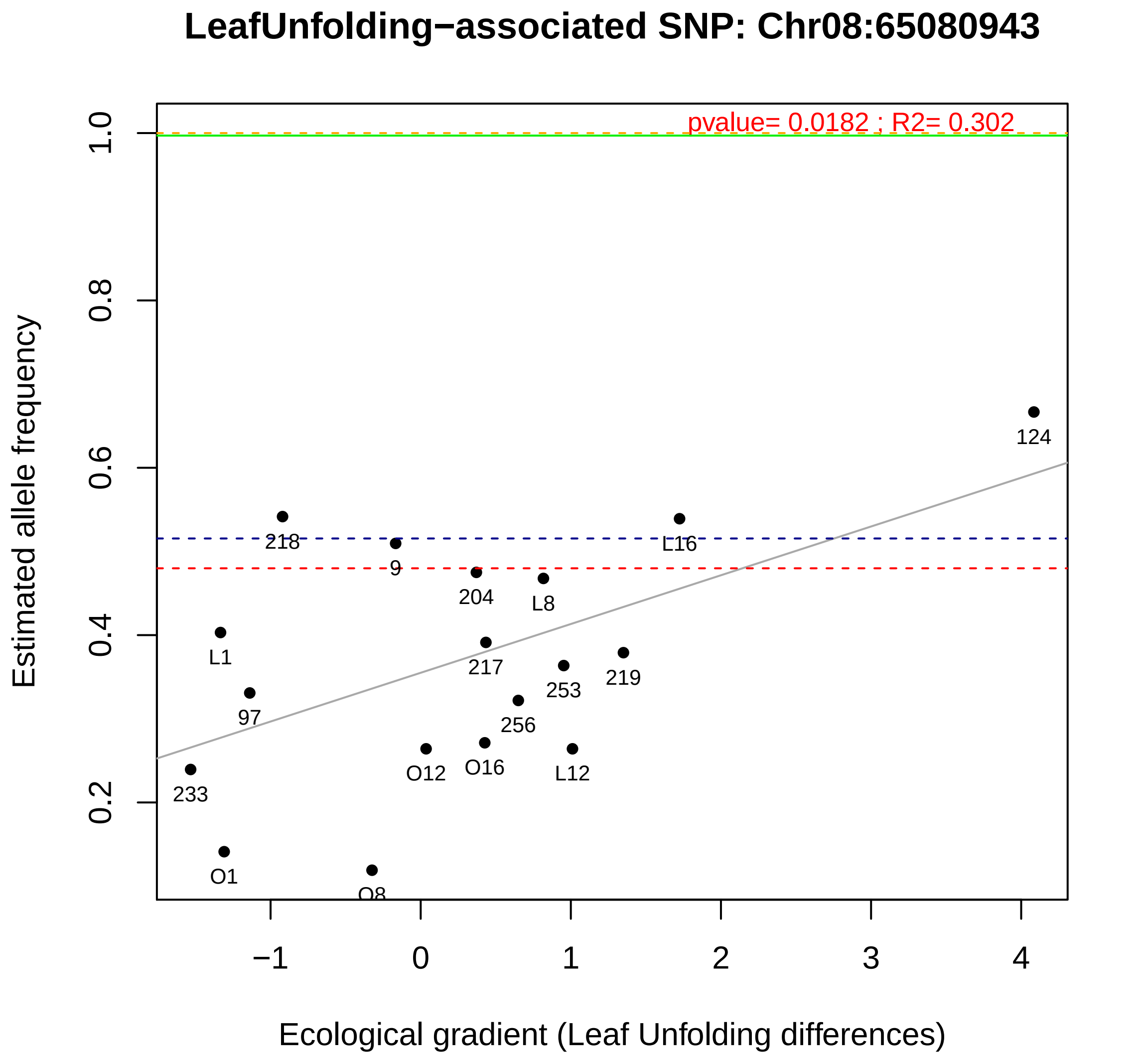

LeafUnfolding−associated SNP: Chr08:65080943
pvalue= 0.0182 ; R2= 0.302
Estimated allele frequency
1.0
0.8
0.6
0.4
0.2
124
218
L16
9
204
L8
L1
217
219
253
97
256
O12
O16
L12
233
O1
O8
−1
0
1
2
3
4
Ecological gradient (Leaf Unfolding differences)

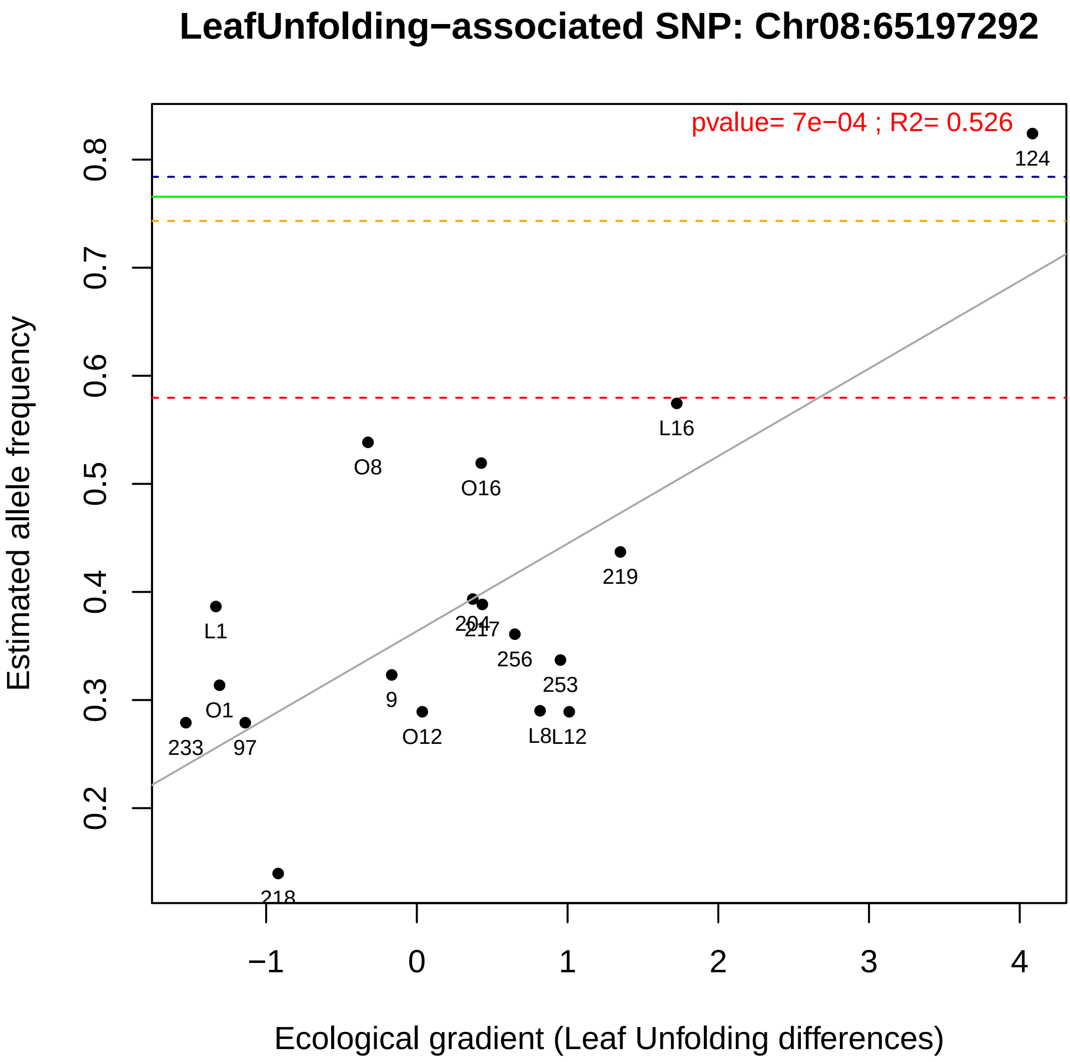

LeafUnfolding−associated SNP: Chr08:65197292
pvalue= 7e−04 ; R2= 0.526
Estimated allele frequency
Ecological gradient (Leaf Unfolding differences)
0.2
0.3
0.4
0.5
0.6
0.7
0.8
−1
0
1
2
3
4
124
L16
O8
O16
219
L1
204
217
256
253
9
O1
233
97
O12
L8
L12
218

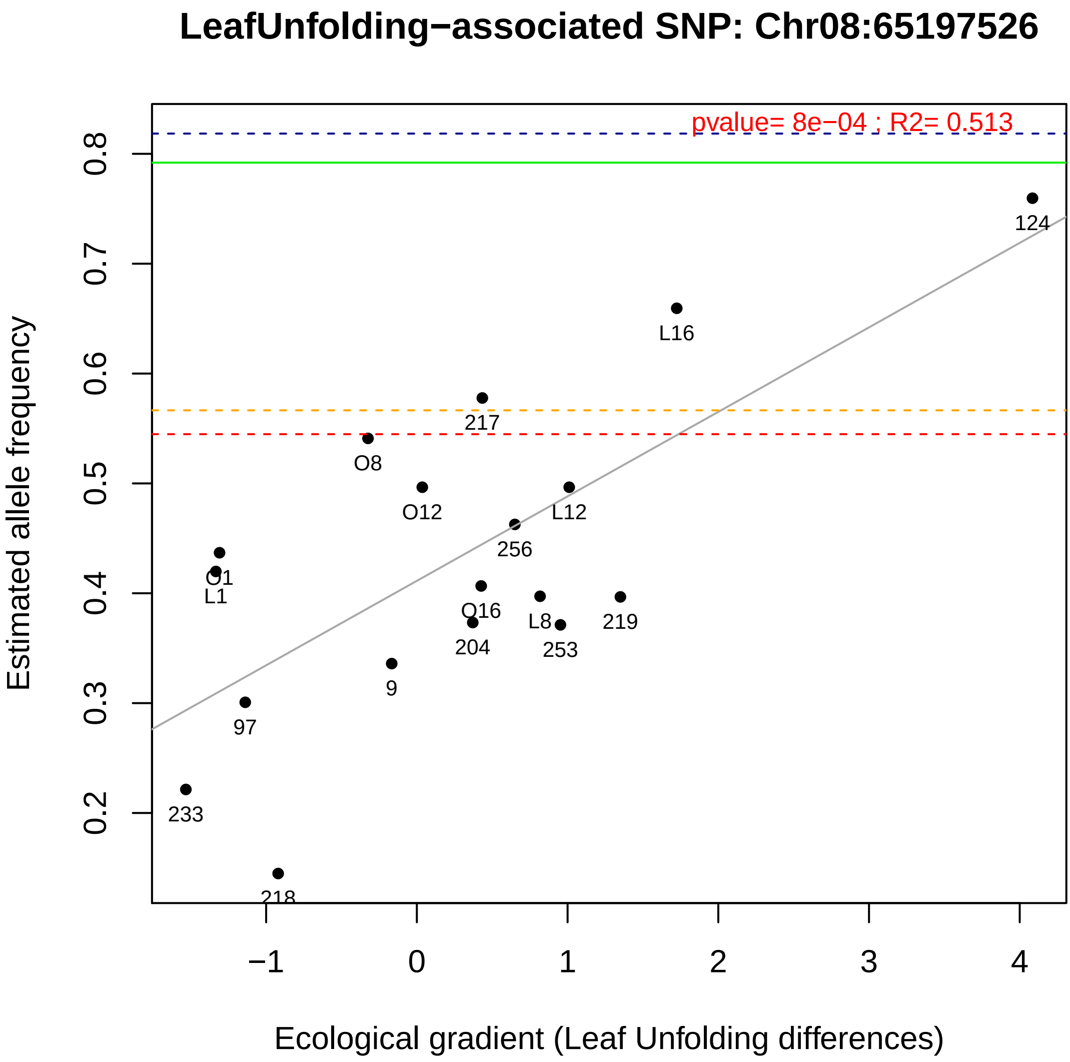

LeafUnfolding−associated SNP: Chr08:65197526
pvalue= 8e−04 ; R2= 0.513
Estimated allele frequency
Ecological gradient (Leaf Unfolding differences)
0.2
0.3
0.4
0.5
0.6
0.7
0.8
−1
0
1
2
3
4
124
L16
217
O8
O12
L12
256
O1
L1
O16
L8
219
204
253
9
97
233
218

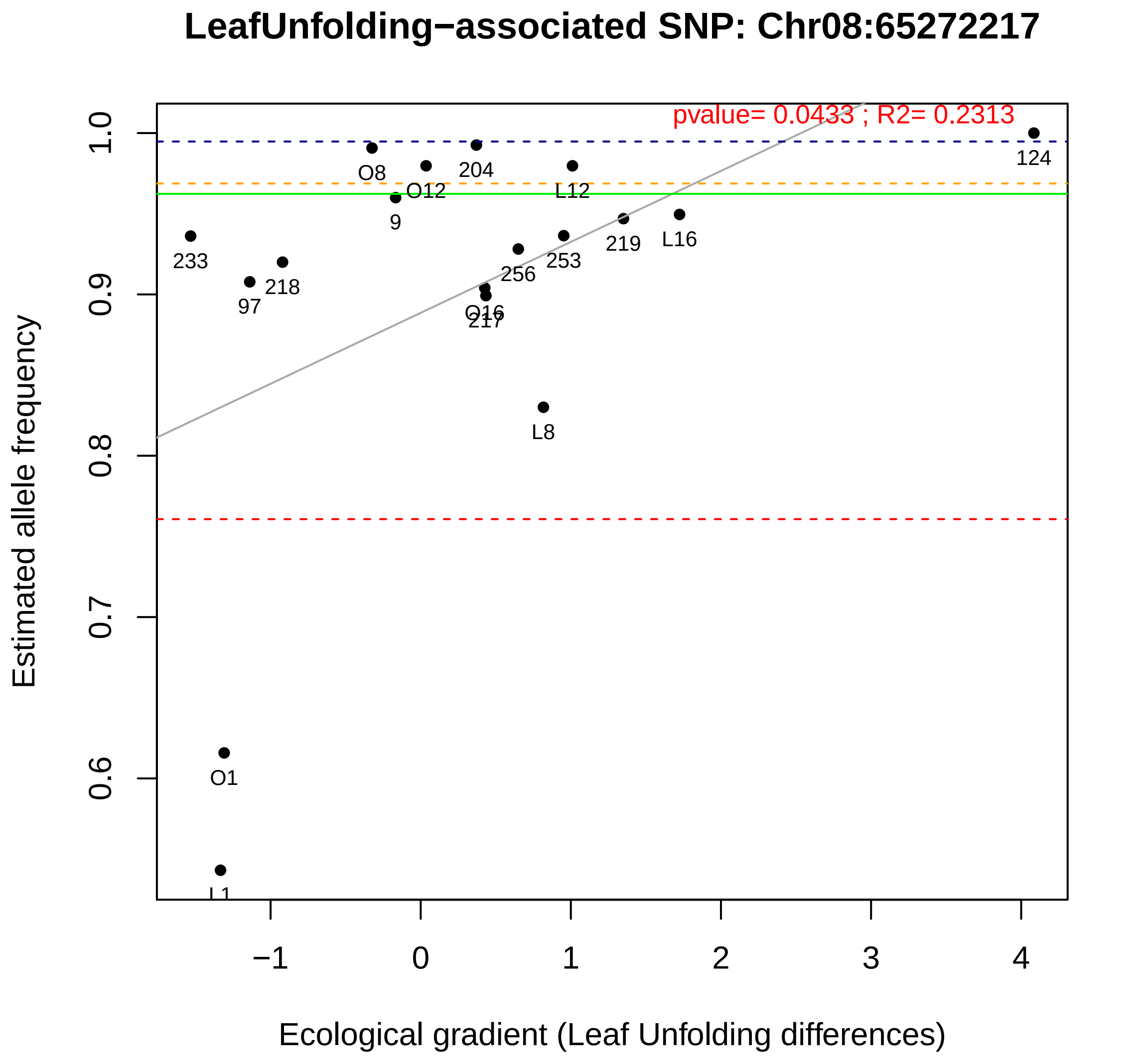
LeafUnfolding−associated SNP: Chr08:65272217
pvalue= 0.0433 ; R2= 0.2313
Estimated allele frequency
Ecological gradient (Leaf Unfolding differences)
1.0
0.9
0.8
0.7
0.6
−1
0
1
2
3
4
O8
204
O12
L12
9
124
219
L16
233
253
256
218
97
O16
217
L8
O1
L1

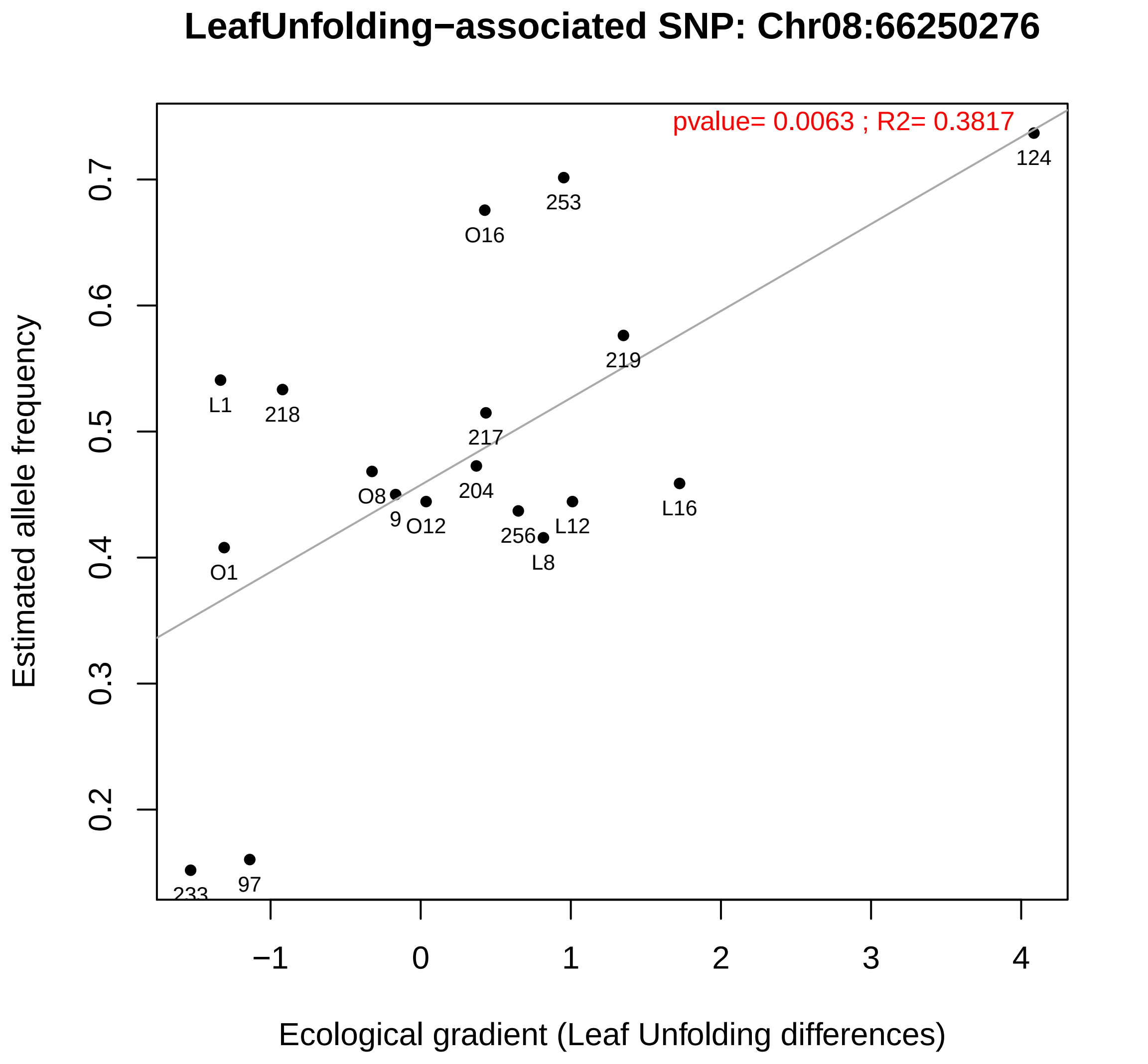

LeafUnfolding−associated SNP: Chr08:66250276
pvalue= 0.0063 ; R2= 0.3817
Estimated allele frequency
Ecological gradient (Leaf Unfolding differences)
0.2
0.3
0.4
0.5
0.6
0.7
−1
0
1
2
3
4
124
253
O16
219
L1
218
217
204
O8
9
O12
256
L12
L8
L16
O1
233
97

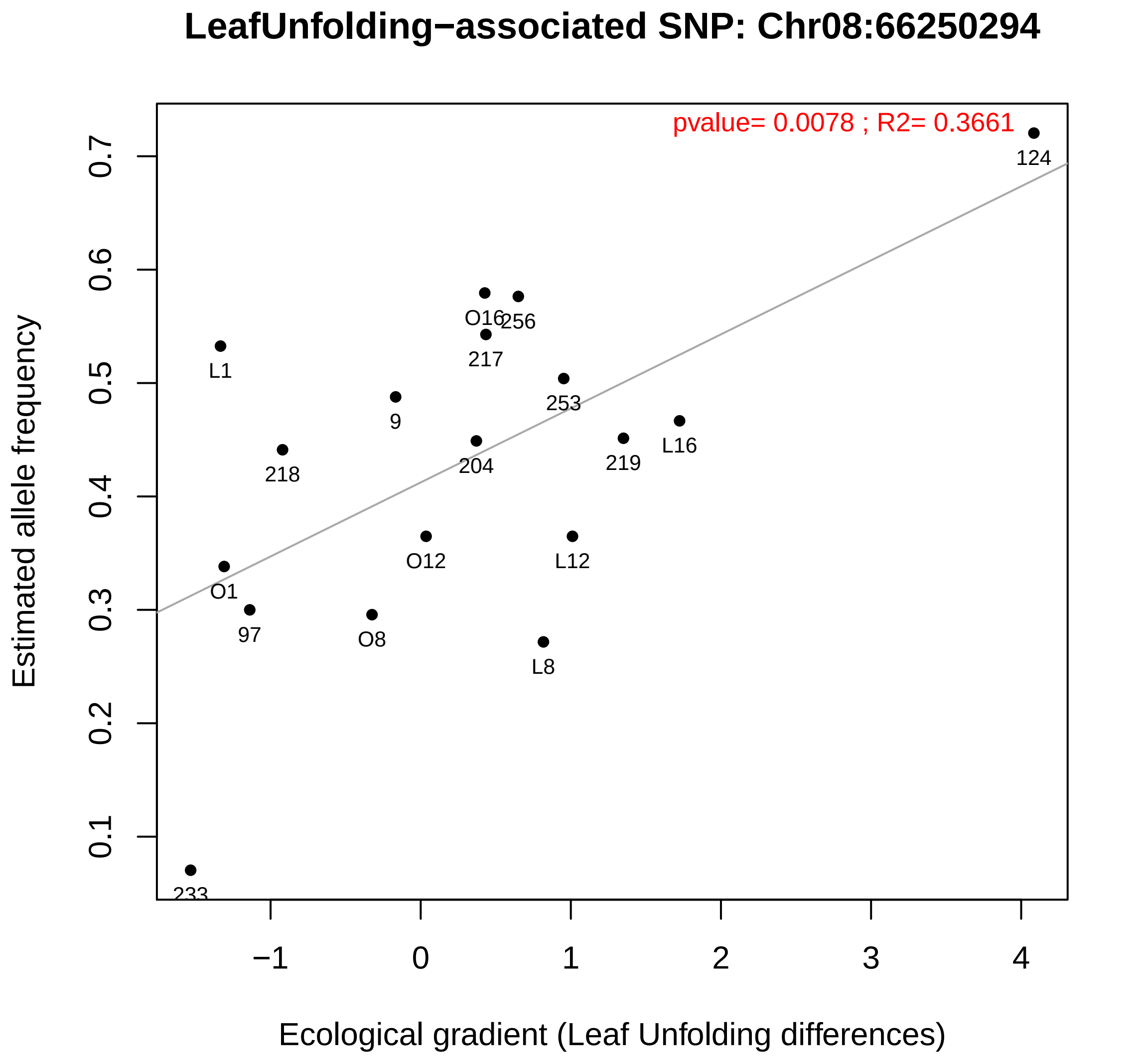

LeafUnfolding−associated SNP: Chr08:66250294
pvalue= 0.0078 ; R2= 0.3661
Estimated allele frequency
Ecological gradient (Leaf Unfolding differences)
0.1
0.2
0.3
0.4
0.5
0.6
0.7
−1
0
1
2
3
4
124
O16
256
217
L1
253
9
204
218
219
L16
O12
L12
O1
97
O8
L8
233

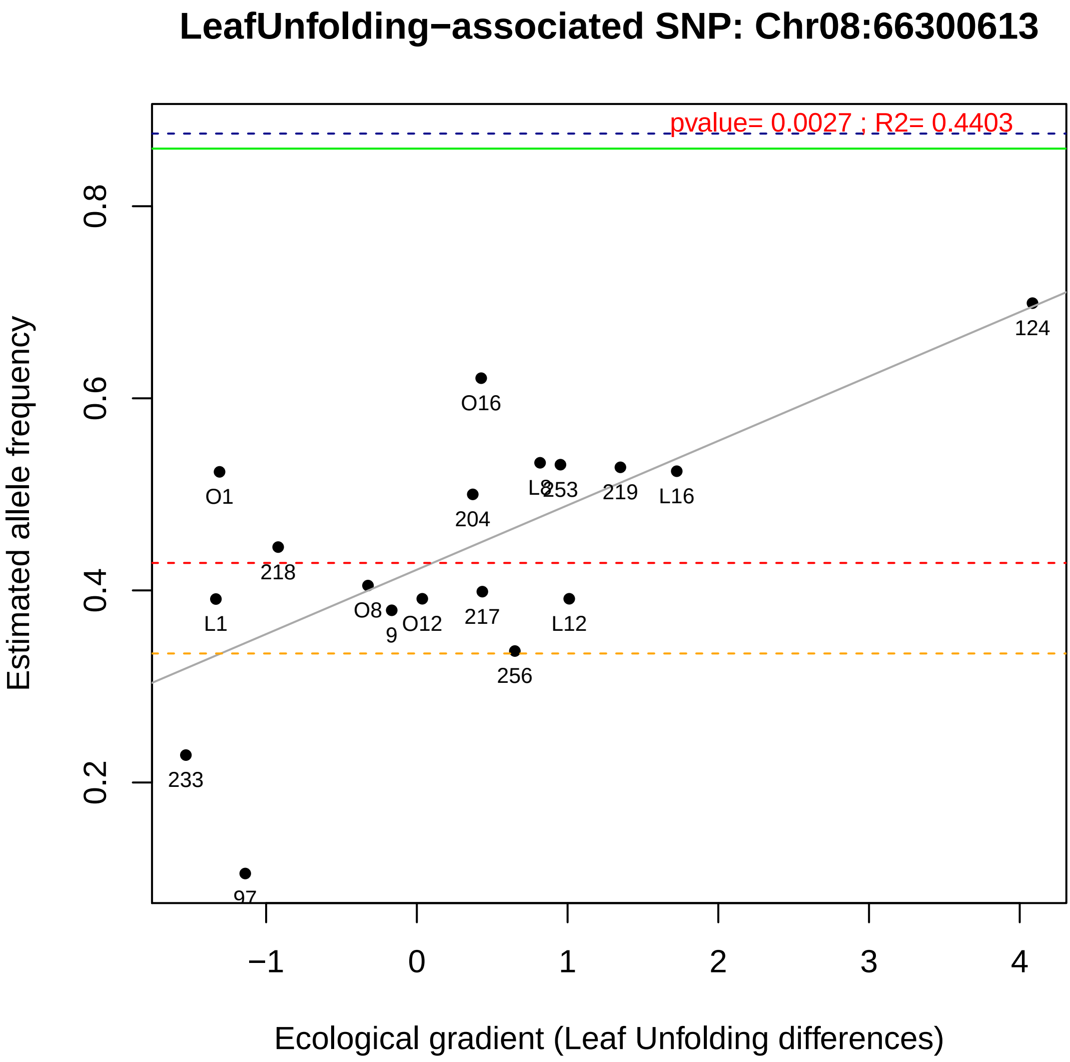

LeafUnfolding−associated SNP: Chr08:66300613
pvalue= 0.0027 ; R2= 0.4403
Estimated allele frequency
Ecological gradient (Leaf Unfolding differences)
0.2
0.4
0.6
0.8
−1
0
1
2
3
4
124
O16
O1
204
L8
253
219
L16
218
L1
O8
9
O12
217
L12
256
233
97

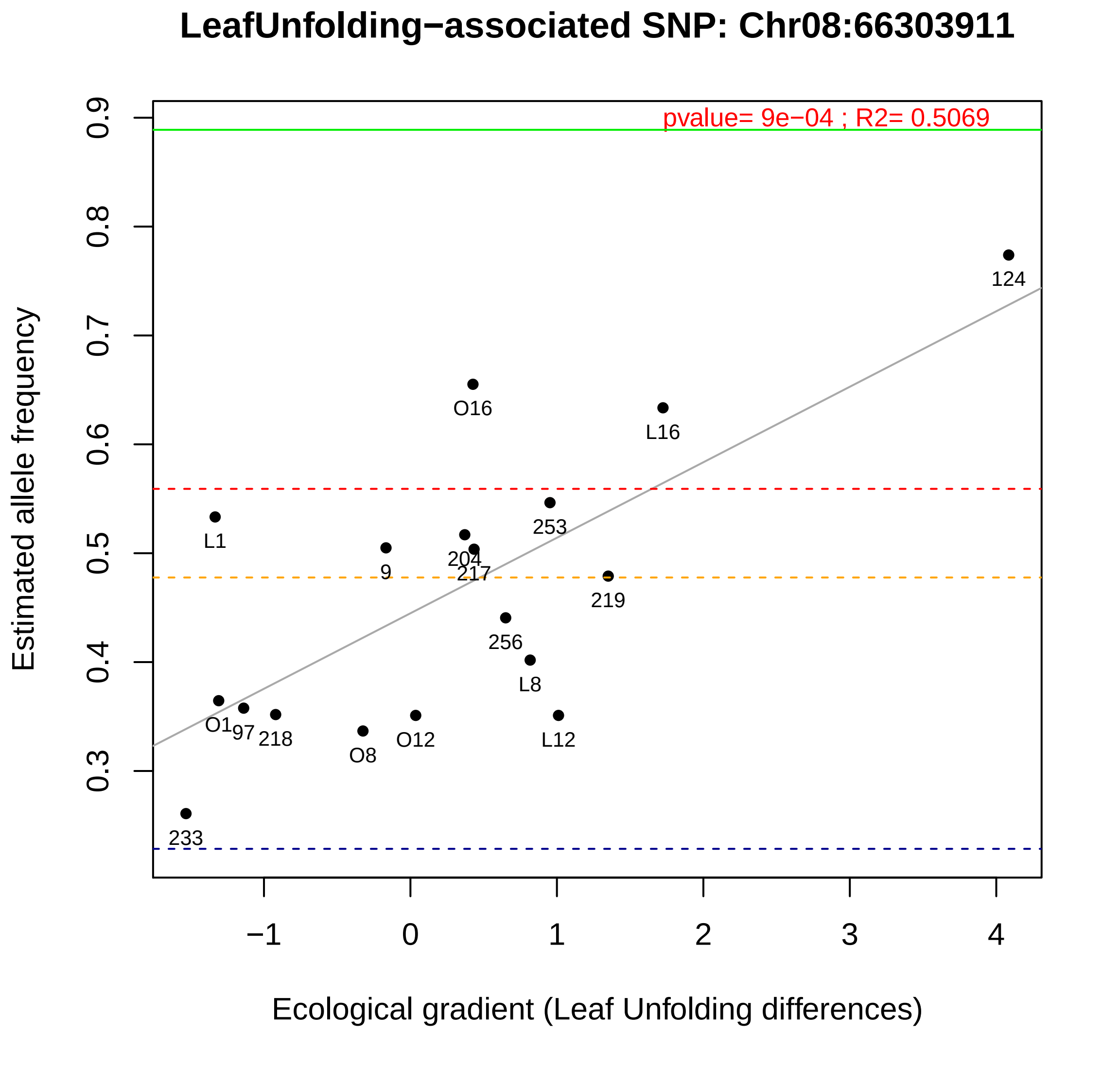

LeafUnfolding−associated SNP: Chr08:66303911
pvalue= 9e−04 ; R2= 0.5069
Estimated allele frequency
Ecological gradient (Leaf Unfolding differences)
0.3
0.4
0.5
0.6
0.7
0.8
0.9
−1
0
1
2
3
4
124
O16
L16
253
L1
9
204
217
219
256
L8
O1
97
218
O8
O12
L12
233

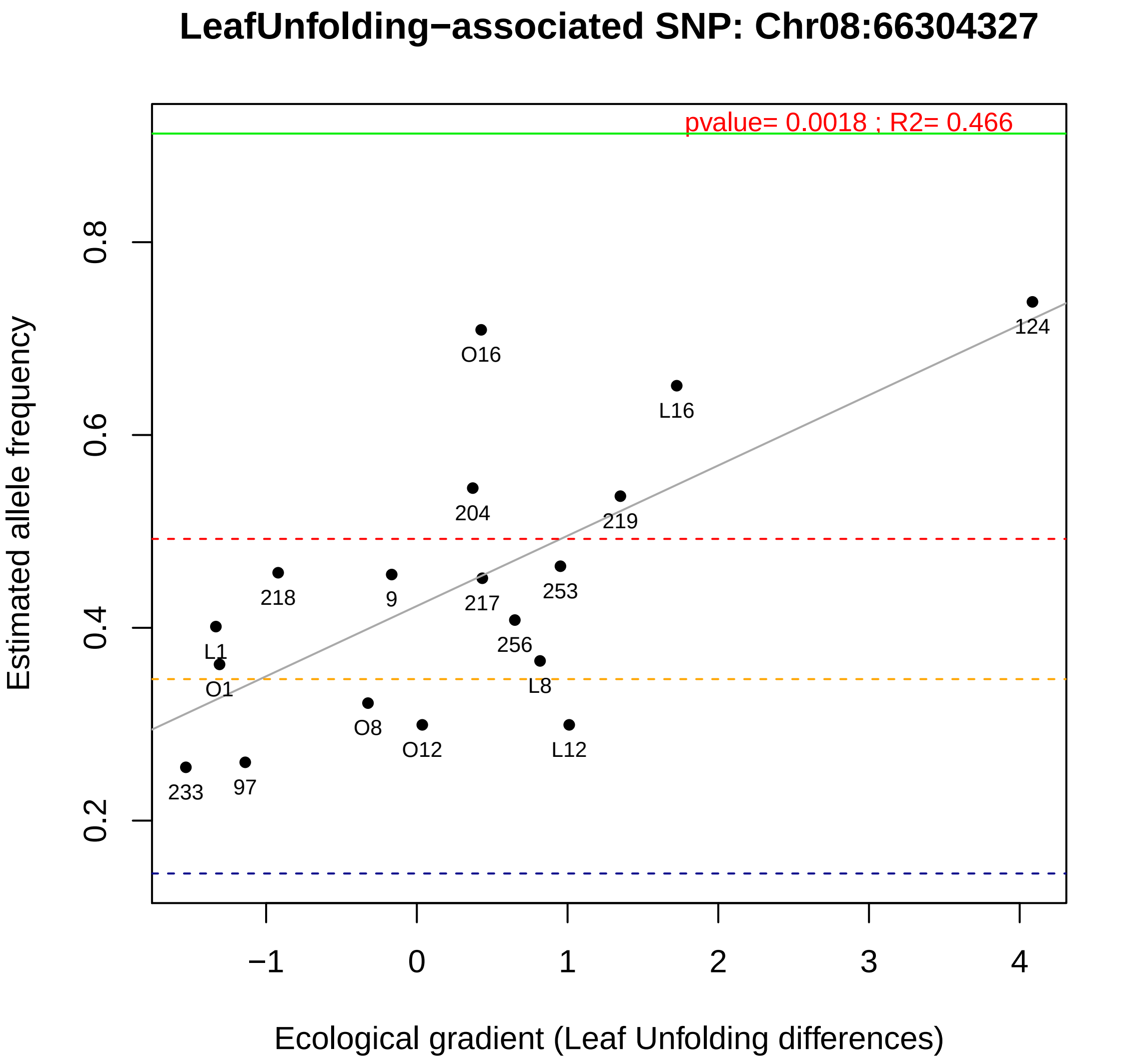

LeafUnfolding−associated SNP: Chr08:66304327
pvalue= 0.0018 ; R2= 0.466
Estimated allele frequency
0.2
0.4
0.6
0.8
−1
0
1
2
3
4
Ecological gradient (Leaf Unfolding differences)
124
O16
L16
204
219
218
9
217
253
256
L1
O1
L8
O8
O12
L12
233
97

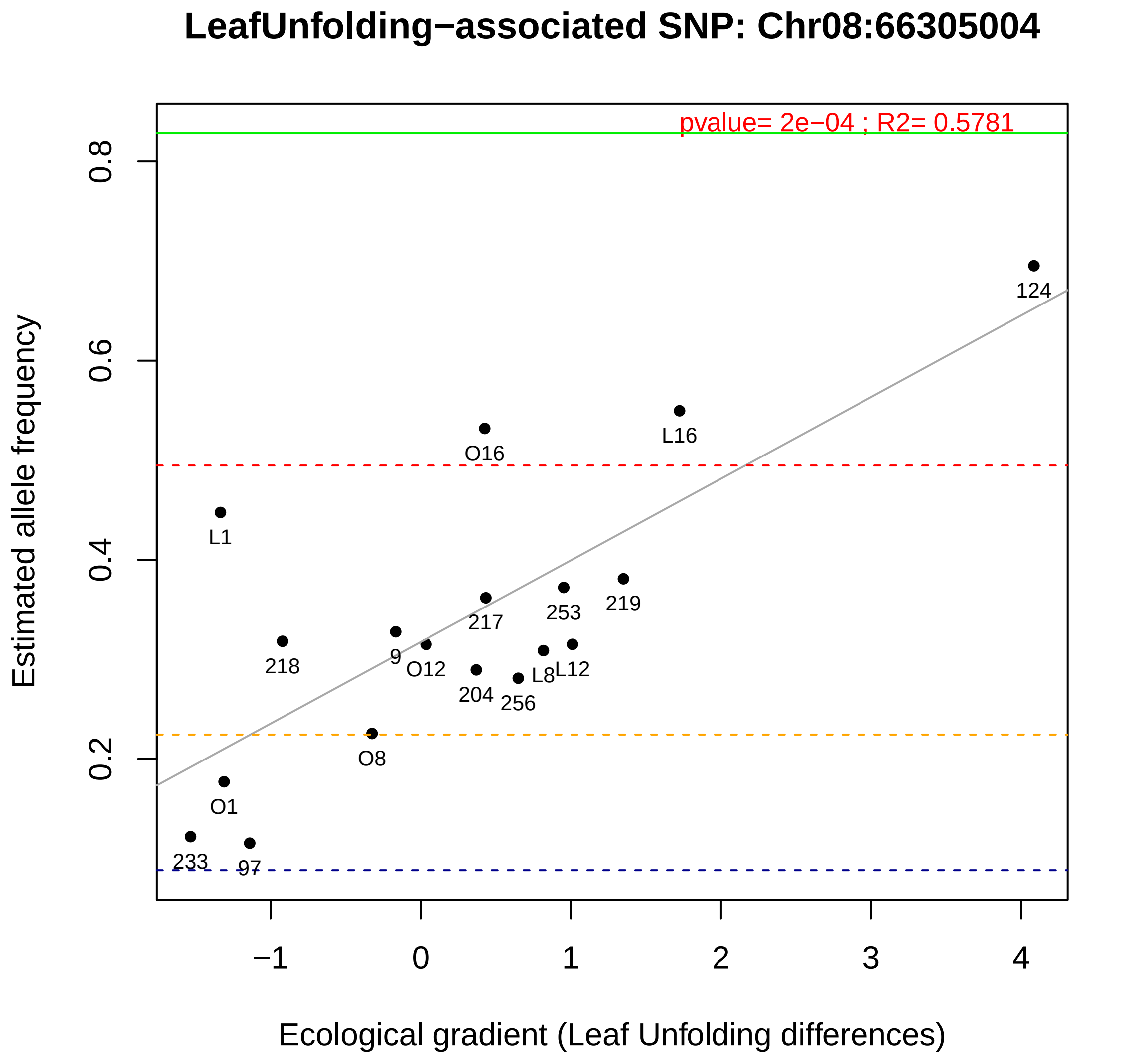
LeafUnfolding−associated SNP: Chr08:66305004
pvalue= 2e−04 ; R2= 0.5781
Estimated allele frequency
0.2
0.4
0.6
0.8
Ecological gradient (Leaf Unfolding differences)
−1
0
1
2
3
4
124
L16
O16
L1
217
253
219
218
9
O12
204
256
L8
L12
O8
O1
233
97

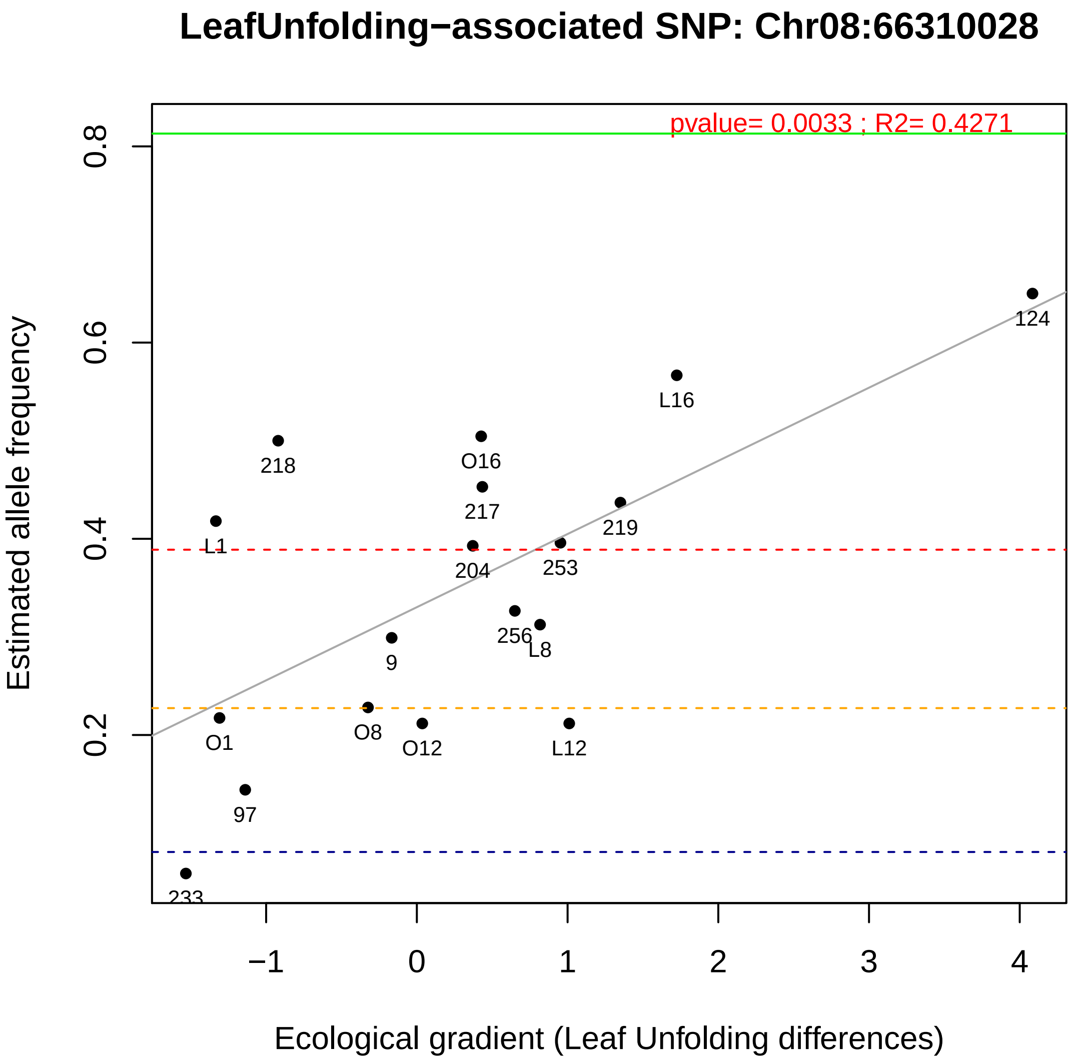
LeafUnfolding−associated SNP: Chr08:66310028
pvalue= 0.0033 ; R2= 0.4271
Estimated allele frequency
0.2
0.4
0.6
0.8
Ecological gradient (Leaf Unfolding differences)
−1
0
1
2
3
4
124
L16
218
O16
217
219
L1
204
253
256
L8
9
O1
O8
O12
L12
97
233

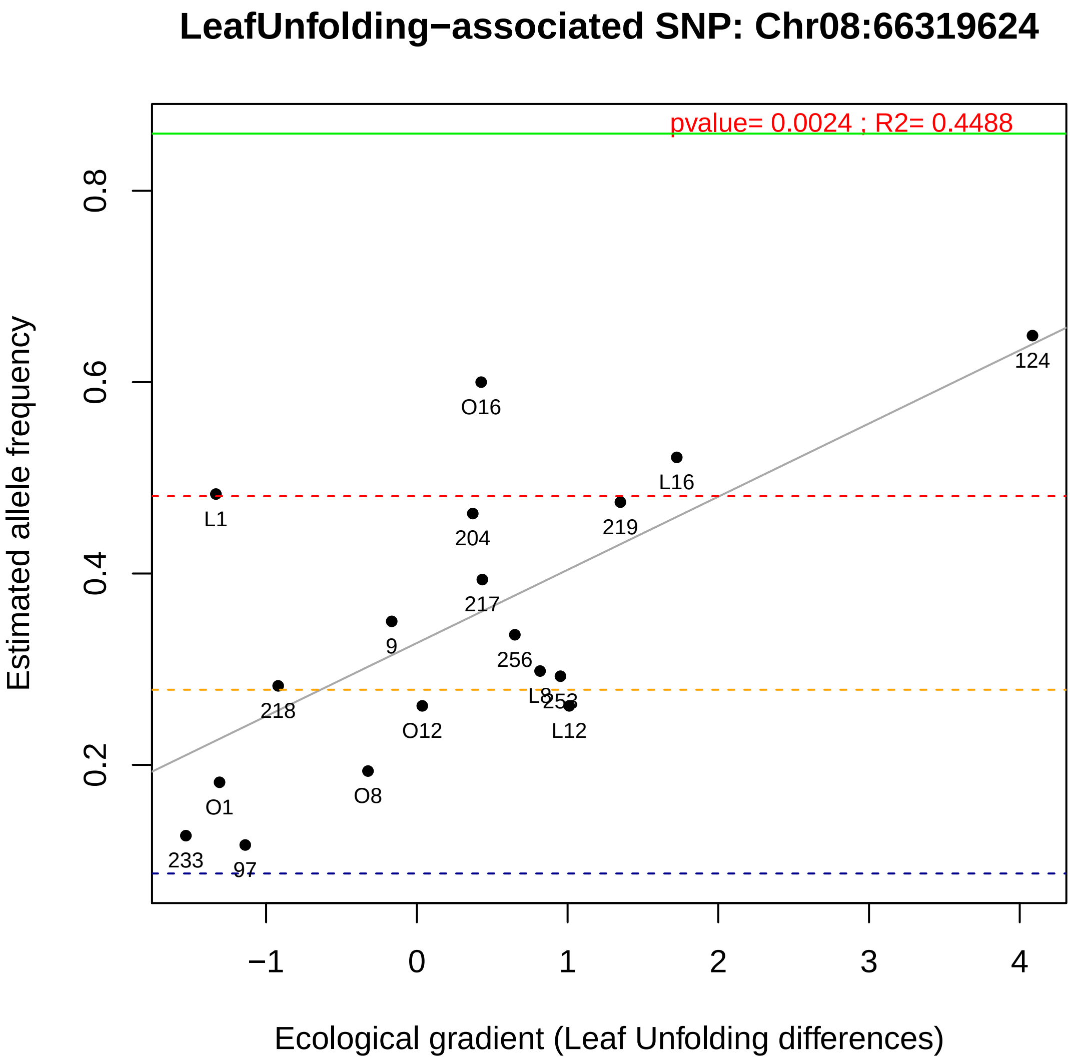

LeafUnfolding−associated SNP: Chr08:66319624
pvalue= 0.0024 ; R2= 0.4488
Estimated allele frequency
0.2
0.4
0.6
0.8
Ecological gradient (Leaf Unfolding differences)
−1
0
1
2
3
4
124
O16
L16
L1
204
219
217
9
256
L8
253
218
O12
L12
O1
O8
233
97

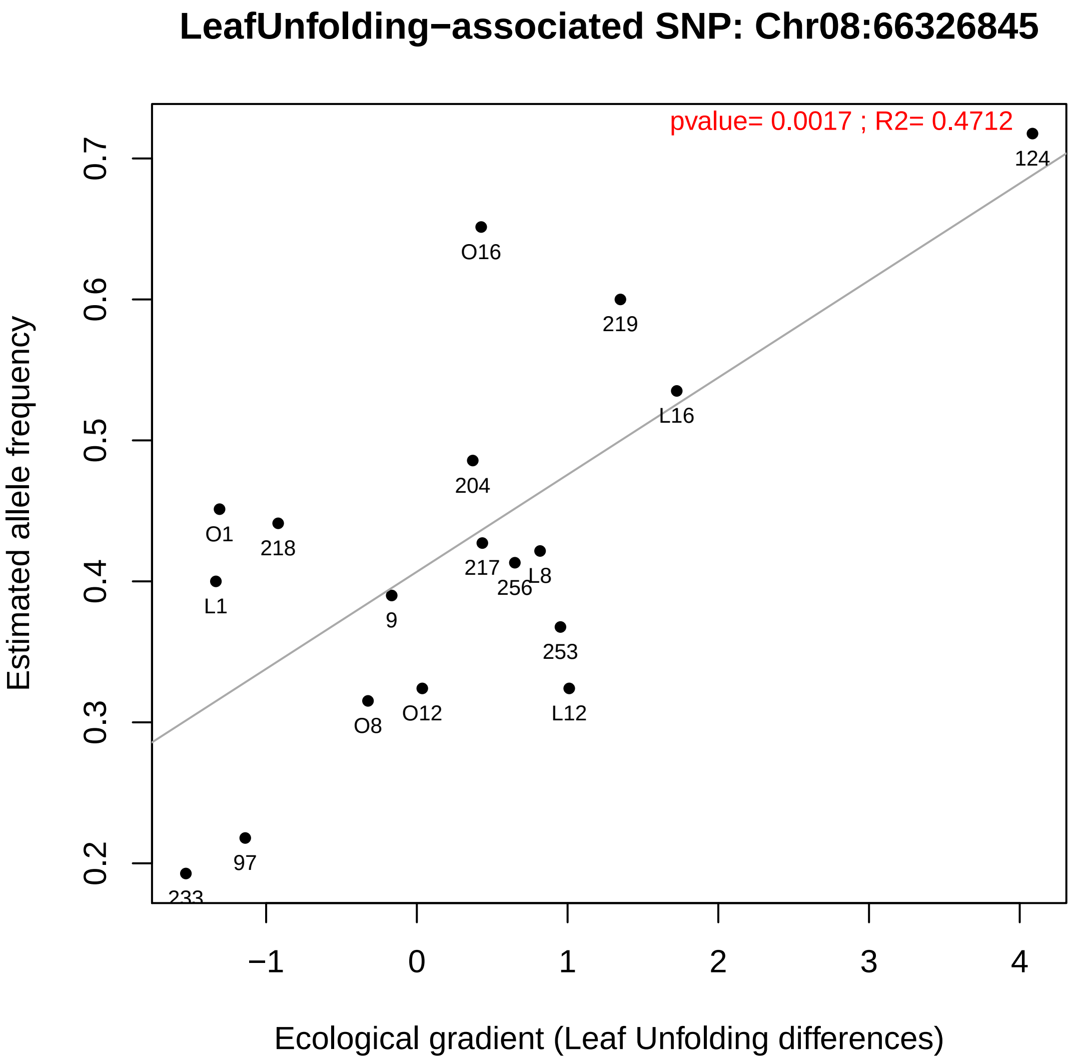

LeafUnfolding−associated SNP: Chr08:66326845
pvalue= 0.0017 ; R2= 0.4712
Estimated allele frequency
Ecological gradient (Leaf Unfolding differences)
0.2
0.3
0.4
0.5
0.6
0.7
−1
0
1
2
3
4
124
O16
219
L16
204
O1
218
217
256
L8
L1
9
253
O8
O12
L12
97
233

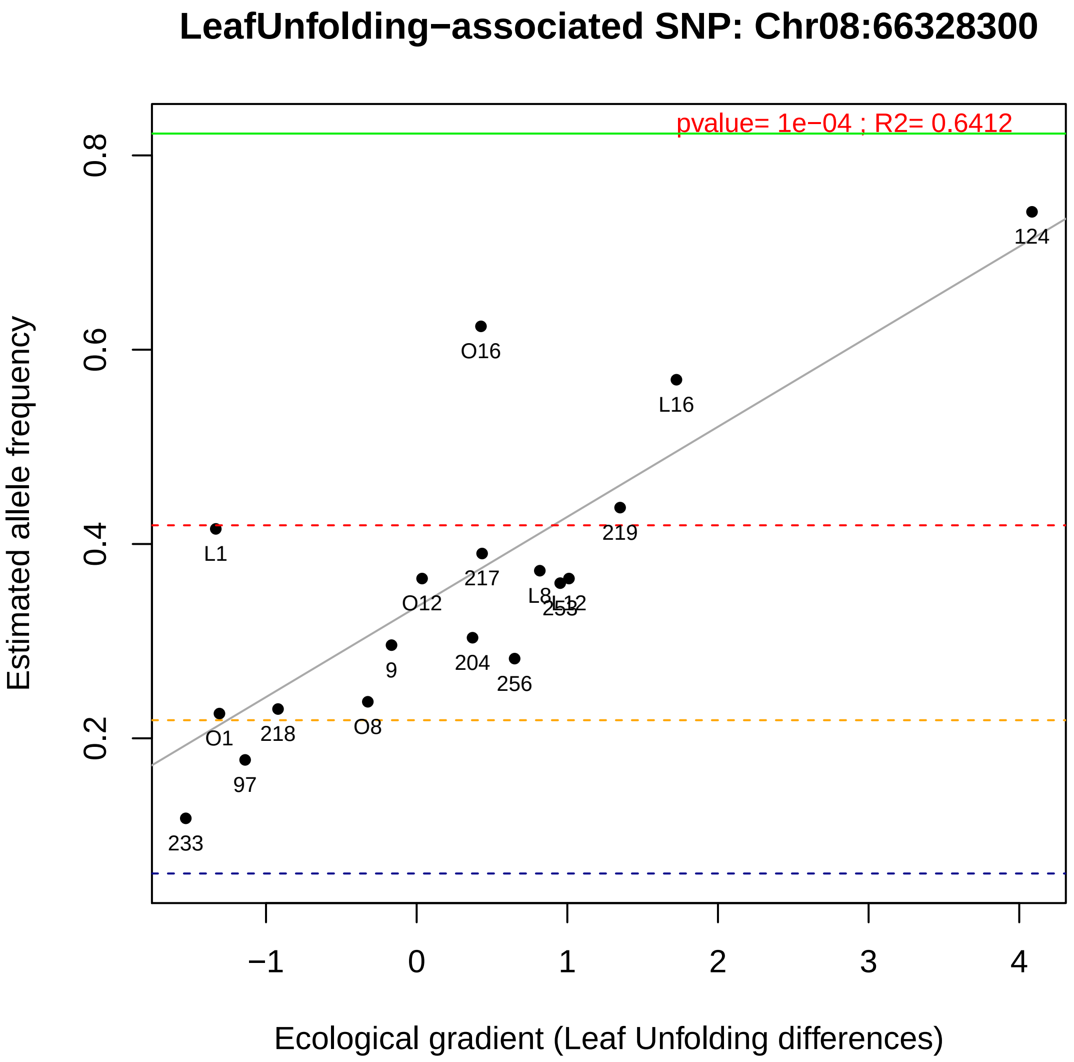

LeafUnfolding−associated SNP: Chr08:66328300
pvalue= 1e−04 ; R2= 0.6412
Estimated allele frequency
0.2
0.4
0.6
0.8
Ecological gradient (Leaf Unfolding differences)
−1
0
1
2
3
4
124
O16
L16
219
L1
217
O12
L8
9
204
256
O1
218
O8
97
233

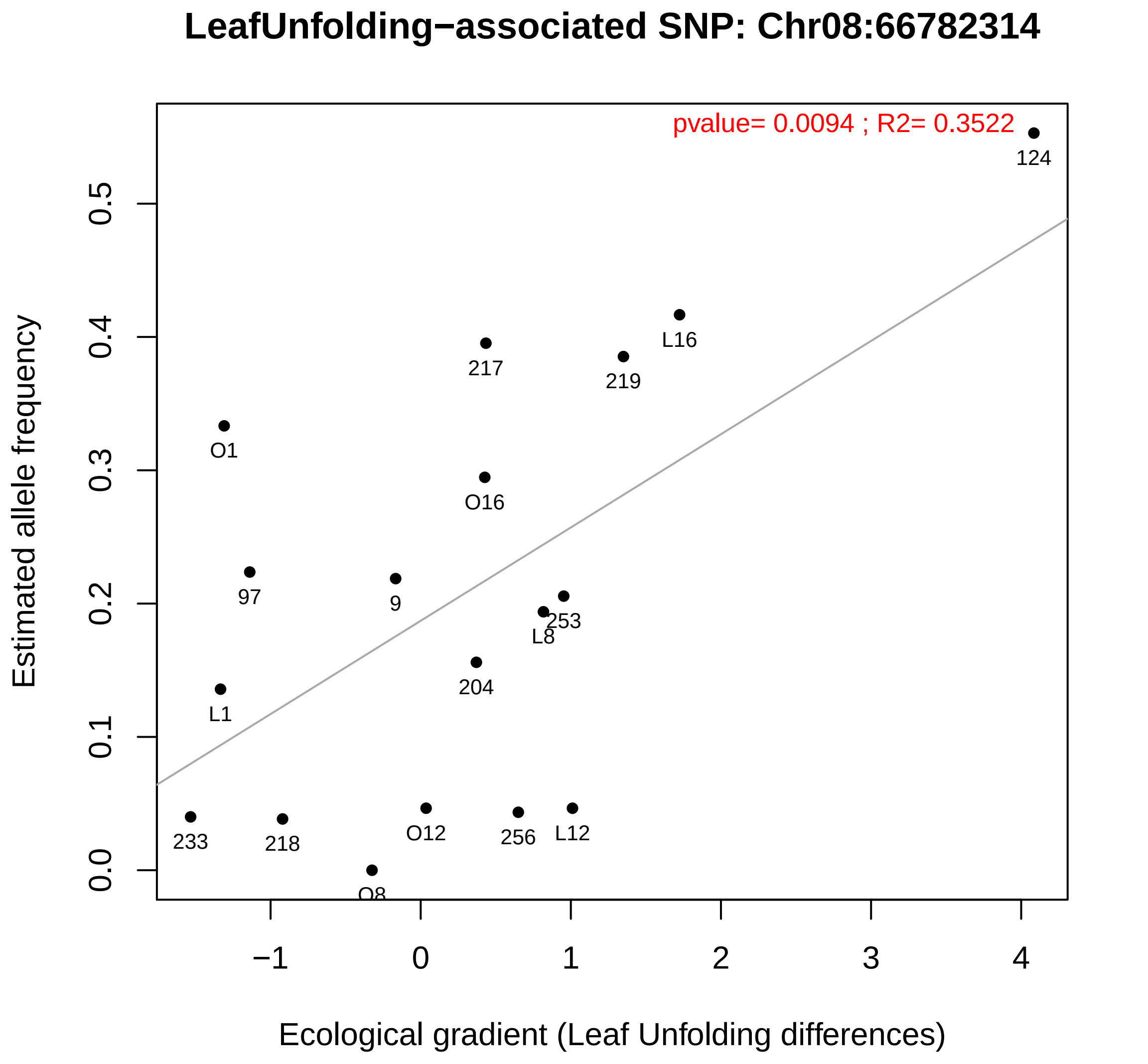
LeafUnfolding−associated SNP: Chr08:66782314
pvalue= 0.0094 ; R2= 0.3522
124
L16
217
219
O1
O16
97
9
253
L8
204
L1
233
218
O12
256
L12
O8
0.0
0.1
0.2
0.3
0.4
0.5
−1
0
1
2
3
4
Estimated allele frequency
Ecological gradient (Leaf Unfolding differences)

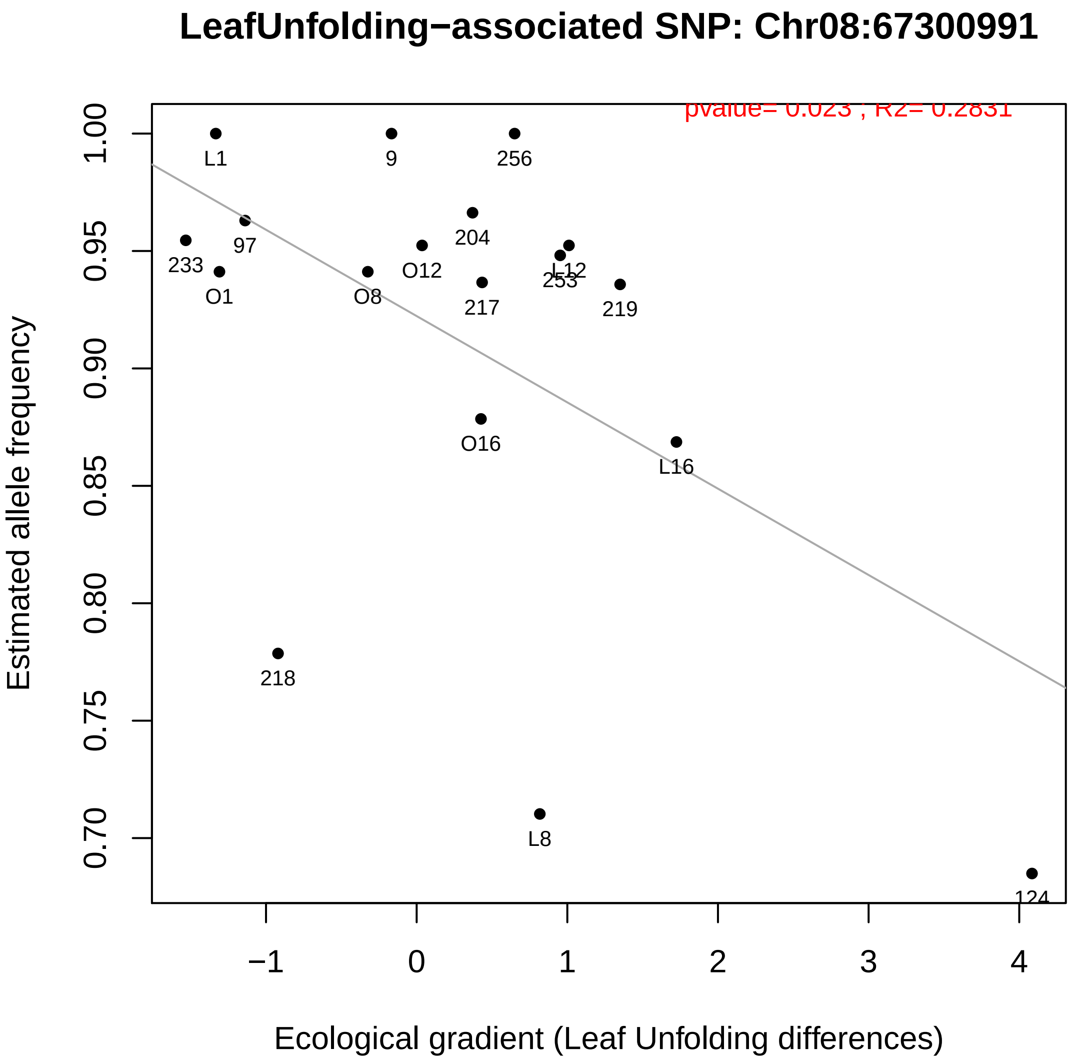
LeafUnfolding−associated SNP: Chr08:67300991
pvalue= 0.023 ; R2= 0.2831
Estimated allele frequency
1.00
0.95
0.90
0.85
0.80
0.75
0.70
−1
0
1
2
3
4
Ecological gradient (Leaf Unfolding differences)
L1
9
256
204
97
233
O12
O1
O8
217
L12
253
219
O16
L16
218
L8
124

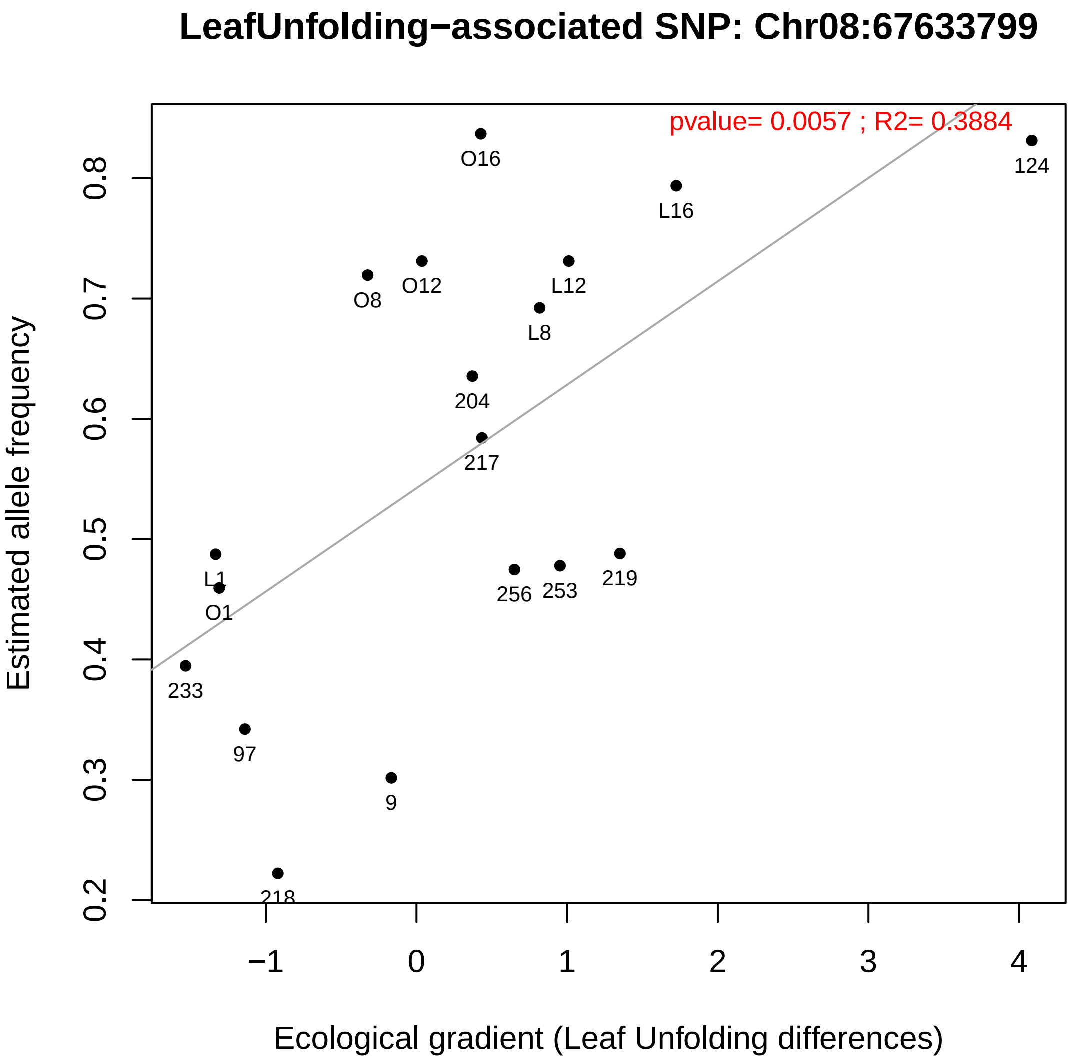

LeafUnfolding−associated SNP: Chr08:67633799
pvalue= 0.0057 ; R2= 0.3884
O16
124
L16
O8
O12
L12
L8
204
217
L1
O1
256
253
219
233
97
9
218
0.2
0.3
0.4
0.5
0.6
0.7
0.8
−1
0
1
2
3
4
Estimated allele frequency
Ecological gradient (Leaf Unfolding differences)

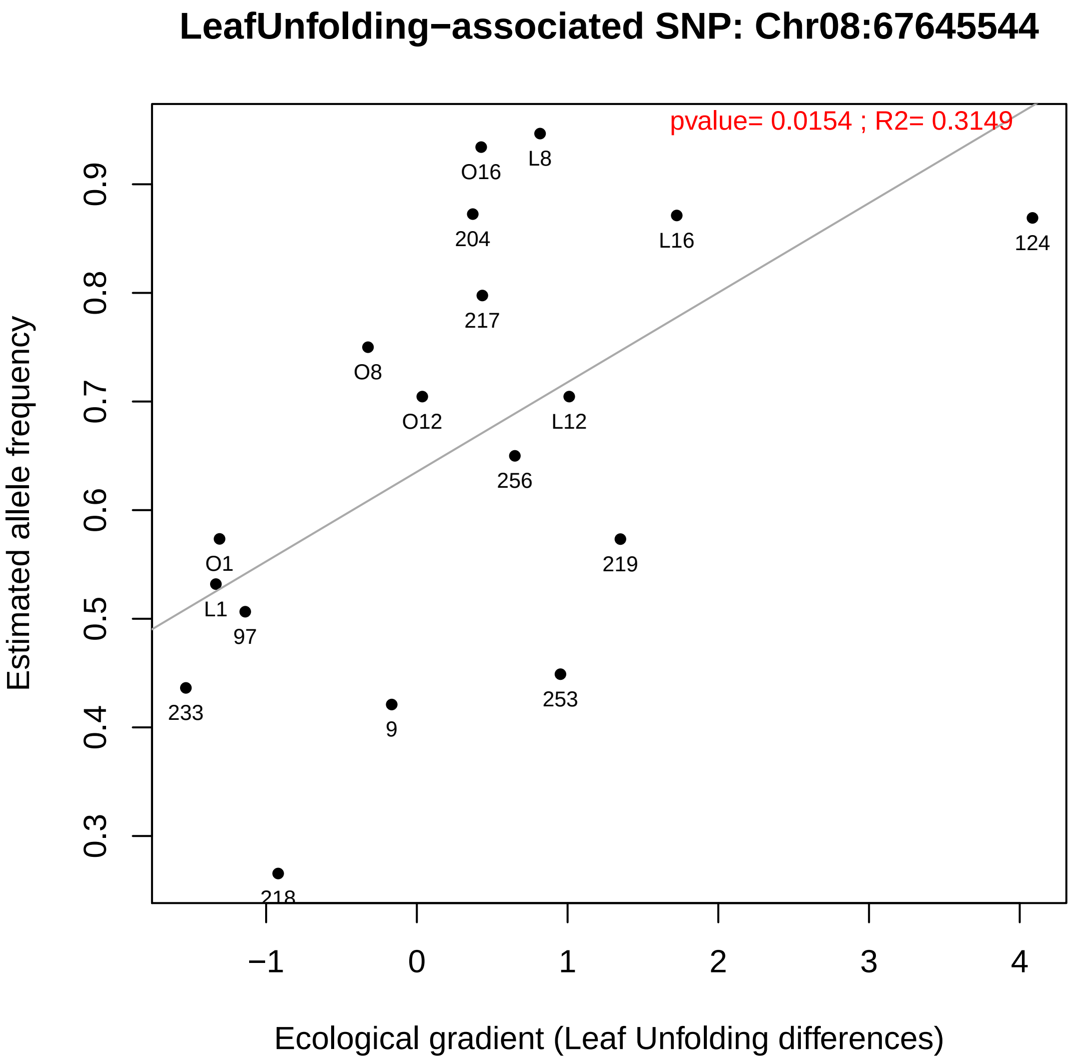
LeafUnfolding−associated SNP: Chr08:67645544
pvalue= 0.0154 ; R2= 0.3149
Estimated allele frequency
0.3
0.4
0.5
0.6
0.7
0.8
0.9
−1
0
1
2
3
4
Ecological gradient (Leaf Unfolding differences)
O16
L8
204
L16
124
217
O8
O12
L12
256
O1
L1
97
219
233
9
253
218

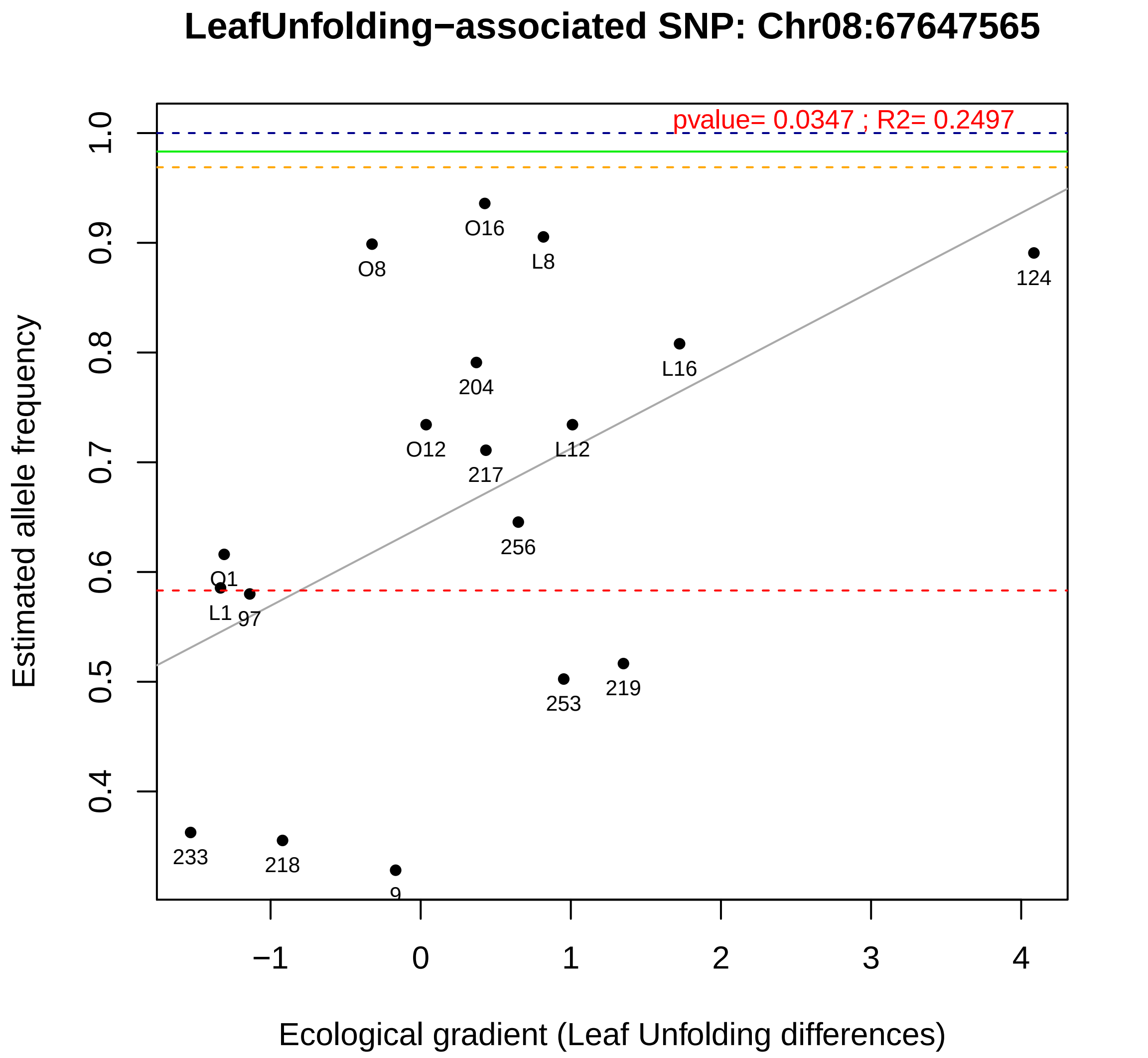
LeafUnfolding−associated SNP: Chr08:67647565
pvalue= 0.0347 ; R2= 0.2497
Estimated allele frequency
Ecological gradient (Leaf Unfolding differences)
1.0
0.9
0.8
0.7
0.6
0.5
0.4
−1
0
1
2
3
4
O16
O8
L8
124
L16
204
O12
217
L12
256
O1
L1
97
253
219
233
218
9

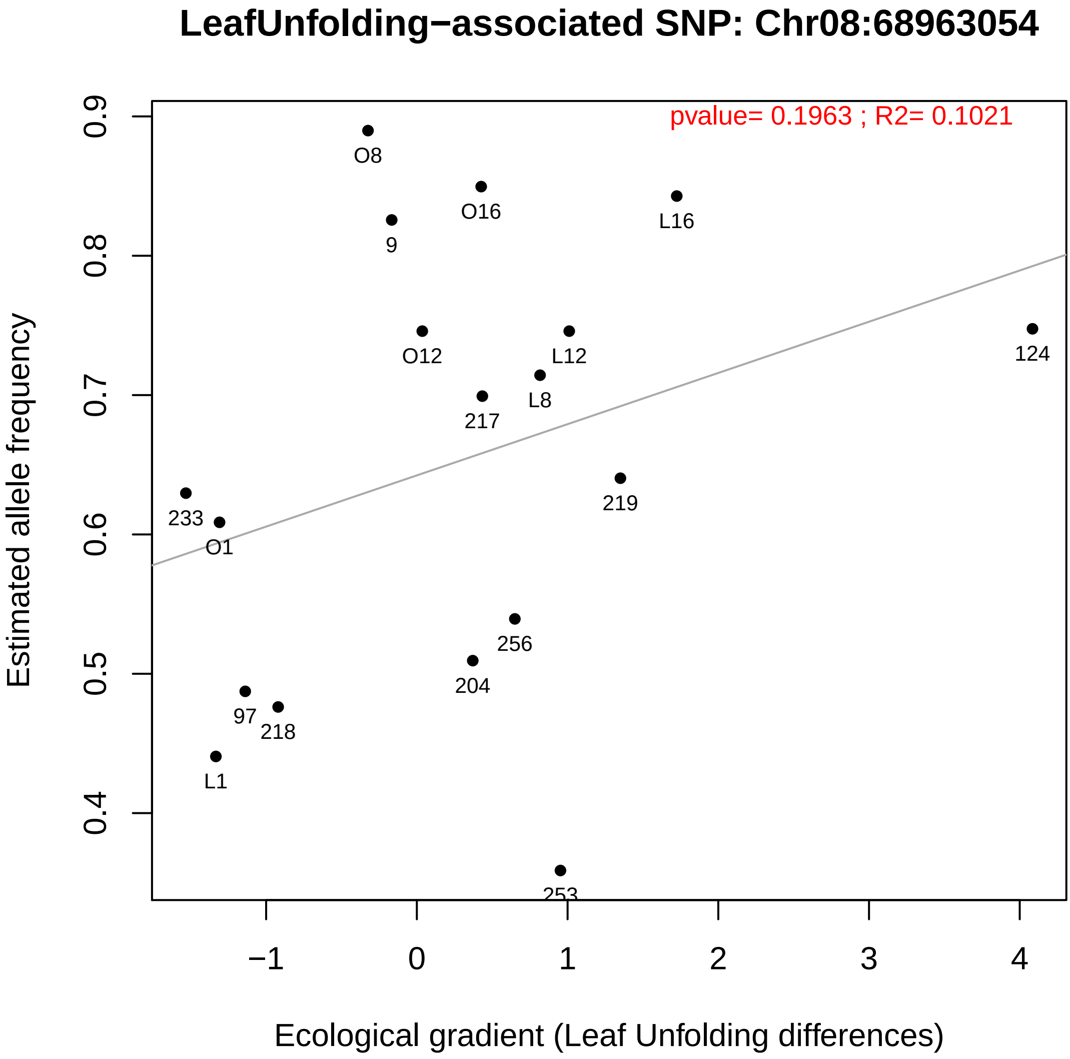
LeafUnfolding−associated SNP: Chr08:68963054
pvalue= 0.1963 ; R2= 0.1021
Estimated allele frequency
Ecological gradient (Leaf Unfolding differences)
0.4
0.5
0.6
0.7
0.8
0.9
−1
0
1
2
3
4
O8
O16
L16
9
O12
L12
124
L8
217
219
233
O1
256
204
97
218
L1
253

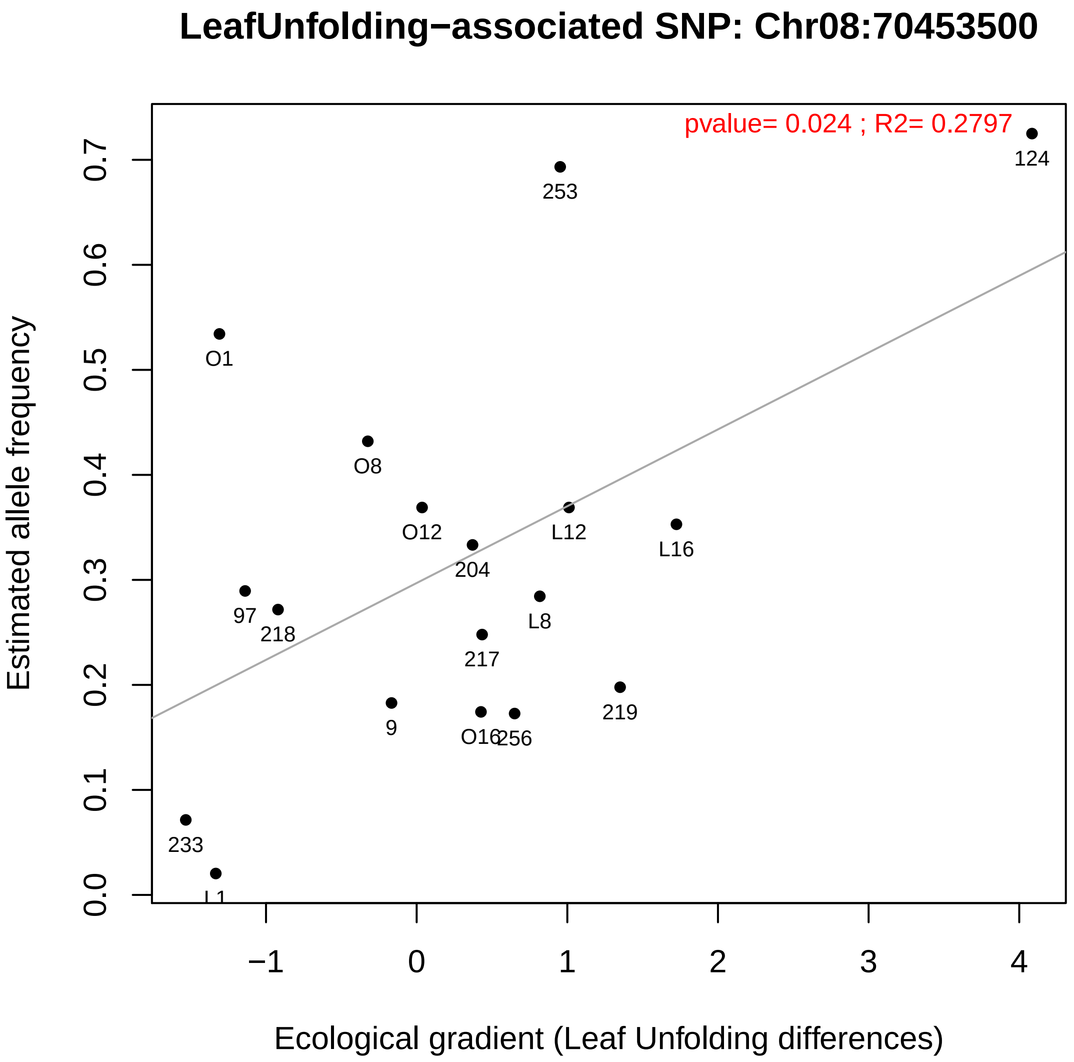

LeafUnfolding−associated SNP: Chr08:70453500
pvalue= 0.024 ; R2= 0.2797
124
253
O1
O8
O12
L12
L16
204
97
218
L8
217
219
9
O16
256
233
L1
Estimated allele frequency
0.0
0.1
0.2
0.3
0.4
0.5
0.6
0.7
−1
0
1
2
3
4
Ecological gradient (Leaf Unfolding differences)

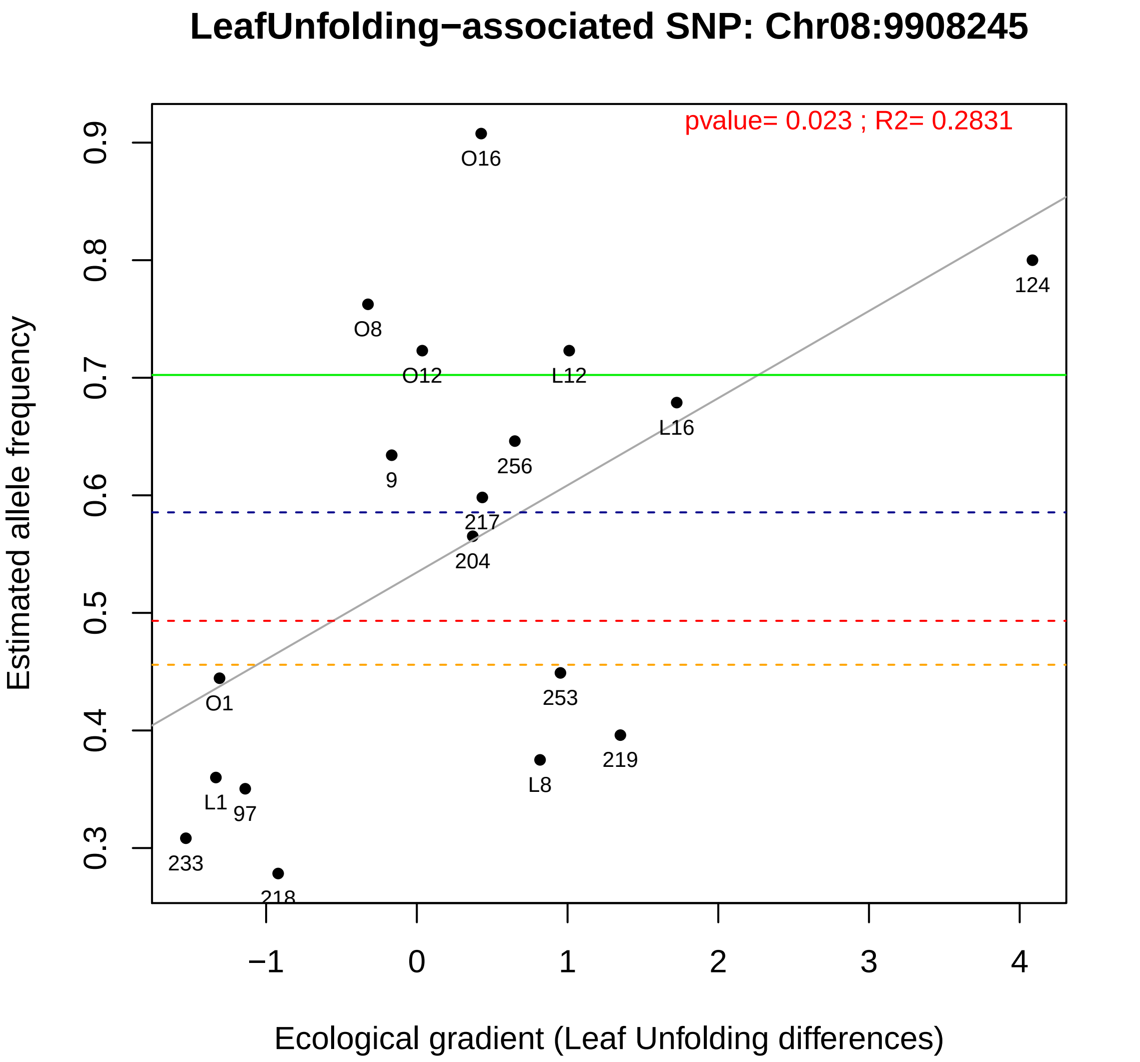
LeafUnfolding−associated SNP: Chr08:9908245
pvalue= 0.023 ; R2= 0.2831
Estimated allele frequency
Ecological gradient (Leaf Unfolding differences)
0.3
0.4
0.5
0.6
0.7
0.8
0.9
−1
0
1
2
3
4
O16
124
O8
O12
L12
L16
256
9
217
204
O1
253
219
L8
L1
97
233
218

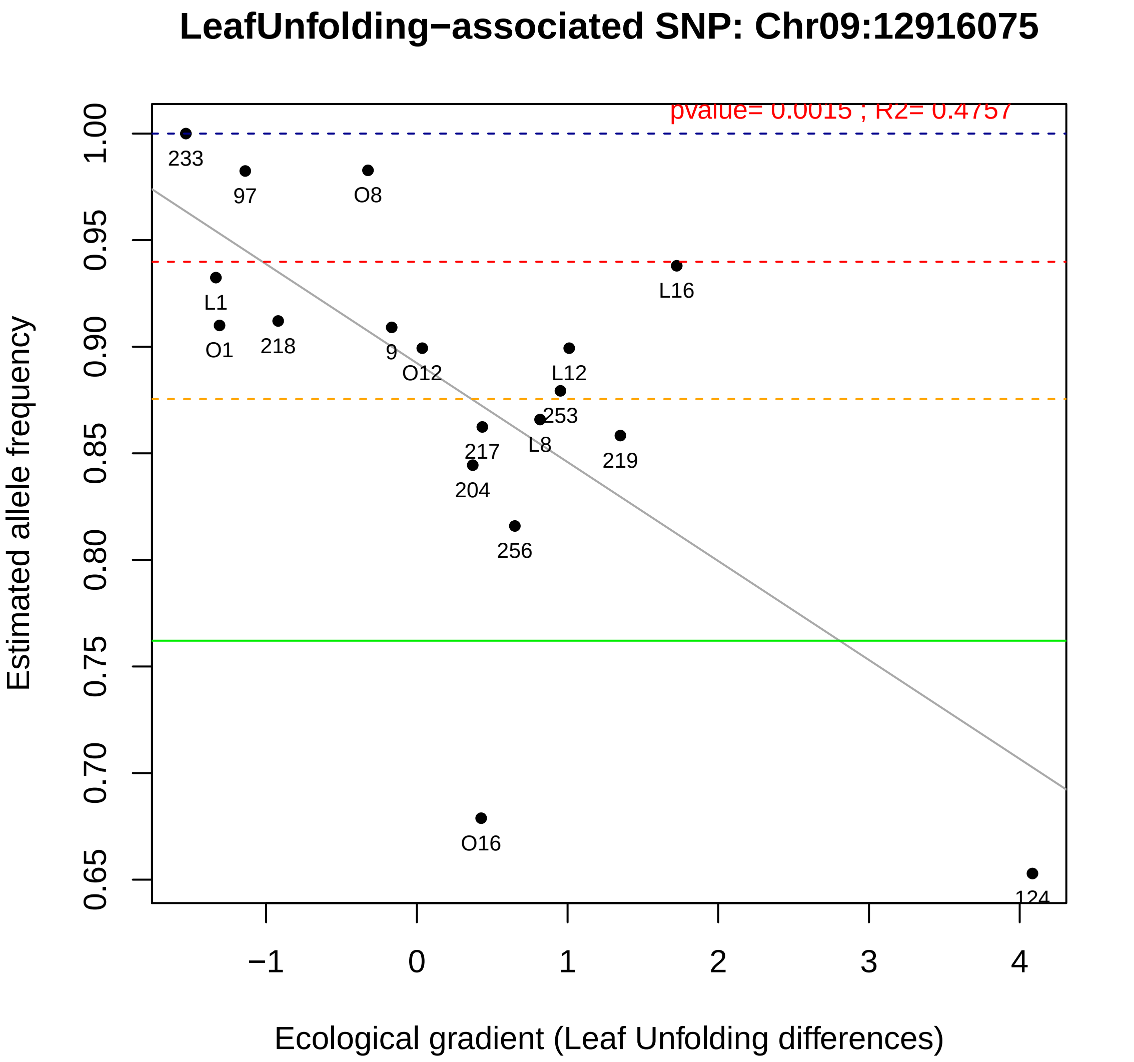
LeafUnfolding−associated SNP: Chr09:12916075
pvalue= 0.0015 ; R2= 0.4757
Estimated allele frequency
0.65
0.70
0.75
0.80
0.85
0.90
0.95
1.00
Ecological gradient (Leaf Unfolding differences)
−1
0
1
2
3
4
233
97
O8
L1
O1
218
9
O12
L12
253
L8
217
204
219
L16
256
O16
124

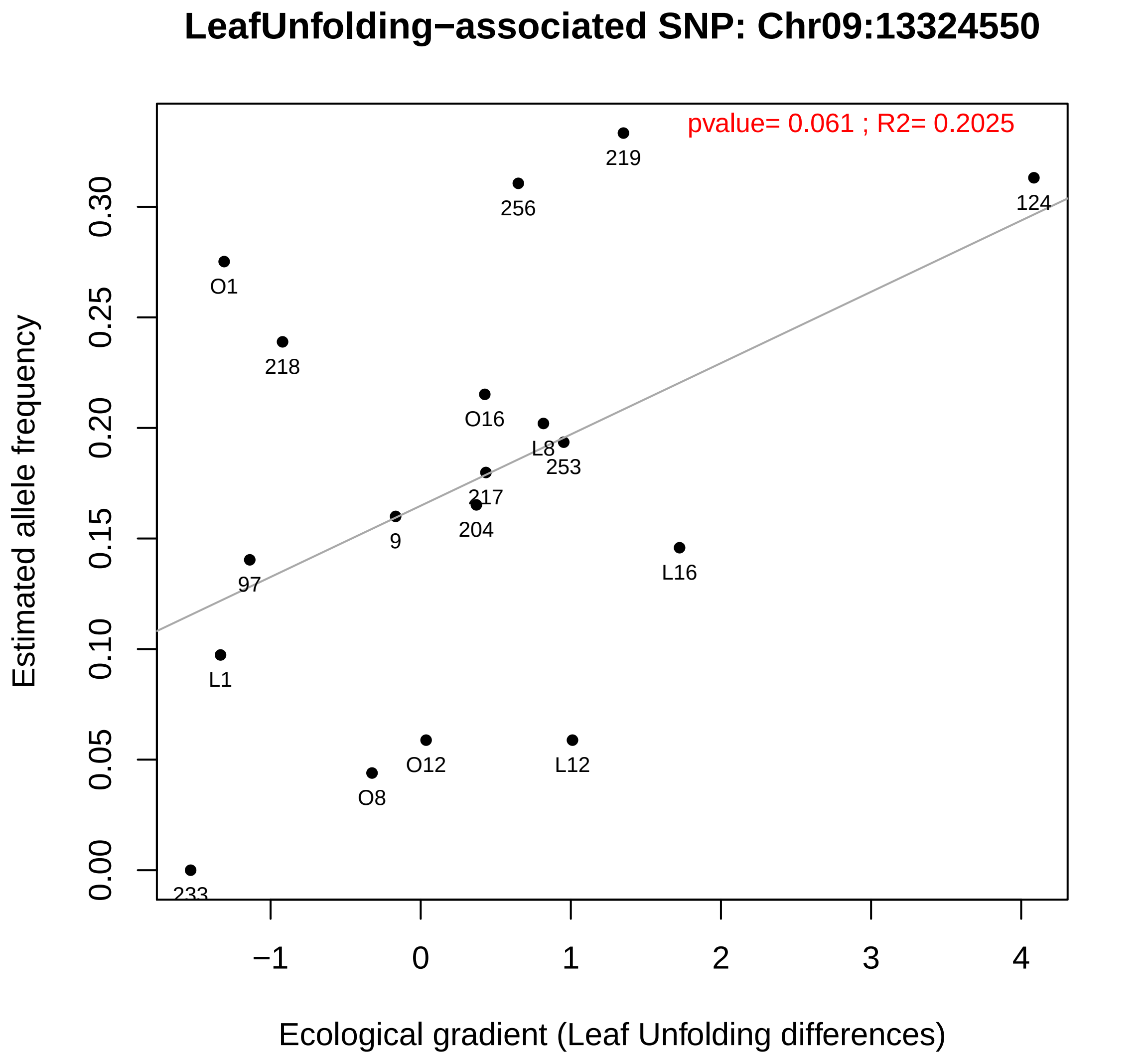

LeafUnfolding−associated SNP: Chr09:13324550
pvalue= 0.061 ; R2= 0.2025
Estimated allele frequency
Ecological gradient (Leaf Unfolding differences)
0.00
0.05
0.10
0.15
0.20
0.25
0.30
−1
0
1
2
3
4
219
256
124
O1
218
O16
L8
253
217
204
9
L16
97
L1
O12
L12
O8
233

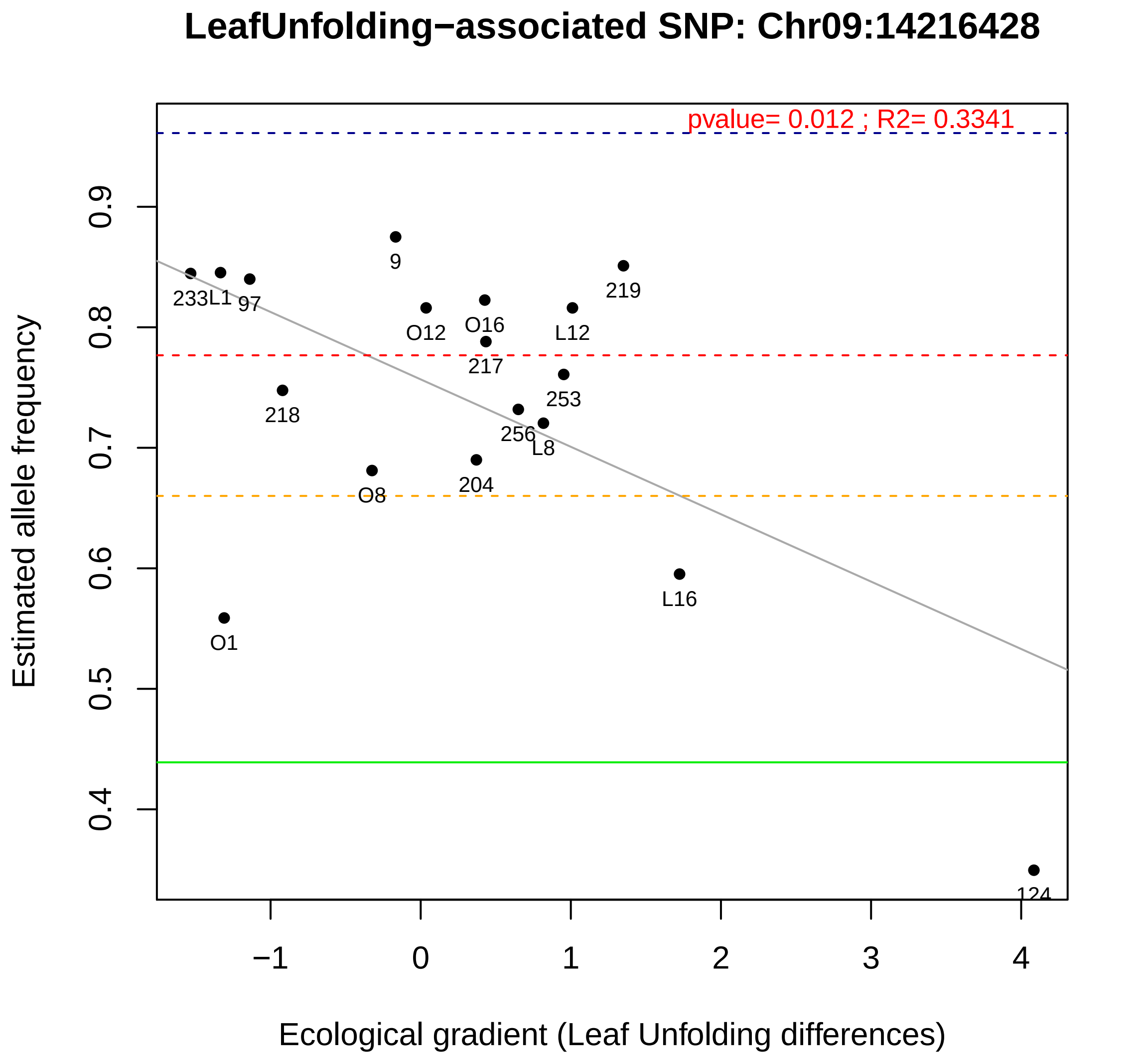
LeafUnfolding−associated SNP: Chr09:14216428
pvalue= 0.012 ; R2= 0.3341
Estimated allele frequency
0.4
0.5
0.6
0.7
0.8
0.9
−1
0
1
2
3
4
Ecological gradient (Leaf Unfolding differences)
9
219
233
L1
97
O12
O16
L12
217
253
218
256
L8
204
O8
L16
O1
124

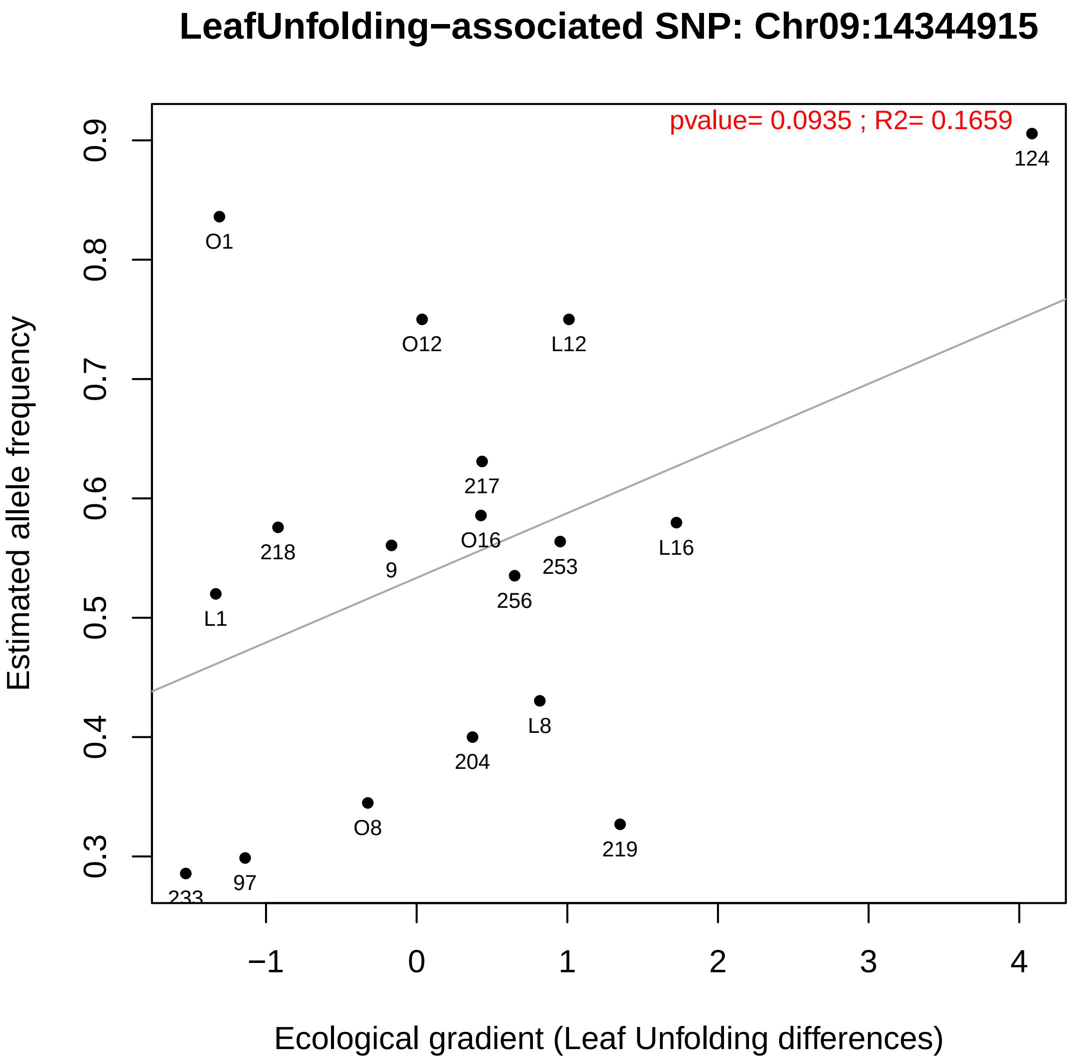
LeafUnfolding−associated SNP: Chr09:14344915
pvalue= 0.0935 ; R2= 0.1659
Estimated allele frequency
0.3
0.4
0.5
0.6
0.7
0.8
0.9
−1
0
1
2
3
4
Ecological gradient (Leaf Unfolding differences)
124
O1
O12
L12
217
O16
218
9
253
L16
256
L1
L8
204
O8
219
97
233

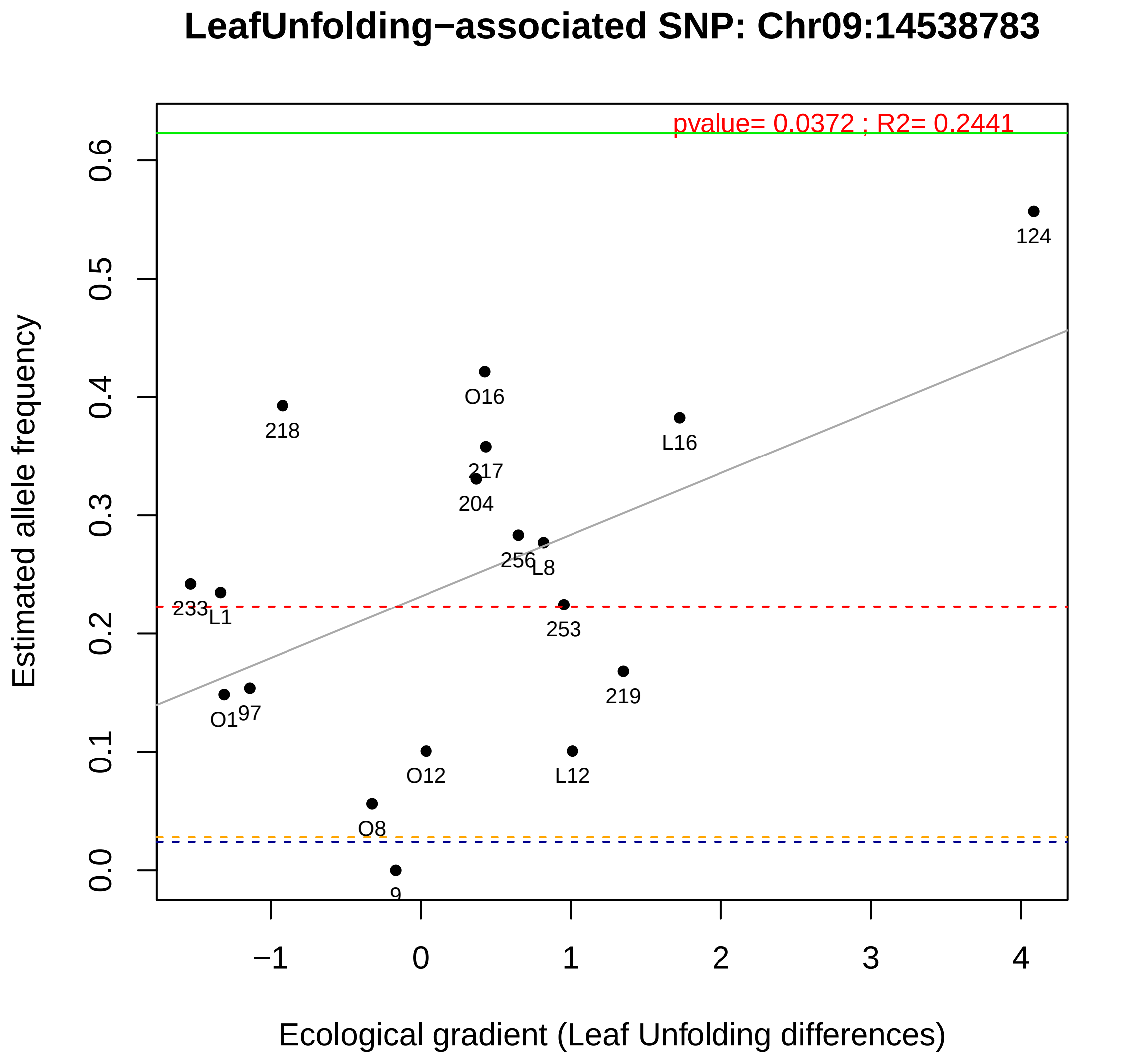

LeafUnfolding−associated SNP: Chr09:14538783
pvalue= 0.0372 ; R2= 0.2441
Estimated allele frequency
0.0
0.1
0.2
0.3
0.4
0.5
0.6
−1
0
1
2
3
4
Ecological gradient (Leaf Unfolding differences)
124
O16
218
L16
217
204
256
L8
233
L1
253
219
O1
97
O12
L12
O8
9

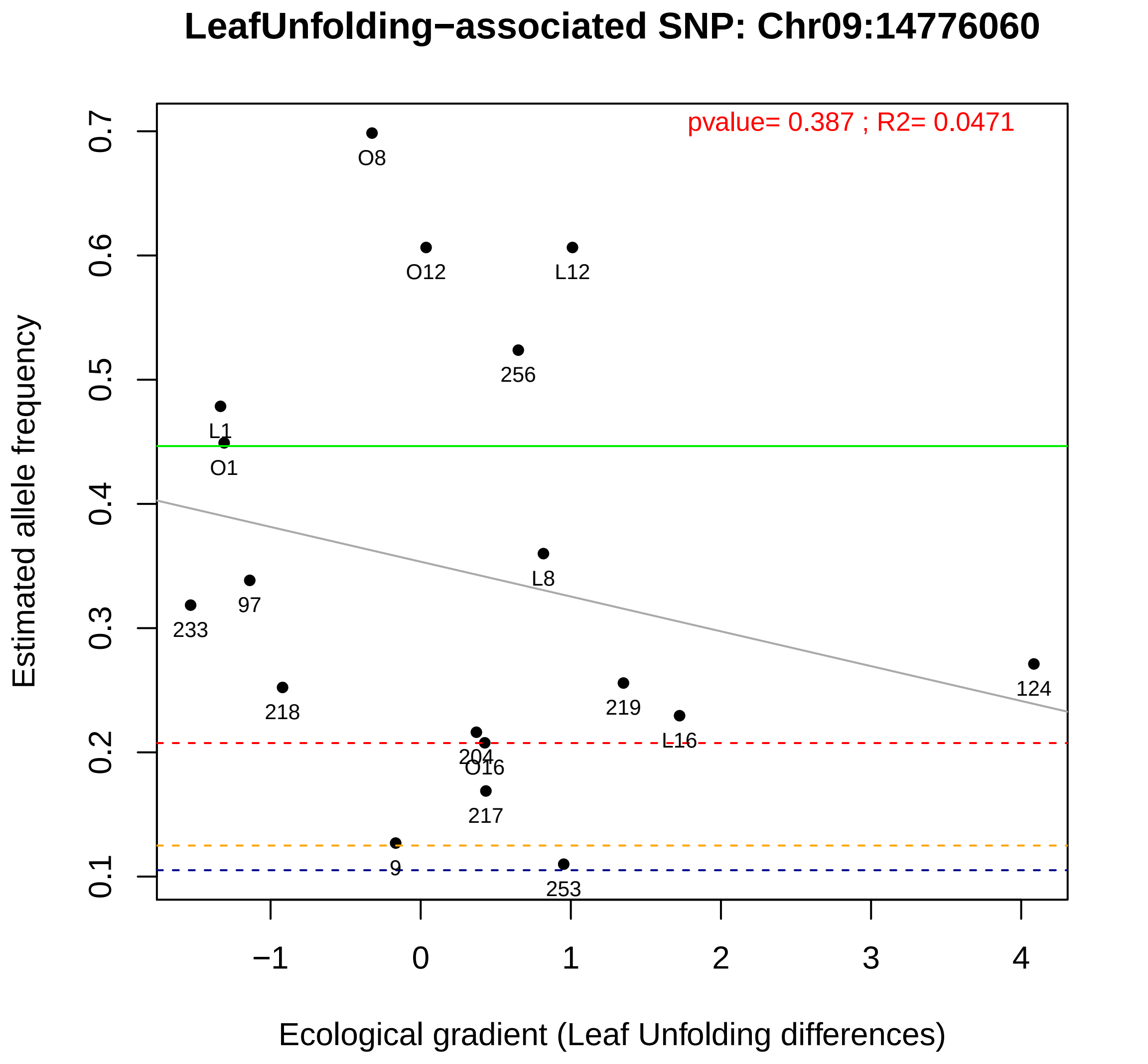
LeafUnfolding−associated SNP: Chr09:14776060
pvalue= 0.387 ; R2= 0.0471
Estimated allele frequency
Ecological gradient (Leaf Unfolding differences)
0.1
0.2
0.3
0.4
0.5
0.6
0.7
−1
0
1
2
3
4
O8
O12
L12
256
L1
O1
L8
97
233
218
124
219
L16
204
O16
217
9
253

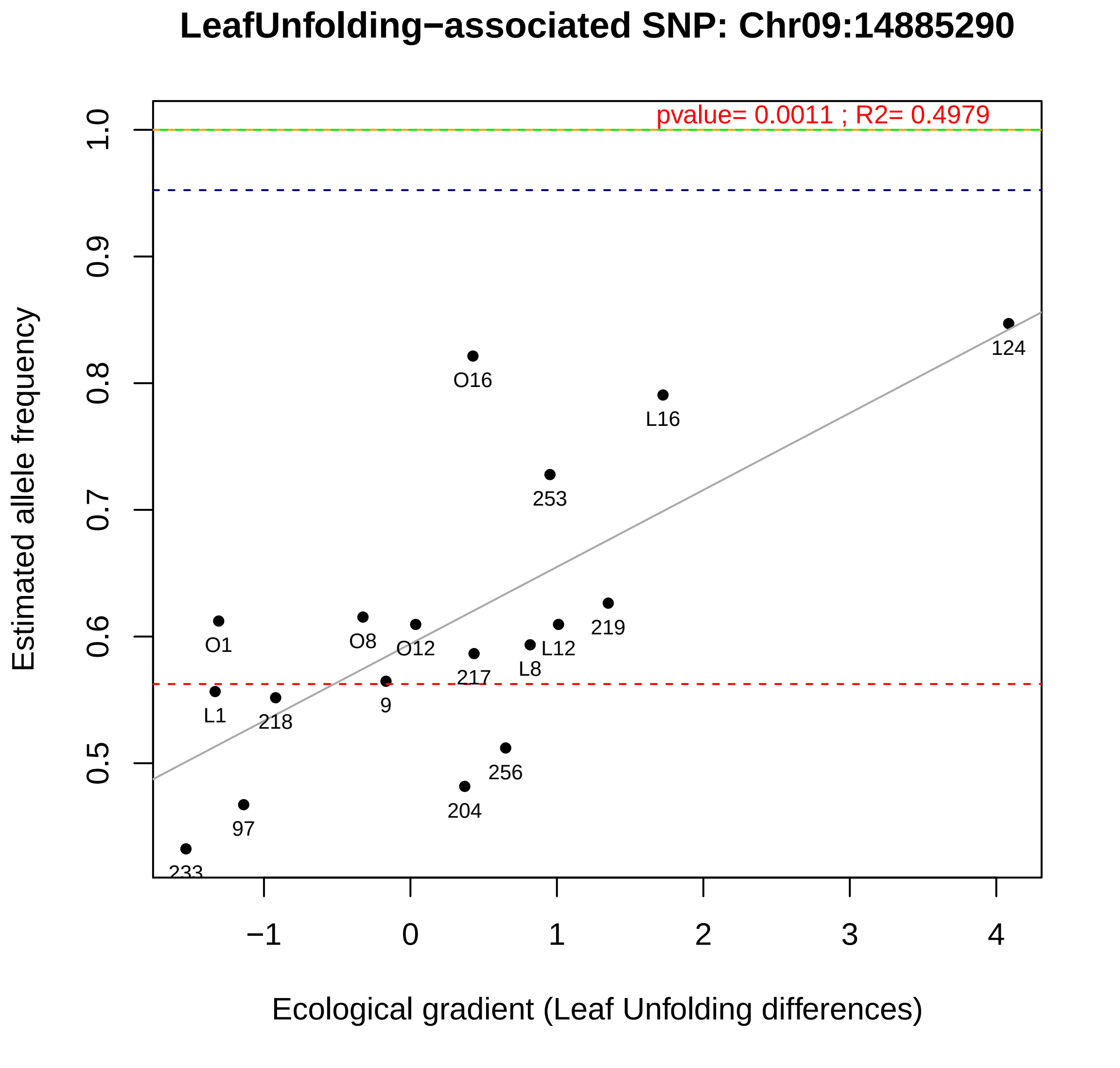

LeafUnfolding−associated SNP: Chr09:14885290
pvalue= 0.0011 ; R2= 0.4979
Estimated allele frequency
1.0
0.9
0.8
0.7
0.6
0.5
124
O16
L16
253
219
O1
O8
O12
L12
217
L8
9
L1
218
256
204
97
233
−1
0
1
2
3
4
Ecological gradient (Leaf Unfolding differences)

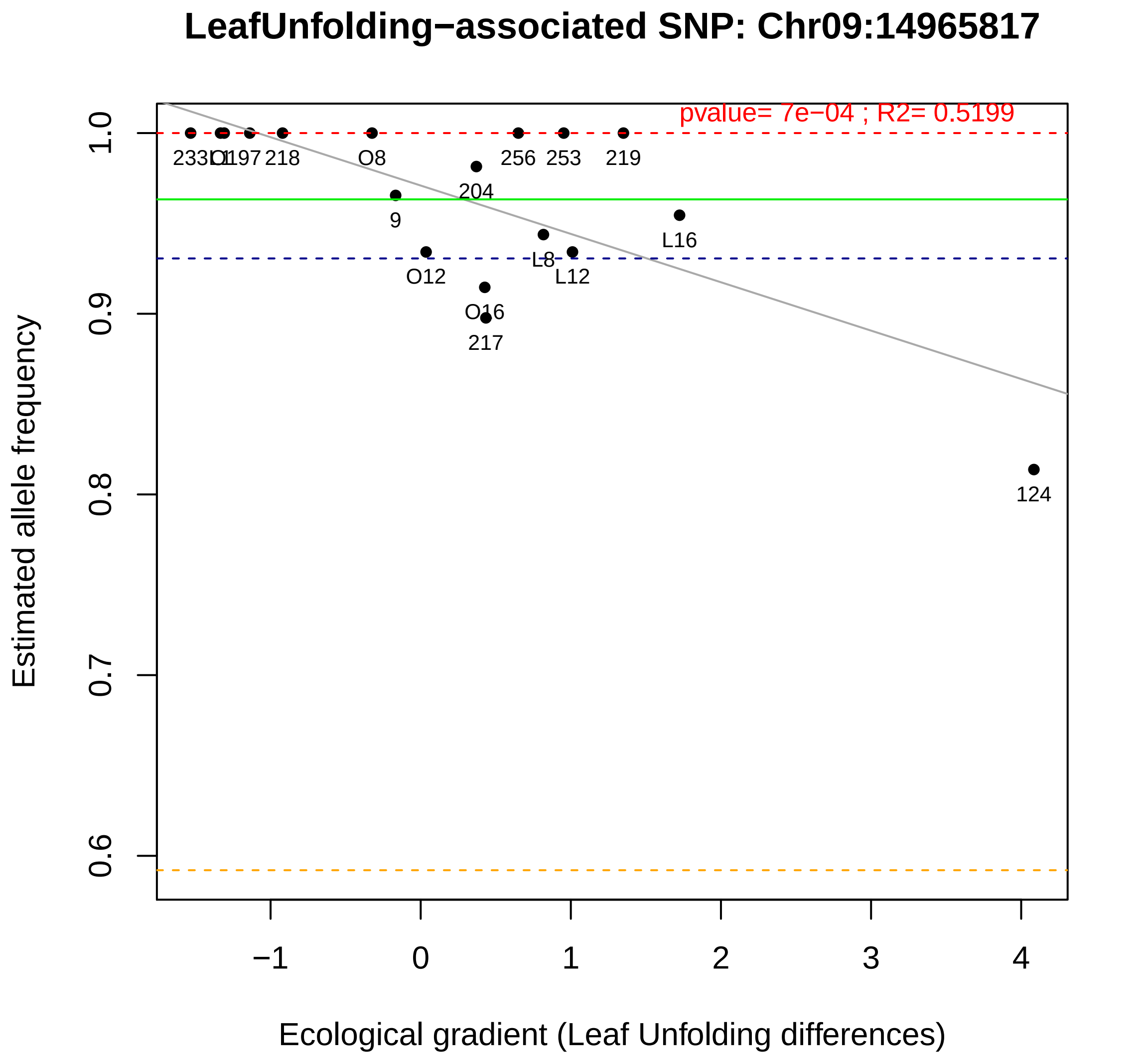

LeafUnfolding−associated SNP: Chr09:14965817
pvalue= 7e−04 ; R2= 0.5199
Estimated allele frequency
1.0
0.9
0.8
0.7
0.6
−1
0
1
2
3
4
Ecological gradient (Leaf Unfolding differences)
233
O1
197
218
O8
256
253
219
204
9
L16
L8
O12
L12
O16
217
124

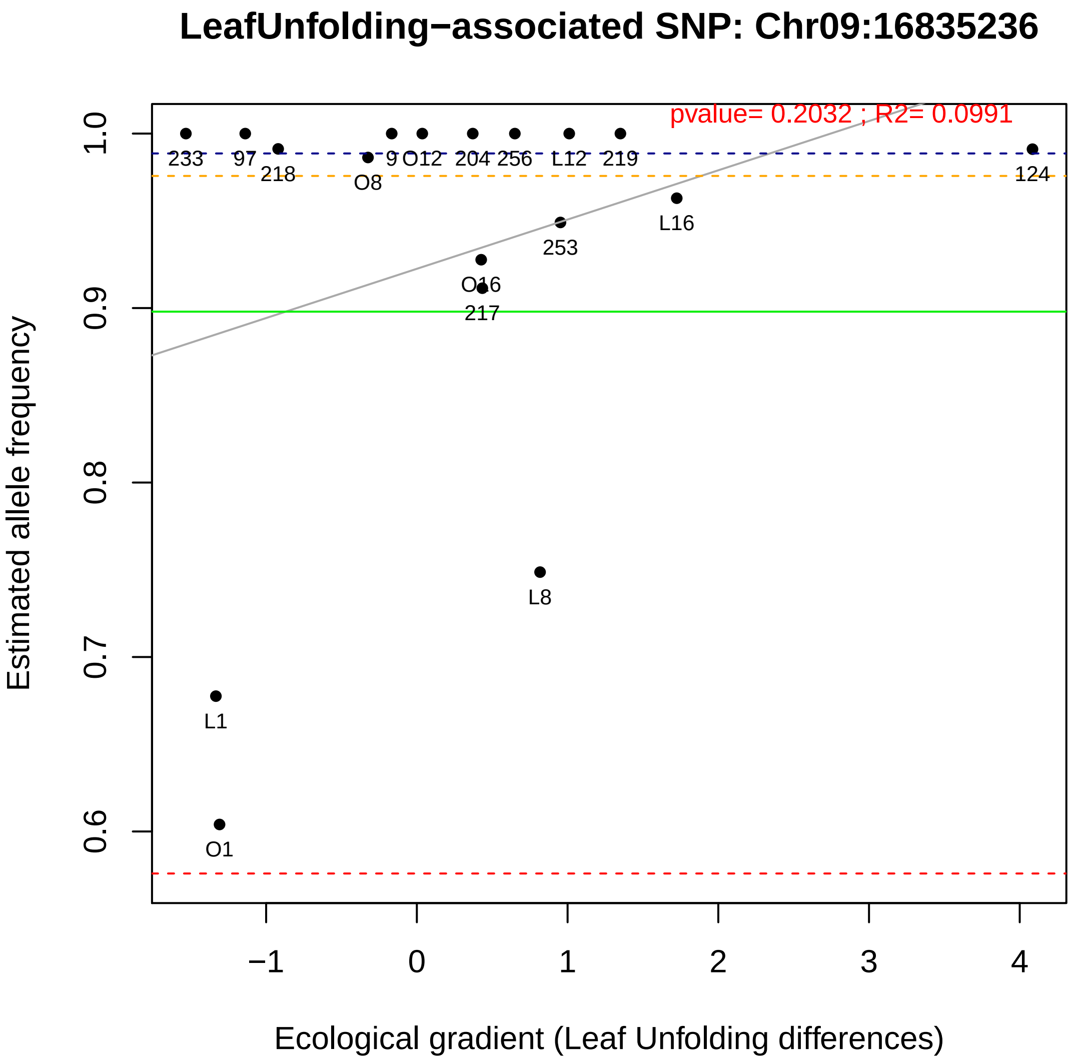

LeafUnfolding−associated SNP: Chr09:16835236
pvalue= 0.2032 ; R2= 0.0991
Estimated allele frequency
Ecological gradient (Leaf Unfolding differences)
1.0
0.9
0.8
0.7
0.6
−1
0
1
2
3
4
233
97
218
O8
9
O12
204
256
L12
219
124
L16
253
O16
217
L8
L1
O1

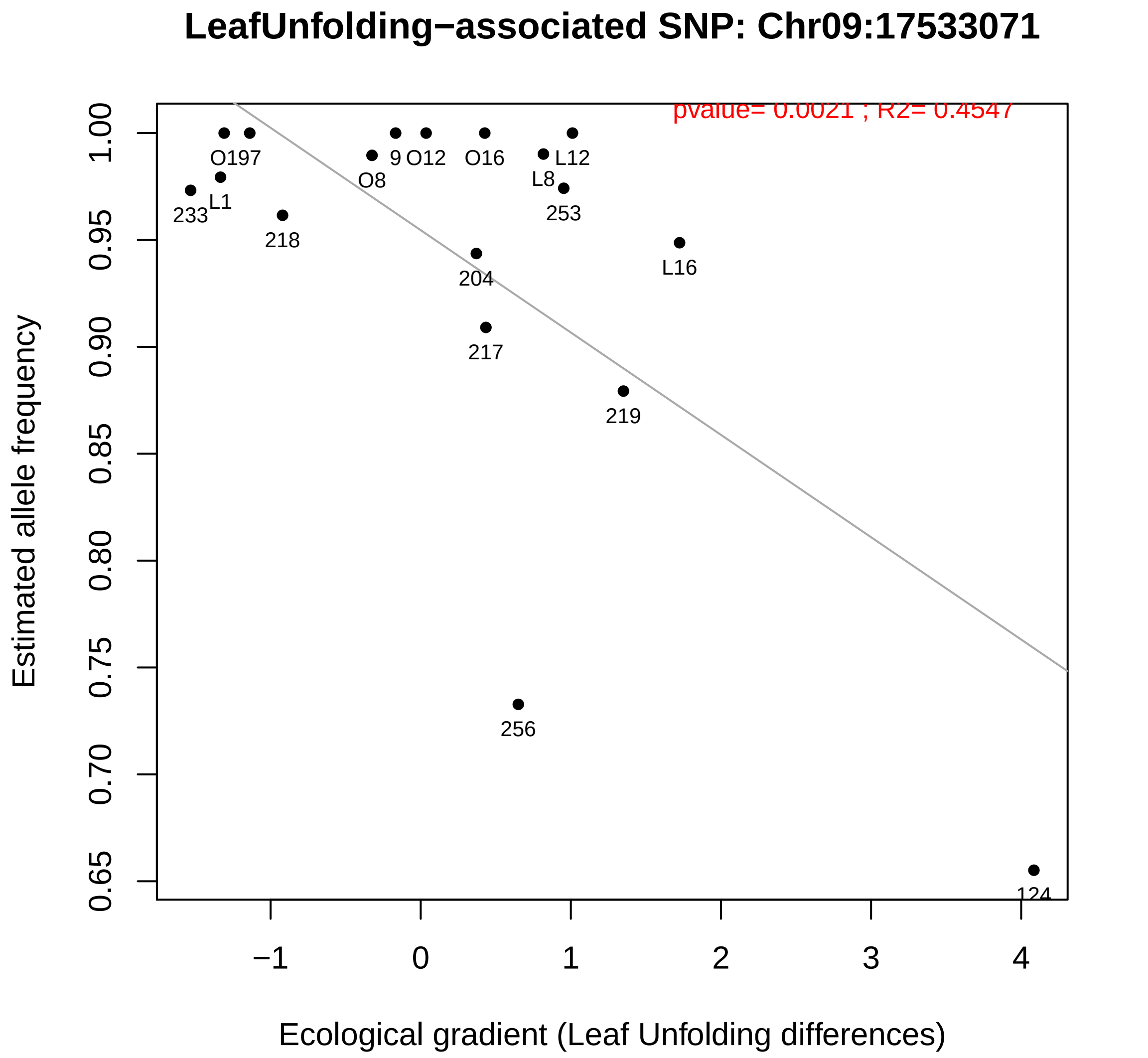

LeafUnfolding−associated SNP: Chr09:17533071
pvalue= 0.0021 ; R2= 0.4547
Estimated allele frequency
1.00
0.95
0.90
0.85
0.80
0.75
0.70
0.65
−1
0
1
2
3
4
Ecological gradient (Leaf Unfolding differences)
O197
L1
233
218
O8
9
O12
O16
L8
L12
253
204
L16
217
219
256
124

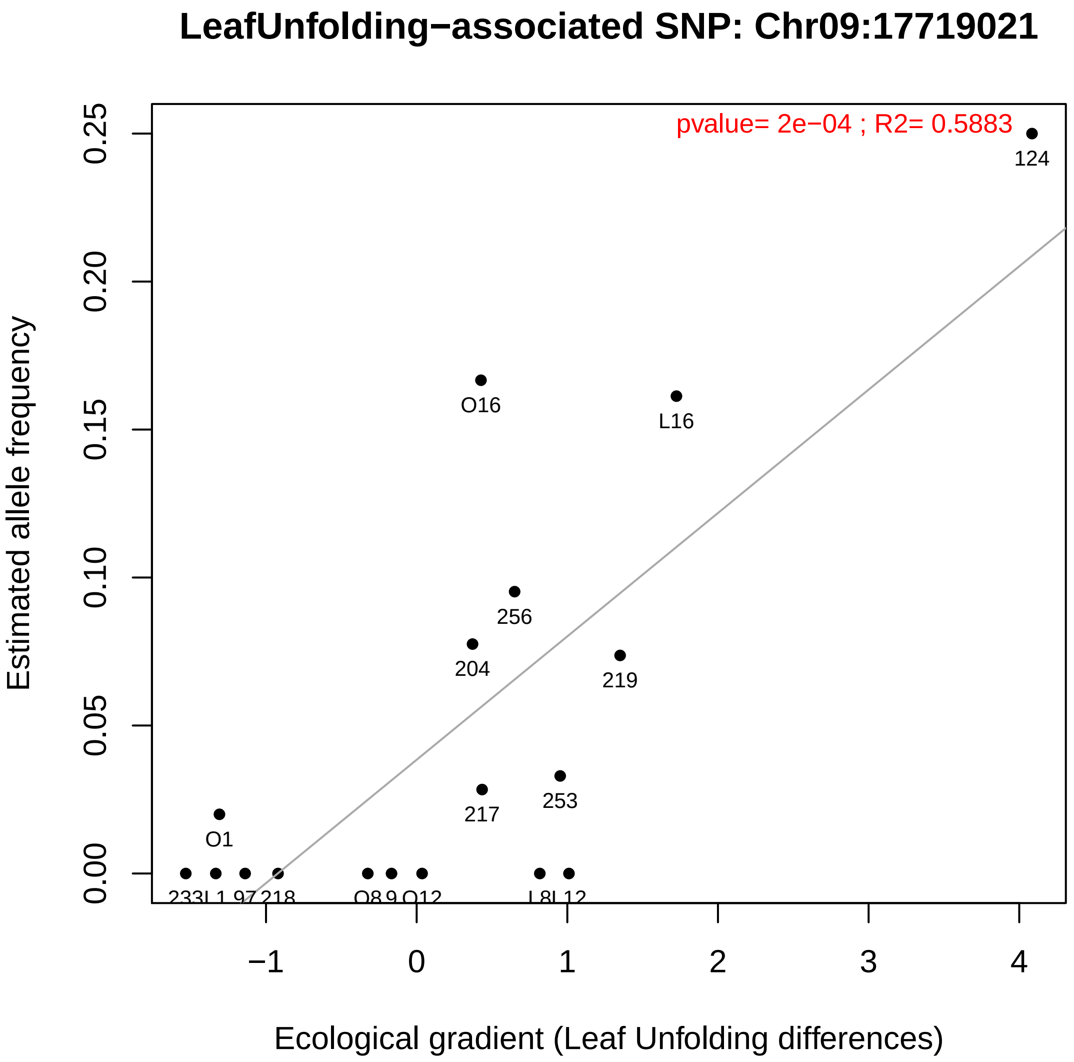
LeafUnfolding−associated SNP: Chr09:17719021
pvalue= 2e−04 ; R2= 0.5883
Estimated allele frequency
Ecological gradient (Leaf Unfolding differences)
0.00
0.05
0.10
0.15
0.20
0.25
−1
0
1
2
3
4
124
O16
L16
256
204
219
253
217
O1

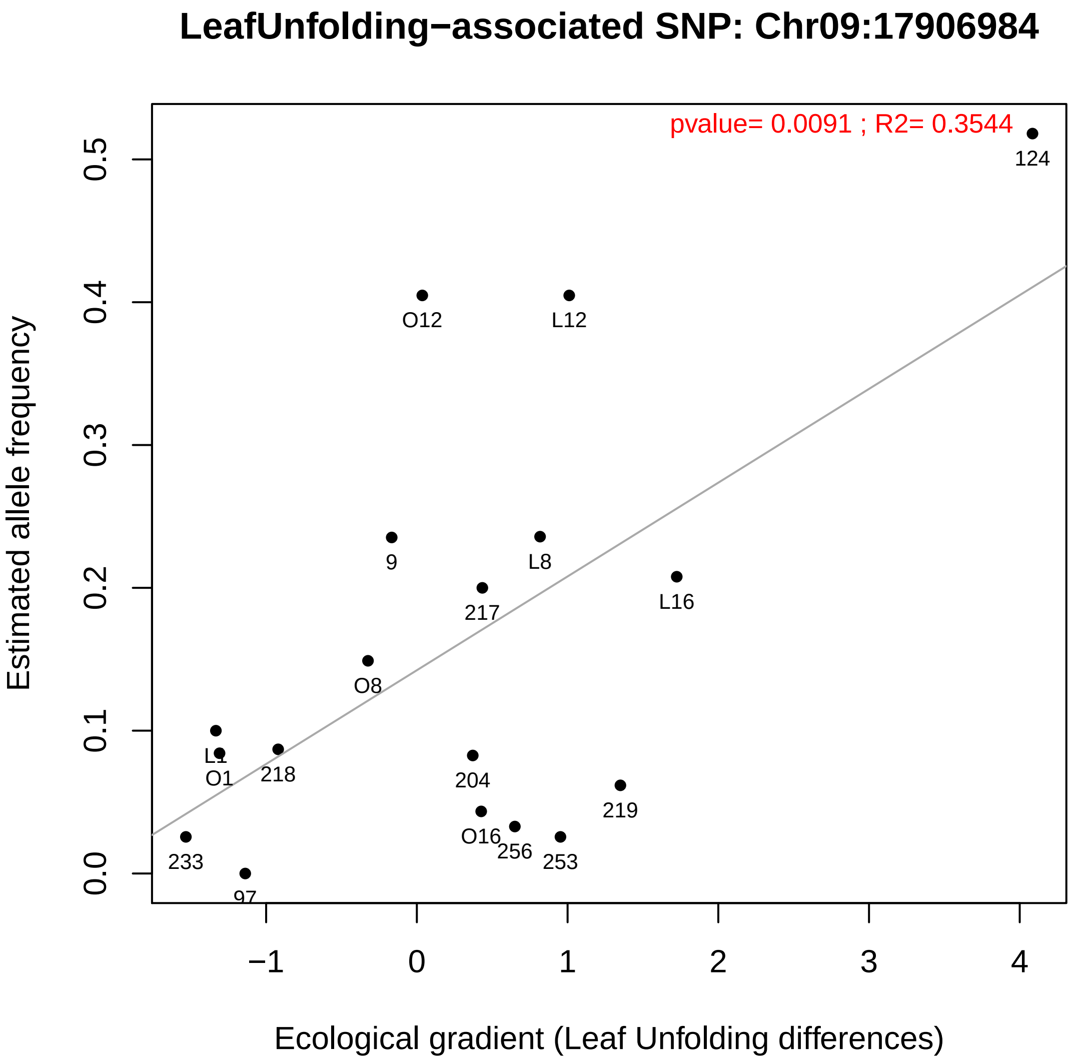

LeafUnfolding−associated SNP: Chr09:17906984
pvalue= 0.0091 ; R2= 0.3544
Estimated allele frequency
0.0
0.1
0.2
0.3
0.4
0.5
−1
0
1
2
3
4
Ecological gradient (Leaf Unfolding differences)
124
O12
L12
9
L8
217
L16
O8
L1
O1
218
204
219
O16
256
253
233
97

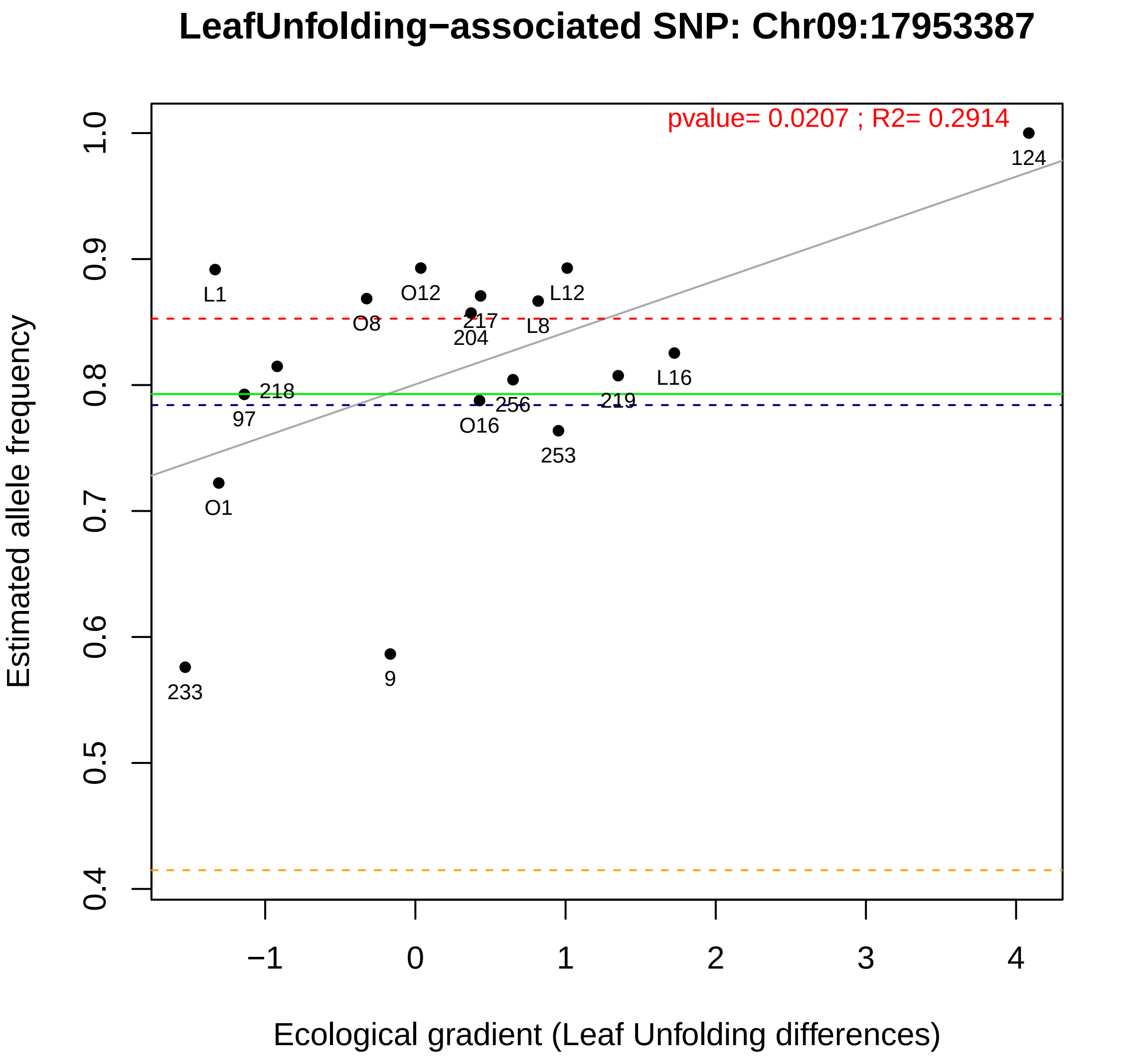

LeafUnfolding−associated SNP: Chr09:17953387
pvalue= 0.0207 ; R2= 0.2914
Estimated allele frequency
Ecological gradient (Leaf Unfolding differences)
0.4
0.5
0.6
0.7
0.8
0.9
1.0
−1
0
1
2
3
4
124
L1
O12
L12
O8
217
204
L8
218
97
256
219
L16
O16
253
O1
233
9

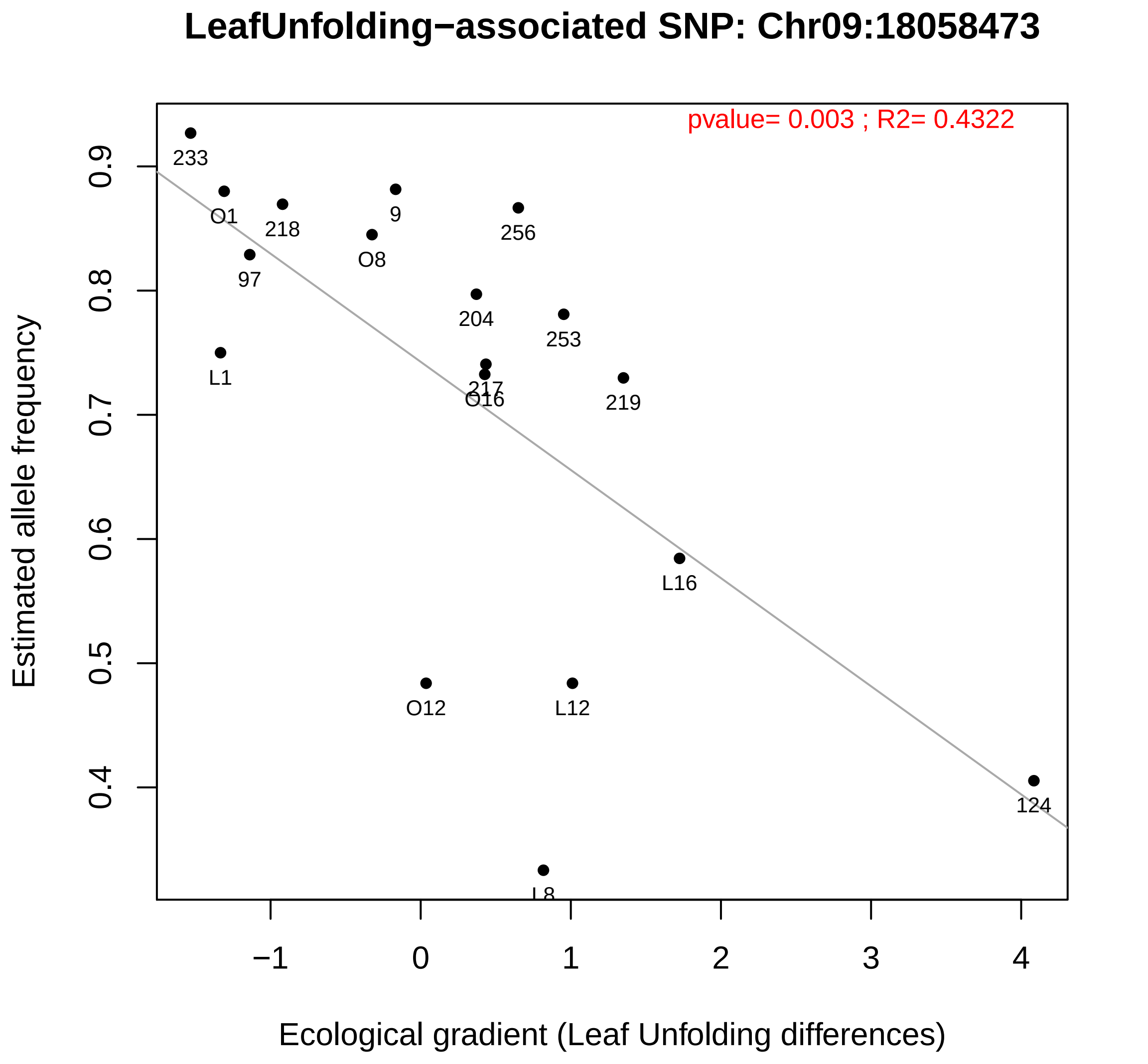
LeafUnfolding−associated SNP: Chr09:18058473
pvalue= 0.003 ; R2= 0.4322
Estimated allele frequency
Ecological gradient (Leaf Unfolding differences)
0.4
0.5
0.6
0.7
0.8
0.9
−1
0
1
2
3
4
233
O1
218
9
256
O8
97
204
253
L1
217
O16
219
L16
O12
L12
124
L8

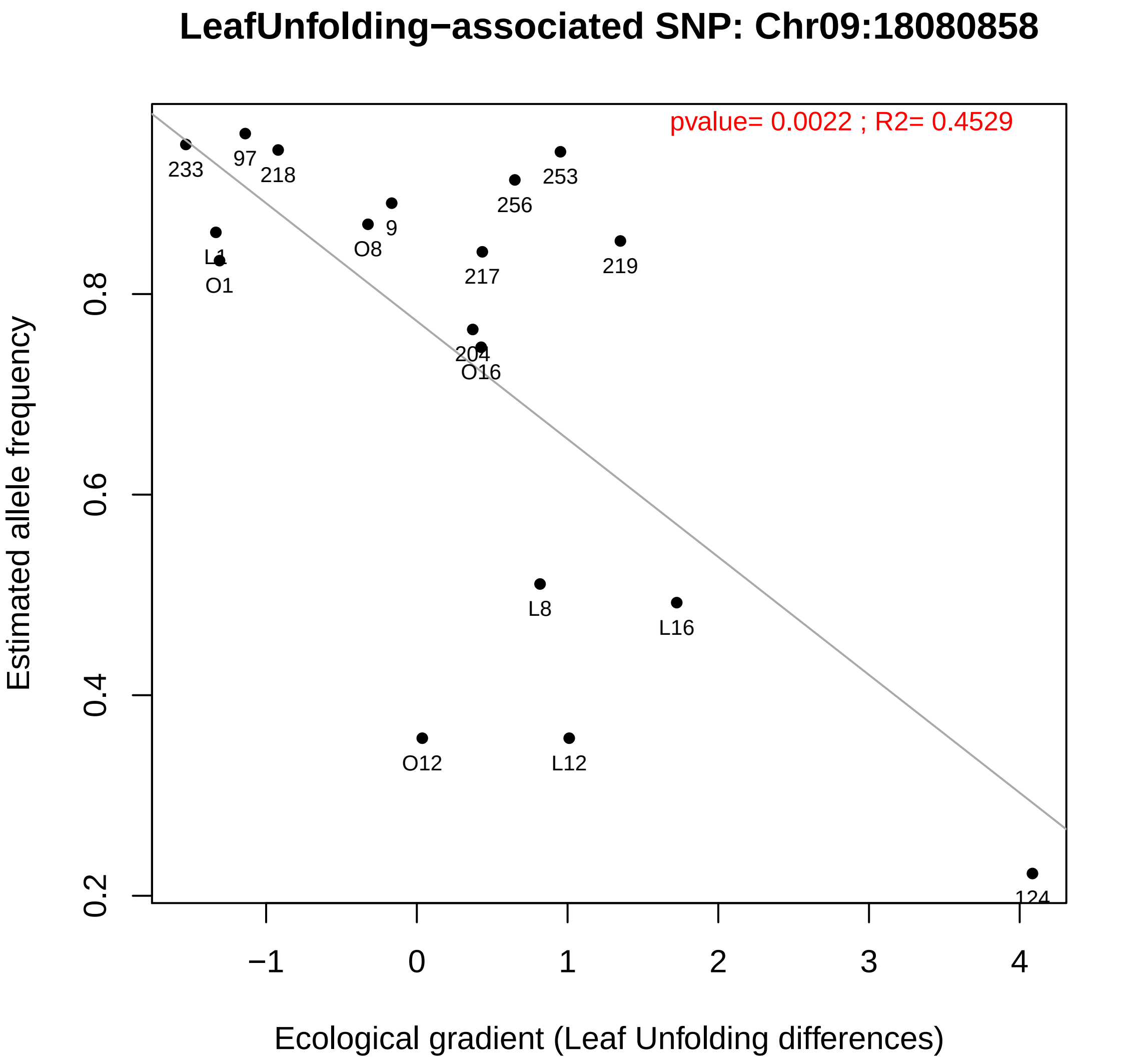
LeafUnfolding−associated SNP: Chr09:18080858
pvalue= 0.0022 ; R2= 0.4529
233
97
218
253
256
9
O8
L1
O1
217
219
204
O16
L8
L16
O12
L12
124
Estimated allele frequency
0.2
0.4
0.6
0.8
−1
0
1
2
3
4
Ecological gradient (Leaf Unfolding differences)

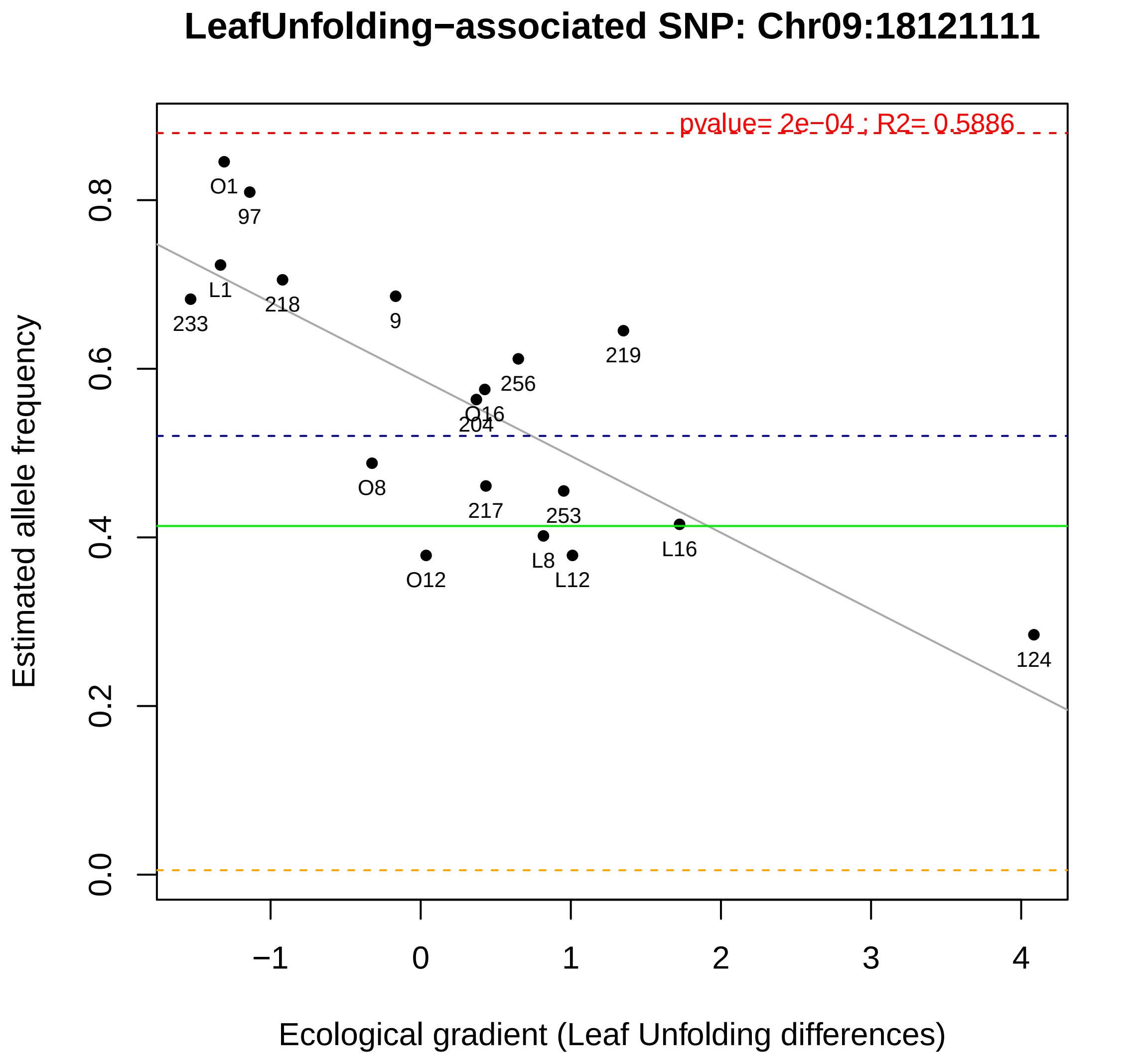
LeafUnfolding−associated SNP: Chr09:18121111
pvalue= 2e−04 ; R2= 0.5886
Estimated allele frequency
Ecological gradient (Leaf Unfolding differences)
0.0
0.2
0.4
0.6
0.8
−1
0
1
2
3
4
O1
97
L1
218
233
9
219
256
O16
204
O8
217
253
L16
L8
L12
O12
124

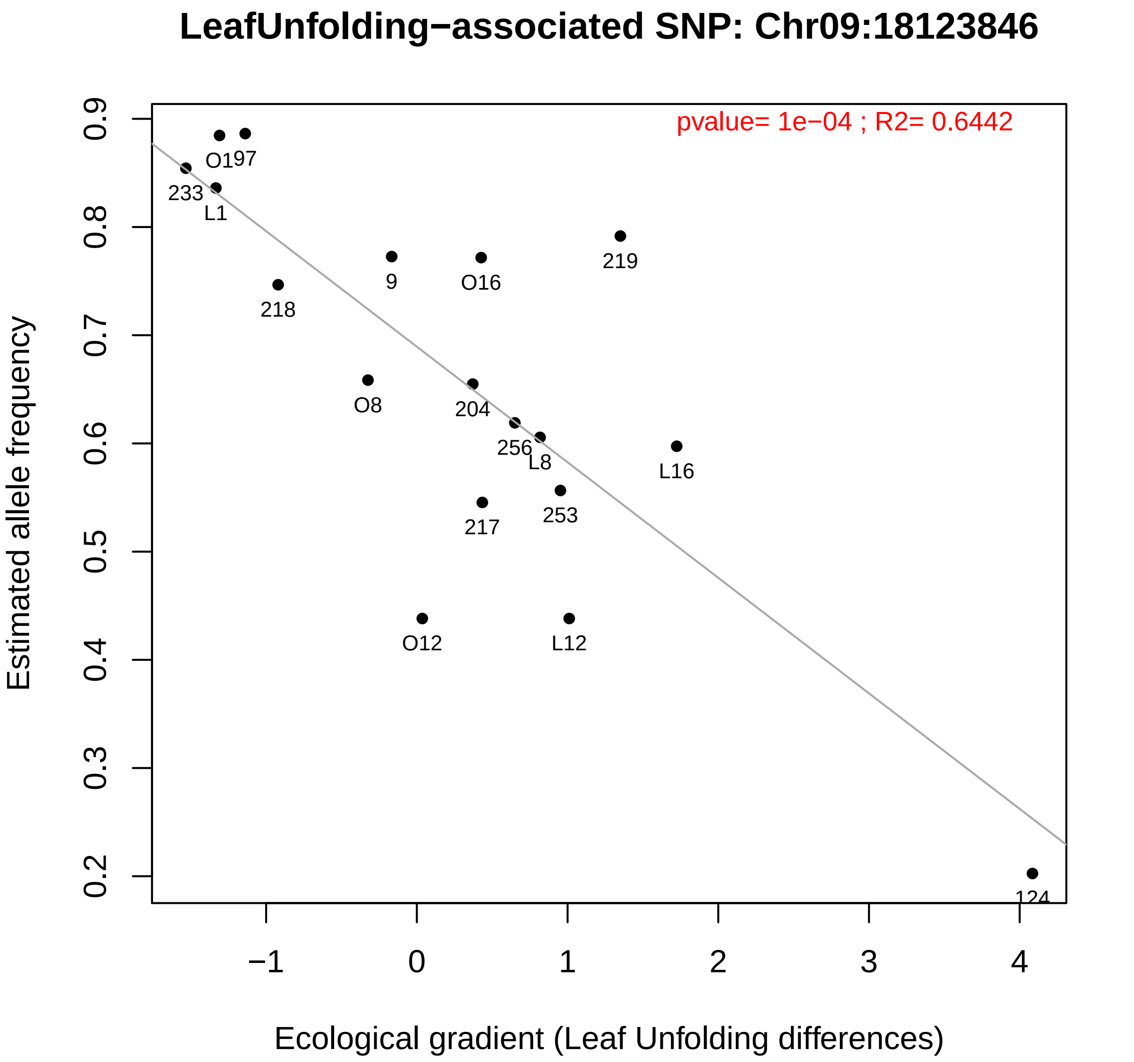

LeafUnfolding−associated SNP: Chr09:18123846
pvalue= 1e−04 ; R2= 0.6442
Estimated allele frequency
Ecological gradient (Leaf Unfolding differences)
0.2
0.3
0.4
0.5
0.6
0.7
0.8
0.9
−1
0
1
2
3
4
O1
97
233
L1
218
9
O16
219
O8
204
256
L8
L16
217
253
O12
L12
124

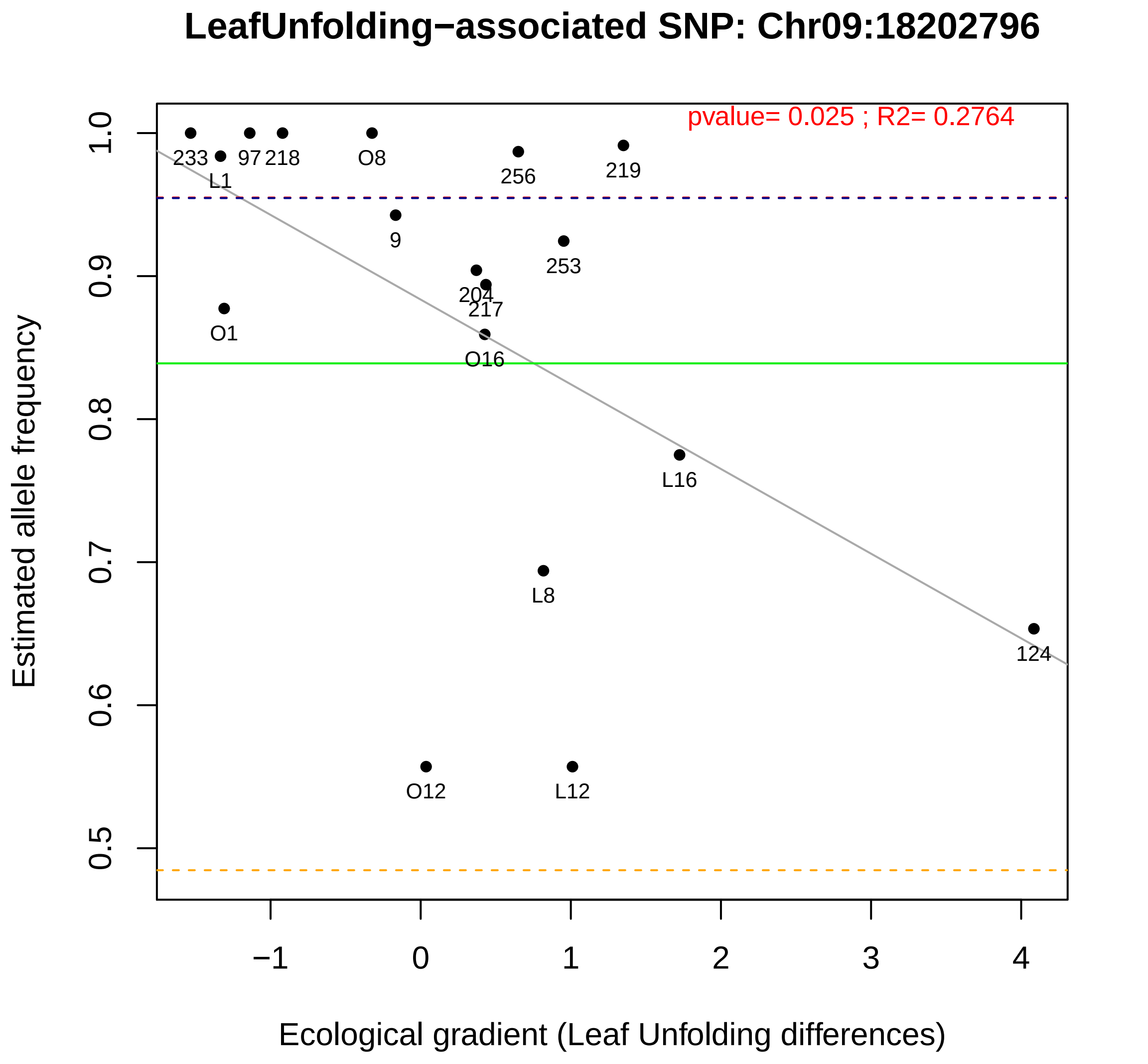

LeafUnfolding−associated SNP: Chr09:18202796
pvalue= 0.025 ; R2= 0.2764
Estimated allele frequency
Ecological gradient (Leaf Unfolding differences)
1.0
0.9
0.8
0.7
0.6
0.5
−1
0
1
2
3
4
233
L1
97
218
O8
256
219
9
253
204
217
O1
O16
L16
L8
124
O12
L12

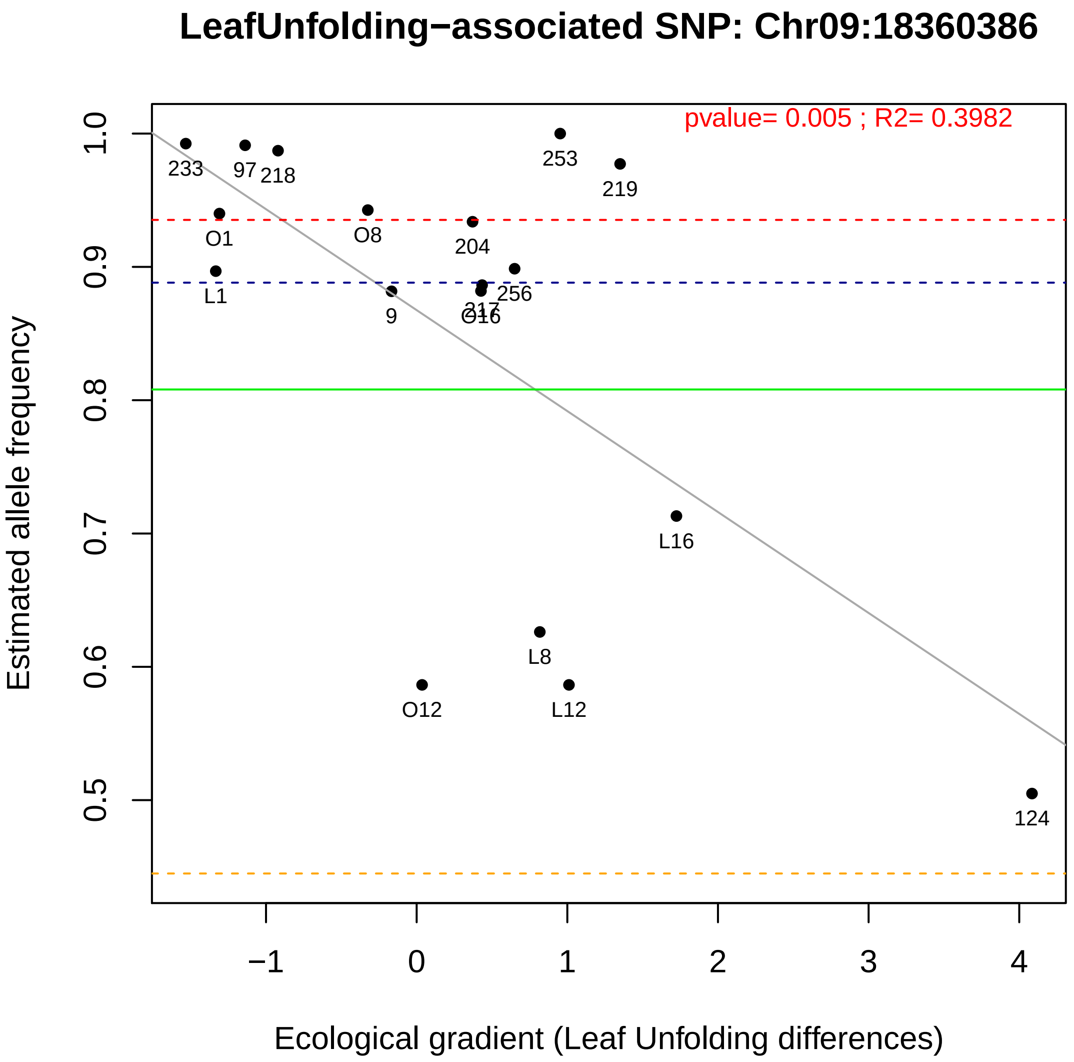

LeafUnfolding−associated SNP: Chr09:18360386
pvalue= 0.005 ; R2= 0.3982
Estimated allele frequency
Ecological gradient (Leaf Unfolding differences)
0.5
0.6
0.7
0.8
0.9
1.0
−1
0
1
2
3
4
233
97
218
253
219
O1
O8
204
L1
9
256
217
O16
L16
L8
O12
L12
124

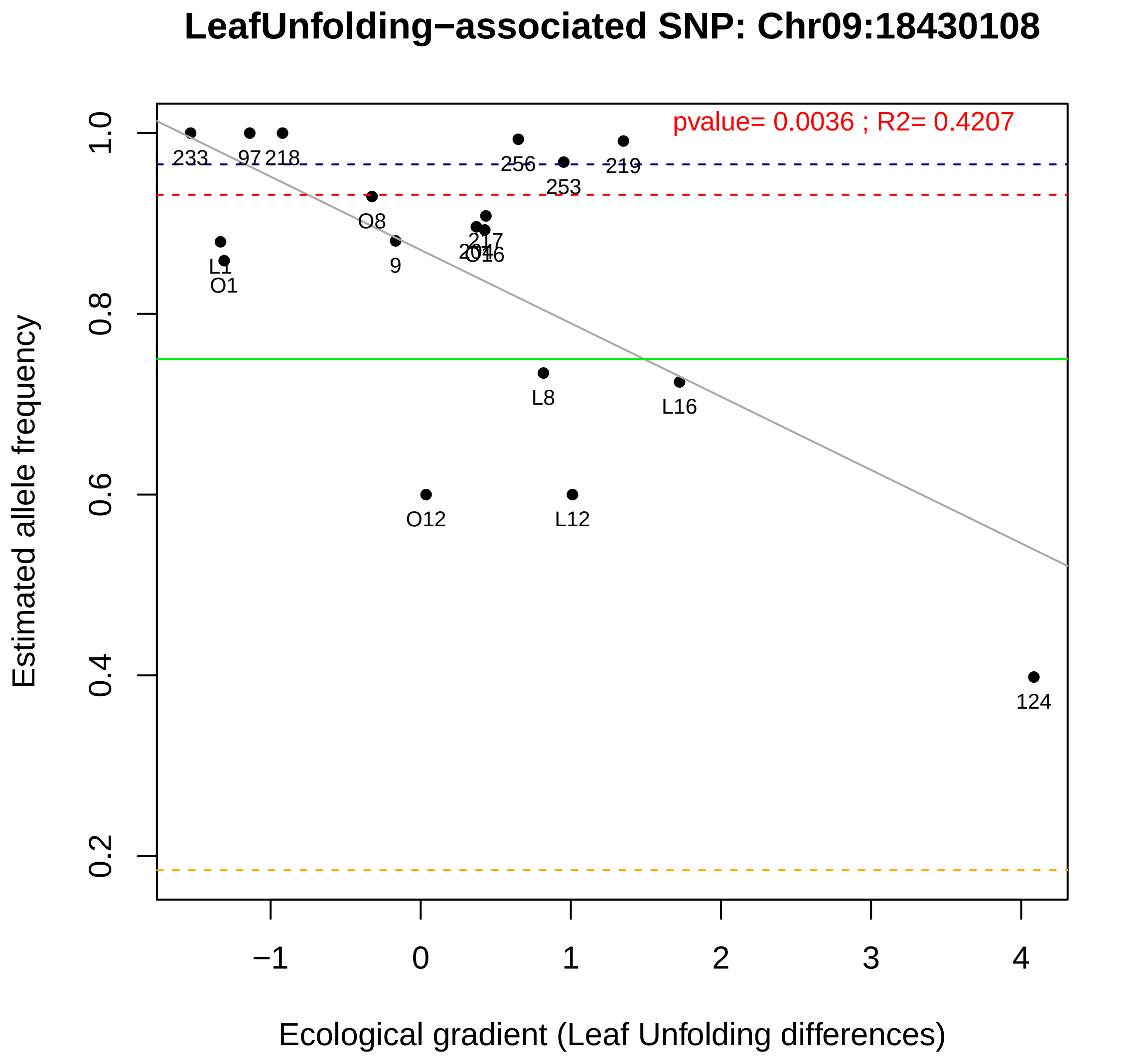
LeafUnfolding−associated SNP: Chr09:18430108
pvalue= 0.0036 ; R2= 0.4207
Estimated allele frequency
Ecological gradient (Leaf Unfolding differences)
1.0
0.8
0.6
0.4
0.2
−1
0
1
2
3
4
233
97
218
256
219
253
O8
9
217
204
O16
L1
O1
L8
L16
O12
L12
124

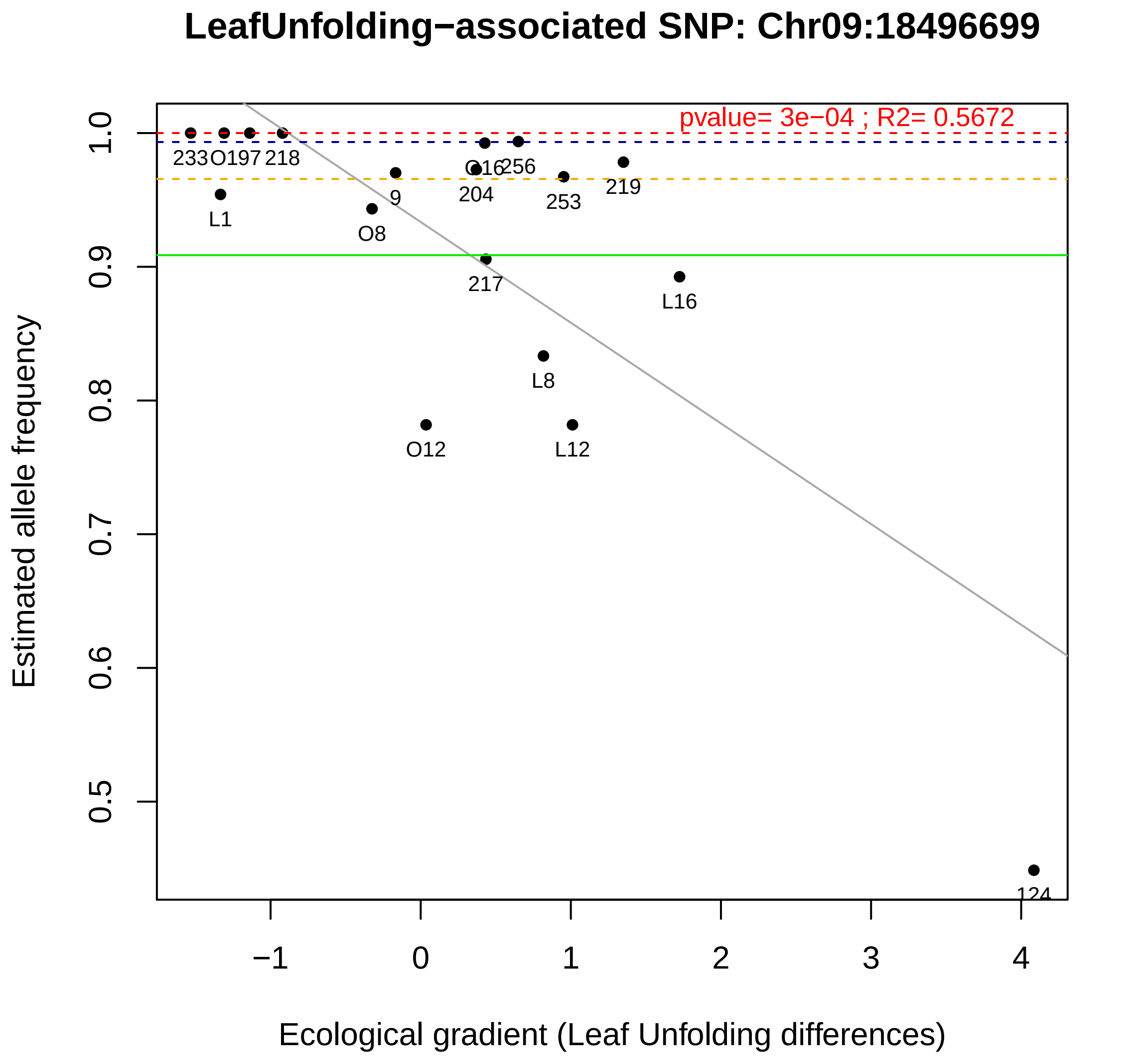
LeafUnfolding−associated SNP: Chr09:18496699
pvalue= 3e−04 ; R2= 0.5672
Estimated allele frequency
0.5
0.6
0.7
0.8
0.9
1.0
−1
0
1
2
3
4
Ecological gradient (Leaf Unfolding differences)
233
O197
218
L1
O8
9
O16
204
256
253
219
217
L16
L8
O12
L12

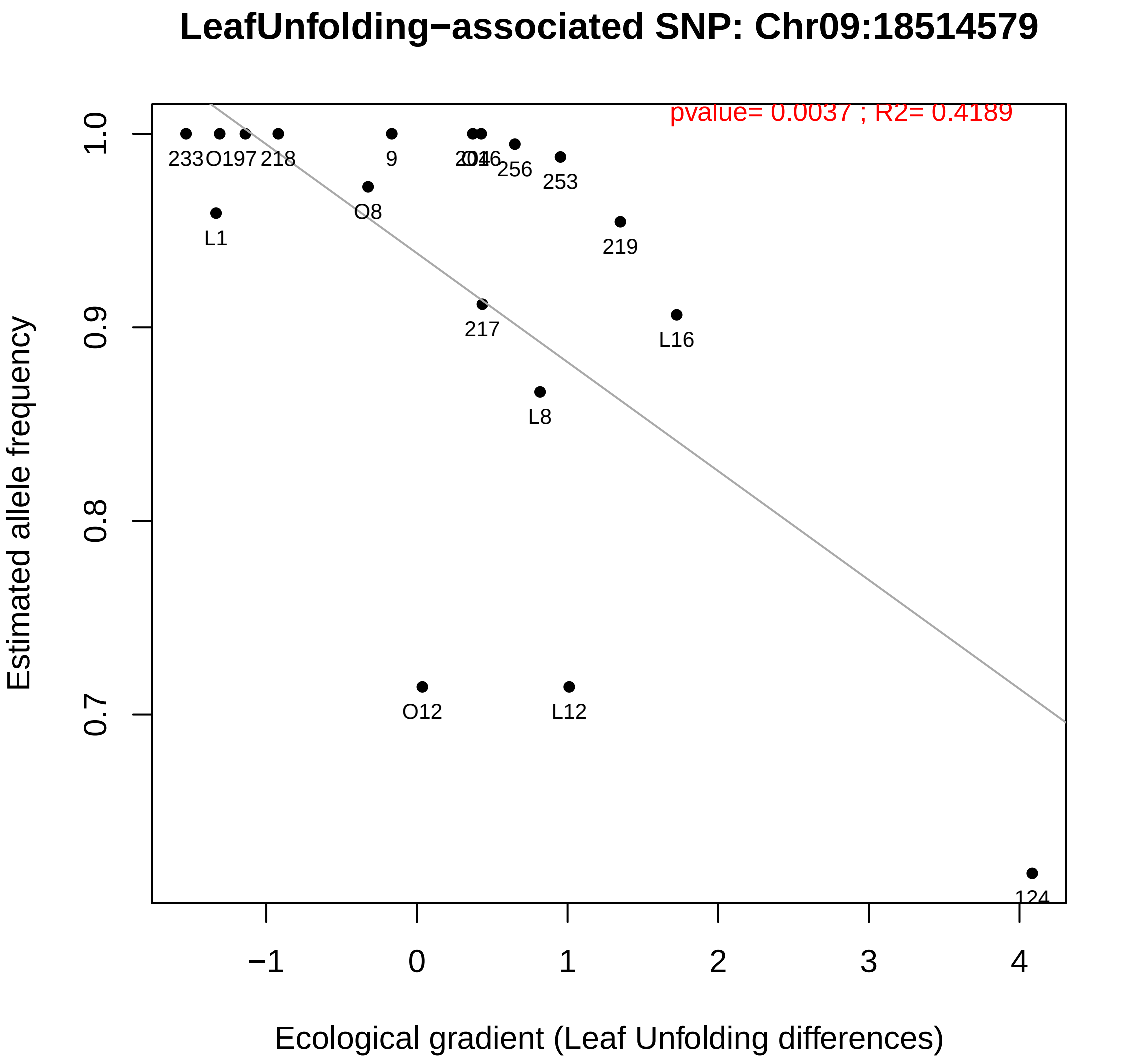
LeafUnfolding−associated SNP: Chr09:18514579
pvalue= 0.0037 ; R2= 0.4189
Estimated allele frequency
Ecological gradient (Leaf Unfolding differences)
1.0
0.9
0.8
0.7
−1
0
1
2
3
4
233
O197
218
9
256
253
L1
O8
219
217
L16
L8
O12
L12
124

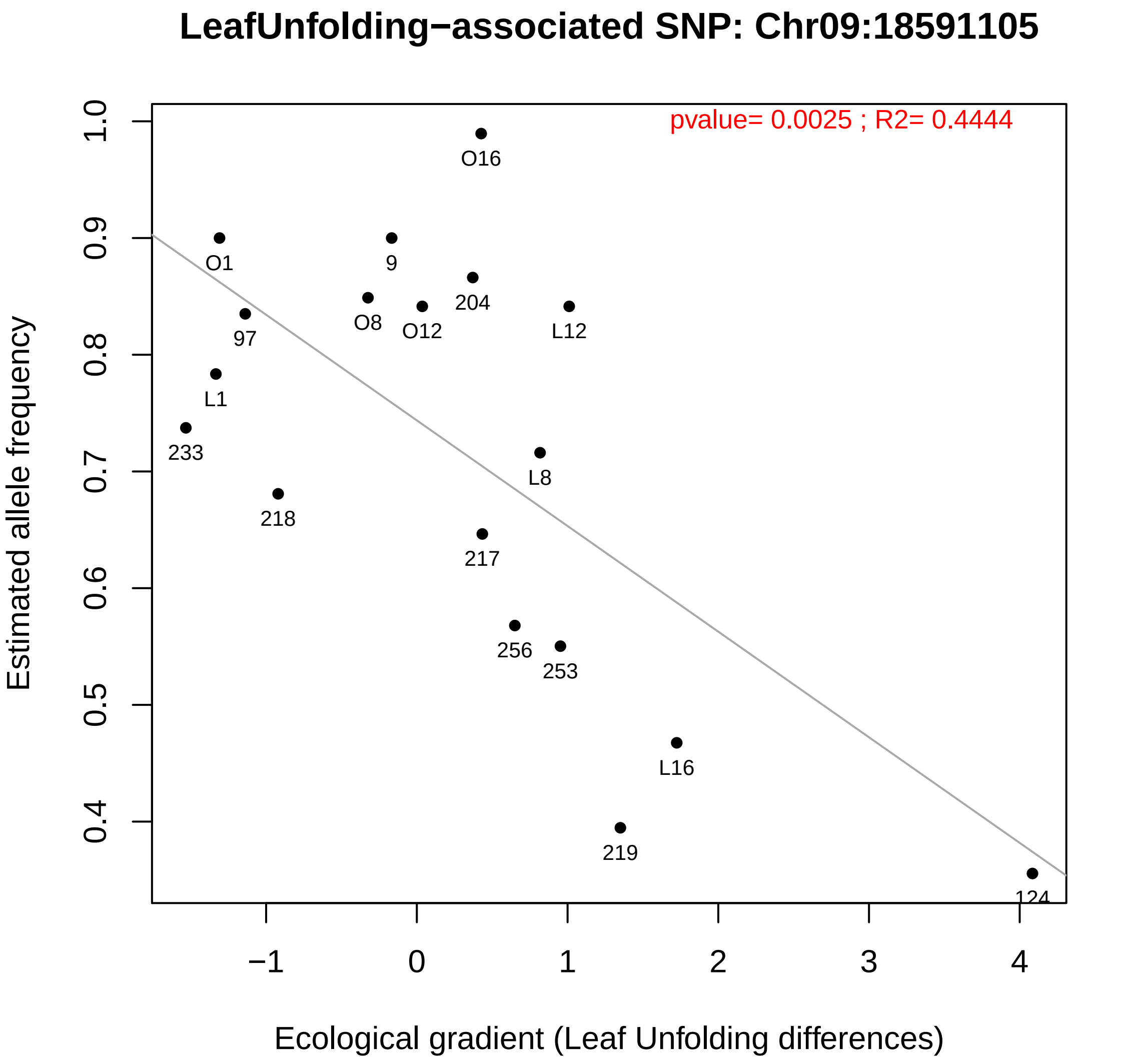
LeafUnfolding−associated SNP: Chr09:18591105
pvalue= 0.0025 ; R2= 0.4444
Estimated allele frequency
0.4
0.5
0.6
0.7
0.8
0.9
1.0
−1
0
1
2
3
4
Ecological gradient (Leaf Unfolding differences)
O16
O1
9
204
O8
O12
L12
97
L1
233
L8
218
217
256
253
L16
219
124

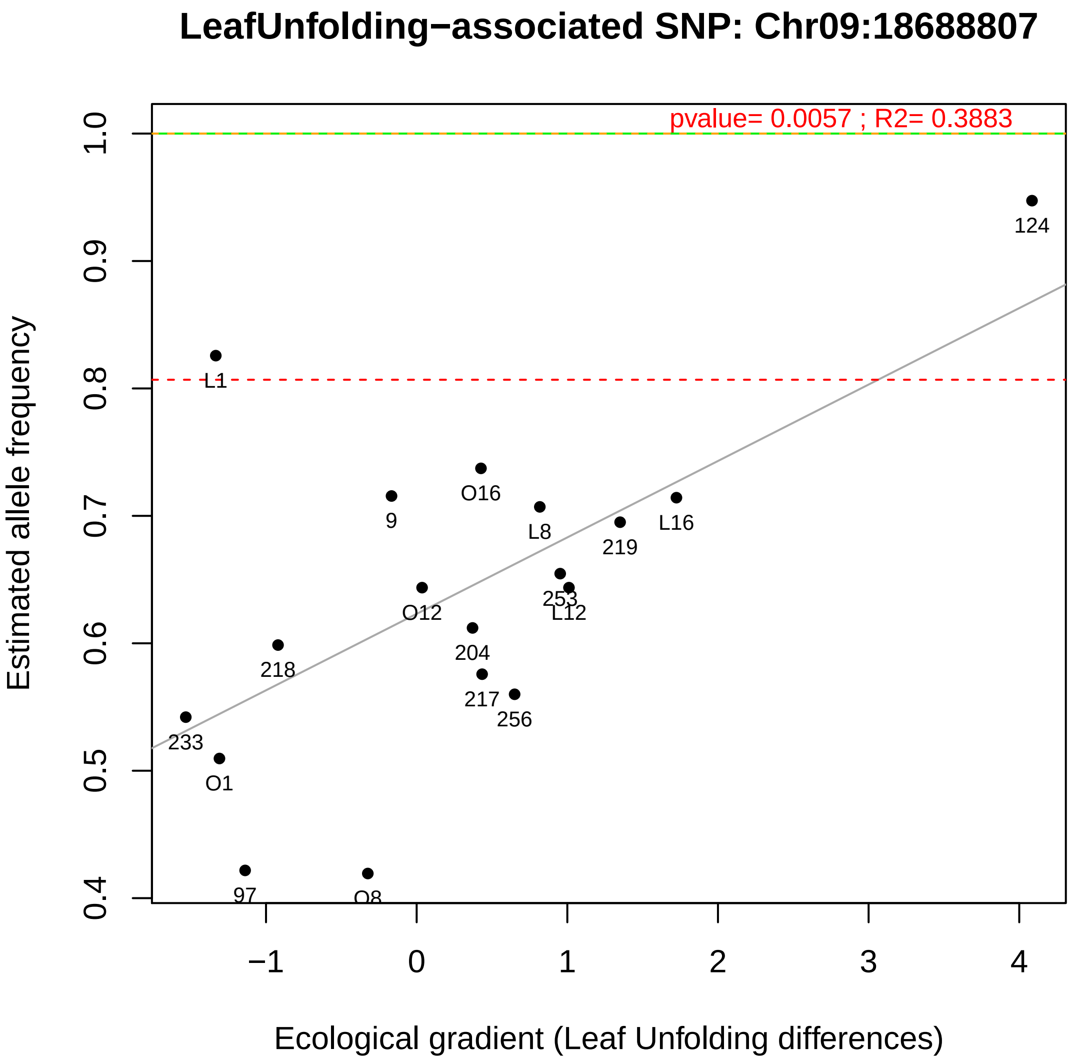

LeafUnfolding−associated SNP: Chr09:18688807
pvalue= 0.0057 ; R2= 0.3883
Estimated allele frequency
Ecological gradient (Leaf Unfolding differences)
0.4
0.5
0.6
0.7
0.8
0.9
1.0
−1
0
1
2
3
4
124
L1
O16
9
L8
219
L16
253
L12
O12
204
218
217
256
233
O1
97
O8

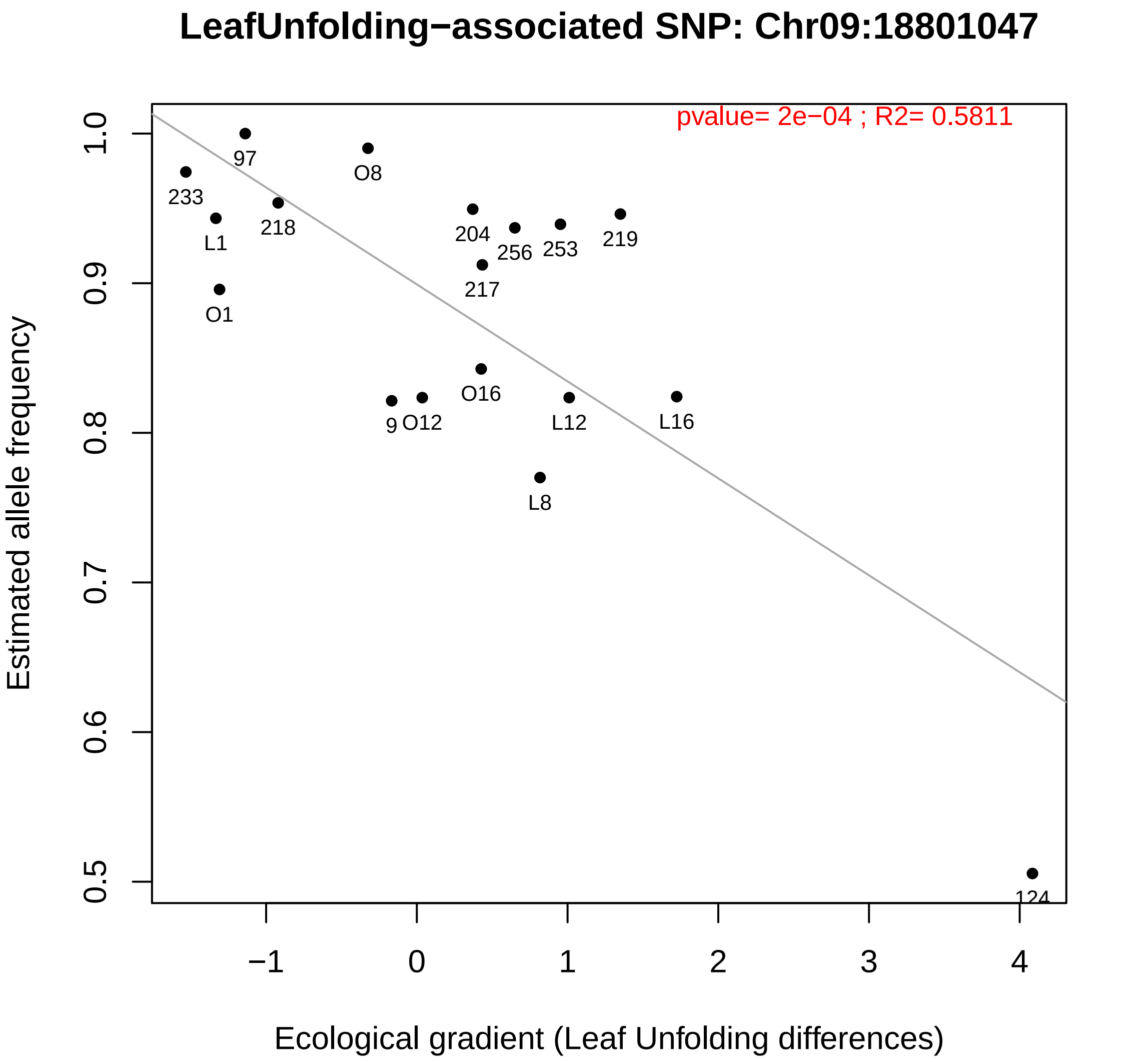

LeafUnfolding−associated SNP: Chr09:18801047
pvalue= 2e−04 ; R2= 0.5811
Estimated allele frequency
Ecological gradient (Leaf Unfolding differences)
0.5
0.6
0.7
0.8
0.9
1.0
−1
0
1
2
3
4
97
233
O8
L1
218
204
256
253
219
217
O1
O16
9
O12
L12
L16
L8
124

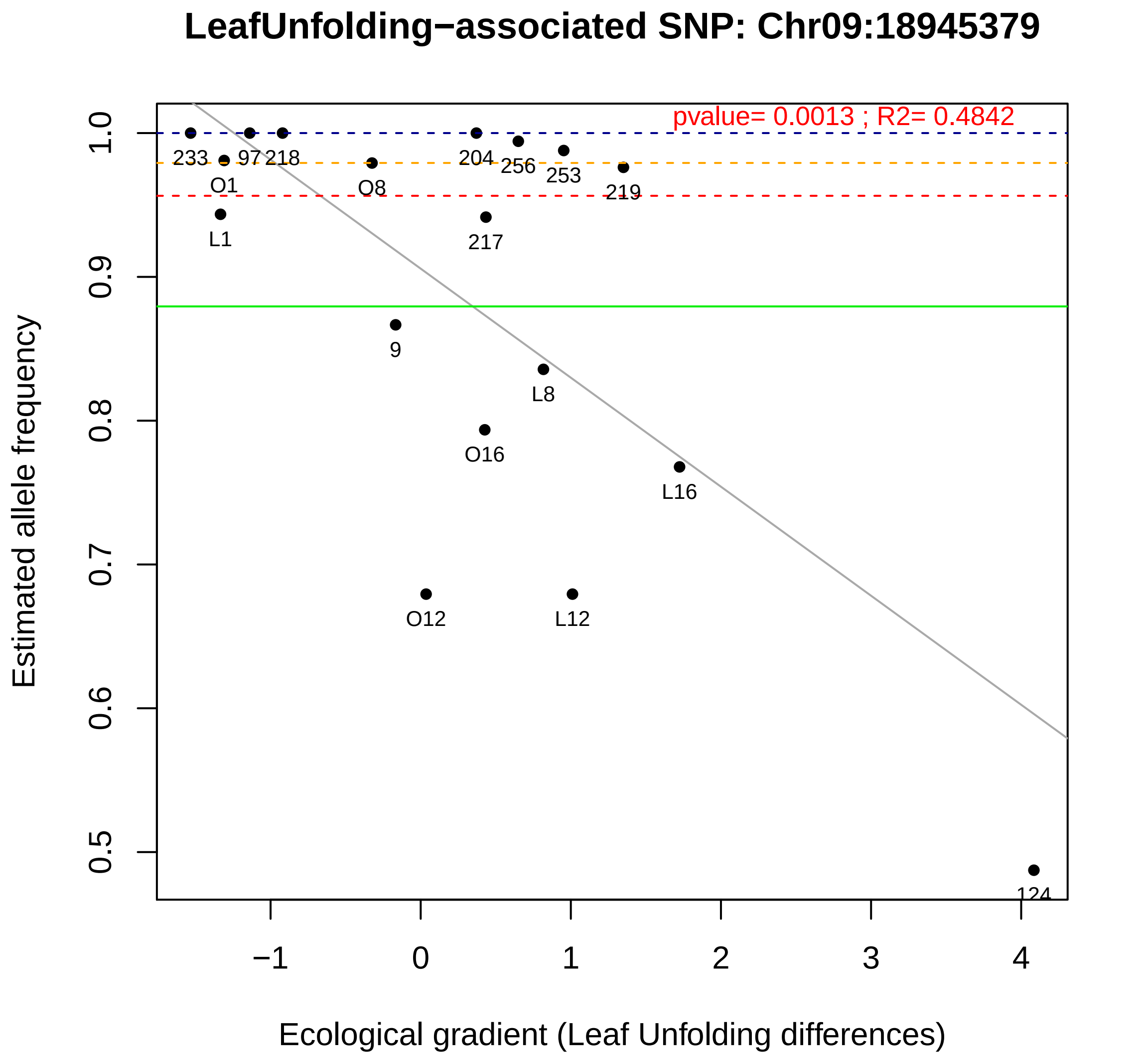
LeafUnfolding−associated SNP: Chr09:18945379
pvalue= 0.0013 ; R2= 0.4842
Estimated allele frequency
Ecological gradient (Leaf Unfolding differences)
0.5
0.6
0.7
0.8
0.9
1.0
−1
0
1
2
3
4
233
97
218
O1
L1
O8
204
256
253
219
217
9
L8
O16
L16
O12
L12
124

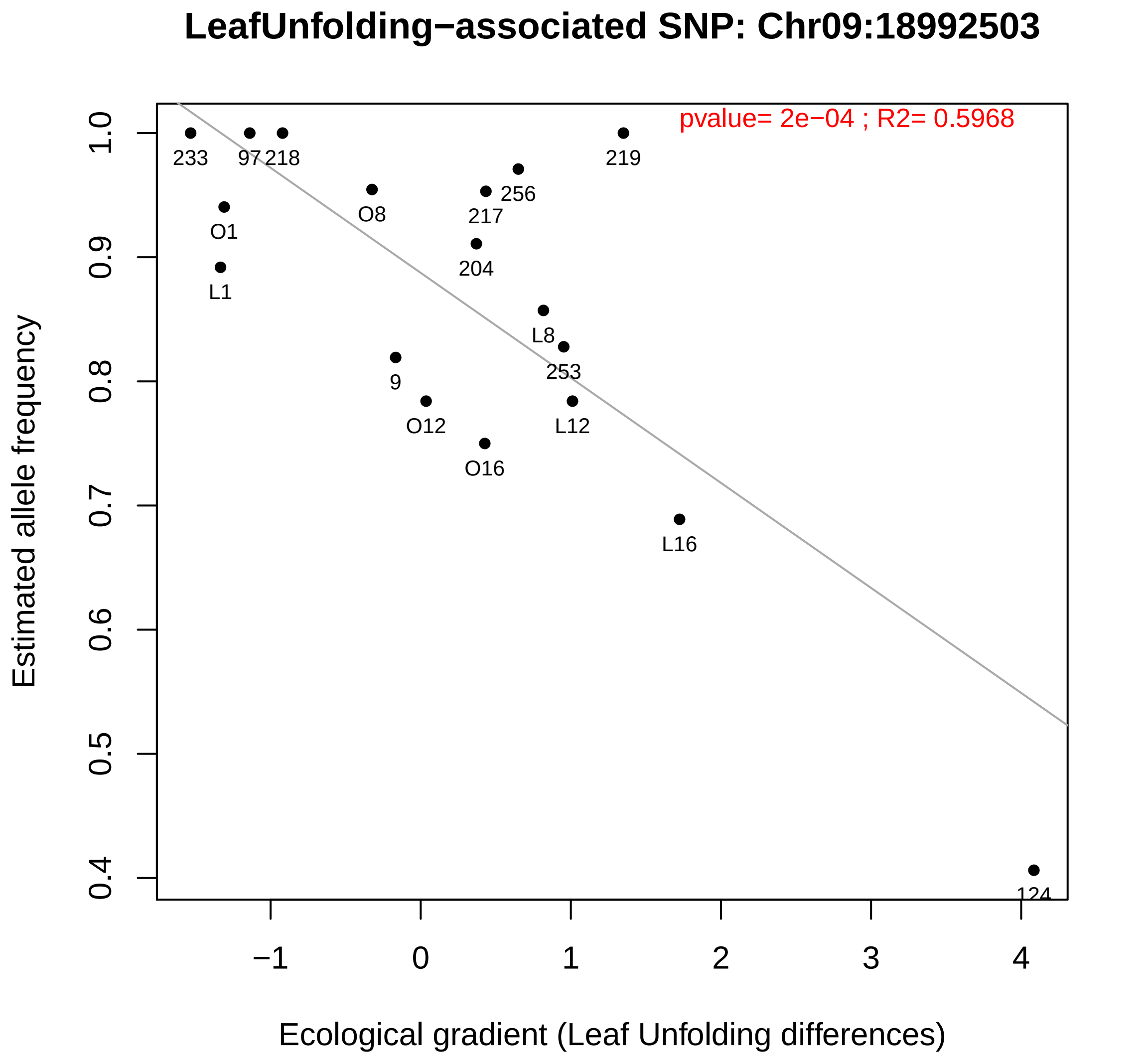
LeafUnfolding−associated SNP: Chr09:18992503
pvalue= 2e−04 ; R2= 0.5968
Estimated allele frequency
Ecological gradient (Leaf Unfolding differences)
1.0
0.9
0.8
0.7
0.6
0.5
0.4
−1
0
1
2
3
4
233
97
218
219
O1
L1
O8
256
217
204
L8
253
9
O12
L12
O16
L16
124

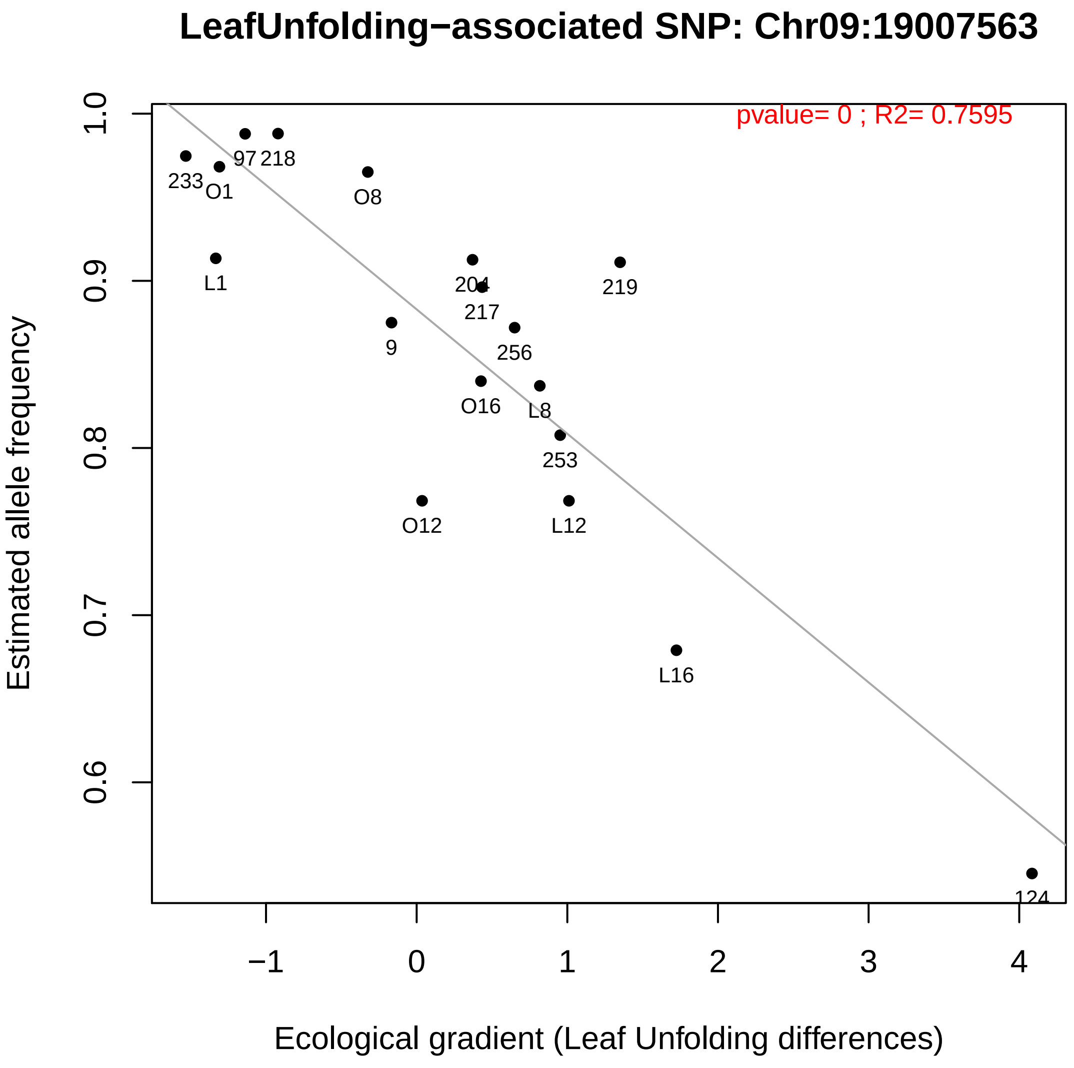

LeafUnfolding−associated SNP: Chr09:19007563
pvalue= 0 ; R2= 0.7595
Estimated allele frequency
Ecological gradient (Leaf Unfolding differences)
1.0
0.9
0.8
0.7
0.6
−1
0
1
2
3
4
233
97
218
O1
O8
L1
204
217
219
9
256
O16
L8
253
O12
L12
L16
124

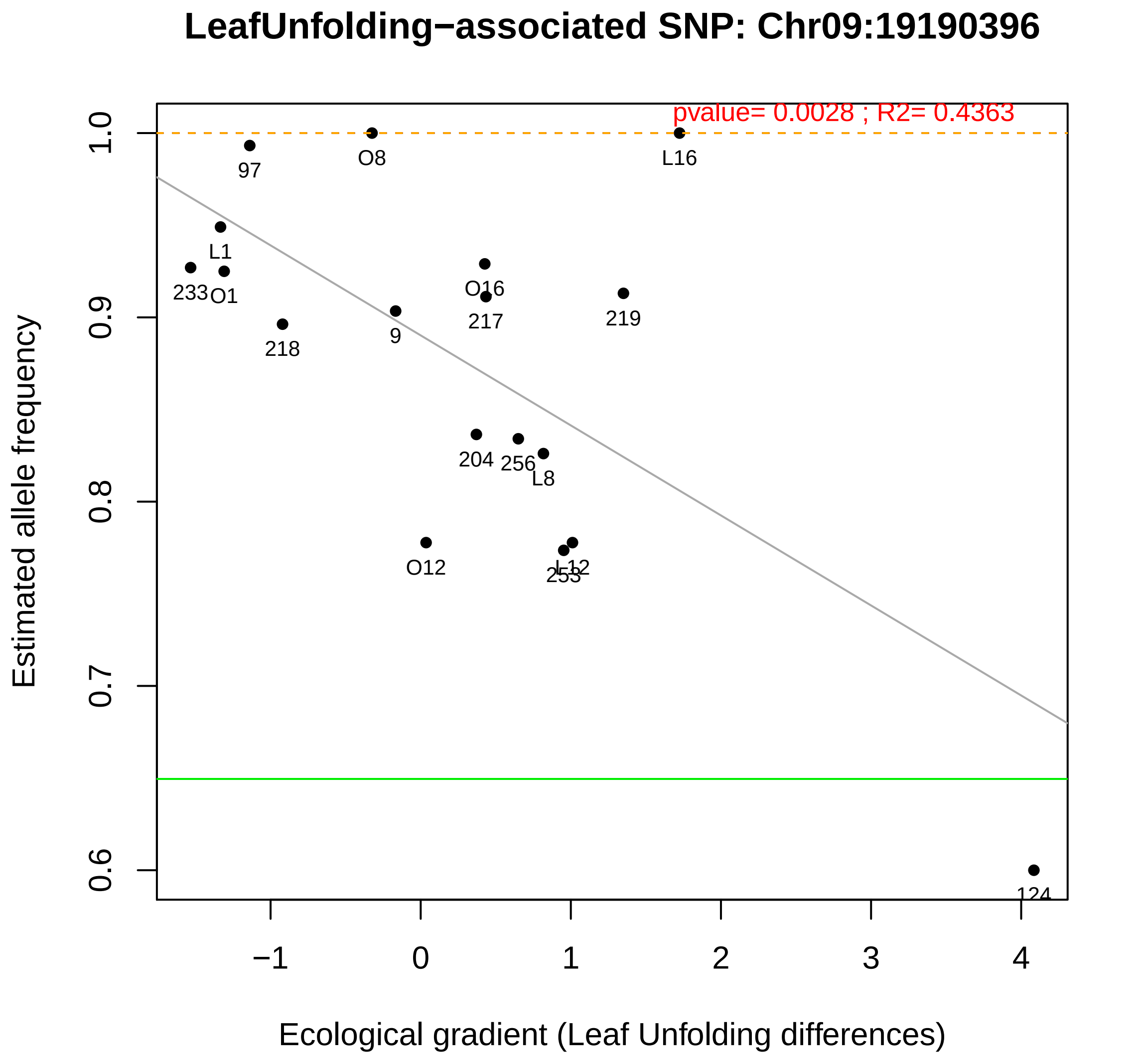
LeafUnfolding−associated SNP: Chr09:19190396
pvalue= 0.0028 ; R2= 0.4363
Estimated allele frequency
Ecological gradient (Leaf Unfolding differences)
1.0
0.9
0.8
0.7
0.6
−1
0
1
2
3
4
97
O8
L16
L1
233
O1
218
9
O16
217
219
204
256
L8
O12
L12
253
124

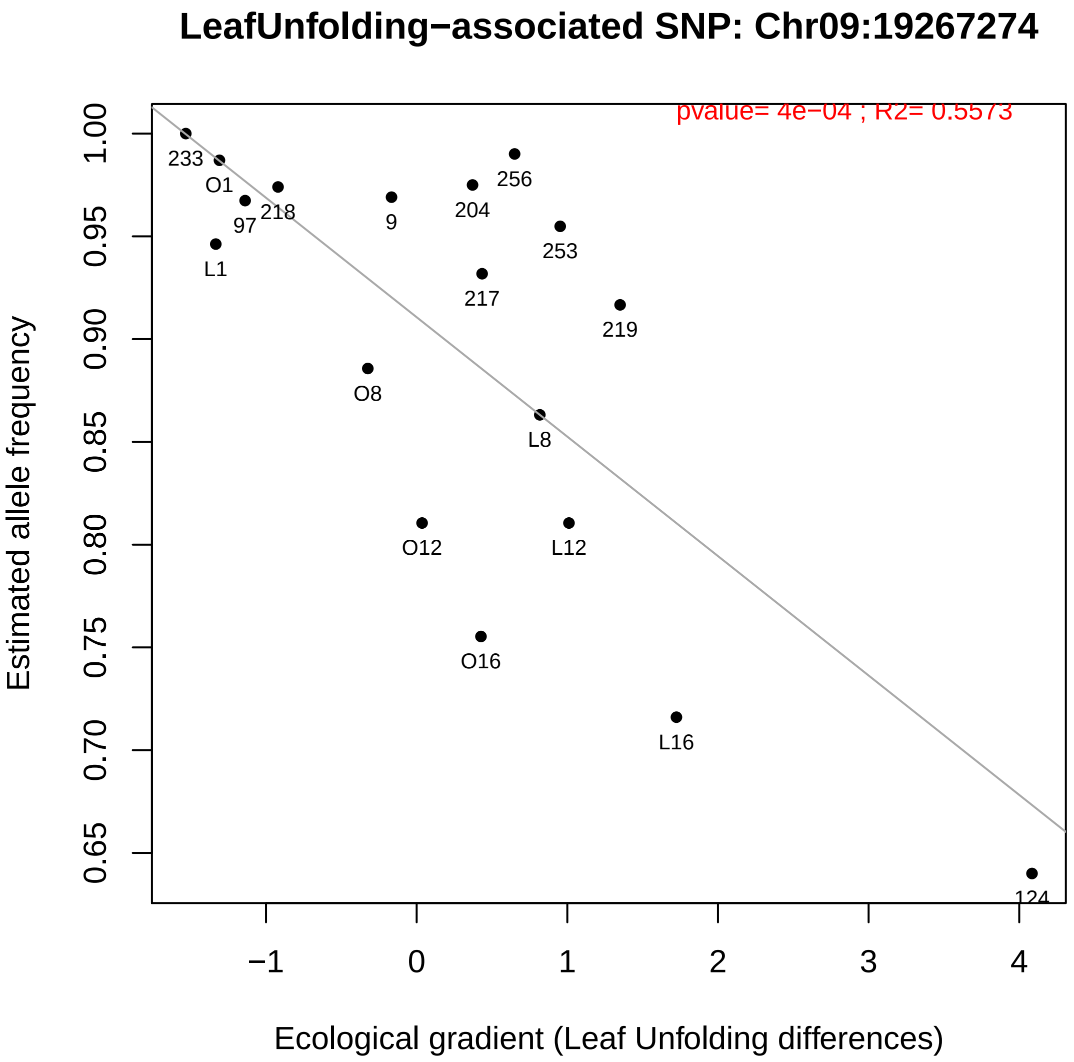

LeafUnfolding−associated SNP: Chr09:19267274
pvalue= 4e−04 ; R2= 0.5573
Estimated allele frequency
1.00
0.95
0.90
0.85
0.80
0.75
0.70
0.65
−1
0
1
2
3
4
Ecological gradient (Leaf Unfolding differences)
233
O1
218
97
L1
9
204
256
253
217
219
O8
L8
O12
L12
O16
L16
124

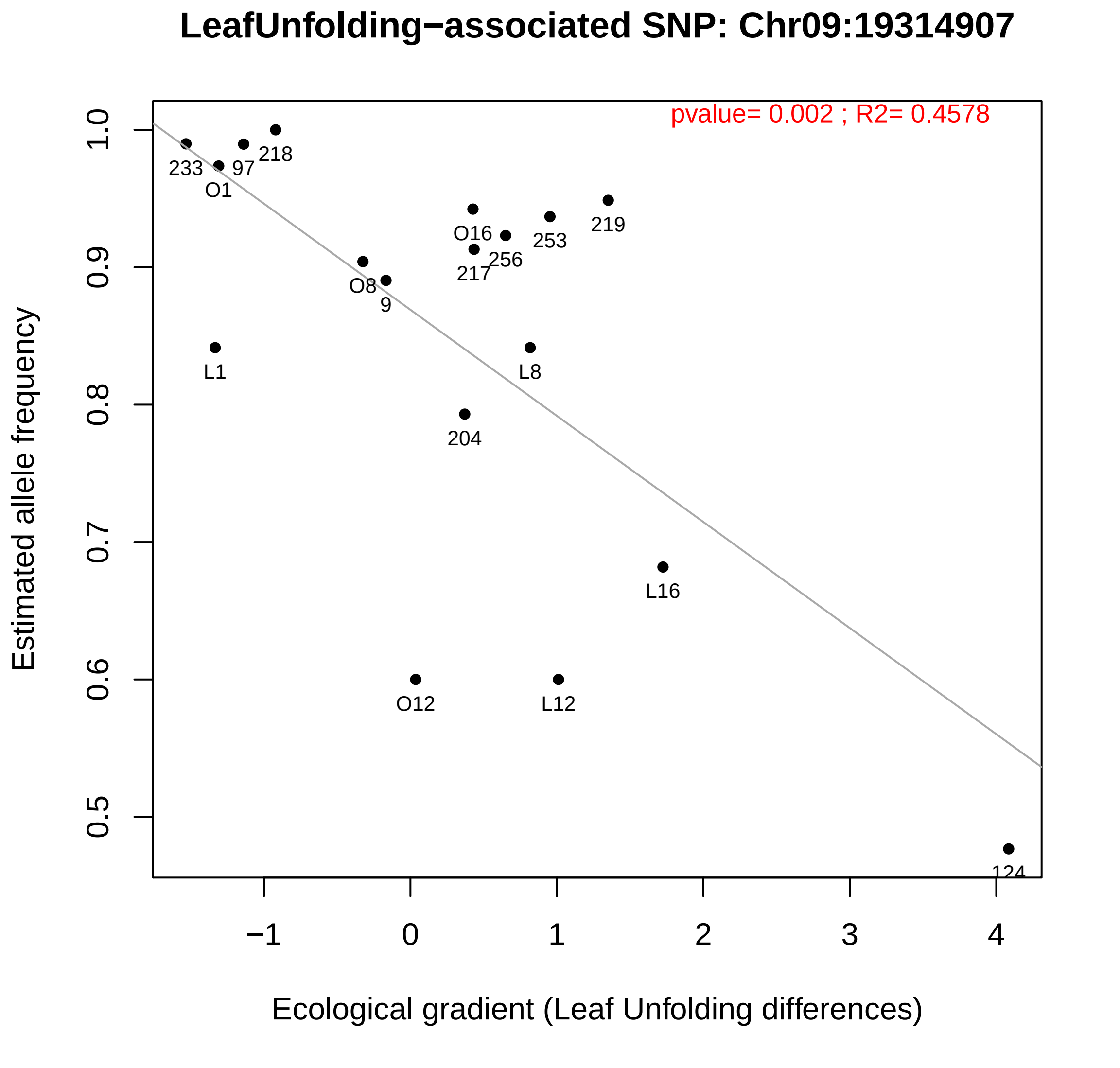

LeafUnfolding−associated SNP: Chr09:19314907
pvalue= 0.002 ; R2= 0.4578
Estimated allele frequency
0.5
0.6
0.7
0.8
0.9
1.0
−1
0
1
2
3
4
Ecological gradient (Leaf Unfolding differences)
233
O1
97
218
O16
256
217
253
219
O8
9
L1
L8
204
L16
O12
L12
124

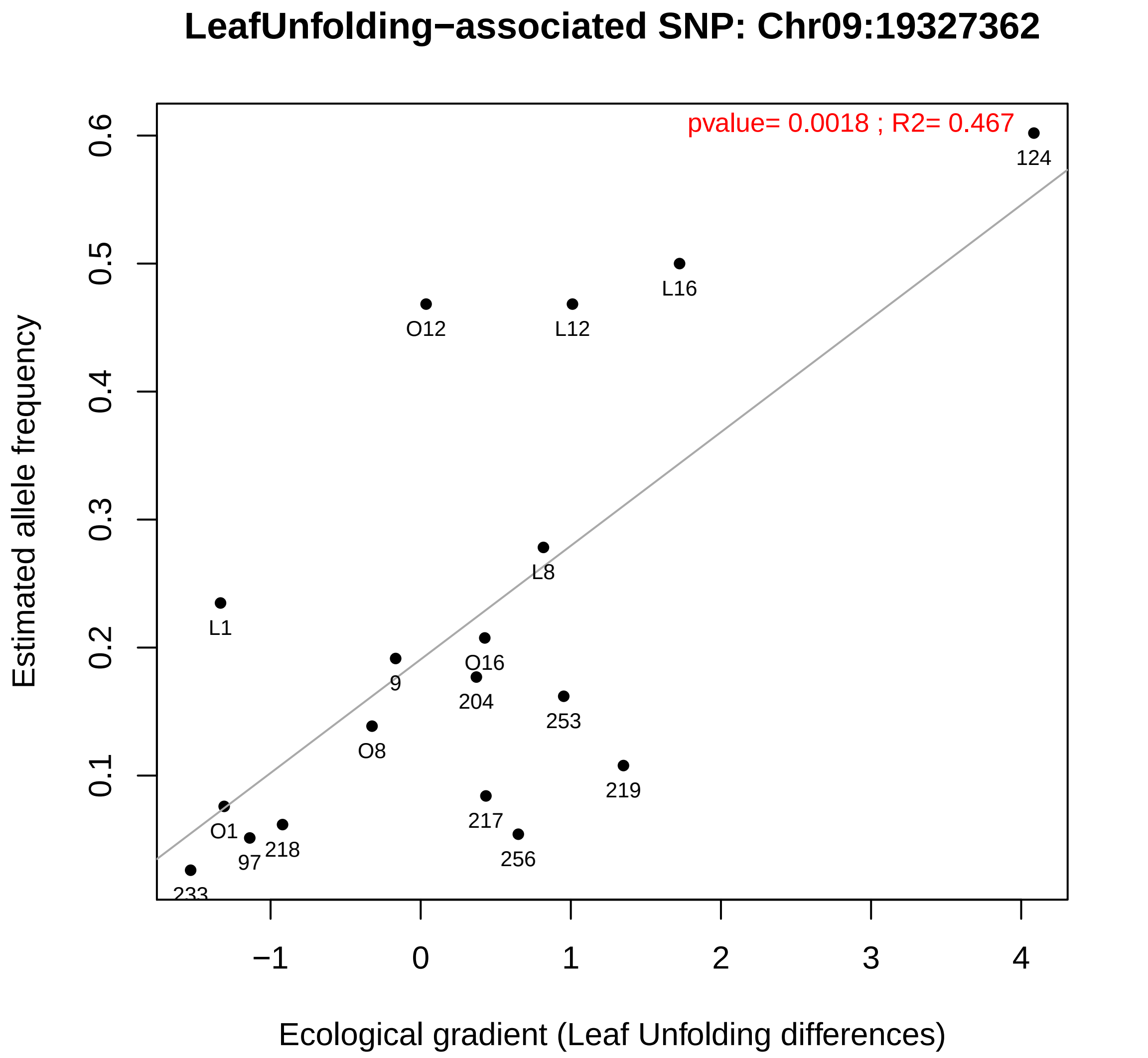
LeafUnfolding−associated SNP: Chr09:19327362
pvalue= 0.0018 ; R2= 0.467
Estimated allele frequency
Ecological gradient (Leaf Unfolding differences)
0.1
0.2
0.3
0.4
0.5
0.6
−1
0
1
2
3
4
124
L16
O12
L12
L8
L1
O16
9
204
253
O8
219
217
O1
218
97
256
233

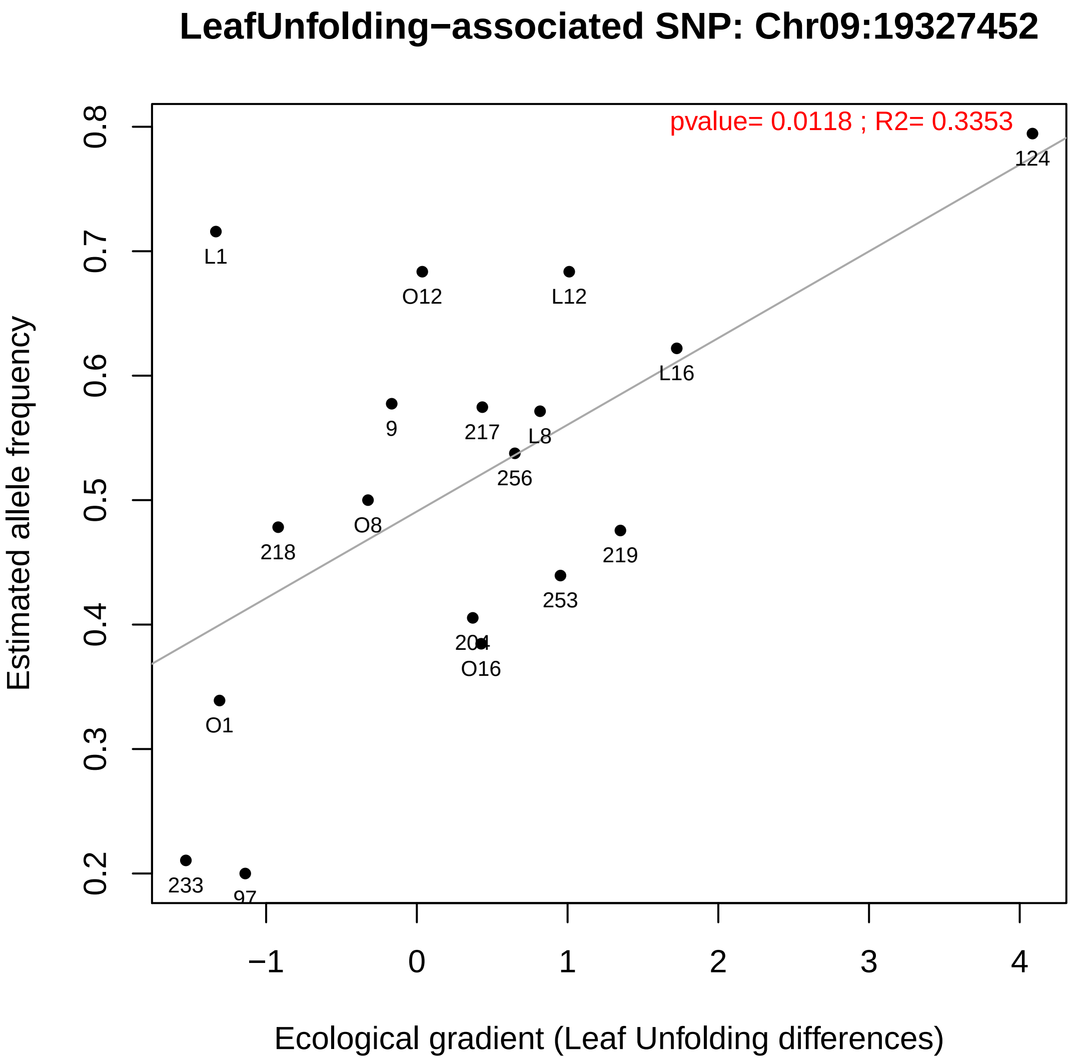

LeafUnfolding−associated SNP: Chr09:19327452
pvalue= 0.0118 ; R2= 0.3353
124
L1
O12
L12
L16
9
217
L8
256
O8
218
219
253
204
O16
O1
233
97
0.2
0.3
0.4
0.5
0.6
0.7
0.8
−1
0
1
2
3
4
Estimated allele frequency
Ecological gradient (Leaf Unfolding differences)

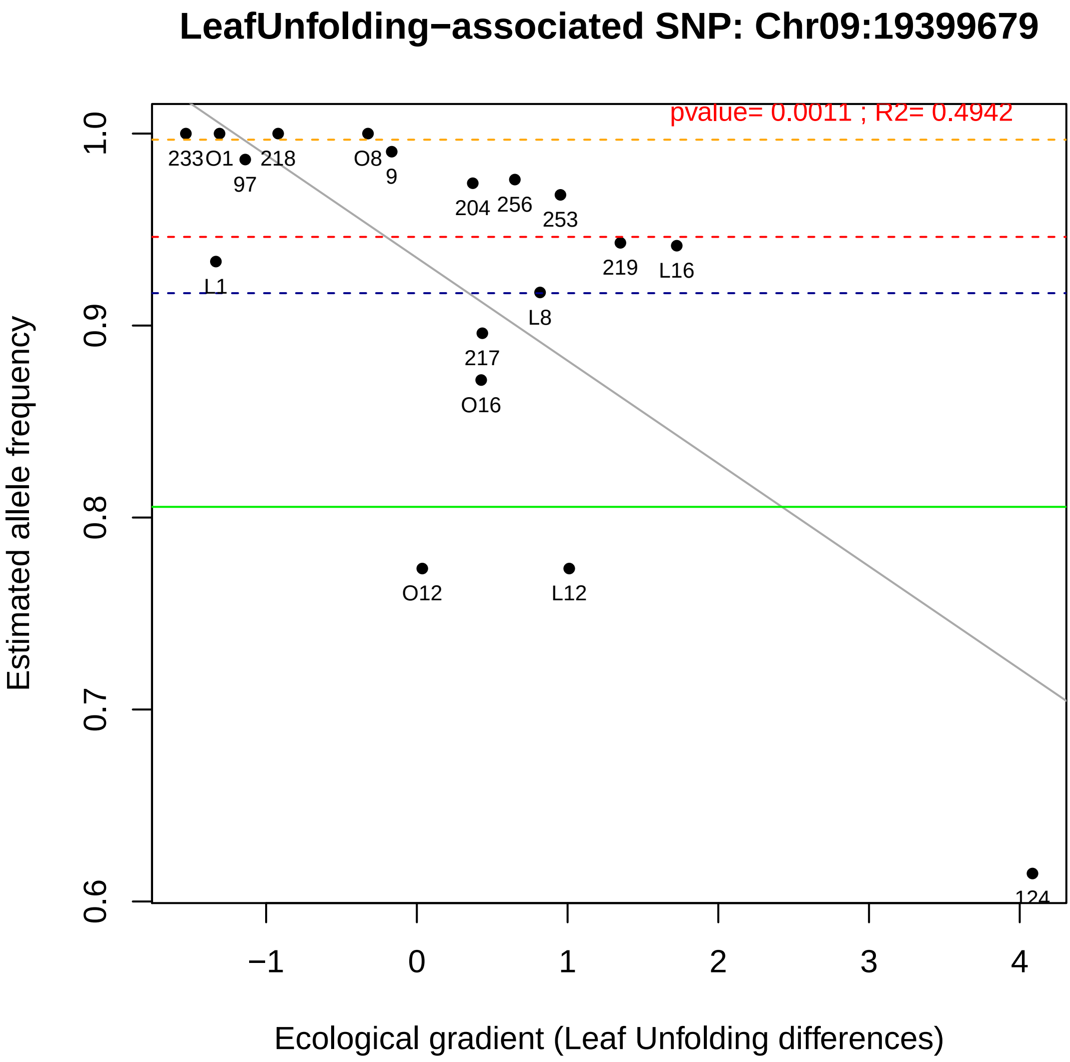
LeafUnfolding−associated SNP: Chr09:19399679
pvalue= 0.0011 ; R2= 0.4942
Estimated allele frequency
Ecological gradient (Leaf Unfolding differences)
0.6
0.7
0.8
0.9
1.0
−1
0
1
2
3
4
233
O1
97
218
O8
9
204
256
253
L1
219
L16
L8
217
O16
O12
L12
124

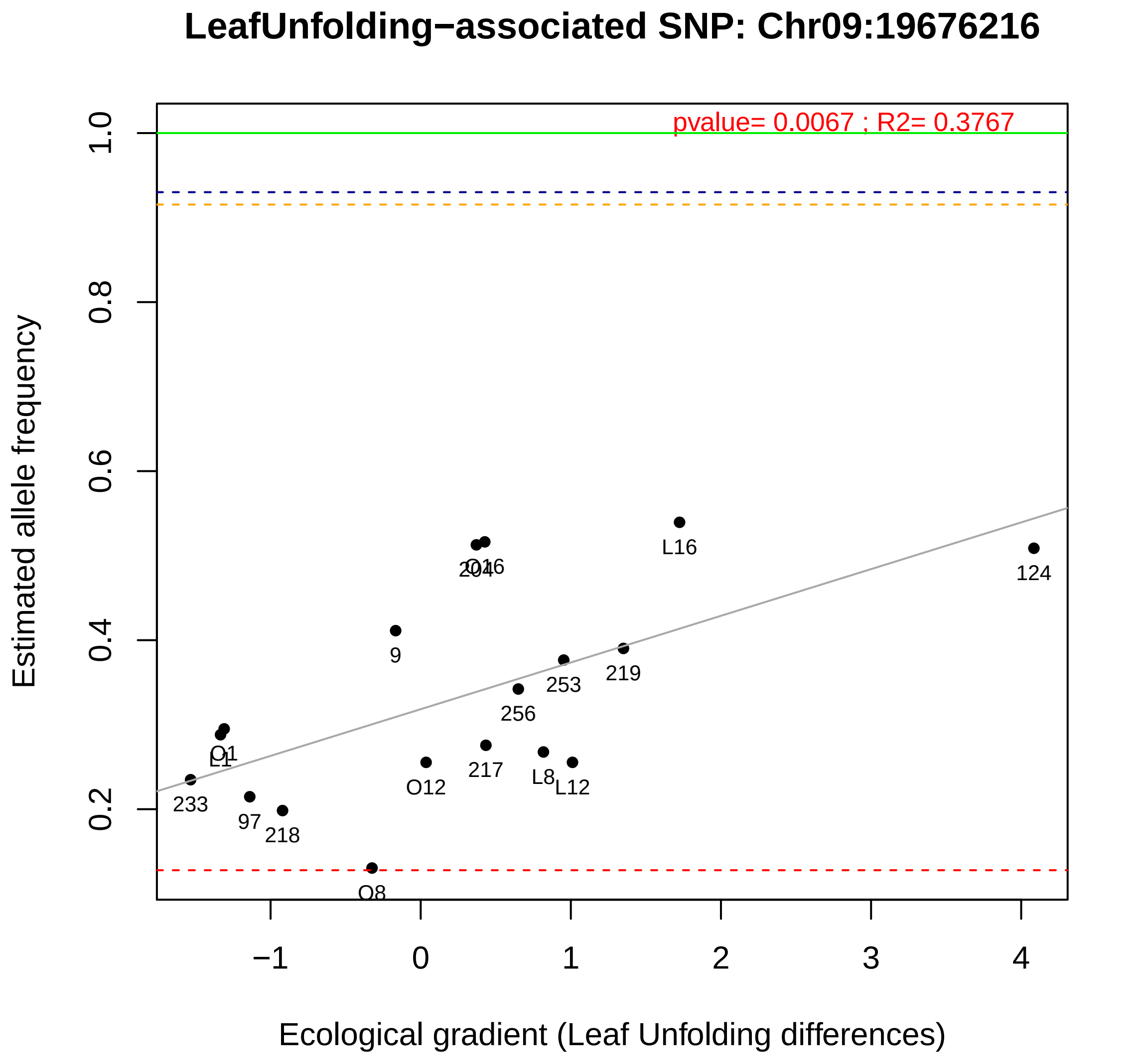

LeafUnfolding−associated SNP: Chr09:19676216
pvalue= 0.0067 ; R2= 0.3767
Estimated allele frequency
Ecological gradient (Leaf Unfolding differences)
1.0
0.8
0.6
0.4
0.2
−1
0
1
2
3
4
L16
124
204
O16
9
253
219
256
O1
L1
233
97
218
O12
217
L8
L12
O8

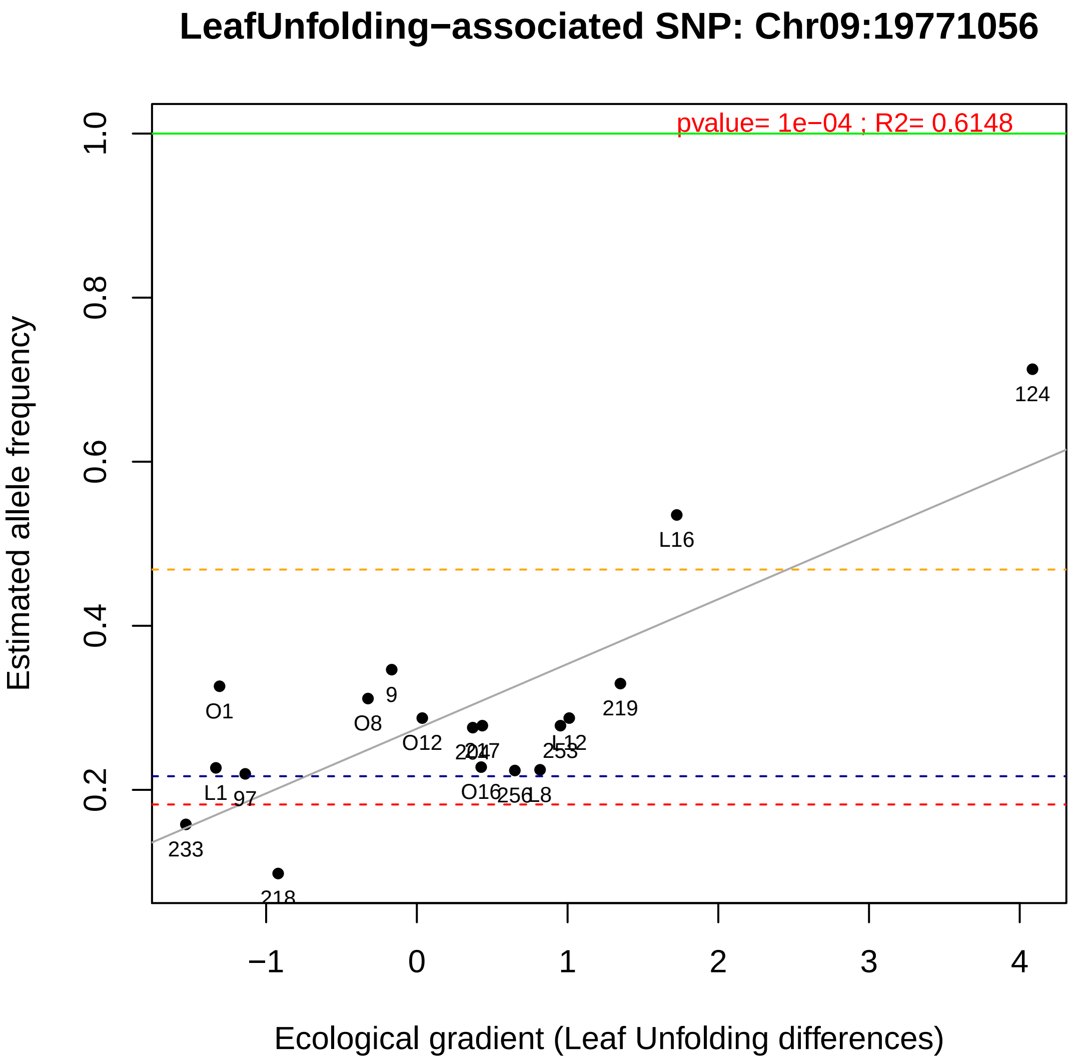
LeafUnfolding−associated SNP: Chr09:19771056
pvalue= 1e−04 ; R2= 0.6148
Estimated allele frequency
0.2
0.4
0.6
0.8
1.0
−1
0
1
2
3
4
Ecological gradient (Leaf Unfolding differences)
124
L16
219
9
O1
O8
O12
L12
253
O16
L8
L1
97
233
218

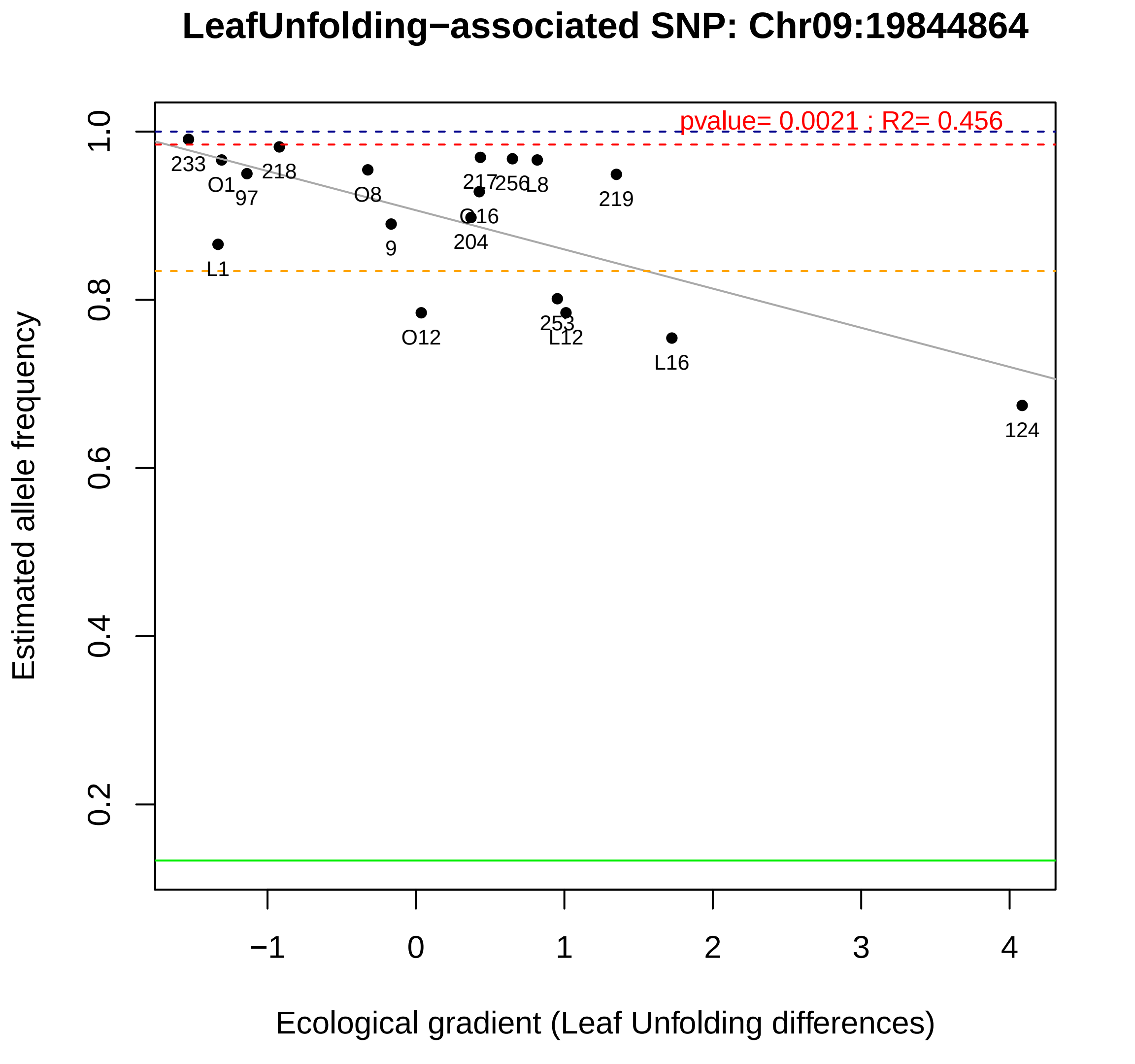

LeafUnfolding−associated SNP: Chr09:19844864
pvalue= 0.0021 ; R2= 0.456
Estimated allele frequency
Ecological gradient (Leaf Unfolding differences)
0.2
0.4
0.6
0.8
1.0
−1
0
1
2
3
4
233
O1
97
218
L1
O8
9
O12
217
256
L8
O16
204
219
253
L12
L16
124

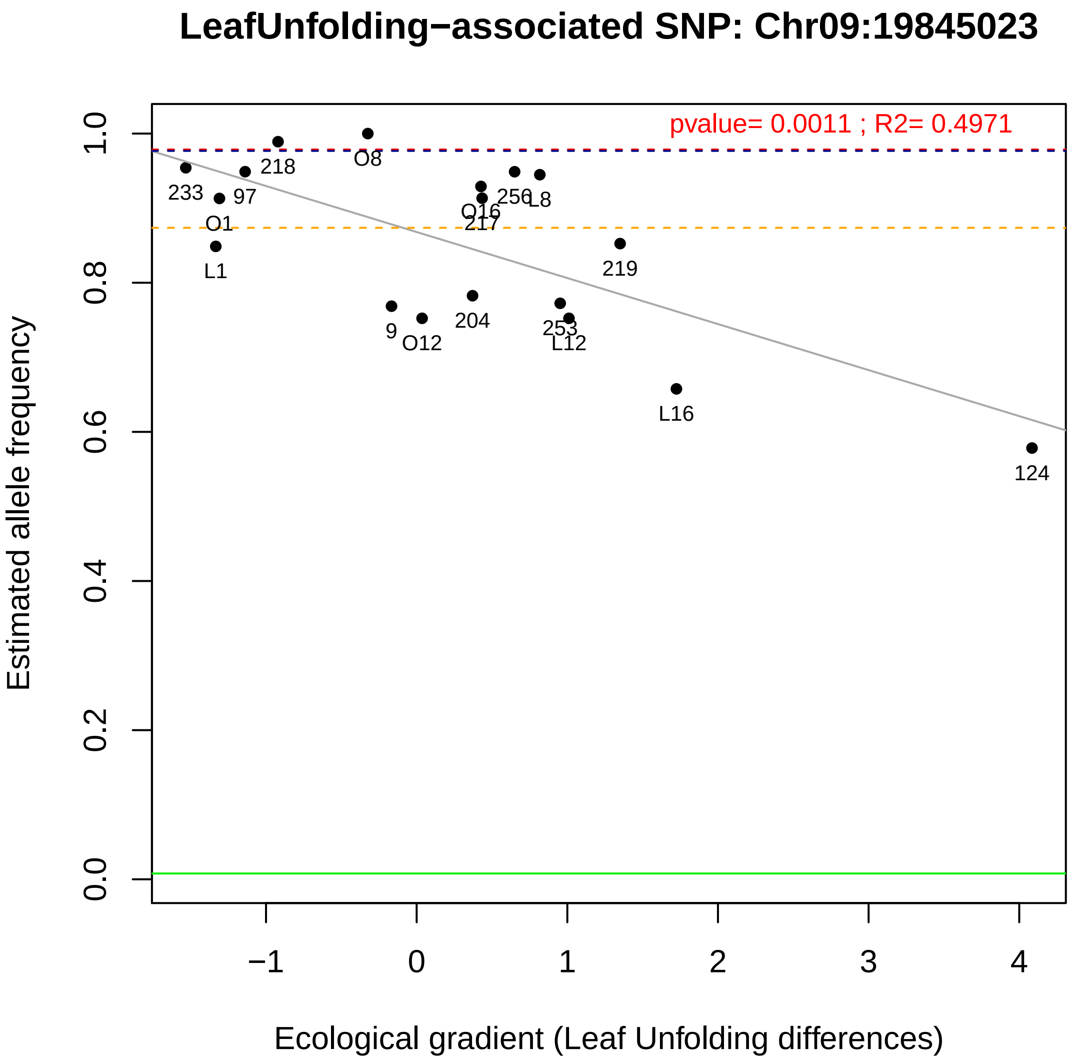
LeafUnfolding−associated SNP: Chr09:19845023
pvalue= 0.0011 ; R2= 0.4971
Estimated allele frequency
Ecological gradient (Leaf Unfolding differences)
0.0
0.2
0.4
0.6
0.8
1.0
−1
0
1
2
3
4
233
218
97
O8
O1
L1
256
L8
O16
217
219
9
O12
204
253
L12
L16
124

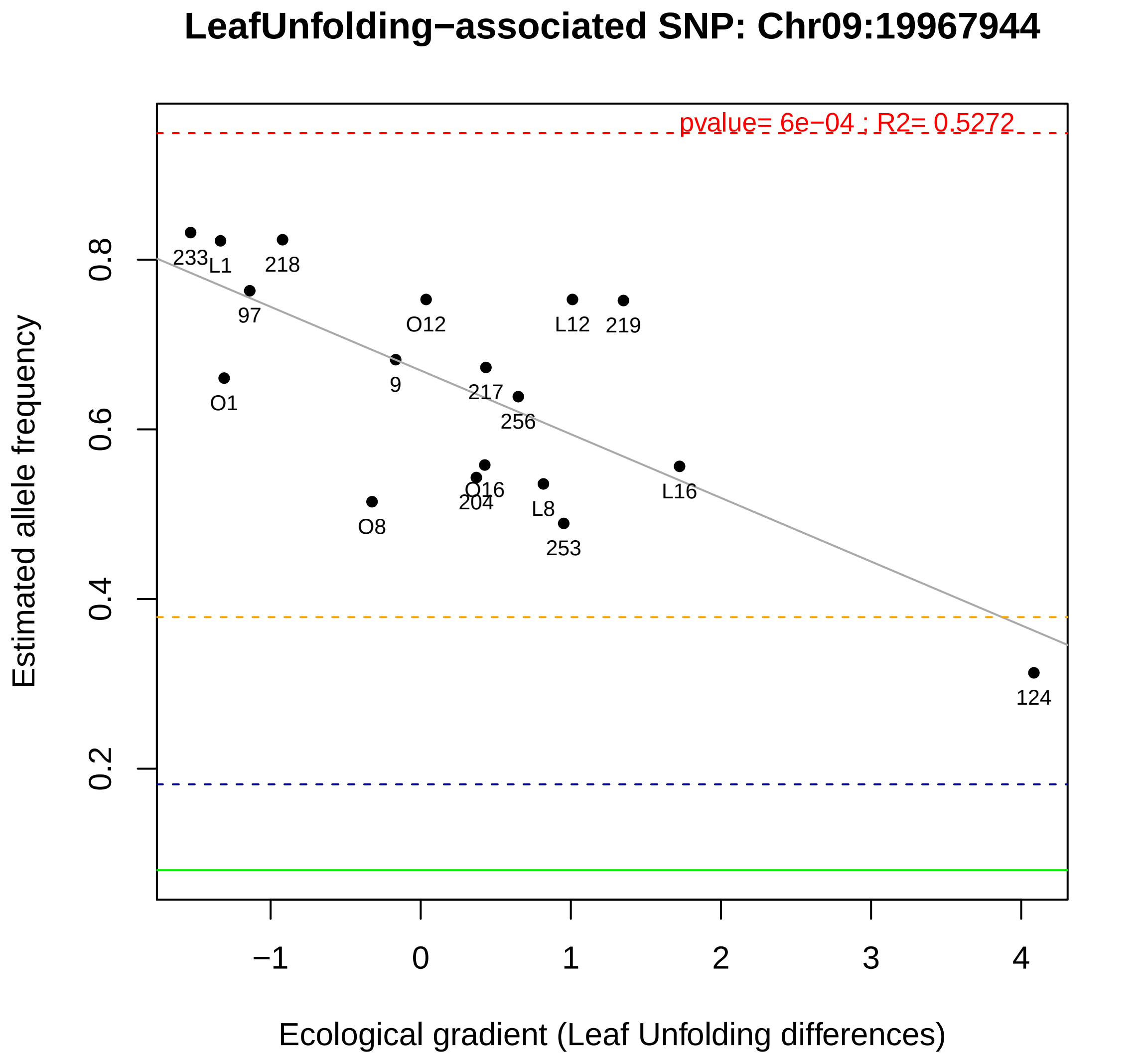

LeafUnfolding−associated SNP: Chr09:19967944
pvalue= 6e−04 ; R2= 0.5272
Estimated allele frequency
0.2
0.4
0.6
0.8
−1
0
1
2
3
4
Ecological gradient (Leaf Unfolding differences)
233
L1
218
97
O12
L12
219
9
217
O1
256
O16
204
L8
L16
O8
253
124

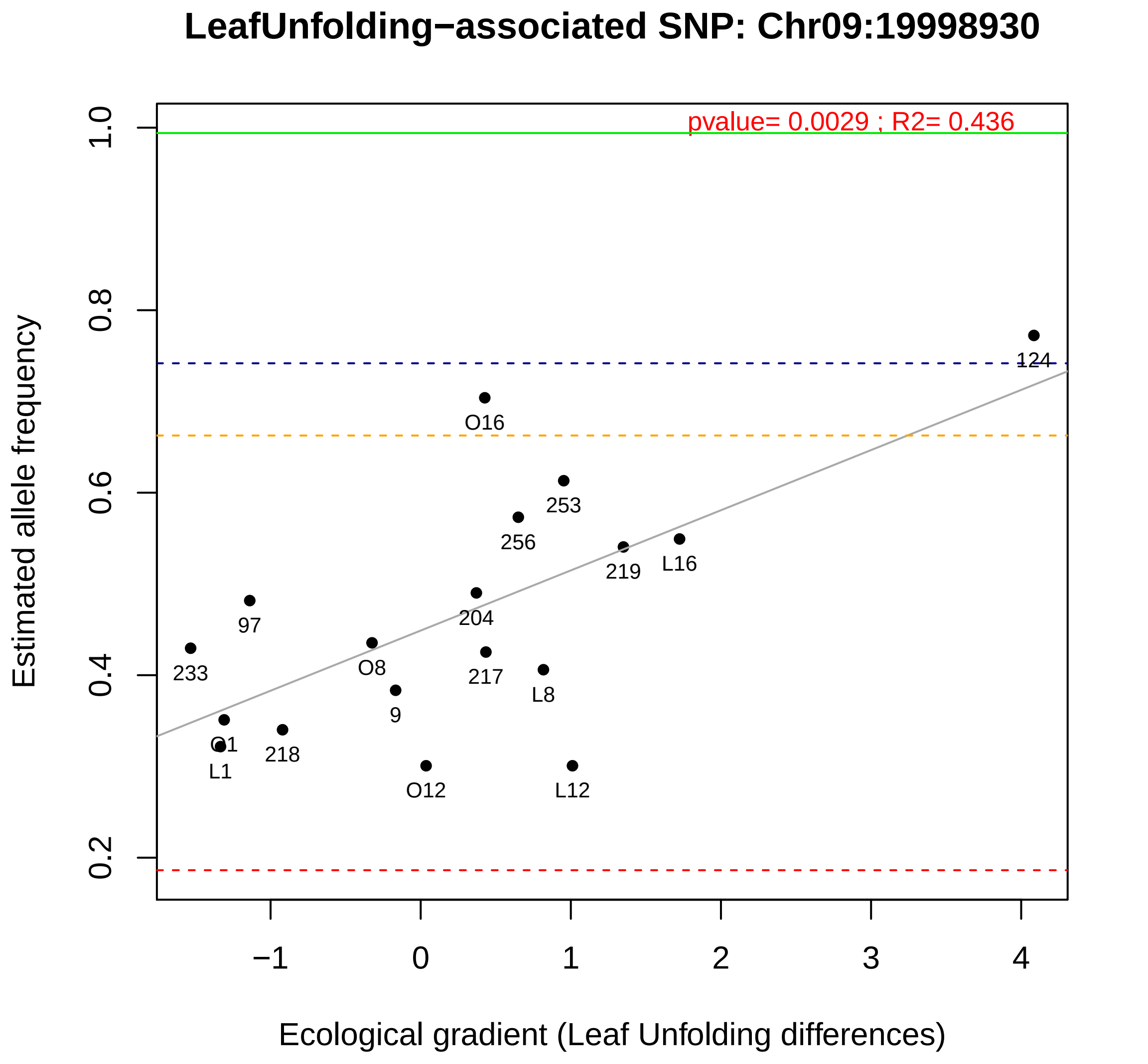

LeafUnfolding−associated SNP: Chr09:19998930
pvalue= 0.0029 ; R2= 0.436
Estimated allele frequency
0.2
0.4
0.6
0.8
1.0
Ecological gradient (Leaf Unfolding differences)
−1
0
1
2
3
4
124
O16
253
256
219
L16
204
97
233
O8
217
L8
9
O1
L1
218
O12
L12

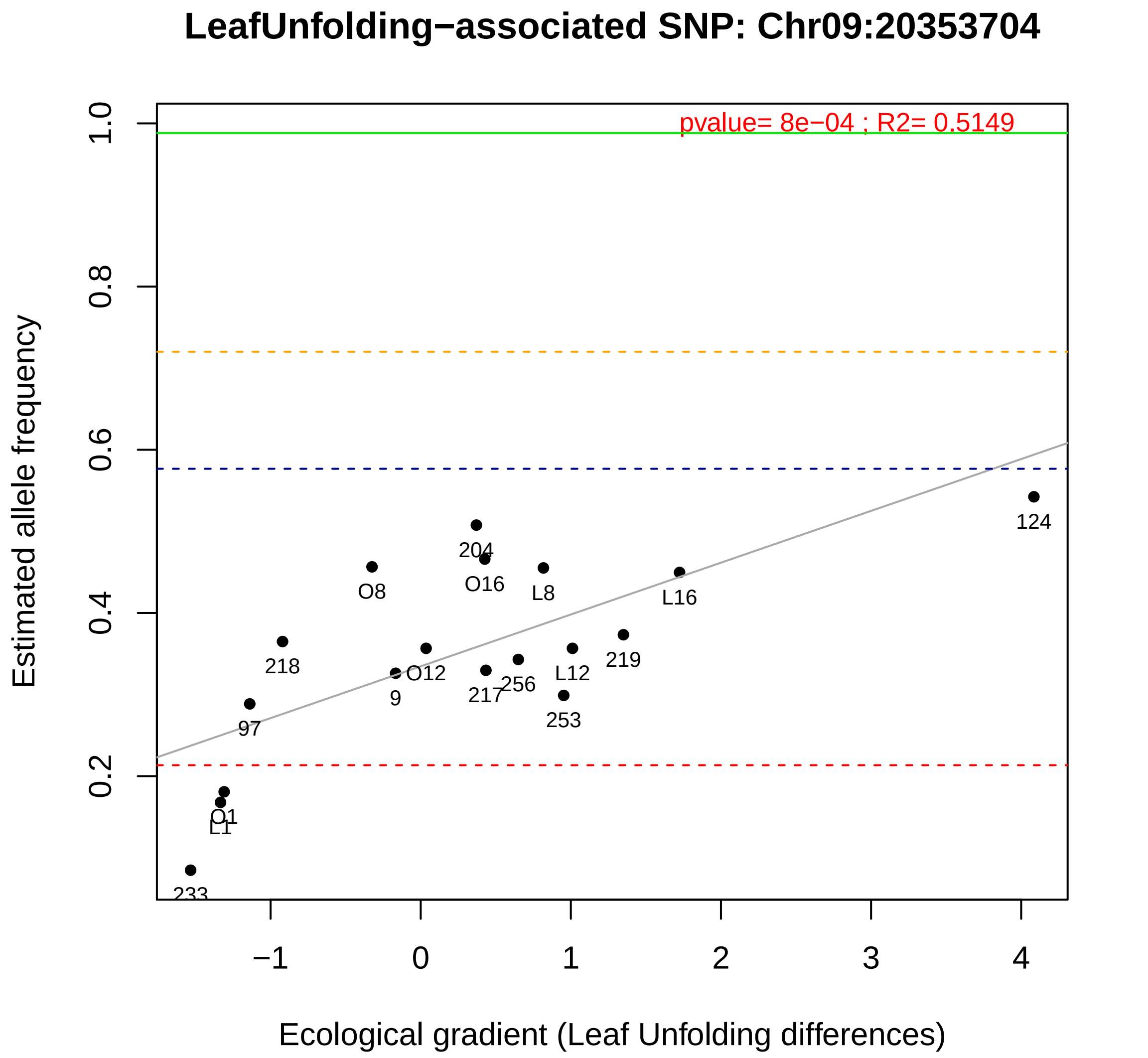
LeafUnfolding−associated SNP: Chr09:20353704
pvalue= 8e−04 ; R2= 0.5149
Estimated allele frequency
1.0
0.8
0.6
0.4
0.2
−1
0
1
2
3
4
Ecological gradient (Leaf Unfolding differences)
124
204
O16
O8
L8
L16
218
O12
9
217
256
L12
219
253
97
O1
L1
233

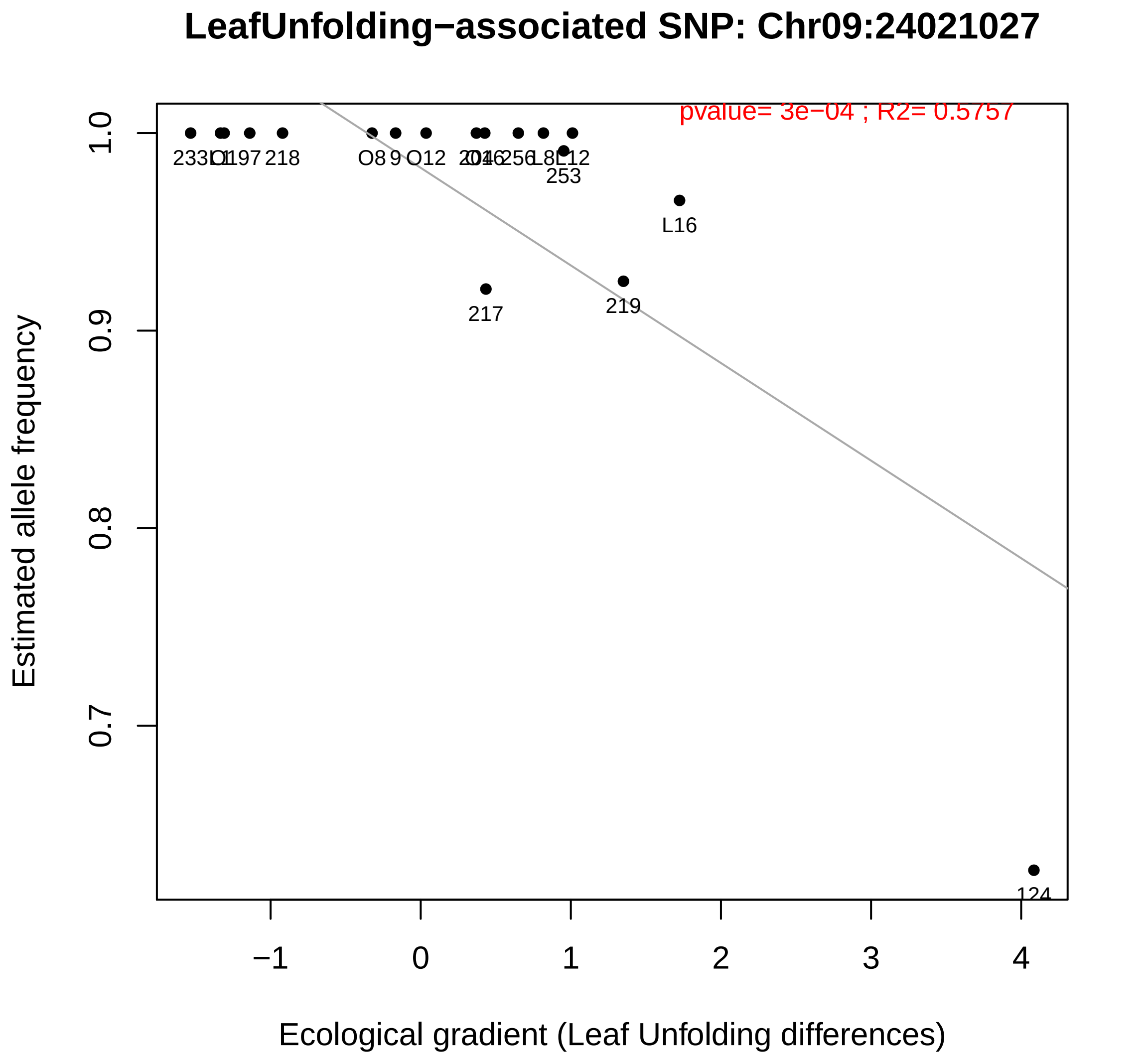

LeafUnfolding−associated SNP: Chr09:24021027
pvalue= 3e−04 ; R2= 0.5757
Estimated allele frequency
Ecological gradient (Leaf Unfolding differences)
1.0
0.9
0.8
0.7
−1
0
1
2
3
4
233
O1
197
218
O8
9
O12
L8
L12
253
L16
217
219
124

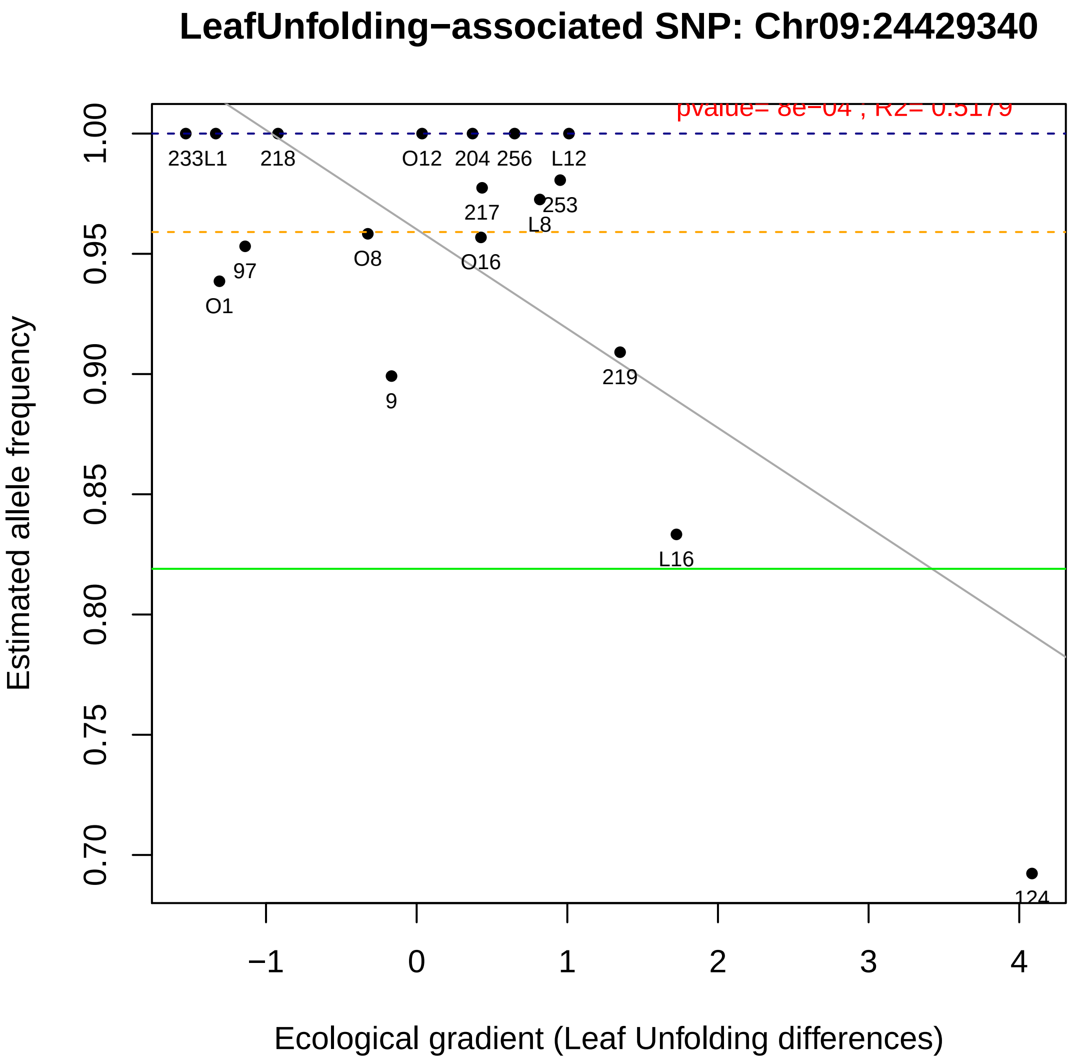

LeafUnfolding−associated SNP: Chr09:24429340
pvalue= 8e−04 ; R2= 0.5179
233
L1
218
O12
204
256
L12
217
253
L8
97
O8
O16
O1
9
219
L16
124
Estimated allele frequency
1.00
0.95
0.90
0.85
0.80
0.75
0.70
−1
0
1
2
3
4
Ecological gradient (Leaf Unfolding differences)

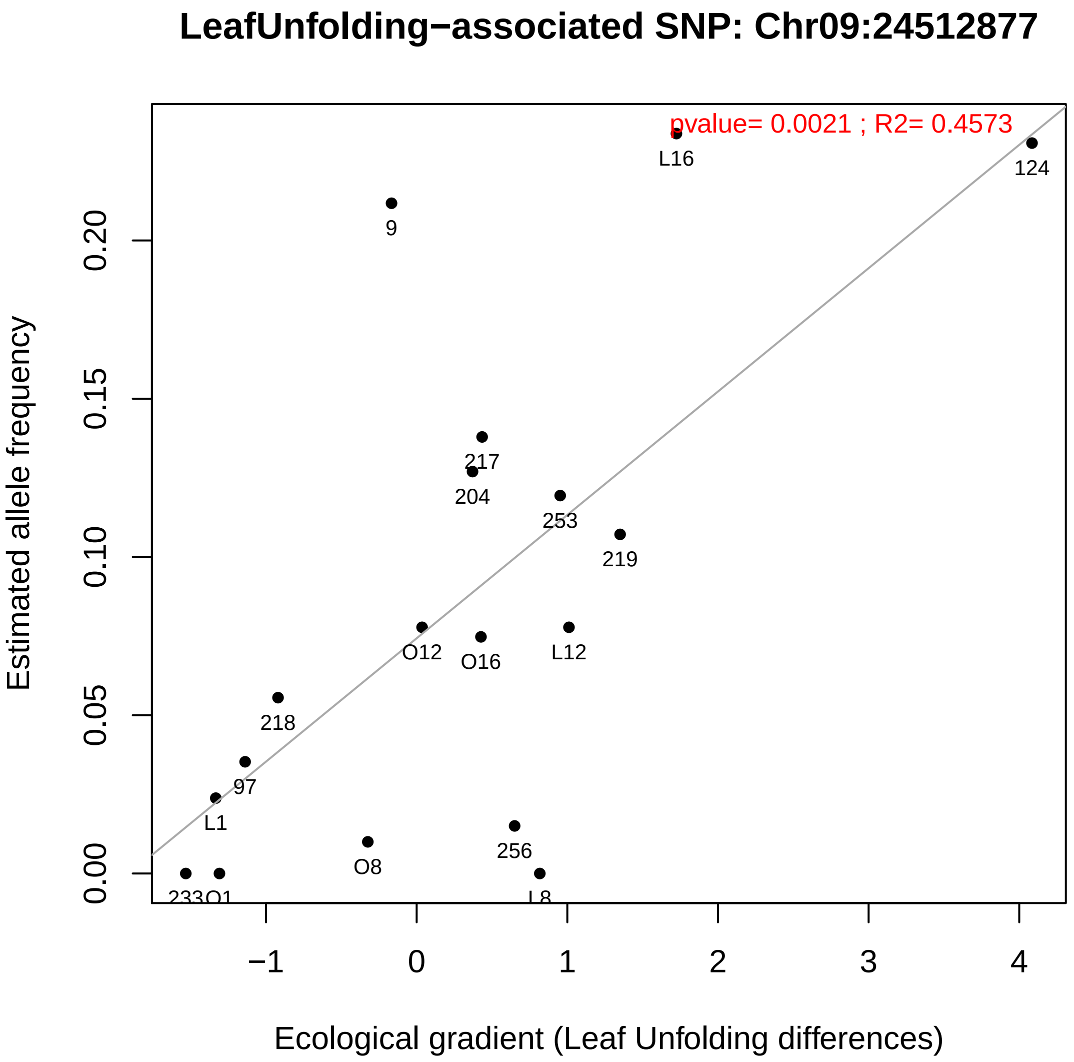

LeafUnfolding−associated SNP: Chr09:24512877
pvalue= 0.0021 ; R2= 0.4573
Estimated allele frequency
0.00
0.05
0.10
0.15
0.20
Ecological gradient (Leaf Unfolding differences)
−1
0
1
2
3
4
L16
124
9
217
204
253
219
O12
O16
L12
218
97
L1
O8
256

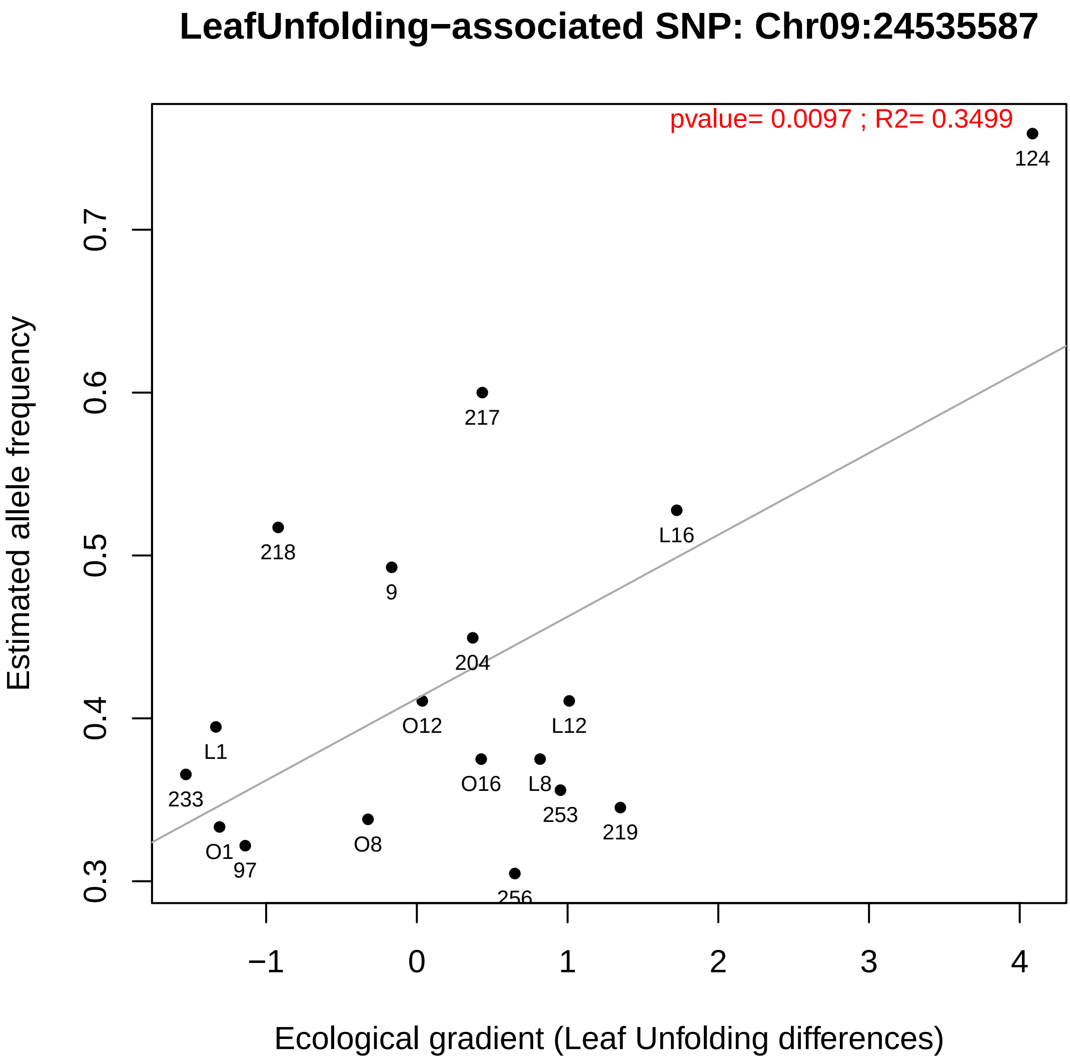
LeafUnfolding−associated SNP: Chr09:24535587
pvalue= 0.0097 ; R2= 0.3499
Estimated allele frequency
Ecological gradient (Leaf Unfolding differences)
0.3
0.4
0.5
0.6
0.7
−1
0
1
2
3
4
124
217
L16
218
9
204
O12
L12
L1
O16
L8
233
253
219
O1
97
O8
256

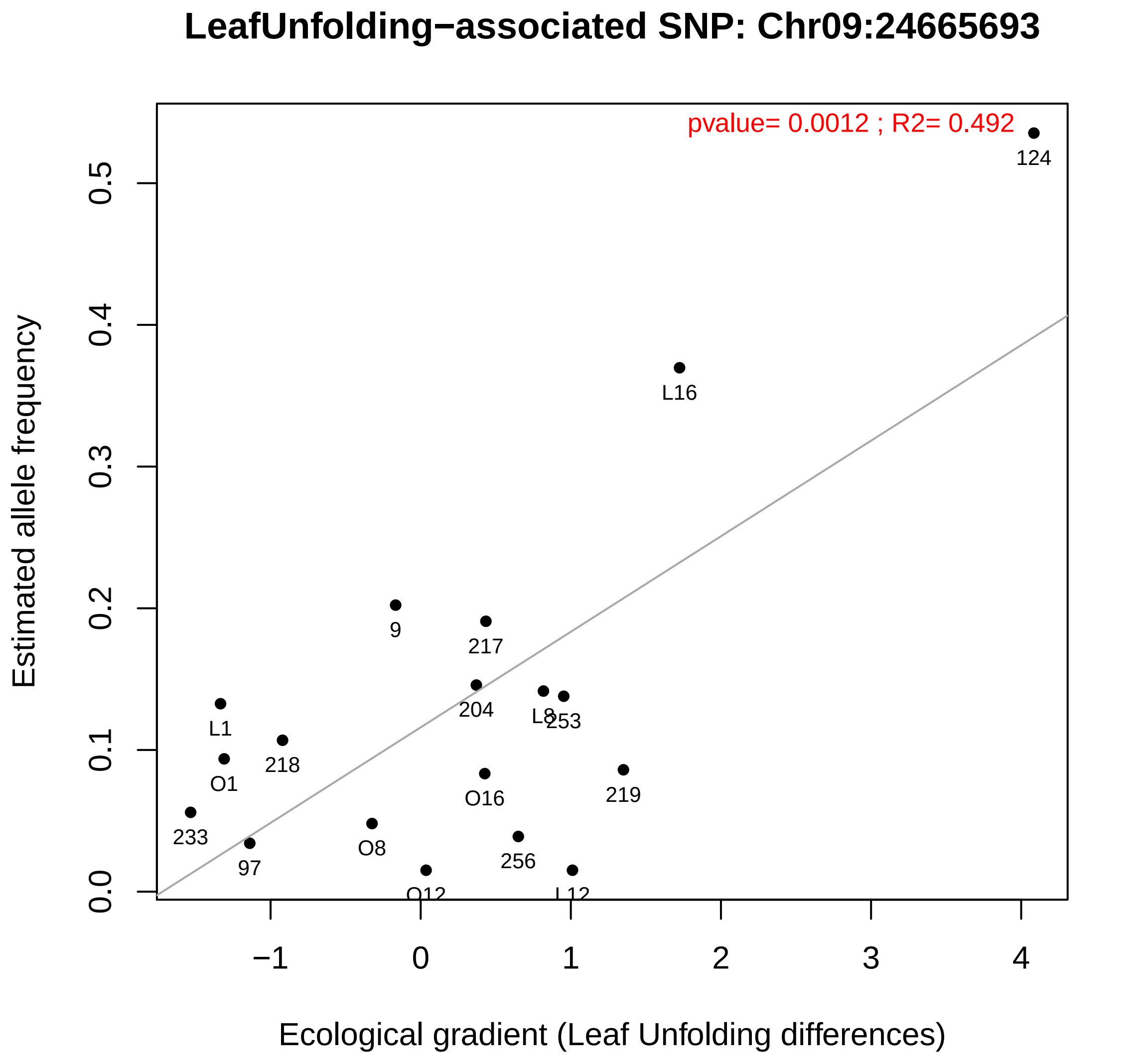
LeafUnfolding−associated SNP: Chr09:24665693
pvalue= 0.0012 ; R2= 0.492
124
L16
9
217
204
L8
253
L1
218
O1
O16
219
233
97
O8
256
O12
L12
0.0
0.1
0.2
0.3
0.4
0.5
−1
0
1
2
3
4
Estimated allele frequency
Ecological gradient (Leaf Unfolding differences)

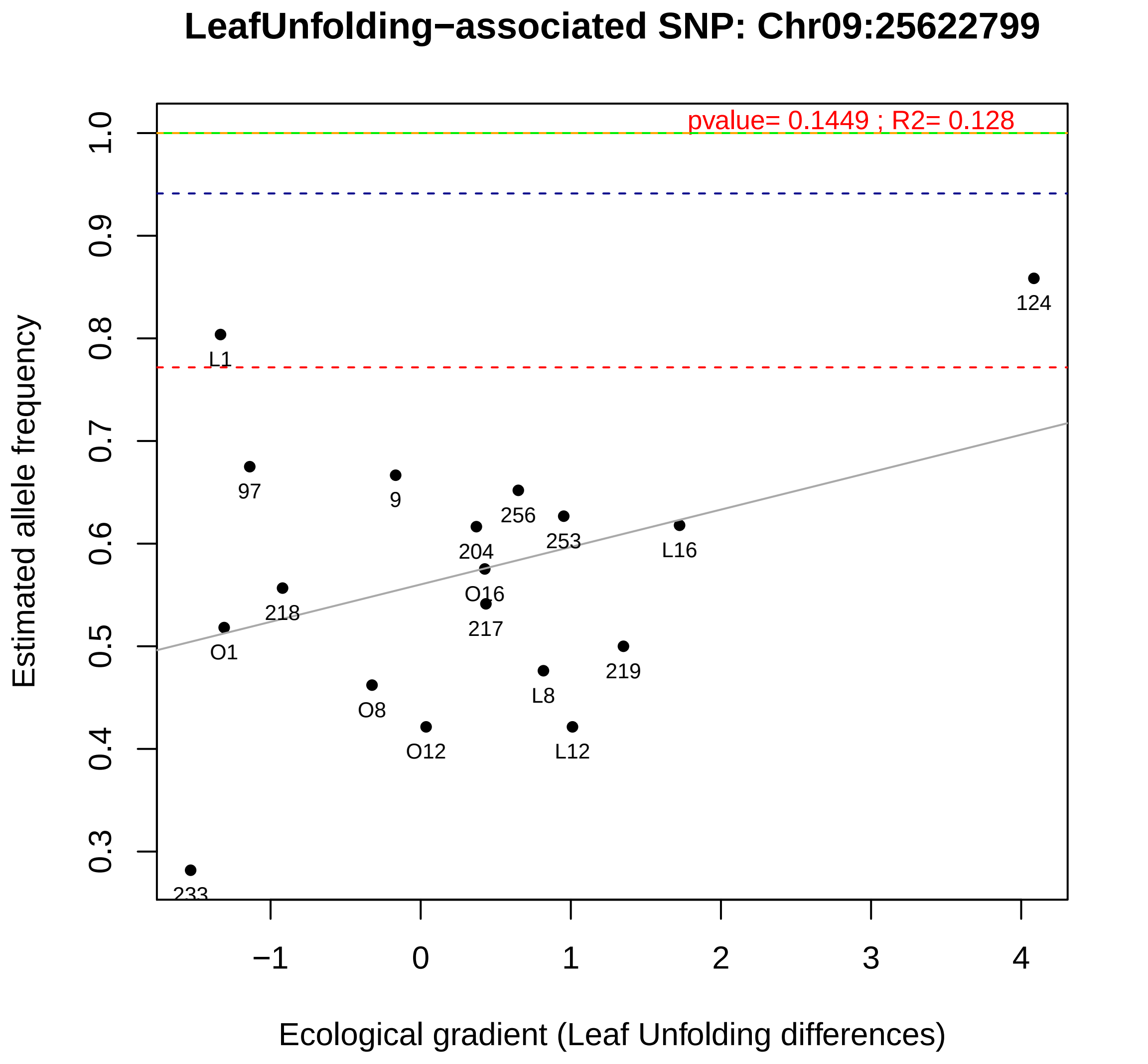

LeafUnfolding−associated SNP: Chr09:25622799
pvalue= 0.1449 ; R2= 0.128
Estimated allele frequency
Ecological gradient (Leaf Unfolding differences)
0.3
0.4
0.5
0.6
0.7
0.8
0.9
1.0
−1
0
1
2
3
4
124
L1
97
9
256
253
204
L16
O16
218
217
O1
219
L8
O8
O12
L12
233

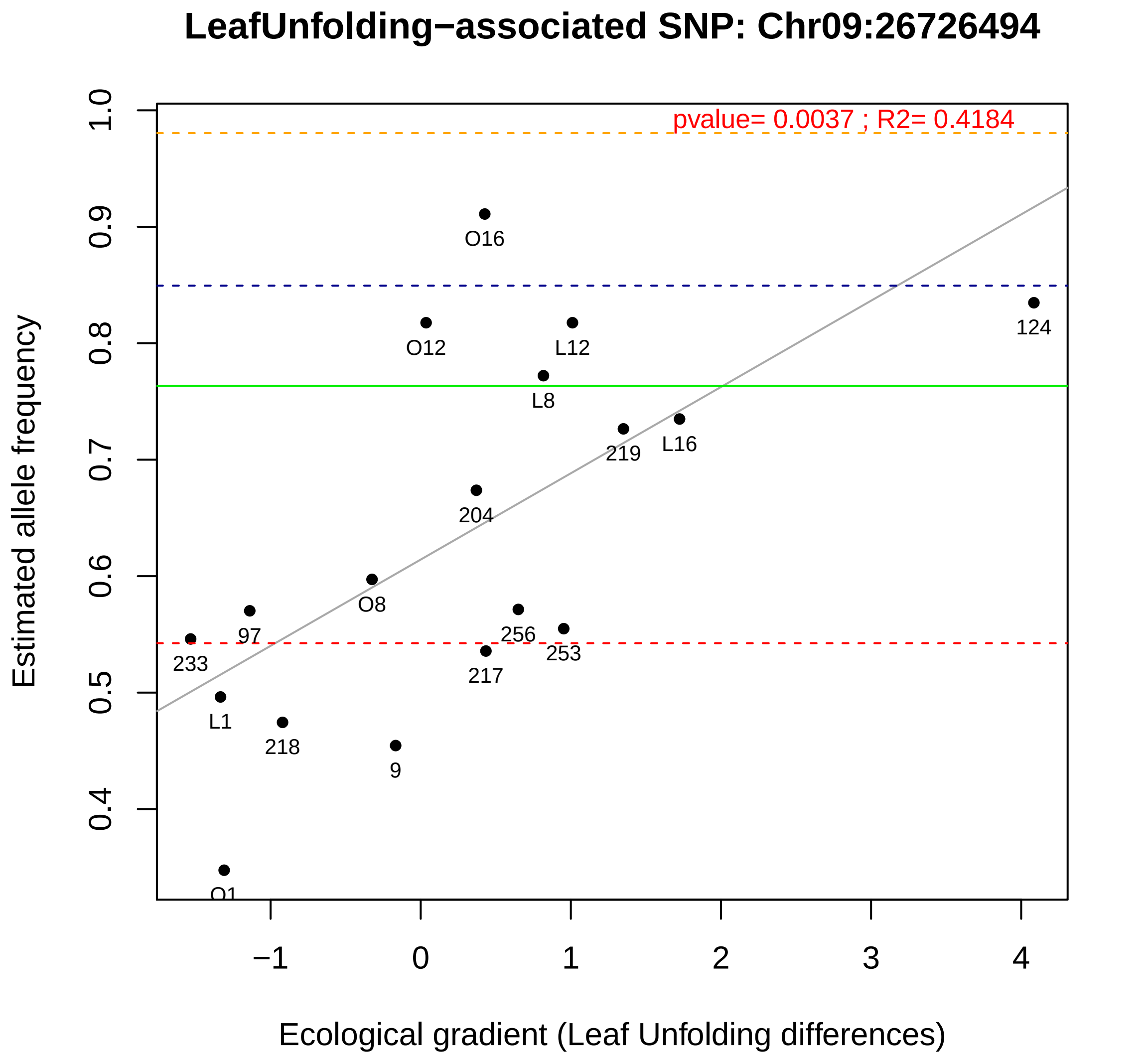
LeafUnfolding−associated SNP: Chr09:26726494
pvalue= 0.0037 ; R2= 0.4184
Estimated allele frequency
Ecological gradient (Leaf Unfolding differences)
O16
O12
L12
124
L8
219
L16
204
O8
97
256
253
233
217
L1
218
9
O1
1.0
0.9
0.8
0.7
0.6
0.5
0.4
−1
0
1
2
3
4

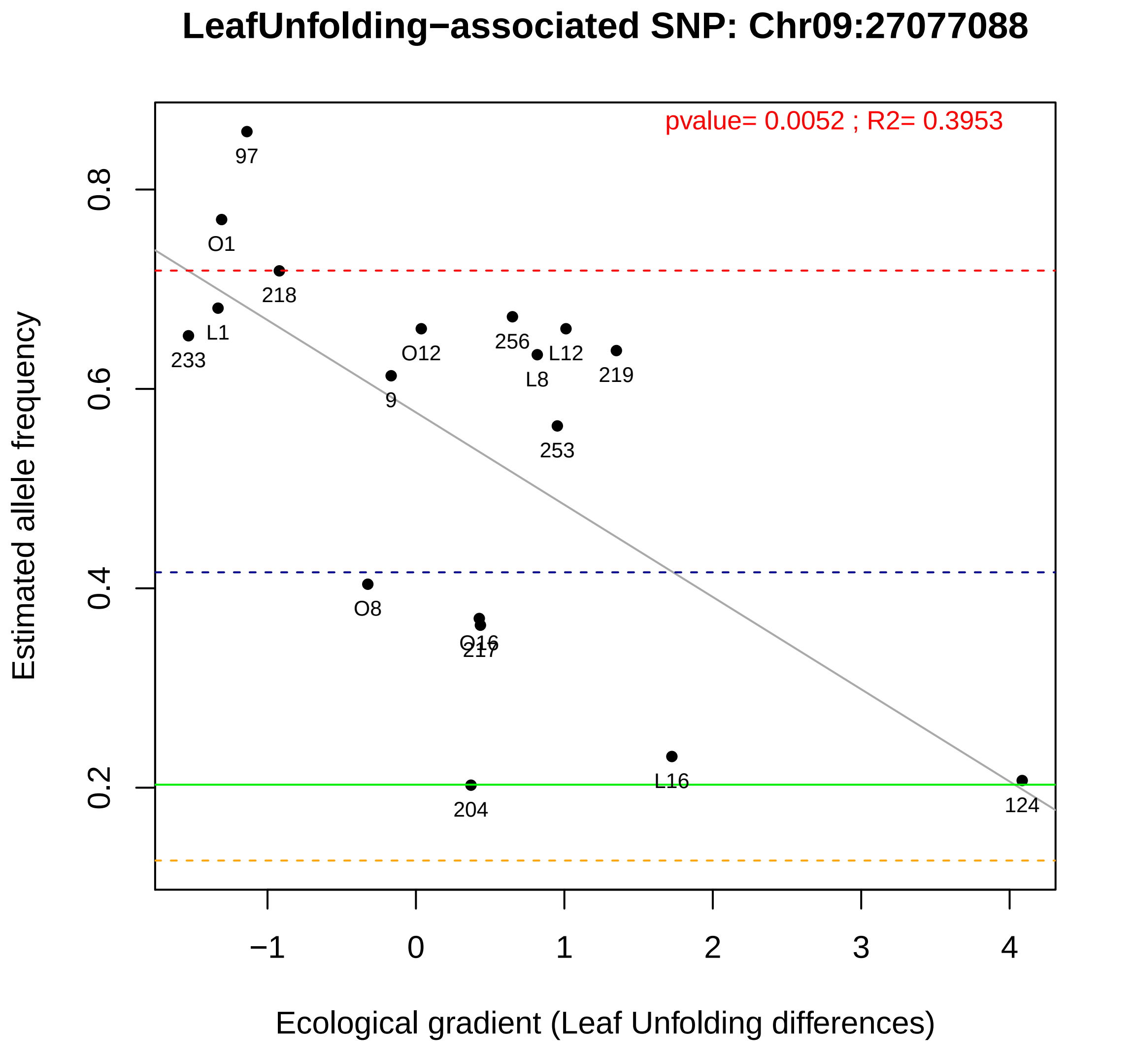
LeafUnfolding−associated SNP: Chr09:27077088
pvalue= 0.0052 ; R2= 0.3953
97
O1
218
L1
233
O12
9
256
L8
L12
219
253
O8
O16
217
204
L16
124
0.2
0.4
0.6
0.8
−1
0
1
2
3
4
Estimated allele frequency
Ecological gradient (Leaf Unfolding differences)

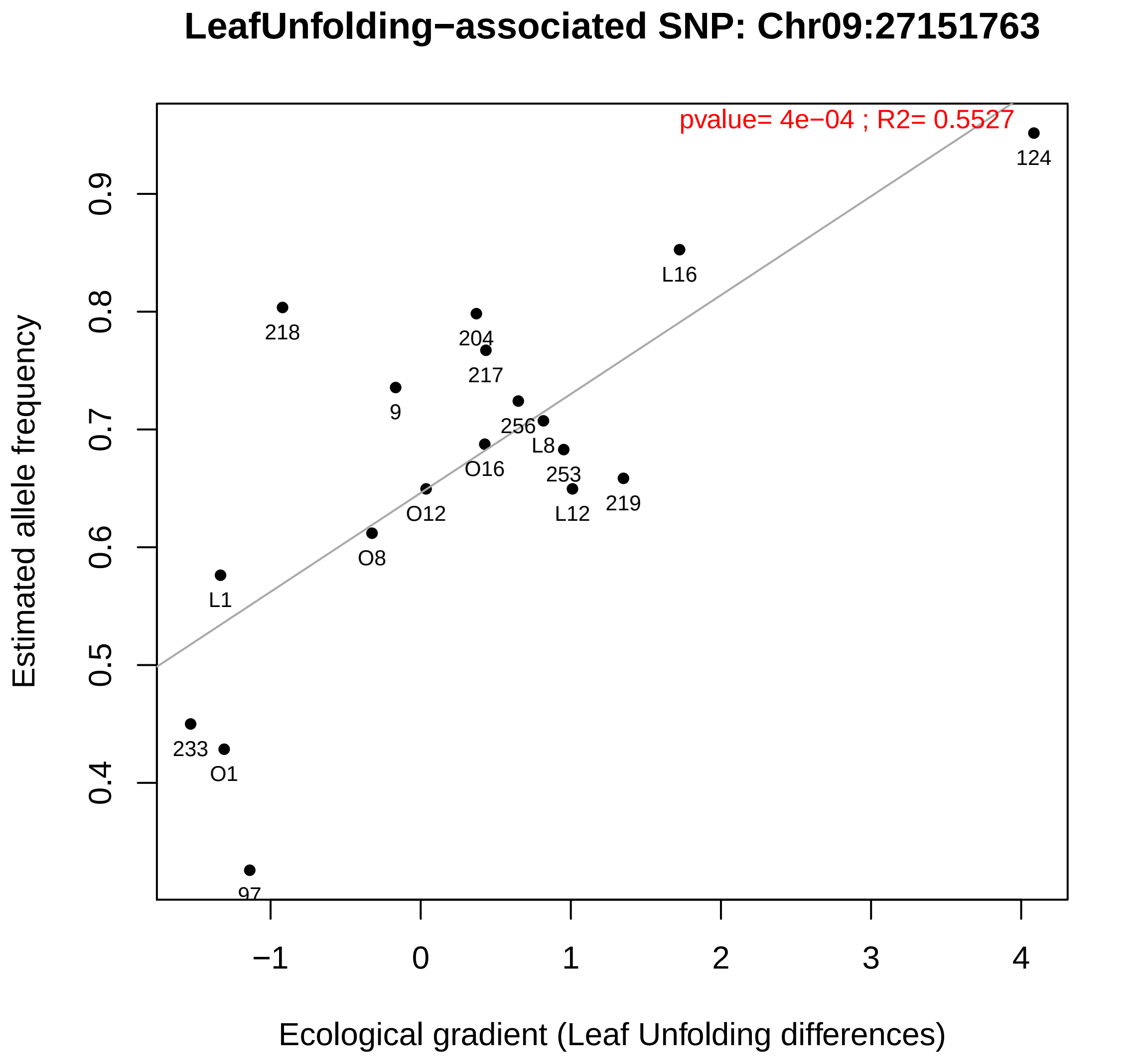
LeafUnfolding−associated SNP: Chr09:27151763
pvalue= 4e−04 ; R2= 0.5527
Estimated allele frequency
Ecological gradient (Leaf Unfolding differences)
0.4
0.5
0.6
0.7
0.8
0.9
−1
0
1
2
3
4
124
L16
218
204
217
9
256
L8
O16
253
219
O12
L12
O8
L1
233
O1
97

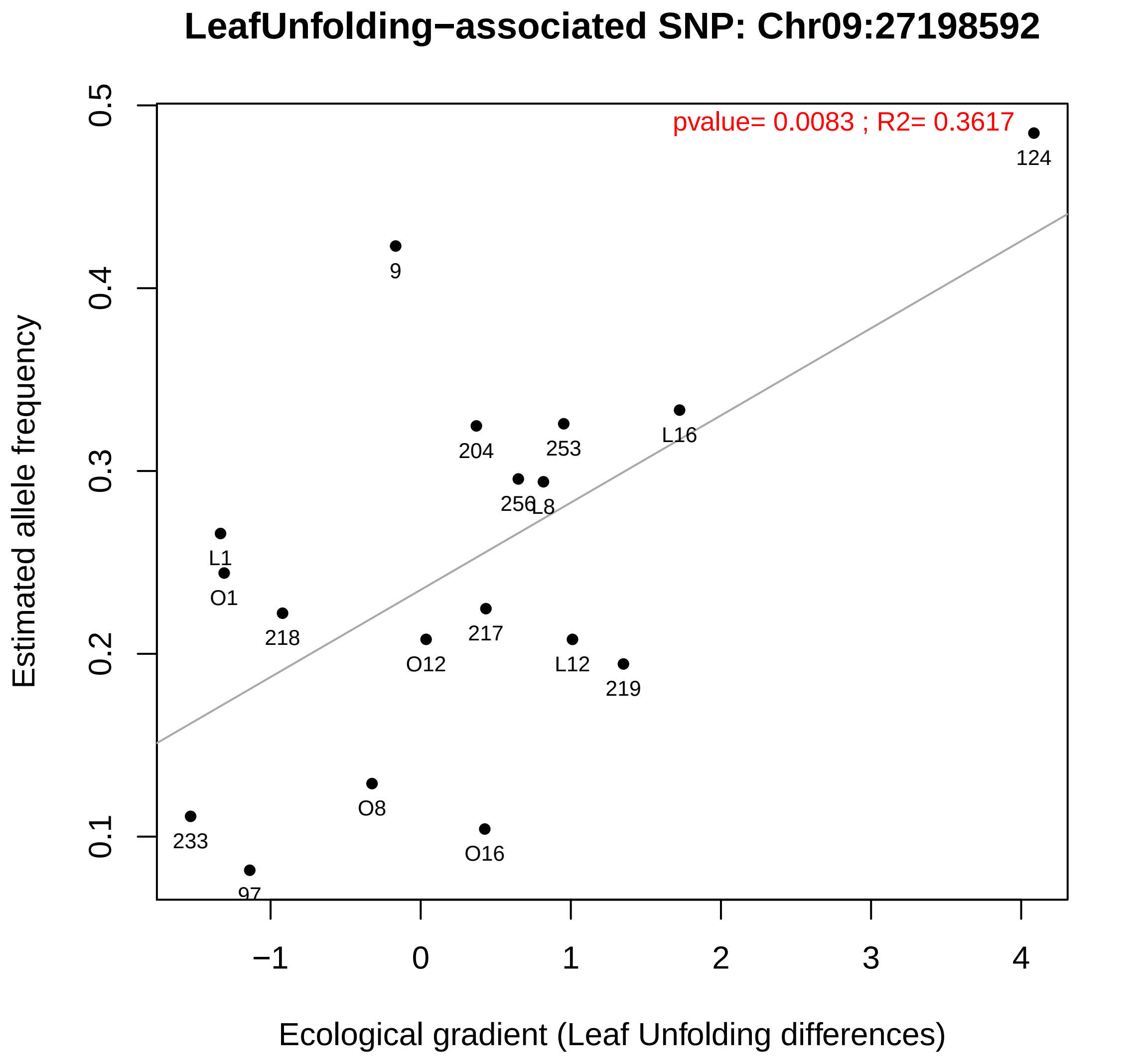

LeafUnfolding−associated SNP: Chr09:27198592
pvalue= 0.0083 ; R2= 0.3617
Estimated allele frequency
Ecological gradient (Leaf Unfolding differences)
0.1
0.2
0.3
0.4
0.5
−1
0
1
2
3
4
124
9
204
253
L16
256
L8
L1
O1
218
O12
217
L12
219
O8
233
O16
97

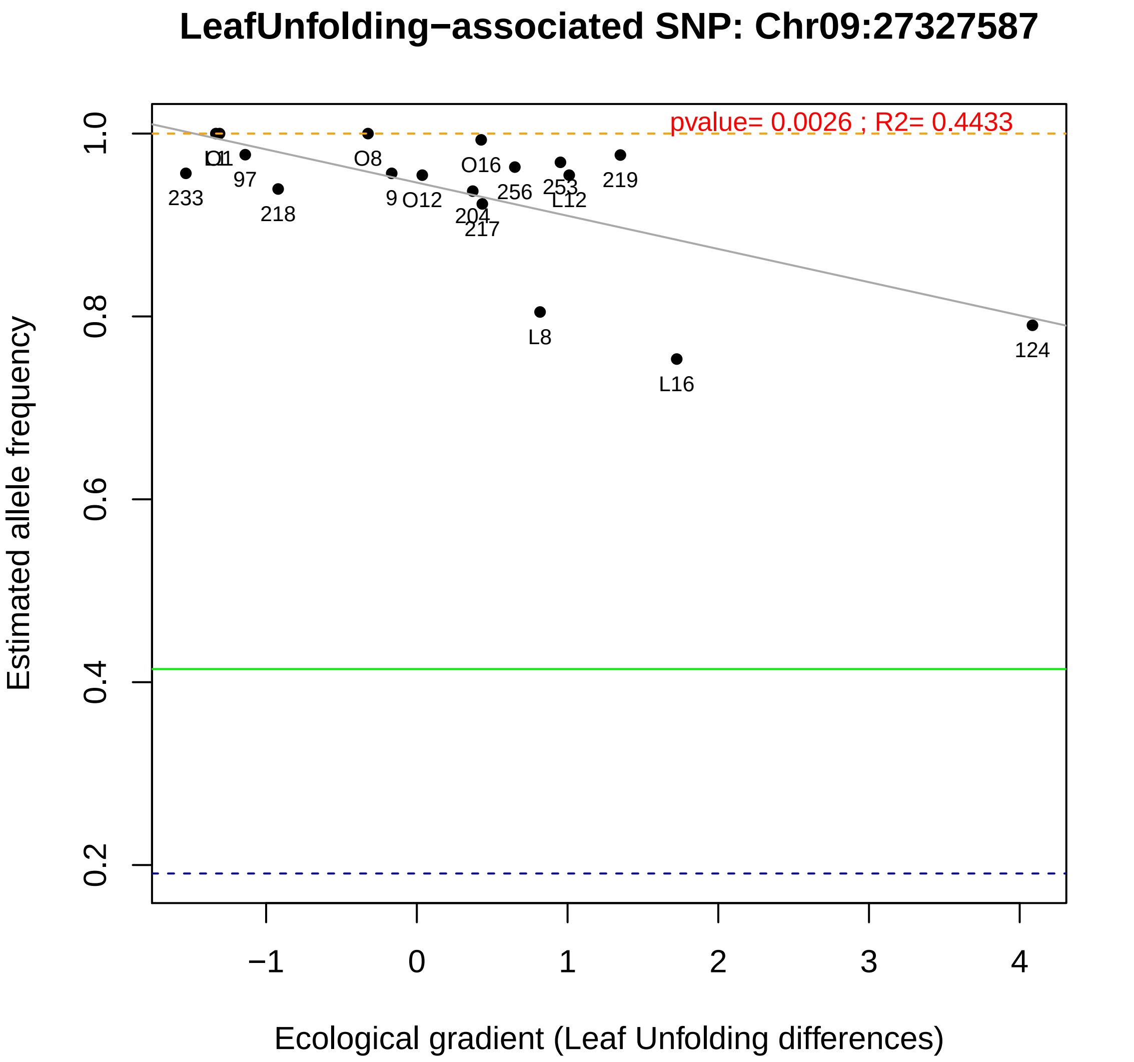
LeafUnfolding−associated SNP: Chr09:27327587
pvalue= 0.0026 ; R2= 0.4433
Estimated allele frequency
Ecological gradient (Leaf Unfolding differences)
0.2
0.4
0.6
0.8
1.0
−1
0
1
2
3
4
233
97
218
O8
9
O12
O16
204
217
256
253
L12
219
L8
L16
124

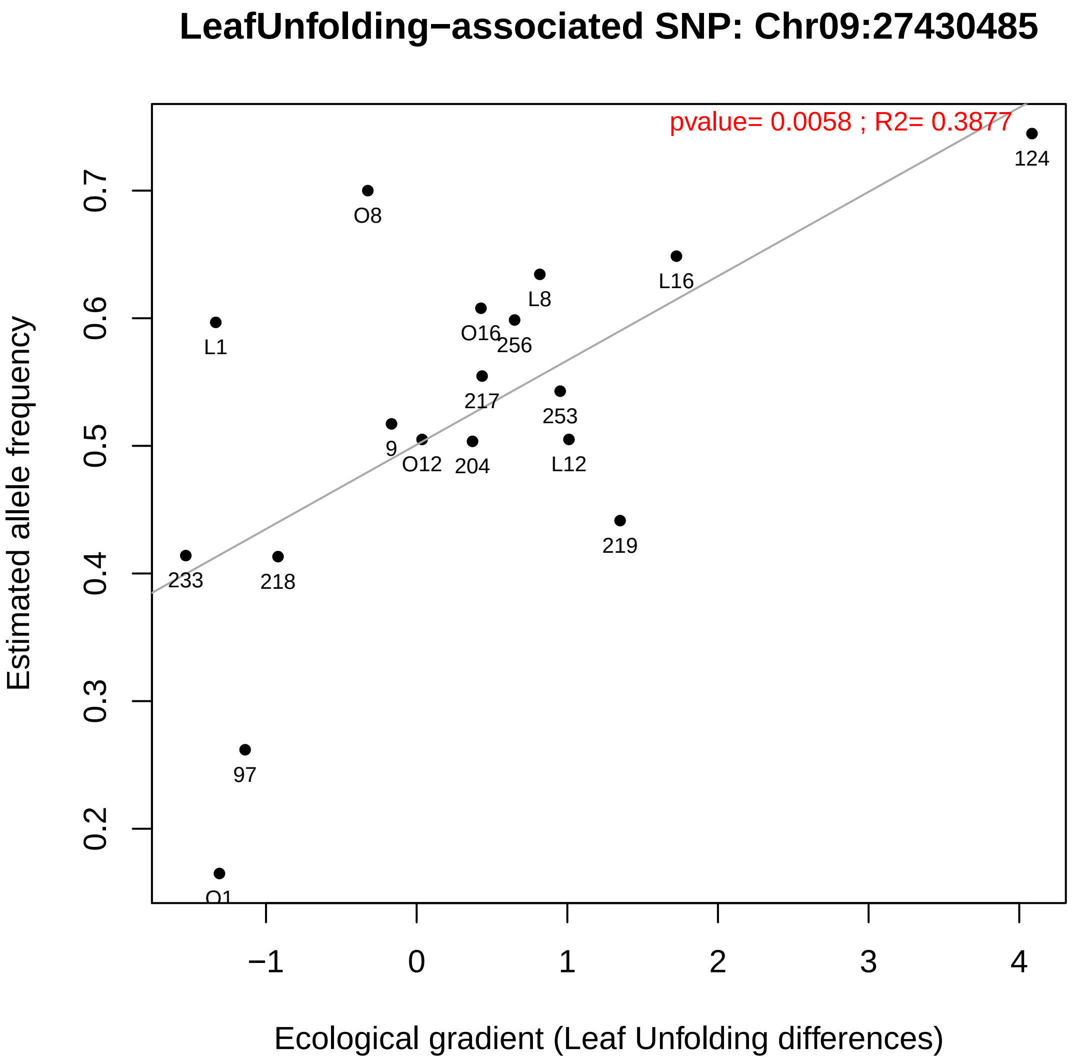

LeafUnfolding−associated SNP: Chr09:27430485
pvalue= 0.0058 ; R2= 0.3877
Estimated allele frequency
0.2
0.3
0.4
0.5
0.6
0.7
−1
0
1
2
3
4
Ecological gradient (Leaf Unfolding differences)
124
O8
L16
L8
O16
256
L1
217
253
9
O12
204
L12
219
233
218
97

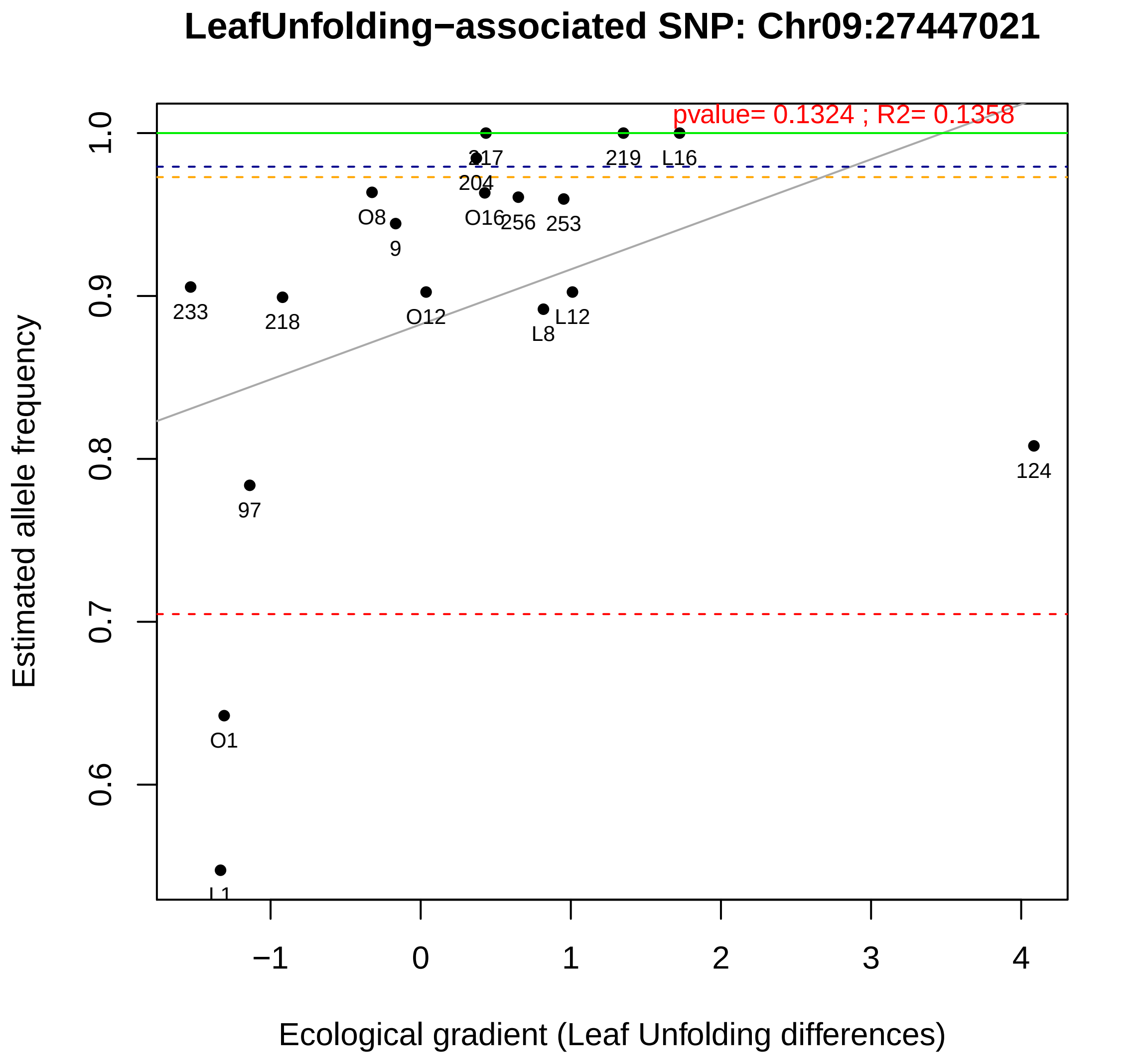
LeafUnfolding−associated SNP: Chr09:27447021
pvalue= 0.1324 ; R2= 0.1358
Estimated allele frequency
1.0
0.9
0.8
0.7
0.6
217
204
219
L16
O8
9
O16
256
253
233
218
O12
L8
L12
124
97
O1
L1
−1
0
1
2
3
4
Ecological gradient (Leaf Unfolding differences)

**LeafUnfolding−associated SNP: Chr09:27516391**

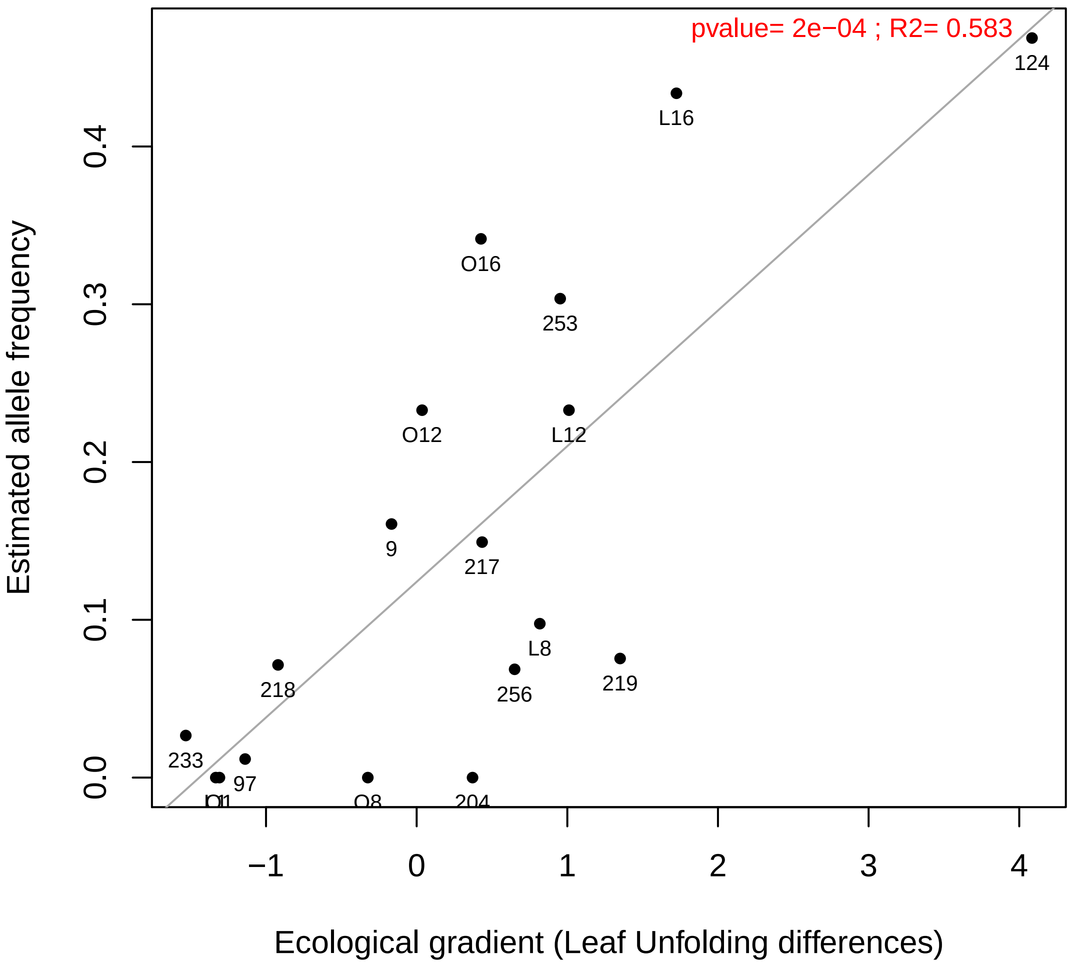

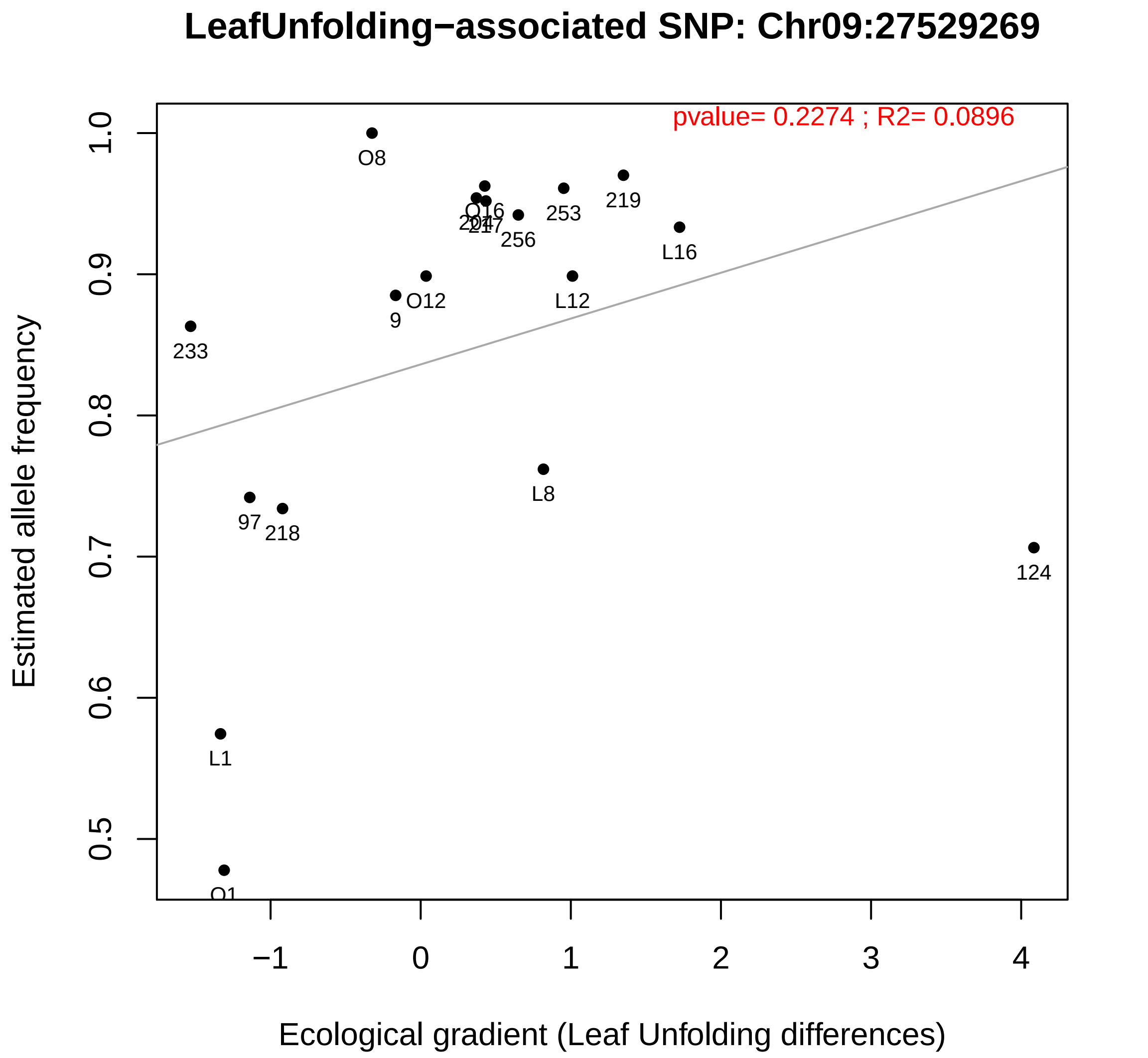

LeafUnfolding−associated SNP: Chr09:27529269
pvalue= 0.2274 ; R2= 0.0896
Estimated allele frequency
Ecological gradient (Leaf Unfolding differences)
1.0
0.9
0.8
0.7
0.6
0.5
−1
0
1
2
3
4
O8
219
253
256
L16
O12
L12
9
233
L8
97
218
124
L1

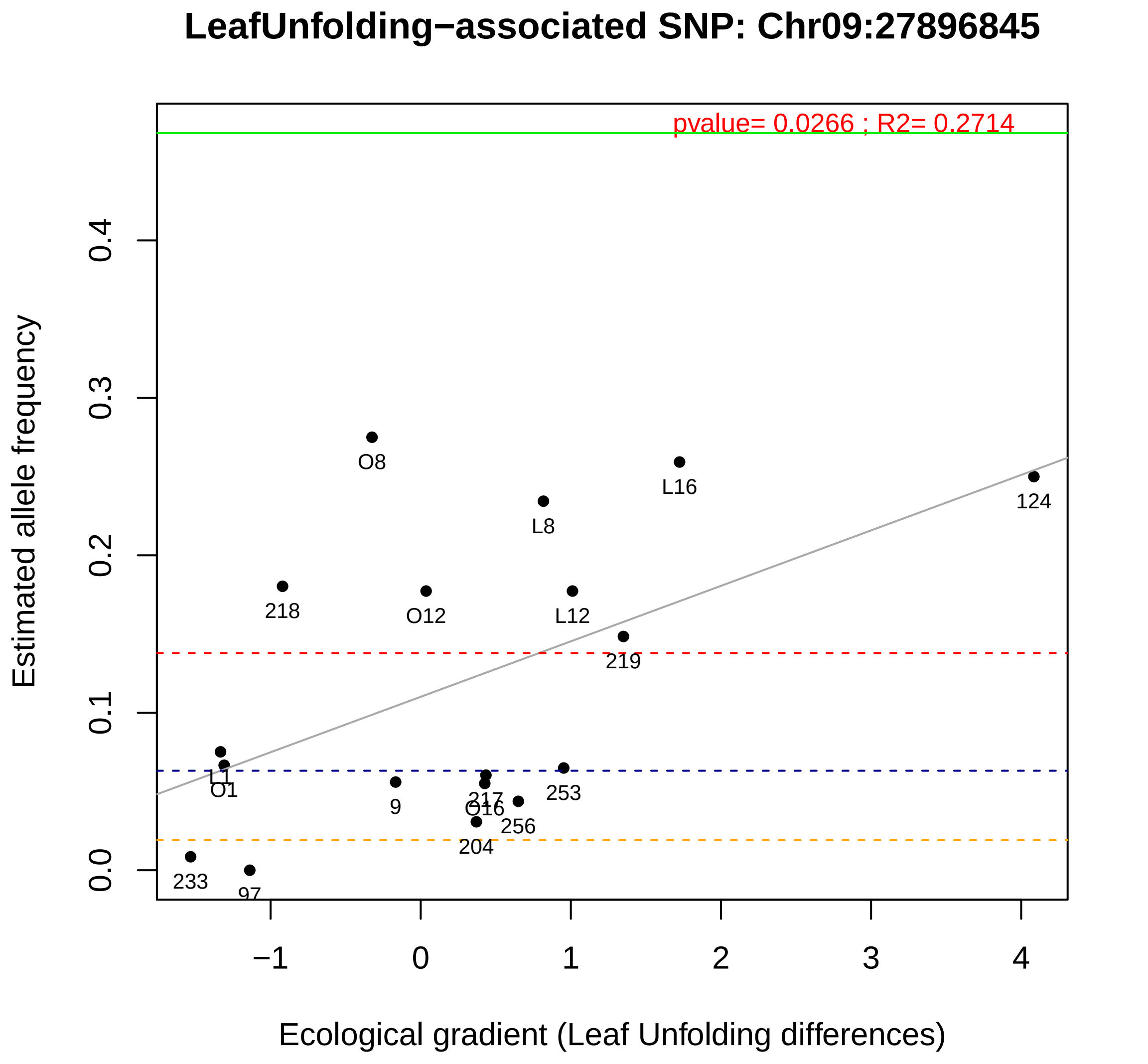
LeafUnfolding−associated SNP: Chr09:27896845
pvalue= 0.0266 ; R2= 0.2714
Estimated allele frequency
0.0
0.1
0.2
0.3
0.4
−1
0
1
2
3
4
Ecological gradient (Leaf Unfolding differences)
O8
L16
124
L8
218
O12
L12
219
L1
O1
9
217
O16
256
253
204
233
97

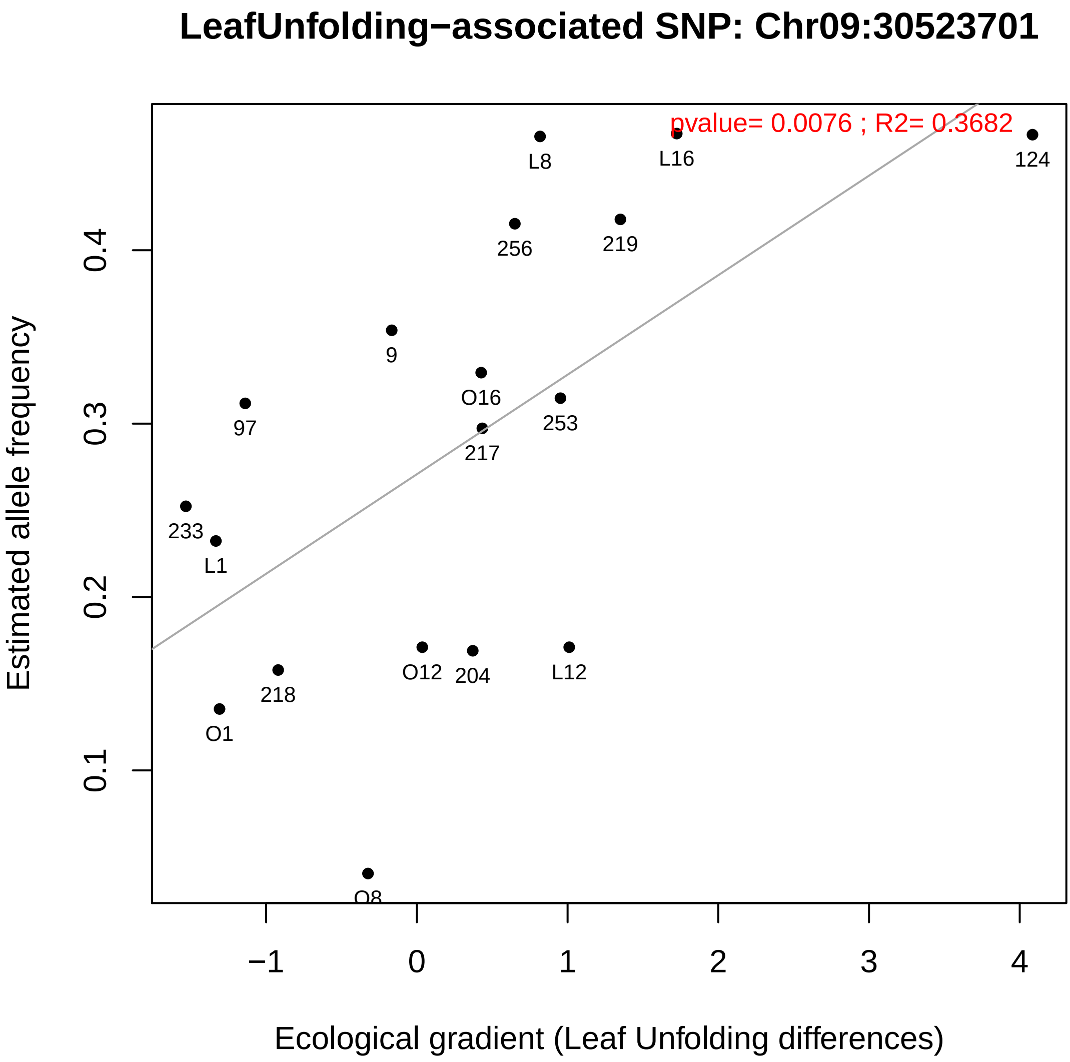
LeafUnfolding−associated SNP: Chr09:30523701
pvalue= 0.0076 ; R2= 0.3682
L8
L16
124
256
219
9
O16
97
253
217
233
L1
O12
204
L12
218
O1
O8
0.1
0.2
0.3
0.4
−1
0
1
2
3
4
Estimated allele frequency
Ecological gradient (Leaf Unfolding differences)

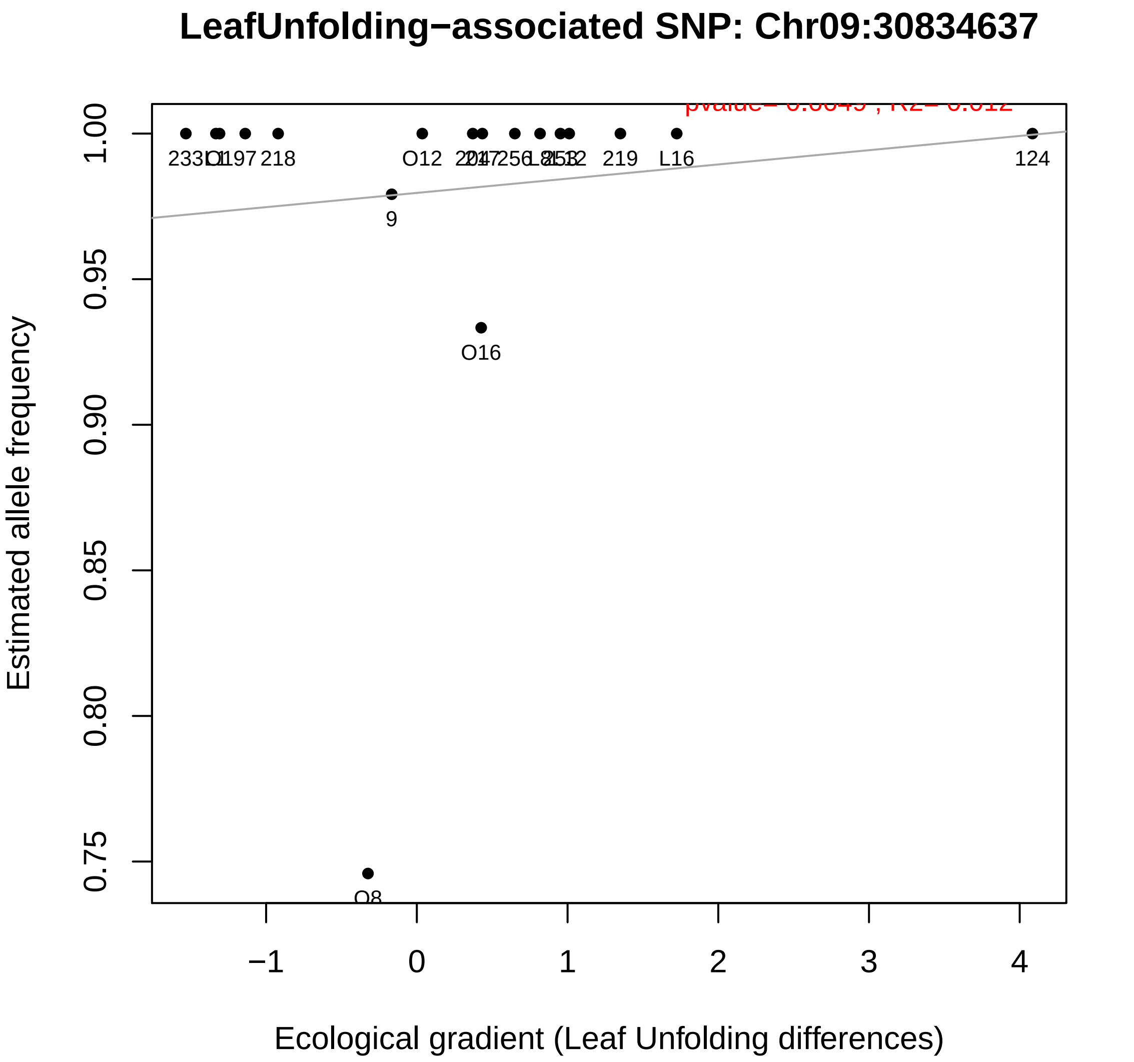

LeafUnfolding−associated SNP: Chr09:30834637
Estimated allele frequency
Ecological gradient (Leaf Unfolding differences)
0.75
0.80
0.85
0.90
0.95
1.00
−1
0
1
2
3
4
9
O16
O12
218
219
L16
124

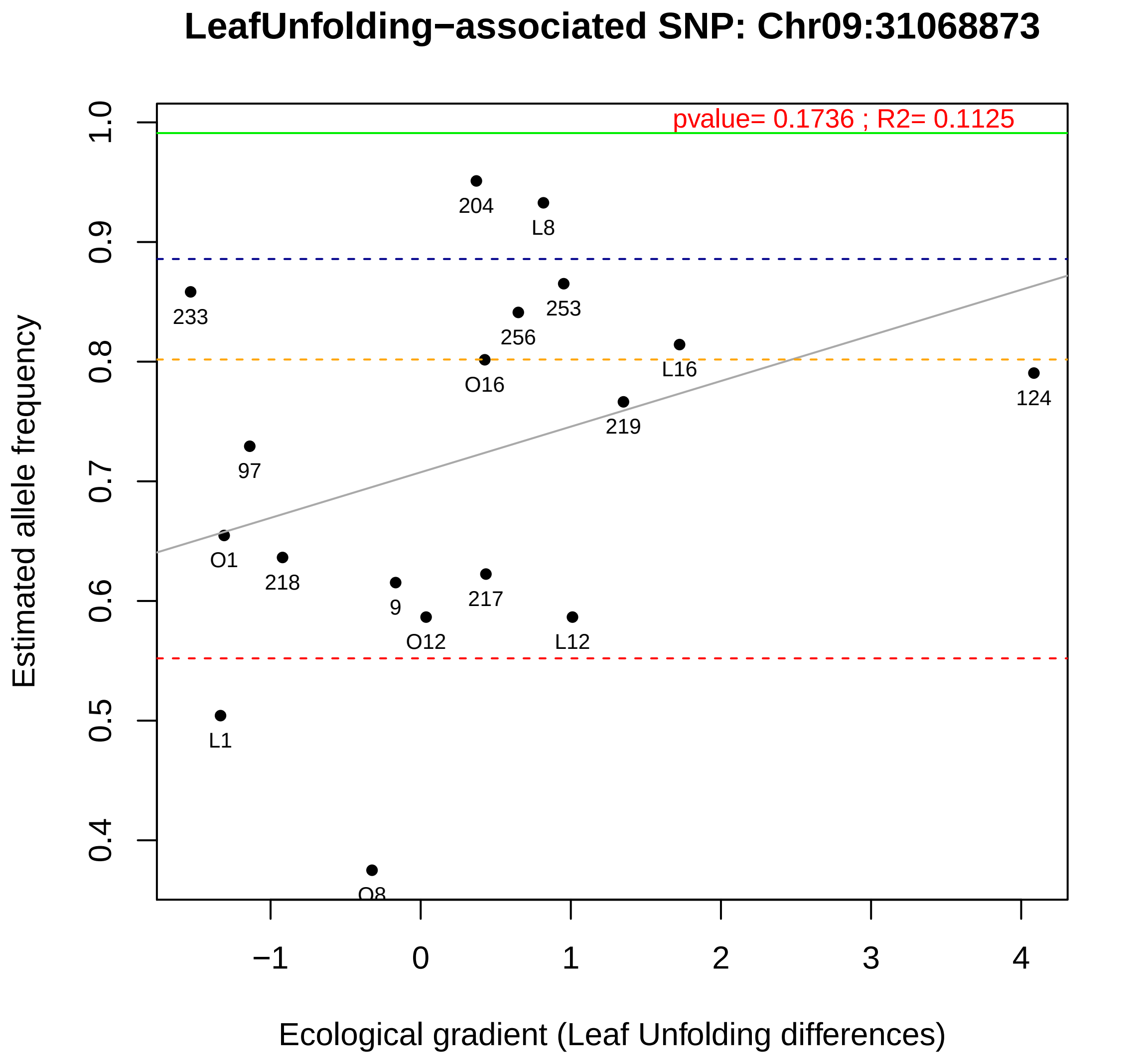
LeafUnfolding−associated SNP: Chr09:31068873
pvalue= 0.1736 ; R2= 0.1125
Estimated allele frequency
Ecological gradient (Leaf Unfolding differences)
1.0
0.9
0.8
0.7
0.6
0.5
0.4
−1
0
1
2
3
4
204
L8
233
253
256
O16
L16
124
219
97
O1
218
9
217
O12
L12
L1
O8

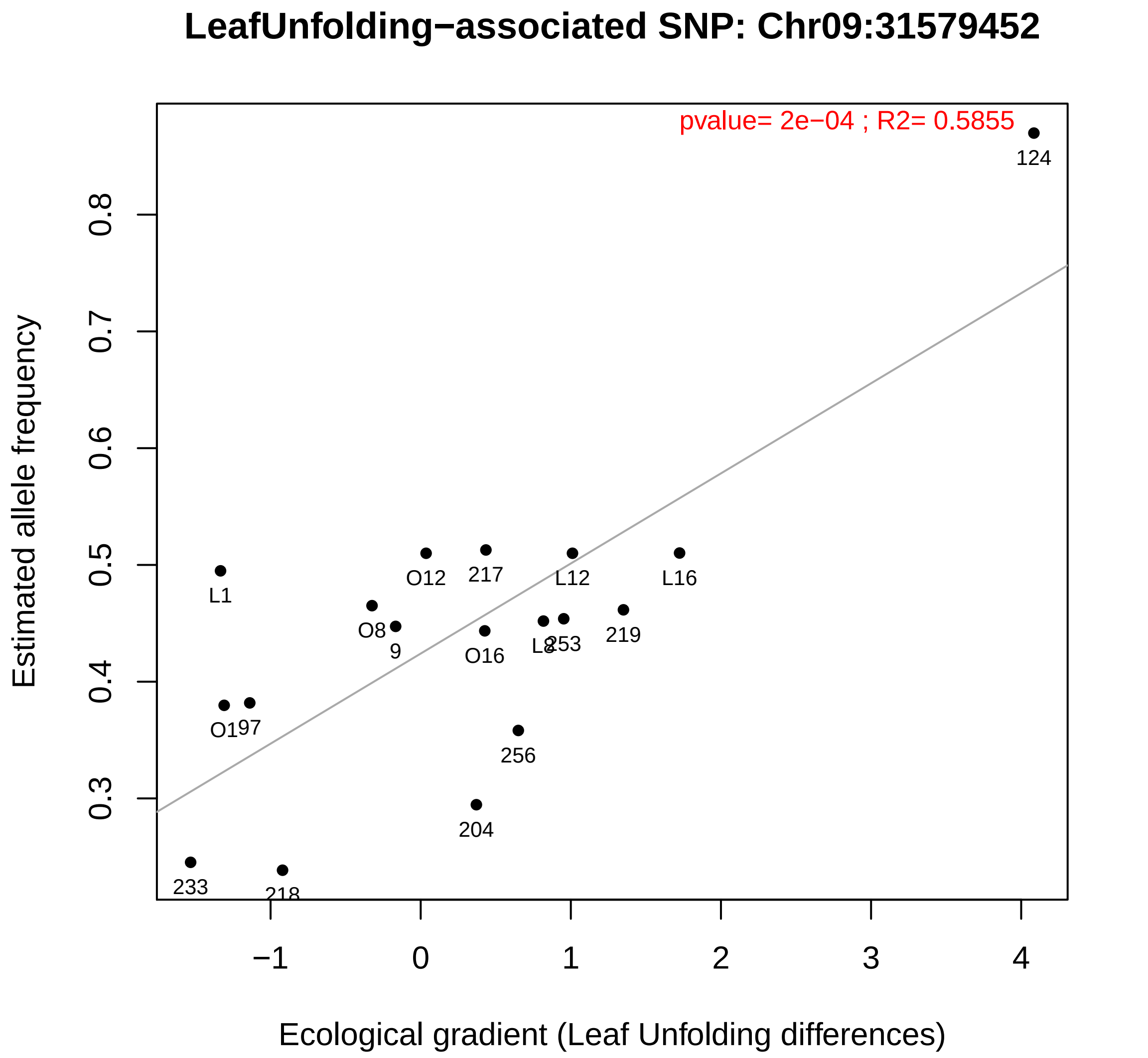

LeafUnfolding−associated SNP: Chr09:31579452
pvalue= 2e−04 ; R2= 0.5855
Estimated allele frequency
Ecological gradient (Leaf Unfolding differences)
0.3
0.4
0.5
0.6
0.7
0.8
−1
0
1
2
3
4
124
L1
O12
217
L12
L16
O8
9
O16
253
219
197
256
204
233
218

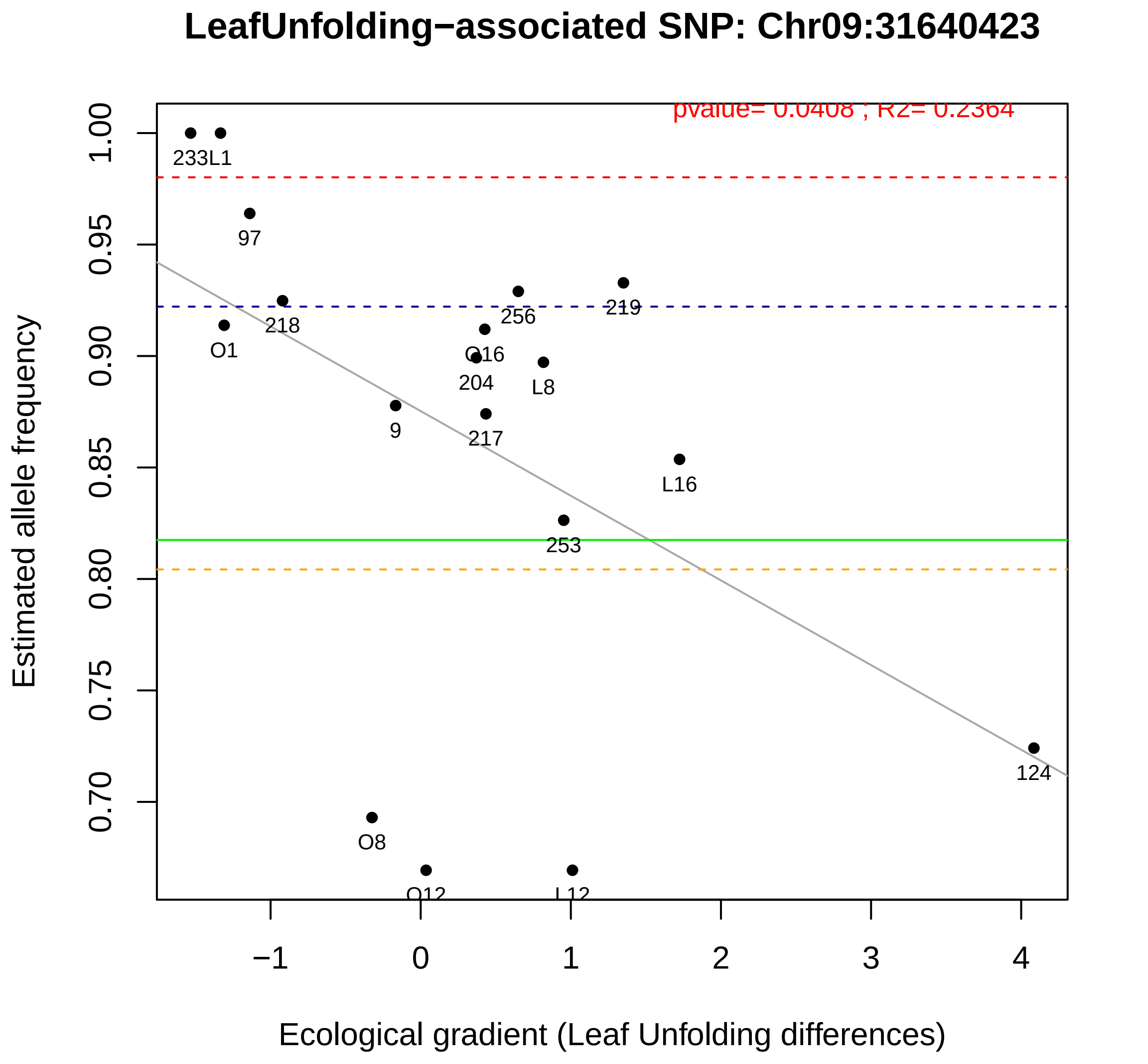
LeafUnfolding−associated SNP: Chr09:31640423
pvalue= 0.0408 ; R2= 0.2364
Estimated allele frequency
0.70
0.75
0.80
0.85
0.90
0.95
1.00
−1
0
1
2
3
4
Ecological gradient (Leaf Unfolding differences)
233
L1
97
218
O1
256
219
O16
204
L8
9
217
L16
253
124
O8
O12
L12

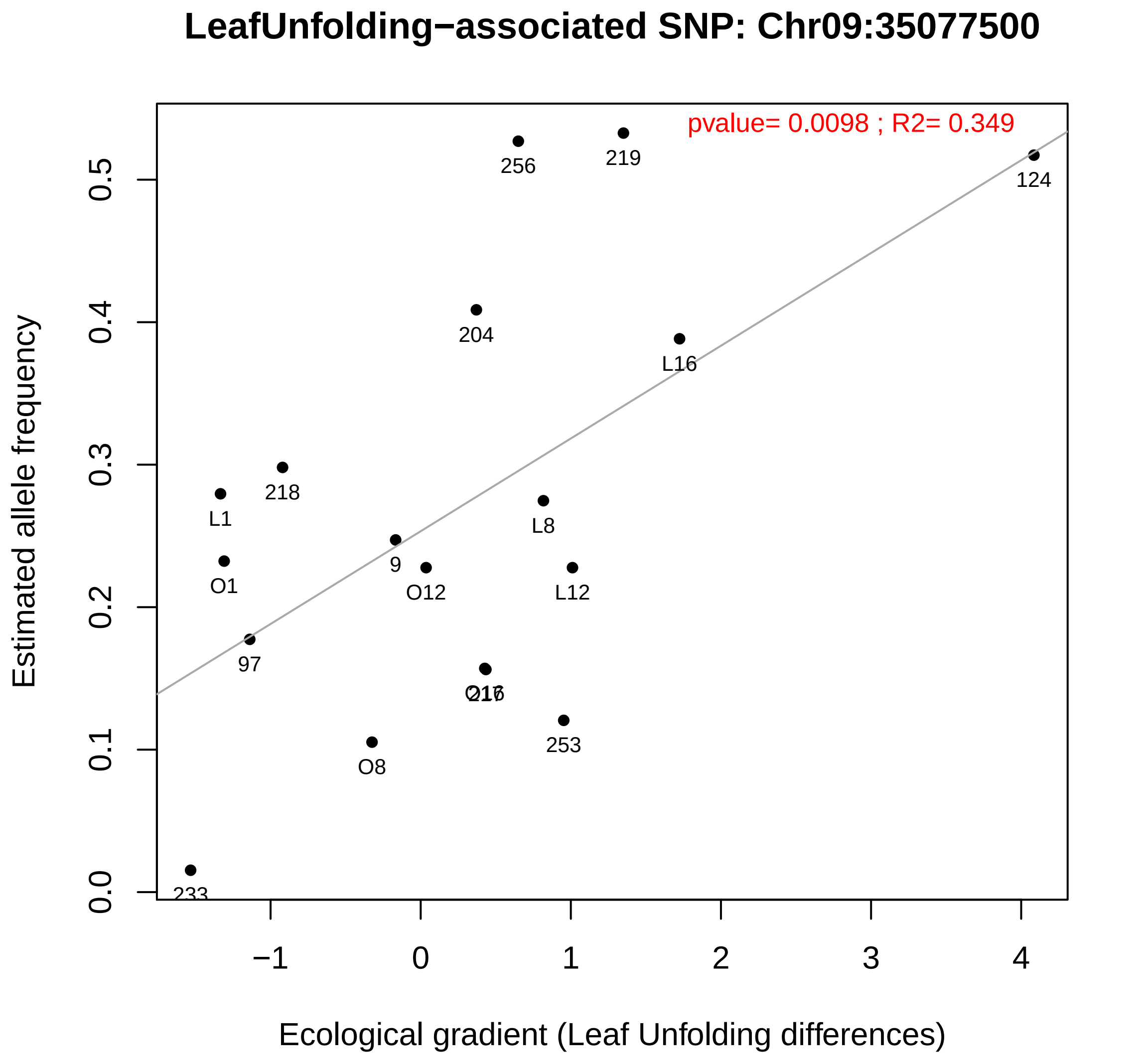
LeafUnfolding−associated SNP: Chr09:35077500
pvalue= 0.0098 ; R2= 0.349
Estimated allele frequency
Ecological gradient (Leaf Unfolding differences)
0.0
0.1
0.2
0.3
0.4
0.5
−1
0
1
2
3
4
256
219
124
204
L16
218
L1
L8
9
O1
O12
L12
97
O17
216
253
O8
233

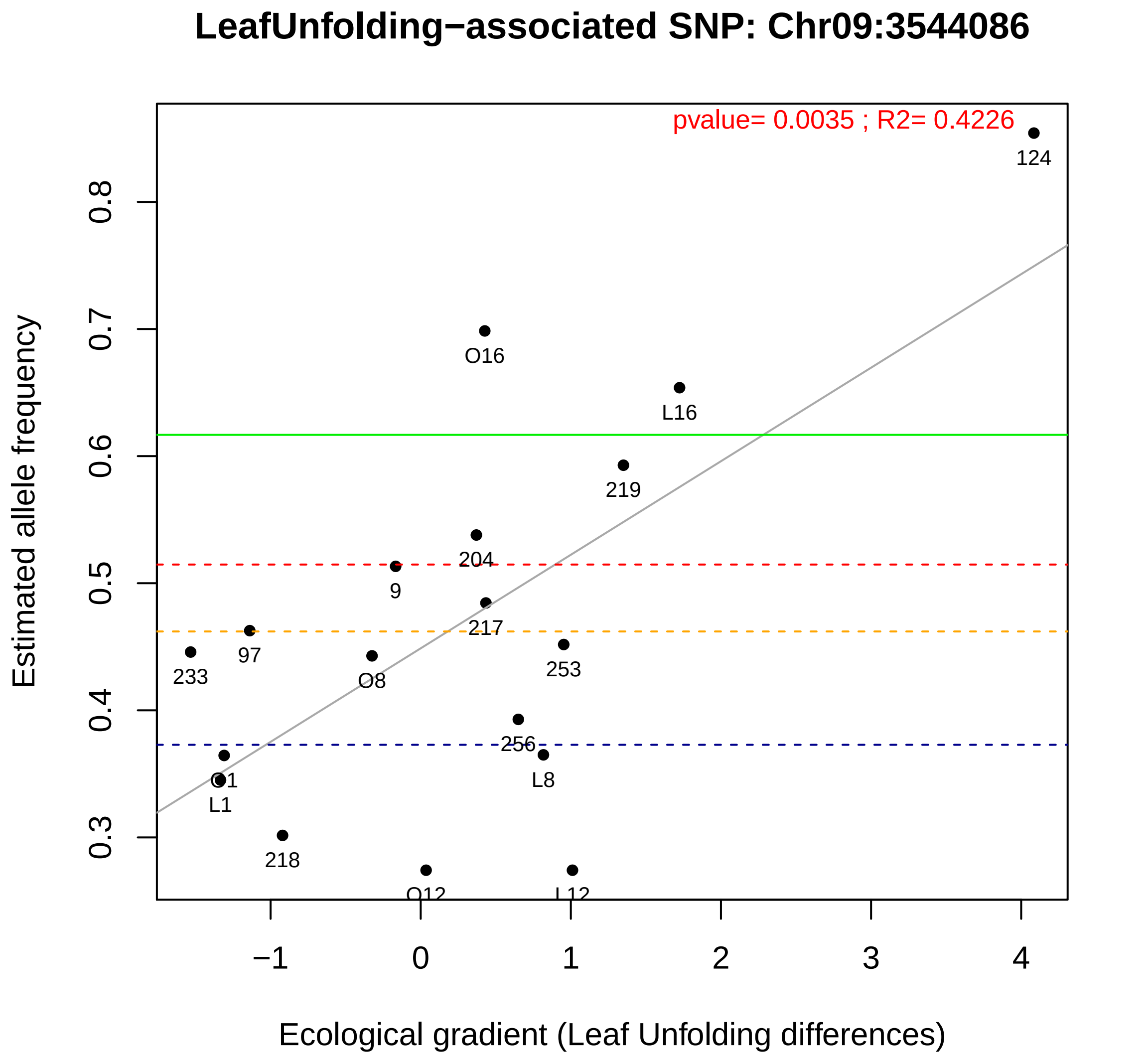

LeafUnfolding−associated SNP: Chr09:3544086
pvalue= 0.0035 ; R2= 0.4226
124
O16
L16
219
204
9
217
97
233
O8
253
256
L8
O1
L1
218
O12
L12
0.3
0.4
0.5
0.6
0.7
0.8
−1
0
1
2
3
4
Estimated allele frequency
Ecological gradient (Leaf Unfolding differences)

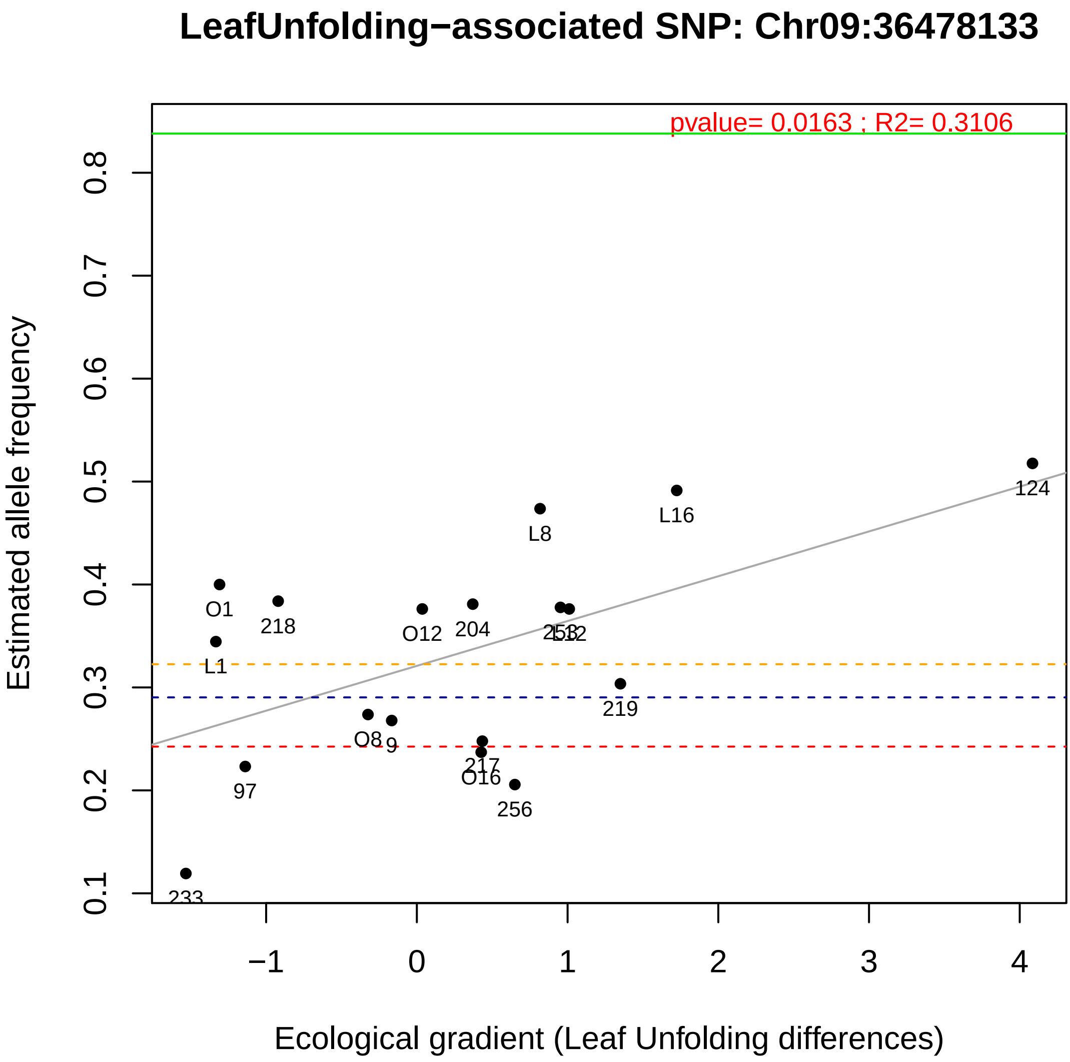

LeafUnfolding−associated SNP: Chr09:36478133
pvalue= 0.0163 ; R2= 0.3106
Estimated allele frequency
0.1
0.2
0.3
0.4
0.5
0.6
0.7
0.8
−1
0
1
2
3
4
Ecological gradient (Leaf Unfolding differences)
124
L16
L8
O1
218
O12
204
L1
219
O8
9
217
O16
256
97
233

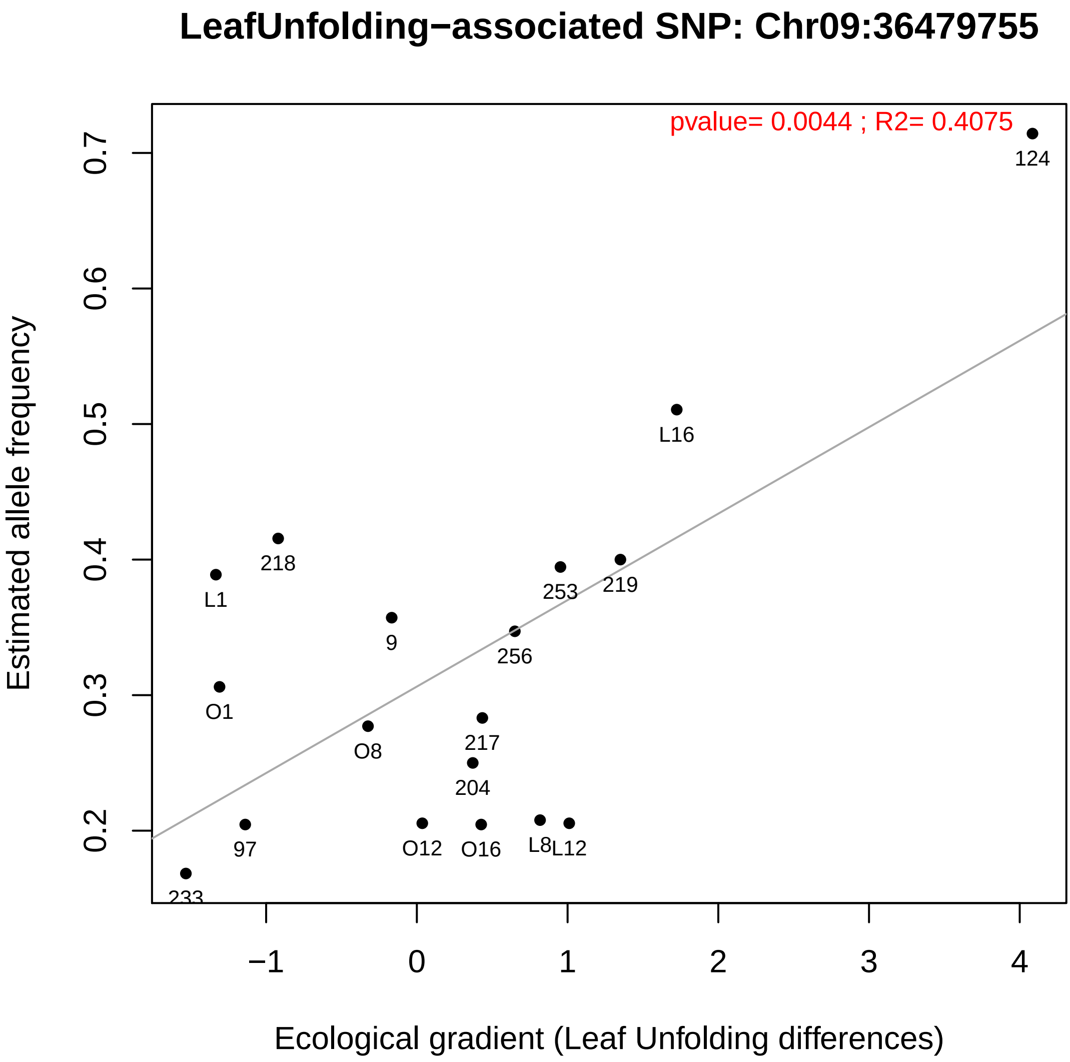
LeafUnfolding−associated SNP: Chr09:36479755
pvalue= 0.0044 ; R2= 0.4075
Estimated allele frequency
0.2
0.3
0.4
0.5
0.6
0.7
−1
0
1
2
3
4
Ecological gradient (Leaf Unfolding differences)
124
L16
218
L1
253
219
9
256
O1
O8
217
204
97
O12
O16
L8
L12
233

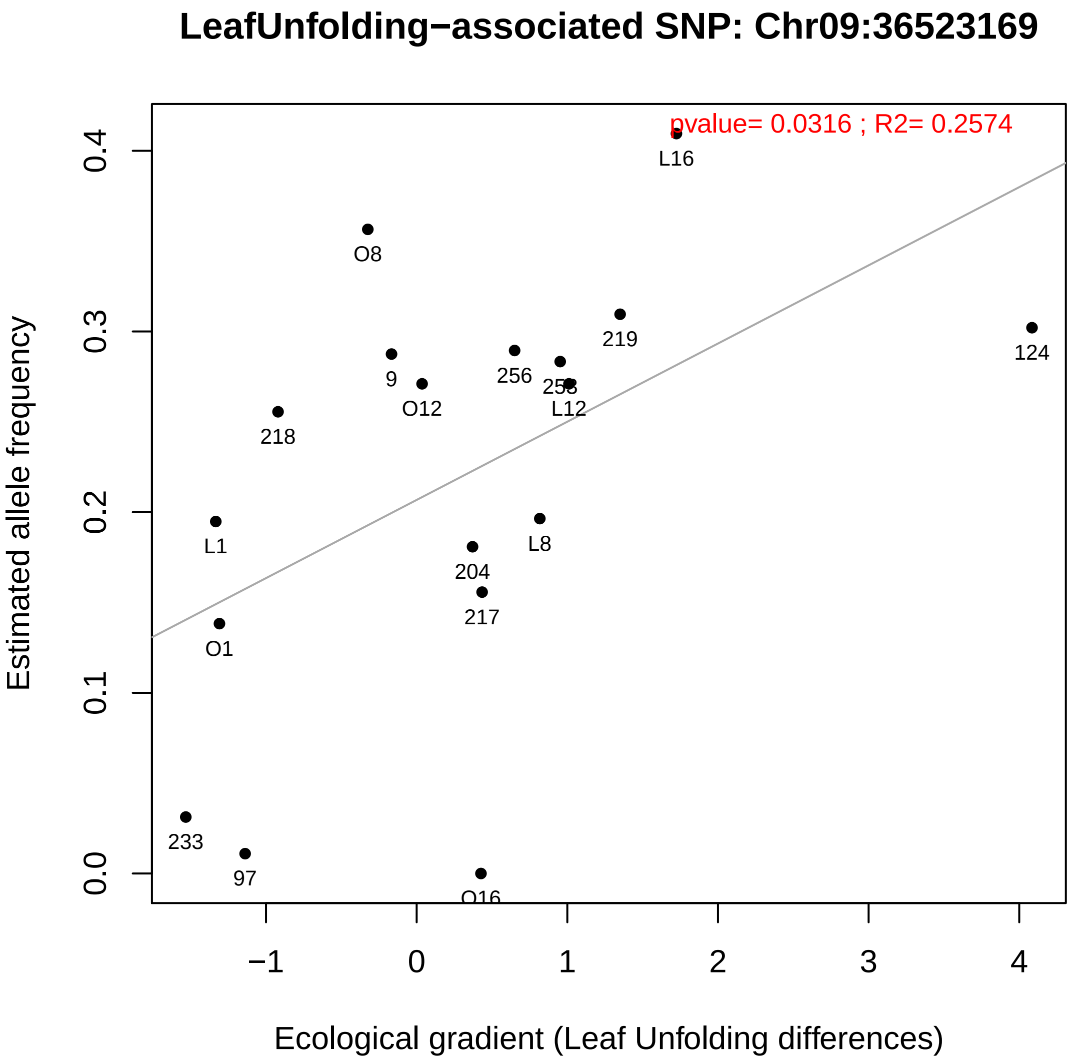
LeafUnfolding−associated SNP: Chr09:36523169
pvalue= 0.0316 ; R2= 0.2574
Estimated allele frequency
Ecological gradient (Leaf Unfolding differences)
0.0
0.1
0.2
0.3
0.4
−1
0
1
2
3
4
L16
O8
219
124
9
256
O12
L12
218
L1
L8
204
217
O1
233
97
O16

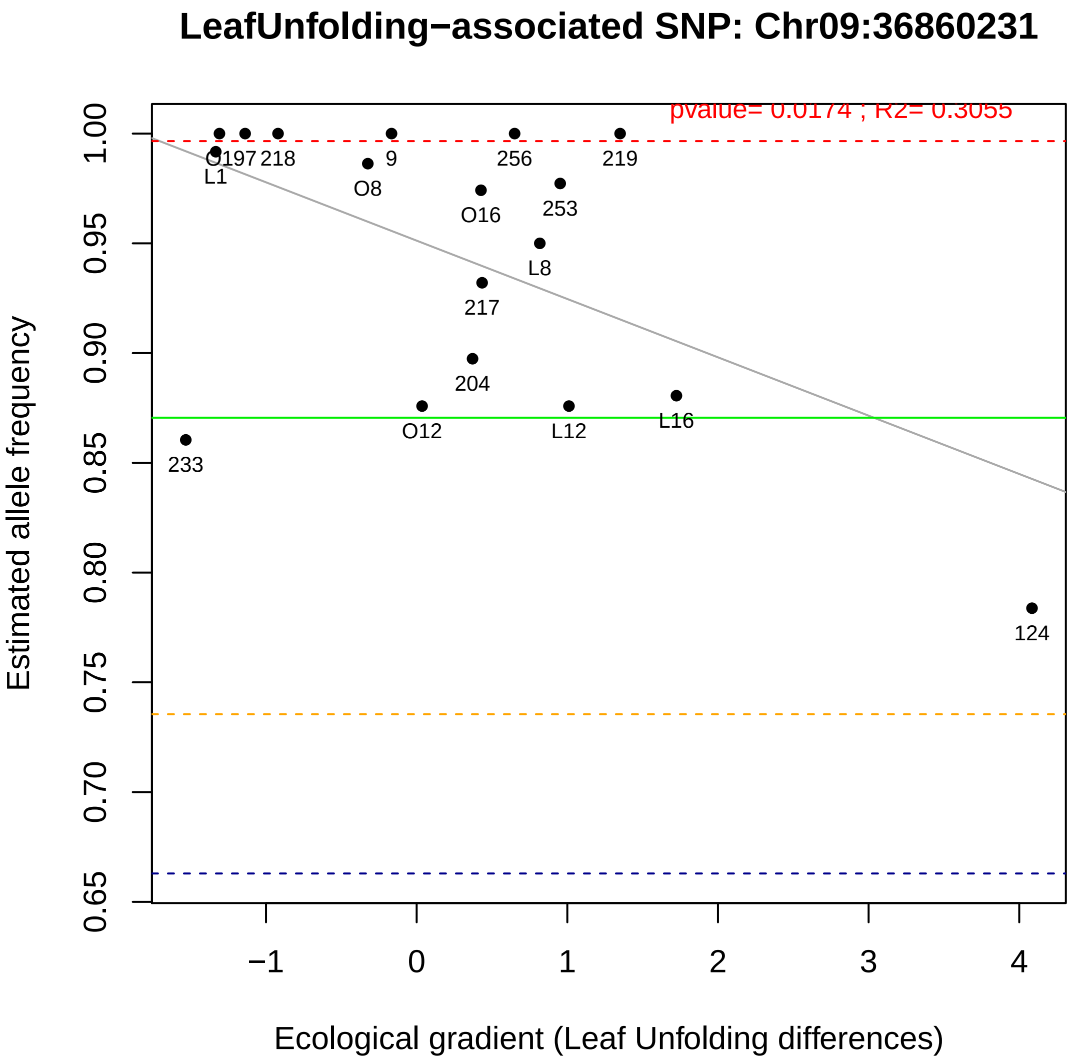

LeafUnfolding−associated SNP: Chr09:36860231
pvalue= 0.0174 ; R2= 0.3055
Estimated allele frequency
Ecological gradient (Leaf Unfolding differences)
1.00
0.95
0.90
0.85
0.80
0.75
0.70
0.65
−1
0
1
2
3
4
O197
218
9
256
219
L1
O8
O16
253
L8
217
204
O12
L12
L16
233
124

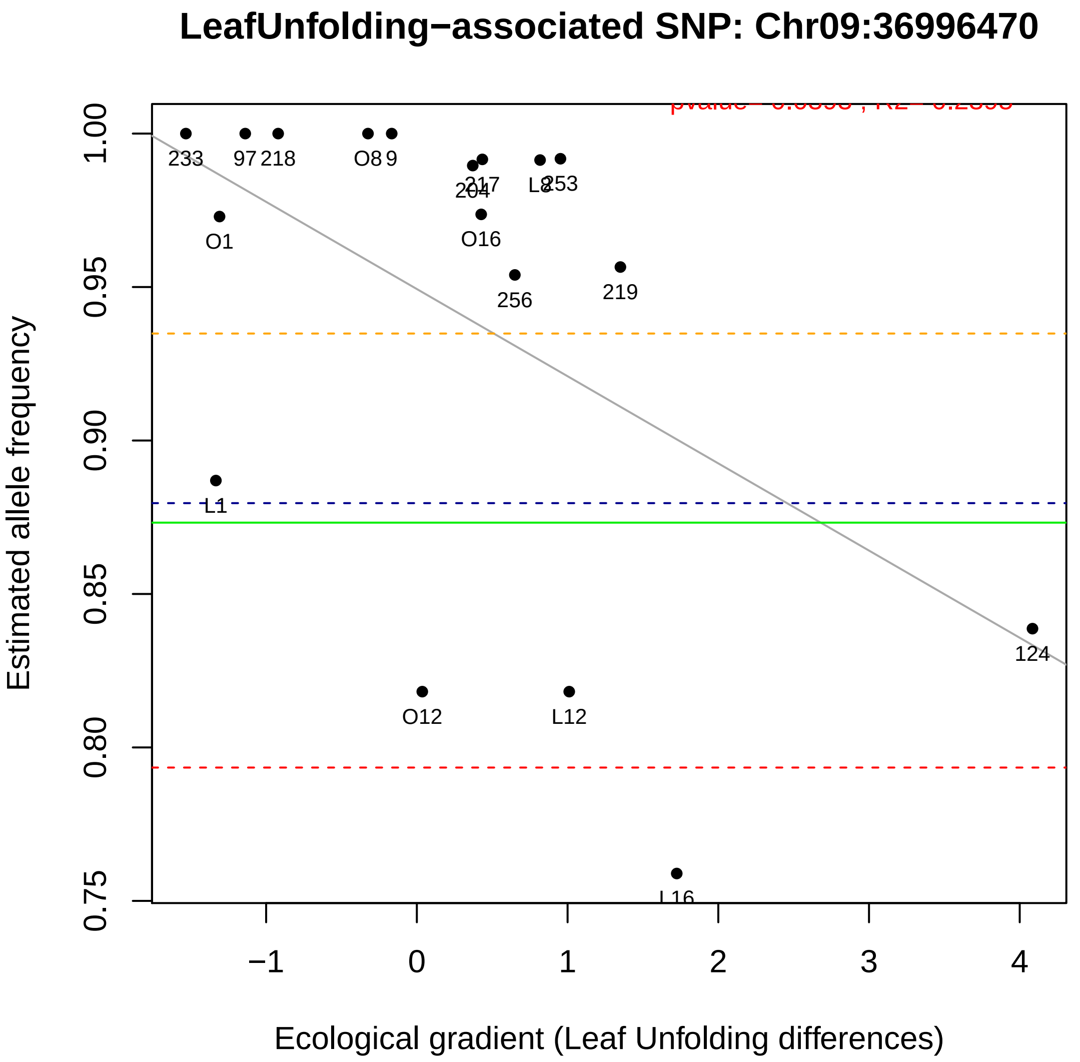
LeafUnfolding−associated SNP: Chr09:36996470
Estimated allele frequency
0.75
0.80
0.85
0.90
0.95
1.00
−1
0
1
2
3
4
Ecological gradient (Leaf Unfolding differences)
233
97
218
O8
9
204
217
L8
253
O1
O16
256
219
L1
124
O12
L12
L16

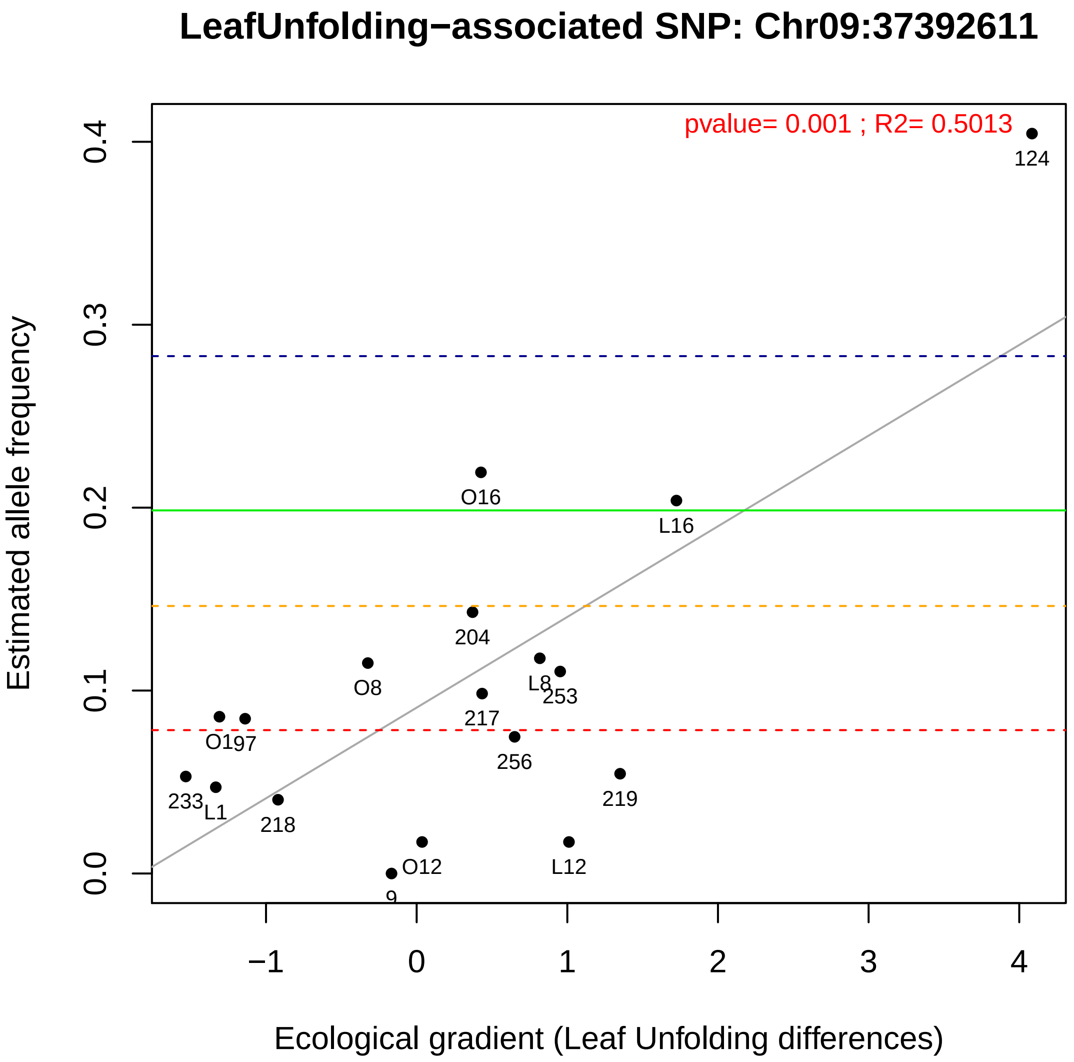

LeafUnfolding−associated SNP: Chr09:37392611
pvalue= 0.001 ; R2= 0.5013
Estimated allele frequency
0.0
0.1
0.2
0.3
0.4
−1
0
1
2
3
4
Ecological gradient (Leaf Unfolding differences)
124
O16
L16
204
O8
L8
253
217
256
O1
197
233
L1
218
219
O12
L12
9

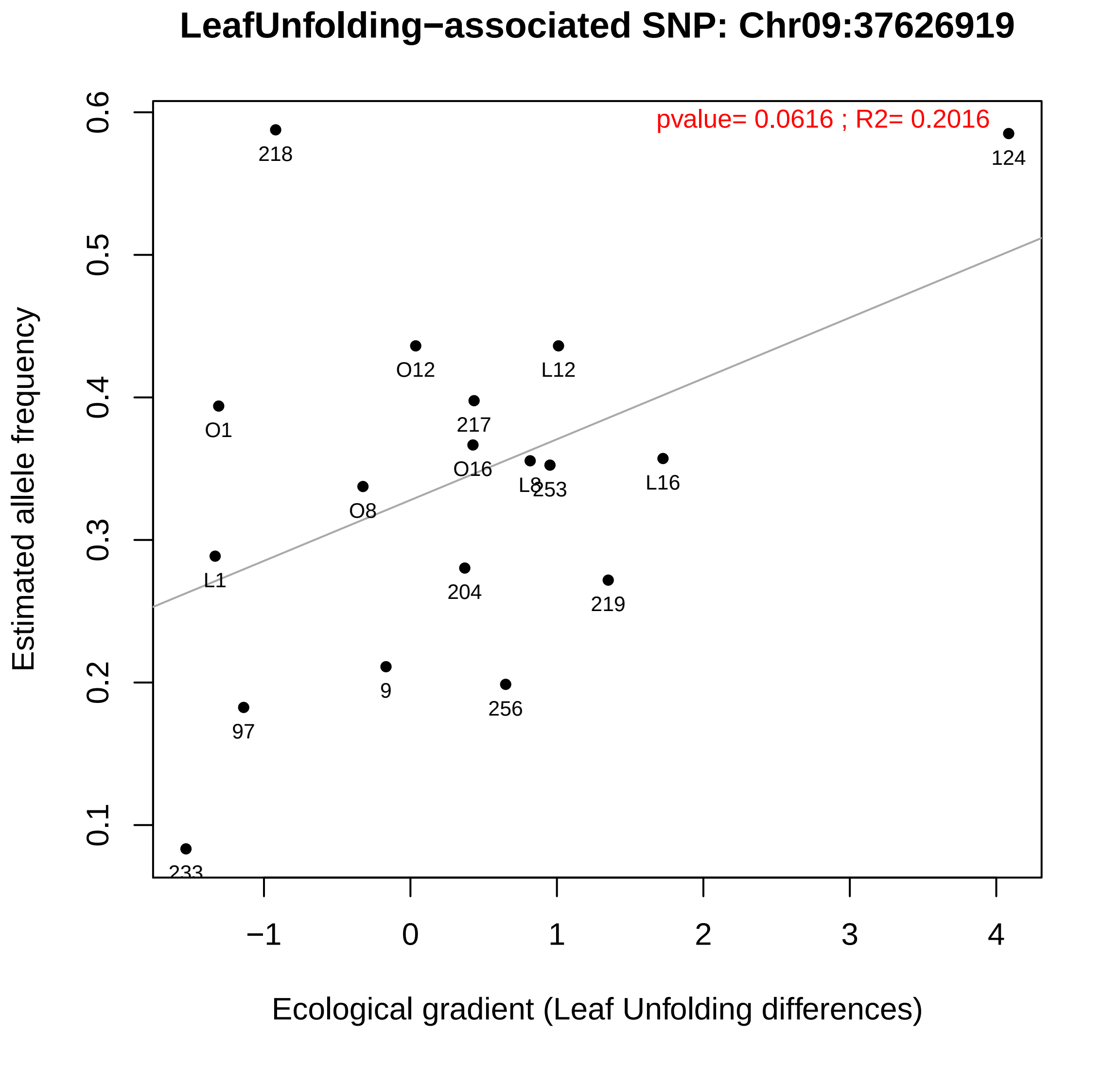

LeafUnfolding−associated SNP: Chr09:37626919
pvalue= 0.0616 ; R2= 0.2016
Estimated allele frequency
0.1
0.2
0.3
0.4
0.5
0.6
−1
0
1
2
3
4
Ecological gradient (Leaf Unfolding differences)
218
124
O12
L12
O1
217
O16
L8
253
L16
O8
L1
204
219
9
256
97
233

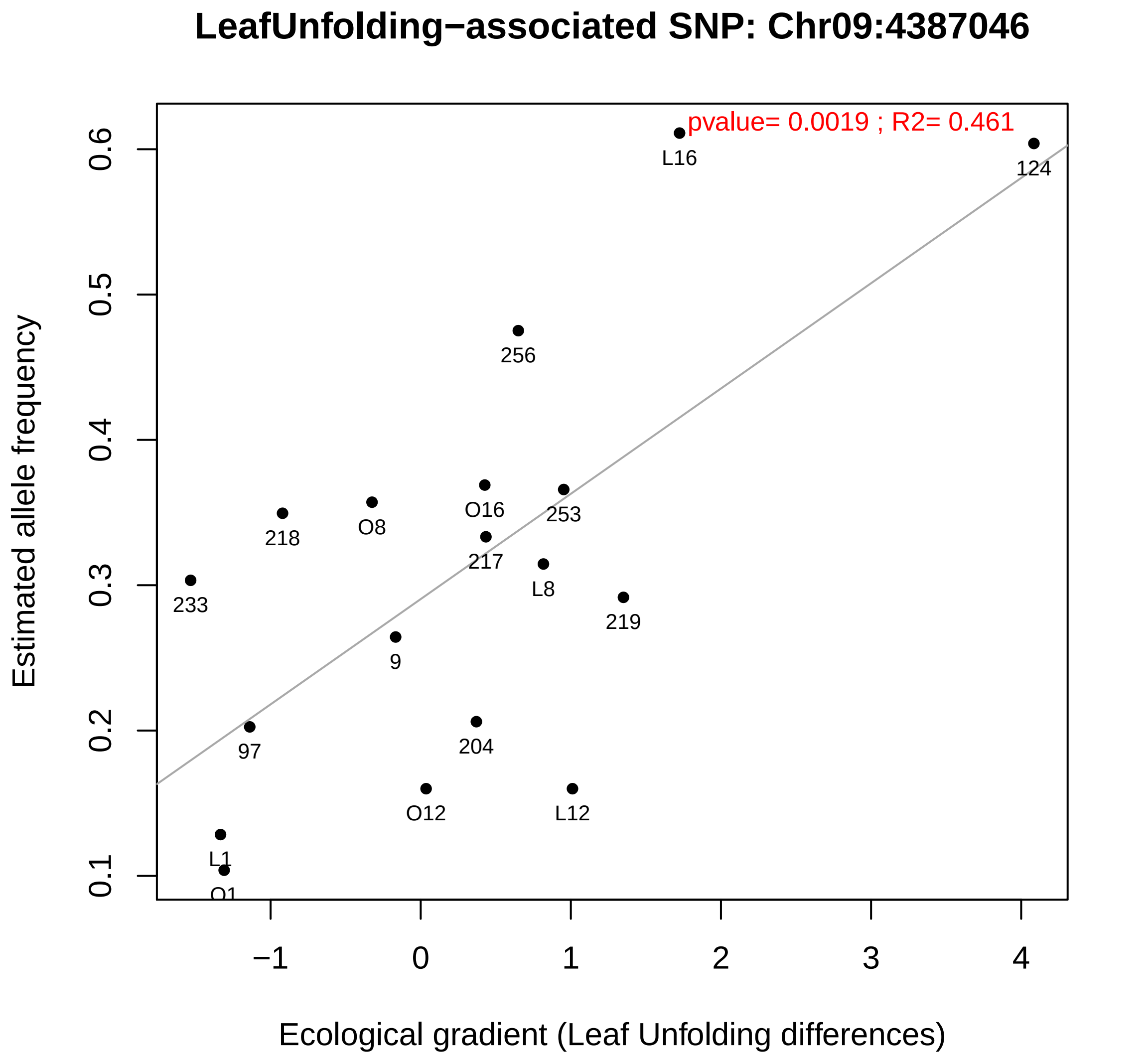
LeafUnfolding−associated SNP: Chr09:4387046
pvalue= 0.0019 ; R2= 0.461
Estimated allele frequency
Ecological gradient (Leaf Unfolding differences)
0.1
0.2
0.3
0.4
0.5
0.6
−1
0
1
2
3
4
L16
124
256
O16
253
218
O8
217
L8
233
219
9
97
204
O12
L12
L1
O1

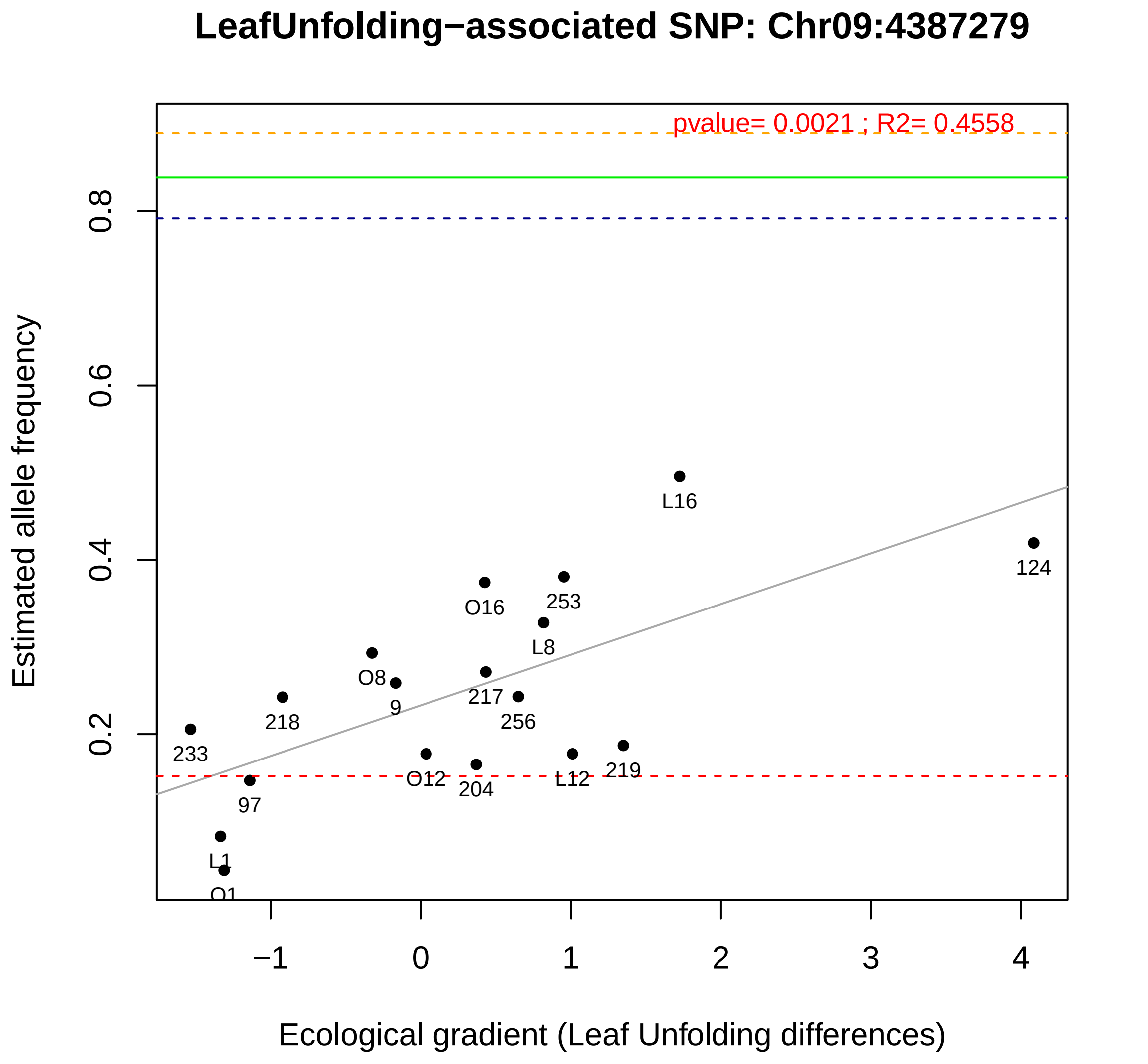

LeafUnfolding−associated SNP: Chr09:4387279
pvalue= 0.0021 ; R2= 0.4558
Estimated allele frequency
0.2
0.4
0.6
0.8
−1
0
1
2
3
4
Ecological gradient (Leaf Unfolding differences)
L16
124
O16
253
L8
O8
9
217
256
218
233
97
O12
204
L12
219
L1
O1

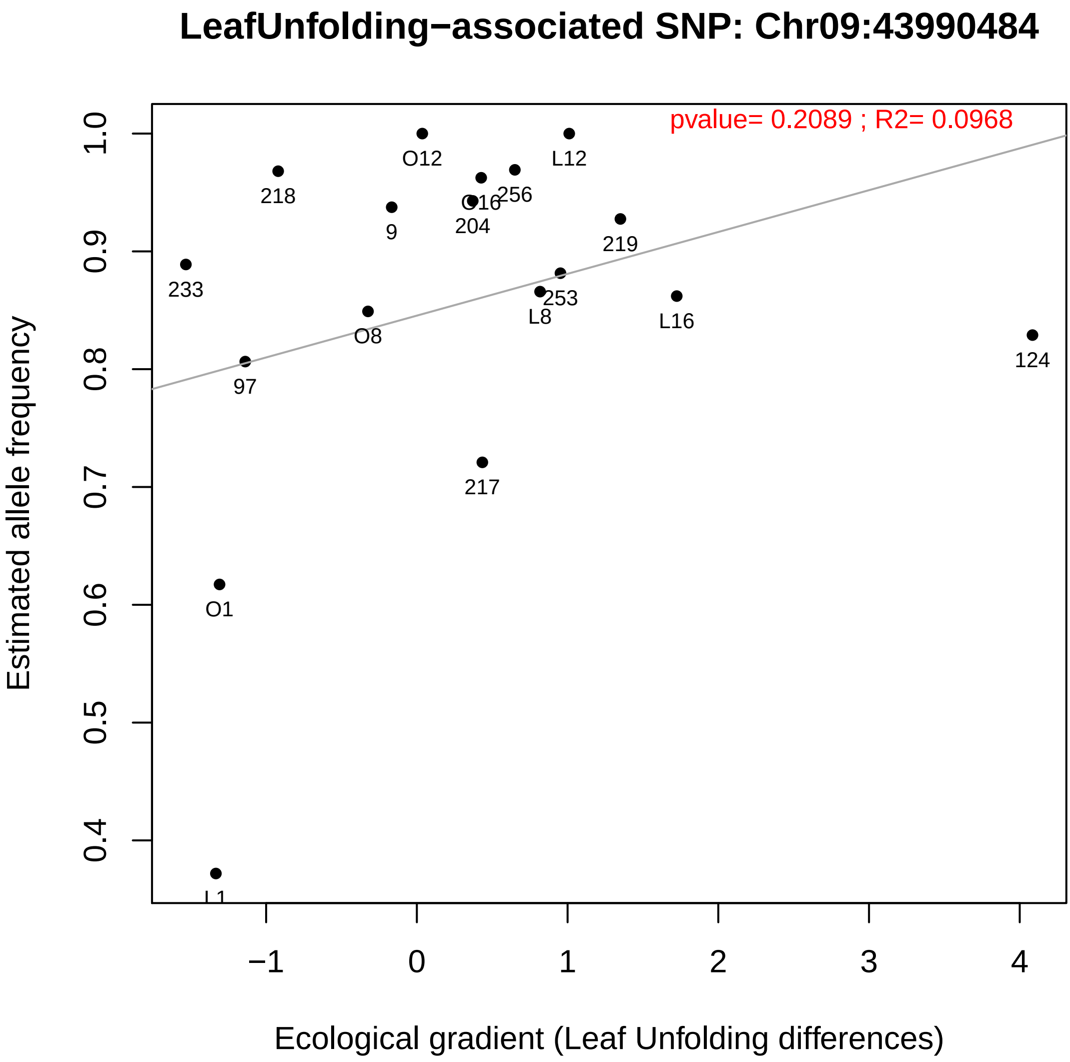
LeafUnfolding−associated SNP: Chr09:43990484
pvalue= 0.2089 ; R2= 0.0968
Estimated allele frequency
Ecological gradient (Leaf Unfolding differences)
0.4
0.5
0.6
0.7
0.8
0.9
1.0
−1
0
1
2
3
4
O12
L12
218
256
O16
9
204
219
233
253
L8
L16
O8
124
97
217
O1

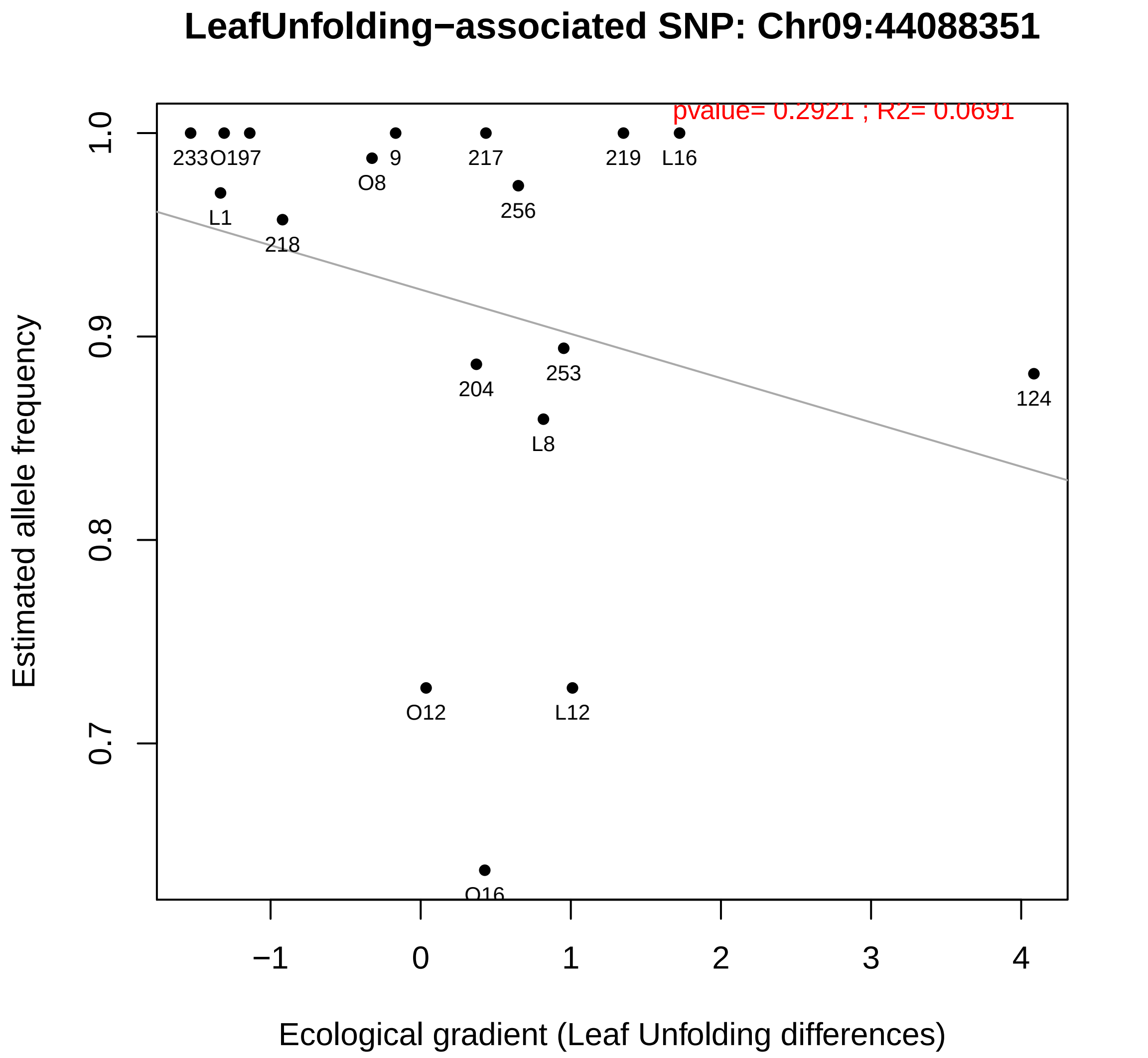
LeafUnfolding−associated SNP: Chr09:44088351
pvalue= 0.2921 ; R2= 0.0691
Estimated allele frequency
1.0
0.9
0.8
0.7
−1
0
1
2
3
4
Ecological gradient (Leaf Unfolding differences)
233
O197
L1
218
O8
9
217
256
219
L16
204
253
124
L8
O12
L12
O16

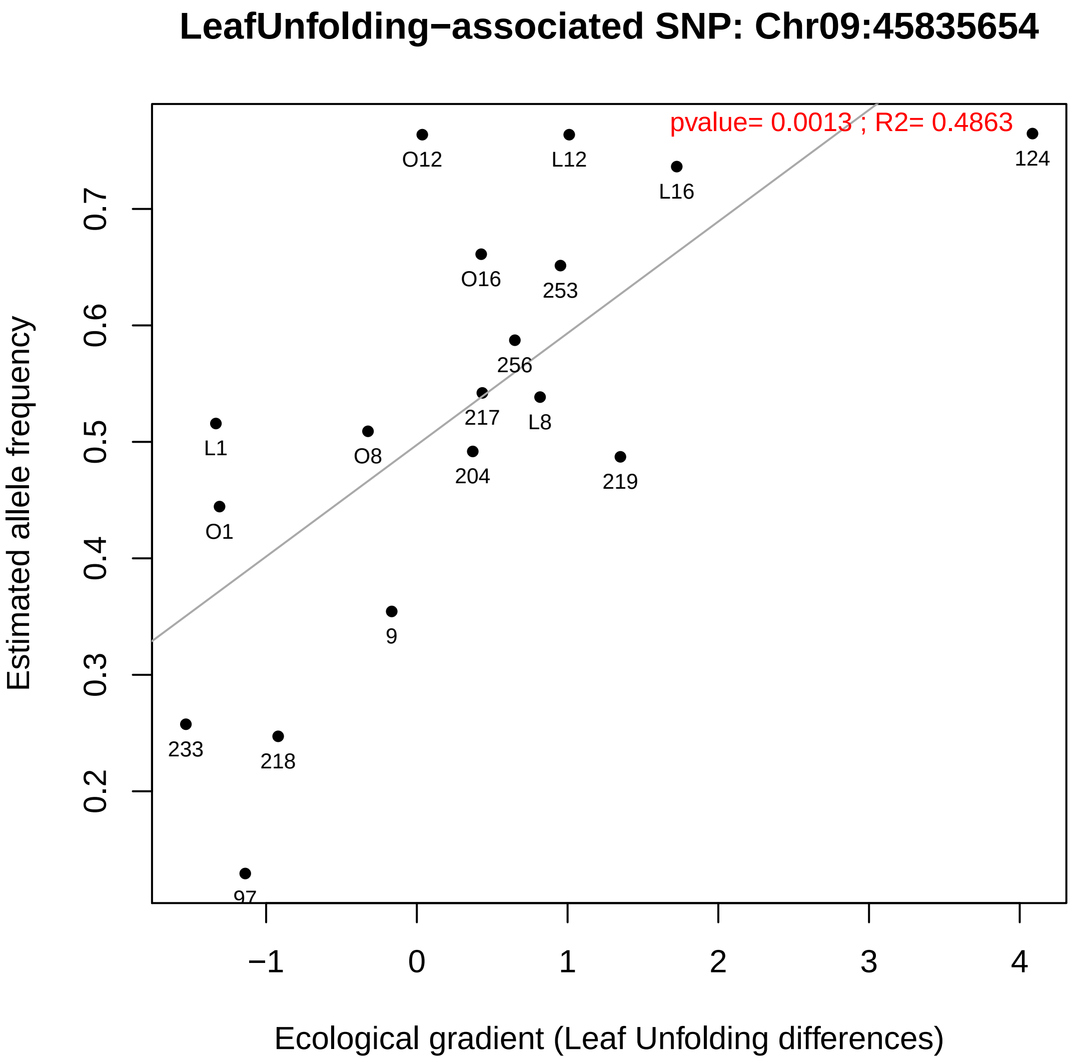
LeafUnfolding−associated SNP: Chr09:45835654
pvalue= 0.0013 ; R2= 0.4863
Estimated allele frequency
Ecological gradient (Leaf Unfolding differences)
0.2
0.3
0.4
0.5
0.6
0.7
−1
0
1
2
3
4
O12
L12
124
L16
O16
253
256
217
L8
L1
O8
204
219
O1
9
233
218
97

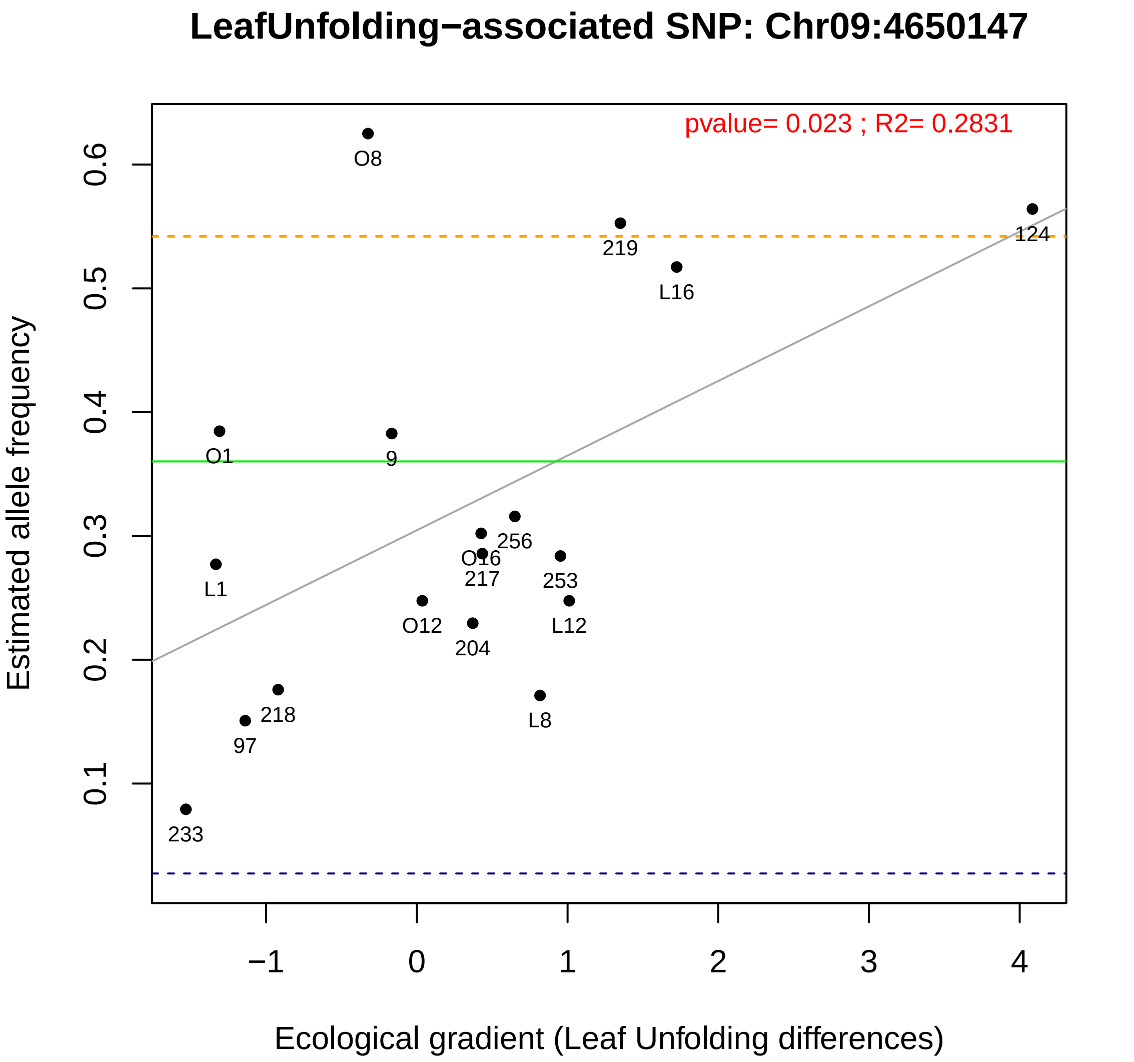
LeafUnfolding−associated SNP: Chr09:4650147
pvalue= 0.023 ; R2= 0.2831
Estimated allele frequency
Ecological gradient (Leaf Unfolding differences)
0.1
0.2
0.3
0.4
0.5
0.6
−1
0
1
2
3
4
O8
219
124
L16
O1
9
256
O16
217
253
L1
O12
L12
204
218
97
L8
233

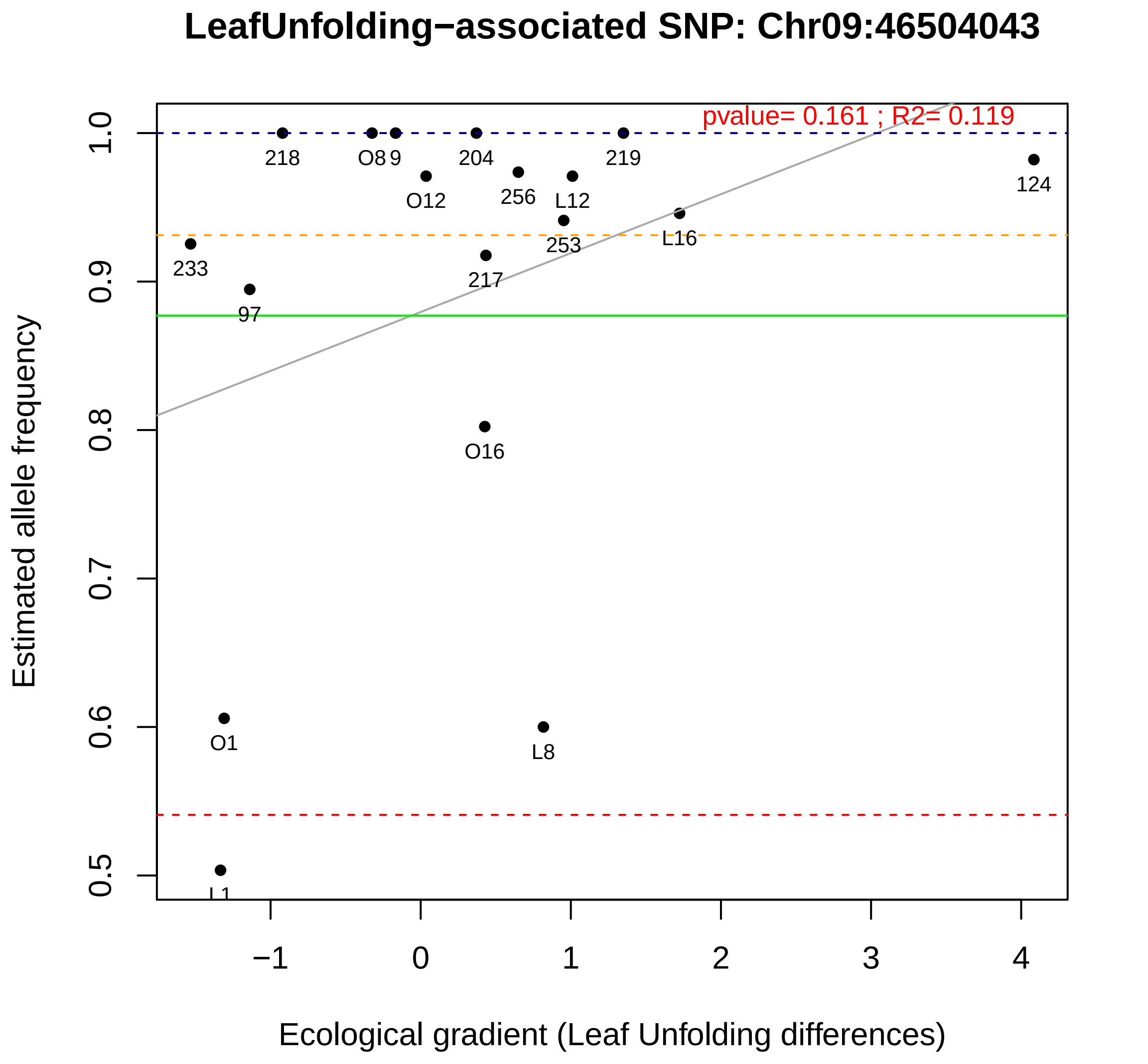

LeafUnfolding−associated SNP: Chr09:46504043
pvalue= 0.161 ; R2= 0.119
Estimated allele frequency
Ecological gradient (Leaf Unfolding differences)
1.0
0.9
0.8
0.7
0.6
0.5
−1
0
1
2
3
4
218
O8
9
204
219
124
O12
256
L12
253
L16
233
217
97
O16
O1
L8
L1

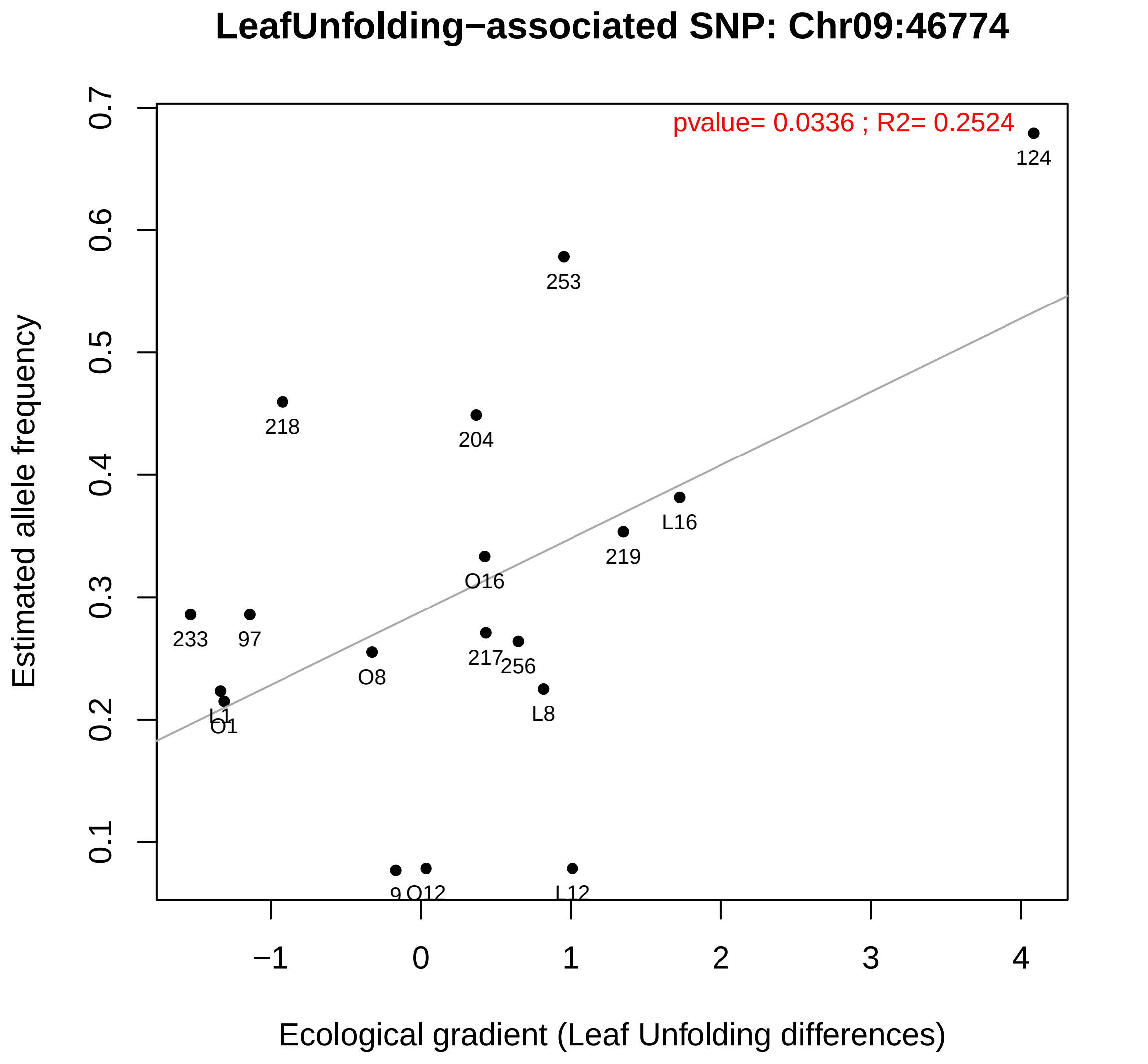
LeafUnfolding−associated SNP: Chr09:46774
pvalue= 0.0336 ; R2= 0.2524
Estimated allele frequency
Ecological gradient (Leaf Unfolding differences)
0.1
0.2
0.3
0.4
0.5
0.6
0.7
−1
0
1
2
3
4
124
253
218
204
L16
219
O16
233
97
O8
217
256
L8
L1
O1
9
O12
L12

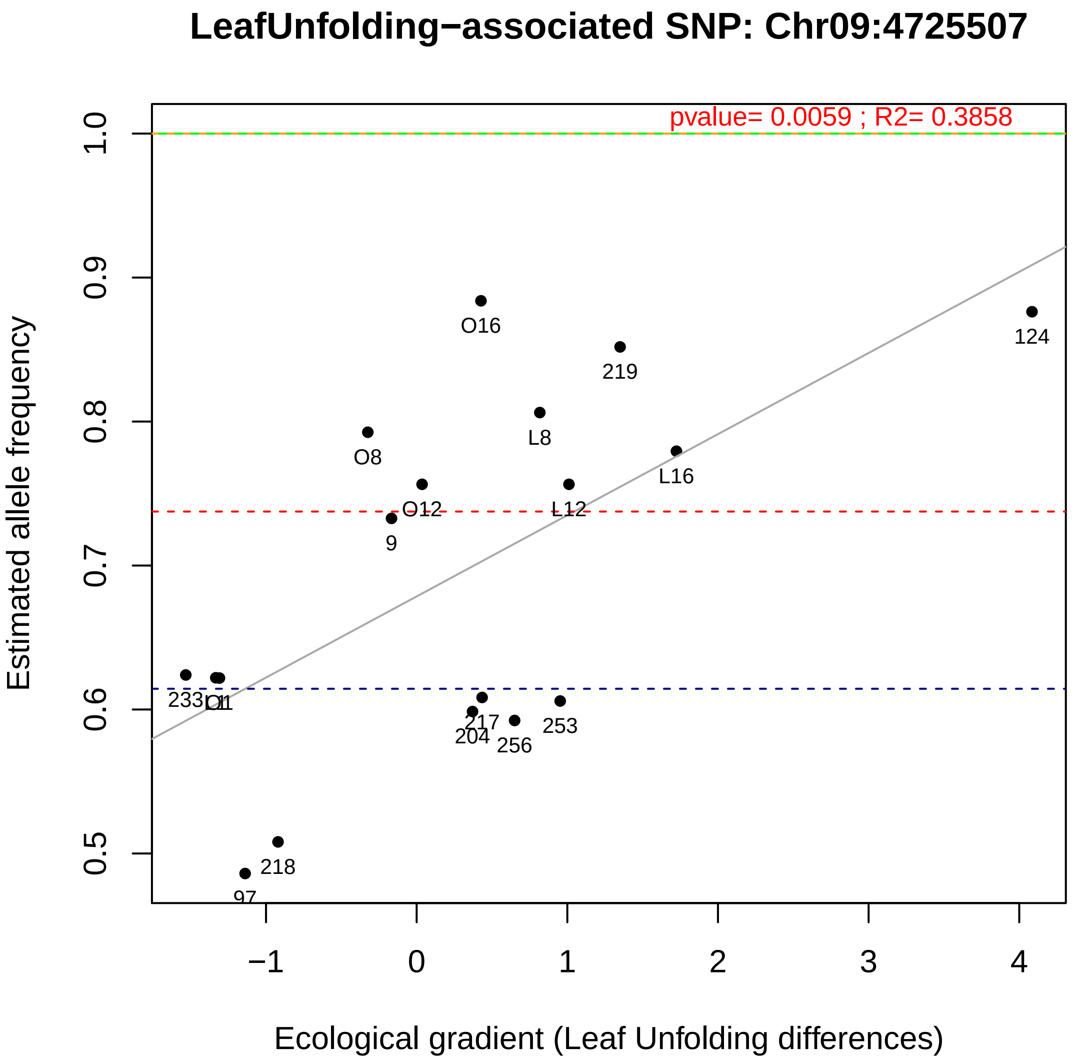

LeafUnfolding−associated SNP: Chr09:4725507
pvalue= 0.0059 ; R2= 0.3858
Estimated allele frequency
Ecological gradient (Leaf Unfolding differences)
0.5
0.6
0.7
0.8
0.9
1.0
−1
0
1
2
3
4
O16
124
219
L8
O8
L16
O12
L12
9
233
O1
217
204
256
253
218
97

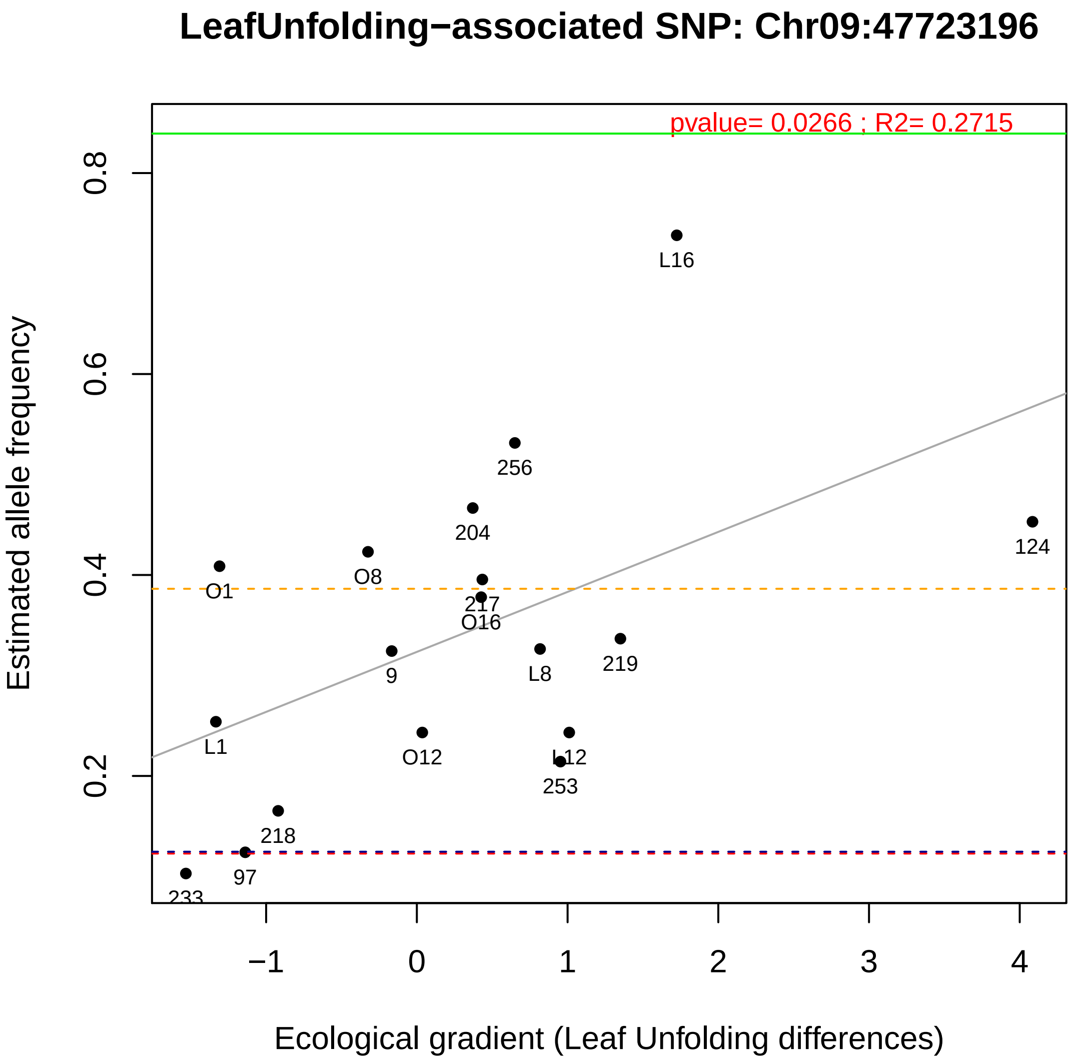
LeafUnfolding−associated SNP: Chr09:47723196
pvalue= 0.0266 ; R2= 0.2715
Estimated allele frequency
0.2
0.4
0.6
0.8
−1
0
1
2
3
4
Ecological gradient (Leaf Unfolding differences)
L16
256
204
124
O8
O1
217
O16
9
L8
219
L1
O12
L12
253
218
97
233

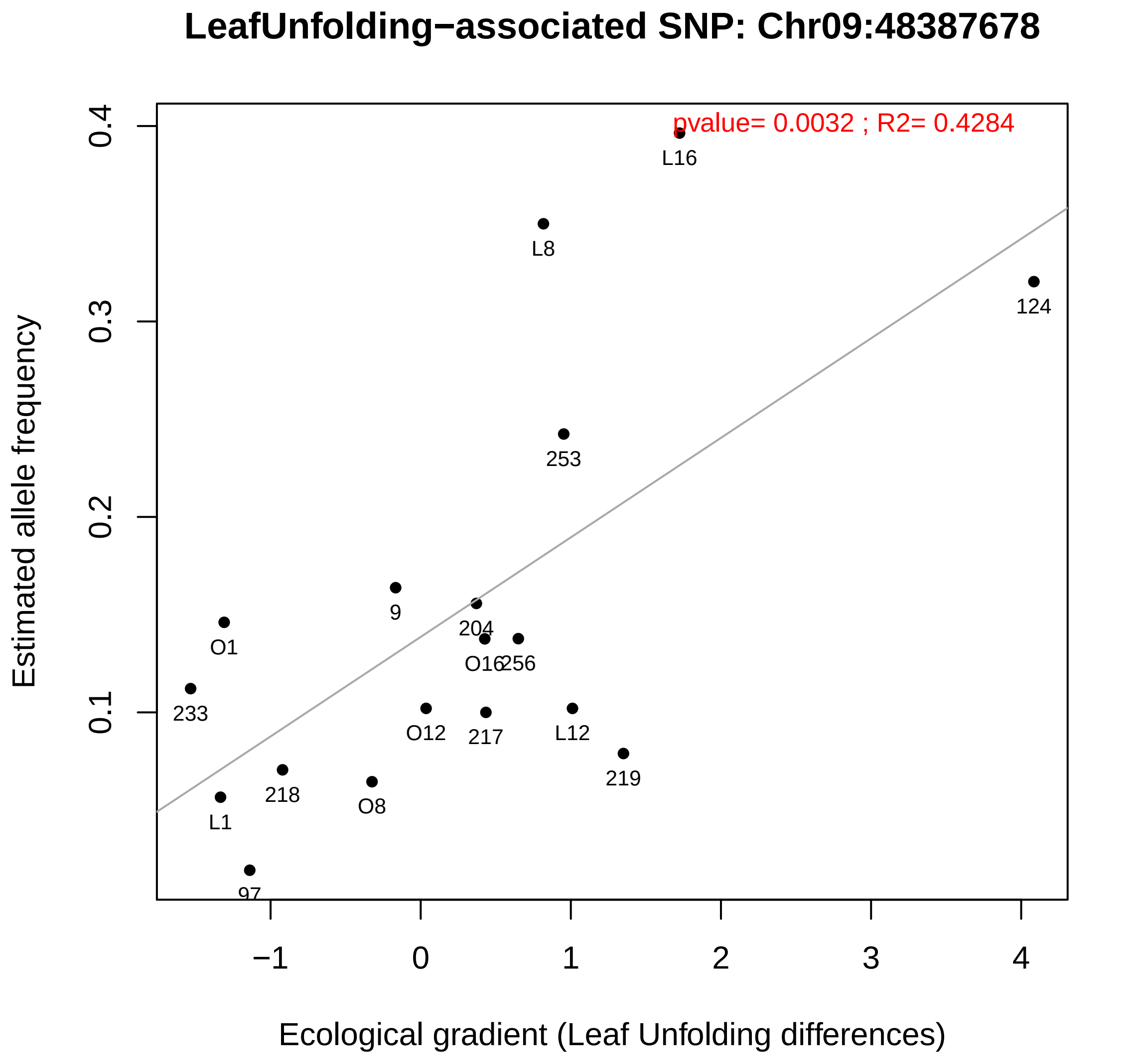

LeafUnfolding−associated SNP: Chr09:48387678
pvalue= 0.0032 ; R2= 0.4284
Estimated allele frequency
0.1
0.2
0.3
0.4
Ecological gradient (Leaf Unfolding differences)
−1
0
1
2
3
4
L16
L8
124
253
9
204
O1
O16
256
233
O12
217
L12
219
218
O8
L1
97

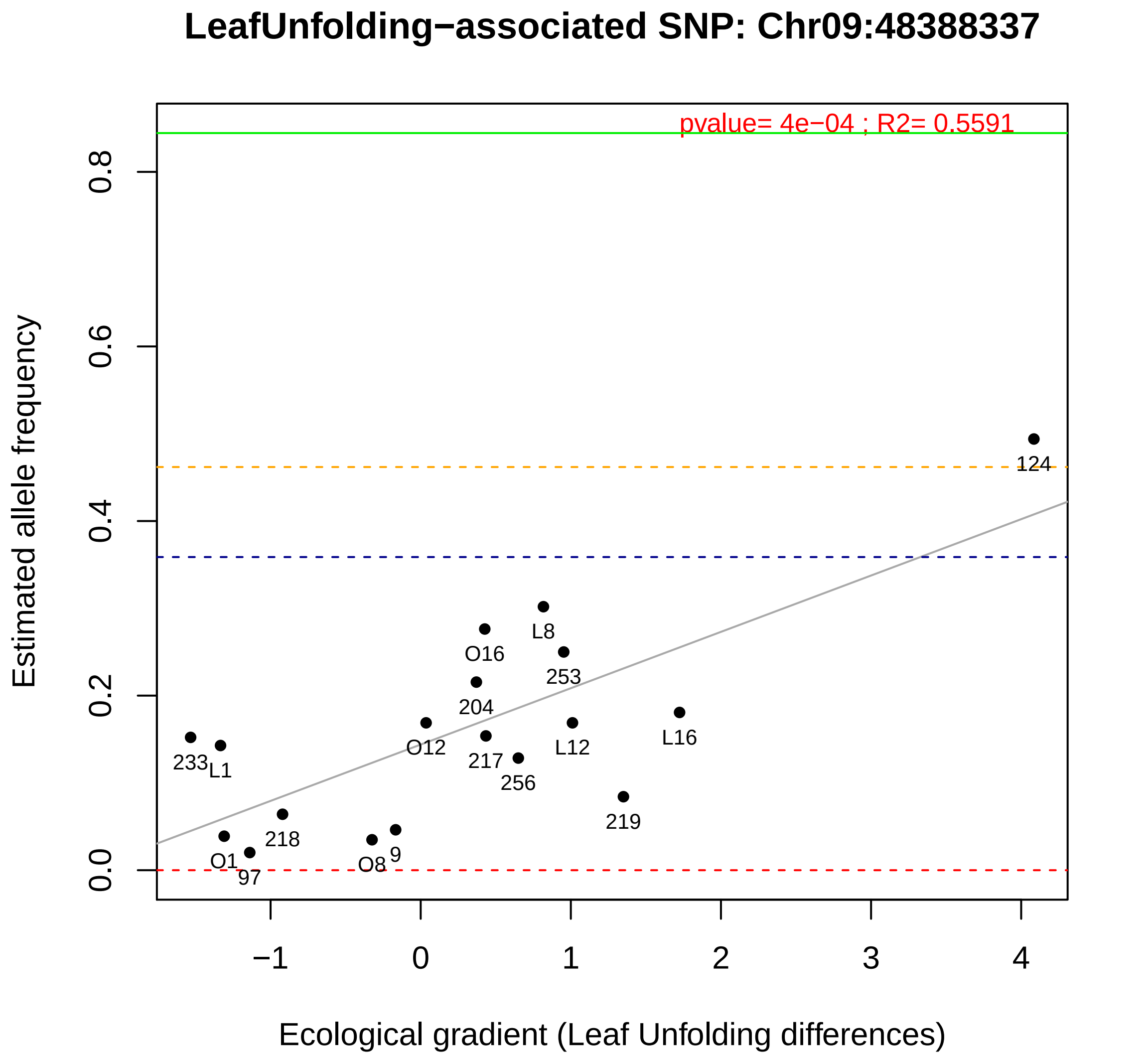

LeafUnfolding−associated SNP: Chr09:48388337
pvalue= 4e−04 ; R2= 0.5591
Estimated allele frequency
0.0
0.2
0.4
0.6
0.8
−1
0
1
2
3
4
Ecological gradient (Leaf Unfolding differences)
124
L8
O16
253
204
O12
217
L12
L16
256
233
L1
218
219
O1
97
O8
9

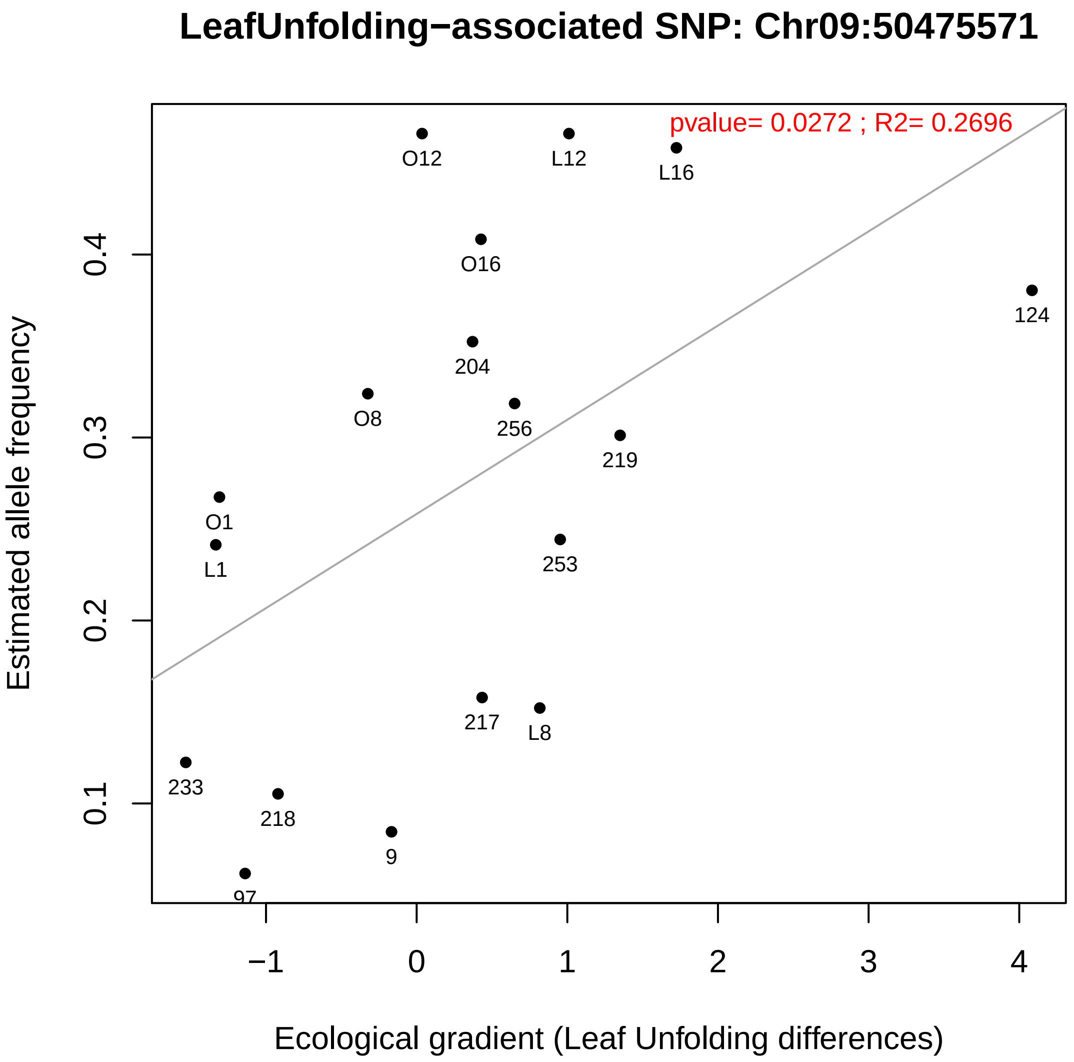
LeafUnfolding−associated SNP: Chr09:50475571
pvalue= 0.0272 ; R2= 0.2696
O12
L12
L16
O16
124
204
O8
256
219
O1
L1
253
217
L8
233
218
9
97
0.1
0.2
0.3
0.4
−1
0
1
2
3
4
Estimated allele frequency
Ecological gradient (Leaf Unfolding differences)

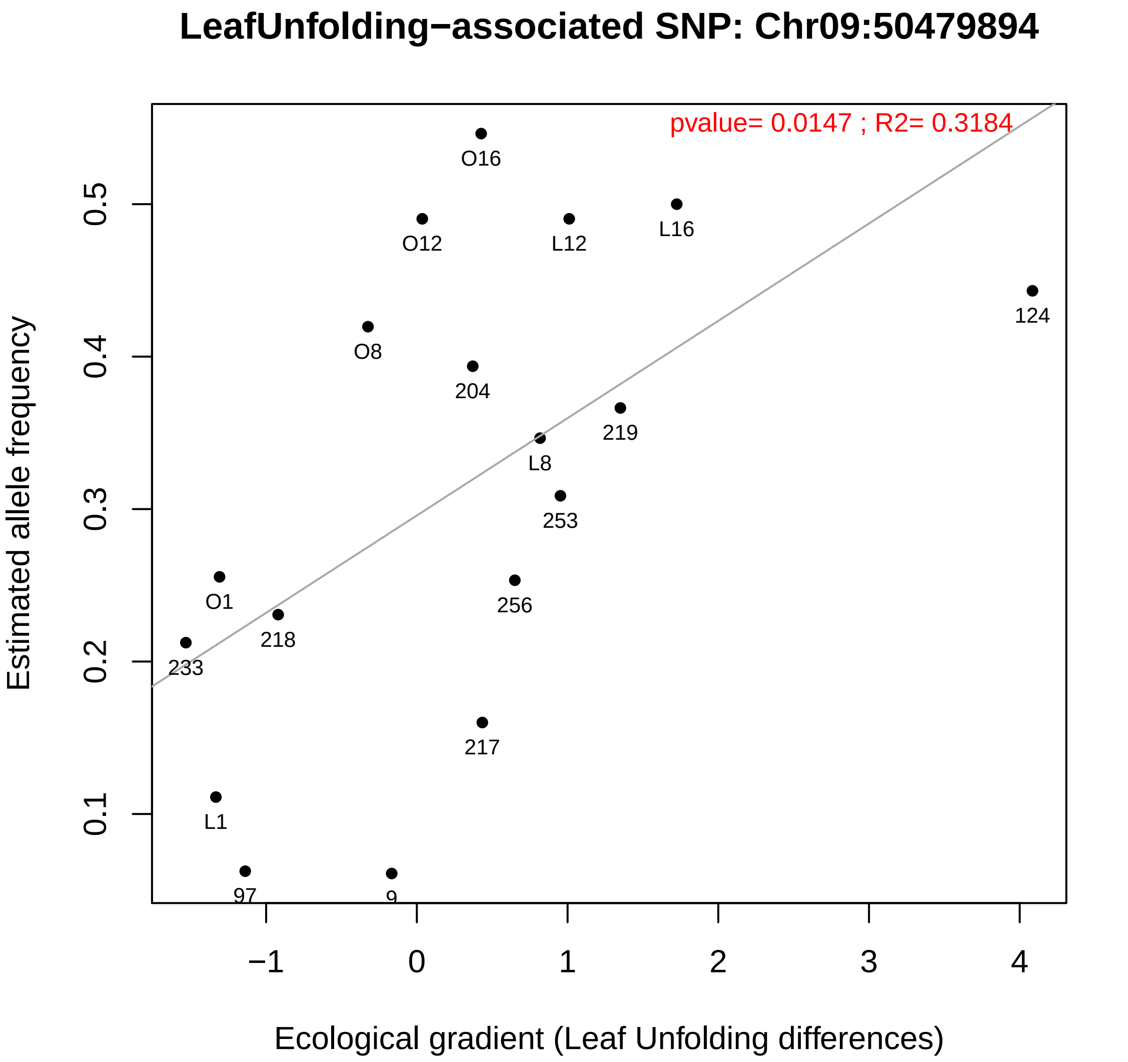

LeafUnfolding−associated SNP: Chr09:50479894
pvalue= 0.0147 ; R2= 0.3184
O16
O12
L12
L16
124
O8
204
219
L8
253
O1
256
218
233
217
L1
97
9
0.1
0.2
0.3
0.4
0.5
−1
0
1
2
3
4
Estimated allele frequency
Ecological gradient (Leaf Unfolding differences)

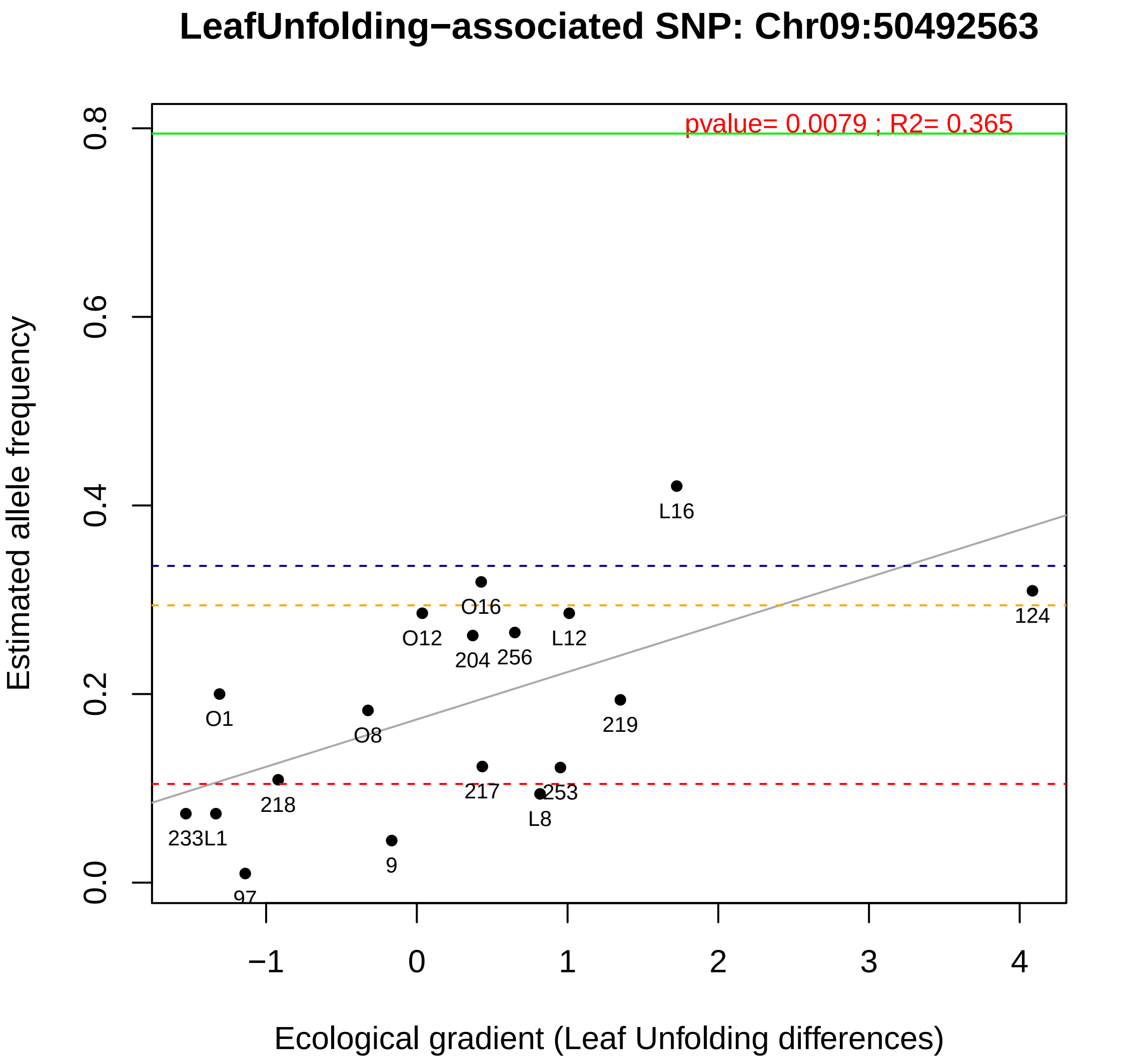

LeafUnfolding−associated SNP: Chr09:50492563
pvalue= 0.0079 ; R2= 0.365
Estimated allele frequency
0.0
0.2
0.4
0.6
0.8
−1
0
1
2
3
4
Ecological gradient (Leaf Unfolding differences)
L16
O16
124
O12
L12
204
256
O1
O8
219
218
217
253
L8
233
L1
9
97

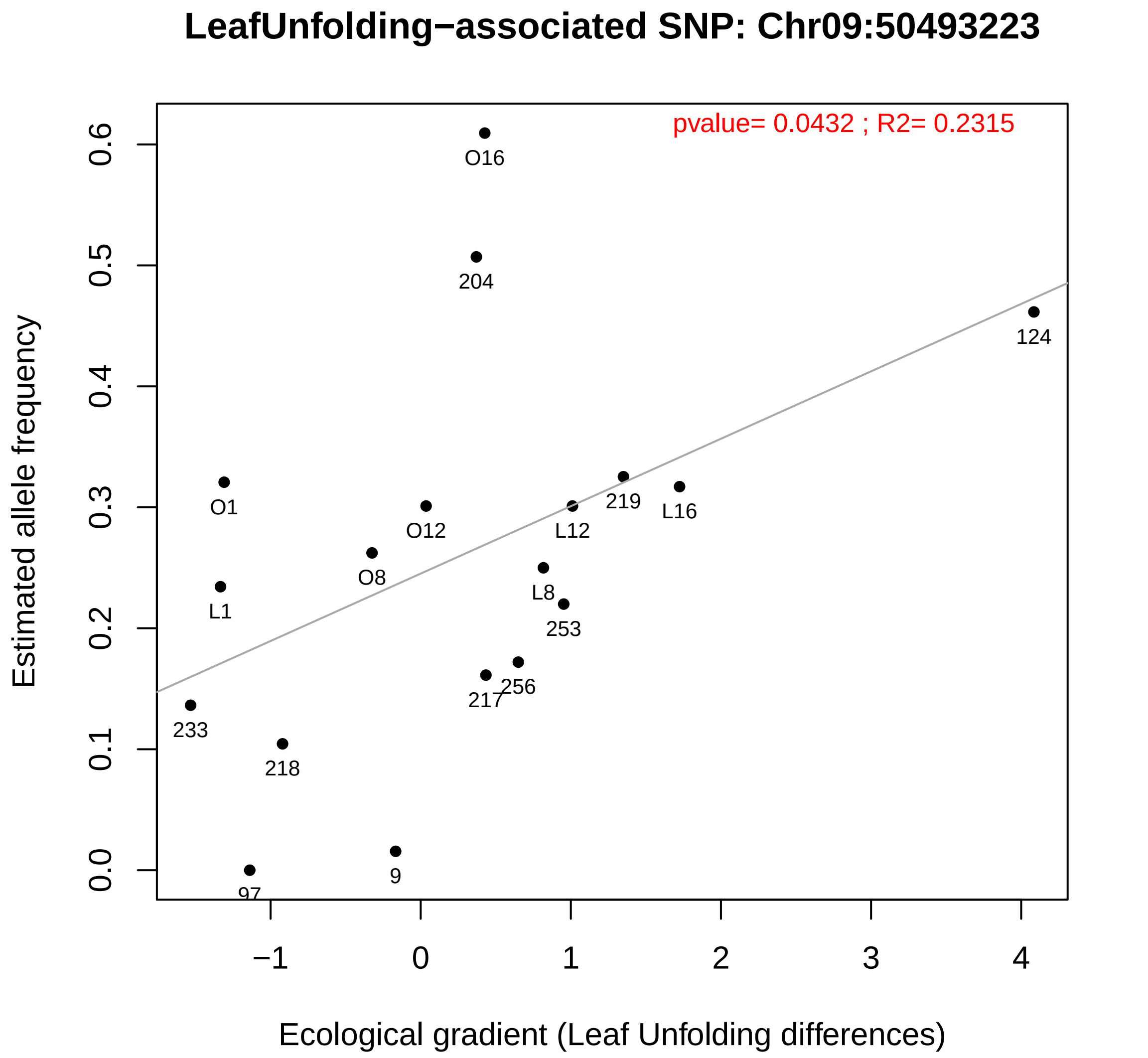
LeafUnfolding−associated SNP: Chr09:50493223
pvalue= 0.0432 ; R2= 0.2315
Estimated allele frequency
0.0
0.1
0.2
0.3
0.4
0.5
0.6
O16
204
124
219
L16
O1
O12
L12
O8
L8
L1
253
256
217
233
218
9
97
−1
0
1
2
3
4
Ecological gradient (Leaf Unfolding differences)

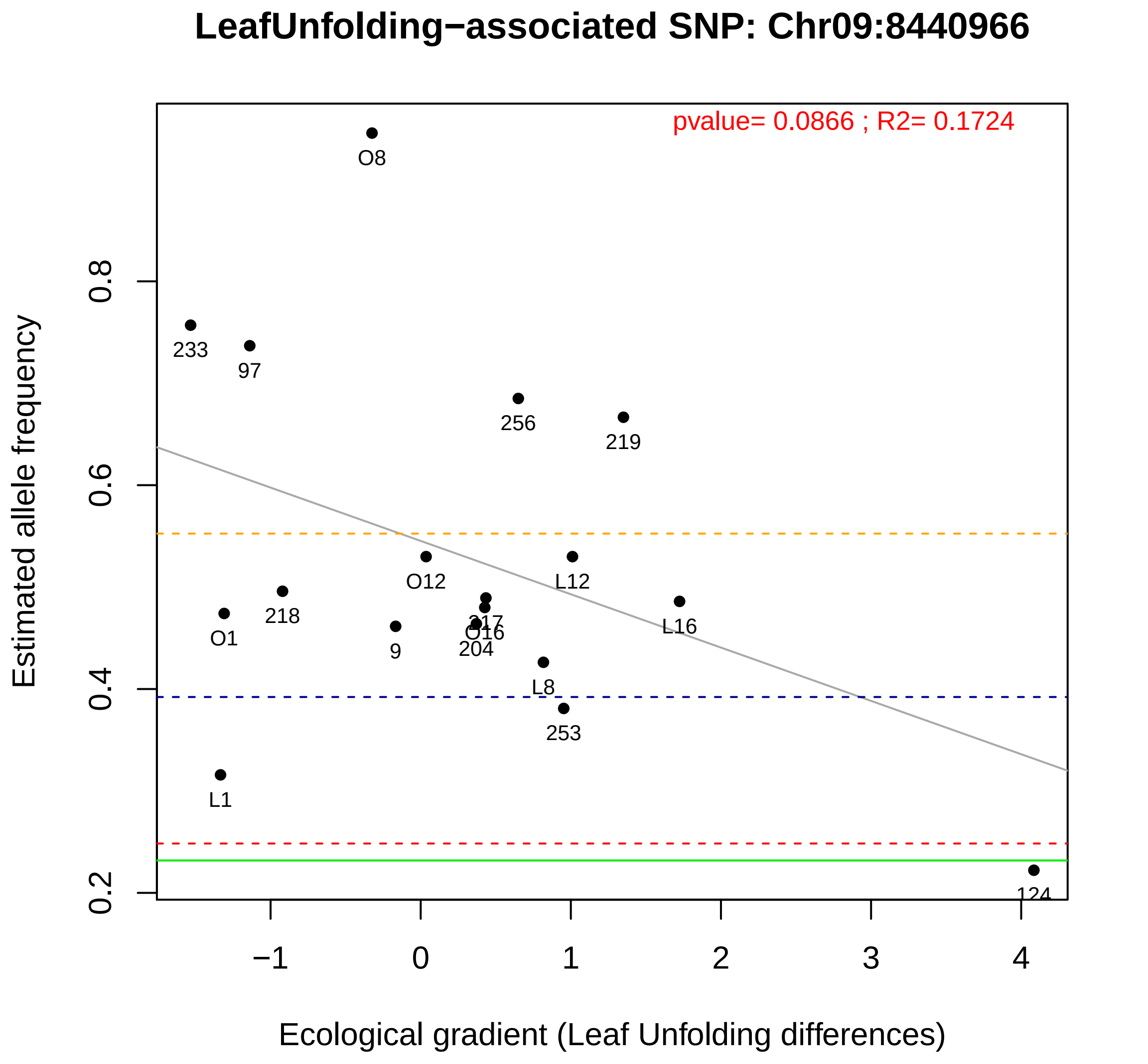

LeafUnfolding−associated SNP: Chr09:8440966
pvalue= 0.0866 ; R2= 0.1724
Estimated allele frequency
0.2
0.4
0.6
0.8
−1
0
1
2
3
4
Ecological gradient (Leaf Unfolding differences)
O8
233
97
256
219
O12
L12
218
O1
9
217
O16
204
L16
L8
253
L1
124

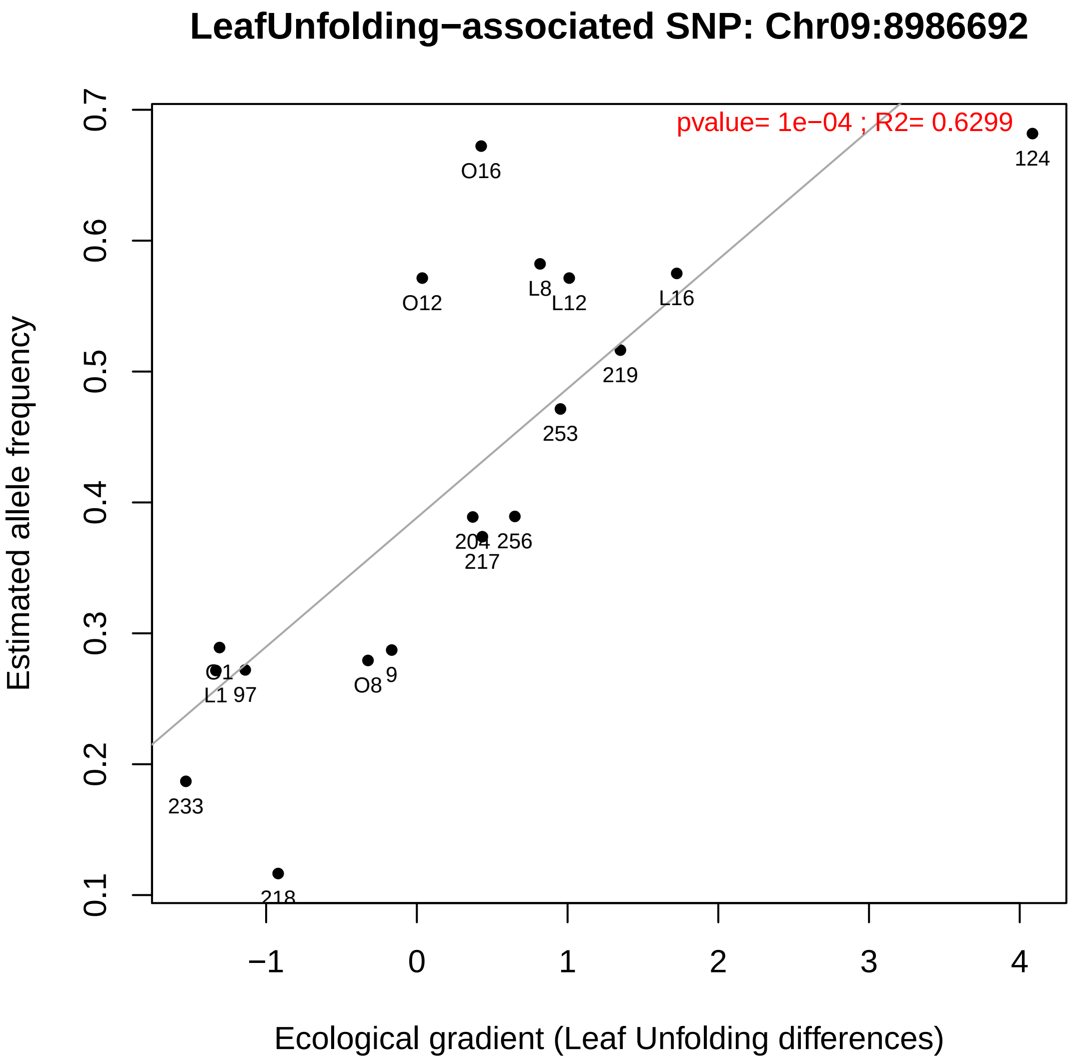
LeafUnfolding−associated SNP: Chr09:8986692
pvalue= 1e−04 ; R2= 0.6299
Estimated allele frequency
Ecological gradient (Leaf Unfolding differences)
0.1
0.2
0.3
0.4
0.5
0.6
0.7
−1
0
1
2
3
4
124
O16
O12
L8
L12
L16
219
253
204
217
256
O1
L1
97
O8
9
233
218

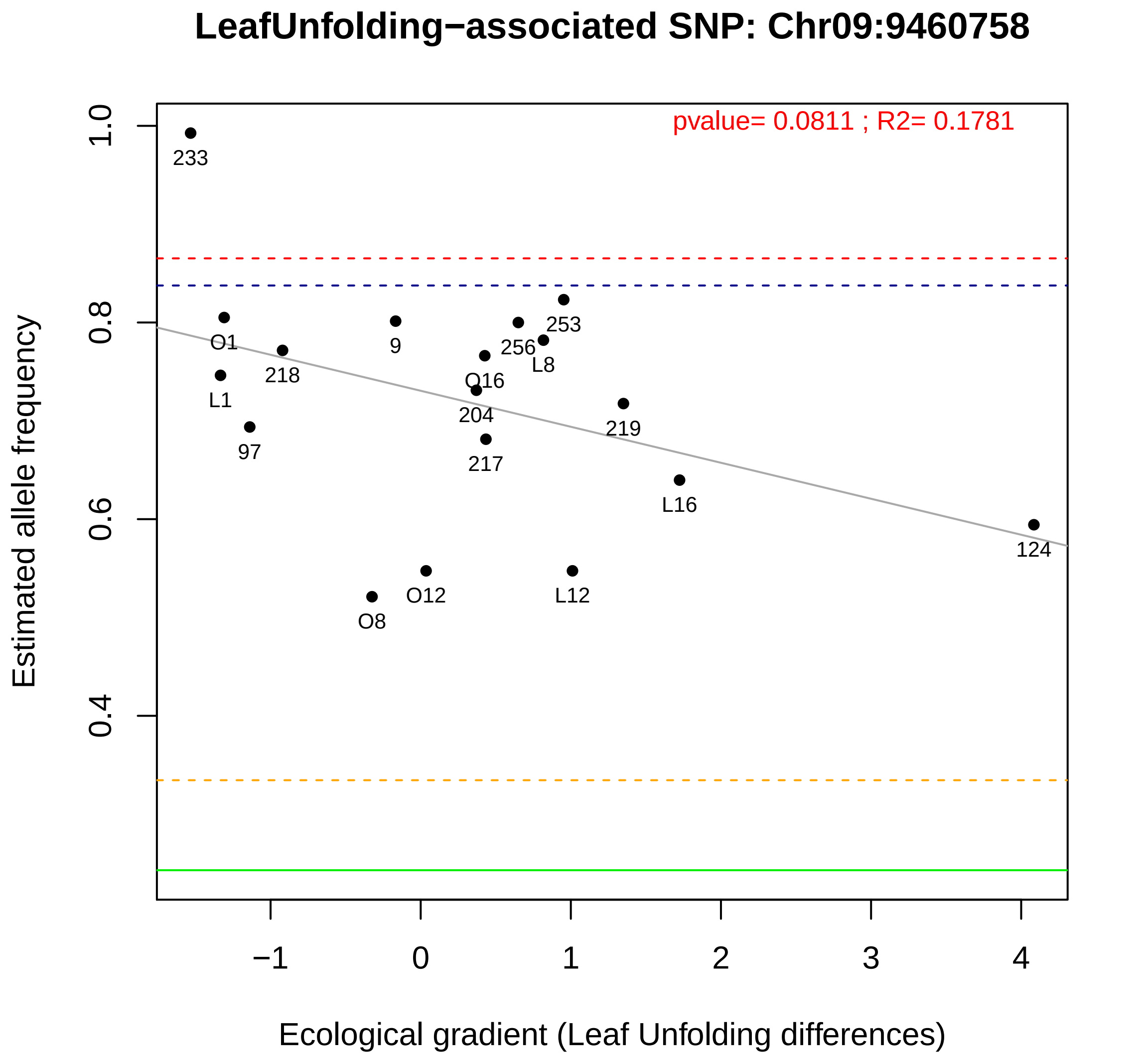
LeafUnfolding−associated SNP: Chr09:9460758
pvalue= 0.0811 ; R2= 0.1781
Estimated allele frequency
Ecological gradient (Leaf Unfolding differences)
0.4
0.6
0.8
1.0
−1
0
1
2
3
4
233
O1
L1
97
218
9
O16
204
217
256
L8
253
219
L16
124
O8
O12
L12

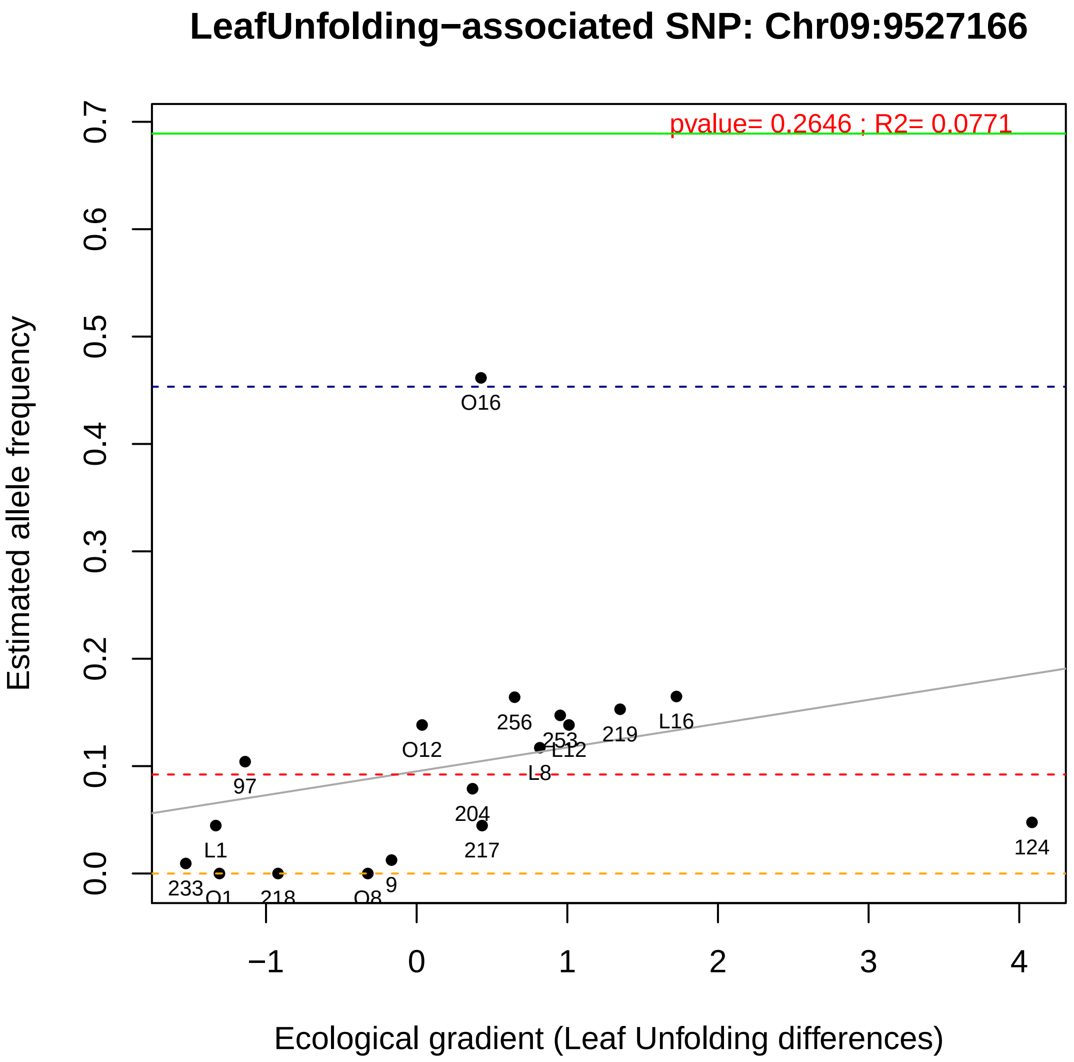
LeafUnfolding−associated SNP: Chr09:9527166
pvalue= 0.2646 ; R2= 0.0771
Estimated allele frequency
Ecological gradient (Leaf Unfolding differences)
0.0
0.1
0.2
0.3
0.4
0.5
0.6
0.7
−1
0
1
2
3
4
O16
256
253
L12
219
L16
O12
L8
97
204
217
L1
124
233
O1
218
O8
9

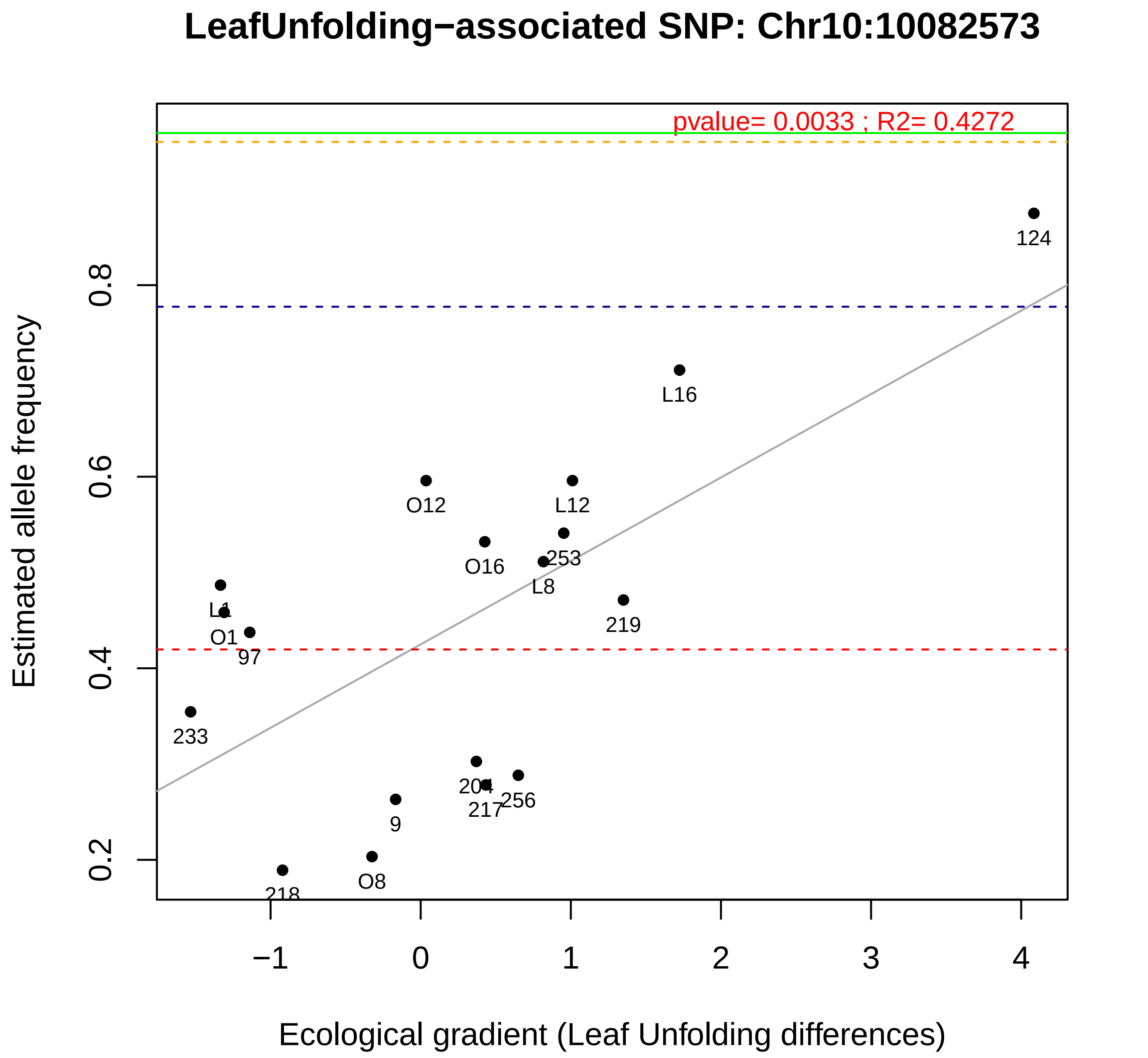

LeafUnfolding−associated SNP: Chr10:10082573
pvalue= 0.0033 ; R2= 0.4272
Estimated allele frequency
Ecological gradient (Leaf Unfolding differences)
0.2
0.4
0.6
0.8
−1
0
1
2
3
4
124
L16
O12
L12
253
O16
L8
219
L1
O1
97
233
204
217
256
9
O8
218

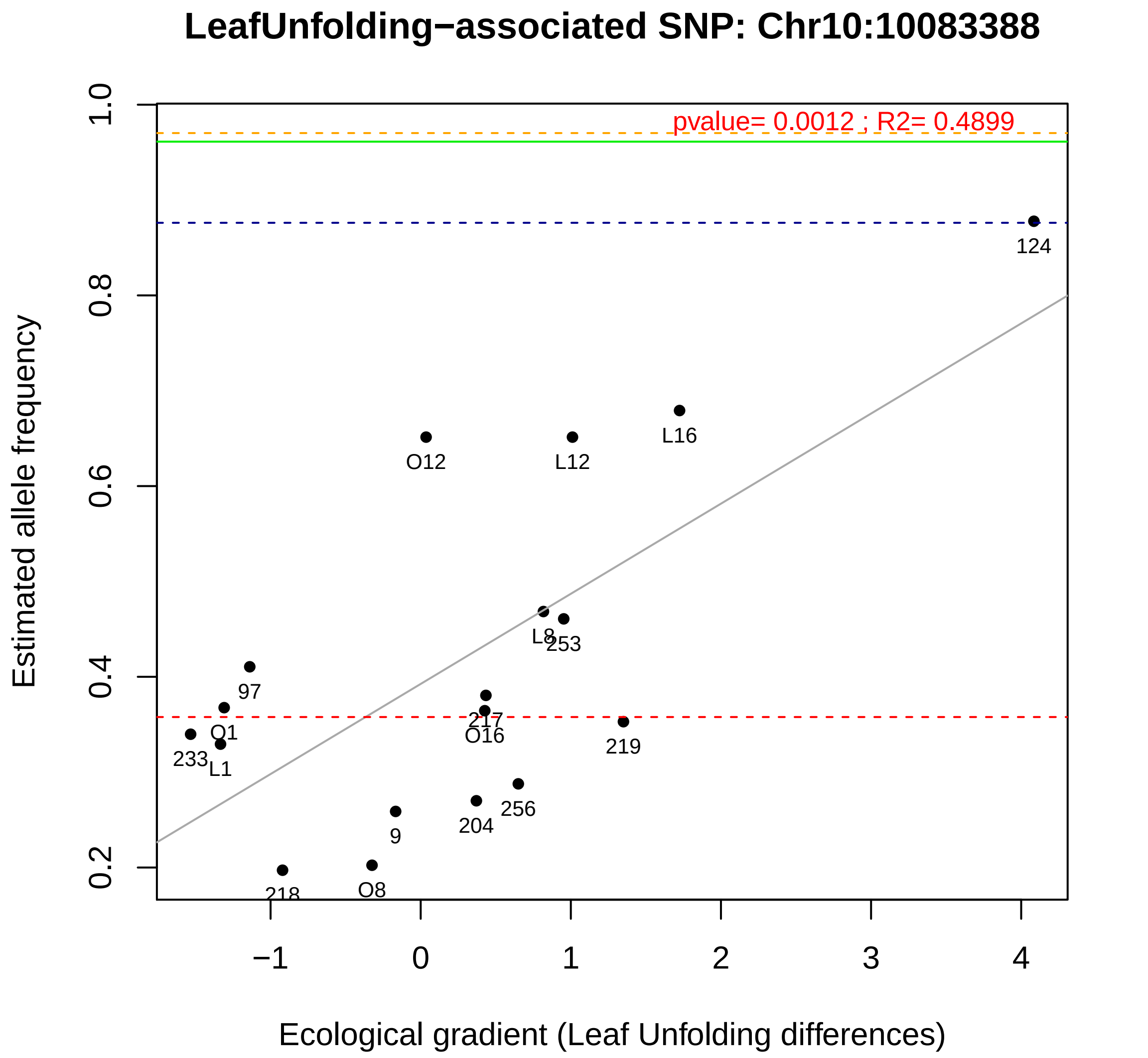
LeafUnfolding−associated SNP: Chr10:10083388
pvalue= 0.0012 ; R2= 0.4899
Estimated allele frequency
0.2
0.4
0.6
0.8
1.0
−1
0
1
2
3
4
Ecological gradient (Leaf Unfolding differences)
124
L16
O12
L12
L8
253
97
217
O16
219
Q1
233
L1
256
204
9
218
O8

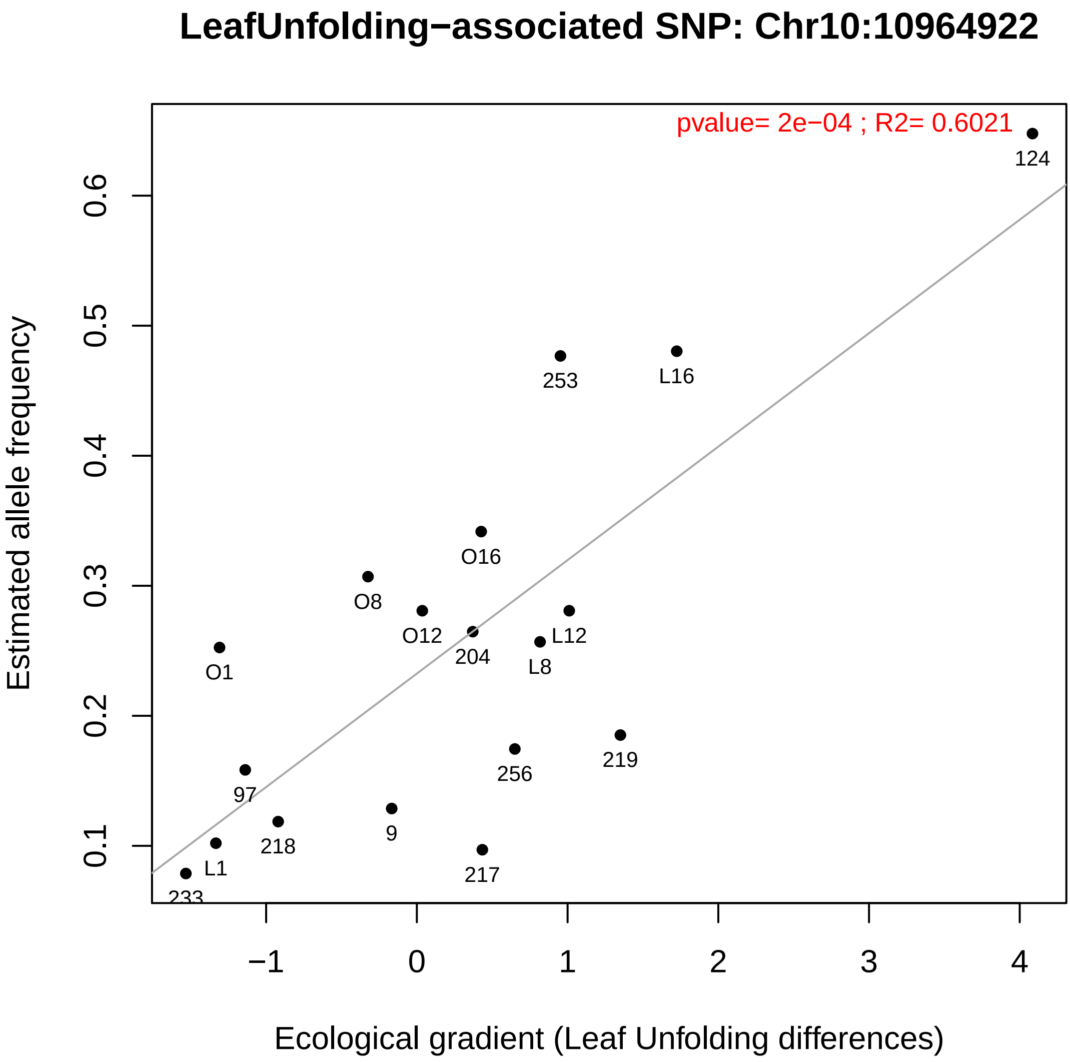

LeafUnfolding−associated SNP: Chr10:10964922
pvalue= 2e−04 ; R2= 0.6021
Estimated allele frequency
Ecological gradient (Leaf Unfolding differences)
0.1
0.2
0.3
0.4
0.5
0.6
−1
0
1
2
3
4
124
253
L16
O16
O8
O12
204
L12
L8
O1
256
219
97
218
9
217
L1
233

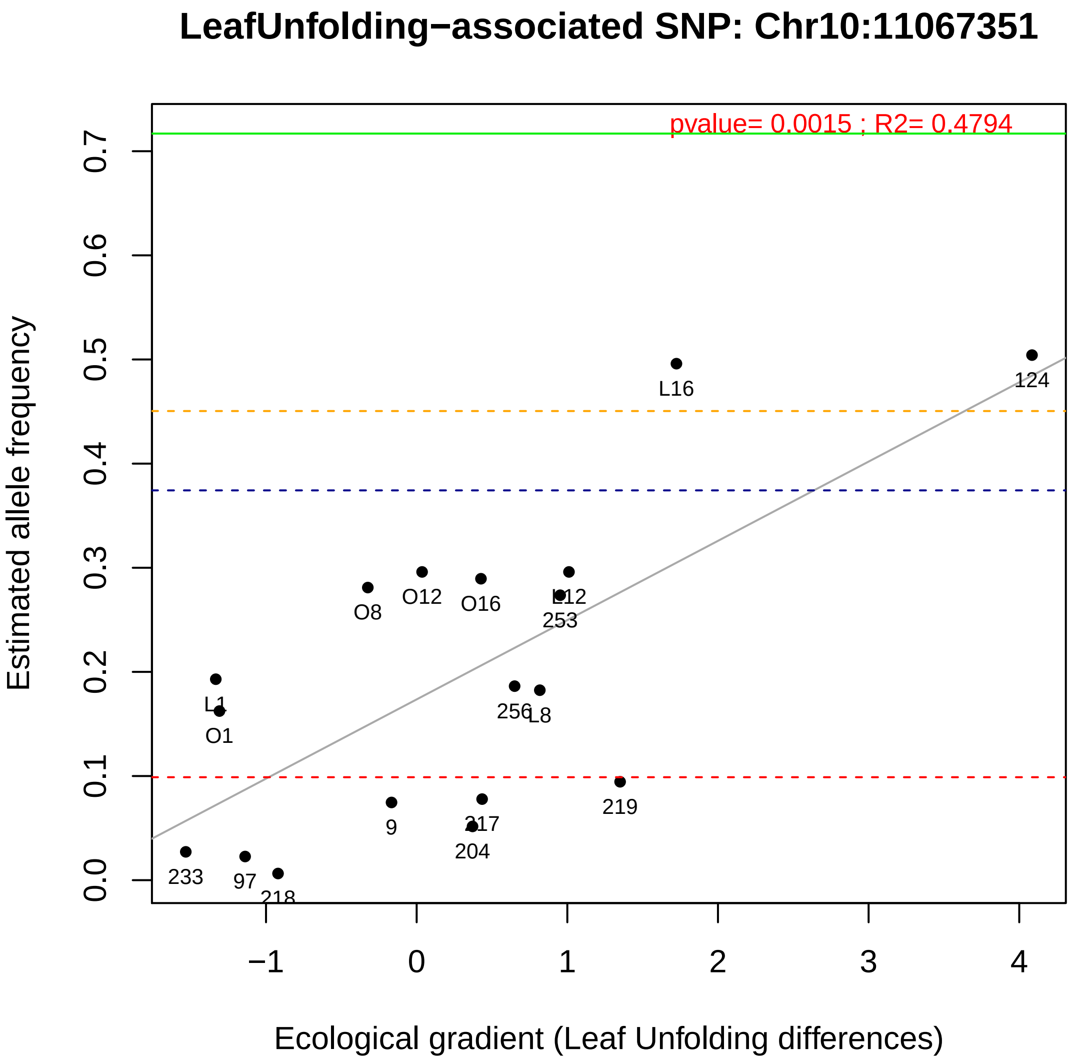
LeafUnfolding−associated SNP: Chr10:11067351
pvalue= 0.0015 ; R2= 0.4794
Estimated allele frequency
Ecological gradient (Leaf Unfolding differences)
0.0
0.1
0.2
0.3
0.4
0.5
0.6
0.7
−1
0
1
2
3
4
L16
124
O8
O12
O16
L12
253
L1
O1
256
L8
219
9
217
204
233
97
218

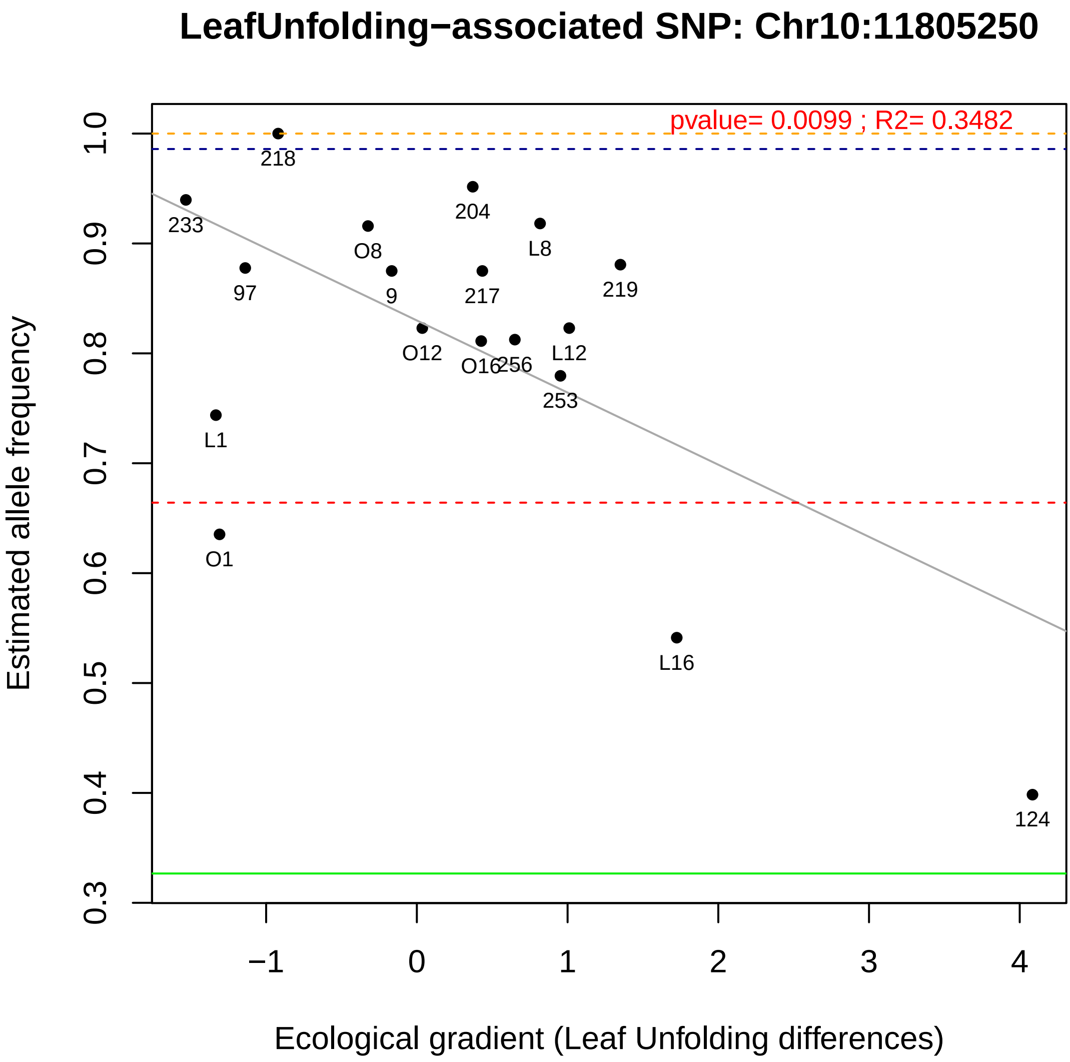
LeafUnfolding−associated SNP: Chr10:11805250
pvalue= 0.0099 ; R2= 0.3482
Estimated allele frequency
Ecological gradient (Leaf Unfolding differences)
1.0
0.9
0.8
0.7
0.6
0.5
0.4
0.3
−1
0
1
2
3
4
218
233
204
O8
L8
97
9
217
219
O12
L12
O16
256
253
L1
O1
L16
124

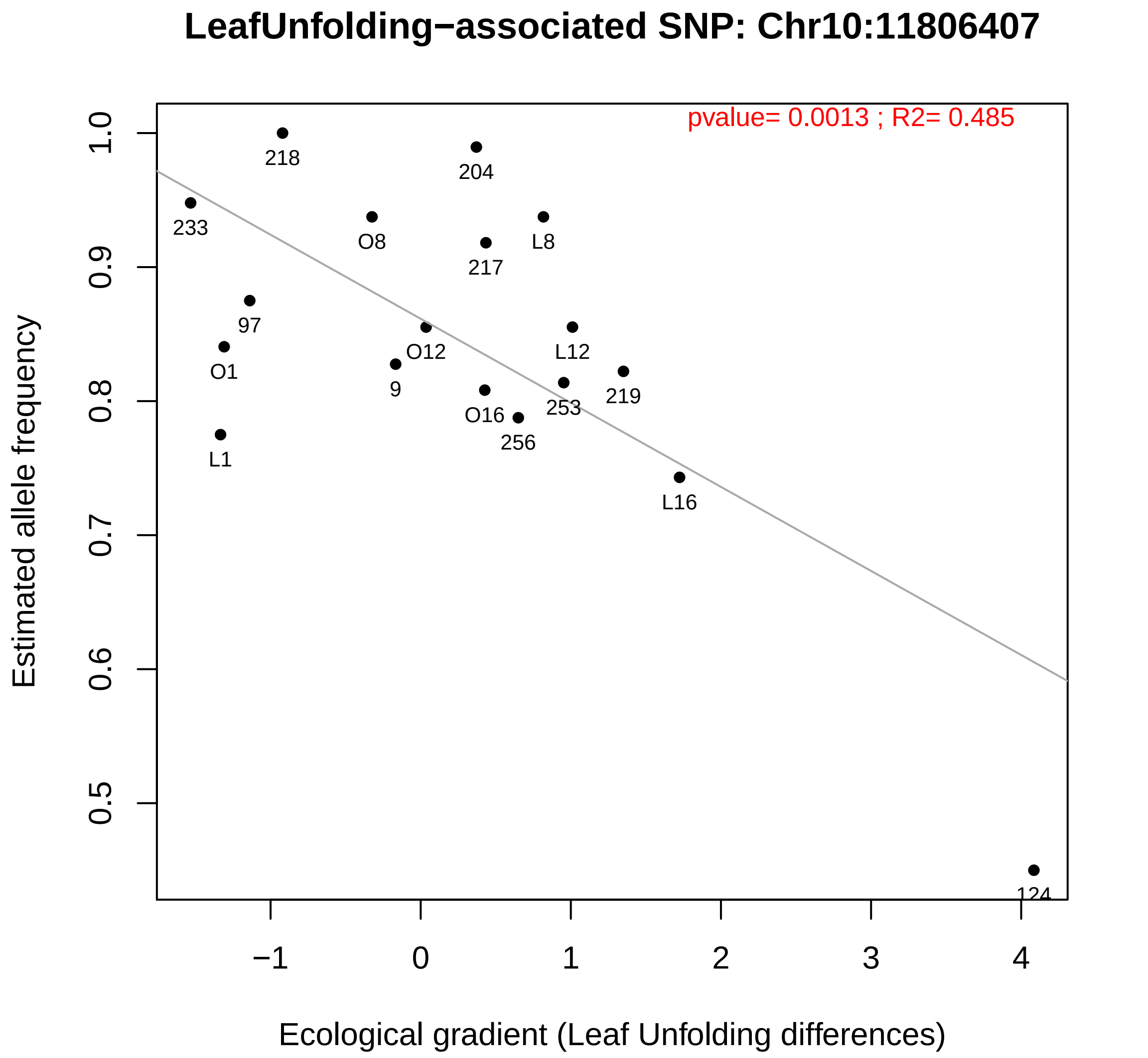

LeafUnfolding−associated SNP: Chr10:11806407
pvalue= 0.0013 ; R2= 0.485
Estimated allele frequency
1.0
0.9
0.8
0.7
0.6
0.5
218
204
233
O8
L8
217
97
O12
L12
O1
9
219
253
O16
256
L1
L16
124
−1
0
1
2
3
4
Ecological gradient (Leaf Unfolding differences)

**LeafUnfolding−associated SNP: Chr10:13071652**

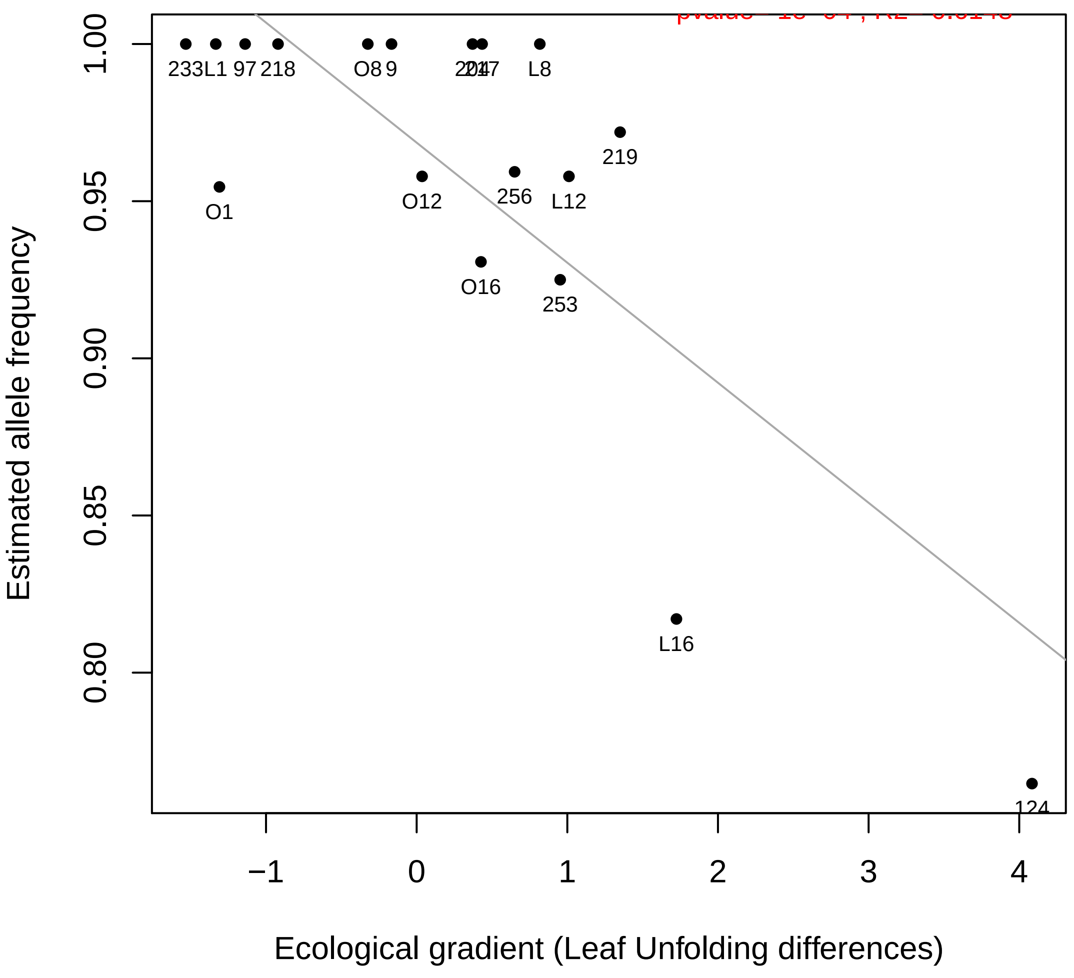

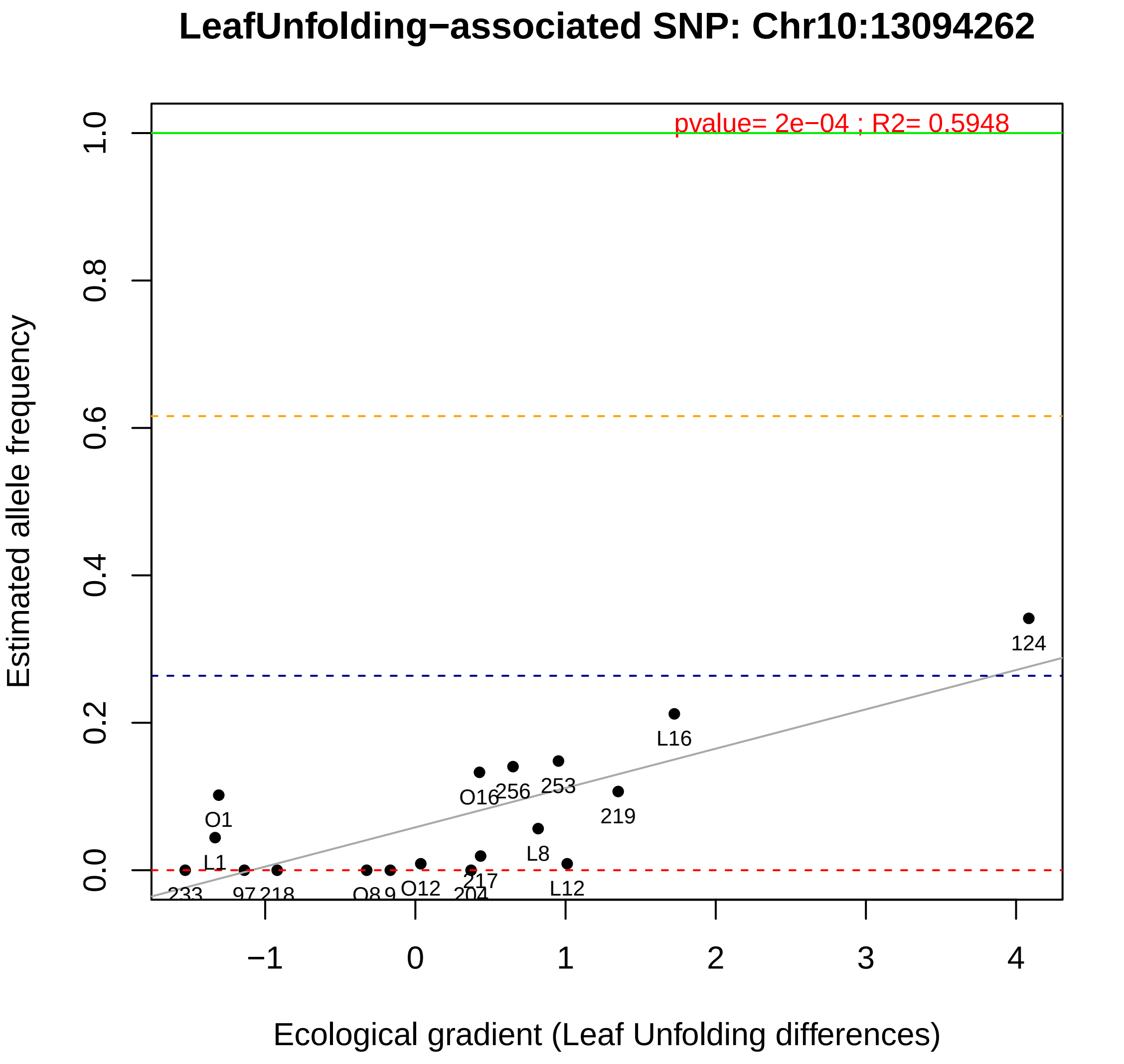

LeafUnfolding−associated SNP: Chr10:13094262
pvalue= 2e−04 ; R2= 0.5948
Estimated allele frequency
0.0
0.2
0.4
0.6
0.8
1.0
Ecological gradient (Leaf Unfolding differences)
−1
0
1
2
3
4
124
L16
O16
256
253
219
O1
L1
L8
L12
217
O12
204
233
97
218
O8
9

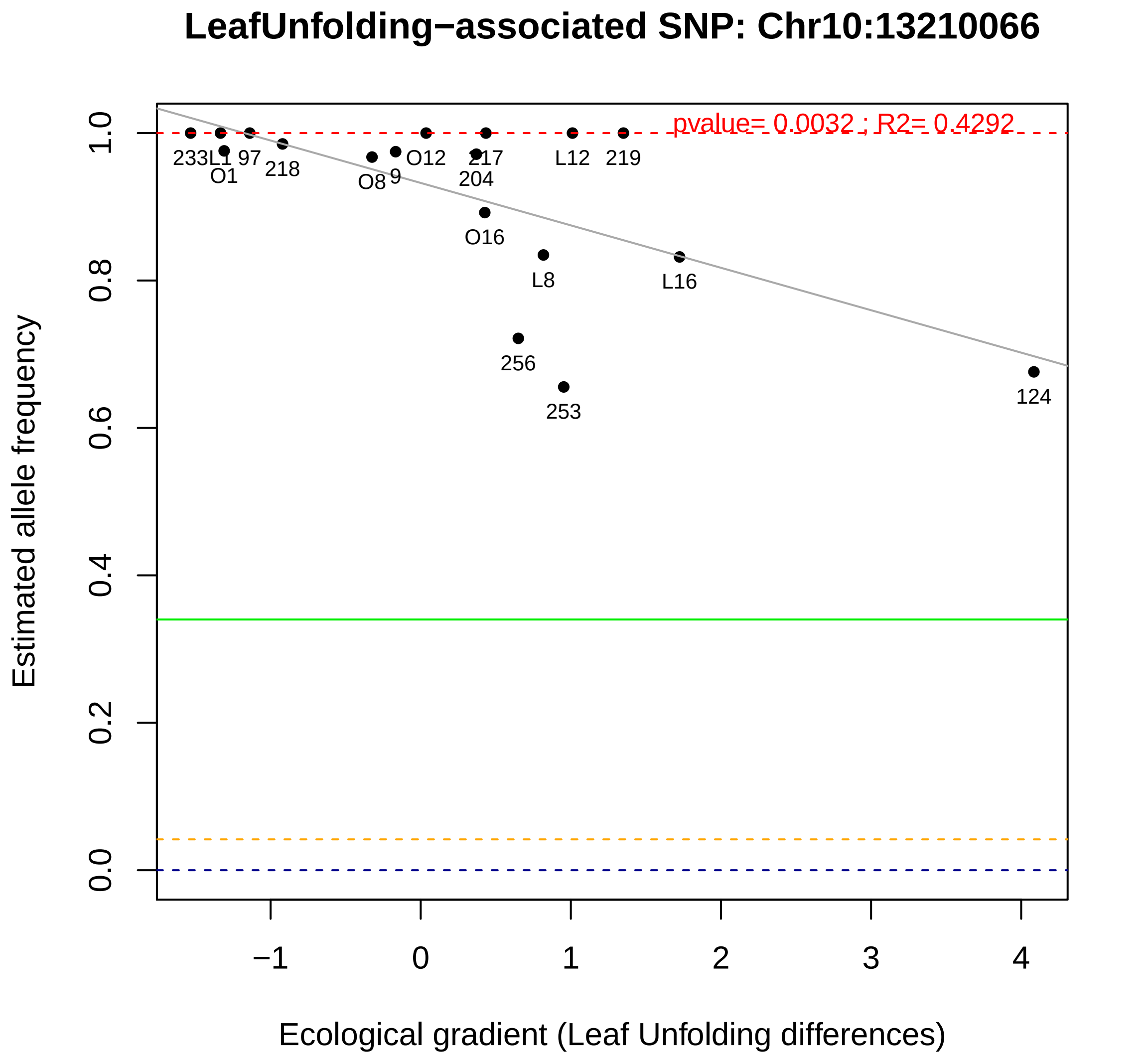

LeafUnfolding−associated SNP: Chr10:13210066
pvalue= 0.0032 ; R2= 0.4292
233
L1
97
O1
218
O8
9
O12
217
204
L12
219
O16
L8
L16
256
253
124
1.0
0.8
0.6
0.4
0.2
0.0
−1
0
1
2
3
4
Estimated allele frequency
Ecological gradient (Leaf Unfolding differences)

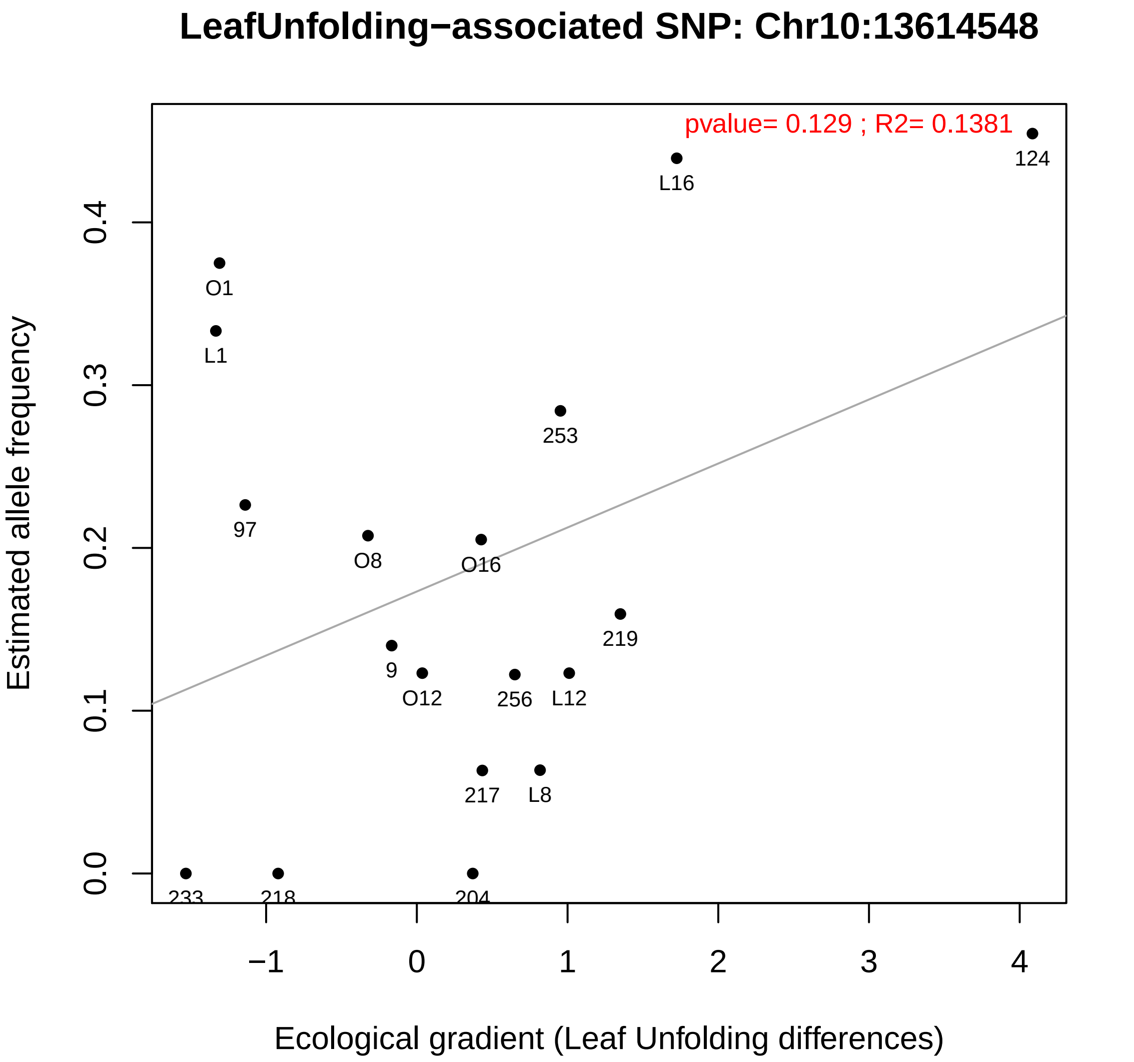
LeafUnfolding−associated SNP: Chr10:13614548
pvalue= 0.129 ; R2= 0.1381
124
L16
O1
L1
253
97
O8
O16
219
9
O12
256
L12
217
L8
233
218
204
0.0
0.1
0.2
0.3
0.4
Estimated allele frequency
−1
0
1
2
3
4
Ecological gradient (Leaf Unfolding differences)

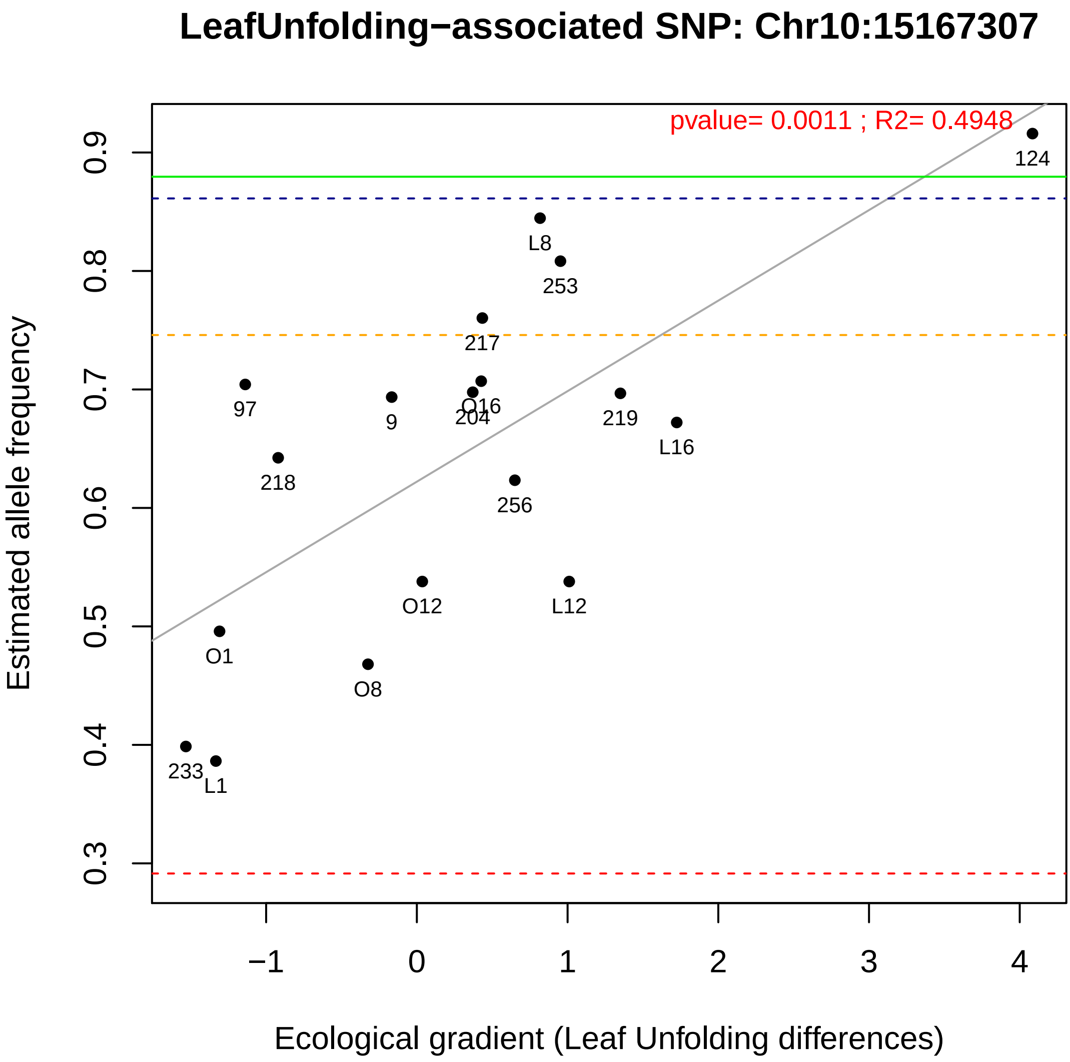
LeafUnfolding−associated SNP: Chr10:15167307
pvalue= 0.0011 ; R2= 0.4948
Estimated allele frequency
Ecological gradient (Leaf Unfolding differences)
0.3
0.4
0.5
0.6
0.7
0.8
0.9
−1
0
1
2
3
4
124
L8
253
217
97
9
O16
204
219
L16
218
256
O12
L12
O1
O8
233
L1

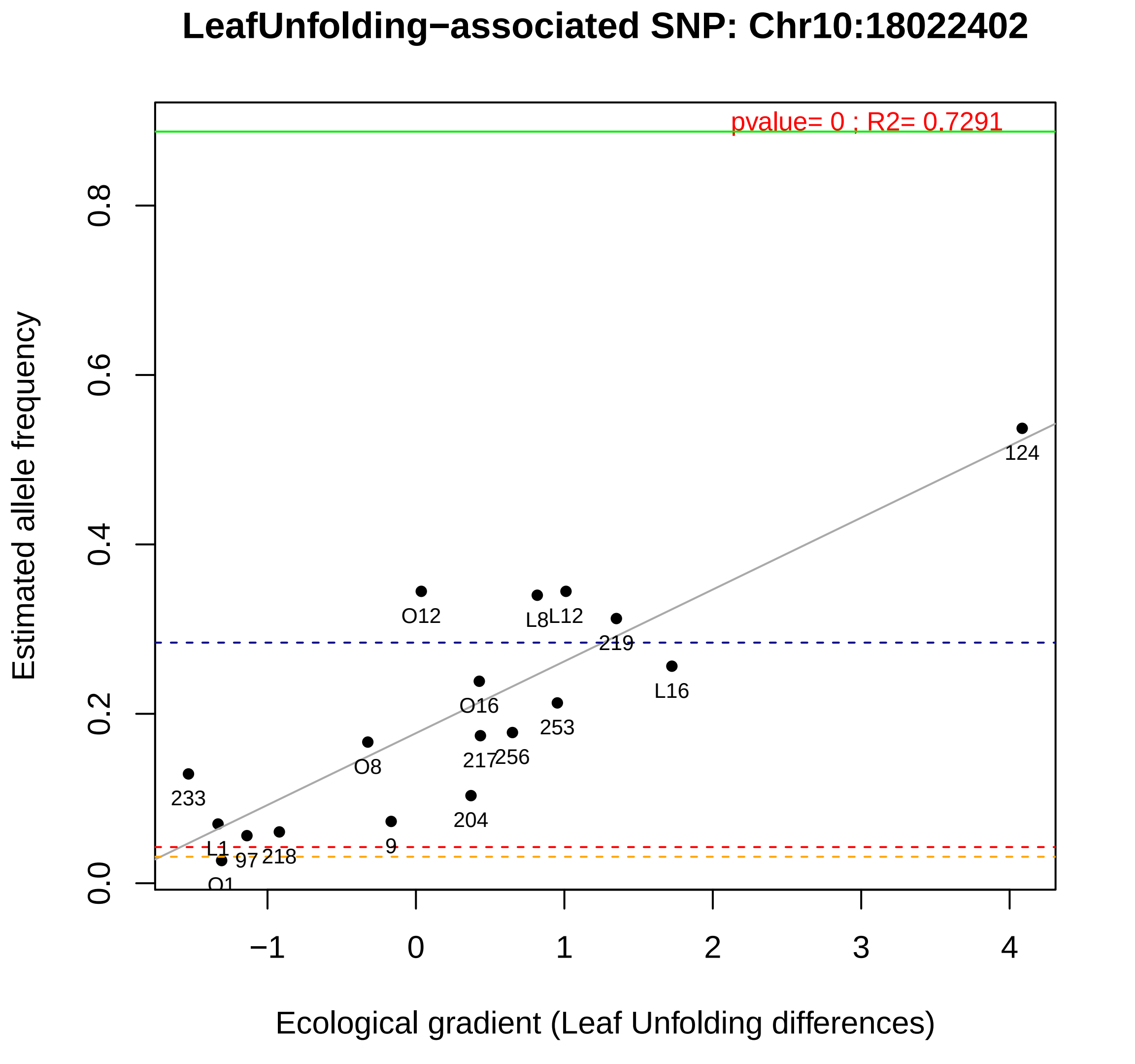

LeafUnfolding−associated SNP: Chr10:18022402
pvalue= 0 ; R2= 0.7291
Estimated allele frequency
0.0
0.2
0.4
0.6
0.8
Ecological gradient (Leaf Unfolding differences)
−1
0
1
2
3
4
124
O12
L8
L12
219
L16
O16
253
217
256
O8
233
204
L1
97
218
9
O1

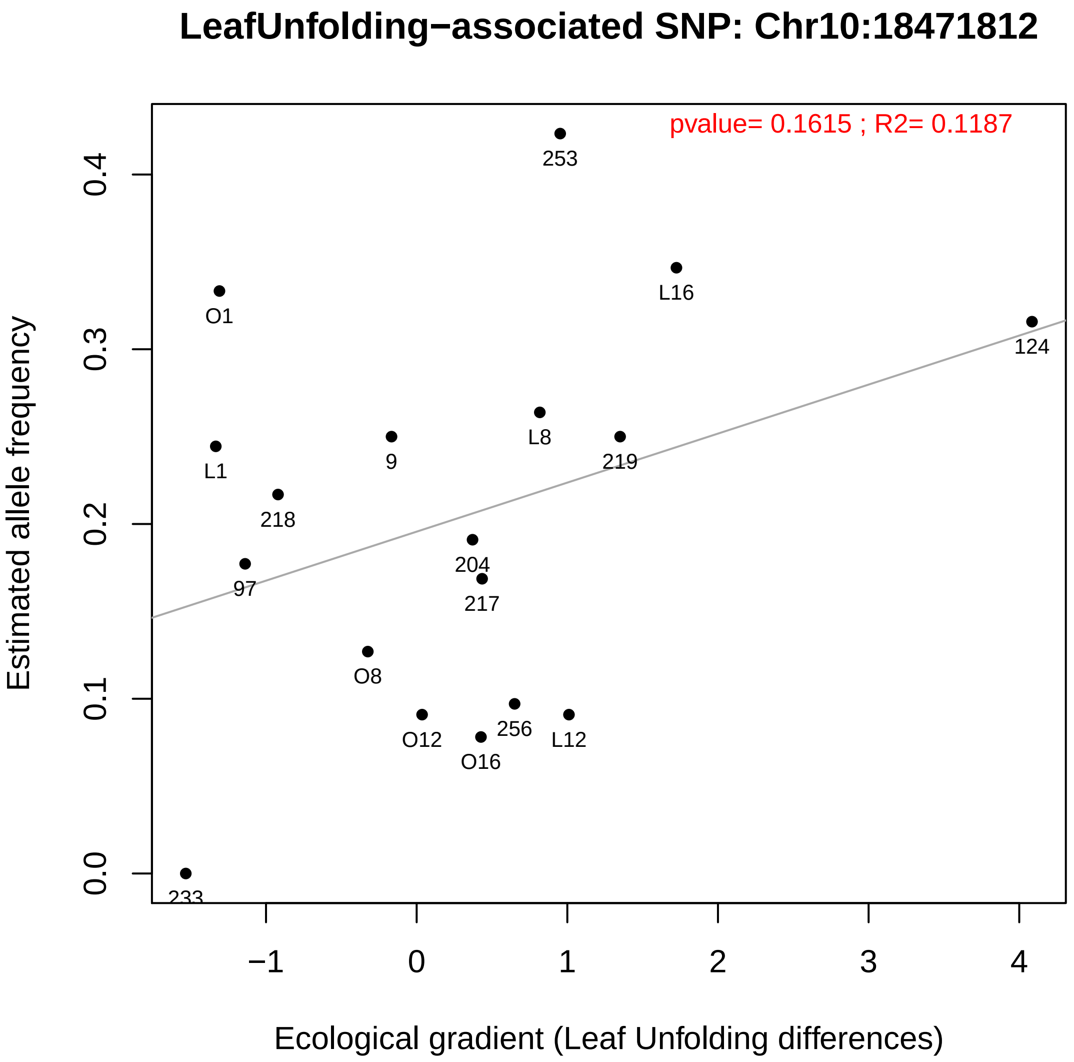

LeafUnfolding−associated SNP: Chr10:18471812
pvalue= 0.1615 ; R2= 0.1187
Estimated allele frequency
Ecological gradient (Leaf Unfolding differences)
0.0
0.1
0.2
0.3
0.4
−1
0
1
2
3
4
253
L16
O1
124
L8
L1
9
219
218
204
97
217
O8
256
O12
L12
O16
233

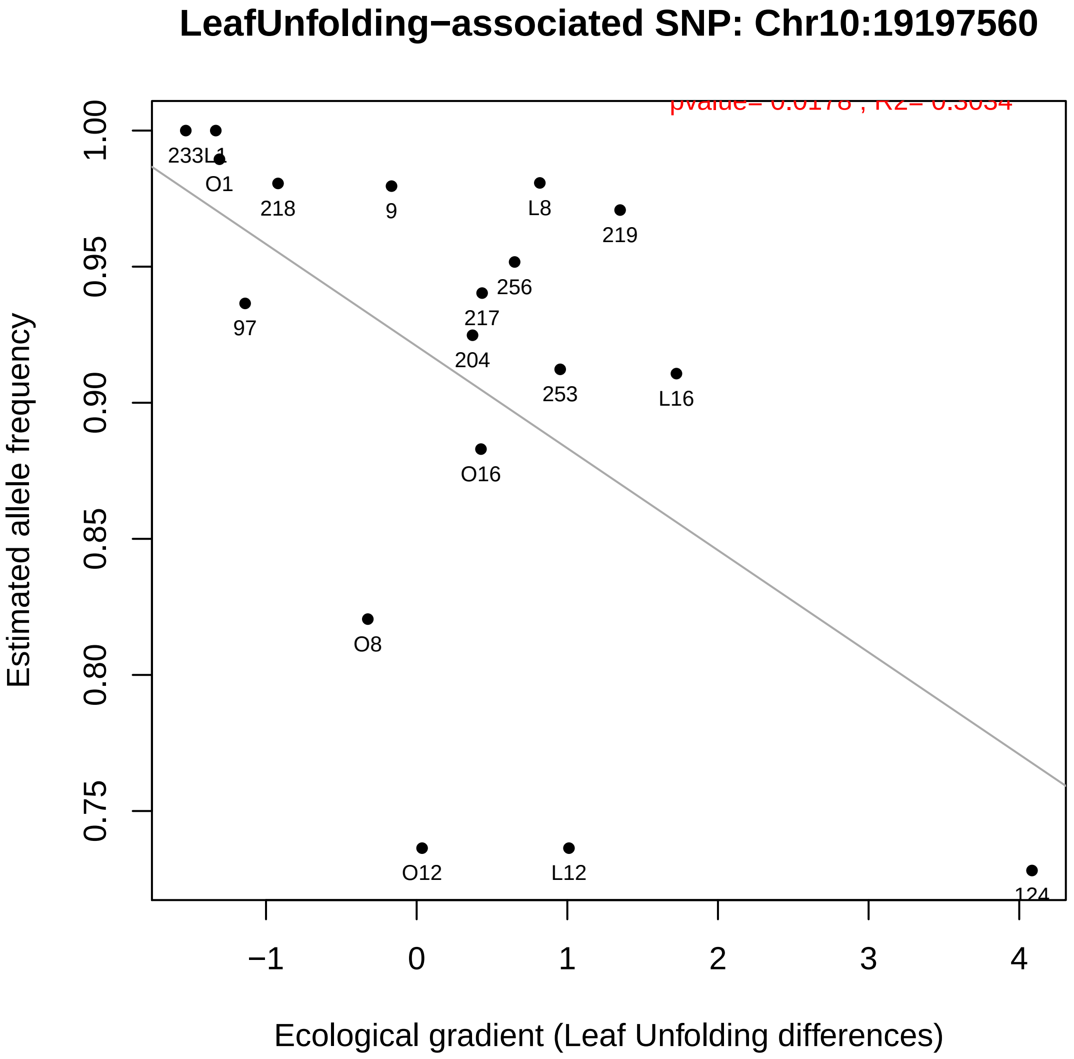
LeafUnfolding−associated SNP: Chr10:19197560
pvalue= 0.0178 ; R2= 0.3034
Estimated allele frequency
Ecological gradient (Leaf Unfolding differences)
1.00
0.95
0.90
0.85
0.80
0.75
−1
0
1
2
3
4
233
L1
O1
218
9
L8
219
256
217
97
204
253
L16
O16
O8
O12
L12
124

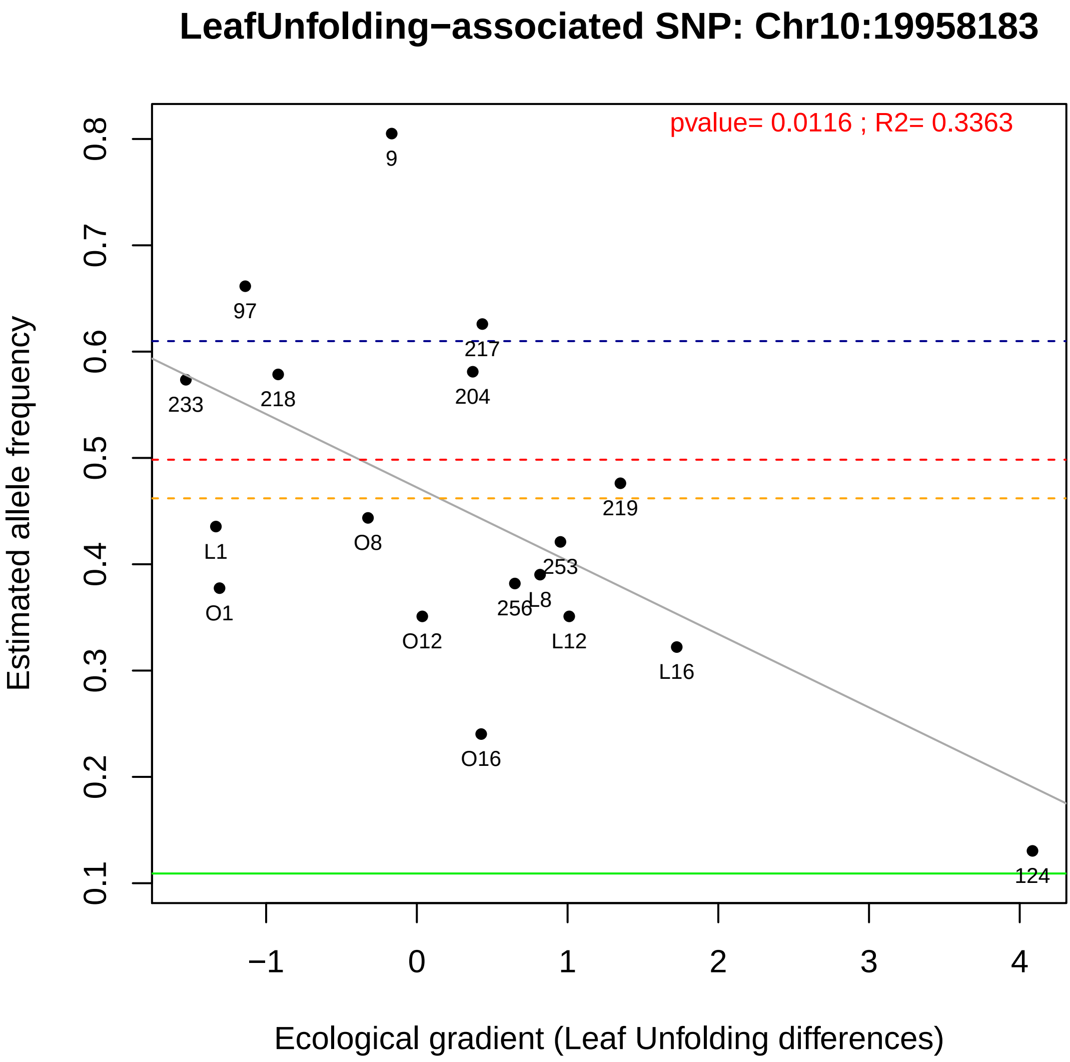

LeafUnfolding−associated SNP: Chr10:19958183
pvalue= 0.0116 ; R2= 0.3363
9
97
217
233
218
204
219
L1
O8
253
O1
256
L8
O12
L12
L16
O16
124
0.1
0.2
0.3
0.4
0.5
0.6
0.7
0.8
−1
0
1
2
3
4
Estimated allele frequency
Ecological gradient (Leaf Unfolding differences)

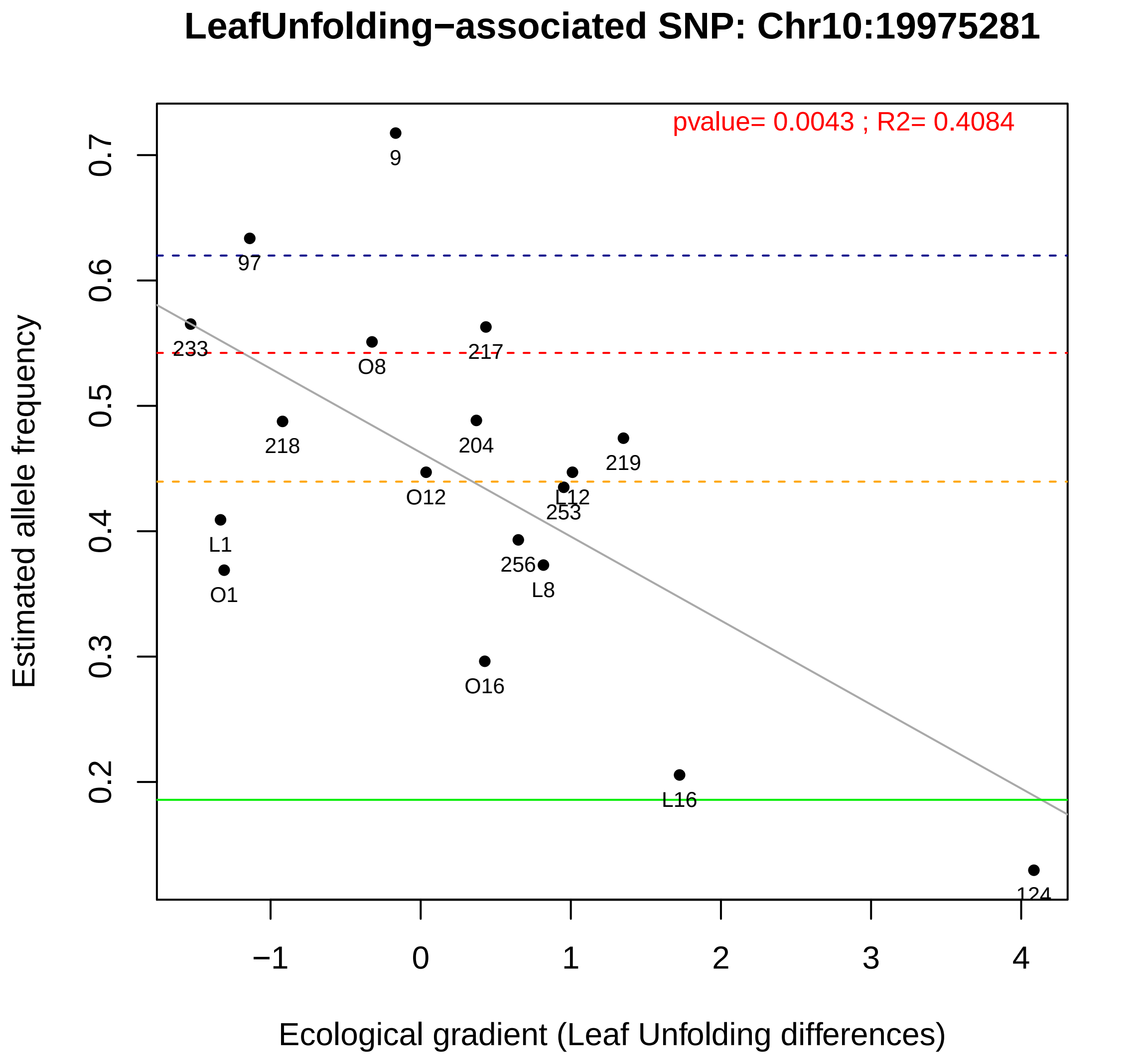

LeafUnfolding−associated SNP: Chr10:19975281
pvalue= 0.0043 ; R2= 0.4084
Estimated allele frequency
0.2
0.3
0.4
0.5
0.6
0.7
−1
0
1
2
3
4
Ecological gradient (Leaf Unfolding differences)
9
97
233
O8
217
218
204
219
O12
L12
253
L1
256
L8
O1
O16
L16
124

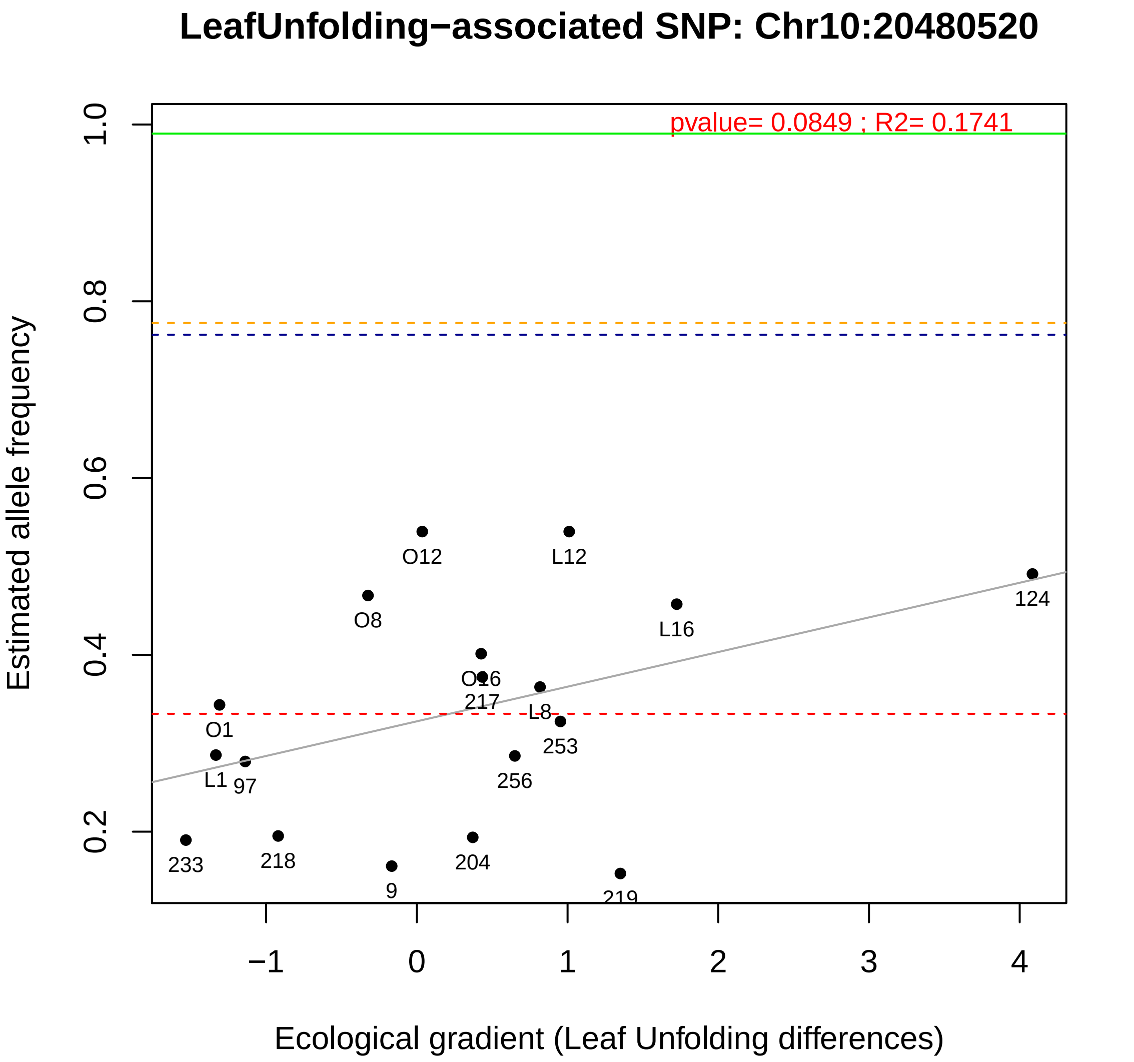
LeafUnfolding−associated SNP: Chr10:20480520
pvalue= 0.0849 ; R2= 0.1741
Estimated allele frequency
Ecological gradient (Leaf Unfolding differences)
1.0
0.8
0.6
0.4
0.2
−1
0
1
2
3
4
O12
L12
124
O8
L16
O16
217
L8
253
O1
L1
97
256
233
218
9
204
219

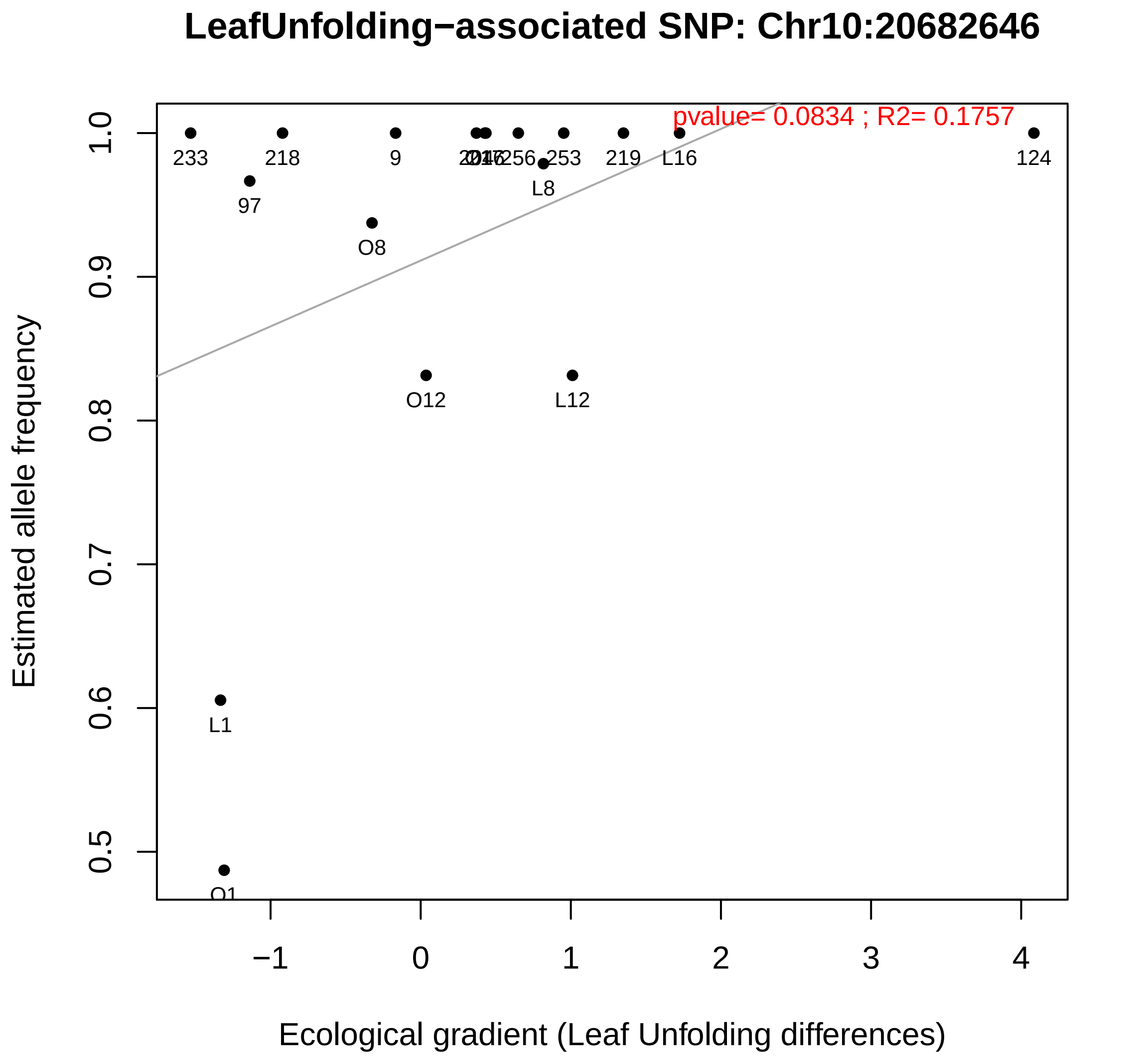
LeafUnfolding−associated SNP: Chr10:20682646
pvalue= 0.0834 ; R2= 0.1757
Estimated allele frequency
Ecological gradient (Leaf Unfolding differences)
233
218
9
256
253
219
L16
124
97
L8
O8
O12
L12
L1
O1
0.5
0.6
0.7
0.8
0.9
1.0
−1
0
1
2
3
4

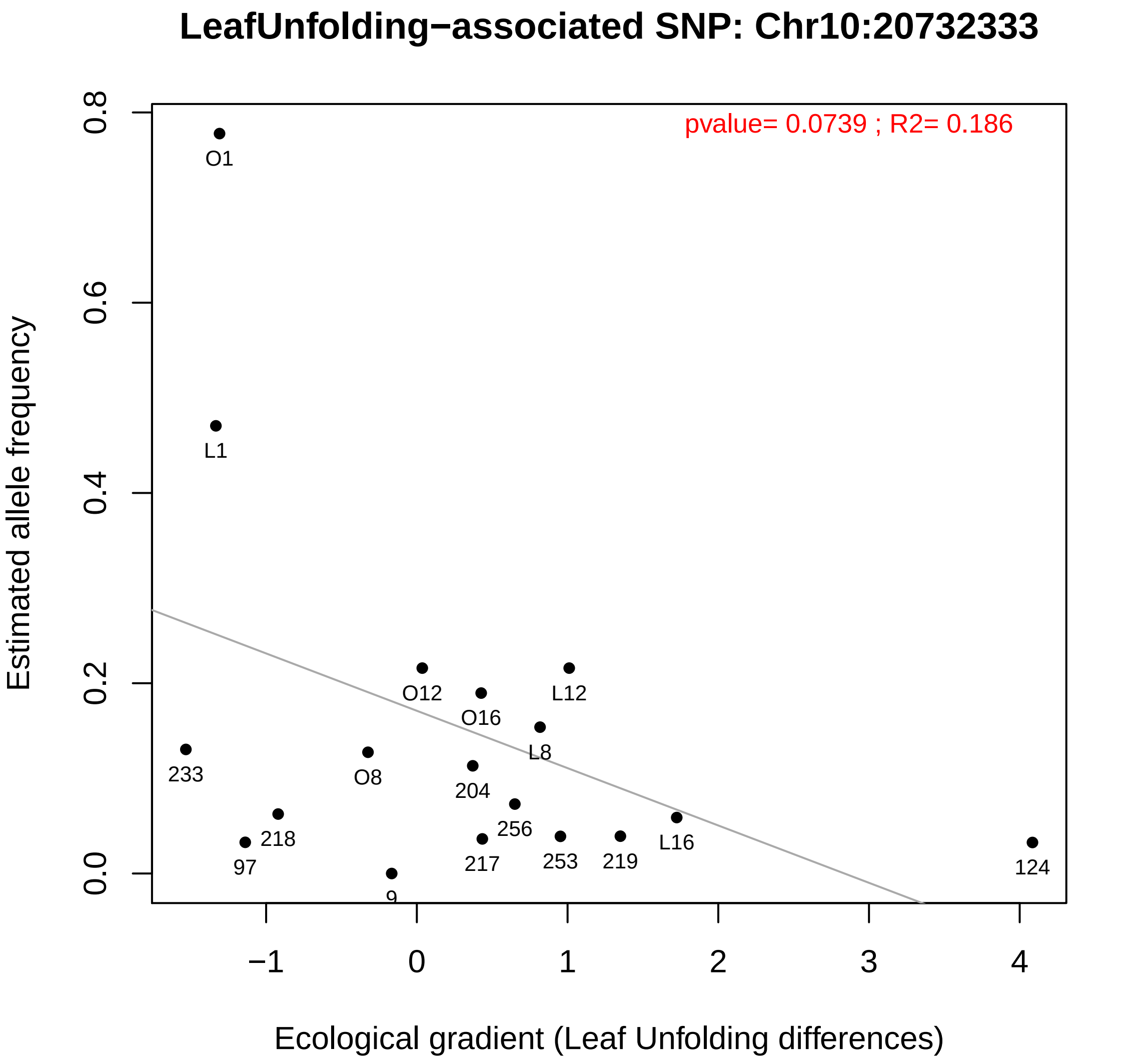

LeafUnfolding−associated SNP: Chr10:20732333
pvalue= 0.0739 ; R2= 0.186
Estimated allele frequency
0.0
0.2
0.4
0.6
0.8
O1
L1
O12
L12
O16
L8
233
O8
204
256
218
97
217
253
219
L16
124
9
−1
0
1
2
3
4
Ecological gradient (Leaf Unfolding differences)

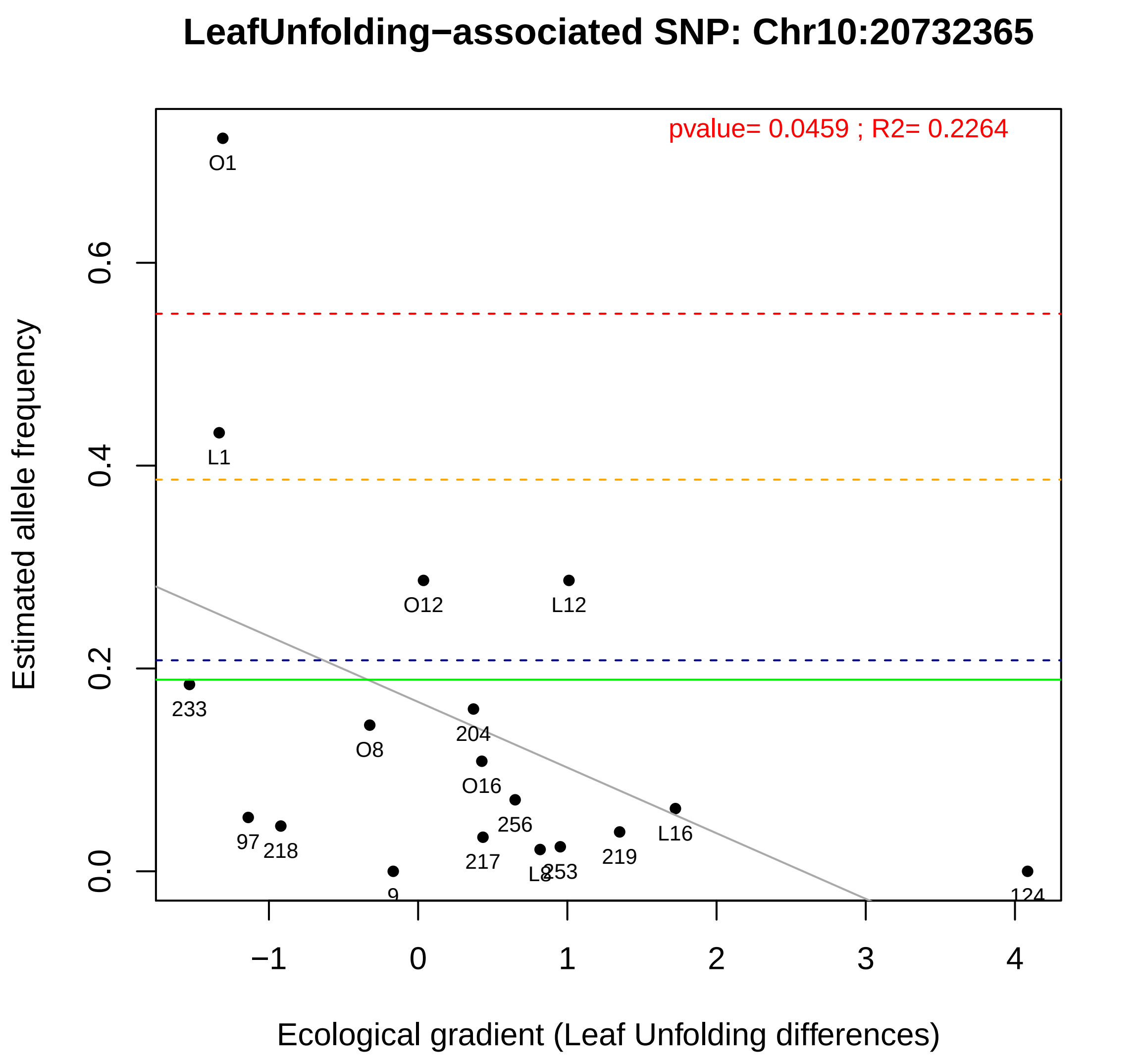
LeafUnfolding−associated SNP: Chr10:20732365
pvalue= 0.0459 ; R2= 0.2264
Estimated allele frequency
0.0
0.2
0.4
0.6
Ecological gradient (Leaf Unfolding differences)
−1
0
1
2
3
4
O1
L1
O12
L12
233
O8
204
O16
256
L16
97
218
217
219
9
L8
253
124

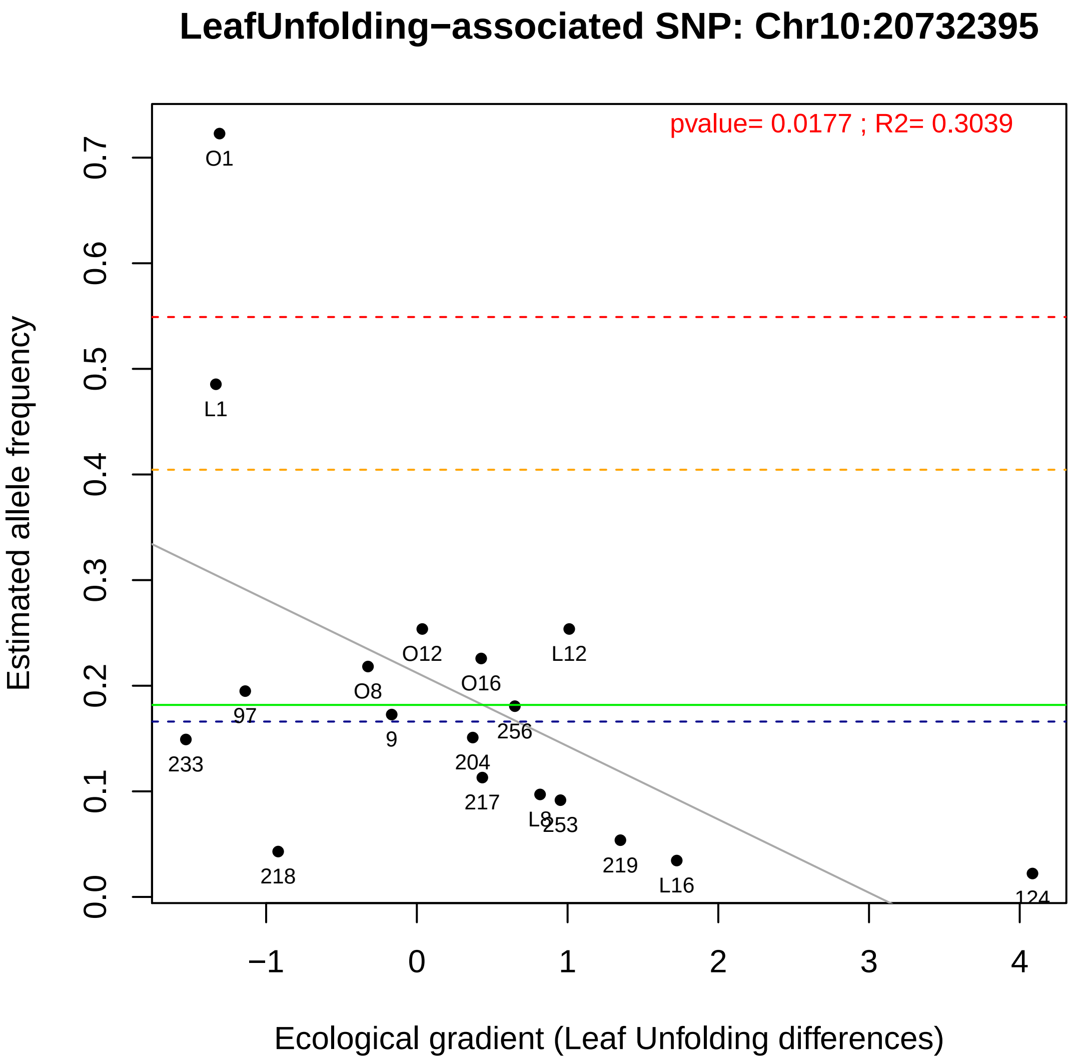

LeafUnfolding−associated SNP: Chr10:20732395
pvalue= 0.0177 ; R2= 0.3039
Estimated allele frequency
0.0
0.1
0.2
0.3
0.4
0.5
0.6
0.7
−1
0
1
2
3
4
Ecological gradient (Leaf Unfolding differences)
O1
L1
O12
L12
O8
O16
97
9
256
233
204
217
L8
253
218
219
L16
124

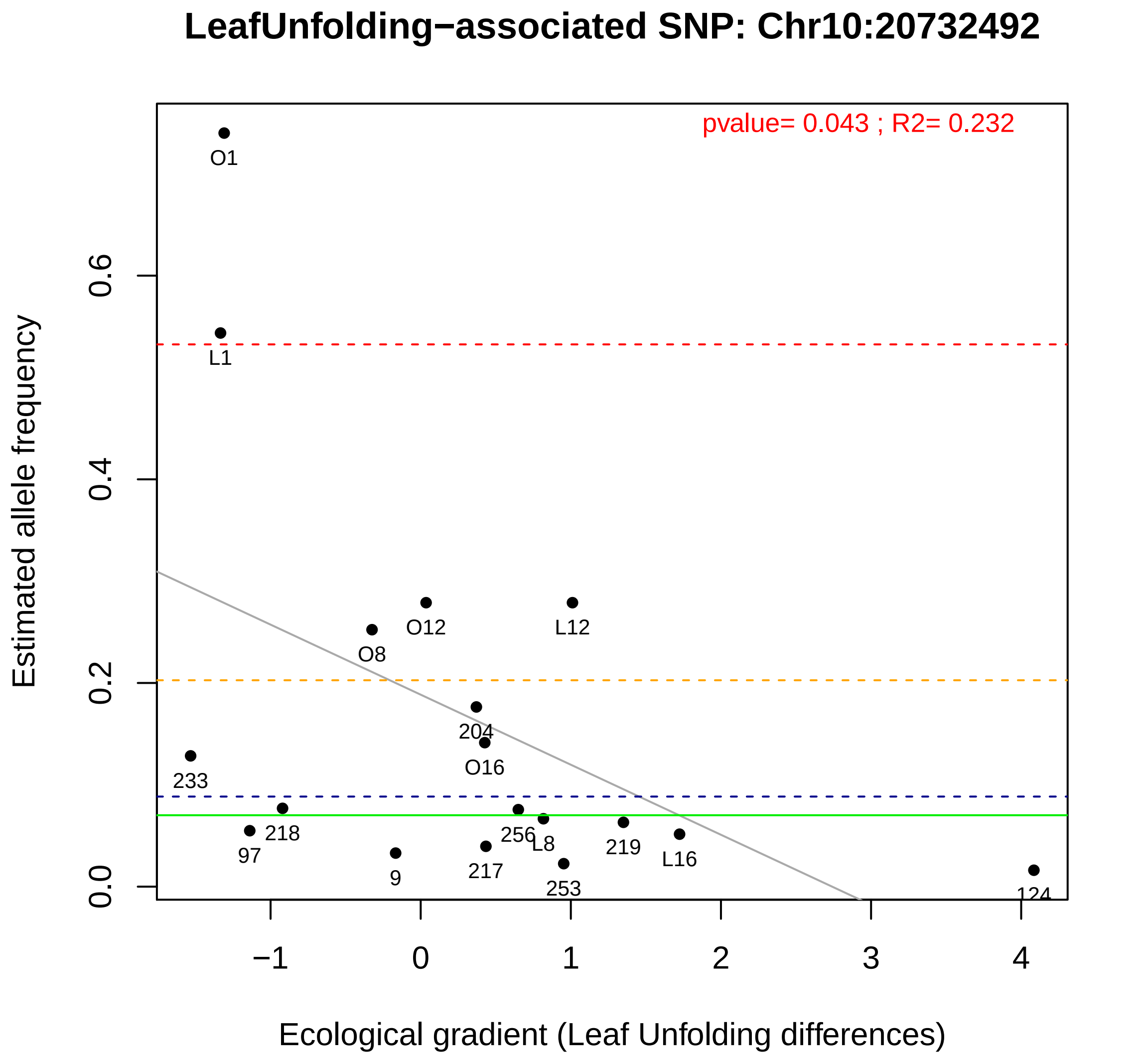
LeafUnfolding−associated SNP: Chr10:20732492
pvalue= 0.043 ; R2= 0.232
Estimated allele frequency
0.0
0.2
0.4
0.6
Ecological gradient (Leaf Unfolding differences)
−1
0
1
2
3
4
O1
L1
O12
L12
O8
204
O16
233
218
97
9
217
256
L8
253
219
L16
124

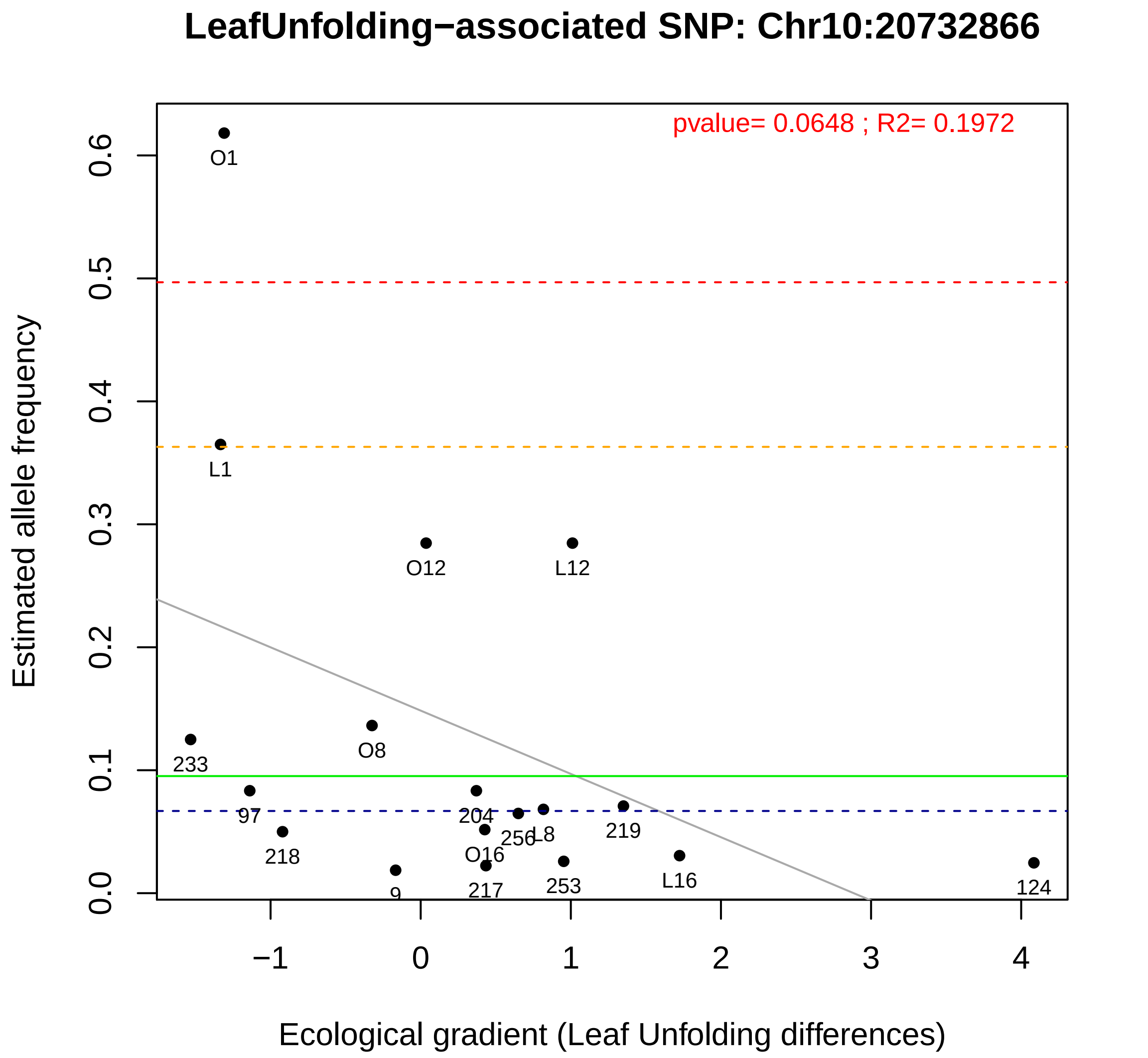
LeafUnfolding−associated SNP: Chr10:20732866
pvalue= 0.0648 ; R2= 0.1972
Estimated allele frequency
0.0
0.1
0.2
0.3
0.4
0.5
0.6
−1
0
1
2
3
4
Ecological gradient (Leaf Unfolding differences)
O1
L1
O12
L12
233
O8
97
204
218
256
L8
219
O16
9
217
253
L16
124

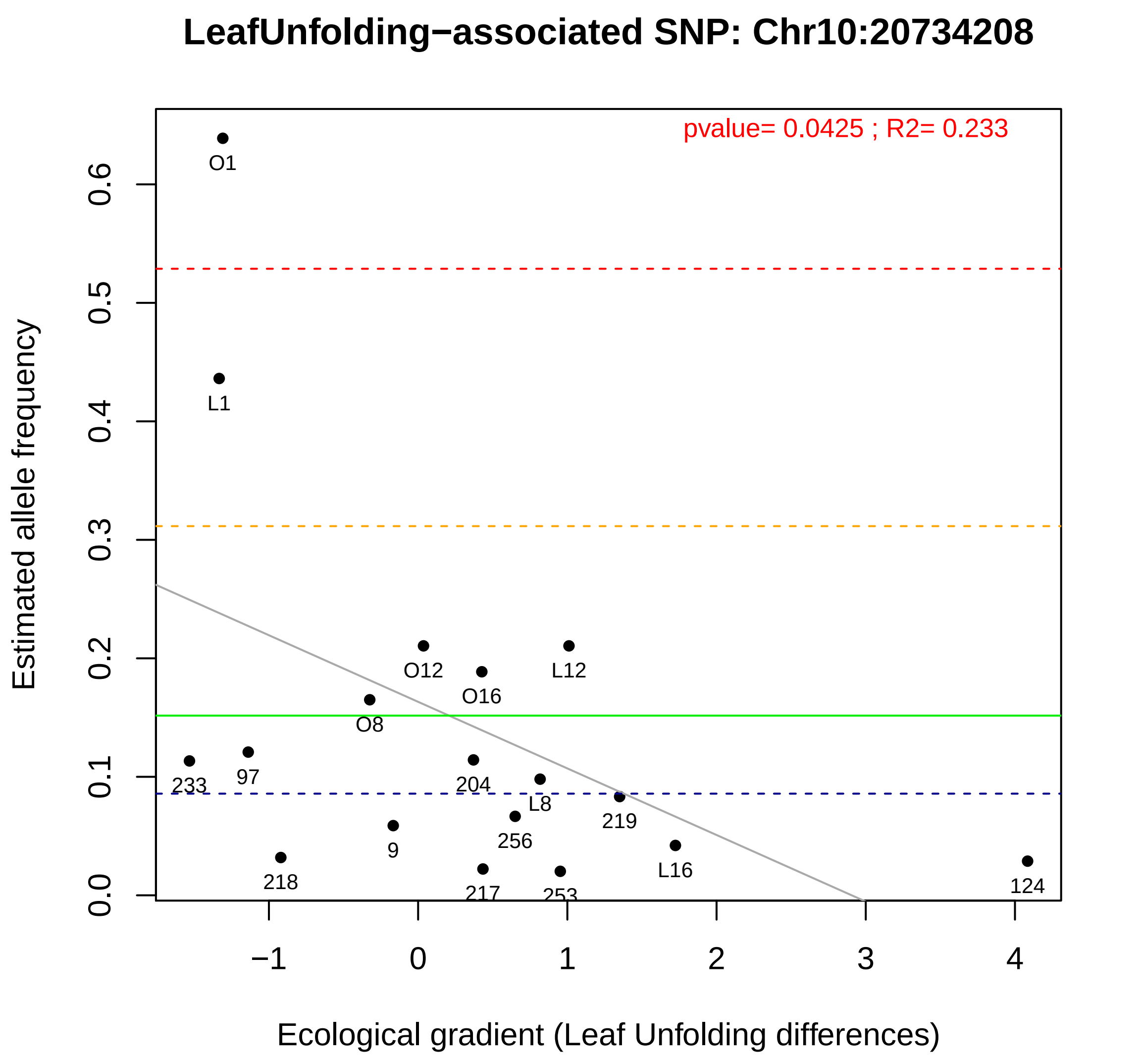

LeafUnfolding−associated SNP: Chr10:20734208
pvalue= 0.0425 ; R2= 0.233
Estimated allele frequency
0.0
0.1
0.2
0.3
0.4
0.5
0.6
−1
0
1
2
3
4
Ecological gradient (Leaf Unfolding differences)
O1
L1
O12
L12
O16
O8
233
97
204
L8
219
256
9
218
L16
217
253
124

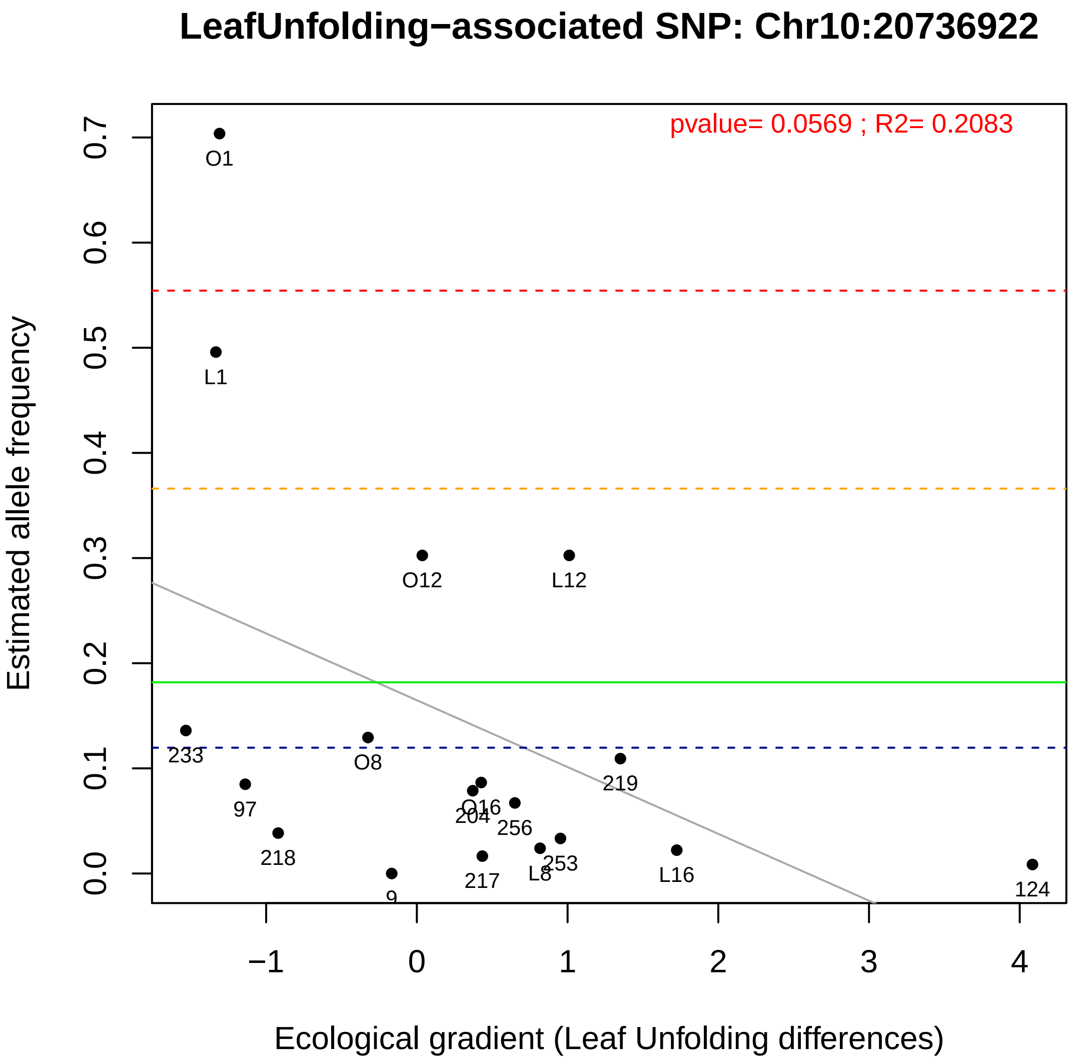

LeafUnfolding−associated SNP: Chr10:20736922
pvalue= 0.0569 ; R2= 0.2083
Estimated allele frequency
0.0
0.1
0.2
0.3
0.4
0.5
0.6
0.7
Ecological gradient (Leaf Unfolding differences)
−1
0
1
2
3
4
O1
L1
O12
L12
233
O8
97
218
9
219
O16
204
256
217
L8
253
L16
124

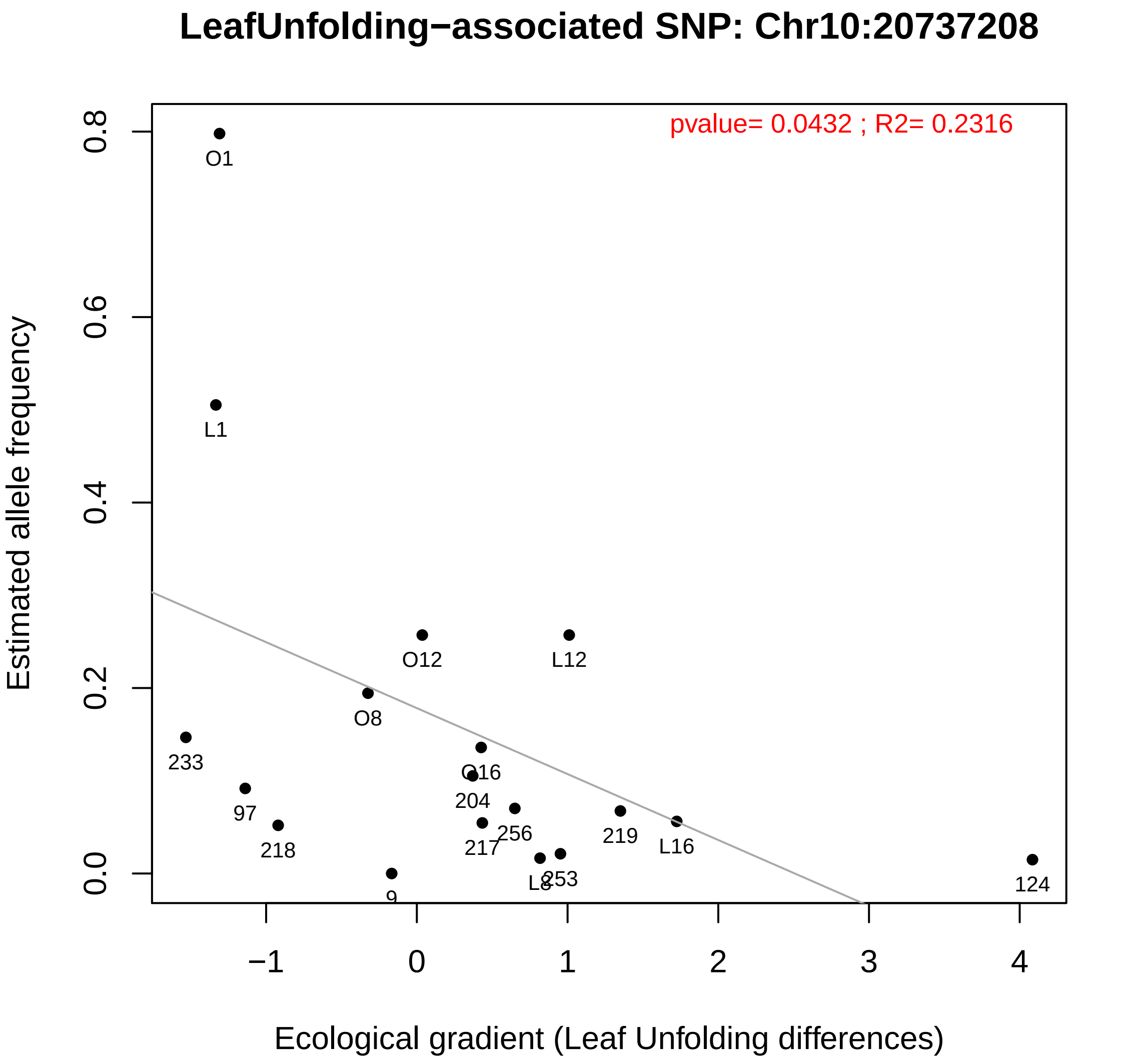
LeafUnfolding−associated SNP: Chr10:20737208
pvalue= 0.0432 ; R2= 0.2316
Estimated allele frequency
0.0
0.2
0.4
0.6
0.8
−1
0
1
2
3
4
Ecological gradient (Leaf Unfolding differences)
O1
L1
O12
L12
O8
233
O16
204
97
218
256
217
219
L16
L8
253
9
124

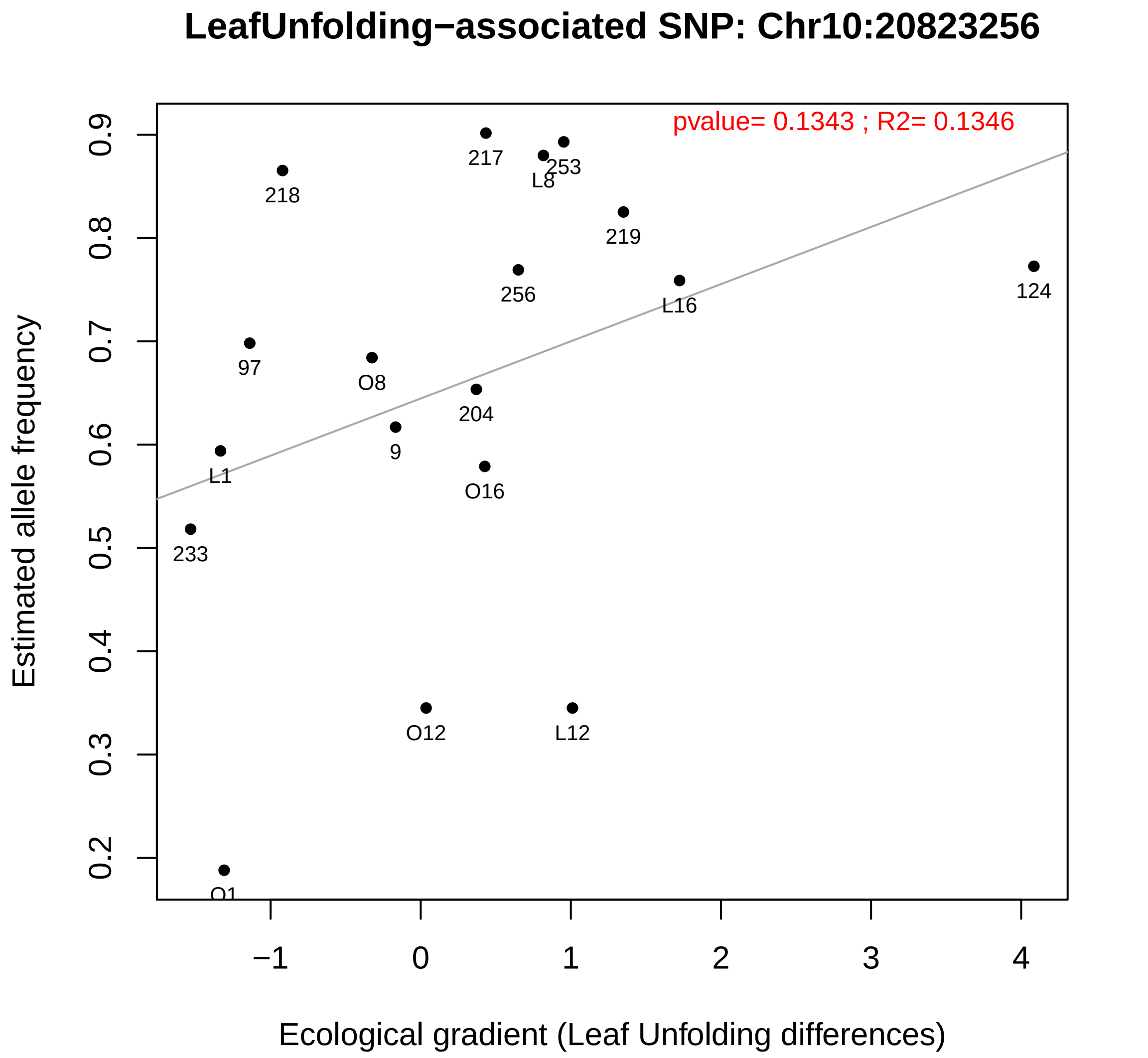
LeafUnfolding−associated SNP: Chr10:20823256
pvalue= 0.1343 ; R2= 0.1346
Estimated allele frequency
Ecological gradient (Leaf Unfolding differences)
0.2
0.3
0.4
0.5
0.6
0.7
0.8
0.9
−1
0
1
2
3
4
217
253
L8
218
219
256
L16
124
97
O8
204
9
L1
O16
233
O12
L12
O1

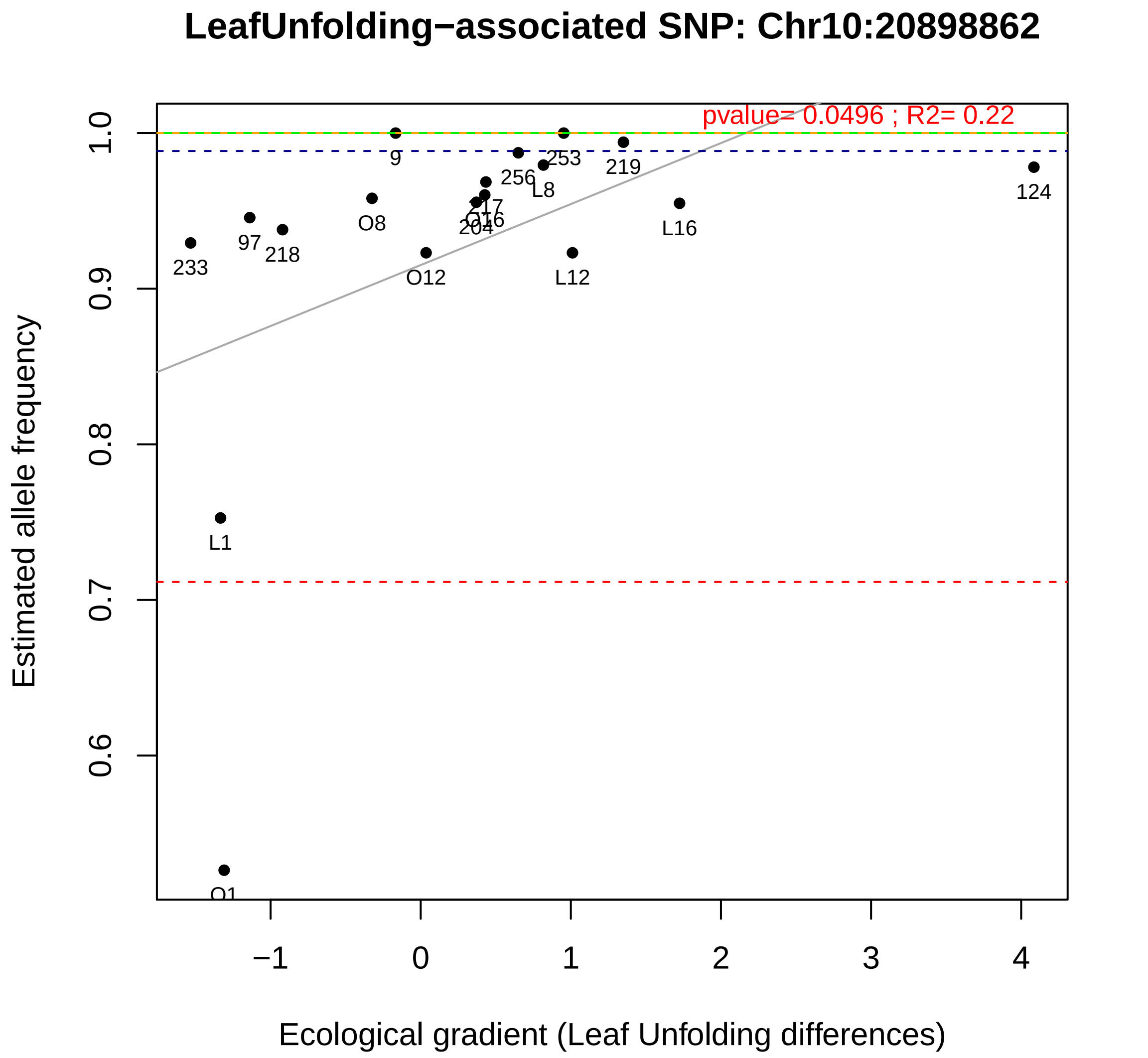
LeafUnfolding−associated SNP: Chr10:20898862
pvalue= 0.0496 ; R2= 0.22
Estimated allele frequency
Ecological gradient (Leaf Unfolding differences)
1.0
0.9
0.8
0.7
0.6
−1
0
1
2
3
4
9
253
256
L8
219
124
O8
97
218
233
O12
L12
L16
204
O16
L1
O1

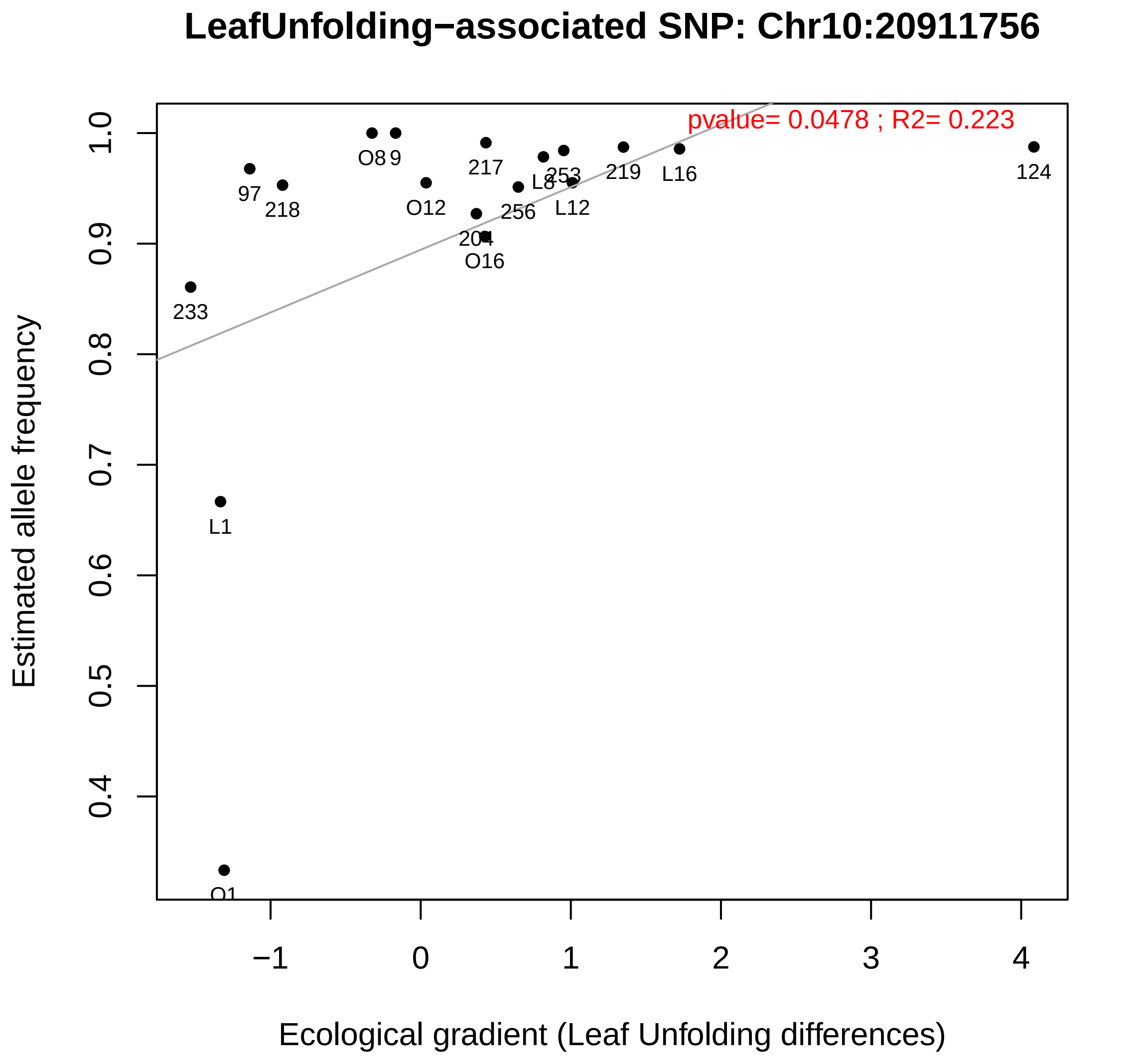
LeafUnfolding−associated SNP: Chr10:20911756
pvalue= 0.0478 ; R2= 0.223
Estimated allele frequency
Ecological gradient (Leaf Unfolding differences)
0.4
0.5
0.6
0.7
0.8
0.9
1.0
−1
0
1
2
3
4
O8
9
217
97
218
O12
256
L8
253
L12
219
L16
124
204
O16
233
L1
O1

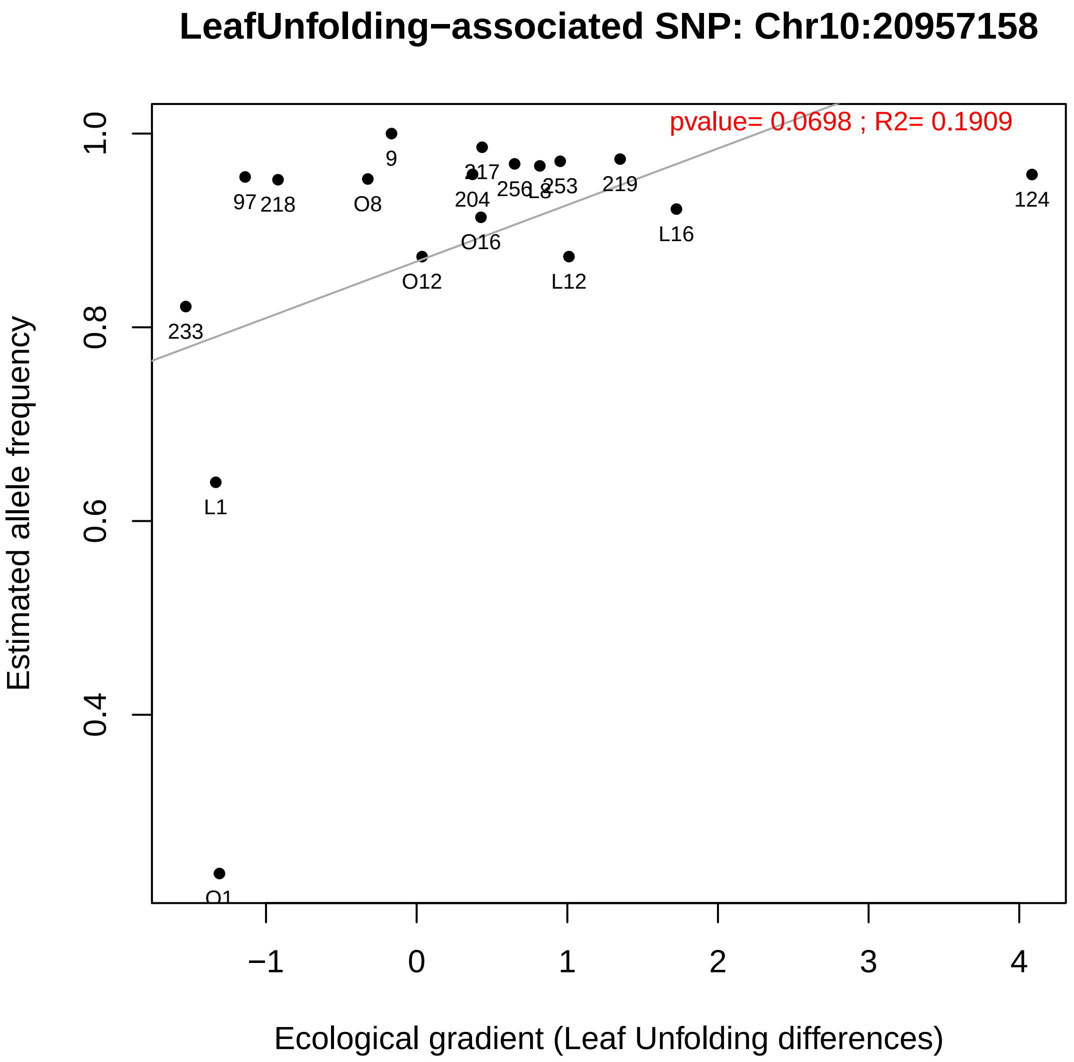
LeafUnfolding−associated SNP: Chr10:20957158
pvalue= 0.0698 ; R2= 0.1909
Estimated allele frequency
Ecological gradient (Leaf Unfolding differences)
1.0
0.8
0.6
0.4
−1
0
1
2
3
4
9
97
218
O8
217
204
256
L8
253
219
124
O16
L16
O12
L12
233
L1
O1

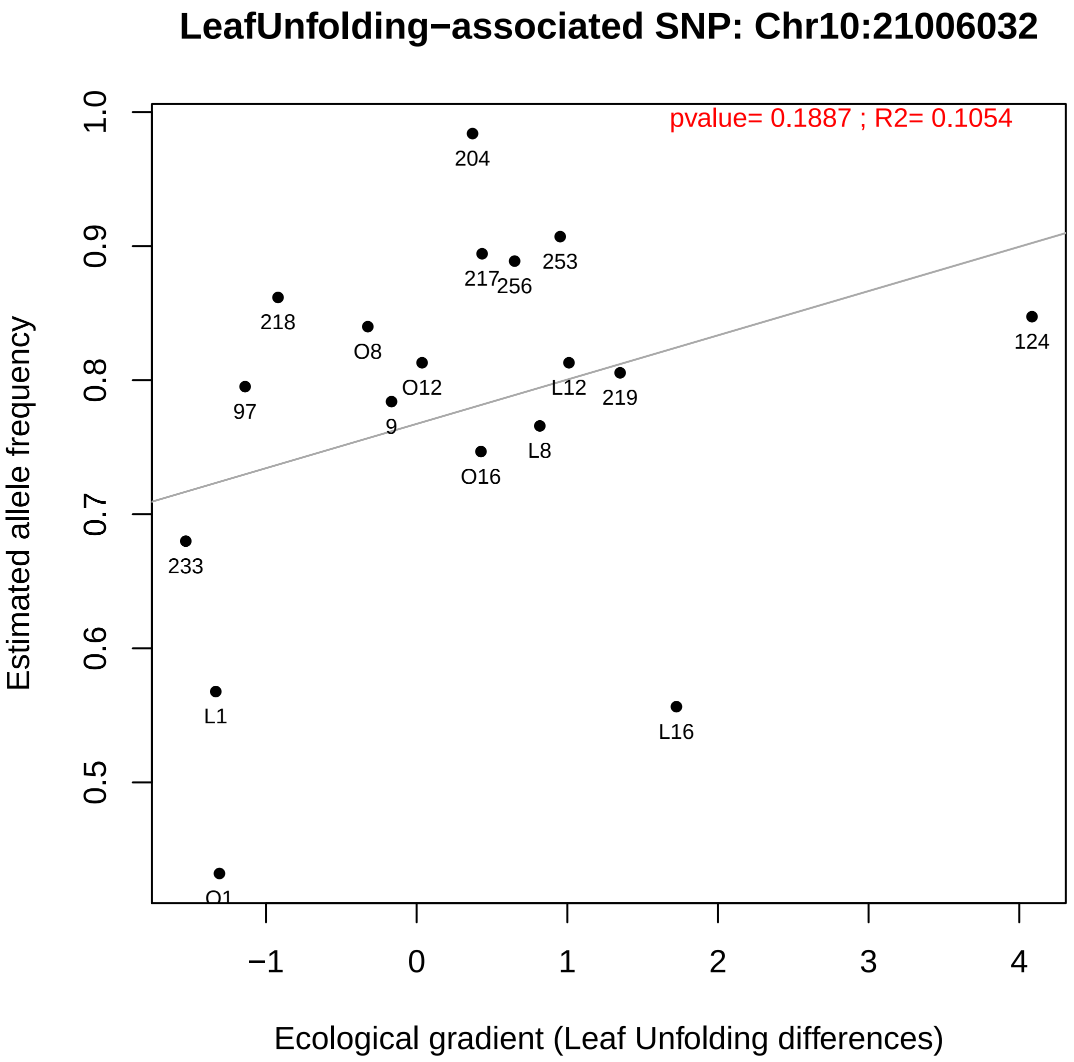
LeafUnfolding−associated SNP: Chr10:21006032
pvalue= 0.1887 ; R2= 0.1054
Estimated allele frequency
Ecological gradient (Leaf Unfolding differences)
1.0
0.9
0.8
0.7
0.6
0.5
−1
0
1
2
3
4
204
253
217
256
218
O8
124
O12
L12
219
97
9
L8
O16
233
L1
L16
O1

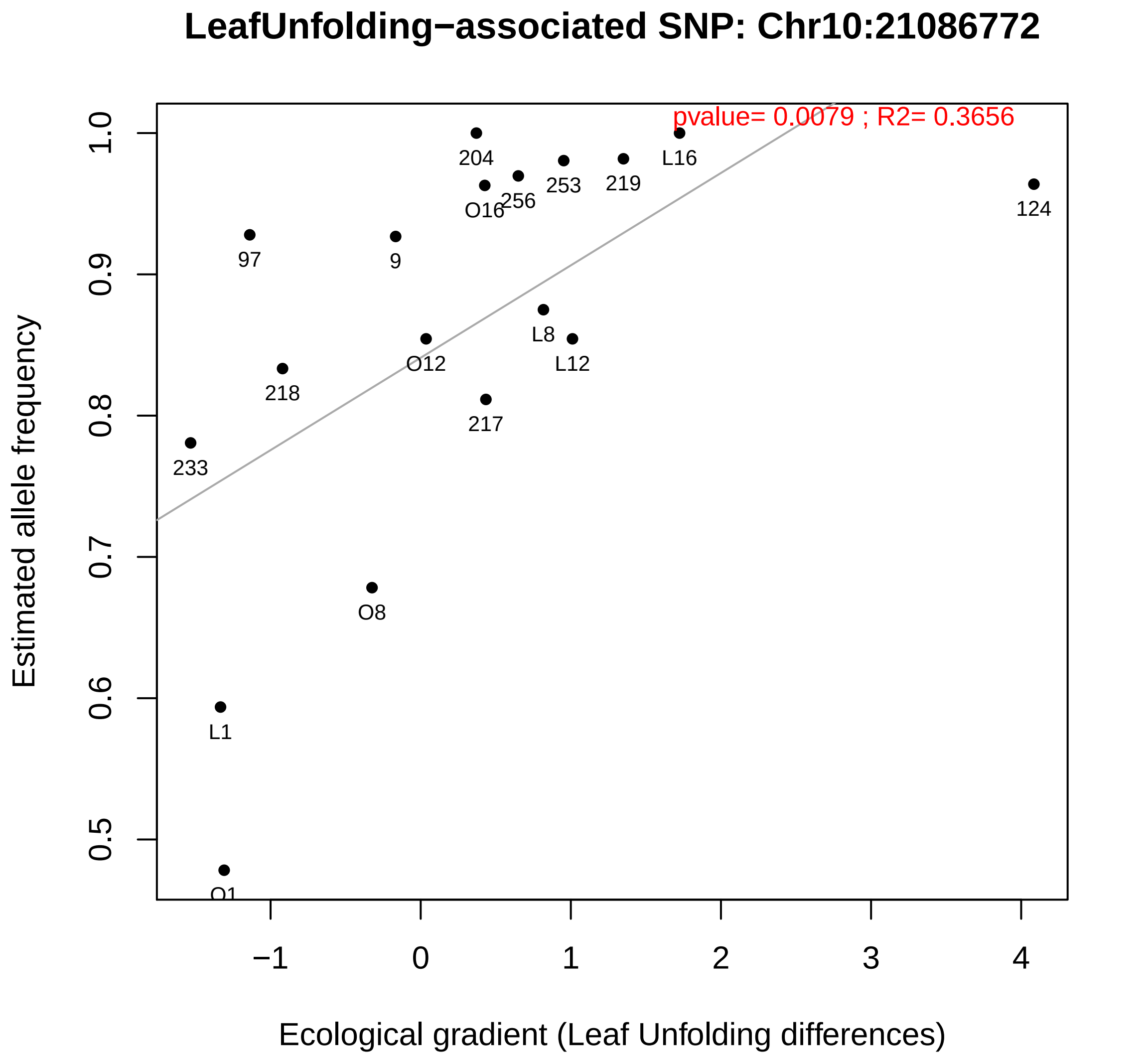
LeafUnfolding−associated SNP: Chr10:21086772
pvalue= 0.0079 ; R2= 0.3656
Estimated allele frequency
Ecological gradient (Leaf Unfolding differences)
1.0
0.9
0.8
0.7
0.6
0.5
−1
0
1
2
3
4
204
L16
253
219
256
O16
124
97
9
L8
L12
O12
218
217
233
O8
L1
O1

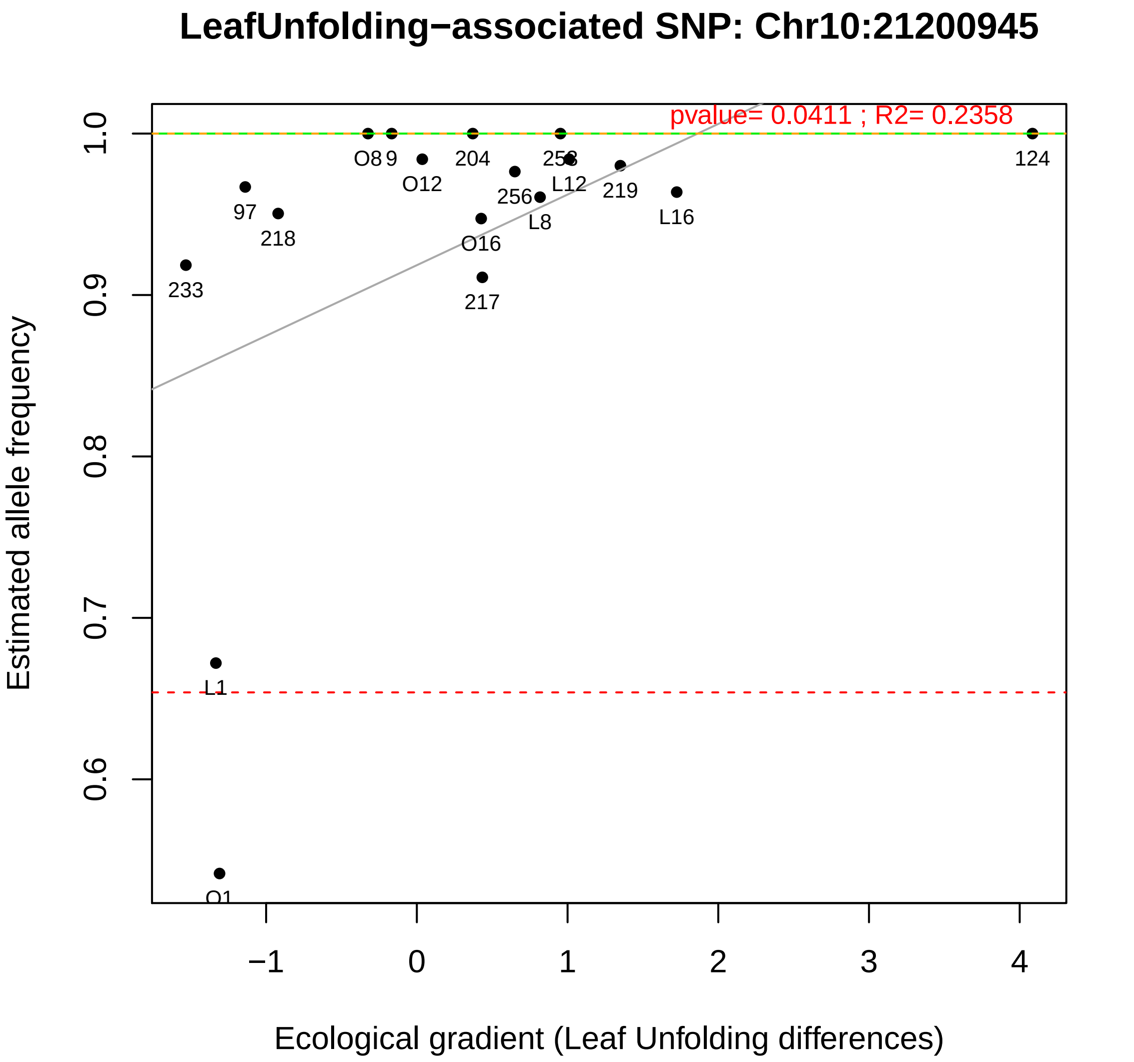

LeafUnfolding−associated SNP: Chr10:21200945
pvalue= 0.0411 ; R2= 0.2358
Estimated allele frequency
Ecological gradient (Leaf Unfolding differences)
1.0
0.9
0.8
0.7
0.6
−1
0
1
2
3
4
O8
9
204
258
124
O12
L12
256
219
97
218
L8
L16
O16
233
217
L1
O1

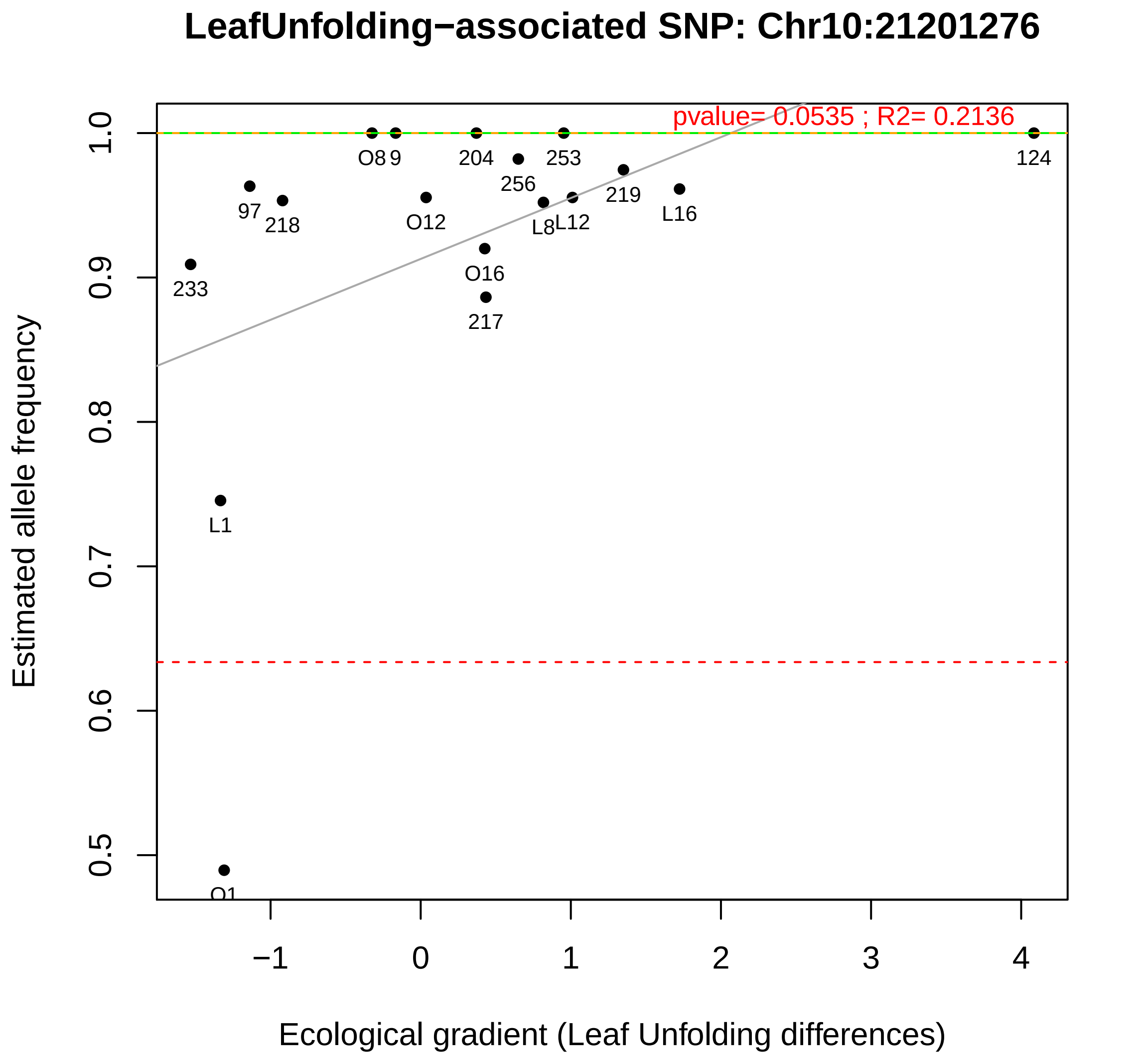
LeafUnfolding−associated SNP: Chr10:21201276
pvalue= 0.0535 ; R2= 0.2136
Estimated allele frequency
Ecological gradient (Leaf Unfolding differences)
1.0
0.9
0.8
0.7
0.6
0.5
−1
0
1
2
3
4
O8
9
204
253
124
256
219
97
218
O12
L8
L12
L16
233
O16
217
L1
O1

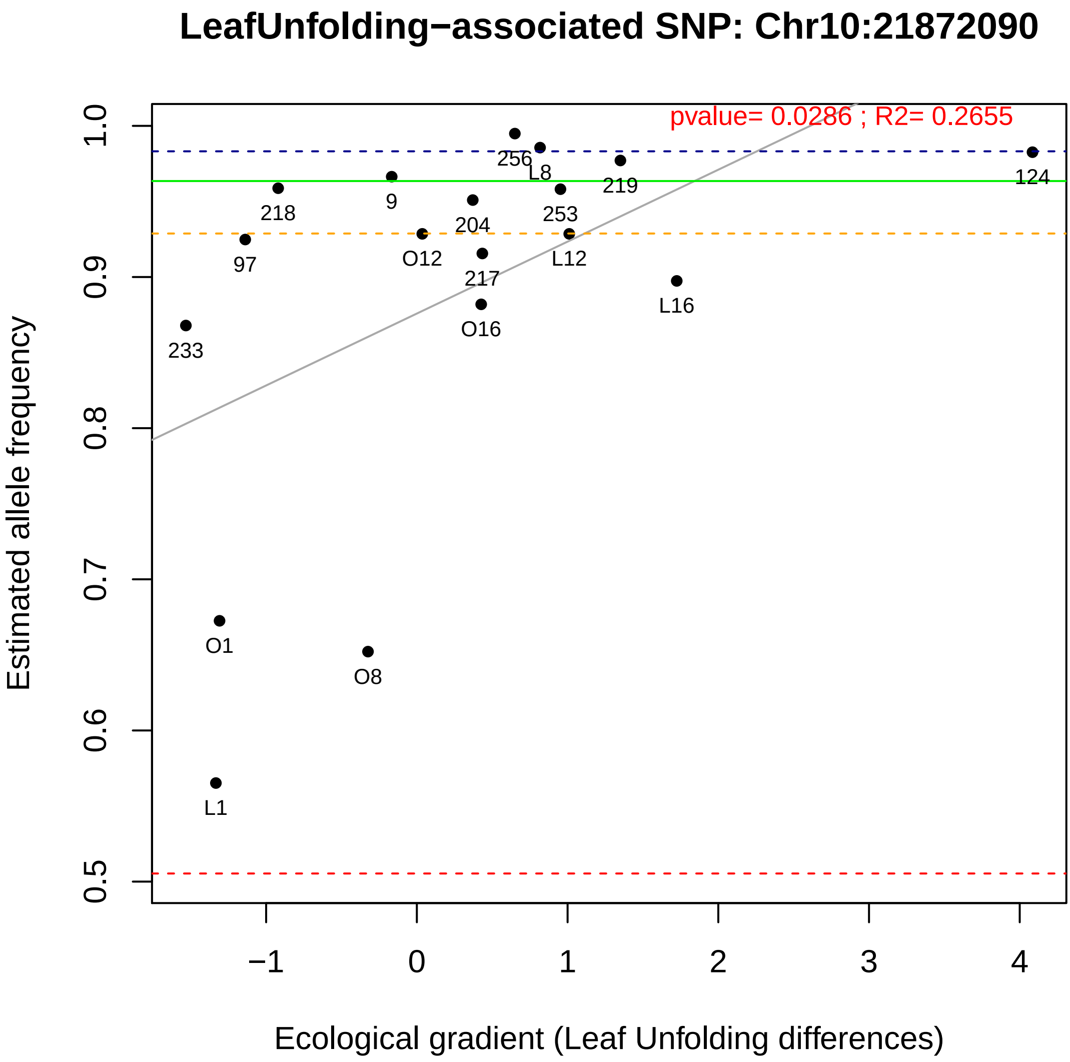
LeafUnfolding−associated SNP: Chr10:21872090
pvalue= 0.0286 ; R2= 0.2655
Estimated allele frequency
Ecological gradient (Leaf Unfolding differences)
1.0
0.9
0.8
0.7
0.6
0.5
−1
0
1
2
3
4
256
L8
219
124
218
9
204
253
97
O12
L12
217
L16
O16
233
O1
O8
L1

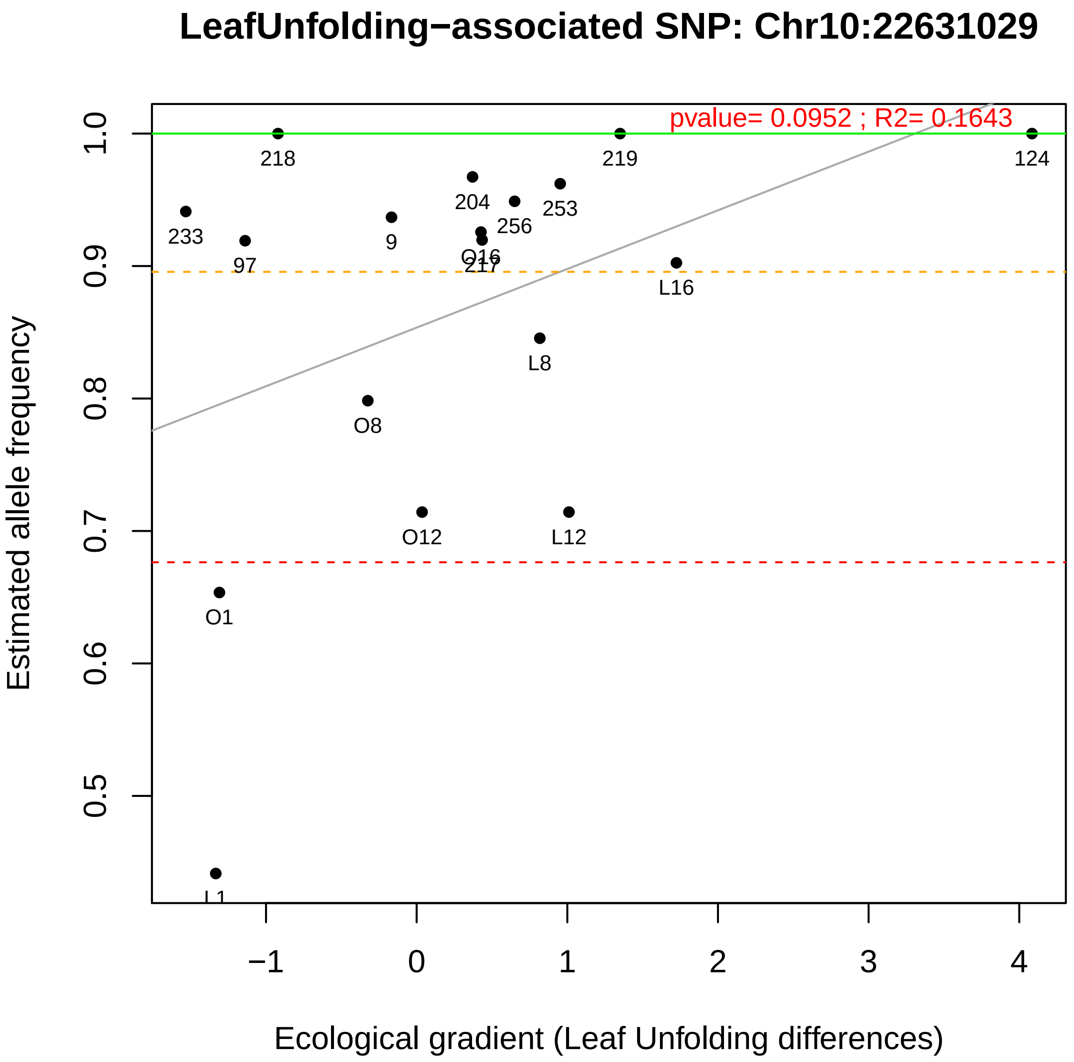

LeafUnfolding−associated SNP: Chr10:22631029
pvalue= 0.0952 ; R2= 0.1643
218
219
124
204
253
256
233
9
97
O16
217
L16
L8
O8
O12
L12
O1
L1
1.0
0.9
0.8
0.7
0.6
0.5
−1
0
1
2
3
4
Estimated allele frequency
Ecological gradient (Leaf Unfolding differences)

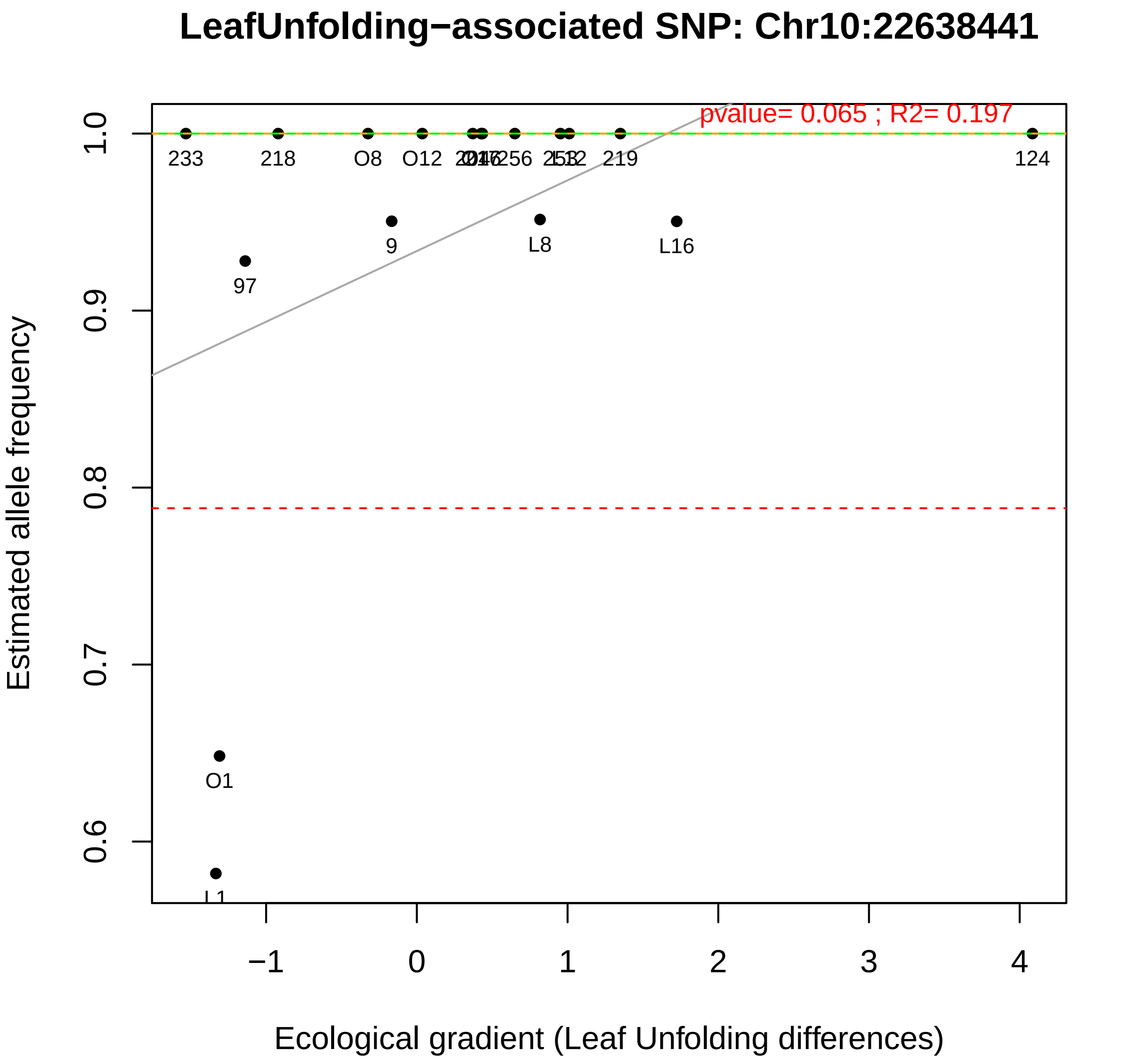

LeafUnfolding−associated SNP: Chr10:22638441
pvalue= 0.065 ; R2= 0.197
Estimated allele frequency
Ecological gradient (Leaf Unfolding differences)
0.6
0.7
0.8
0.9
1.0
−1
0
1
2
3
4
233
218
O8
O12
219
124
97
9
L8
L16
O1

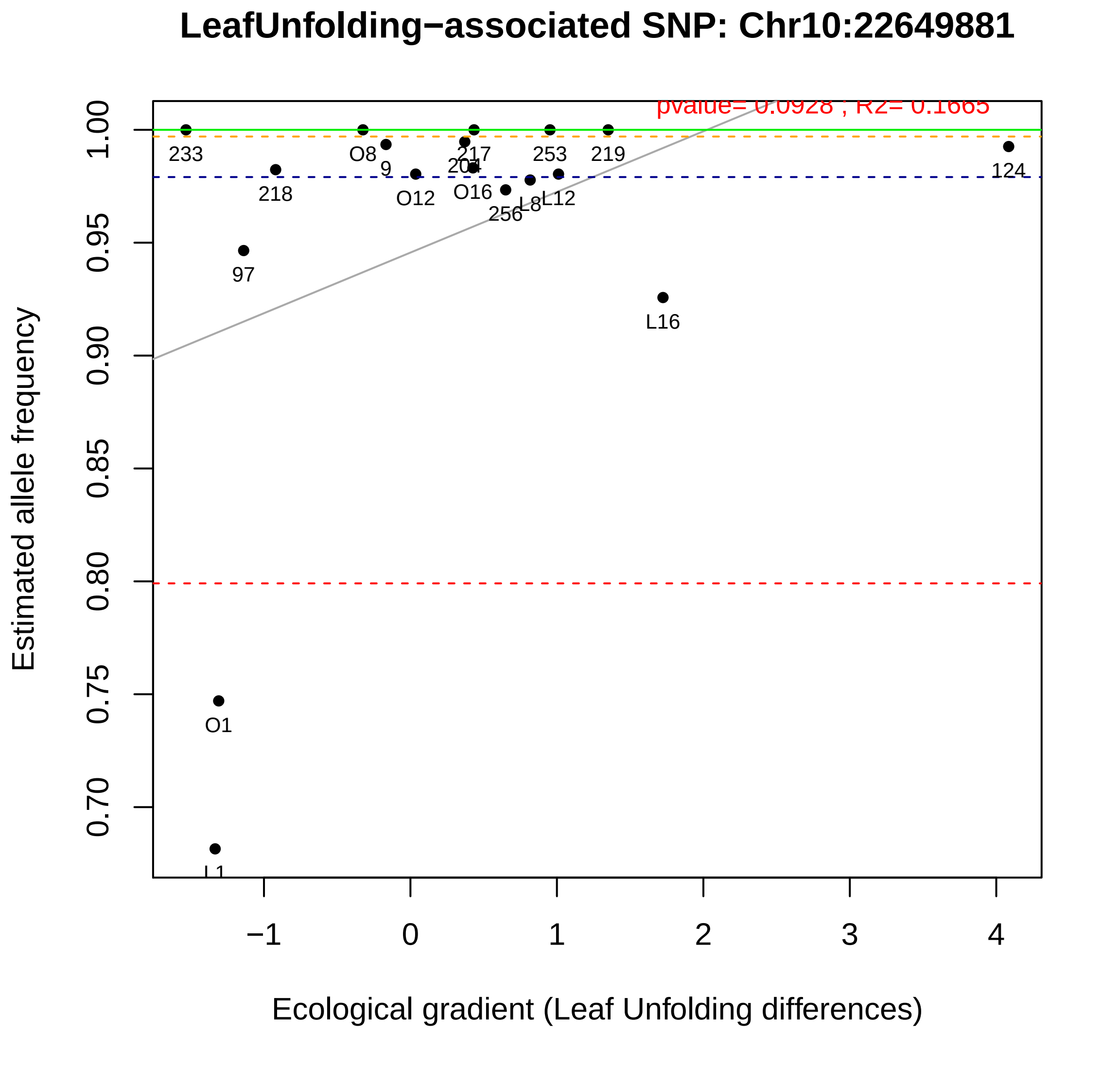

LeafUnfolding−associated SNP: Chr10:22649881
pvalue= 0.0928 ; R2= 0.1665
Estimated allele frequency
Ecological gradient (Leaf Unfolding differences)
1.00
0.95
0.90
0.85
0.80
0.75
0.70
−1
0
1
2
3
4
233
O8
9
217
204
253
219
124
218
O12
O16
256
L8
L12
97
L16
O1
L1

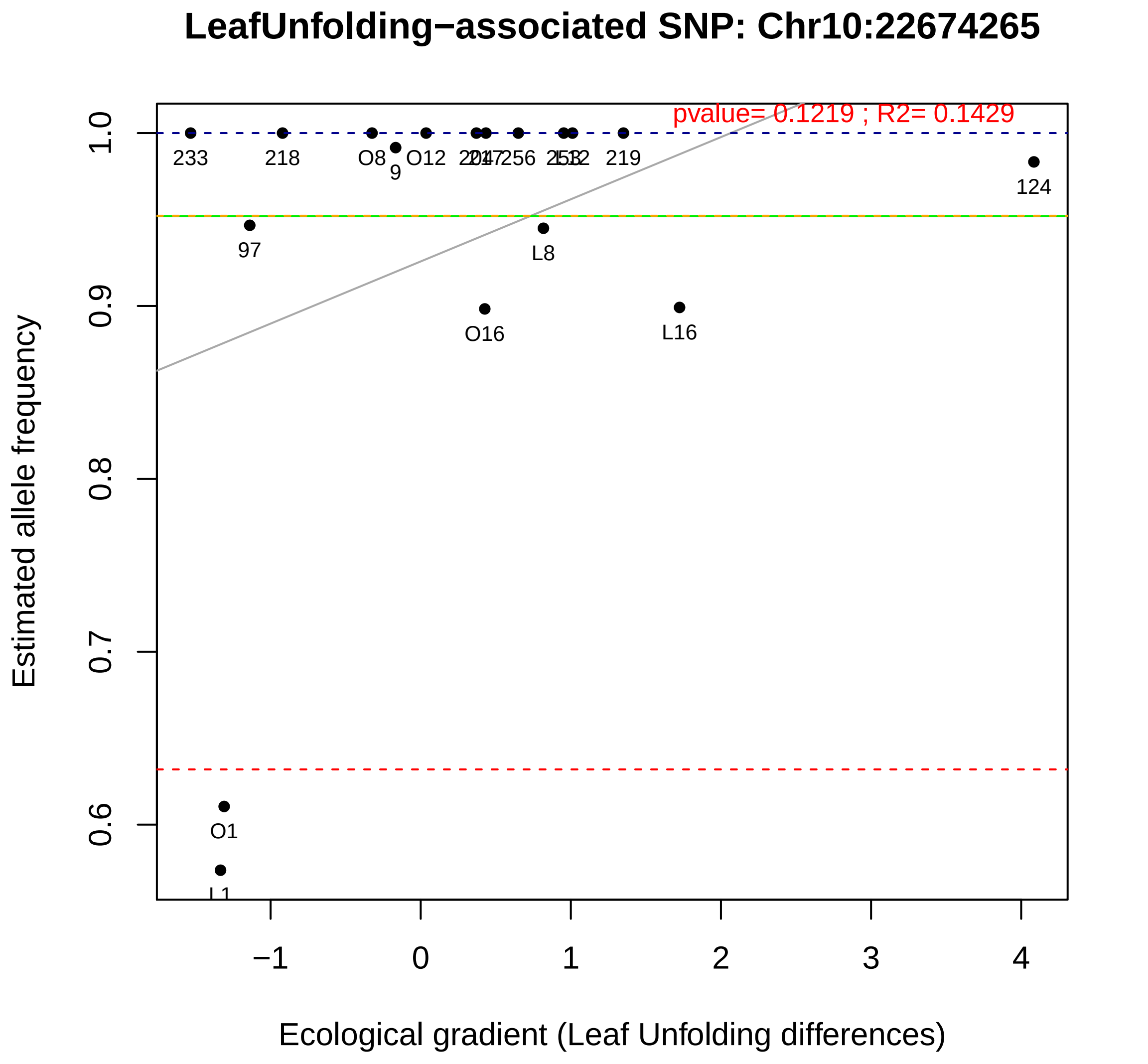

LeafUnfolding−associated SNP: Chr10:22674265
pvalue= 0.1219 ; R2= 0.1429
Estimated allele frequency
Ecological gradient (Leaf Unfolding differences)
1.0
0.9
0.8
0.7
0.6
−1
0
1
2
3
4
233
218
O8
9
O12
219
256
124
97
L8
O16
L16
O1

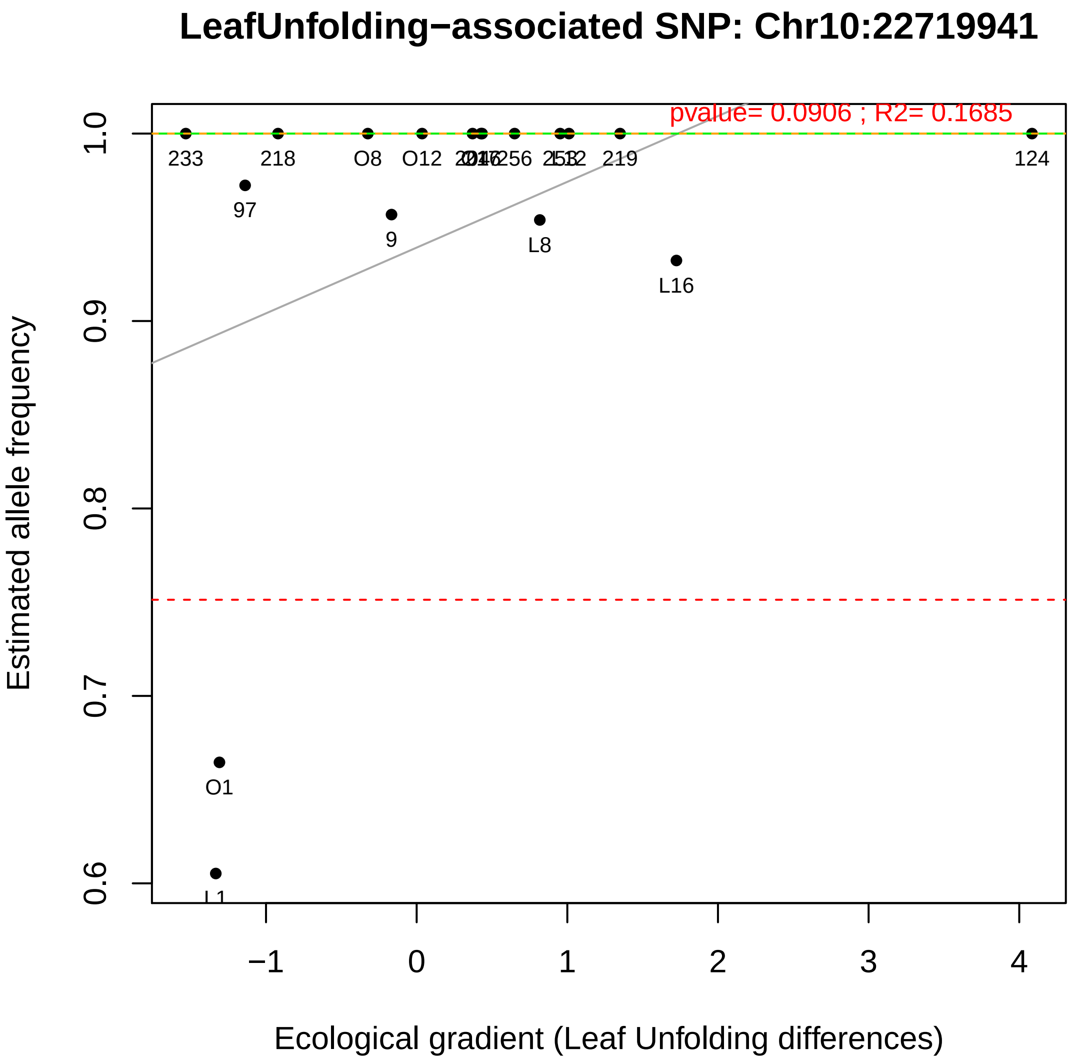

LeafUnfolding−associated SNP: Chr10:22719941
pvalue= 0.0906 ; R2= 0.1685
233
218
O8
O12
256
219
124
97
9
L8
L16
O1
Estimated allele frequency
0.6
0.7
0.8
0.9
1.0
−1
0
1
2
3
4
Ecological gradient (Leaf Unfolding differences)

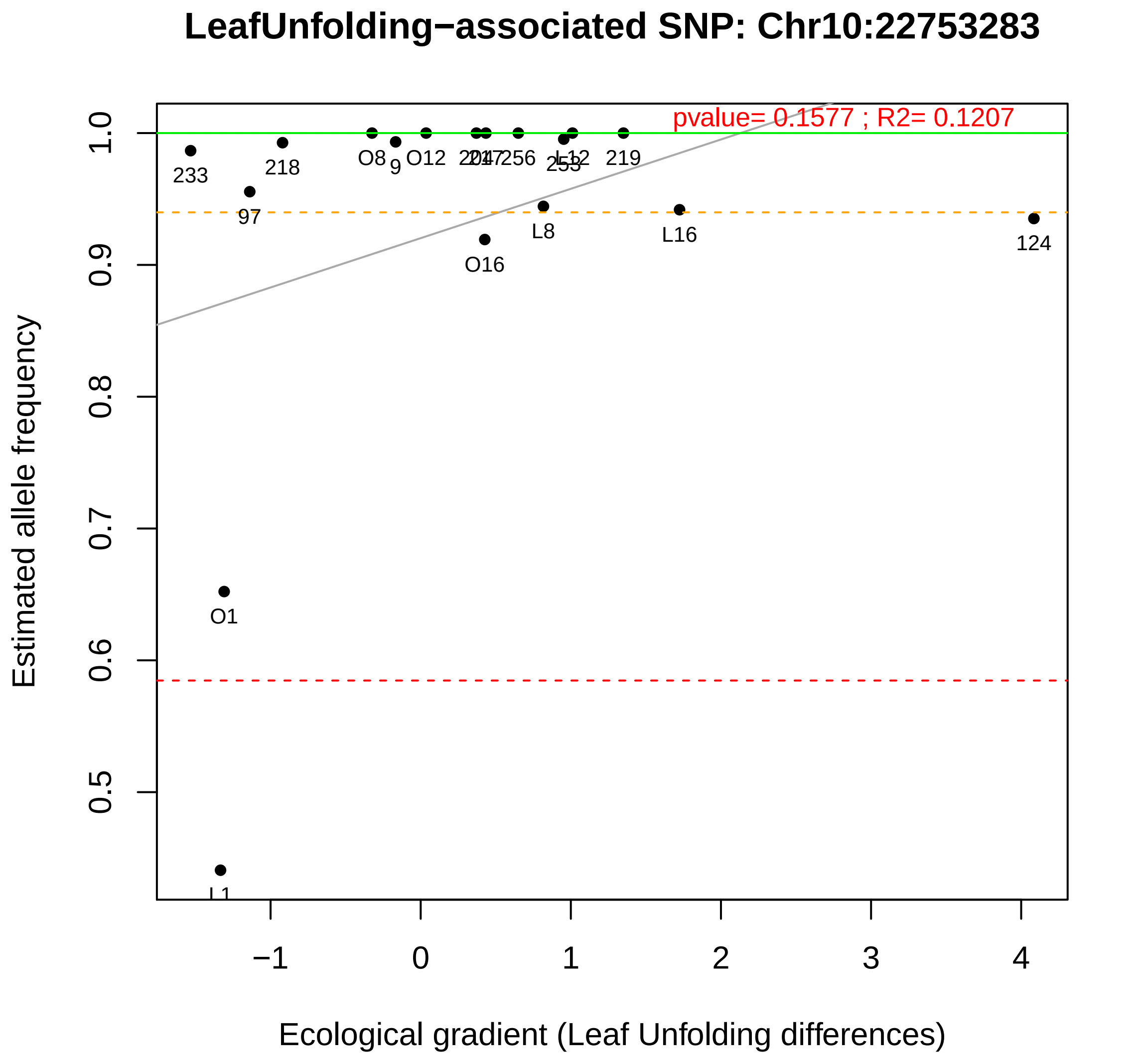

LeafUnfolding−associated SNP: Chr10:22753283
pvalue= 0.1577 ; R2= 0.1207
Estimated allele frequency
Ecological gradient (Leaf Unfolding differences)
1.0
0.9
0.8
0.7
0.6
0.5
−1
0
1
2
3
4
233
97
218
O8
9
O12
256
L12
253
219
L8
O16
L16
124
O1
L1

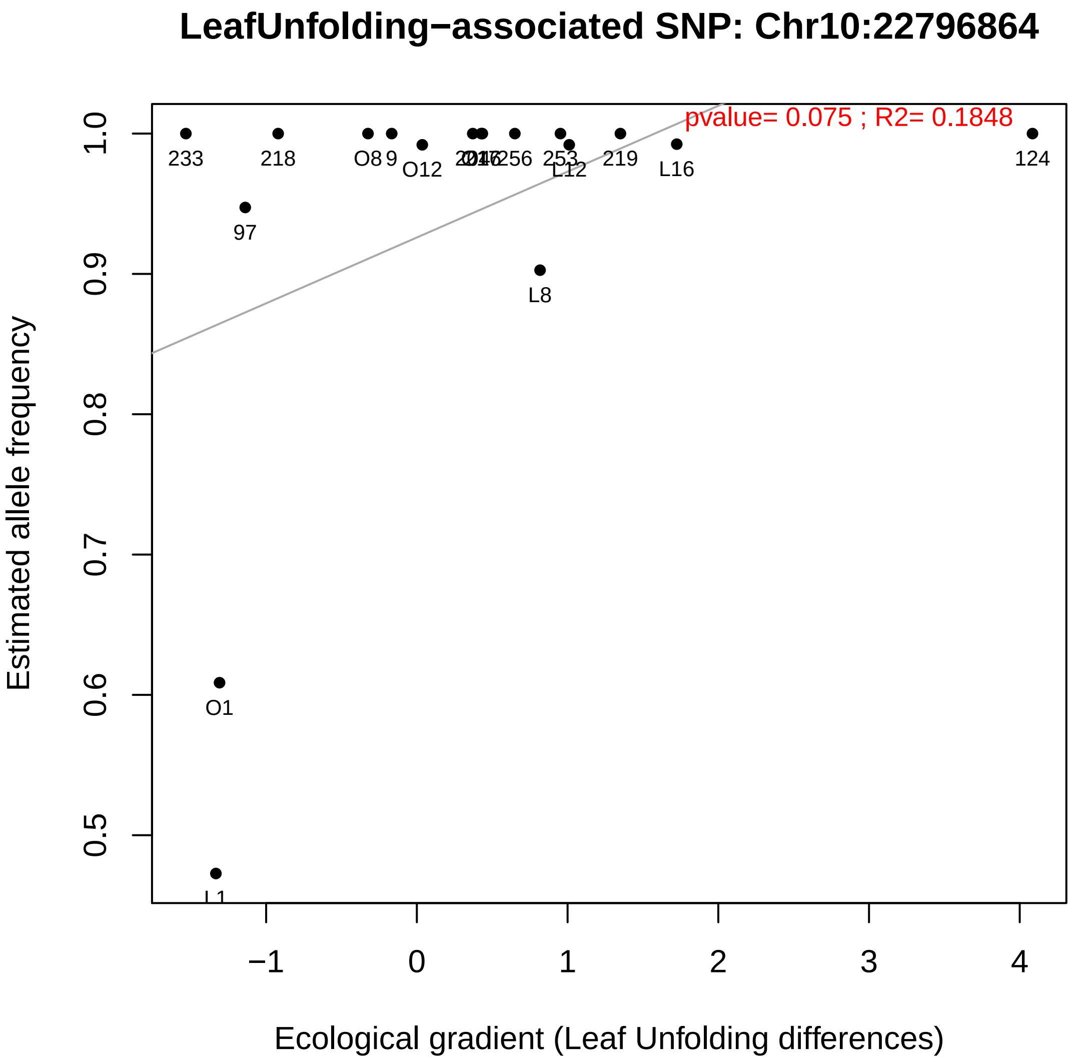

LeafUnfolding−associated SNP: Chr10:22796864
pvalue= 0.075 ; R2= 0.1848
Estimated allele frequency
Ecological gradient (Leaf Unfolding differences)
233
218
O8
9
O12
256
253
L12
219
L16
124
97
L8
O1

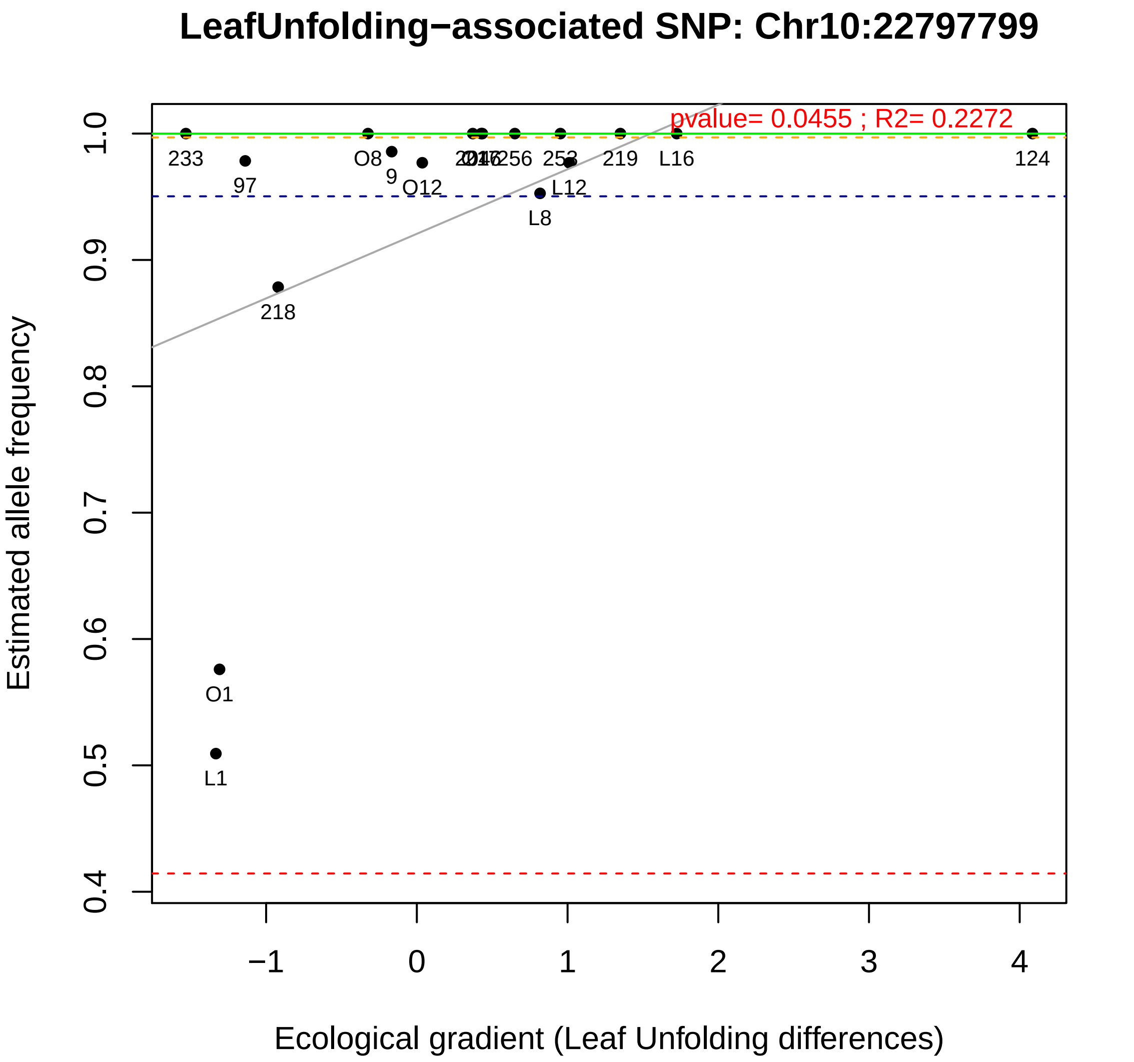

LeafUnfolding−associated SNP: Chr10:22797799
pvalue= 0.0455 ; R2= 0.2272
Estimated allele frequency
Ecological gradient (Leaf Unfolding differences)
1.0
0.9
0.8
0.7
0.6
0.5
0.4
−1
0
1
2
3
4
233
97
218
O8
9
O12
256
253
L12
L8
219
L16
124
O1
L1

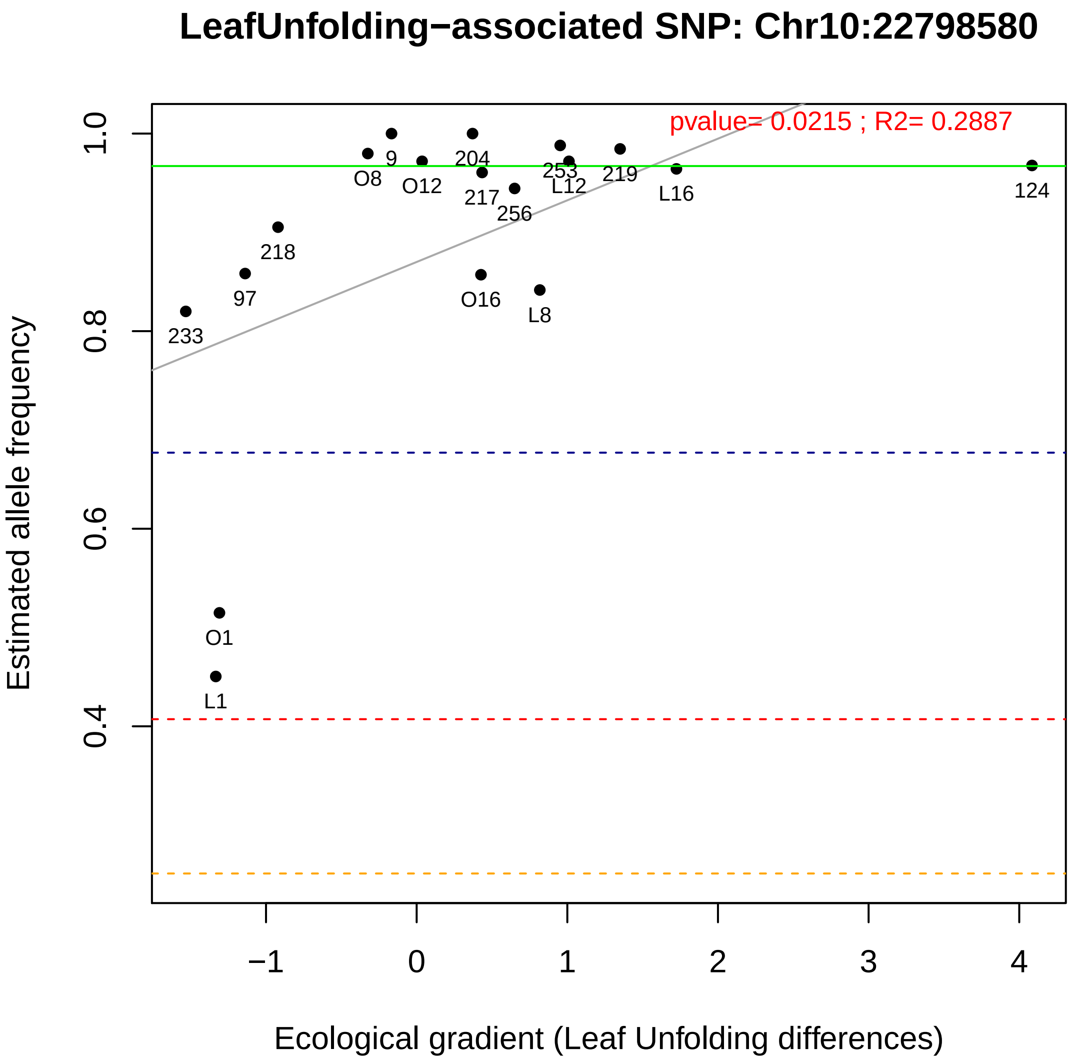
LeafUnfolding−associated SNP: Chr10:22798580
pvalue= 0.0215 ; R2= 0.2887
Estimated allele frequency
Ecological gradient (Leaf Unfolding differences)
1.0
0.8
0.6
0.4
−1
0
1
2
3
4
O8
9
O12
204
217
256
253
L12
219
L16
124
218
97
233
O16
L8
O1
L1

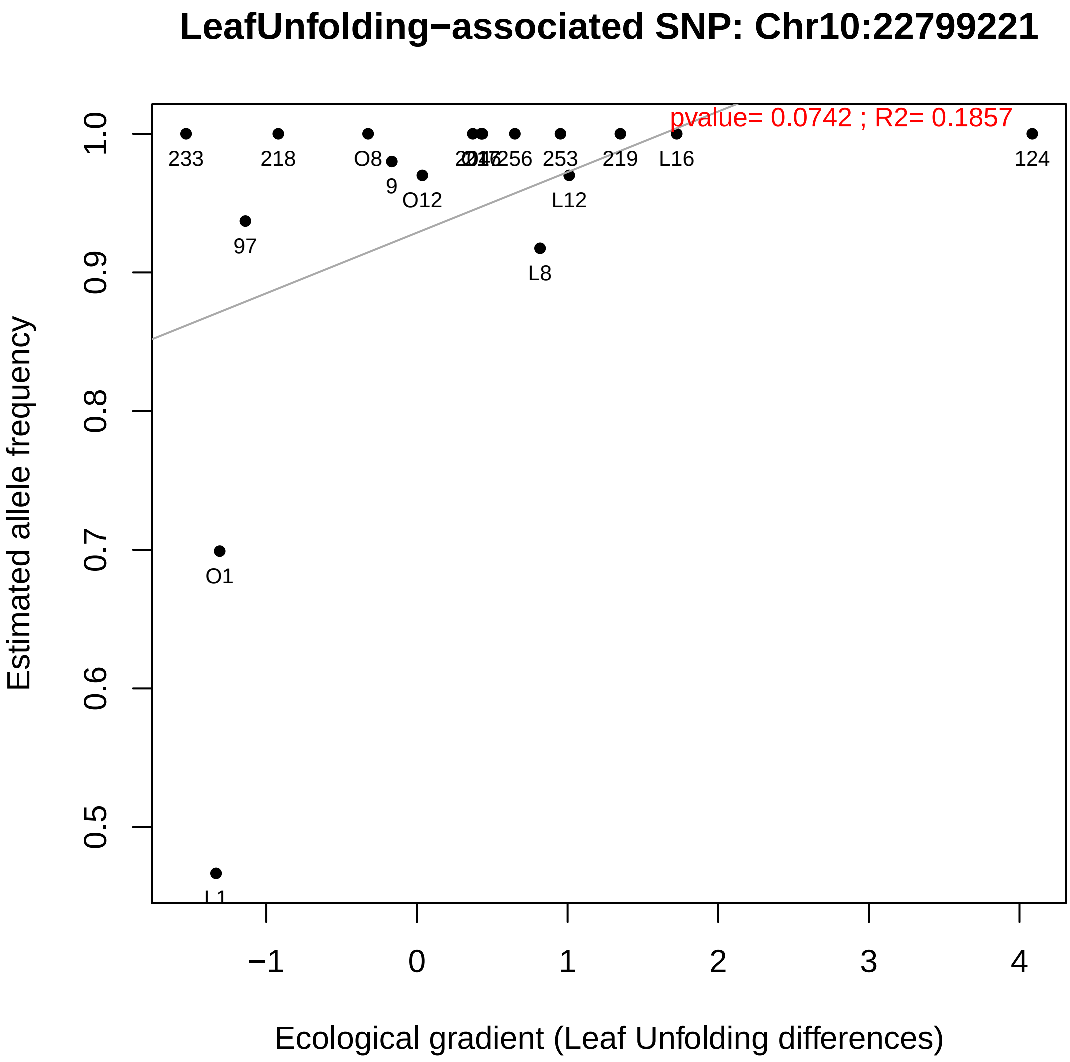

LeafUnfolding−associated SNP: Chr10:22799221
pvalue= 0.0742 ; R2= 0.1857
Estimated allele frequency
Ecological gradient (Leaf Unfolding differences)
1.0
0.9
0.8
0.7
0.6
0.5
−1
0
1
2
3
4
233
218
O8
9
O12
256
253
L12
219
L16
124
97
L8
O1

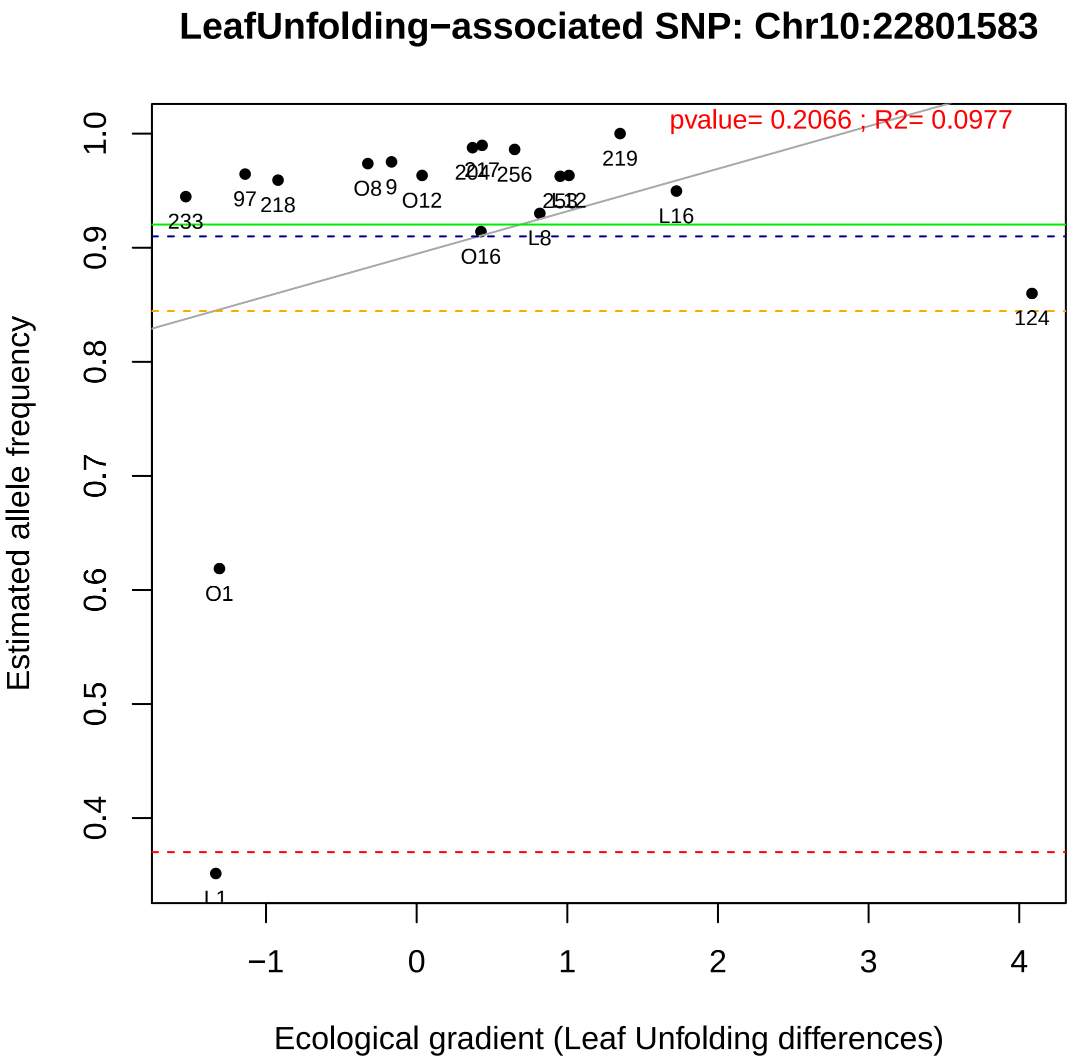
LeafUnfolding−associated SNP: Chr10:22801583
pvalue= 0.2066 ; R2= 0.0977
Estimated allele frequency
Ecological gradient (Leaf Unfolding differences)
1.0
0.9
0.8
0.7
0.6
0.5
0.4
−1
0
1
2
3
4
233
97
218
O8
9
O12
204
217
256
219
L16
O16
L8
124
O1

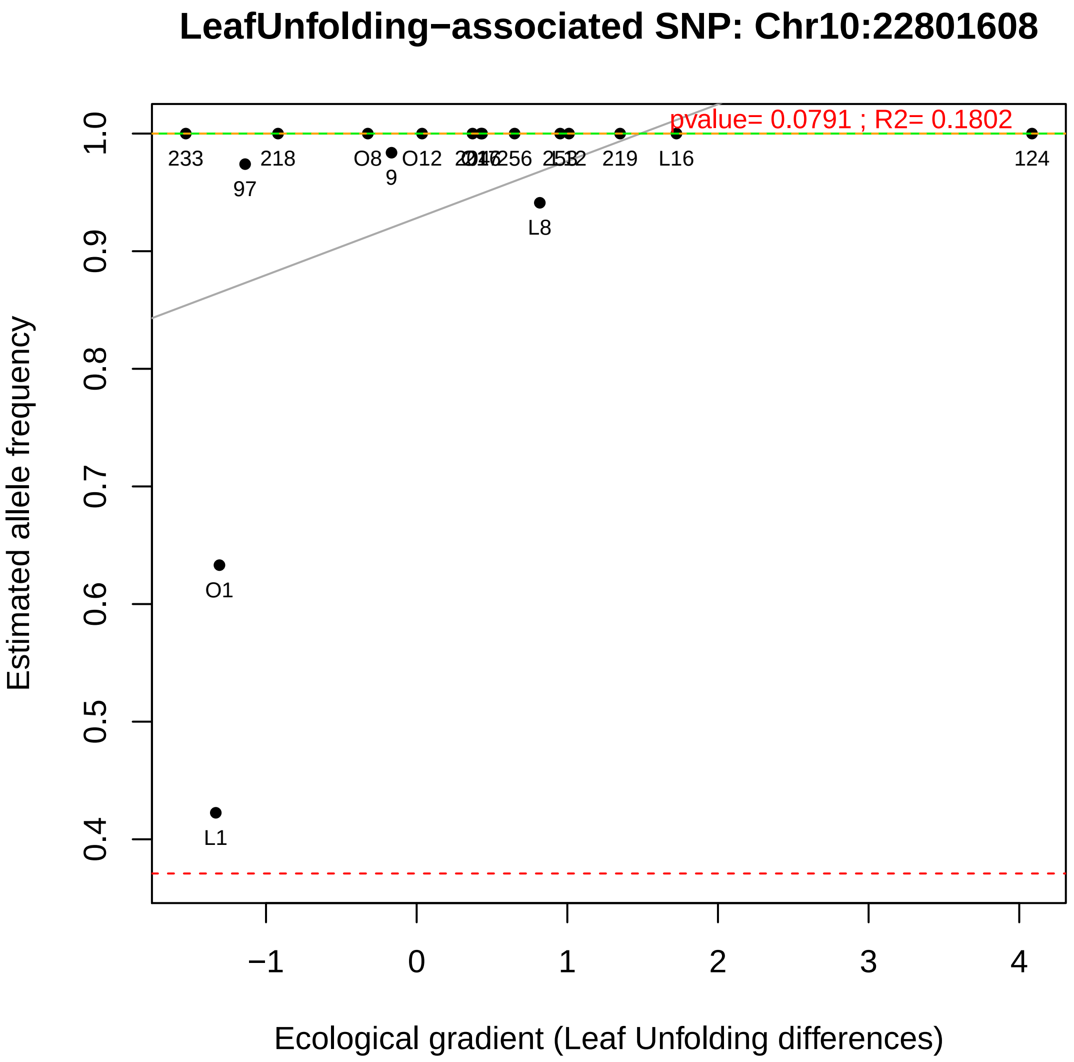

LeafUnfolding−associated SNP: Chr10:22801608
pvalue= 0.0791 ; R2= 0.1802
Estimated allele frequency
Ecological gradient (Leaf Unfolding differences)
233
97
218
O8
9
O12
L8
219
L16
124
O1
L1

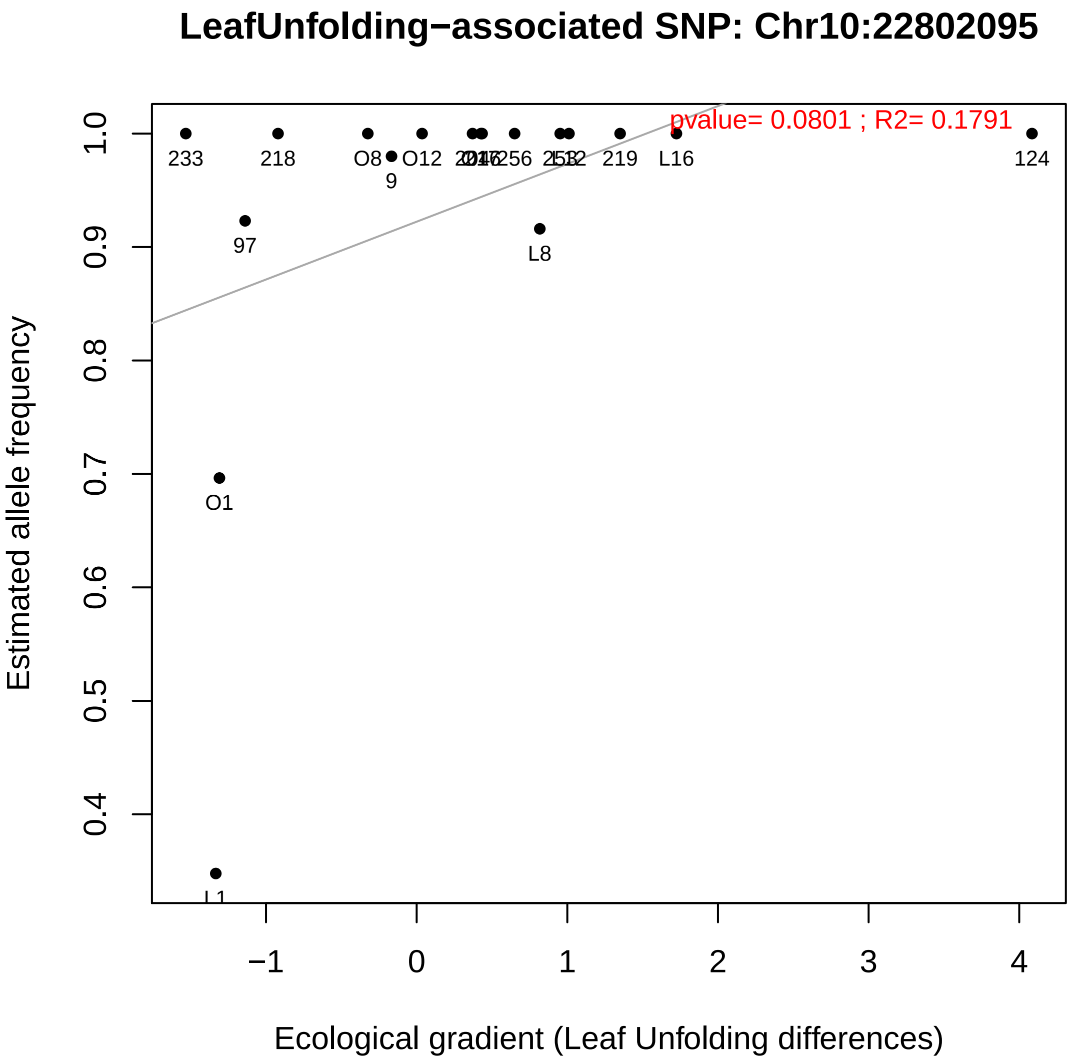
LeafUnfolding−associated SNP: Chr10:22802095
pvalue= 0.0801 ; R2= 0.1791
233
218
O8
9
O12
219
L16
124
97
L8
O1
Estimated allele frequency
Ecological gradient (Leaf Unfolding differences)
1.0
0.9
0.8
0.7
0.6
0.5
0.4
−1
0
1
2
3
4

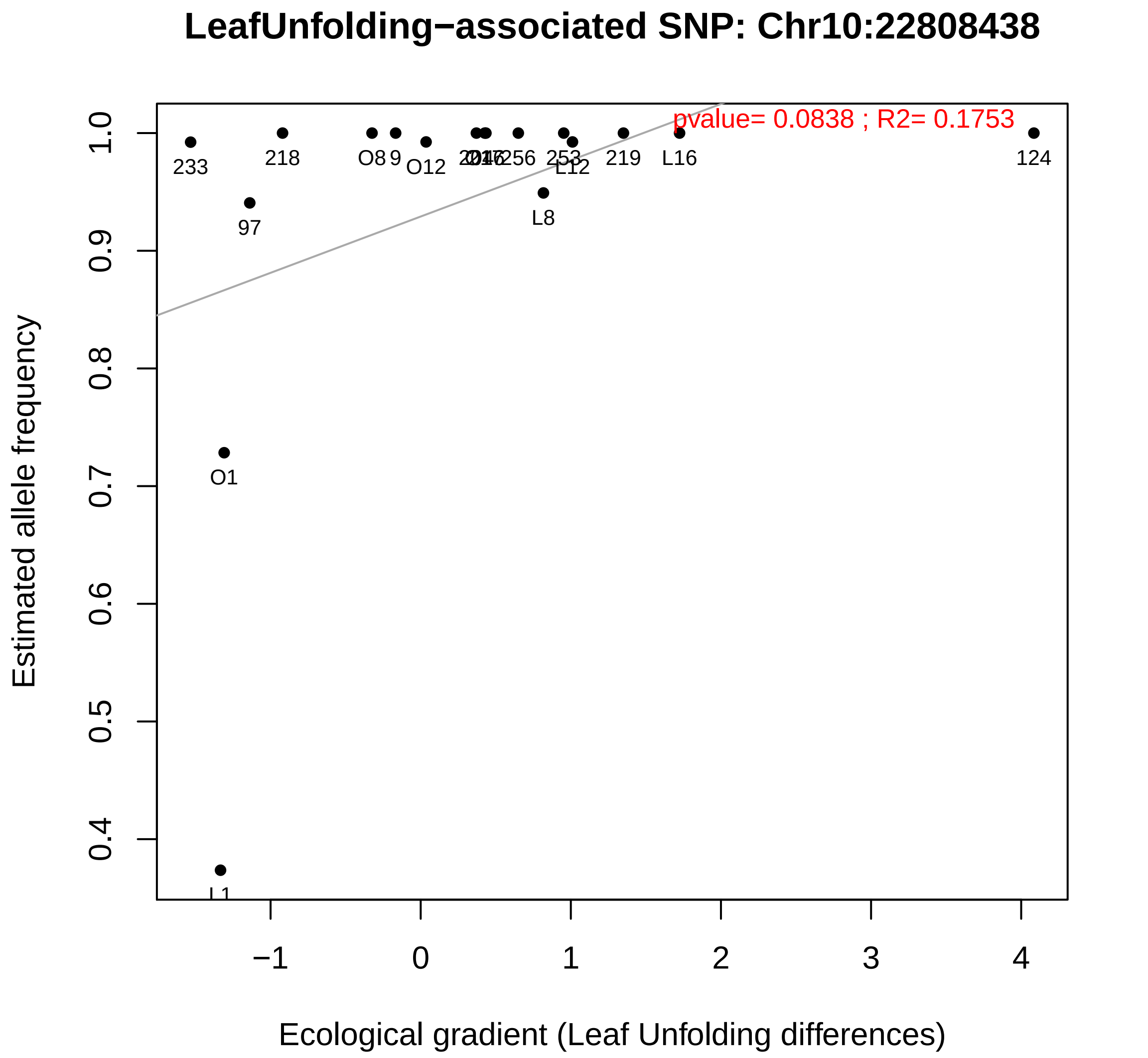

LeafUnfolding−associated SNP: Chr10:22808438
pvalue= 0.0838 ; R2= 0.1753
Estimated allele frequency
Ecological gradient (Leaf Unfolding differences)
1.0
0.9
0.8
0.7
0.6
0.5
0.4
−1
0
1
2
3
4
233
218
O8
9
O12
256
253
L12
219
L16
124
97
L8
O1
L1

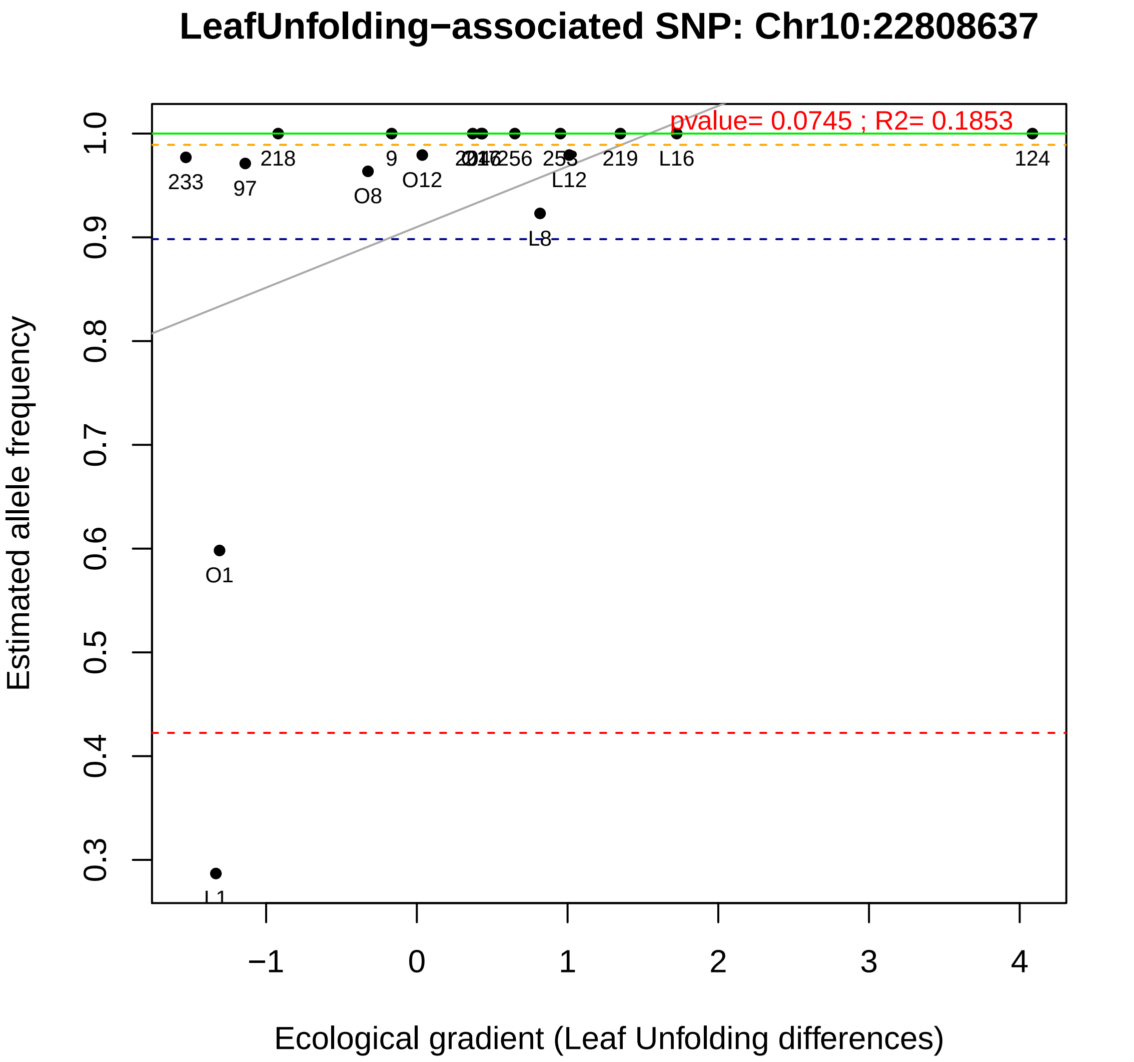

LeafUnfolding−associated SNP: Chr10:22808637
pvalue= 0.0745 ; R2= 0.1853
Estimated allele frequency
Ecological gradient (Leaf Unfolding differences)
1.0
0.9
0.8
0.7
0.6
0.5
0.4
0.3
−1
0
1
2
3
4
233
97
218
O8
9
O12
L8
L12
219
L16
124
O1
L1

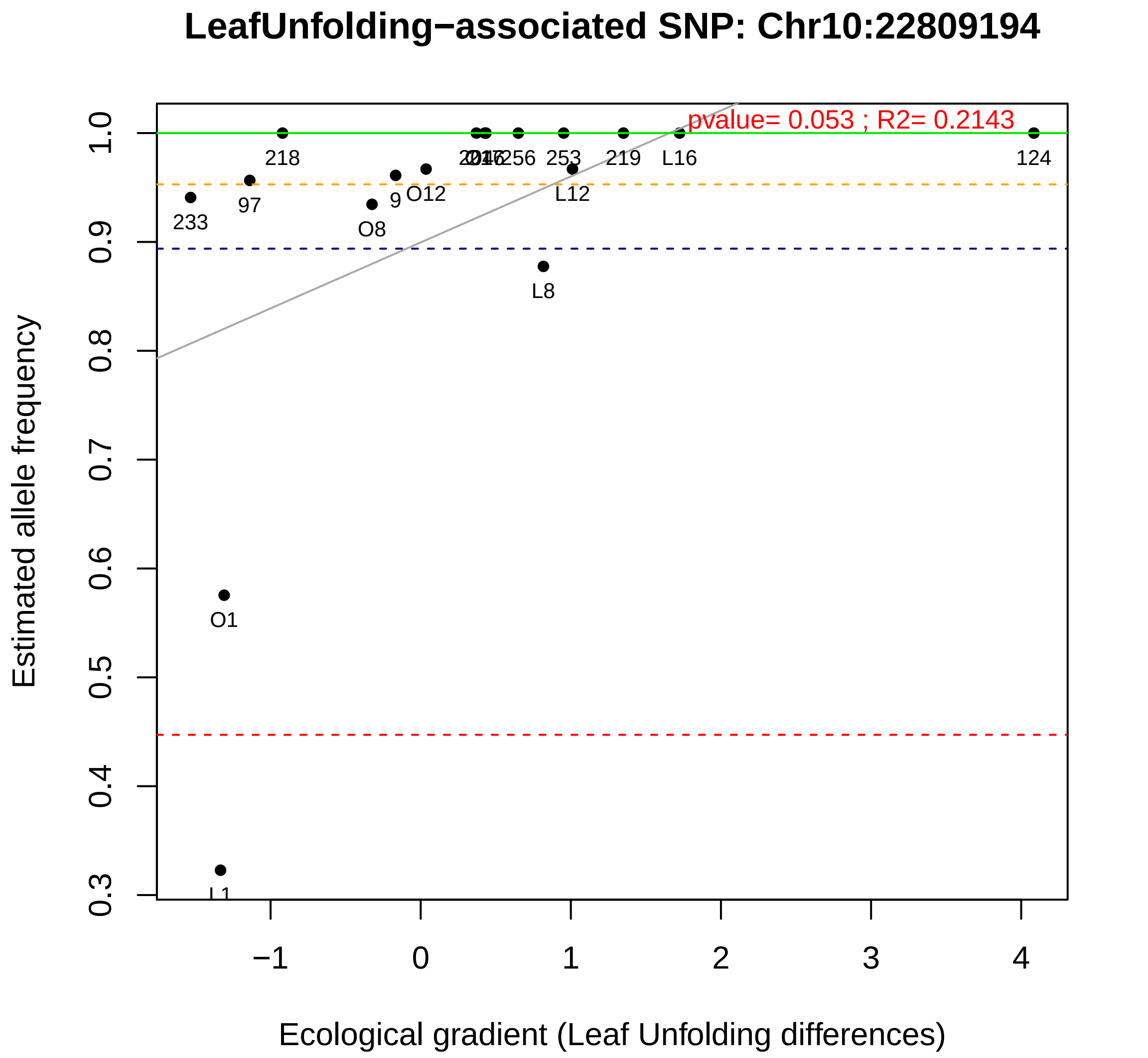

LeafUnfolding−associated SNP: Chr10:22809194
pvalue= 0.053 ; R2= 0.2143
Estimated allele frequency
Ecological gradient (Leaf Unfolding differences)
1.0
0.9
0.8
0.7
0.6
0.5
0.4
0.3
−1
0
1
2
3
4
218
233
97
O8
9
O12
256
253
219
L16
124
L12
L8
O1

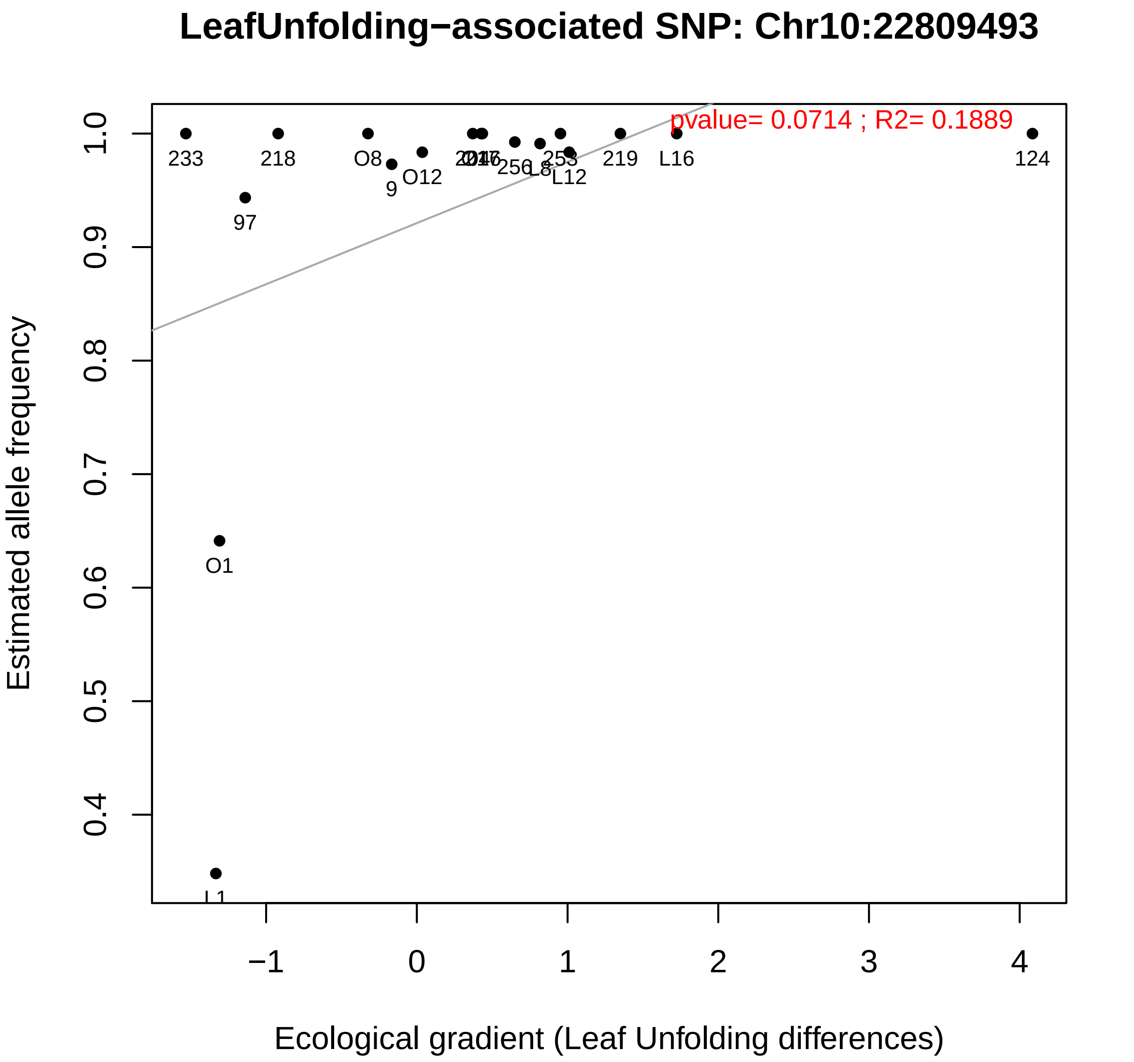

LeafUnfolding−associated SNP: Chr10:22809493
pvalue= 0.0714 ; R2= 0.1889
Estimated allele frequency
Ecological gradient (Leaf Unfolding differences)
0.4
0.5
0.6
0.7
0.8
0.9
1.0
−1
0
1
2
3
4
233
218
O8
9
O12
97
O1
219
L16
124
256
L8
253
L12
L1

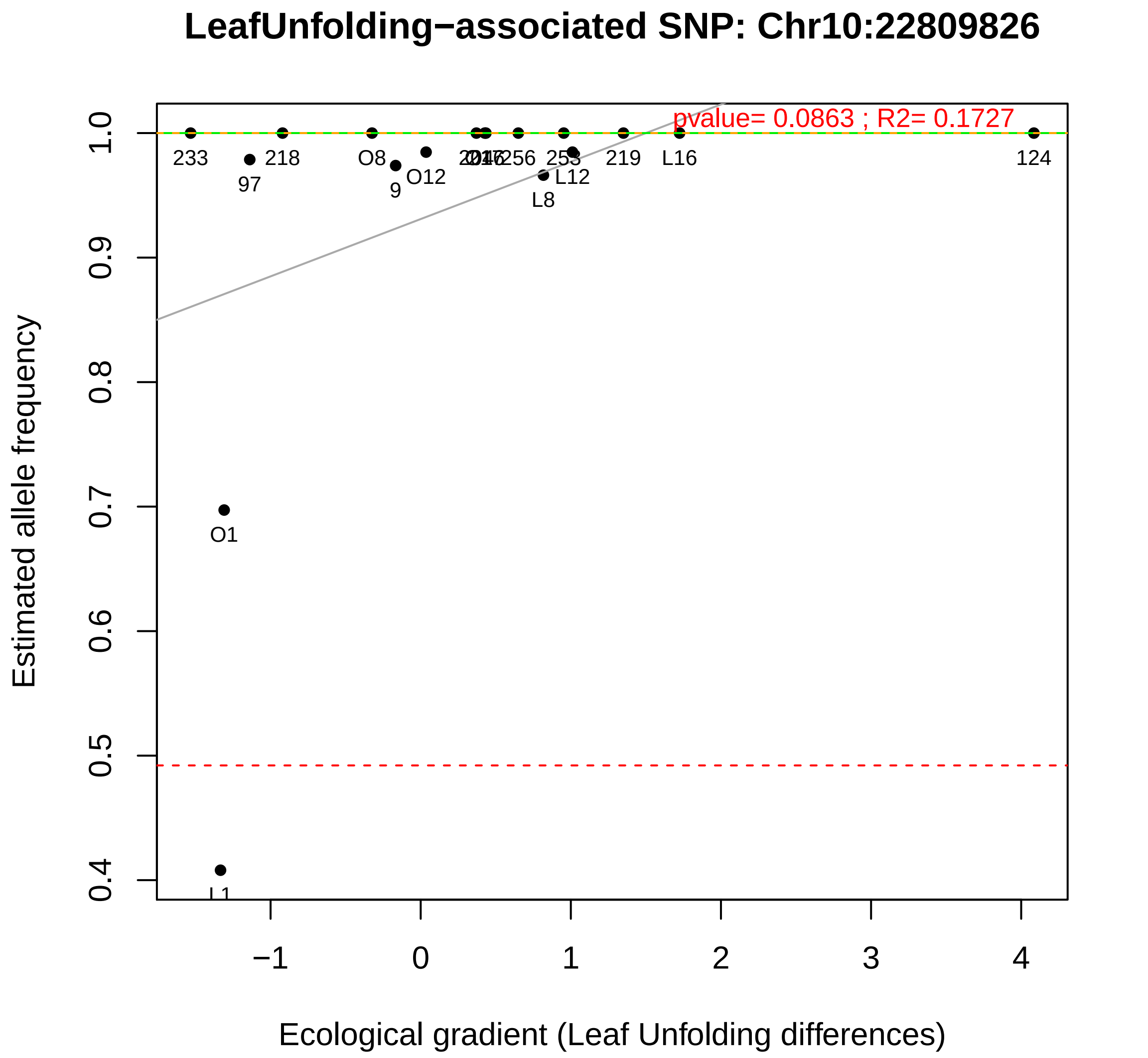

LeafUnfolding−associated SNP: Chr10:22809826
pvalue= 0.0863 ; R2= 0.1727
Estimated allele frequency
Ecological gradient (Leaf Unfolding differences)
233
97
218
O8
9
O12
253
L12
L8
219
L16
124
O1
0.4
0.5
0.6
0.7
0.8
0.9
1.0
−1
0
1
2
3
4

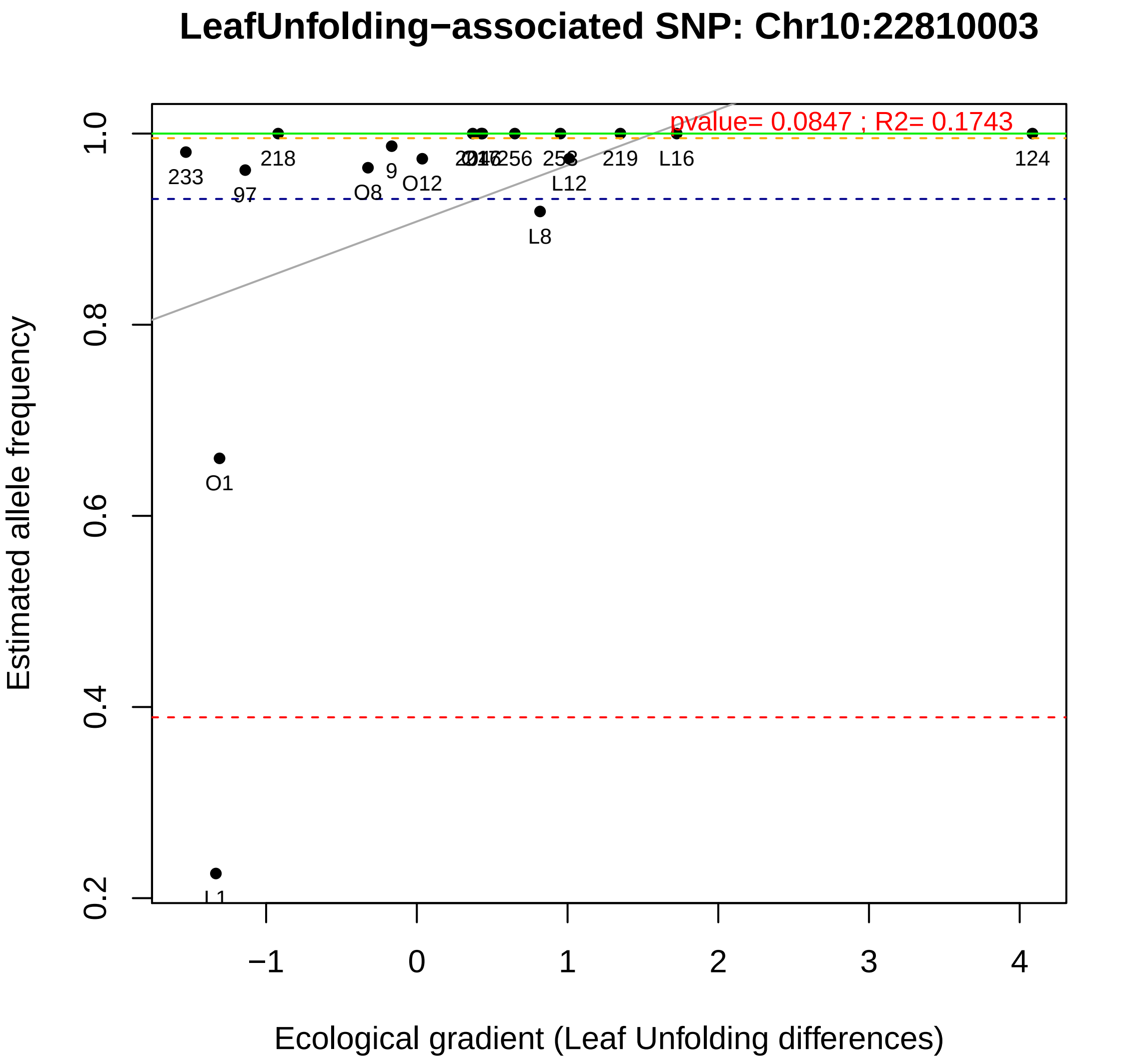
LeafUnfolding−associated SNP: Chr10:22810003
pvalue= 0.0847 ; R2= 0.1743
Estimated allele frequency
Ecological gradient (Leaf Unfolding differences)
0.2
0.4
0.6
0.8
1.0
−1
0
1
2
3
4
233
97
218
O8
9
O12
256
219
L16
124
L12
L8
O1

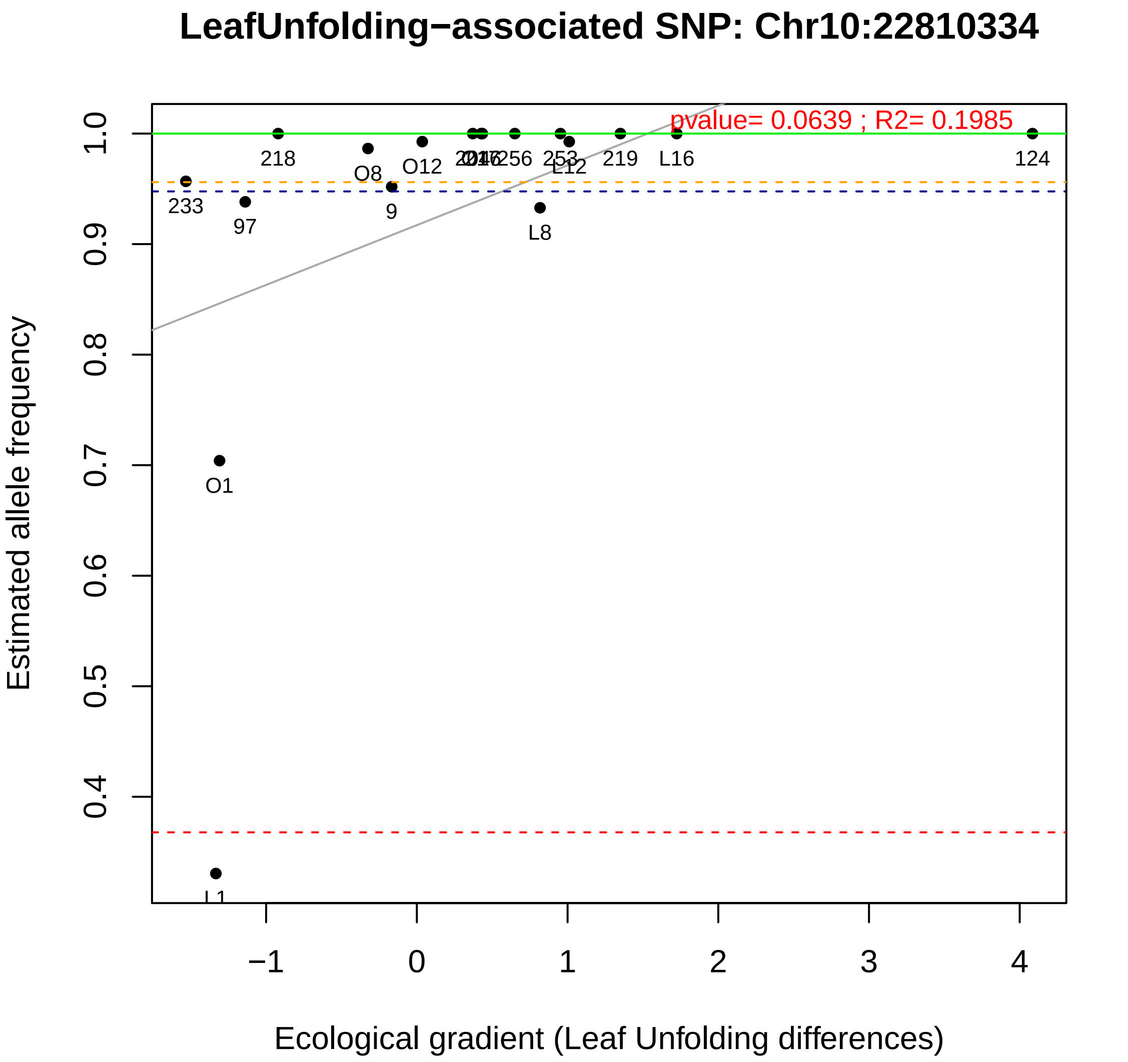

LeafUnfolding−associated SNP: Chr10:22810334
pvalue= 0.0639 ; R2= 0.1985
Estimated allele frequency
Ecological gradient (Leaf Unfolding differences)
1.0
0.9
0.8
0.7
0.6
0.5
0.4
−1
0
1
2
3
4
218
O8
O12
256
253
L12
219
L16
124
233
97
9
L8
O1
L1

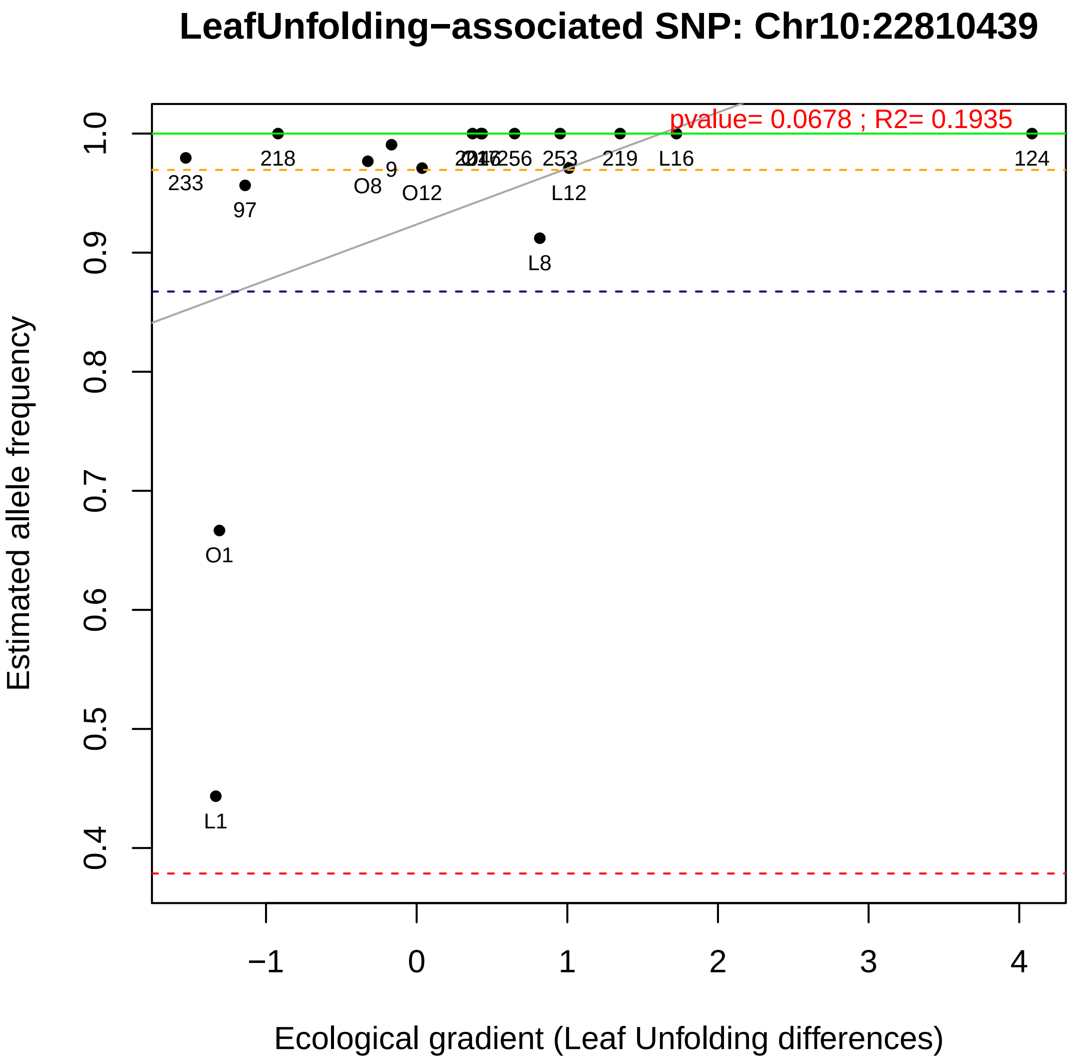

LeafUnfolding−associated SNP: Chr10:22810439
pvalue= 0.0678 ; R2= 0.1935
Estimated allele frequency
Ecological gradient (Leaf Unfolding differences)
1.0
0.9
0.8
0.7
0.6
0.5
0.4
−1
0
1
2
3
4
233
97
218
O8
9
O12
256
253
L12
219
L16
124
L8
O1
L1

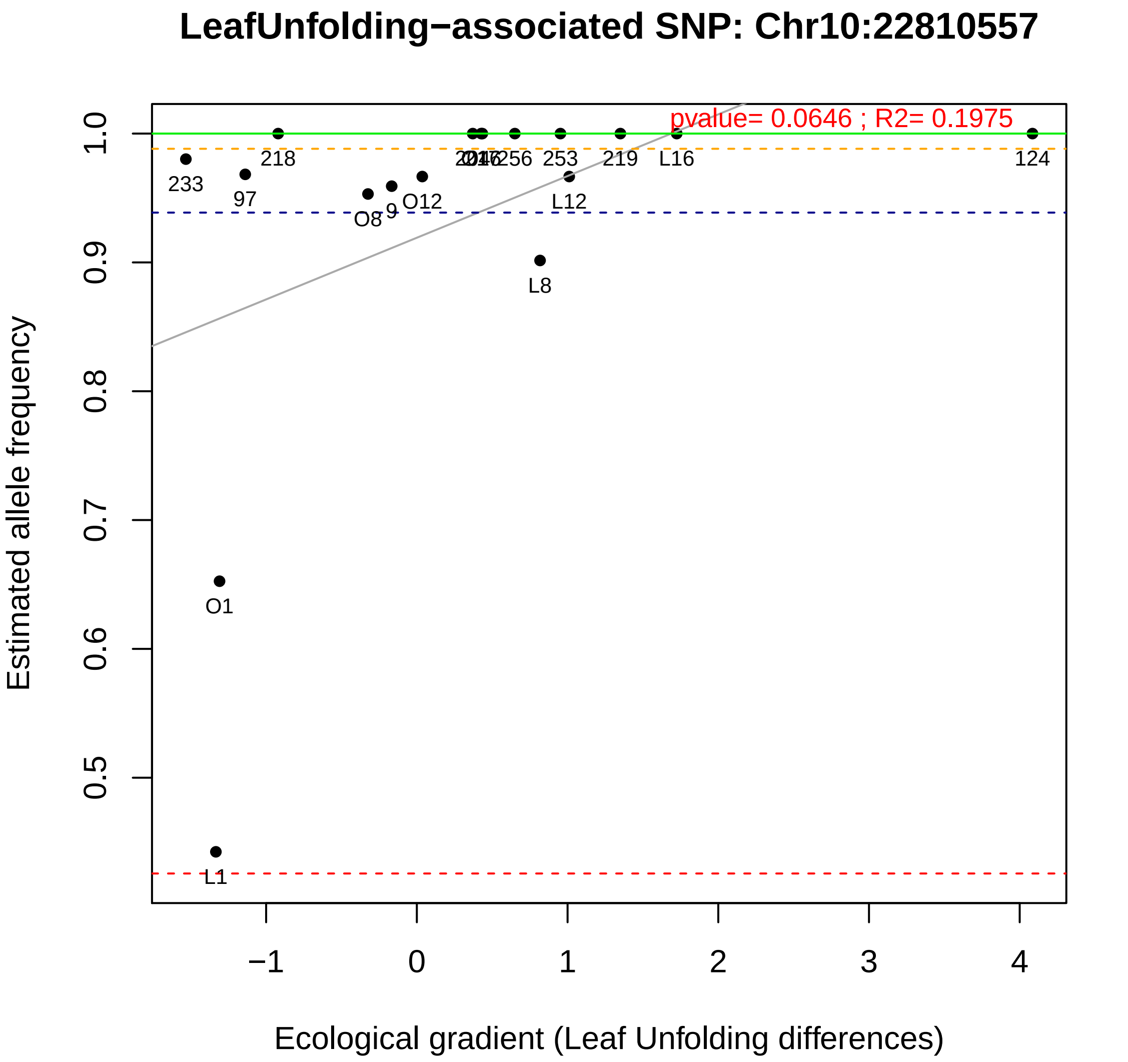
LeafUnfolding−associated SNP: Chr10:22810557
pvalue= 0.0646 ; R2= 0.1975
Estimated allele frequency
Ecological gradient (Leaf Unfolding differences)
1.0
0.9
0.8
0.7
0.6
0.5
−1
0
1
2
3
4
218
233
97
O8
9
O12
256
253
219
L16
124
L12
L8
O1
L1

**LeafUnfolding−associated SNP: Chr10:22810590**

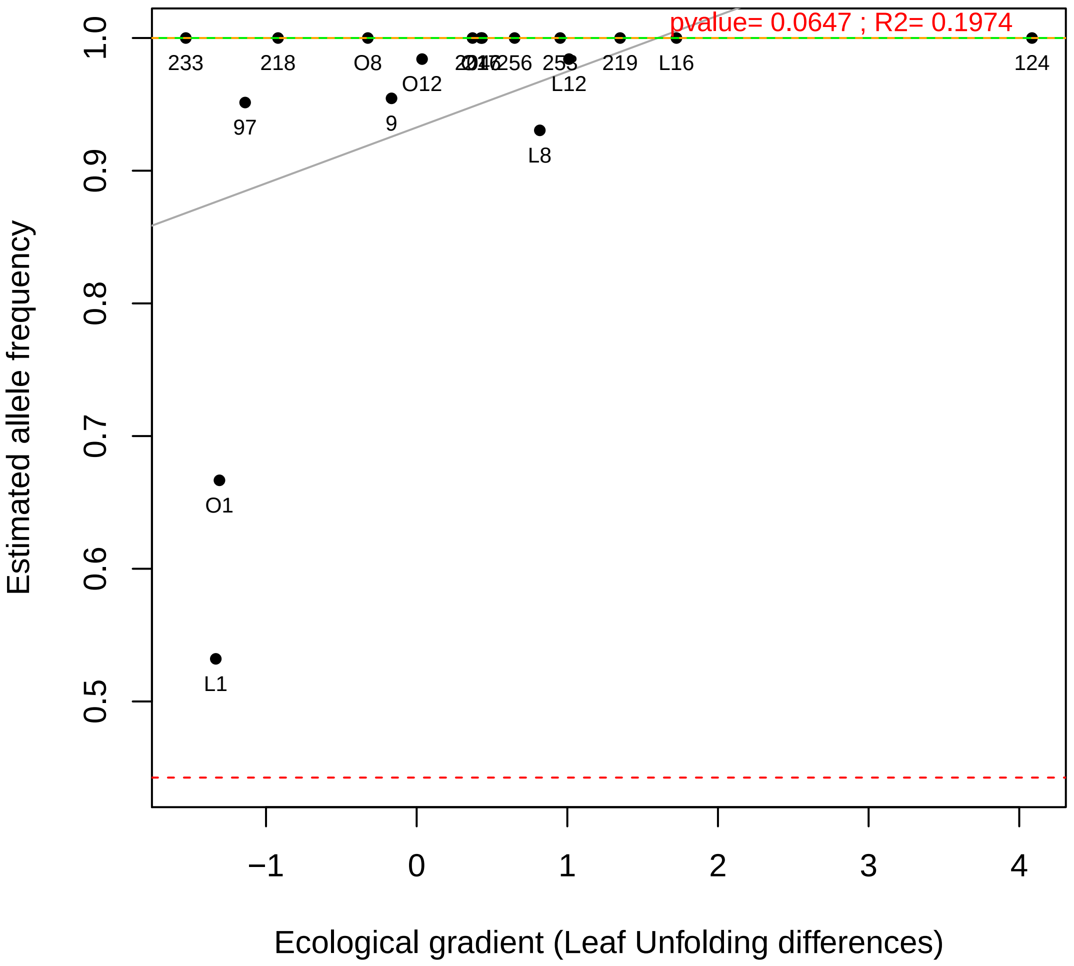

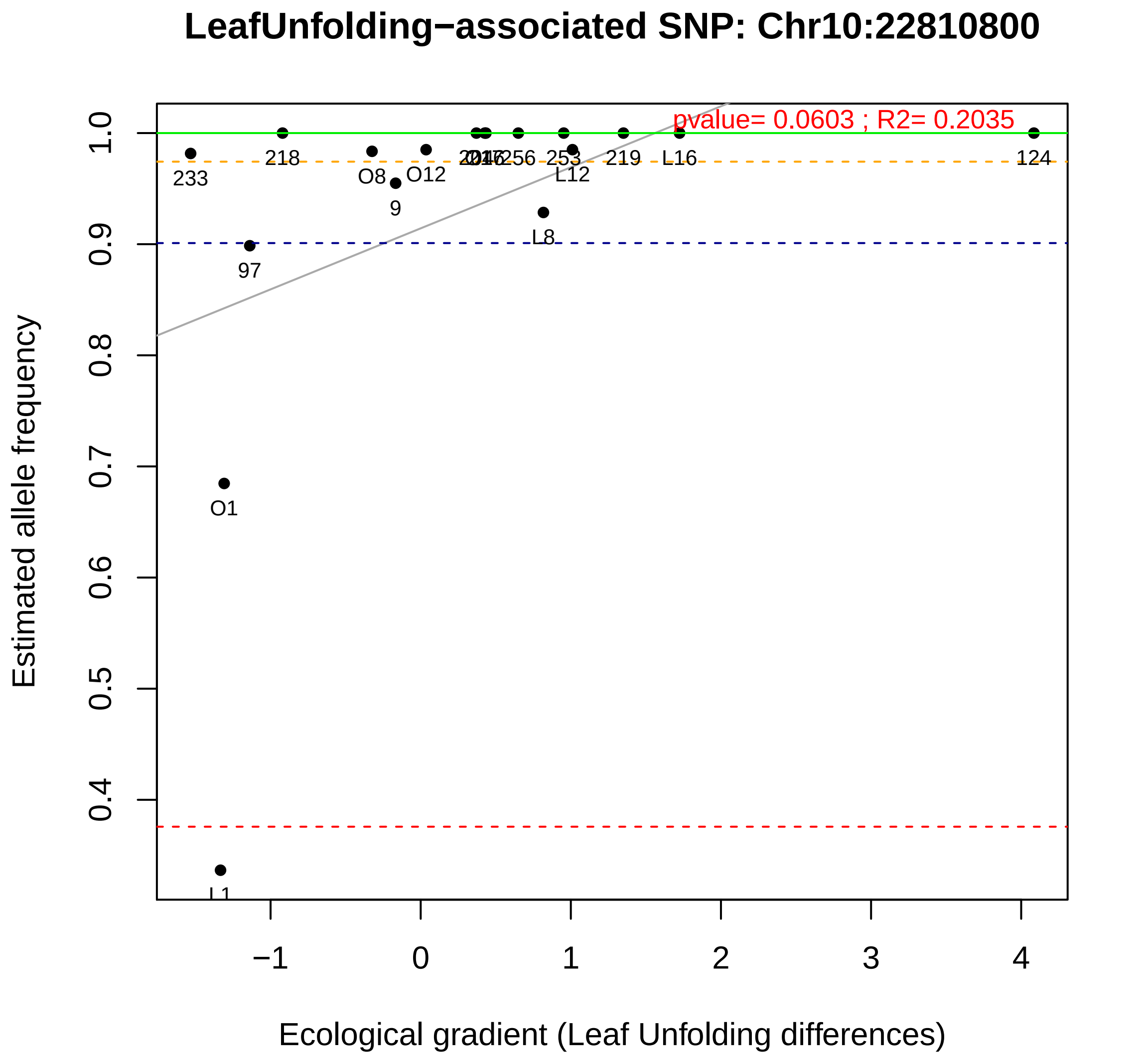
LeafUnfolding−associated SNP: Chr10:22810800
pvalue= 0.0603 ; R2= 0.2035
Estimated allele frequency
Ecological gradient (Leaf Unfolding differences)
1.0
0.9
0.8
0.7
0.6
0.5
0.4
−1
0
1
2
3
4
233
218
O8
O12
9
97
O1
L8
L12
253
219
L16
124
256
L1

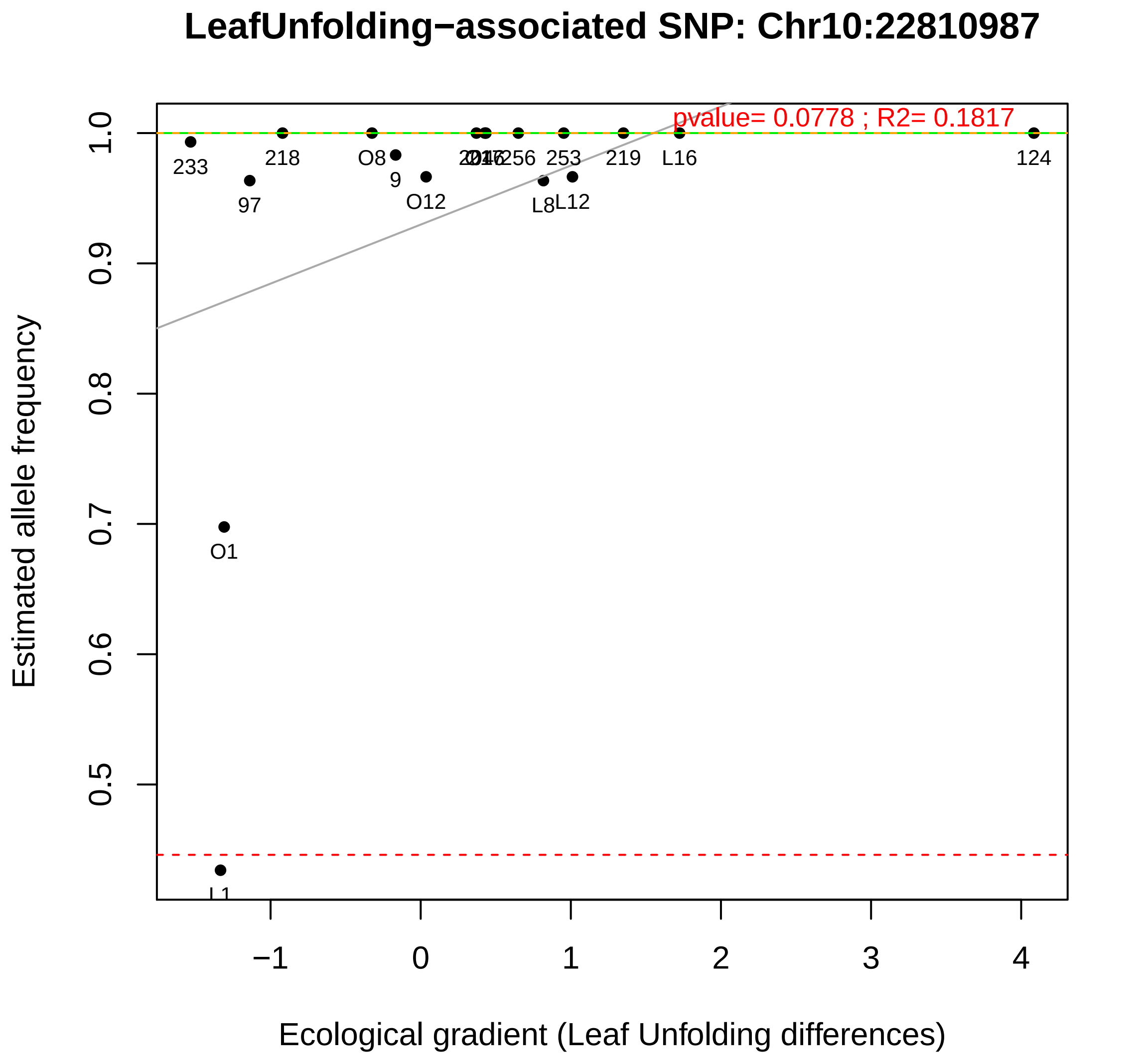

LeafUnfolding−associated SNP: Chr10:22810987
pvalue= 0.0778 ; R2= 0.1817
Estimated allele frequency
Ecological gradient (Leaf Unfolding differences)
0.5
0.6
0.7
0.8
0.9
1.0
−1
0
1
2
3
4
233
218
O8
9
97
O12
256
253
219
L16
124
L8
L12
O1

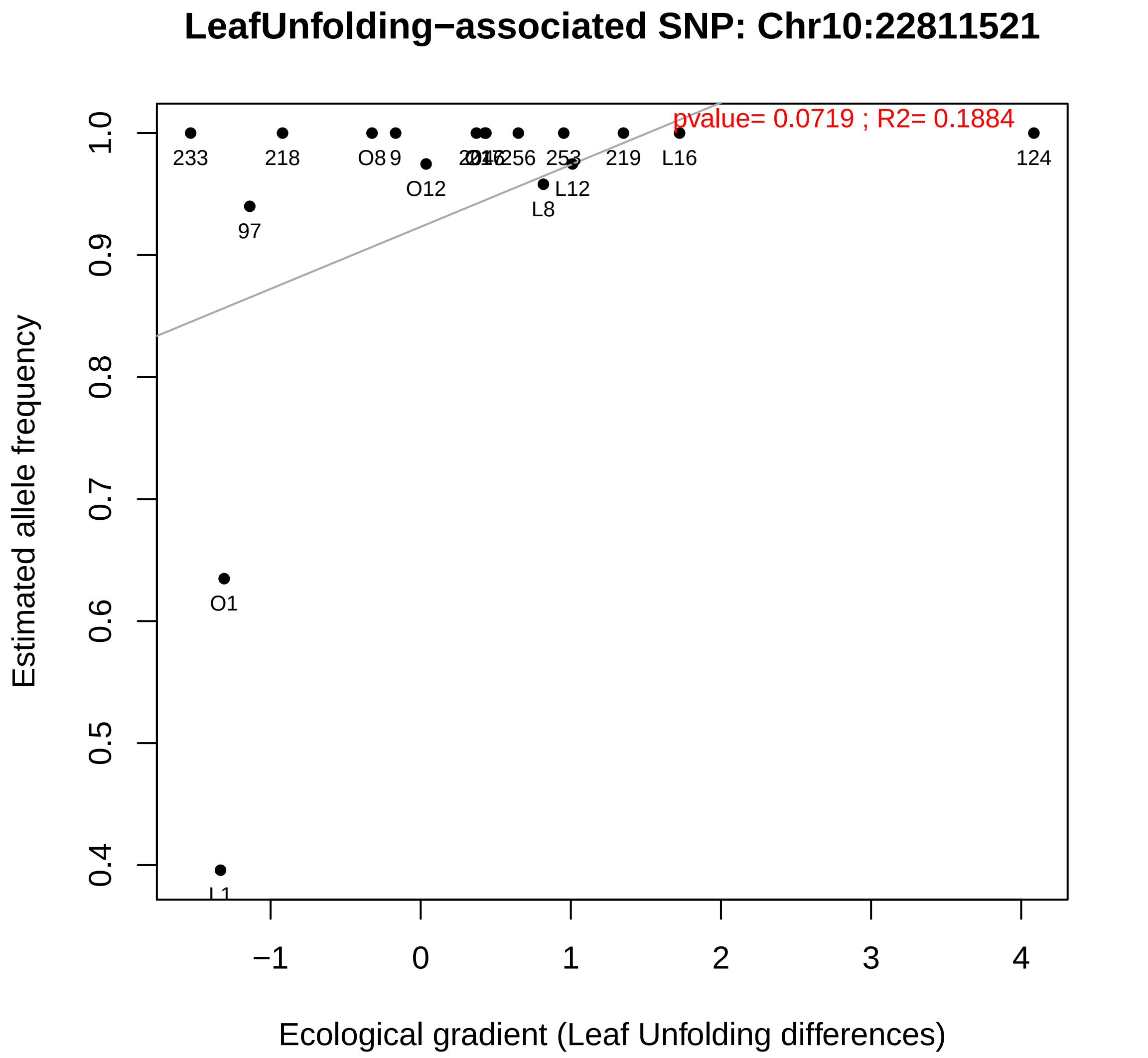
LeafUnfolding−associated SNP: Chr10:22811521
pvalue= 0.0719 ; R2= 0.1884
Estimated allele frequency
Ecological gradient (Leaf Unfolding differences)
233
218
O8
9
O12
256
253
L12
L8
219
L16
124
97
O1
0.4
0.5
0.6
0.7
0.8
0.9
1.0
−1
0
1
2
3
4

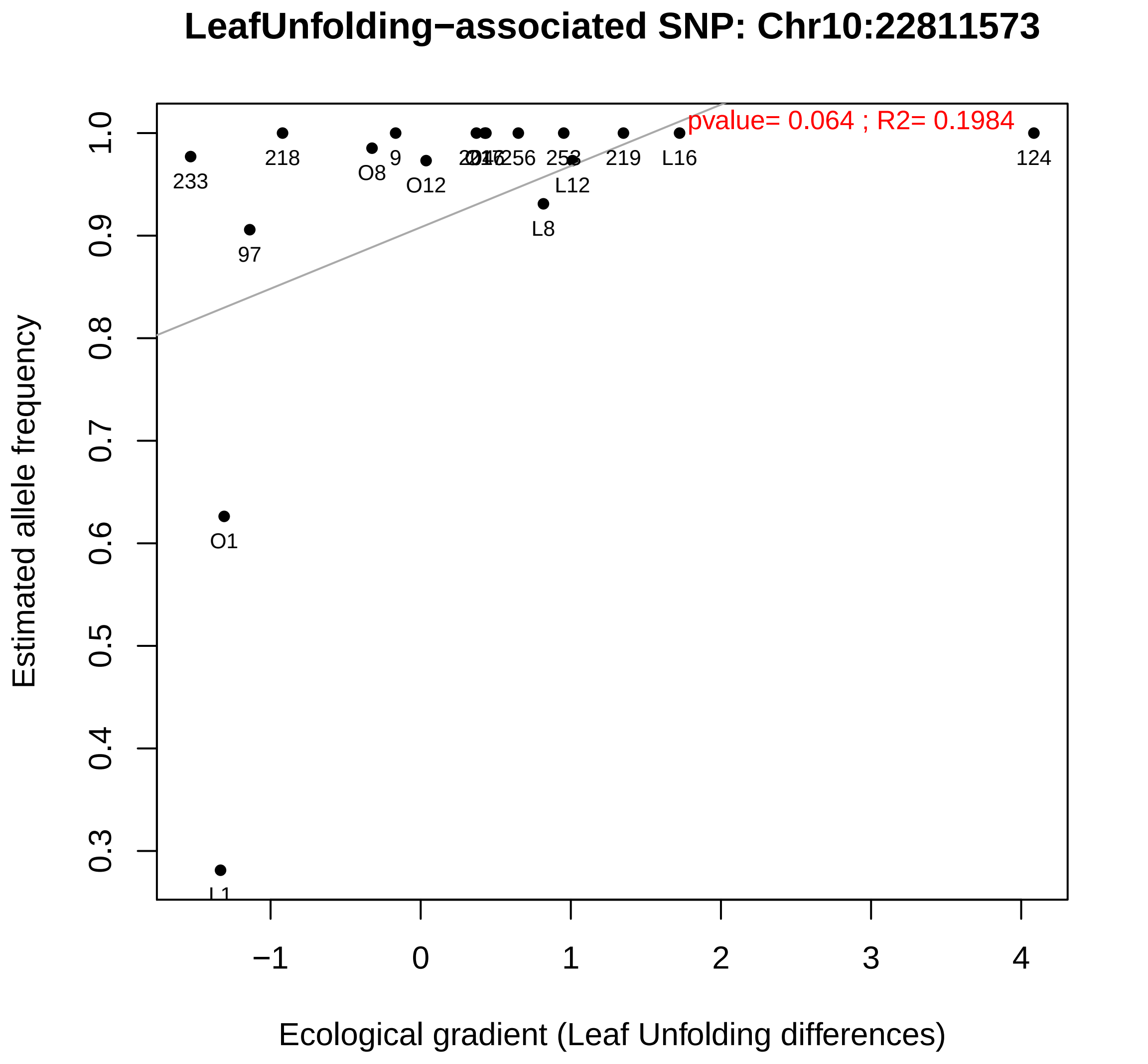

LeafUnfolding−associated SNP: Chr10:22811573
pvalue= 0.064 ; R2= 0.1984
Estimated allele frequency
Ecological gradient (Leaf Unfolding differences)
1.0
0.9
0.8
0.7
0.6
0.5
0.4
0.3
−1
0
1
2
3
4
233
218
O8
9
O12
97
L8
L12
219
L16
124
O1
L1

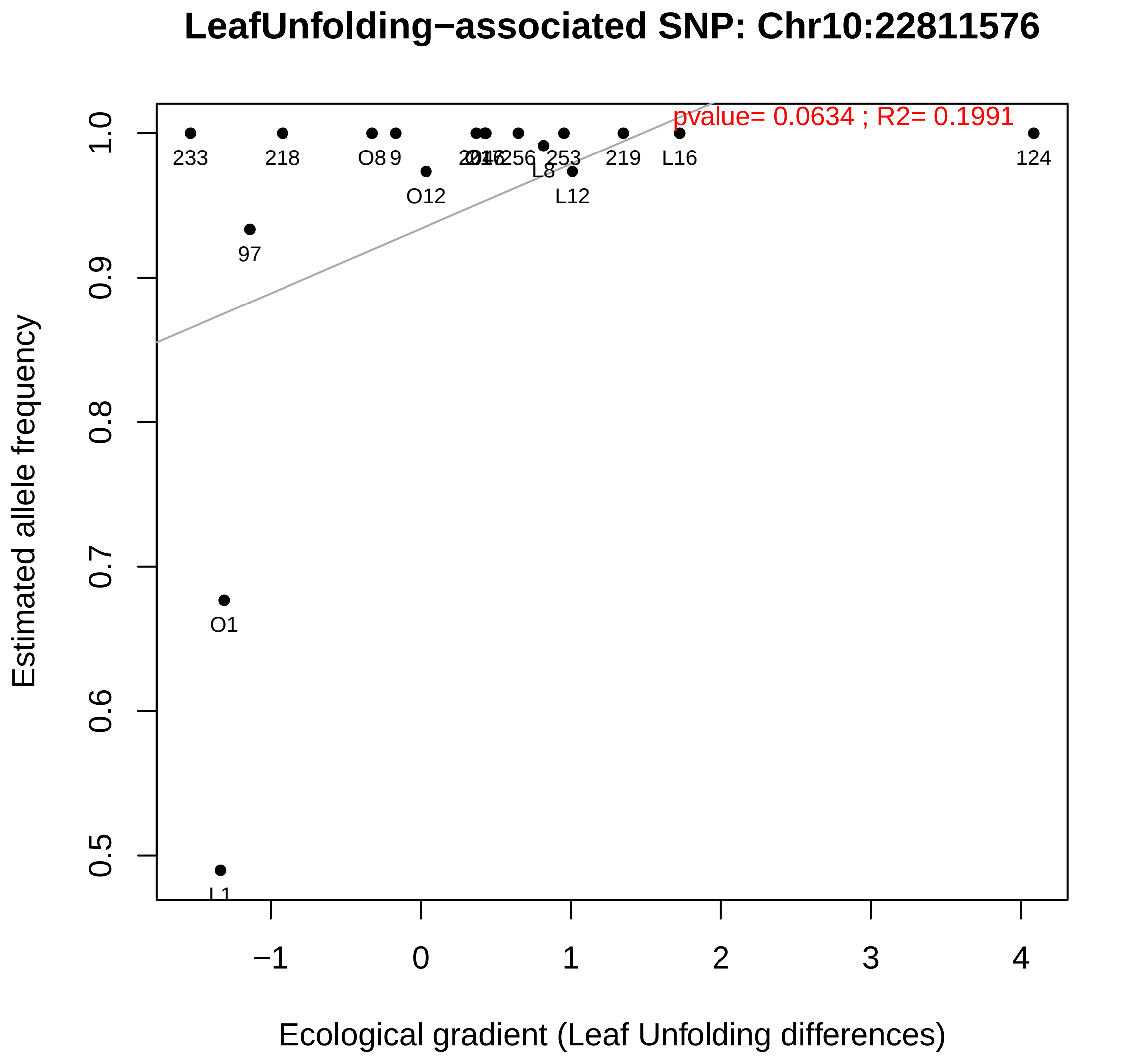
LeafUnfolding−associated SNP: Chr10:22811576
pvalue= 0.0634 ; R2= 0.1991
Estimated allele frequency
Ecological gradient (Leaf Unfolding differences)
0.5
0.6
0.7
0.8
0.9
1.0
−1
0
1
2
3
4
233
218
O8
9
O12
256
253
L8
L12
219
L16
124
97
O1
L1

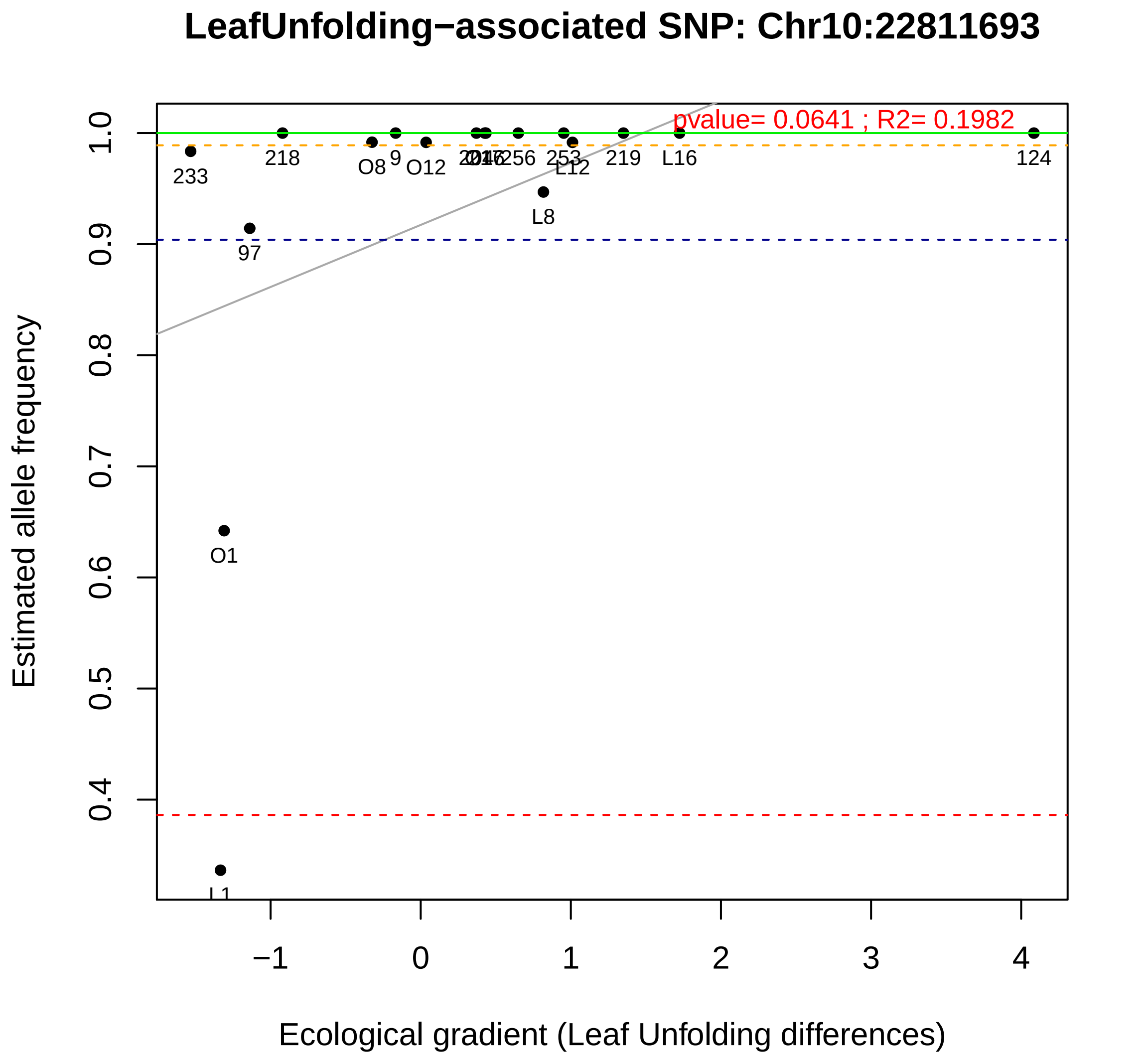

LeafUnfolding−associated SNP: Chr10:22811693
pvalue= 0.0641 ; R2= 0.1982
Estimated allele frequency
Ecological gradient (Leaf Unfolding differences)
1.0
0.9
0.8
0.7
0.6
0.5
0.4
−1
0
1
2
3
4
233
218
O8
9
O12
256
253
L12
219
L16
124
97
L8
O1

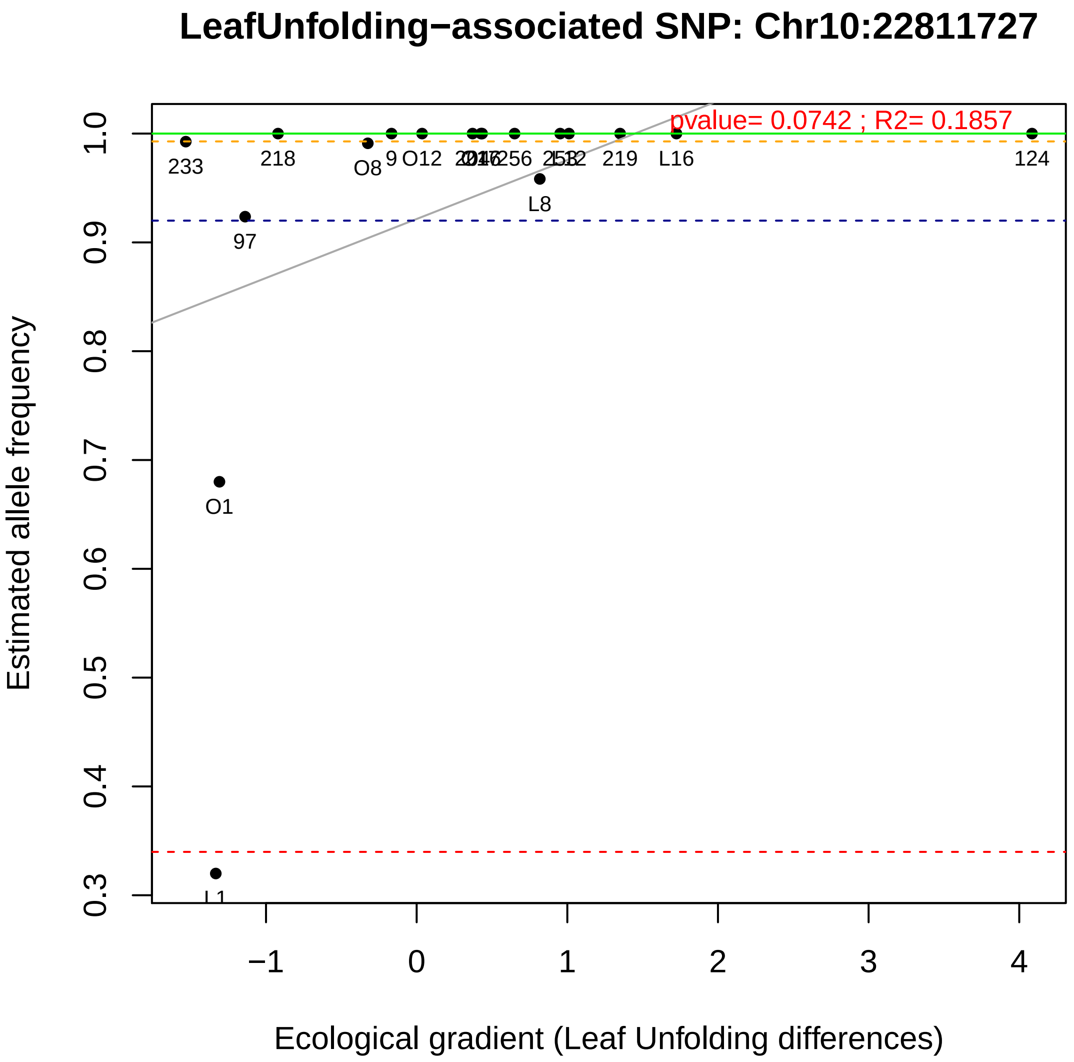

LeafUnfolding−associated SNP: Chr10:22811727
pvalue= 0.0742 ; R2= 0.1857
Estimated allele frequency
Ecological gradient (Leaf Unfolding differences)
1.0
0.9
0.8
0.7
0.6
0.5
0.4
0.3
−1
0
1
2
3
4
233
218
O8
9
O12
256
219
L16
124
L8
97
O1

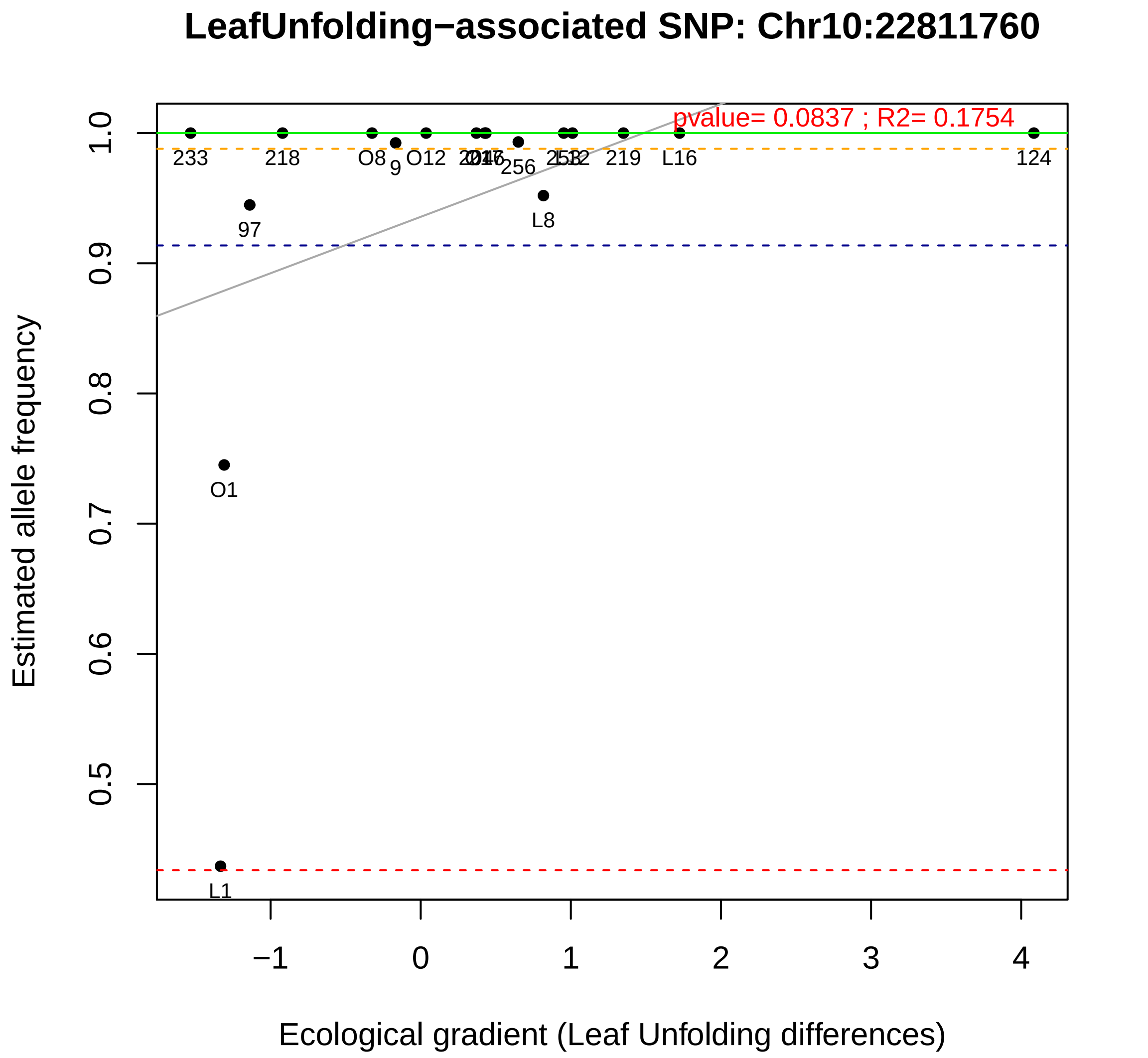
LeafUnfolding−associated SNP: Chr10:22811760
pvalue= 0.0837 ; R2= 0.1754
Estimated allele frequency
Ecological gradient (Leaf Unfolding differences)
1.0
0.9
0.8
0.7
0.6
0.5
−1
0
1
2
3
4
233
218
O8
9
O12
256
219
L16
124
97
L8
O1
L1

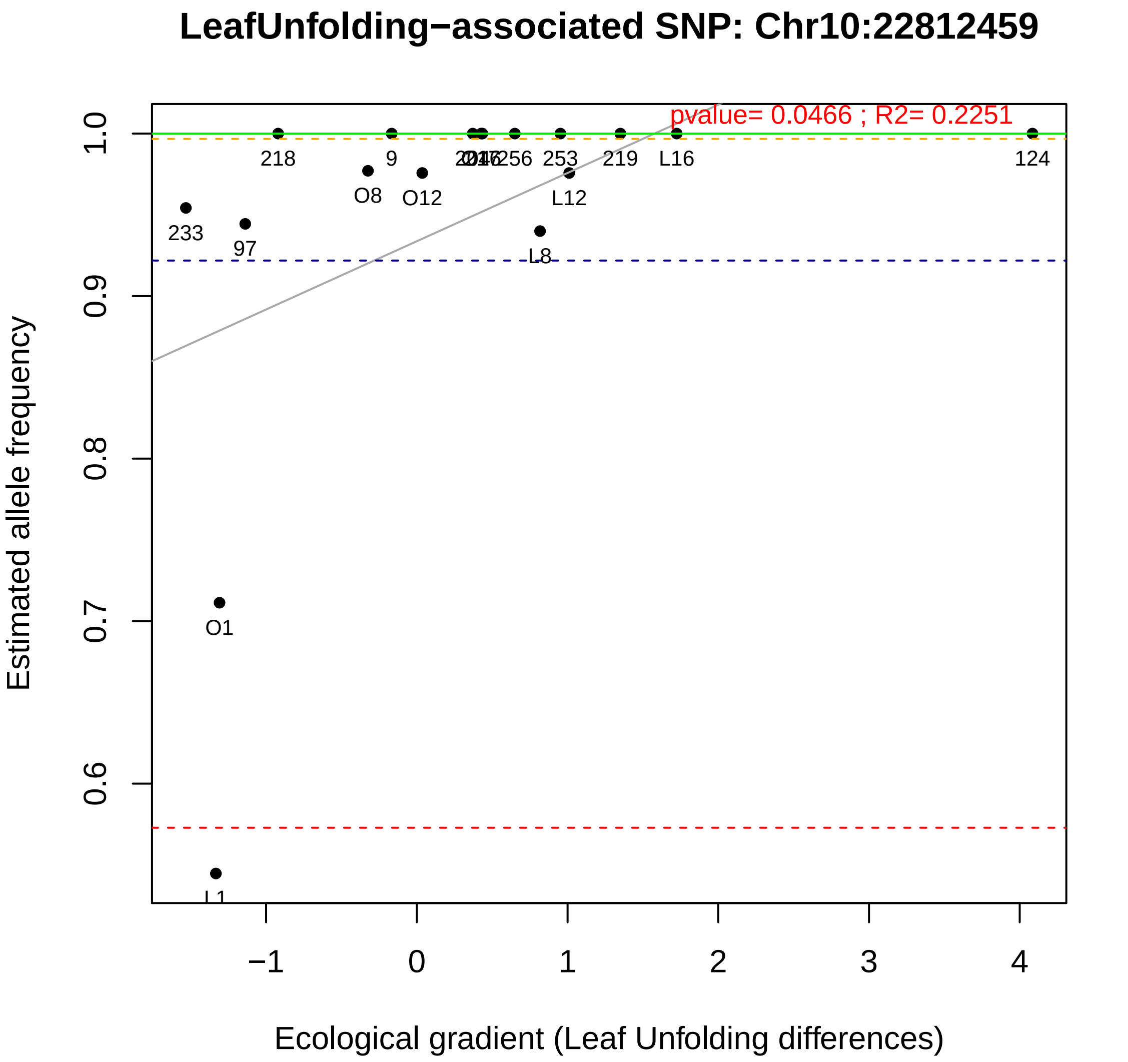
LeafUnfolding−associated SNP: Chr10:22812459
pvalue= 0.0466 ; R2= 0.2251
Estimated allele frequency
Ecological gradient (Leaf Unfolding differences)
0.6
0.7
0.8
0.9
1.0
−1
0
1
2
3
4
218
9
256
253
219
L16
124
O8
O12
L12
233
97
L8
O1
L1

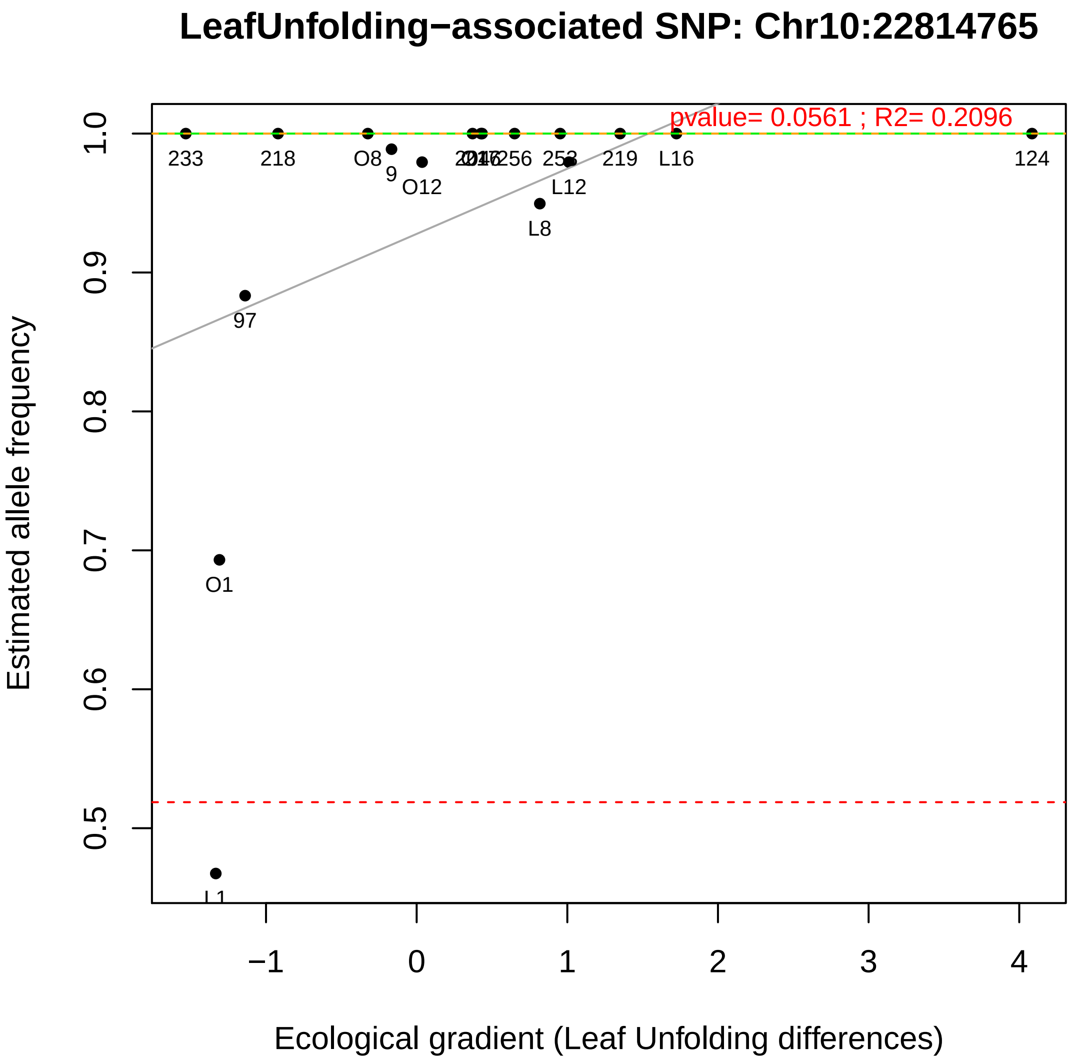
LeafUnfolding−associated SNP: Chr10:22814765
pvalue= 0.0561 ; R2= 0.2096
Estimated allele frequency
Ecological gradient (Leaf Unfolding differences)
1.0
0.9
0.8
0.7
0.6
0.5
−1
0
1
2
3
4
233
218
O8
9
O12
256
219
L16
124
L12
L8
97
O1

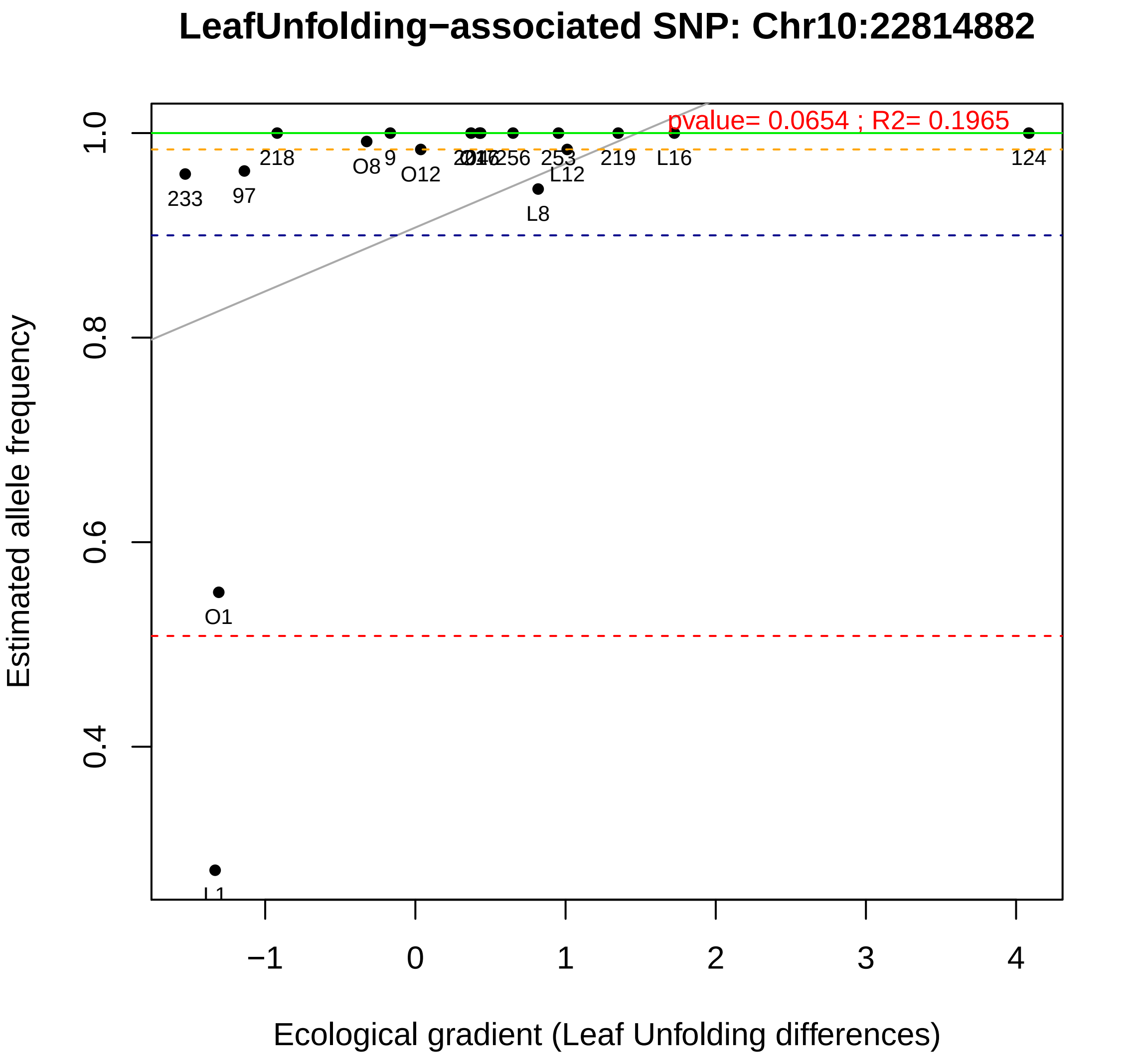

LeafUnfolding−associated SNP: Chr10:22814882
pvalue= 0.0654 ; R2= 0.1965
1.0
0.8
0.6
0.4
Estimated allele frequency
218
233
97
O8
9
O12
256
253
L12
L8
219
L16
124
O1
L1
−1
0
1
2
3
4
Ecological gradient (Leaf Unfolding differences)

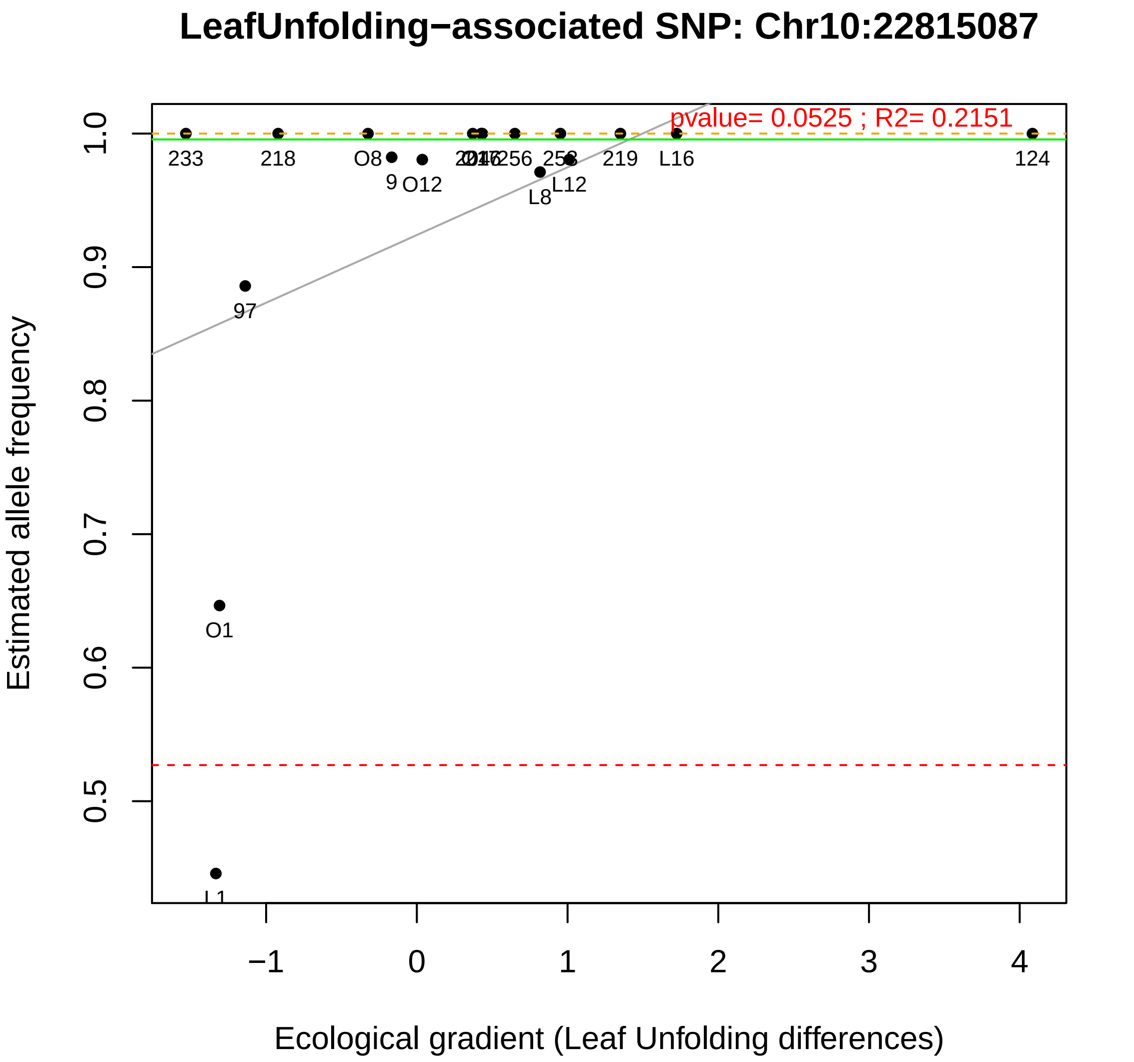

LeafUnfolding−associated SNP: Chr10:22815087
pvalue= 0.0525 ; R2= 0.2151
Estimated allele frequency
Ecological gradient (Leaf Unfolding differences)
1.0
0.9
0.8
0.7
0.6
0.5
−1
0
1
2
3
4
233
218
O8
9
O12
256
L12
L8
219
L16
124
97
O1

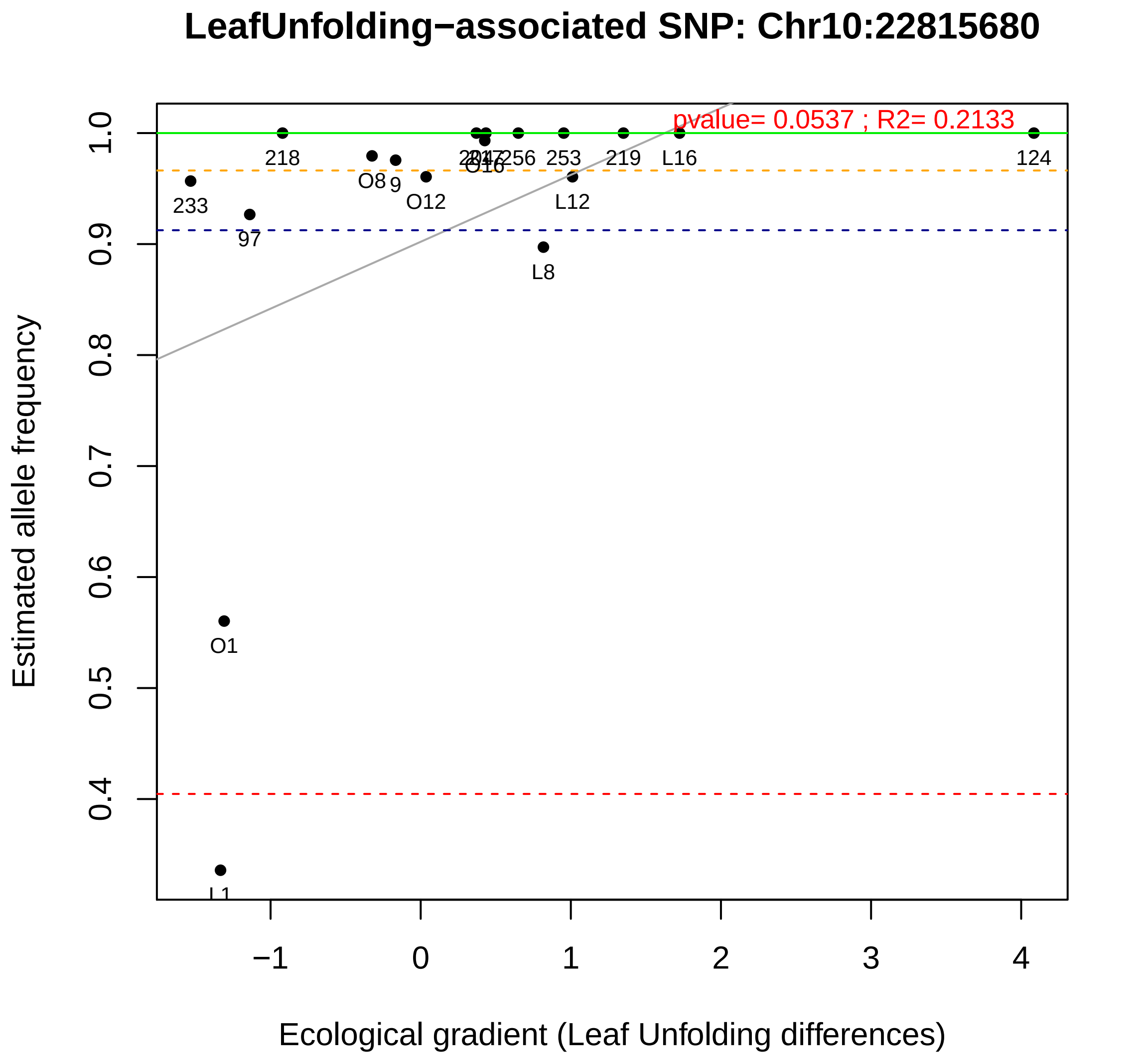

LeafUnfolding−associated SNP: Chr10:22815680
pvalue= 0.0537 ; R2= 0.2133
Estimated allele frequency
Ecological gradient (Leaf Unfolding differences)
1.0
0.9
0.8
0.7
0.6
0.5
0.4
−1
0
1
2
3
4
218
233
97
O8
9
O12
O16
256
253
219
L16
124
L12
L8
O1

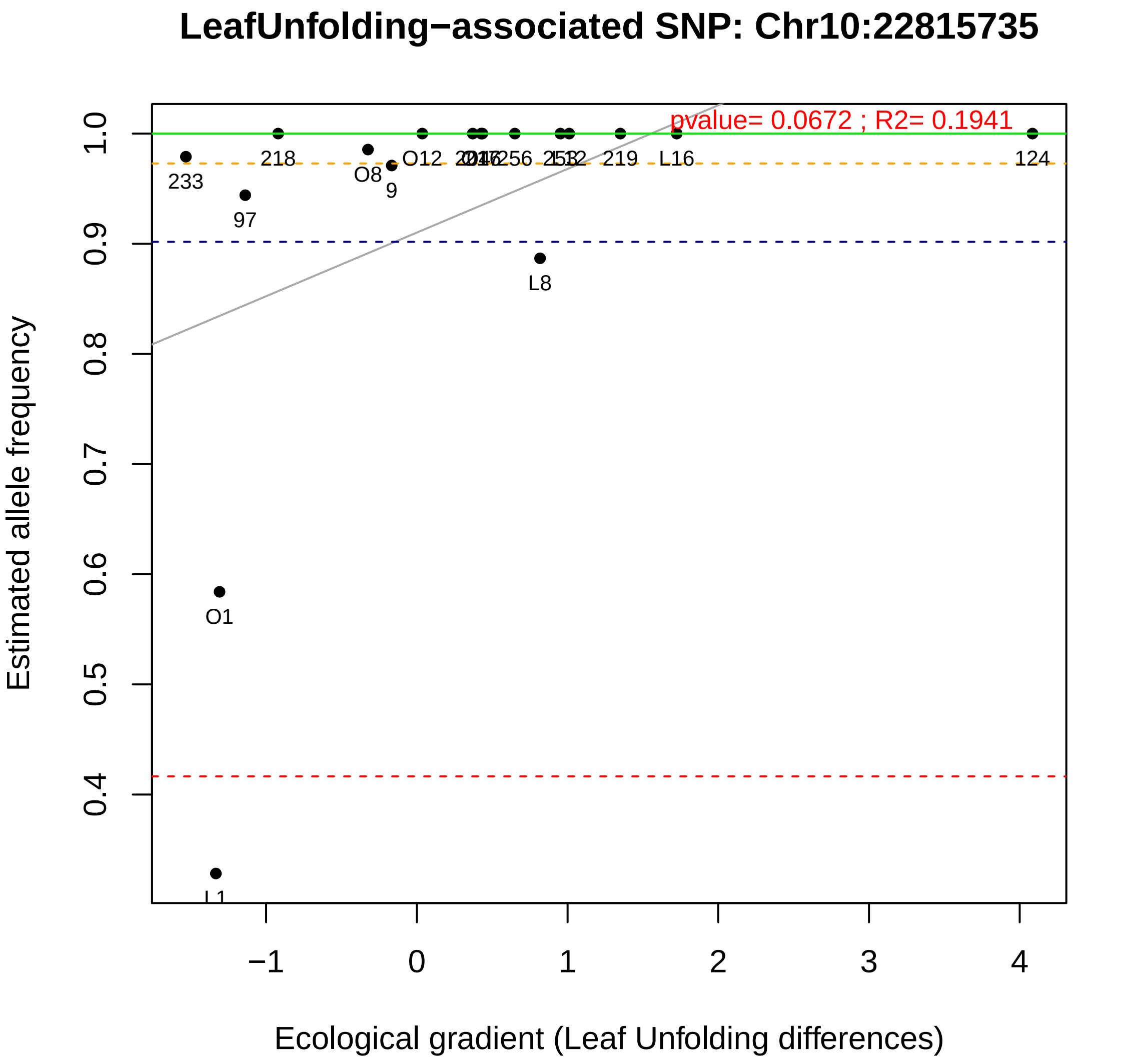

LeafUnfolding−associated SNP: Chr10:22815735
pvalue= 0.0672 ; R2= 0.1941
Estimated allele frequency
Ecological gradient (Leaf Unfolding differences)
1.0
0.9
0.8
0.7
0.6
0.5
0.4
−1
0
1
2
3
4
233
218
97
O8
9
O12
L8
219
L16
124
O1

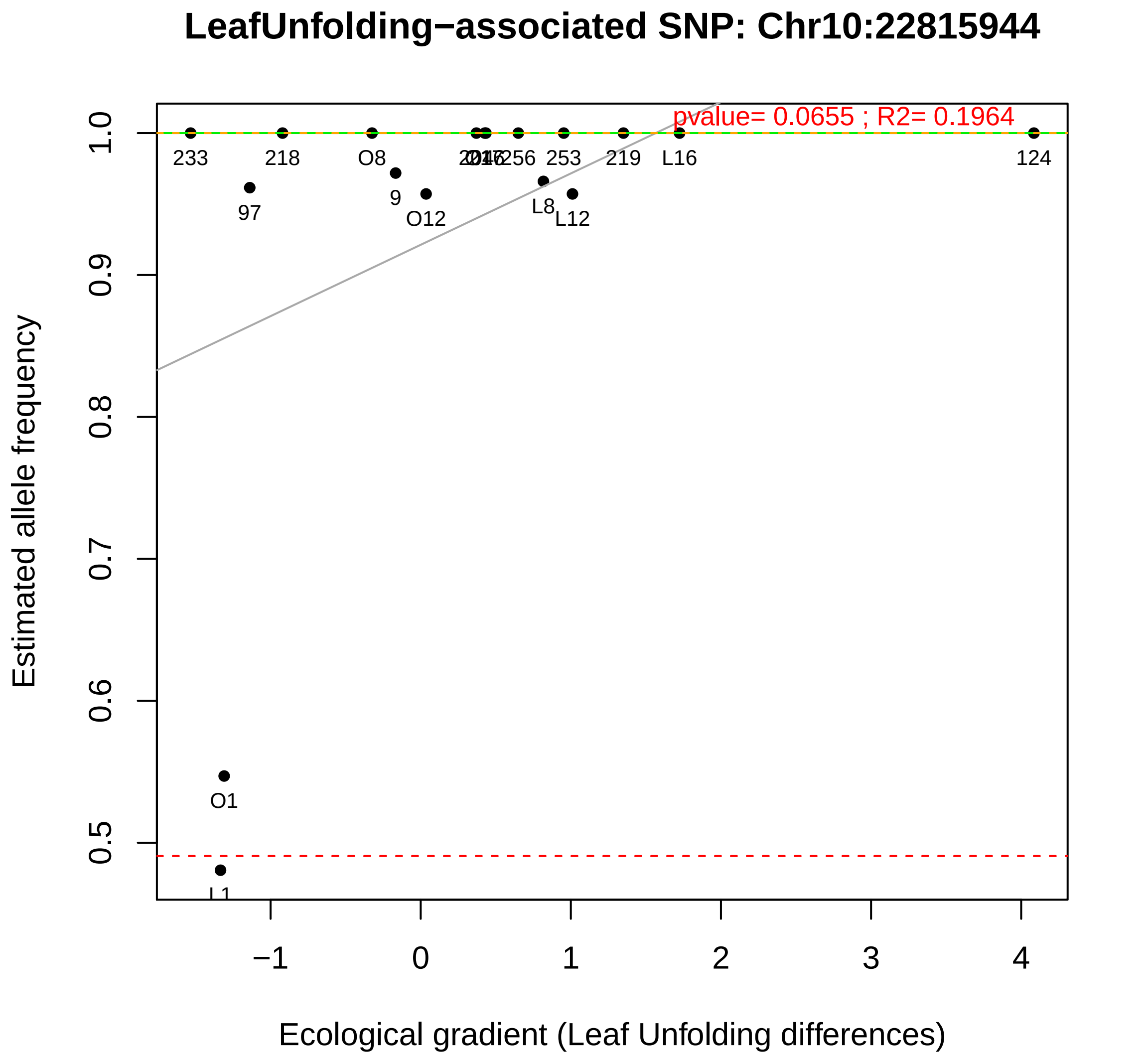
LeafUnfolding−associated SNP: Chr10:22815944
pvalue= 0.0655 ; R2= 0.1964
Estimated allele frequency
Ecological gradient (Leaf Unfolding differences)
233
218
O8
256
253
219
L16
124
97
9
O12
L8
L12
O1
L1

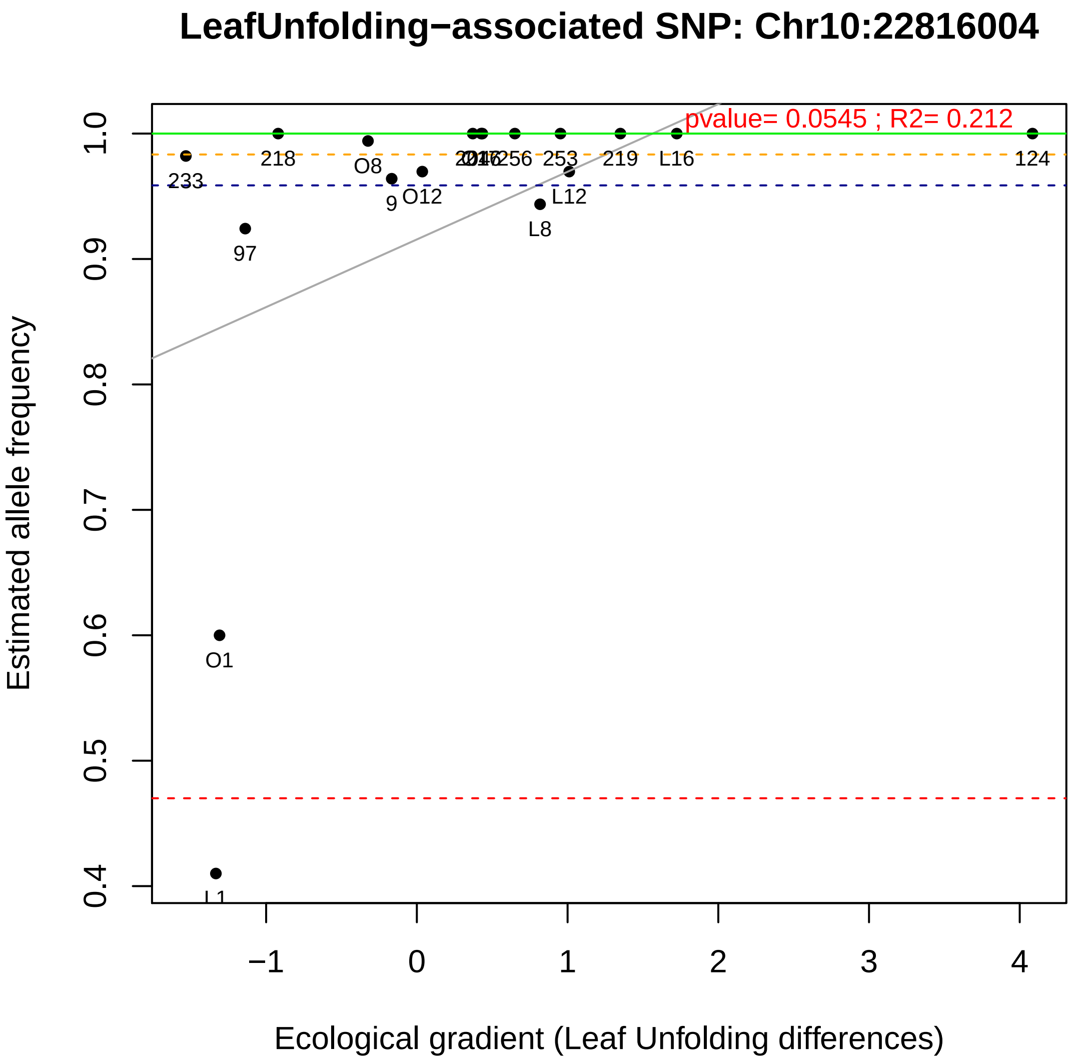

LeafUnfolding−associated SNP: Chr10:22816004
pvalue= 0.0545 ; R2= 0.212
Estimated allele frequency
Ecological gradient (Leaf Unfolding differences)
1.0
0.9
0.8
0.7
0.6
0.5
0.4
−1
0
1
2
3
4
233
218
O8
9
O12
L12
L8
97
256
253
219
L16
124
O1
L1

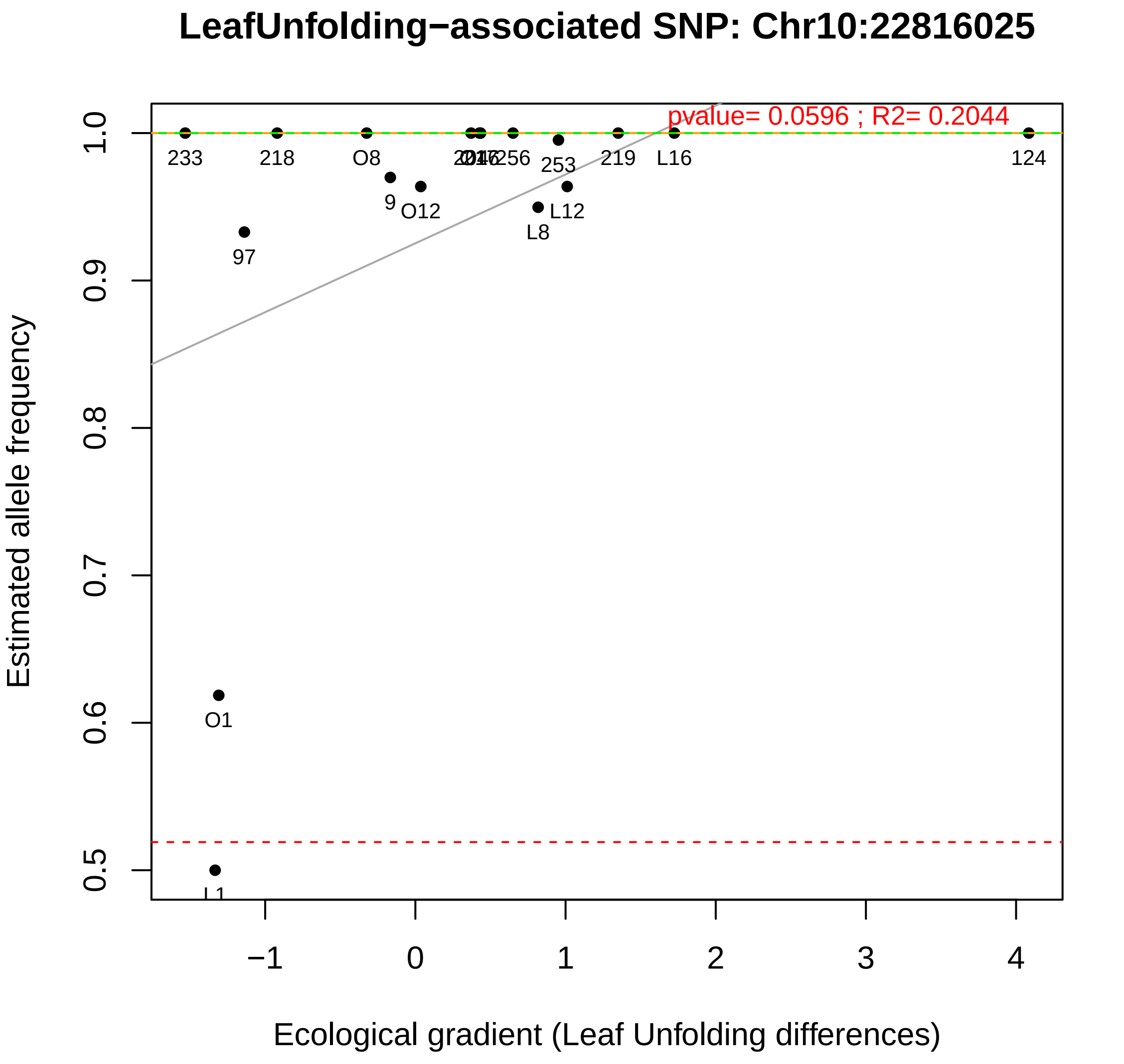
LeafUnfolding−associated SNP: Chr10:22816025
pvalue= 0.0596 ; R2= 0.2044
233
218
O8
256
253
219
L16
124
9
O12
L8
L12
97
O1
1.0
0.9
0.8
0.7
0.6
0.5
−1
0
1
2
3
4
Estimated allele frequency
Ecological gradient (Leaf Unfolding differences)

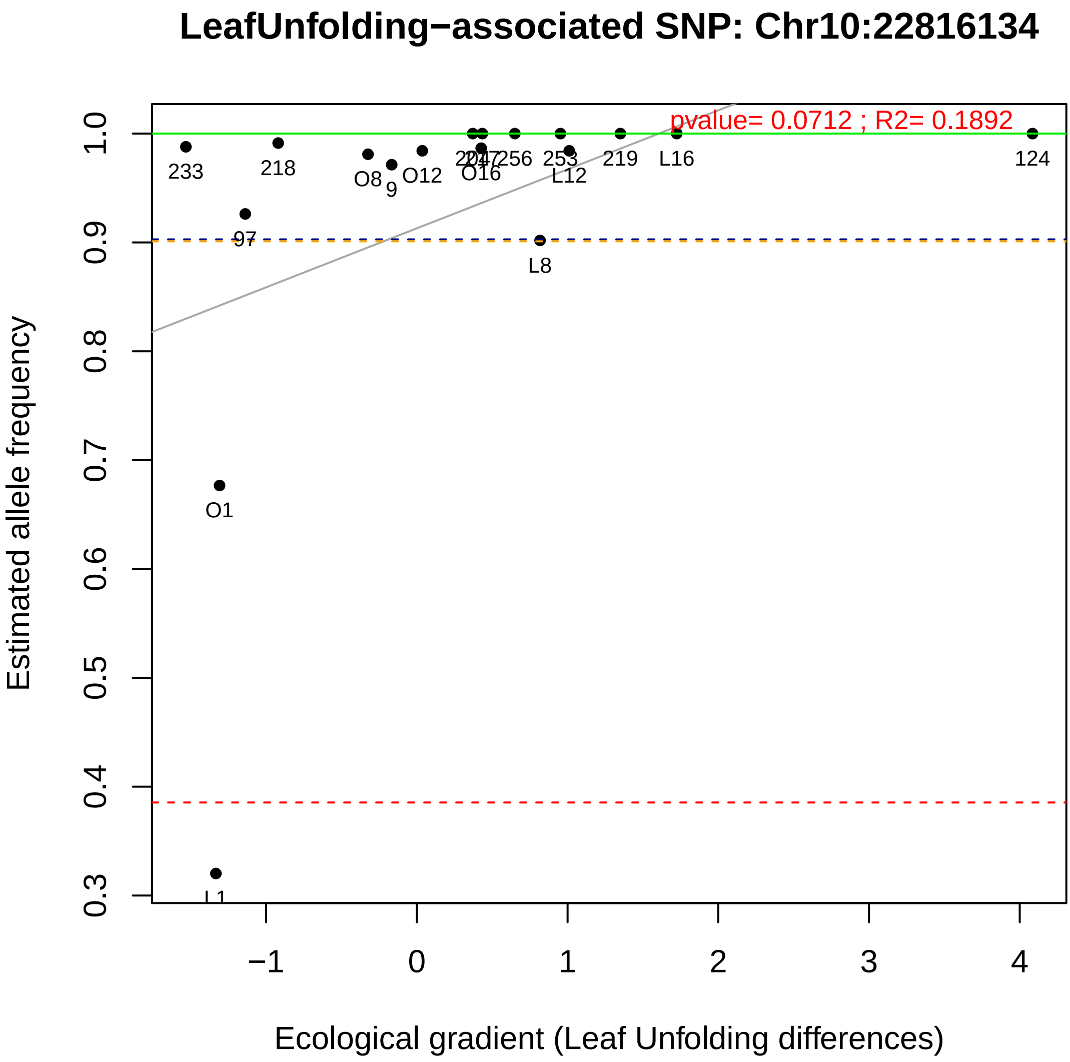

LeafUnfolding−associated SNP: Chr10:22816134
pvalue= 0.0712 ; R2= 0.1892
Estimated allele frequency
Ecological gradient (Leaf Unfolding differences)
233
218
97
O8
9
O12
O16
256
253
L12
219
L16
124
L8
O1

#### LeafUnfolding−associated SNP: Chr10:22816503

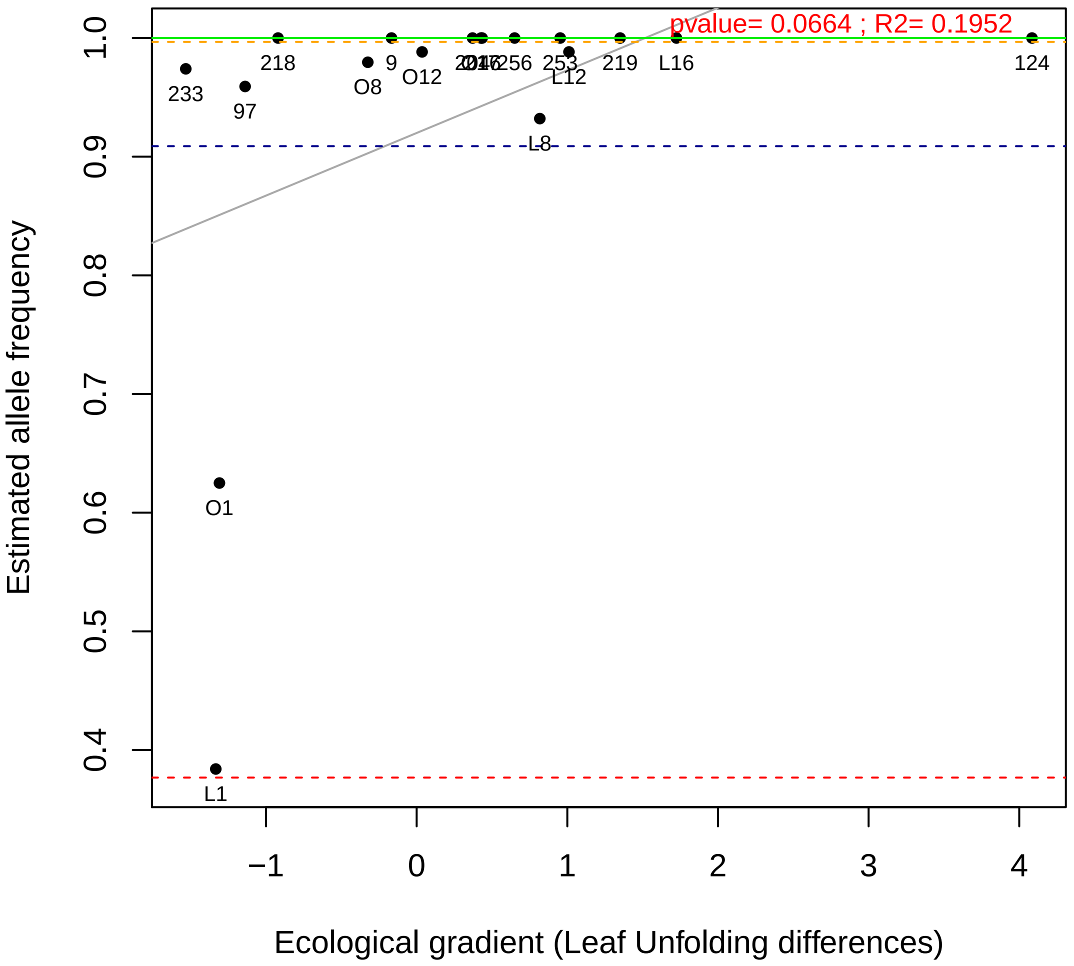

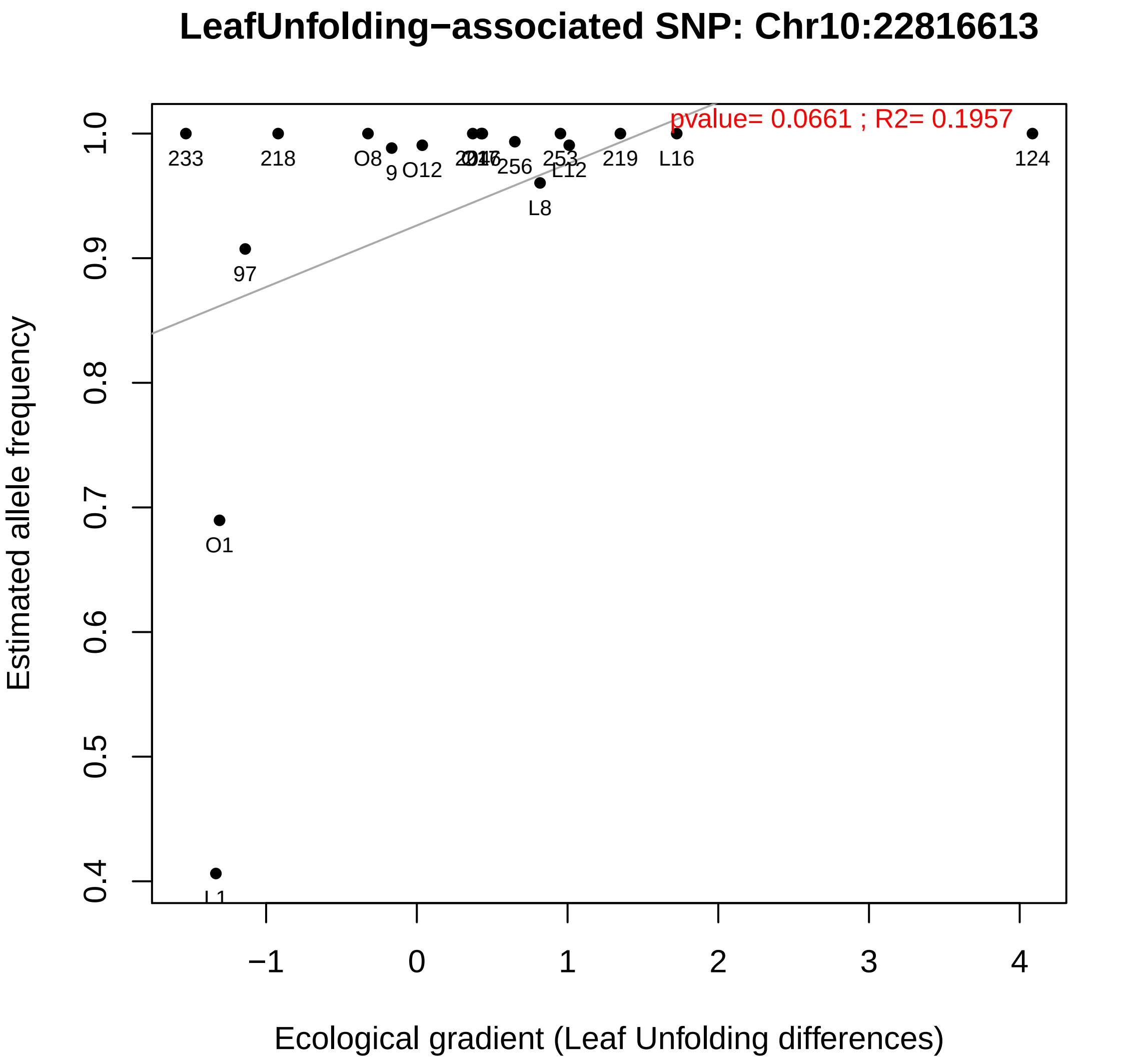

LeafUnfolding−associated SNP: Chr10:22816613
pvalue= 0.0661 ; R2= 0.1957
Estimated allele frequency
Ecological gradient (Leaf Unfolding differences)
1.0
0.9
0.8
0.7
0.6
0.5
0.4
−1
0
1
2
3
4
233
218
O8
9
O12
256
253
L12
219
L16
124
L8
97
O1
L1

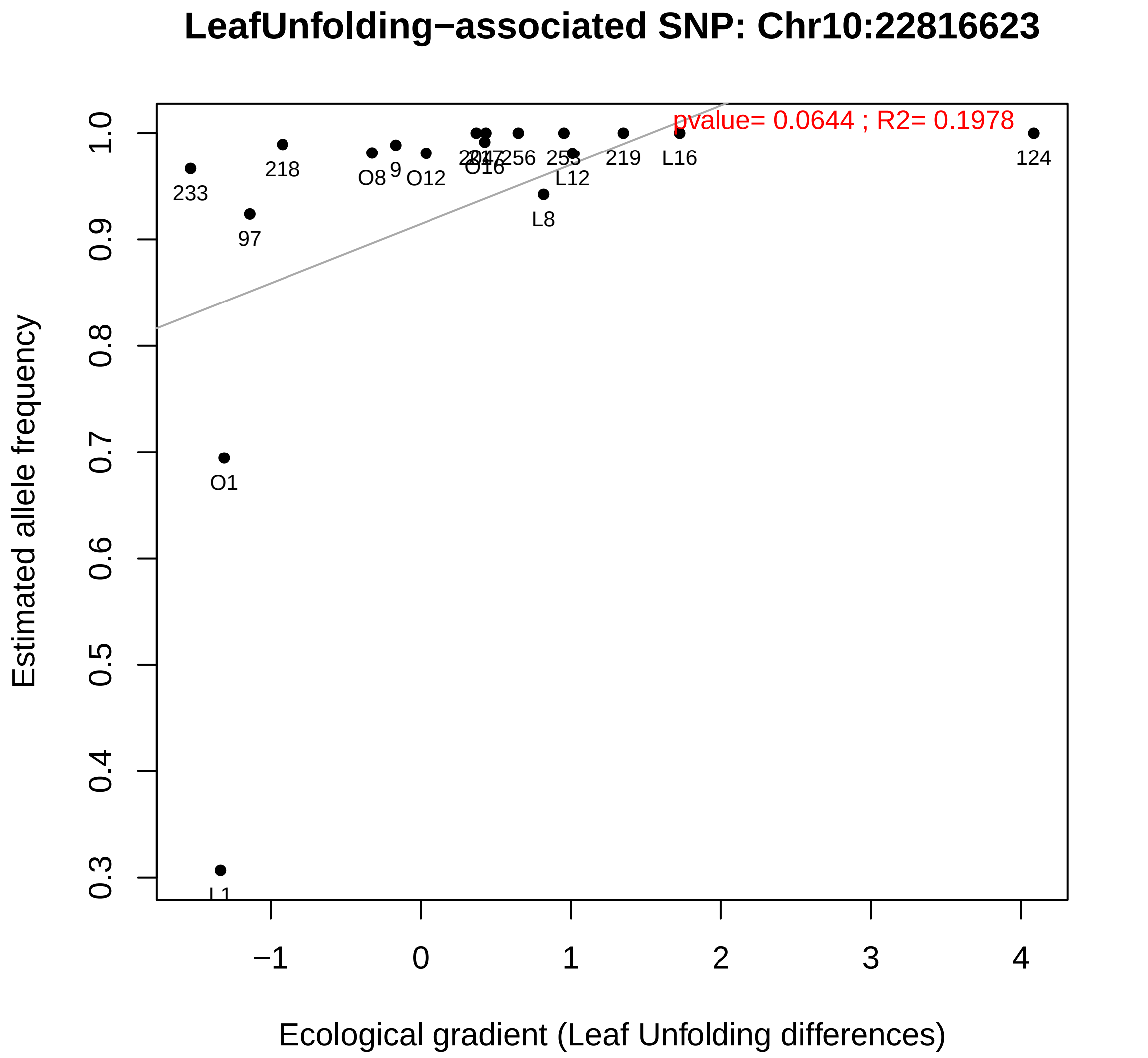
LeafUnfolding−associated SNP: Chr10:22816623
pvalue= 0.0644 ; R2= 0.1978
Estimated allele frequency
Ecological gradient (Leaf Unfolding differences)
1.0
0.9
0.8
0.7
0.6
0.5
0.4
0.3
−1
0
1
2
3
4
233
97
218
O8
9
O12
O16
256
253
L12
L8
219
L16
124
O1

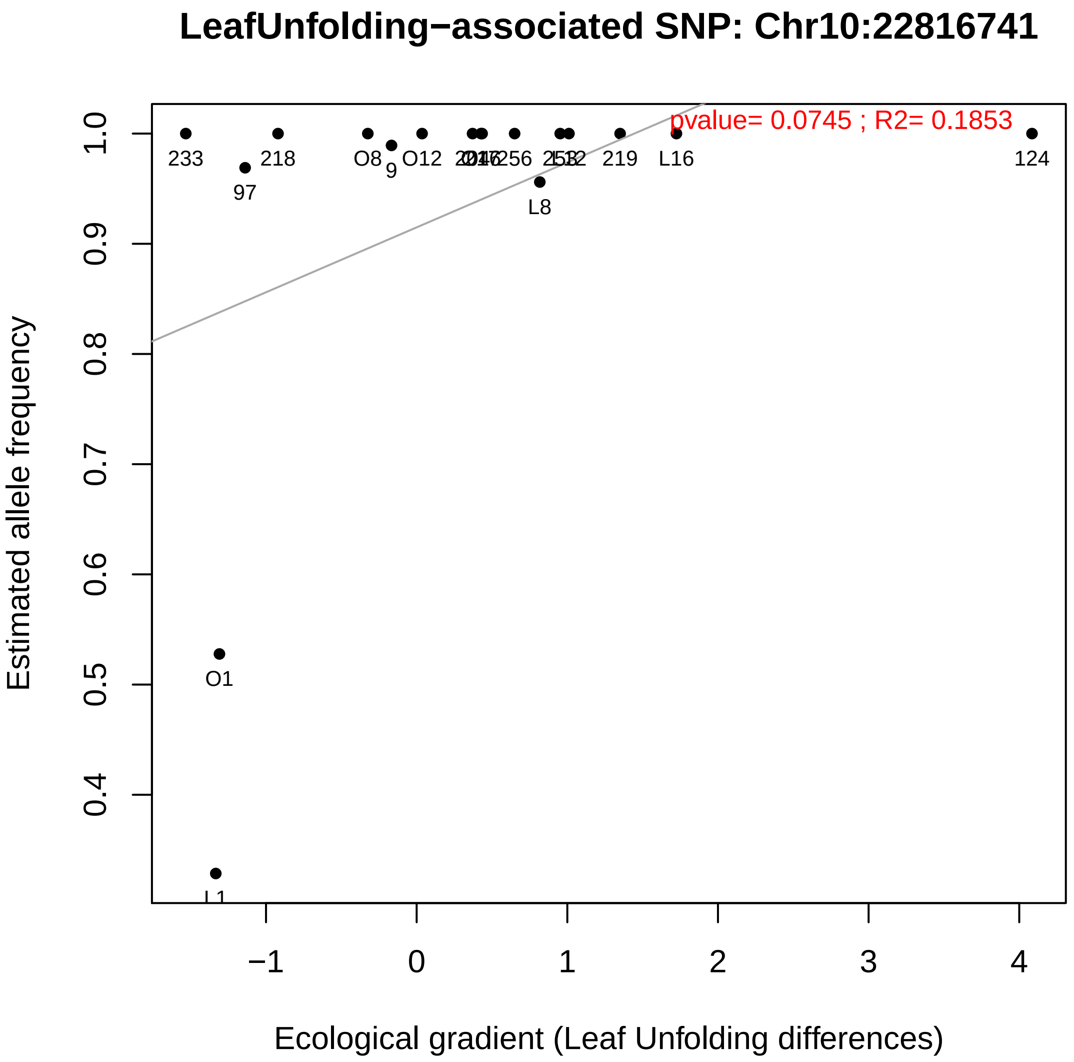
LeafUnfolding−associated SNP: Chr10:22816741
pvalue= 0.0745 ; R2= 0.1853
Estimated allele frequency
Ecological gradient (Leaf Unfolding differences)
233
97
218
O8
9
O12
256
L8
219
L16
124
O1
L1
1.0
0.9
0.8
0.7
0.6
0.5
0.4
−1
0
1
2
3
4

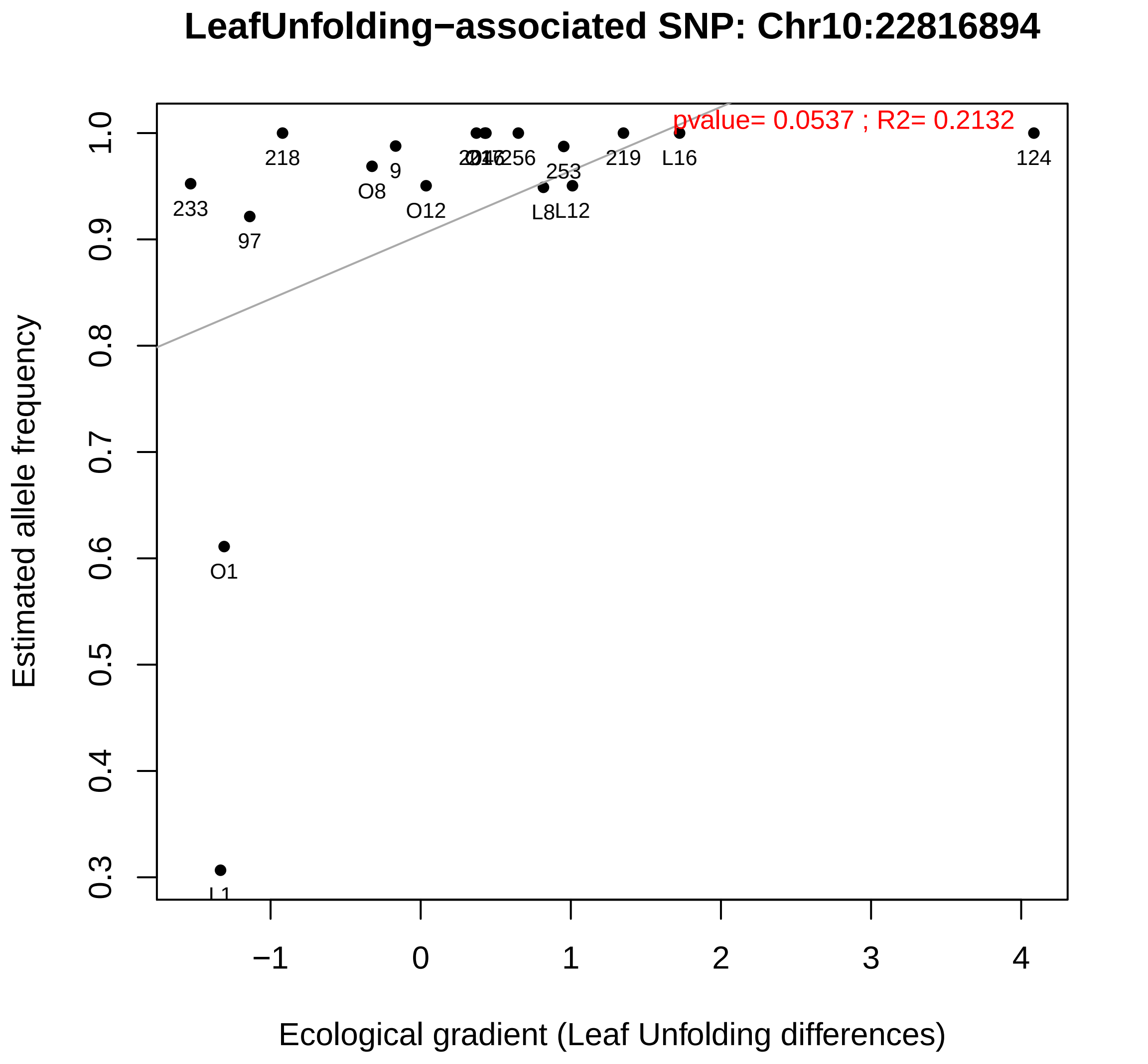

LeafUnfolding−associated SNP: Chr10:22816894
pvalue= 0.0537 ; R2= 0.2132
Estimated allele frequency
Ecological gradient (Leaf Unfolding differences)
1.0
0.9
0.8
0.7
0.6
0.5
0.4
0.3
−1
0
1
2
3
4
218
233
97
O8
9
O12
256
253
L8
L12
219
L16
124
O1

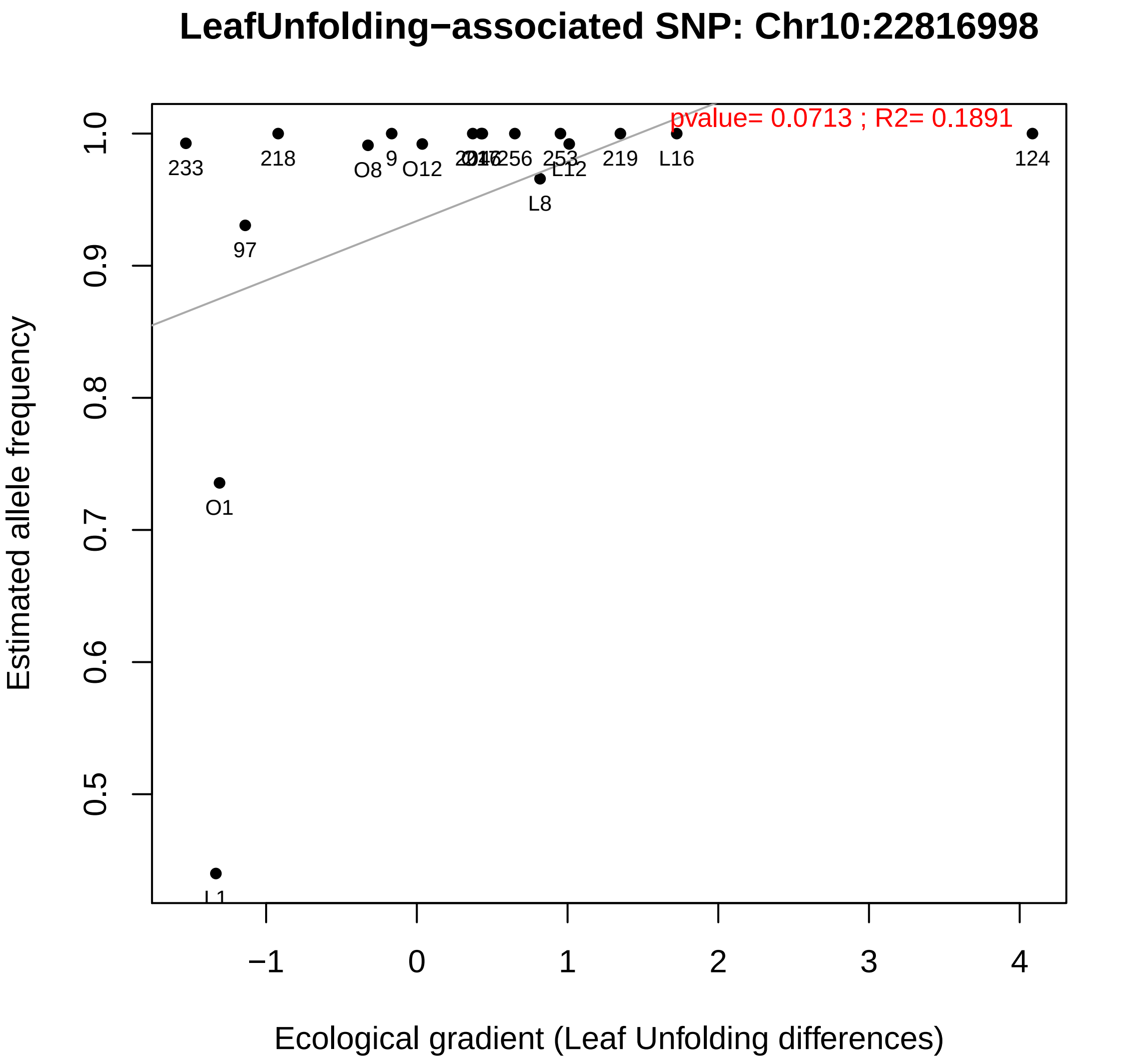
LeafUnfolding−associated SNP: Chr10:22816998
pvalue= 0.0713 ; R2= 0.1891
Estimated allele frequency
Ecological gradient (Leaf Unfolding differences)
1.0
0.9
0.8
0.7
0.6
0.5
−1
0
1
2
3
4
233
218
O8
9
O12
256
253
L12
219
L16
124
L8
97
O1
L1

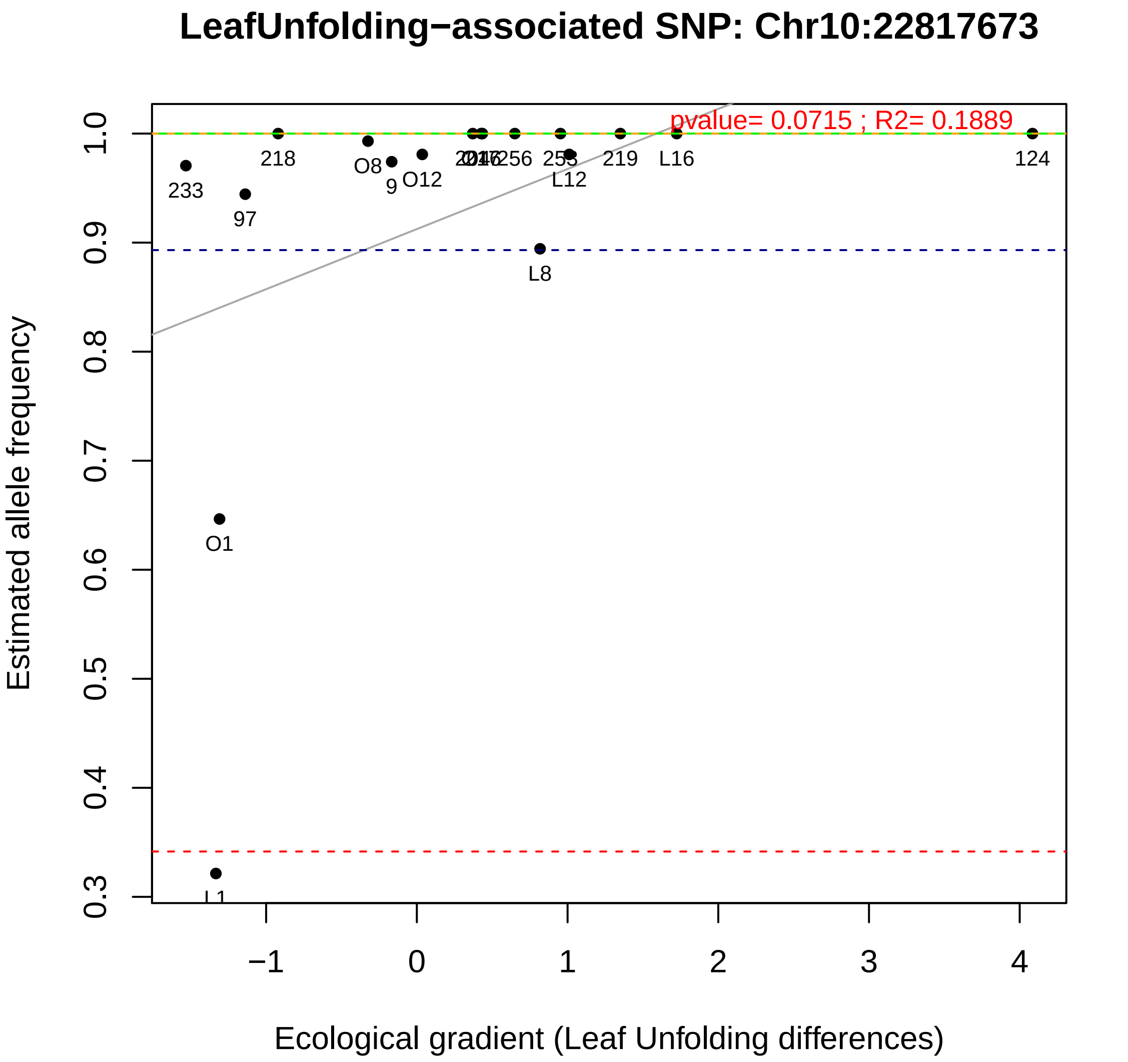

LeafUnfolding−associated SNP: Chr10:22817673
pvalue= 0.0715 ; R2= 0.1889
Estimated allele frequency
Ecological gradient (Leaf Unfolding differences)
1.0
0.9
0.8
0.7
0.6
0.5
0.4
0.3
−1
0
1
2
3
4
218
233
97
O8
9
O12
256
253
L12
219
L16
124
L8
O1
L1

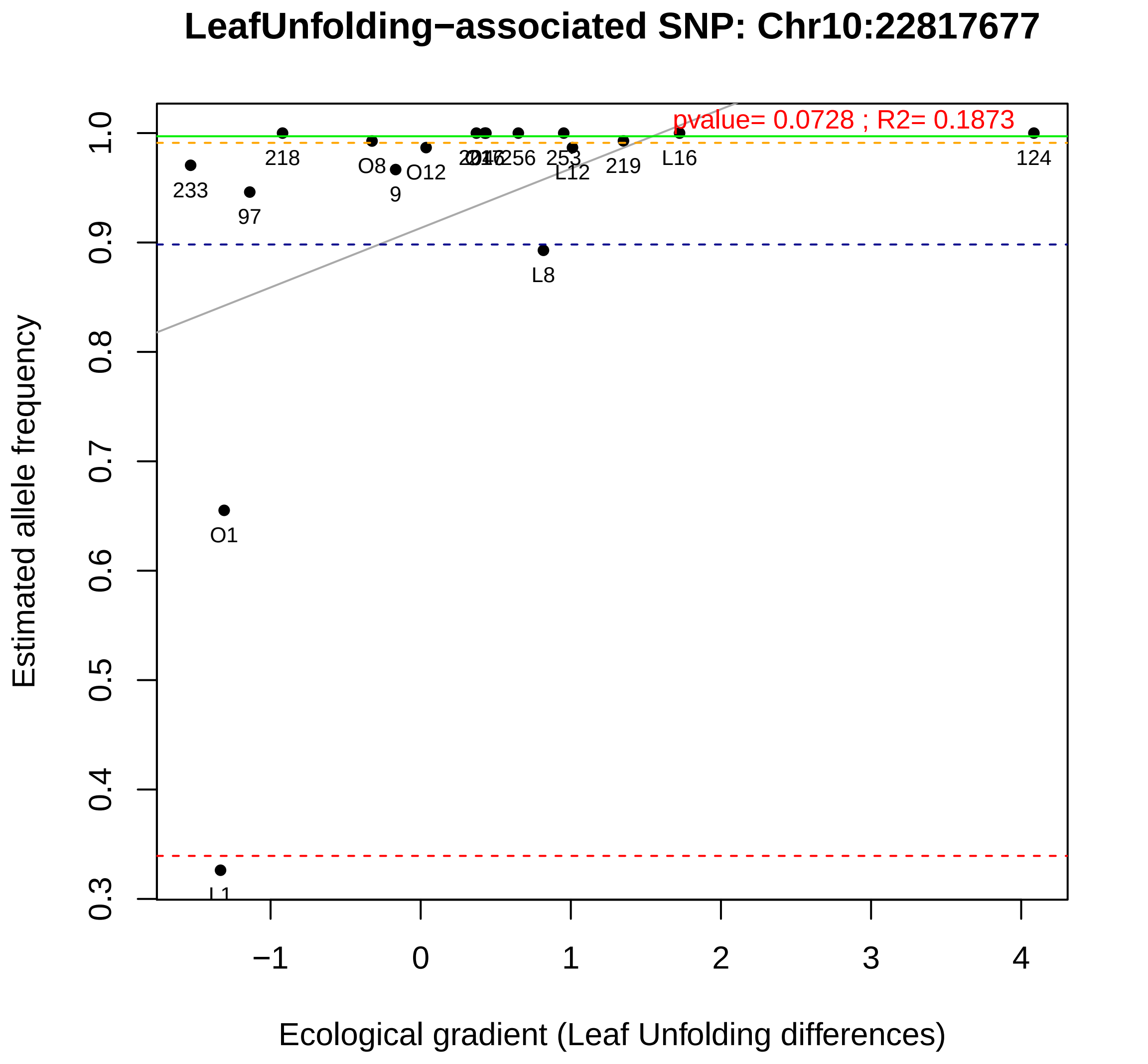
LeafUnfolding−associated SNP: Chr10:22817677
pvalue= 0.0728 ; R2= 0.1873
Estimated allele frequency
Ecological gradient (Leaf Unfolding differences)
1.0
0.9
0.8
0.7
0.6
0.5
0.4
0.3
−1
0
1
2
3
4
218
O8
O12
9
233
97
256
253
L12
219
L16
124
L8
O1

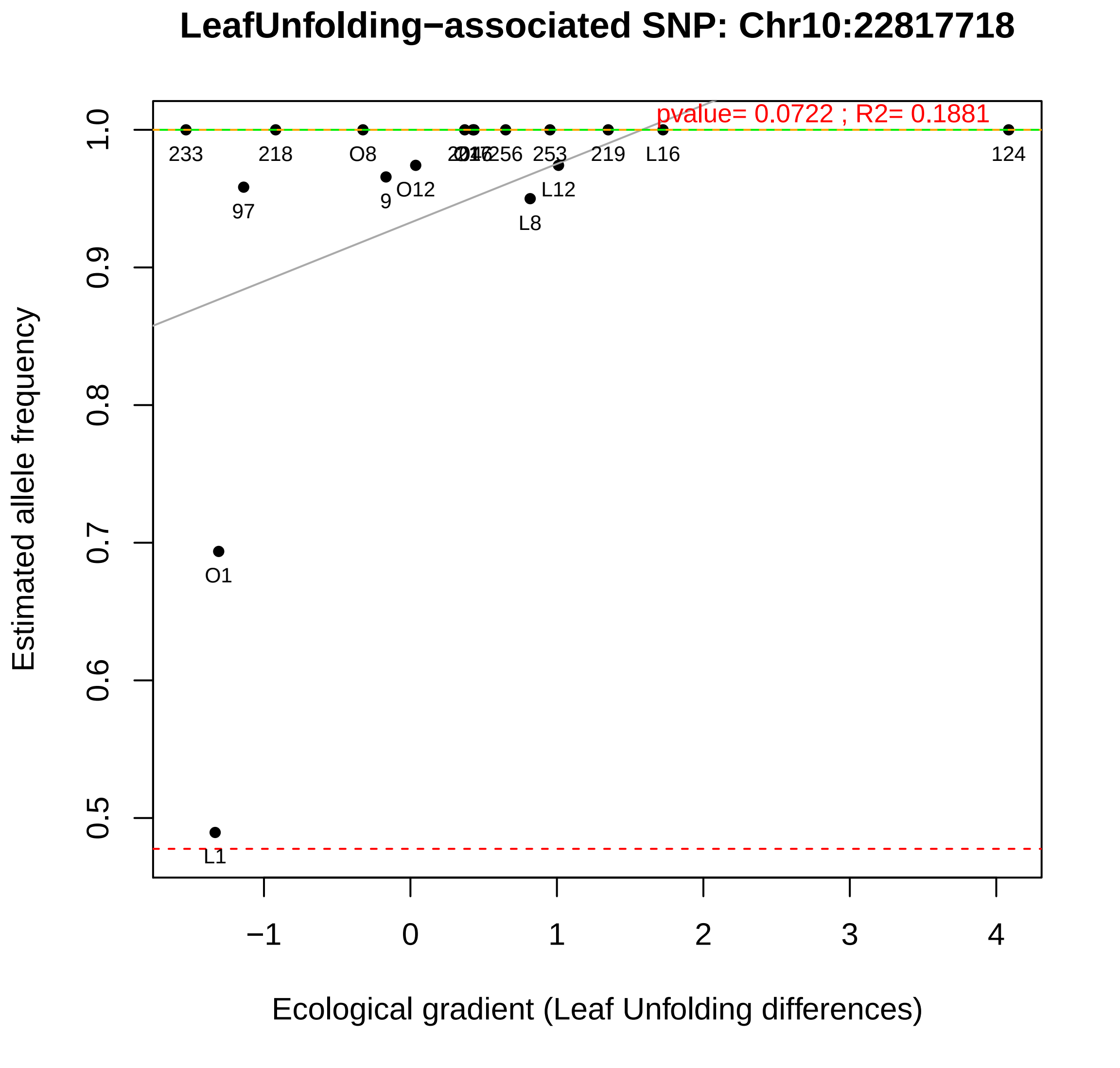

LeafUnfolding−associated SNP: Chr10:22817718
pvalue= 0.0722 ; R2= 0.1881
233
218
O8
256
253
219
L16
124
97
9
O12
L12
L8
O1
L1
Estimated allele frequency
Ecological gradient (Leaf Unfolding differences)

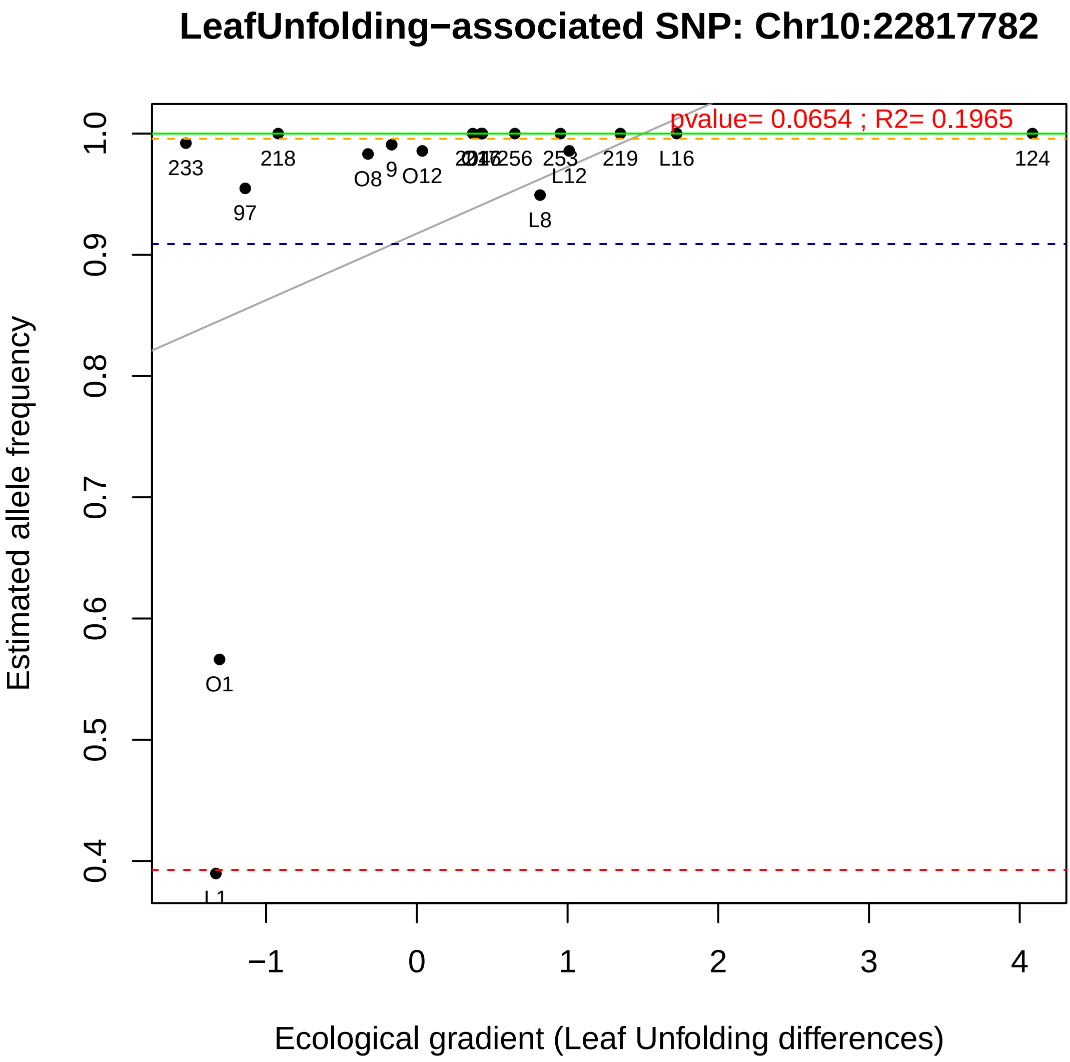

LeafUnfolding−associated SNP: Chr10:22817782
pvalue= 0.0654 ; R2= 0.1965
Estimated allele frequency
Ecological gradient (Leaf Unfolding differences)
1.0
0.9
0.8
0.7
0.6
0.5
0.4
−1
0
1
2
3
4
233
218
97
O8
9
O12
256
253
L12
L8
219
L16
124
O1

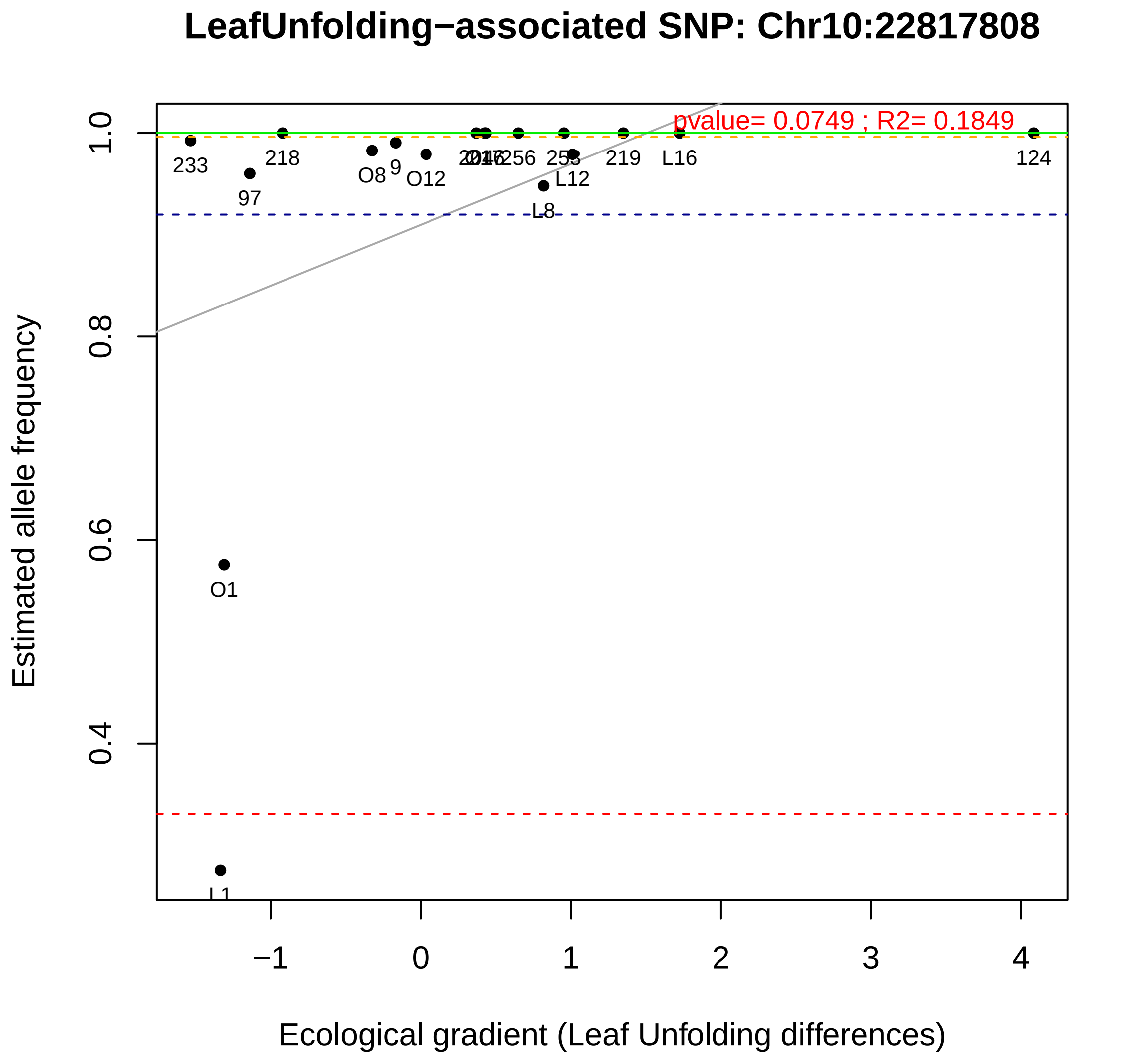

LeafUnfolding−associated SNP: Chr10:22817808
pvalue= 0.0749 ; R2= 0.1849
Estimated allele frequency
Ecological gradient (Leaf Unfolding differences)
1.0
0.8
0.6
0.4
−1
0
1
2
3
4
233
218
97
O8
9
O12
256
253
L12
L8
219
L16
124
O1

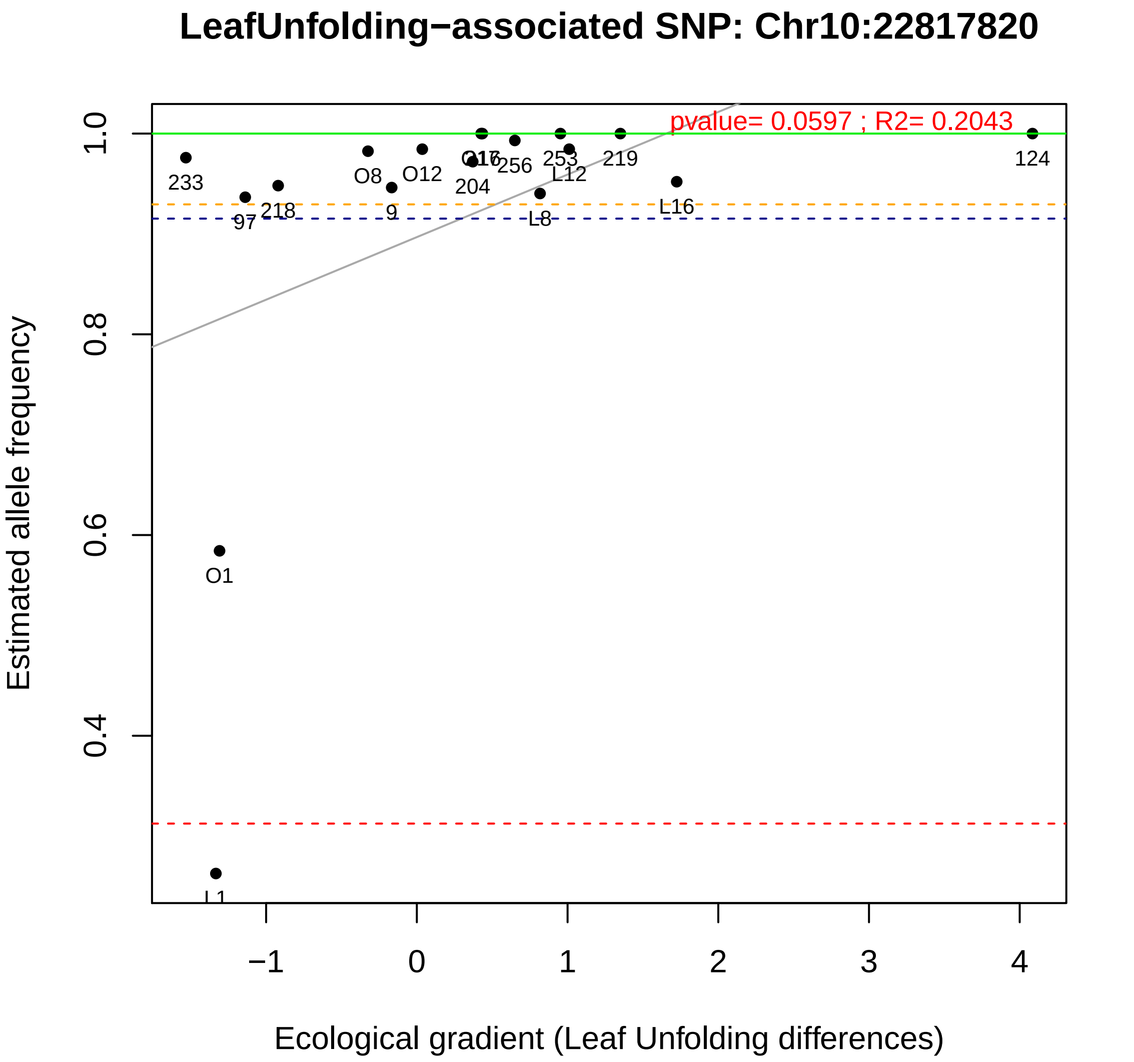
LeafUnfolding−associated SNP: Chr10:22817820
pvalue= 0.0597 ; R2= 0.2043
Estimated allele frequency
Ecological gradient (Leaf Unfolding differences)
1.0
0.8
0.6
0.4
−1
0
1
2
3
4
233
97
218
O8
9
O12
204
256
253
L12
L8
219
L16
124
O1
L1

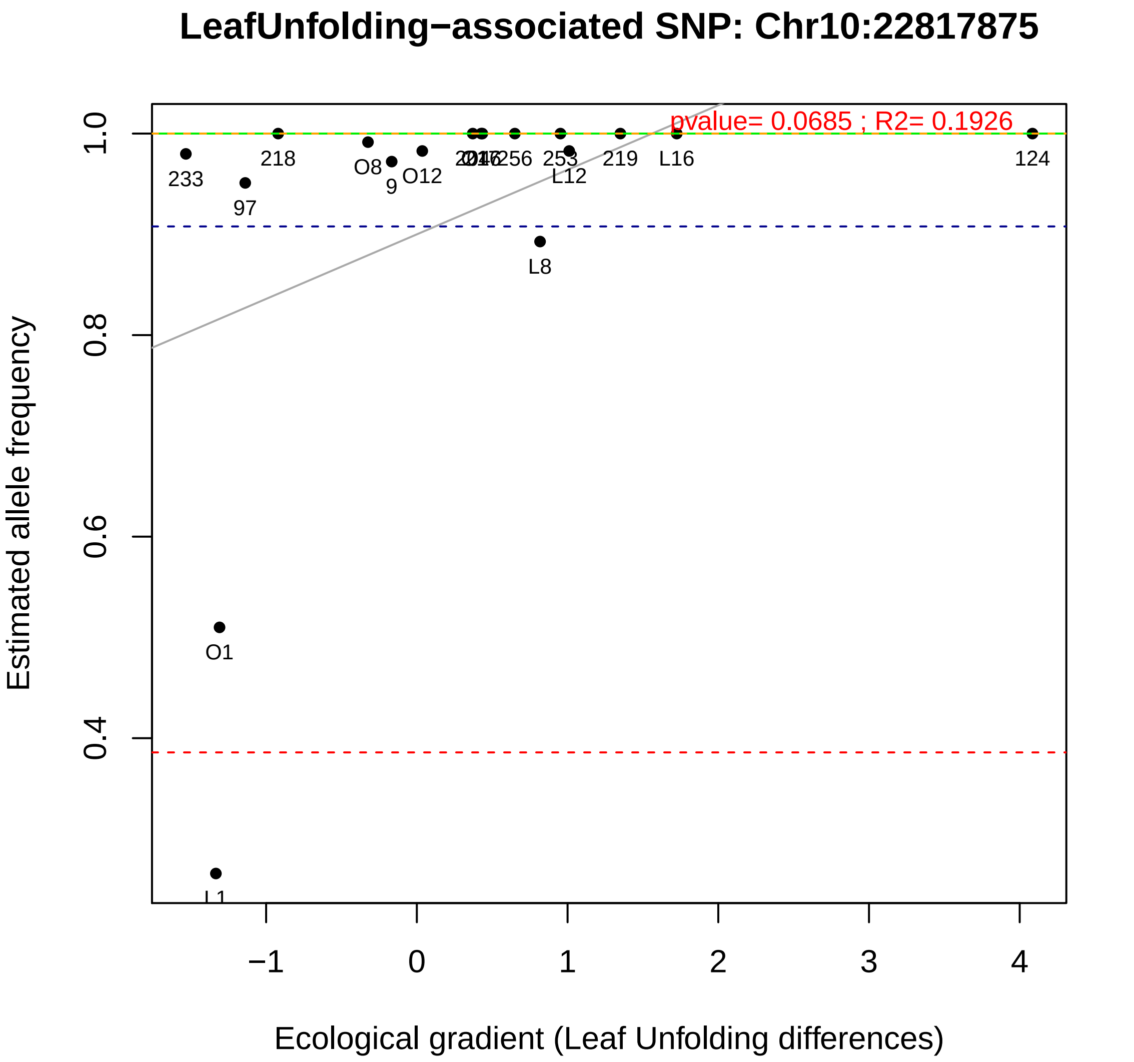
LeafUnfolding−associated SNP: Chr10:22817875
pvalue= 0.0685 ; R2= 0.1926
Estimated allele frequency
Ecological gradient (Leaf Unfolding differences)
1.0
0.8
0.6
0.4
−1
0
1
2
3
4
233
97
218
O8
9
O12
256
253
L12
219
L16
124
L8
O1

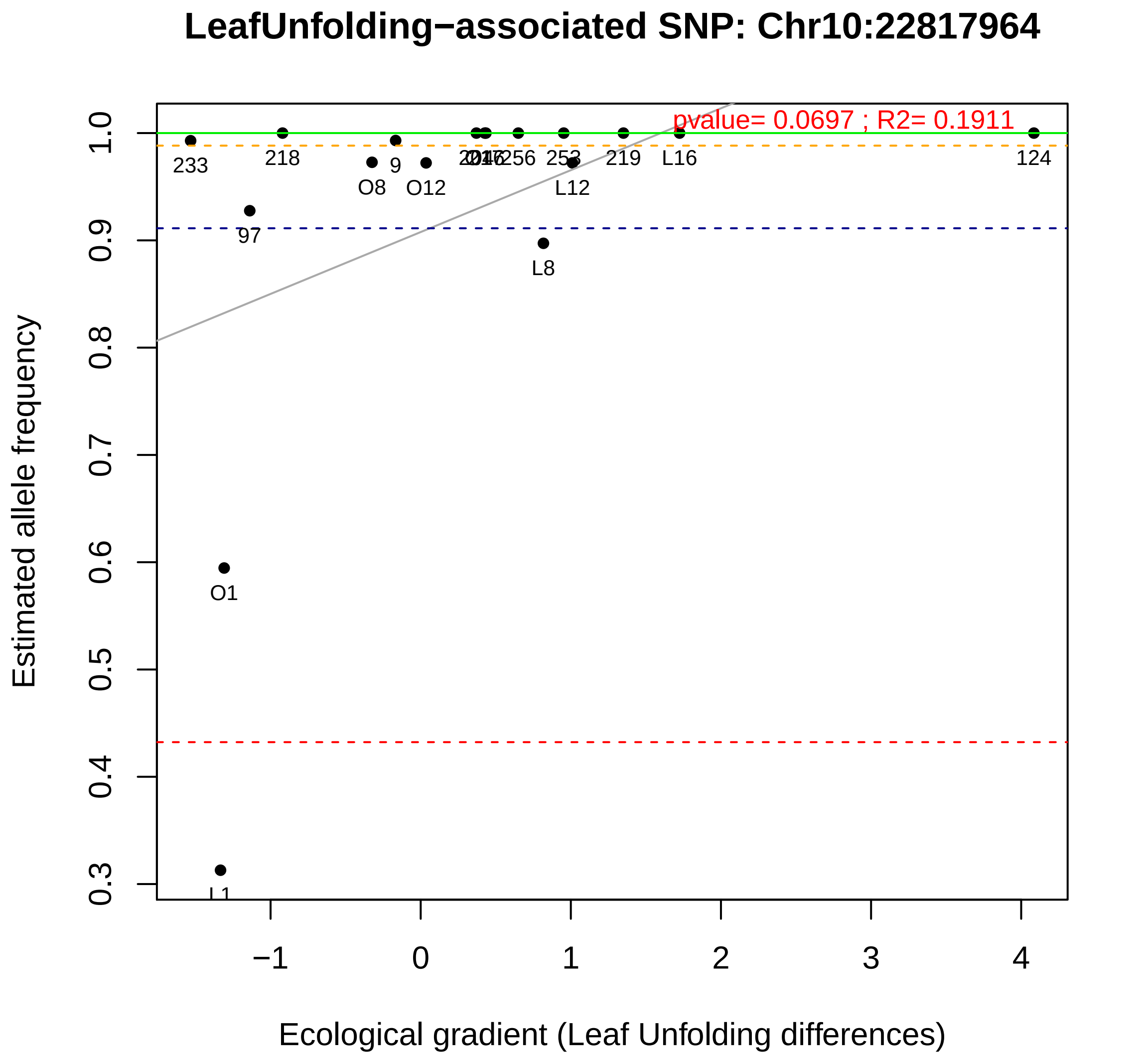

LeafUnfolding−associated SNP: Chr10:22817964
pvalue= 0.0697 ; R2= 0.1911
Estimated allele frequency
Ecological gradient (Leaf Unfolding differences)
0.3
0.4
0.5
0.6
0.7
0.8
0.9
1.0
−1
0
1
2
3
4
233
218
97
O8
9
O12
256
253
L12
219
L16
124
L8
O1

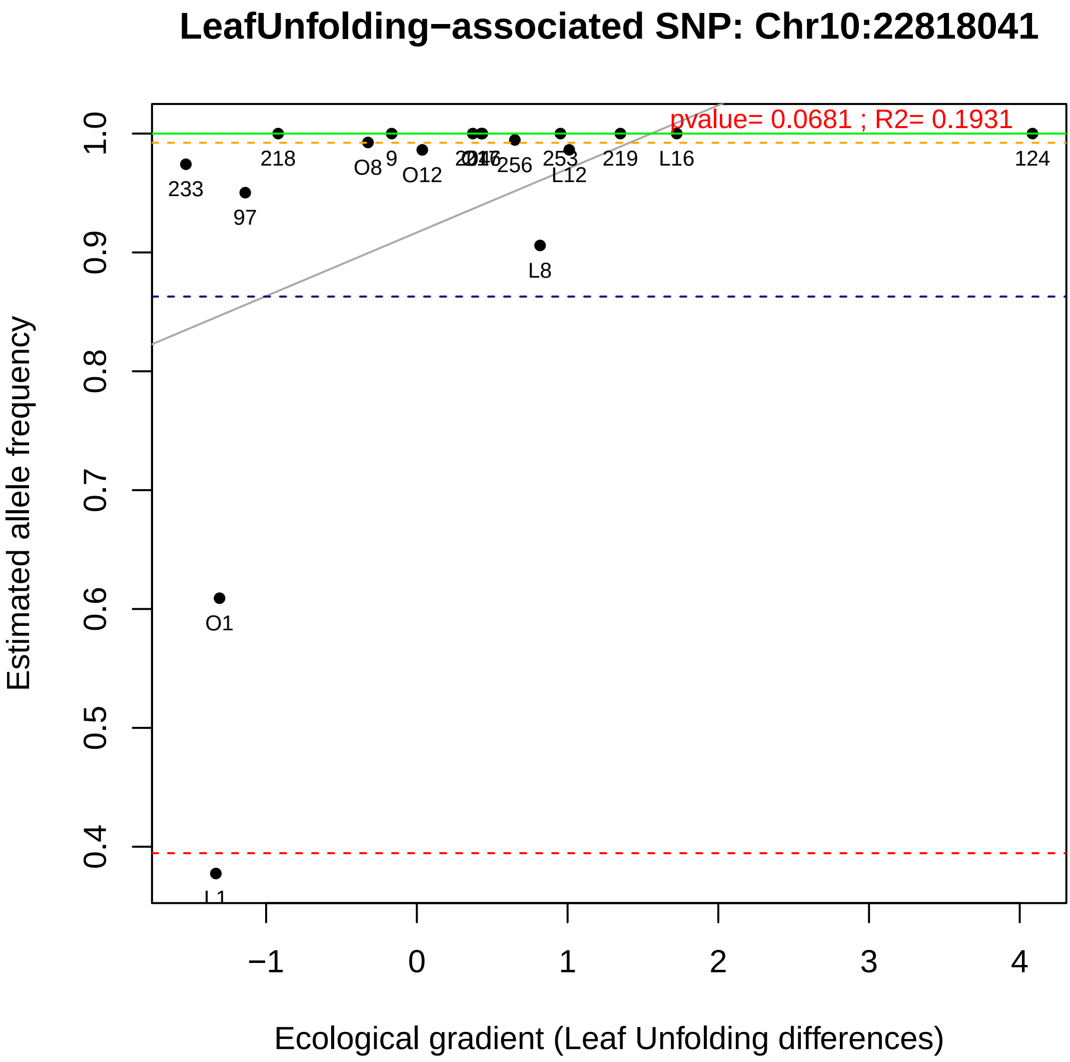

LeafUnfolding−associated SNP: Chr10:22818041
pvalue= 0.0681 ; R2= 0.1931
Estimated allele frequency
Ecological gradient (Leaf Unfolding differences)
1.0
0.9
0.8
0.7
0.6
0.5
0.4
−1
0
1
2
3
4
218
233
97
O8
9
O12
256
253
L12
219
L16
124
L8
O1

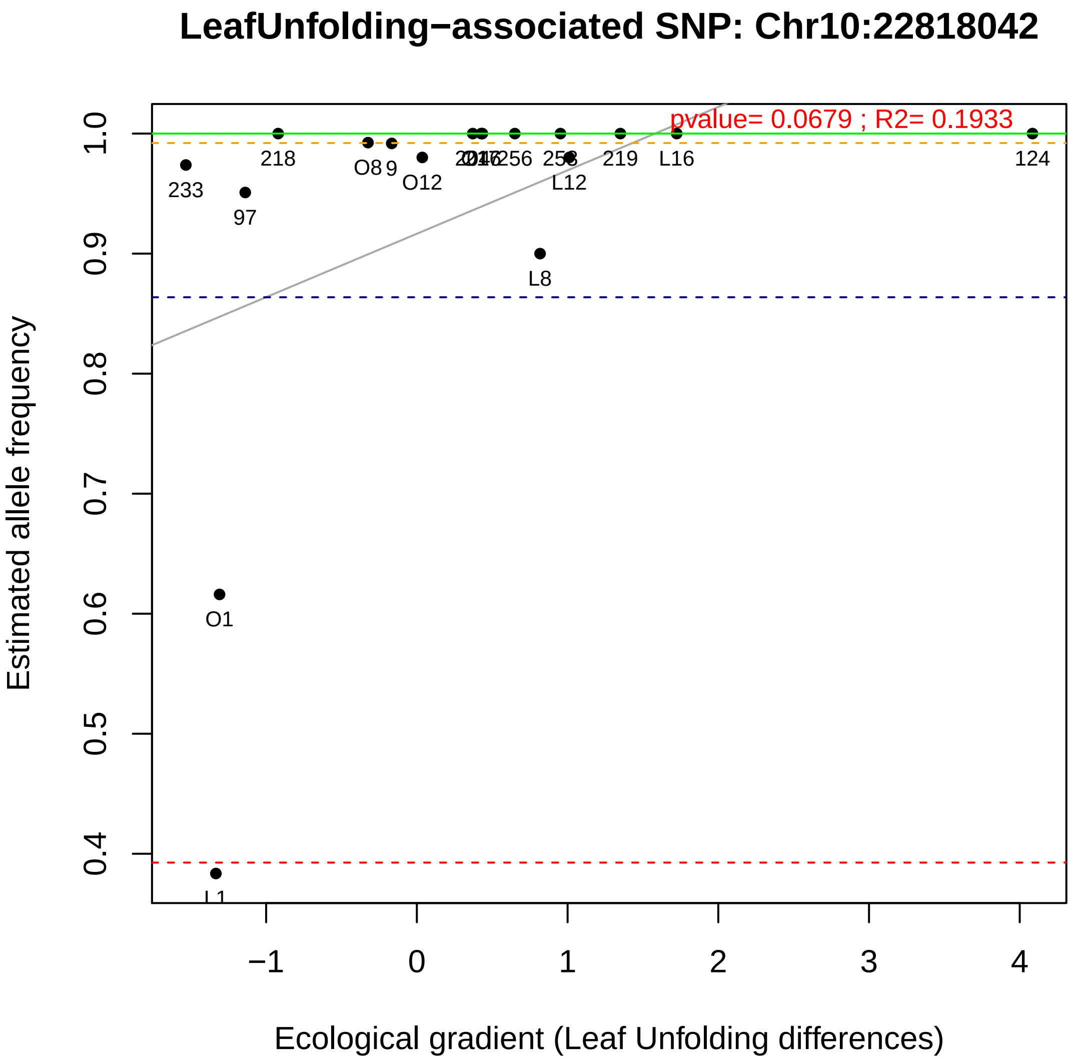

LeafUnfolding−associated SNP: Chr10:22818042
pvalue= 0.0679 ; R2= 0.1933
Estimated allele frequency
Ecological gradient (Leaf Unfolding differences)
1.0
0.9
0.8
0.7
0.6
0.5
0.4
−1
0
1
2
3
4
218
233
97
O8
9
O12
256
219
L16
124
L12
L8
O1

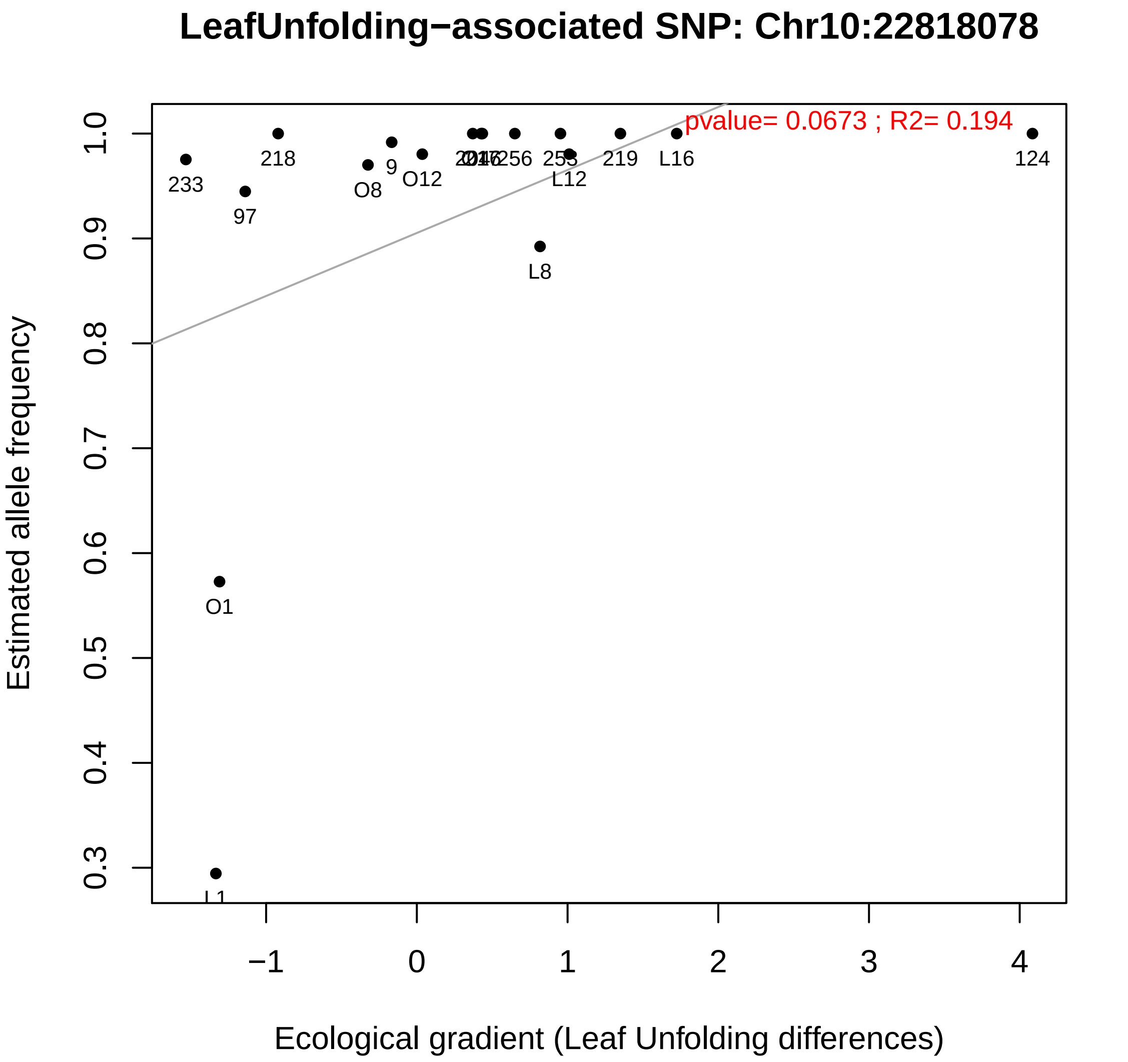

LeafUnfolding−associated SNP: Chr10:22818078
pvalue= 0.0673 ; R2= 0.194
Estimated allele frequency
Ecological gradient (Leaf Unfolding differences)
1.0
0.9
0.8
0.7
0.6
0.5
0.4
0.3
−1
0
1
2
3
4
233
97
218
O8
9
O12
256
253
L12
219
L16
124
L8
O1

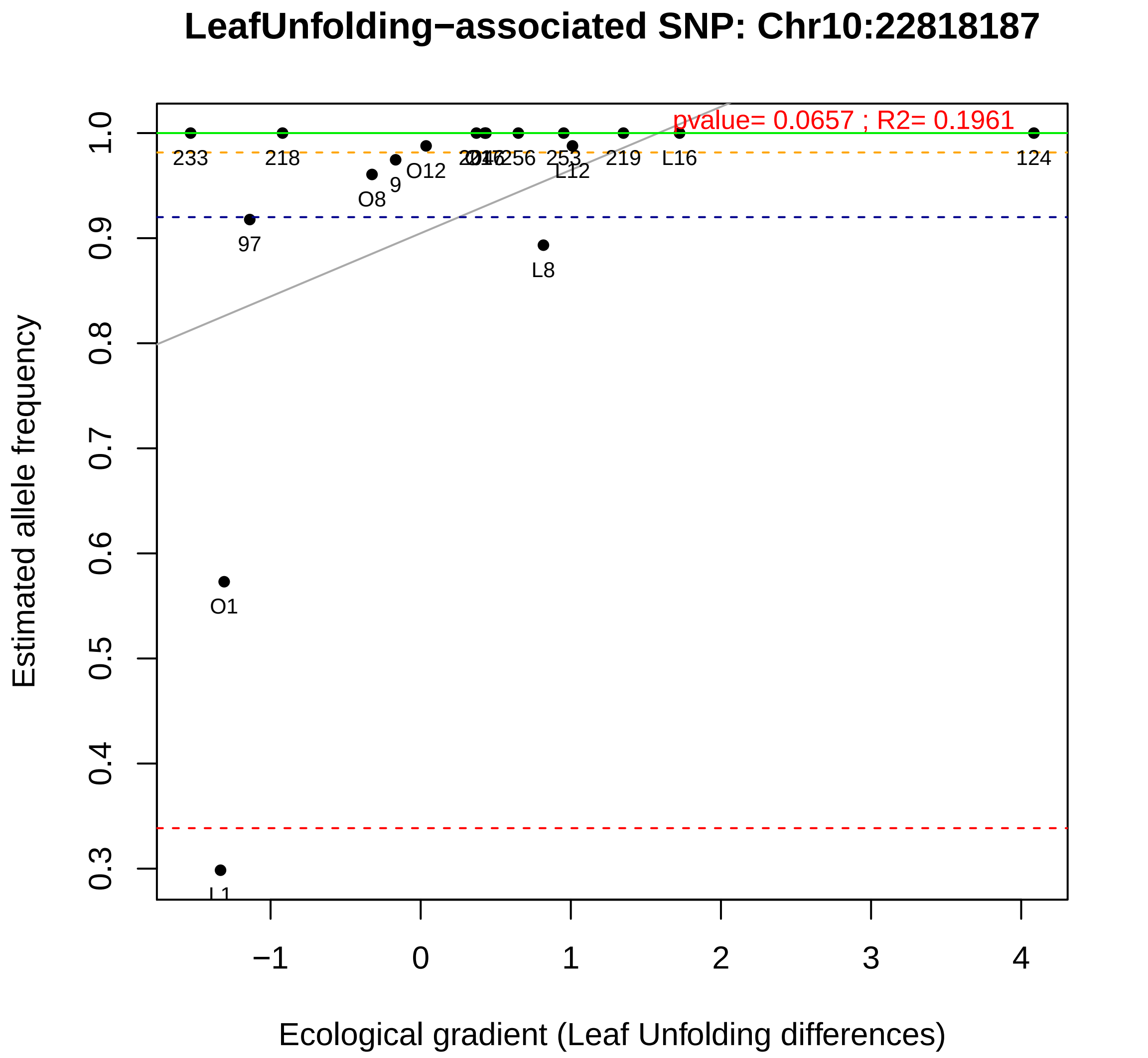
LeafUnfolding−associated SNP: Chr10:22818187
pvalue= 0.0657 ; R2= 0.1961
Estimated allele frequency
Ecological gradient (Leaf Unfolding differences)
1.0
0.9
0.8
0.7
0.6
0.5
0.4
0.3
−1
0
1
2
3
4
233
218
97
O8
9
O12
219
L16
124
L8
O1
L12
253
256

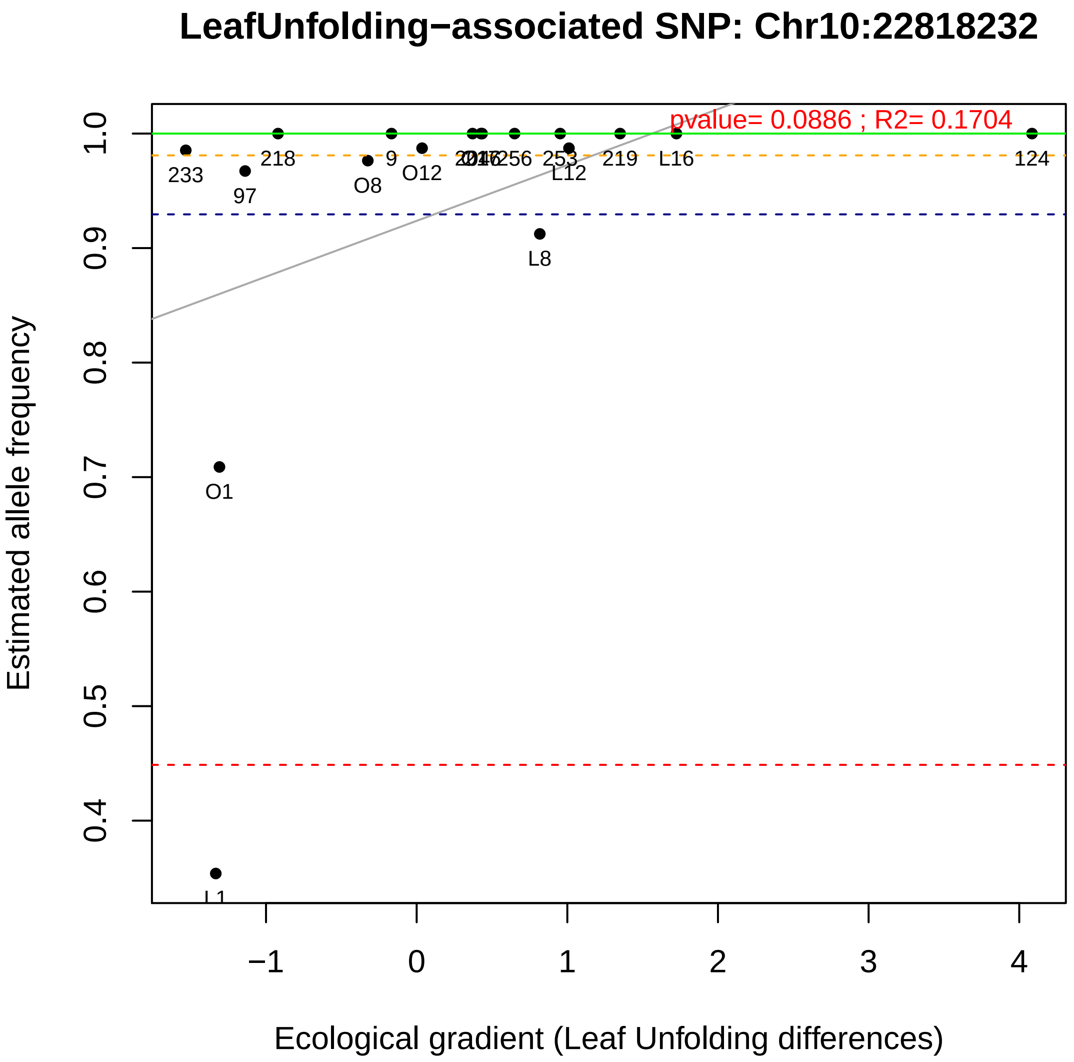
LeafUnfolding−associated SNP: Chr10:22818232
pvalue= 0.0886 ; R2= 0.1704
Estimated allele frequency
Ecological gradient (Leaf Unfolding differences)
0.4
0.5
0.6
0.7
0.8
0.9
1.0
−1
0
1
2
3
4
233
97
218
O8
9
O12
L8
253
L12
219
L16
124
O1
L1

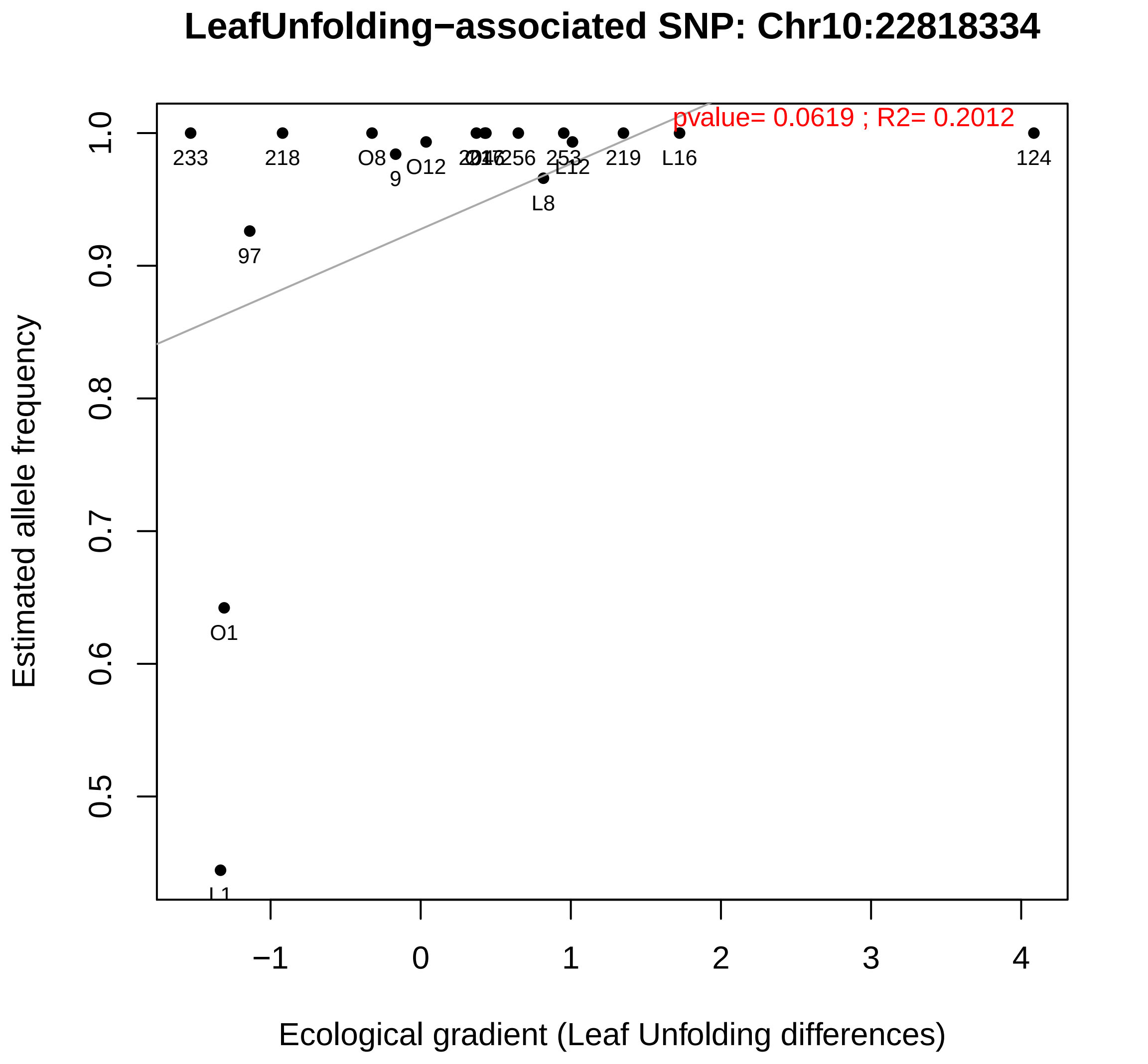

LeafUnfolding−associated SNP: Chr10:22818334
pvalue= 0.0619 ; R2= 0.2012
Estimated allele frequency
Ecological gradient (Leaf Unfolding differences)
233
218
O8
9
O12
256
253
L12
219
L16
124
L8
97
O1
L1

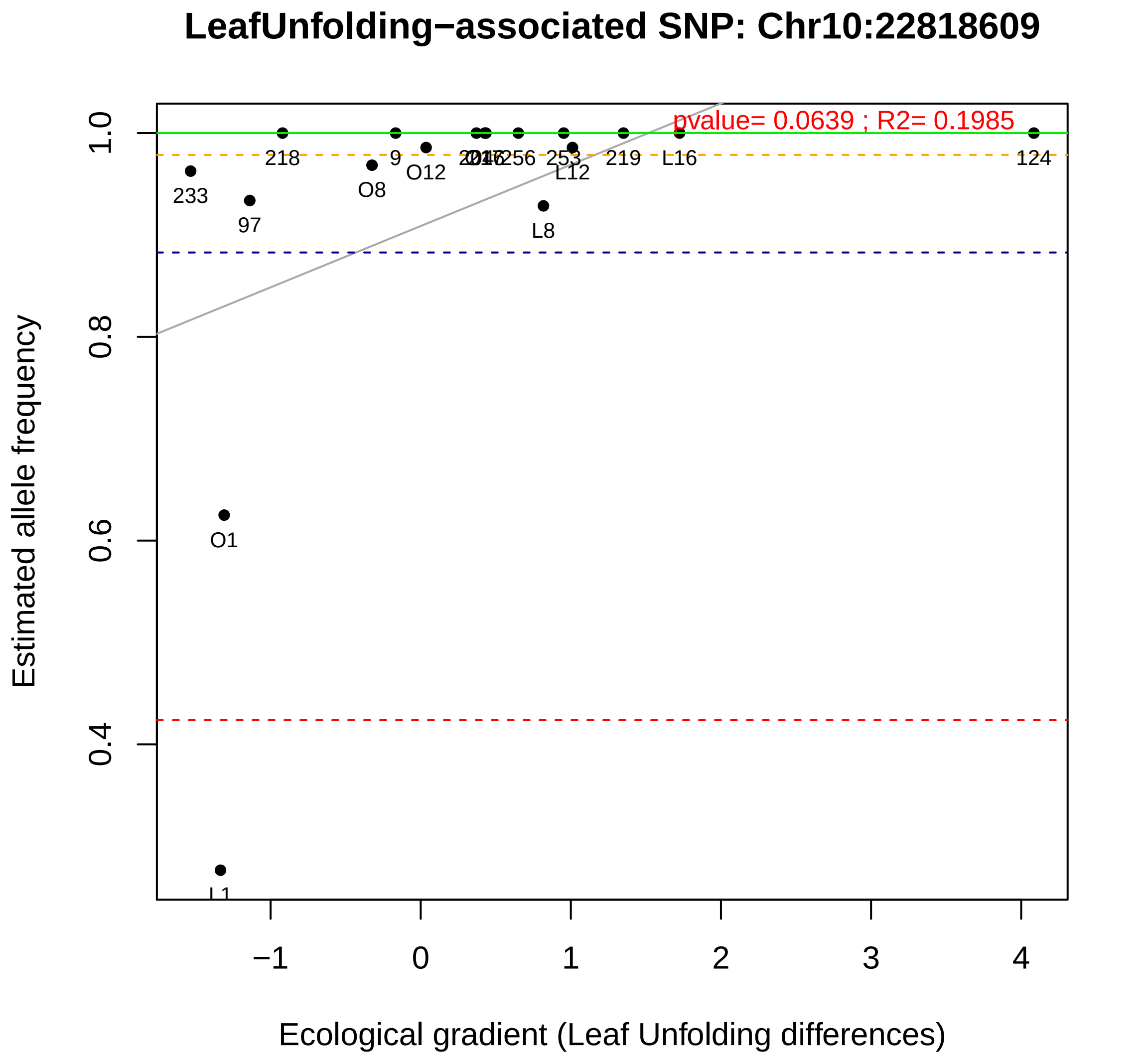

LeafUnfolding−associated SNP: Chr10:22818609
pvalue= 0.0639 ; R2= 0.1985
Estimated allele frequency
Ecological gradient (Leaf Unfolding differences)
1.0
0.8
0.6
0.4
−1
0
1
2
3
4
218
9
O12
256
253
L12
219
L16
124
233
O8
97
L8
O1
L1

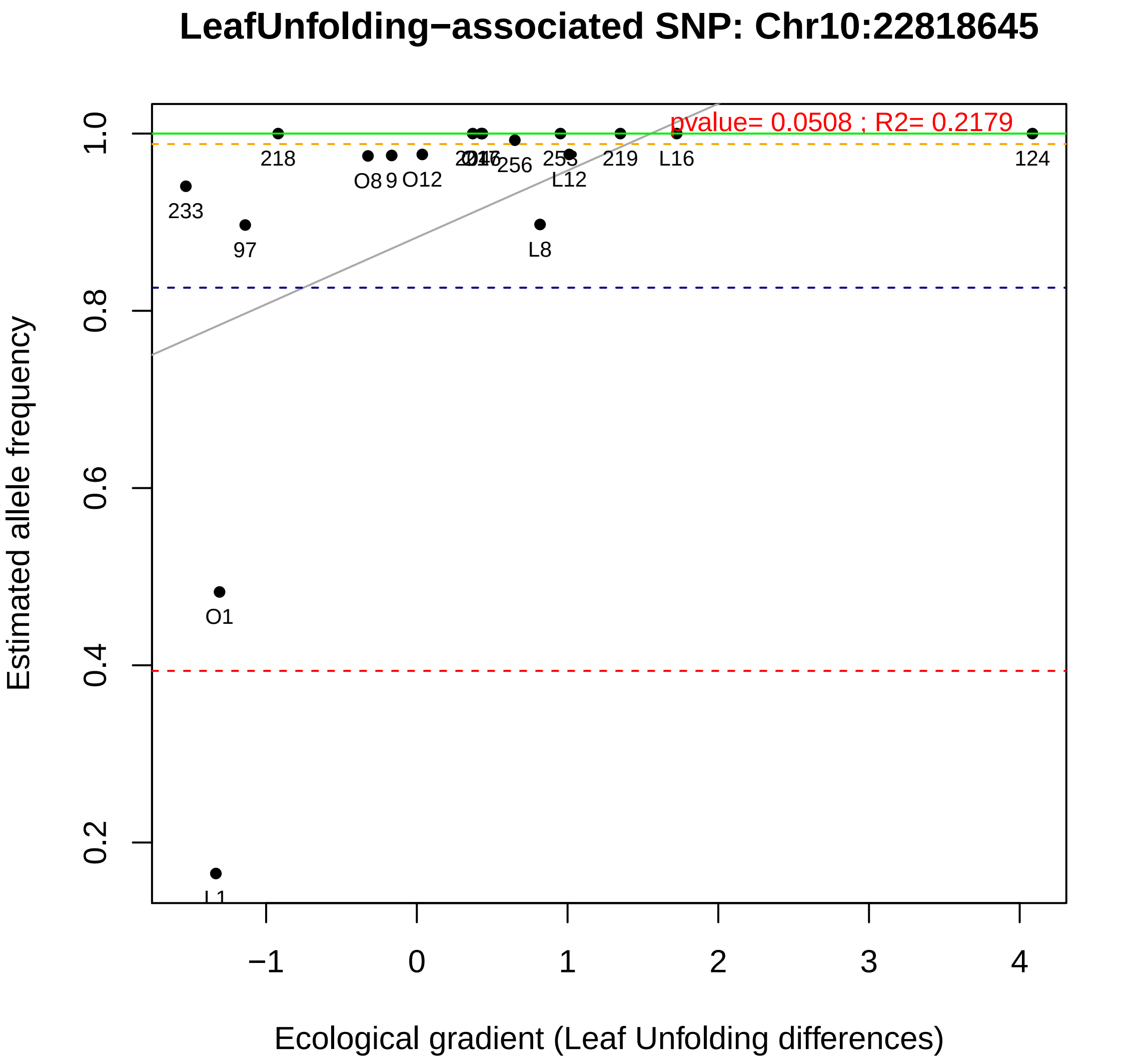

LeafUnfolding−associated SNP: Chr10:22818645
pvalue= 0.0508 ; R2= 0.2179
Estimated allele frequency
Ecological gradient (Leaf Unfolding differences)
0.2
0.4
0.6
0.8
1.0
−1
0
1
2
3
4
218
O8
9
O12
256
L12
219
L16
124
233
97
L8
O1

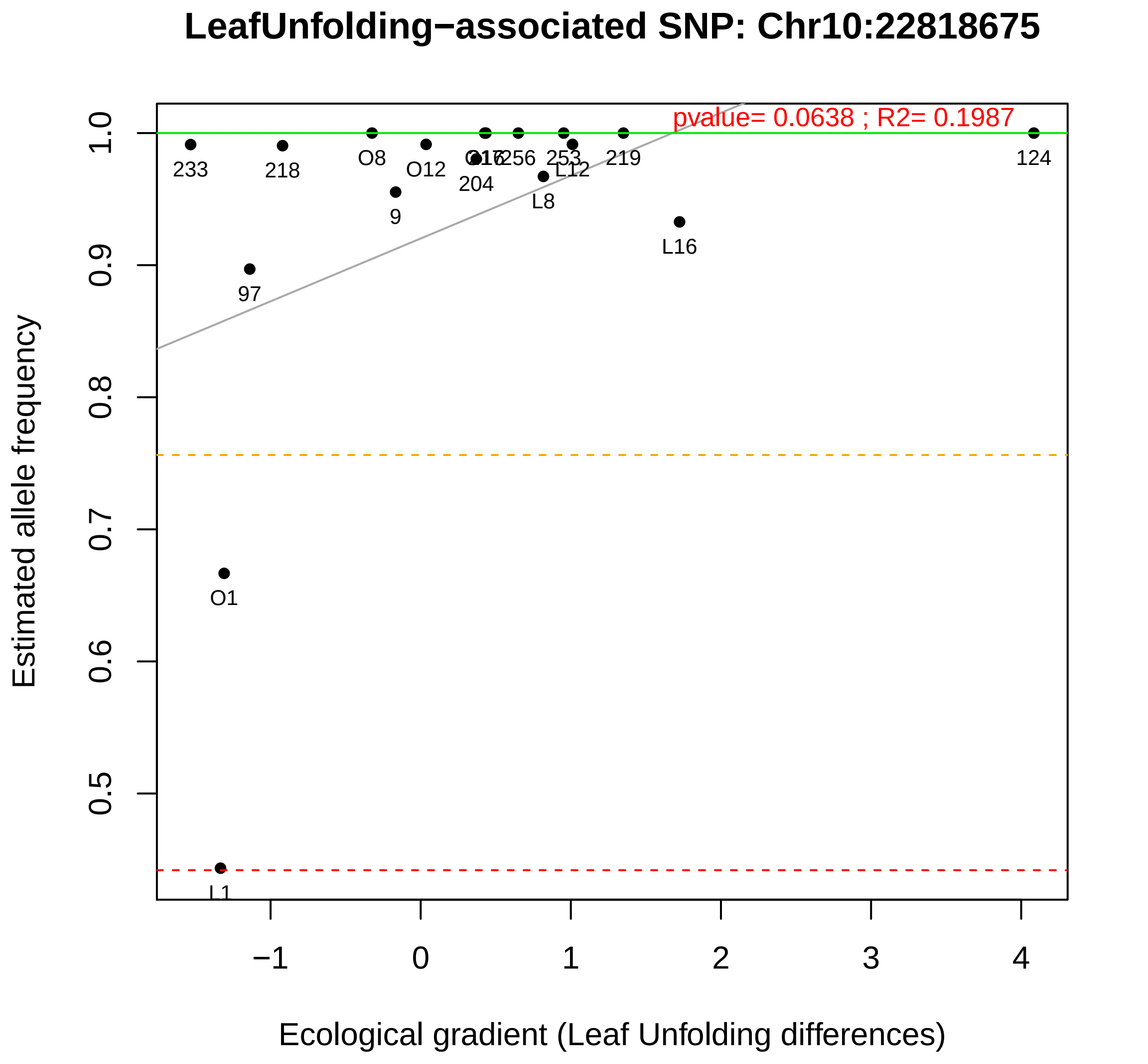
LeafUnfolding−associated SNP: Chr10:22818675
pvalue= 0.0638 ; R2= 0.1987
Estimated allele frequency
Ecological gradient (Leaf Unfolding differences)
1.0
0.9
0.8
0.7
0.6
0.5
−1
0
1
2
3
4
233
218
O8
O12
256
253
219
124
204
L12
L8
9
L16
97
O1
L1

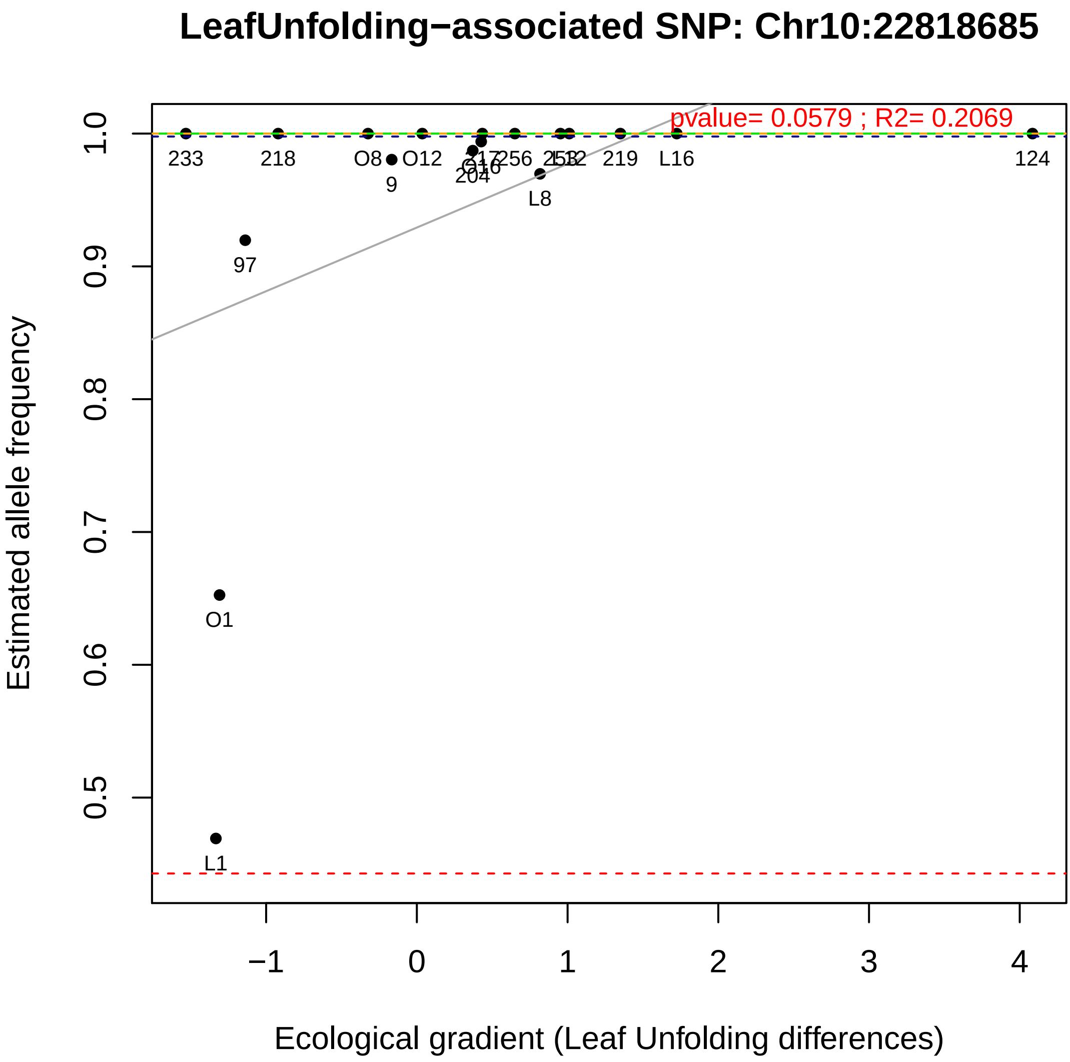

LeafUnfolding−associated SNP: Chr10:22818685
pvalue= 0.0579 ; R2= 0.2069
233
218
O8
O12
9
217
256
204
O16
251
L12
219
L16
124
L8
97
O1
L1
1.0
0.9
0.8
0.7
0.6
0.5
−1
0
1
2
3
4
Estimated allele frequency
Ecological gradient (Leaf Unfolding differences)

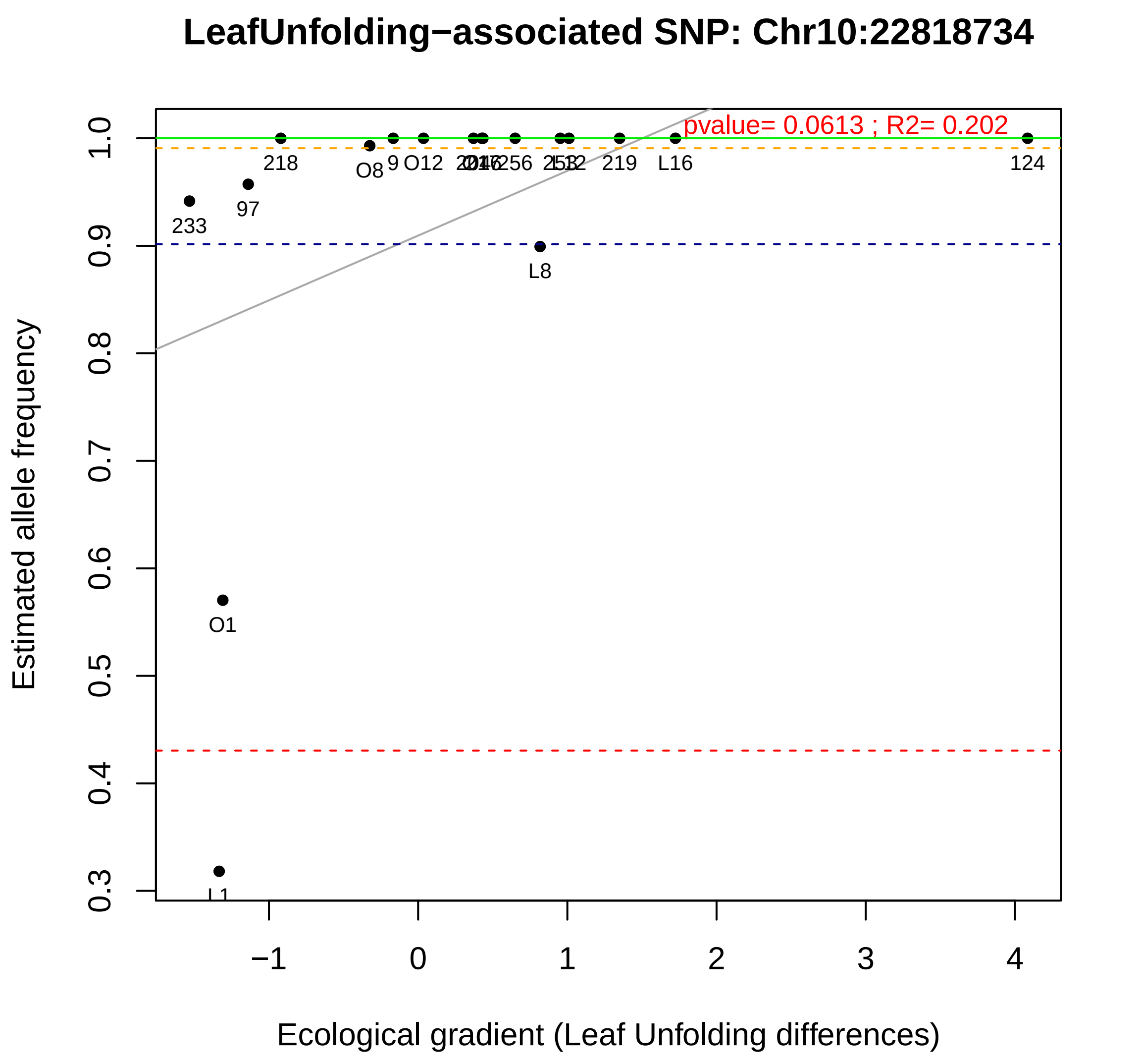
LeafUnfolding−associated SNP: Chr10:22818734
pvalue= 0.0613 ; R2= 0.202
Estimated allele frequency
Ecological gradient (Leaf Unfolding differences)
1.0
0.9
0.8
0.7
0.6
0.5
0.4
0.3
−1
0
1
2
3
4
218
O8
9
O12
256
219
L16
124
233
97
L8
O1

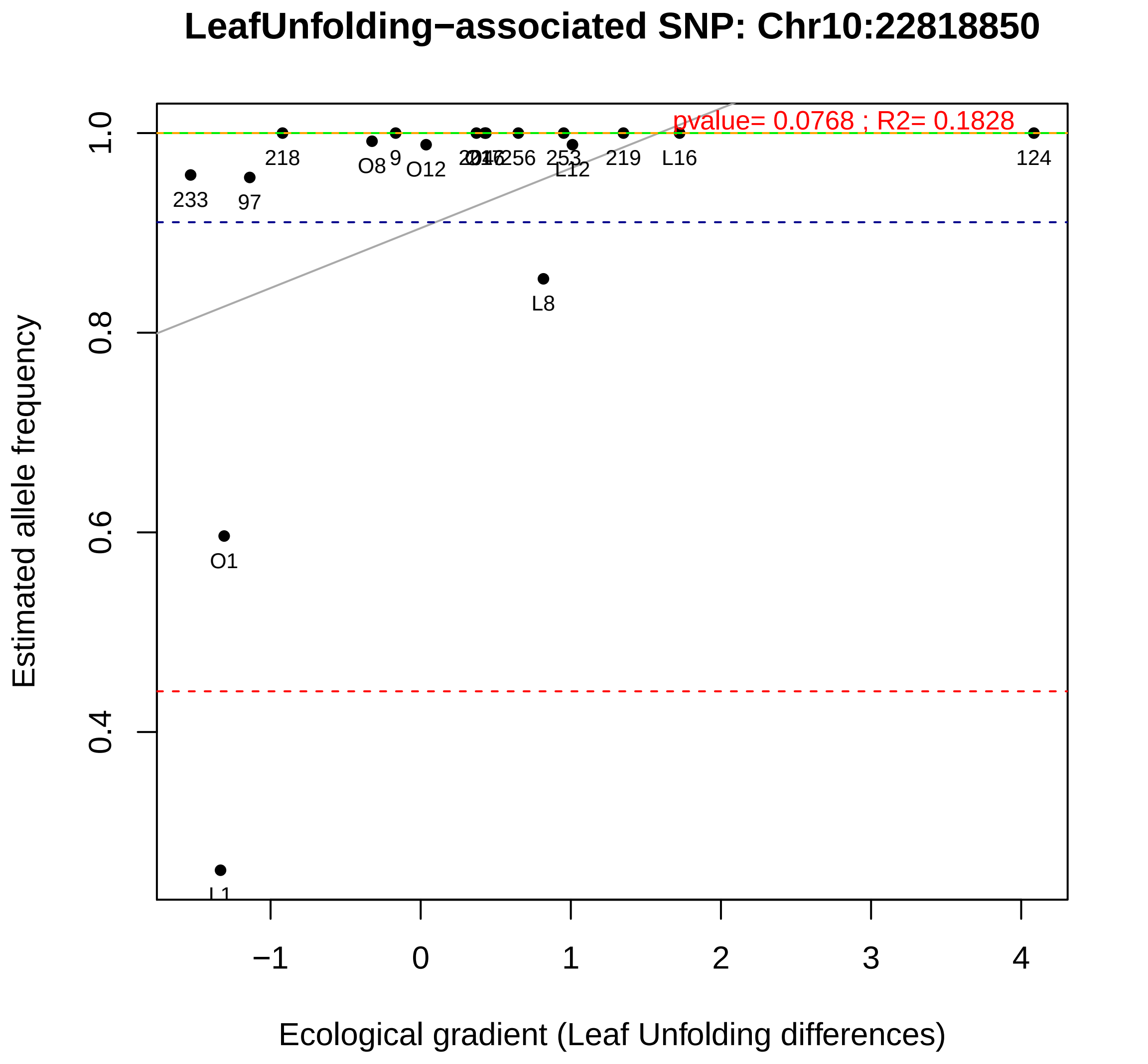
LeafUnfolding−associated SNP: Chr10:22818850
pvalue= 0.0768 ; R2= 0.1828
Estimated allele frequency
Ecological gradient (Leaf Unfolding differences)
1.0
0.8
0.6
0.4
−1
0
1
2
3
4
218
O8
9
O12
256
253
L12
219
L16
124
233
97
L8
O1
L1

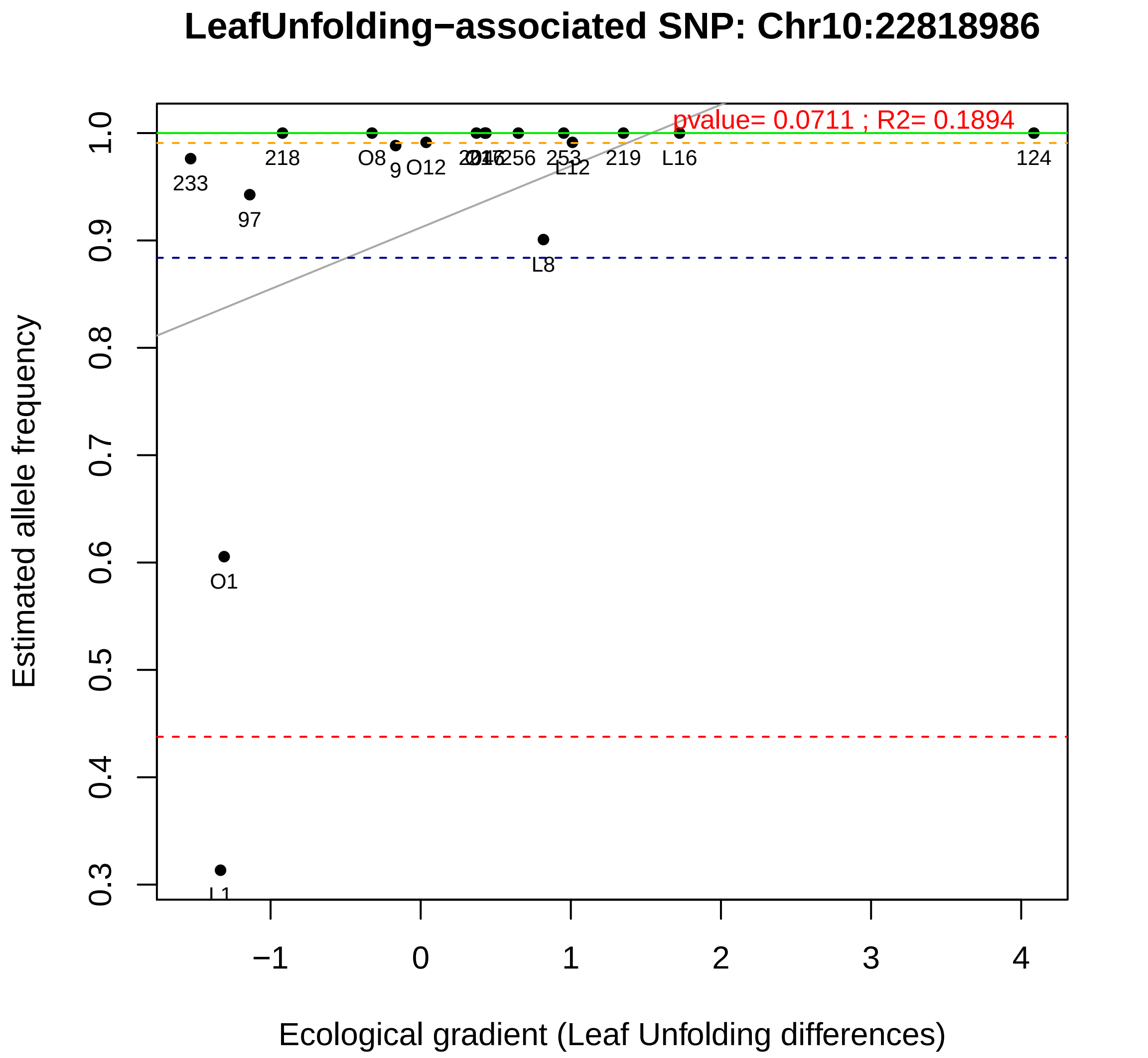

LeafUnfolding−associated SNP: Chr10:22818986
pvalue= 0.0711 ; R2= 0.1894
Estimated allele frequency
Ecological gradient (Leaf Unfolding differences)
0.3
0.4
0.5
0.6
0.7
0.8
0.9
1.0
−1
0
1
2
3
4
233
218
O8
9
O12
256
253
L12
219
L16
124
97
L8
O1
L1

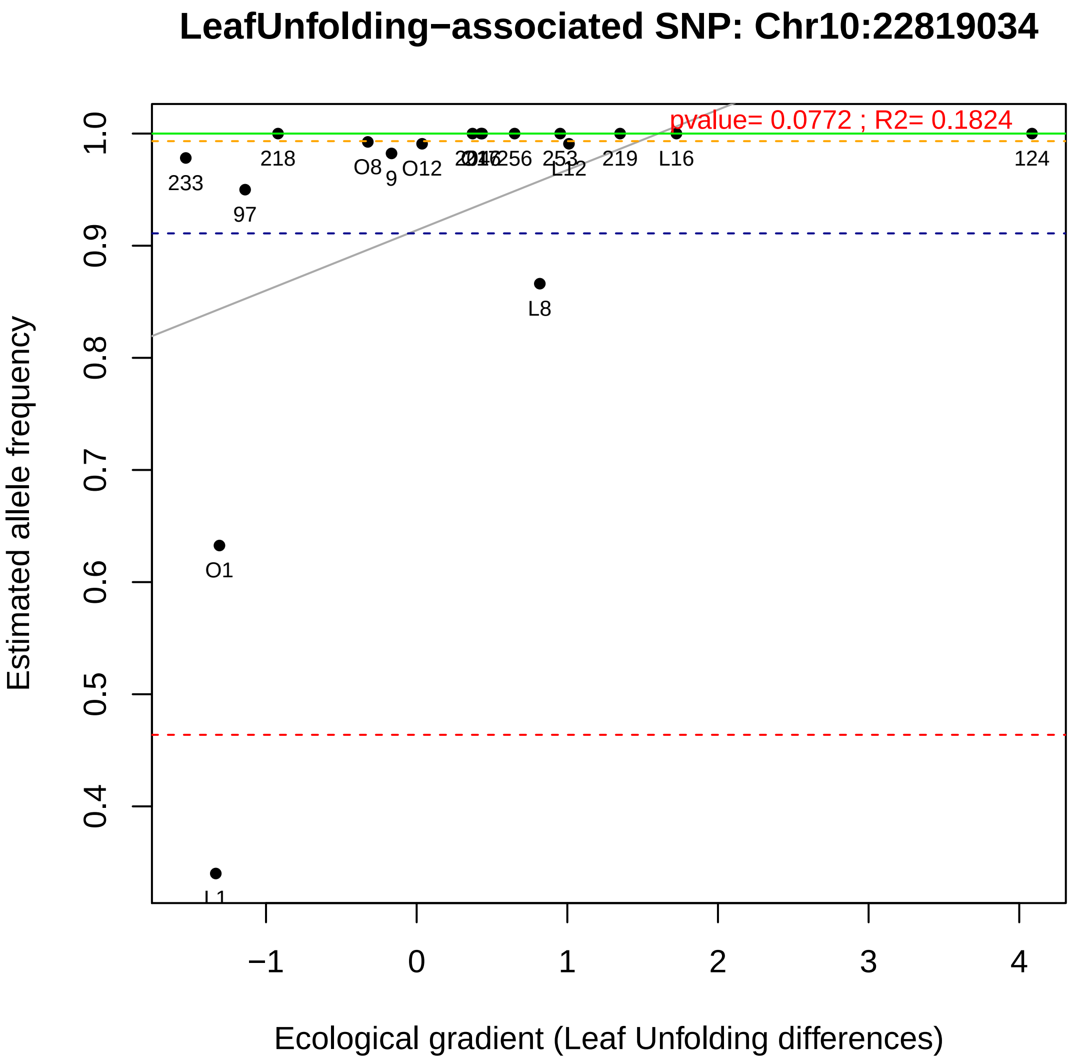
LeafUnfolding−associated SNP: Chr10:22819034
pvalue= 0.0772 ; R2= 0.1824
Estimated allele frequency
Ecological gradient (Leaf Unfolding differences)
1.0
0.9
0.8
0.7
0.6
0.5
0.4
−1
0
1
2
3
4
233
218
O8
9
O12
256
253
L12
219
L16
124
97
L8
O1
L1

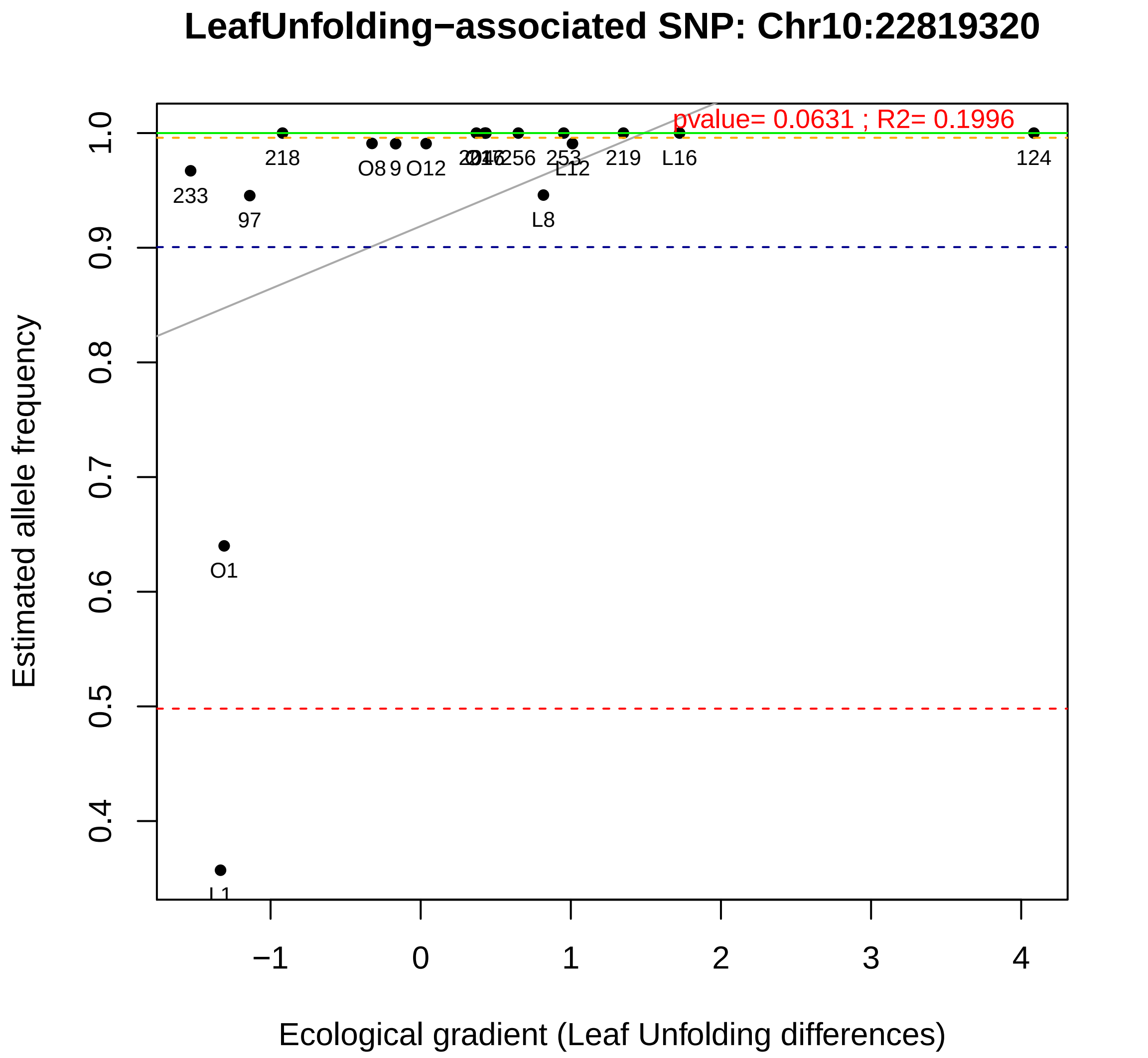

LeafUnfolding−associated SNP: Chr10:22819320
pvalue= 0.0631 ; R2= 0.1996
Estimated allele frequency
Ecological gradient (Leaf Unfolding differences)
1.0
0.9
0.8
0.7
0.6
0.5
0.4
−1
0
1
2
3
4
218
233
97
O8
9
O12
256
253
L12
L8
219
L16
124
O1
L1

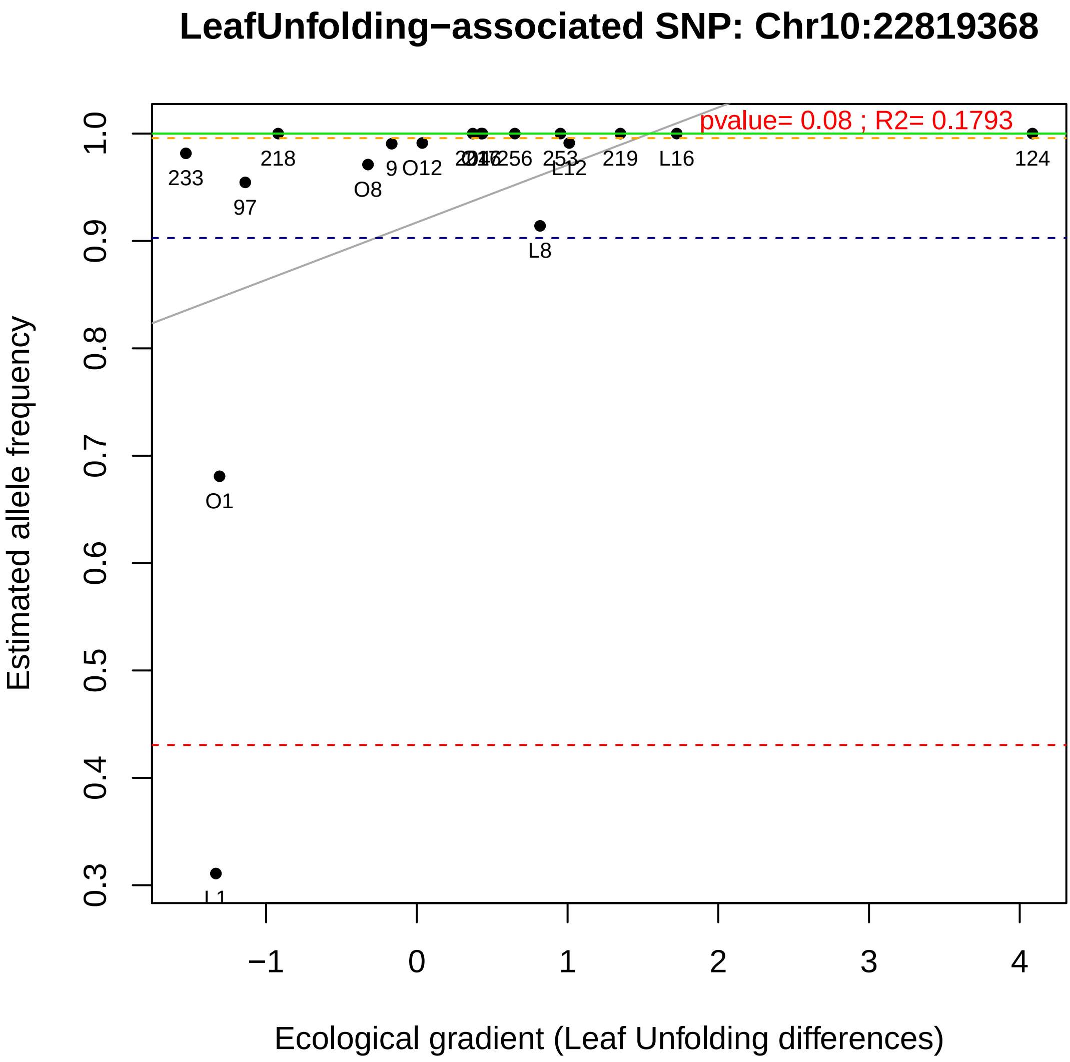

LeafUnfolding−associated SNP: Chr10:22819368
pvalue= 0.08 ; R2= 0.1793
Estimated allele frequency
Ecological gradient (Leaf Unfolding differences)
0.3
0.4
0.5
0.6
0.7
0.8
0.9
1.0
−1
0
1
2
3
4
233
218
97
O8
9
O12
256
253
L12
219
L16
124
L8
O1

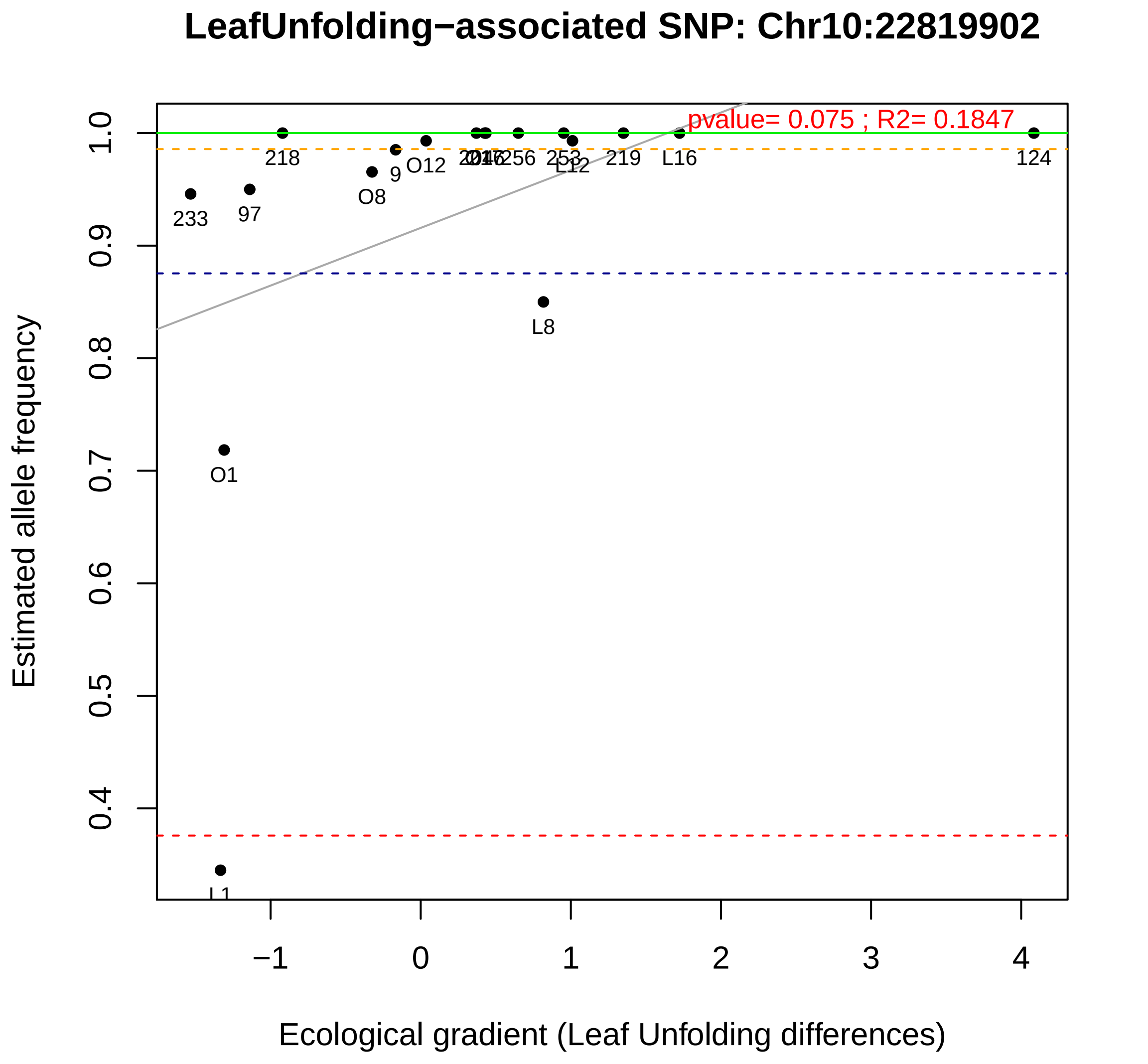
LeafUnfolding−associated SNP: Chr10:22819902
pvalue= 0.075 ; R2= 0.1847
Estimated allele frequency
Ecological gradient (Leaf Unfolding differences)
218
O12
9
O8
233
97
256
253
L12
219
L16
124
L8
O1
L1
1.0
0.9
0.8
0.7
0.6
0.5
0.4
−1
0
1
2
3
4

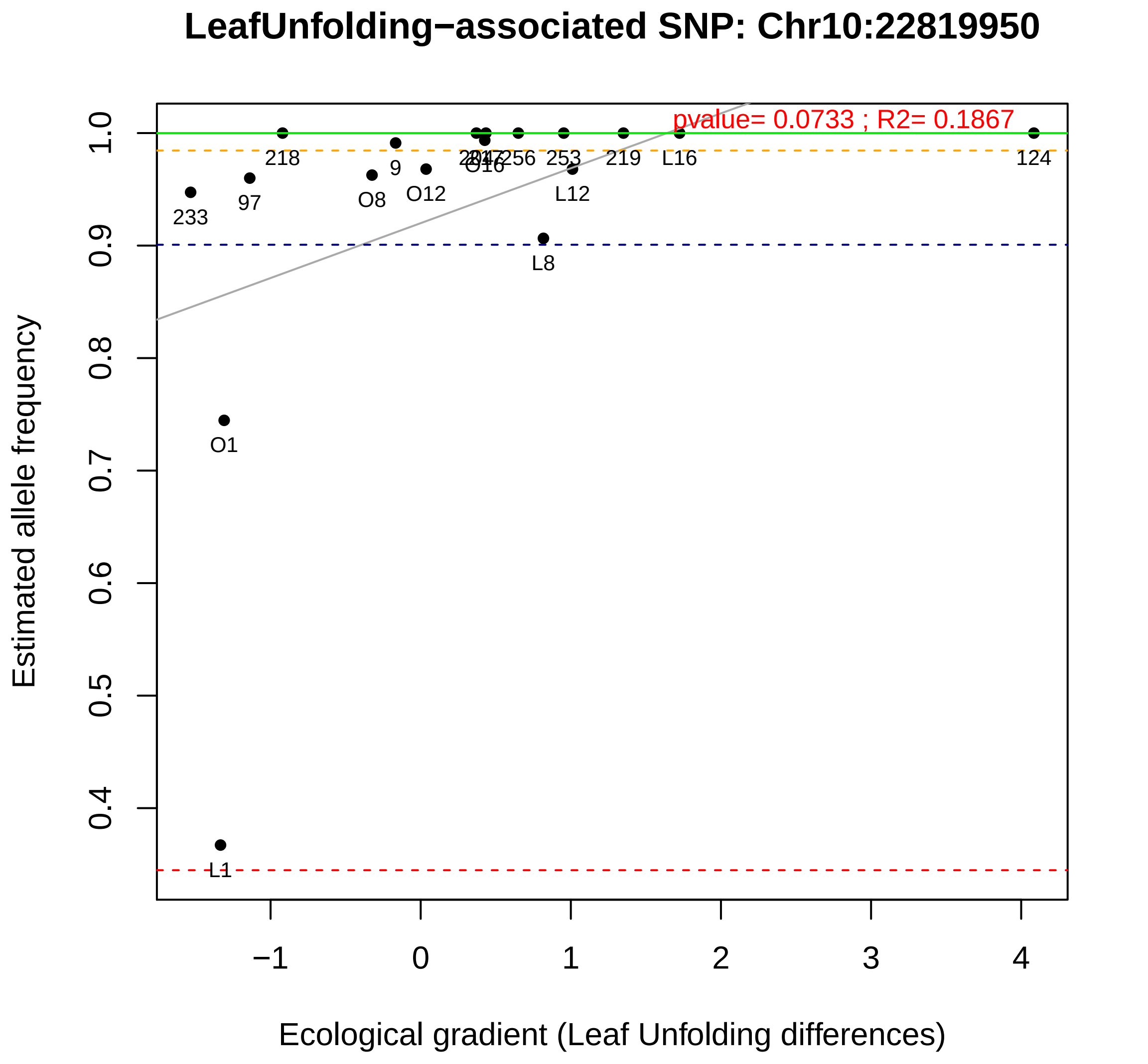

LeafUnfolding−associated SNP: Chr10:22819950
pvalue= 0.0733 ; R2= 0.1867
Estimated allele frequency
Ecological gradient (Leaf Unfolding differences)
1.0
0.9
0.8
0.7
0.6
0.5
0.4
−1
0
1
2
3
4
218
9
256
253
219
L16
124
233
97
O8
O12
O16
L12
L8
O1
L1

**LeafUnfolding−associated SNP: Chr10:22820059**

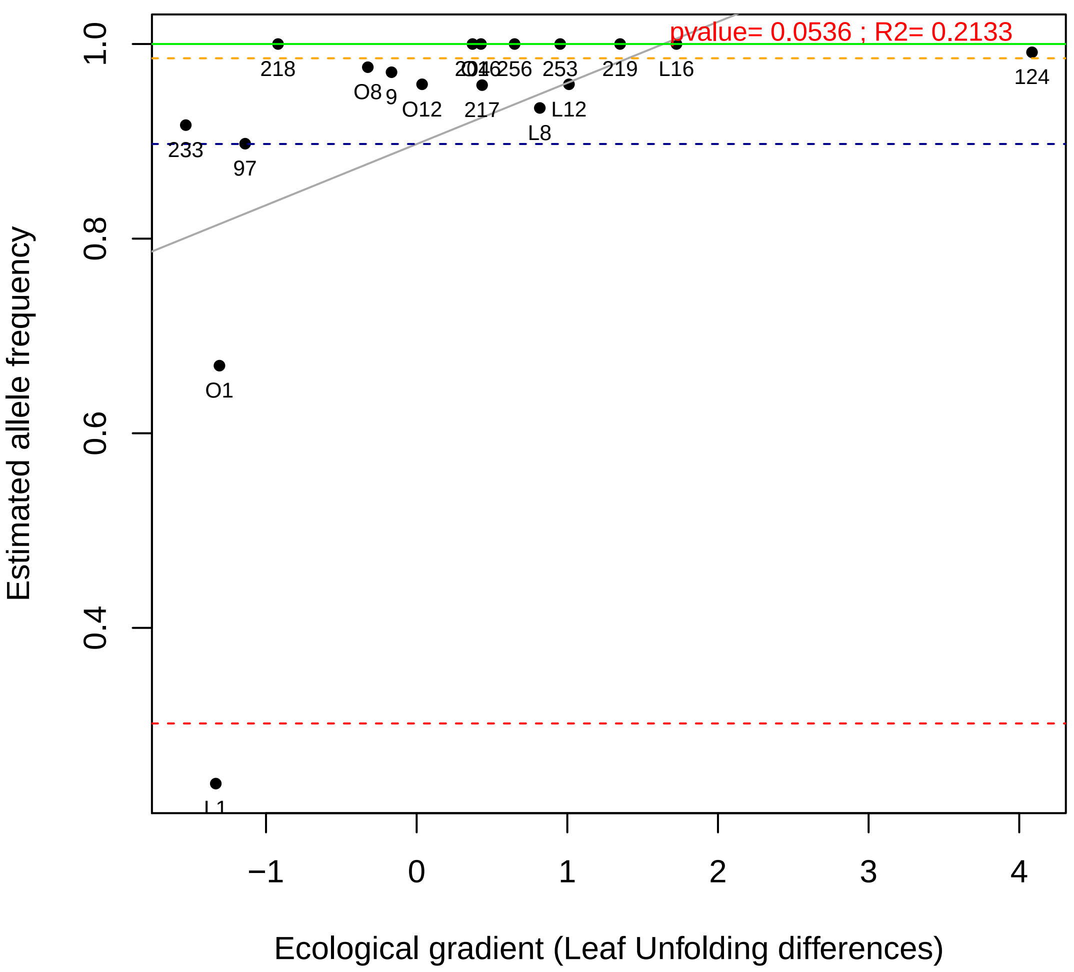

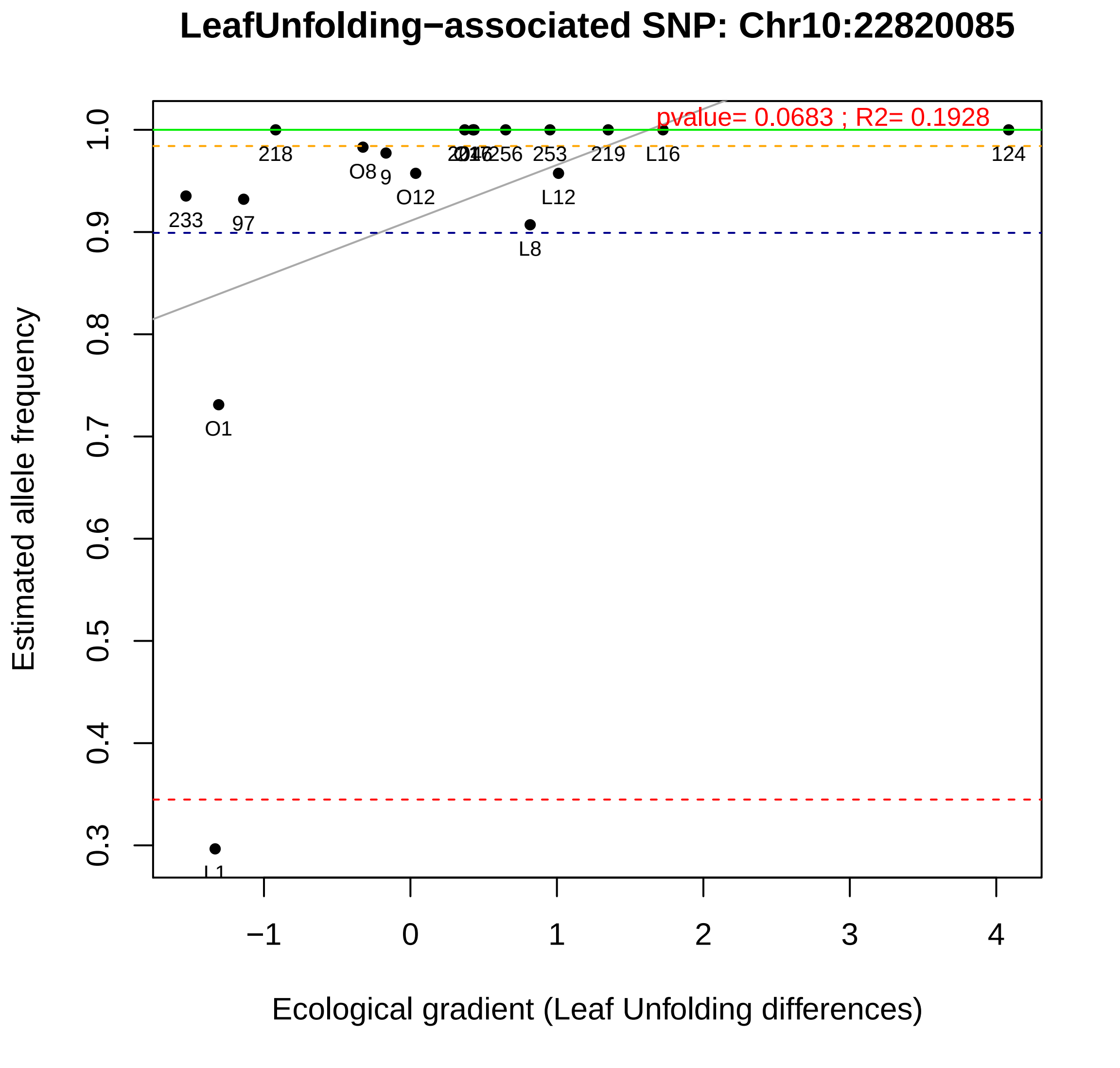

LeafUnfolding−associated SNP: Chr10:22820085
pvalue= 0.0683 ; R2= 0.1928
Estimated allele frequency
Ecological gradient (Leaf Unfolding differences)
1.0
0.9
0.8
0.7
0.6
0.5
0.4
0.3
−1
0
1
2
3
4
218
O8
9
O12
256
253
219
L16
124
L12
L8
233
97
O1

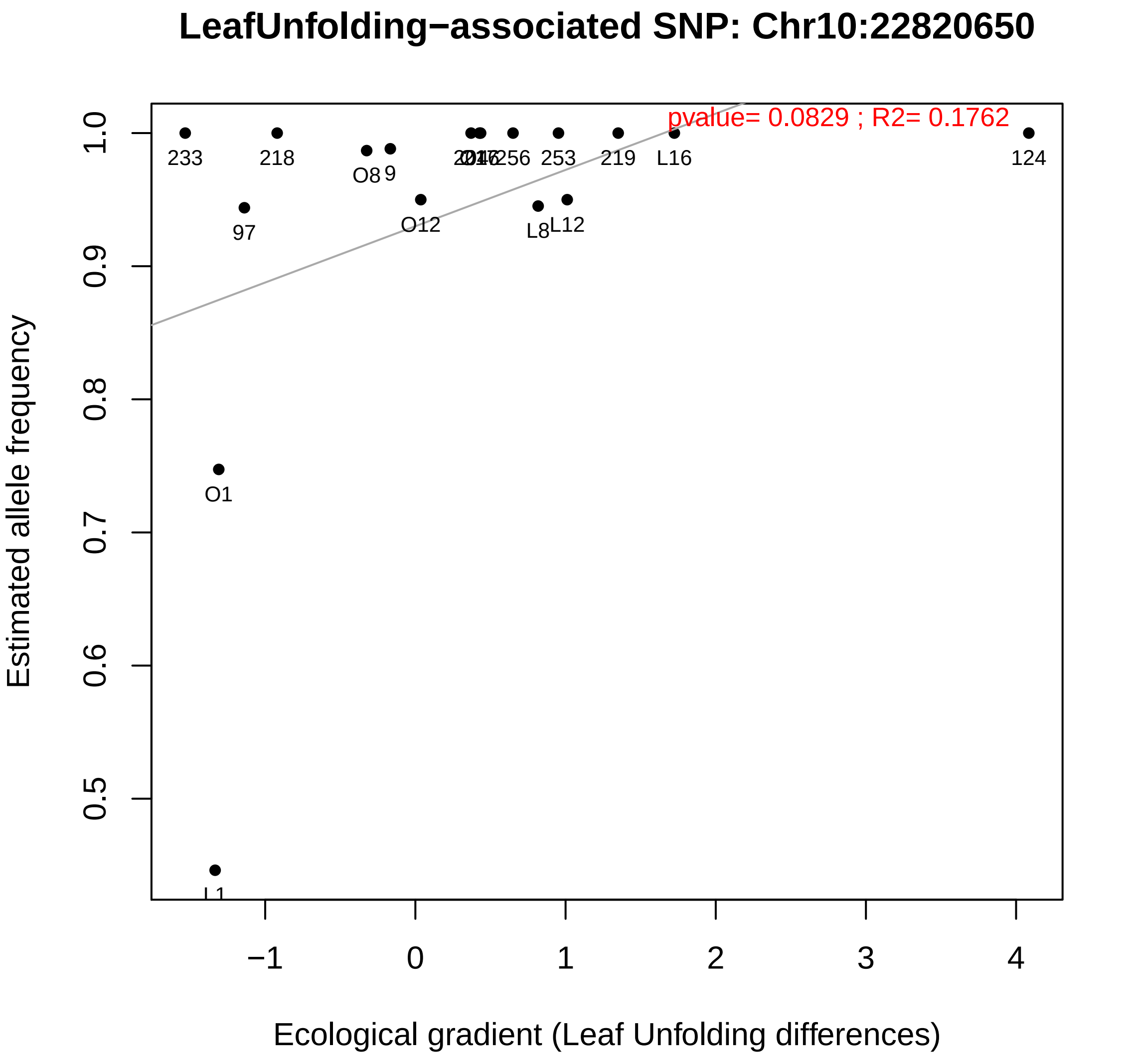
LeafUnfolding−associated SNP: Chr10:22820650
pvalue= 0.0829 ; R2= 0.1762
Estimated allele frequency
Ecological gradient (Leaf Unfolding differences)
1.0
0.9
0.8
0.7
0.6
0.5
−1
0
1
2
3
4
233
218
O8
9
256
253
219
L16
124
97
O12
L8
L12
O1
L1

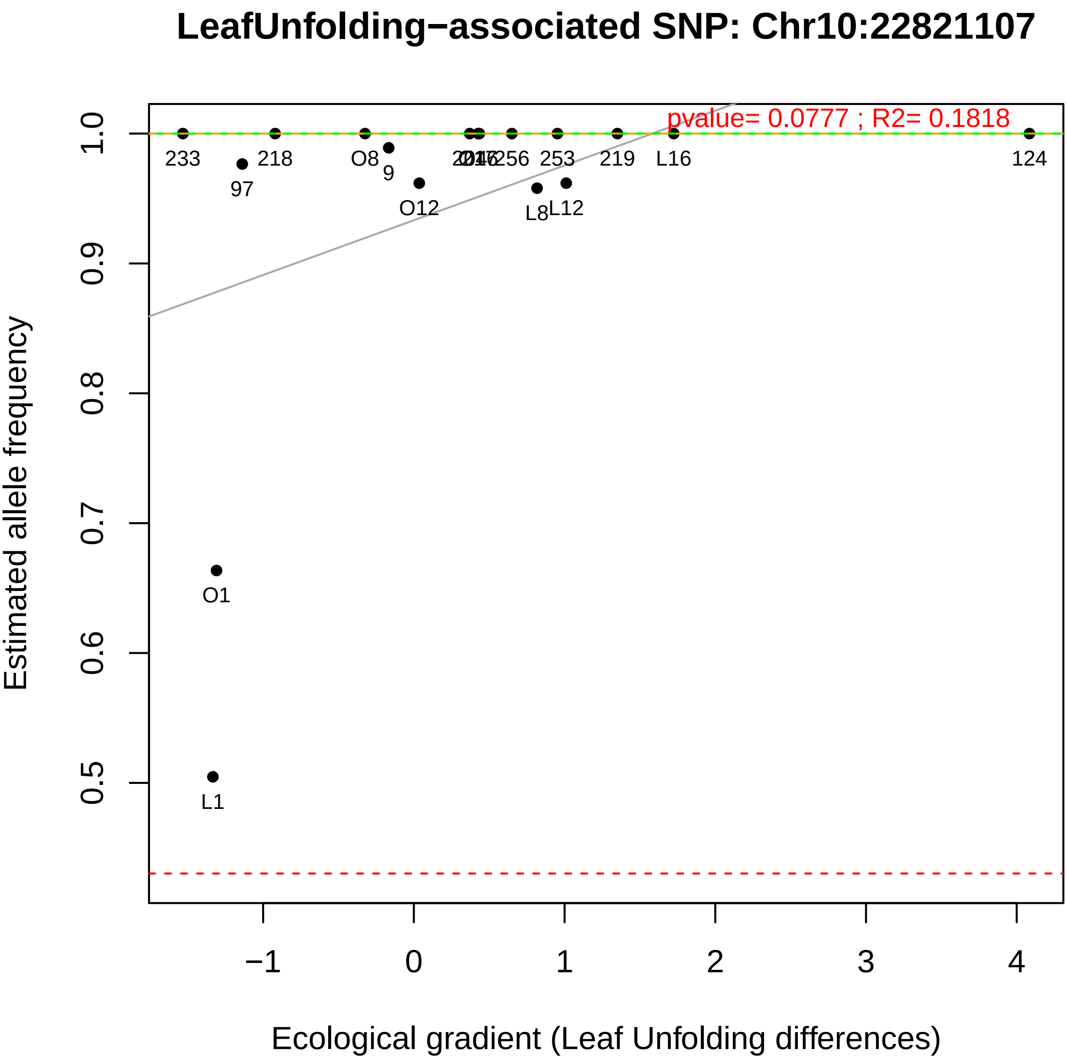
LeafUnfolding−associated SNP: Chr10:22821107
pvalue= 0.0777 ; R2= 0.1818
Estimated allele frequency
Ecological gradient (Leaf Unfolding differences)
1.0
0.9
0.8
0.7
0.6
0.5
−1
0
1
2
3
4
233
97
218
O8
9
O12
256
253
L8
L12
219
L16
124
O1
L1

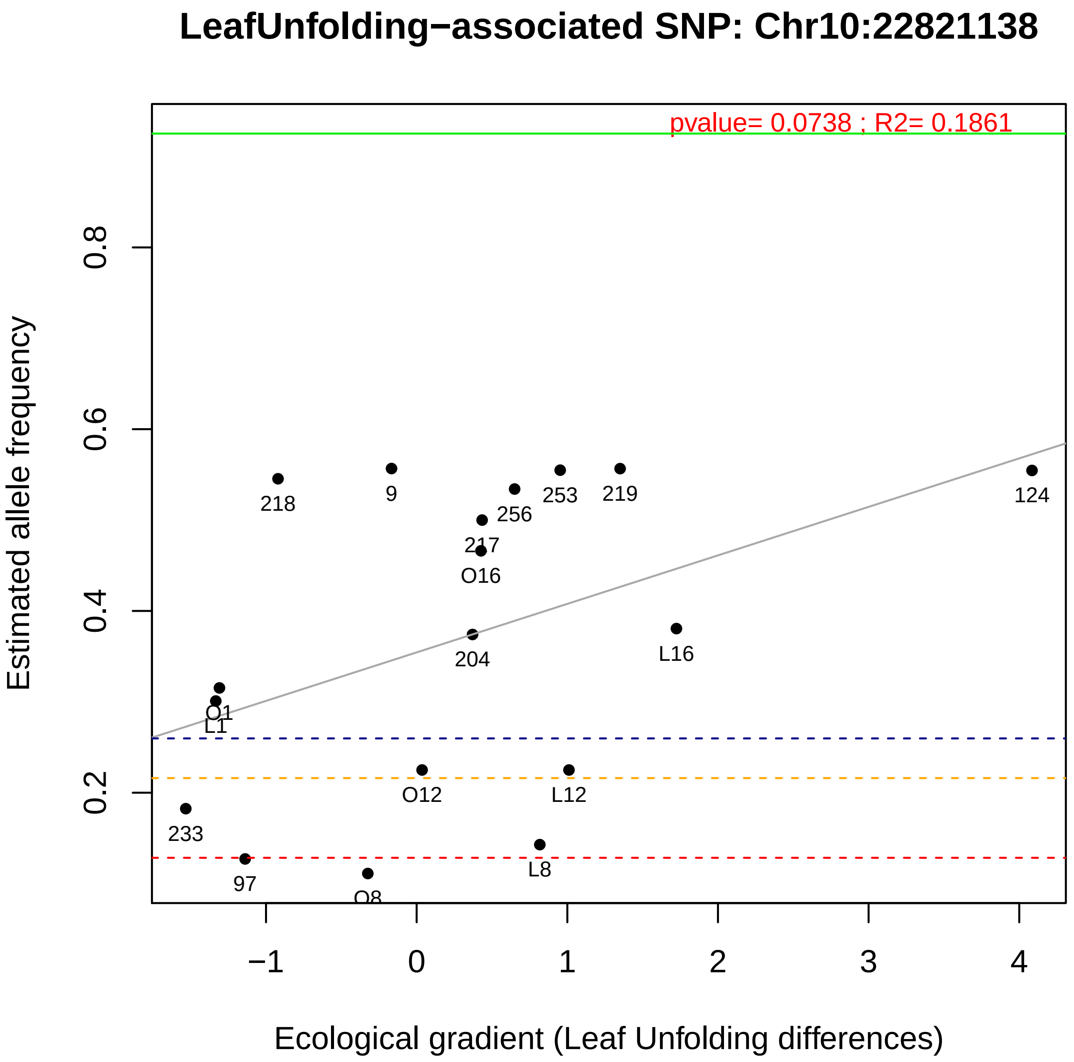
LeafUnfolding−associated SNP: Chr10:22821138
pvalue= 0.0738 ; R2= 0.1861
Estimated allele frequency
0.2
0.4
0.6
0.8
Ecological gradient (Leaf Unfolding differences)
−1
0
1
2
3
4
218
9
256
253
219
124
217
O16
204
L16
O1
L1
O12
L12
233
97
O8
L8

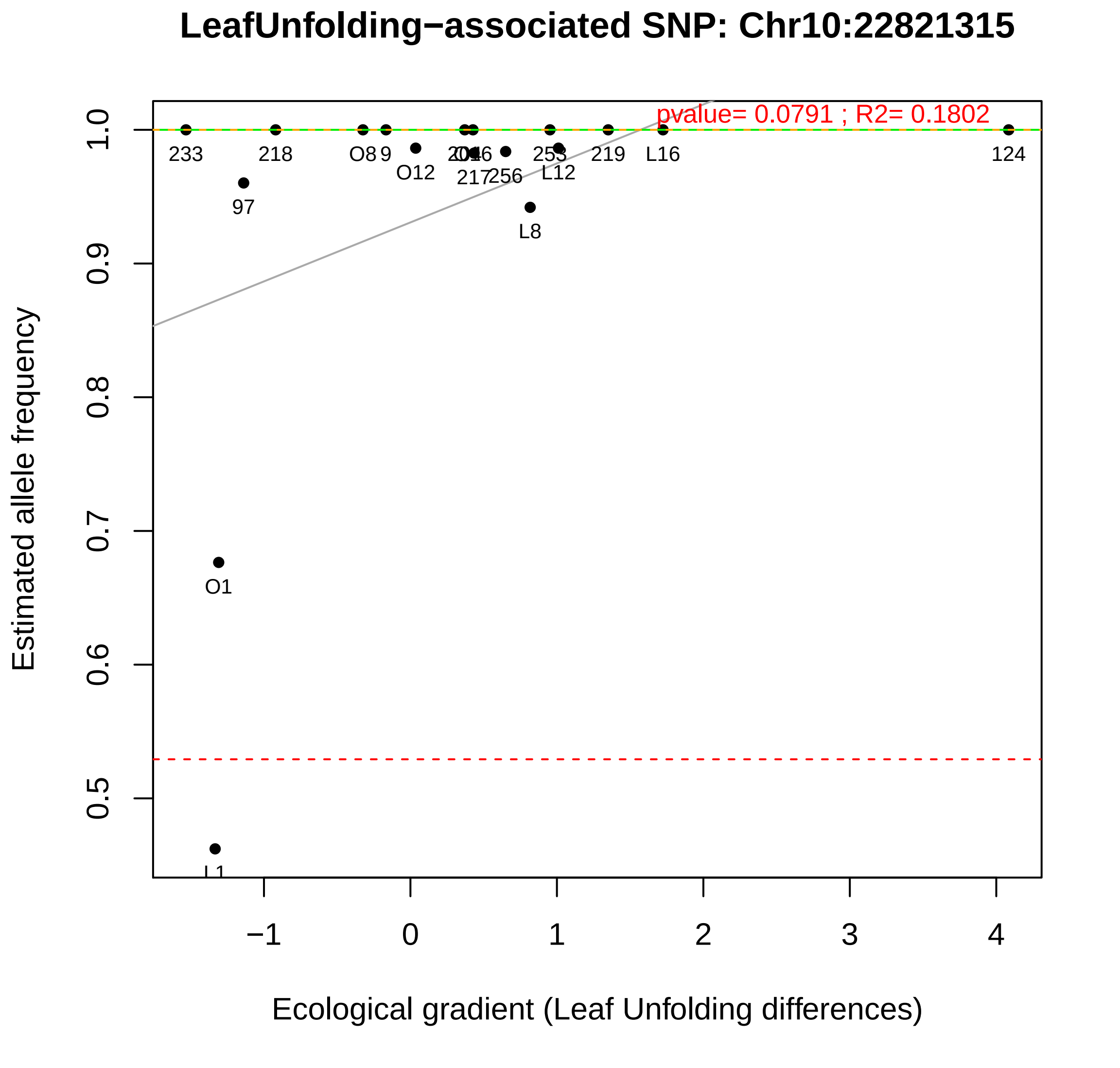

LeafUnfolding−associated SNP: Chr10:22821315
pvalue= 0.0791 ; R2= 0.1802
Estimated allele frequency
Ecological gradient (Leaf Unfolding differences)
0.5
0.6
0.7
0.8
0.9
1.0
−1
0
1
2
3
4
233
218
O8
9
O12
217
256
253
L12
219
L16
124
97
L8
O1

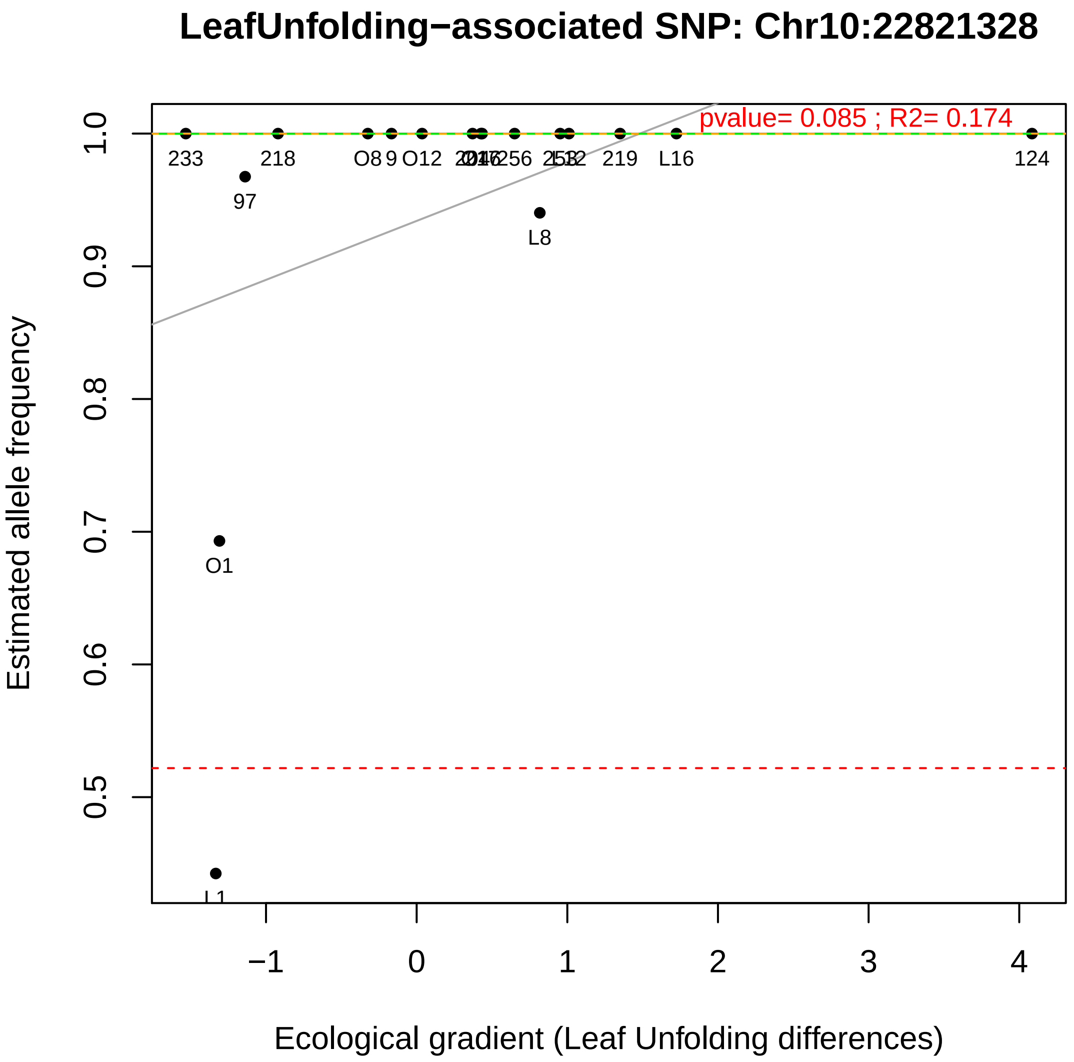
LeafUnfolding−associated SNP: Chr10:22821328
pvalue= 0.085 ; R2= 0.174
Estimated allele frequency
Ecological gradient (Leaf Unfolding differences)
1.0
0.9
0.8
0.7
0.6
0.5
−1
0
1
2
3
4
233
218
O8
9
O12
219
L16
124
97
L8
O1

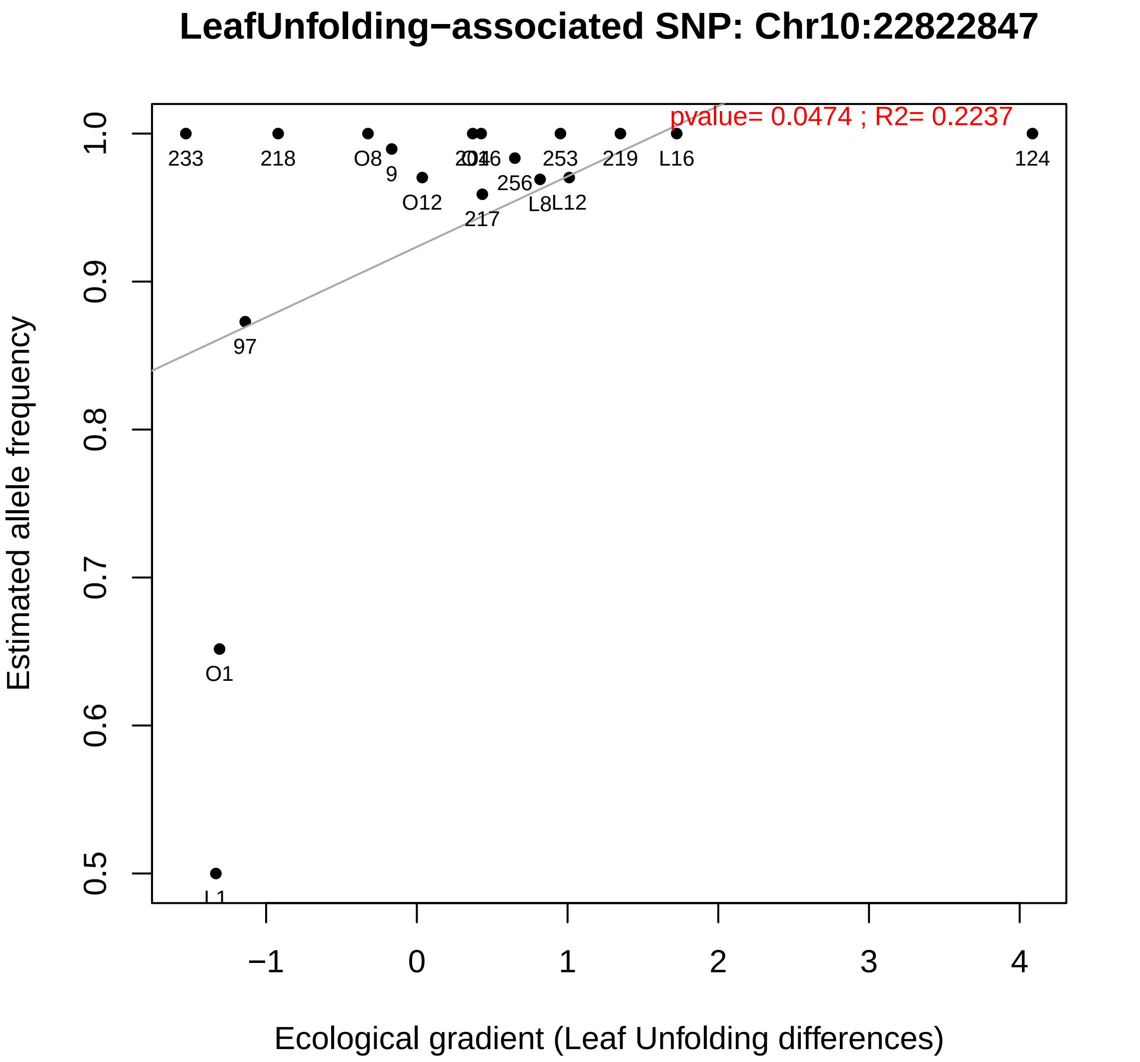

LeafUnfolding−associated SNP: Chr10:22822847
pvalue= 0.0474 ; R2= 0.2237
Estimated allele frequency
Ecological gradient (Leaf Unfolding differences)
1.0
0.9
0.8
0.7
0.6
0.5
−1
0
1
2
3
4
233
218
O8
9
O12
217
256
L8
L12
253
219
L16
124
97
O1

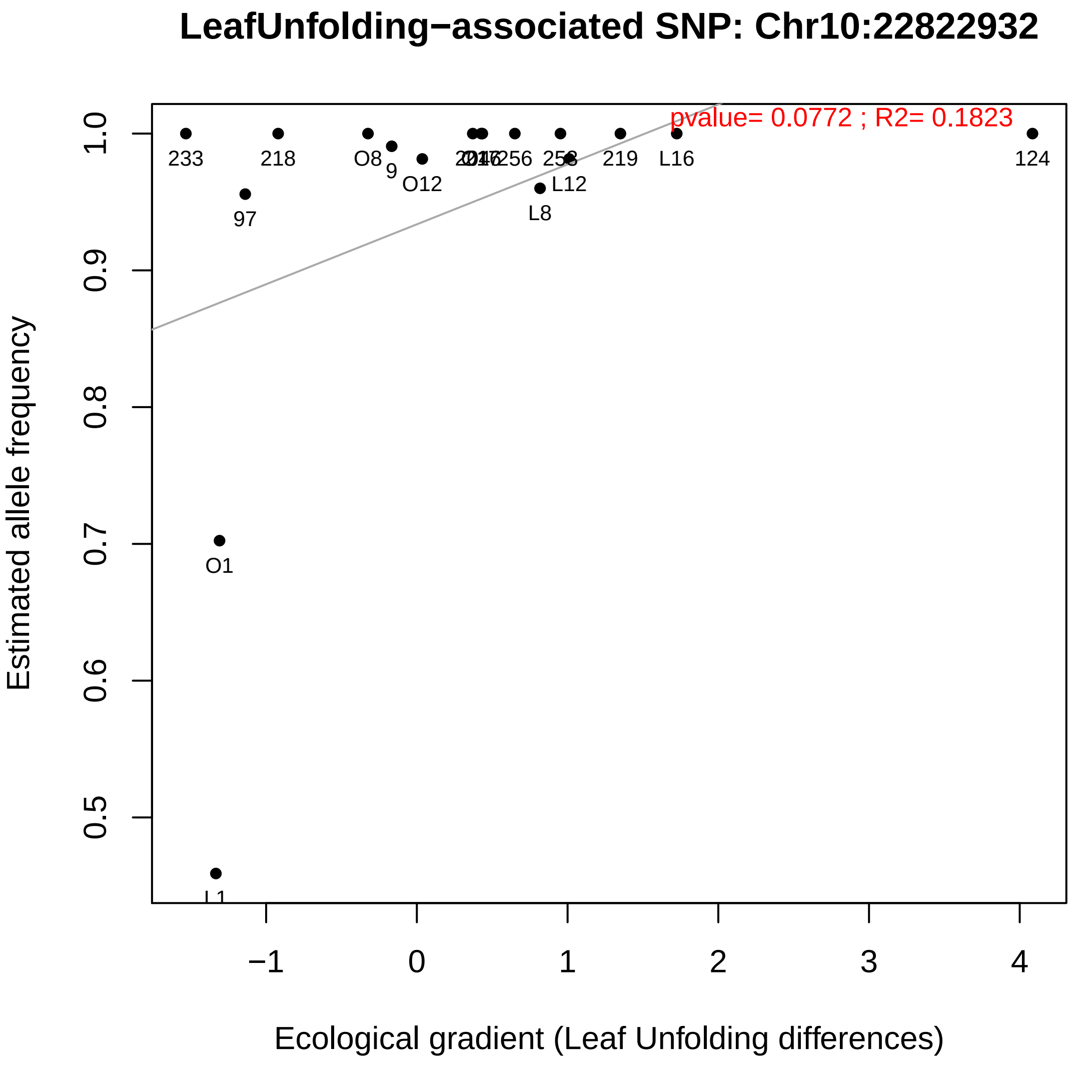
LeafUnfolding−associated SNP: Chr10:22822932
pvalue= 0.0772 ; R2= 0.1823
233
218
O8
9
O12
256
219
L16
124
97
L12
L8
O1
−1
0
1
2
3
4
0.5
0.6
0.7
0.8
0.9
1.0
Estimated allele frequency
Ecological gradient (Leaf Unfolding differences)

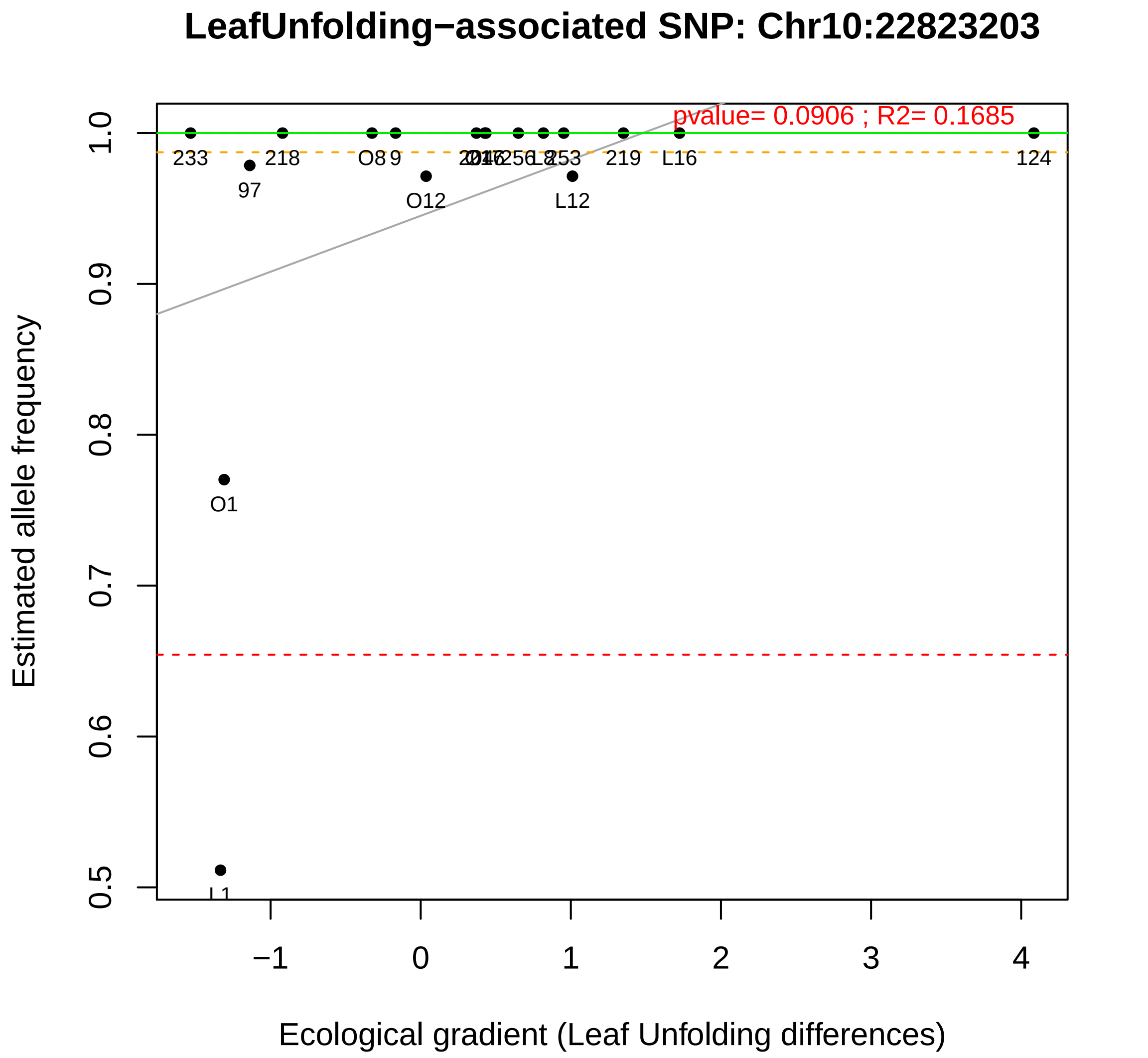
LeafUnfolding−associated SNP: Chr10:22823203
pvalue= 0.0906 ; R2= 0.1685
Estimated allele frequency
Ecological gradient (Leaf Unfolding differences)
233
218
O8
9
219
L16
124
97
O12
L12
O1
L1
1.0
0.9
0.8
0.7
0.6
0.5
−1
0
1
2
3
4

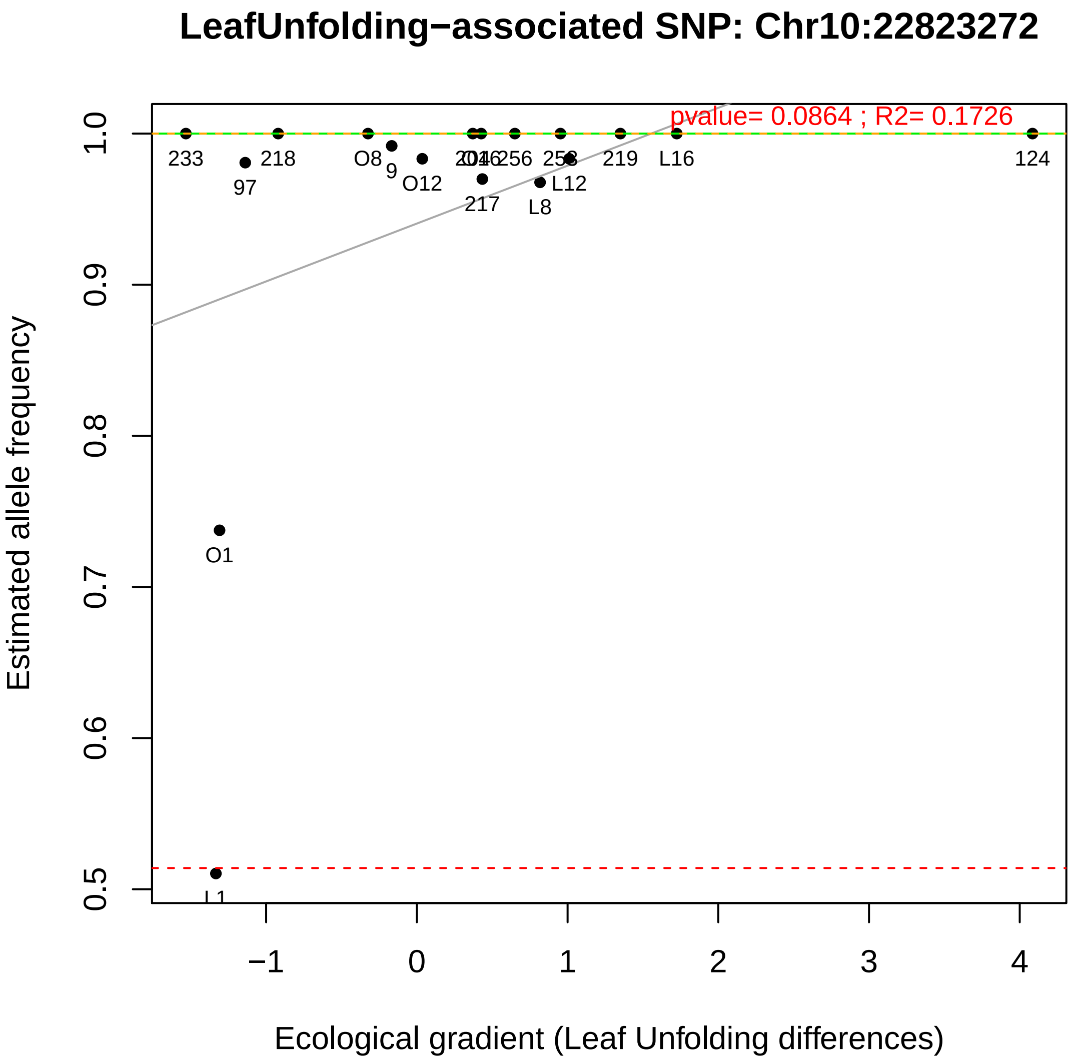

LeafUnfolding−associated SNP: Chr10:22823272
pvalue= 0.0864 ; R2= 0.1726
Estimated allele frequency
Ecological gradient (Leaf Unfolding differences)
1.0
0.9
0.8
0.7
0.6
0.5
−1
0
1
2
3
4
233
97
218
O8
9
O12
217
256
L8
L12
219
L16
124
O1

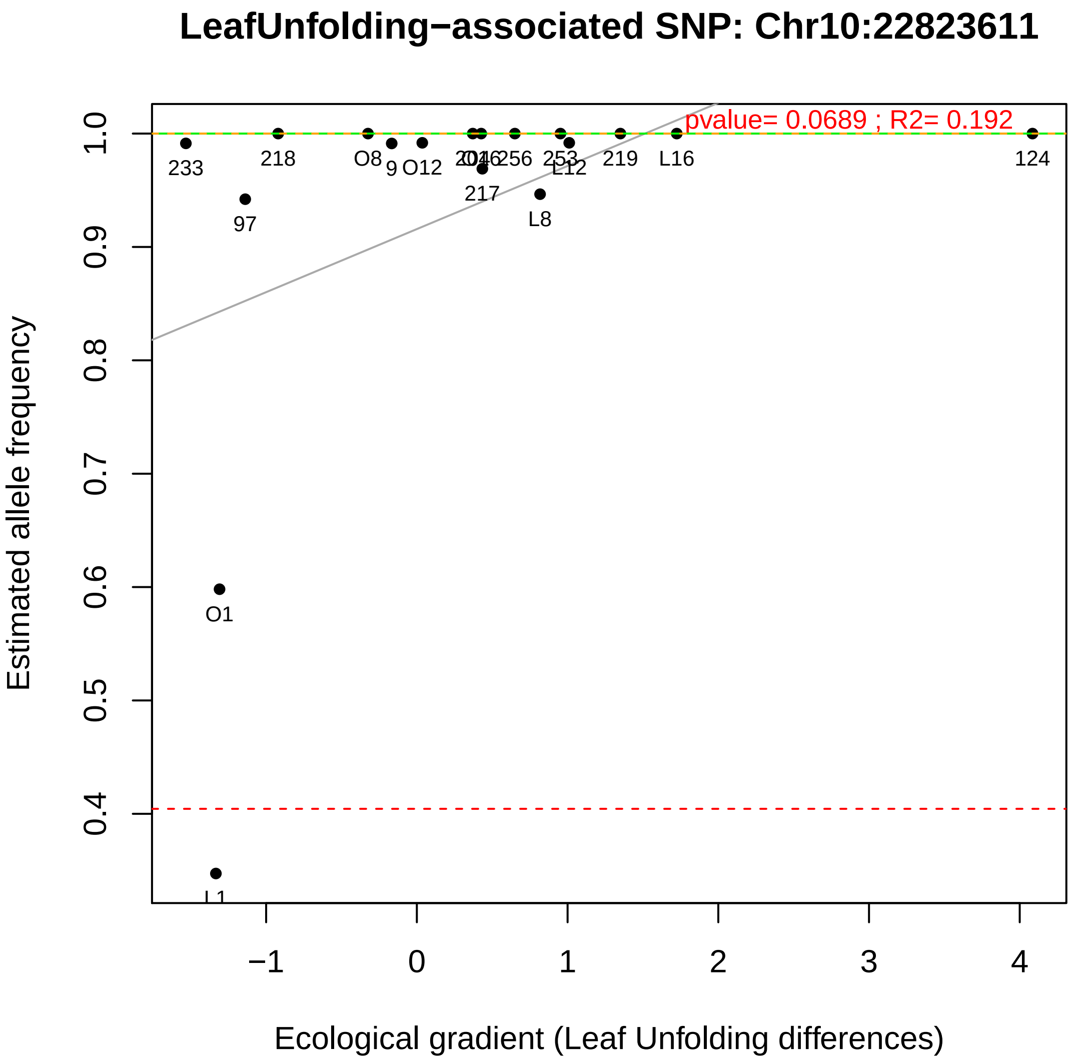
LeafUnfolding−associated SNP: Chr10:22823611
pvalue= 0.0689 ; R2= 0.192
Estimated allele frequency
0.4
0.5
0.6
0.7
0.8
0.9
1.0
−1
0
1
2
3
4
Ecological gradient (Leaf Unfolding differences)
233
218
O8
9
O12
217
L8
253
L12
219
L16
124
97
O1

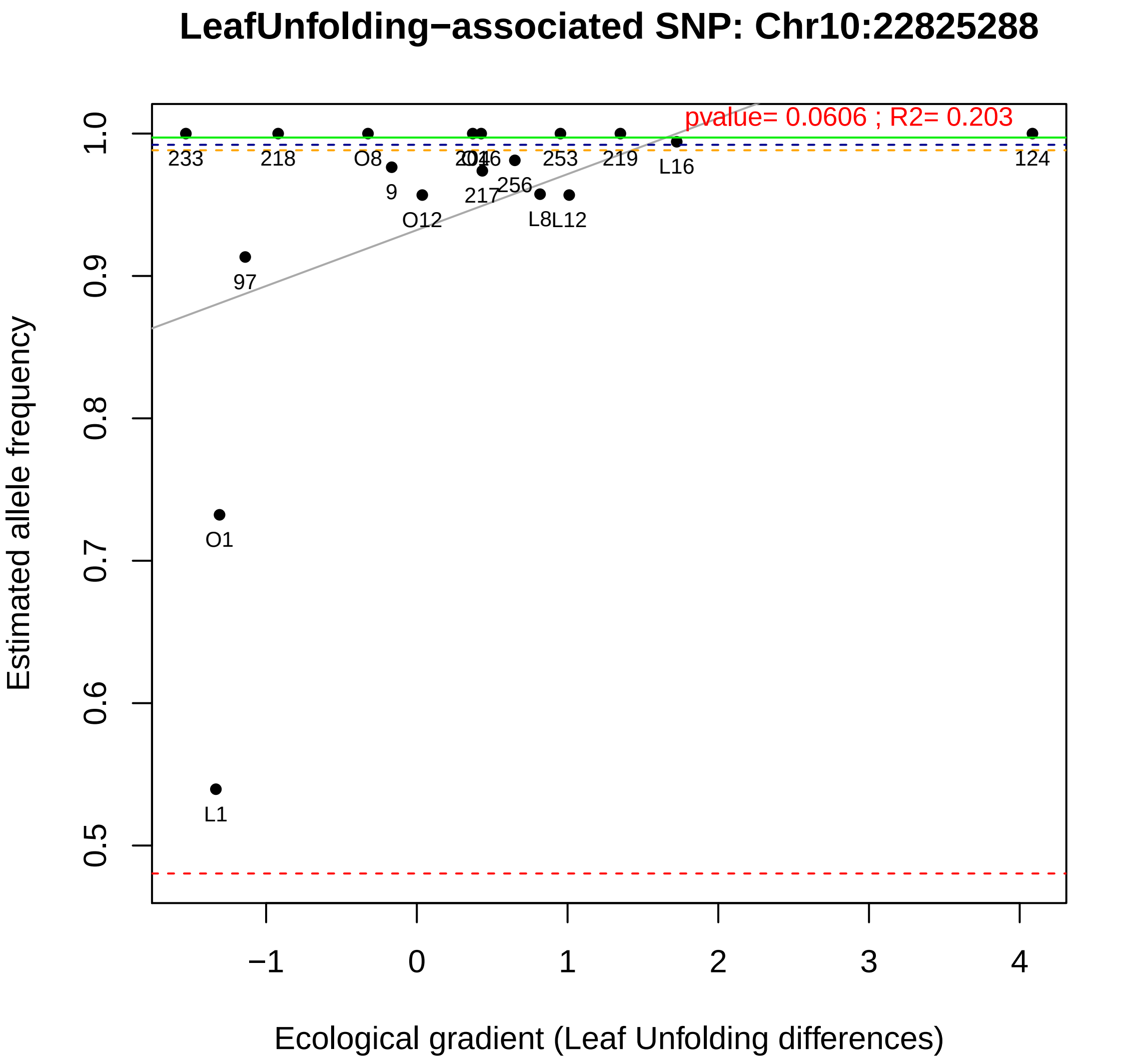

LeafUnfolding−associated SNP: Chr10:22825288
pvalue= 0.0606 ; R2= 0.203
Estimated allele frequency
Ecological gradient (Leaf Unfolding differences)
0.5
0.6
0.7
0.8
0.9
1.0
−1
0
1
2
3
4
233
218
O8
9
O12
217
256
253
219
L16
124
L8
L12
97
O1
L1

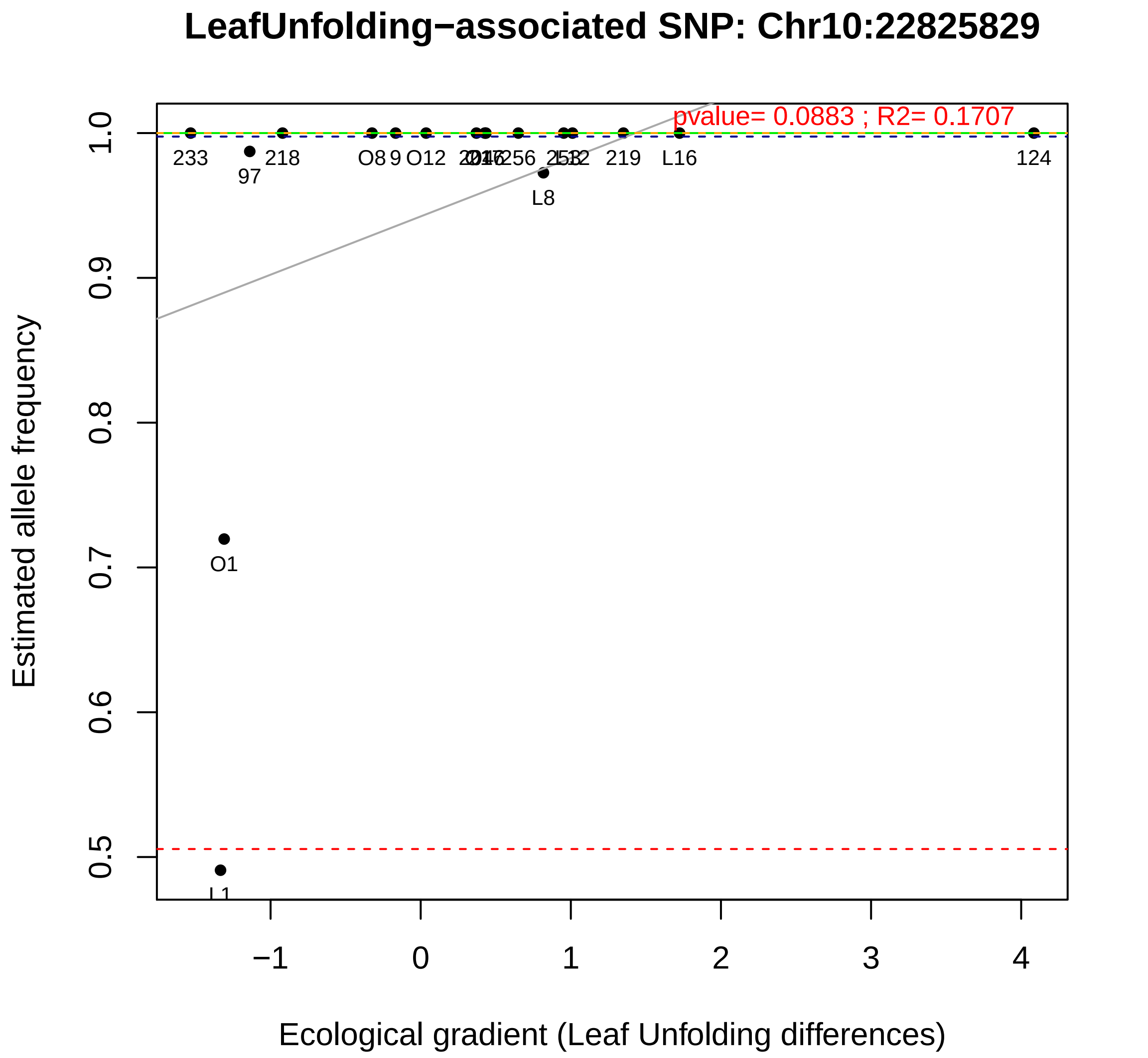

LeafUnfolding−associated SNP: Chr10:22825829
pvalue= 0.0883 ; R2= 0.1707
Estimated allele frequency
Ecological gradient (Leaf Unfolding differences)
0.5
0.6
0.7
0.8
0.9
1.0
−1
0
1
2
3
4
233
97
218
O8
9
O12
L16
219
124
L8
O1

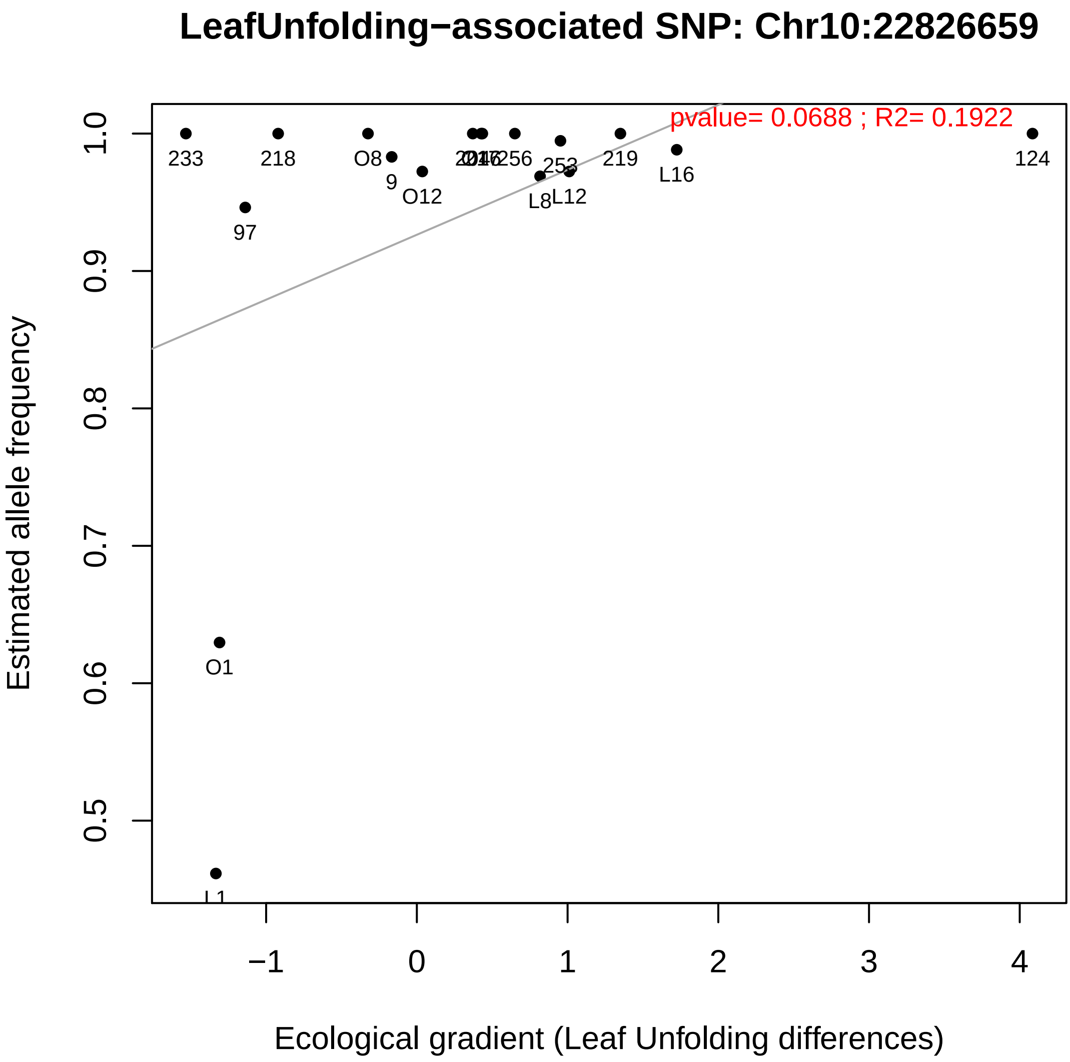
LeafUnfolding−associated SNP: Chr10:22826659
pvalue= 0.0688 ; R2= 0.1922
Estimated allele frequency
Ecological gradient (Leaf Unfolding differences)
1.0
0.9
0.8
0.7
0.6
0.5
−1
0
1
2
3
4
233
218
O8
9
O12
256
253
L8
L12
219
L16
124
97
O1

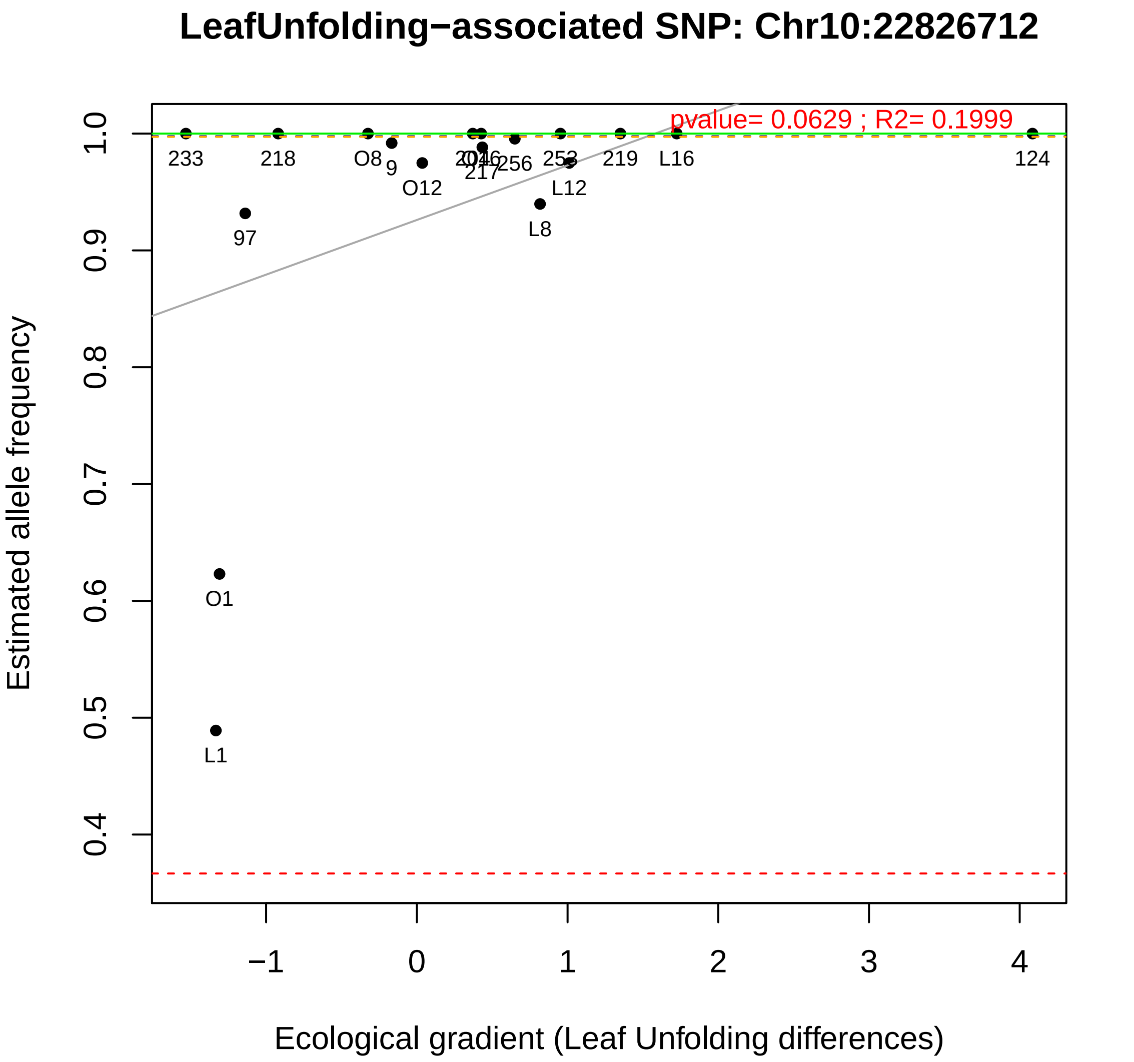

LeafUnfolding−associated SNP: Chr10:22826712
pvalue= 0.0629 ; R2= 0.1999
Estimated allele frequency
Ecological gradient (Leaf Unfolding differences)
1.0
0.9
0.8
0.7
0.6
0.5
0.4
−1
0
1
2
3
4
233
218
O8
9
O12
204
O16
217
256
253
L12
219
L16
124
97
L8
O1
L1

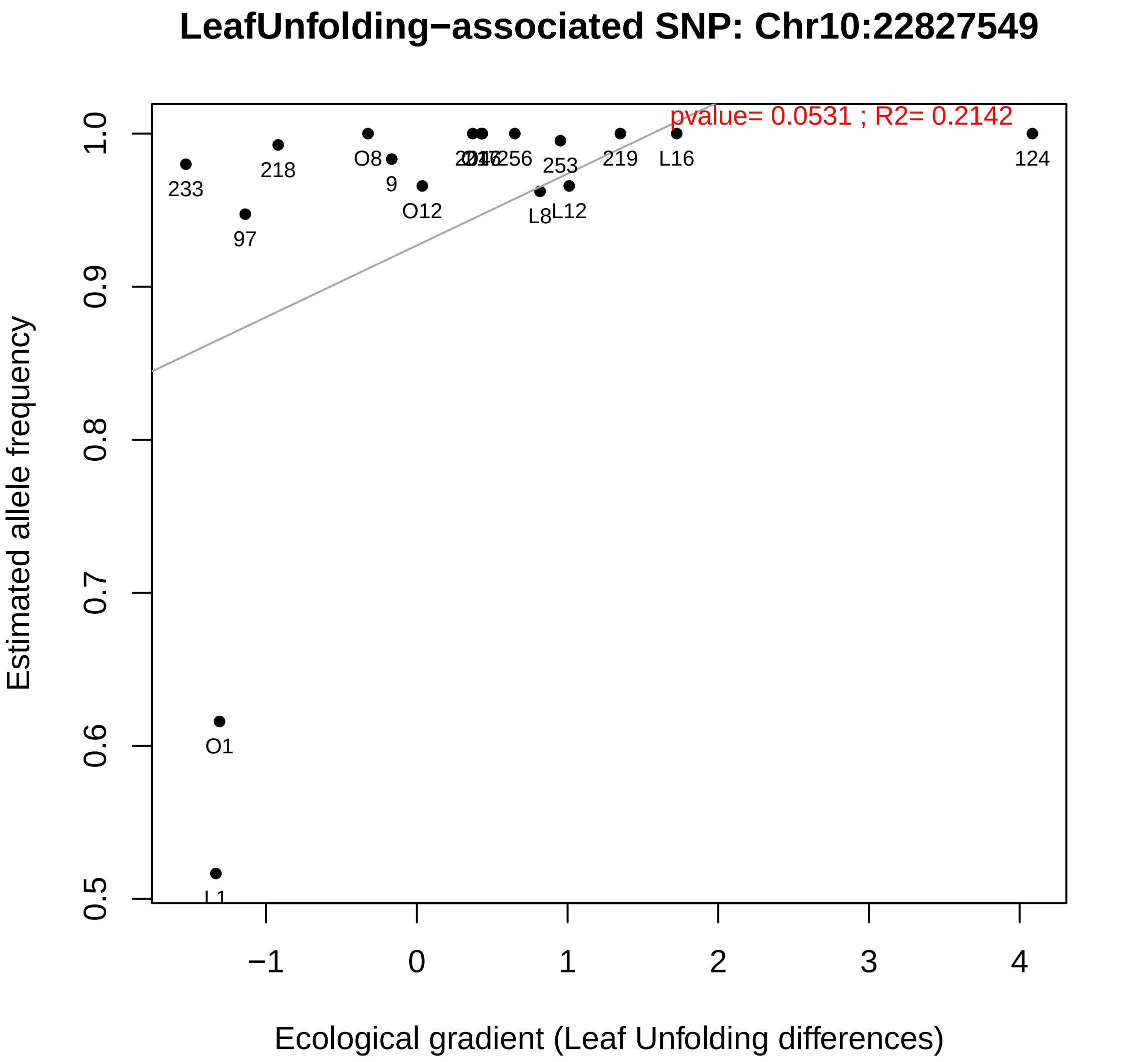
LeafUnfolding−associated SNP: Chr10:22827549
pvalue= 0.0531 ; R2= 0.2142
Estimated allele frequency
Ecological gradient (Leaf Unfolding differences)
1.0
0.9
0.8
0.7
0.6
0.5
−1
0
1
2
3
4
233
97
218
O8
9
O12
256
253
219
L16
124
L8
L12
O1

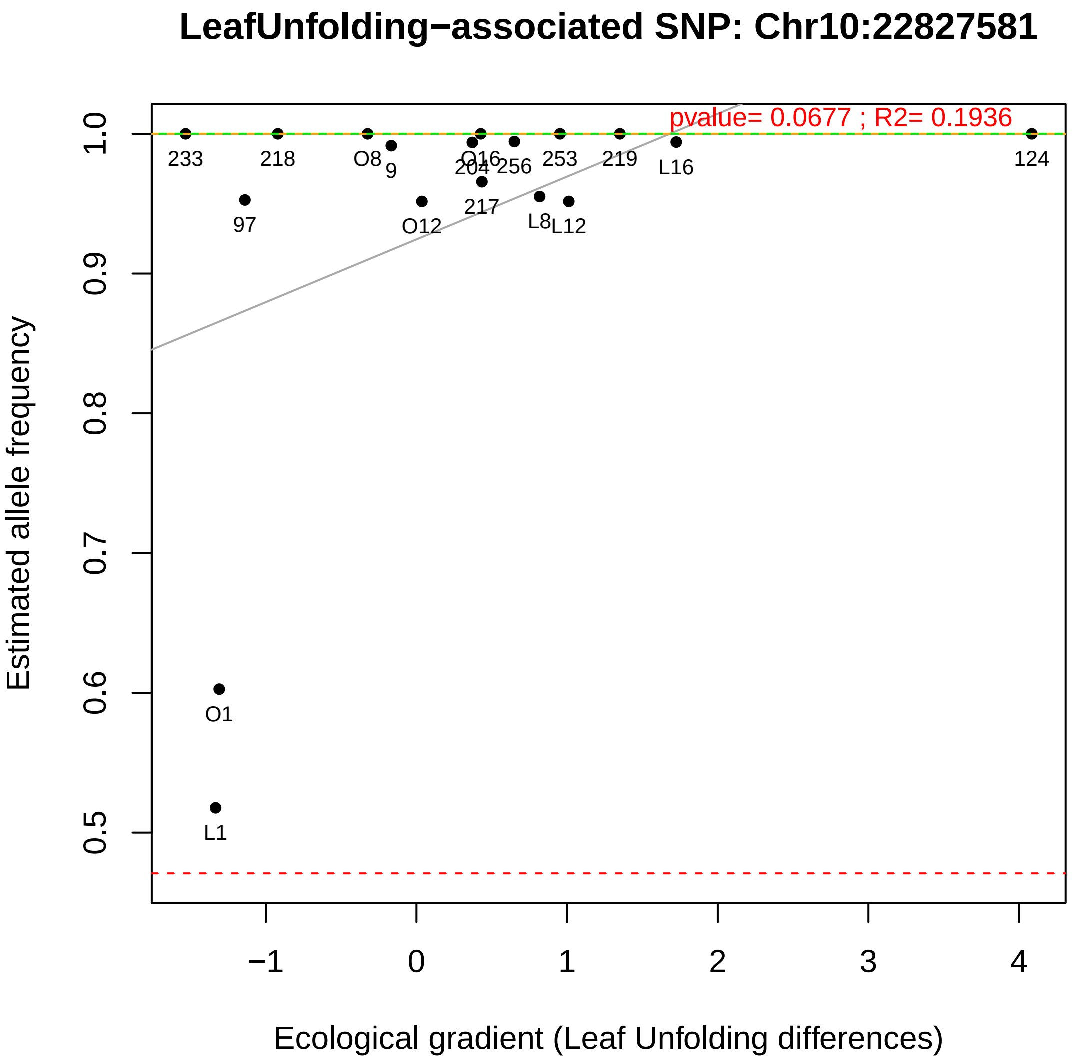
LeafUnfolding−associated SNP: Chr10:22827581
pvalue= 0.0677 ; R2= 0.1936
233
218
O8
9
O16
204
256
253
219
L16
124
97
O12
217
L8
L12
O1
L1
Estimated allele frequency
1.0
0.9
0.8
0.7
0.6
0.5
−1
0
1
2
3
4
Ecological gradient (Leaf Unfolding differences)

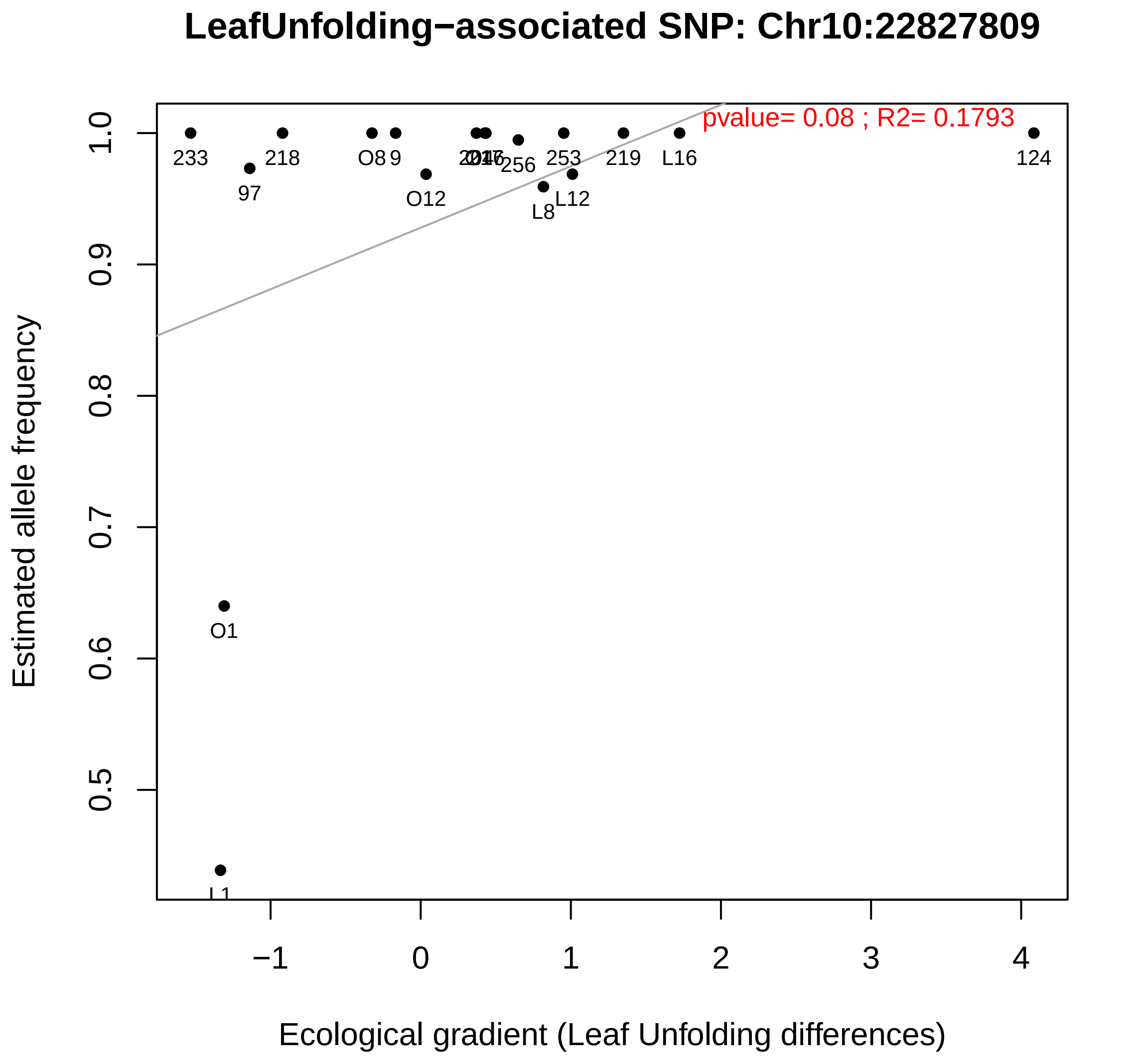
LeafUnfolding−associated SNP: Chr10:22827809
pvalue= 0.08 ; R2= 0.1793
Estimated allele frequency
Ecological gradient (Leaf Unfolding differences)
1.0
0.9
0.8
0.7
0.6
0.5
−1
0
1
2
3
4
233
97
218
O8
9
O12
256
253
L8
L12
219
L16
124
O1
L1

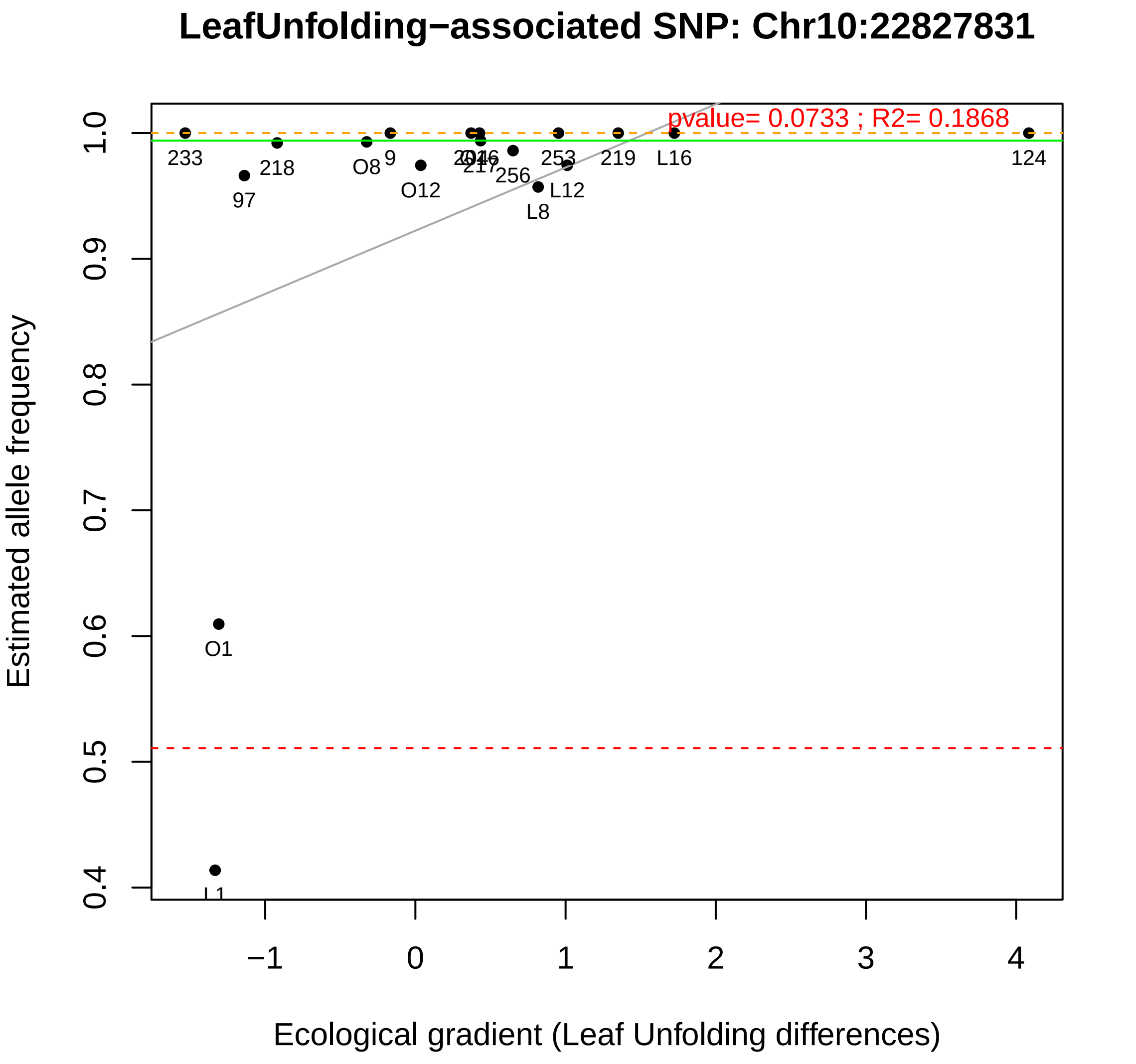

LeafUnfolding−associated SNP: Chr10:22827831
pvalue= 0.0733 ; R2= 0.1868
Estimated allele frequency
Ecological gradient (Leaf Unfolding differences)
1.0
0.9
0.8
0.7
0.6
0.5
0.4
−1
0
1
2
3
4
233
218
97
O8
9
O12
256
253
L12
L8
219
L16
124
O1

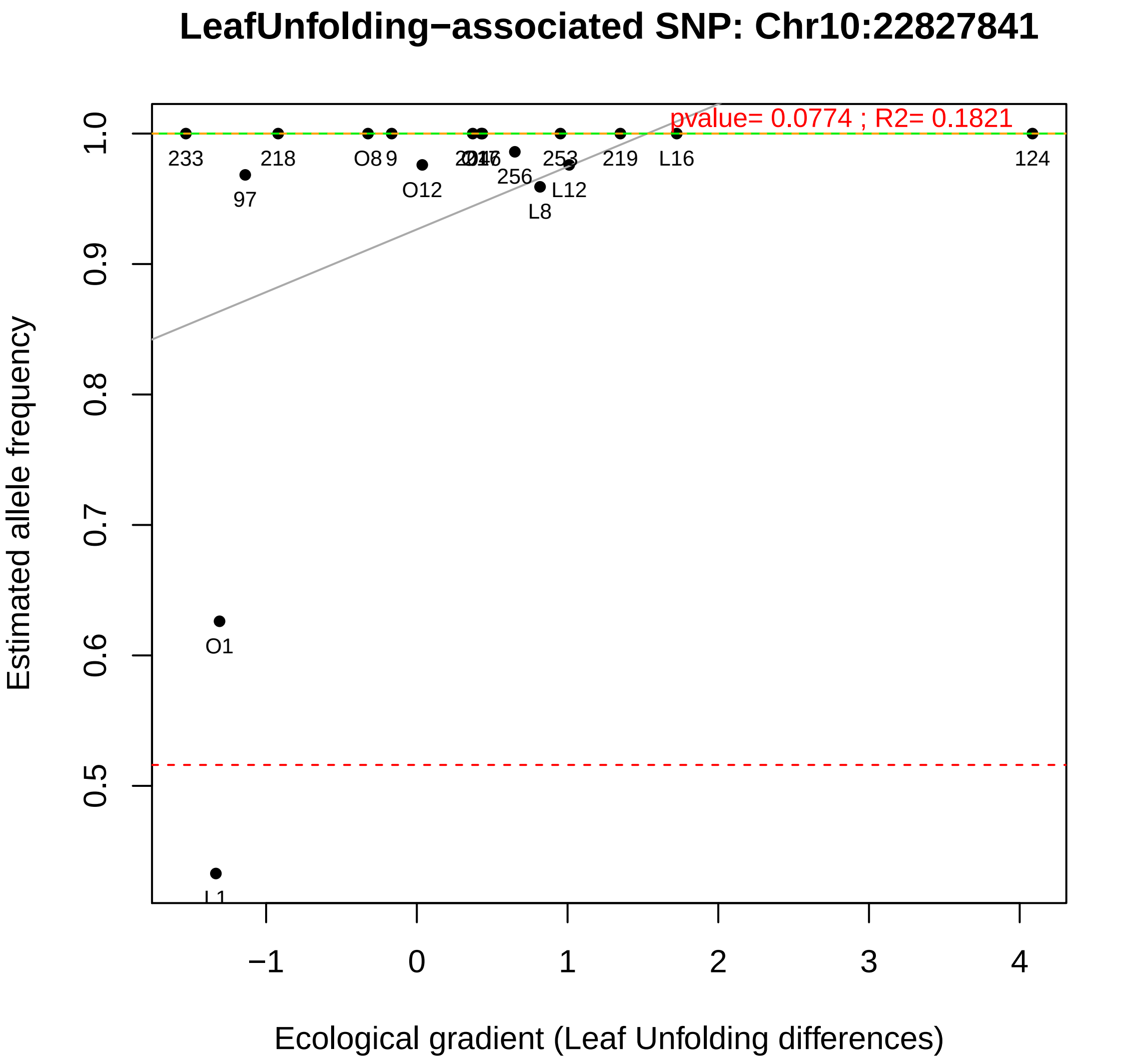
LeafUnfolding−associated SNP: Chr10:22827841
pvalue= 0.0774 ; R2= 0.1821
Estimated allele frequency
Ecological gradient (Leaf Unfolding differences)
1.0
0.9
0.8
0.7
0.6
0.5
−1
0
1
2
3
4
233
218
O8
9
97
O12
256
253
L12
L8
219
L16
124
O1
L1

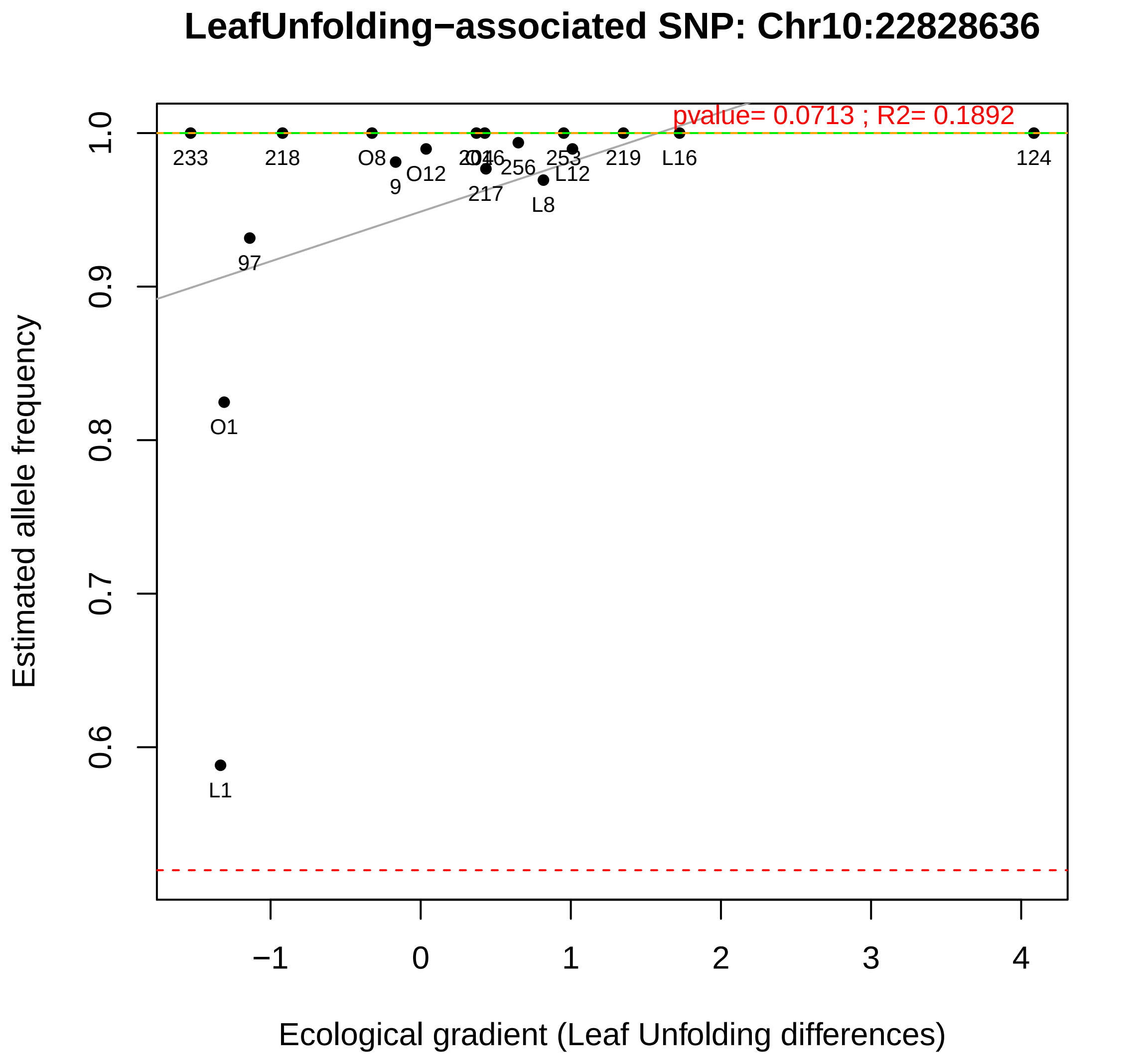

LeafUnfolding−associated SNP: Chr10:22828636
pvalue= 0.0713 ; R2= 0.1892
233
218
O8
9
O12
204
O16
217
256
253
L12
L8
219
L16
124
97
O1
L1
1.0
0.9
0.8
0.7
0.6
−1
0
1
2
3
4
Estimated allele frequency
Ecological gradient (Leaf Unfolding differences)

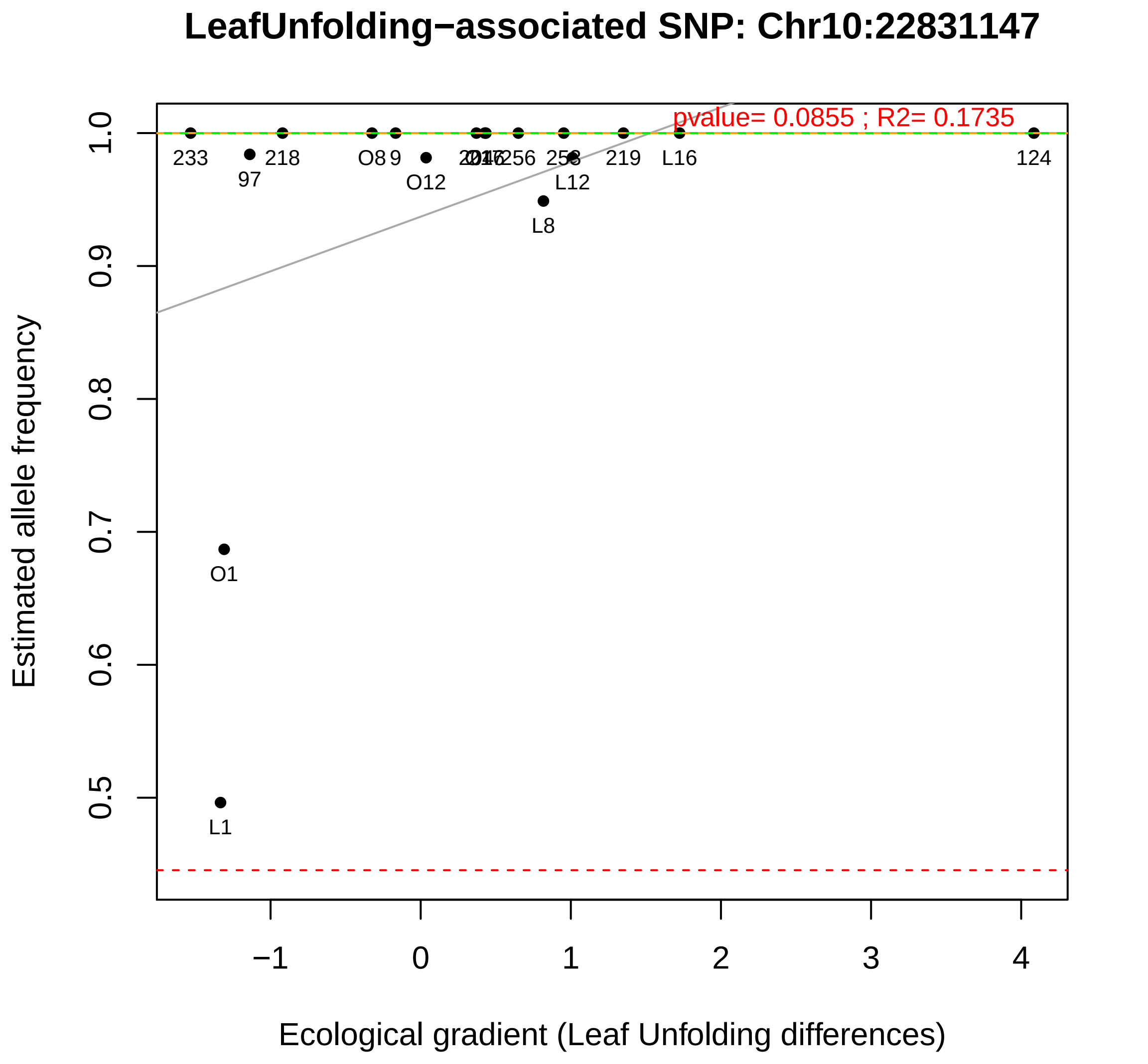

LeafUnfolding−associated SNP: Chr10:22831147
pvalue= 0.0855 ; R2= 0.1735
Estimated allele frequency
Ecological gradient (Leaf Unfolding differences)
1.0
0.9
0.8
0.7
0.6
0.5
−1
0
1
2
3
4
233
97
218
O8
9
O12
L8
L12
219
L16
124
O1
L1

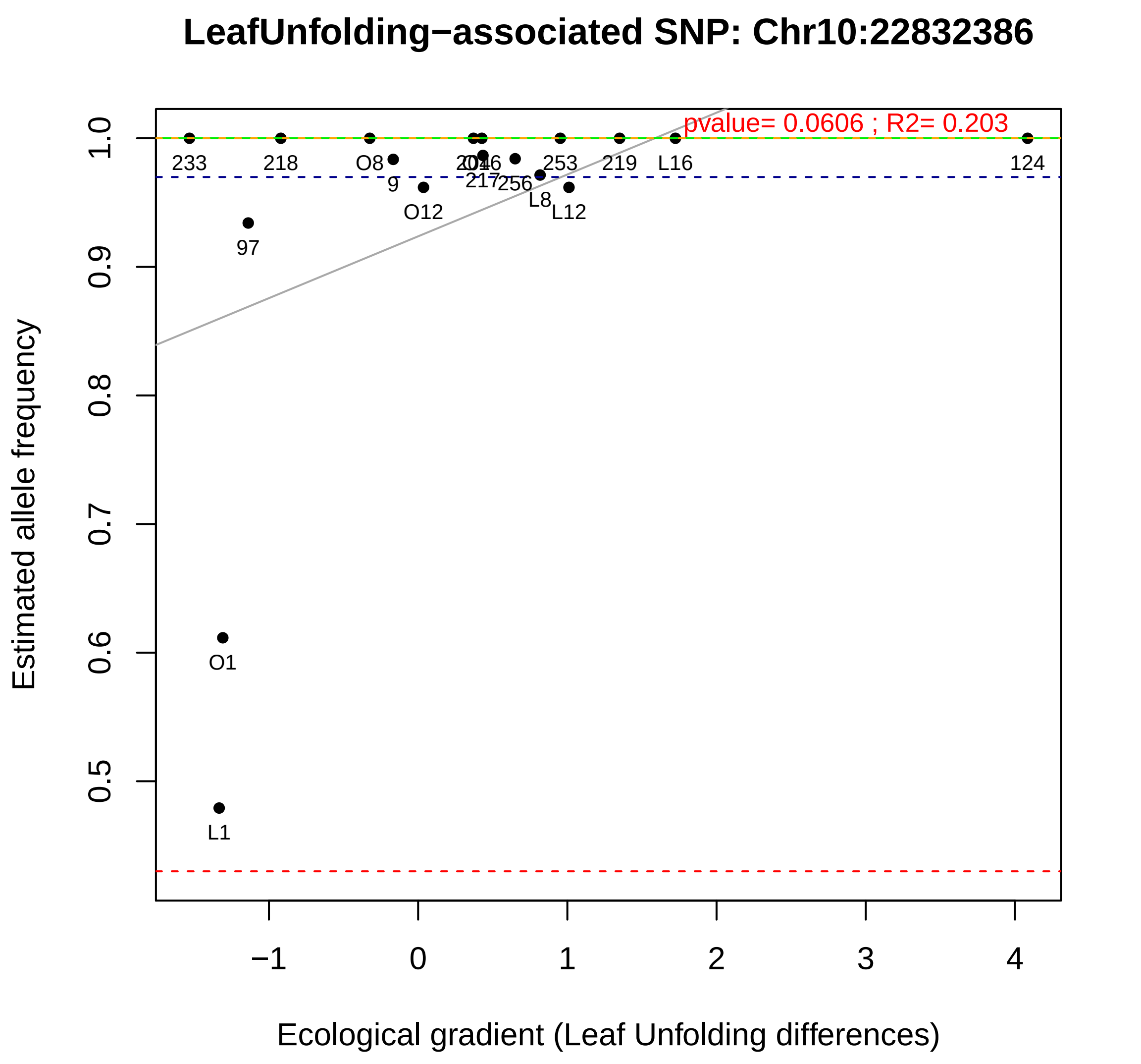

LeafUnfolding−associated SNP: Chr10:22832386
pvalue= 0.0606 ; R2= 0.203
Estimated allele frequency
Ecological gradient (Leaf Unfolding differences)
233
218
O8
9
O12
97
O1
L1
204
O46
217
256
253
L8
L12
219
L16
124
1.0
0.9
0.8
0.7
0.6
0.5
−1
0
1
2
3
4

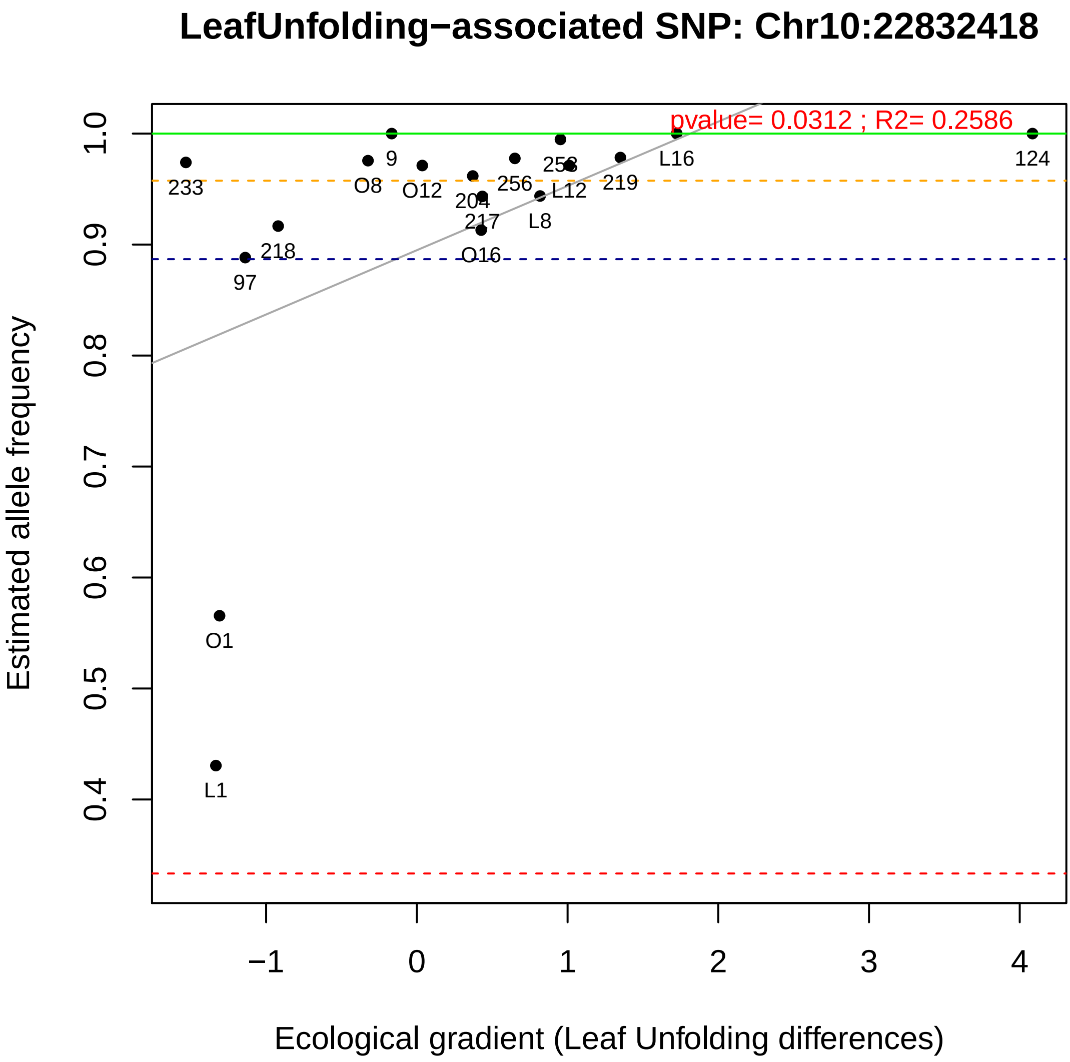
LeafUnfolding−associated SNP: Chr10:22832418
pvalue= 0.0312 ; R2= 0.2586
Estimated allele frequency
Ecological gradient (Leaf Unfolding differences)
0.4
0.5
0.6
0.7
0.8
0.9
1.0
−1
0
1
2
3
4
233
9
O8
O12
204
256
253
L12
219
L16
124
217
L8
218
O16
97
O1
L1

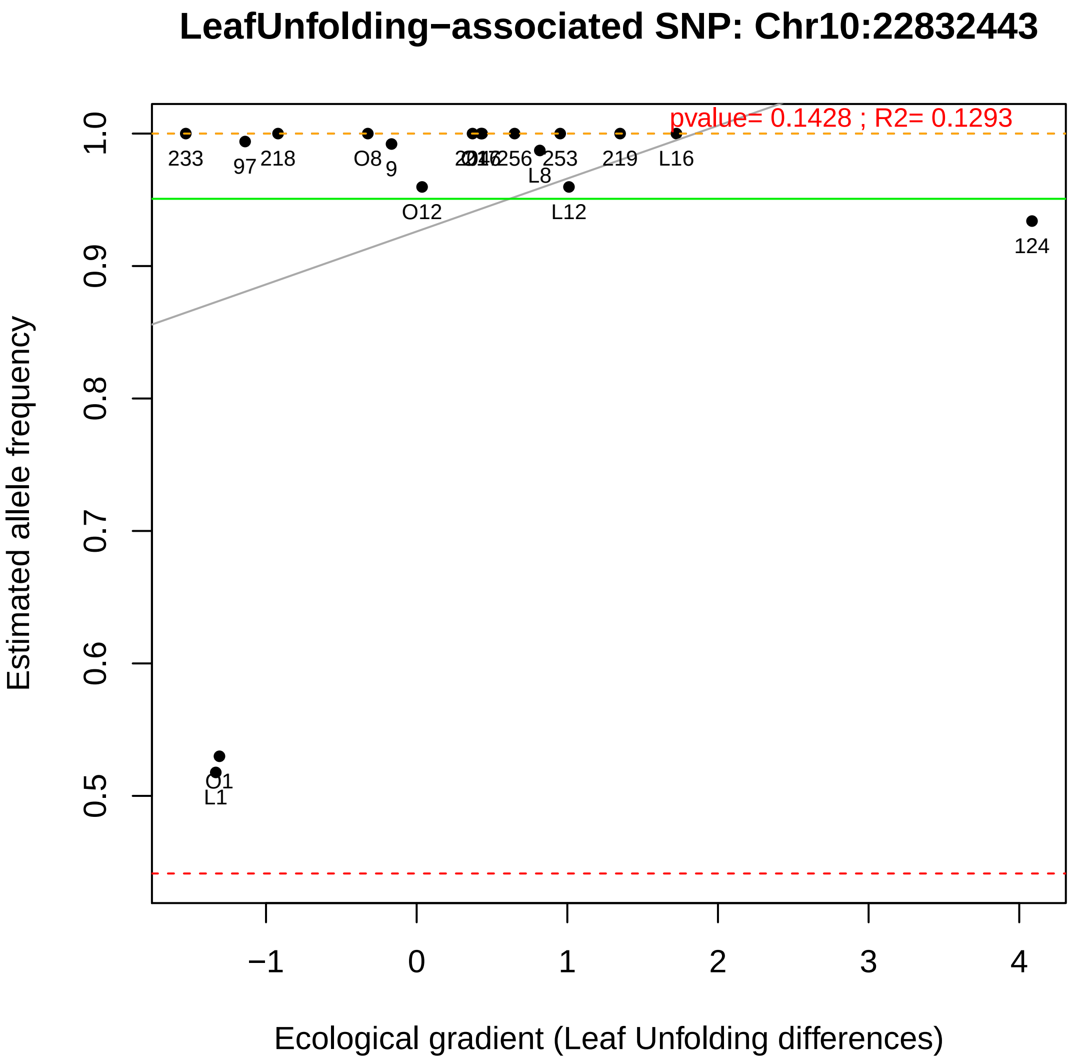
LeafUnfolding−associated SNP: Chr10:22832443
pvalue= 0.1428 ; R2= 0.1293
233
97
218
O8
9
O12
256
L8
253
L12
219
L16
124
O1
L1
1.0
0.9
0.8
0.7
0.6
0.5
−1
0
1
2
3
4
Estimated allele frequency
Ecological gradient (Leaf Unfolding differences)

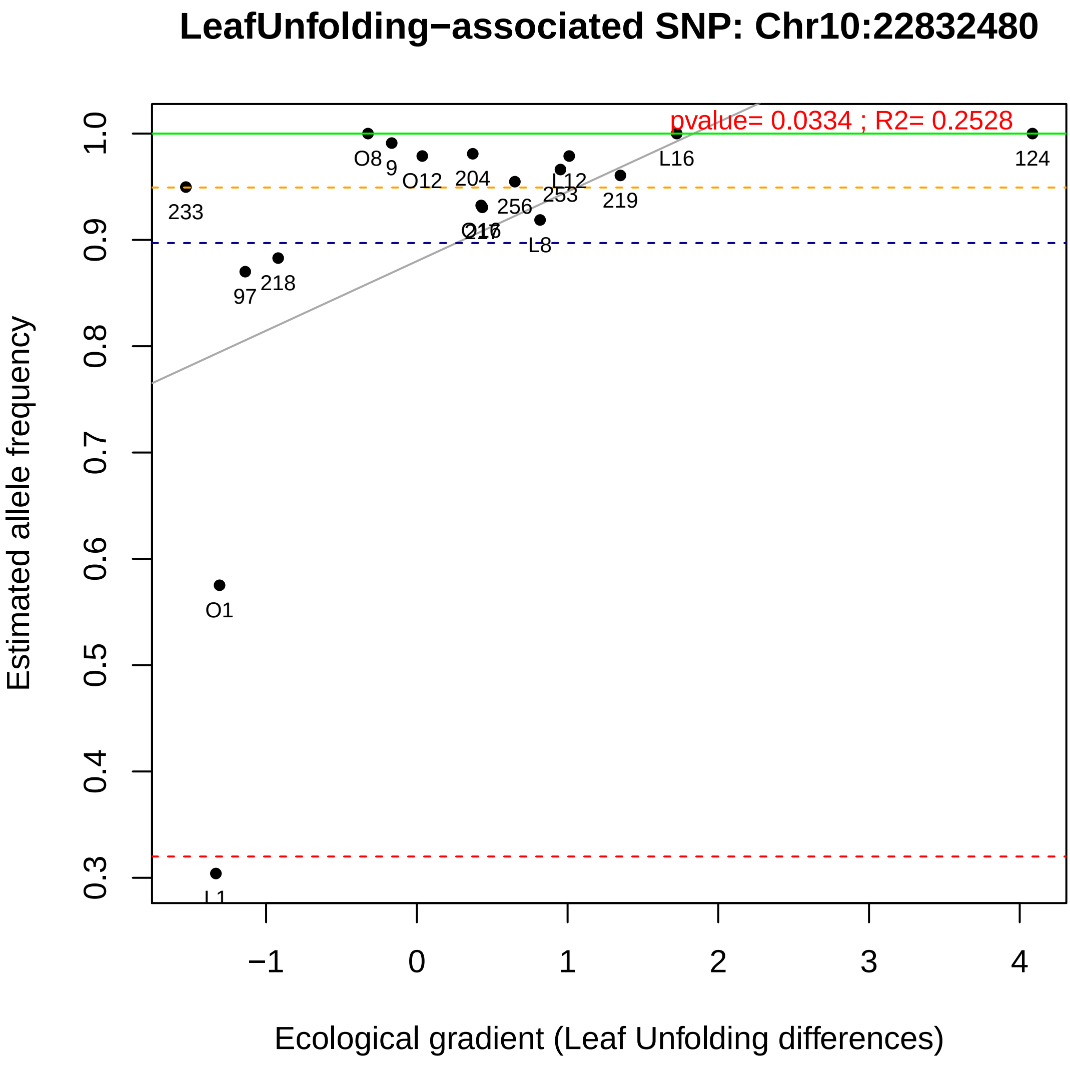

LeafUnfolding−associated SNP: Chr10:22832480
pvalue= 0.0334 ; R2= 0.2528
Estimated allele frequency
Ecological gradient (Leaf Unfolding differences)
1.0
0.9
0.8
0.7
0.6
0.5
0.4
0.3
−1
0
1
2
3
4
O8
9
O12
204
256
L12
253
219
L16
124
233
L8
97
218
O1
L1

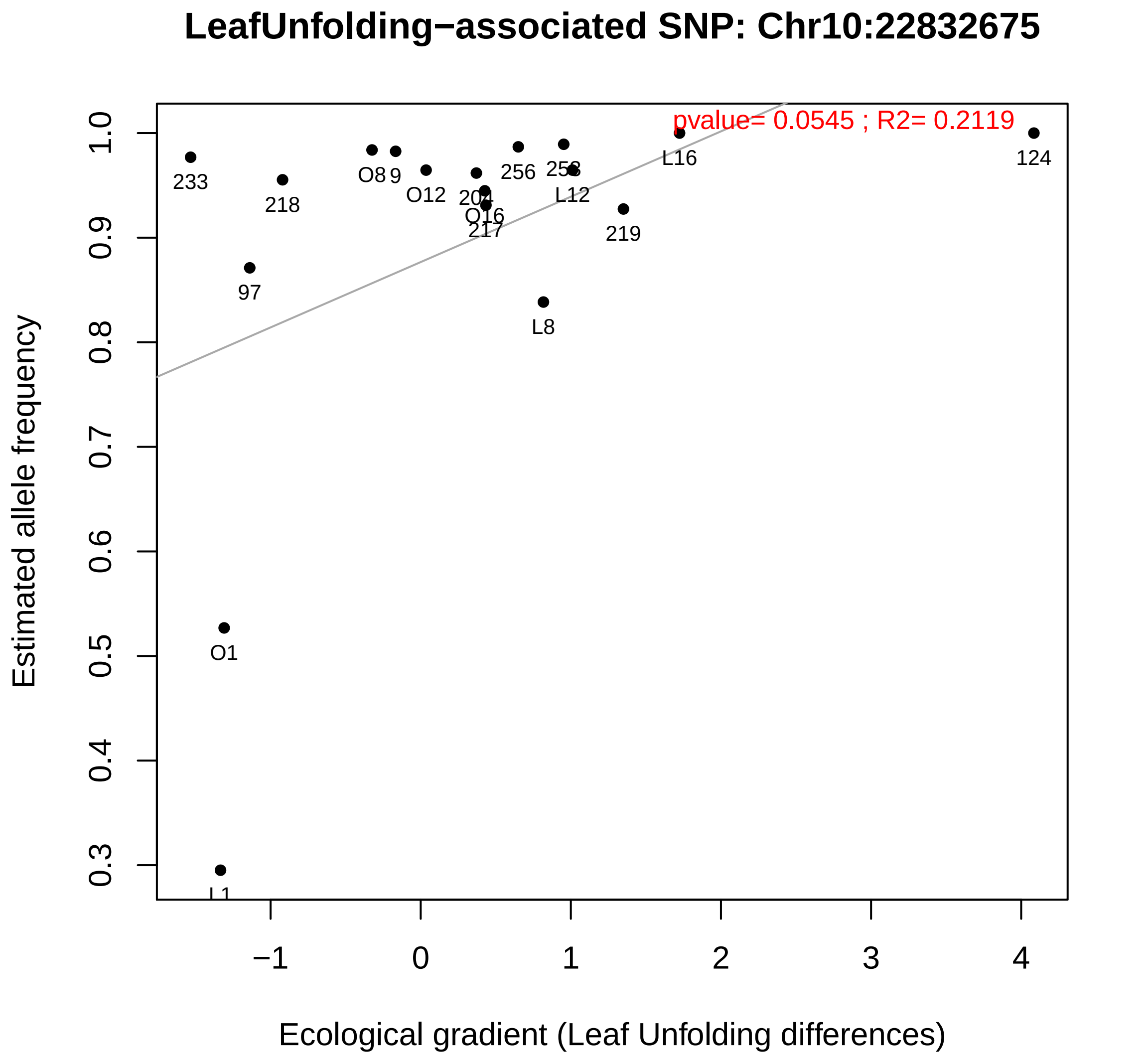
LeafUnfolding−associated SNP: Chr10:22832675
pvalue= 0.0545 ; R2= 0.2119
Estimated allele frequency
Ecological gradient (Leaf Unfolding differences)
233
218
97
O1
L1
O8
9
O12
204
O16
217
256
253
L12
L8
219
L16
124
1.0
0.9
0.8
0.7
0.6
0.5
0.4
0.3
−1
0
1
2
3
4

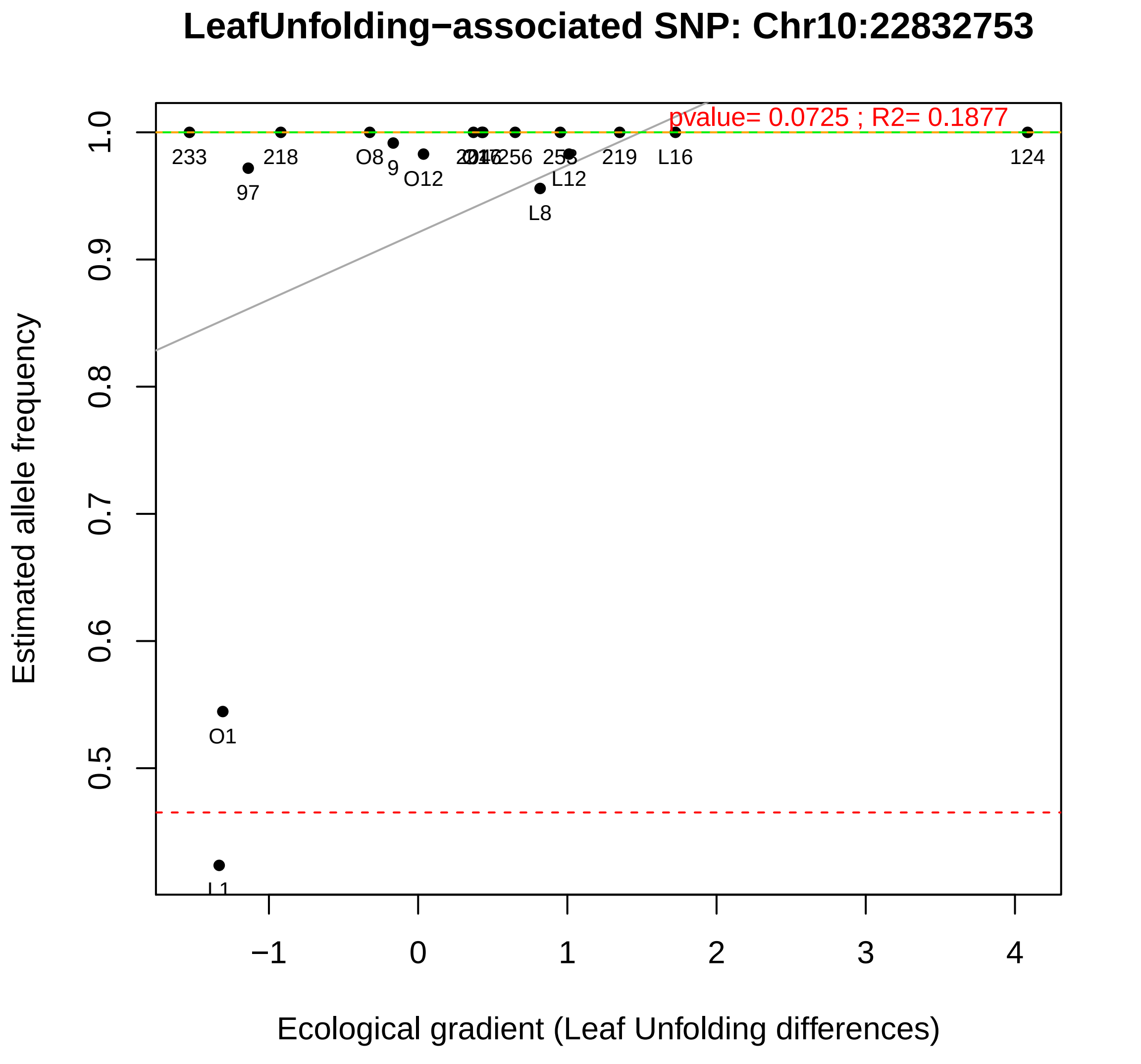

LeafUnfolding−associated SNP: Chr10:22832753
pvalue= 0.0725 ; R2= 0.1877
Estimated allele frequency
Ecological gradient (Leaf Unfolding differences)
1.0
0.9
0.8
0.7
0.6
0.5
−1
0
1
2
3
4
233
218
O8
9
O12
97
256
L12
L8
219
L16
124
O1

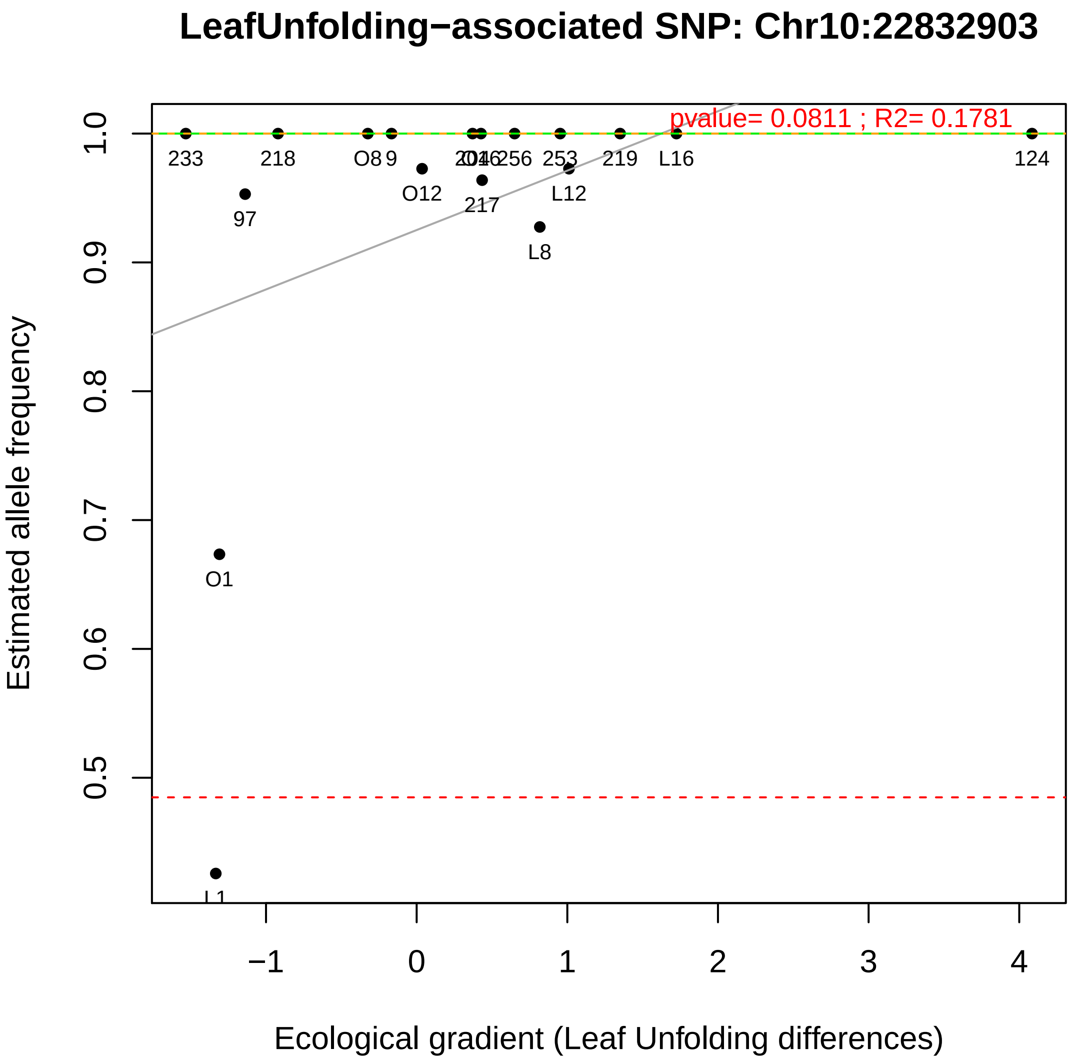
LeafUnfolding−associated SNP: Chr10:22832903
pvalue= 0.0811 ; R2= 0.1781
Estimated allele frequency
1.0
0.9
0.8
0.7
0.6
0.5
233
218
O8
9
204
O16
256
253
219
L16
124
97
O12
217
L12
L8
O1
L1
−1
0
1
2
3
4
Ecological gradient (Leaf Unfolding differences)

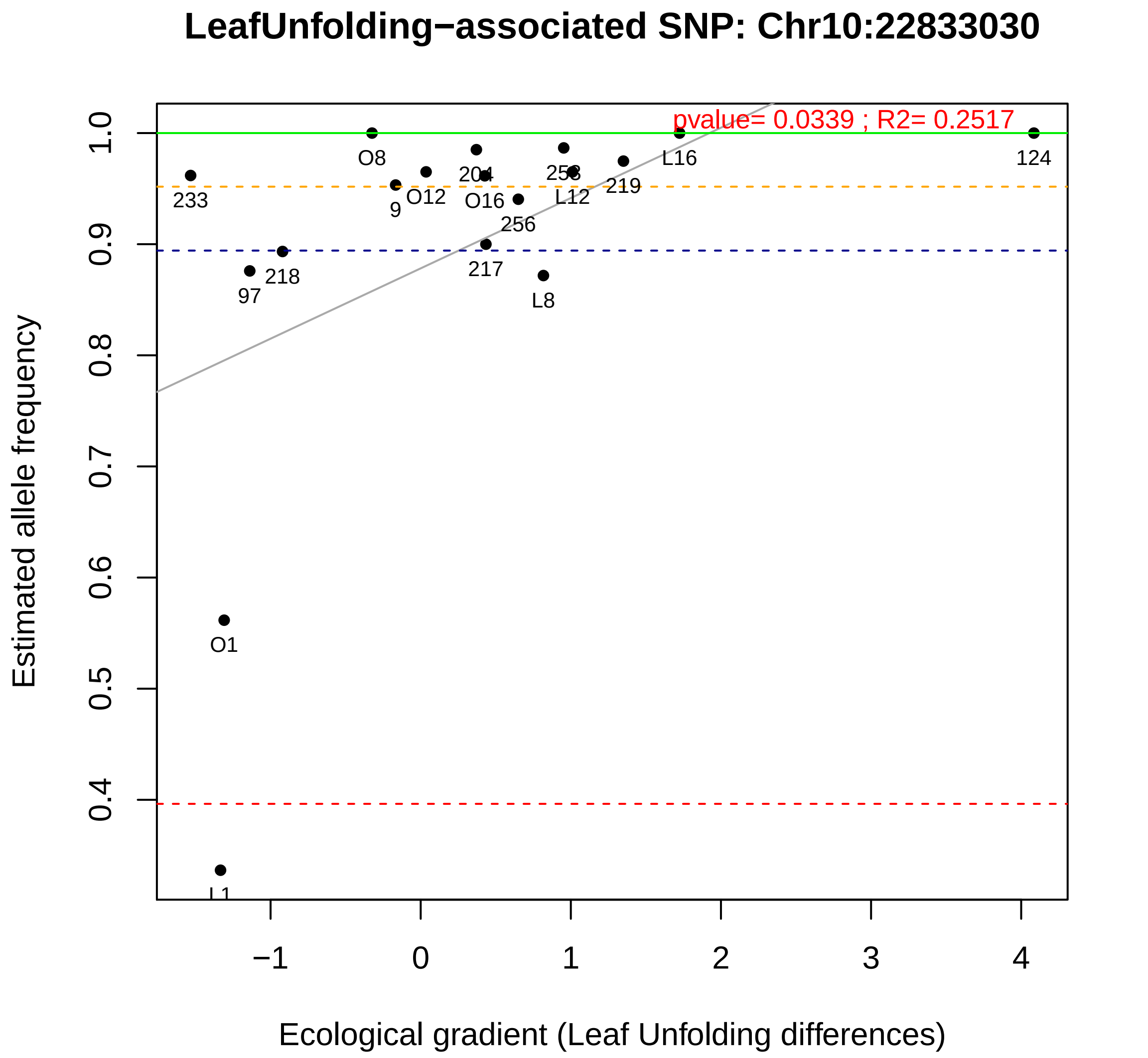

LeafUnfolding−associated SNP: Chr10:22833030
pvalue= 0.0339 ; R2= 0.2517
Estimated allele frequency
Ecological gradient (Leaf Unfolding differences)
0.4
0.5
0.6
0.7
0.8
0.9
1.0
−1
0
1
2
3
4
O8
233
9
O12
204
O16
256
253
L12
219
L16
124
218
97
217
L8
O1

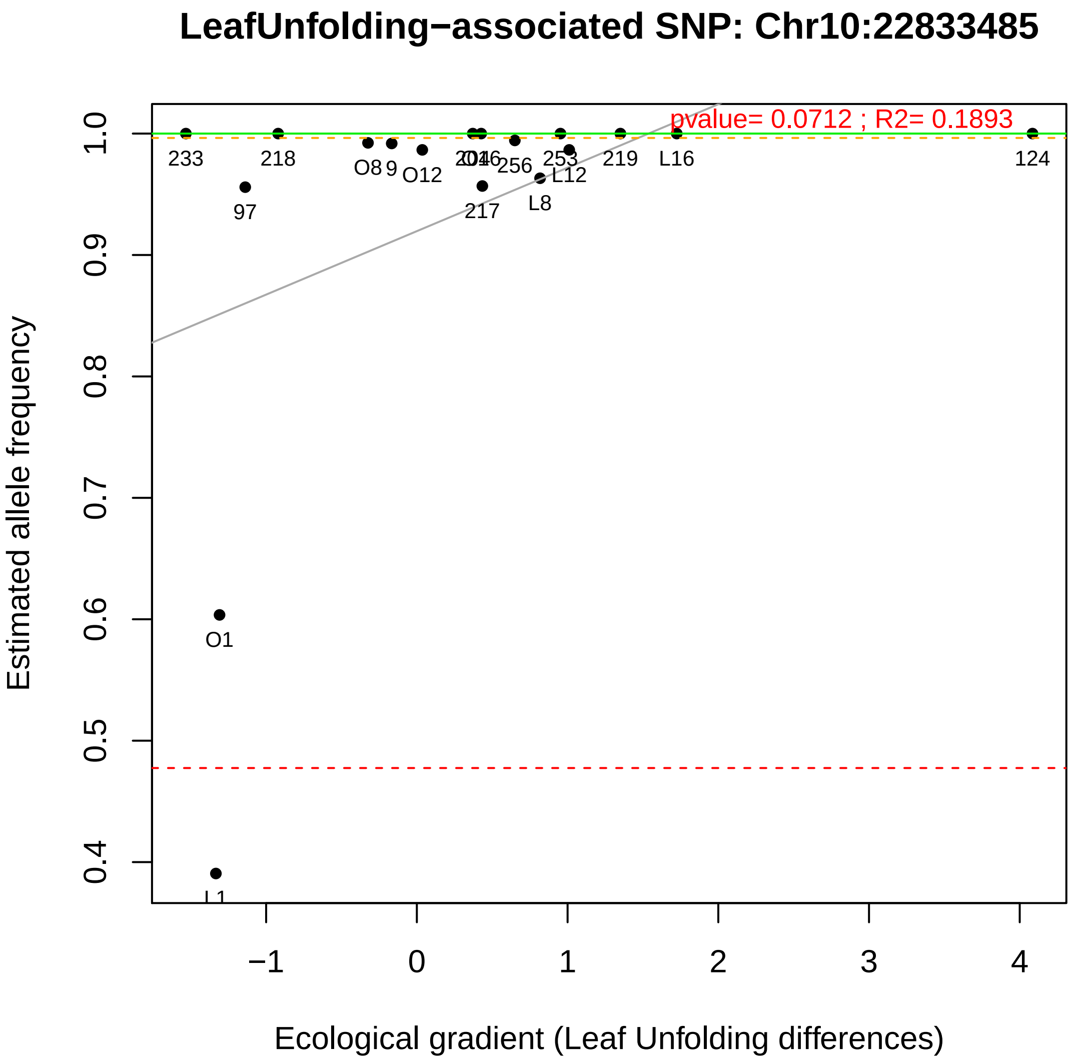
LeafUnfolding−associated SNP: Chr10:22833485
pvalue= 0.0712 ; R2= 0.1893
Estimated allele frequency
Ecological gradient (Leaf Unfolding differences)
1.0
0.9
0.8
0.7
0.6
0.5
0.4
−1
0
1
2
3
4
233
218
O8
9
O12
204
O16
256
253
L12
219
L16
124
97
217
L8
O1
L1

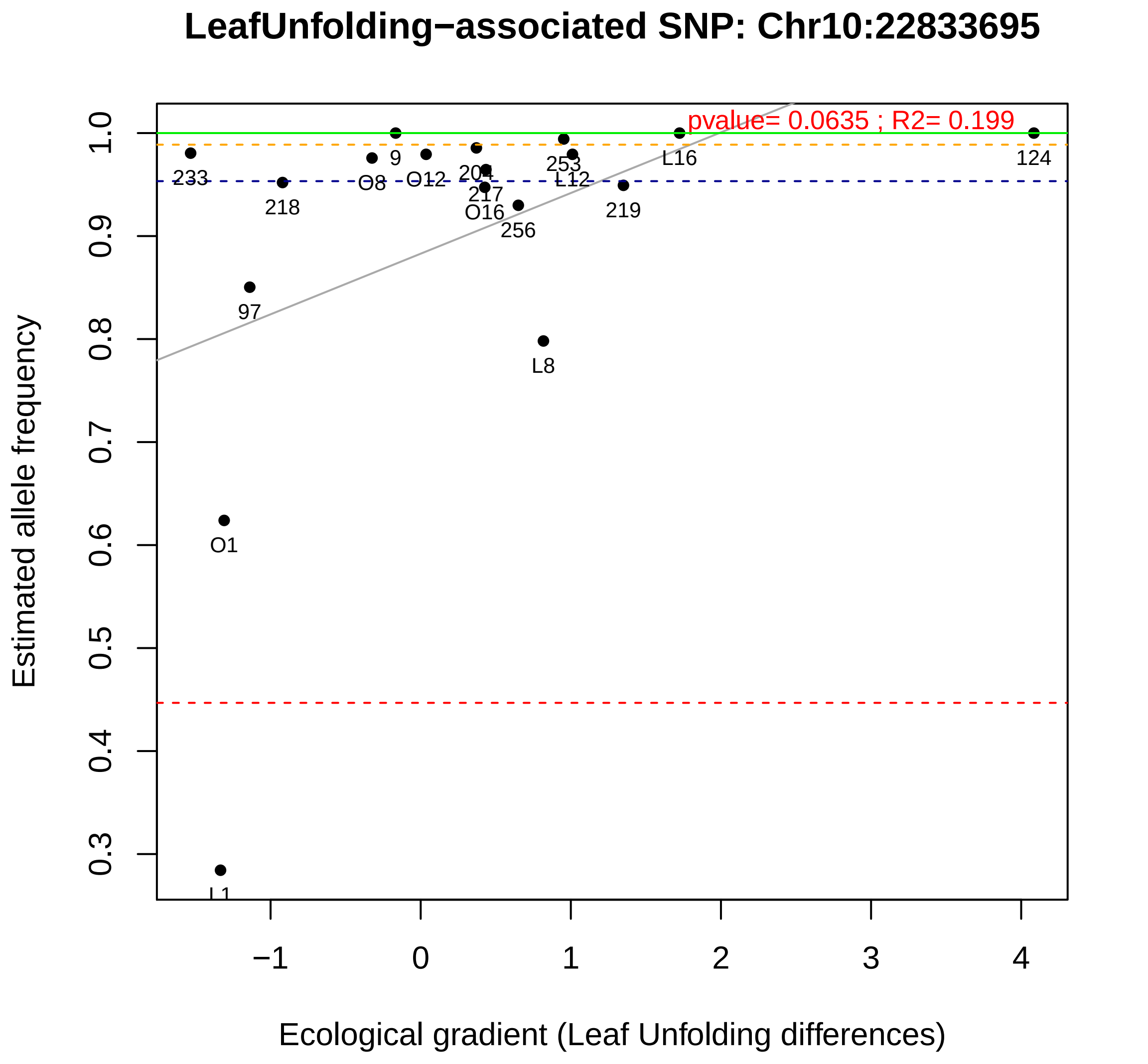

LeafUnfolding−associated SNP: Chr10:22833695
pvalue= 0.0635 ; R2= 0.199
Estimated allele frequency
Ecological gradient (Leaf Unfolding differences)
0.3
0.4
0.5
0.6
0.7
0.8
0.9
1.0
−1
0
1
2
3
4
233
218
97
O1
L1
O8
9
O12
204
217
O16
256
L8
253
L12
219
L16
124

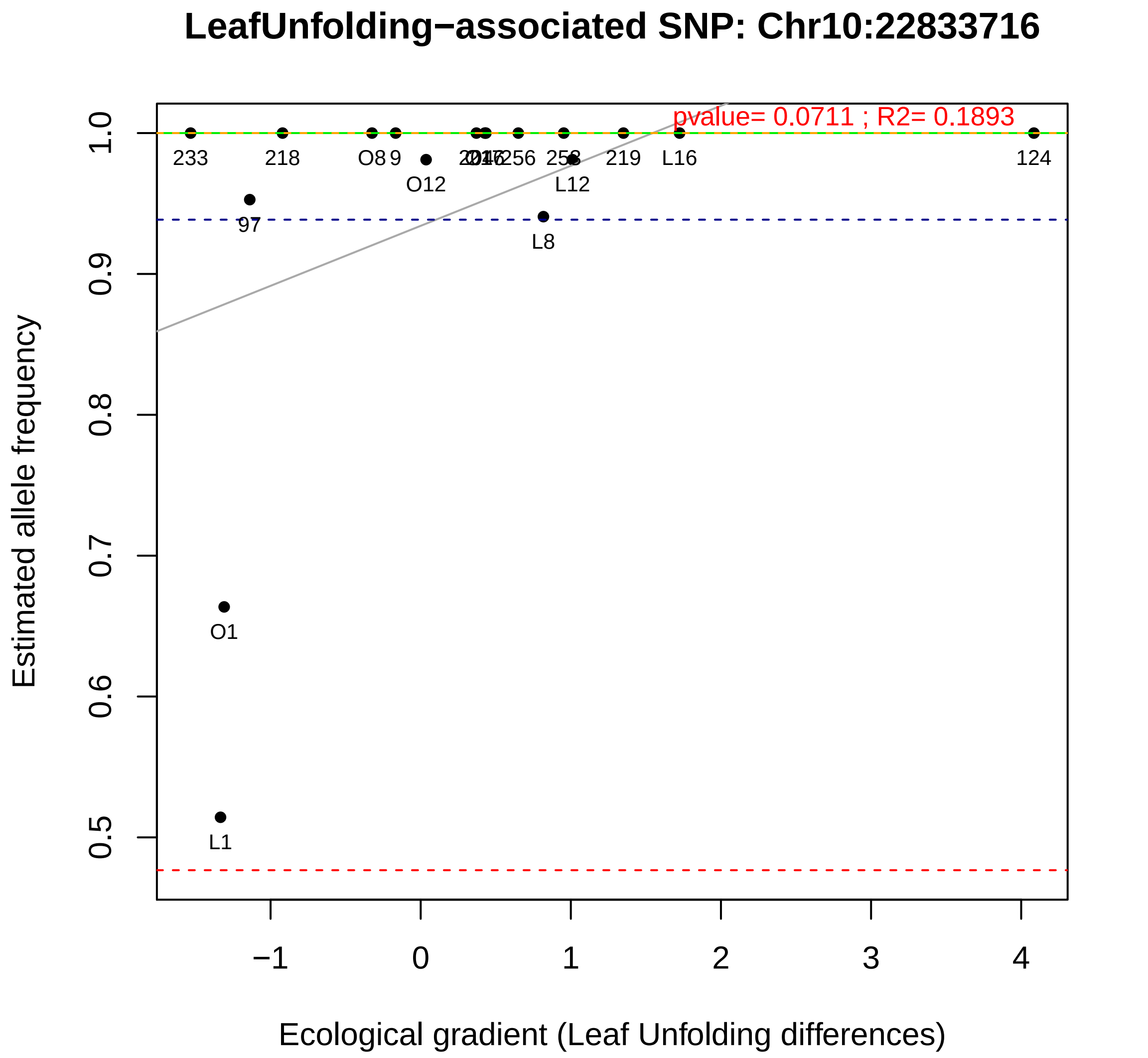
LeafUnfolding−associated SNP: Chr10:22833716
pvalue= 0.0711 ; R2= 0.1893
Estimated allele frequency
Ecological gradient (Leaf Unfolding differences)
1.0
0.9
0.8
0.7
0.6
0.5
−1
0
1
2
3
4
233
218
O8
9
256
219
L16
124
O12
L12
97
L8
O1
L1

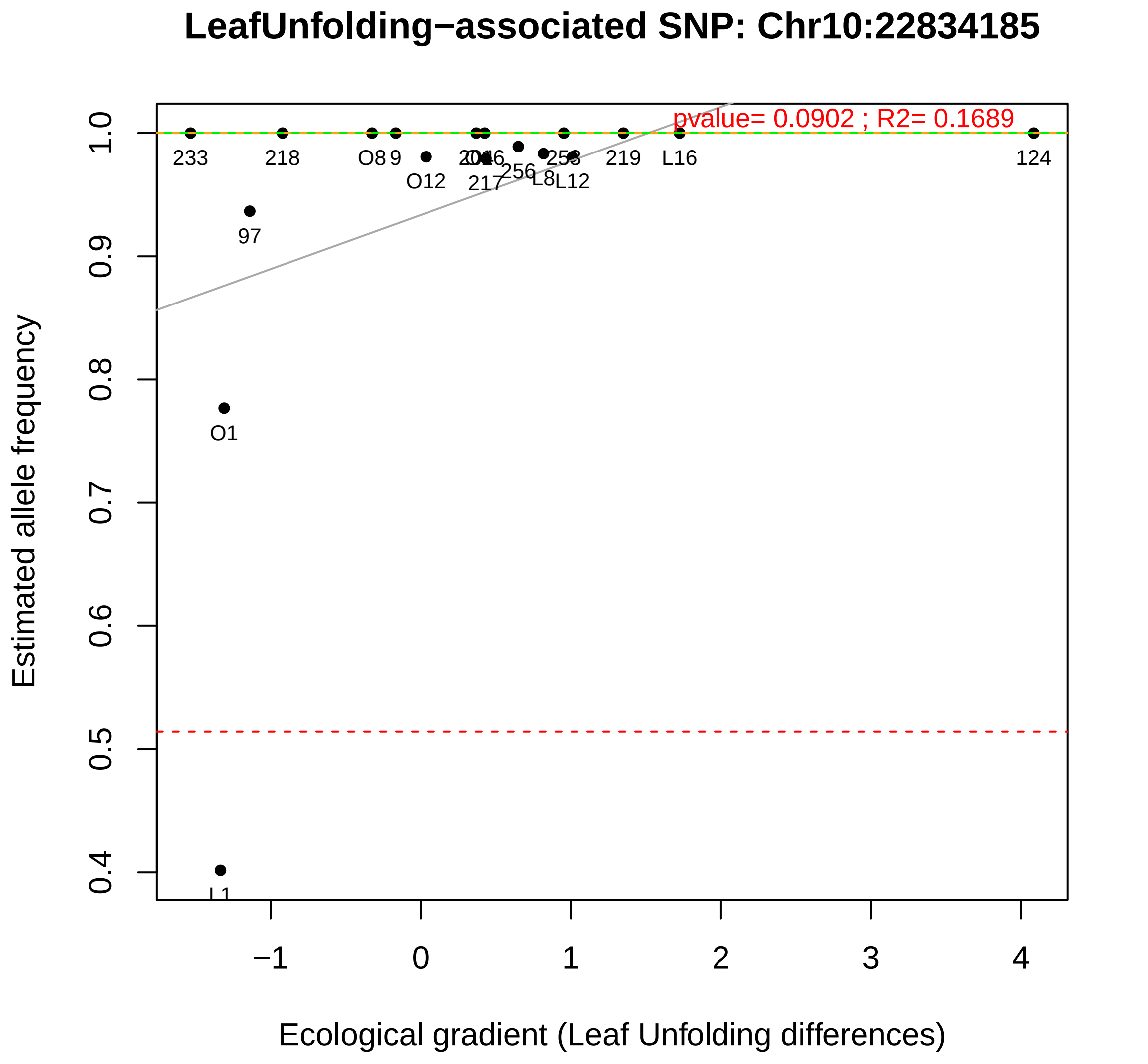
LeafUnfolding−associated SNP: Chr10:22834185
pvalue= 0.0902 ; R2= 0.1689
Estimated allele frequency
Ecological gradient (Leaf Unfolding differences)
0.4
0.5
0.6
0.7
0.8
0.9
1.0
−1
0
1
2
3
4
233
218
O8
9
O12
217
256
L8
L12
219
L16
124
97
O1

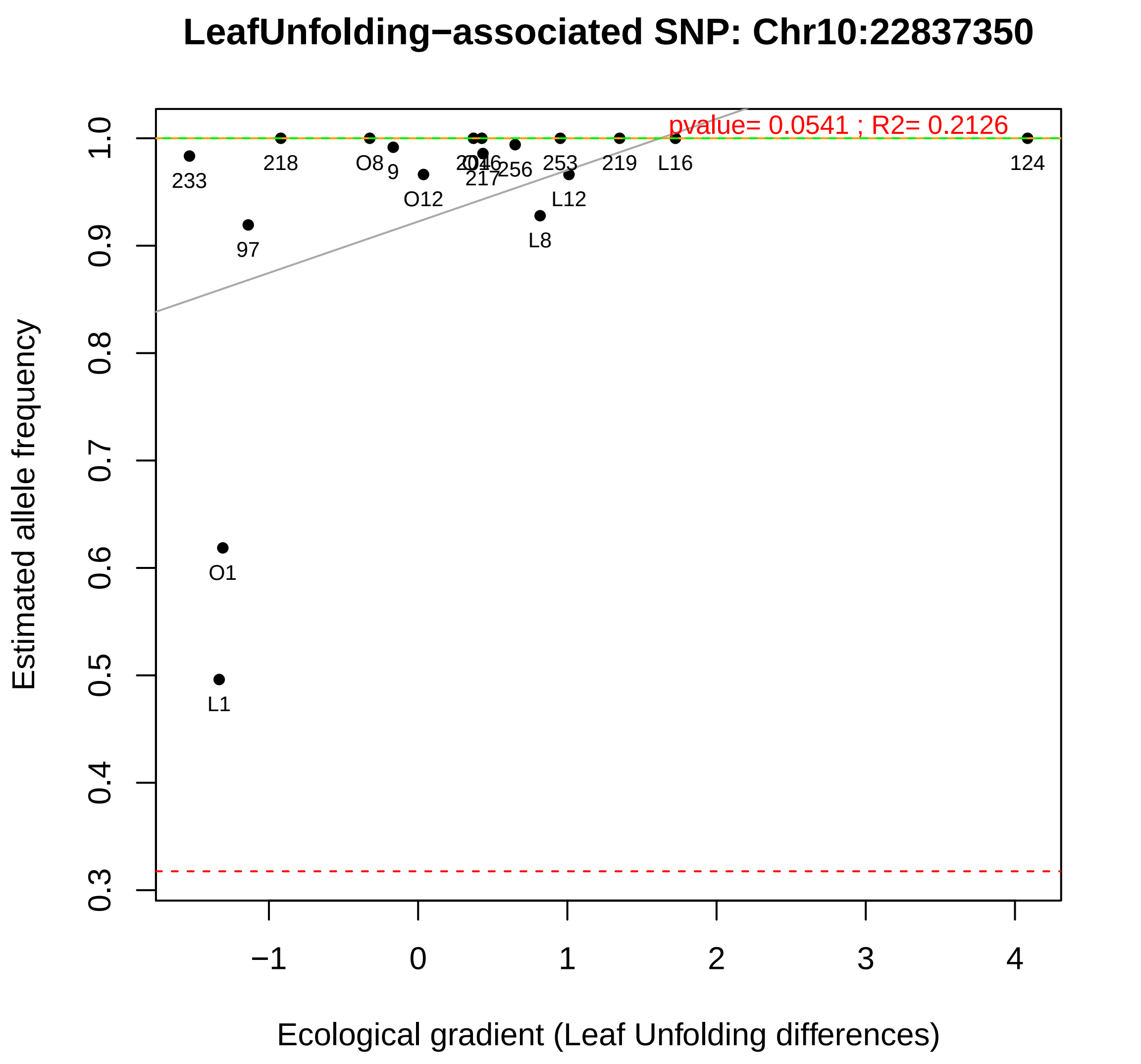
LeafUnfolding−associated SNP: Chr10:22837350
pvalue= 0.0541 ; R2= 0.2126
Estimated allele frequency
Ecological gradient (Leaf Unfolding differences)
0.3
0.4
0.5
0.6
0.7
0.8
0.9
1.0
−1
0
1
2
3
4
233
218
O8
9
O12
217
256
253
L12
219
L16
124
97
L8
O1
L1

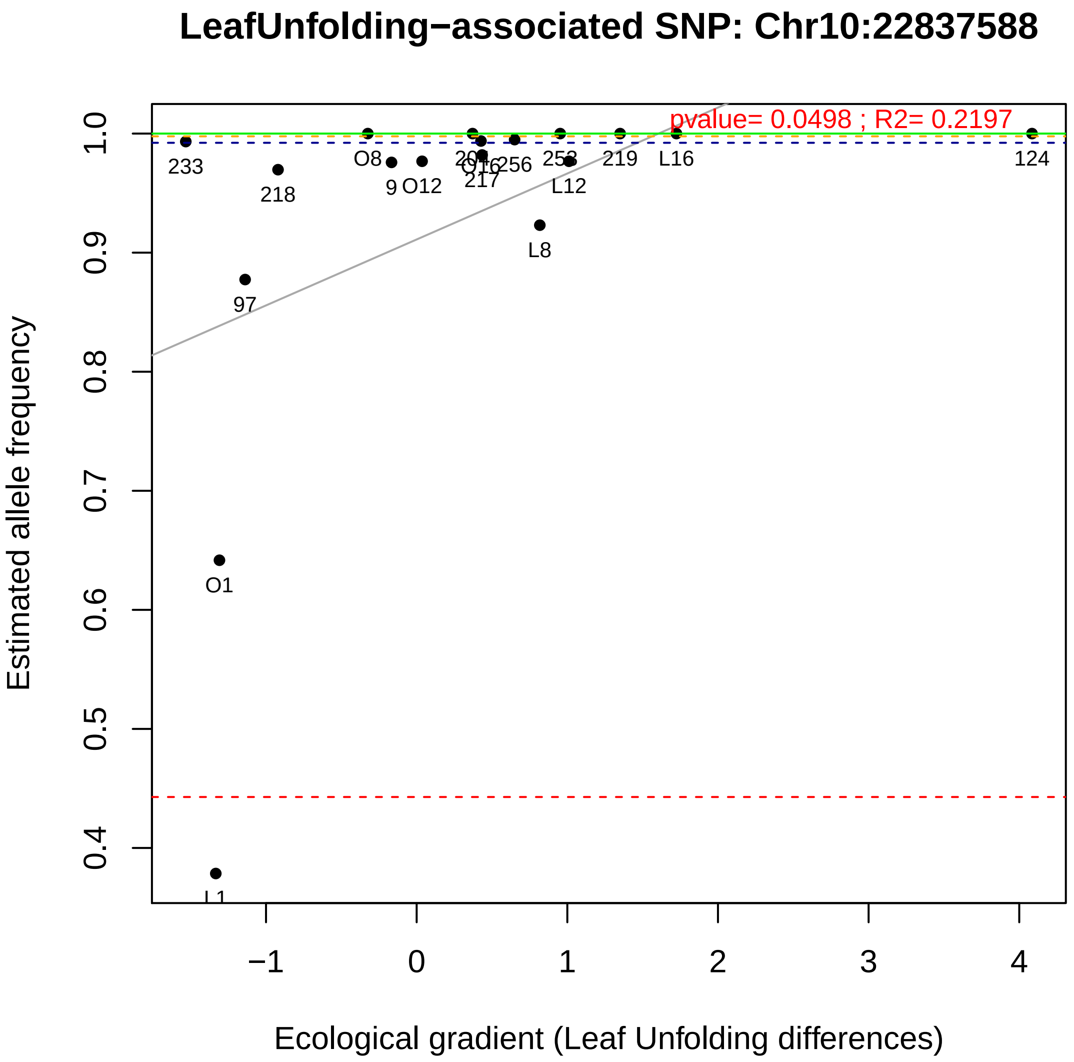
LeafUnfolding−associated SNP: Chr10:22837588
pvalue= 0.0498 ; R2= 0.2197
Estimated allele frequency
Ecological gradient (Leaf Unfolding differences)
0.4
0.5
0.6
0.7
0.8
0.9
1.0
−1
0
1
2
3
4
233
218
97
O1
O8
9
O12
217
256
L8
L12
219
L16
124

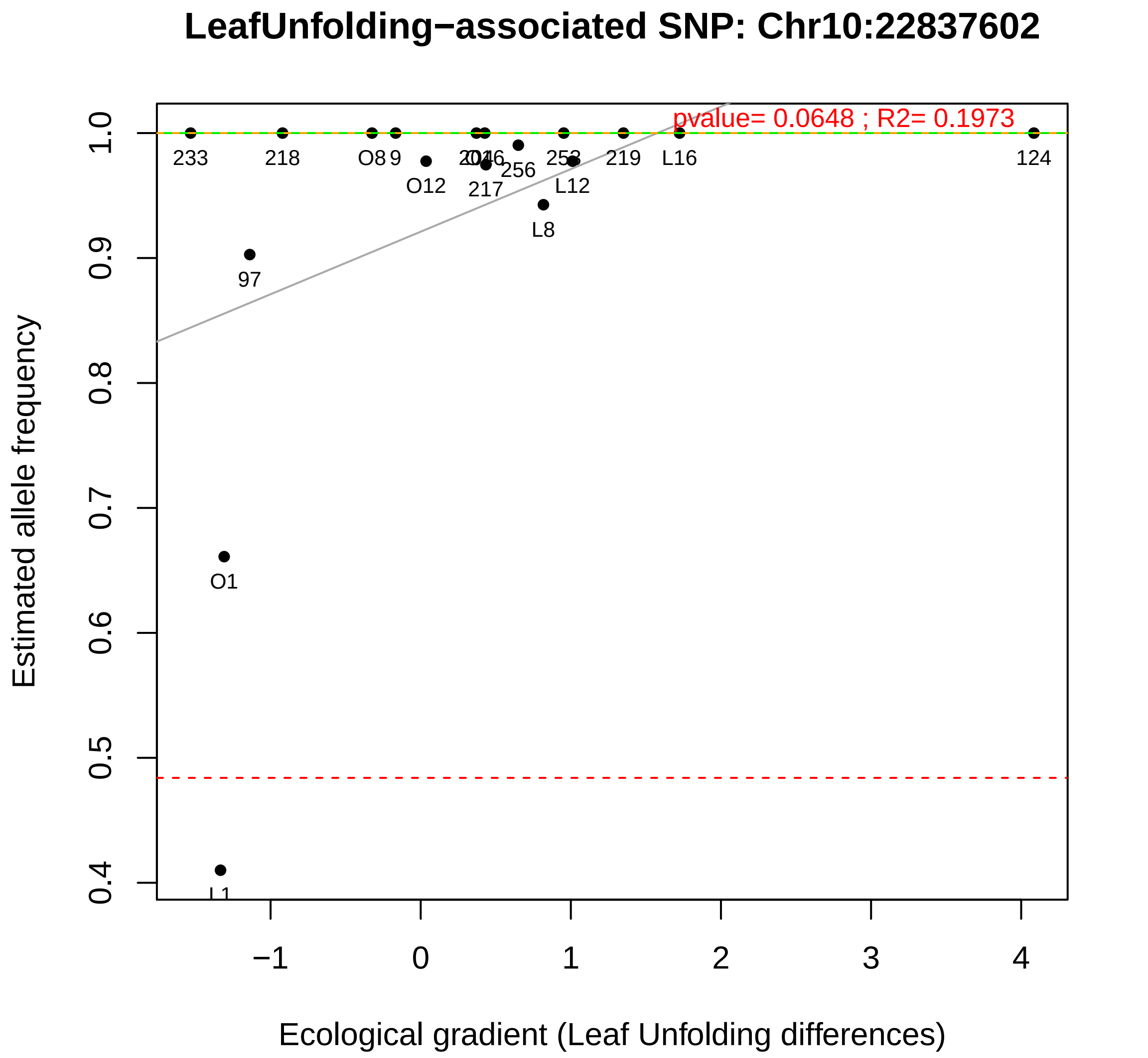

LeafUnfolding−associated SNP: Chr10:22837602
pvalue= 0.0648 ; R2= 0.1973
Estimated allele frequency
Ecological gradient (Leaf Unfolding differences)
0.4
0.5
0.6
0.7
0.8
0.9
1.0
−1
0
1
2
3
4
233
218
O8
9
O12
217
256
L12
L8
219
L16
124
97
O1

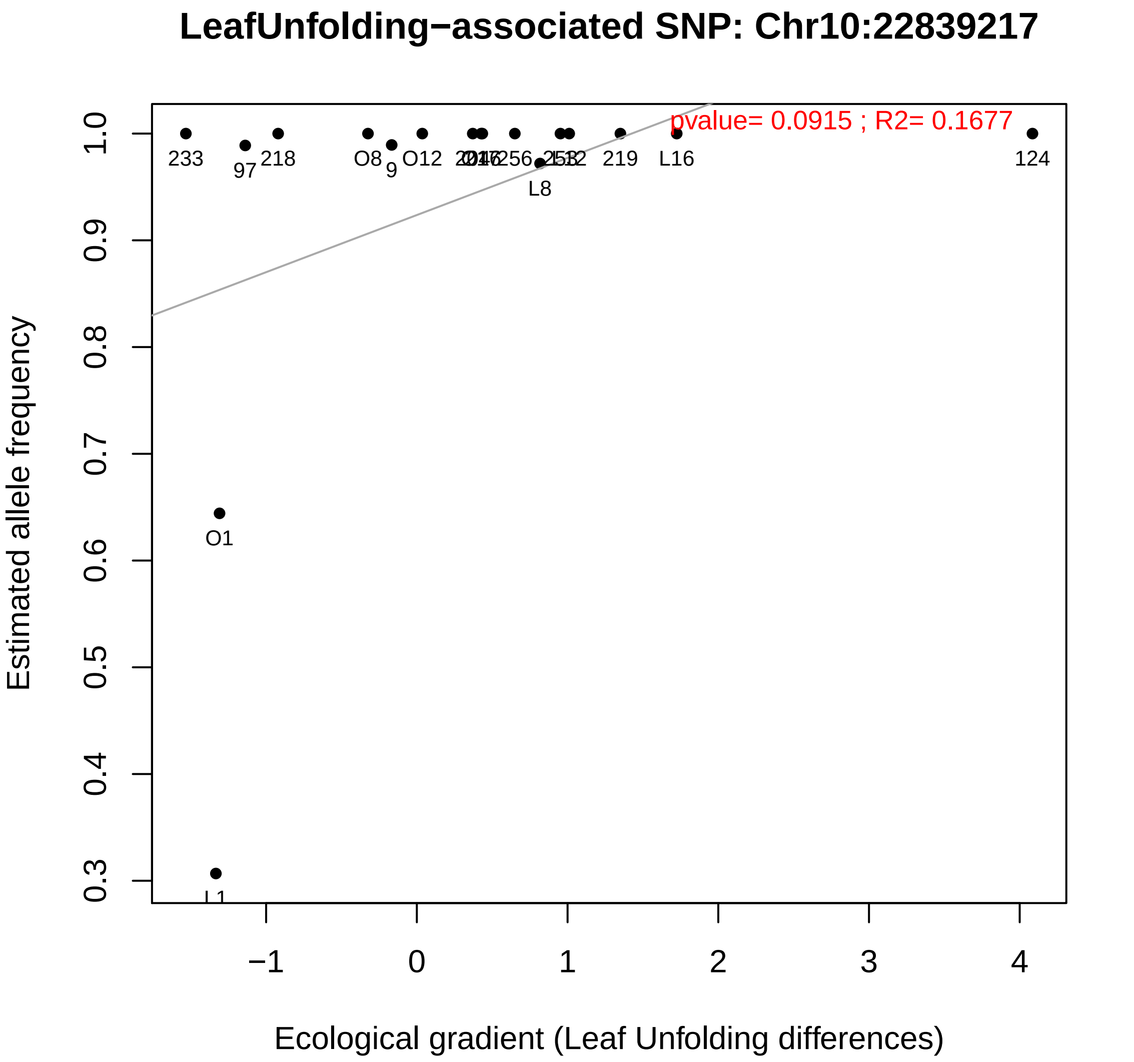
LeafUnfolding−associated SNP: Chr10:22839217
pvalue= 0.0915 ; R2= 0.1677
Estimated allele frequency
Ecological gradient (Leaf Unfolding differences)
1.0
0.9
0.8
0.7
0.6
0.5
0.4
0.3
−1
0
1
2
3
4
233
97
218
O8
9
O12
L8
219
L16
124
O1

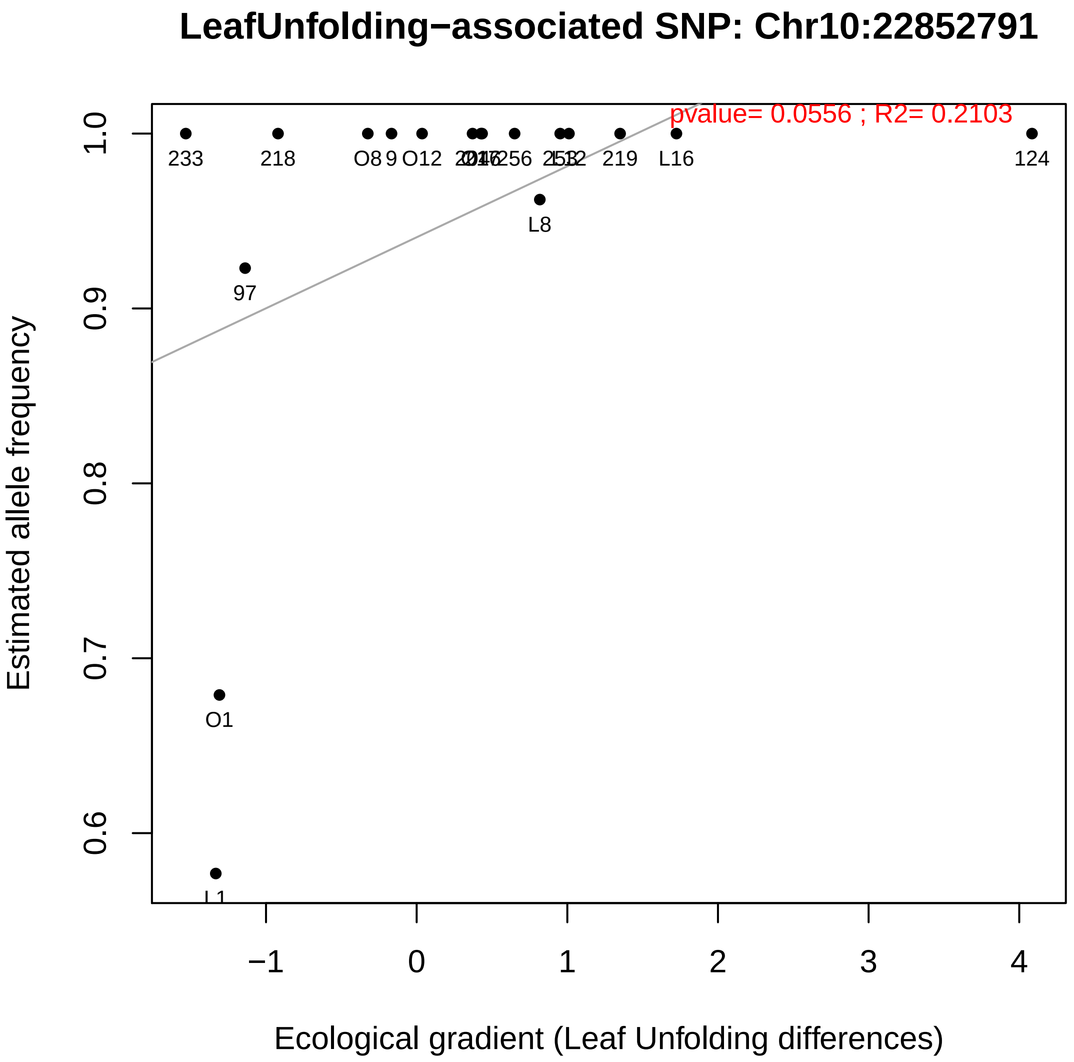

LeafUnfolding−associated SNP: Chr10:22852791
pvalue= 0.0556 ; R2= 0.2103
Estimated allele frequency
Ecological gradient (Leaf Unfolding differences)
1.0
0.9
0.8
0.7
0.6
−1
0
1
2
3
4
233
218
O8
9
O12
219
L16
124
L8
97
O1

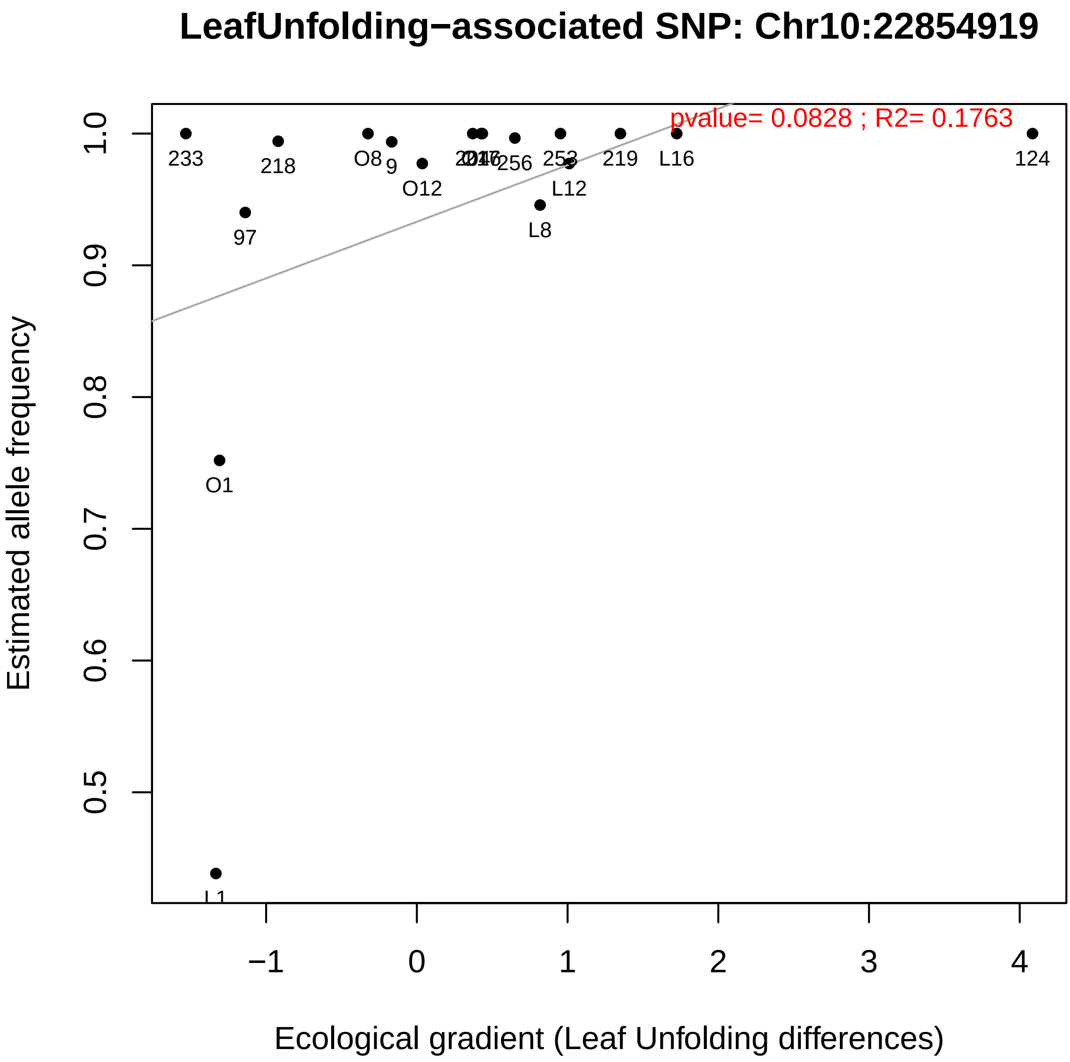
LeafUnfolding−associated SNP: Chr10:22854919
pvalue= 0.0828 ; R2= 0.1763
Estimated allele frequency
1.0
0.9
0.8
0.7
0.6
0.5
−1
0
1
2
3
4
Ecological gradient (Leaf Unfolding differences)
233
218
97
O8
9
O12
256
253
L12
L8
219
L16
124
O1

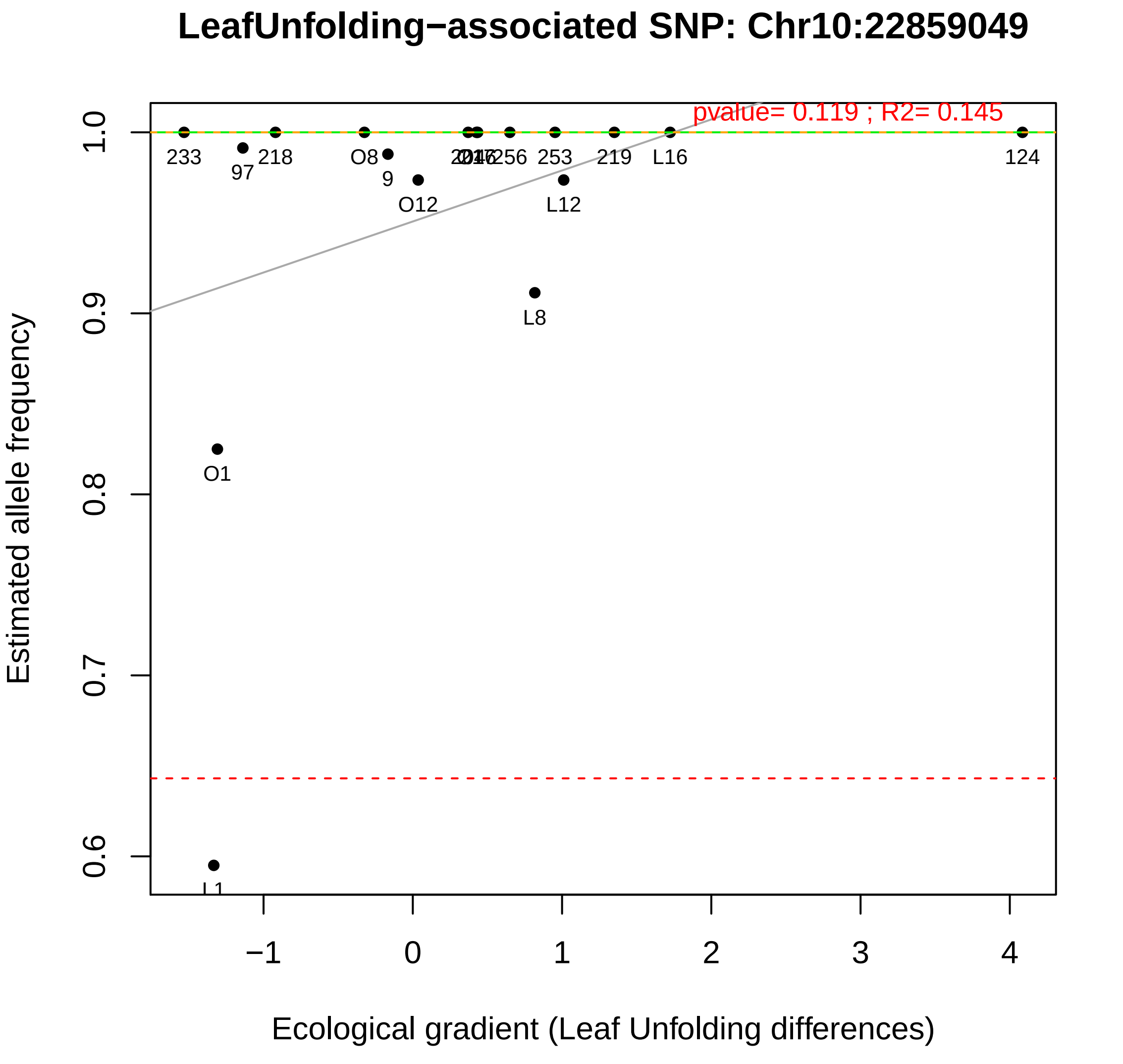
LeafUnfolding−associated SNP: Chr10:22859049
pvalue= 0.119 ; R2= 0.145
Estimated allele frequency
Ecological gradient (Leaf Unfolding differences)
1.0
0.9
0.8
0.7
0.6
−1
0
1
2
3
4
233
97
218
O8
9
O12
256
253
L12
219
L16
124
L8
O1

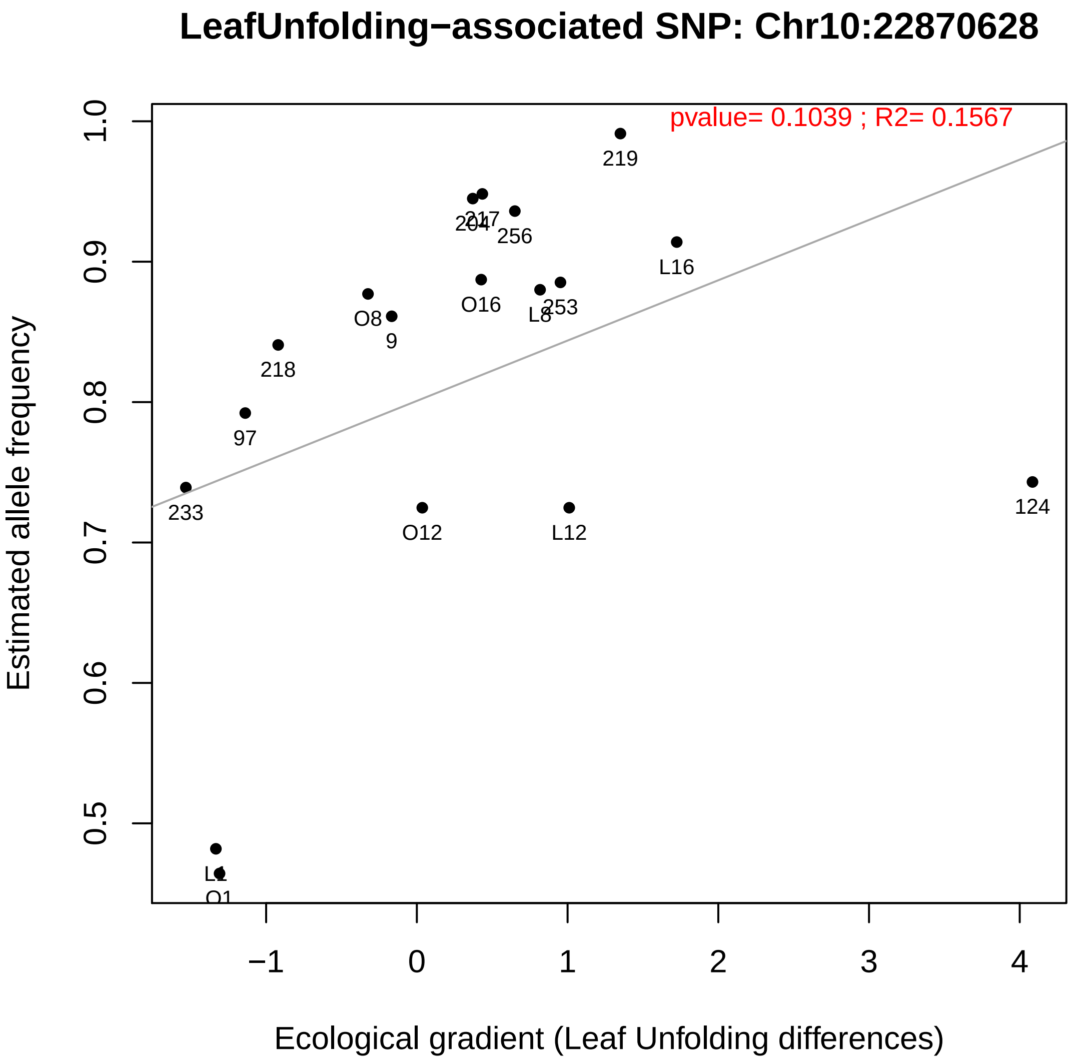
LeafUnfolding−associated SNP: Chr10:22870628
pvalue= 0.1039 ; R2= 0.1567
Estimated allele frequency
Ecological gradient (Leaf Unfolding differences)
0.5
0.6
0.7
0.8
0.9
1.0
−1
0
1
2
3
4
219
217
204
256
L16
O16
253
L8
O8
9
218
97
233
O12
L12
124
L1
O1

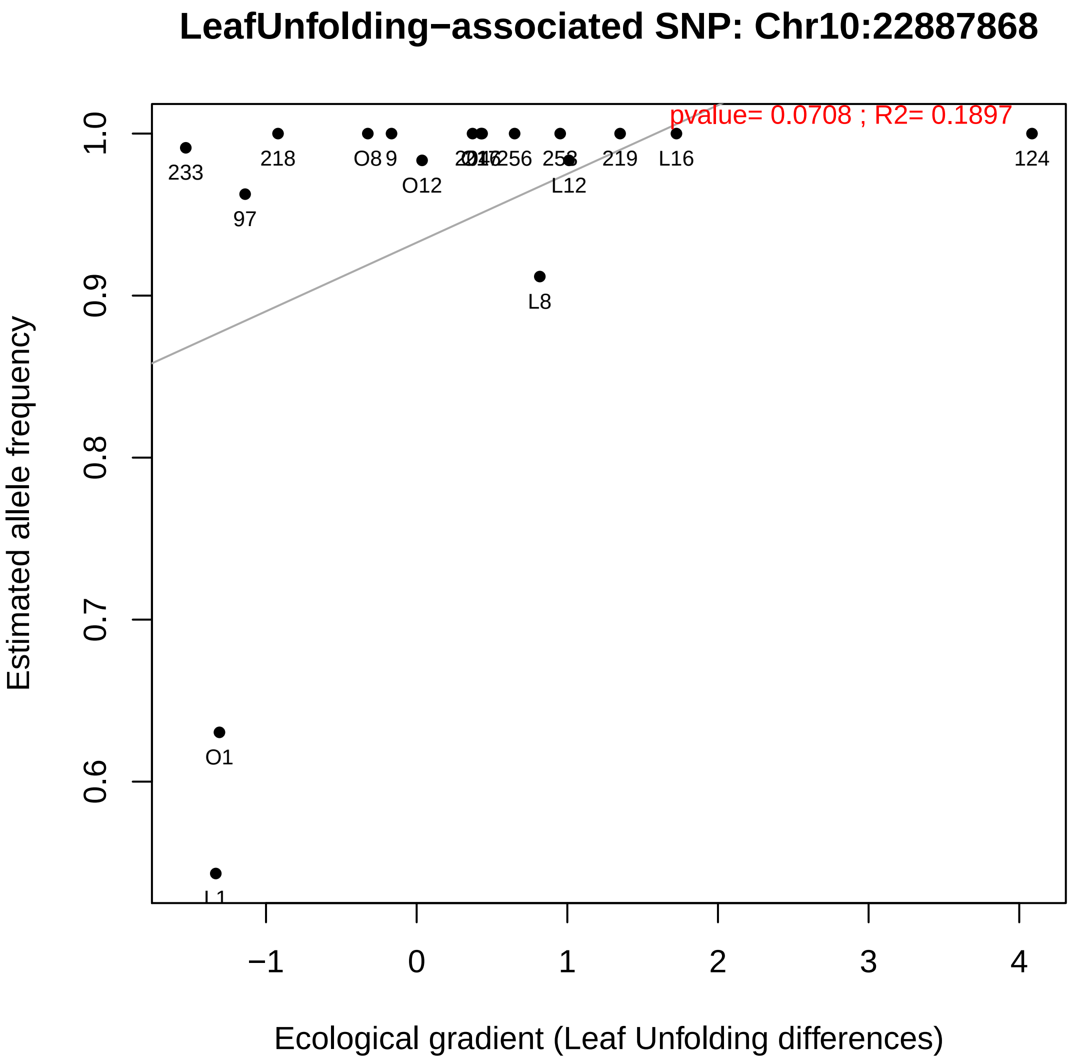
LeafUnfolding−associated SNP: Chr10:22887868
pvalue= 0.0708 ; R2= 0.1897
Estimated allele frequency
Ecological gradient (Leaf Unfolding differences)
0.6
0.7
0.8
0.9
1.0
−1
0
1
2
3
4
233
218
O8
9
O12
256
219
L16
124
97
L12
L8
O1

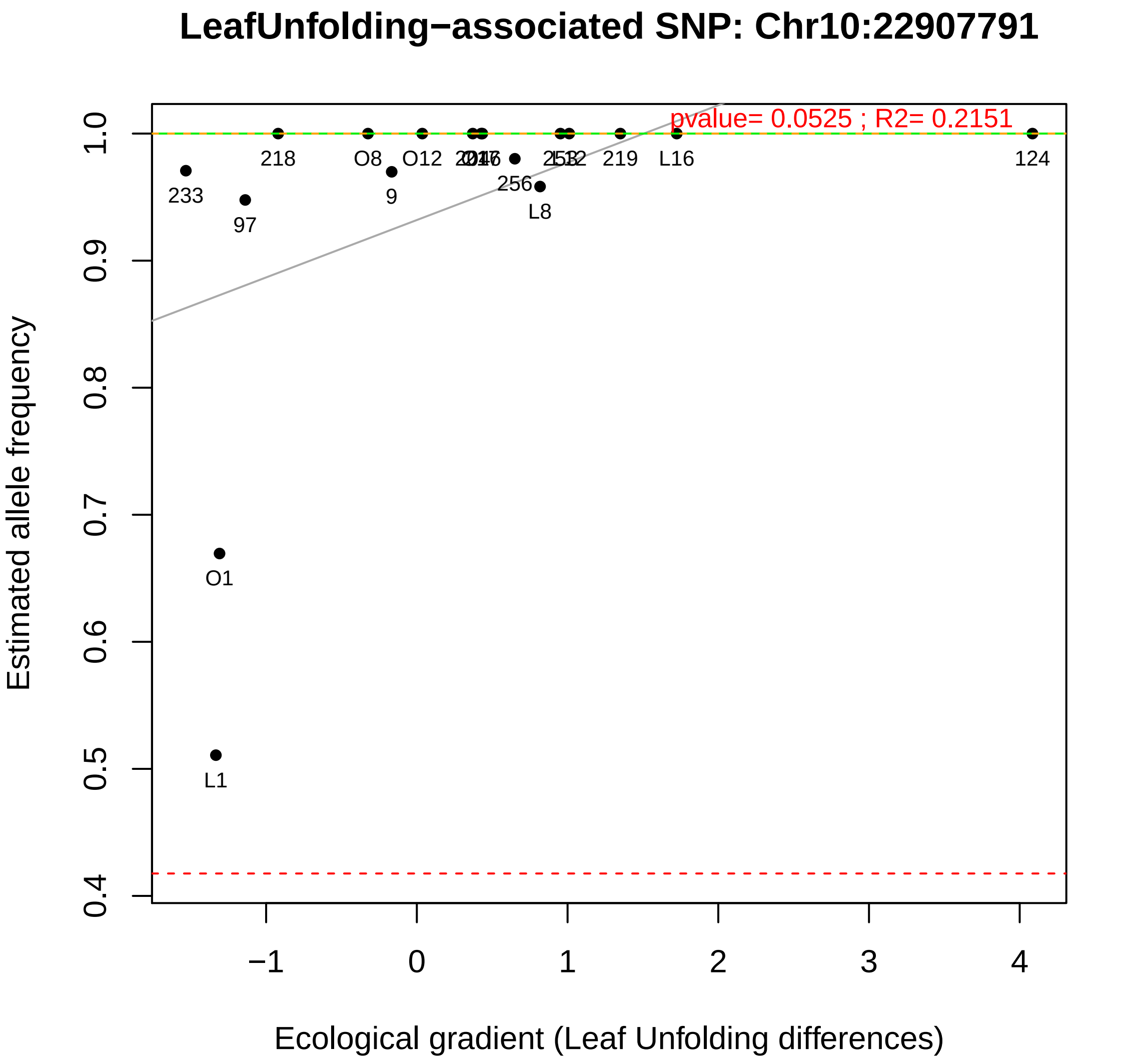
LeafUnfolding−associated SNP: Chr10:22907791
pvalue= 0.0525 ; R2= 0.2151
Estimated allele frequency
0.4
0.5
0.6
0.7
0.8
0.9
1.0
Ecological gradient (Leaf Unfolding differences)
−1
0
1
2
3
4
233
97
218
O8
9
O12
256
L8
219
L16
124
O1
L1

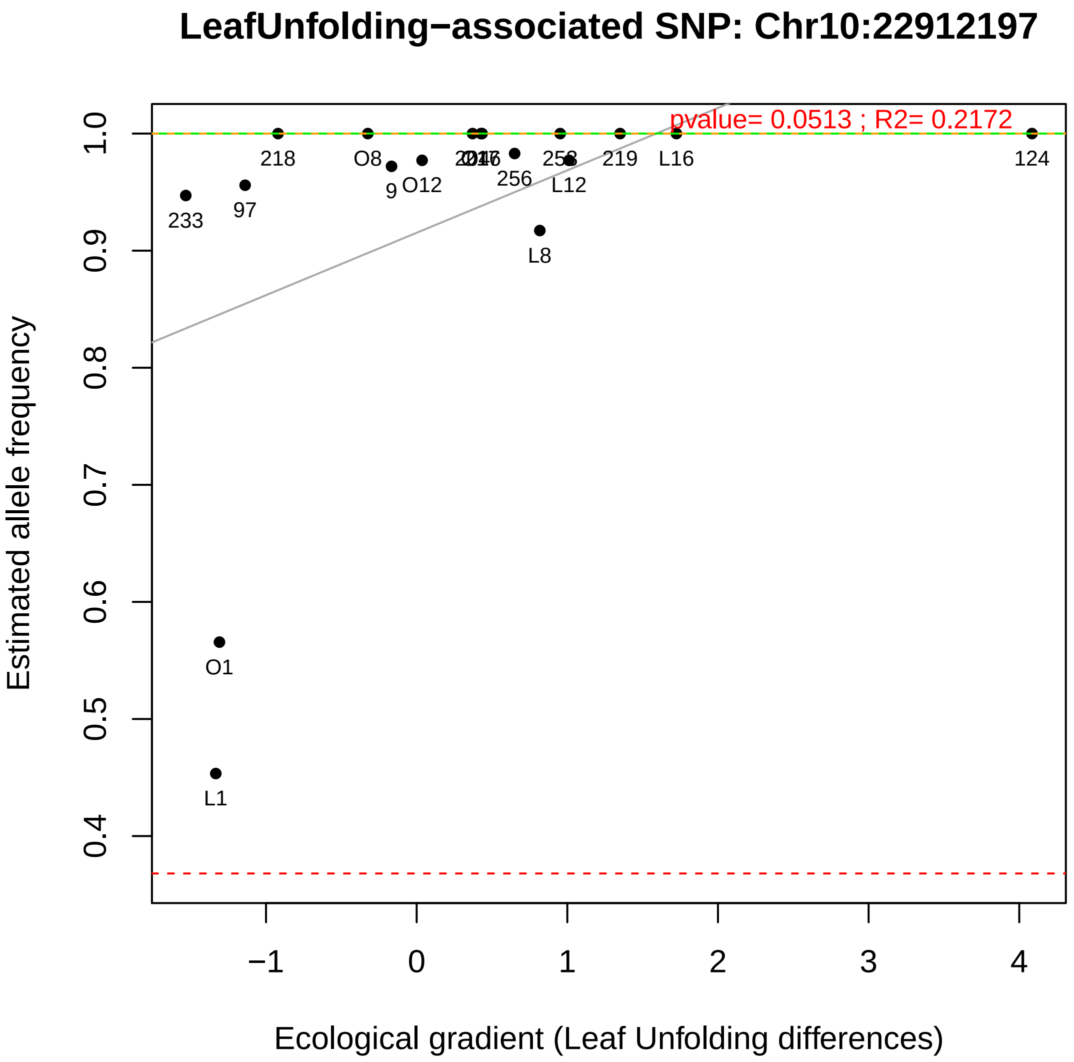
LeafUnfolding−associated SNP: Chr10:22912197
pvalue= 0.0513 ; R2= 0.2172
Estimated allele frequency
Ecological gradient (Leaf Unfolding differences)
0.4
0.5
0.6
0.7
0.8
0.9
1.0
−1
0
1
2
3
4
218
O8
256
219
L16
124
233
97
9
O12
L12
L8
O1
L1

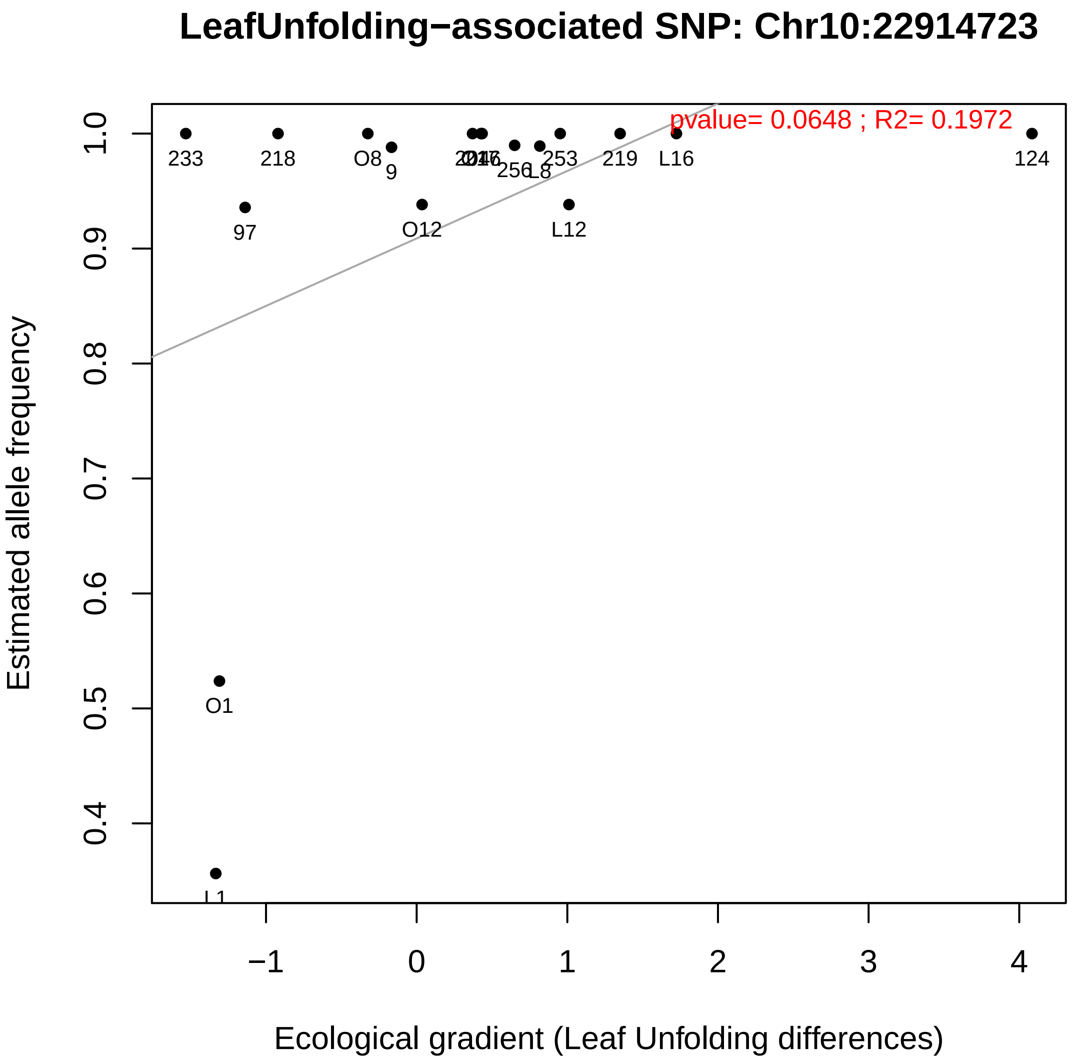

LeafUnfolding−associated SNP: Chr10:22914723
pvalue= 0.0648 ; R2= 0.1972
Estimated allele frequency
Ecological gradient (Leaf Unfolding differences)
1.0
0.9
0.8
0.7
0.6
0.5
0.4
−1
0
1
2
3
4
233
218
O8
9
256
L8
253
219
L16
124
97
O12
L12
O1

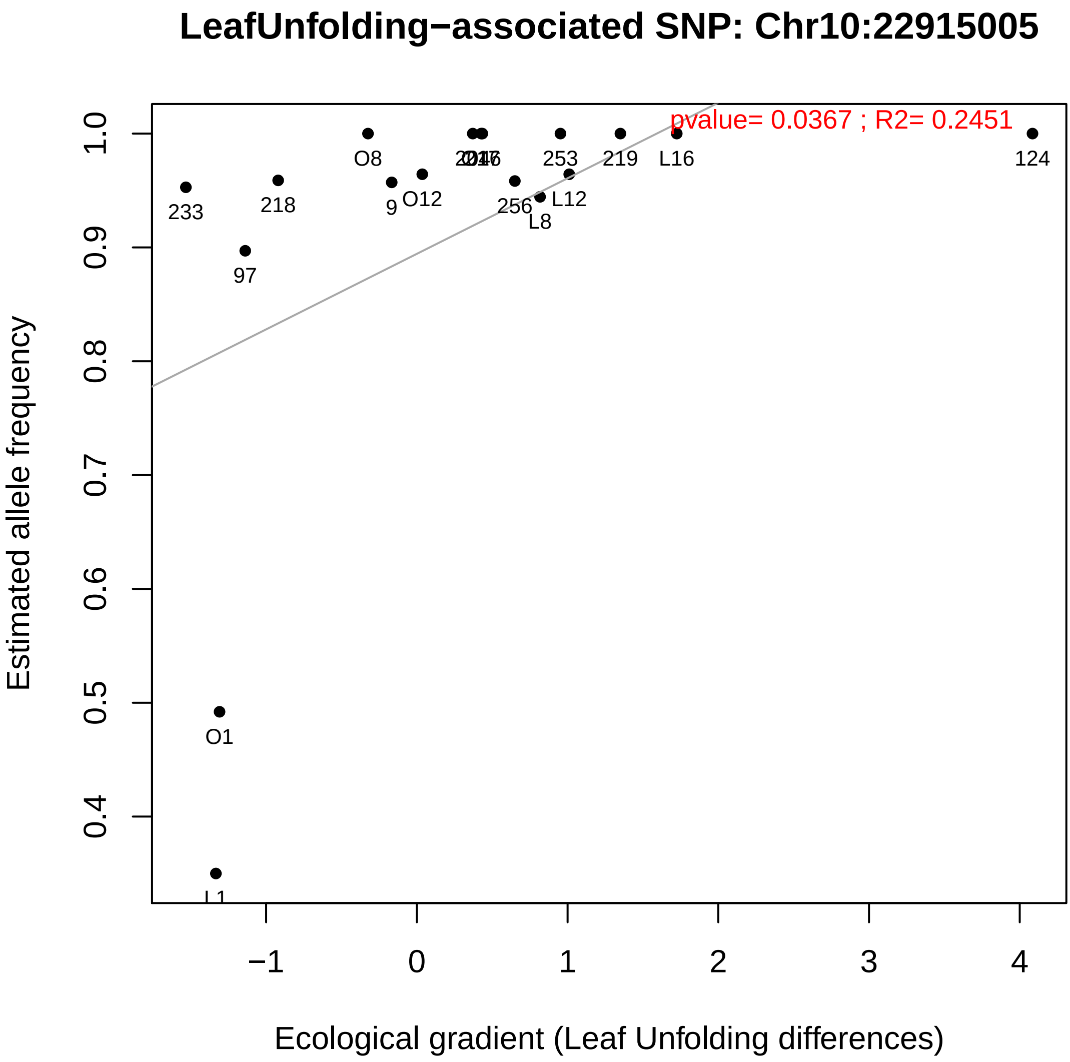

LeafUnfolding−associated SNP: Chr10:22915005
pvalue= 0.0367 ; R2= 0.2451
Estimated allele frequency
Ecological gradient (Leaf Unfolding differences)
1.0
0.9
0.8
0.7
0.6
0.5
0.4
−1
0
1
2
3
4
233
97
218
O8
9
O12
256
L8
L12
253
219
L16
124
O1
L1

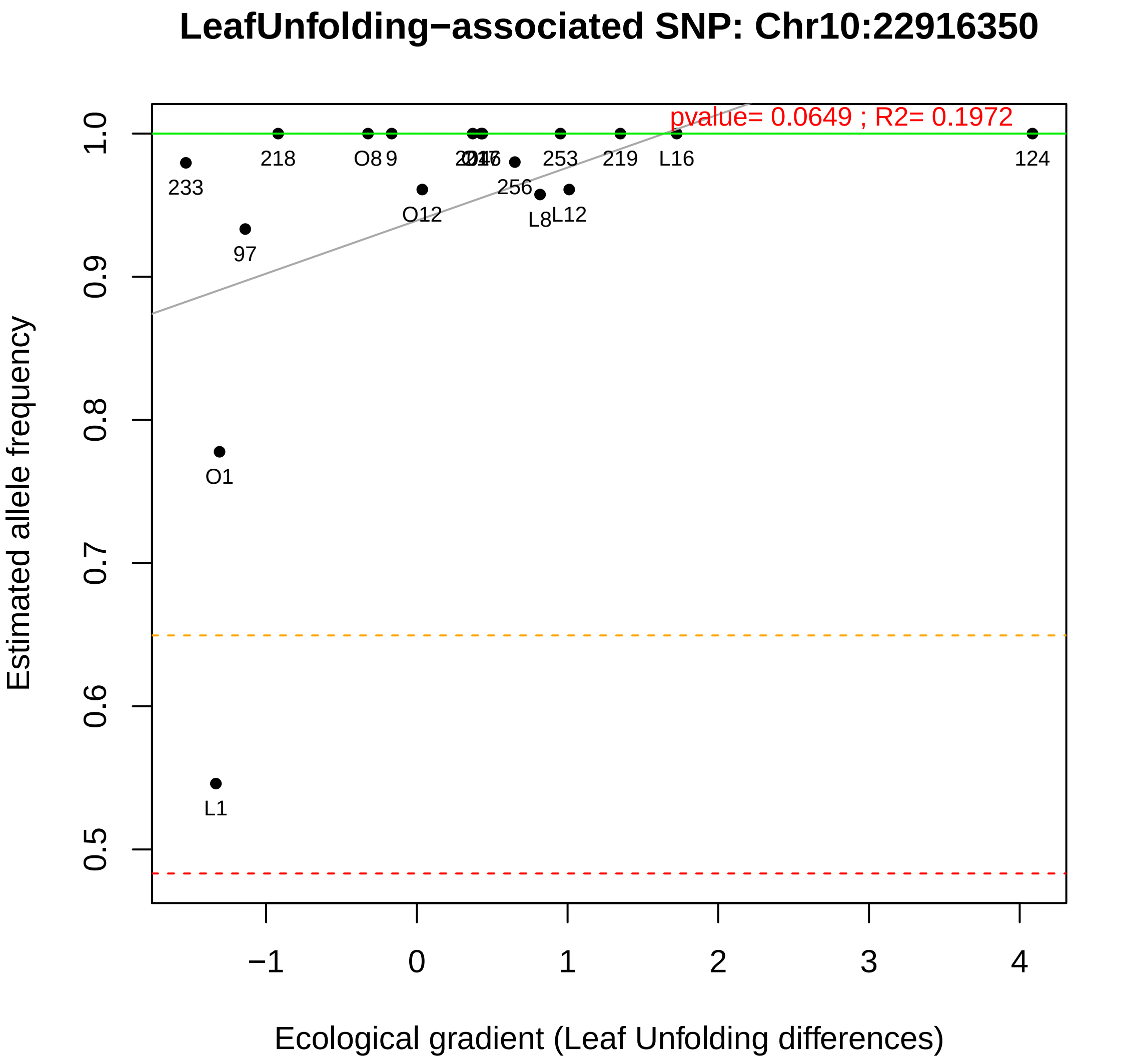
LeafUnfolding−associated SNP: Chr10:22916350
pvalue= 0.0649 ; R2= 0.1972
Estimated allele frequency
Ecological gradient (Leaf Unfolding differences)
1.0
0.9
0.8
0.7
0.6
0.5
−1
0
1
2
3
4
233
218
O8
9
256
253
219
L16
124
O12
L8
L12
97
O1
L1

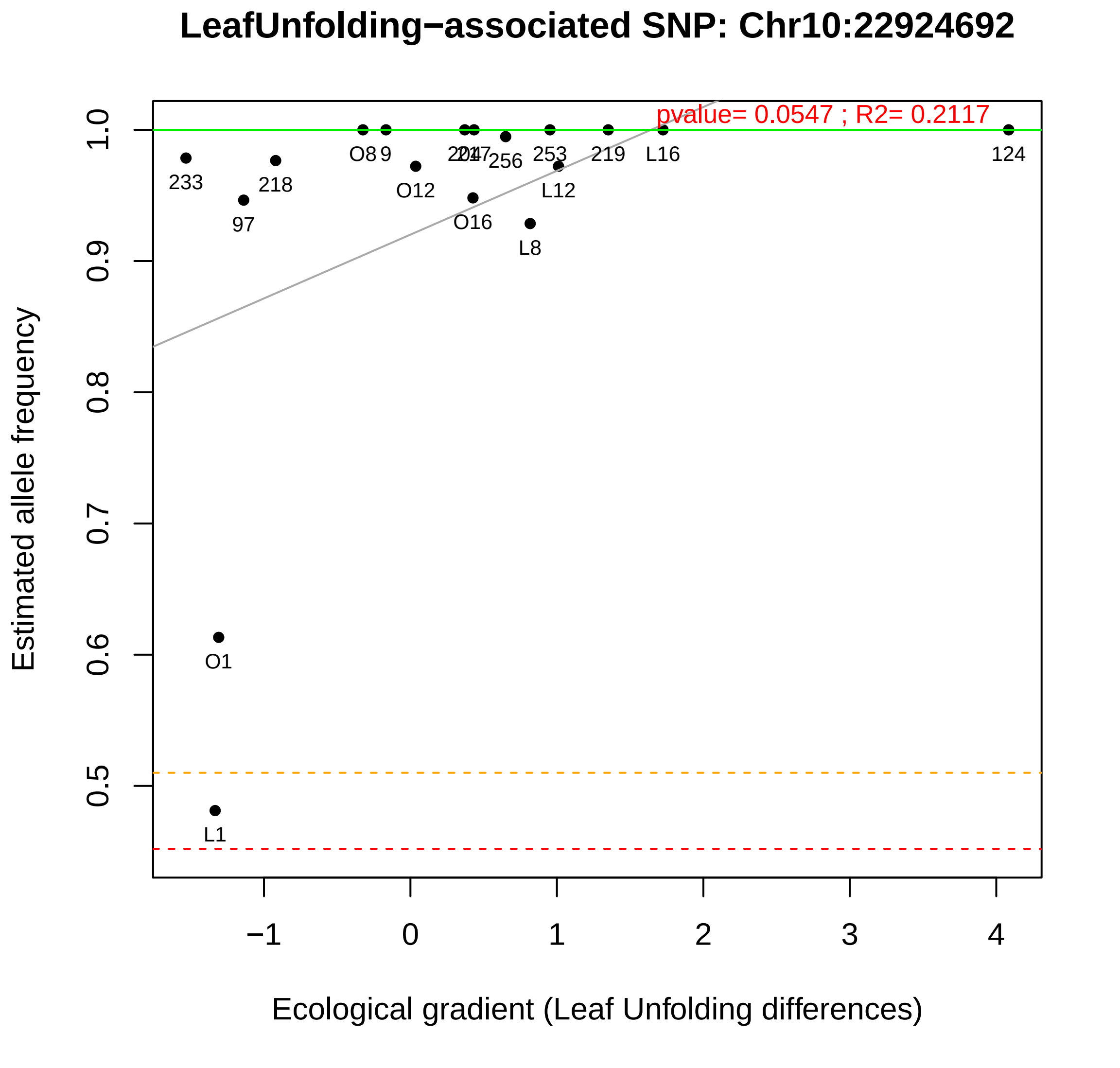
LeafUnfolding−associated SNP: Chr10:22924692
pvalue= 0.0547 ; R2= 0.2117
Estimated allele frequency
Ecological gradient (Leaf Unfolding differences)
1.0
0.9
0.8
0.7
0.6
0.5
−1
0
1
2
3
4
233
97
218
O8
9
O12
O16
256
253
L12
L8
219
L16
124
O1
L1

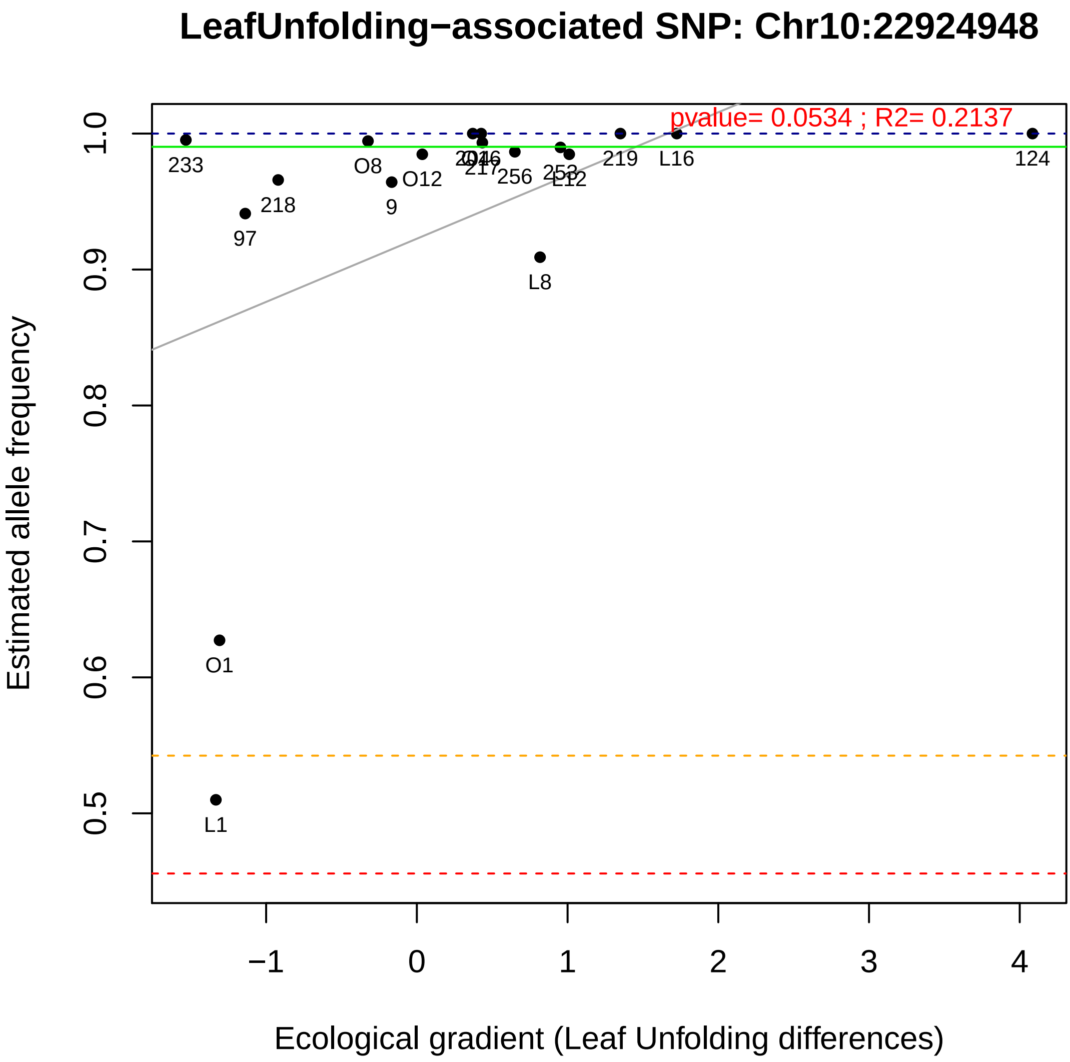

LeafUnfolding−associated SNP: Chr10:22924948
pvalue= 0.0534 ; R2= 0.2137
Estimated allele frequency
Ecological gradient (Leaf Unfolding differences)
0.5
0.6
0.7
0.8
0.9
1.0
−1
0
1
2
3
4
233
218
97
O8
O12
9
217
256
L12
L8
219
L16
124
O1
L1

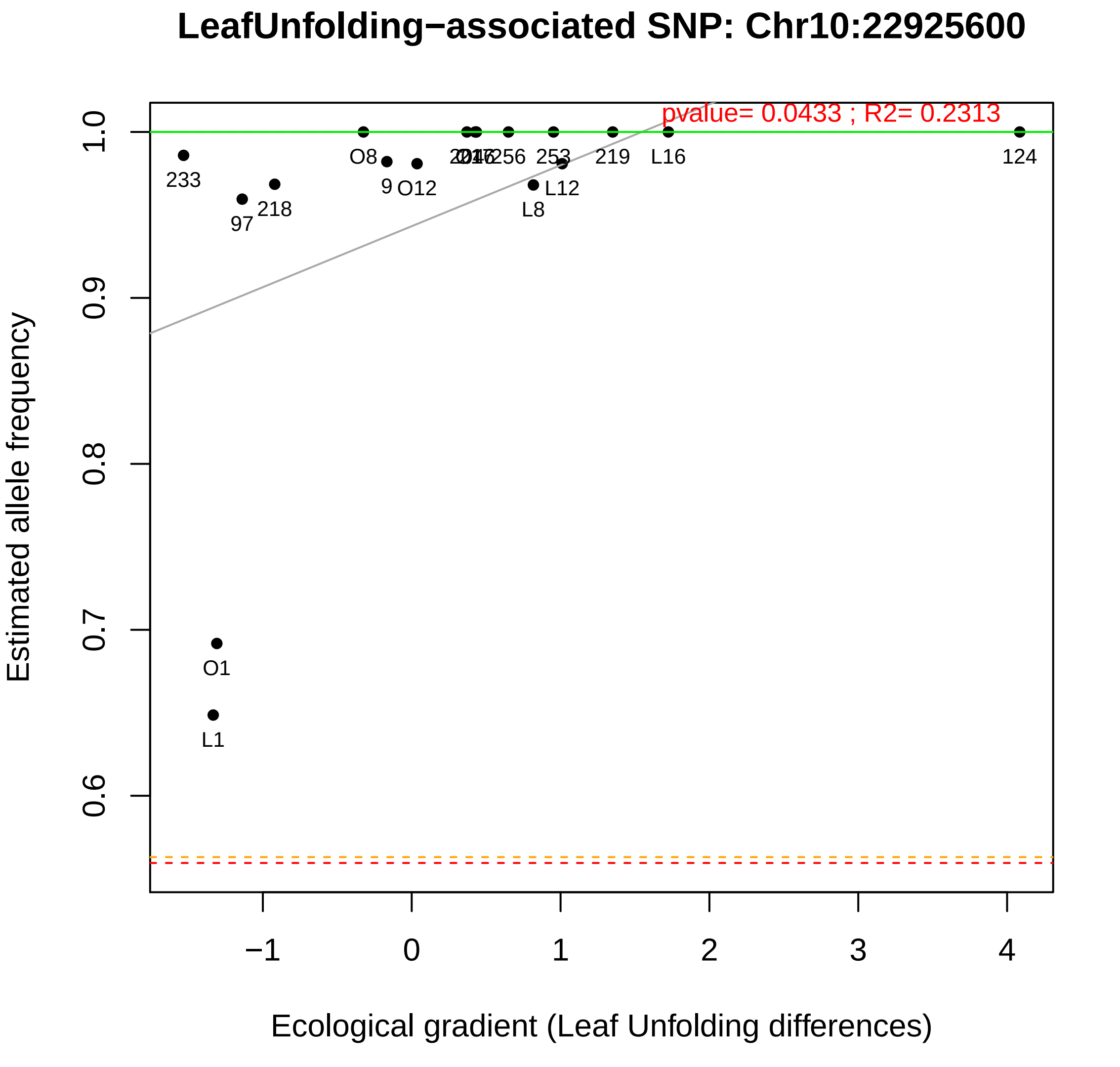
LeafUnfolding−associated SNP: Chr10:22925600
pvalue= 0.0433 ; R2= 0.2313
Estimated allele frequency
Ecological gradient (Leaf Unfolding differences)
1.0
0.9
0.8
0.7
0.6
−1
0
1
2
3
4
233
97
218
O8
9
O12
256
253
L12
L8
219
L16
124
O1
L1

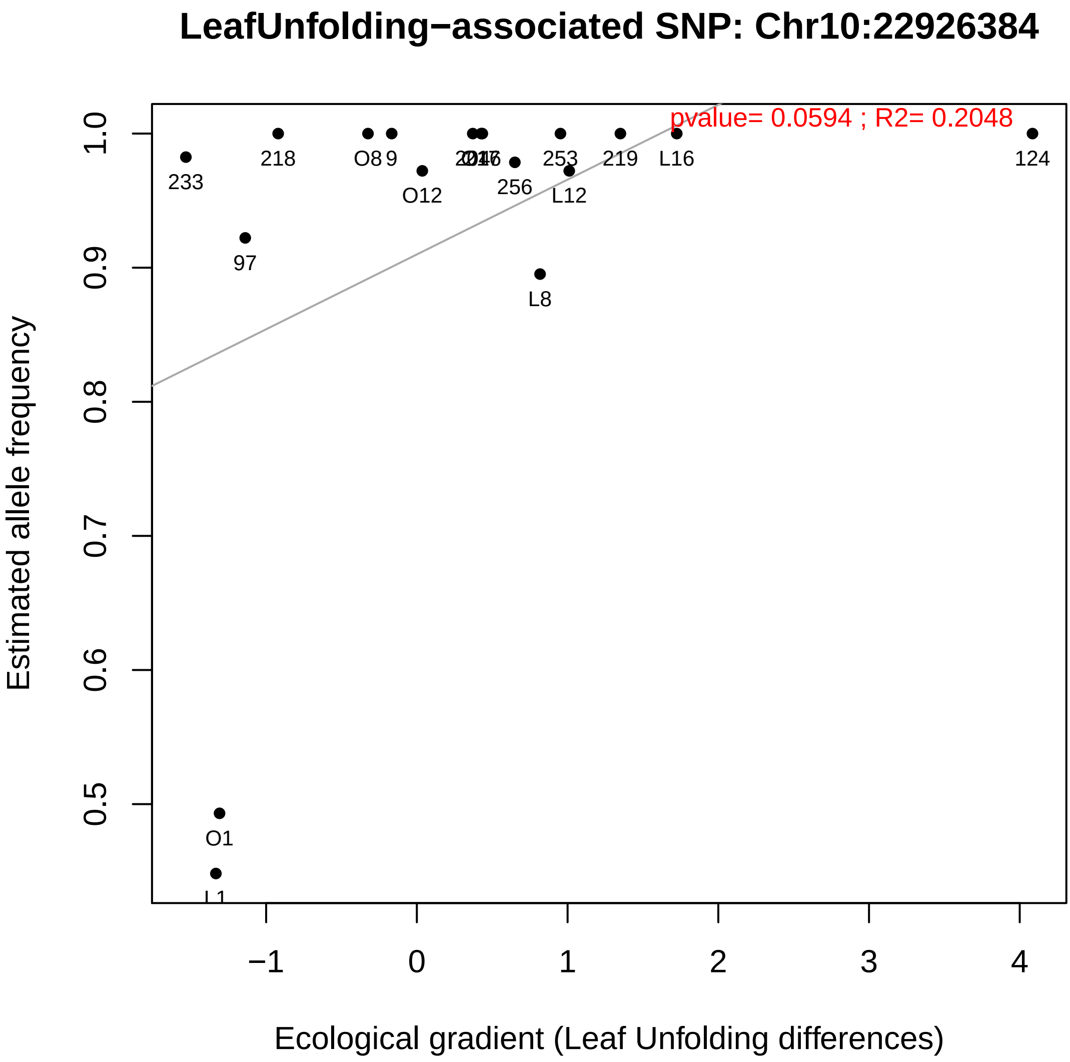
LeafUnfolding−associated SNP: Chr10:22926384
pvalue= 0.0594 ; R2= 0.2048
Estimated allele frequency
Ecological gradient (Leaf Unfolding differences)
1.0
0.9
0.8
0.7
0.6
0.5
−1
0
1
2
3
4
233
218
O8
9
O12
256
253
L12
219
L16
124
97
L8
O1
L1

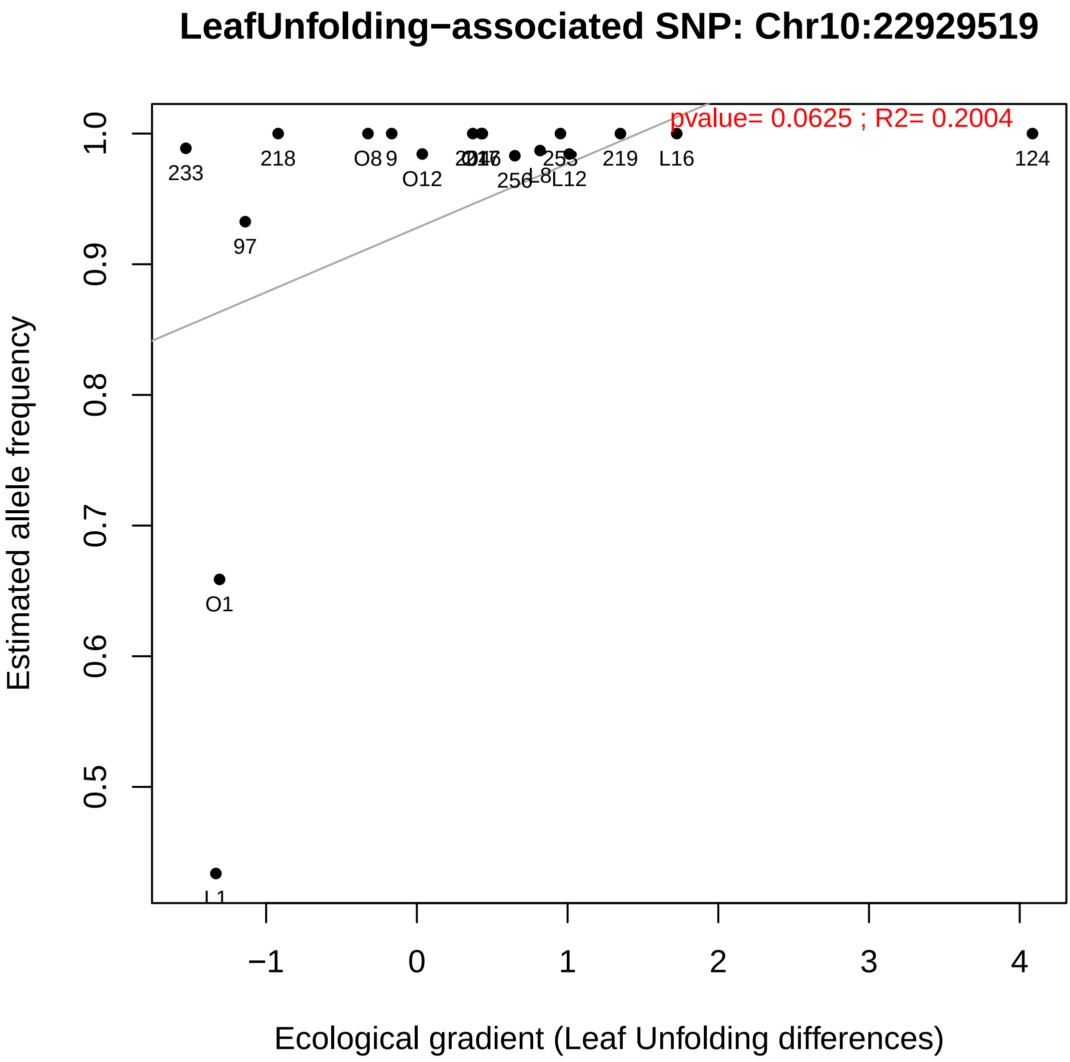
LeafUnfolding−associated SNP: Chr10:22929519
pvalue= 0.0625 ; R2= 0.2004
Estimated allele frequency
Ecological gradient (Leaf Unfolding differences)
1.0
0.9
0.8
0.7
0.6
0.5
−1
0
1
2
3
4
233
218
O8
9
O12
L8
256
253
L12
219
L16
124
97
O1
L1

### LeafUnfolding−associated SNP: Chr10:22929615

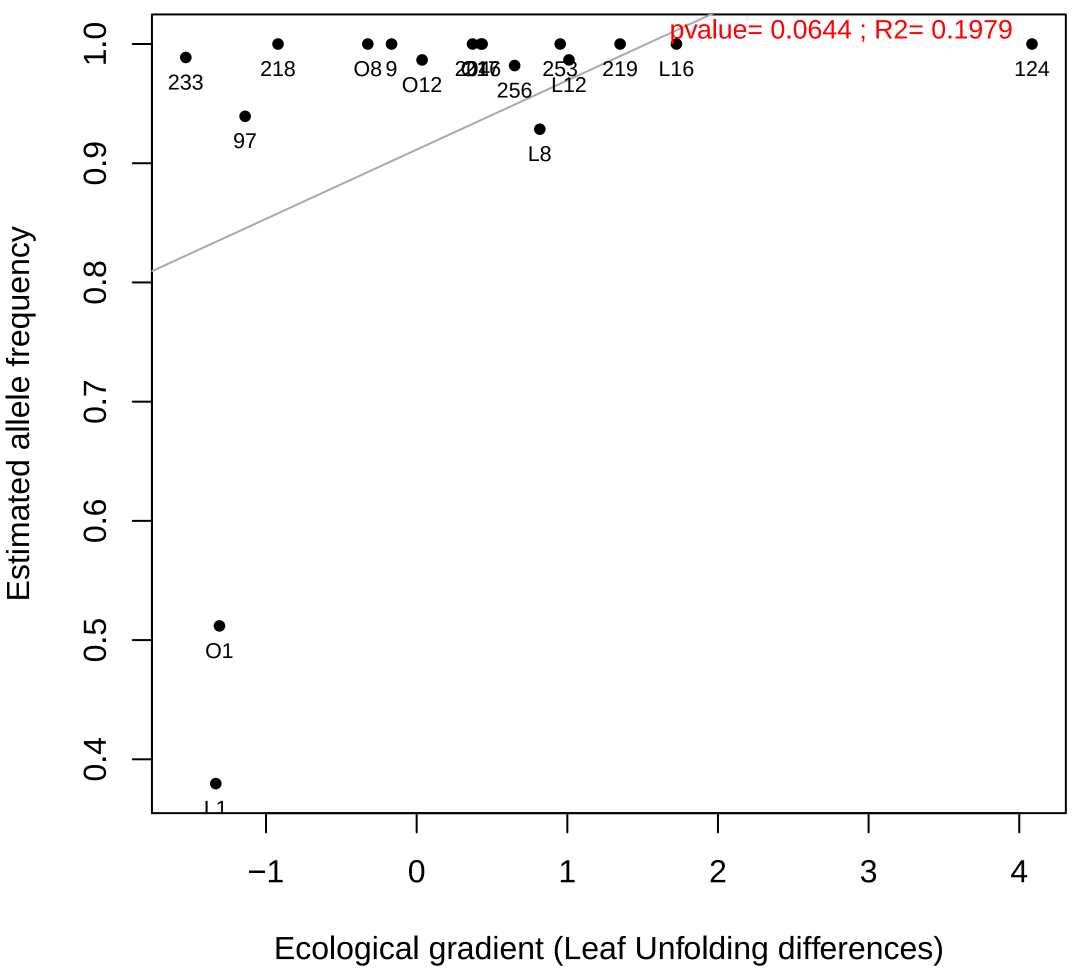

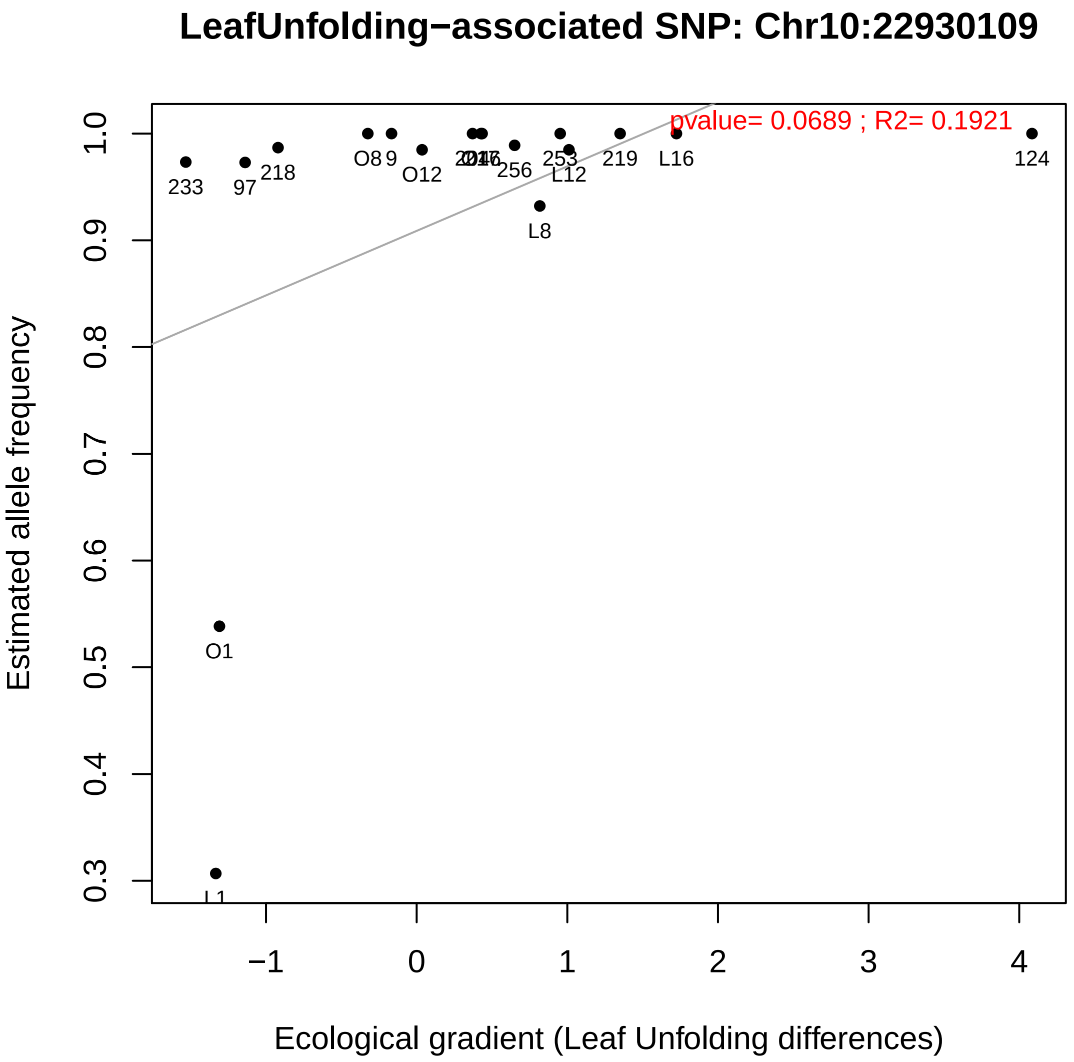
LeafUnfolding−associated SNP: Chr10:22930109
pvalue= 0.0689 ; R2= 0.1921
Estimated allele frequency
Ecological gradient (Leaf Unfolding differences)
0.3
0.4
0.5
0.6
0.7
0.8
0.9
1.0
−1
0
1
2
3
4
233
97
218
O8
9
O12
256
253
L12
219
L16
124
L8
O1

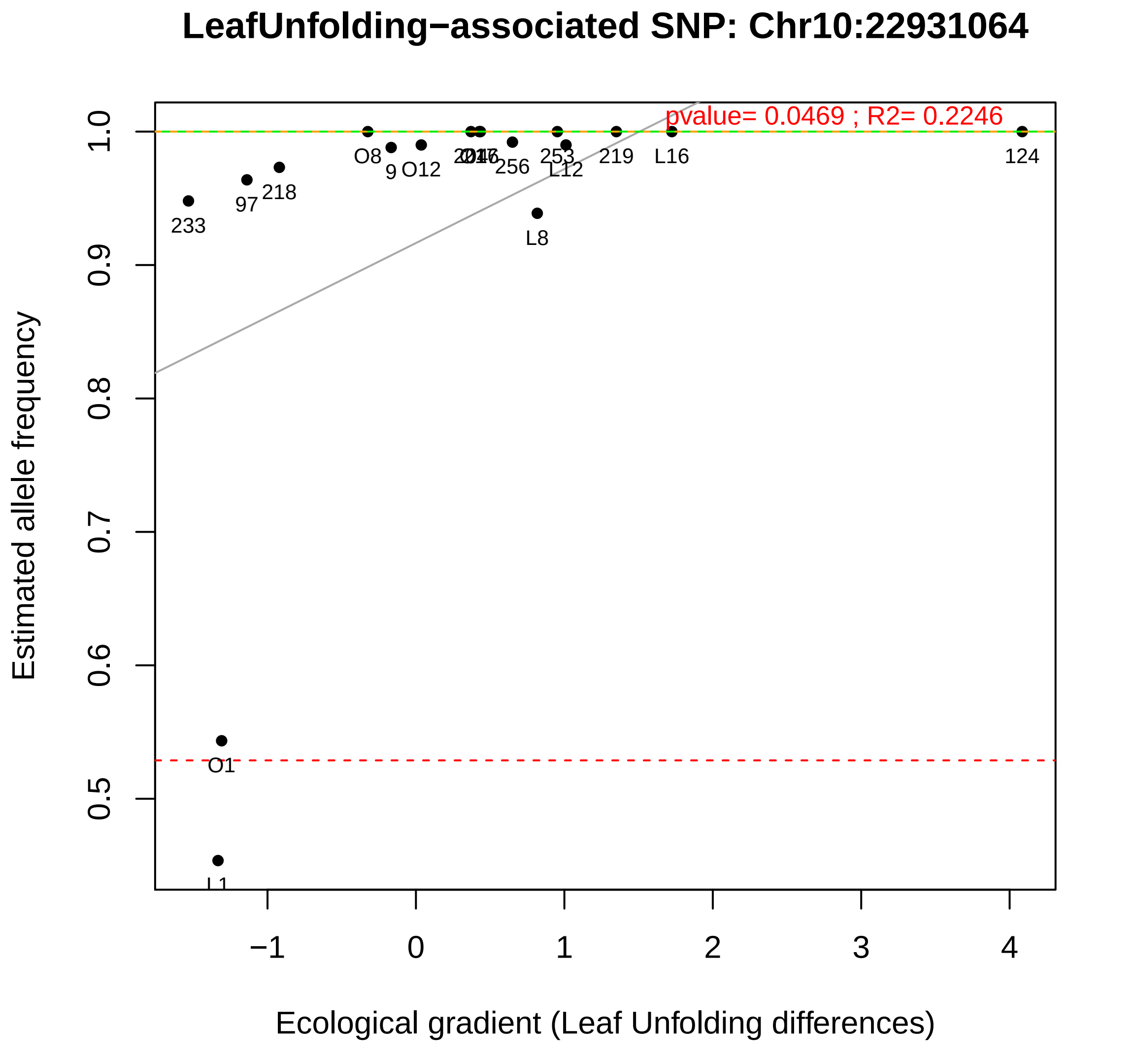

LeafUnfolding−associated SNP: Chr10:22931064
pvalue= 0.0469 ; R2= 0.2246
Estimated allele frequency
Ecological gradient (Leaf Unfolding differences)
233
97
218
O8
9
O12
256
253
L12
219
L16
124
L8
O1

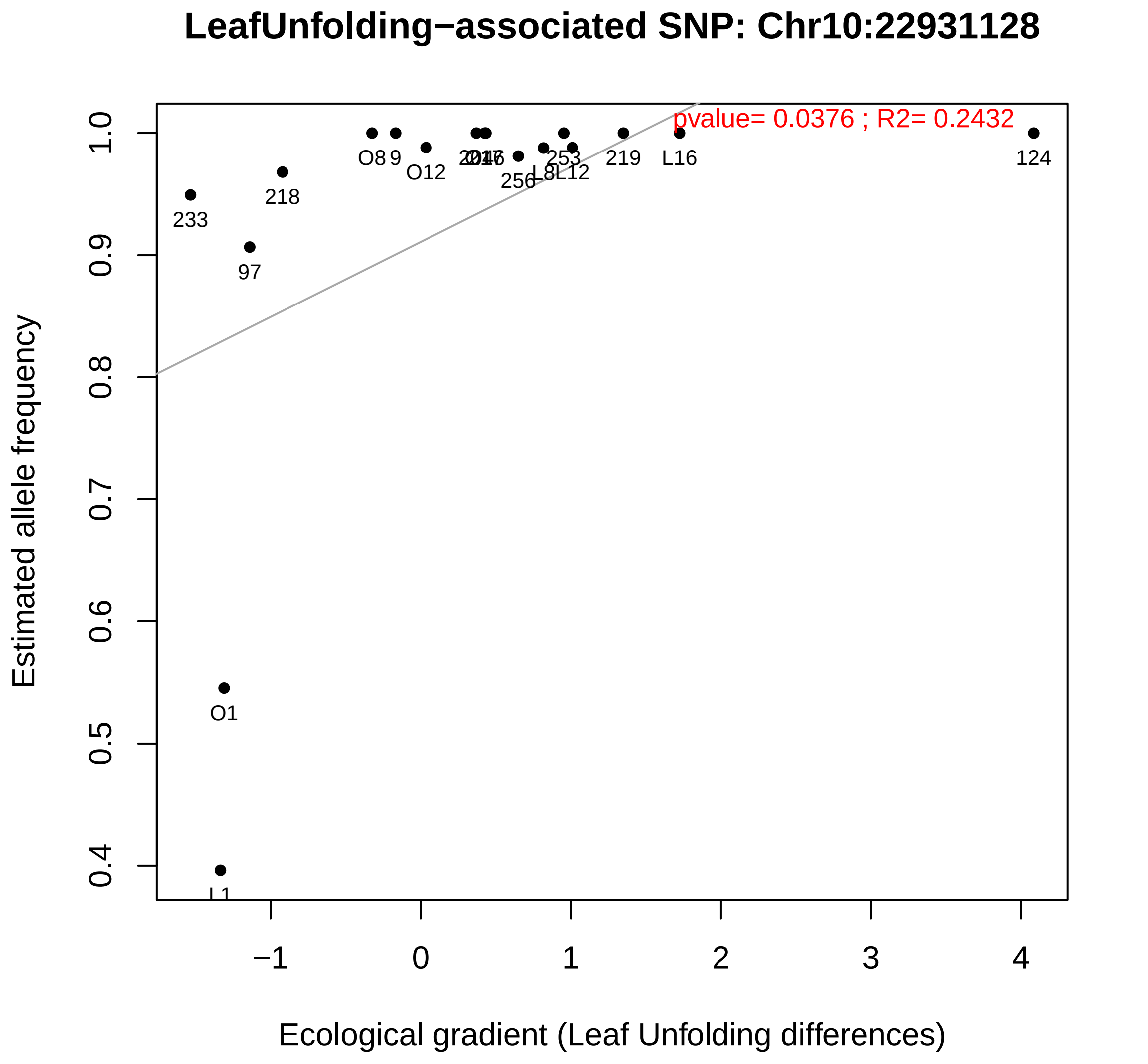

LeafUnfolding−associated SNP: Chr10:22931128
pvalue= 0.0376 ; R2= 0.2432
Estimated allele frequency
Ecological gradient (Leaf Unfolding differences)
1.0
0.9
0.8
0.7
0.6
0.5
0.4
−1
0
1
2
3
4
233
97
218
O8
9
O12
256
L8
253
L12
219
L16
124
O1

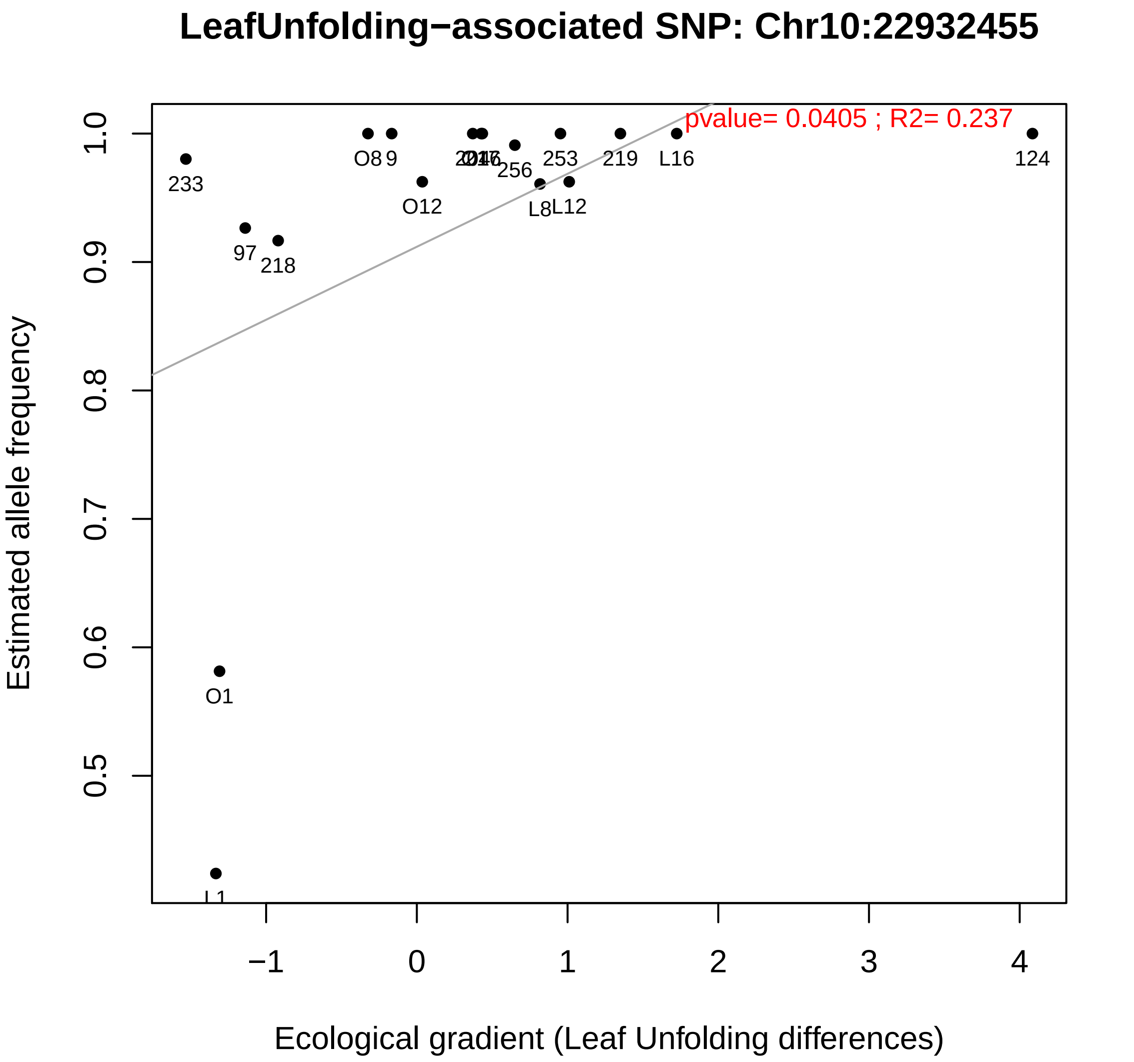
LeafUnfolding−associated SNP: Chr10:22932455
pvalue= 0.0405 ; R2= 0.237
Estimated allele frequency
Ecological gradient (Leaf Unfolding differences)
0.5
0.6
0.7
0.8
0.9
1.0
−1
0
1
2
3
4
233
97
218
O8
9
O12
256
253
L8
L12
219
L16
124
O1

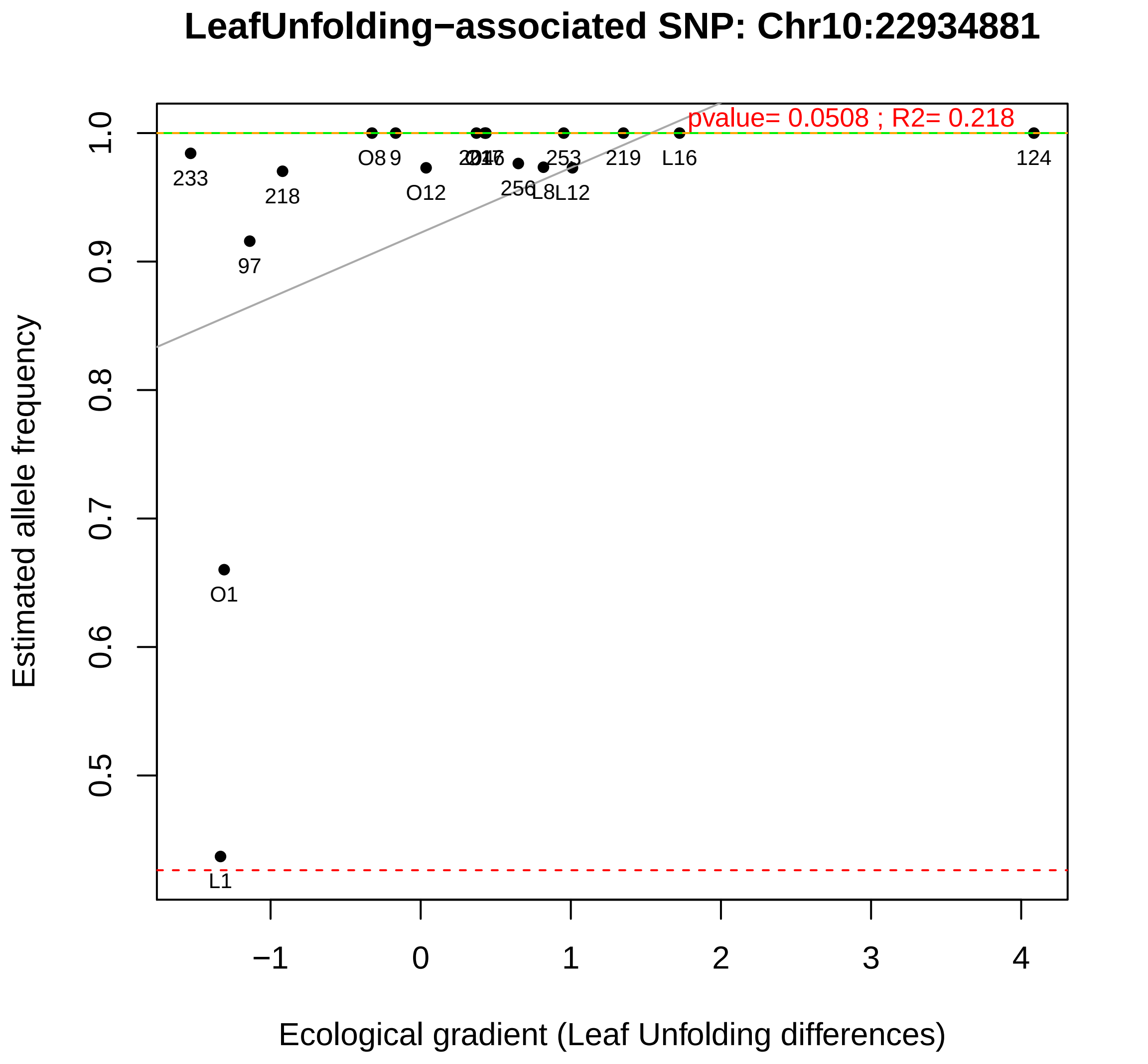

LeafUnfolding−associated SNP: Chr10:22934881
pvalue= 0.0508 ; R2= 0.218
Estimated allele frequency
Ecological gradient (Leaf Unfolding differences)
1.0
0.9
0.8
0.7
0.6
0.5
−1
0
1
2
3
4
233
218
97
O1
L1
O8
9
O12
256
L8
L12
253
219
L16
124

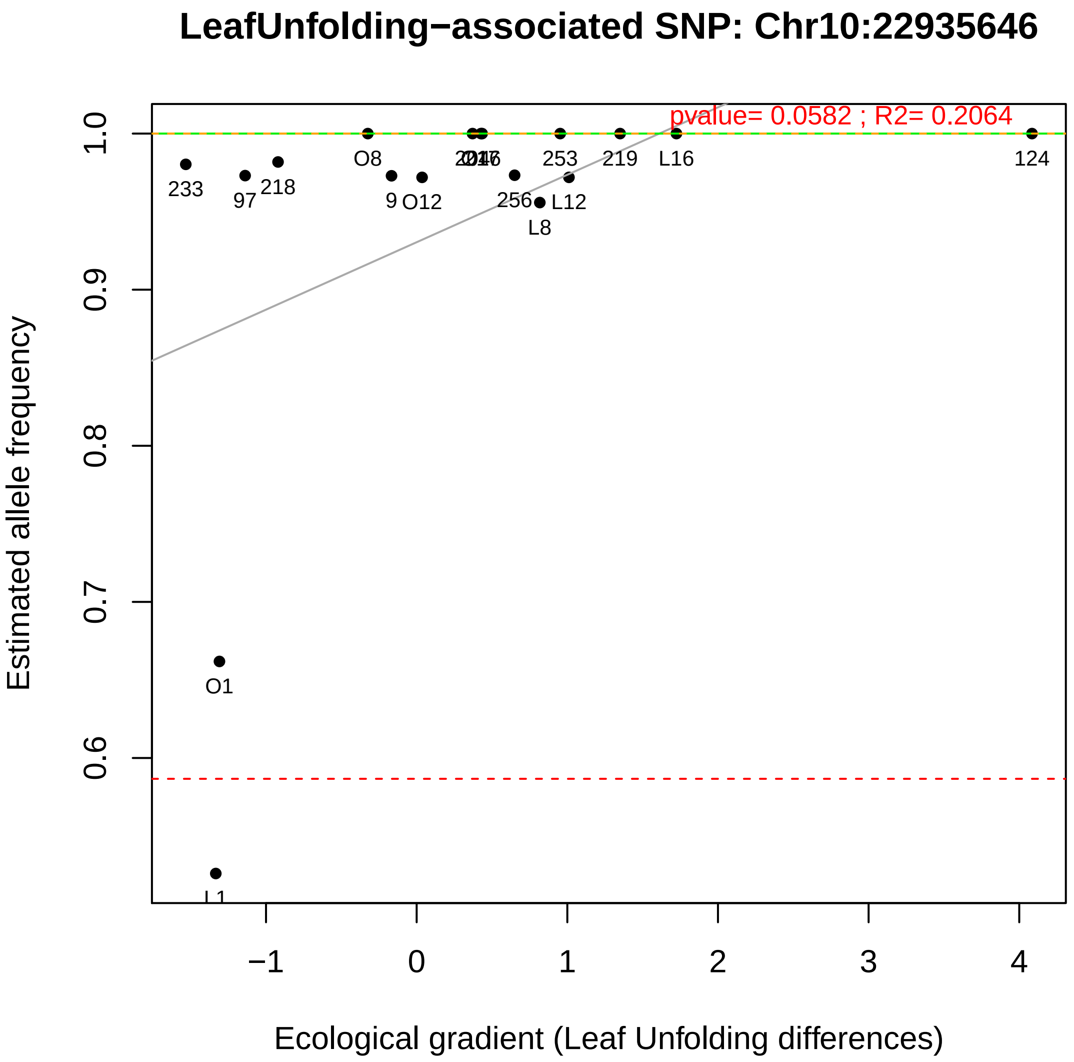

LeafUnfolding−associated SNP: Chr10:22935646
pvalue= 0.0582 ; R2= 0.2064
Estimated allele frequency
Ecological gradient (Leaf Unfolding differences)
1.0
0.9
0.8
0.7
0.6
−1
0
1
2
3
4
233
97
218
O8
9
O12
256
L8
L12
253
219
L16
124
O1
L1

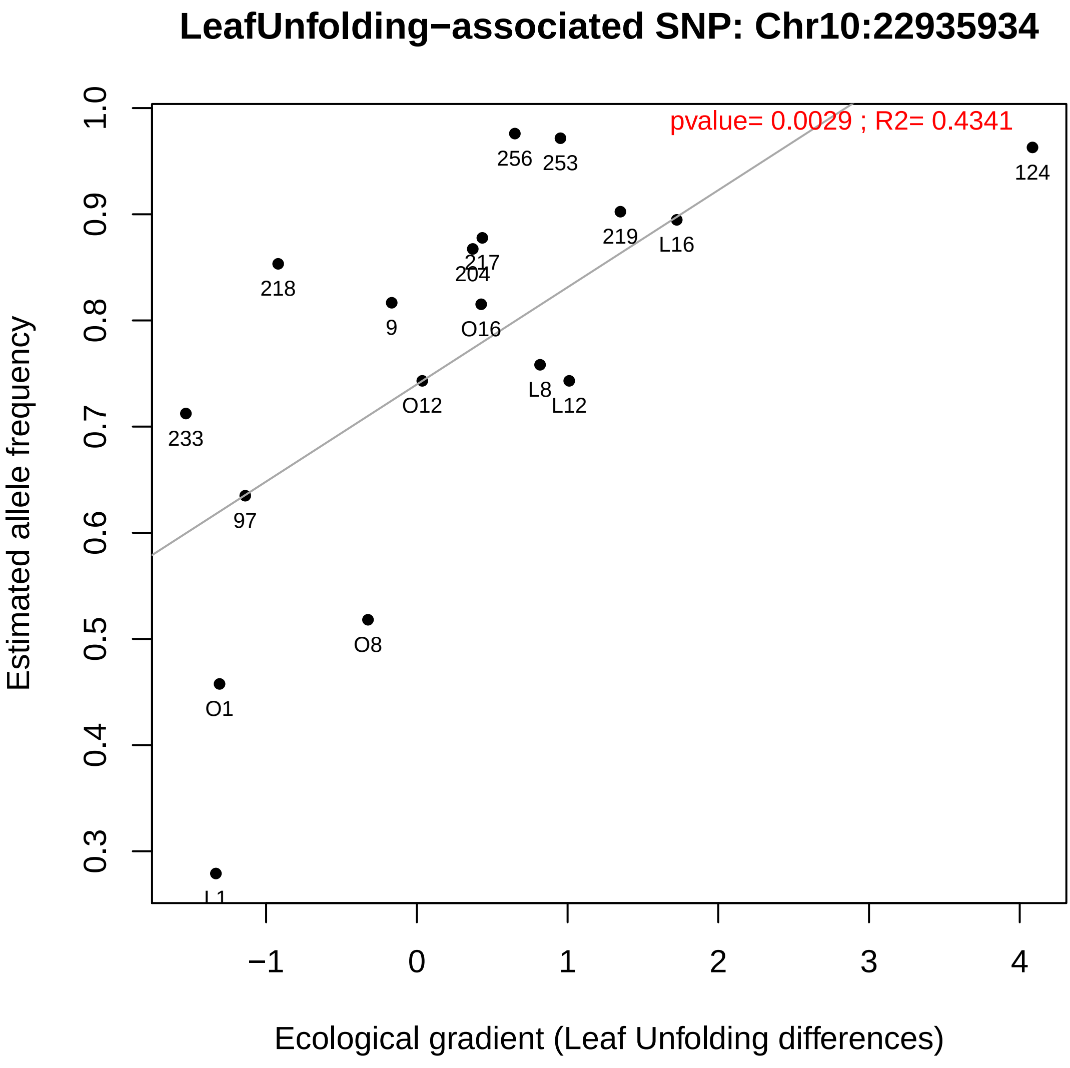
LeafUnfolding−associated SNP: Chr10:22935934
pvalue= 0.0029 ; R2= 0.4341
Estimated allele frequency
Ecological gradient (Leaf Unfolding differences)
0.3
0.4
0.5
0.6
0.7
0.8
0.9
1.0
−1
0
1
2
3
4
256
253
124
219
L16
217
204
218
9
O16
O12
L8
L12
233
97
O8
O1
L1

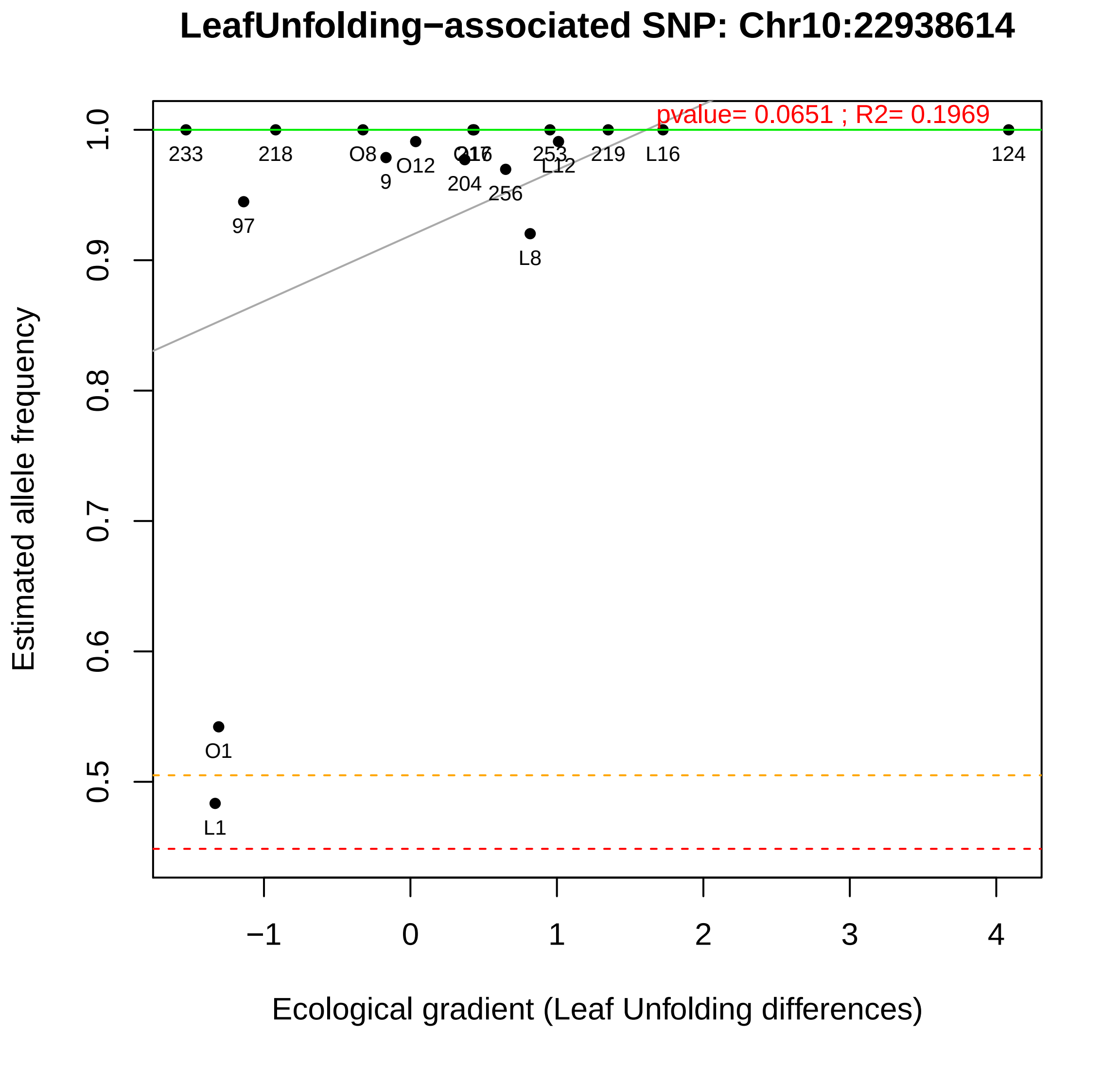

LeafUnfolding−associated SNP: Chr10:22938614
pvalue= 0.0651 ; R2= 0.1969
233
218
O8
9
O12
204
256
253
L12
219
L16
124
97
L8
O1
L1
1.0
0.9
0.8
0.7
0.6
0.5
−1
0
1
2
3
4
Estimated allele frequency
Ecological gradient (Leaf Unfolding differences)

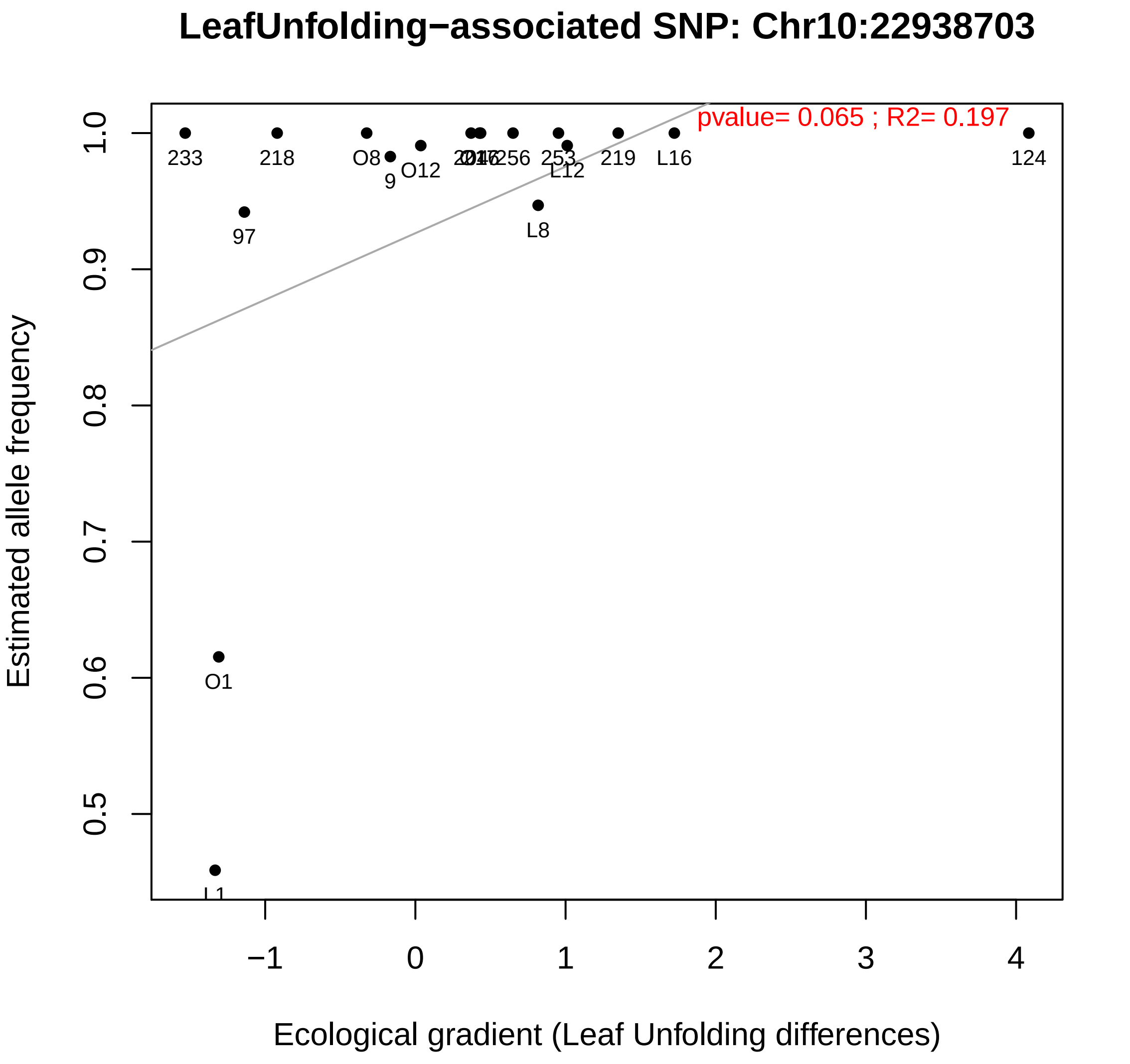

LeafUnfolding−associated SNP: Chr10:22938703
pvalue= 0.065 ; R2= 0.197
Estimated allele frequency
Ecological gradient (Leaf Unfolding differences)
1.0
0.9
0.8
0.7
0.6
0.5
−1
0
1
2
3
4
233
218
O8
9
O12
256
253
L12
219
L16
124
97
L8
O1

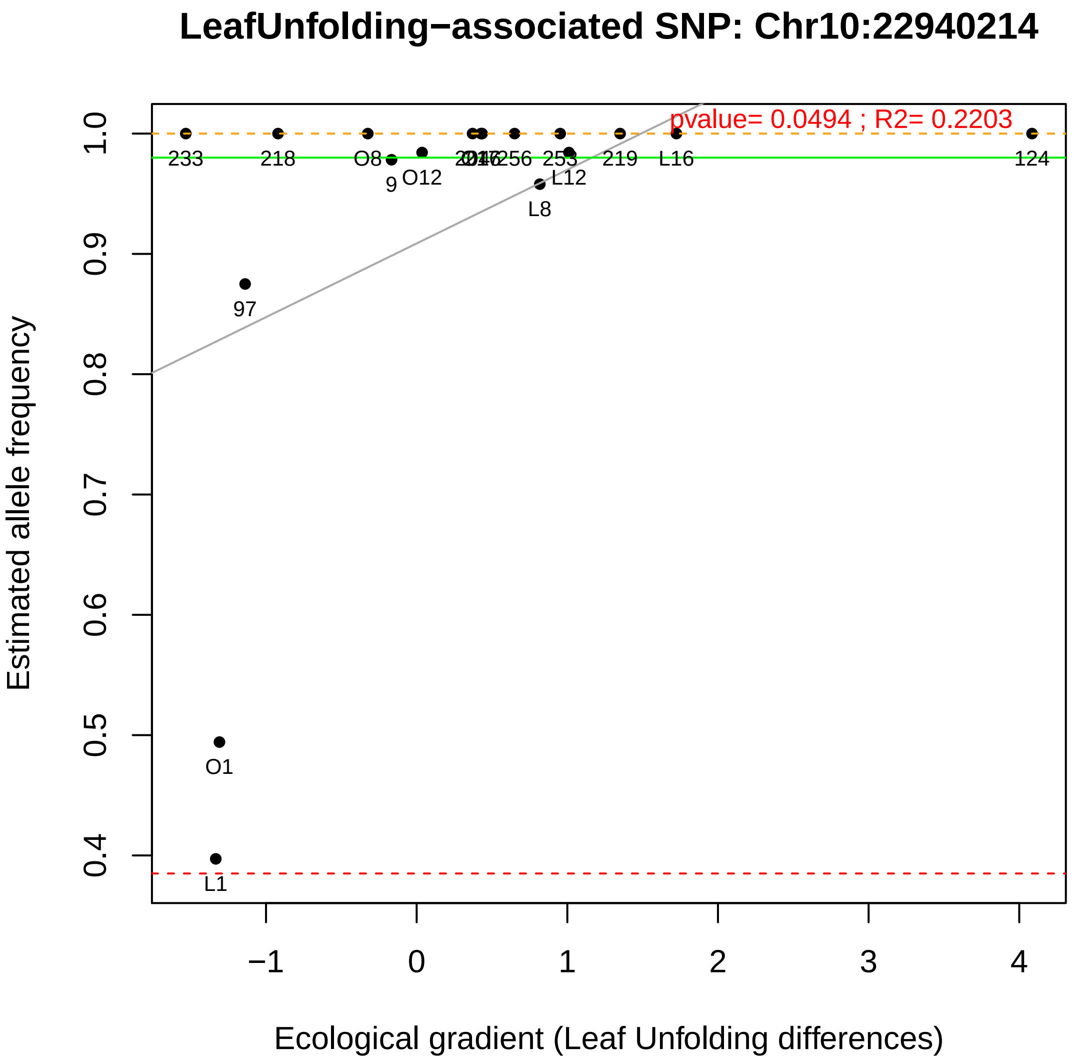
LeafUnfolding−associated SNP: Chr10:22940214
pvalue= 0.0494 ; R2= 0.2203
Estimated allele frequency
Ecological gradient (Leaf Unfolding differences)
0.4
0.5
0.6
0.7
0.8
0.9
1.0
−1
0
1
2
3
4
233
218
O8
9
O12
256
253
L12
L8
219
L16
124
97
O1
L1

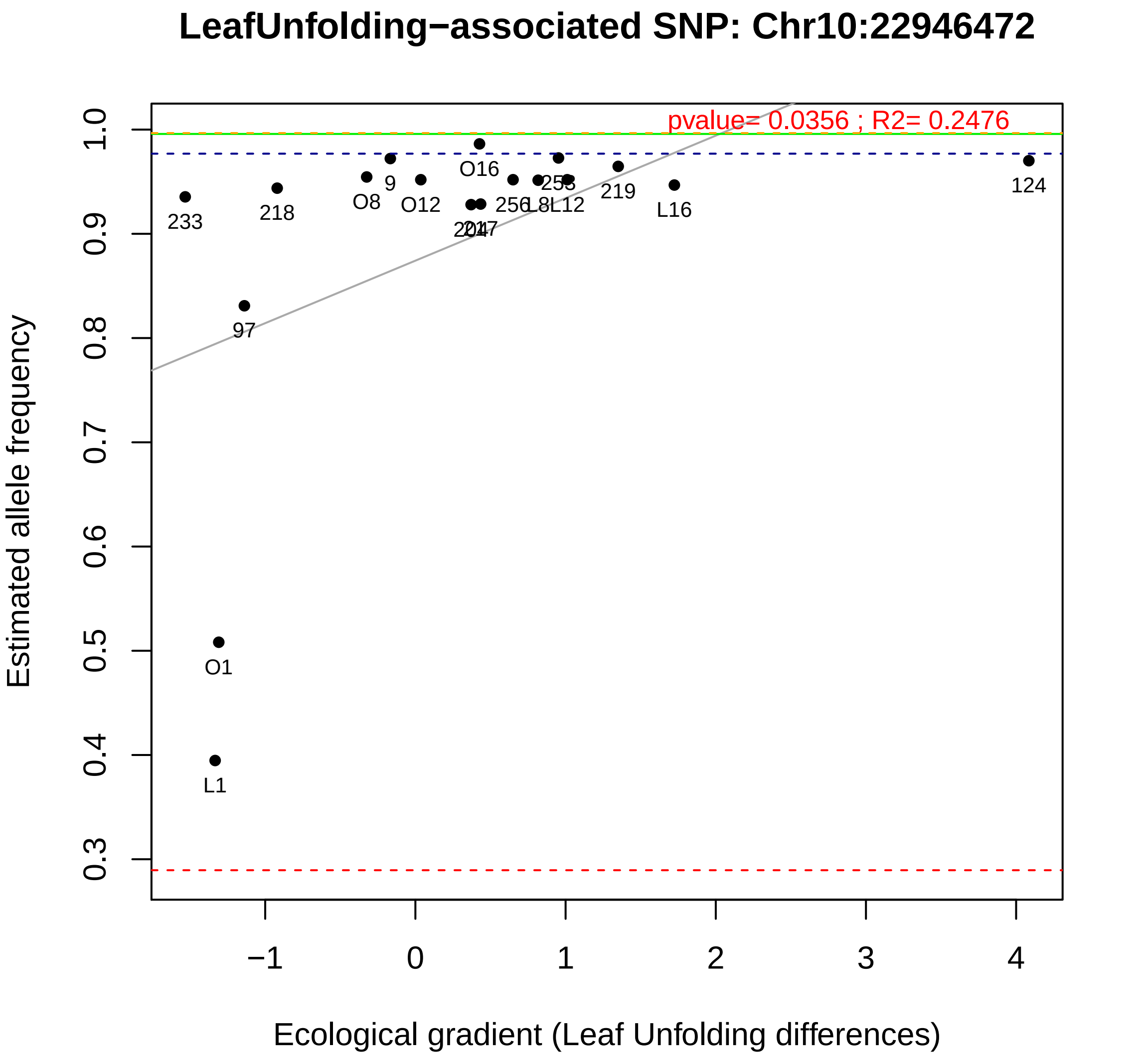
LeafUnfolding−associated SNP: Chr10:22946472
pvalue= 0.0356 ; R2= 0.2476
Estimated allele frequency
Ecological gradient (Leaf Unfolding differences)
233
218
97
O1
L1
O8
9
O12
O16
256
L8
253
L12
219
L16
124
−1
0
1
2
3
4
0.3
0.4
0.5
0.6
0.7
0.8
0.9
1.0

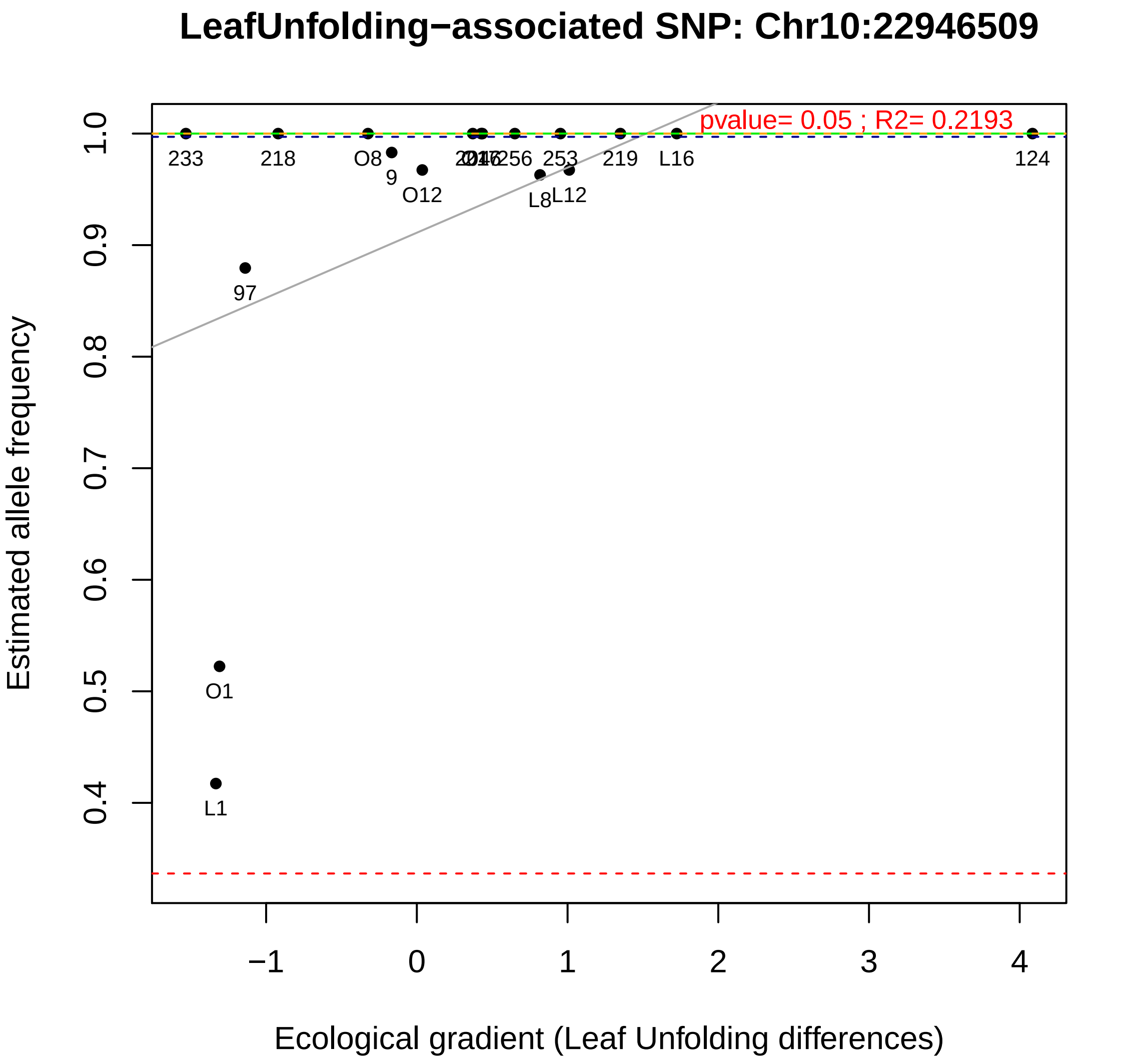

LeafUnfolding−associated SNP: Chr10:22946509
pvalue= 0.05 ; R2= 0.2193
Estimated allele frequency
Ecological gradient (Leaf Unfolding differences)
233
218
O8
9
O12
256
253
219
L16
124
L8
L12
97
O1
L1

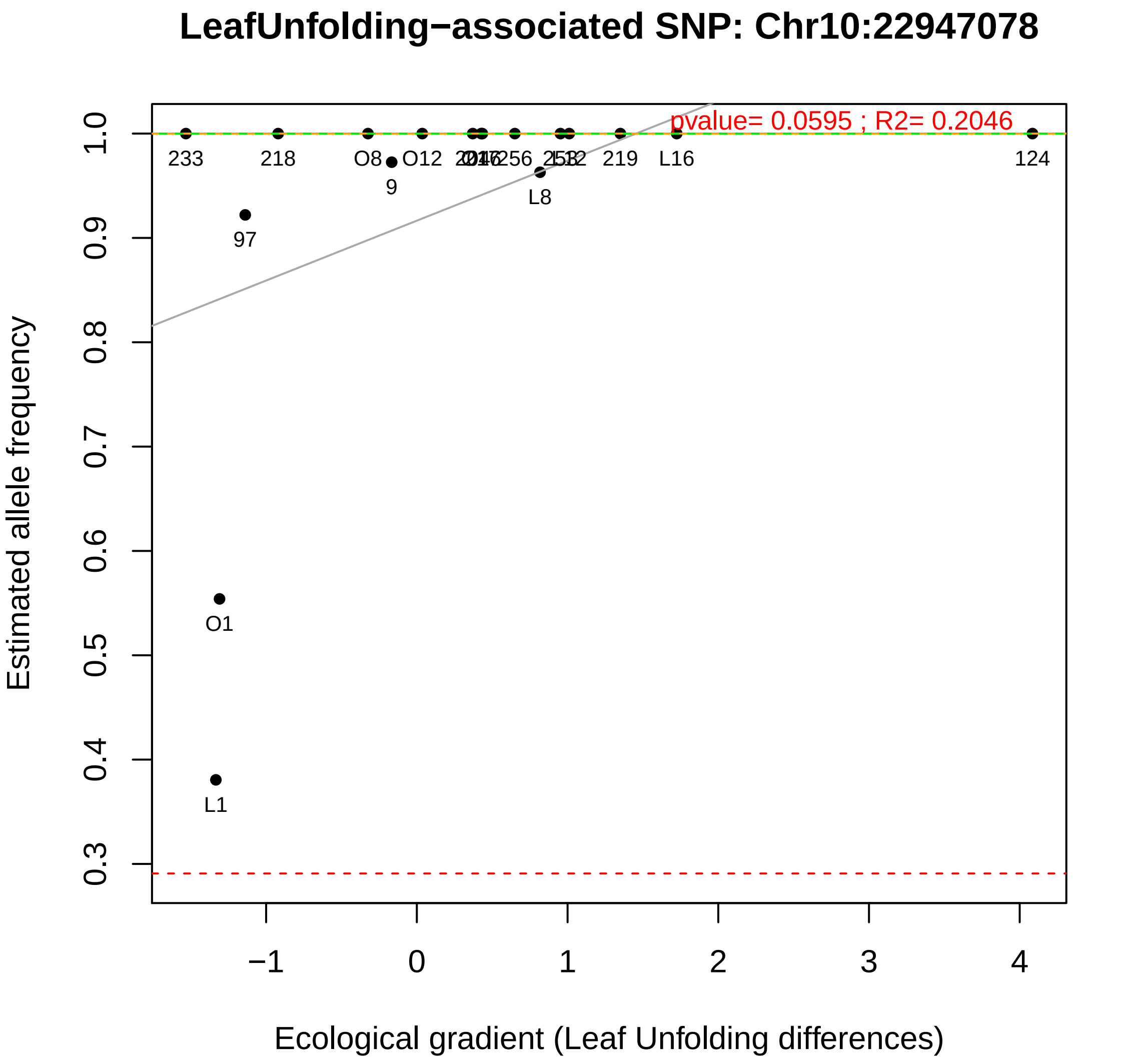

LeafUnfolding−associated SNP: Chr10:22947078
pvalue= 0.0595 ; R2= 0.2046
Estimated allele frequency
Ecological gradient (Leaf Unfolding differences)
233
218
O8
O12
219
L16
124
9
L8
97
O1
L1
1.0
0.9
0.8
0.7
0.6
0.5
0.4
0.3
−1
0
1
2
3
4

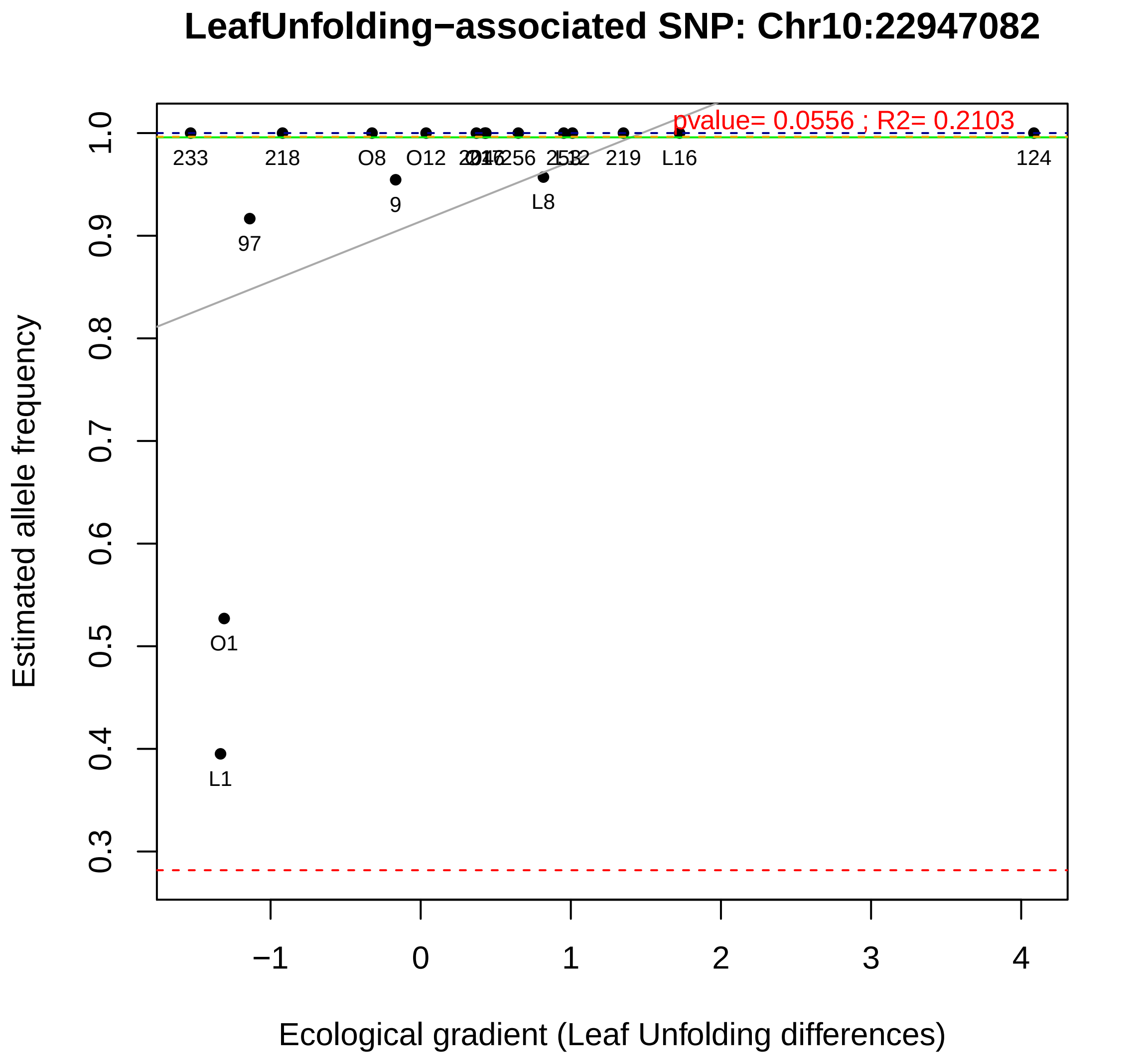
LeafUnfolding−associated SNP: Chr10:22947082
pvalue= 0.0556 ; R2= 0.2103
Estimated allele frequency
Ecological gradient (Leaf Unfolding differences)
233
218
O8
O12
219
L16
124
9
L8
97
O1
L1
1.0
0.9
0.8
0.7
0.6
0.5
0.4
0.3
−1
0
1
2
3
4

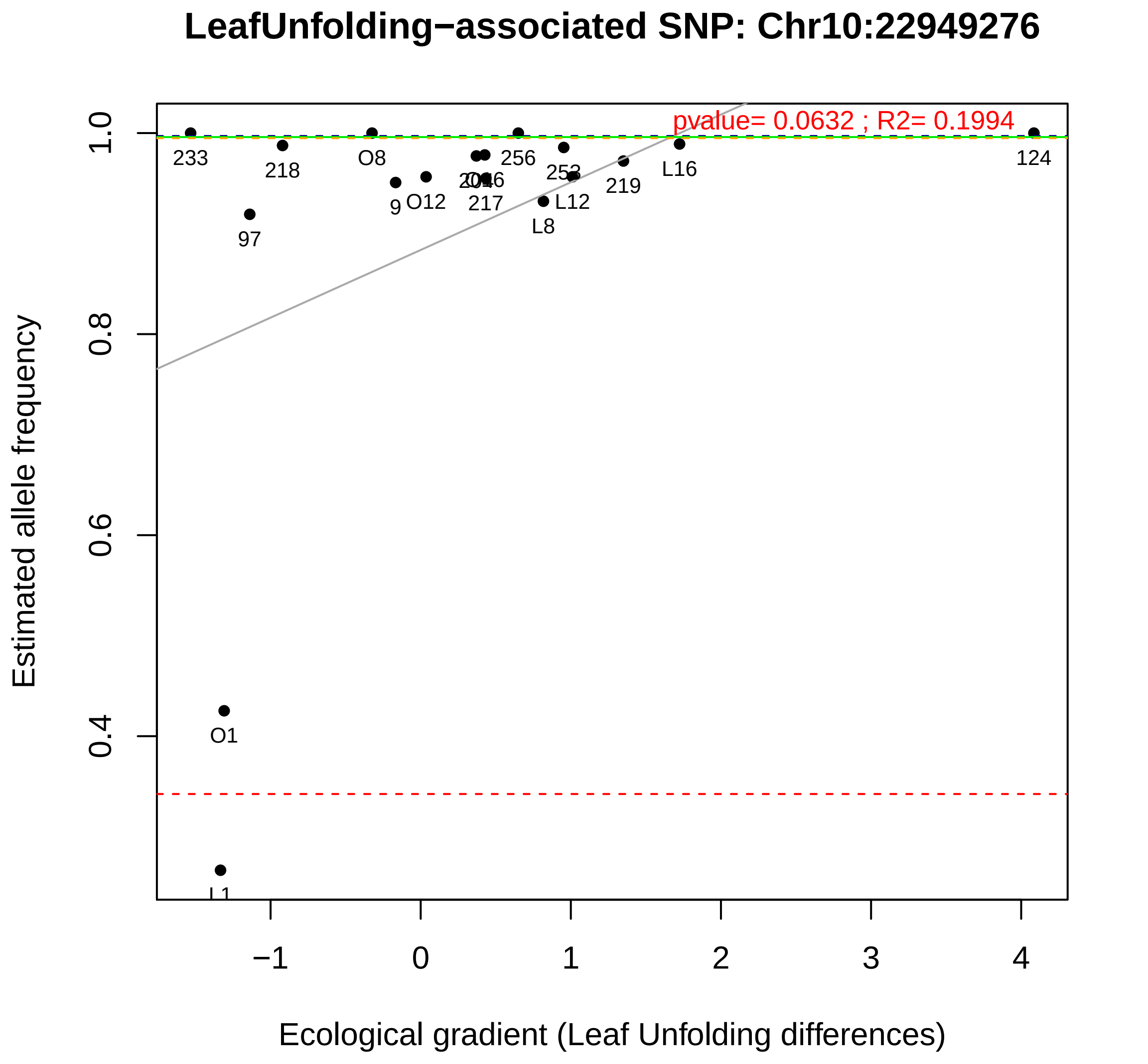

LeafUnfolding−associated SNP: Chr10:22949276
pvalue= 0.0632 ; R2= 0.1994
Estimated allele frequency
Ecological gradient (Leaf Unfolding differences)
1.0
0.8
0.6
0.4
−1
0
1
2
3
4
233
218
97
O8
9
O12
217
256
253
L12
L8
219
L16
124
O1
L1

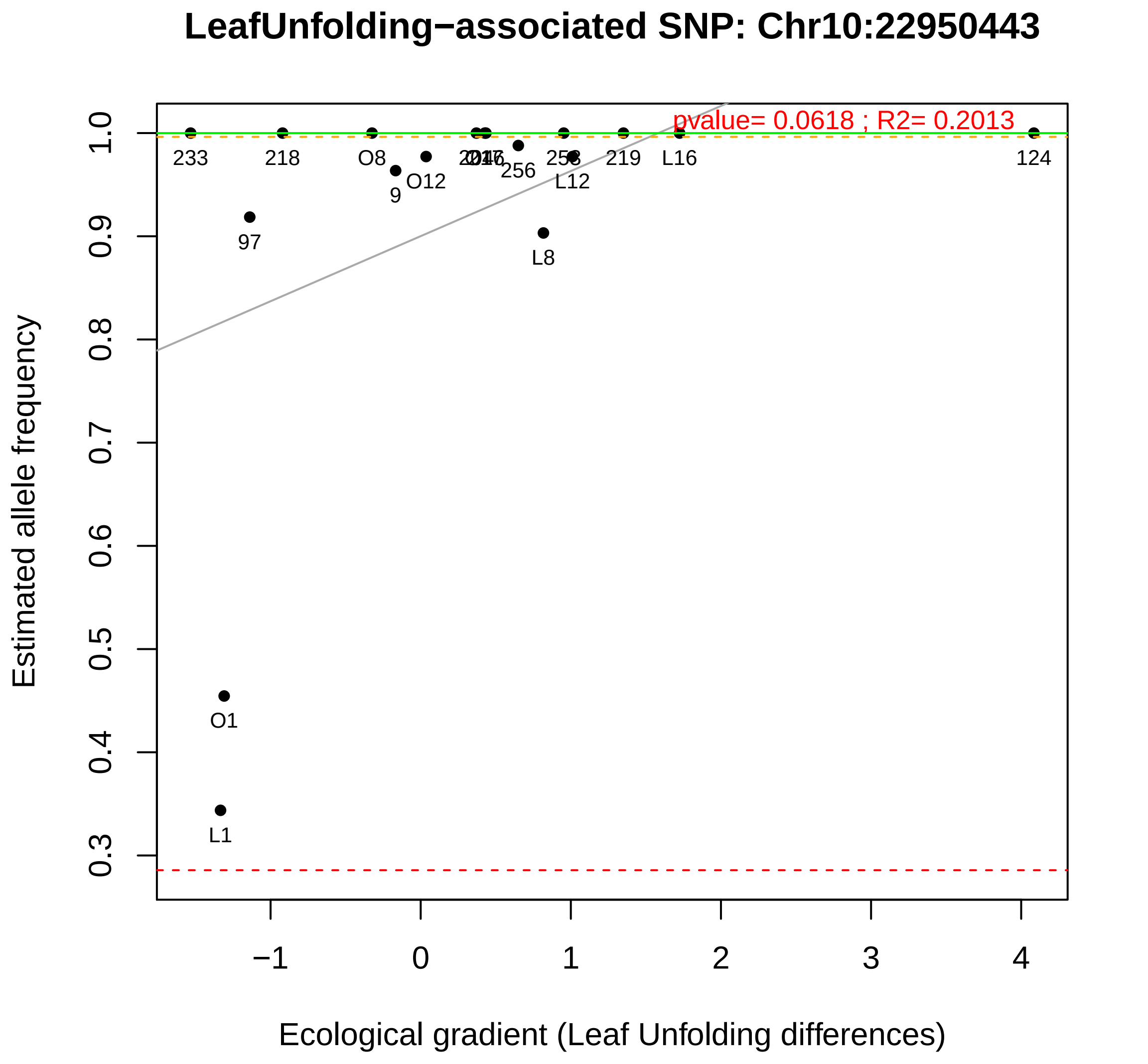

LeafUnfolding−associated SNP: Chr10:22950443
pvalue= 0.0618 ; R2= 0.2013
Estimated allele frequency
Ecological gradient (Leaf Unfolding differences)
1.0
0.9
0.8
0.7
0.6
0.5
0.4
0.3
−1
0
1
2
3
4
233
218
O8
9
O12
256
L12
219
L16
124
97
L8
O1
L1

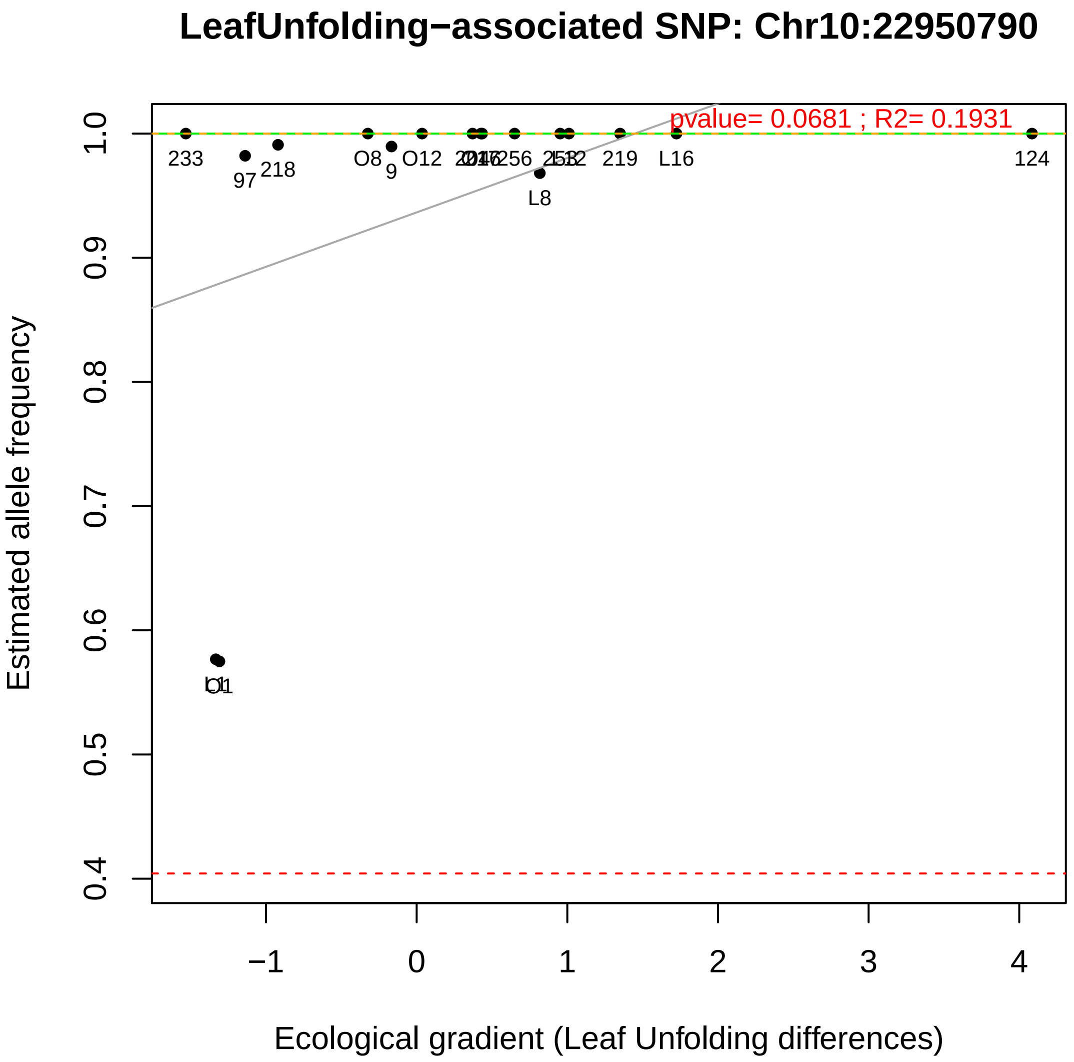
LeafUnfolding−associated SNP: Chr10:22950790
pvalue= 0.0681 ; R2= 0.1931
Estimated allele frequency
Ecological gradient (Leaf Unfolding differences)
1.0
0.9
0.8
0.7
0.6
0.5
0.4
−1
0
1
2
3
4
233
97
218
O8
9
O12
256
L8
219
L16
124

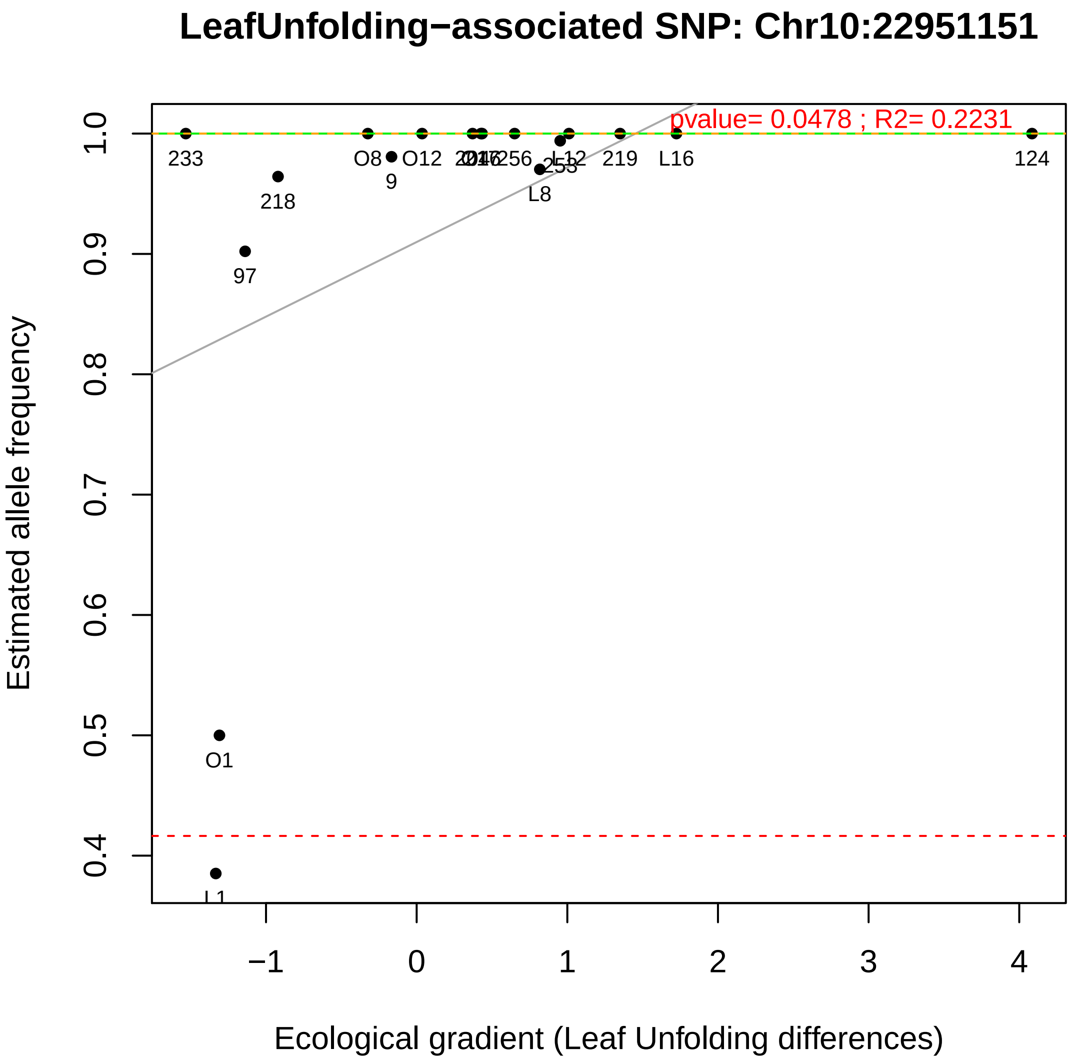

LeafUnfolding−associated SNP: Chr10:22951151
pvalue= 0.0478 ; R2= 0.2231
233
O8
9
O12
256
L12
253
L8
219
L16
124
218
97
O1
L1
1.0
0.9
0.8
0.7
0.6
0.5
0.4
−1
0
1
2
3
4
Estimated allele frequency
Ecological gradient (Leaf Unfolding differences)

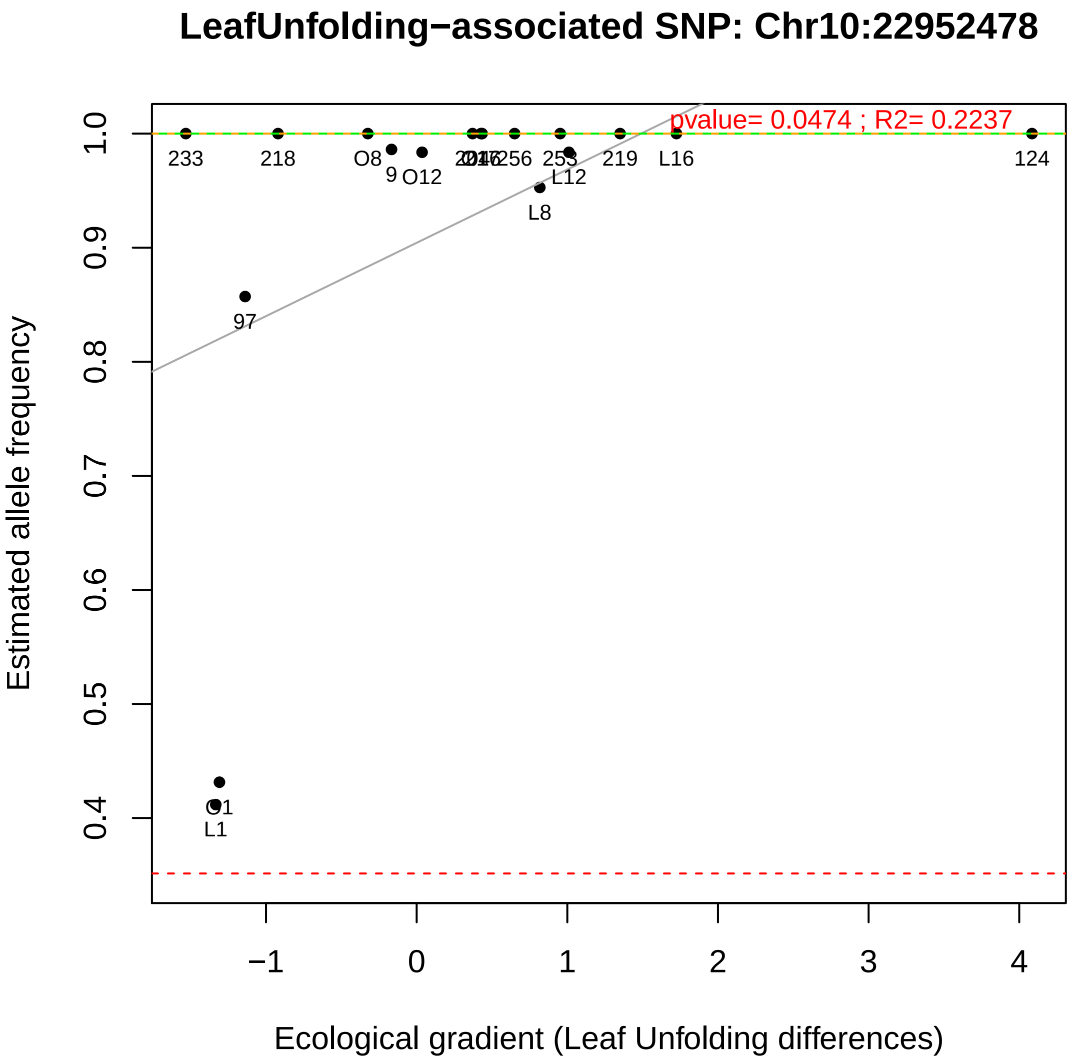

LeafUnfolding−associated SNP: Chr10:22952478
pvalue= 0.0474 ; R2= 0.2237
Estimated allele frequency
Ecological gradient (Leaf Unfolding differences)
0.4
0.5
0.6
0.7
0.8
0.9
1.0
−1
0
1
2
3
4
233
218
O8
9
O12
256
253
L12
L8
219
L16
124
97
L1

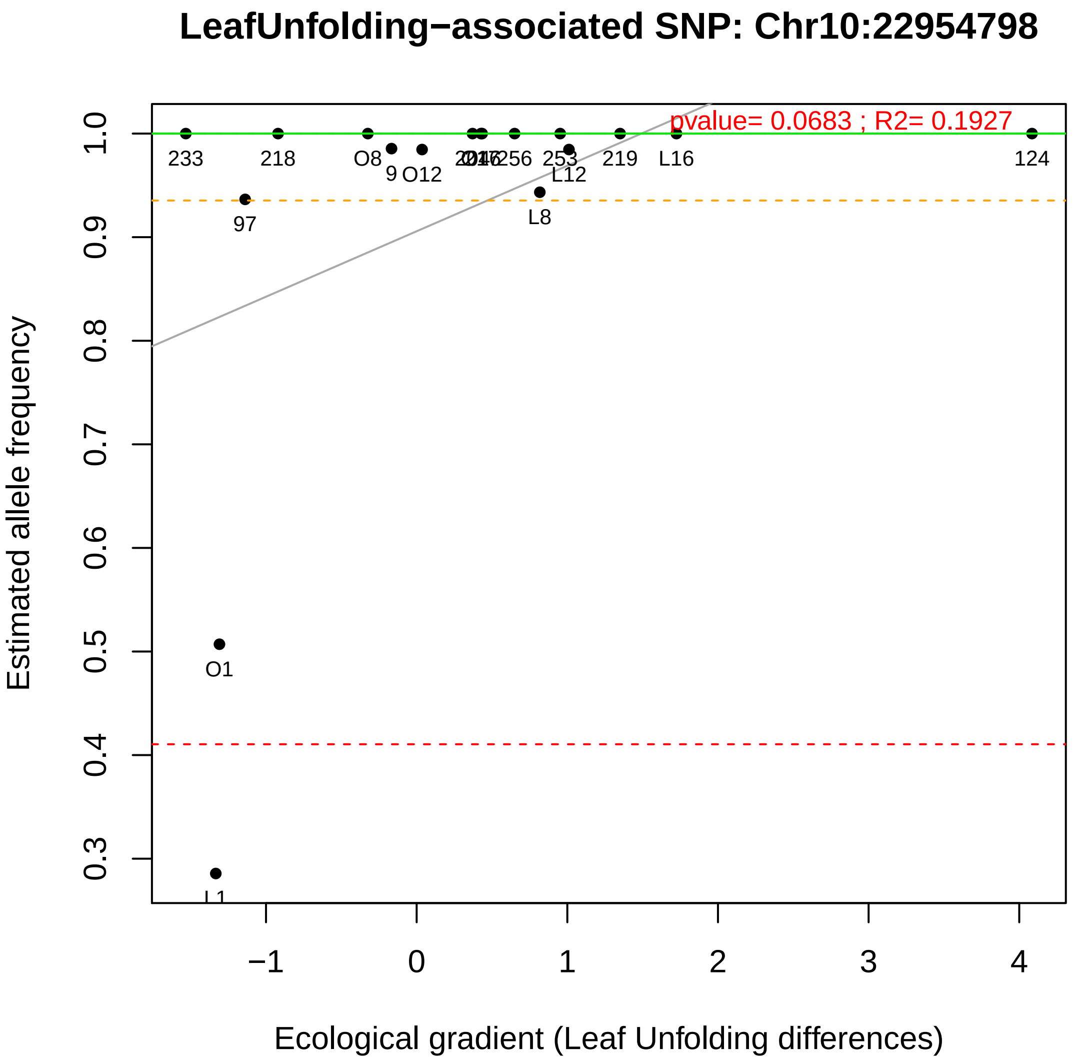
LeafUnfolding−associated SNP: Chr10:22954798
pvalue= 0.0683 ; R2= 0.1927
Estimated allele frequency
Ecological gradient (Leaf Unfolding differences)
0.3
0.4
0.5
0.6
0.7
0.8
0.9
1.0
−1
0
1
2
3
4
233
218
O8
9
O12
256
253
L12
219
L16
124
97
L8
O1

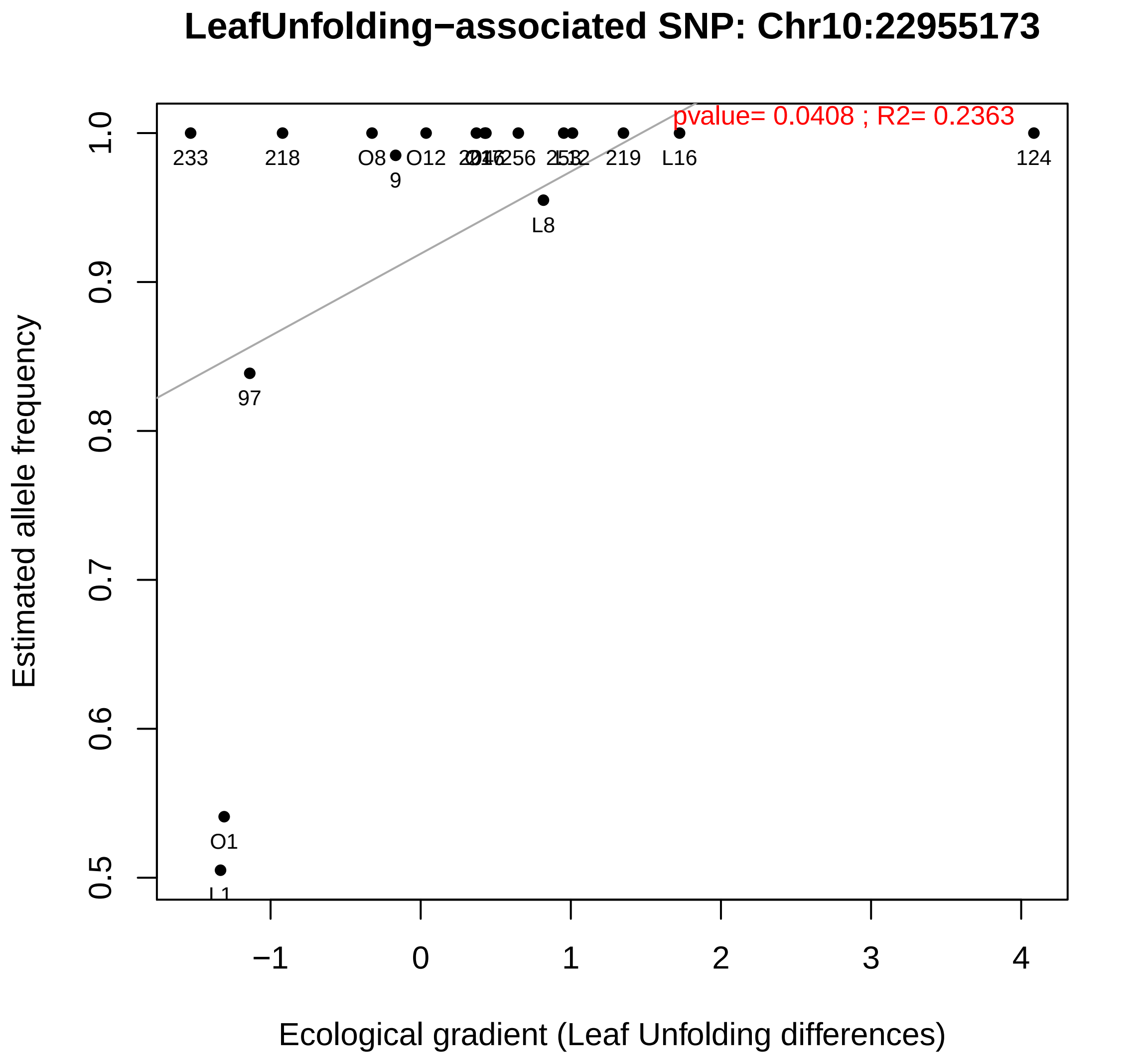

LeafUnfolding−associated SNP: Chr10:22955173
pvalue= 0.0408 ; R2= 0.2363
233
218
O8
9
O12
L16
219
124
L8
97
O1
L1
1.0
0.9
0.8
0.7
0.6
0.5
−1
0
1
2
3
4
Estimated allele frequency
Ecological gradient (Leaf Unfolding differences)

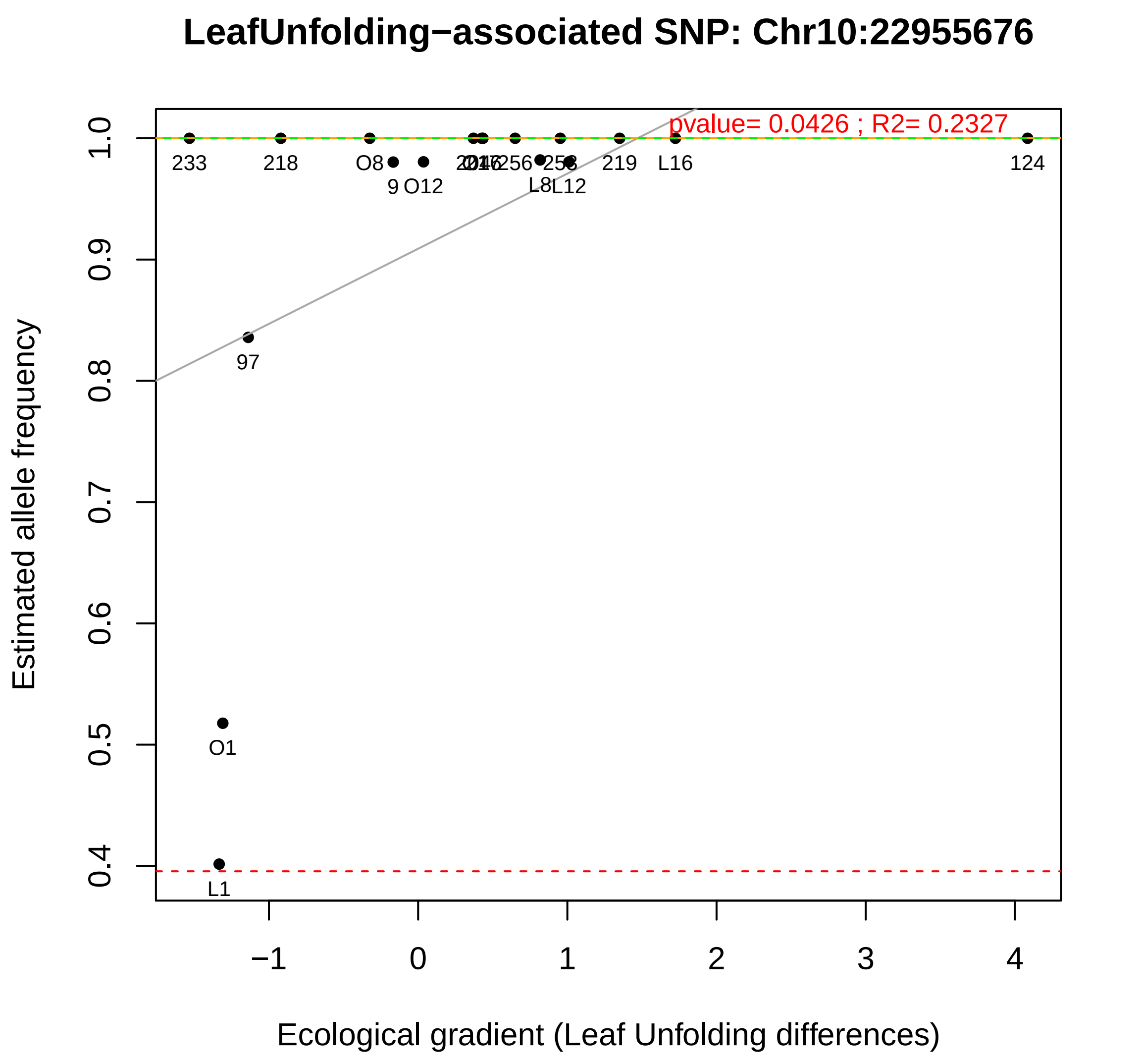
LeafUnfolding−associated SNP: Chr10:22955676
pvalue= 0.0426 ; R2= 0.2327
Estimated allele frequency
Ecological gradient (Leaf Unfolding differences)
1.0
0.9
0.8
0.7
0.6
0.5
0.4
−1
0
1
2
3
4
233
218
O8
9
O12
256
L8
L12
219
L16
124
97
O1
L1

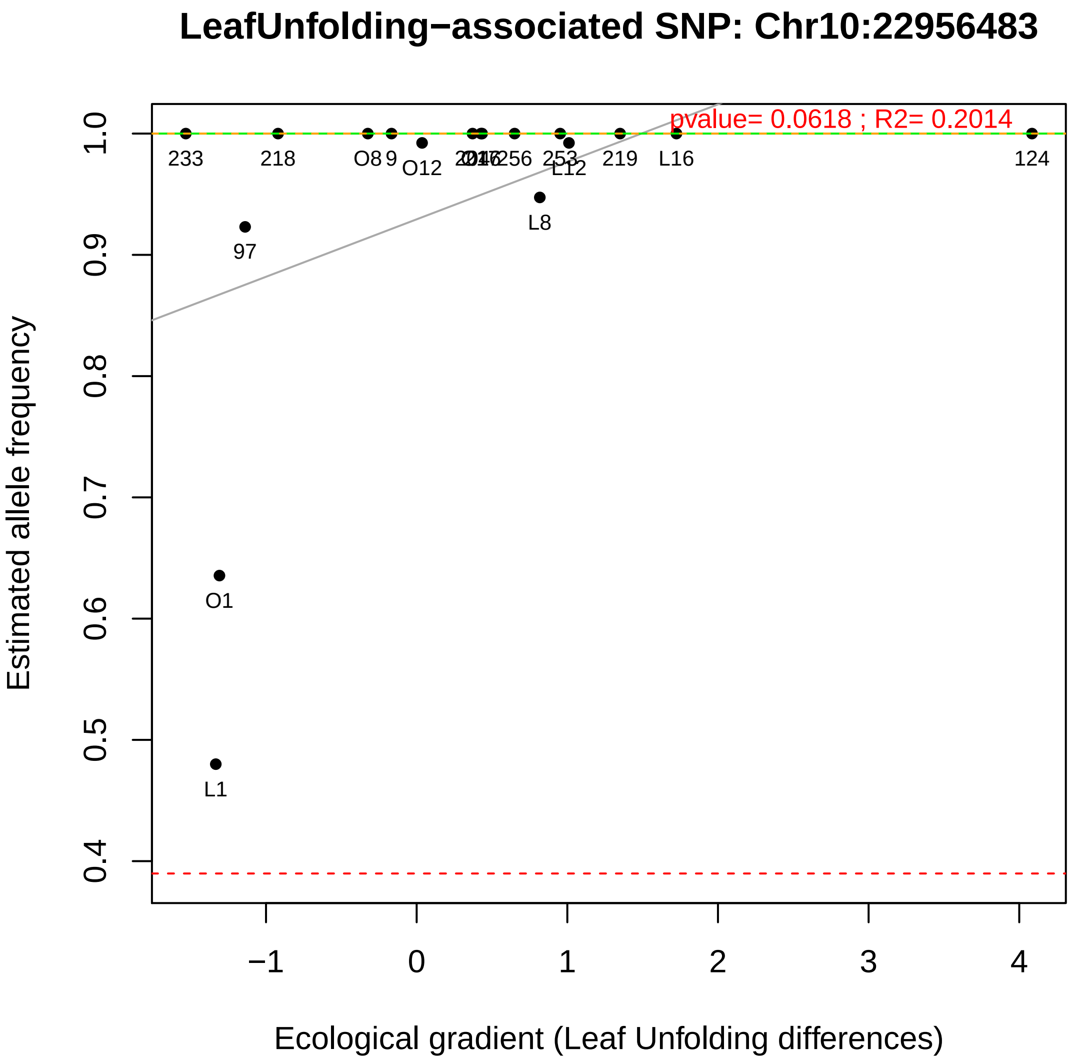

LeafUnfolding−associated SNP: Chr10:22956483
pvalue= 0.0618 ; R2= 0.2014
Estimated allele frequency
Ecological gradient (Leaf Unfolding differences)
0.4
0.5
0.6
0.7
0.8
0.9
1.0
−1
0
1
2
3
4
233
218
O8
9
O12
256
253
L12
219
L16
124
L8
97
O1
L1

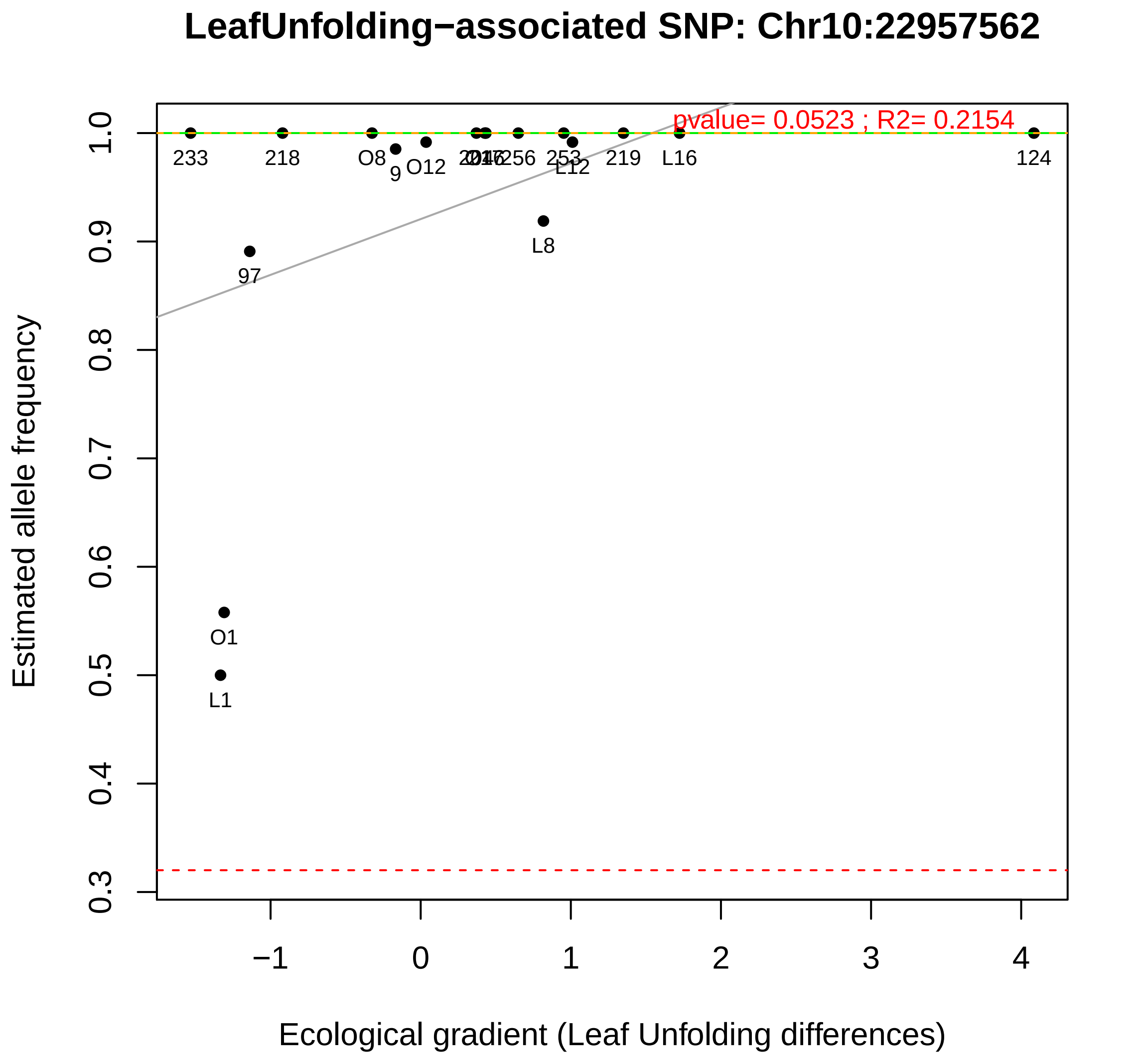
LeafUnfolding−associated SNP: Chr10:22957562
pvalue= 0.0523 ; R2= 0.2154
233
218
O8
9
O12
256
253
L12
219
L16
124
97
L8
O1
L1
Estimated allele frequency
1.0
0.9
0.8
0.7
0.6
0.5
0.4
0.3
−1
0
1
2
3
4
Ecological gradient (Leaf Unfolding differences)

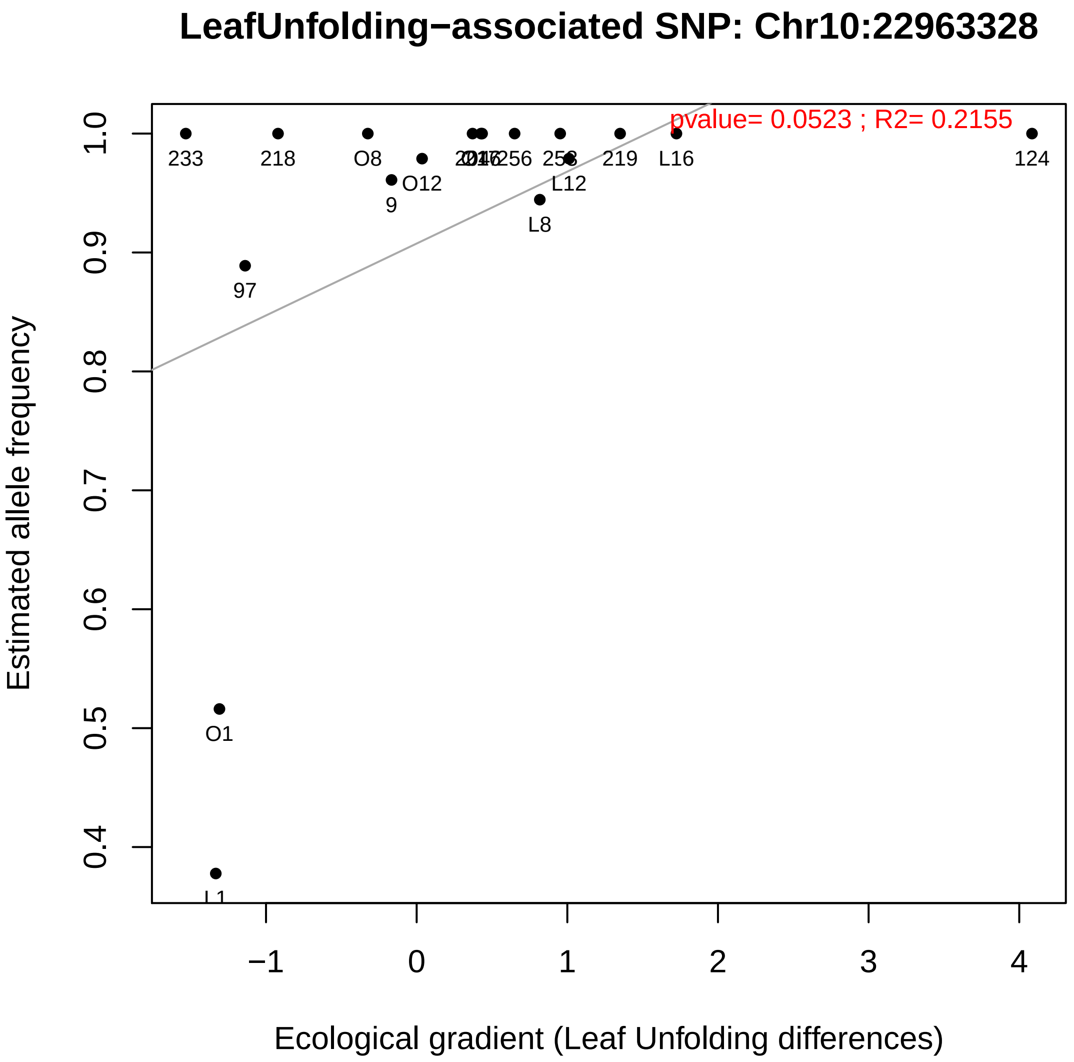

LeafUnfolding−associated SNP: Chr10:22963328
pvalue= 0.0523 ; R2= 0.2155
Estimated allele frequency
Ecological gradient (Leaf Unfolding differences)
233
218
O8
O12
9
97
O1
L8
L12
219
L16
124

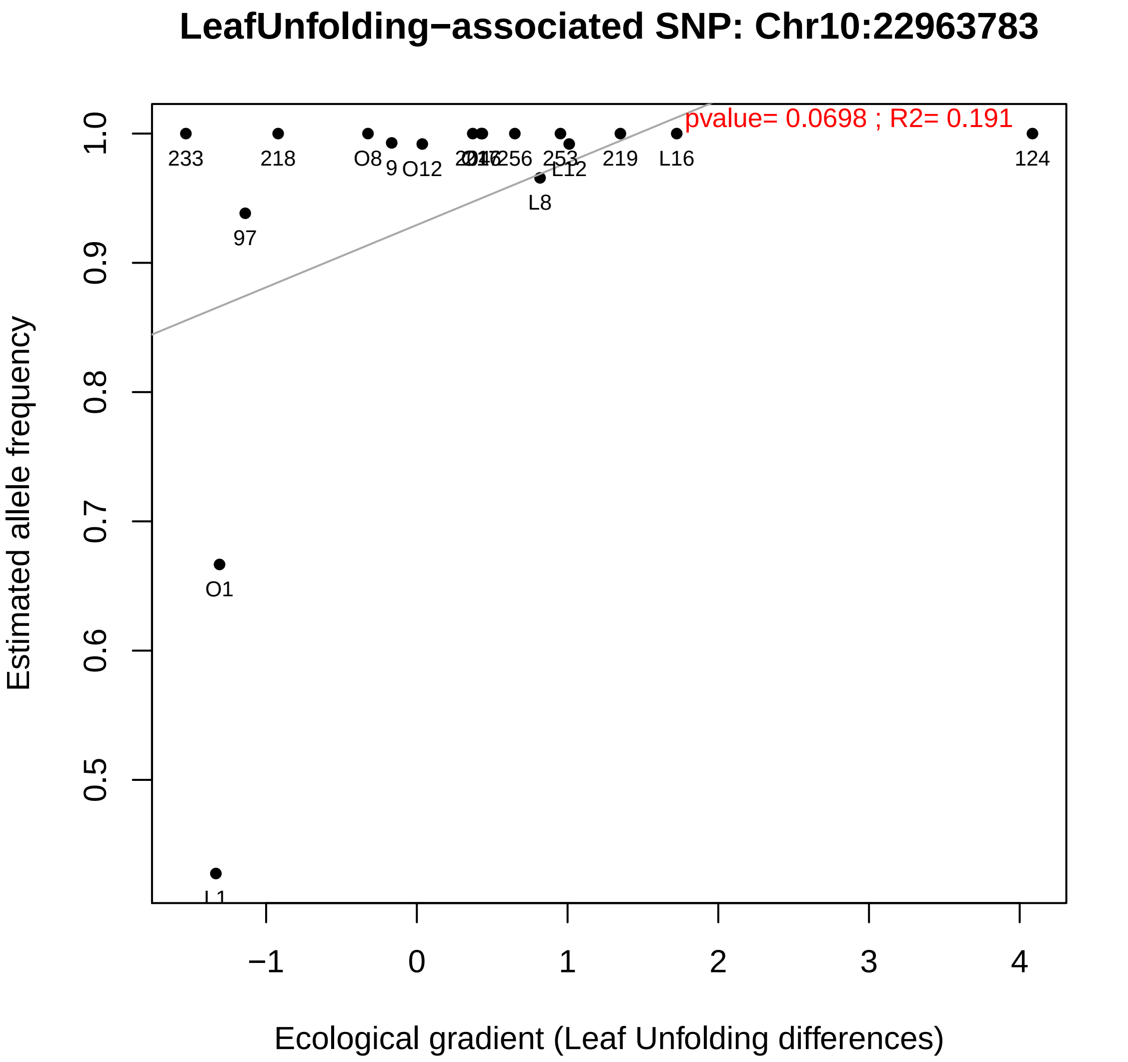
LeafUnfolding−associated SNP: Chr10:22963783
pvalue= 0.0698 ; R2= 0.191
Estimated allele frequency
Ecological gradient (Leaf Unfolding differences)
233
218
O8
9
O12
256
253
L12
219
L16
124
L8
97
O1
1.0
0.9
0.8
0.7
0.6
0.5
−1
0
1
2
3
4

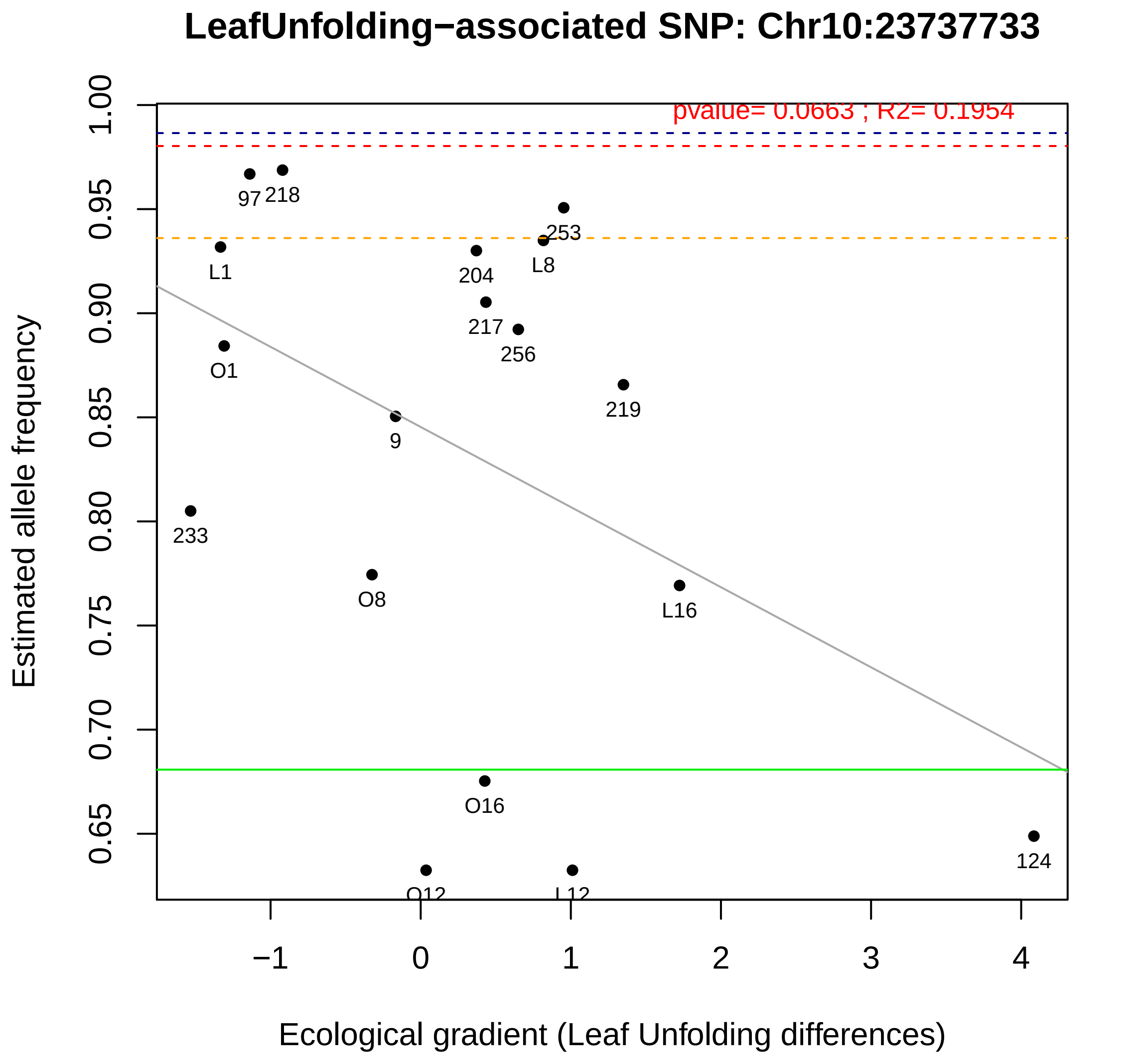

LeafUnfolding−associated SNP: Chr10:23737733
pvalue= 0.0663 ; R2= 0.1954
Estimated allele frequency
Ecological gradient (Leaf Unfolding differences)
1.00
0.95
0.90
0.85
0.80
0.75
0.70
0.65
−1
0
1
2
3
4
97
218
L1
204
253
L8
217
256
O1
219
9
233
O8
L16
O16
124
O12
L12

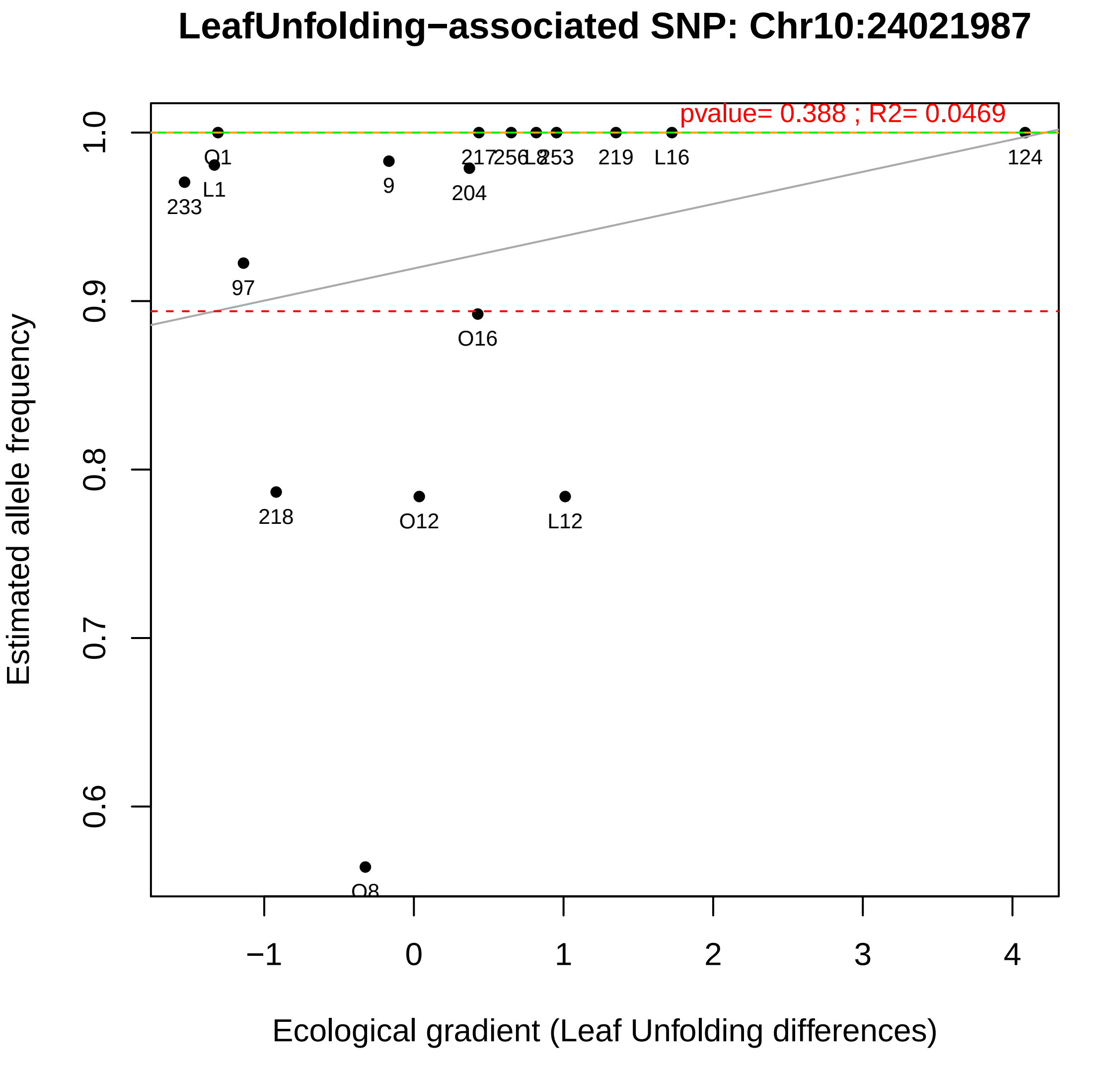

LeafUnfolding−associated SNP: Chr10:24021987
pvalue= 0.388 ; R2= 0.0469
Estimated allele frequency
Ecological gradient (Leaf Unfolding differences)
1.0
0.9
0.8
0.7
0.6
−1
0
1
2
3
4
O1
L1
233
97
9
204
217
256
253
219
L16
124
O16
218
O12
L12
O8

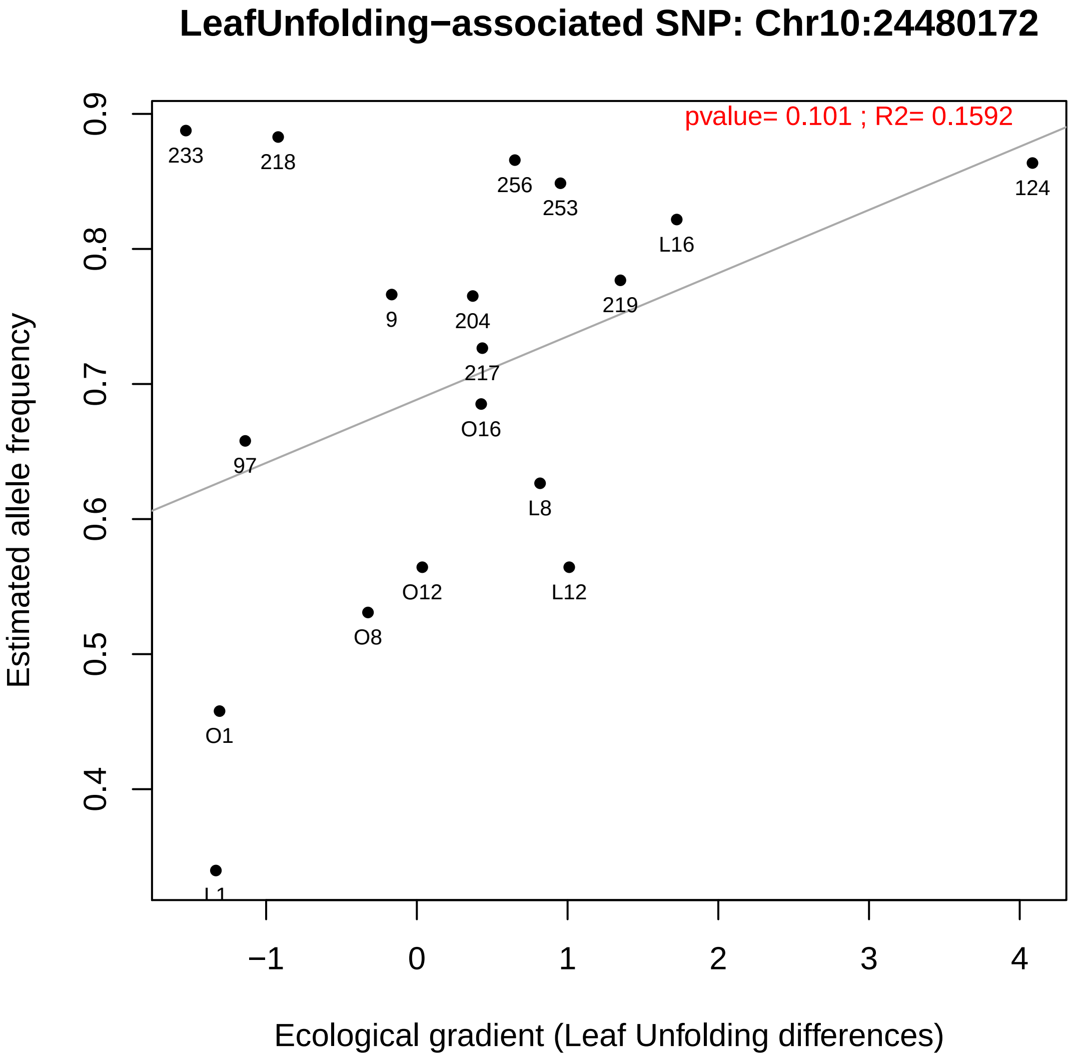
LeafUnfolding−associated SNP: Chr10:24480172
pvalue= 0.101 ; R2= 0.1592
Estimated allele frequency
0.4
0.5
0.6
0.7
0.8
0.9
−1
0
1
2
3
4
Ecological gradient (Leaf Unfolding differences)
233
218
256
253
124
L16
9
204
219
217
O16
97
L8
O12
L12
O8
O1
L1

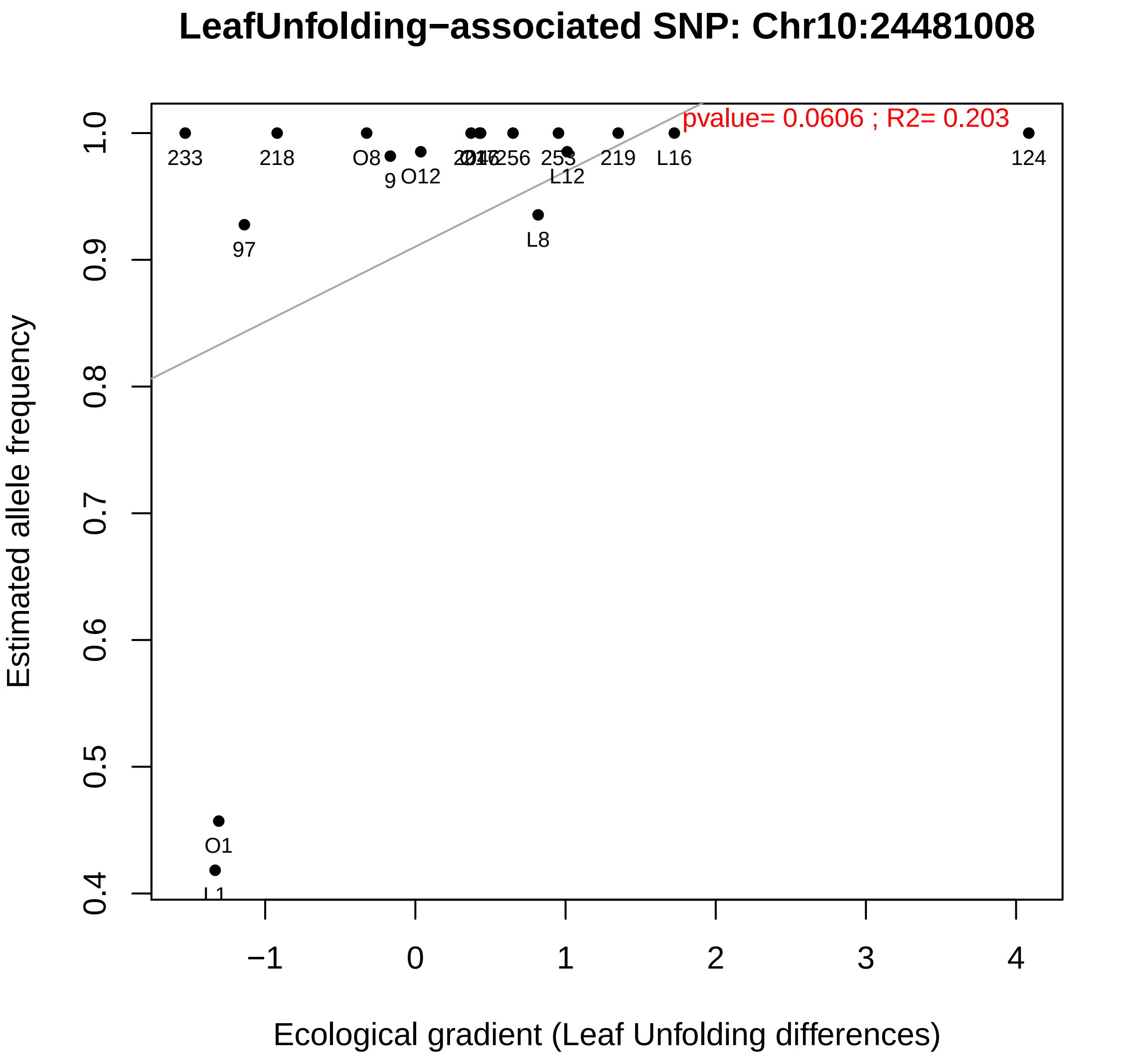

LeafUnfolding−associated SNP: Chr10:24481008
pvalue= 0.0606 ; R2= 0.203
Estimated allele frequency
Ecological gradient (Leaf Unfolding differences)
1.0
0.9
0.8
0.7
0.6
0.5
0.4
−1
0
1
2
3
4
233
218
O8
9
O12
256
253
L12
219
L16
124
97
L8
O1
L1

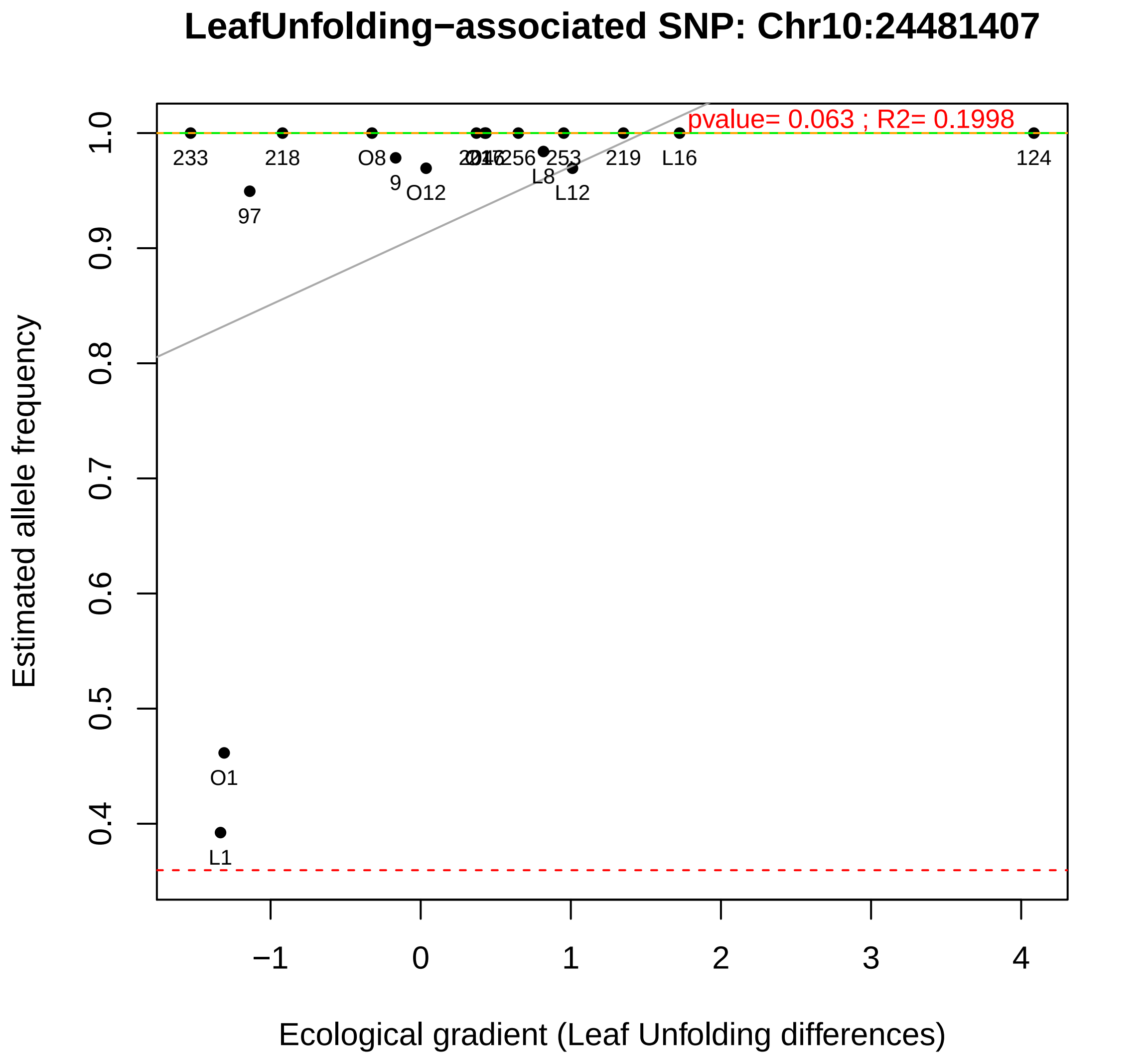

LeafUnfolding−associated SNP: Chr10:24481407
pvalue= 0.063 ; R2= 0.1998
Estimated allele frequency
Ecological gradient (Leaf Unfolding differences)
0.4
0.5
0.6
0.7
0.8
0.9
1.0
−1
0
1
2
3
4
233
218
O8
9
O12
97
253
L8
L12
219
L16
124
O1
L1

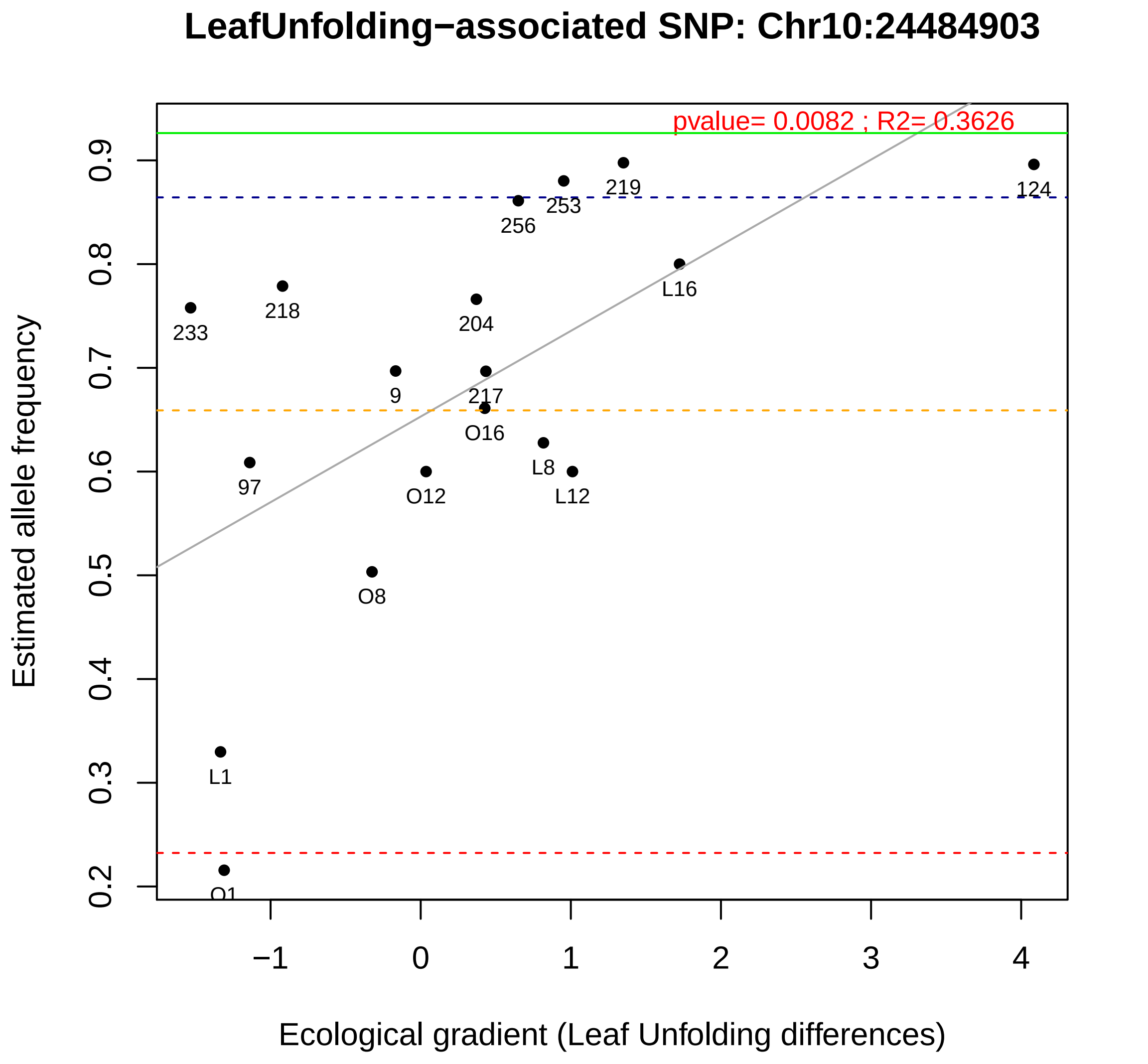

LeafUnfolding−associated SNP: Chr10:24484903
pvalue= 0.0082 ; R2= 0.3626
Estimated allele frequency
Ecological gradient (Leaf Unfolding differences)
0.2
0.3
0.4
0.5
0.6
0.7
0.8
0.9
−1
0
1
2
3
4
233
218
97
L1
O1
O8
9
O12
204
217
O16
256
L8
L12
253
219
L16
124

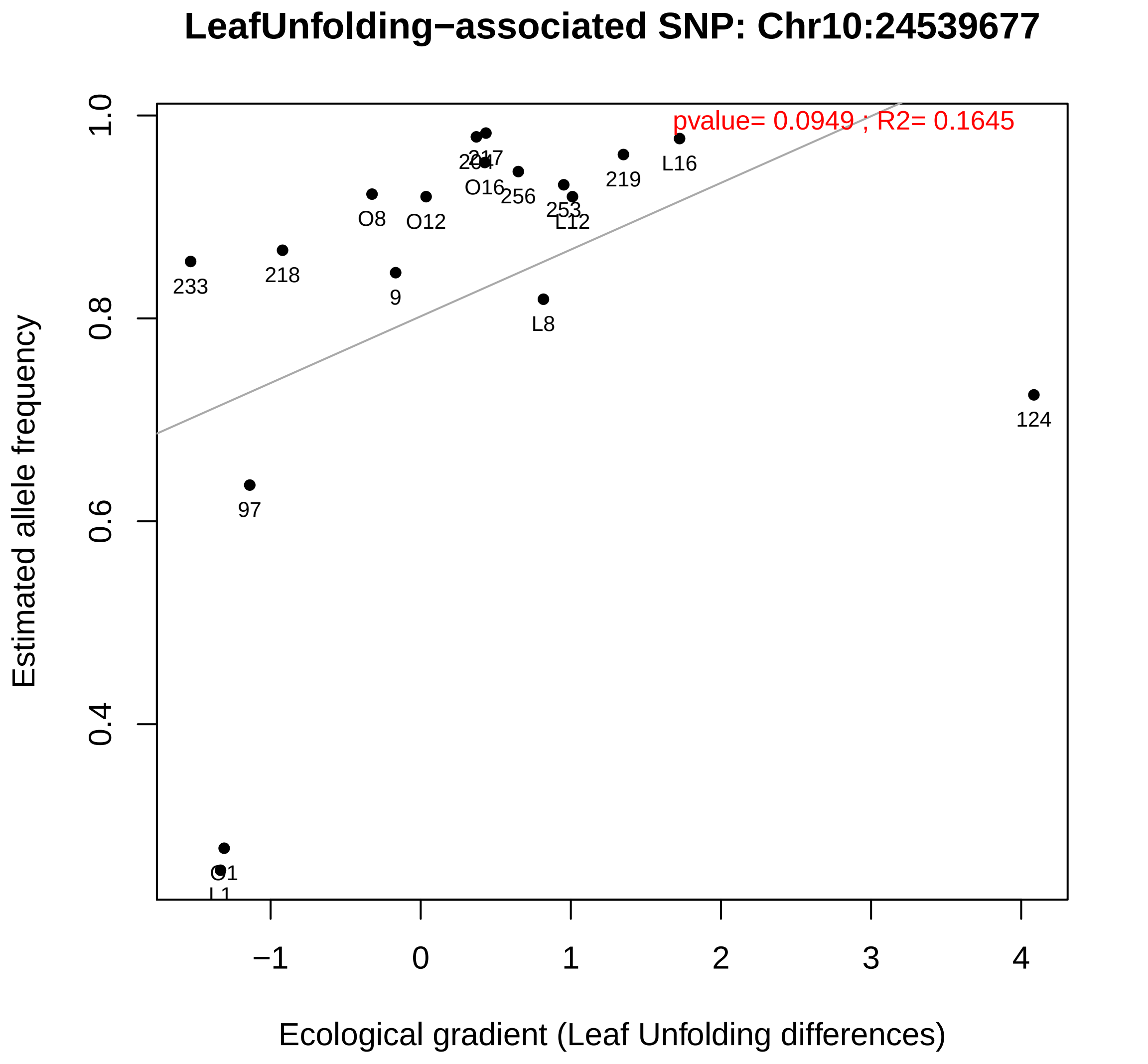

LeafUnfolding−associated SNP: Chr10:24539677
pvalue= 0.0949 ; R2= 0.1645
Estimated allele frequency
Ecological gradient (Leaf Unfolding differences)
1.0
0.8
0.6
0.4
−1
0
1
2
3
4
233
218
97
O8
O12
9
217
204
O16
256
253
L12
L8
219
L16
124
O1
L1

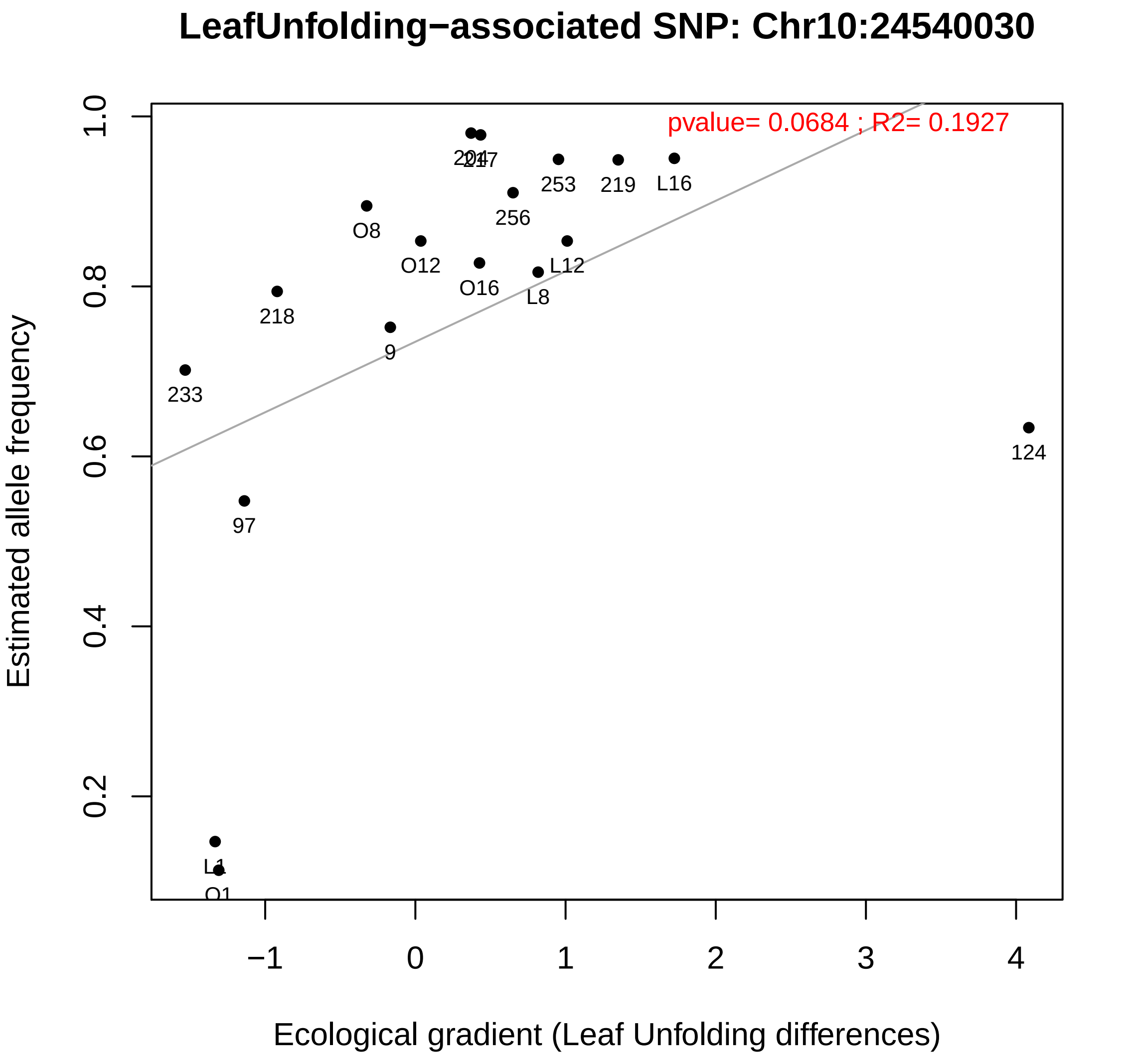
LeafUnfolding−associated SNP: Chr10:24540030
pvalue= 0.0684 ; R2= 0.1927
Estimated allele frequency
Ecological gradient (Leaf Unfolding differences)
1.0
0.8
0.6
0.4
0.2
−1
0
1
2
3
4
253
219
L16
256
O8
O12
L12
O16
L8
218
9
233
124
97
O1

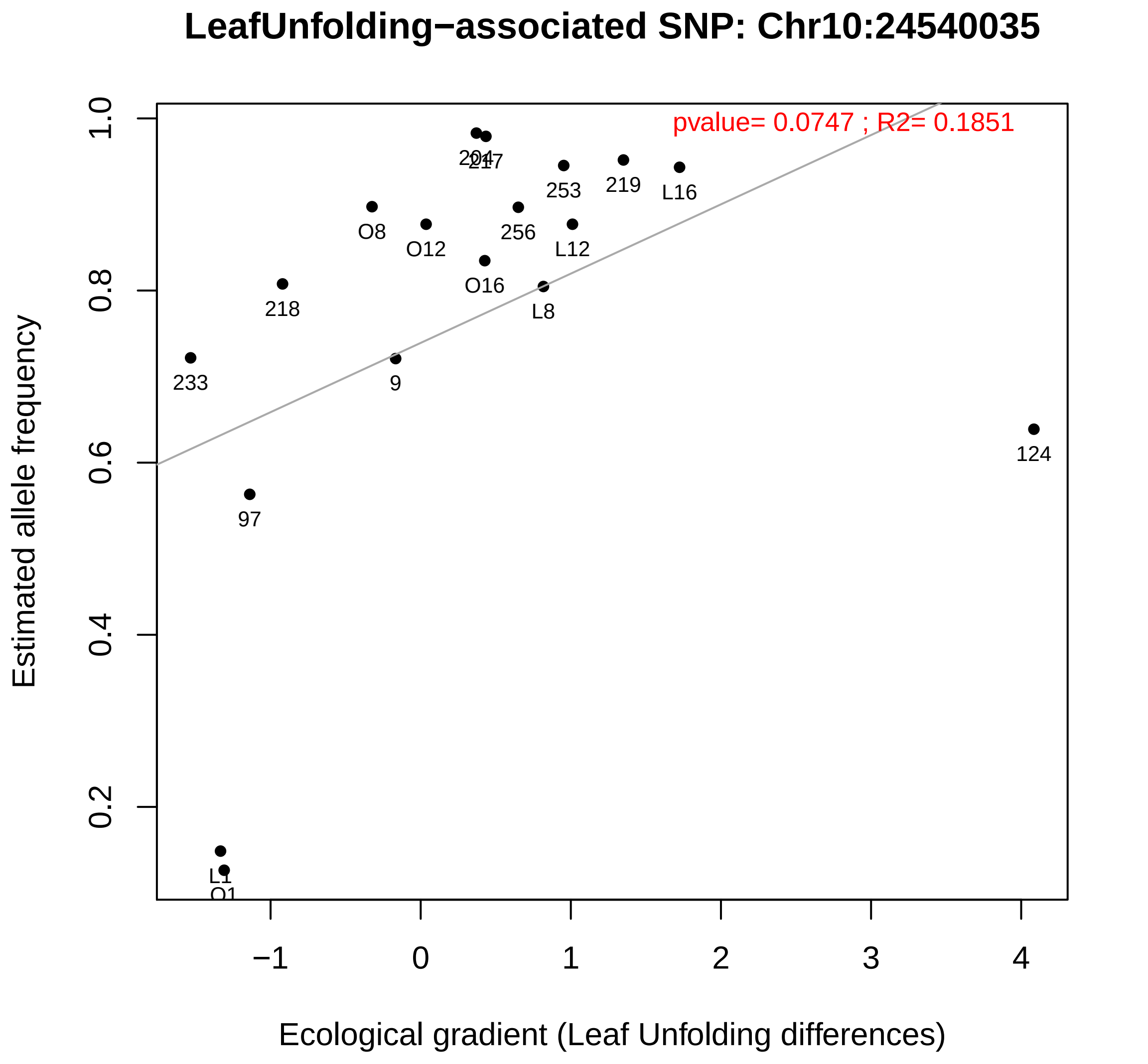

LeafUnfolding−associated SNP: Chr10:24540035
pvalue= 0.0747 ; R2= 0.1851
Estimated allele frequency
Ecological gradient (Leaf Unfolding differences)
0.2
0.4
0.6
0.8
1.0
−1
0
1
2
3
4
204
217
253
219
L16
O8
256
O12
L12
O16
218
L8
233
9
124
97
L1
O1

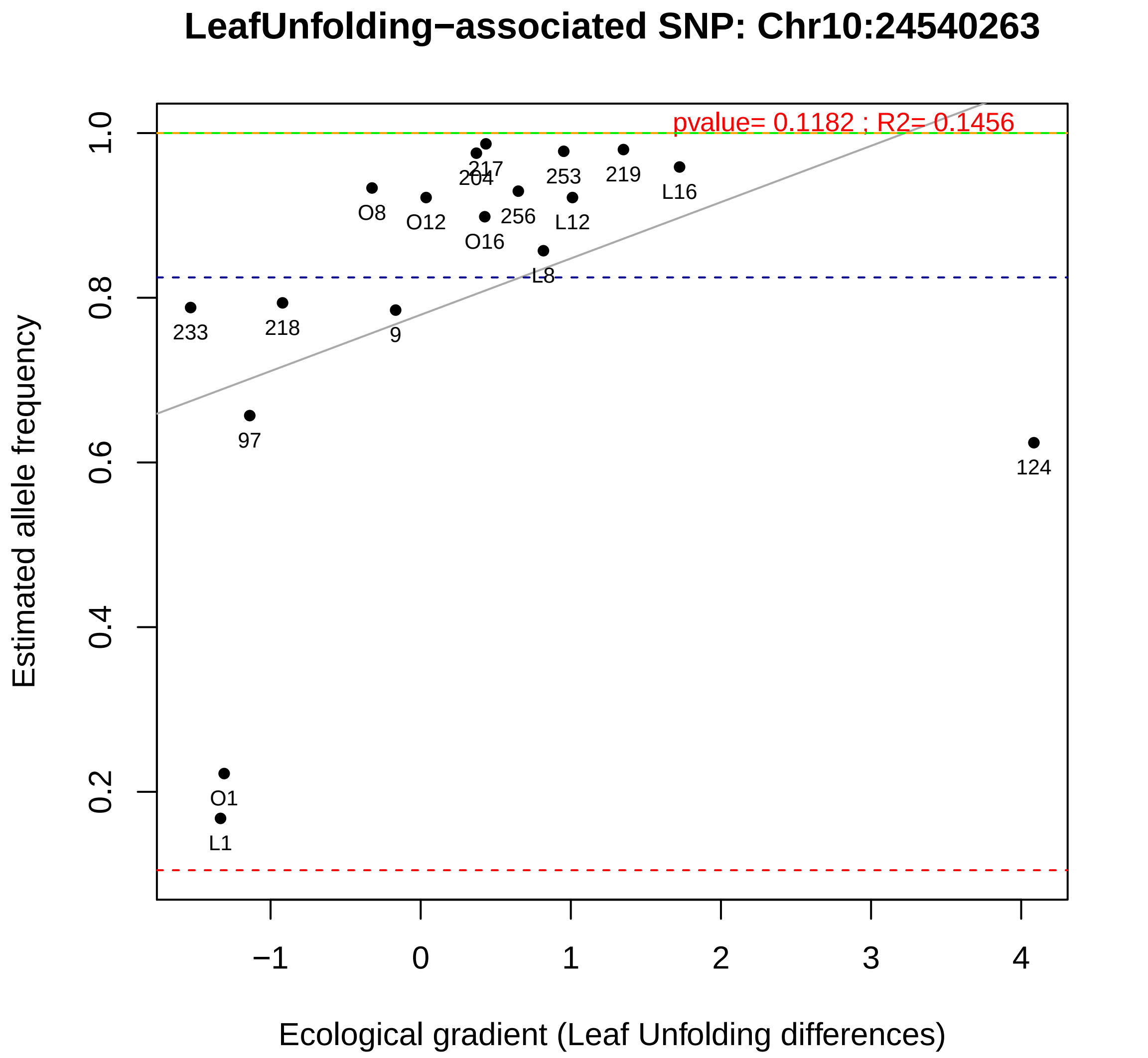
LeafUnfolding−associated SNP: Chr10:24540263
pvalue= 0.1182 ; R2= 0.1456
Estimated allele frequency
1.0
0.8
0.6
0.4
0.2
217
204
253
219
L16
O8
O12
256
L12
O16
L8
233
218
9
97
124
O1
L1
−1
0
1
2
3
4
Ecological gradient (Leaf Unfolding differences)

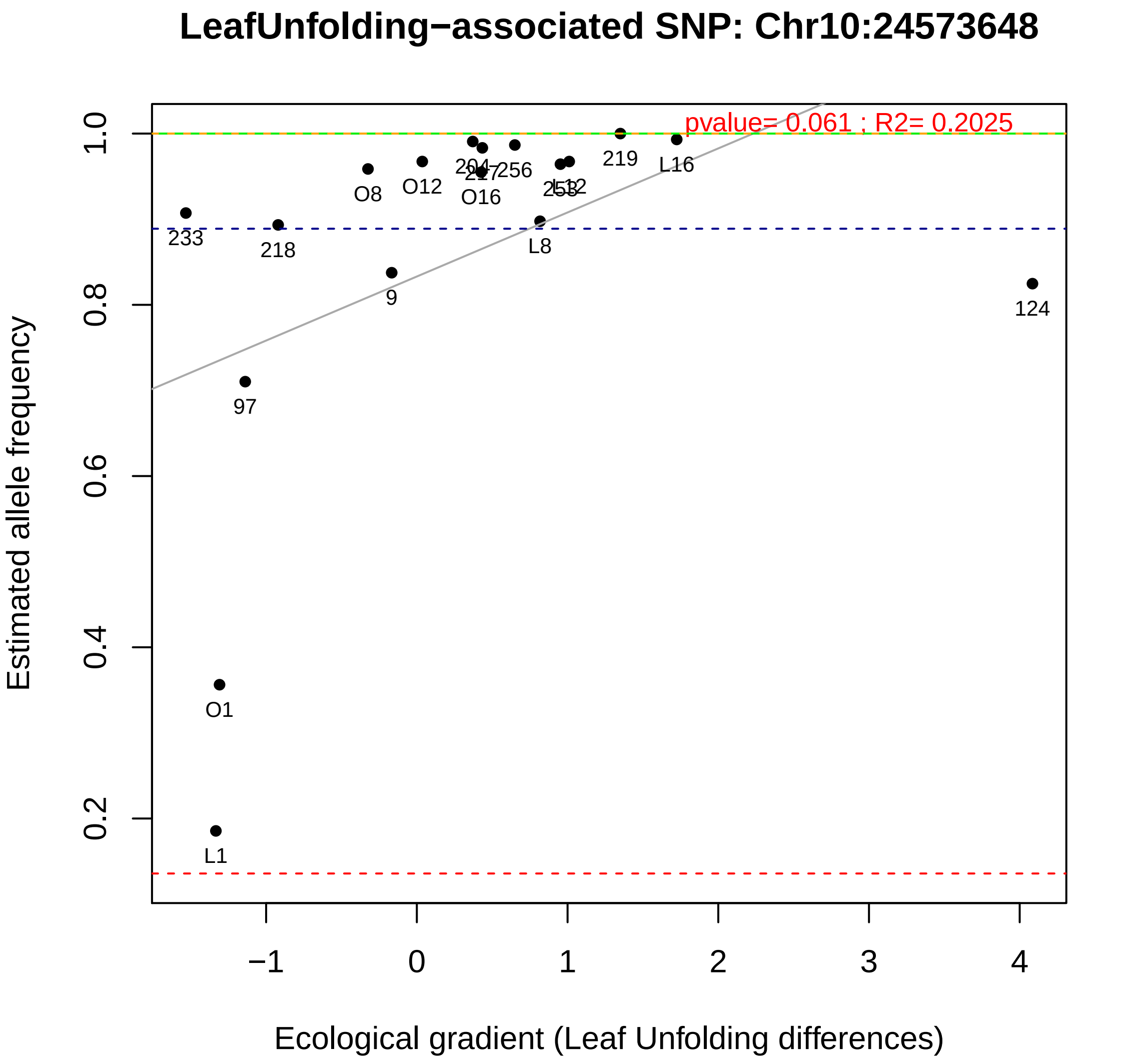
LeafUnfolding−associated SNP: Chr10:24573648
pvalue= 0.061 ; R2= 0.2025
Estimated allele frequency
Ecological gradient (Leaf Unfolding differences)
1.0
0.8
0.6
0.4
0.2
−1
0
1
2
3
4
233
218
97
O1
L1
O8
9
O12
204
217
O16
256
253
L12
L8
219
L16
124

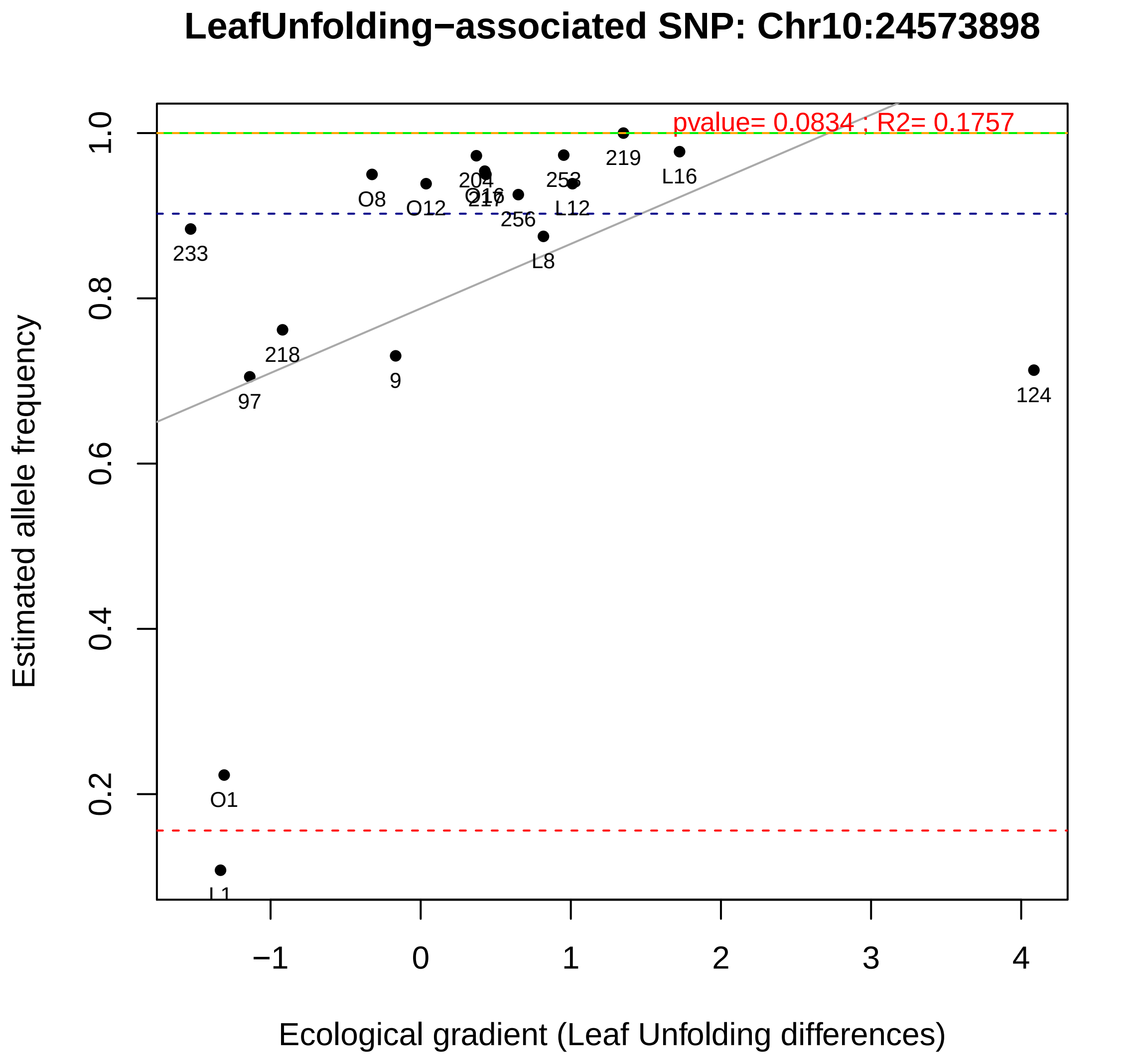

LeafUnfolding−associated SNP: Chr10:24573898
pvalue= 0.0834 ; R2= 0.1757
Estimated allele frequency
Ecological gradient (Leaf Unfolding differences)
1.0
0.8
0.6
0.4
0.2
−1
0
1
2
3
4
219
L16
204
253
O8
O12
O16
217
256
L12
233
L8
218
97
9
124
O1
L1

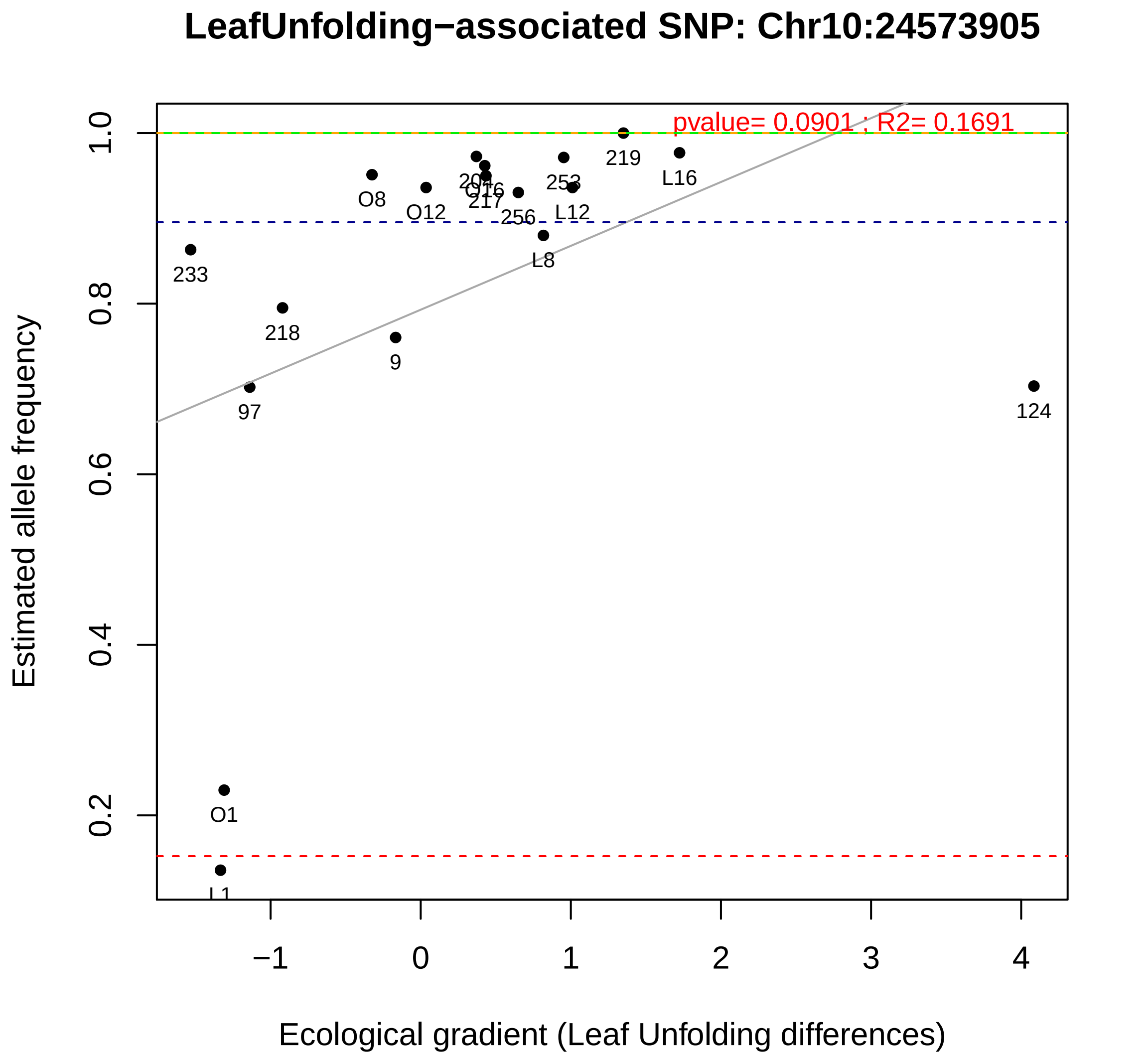

LeafUnfolding−associated SNP: Chr10:24573905
pvalue= 0.0901 ; R2= 0.1691
Estimated allele frequency
Ecological gradient (Leaf Unfolding differences)
0.2
0.4
0.6
0.8
1.0
−1
0
1
2
3
4
219
L16
253
O8
O12
217
256
L12
L8
233
218
9
97
124
O1

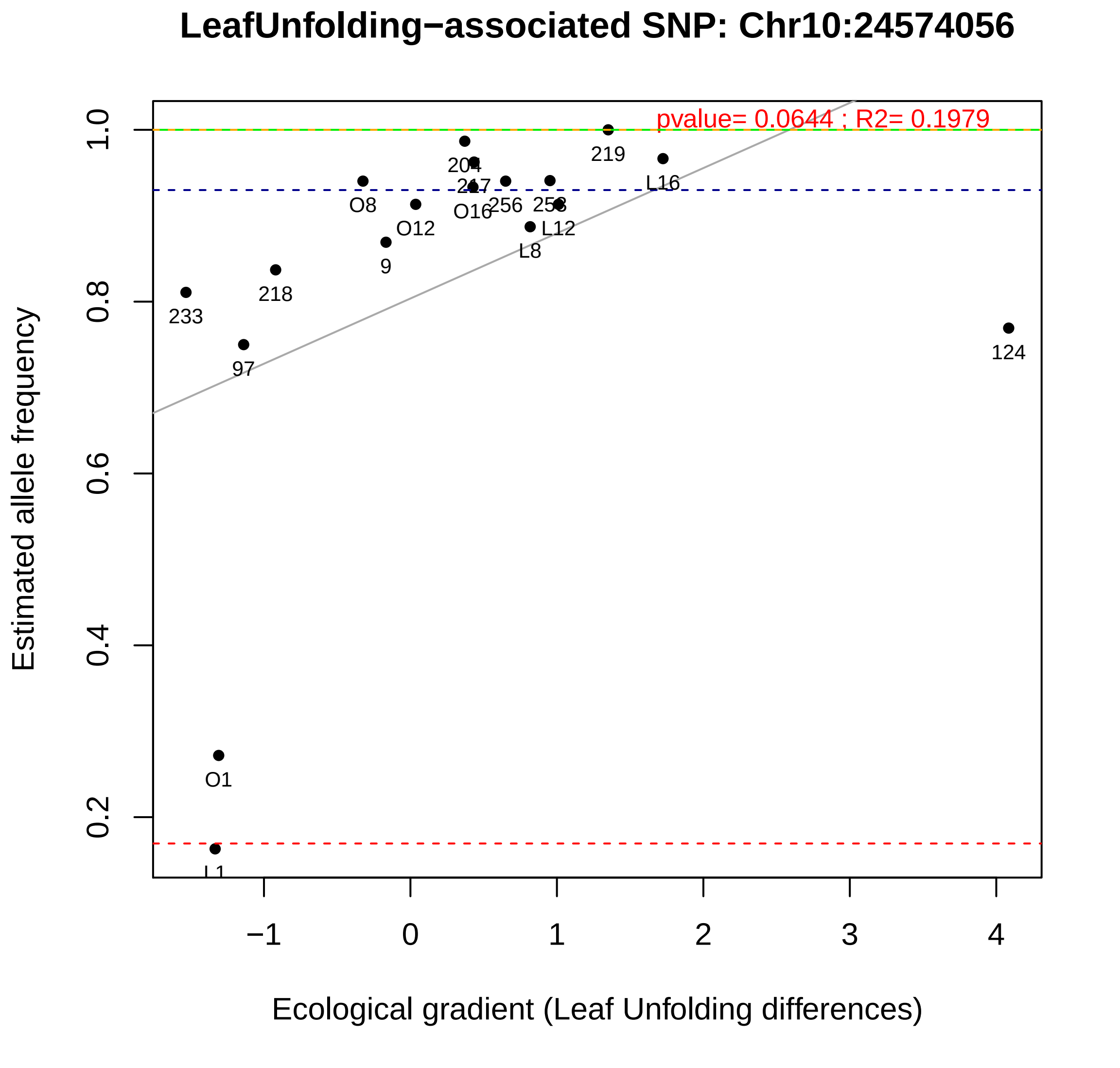

LeafUnfolding−associated SNP: Chr10:24574056
pvalue= 0.0644 ; R2= 0.1979
Estimated allele frequency
Ecological gradient (Leaf Unfolding differences)
0.2
0.4
0.6
0.8
1.0
−1
0
1
2
3
4
219
L16
204
217
O16
256
253
L12
L8
O8
O12
9
218
233
97
124
O1
L1

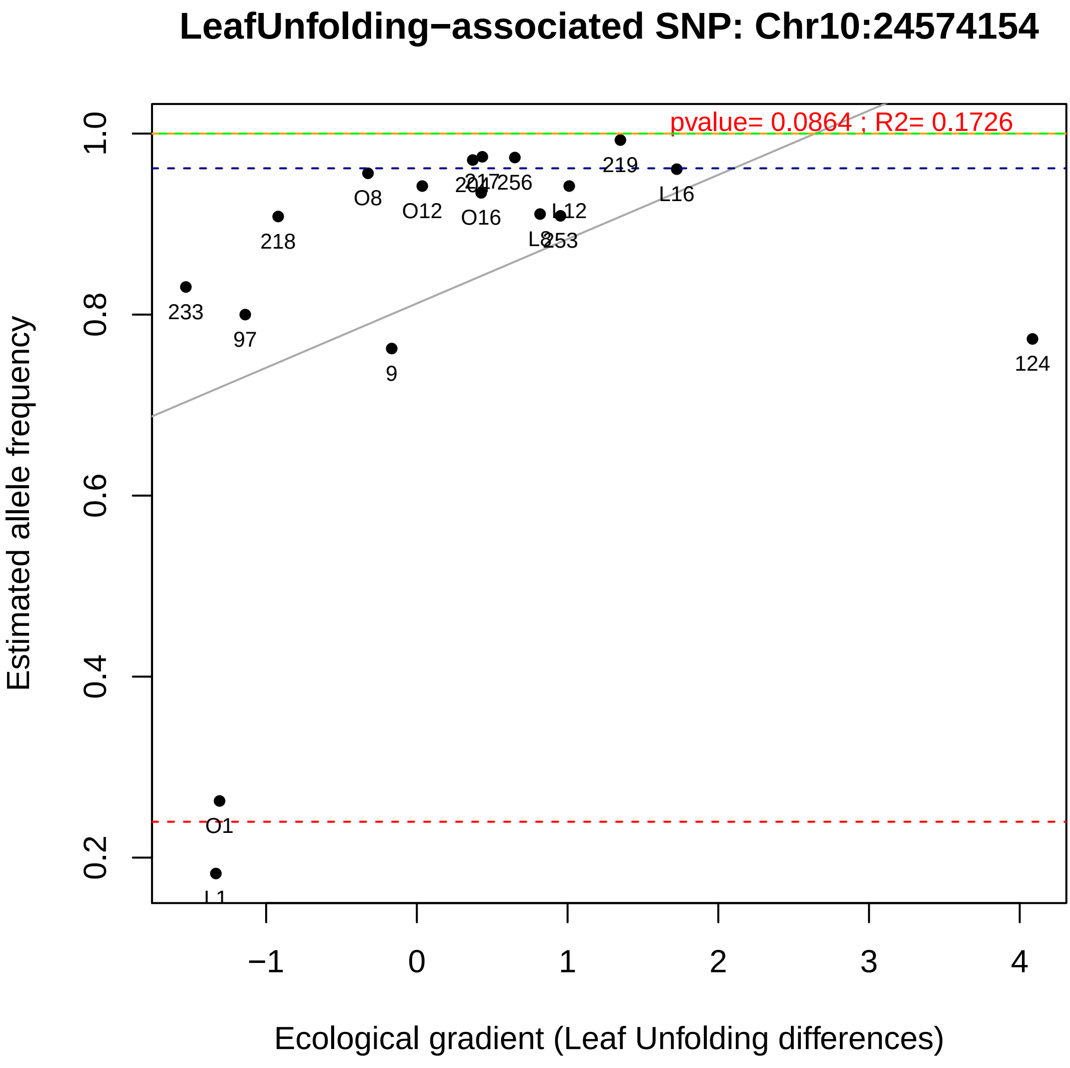

LeafUnfolding−associated SNP: Chr10:24574154
pvalue= 0.0864 ; R2= 0.1726
Estimated allele frequency
Ecological gradient (Leaf Unfolding differences)
1.0
0.8
0.6
0.4
0.2
−1
0
1
2
3
4
219
L16
O8
O12
O16
256
218
L12
253
233
97
9
124
O1

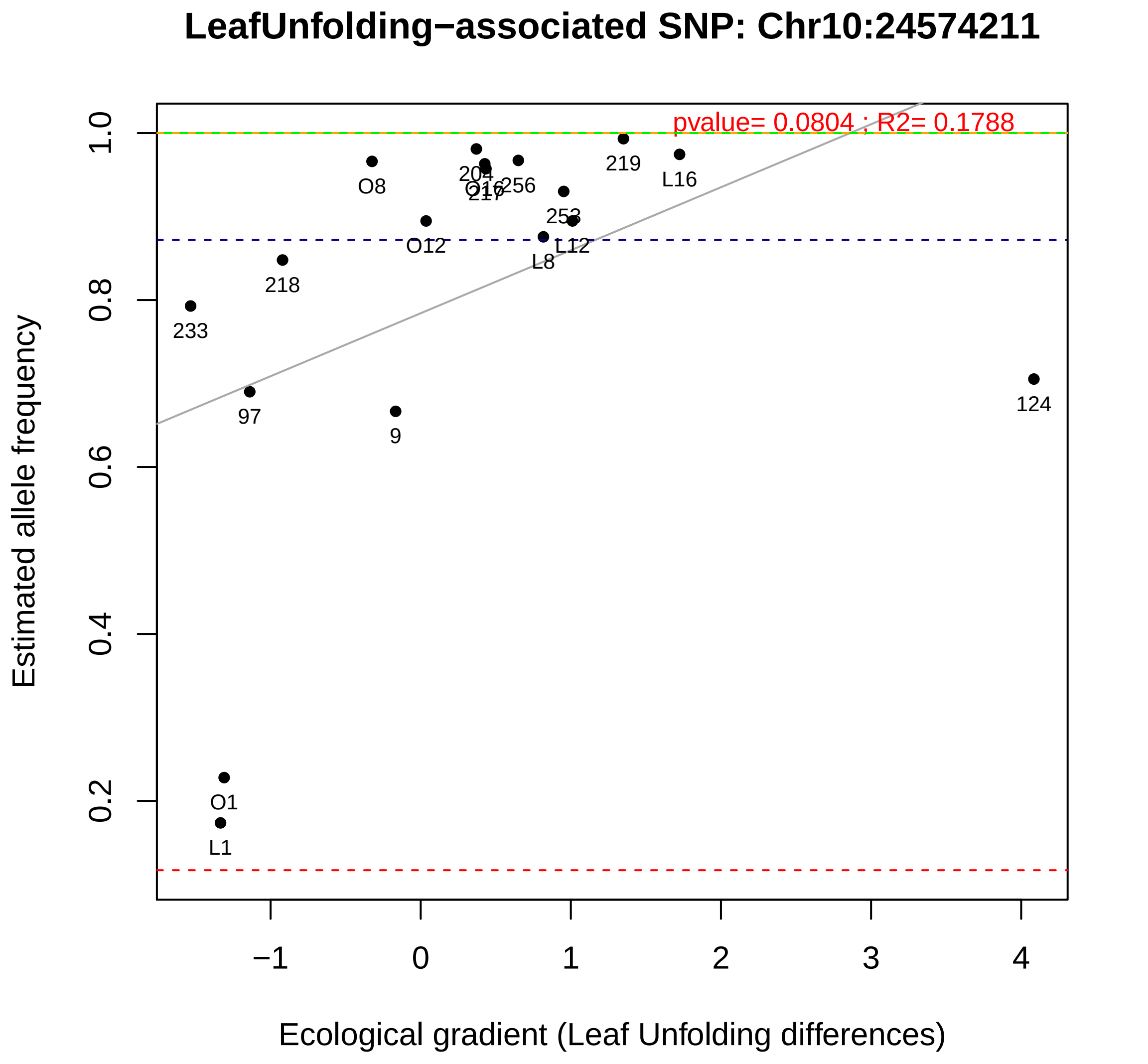

LeafUnfolding−associated SNP: Chr10:24574211
pvalue= 0.0804 ; R2= 0.1788
Estimated allele frequency
Ecological gradient (Leaf Unfolding differences)
0.2
0.4
0.6
0.8
1.0
−1
0
1
2
3
4
O8
204
O16
217
256
219
L16
253
L12
O12
L8
218
233
97
9
124
O1
L1

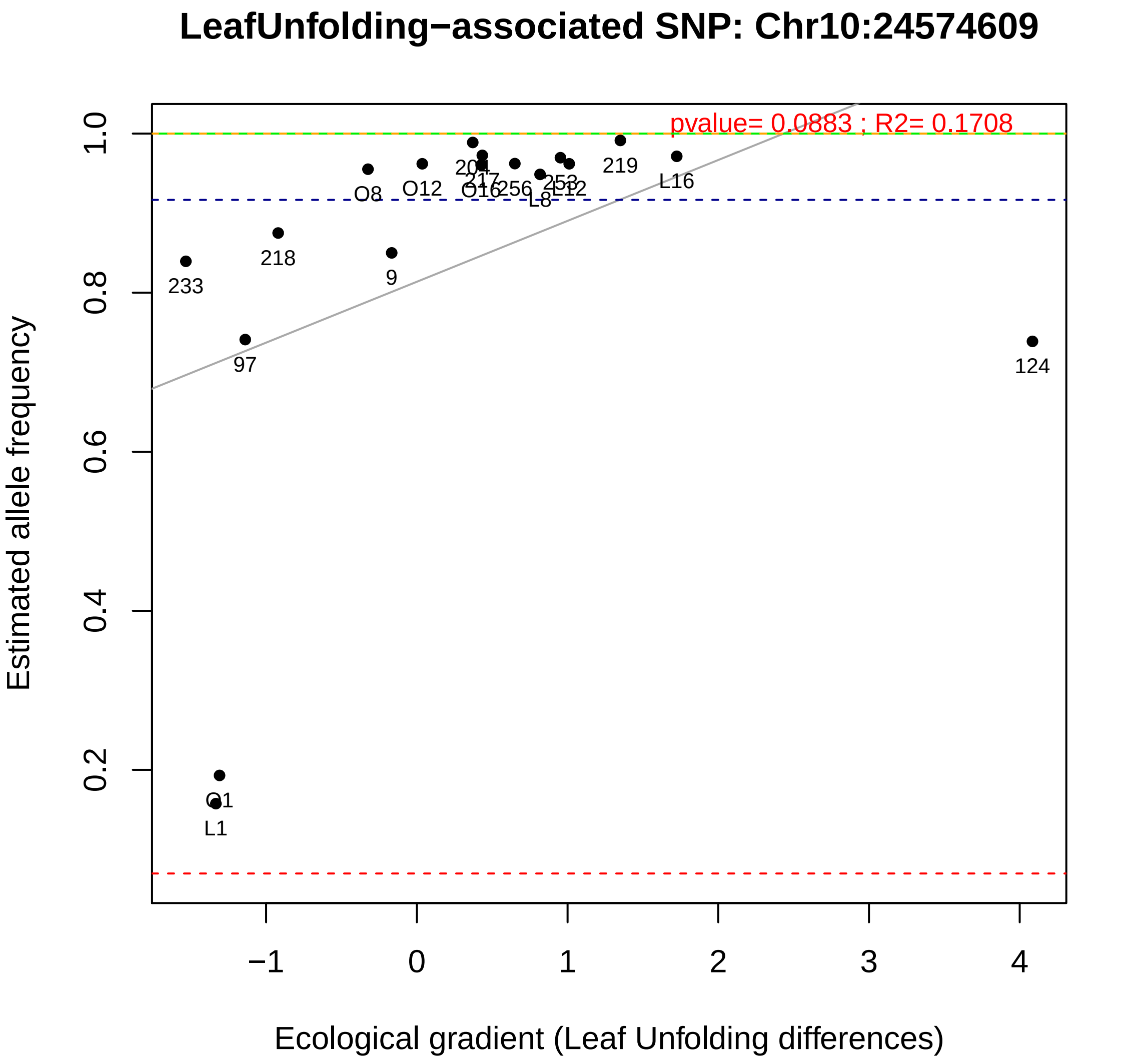
LeafUnfolding−associated SNP: Chr10:24574609
pvalue= 0.0883 ; R2= 0.1708
Estimated allele frequency
0.2
0.4
0.6
0.8
1.0
−1
0
1
2
3
4
Ecological gradient (Leaf Unfolding differences)
233
97
218
O8
9
O12
204
217
O16
256
L8
253
L12
219
L16
124
O1
L1

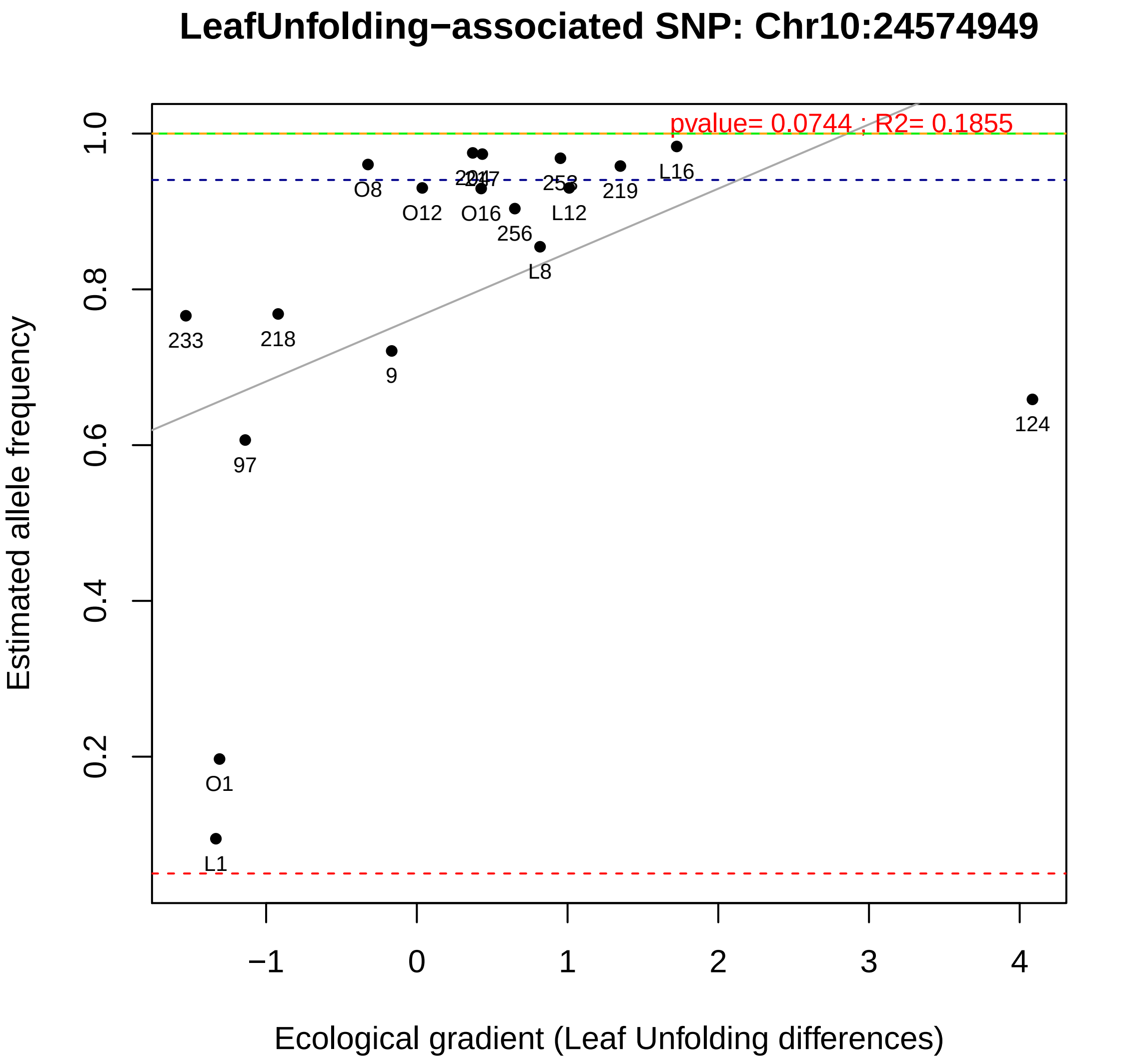
LeafUnfolding−associated SNP: Chr10:24574949
pvalue= 0.0744 ; R2= 0.1855
Estimated allele frequency
Ecological gradient (Leaf Unfolding differences)
1.0
0.8
0.6
0.4
0.2
−1
0
1
2
3
4
O8
O12
O16
256
L12
219
L16
L8
233
218
9
97
124
O1
L1

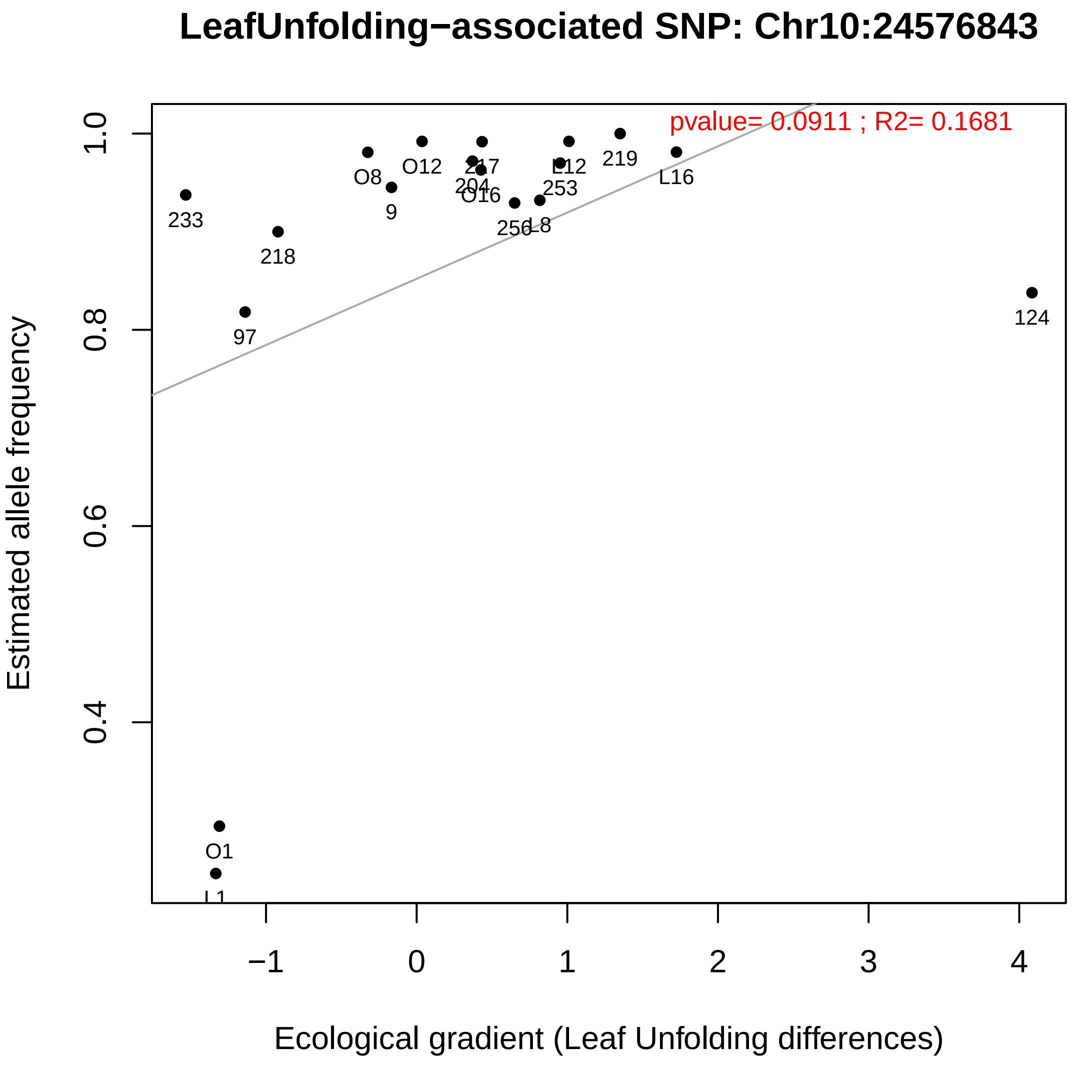
LeafUnfolding−associated SNP: Chr10:24576843
pvalue= 0.0911 ; R2= 0.1681
Estimated allele frequency
0.4
0.6
0.8
1.0
−1
0
1
2
3
4
Ecological gradient (Leaf Unfolding differences)
233
218
97
O8
9
O12
217
204
O16
256
L8
L12
253
219
L16
124
O1
L1

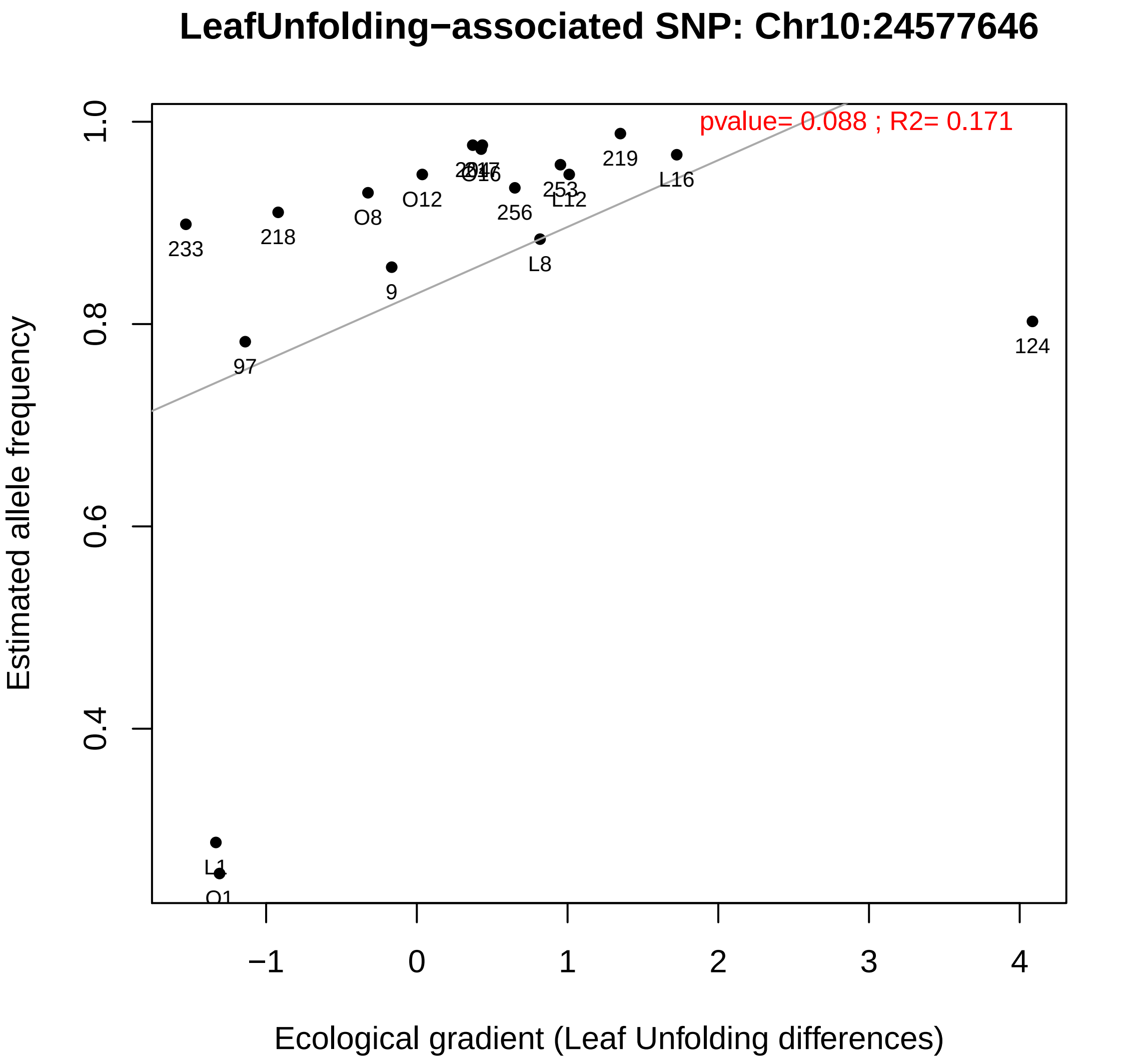

LeafUnfolding−associated SNP: Chr10:24577646
pvalue= 0.088 ; R2= 0.171
Estimated allele frequency
Ecological gradient (Leaf Unfolding differences)
1.0
0.8
0.6
0.4
−1
0
1
2
3
4
233
218
97
O8
9
O12
256
253
L12
L8
219
L16
124
L1
O1

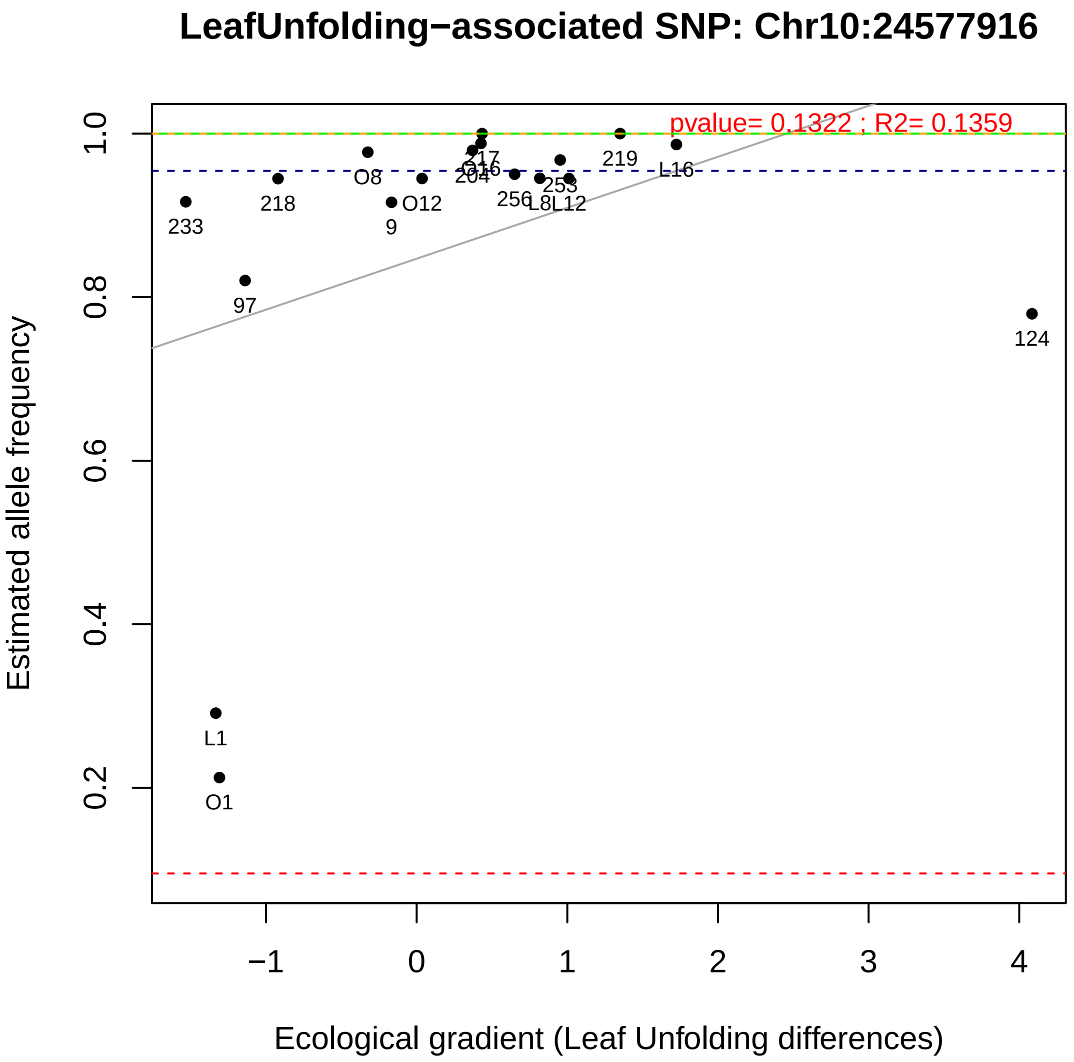

LeafUnfolding−associated SNP: Chr10:24577916
pvalue= 0.1322 ; R2= 0.1359
Estimated allele frequency
Ecological gradient (Leaf Unfolding differences)
1.0
0.8
0.6
0.4
0.2
−1
0
1
2
3
4
233
218
97
O8
9
O12
217
O16
204
256
L8
253
L12
219
L16
124
L1
O1

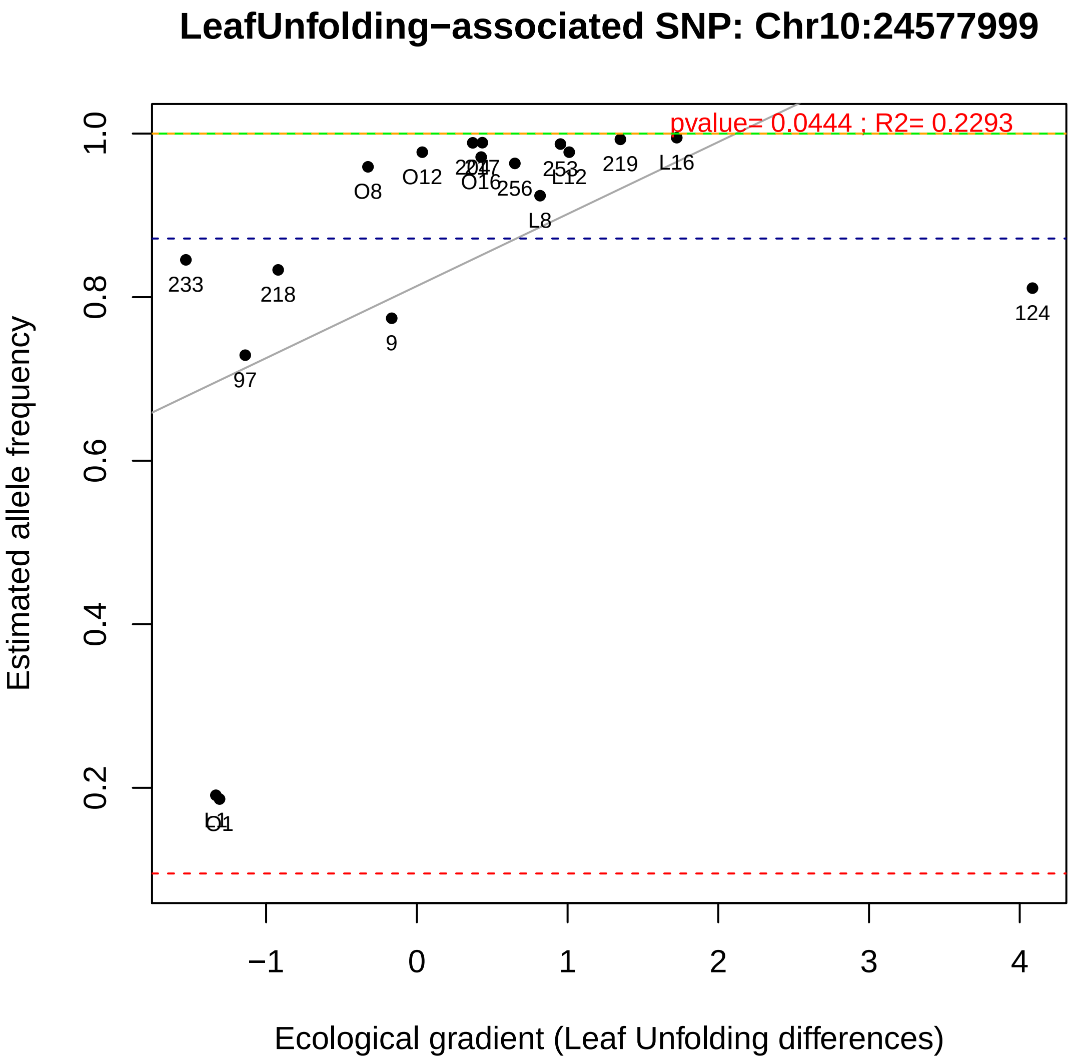
LeafUnfolding−associated SNP: Chr10:24577999
pvalue= 0.0444 ; R2= 0.2293
Estimated allele frequency
Ecological gradient (Leaf Unfolding differences)
0.2
0.4
0.6
0.8
1.0
−1
0
1
2
3
4
233
218
97
9
O8
O12
O16
256
L8
253
L12
219
L16
124

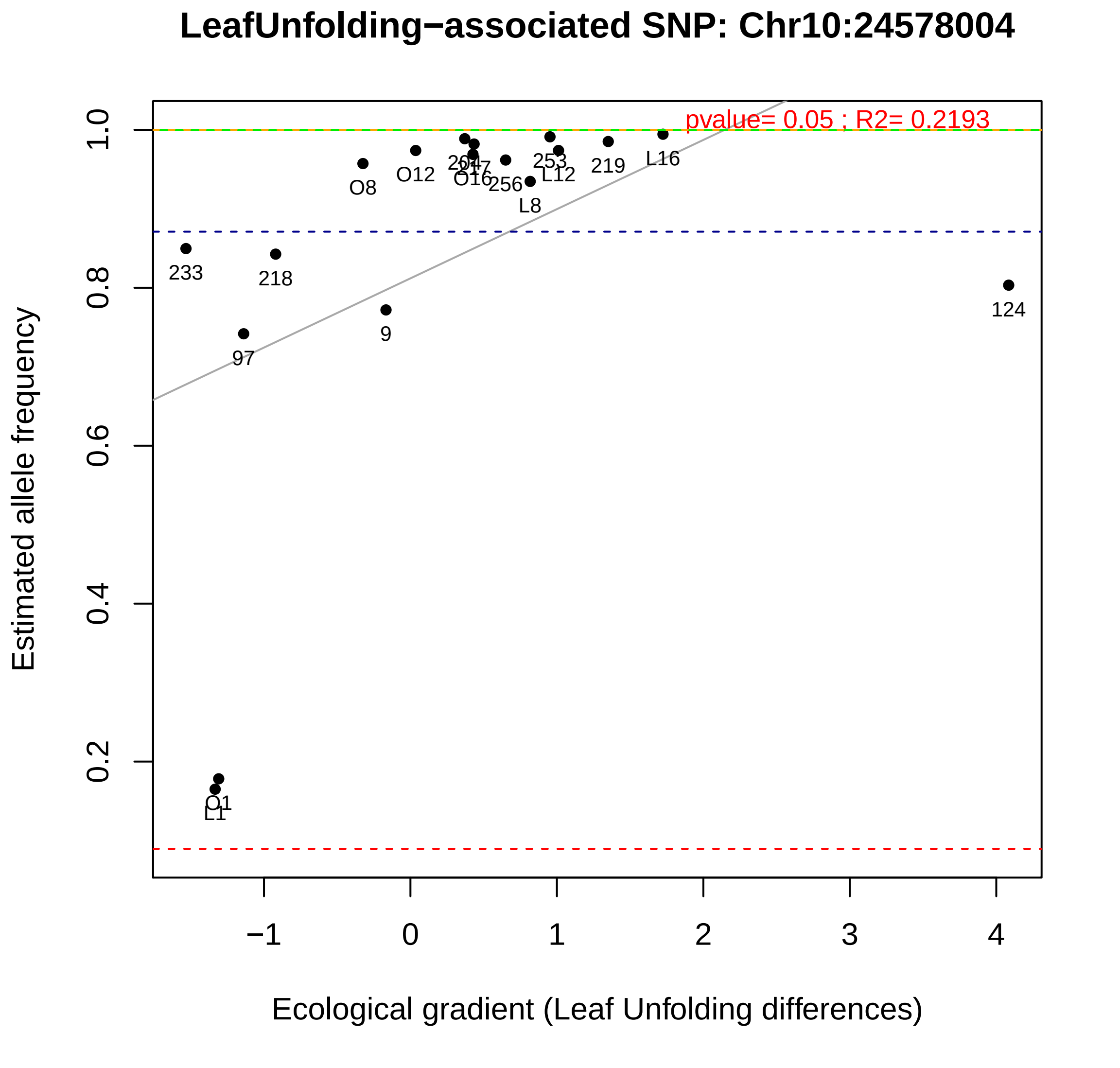

LeafUnfolding−associated SNP: Chr10:24578004
pvalue= 0.05 ; R2= 0.2193
Estimated allele frequency
Ecological gradient (Leaf Unfolding differences)
0.2
0.4
0.6
0.8
1.0
−1
0
1
2
3
4
233
218
97
9
O8
O12
O16
256
L8
253
L12
219
L16
124
O1
L1

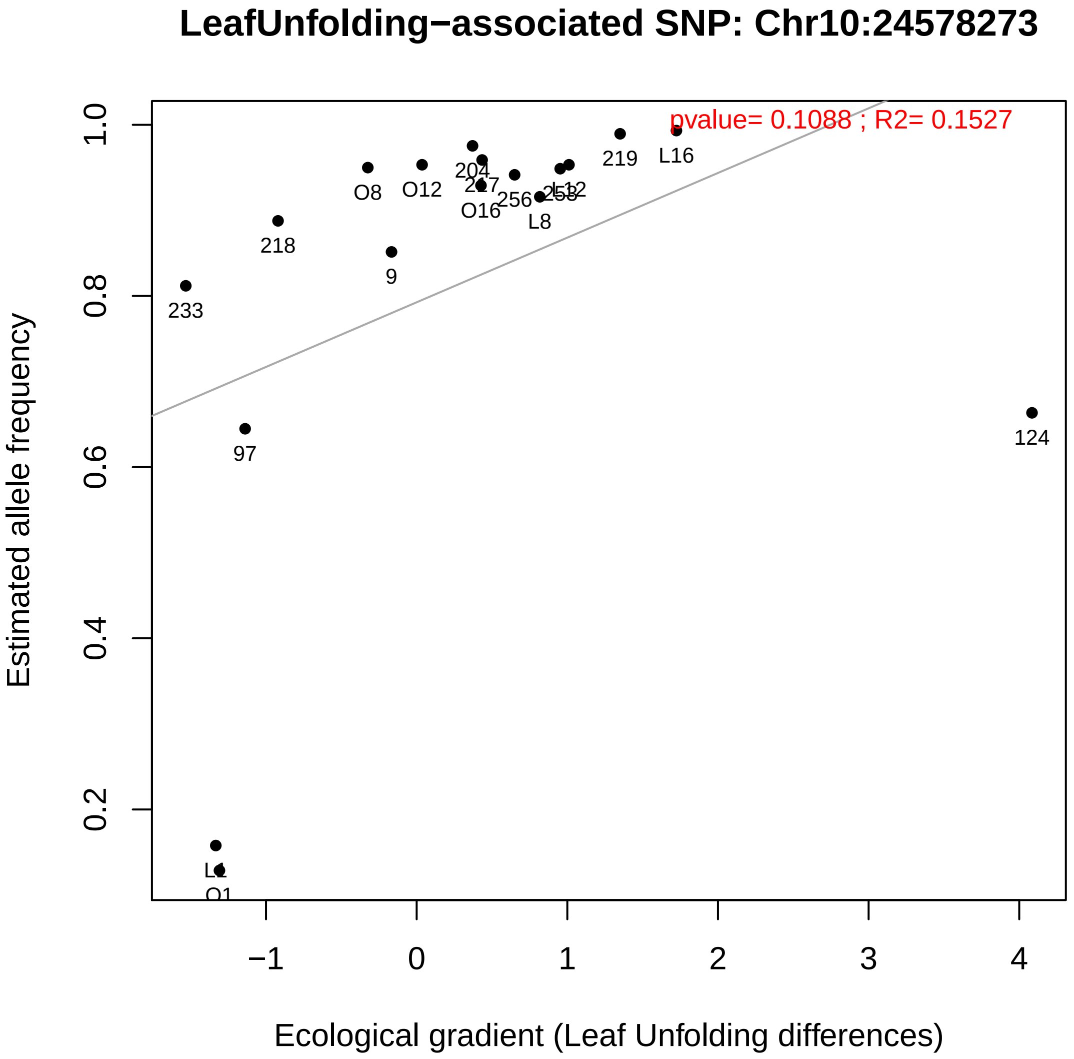
LeafUnfolding−associated SNP: Chr10:24578273
pvalue= 0.1088 ; R2= 0.1527
Estimated allele frequency
Ecological gradient (Leaf Unfolding differences)
1.0
0.8
0.6
0.4
0.2
−1
0
1
2
3
4
219
L16
204
O8
O12
O16
256
L8
218
9
233
97
124

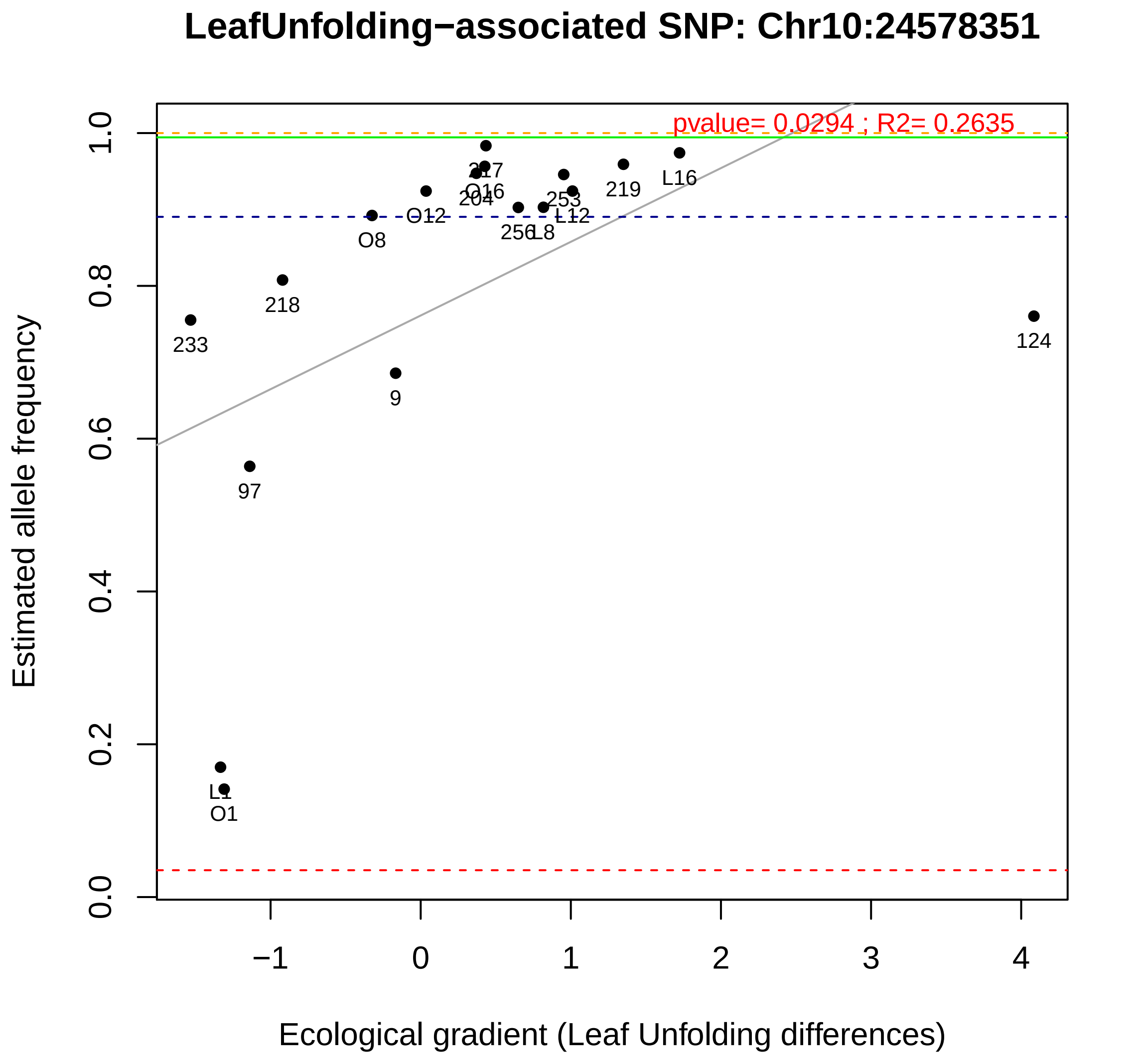

LeafUnfolding−associated SNP: Chr10:24578351
pvalue= 0.0294 ; R2= 0.2635
Estimated allele frequency
Ecological gradient (Leaf Unfolding differences)
0.0
0.2
0.4
0.6
0.8
1.0
−1
0
1
2
3
4
233
218
97
L1
O1
O8
9
O12
217
O16
204
256
L8
253
L12
219
L16
124

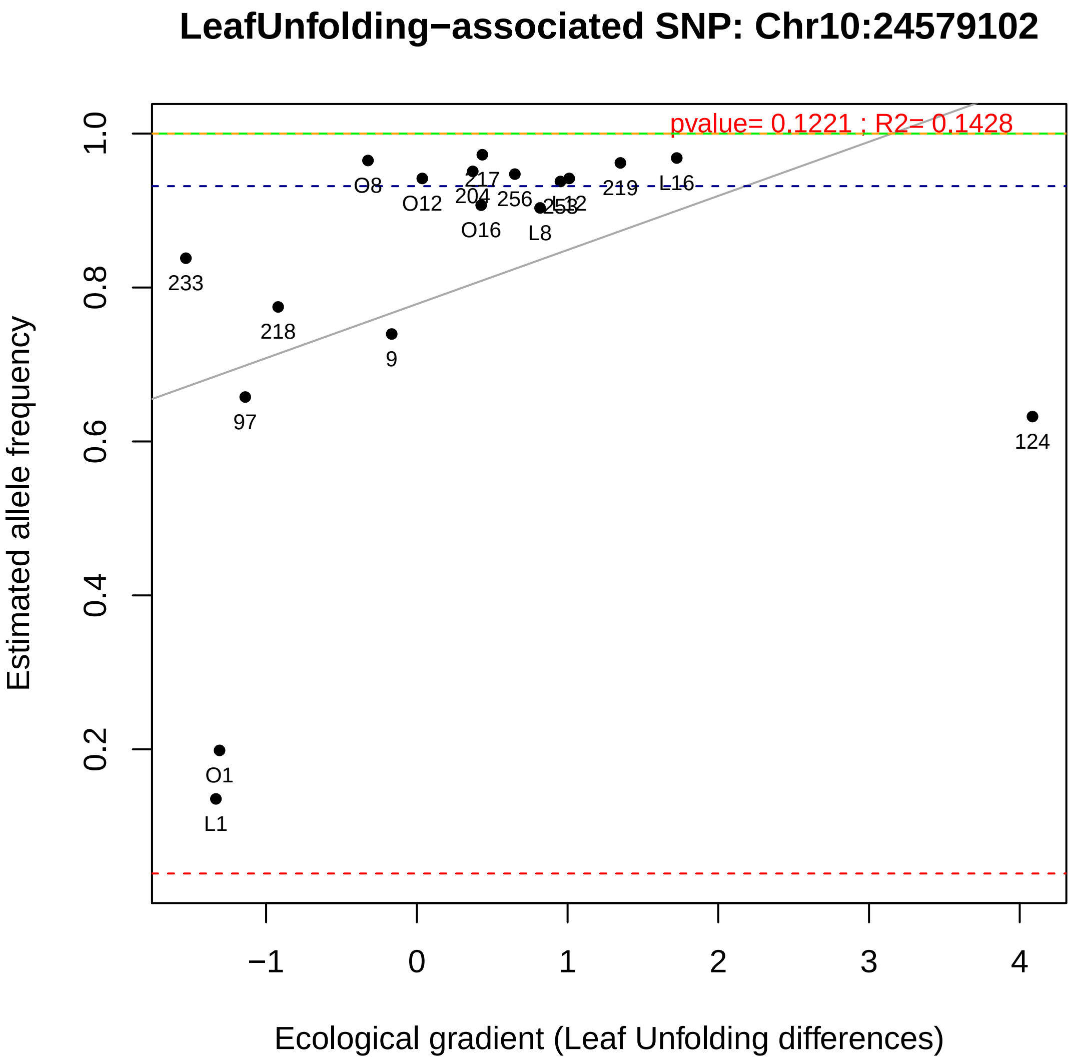

LeafUnfolding−associated SNP: Chr10:24579102
pvalue= 0.1221 ; R2= 0.1428
Estimated allele frequency
Ecological gradient (Leaf Unfolding differences)
0.2
0.4
0.6
0.8
1.0
−1
0
1
2
3
4
O8
O12
217
204
256
O16
L8
L12
253
219
L16
233
218
9
97
124
O1
L1

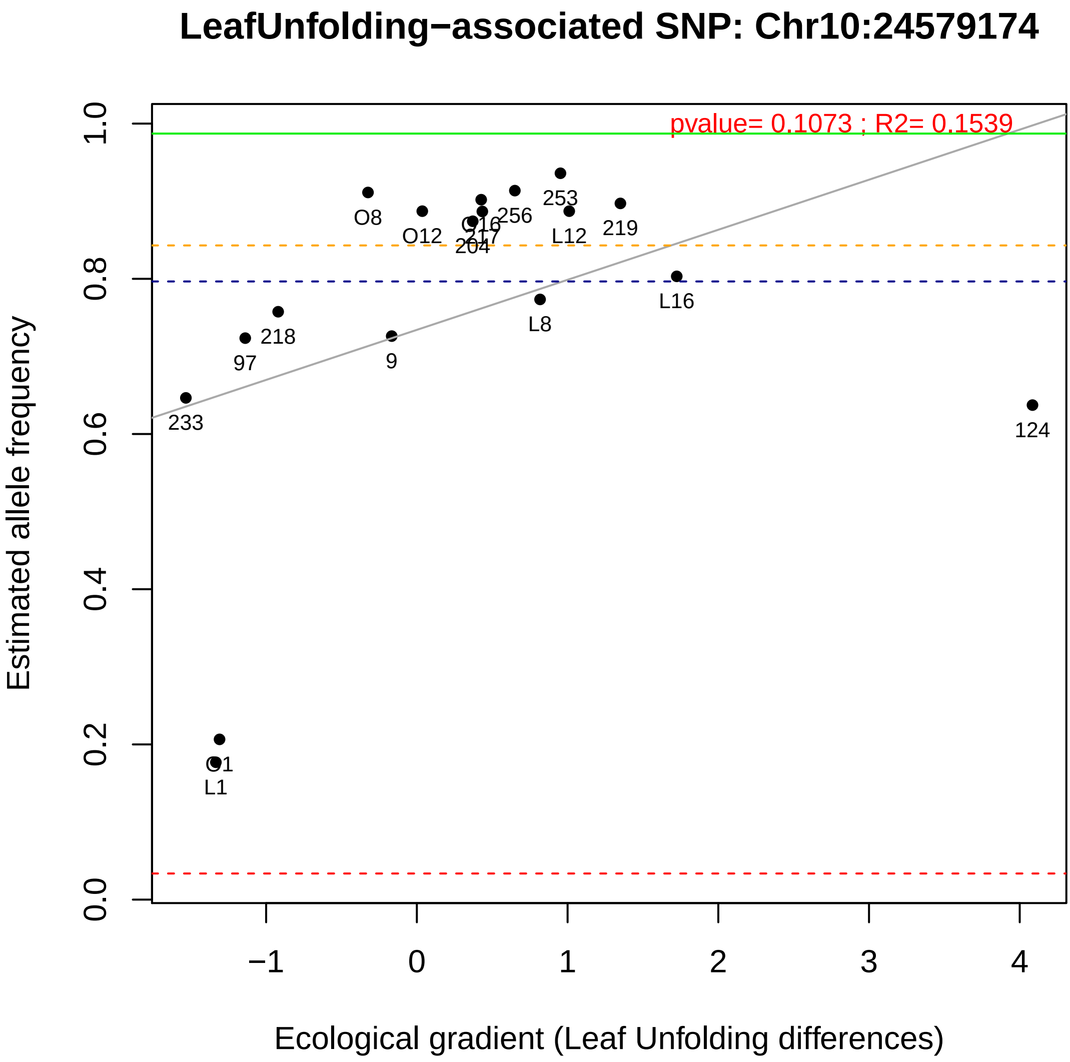
LeafUnfolding−associated SNP: Chr10:24579174
pvalue= 0.1073 ; R2= 0.1539
Estimated allele frequency
Ecological gradient (Leaf Unfolding differences)
0.0
0.2
0.4
0.6
0.8
1.0
−1
0
1
2
3
4
O8
O12
204
217
256
253
L12
219
L16
L8
218
97
9
233
124
O1
L1

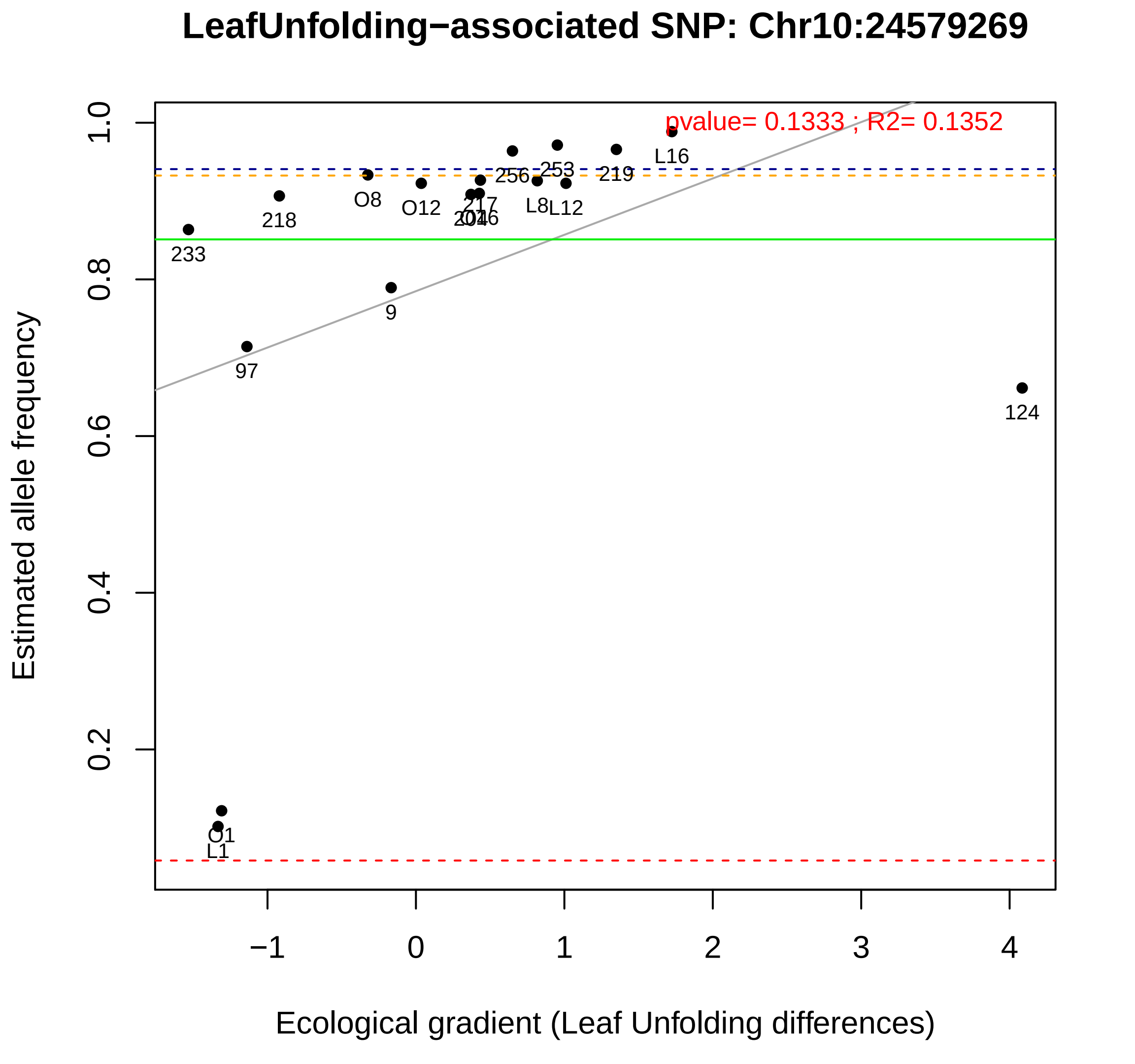

LeafUnfolding−associated SNP: Chr10:24579269
pvalue= 0.1333 ; R2= 0.1352
Estimated allele frequency
Ecological gradient (Leaf Unfolding differences)
0.2
0.4
0.6
0.8
1.0
−1
0
1
2
3
4
233
218
O8
O12
217
L8
L12
256
253
219
L16
9
97
124
O1
L1

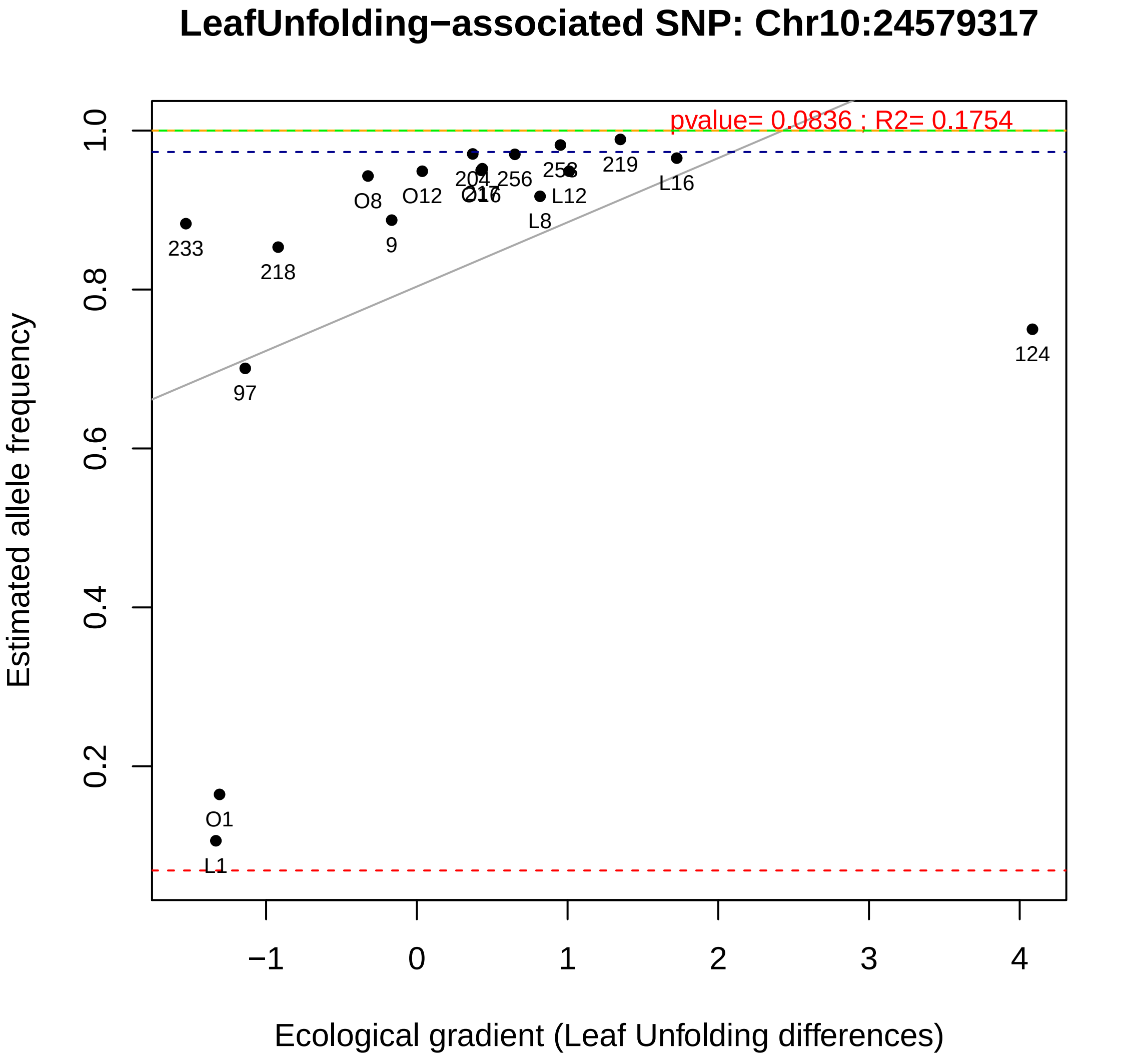

LeafUnfolding−associated SNP: Chr10:24579317
pvalue= 0.0836 ; R2= 0.1754
Estimated allele frequency
Ecological gradient (Leaf Unfolding differences)
0.2
0.4
0.6
0.8
1.0
−1
0
1
2
3
4
233
218
97
O8
9
O12
204
256
L12
L8
219
L16
124
O1
L1

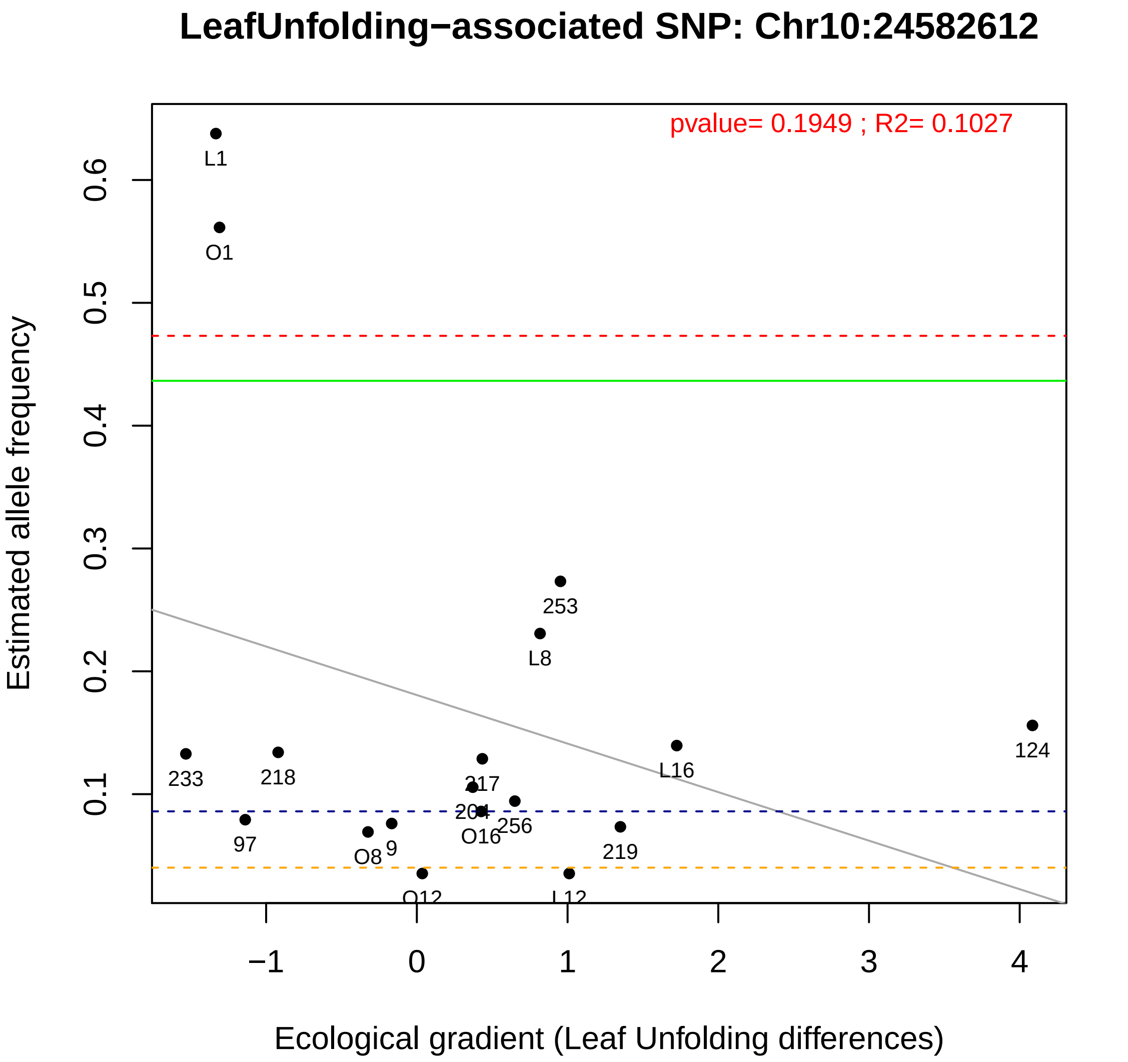

LeafUnfolding−associated SNP: Chr10:24582612
pvalue= 0.1949 ; R2= 0.1027
Estimated allele frequency
0.1
0.2
0.3
0.4
0.5
0.6
−1
0
1
2
3
4
Ecological gradient (Leaf Unfolding differences)
L1
O1
253
L8
233
218
97
O8
9
217
204
O16
256
219
L16
124

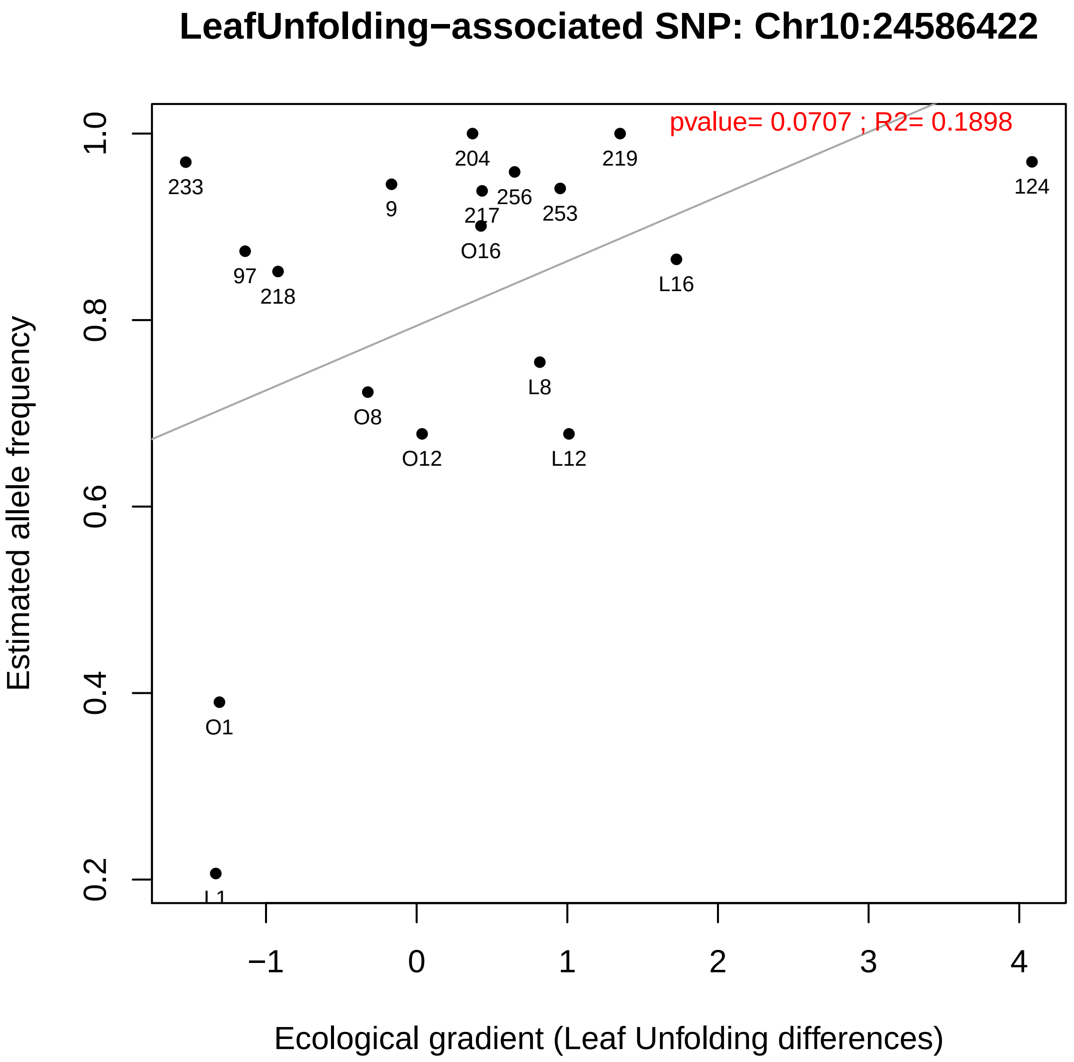

LeafUnfolding−associated SNP: Chr10:24586422
pvalue= 0.0707 ; R2= 0.1898
Estimated allele frequency
Ecological gradient (Leaf Unfolding differences)
0.2
0.4
0.6
0.8
1.0
−1
0
1
2
3
4
233
97
218
O1
9
O8
O12
204
217
O16
256
253
219
L8
L12
L16
124

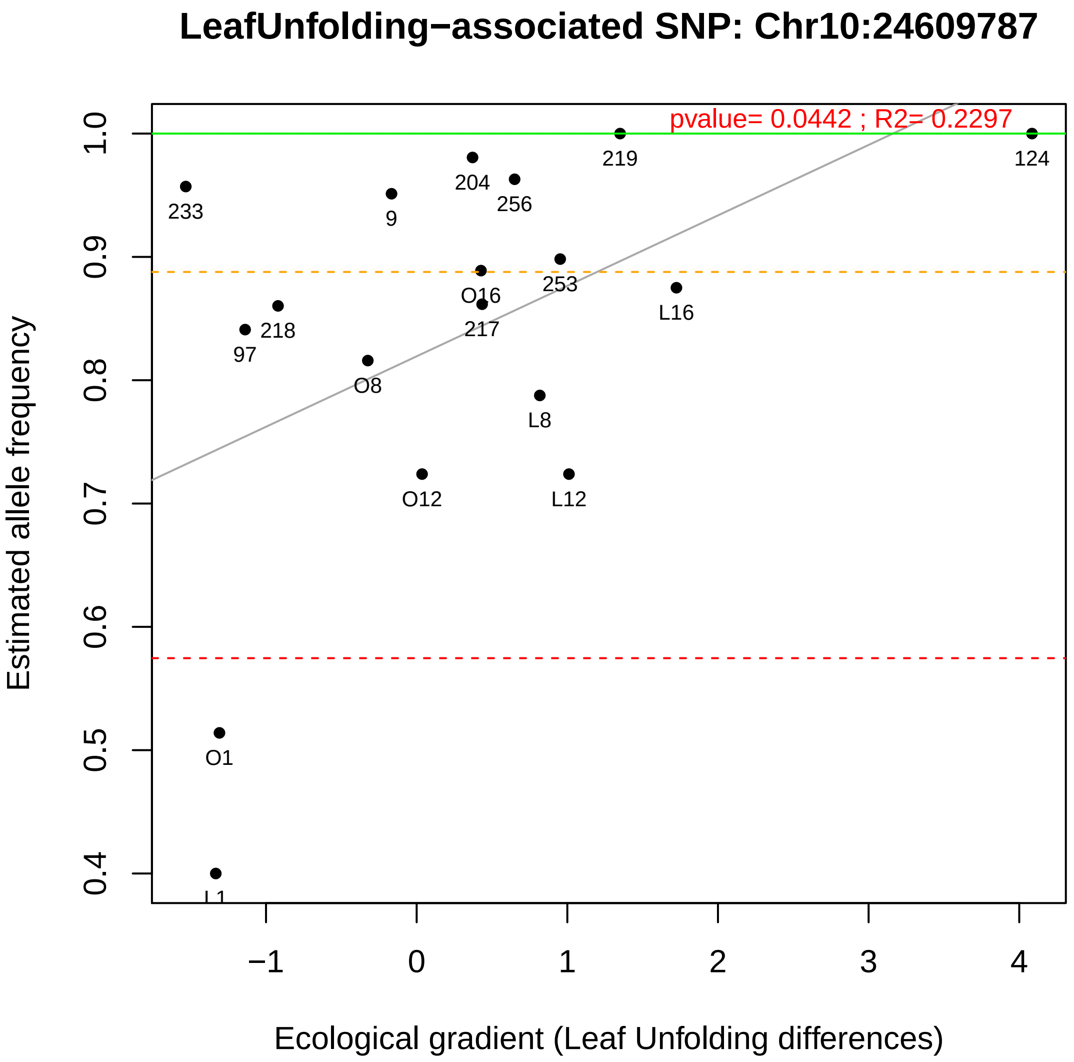

LeafUnfolding−associated SNP: Chr10:24609787
pvalue= 0.0442 ; R2= 0.2297
Estimated allele frequency
1.0
0.9
0.8
0.7
0.6
0.5
0.4
−1
0
1
2
3
4
Ecological gradient (Leaf Unfolding differences)
219
124
204
256
233
9
253
O16
217
L16
218
97
O8
L8
O12
L12
O1

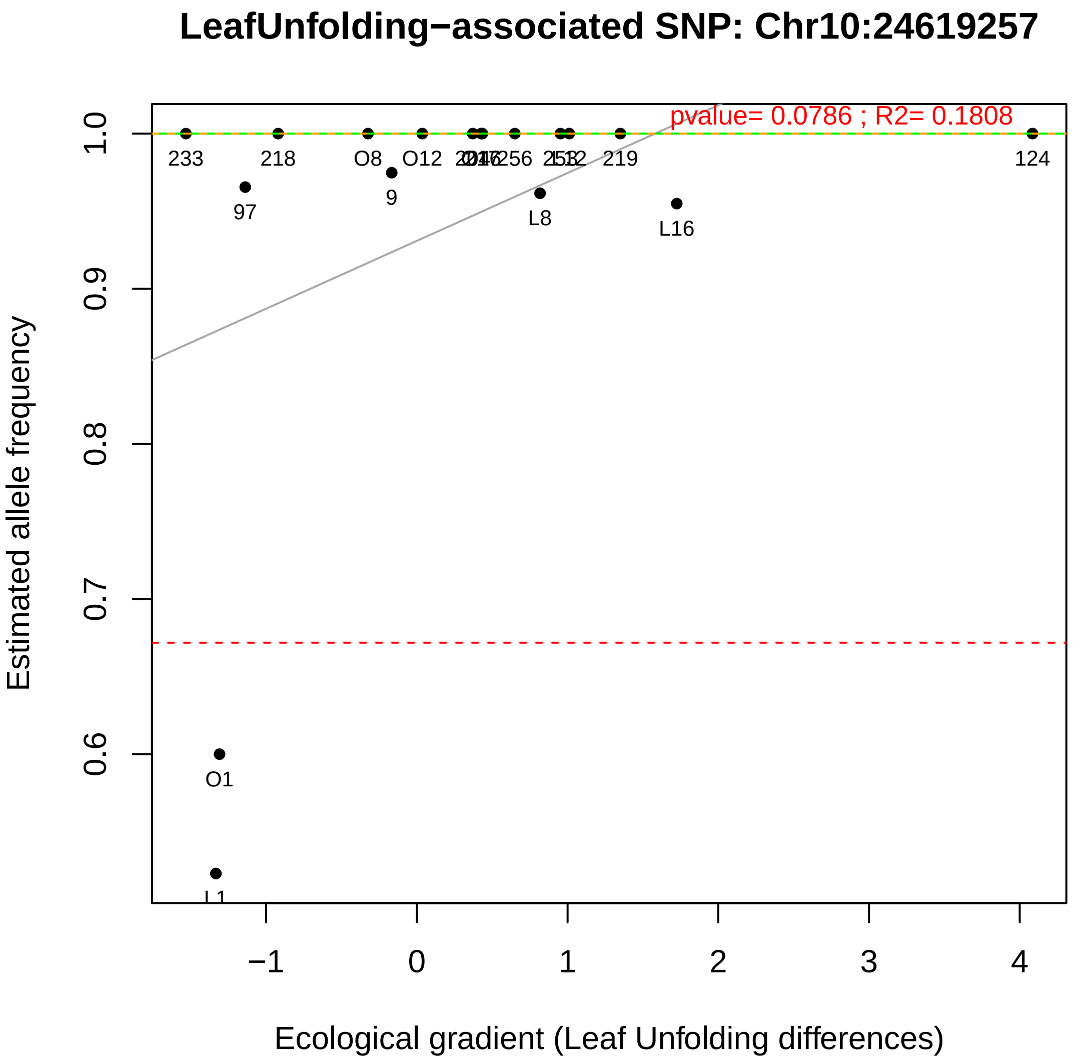
LeafUnfolding−associated SNP: Chr10:24619257
pvalue= 0.0786 ; R2= 0.1808
233
218
O8
O12
219
124
97
9
L8
L16
O1
Estimated allele frequency
1.0
0.9
0.8
0.7
0.6
−1
0
1
2
3
4
Ecological gradient (Leaf Unfolding differences)

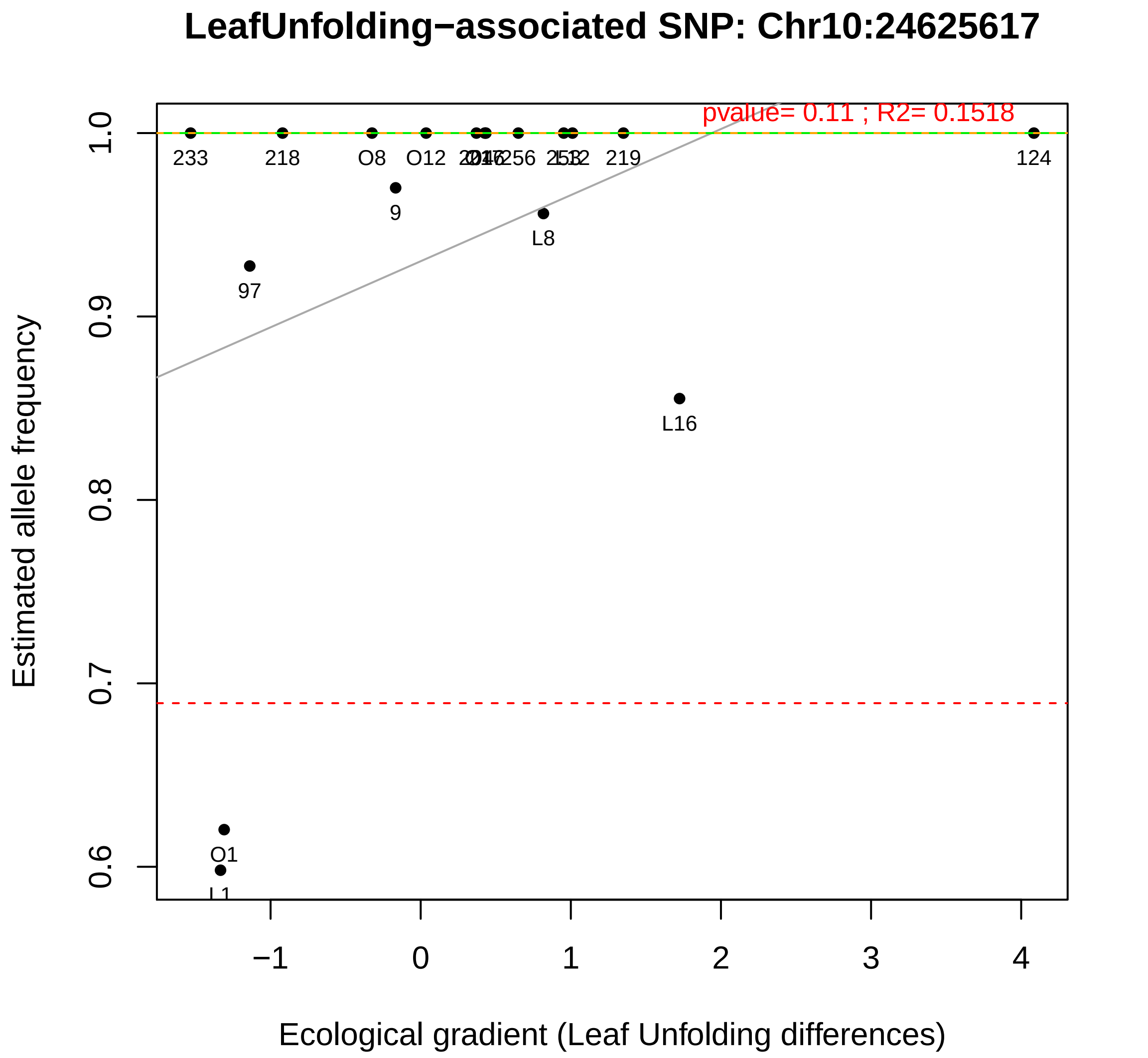

LeafUnfolding−associated SNP: Chr10:24625617
pvalue= 0.11 ; R2= 0.1518
233
218
O8
O12
219
124
9
L8
97
L16
O1
1.0
0.9
0.8
0.7
0.6
−1
0
1
2
3
4
Estimated allele frequency
Ecological gradient (Leaf Unfolding differences)

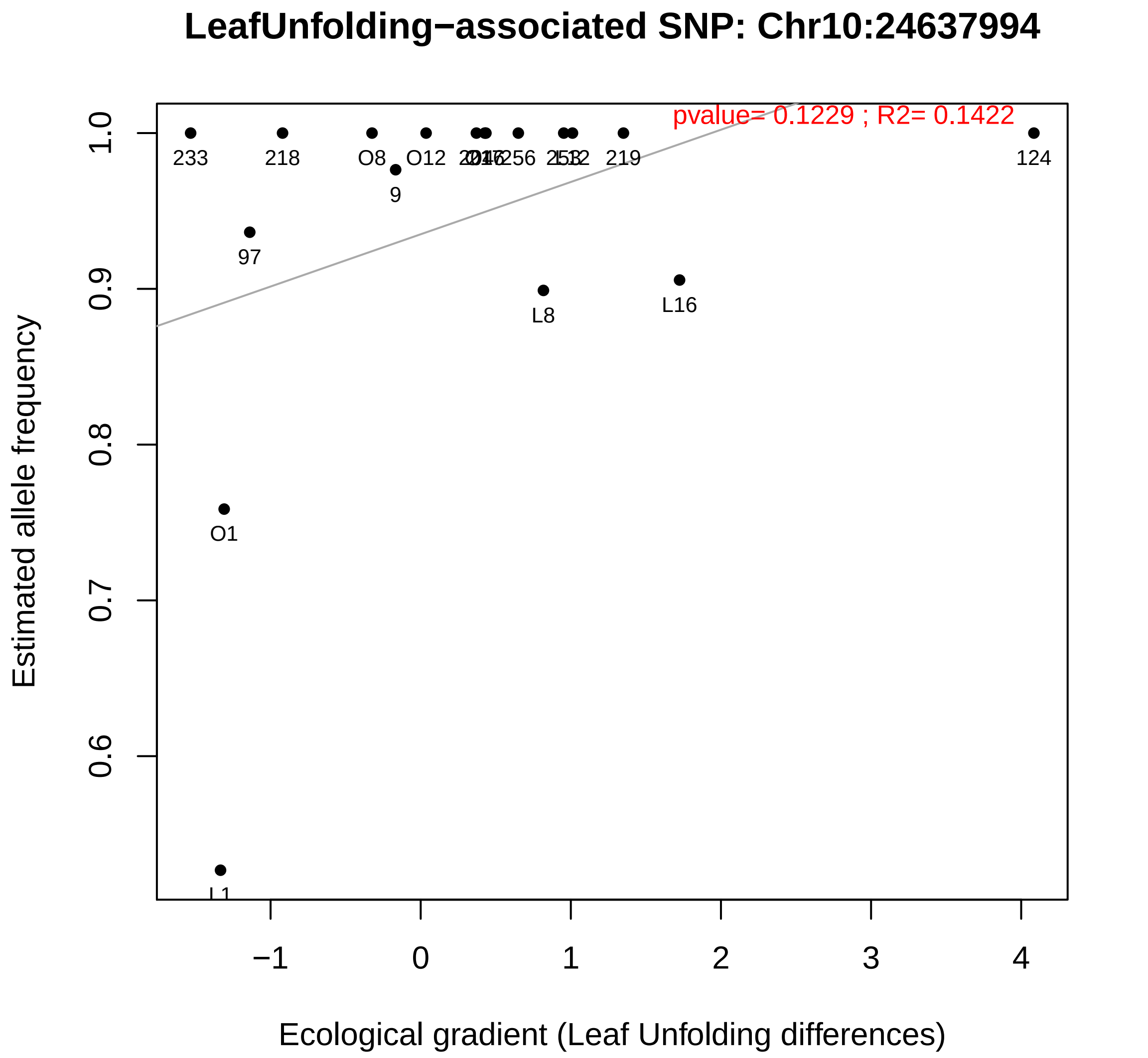
LeafUnfolding−associated SNP: Chr10:24637994
pvalue= 0.1229 ; R2= 0.1422
Estimated allele frequency
Ecological gradient (Leaf Unfolding differences)
233
218
O8
O12
219
124
9
97
L8
L16
O1
0.6
0.7
0.8
0.9
1.0
−1
0
1
2
3
4

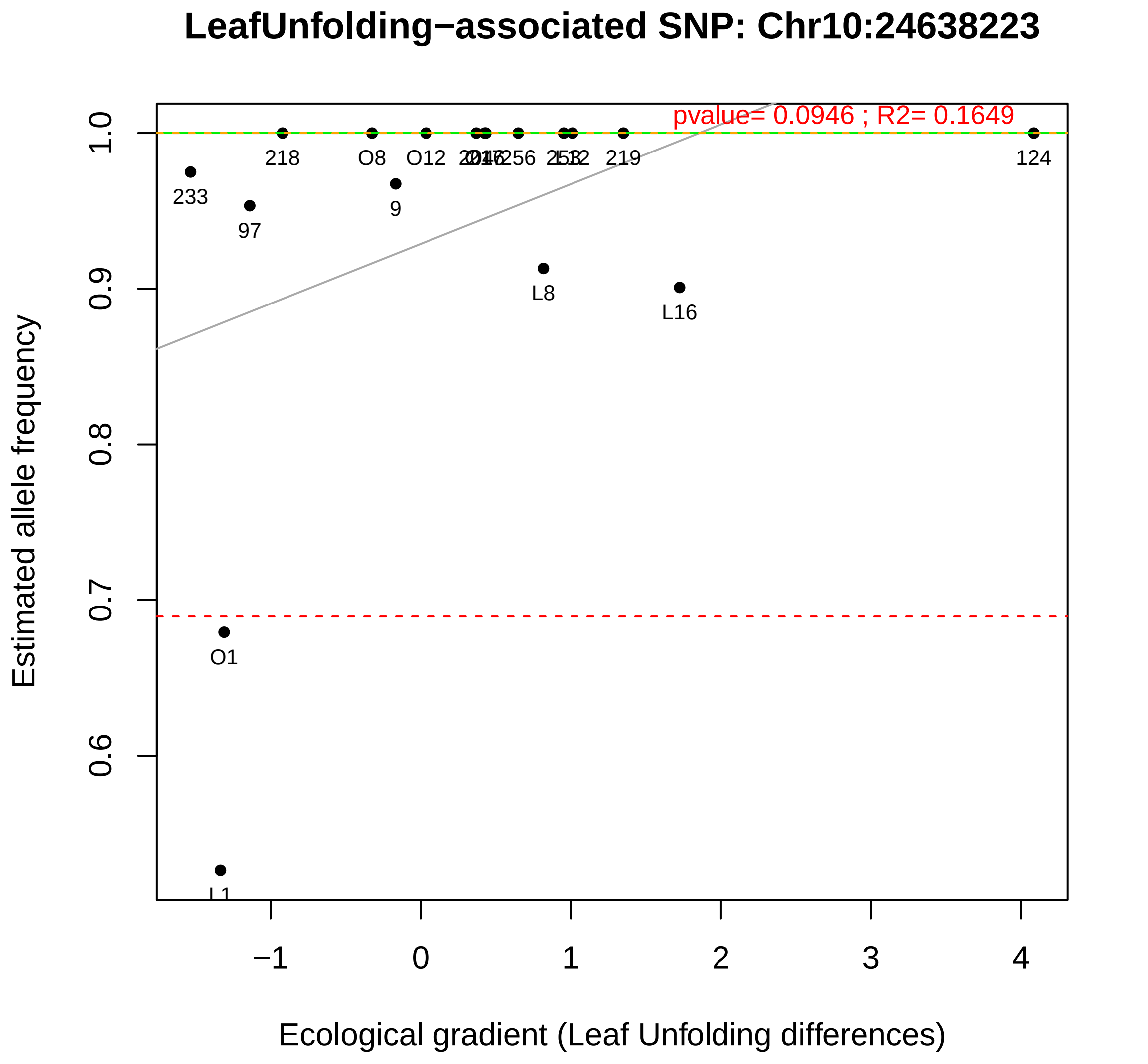

LeafUnfolding−associated SNP: Chr10:24638223
pvalue= 0.0946 ; R2= 0.1649
Estimated allele frequency
Ecological gradient (Leaf Unfolding differences)
1.0
0.9
0.8
0.7
0.6
−1
0
1
2
3
4
218
O8
O12
219
124
233
97
9
L8
L16
O1

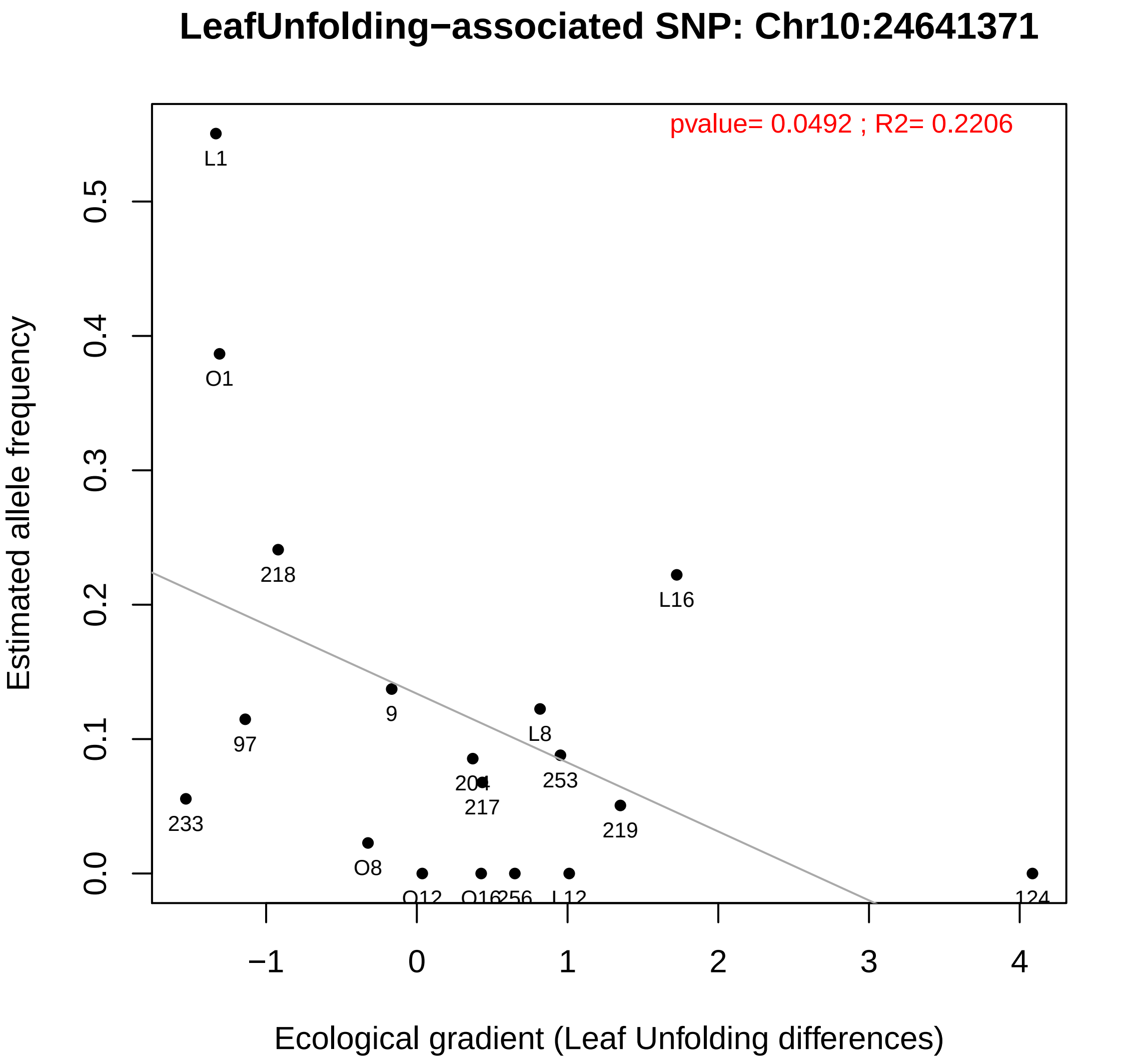
LeafUnfolding−associated SNP: Chr10:24641371
pvalue= 0.0492 ; R2= 0.2206
Estimated allele frequency
0.0
0.1
0.2
0.3
0.4
0.5
−1
0
1
2
3
4
Ecological gradient (Leaf Unfolding differences)
L1
O1
218
L16
9
97
L8
204
217
253
233
219
O8
124

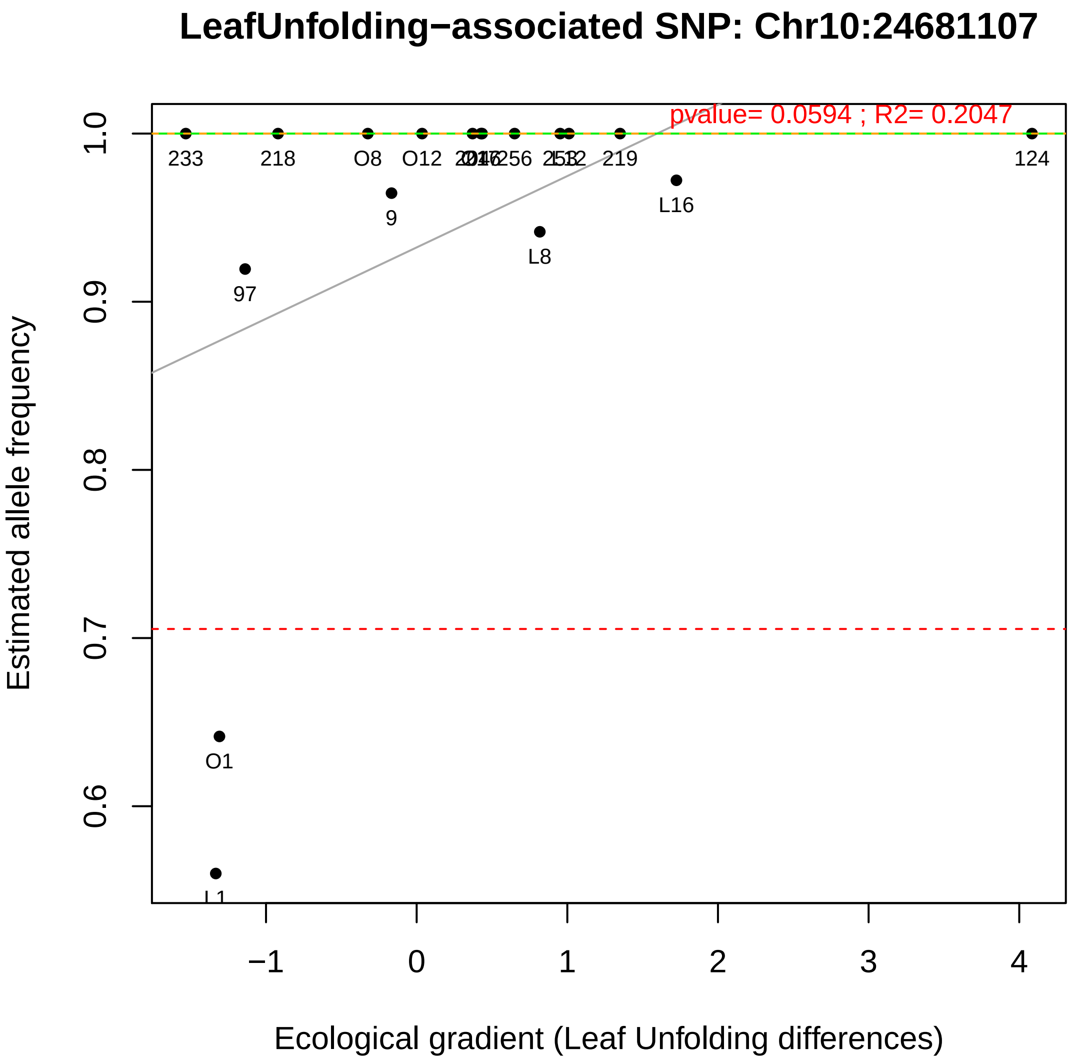
LeafUnfolding−associated SNP: Chr10:24681107
pvalue= 0.0594 ; R2= 0.2047
Estimated allele frequency
Ecological gradient (Leaf Unfolding differences)
1.0
0.9
0.8
0.7
0.6
−1
0
1
2
3
4
233
218
O8
O12
219
124
9
97
L8
L16
O1

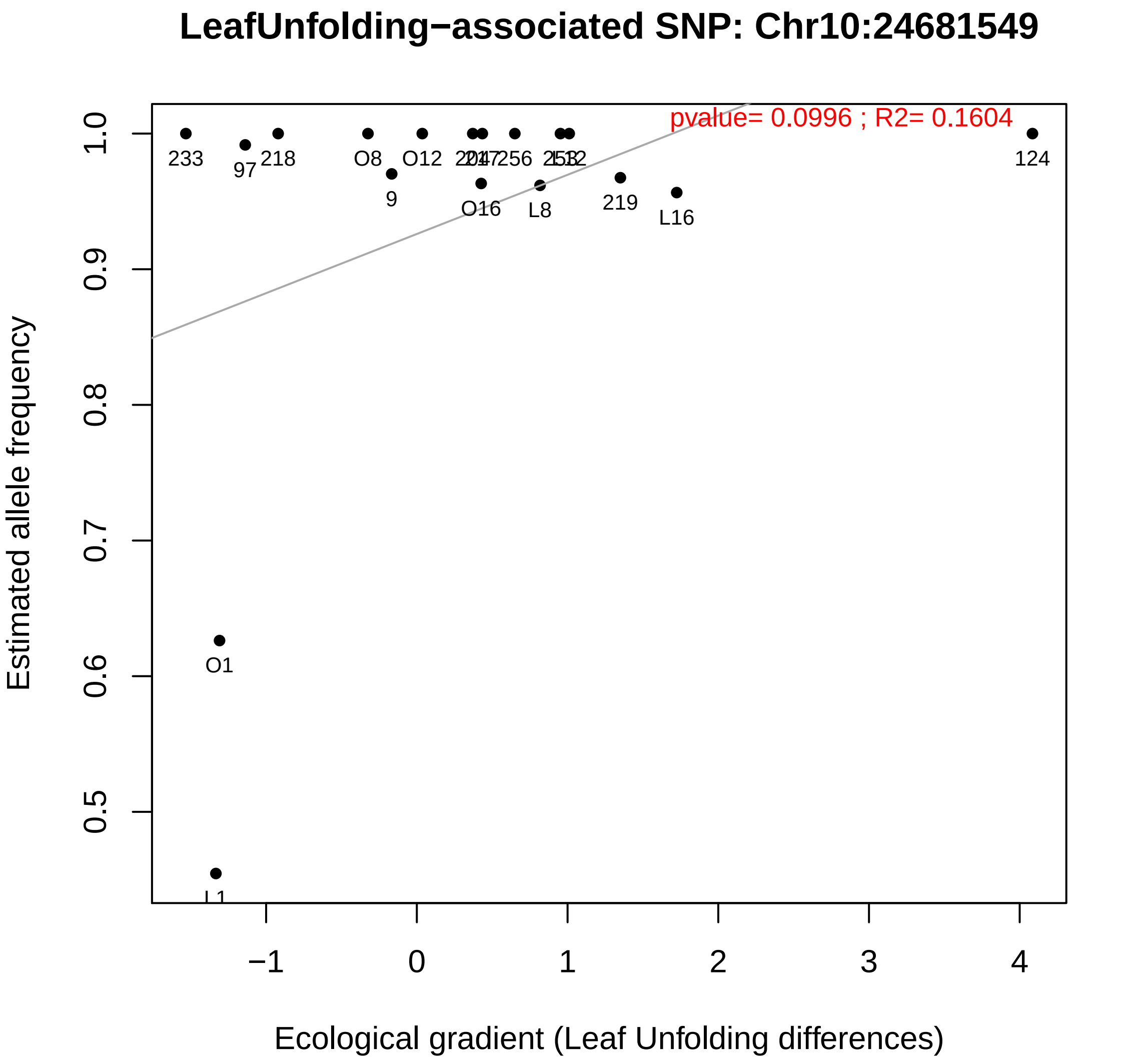

LeafUnfolding−associated SNP: Chr10:24681549
pvalue= 0.0996 ; R2= 0.1604
Estimated allele frequency
Ecological gradient (Leaf Unfolding differences)
0.5
0.6
0.7
0.8
0.9
1.0
−1
0
1
2
3
4
233
97
218
O8
9
O12
O16
256
L8
219
L16
124
O1

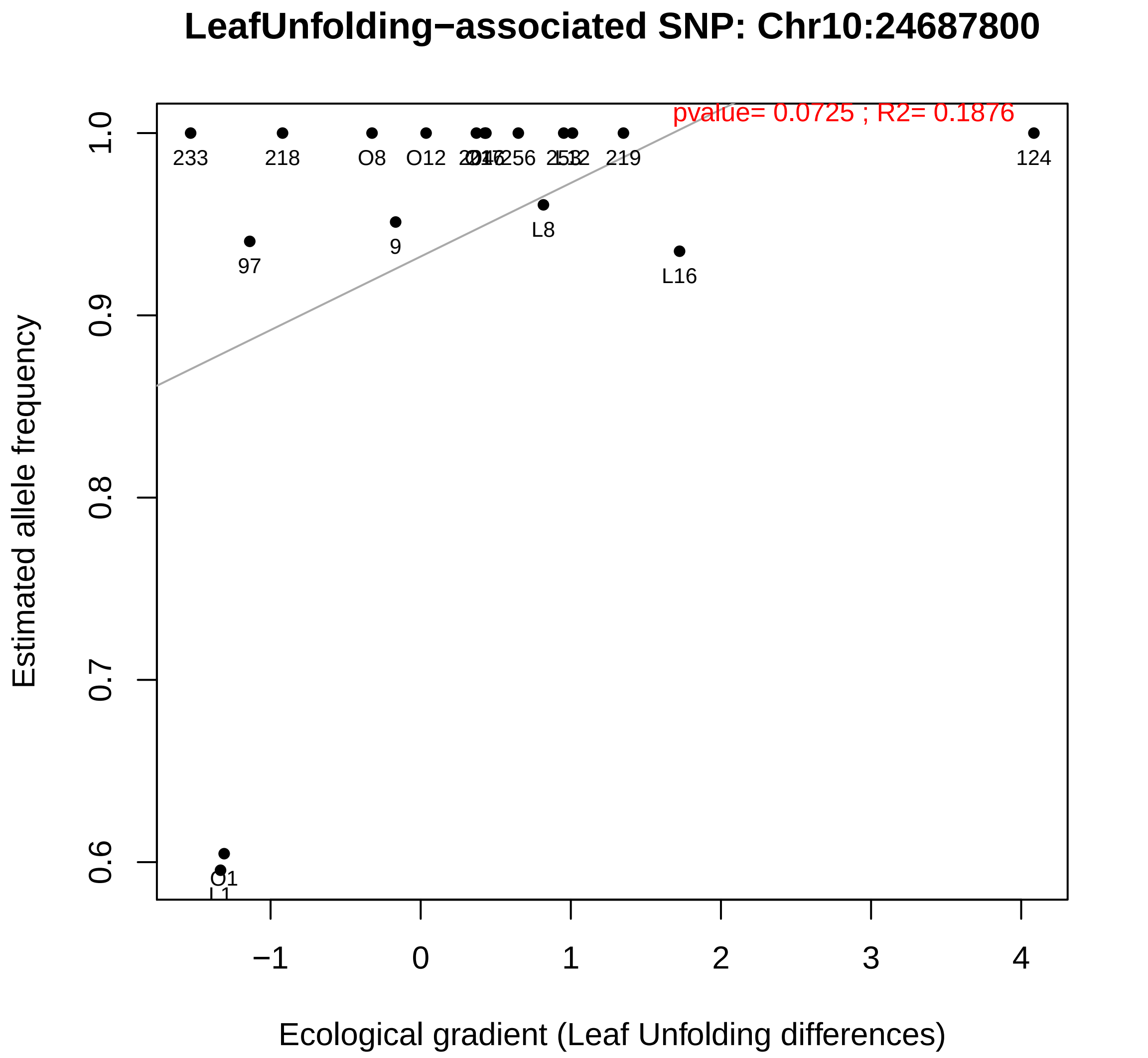
LeafUnfolding−associated SNP: Chr10:24687800
pvalue= 0.0725 ; R2= 0.1876
Estimated allele frequency
Ecological gradient (Leaf Unfolding differences)
233
218
O8
O12
256
219
124
97
9
L8
L16
O1
1.0
0.9
0.8
0.7
0.6
−1
0
1
2
3
4

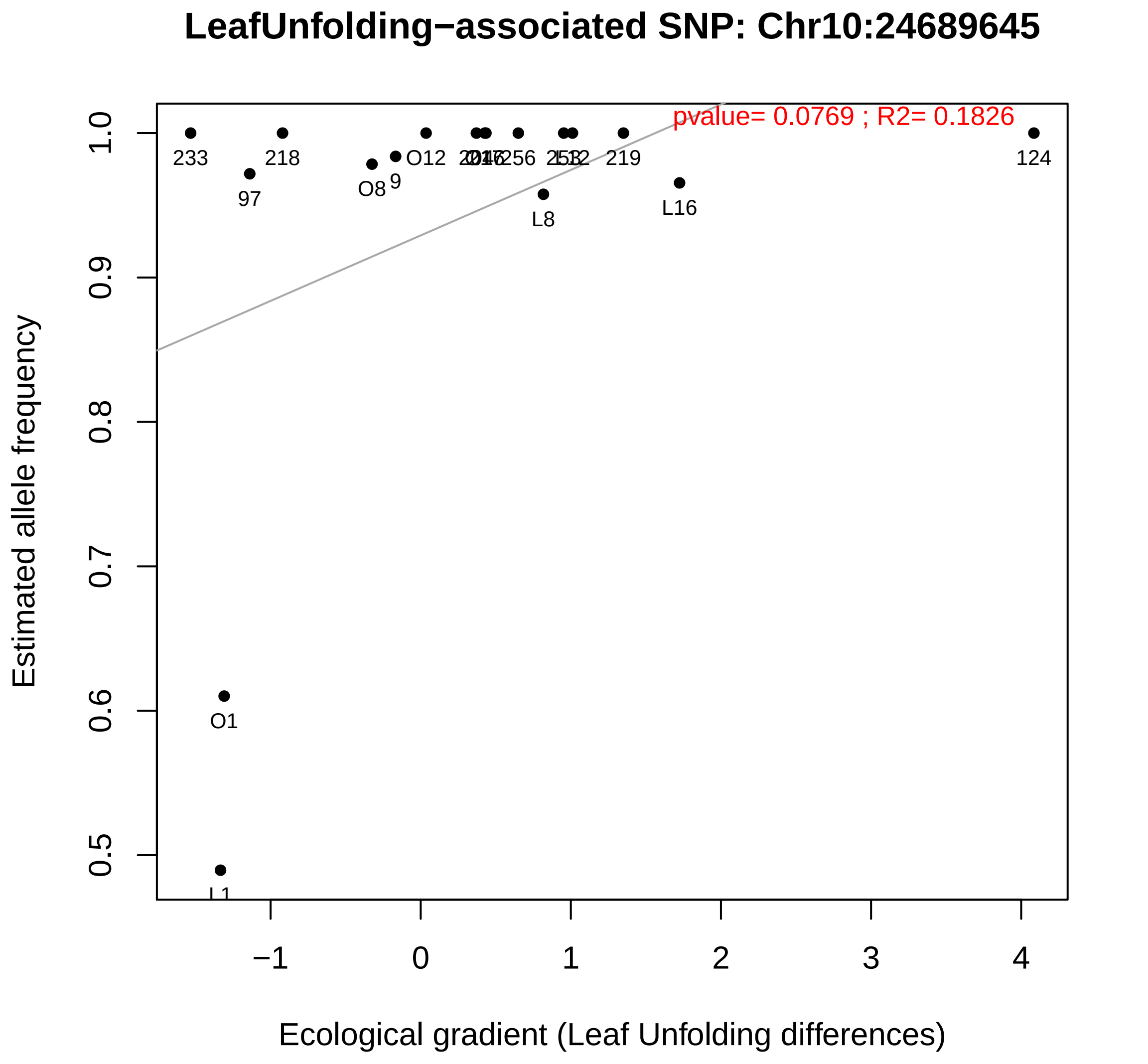

LeafUnfolding−associated SNP: Chr10:24689645
pvalue= 0.0769 ; R2= 0.1826
Estimated allele frequency
Ecological gradient (Leaf Unfolding differences)
1.0
0.9
0.8
0.7
0.6
0.5
−1
0
1
2
3
4
233
218
97
O8
9
O12
256
219
L8
L16
124
O1

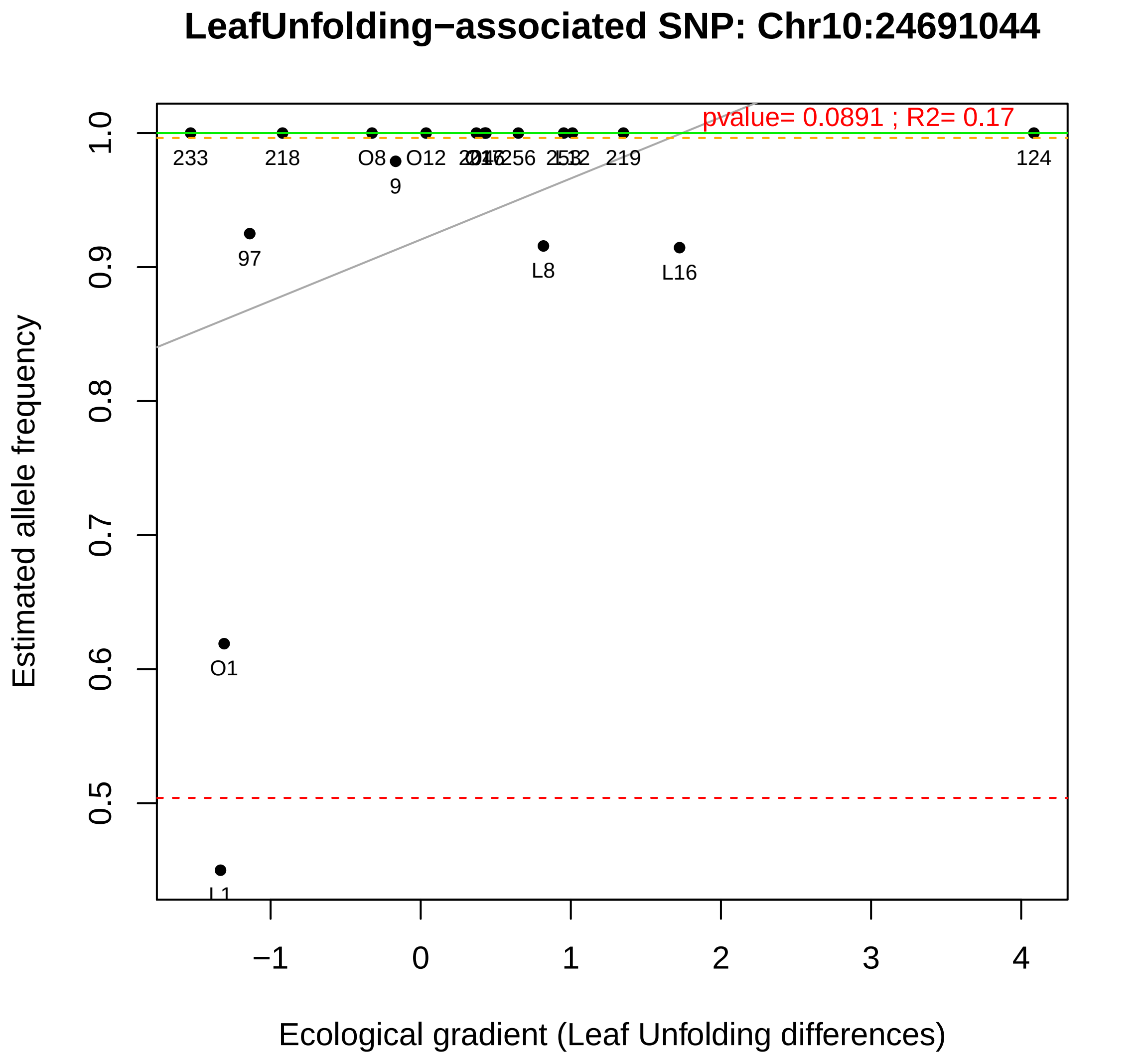
LeafUnfolding−associated SNP: Chr10:24691044
pvalue= 0.0891 ; R2= 0.17
233
218
O8
9
O12
219
124
97
L8
L16
O1
1.0
0.9
0.8
0.7
0.6
0.5
−1
0
1
2
3
4
Estimated allele frequency
Ecological gradient (Leaf Unfolding differences)

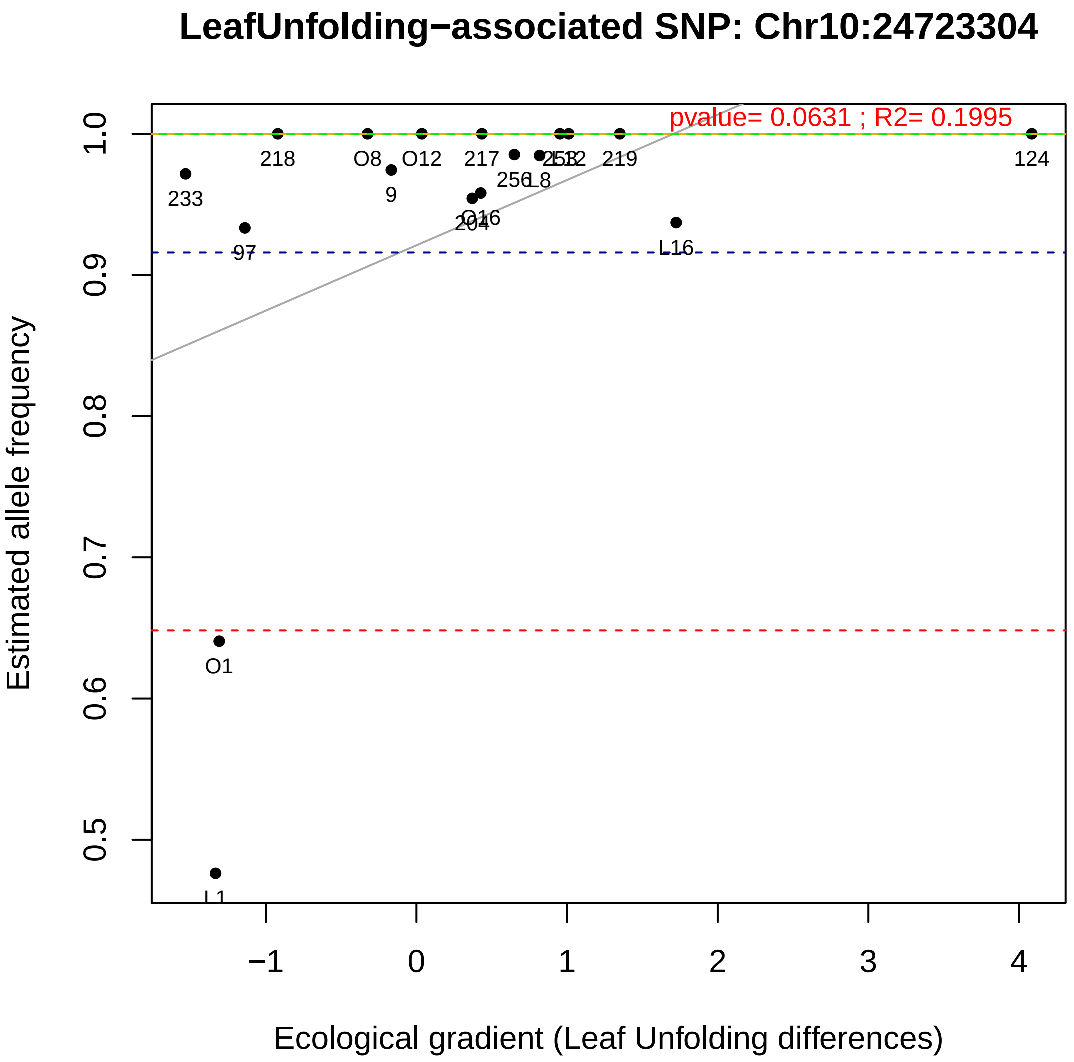

LeafUnfolding−associated SNP: Chr10:24723304
pvalue= 0.0631 ; R2= 0.1995
Estimated allele frequency
Ecological gradient (Leaf Unfolding differences)
1.0
0.9
0.8
0.7
0.6
0.5
−1
0
1
2
3
4
218
O8
O12
217
219
124
233
9
256
L8
97
L16
O1
204
O16

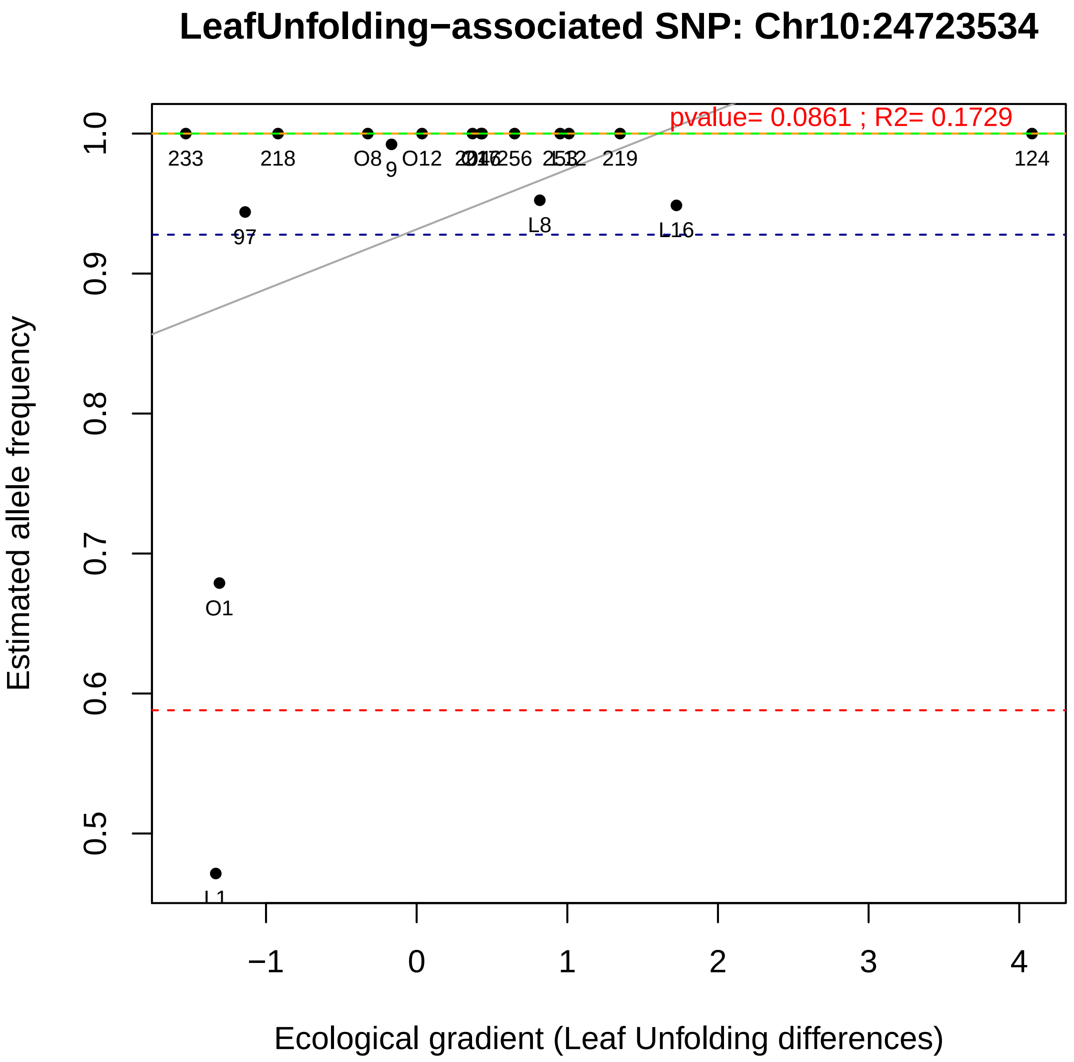

LeafUnfolding−associated SNP: Chr10:24723534
pvalue= 0.0861 ; R2= 0.1729
Estimated allele frequency
Ecological gradient (Leaf Unfolding differences)
1.0
0.9
0.8
0.7
0.6
0.5
−1
0
1
2
3
4
233
218
O8
9
O12
256
219
124
97
L8
L16
O1

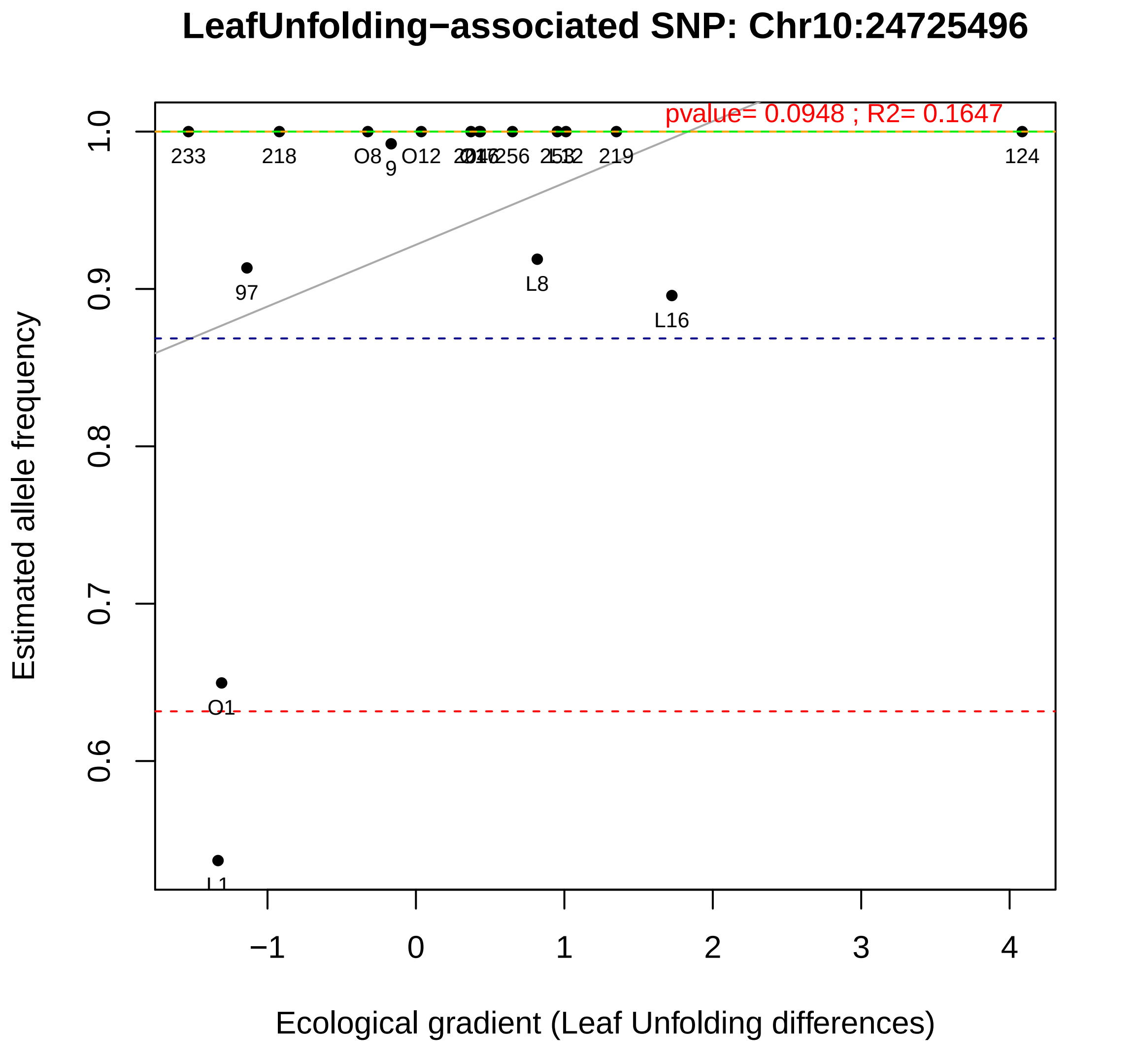

LeafUnfolding−associated SNP: Chr10:24725496
pvalue= 0.0948 ; R2= 0.1647
Estimated allele frequency
Ecological gradient (Leaf Unfolding differences)
233
218
O8
9
O12
256
219
124
97
L8
L16
O1
L1
1.0
0.9
0.8
0.7
0.6
−1
0
1
2
3
4

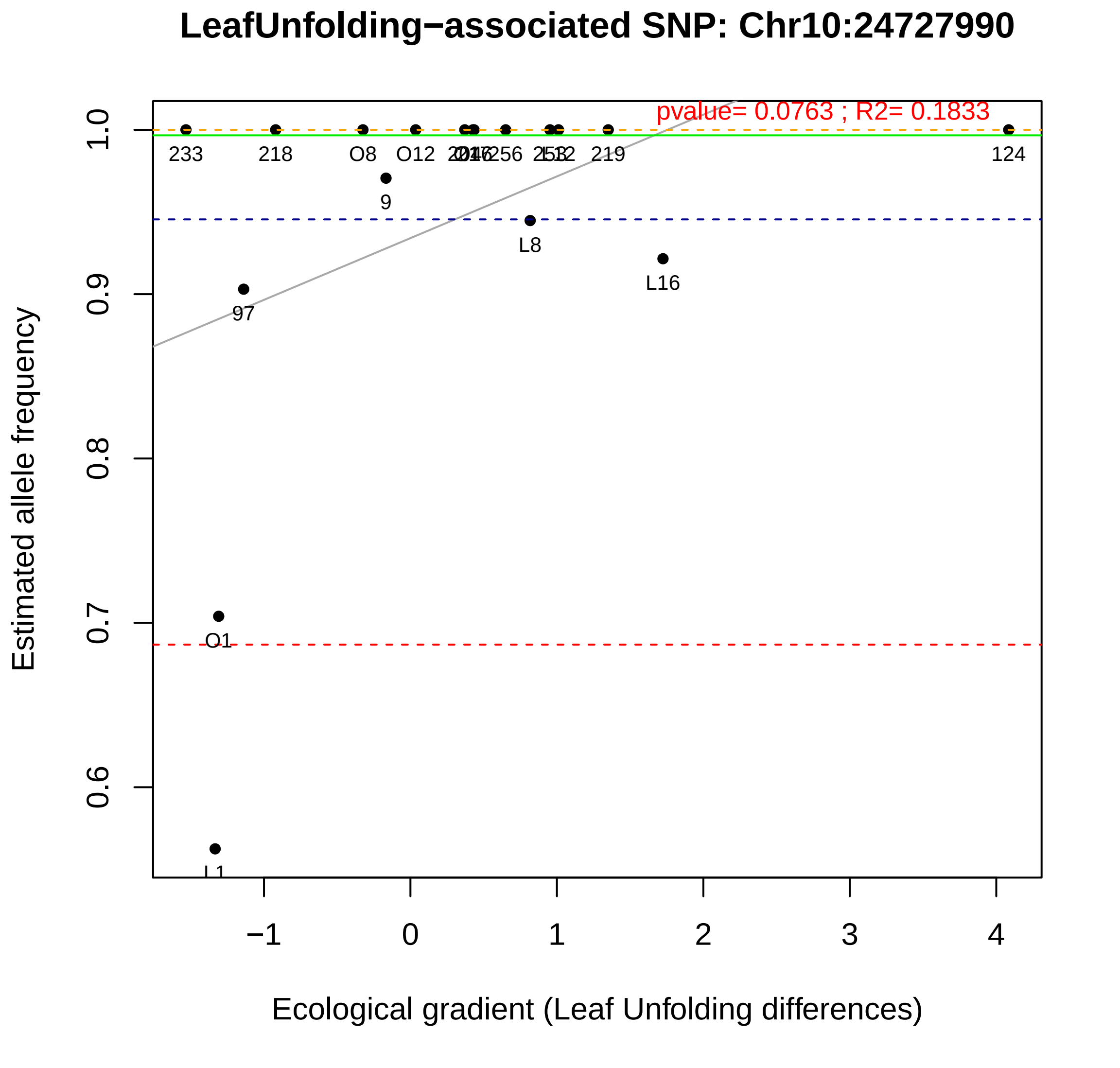

LeafUnfolding−associated SNP: Chr10:24727990
pvalue= 0.0763 ; R2= 0.1833
Estimated allele frequency
Ecological gradient (Leaf Unfolding differences)
1.0
0.9
0.8
0.7
0.6
−1
0
1
2
3
4
233
218
O8
O12
256
219
124
9
L8
L16
97
O1

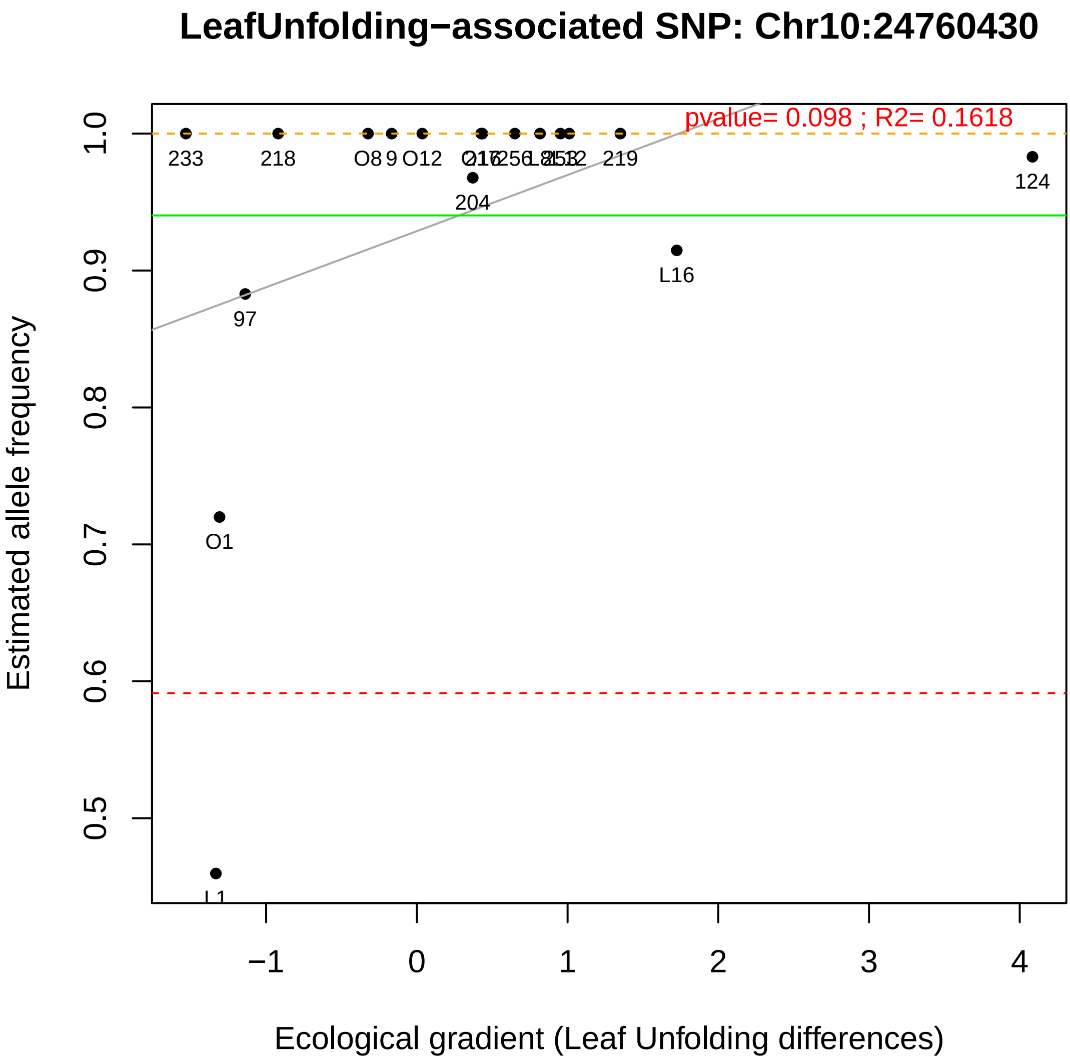
LeafUnfolding−associated SNP: Chr10:24760430
pvalue= 0.098 ; R2= 0.1618
Estimated allele frequency
Ecological gradient (Leaf Unfolding differences)
233
218
O8
9
O12
219
204
124
L16
97
O1
0.5
0.6
0.7
0.8
0.9
1.0
−1
0
1
2
3
4

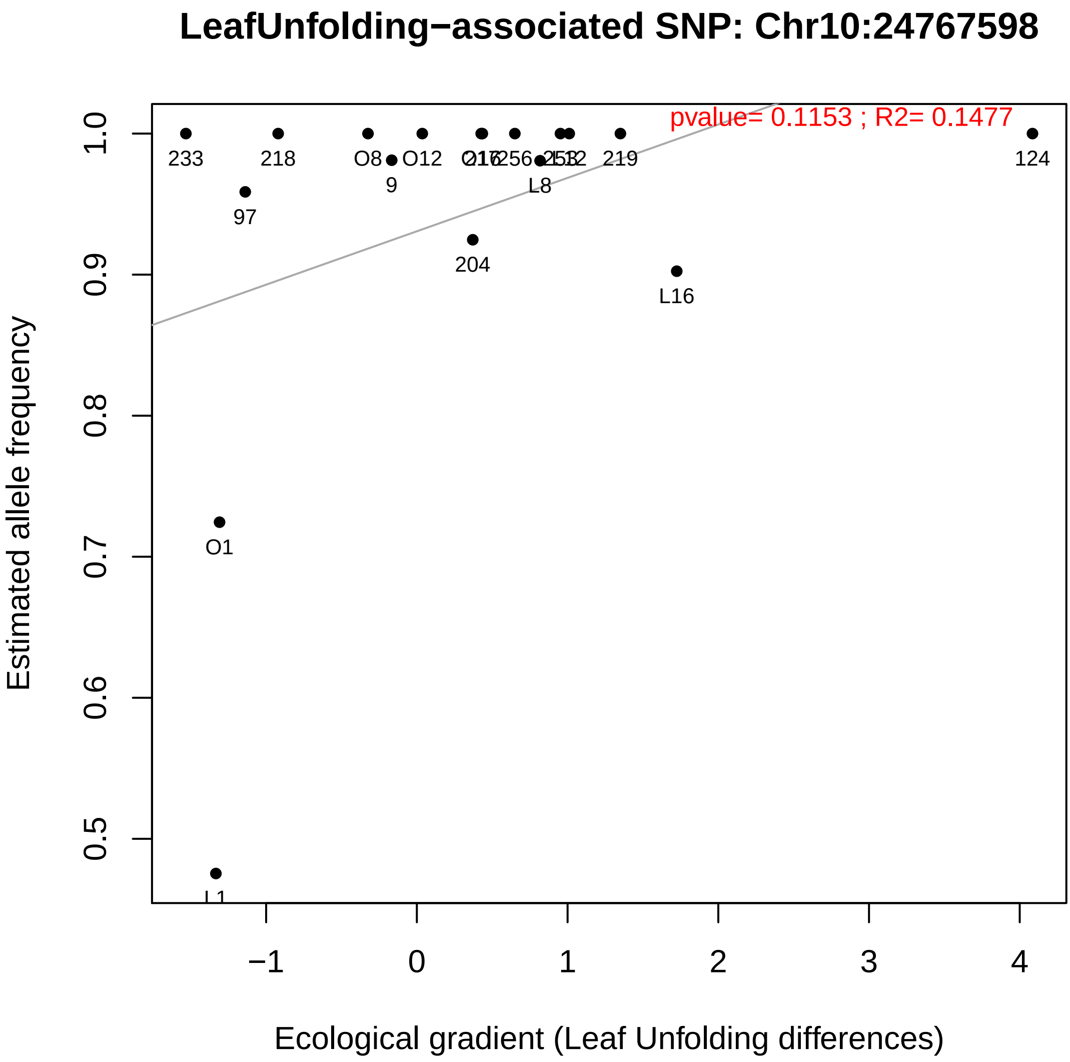

LeafUnfolding−associated SNP: Chr10:24767598
pvalue= 0.1153 ; R2= 0.1477
233
218
O8
O12
9
204
219
124
97
L8
L16
O1
L1
Estimated allele frequency
Ecological gradient (Leaf Unfolding differences)
1.0
0.9
0.8
0.7
0.6
0.5
−1
0
1
2
3
4

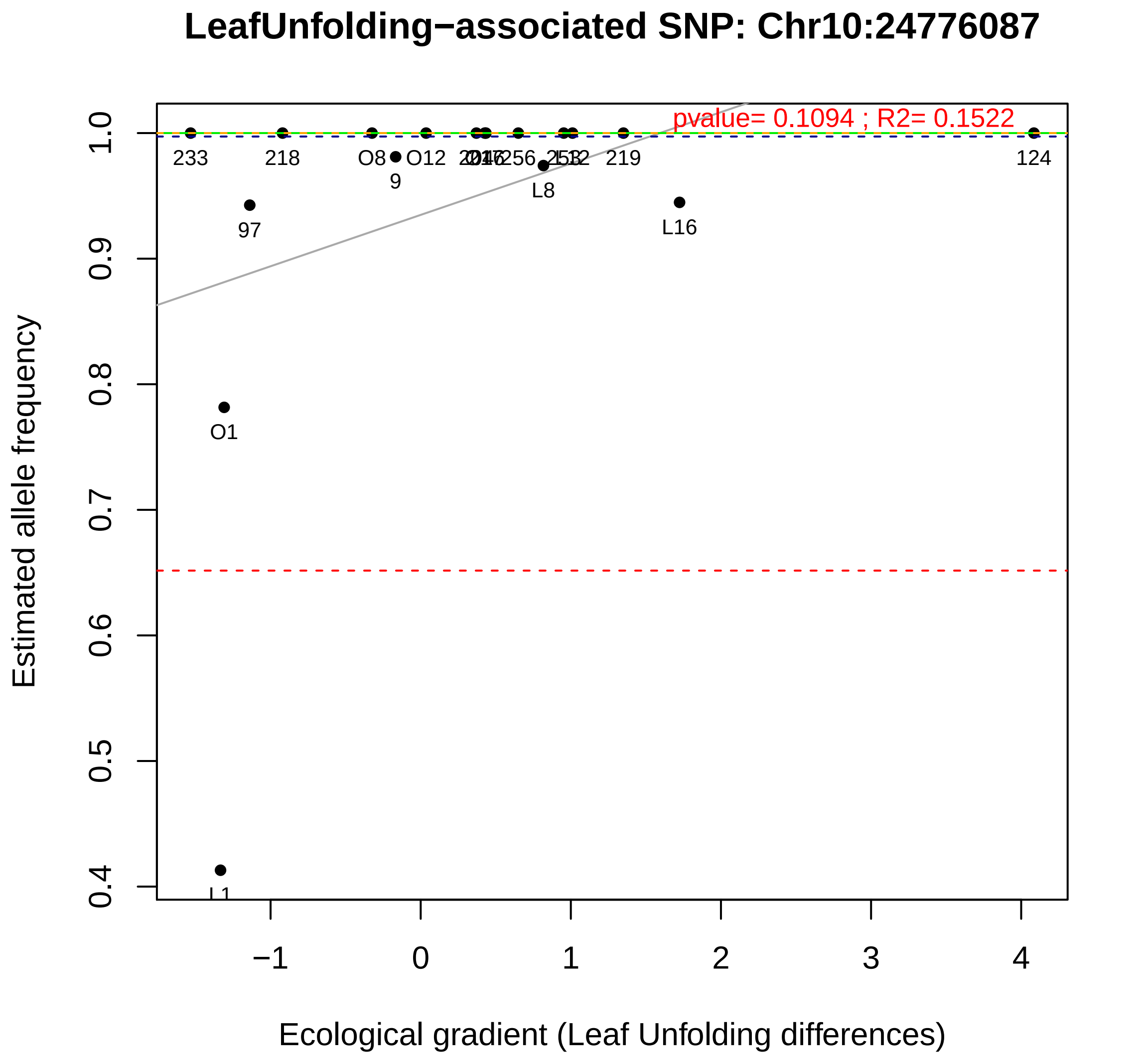

LeafUnfolding−associated SNP: Chr10:24776087
pvalue= 0.1094 ; R2= 0.1522
233
218
O8
O12
219
124
9
L8
97
L16
O1
Estimated allele frequency
0.4
0.5
0.6
0.7
0.8
0.9
1.0
−1
0
1
2
3
4
Ecological gradient (Leaf Unfolding differences)

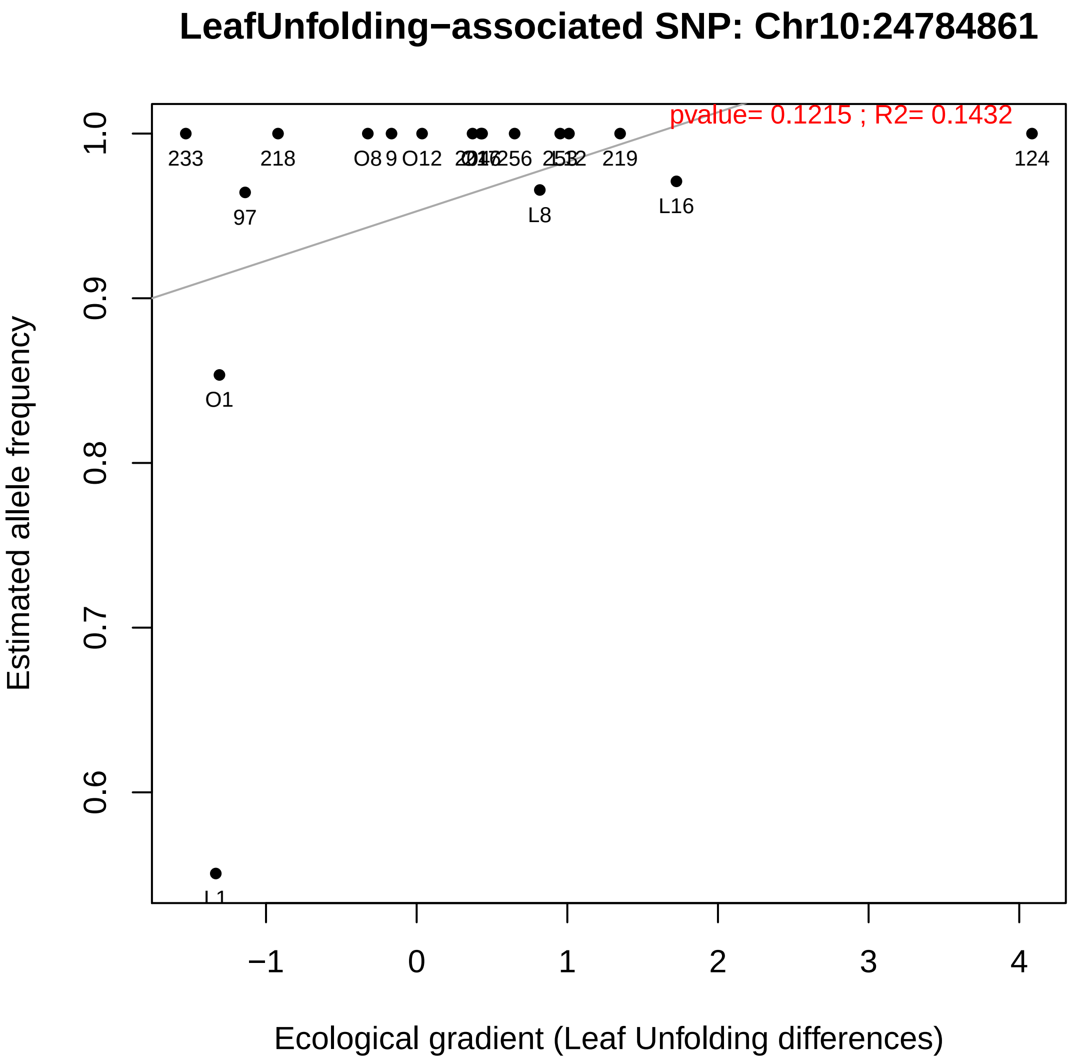

LeafUnfolding−associated SNP: Chr10:24784861
pvalue= 0.1215 ; R2= 0.1432
233
218
O8
9
O12
219
124
97
L8
L16
O1
1.0
0.9
0.8
0.7
0.6
−1
0
1
2
3
4
Estimated allele frequency
Ecological gradient (Leaf Unfolding differences)

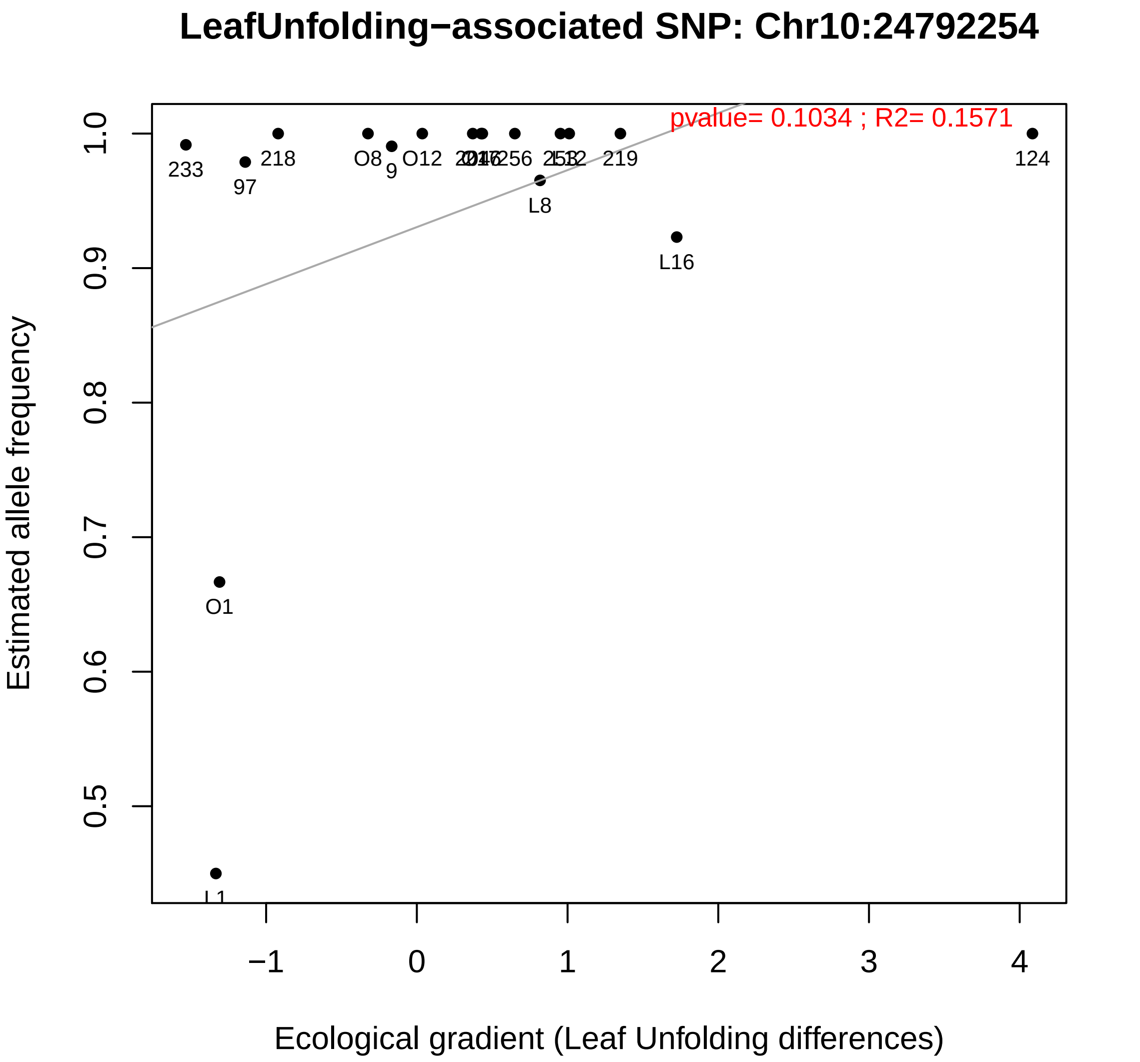

LeafUnfolding−associated SNP: Chr10:24792254
pvalue= 0.1034 ; R2= 0.1571
Estimated allele frequency
Ecological gradient (Leaf Unfolding differences)
1.0
0.9
0.8
0.7
0.6
0.5
−1
0
1
2
3
4
233
97
218
O8
9
O12
256
219
124
L8
L16
O1

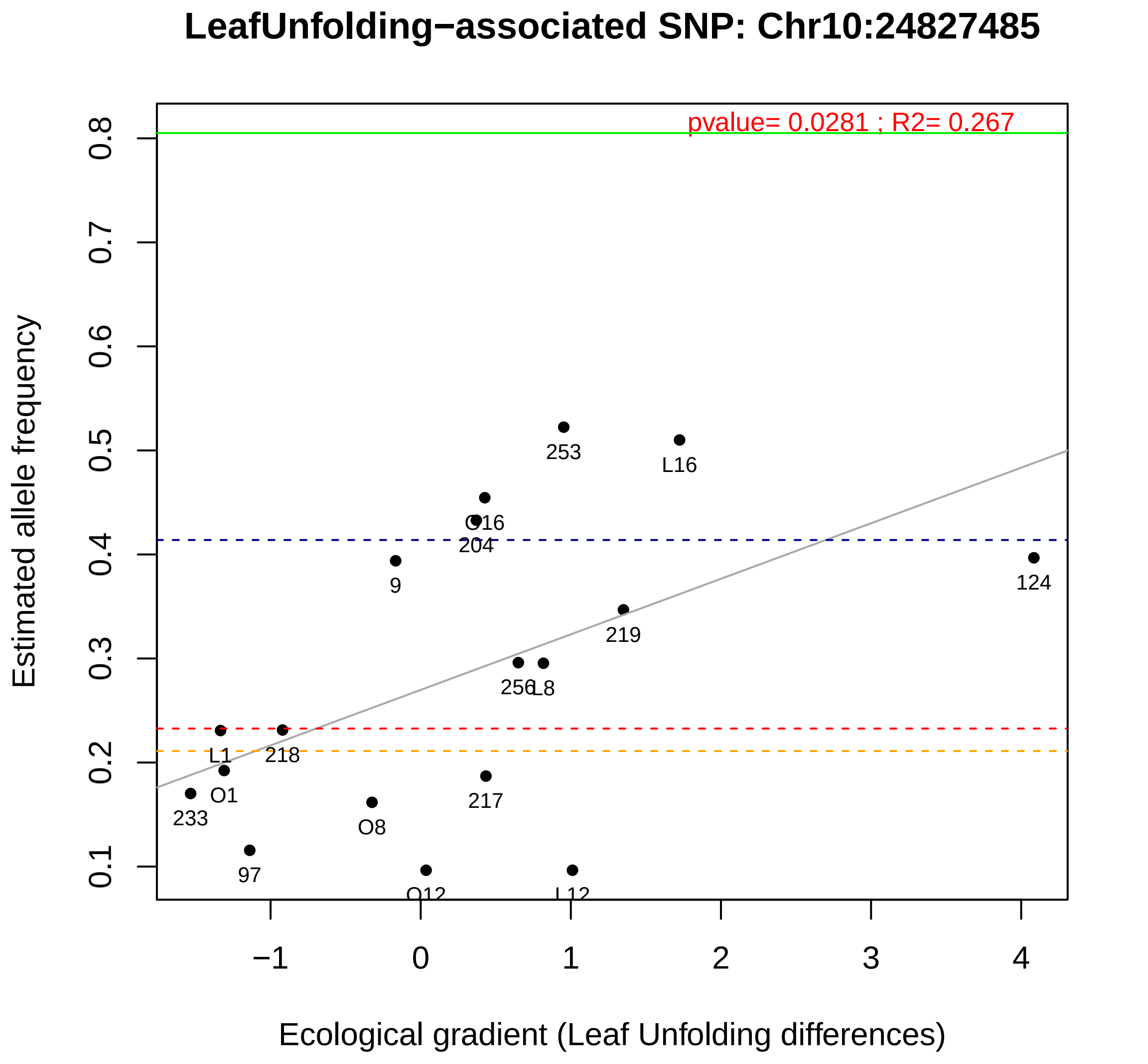
LeafUnfolding−associated SNP: Chr10:24827485
pvalue= 0.0281 ; R2= 0.267
Estimated allele frequency
0.1
0.2
0.3
0.4
0.5
0.6
0.7
0.8
−1
0
1
2
3
4
Ecological gradient (Leaf Unfolding differences)
253
L16
O16
204
9
124
219
256
L8
L1
218
O1
233
O8
217
97
O12
L12

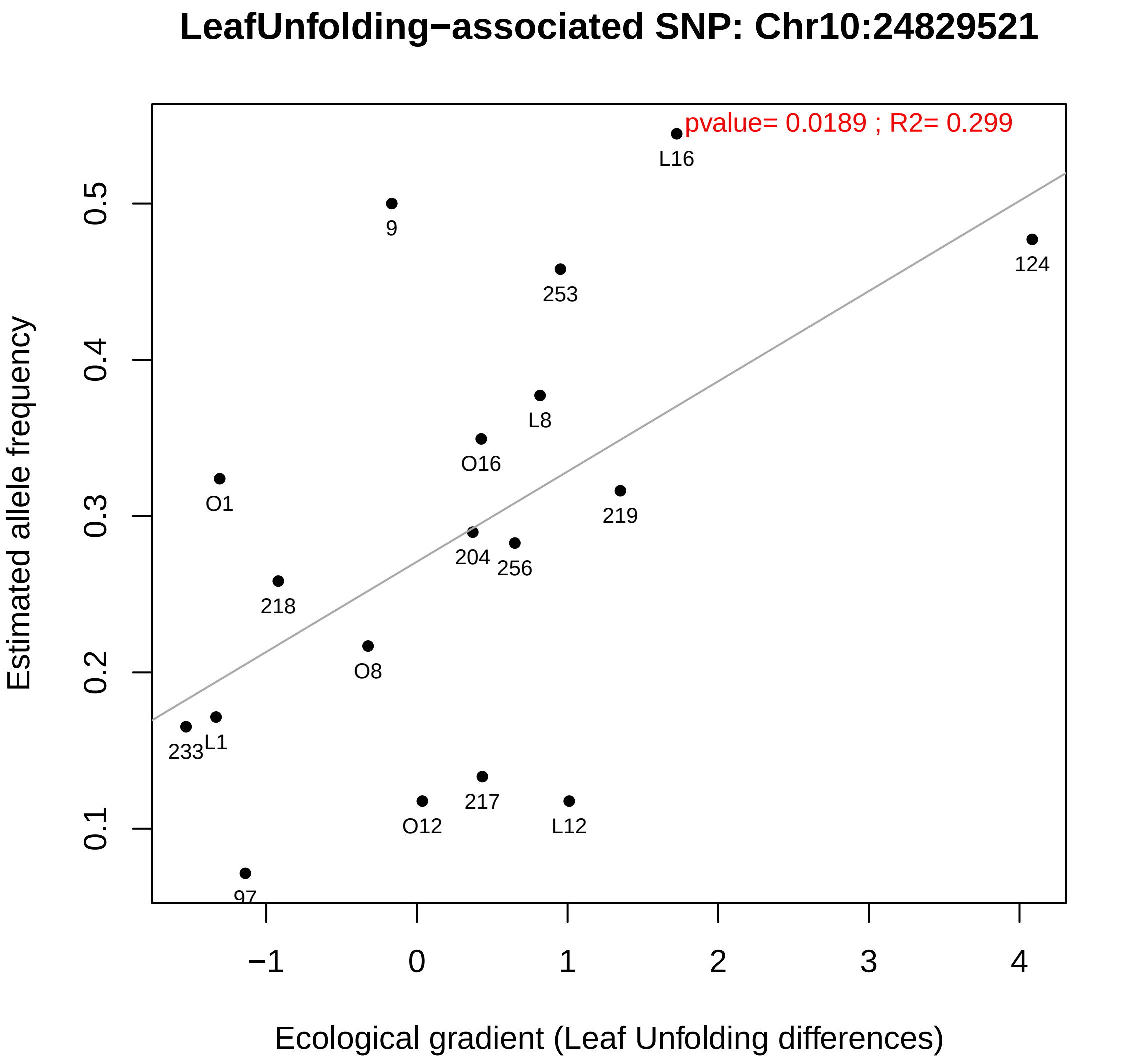
LeafUnfolding−associated SNP: Chr10:24829521
pvalue= 0.0189 ; R2= 0.299
Estimated allele frequency
0.1
0.2
0.3
0.4
0.5
−1
0
1
2
3
4
Ecological gradient (Leaf Unfolding differences)
L16
9
124
253
L8
O16
O1
219
204
256
218
O8
233
L1
217
O12
L12
97

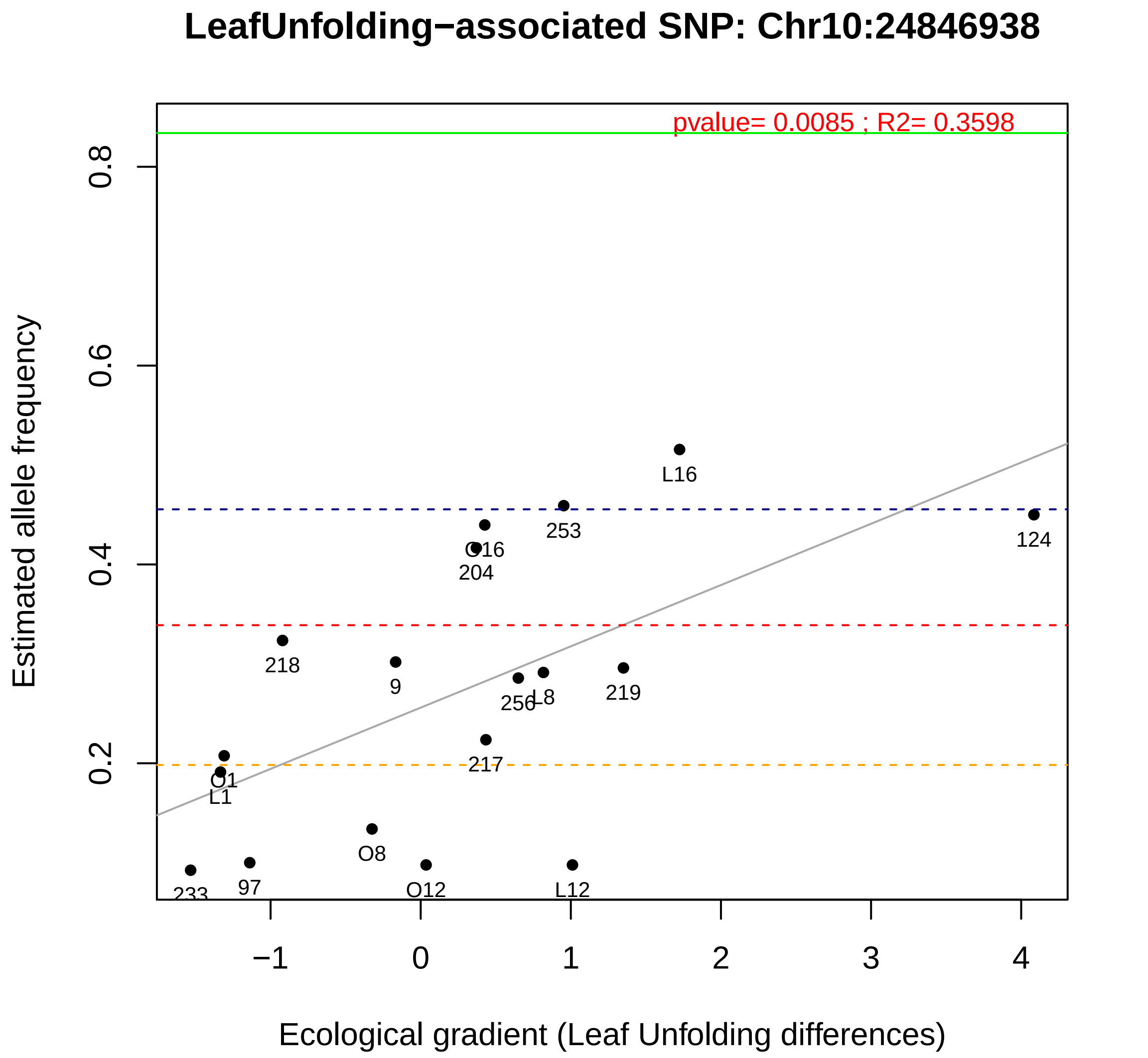

LeafUnfolding−associated SNP: Chr10:24846938
pvalue= 0.0085 ; R2= 0.3598
Estimated allele frequency
Ecological gradient (Leaf Unfolding differences)
0.2
0.4
0.6
0.8
−1
0
1
2
3
4
L16
253
124
O16
204
218
9
256
L8
219
217
O1
L1
O8
233
97
O12
L12

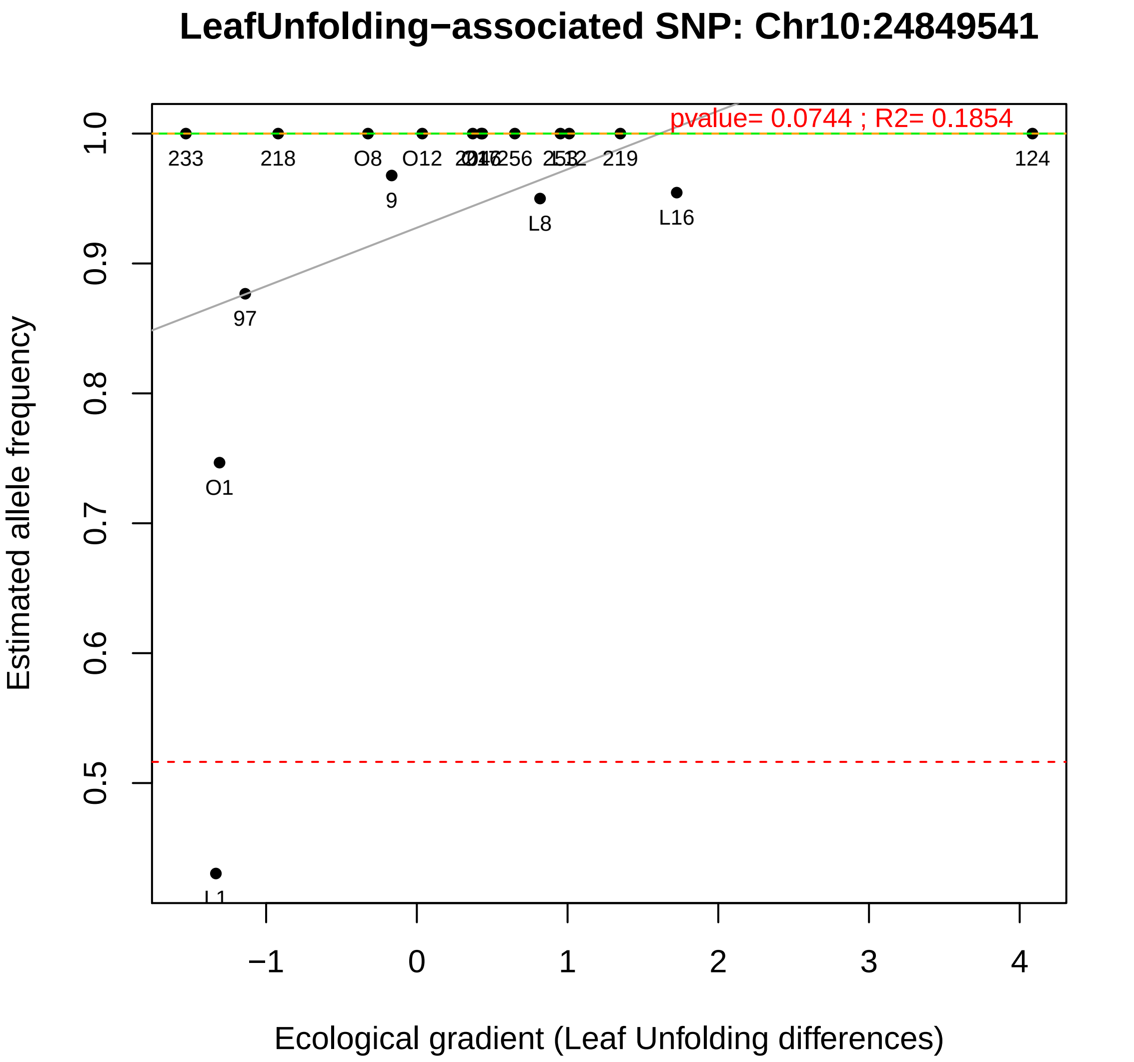

LeafUnfolding−associated SNP: Chr10:24849541
pvalue= 0.0744 ; R2= 0.1854
Estimated allele frequency
Ecological gradient (Leaf Unfolding differences)
233
218
O8
O12
219
124
9
L8
L16
97
O1
1.0
0.9
0.8
0.7
0.6
0.5
−1
0
1
2
3
4

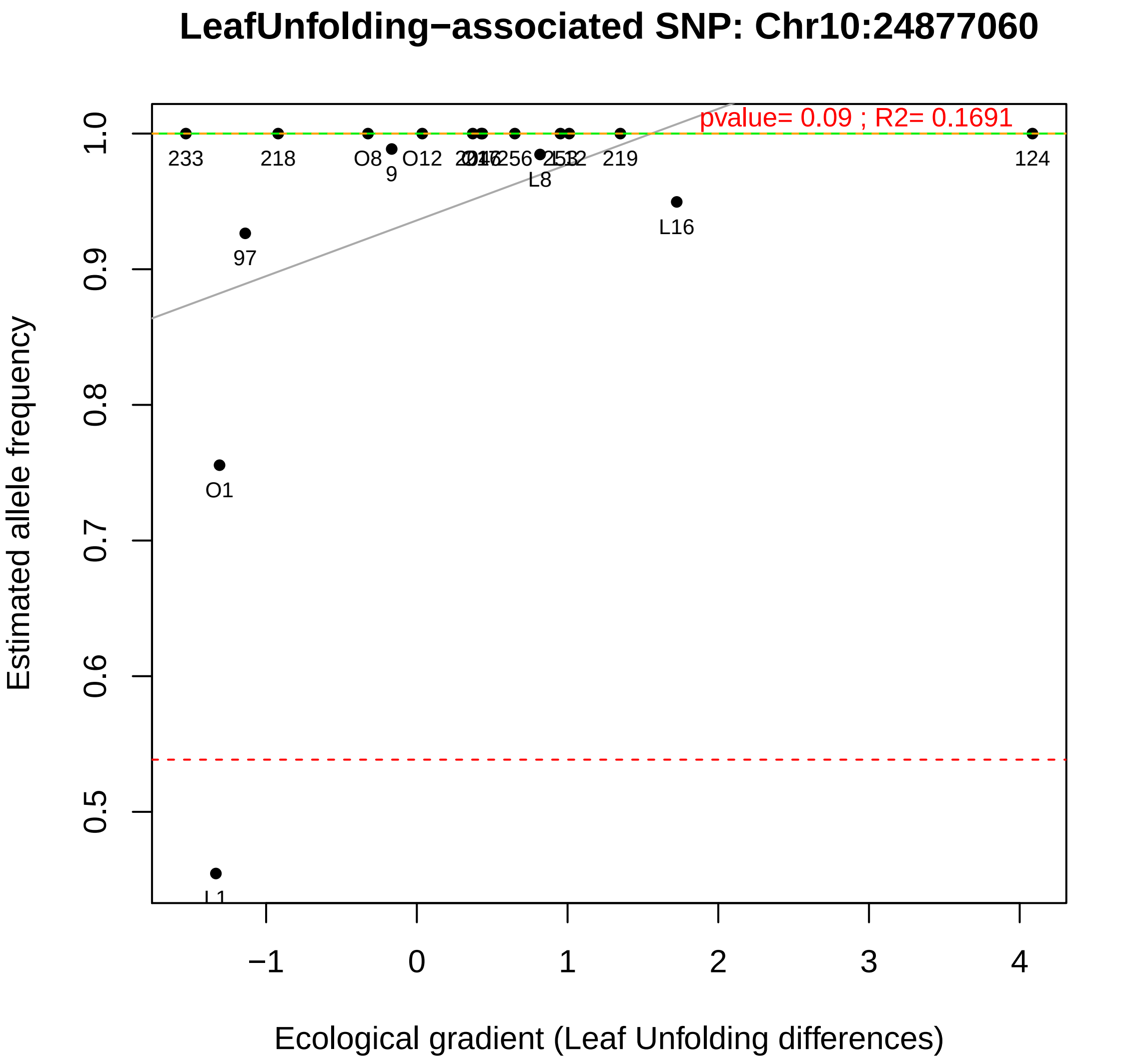

LeafUnfolding−associated SNP: Chr10:24877060
pvalue= 0.09 ; R2= 0.1691
Estimated allele frequency
Ecological gradient (Leaf Unfolding differences)
233
218
O8
9
O12
219
124
L16
97
O1
L8
L1

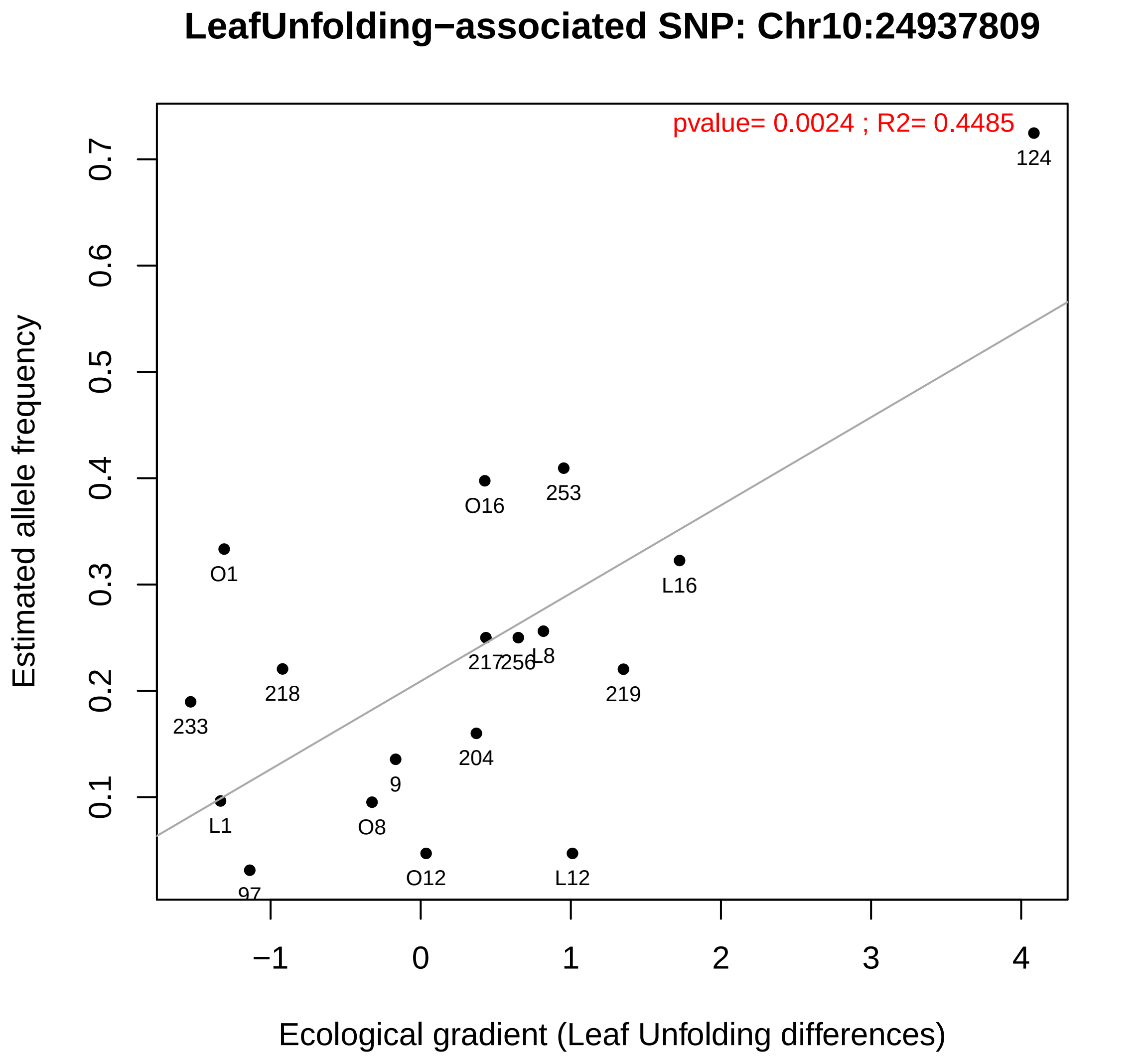

LeafUnfolding−associated SNP: Chr10:24937809
pvalue= 0.0024 ; R2= 0.4485
Estimated allele frequency
Ecological gradient (Leaf Unfolding differences)
0.1
0.2
0.3
0.4
0.5
0.6
0.7
−1
0
1
2
3
4
124
O16
253
O1
L16
217
256
L8
218
219
233
204
9
L1
O8
O12
L12
97

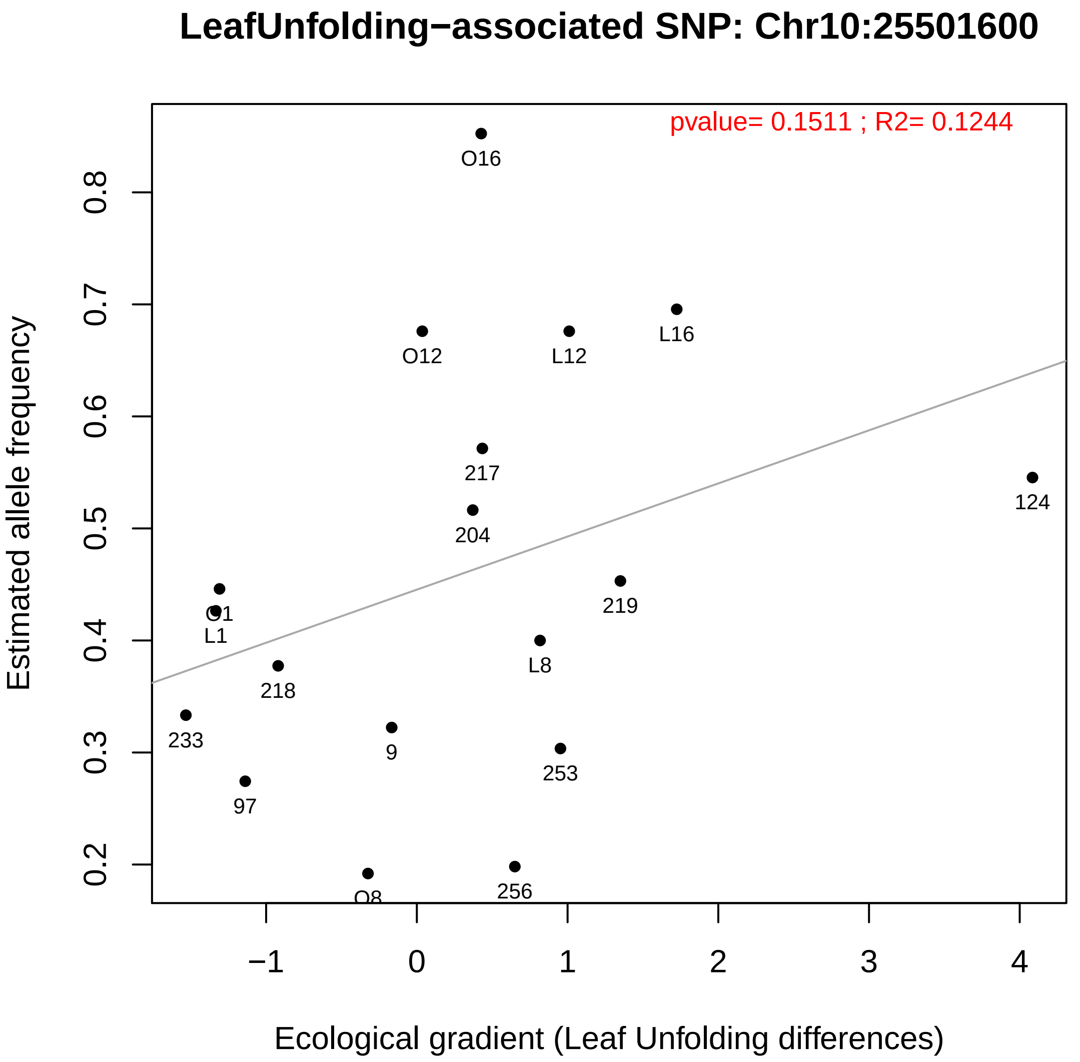
LeafUnfolding−associated SNP: Chr10:25501600
pvalue= 0.1511 ; R2= 0.1244
O16
O12
L12
L16
217
124
204
219
O1
L1
L8
218
233
9
253
97
O8
256
Estimated allele frequency
0.2
0.3
0.4
0.5
0.6
0.7
0.8
−1
0
1
2
3
4
Ecological gradient (Leaf Unfolding differences)

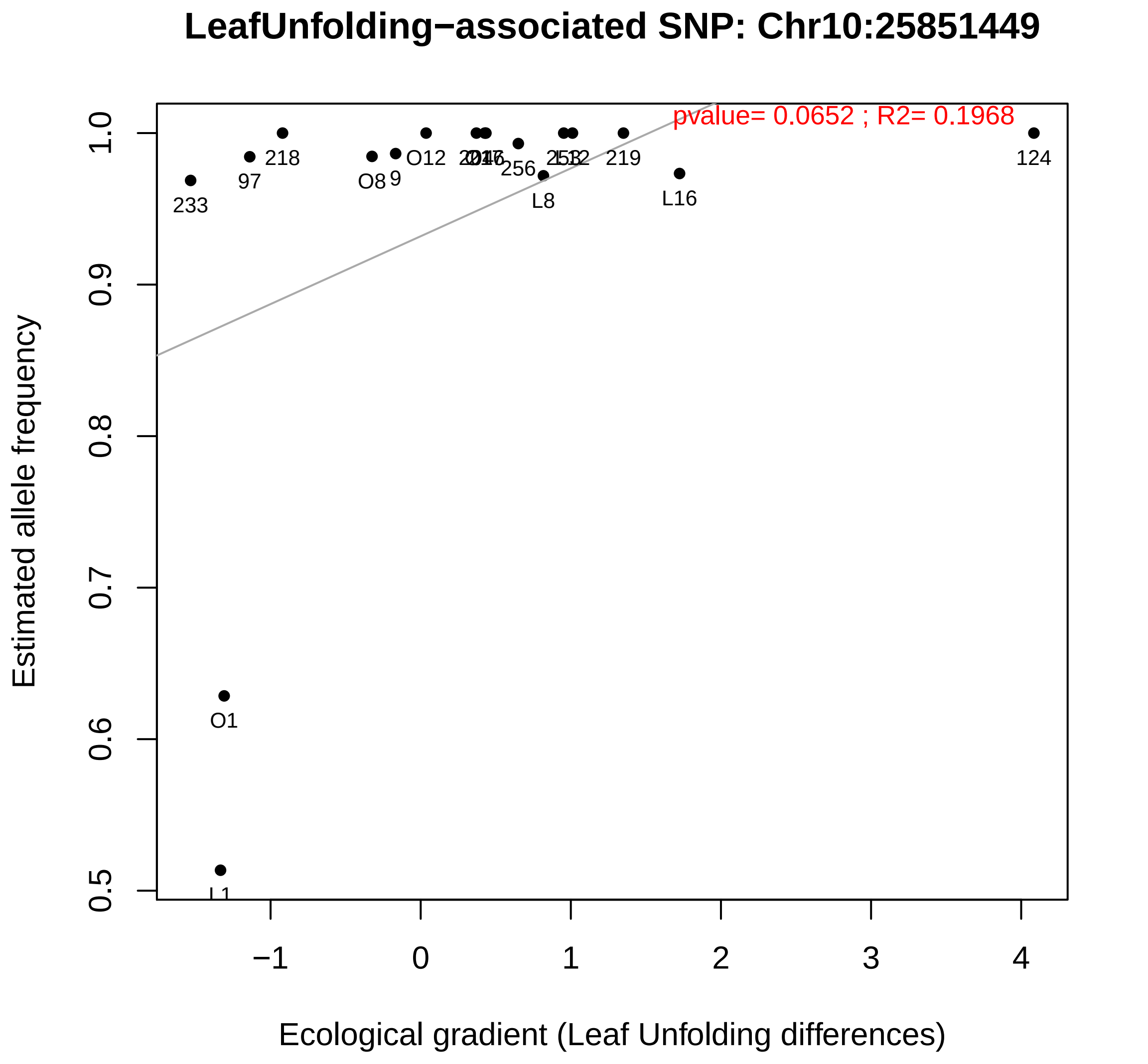
LeafUnfolding−associated SNP: Chr10:25851449
pvalue= 0.0652 ; R2= 0.1968
Estimated allele frequency
Ecological gradient (Leaf Unfolding differences)
1.0
0.9
0.8
0.7
0.6
0.5
−1
0
1
2
3
4
233
97
218
O8
9
O12
256
L8
219
L16
124
O1
L1

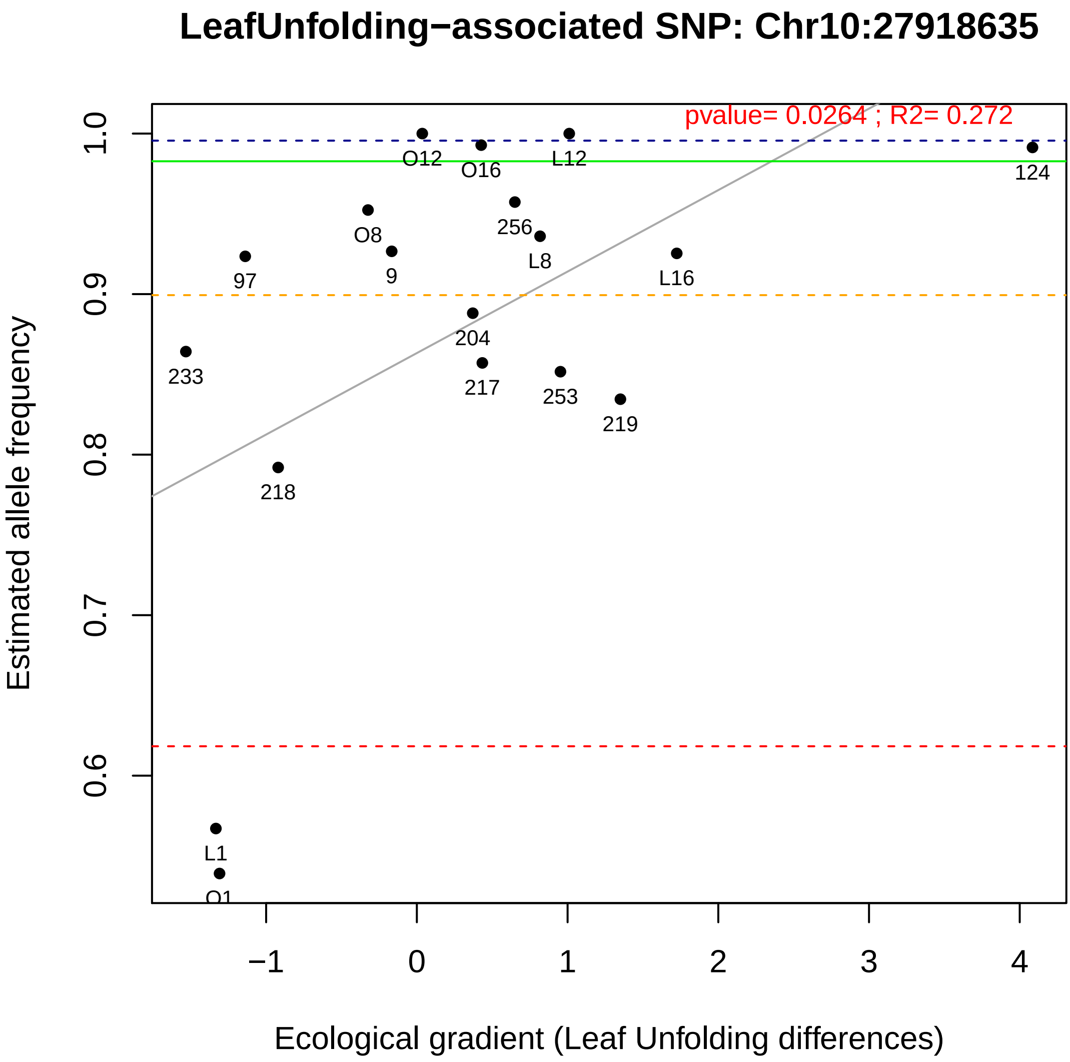

LeafUnfolding−associated SNP: Chr10:27918635
pvalue= 0.0264 ; R2= 0.272
Estimated allele frequency
Ecological gradient (Leaf Unfolding differences)
1.0
0.9
0.8
0.7
0.6
−1
0
1
2
3
4
O12
O16
L12
124
O8
256
L8
97
9
L16
204
233
217
253
219
218
L1
O1

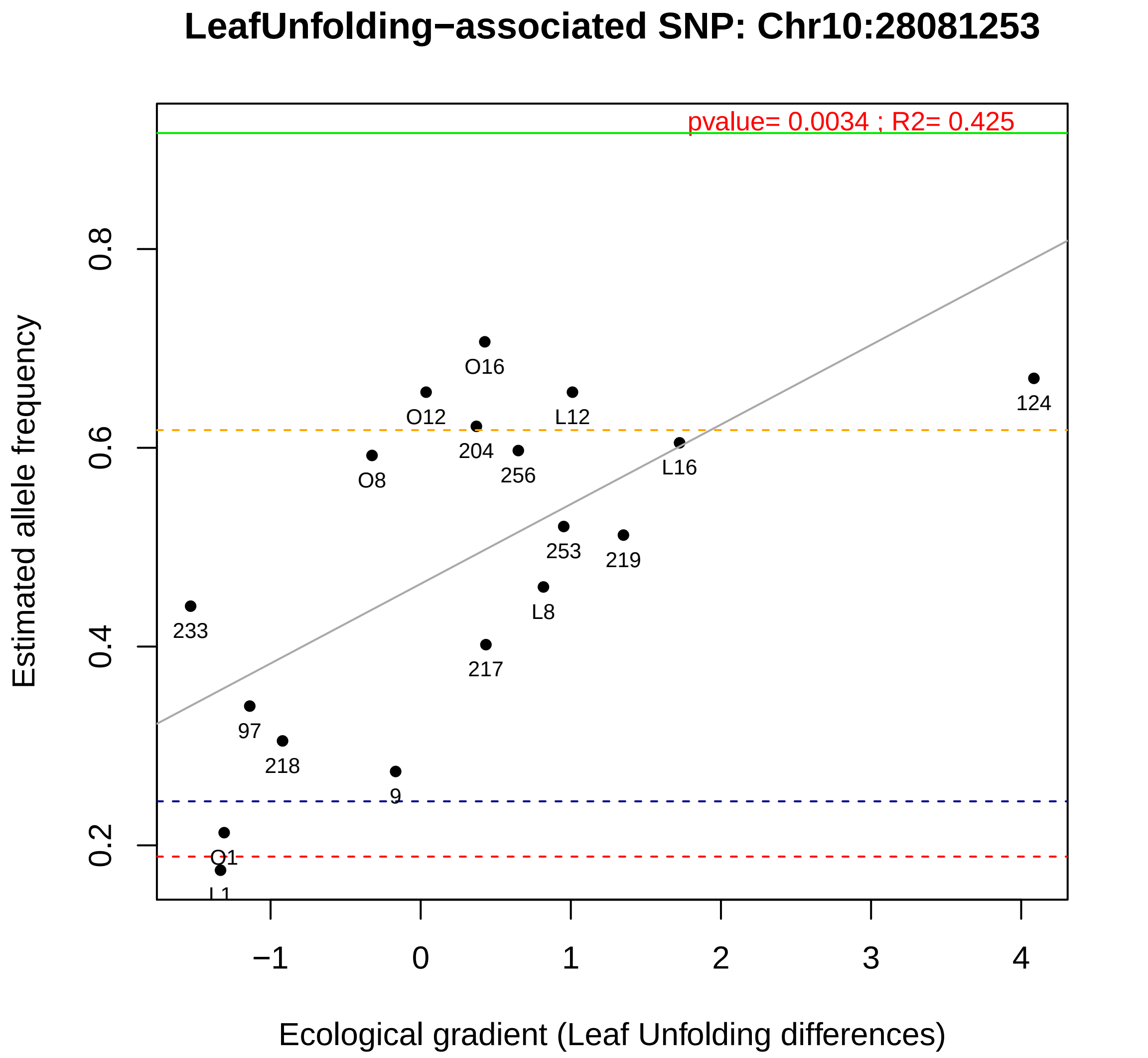

LeafUnfolding−associated SNP: Chr10:28081253
pvalue= 0.0034 ; R2= 0.425
Estimated allele frequency
0.2
0.4
0.6
0.8
−1
0
1
2
3
4
Ecological gradient (Leaf Unfolding differences)
O16
O12
L12
124
204
256
L16
O8
253
219
L8
233
217
97
218
9
O1
L1

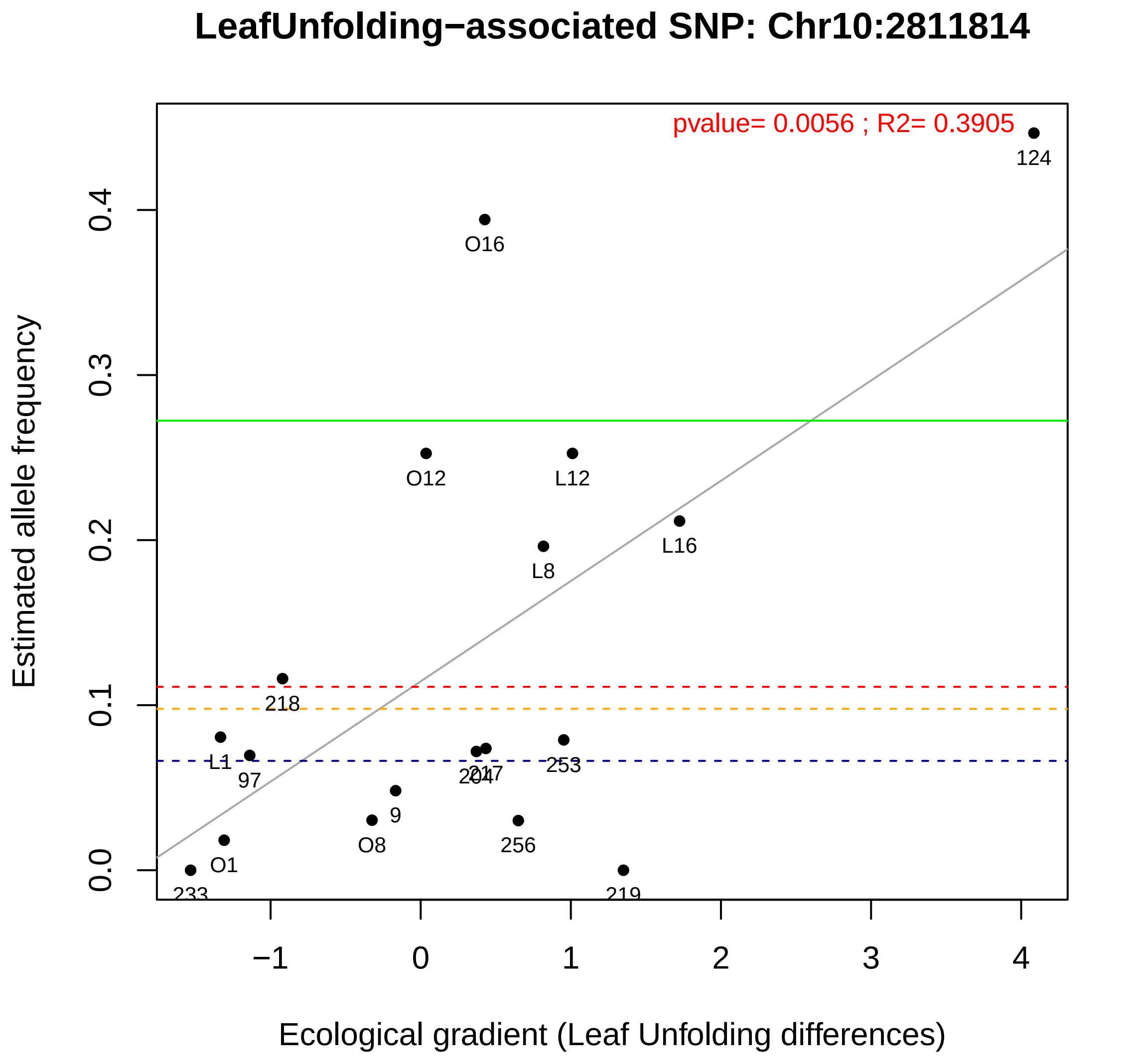
LeafUnfolding−associated SNP: Chr10:2811814
pvalue= 0.0056 ; R2= 0.3905
Estimated allele frequency
Ecological gradient (Leaf Unfolding differences)
0.0
0.1
0.2
0.3
0.4
−1
0
1
2
3
4
124
O16
O12
L12
L16
L8
218
L1
97
204
217
253
9
O8
256
O1
233
219

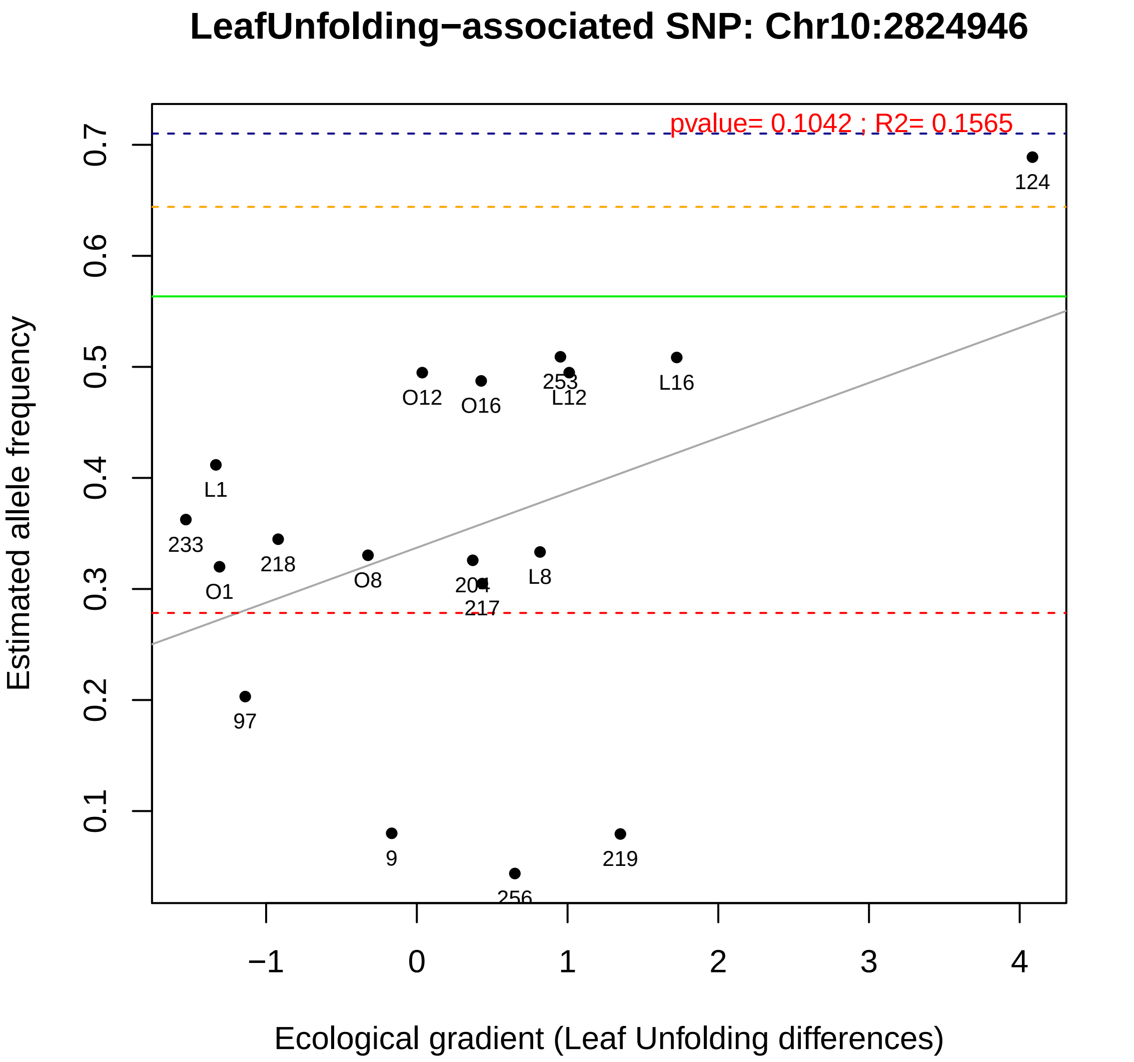

LeafUnfolding−associated SNP: Chr10:2824946
pvalue= 0.1042 ; R2= 0.1565
Estimated allele frequency
Ecological gradient (Leaf Unfolding differences)
0.1
0.2
0.3
0.4
0.5
0.6
0.7
−1
0
1
2
3
4
124
O12
O16
253
L12
L16
L1
233
218
O1
O8
204
217
L8
97
9
219
256

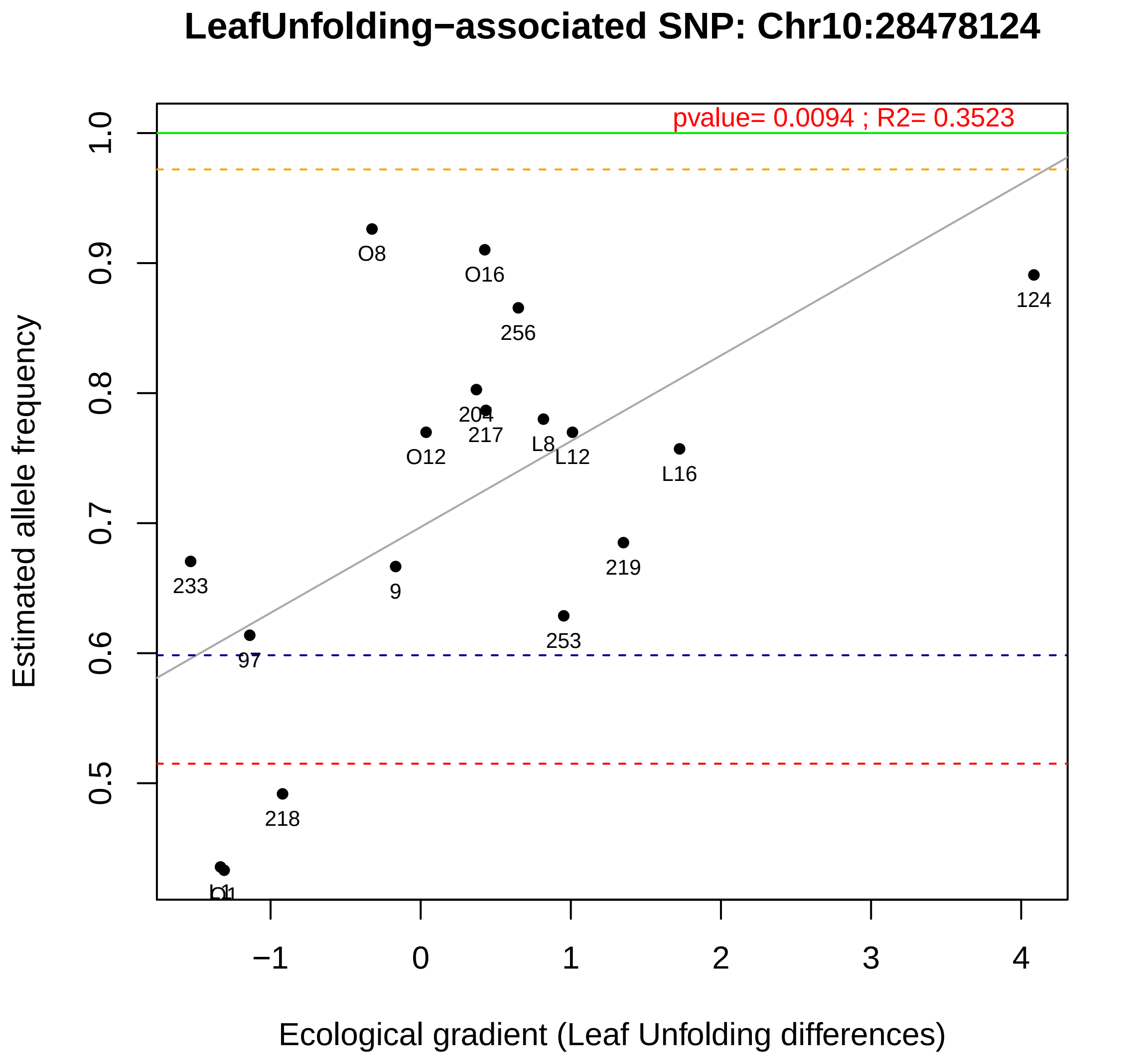
LeafUnfolding−associated SNP: Chr10:28478124
pvalue= 0.0094 ; R2= 0.3523
Estimated allele frequency
Ecological gradient (Leaf Unfolding differences)
1.0
0.9
0.8
0.7
0.6
0.5
−1
0
1
2
3
4
O8
O16
124
256
204
217
L8
L12
O12
L16
219
233
9
253
97
218

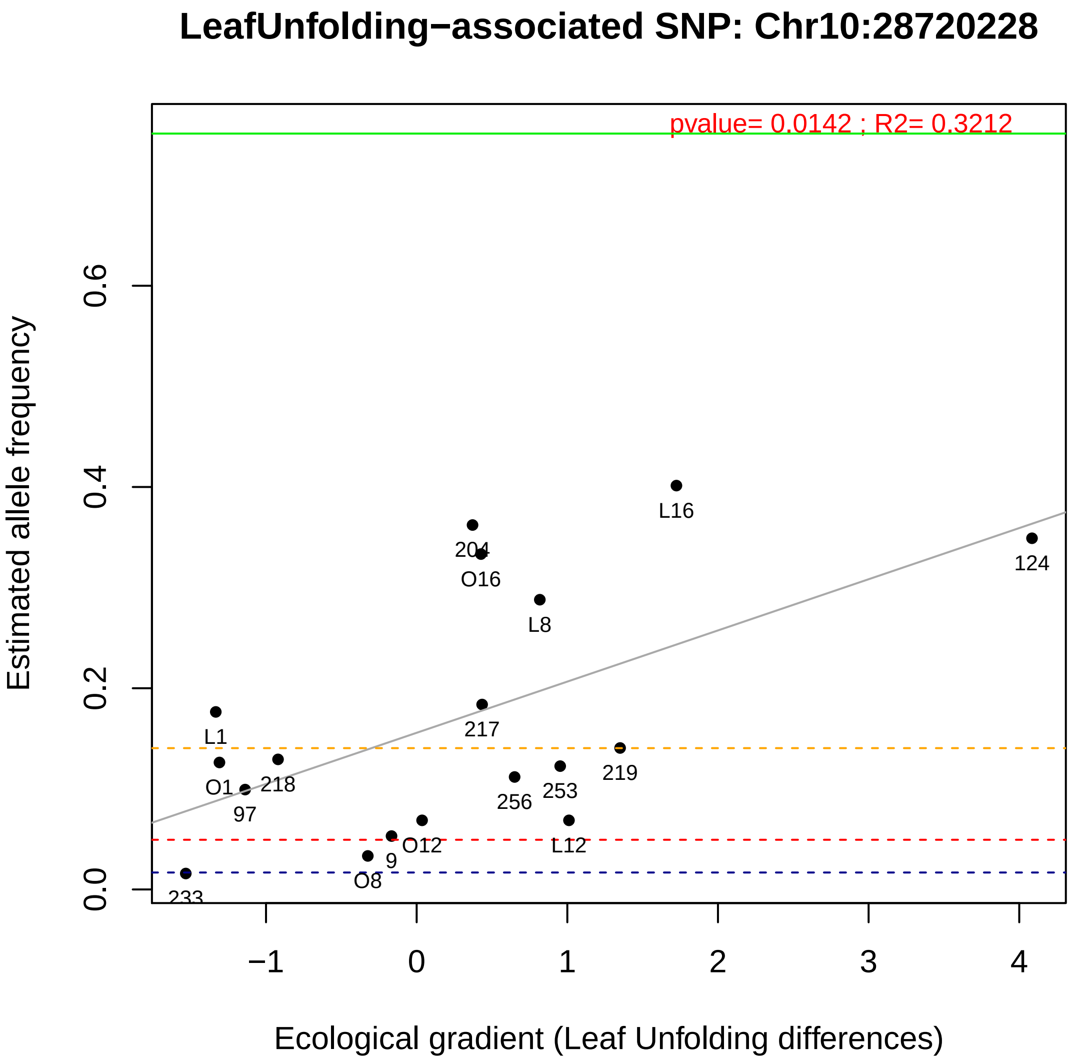

LeafUnfolding−associated SNP: Chr10:28720228
pvalue= 0.0142 ; R2= 0.3212
Estimated allele frequency
0.0
0.2
0.4
0.6
−1
0
1
2
3
4
Ecological gradient (Leaf Unfolding differences)
L16
204
O16
124
L8
217
L1
219
O1
218
253
97
256
O12
L12
9
O8
233

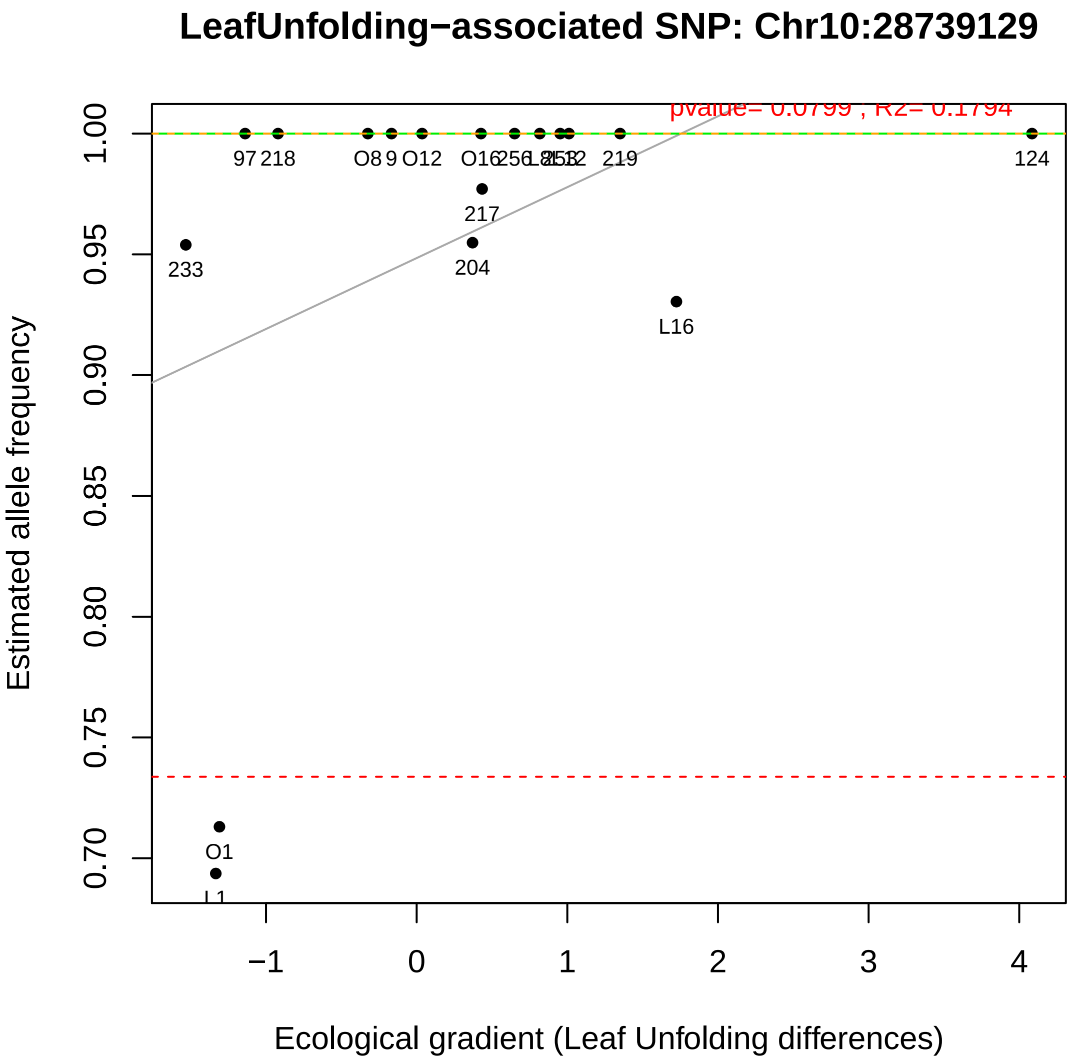

LeafUnfolding−associated SNP: Chr10:28739129
pvalue= 0.0799 ; R2= 0.1794
Estimated allele frequency
Ecological gradient (Leaf Unfolding differences)
1.00
0.95
0.90
0.85
0.80
0.75
0.70
−1
0
1
2
3
4
97
218
O8
9
O12
O16
256
219
124
217
233
204
L16
O1

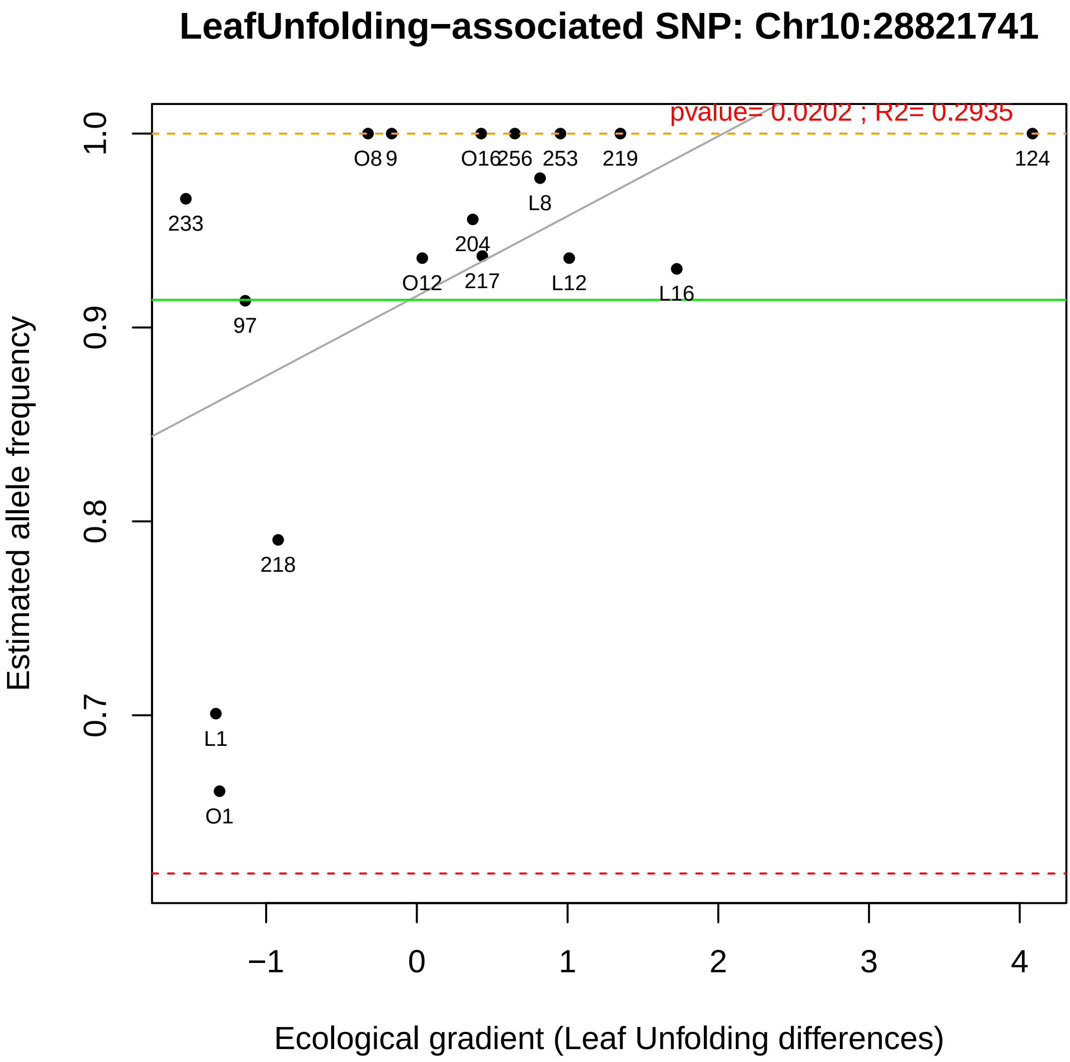
LeafUnfolding−associated SNP: Chr10:28821741
pvalue= 0.0202 ; R2= 0.2935
Estimated allele frequency
Ecological gradient (Leaf Unfolding differences)
1.0
0.9
0.8
0.7
−1
0
1
2
3
4
O8
9
O16
256
253
219
124
L8
233
204
O12
217
L12
L16
97
218
L1
O1

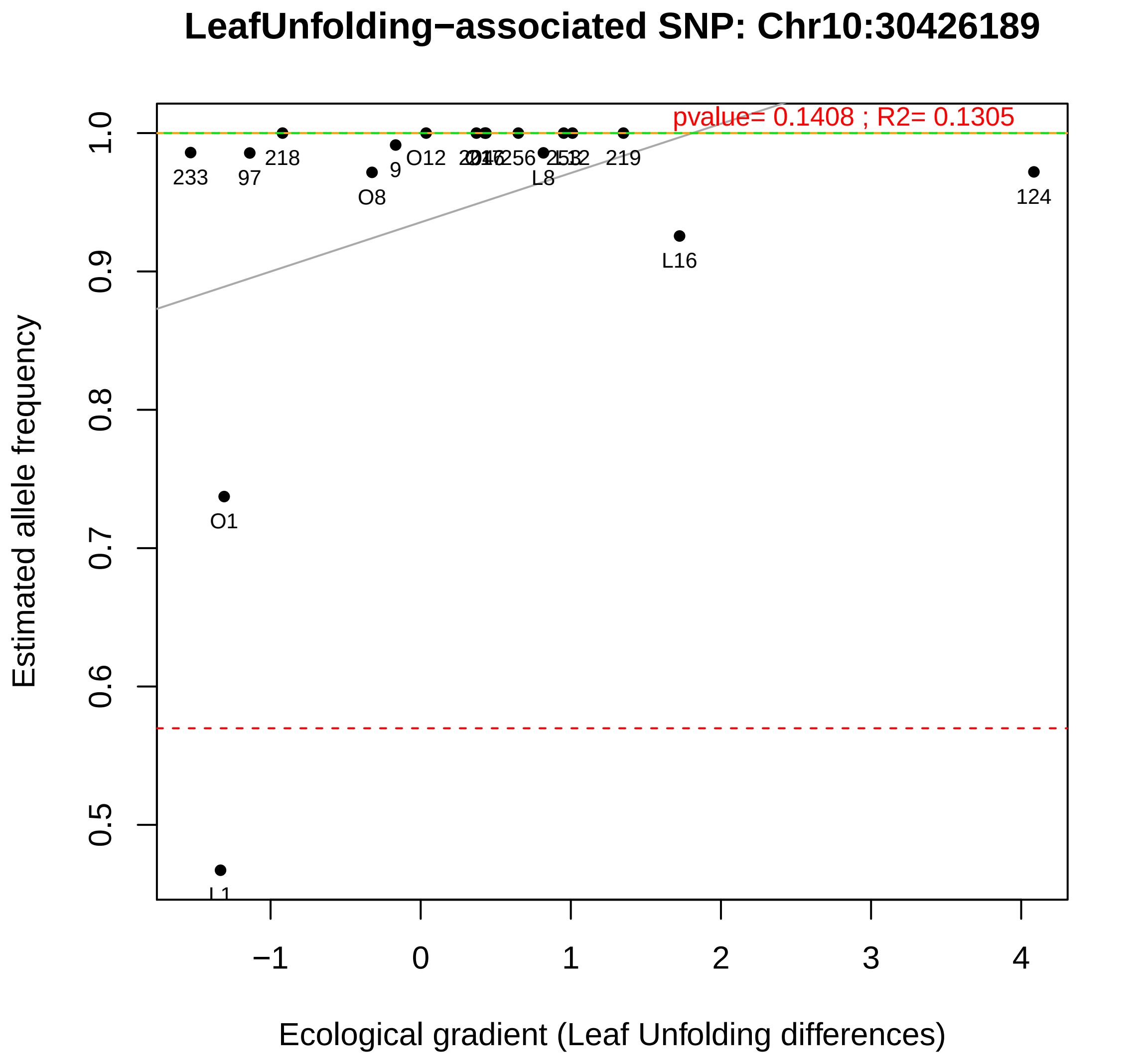
LeafUnfolding−associated SNP: Chr10:30426189
pvalue= 0.1408 ; R2= 0.1305
Estimated allele frequency
Ecological gradient (Leaf Unfolding differences)
1.0
0.9
0.8
0.7
0.6
0.5
−1
0
1
2
3
4
233
97
218
O8
9
O12
219
L8
124
L16
O1

#### LeafUnfolding−associated SNP: Chr10:30429220

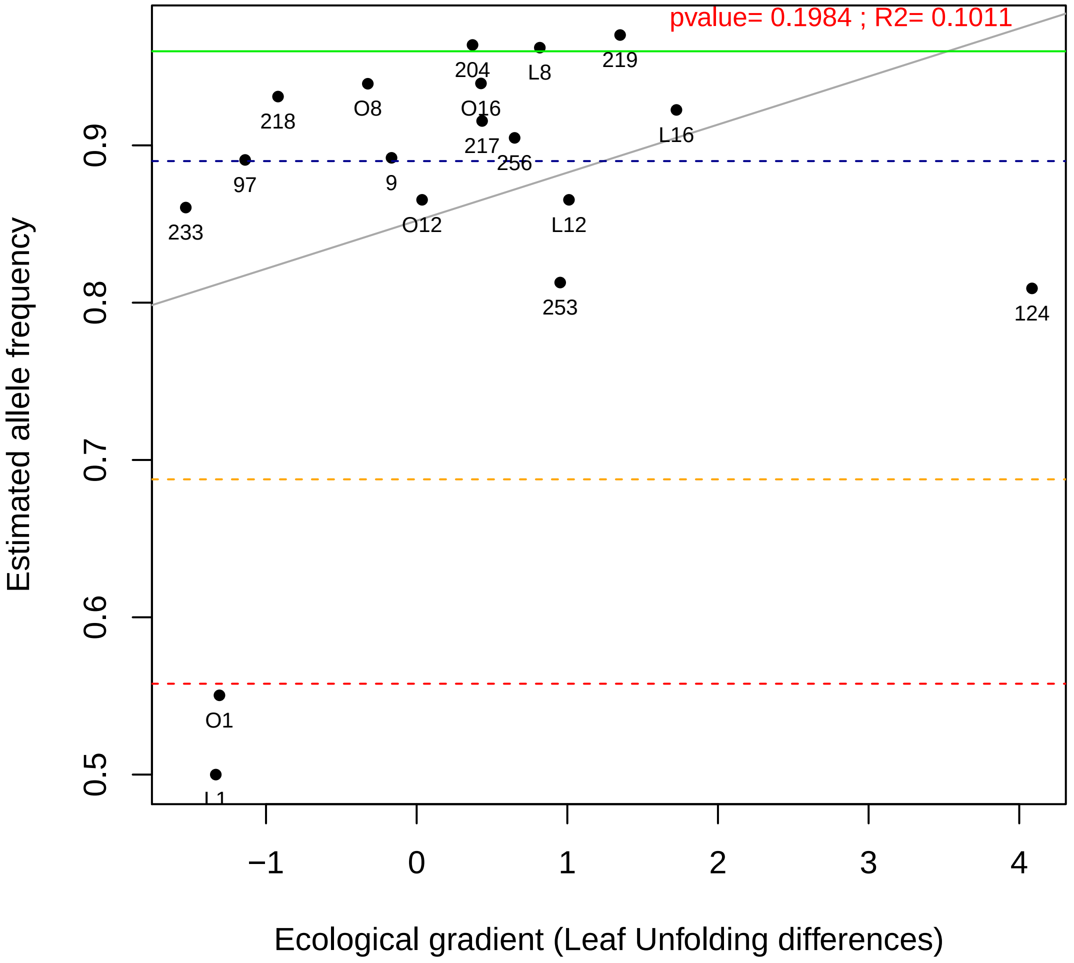

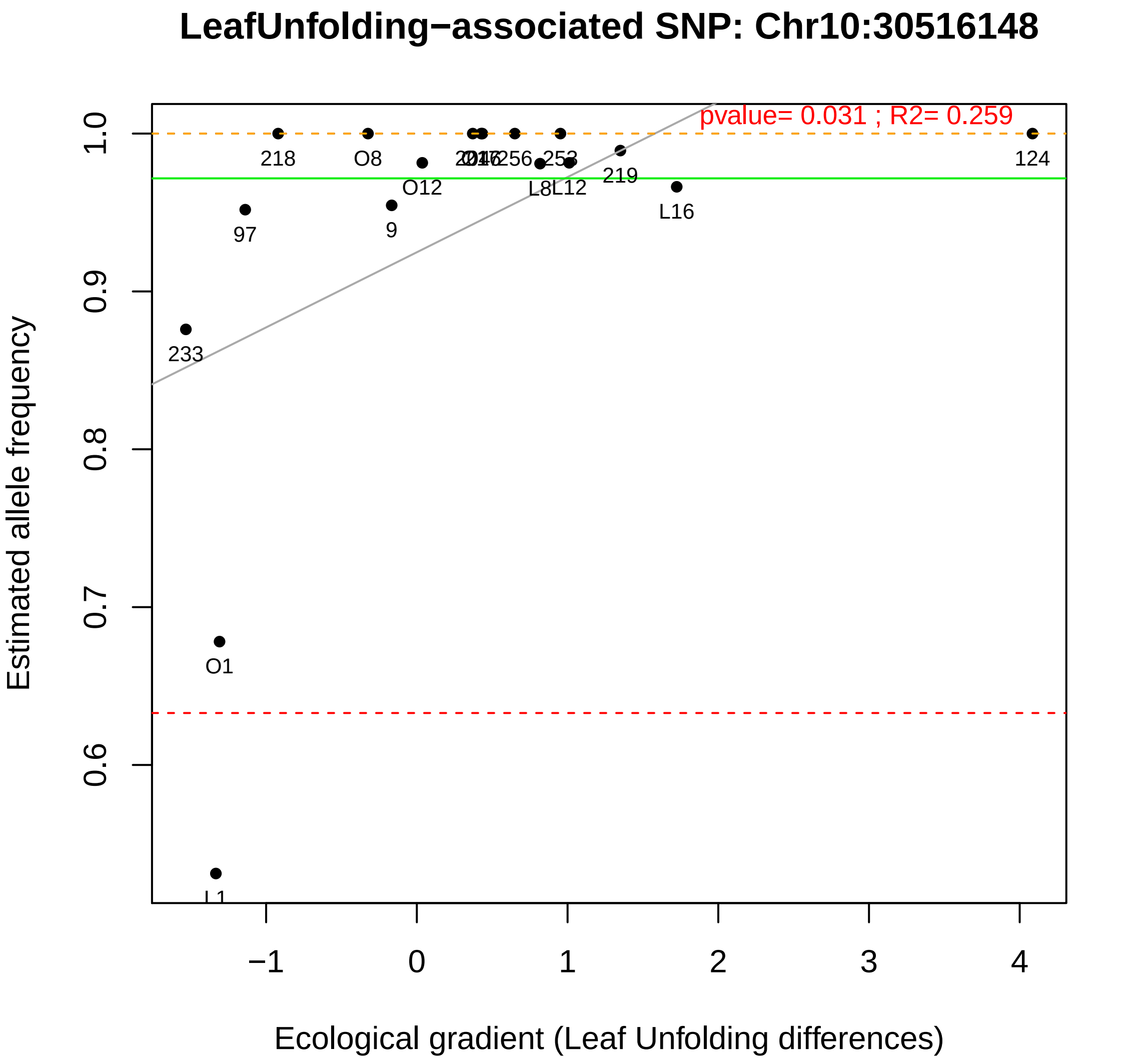

LeafUnfolding−associated SNP: Chr10:30516148
pvalue= 0.031 ; R2= 0.259
218
O8
O12
253
219
124
L8
L12
L16
97
9
233
O1
1.0
0.9
0.8
0.7
0.6
−1
0
1
2
3
4
Estimated allele frequency
Ecological gradient (Leaf Unfolding differences)

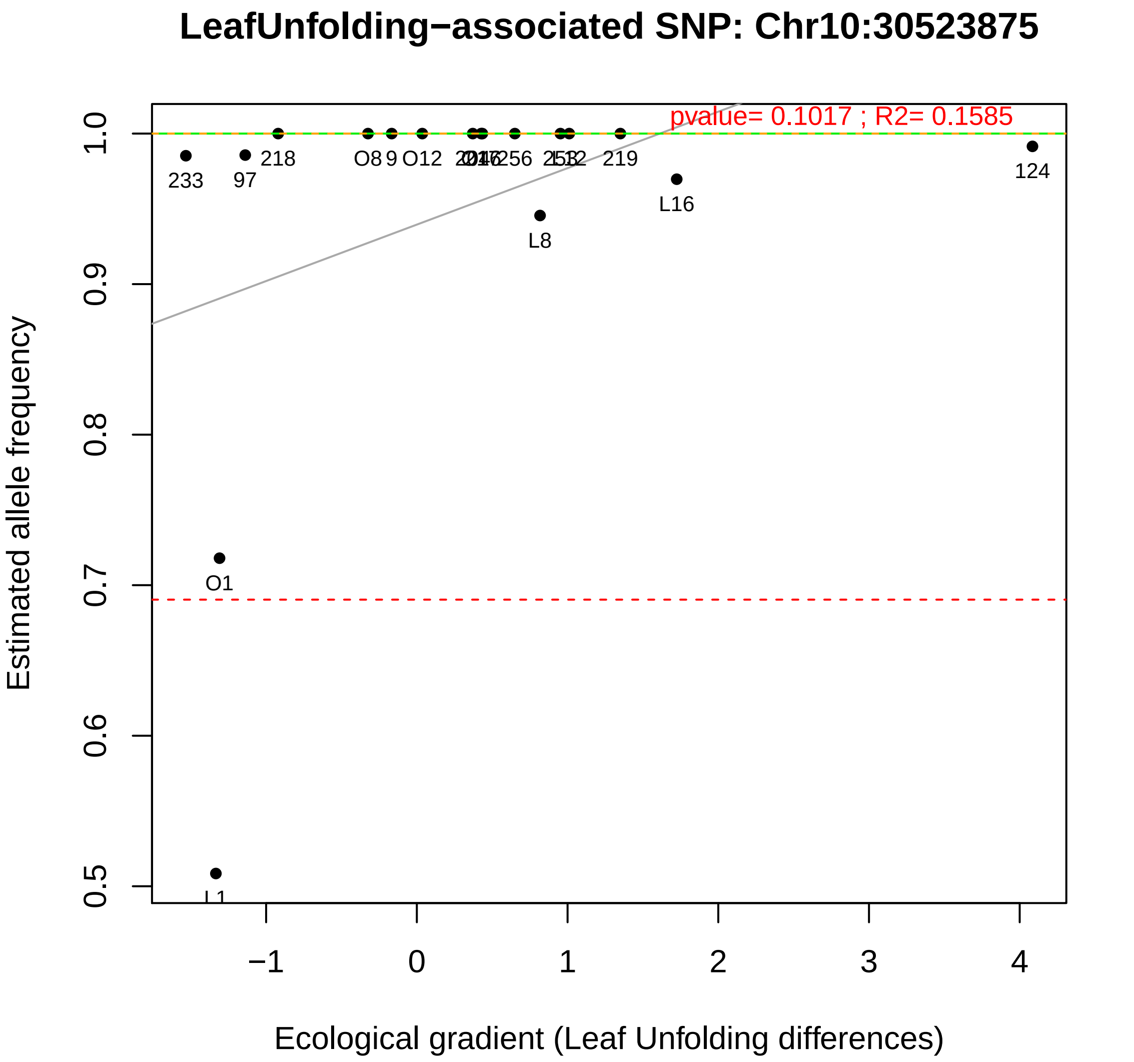

LeafUnfolding−associated SNP: Chr10:30523875
pvalue= 0.1017 ; R2= 0.1585
Estimated allele frequency
Ecological gradient (Leaf Unfolding differences)
233
97
218
O8
9
O12
219
L8
L16
124
O1
0.5
0.6
0.7
0.8
0.9
1.0
−1
0
1
2
3
4

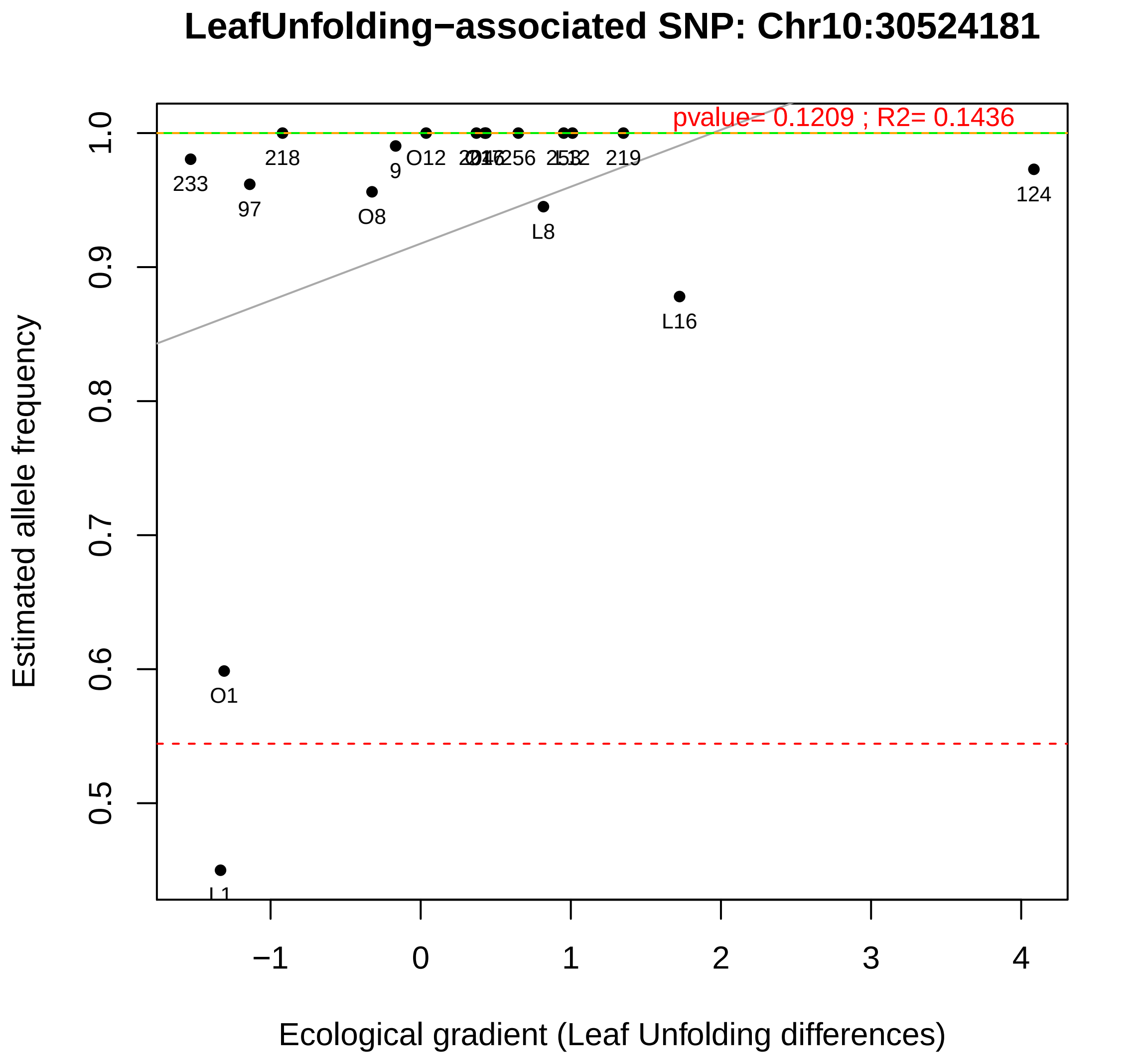

LeafUnfolding−associated SNP: Chr10:30524181
pvalue= 0.1209 ; R2= 0.1436
Estimated allele frequency
Ecological gradient (Leaf Unfolding differences)
1.0
0.9
0.8
0.7
0.6
0.5
−1
0
1
2
3
4
233
218
97
O8
9
O12
L8
219
124
L16
O1
L1

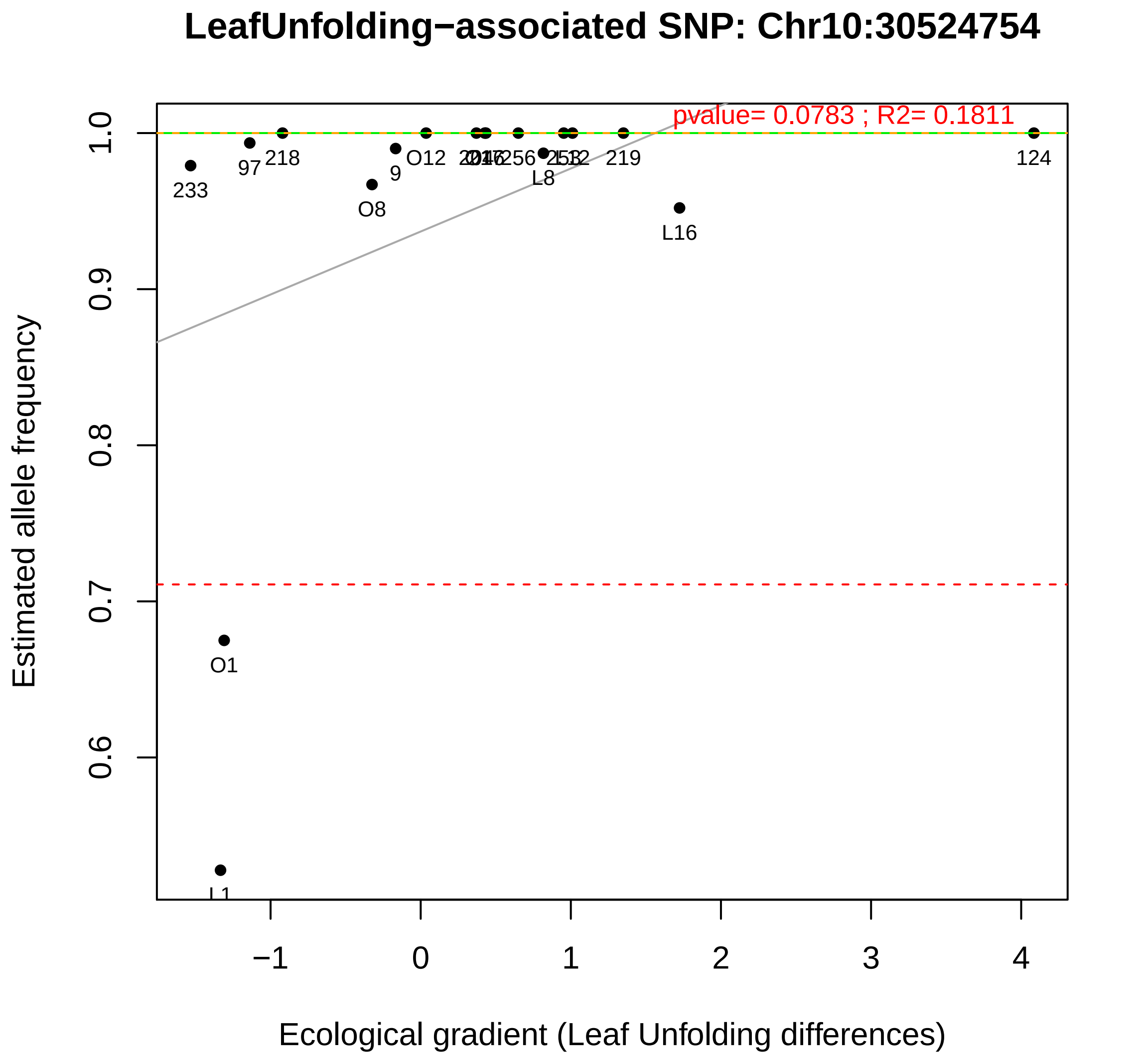

LeafUnfolding−associated SNP: Chr10:30524754
pvalue= 0.0783 ; R2= 0.1811
Estimated allele frequency
Ecological gradient (Leaf Unfolding differences)
1.0
0.9
0.8
0.7
0.6
−1
0
1
2
3
4
233
97
218
O8
9
O12
L8
219
L16
124
O1

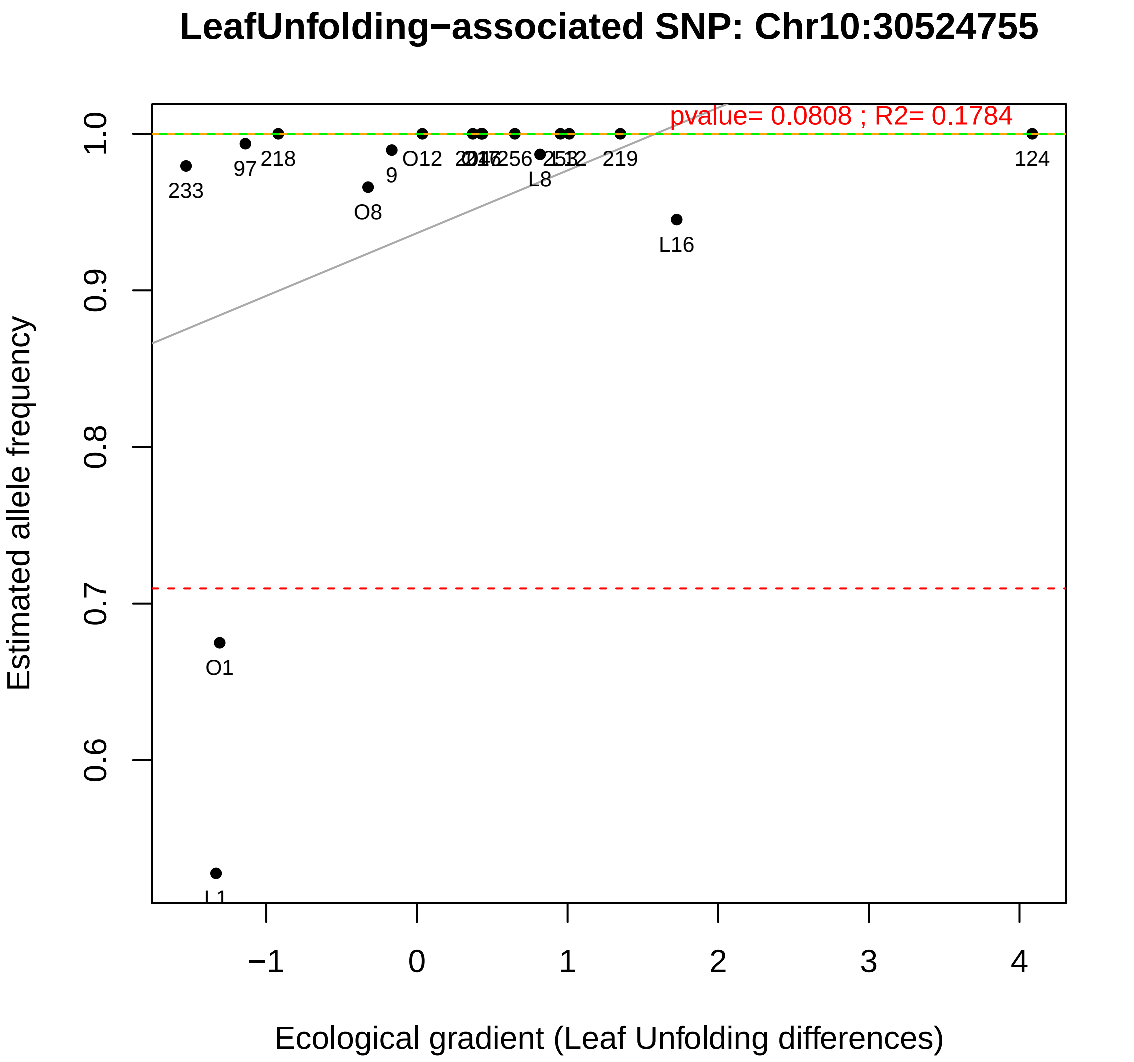

LeafUnfolding−associated SNP: Chr10:30524755
pvalue= 0.0808 ; R2= 0.1784
Estimated allele frequency
Ecological gradient (Leaf Unfolding differences)
1.0
0.9
0.8
0.7
0.6
−1
0
1
2
3
4
233
97
218
O8
9
O12
L8
219
124
L16
O1

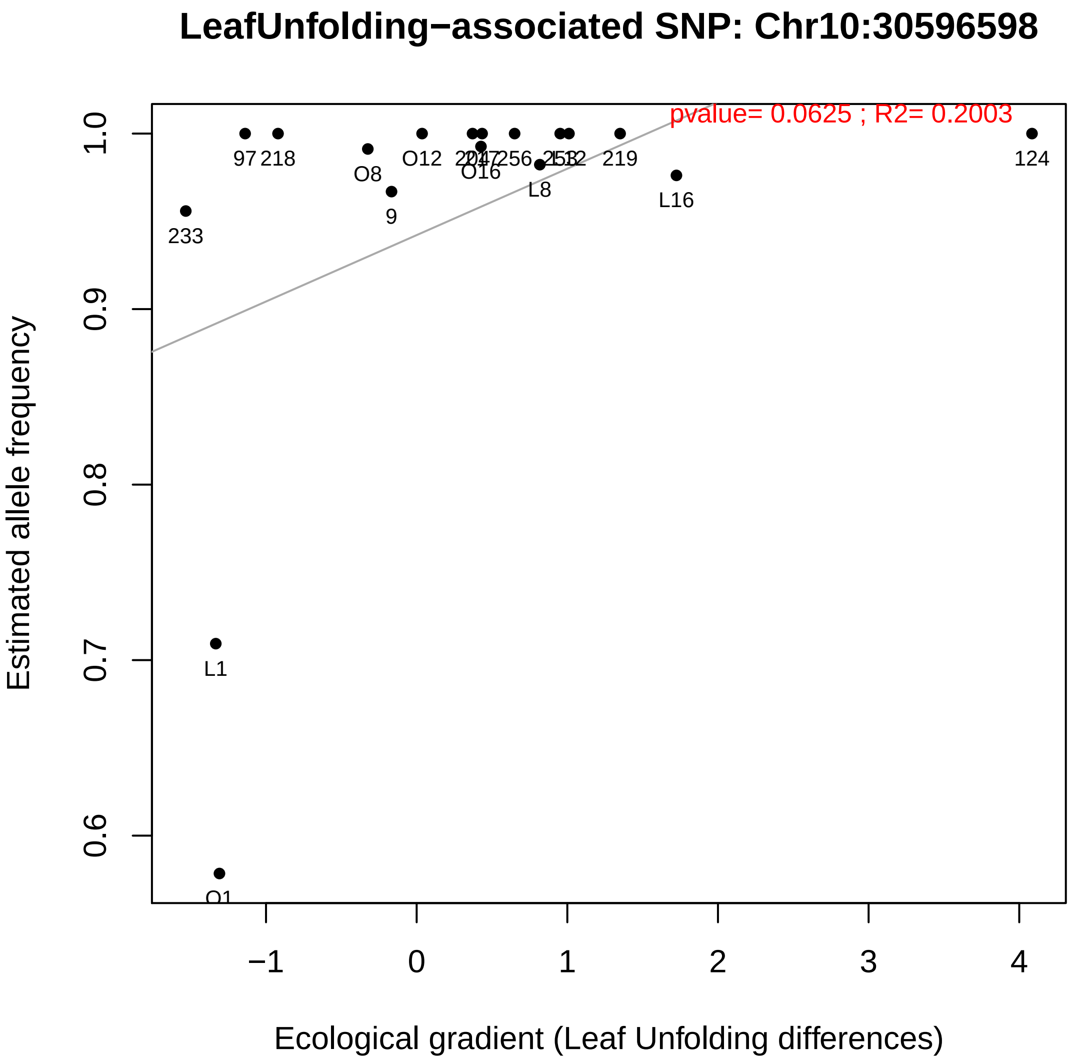
LeafUnfolding−associated SNP: Chr10:30596598
pvalue= 0.0625 ; R2= 0.2003
Estimated allele frequency
0.6
0.7
0.8
0.9
1.0
−1
0
1
2
3
4
Ecological gradient (Leaf Unfolding differences)
97
218
O8
O12
O16
256
L8
219
L16
124
233
9
L1
O1

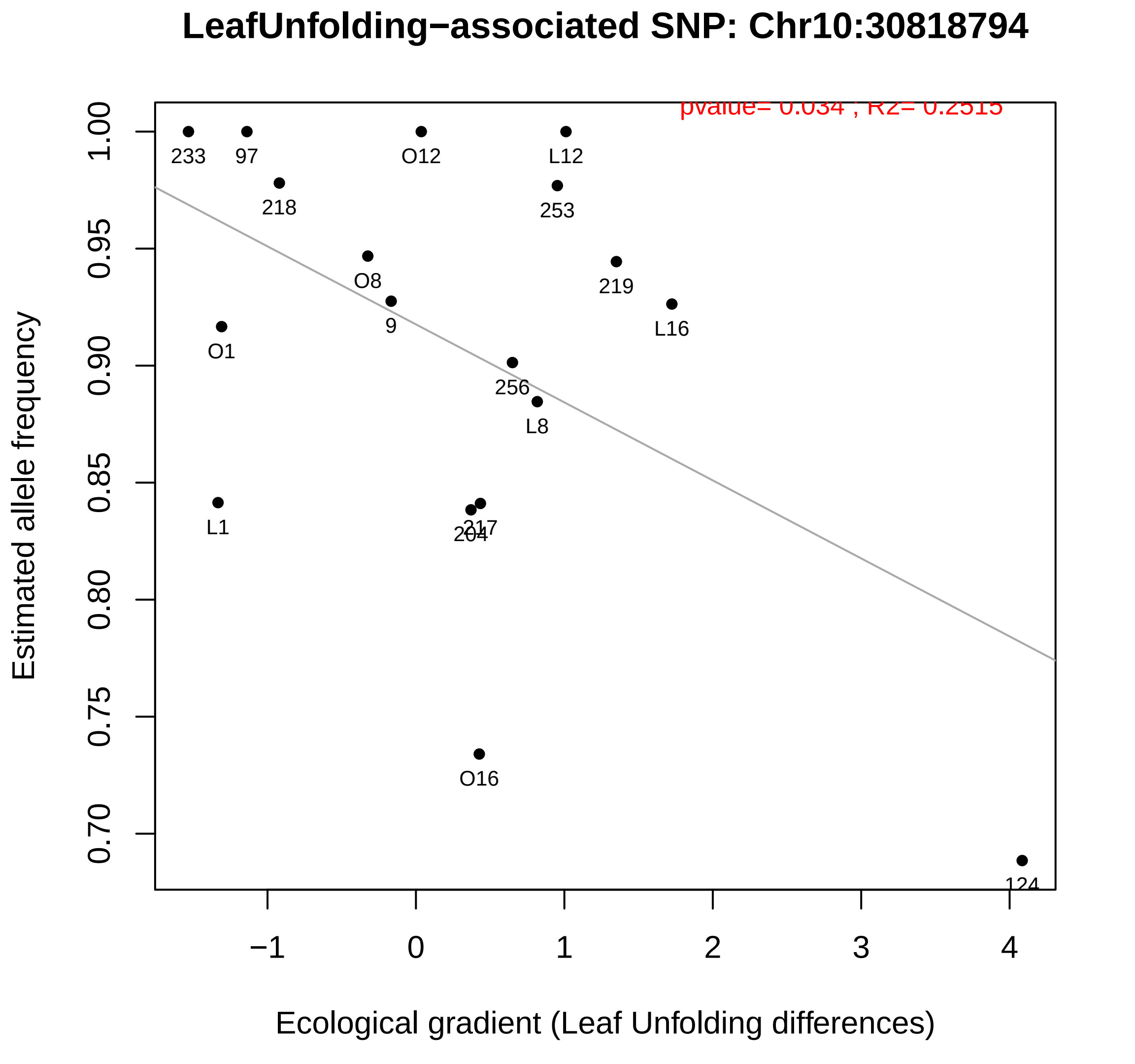
LeafUnfolding−associated SNP: Chr10:30818794
pvalue= 0.034 ; R2= 0.2515
Estimated allele frequency
Ecological gradient (Leaf Unfolding differences)
0.70
0.75
0.80
0.85
0.90
0.95
1.00
−1
0
1
2
3
4
233
97
O12
L12
218
253
O8
219
9
L16
O1
256
L8
L1
217
204
O16
124

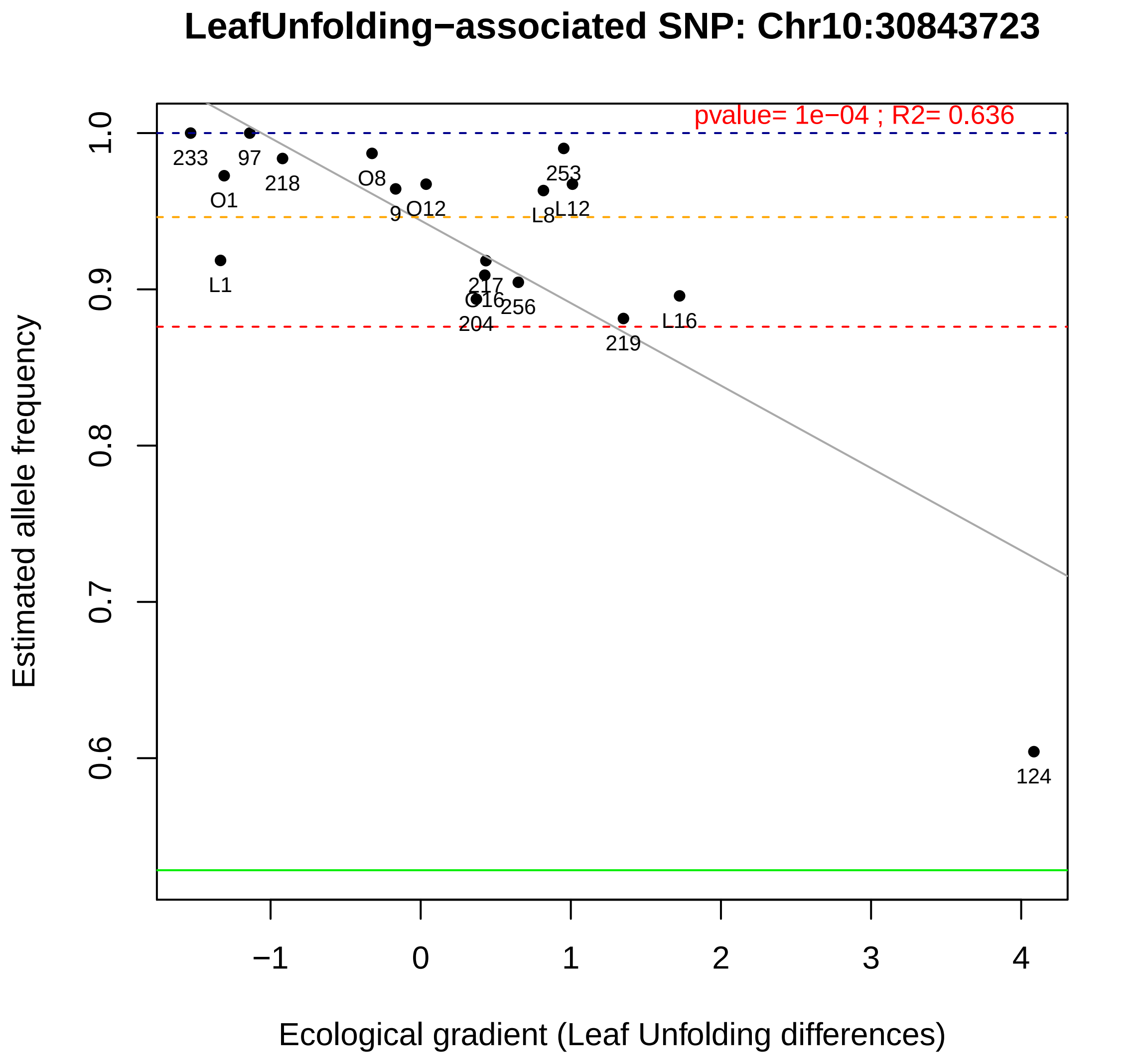
LeafUnfolding−associated SNP: Chr10:30843723
pvalue= 1e−04 ; R2= 0.636
Estimated allele frequency
Ecological gradient (Leaf Unfolding differences)
1.0
0.9
0.8
0.7
0.6
−1
0
1
2
3
4
233
97
218
O1
L1
O8
9
O12
253
L8
L12
217
O16
256
204
219
L16
124

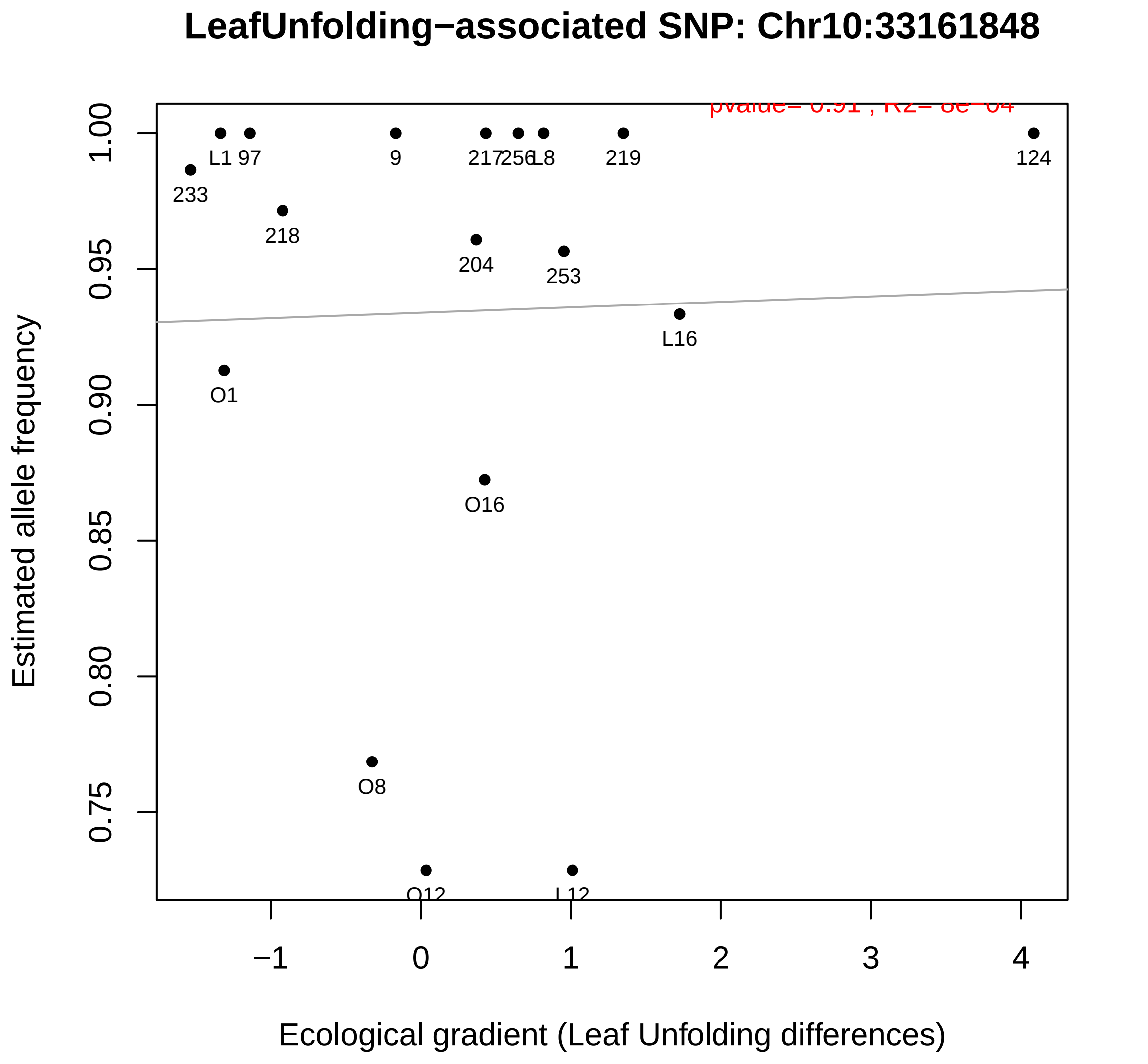
LeafUnfolding−associated SNP: Chr10:33161848
pvalue= 0.91 ; R2= 8e−04
Estimated allele frequency
Ecological gradient (Leaf Unfolding differences)
0.75
0.80
0.85
0.90
0.95
1.00
−1
0
1
2
3
4
L1
97
9
217
256
L8
219
124
233
218
204
253
L16
O1
O16
O8
O12
L12

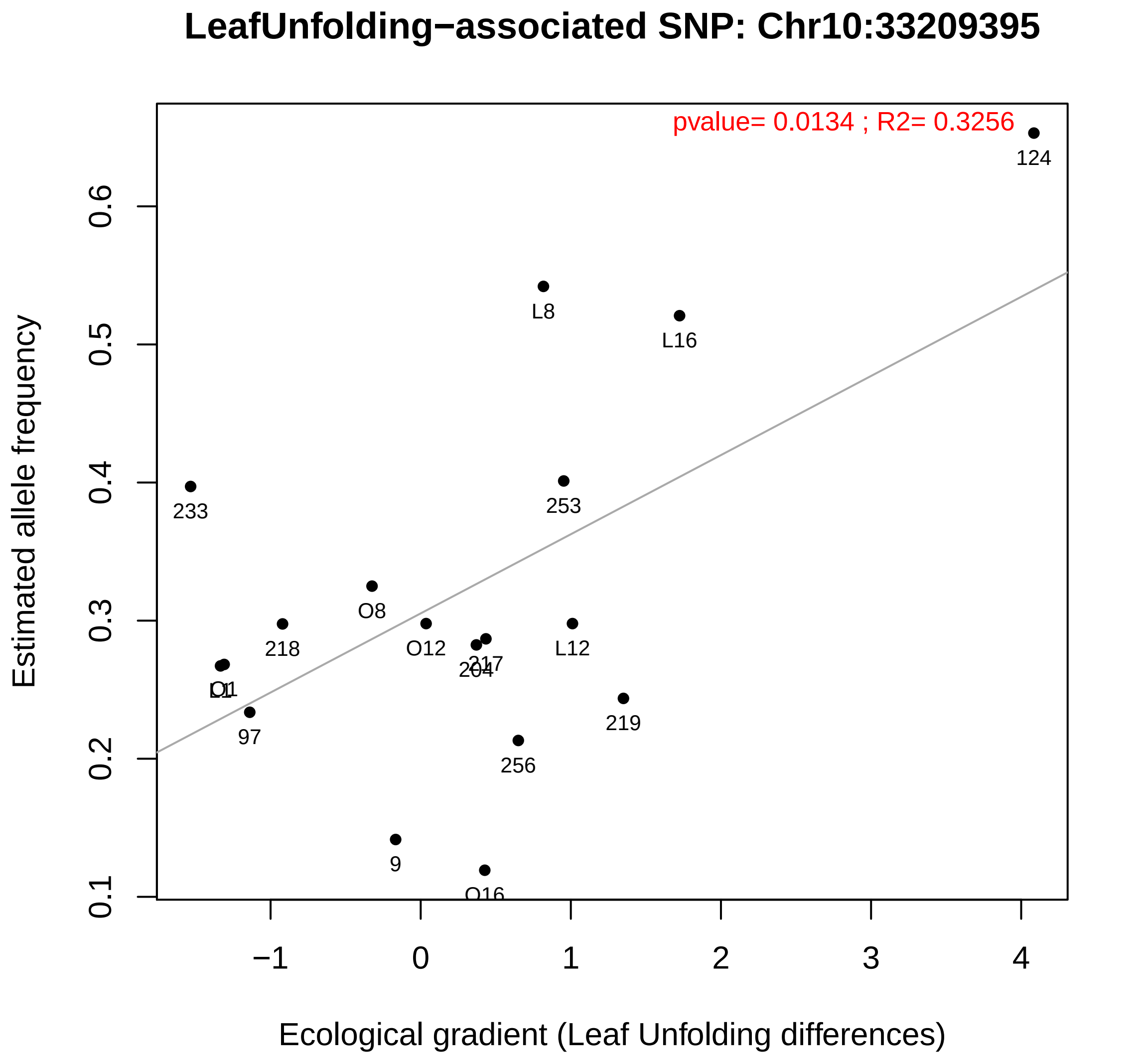

LeafUnfolding−associated SNP: Chr10:33209395
pvalue= 0.0134 ; R2= 0.3256
Estimated allele frequency
Ecological gradient (Leaf Unfolding differences)
0.1
0.2
0.3
0.4
0.5
0.6
−1
0
1
2
3
4
124
L8
L16
233
253
O8
218
O12
L12
217
204
219
97
256
9
O16

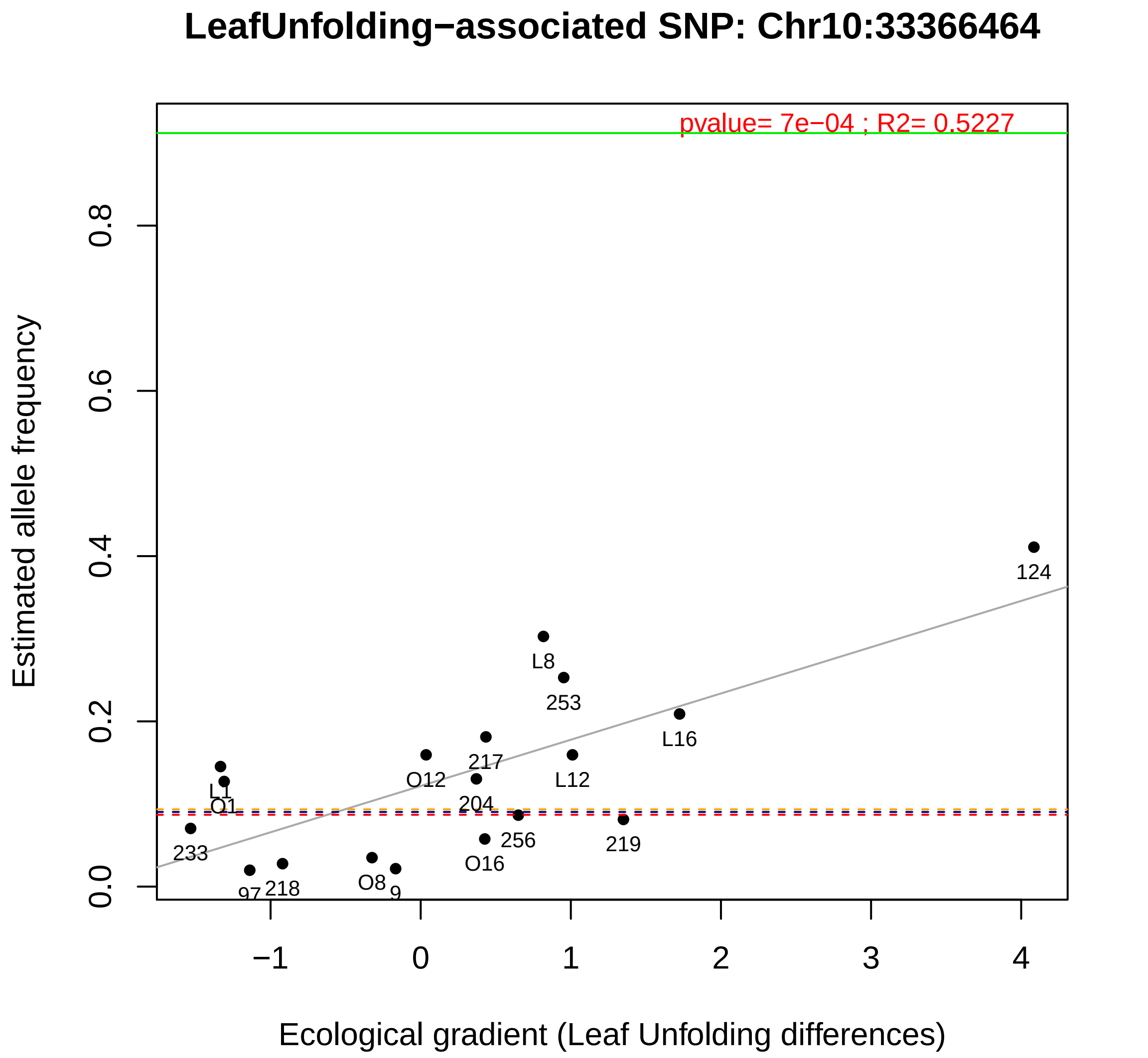
LeafUnfolding−associated SNP: Chr10:33366464
pvalue= 7e−04 ; R2= 0.5227
Estimated allele frequency
0.0
0.2
0.4
0.6
0.8
−1
0
1
2
3
4
Ecological gradient (Leaf Unfolding differences)
124
L8
253
L16
217
L12
O12
204
L1
O1
233
256
219
O16
97
218
O8
9

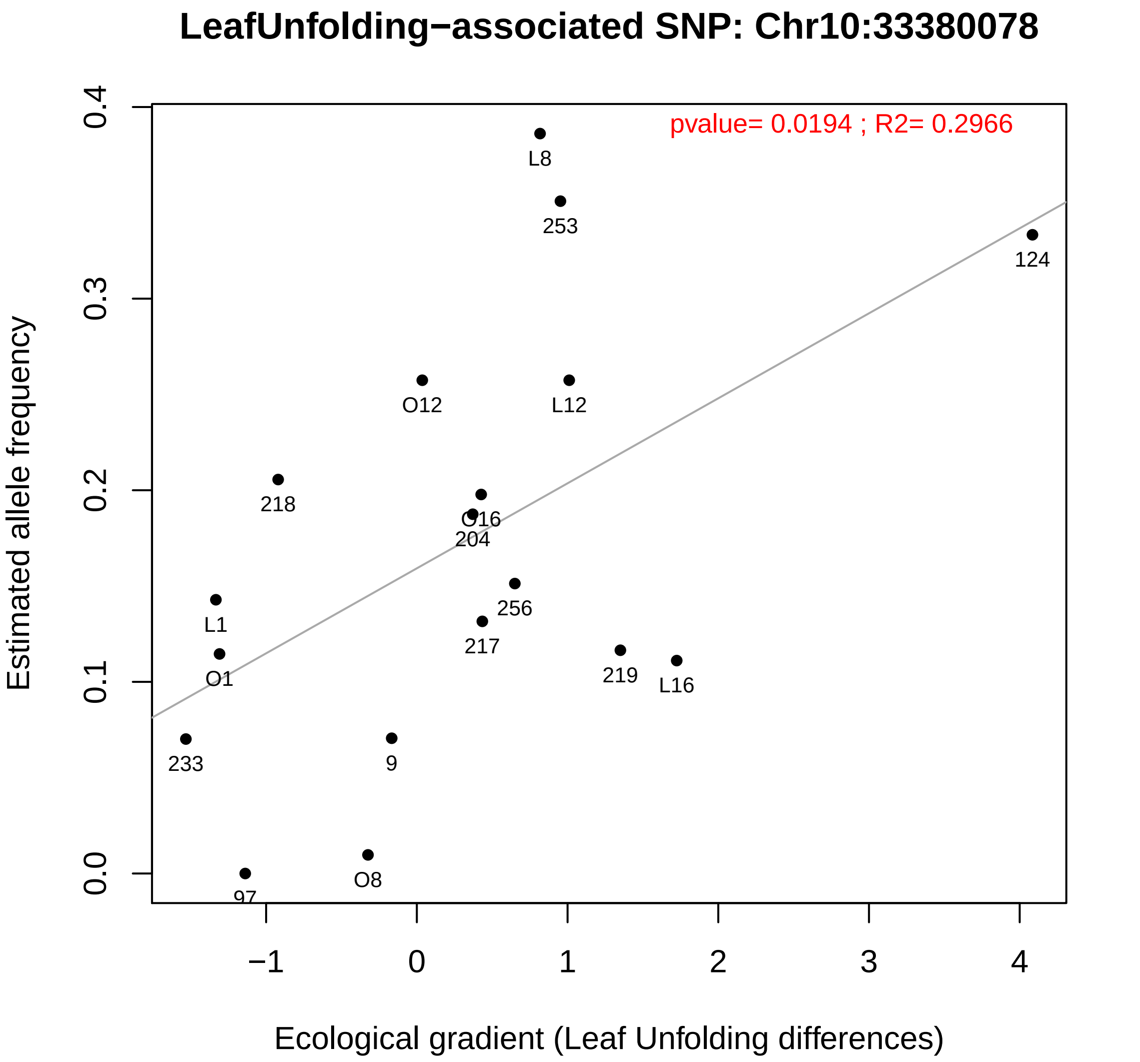
LeafUnfolding−associated SNP: Chr10:33380078
pvalue= 0.0194 ; R2= 0.2966
L8
253
124
O12
L12
218
O16
204
256
L1
217
O1
219
L16
233
9
O8
97
0.0
0.1
0.2
0.3
0.4
−1
0
1
2
3
4
Estimated allele frequency
Ecological gradient (Leaf Unfolding differences)

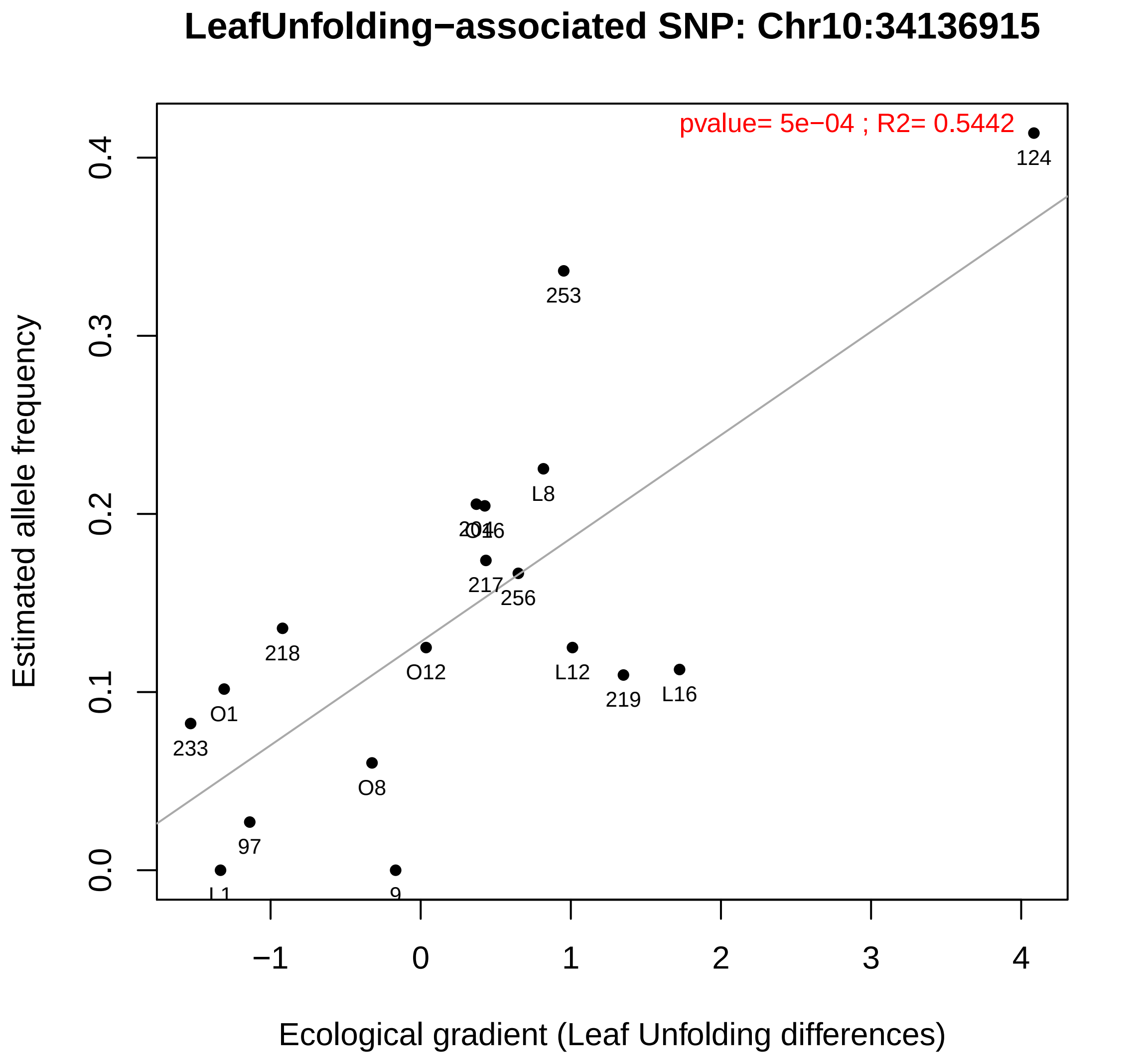
LeafUnfolding−associated SNP: Chr10:34136915
pvalue= 5e−04 ; R2= 0.5442
Estimated allele frequency
Ecological gradient (Leaf Unfolding differences)
0.0
0.1
0.2
0.3
0.4
−1
0
1
2
3
4
124
253
L8
O16
217
256
218
O12
L12
219
L16
O1
233
O8
97
L1
9

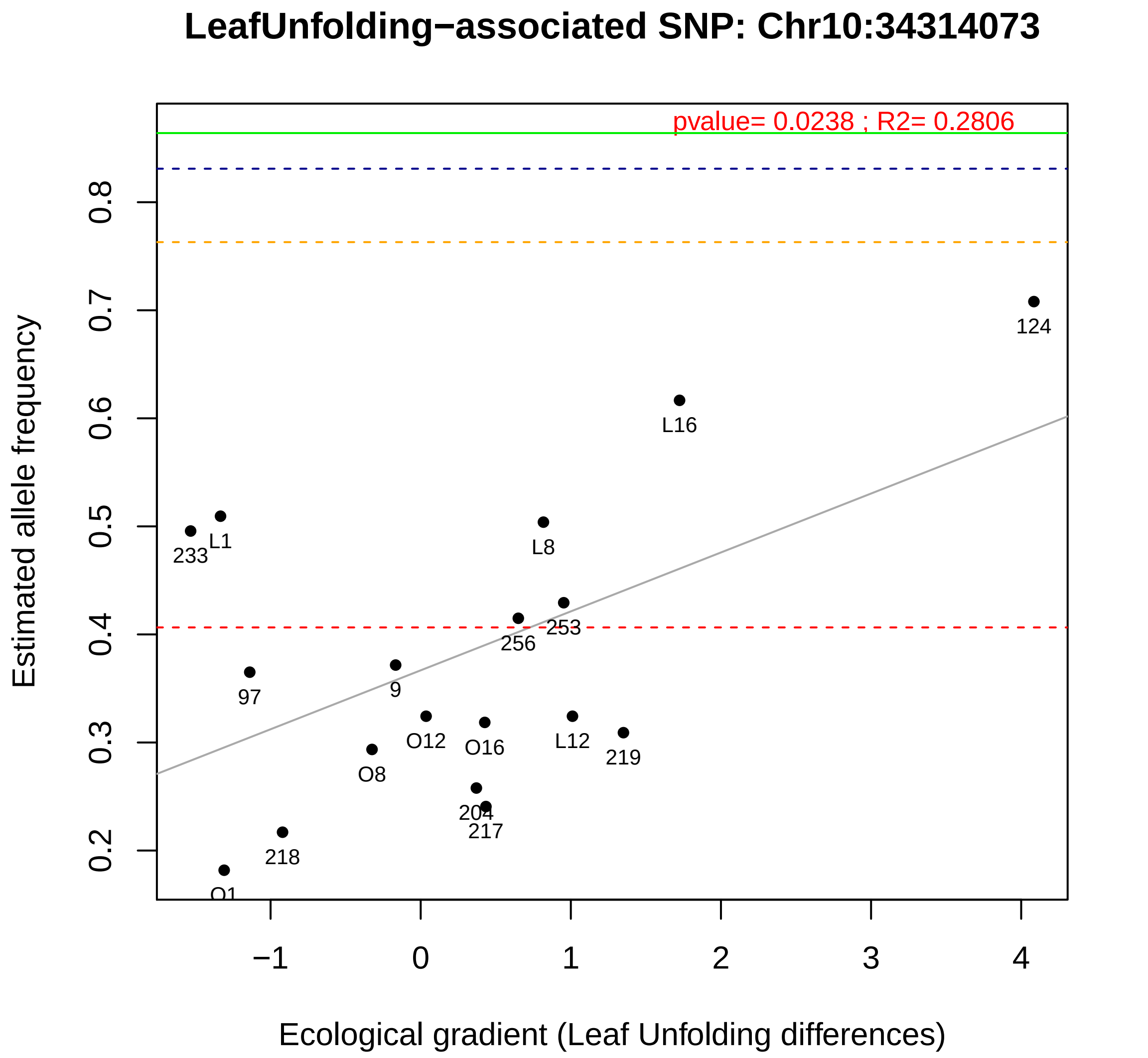

LeafUnfolding−associated SNP: Chr10:34314073
pvalue= 0.0238 ; R2= 0.2806
Estimated allele frequency
0.2
0.3
0.4
0.5
0.6
0.7
0.8
−1
0
1
2
3
4
Ecological gradient (Leaf Unfolding differences)
124
L16
233
L1
L8
253
256
97
9
O12
O16
L12
219
O8
204
217
218
O1

LeafUnfolding−associated SNP: Chr10:34687730

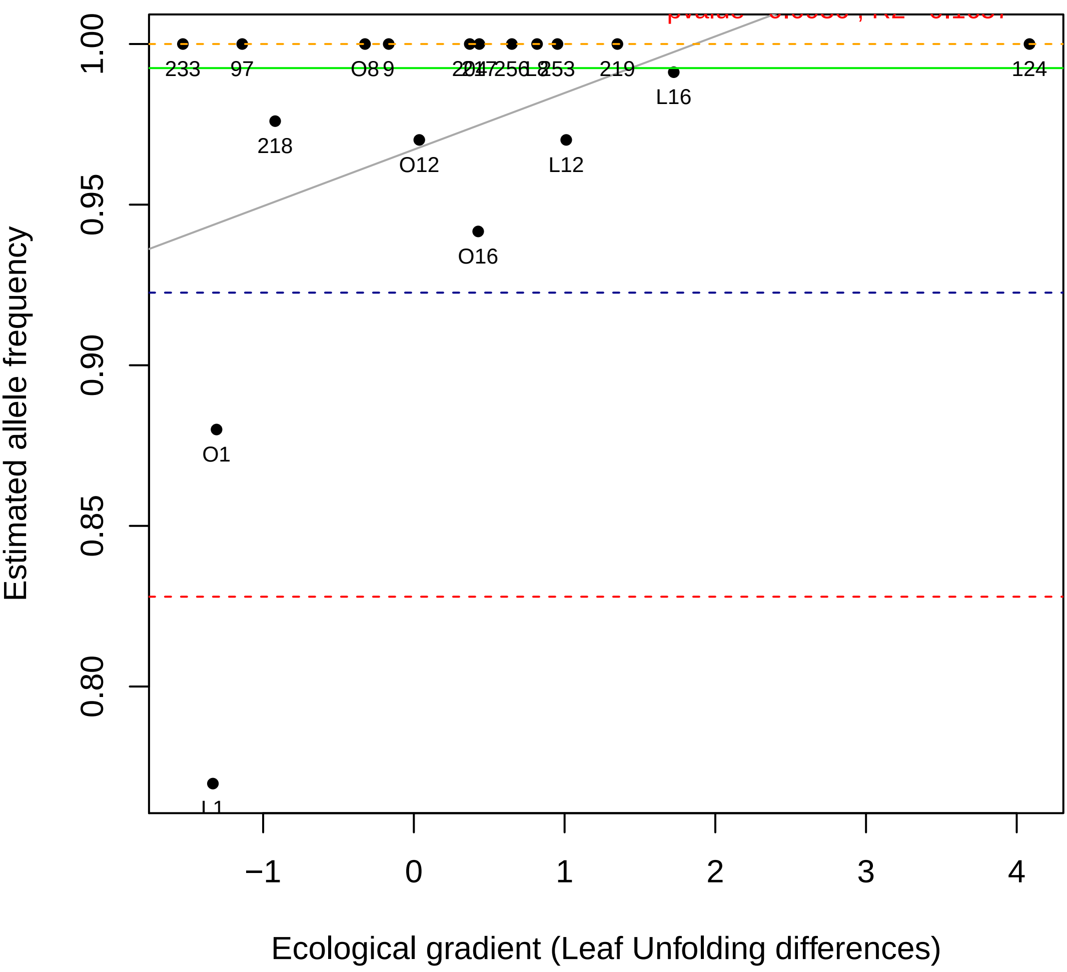

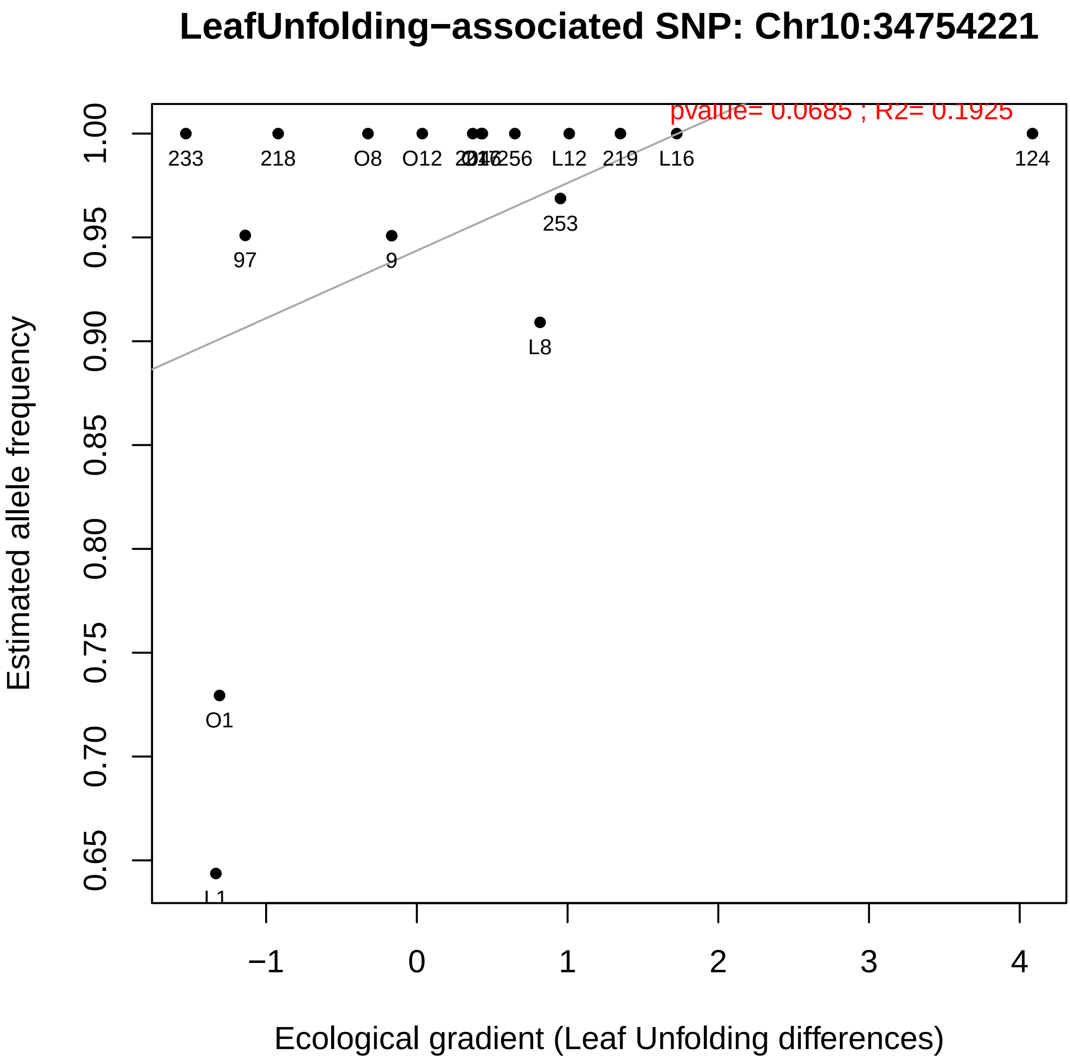
LeafUnfolding−associated SNP: Chr10:34754221
pvalue= 0.0685 ; R2= 0.1925
Estimated allele frequency
1.00
0.95
0.90
0.85
0.80
0.75
0.70
0.65
233
218
O8
O12
256
L12
219
L16
124
97
9
253
L8
O1
−1
0
1
2
3
4
Ecological gradient (Leaf Unfolding differences)

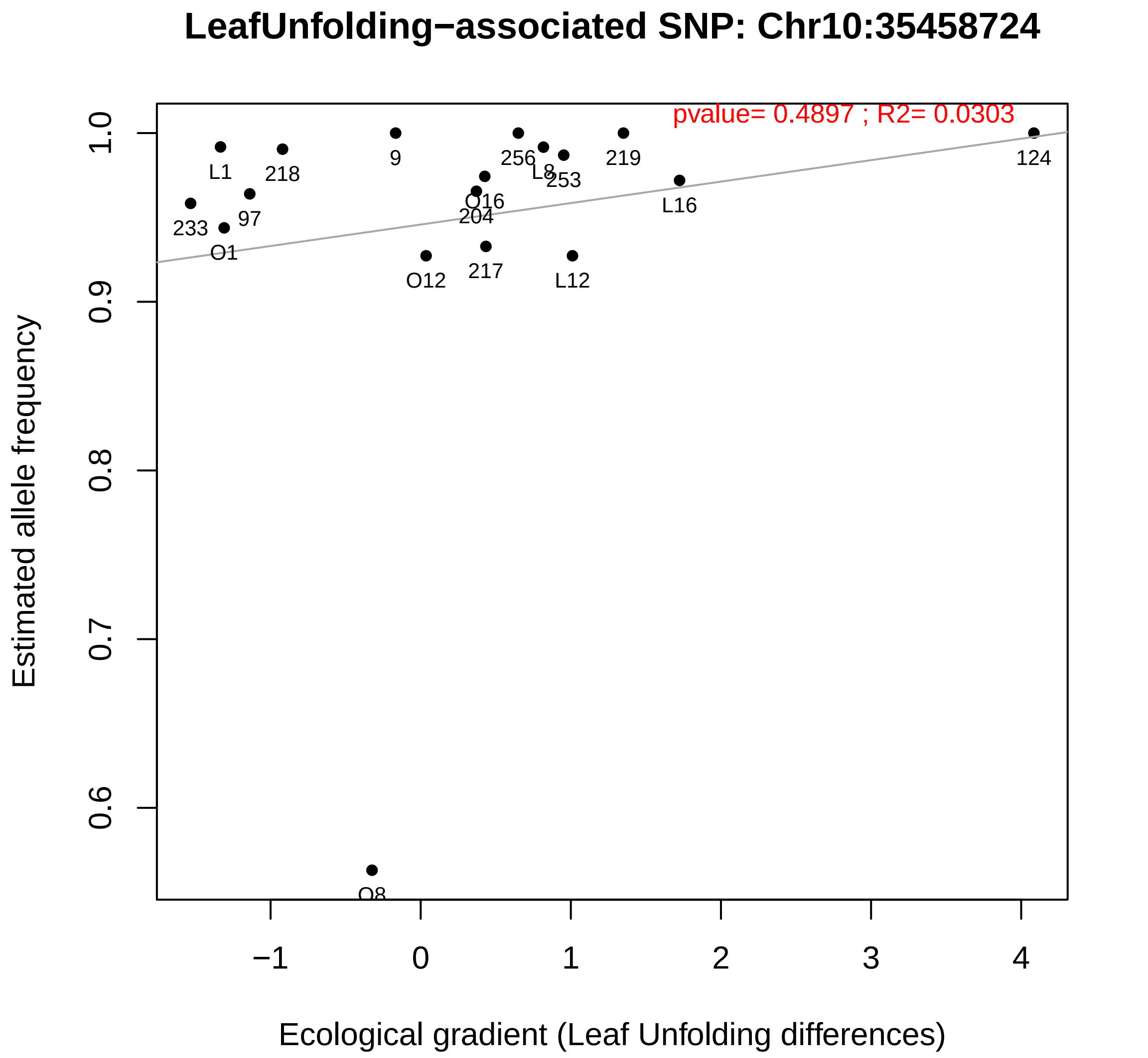
LeafUnfolding−associated SNP: Chr10:35458724
pvalue= 0.4897 ; R2= 0.0303
Estimated allele frequency
Ecological gradient (Leaf Unfolding differences)
0.6
0.7
0.8
0.9
1.0
−1
0
1
2
3
4
L1
218
233
97
O1
9
256
L8
253
219
124
O16
204
L16
O12
217
L12
O8

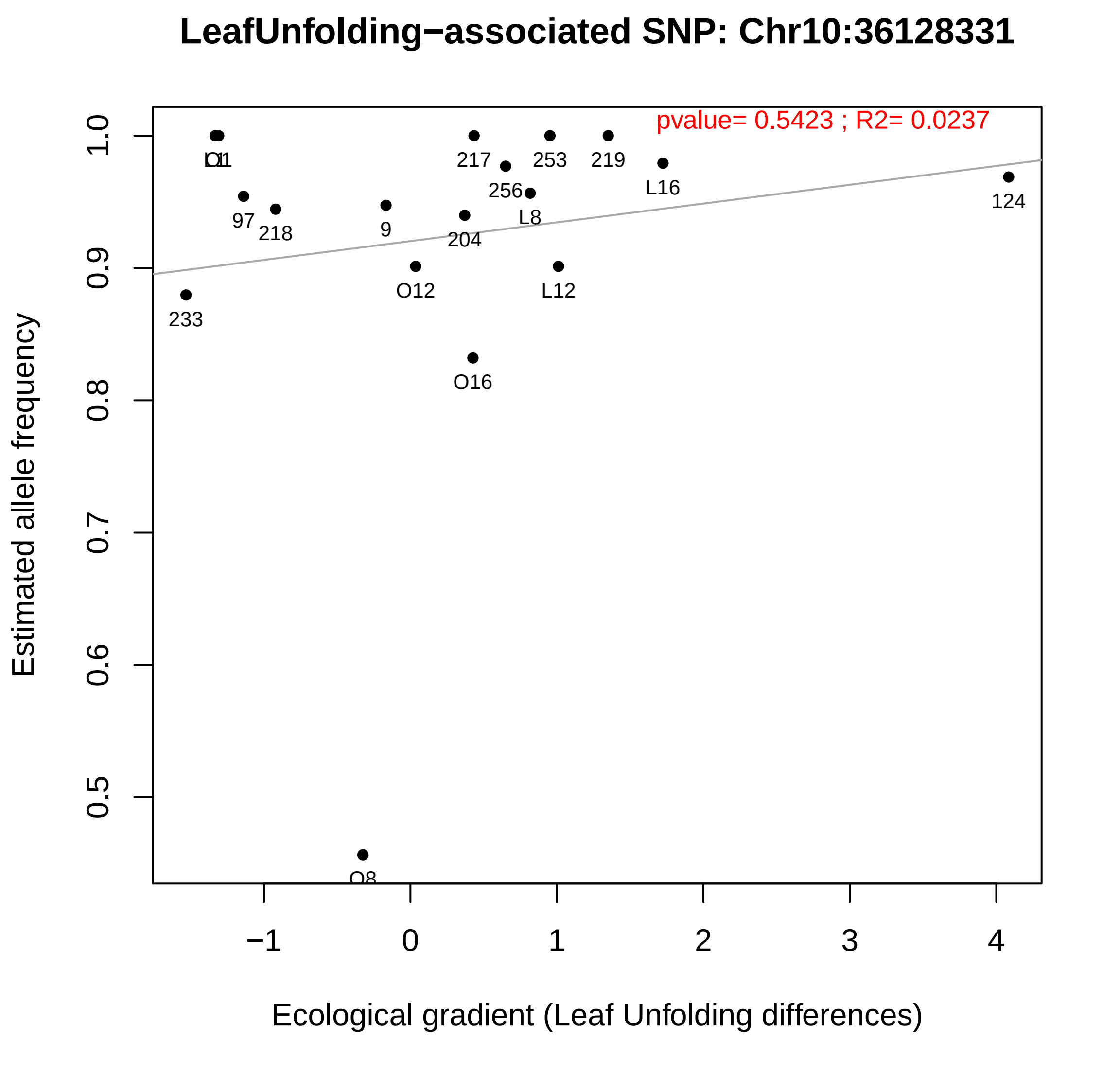

LeafUnfolding−associated SNP: Chr10:36128331
pvalue= 0.5423 ; R2= 0.0237
Estimated allele frequency
Ecological gradient (Leaf Unfolding differences)
1.0
0.9
0.8
0.7
0.6
0.5
−1
0
1
2
3
4
217
253
219
256
L16
124
97
218
9
204
L8
O12
L12
233
O16

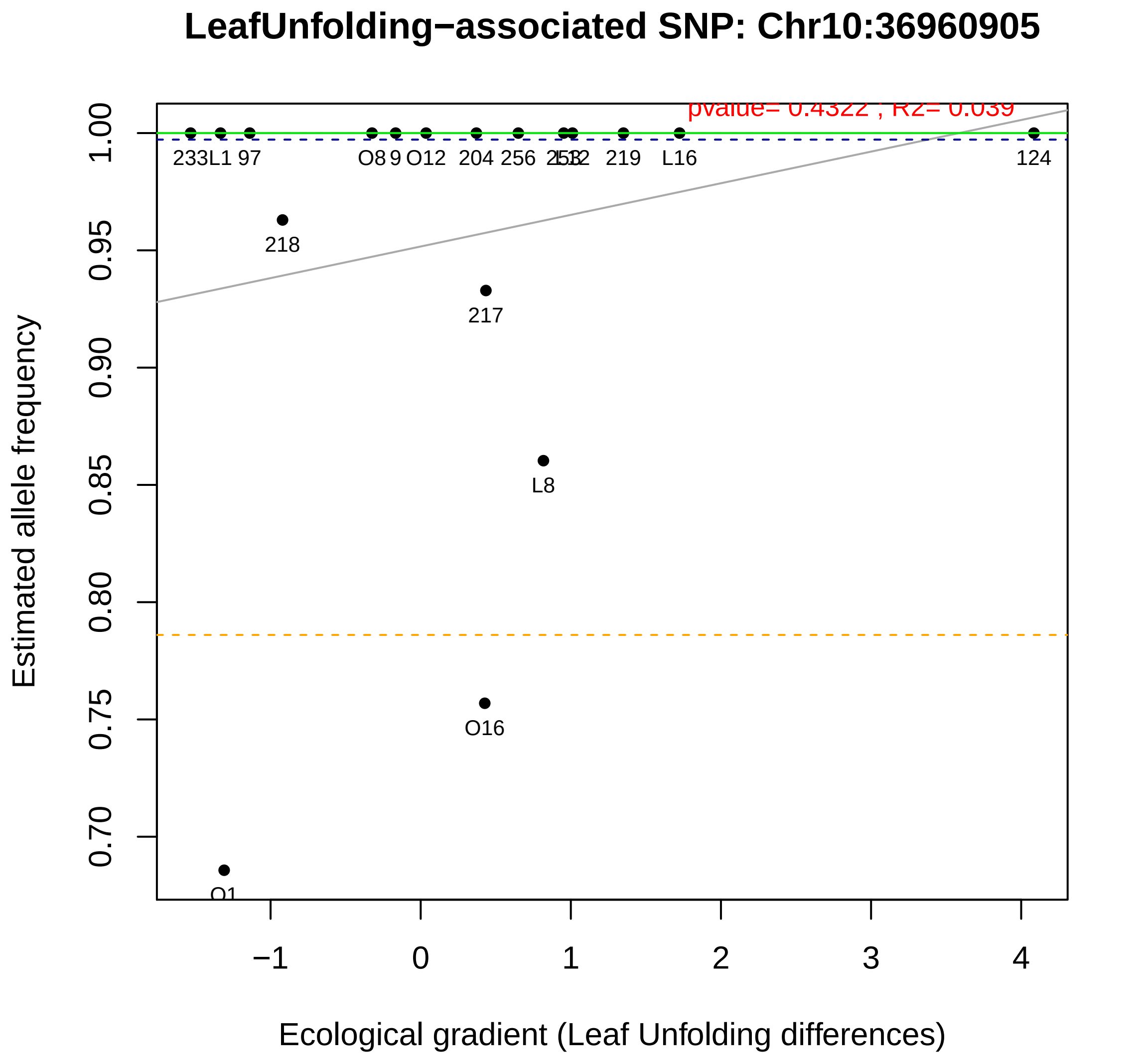

LeafUnfolding−associated SNP: Chr10:36960905
pvalue= 0.4322 ; R2= 0.039
Estimated allele frequency
Ecological gradient (Leaf Unfolding differences)
1.00
0.95
0.90
0.85
0.80
0.75
0.70
−1
0
1
2
3
4
233
L1
97
O8
9
O12
204
256
219
L16
124
218
217
L8
O16
O1

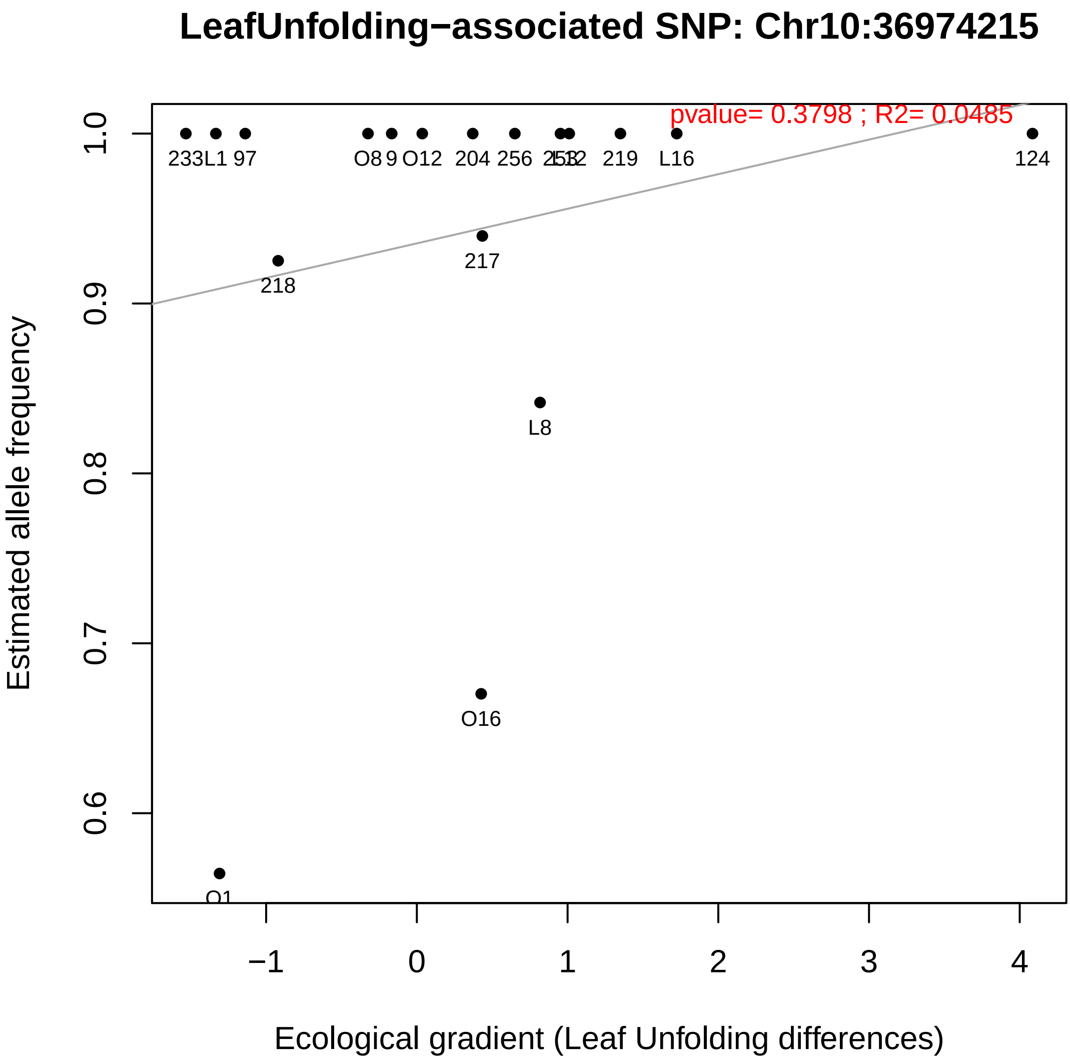

LeafUnfolding−associated SNP: Chr10:36974215
pvalue= 0.3798 ; R2= 0.0485
Estimated allele frequency
Ecological gradient (Leaf Unfolding differences)
1.0
0.9
0.8
0.7
0.6
−1
0
1
2
3
4
233
L1
97
O8
9
O12
204
256
219
L16
124
218
217
L8
O16
O1

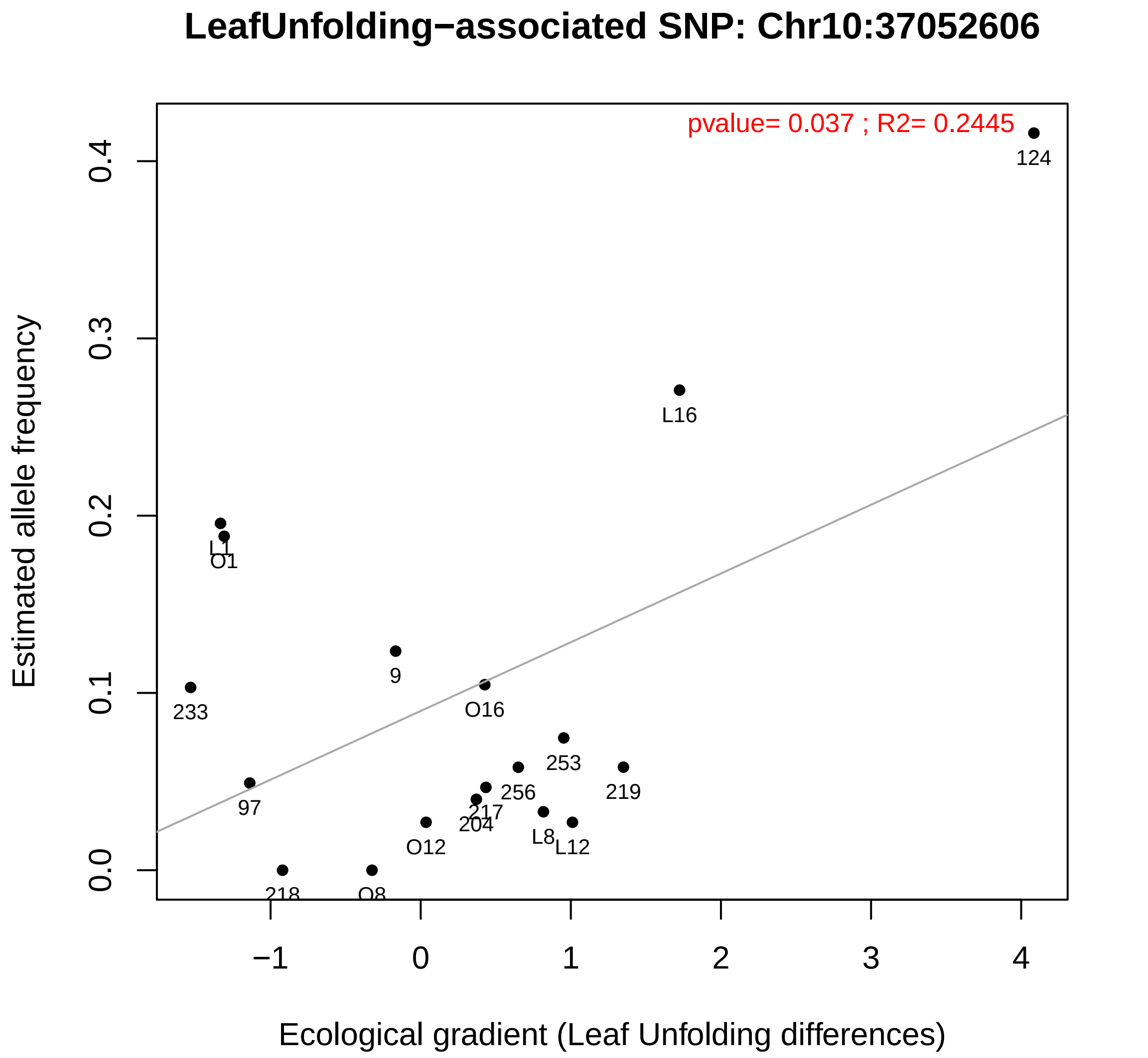

LeafUnfolding−associated SNP: Chr10:37052606
pvalue= 0.037 ; R2= 0.2445
Estimated allele frequency
Ecological gradient (Leaf Unfolding differences)
0.0
0.1
0.2
0.3
0.4
−1
0
1
2
3
4
124
L16
L1
O1
9
233
O16
253
256
219
97
217
204
L8
L12
O12
218
O8

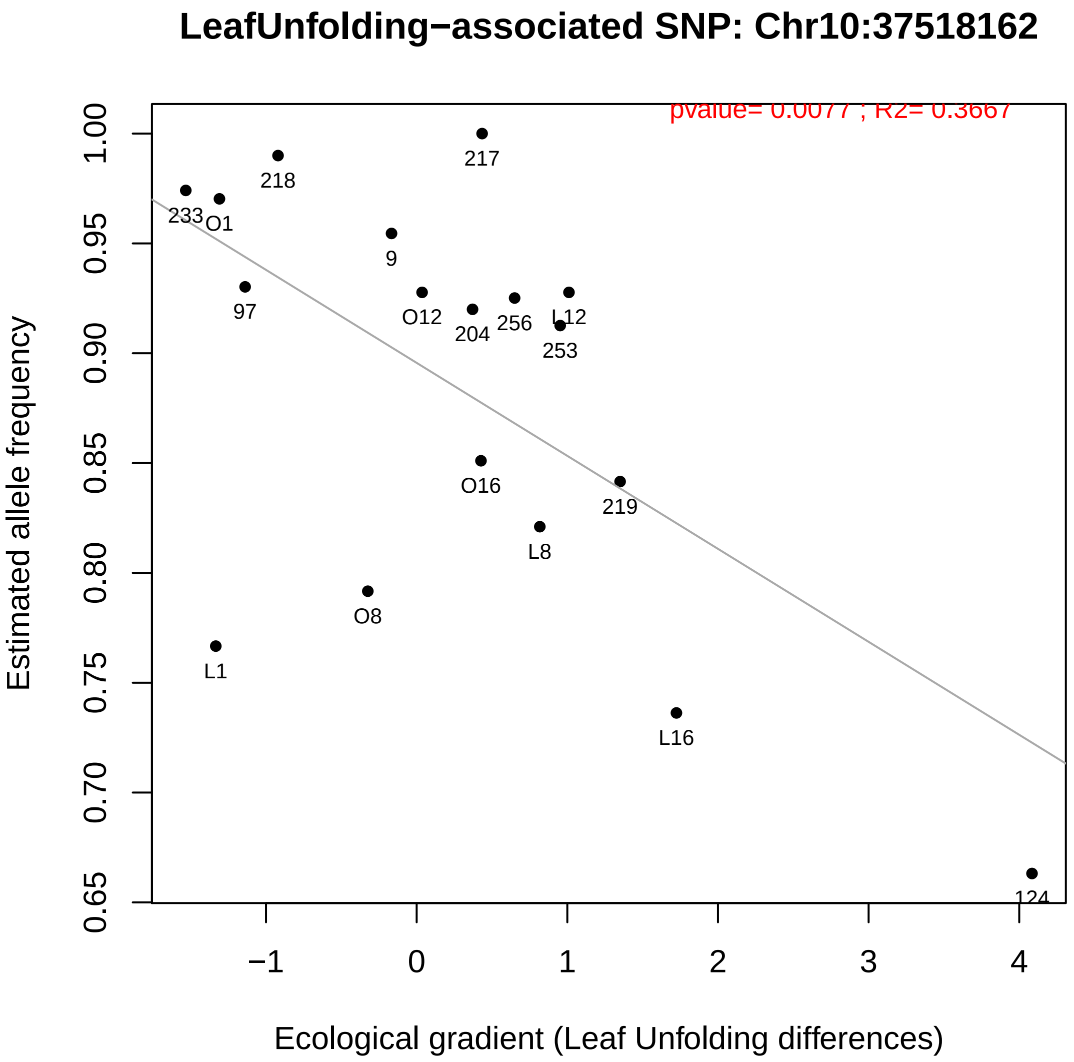

LeafUnfolding−associated SNP: Chr10:37518162
pvalue= 0.0077 ; R2= 0.3667
Estimated allele frequency
1.00
0.95
0.90
0.85
0.80
0.75
0.70
0.65
−1
0
1
2
3
4
Ecological gradient (Leaf Unfolding differences)
233
O1
218
217
9
97
O12
204
256
L12
253
O16
219
L8
O8
L1
L16
124

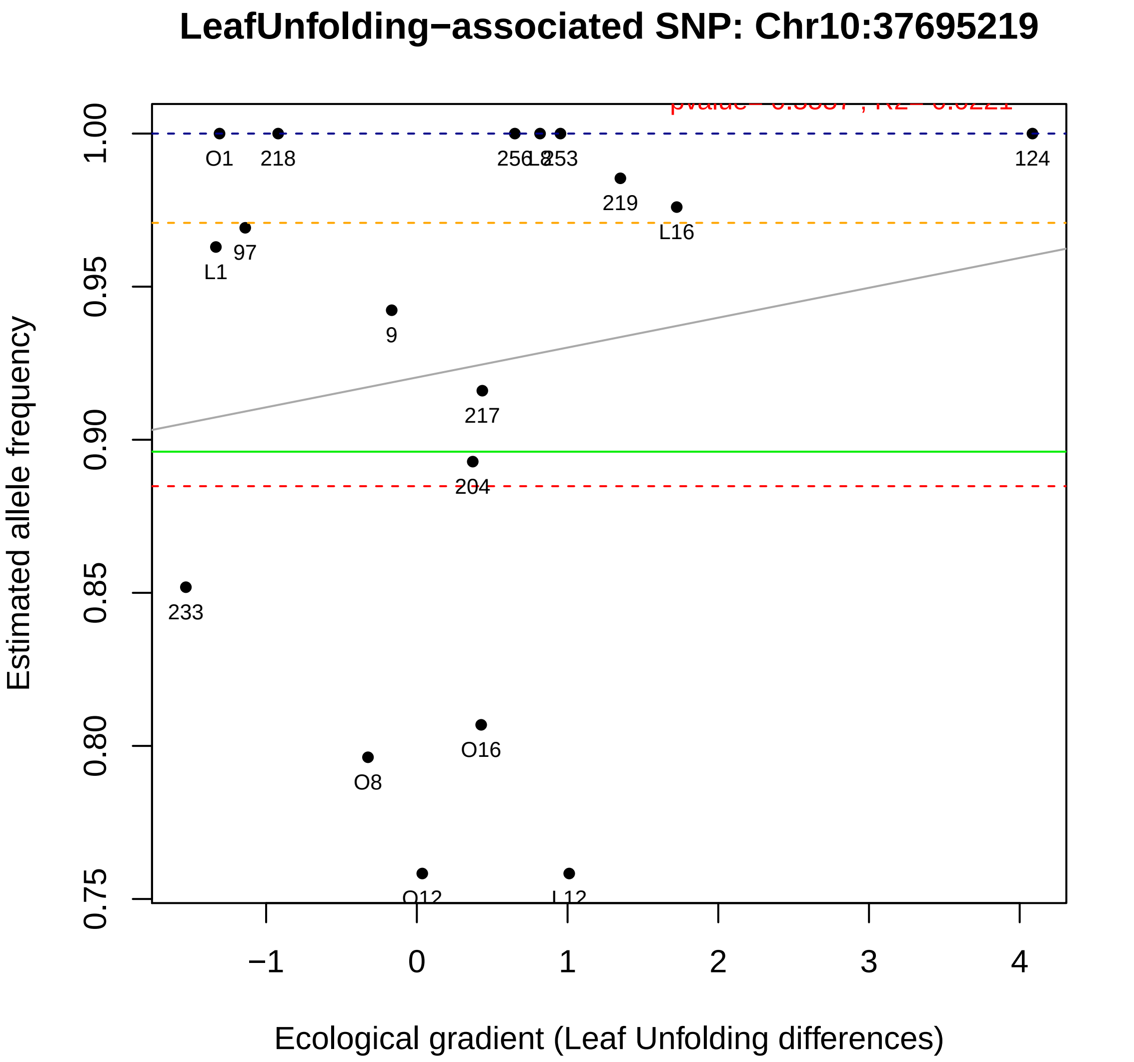
LeafUnfolding−associated SNP: Chr10:37695219
Estimated allele frequency
Ecological gradient (Leaf Unfolding differences)
1.00
0.95
0.90
0.85
0.80
0.75
−1
0
1
2
3
4
O1
218
256
L8
253
124
219
L16
97
L1
9
217
204
233
O16
O8

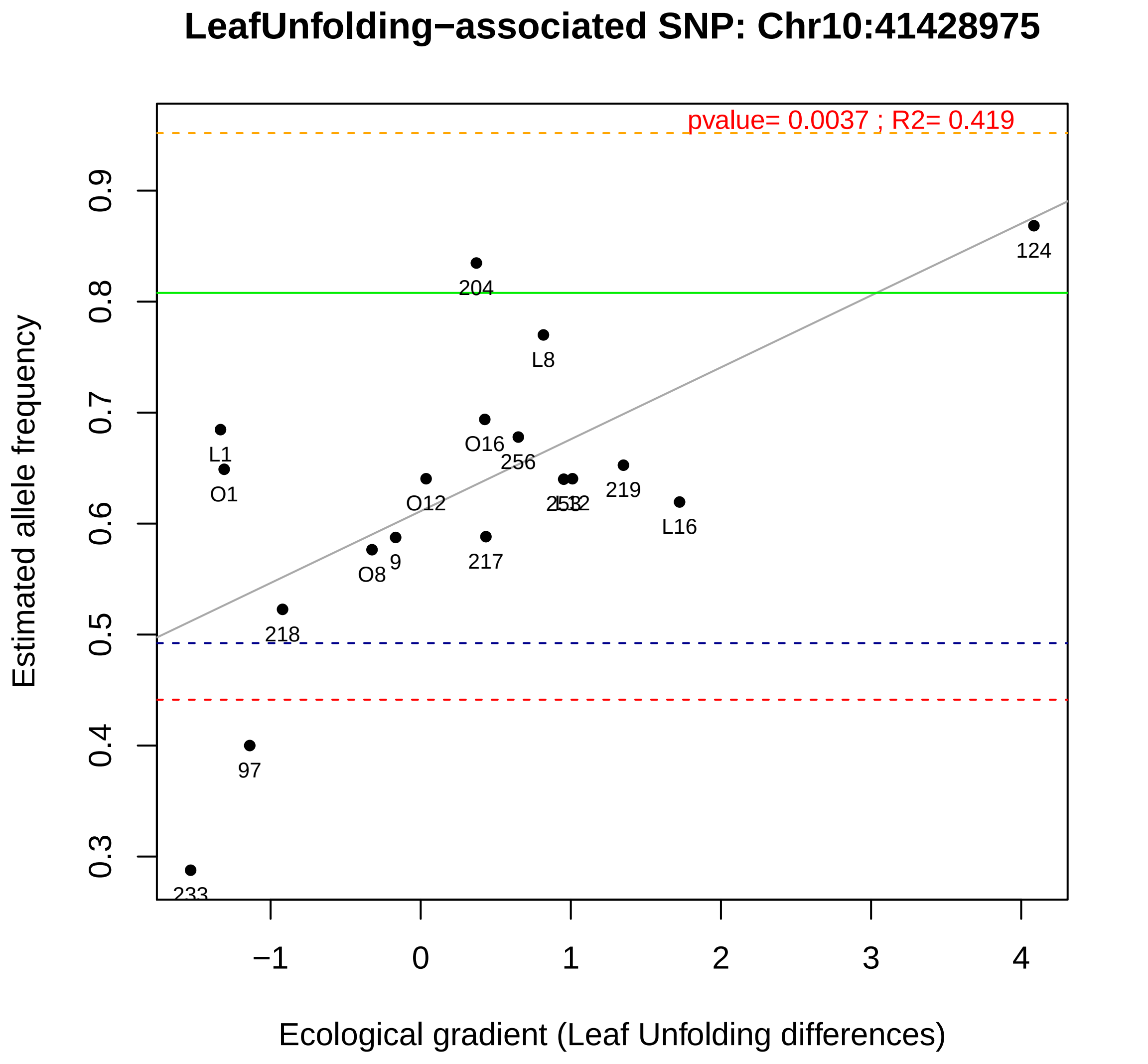

LeafUnfolding−associated SNP: Chr10:41428975
pvalue= 0.0037 ; R2= 0.419
Estimated allele frequency
Ecological gradient (Leaf Unfolding differences)
0.3
0.4
0.5
0.6
0.7
0.8
0.9
−1
0
1
2
3
4
124
204
L8
L1
O1
O16
256
O12
219
L16
9
O8
217
218
97
233

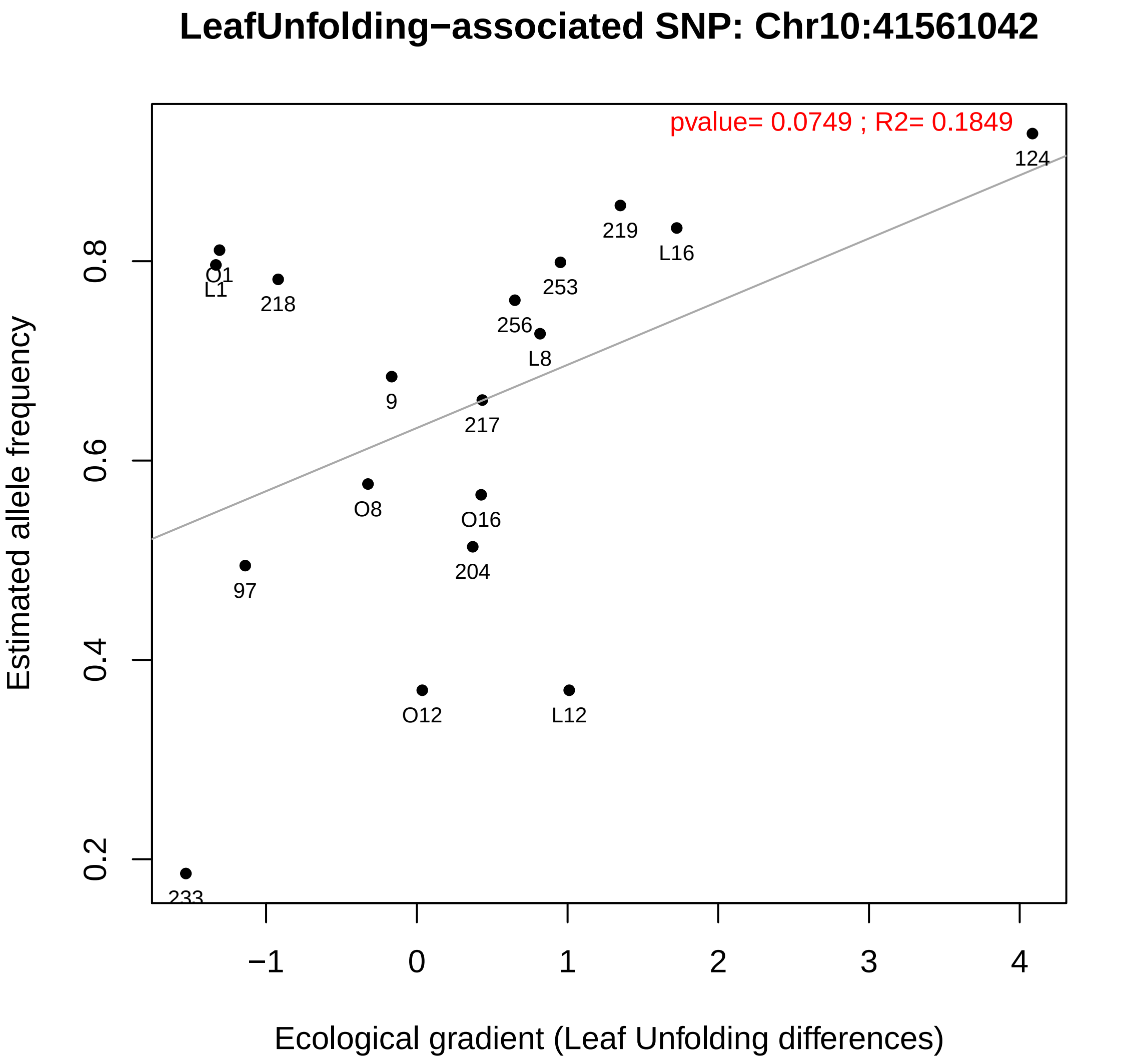

LeafUnfolding−associated SNP: Chr10:41561042
pvalue= 0.0749 ; R2= 0.1849
Estimated allele frequency
Ecological gradient (Leaf Unfolding differences)
0.2
0.4
0.6
0.8
−1
0
1
2
3
4
124
219
L16
253
O1
L1
218
256
L8
9
217
O8
O16
204
97
O12
L12
233

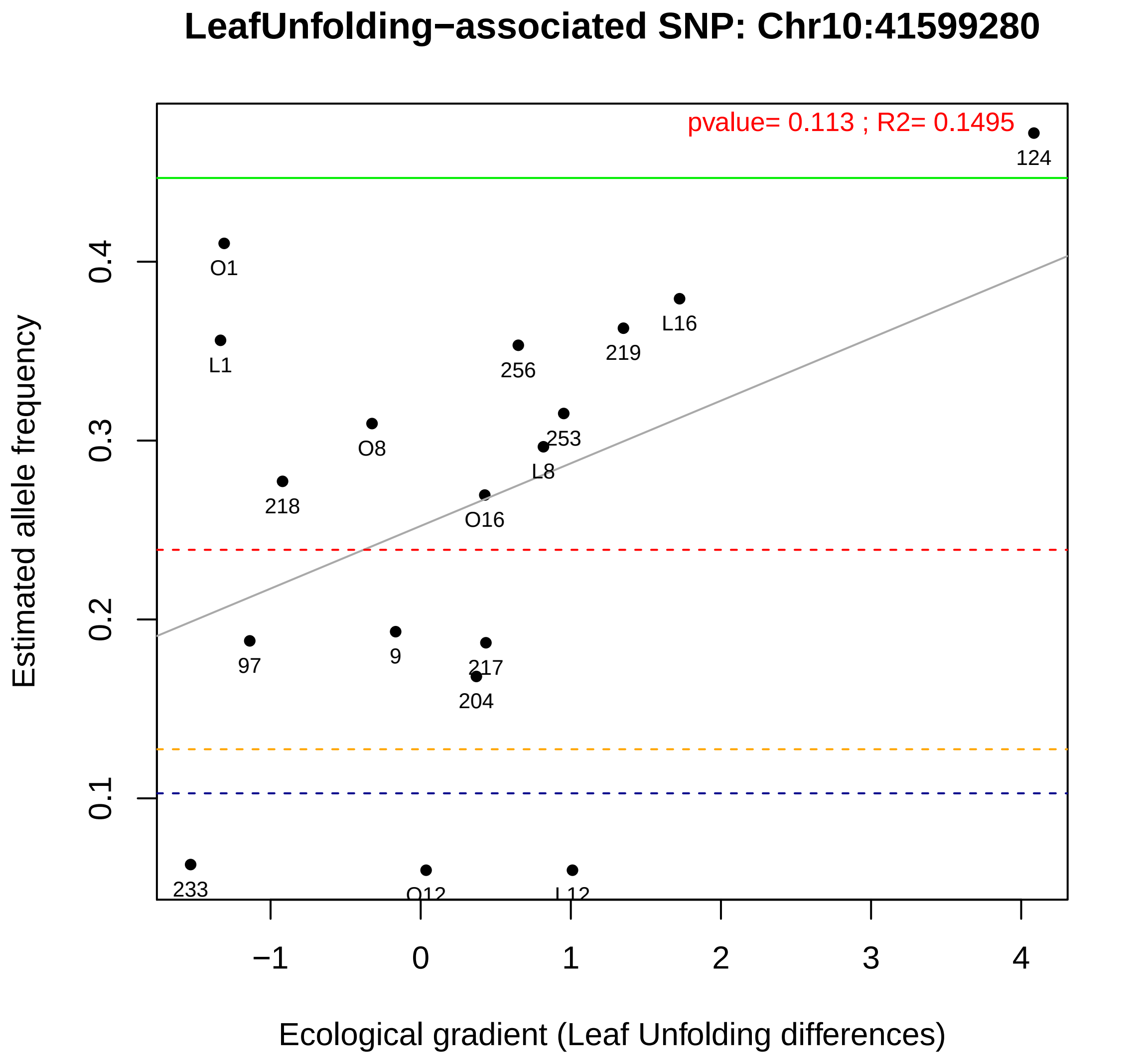
LeafUnfolding−associated SNP: Chr10:41599280
pvalue= 0.113 ; R2= 0.1495
Estimated allele frequency
0.1
0.2
0.3
0.4
−1
0
1
2
3
4
Ecological gradient (Leaf Unfolding differences)
124
O1
L16
L1
219
256
253
O8
L8
218
O16
97
9
217
204
233
O12
L12

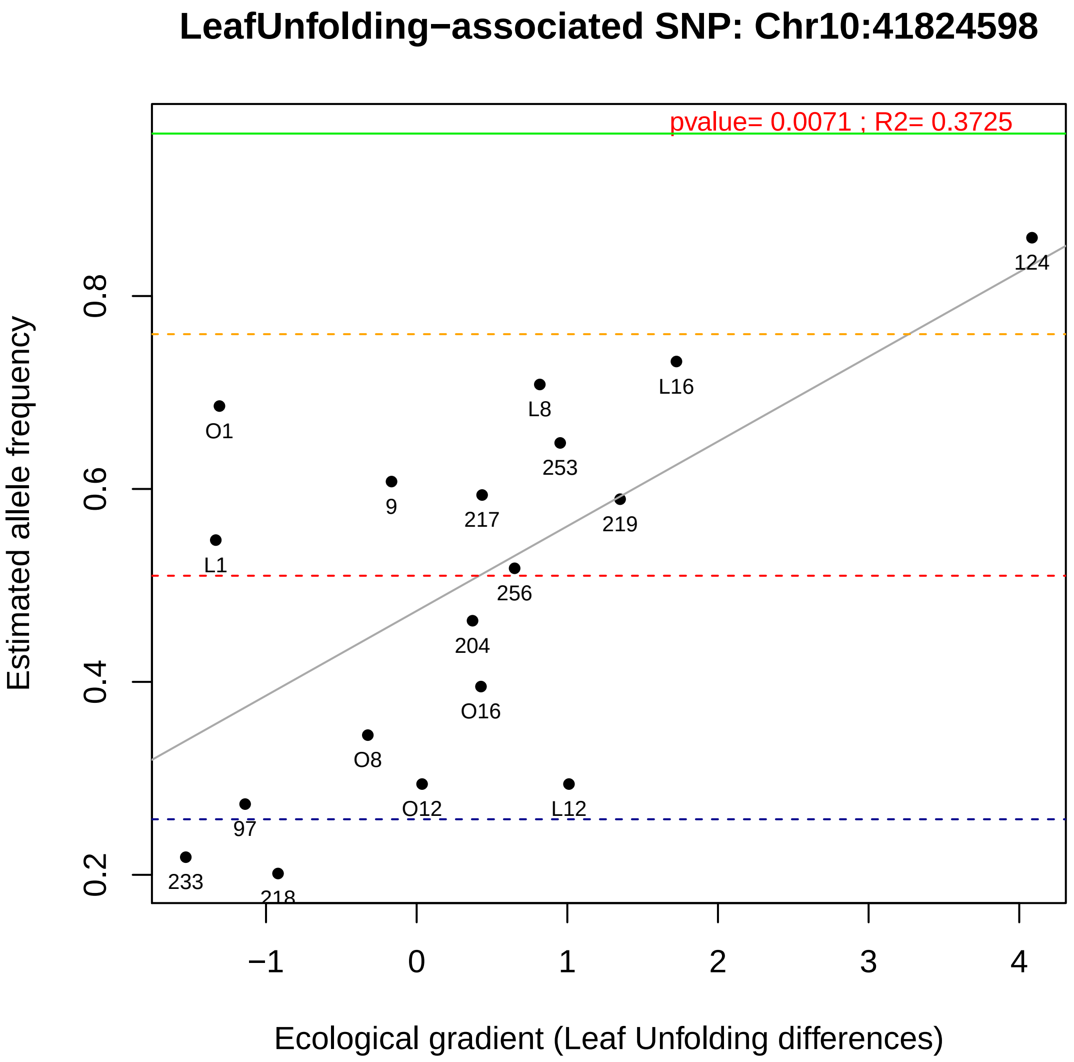
LeafUnfolding−associated SNP: Chr10:41824598
pvalue= 0.0071 ; R2= 0.3725
Estimated allele frequency
0.2
0.4
0.6
0.8
−1
0
1
2
3
4
Ecological gradient (Leaf Unfolding differences)
124
L16
L8
O1
253
9
217
219
L1
256
204
O16
O8
O12
L12
97
233
218

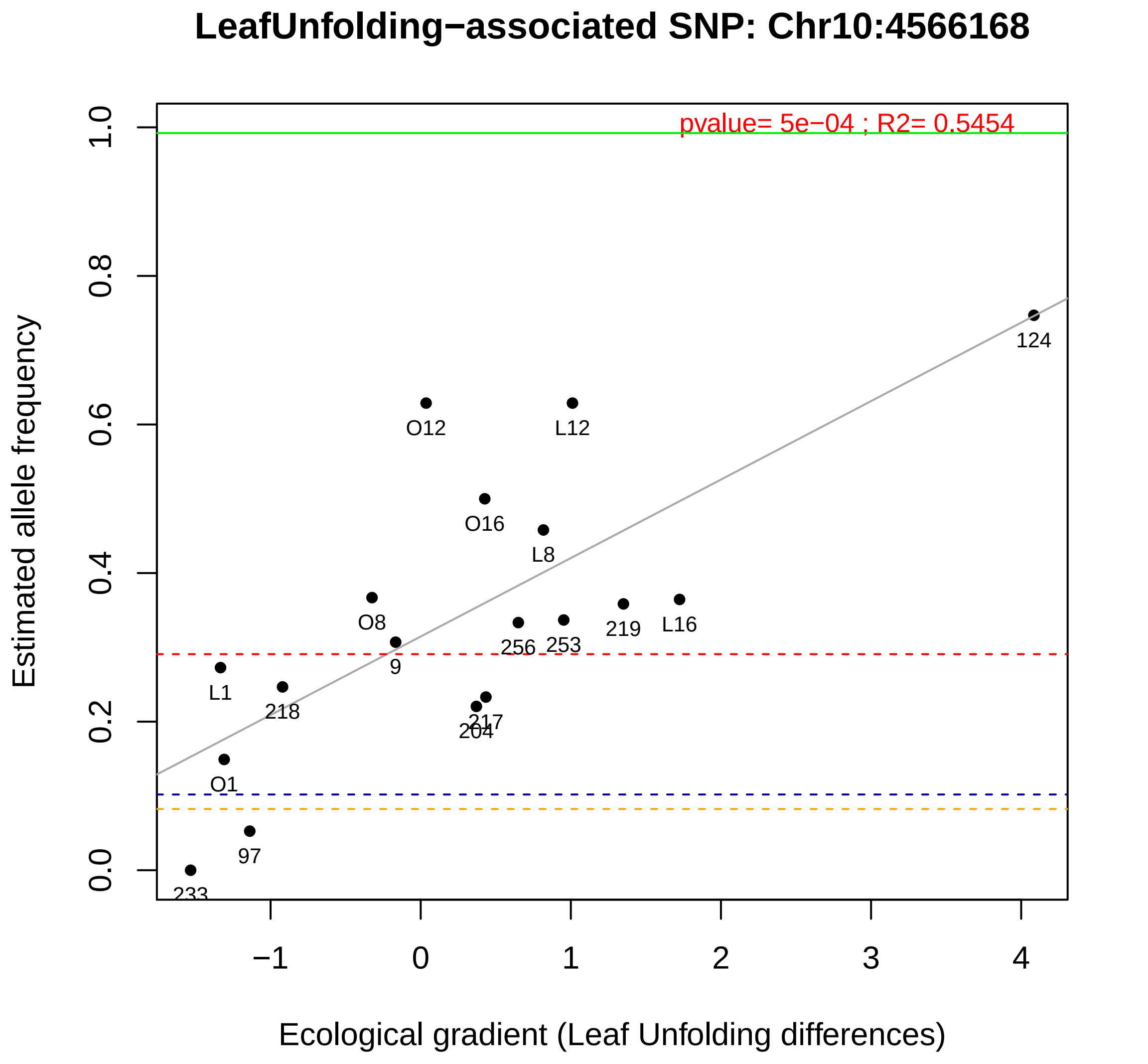

LeafUnfolding−associated SNP: Chr10:4566168
pvalue= 5e−04 ; R2= 0.5454
Estimated allele frequency
0.0
0.2
0.4
0.6
0.8
1.0
Ecological gradient (Leaf Unfolding differences)
−1
0
1
2
3
4
124
O12
L12
O16
L8
O8
9
256
253
219
L16
L1
218
217
204
O1
97
233

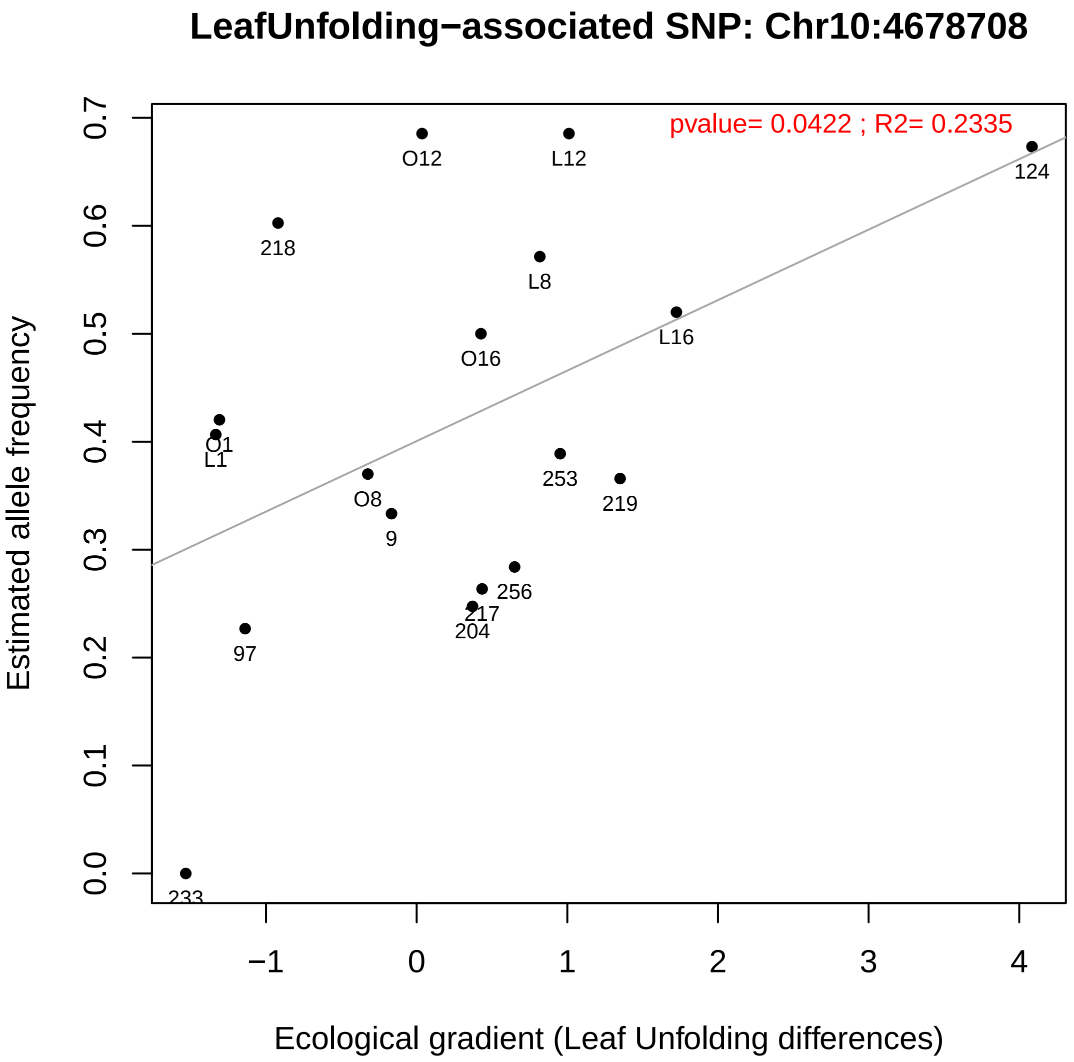
LeafUnfolding−associated SNP: Chr10:4678708
pvalue= 0.0422 ; R2= 0.2335
Estimated allele frequency
Ecological gradient (Leaf Unfolding differences)
0.0
0.1
0.2
0.3
0.4
0.5
0.6
0.7
−1
0
1
2
3
4
O12
L12
124
218
L8
L16
O16
O1
L1
253
219
O8
9
256
217
204
97
233

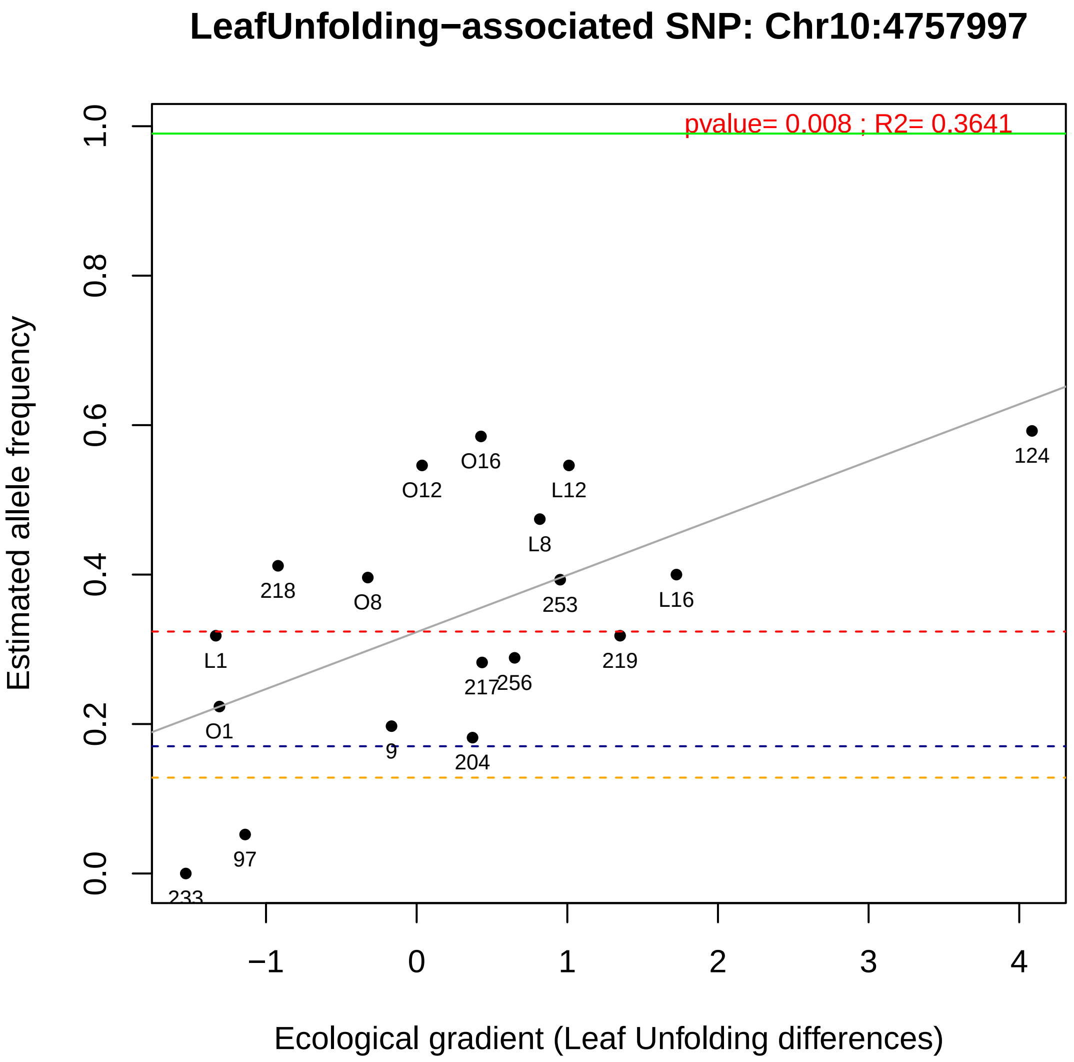
LeafUnfolding−associated SNP: Chr10:4757997
pvalue= 0.008 ; R2= 0.3641
Estimated allele frequency
Ecological gradient (Leaf Unfolding differences)
0.0
0.2
0.4
0.6
0.8
1.0
−1
0
1
2
3
4
O16
O12
L12
L8
218
O8
253
L16
L1
219
217
256
O1
9
204
97
233
124

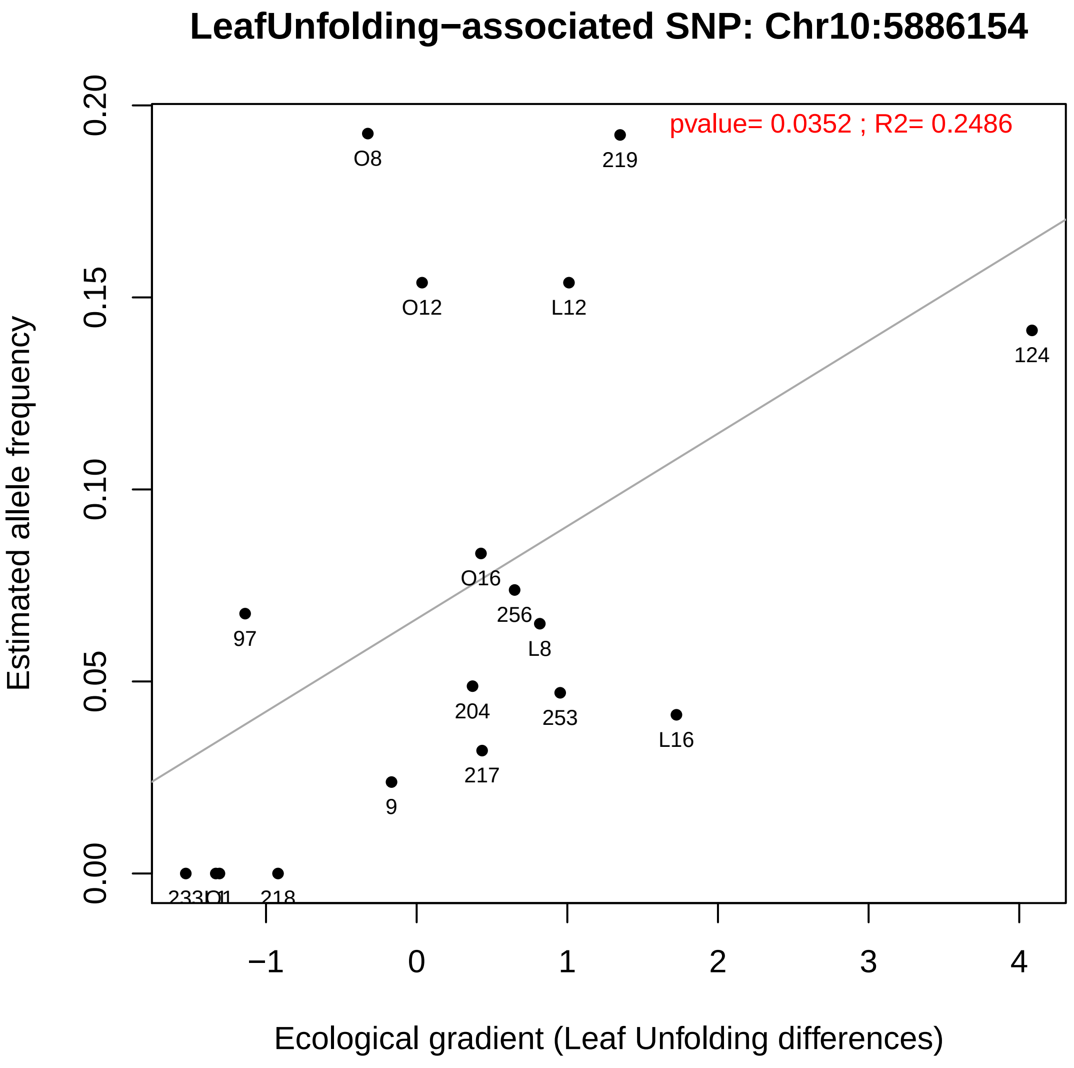
LeafUnfolding−associated SNP: Chr10:5886154
pvalue= 0.0352 ; R2= 0.2486
Estimated allele frequency
Ecological gradient (Leaf Unfolding differences)
0.00
0.05
0.10
0.15
0.20
−1
0
1
2
3
4
O8
219
O12
L12
124
O16
256
L8
97
204
253
L16
217
9
233
O1
218

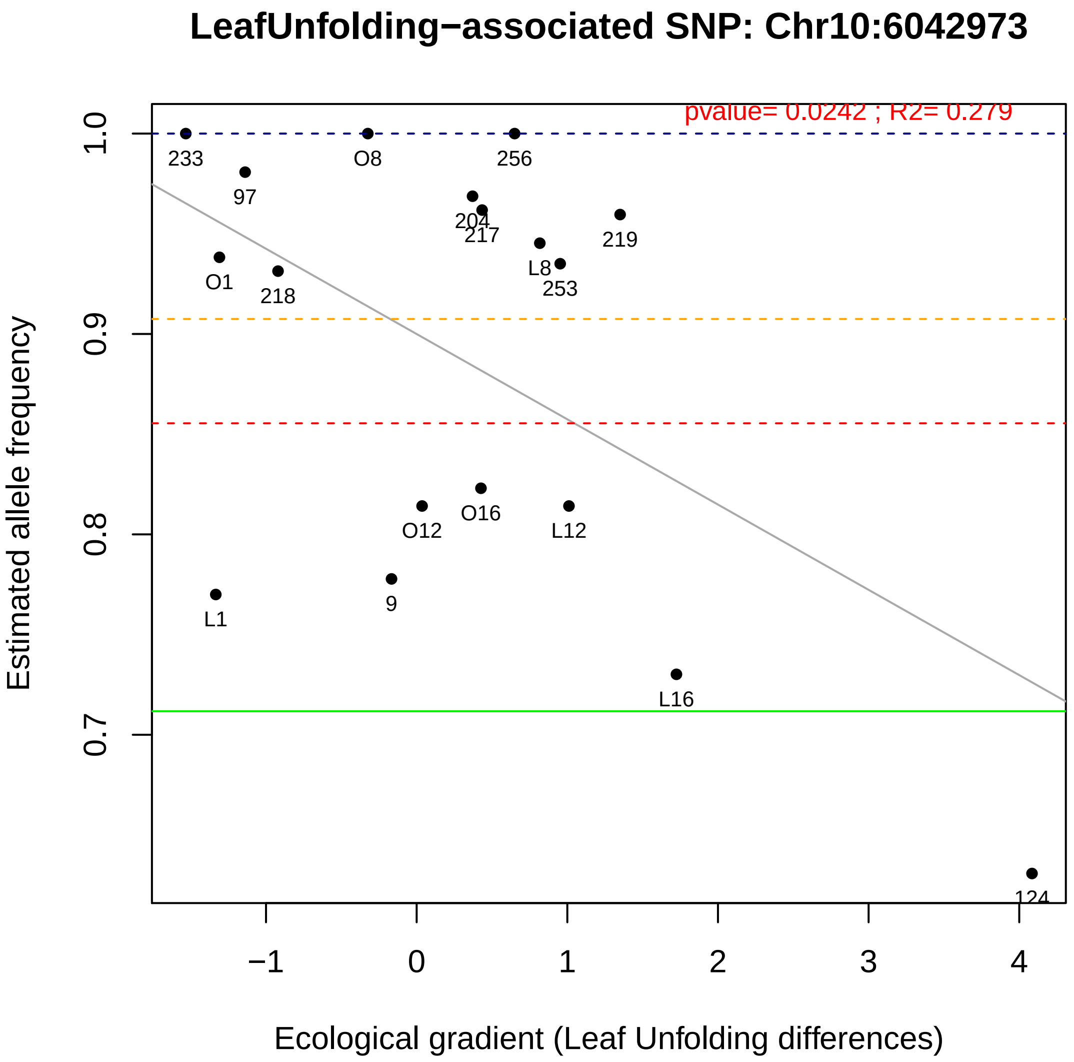
LeafUnfolding−associated SNP: Chr10:6042973
pvalue= 0.0242 ; R2= 0.279
Estimated allele frequency
1.0
0.9
0.8
0.7
233
97
O8
256
204
217
219
O1
218
L8
253
O16
O12
L12
L1
9
L16
124
−1
0
1
2
3
4
Ecological gradient (Leaf Unfolding differences)

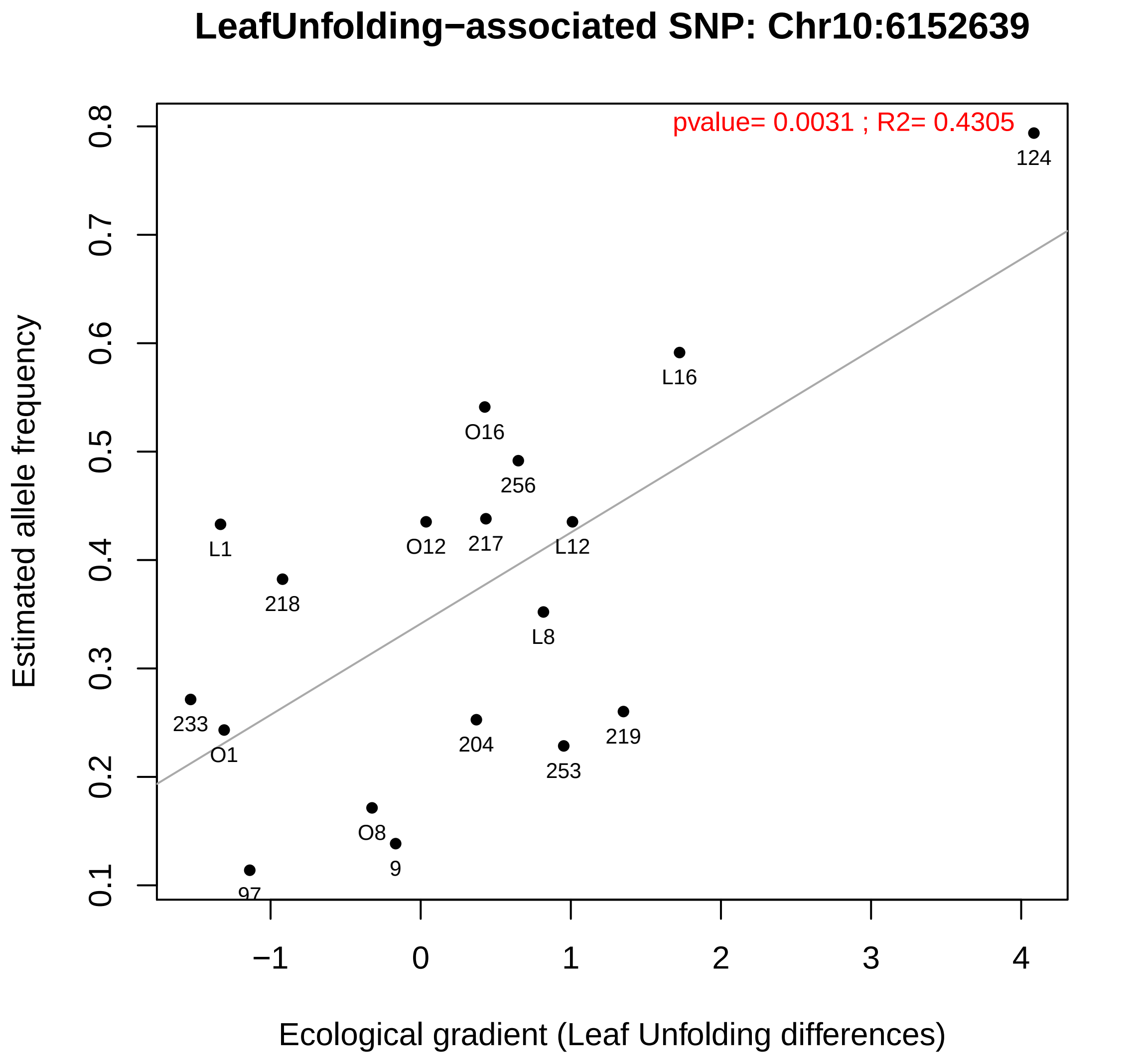

LeafUnfolding−associated SNP: Chr10:6152639
pvalue= 0.0031 ; R2= 0.4305
Estimated allele frequency
Ecological gradient (Leaf Unfolding differences)
0.1
0.2
0.3
0.4
0.5
0.6
0.7
0.8
−1
0
1
2
3
4
124
L16
O16
256
L1
O12
217
L12
218
L8
233
O1
204
219
253
O8
9
97

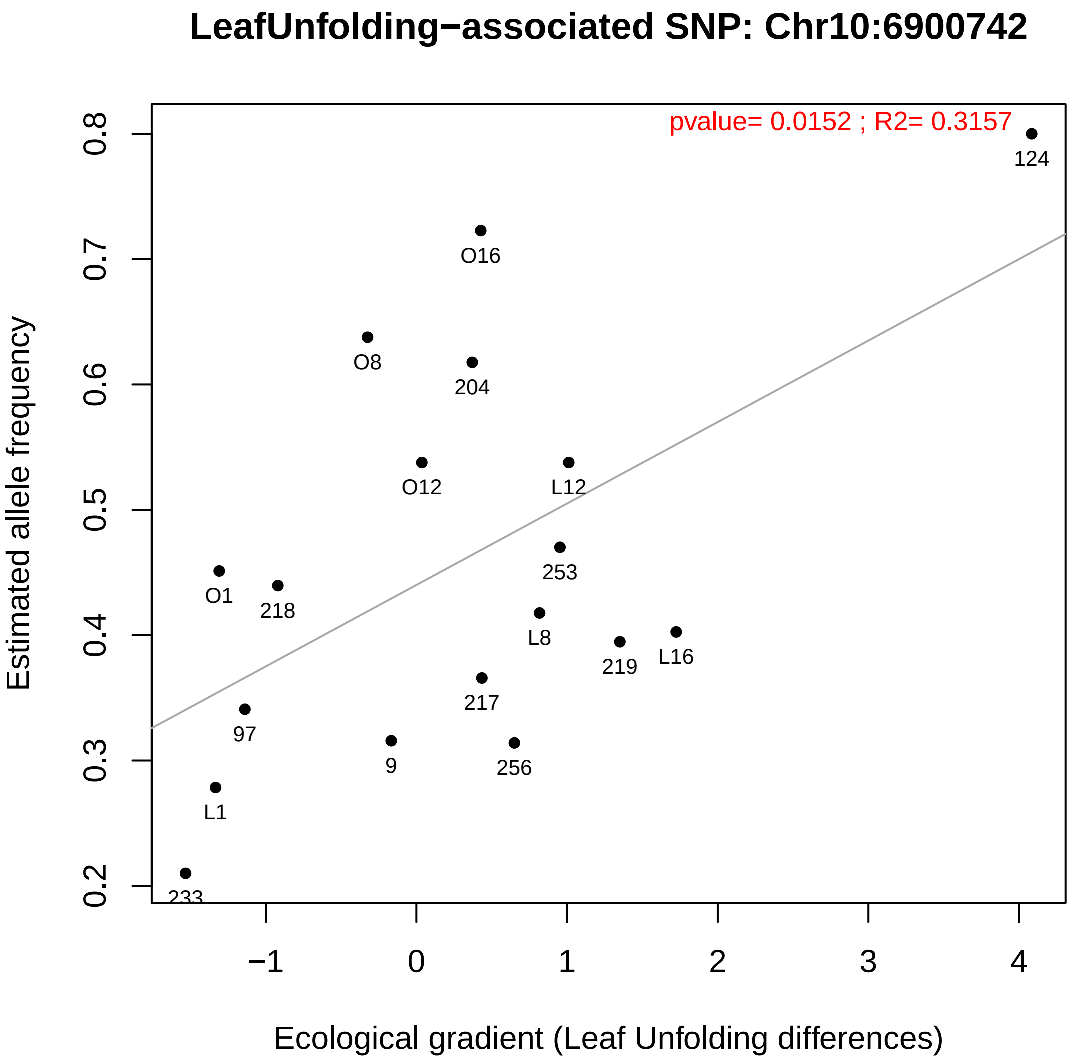
LeafUnfolding−associated SNP: Chr10:6900742
pvalue= 0.0152 ; R2= 0.3157
Estimated allele frequency
0.2
0.3
0.4
0.5
0.6
0.7
0.8
Ecological gradient (Leaf Unfolding differences)
−1
0
1
2
3
4
124
O16
O8
204
O12
L12
253
O1
218
L8
219
L16
217
97
9
256
L1
233

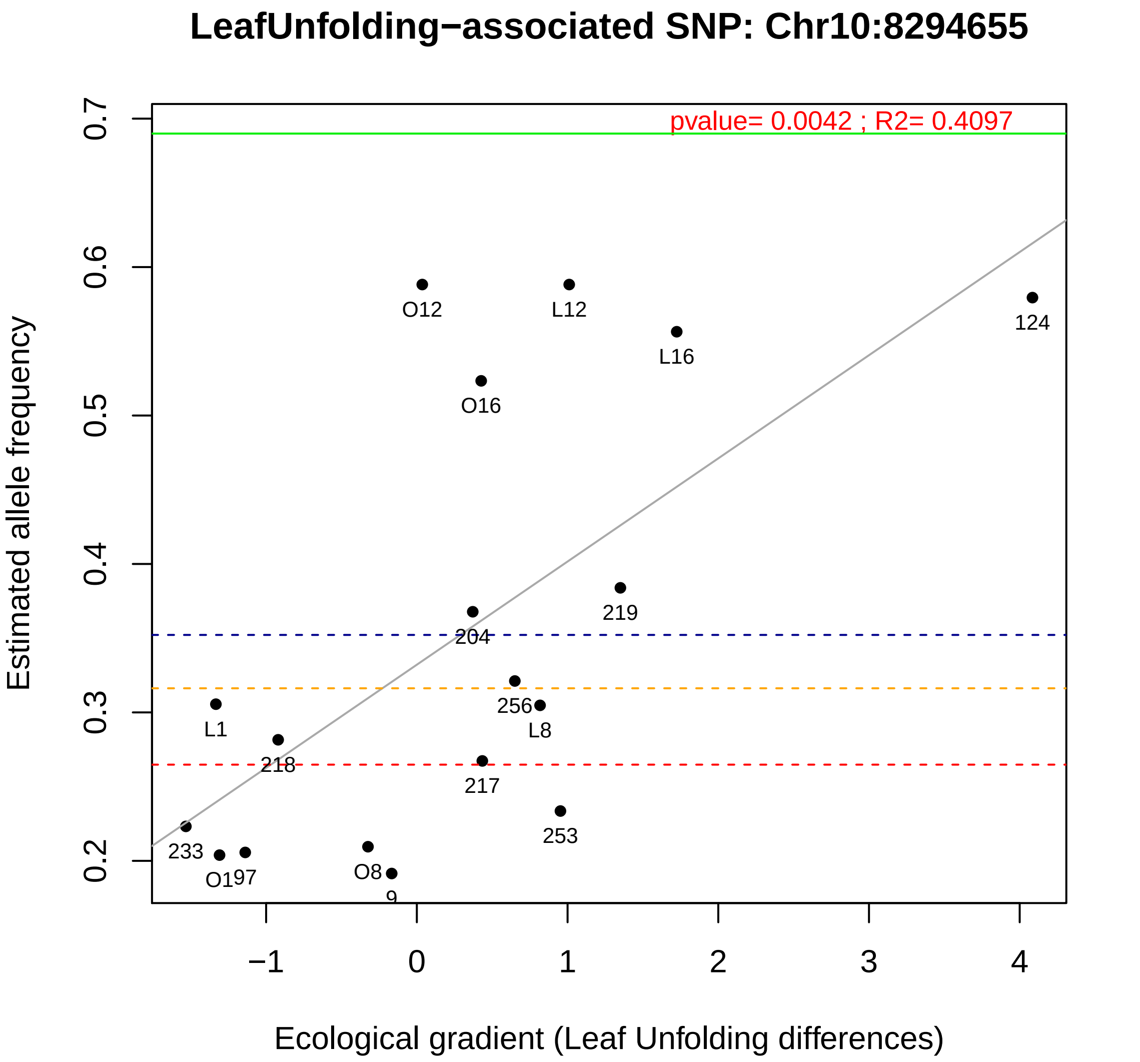
LeafUnfolding−associated SNP: Chr10:8294655
pvalue= 0.0042 ; R2= 0.4097
Estimated allele frequency
0.2
0.3
0.4
0.5
0.6
0.7
−1
0
1
2
3
4
Ecological gradient (Leaf Unfolding differences)
O12
L12
124
L16
O16
219
204
256
L8
L1
218
217
253
233
O197
O8
9

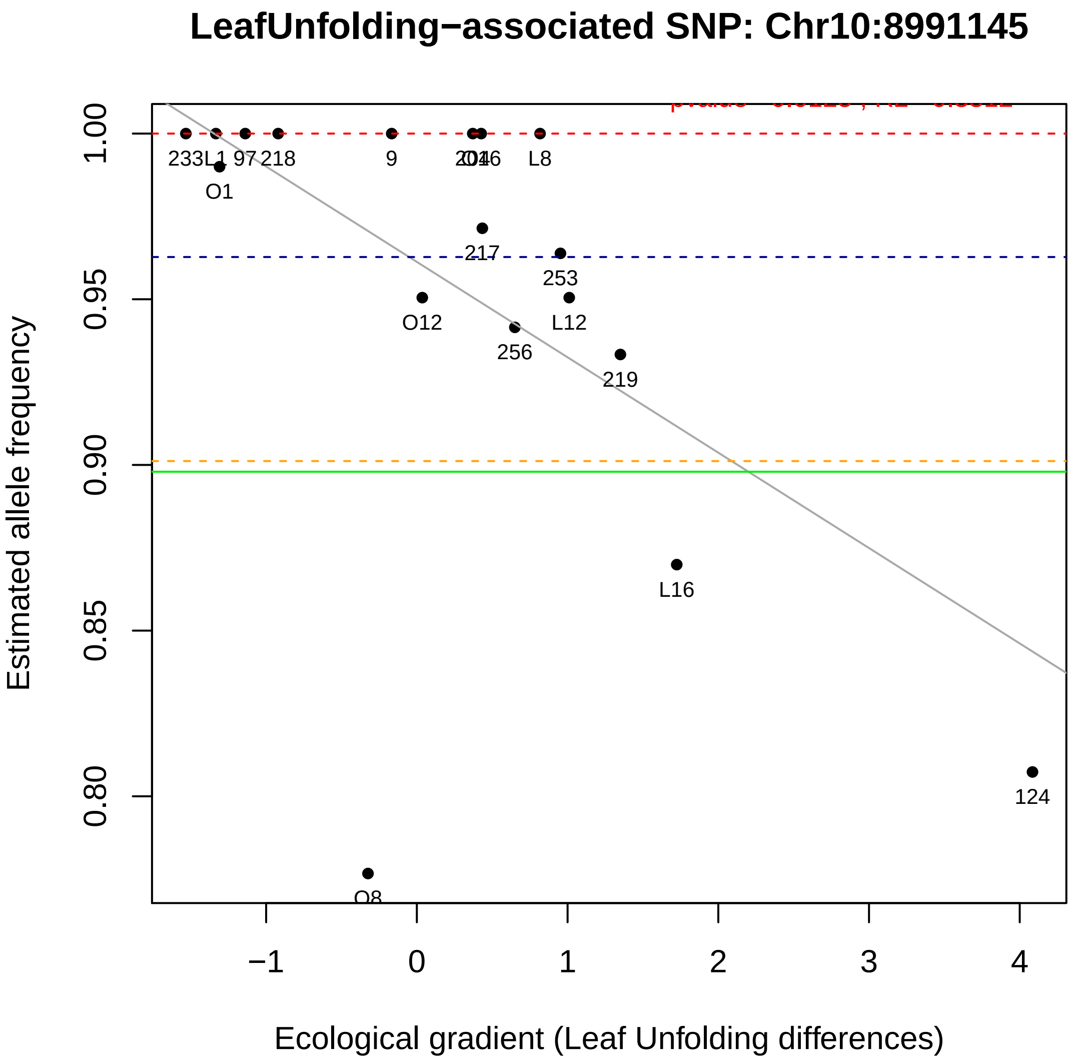

LeafUnfolding−associated SNP: Chr10:8991145
Estimated allele frequency
Ecological gradient (Leaf Unfolding differences)
1.00
0.95
0.90
0.85
0.80
−1
0
1
2
3
4
233
L1
97
218
9
L8
O1
217
253
O12
L12
256
219
L16
124

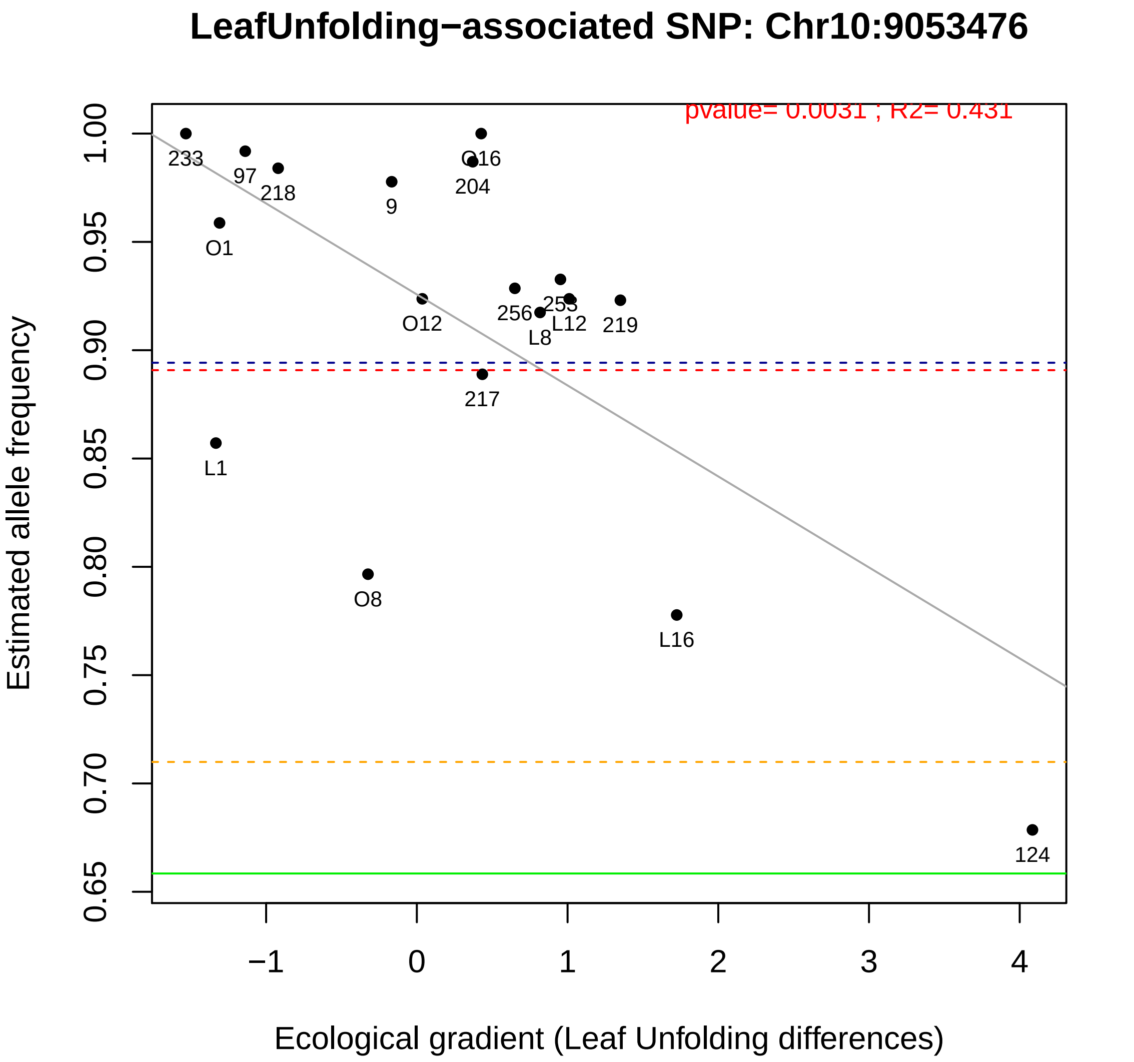

LeafUnfolding−associated SNP: Chr10:9053476
pvalue= 0.0031 ; R2= 0.431
Estimated allele frequency
Ecological gradient (Leaf Unfolding differences)
0.65
0.70
0.75
0.80
0.85
0.90
0.95
1.00
−1
0
1
2
3
4
233
97
218
9
O16
204
O1
O12
256
253
L12
219
L8
217
L1
O8
L16
124

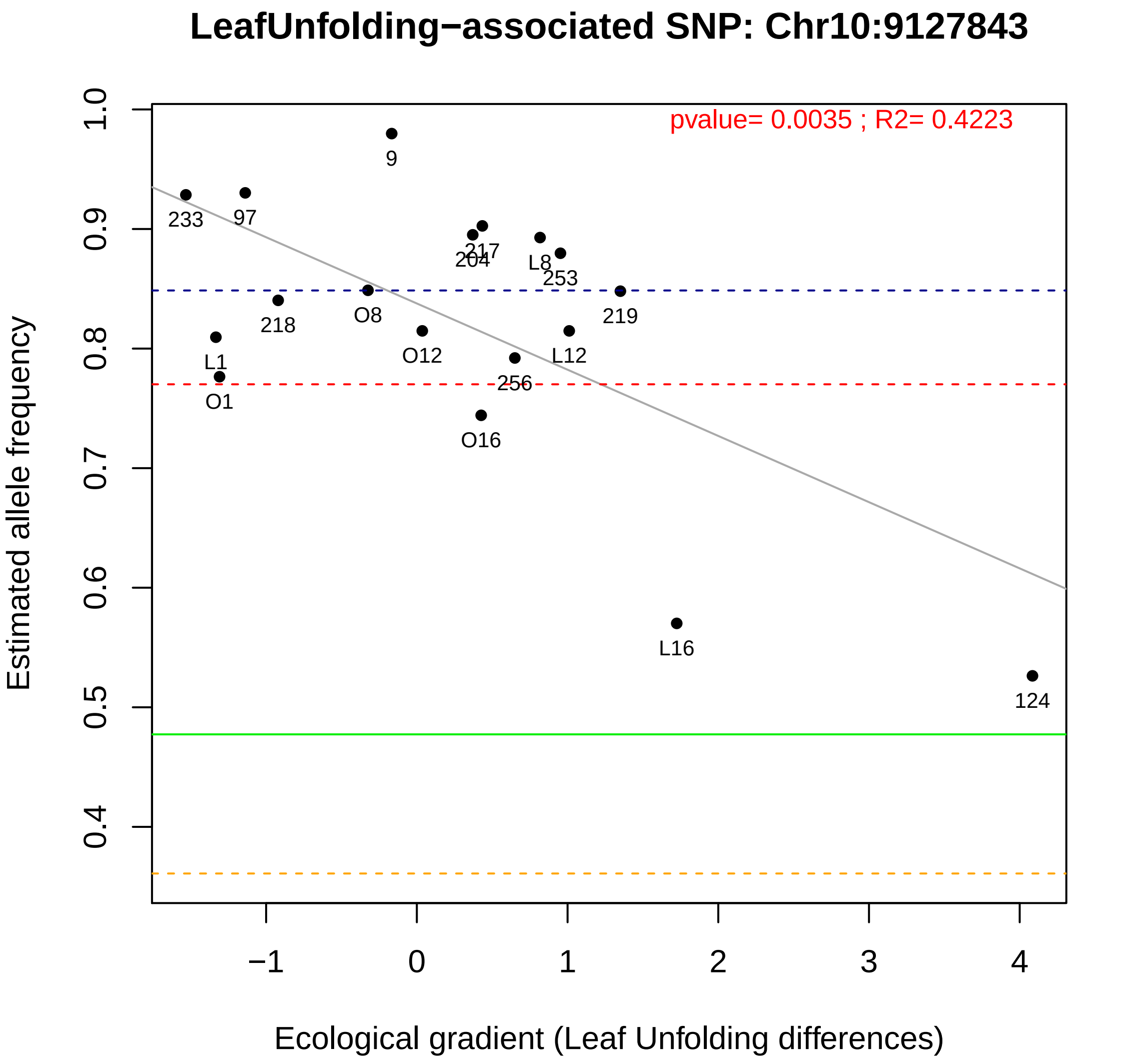
LeafUnfolding−associated SNP: Chr10:9127843
pvalue= 0.0035 ; R2= 0.4223
Estimated allele frequency
Ecological gradient (Leaf Unfolding differences)
9
233
97
217
204
L8
253
219
218
O8
L1
O12
L12
256
O1
O16
L16
124
0.4
0.5
0.6
0.7
0.8
0.9
1.0
−1
0
1
2
3
4

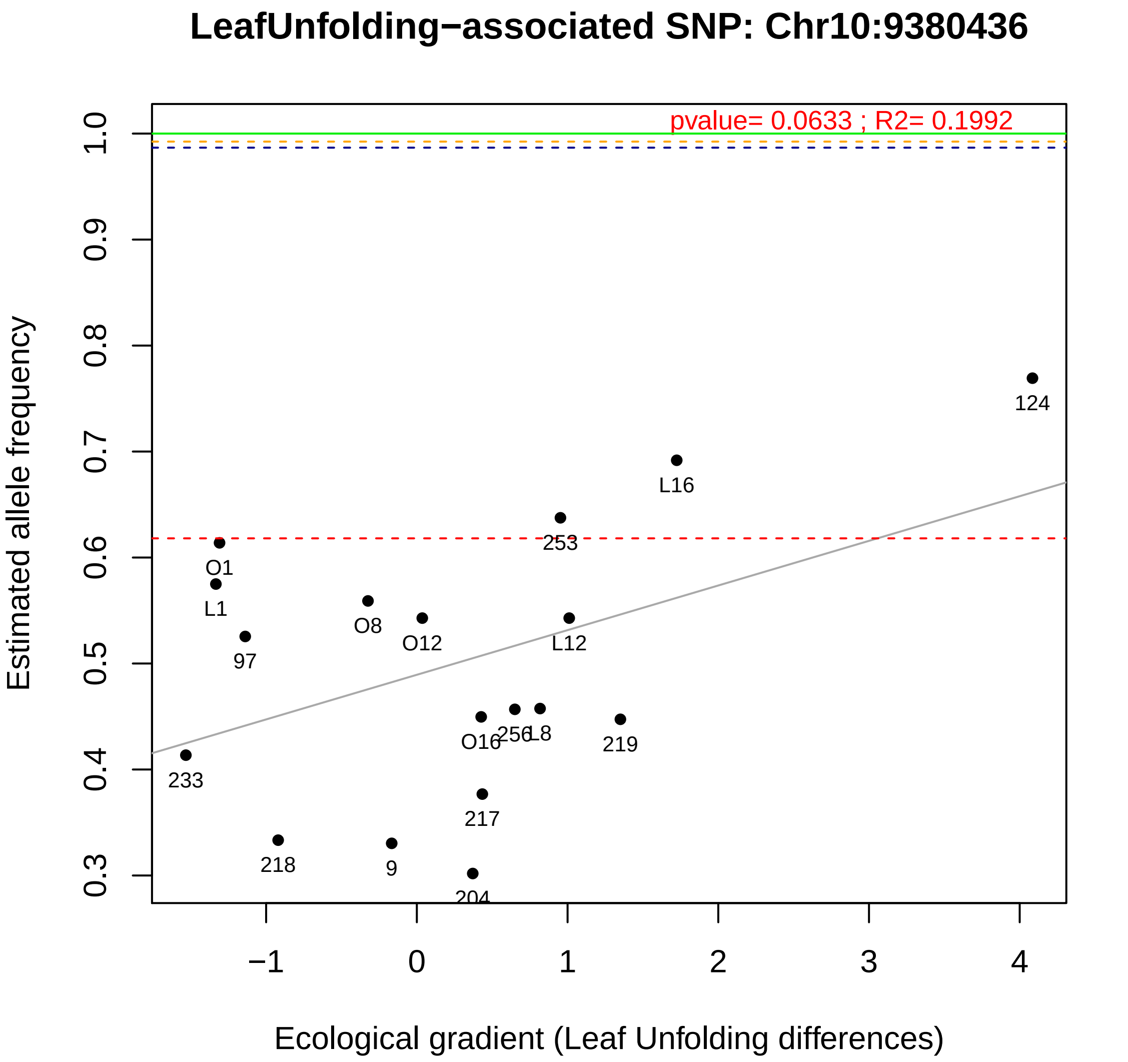

LeafUnfolding−associated SNP: Chr10:9380436
pvalue= 0.0633 ; R2= 0.1992
Estimated allele frequency
0.3
0.4
0.5
0.6
0.7
0.8
0.9
1.0
Ecological gradient (Leaf Unfolding differences)
−1
0
1
2
3
4
124
L16
253
O1
L1
O8
O12
L12
97
O16
256
L8
219
233
217
218
9
204

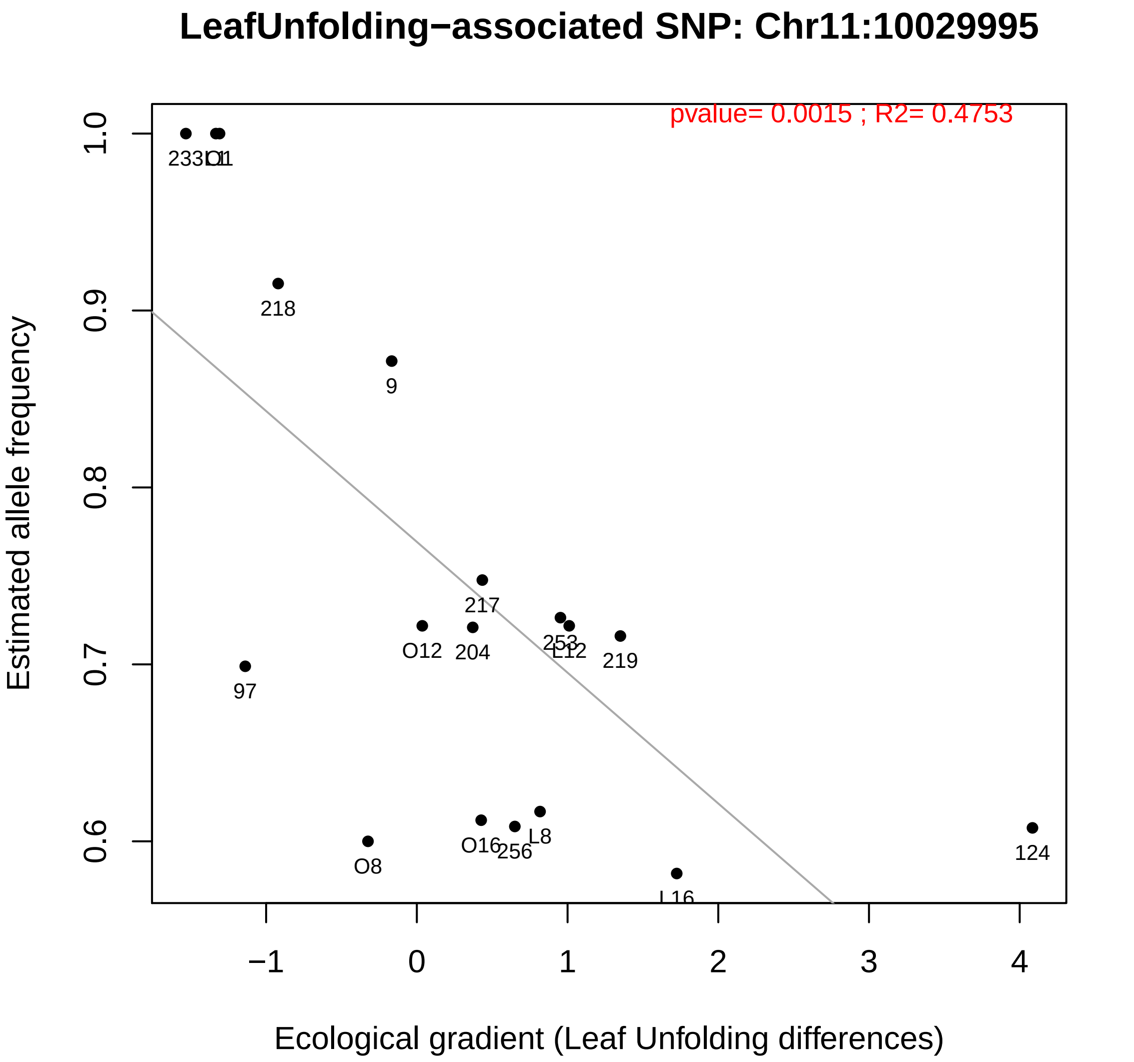

LeafUnfolding−associated SNP: Chr11:10029995
pvalue= 0.0015 ; R2= 0.4753
Estimated allele frequency
Ecological gradient (Leaf Unfolding differences)
0.6
0.7
0.8
0.9
1.0
−1
0
1
2
3
4
218
9
217
O12
204
253
L12
219
97
O16
256
L8
O8
124

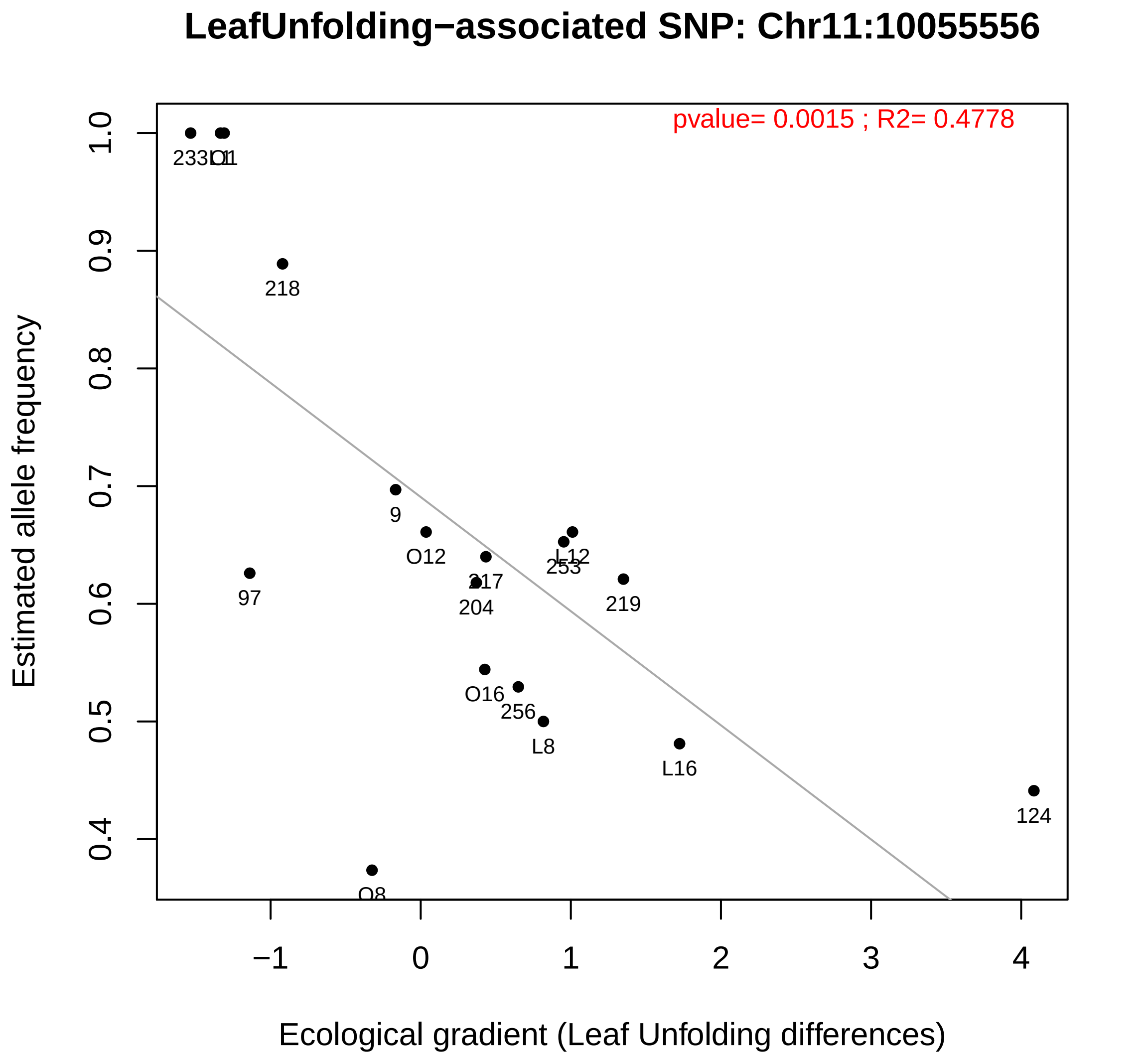

LeafUnfolding−associated SNP: Chr11:10055556
pvalue= 0.0015 ; R2= 0.4778
Estimated allele frequency
Ecological gradient (Leaf Unfolding differences)
1.0
0.9
0.8
0.7
0.6
0.5
0.4
−1
0
1
2
3
4
233
O1
218
9
O12
97
217
204
L12
253
219
O16
256
L8
L16
124
O8

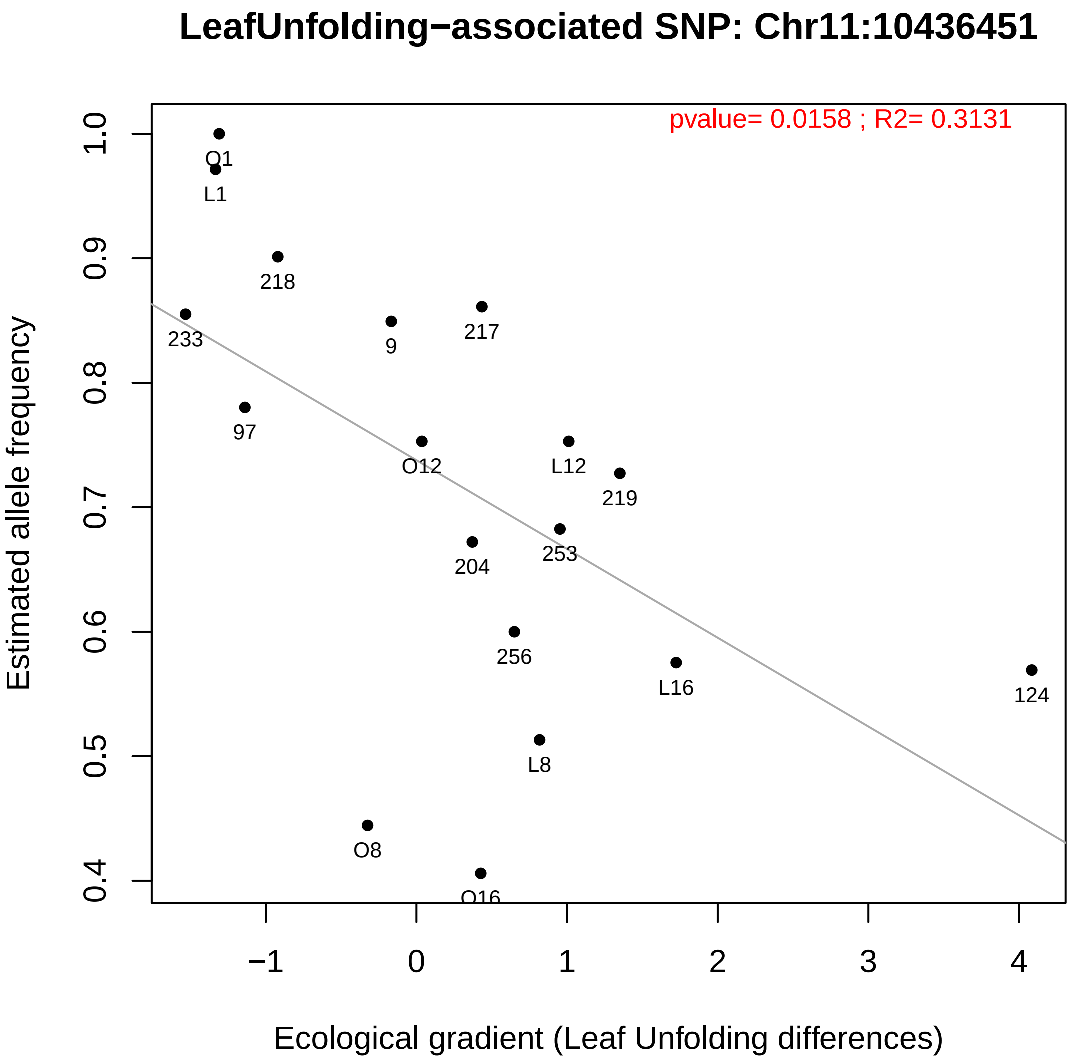
LeafUnfolding−associated SNP: Chr11:10436451
pvalue= 0.0158 ; R2= 0.3131
Estimated allele frequency
Ecological gradient (Leaf Unfolding differences)
1.0
0.9
0.8
0.7
0.6
0.5
0.4
−1
0
1
2
3
4
O1
L1
218
233
9
217
97
O12
L12
219
253
204
256
L16
124
L8
O8
O16

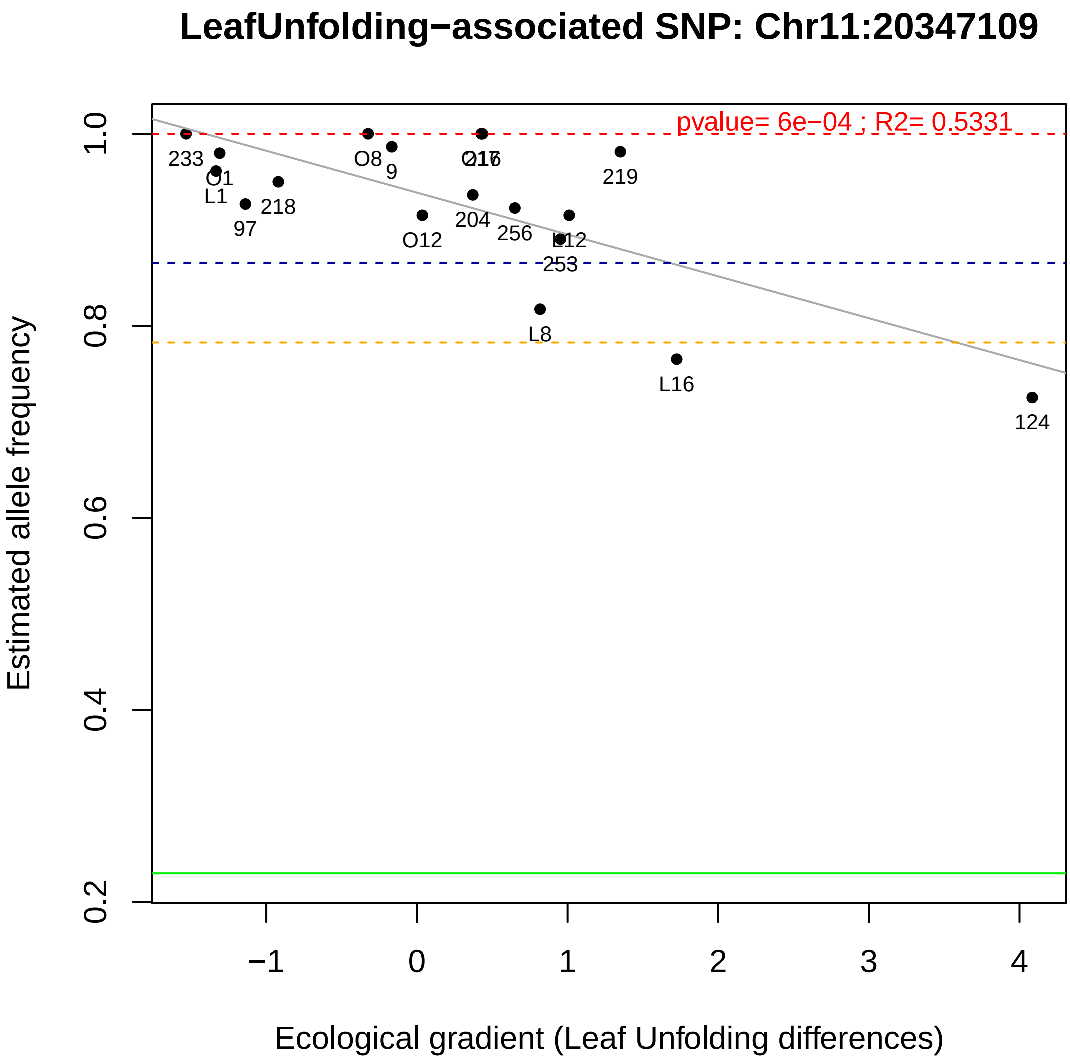
LeafUnfolding−associated SNP: Chr11:20347109
pvalue= 6e−04 ; R2= 0.5331
Estimated allele frequency
Ecological gradient (Leaf Unfolding differences)
233
O1
L1
97
218
O8
9
O12
204
256
219
L8
253
L16
124
1.0
0.8
0.6
0.4
0.2
−1
0
1
2
3
4

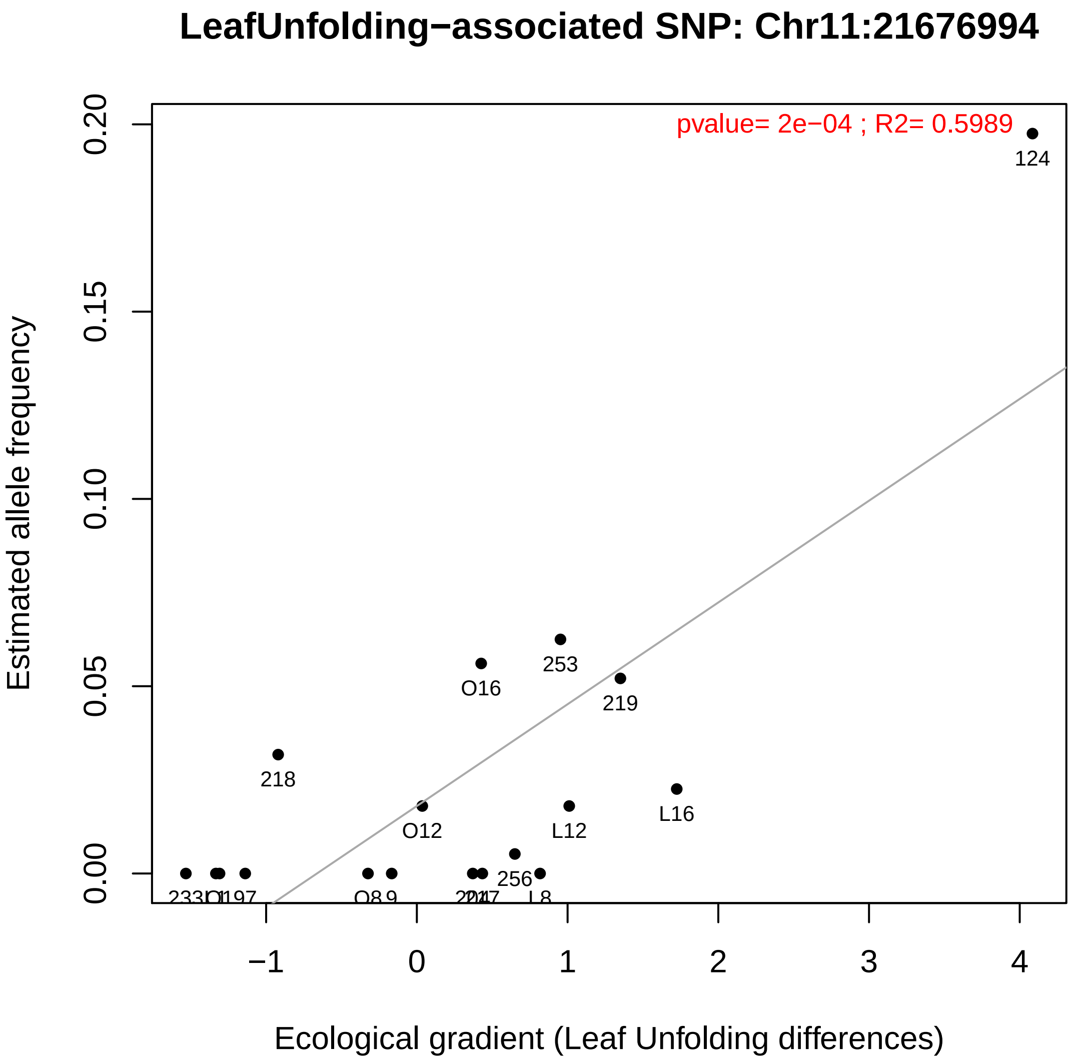
LeafUnfolding−associated SNP: Chr11:21676994
pvalue= 2e−04 ; R2= 0.5989
124
253
O16
219
218
O12
L12
L16
256
Estimated allele frequency
0.00
0.05
0.10
0.15
0.20
−1
0
1
2
3
4
Ecological gradient (Leaf Unfolding differences)

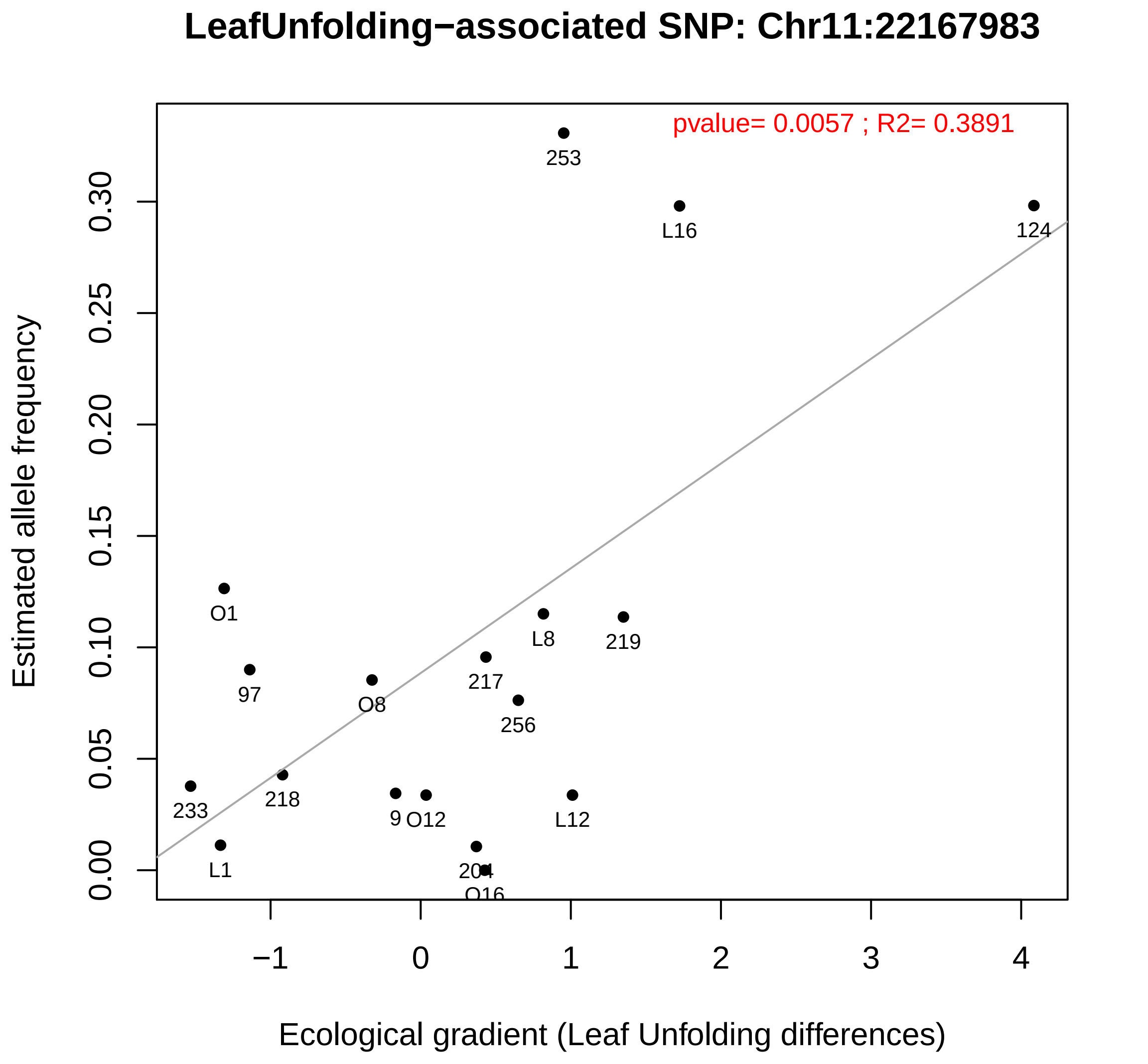
LeafUnfolding−associated SNP: Chr11:22167983
pvalue= 0.0057 ; R2= 0.3891
Estimated allele frequency
0.00
0.05
0.10
0.15
0.20
0.25
0.30
−1
0
1
2
3
4
Ecological gradient (Leaf Unfolding differences)
253
L16
124
O1
L8
219
97
O8
217
256
233
218
9
O12
L12
L1
204
O16

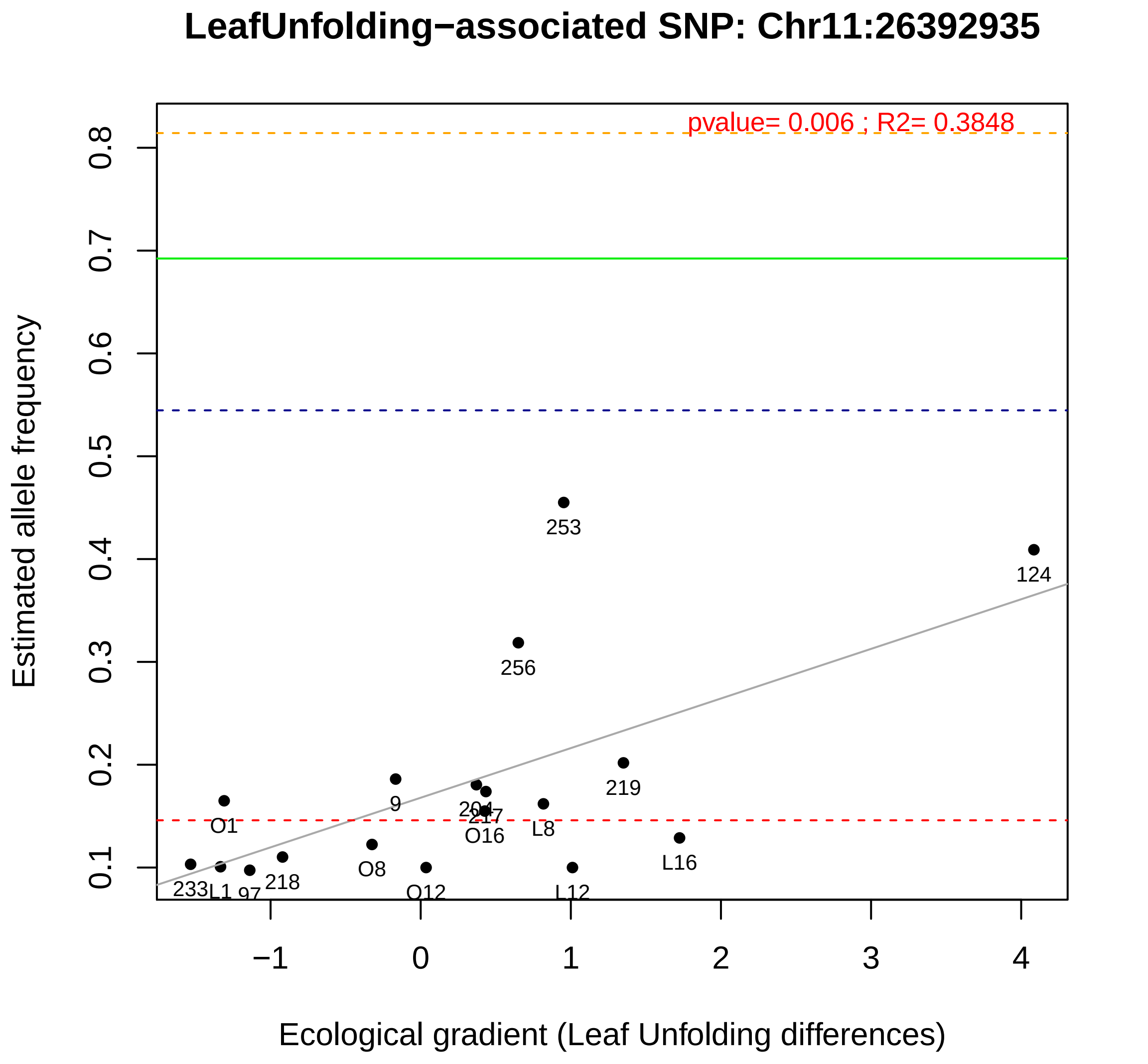

LeafUnfolding−associated SNP: Chr11:26392935
pvalue= 0.006 ; R2= 0.3848
Estimated allele frequency
Ecological gradient (Leaf Unfolding differences)
0.1
0.2
0.3
0.4
0.5
0.6
0.7
0.8
−1
0
1
2
3
4
253
124
256
219
9
O1
204
217
O16
L8
L16
O8
233
L1
97
218
O12
L12

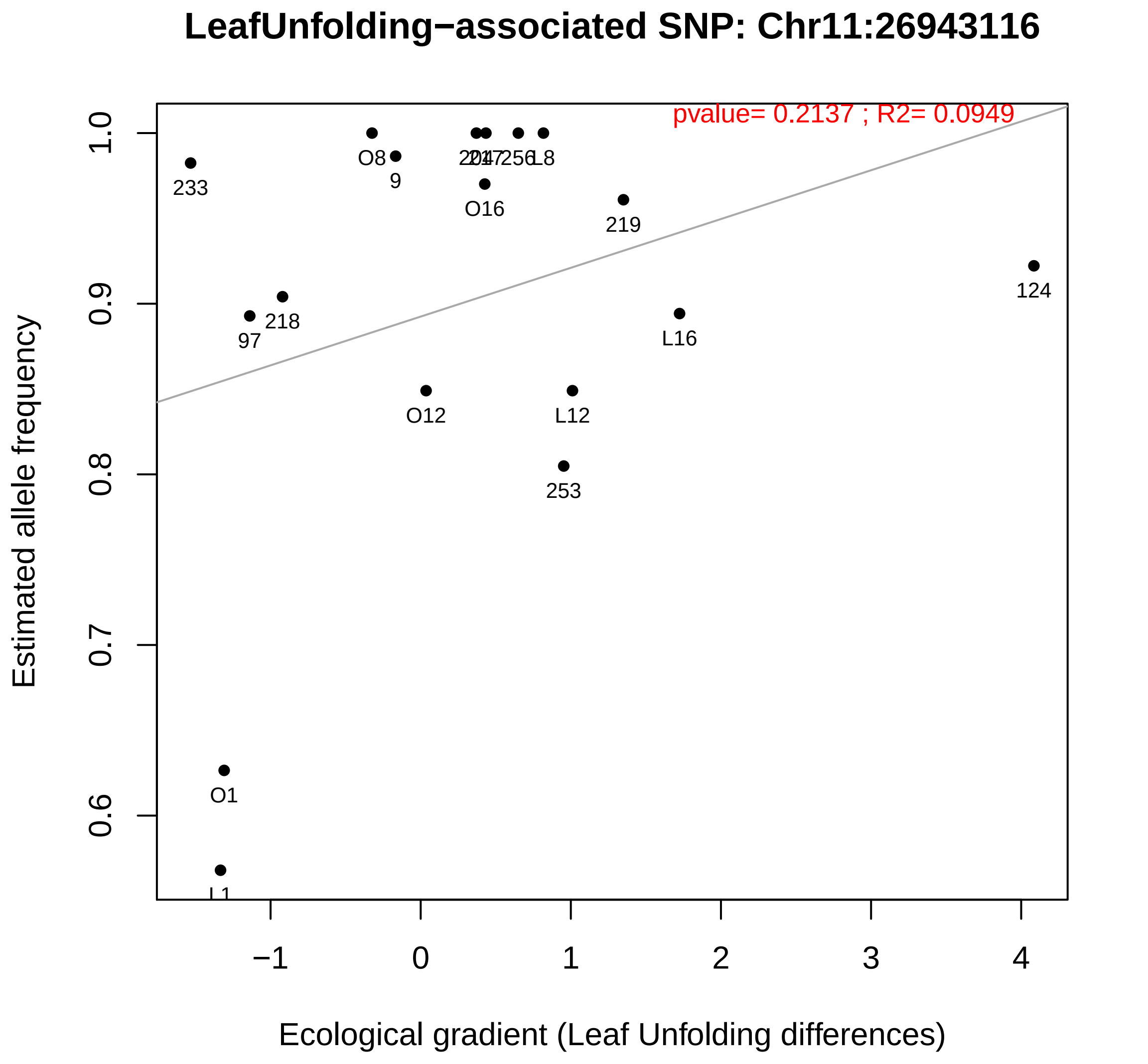
LeafUnfolding−associated SNP: Chr11:26943116
pvalue= 0.2137 ; R2= 0.0949
Estimated allele frequency
Ecological gradient (Leaf Unfolding differences)
0.6
0.7
0.8
0.9
1.0
−1
0
1
2
3
4
233
O8
9
O16
219
124
218
97
L16
O12
L12
253
O1
L8

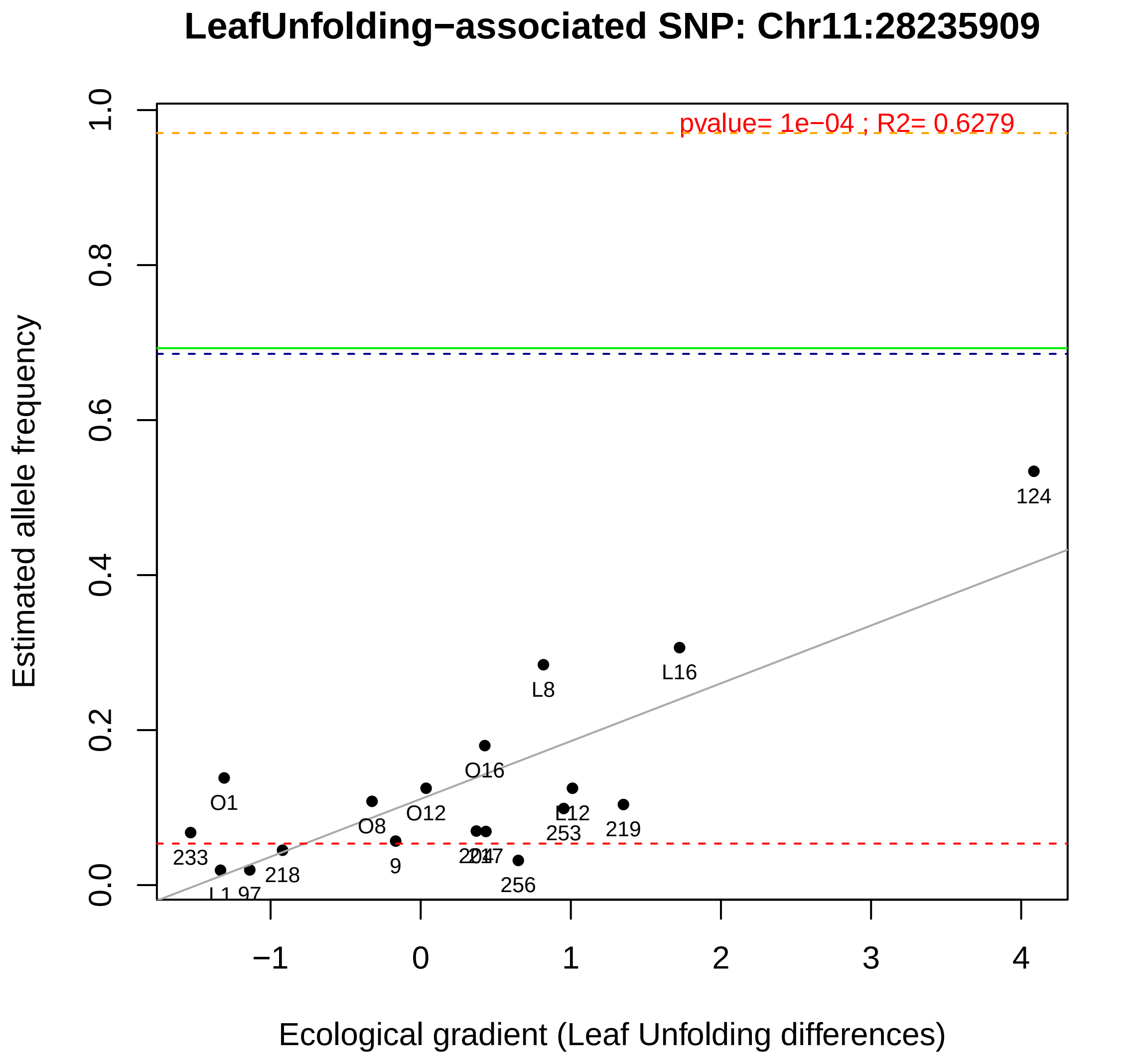
LeafUnfolding−associated SNP: Chr11:28235909
pvalue= 1e−04 ; R2= 0.6279
Estimated allele frequency
0.0
0.2
0.4
0.6
0.8
1.0
Ecological gradient (Leaf Unfolding differences)
−1
0
1
2
3
4
124
L16
L8
O16
O1
O8
O12
L12
253
219
233
218
9
256

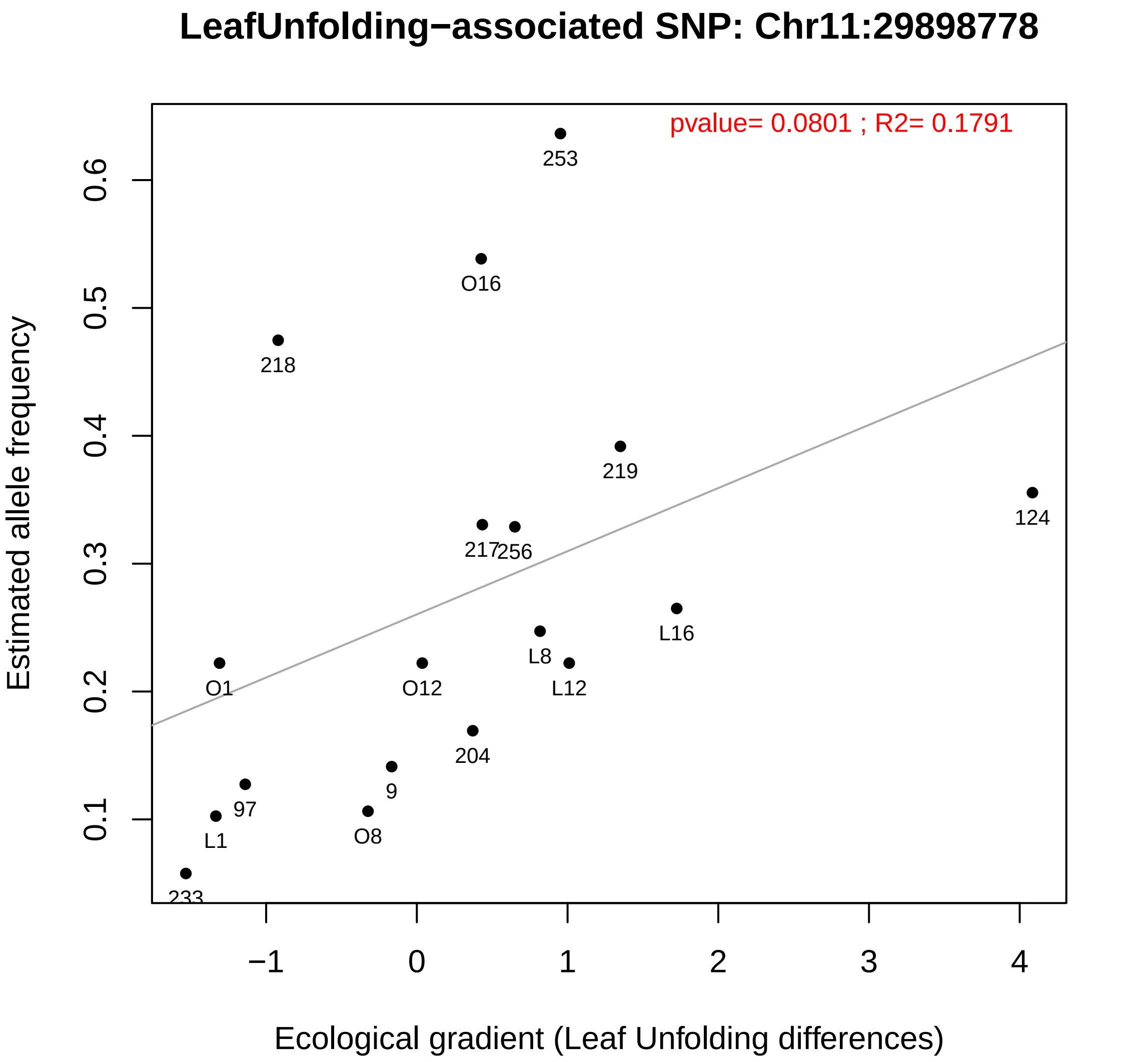
LeafUnfolding−associated SNP: Chr11:29898778
pvalue= 0.0801 ; R2= 0.1791
Estimated allele frequency
Ecological gradient (Leaf Unfolding differences)
253
O16
218
219
124
217
256
L16
L8
L12
O1
O12
204
9
97
L1
O8
233
0.1
0.2
0.3
0.4
0.5
0.6
−1
0
1
2
3
4

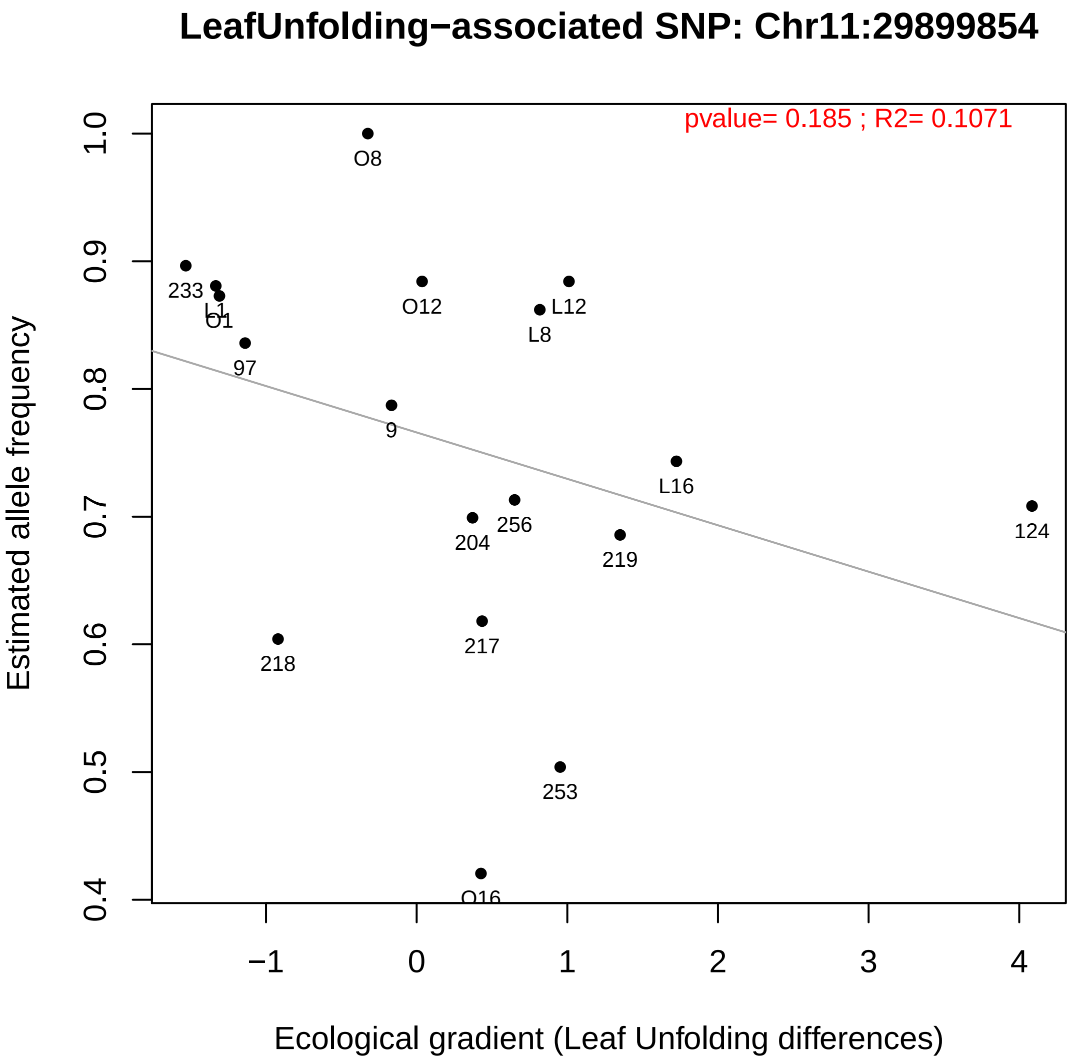

LeafUnfolding−associated SNP: Chr11:29899854
pvalue= 0.185 ; R2= 0.1071
Estimated allele frequency
Ecological gradient (Leaf Unfolding differences)
1.0
0.9
0.8
0.7
0.6
0.5
0.4
−1
0
1
2
3
4
O8
233
L1
O1
97
O12
L12
L8
9
L16
256
204
219
124
218
217
253
O16

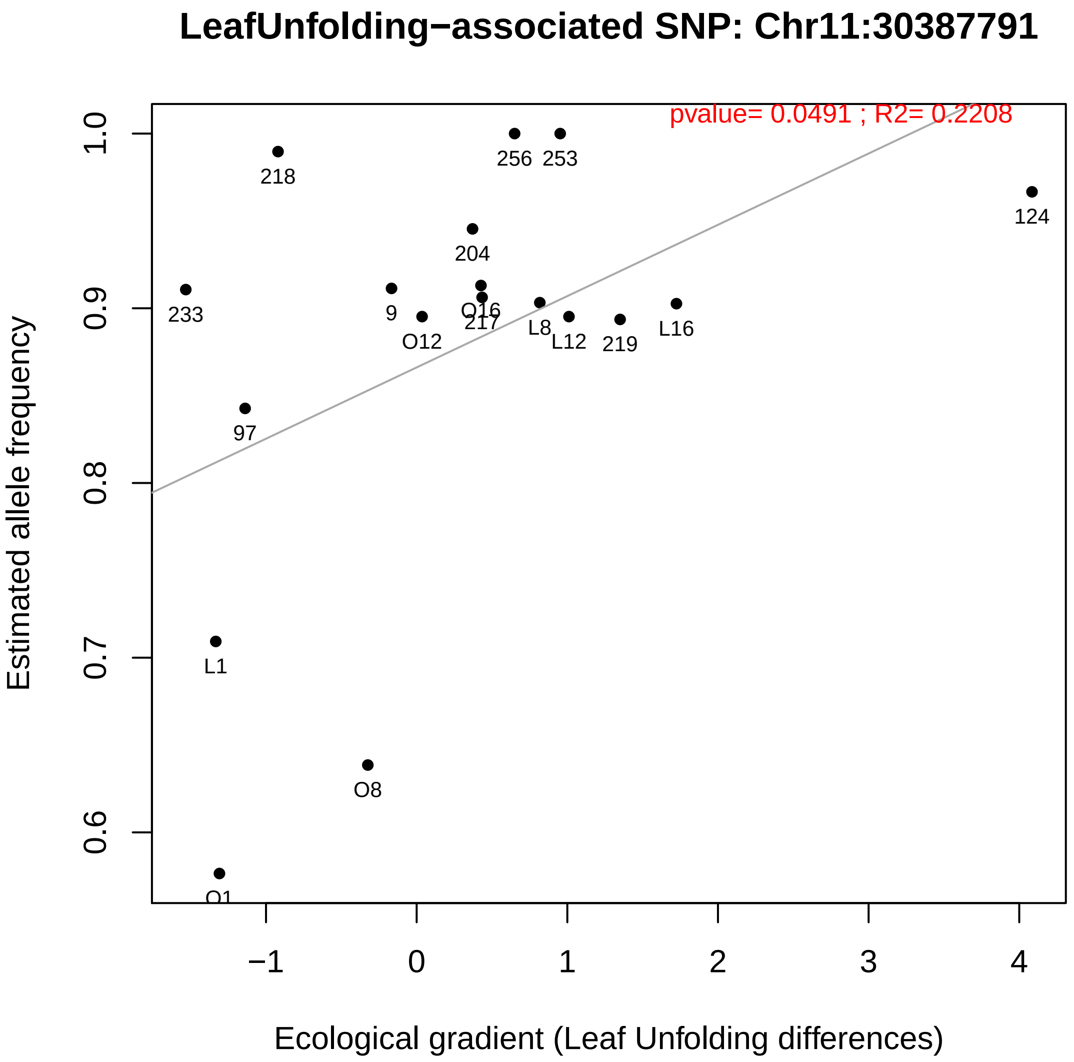
LeafUnfolding−associated SNP: Chr11:30387791
pvalue= 0.0491 ; R2= 0.2208
Estimated allele frequency
Ecological gradient (Leaf Unfolding differences)
0.6
0.7
0.8
0.9
1.0
−1
0
1
2
3
4
218
256
253
124
204
233
9
O16
217
O12
L8
L12
219
L16
97
L1
O8
O1

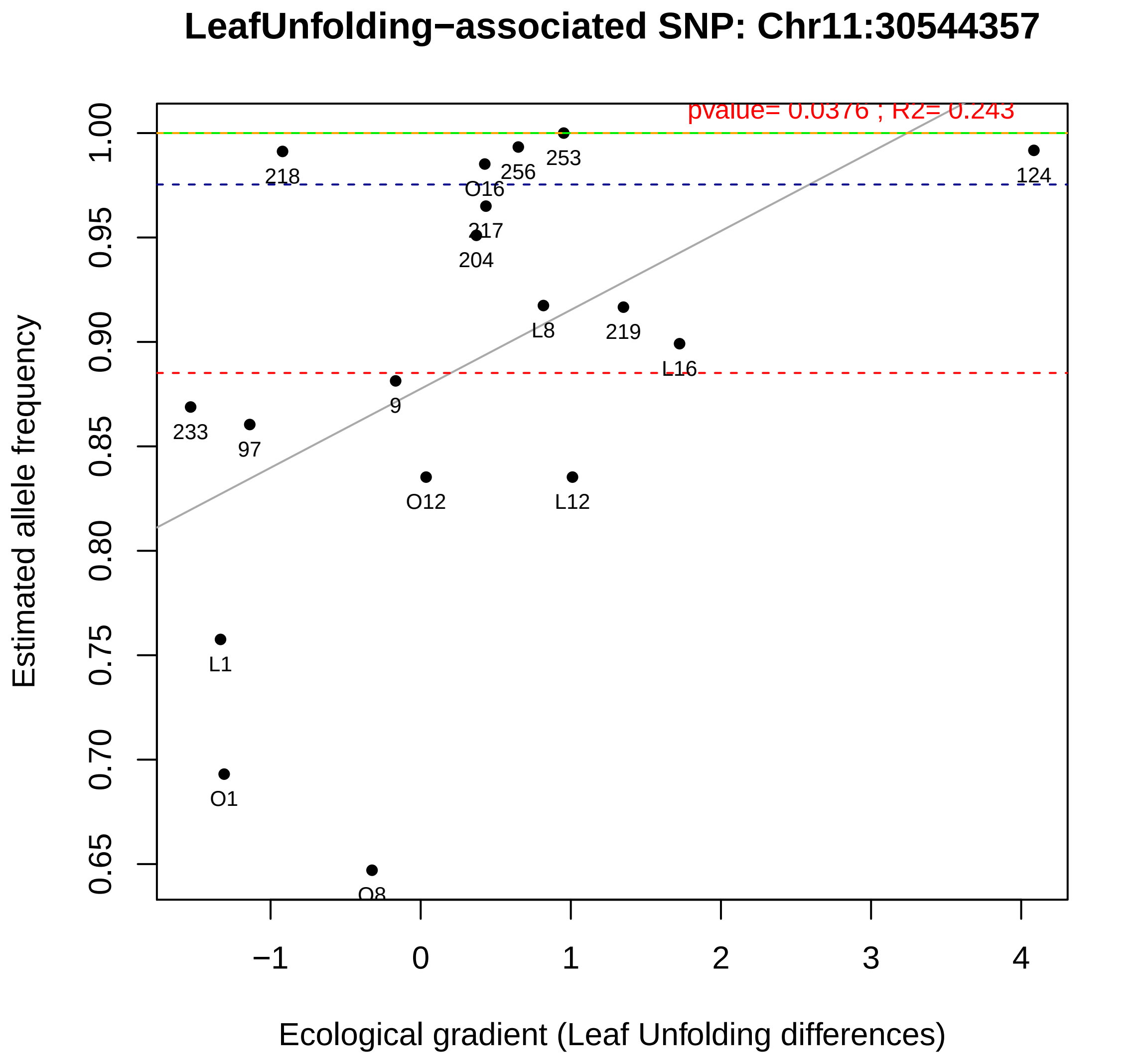
LeafUnfolding−associated SNP: Chr11:30544357
pvalue= 0.0376 ; R2= 0.243
Estimated allele frequency
Ecological gradient (Leaf Unfolding differences)
1.00
0.95
0.90
0.85
0.80
0.75
0.70
0.65
−1
0
1
2
3
4
218
256
253
124
O16
217
204
L8
219
L16
9
233
97
O12
L12
L1
O1
O8

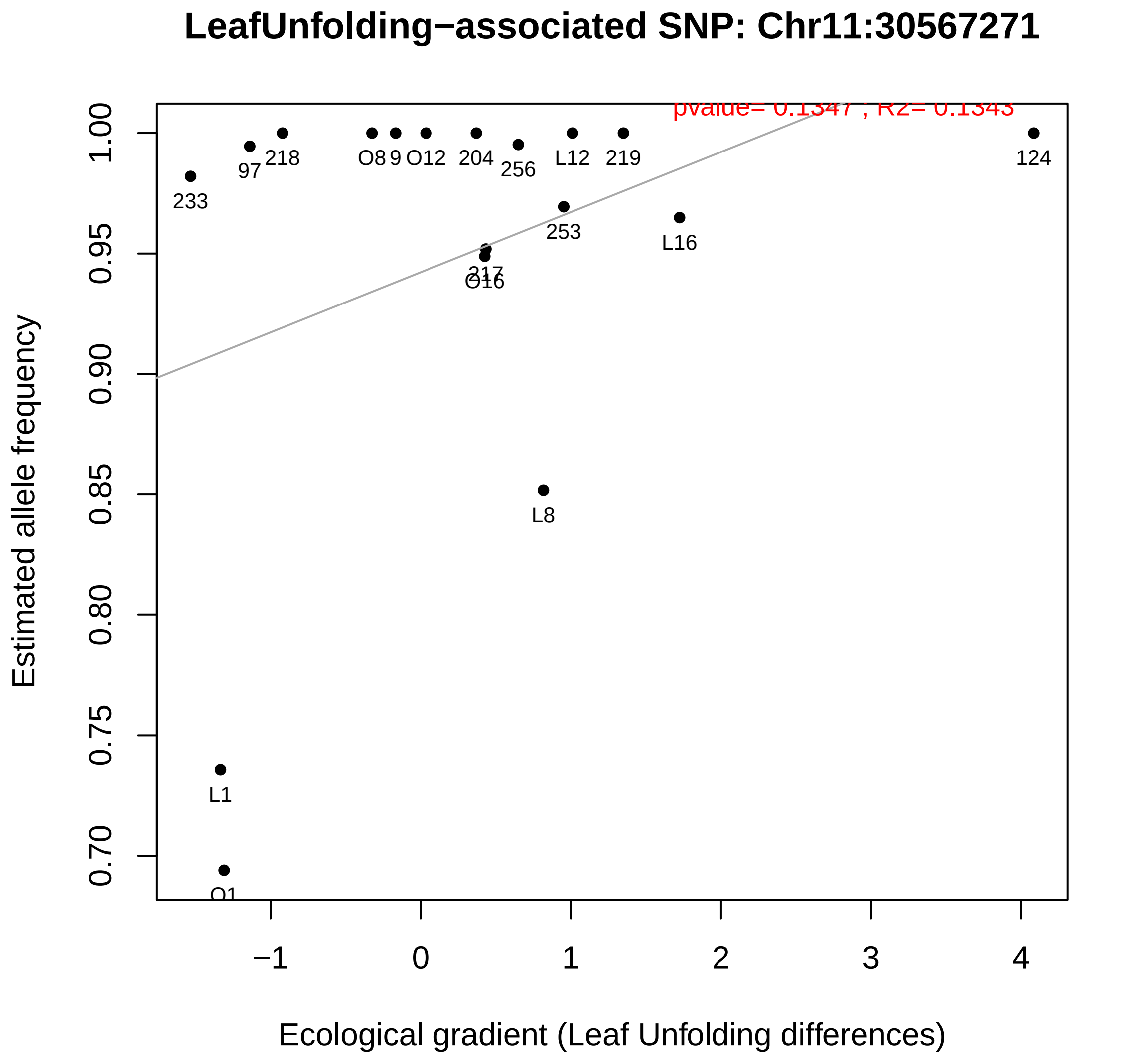
LeafUnfolding−associated SNP: Chr11:30567271
pvalue= 0.1347 ; R2= 0.1343
Estimated allele frequency
Ecological gradient (Leaf Unfolding differences)
1.00
0.95
0.90
0.85
0.80
0.75
0.70
−1
0
1
2
3
4
233
97
218
O8
9
O12
204
256
L12
219
124
253
L16
217
O16
L8
L1
O1

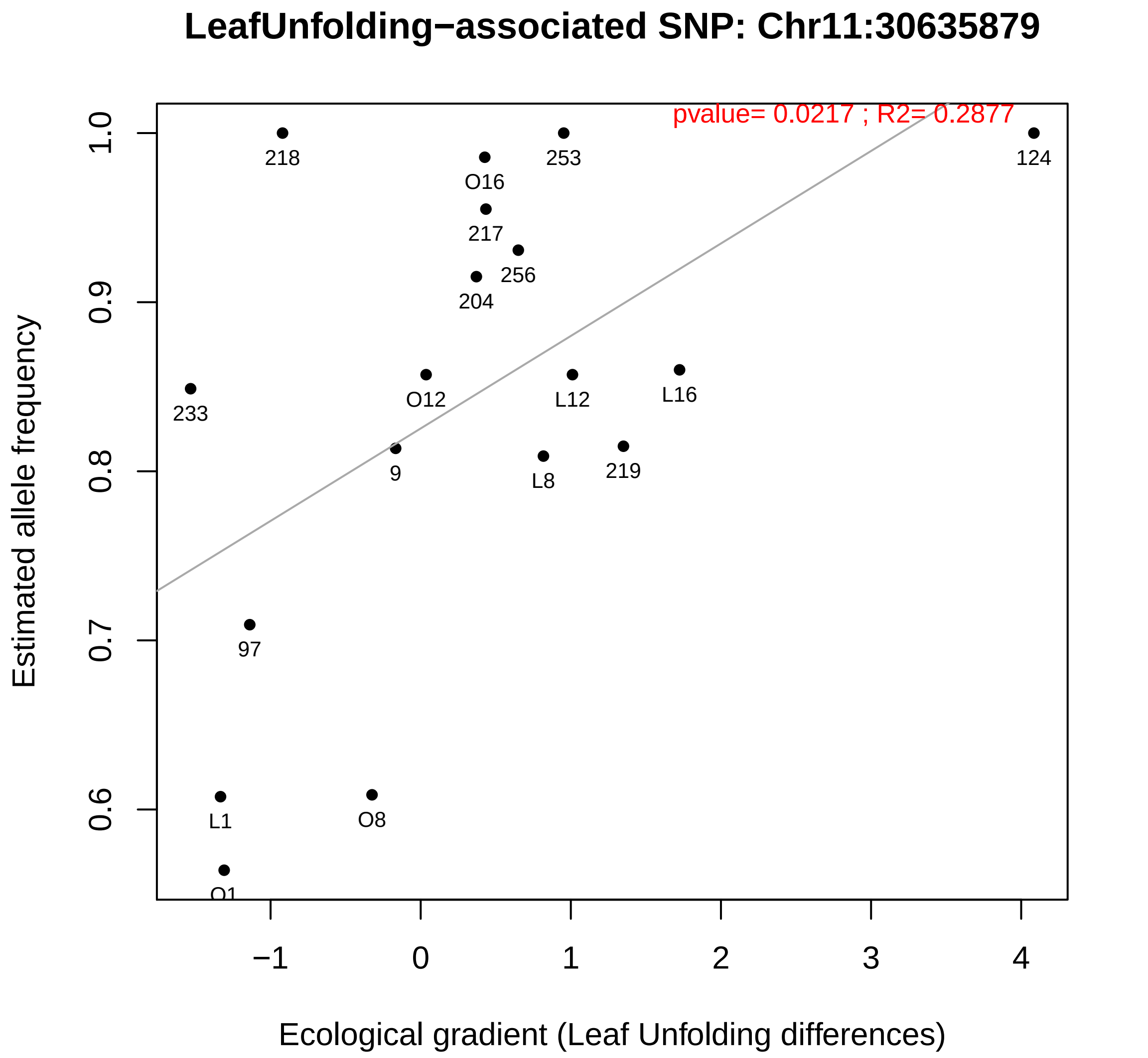

LeafUnfolding−associated SNP: Chr11:30635879
pvalue= 0.0217 ; R2= 0.2877
Estimated allele frequency
Ecological gradient (Leaf Unfolding differences)
1.0
0.9
0.8
0.7
0.6
−1
0
1
2
3
4
218
O16
253
124
217
256
204
233
O12
L12
L16
9
L8
219
97
L1
O8
O1

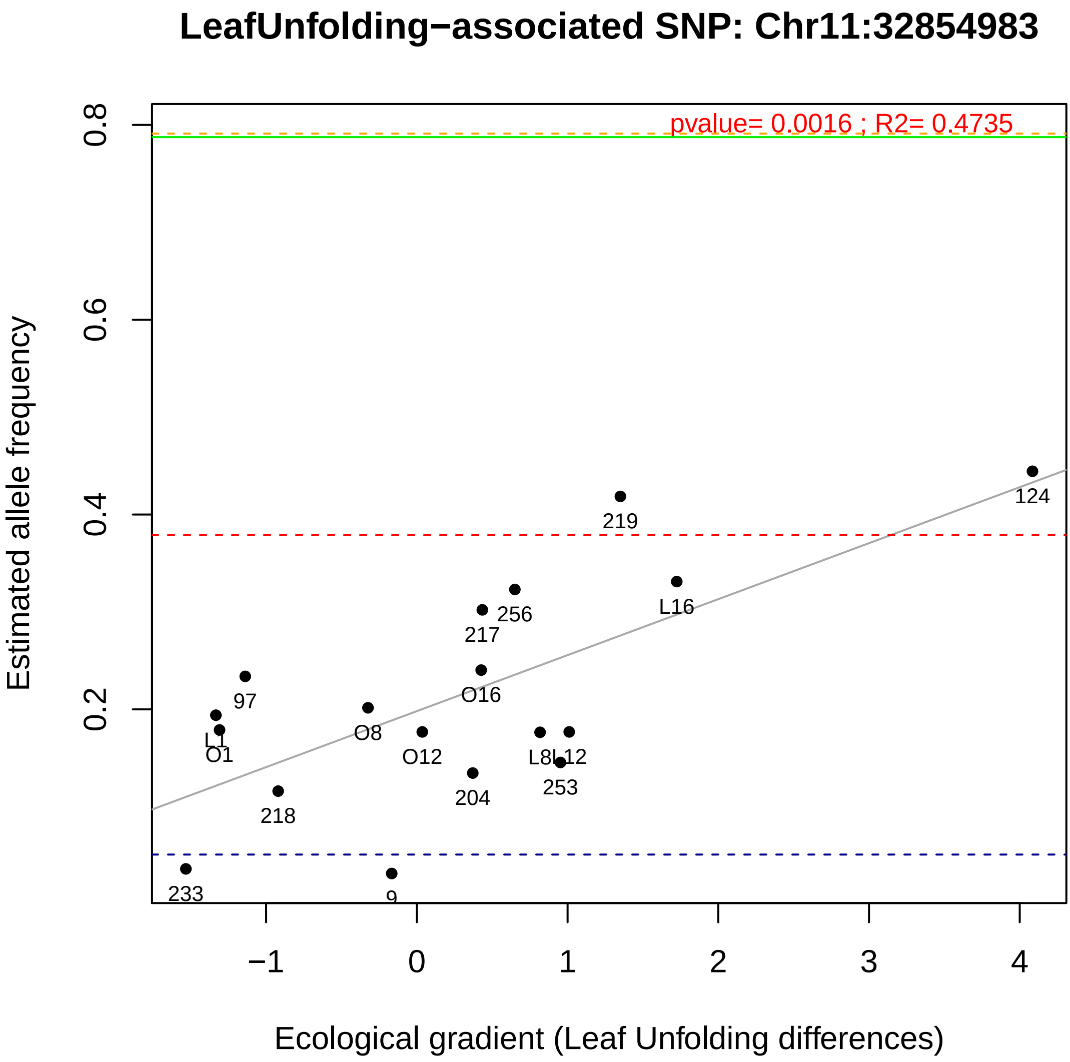

LeafUnfolding−associated SNP: Chr11:32854983
pvalue= 0.0016 ; R2= 0.4735
Estimated allele frequency
0.2
0.4
0.6
0.8
−1
0
1
2
3
4
Ecological gradient (Leaf Unfolding differences)
124
219
L16
256
217
O16
97
L1
O1
O8
O12
L8
L12
253
204
218
233
9

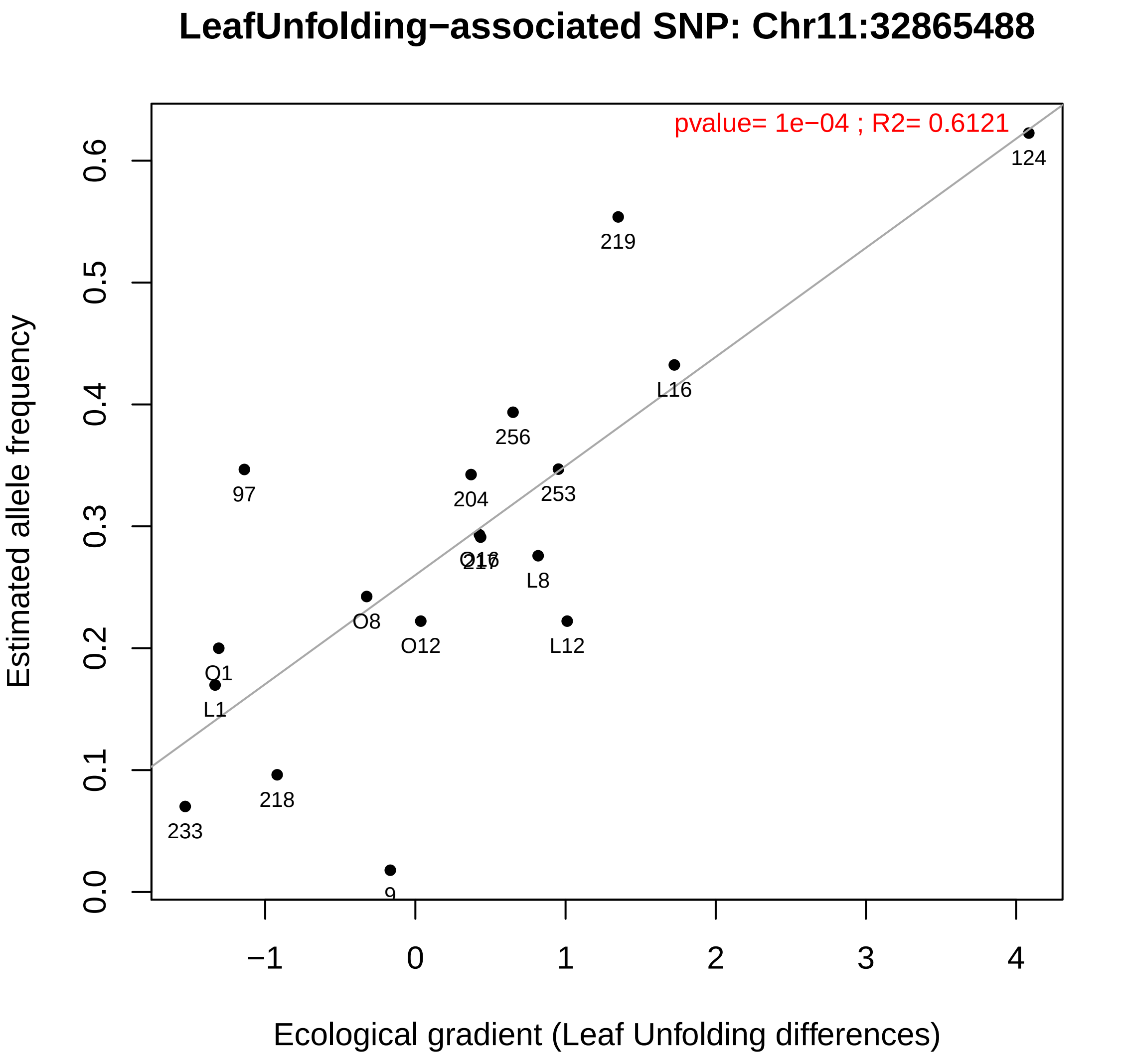

LeafUnfolding−associated SNP: Chr11:32865488
pvalue= 1e−04 ; R2= 0.6121
Estimated allele frequency
Ecological gradient (Leaf Unfolding differences)
0.0
0.1
0.2
0.3
0.4
0.5
0.6
−1
0
1
2
3
4
124
219
L16
256
97
204
253
L8
O8
O12
L12
O1
L1
218
233
9

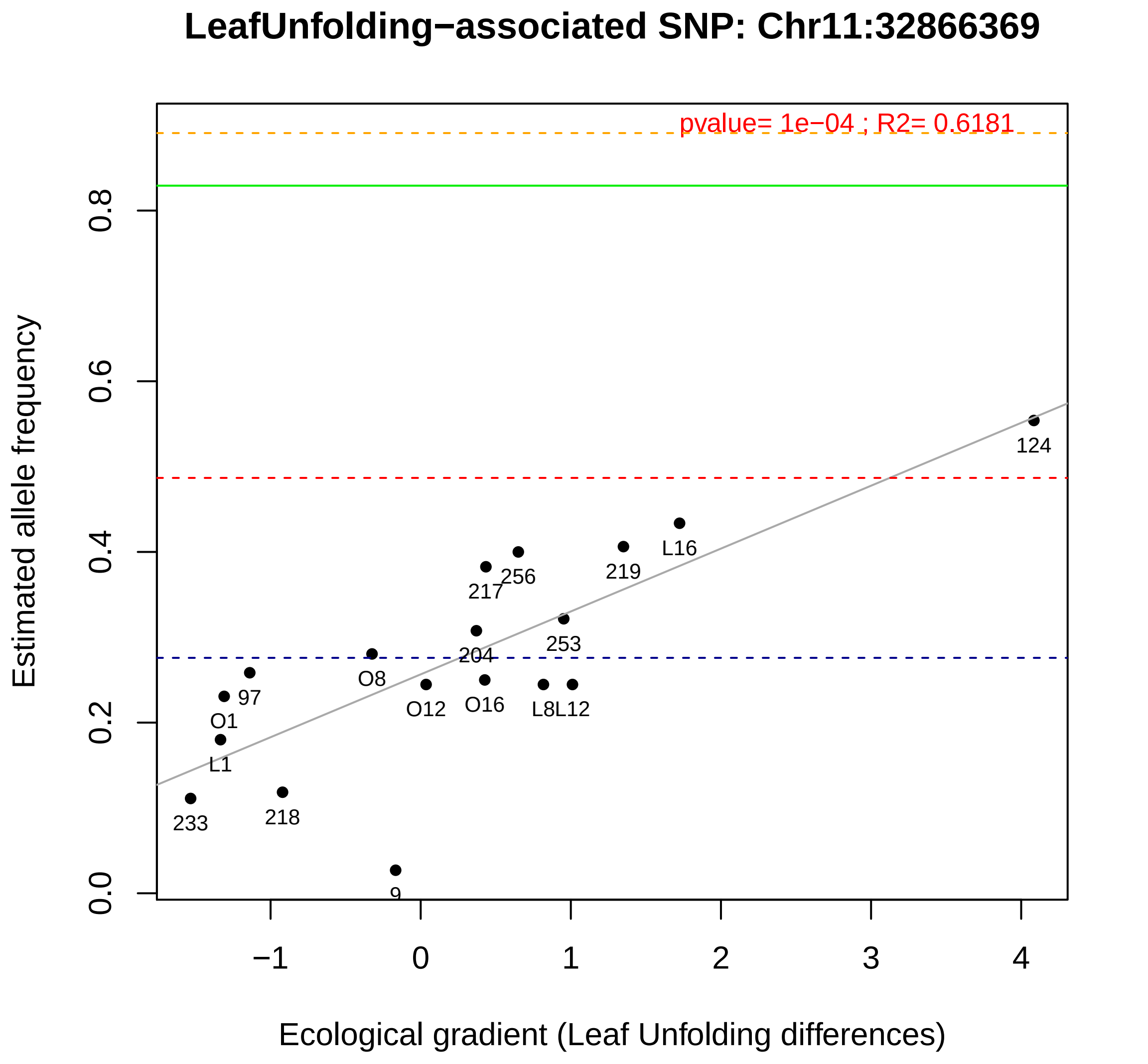

LeafUnfolding−associated SNP: Chr11:32866369
pvalue= 1e−04 ; R2= 0.6181
Estimated allele frequency
0.0
0.2
0.4
0.6
0.8
−1
0
1
2
3
4
Ecological gradient (Leaf Unfolding differences)
124
L16
219
256
217
253
204
O8
97
O1
O12
O16
L8
L12
L1
233
218
9

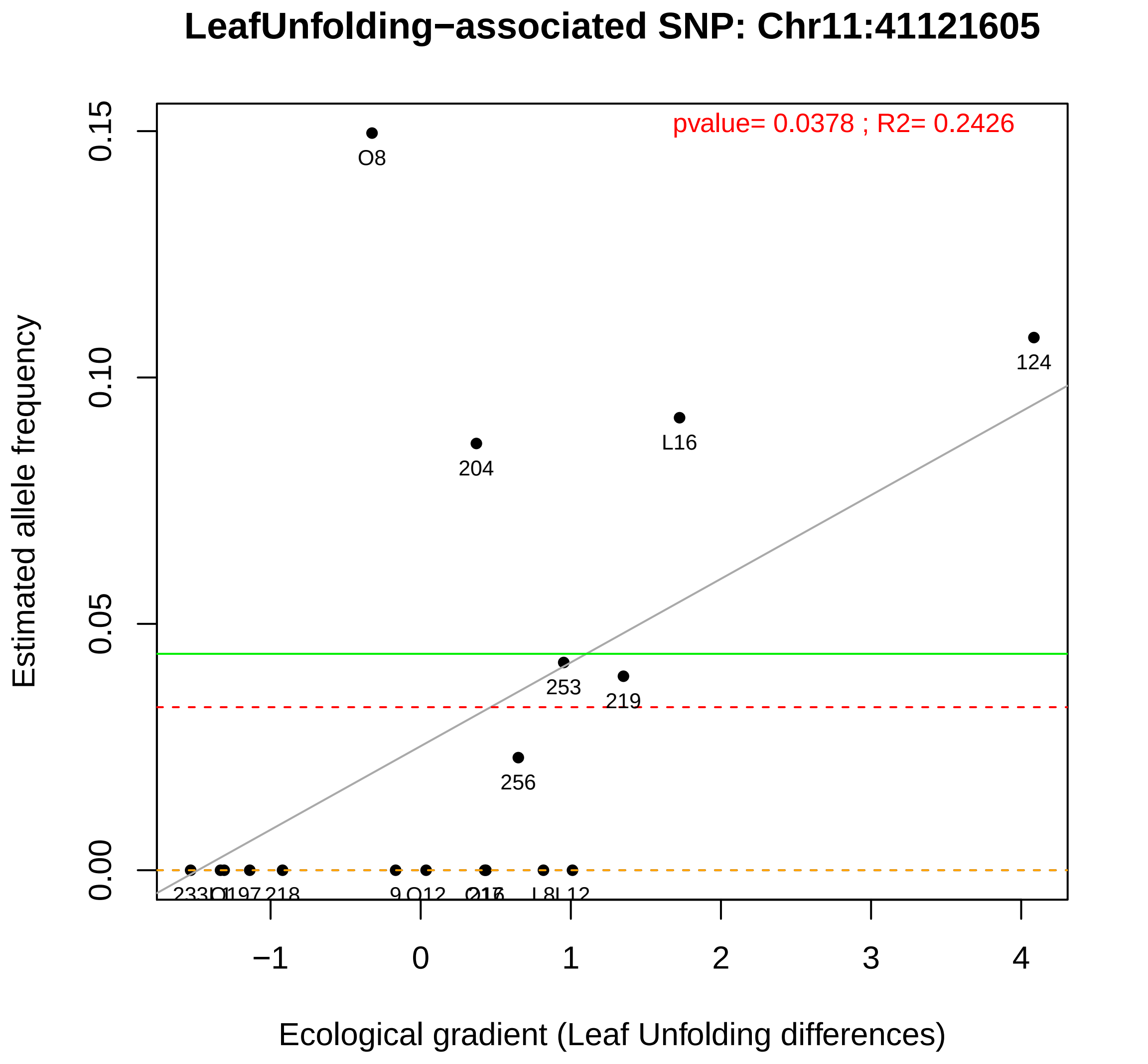

LeafUnfolding−associated SNP: Chr11:41121605
pvalue= 0.0378 ; R2= 0.2426
Estimated allele frequency
Ecological gradient (Leaf Unfolding differences)
0.00
0.05
0.10
0.15
−1
0
1
2
3
4
O8
124
L16
204
253
219
256
218
L12

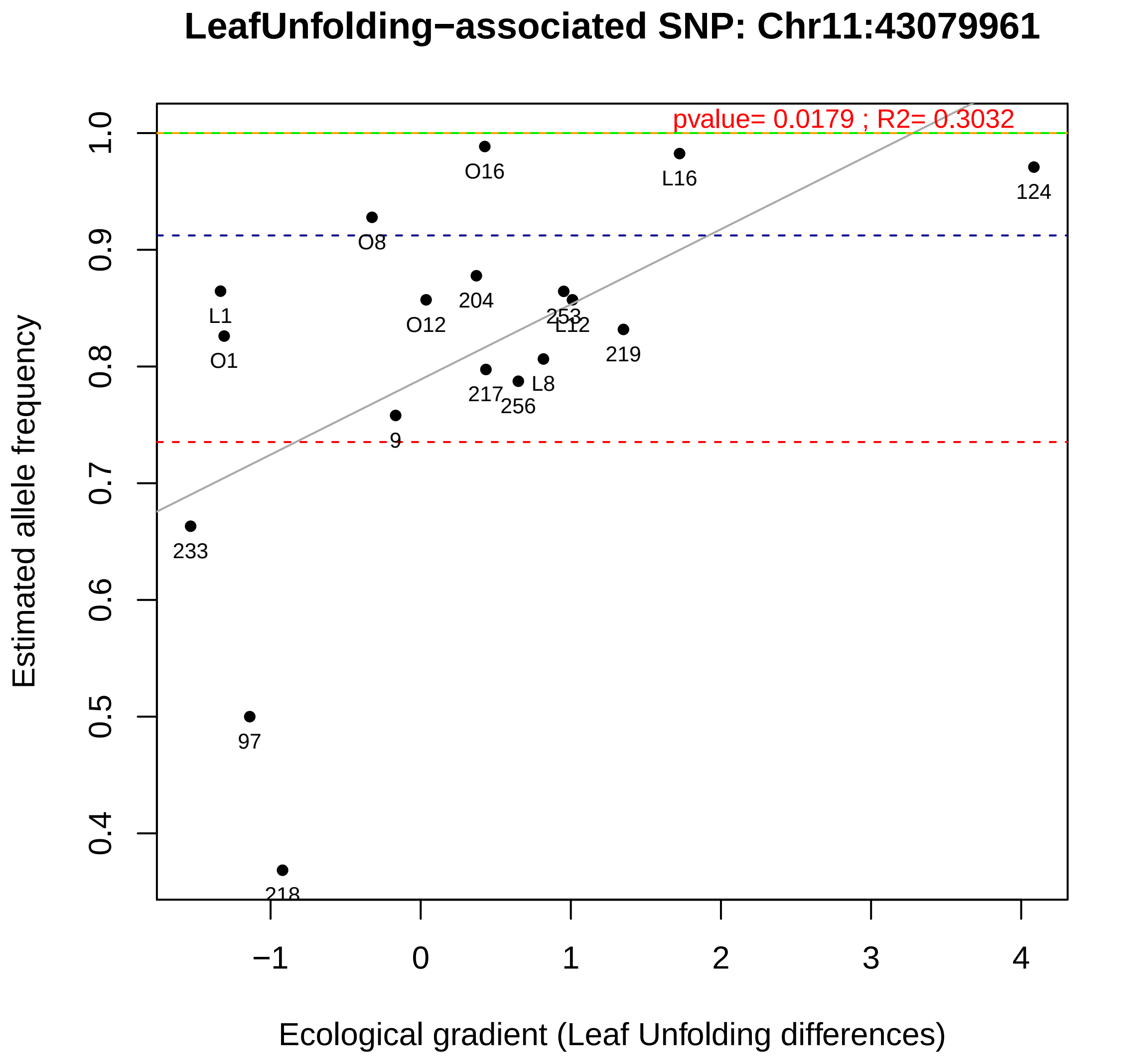

LeafUnfolding−associated SNP: Chr11:43079961
pvalue= 0.0179 ; R2= 0.3032
Estimated allele frequency
Ecological gradient (Leaf Unfolding differences)
0.4
0.5
0.6
0.7
0.8
0.9
1.0
−1
0
1
2
3
4
O16
L16
124
O8
204
L1
O12
253
L12
219
O1
217
L8
256
9
233
97
218

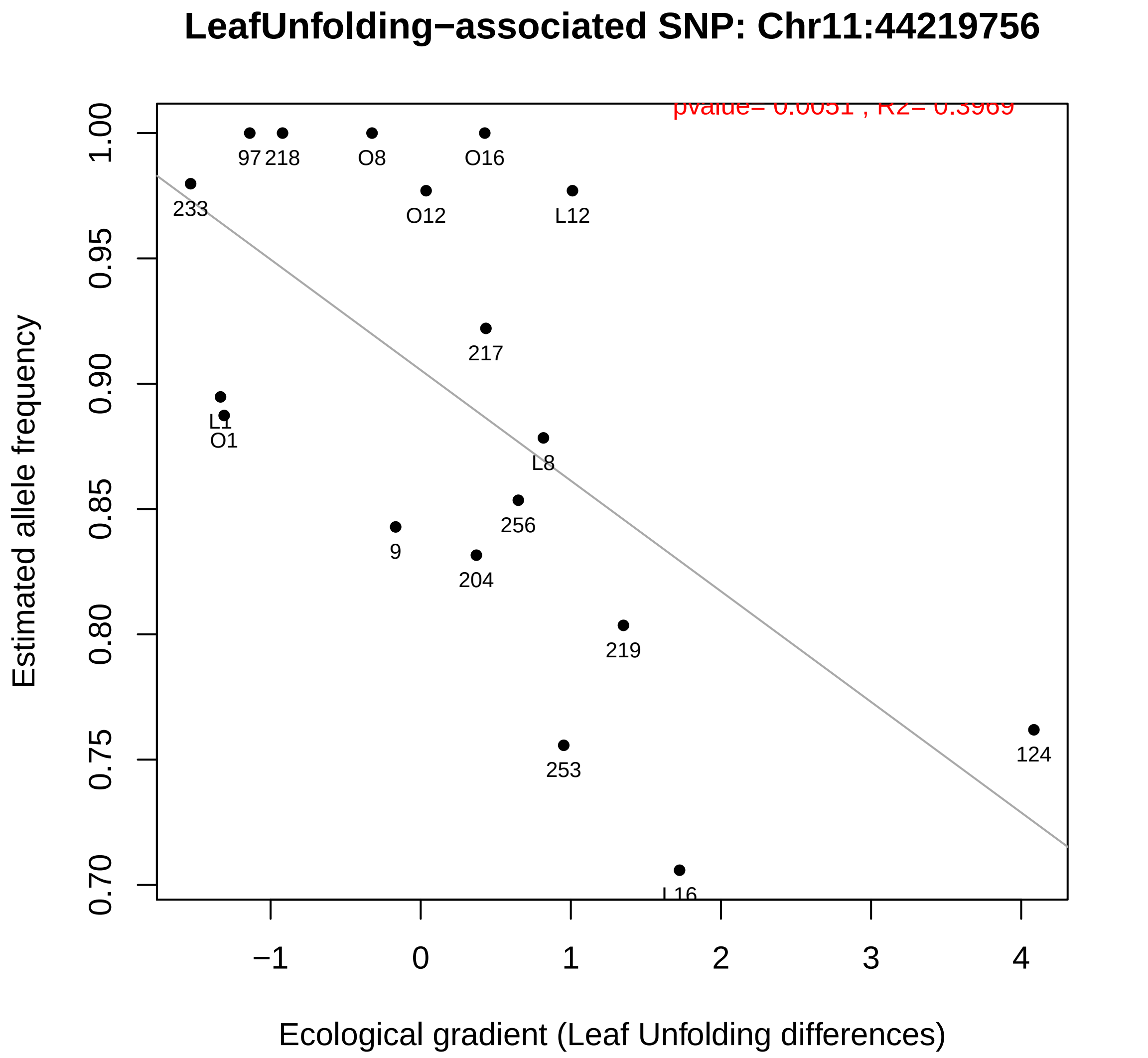

LeafUnfolding−associated SNP: Chr11:44219756
pvalue= 0.0051 , R2= 0.3969
Estimated allele frequency
1.00
0.95
0.90
0.85
0.80
0.75
0.70
−1
0
1
2
3
4
Ecological gradient (Leaf Unfolding differences)
97
218
O8
O16
233
O12
L12
217
L1
O1
L8
256
9
204
219
253
124
L16

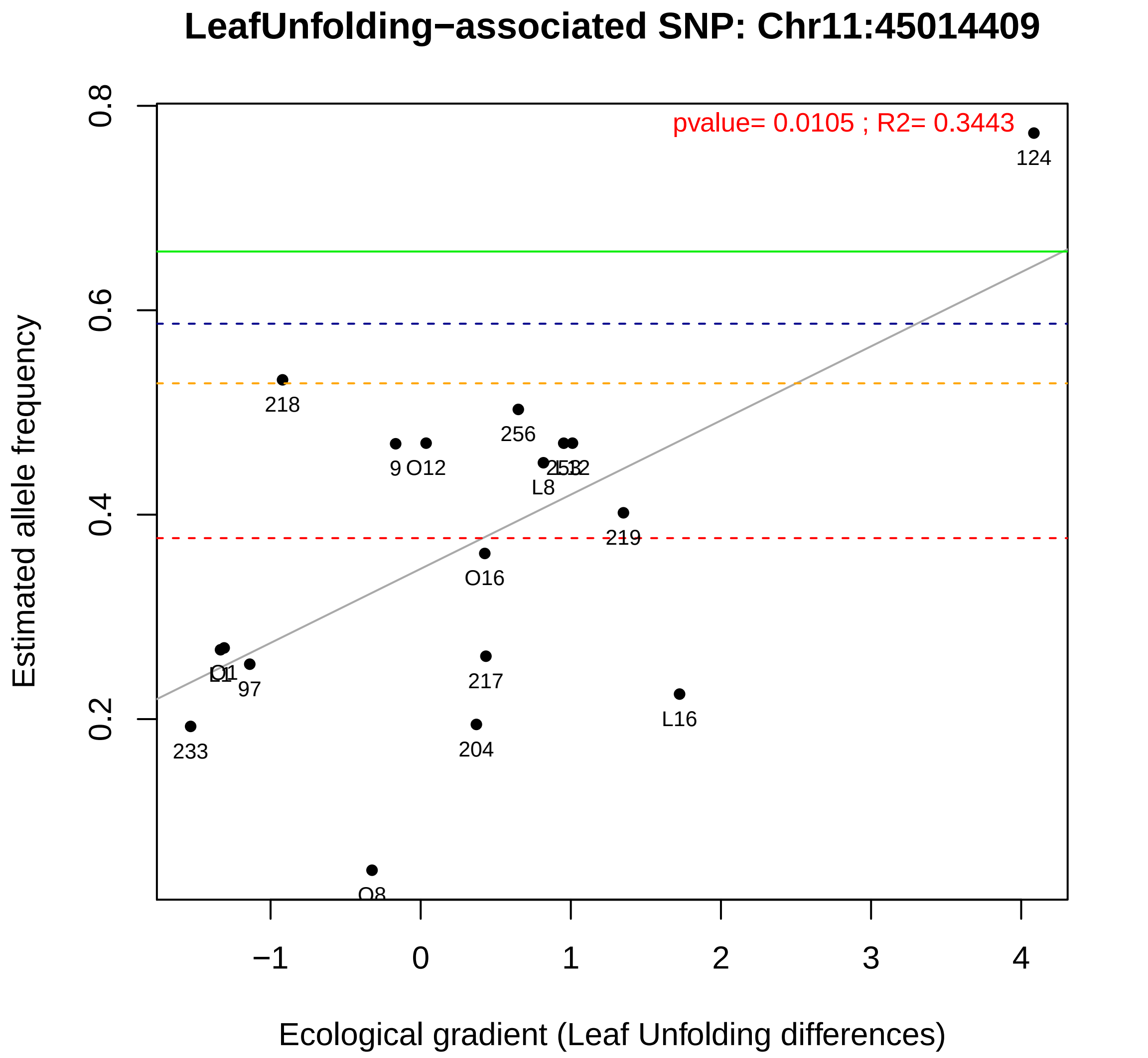
LeafUnfolding−associated SNP: Chr11:45014409
pvalue= 0.0105 ; R2= 0.3443
Estimated allele frequency
0.8
0.6
0.4
0.2
Ecological gradient (Leaf Unfolding differences)
−1
0
1
2
3
4
124
218
256
9
O12
L8
219
O16
217
L16
97
233
204
O8

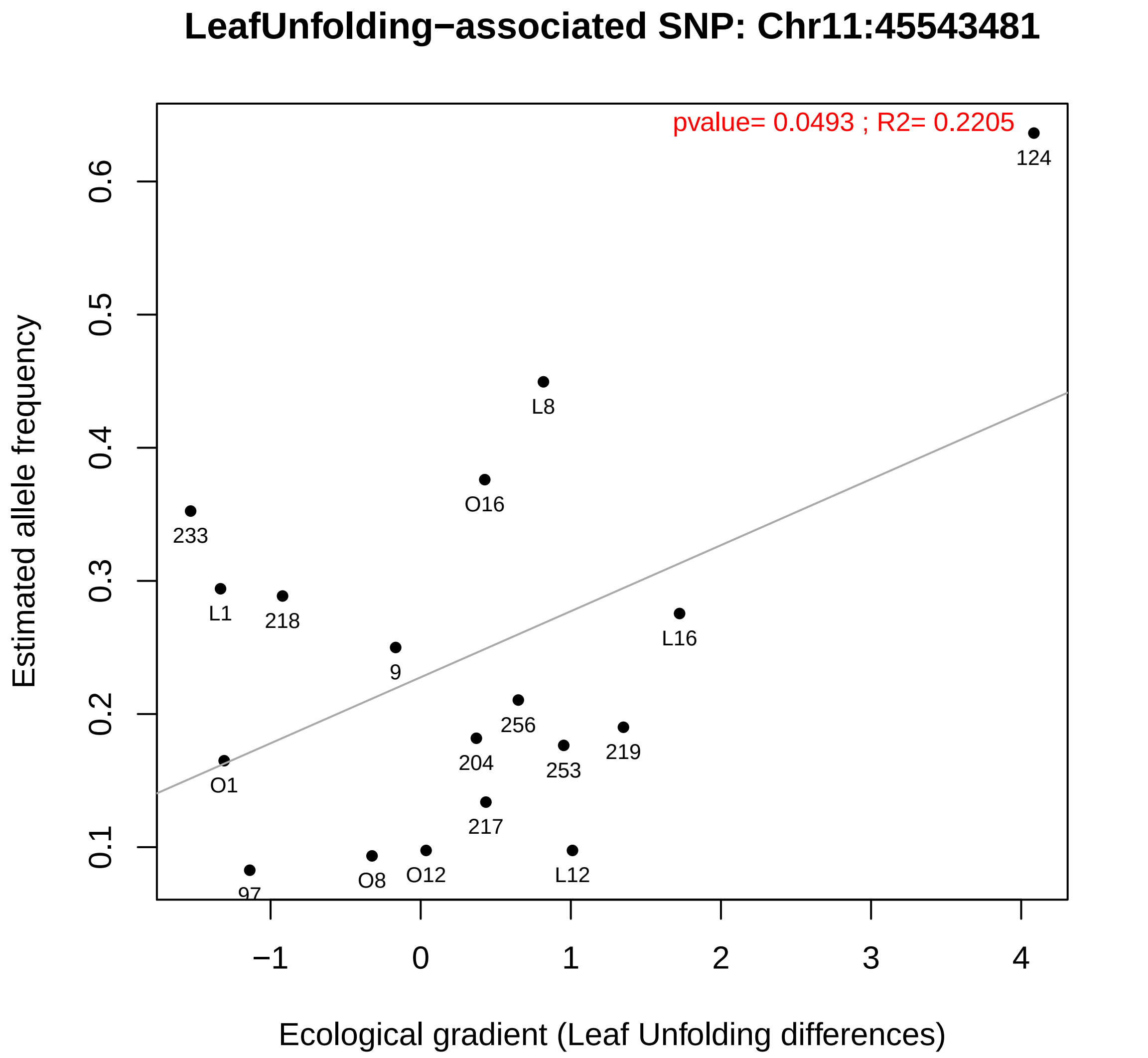
LeafUnfolding−associated SNP: Chr11:45543481
pvalue= 0.0493 ; R2= 0.2205
Estimated allele frequency
Ecological gradient (Leaf Unfolding differences)
0.1
0.2
0.3
0.4
0.5
0.6
−1
0
1
2
3
4
124
L8
O16
233
L1
218
L16
9
256
204
253
219
O1
217
97
O8
O12
L12

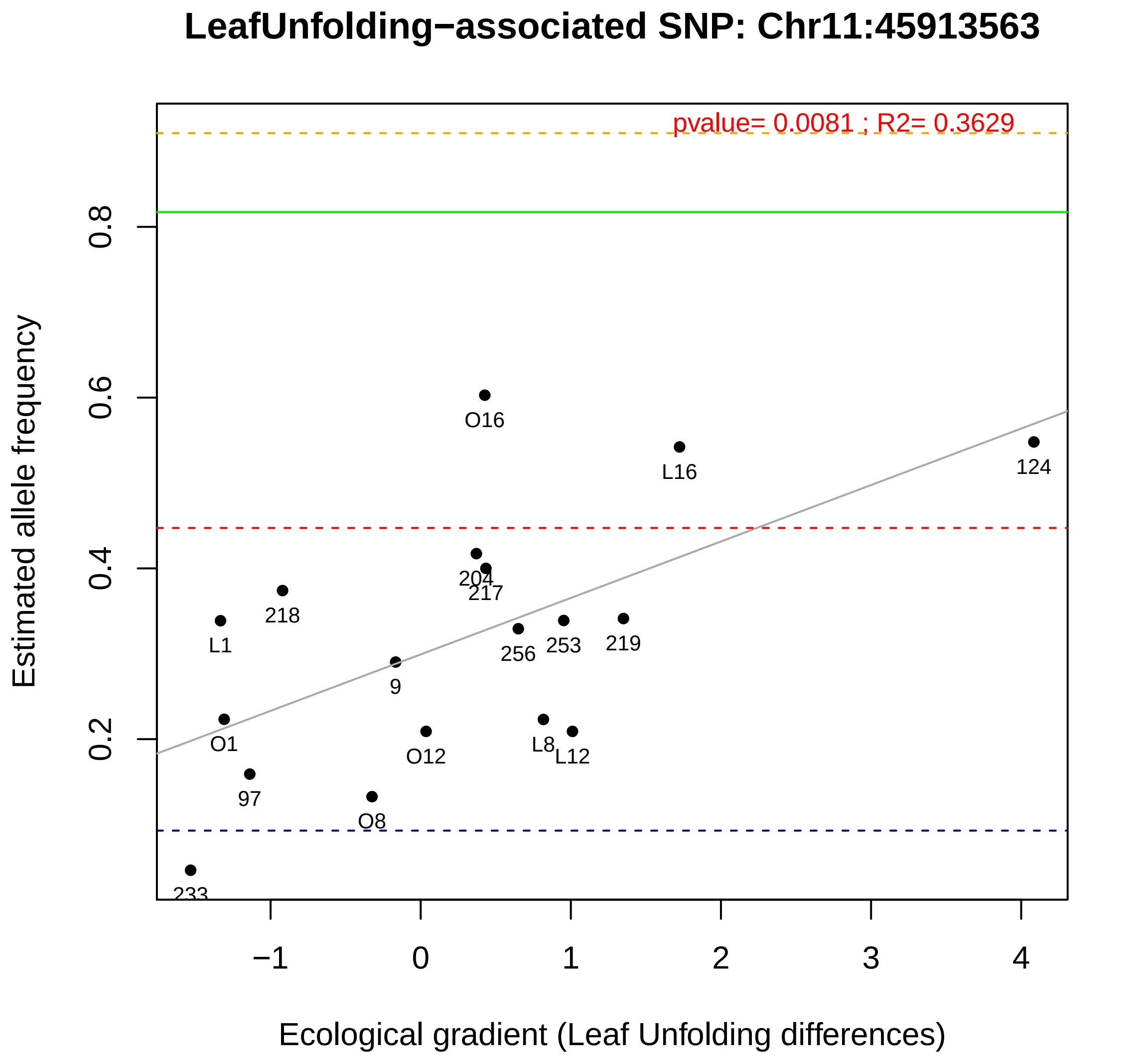

LeafUnfolding−associated SNP: Chr11:45913563
pvalue= 0.0081 ; R2= 0.3629
Estimated allele frequency
0.2
0.4
0.6
0.8
−1
0
1
2
3
4
Ecological gradient (Leaf Unfolding differences)
O16
L16
124
204
217
218
L1
256
253
219
9
O1
O12
L8
L12
97
O8
233

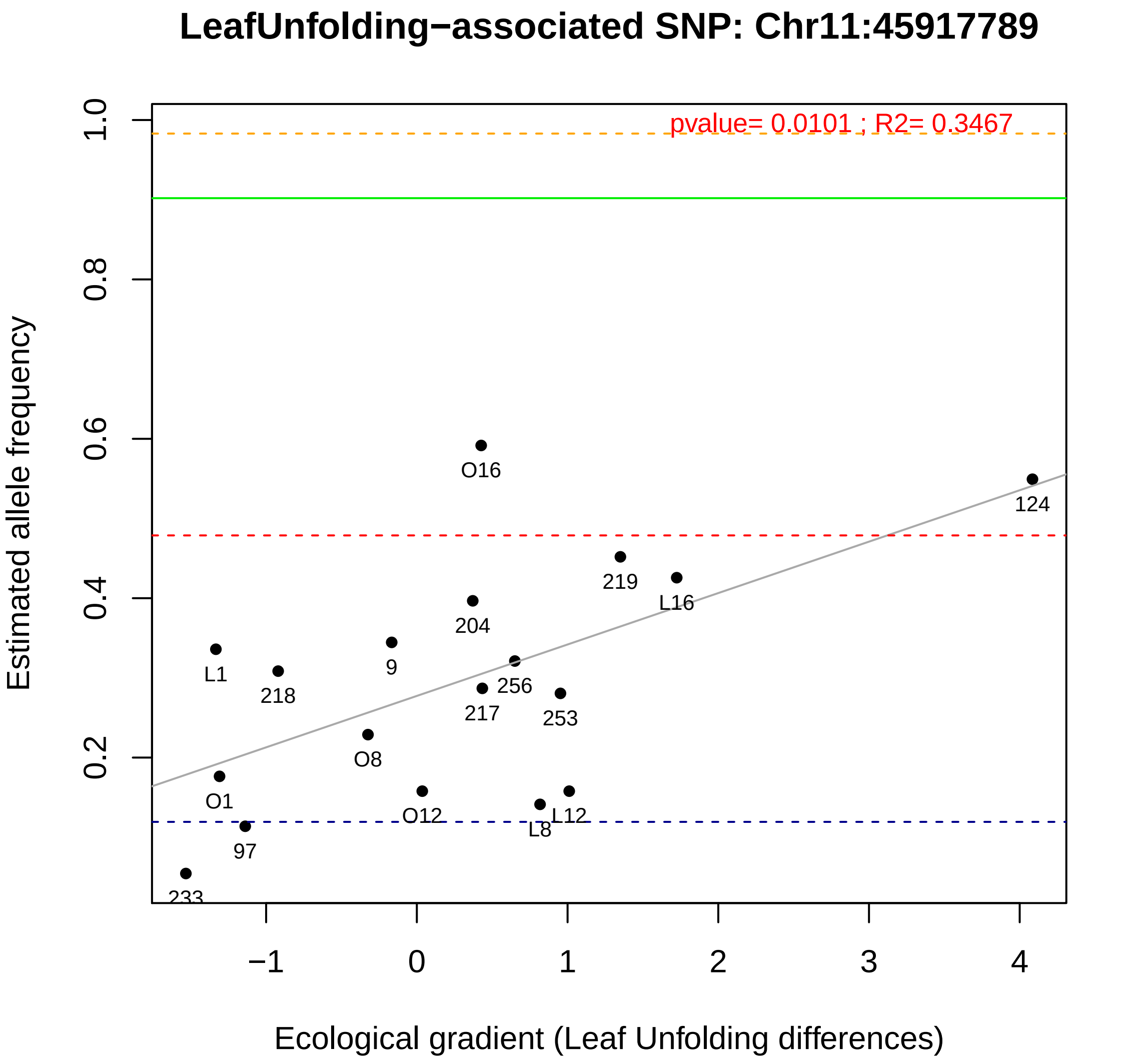
LeafUnfolding−associated SNP: Chr11:45917789
pvalue= 0.0101 ; R2= 0.3467
Estimated allele frequency
0.2
0.4
0.6
0.8
1.0
−1
0
1
2
3
4
Ecological gradient (Leaf Unfolding differences)
O16
124
219
L16
204
L1
218
9
256
217
253
O8
O1
O12
L8
L12
97
233

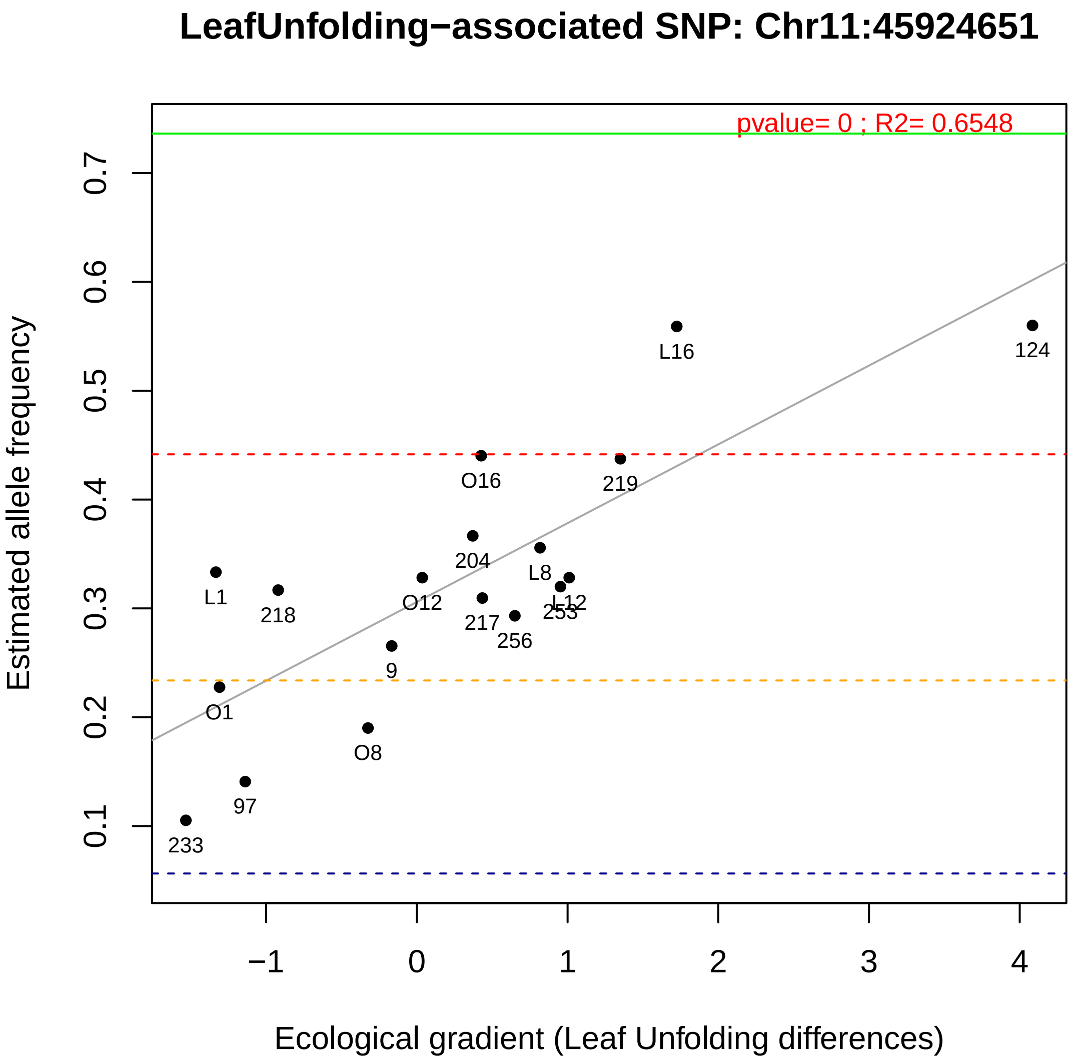
LeafUnfolding−associated SNP: Chr11:45924651
pvalue= 0 ; R2= 0.6548
Estimated allele frequency
Ecological gradient (Leaf Unfolding differences)
0.1
0.2
0.3
0.4
0.5
0.6
0.7
−1
0
1
2
3
4
L16
124
O16
219
204
L8
L1
218
O12
L12
253
217
256
9
O1
O8
97
233

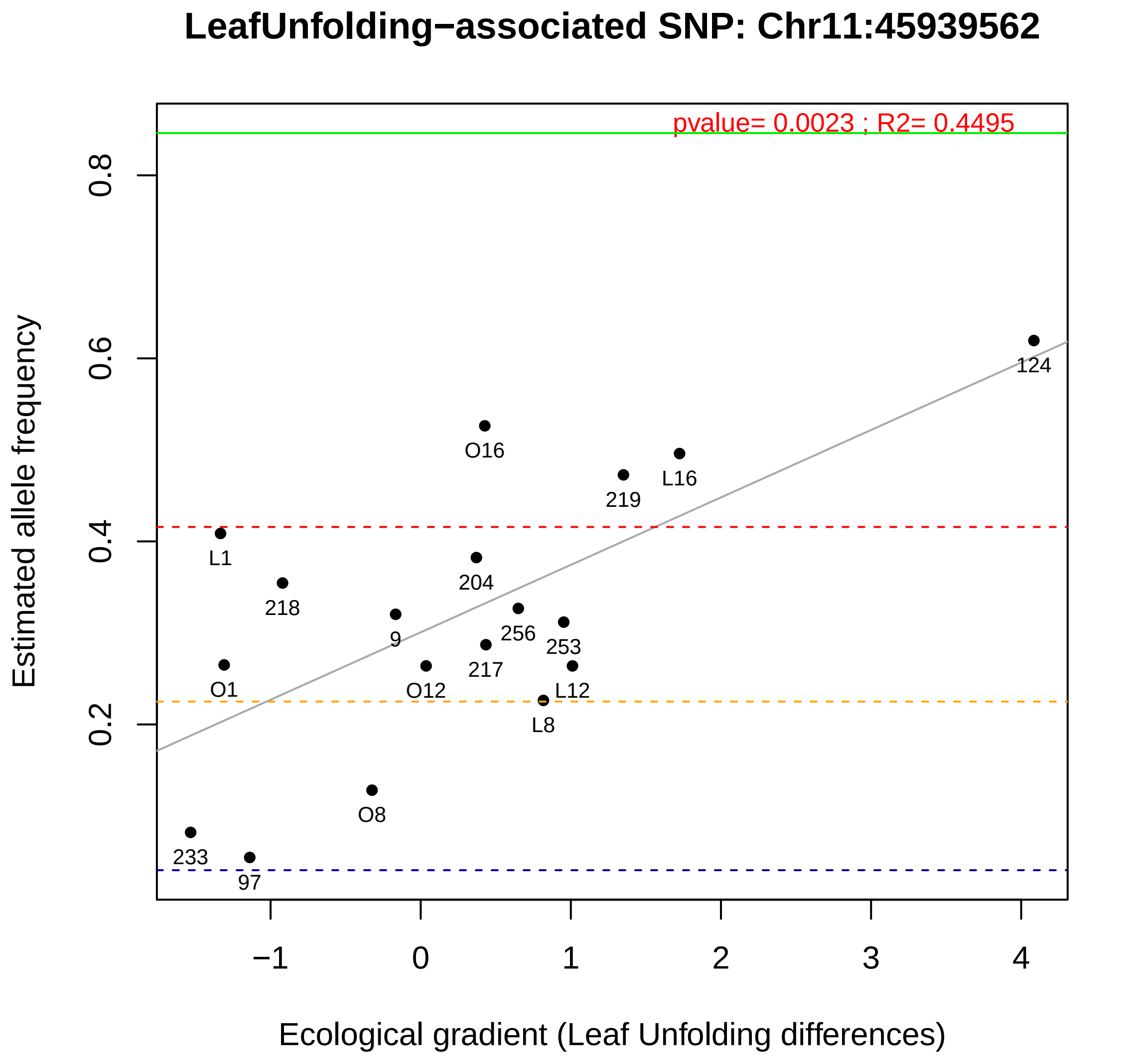
LeafUnfolding−associated SNP: Chr11:45939562
pvalue= 0.0023 ; R2= 0.4495
Estimated allele frequency
0.2
0.4
0.6
0.8
−1
0
1
2
3
4
Ecological gradient (Leaf Unfolding differences)
124
O16
L16
219
L1
218
204
9
256
253
217
O1
O12
L12
L8
O8
233
97

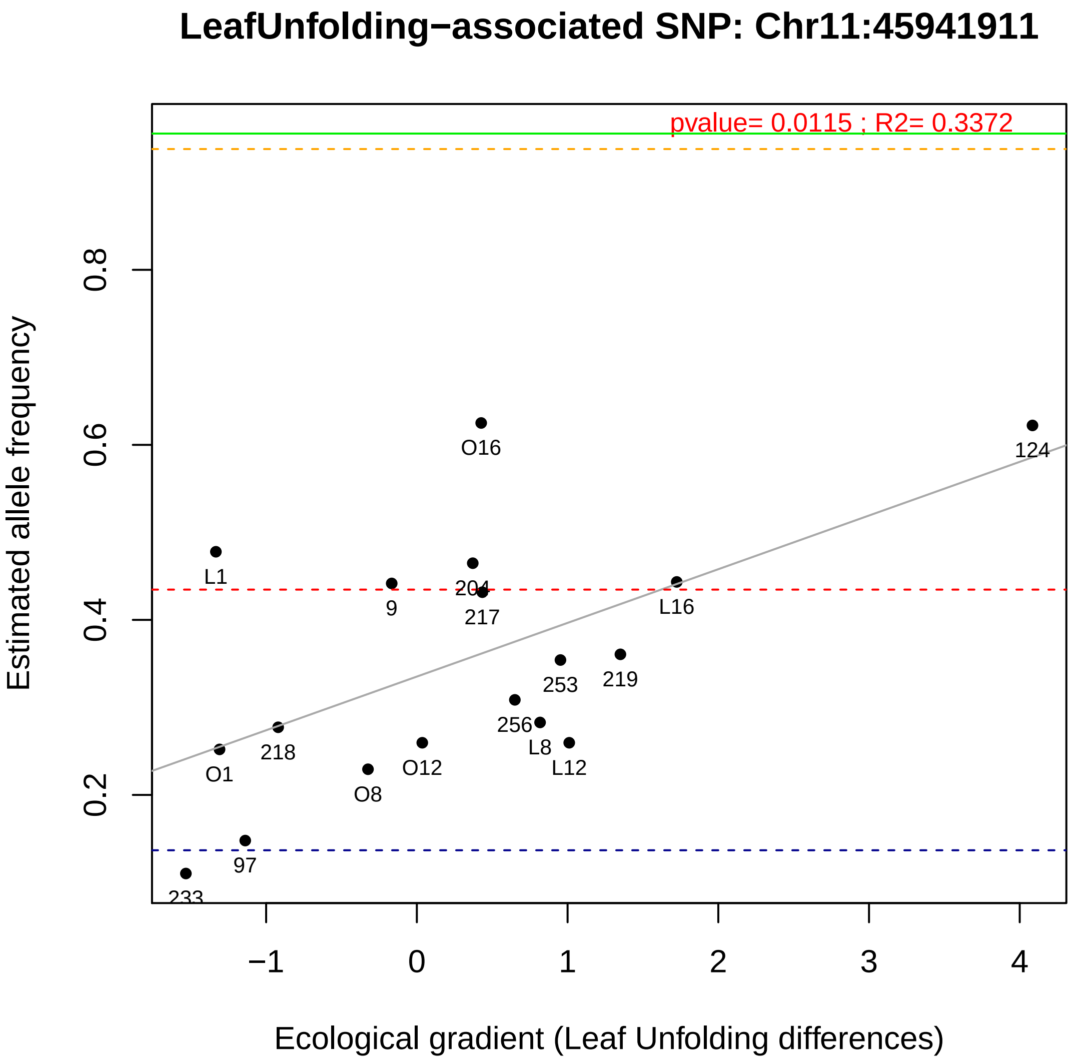
LeafUnfolding−associated SNP: Chr11:45941911
pvalue= 0.0115 ; R2= 0.3372
Estimated allele frequency
0.2
0.4
0.6
0.8
−1
0
1
2
3
4
Ecological gradient (Leaf Unfolding differences)
O16
124
L1
9
204
217
L16
253
219
256
L8
L12
218
O1
O12
O8
97
233

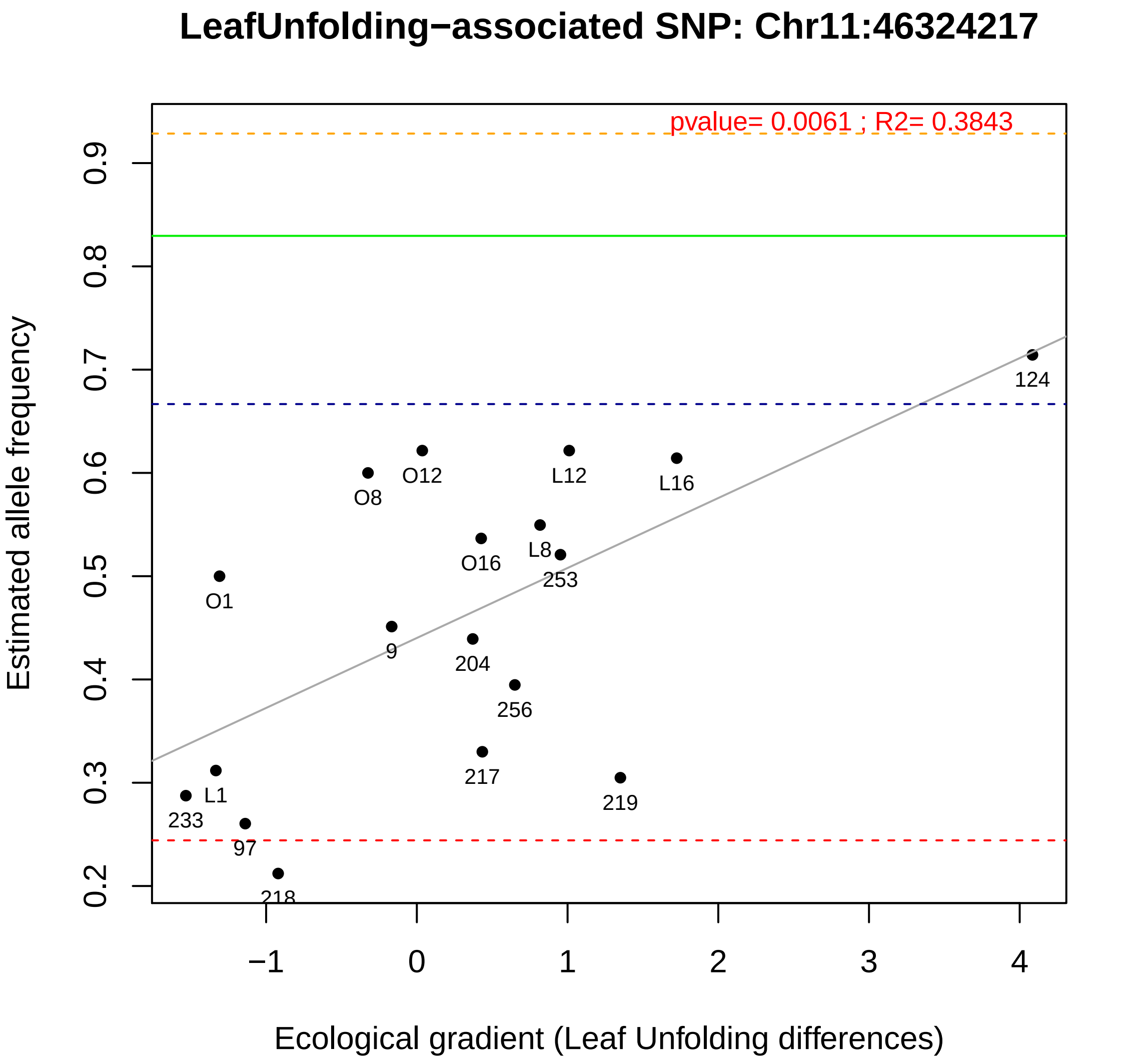

LeafUnfolding−associated SNP: Chr11:46324217
pvalue= 0.0061 ; R2= 0.3843
Estimated allele frequency
0.2
0.3
0.4
0.5
0.6
0.7
0.8
0.9
−1
0
1
2
3
4
Ecological gradient (Leaf Unfolding differences)
124
O12
L12
L16
O8
L8
O16
253
O1
9
204
256
217
219
L1
233
97
218

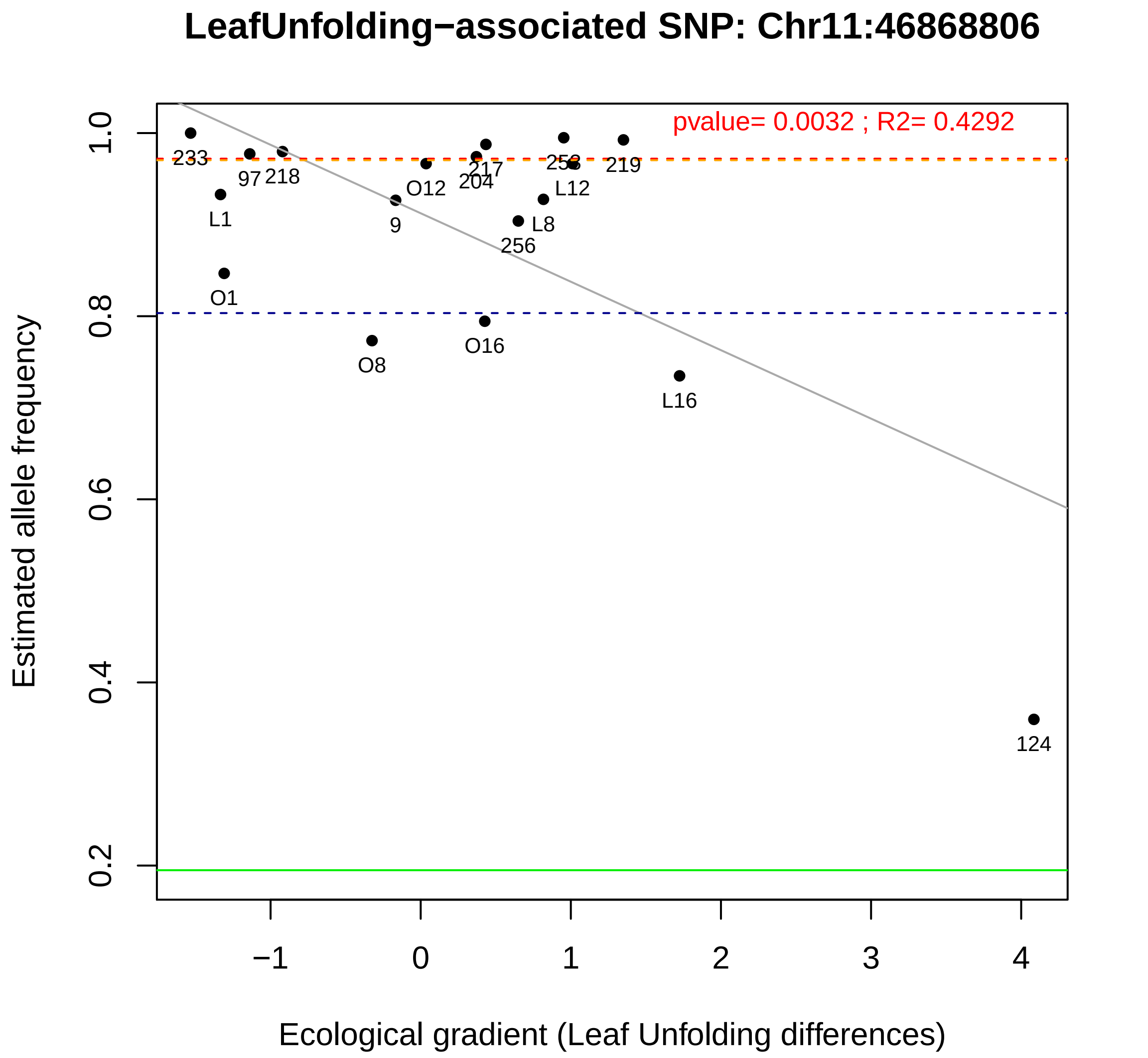

LeafUnfolding−associated SNP: Chr11:46868806
pvalue= 0.0032 ; R2= 0.4292
Estimated allele frequency
Ecological gradient (Leaf Unfolding differences)
233
97
218
L1
O1
O12
9
217
204
256
L8
L12
219
O8
O16
L16
124
0.2
0.4
0.6
0.8
1.0
−1
0
1
2
3
4

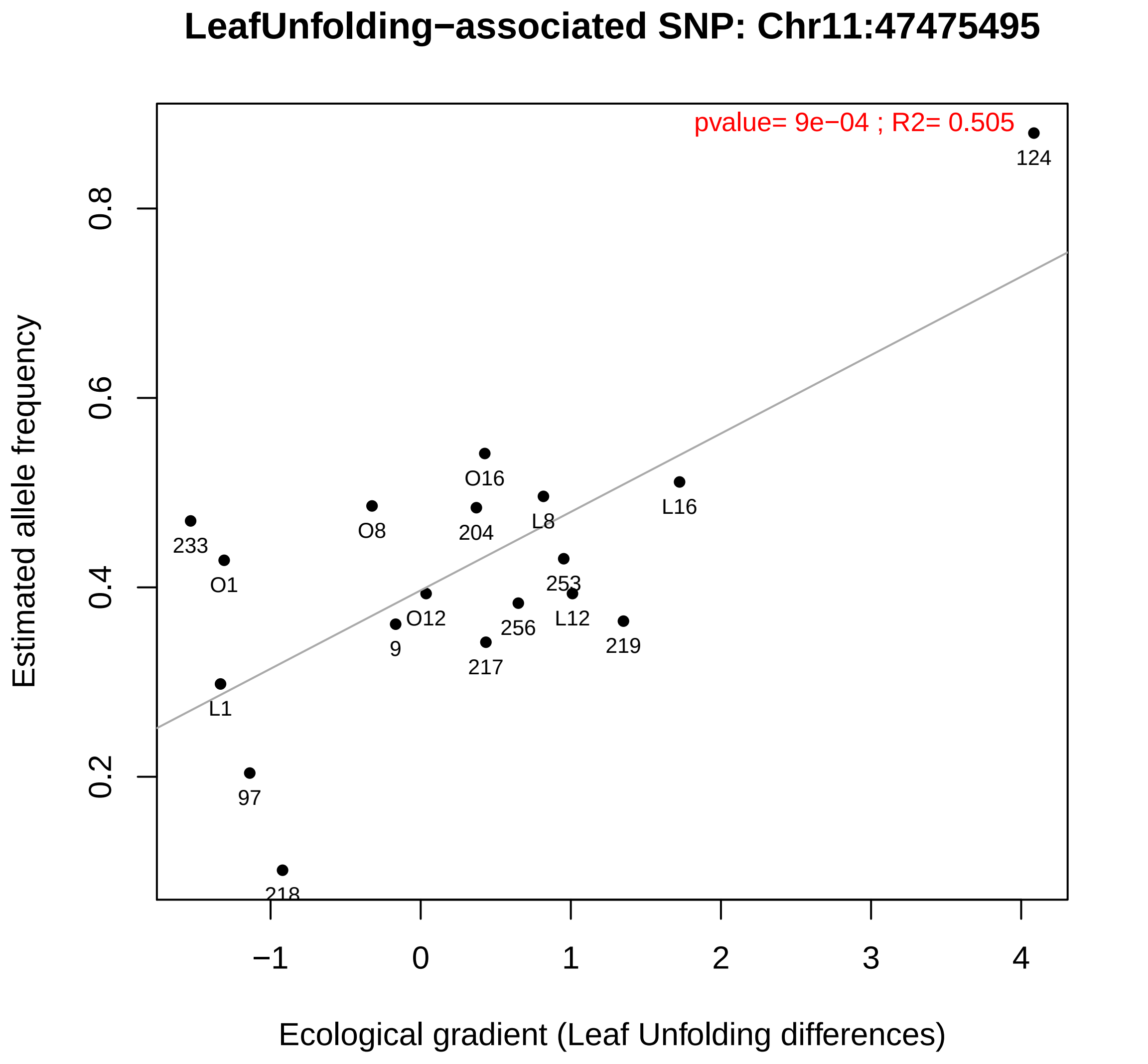
LeafUnfolding−associated SNP: Chr11:47475495
pvalue= 9e−04 ; R2= 0.505
Estimated allele frequency
0.2
0.4
0.6
0.8
−1
0
1
2
3
4
Ecological gradient (Leaf Unfolding differences)
124
O16
L16
L8
204
O8
233
O1
253
L12
O12
256
219
9
217
L1
97
218

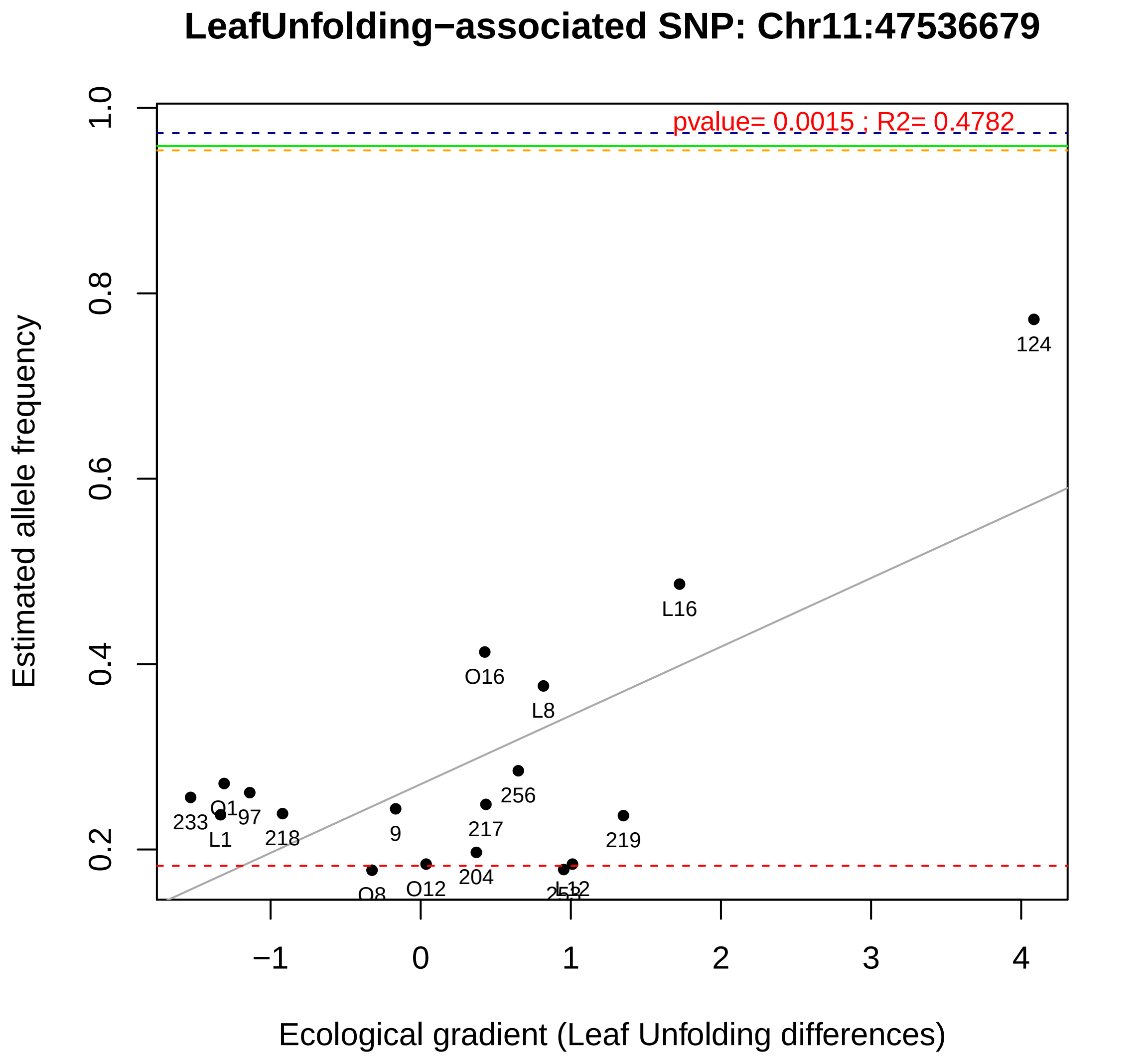

LeafUnfolding−associated SNP: Chr11:47536679
pvalue= 0.0015 ; R2= 0.4782
Estimated allele frequency
0.2
0.4
0.6
0.8
1.0
−1
0
1
2
3
4
Ecological gradient (Leaf Unfolding differences)
124
L16
O16
L8
256
217
219
233
O1
97
L1
218
9
204
O12
O8
L12

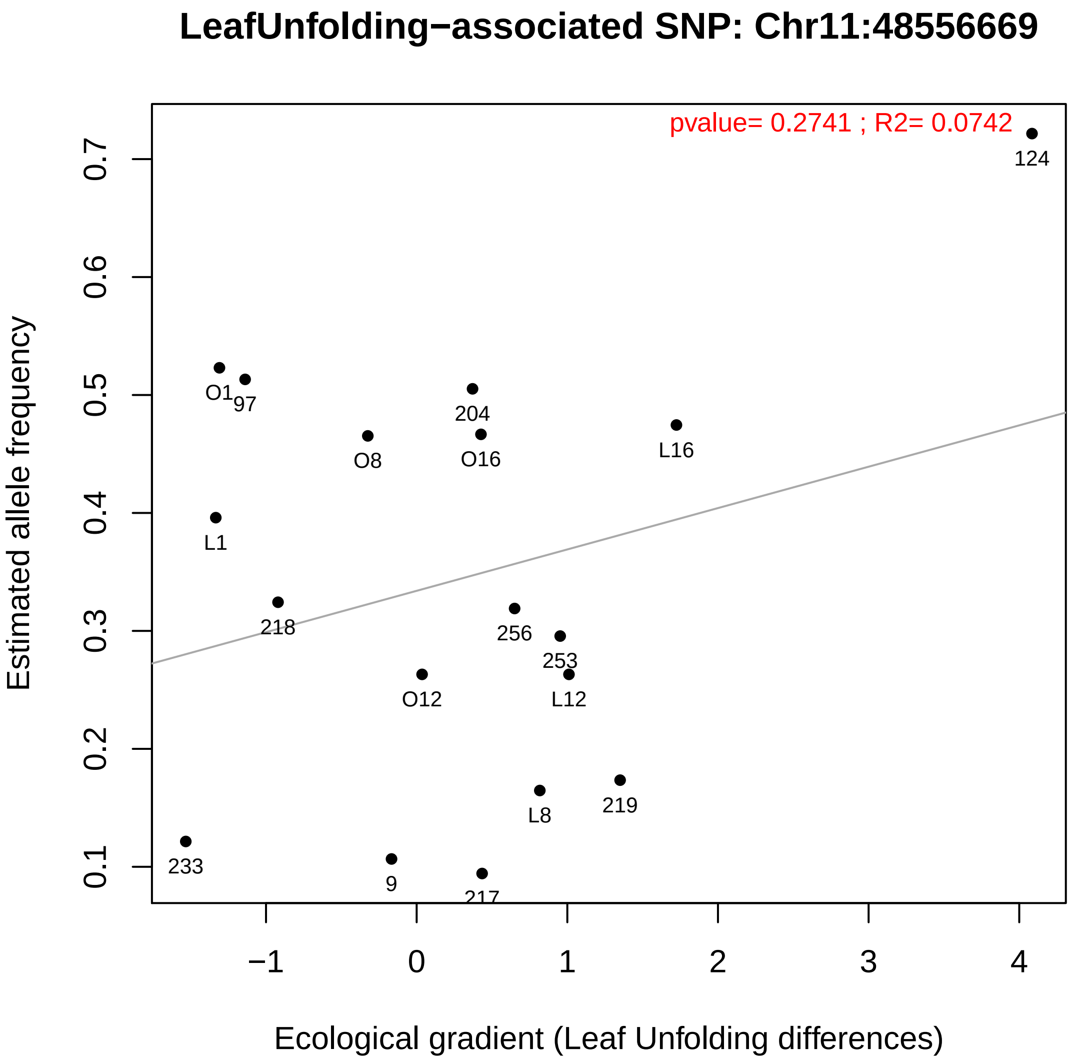
LeafUnfolding−associated SNP: Chr11:48556669
pvalue= 0.2741 ; R2= 0.0742
Estimated allele frequency
Ecological gradient (Leaf Unfolding differences)
0.1
0.2
0.3
0.4
0.5
0.6
0.7
−1
0
1
2
3
4
124
O1
97
204
O16
O8
L16
L1
218
256
253
O12
L12
L8
219
233
9
217

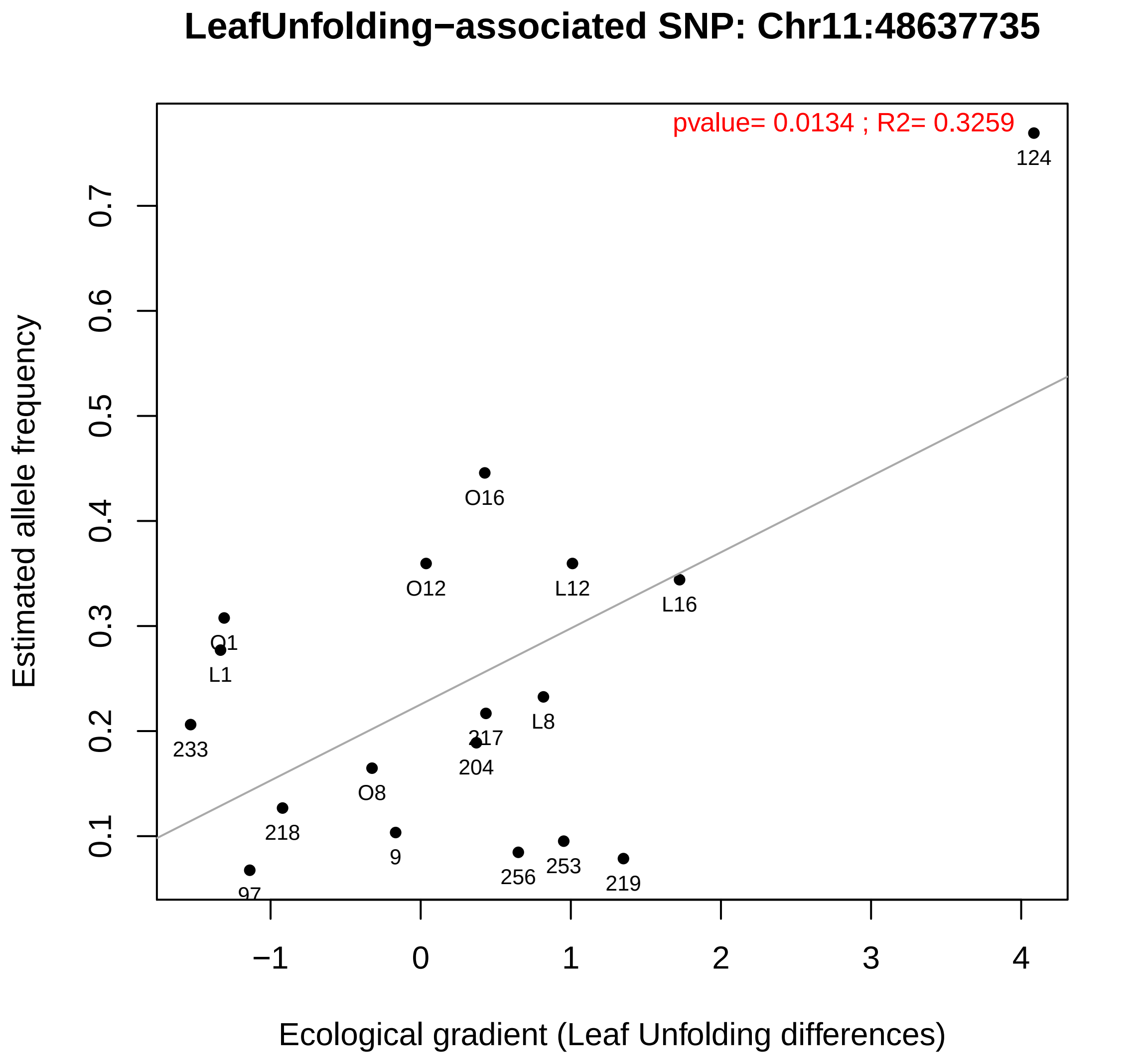

LeafUnfolding−associated SNP: Chr11:48637735
pvalue= 0.0134 ; R2= 0.3259
Estimated allele frequency
0.1
0.2
0.3
0.4
0.5
0.6
0.7
−1
0
1
2
3
4
Ecological gradient (Leaf Unfolding differences)
124
O16
O12
L12
L16
O1
L1
233
L8
217
204
O8
218
9
256
253
219
97

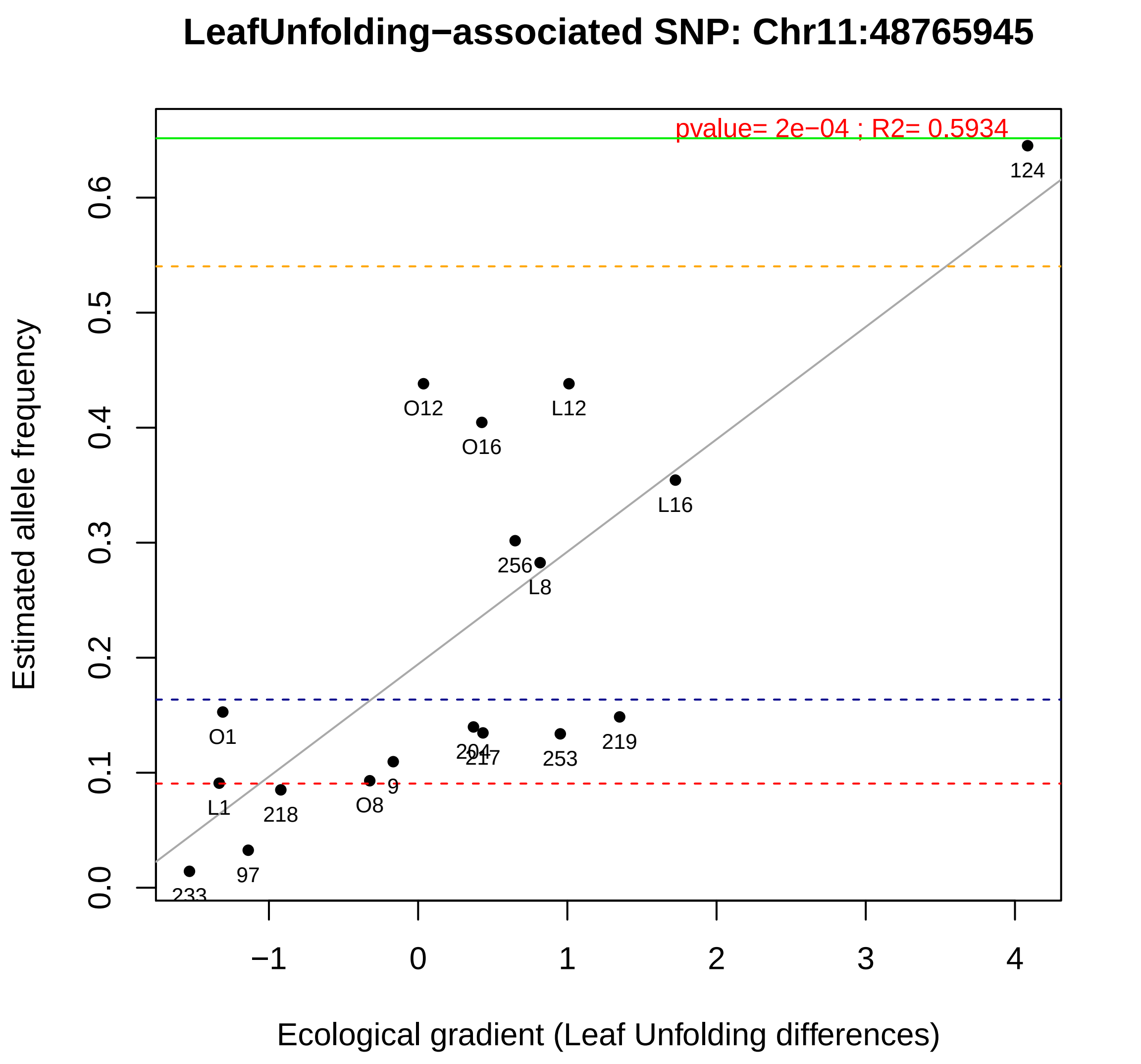
LeafUnfolding−associated SNP: Chr11:48765945
pvalue= 2e−04 ; R2= 0.5934
Estimated allele frequency
Ecological gradient (Leaf Unfolding differences)
0.0
0.1
0.2
0.3
0.4
0.5
0.6
−1
0
1
2
3
4
124
O12
L12
O16
L16
256
L8
O1
204
217
253
219
L1
218
O8
9
97
233

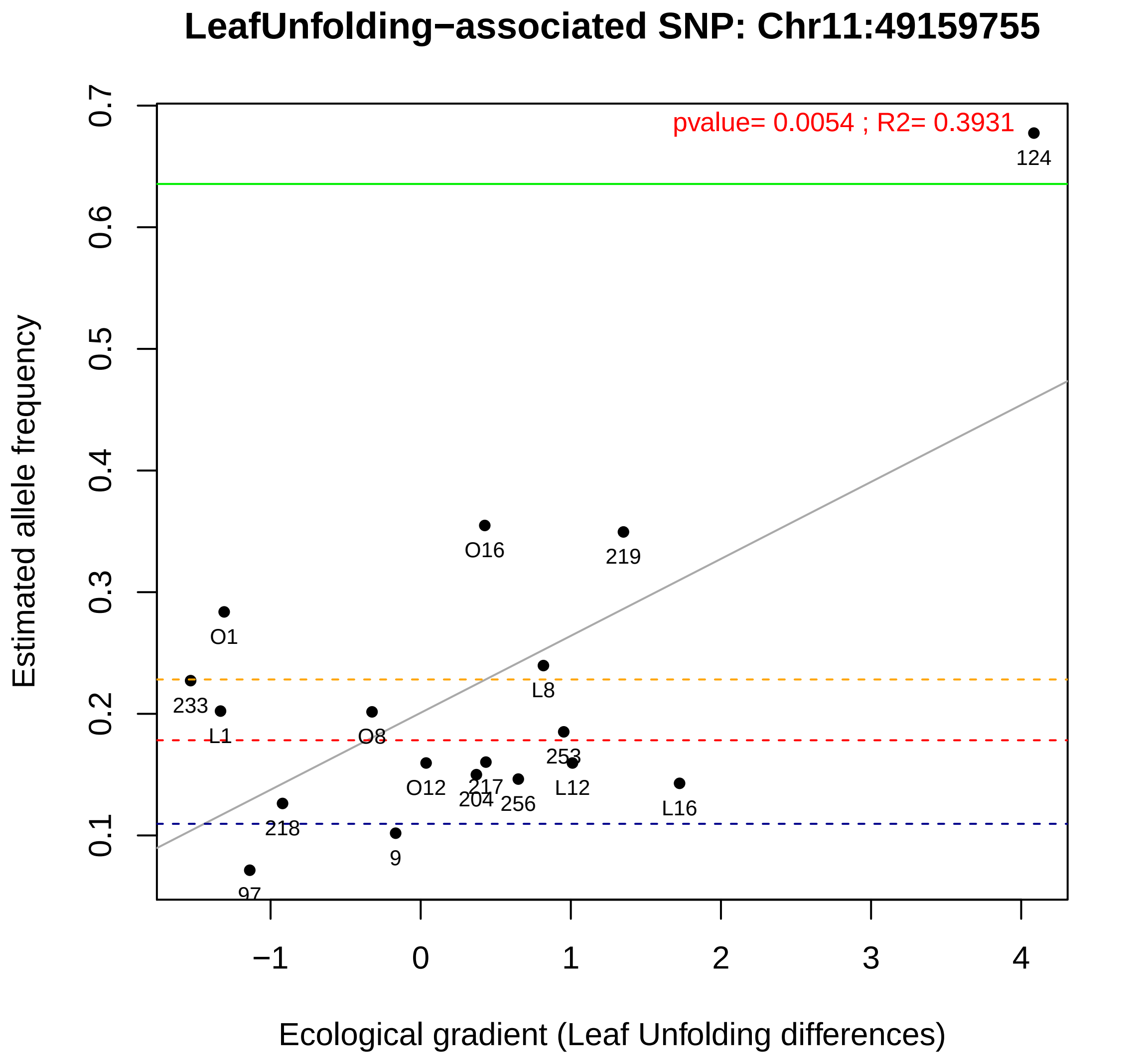

LeafUnfolding−associated SNP: Chr11:49159755
pvalue= 0.0054 ; R2= 0.3931
Estimated allele frequency
0.1
0.2
0.3
0.4
0.5
0.6
0.7
−1
0
1
2
3
4
Ecological gradient (Leaf Unfolding differences)
124
O16
219
O1
233
L1
O8
L8
253
O12
217
204
256
L12
L16
218
9
97

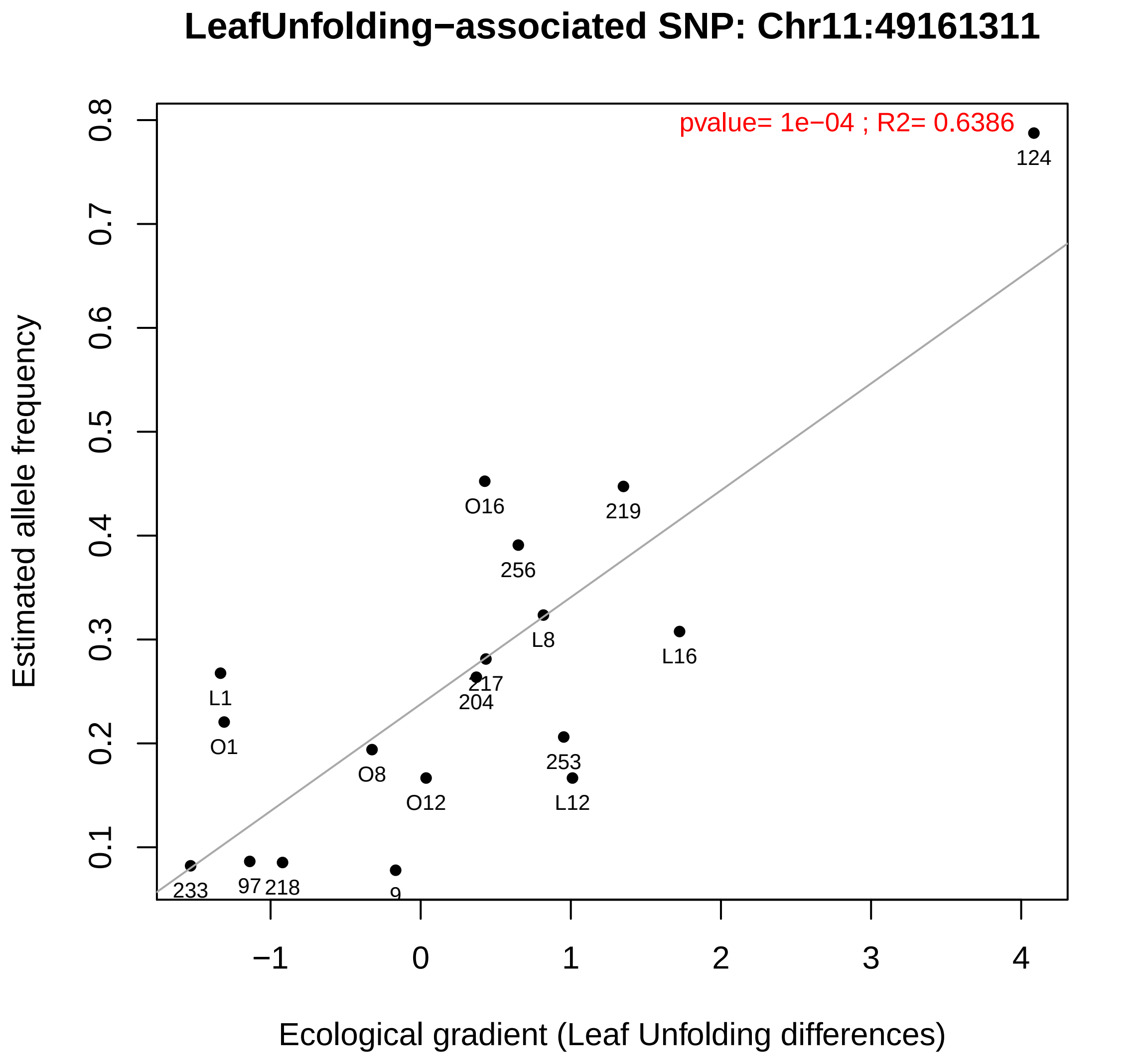

LeafUnfolding−associated SNP: Chr11:49161311
pvalue= 1e−04 ; R2= 0.6386
Estimated allele frequency
Ecological gradient (Leaf Unfolding differences)
0.1
0.2
0.3
0.4
0.5
0.6
0.7
0.8
−1
0
1
2
3
4
124
O16
219
256
L8
L16
217
204
L1
O1
O8
O12
253
L12
233
97
218
9

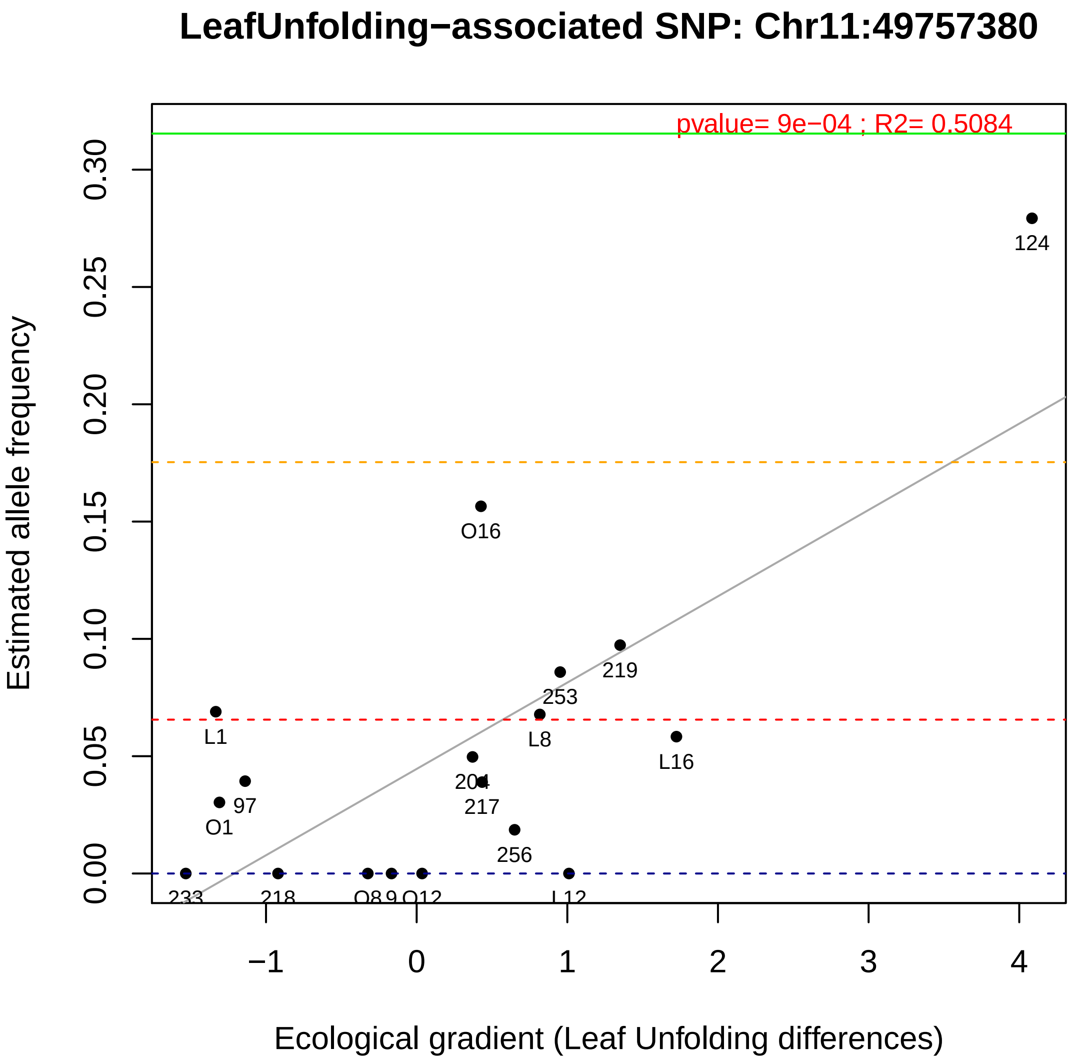

LeafUnfolding−associated SNP: Chr11:49757380
pvalue= 9e−04 ; R2= 0.5084
Estimated allele frequency
0.00
0.05
0.10
0.15
0.20
0.25
0.30
−1
0
1
2
3
4
Ecological gradient (Leaf Unfolding differences)
124
O16
219
253
L1
L8
L16
97
O1
204
217
256
233
218
O8
9
O12
L12

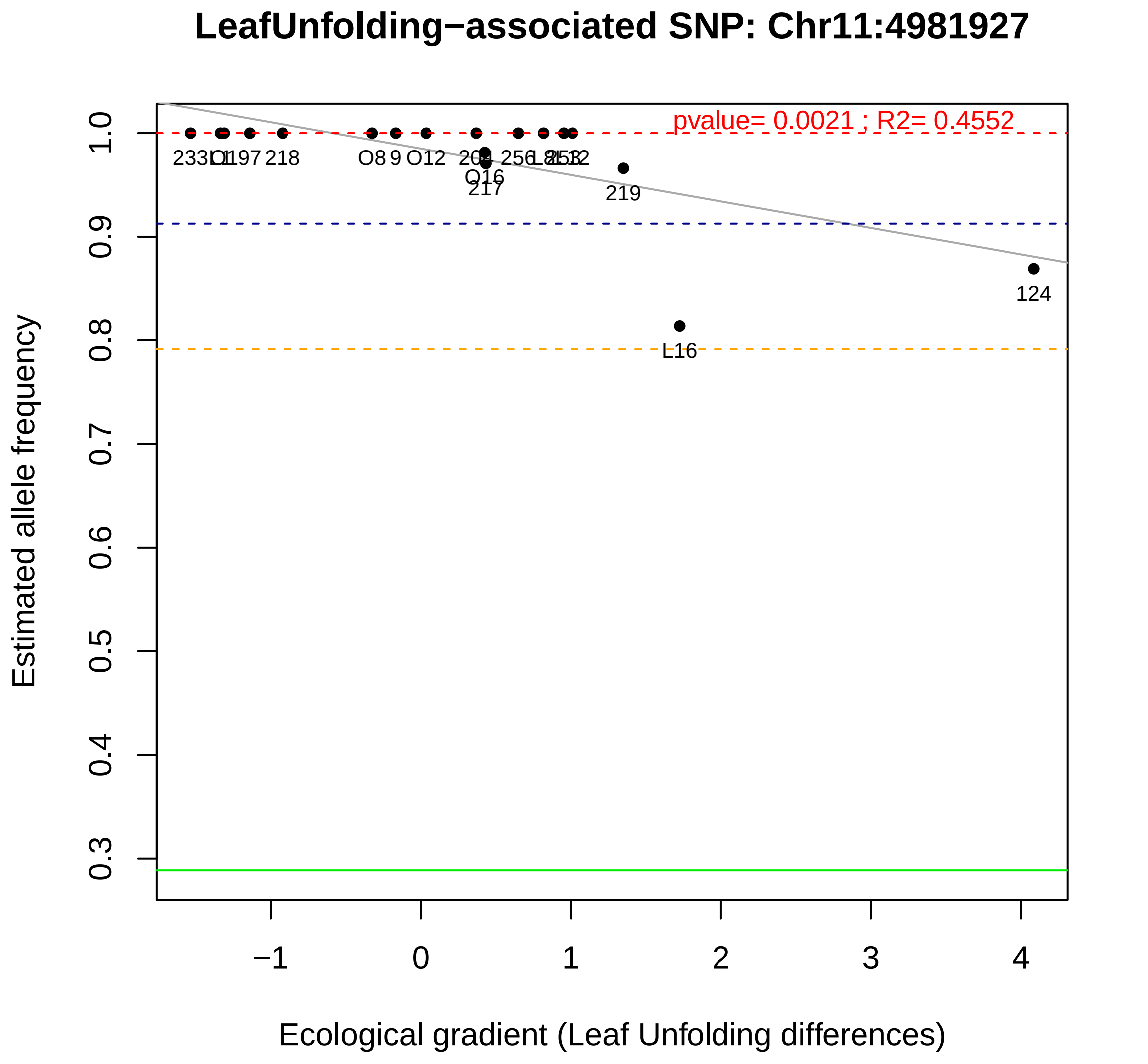
LeafUnfolding−associated SNP: Chr11:4981927
pvalue= 0.0021 ; R2= 0.4552
Estimated allele frequency
Ecological gradient (Leaf Unfolding differences)
1.0
0.9
0.8
0.7
0.6
0.5
0.4
0.3
−1
0
1
2
3
4
218
O8
9
O12
Q16
217
219
124
L16

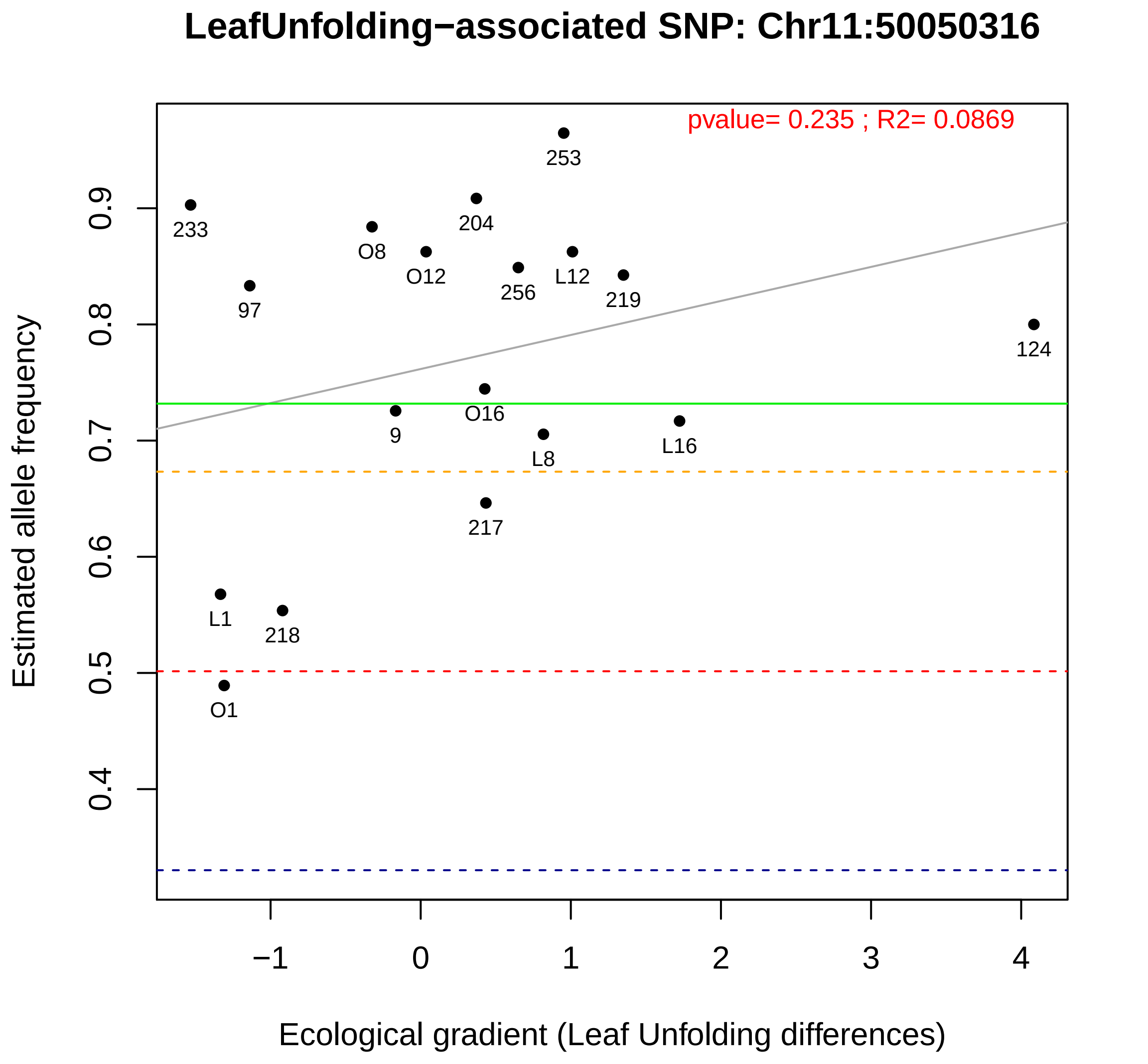
LeafUnfolding−associated SNP: Chr11:50050316
pvalue= 0.235 ; R2= 0.0869
Estimated allele frequency
Ecological gradient (Leaf Unfolding differences)
0.4
0.5
0.6
0.7
0.8
0.9
−1
0
1
2
3
4
253
233
204
O8
O12
256
L12
219
97
124
O16
9
L8
L16
217
L1
218
O1

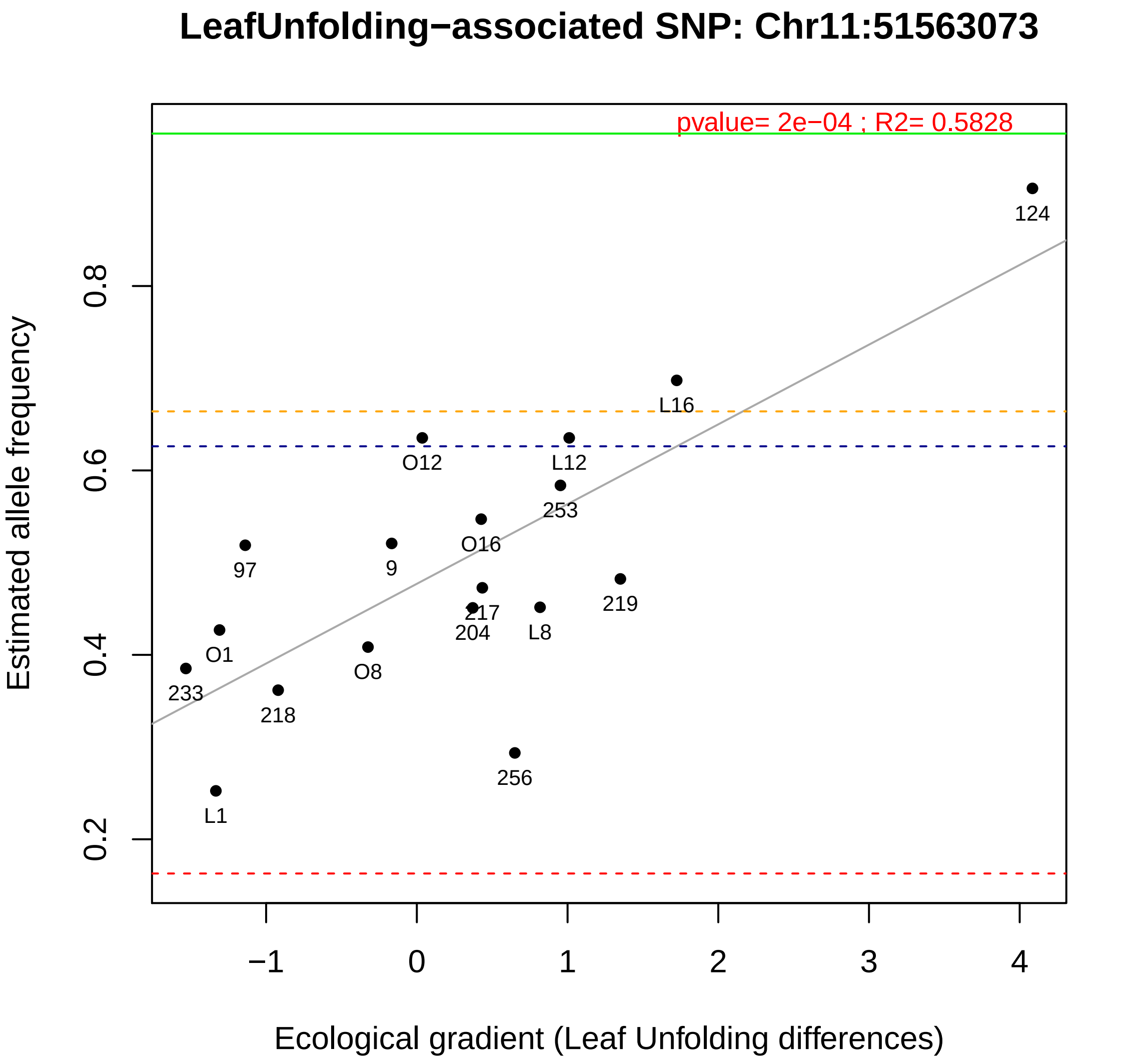

LeafUnfolding−associated SNP: Chr11:51563073
pvalue= 2e−04 ; R2= 0.5828
Estimated allele frequency
0.2
0.4
0.6
0.8
−1
0
1
2
3
4
Ecological gradient (Leaf Unfolding differences)
124
L16
O12
L12
253
O16
97
9
217
204
L8
219
O1
O8
233
218
256
L1

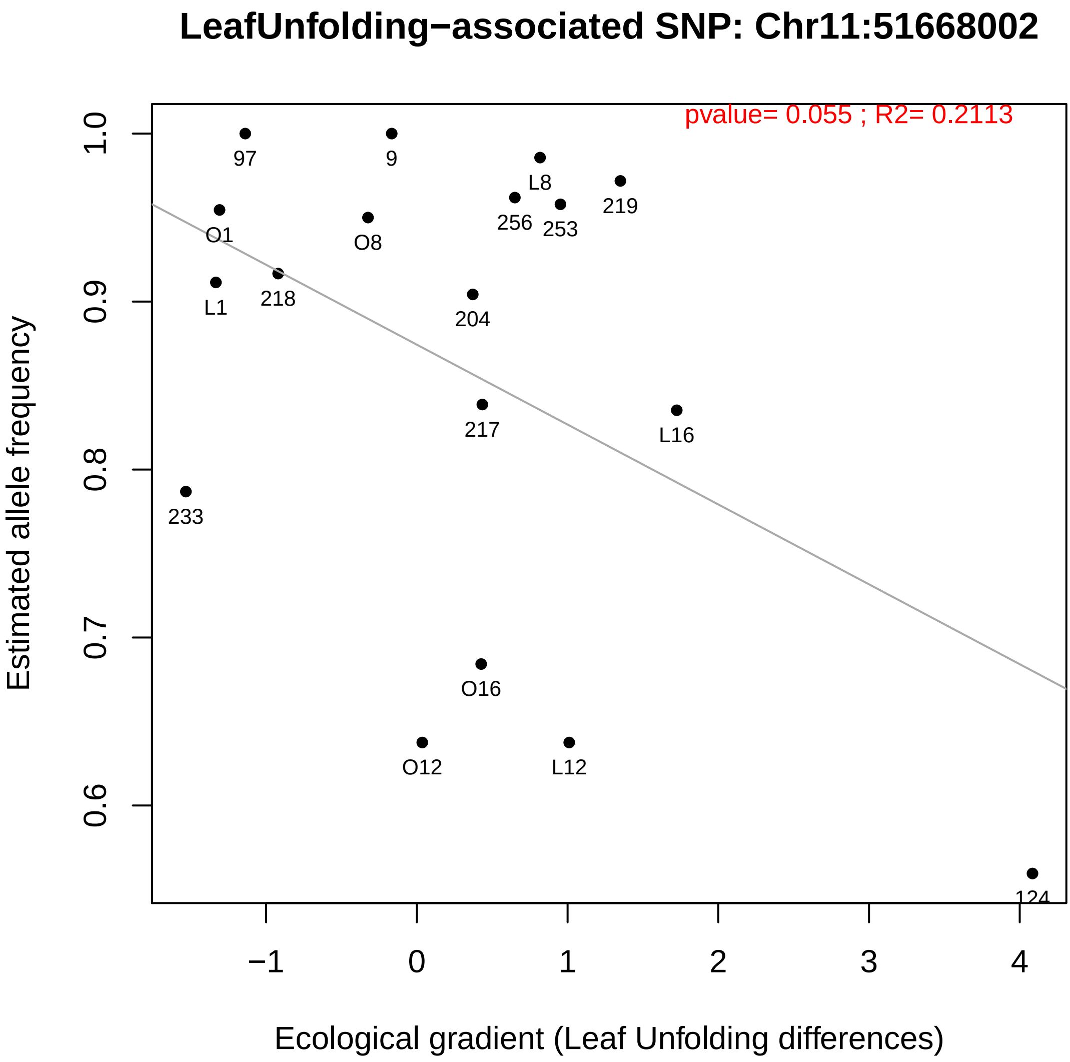
LeafUnfolding−associated SNP: Chr11:51668002
pvalue= 0.055 ; R2= 0.2113
Estimated allele frequency
Ecological gradient (Leaf Unfolding differences)
1.0
0.9
0.8
0.7
0.6
−1
0
1
2
3
4
97
9
L8
219
256
253
O1
O8
L1
218
204
217
L16
233
O16
O12
L12
124

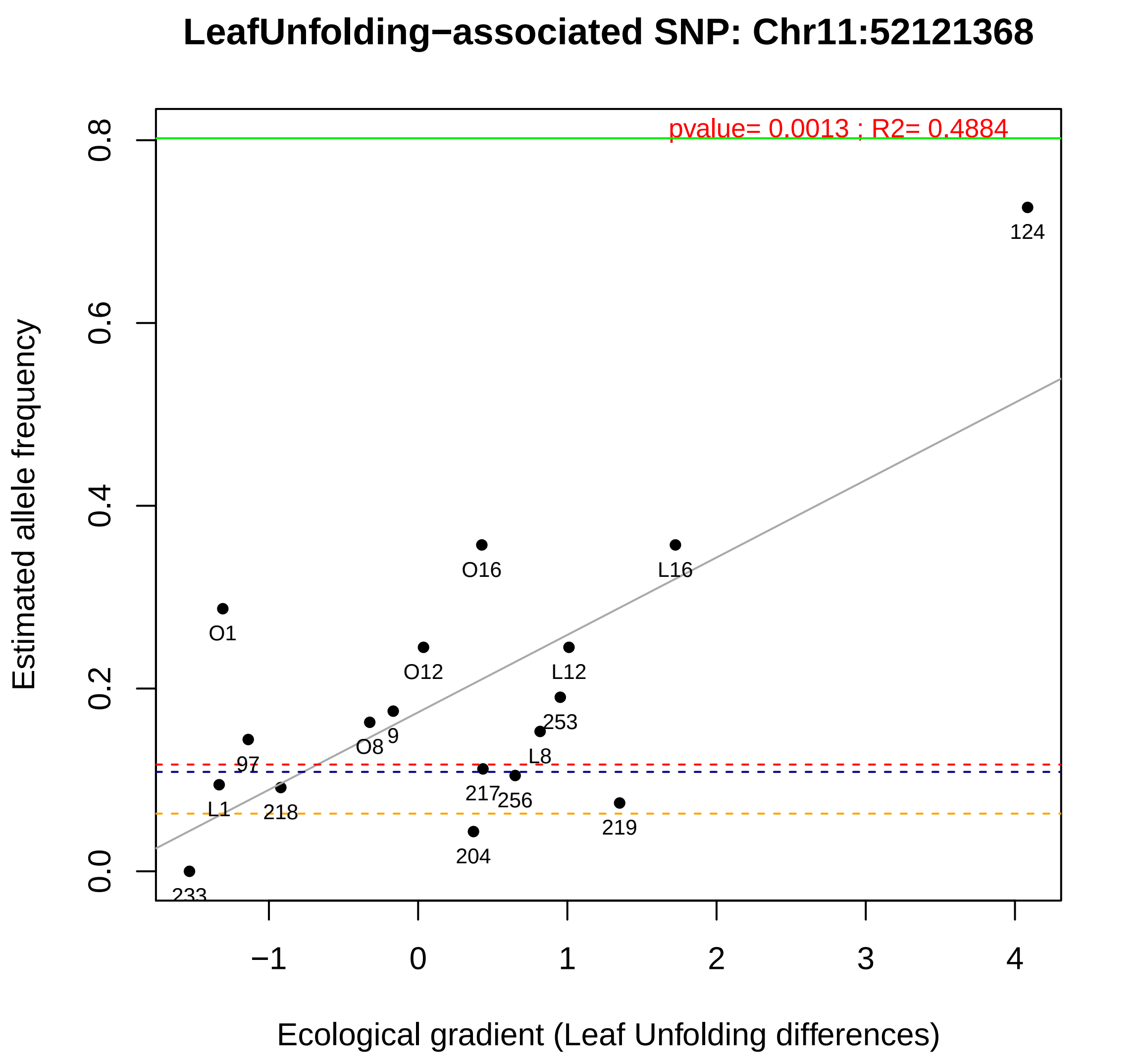

LeafUnfolding−associated SNP: Chr11:52121368
pvalue= 0.0013 ; R2= 0.4884
Estimated allele frequency
0.0
0.2
0.4
0.6
0.8
−1
0
1
2
3
4
Ecological gradient (Leaf Unfolding differences)
124
O16
L16
O1
O12
L12
253
9
O8
L8
97
217
256
L1
218
219
204
233

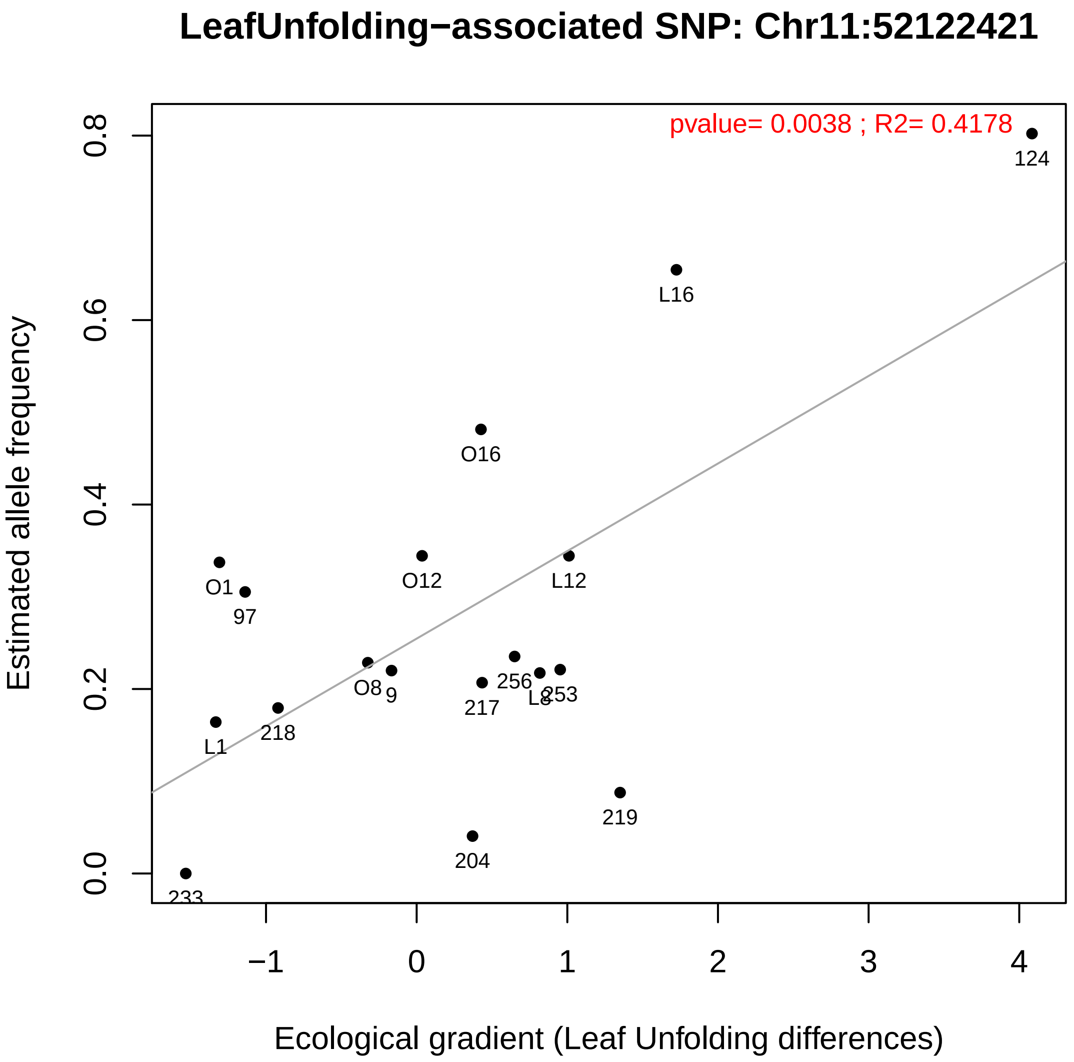
LeafUnfolding−associated SNP: Chr11:52122421
pvalue= 0.0038 ; R2= 0.4178
Estimated allele frequency
0.0
0.2
0.4
0.6
0.8
Ecological gradient (Leaf Unfolding differences)
−1
0
1
2
3
4
124
L16
O16
O12
L12
O1
97
O8
9
256
253
217
218
L1
219
204
233

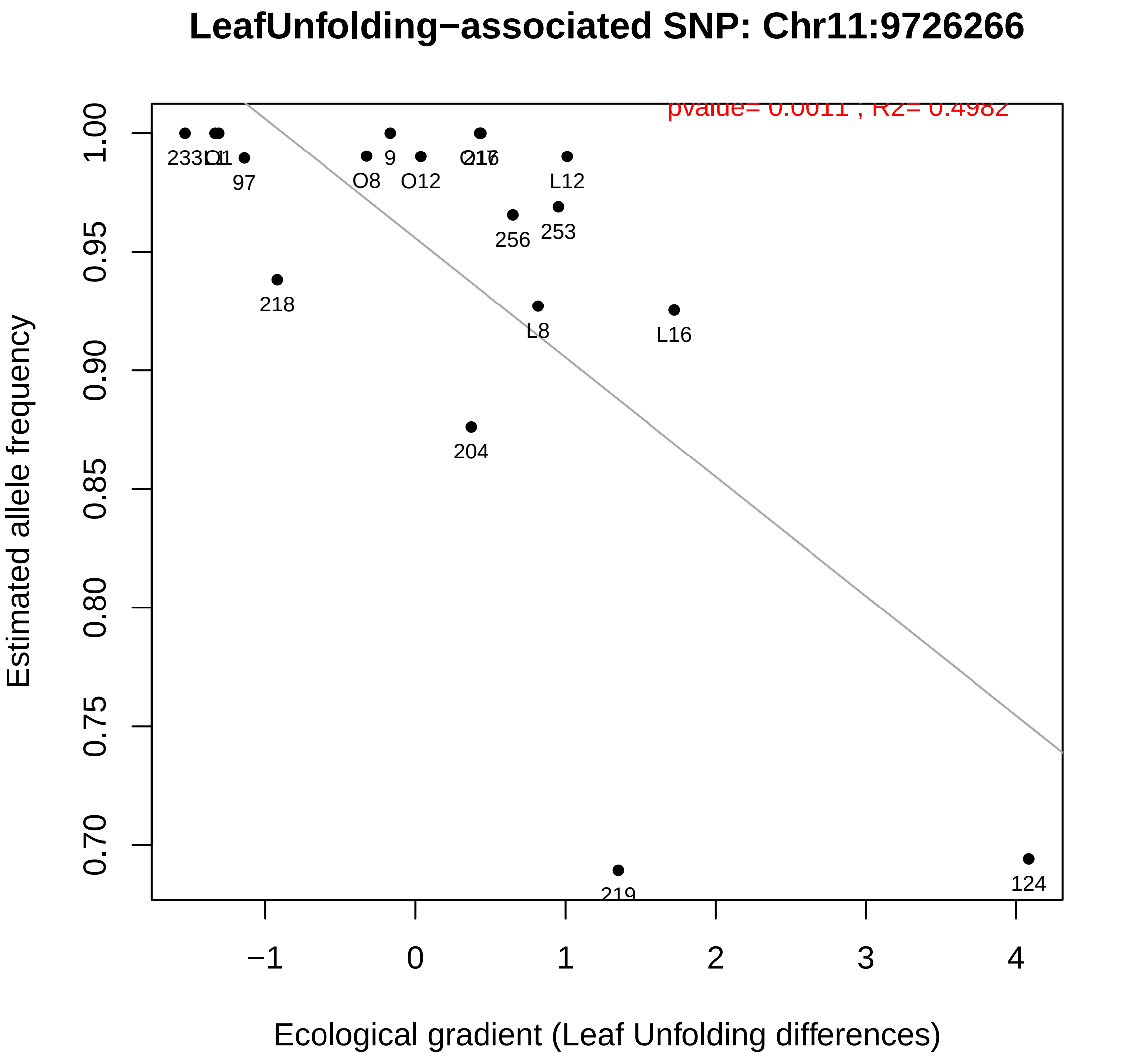

LeafUnfolding−associated SNP: Chr11:9726266
pvalue= 0.0011 ; R2= 0.4982
Estimated allele frequency
Ecological gradient (Leaf Unfolding differences)
1.00
0.95
0.90
0.85
0.80
0.75
0.70
−1
0
1
2
3
4
97
O8
9
O12
L12
256
253
218
L8
L16
204
219
124

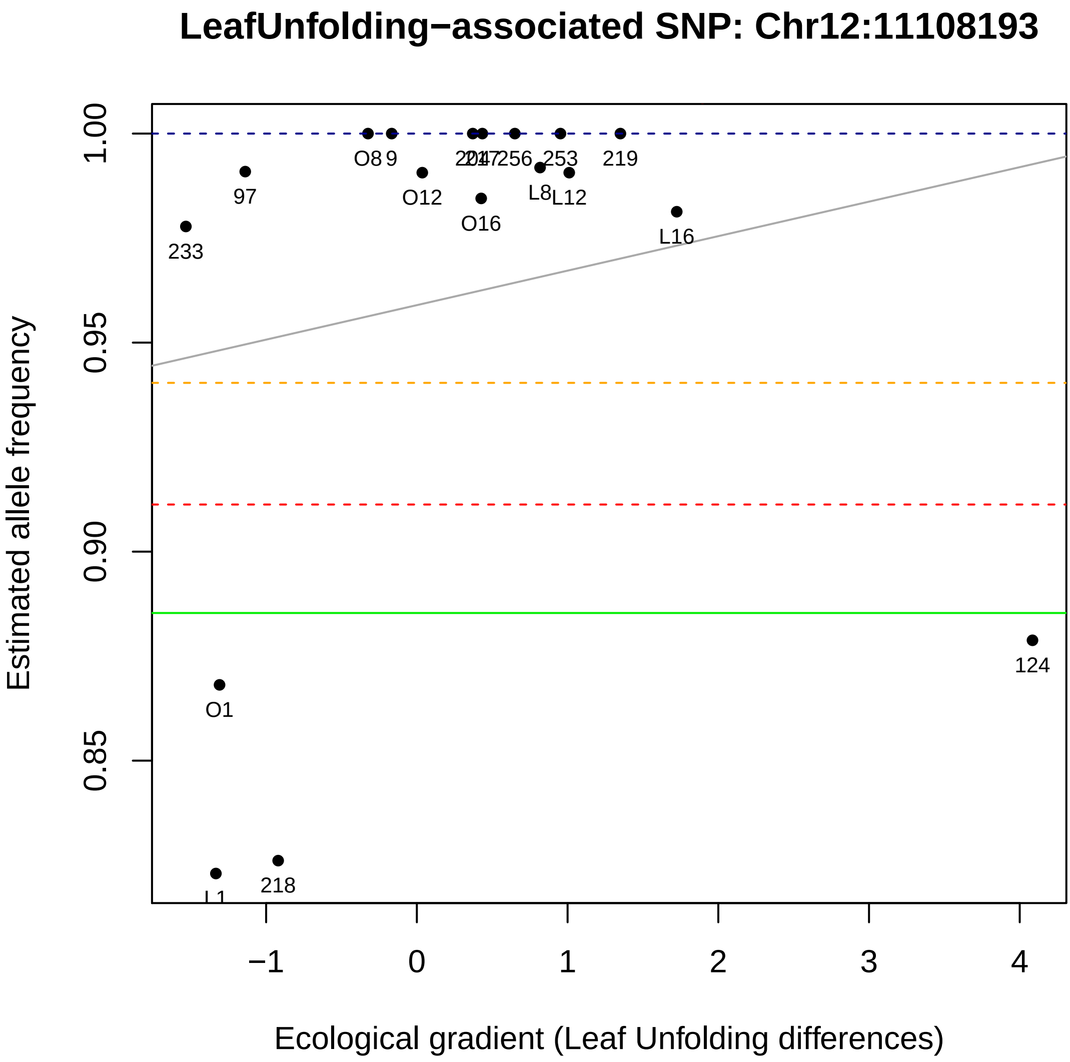

LeafUnfolding−associated SNP: Chr12:11108193
Estimated allele frequency
Ecological gradient (Leaf Unfolding differences)
1.00
0.95
0.90
0.85
−1
0
1
2
3
4
O8
9
O12
O16
256
253
219
L8
L12
L16
97
233
124
O1
218

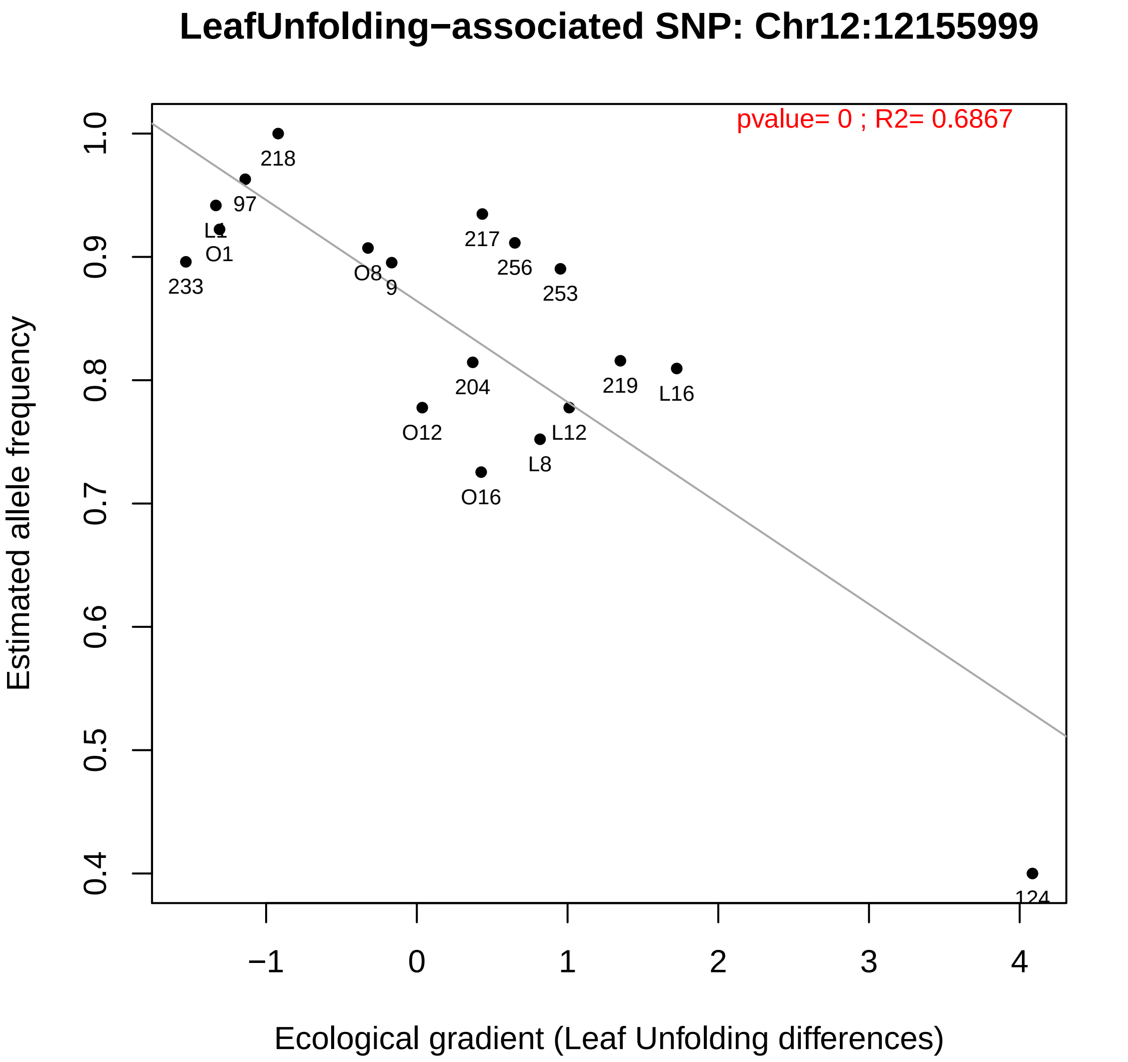
LeafUnfolding−associated SNP: Chr12:12155999
pvalue= 0 ; R2= 0.6867
Estimated allele frequency
Ecological gradient (Leaf Unfolding differences)
1.0
0.9
0.8
0.7
0.6
0.5
0.4
−1
0
1
2
3
4
218
97
L1
O1
233
O8
9
217
256
253
204
219
L16
O12
L12
L8
O16
124

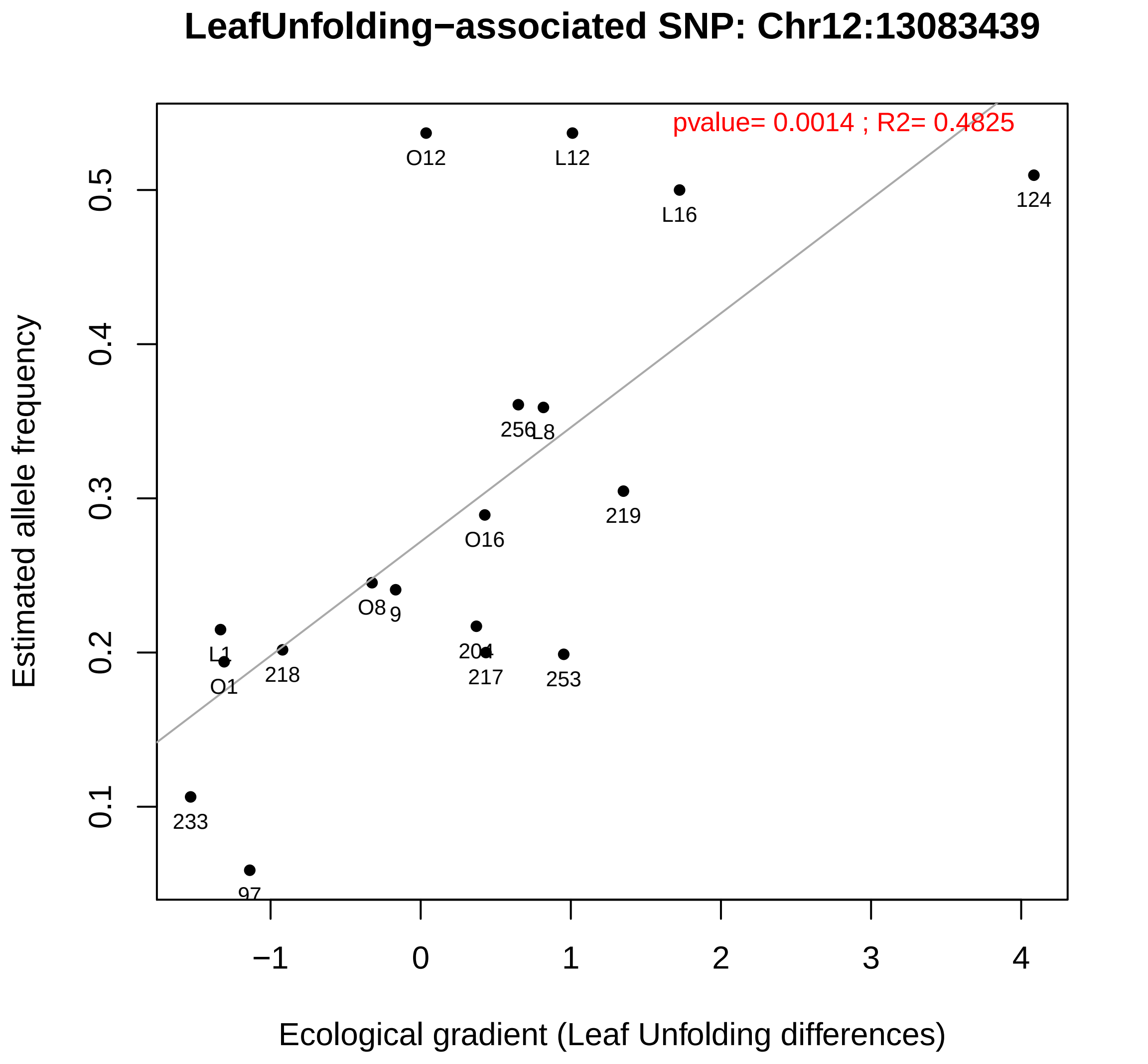

LeafUnfolding−associated SNP: Chr12:13083439
pvalue= 0.0014 ; R2= 0.4825
Estimated allele frequency
0.1
0.2
0.3
0.4
0.5
−1
0
1
2
3
4
Ecological gradient (Leaf Unfolding differences)
O12
L12
L16
124
256
L8
219
O16
O8
9
204
217
253
L1
O1
218
233
97

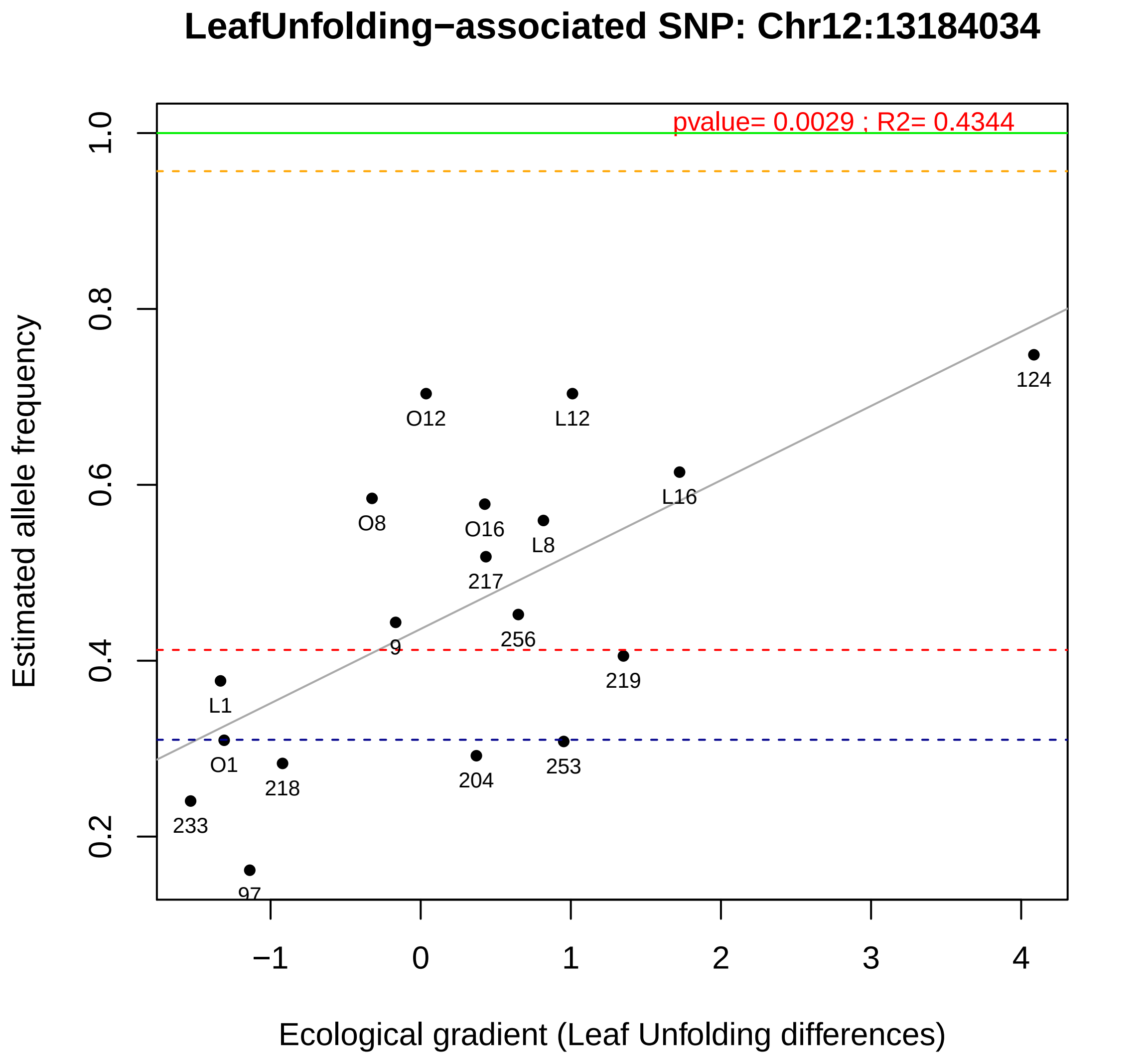
LeafUnfolding−associated SNP: Chr12:13184034
pvalue= 0.0029 ; R2= 0.4344
Estimated allele frequency
0.2
0.4
0.6
0.8
1.0
−1
0
1
2
3
4
Ecological gradient (Leaf Unfolding differences)
124
O12
L12
L16
O8
O16
L8
217
256
9
219
L1
O1
218
204
253
233
97

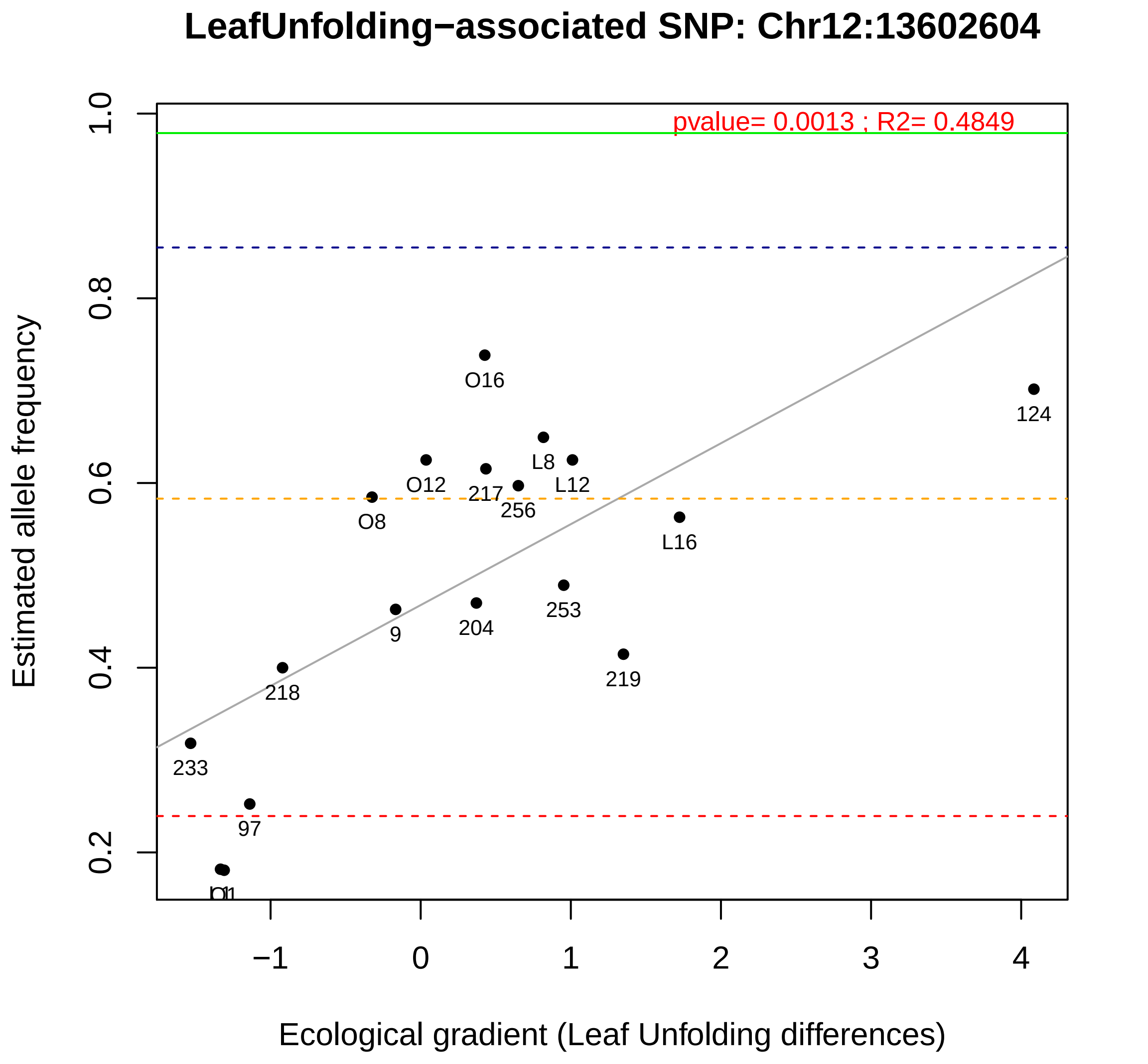
LeafUnfolding−associated SNP: Chr12:13602604
pvalue= 0.0013 ; R2= 0.4849
Estimated allele frequency
0.2
0.4
0.6
0.8
1.0
−1
0
1
2
3
4
Ecological gradient (Leaf Unfolding differences)
O16
124
L8
O12
L12
217
256
O8
L16
253
9
204
219
218
233
97

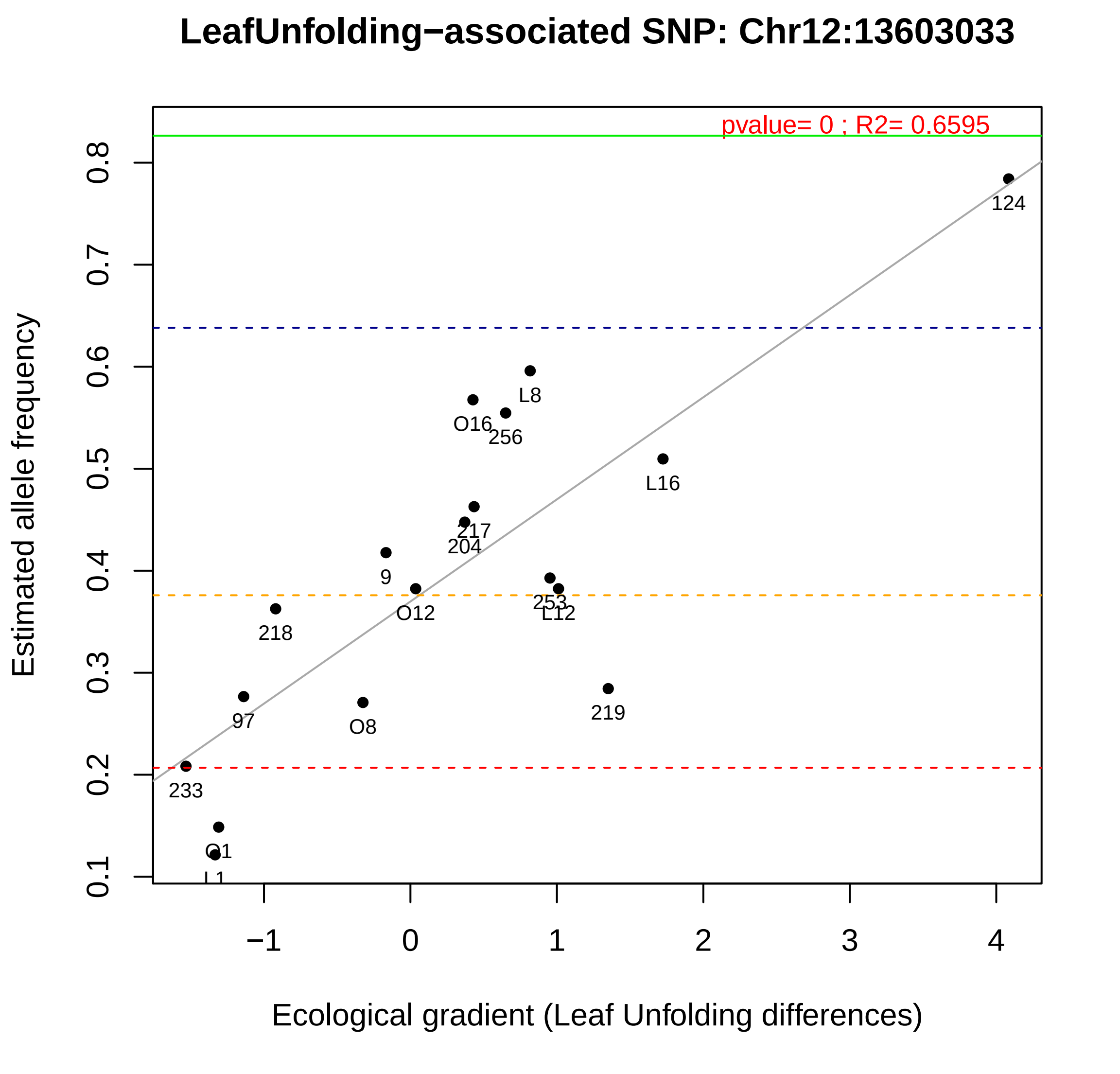

LeafUnfolding−associated SNP: Chr12:13603033
pvalue= 0 ; R2= 0.6595
Estimated allele frequency
Ecological gradient (Leaf Unfolding differences)
0.1
0.2
0.3
0.4
0.5
0.6
0.7
0.8
−1
0
1
2
3
4
124
L8
O16
256
L16
217
204
9
O12
253
L12
218
97
O8
219
233
O1
L1

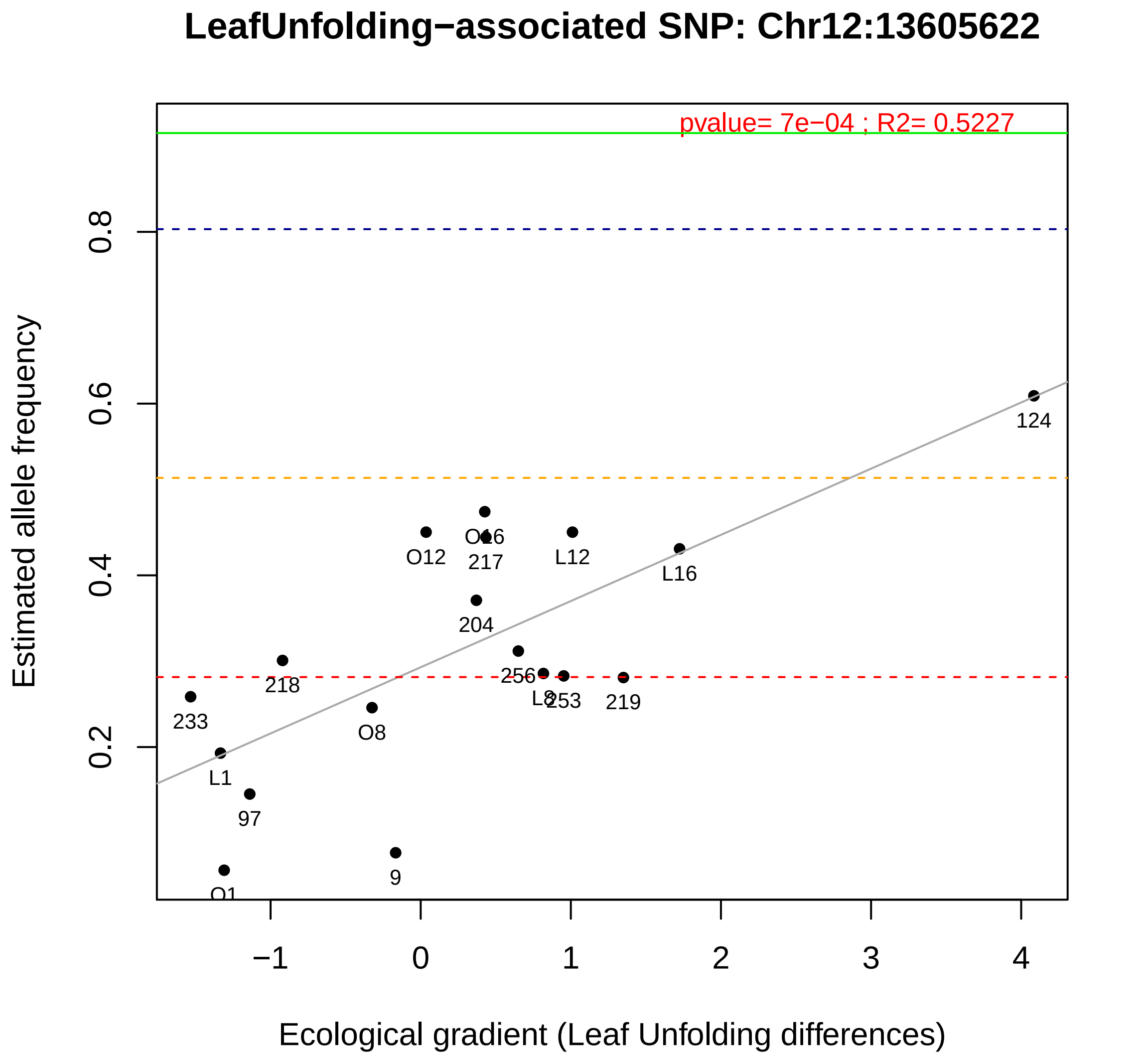

LeafUnfolding−associated SNP: Chr12:13605622
pvalue= 7e−04 ; R2= 0.5227
Estimated allele frequency
0.2
0.4
0.6
0.8
−1
0
1
2
3
4
Ecological gradient (Leaf Unfolding differences)
124
O16
217
O12
L12
L16
204
256
L8
253
219
218
233
O8
L1
97
9
O1

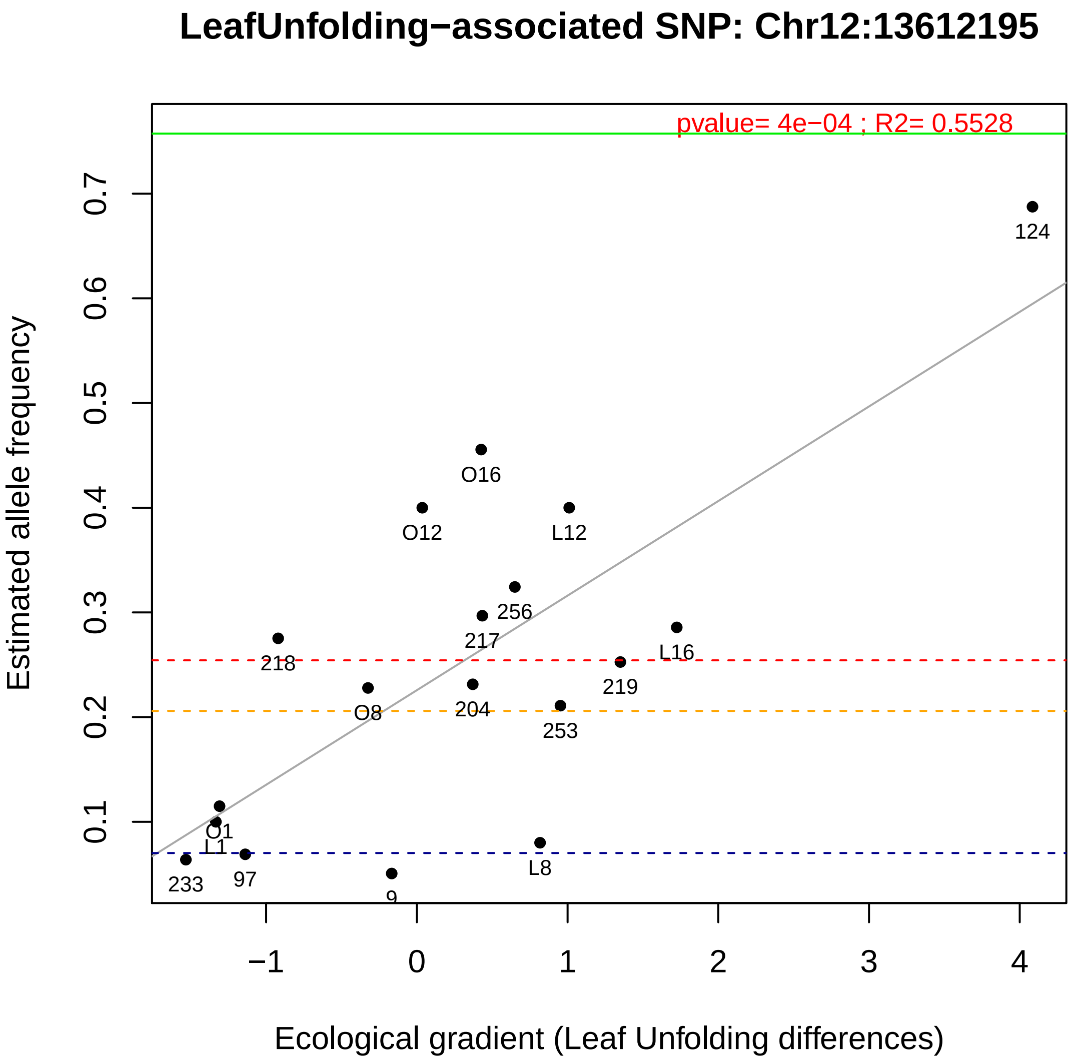

LeafUnfolding−associated SNP: Chr12:13612195
pvalue= 4e−04 ; R2= 0.5528
Estimated allele frequency
0.1
0.2
0.3
0.4
0.5
0.6
0.7
−1
0
1
2
3
4
Ecological gradient (Leaf Unfolding differences)
124
O16
O12
L12
256
217
218
L16
219
O8
204
253
O1
L1
233
97
9
L8

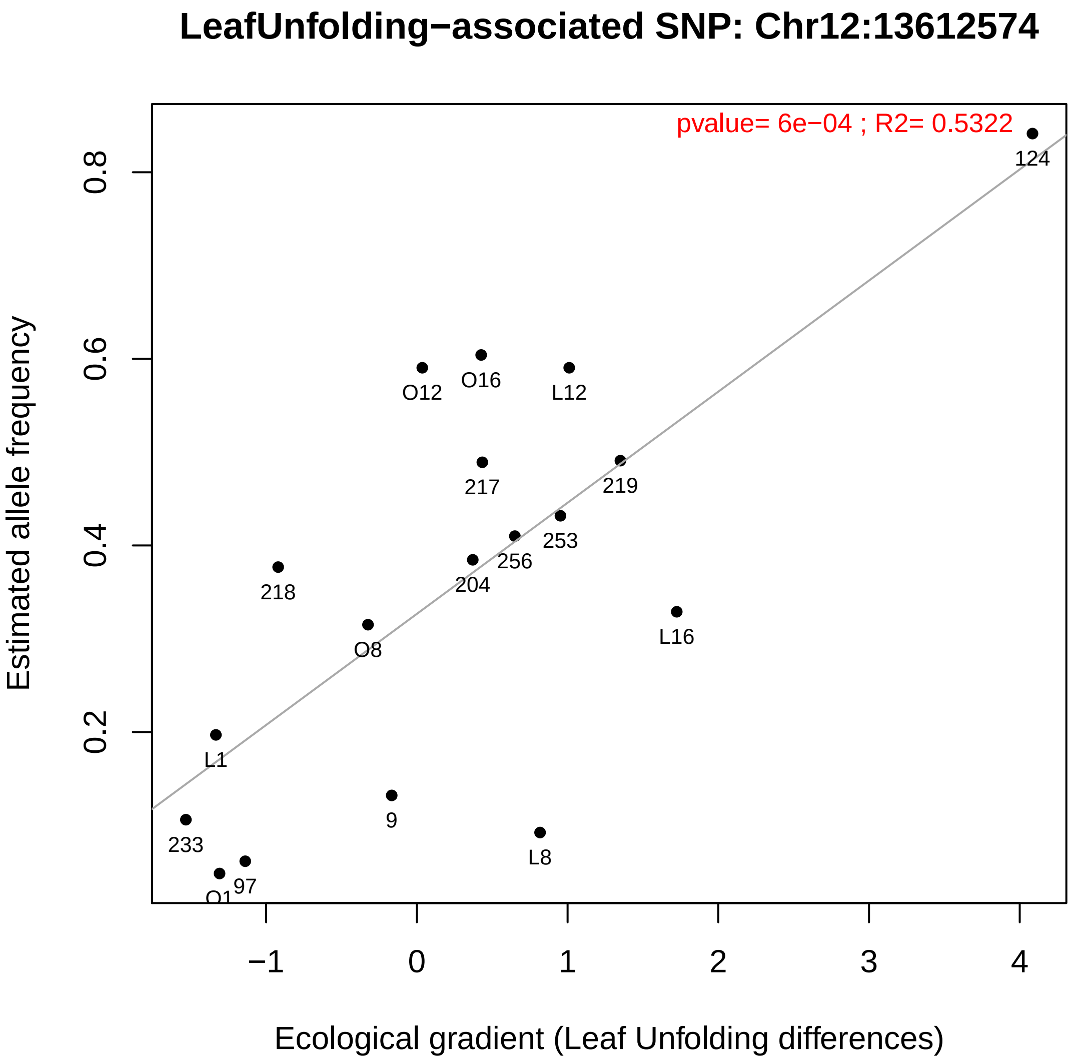

LeafUnfolding−associated SNP: Chr12:13612574
pvalue= 6e−04 ; R2= 0.5322
Estimated allele frequency
0.2
0.4
0.6
0.8
−1
0
1
2
3
4
Ecological gradient (Leaf Unfolding differences)
124
O12
O16
L12
217
219
253
256
204
218
O8
L16
L1
9
233
L8
97

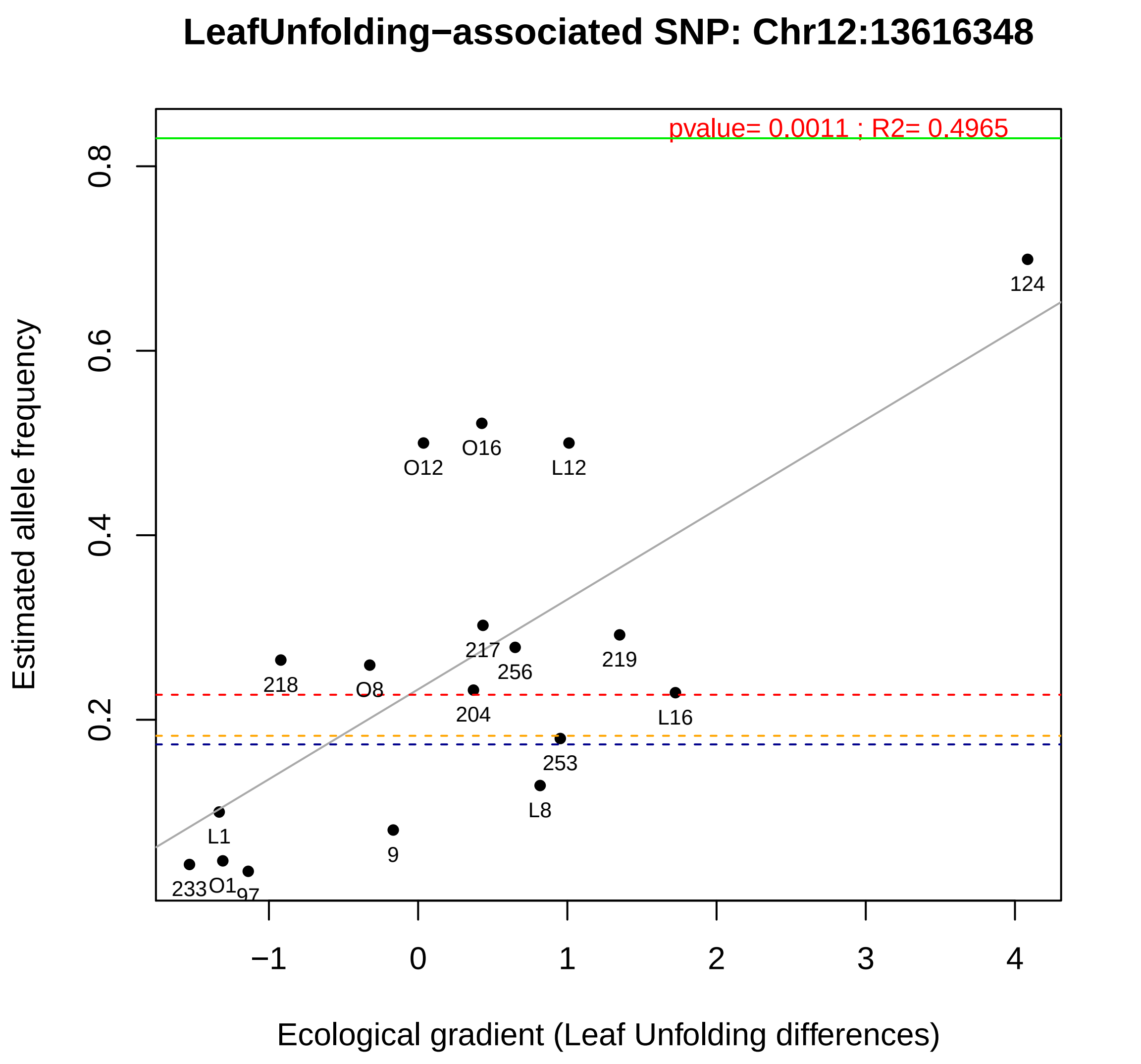

LeafUnfolding−associated SNP: Chr12:13616348
pvalue= 0.0011 ; R2= 0.4965
Estimated allele frequency
0.2
0.4
0.6
0.8
124
O16
O12
L12
217
256
219
218
O8
204
L16
253
L8
L1
9
233
O1
97
−1
0
1
2
3
4
Ecological gradient (Leaf Unfolding differences)

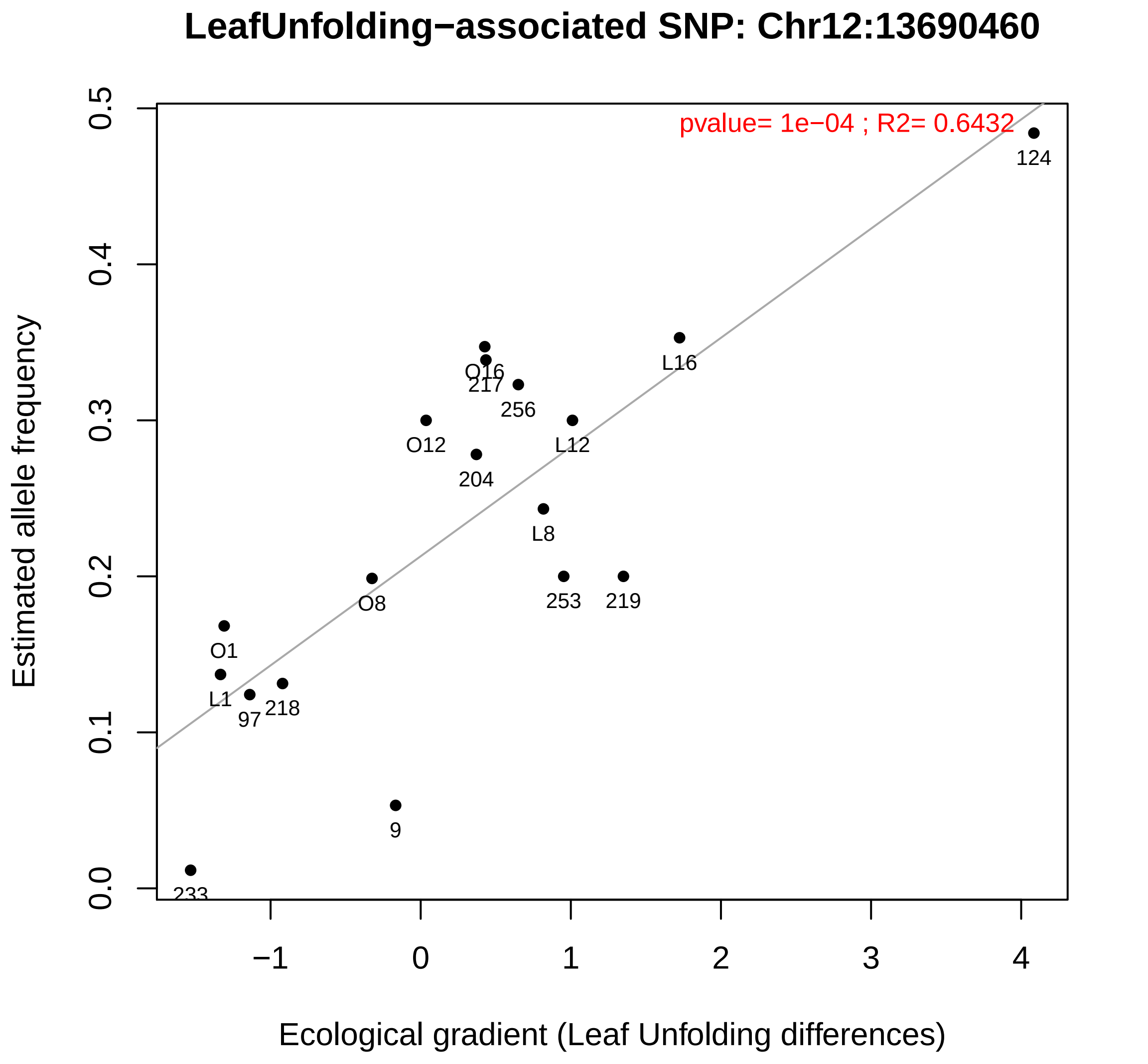
LeafUnfolding−associated SNP: Chr12:13690460
pvalue= 1e−04 ; R2= 0.6432
Estimated allele frequency
Ecological gradient (Leaf Unfolding differences)
0.0
0.1
0.2
0.3
0.4
0.5
−1
0
1
2
3
4
124
L16
O16
217
256
O12
L12
204
L8
O8
253
219
O1
L1
97
218
9
233

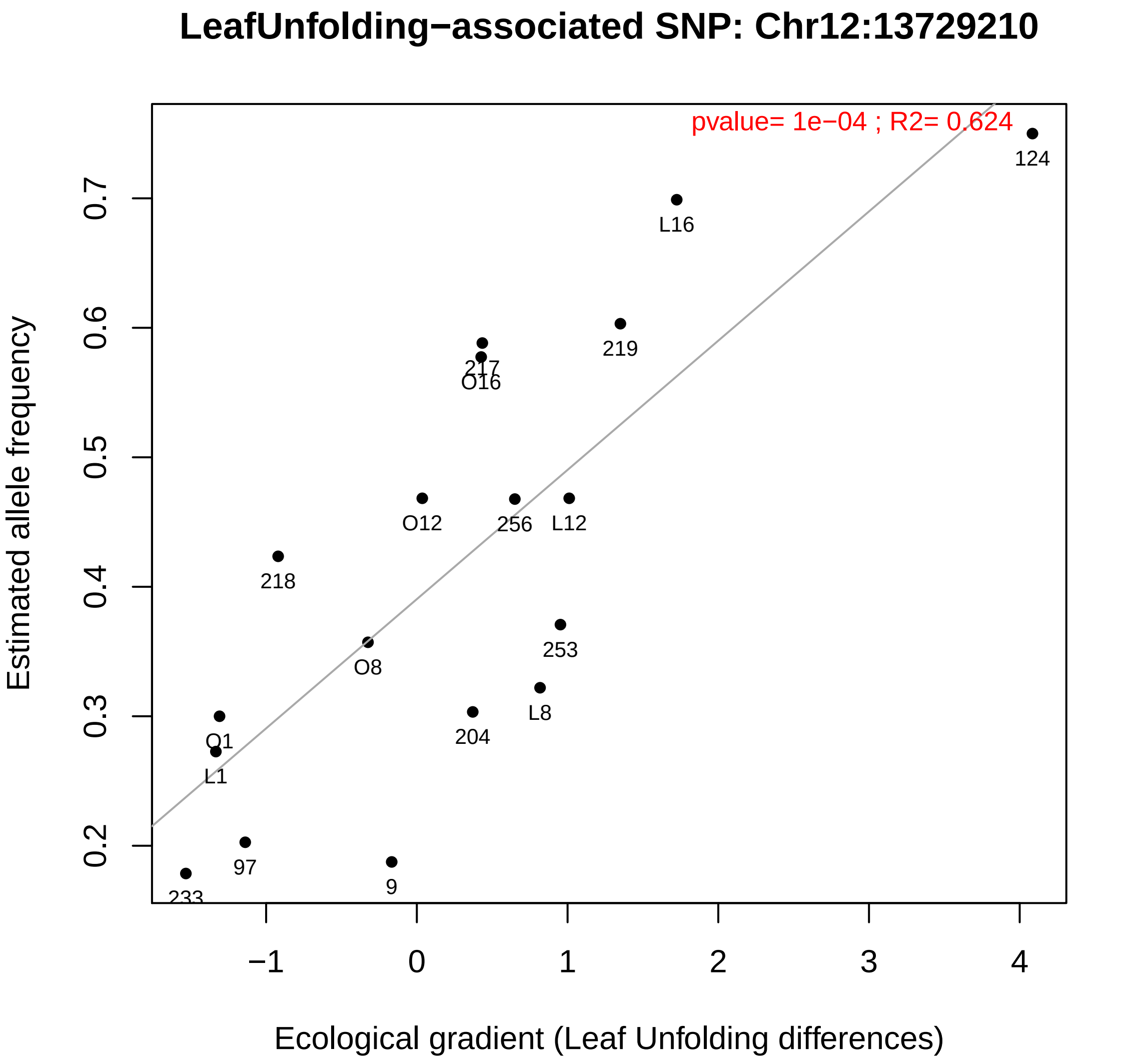

LeafUnfolding−associated SNP: Chr12:13729210
pvalue= 1e−04 ; R2= 0.624
124
L16
219
217
O16
O12
256
L12
218
O8
253
L8
204
O1
L1
97
9
233
0.2
0.3
0.4
0.5
0.6
0.7
−1
0
1
2
3
4
Estimated allele frequency
Ecological gradient (Leaf Unfolding differences)

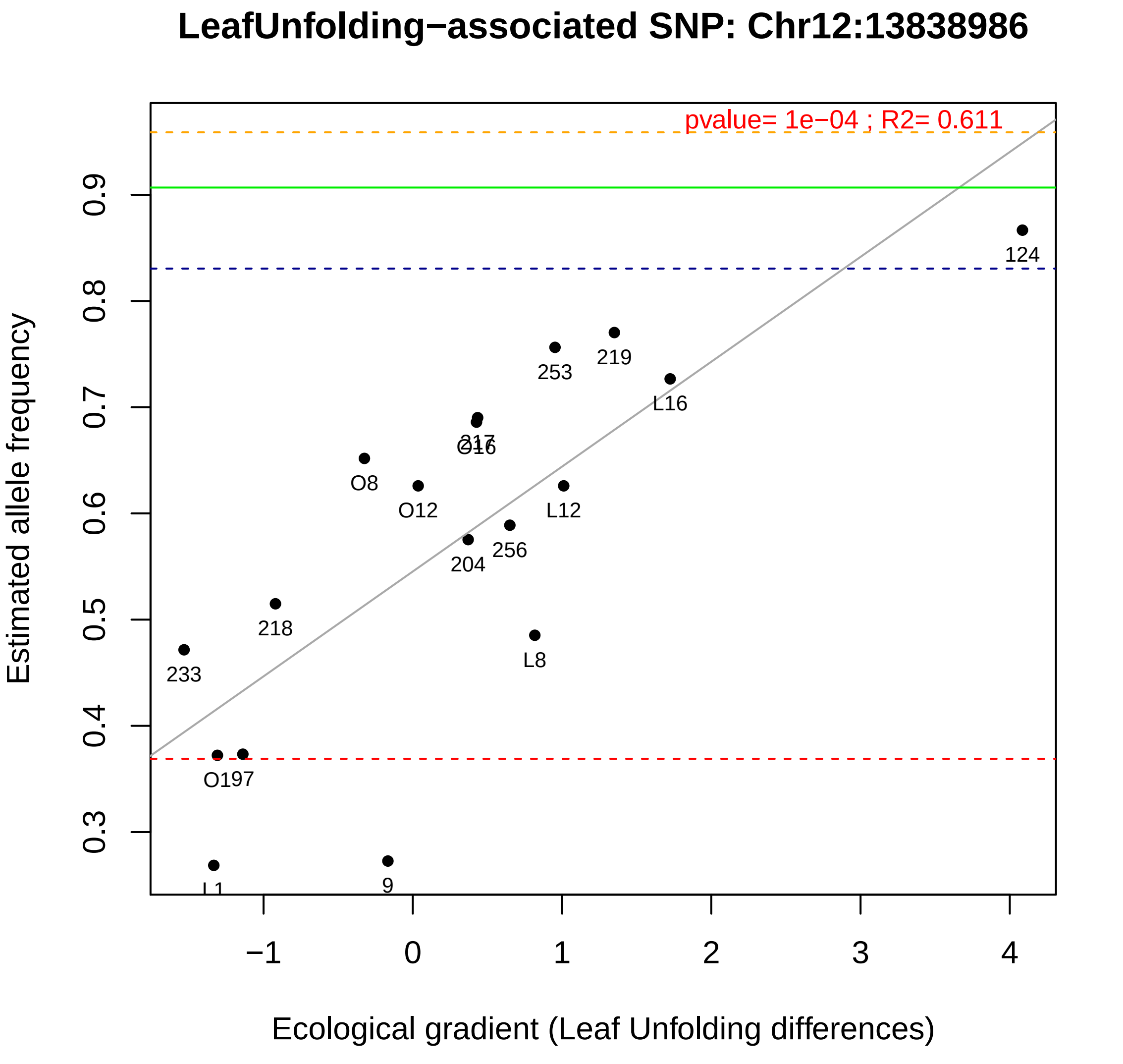

LeafUnfolding−associated SNP: Chr12:13838986
pvalue= 1e−04 ; R2= 0.611
Estimated allele frequency
Ecological gradient (Leaf Unfolding differences)
0.3
0.4
0.5
0.6
0.7
0.8
0.9
−1
0
1
2
3
4
124
219
253
L16
217
O16
O8
O12
L12
256
204
218
233
L8
O197
9

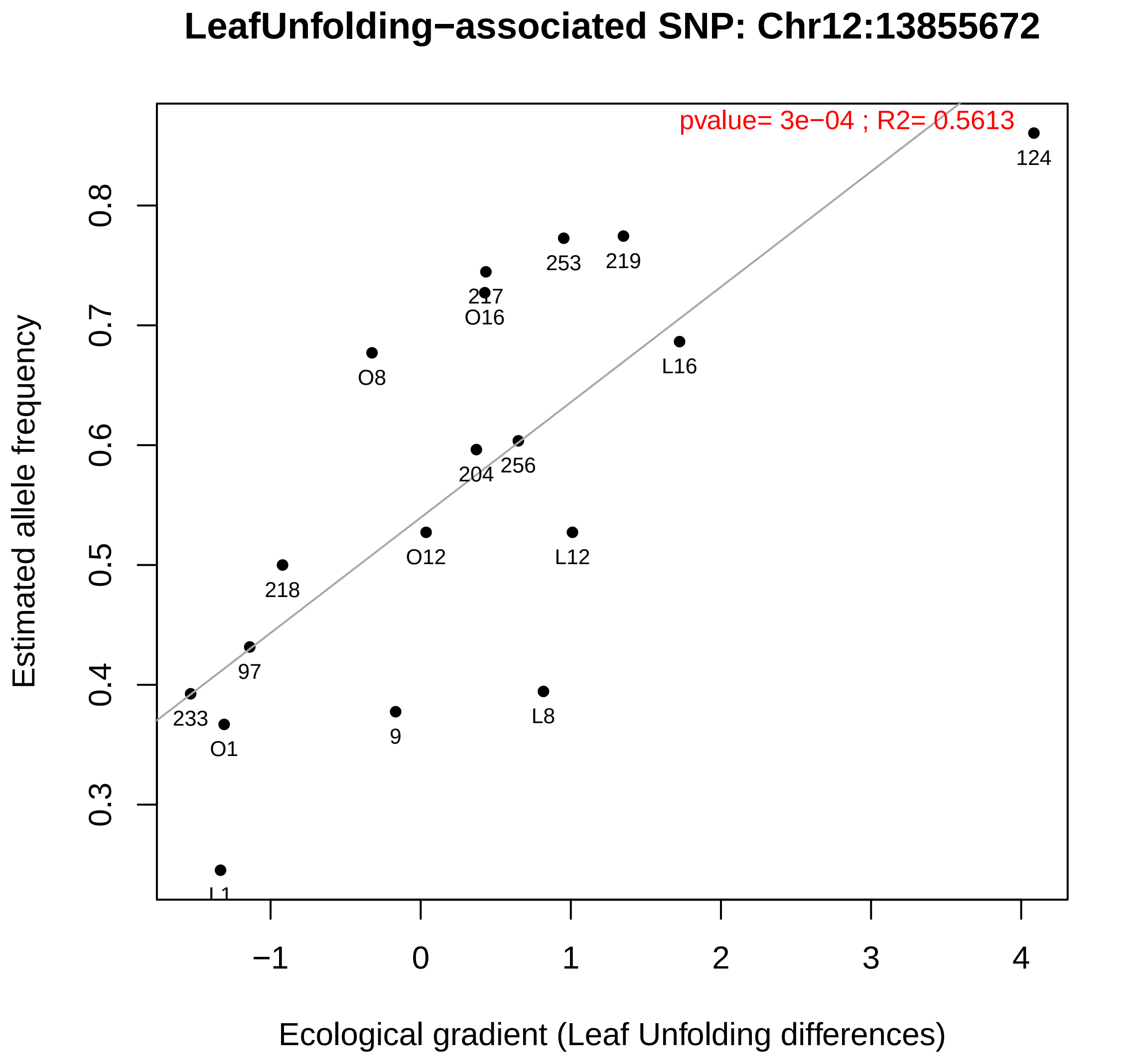
LeafUnfolding−associated SNP: Chr12:13855672
pvalue= 3e−04 ; R2= 0.5613
Estimated allele frequency
Ecological gradient (Leaf Unfolding differences)
0.3
0.4
0.5
0.6
0.7
0.8
−1
0
1
2
3
4
124
253
219
217
O16
O8
L16
204
256
O12
L12
218
97
233
O1
9
L8
L1

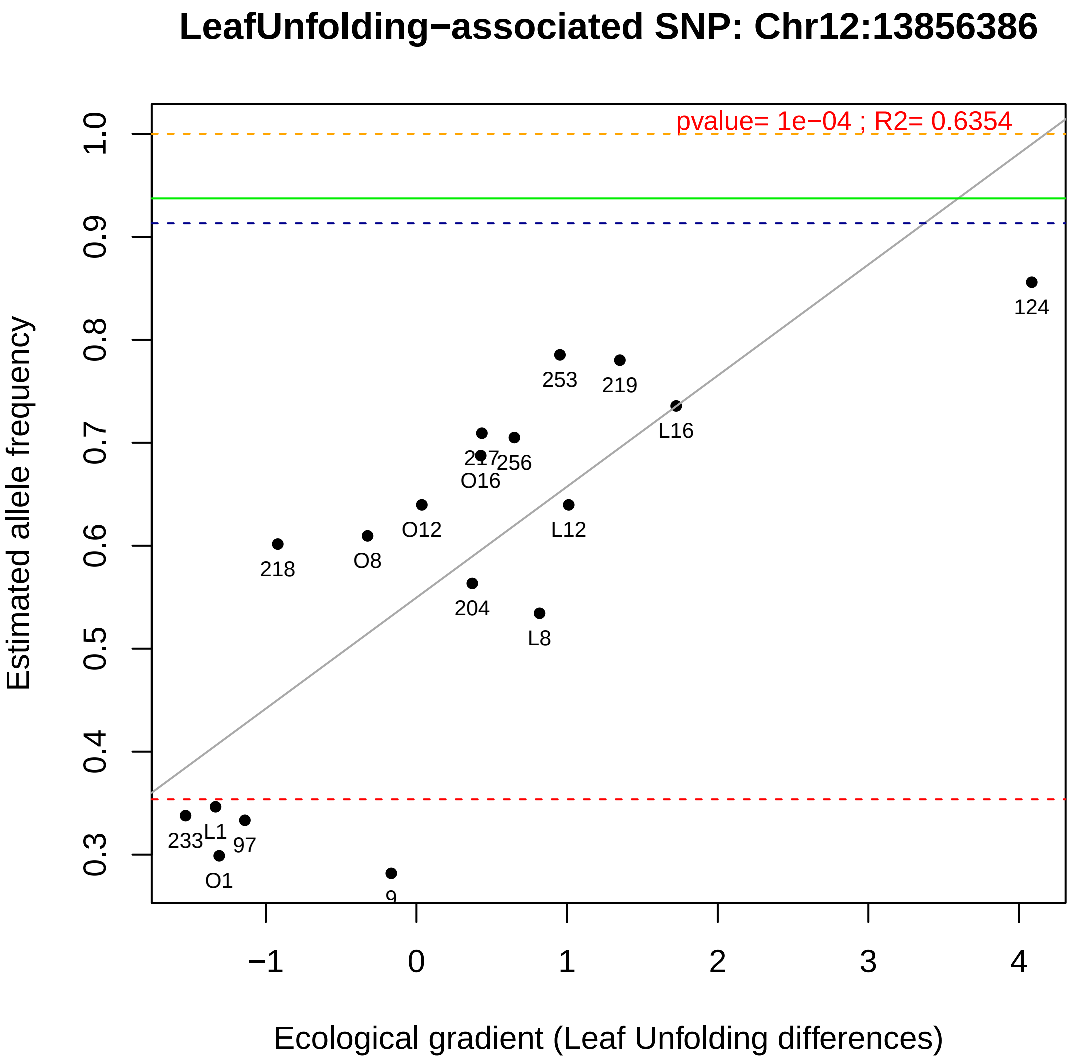
LeafUnfolding−associated SNP: Chr12:13856386
pvalue= 1e−04 ; R2= 0.6354
Estimated allele frequency
0.3
0.4
0.5
0.6
0.7
0.8
0.9
1.0
−1
0
1
2
3
4
Ecological gradient (Leaf Unfolding differences)
124
253
219
L16
217
256
O16
O12
L12
218
O8
204
L8
233
L1
97
O1
9

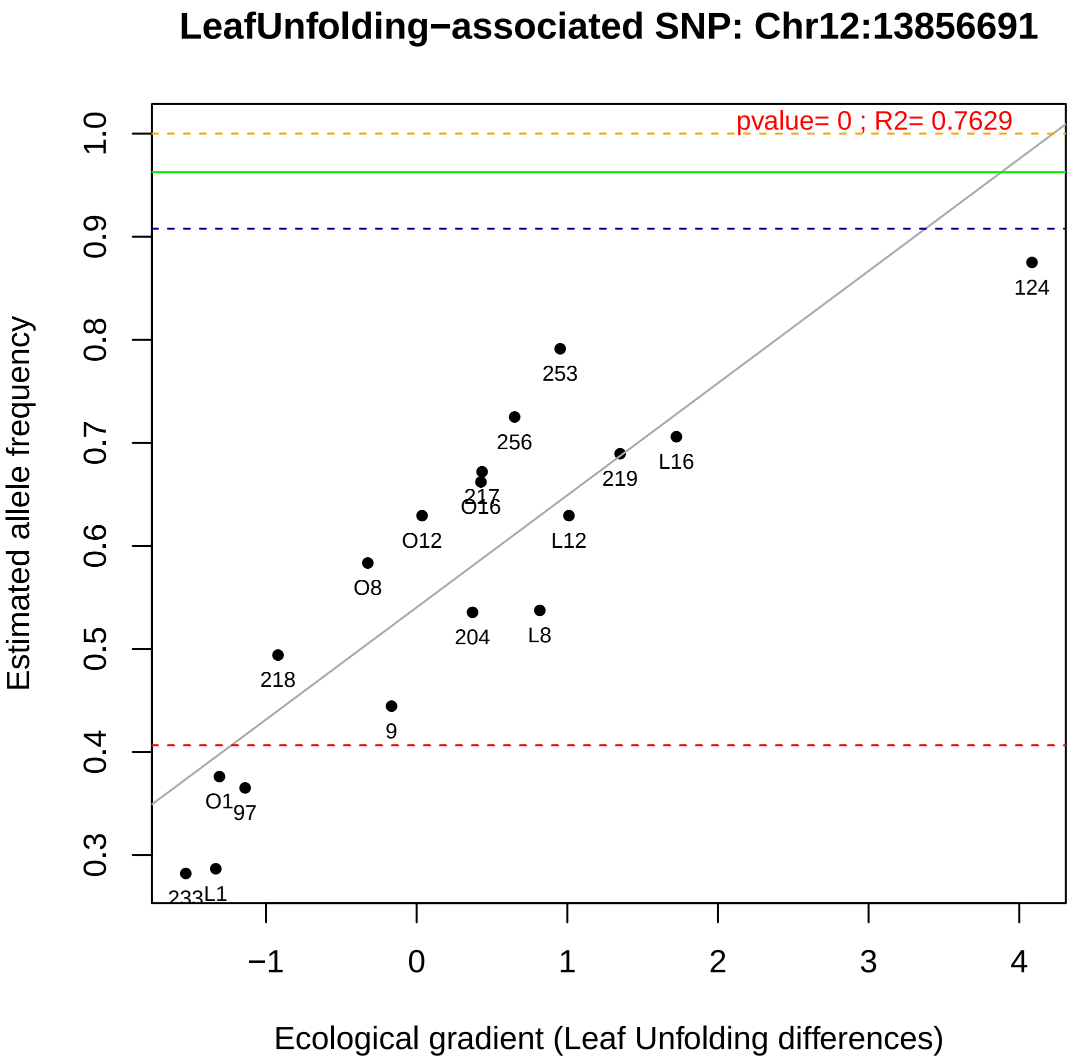
LeafUnfolding−associated SNP: Chr12:13856691
pvalue= 0 ; R2= 0.7629
Estimated allele frequency
Ecological gradient (Leaf Unfolding differences)
0.3
0.4
0.5
0.6
0.7
0.8
0.9
1.0
−1
0
1
2
3
4
124
253
256
L16
219
217
O16
O12
L12
O8
204
L8
218
9
O1
97
233
L1

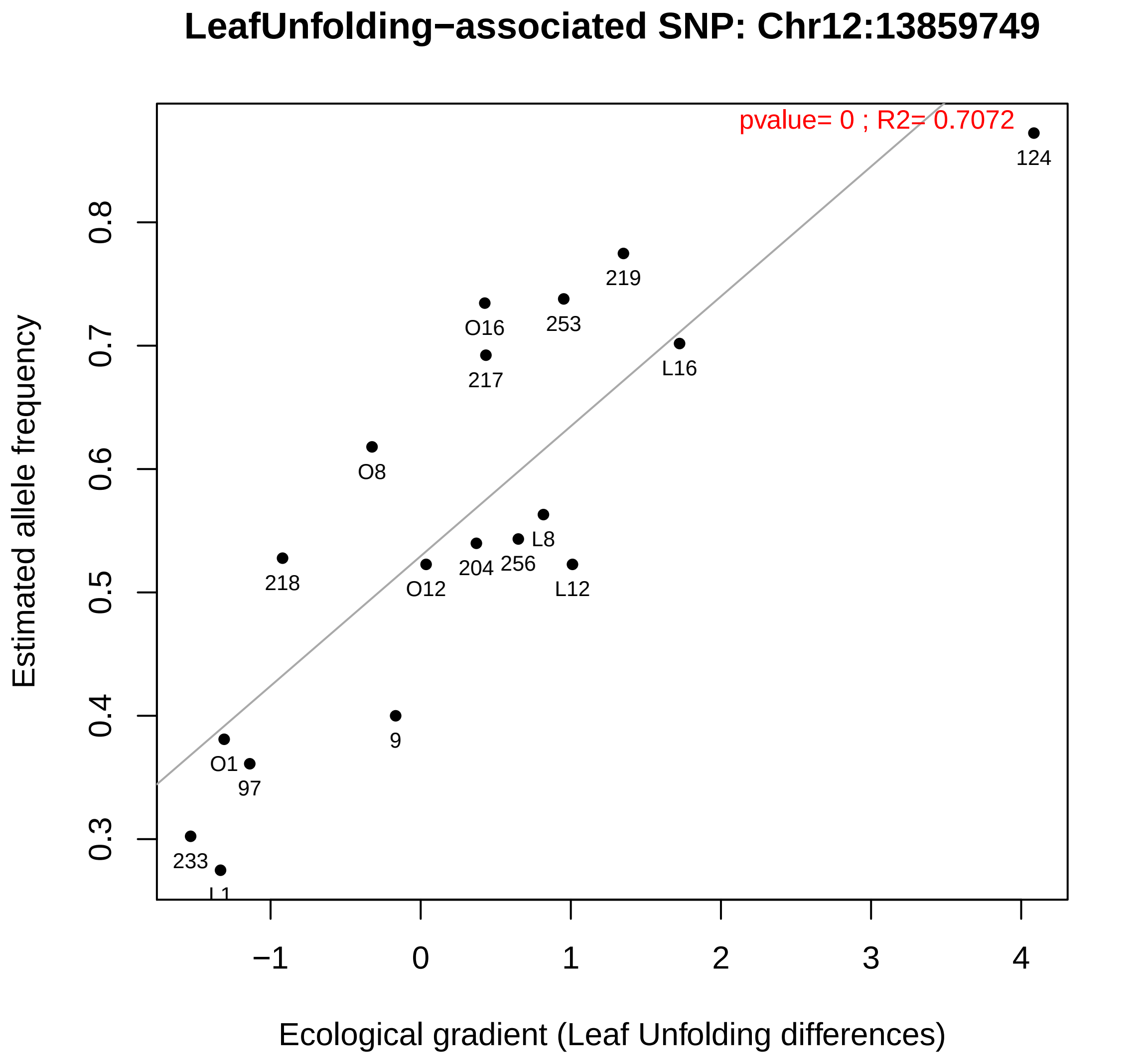

LeafUnfolding−associated SNP: Chr12:13859749
pvalue= 0 ; R2= 0.7072
124
219
O16
253
L16
217
O8
L8
218
204
256
O12
L12
9
O1
97
233
L1
0.3
0.4
0.5
0.6
0.7
0.8
Estimated allele frequency
−1
0
1
2
3
4
Ecological gradient (Leaf Unfolding differences)

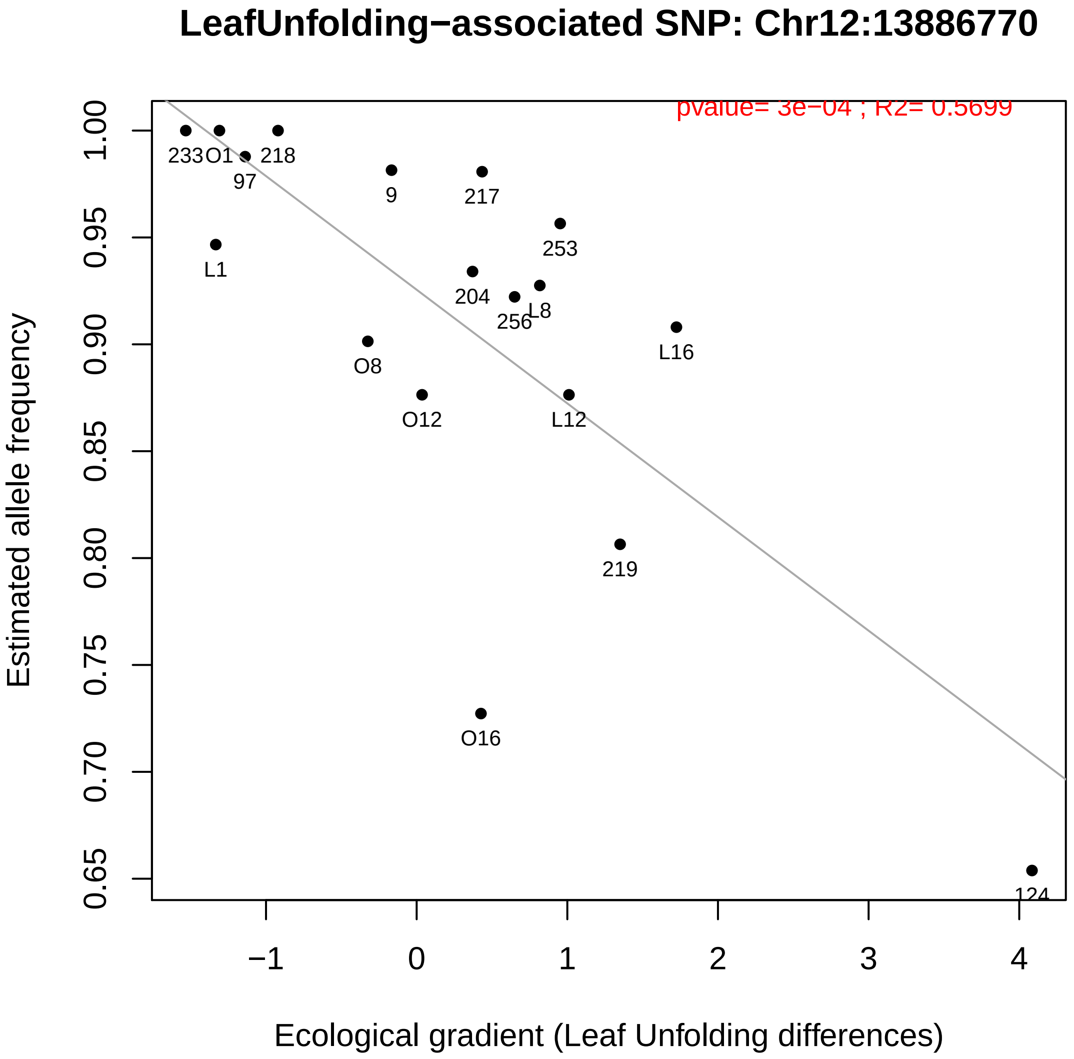
LeafUnfolding−associated SNP: Chr12:13886770
pvalue= 3e−04 ; R2= 0.5699
Estimated allele frequency
Ecological gradient (Leaf Unfolding differences)
0.65
0.70
0.75
0.80
0.85
0.90
0.95
1.00
−1
0
1
2
3
4
233
O1
97
218
9
217
253
L1
204
256
L8
L16
O8
O12
L12
219
O16
124

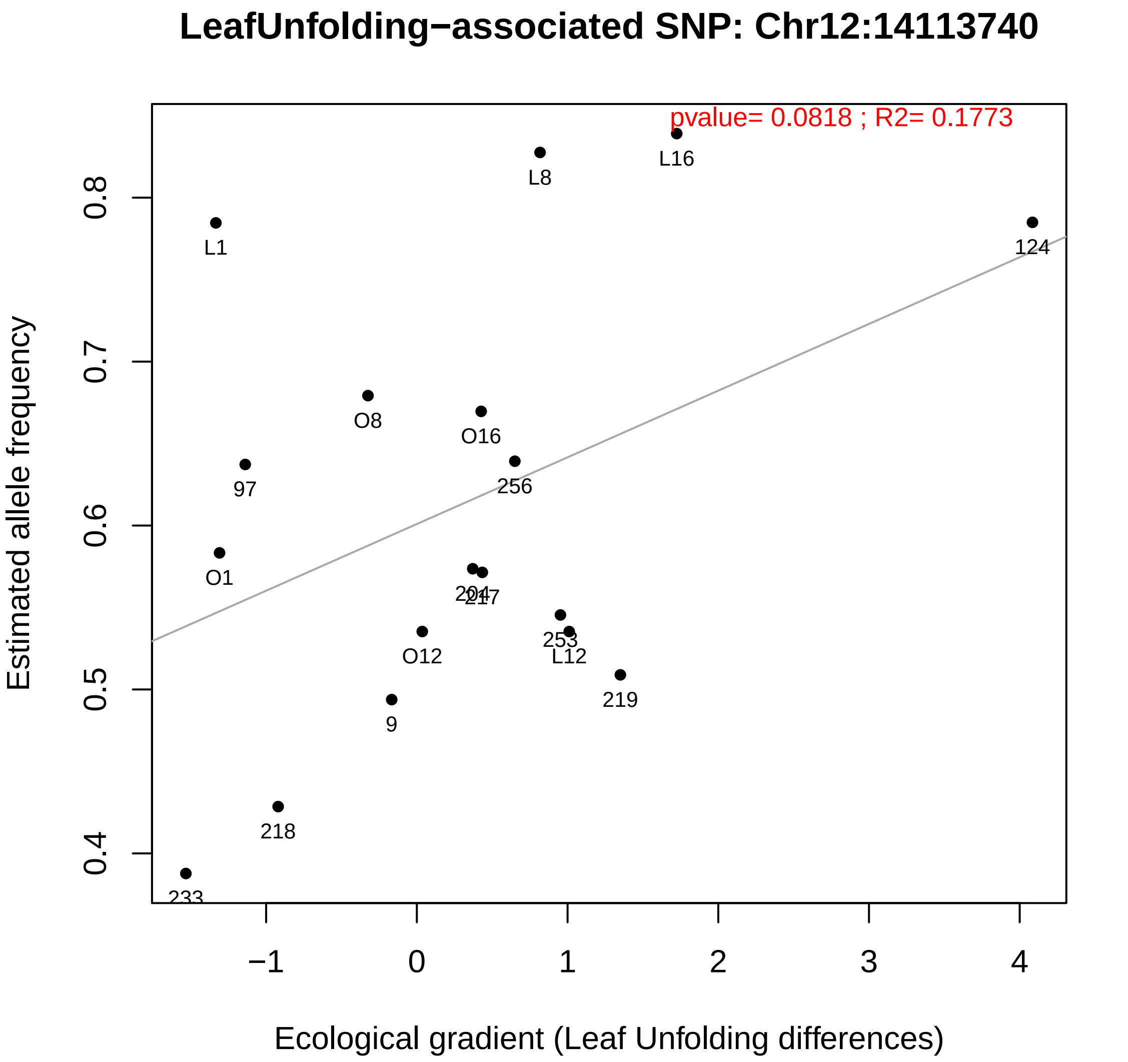

LeafUnfolding−associated SNP: Chr12:14113740
pvalue= 0.0818 ; R2= 0.1773
Estimated allele frequency
0.4
0.5
0.6
0.7
0.8
−1
0
1
2
3
4
Ecological gradient (Leaf Unfolding differences)
L16
L8
L1
124
O8
O16
97
256
O1
204
217
253
L12
O12
219
9
218
233

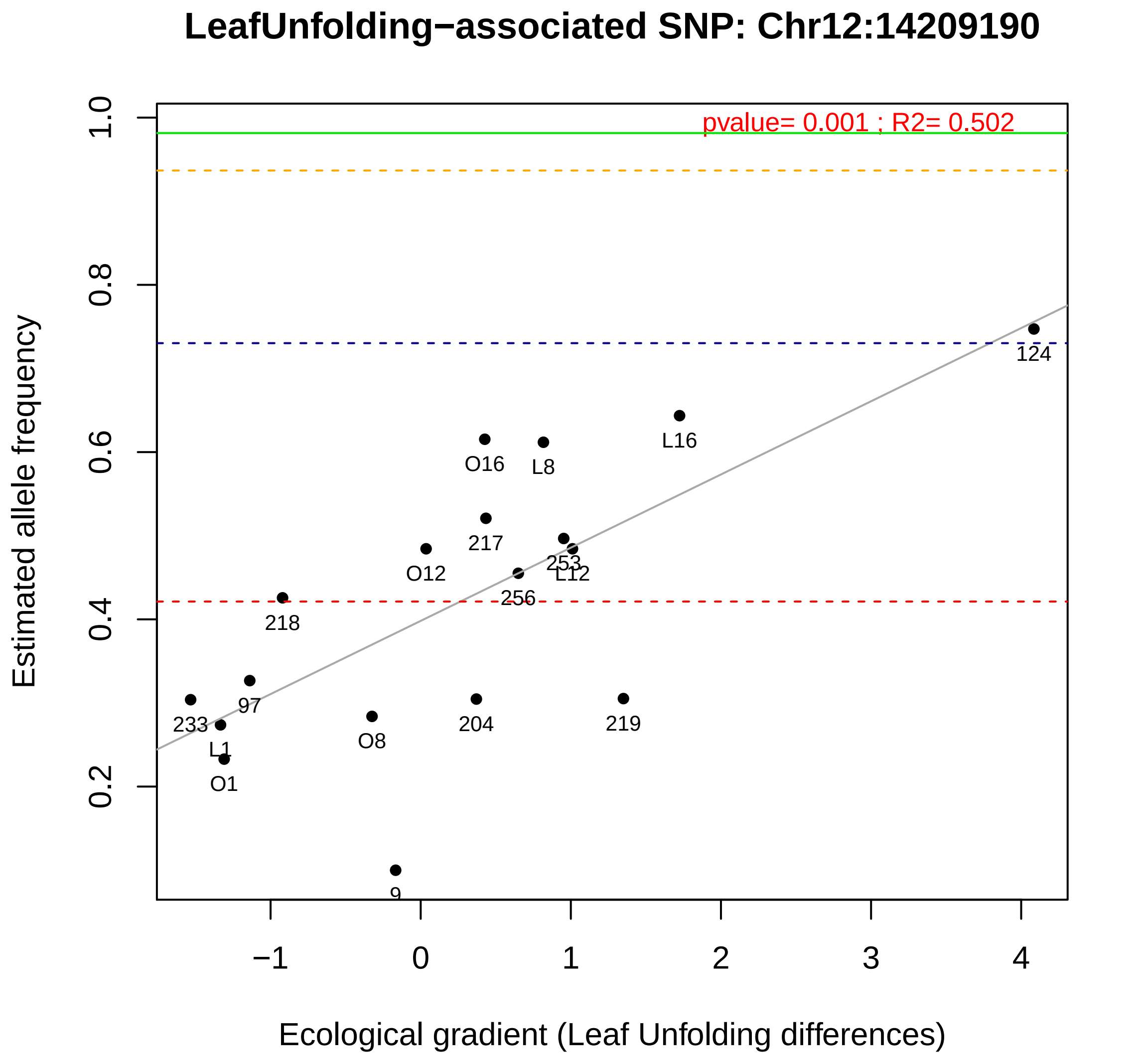

LeafUnfolding−associated SNP: Chr12:14209190
pvalue= 0.001 ; R2= 0.502
Estimated allele frequency
0.2
0.4
0.6
0.8
1.0
−1
0
1
2
3
4
Ecological gradient (Leaf Unfolding differences)
124
L16
O16
L8
217
253
L12
O12
256
218
97
233
L1
O1
O8
204
219
9

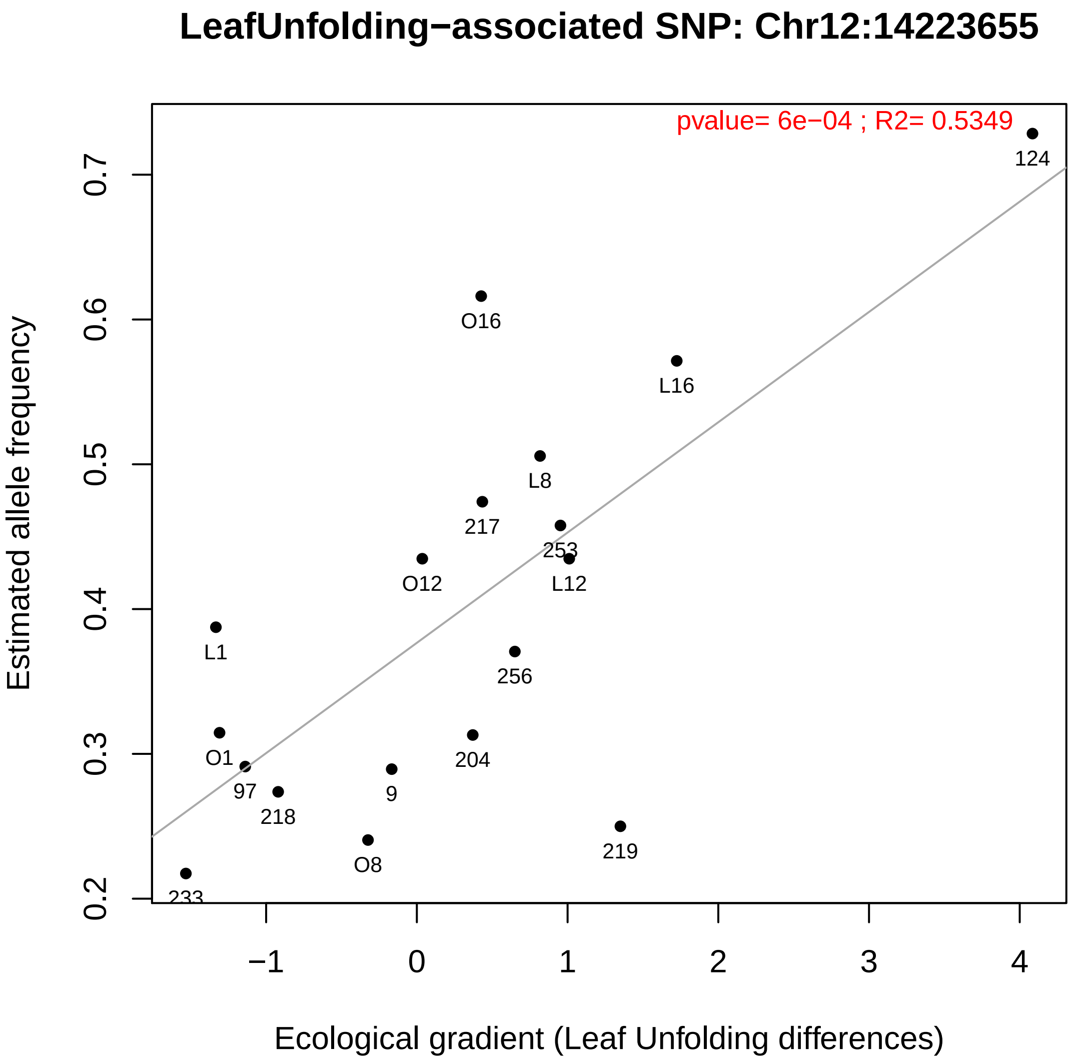

LeafUnfolding−associated SNP: Chr12:14223655
pvalue= 6e−04 ; R2= 0.5349
Estimated allele frequency
Ecological gradient (Leaf Unfolding differences)
0.2
0.3
0.4
0.5
0.6
0.7
−1
0
1
2
3
4
124
O16
L16
L8
217
253
L12
O12
L1
256
O1
97
218
9
204
O8
219
233

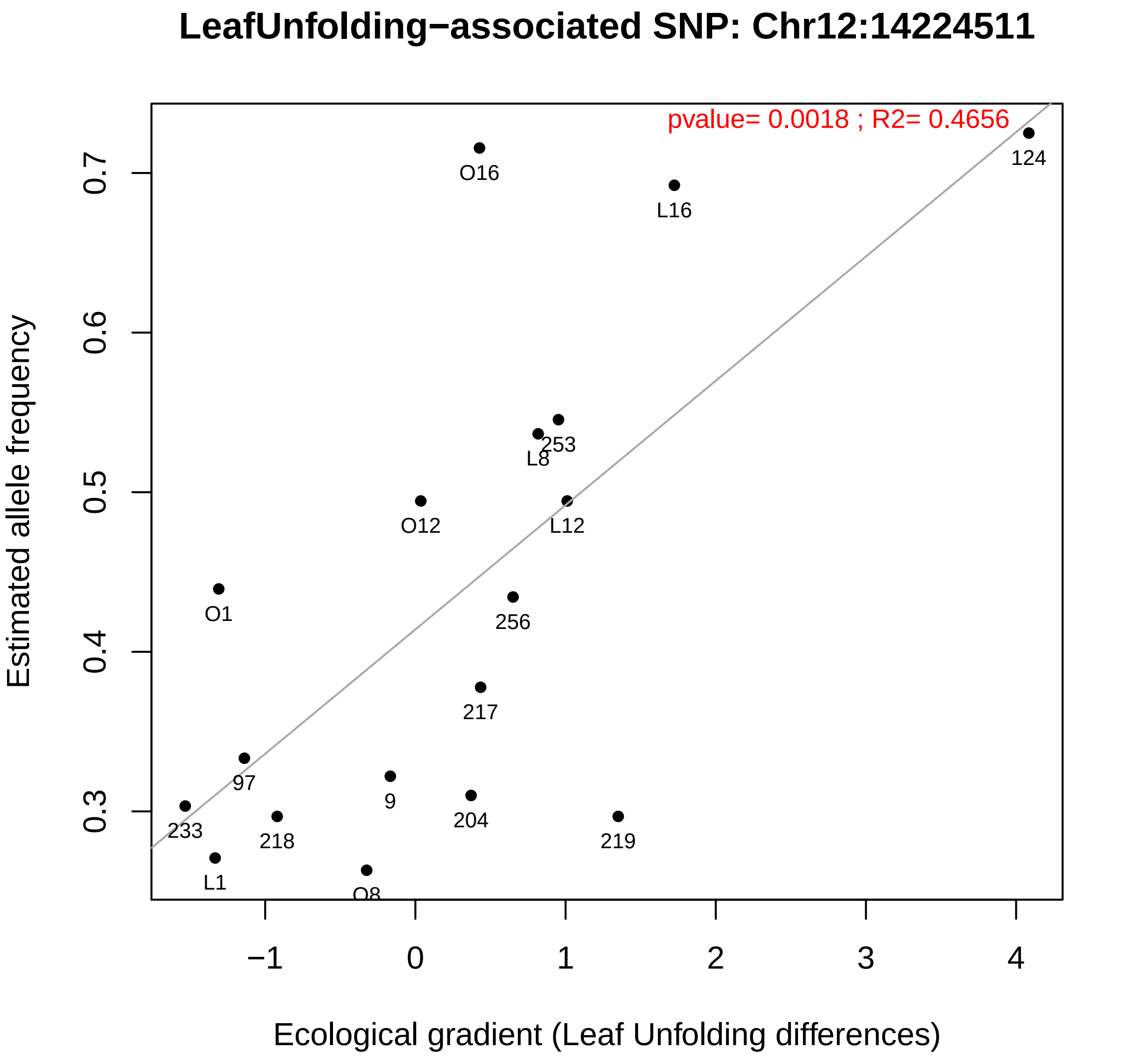
LeafUnfolding−associated SNP: Chr12:14224511
pvalue= 0.0018 ; R2= 0.4656
Estimated allele frequency
Ecological gradient (Leaf Unfolding differences)
0.3
0.4
0.5
0.6
0.7
−1
0
1
2
3
4
O16
124
L16
253
L8
O12
L12
O1
256
217
97
9
204
233
218
219
L1
O8

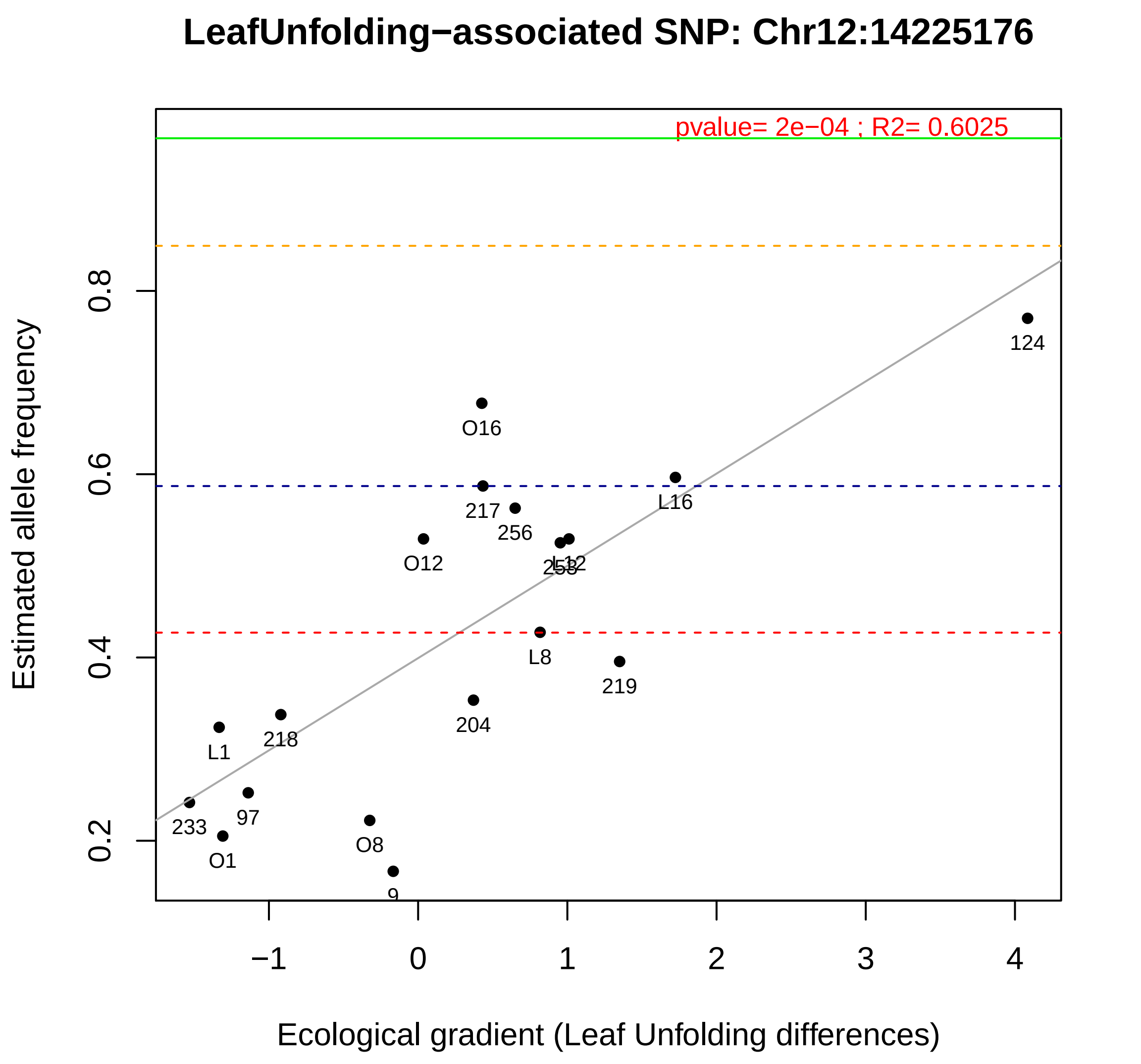

LeafUnfolding−associated SNP: Chr12:14225176
pvalue= 2e−04 ; R2= 0.6025
Estimated allele frequency
0.2
0.4
0.6
0.8
−1
0
1
2
3
4
Ecological gradient (Leaf Unfolding differences)
124
O16
217
L16
256
O12
L8
219
204
218
L1
97
233
O1
O8
9

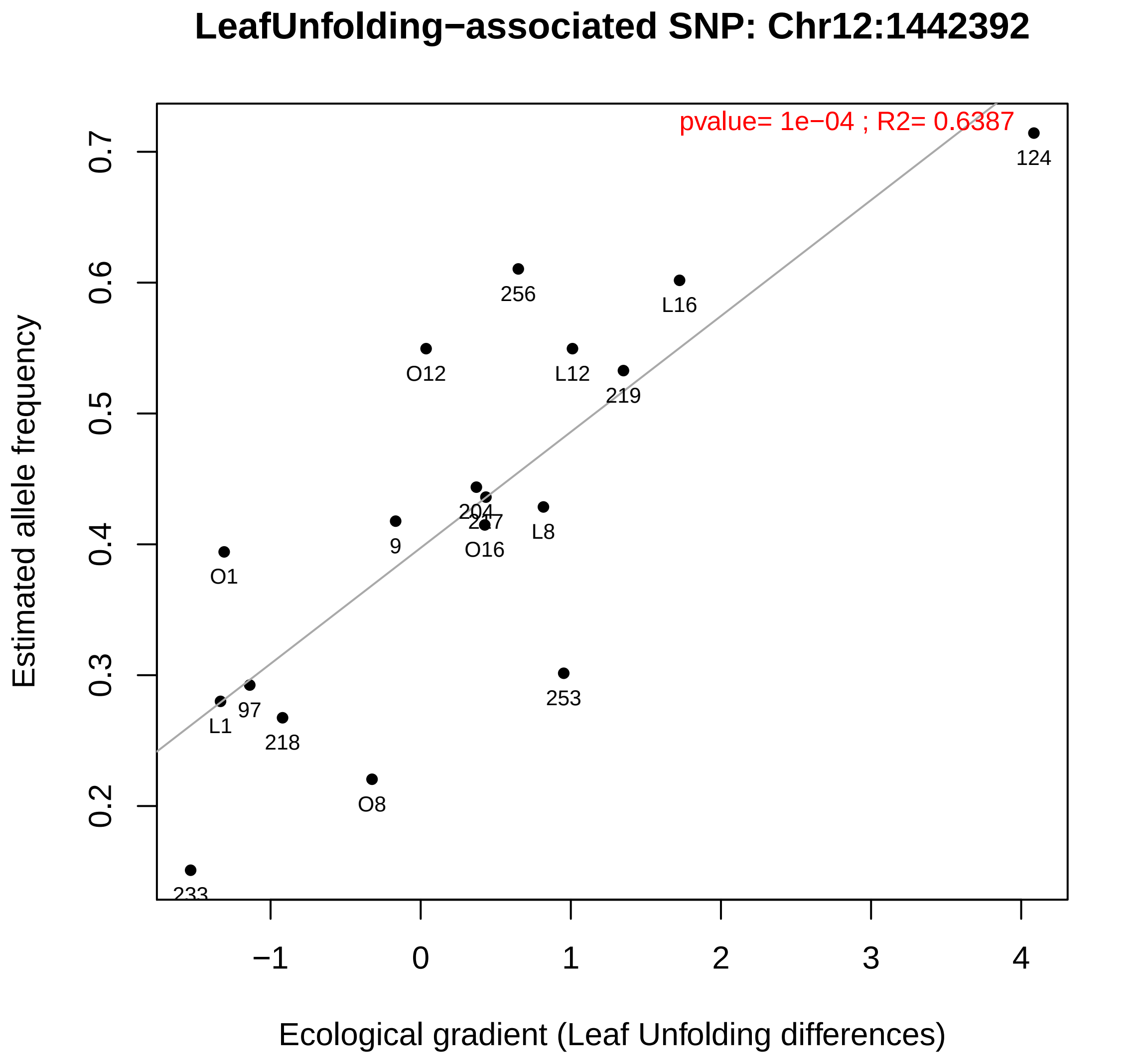
LeafUnfolding−associated SNP: Chr12:1442392
pvalue= 1e−04 ; R2= 0.6387
Estimated allele frequency
Ecological gradient (Leaf Unfolding differences)
0.2
0.3
0.4
0.5
0.6
0.7
−1
0
1
2
3
4
124
256
L16
O12
L12
219
204
217
O16
L8
9
O1
253
97
L1
218
O8
233

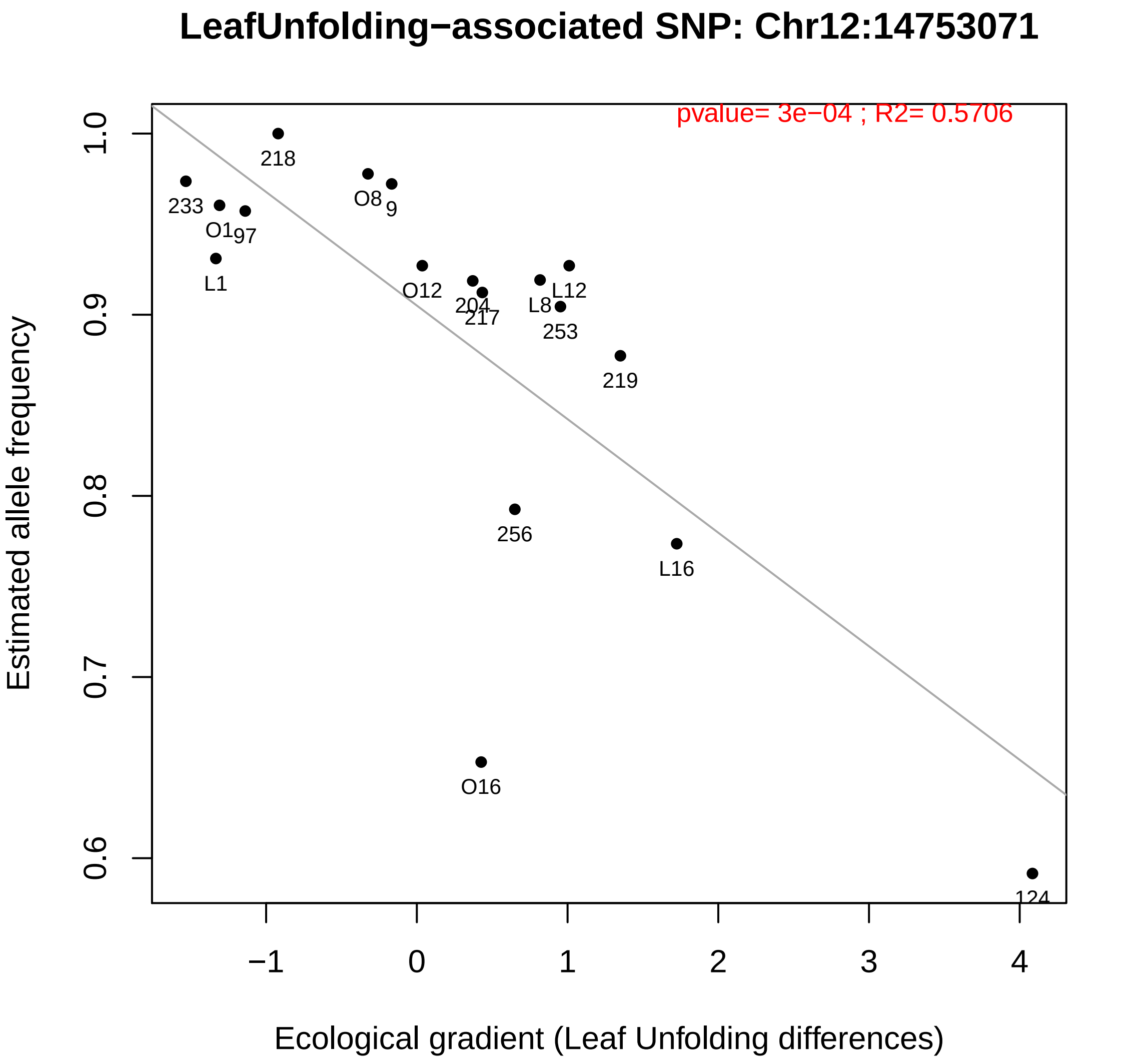
LeafUnfolding−associated SNP: Chr12:14753071
pvalue= 3e−04 ; R2= 0.5706
Estimated allele frequency
Ecological gradient (Leaf Unfolding differences)
0.6
0.7
0.8
0.9
1.0
−1
0
1
2
3
4
218
233
O1
97
L1
O8
9
O12
204
217
L8
L12
253
219
256
L16
O16
124

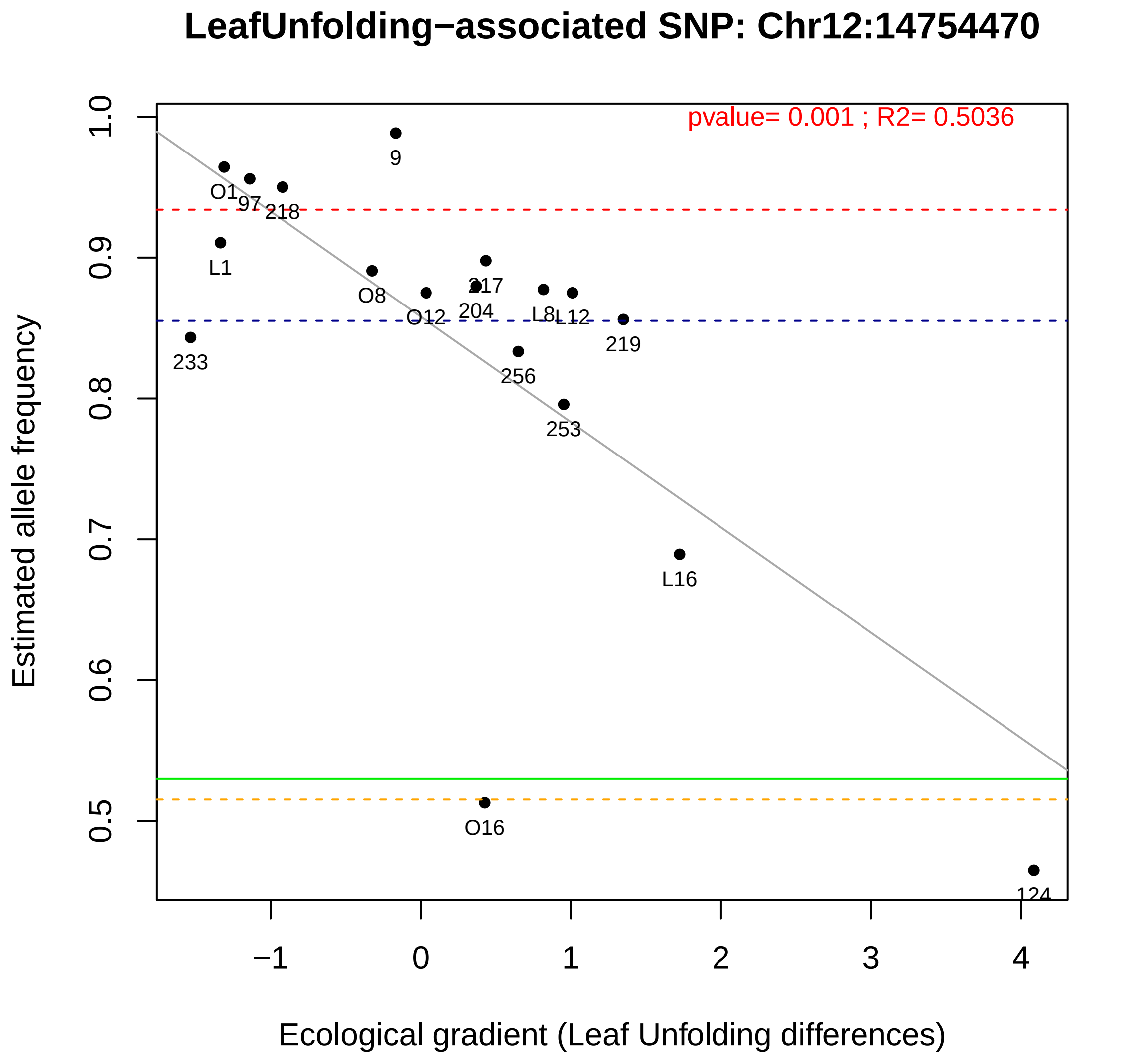
LeafUnfolding−associated SNP: Chr12:14754470
pvalue= 0.001 ; R2= 0.5036
Estimated allele frequency
Ecological gradient (Leaf Unfolding differences)
1.0
0.9
0.8
0.7
0.6
0.5
−1
0
1
2
3
4
9
O1
97
218
L1
O8
O12
217
204
L8
L12
219
233
256
253
L16
O16
124

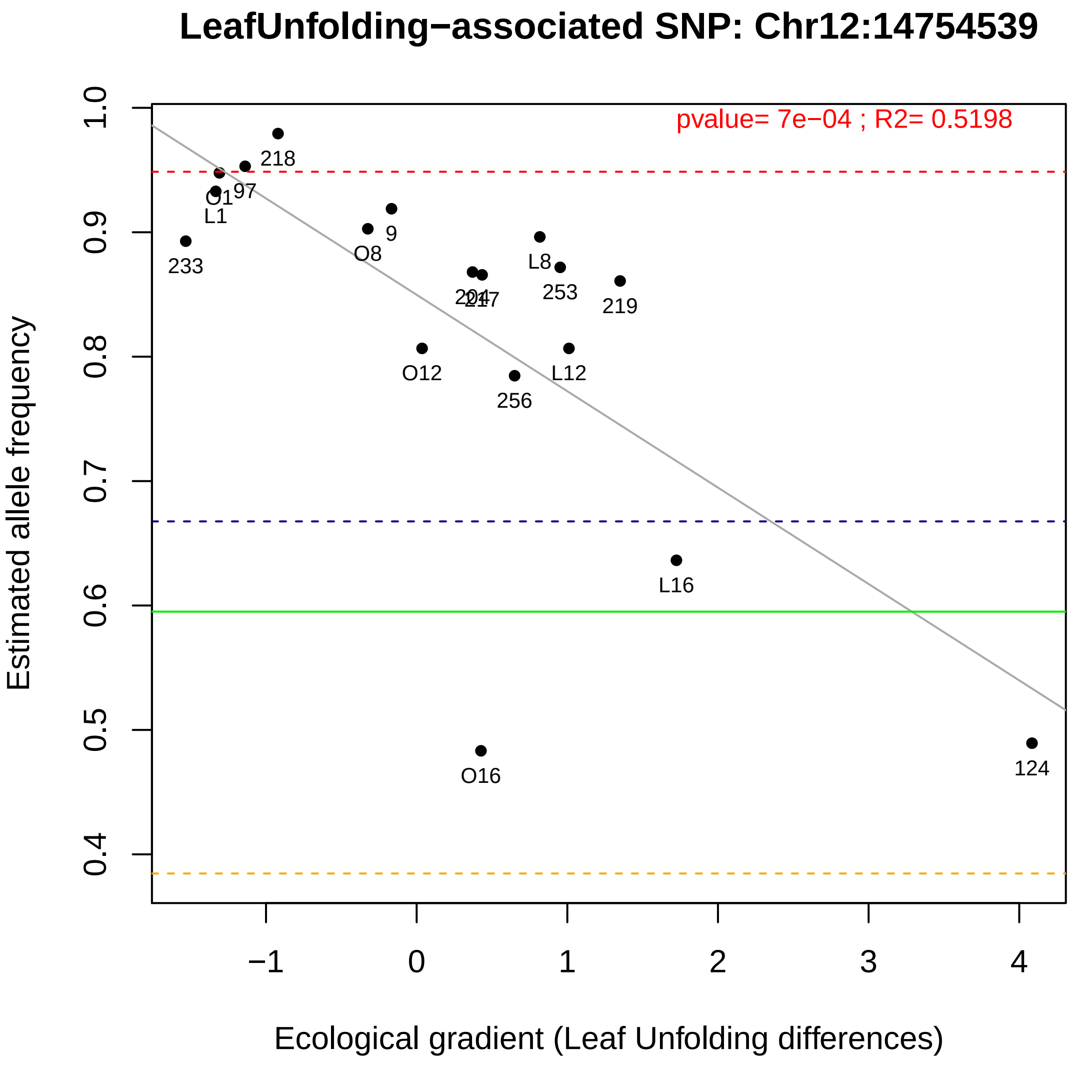

LeafUnfolding−associated SNP: Chr12:14754539
pvalue= 7e−04 ; R2= 0.5198
Estimated allele frequency
Ecological gradient (Leaf Unfolding differences)
1.0
0.9
0.8
0.7
0.6
0.5
0.4
−1
0
1
2
3
4
218
97
O1
L1
233
O8
9
L8
253
204
217
219
O12
256
L12
L16
O16
124

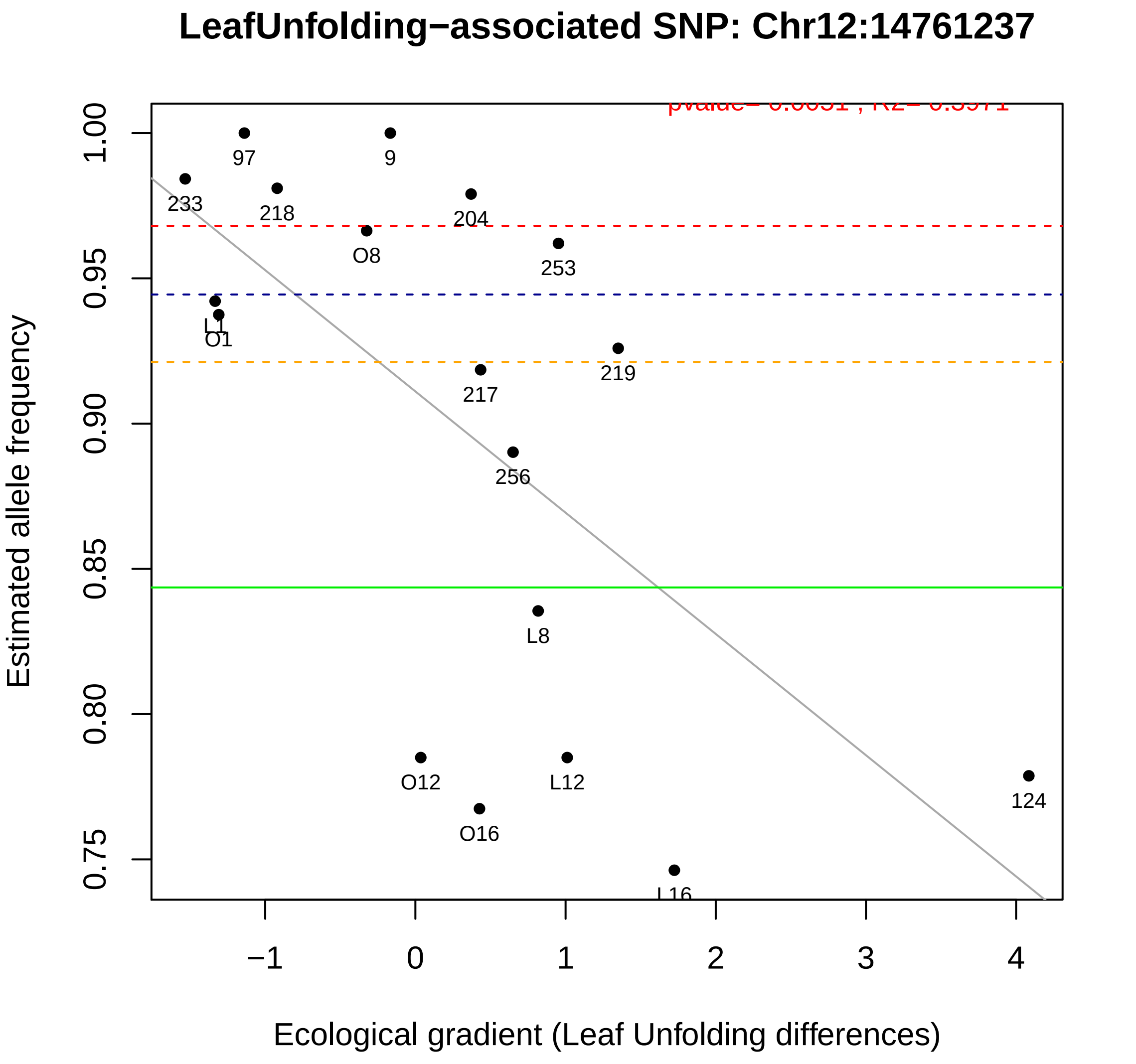

LeafUnfolding−associated SNP: Chr12:14761237
pvalue= 0.0031 ; R2= 0.3971
Estimated allele frequency
Ecological gradient (Leaf Unfolding differences)
1.00
0.95
0.90
0.85
0.80
0.75
−1
0
1
2
3
4
97
9
233
218
204
O8
253
L1
O1
219
217
256
L8
O12
L12
124
O16
L16

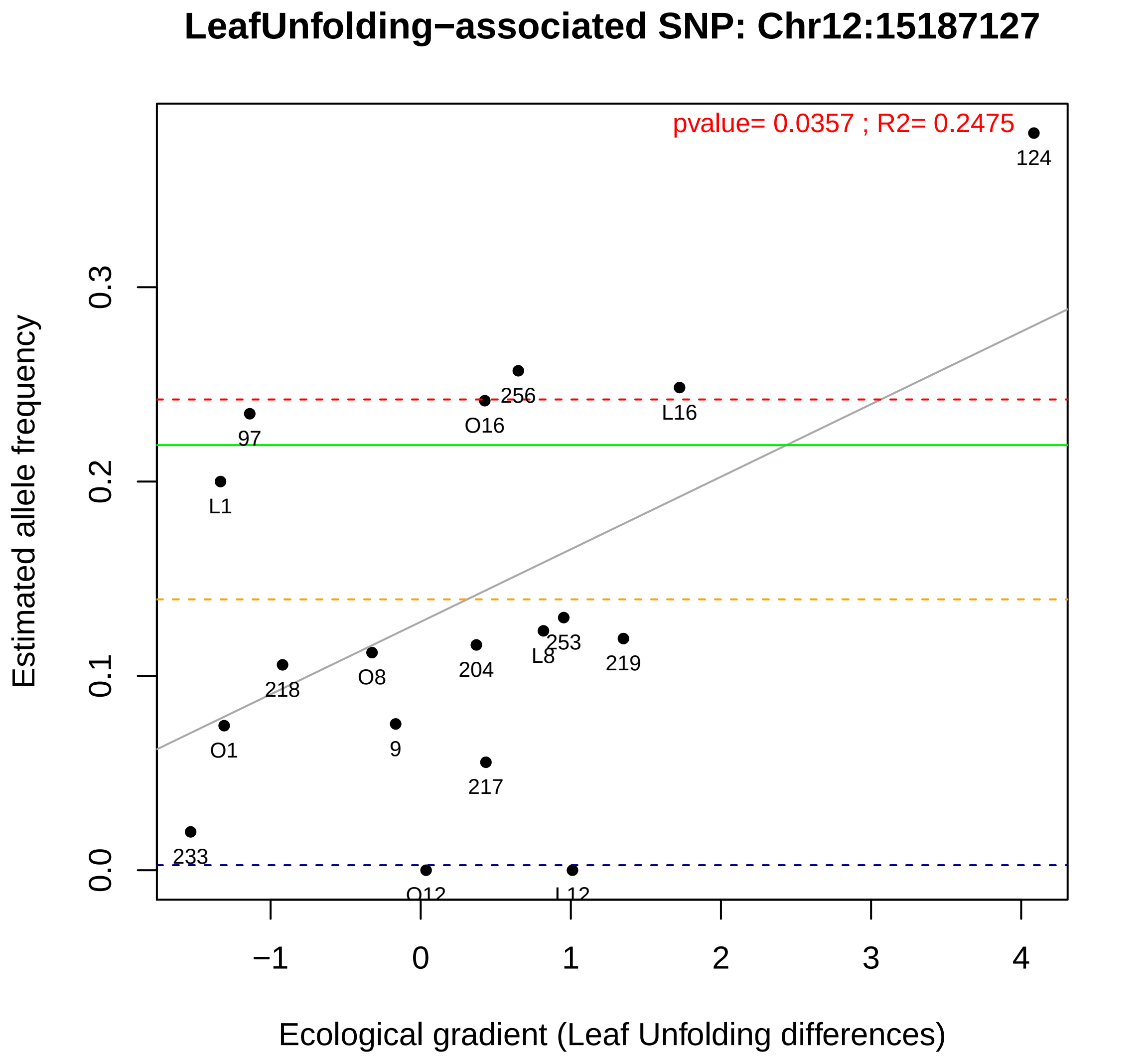
LeafUnfolding−associated SNP: Chr12:15187127
pvalue= 0.0357 ; R2= 0.2475
Estimated allele frequency
Ecological gradient (Leaf Unfolding differences)
0.0
0.1
0.2
0.3
−1
0
1
2
3
4
124
256
O16
L16
97
L1
253
L8
219
204
O8
218
O1
9
217
233
O12
L12

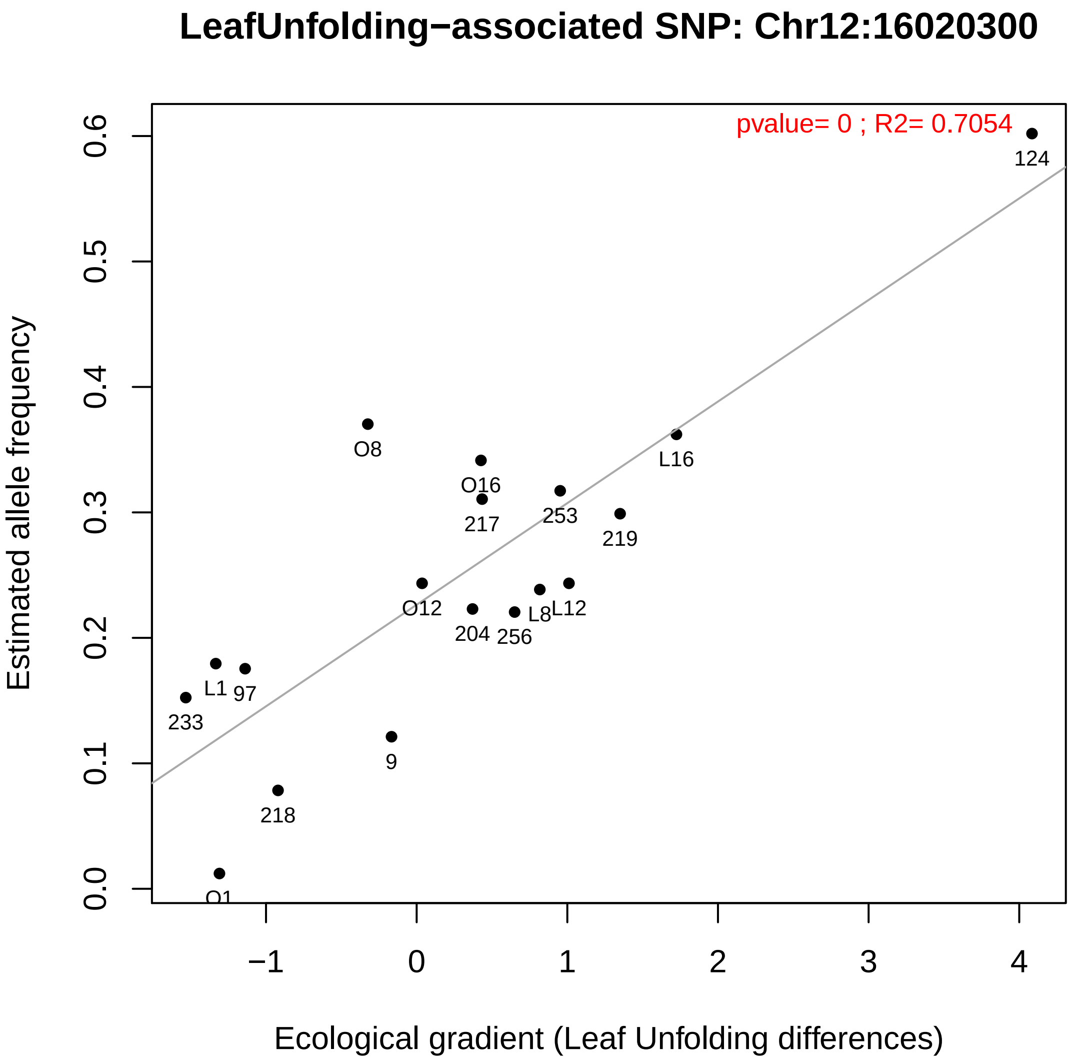
LeafUnfolding−associated SNP: Chr12:16020300
pvalue= 0 ; R2= 0.7054
Estimated allele frequency
0.0
0.1
0.2
0.3
0.4
0.5
0.6
−1
0
1
2
3
4
Ecological gradient (Leaf Unfolding differences)
124
L16
O8
O16
217
253
219
O12
204
256
L8
L12
L1
97
233
9
218
O1

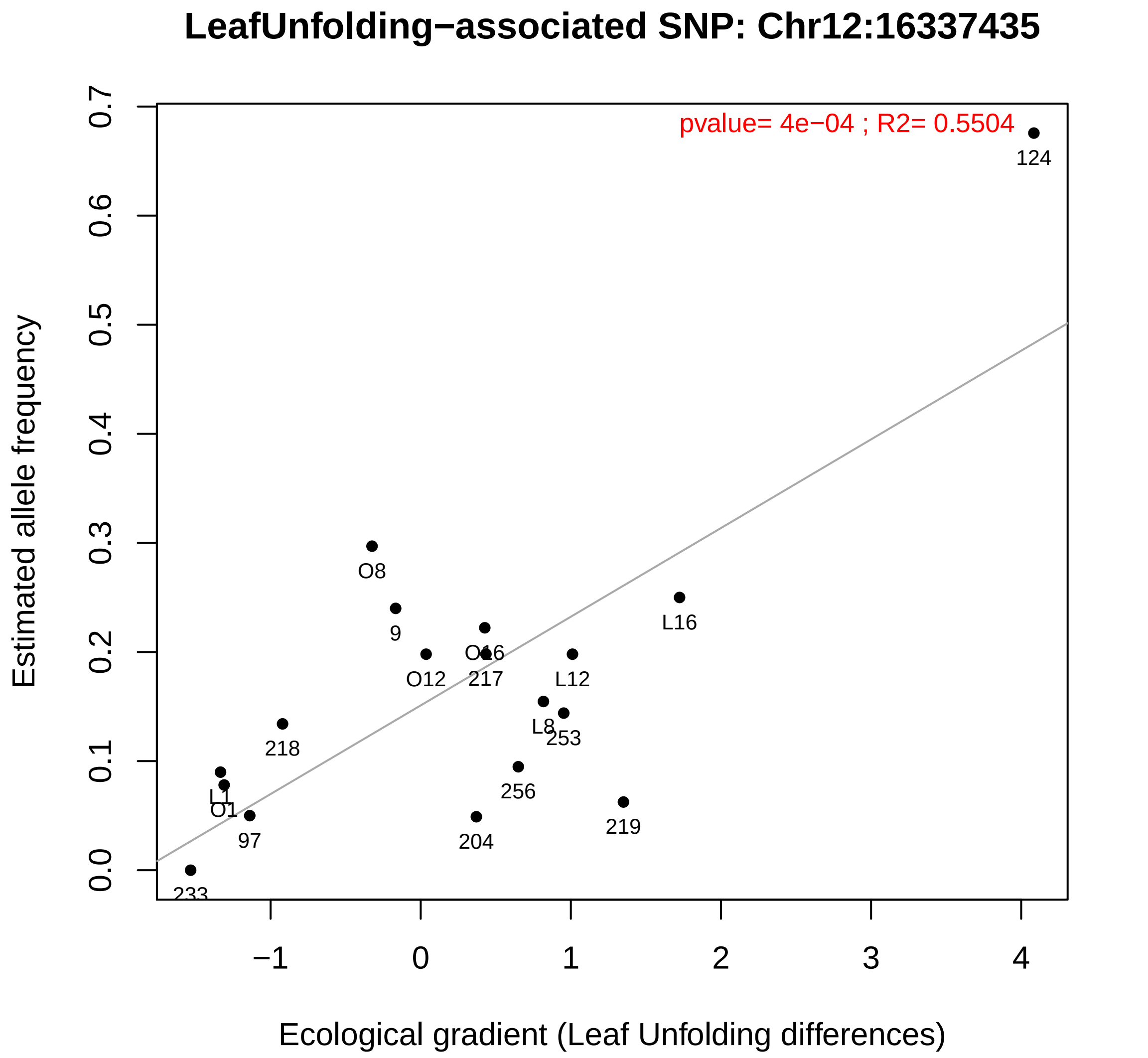
LeafUnfolding−associated SNP: Chr12:16337435
pvalue= 4e−04 ; R2= 0.5504
Estimated allele frequency
0.0
0.1
0.2
0.3
0.4
0.5
0.6
0.7
Ecological gradient (Leaf Unfolding differences)
−1
0
1
2
3
4
124
O8
9
L16
O16
217
O12
L12
L8
253
218
256
L1
O1
97
204
219
233

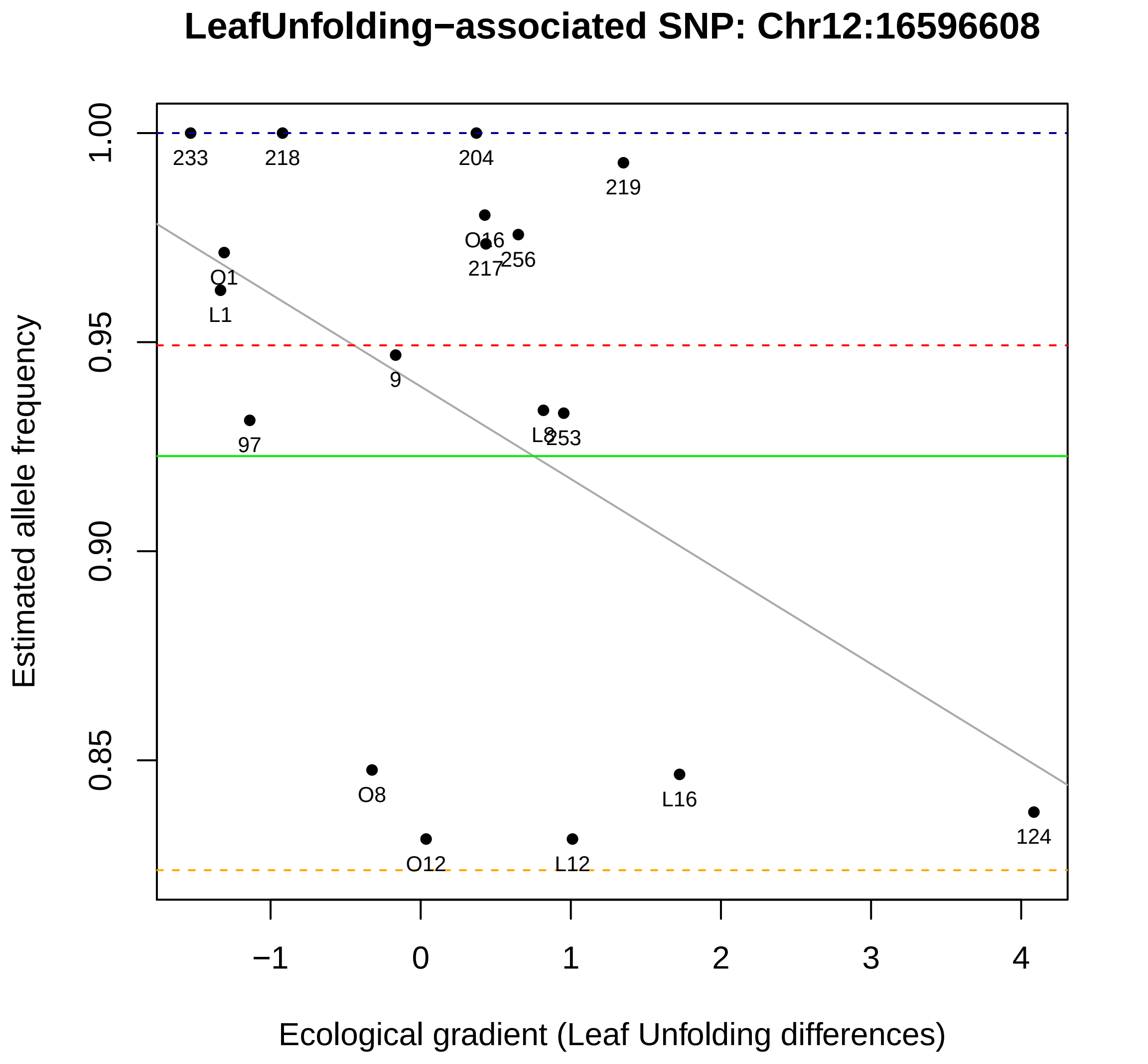
LeafUnfolding−associated SNP: Chr12:16596608
Estimated allele frequency
Ecological gradient (Leaf Unfolding differences)
1.00
0.95
0.90
0.85
−1
0
1
2
3
4
233
218
204
219
O16
217
256
O1
L1
9
97
L8
253
O8
L16
124
O12
L12

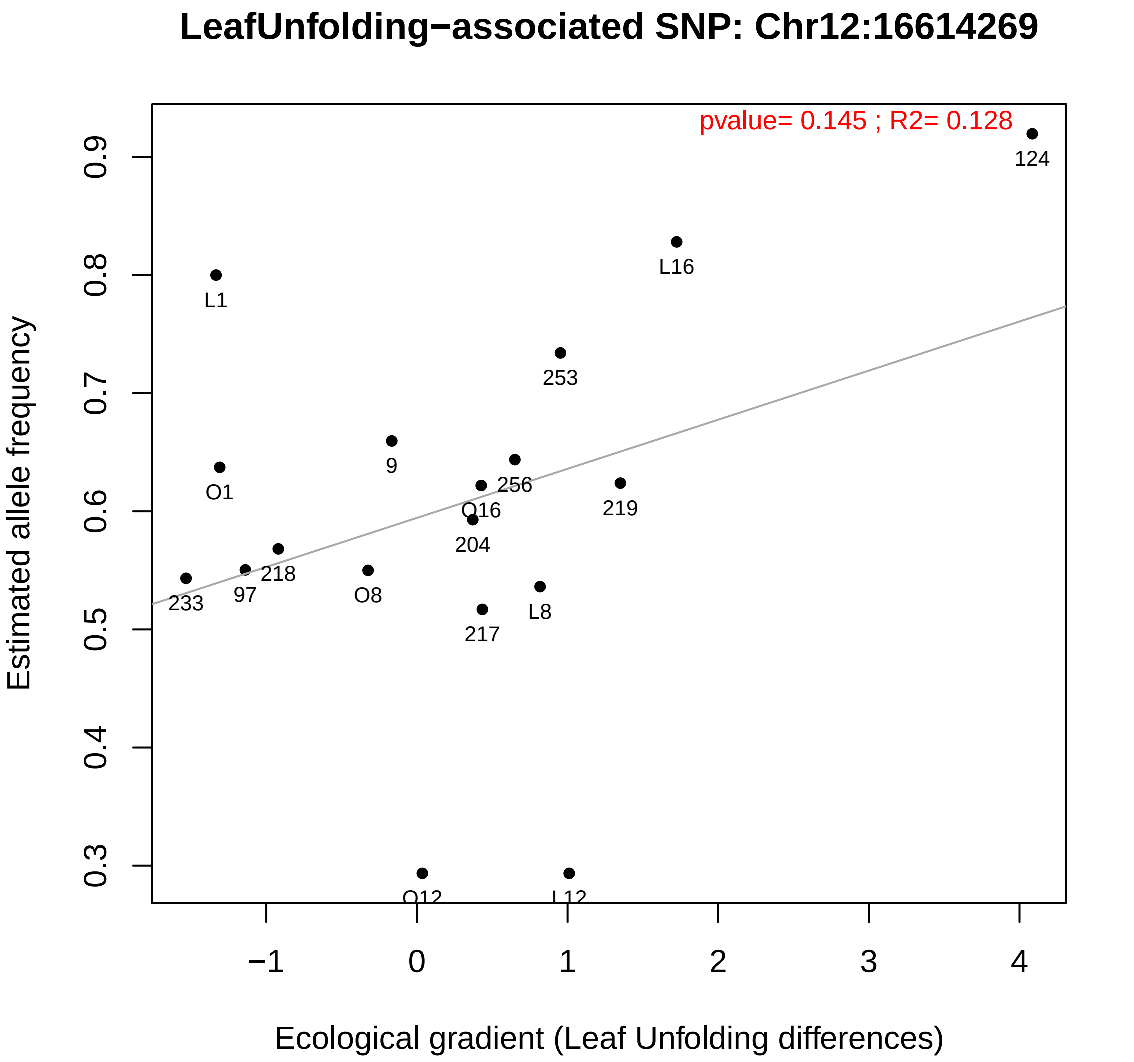

LeafUnfolding−associated SNP: Chr12:16614269
pvalue= 0.145 ; R2= 0.128
124
L16
L1
253
9
O1
256
Q16
219
204
218
97
233
O8
L8
217
O12
L12
0.3
0.4
0.5
0.6
0.7
0.8
0.9
−1
0
1
2
3
4
Estimated allele frequency
Ecological gradient (Leaf Unfolding differences)

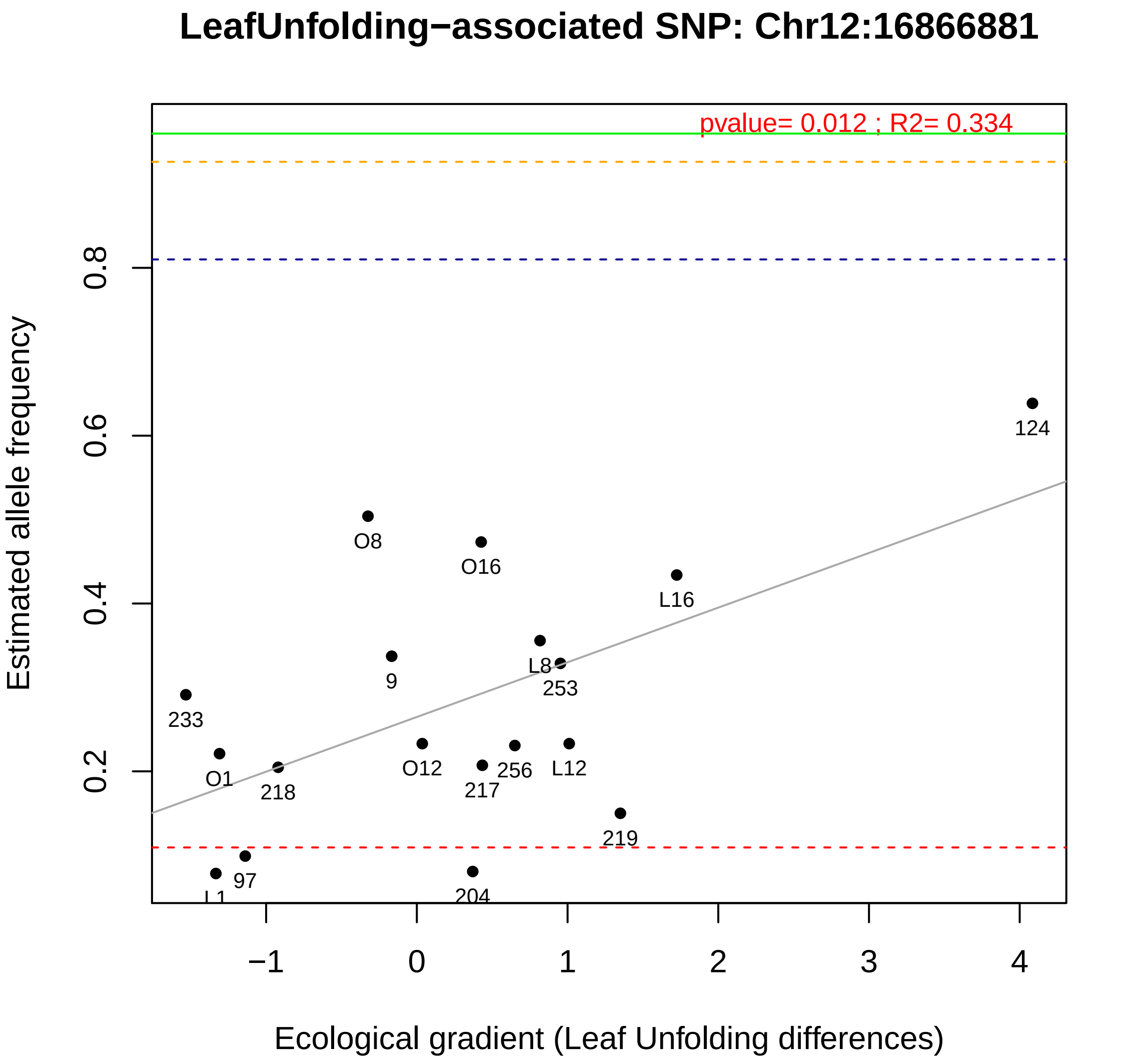
LeafUnfolding−associated SNP: Chr12:16866881
pvalue= 0.012 ; R2= 0.334
Estimated allele frequency
0.2
0.4
0.6
0.8
−1
0
1
2
3
4
Ecological gradient (Leaf Unfolding differences)
124
O8
O16
L16
L8
253
9
233
O1
218
O12
217
256
L12
219
97
204

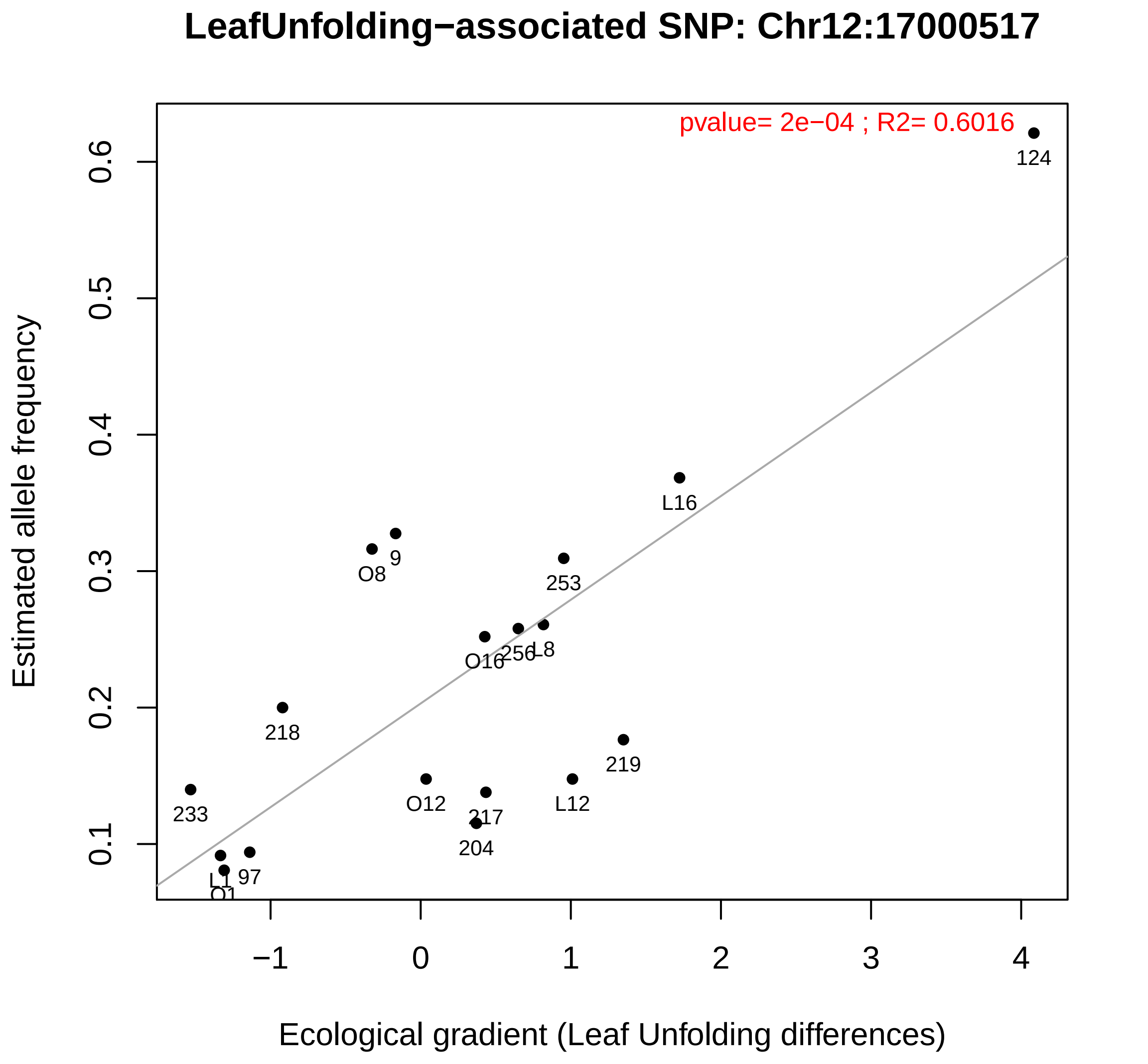
LeafUnfolding−associated SNP: Chr12:17000517
pvalue= 2e−04 ; R2= 0.6016
Estimated allele frequency
Ecological gradient (Leaf Unfolding differences)
0.1
0.2
0.3
0.4
0.5
0.6
−1
0
1
2
3
4
124
L16
9
O8
253
O16
256
L8
218
219
233
O12
217
L12
204
L1
97
O1

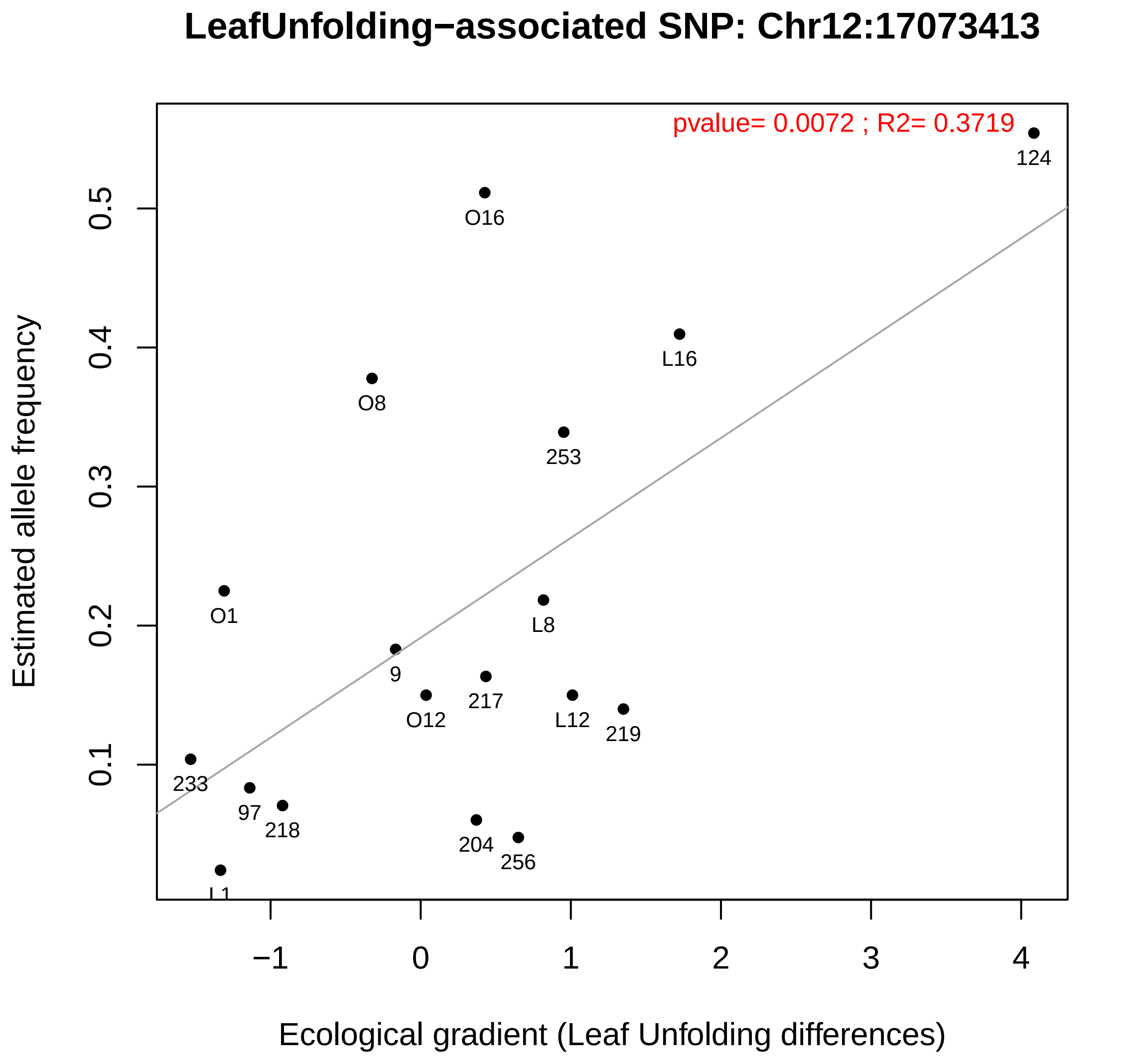

LeafUnfolding−associated SNP: Chr12:17073413
pvalue= 0.0072 ; R2= 0.3719
124
O16
L16
O8
253
O1
L8
9
217
O12
L12
219
233
97
218
204
256
L1
0.1
0.2
0.3
0.4
0.5
−1
0
1
2
3
4
Estimated allele frequency
Ecological gradient (Leaf Unfolding differences)

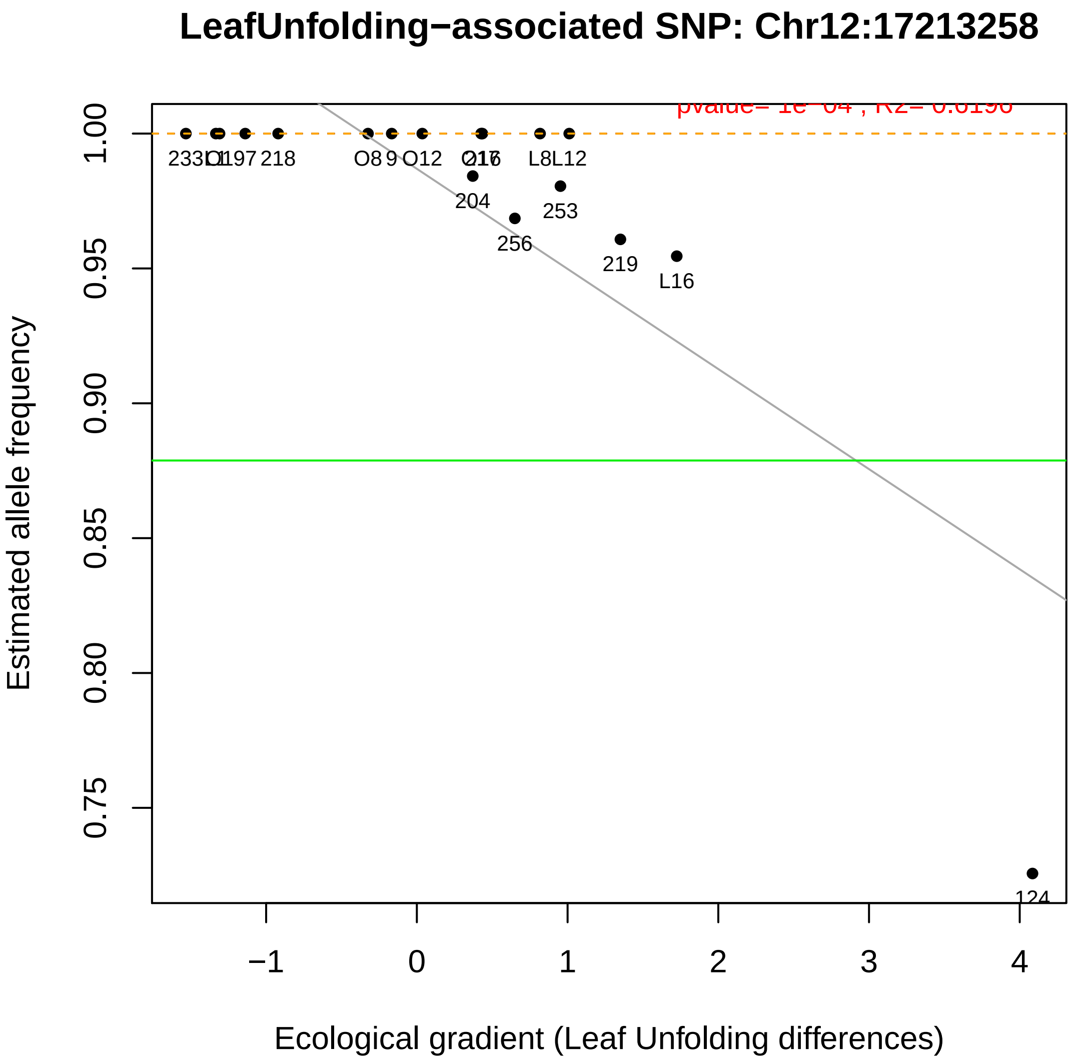

LeafUnfolding−associated SNP: Chr12:17213258
pvalue= 1e−04 , R2= 0.6196
Estimated allele frequency
Ecological gradient (Leaf Unfolding differences)
1.00
0.95
0.90
0.85
0.80
0.75
−1
0
1
2
3
4
233
O1
197
218
O8
9
O12
O17
216
L8
L12
204
253
256
219
L16
124

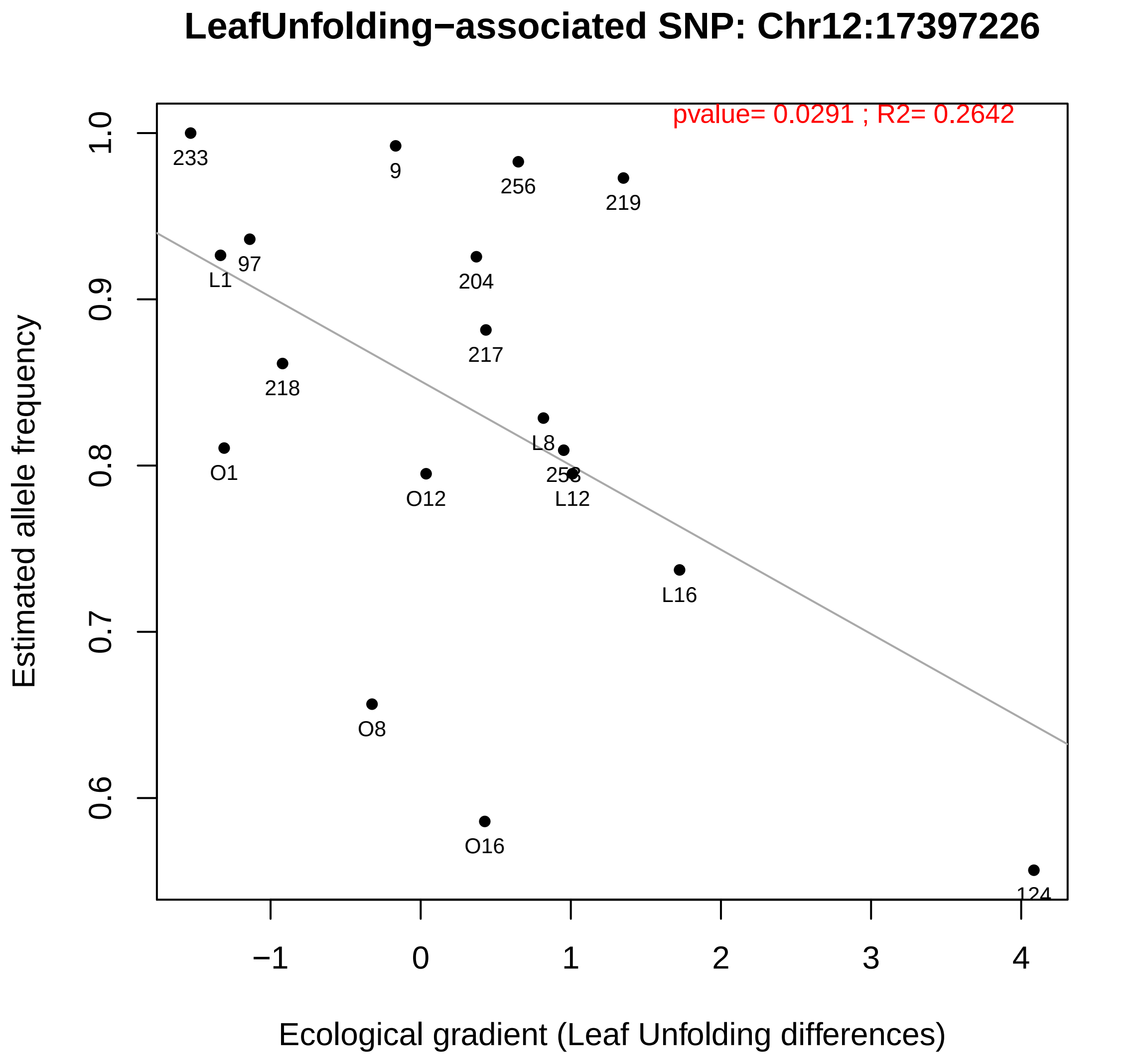

LeafUnfolding−associated SNP: Chr12:17397226
pvalue= 0.0291 ; R2= 0.2642
Estimated allele frequency
Ecological gradient (Leaf Unfolding differences)
0.6
0.7
0.8
0.9
1.0
−1
0
1
2
3
4
233
9
256
219
97
L1
204
217
218
L8
O1
O12
25
L12
L16
O8
O16
124

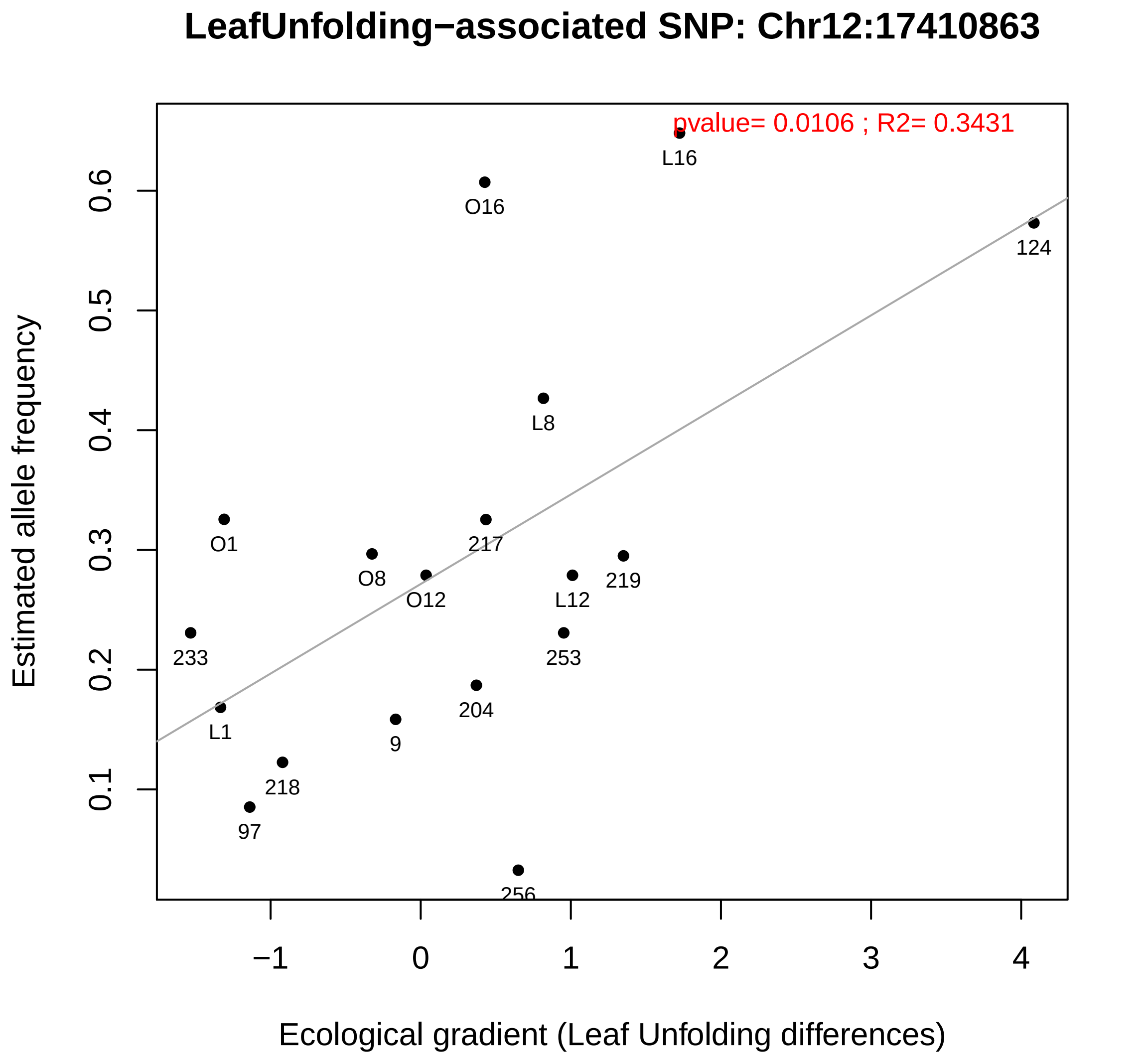
LeafUnfolding−associated SNP: Chr12:17410863
pvalue= 0.0106 ; R2= 0.3431
Estimated allele frequency
0.1
0.2
0.3
0.4
0.5
0.6
−1
0
1
2
3
4
Ecological gradient (Leaf Unfolding differences)
L16
O16
124
L8
O1
217
O8
O12
L12
219
233
253
L1
204
9
218
97
256

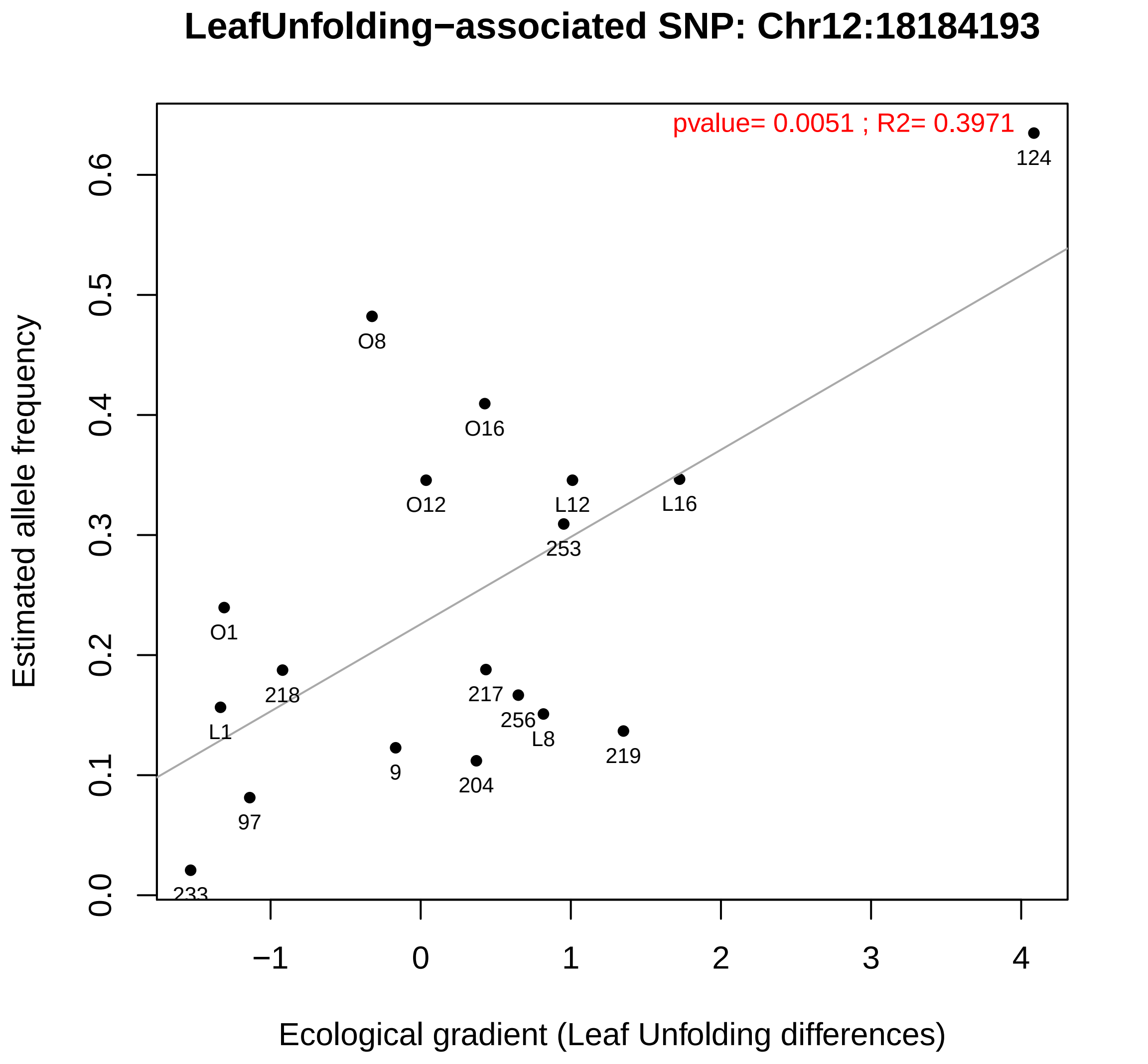
LeafUnfolding−associated SNP: Chr12:18184193
pvalue= 0.0051 ; R2= 0.3971
Estimated allele frequency
Ecological gradient (Leaf Unfolding differences)
0.0
0.1
0.2
0.3
0.4
0.5
0.6
−1
0
1
2
3
4
124
O8
O16
O12
L12
L16
253
O1
218
L1
217
256
L8
219
9
204
97
233

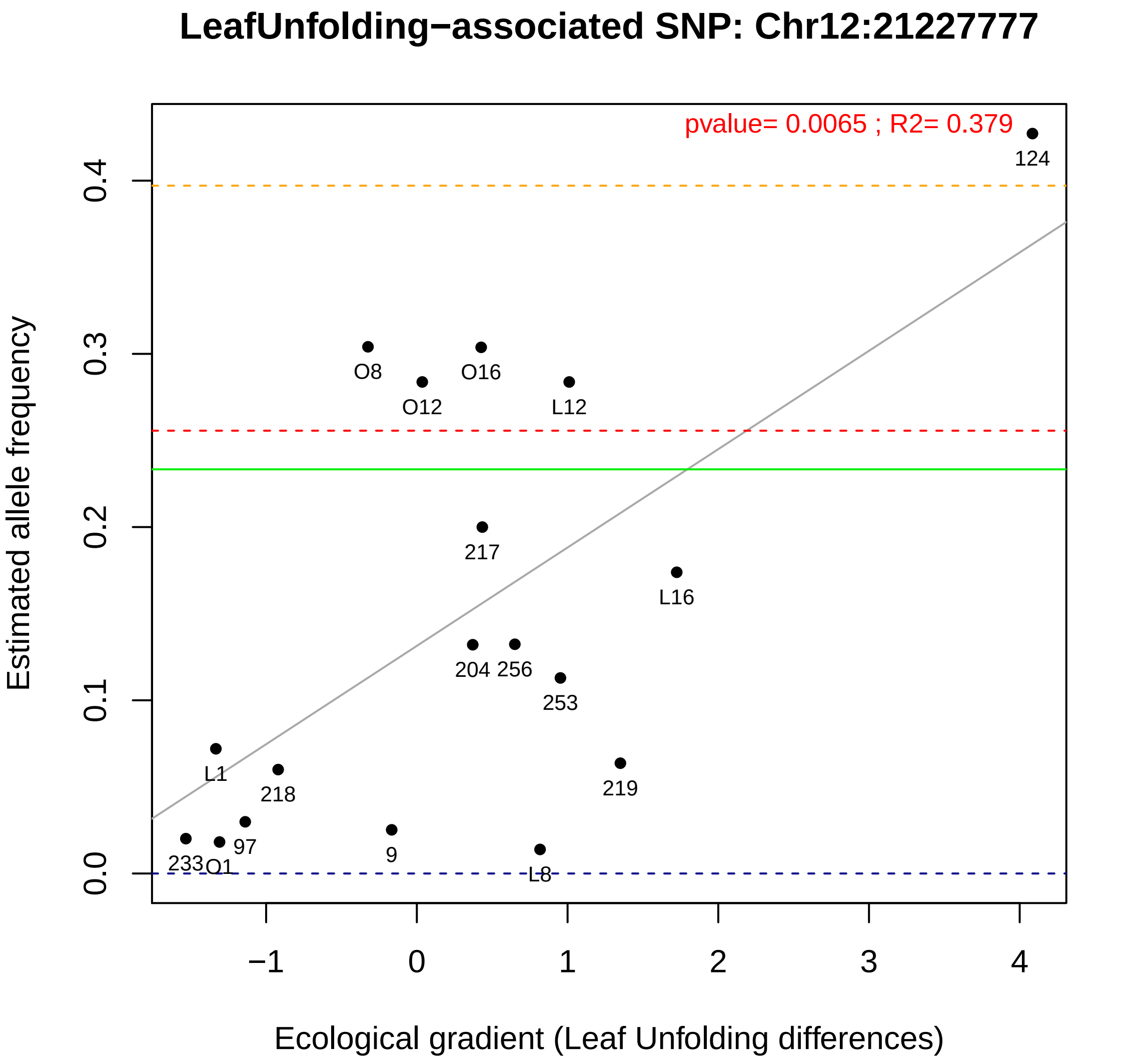

LeafUnfolding−associated SNP: Chr12:21227777
pvalue= 0.0065 ; R2= 0.379
124
O8
O16
O12
L12
217
L16
204
256
253
L1
218
219
97
9
233
O1
L8
0.0
0.1
0.2
0.3
0.4
−1
0
1
2
3
4
Estimated allele frequency
Ecological gradient (Leaf Unfolding differences)

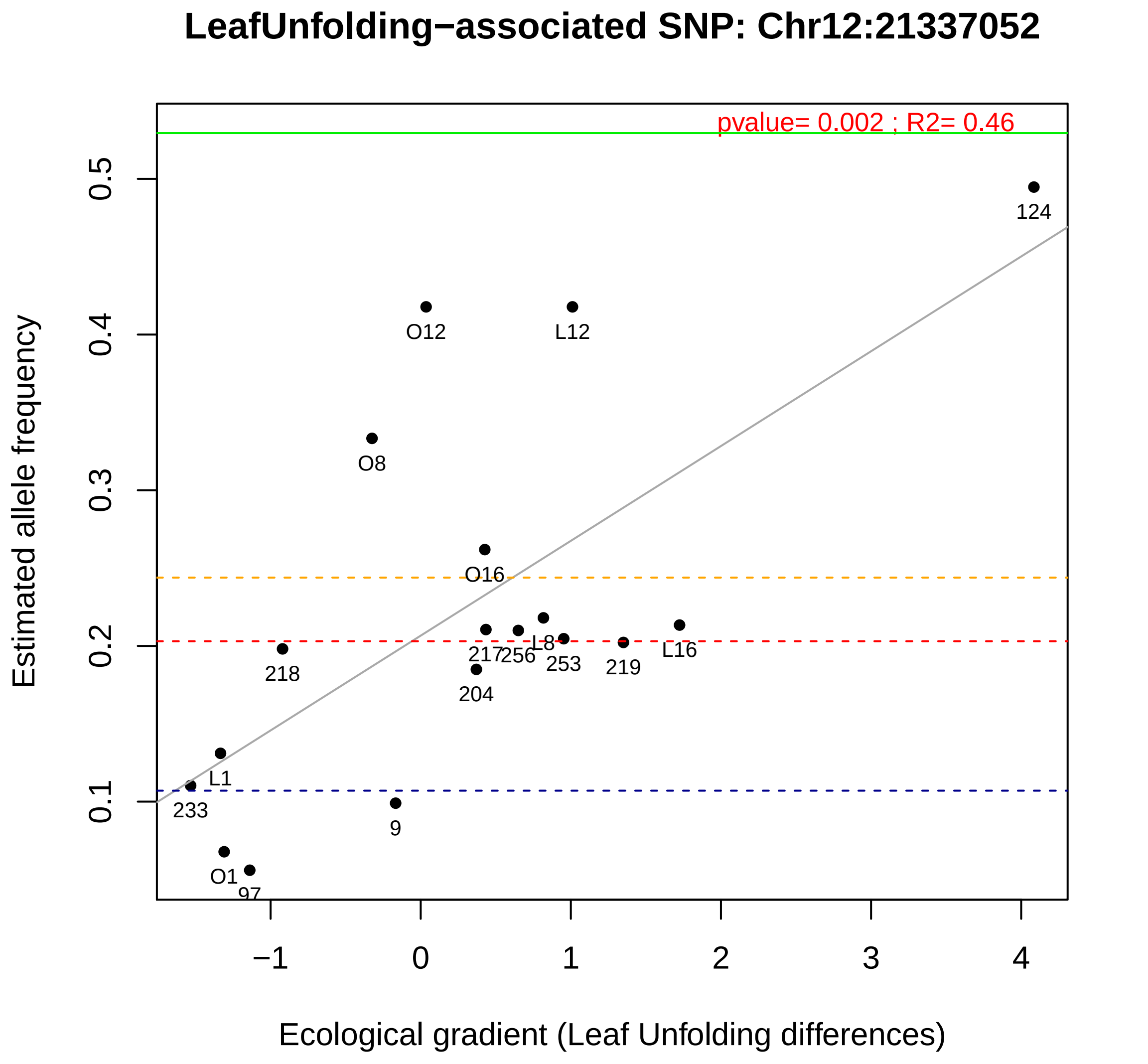
LeafUnfolding−associated SNP: Chr12:21337052
pvalue= 0.002 ; R2= 0.46
Estimated allele frequency
0.1
0.2
0.3
0.4
0.5
Ecological gradient (Leaf Unfolding differences)
−1
0
1
2
3
4
124
O12
L12
O8
O16
217
256
L8
253
219
L16
218
204
L1
233
9
O1
97

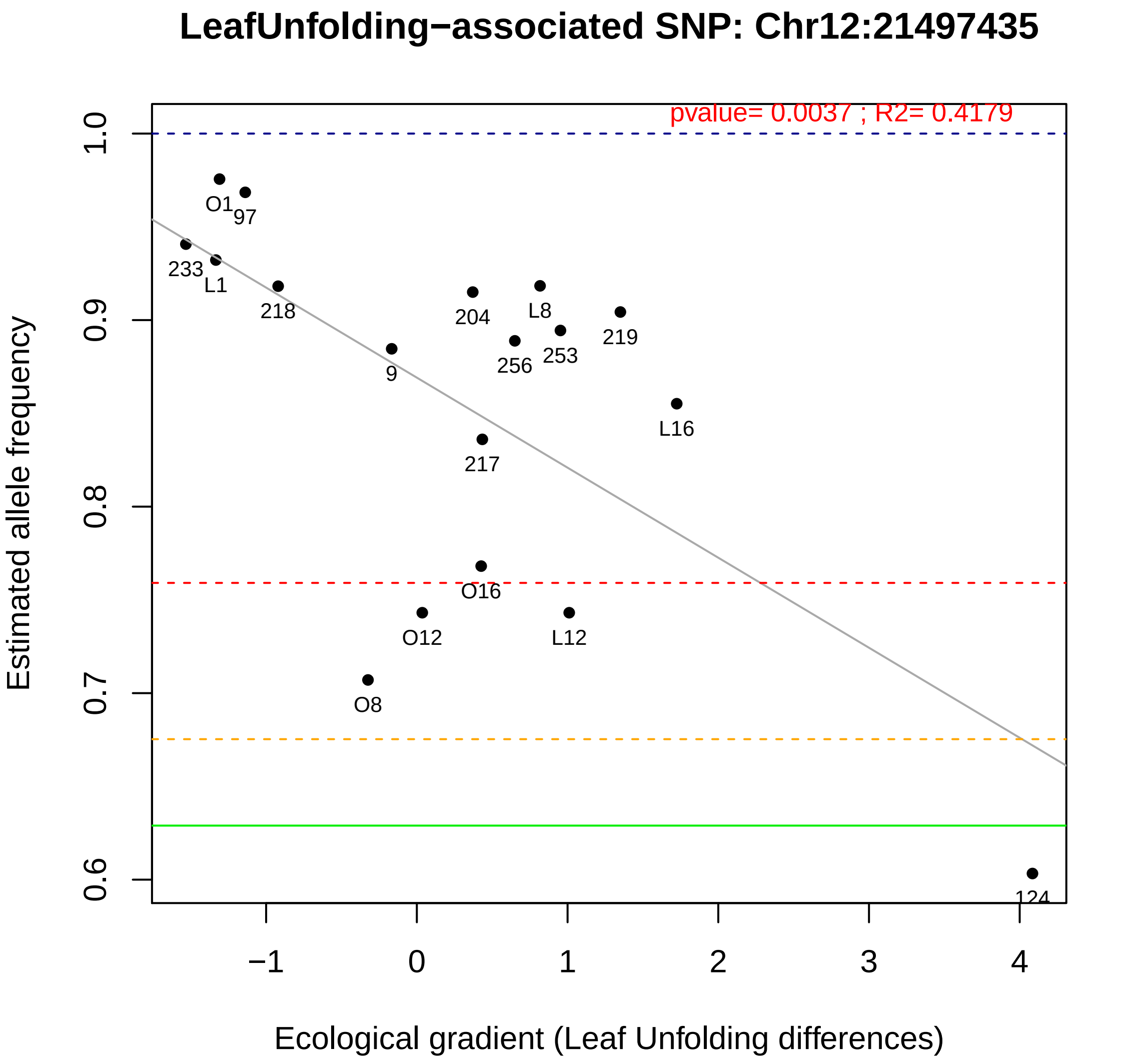
LeafUnfolding−associated SNP: Chr12:21497435
pvalue= 0.0037 ; R2= 0.4179
Estimated allele frequency
1.0
0.9
0.8
0.7
0.6
O1
97
233
L1
218
204
L8
219
253
256
9
L16
217
O16
O12
L12
O8
124
−1
0
1
2
3
4
Ecological gradient (Leaf Unfolding differences)

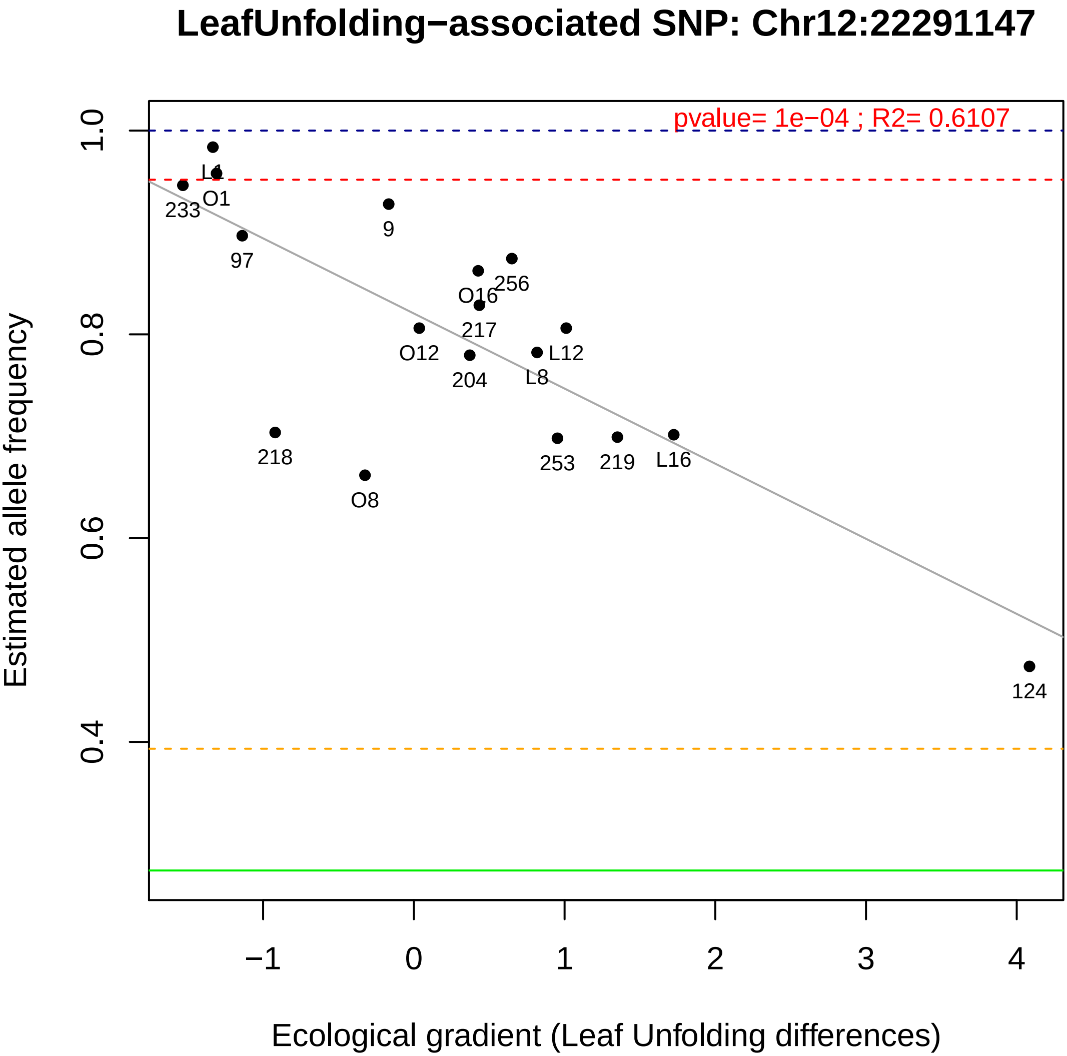

LeafUnfolding−associated SNP: Chr12:22291147
pvalue= 1e−04 ; R2= 0.6107
Estimated allele frequency
Ecological gradient (Leaf Unfolding differences)
1.0
0.8
0.6
0.4
−1
0
1
2
3
4
L1
O1
233
97
9
256
O16
217
O12
L12
204
L8
218
O8
253
219
L16
124

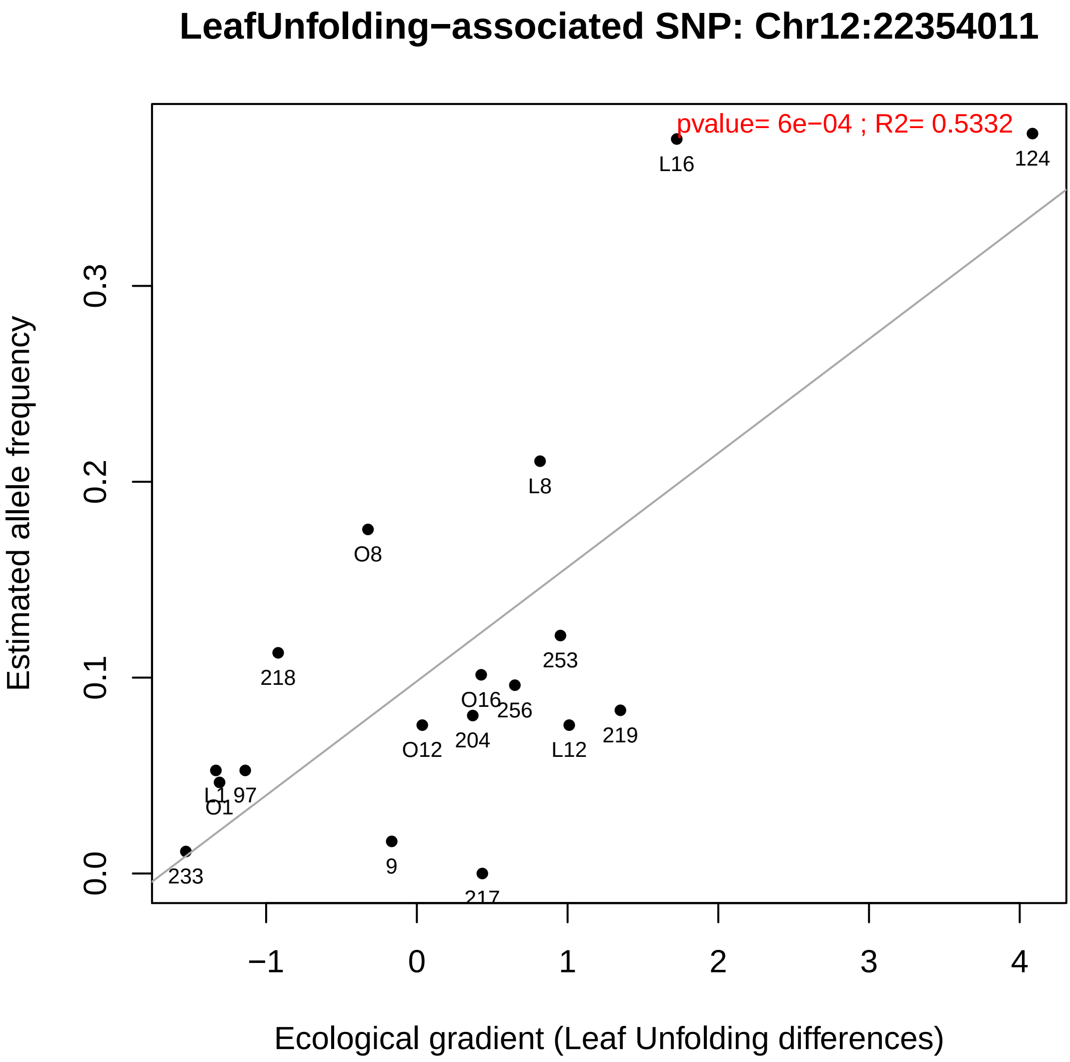
LeafUnfolding−associated SNP: Chr12:22354011
pvalue= 6e−04 ; R2= 0.5332
Estimated allele frequency
Ecological gradient (Leaf Unfolding differences)
0.0
0.1
0.2
0.3
−1
0
1
2
3
4
L16
124
L8
O8
253
218
O16
256
204
O12
L12
219
L1
97
O1
233
9
217

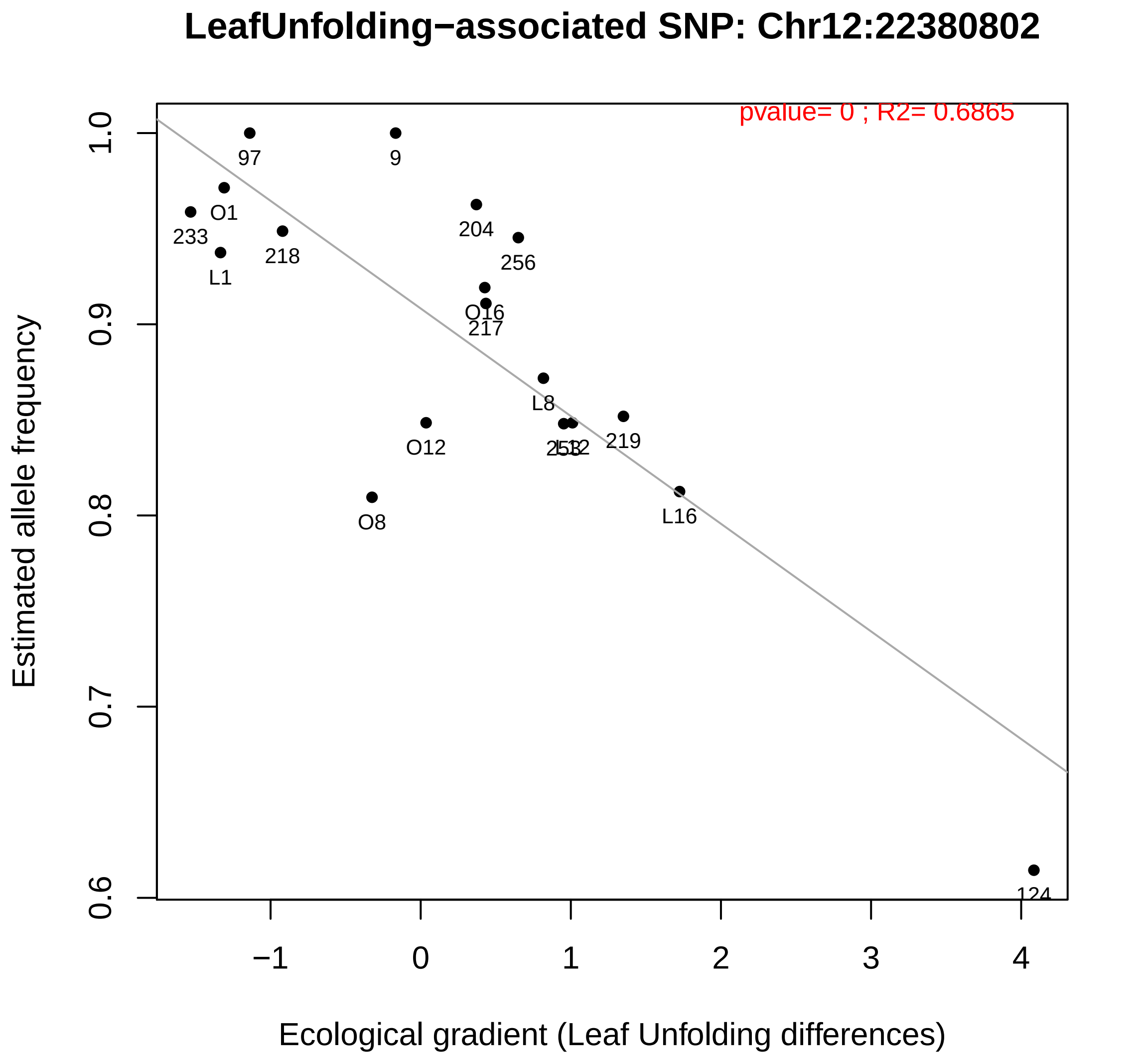
LeafUnfolding−associated SNP: Chr12:22380802
pvalue= 0 ; R2= 0.6865
Estimated allele frequency
Ecological gradient (Leaf Unfolding differences)
1.0
0.9
0.8
0.7
0.6
−1
0
1
2
3
4
97
9
O1
233
218
L1
204
256
O16
217
L8
O12
253
L12
219
O8
L16
124

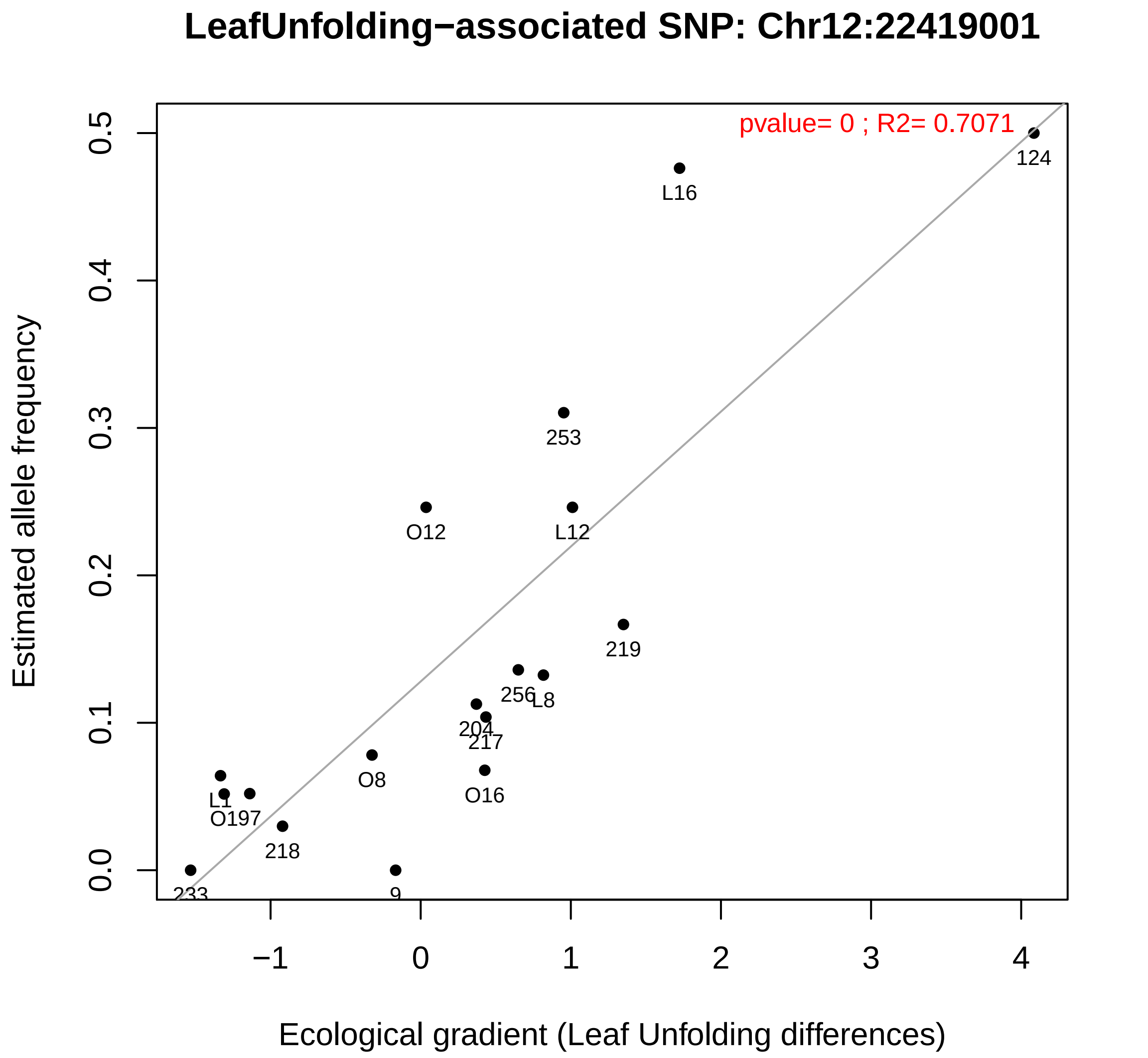

LeafUnfolding−associated SNP: Chr12:22419001
pvalue= 0 ; R2= 0.7071
Estimated allele frequency
Ecological gradient (Leaf Unfolding differences)
0.0
0.1
0.2
0.3
0.4
0.5
−1
0
1
2
3
4
124
L16
253
O12
L12
219
256
L8
204
217
O8
O16
L1
O197
218
233
9

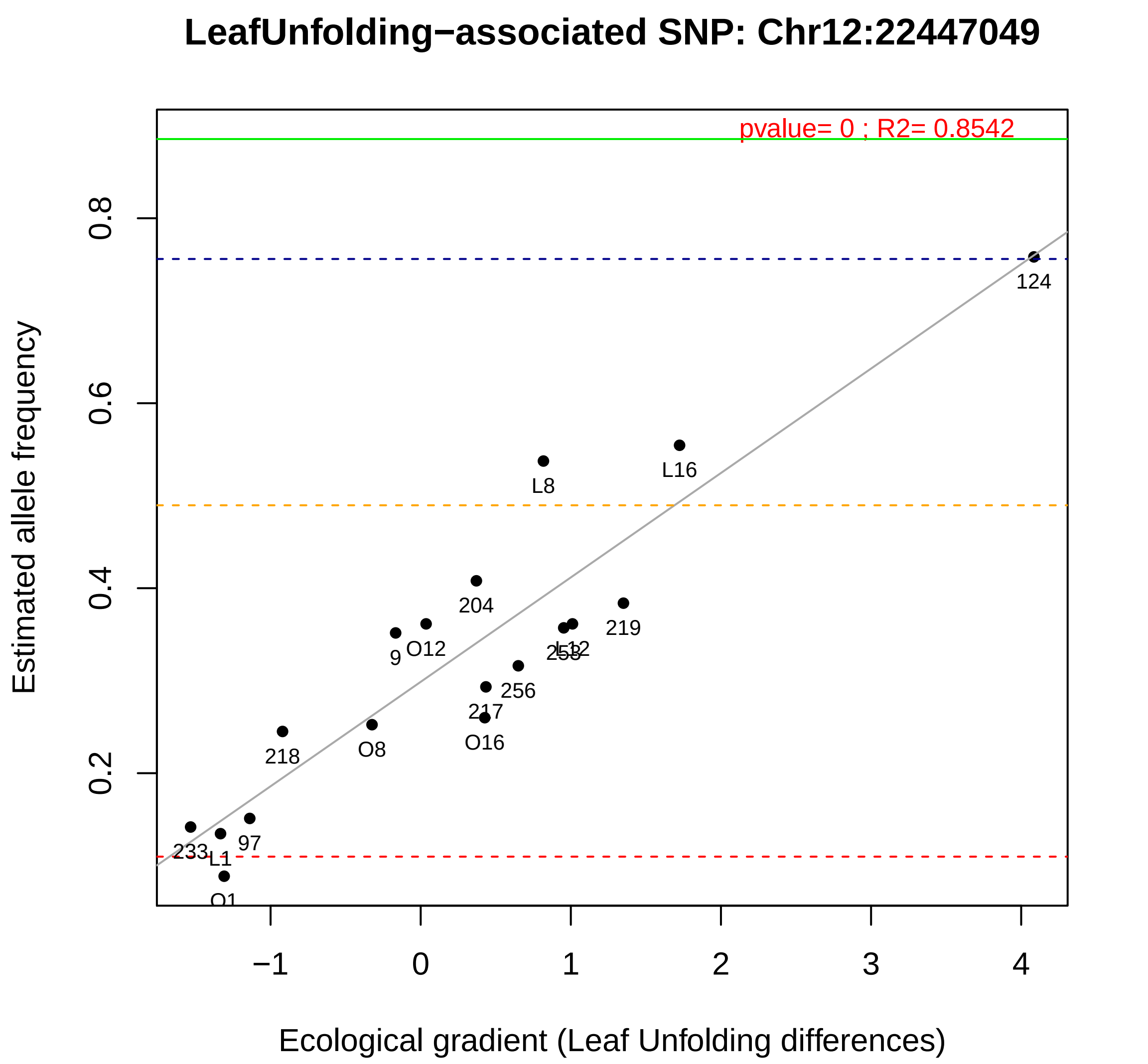

LeafUnfolding−associated SNP: Chr12:22447049
pvalue= 0 ; R2= 0.8542
Estimated allele frequency
0.2
0.4
0.6
0.8
−1
0
1
2
3
4
Ecological gradient (Leaf Unfolding differences)
124
L16
L8
204
219
9
O12
256
217
O16
218
O8
233
L1
97

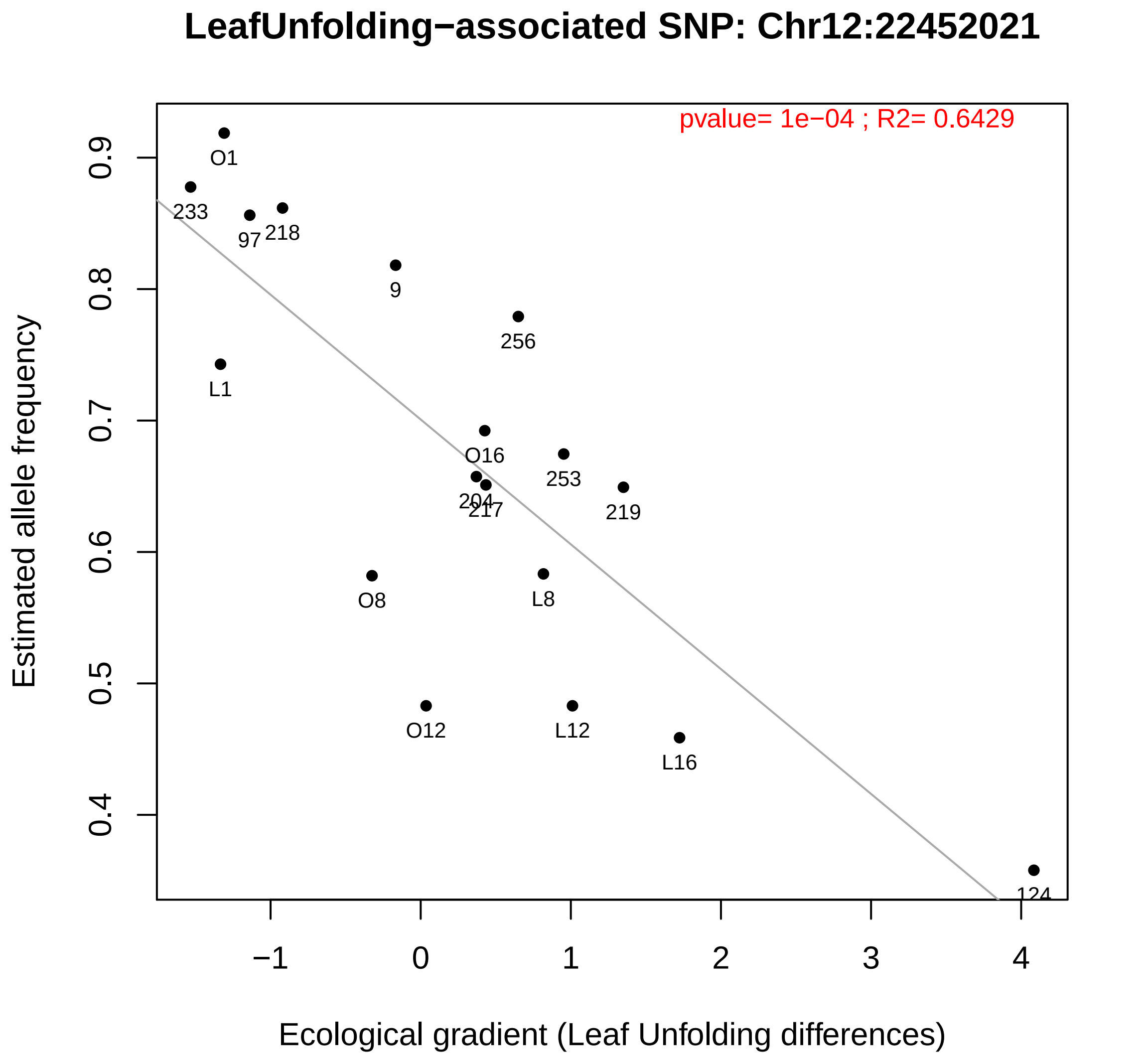
LeafUnfolding−associated SNP: Chr12:22452021
pvalue= 1e−04 ; R2= 0.6429
Estimated allele frequency
Ecological gradient (Leaf Unfolding differences)
0.4
0.5
0.6
0.7
0.8
0.9
−1
0
1
2
3
4
O1
233
97
218
9
256
L1
O16
253
204
217
219
O8
L8
O12
L12
L16
124

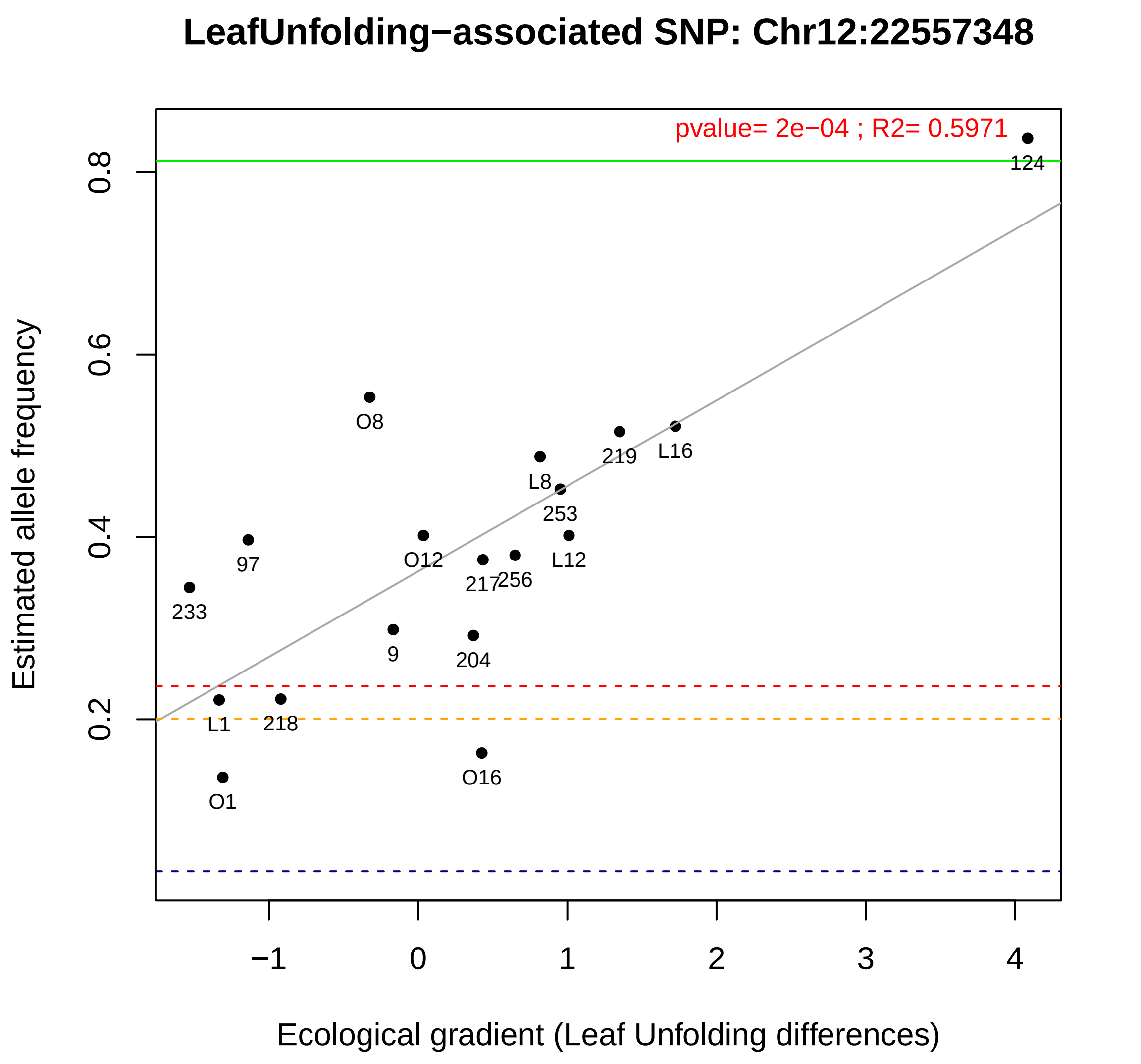
LeafUnfolding−associated SNP: Chr12:22557348
pvalue= 2e−04 ; R2= 0.5971
Estimated allele frequency
0.2
0.4
0.6
0.8
−1
0
1
2
3
4
Ecological gradient (Leaf Unfolding differences)
124
O8
219
L16
L8
253
97
O12
L12
217
256
233
9
204
L1
218
O16
O1

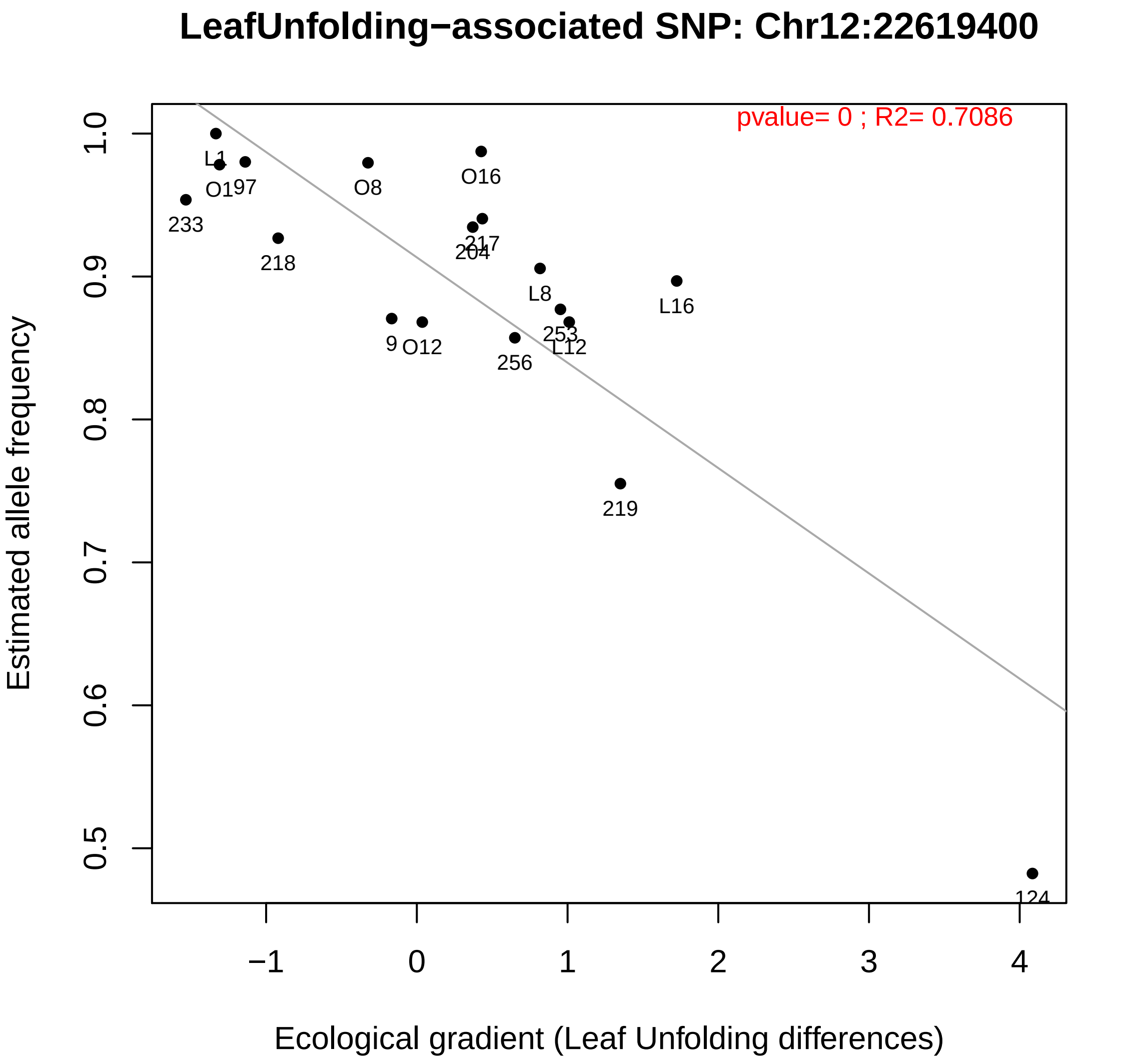

LeafUnfolding−associated SNP: Chr12:22619400
pvalue= 0 ; R2= 0.7086
Estimated allele frequency
Ecological gradient (Leaf Unfolding differences)
1.0
0.9
0.8
0.7
0.6
0.5
−1
0
1
2
3
4
L1
O1
97
233
218
O8
O16
217
204
L8
L16
9
O12
256
253
L12
219
124

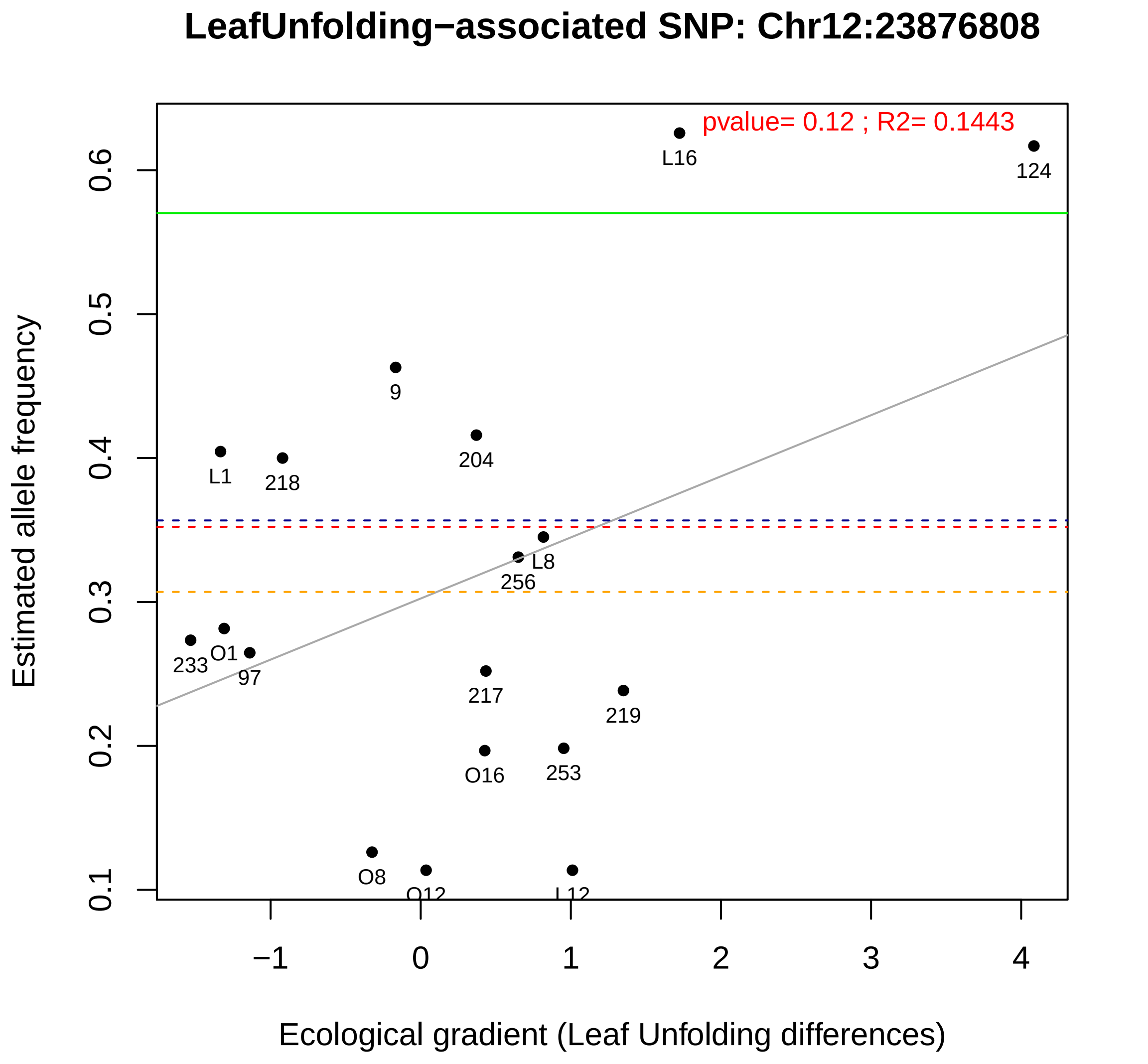

LeafUnfolding−associated SNP: Chr12:23876808
pvalue= 0.12 ; R2= 0.1443
Estimated allele frequency
Ecological gradient (Leaf Unfolding differences)
0.1
0.2
0.3
0.4
0.5
0.6
−1
0
1
2
3
4
L16
124
9
204
L1
218
L8
256
233
O1
97
217
219
O16
253
O8
O12
L12

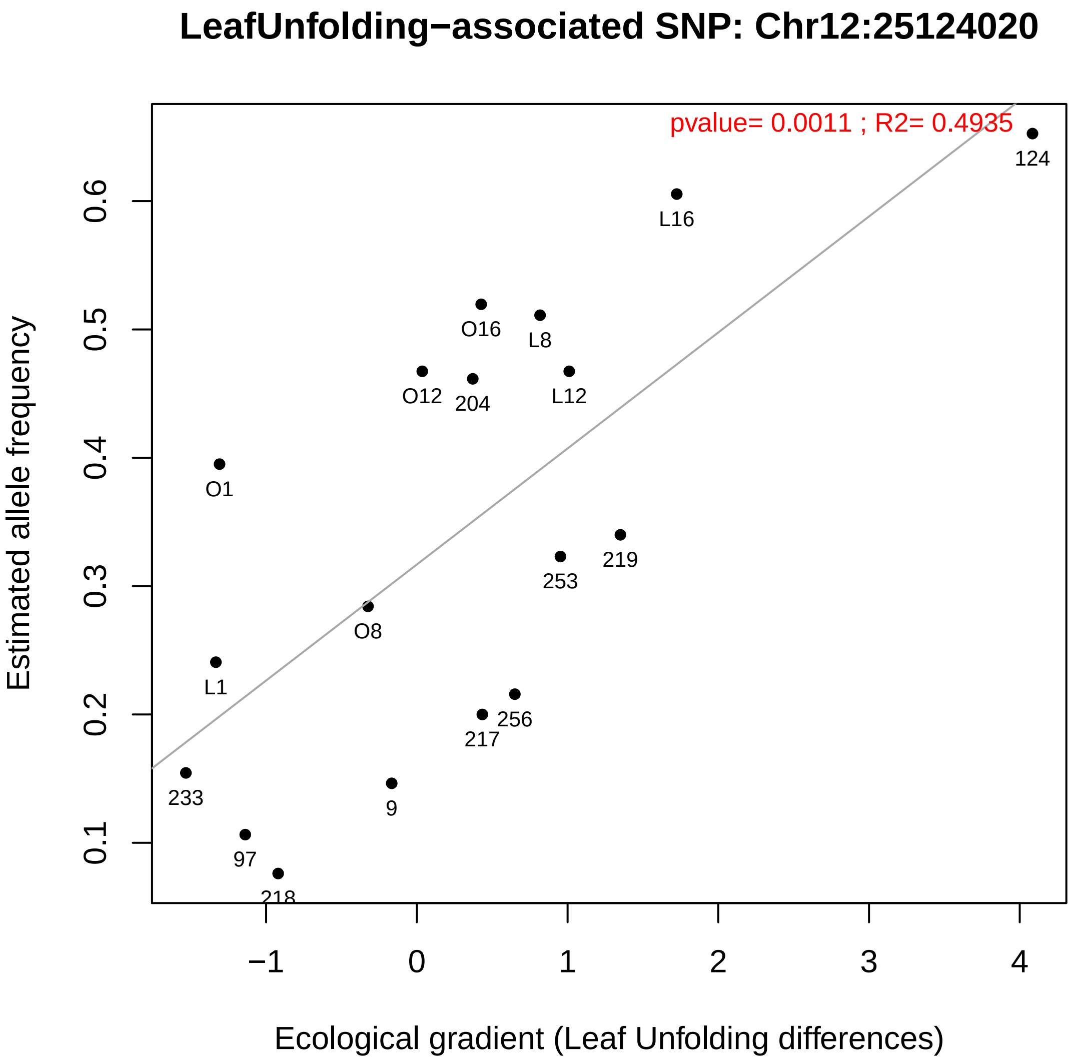
LeafUnfolding−associated SNP: Chr12:25124020
pvalue= 0.0011 ; R2= 0.4935
Estimated allele frequency
Ecological gradient (Leaf Unfolding differences)
0.1
0.2
0.3
0.4
0.5
0.6
−1
0
1
2
3
4
124
L16
O16
L8
O12
204
L12
O1
219
253
O8
L1
256
217
233
9
97
218

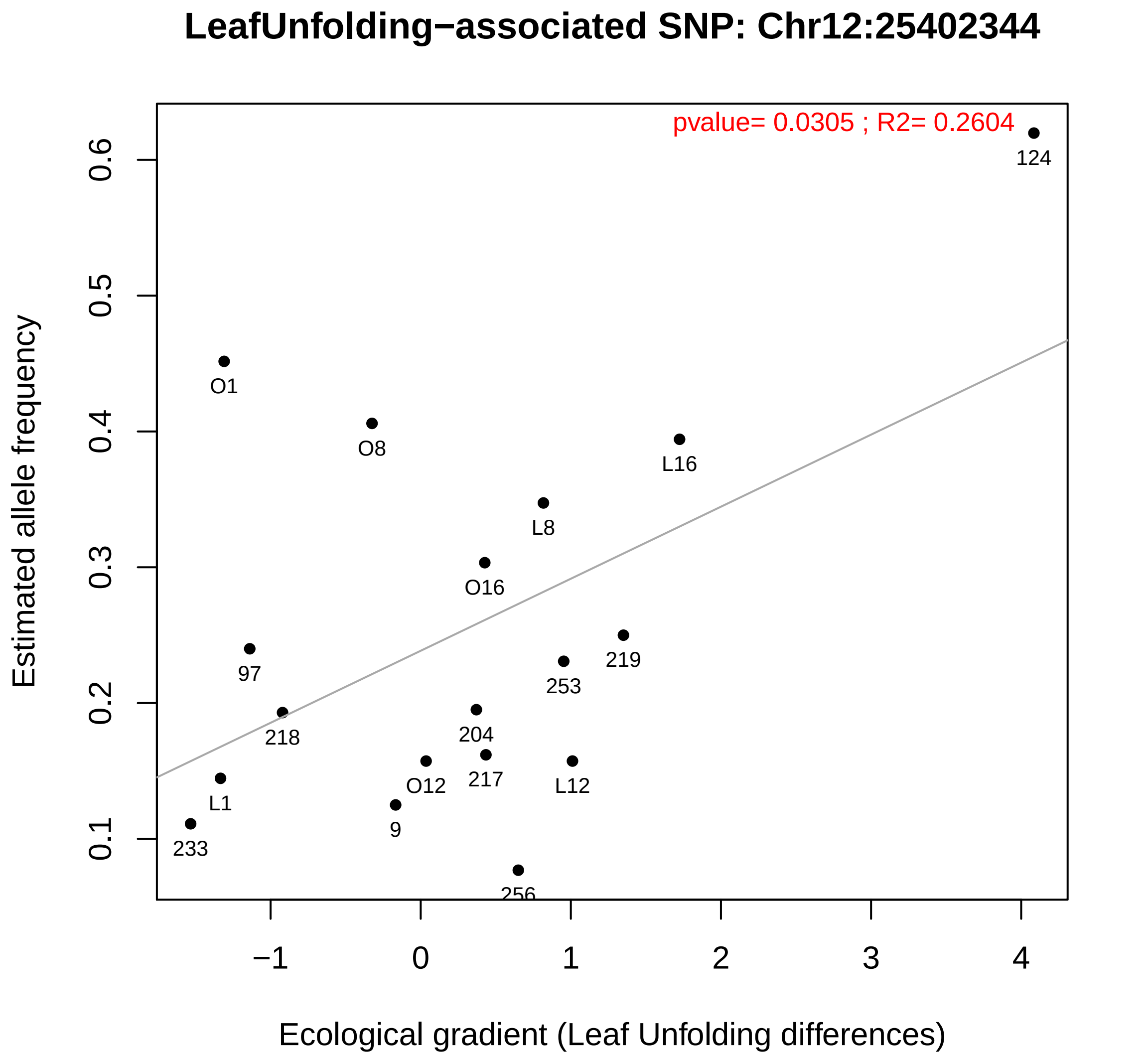
LeafUnfolding−associated SNP: Chr12:25402344
pvalue= 0.0305 ; R2= 0.2604
Estimated allele frequency
0.1
0.2
0.3
0.4
0.5
0.6
Ecological gradient (Leaf Unfolding differences)
−1
0
1
2
3
4
124
O1
O8
L16
L8
O16
97
219
253
218
204
L1
O12
217
L12
233
9
256

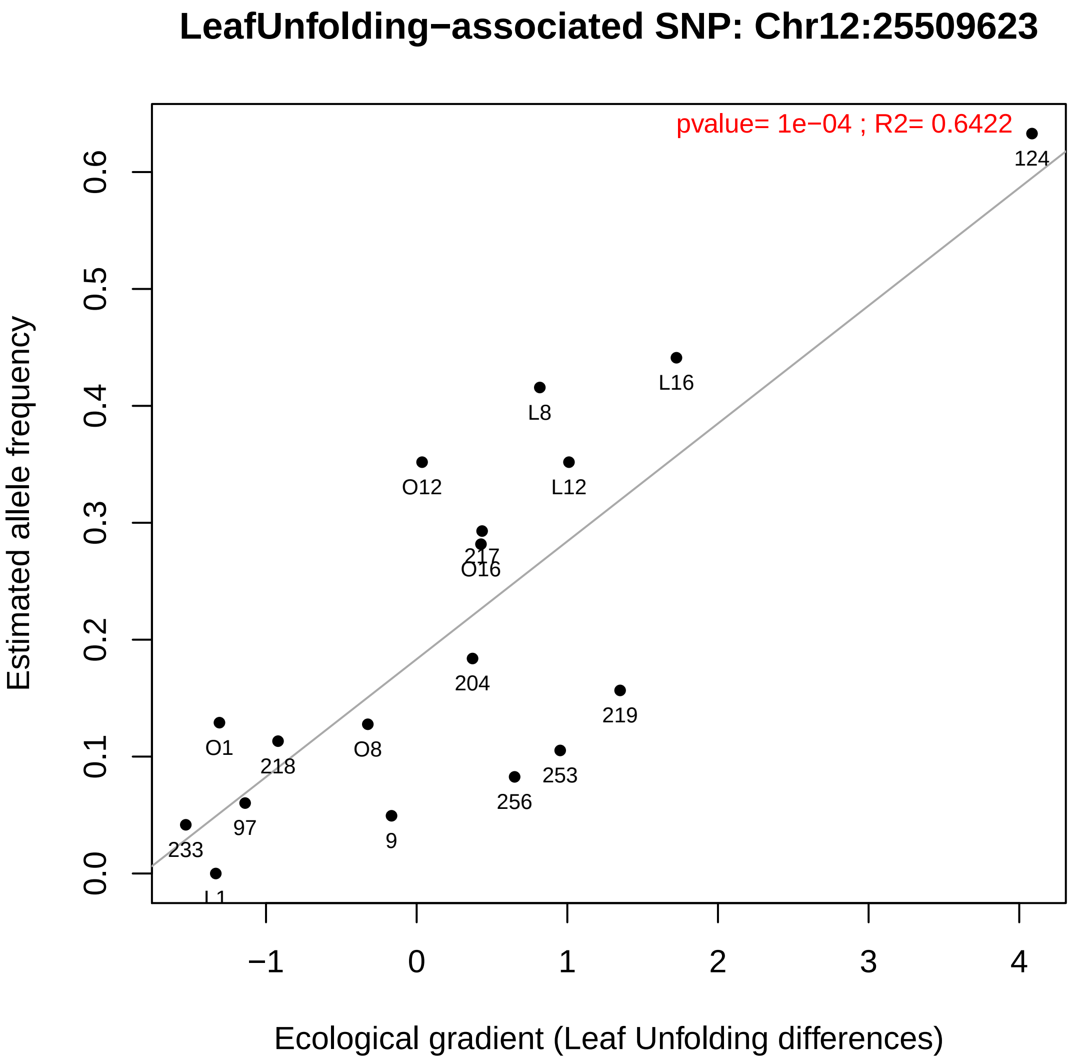

LeafUnfolding−associated SNP: Chr12:25509623
pvalue= 1e−04 ; R2= 0.6422
Estimated allele frequency
0.0
0.1
0.2
0.3
0.4
0.5
0.6
−1
0
1
2
3
4
Ecological gradient (Leaf Unfolding differences)
124
L16
L8
O12
L12
217
O16
204
219
O1
218
O8
253
256
97
9
233
L1

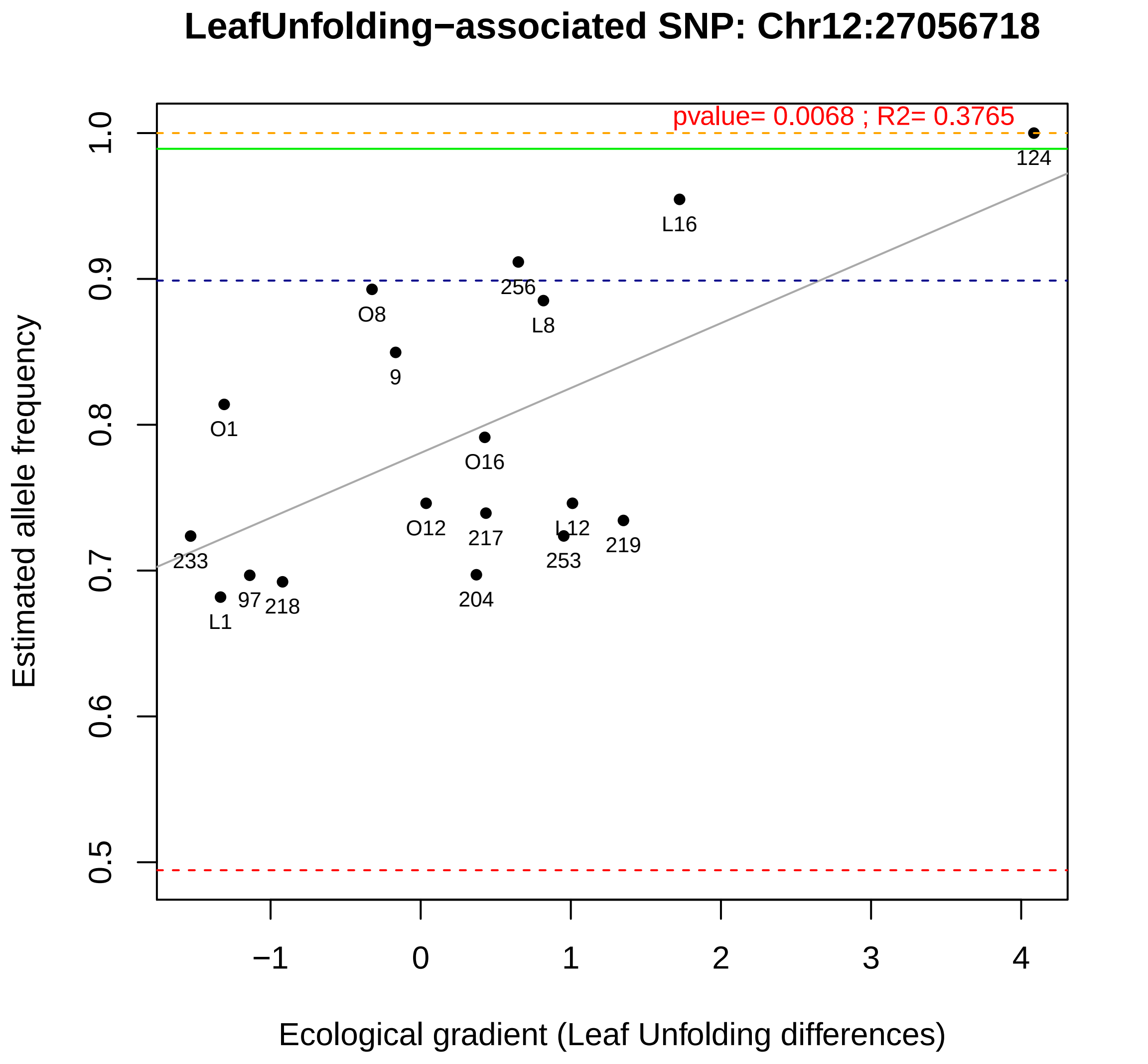
LeafUnfolding−associated SNP: Chr12:27056718
pvalue= 0.0068 ; R2= 0.3765
Estimated allele frequency
Ecological gradient (Leaf Unfolding differences)
1.0
0.9
0.8
0.7
0.6
0.5
−1
0
1
2
3
4
124
L16
256
O8
L8
9
O1
O16
O12
217
L12
219
253
233
97
218
L1
204

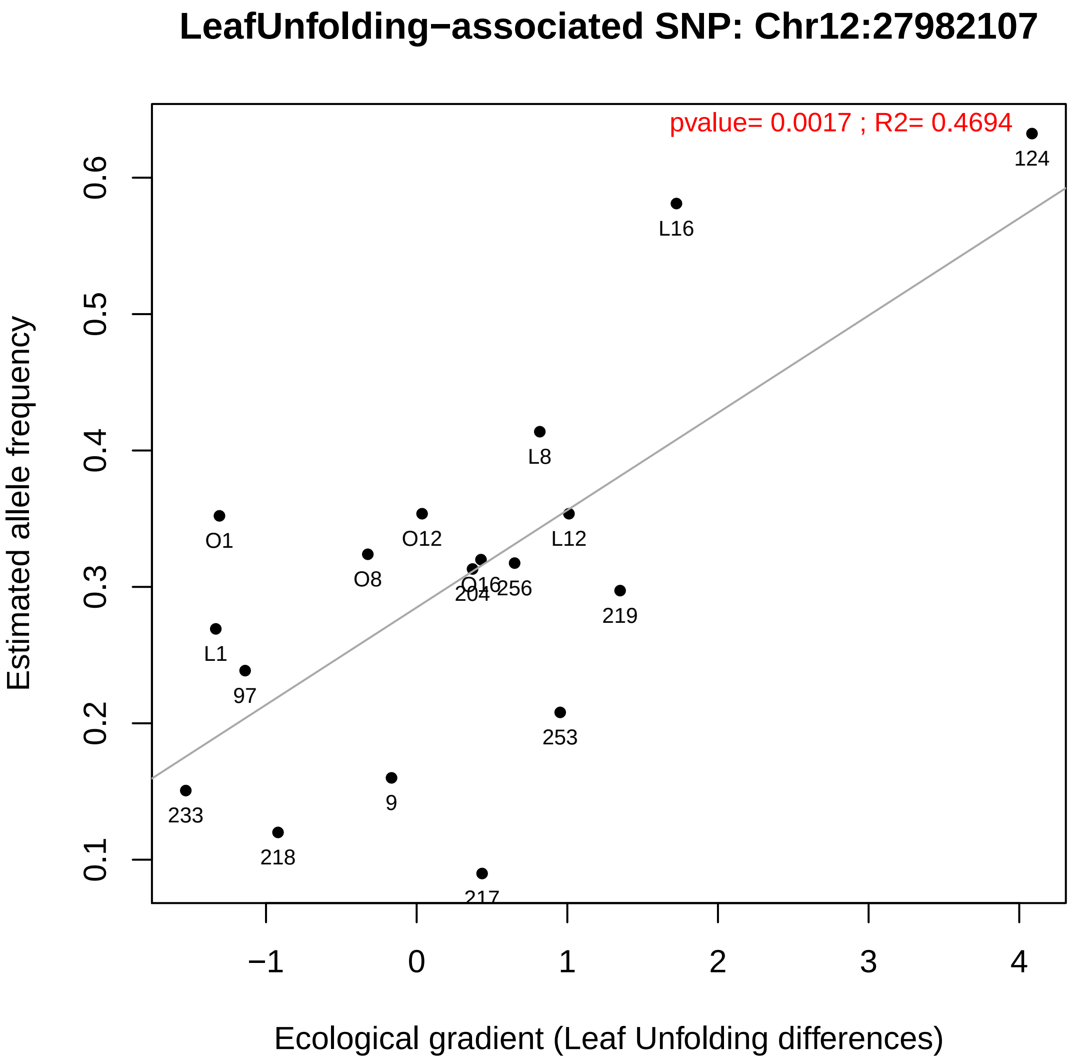

LeafUnfolding−associated SNP: Chr12:27982107
pvalue= 0.0017 ; R2= 0.4694
Estimated allele frequency
Ecological gradient (Leaf Unfolding differences)
0.1
0.2
0.3
0.4
0.5
0.6
−1
0
1
2
3
4
124
L16
L8
O1
O12
L12
O8
O16
204
256
219
L1
97
253
233
9
218
217

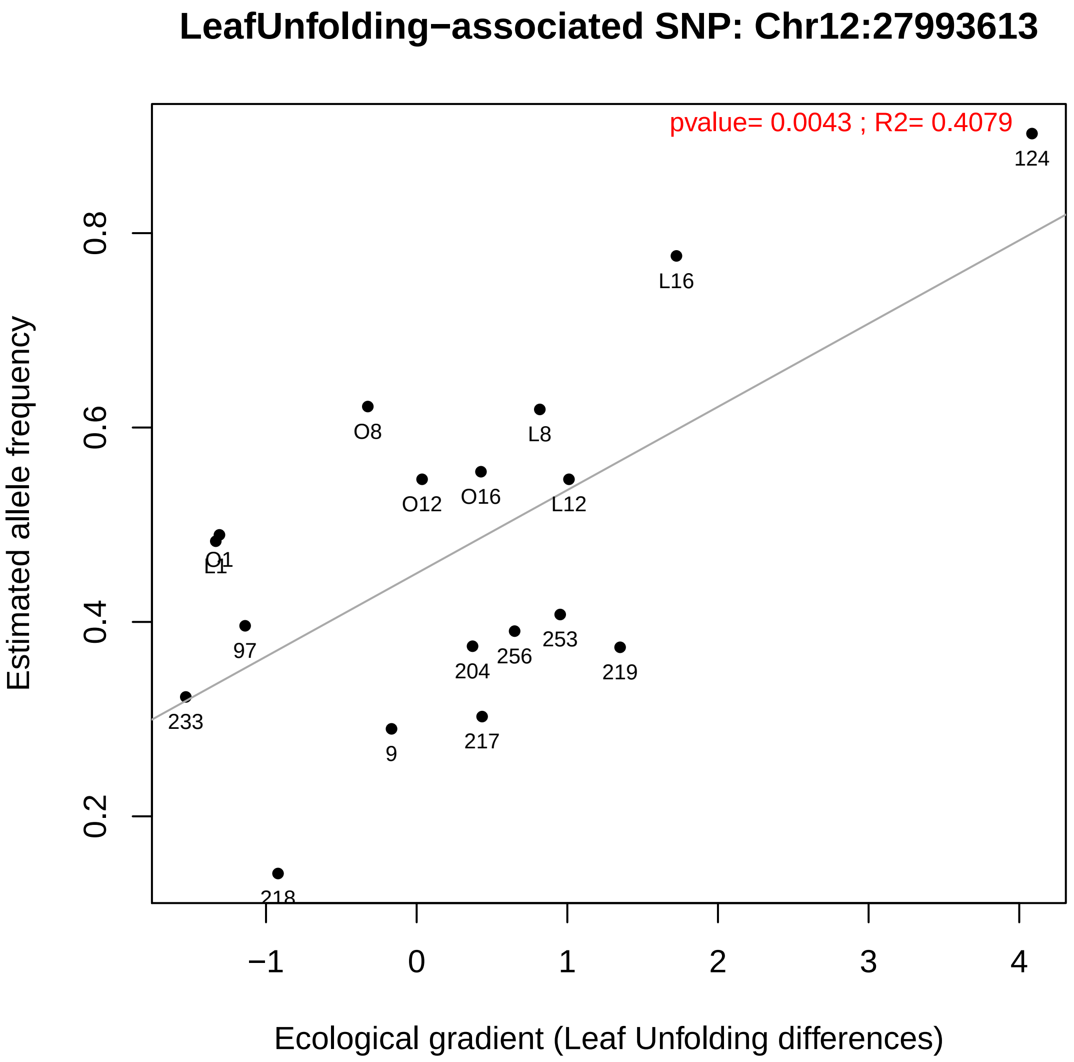
LeafUnfolding−associated SNP: Chr12:27993613
pvalue= 0.0043 ; R2= 0.4079
Estimated allele frequency
0.2
0.4
0.6
0.8
−1
0
1
2
3
4
Ecological gradient (Leaf Unfolding differences)
124
L16
O8
L8
O12
O16
L12
O1
L1
97
233
204
256
253
219
9
217
218

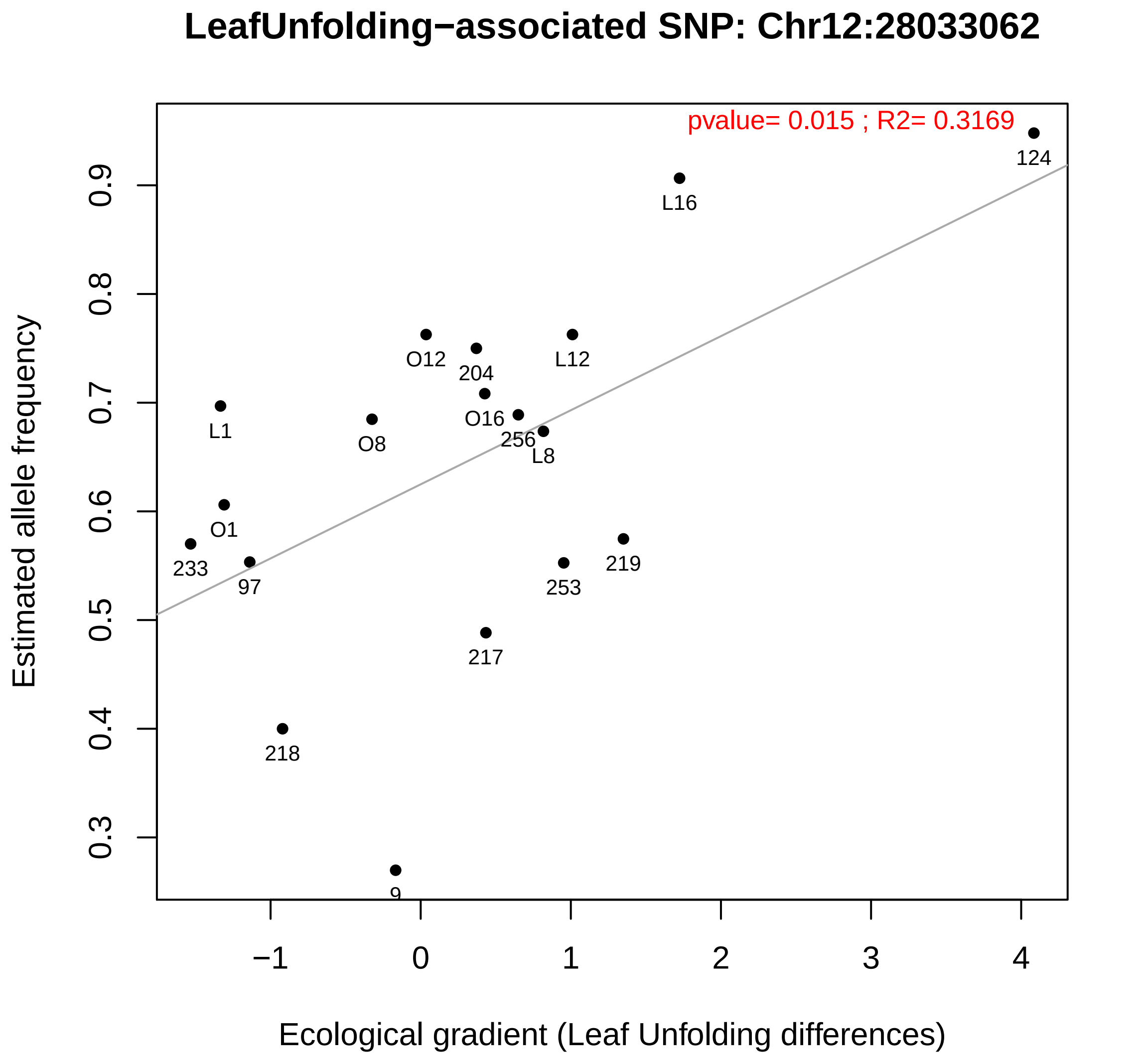

LeafUnfolding−associated SNP: Chr12:28033062
pvalue= 0.015 ; R2= 0.3169
Estimated allele frequency
Ecological gradient (Leaf Unfolding differences)
0.3
0.4
0.5
0.6
0.7
0.8
0.9
−1
0
1
2
3
4
124
L16
O12
204
L12
L1
O8
O16
256
L8
O1
233
97
219
253
217
218
9

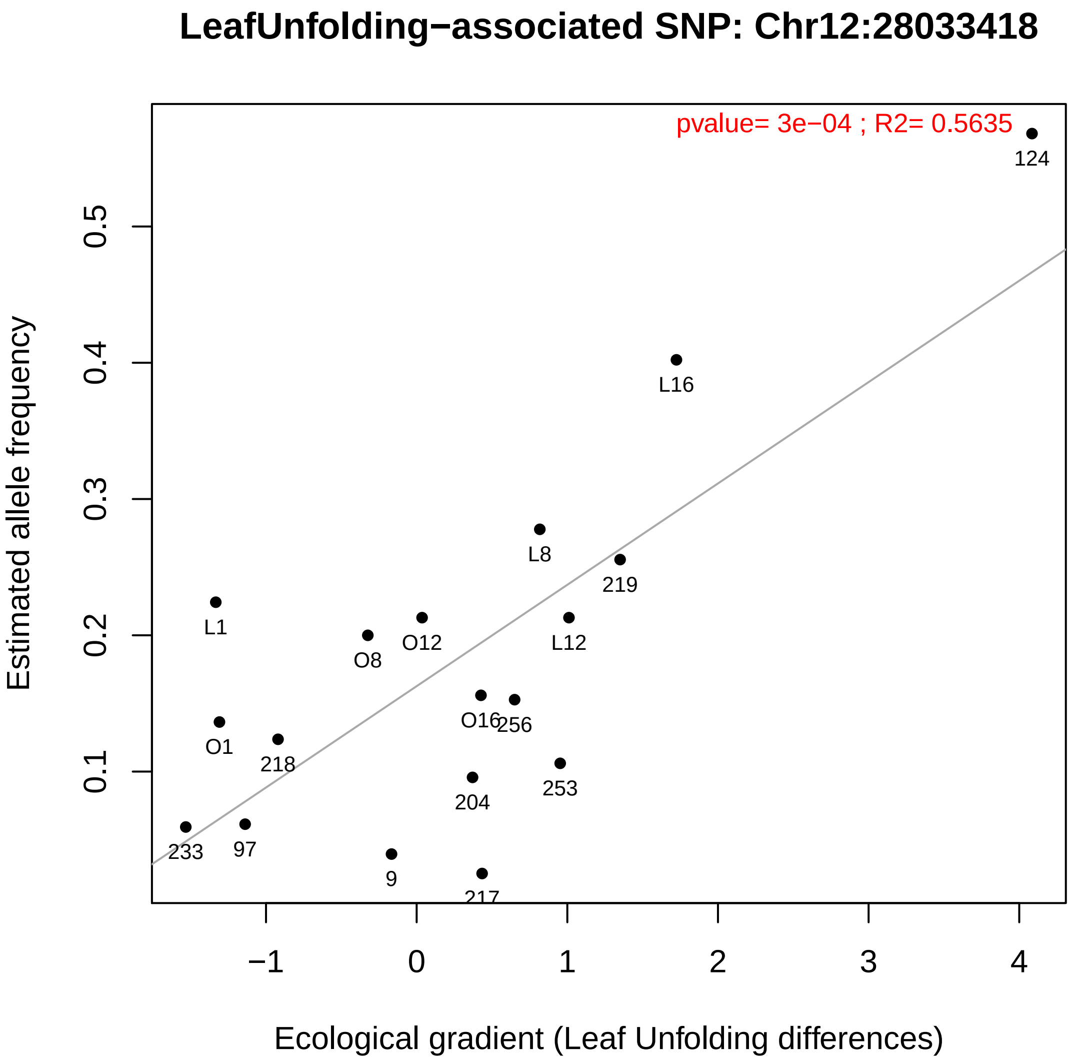
LeafUnfolding−associated SNP: Chr12:28033418
pvalue= 3e−04 ; R2= 0.5635
Estimated allele frequency
Ecological gradient (Leaf Unfolding differences)
0.1
0.2
0.3
0.4
0.5
−1
0
1
2
3
4
124
L16
L8
219
L1
O12
L12
O8
O16
256
O1
218
253
204
233
97
9
217

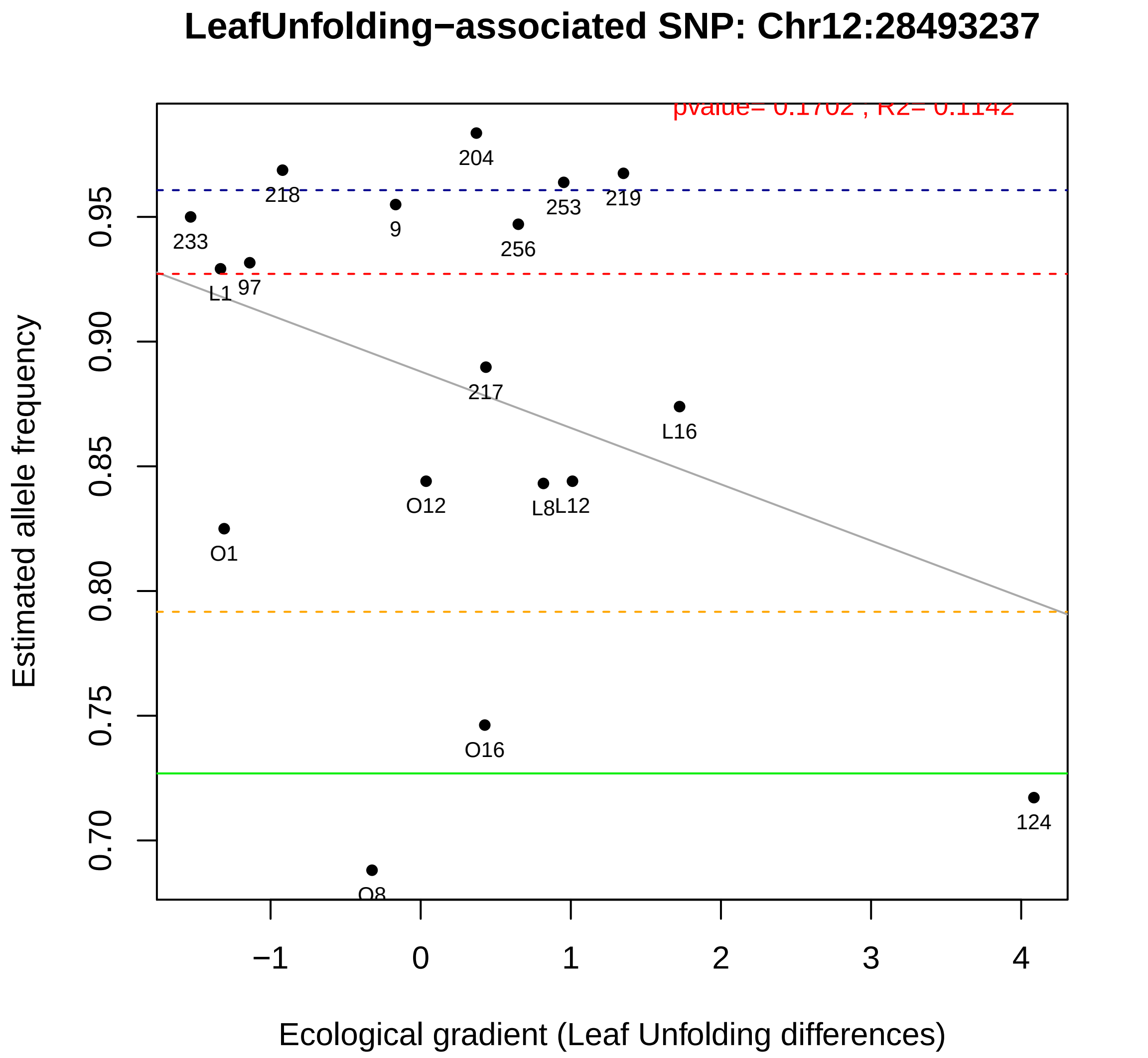
LeafUnfolding−associated SNP: Chr12:28493237
pvalue= 0.1702 , R2= 0.1142
Estimated allele frequency
0.70
0.75
0.80
0.85
0.90
0.95
−1
0
1
2
3
4
Ecological gradient (Leaf Unfolding differences)
204
218
253
219
9
233
256
L1
97
217
L16
O12
L8
L12
O1
O16
124
O8

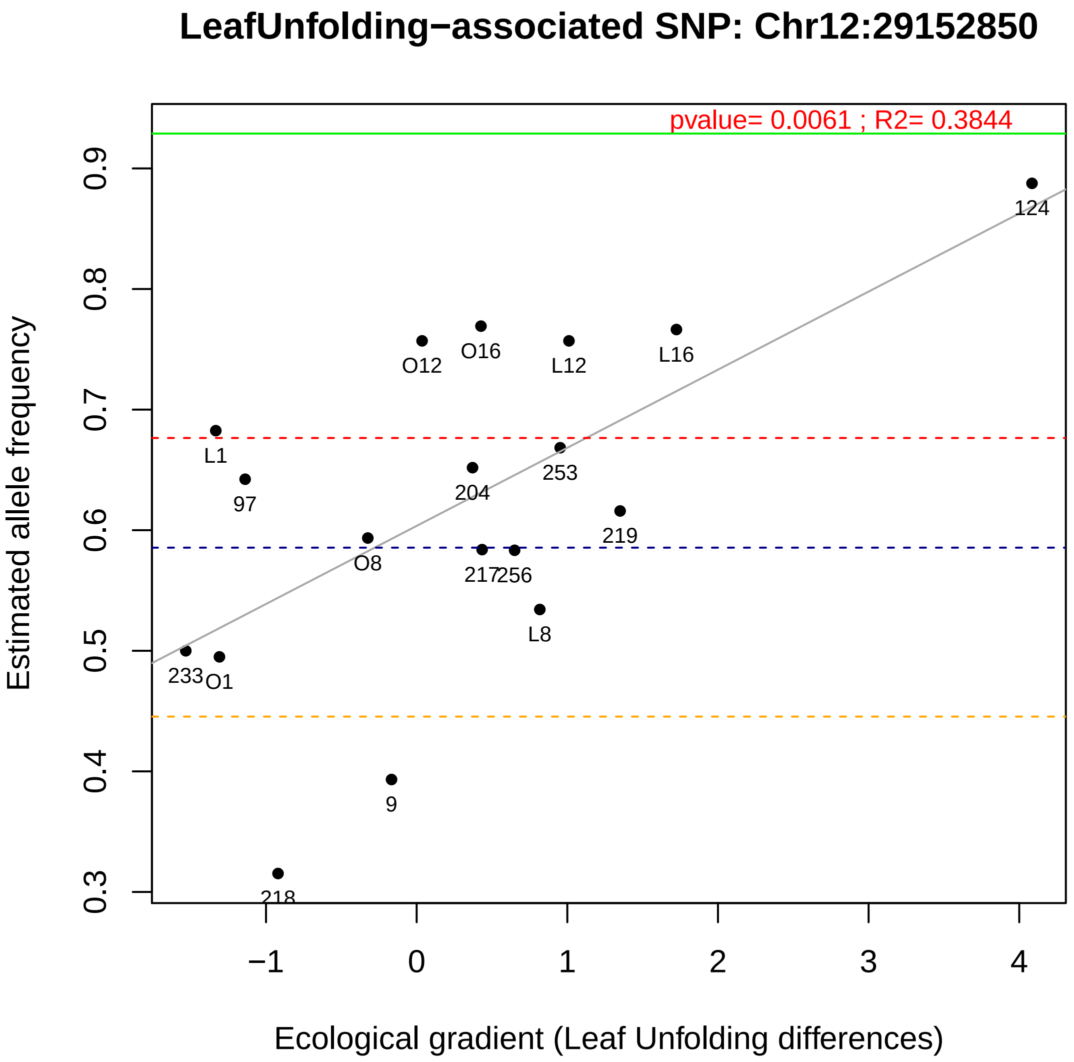

LeafUnfolding−associated SNP: Chr12:29152850
pvalue= 0.0061 ; R2= 0.3844
Estimated allele frequency
0.3
0.4
0.5
0.6
0.7
0.8
0.9
−1
0
1
2
3
4
Ecological gradient (Leaf Unfolding differences)
124
O12
O16
L12
L16
L1
97
204
253
219
O8
217
256
L8
233
O1
9
218

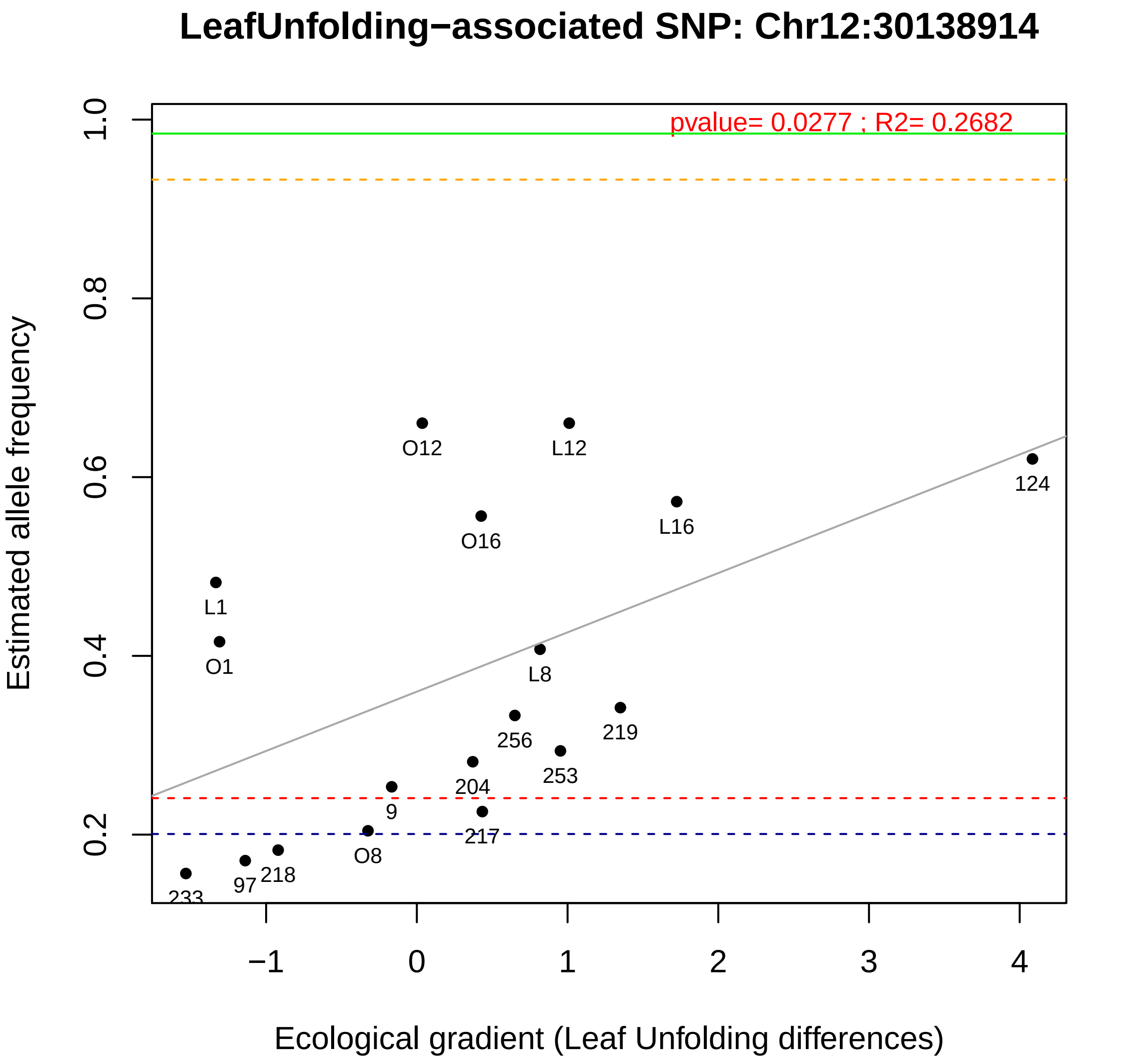
LeafUnfolding−associated SNP: Chr12:30138914
pvalue= 0.0277 ; R2= 0.2682
Estimated allele frequency
1.0
0.8
0.6
0.4
0.2
O12
L12
124
O16
L16
L1
O1
L8
256
219
253
204
9
217
O8
233
97
218
−1
0
1
2
3
4
Ecological gradient (Leaf Unfolding differences)

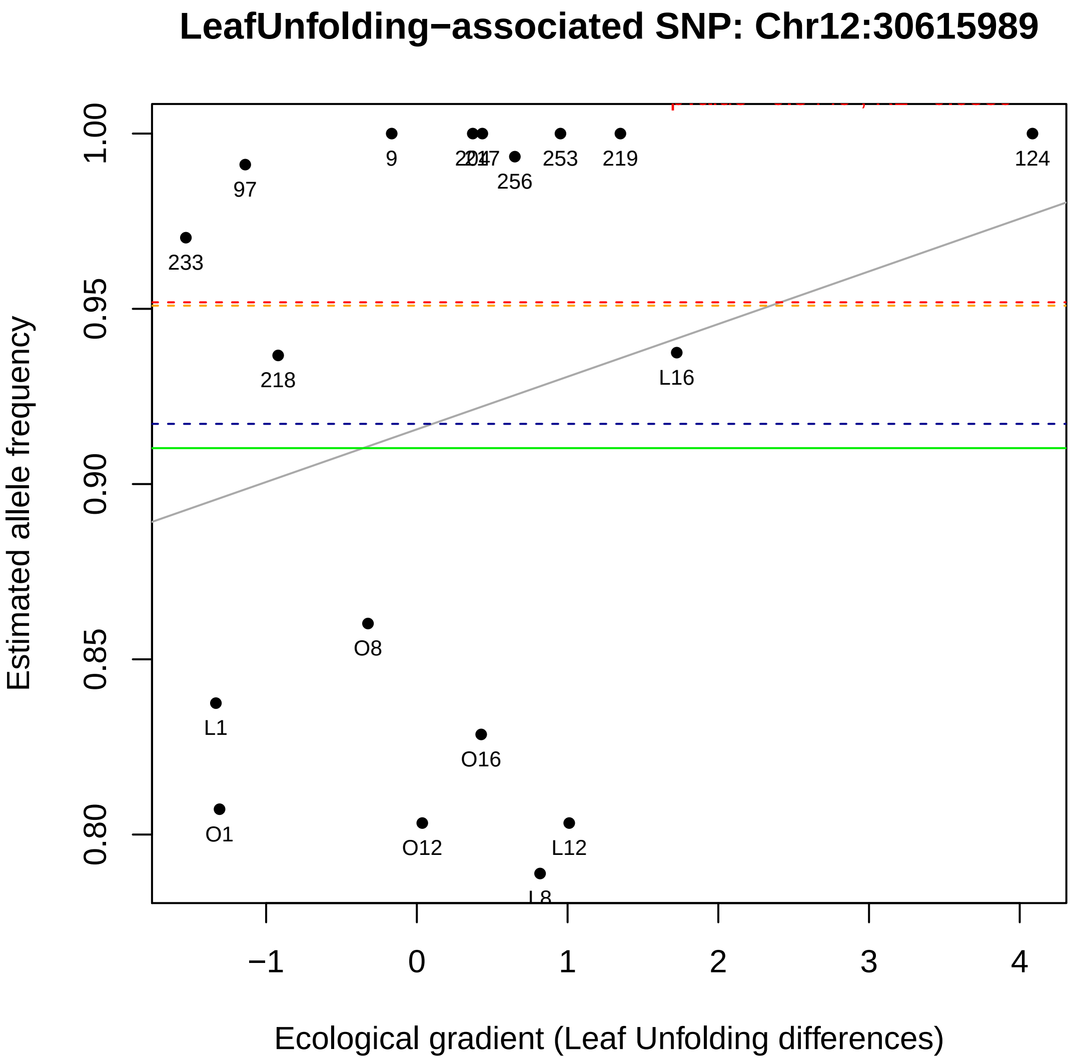

LeafUnfolding−associated SNP: Chr12:30615989
Estimated allele frequency
Ecological gradient (Leaf Unfolding differences)
0.80
0.85
0.90
0.95
1.00
−1
0
1
2
3
4
9
204
217
256
253
219
124
97
233
218
L16
O8
L1
O16
O1
O12
L12

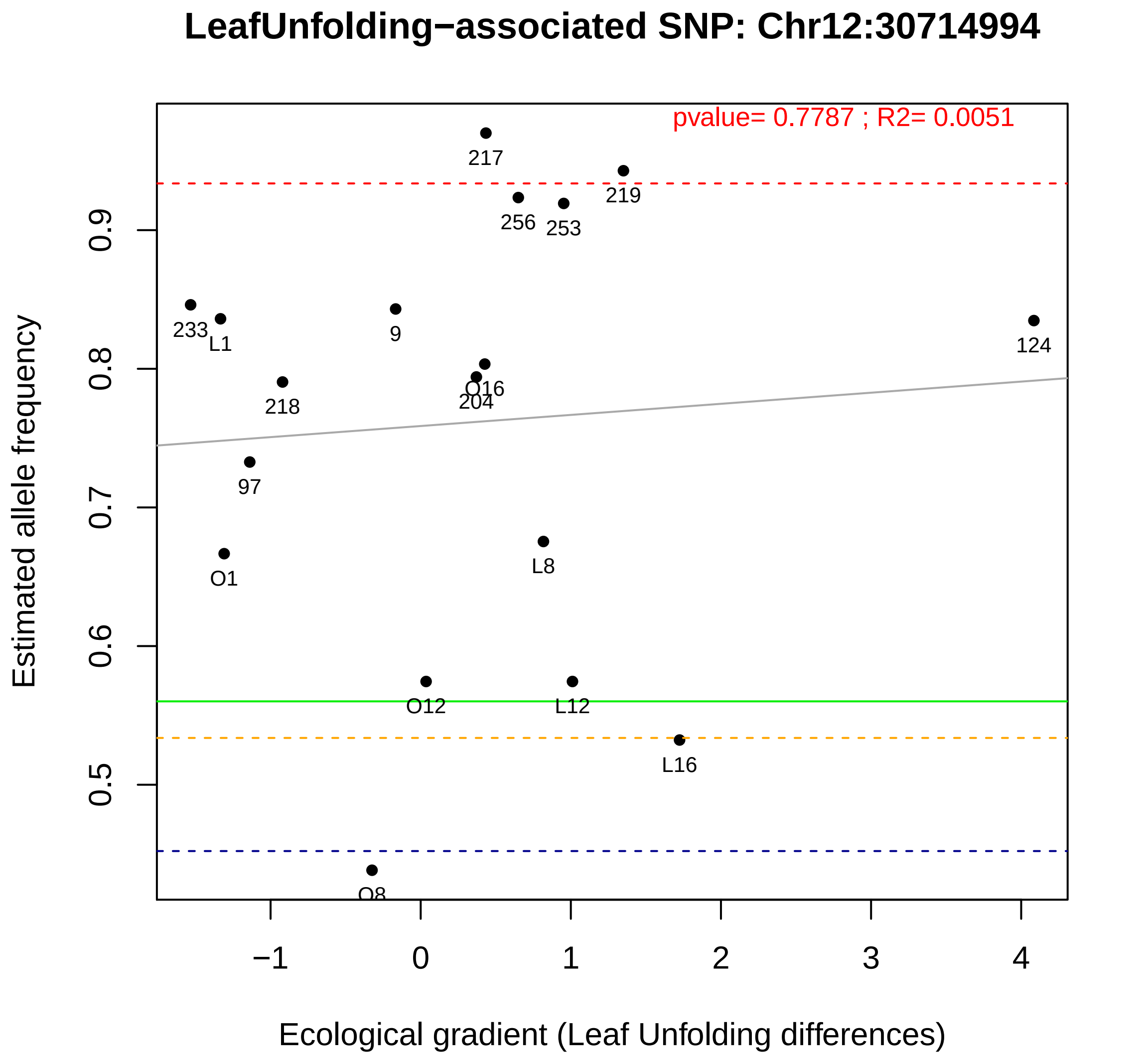

LeafUnfolding−associated SNP: Chr12:30714994
pvalue= 0.7787 ; R2= 0.0051
Estimated allele frequency
0.5
0.6
0.7
0.8
0.9
−1
0
1
2
3
4
Ecological gradient (Leaf Unfolding differences)
217
219
256
253
233
L1
9
124
218
O16
204
97
O1
L8
O12
L12
L16
O8

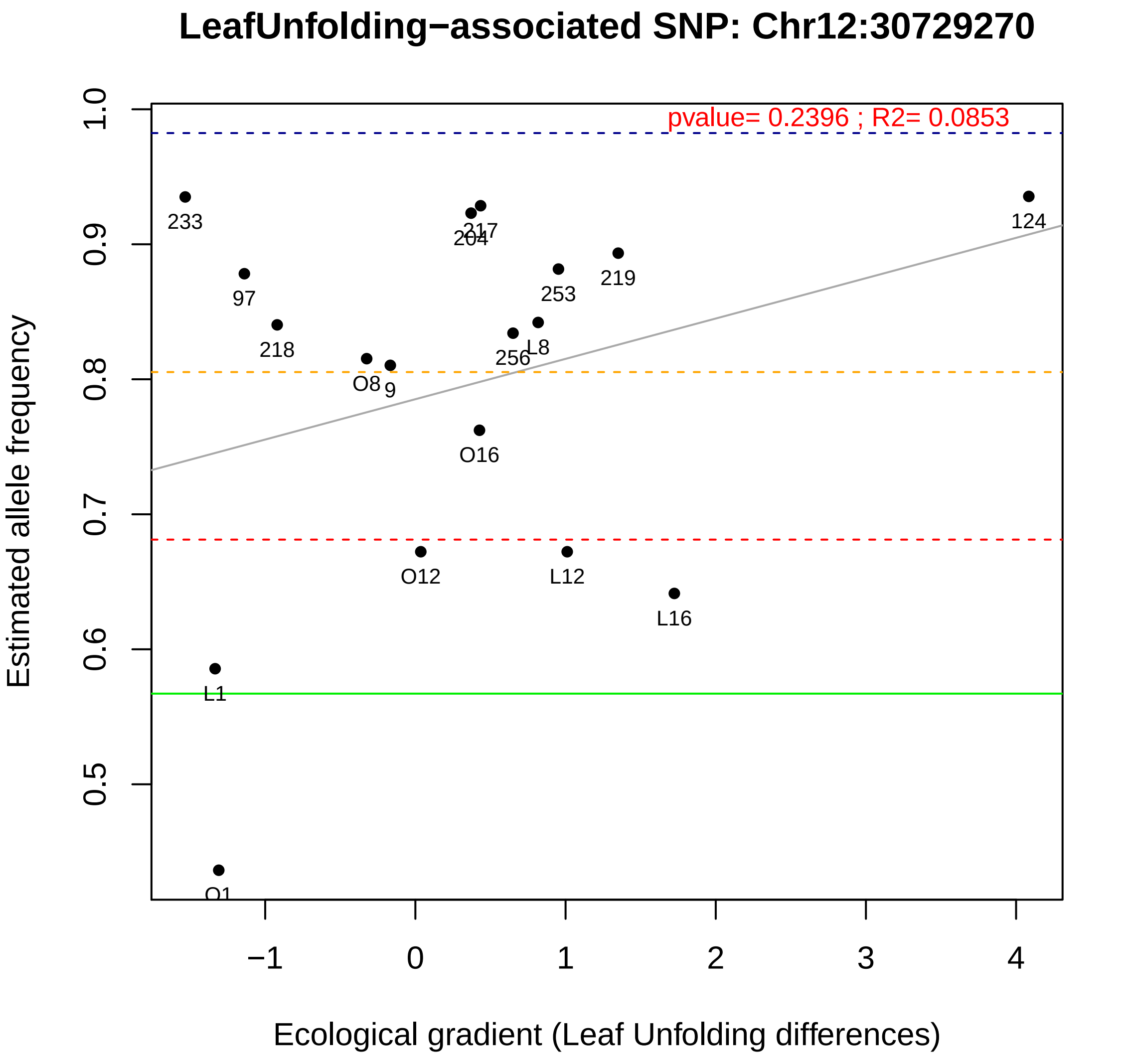
LeafUnfolding−associated SNP: Chr12:30729270
pvalue= 0.2396 ; R2= 0.0853
Estimated allele frequency
Ecological gradient (Leaf Unfolding differences)
1.0
0.9
0.8
0.7
0.6
0.5
−1
0
1
2
3
4
233
97
218
O8
9
204
217
O16
256
L8
253
219
124
O12
L12
L16
L1
O1

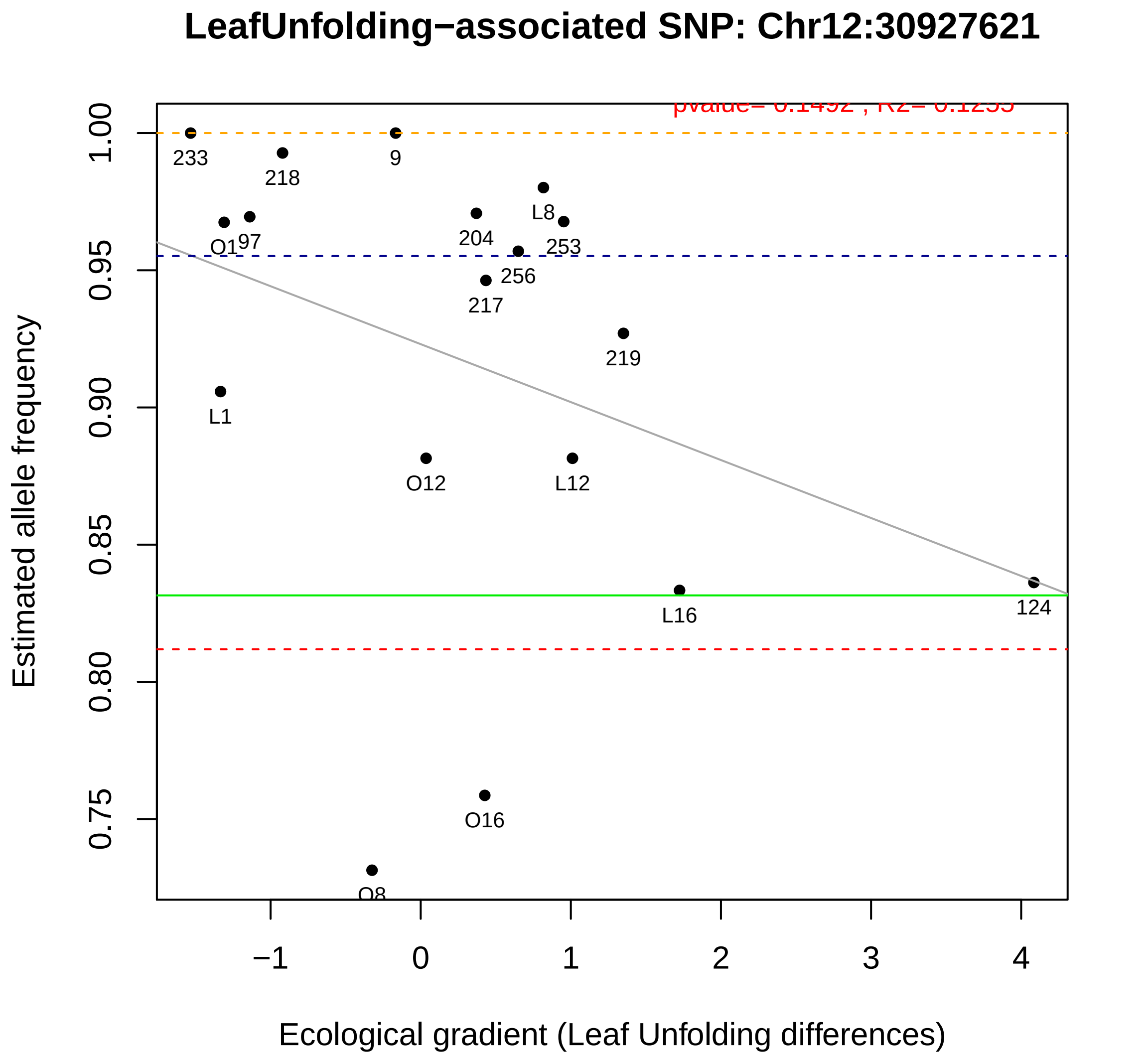

LeafUnfolding−associated SNP: Chr12:30927621
pvalue= 0.1492 ; R2= 0.1235
Estimated allele frequency
Ecological gradient (Leaf Unfolding differences)
1.00
0.95
0.90
0.85
0.80
0.75
−1
0
1
2
3
4
233
218
9
L8
204
253
O1
97
256
217
219
L1
O12
L12
L16
124
O16
O8

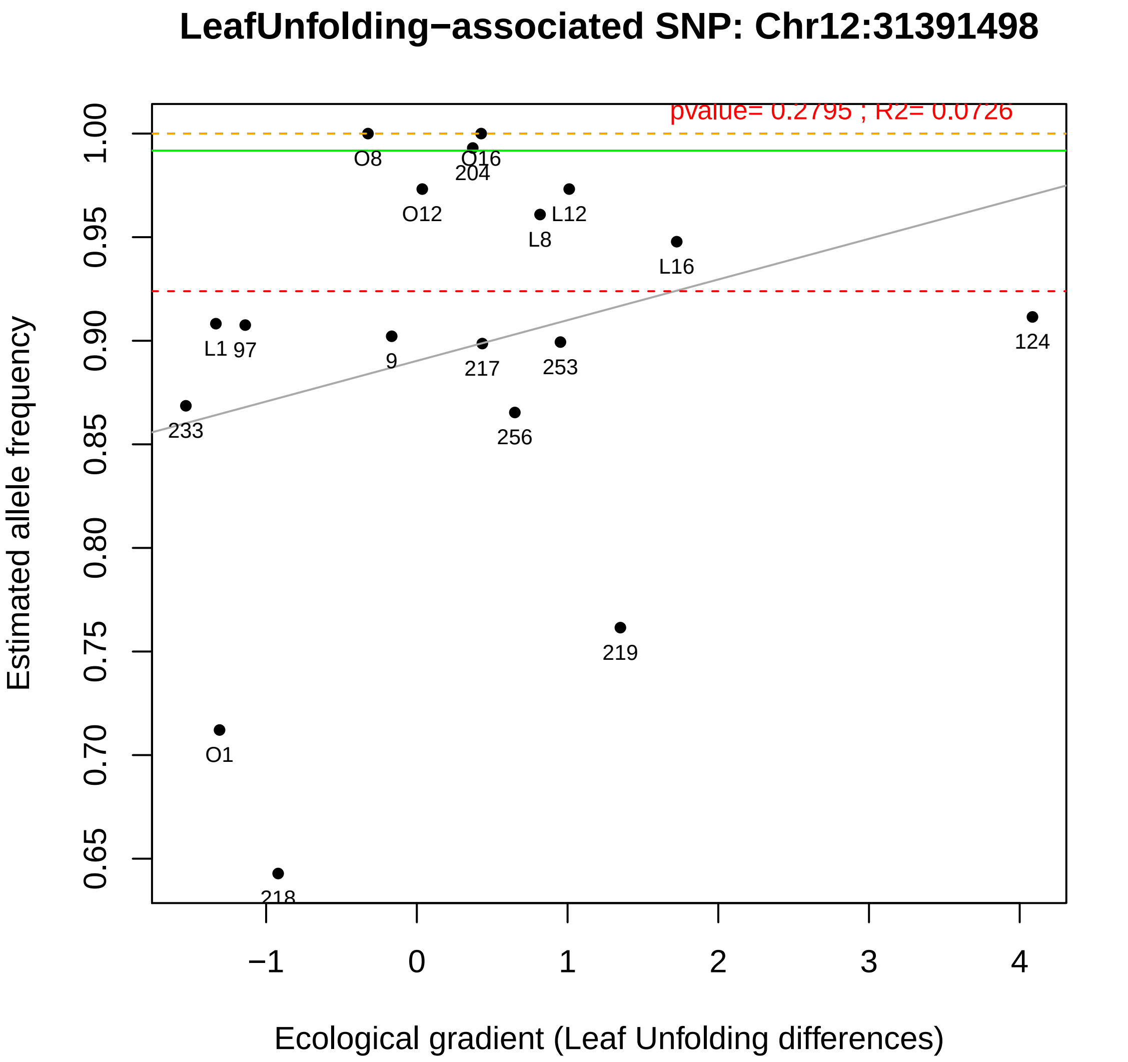
LeafUnfolding−associated SNP: Chr12:31391498
pvalue= 0.2795 ; R2= 0.0726
Estimated allele frequency
Ecological gradient (Leaf Unfolding differences)
0.65
0.70
0.75
0.80
0.85
0.90
0.95
1.00
−1
0
1
2
3
4
O8
O16
204
O12
L12
L8
L16
L1
97
9
217
253
124
233
256
219
O1
218

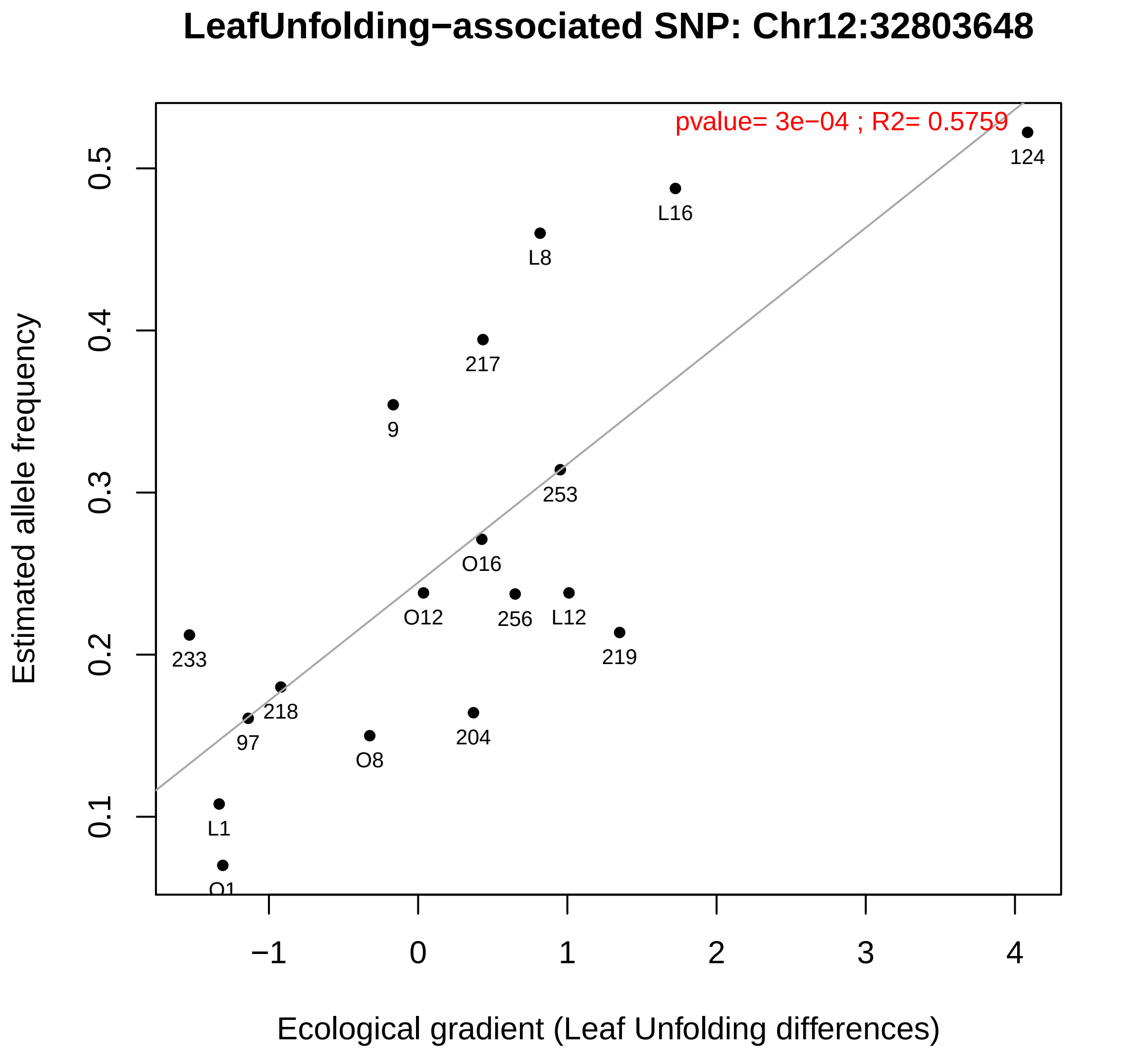
LeafUnfolding−associated SNP: Chr12:32803648
pvalue= 3e−04 ; R2= 0.5759
Estimated allele frequency
0.1
0.2
0.3
0.4
0.5
Ecological gradient (Leaf Unfolding differences)
−1
0
1
2
3
4
124
L16
L8
217
9
253
O16
O12
256
L12
219
233
218
97
O8
204
L1
O1

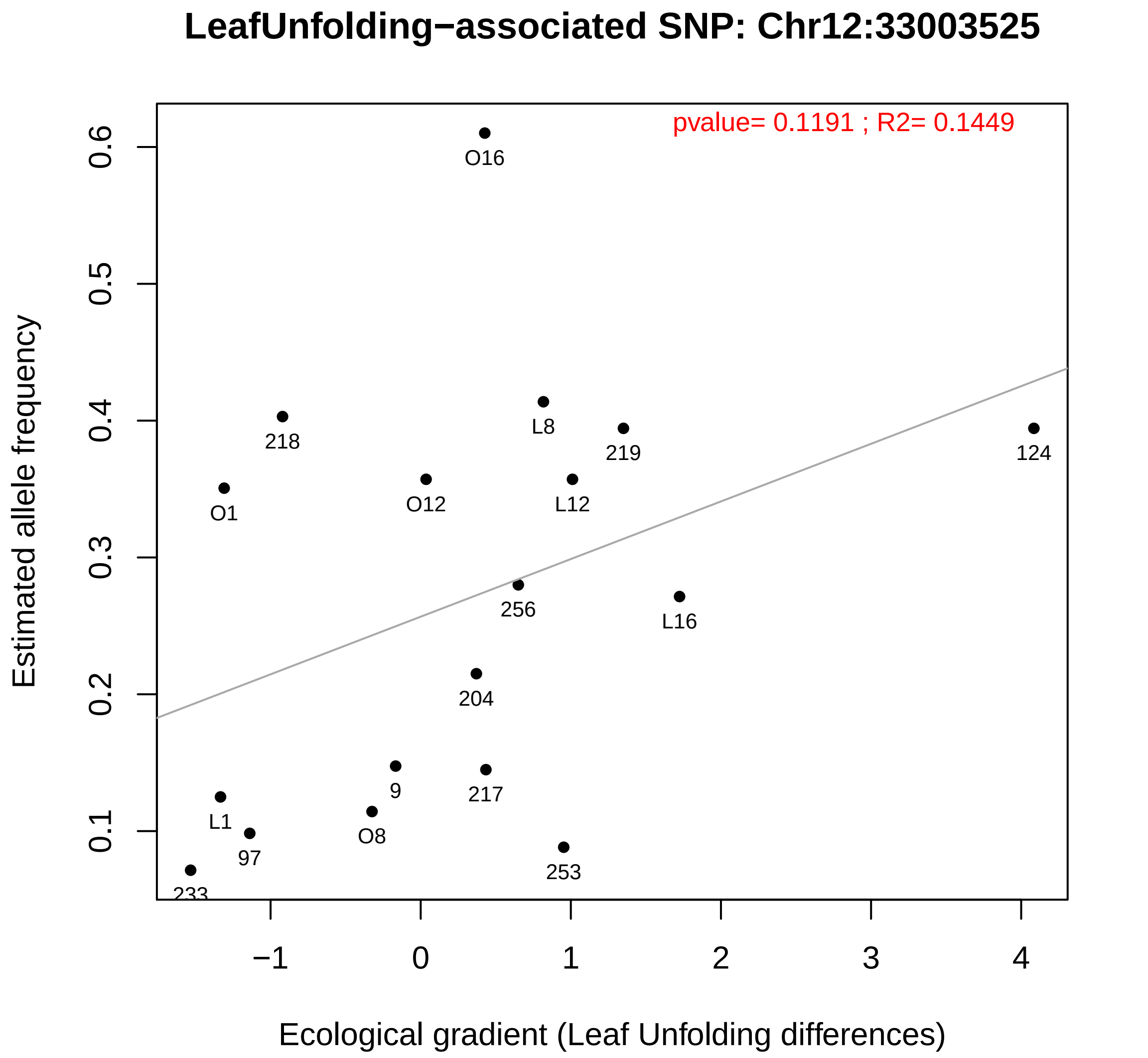
LeafUnfolding−associated SNP: Chr12:33003525
pvalue= 0.1191 ; R2= 0.1449
Estimated allele frequency
0.1
0.2
0.3
0.4
0.5
0.6
−1
0
1
2
3
4
Ecological gradient (Leaf Unfolding differences)
O16
L8
218
219
124
O1
O12
L12
256
L16
204
9
217
L1
O8
97
253
233

**LeafUnfolding−associated SNP: Chr12:33101671**

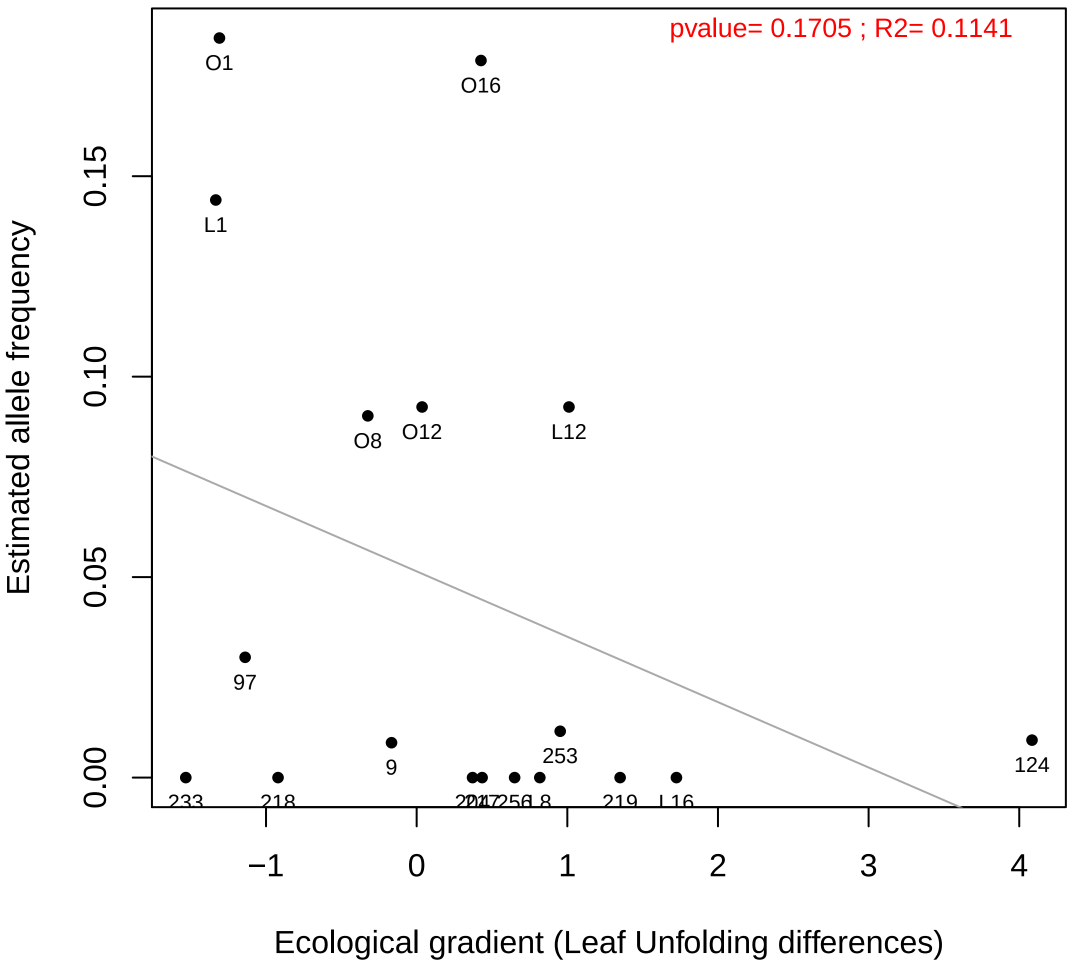

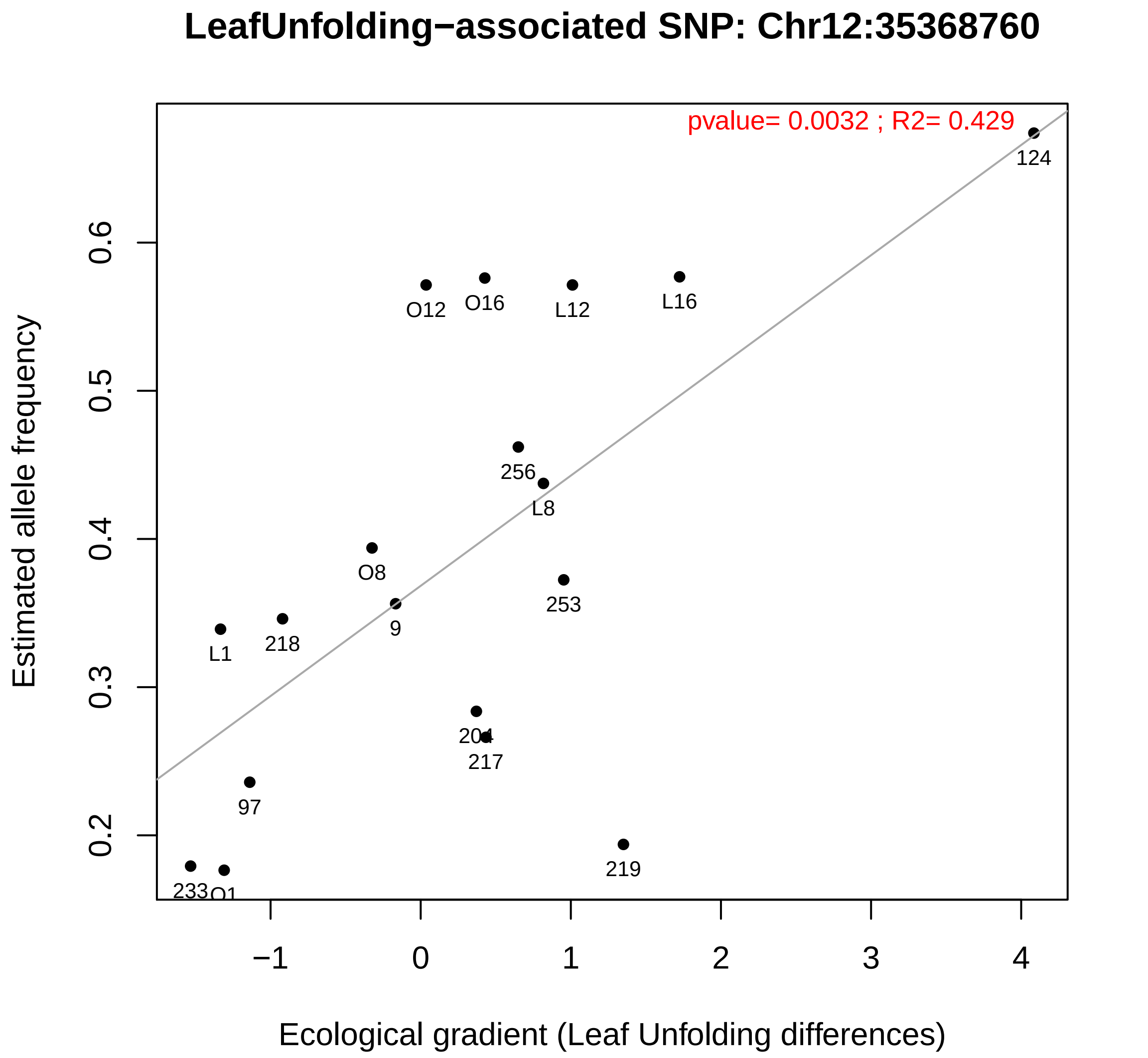

LeafUnfolding−associated SNP: Chr12:35368760
pvalue= 0.0032 ; R2= 0.429
Estimated allele frequency
Ecological gradient (Leaf Unfolding differences)
0.2
0.3
0.4
0.5
0.6
−1
0
1
2
3
4
124
O12
O16
L12
L16
256
L8
O8
253
9
L1
218
204
217
97
219
233
O1

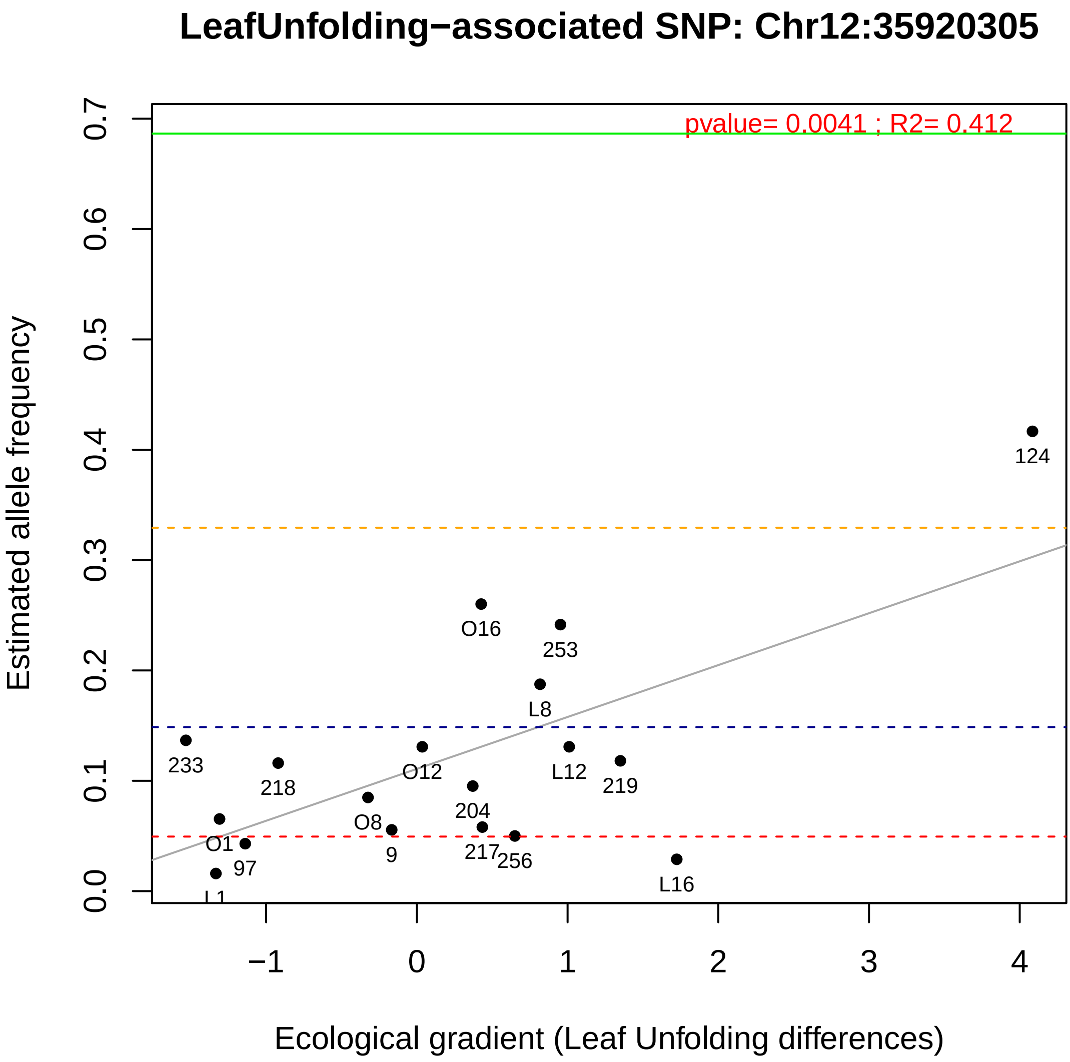

LeafUnfolding−associated SNP: Chr12:35920305
pvalue= 0.0041 ; R2= 0.412
Estimated allele frequency
Ecological gradient (Leaf Unfolding differences)
0.0
0.1
0.2
0.3
0.4
0.5
0.6
0.7
−1
0
1
2
3
4
124
O16
253
L8
233
218
O12
L12
219
204
O8
9
O1
97
217
256
L16
L1

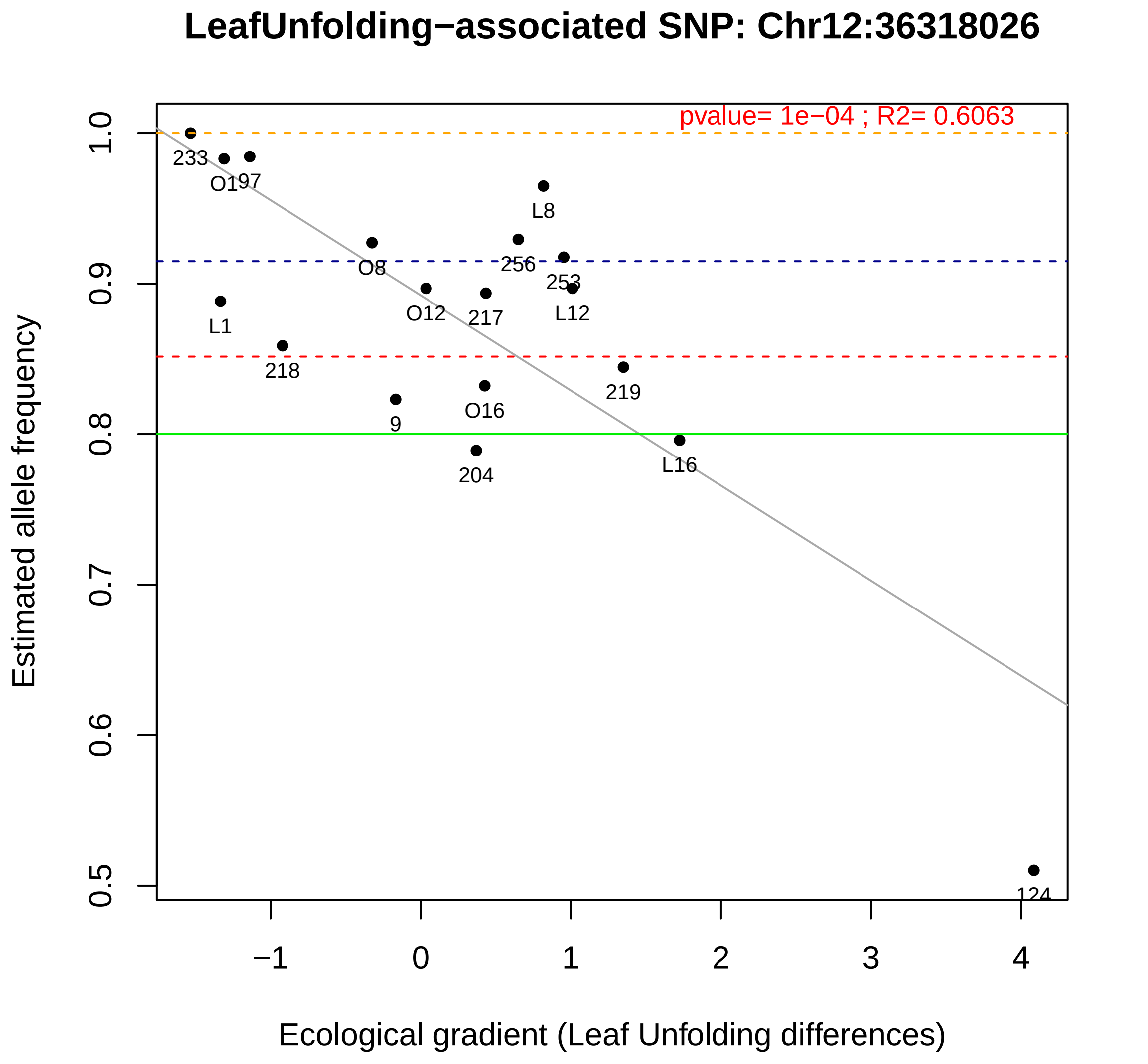
LeafUnfolding−associated SNP: Chr12:36318026
pvalue= 1e−04 ; R2= 0.6063
Estimated allele frequency
Ecological gradient (Leaf Unfolding differences)
0.5
0.6
0.7
0.8
0.9
1.0
−1
0
1
2
3
4
233
O1
97
L8
O8
256
253
L12
O12
217
L1
218
219
O16
9
204
L16
124

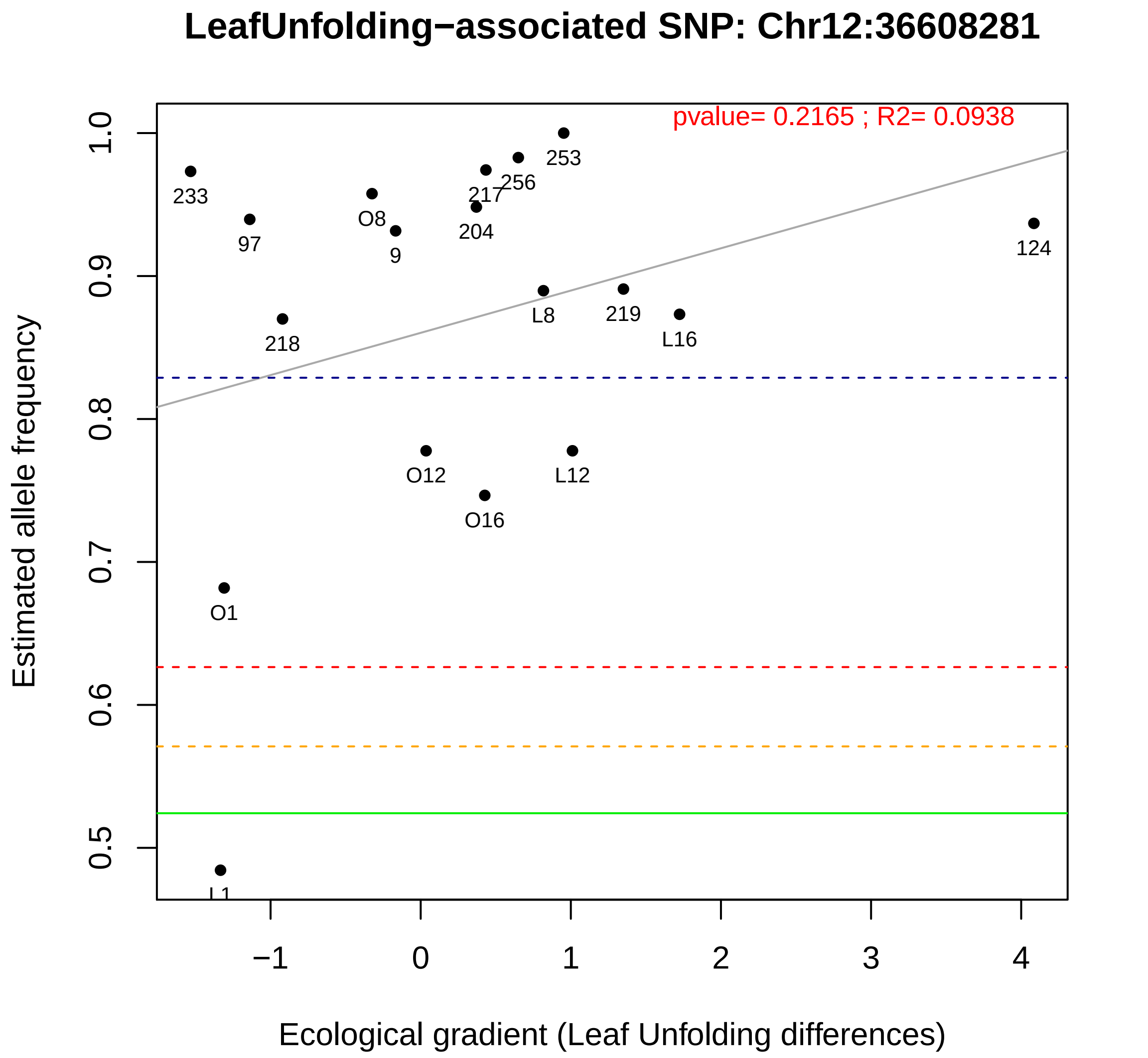
LeafUnfolding−associated SNP: Chr12:36608281
pvalue= 0.2165 ; R2= 0.0938
Estimated allele frequency
Ecological gradient (Leaf Unfolding differences)
0.5
0.6
0.7
0.8
0.9
1.0
−1
0
1
2
3
4
233
97
218
O8
9
217
256
204
253
124
L8
219
L16
O12
O16
L12
O1
L1

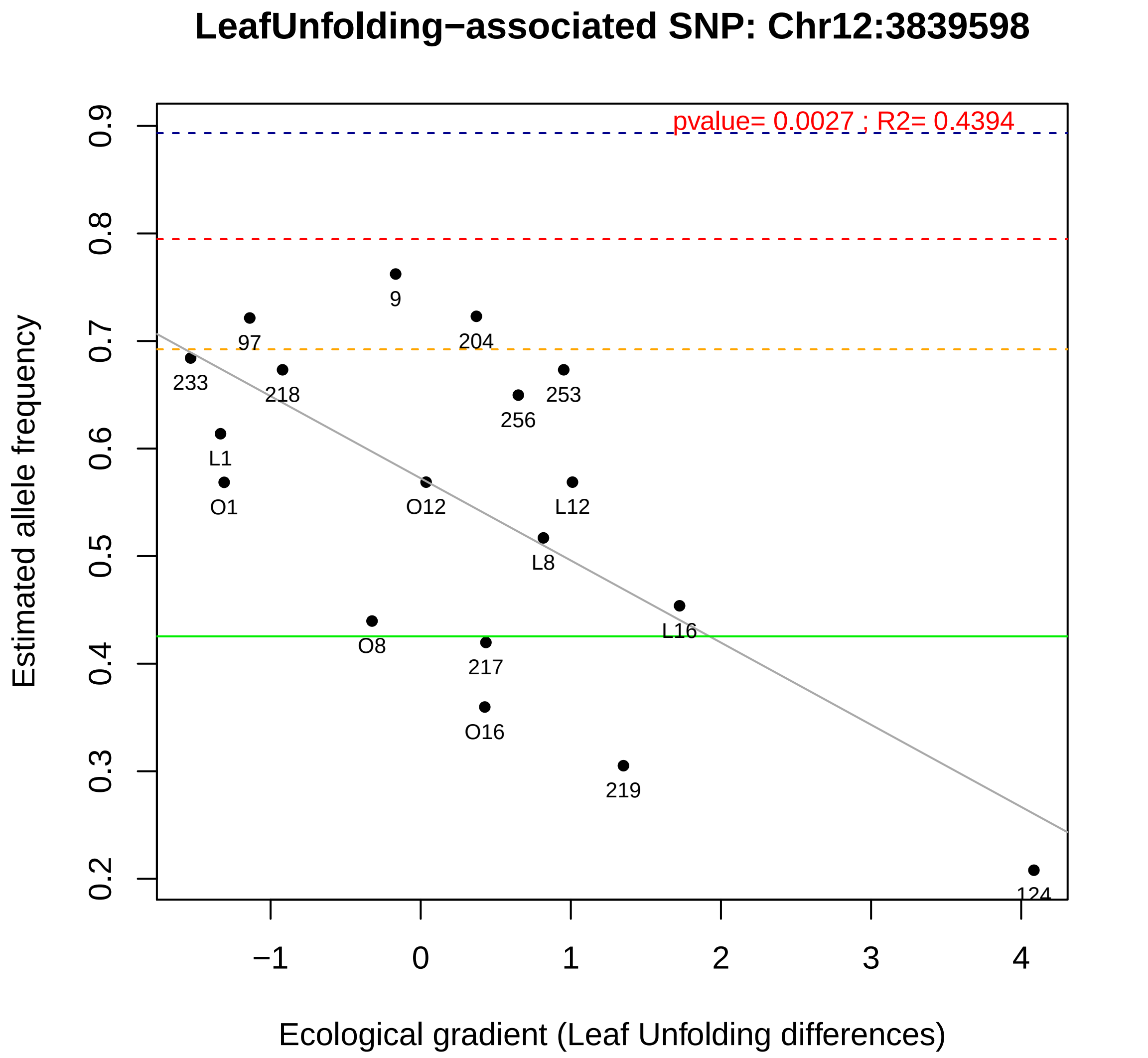

LeafUnfolding−associated SNP: Chr12:3839598
pvalue= 0.0027 ; R2= 0.4394
Estimated allele frequency
0.2
0.3
0.4
0.5
0.6
0.7
0.8
0.9
−1
0
1
2
3
4
Ecological gradient (Leaf Unfolding differences)
9
97
204
233
218
253
256
L1
O1
O12
L12
L8
L16
O8
217
O16
219
124

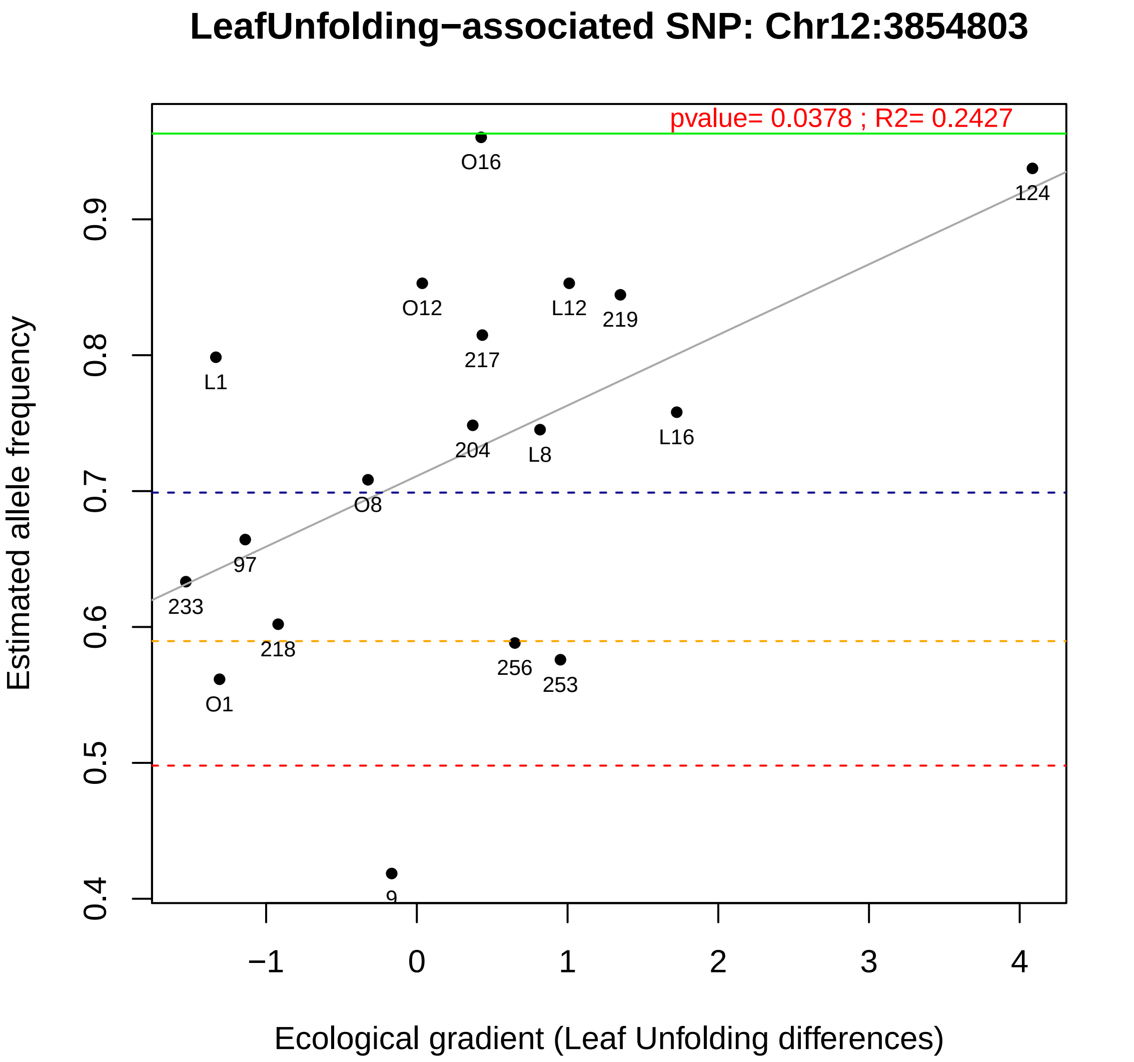
LeafUnfolding−associated SNP: Chr12:3854803
pvalue= 0.0378 ; R2= 0.2427
Estimated allele frequency
Ecological gradient (Leaf Unfolding differences)
0.4
0.5
0.6
0.7
0.8
0.9
−1
0
1
2
3
4
O16
124
O12
L12
219
217
L1
204
L8
L16
O8
97
233
218
O1
256
253
9

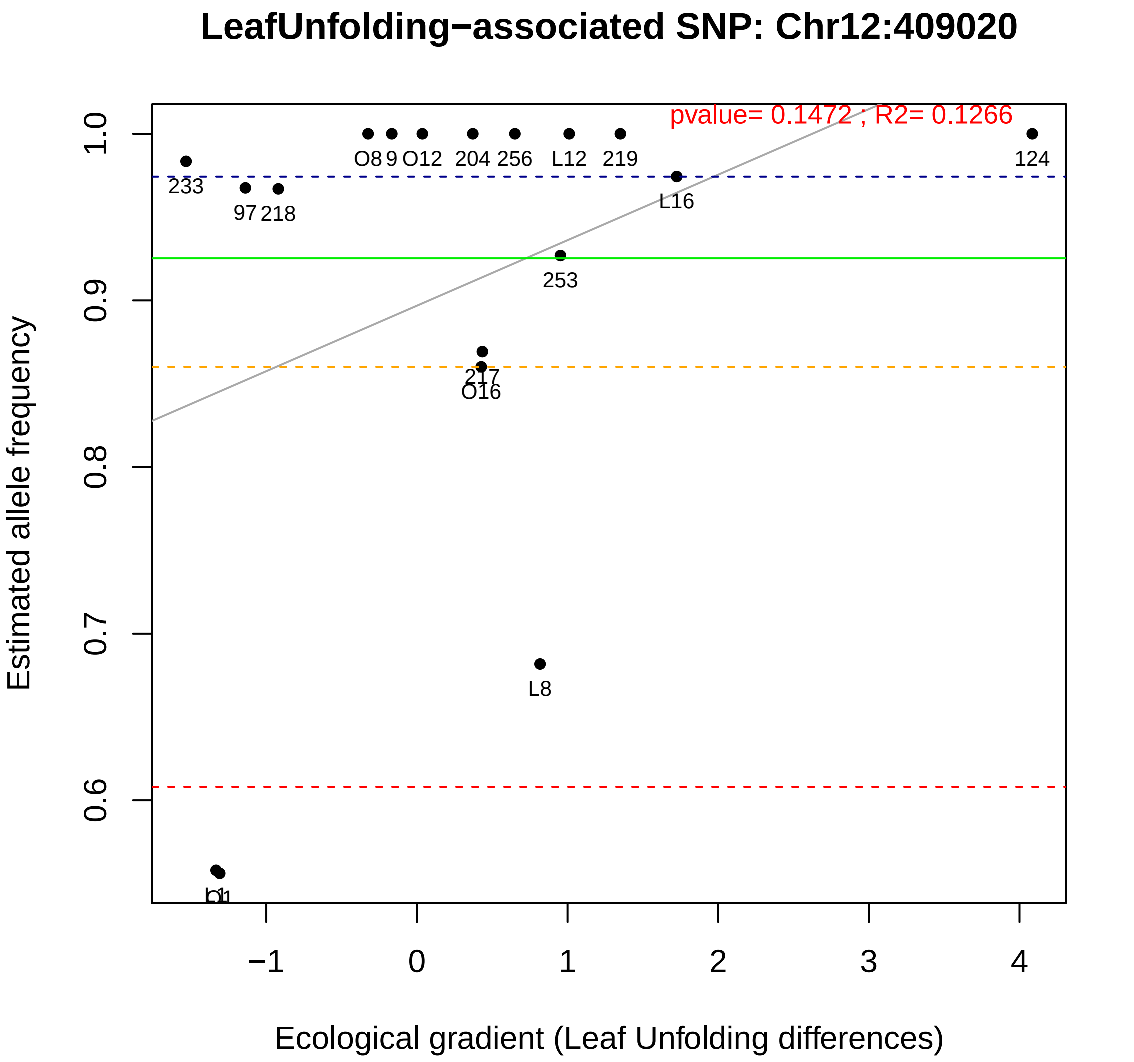

LeafUnfolding−associated SNP: Chr12:409020
pvalue= 0.1472 ; R2= 0.1266
Estimated allele frequency
Ecological gradient (Leaf Unfolding differences)
1.0
0.9
0.8
0.7
0.6
−1
0
1
2
3
4
233
97
218
O8
9
O12
204
256
L12
219
124
L16
253
217
O16
L8

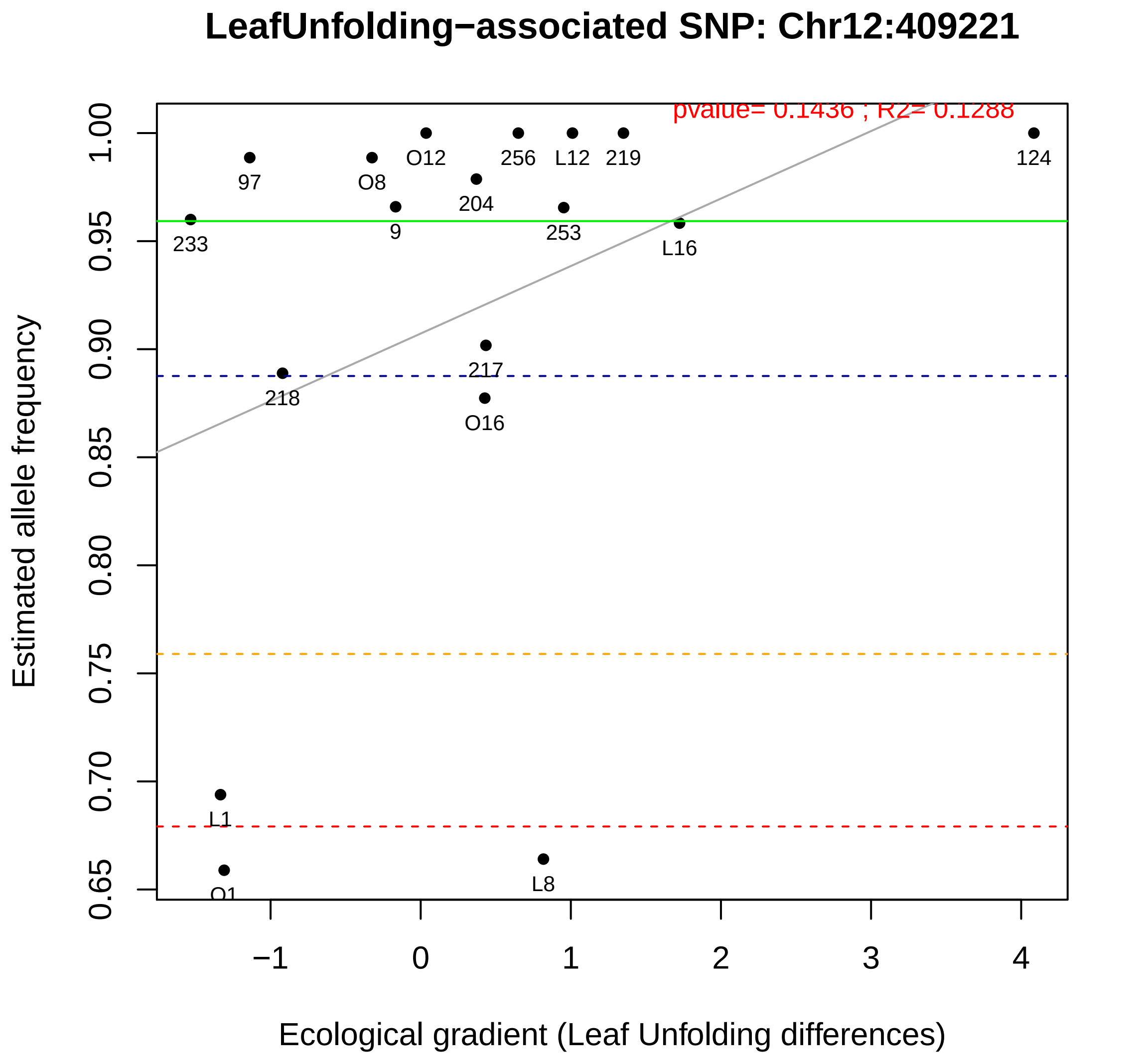
LeafUnfolding−associated SNP: Chr12:409221
pvalue= 0.1436 ; R2= 0.1288
Estimated allele frequency
Ecological gradient (Leaf Unfolding differences)
1.00
0.95
0.90
0.85
0.80
0.75
0.70
0.65
−1
0
1
2
3
4
97
O8
O12
256
L12
219
124
204
9
253
233
L16
217
218
O16
L1
O1
L8

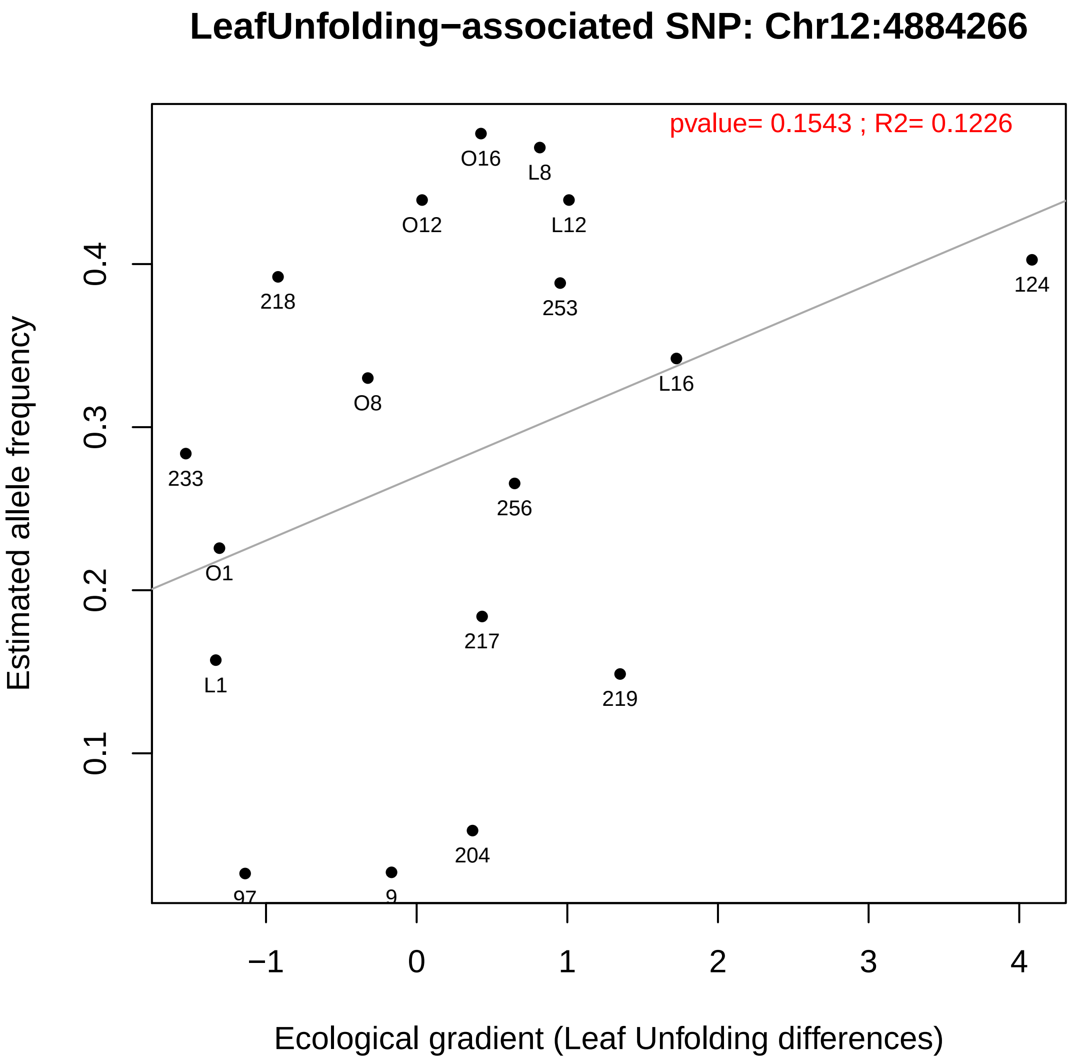
LeafUnfolding−associated SNP: Chr12:4884266
pvalue= 0.1543 ; R2= 0.1226
Estimated allele frequency
0.1
0.2
0.3
0.4
Ecological gradient (Leaf Unfolding differences)
−1
0
1
2
3
4
O16
L8
O12
L12
218
253
124
L16
O8
233
256
O1
217
L1
219
204
97
9

**LeafUnfolding−associated SNP: Chr12:6712422**

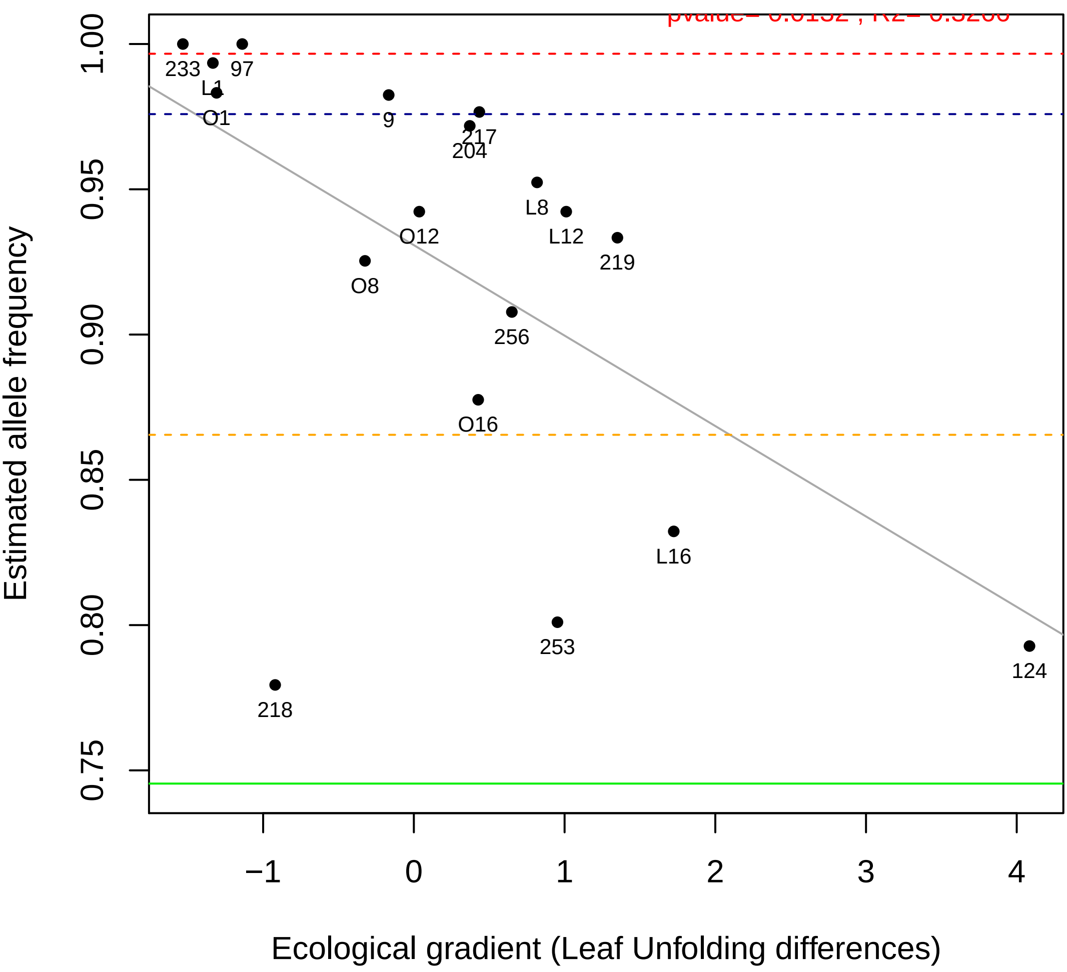

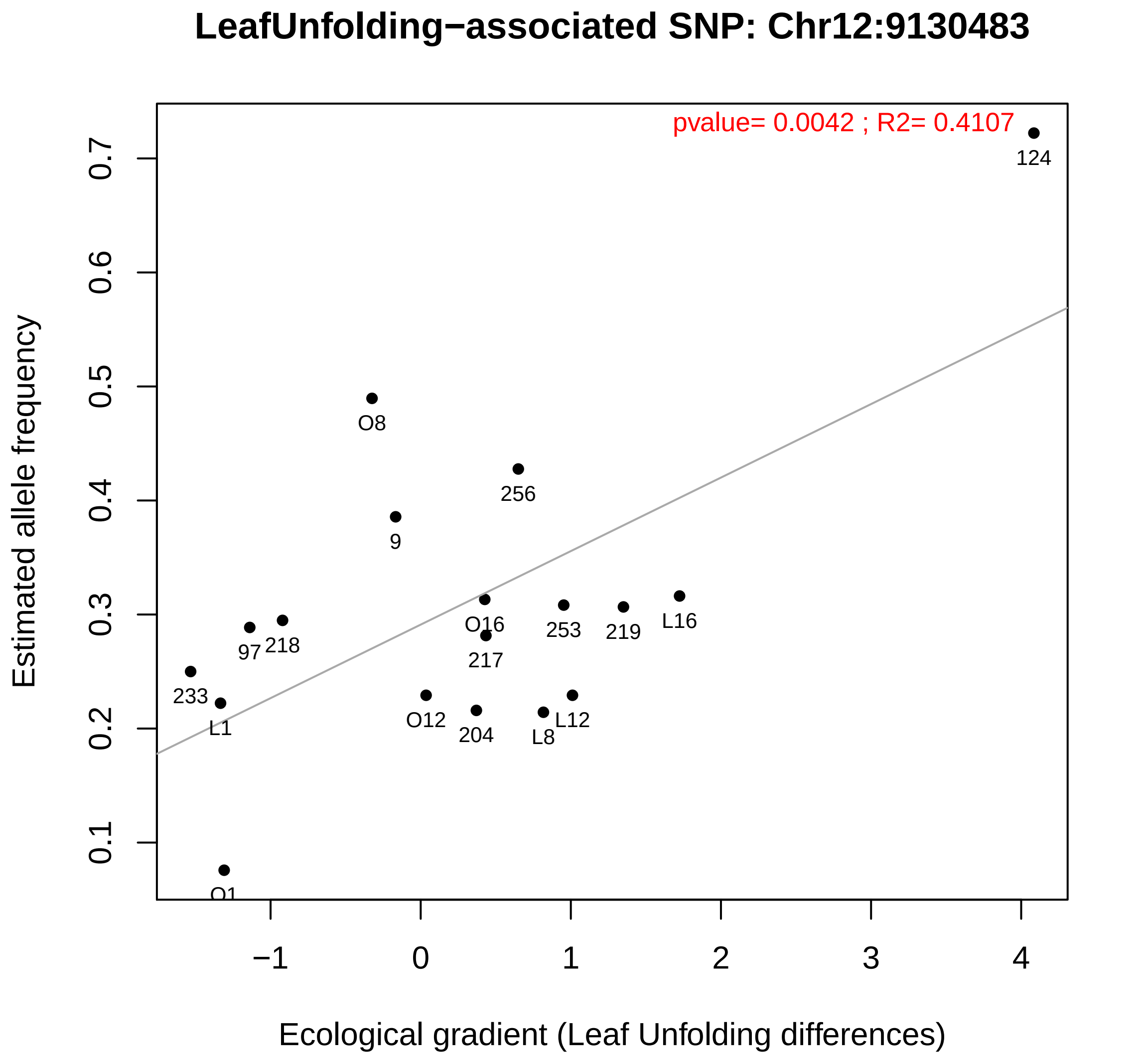
LeafUnfolding−associated SNP: Chr12:9130483
pvalue= 0.0042 ; R2= 0.4107
Estimated allele frequency
0.1
0.2
0.3
0.4
0.5
0.6
0.7
−1
0
1
2
3
4
Ecological gradient (Leaf Unfolding differences)
124
O8
256
9
O16
217
253
219
L16
97
218
233
L1
O12
204
L8
L12
O1

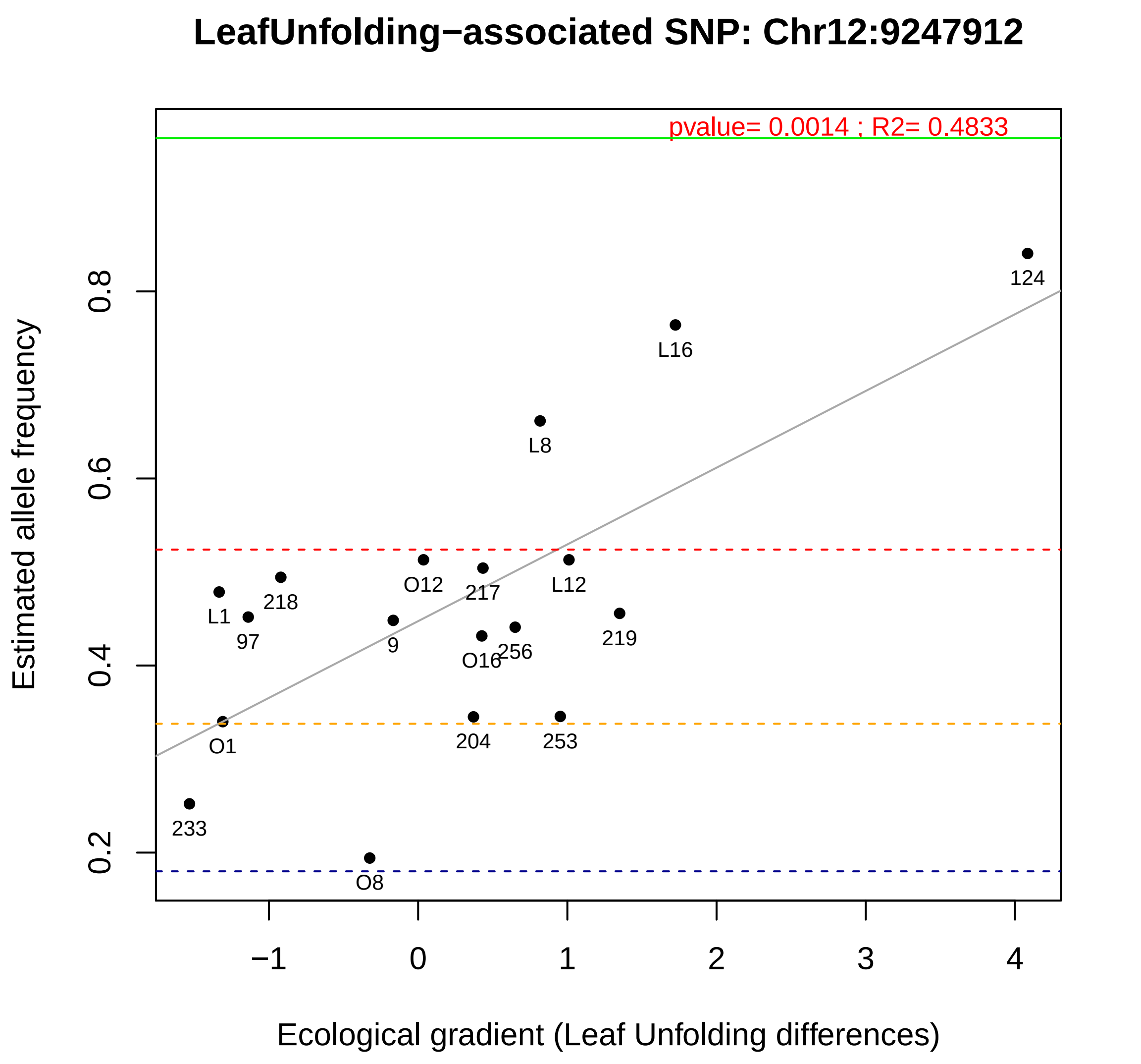

LeafUnfolding−associated SNP: Chr12:9247912
pvalue= 0.0014 ; R2= 0.4833
Estimated allele frequency
0.2
0.4
0.6
0.8
−1
0
1
2
3
4
Ecological gradient (Leaf Unfolding differences)
124
L16
L8
O12
217
L12
218
L1
97
9
O16
256
219
O1
204
253
233
O8

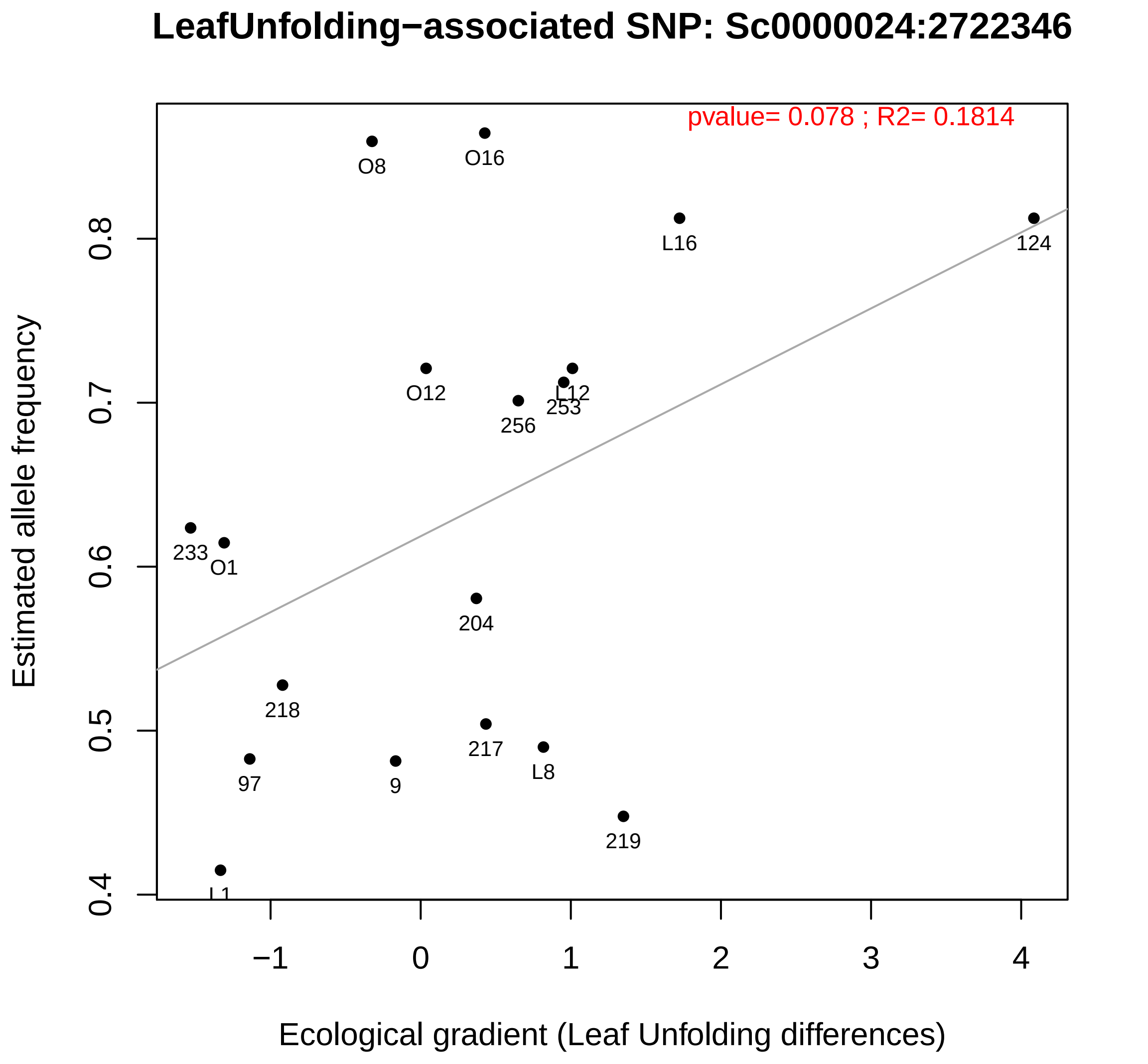
LeafUnfolding−associated SNP: Sc0000024:2722346
pvalue= 0.078 ; R2= 0.1814
Estimated allele frequency
Ecological gradient (Leaf Unfolding differences)
0.4
0.5
0.6
0.7
0.8
−1
0
1
2
3
4
O8
O16
L16
124
O12
L12
253
256
233
O1
204
218
97
9
217
L8
219

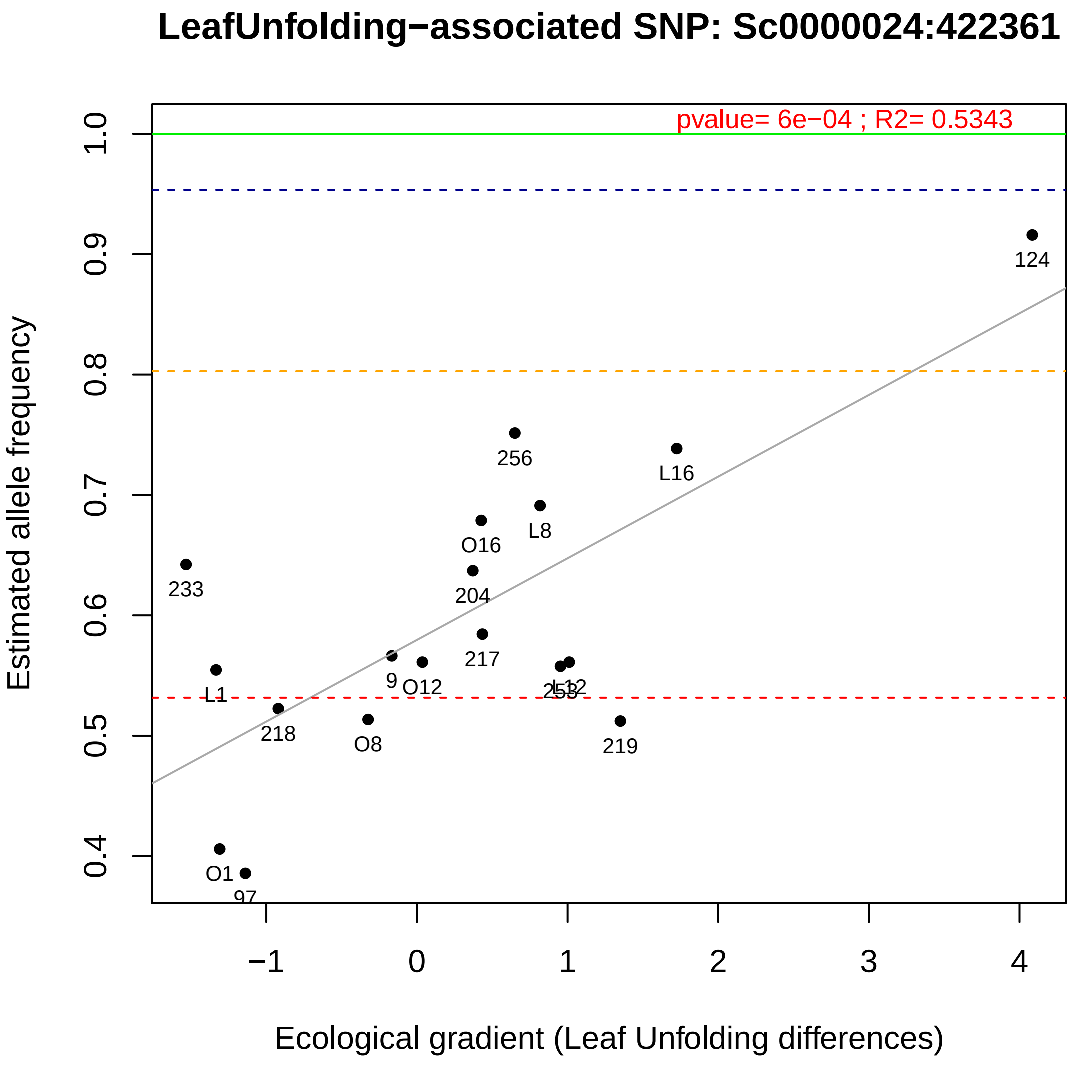

LeafUnfolding−associated SNP: Sc0000024:422361
pvalue= 6e−04 ; R2= 0.5343
Estimated allele frequency
Ecological gradient (Leaf Unfolding differences)
0.4
0.5
0.6
0.7
0.8
0.9
1.0
−1
0
1
2
3
4
124
256
L16
L8
O16
233
204
217
9
O12
L1
218
O8
219
O1

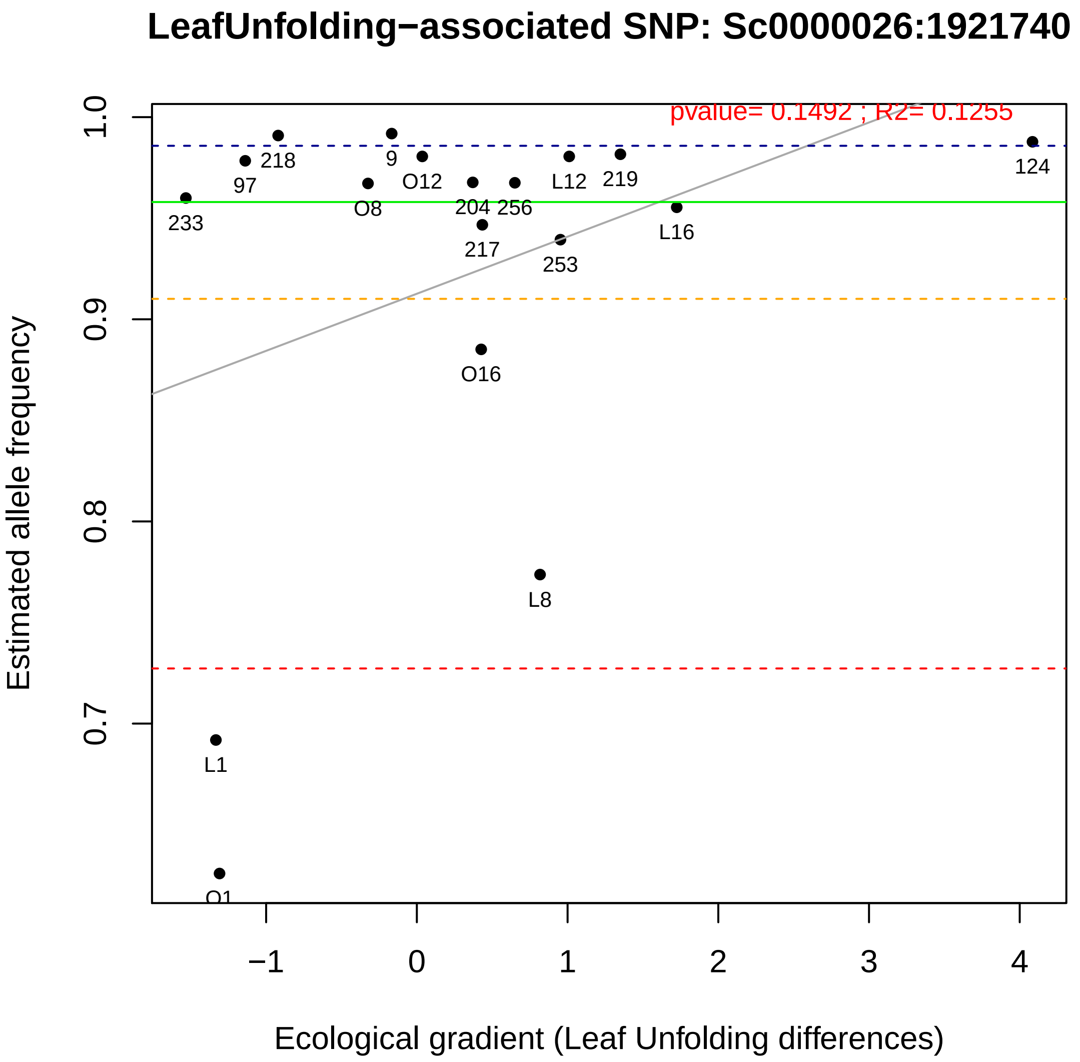

LeafUnfolding−associated SNP: Sc0000026:1921740
pvalue= 0.1492 ; R2= 0.1255
Estimated allele frequency
Ecological gradient (Leaf Unfolding differences)
1.0
0.9
0.8
0.7
−1
0
1
2
3
4
233
97
218
O8
9
O12
204
256
217
L12
219
253
L16
124
O16
L8
L1
O1

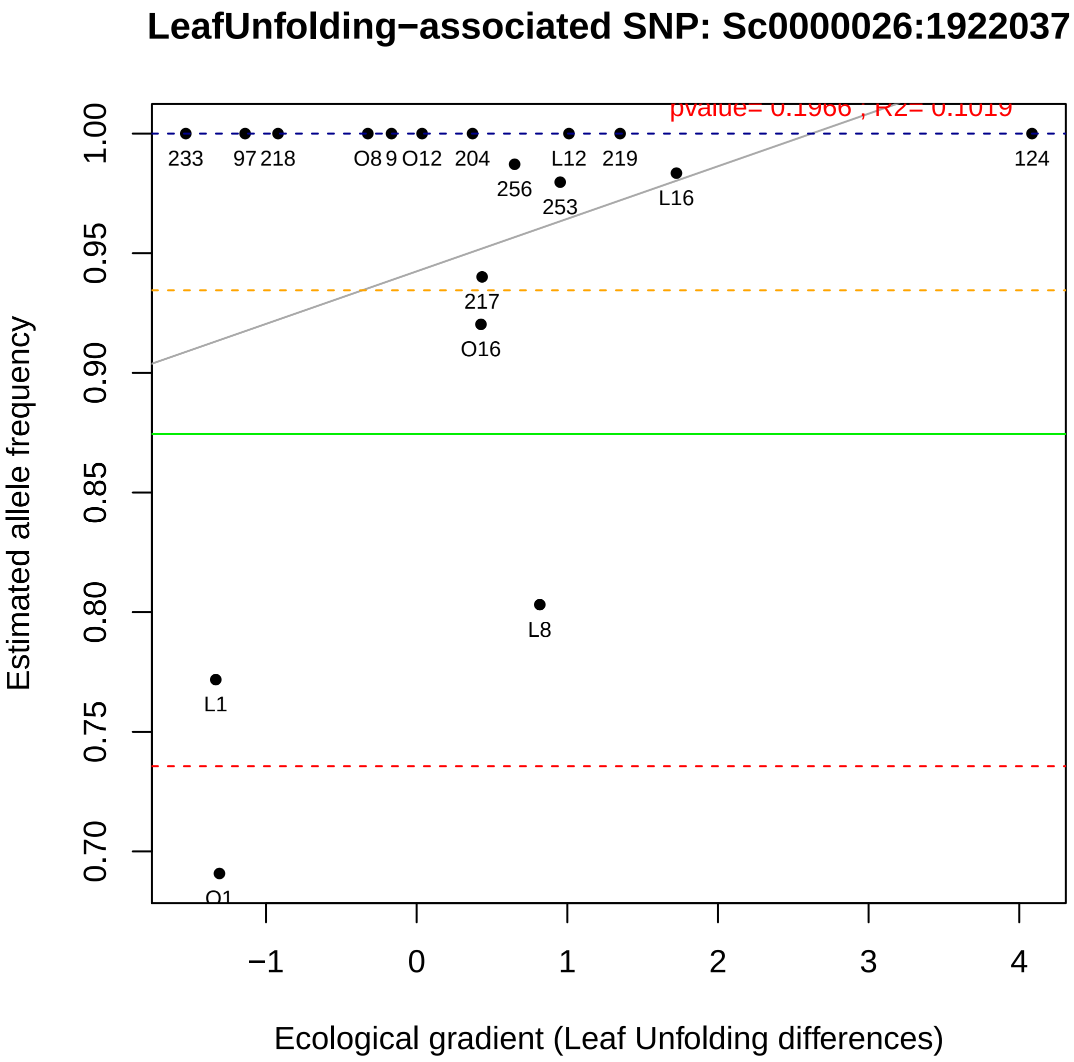
LeafUnfolding−associated SNP: Sc0000026:1922037
pvalue= 0.1966 ; R2= 0.1019
Estimated allele frequency
Ecological gradient (Leaf Unfolding differences)
1.00
0.95
0.90
0.85
0.80
0.75
0.70
−1
0
1
2
3
4
233
97
218
O8
9
O12
204
L12
219
124
256
253
L16
217
O16
L8
L1
O1

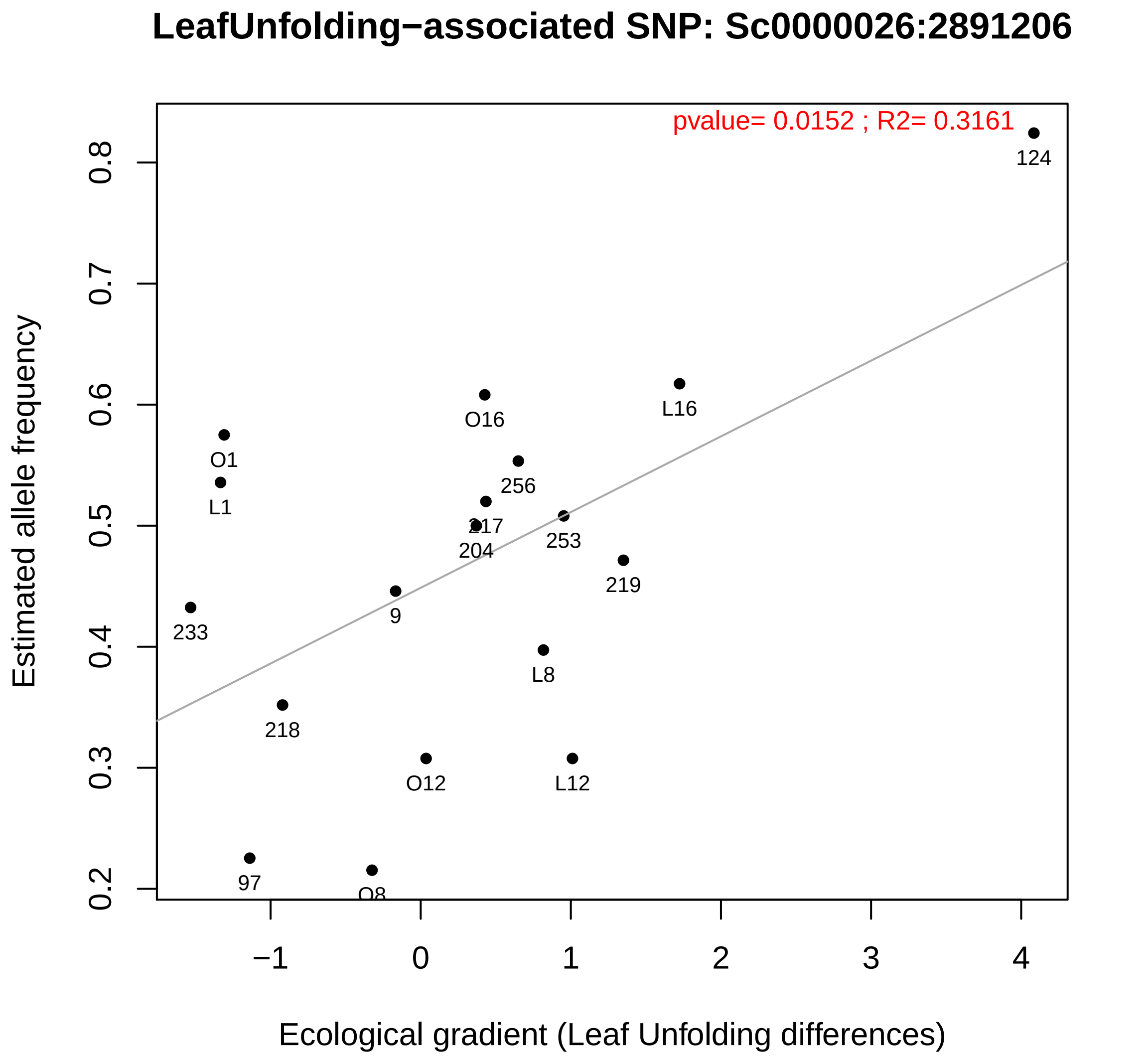
LeafUnfolding−associated SNP: Sc0000026:2891206
pvalue= 0.0152 ; R2= 0.3161
Estimated allele frequency
0.2
0.3
0.4
0.5
0.6
0.7
0.8
−1
0
1
2
3
4
Ecological gradient (Leaf Unfolding differences)
124
L16
O16
O1
L1
256
217
204
253
219
9
233
L8
218
O12
L12
97
O8

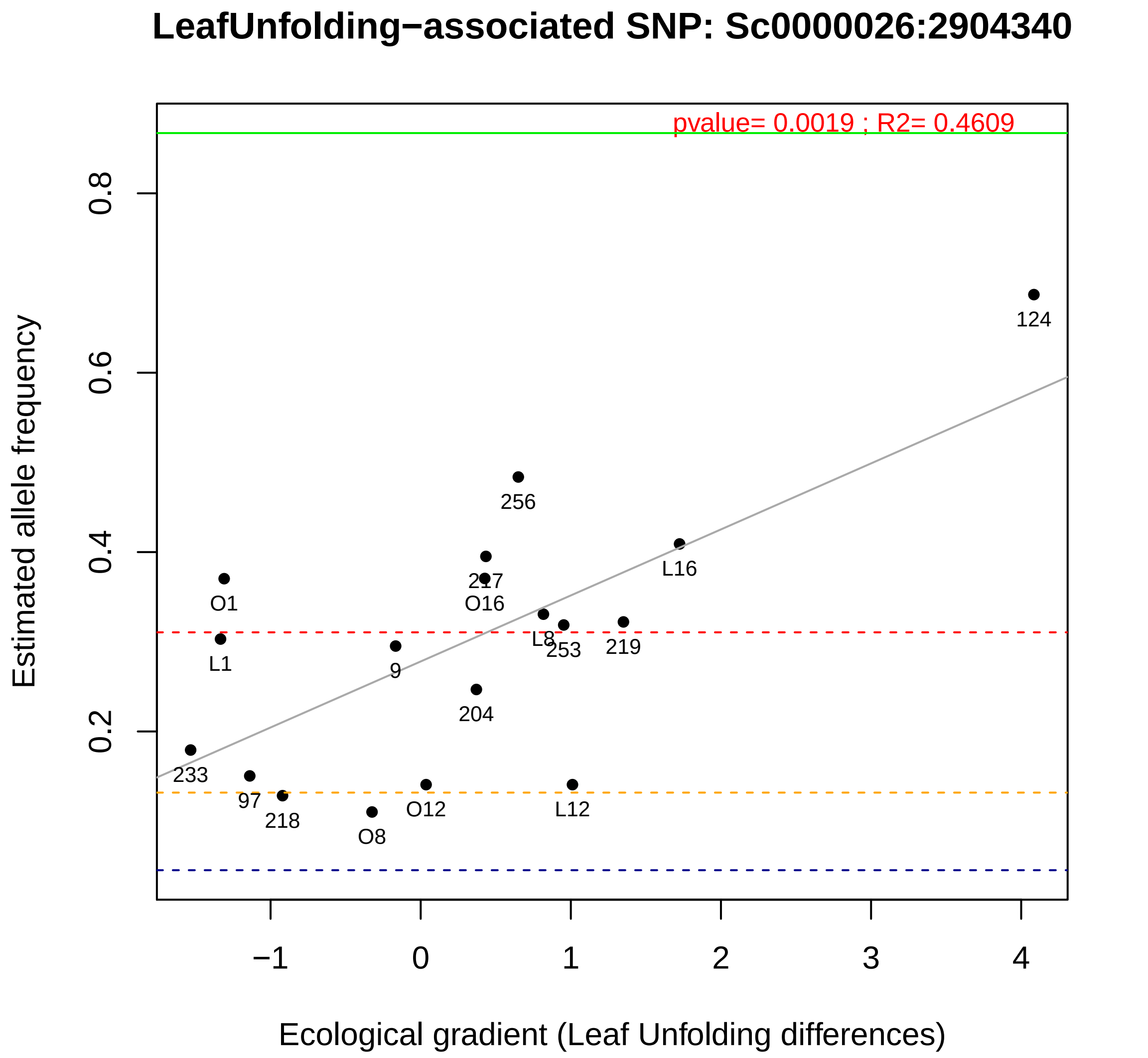

LeafUnfolding−associated SNP: Sc0000026:2904340
pvalue= 0.0019 ; R2= 0.4609
Estimated allele frequency
0.2
0.4
0.6
0.8
−1
0
1
2
3
4
Ecological gradient (Leaf Unfolding differences)
124
256
L16
217
O16
O1
L8
253
219
L1
9
204
233
97
218
O8
O12
L12

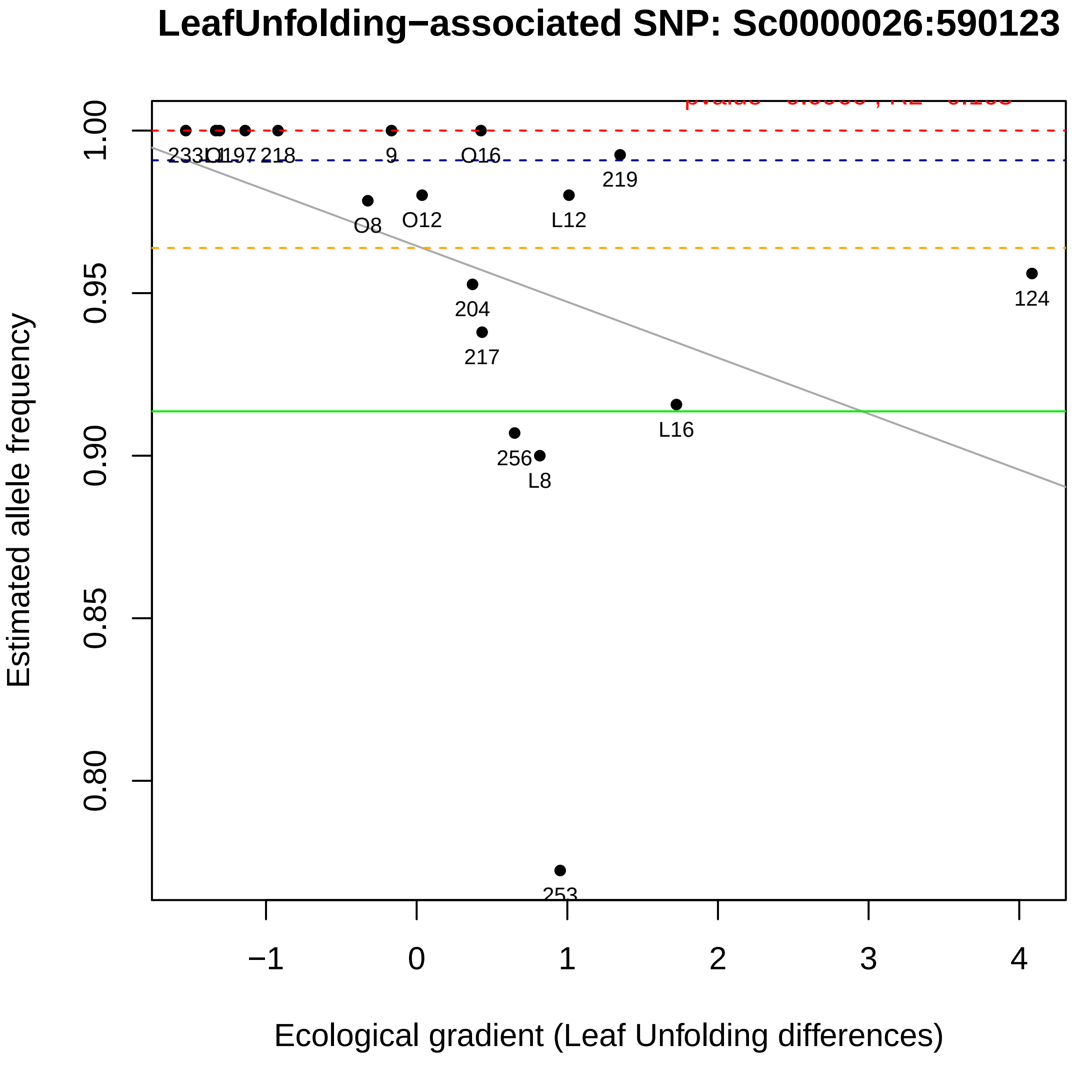
LeafUnfolding−associated SNP: Sc0000026:590123
Estimated allele frequency
Ecological gradient (Leaf Unfolding differences)
1.00
0.95
0.90
0.85
0.80
−1
0
1
2
3
4
233
O1
197
218
9
O16
219
O8
O12
L12
204
217
124
L16
256
L8
253

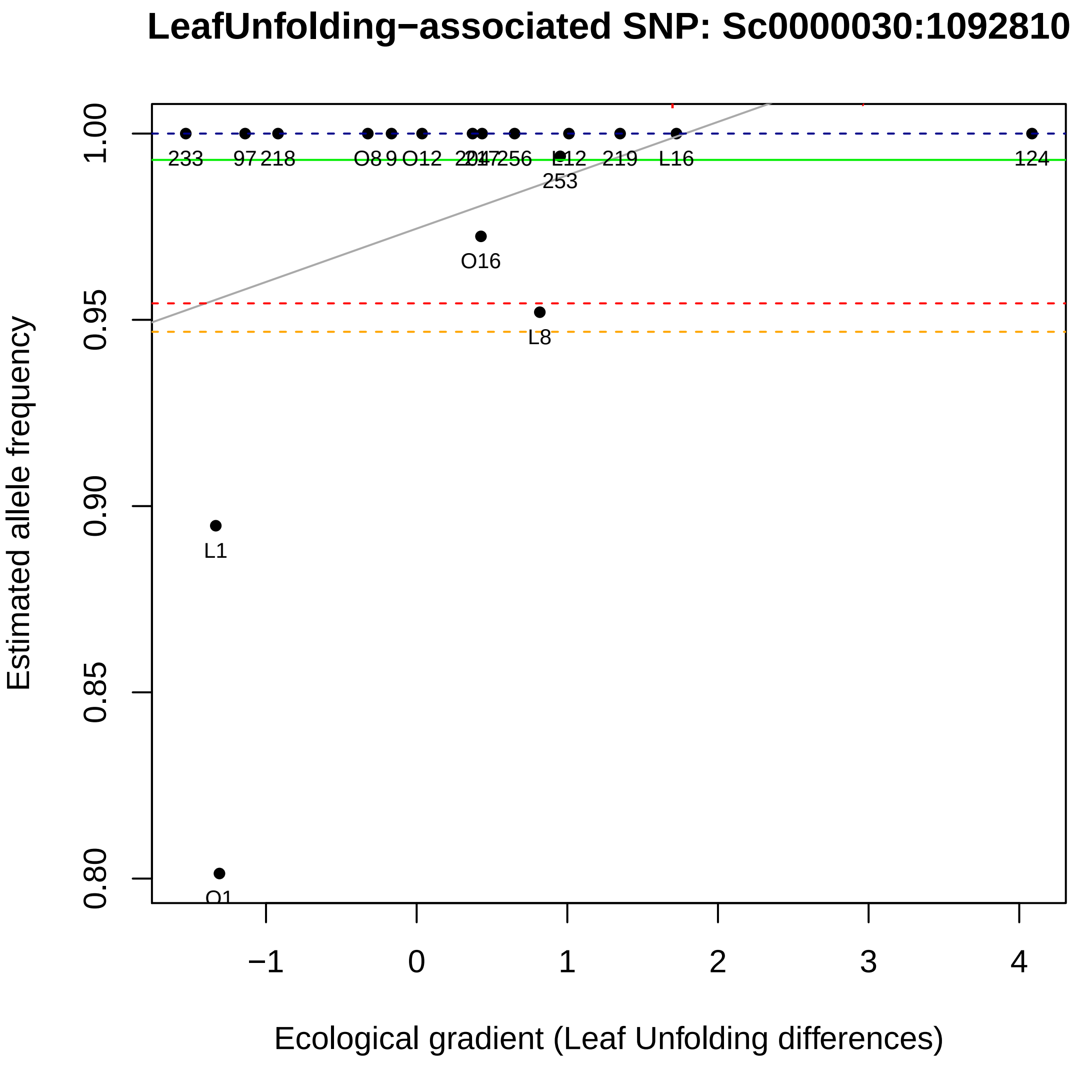

LeafUnfolding−associated SNP: Sc0000030:1092810
Estimated allele frequency
Ecological gradient (Leaf Unfolding differences)
1.00
0.95
0.90
0.85
0.80
−1
0
1
2
3
4
233
97
218
O8
9
O12
256
L12
253
219
L16
124
O16
L8
L1

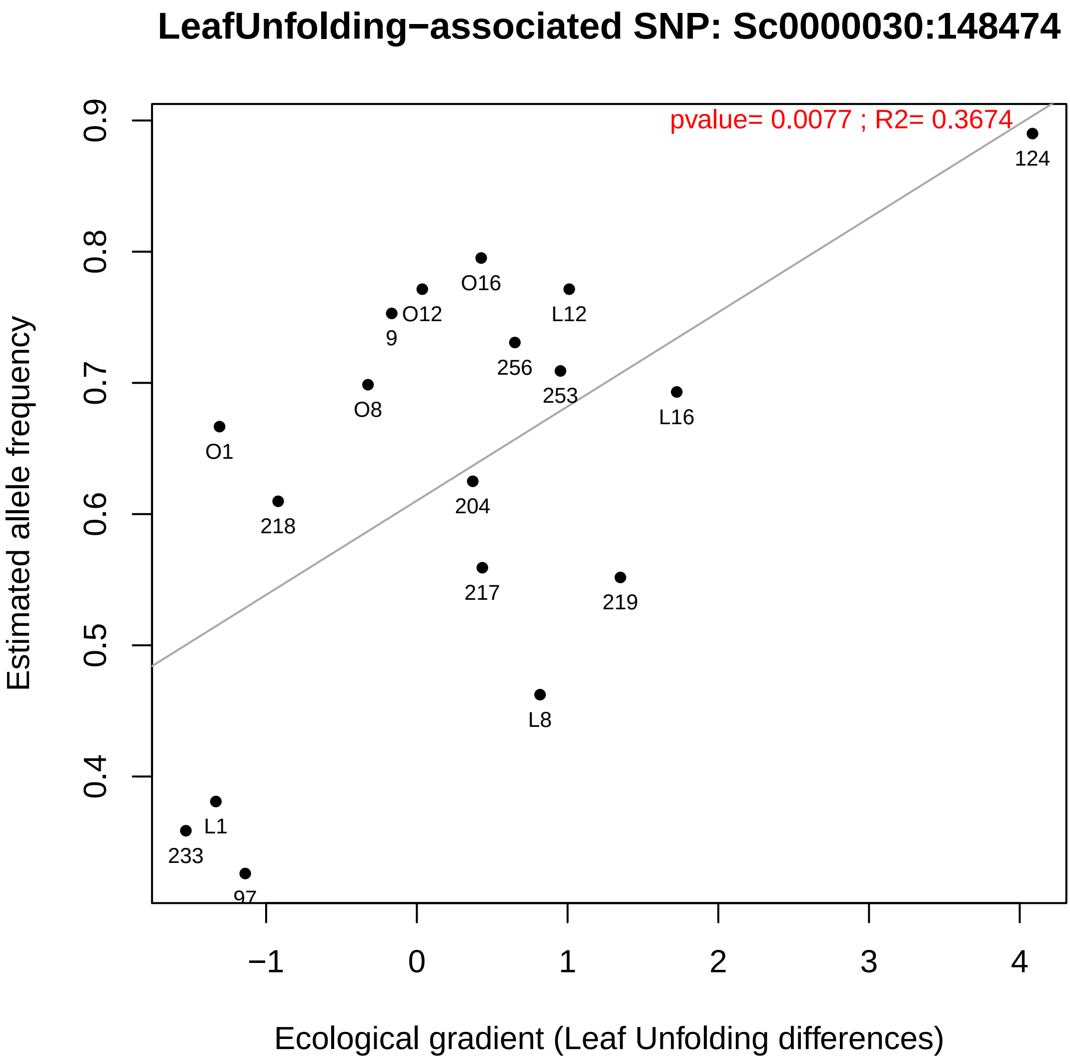

LeafUnfolding−associated SNP: Sc0000030:148474
pvalue= 0.0077 ; R2= 0.3674
124
O16
O12
9
L12
256
253
O8
L16
O1
204
218
217
219
L8
L1
233
97
0.9
0.8
0.7
0.6
0.5
0.4
−1
0
1
2
3
4
Estimated allele frequency
Ecological gradient (Leaf Unfolding differences)

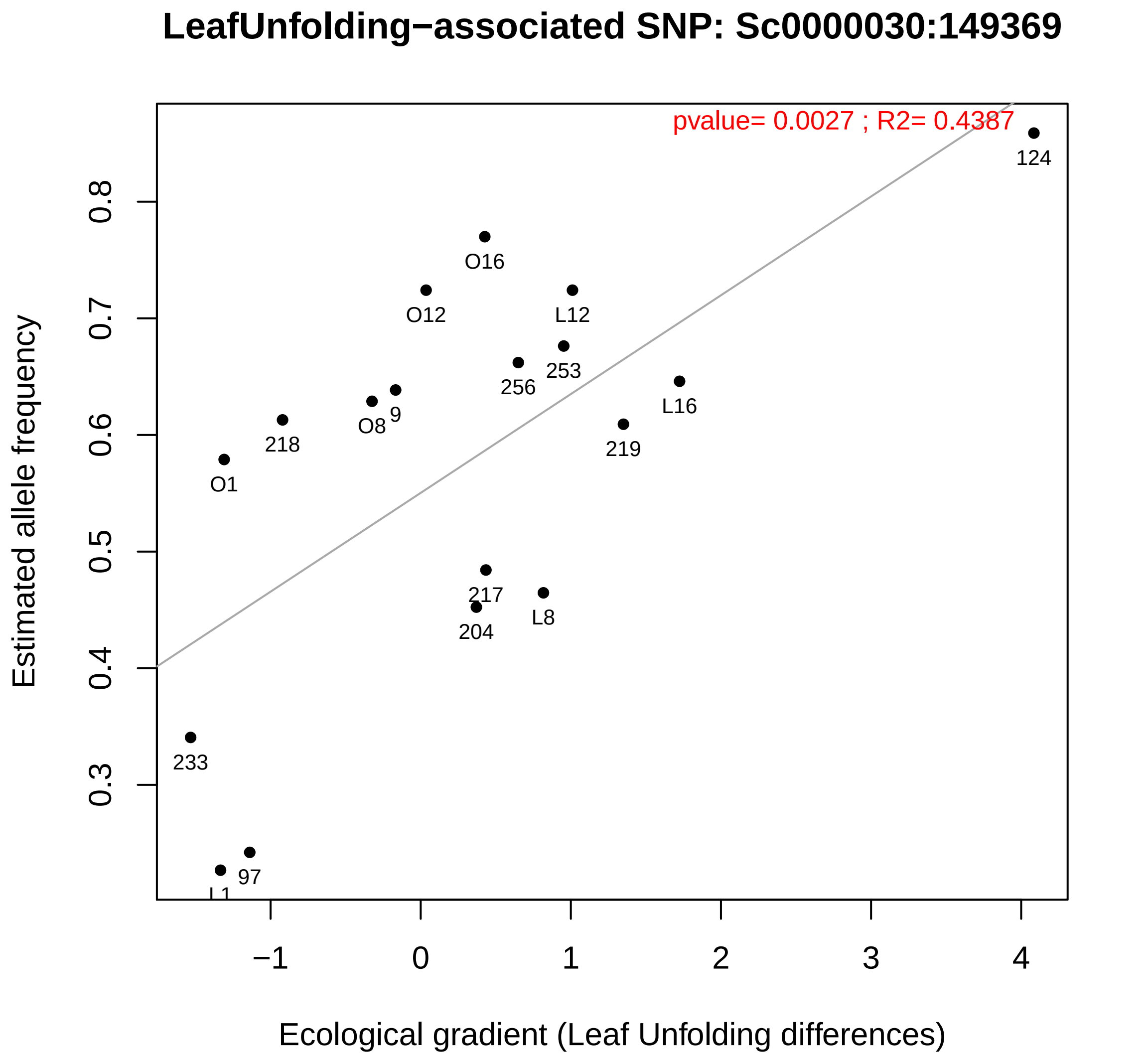

LeafUnfolding−associated SNP: Sc0000030:149369
pvalue= 0.0027 ; R2= 0.4387
Estimated allele frequency
Ecological gradient (Leaf Unfolding differences)
0.3
0.4
0.5
0.6
0.7
0.8
−1
0
1
2
3
4
124
O16
O12
L12
253
256
L16
9
O8
218
219
O1
217
L8
204
233
97
L1

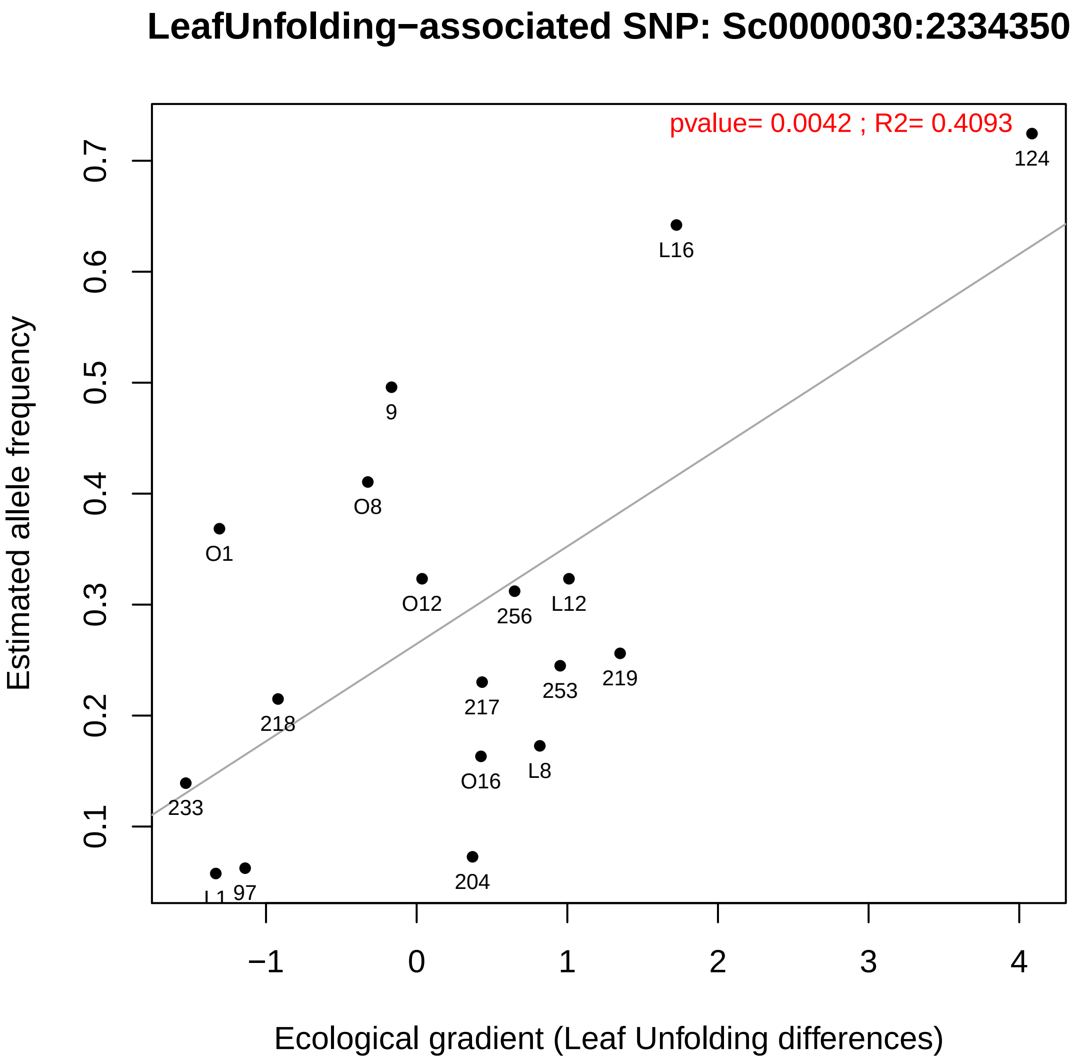

LeafUnfolding−associated SNP: Sc0000030:2334350
pvalue= 0.0042 ; R2= 0.4093
Estimated allele frequency
0.1
0.2
0.3
0.4
0.5
0.6
0.7
Ecological gradient (Leaf Unfolding differences)
−1
0
1
2
3
4
124
L16
9
O8
O1
O12
256
L12
219
253
217
218
233
O16
L8
204
97

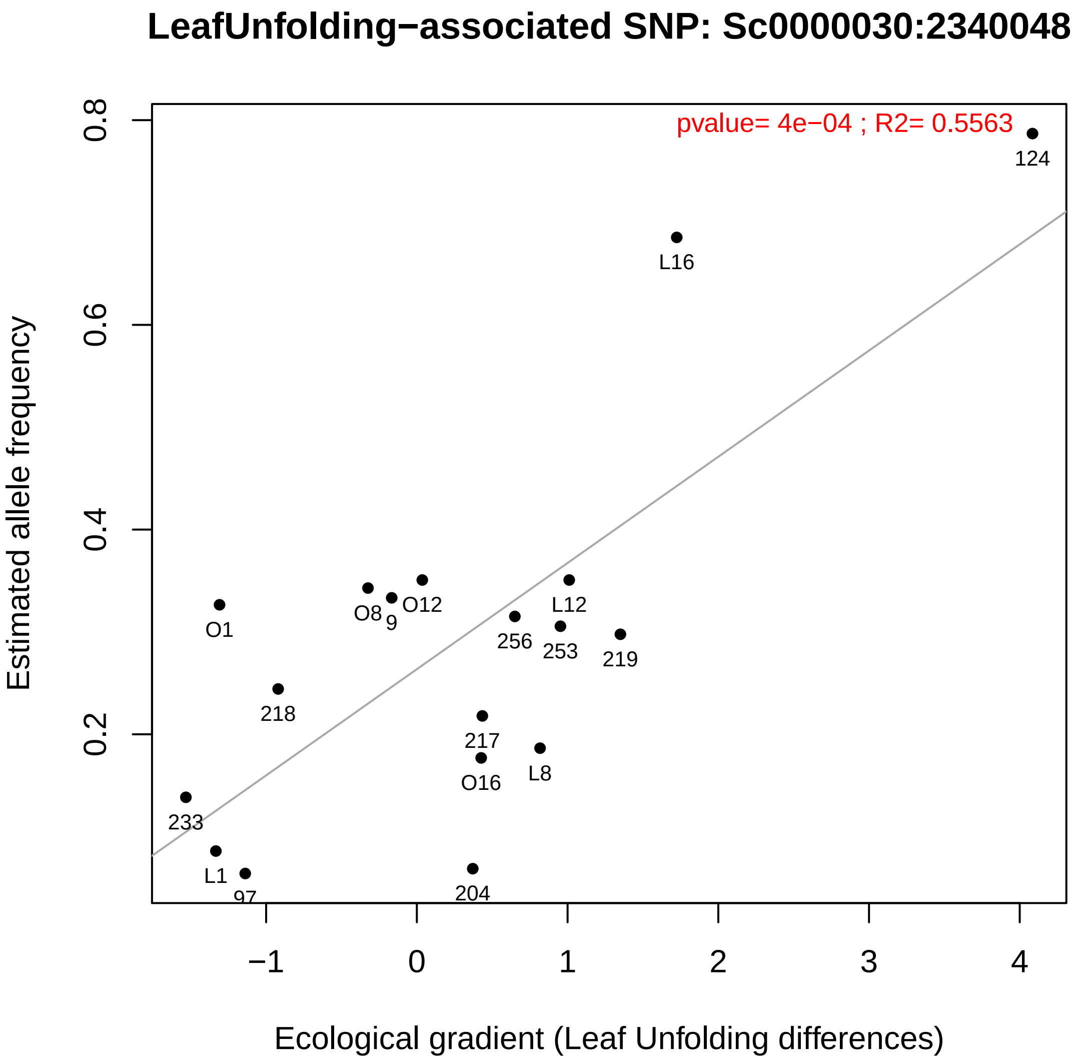

LeafUnfolding−associated SNP: Sc0000030:2340048
pvalue= 4e−04 ; R2= 0.5563
124
L16
L12
O1
O8
9
O12
256
253
219
218
217
L8
O16
233
L1
97
204
Estimated allele frequency
0.2
0.4
0.6
0.8
−1
0
1
2
3
4
Ecological gradient (Leaf Unfolding differences)

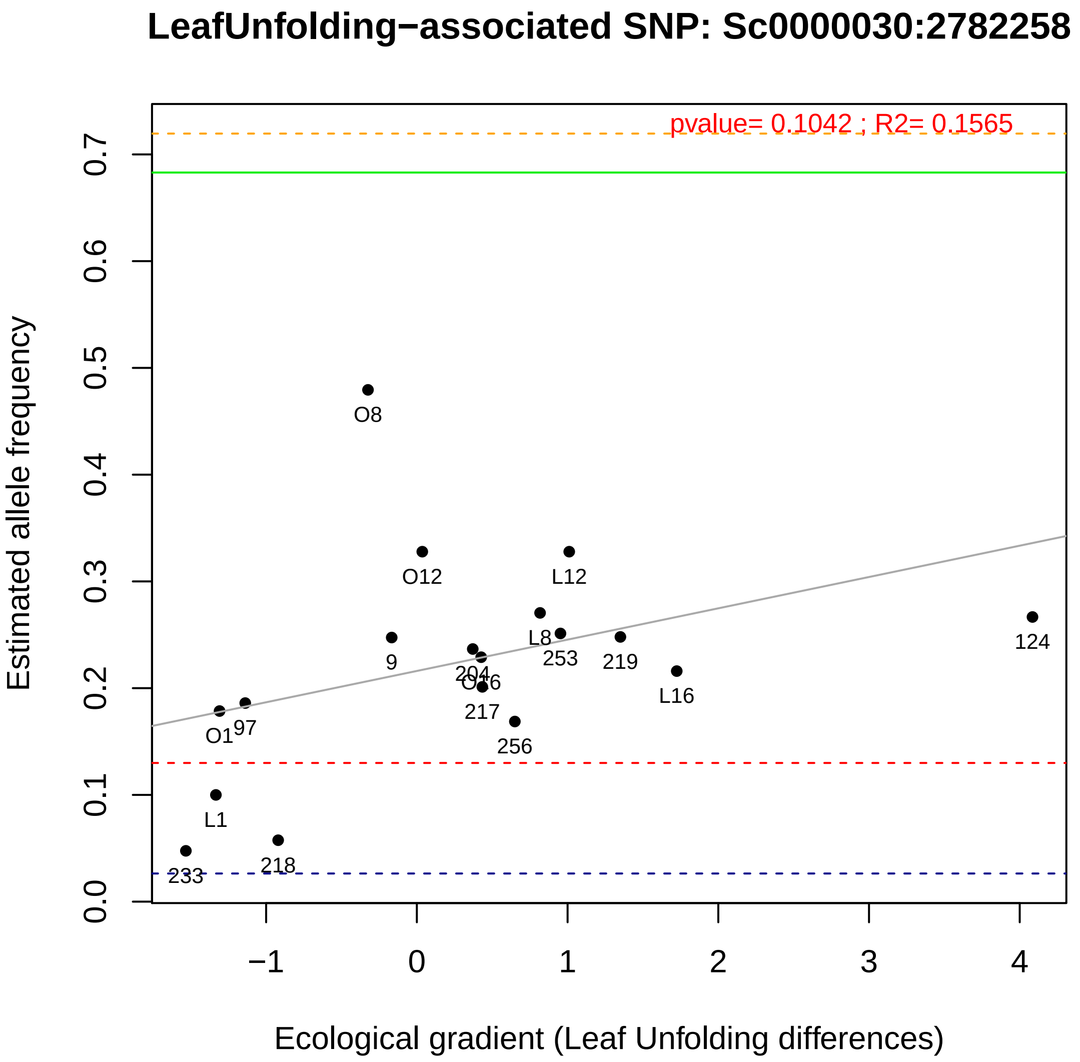

LeafUnfolding−associated SNP: Sc0000030:2782258
pvalue= 0.1042 ; R2= 0.1565
Estimated allele frequency
0.0
0.1
0.2
0.3
0.4
0.5
0.6
0.7
−1
0
1
2
3
4
Ecological gradient (Leaf Unfolding differences)
O8
O12
L12
L8
253
219
124
9
204
O16
217
L16
256
O1
97
L1
218
233

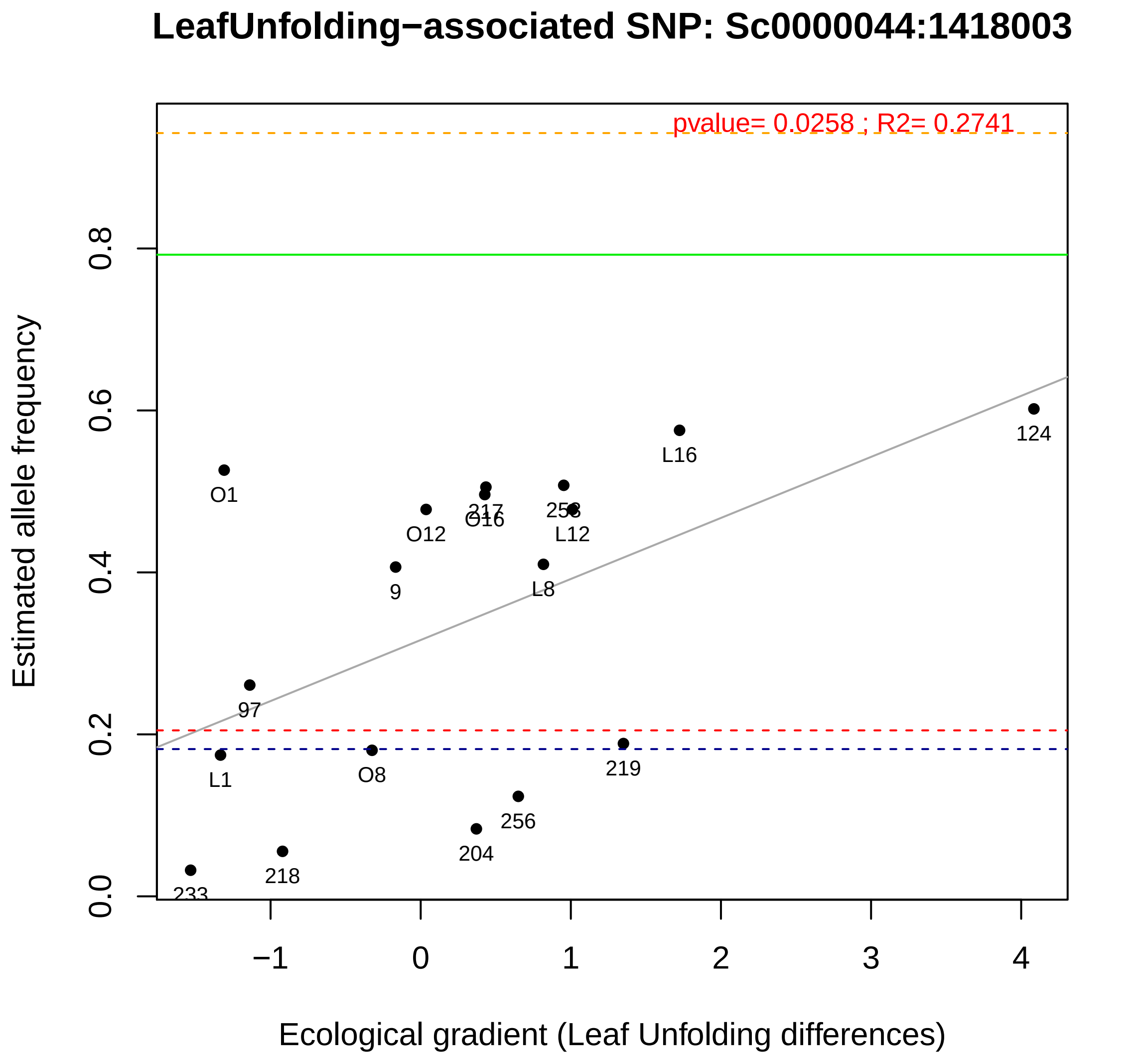
LeafUnfolding−associated SNP: Sc0000044:1418003
pvalue= 0.0258 ; R2= 0.2741
Estimated allele frequency
0.0
0.2
0.4
0.6
0.8
−1
0
1
2
3
4
Ecological gradient (Leaf Unfolding differences)
O1
O12
217
O16
253
L12
L16
124
9
L8
97
L1
O8
219
256
204
218
233

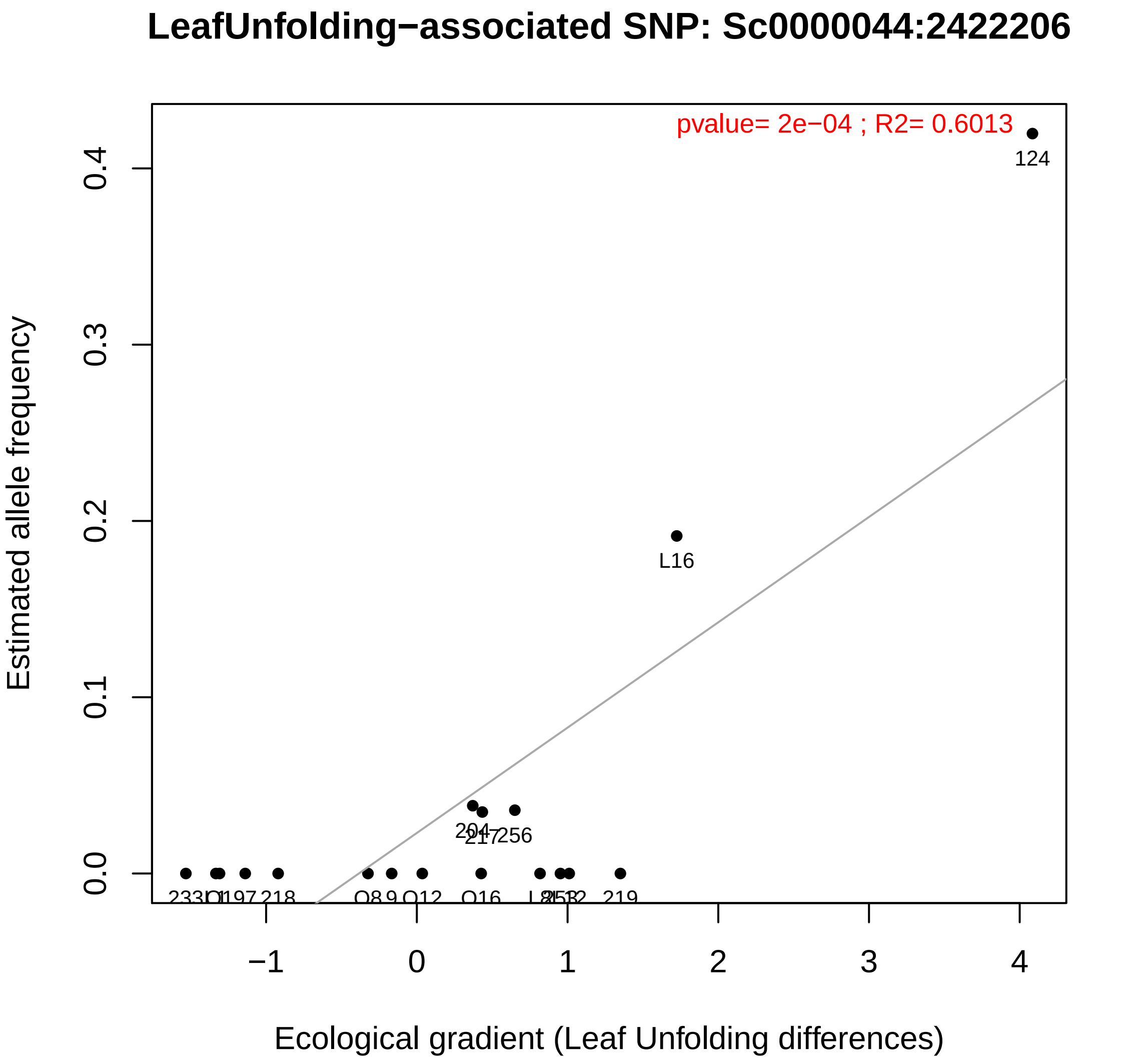
LeafUnfolding−associated SNP: Sc0000044:2422206
pvalue= 2e−04 ; R2= 0.6013
Estimated allele frequency
Ecological gradient (Leaf Unfolding differences)
0.0
0.1
0.2
0.3
0.4
−1
0
1
2
3
4
124
L16
204
217
256
218
O8
9
O12
O16
219

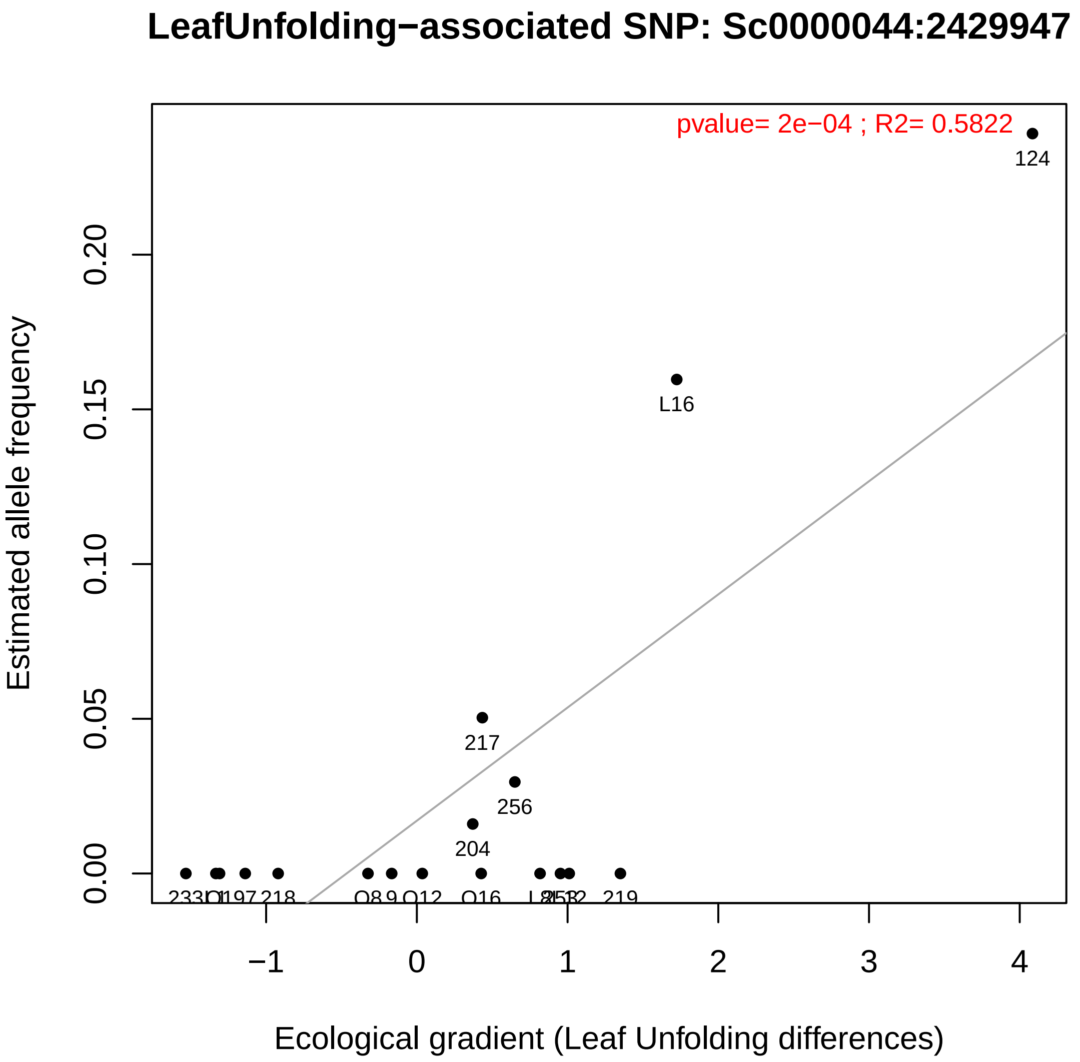

LeafUnfolding−associated SNP: Sc0000044:2429947
pvalue= 2e−04 ; R2= 0.5822
124
L16
217
256
204
218
O8
9
O12
O16
219
0.00
0.05
0.10
0.15
0.20
−1
0
1
2
3
4
Estimated allele frequency
Ecological gradient (Leaf Unfolding differences)

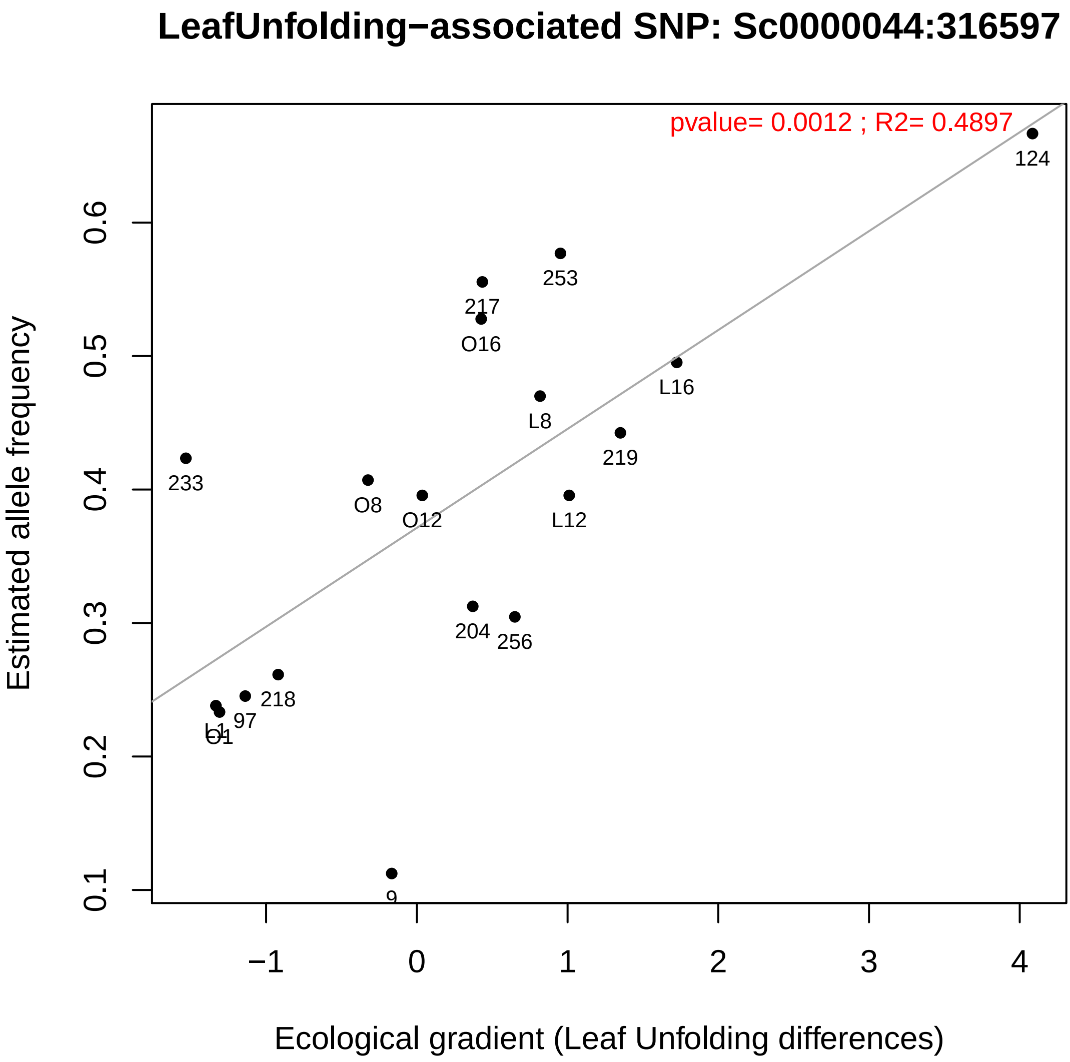
LeafUnfolding−associated SNP: Sc0000044:316597
pvalue= 0.0012 ; R2= 0.4897
124
253
217
O16
L16
L8
219
233
O8
O12
L12
204
256
218
97
L1
O1
9
0.1
0.2
0.3
0.4
0.5
0.6
Estimated allele frequency
−1
0
1
2
3
4
Ecological gradient (Leaf Unfolding differences)

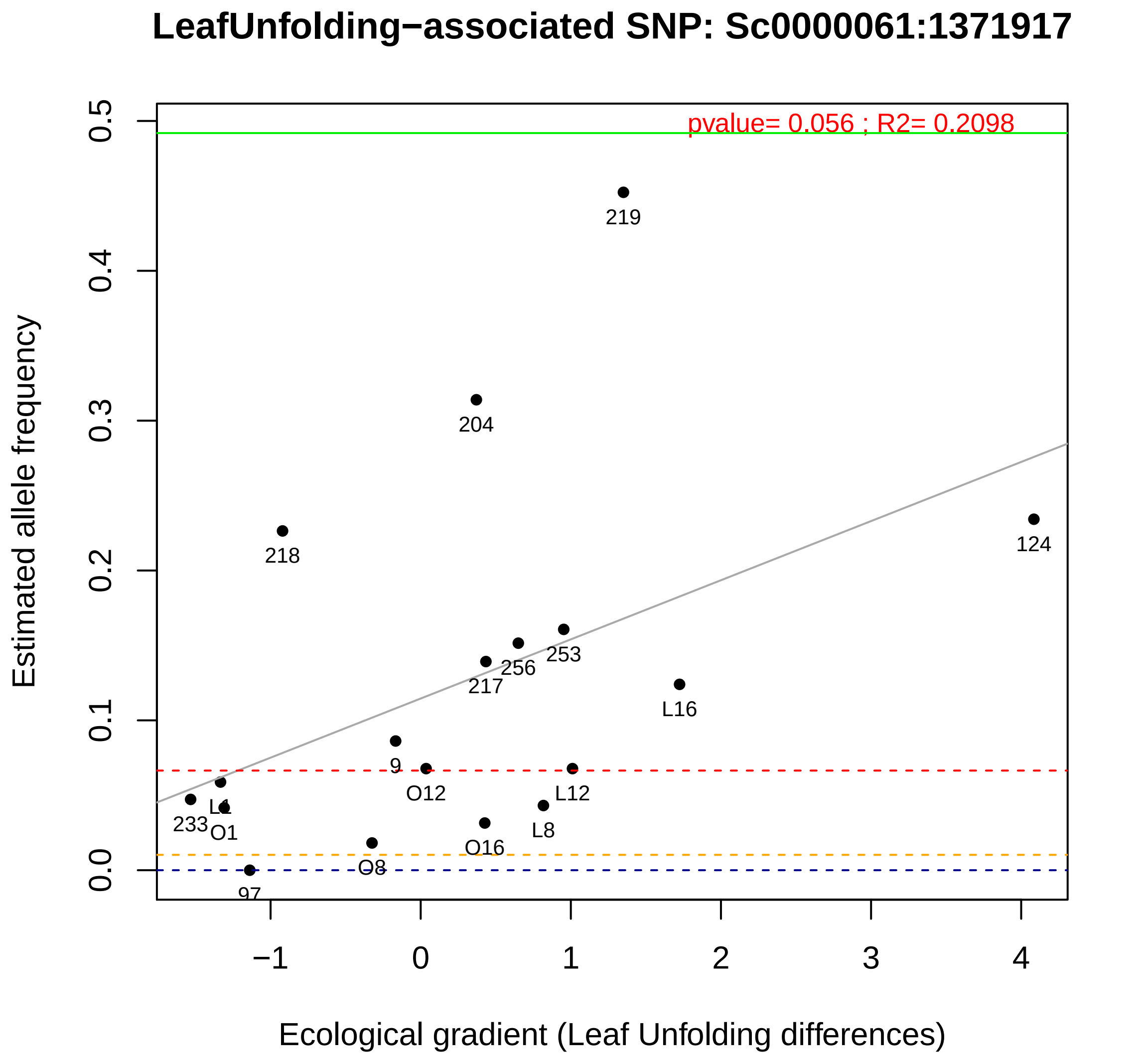
LeafUnfolding−associated SNP: Sc0000061:1371917
pvalue= 0.056 ; R2= 0.2098
Estimated allele frequency
Ecological gradient (Leaf Unfolding differences)
0.0
0.1
0.2
0.3
0.4
0.5
−1
0
1
2
3
4
219
204
218
124
253
256
217
L16
9
O12
L12
L1
233
O1
L8
O16
O8
97

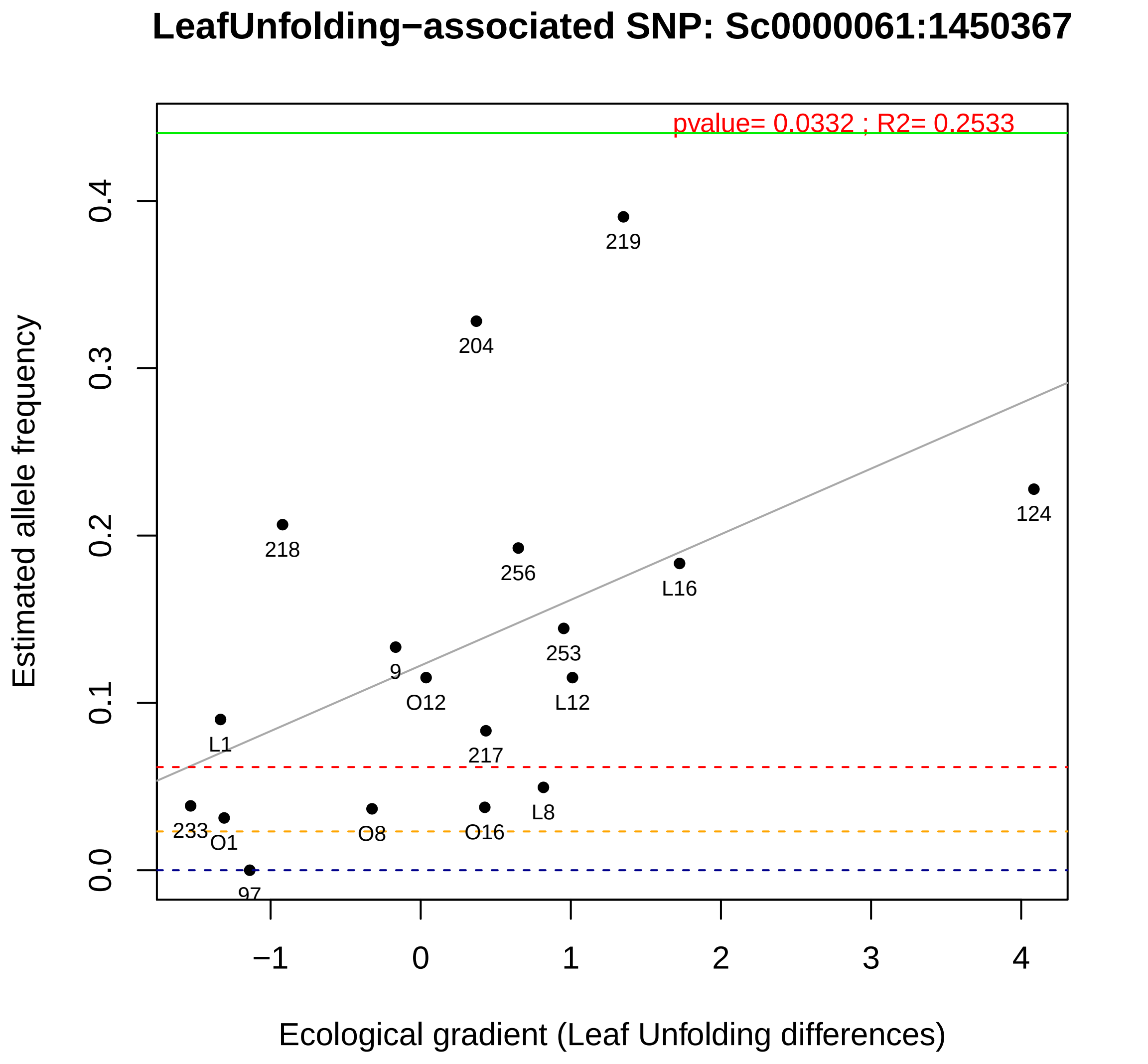
LeafUnfolding−associated SNP: Sc0000061:1450367
pvalue= 0.0332 ; R2= 0.2533
Estimated allele frequency
0.0
0.1
0.2
0.3
0.4
219
204
124
218
256
L16
253
9
O12
L12
L1
217
L8
233
O1
O8
O16
97
−1
0
1
2
3
4
Ecological gradient (Leaf Unfolding differences)

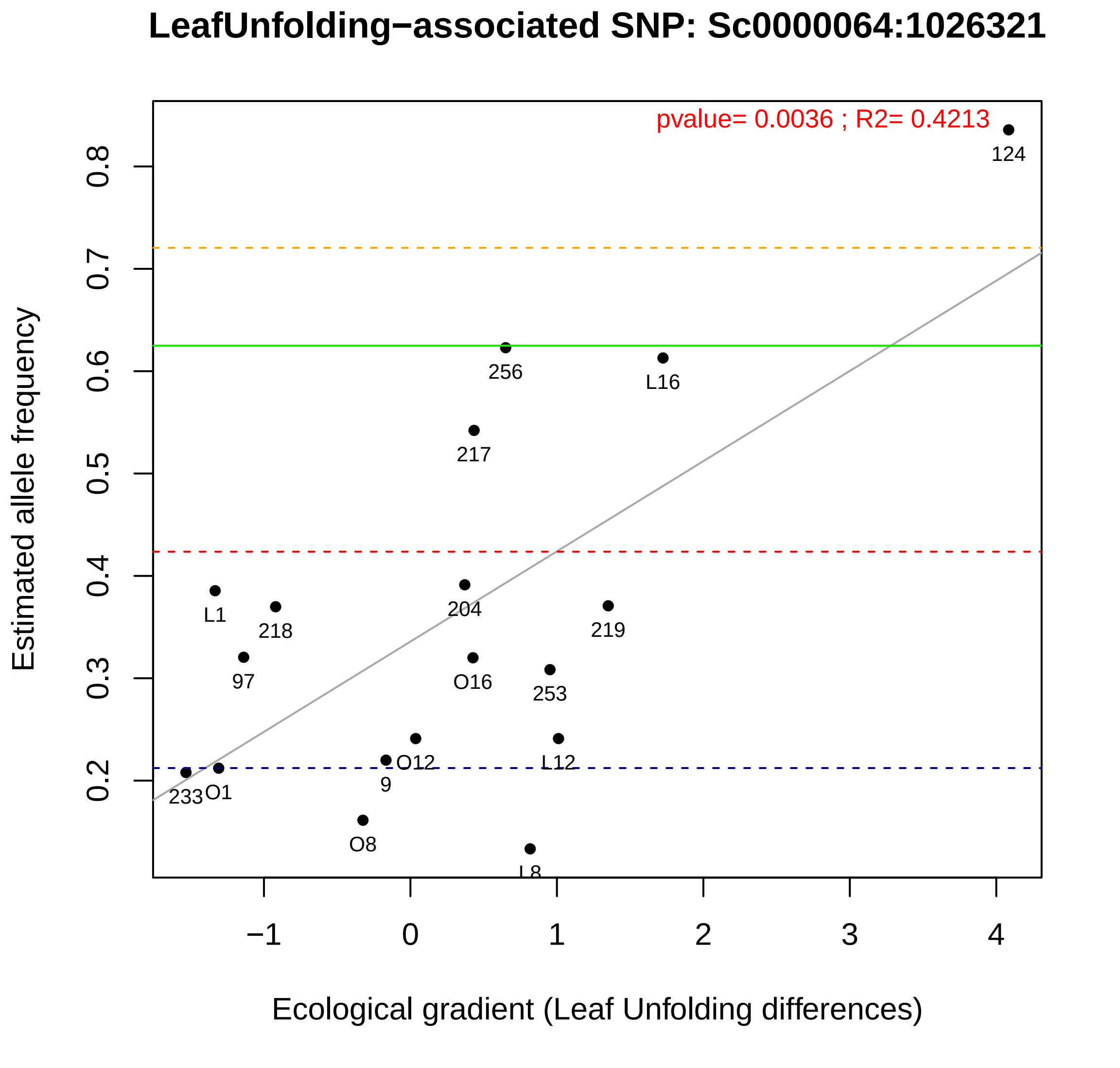
LeafUnfolding−associated SNP: Sc0000064:1026321
pvalue= 0.0036 ; R2= 0.4213
Estimated allele frequency
Ecological gradient (Leaf Unfolding differences)
0.2
0.3
0.4
0.5
0.6
0.7
0.8
−1
0
1
2
3
4
124
256
L16
217
204
L1
218
219
97
O16
253
O12
L12
9
233
O1
O8

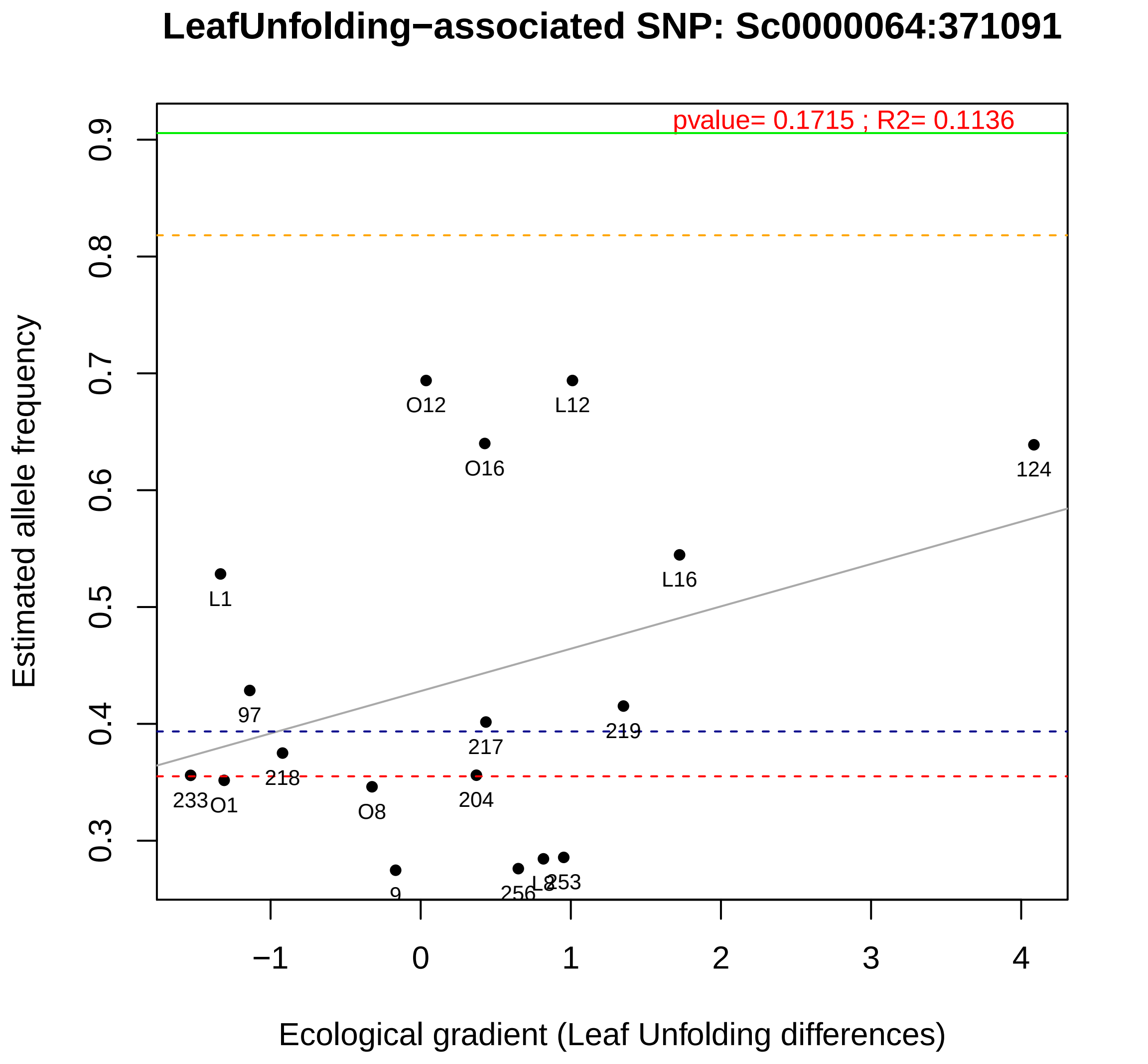
LeafUnfolding−associated SNP: Sc0000064:371091
pvalue= 0.1715 ; R2= 0.1136
Estimated allele frequency
Ecological gradient (Leaf Unfolding differences)
0.3
0.4
0.5
0.6
0.7
0.8
0.9
−1
0
1
2
3
4
O12
L12
O16
124
L16
L1
97
217
219
218
233
O1
O8
204
9
256
L8
253

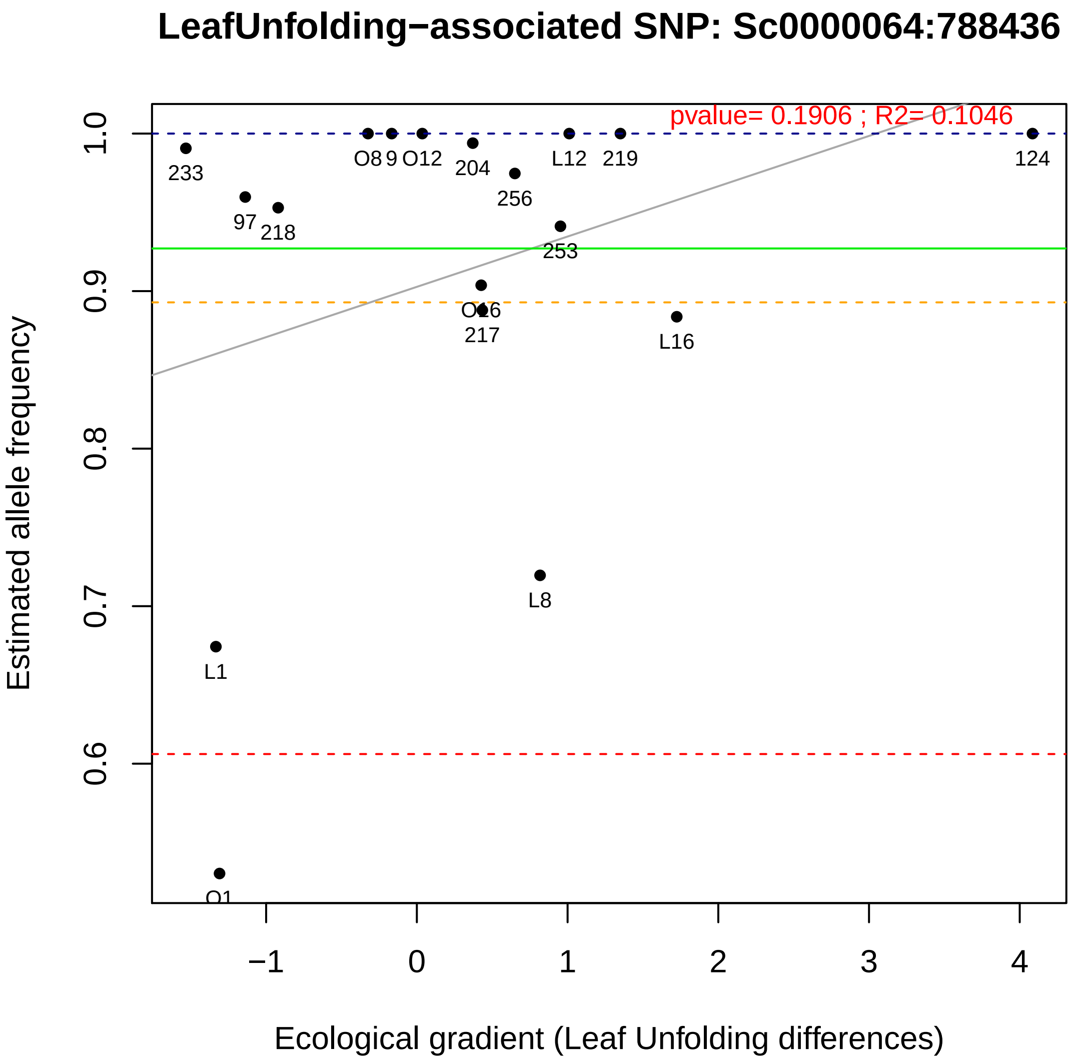
LeafUnfolding−associated SNP: Sc0000064:788436
pvalue= 0.1906 ; R2= 0.1046
Estimated allele frequency
Ecological gradient (Leaf Unfolding differences)
1.0
0.9
0.8
0.7
0.6
−1
0
1
2
3
4
233
97
218
O8
9
O12
204
256
L12
219
124
253
O26
217
L16
L8
L1
O1

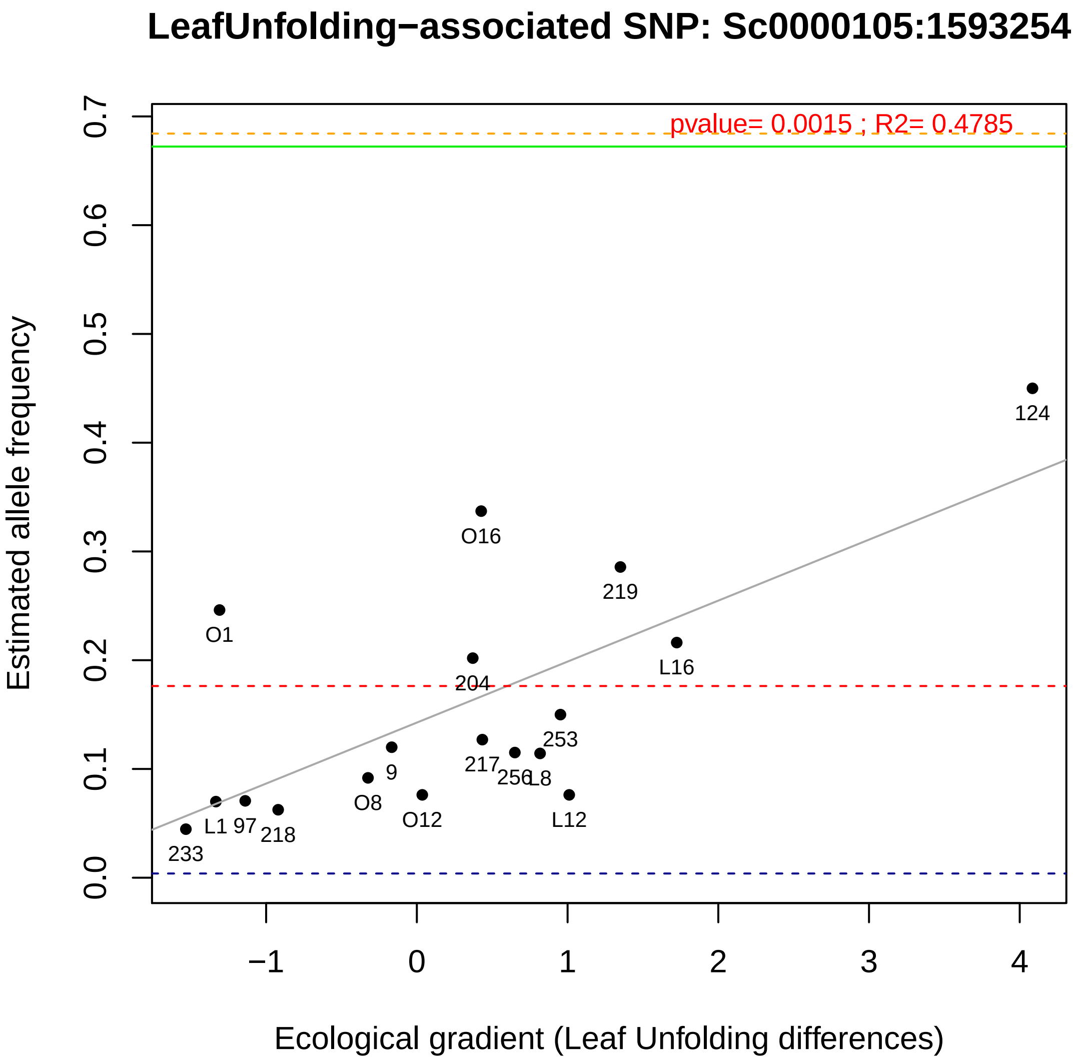
LeafUnfolding−associated SNP: Sc0000105:1593254
pvalue= 0.0015 ; R2= 0.4785
Estimated allele frequency
Ecological gradient (Leaf Unfolding differences)
0.0
0.1
0.2
0.3
0.4
0.5
0.6
0.7
−1
0
1
2
3
4
124
O16
219
O1
L16
204
253
9
217
256
L8
O8
O12
L12
L1
97
218
233

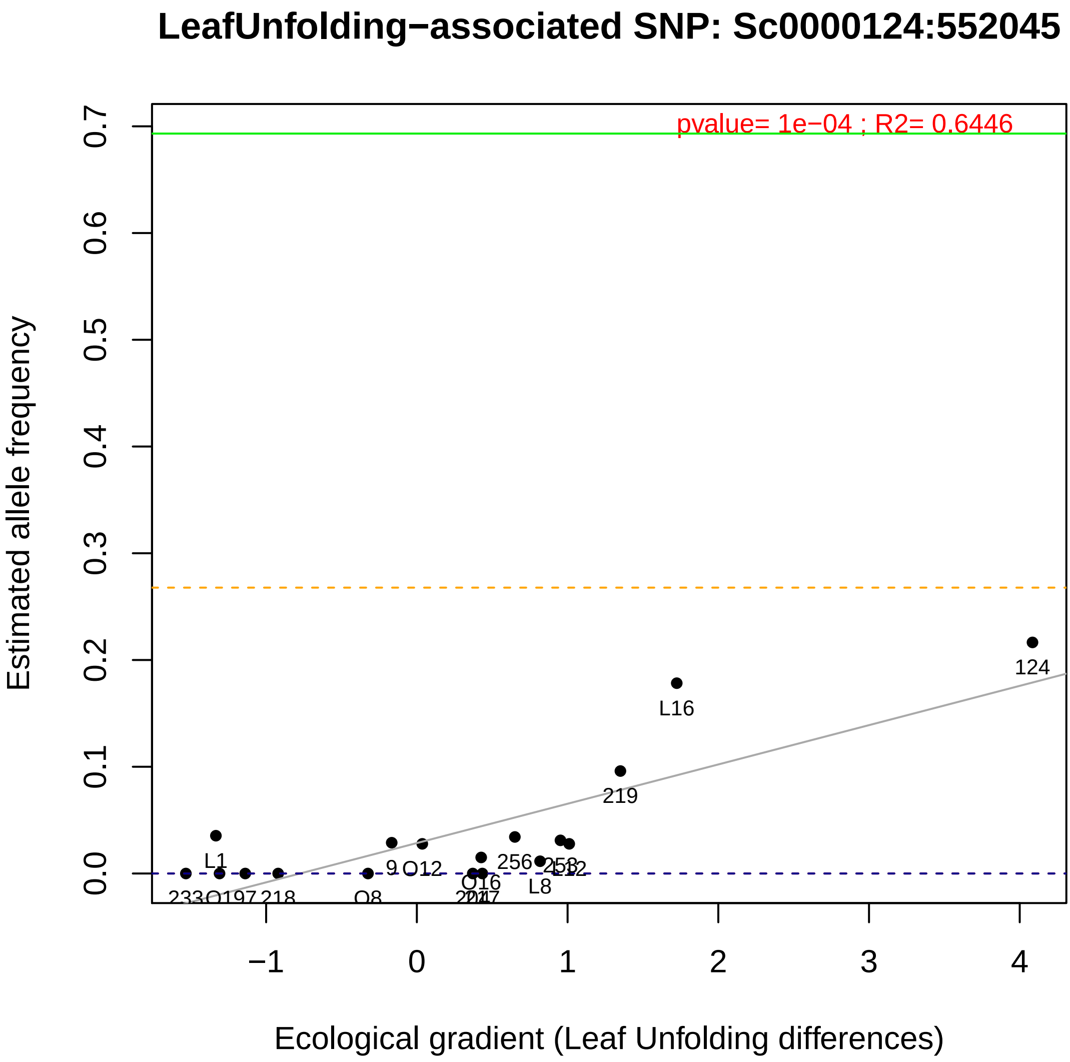

LeafUnfolding−associated SNP: Sc0000124:552045
pvalue= 1e−04 ; R2= 0.6446
Estimated allele frequency
Ecological gradient (Leaf Unfolding differences)
0.0
0.1
0.2
0.3
0.4
0.5
0.6
0.7
−1
0
1
2
3
4
124
L16
219
256
L8
L1
9
O12
O16
O8
218

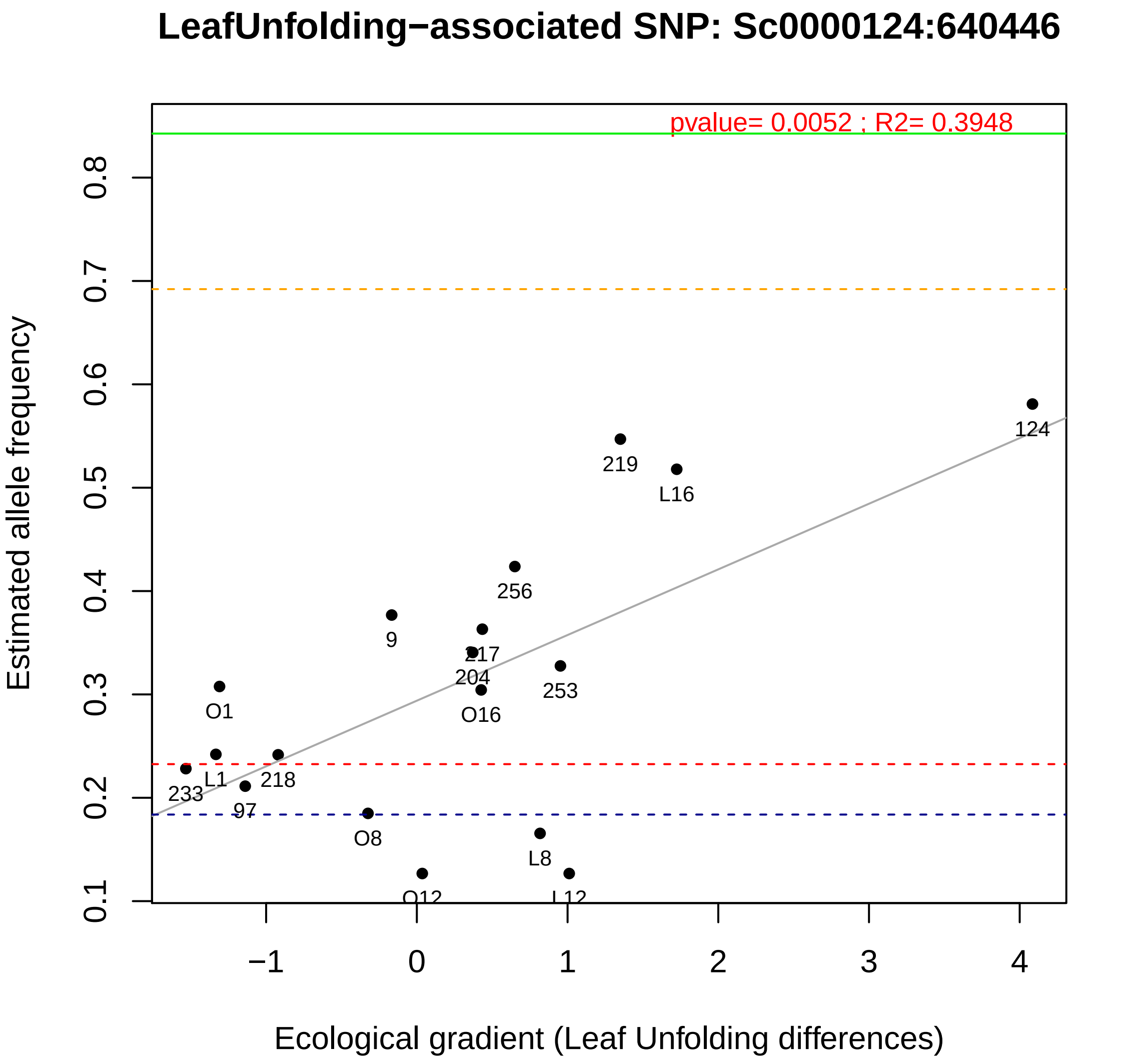
LeafUnfolding−associated SNP: Sc0000124:640446
pvalue= 0.0052 ; R2= 0.3948
Estimated allele frequency
0.1
0.2
0.3
0.4
0.5
0.6
0.7
0.8
−1
0
1
2
3
4
Ecological gradient (Leaf Unfolding differences)
124
219
L16
256
9
217
204
253
O16
O1
L1
218
233
97
O8
L8
O12
L12

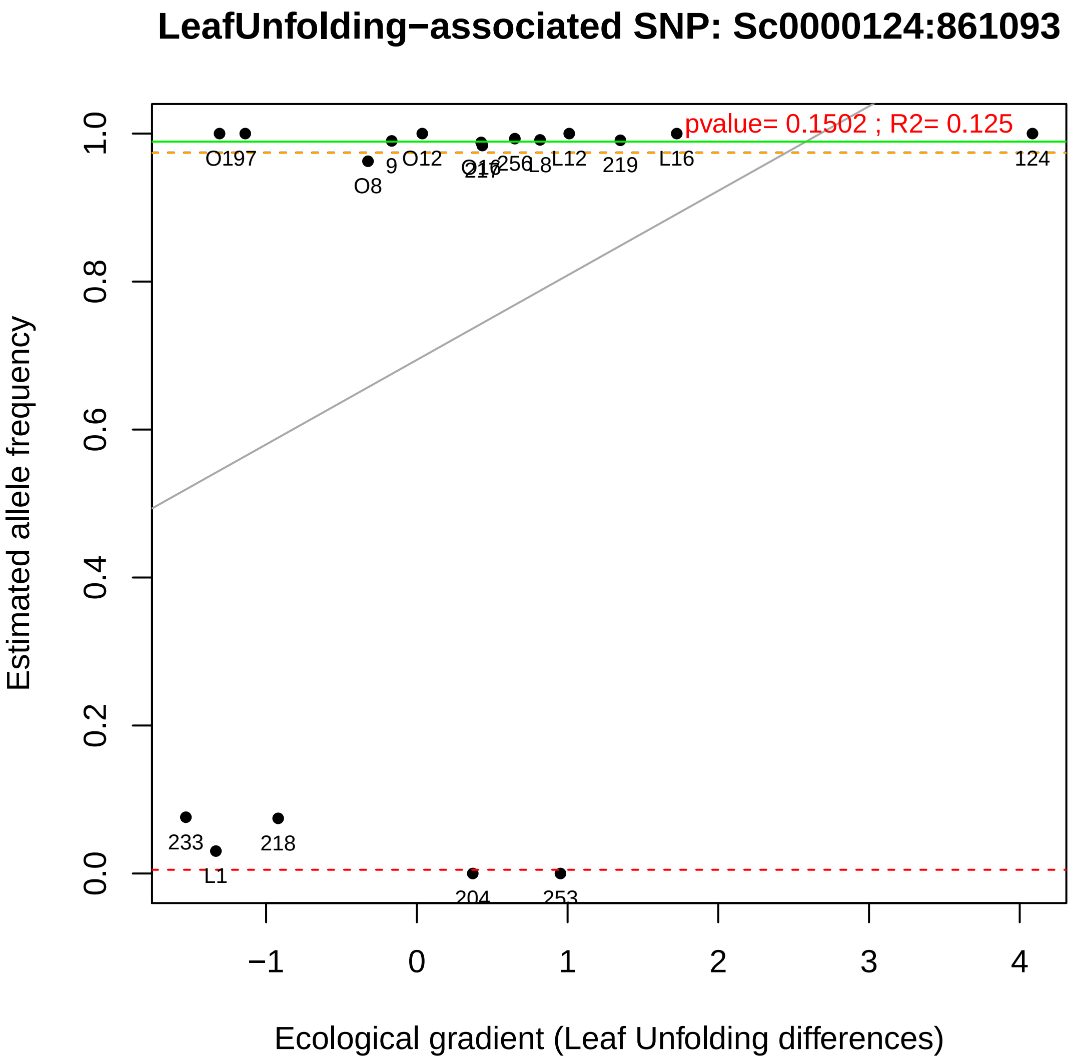
LeafUnfolding−associated SNP: Sc0000124:861093
pvalue= 0.1502 ; R2= 0.125
Estimated allele frequency
Ecological gradient (Leaf Unfolding differences)
0.0
0.2
0.4
0.6
0.8
1.0
−1
0
1
2
3
4
O19
197
O8
9
O12
256
L8
L12
219
L16
124
233
L1
218
204
253

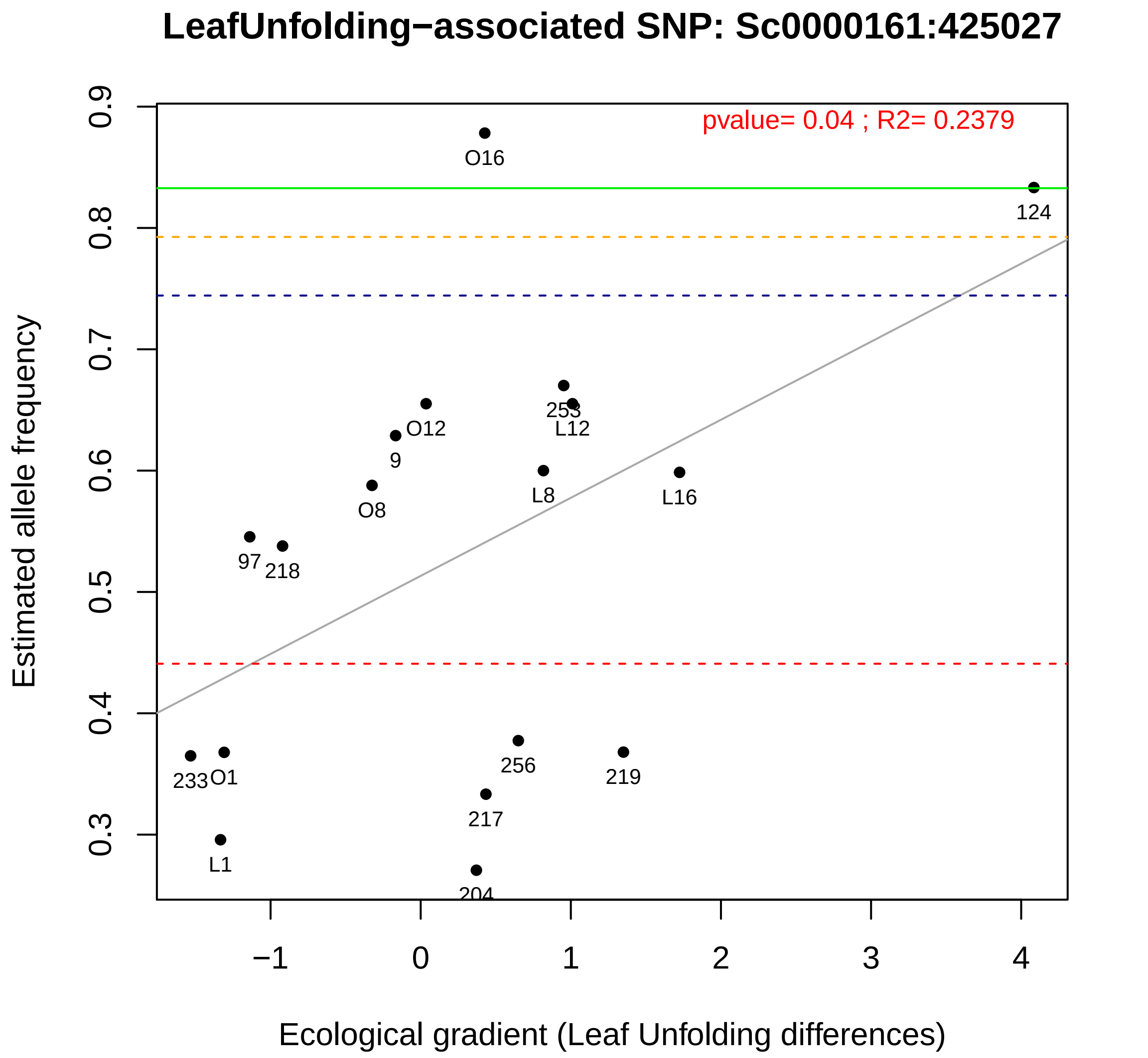
LeafUnfolding−associated SNP: Sc0000161:425027
pvalue= 0.04 ; R2= 0.2379
Estimated allele frequency
Ecological gradient (Leaf Unfolding differences)
0.3
0.4
0.5
0.6
0.7
0.8
0.9
−1
0
1
2
3
4
O16
124
253
L12
O12
9
L8
L16
O8
97
218
256
219
233
O1
217
L1
204

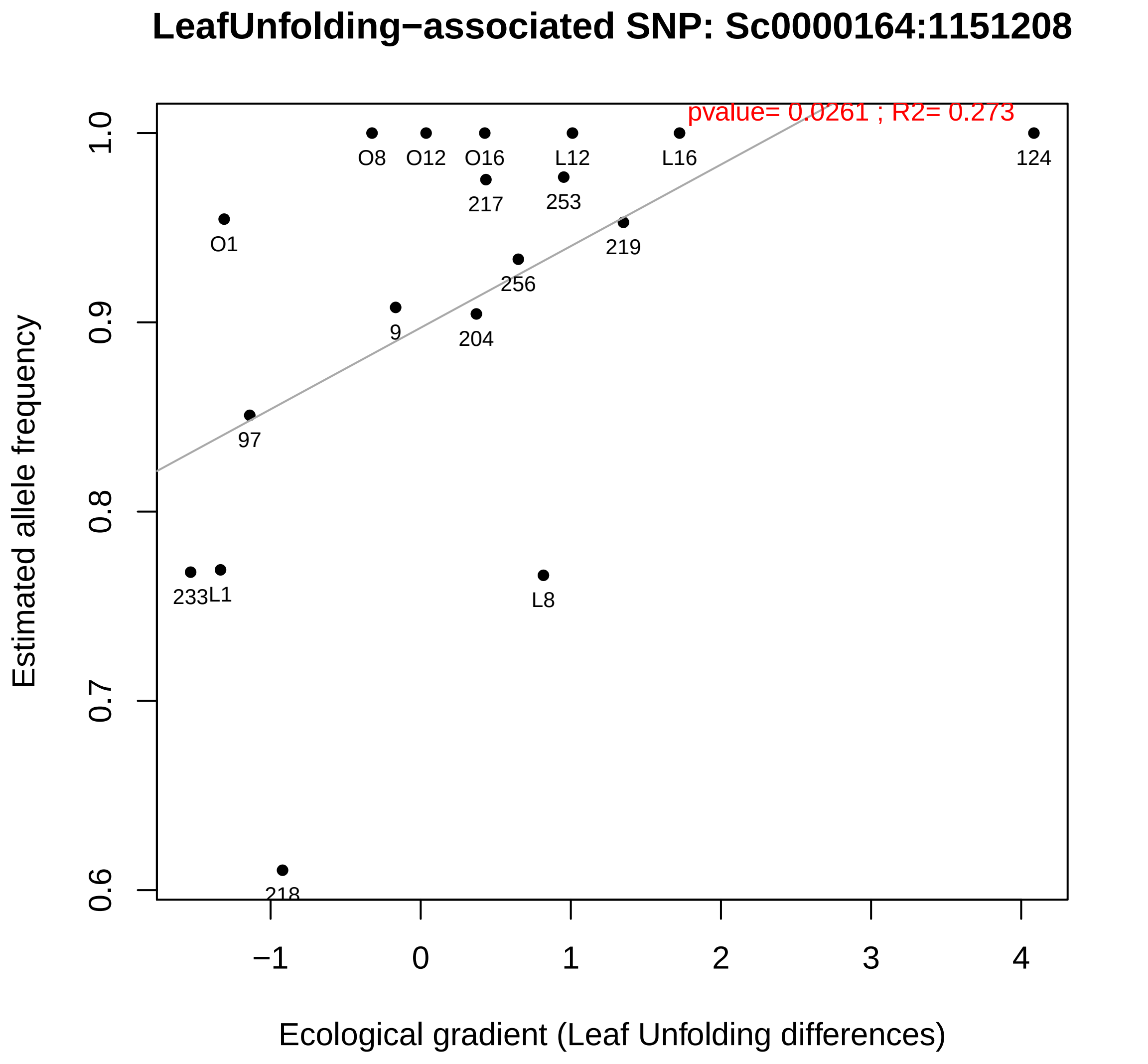

LeafUnfolding−associated SNP: Sc0000164:1151208
pvalue= 0.0261 ; R2= 0.273
Estimated allele frequency
Ecological gradient (Leaf Unfolding differences)
0.6
0.7
0.8
0.9
1.0
−1
0
1
2
3
4
O8
O12
O16
L12
L16
124
217
253
O1
219
256
9
204
97
233
L1
L8
218

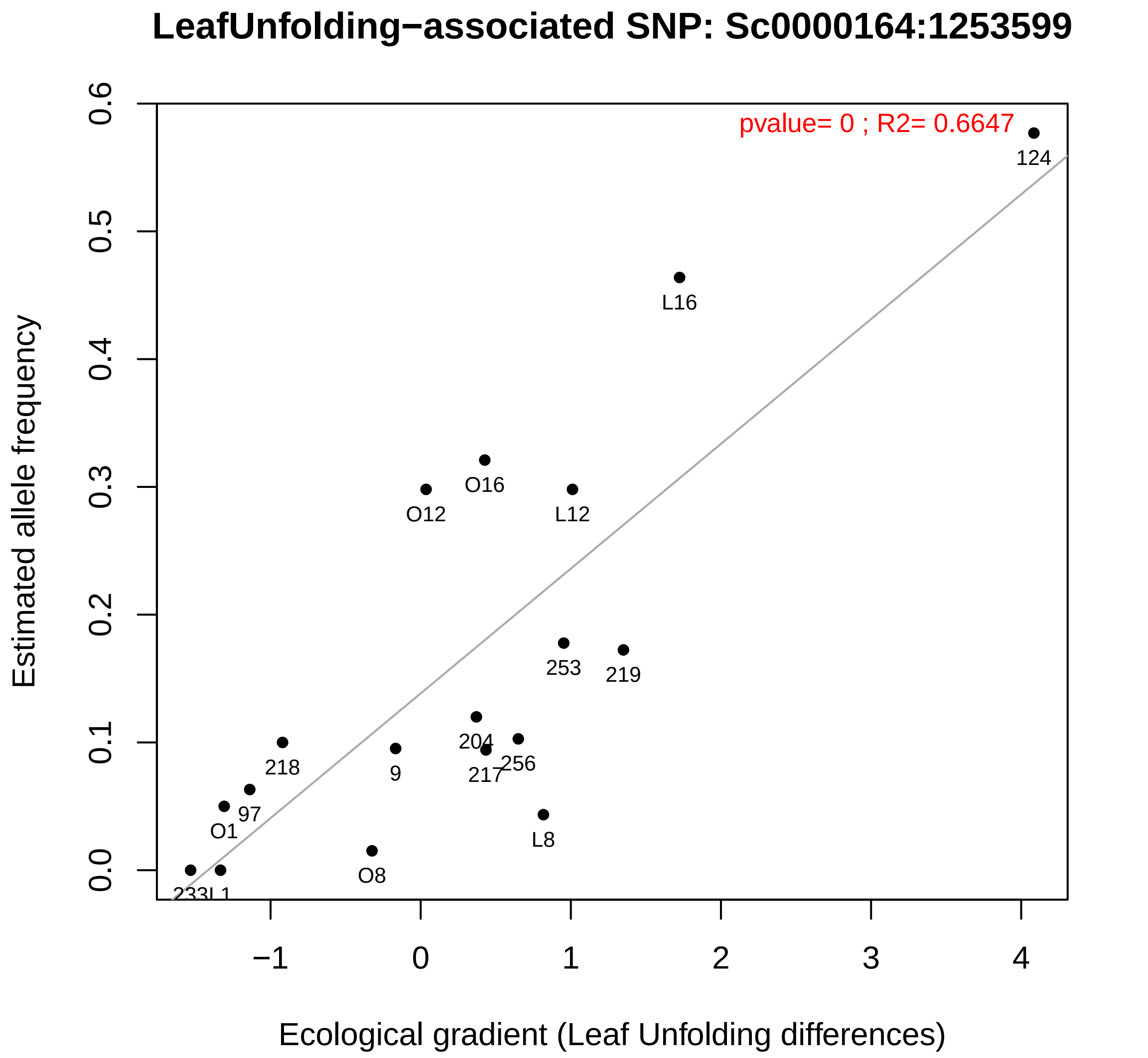
LeafUnfolding−associated SNP: Sc0000164:1253599
pvalue= 0 ; R2= 0.6647
Estimated allele frequency
Ecological gradient (Leaf Unfolding differences)
0.0
0.1
0.2
0.3
0.4
0.5
0.6
−1
0
1
2
3
4
124
L16
O16
O12
L12
253
219
204
217
256
218
9
97
O1
L8
O8
233
L1

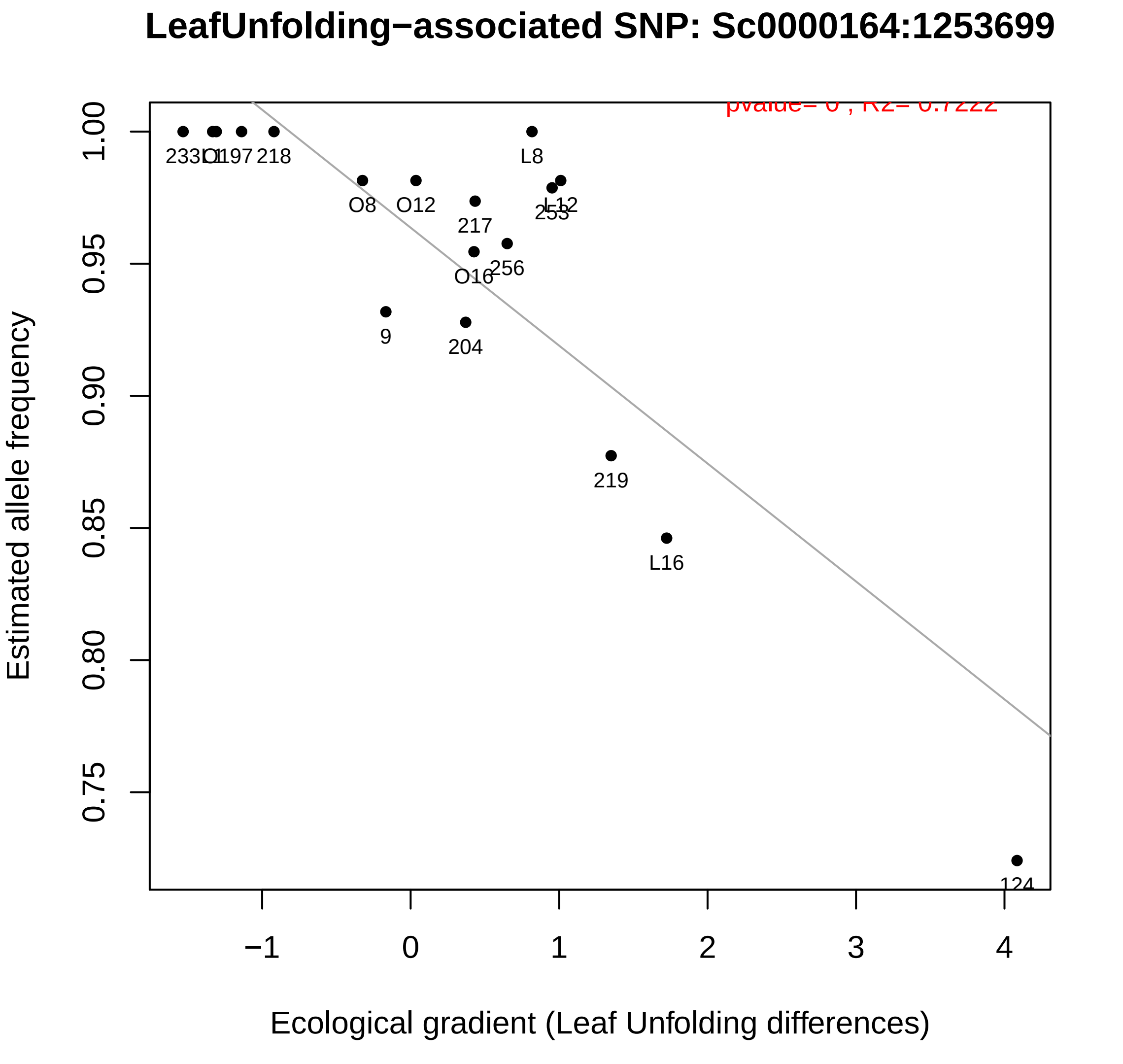

LeafUnfolding−associated SNP: Sc0000164:1253699
pvalue= 0 , R2= 0.7222
Estimated allele frequency
1.00
0.95
0.90
0.85
0.80
0.75
−1
0
1
2
3
4
Ecological gradient (Leaf Unfolding differences)
218
L8
O8
O12
217
256
O16
9
204
219
L16
124

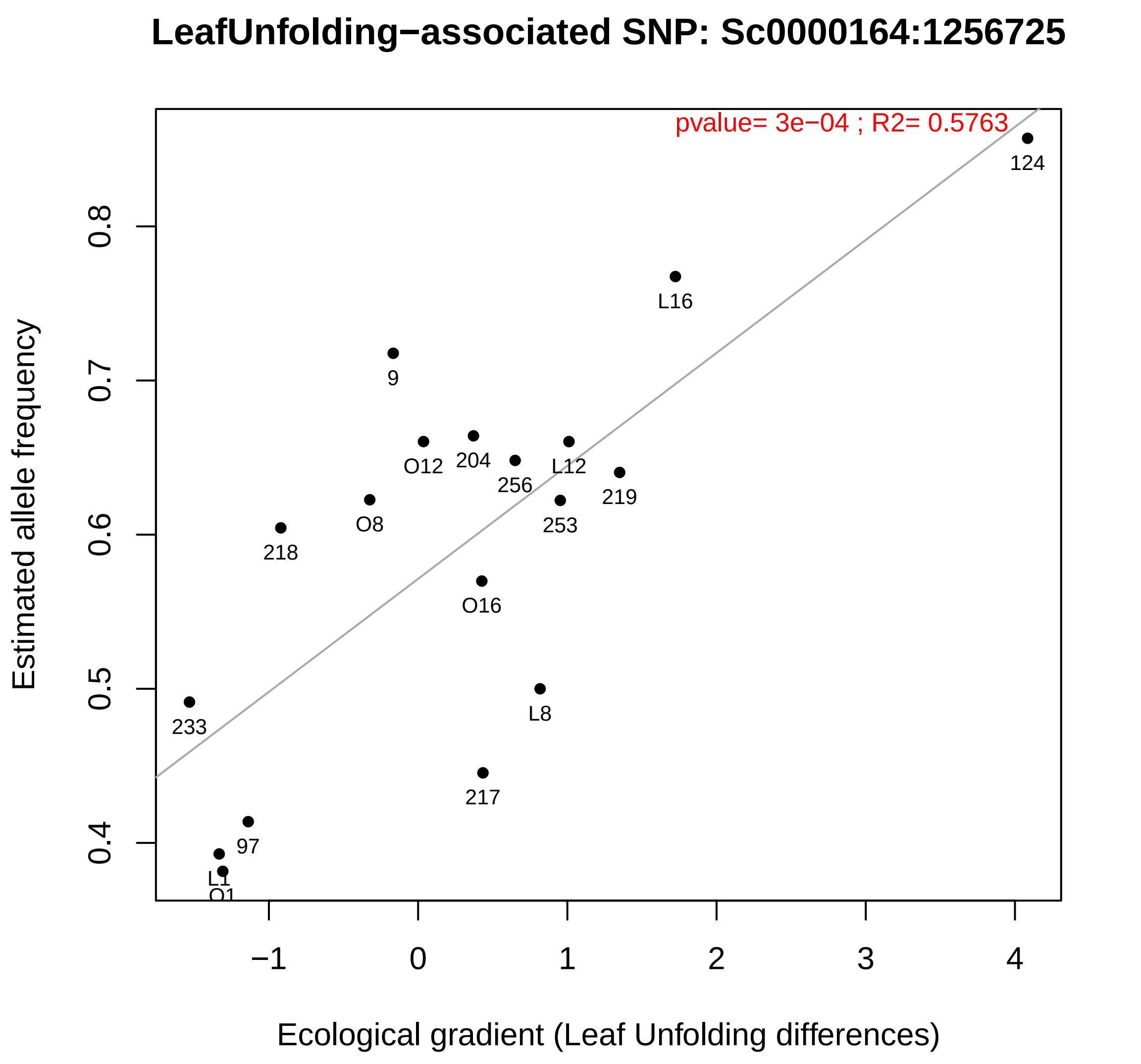

LeafUnfolding−associated SNP: Sc0000164:1256725
pvalue= 3e−04 ; R2= 0.5763
Estimated allele frequency
Ecological gradient (Leaf Unfolding differences)
0.4
0.5
0.6
0.7
0.8
−1
0
1
2
3
4
124
L16
9
O12
204
256
L12
219
253
O8
218
O16
233
L8
217
97
L1
O1

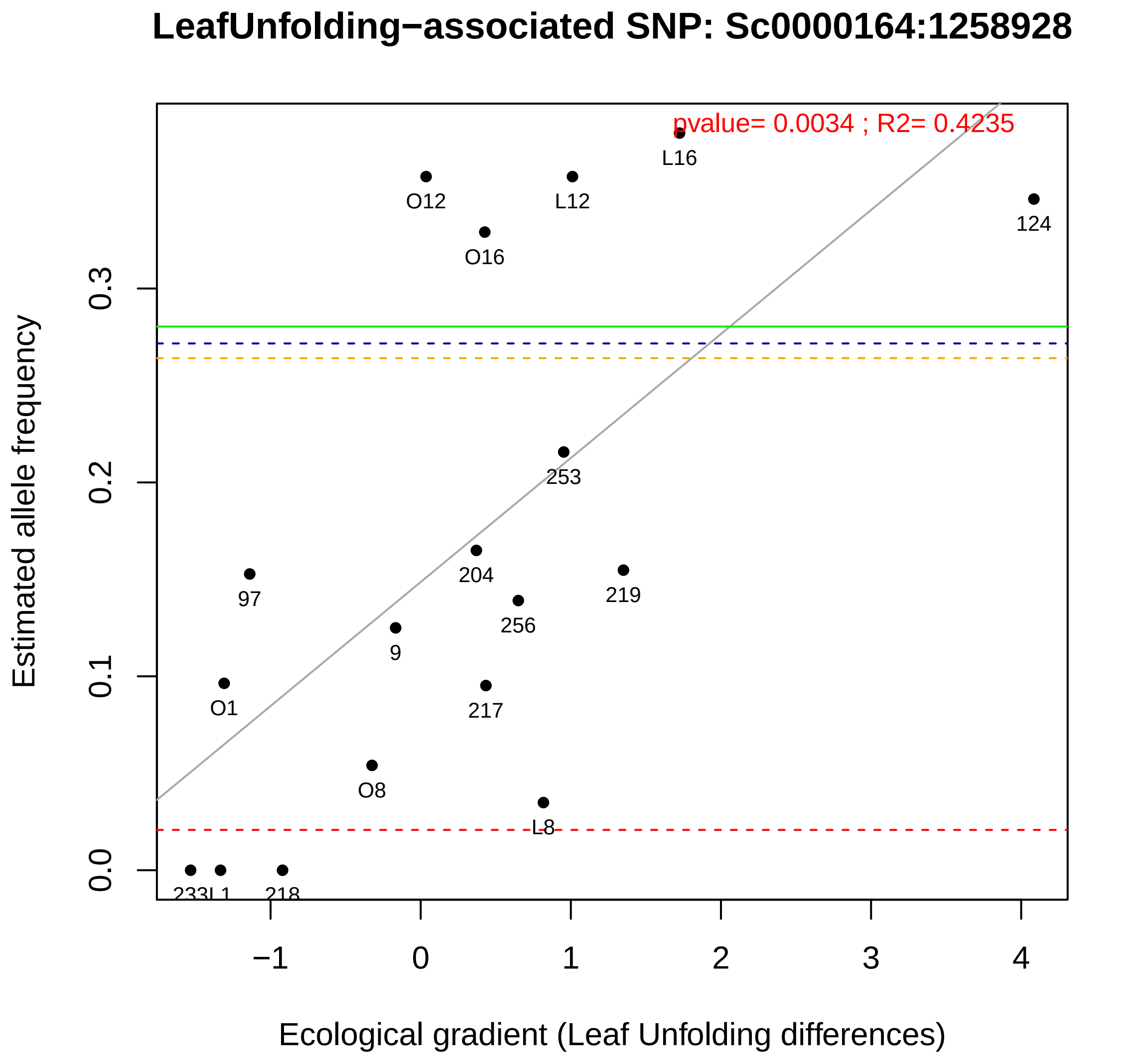

LeafUnfolding−associated SNP: Sc0000164:1258928
pvalue= 0.0034 ; R2= 0.4235
Estimated allele frequency
Ecological gradient (Leaf Unfolding differences)
0.0
0.1
0.2
0.3
−1
0
1
2
3
4
L16
O12
L12
124
O16
253
204
219
97
256
9
O1
217
O8
L8
233
L1
218

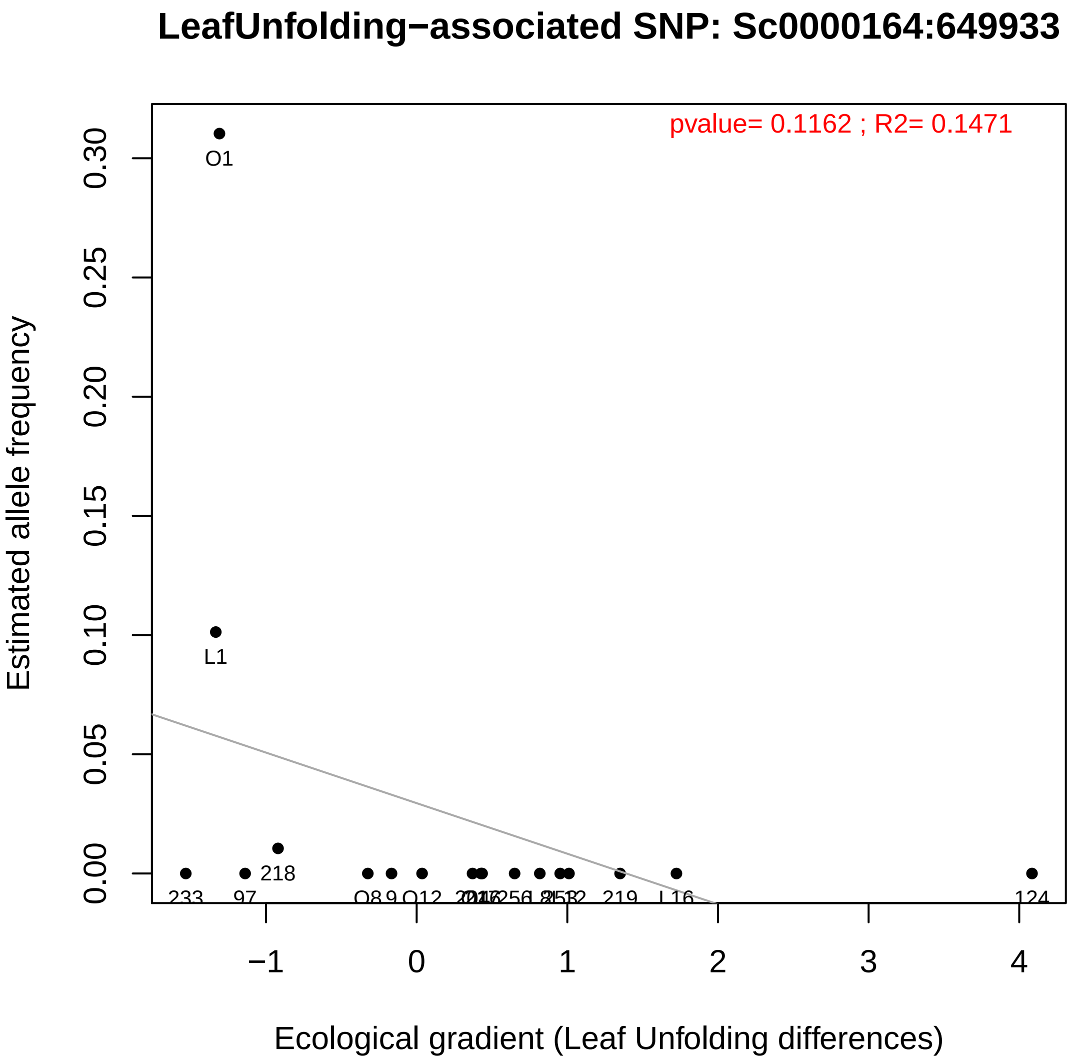
LeafUnfolding−associated SNP: Sc0000164:649933
pvalue= 0.1162 ; R2= 0.1471
Estimated allele frequency
0.00
0.05
0.10
0.15
0.20
0.25
0.30
Ecological gradient (Leaf Unfolding differences)
−1
0
1
2
3
4
O1
L1
233
97
218
O8
9
O12
219
L16
124

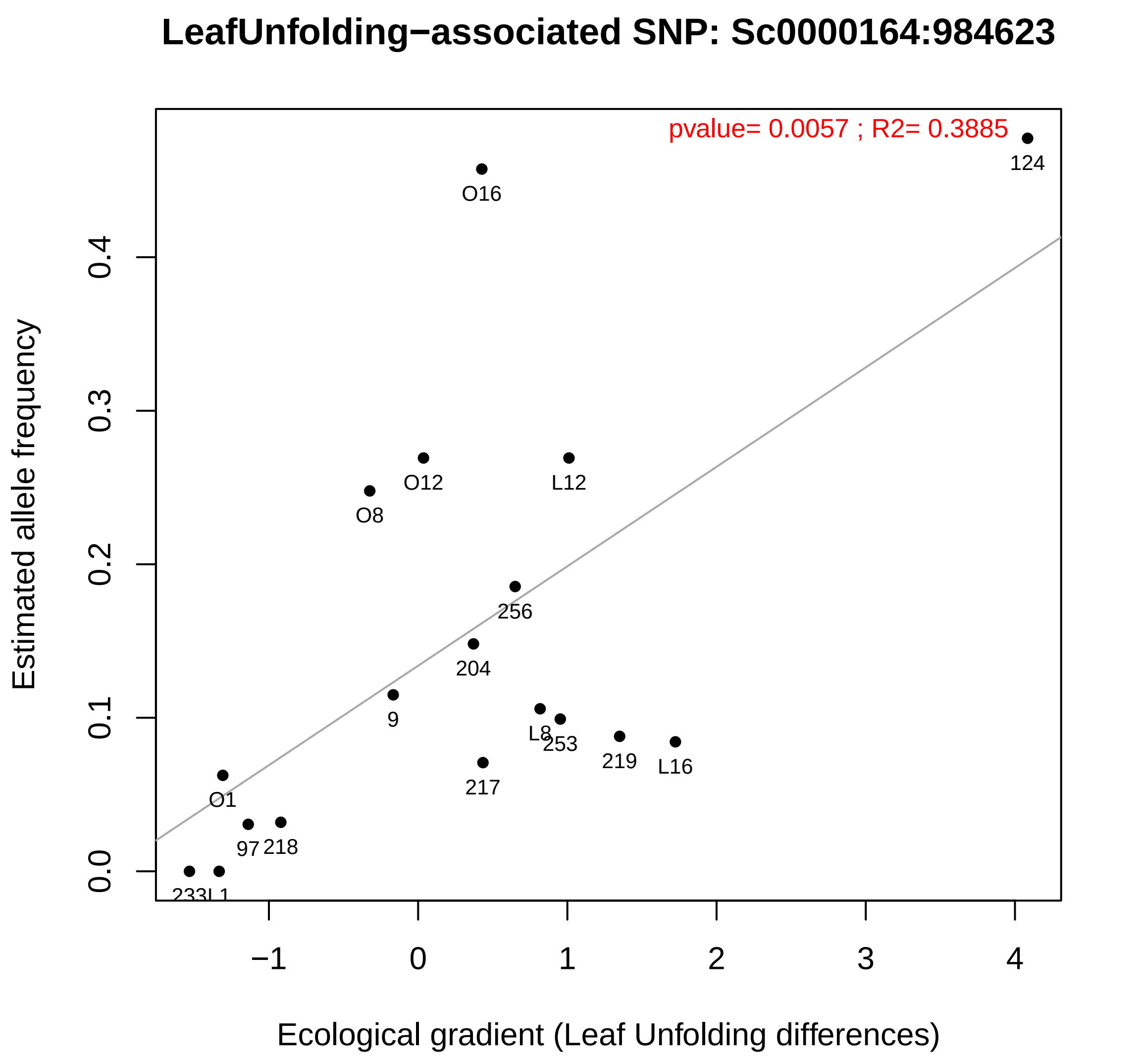

LeafUnfolding−associated SNP: Sc0000164:984623
pvalue= 0.0057 ; R2= 0.3885
Estimated allele frequency
Ecological gradient (Leaf Unfolding differences)
0.0
0.1
0.2
0.3
0.4
−1
0
1
2
3
4
124
O16
O12
L12
O8
256
204
9
L8
253
219
L16
217
O1
97
218
233
L1

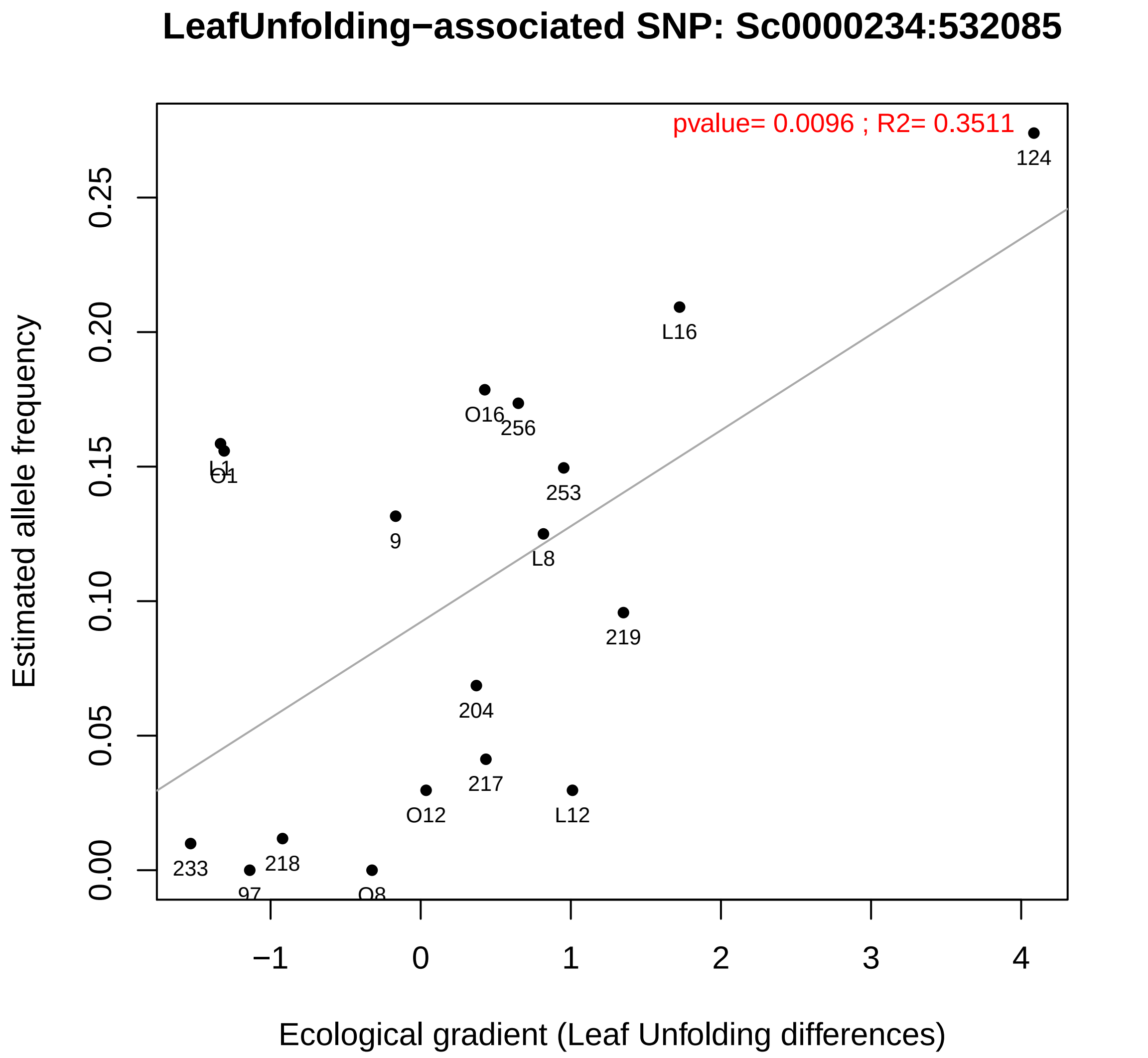

LeafUnfolding−associated SNP: Sc0000234:532085
pvalue= 0.0096 ; R2= 0.3511
Estimated allele frequency
0.00
0.05
0.10
0.15
0.20
0.25
−1
0
1
2
3
4
Ecological gradient (Leaf Unfolding differences)
124
L16
O16
256
L1
O1
253
9
L8
219
204
217
O12
L12
233
218
97
O8

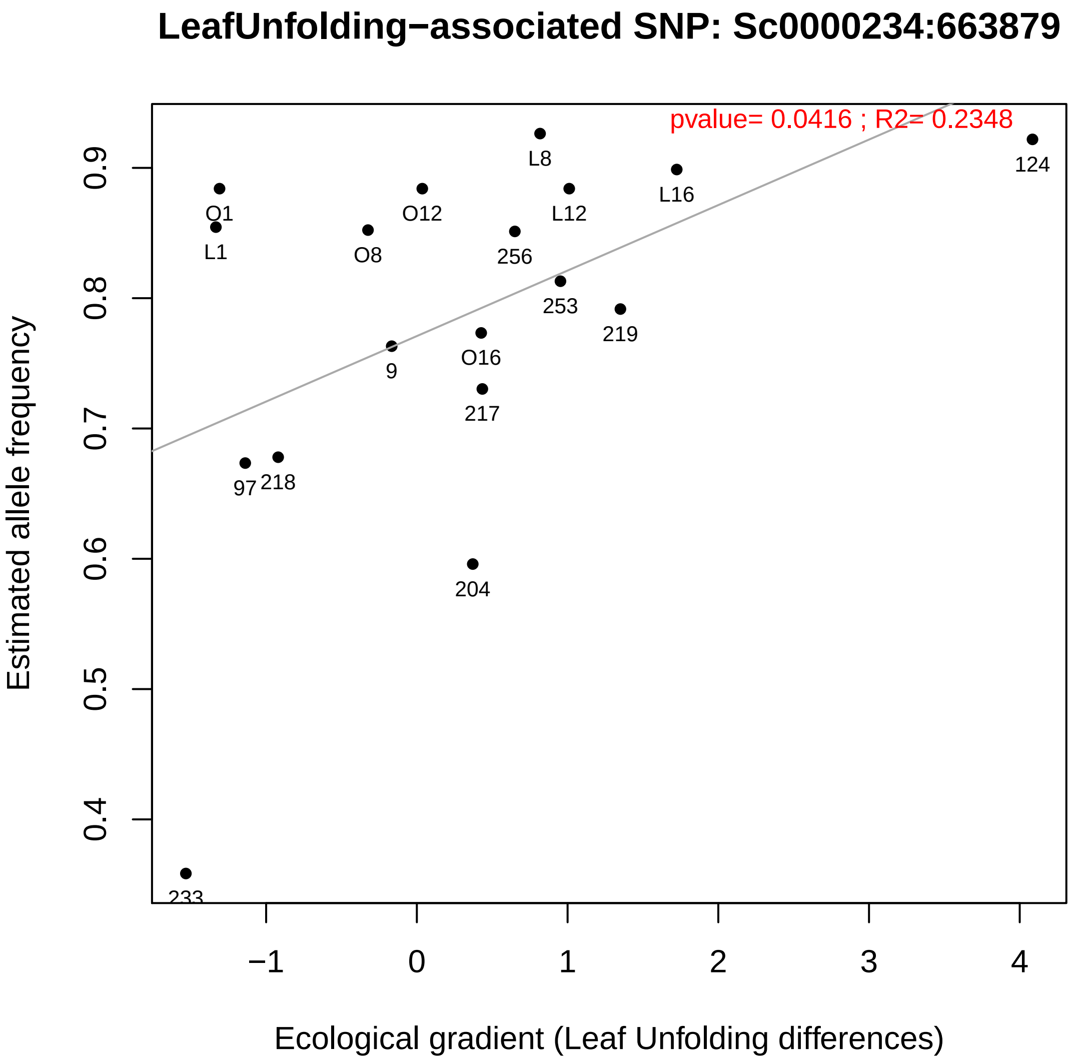
LeafUnfolding−associated SNP: Sc0000234:663879
pvalue= 0.0416 ; R2= 0.2348
Estimated allele frequency
0.4
0.5
0.6
0.7
0.8
0.9
−1
0
1
2
3
4
Ecological gradient (Leaf Unfolding differences)
L8
124
O1
O12
L12
L16
L1
O8
256
253
219
9
O16
217
97
218
204
233

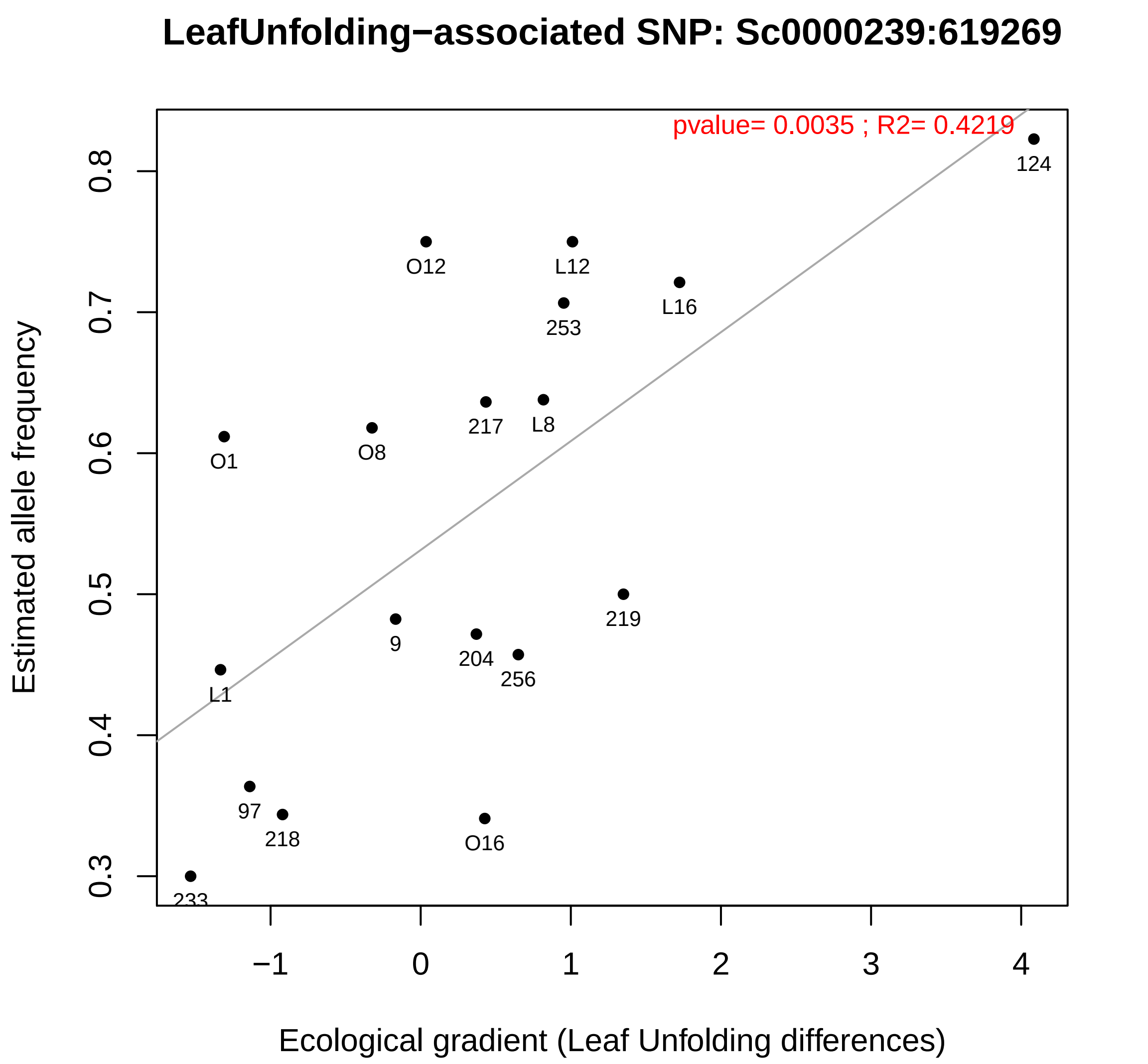
LeafUnfolding−associated SNP: Sc0000239:619269
pvalue= 0.0035 ; R2= 0.4219
Estimated allele frequency
0.3
0.4
0.5
0.6
0.7
0.8
−1
0
1
2
3
4
Ecological gradient (Leaf Unfolding differences)
124
O12
L12
L16
253
217
L8
O8
O1
219
9
204
256
L1
97
218
O16

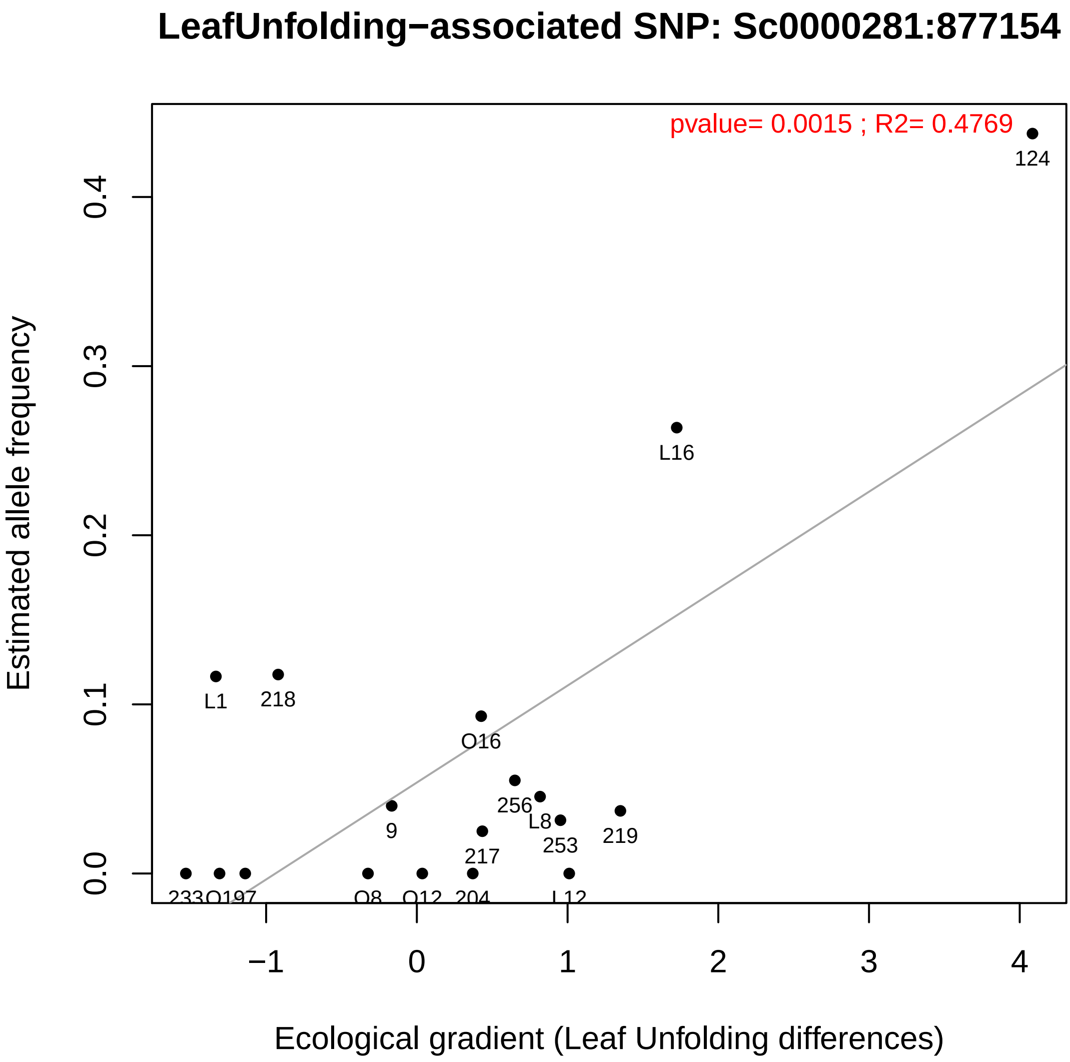

LeafUnfolding−associated SNP: Sc0000281:877154
pvalue= 0.0015 ; R2= 0.4769
Estimated allele frequency
Ecological gradient (Leaf Unfolding differences)
0.0
0.1
0.2
0.3
0.4
−1
0
1
2
3
4
124
L16
L1
218
O16
256
L8
9
217
253
219

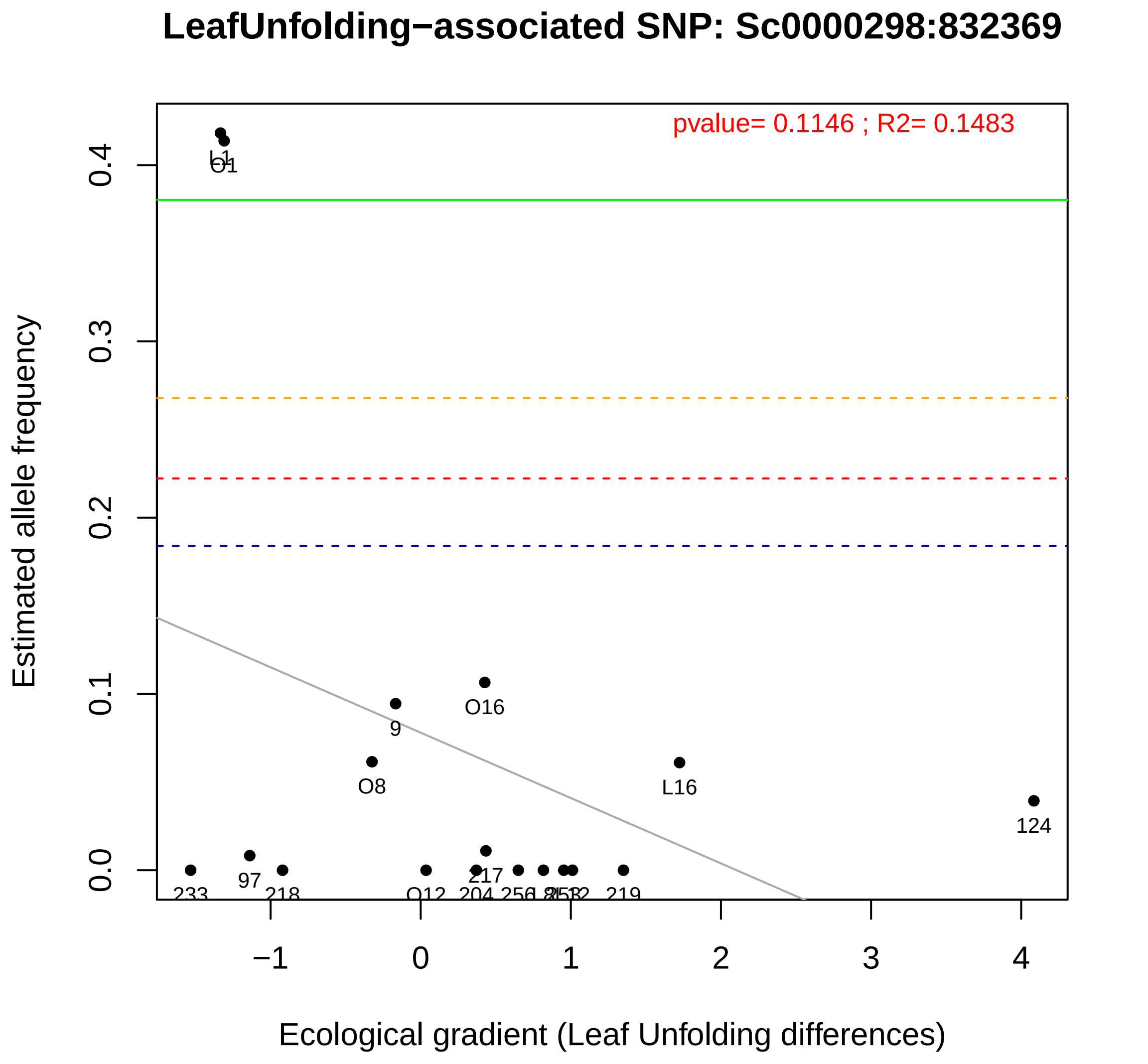

LeafUnfolding−associated SNP: Sc0000298:832369
pvalue= 0.1146 ; R2= 0.1483
Estimated allele frequency
Ecological gradient (Leaf Unfolding differences)
0.0
0.1
0.2
0.3
0.4
−1
0
1
2
3
4
L1
O1
O16
9
O8
L16
124
97
217
233
218
O12
204
219

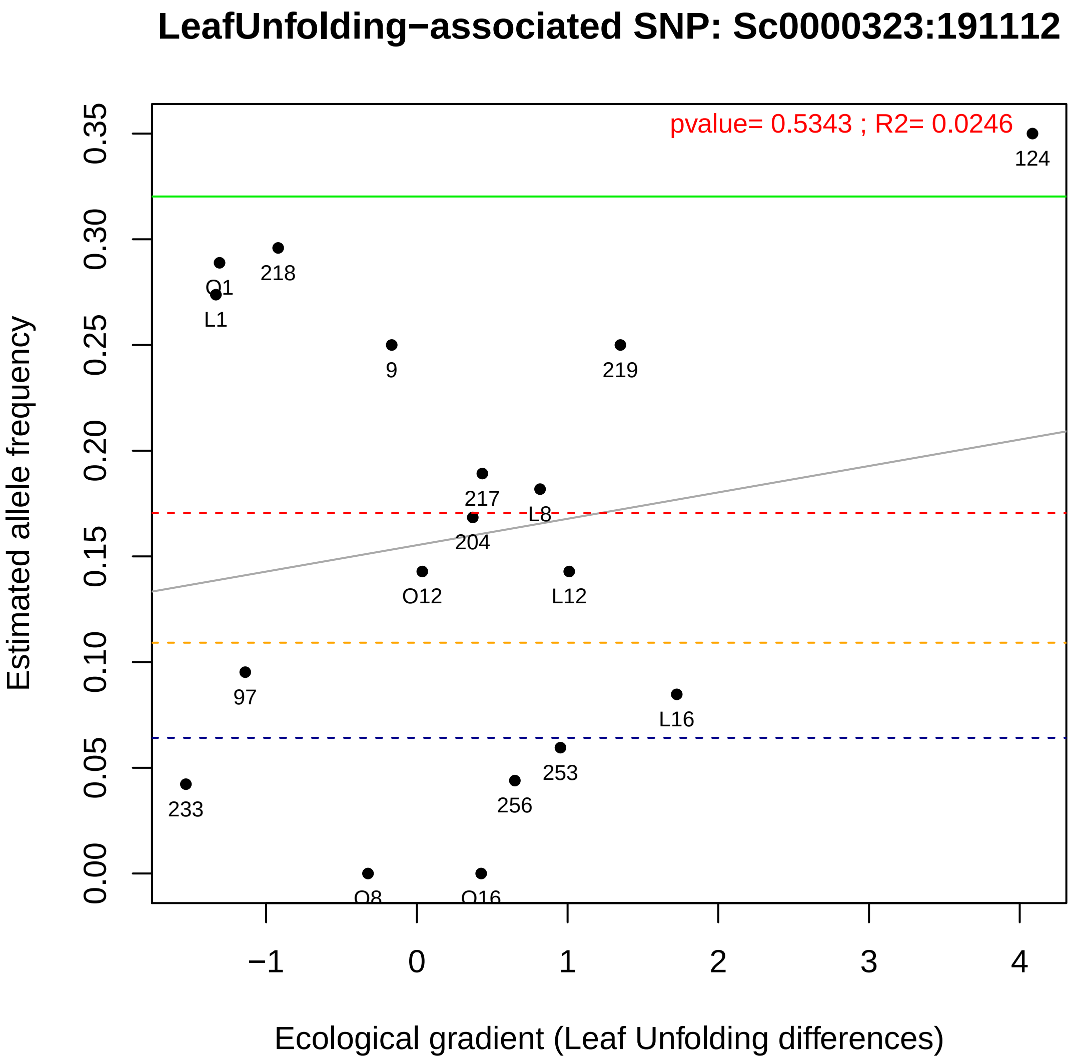
LeafUnfolding−associated SNP: Sc0000323:191112
pvalue= 0.5343 ; R2= 0.0246
Estimated allele frequency
Ecological gradient (Leaf Unfolding differences)
0.00
0.05
0.10
0.15
0.20
0.25
0.30
0.35
−1
0
1
2
3
4
124
218
O1
L1
9
219
217
L8
204
O12
L12
97
L16
253
256
233
O8
O16

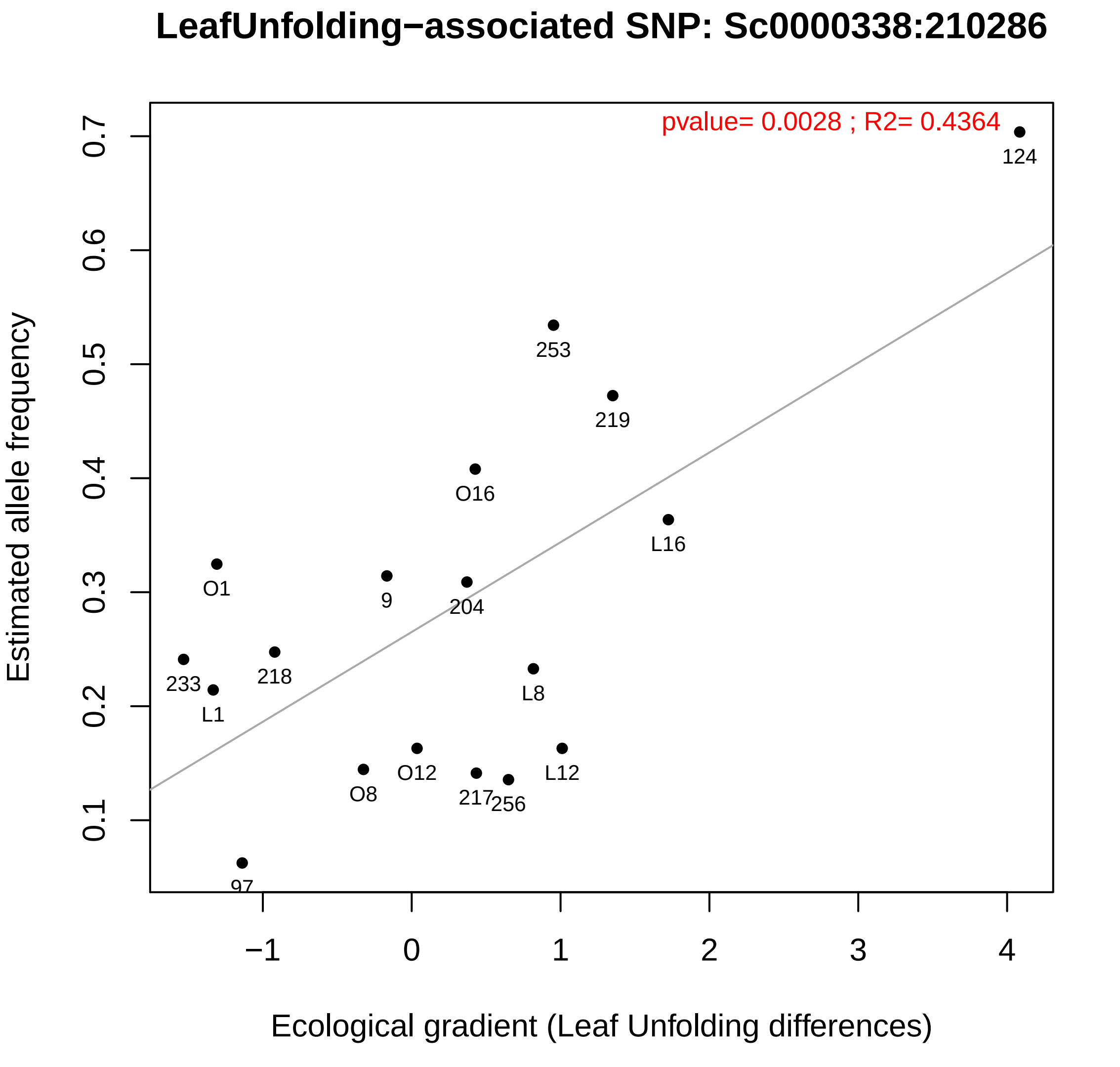

LeafUnfolding−associated SNP: Sc0000338:210286
pvalue= 0.0028 ; R2= 0.4364
Estimated allele frequency
0.1
0.2
0.3
0.4
0.5
0.6
0.7
−1
0
1
2
3
4
Ecological gradient (Leaf Unfolding differences)
124
253
219
O16
L16
O1
9
204
233
218
L1
L8
O12
L12
O8
217
256
97

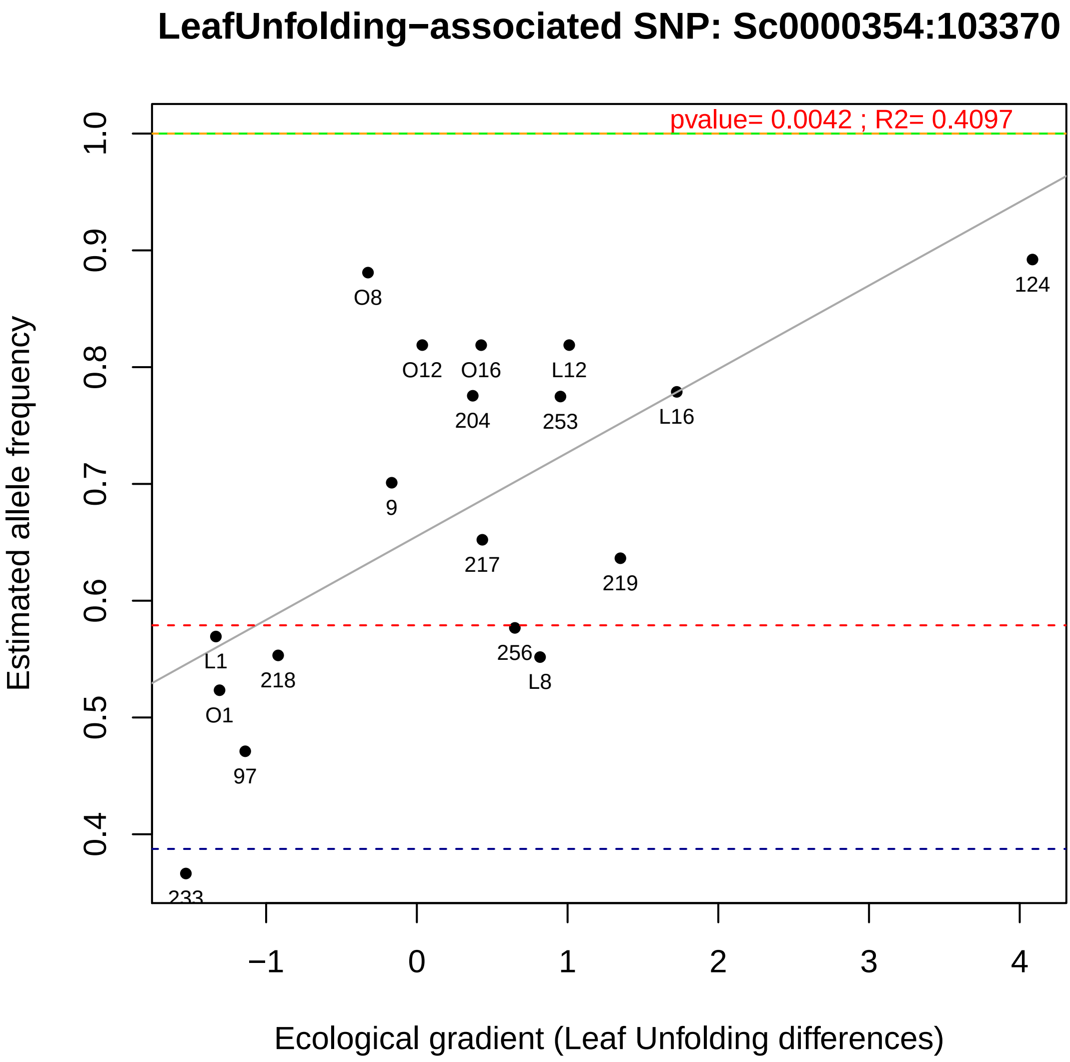
LeafUnfolding−associated SNP: Sc0000354:103370
pvalue= 0.0042 ; R2= 0.4097
Estimated allele frequency
0.4
0.5
0.6
0.7
0.8
0.9
1.0
−1
0
1
2
3
4
Ecological gradient (Leaf Unfolding differences)
O8
O12
O16
L12
124
204
253
L16
9
217
219
L1
218
256
L8
O1
97
233

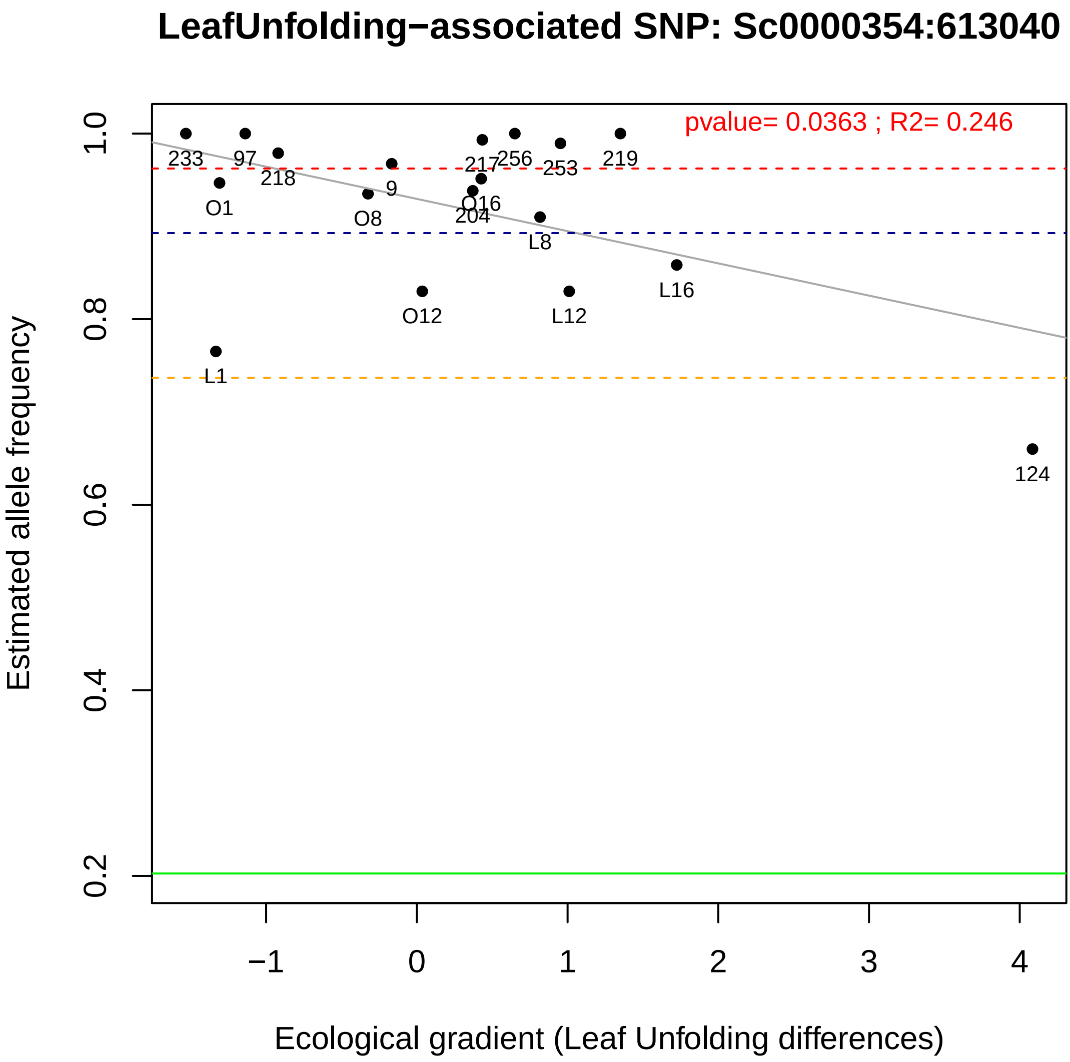
LeafUnfolding−associated SNP: Sc0000354:613040
pvalue= 0.0363 ; R2= 0.246
Estimated allele frequency
0.2
0.4
0.6
0.8
1.0
−1
0
1
2
3
4
Ecological gradient (Leaf Unfolding differences)
233
97
218
O1
L1
O8
9
O12
217
256
O16
204
253
L8
L12
219
L16
124

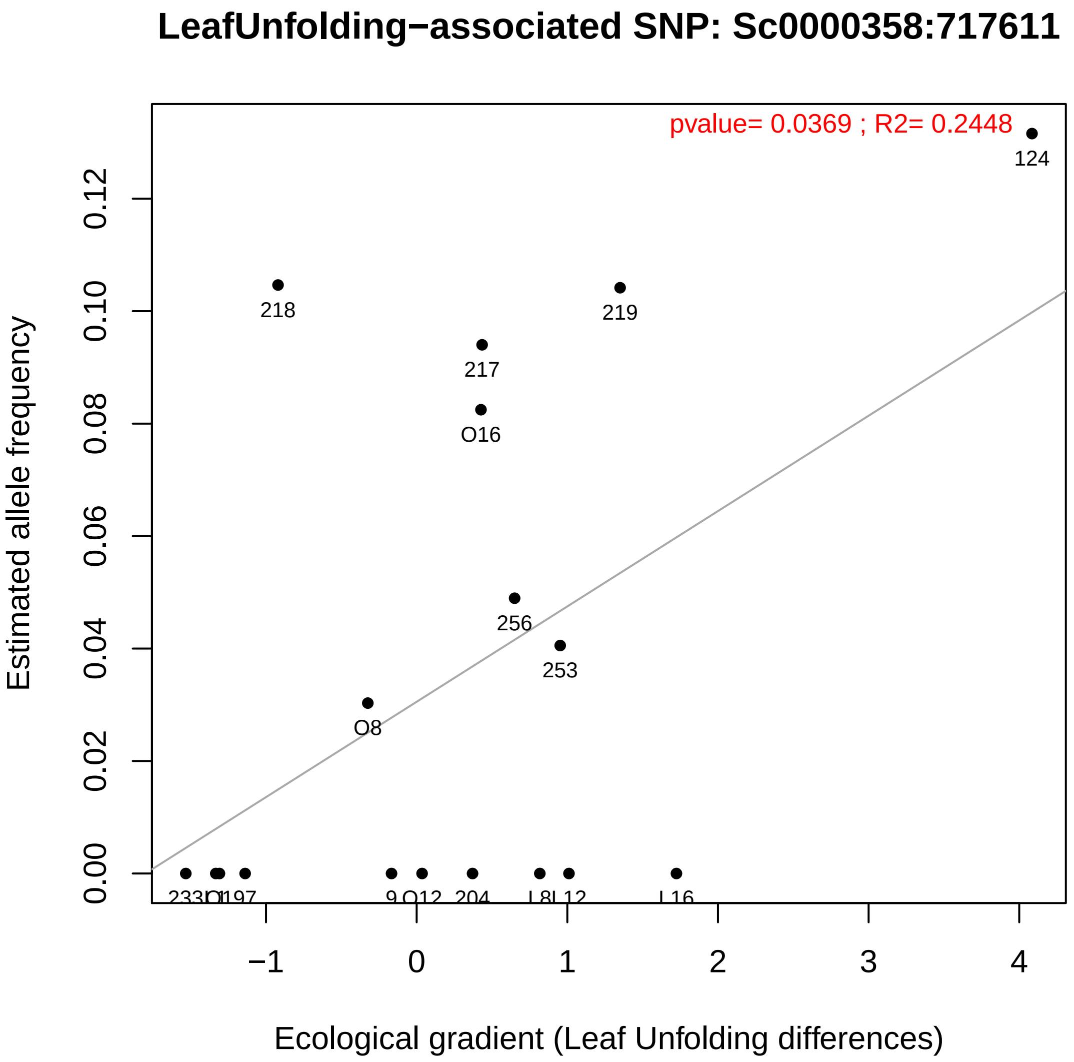
LeafUnfolding−associated SNP: Sc0000358:717611
pvalue= 0.0369 ; R2= 0.2448
Estimated allele frequency
0.00
0.02
0.04
0.06
0.08
0.10
0.12
Ecological gradient (Leaf Unfolding differences)
−1
0
1
2
3
4
124
218
219
217
O16
256
253
O8
204
L16

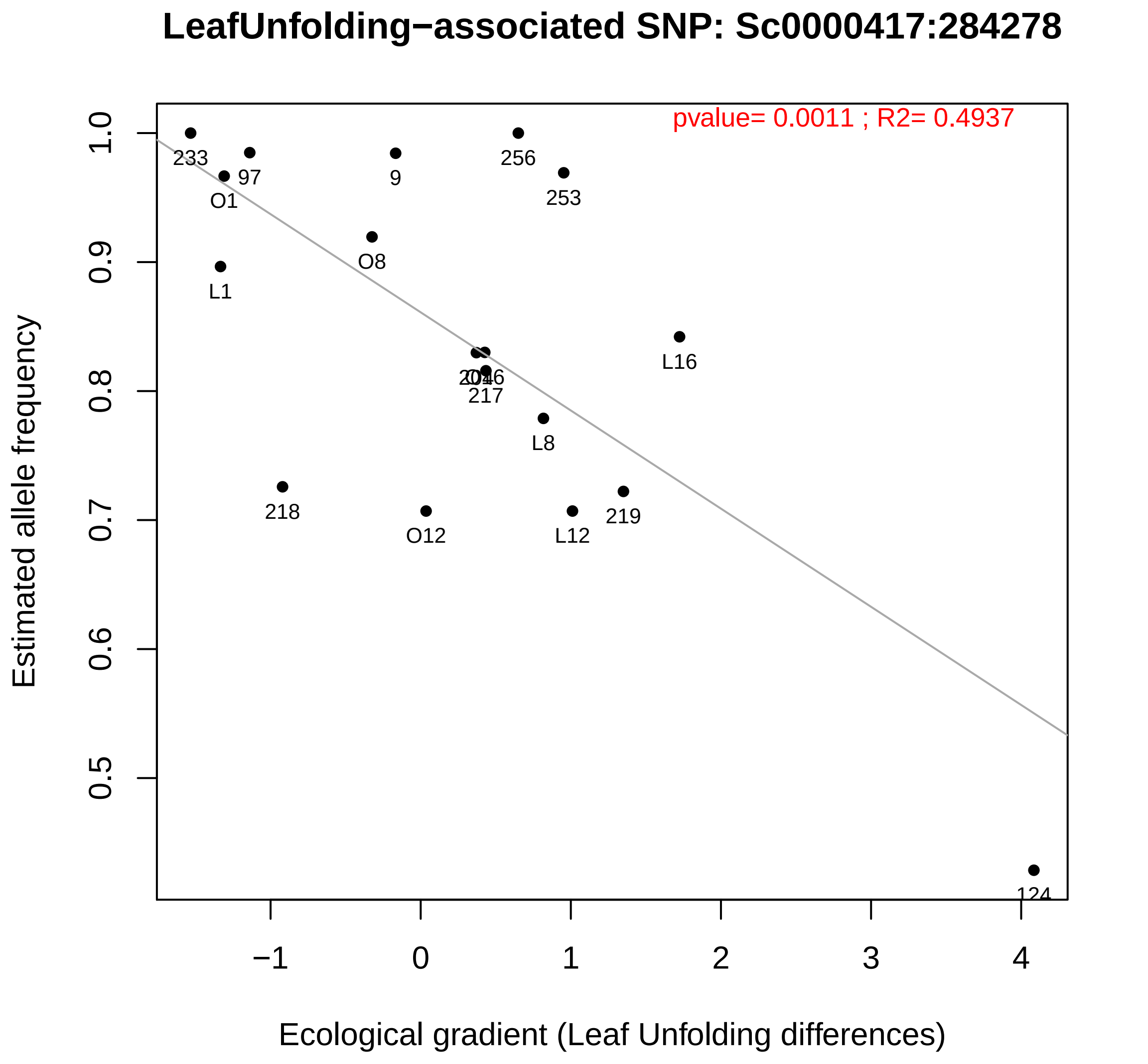

LeafUnfolding−associated SNP: Sc0000417:284278
pvalue= 0.0011 ; R2= 0.4937
Estimated allele frequency
Ecological gradient (Leaf Unfolding differences)
1.0
0.9
0.8
0.7
0.6
0.5
−1
0
1
2
3
4
233
97
O1
9
256
253
O8
L1
L16
217
L8
218
O12
L12
219
124

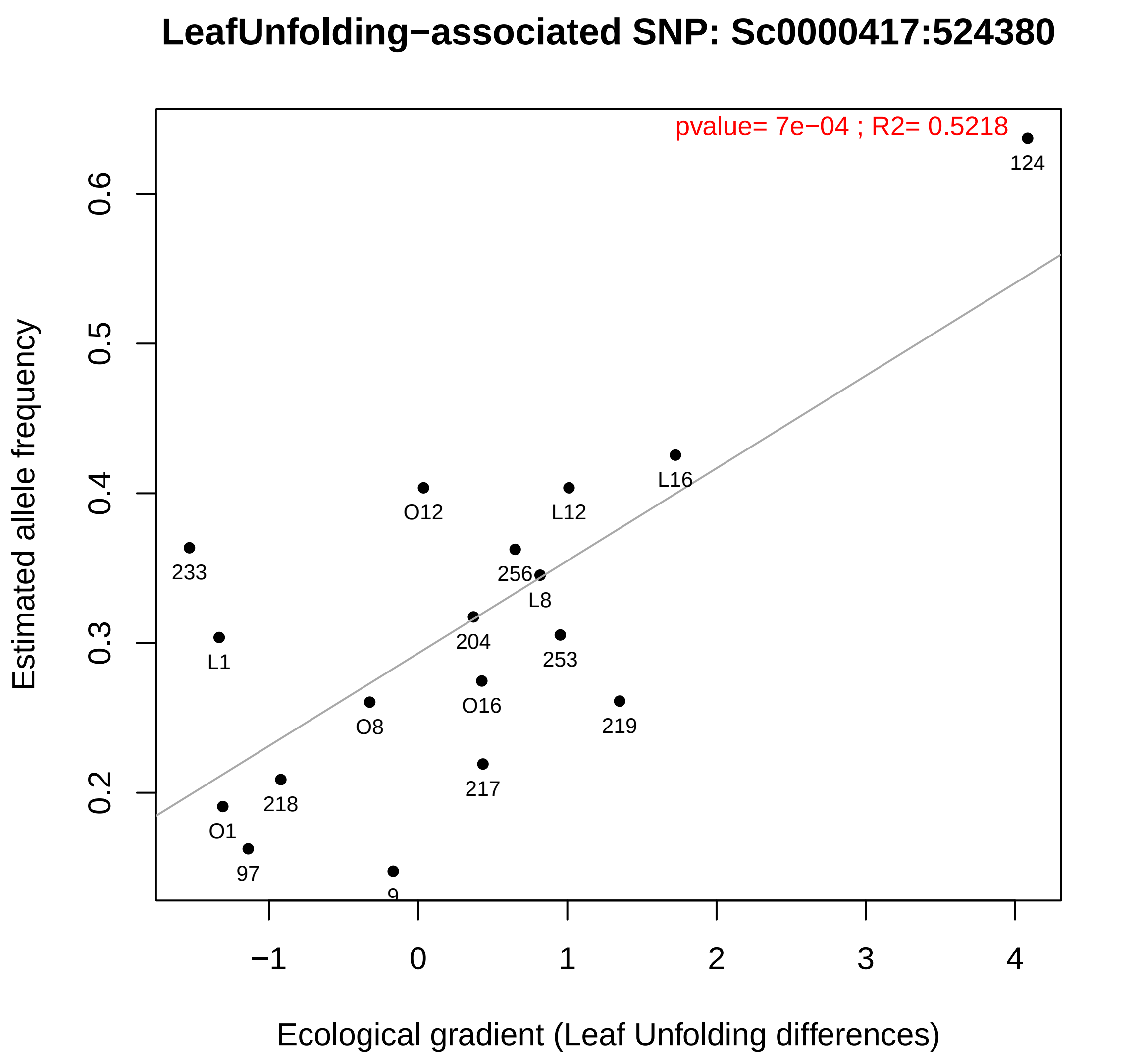

LeafUnfolding−associated SNP: Sc0000417:524380
pvalue= 7e−04 ; R2= 0.5218
Estimated allele frequency
Ecological gradient (Leaf Unfolding differences)
0.2
0.3
0.4
0.5
0.6
−1
0
1
2
3
4
124
L16
O12
L12
233
256
L8
204
253
L1
O16
O8
219
217
218
O1
97
9

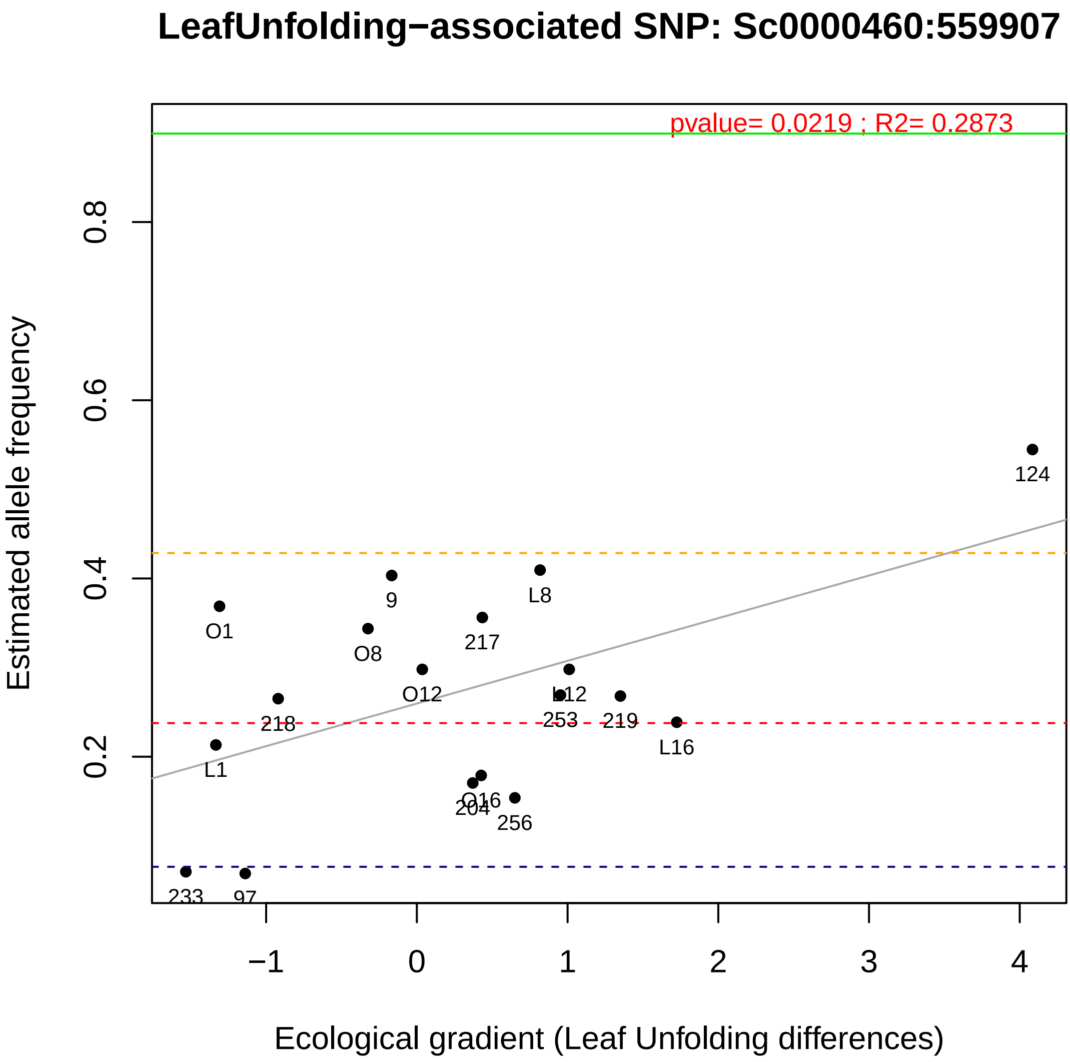
LeafUnfolding−associated SNP: Sc0000460:559907
pvalue= 0.0219 ; R2= 0.2873
Estimated allele frequency
0.2
0.4
0.6
0.8
−1
0
1
2
3
4
Ecological gradient (Leaf Unfolding differences)
124
L8
9
O1
O8
217
O12
L12
253
219
L16
218
L1
O16
204
256
233
97

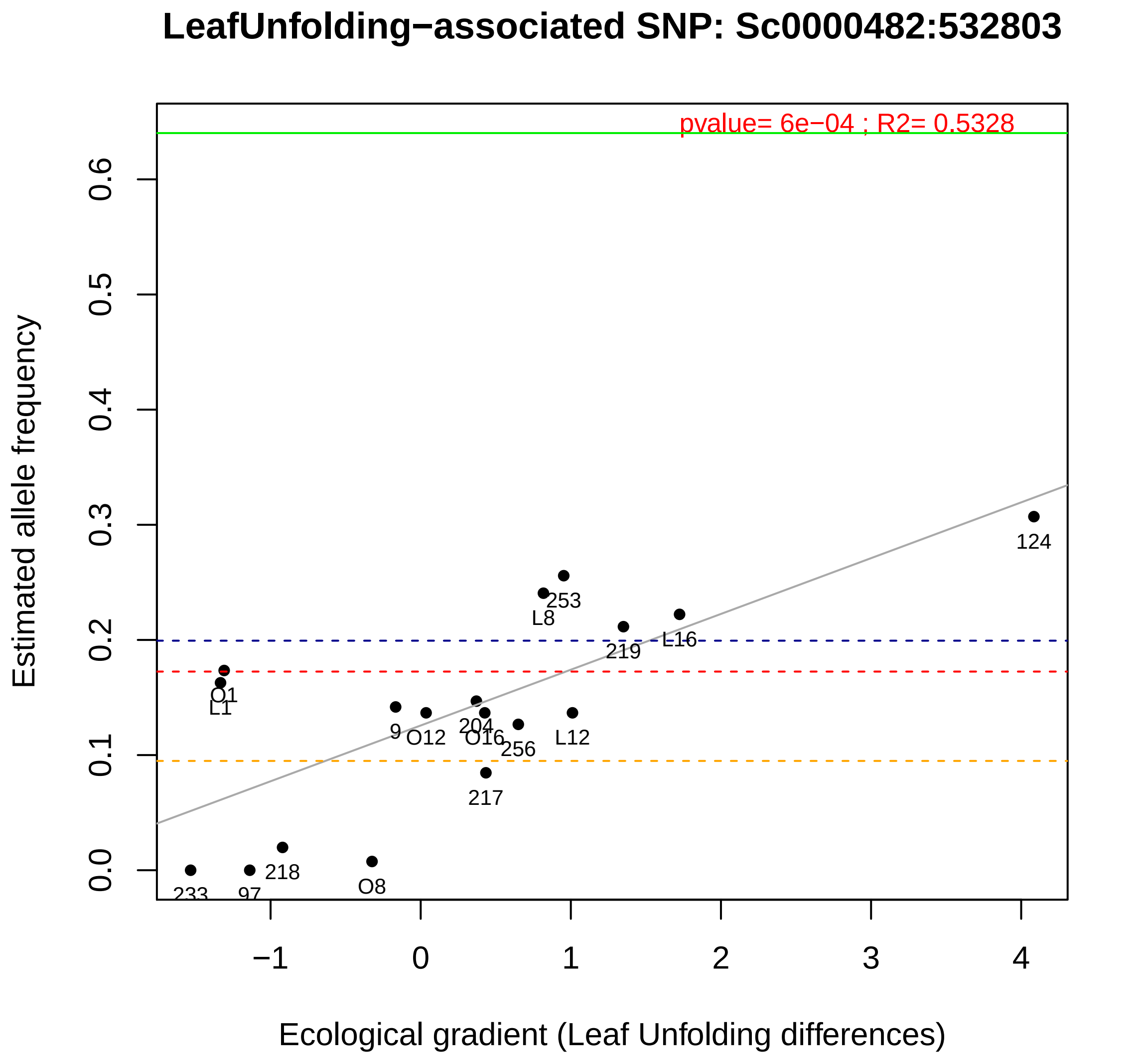
LeafUnfolding−associated SNP: Sc0000482:532803
pvalue= 6e−04 ; R2= 0.5328
Estimated allele frequency
0.0
0.1
0.2
0.3
0.4
0.5
0.6
−1
0
1
2
3
4
Ecological gradient (Leaf Unfolding differences)
124
253
L8
219
L16
O1
L1
9
O12
204
O16
256
L12
217
218
233
97
O8

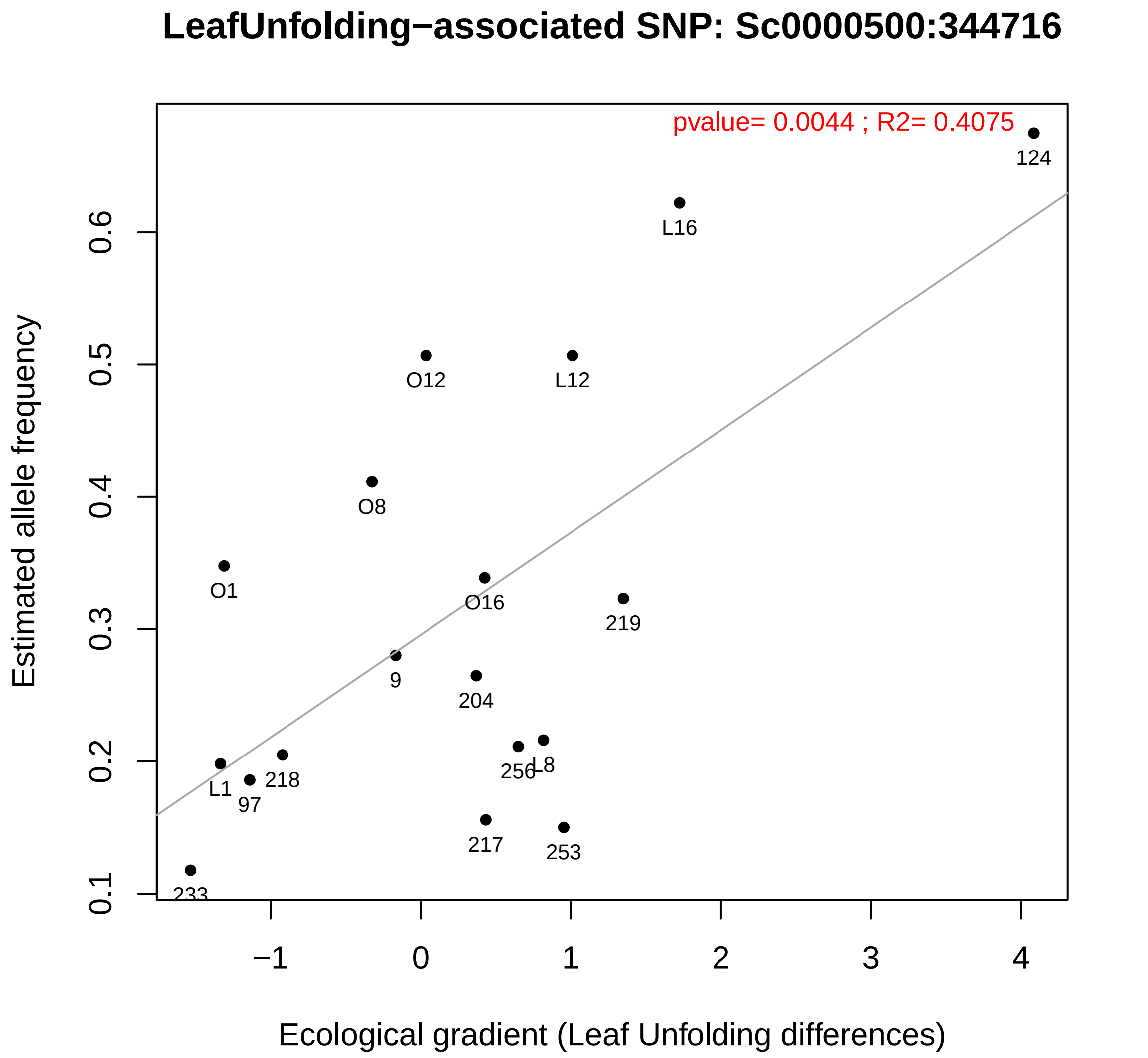

LeafUnfolding−associated SNP: Sc0000500:344716
pvalue= 0.0044 ; R2= 0.4075
124
L16
O12
L12
O8
O1
O16
219
9
204
256
L8
L1
97
218
217
253
233
0.1
0.2
0.3
0.4
0.5
0.6
−1
0
1
2
3
4
Estimated allele frequency
Ecological gradient (Leaf Unfolding differences)

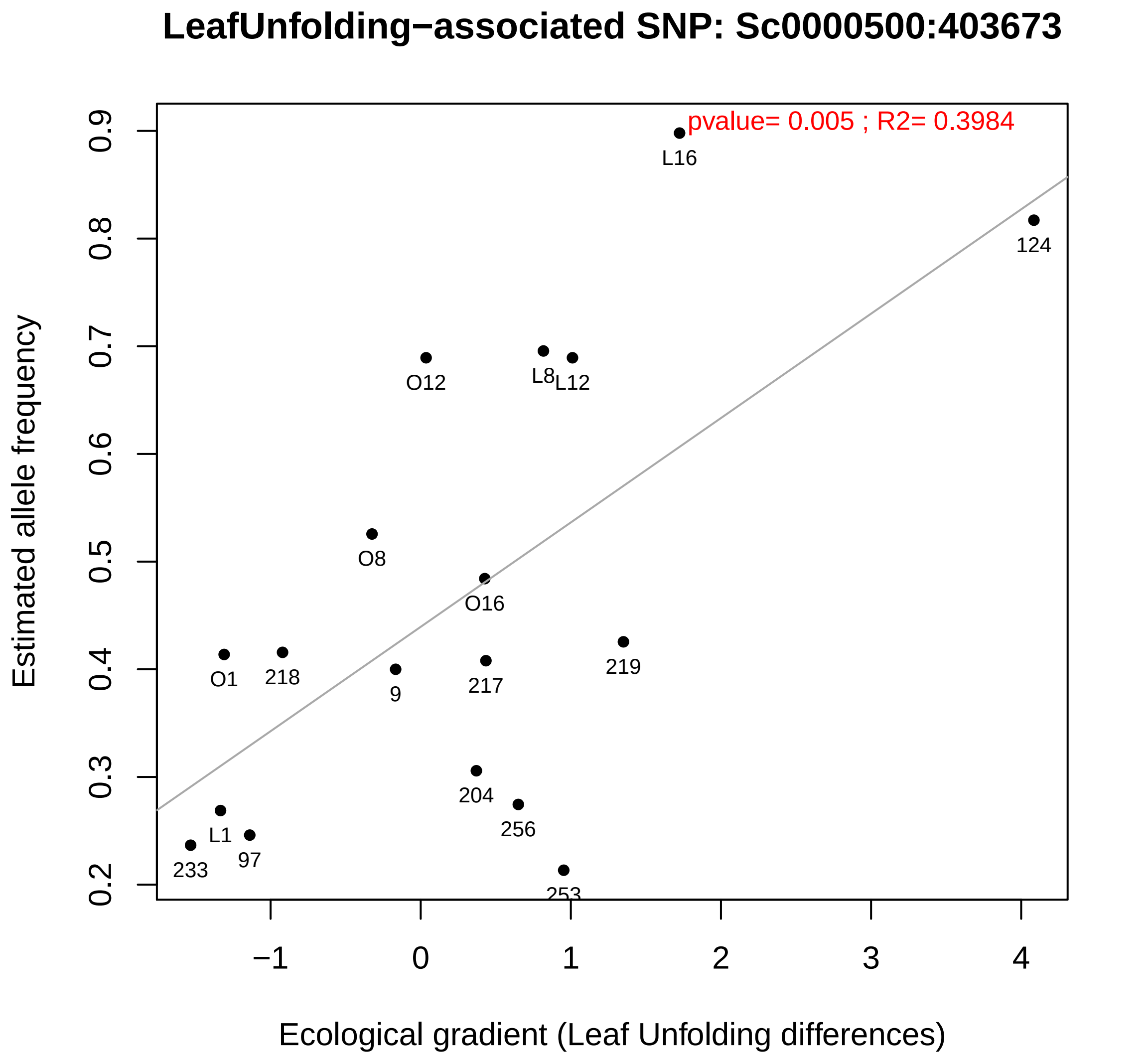
LeafUnfolding−associated SNP: Sc0000500:403673
pvalue= 0.005 ; R2= 0.3984
Estimated allele frequency
Ecological gradient (Leaf Unfolding differences)
0.2
0.3
0.4
0.5
0.6
0.7
0.8
0.9
−1
0
1
2
3
4
L16
124
O12
L8
L12
O8
O16
O1
218
9
217
219
204
256
L1
97
233
253

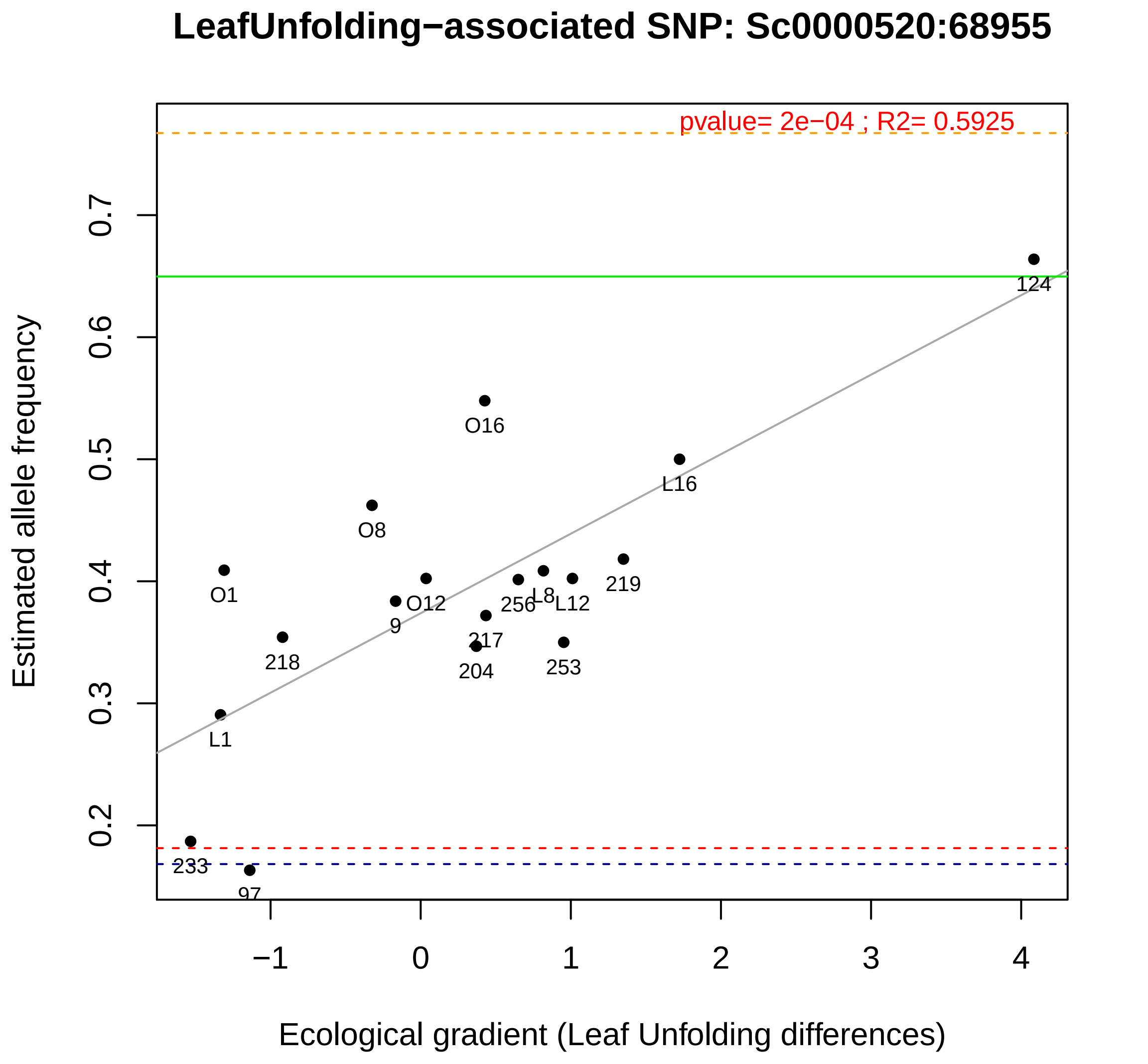
LeafUnfolding−associated SNP: Sc0000520:68955
pvalue= 2e−04 ; R2= 0.5925
Estimated allele frequency
0.2
0.3
0.4
0.5
0.6
0.7
124
O16
L16
O8
O1
O12
9
256
L8
L12
219
217
204
253
218
L1
233
97
−1
0
1
2
3
4
Ecological gradient (Leaf Unfolding differences)

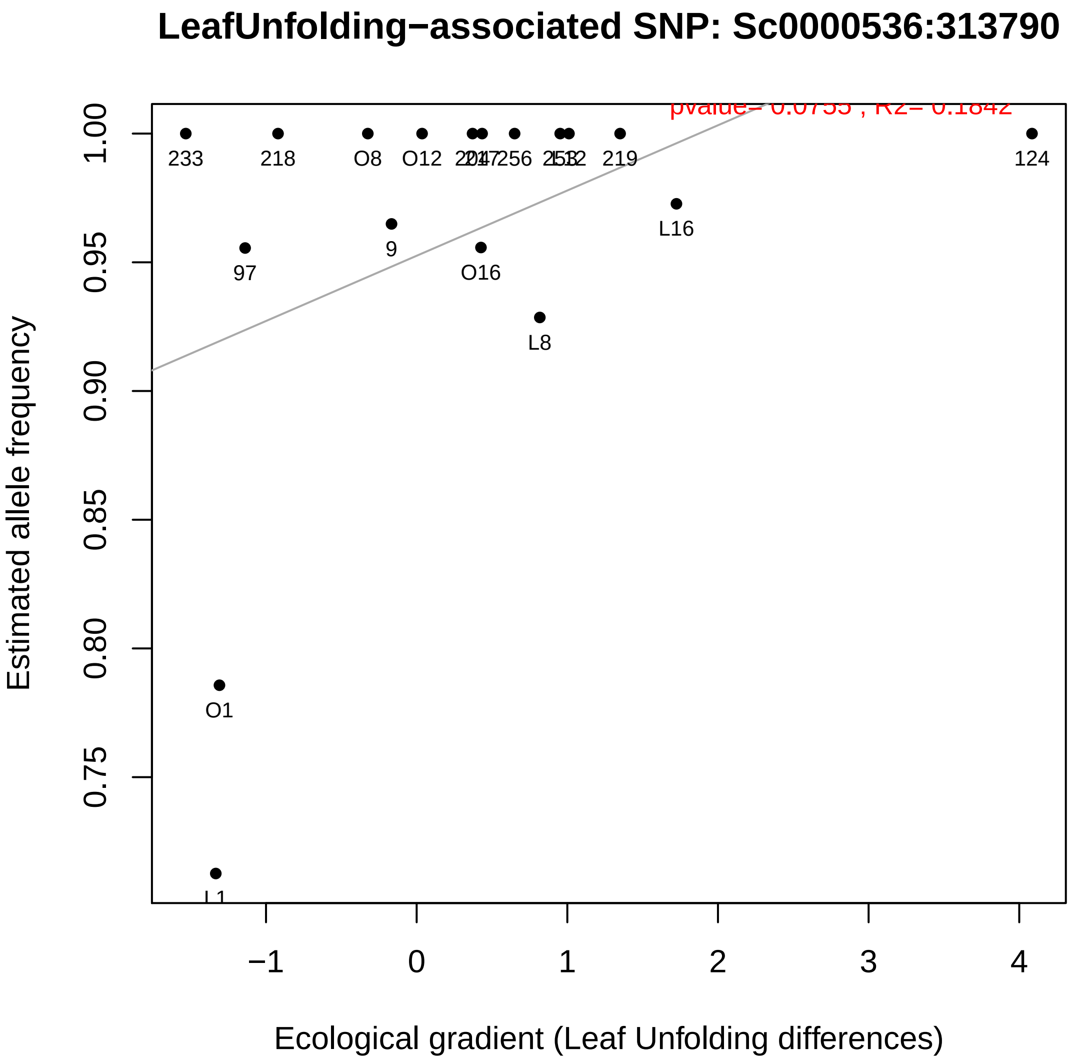

LeafUnfolding−associated SNP: Sc0000536:313790
pvalue= 0.0755 , R2= 0.1842
Estimated allele frequency
Ecological gradient (Leaf Unfolding differences)
1.00
0.95
0.90
0.85
0.80
0.75
−1
0
1
2
3
4
233
218
O8
O12
256
219
124
97
9
O16
L16
L8
O1

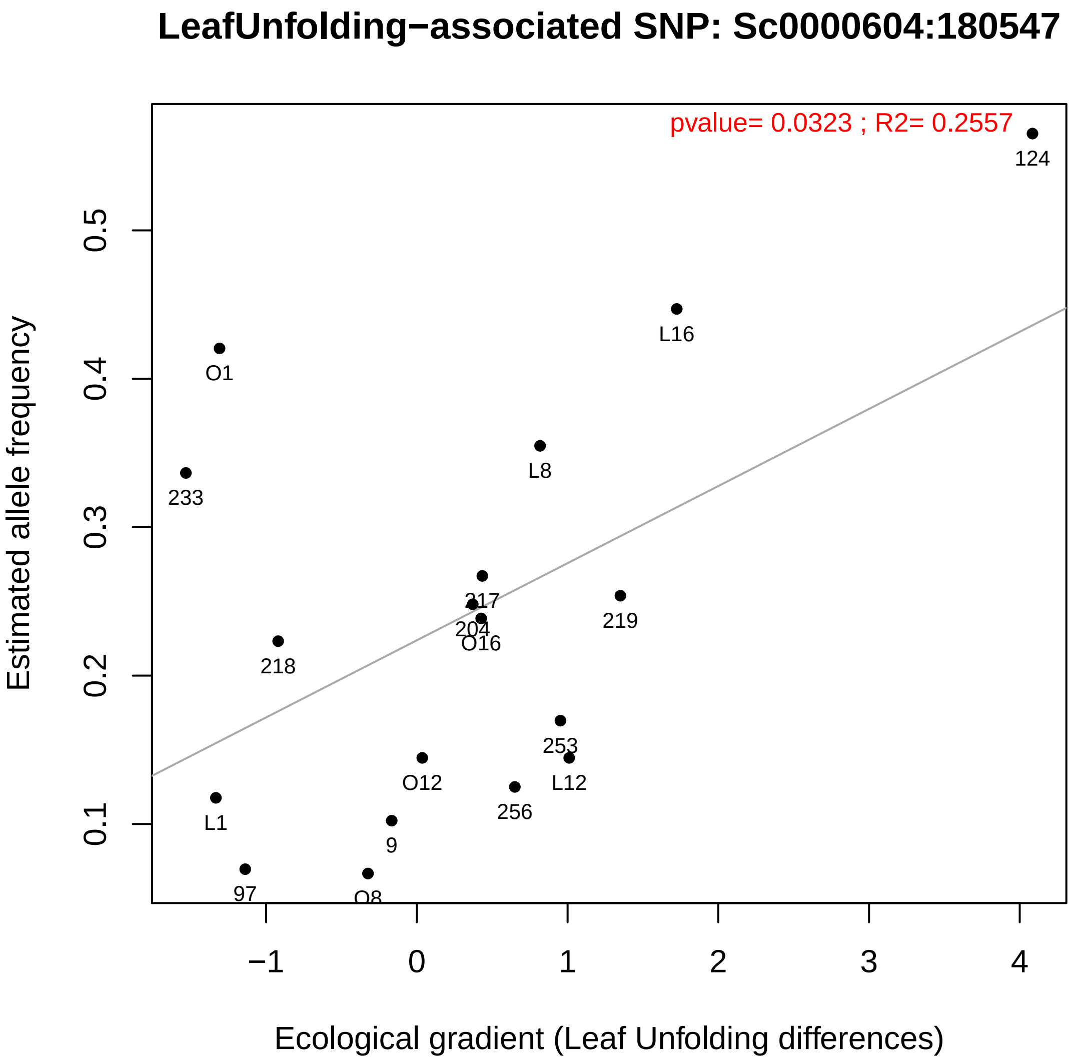

LeafUnfolding−associated SNP: Sc0000604:180547
pvalue= 0.0323 ; R2= 0.2557
Estimated allele frequency
Ecological gradient (Leaf Unfolding differences)
0.1
0.2
0.3
0.4
0.5
−1
0
1
2
3
4
124
L16
O1
L8
233
217
204
O16
219
218
253
L12
O12
256
L1
9
97
O8

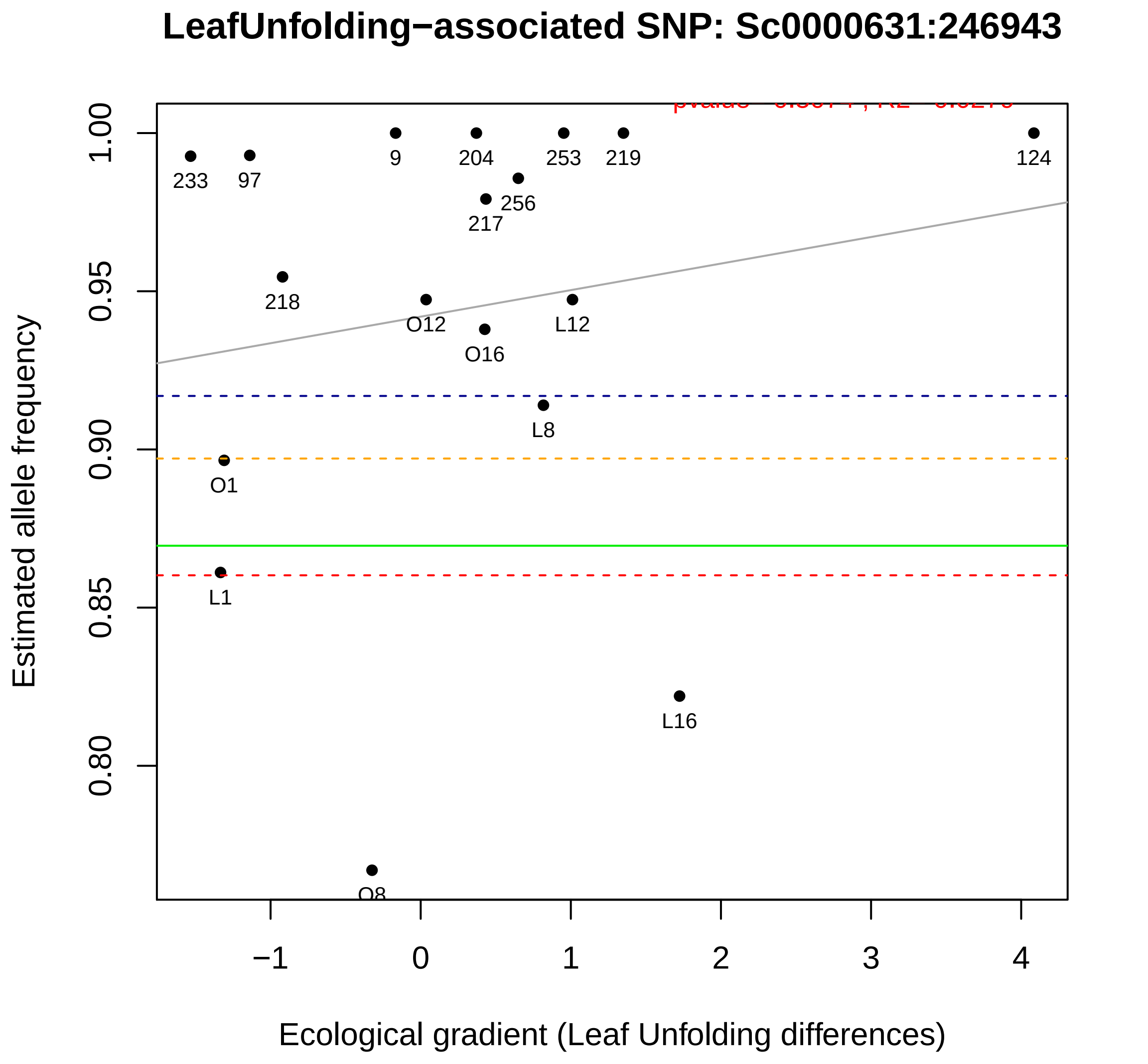

LeafUnfolding−associated SNP: Sc0000631:246943
Estimated allele frequency
Ecological gradient (Leaf Unfolding differences)
1.00
0.95
0.90
0.85
0.80
−1
0
1
2
3
4
233
97
9
204
253
219
124
256
217
218
O12
L12
O16
L8
O1
L1
L16
O8

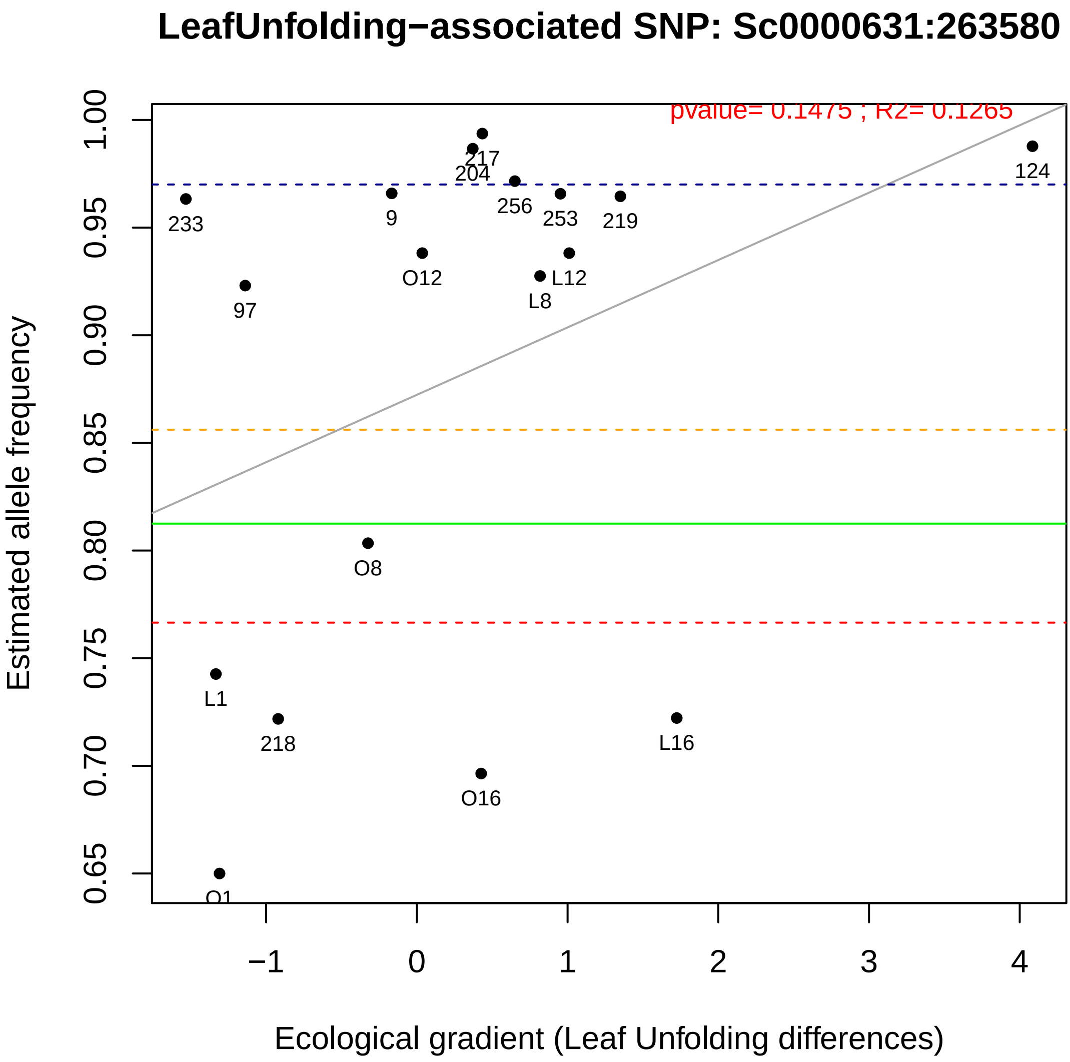
LeafUnfolding−associated SNP: Sc0000631:263580
pvalue= 0.1475 ; R2= 0.1265
Estimated allele frequency
0.65
0.70
0.75
0.80
0.85
0.90
0.95
1.00
−1
0
1
2
3
4
Ecological gradient (Leaf Unfolding differences)
233
97
9
O12
217
204
256
253
219
L12
L8
124
O8
L1
218
O16
L16
O1

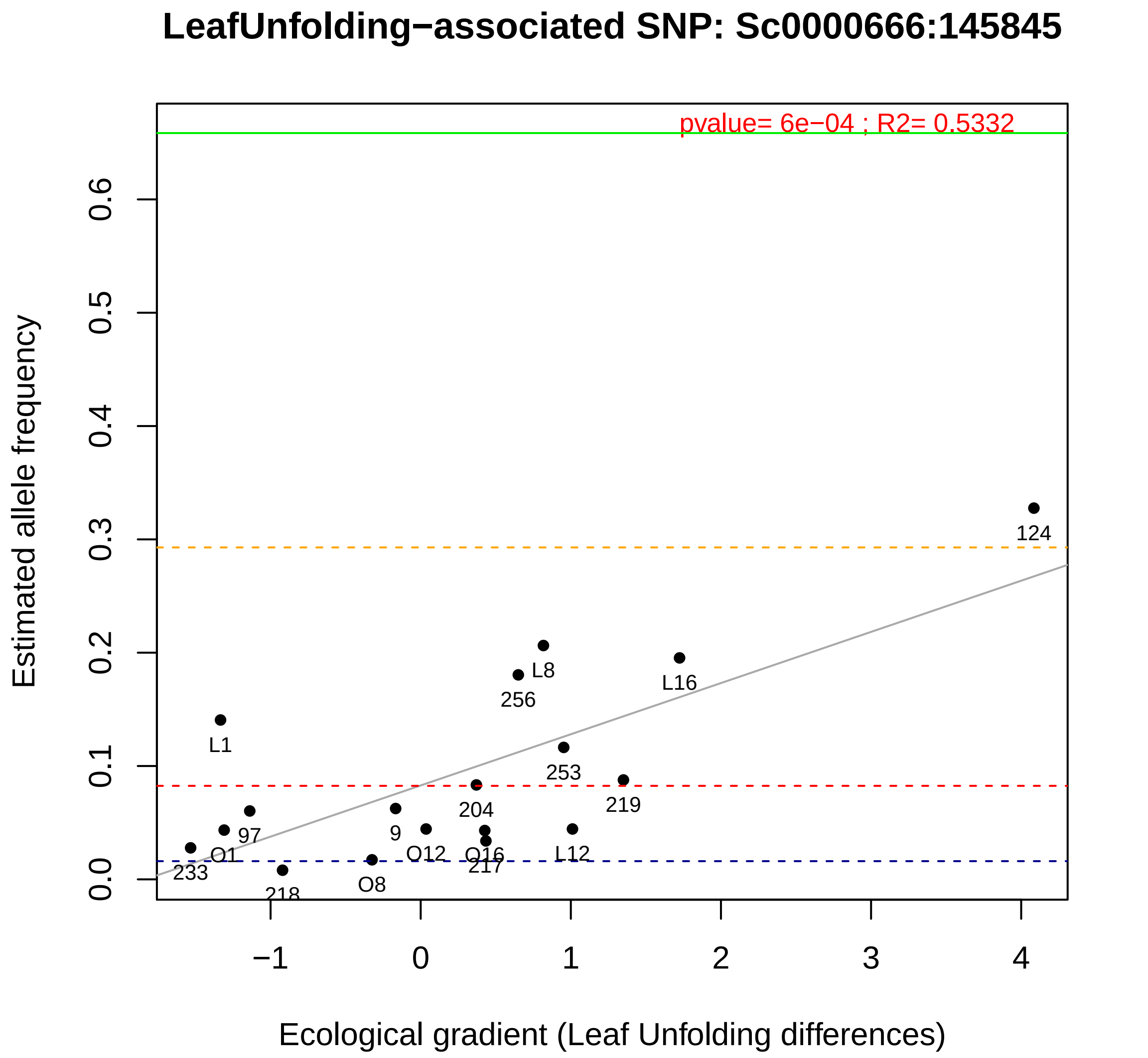

LeafUnfolding−associated SNP: Sc0000666:145845
pvalue= 6e−04 ; R2= 0.5332
Estimated allele frequency
0.0
0.1
0.2
0.3
0.4
0.5
0.6
−1
0
1
2
3
4
Ecological gradient (Leaf Unfolding differences)
124
L8
L16
256
L1
253
204
219
97
9
O1
O12
Q16
217
L12
233
218
O8

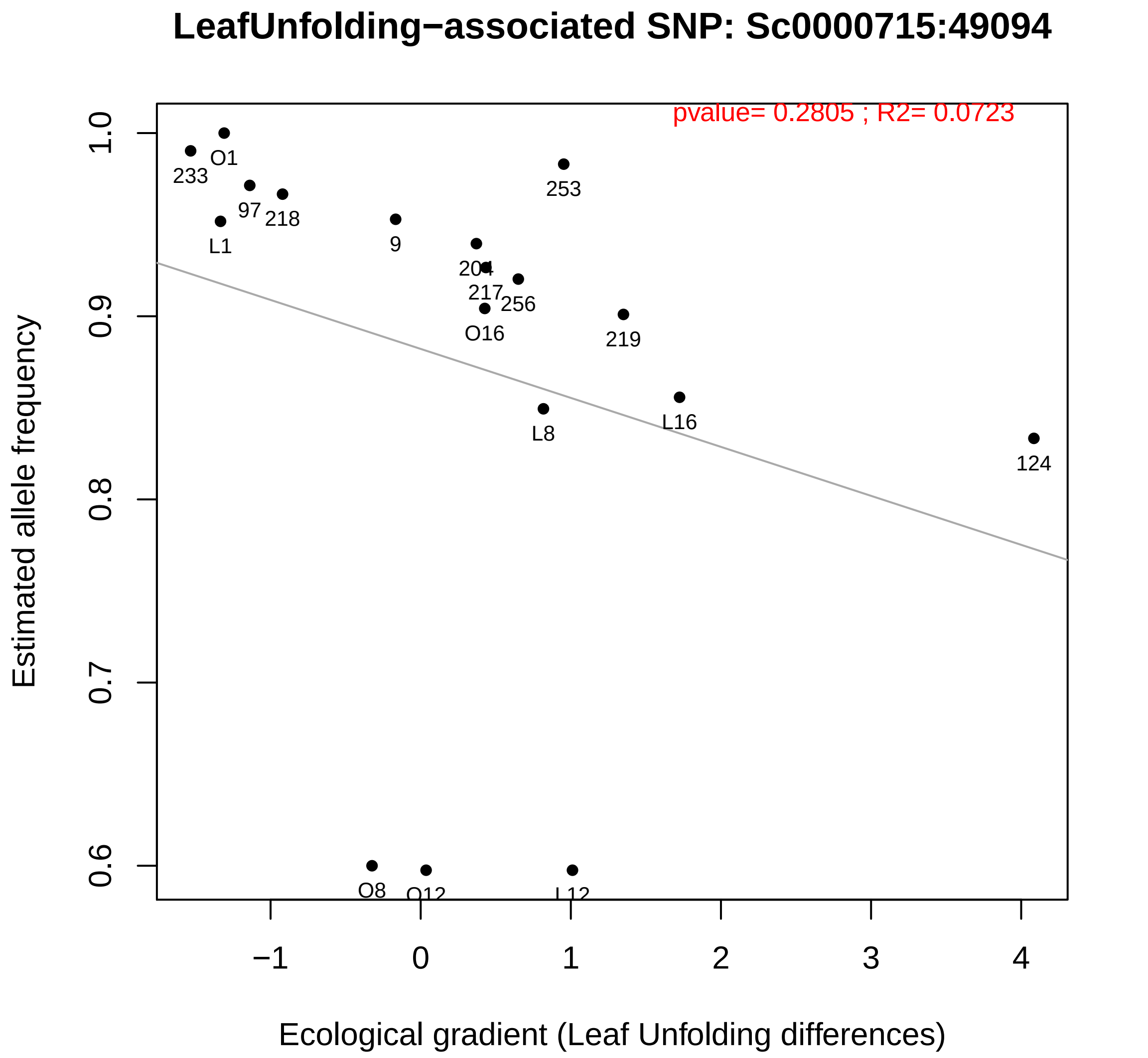
LeafUnfolding−associated SNP: Sc0000715:49094
pvalue= 0.2805 ; R2= 0.0723
Estimated allele frequency
Ecological gradient (Leaf Unfolding differences)
1.0
0.9
0.8
0.7
0.6
−1
0
1
2
3
4
233
O1
97
218
L1
9
253
204
217
256
O16
219
L16
L8
124
O8
O12
L12

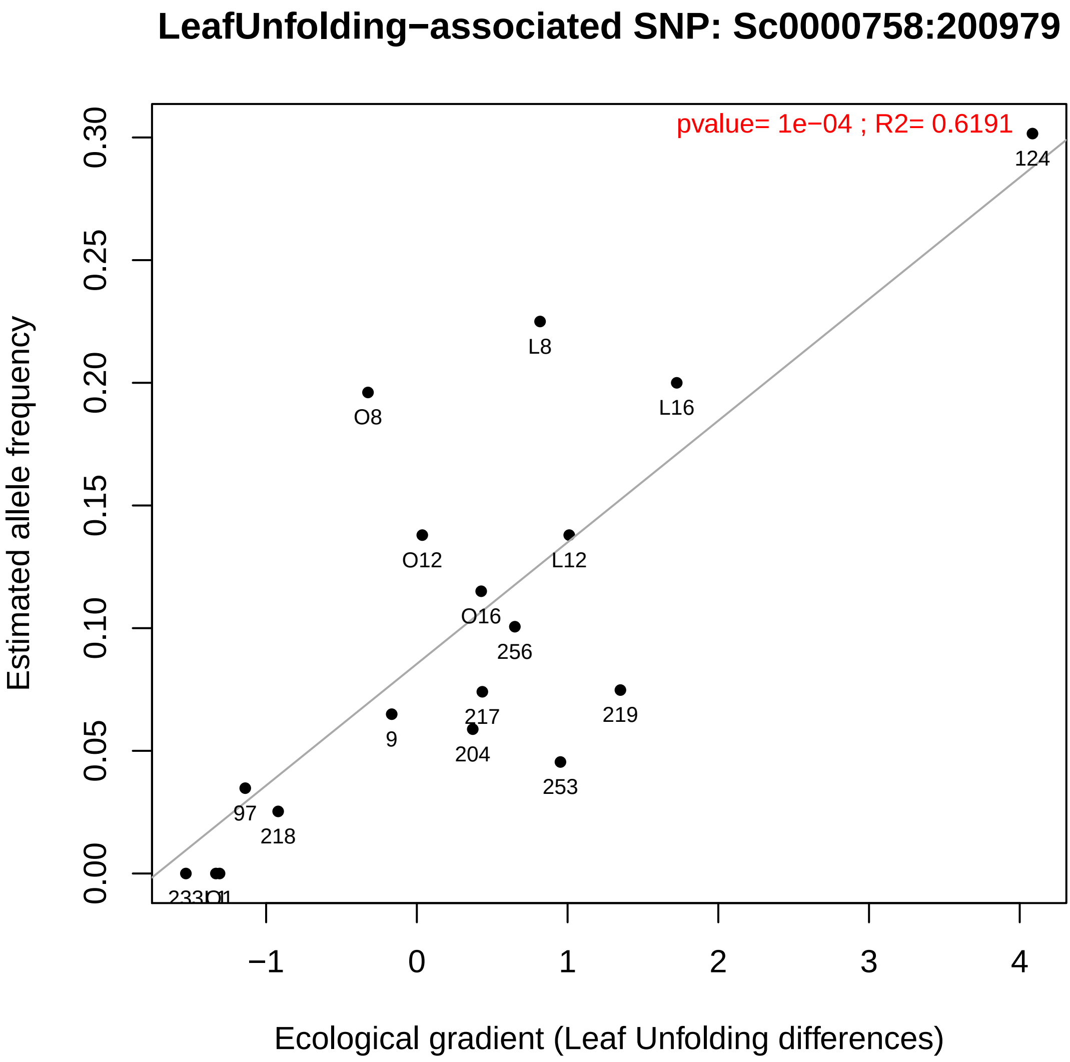

LeafUnfolding−associated SNP: Sc0000758:200979
pvalue= 1e−04 ; R2= 0.6191
Estimated allele frequency
0.00
0.05
0.10
0.15
0.20
0.25
0.30
−1
0
1
2
3
4
Ecological gradient (Leaf Unfolding differences)
124
L8
L16
O8
O12
L12
O16
256
217
219
9
204
253
97
218

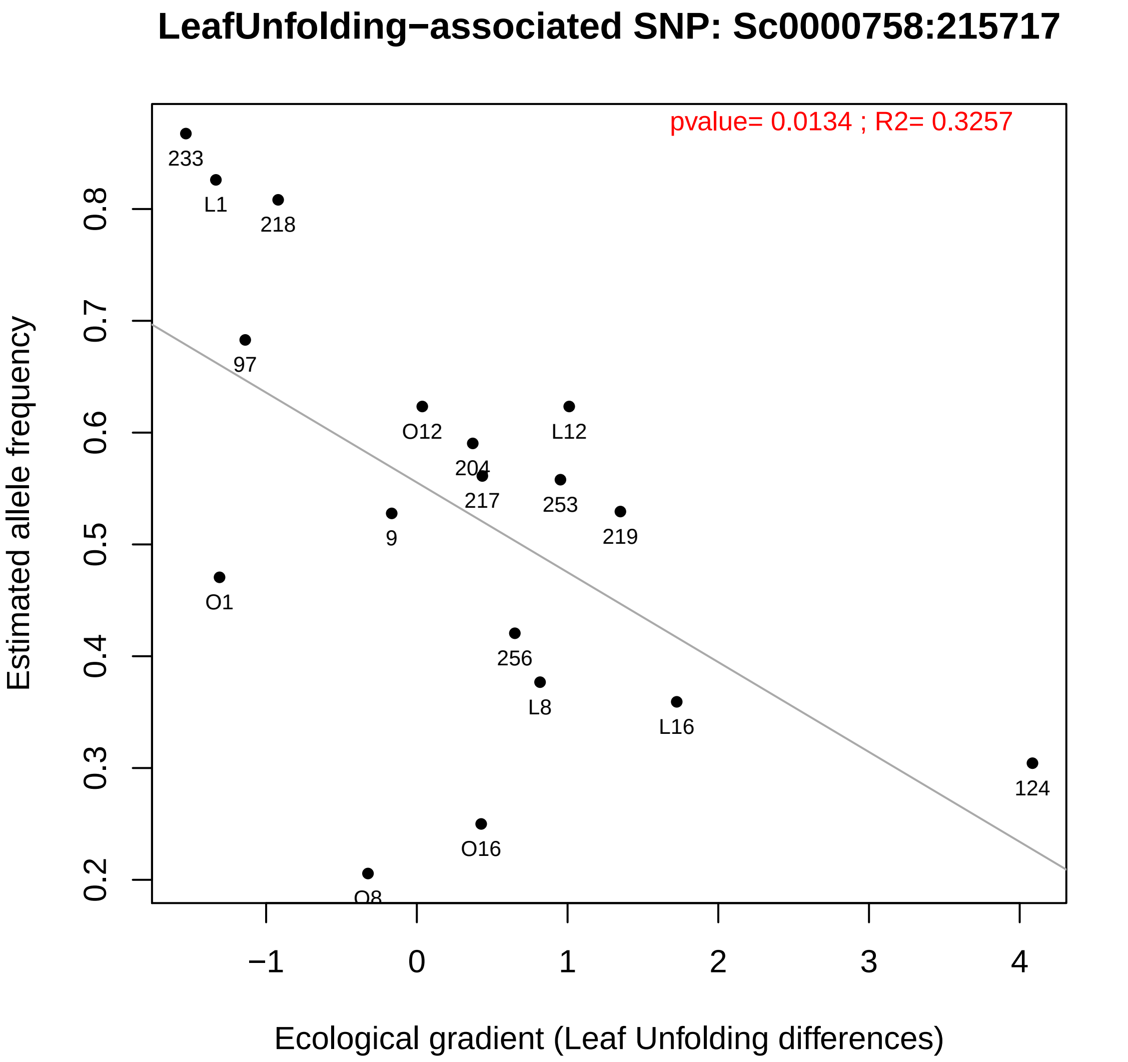

LeafUnfolding−associated SNP: Sc0000758:215717
pvalue= 0.0134 ; R2= 0.3257
233
L1
218
97
O12
L12
204
217
253
9
219
O1
256
L8
L16
124
O16
O8
0.2
0.3
0.4
0.5
0.6
0.7
0.8
−1
0
1
2
3
4
Estimated allele frequency
Ecological gradient (Leaf Unfolding differences)

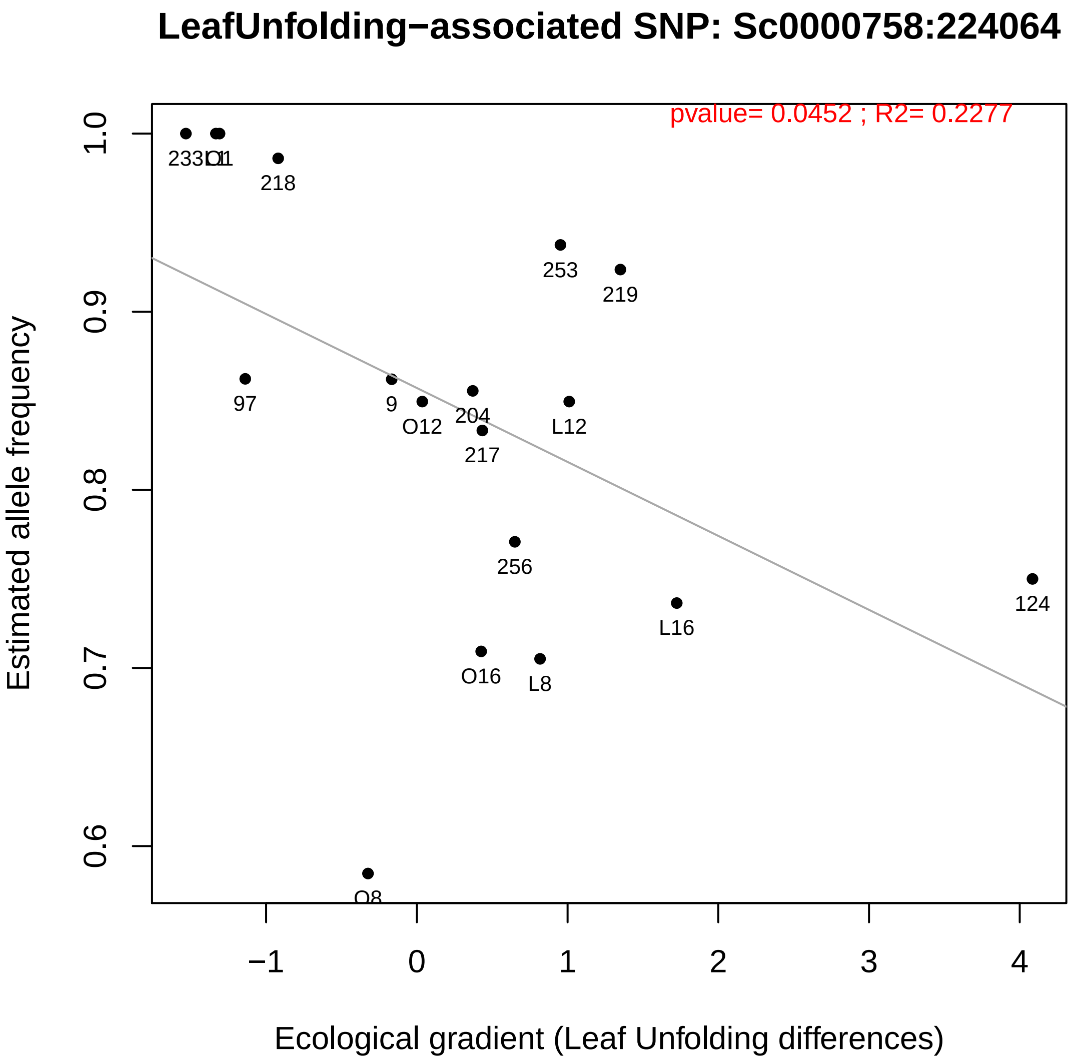

LeafUnfolding−associated SNP: Sc0000758:224064
pvalue= 0.0452 ; R2= 0.2277
Estimated allele frequency
Ecological gradient (Leaf Unfolding differences)
0.6
0.7
0.8
0.9
1.0
−1
0
1
2
3
4
218
253
219
97
9
O12
204
217
L12
256
124
L16
O16
L8

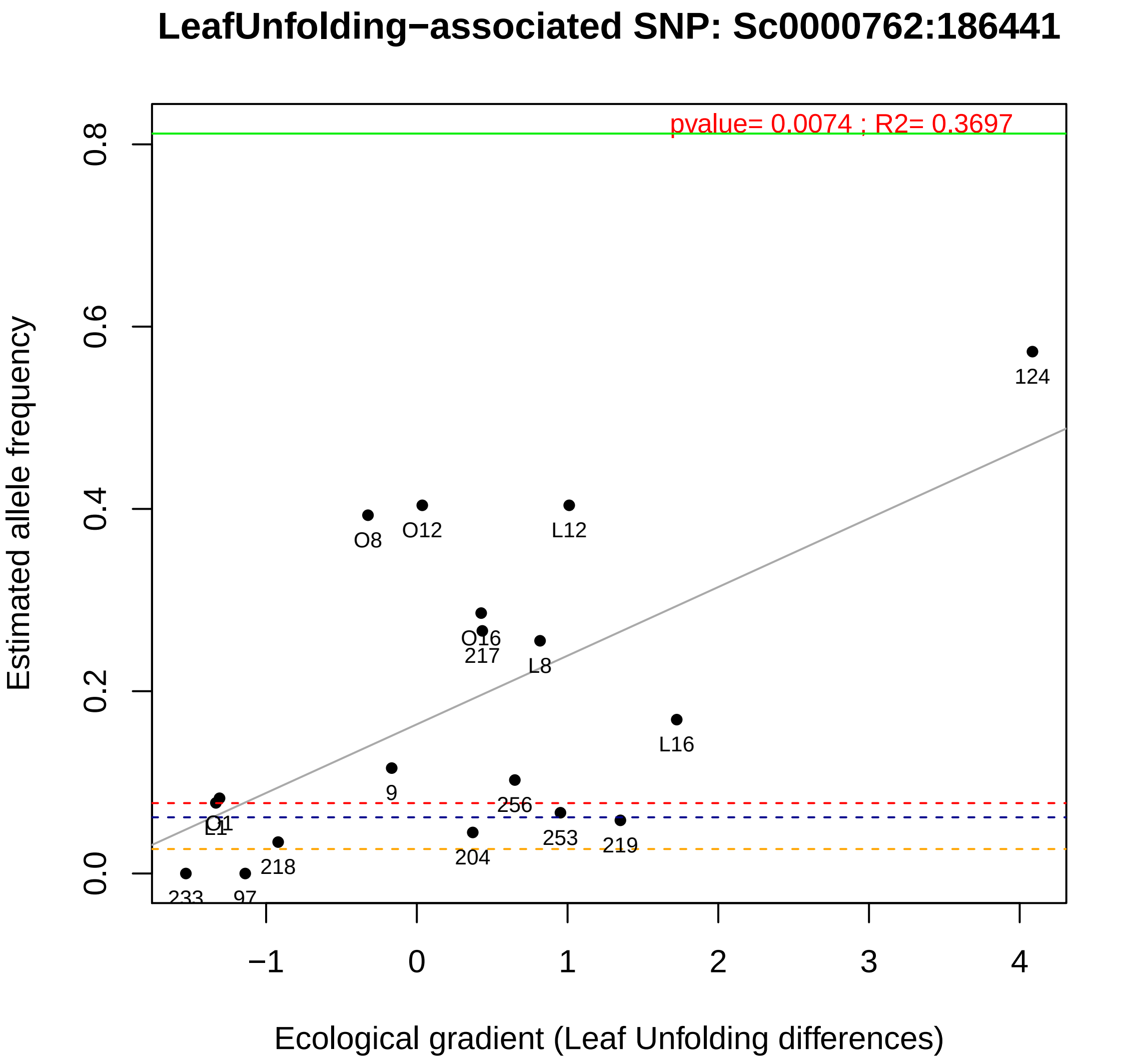
LeafUnfolding−associated SNP: Sc0000762:186441
pvalue= 0.0074 ; R2= 0.3697
Estimated allele frequency
0.0
0.2
0.4
0.6
0.8
Ecological gradient (Leaf Unfolding differences)
−1
0
1
2
3
4
124
O8
O12
L12
O16
217
L8
L16
9
256
O1
L1
253
219
204
218
233
97

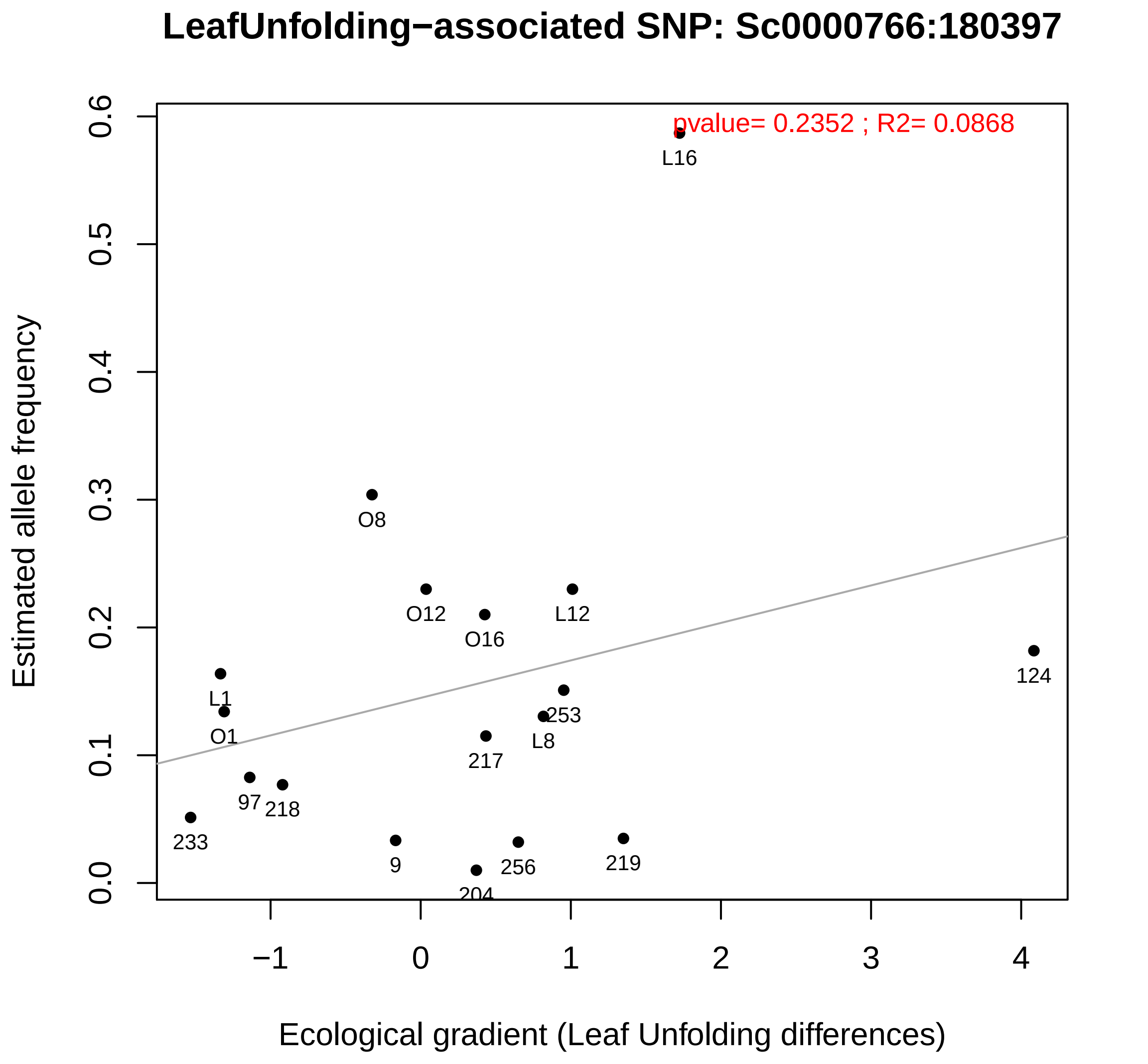
LeafUnfolding−associated SNP: Sc0000766:180397
pvalue= 0.2352 ; R2= 0.0868
Estimated allele frequency
Ecological gradient (Leaf Unfolding differences)
0.0
0.1
0.2
0.3
0.4
0.5
0.6
−1
0
1
2
3
4
L16
O8
O12
O16
L12
124
L1
O1
253
L8
217
97
218
233
9
204
256
219

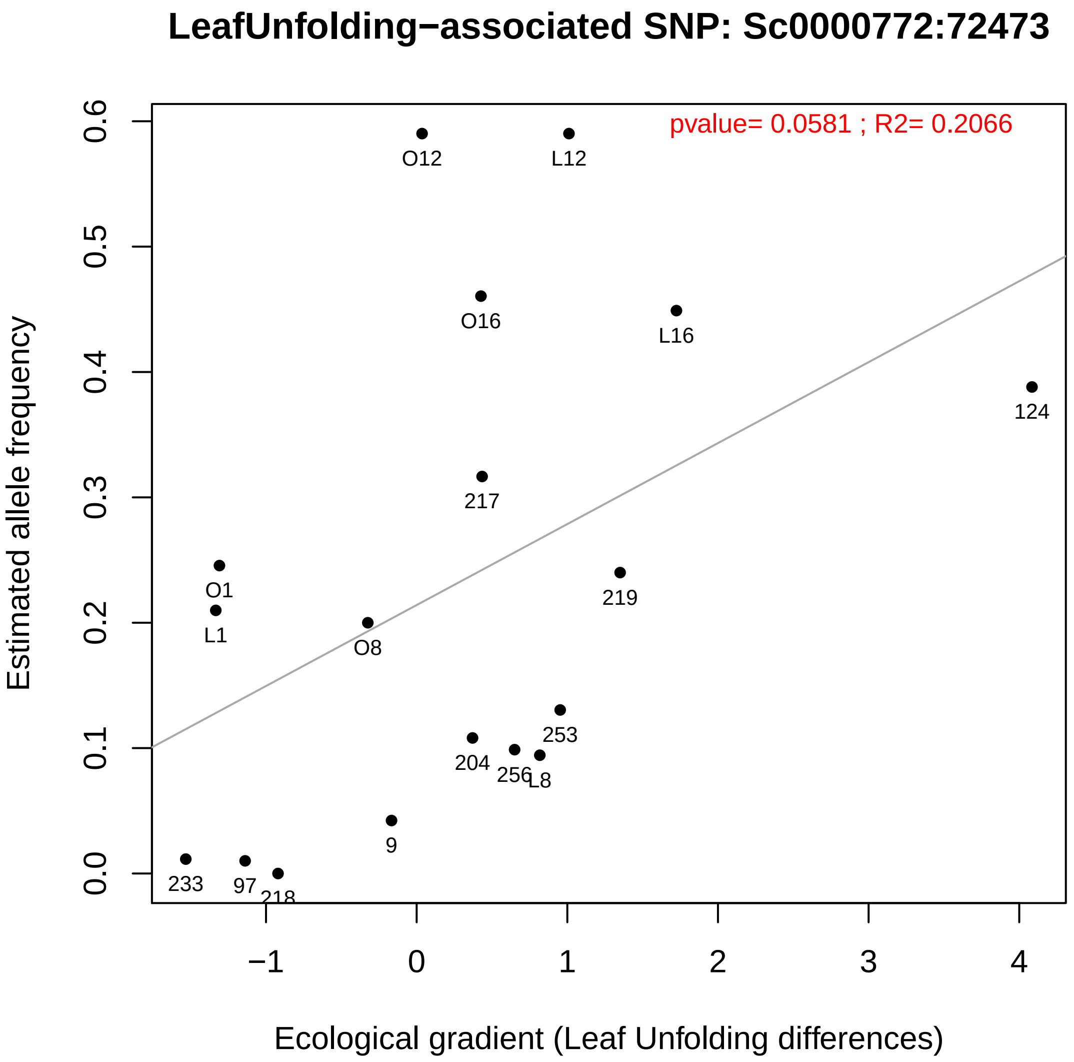
LeafUnfolding−associated SNP: Sc0000772:72473
pvalue= 0.0581 ; R2= 0.2066
Estimated allele frequency
Ecological gradient (Leaf Unfolding differences)
O12
L12
O16
L16
124
217
O1
L1
O8
219
253
204
256
L8
9
233
97
218
0.0
0.1
0.2
0.3
0.4
0.5
0.6
−1
0
1
2
3
4

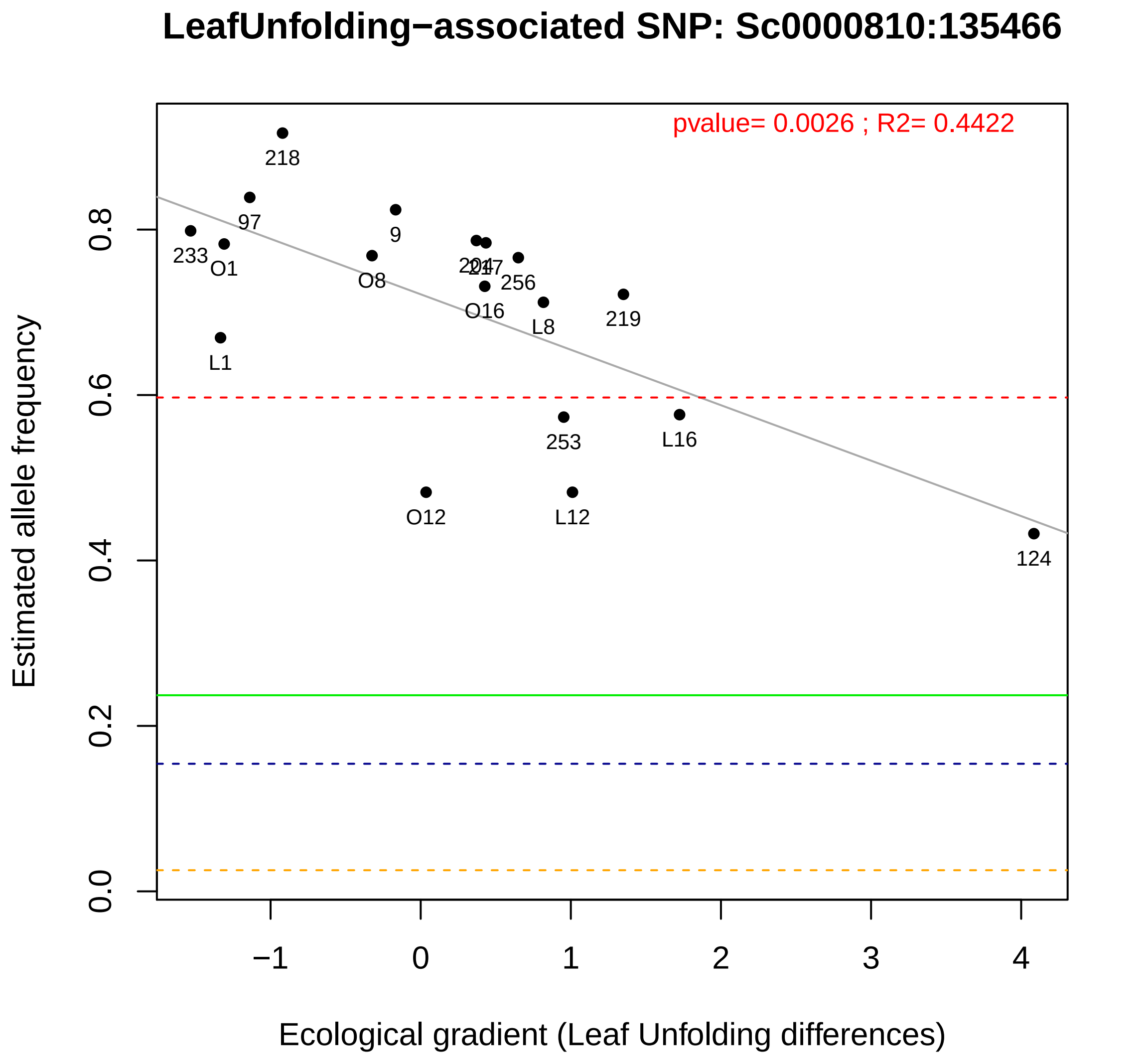
LeafUnfolding−associated SNP: Sc0000810:135466
pvalue= 0.0026 ; R2= 0.4422
Estimated allele frequency
Ecological gradient (Leaf Unfolding differences)
0.0
0.2
0.4
0.6
0.8
−1
0
1
2
3
4
218
97
233
O1
9
O8
204
217
256
O16
L8
219
L1
253
L16
O12
L12
124

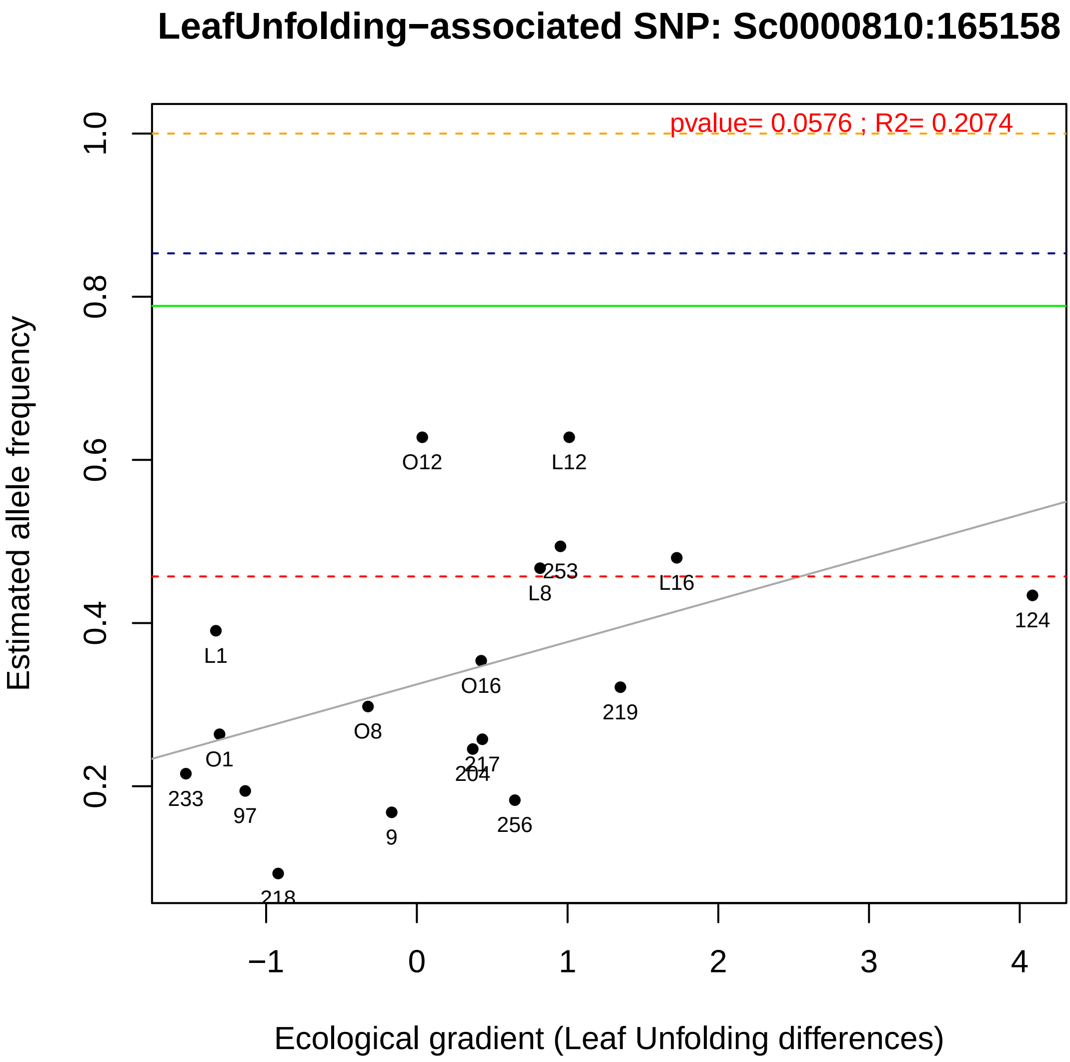
LeafUnfolding−associated SNP: Sc0000810:165158
pvalue= 0.0576 ; R2= 0.2074
Estimated allele frequency
0.2
0.4
0.6
0.8
1.0
−1
0
1
2
3
4
Ecological gradient (Leaf Unfolding differences)
O12
L12
253
L8
L16
124
L1
O16
219
O8
O1
217
204
233
97
9
256
218

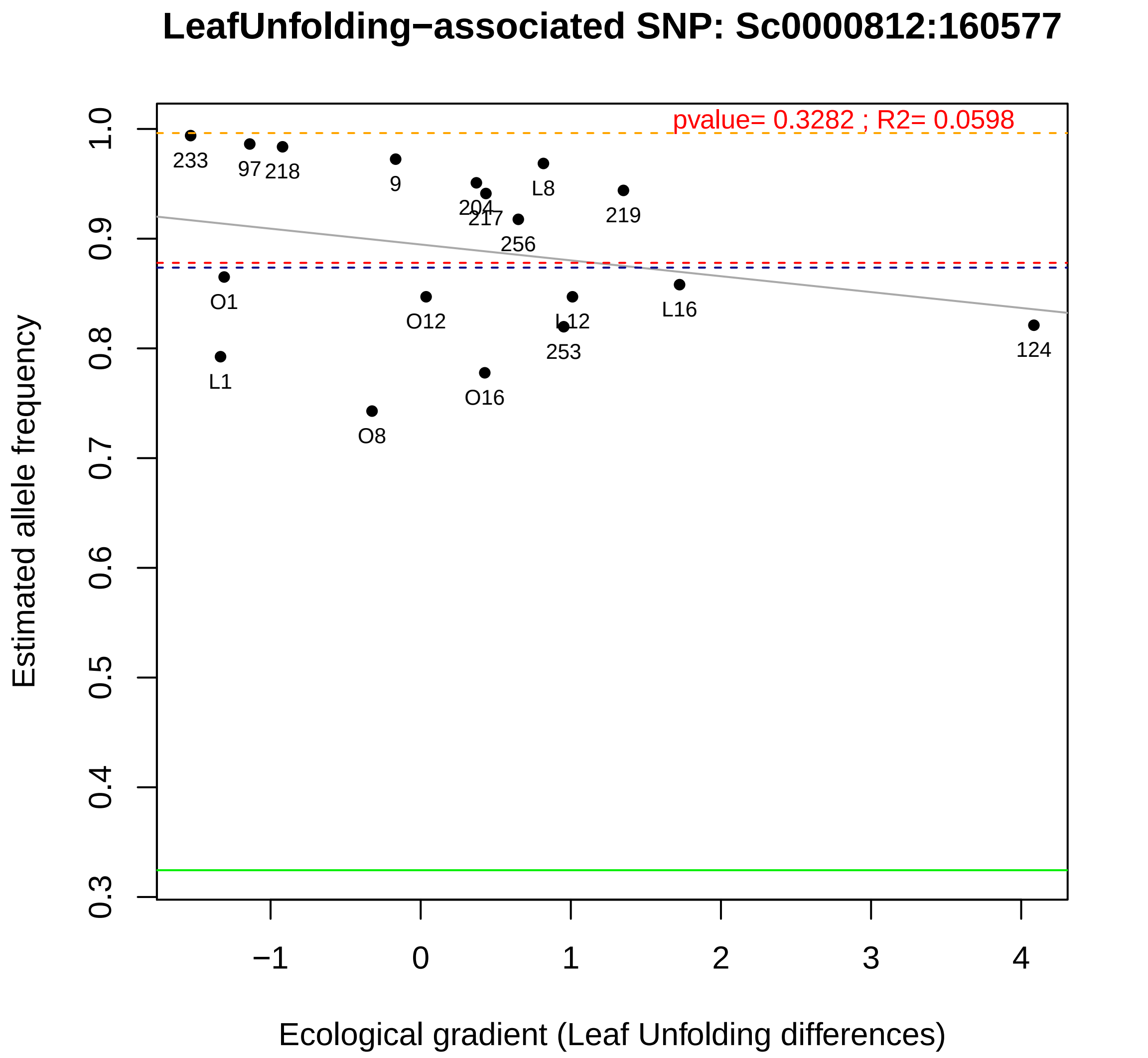

LeafUnfolding−associated SNP: Sc0000812:160577
pvalue= 0.3282 ; R2= 0.0598
Estimated allele frequency
Ecological gradient (Leaf Unfolding differences)
1.0
0.9
0.8
0.7
0.6
0.5
0.4
0.3
−1
0
1
2
3
4
233
97
218
9
204
217
L8
219
256
O1
O12
L12
253
L16
124
L1
O16
O8

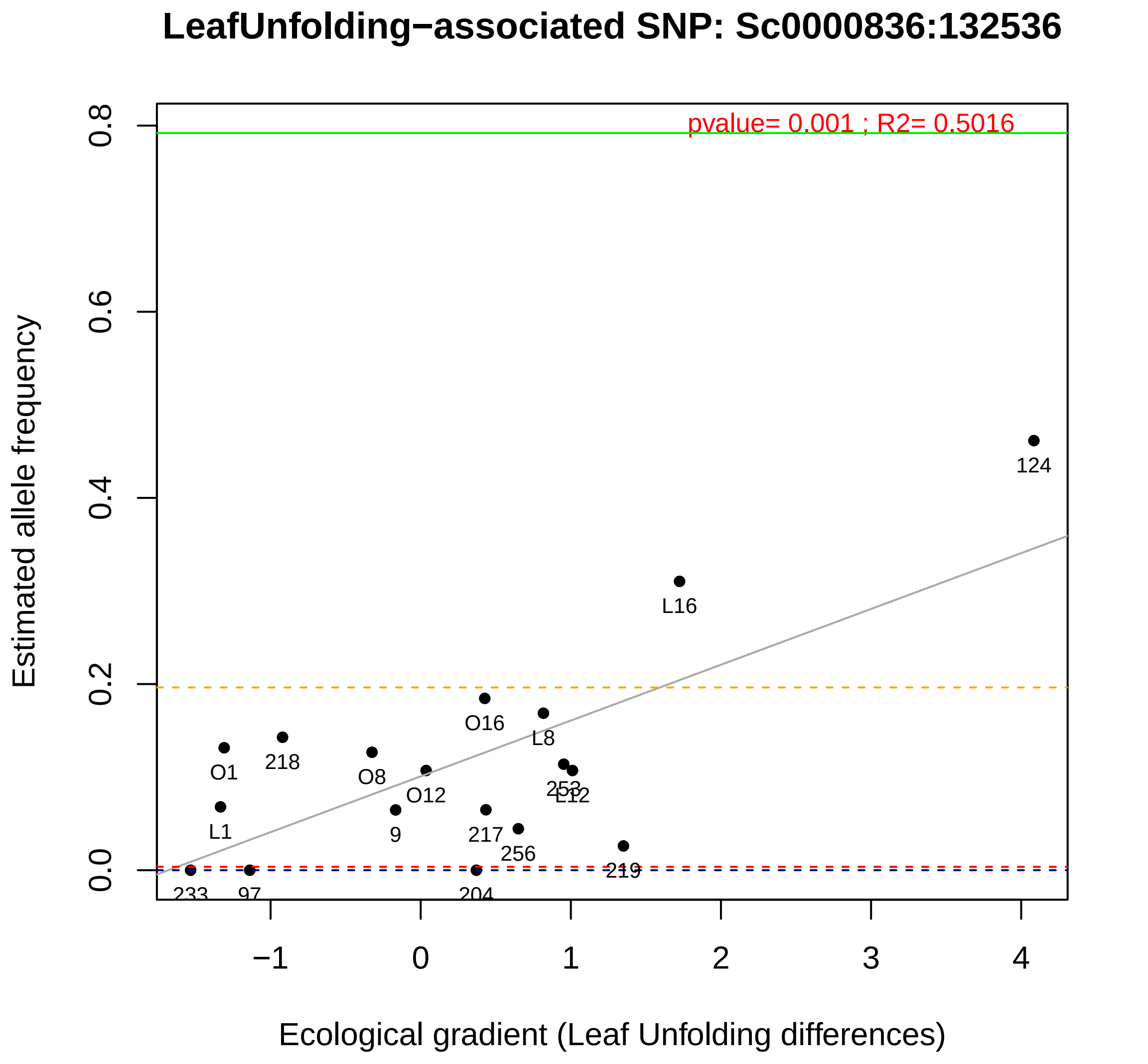

LeafUnfolding−associated SNP: Sc0000836:132536
pvalue= 0.001 ; R2= 0.5016
Estimated allele frequency
0.0
0.2
0.4
0.6
0.8
−1
0
1
2
3
4
Ecological gradient (Leaf Unfolding differences)
124
L16
O16
L8
O1
218
O8
O12
253
L12
L1
9
217
256
219
233
97
204

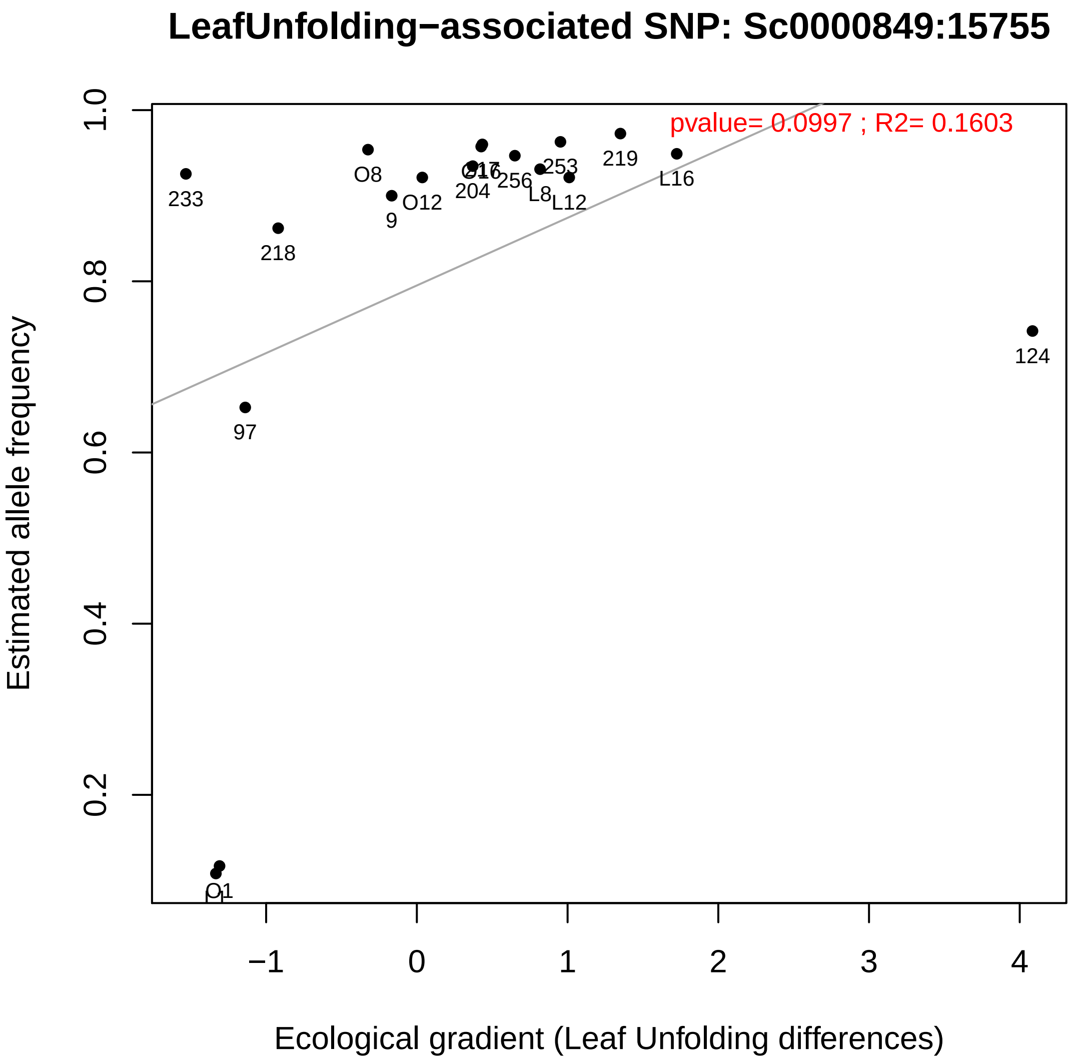

LeafUnfolding−associated SNP: Sc0000849:15755
pvalue= 0.0997 ; R2= 0.1603
Estimated allele frequency
Ecological gradient (Leaf Unfolding differences)
0.2
0.4
0.6
0.8
1.0
−1
0
1
2
3
4
233
218
97
O8
9
O12
204
256
253
L8
L12
219
L16
124
O1

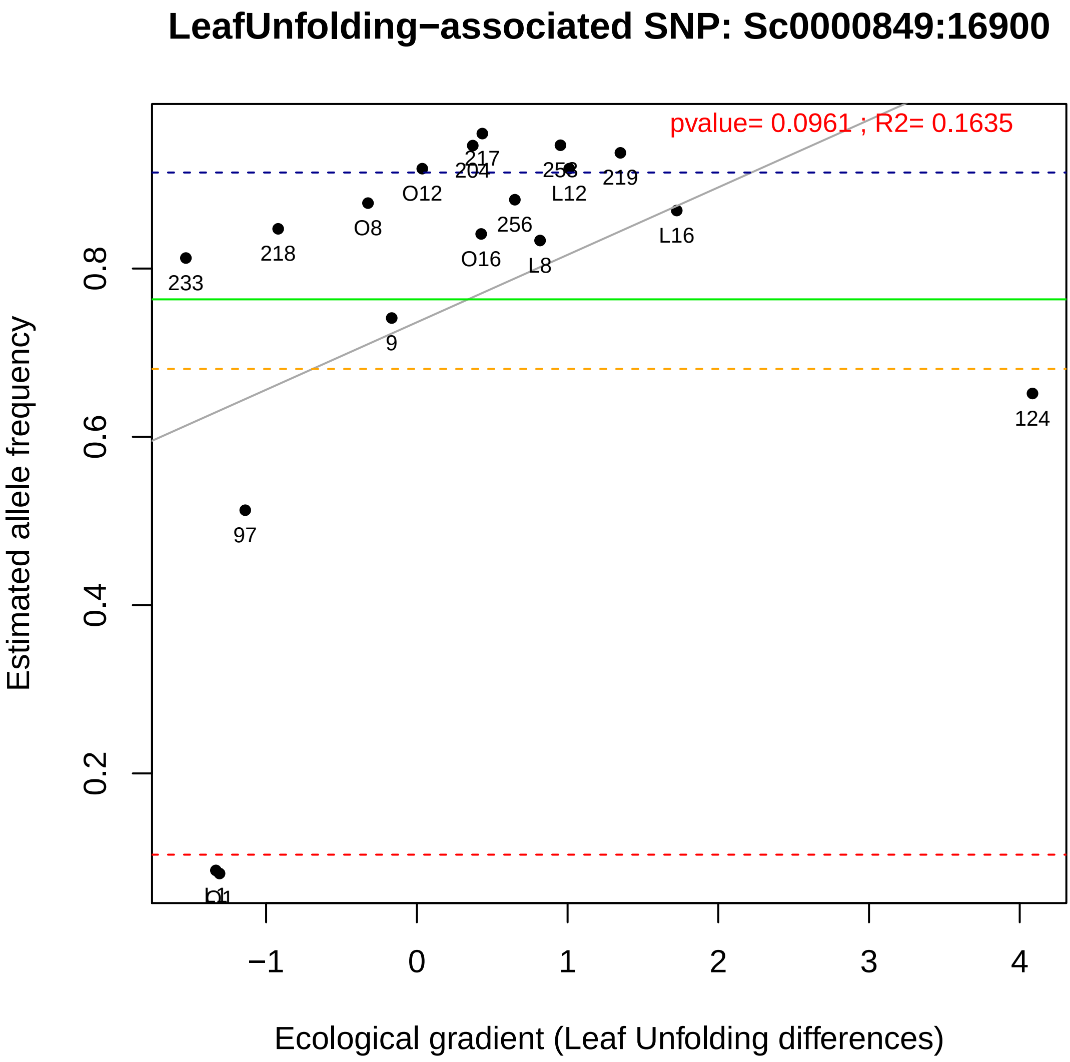

LeafUnfolding−associated SNP: Sc0000849:16900
pvalue= 0.0961 ; R2= 0.1635
Estimated allele frequency
0.2
0.4
0.6
0.8
−1
0
1
2
3
4
Ecological gradient (Leaf Unfolding differences)
233
218
O8
O12
204
217
256
O16
L8
L12
219
L16
9
97
124

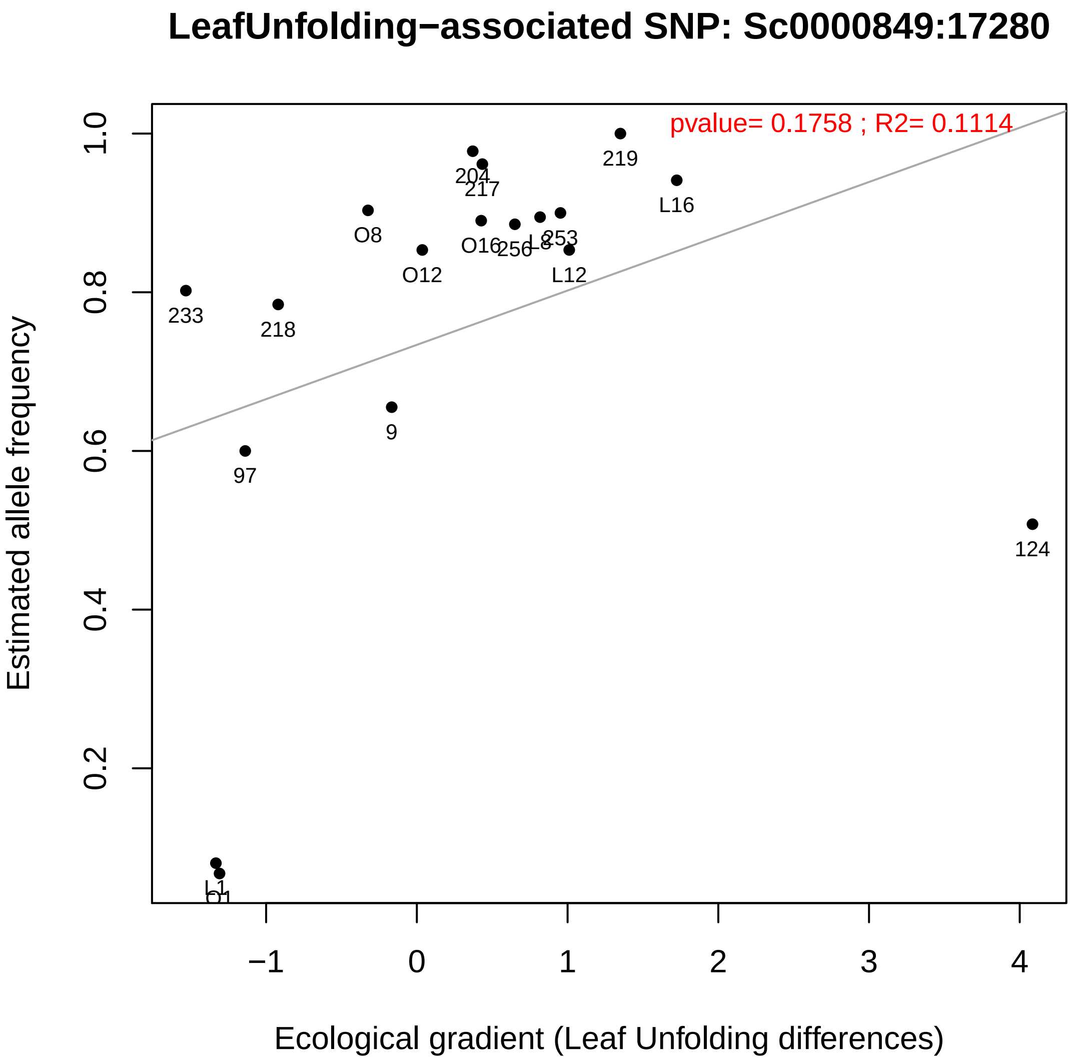
LeafUnfolding−associated SNP: Sc0000849:17280
pvalue= 0.1758 ; R2= 0.1114
Estimated allele frequency
Ecological gradient (Leaf Unfolding differences)
1.0
0.8
0.6
0.4
0.2
−1
0
1
2
3
4
219
204
217
L16
O8
O16
256
L8
253
L12
O12
233
218
9
97
124
L1
O1

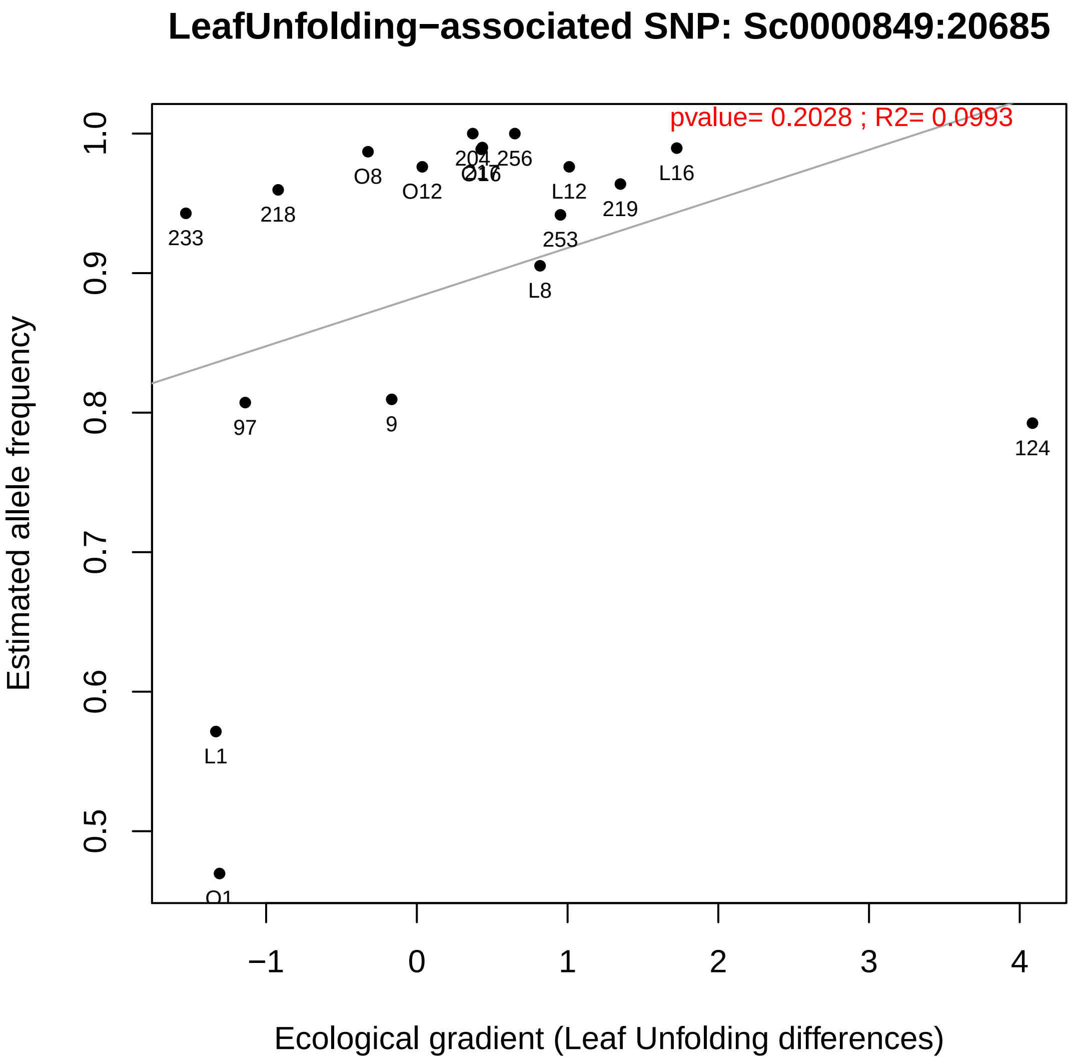
LeafUnfolding−associated SNP: Sc0000849:20685
pvalue= 0.2028 ; R2= 0.0993
Estimated allele frequency
Ecological gradient (Leaf Unfolding differences)
1.0
0.9
0.8
0.7
0.6
0.5
−1
0
1
2
3
4
233
218
O8
O12
204
256
L12
L16
219
253
L8
97
9
124
L1
O1

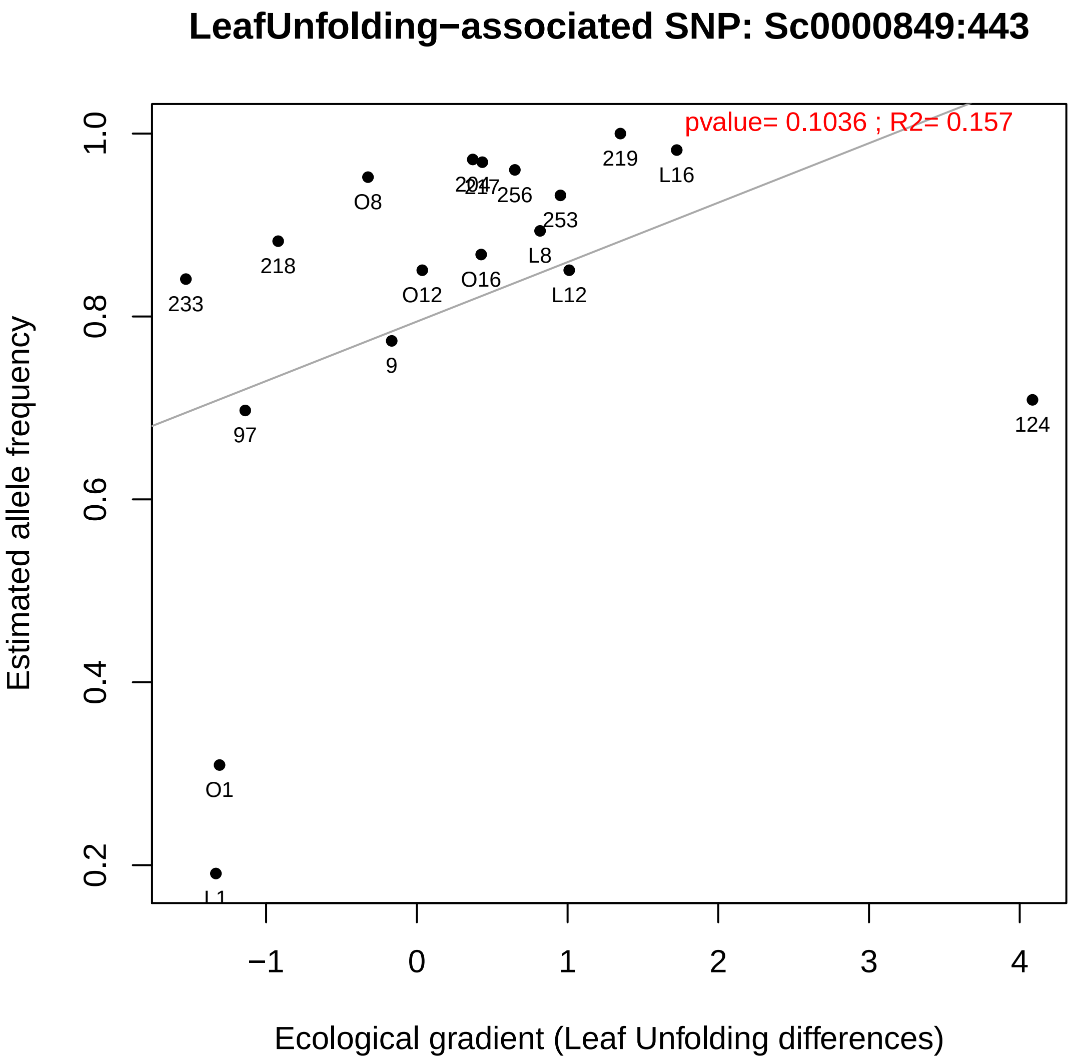

LeafUnfolding−associated SNP: Sc0000849:443
pvalue= 0.1036 ; R2= 0.157
Estimated allele frequency
Ecological gradient (Leaf Unfolding differences)
0.2
0.4
0.6
0.8
1.0
−1
0
1
2
3
4
219
L16
O8
256
253
L8
218
O16
O12
L12
233
9
97
124
O1

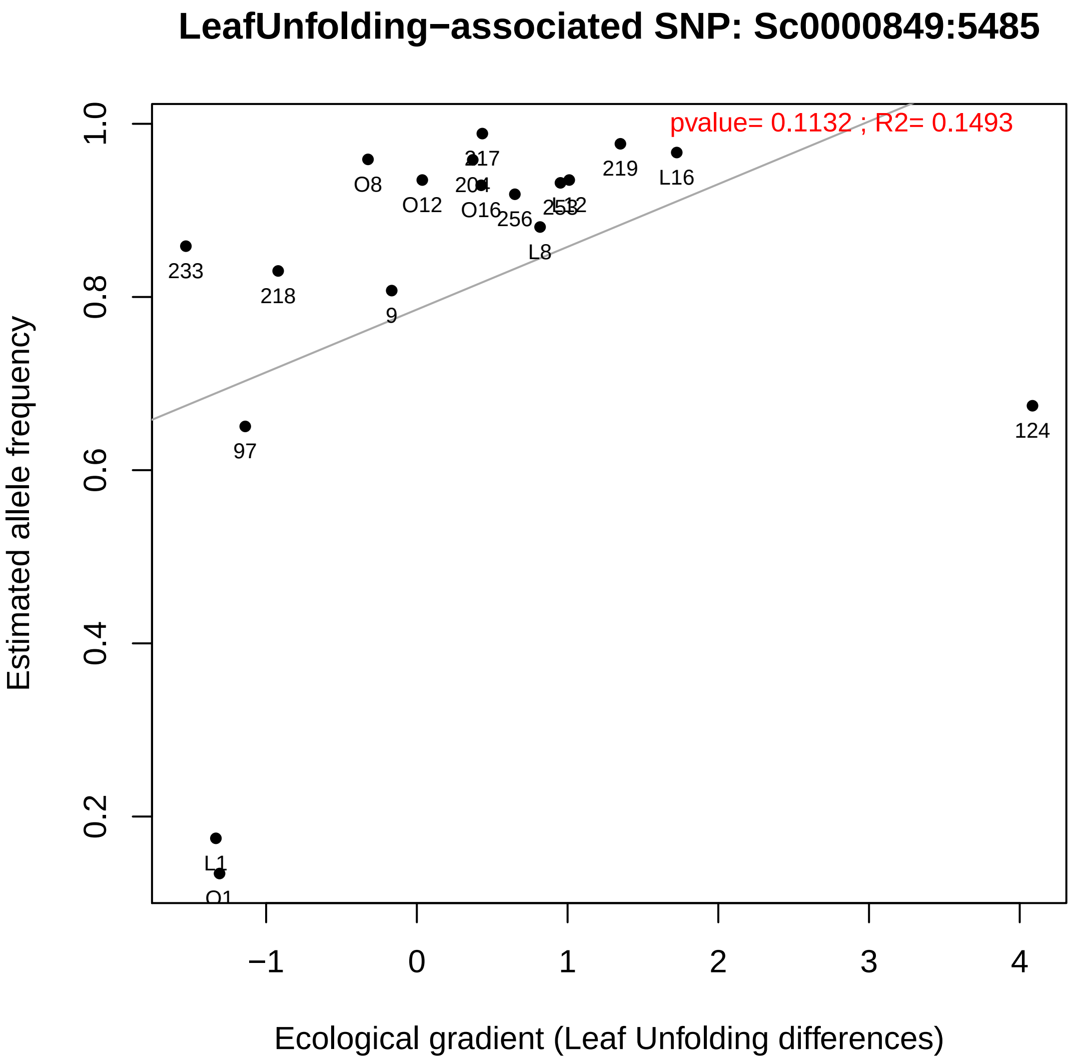

LeafUnfolding−associated SNP: Sc0000849:5485
pvalue= 0.1132 ; R2= 0.1493
Estimated allele frequency
Ecological gradient (Leaf Unfolding differences)
0.2
0.4
0.6
0.8
1.0
−1
0
1
2
3
4
233
218
97
9
O8
O12
217
204
O16
256
253
L12
L8
219
L16
124
L1
O1

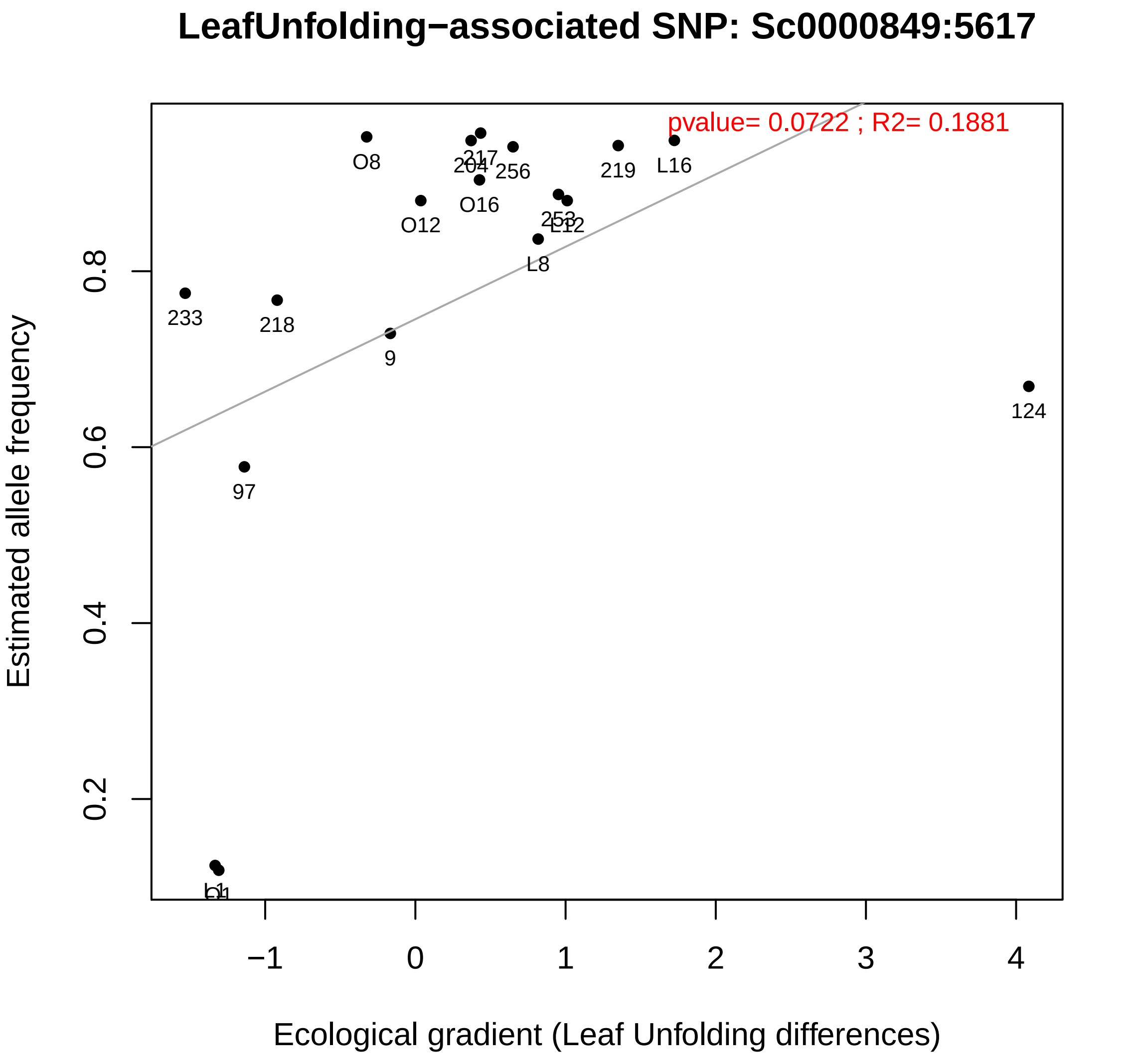
LeafUnfolding−associated SNP: Sc0000849:5617
pvalue= 0.0722 ; R2= 0.1881
Estimated allele frequency
0.2
0.4
0.6
0.8
Ecological gradient (Leaf Unfolding differences)
−1
0
1
2
3
4
O8
217
204
256
219
L16
O16
O12
253
L12
L8
233
218
9
124
97

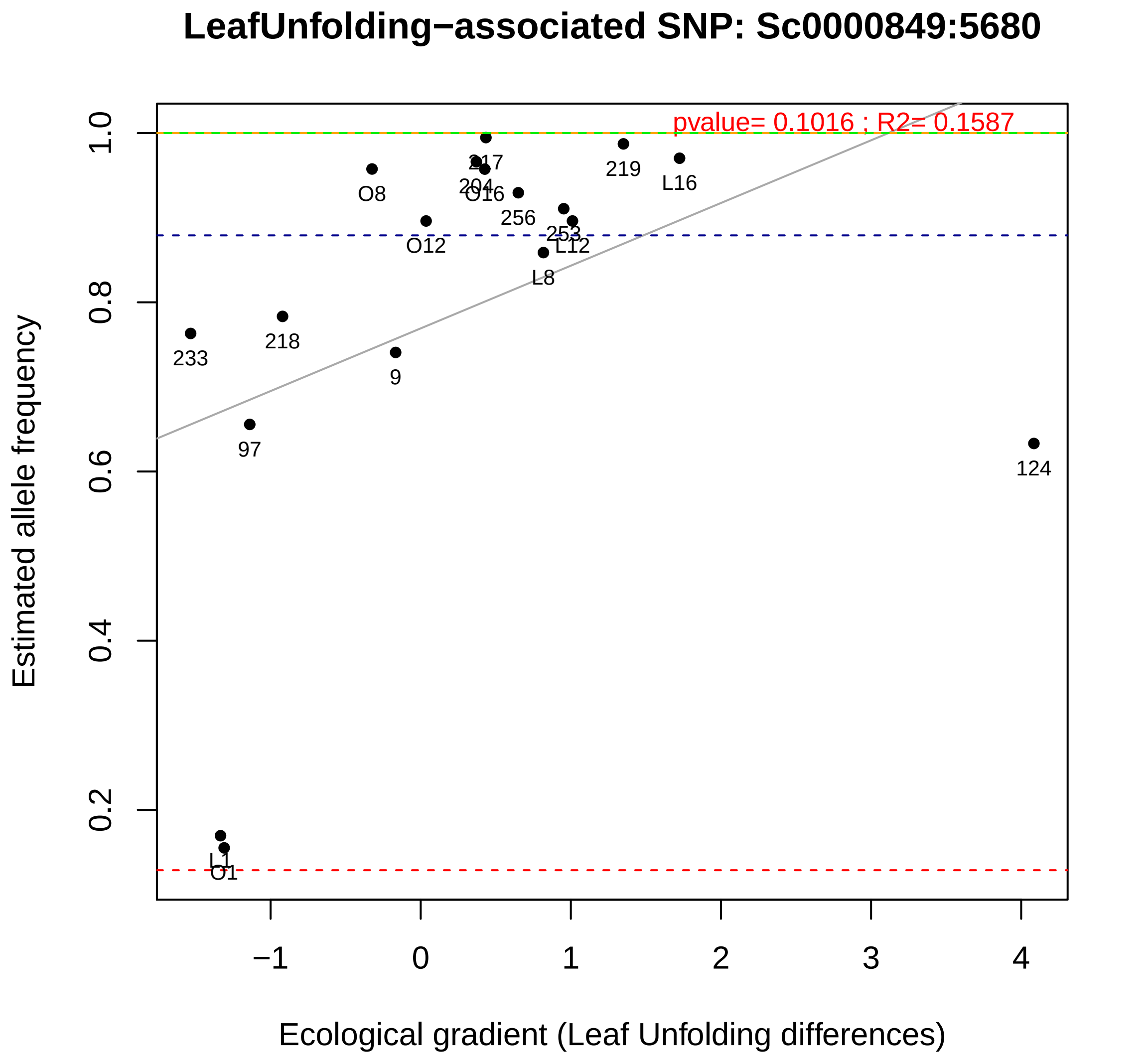

LeafUnfolding−associated SNP: Sc0000849:5680
pvalue= 0.1016 ; R2= 0.1587
Estimated allele frequency
Ecological gradient (Leaf Unfolding differences)
0.2
0.4
0.6
0.8
1.0
−1
0
1
2
3
4
217
204
O16
219
L16
O8
256
253
L12
O12
L8
218
233
9
97
124
L1
O1

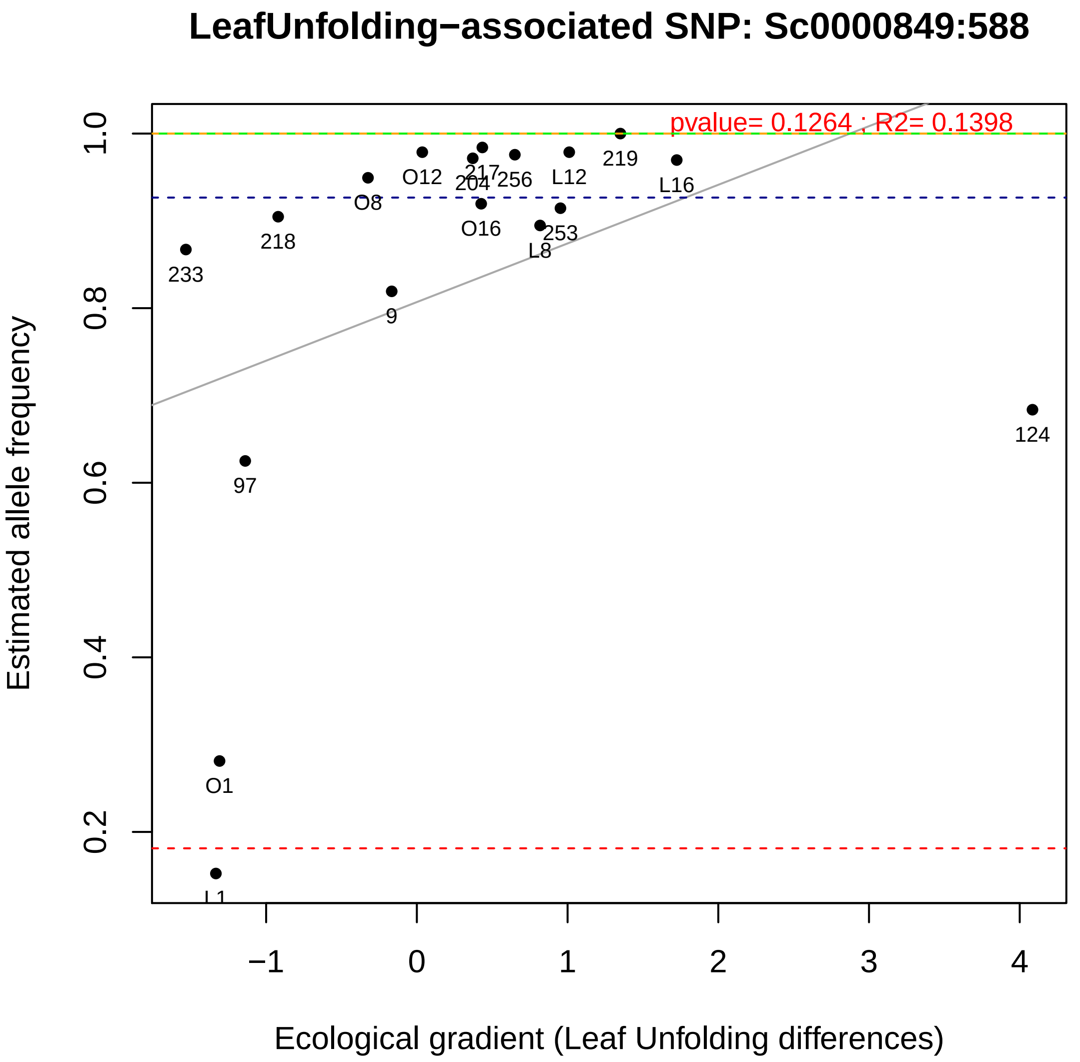

LeafUnfolding−associated SNP: Sc0000849:588
pvalue= 0.1264 ; R2= 0.1398
Estimated allele frequency
Ecological gradient (Leaf Unfolding differences)
1.0
0.8
0.6
0.4
0.2
−1
0
1
2
3
4
219
O12
217
204
256
L12
L16
O8
O16
253
L8
218
233
9
124
97
O1

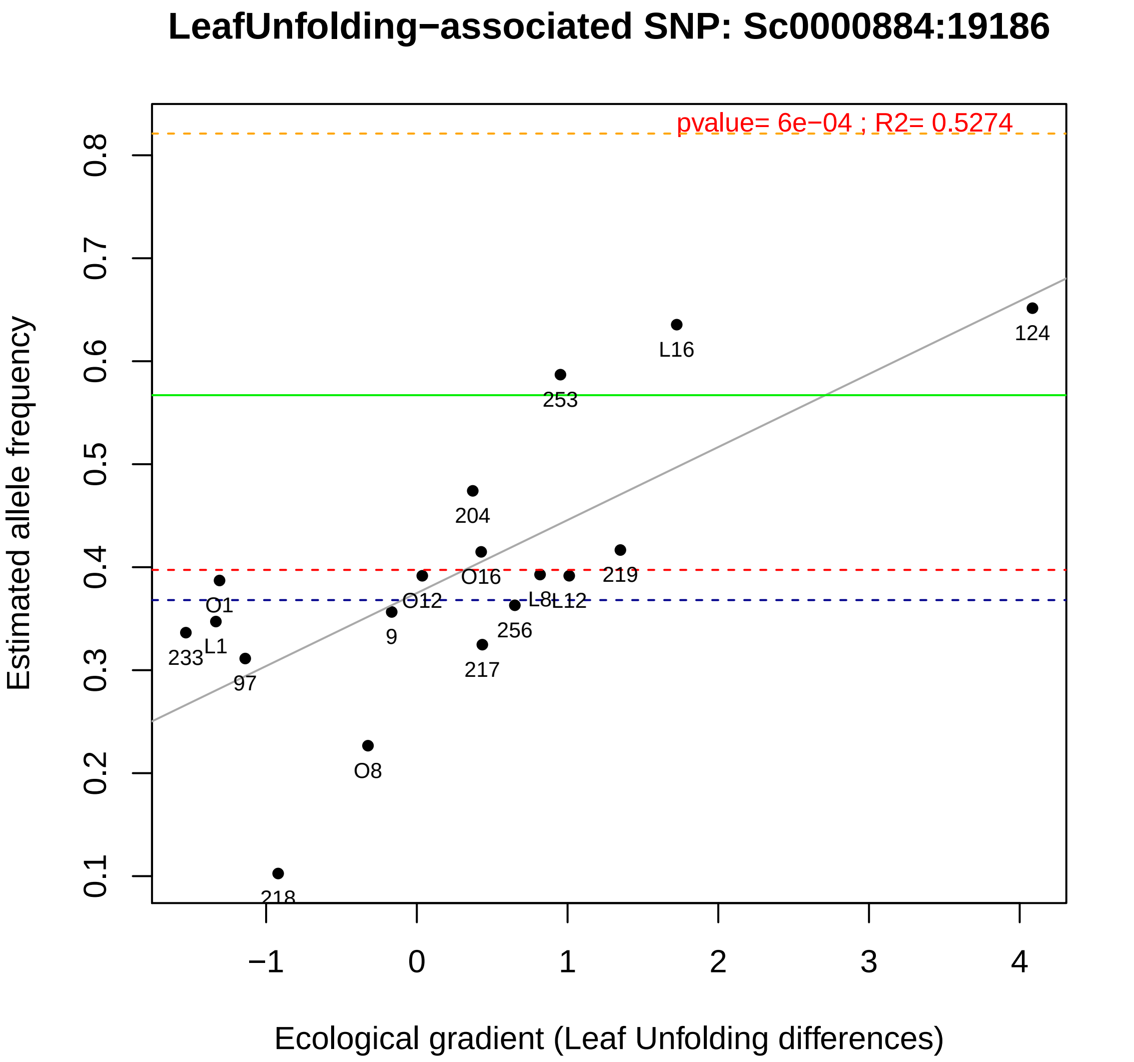

LeafUnfolding−associated SNP: Sc0000884:19186
pvalue= 6e−04 ; R2= 0.5274
Estimated allele frequency
0.1
0.2
0.3
0.4
0.5
0.6
0.7
0.8
−1
0
1
2
3
4
Ecological gradient (Leaf Unfolding differences)
124
L16
253
204
O16
219
L8
L12
O12
O1
L1
233
97
9
256
217
O8
218

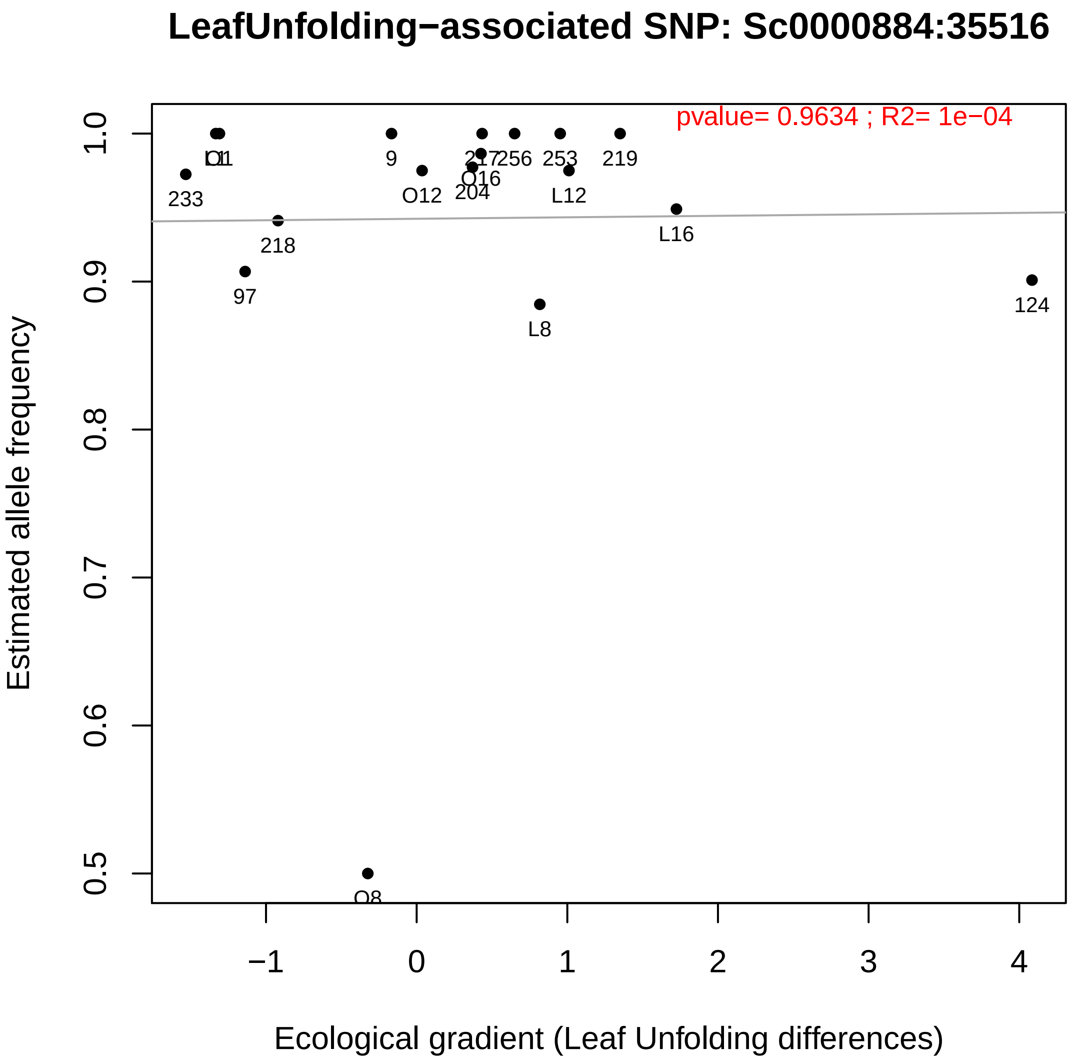

LeafUnfolding−associated SNP: Sc0000884:35516
pvalue= 0.9634 ; R2= 1e−04
Estimated allele frequency
Ecological gradient (Leaf Unfolding differences)
0.5
0.6
0.7
0.8
0.9
1.0
−1
0
1
2
3
4
233
97
218
9
O12
204
O16
L12
L16
L8
124
256
253
219
O8

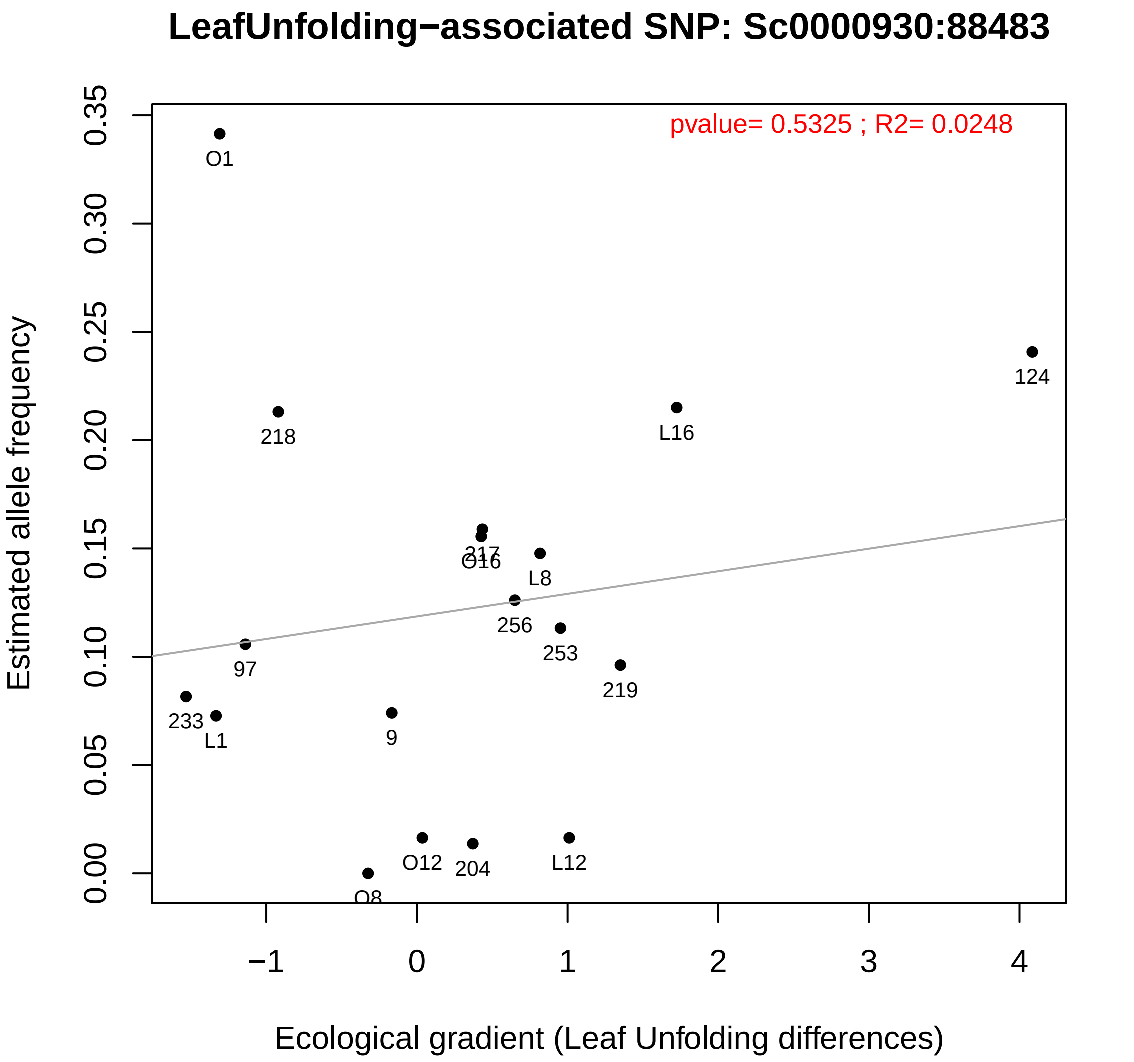

LeafUnfolding−associated SNP: Sc0000930:88483
pvalue= 0.5325 ; R2= 0.0248
Estimated allele frequency
Ecological gradient (Leaf Unfolding differences)
0.00
0.05
0.10
0.15
0.20
0.25
0.30
0.35
−1
0
1
2
3
4
O1
218
124
L16
217
O16
L8
256
253
97
219
233
L1
9
O12
204
L12
O8

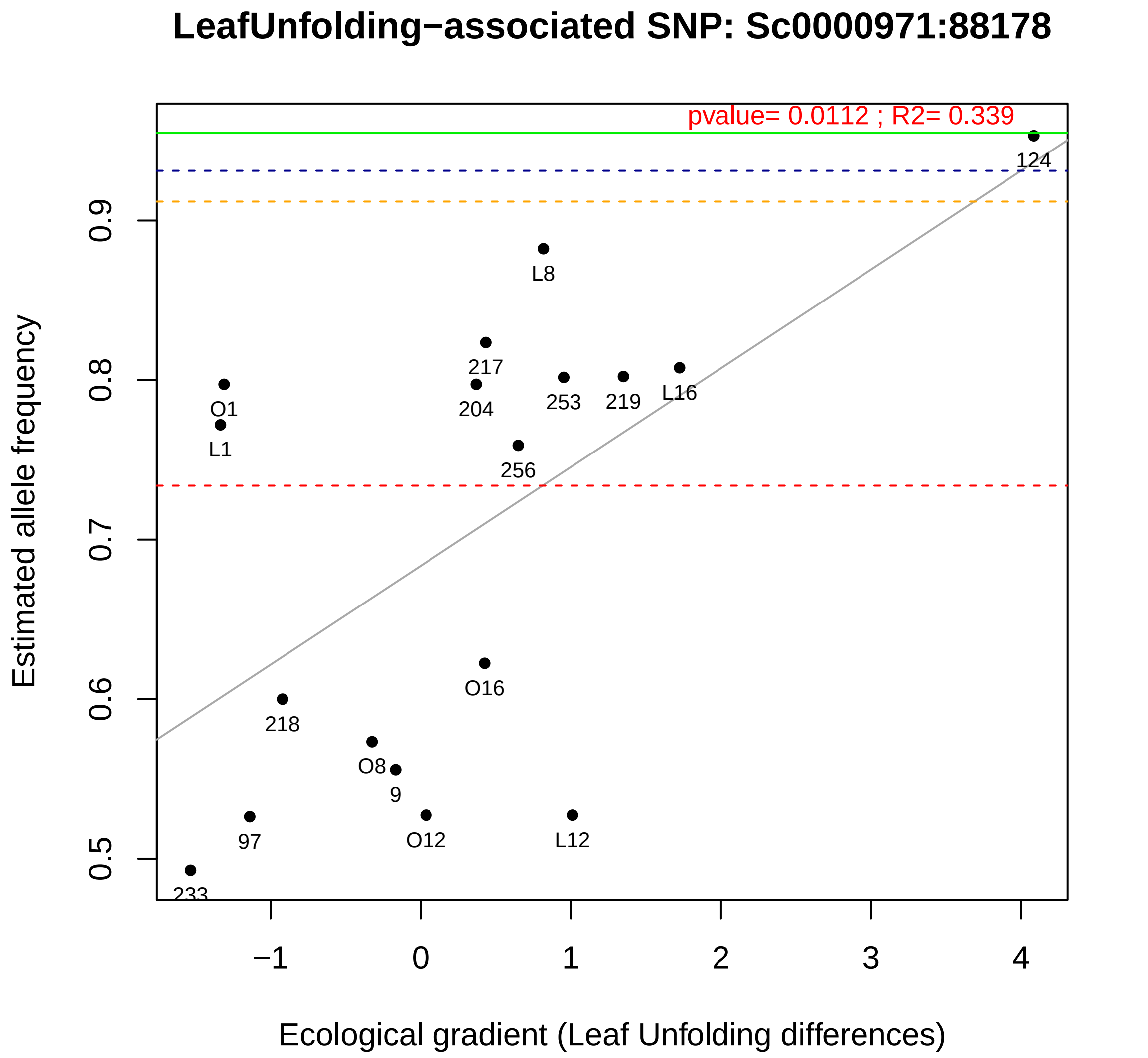

LeafUnfolding−associated SNP: Sc0000971:88178
pvalue= 0.0112 ; R2= 0.339
Estimated allele frequency
Ecological gradient (Leaf Unfolding differences)
0.5
0.6
0.7
0.8
0.9
−1
0
1
2
3
4
124
L8
217
253
219
L16
O1
204
L1
256
O16
218
O8
9
97
O12
L12
233

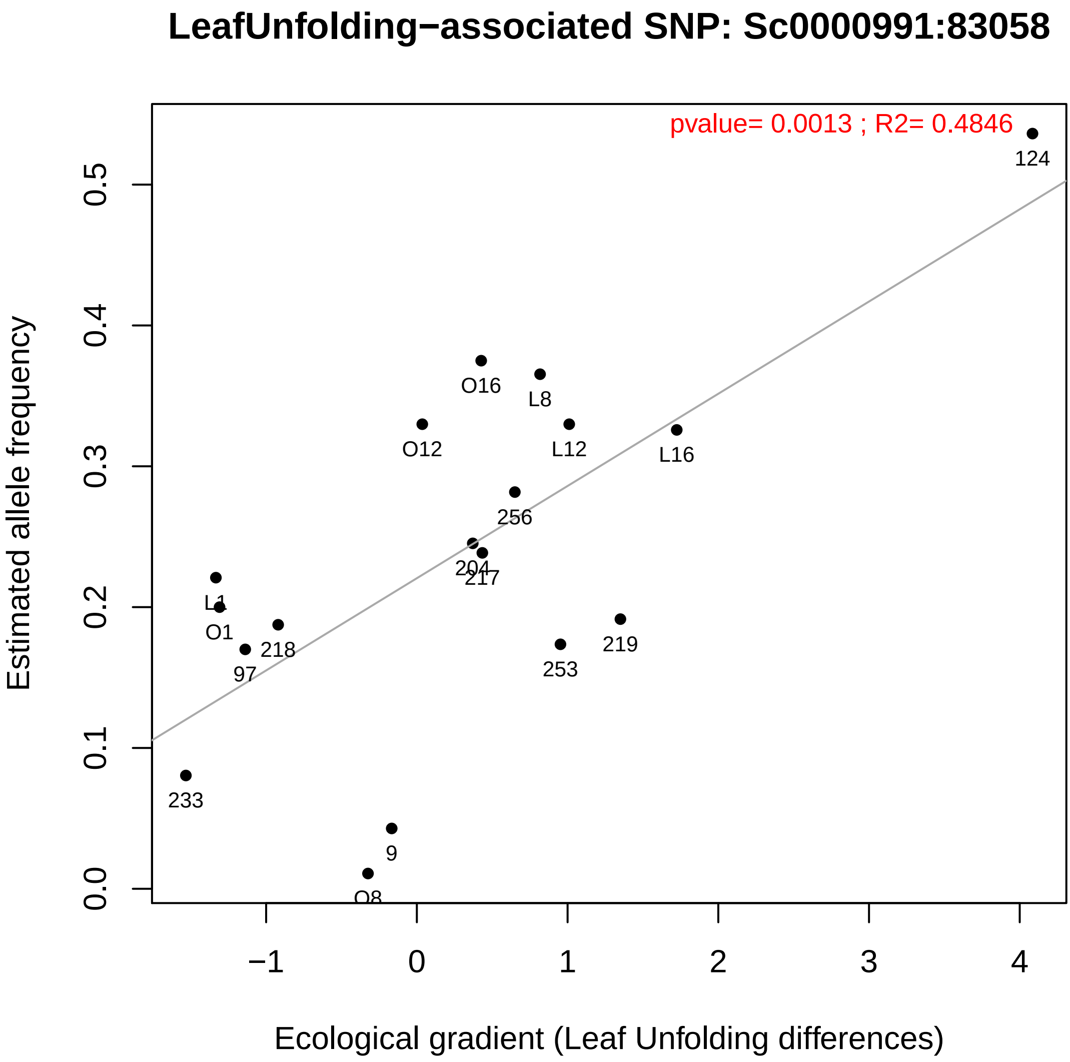
LeafUnfolding−associated SNP: Sc0000991:83058
pvalue= 0.0013 ; R2= 0.4846
Estimated allele frequency
Ecological gradient (Leaf Unfolding differences)
0.0
0.1
0.2
0.3
0.4
0.5
−1
0
1
2
3
4
124
O16
L8
O12
L12
L16
256
204
217
L1
O1
218
97
219
253
233
9
O8

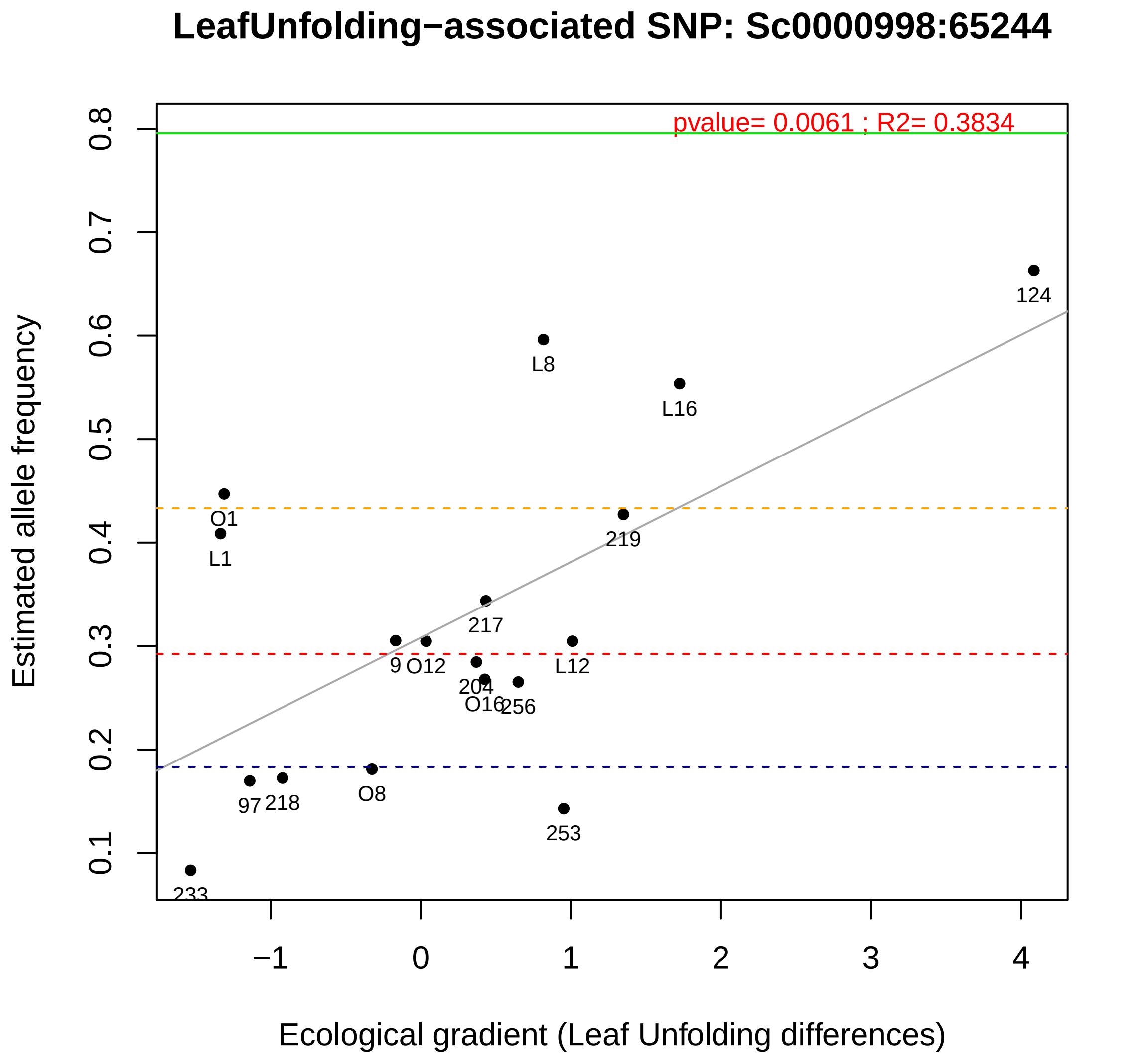

LeafUnfolding−associated SNP: Sc0000998:65244
pvalue= 0.0061 ; R2= 0.3834
Estimated allele frequency
0.1
0.2
0.3
0.4
0.5
0.6
0.7
0.8
−1
0
1
2
3
4
Ecological gradient (Leaf Unfolding differences)
124
L8
L16
O1
L1
219
217
9
O12
L12
204
O16
256
97
218
O8
253
233

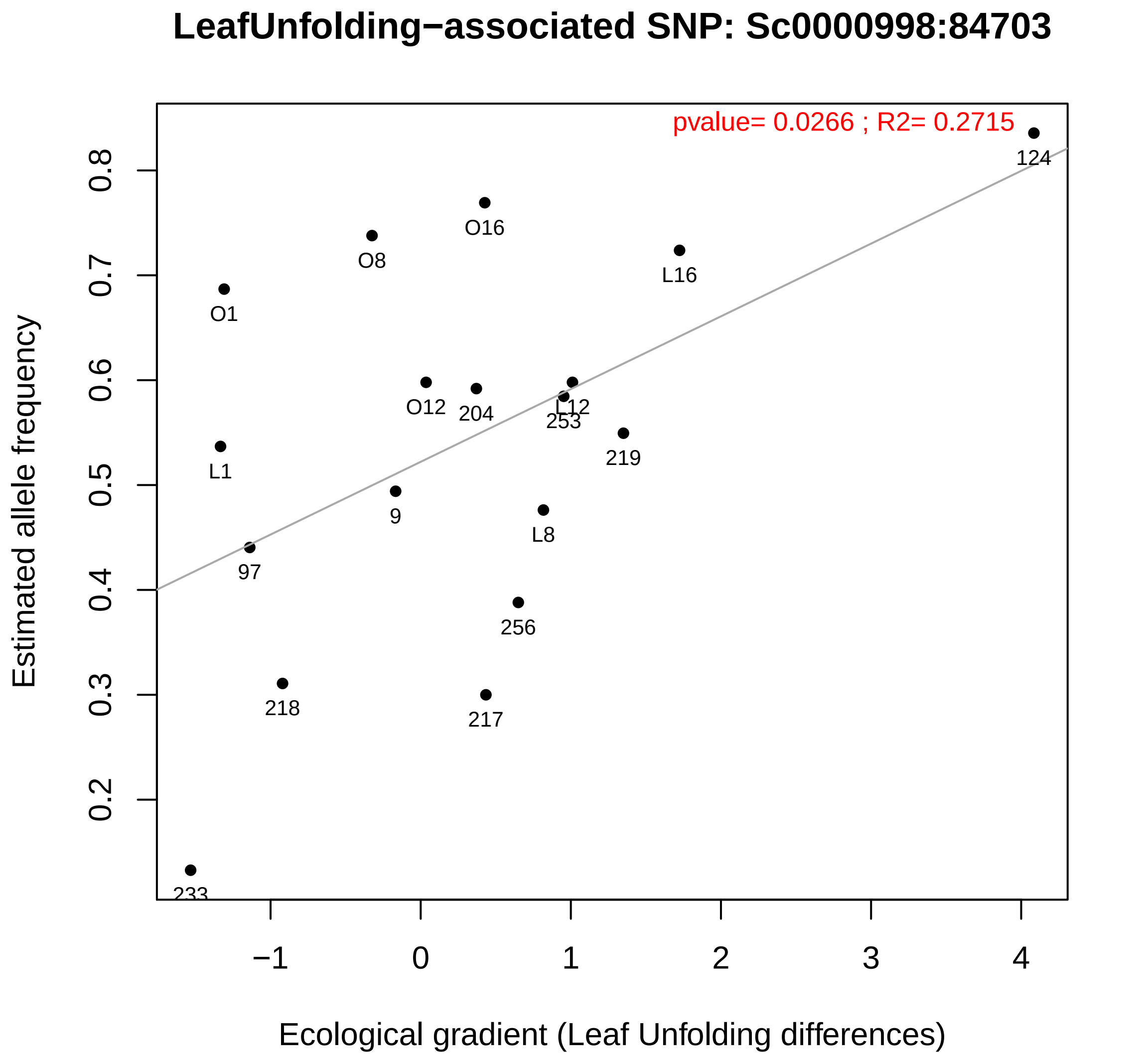
LeafUnfolding−associated SNP: Sc0000998:84703
pvalue= 0.0266 ; R2= 0.2715
Estimated allele frequency
Ecological gradient (Leaf Unfolding differences)
0.2
0.3
0.4
0.5
0.6
0.7
0.8
−1
0
1
2
3
4
124
O16
O8
L16
O1
O12
204
L12
253
219
L1
9
L8
97
256
218
217
233

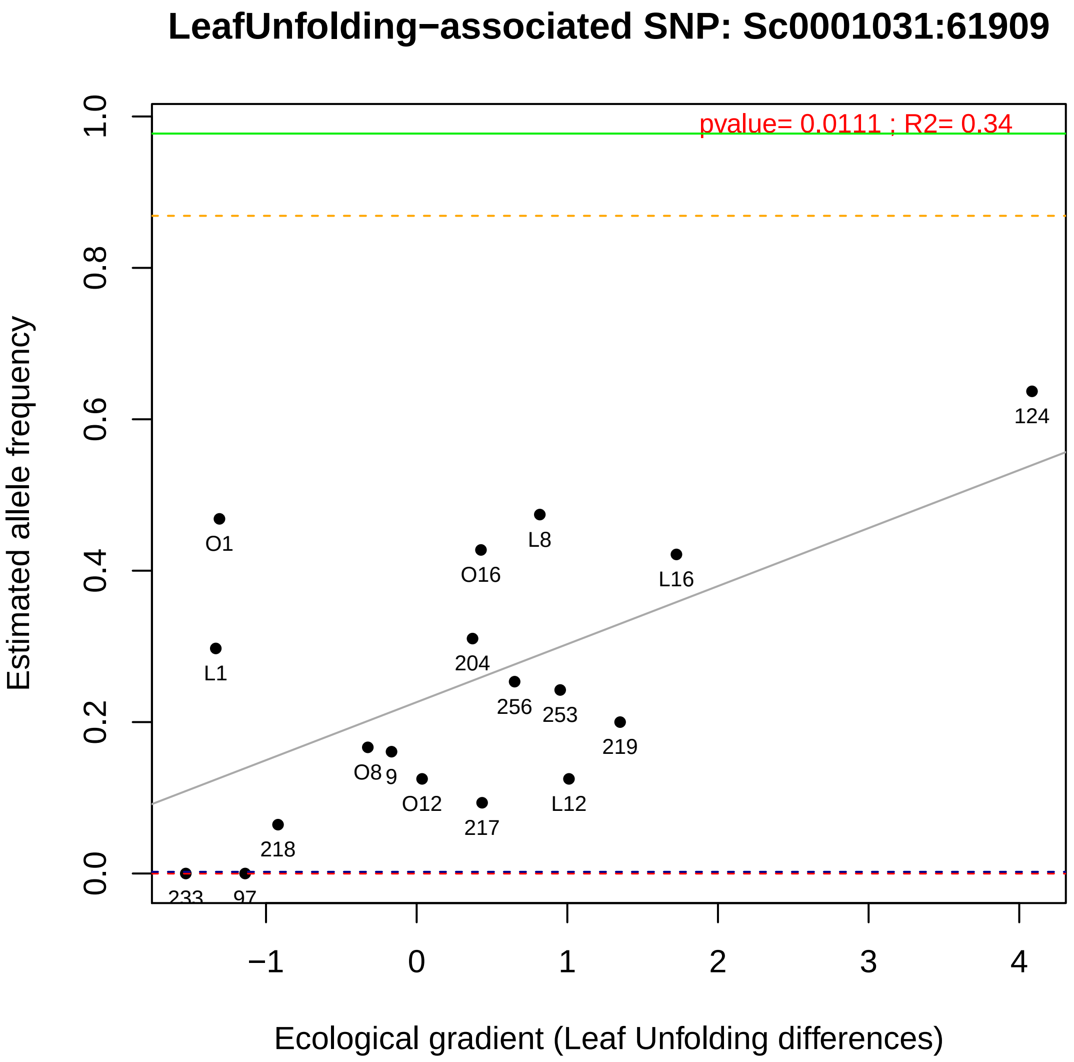
LeafUnfolding−associated SNP: Sc0001031:61909
pvalue= 0.0111 ; R2= 0.34
Estimated allele frequency
0.0
0.2
0.4
0.6
0.8
1.0
−1
0
1
2
3
4
Ecological gradient (Leaf Unfolding differences)
O1
L1
218
233
97
O8
9
O12
217
204
O16
L8
256
253
L12
219
L16
124

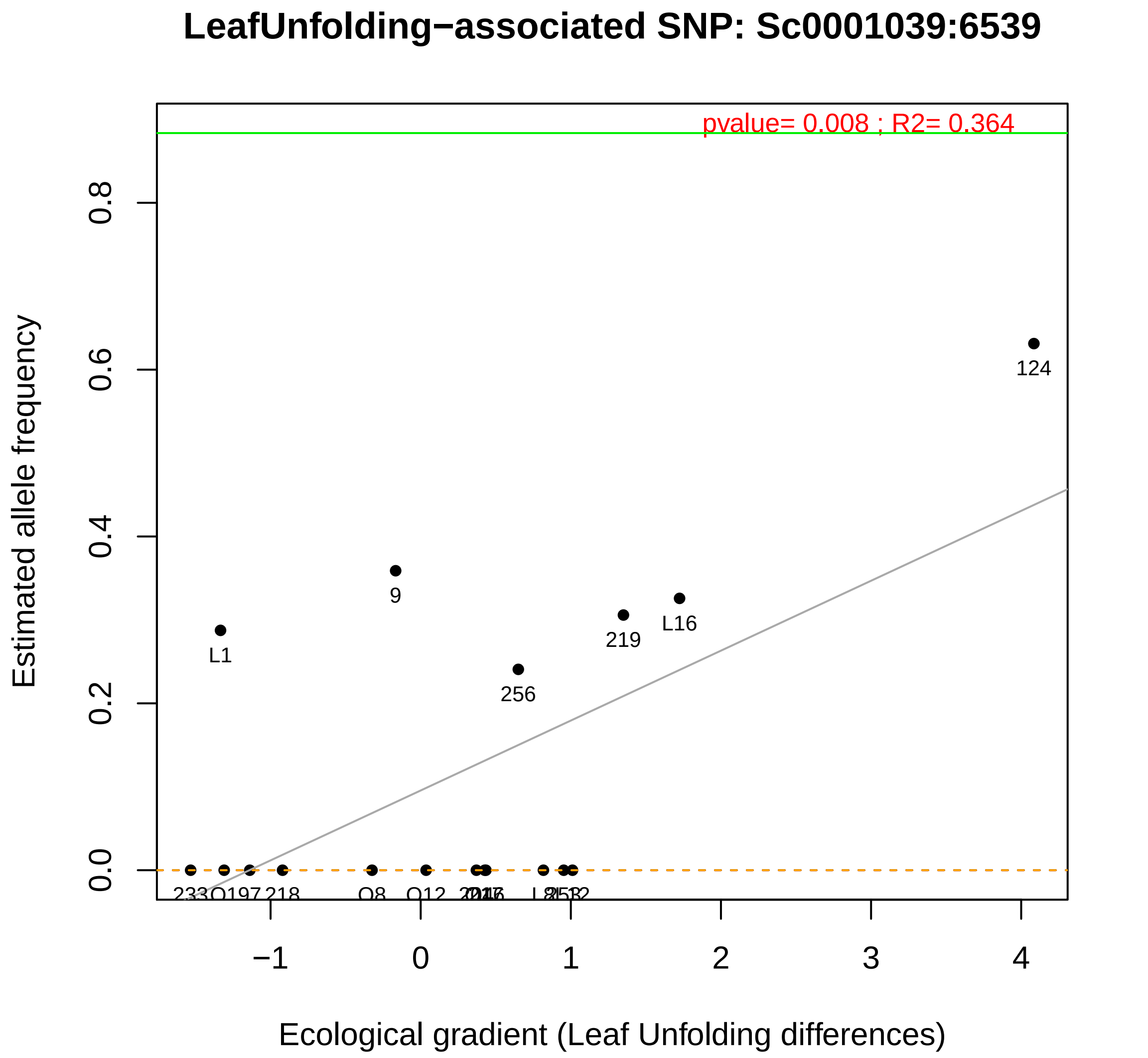

LeafUnfolding−associated SNP: Sc0001039:6539
pvalue= 0.008 ; R2= 0.364
Estimated allele frequency
0.0
0.2
0.4
0.6
0.8
124
9
L1
L16
219
256
−1
0
1
2
3
4
Ecological gradient (Leaf Unfolding differences)

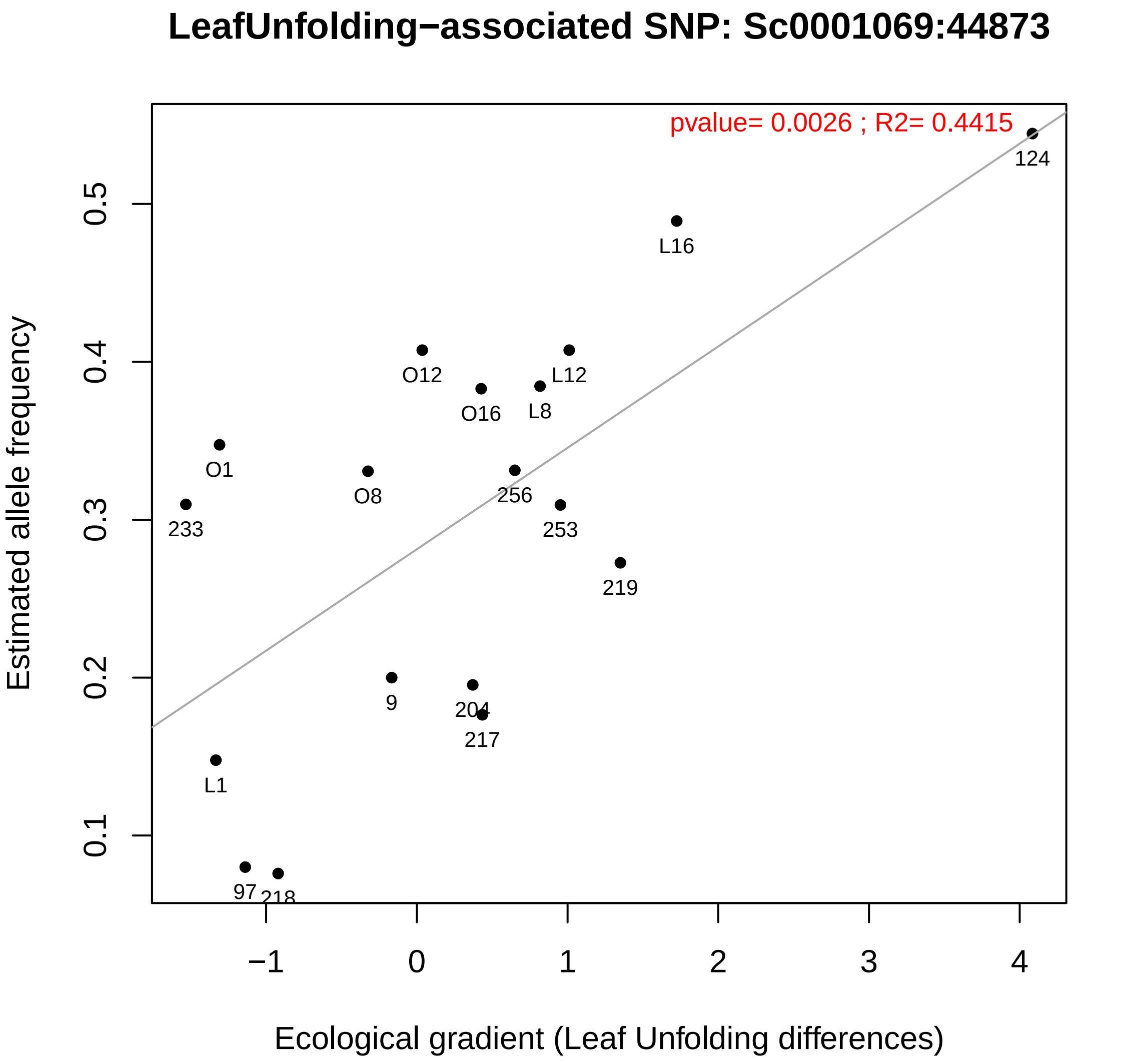
LeafUnfolding−associated SNP: Sc0001069:44873
pvalue= 0.0026 ; R2= 0.4415
Estimated allele frequency
0.1
0.2
0.3
0.4
0.5
−1
0
1
2
3
4
Ecological gradient (Leaf Unfolding differences)
124
L16
O12
L12
O16
L8
O1
O8
256
233
253
219
9
204
217
L1
97
218

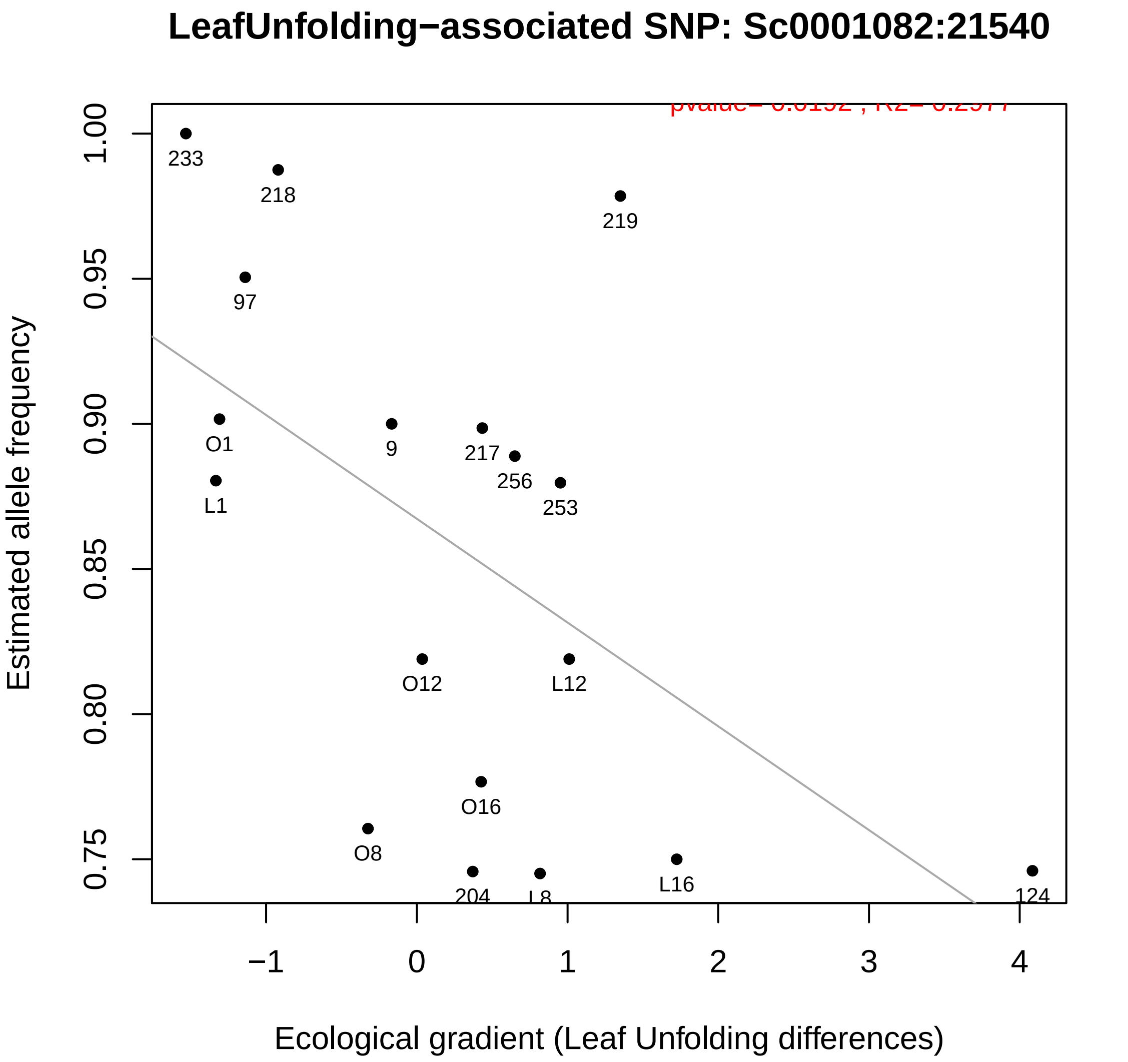

LeafUnfolding−associated SNP: Sc0001082:21540
Estimated allele frequency
1.00
0.95
0.90
0.85
0.80
0.75
−1
0
1
2
3
4
Ecological gradient (Leaf Unfolding differences)
233
218
219
97
O1
9
217
256
253
L1
O12
L12
O16
O8
204
L8
L16
124

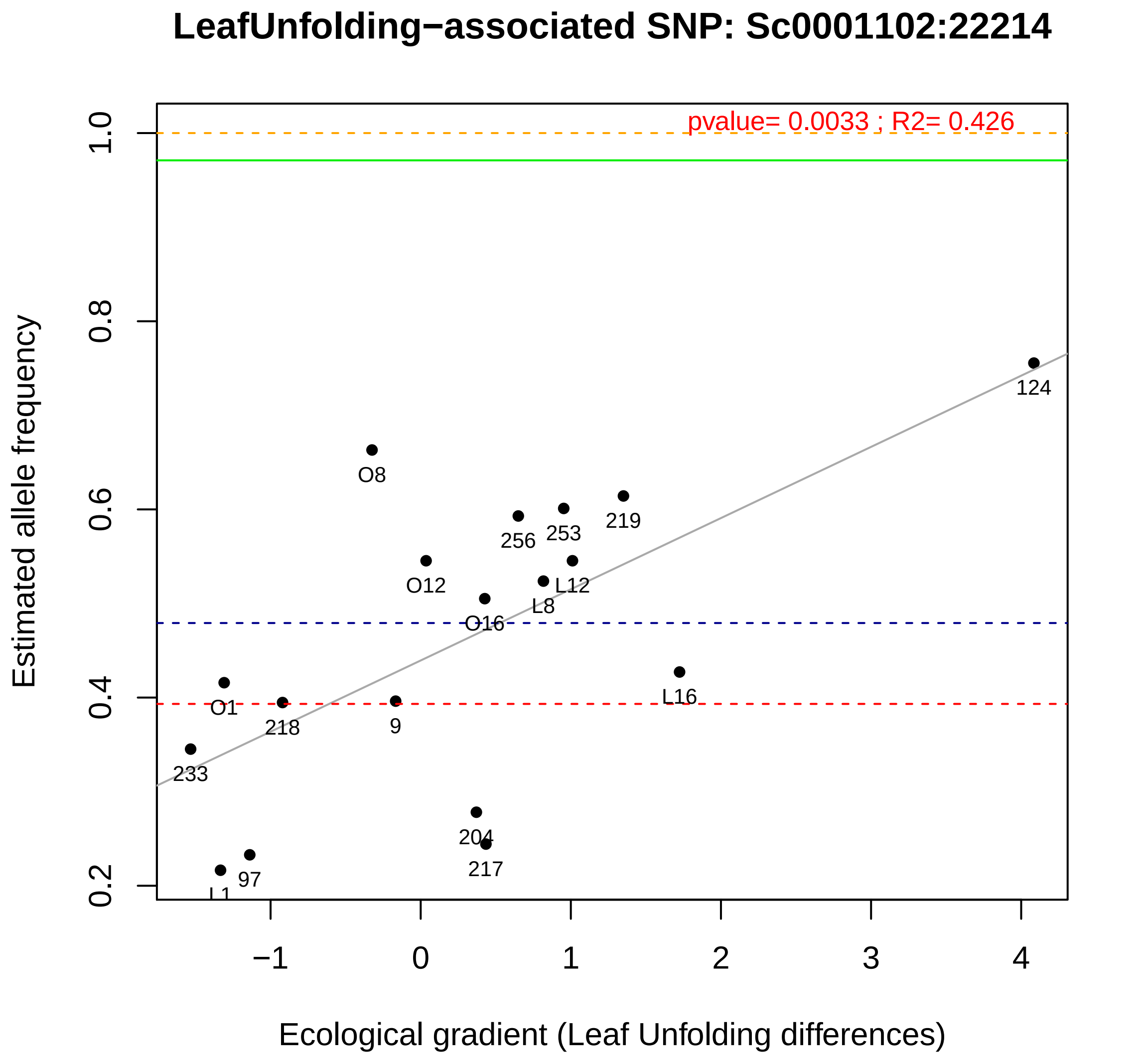
LeafUnfolding−associated SNP: Sc0001102:22214
pvalue= 0.0033 ; R2= 0.426
Estimated allele frequency
Ecological gradient (Leaf Unfolding differences)
0.2
0.4
0.6
0.8
1.0
−1
0
1
2
3
4
124
O8
219
253
256
O12
L12
L8
O16
L16
O1
218
9
233
204
217
97
L1

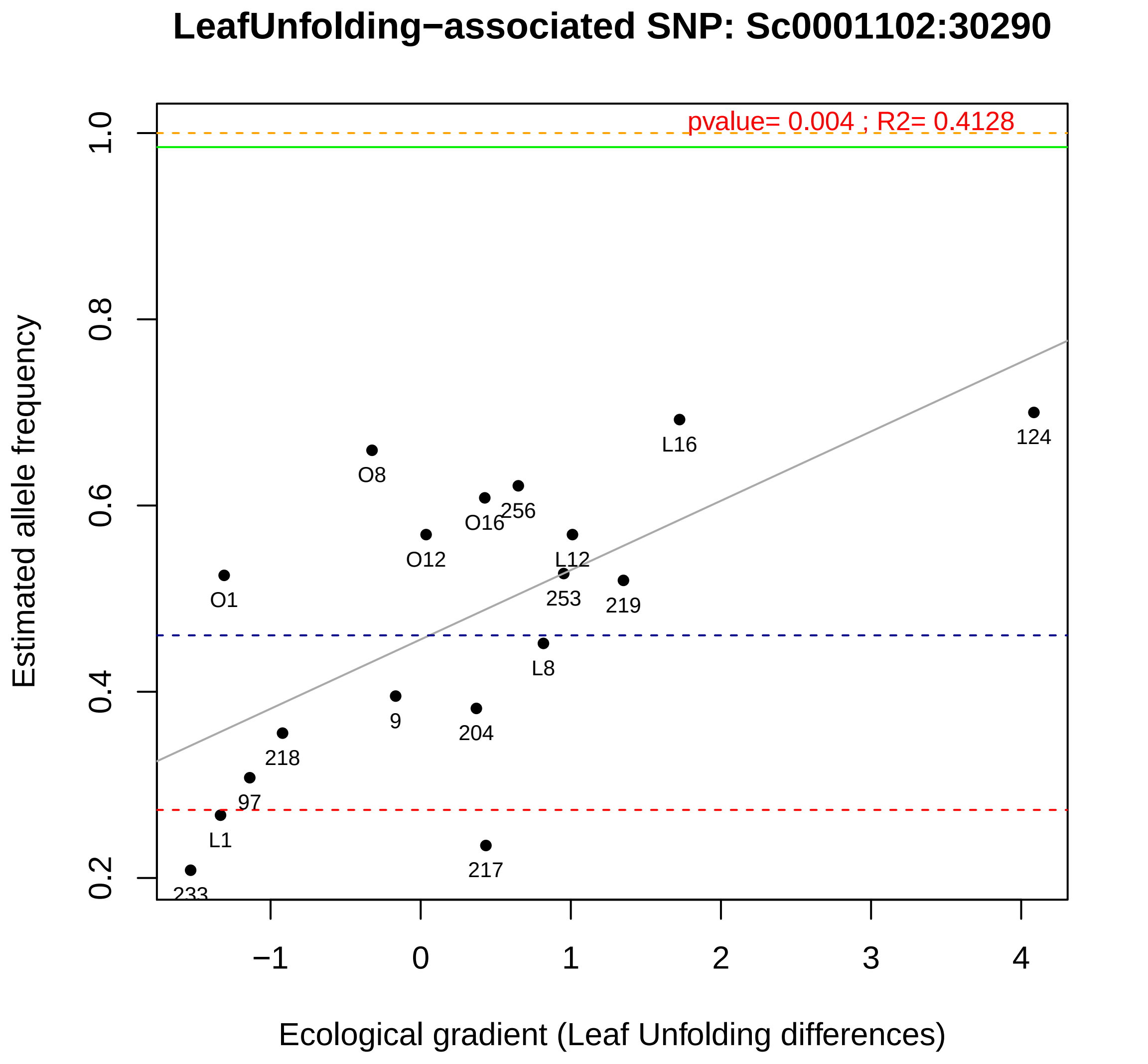
LeafUnfolding−associated SNP: Sc0001102:30290
pvalue= 0.004 ; R2= 0.4128
Estimated allele frequency
Ecological gradient (Leaf Unfolding differences)
0.2
0.4
0.6
0.8
1.0
−1
0
1
2
3
4
233
L1
97
218
O1
9
O8
O12
O16
256
204
217
L8
253
L12
219
L16
124

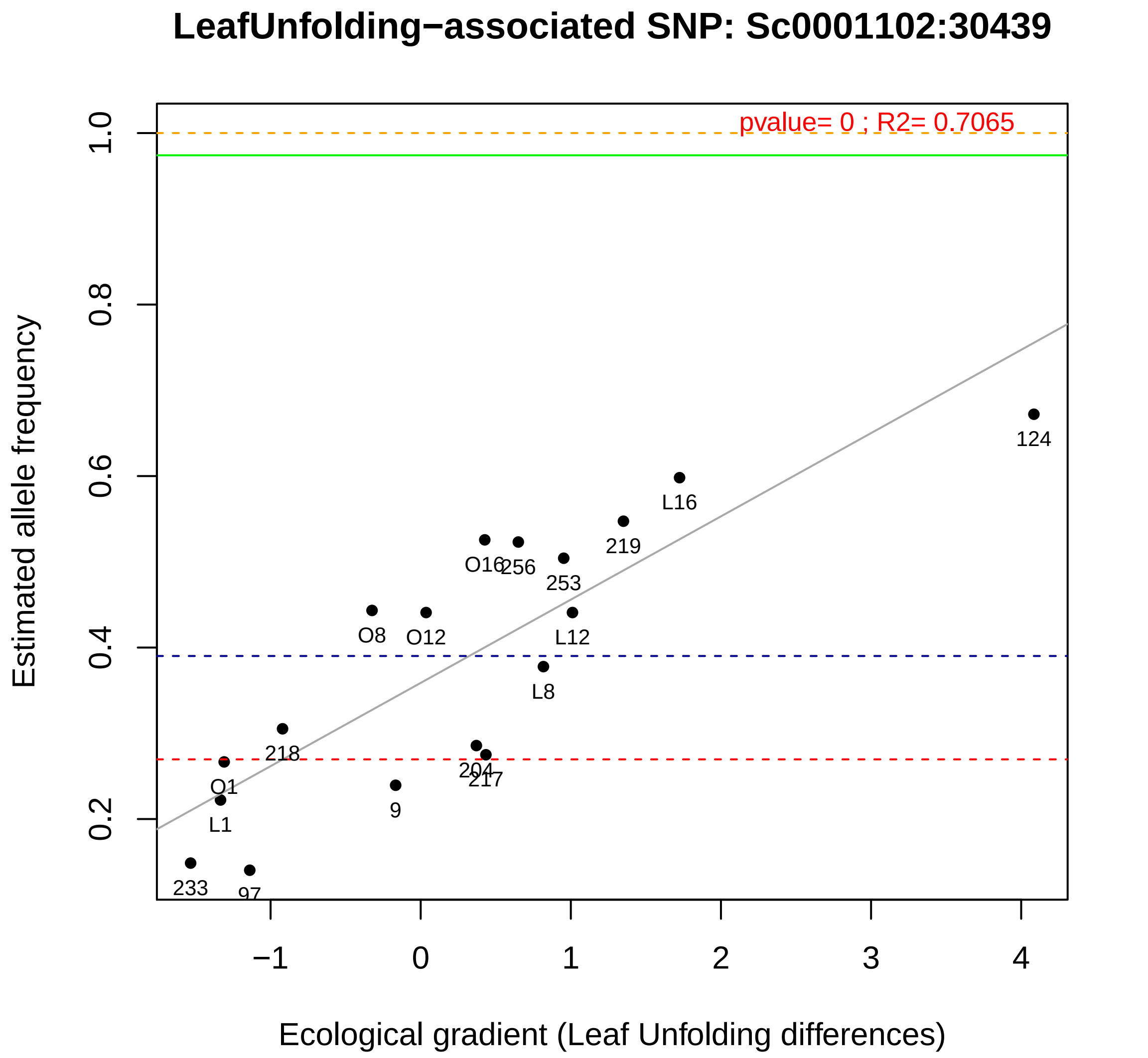

LeafUnfolding−associated SNP: Sc0001102:30439
pvalue= 0 ; R2= 0.7065
Estimated allele frequency
Ecological gradient (Leaf Unfolding differences)
0.2
0.4
0.6
0.8
1.0
−1
0
1
2
3
4
124
L16
219
O16
256
253
O8
O12
L12
L8
218
204
217
O1
L1
9
233
97

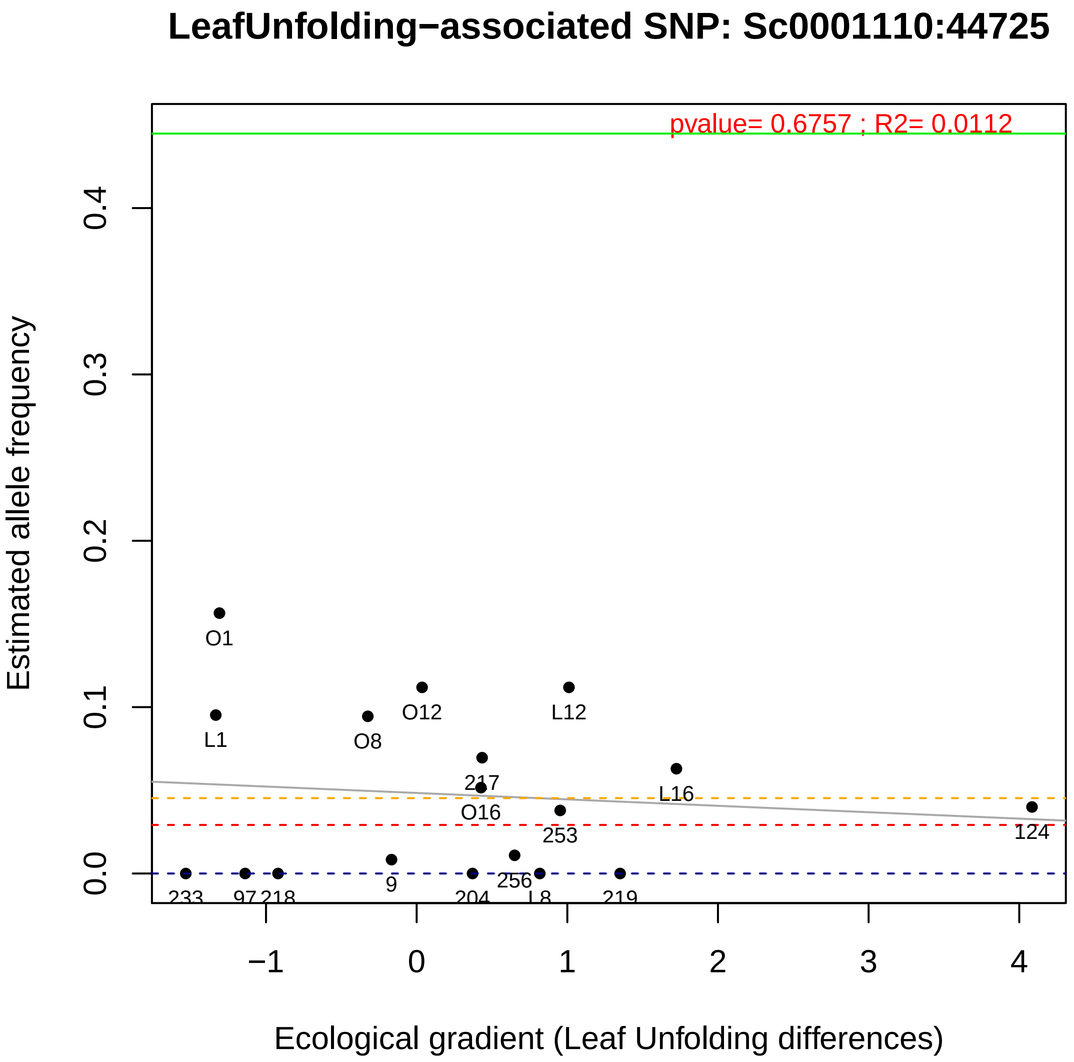

LeafUnfolding−associated SNP: Sc0001110:44725
pvalue= 0.6757 ; R2= 0.0112
Estimated allele frequency
0.0
0.1
0.2
0.3
0.4
−1
0
1
2
3
4
Ecological gradient (Leaf Unfolding differences)
O1
L1
O8
O12
L12
217
O16
L16
253
124
9
256
233
97
218
204
L8
219

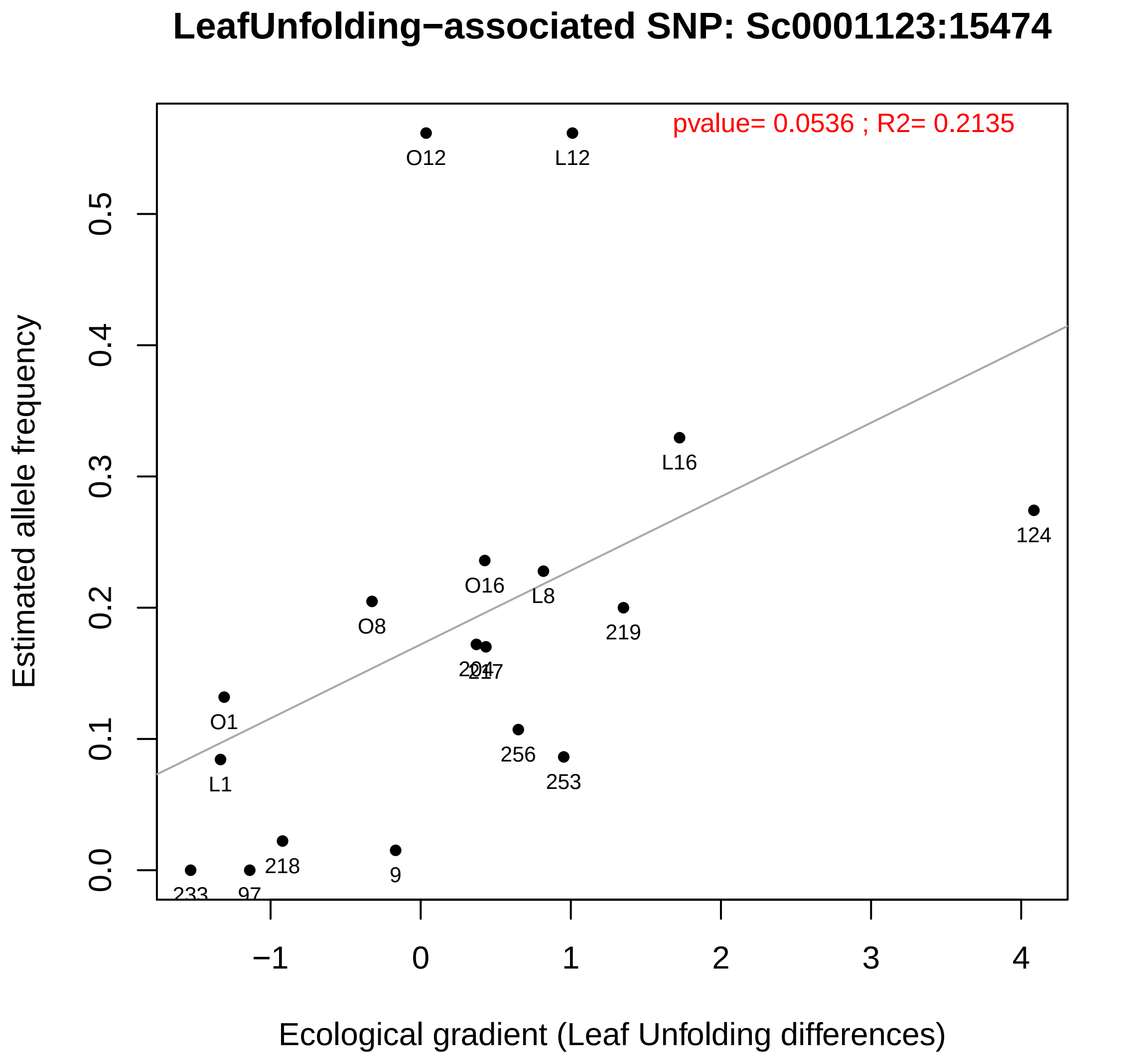
LeafUnfolding−associated SNP: Sc0001123:15474
pvalue= 0.0536 ; R2= 0.2135
Estimated allele frequency
Ecological gradient (Leaf Unfolding differences)
0.0
0.1
0.2
0.3
0.4
0.5
−1
0
1
2
3
4
O12
L12
L16
124
O16
L8
O8
219
O1
256
L1
253
218
9
233
97

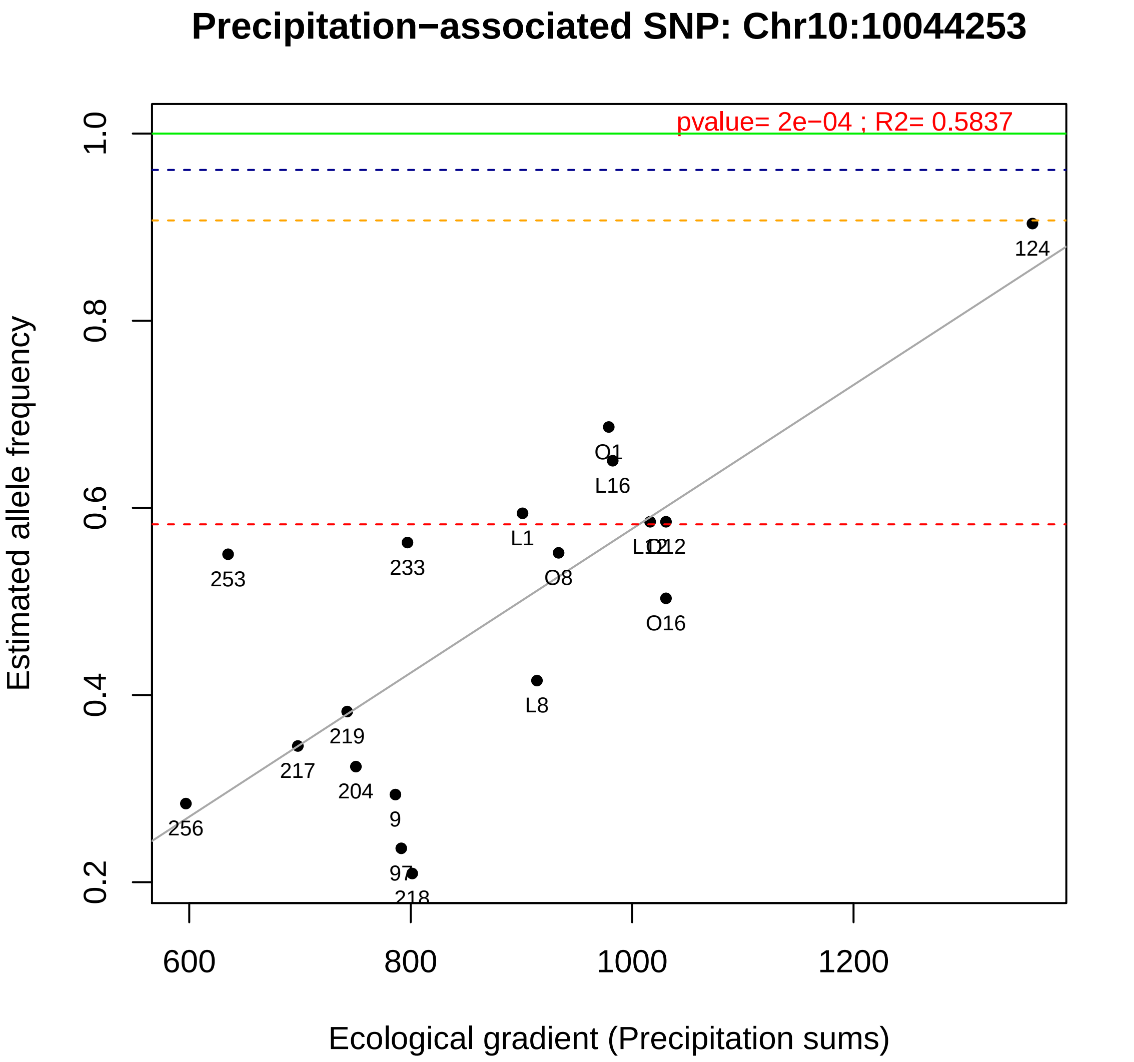
Precipitation−associated SNP: Chr10:10044253
pvalue= 2e−04 ; R2= 0.5837
Estimated allele frequency
Ecological gradient (Precipitation sums)
0.2
0.4
0.6
0.8
1.0
600
800
1000
1200
124
O1
L16
L1
L12
O12
O8
O16
L8
253
233
219
217
204
256
9
97
218

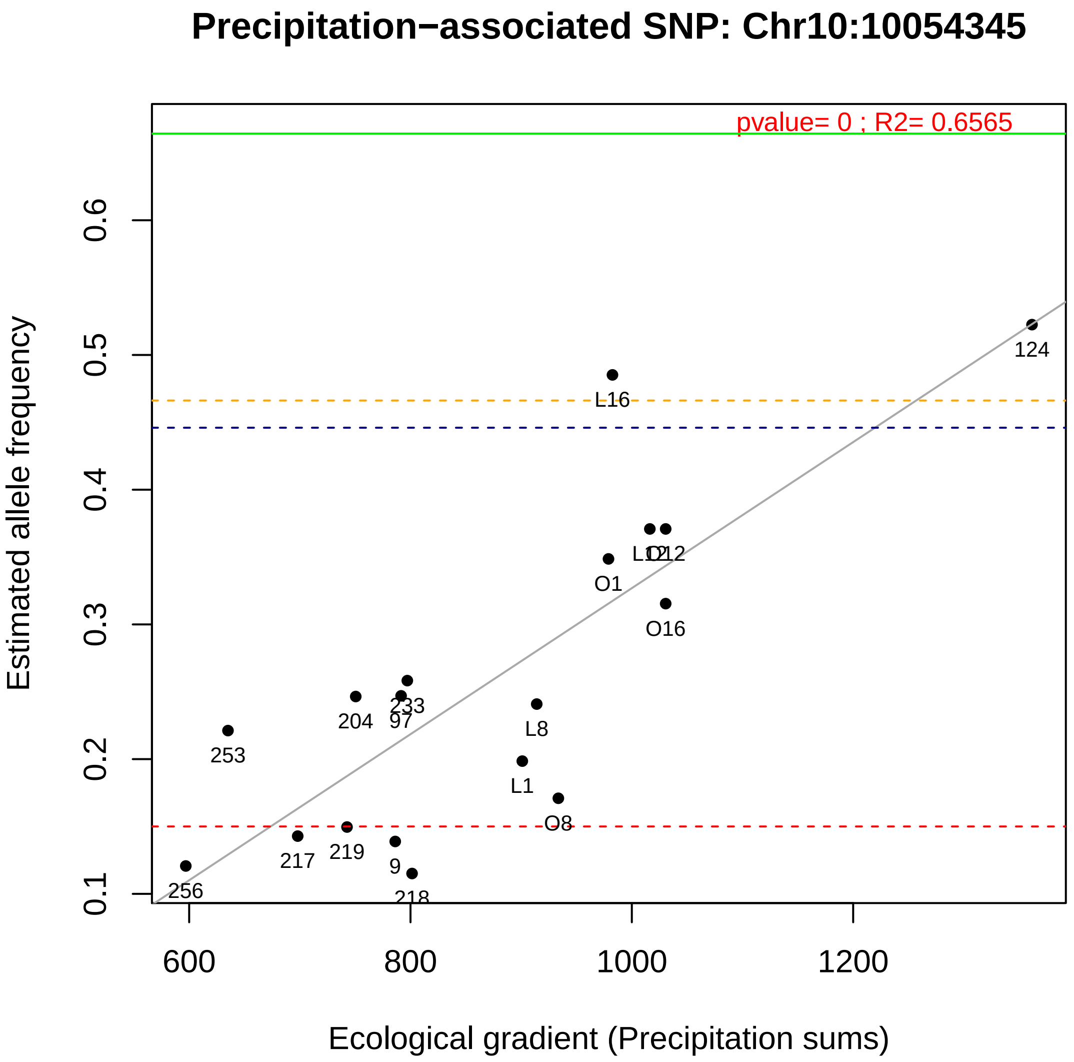

Precipitation−associated SNP: Chr10:10054345
pvalue= 0 ; R2= 0.6565
Estimated allele frequency
Ecological gradient (Precipitation sums)
0.1
0.2
0.3
0.4
0.5
0.6
600
800
1000
1200
124
L16
L12
O12
O1
O16
233
204
97
L8
253
L1
O8
217
219
9
256
218

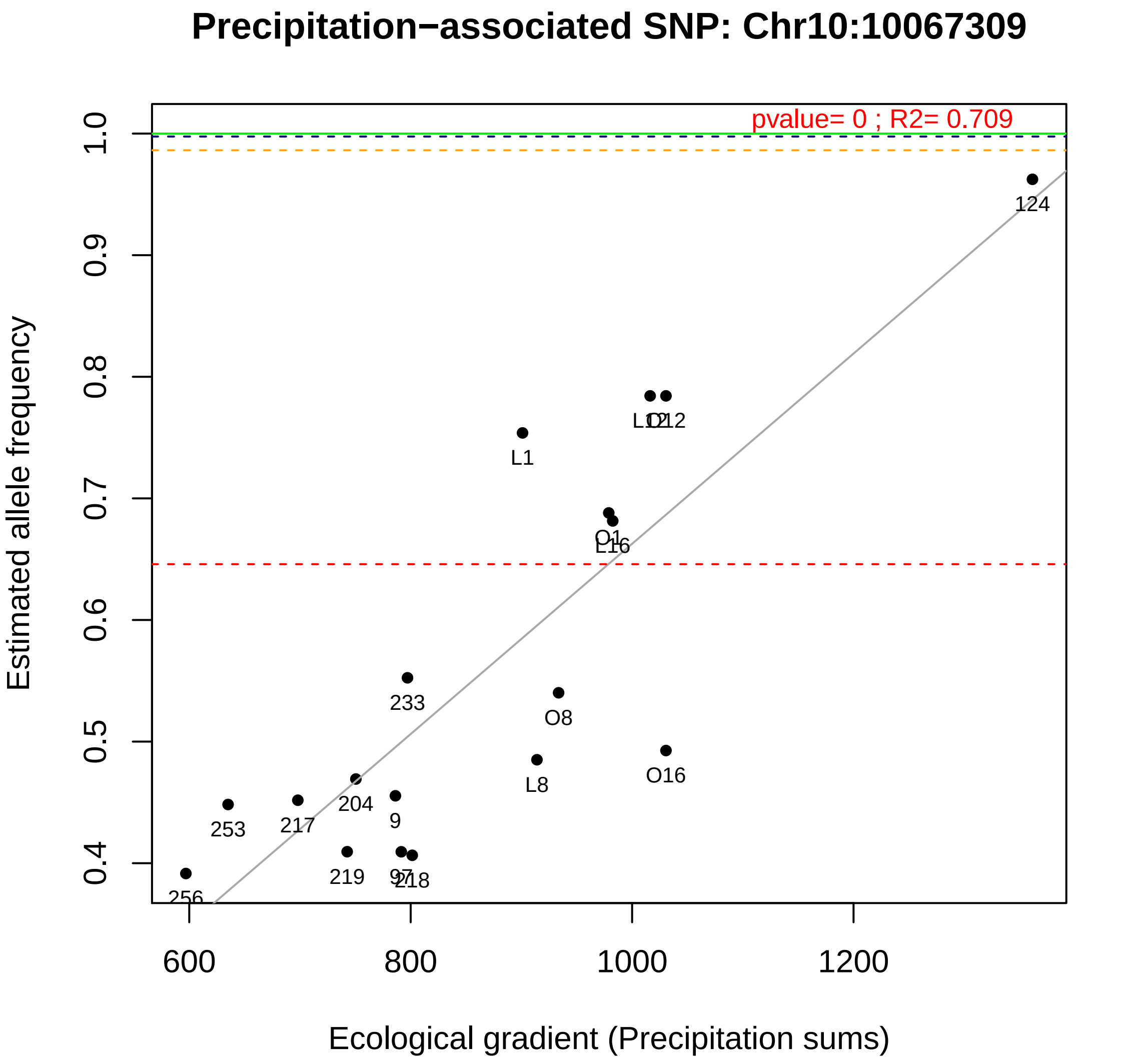

Precipitation−associated SNP: Chr10:10067309
pvalue= 0 ; R2= 0.709
Estimated allele frequency
Ecological gradient (Precipitation sums)
0.4
0.5
0.6
0.7
0.8
0.9
1.0
600
800
1000
1200
124
L12
O12
L1
O1
L16
233
O8
O16
L8
204
9
253
217
219
97
218
256

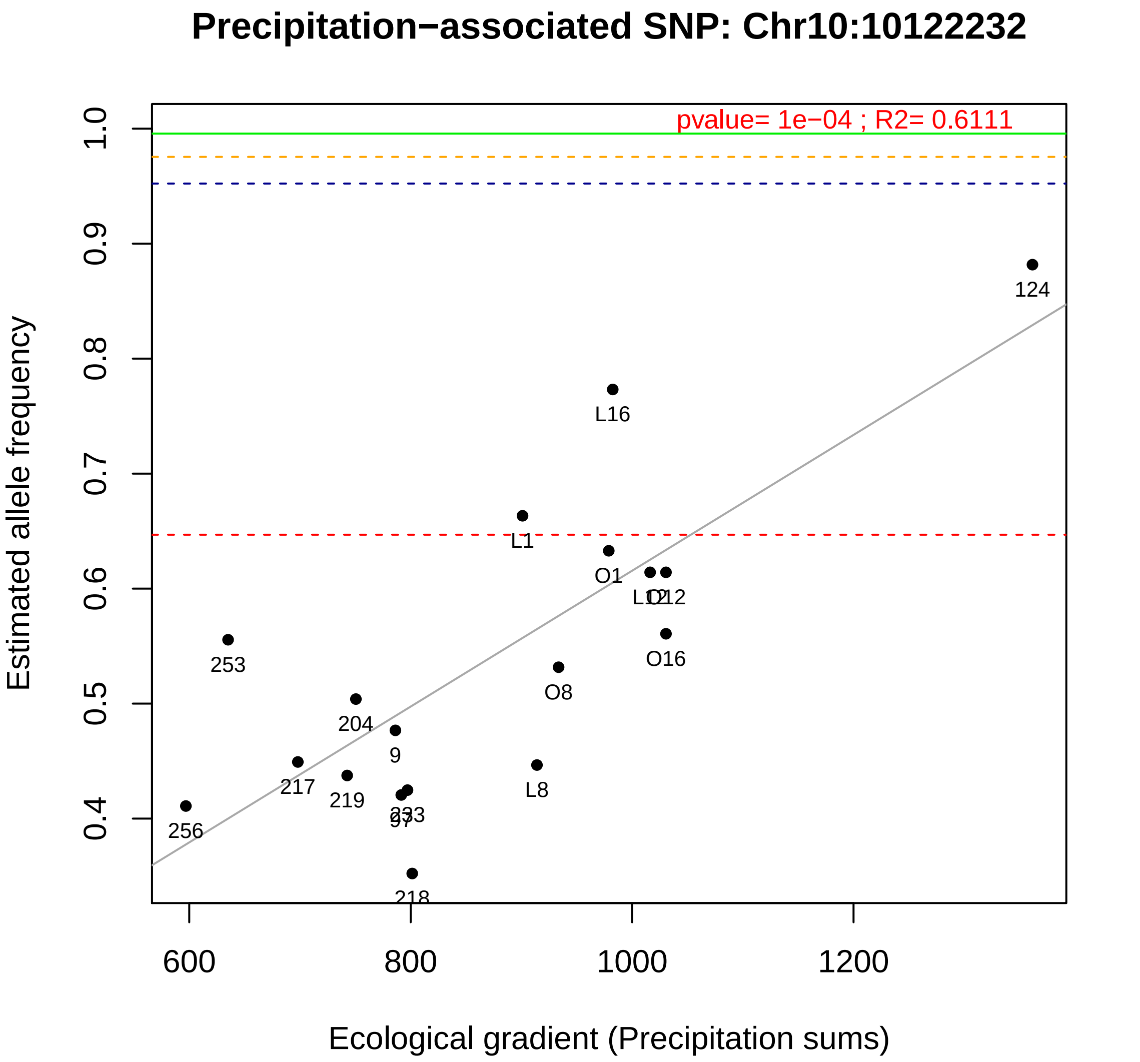
Precipitation−associated SNP: Chr10:10122232
pvalue= 1e−04 ; R2= 0.6111
Estimated allele frequency
Ecological gradient (Precipitation sums)
0.4
0.5
0.6
0.7
0.8
0.9
1.0
600
800
1000
1200
124
L16
L1
O1
O16
O8
253
204
9
217
219
L8
256
218

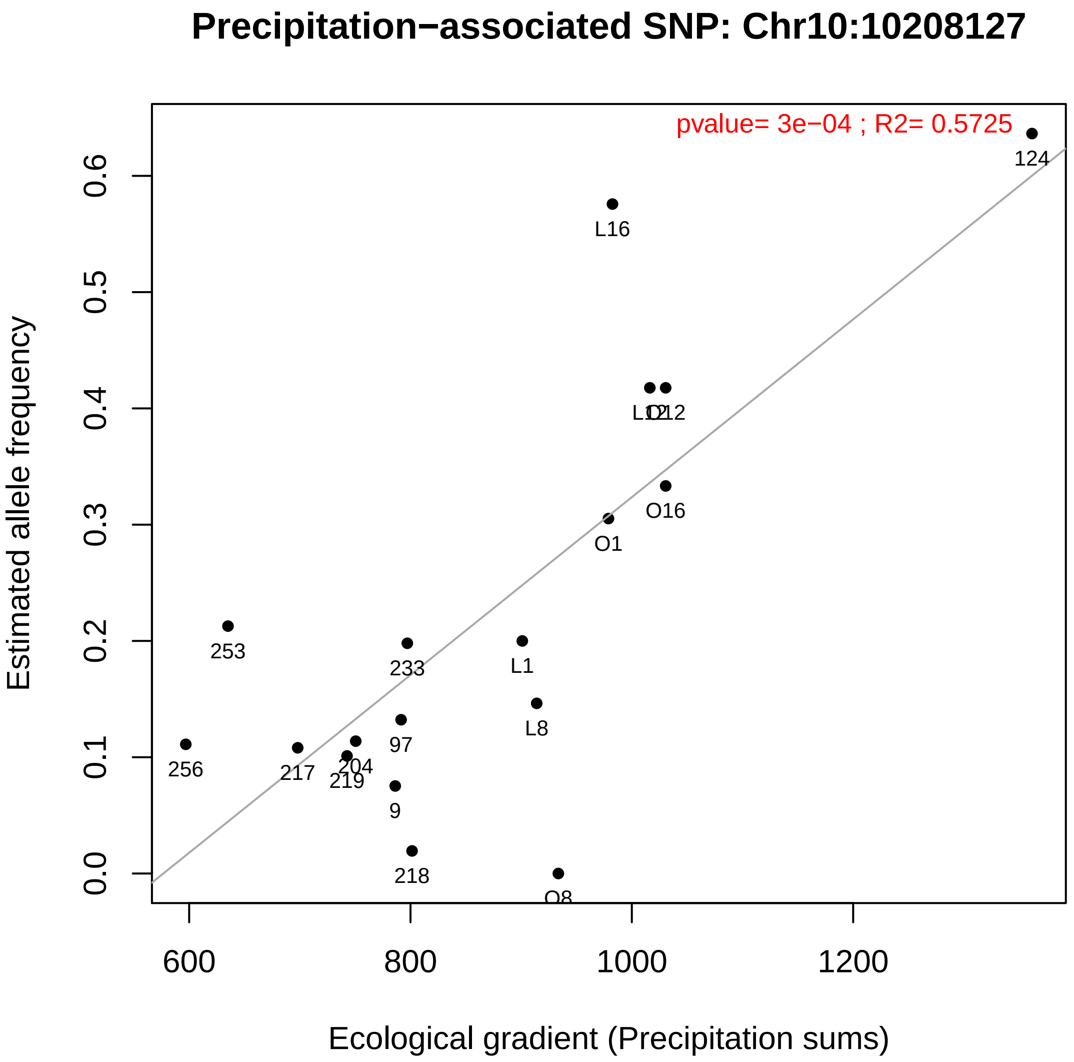

Precipitation−associated SNP: Chr10:10208127
pvalue= 3e−04 ; R2= 0.5725
124
L16
L12
O12
O16
O1
253
233
L1
L8
97
256
217
204
219
9
218
O8
0.0
0.1
0.2
0.3
0.4
0.5
0.6
600
800
1000
1200
Estimated allele frequency
Ecological gradient (Precipitation sums)

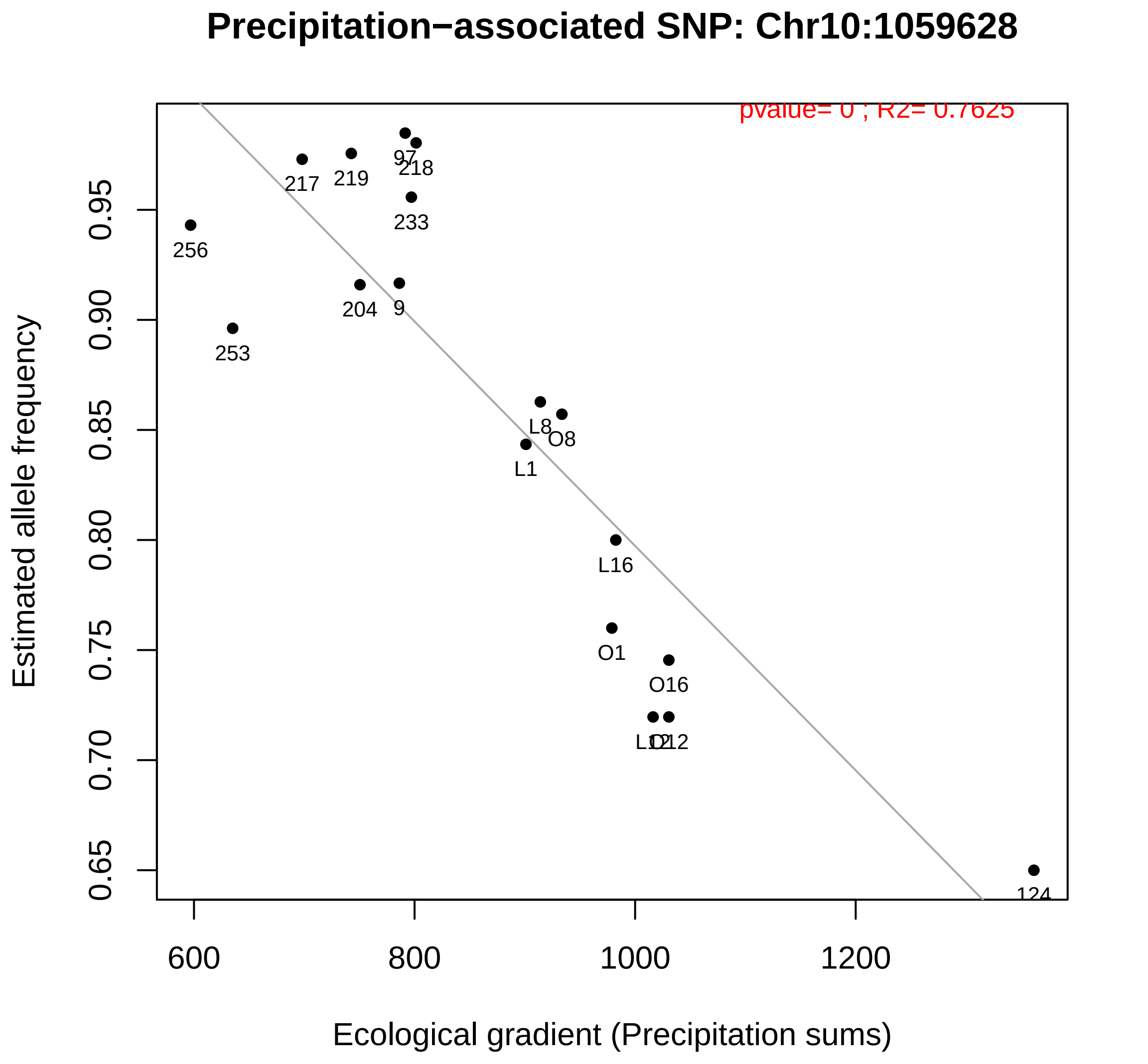
Precipitation−associated SNP: Chr10:1059628
pvalue= 0 ; R2= 0.7625
Estimated allele frequency
0.65
0.70
0.75
0.80
0.85
0.90
0.95
600
800
1000
1200
Ecological gradient (Precipitation sums)
97
218
217
219
233
256
204
9
253
L8
O8
L1
L16
O1
O16
L12
O12
124

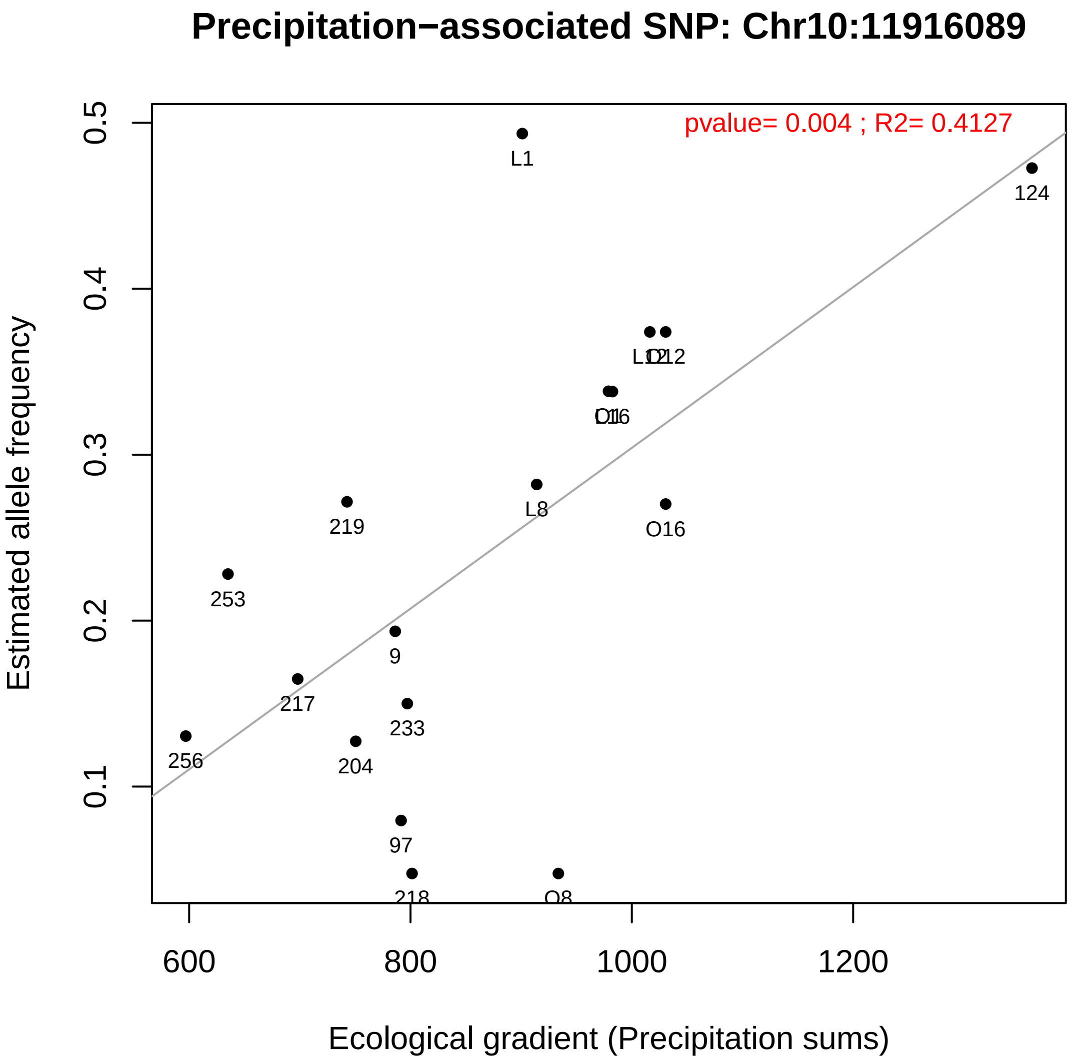
Precipitation−associated SNP: Chr10:11916089
pvalue= 0.004 ; R2= 0.4127
Estimated allele frequency
0.1
0.2
0.3
0.4
0.5
600
800
1000
1200
Ecological gradient (Precipitation sums)
L1
124
L12
O12
O11
L16
L8
O16
219
253
9
217
233
256
204
97
218
O8

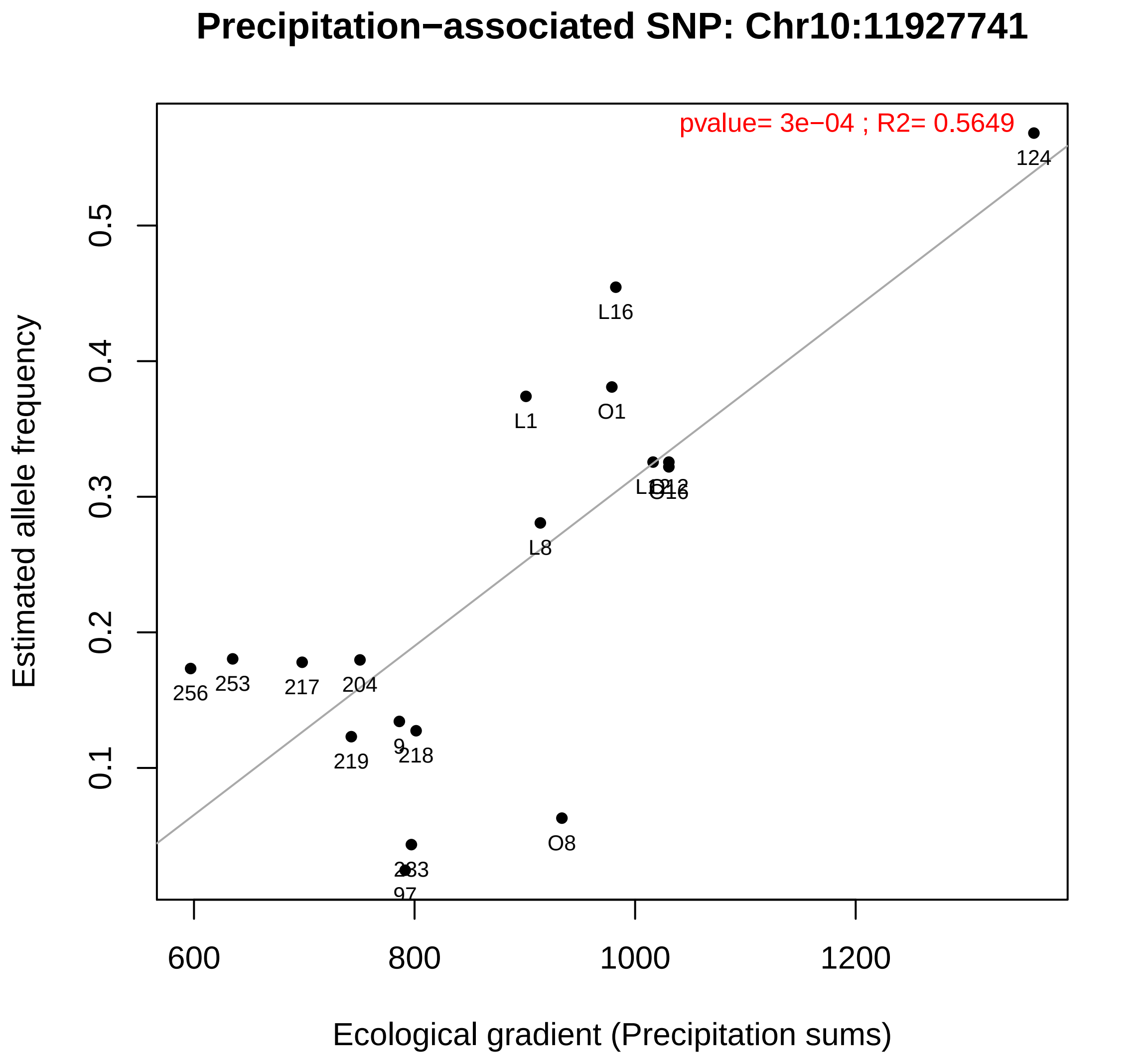

Precipitation−associated SNP: Chr10:11927741
pvalue= 3e−04 ; R2= 0.5649
124
L16
L1
O1
L8
256
253
217
204
219
9
218
O8
97
Estimated allele frequency
0.1
0.2
0.3
0.4
0.5
600
800
1000
1200
Ecological gradient (Precipitation sums)

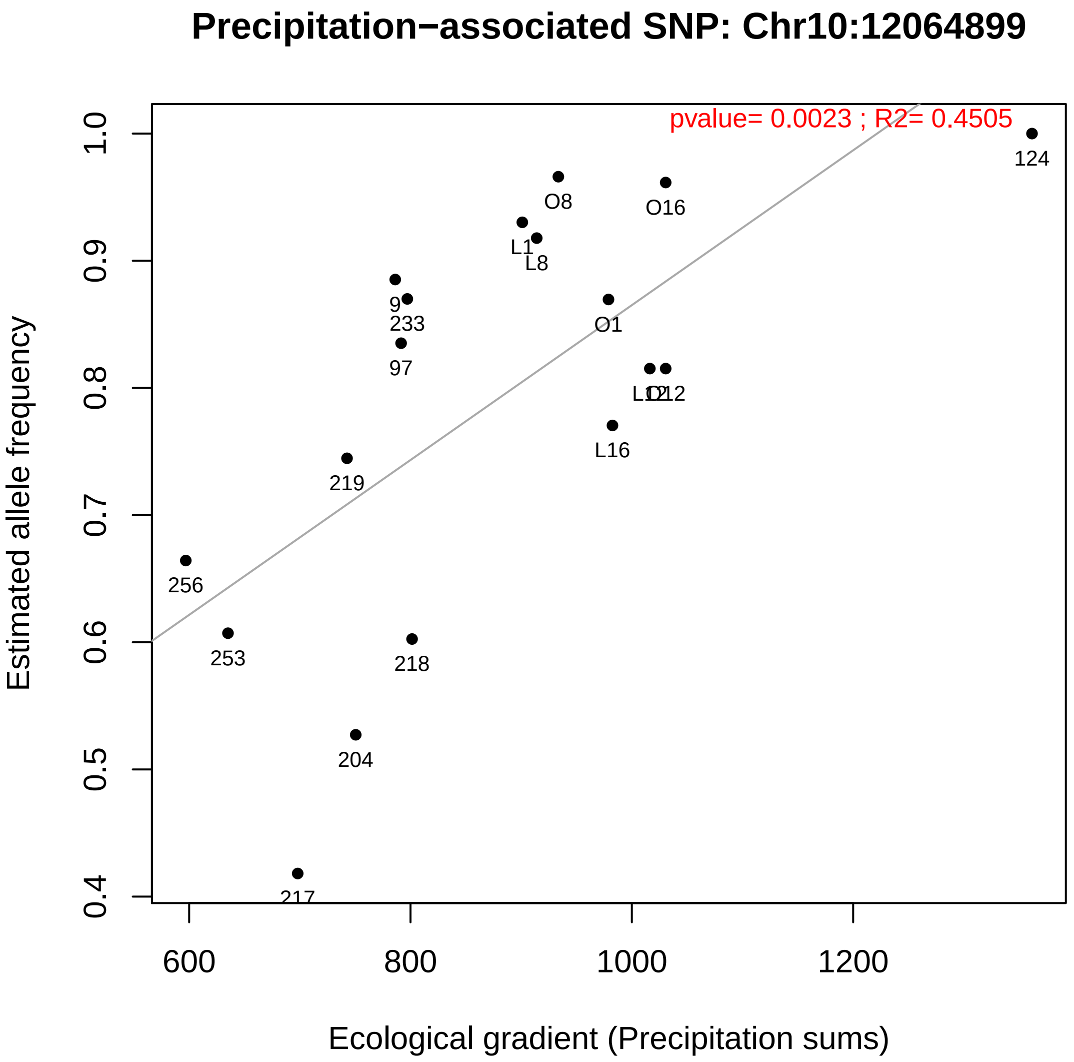
Precipitation−associated SNP: Chr10:12064899
pvalue= 0.0023 ; R2= 0.4505
Estimated allele frequency
Ecological gradient (Precipitation sums)
0.4
0.5
0.6
0.7
0.8
0.9
1.0
600
800
1000
1200
124
O8
O16
L1
L8
9
233
97
O1
L16
219
256
253
218
204
217

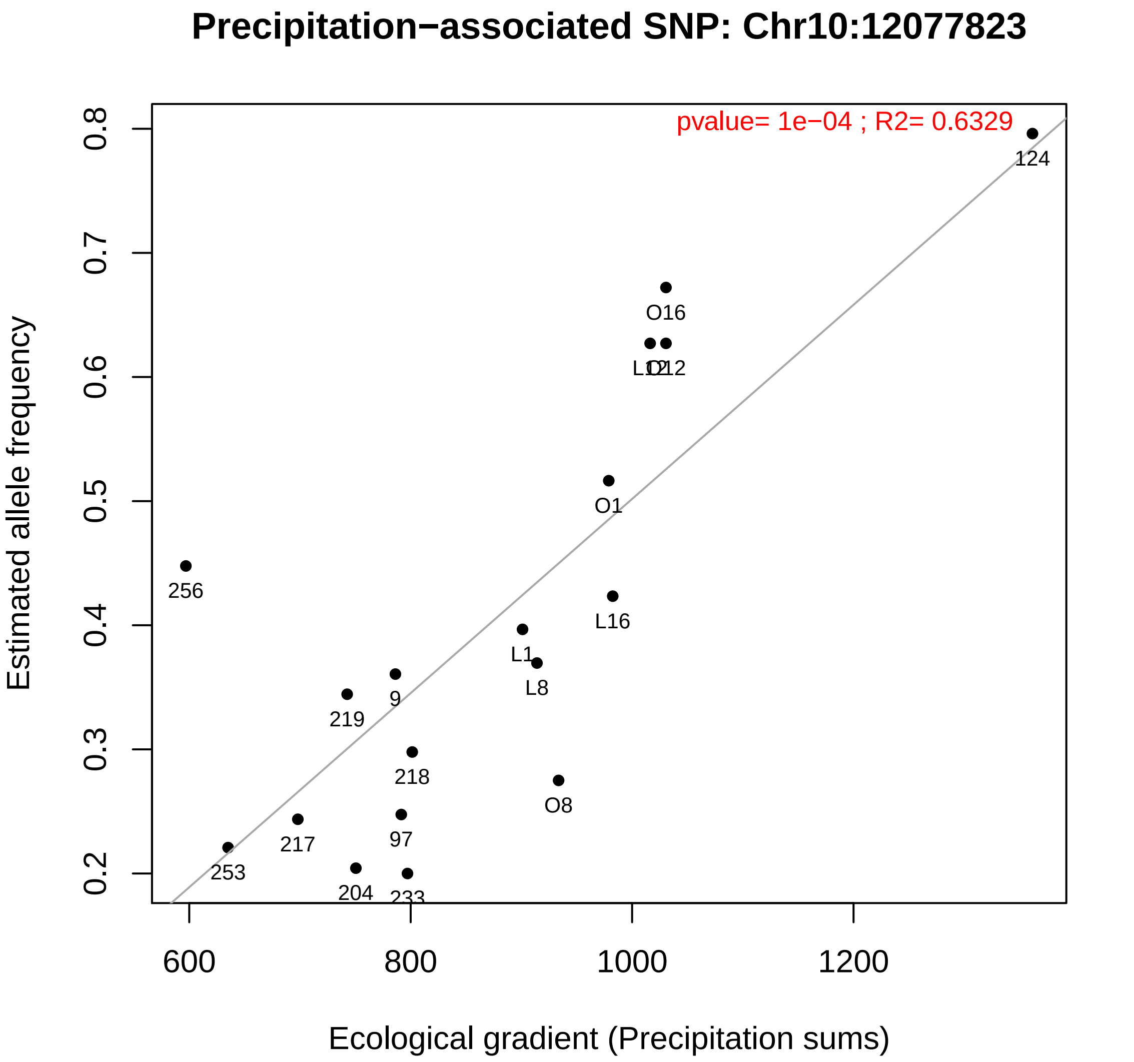
Precipitation−associated SNP: Chr10:12077823
pvalue= 1e−04 ; R2= 0.6329
Estimated allele frequency
Ecological gradient (Precipitation sums)
0.2
0.3
0.4
0.5
0.6
0.7
0.8
600
800
1000
1200
124
O16
O12
O1
256
L16
L1
L8
9
219
218
O8
217
97
253
204
233

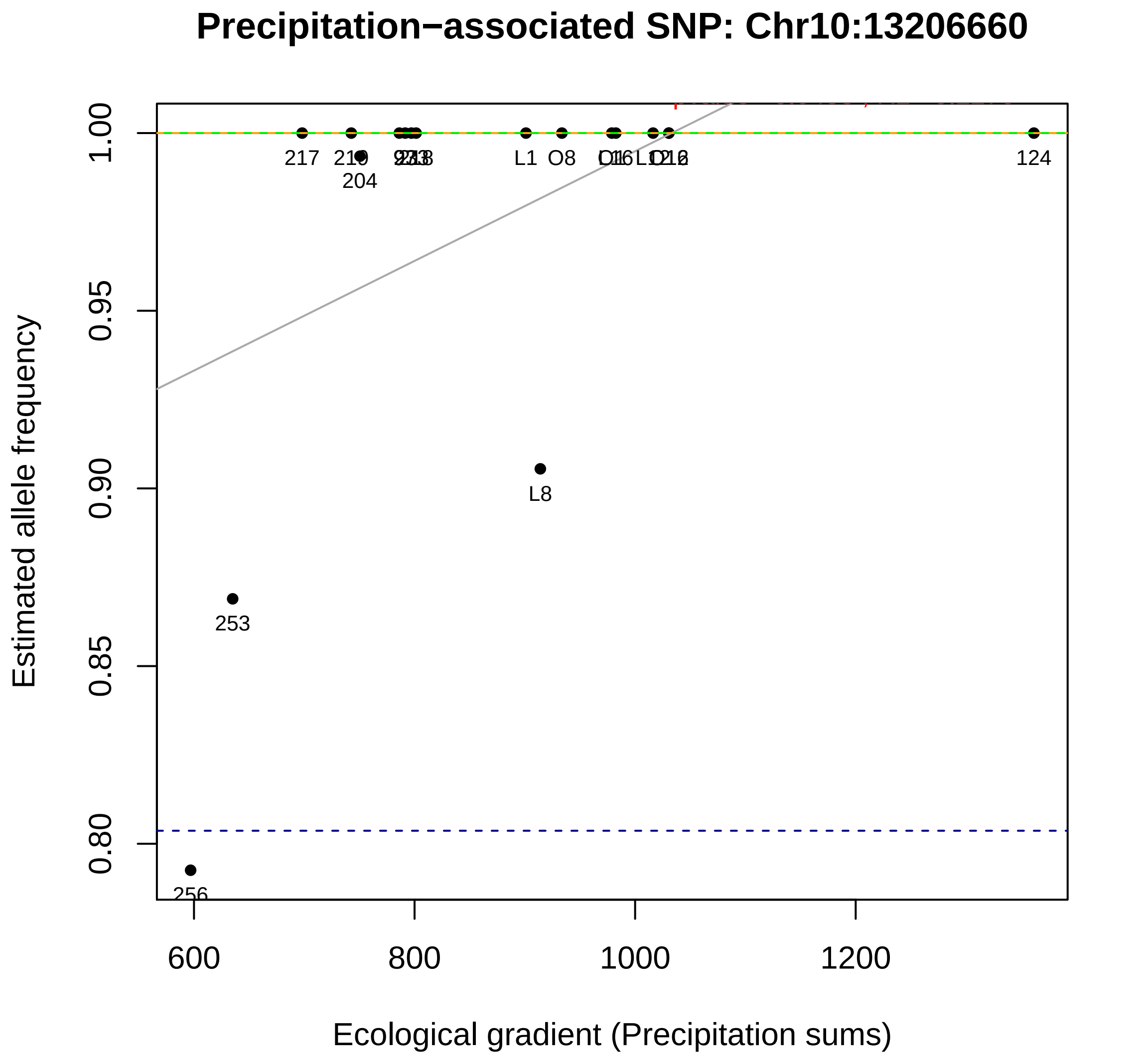

Precipitation−associated SNP: Chr10:13206660
Estimated allele frequency
Ecological gradient (Precipitation sums)
0.80
0.85
0.90
0.95
1.00
600
800
1000
1200
217
204
L1
O8
124
L8
253
256

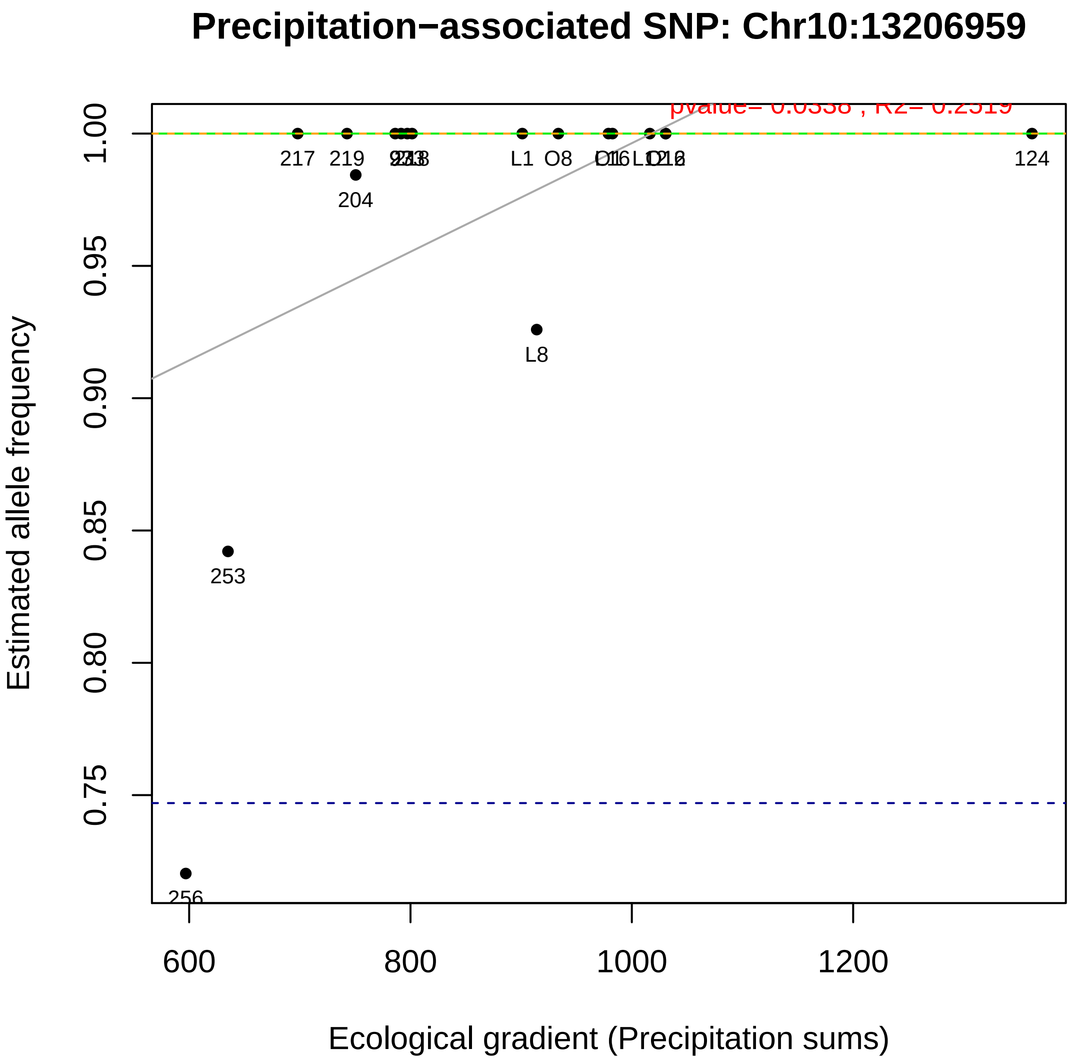
Precipitation−associated SNP: Chr10:13206959
pvalue= 0.0338 , R2= 0.2519
Estimated allele frequency
1.00
0.95
0.90
0.85
0.80
0.75
217
219
204
L1
O8
124
L8
253
256
600
800
1000
1200
Ecological gradient (Precipitation sums)

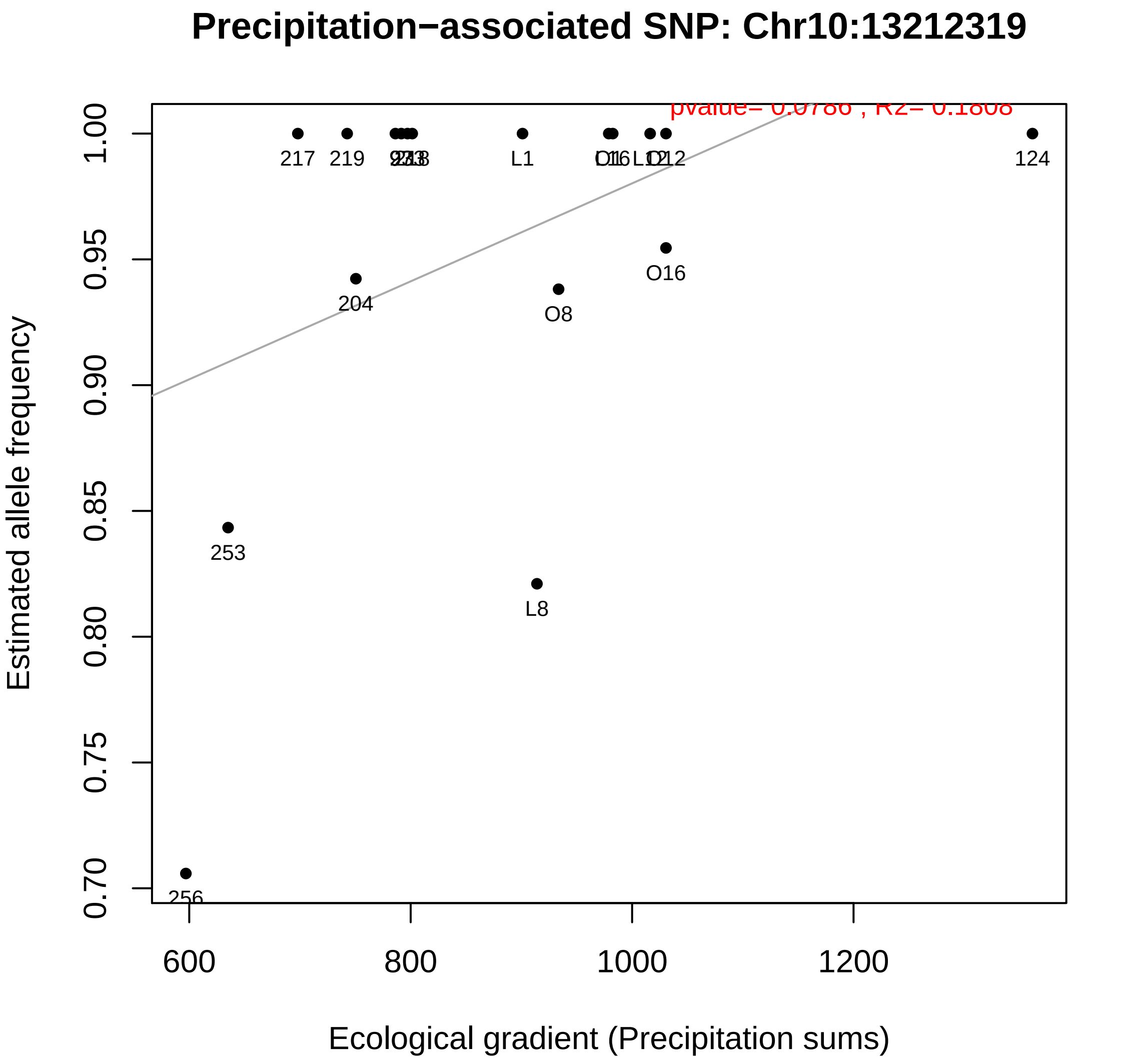
Precipitation−associated SNP: Chr10:13212319
pvalue= 0.0786 , R2= 0.1808
Estimated allele frequency
Ecological gradient (Precipitation sums)
1.00
0.95
0.90
0.85
0.80
0.75
0.70
600
800
1000
1200
217
219
L1
124
204
O8
O16
253
L8
256

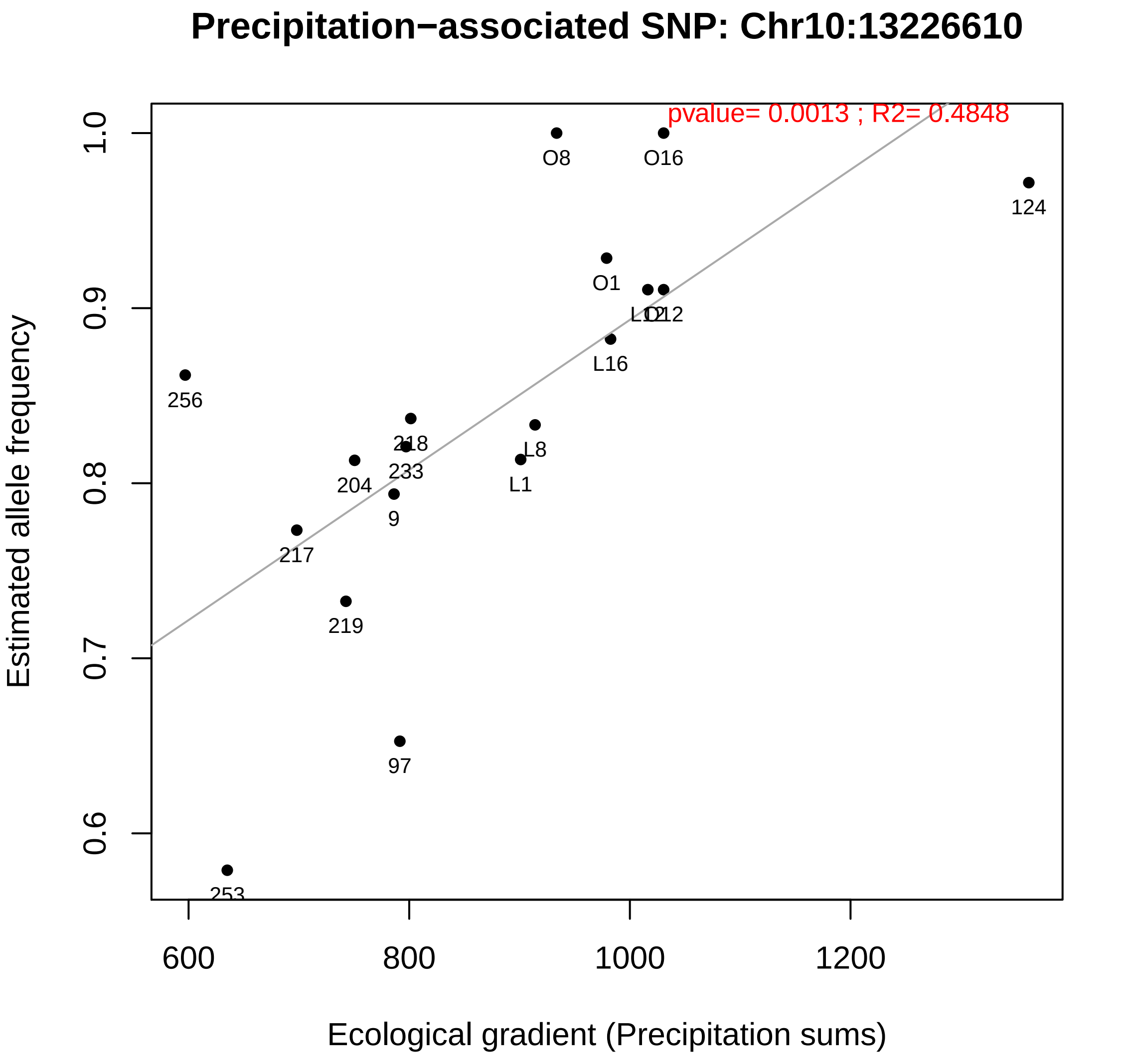

Precipitation−associated SNP: Chr10:13226610
pvalue= 0.0013 ; R2= 0.4848
Estimated allele frequency
Ecological gradient (Precipitation sums)
0.6
0.7
0.8
0.9
1.0
600
800
1000
1200
O8
O16
124
O1
L12
O12
L16
256
218
L8
233
L1
204
9
217
219
97
253

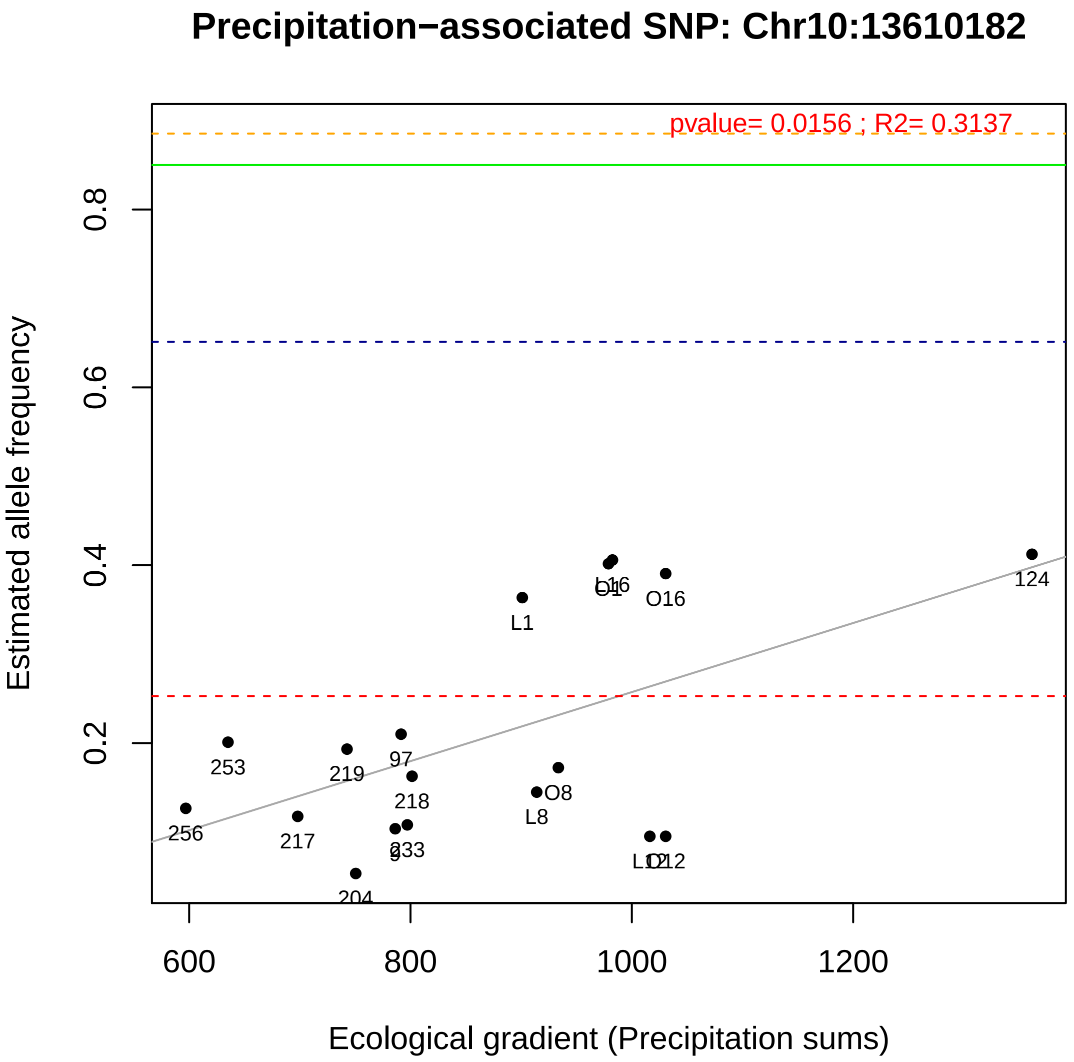
Precipitation−associated SNP: Chr10:13610182
pvalue= 0.0156 ; R2= 0.3137
Estimated allele frequency
0.2
0.4
0.6
0.8
600
800
1000
1200
Ecological gradient (Precipitation sums)
124
L16
O1
O16
L1
253
219
97
218
256
217
233
204
O8
L8
L12
O12

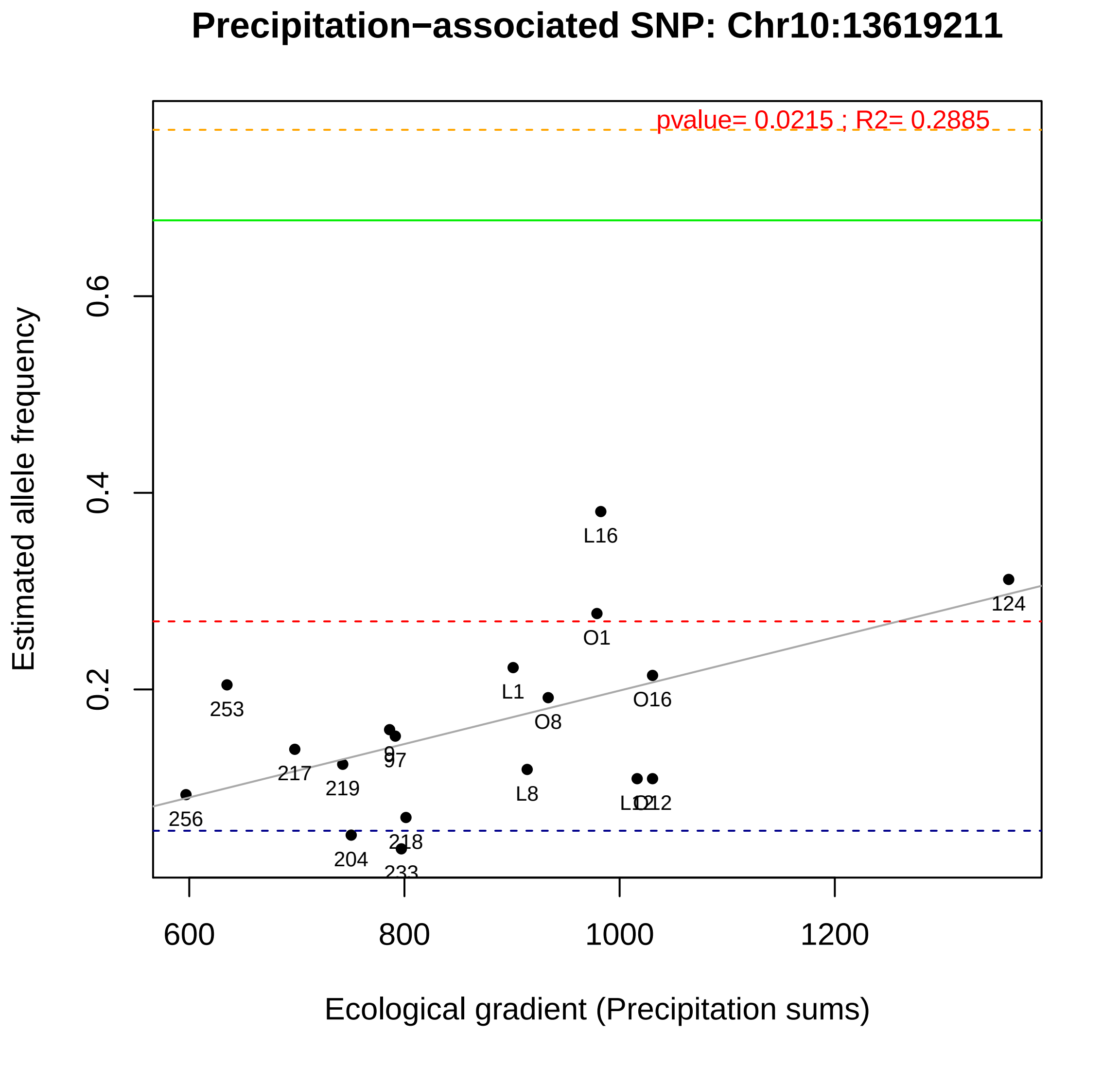
Precipitation−associated SNP: Chr10:13619211
pvalue= 0.0215 ; R2= 0.2885
Estimated allele frequency
0.2
0.4
0.6
600
800
1000
1200
Ecological gradient (Precipitation sums)
L16
124
O1
L1
O16
253
O8
9
97
217
219
L8
L12
O12
256
218
204
233

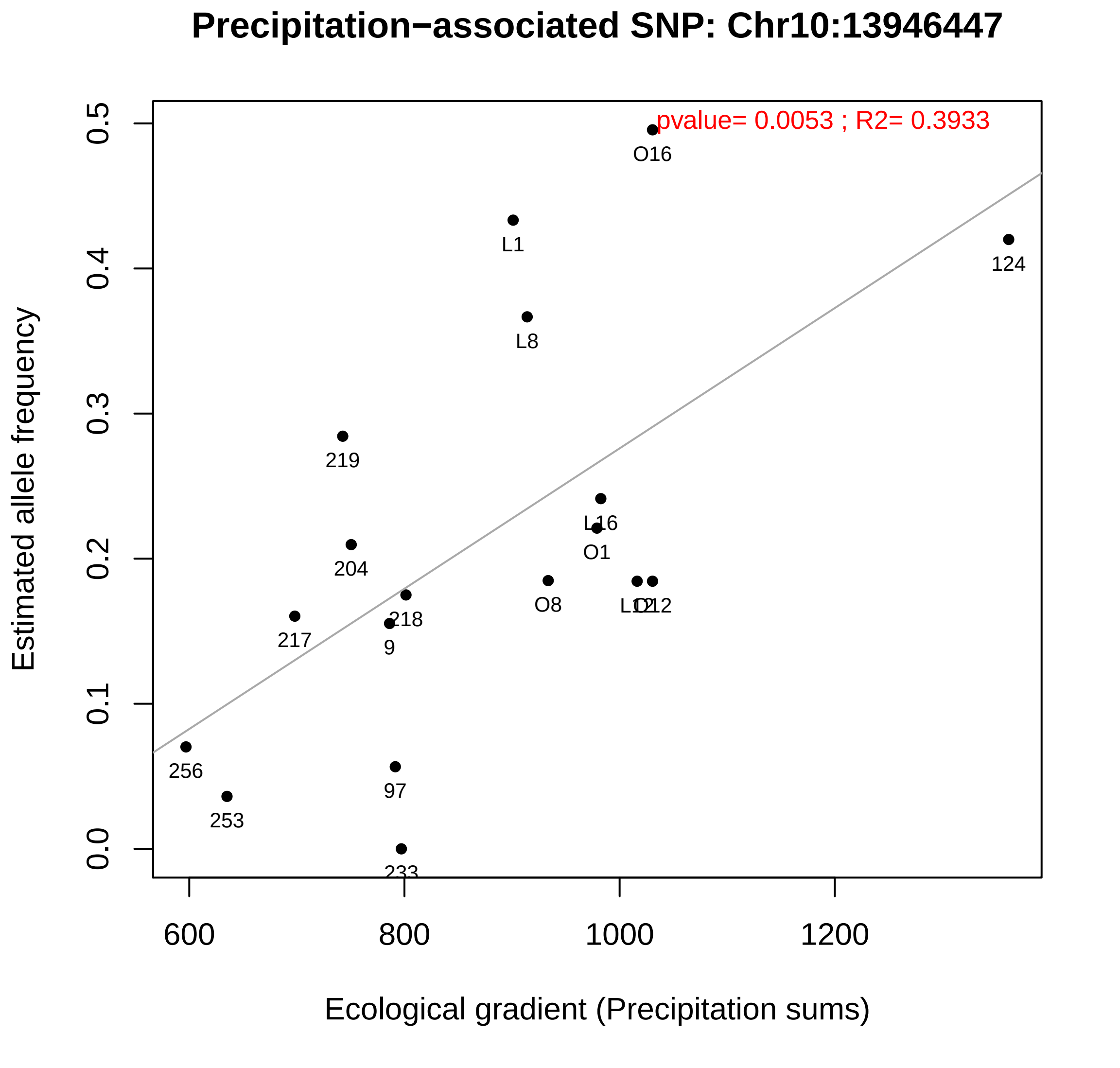

Precipitation−associated SNP: Chr10:13946447
pvalue= 0.0053 ; R2= 0.3933
Estimated allele frequency
0.0
0.1
0.2
0.3
0.4
0.5
600
800
1000
1200
Ecological gradient (Precipitation sums)
O16
L1
124
L8
219
L16
O1
204
O8
L12
O12
218
217
9
256
97
253
233

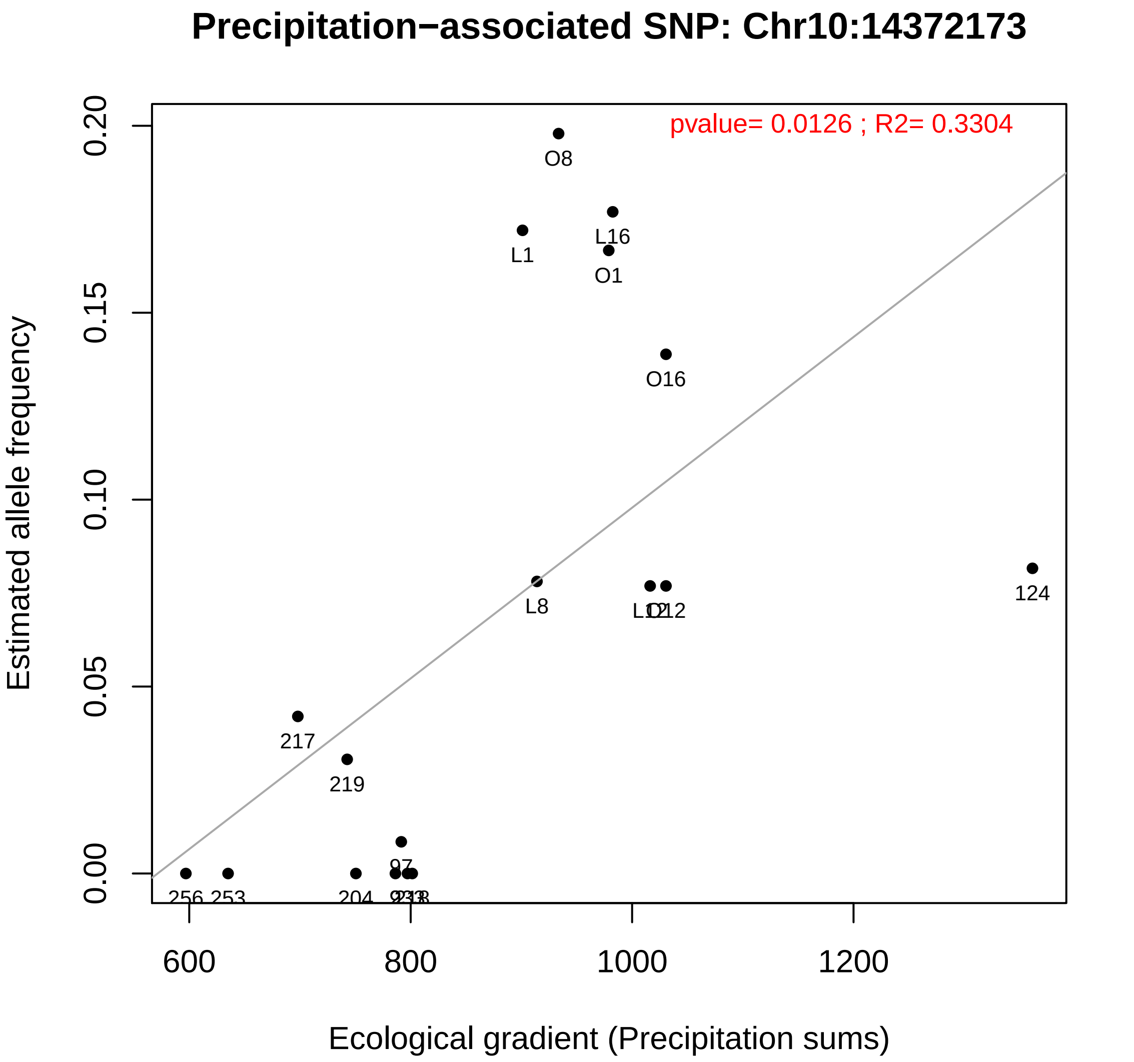
Precipitation−associated SNP: Chr10:14372173
pvalue= 0.0126 ; R2= 0.3304
Estimated allele frequency
Ecological gradient (Precipitation sums)
0.00
0.05
0.10
0.15
0.20
600
800
1000
1200
O8
L16
L1
O1
O16
L8
124
217
219
97
256
253
204

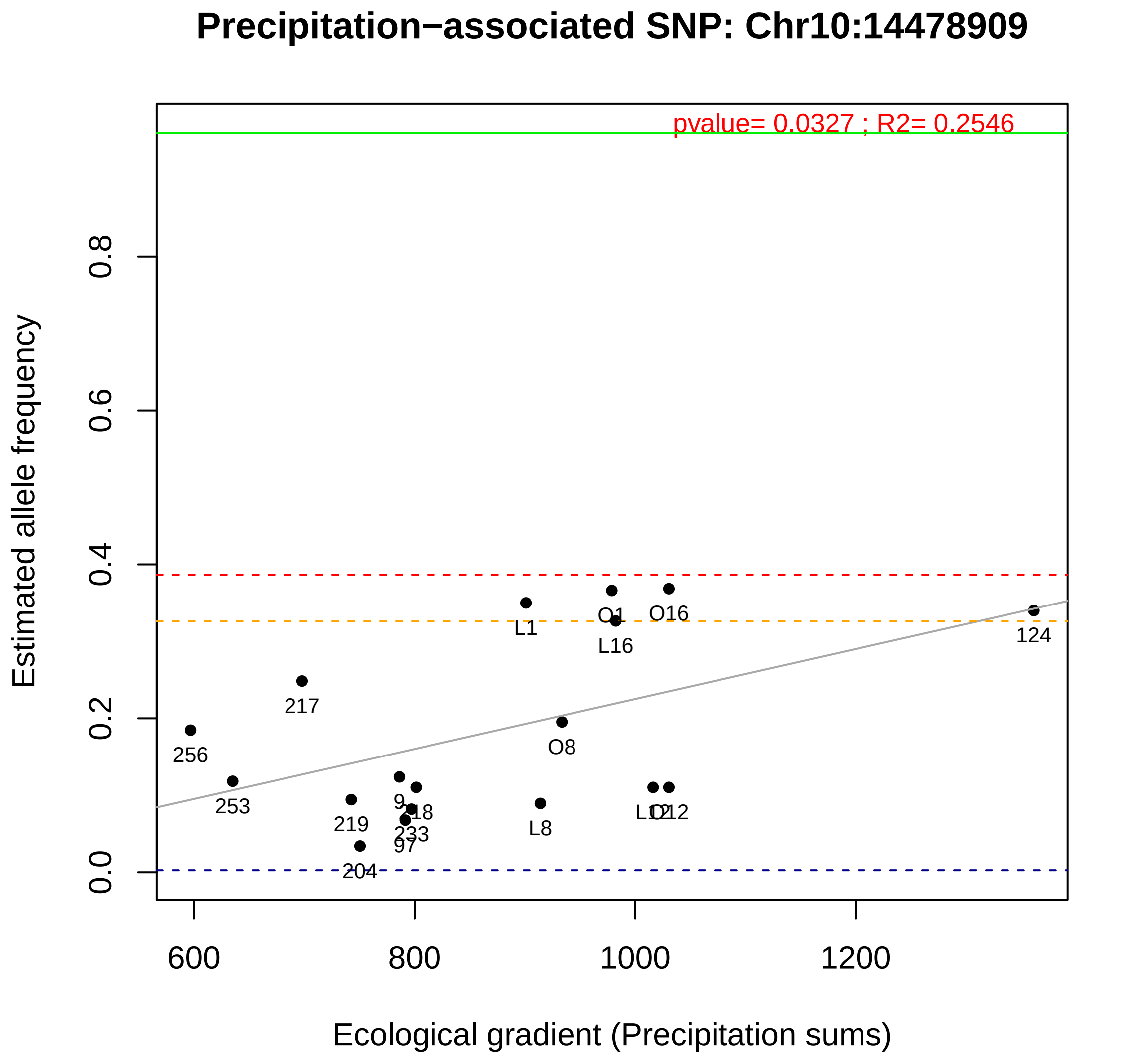

Precipitation−associated SNP: Chr10:14478909
pvalue= 0.0327 ; R2= 0.2546
Estimated allele frequency
0.0
0.2
0.4
0.6
0.8
600
800
1000
1200
Ecological gradient (Precipitation sums)
256
253
217
219
204
9
218
233
97
L1
O8
L8
O1
L16
O16
L12
O12
124

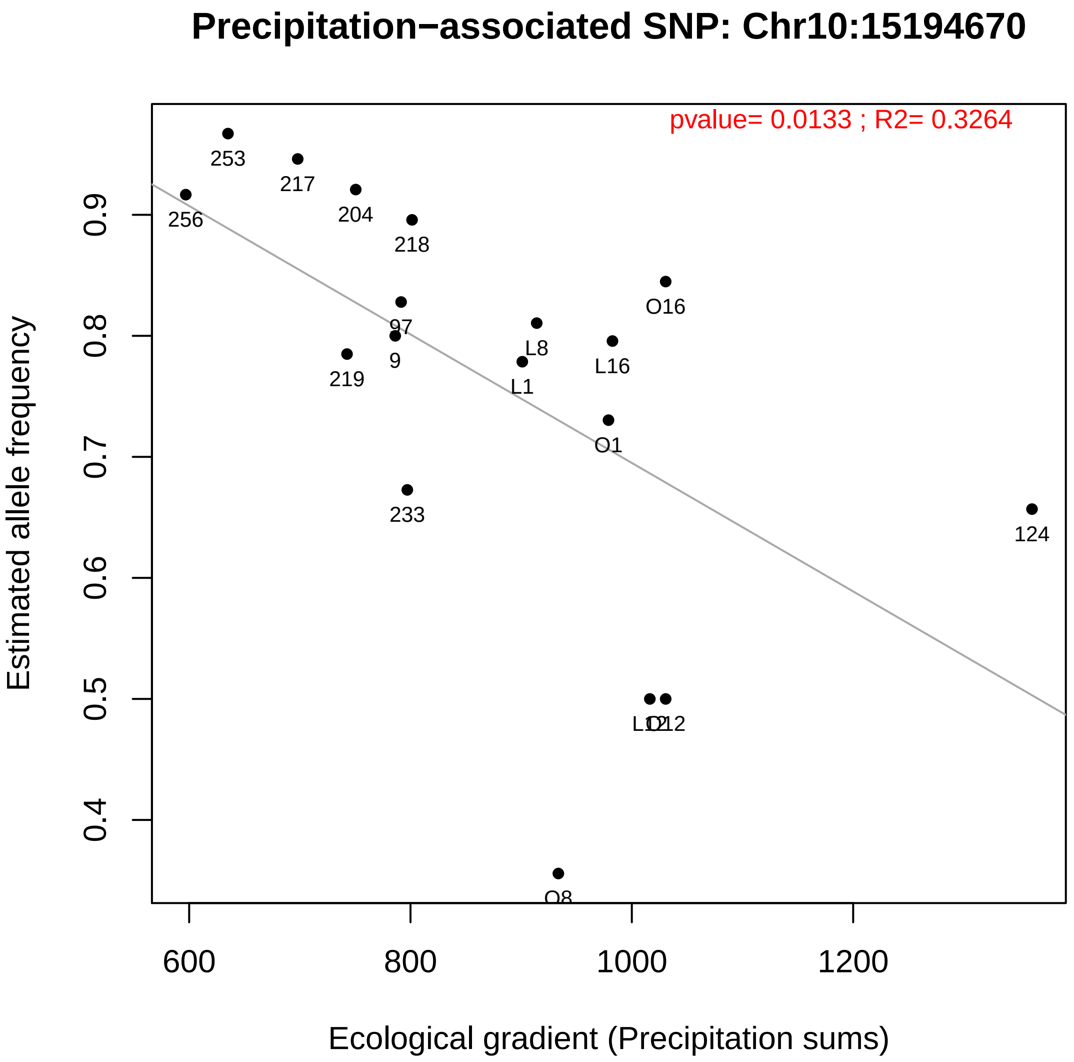

Precipitation−associated SNP: Chr10:15194670
pvalue= 0.0133 ; R2= 0.3264
Estimated allele frequency
Ecological gradient (Precipitation sums)
0.4
0.5
0.6
0.7
0.8
0.9
600
800
1000
1200
253
217
256
204
218
O16
97
L8
9
L16
219
L1
O1
233
124
L12
O12
O8

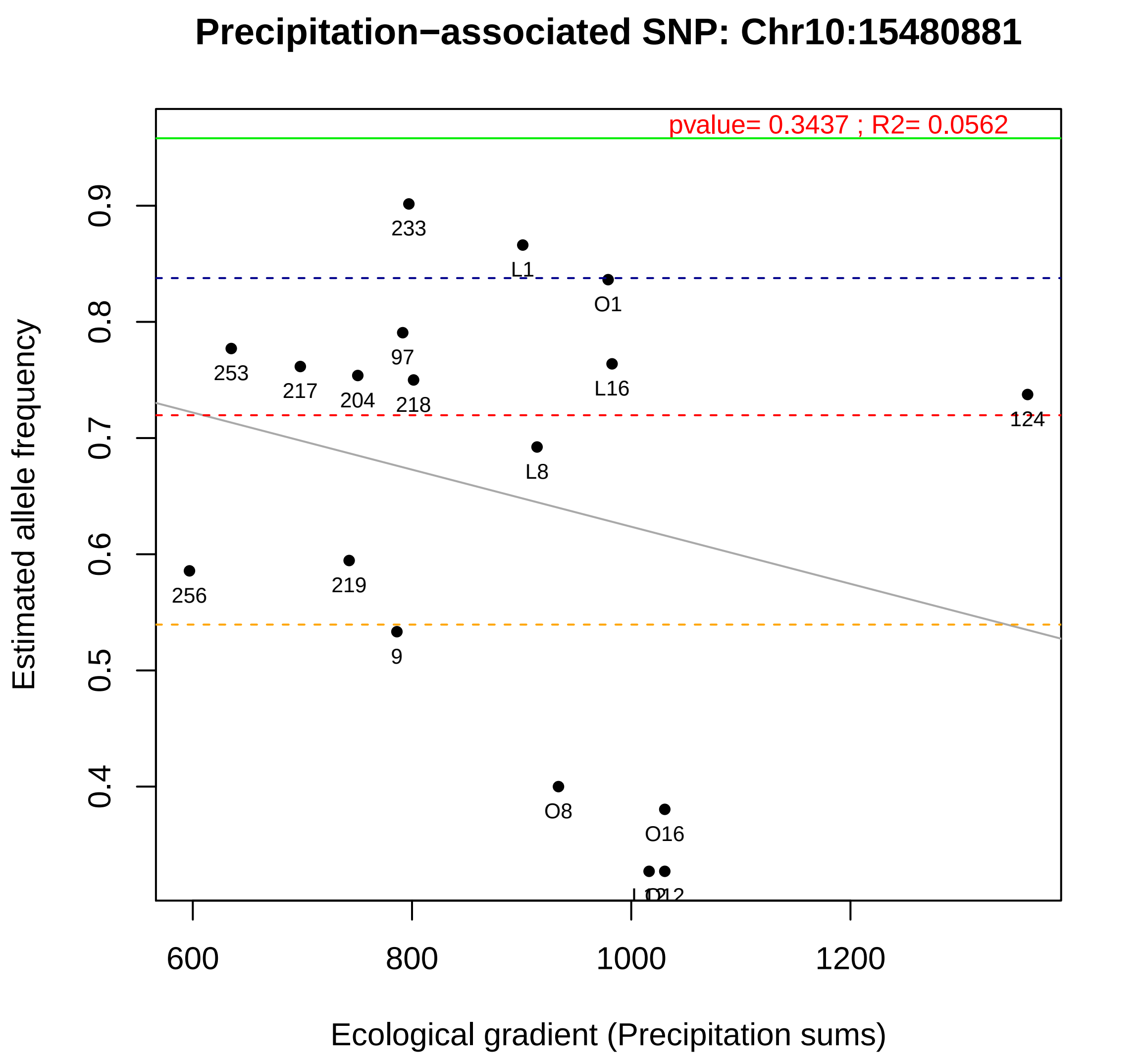

Precipitation−associated SNP: Chr10:15480881
pvalue= 0.3437 ; R2= 0.0562
Estimated allele frequency
0.4
0.5
0.6
0.7
0.8
0.9
600
800
1000
1200
Ecological gradient (Precipitation sums)
233
L1
O1
253
217
204
97
218
L16
124
L8
256
219
9
O8
O16

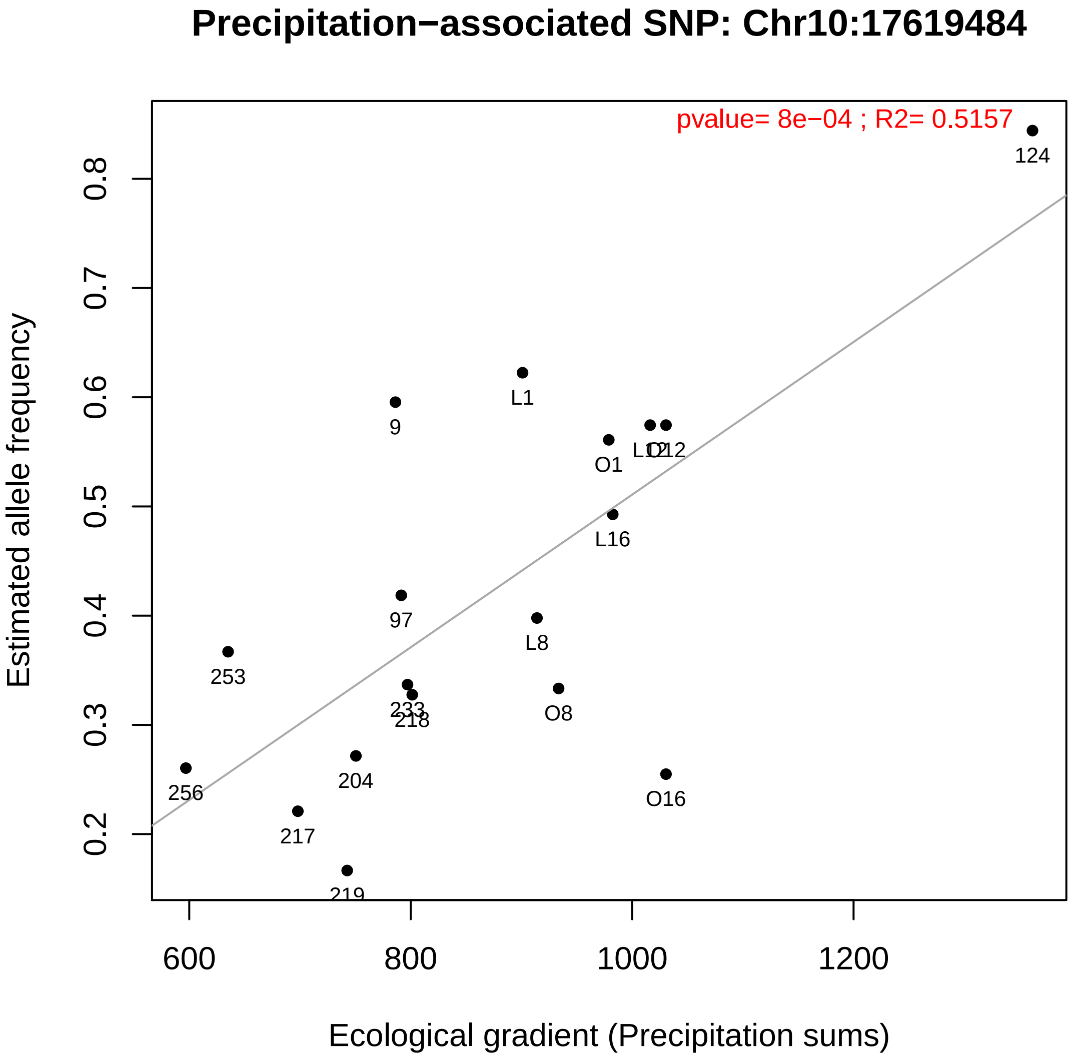
Precipitation−associated SNP: Chr10:17619484
pvalue= 8e−04 ; R2= 0.5157
124
L1
9
L12
O12
O1
L16
97
L8
253
233
218
O8
204
256
O16
217
219
0.2
0.3
0.4
0.5
0.6
0.7
0.8
600
800
1000
1200
Estimated allele frequency
Ecological gradient (Precipitation sums)

#### Precipitation−associated SNP: Chr10:17665851

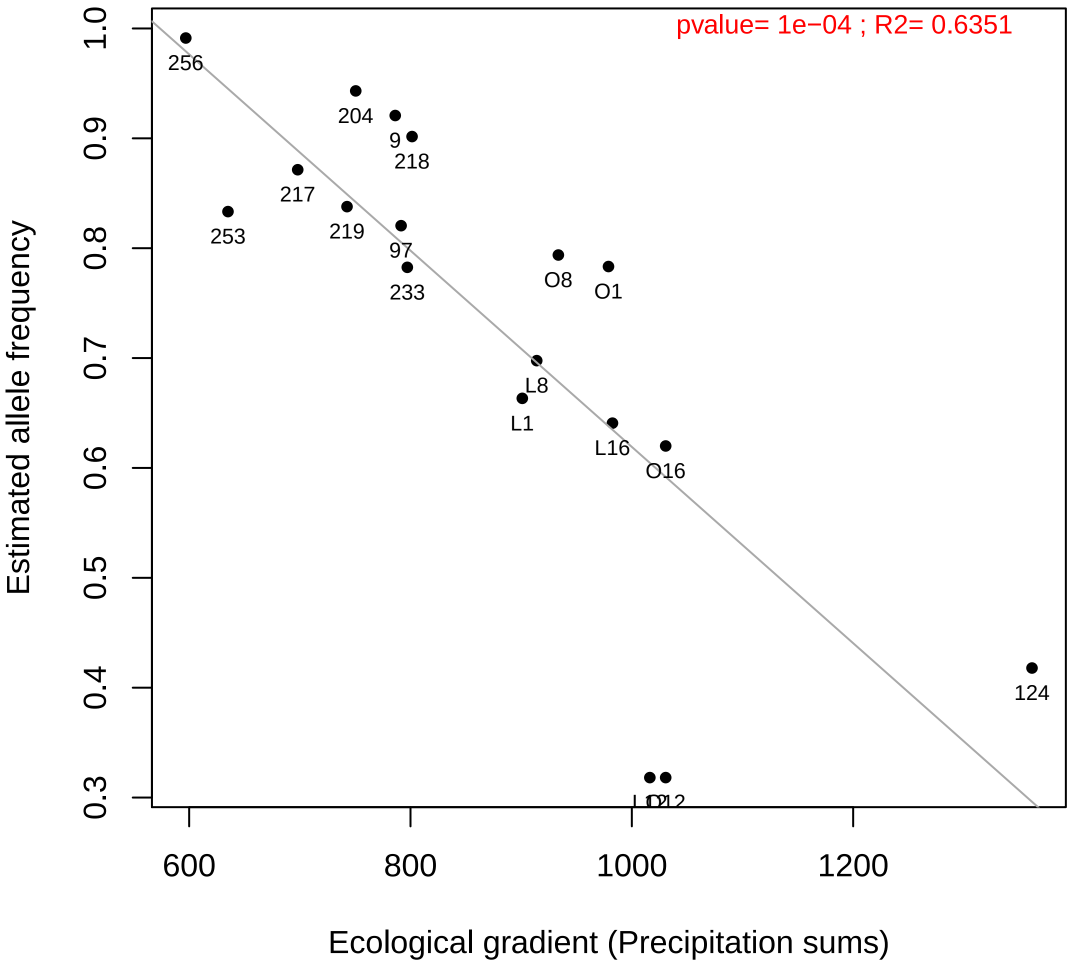

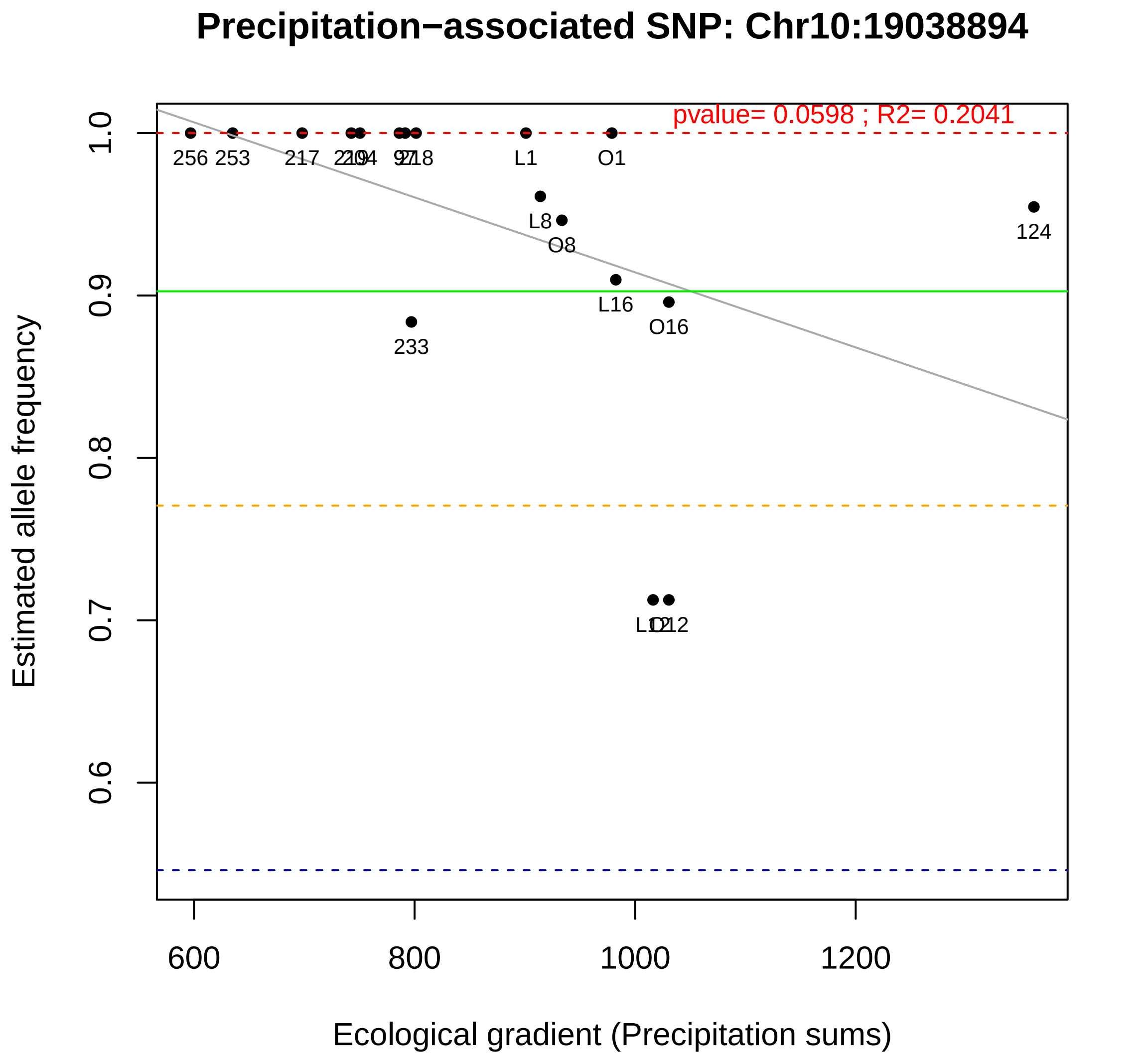
Precipitation−associated SNP: Chr10:19038894
pvalue= 0.0598 ; R2= 0.2041
Estimated allele frequency
1.0
0.9
0.8
0.7
0.6
600
800
1000
1200
Ecological gradient (Precipitation sums)
256
253
217
L1
O1
L8
O8
124
L16
O16
233

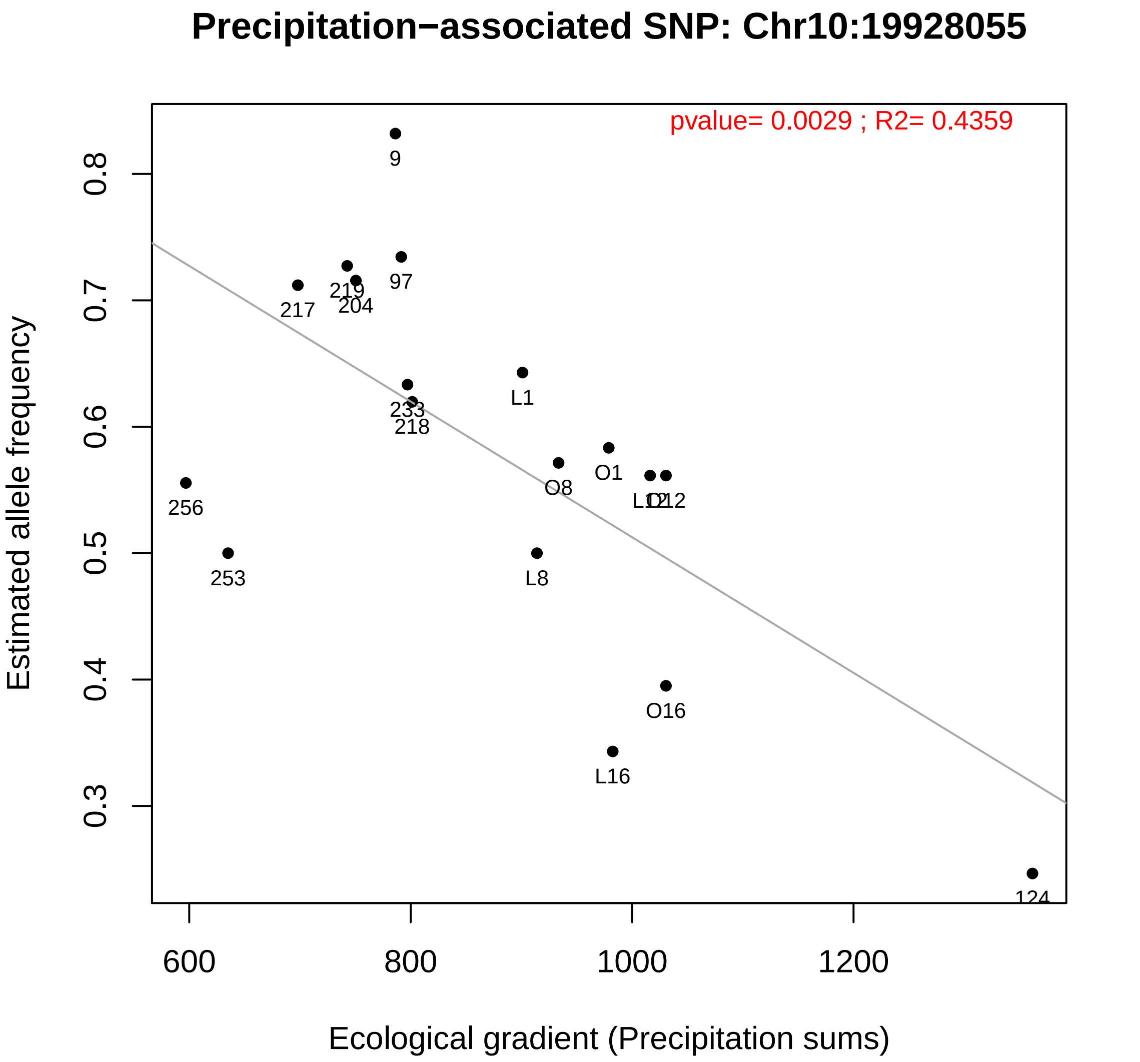

Precipitation−associated SNP: Chr10:19928055
pvalue= 0.0029 ; R2= 0.4359
Estimated allele frequency
Ecological gradient (Precipitation sums)
0.3
0.4
0.5
0.6
0.7
0.8
600
800
1000
1200
9
97
219
204
217
233
218
L1
O1
O8
L12
O12
256
253
L8
O16
L16
124

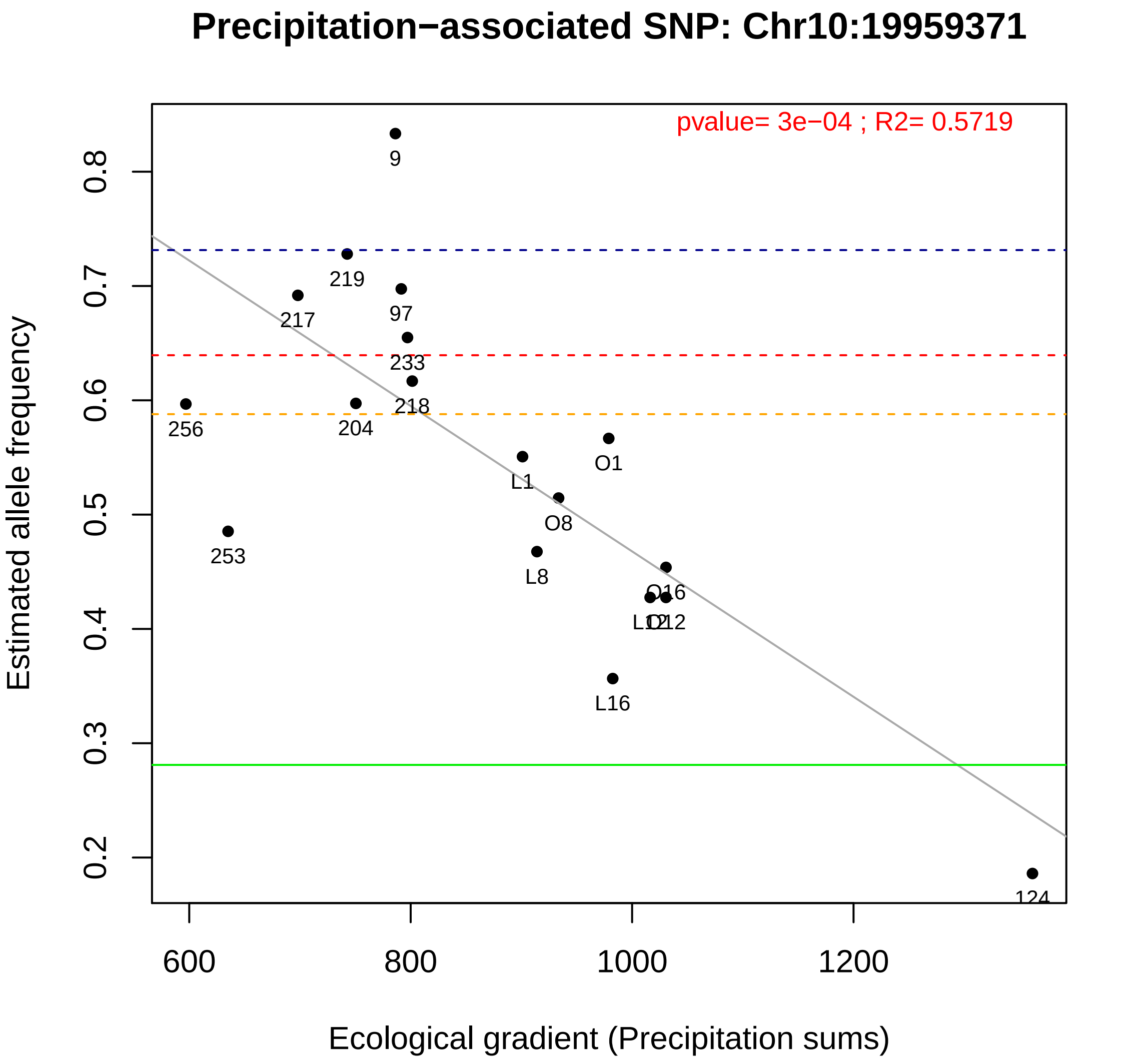

Precipitation−associated SNP: Chr10:19959371
pvalue= 3e−04 ; R2= 0.5719
Estimated allele frequency
0.2
0.3
0.4
0.5
0.6
0.7
0.8
600
800
1000
1200
Ecological gradient (Precipitation sums)
9
219
217
97
233
218
256
204
O1
L1
O8
253
L8
O16
L12
O2
L16
124

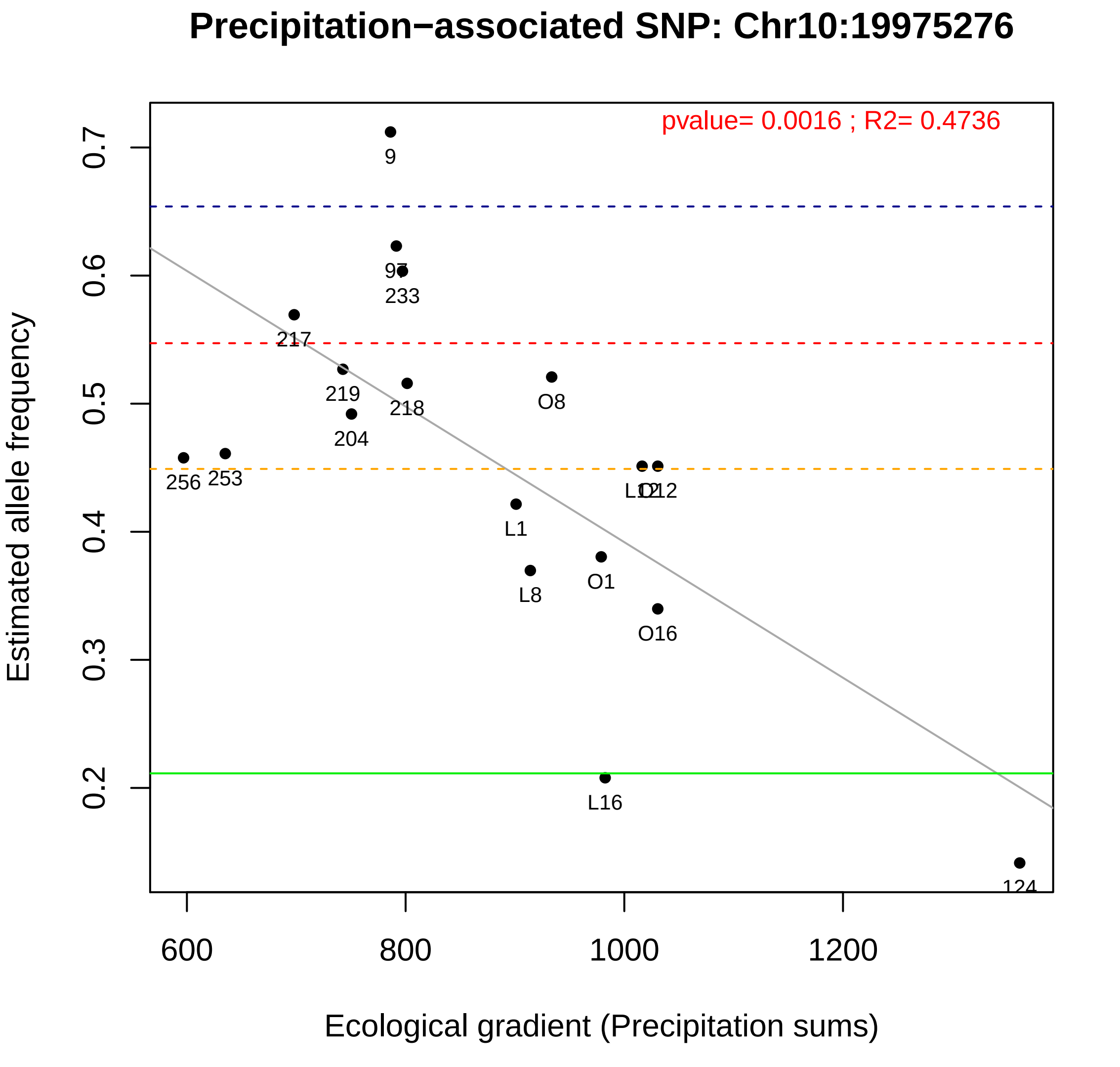

Precipitation−associated SNP: Chr10:19975276
pvalue= 0.0016 ; R2= 0.4736
Estimated allele frequency
0.2
0.3
0.4
0.5
0.6
0.7
600
800
1000
1200
Ecological gradient (Precipitation sums)
9
97
233
217
219
218
204
O8
256
253
L1
L8
O1
O16
L16
124

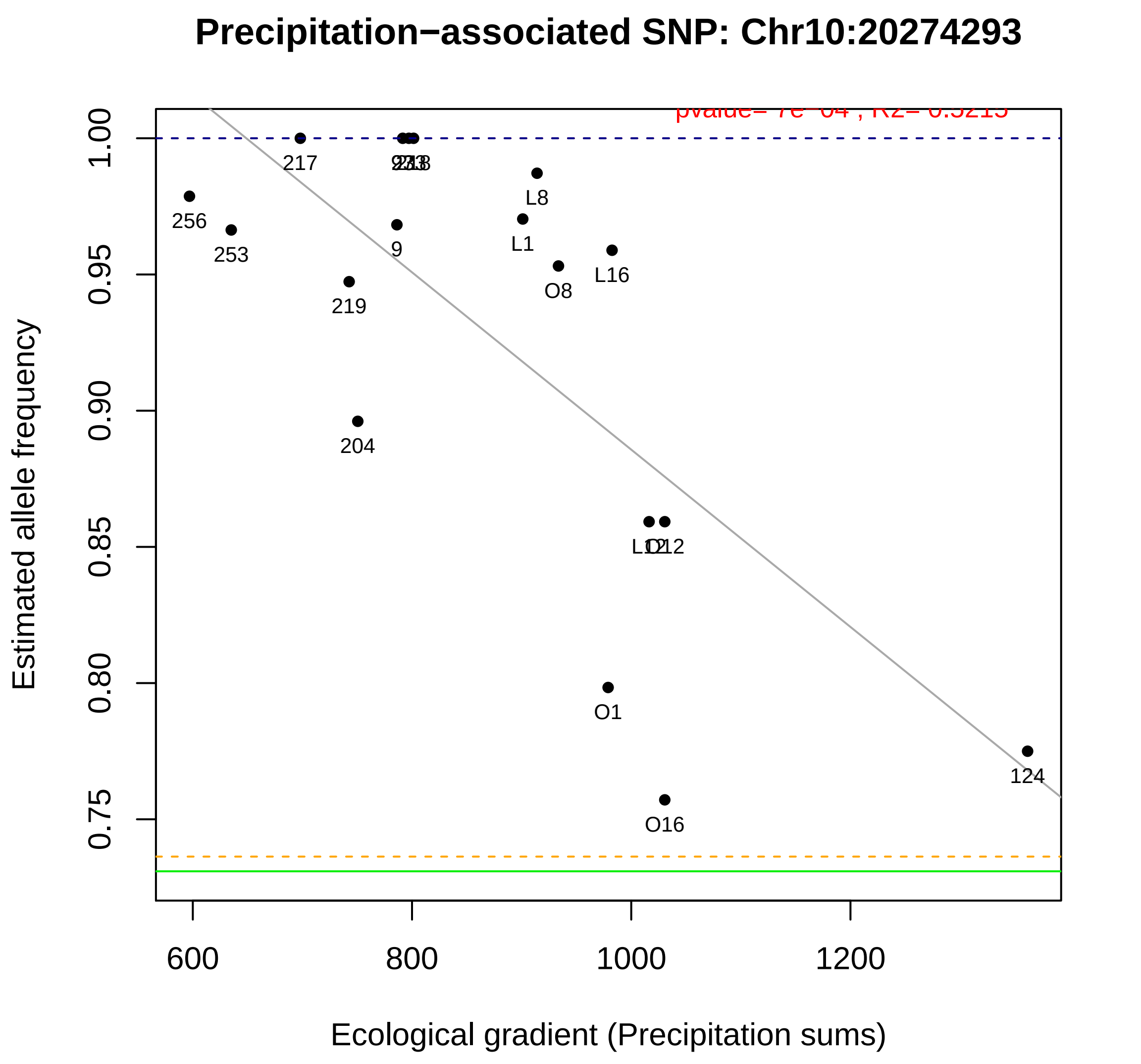
Precipitation−associated SNP: Chr10:20274293
pvalue= 7e−04 ; R2= 0.5215
Estimated allele frequency
1.00
0.95
0.90
0.85
0.80
0.75
600
800
1000
1200
Ecological gradient (Precipitation sums)
217
256
253
219
204
9
L8
L1
O8
L16
O1
O16
124

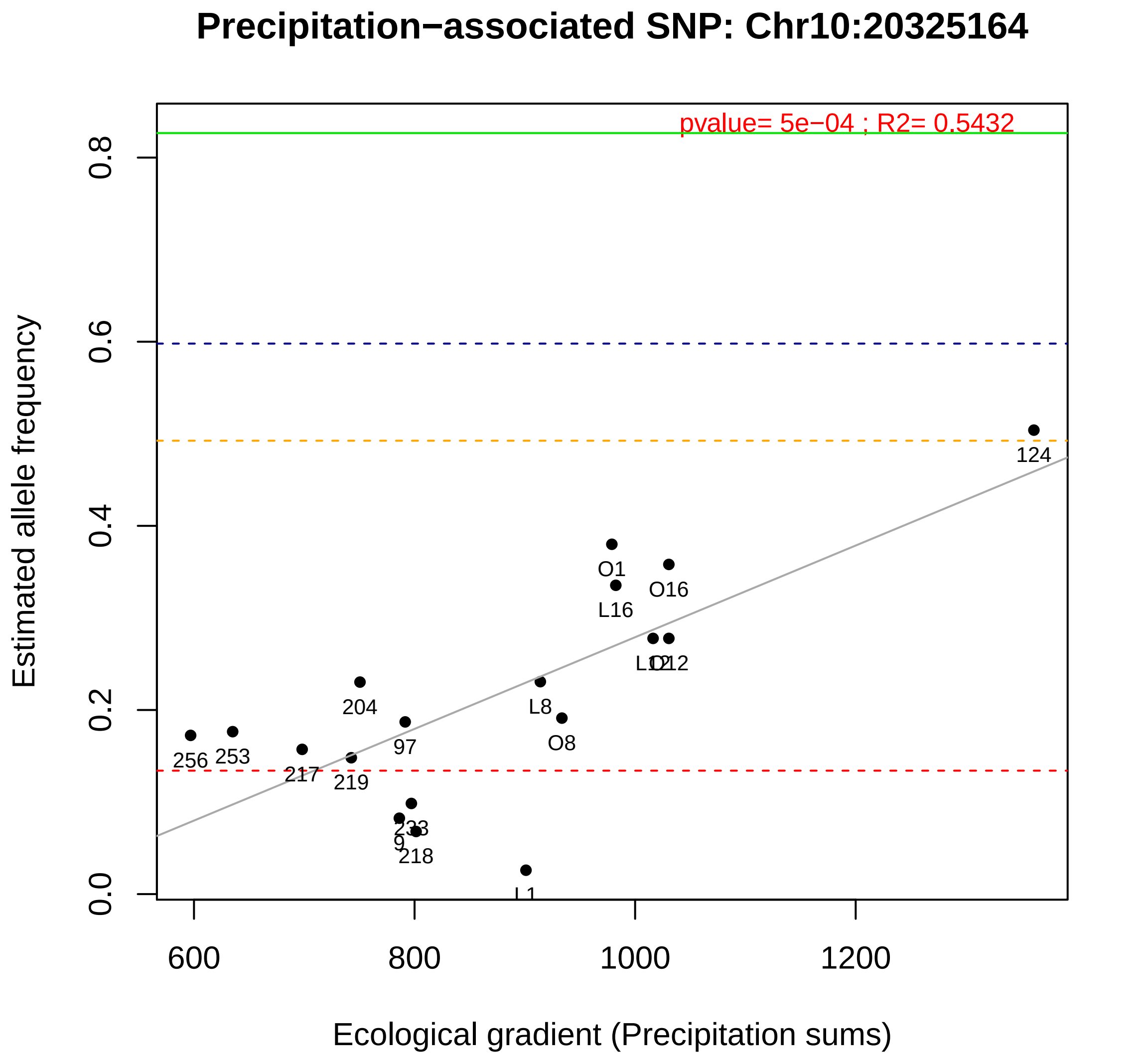

Precipitation−associated SNP: Chr10:20325164
pvalue= 5e−04 ; R2= 0.5432
Estimated allele frequency
Ecological gradient (Precipitation sums)
0.0
0.2
0.4
0.6
0.8
600
800
1000
1200
124
O1
O16
L16
L12
O12
204
L8
O8
97
256
253
217
219
233
9
218
L1

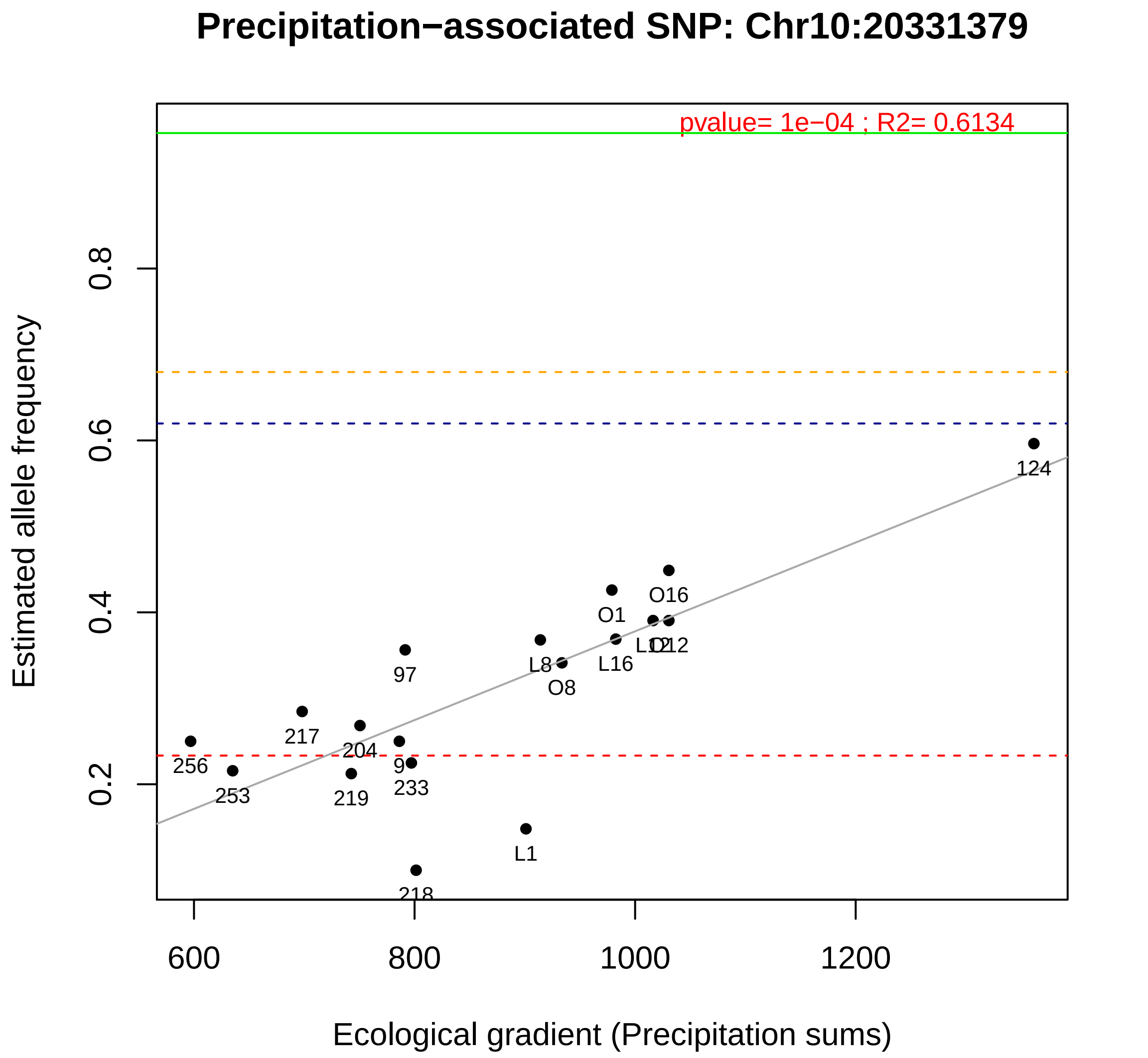
Precipitation−associated SNP: Chr10:20331379
pvalue= 1e−04 ; R2= 0.6134
Estimated allele frequency
0.2
0.4
0.6
0.8
600
800
1000
1200
Ecological gradient (Precipitation sums)
124
O16
O1
L16
L8
O8
97
217
204
256
253
219
233
L1
218

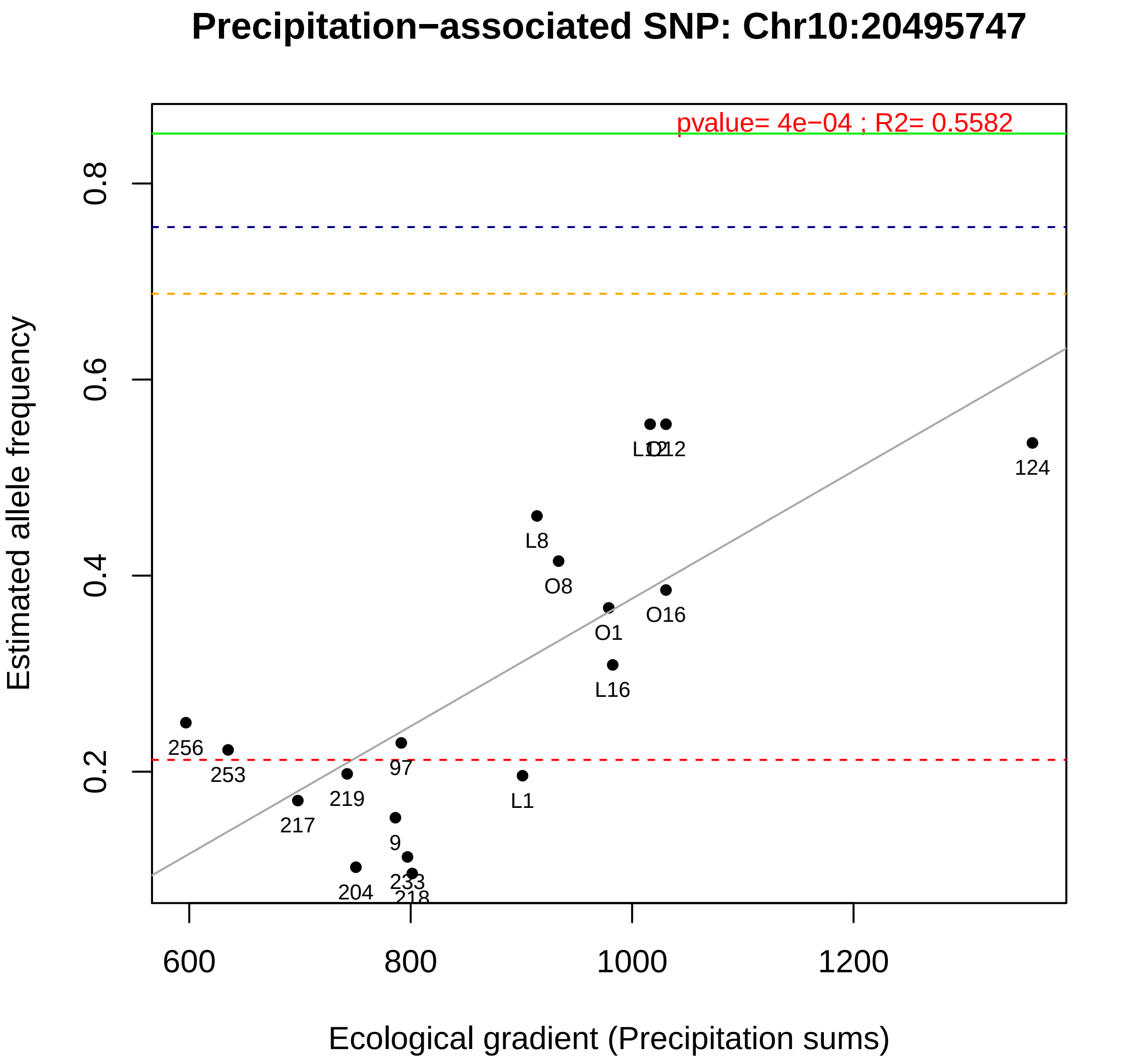
Precipitation−associated SNP: Chr10:20495747
pvalue= 4e−04 ; R2= 0.5582
Estimated allele frequency
0.2
0.4
0.6
0.8
600
800
1000
1200
Ecological gradient (Precipitation sums)
124
L8
O8
O16
O1
L16
256
253
217
219
97
L1
9
204
233
218

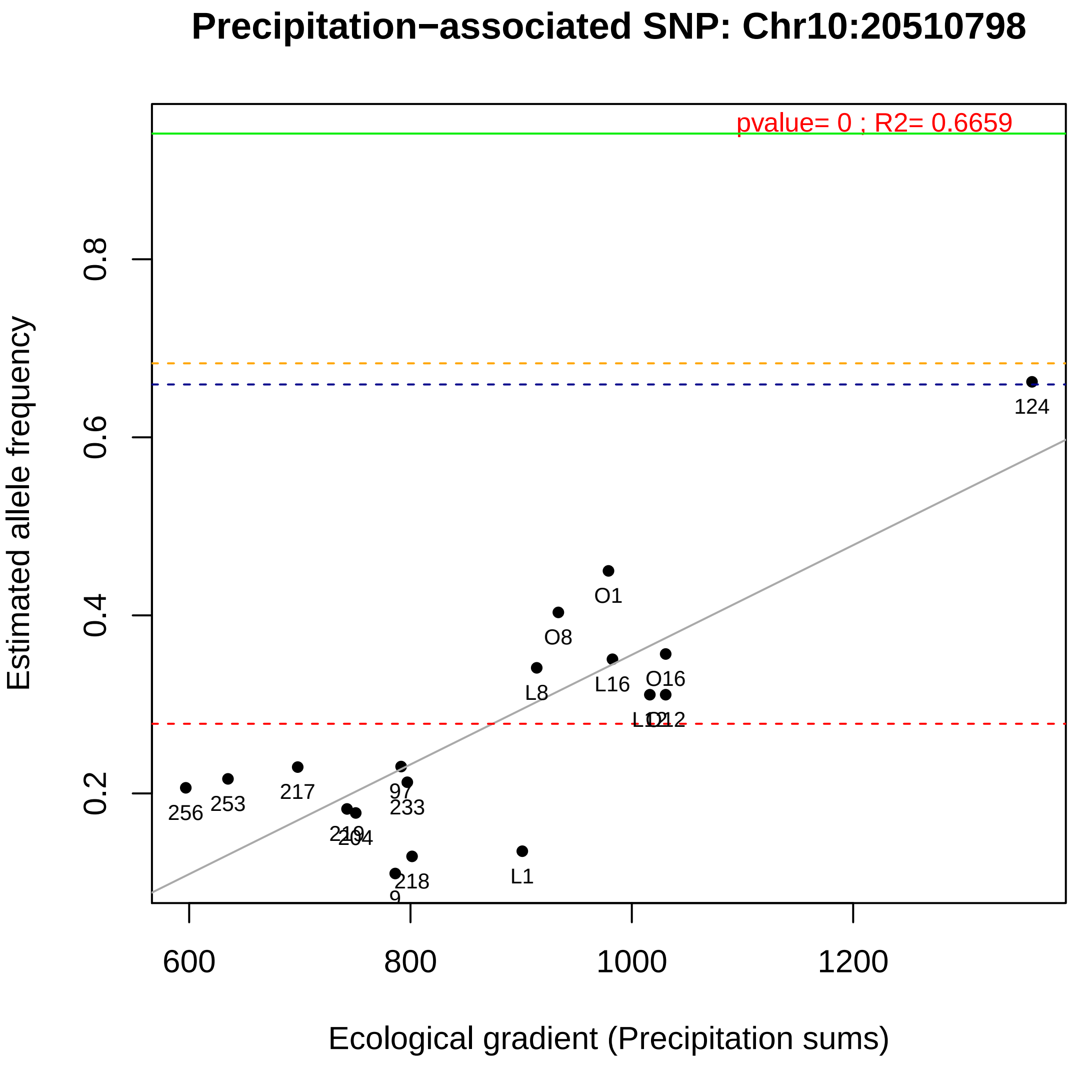
Precipitation−associated SNP: Chr10:20510798
pvalue= 0 ; R2= 0.6659
124
O1
O8
L8
L16
O16
L10
O12
256
253
217
97
233
210
204
218
9
L1
0.2
0.4
0.6
0.8
600
800
1000
1200
Estimated allele frequency
Ecological gradient (Precipitation sums)

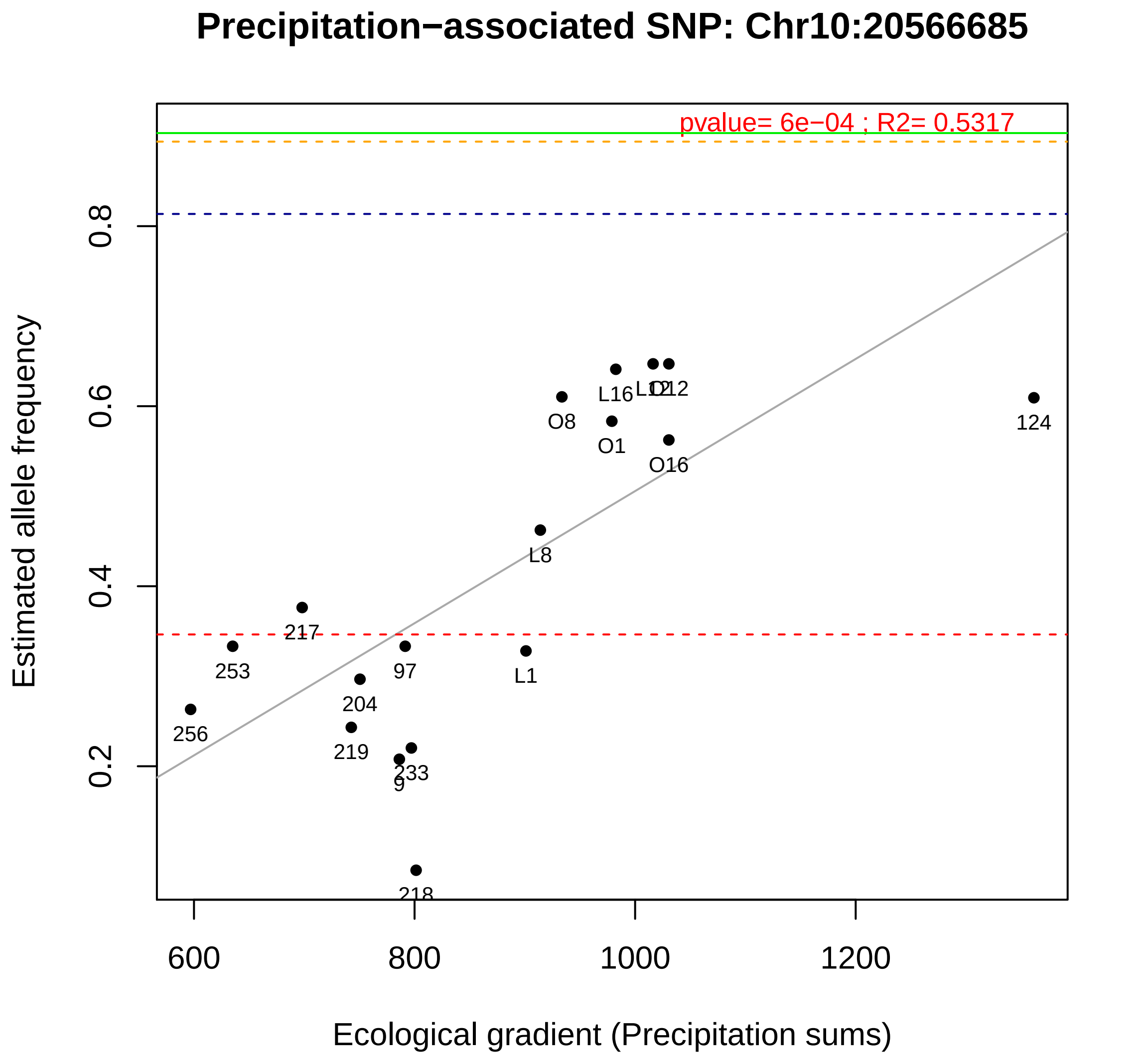

Precipitation−associated SNP: Chr10:20566685
pvalue= 6e−04 ; R2= 0.5317
Estimated allele frequency
0.2
0.4
0.6
0.8
600
800
1000
1200
Ecological gradient (Precipitation sums)
256
253
217
204
219
97
233
9
218
L1
L8
O8
O1
L16
O16
124

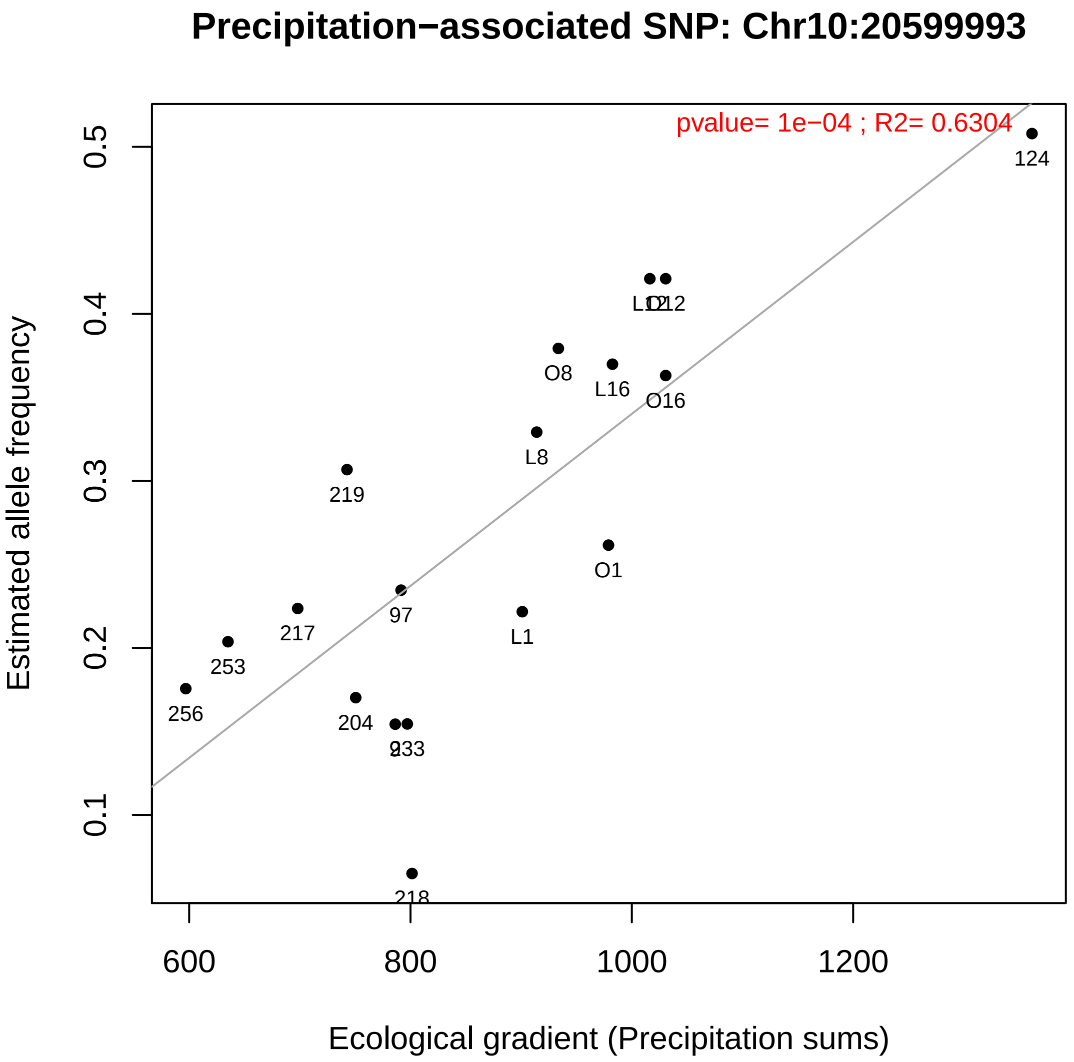
Precipitation−associated SNP: Chr10:20599993
pvalue= 1e−04 ; R2= 0.6304
Estimated allele frequency
0.1
0.2
0.3
0.4
0.5
Ecological gradient (Precipitation sums)
600
800
1000
1200
124
O8
L16
O16
L8
219
O1
97
217
L1
253
256
204
218

#### Precipitation−associated SNP: Chr10:20818183

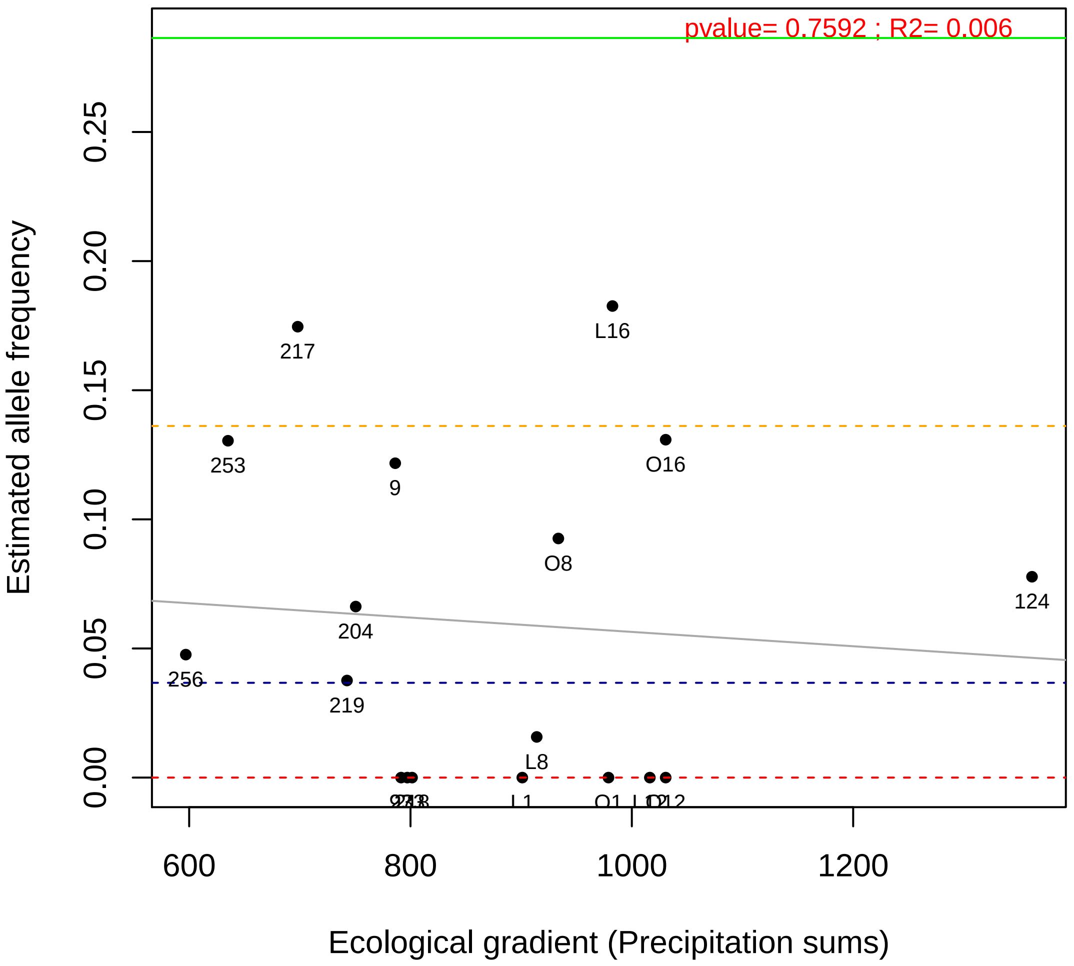

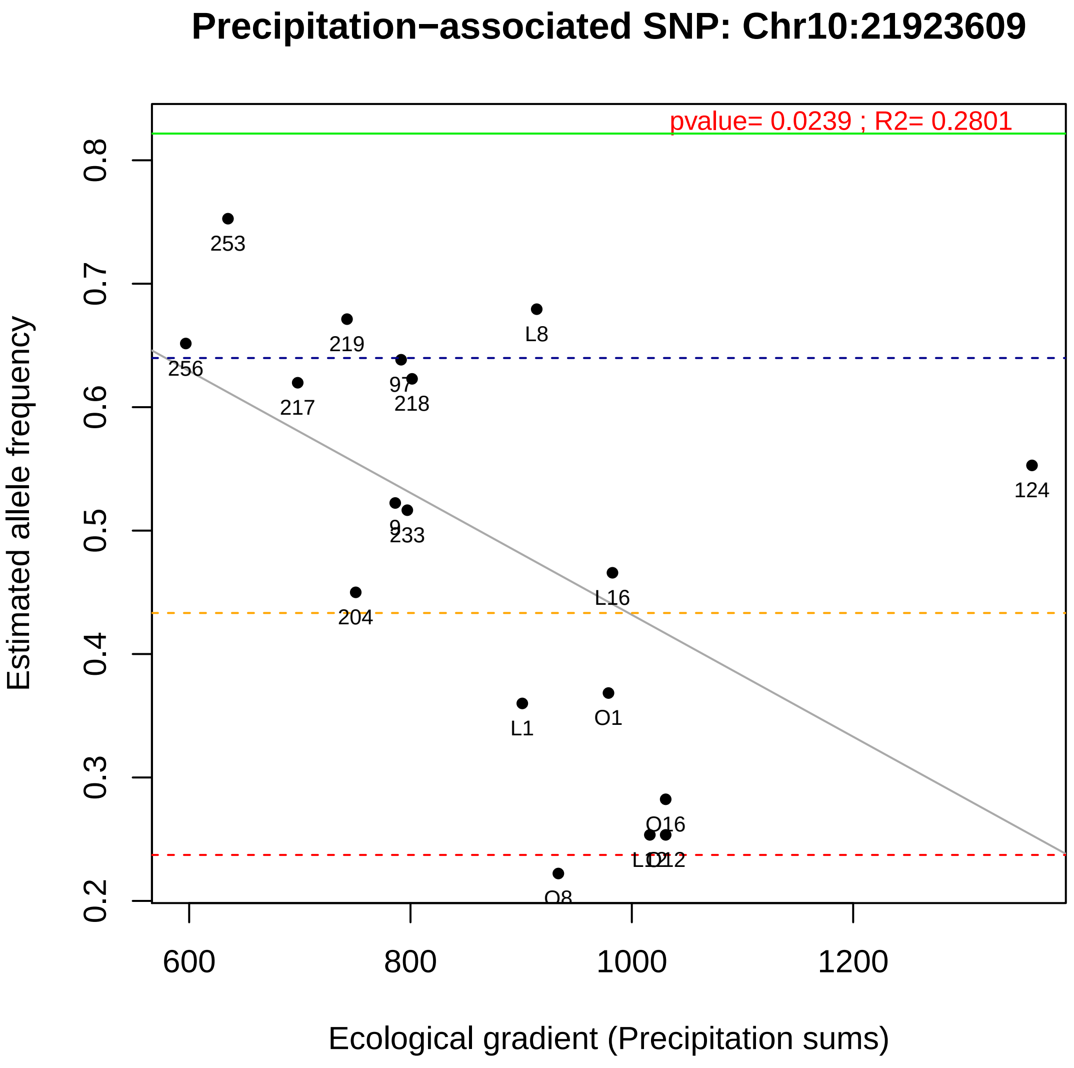

Precipitation−associated SNP: Chr10:21923609
pvalue= 0.0239 ; R2= 0.2801
Estimated allele frequency
0.2
0.3
0.4
0.5
0.6
0.7
0.8
600
800
1000
1200
Ecological gradient (Precipitation sums)
253
219
L8
256
217
97
218
124
9
233
L16
204
L1
O1
O16
O12
O8

### Precipitation−associated SNP: Chr10:22062682

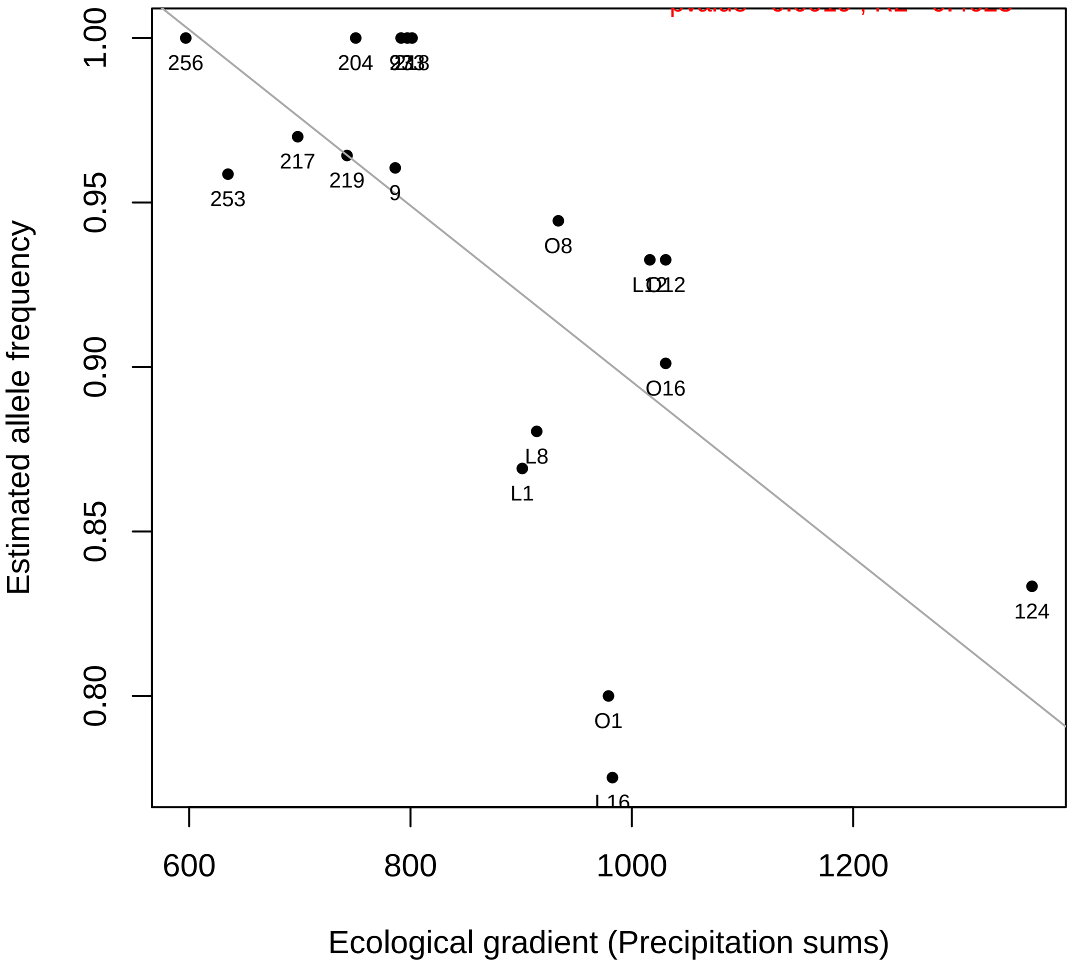

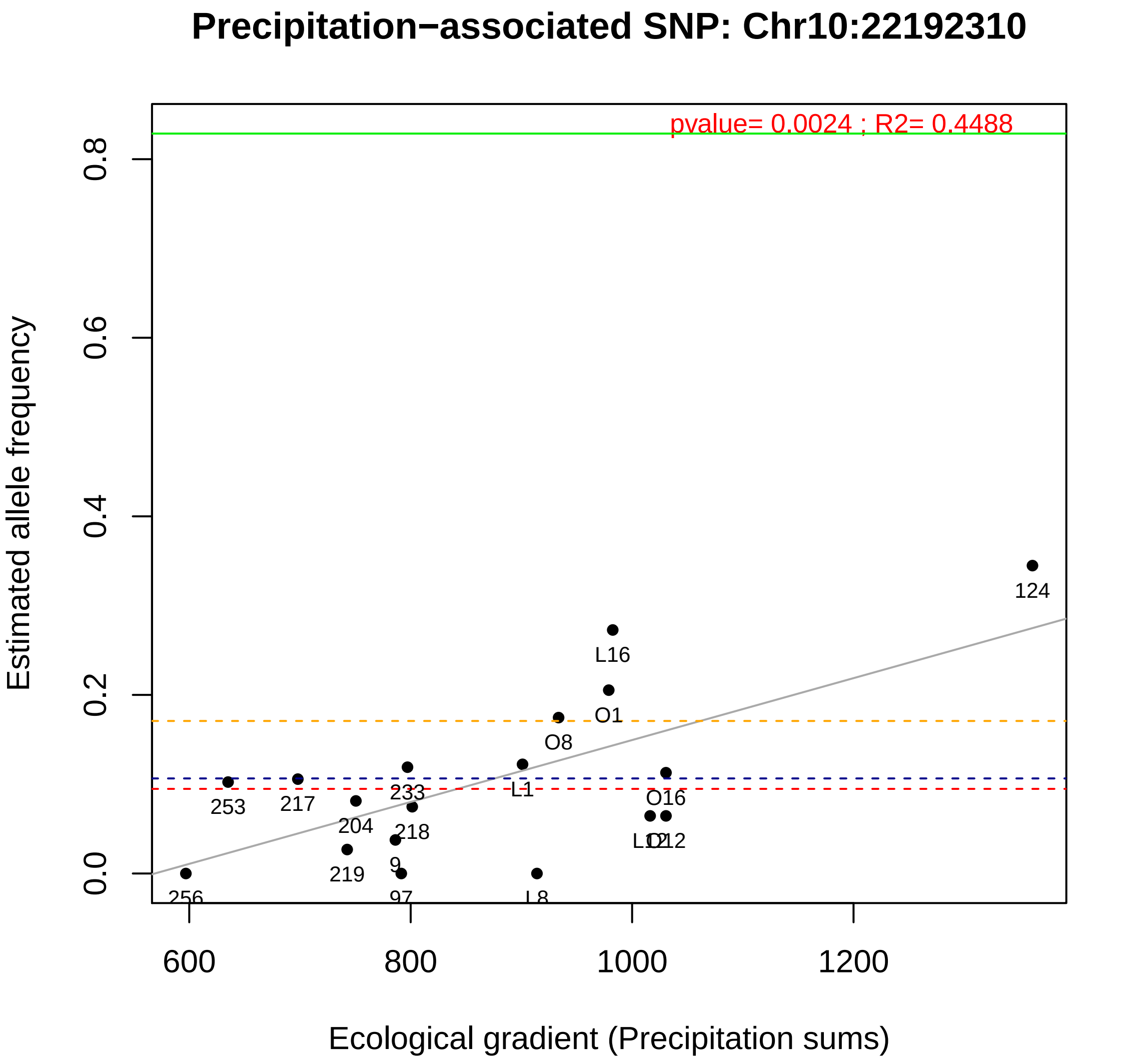

Precipitation−associated SNP: Chr10:22192310
pvalue= 0.0024 ; R2= 0.4488
Estimated allele frequency
0.0
0.2
0.4
0.6
0.8
600
800
1000
1200
Ecological gradient (Precipitation sums)
124
L16
O1
O8
L1
O16
233
253
217
204
218
219
9
97
256
L8

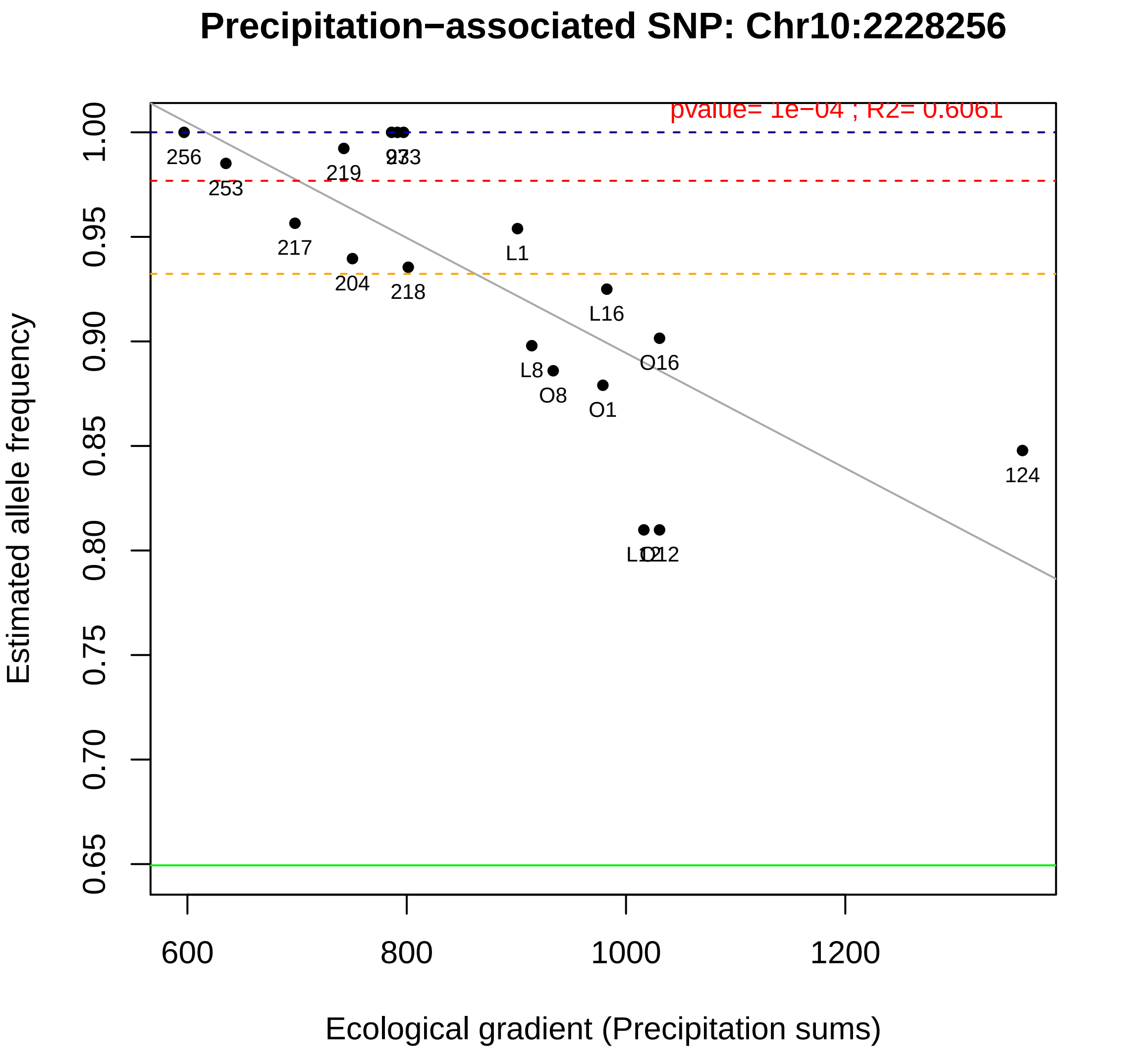
Precipitation−associated SNP: Chr10:2228256
pvalue= 1e−04 ; R2= 0.6061
Estimated allele frequency
Ecological gradient (Precipitation sums)
0.65
0.70
0.75
0.80
0.85
0.90
0.95
1.00
600
800
1000
1200
256
253
219
217
204
218
L1
L16
L8
O8
O16
O1
124

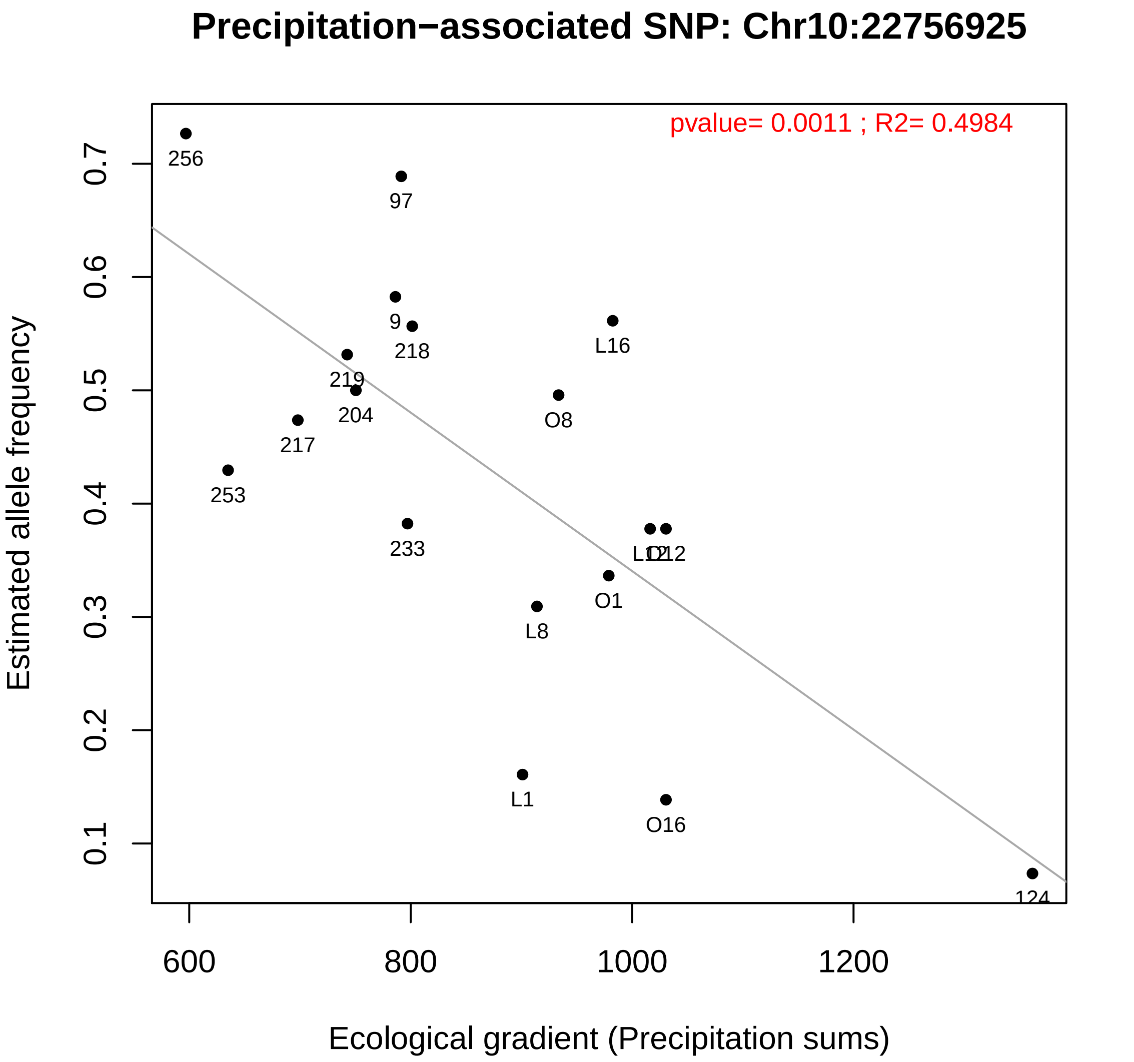

Precipitation−associated SNP: Chr10:22756925
pvalue= 0.0011 ; R2= 0.4984
Estimated allele frequency
0.1
0.2
0.3
0.4
0.5
0.6
0.7
600
800
1000
1200
Ecological gradient (Precipitation sums)
256
97
9
218
219
204
217
253
233
L16
O8
L12
O12
O1
L8
L1
O16
124

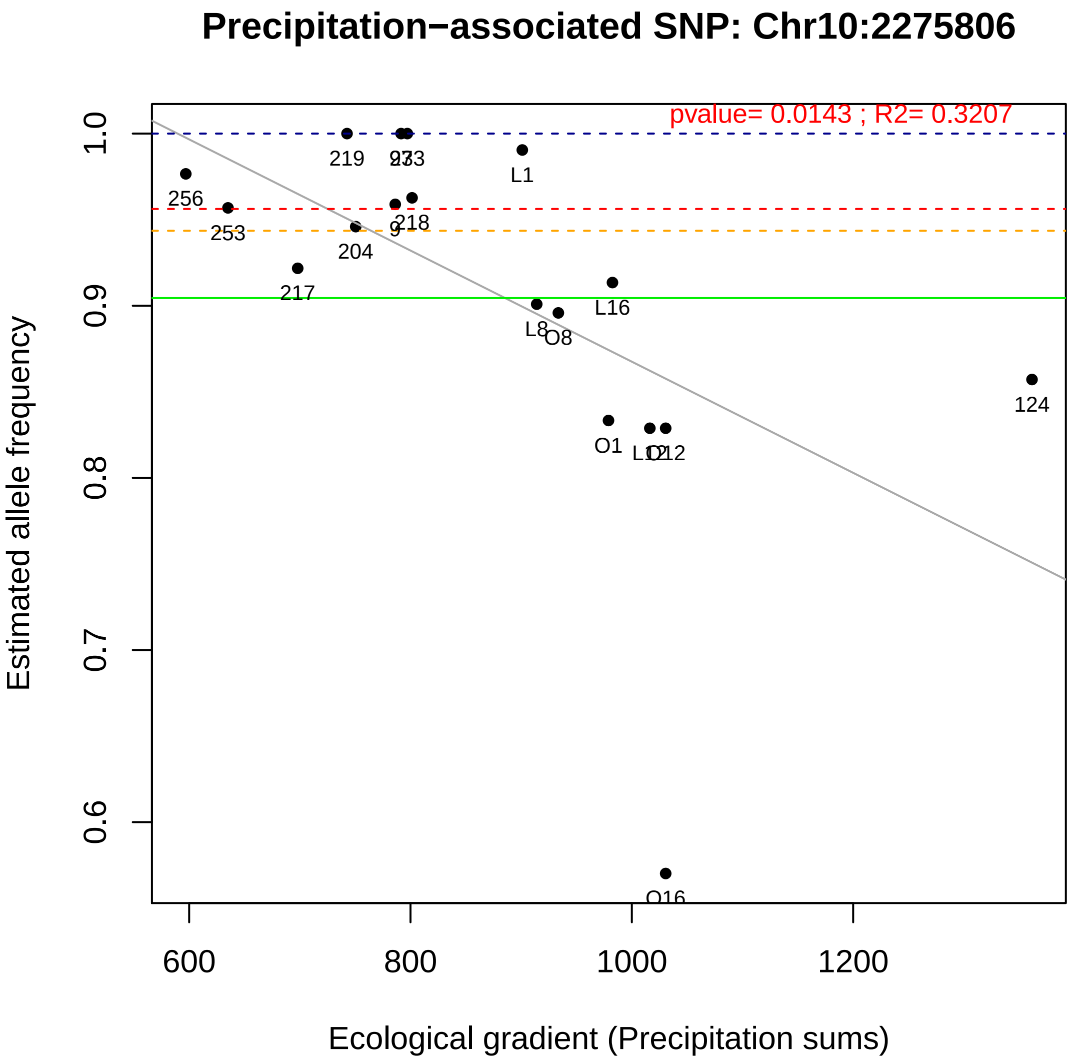

Precipitation−associated SNP: Chr10:2275806
pvalue= 0.0143 ; R2= 0.3207
Estimated allele frequency
Ecological gradient (Precipitation sums)
0.6
0.7
0.8
0.9
1.0
600
800
1000
1200
219
256
253
204
217
218
L1
L16
L8
O8
O1
124
O16

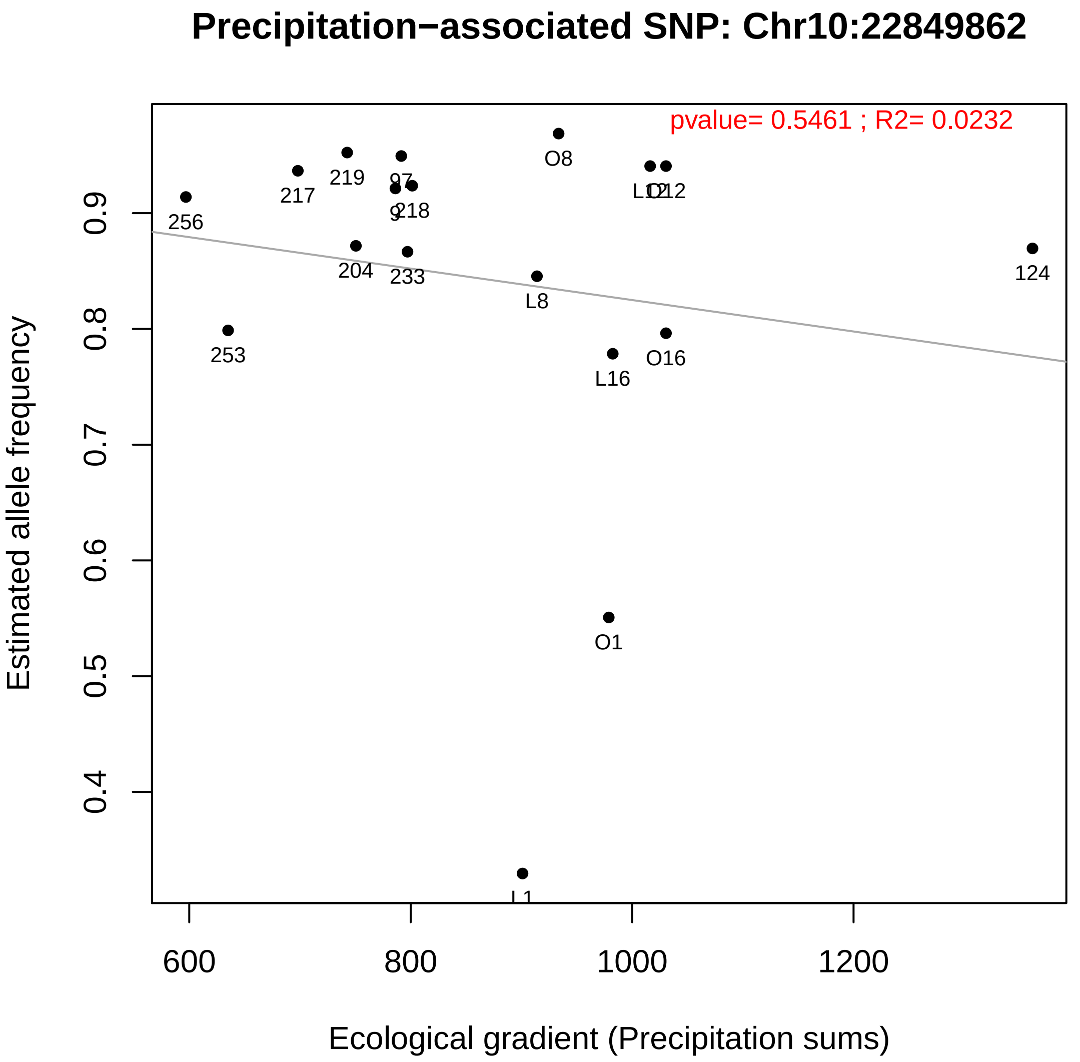

Precipitation−associated SNP: Chr10:22849862
pvalue= 0.5461 ; R2= 0.0232
Estimated allele frequency
Ecological gradient (Precipitation sums)
0.4
0.5
0.6
0.7
0.8
0.9
600
800
1000
1200
256
253
217
219
204
97
218
233
L8
O8
L12
O12
124
L16
O16
O1
L1

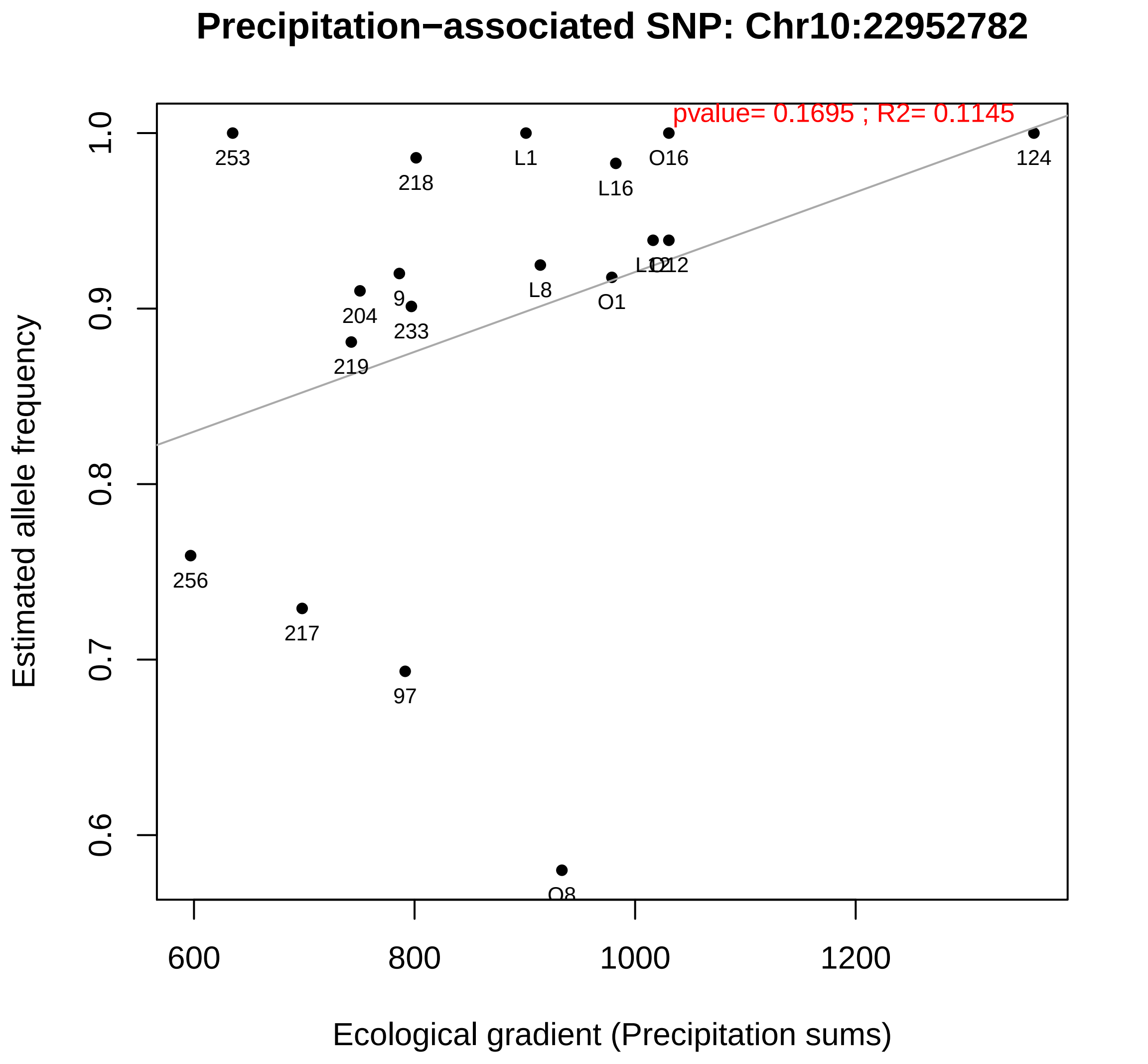

Precipitation−associated SNP: Chr10:22952782
pvalue= 0.1695 ; R2= 0.1145
Estimated allele frequency
Ecological gradient (Precipitation sums)
0.6
0.7
0.8
0.9
1.0
600
800
1000
1200
253
218
L1
L16
O16
124
204
9
233
219
L8
O1
L12
O12
256
217
97
O8

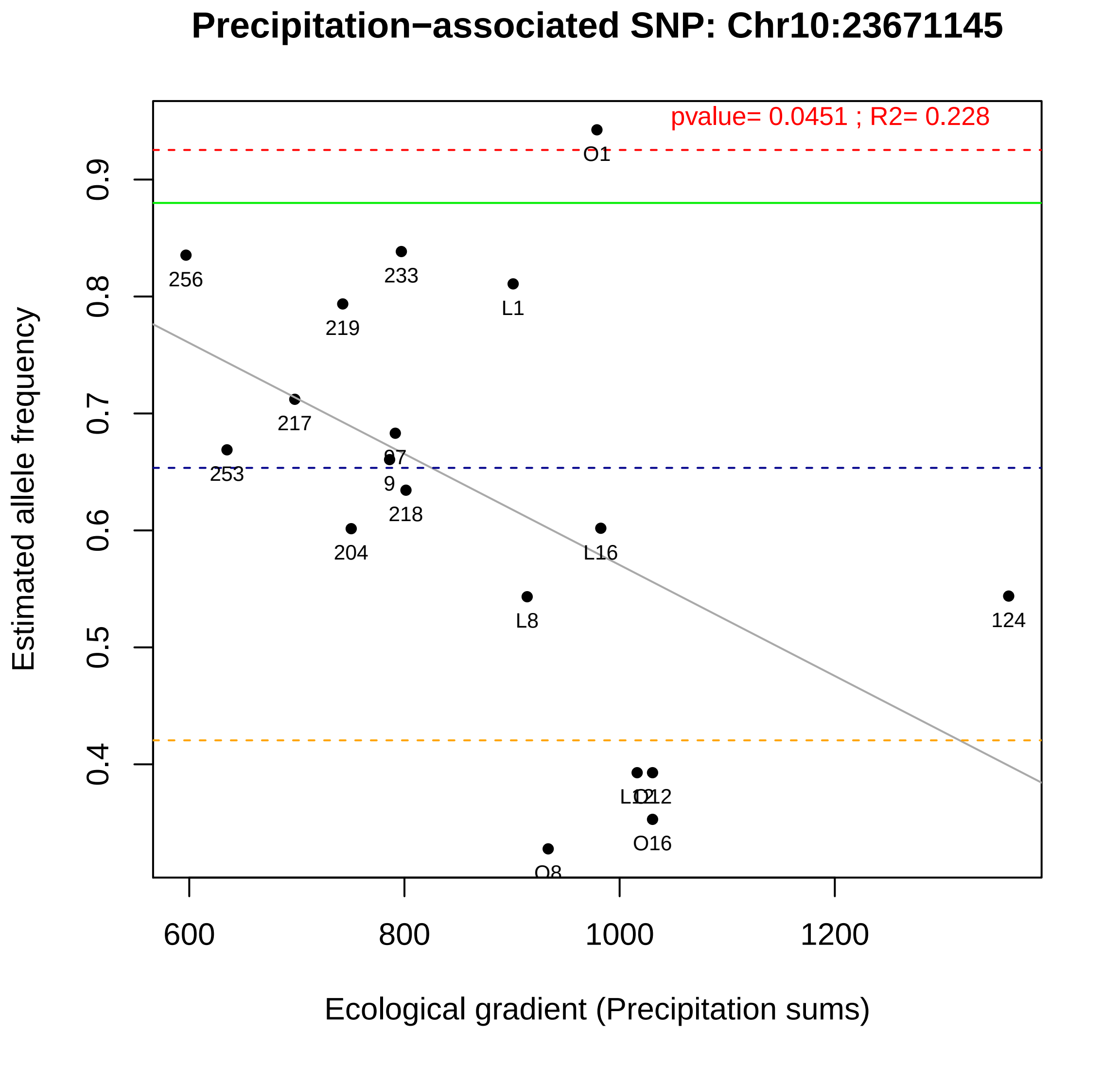

Precipitation−associated SNP: Chr10:23671145
pvalue= 0.0451 ; R2= 0.228
Estimated allele frequency
Ecological gradient (Precipitation sums)
0.4
0.5
0.6
0.7
0.8
0.9
600
800
1000
1200
O1
256
233
L1
219
217
253
97
9
218
204
L16
L8
124
L12
O12
O16
O8

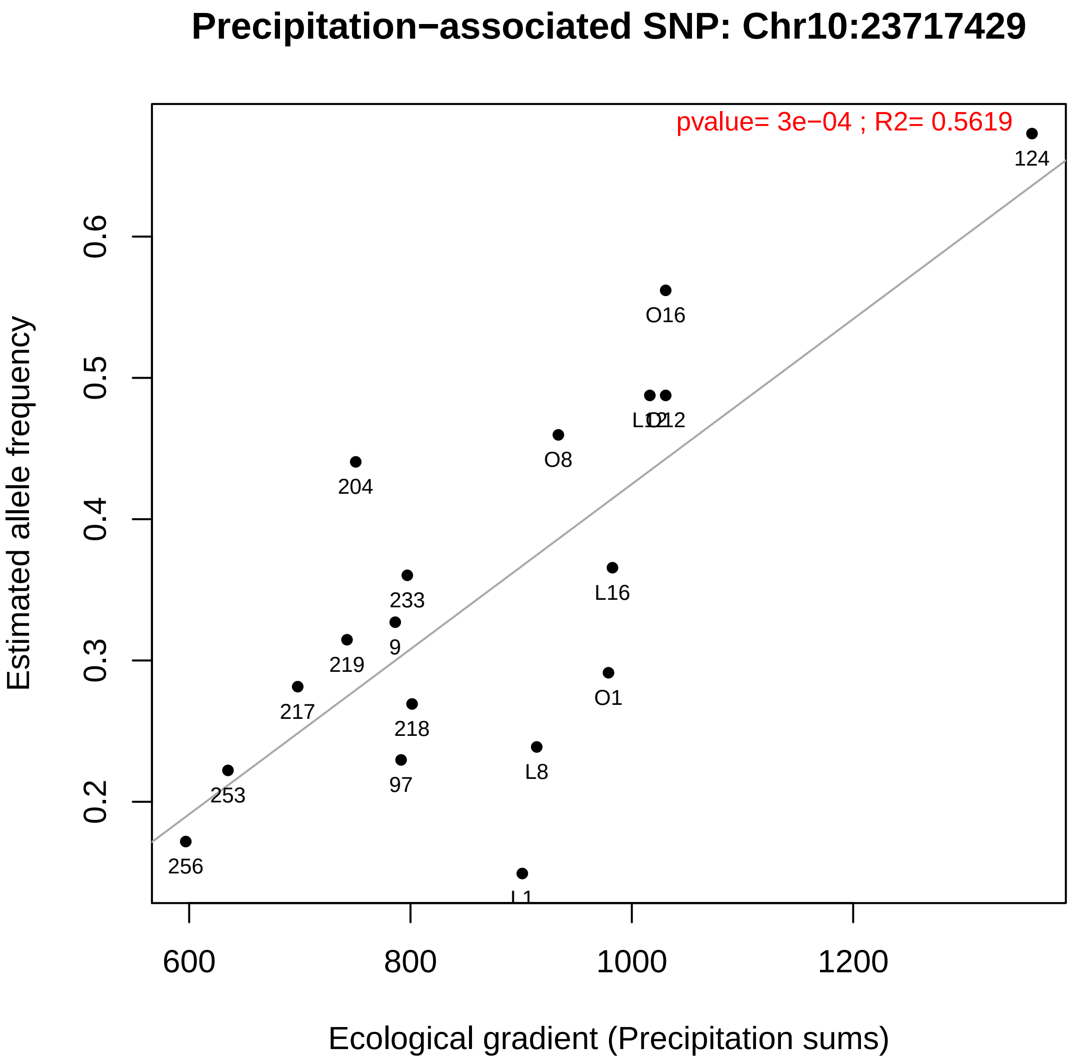

Precipitation−associated SNP: Chr10:23717429
pvalue= 3e−04 ; R2= 0.5619
Estimated allele frequency
0.2
0.3
0.4
0.5
0.6
600
800
1000
1200
Ecological gradient (Precipitation sums)
124
O16
O8
204
233
L16
9
219
217
O1
218
L8
97
253
256

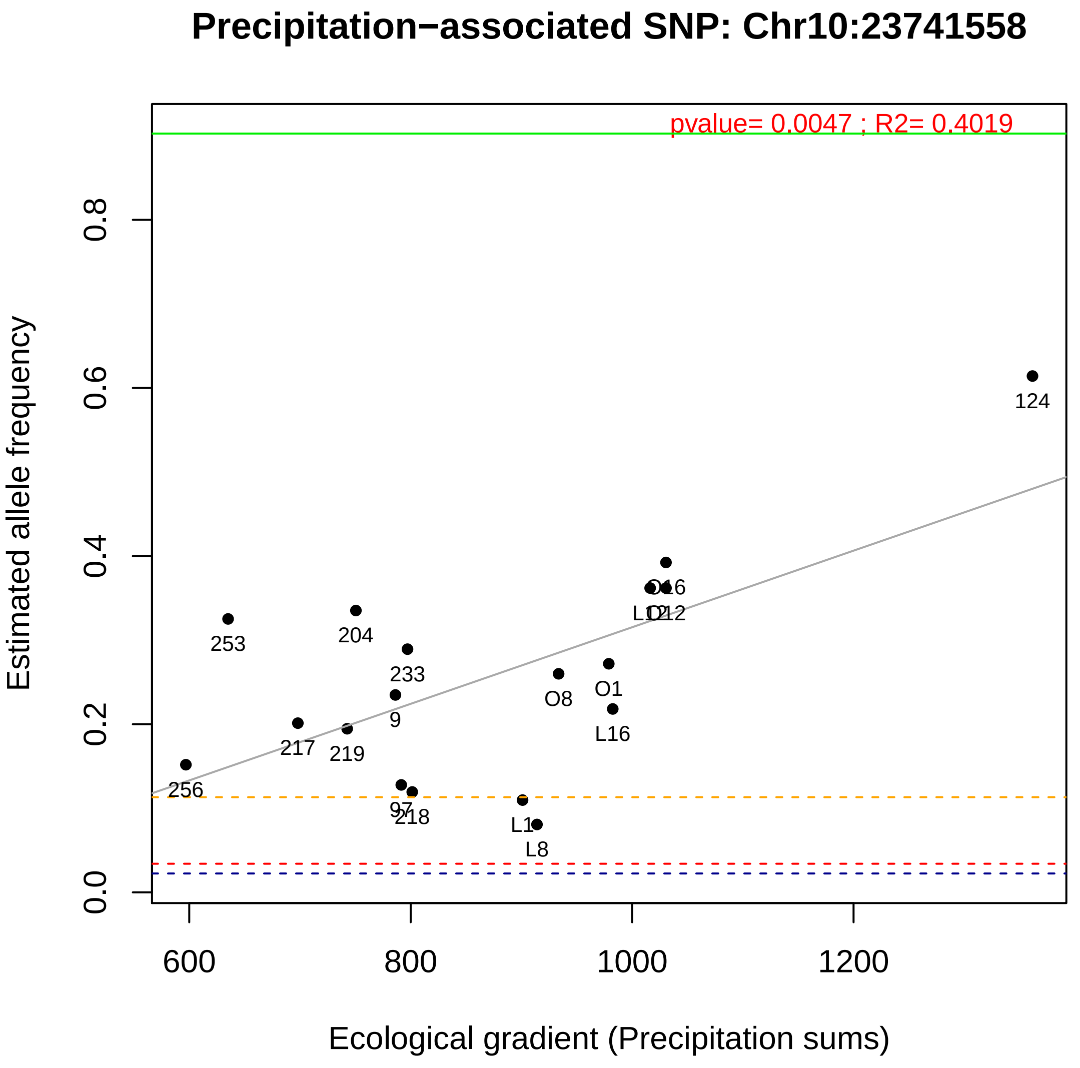
Precipitation−associated SNP: Chr10:23741558
pvalue= 0.0047 ; R2= 0.4019
Estimated allele frequency
Ecological gradient (Precipitation sums)
0.0
0.2
0.4
0.6
0.8
600
800
1000
1200
124
253
204
233
9
O8
O1
L16
217
219
256
97
218
L1
L8

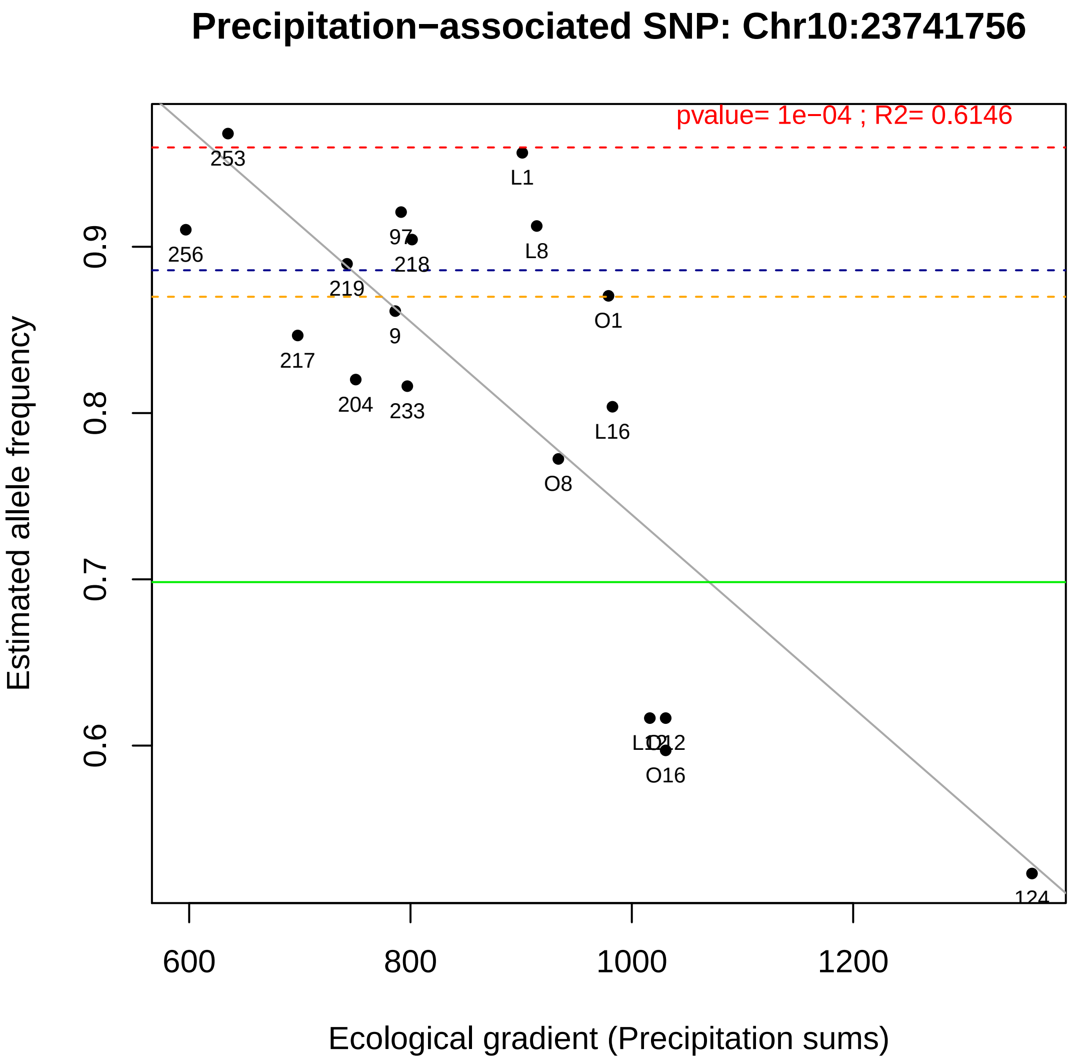
Precipitation−associated SNP: Chr10:23741756
pvalue= 1e−04 ; R2= 0.6146
253
L1
256
97
218
L8
219
O1
9
217
204
233
L16
O8
L10
O12
O16
124
0.9
0.8
0.7
0.6
600
800
1000
1200
Estimated allele frequency
Ecological gradient (Precipitation sums)

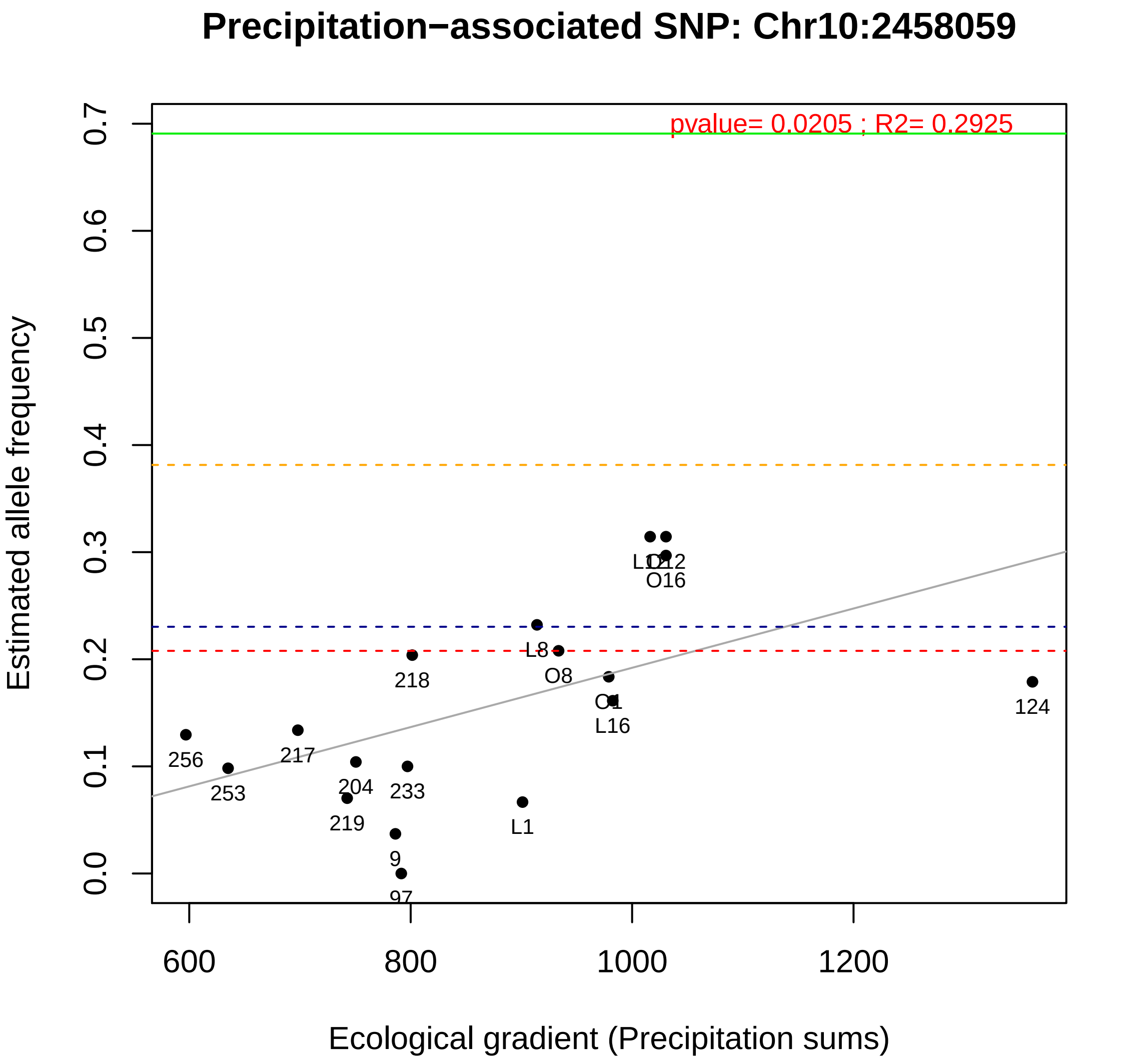

Precipitation−associated SNP: Chr10:2458059
pvalue= 0.0205 ; R2= 0.2925
Estimated allele frequency
0.0
0.1
0.2
0.3
0.4
0.5
0.6
0.7
256
253
217
204
219
233
218
9
97
L1
L8
O8
O1
L16
L12
O12
O16
124
600
800
1000
1200
Ecological gradient (Precipitation sums)

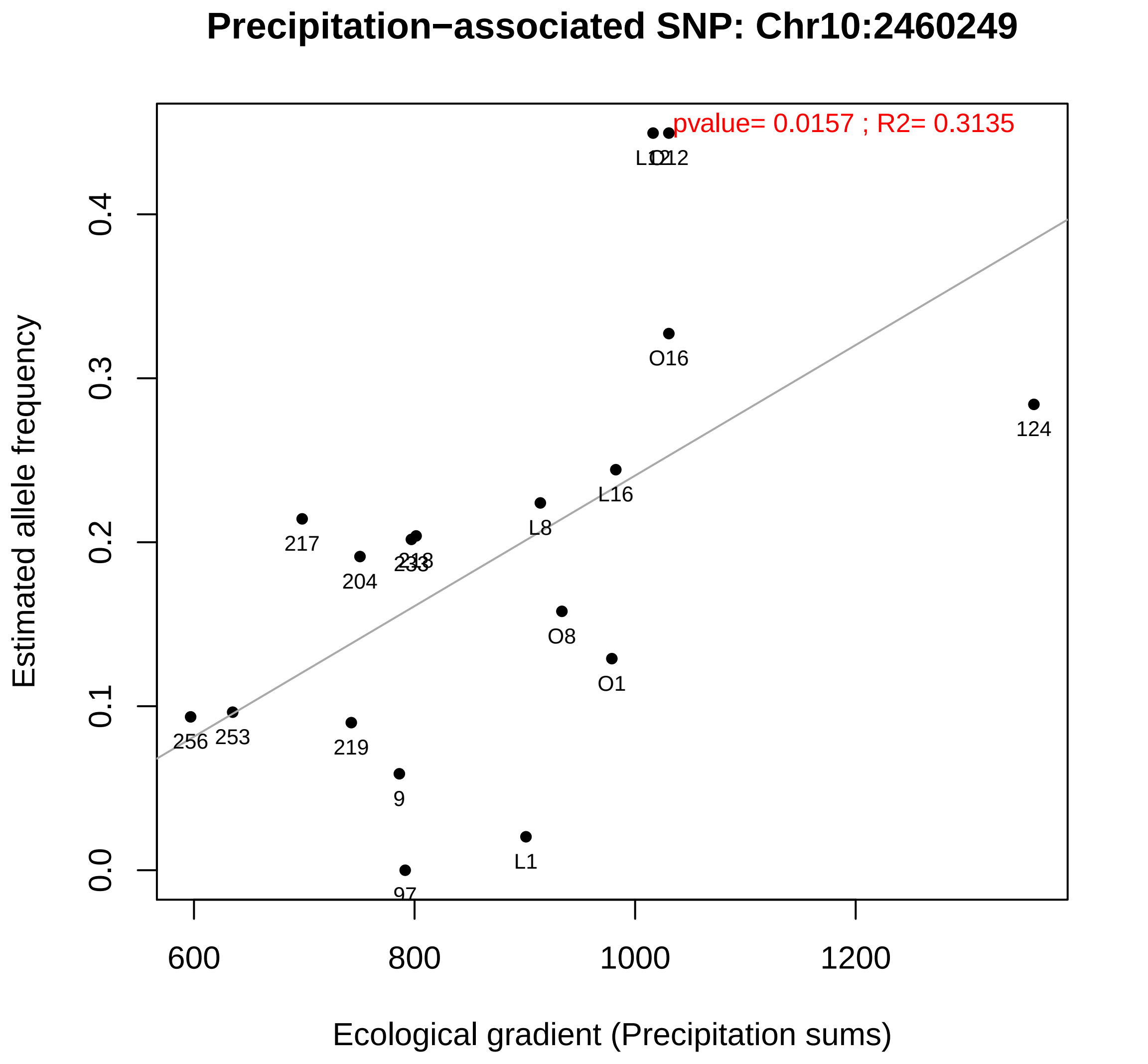

Precipitation−associated SNP: Chr10:2460249
pvalue= 0.0157 ; R2= 0.3135
Estimated allele frequency
Ecological gradient (Precipitation sums)
0.0
0.1
0.2
0.3
0.4
600
800
1000
1200
L12
O12
O16
124
L16
L8
217
204
218
233
O8
O1
256
253
219
9
L1
97

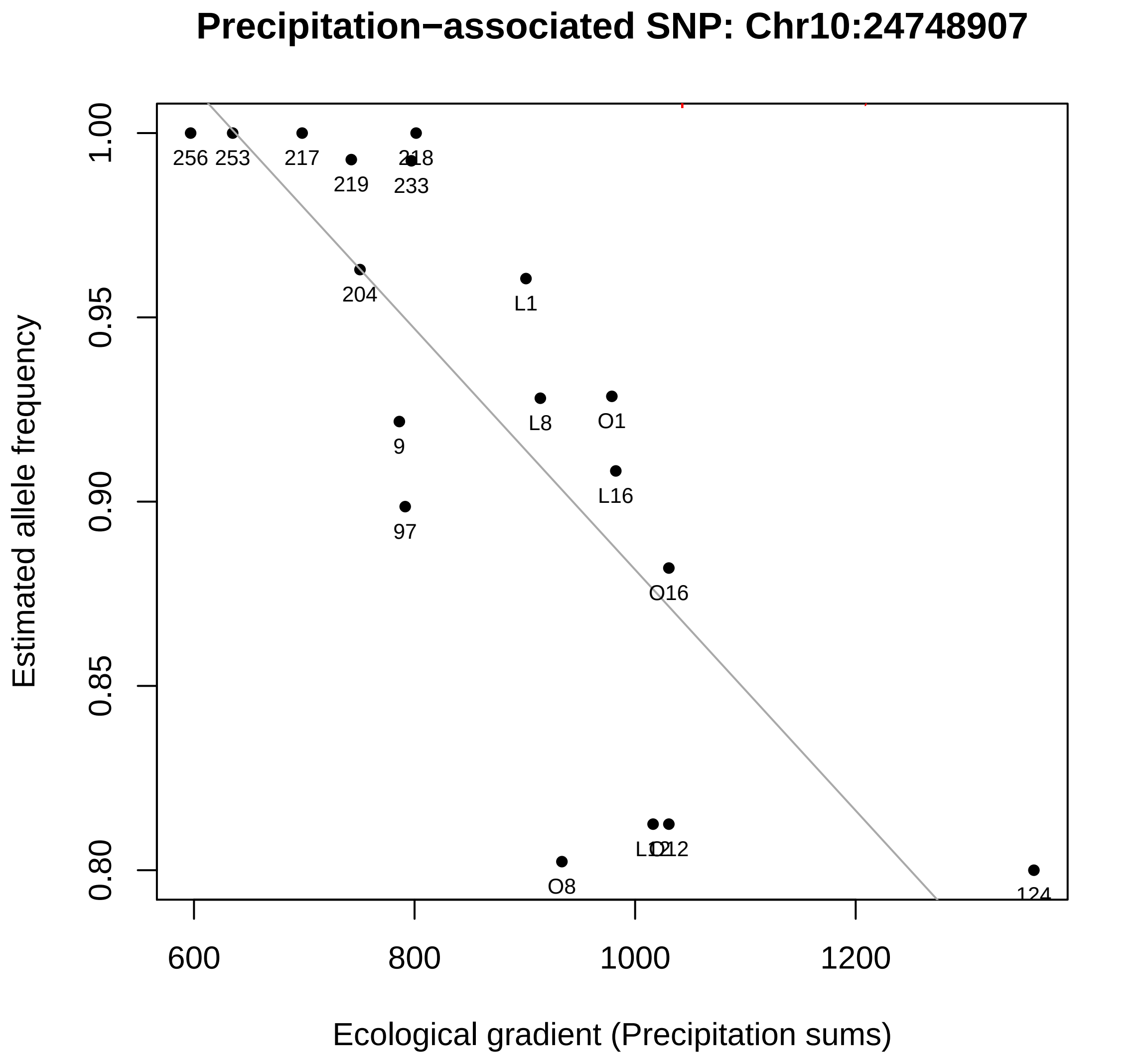
Precipitation−associated SNP: Chr10:24748907
Estimated allele frequency
Ecological gradient (Precipitation sums)
1.00
0.95
0.90
0.85
0.80
600
800
1000
1200
256
253
217
218
219
233
204
L1
L8
O1
9
L16
97
O16
L12
O12
O8
124

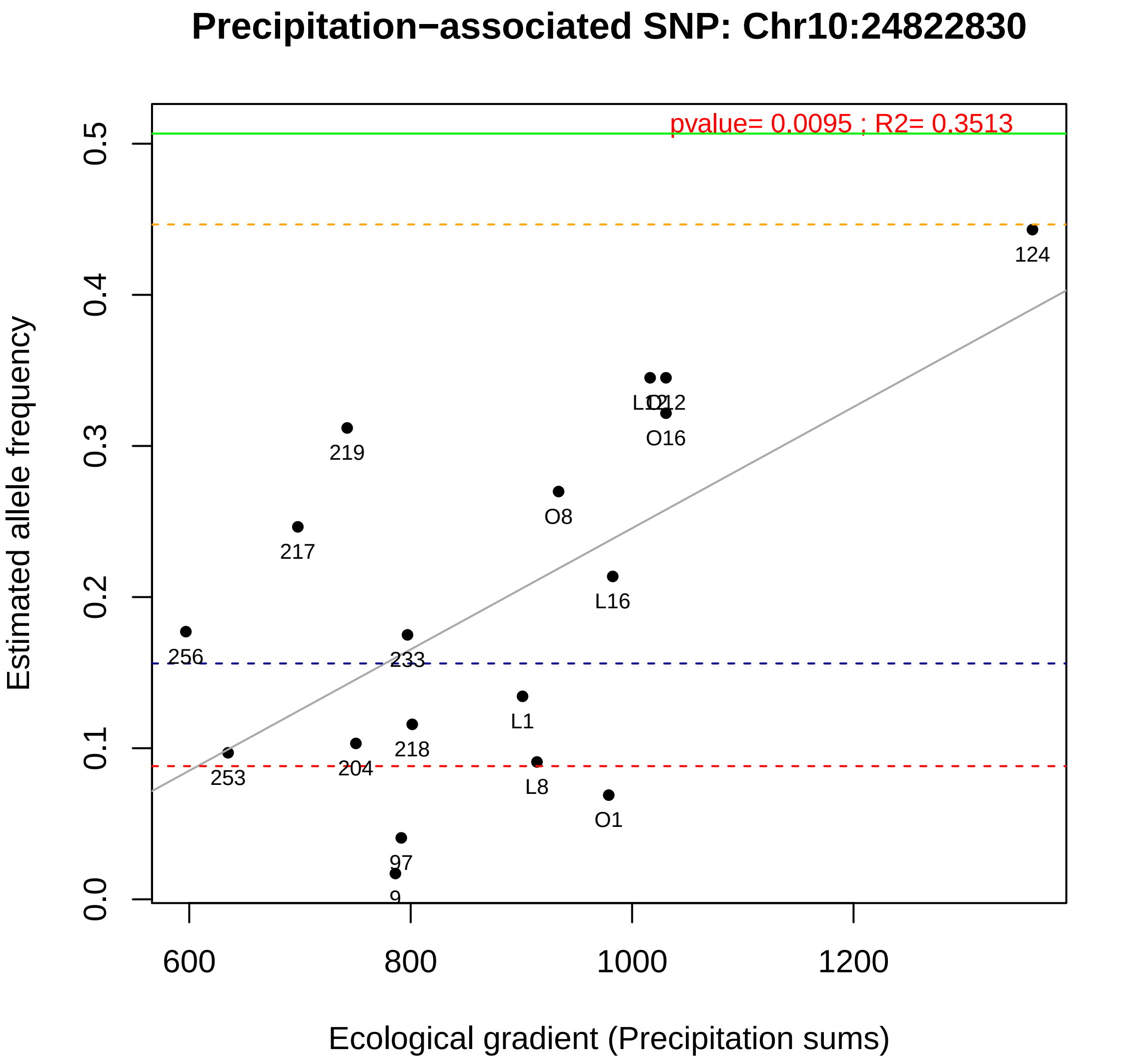
Precipitation−associated SNP: Chr10:24822830
pvalue= 0.0095 ; R2= 0.3513
Estimated allele frequency
Ecological gradient (Precipitation sums)
0.0
0.1
0.2
0.3
0.4
0.5
600
800
1000
1200
124
L12
O12
O16
219
O8
217
L16
256
233
L1
218
204
253
L8
O1
97
9

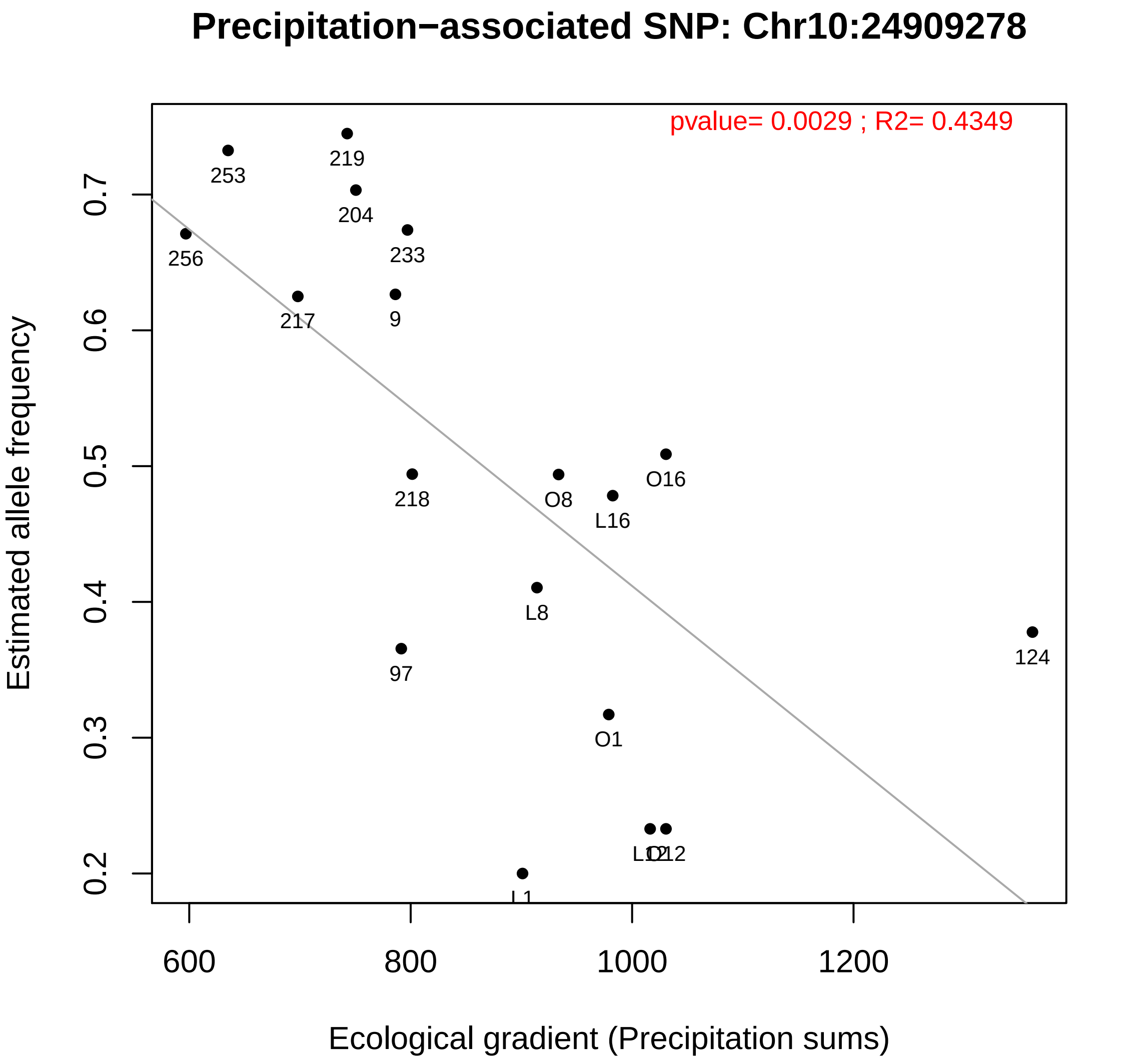
Precipitation−associated SNP: Chr10:24909278
pvalue= 0.0029 ; R2= 0.4349
Estimated allele frequency
0.2
0.3
0.4
0.5
0.6
0.7
600
800
1000
1200
Ecological gradient (Precipitation sums)
219
253
204
256
233
217
9
218
O8
O16
L16
L8
124
97
O1
L12
O12
L1

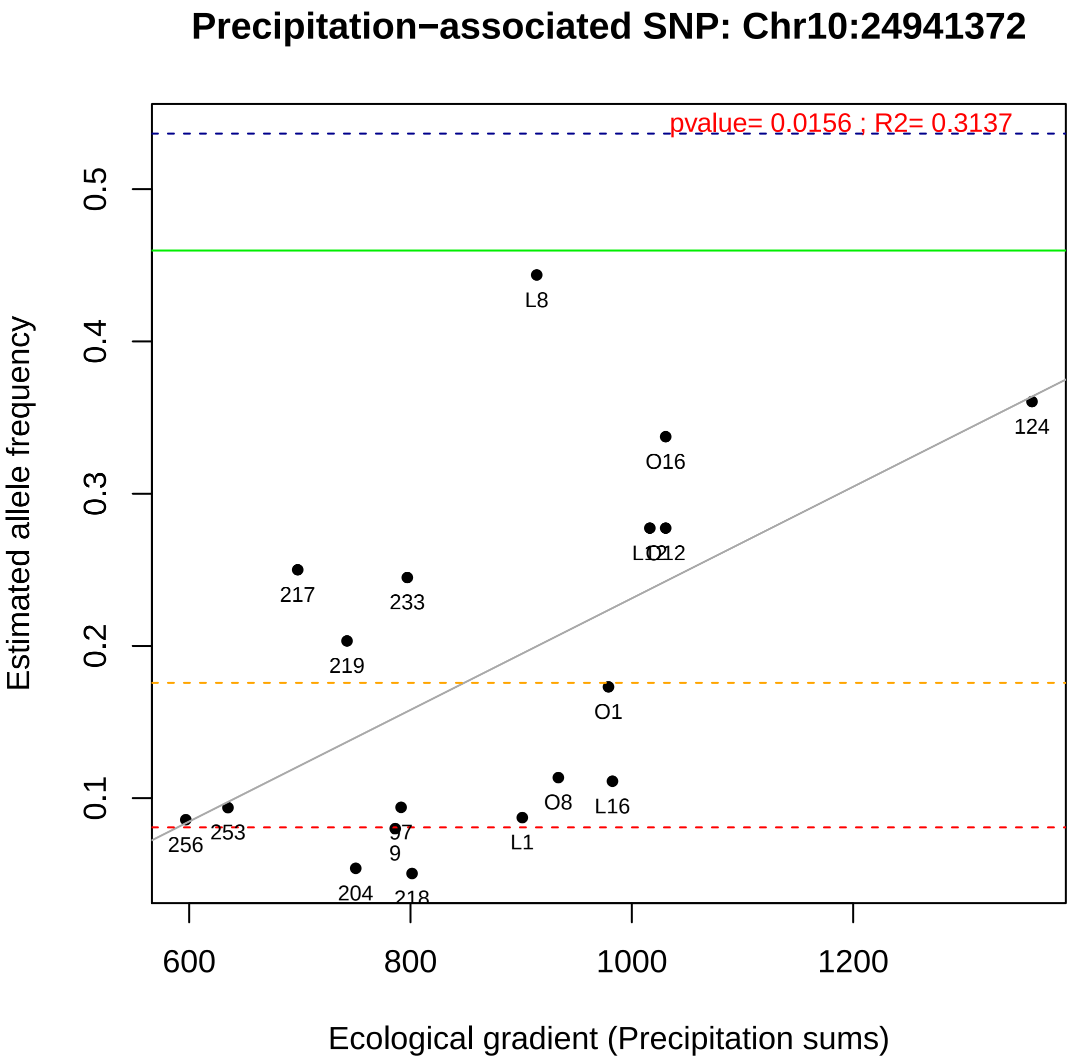

Precipitation−associated SNP: Chr10:24941372
pvalue= 0.0156 ; R2= 0.3137
Estimated allele frequency
0.1
0.2
0.3
0.4
0.5
600
800
1000
1200
Ecological gradient (Precipitation sums)
L8
124
O16
L12
O12
217
233
219
O1
O8
L16
256
253
97
9
L1
204
218

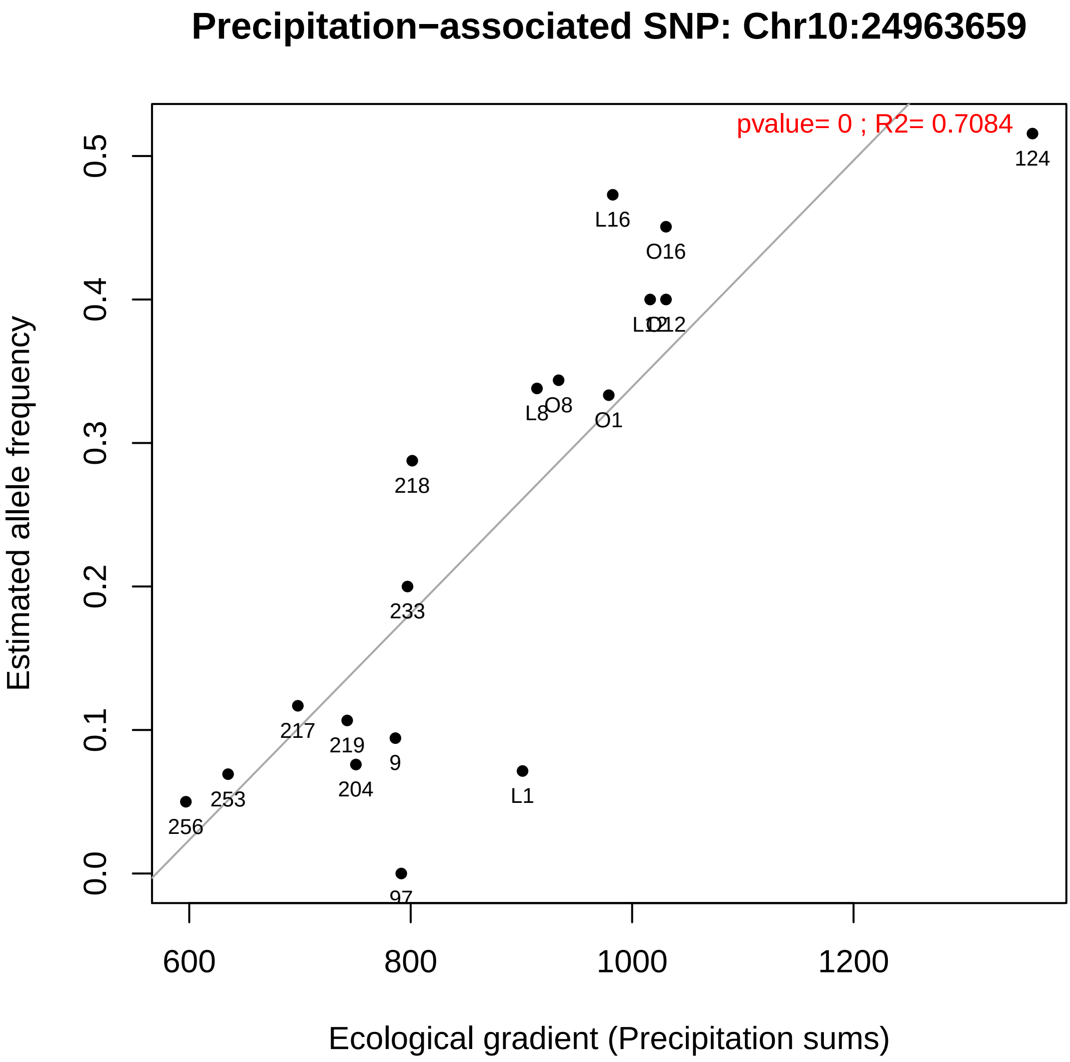

Precipitation−associated SNP: Chr10:24963659
pvalue= 0 ; R2= 0.7084
Estimated allele frequency
Ecological gradient (Precipitation sums)
0.0
0.1
0.2
0.3
0.4
0.5
600
800
1000
1200
124
L16
O16
L12
O12
L8
O8
O1
218
233
217
219
9
204
L1
253
256
97

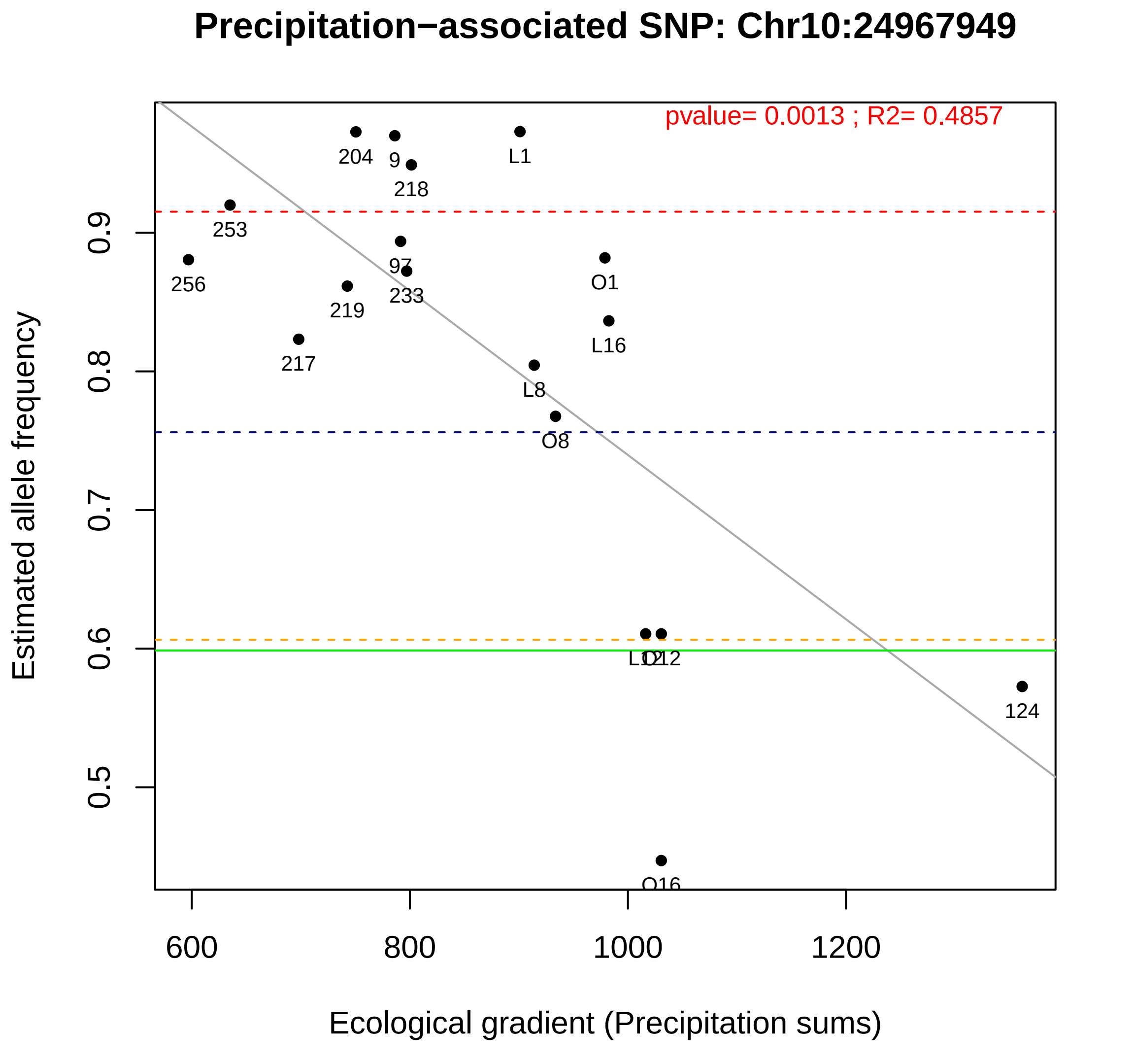

Precipitation−associated SNP: Chr10:24967949
pvalue= 0.0013 ; R2= 0.4857
204
9
218
L1
253
256
97
233
219
O1
L16
217
L8
O8
L12
O12
124
O16
0.5
0.6
0.7
0.8
0.9
600
800
1000
1200
Estimated allele frequency
Ecological gradient (Precipitation sums)

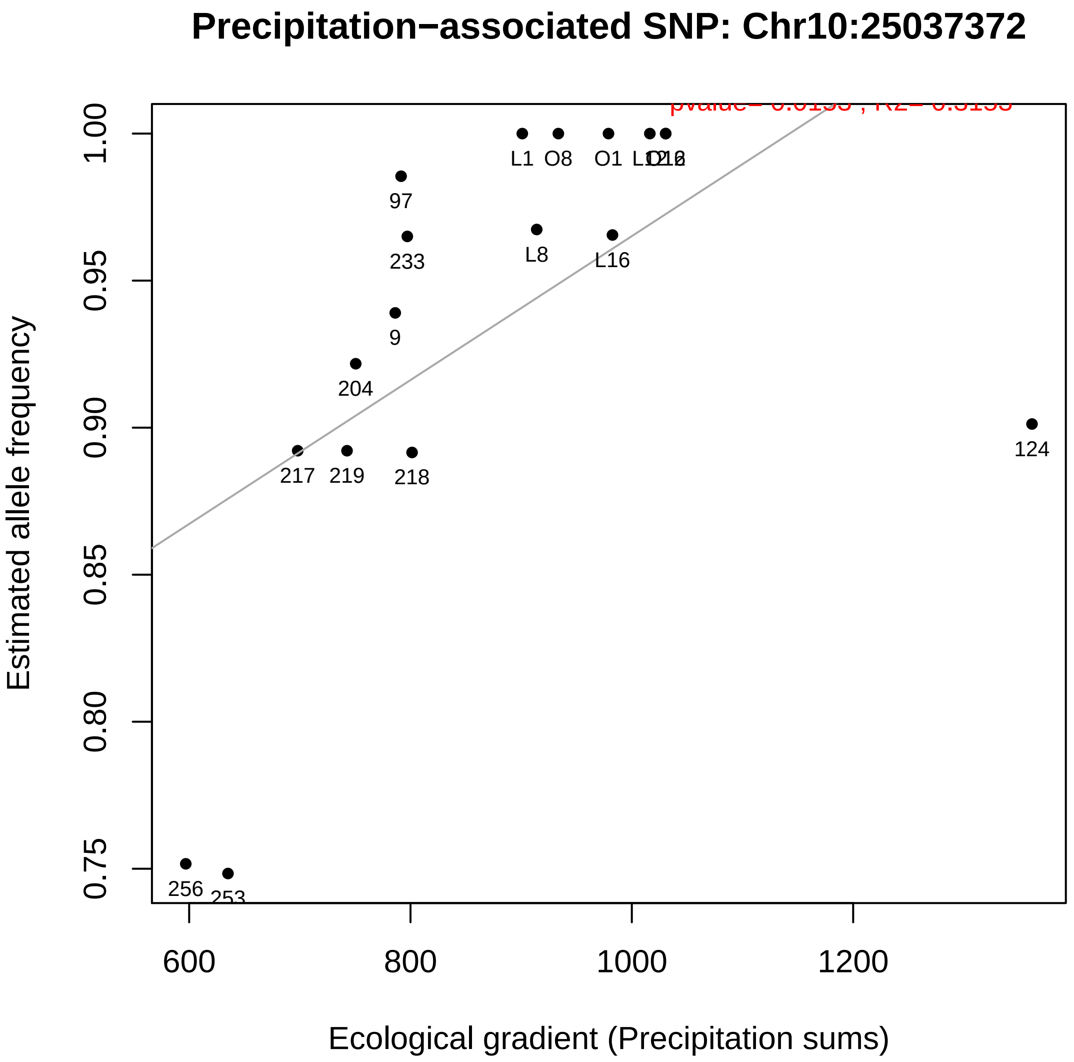
Precipitation−associated SNP: Chr10:25037372
pvalue= ; R2=
Estimated allele frequency
0.75
0.80
0.85
0.90
0.95
1.00
600
800
1000
1200
Ecological gradient (Precipitation sums)
L1
O8
O1
97
233
L8
L16
9
204
217
219
218
124
256
253

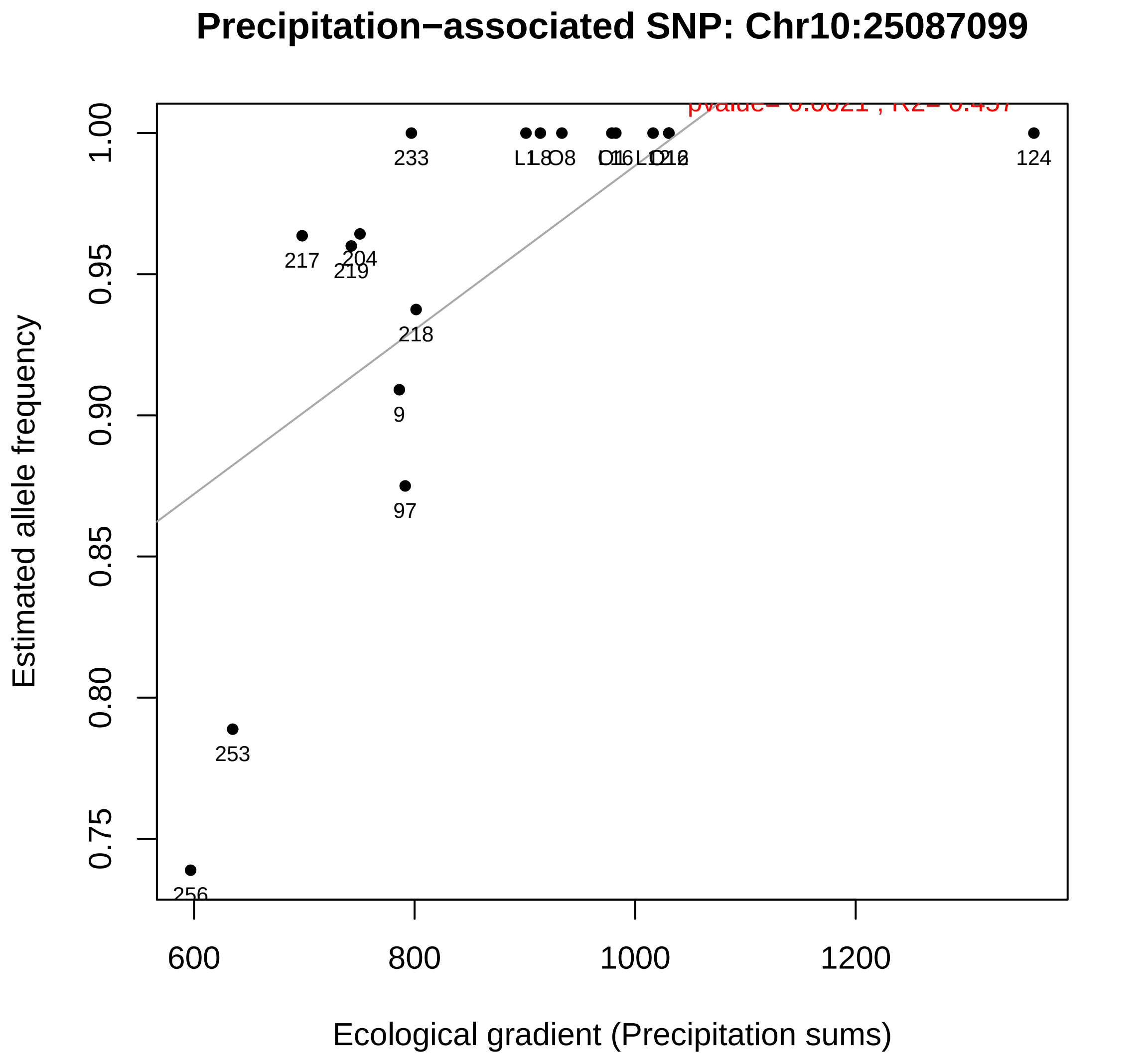

Precipitation−associated SNP: Chr10:25087099
Estimated allele frequency
Ecological gradient (Precipitation sums)
0.75
0.80
0.85
0.90
0.95
1.00
600
800
1000
1200
233
124
217
219
204
218
9
97
253
256

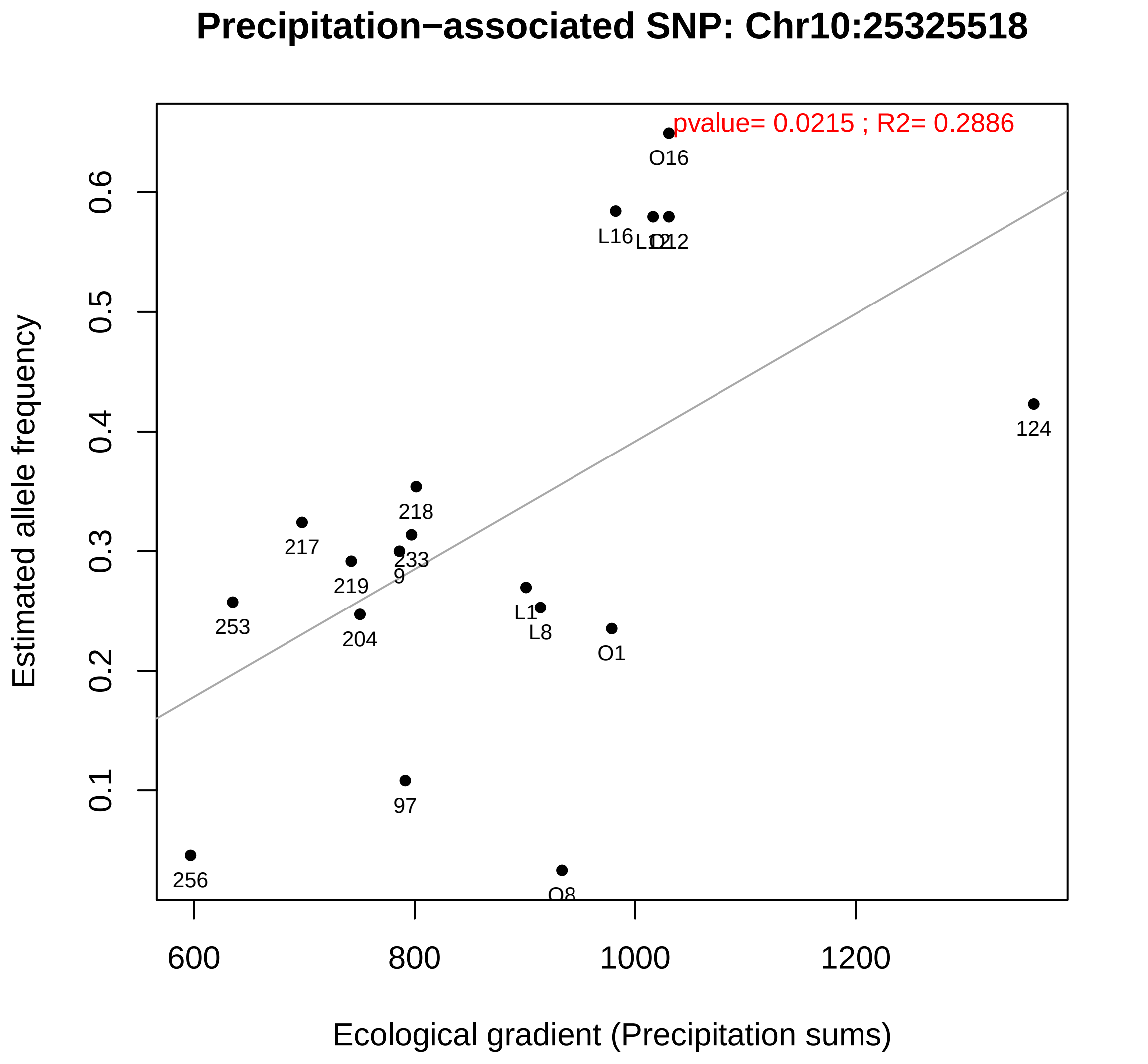

Precipitation−associated SNP: Chr10:25325518
pvalue= 0.0215 ; R2= 0.2886
Estimated allele frequency
0.1
0.2
0.3
0.4
0.5
0.6
600
800
1000
1200
Ecological gradient (Precipitation sums)
O16
L16
L10
O12
124
218
217
233
219
9
253
204
L1
L8
O1
97
256
O8

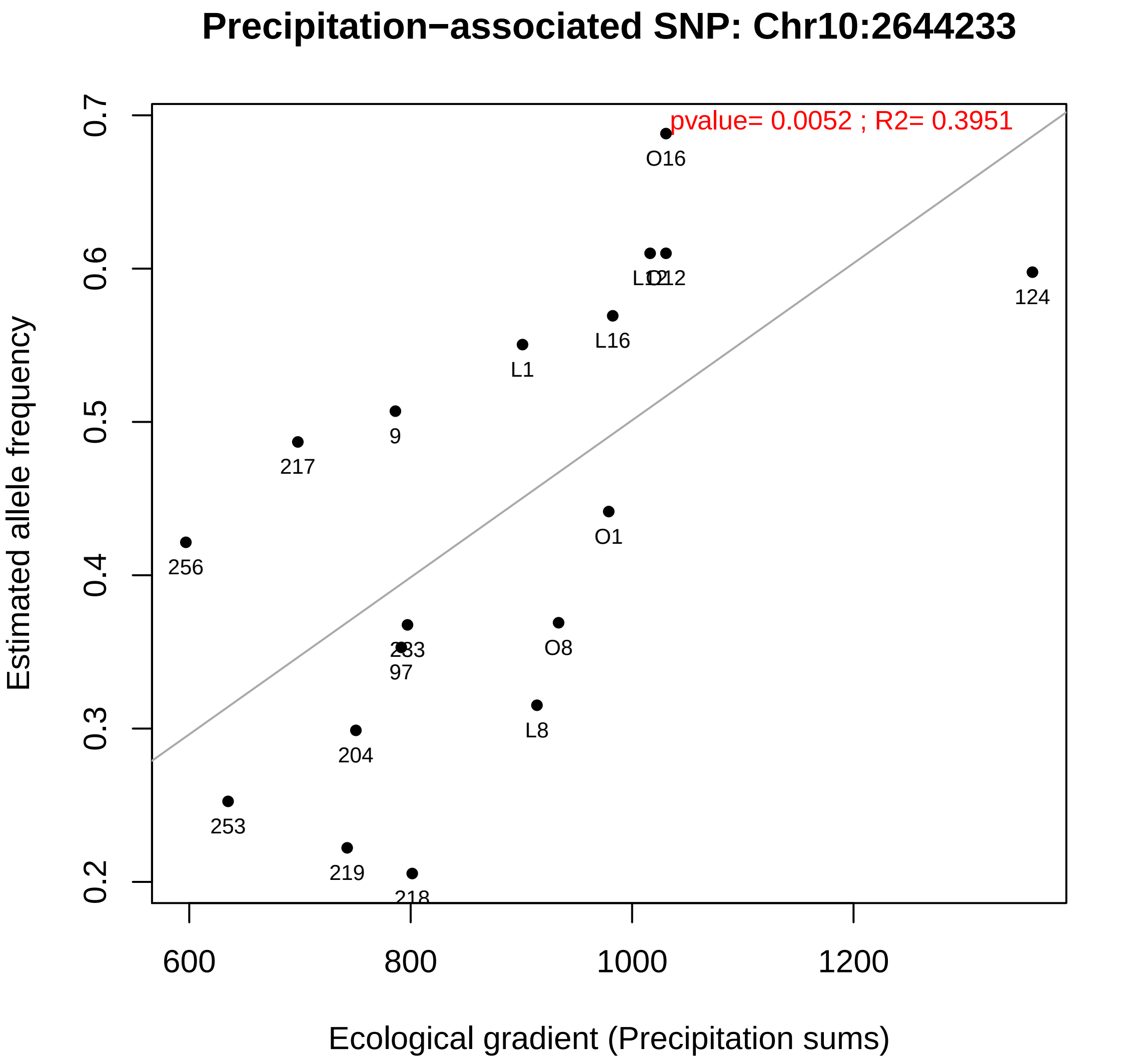

Precipitation−associated SNP: Chr10:2644233
pvalue= 0.0052 ; R2= 0.3951
Estimated allele frequency
Ecological gradient (Precipitation sums)
0.2
0.3
0.4
0.5
0.6
0.7
600
800
1000
1200
O16
124
L16
L1
9
217
O1
256
O8
97
L8
204
253
219
218

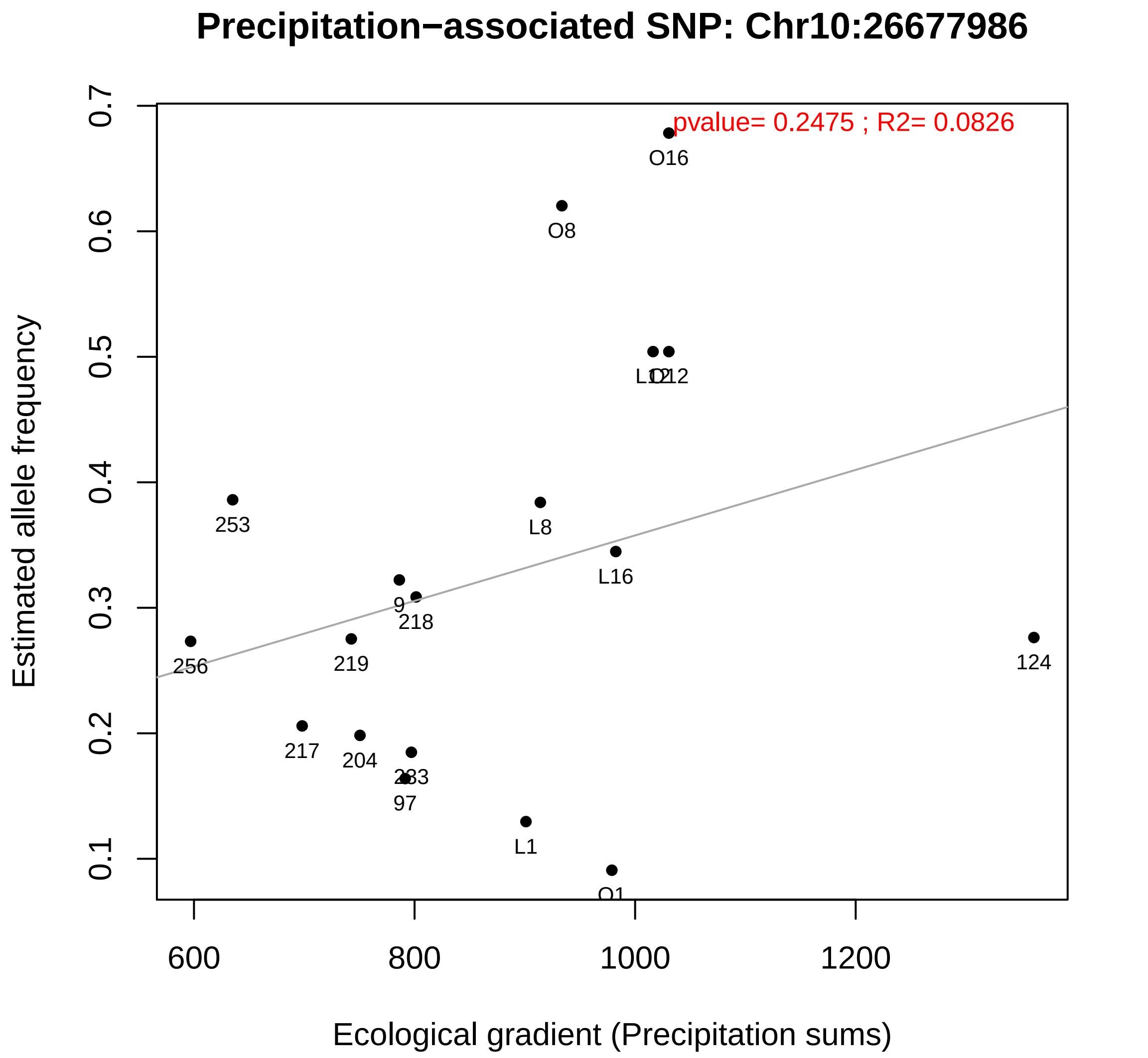

Precipitation−associated SNP: Chr10:26677986
pvalue= 0.2475 ; R2= 0.0826
Estimated allele frequency
0.1
0.2
0.3
0.4
0.5
0.6
0.7
600
800
1000
1200
Ecological gradient (Precipitation sums)
O16
O8
L12
O12
253
L8
L16
9
218
256
219
124
217
204
233
97
L1
O1

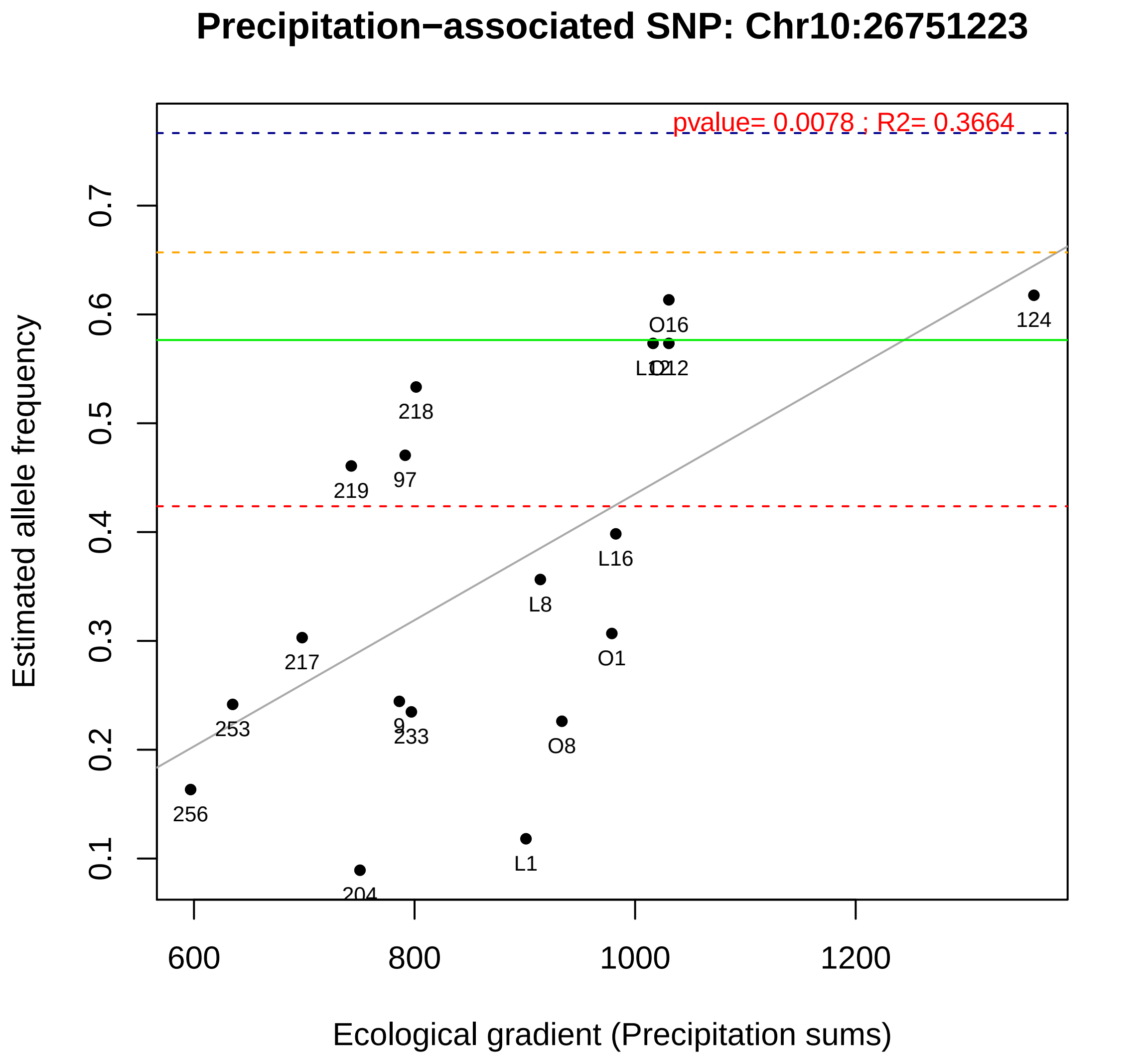

Precipitation−associated SNP: Chr10:26751223
pvalue= 0.0078 ; R2= 0.3664
Estimated allele frequency
0.1
0.2
0.3
0.4
0.5
0.6
0.7
600
800
1000
1200
Ecological gradient (Precipitation sums)
O16
124
L12
O12
218
219
97
L16
L8
217
O1
253
9
233
O8
256
L1
204

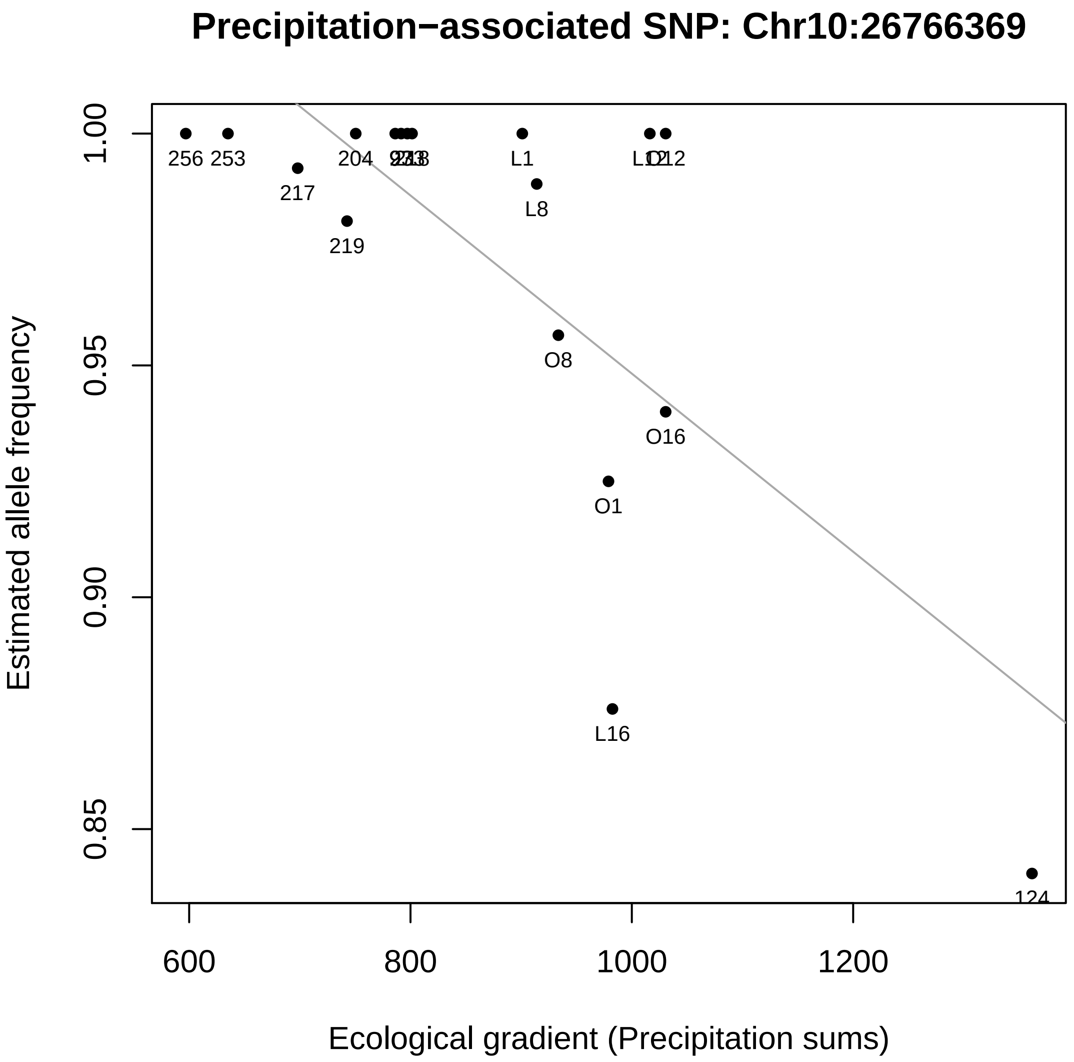

Precipitation−associated SNP: Chr10:26766369
Estimated allele frequency
1.00
0.95
0.90
0.85
256
253
217
204
219
L1
L8
O8
O16
O1
L16
124
600
800
1000
1200
Ecological gradient (Precipitation sums)

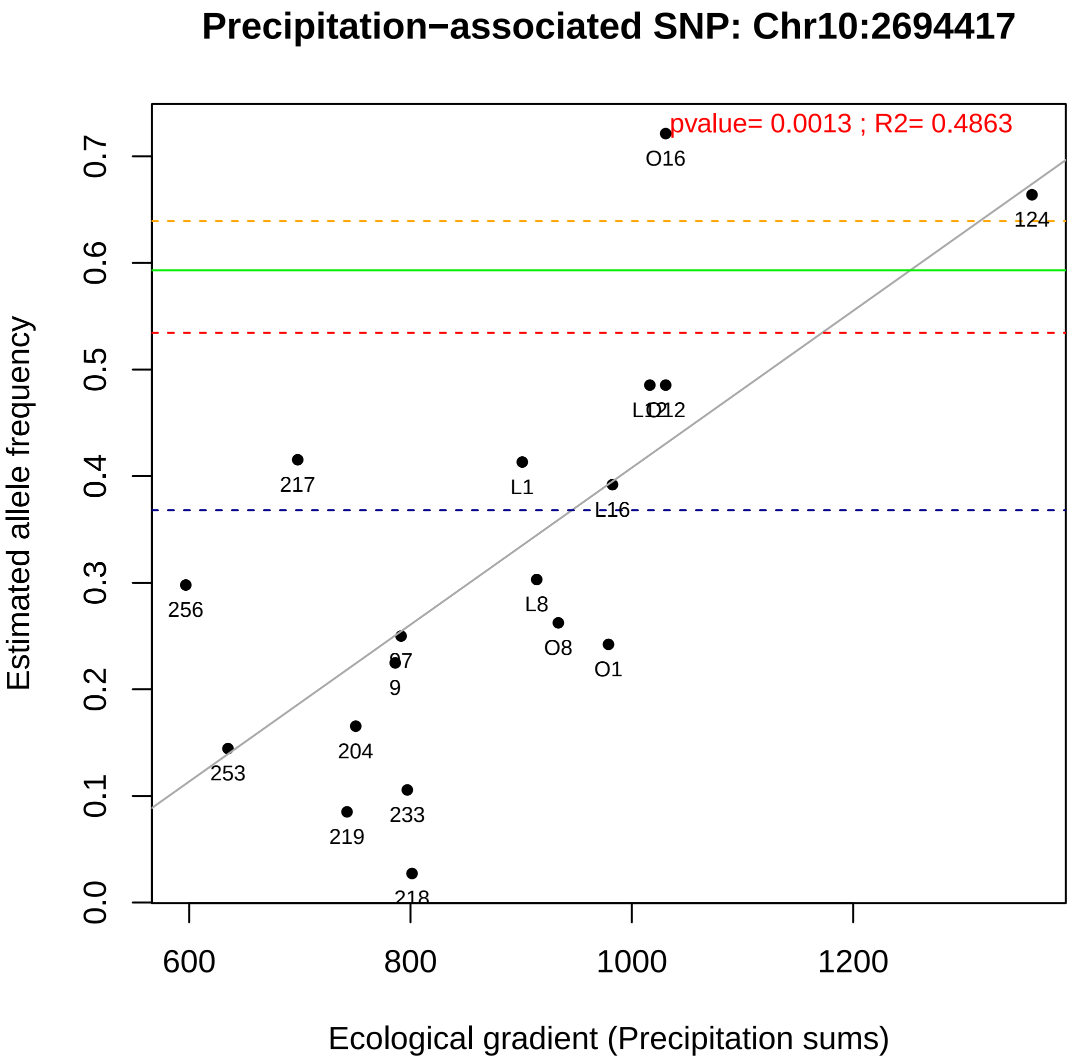

Precipitation−associated SNP: Chr10:2694417
pvalue= 0.0013 ; R2= 0.4863
O16
124
L12
O12
217
L1
L16
256
L8
O8
O1
07
9
204
253
233
219
218
0.0
0.1
0.2
0.3
0.4
0.5
0.6
0.7
600
800
1000
1200
Estimated allele frequency
Ecological gradient (Precipitation sums)

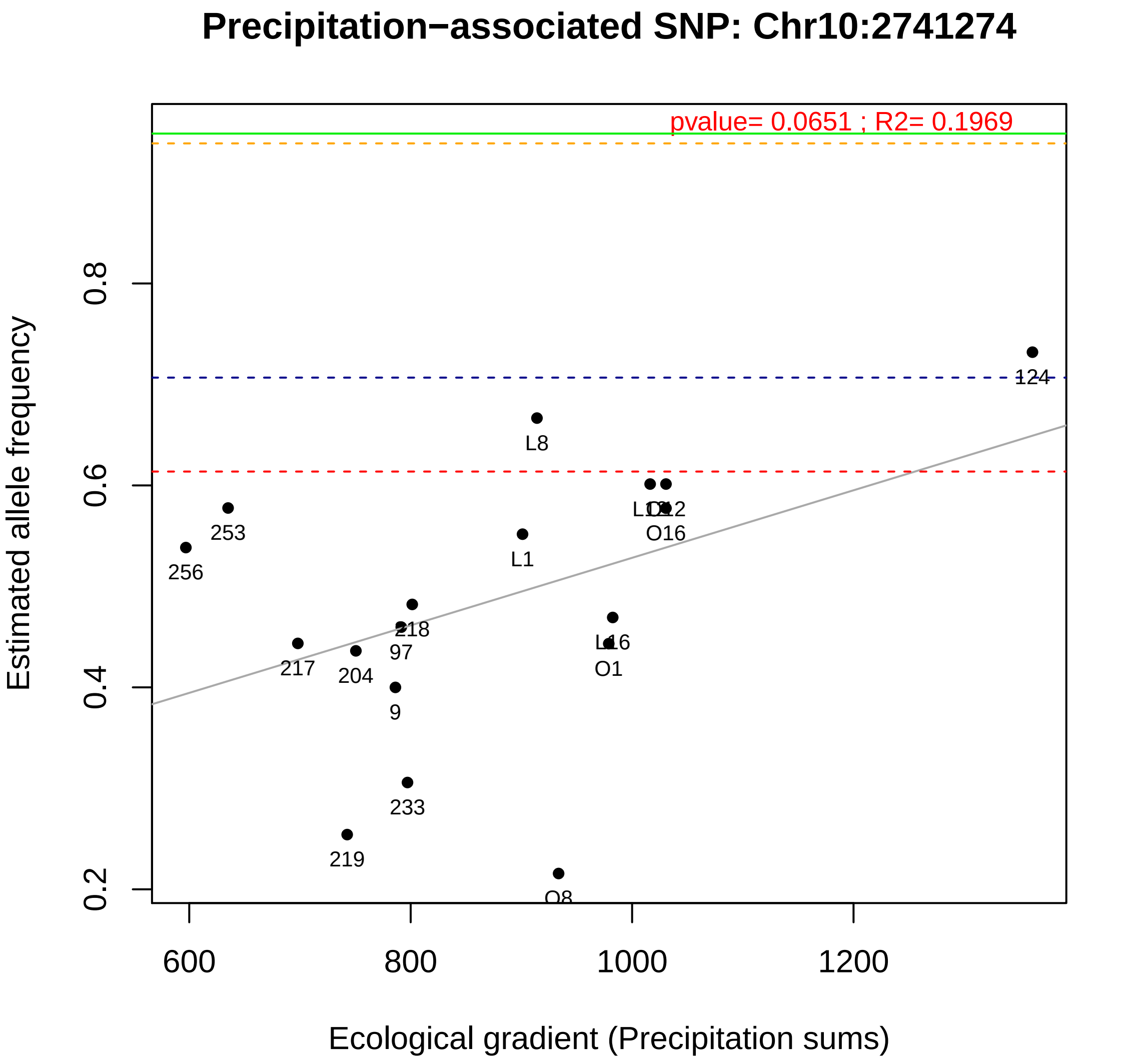
Precipitation−associated SNP: Chr10:2741274
pvalue= 0.0651 ; R2= 0.1969
Estimated allele frequency
0.2
0.4
0.6
0.8
Ecological gradient (Precipitation sums)
600
800
1000
1200
124
L8
253
256
L1
O16
218
97
217
204
9
O1
233
219

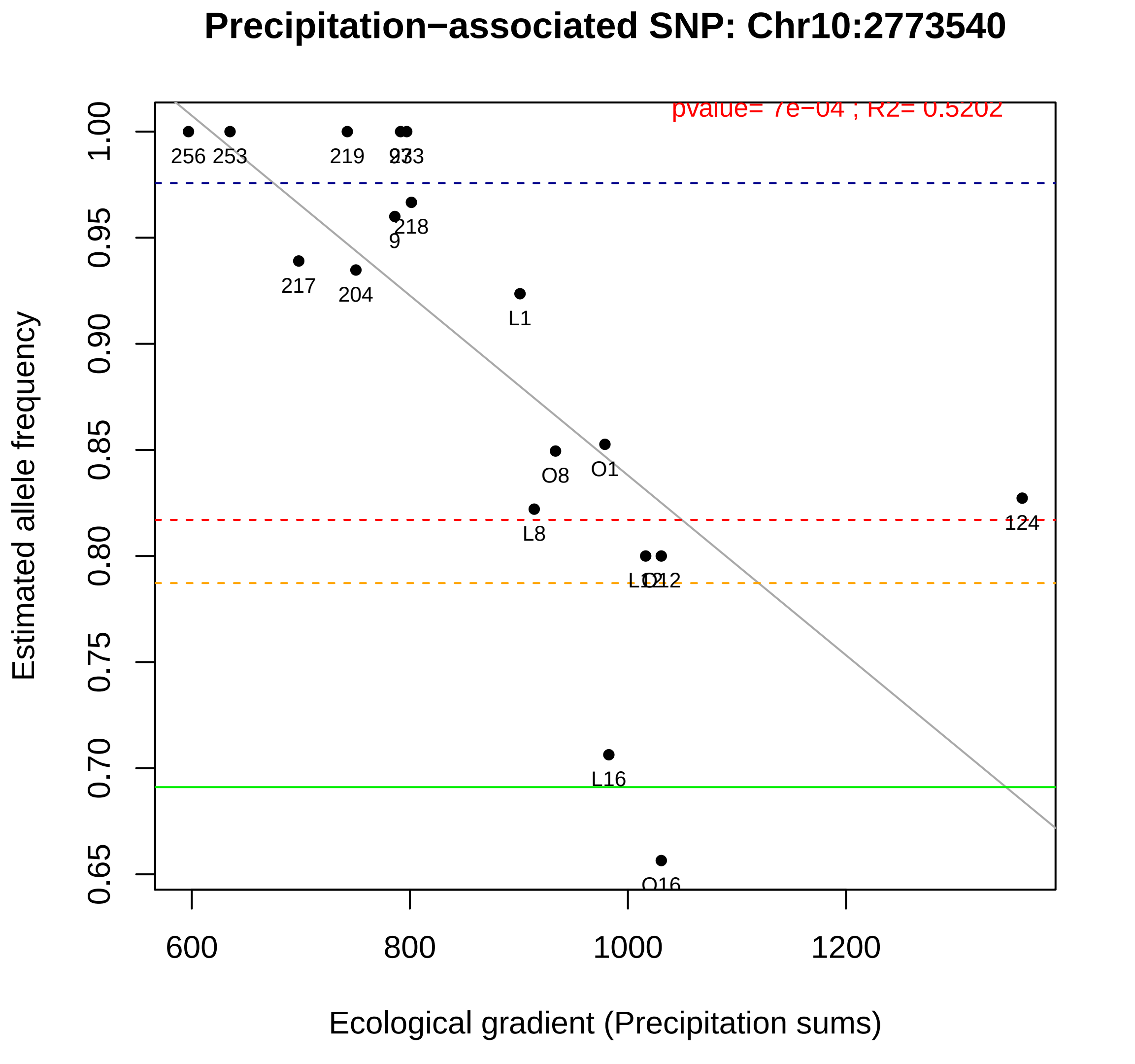

Precipitation−associated SNP: Chr10:2773540
pvalue= 7e−04 ; R2= 0.5202
Estimated allele frequency
Ecological gradient (Precipitation sums)
1.00
0.95
0.90
0.85
0.80
0.75
0.70
0.65
600
800
1000
1200
256
253
219
218
9
217
204
L1
O8
O1
L8
124
L16
O16

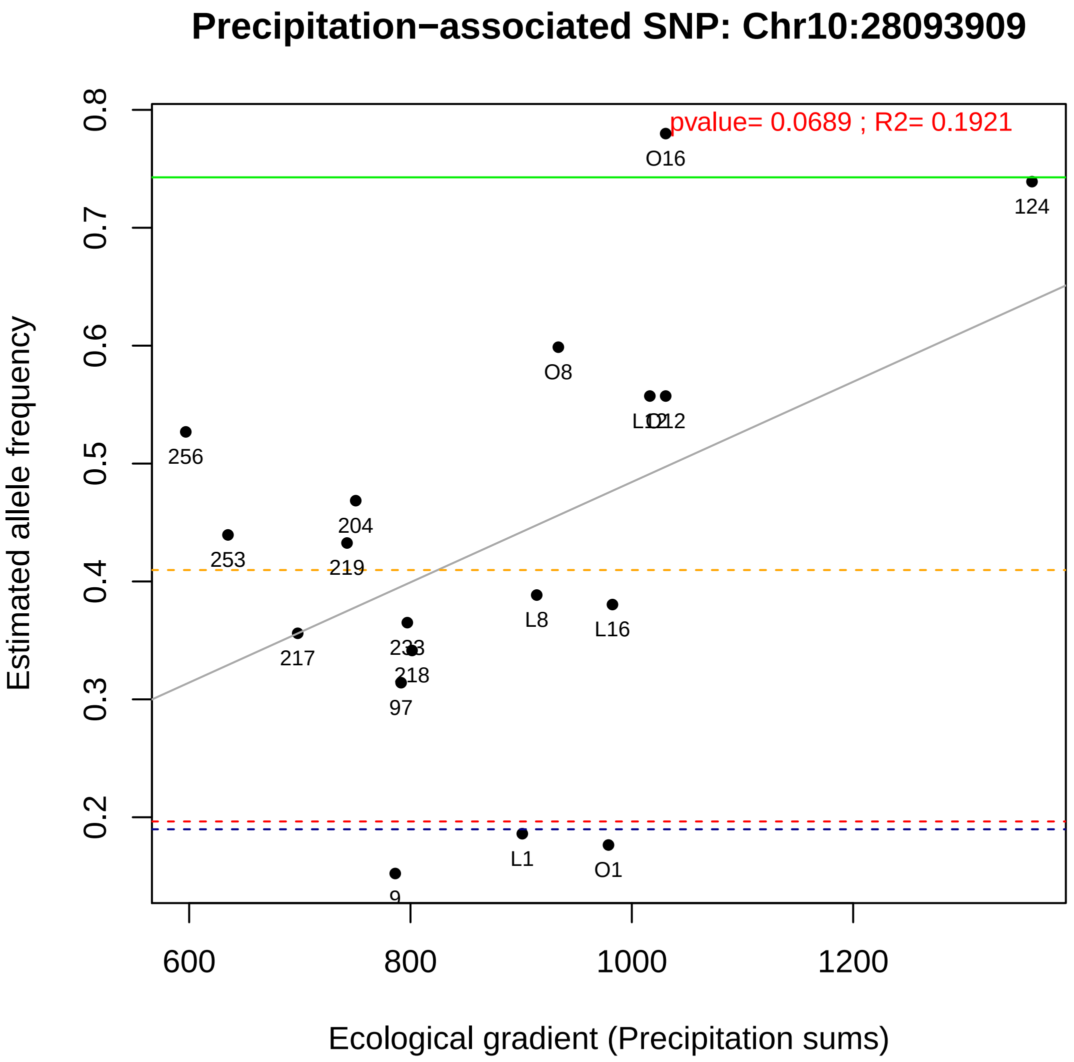

Precipitation−associated SNP: Chr10:28093909
pvalue= 0.0689 ; R2= 0.1921
Estimated allele frequency
0.2
0.3
0.4
0.5
0.6
0.7
0.8
600
800
1000
1200
Ecological gradient (Precipitation sums)
O16
124
O8
L12
O12
256
204
253
219
217
233
218
97
L8
L16
L1
O1
9

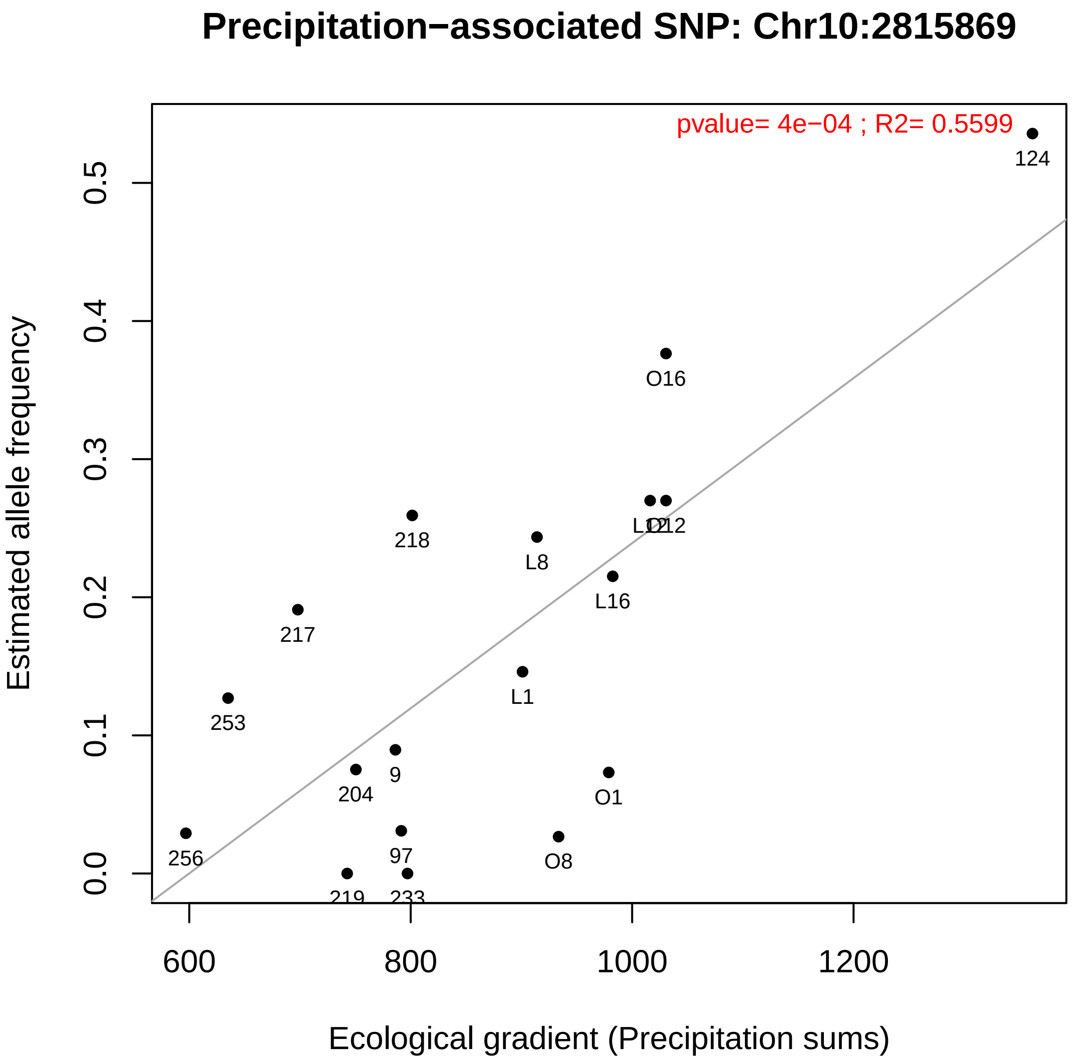

Precipitation−associated SNP: Chr10:2815869
pvalue= 4e−04 ; R2= 0.5599
Estimated allele frequency
Ecological gradient (Precipitation sums)
0.0
0.1
0.2
0.3
0.4
0.5
600
800
1000
1200
124
O16
L12
O12
218
L8
L16
217
L1
253
9
204
O1
256
97
O8
219
233

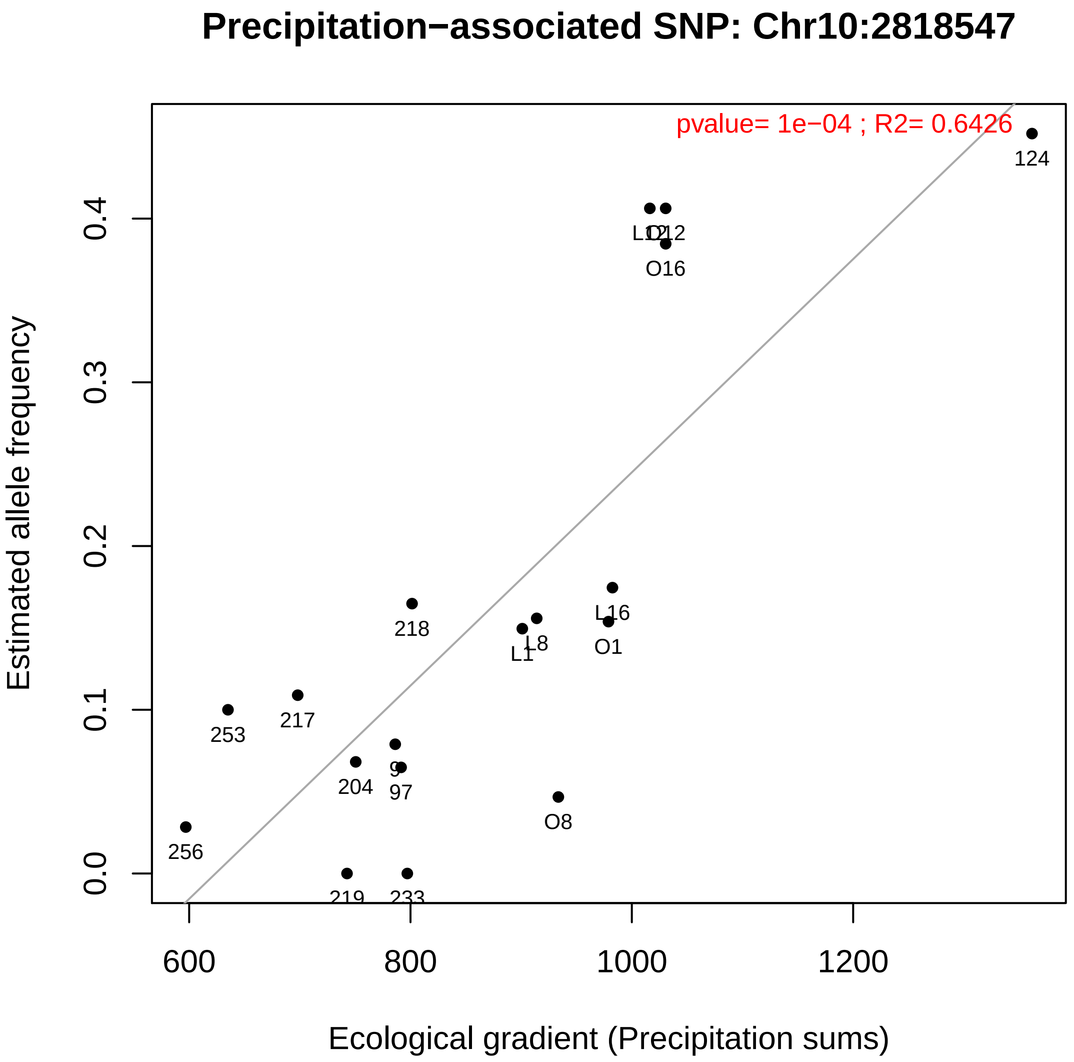
Precipitation−associated SNP: Chr10:2818547
pvalue= 1e−04 ; R2= 0.6426
Estimated allele frequency
Ecological gradient (Precipitation sums)
0.0
0.1
0.2
0.3
0.4
600
800
1000
1200
124
L12
O12
O16
218
L16
L1
L8
O1
253
217
204
97
O8
256
219
233

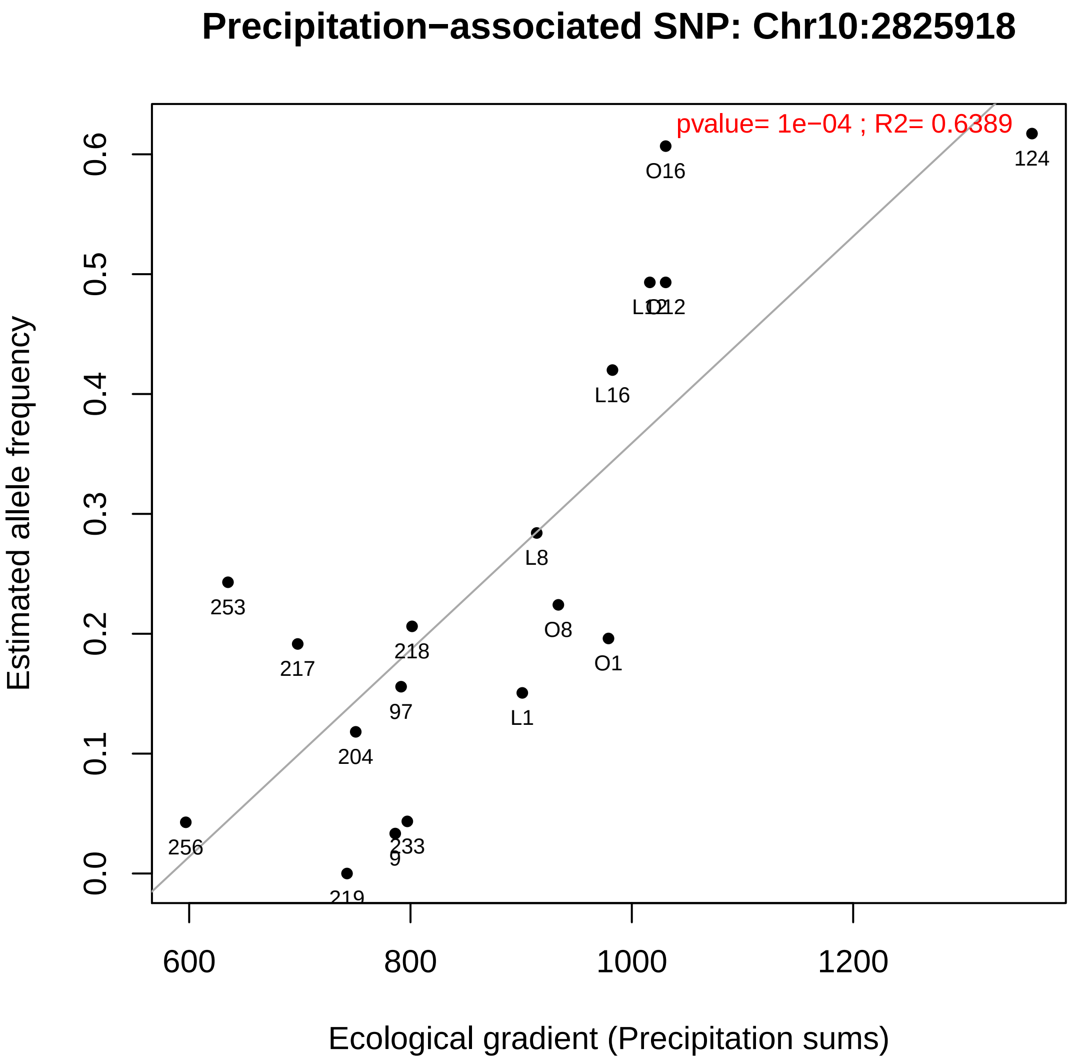
Precipitation−associated SNP: Chr10:2825918
pvalue= 1e−04 ; R2= 0.6389
O16
124
L10
O12
L16
L8
O8
O1
253
217
218
97
L1
204
256
233
9
219
0.0
0.1
0.2
0.3
0.4
0.5
0.6
600
800
1000
1200
Estimated allele frequency
Ecological gradient (Precipitation sums)

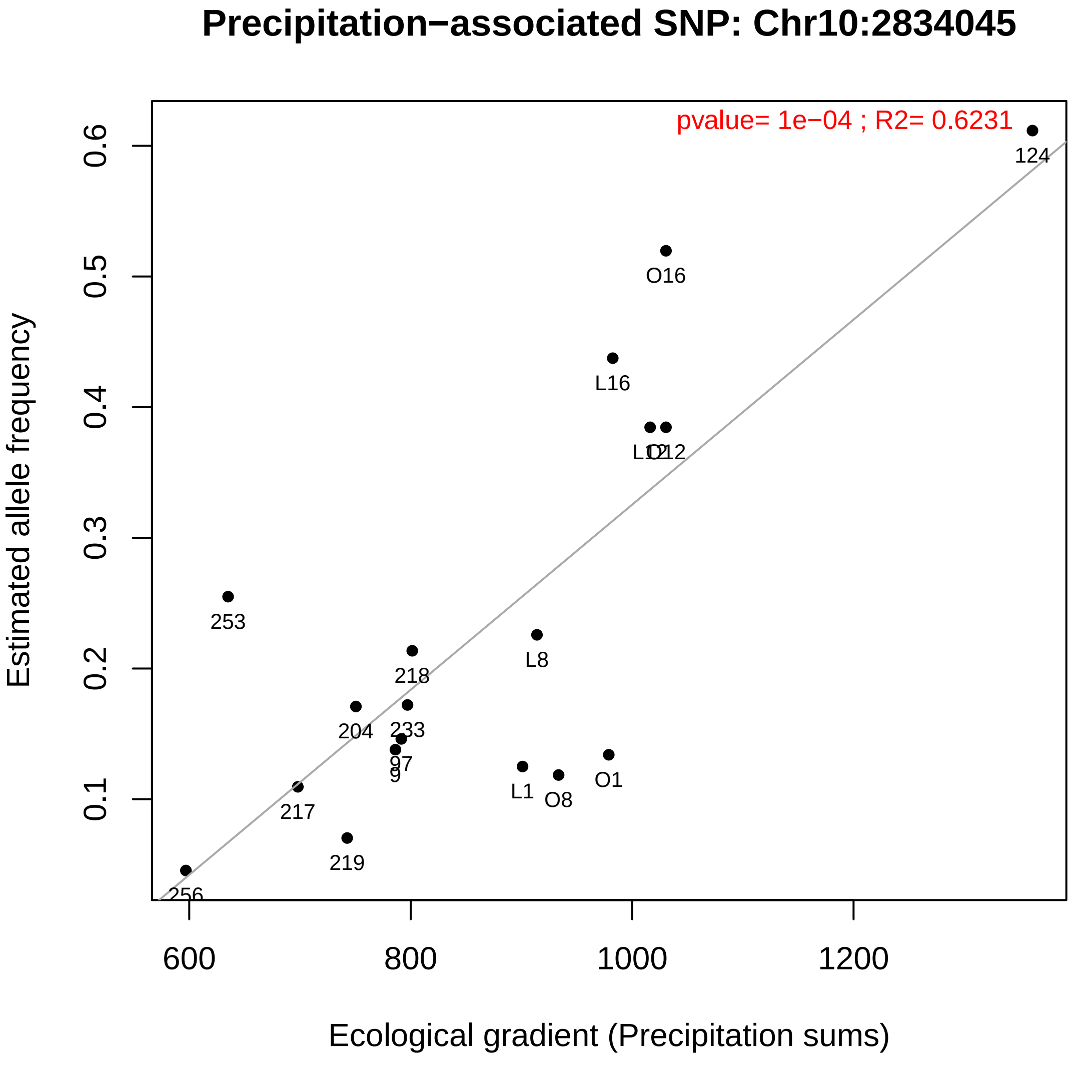
Precipitation−associated SNP: Chr10:2834045
pvalue= 1e−04 ; R2= 0.6231
Estimated allele frequency
0.1
0.2
0.3
0.4
0.5
0.6
Ecological gradient (Precipitation sums)
600
800
1000
1200
124
O16
L16
253
L8
218
204
233
97
9
217
219
256
L1
O8
O1

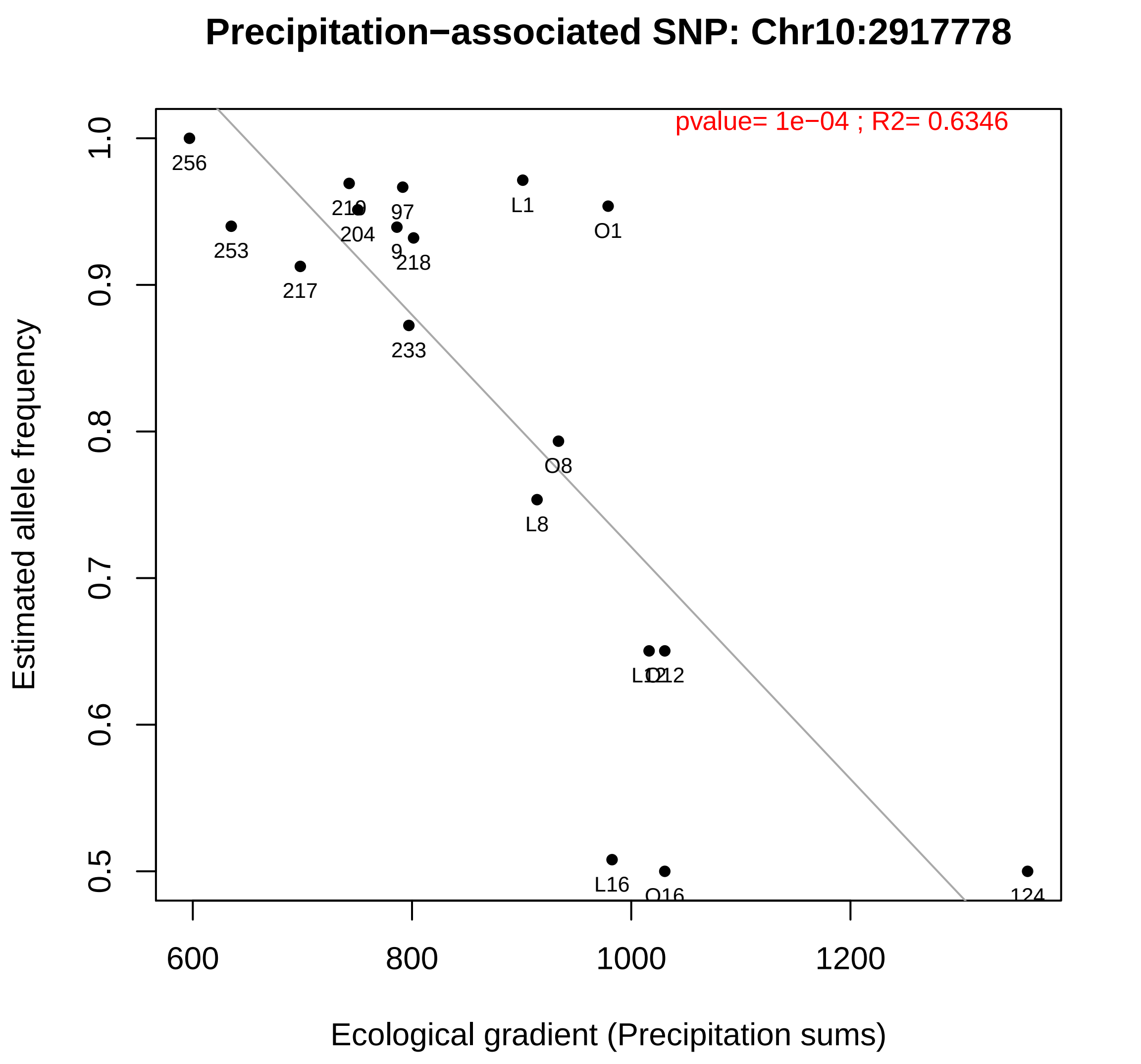
Precipitation−associated SNP: Chr10:2917778
pvalue= 1e−04 ; R2= 0.6346
Estimated allele frequency
Ecological gradient (Precipitation sums)
1.0
0.9
0.8
0.7
0.6
0.5
600
800
1000
1200
256
253
217
210
204
97
9
218
233
L1
O1
O8
L8
L12
O12
L16
O16
124

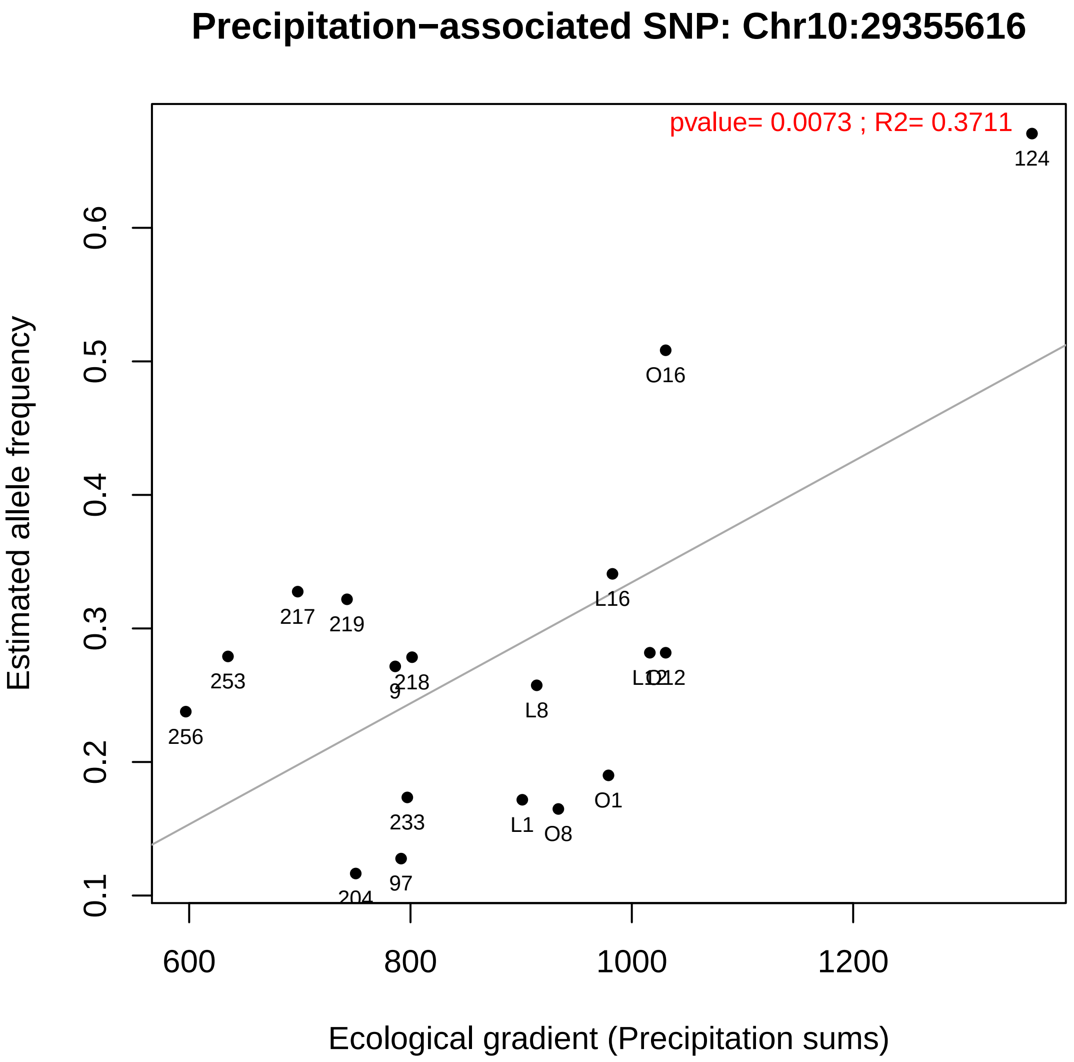
Precipitation−associated SNP: Chr10:29355616
pvalue= 0.0073 ; R2= 0.3711
Estimated allele frequency
Ecological gradient (Precipitation sums)
0.1
0.2
0.3
0.4
0.5
0.6
600
800
1000
1200
124
O16
L16
217
219
253
218
9
L8
L12
O12
256
O1
233
L1
O8
97
204

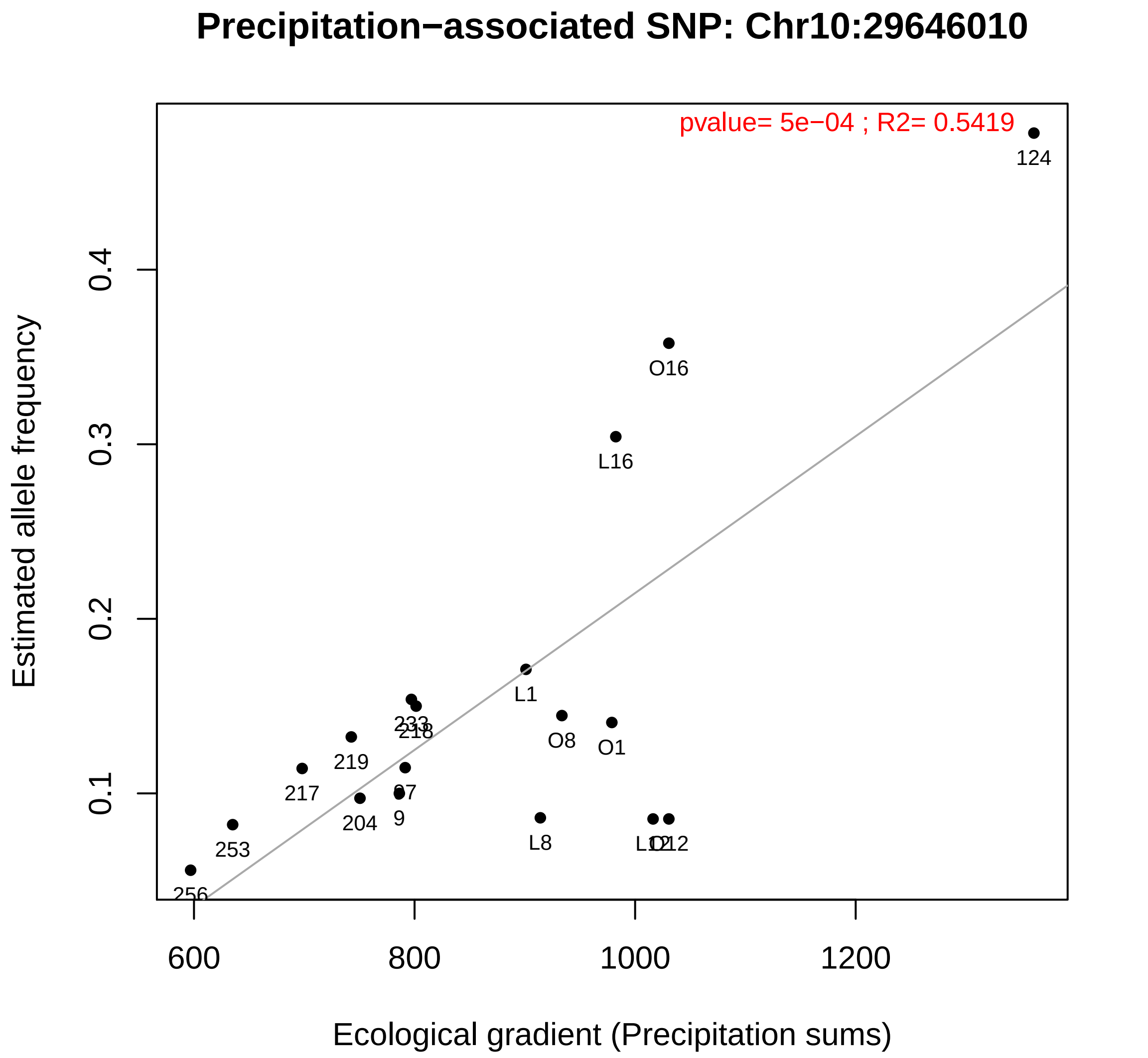

Precipitation−associated SNP: Chr10:29646010
pvalue= 5e−04 ; R2= 0.5419
124
O16
L16
L1
233
218
219
O8
O1
217
97
204
9
L8
L12
O12
253
256
0.1
0.2
0.3
0.4
600
800
1000
1200
Estimated allele frequency
Ecological gradient (Precipitation sums)

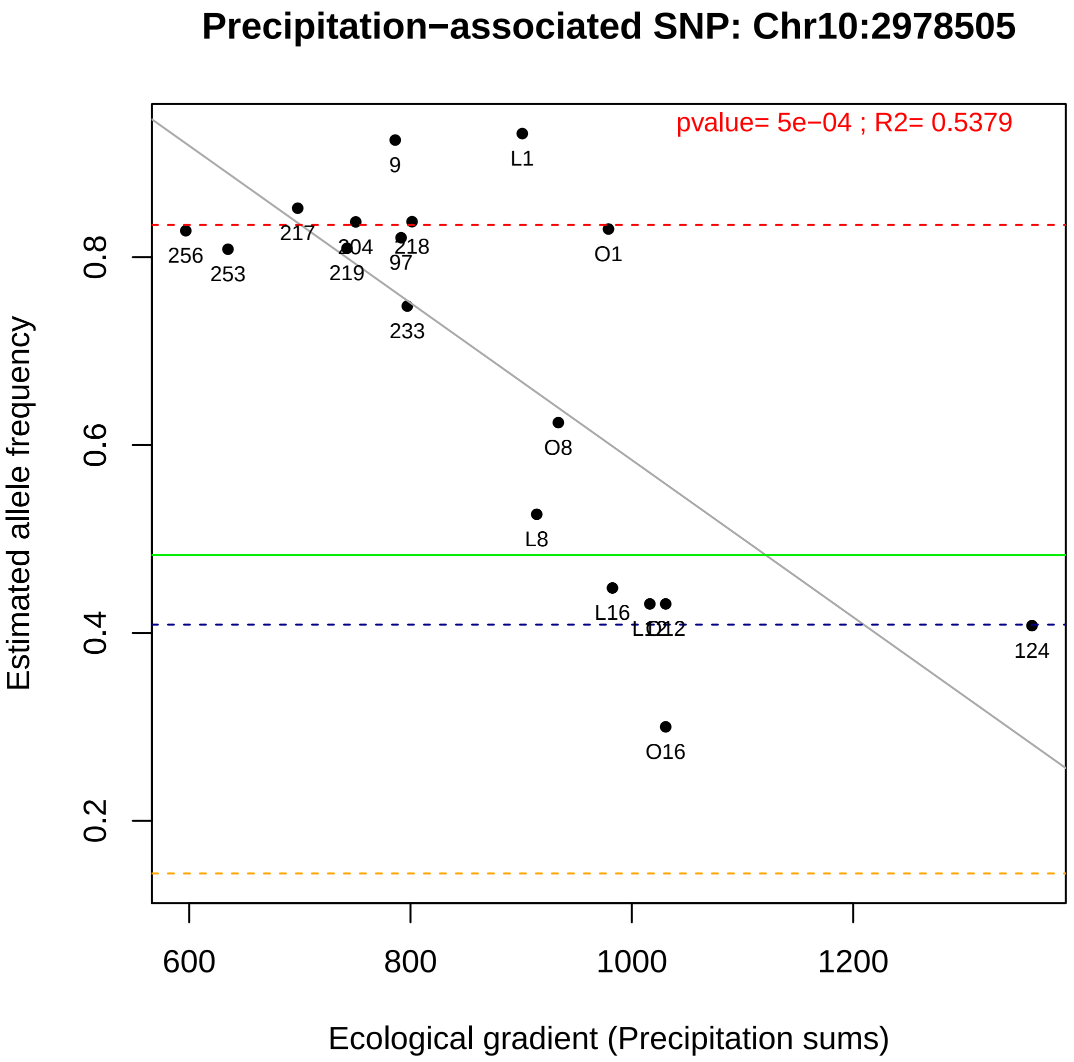

Precipitation−associated SNP: Chr10:2978505
pvalue= 5e−04 ; R2= 0.5379
9
L1
217
256
253
204
219
218
97
O1
233
O8
L8
L16
L12
O12
124
O16
0.2
0.4
0.6
0.8
600
800
1000
1200
Estimated allele frequency
Ecological gradient (Precipitation sums)

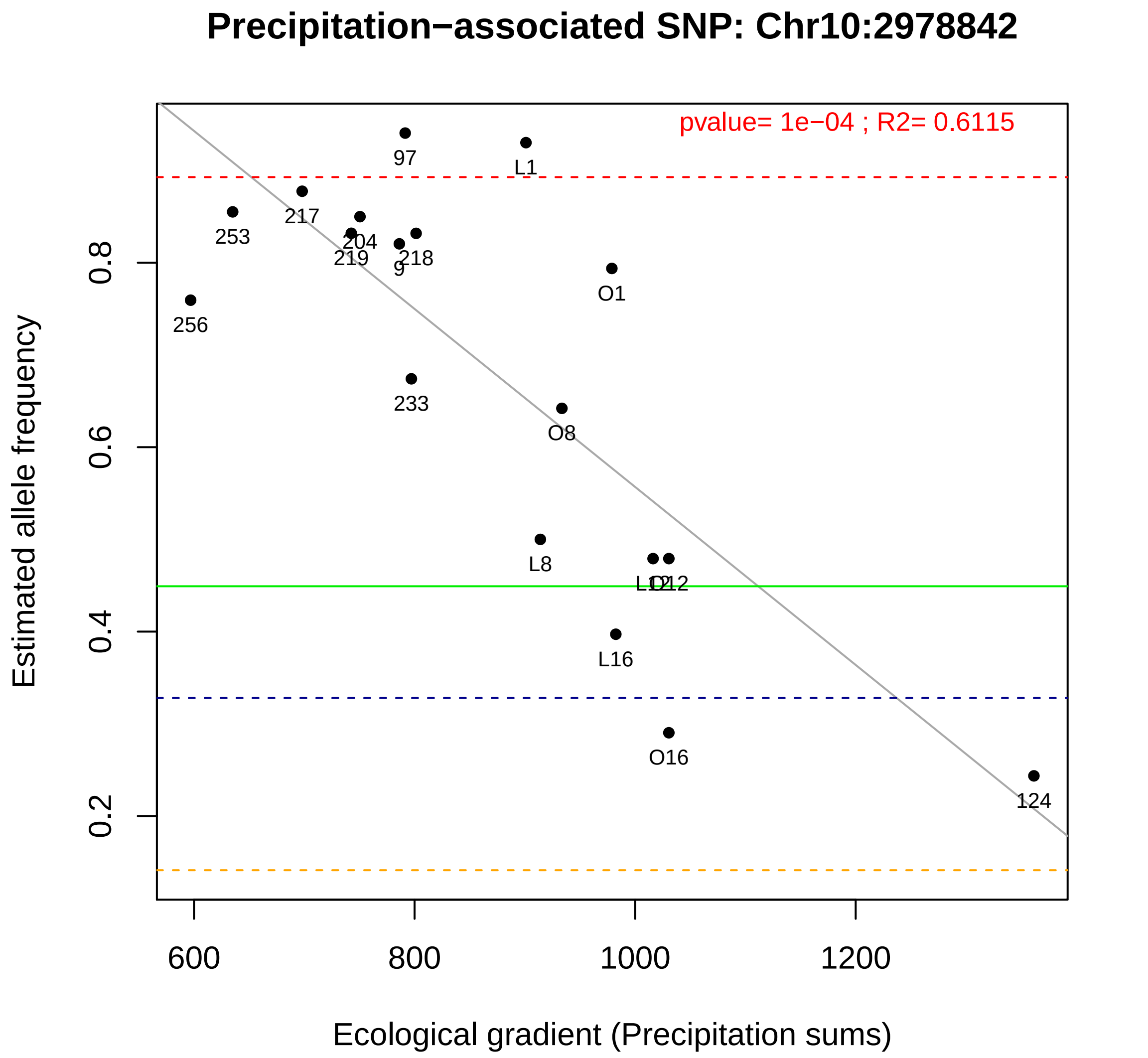

Precipitation−associated SNP: Chr10:2978842
pvalue= 1e−04 ; R2= 0.6115
97
L1
217
253
204
219
218
9
O1
256
233
O8
L8
L10
O12
L16
O16
124
0.2
0.4
0.6
0.8
Estimated allele frequency
600
800
1000
1200
Ecological gradient (Precipitation sums)

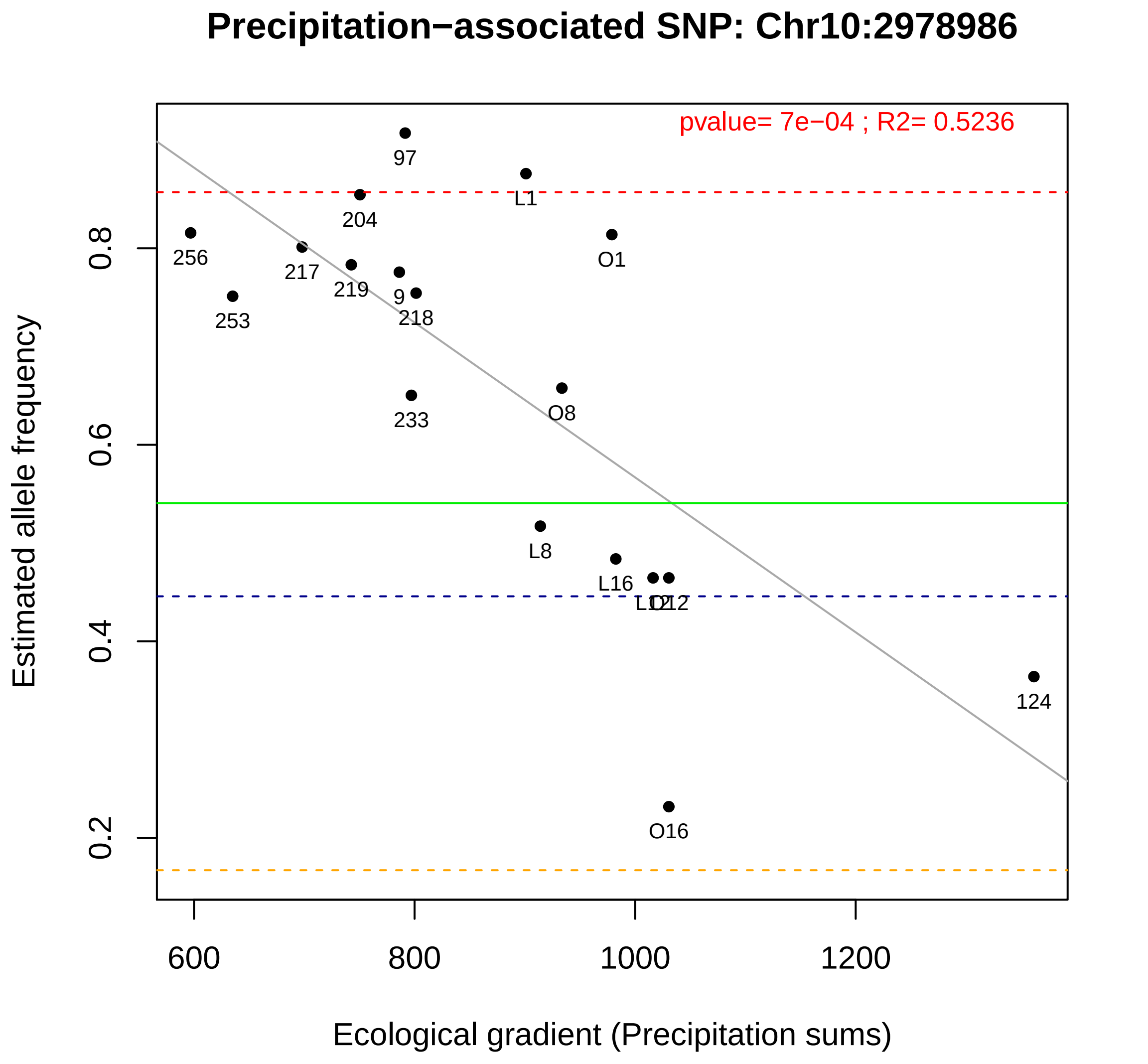
Precipitation−associated SNP: Chr10:2978986
pvalue= 7e−04 ; R2= 0.5236
Estimated allele frequency
0.2
0.4
0.6
0.8
600
800
1000
1200
Ecological gradient (Precipitation sums)
97
L1
204
256
217
O1
219
9
218
253
233
O8
L8
L16
124
O16

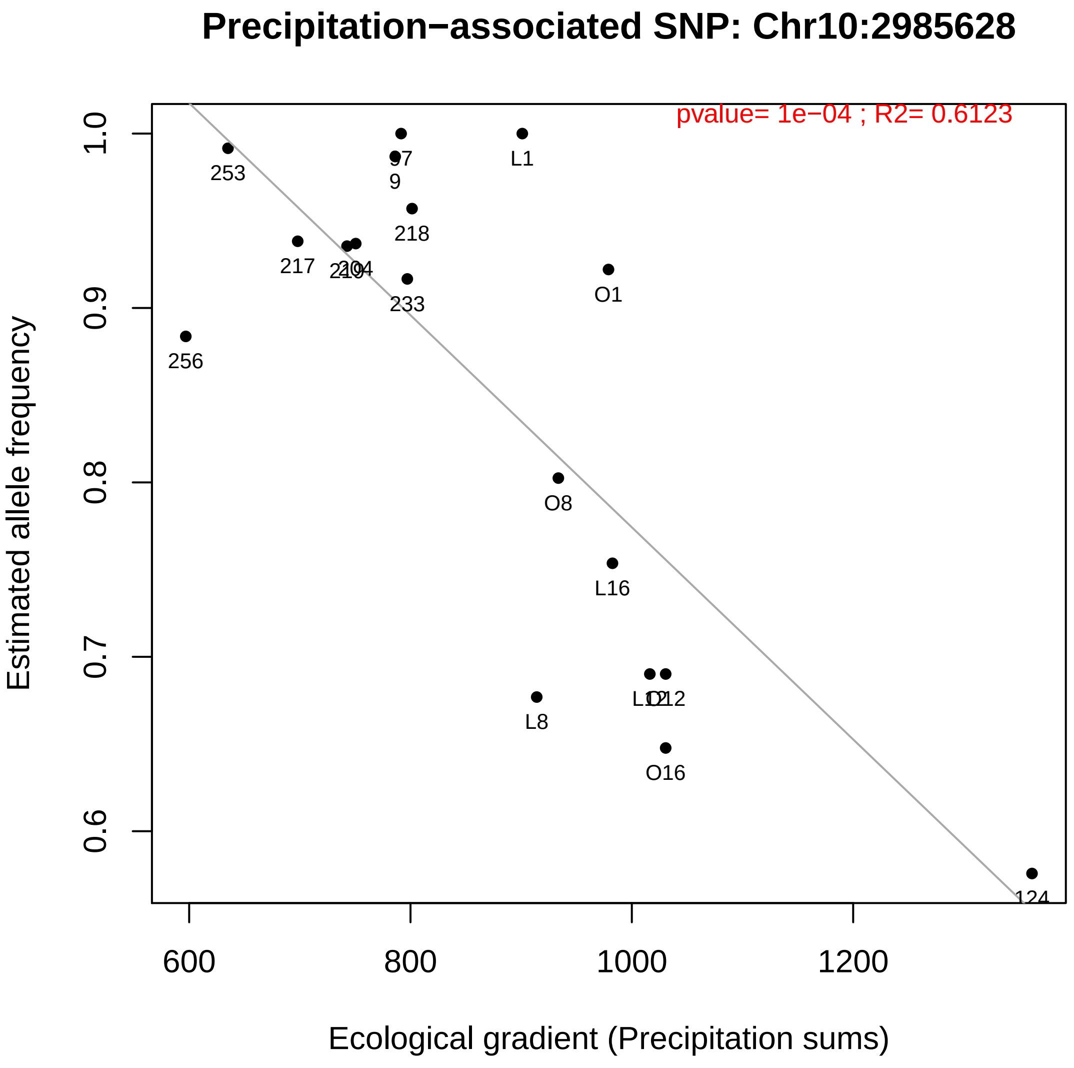
Precipitation−associated SNP: Chr10:2985628
pvalue= 1e−04 ; R2= 0.6123
Estimated allele frequency
Ecological gradient (Precipitation sums)
0.6
0.7
0.8
0.9
1.0
600
800
1000
1200
253
97
9
L1
218
217
204
210
233
O1
256
O8
L16
L8
L12
O12
O16
124

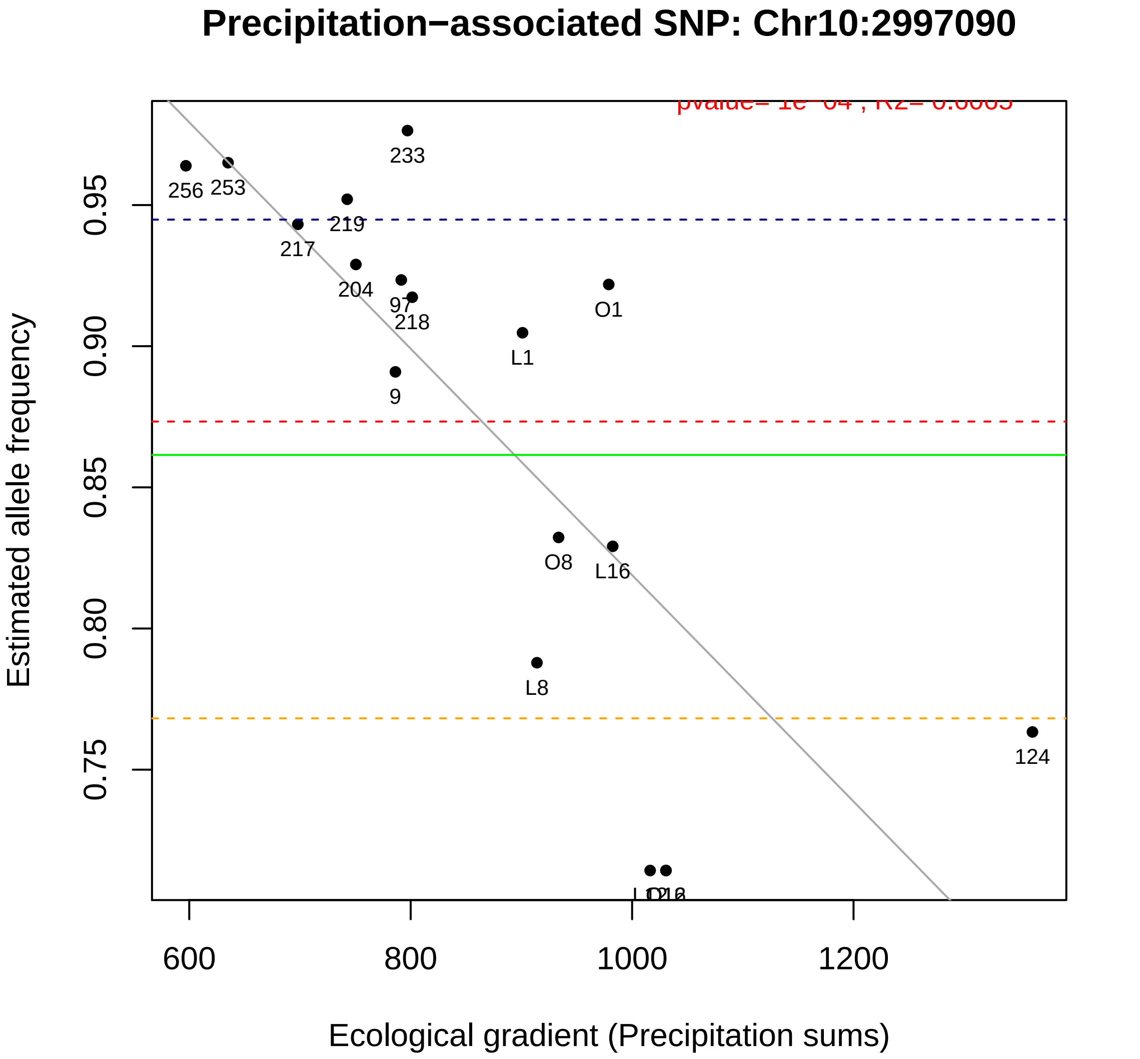
Precipitation−associated SNP: Chr10:2997090
Estimated allele frequency
0.75
0.80
0.85
0.90
0.95
600
800
1000
1200
Ecological gradient (Precipitation sums)
256
253
233
219
217
204
97
218
9
O1
L1
O8
L16
L8
124

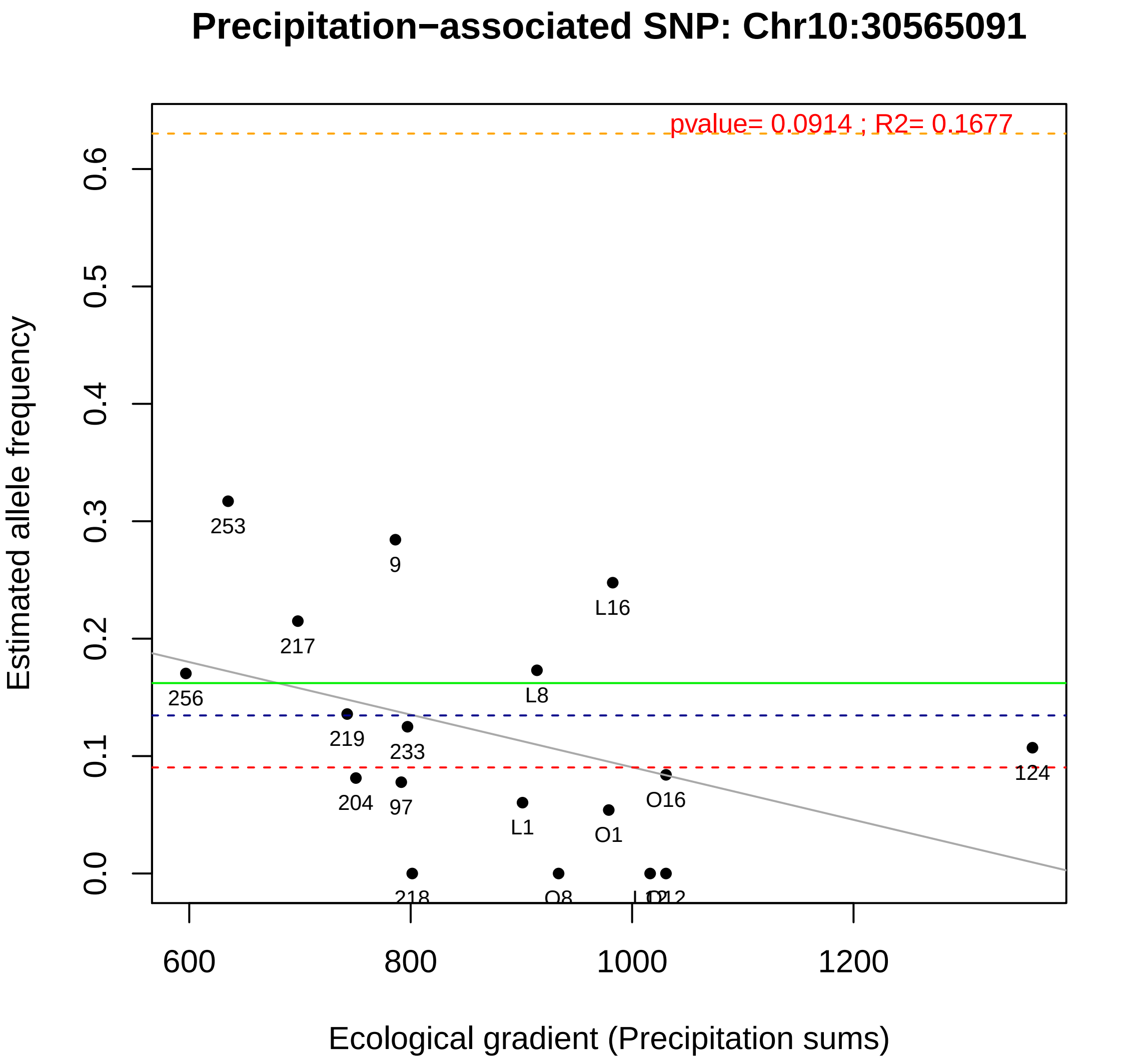
Precipitation−associated SNP: Chr10:30565091
pvalue= 0.0914 ; R2= 0.1677
Estimated allele frequency
0.0
0.1
0.2
0.3
0.4
0.5
0.6
Ecological gradient (Precipitation sums)
600
800
1000
1200
253
9
L16
217
256
L8
219
233
124
204
97
O16
L1
O1
218
O8

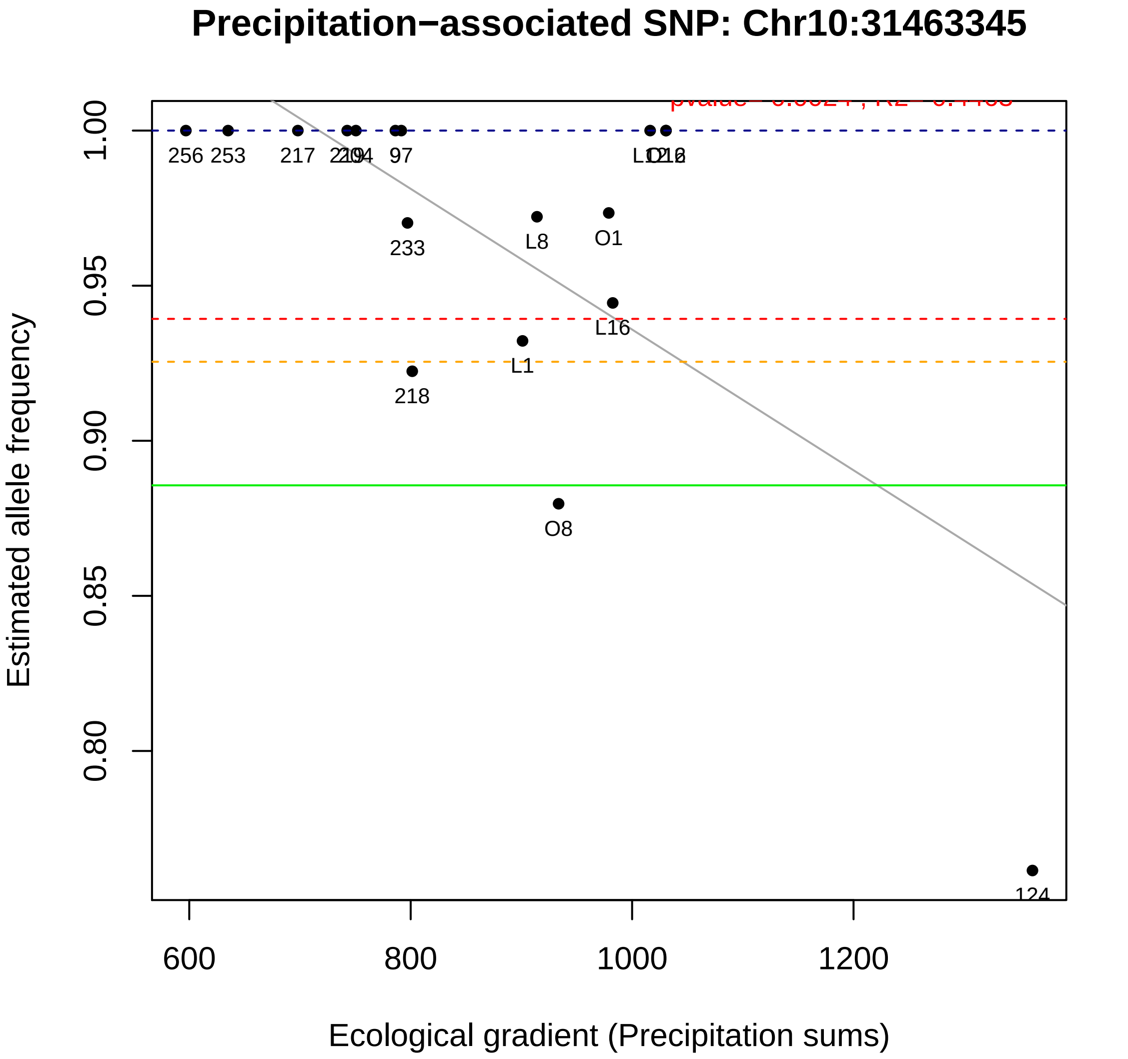

Precipitation−associated SNP: Chr10:31463345
Estimated allele frequency
Ecological gradient (Precipitation sums)
1.00
0.95
0.90
0.85
0.80
600
800
1000
1200
256
253
217
97
233
L8
O1
L16
L1
218
O8
124

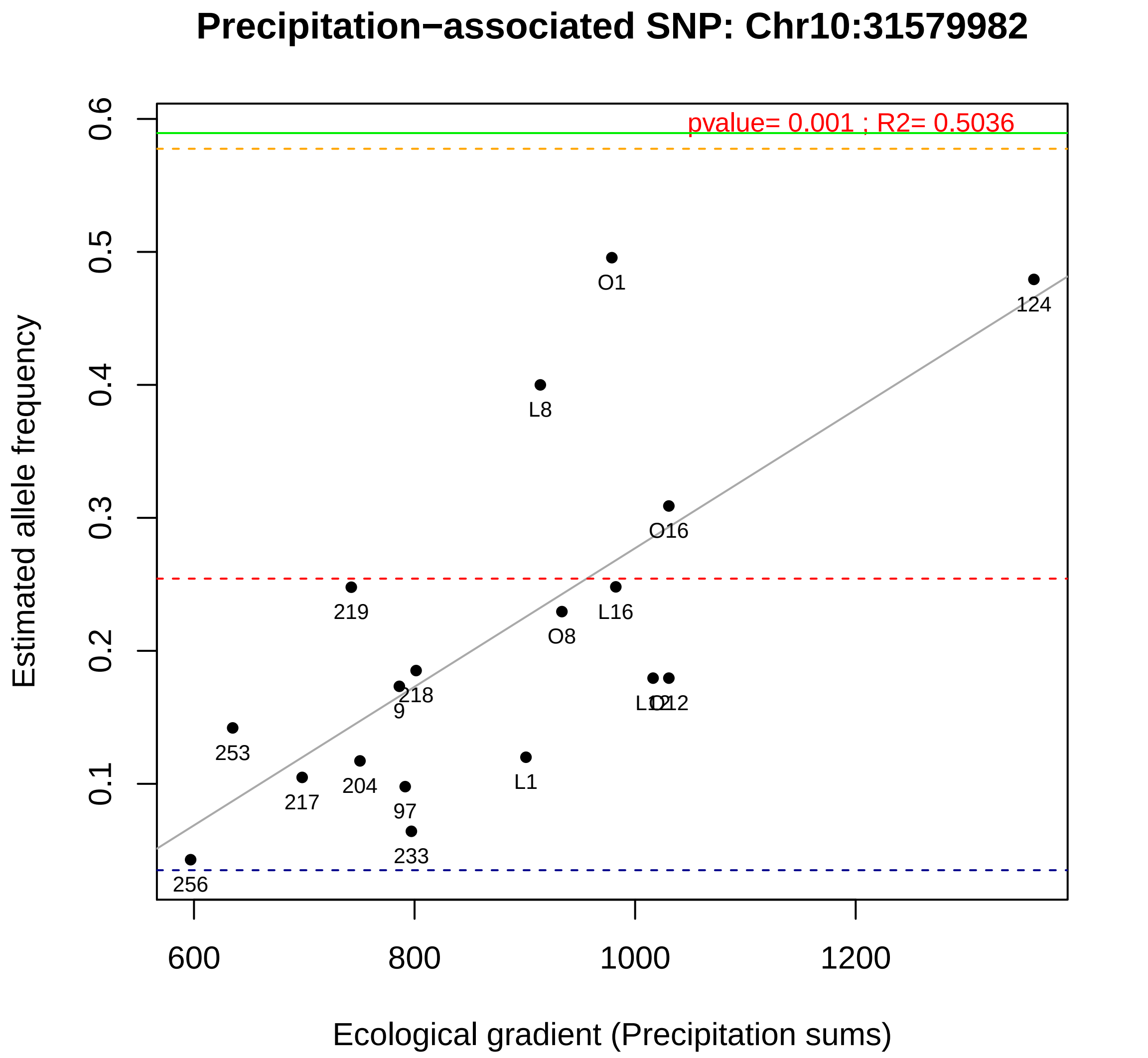
Precipitation−associated SNP: Chr10:31579982
pvalue= 0.001 ; R2= 0.5036
Estimated allele frequency
Ecological gradient (Precipitation sums)
0.1
0.2
0.3
0.4
0.5
0.6
600
800
1000
1200
O1
124
L8
O16
219
L16
O8
218
9
L12
O12
253
204
217
97
L1
233
256

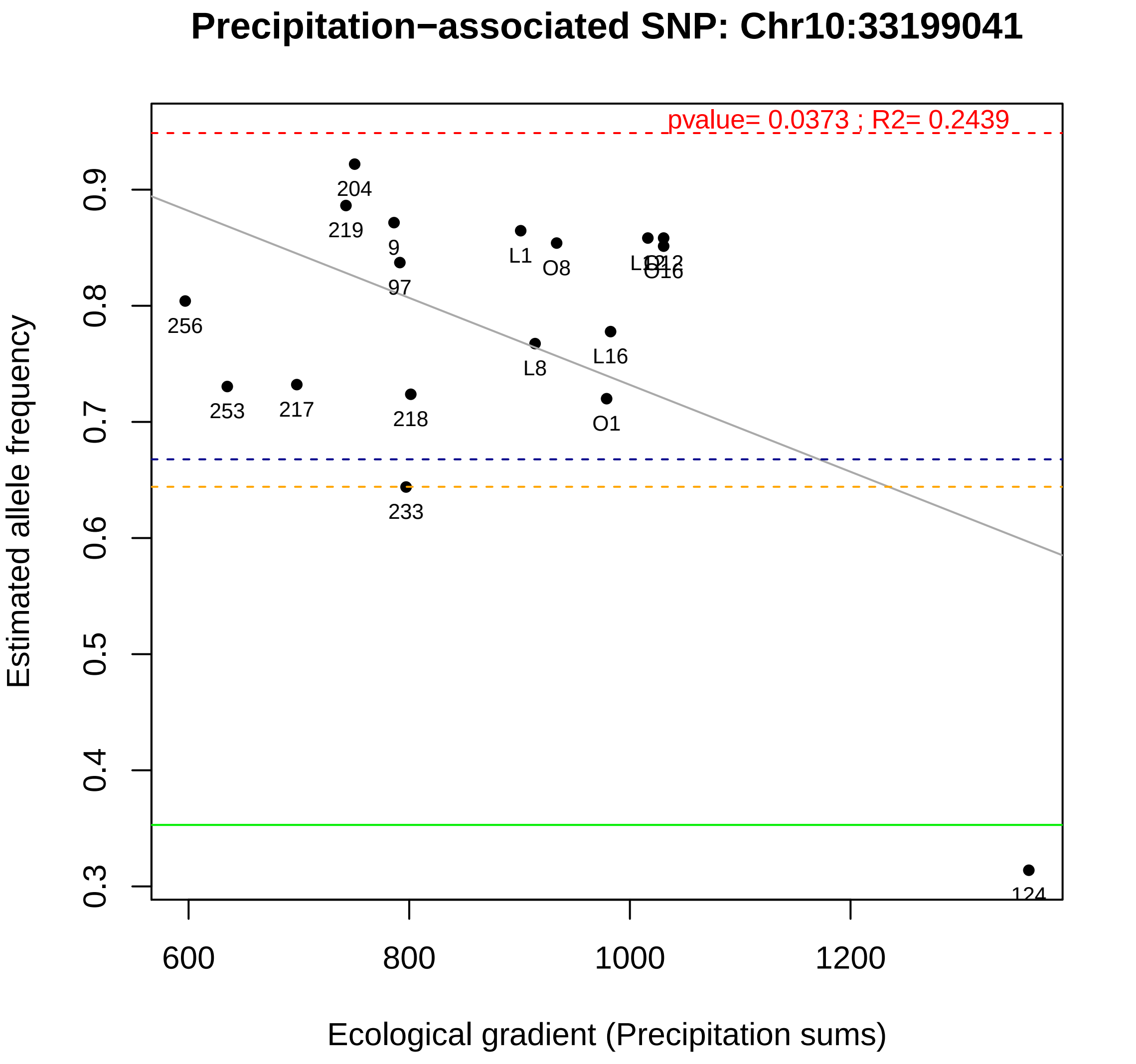
Precipitation−associated SNP: Chr10:33199041
pvalue= 0.0373 ; R2= 0.2439
Estimated allele frequency
0.3
0.4
0.5
0.6
0.7
0.8
0.9
600
800
1000
1200
Ecological gradient (Precipitation sums)
204
219
9
97
256
253
217
218
233
L1
O8
L8
L16
O1
O16
124

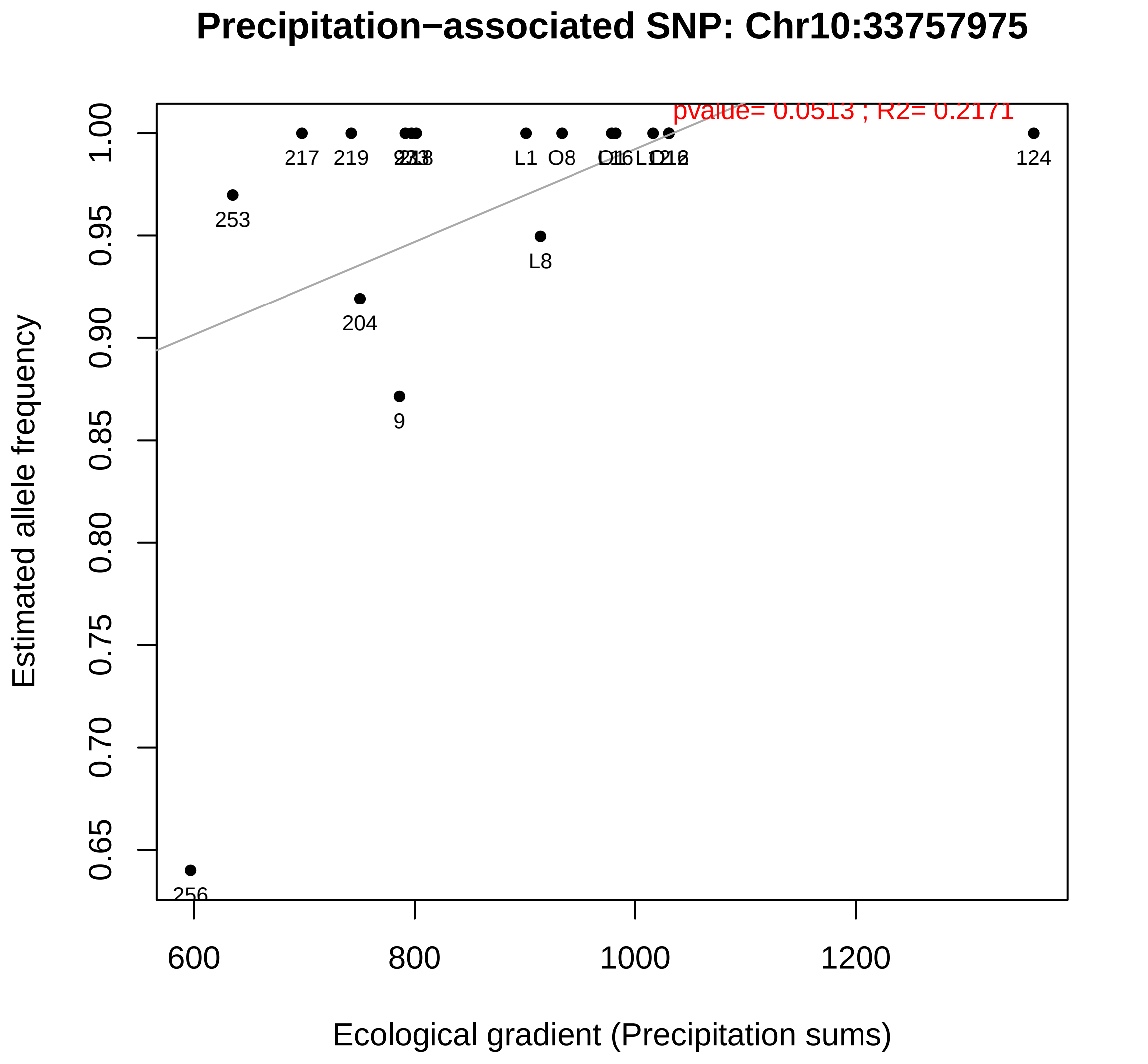

Precipitation−associated SNP: Chr10:33757975
pvalue= 0.0513 ; R2= 0.2171
Estimated allele frequency
Ecological gradient (Precipitation sums)
1.00
0.95
0.90
0.85
0.80
0.75
0.70
0.65
600
800
1000
1200
217
219
L1
O8
124
253
L8
204
9
256

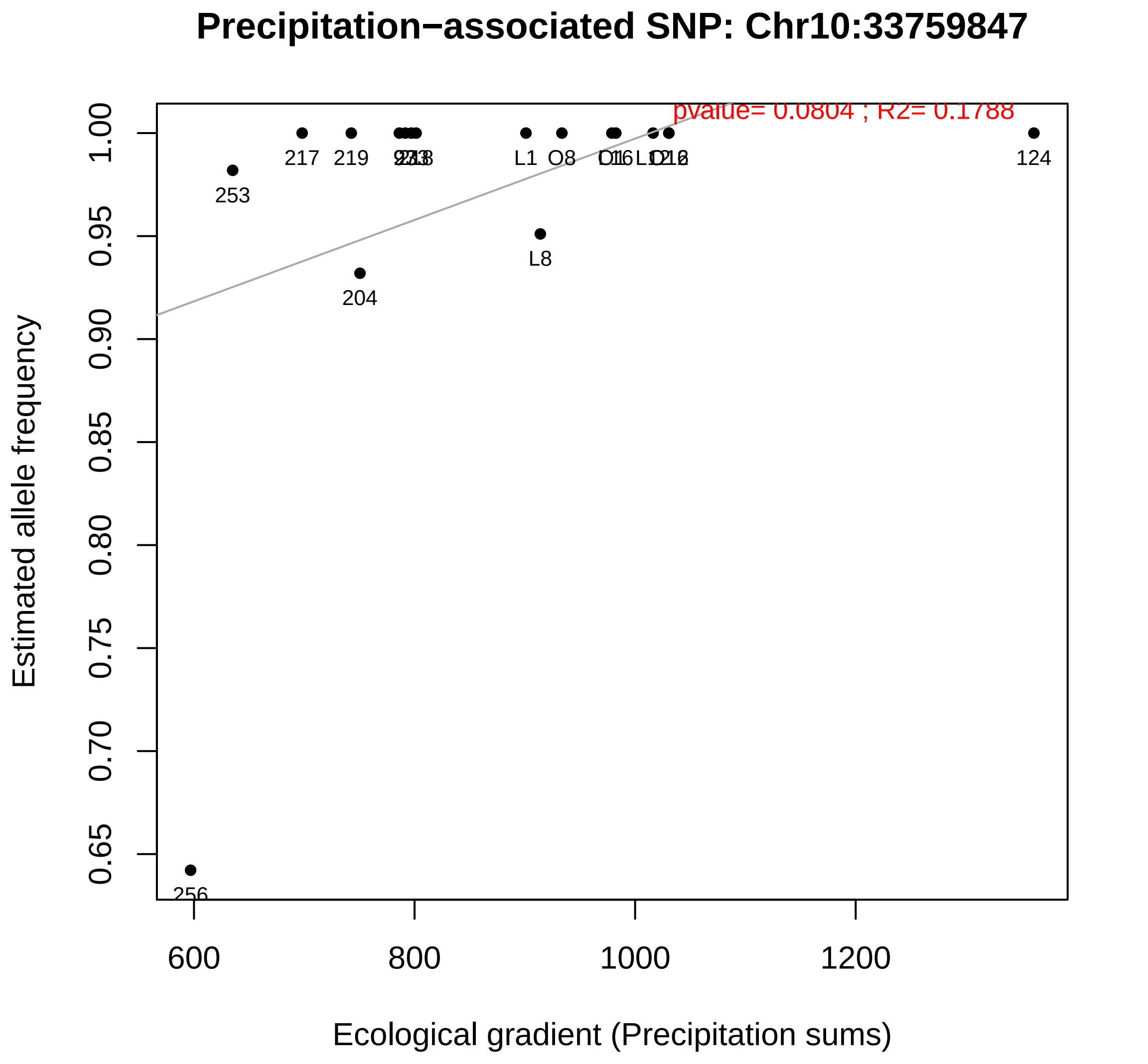

Precipitation−associated SNP: Chr10:33759847
pvalue= 0.0804 ; R2= 0.1788
Estimated allele frequency
1.00
0.95
0.90
0.85
0.80
0.75
0.70
0.65
600
800
1000
1200
Ecological gradient (Precipitation sums)
253
217
219
204
L1
O8
L8
124
256

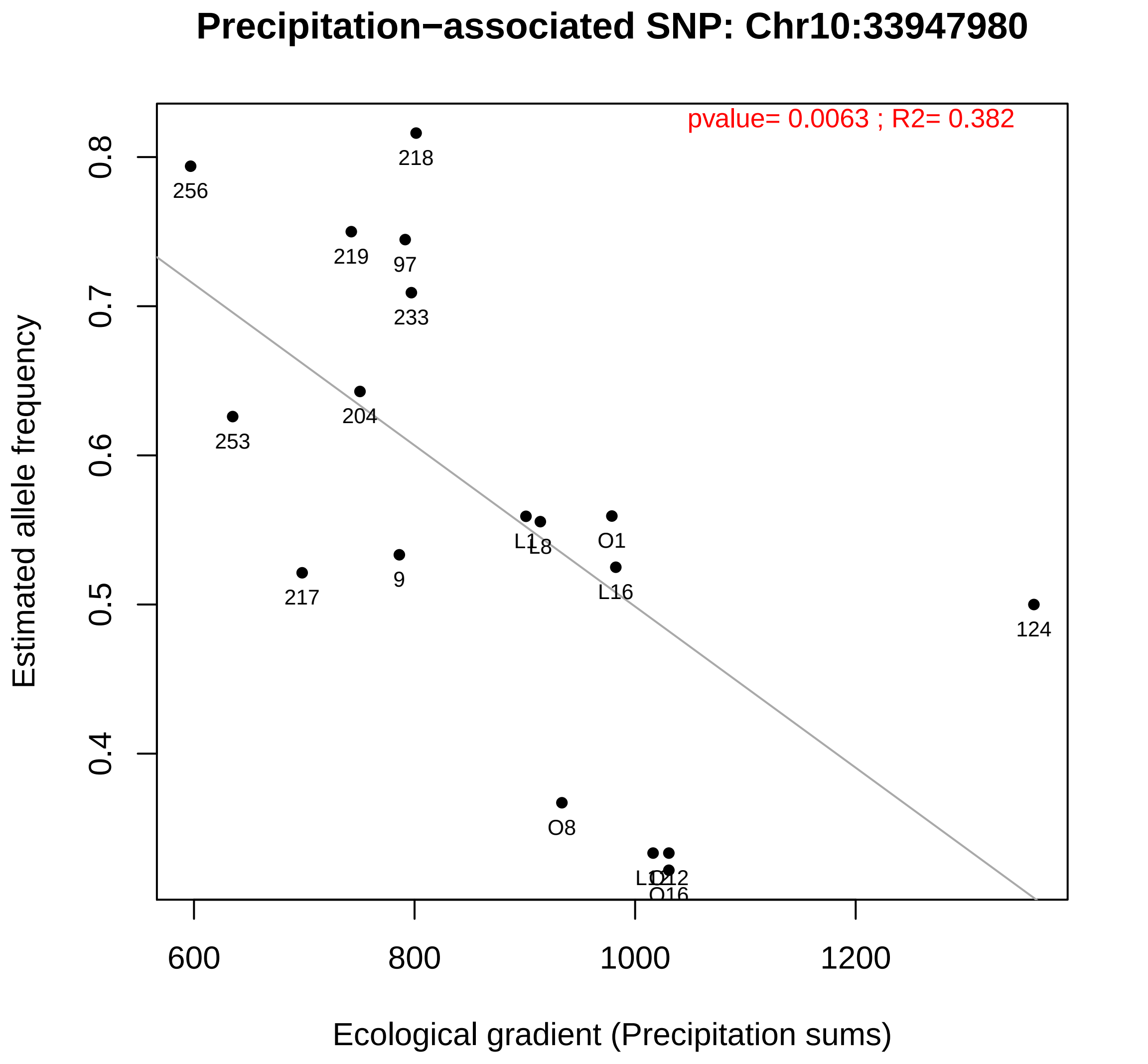

Precipitation−associated SNP: Chr10:33947980
pvalue= 0.0063 ; R2= 0.382
Estimated allele frequency
Ecological gradient (Precipitation sums)
0.4
0.5
0.6
0.7
0.8
600
800
1000
1200
218
256
219
97
233
204
253
L1
L8
O1
L16
9
217
124
O8
O16

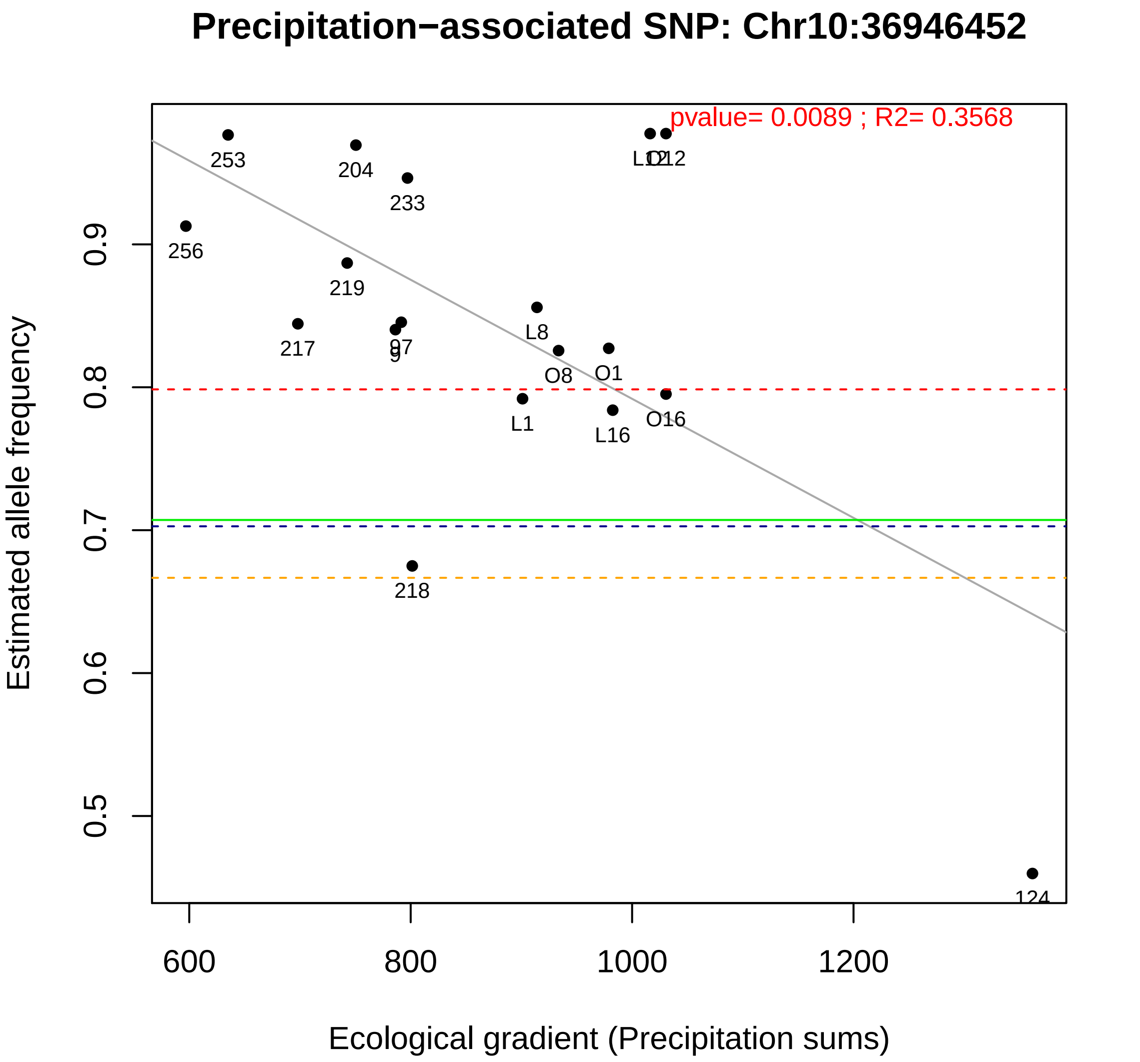

Precipitation−associated SNP: Chr10:36946452
pvalue= 0.0089 ; R2= 0.3568
253
204
233
256
219
217
97
9
L8
O8
O1
L1
L16
O16
218
124
0.5
0.6
0.7
0.8
0.9
600
800
1000
1200
Estimated allele frequency
Ecological gradient (Precipitation sums)

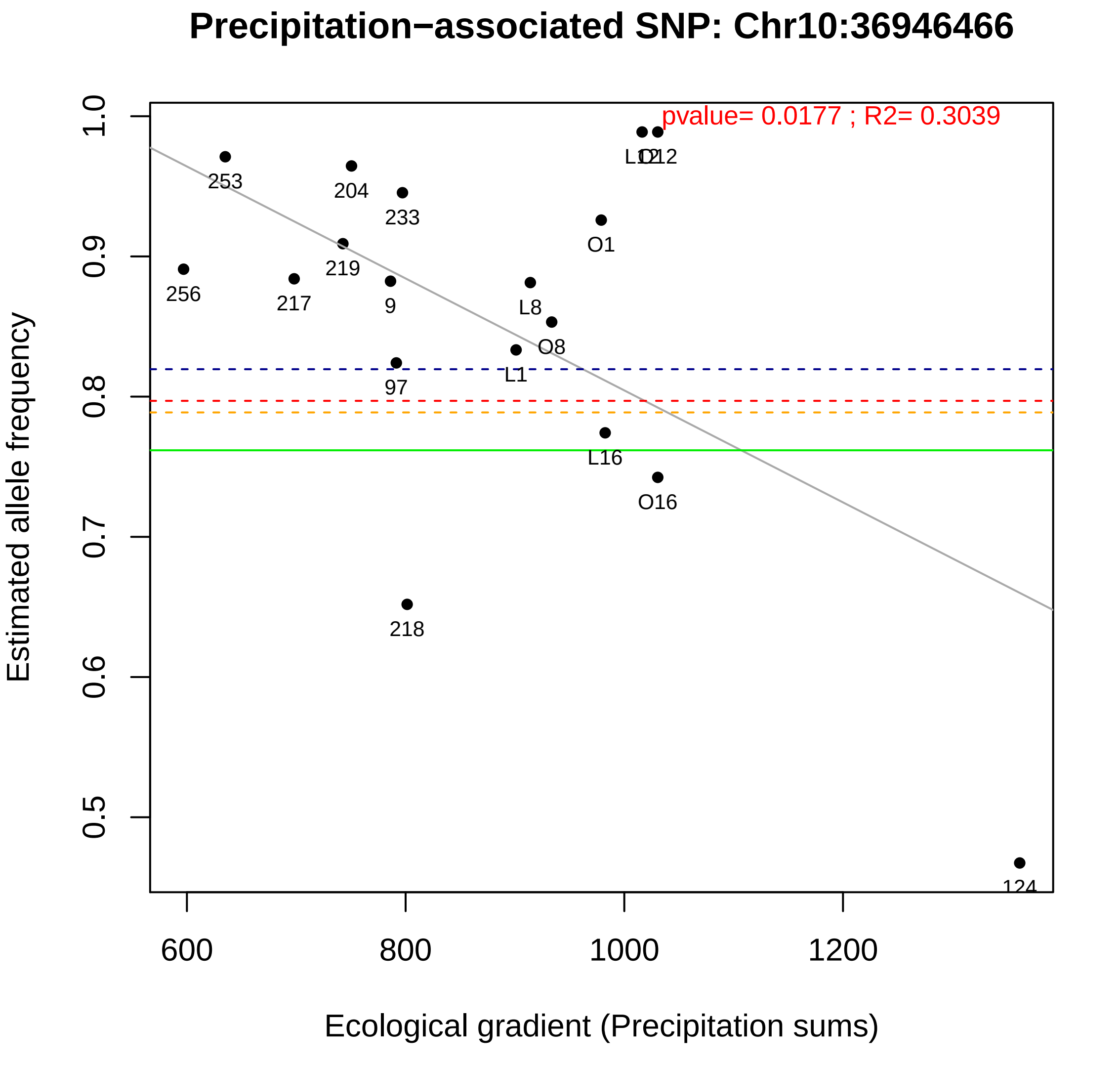
Precipitation−associated SNP: Chr10:36946466
pvalue= 0.0177 ; R2= 0.3039
Estimated allele frequency
Ecological gradient (Precipitation sums)
1.0
0.9
0.8
0.7
0.6
0.5
600
800
1000
1200
253
204
233
L12
O12
O1
219
256
217
9
L8
O8
L1
97
L16
O16
218
124

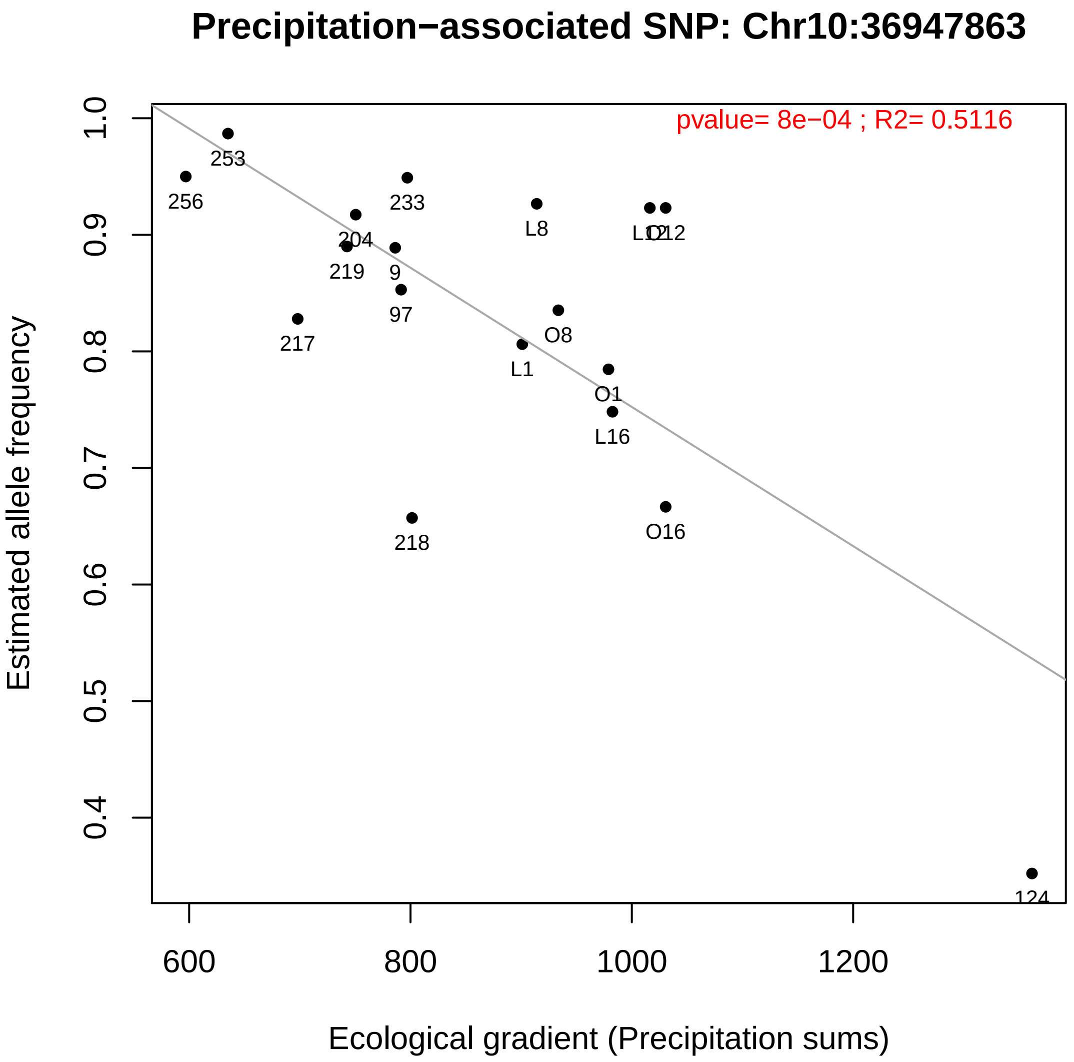

Precipitation−associated SNP: Chr10:36947863
pvalue= 8e−04 ; R2= 0.5116
Estimated allele frequency
Ecological gradient (Precipitation sums)
0.4
0.5
0.6
0.7
0.8
0.9
1.0
600
800
1000
1200
253
256
233
204
219
9
97
217
L8
L10
O12
O8
L1
O1
L16
218
O16
124

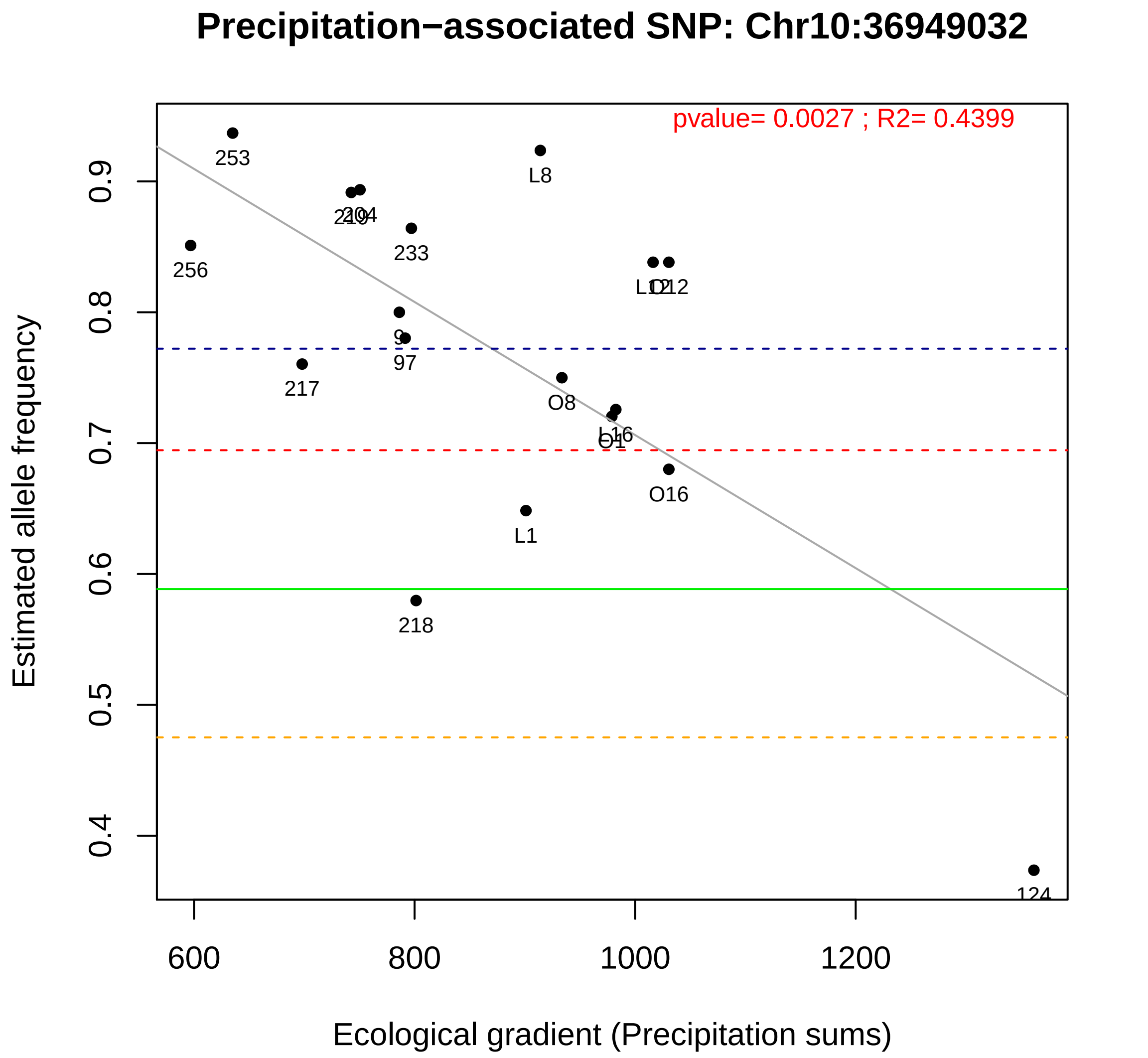

Precipitation−associated SNP: Chr10:36949032
pvalue= 0.0027 ; R2= 0.4399
253
L8
210
204
233
256
L12
O12
9
97
217
O8
L16
O1
O16
L1
218
124
0.4
0.5
0.6
0.7
0.8
0.9
600
800
1000
1200
Estimated allele frequency
Ecological gradient (Precipitation sums)

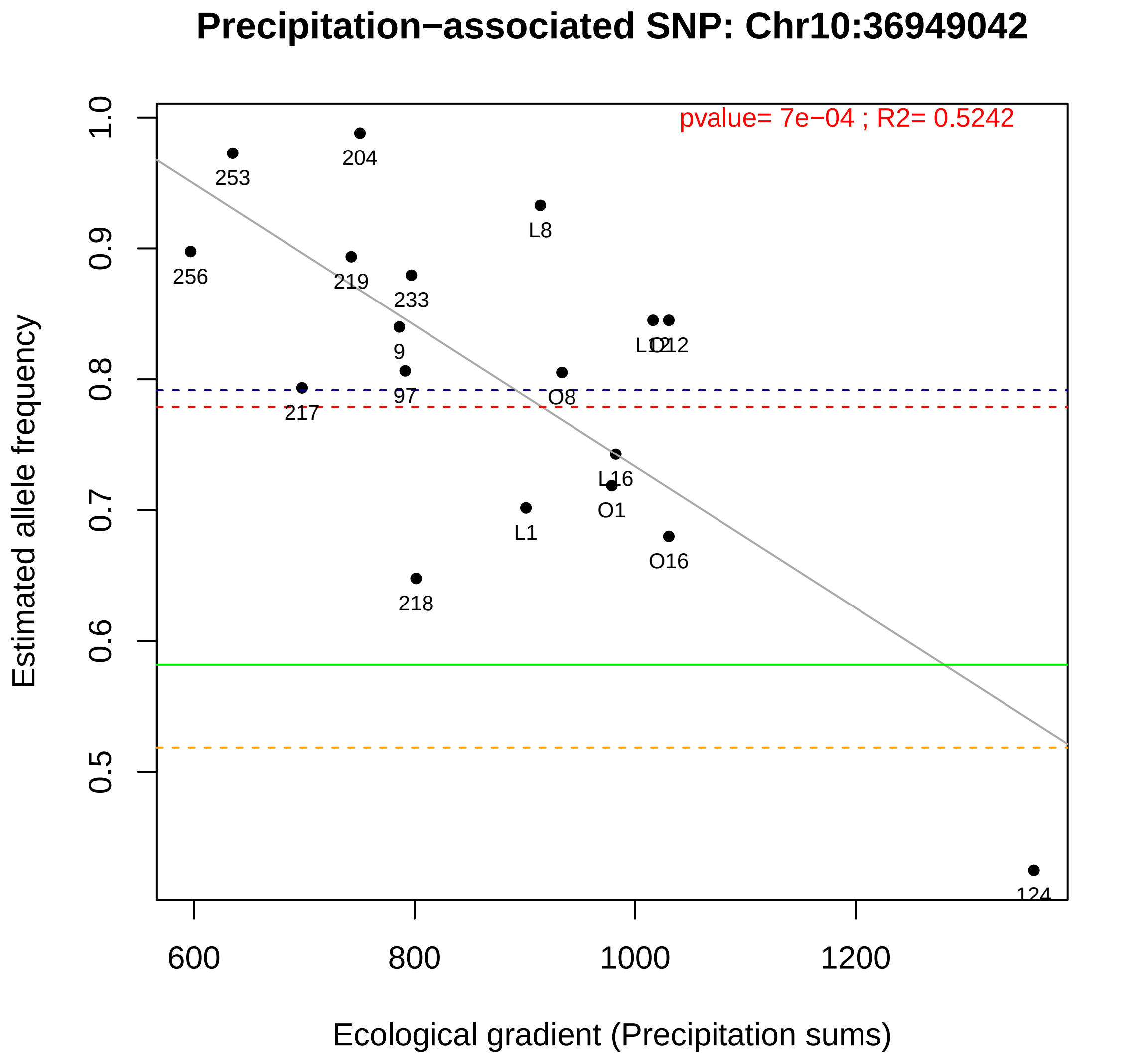

Precipitation−associated SNP: Chr10:36949042
pvalue= 7e−04 ; R2= 0.5242
Estimated allele frequency
Ecological gradient (Precipitation sums)
1.0
0.9
0.8
0.7
0.6
0.5
600
800
1000
1200
204
253
L8
256
219
233
9
97
217
O8
L16
O1
L1
O16
218
124

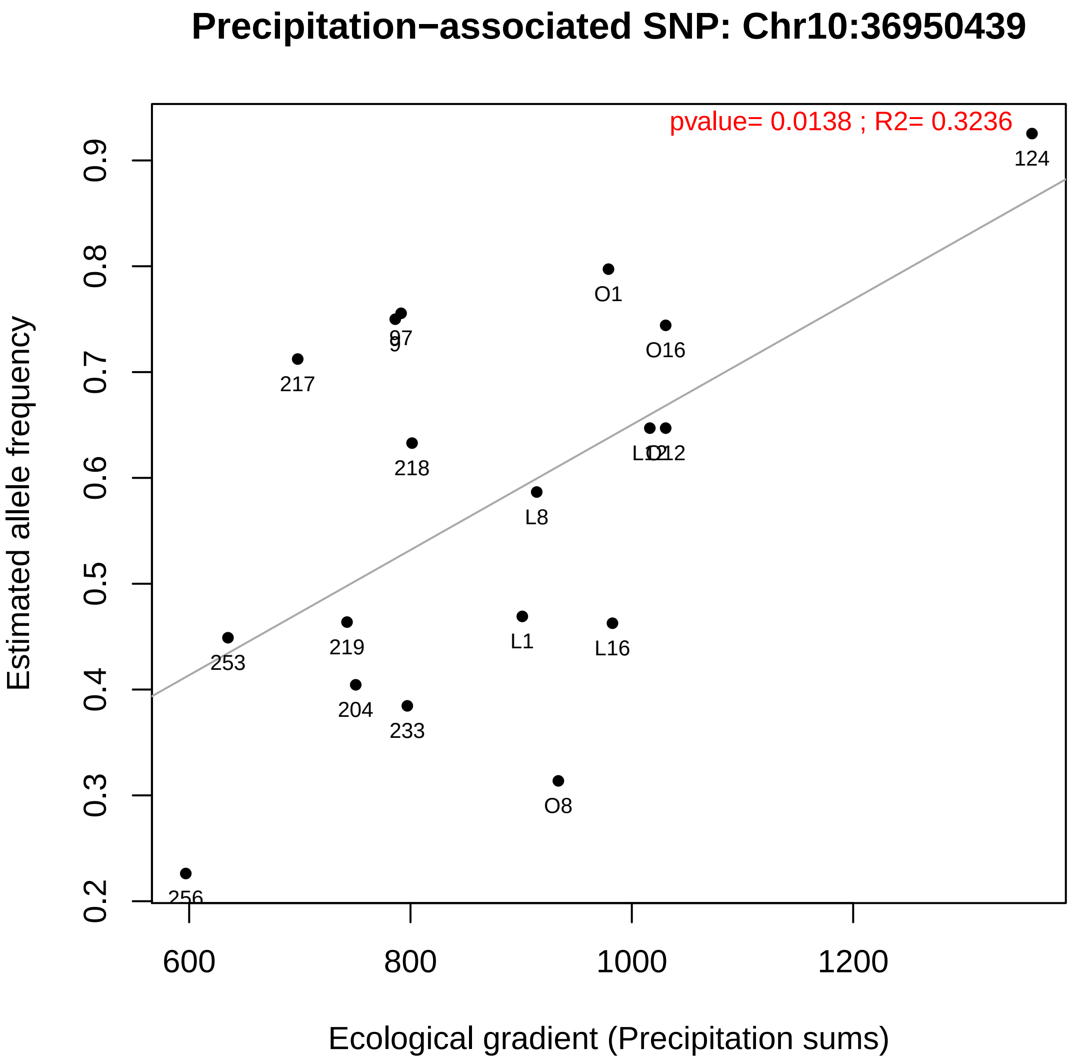
Precipitation−associated SNP: Chr10:36950439
pvalue= 0.0138 ; R2= 0.3236
Estimated allele frequency
Ecological gradient (Precipitation sums)
0.2
0.3
0.4
0.5
0.6
0.7
0.8
0.9
600
800
1000
1200
124
O1
O16
217
218
L8
253
219
L1
L16
204
233
O8
256

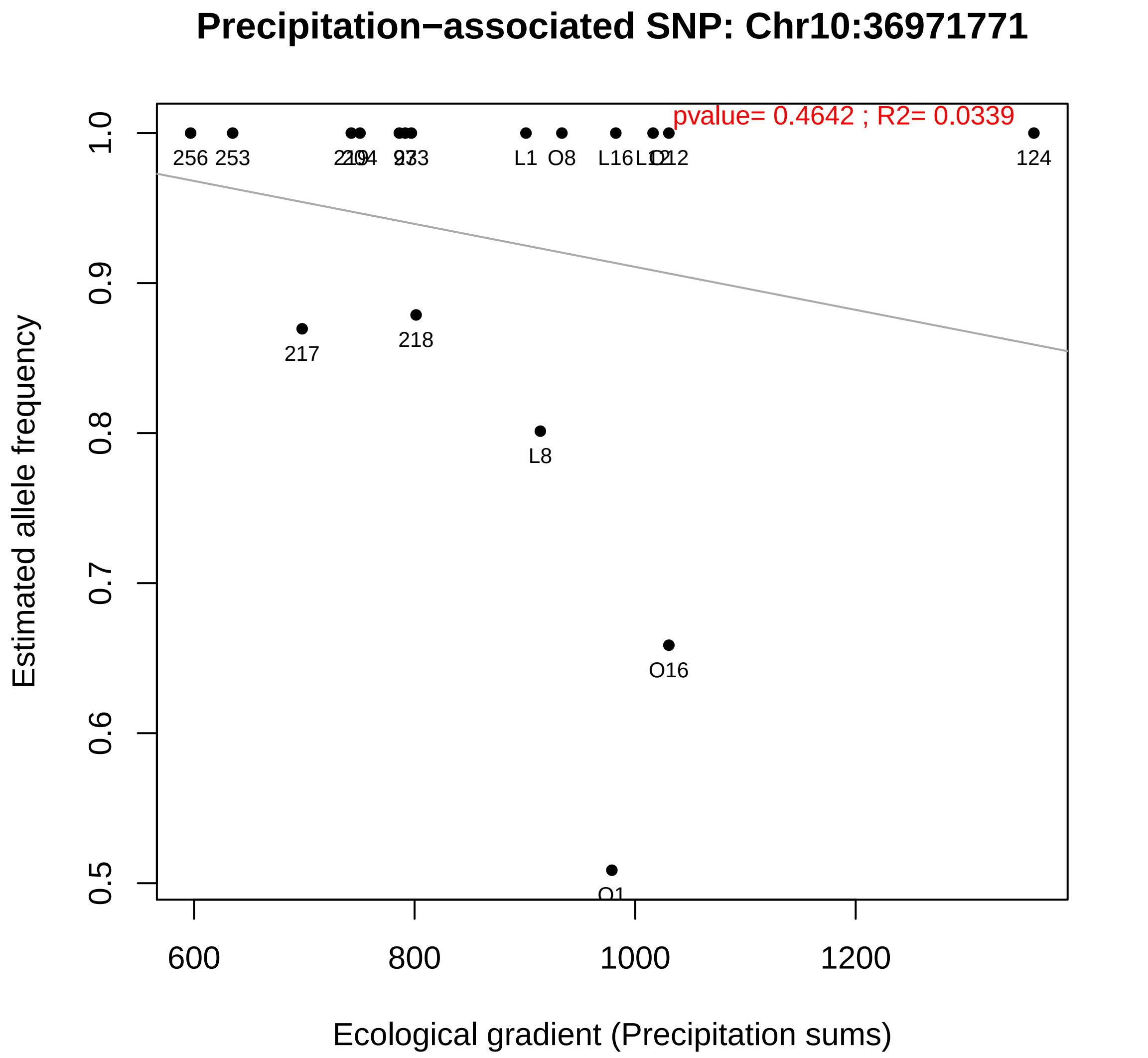

Precipitation−associated SNP: Chr10:36971771
pvalue= 0.4642 ; R2= 0.0339
Estimated allele frequency
Ecological gradient (Precipitation sums)
1.0
0.9
0.8
0.7
0.6
0.5
600
800
1000
1200
256
253
124
217
218
L1
O8
L16
L8
O16

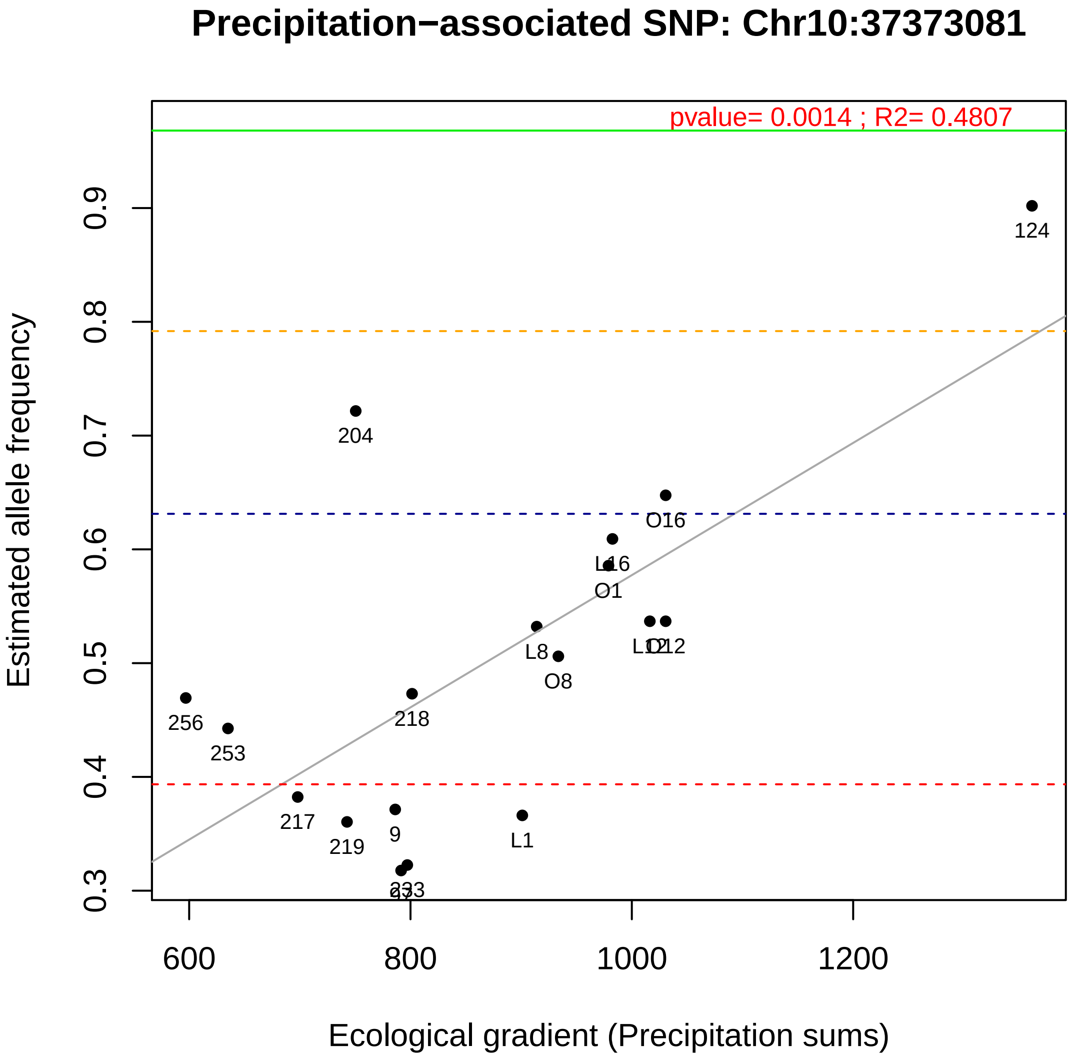
Precipitation−associated SNP: Chr10:37373081
pvalue= 0.0014 ; R2= 0.4807
Estimated allele frequency
0.3
0.4
0.5
0.6
0.7
0.8
0.9
Ecological gradient (Precipitation sums)
600
800
1000
1200
124
204
O16
L16
O1
L8
O8
256
253
218
217
219
9
L1
233

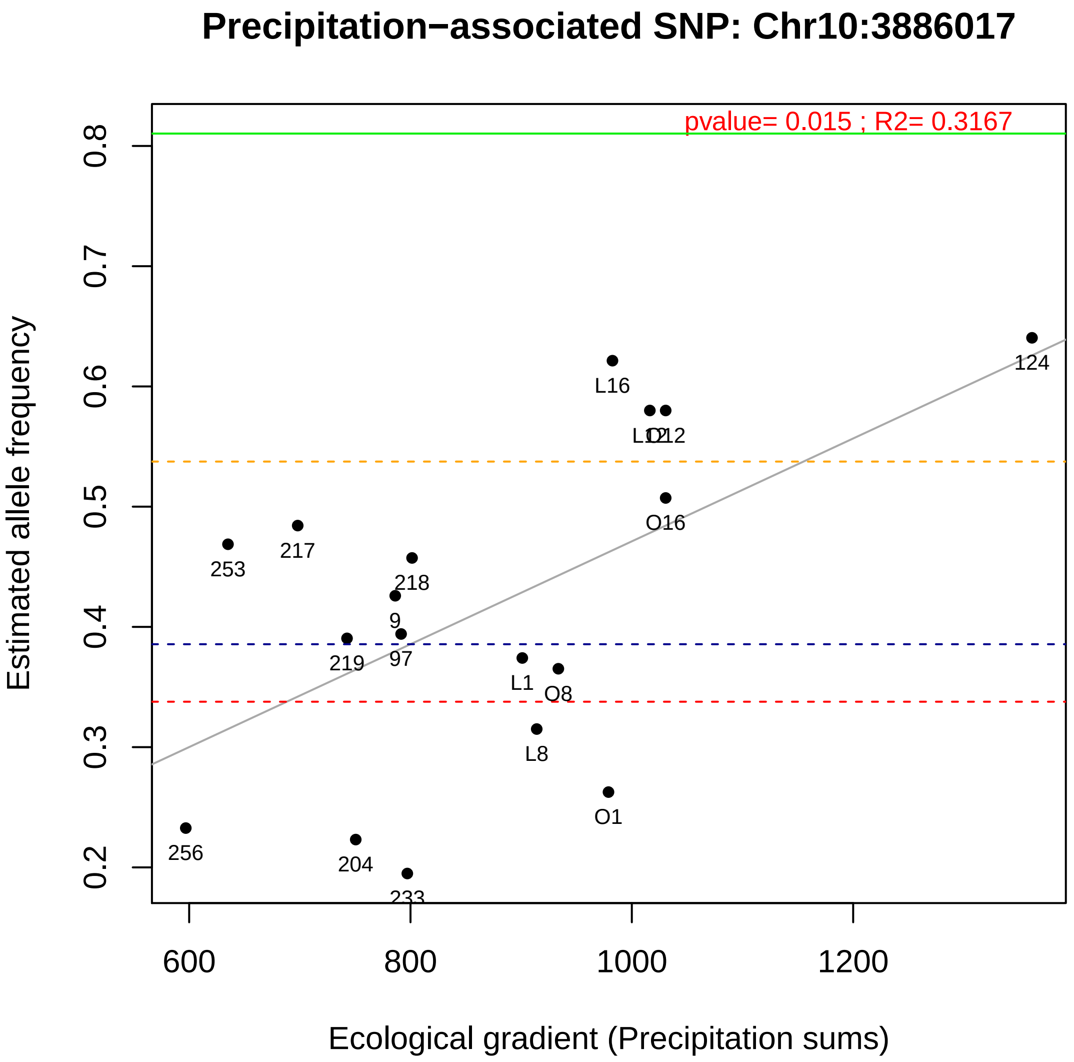

Precipitation−associated SNP: Chr10:3886017
pvalue= 0.015 ; R2= 0.3167
Estimated allele frequency
0.2
0.3
0.4
0.5
0.6
0.7
0.8
600
800
1000
1200
Ecological gradient (Precipitation sums)
124
L16
O12
O16
253
217
218
9
219
97
L1
O8
L8
O1
256
204
233

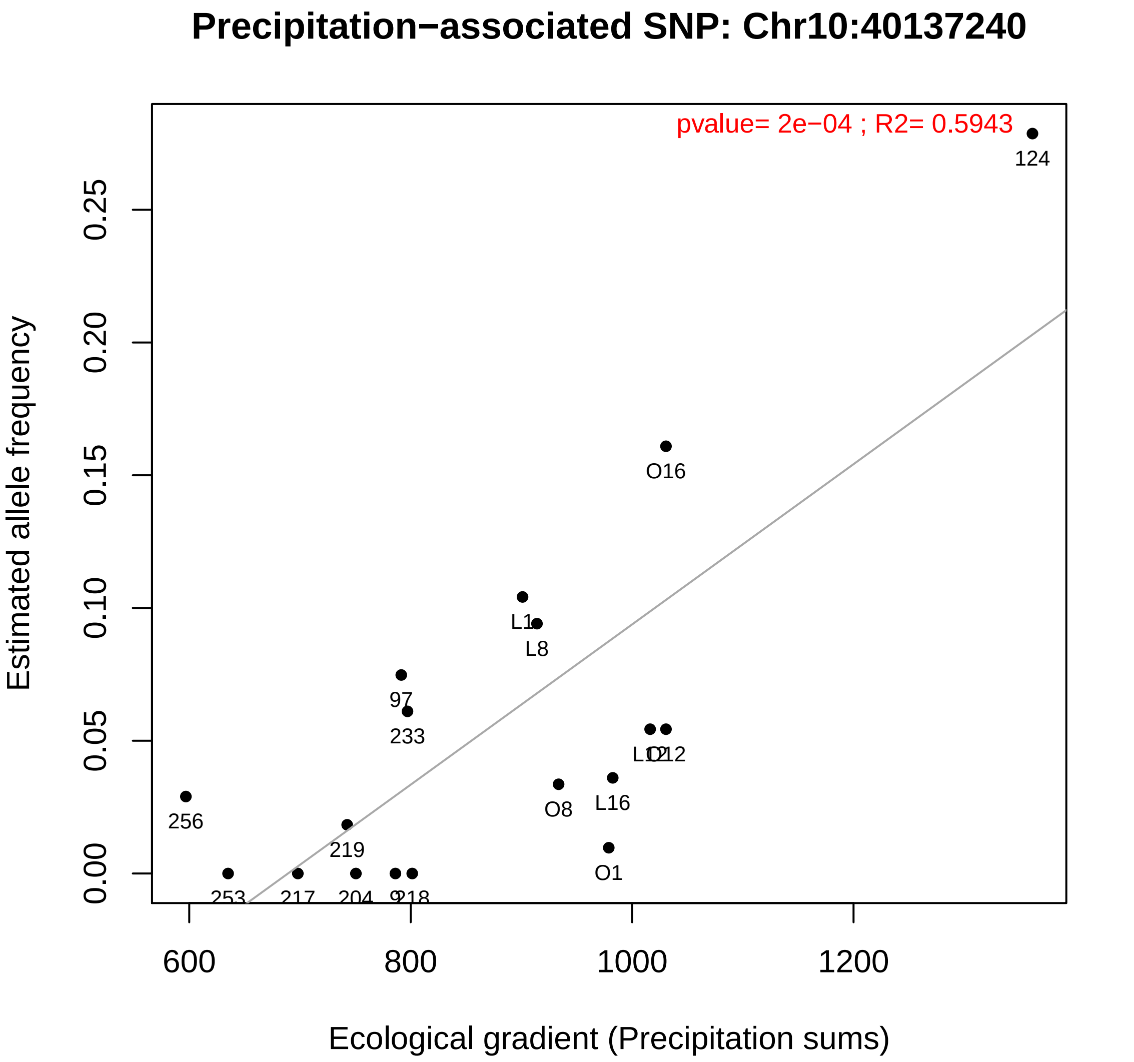
Precipitation−associated SNP: Chr10:40137240
pvalue= 2e−04 ; R2= 0.5943
Estimated allele frequency
Ecological gradient (Precipitation sums)
0.00
0.05
0.10
0.15
0.20
0.25
600
800
1000
1200
124
O16
L1
L8
97
233
L12
O12
O8
L16
256
219
O1
253
217
204
218

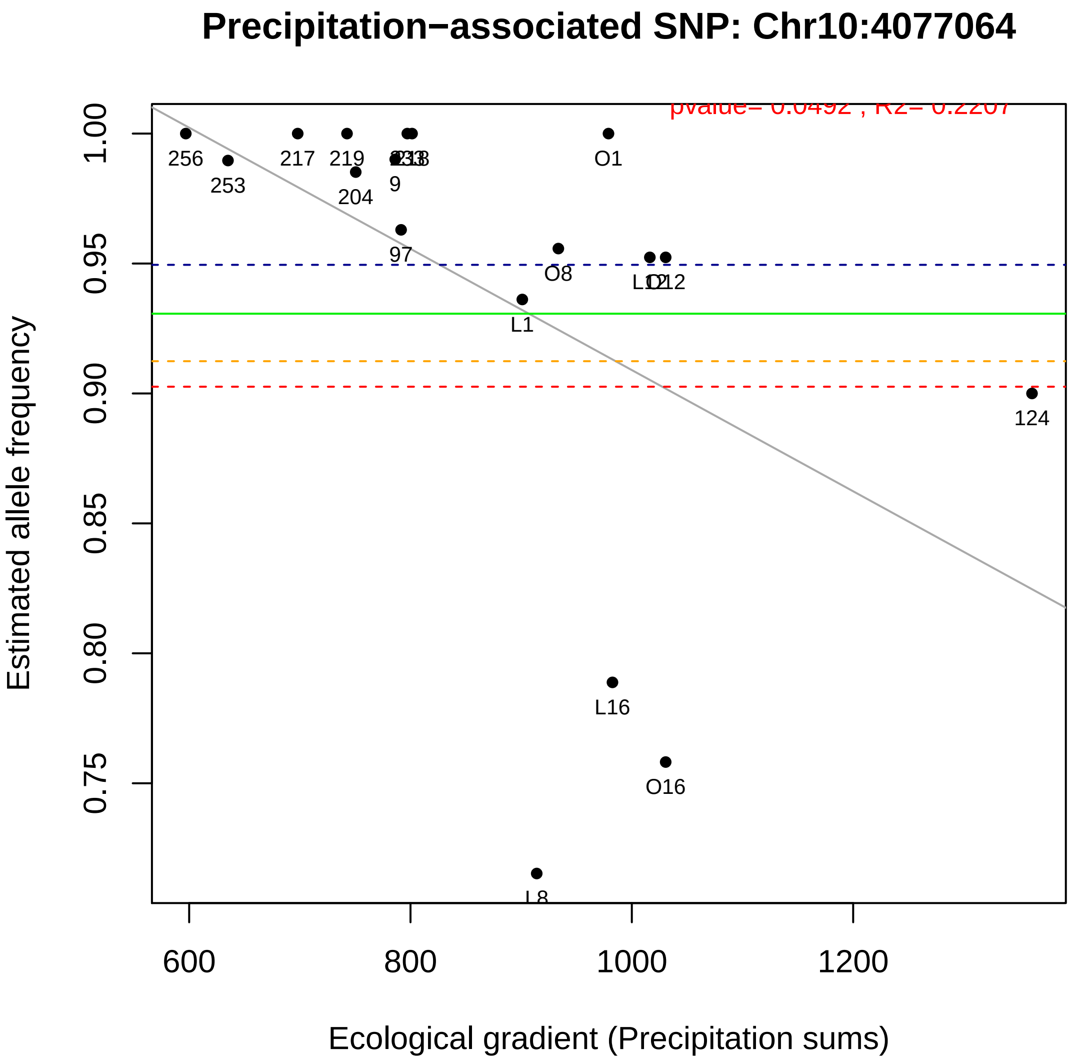
Precipitation−associated SNP: Chr10:4077064
pvalue= 0.0492 , R2= 0.2207
Estimated allele frequency
1.00
0.95
0.90
0.85
0.80
0.75
256
253
217
219
204
9
97
O1
O8
L1
124
L16
O16
L8
600
800
1000
1200
Ecological gradient (Precipitation sums)

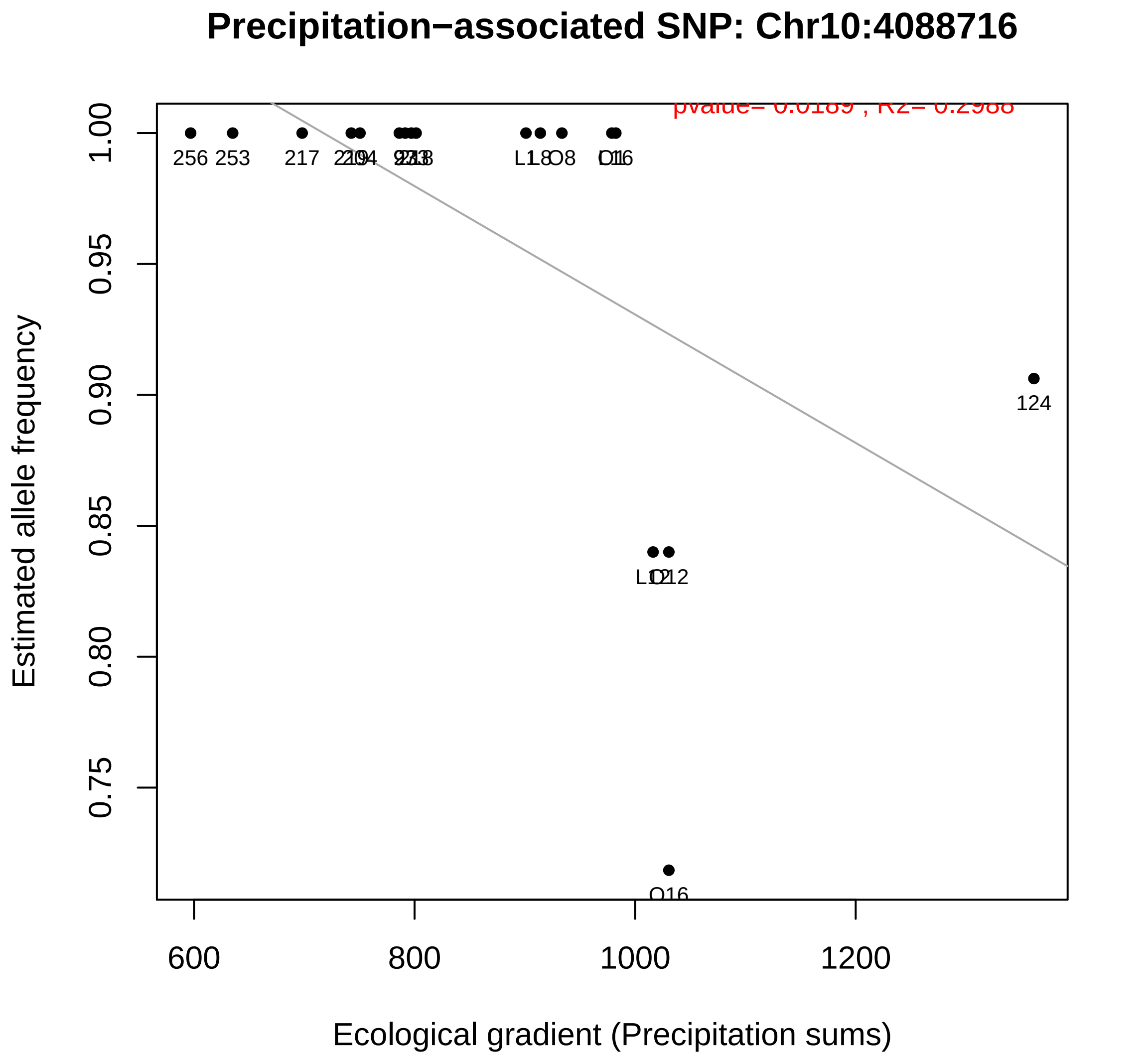

Precipitation−associated SNP: Chr10:4088716
pvalue= 0.0189 , R2= 0.2988
Estimated allele frequency
Ecological gradient (Precipitation sums)
0.75
0.80
0.85
0.90
0.95
1.00
600
800
1000
1200
256
253
217
124
O16

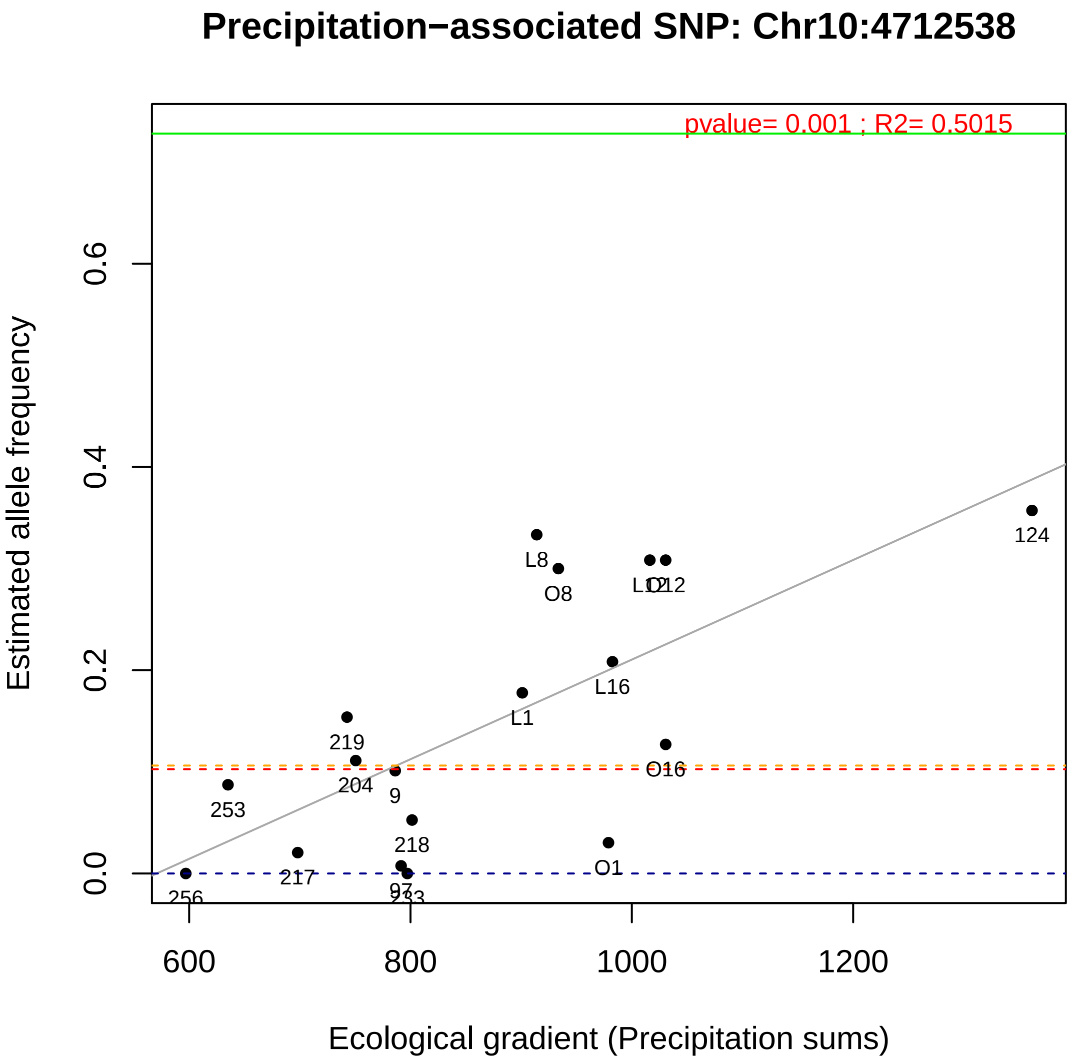

Precipitation−associated SNP: Chr10:4712538
pvalue= 0.001 ; R2= 0.5015
Estimated allele frequency
0.0
0.2
0.4
0.6
600
800
1000
1200
Ecological gradient (Precipitation sums)
124
L8
O8
O12
L16
L1
219
O16
204
9
253
218
217
O1

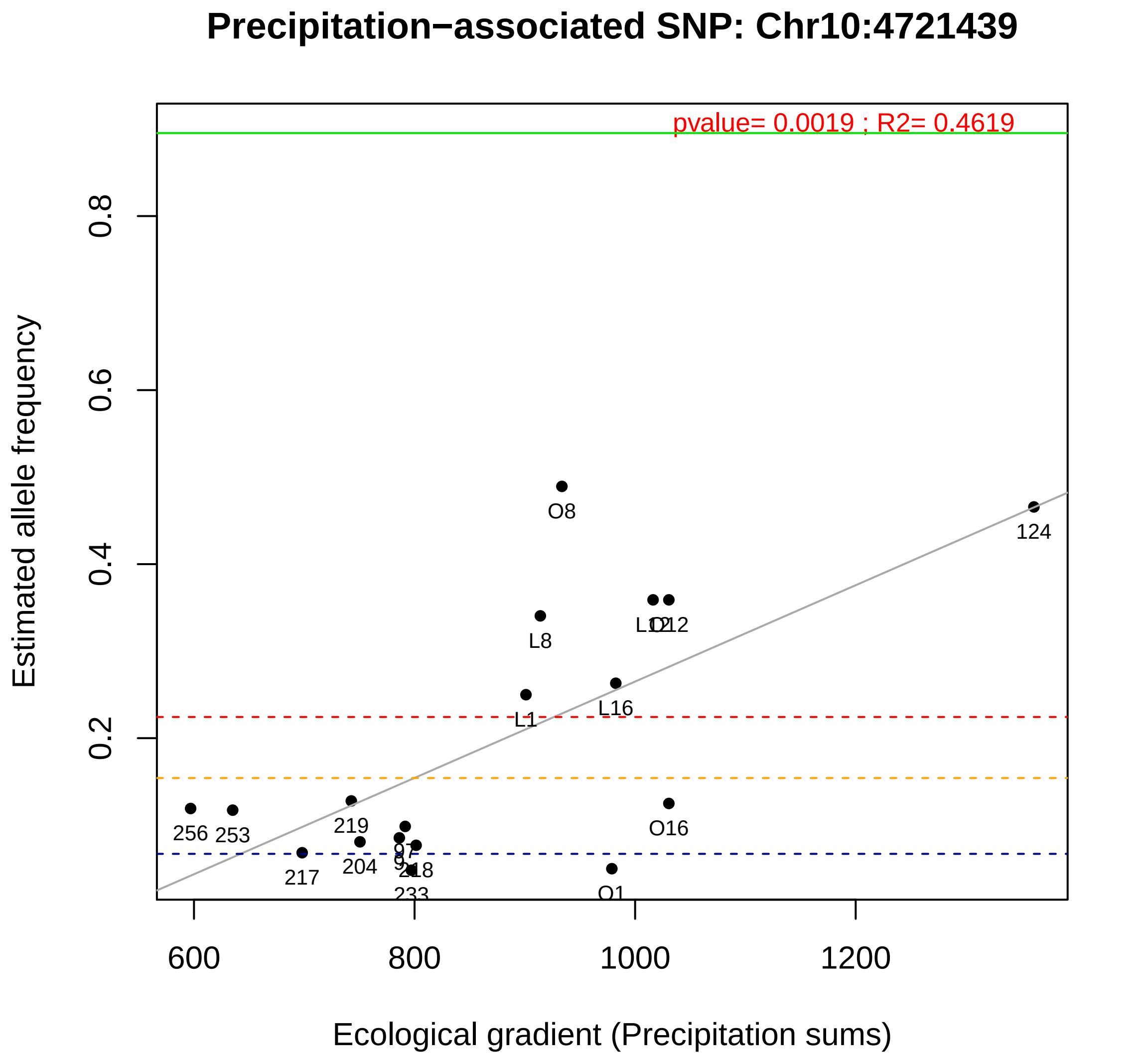

Precipitation−associated SNP: Chr10:4721439
pvalue= 0.0019 ; R2= 0.4619
Estimated allele frequency
0.2
0.4
0.6
0.8
600
800
1000
1200
Ecological gradient (Precipitation sums)
O8
124
L8
L12
O12
L16
L1
256
253
219
204
217
97
218
233
O16
O1

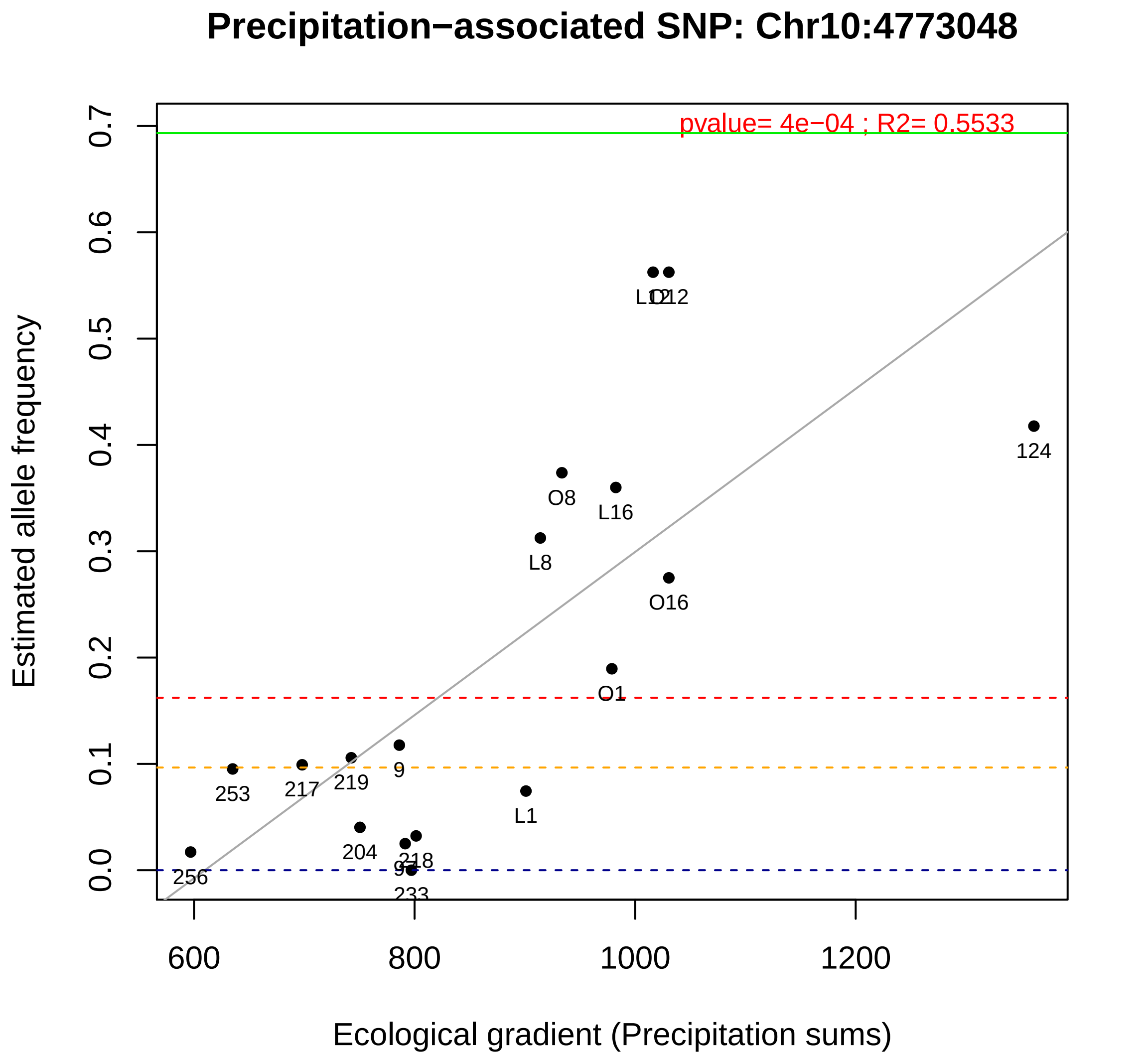

Precipitation−associated SNP: Chr10:4773048
pvalue= 4e−04 ; R2= 0.5533
Estimated allele frequency
Ecological gradient (Precipitation sums)
0.0
0.1
0.2
0.3
0.4
0.5
0.6
0.7
600
800
1000
1200
124
O8
L16
L8
O16
O1
253
217
219
9
L1
204
218
256
233

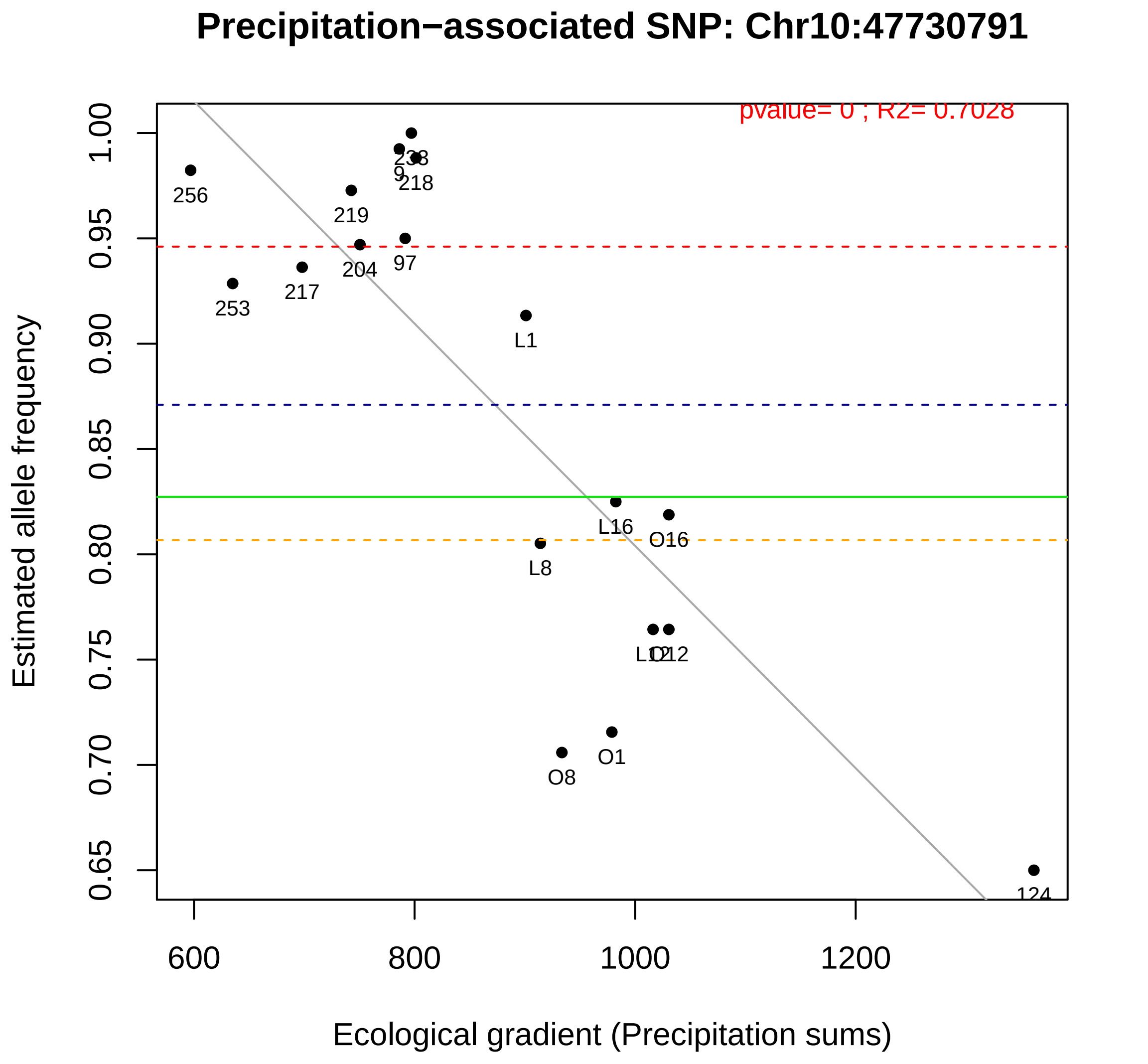

Precipitation−associated SNP: Chr10:47730791
pvalue= 0 ; R2= 0.7028
Estimated allele frequency
Ecological gradient (Precipitation sums)
1.00
0.95
0.90
0.85
0.80
0.75
0.70
0.65
600
800
1000
1200
256
253
217
219
204
238
9
218
97
L1
L16
O16
L8
L12
O12
O1
O8
124

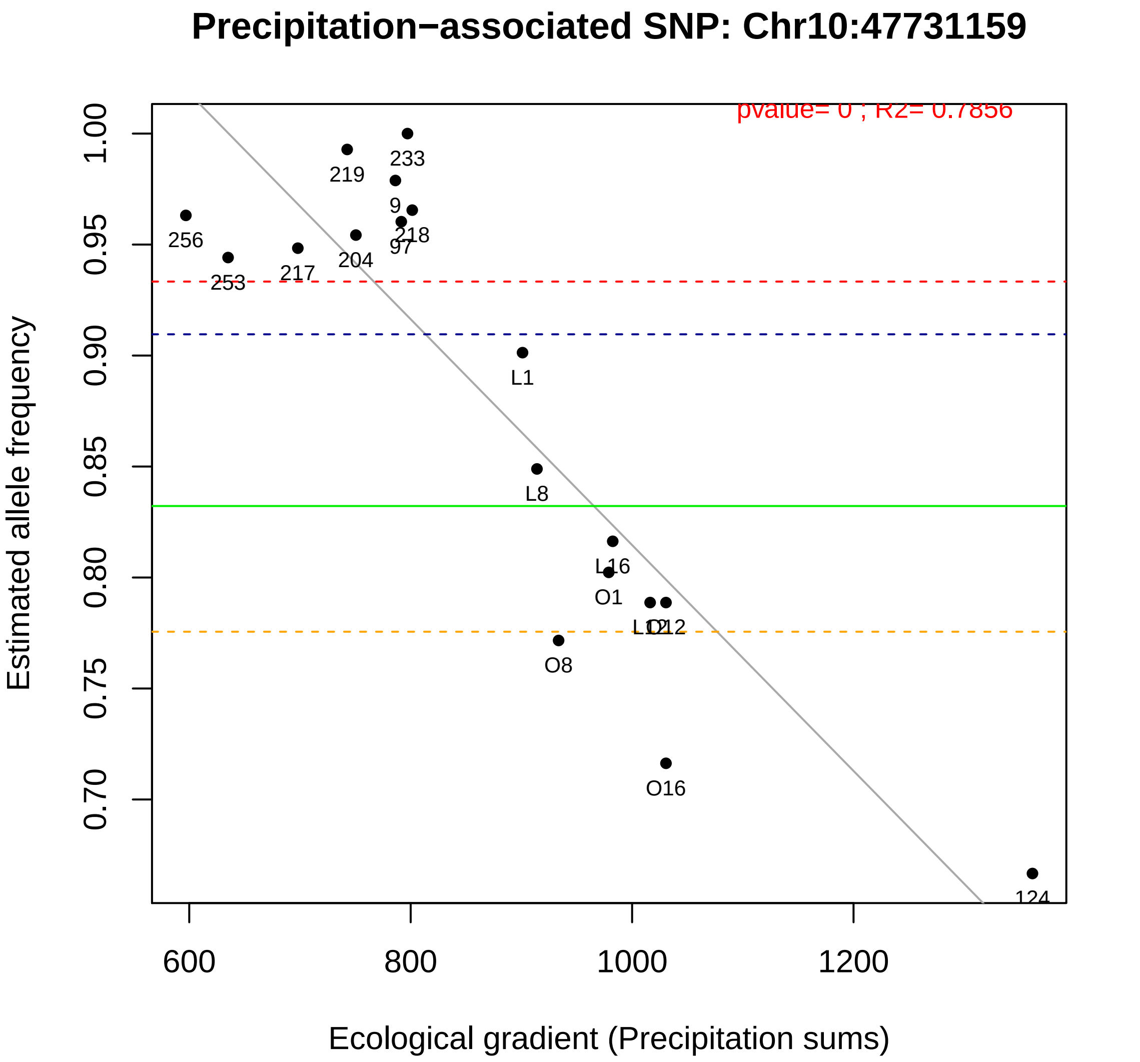
Precipitation−associated SNP: Chr10:47731159
pvalue= 0 ; R2= 0.7856
Estimated allele frequency
1.00
0.95
0.90
0.85
0.80
0.75
0.70
600
800
1000
1200
Ecological gradient (Precipitation sums)
233
219
9
218
97
256
204
253
217
L1
L8
L16
O1
O12
O8
O16
124

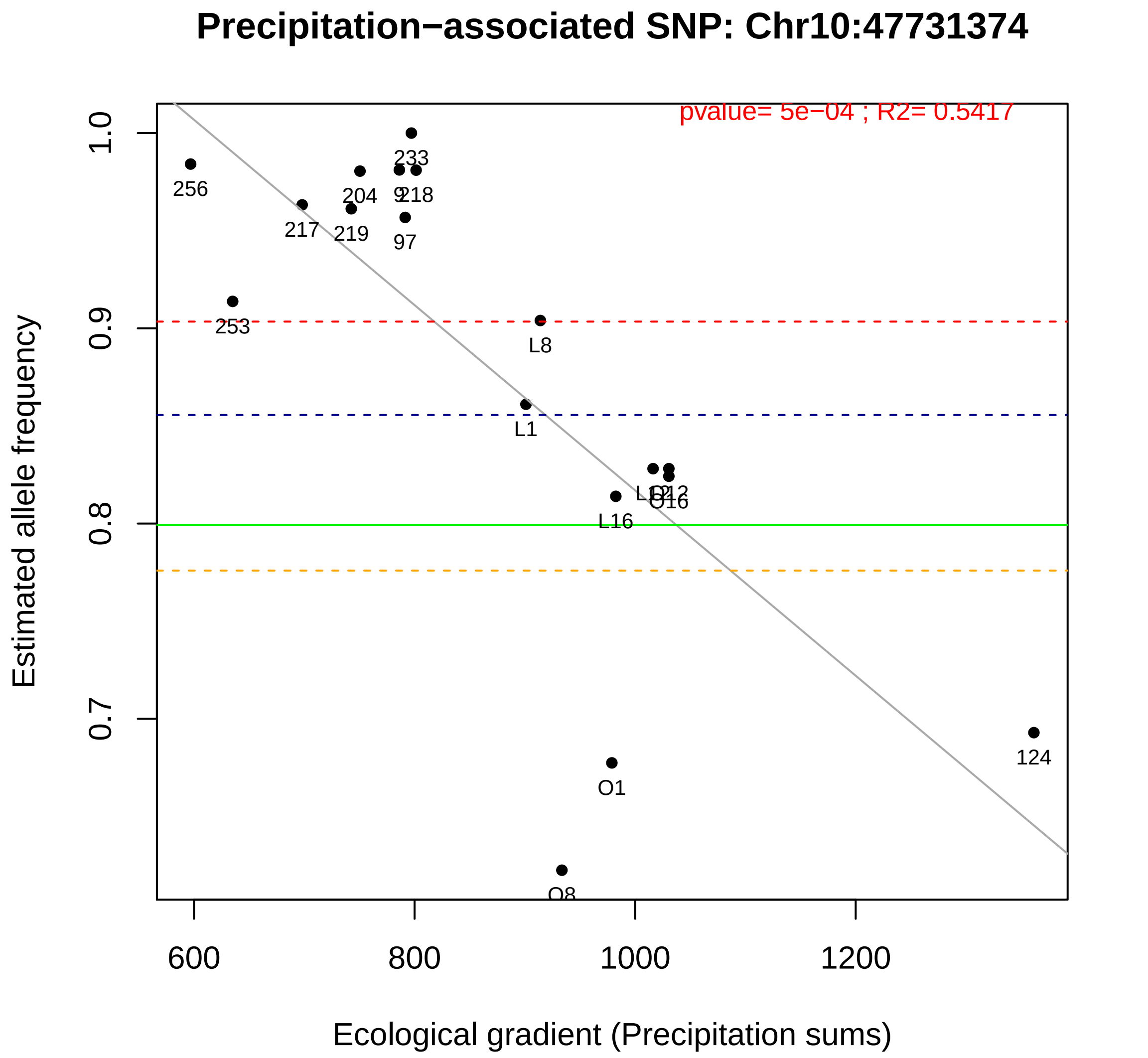

Precipitation−associated SNP: Chr10:47731374
pvalue= 5e−04 ; R2= 0.5417
Estimated allele frequency
Ecological gradient (Precipitation sums)
0.7
0.8
0.9
1.0
600
800
1000
1200
256
253
217
219
204
233
9
218
97
L8
L1
L12
O12
O16
L16
O1
O8
124

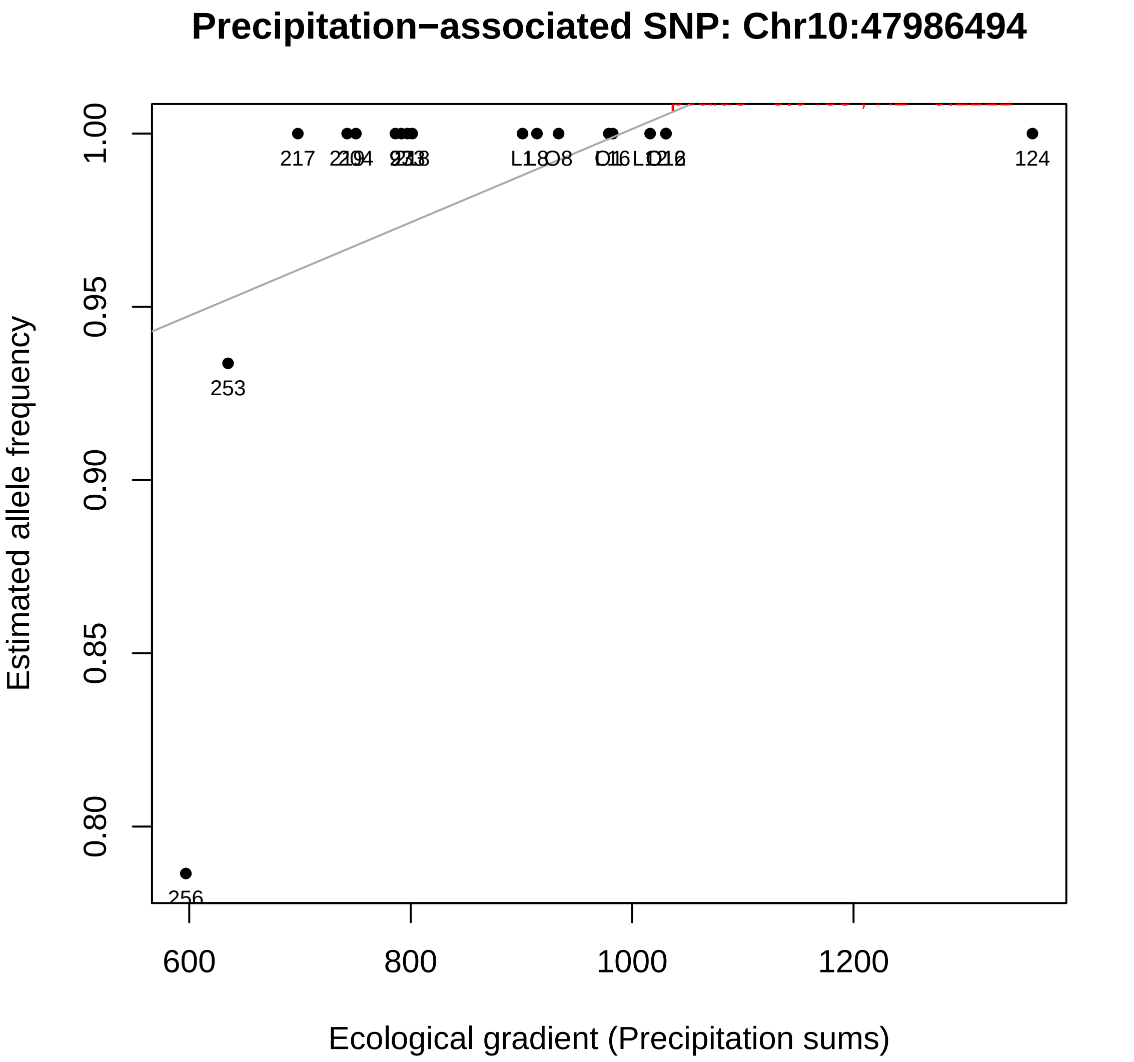
Precipitation−associated SNP: Chr10:47986494
Estimated allele frequency
0.80
0.85
0.90
0.95
1.00
Ecological gradient (Precipitation sums)
600
800
1000
1200
217
124
253
256

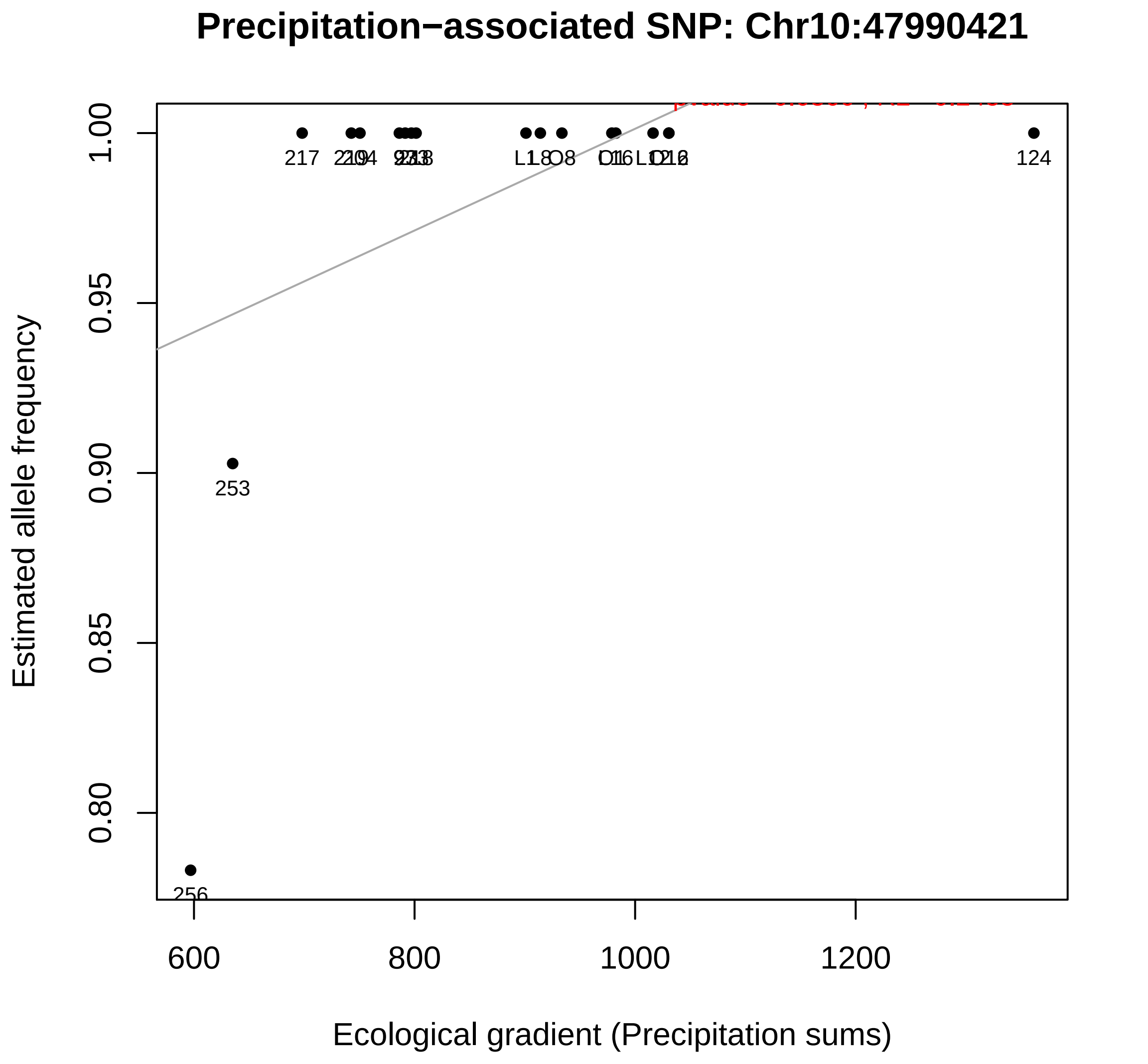

Precipitation−associated SNP: Chr10:47990421
Estimated allele frequency
Ecological gradient (Precipitation sums)
0.80
0.85
0.90
0.95
1.00
600
800
1000
1200
217
124
253
256

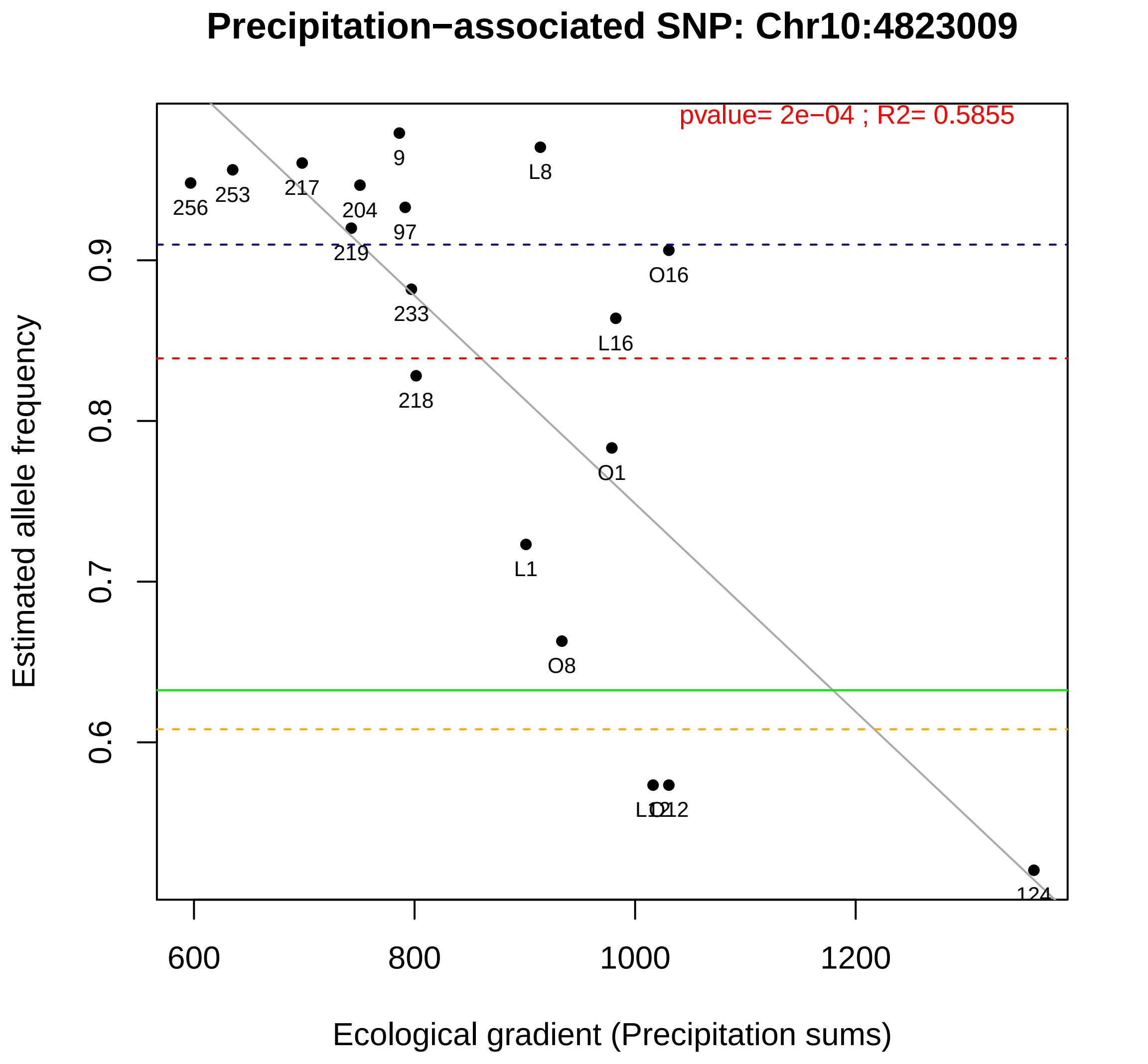
Precipitation−associated SNP: Chr10:4823009
pvalue= 2e−04 ; R2= 0.5855
Estimated allele frequency
0.6
0.7
0.8
0.9
600
800
1000
1200
Ecological gradient (Precipitation sums)
256
253
217
9
L8
204
97
219
O16
233
L16
218
O1
L1
O8
L12
O12
124

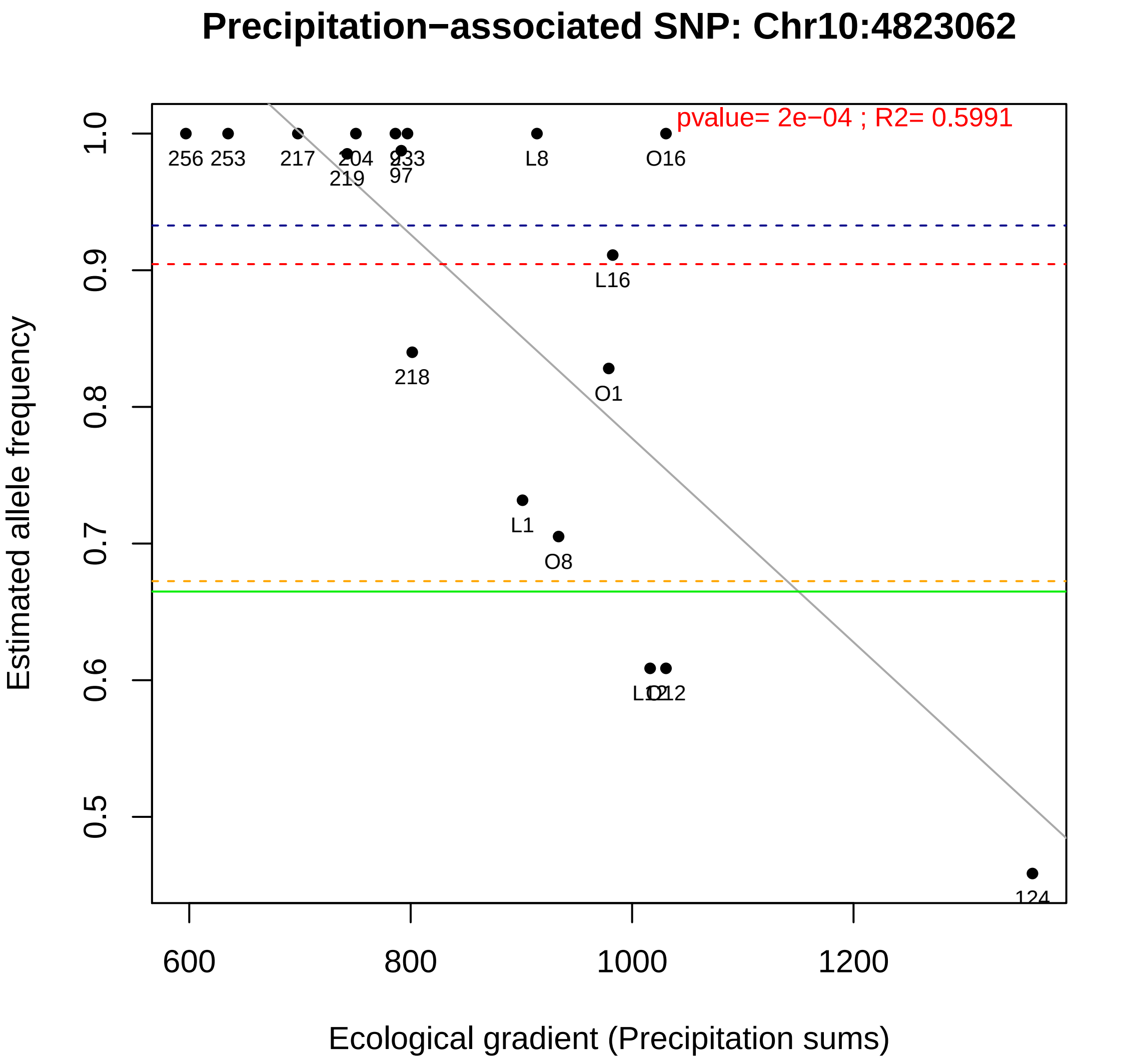
Precipitation−associated SNP: Chr10:4823062
pvalue= 2e−04 ; R2= 0.5991
256
253
217
204
219
233
97
L8
O16
L16
218
O1
L1
O8
124
1.0
0.9
0.8
0.7
0.6
0.5
600
800
1000
1200
Estimated allele frequency
Ecological gradient (Precipitation sums)

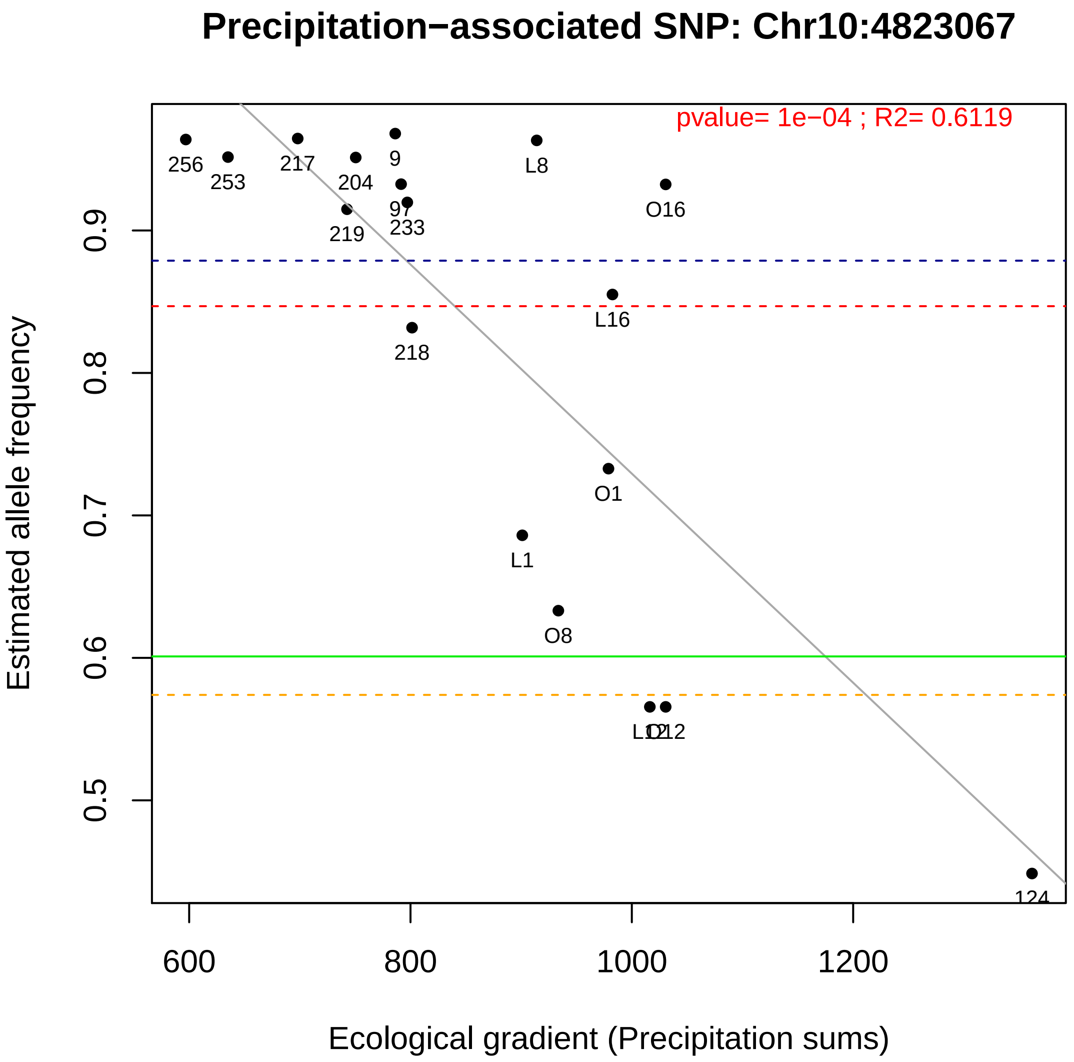

Precipitation−associated SNP: Chr10:4823067
pvalue= 1e−04 ; R2= 0.6119
256
253
217
204
9
97
233
219
L8
O16
L16
218
O1
L1
O8
L10
O12
124
0.5
0.6
0.7
0.8
0.9
600
800
1000
1200
Estimated allele frequency
Ecological gradient (Precipitation sums)

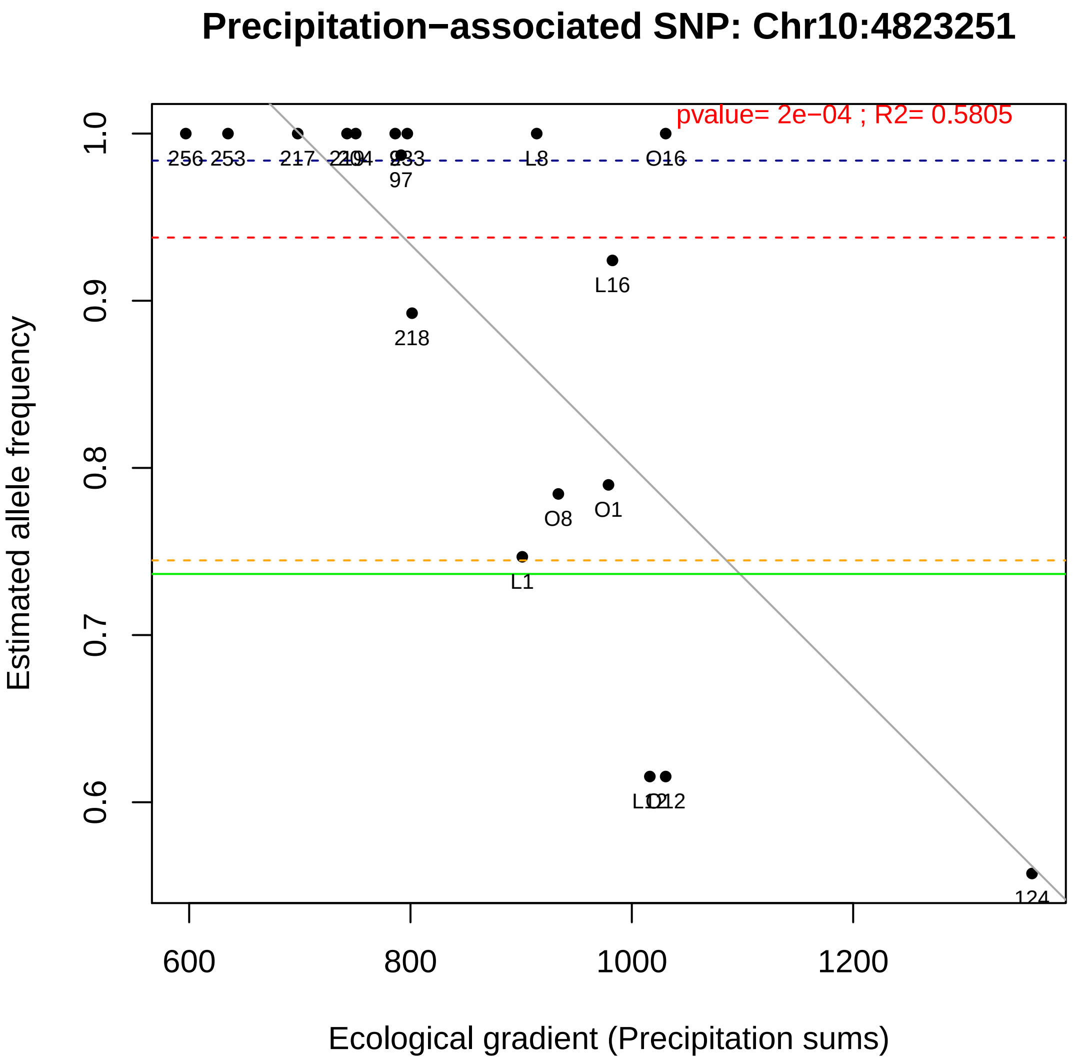

Precipitation−associated SNP: Chr10:4823251
pvalue= 2e−04 ; R2= 0.5805
Estimated allele frequency
Ecological gradient (Precipitation sums)
1.0
0.9
0.8
0.7
0.6
600
800
1000
1200
256
253
217
210
194
93
233
97
L8
O16
L16
218
O8
O1
L1
L12
O12
124

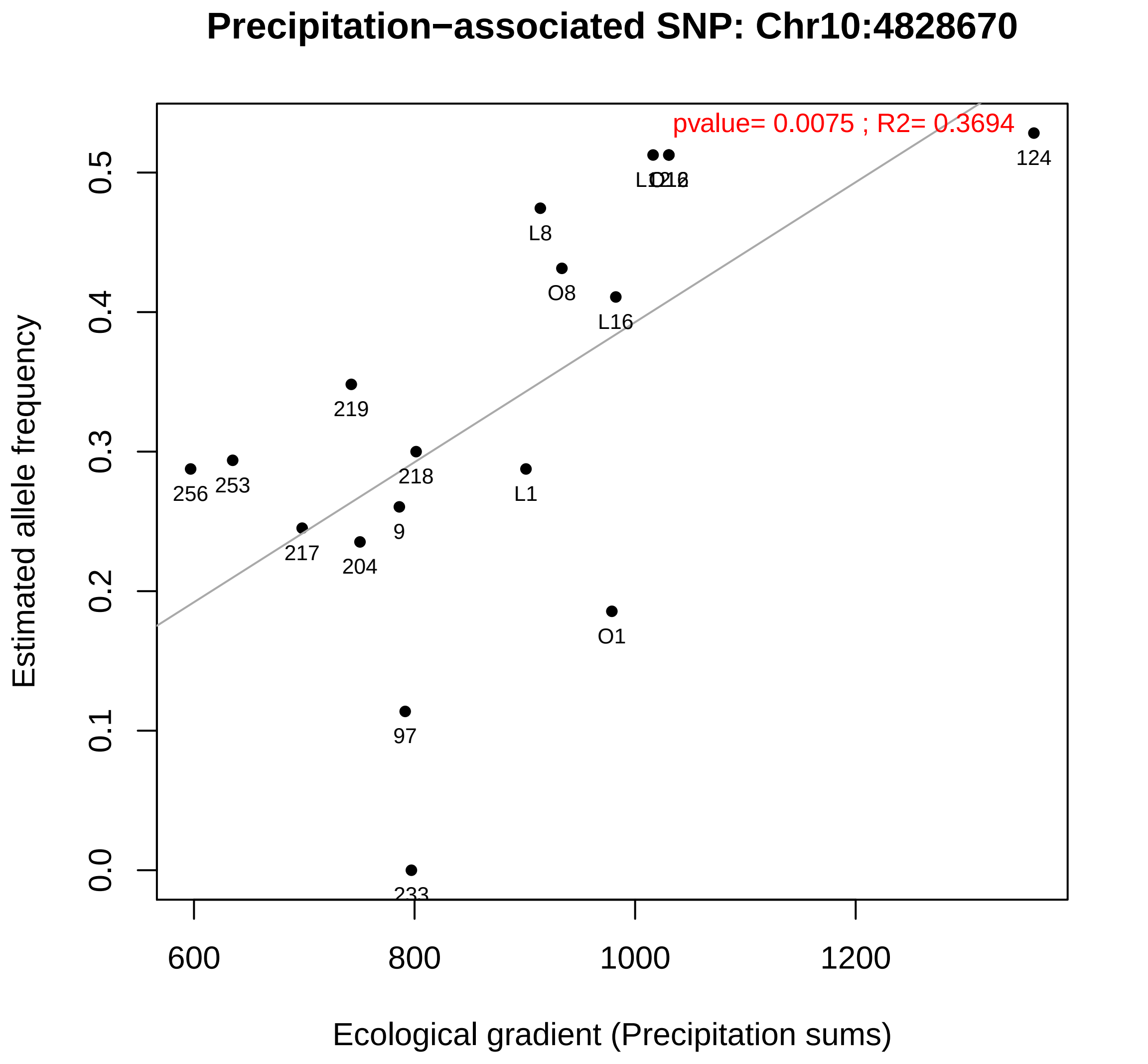
Precipitation−associated SNP: Chr10:4828670
pvalue= 0.0075 ; R2= 0.3694
Estimated allele frequency
Ecological gradient (Precipitation sums)
0.0
0.1
0.2
0.3
0.4
0.5
600
800
1000
1200
124
L8
O8
L16
219
218
256
253
L1
9
217
204
O1
97
233

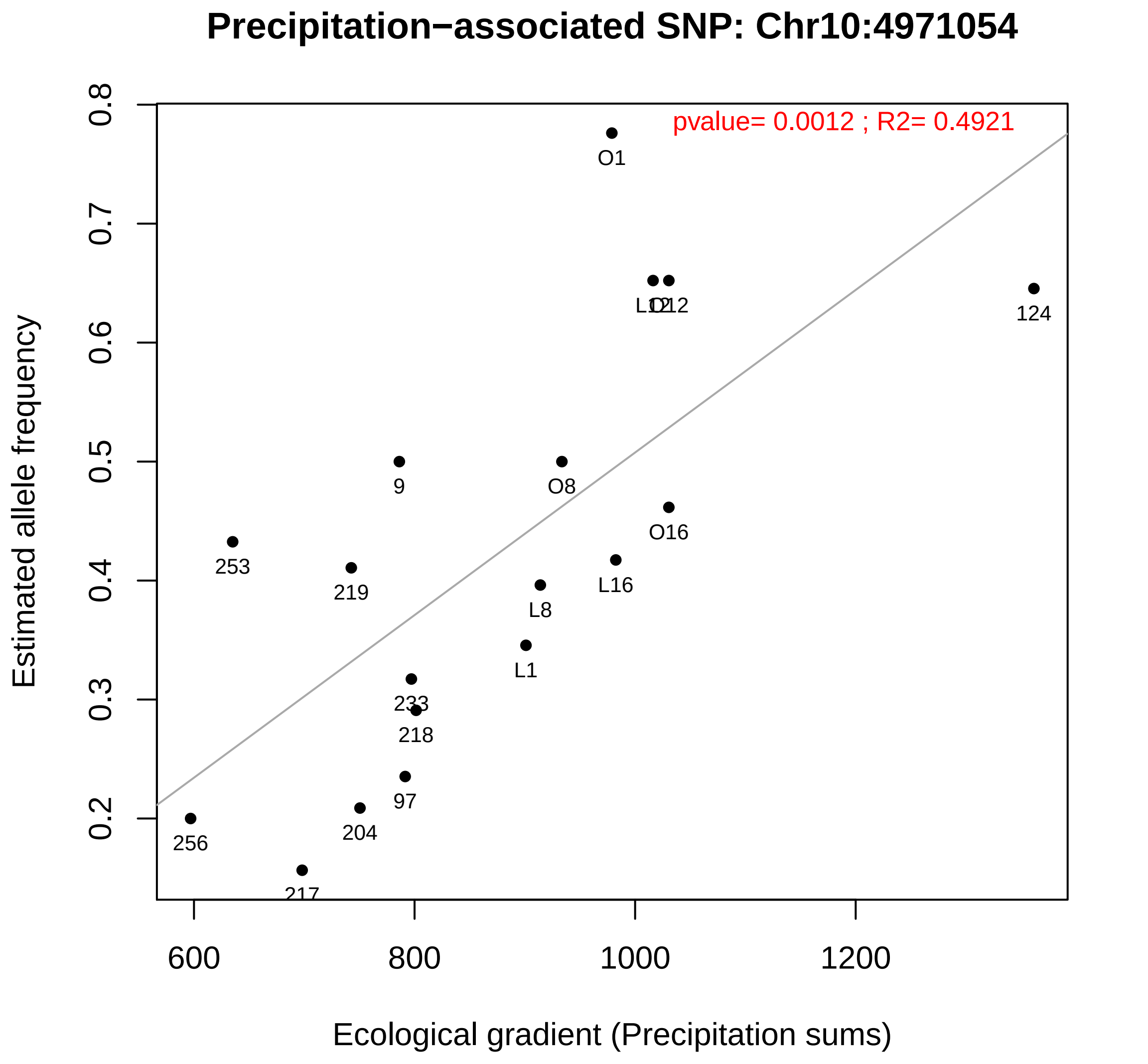
Precipitation−associated SNP: Chr10:4971054
pvalue= 0.0012 ; R2= 0.4921
Estimated allele frequency
0.2
0.3
0.4
0.5
0.6
0.7
0.8
600
800
1000
1200
Ecological gradient (Precipitation sums)
O1
L12
O12
124
9
O8
O16
253
219
L16
L8
L1
233
218
97
204
256
217

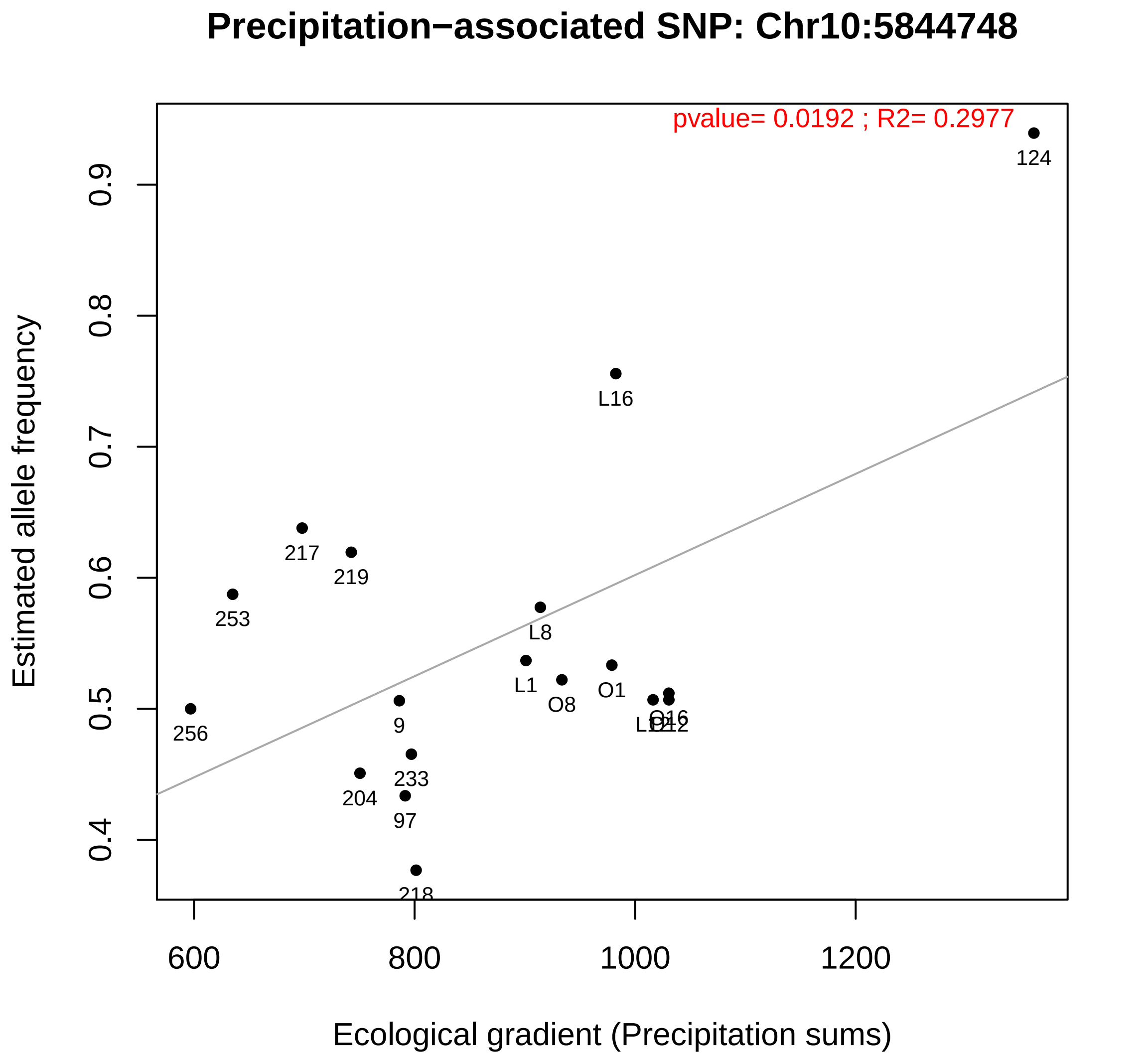
Precipitation−associated SNP: Chr10:5844748
pvalue= 0.0192 ; R2= 0.2977
Estimated allele frequency
0.4
0.5
0.6
0.7
0.8
0.9
600
800
1000
1200
Ecological gradient (Precipitation sums)
124
L16
217
219
253
L8
L1
O8
O1
256
9
233
204
97
218

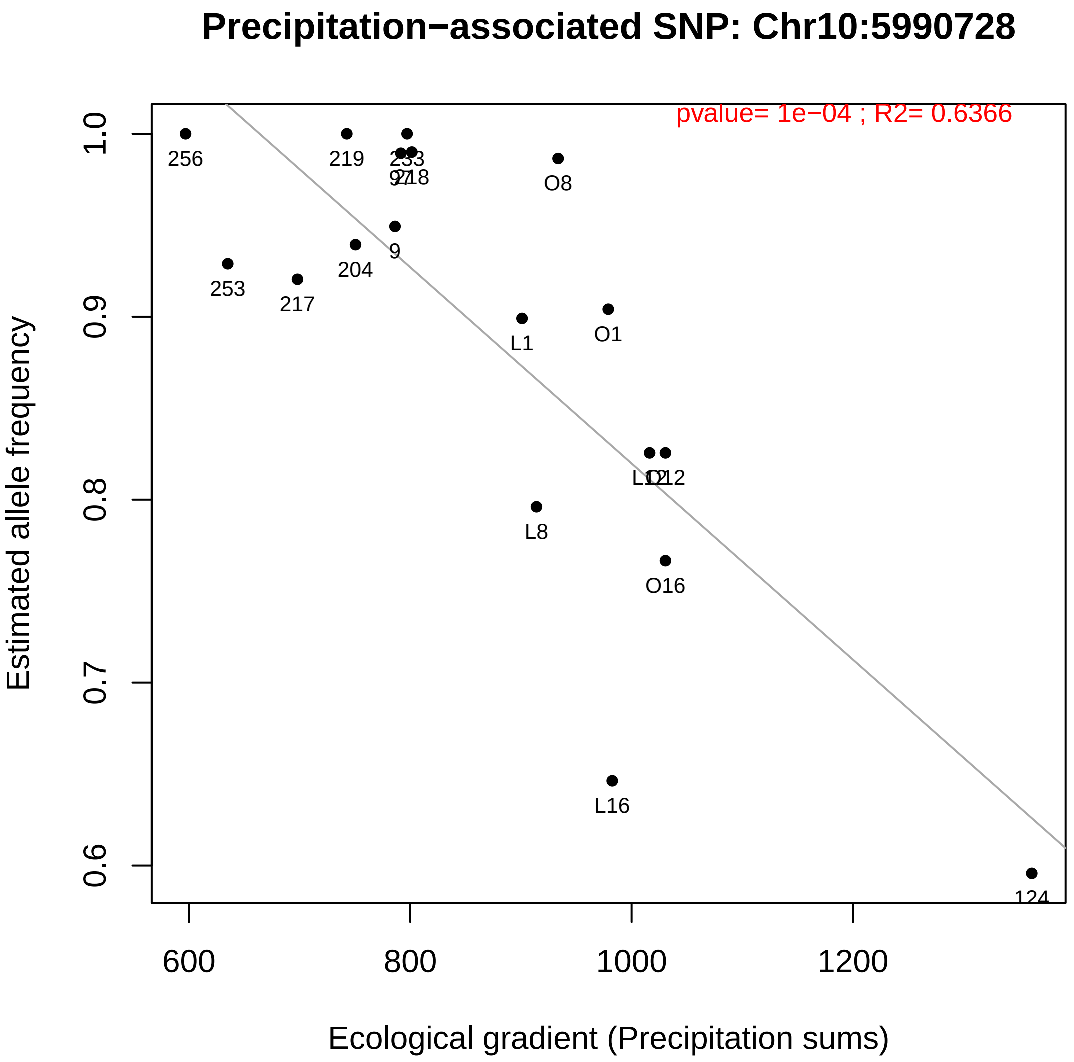

Precipitation−associated SNP: Chr10:5990728
pvalue= 1e−04 ; R2= 0.6366
Estimated allele frequency
Ecological gradient (Precipitation sums)
1.0
0.9
0.8
0.7
0.6
600
800
1000
1200
256
219
233
218
O8
253
217
204
9
L1
O1
O12
L8
O16
L16
124

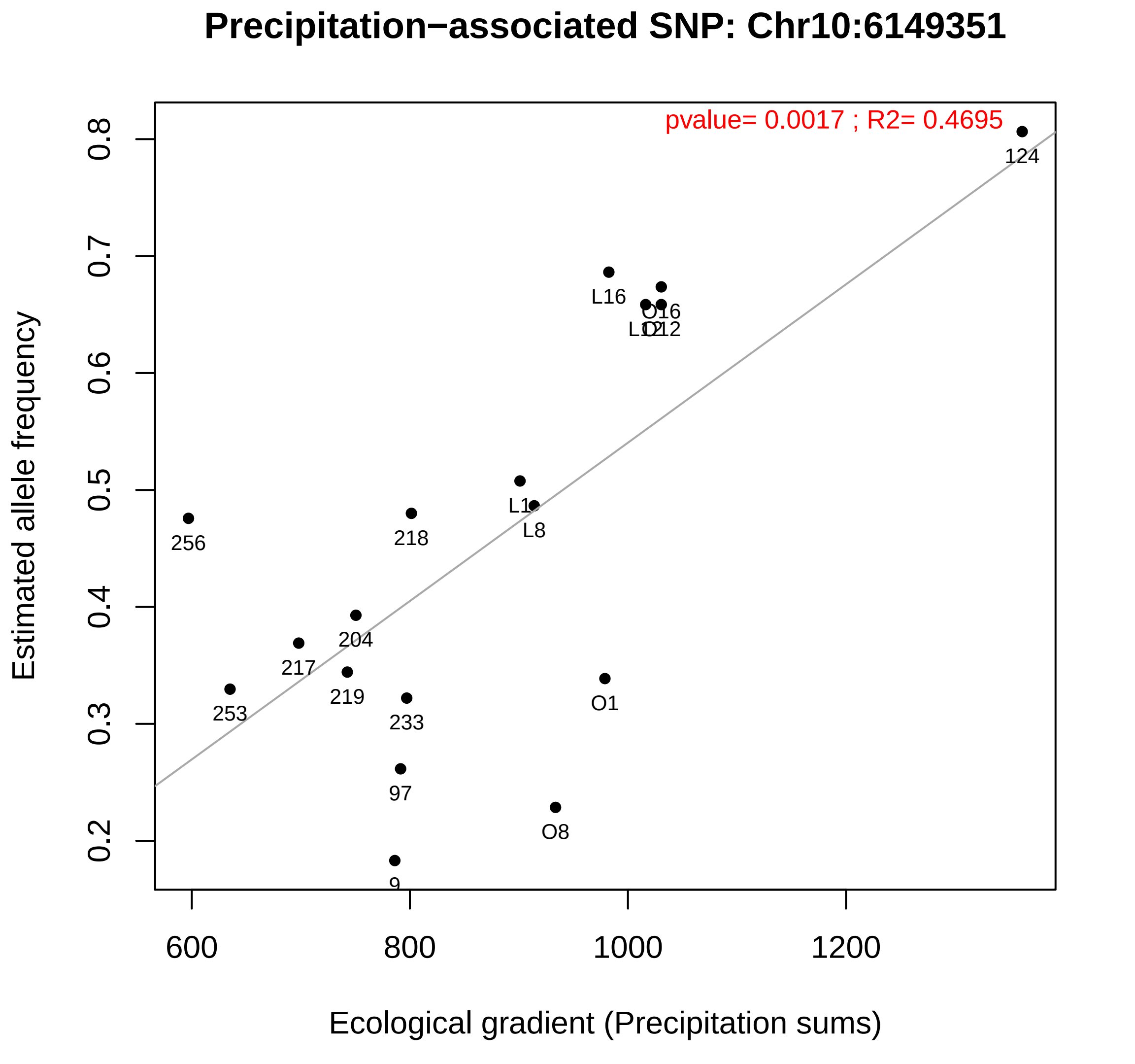

Precipitation−associated SNP: Chr10:6149351
pvalue= 0.0017 ; R2= 0.4695
Estimated allele frequency
0.2
0.3
0.4
0.5
0.6
0.7
0.8
600
800
1000
1200
Ecological gradient (Precipitation sums)
124
L16
O16
L12
O12
L8
256
218
204
217
219
253
233
97
O1
O8
9

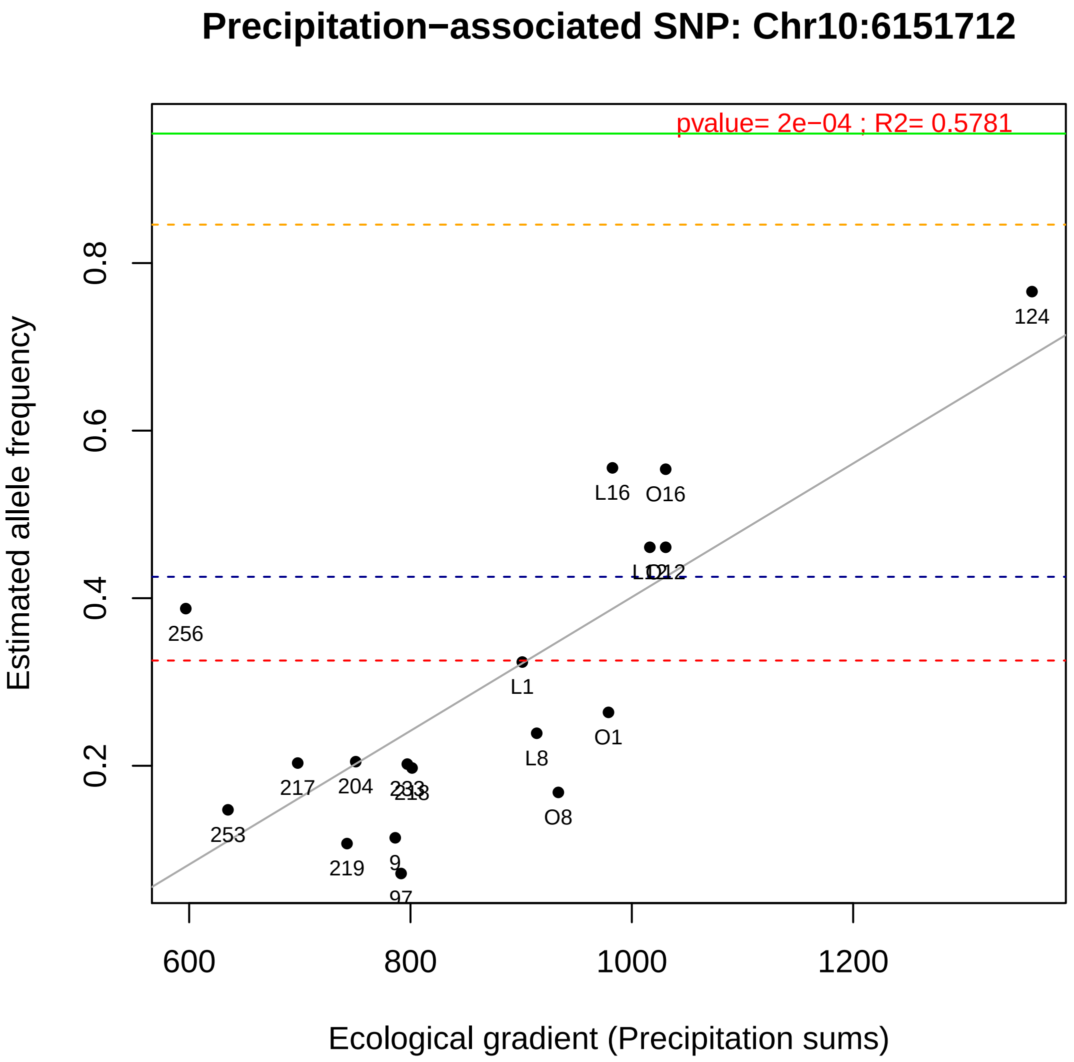

Precipitation−associated SNP: Chr10:6151712
pvalue= 2e−04 ; R2= 0.5781
Estimated allele frequency
0.2
0.4
0.6
0.8
600
800
1000
1200
Ecological gradient (Precipitation sums)
124
L16
O16
256
L1
O1
L8
O8
217
204
253
219
9
97

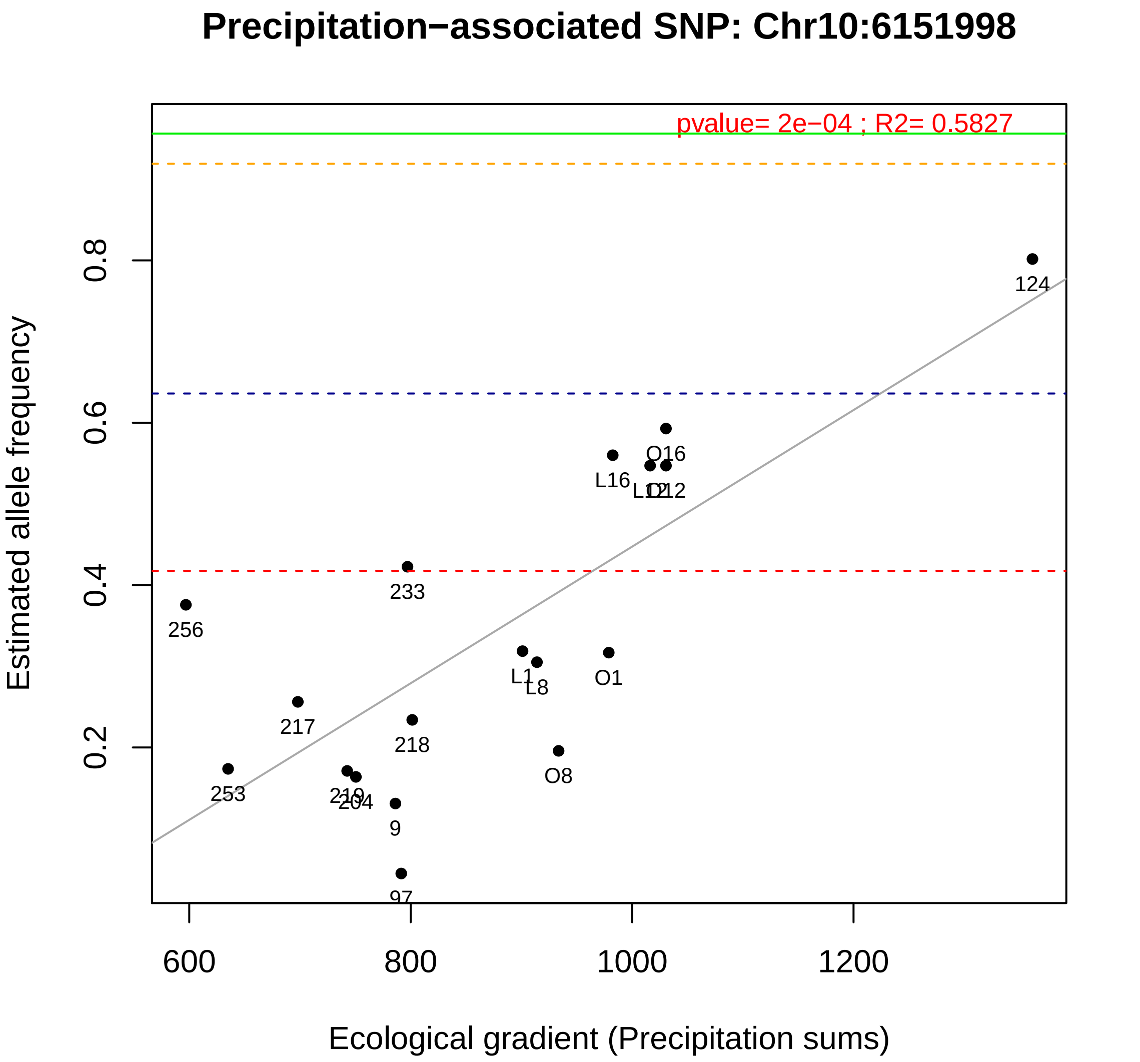

Precipitation−associated SNP: Chr10:6151998
pvalue= 2e−04 ; R2= 0.5827
Estimated allele frequency
0.2
0.4
0.6
0.8
Ecological gradient (Precipitation sums)
600
800
1000
1200
124
O16
L16
L12
O12
233
256
L1
L8
O1
217
218
O8
253
219
204
9
97

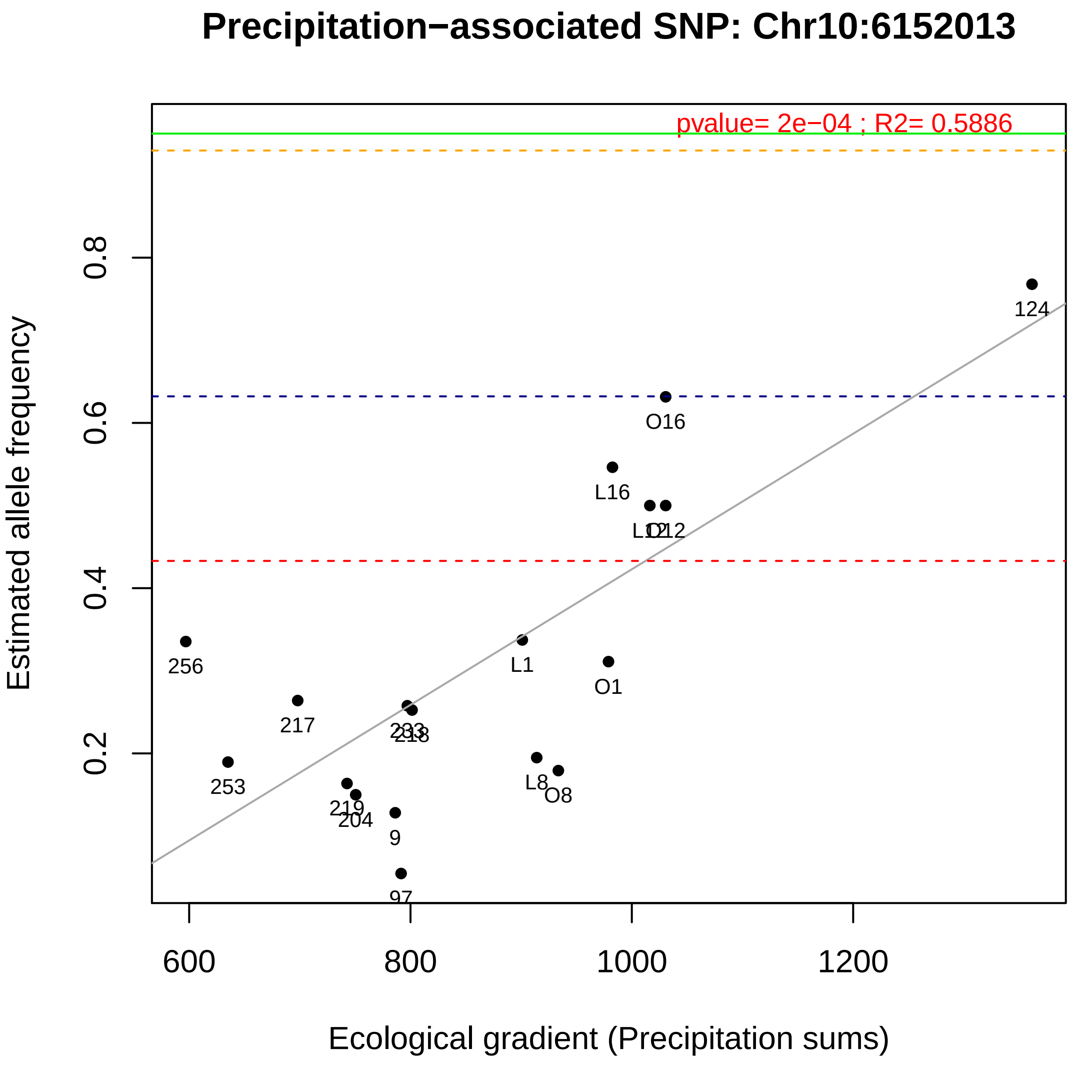

Precipitation−associated SNP: Chr10:6152013
pvalue= 2e−04 ; R2= 0.5886
Estimated allele frequency
0.2
0.4
0.6
0.8
600
800
1000
1200
Ecological gradient (Precipitation sums)
124
O16
L16
L12
O12
256
L1
O1
217
233
218
253
L8
O8
219
204
9
97

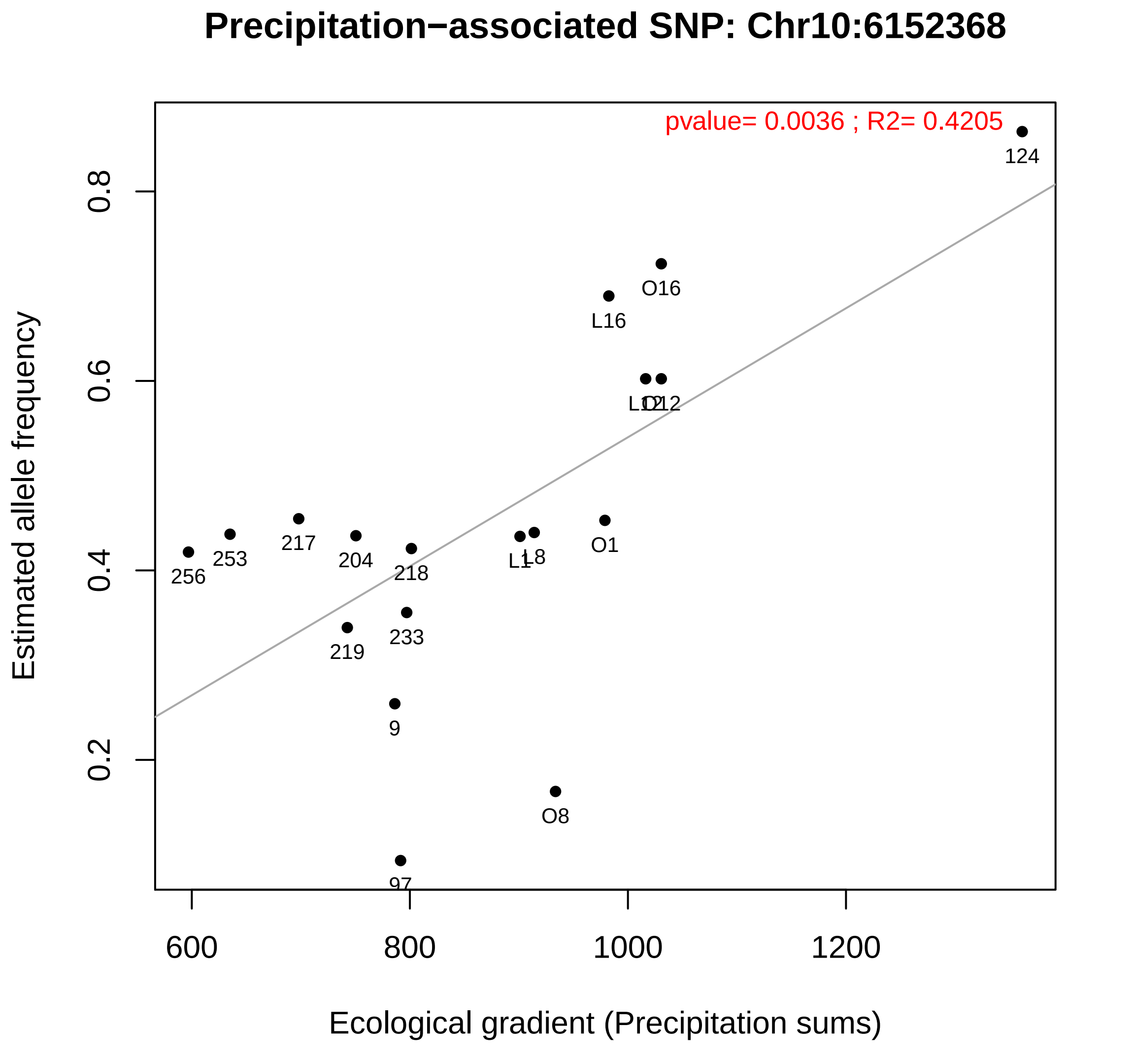
Precipitation−associated SNP: Chr10:6152368
pvalue= 0.0036 ; R2= 0.4205
Estimated allele frequency
Ecological gradient (Precipitation sums)
0.2
0.4
0.6
0.8
600
800
1000
1200
124
O16
L16
L12
O12
256
253
217
204
218
L1
L8
O1
219
233
9
O8
97

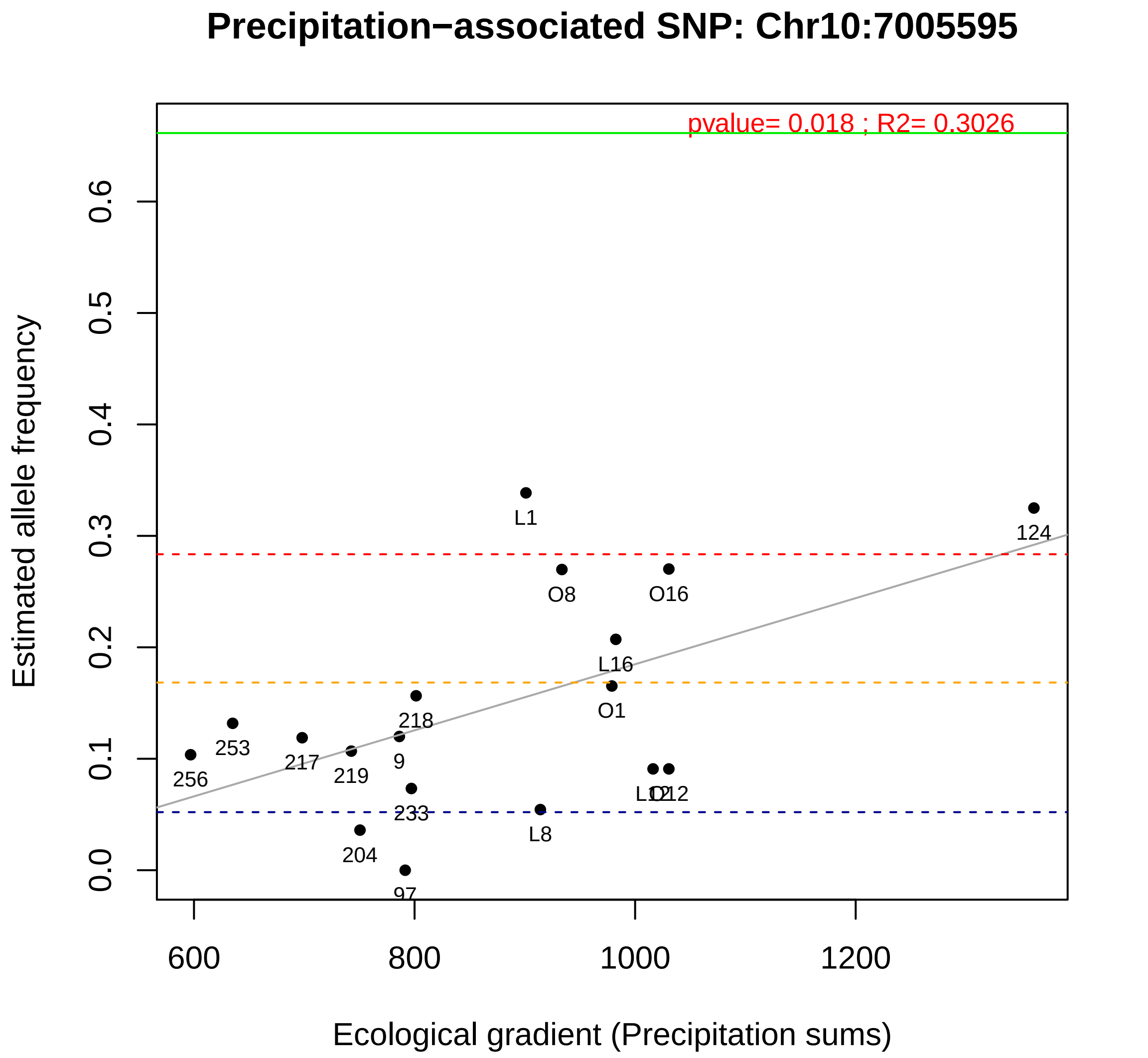

Precipitation−associated SNP: Chr10:7005595
pvalue= 0.018 ; R2= 0.3026
Estimated allele frequency
0.0
0.1
0.2
0.3
0.4
0.5
0.6
600
800
1000
1200
Ecological gradient (Precipitation sums)
L1
124
O8
O16
L16
O1
218
253
217
256
219
9
233
204
97
L8

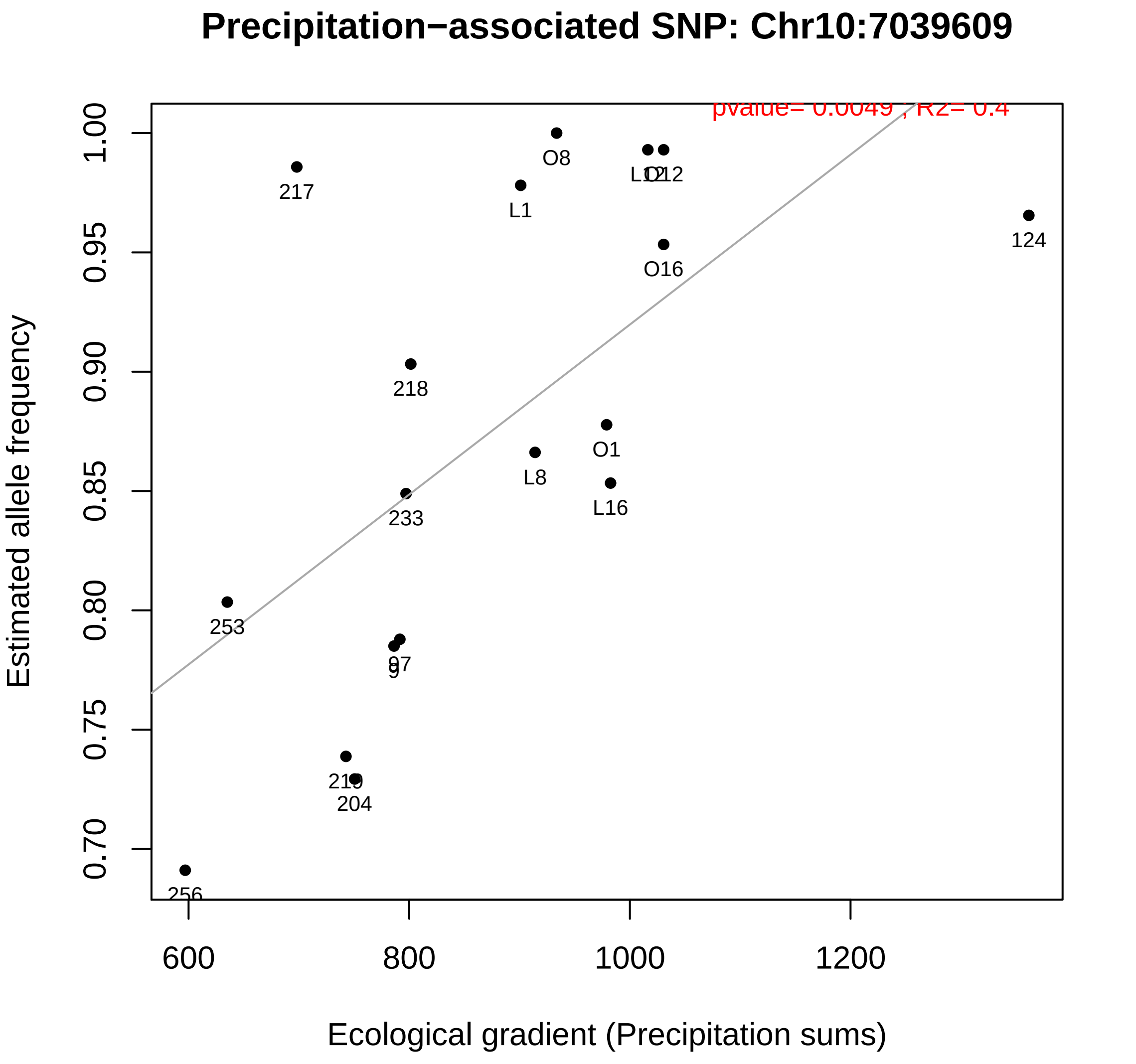

Precipitation−associated SNP: Chr10:7039609
pvalue= 0.0049 ; R2= 0.4
Estimated allele frequency
Ecological gradient (Precipitation sums)
0.70
0.75
0.80
0.85
0.90
0.95
1.00
600
800
1000
1200
O8
217
L1
L12
O12
124
O16
218
O1
L8
L16
233
253
97
9
219
204
256

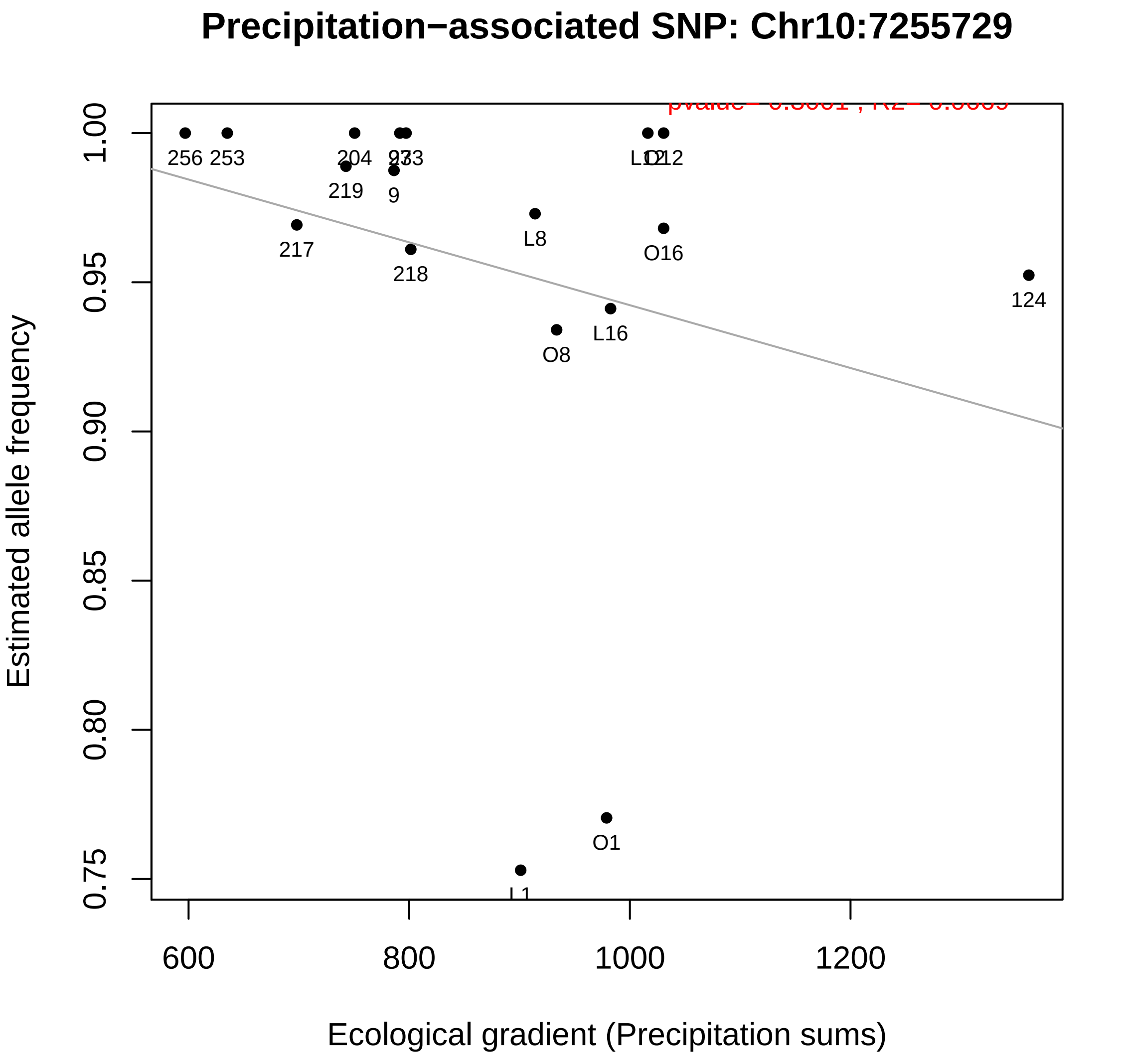
Precipitation−associated SNP: Chr10:7255729
Estimated allele frequency
Ecological gradient (Precipitation sums)
0.75
0.80
0.85
0.90
0.95
1.00
600
800
1000
1200
256
253
204
219
9
217
218
L8
O16
O8
L16
124
O1

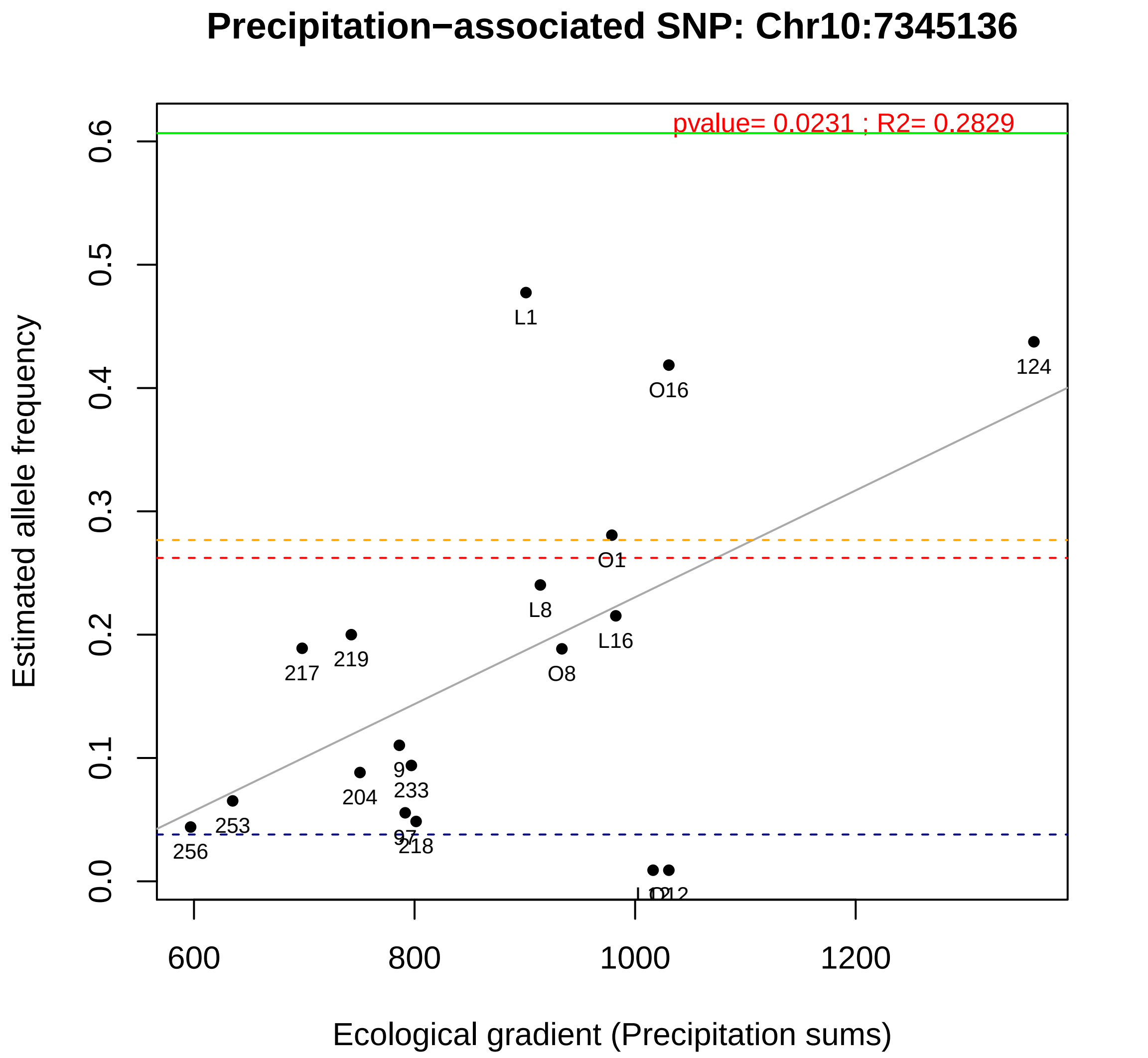

Precipitation−associated SNP: Chr10:7345136
pvalue= 0.0231 ; R2= 0.2829
Estimated allele frequency
Ecological gradient (Precipitation sums)
0.0
0.1
0.2
0.3
0.4
0.5
0.6
600
800
1000
1200
L1
124
O16
O1
L8
L16
O8
217
219
9
233
204
253
256
97
218

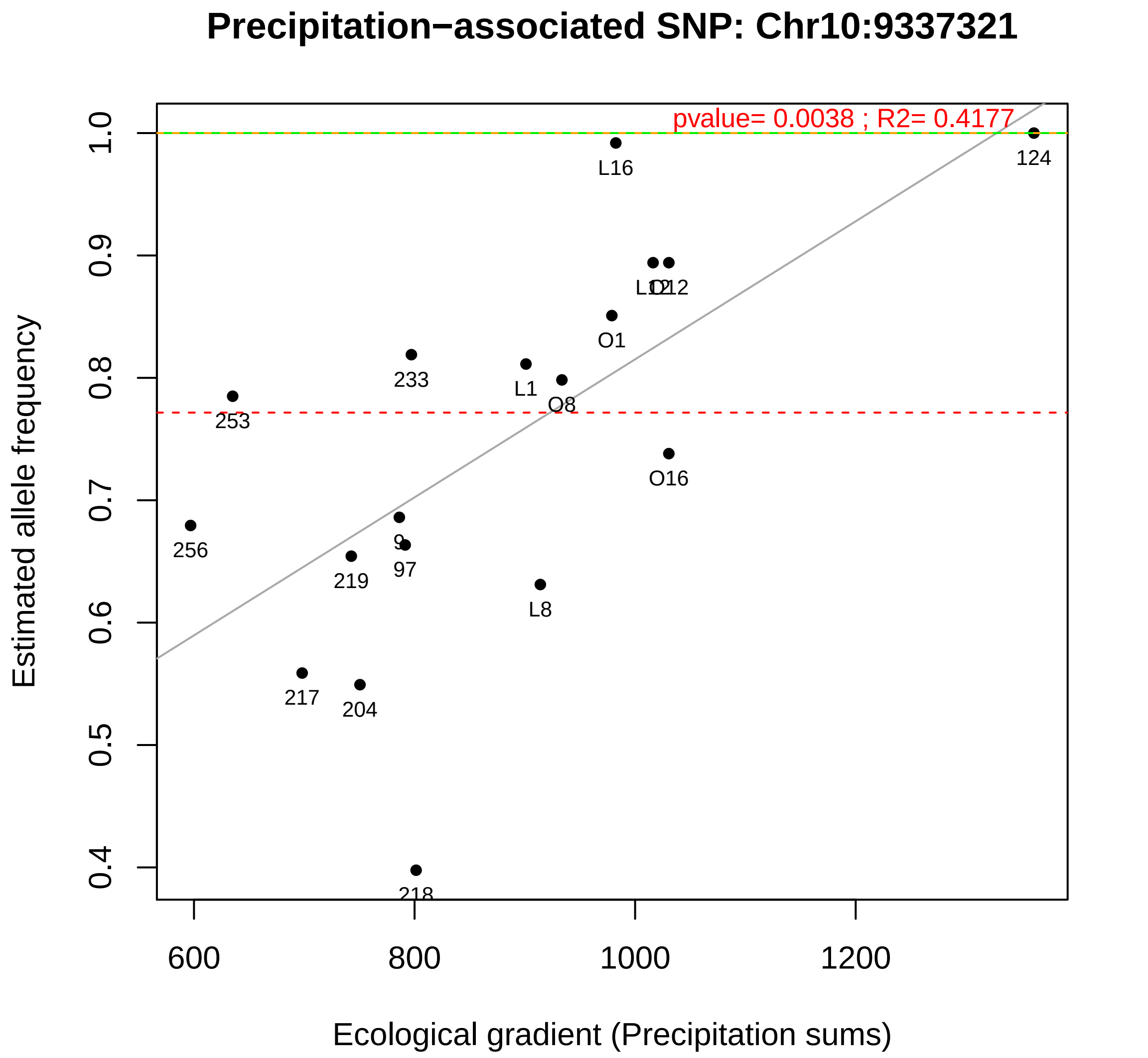
Precipitation−associated SNP: Chr10:9337321
pvalue= 0.0038 ; R2= 0.4177
Estimated allele frequency
Ecological gradient (Precipitation sums)
0.4
0.5
0.6
0.7
0.8
0.9
1.0
600
800
1000
1200
124
L16
O12
O1
233
L1
O8
253
O16
256
219
97
L8
217
204
218

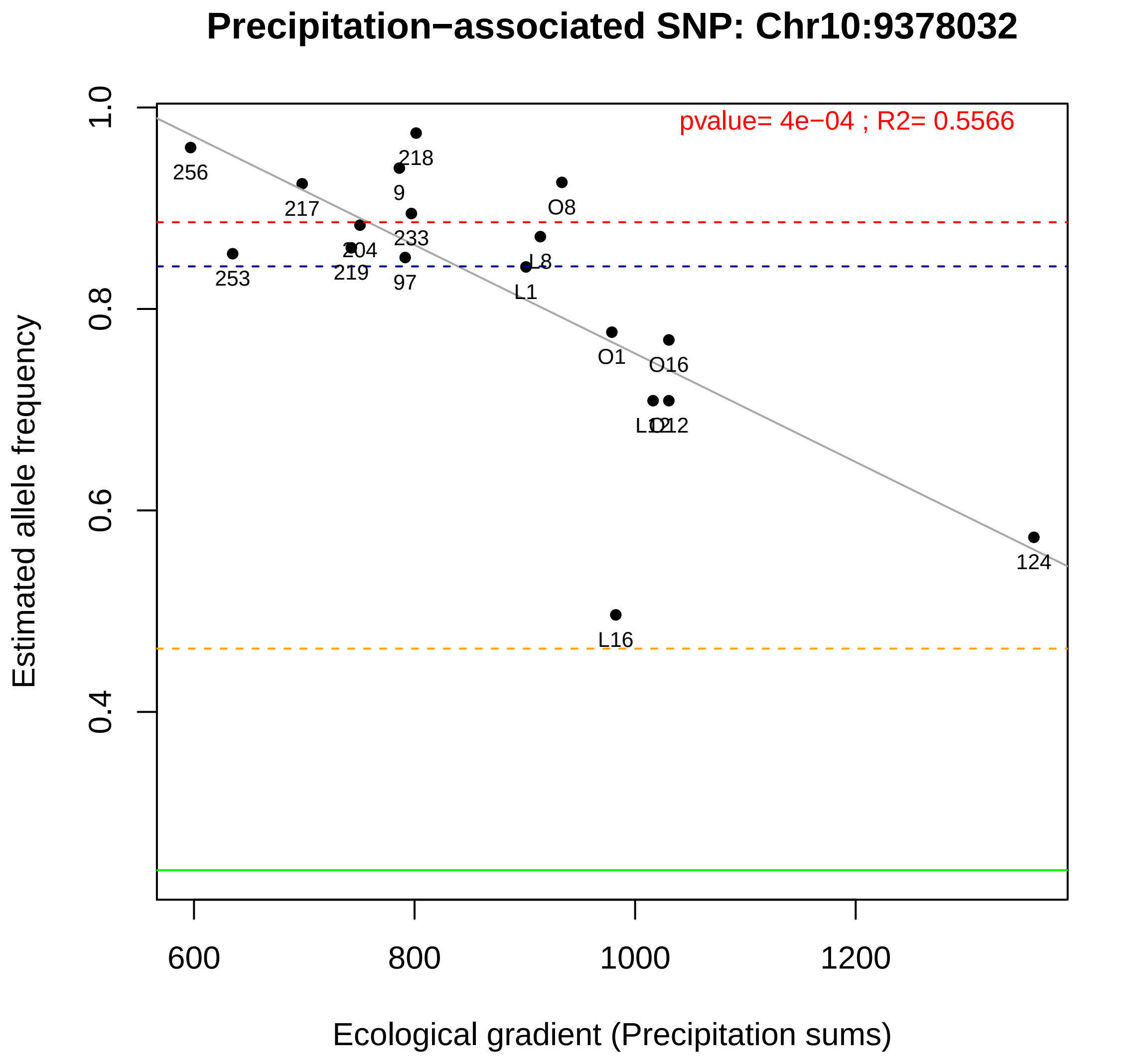

Precipitation−associated SNP: Chr10:9378032
pvalue= 4e−04 ; R2= 0.5566
Estimated allele frequency
Ecological gradient (Precipitation sums)
1.0
0.8
0.6
0.4
600
800
1000
1200
256
218
9
217
O8
233
204
253
219
97
L8
L1
O1
O16
O12
124
L16

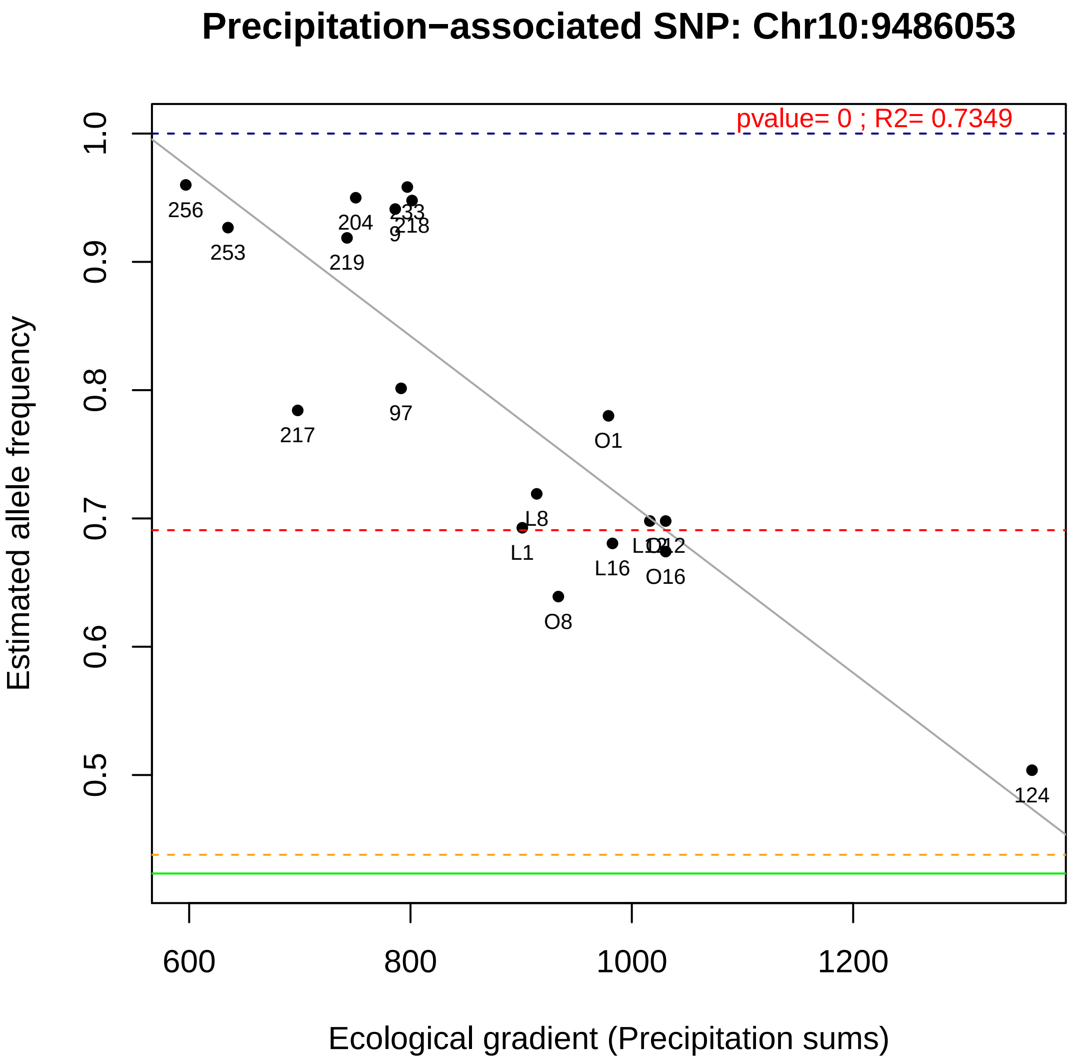

Precipitation−associated SNP: Chr10:9486053
pvalue= 0 ; R2= 0.7349
Estimated allele frequency
1.0
0.9
0.8
0.7
0.6
0.5
256
253
204
219
218
9
217
97
O1
L8
L1
L16
O8
O16
124
600
800
1000
1200
Ecological gradient (Precipitation sums)

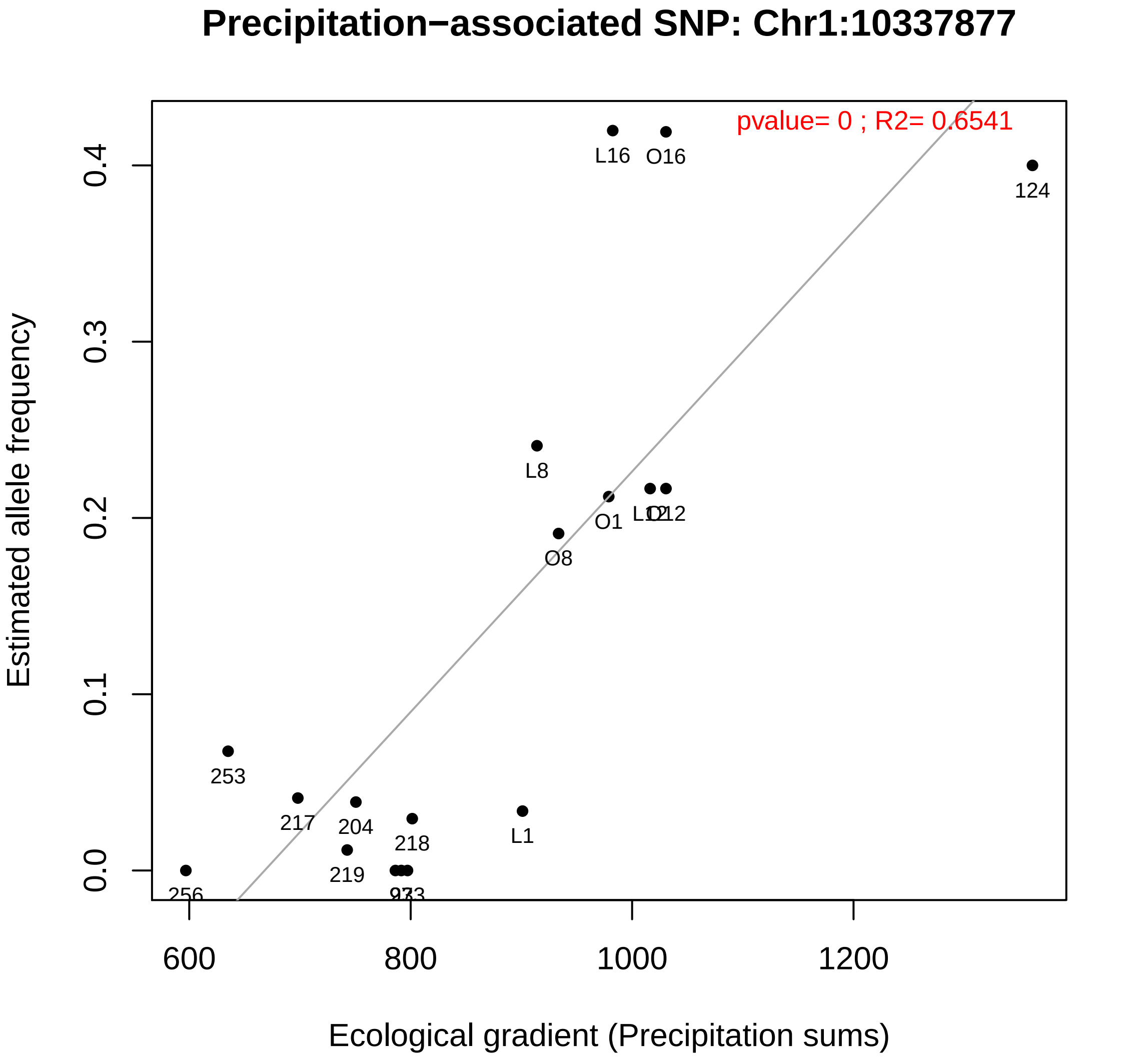
Precipitation−associated SNP: Chr1:10337877
pvalue= 0 ; R2= 0.6541
Estimated allele frequency
Ecological gradient (Precipitation sums)
0.0
0.1
0.2
0.3
0.4
600
800
1000
1200
L16
O16
124
L8
O1
O12
O8
253
217
204
218
L1
219
256

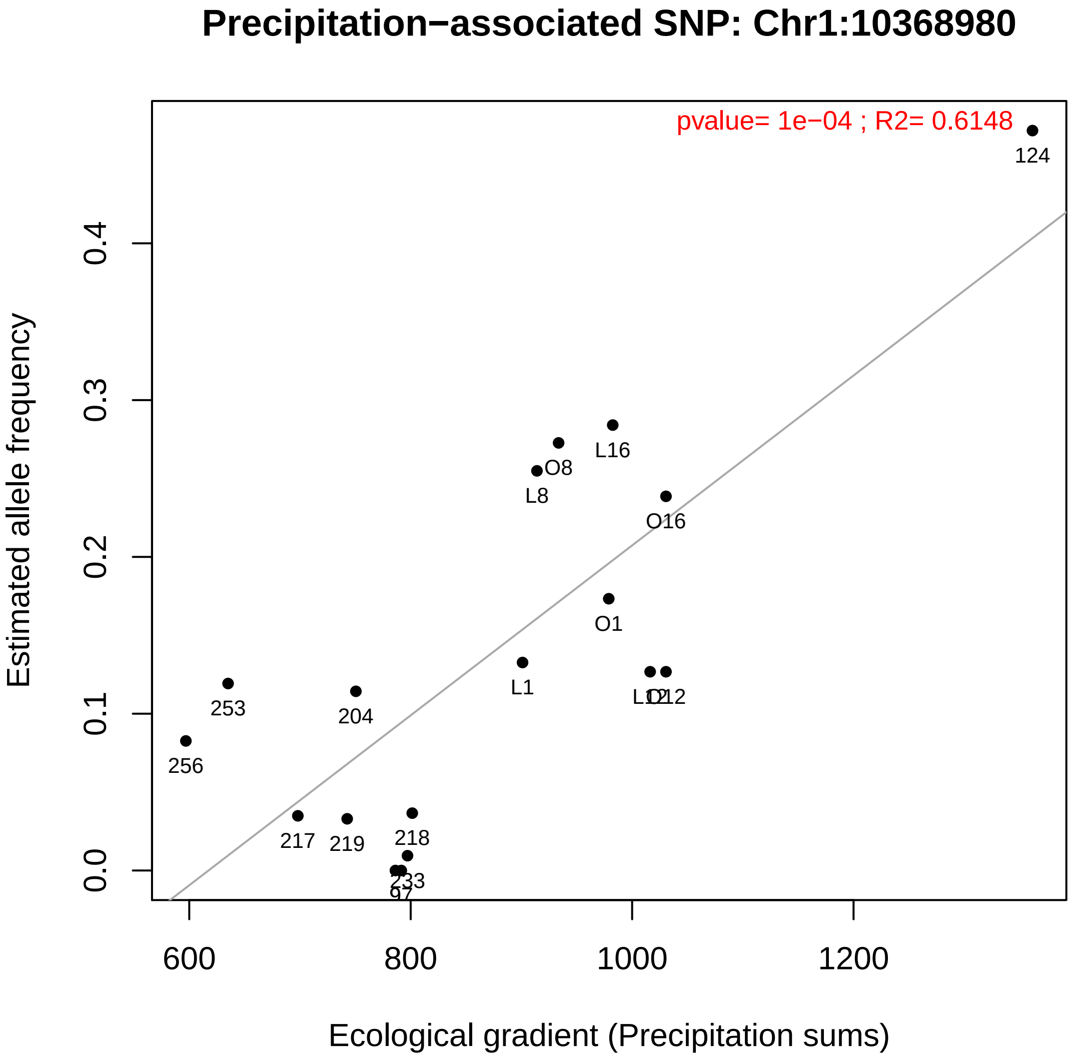

Precipitation−associated SNP: Chr1:10368980
pvalue= 1e−04 ; R2= 0.6148
Estimated allele frequency
Ecological gradient (Precipitation sums)
0.0
0.1
0.2
0.3
0.4
600
800
1000
1200
124
L16
O8
L8
O16
O1
L1
253
204
256
217
219
218
233

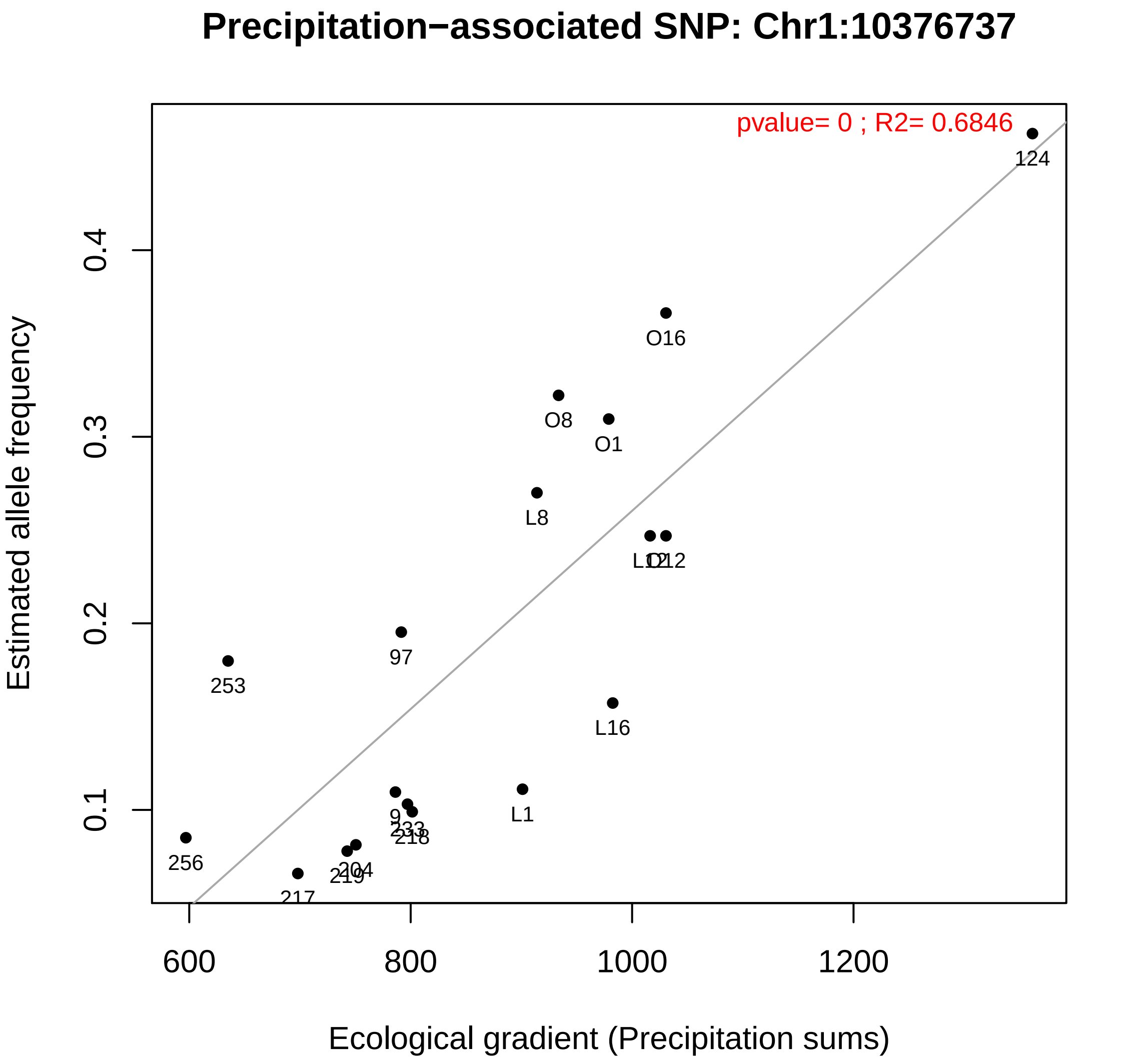
Precipitation−associated SNP: Chr1:10376737
pvalue= 0 ; R2= 0.6846
124
O16
O8
O1
L8
L12
O12
97
253
L16
L1
256
217
219
204
233
218
0.1
0.2
0.3
0.4
600
800
1000
1200
Estimated allele frequency
Ecological gradient (Precipitation sums)

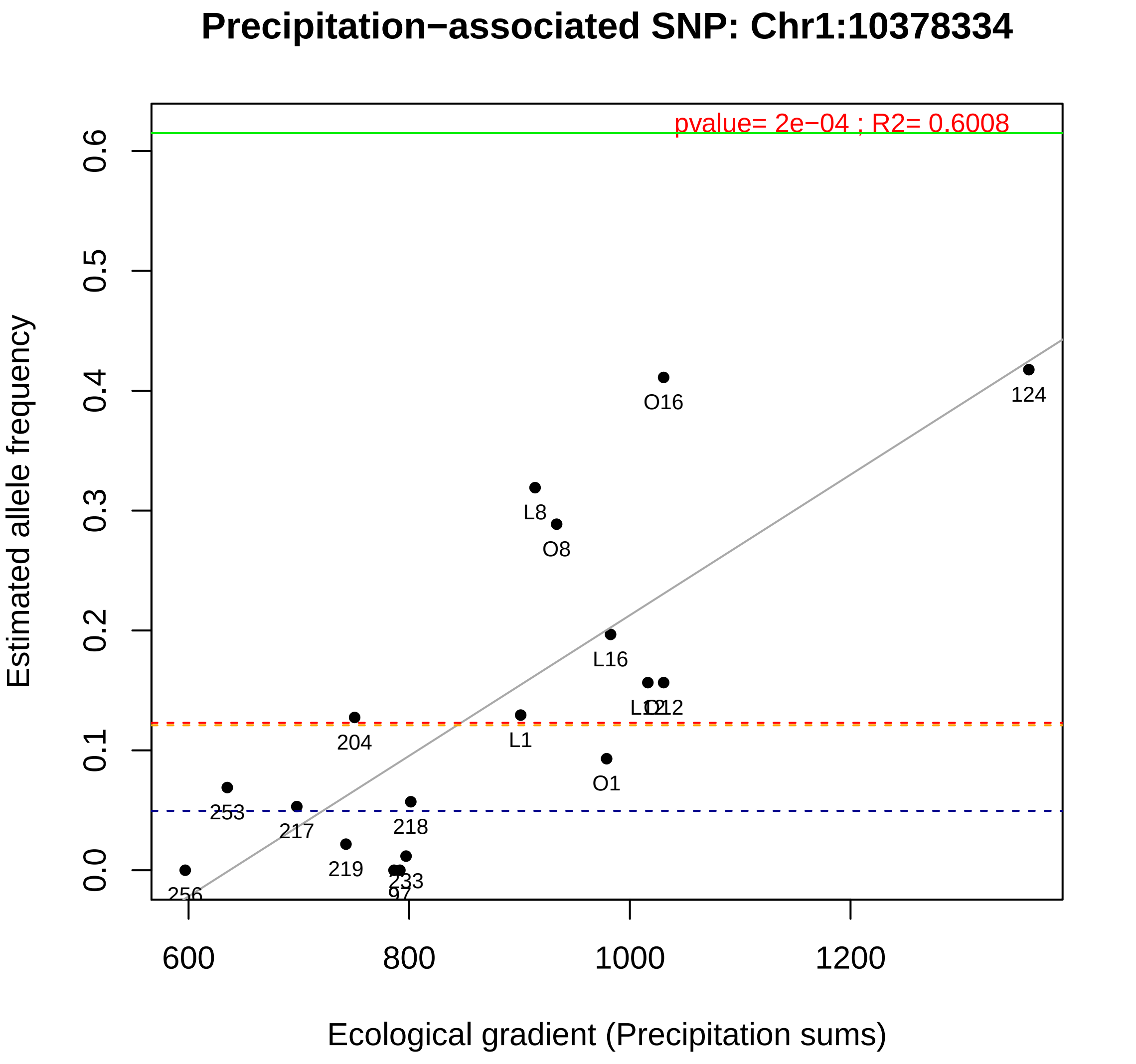
Precipitation−associated SNP: Chr1:10378334
pvalue= 2e−04 ; R2= 0.6008
Estimated allele frequency
0.0
0.1
0.2
0.3
0.4
0.5
0.6
Ecological gradient (Precipitation sums)
600
800
1000
1200
O16
124
L8
O8
L16
204
L1
O1
253
217
218
219
233
256

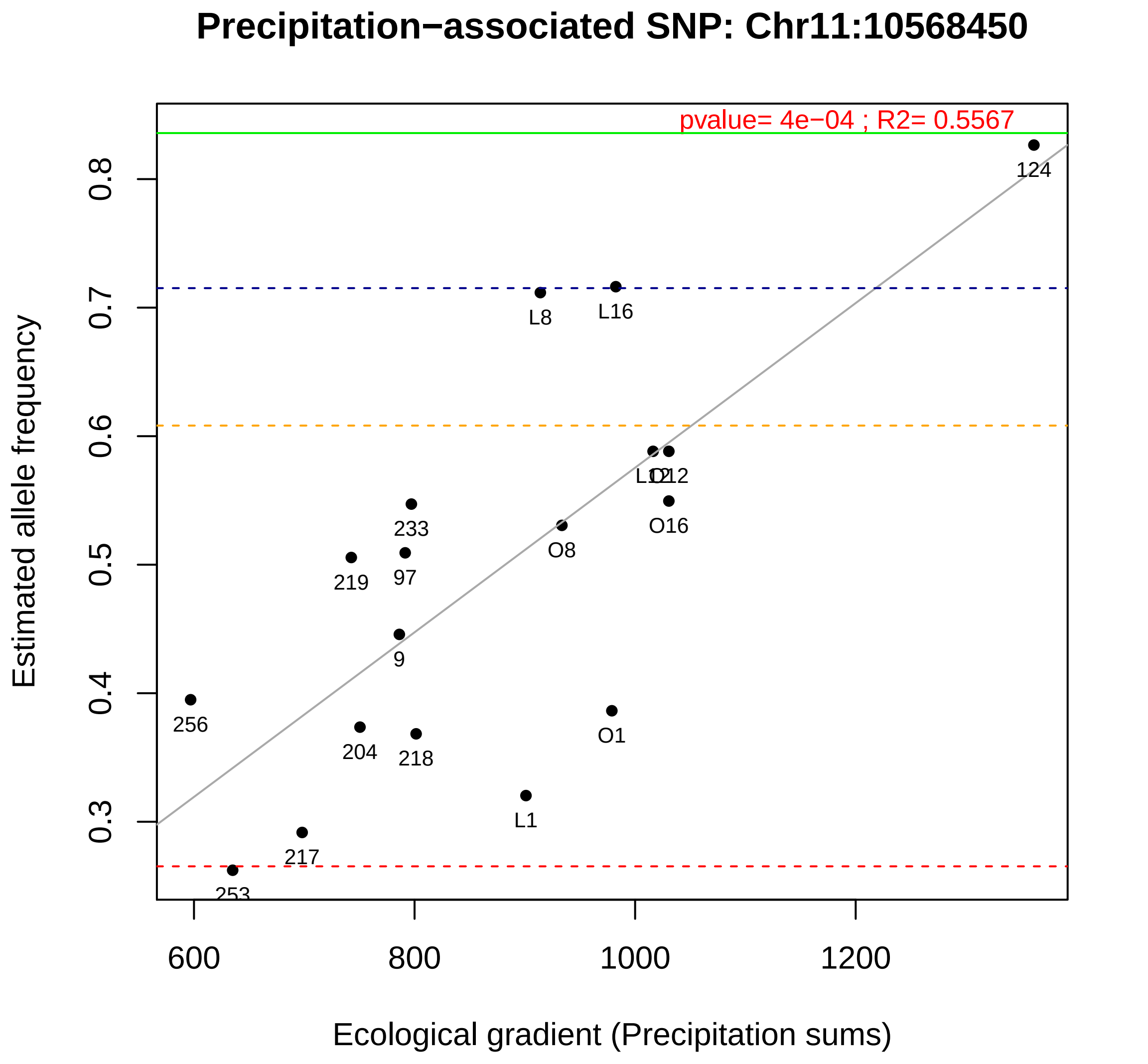
Precipitation−associated SNP: Chr11:10568450
pvalue= 4e−04 ; R2= 0.5567
Estimated allele frequency
0.3
0.4
0.5
0.6
0.7
0.8
600
800
1000
1200
Ecological gradient (Precipitation sums)
124
L8
L16
O12
O16
233
O8
219
97
9
256
204
218
O1
L1
217
253

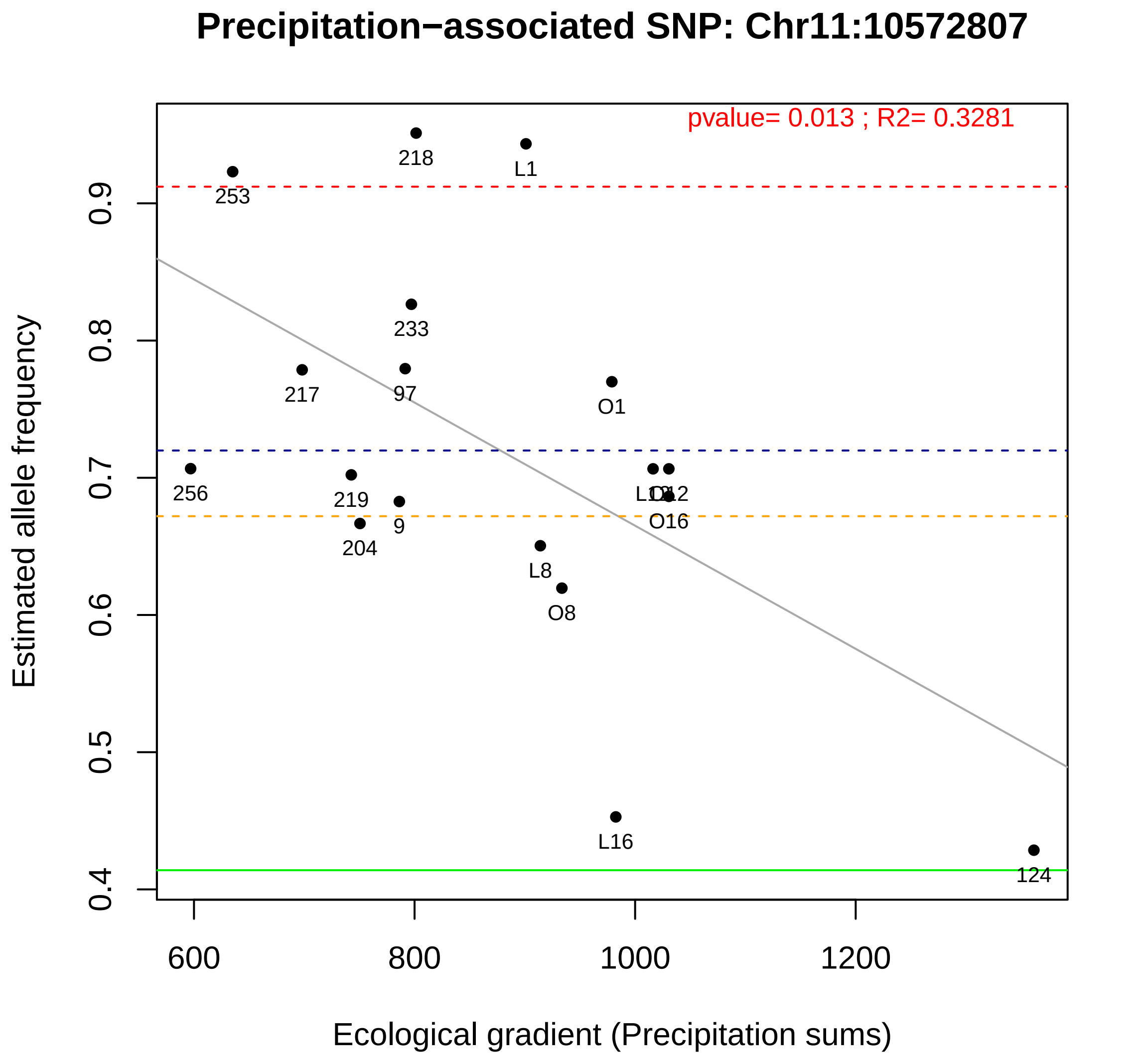
Precipitation−associated SNP: Chr11:10572807
pvalue= 0.013 ; R2= 0.3281
Estimated allele frequency
Ecological gradient (Precipitation sums)
0.4
0.5
0.6
0.7
0.8
0.9
600
800
1000
1200
218
L1
253
233
217
97
O1
256
219
9
204
L12
O2
O16
L8
O8
L16
124

**Precipitation−associated SNP: Chr11:12218834**

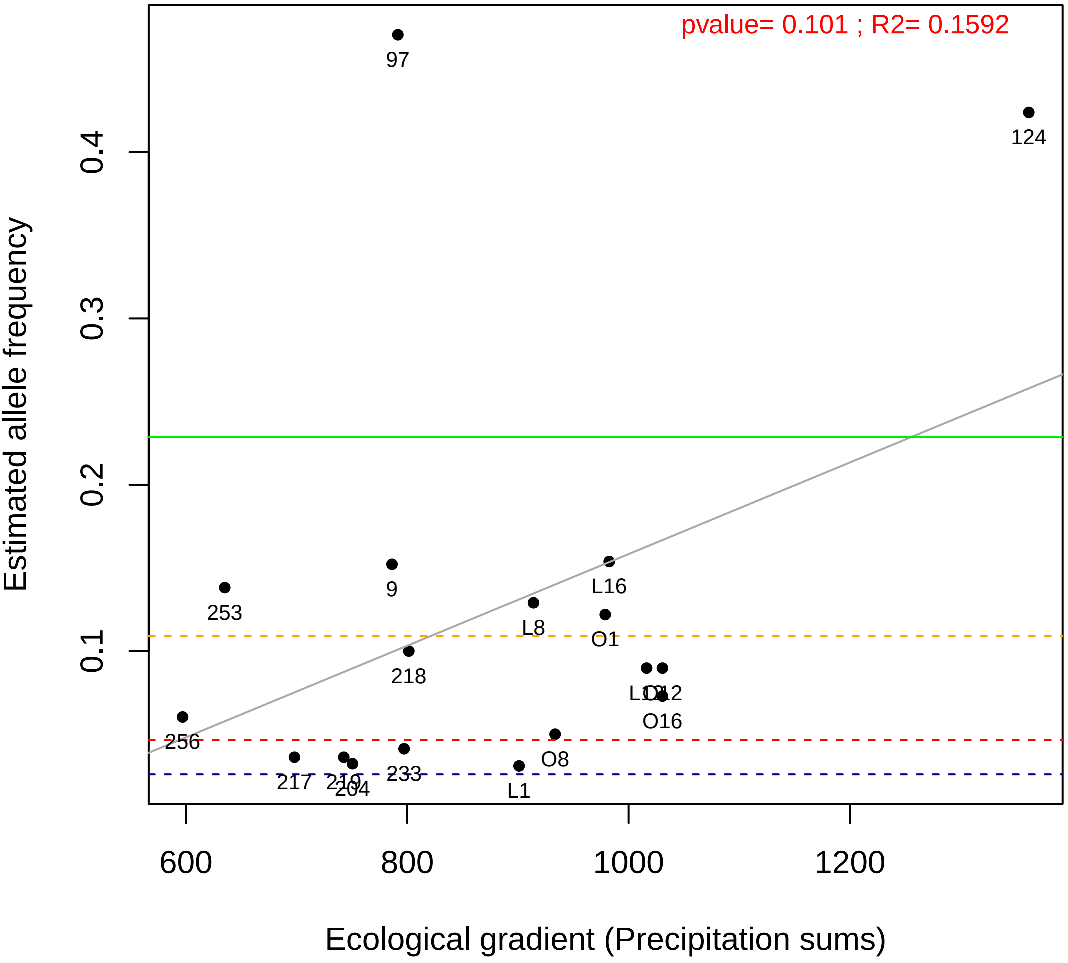

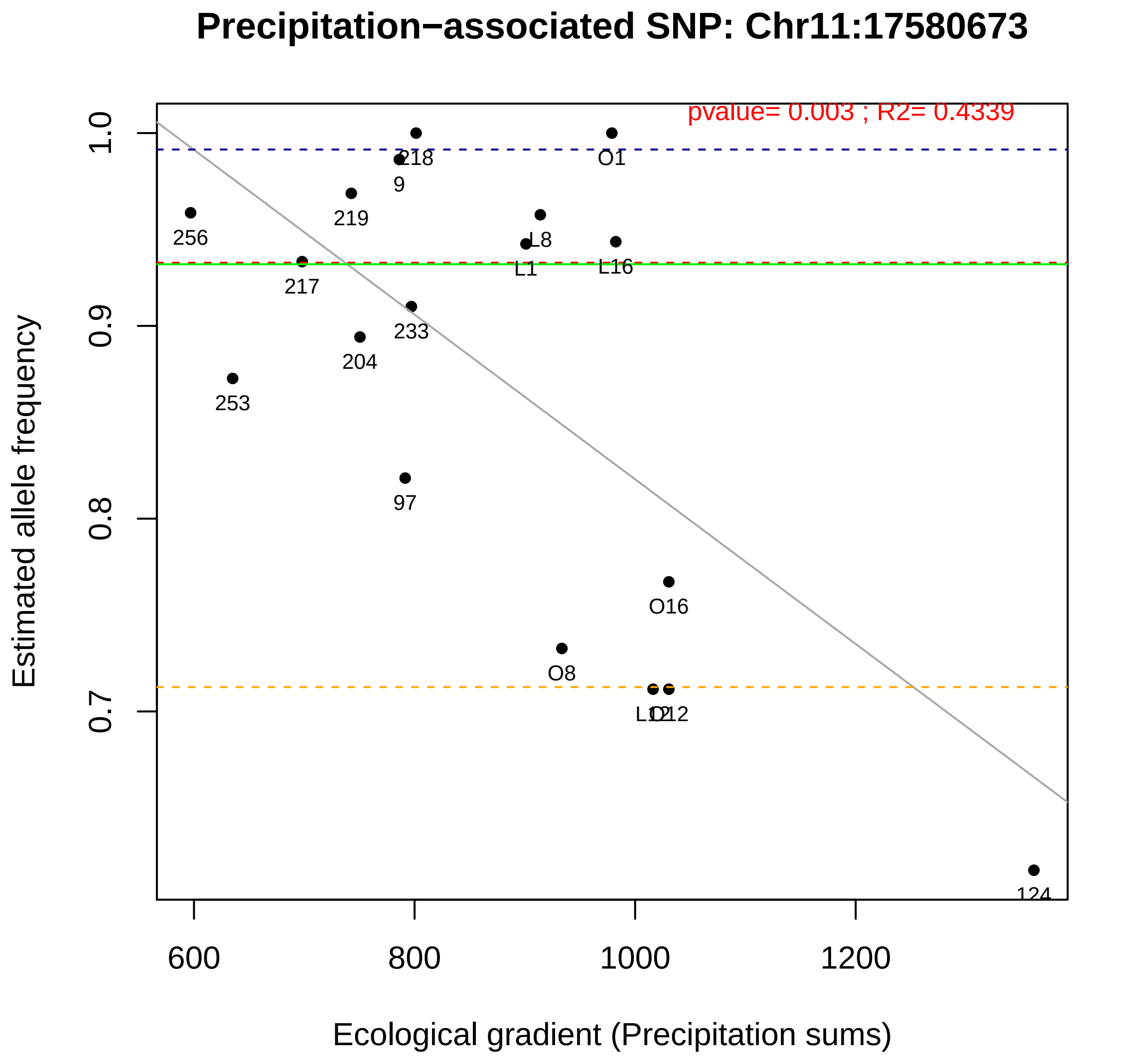
Precipitation−associated SNP: Chr11:17580673
pvalue= 0.003 ; R2= 0.4339
Estimated allele frequency
1.0
0.9
0.8
0.7
600
800
1000
1200
Ecological gradient (Precipitation sums)
218
9
O1
219
256
L8
L1
L16
217
233
204
253
97
O16
O8
L12
O12
124

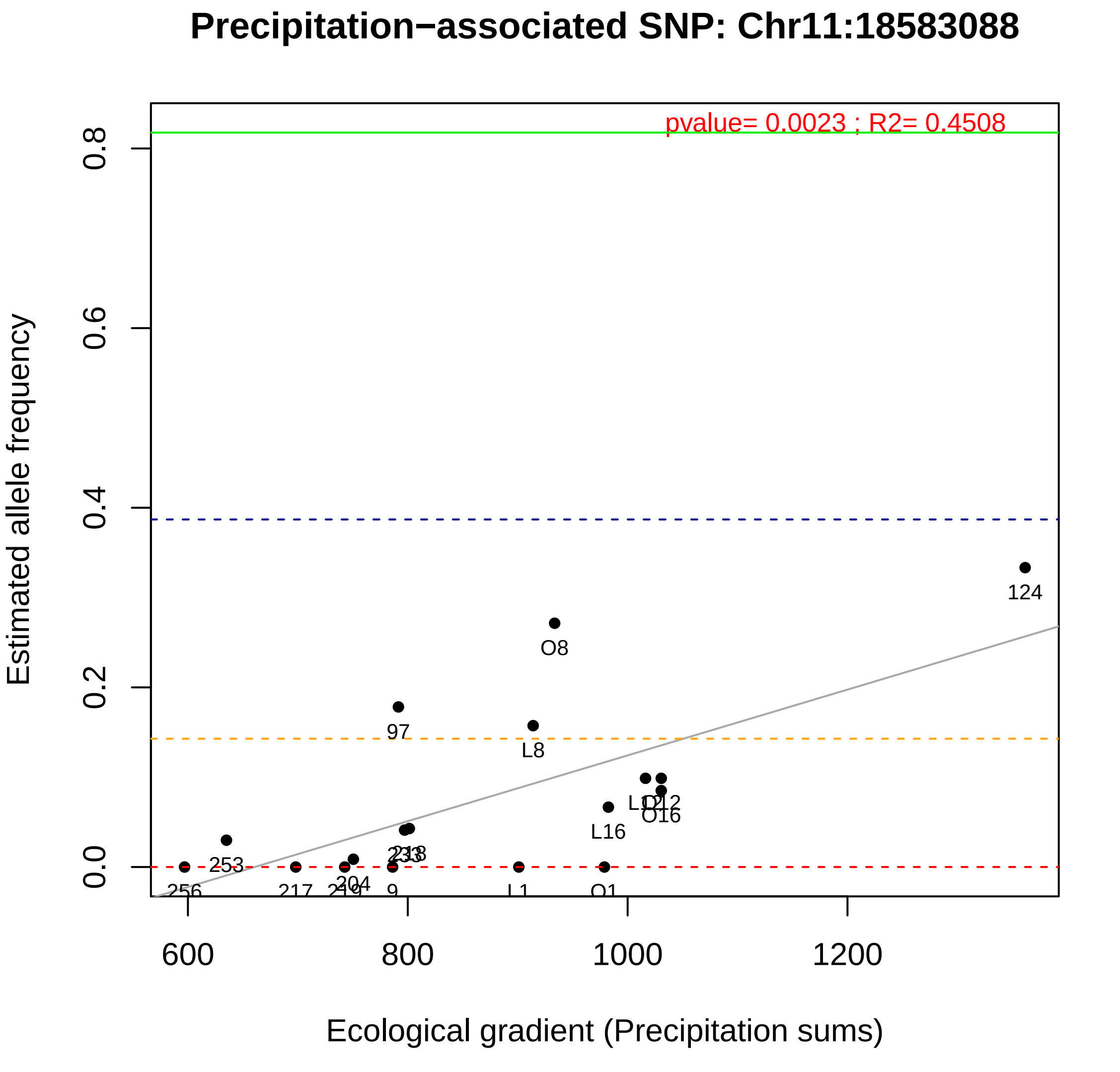

Precipitation−associated SNP: Chr11:18583088
pvalue= 0.0023 ; R2= 0.4508
Estimated allele frequency
0.0
0.2
0.4
0.6
0.8
600
800
1000
1200
Ecological gradient (Precipitation sums)
124
O8
97
L8
L16
O16
253
204

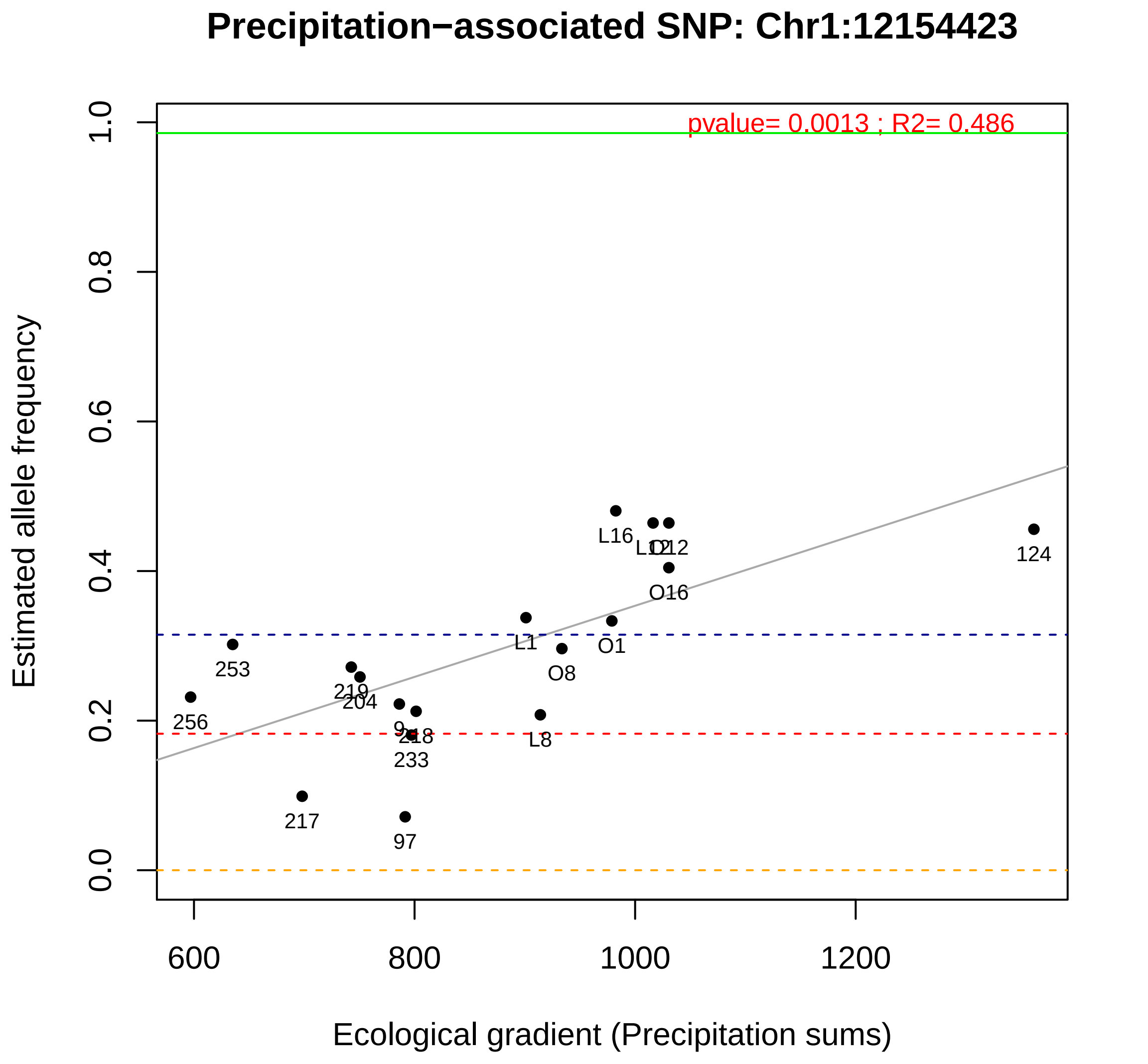
Precipitation−associated SNP: Chr1:12154423
pvalue= 0.0013 ; R2= 0.486
Estimated allele frequency
0.0
0.2
0.4
0.6
0.8
1.0
Ecological gradient (Precipitation sums)
600
800
1000
1200
L16
L12
O12
124
O16
L1
O1
O8
253
219
204
256
9
218
233
L8
217
97

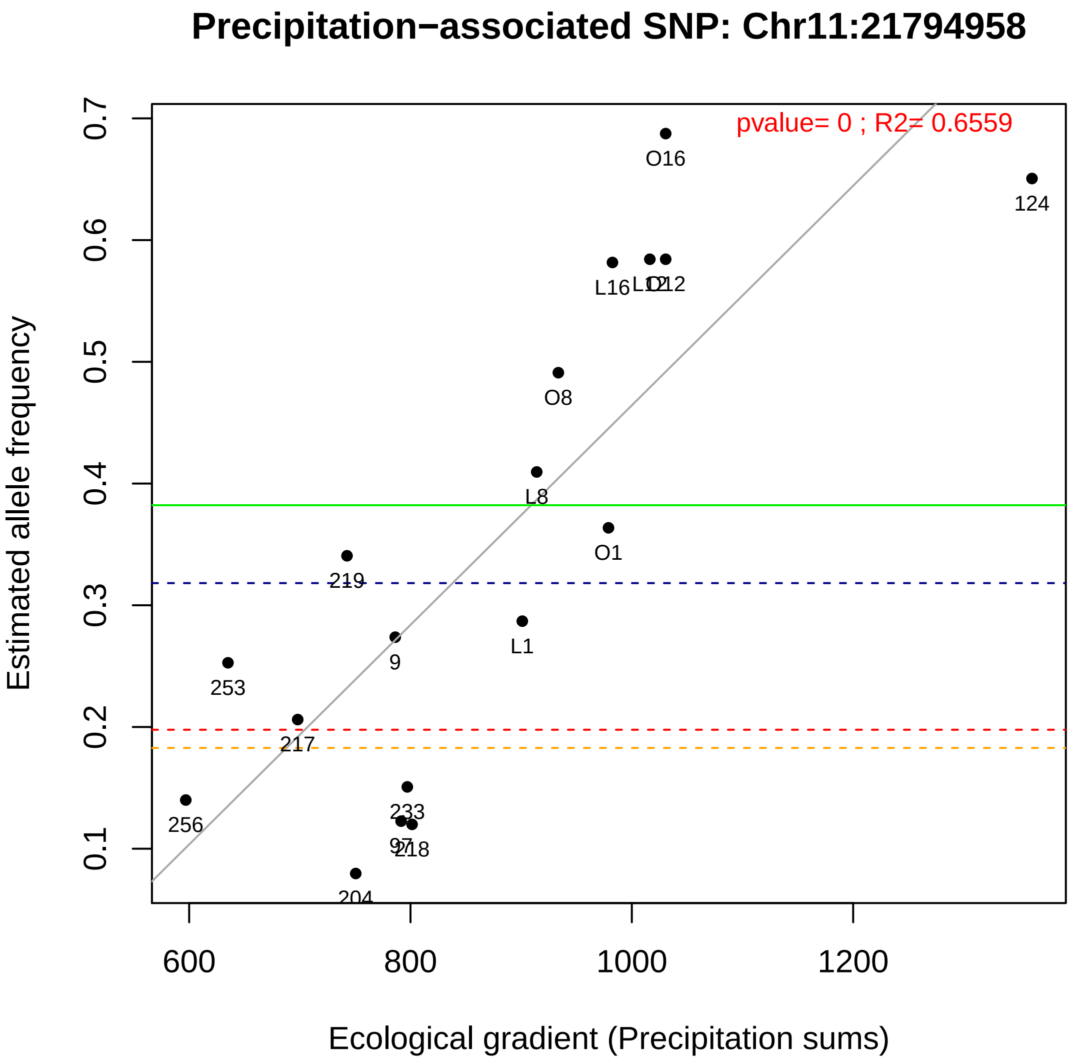

Precipitation−associated SNP: Chr11:21794958
pvalue= 0 ; R2= 0.6559
O16
124
L16
L12
O12
O8
L8
O1
219
9
L1
253
217
256
233
97
218
204
0.1
0.2
0.3
0.4
0.5
0.6
0.7
600
800
1000
1200
Estimated allele frequency
Ecological gradient (Precipitation sums)

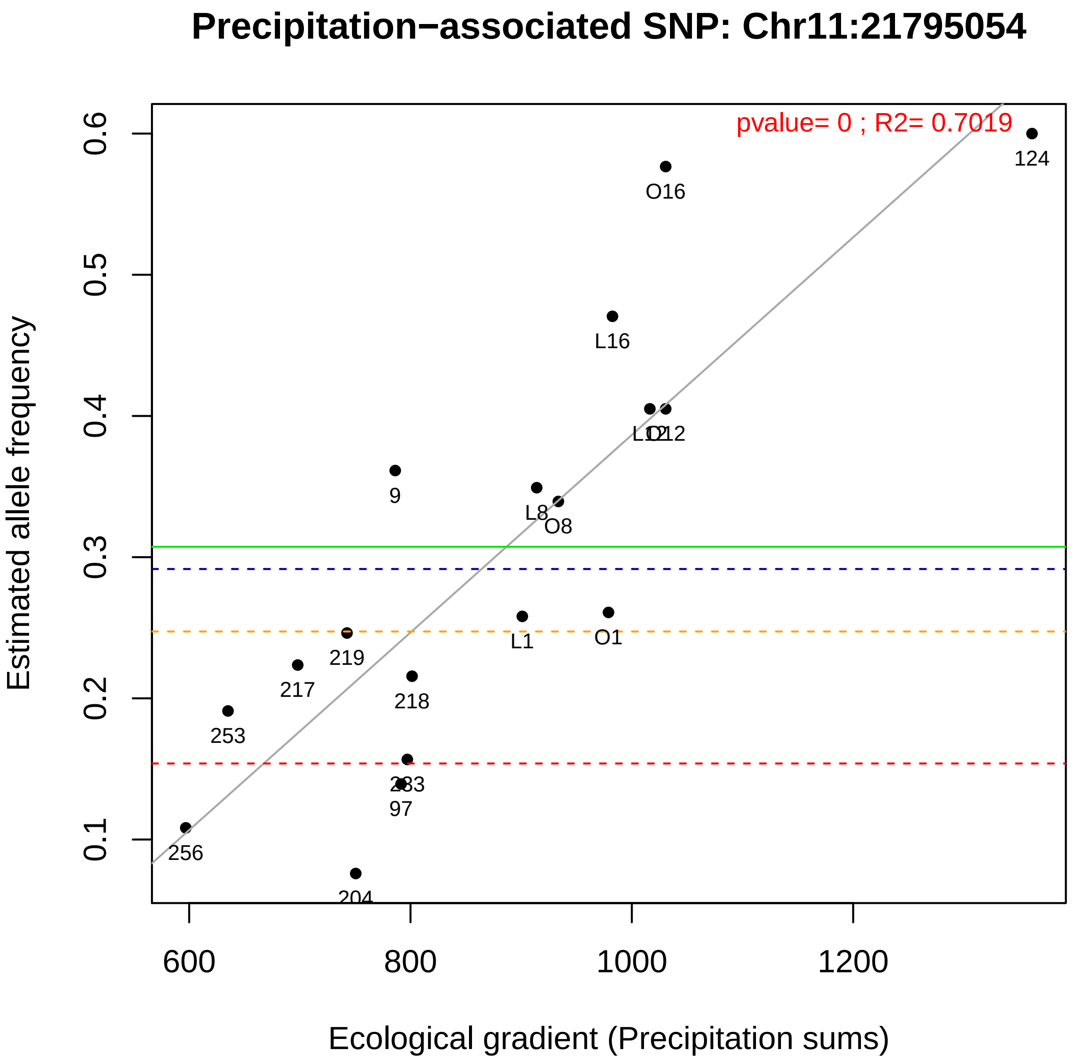

Precipitation−associated SNP: Chr11:21795054
pvalue= 0 ; R2= 0.7019
124
O16
L16
L12
O12
9
L8
O8
L1
O1
219
217
218
253
233
97
256
204
0.1
0.2
0.3
0.4
0.5
0.6
600
800
1000
1200
Estimated allele frequency
Ecological gradient (Precipitation sums)

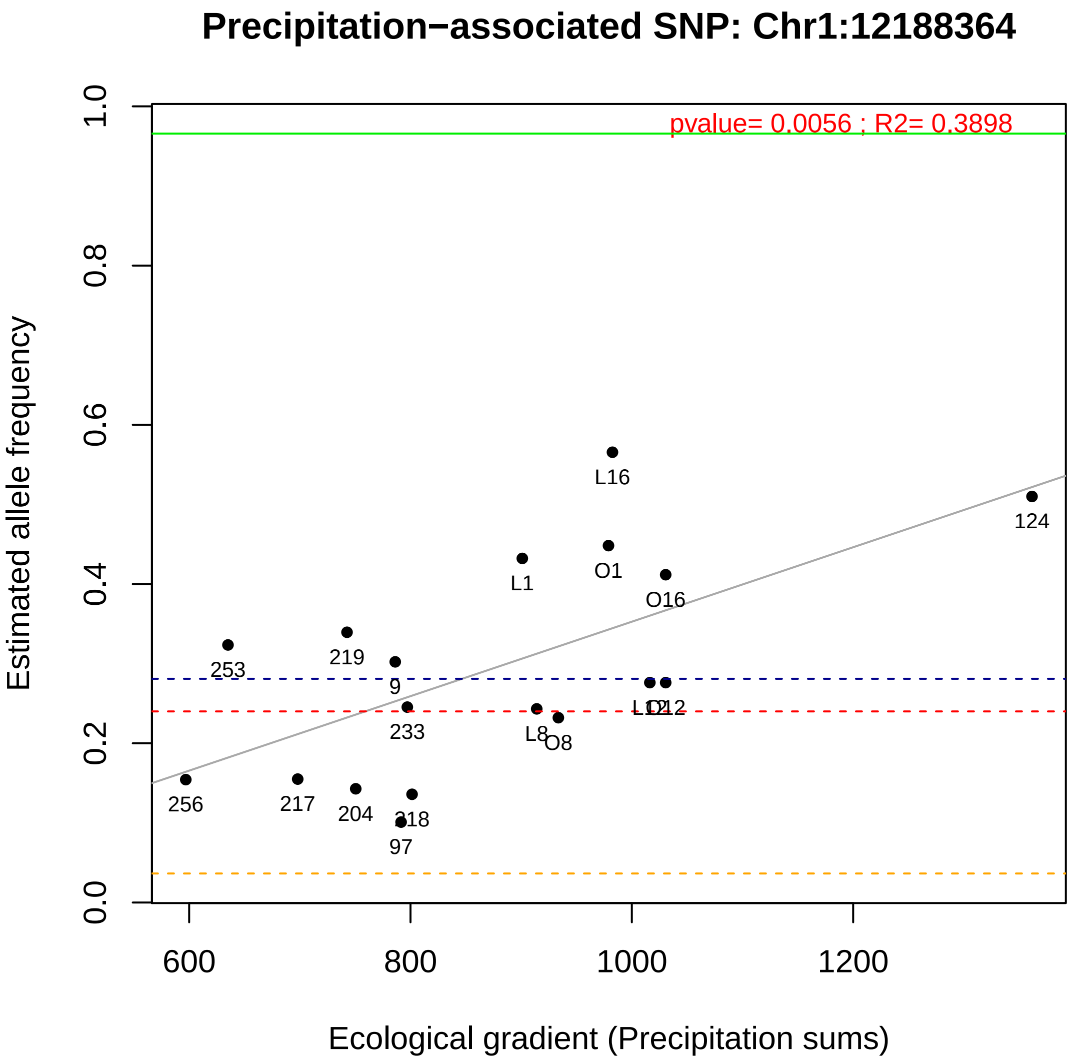

Precipitation−associated SNP: Chr1:12188364
pvalue= 0.0056 ; R2= 0.3898
Estimated allele frequency
0.0
0.2
0.4
0.6
0.8
1.0
600
800
1000
1200
Ecological gradient (Precipitation sums)
L16
124
O1
L1
O16
253
219
9
233
L8
O8
256
217
204
218
97

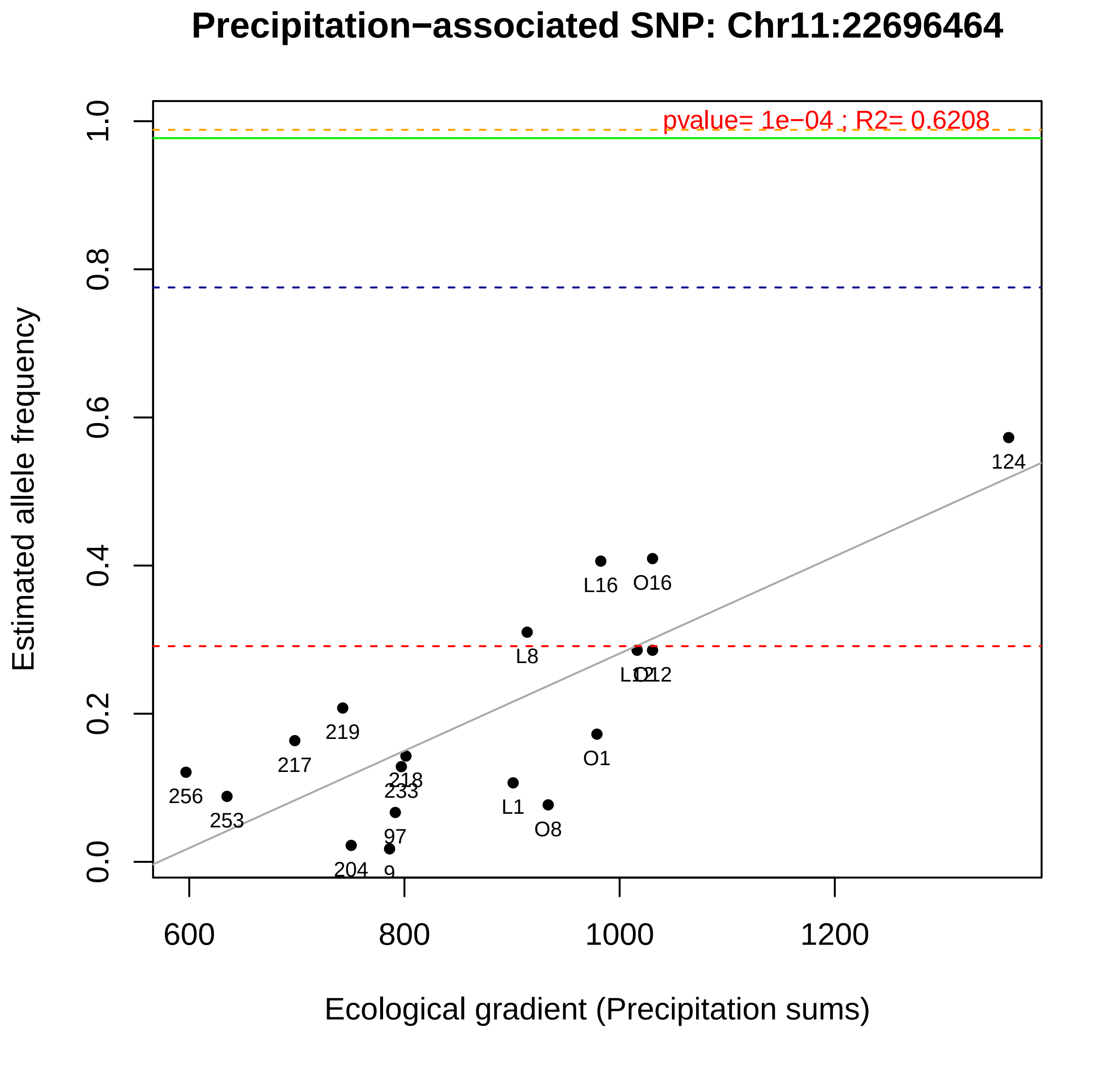
Precipitation−associated SNP: Chr11:22696464
pvalue= 1e−04 ; R2= 0.6208
Estimated allele frequency
0.0
0.2
0.4
0.6
0.8
1.0
600
800
1000
1200
Ecological gradient (Precipitation sums)
124
L16
O16
L8
L12
O12
219
217
O1
218
233
256
253
L1
O8
97
204
9

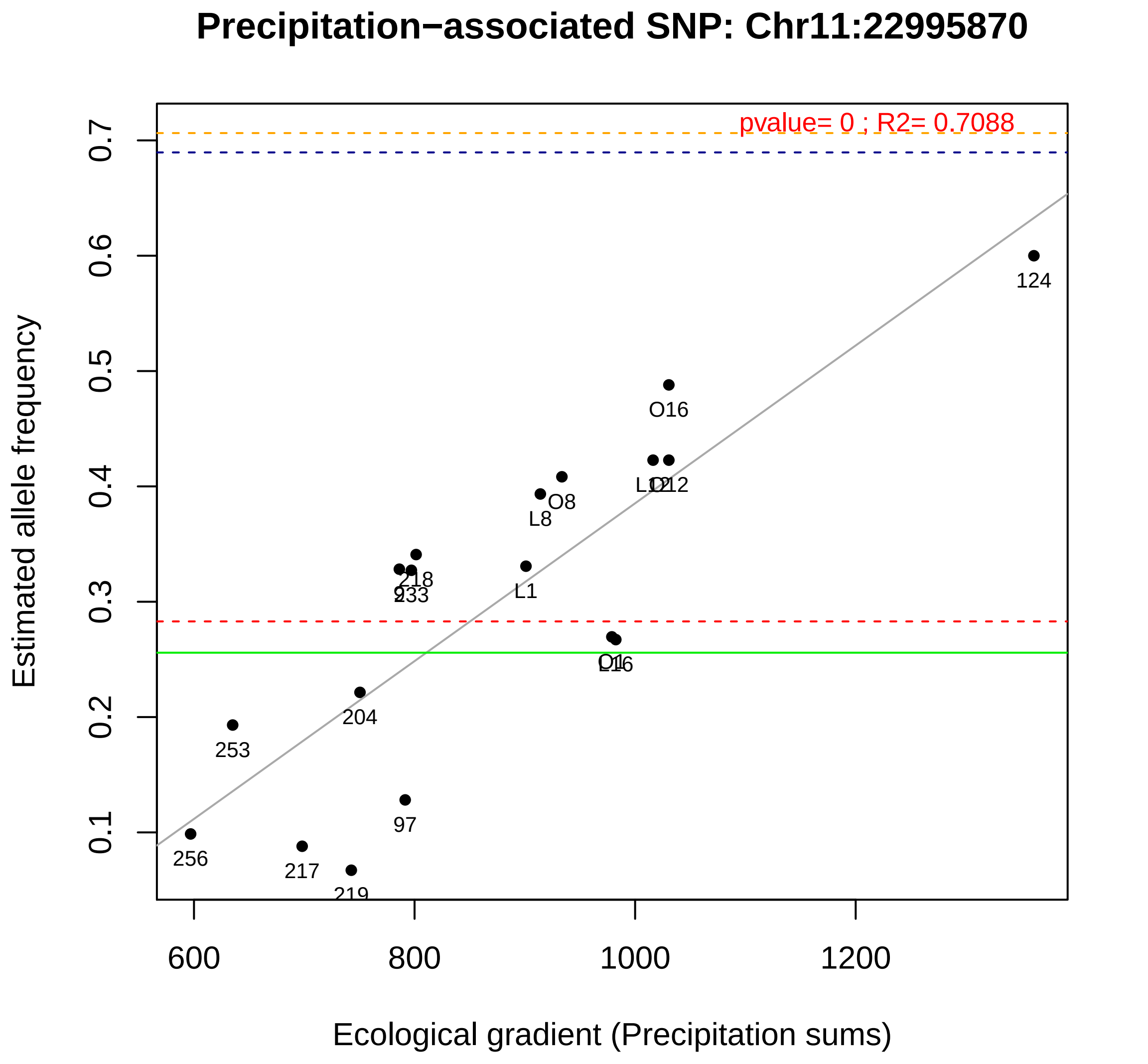
Precipitation−associated SNP: Chr11:22995870
pvalue= 0 ; R2= 0.7088
Estimated allele frequency
0.1
0.2
0.3
0.4
0.5
0.6
0.7
600
800
1000
1200
Ecological gradient (Precipitation sums)
124
O16
L12
O12
O8
L8
L1
218
9
233
O1
L16
204
253
97
256
217
219

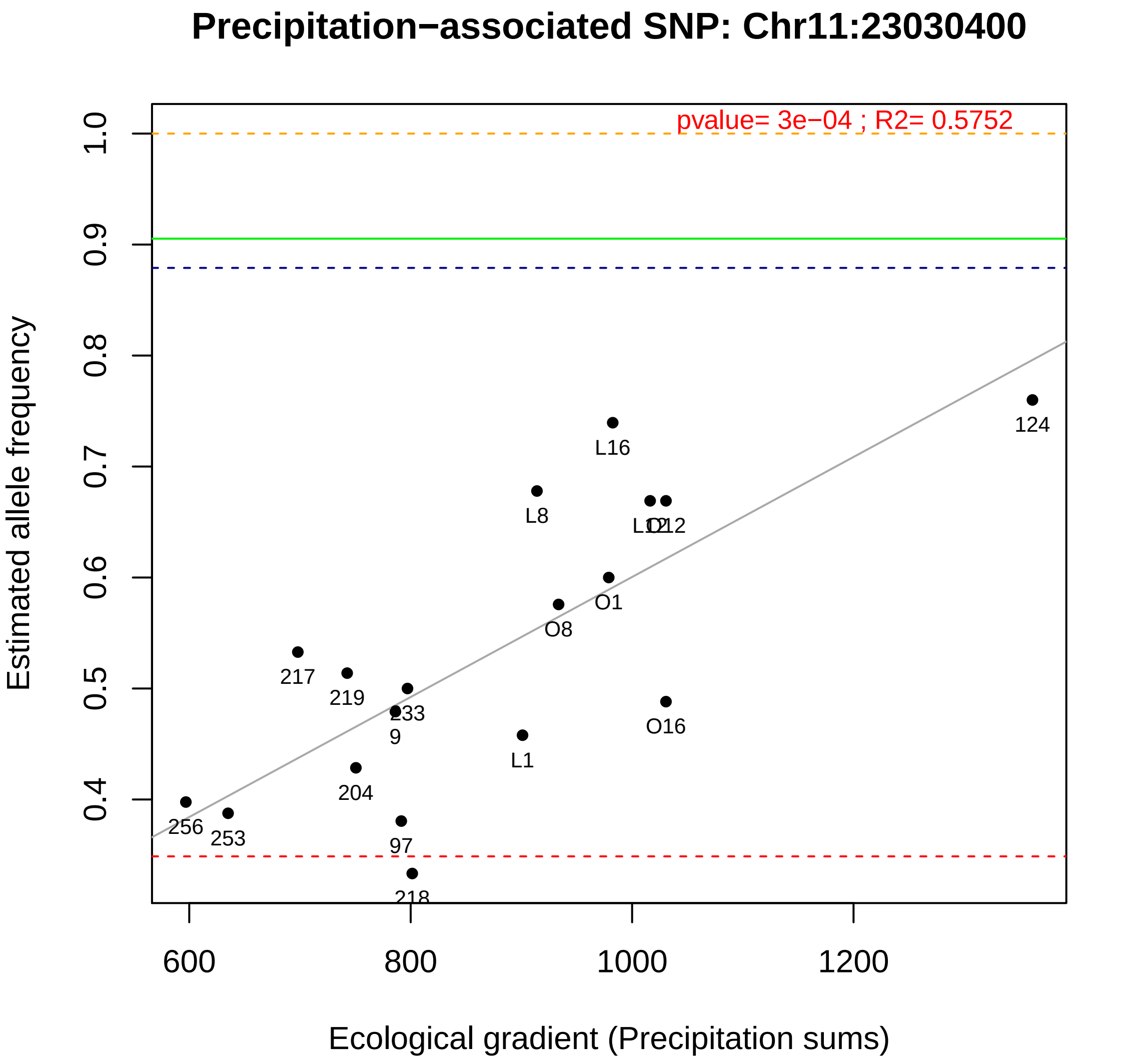

Precipitation−associated SNP: Chr11:23030400
pvalue= 3e−04 ; R2= 0.5752
Estimated allele frequency
0.4
0.5
0.6
0.7
0.8
0.9
1.0
Ecological gradient (Precipitation sums)
600
800
1000
1200
124
L16
L8
L12
O12
O1
O8
217
219
233
9
L1
O16
204
256
253
97
218

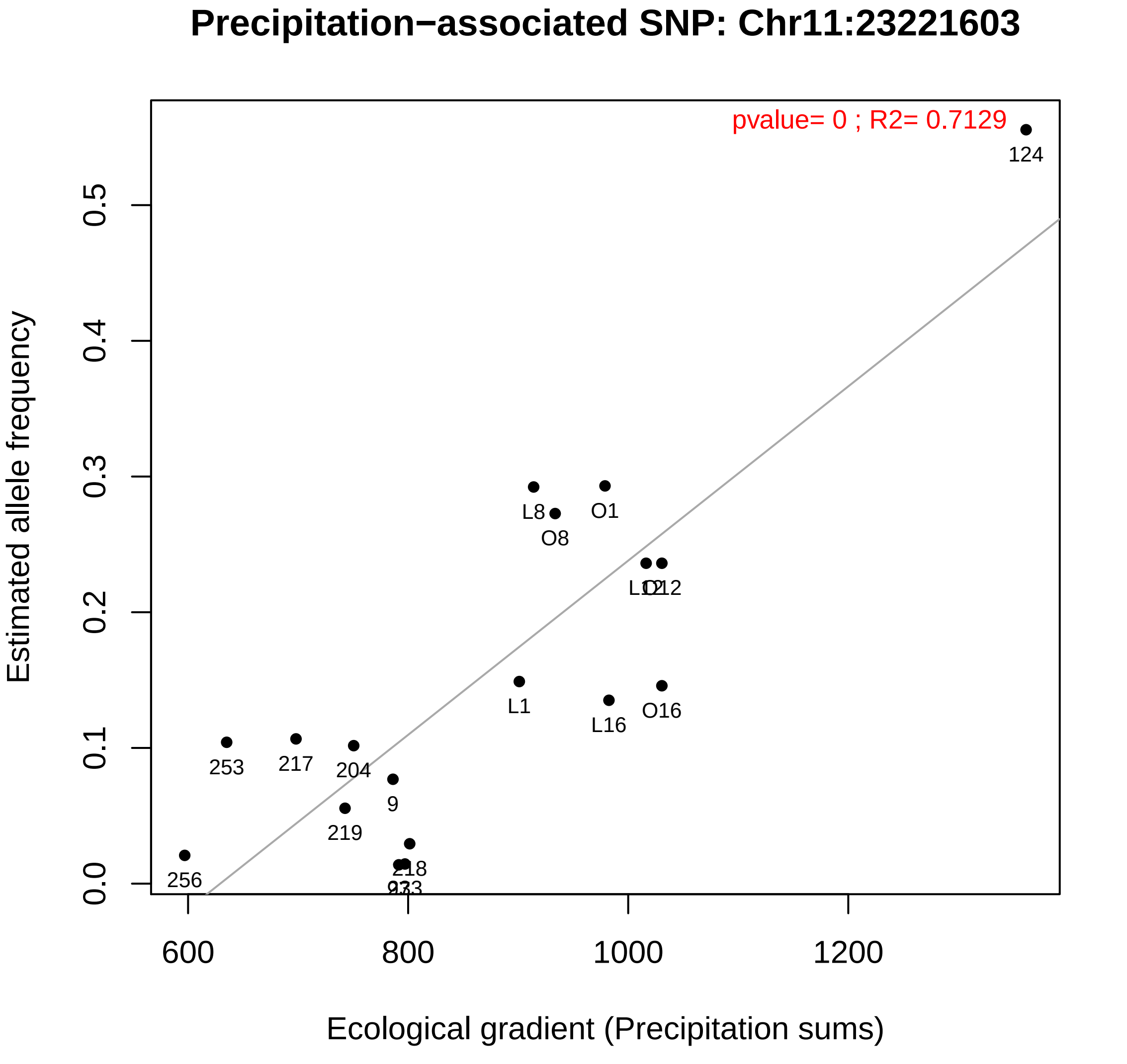

Precipitation−associated SNP: Chr11:23221603
pvalue= 0 ; R2= 0.7129
124
L8
O1
O8
L12
O12
L1
L16
O16
253
217
204
9
219
256
218
0.0
0.1
0.2
0.3
0.4
0.5
600
800
1000
1200
Estimated allele frequency
Ecological gradient (Precipitation sums)

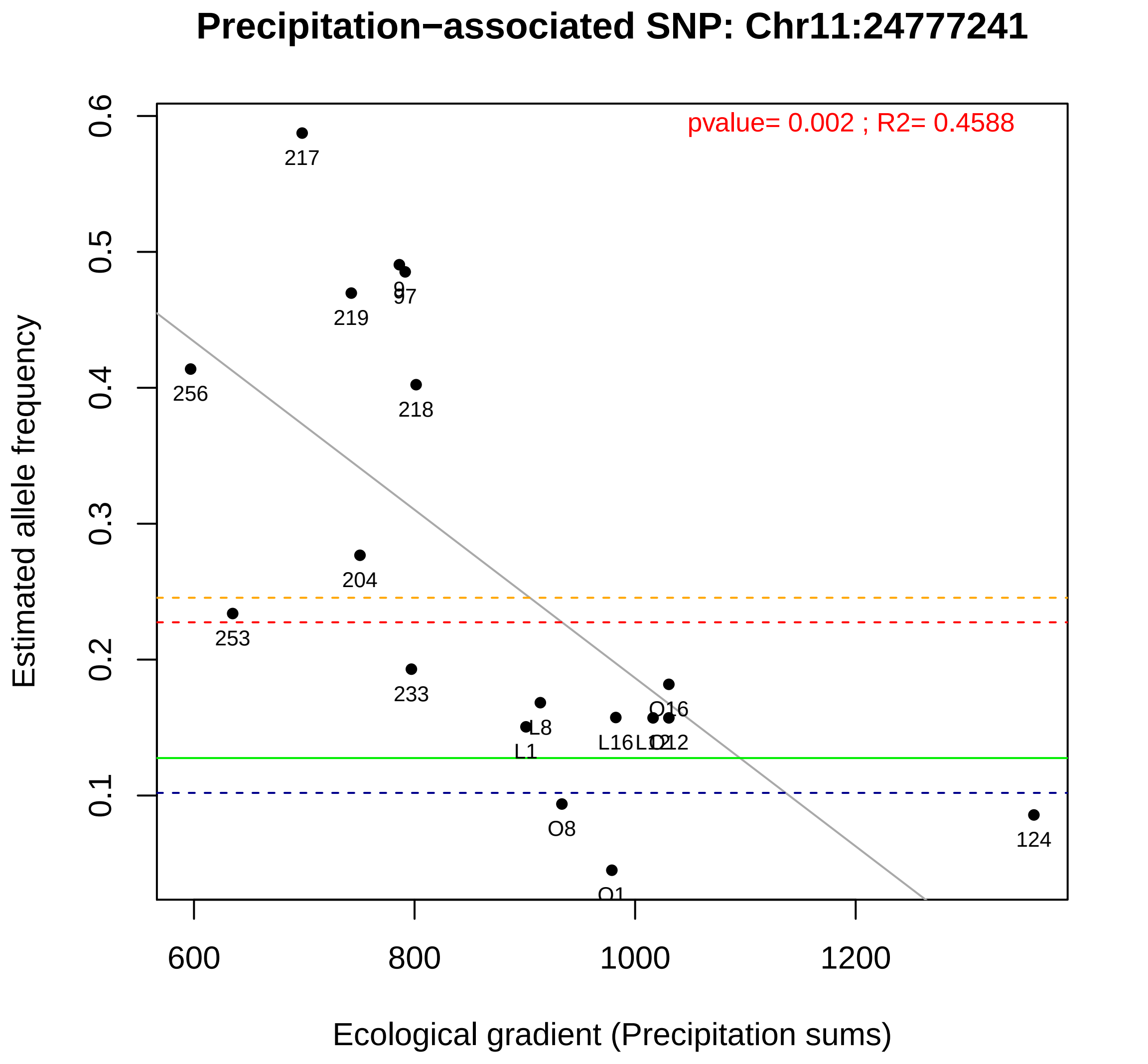

Precipitation−associated SNP: Chr11:24777241
pvalue= 0.002 ; R2= 0.4588
Estimated allele frequency
0.1
0.2
0.3
0.4
0.5
0.6
600
800
1000
1200
Ecological gradient (Precipitation sums)
217
219
9
97
256
218
204
253
233
L8
L1
L16
L12
O12
O16
O8
O1
124

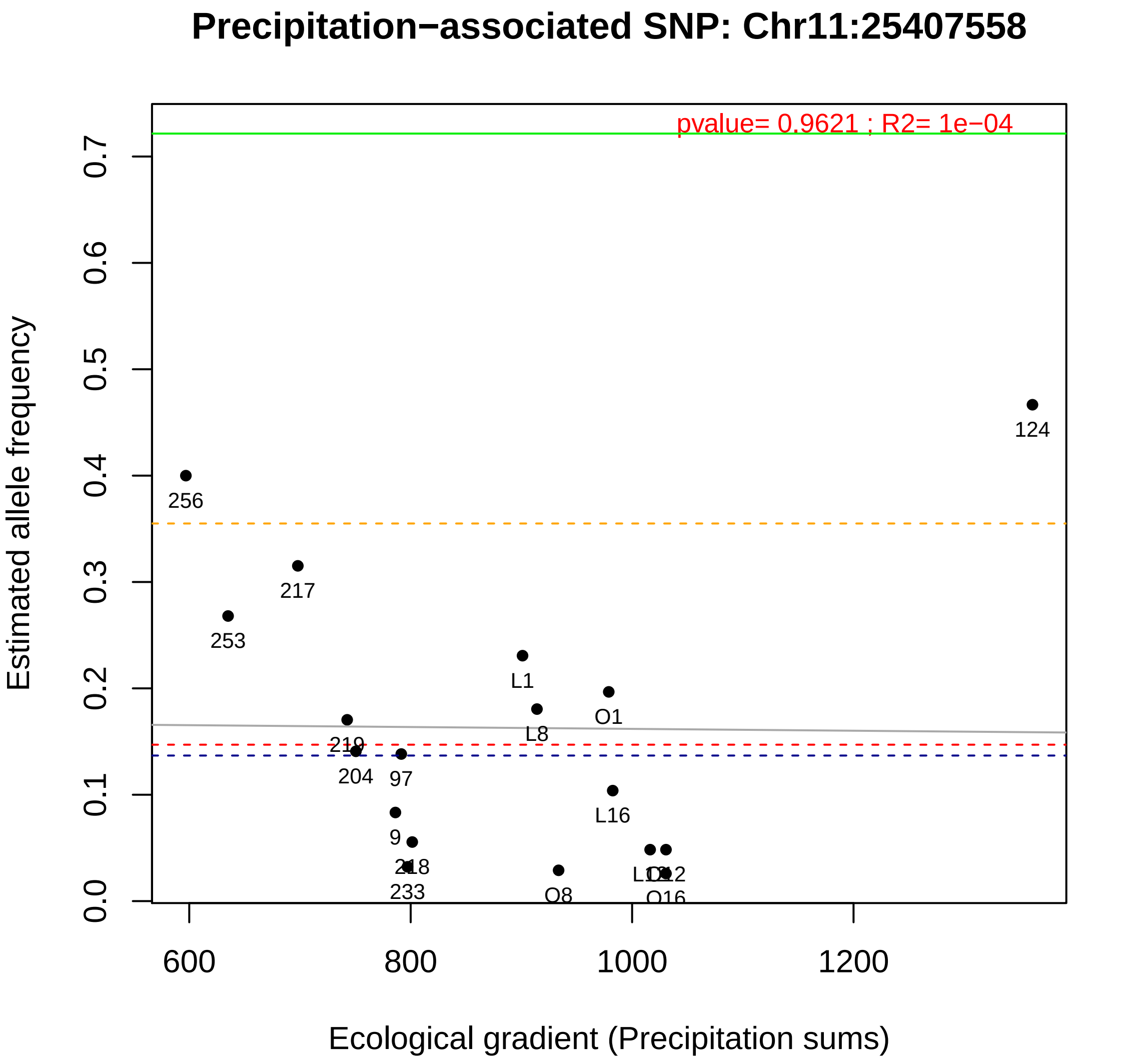

Precipitation−associated SNP: Chr11:25407558
pvalue= 0.9621 ; R2= 1e−04
Estimated allele frequency
0.0
0.1
0.2
0.3
0.4
0.5
0.6
0.7
600
800
1000
1200
Ecological gradient (Precipitation sums)
124
256
217
253
L1
O1
L8
219
204
97
L16
9
218
233
O8
O16

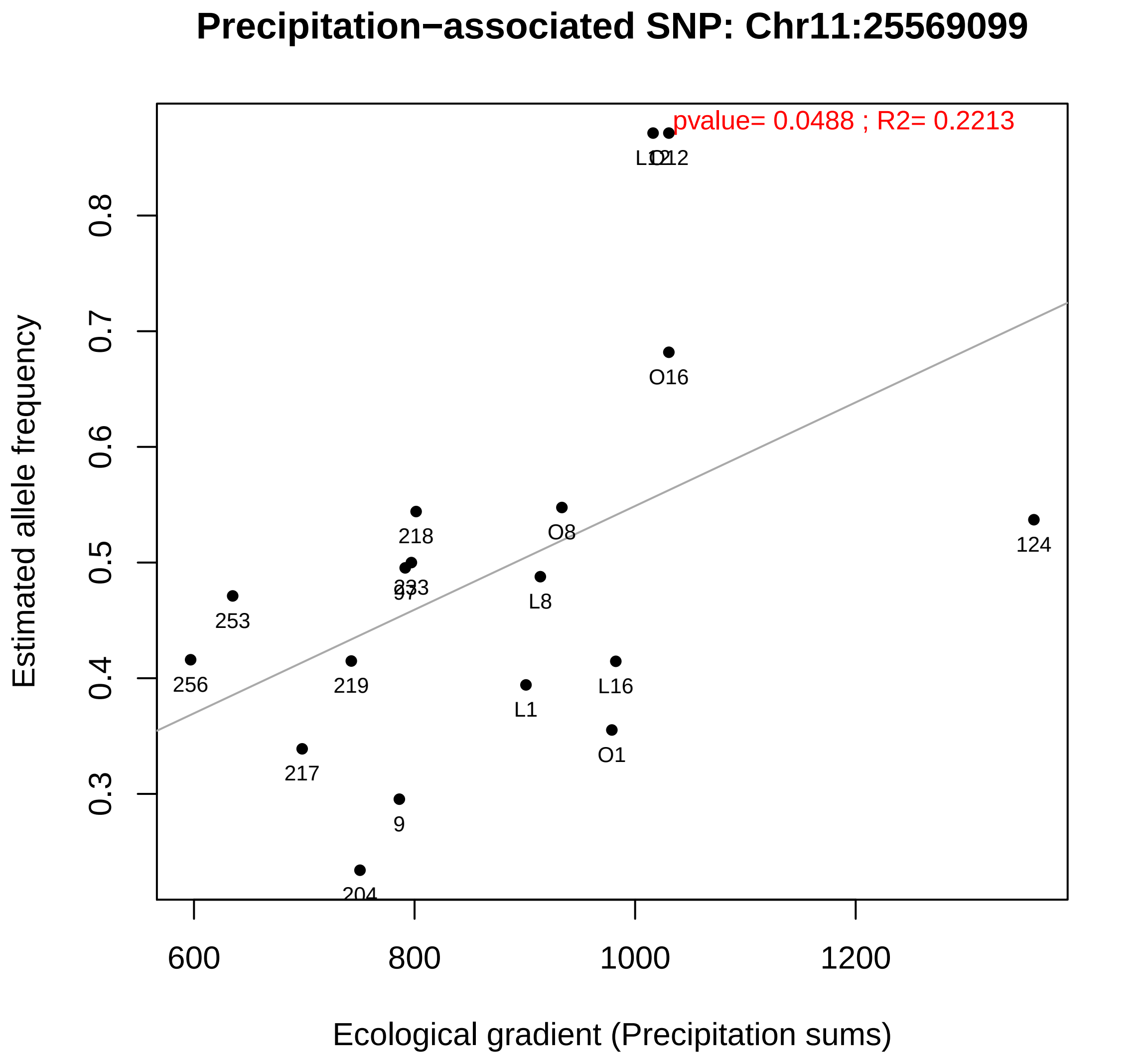
Precipitation−associated SNP: Chr11:25569099
pvalue= 0.0488 ; R2= 0.2213
Estimated allele frequency
0.3
0.4
0.5
0.6
0.7
0.8
600
800
1000
1200
Ecological gradient (Precipitation sums)
O16
218
O8
124
253
L8
256
219
L16
L1
O1
217
9
204

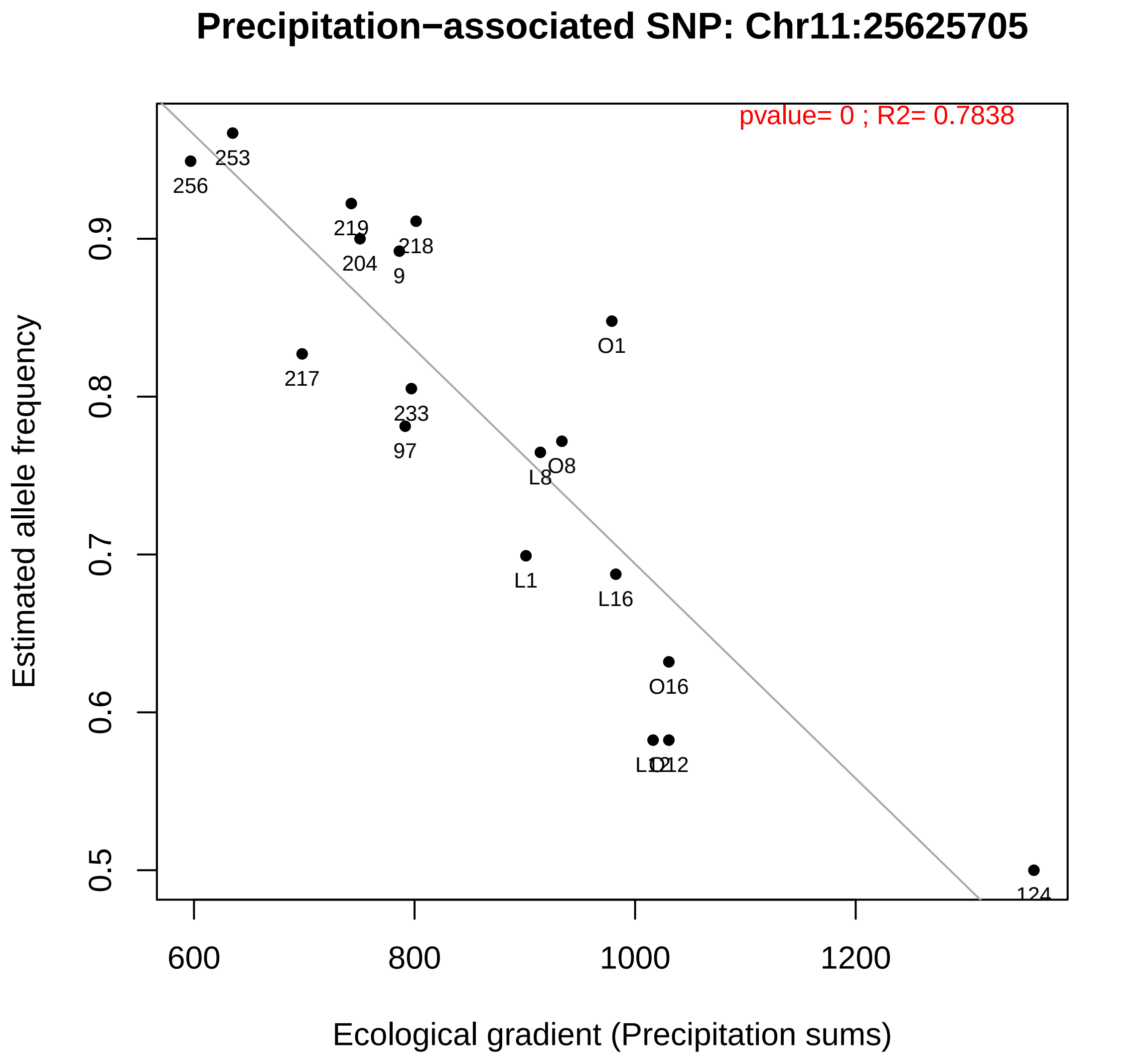

Precipitation−associated SNP: Chr11:25625705
pvalue= 0 ; R2= 0.7838
256
253
219
204
218
9
217
233
97
O1
L8
O8
L1
L16
O16
L12
O12
124
Estimated allele frequency
0.5
0.6
0.7
0.8
0.9
600
800
1000
1200
Ecological gradient (Precipitation sums)

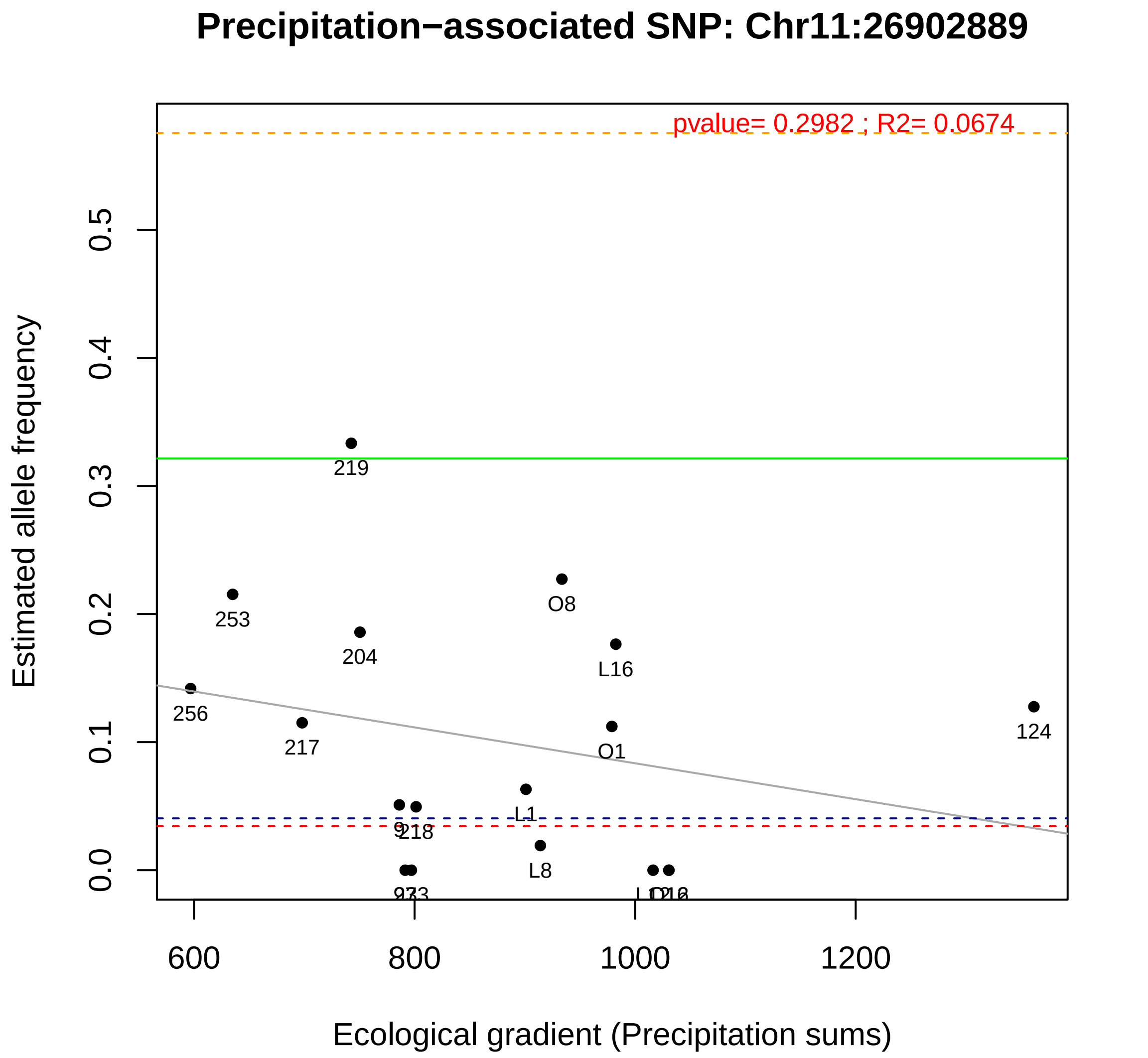

Precipitation−associated SNP: Chr11:26902889
pvalue= 0.2982 ; R2= 0.0674
Estimated allele frequency
0.0
0.1
0.2
0.3
0.4
0.5
600
800
1000
1200
Ecological gradient (Precipitation sums)
219
253
204
O8
L16
256
217
O1
124
L1
218
L8

**Precipitation−associated SNP: Chr11:27002271**

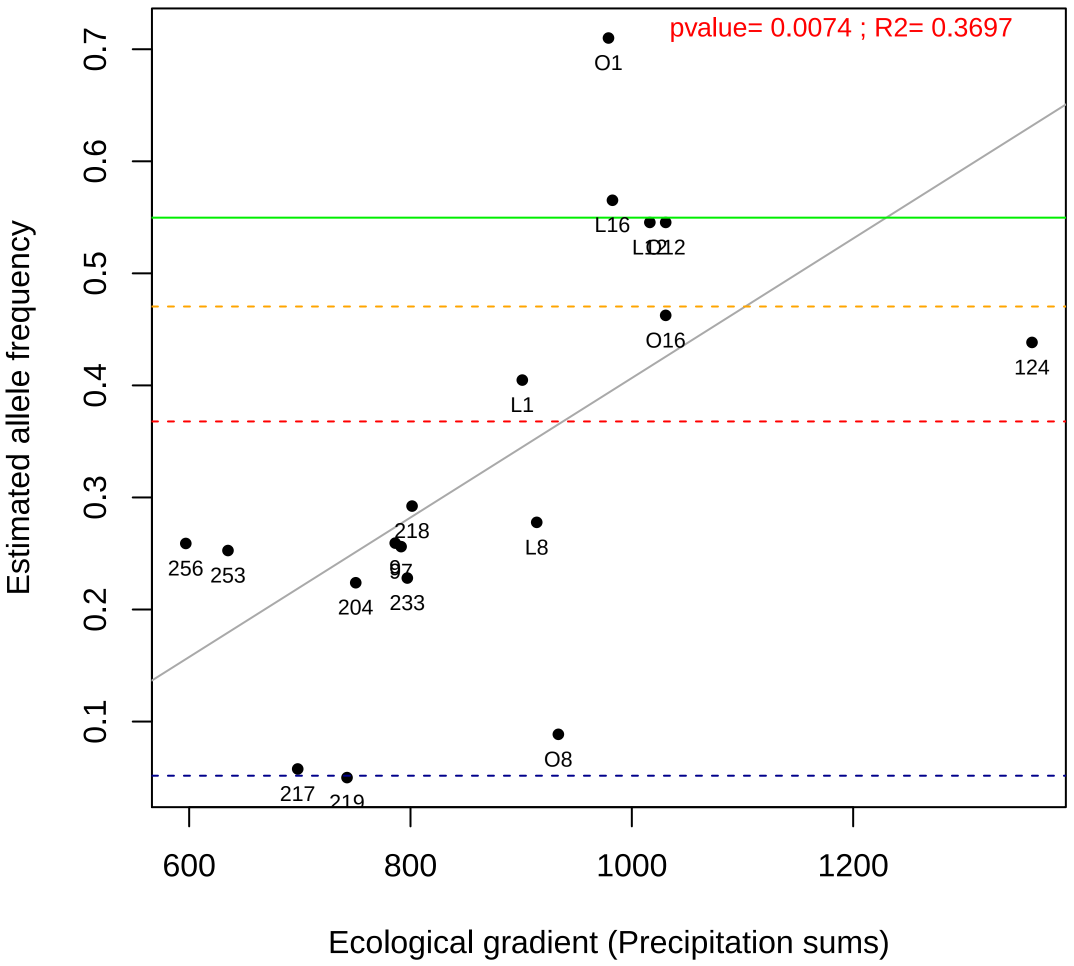

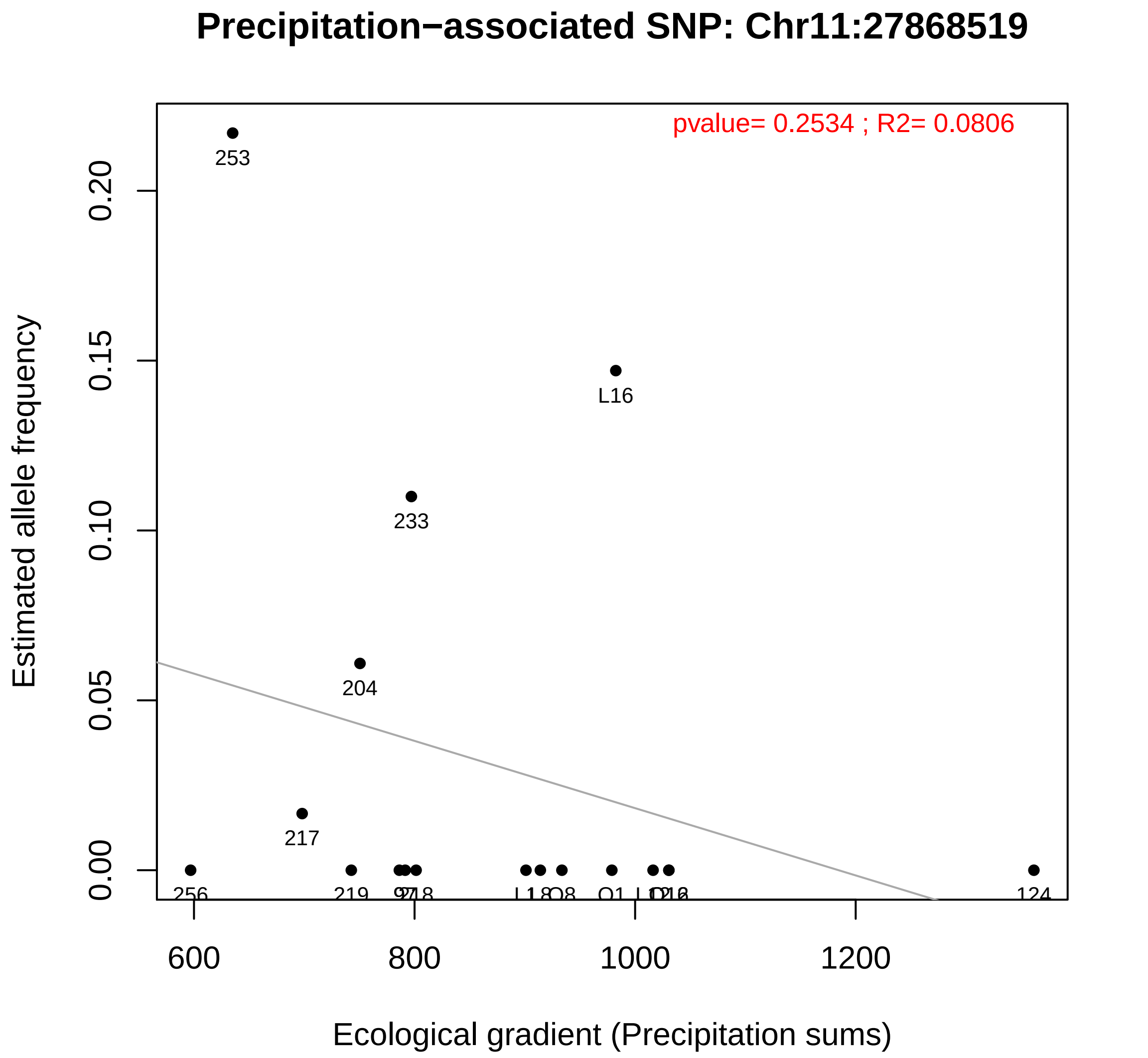

Precipitation−associated SNP: Chr11:27868519
pvalue= 0.2534 ; R2= 0.0806
253
L16
233
204
217
256
219
124
0.00
0.05
0.10
0.15
0.20
600
800
1000
1200
Estimated allele frequency
Ecological gradient (Precipitation sums)

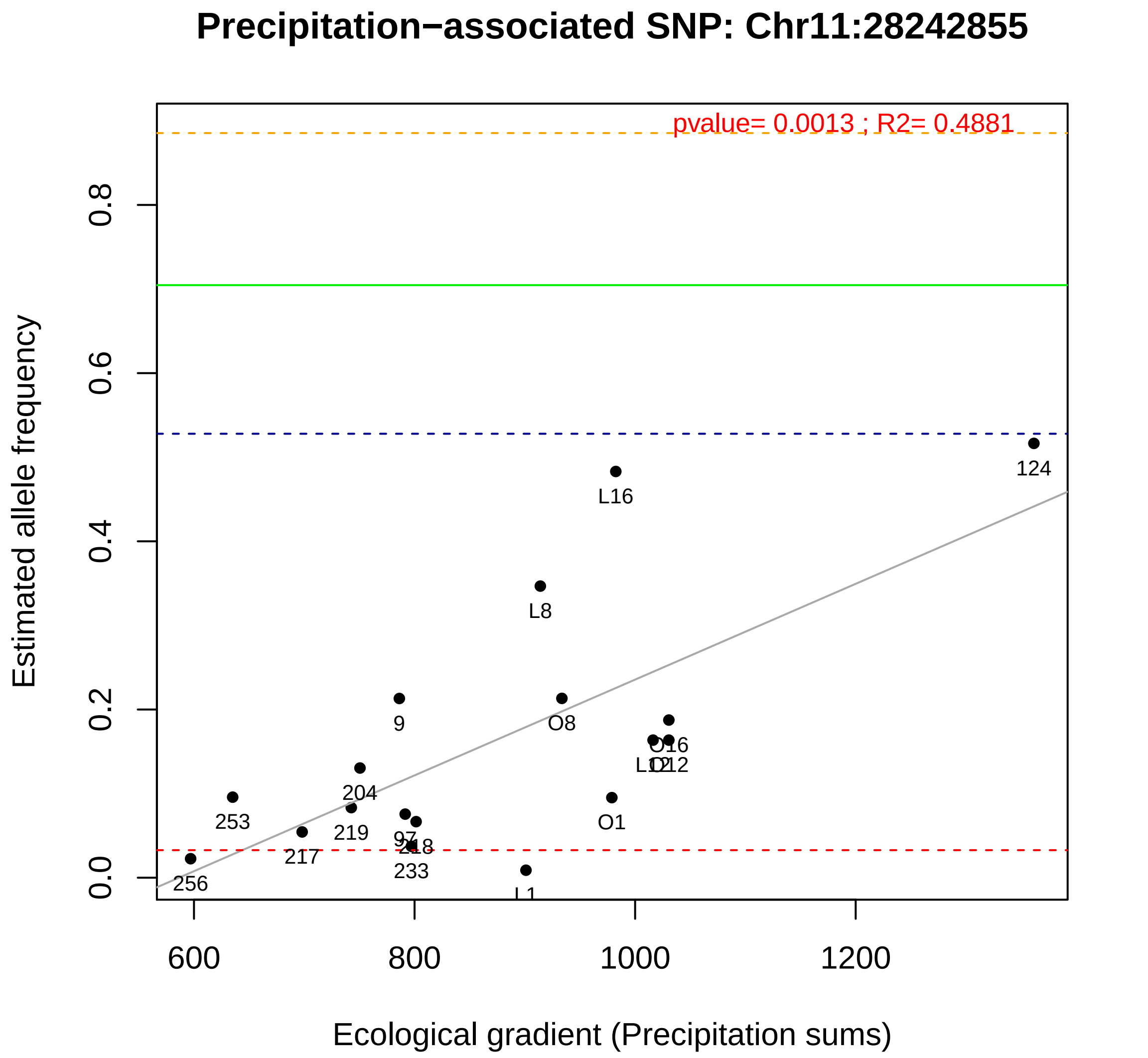
Precipitation−associated SNP: Chr11:28242855
pvalue= 0.0013 ; R2= 0.4881
Estimated allele frequency
0.0
0.2
0.4
0.6
0.8
600
800
1000
1200
Ecological gradient (Precipitation sums)
124
L16
L8
9
O8
O16
O12
204
253
219
O1
217
97
218
233
256

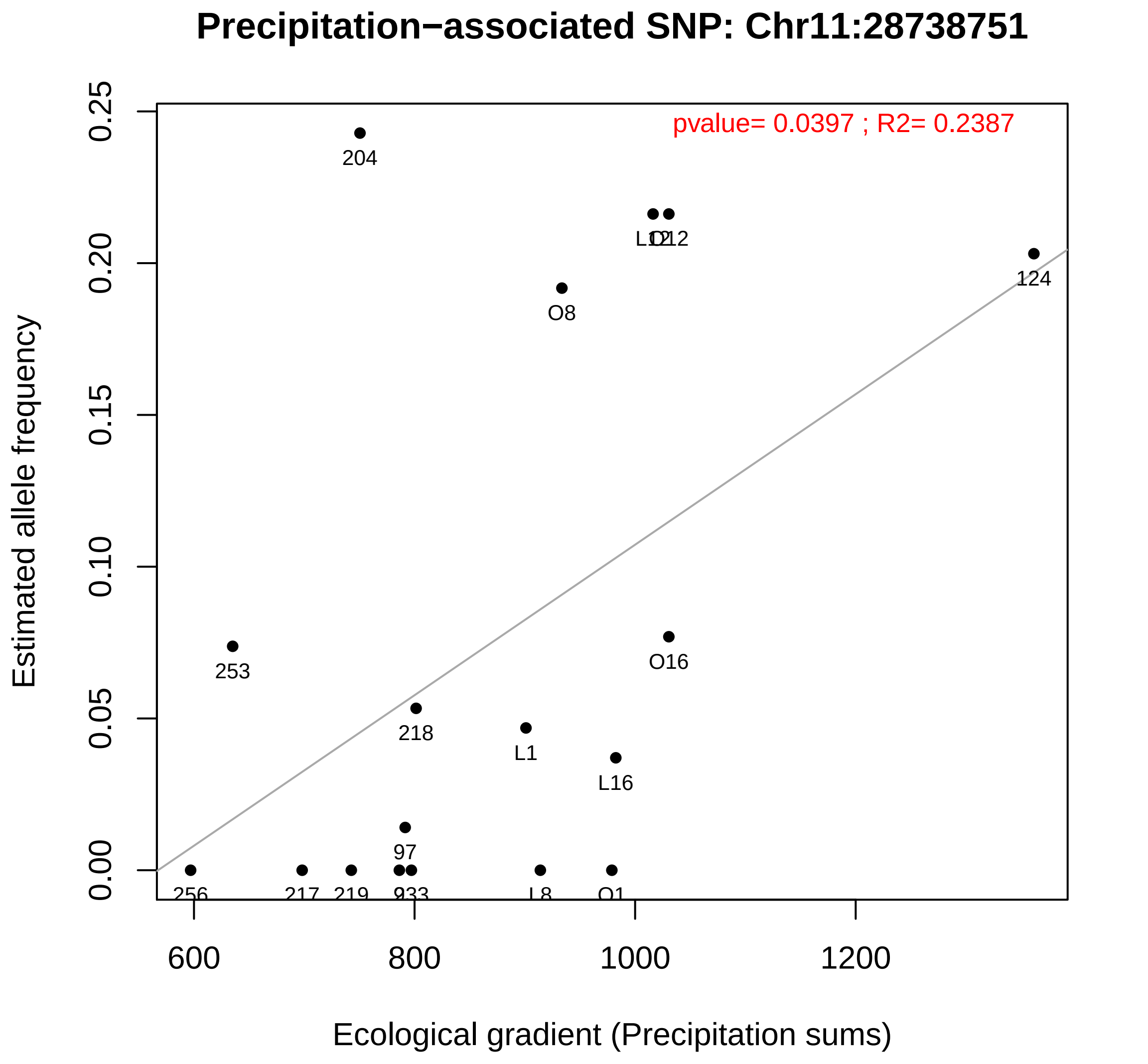

Precipitation−associated SNP: Chr11:28738751
pvalue= 0.0397 ; R2= 0.2387
Estimated allele frequency
0.00
0.05
0.10
0.15
0.20
0.25
600
800
1000
1200
Ecological gradient (Precipitation sums)
204
L12
O12
124
O8
253
O16
218
L1
L16
97
256
217
219
L8
O1

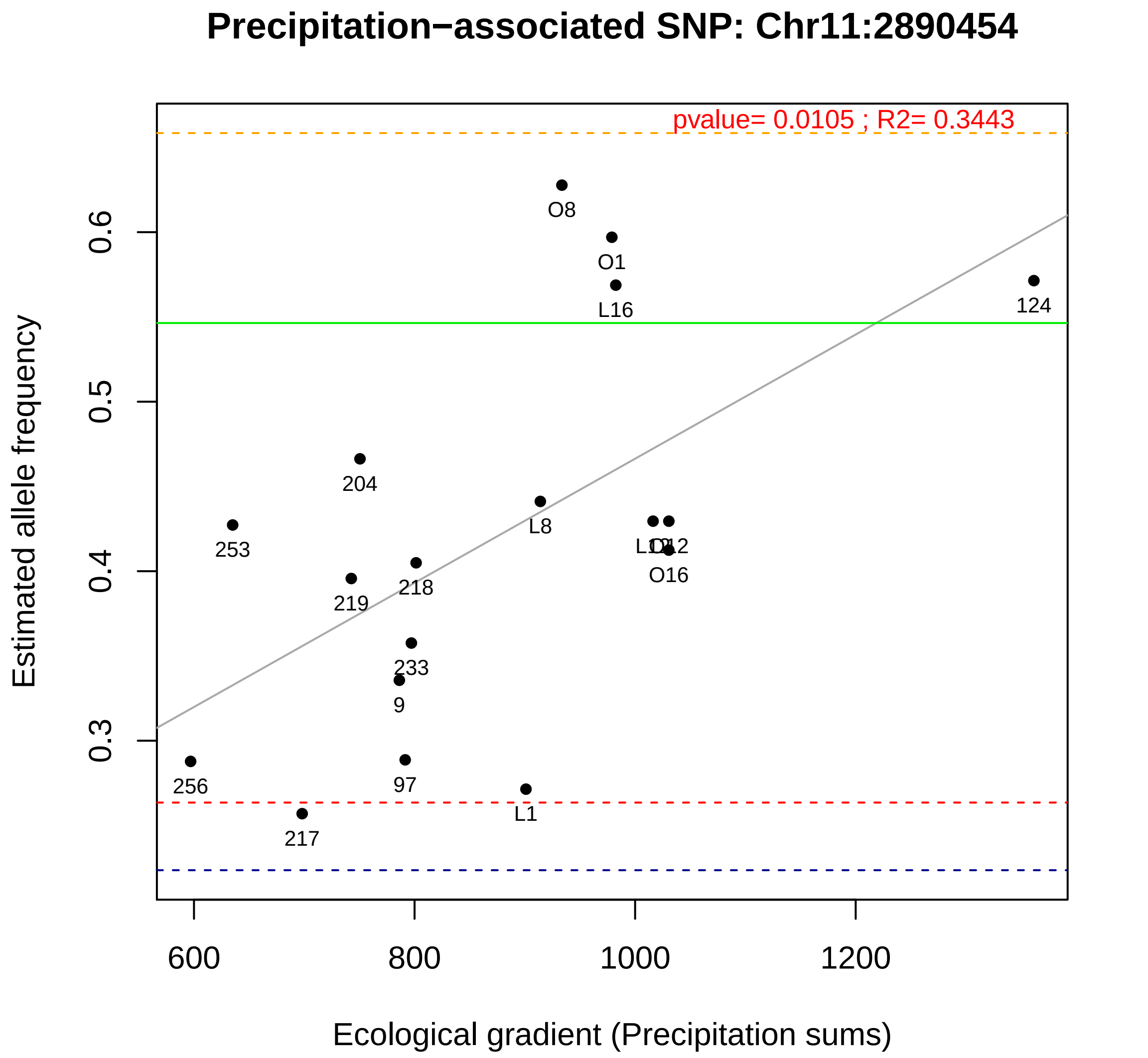

Precipitation−associated SNP: Chr11:2890454
pvalue= 0.0105 ; R2= 0.3443
O8
O1
L16
124
204
L8
253
L12
O12
O16
219
218
233
9
256
97
L1
217
0.3
0.4
0.5
0.6
600
800
1000
1200
Estimated allele frequency
Ecological gradient (Precipitation sums)

Precipitation−associated SNP: Chr11:30007916

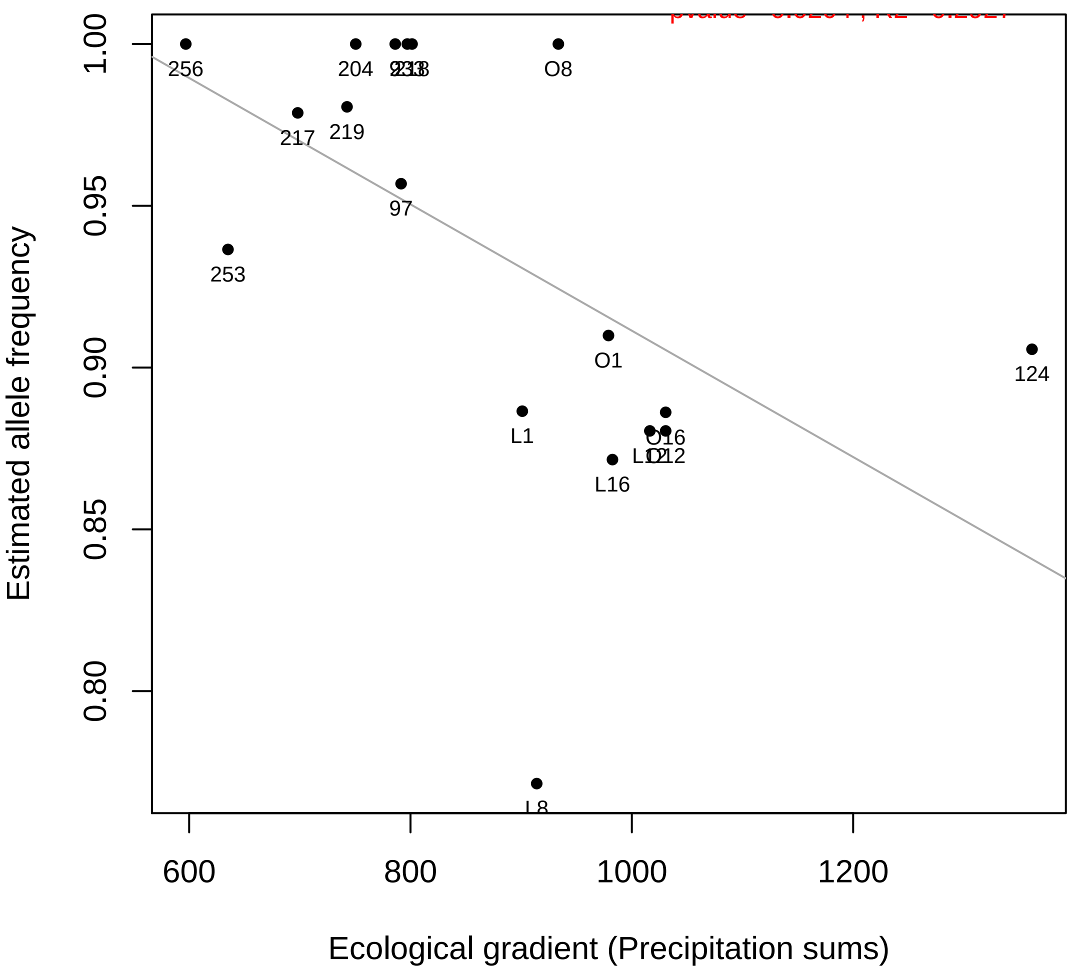

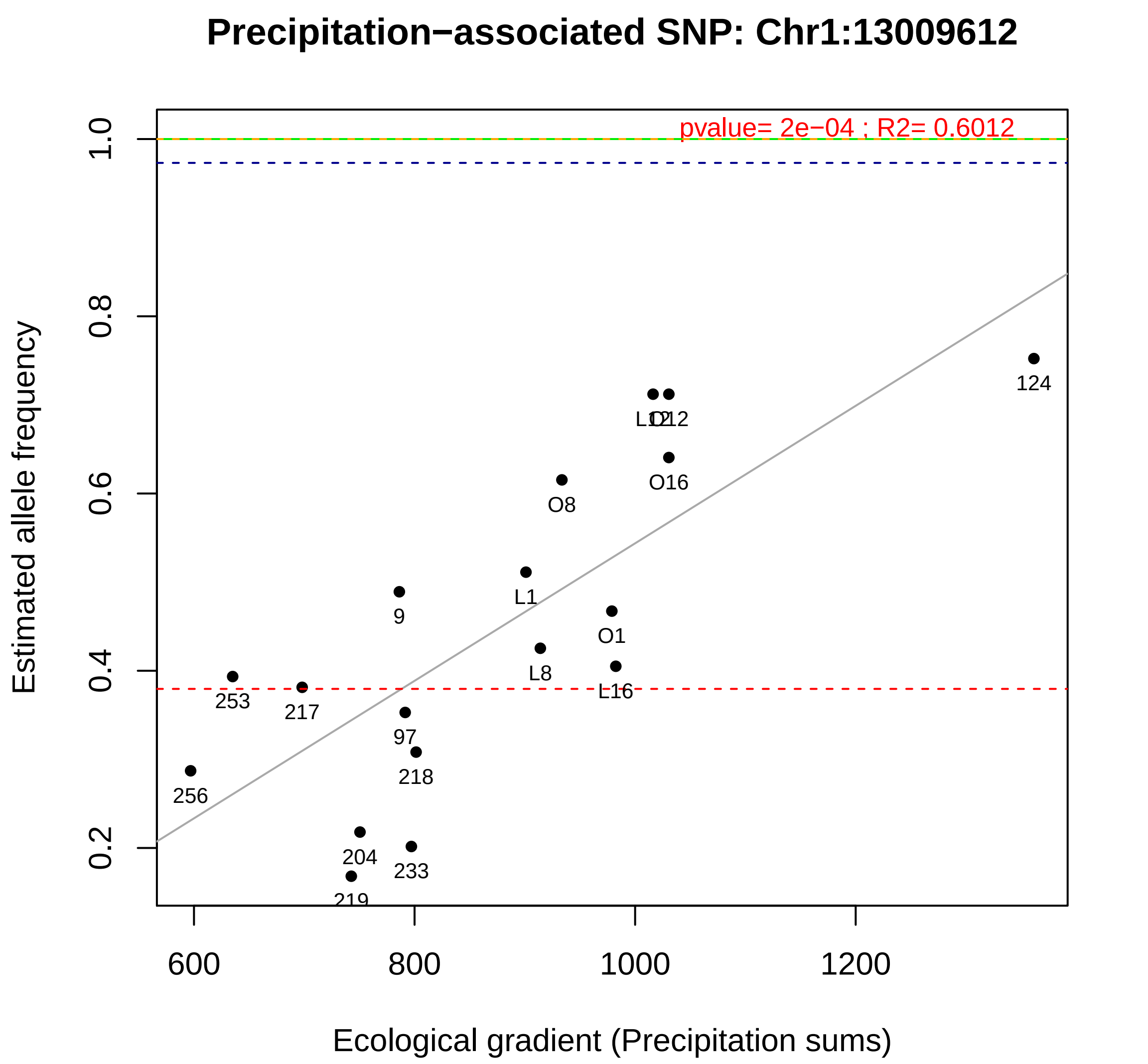

Precipitation−associated SNP: Chr1:13009612
pvalue= 2e−04 ; R2= 0.6012
Estimated allele frequency
0.2
0.4
0.6
0.8
1.0
Ecological gradient (Precipitation sums)
600
800
1000
1200
124
L12
O12
O16
O8
L1
9
O1
L8
L16
253
217
97
218
256
204
233
219

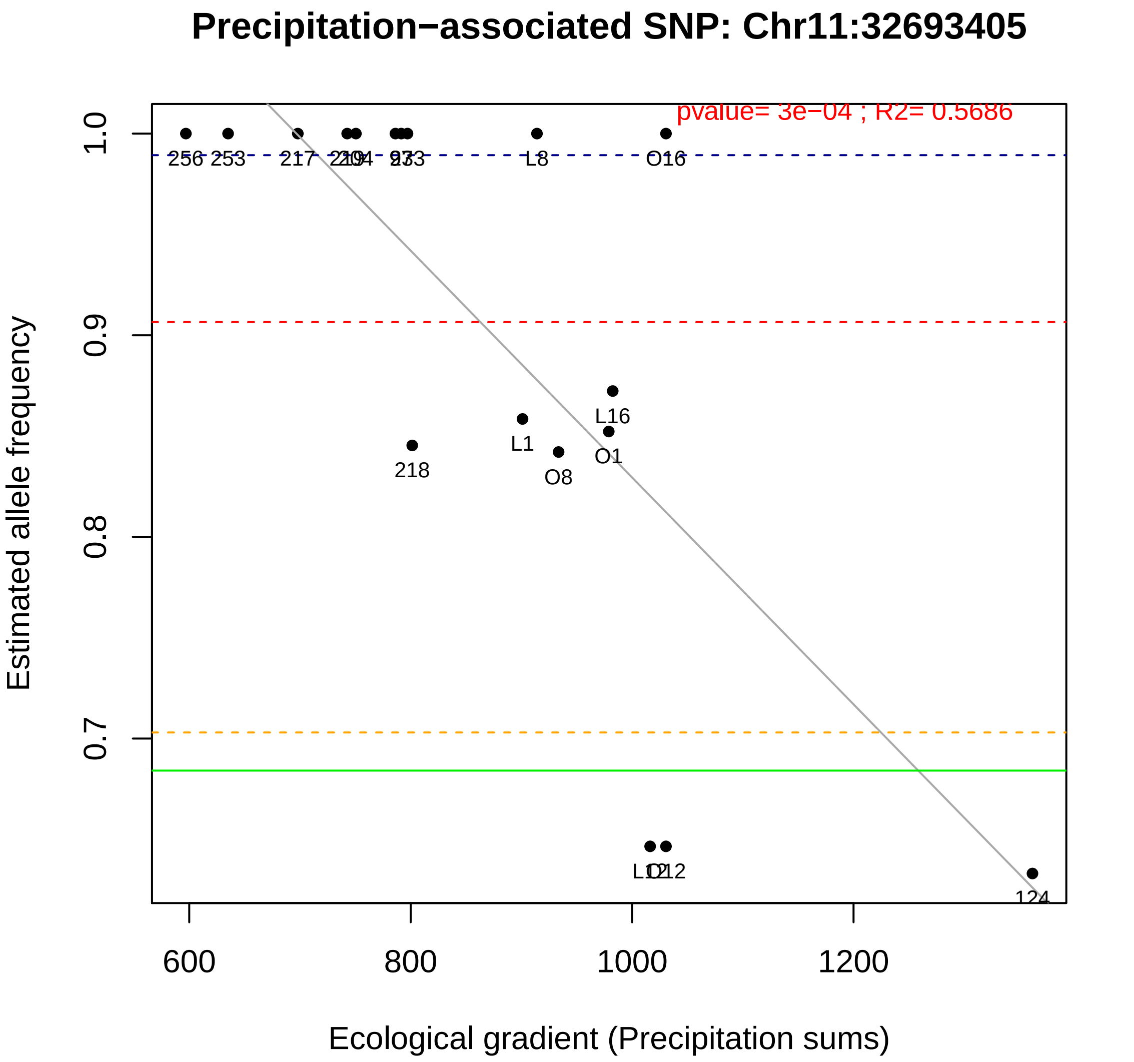

Precipitation−associated SNP: Chr11:32693405
pvalue= 3e−04 ; R2= 0.5686
Estimated allele frequency
Ecological gradient (Precipitation sums)
1.0
0.9
0.8
0.7
600
800
1000
1200
256
253
217
L8
O16
218
L1
O8
L16
O1
124

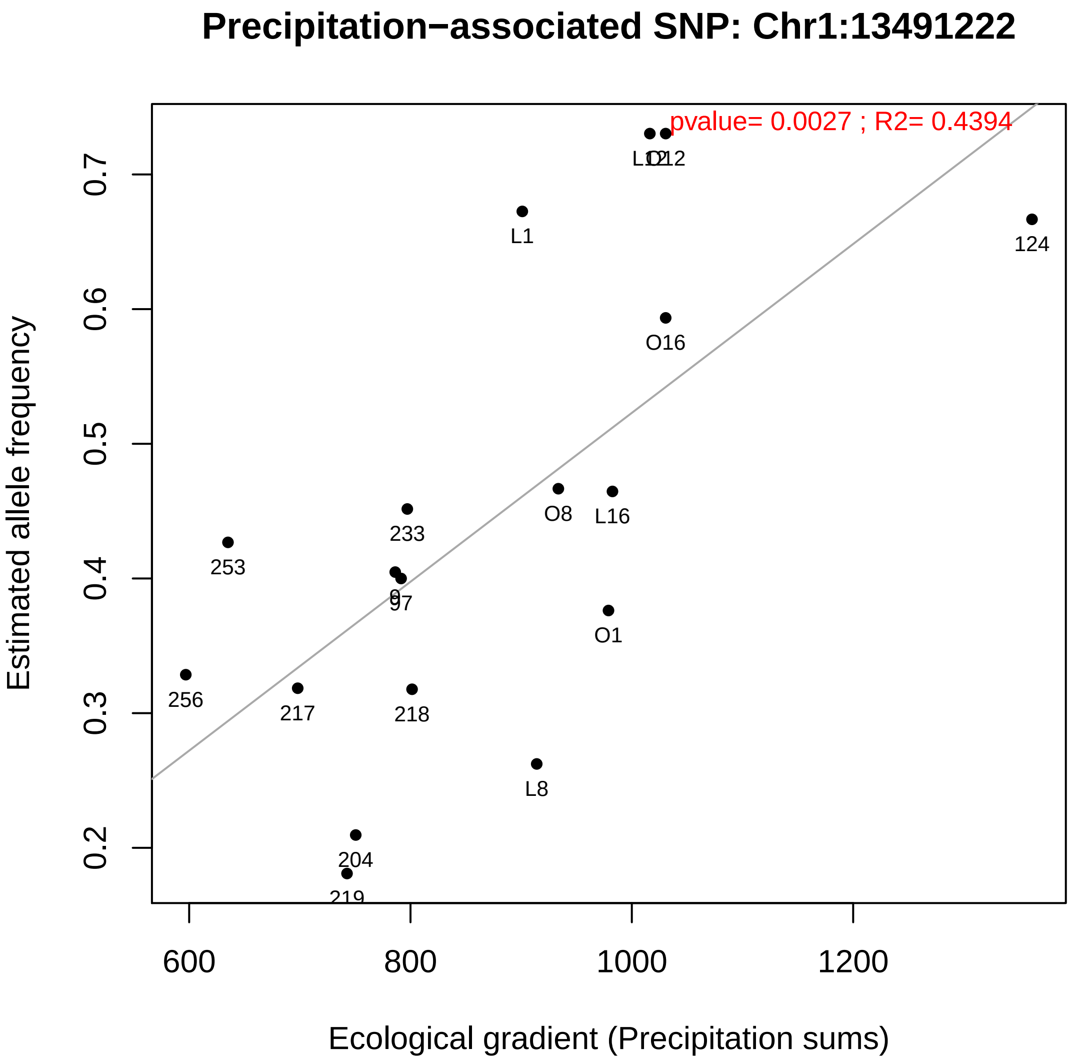

Precipitation−associated SNP: Chr1:13491222
pvalue= 0.0027 ; R2= 0.4394
Estimated allele frequency
0.2
0.3
0.4
0.5
0.6
0.7
600
800
1000
1200
Ecological gradient (Precipitation sums)
L1
124
O16
O8
L16
233
253
97
O1
256
217
218
L8
204
219

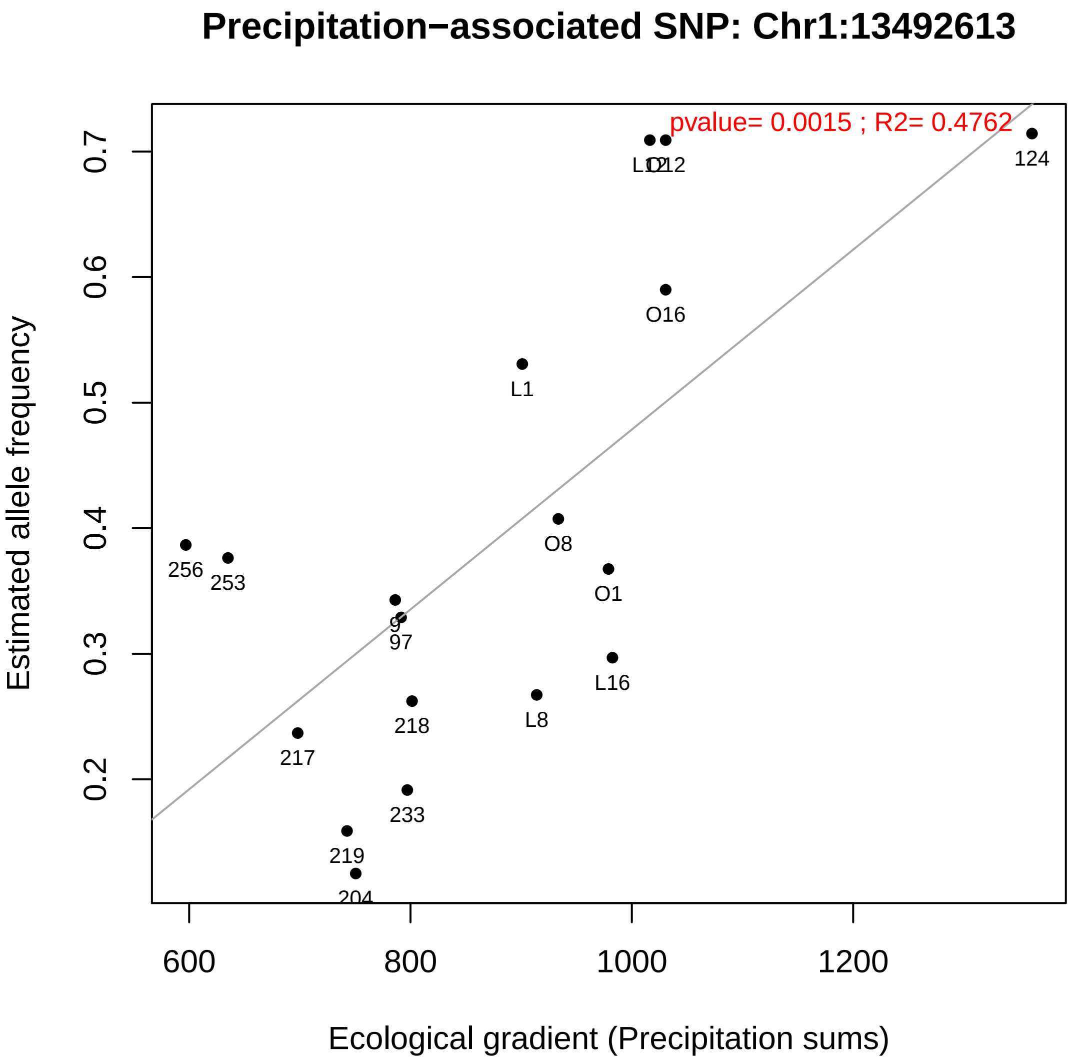
Precipitation−associated SNP: Chr1:13492613
pvalue= 0.0015 ; R2= 0.4762
Estimated allele frequency
0.2
0.3
0.4
0.5
0.6
0.7
600
800
1000
1200
Ecological gradient (Precipitation sums)
124
L12
O12
O16
L1
O8
O1
L16
L8
256
253
9
97
218
217
233
219
204

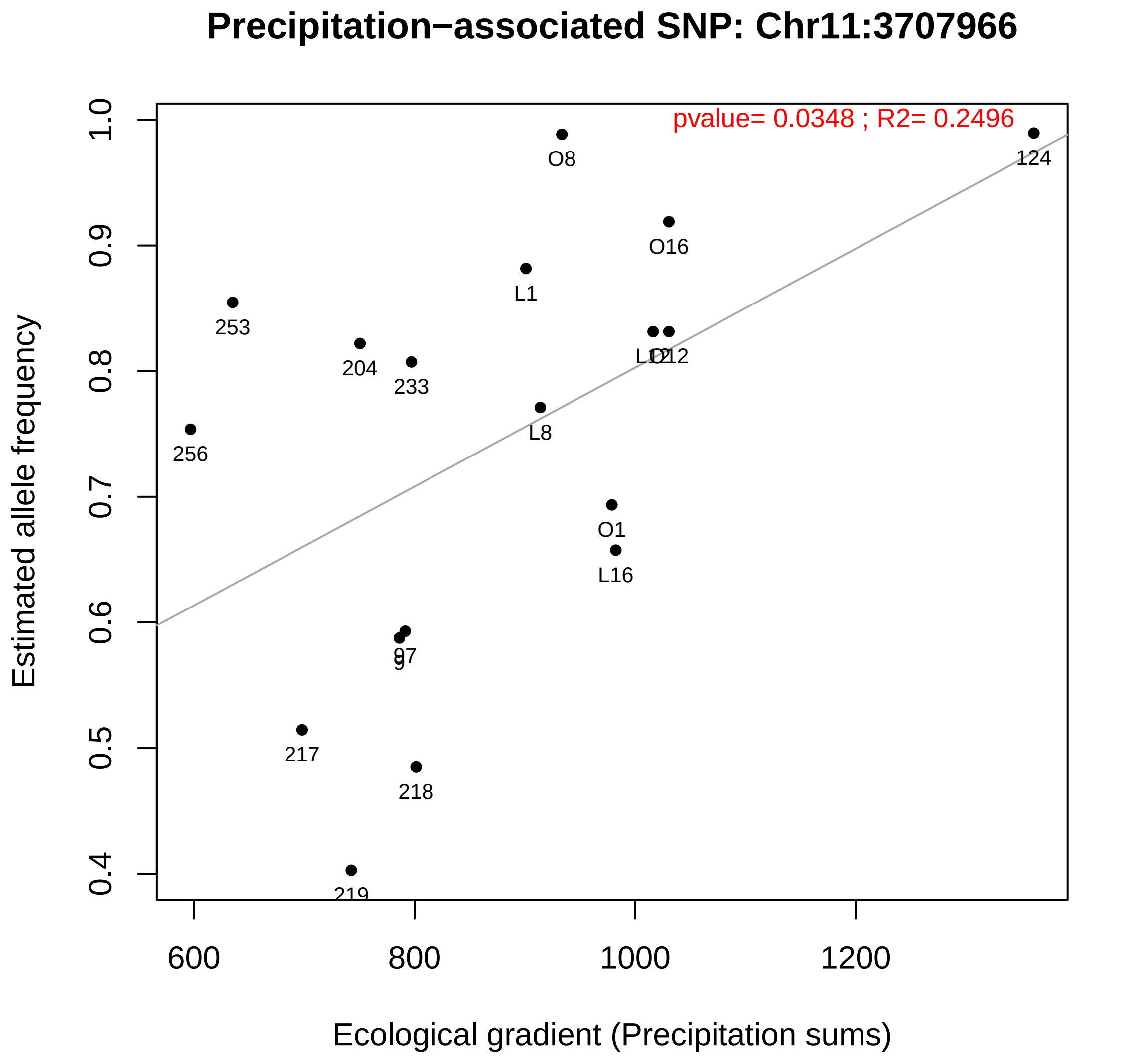
Precipitation−associated SNP: Chr11:3707966
pvalue= 0.0348 ; R2= 0.2496
Estimated allele frequency
Ecological gradient (Precipitation sums)
0.4
0.5
0.6
0.7
0.8
0.9
1.0
600
800
1000
1200
O8
124
O16
L1
253
204
233
L8
256
O1
L16
97
9
217
218
219

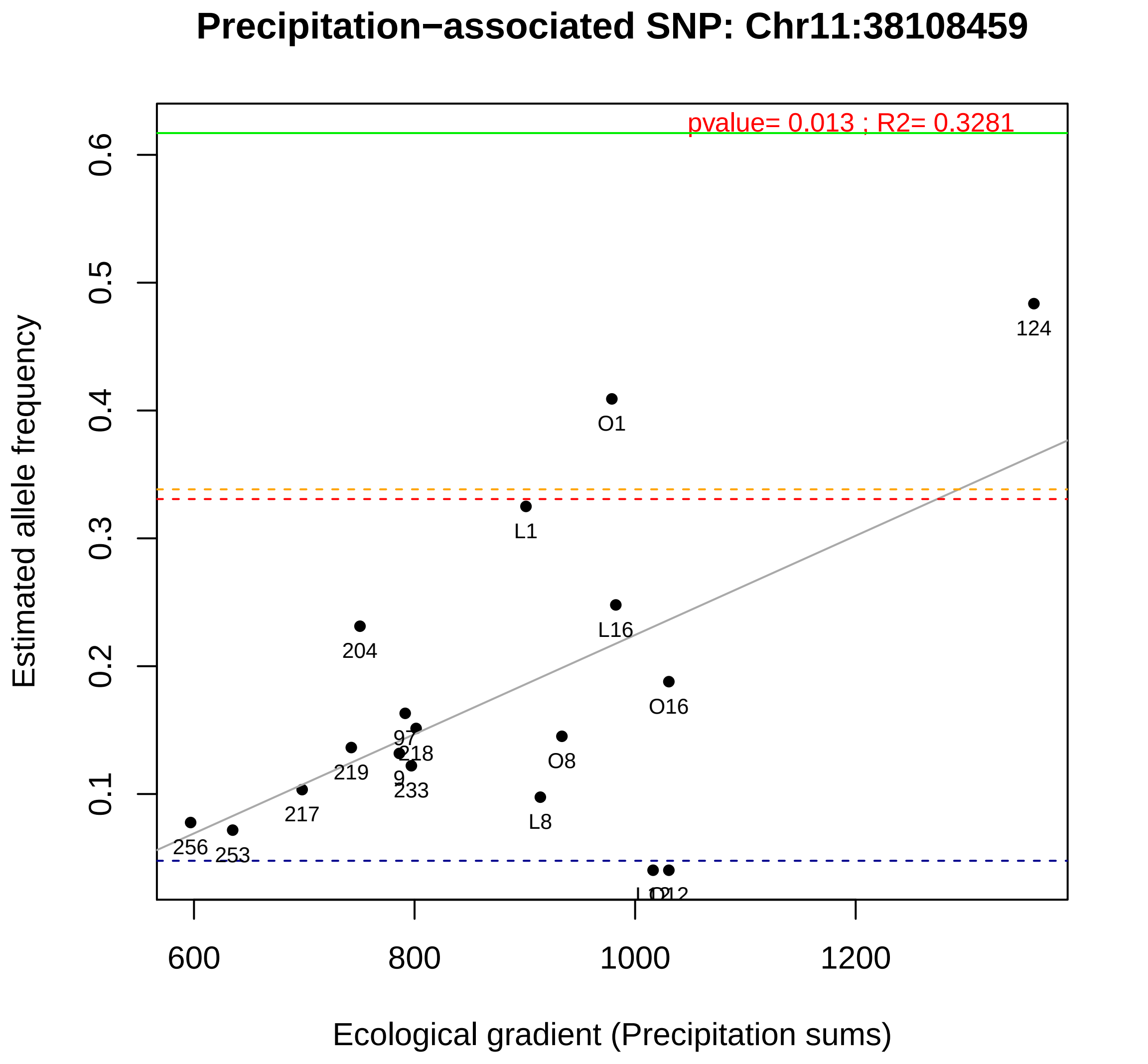
Precipitation−associated SNP: Chr11:38108459
pvalue= 0.013 ; R2= 0.3281
Estimated allele frequency
0.1
0.2
0.3
0.4
0.5
0.6
124
O1
L1
L16
204
O16
97
218
O8
219
9
233
217
L8
256
253
600
800
1000
1200
Ecological gradient (Precipitation sums)

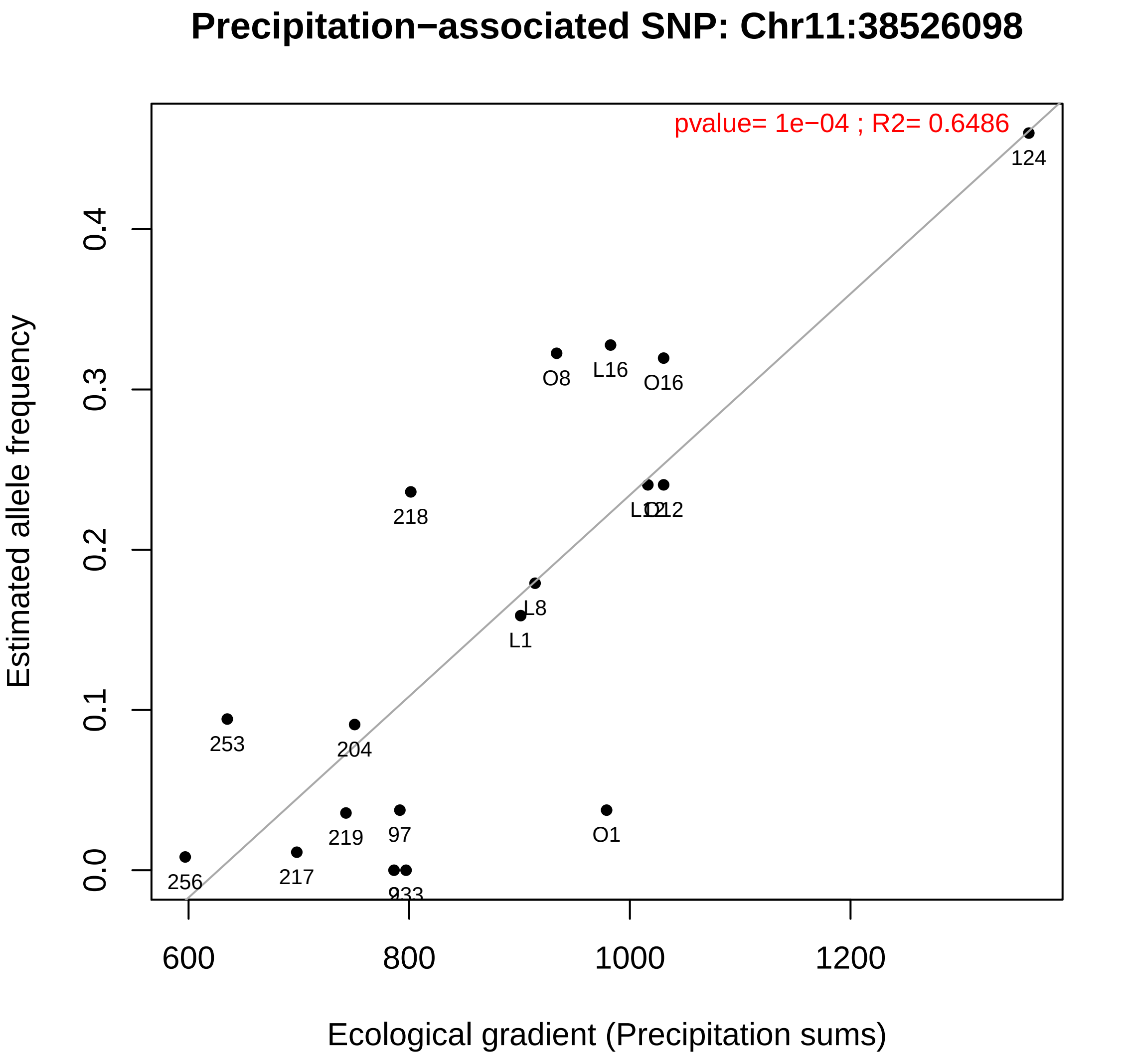

Precipitation−associated SNP: Chr11:38526098
pvalue= 1e−04 ; R2= 0.6486
Estimated allele frequency
Ecological gradient (Precipitation sums)
0.0
0.1
0.2
0.3
0.4
600
800
1000
1200
124
O8
L16
O16
218
L8
L1
253
204
219
97
O1
256
217

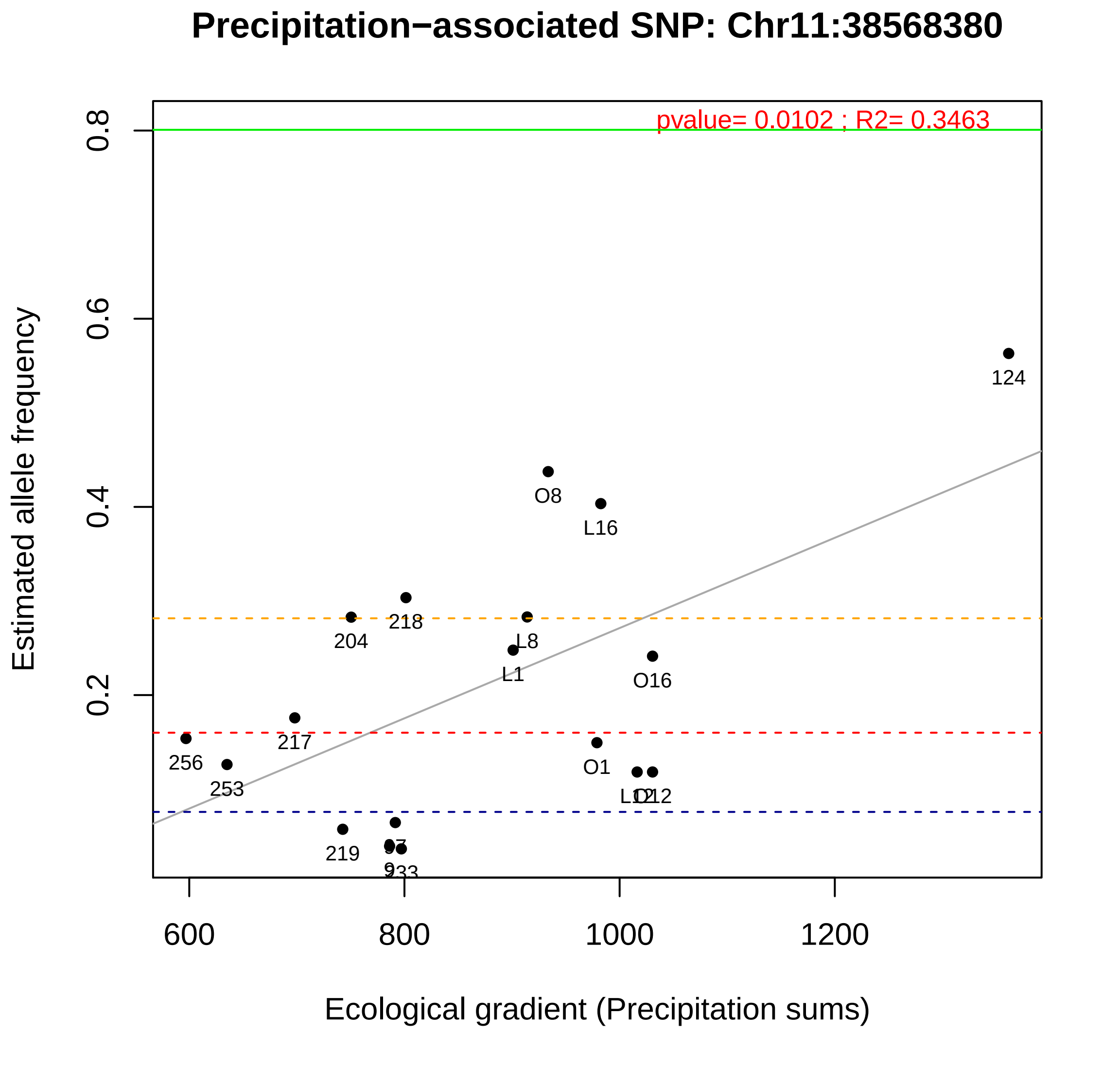

Precipitation−associated SNP: Chr11:38568380
pvalue= 0.0102 ; R2= 0.3463
Estimated allele frequency
0.2
0.4
0.6
0.8
600
800
1000
1200
Ecological gradient (Precipitation sums)
124
O8
L16
218
204
L8
L1
O16
217
256
253
O1
O12
219

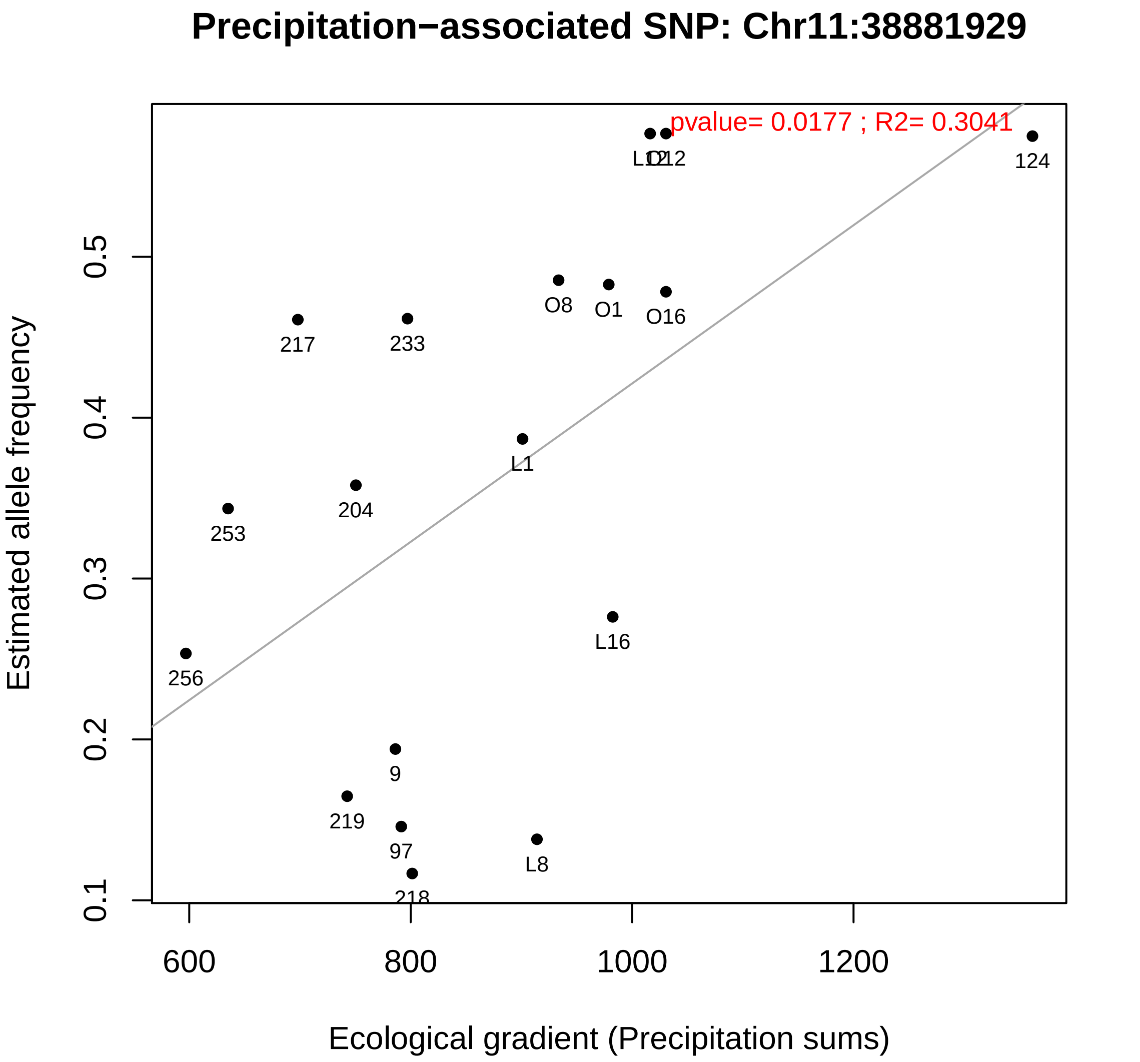
Precipitation−associated SNP: Chr11:38881929
pvalue= 0.0177 ; R2= 0.3041
Estimated allele frequency
0.1
0.2
0.3
0.4
0.5
600
800
1000
1200
Ecological gradient (Precipitation sums)
L10
O12
124
O8
O1
O16
217
233
L1
204
253
L16
256
9
219
97
L8
218

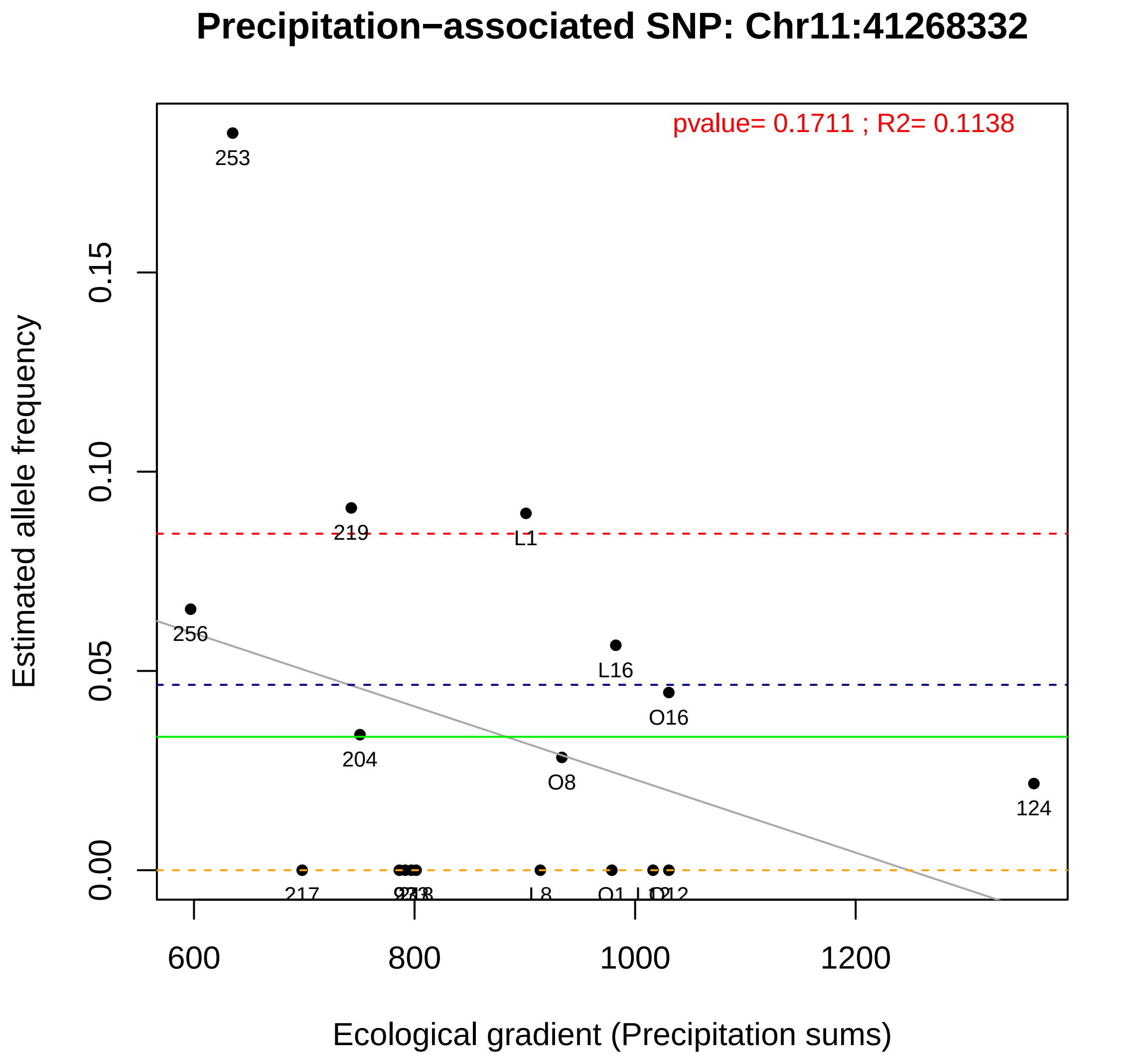

Precipitation−associated SNP: Chr11:41268332
pvalue= 0.1711 ; R2= 0.1138
Estimated allele frequency
Ecological gradient (Precipitation sums)
0.00
0.05
0.10
0.15
600
800
1000
1200
253
219
L1
256
L16
O16
204
O8
124
217
L8

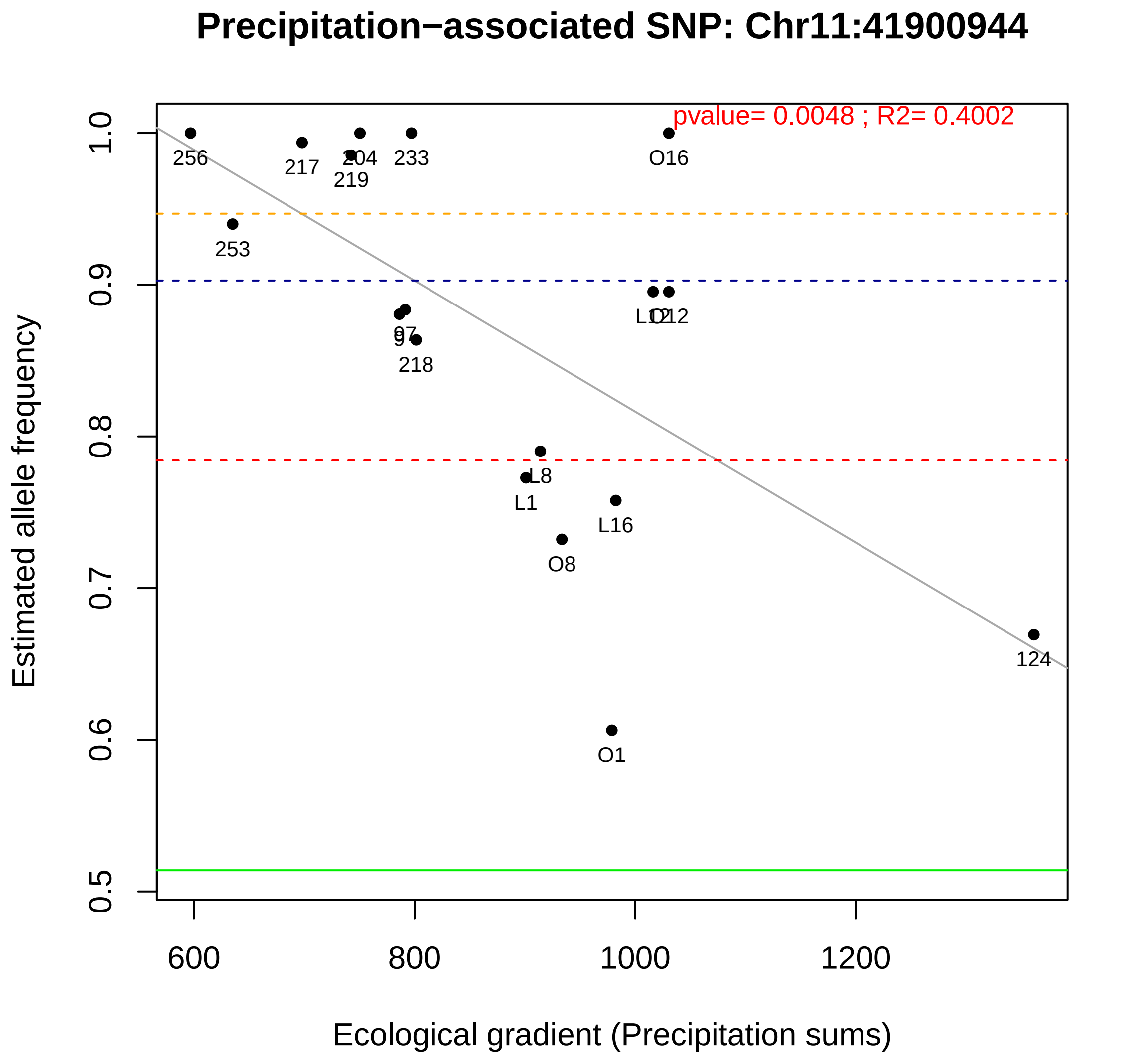
Precipitation−associated SNP: Chr11:41900944
pvalue= 0.0048 ; R2= 0.4002
Estimated allele frequency
Ecological gradient (Precipitation sums)
0.5
0.6
0.7
0.8
0.9
1.0
600
800
1000
1200
256
217
204
219
233
O16
253
97
218
L12
O12
L8
L1
L16
O8
124
O1

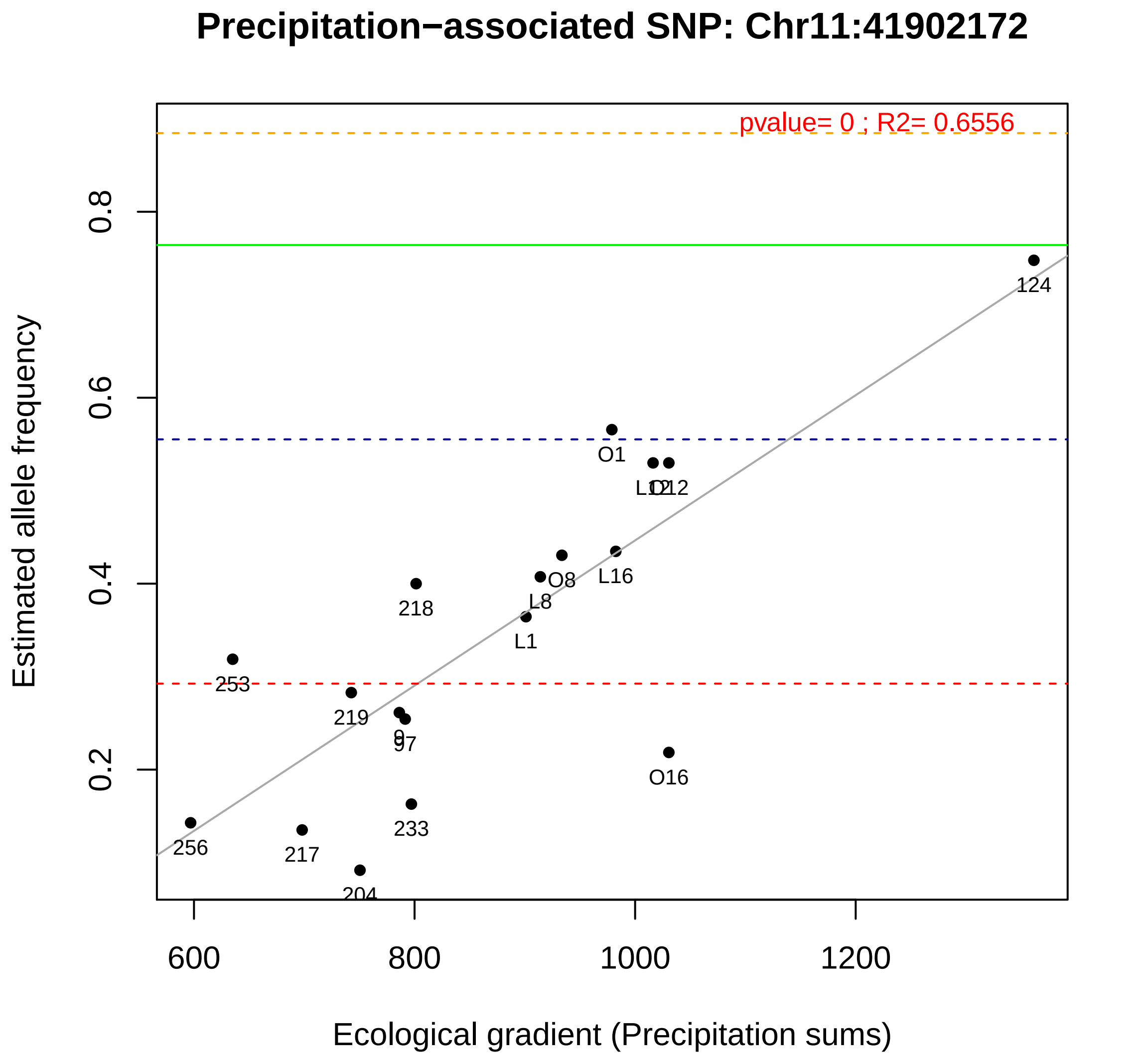

Precipitation−associated SNP: Chr11:41902172
pvalue= 0 ; R2= 0.6556
Estimated allele frequency
0.2
0.4
0.6
0.8
600
800
1000
1200
Ecological gradient (Precipitation sums)
124
O1
L12
O12
O8
L16
L8
218
L1
253
219
9
97
O16
233
256
217
204

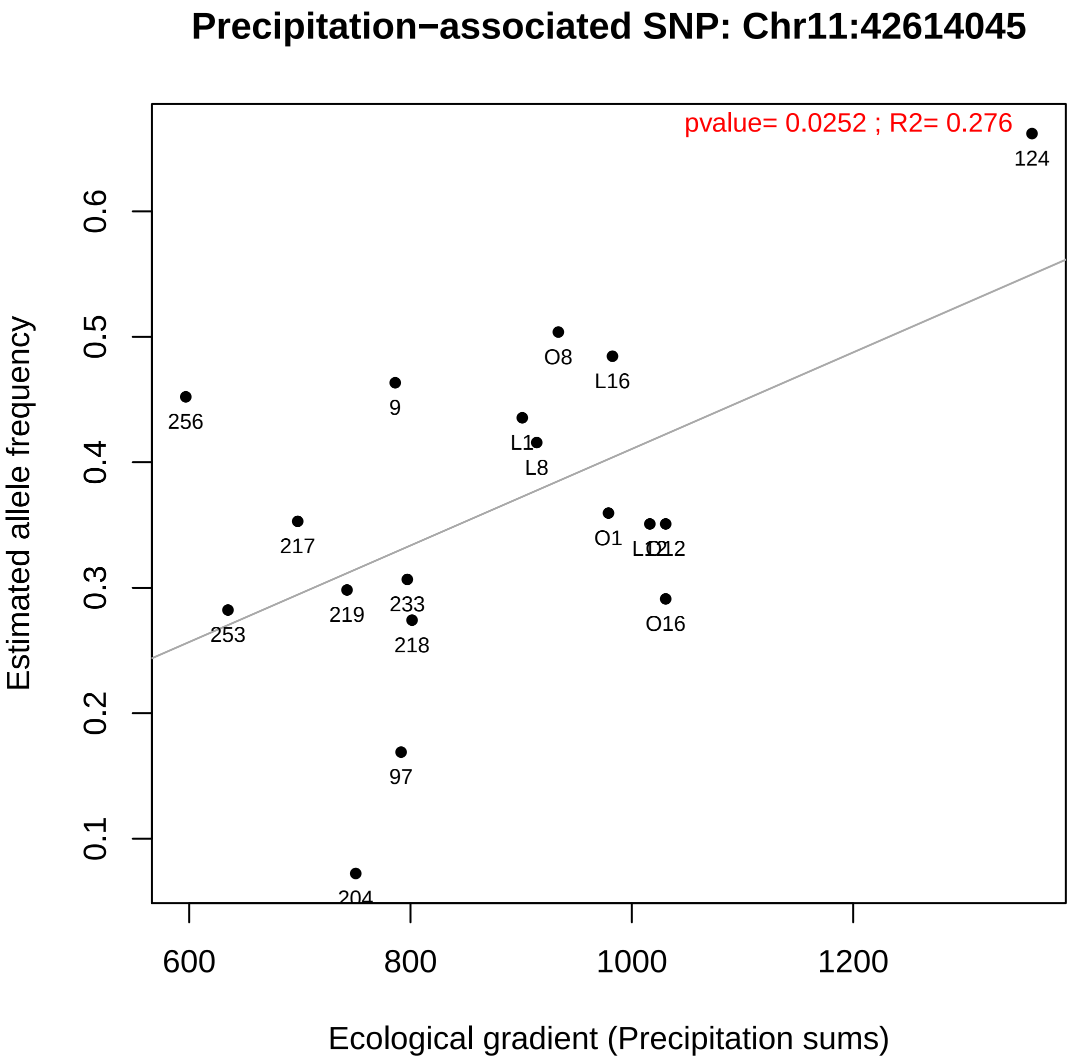

Precipitation−associated SNP: Chr11:42614045
pvalue= 0.0252 ; R2= 0.276
Estimated allele frequency
0.1
0.2
0.3
0.4
0.5
0.6
600
800
1000
1200
Ecological gradient (Precipitation sums)
124
O8
L16
256
9
L1
L8
217
O1
L12
O12
253
219
233
218
O16
97
204

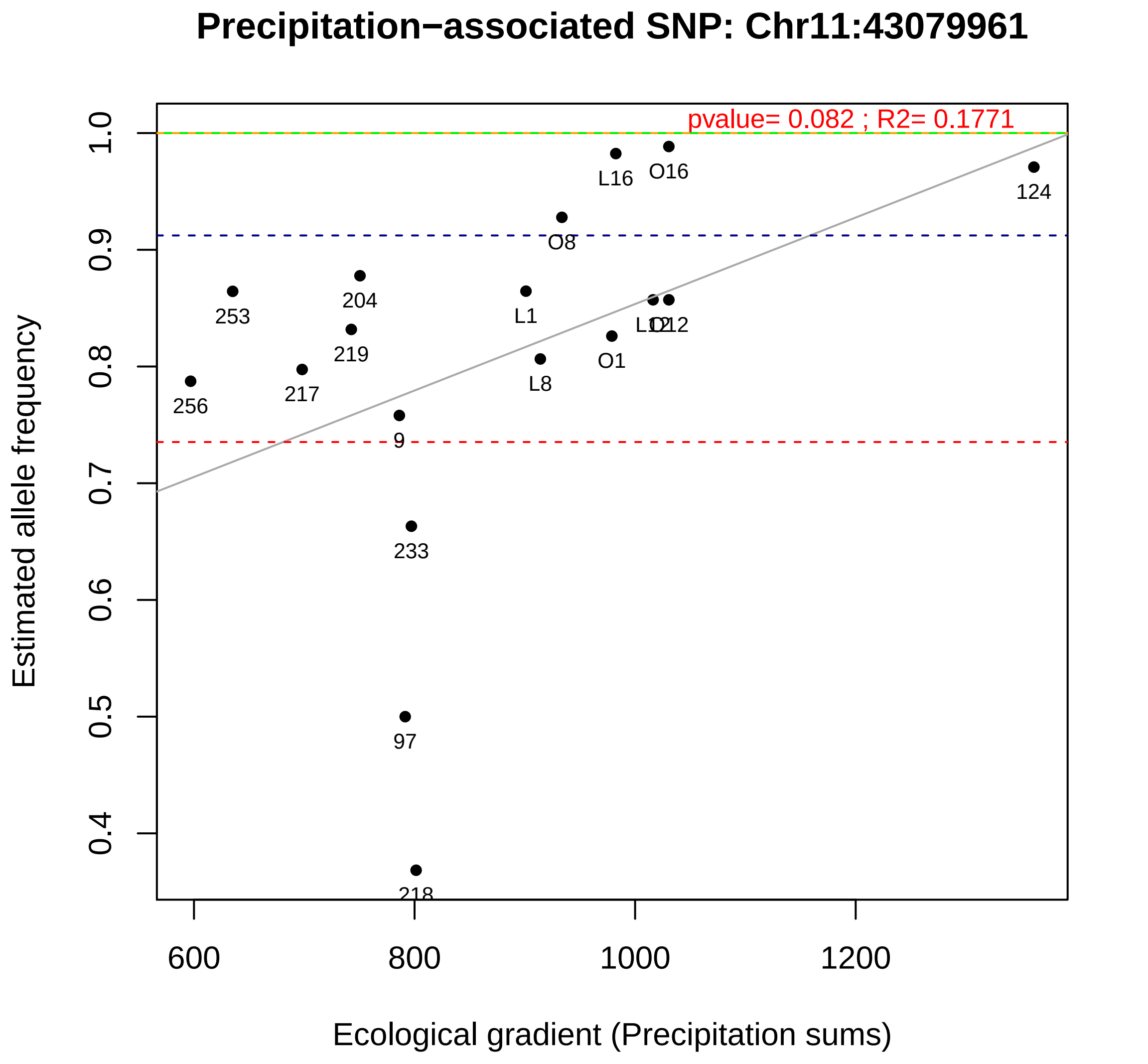
Precipitation−associated SNP: Chr11:43079961
pvalue= 0.082 ; R2= 0.1771
Estimated allele frequency
Ecological gradient (Precipitation sums)
0.4
0.5
0.6
0.7
0.8
0.9
1.0
600
800
1000
1200
L16
O16
124
O8
253
204
L1
L12
O12
219
O1
L8
256
217
9
233
97
218

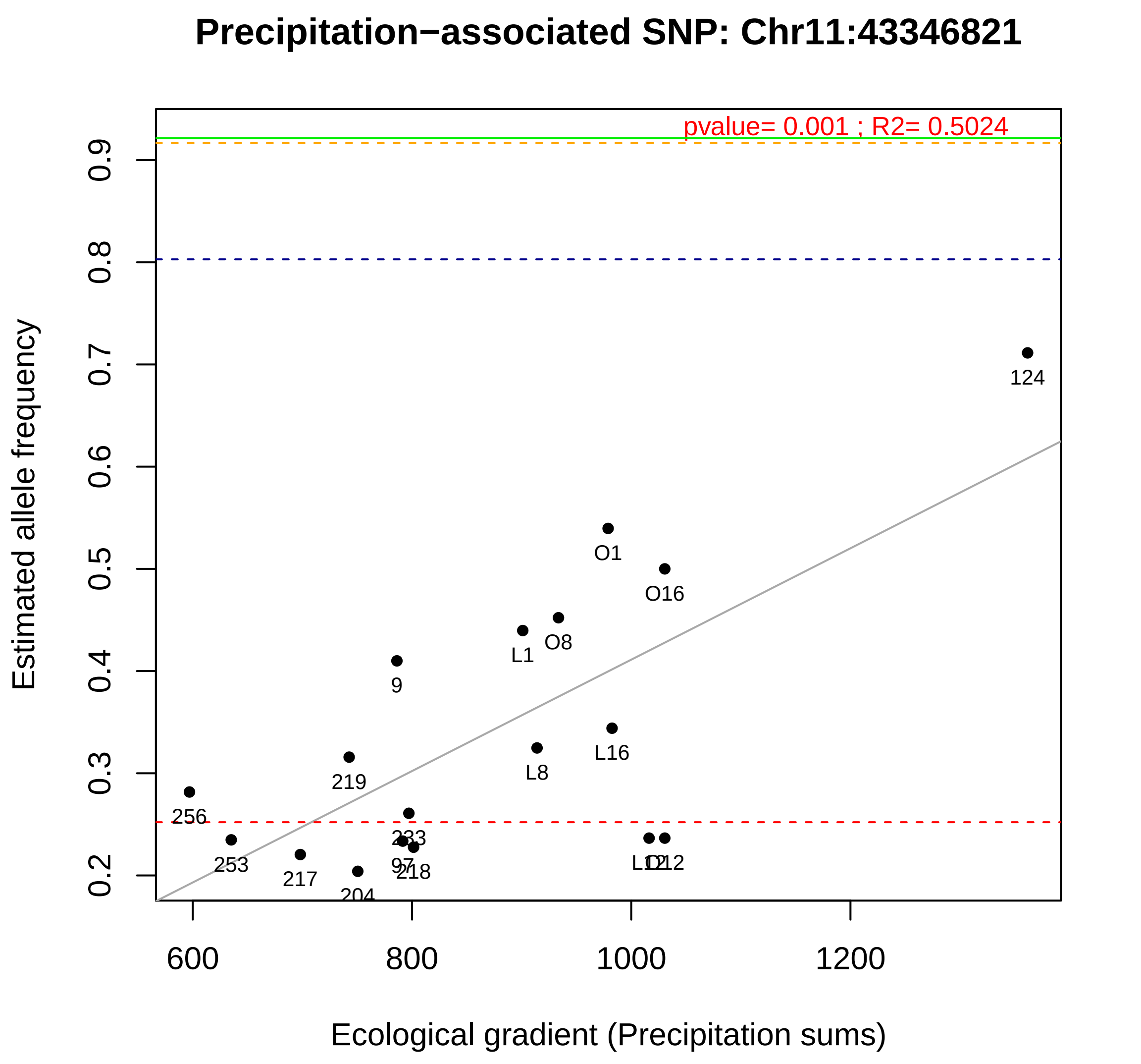
Precipitation−associated SNP: Chr11:43346821
pvalue= 0.001 ; R2= 0.5024
Estimated allele frequency
0.2
0.3
0.4
0.5
0.6
0.7
0.8
0.9
600
800
1000
1200
Ecological gradient (Precipitation sums)
124
O1
O16
O8
L1
9
L16
L8
219
256
233
253
217
97
218
204
L12
O12

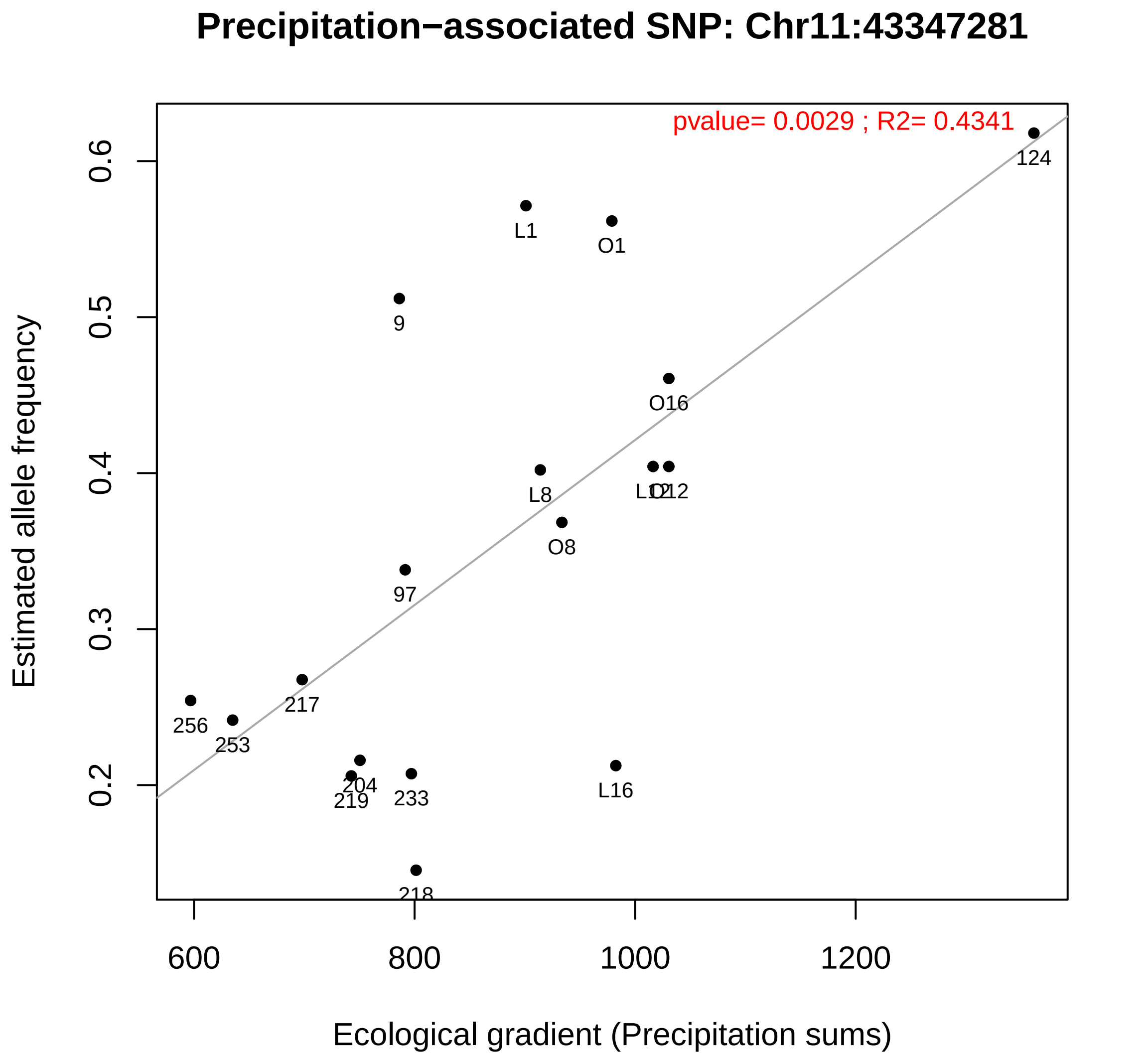

Precipitation−associated SNP: Chr11:43347281
pvalue= 0.0029 ; R2= 0.4341
Estimated allele frequency
0.2
0.3
0.4
0.5
0.6
600
800
1000
1200
Ecological gradient (Precipitation sums)
124
L1
O1
9
O16
L8
O12
O8
97
217
256
253
204
219
233
L16
218

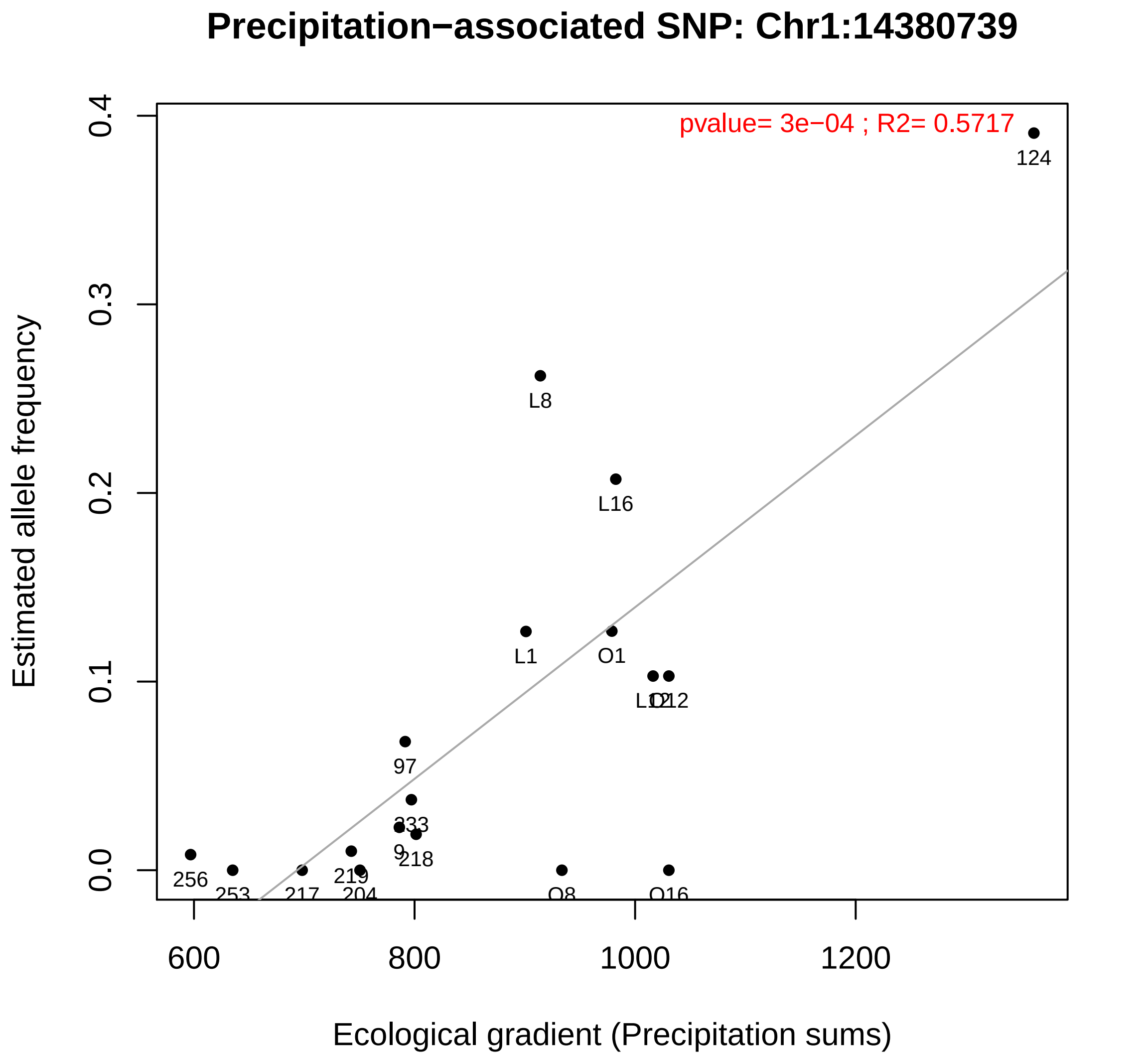

Precipitation−associated SNP: Chr1:14380739
pvalue= 3e−04 ; R2= 0.5717
124
L8
L16
L1
O1
L12
O12
97
233
9
218
256
253
217
219
204
O8
O16
0.0
0.1
0.2
0.3
0.4
600
800
1000
1200
Estimated allele frequency
Ecological gradient (Precipitation sums)

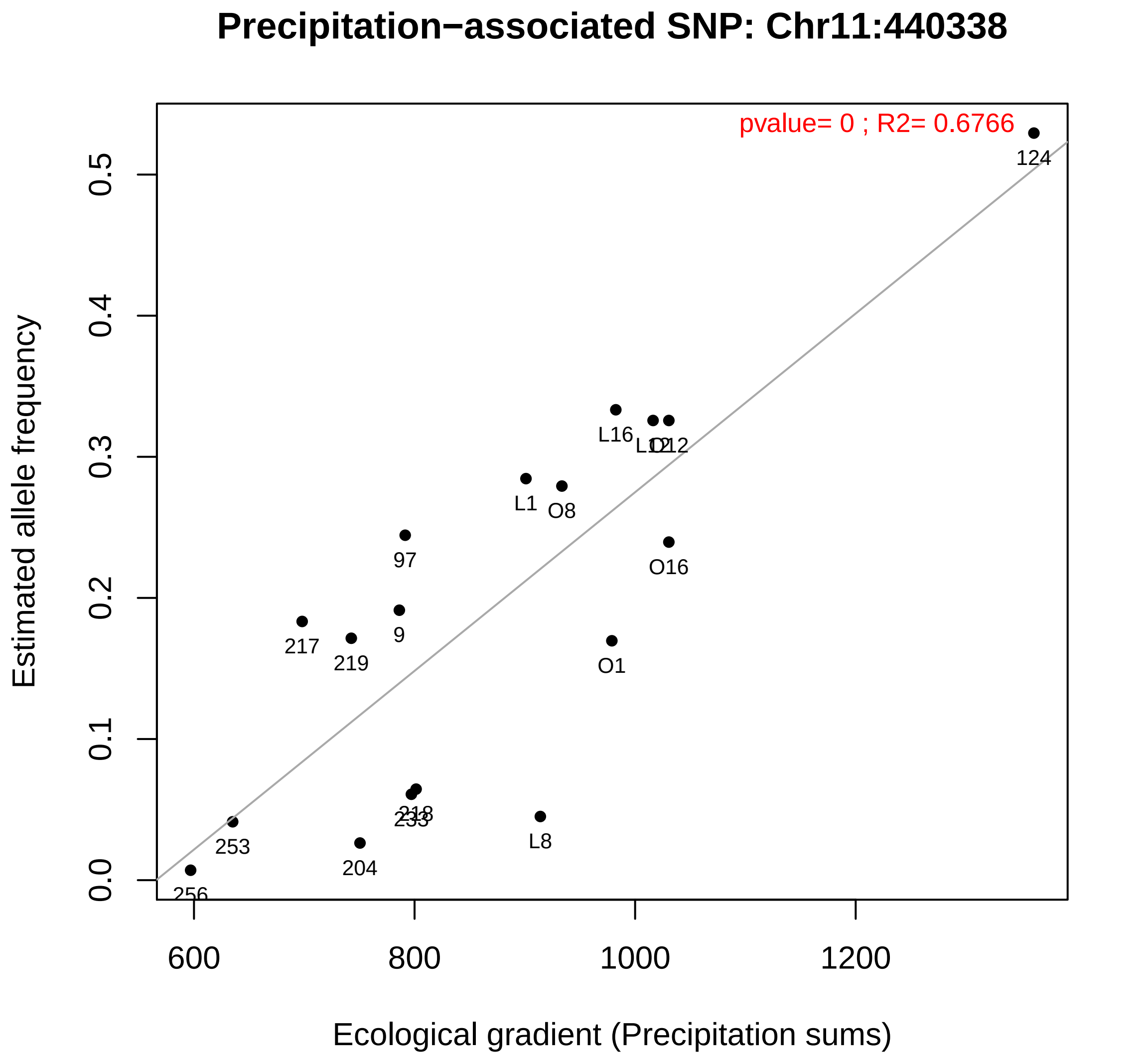

Precipitation−associated SNP: Chr11:440338
pvalue= 0 ; R2= 0.6766
Estimated allele frequency
0.0
0.1
0.2
0.3
0.4
0.5
600
800
1000
1200
Ecological gradient (Precipitation sums)
124
L16
L1
O8
97
O16
217
219
9
O1
253
204
L8

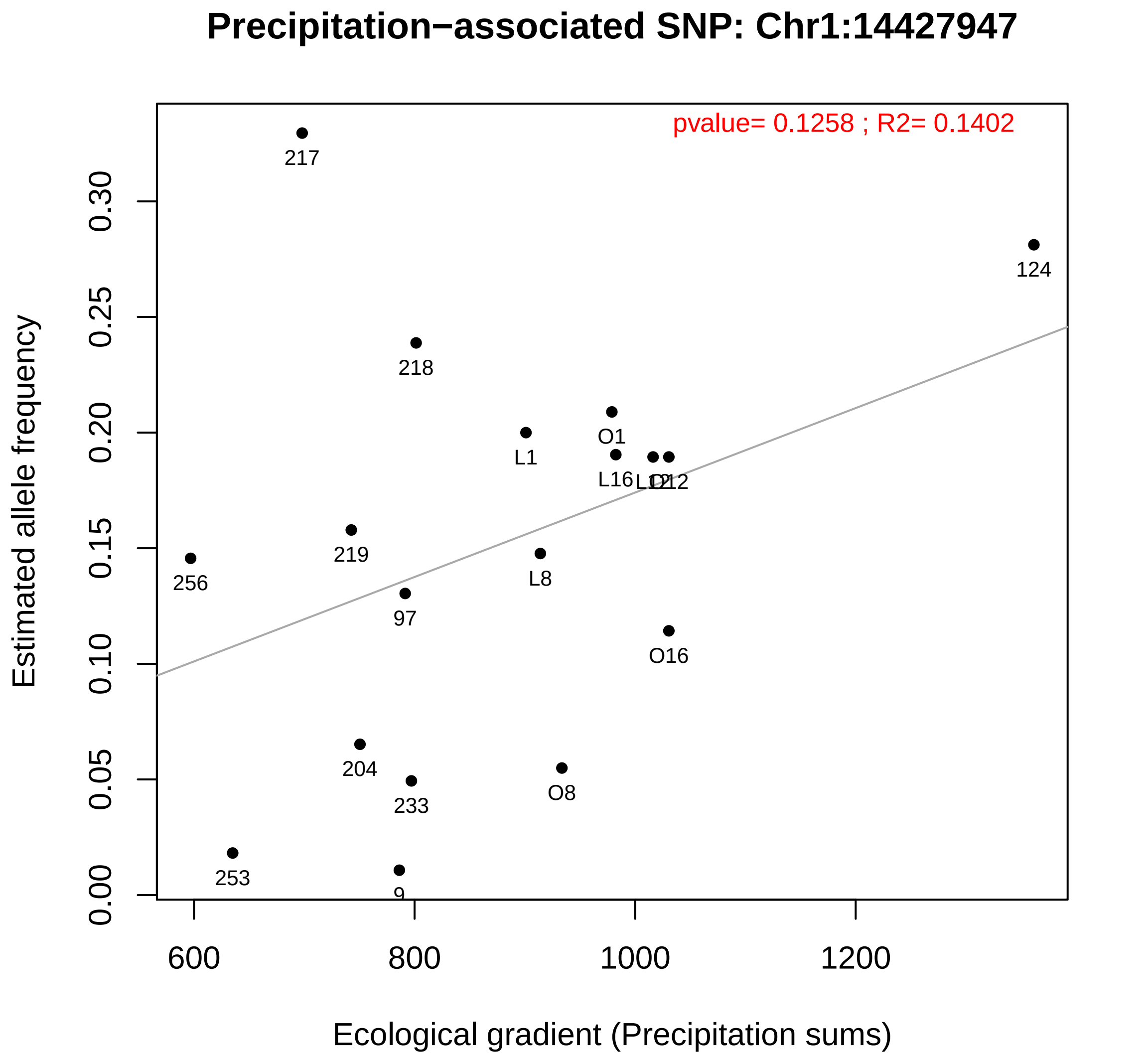
Precipitation−associated SNP: Chr1:14427947
pvalue= 0.1258 ; R2= 0.1402
217
124
218
O1
L1
L16
L12
O12
219
256
L8
97
O16
204
233
O8
253
9
0.00
0.05
0.10
0.15
0.20
0.25
0.30
600
800
1000
1200
Estimated allele frequency
Ecological gradient (Precipitation sums)

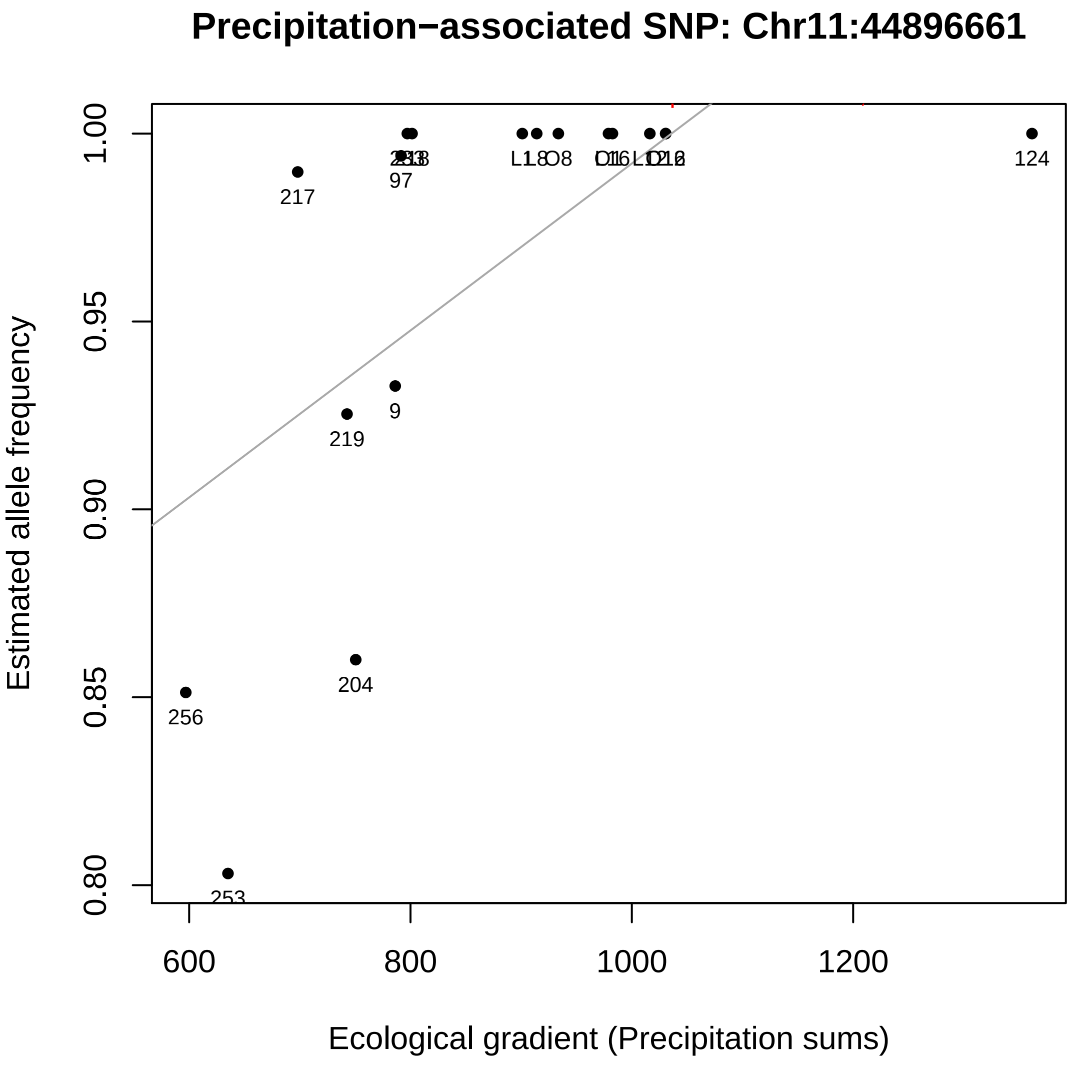
Precipitation−associated SNP: Chr11:44896661
Estimated allele frequency
Ecological gradient (Precipitation sums)
0.80
0.85
0.90
0.95
1.00
600
800
1000
1200
217
97
9
219
204
256
253
124
O8

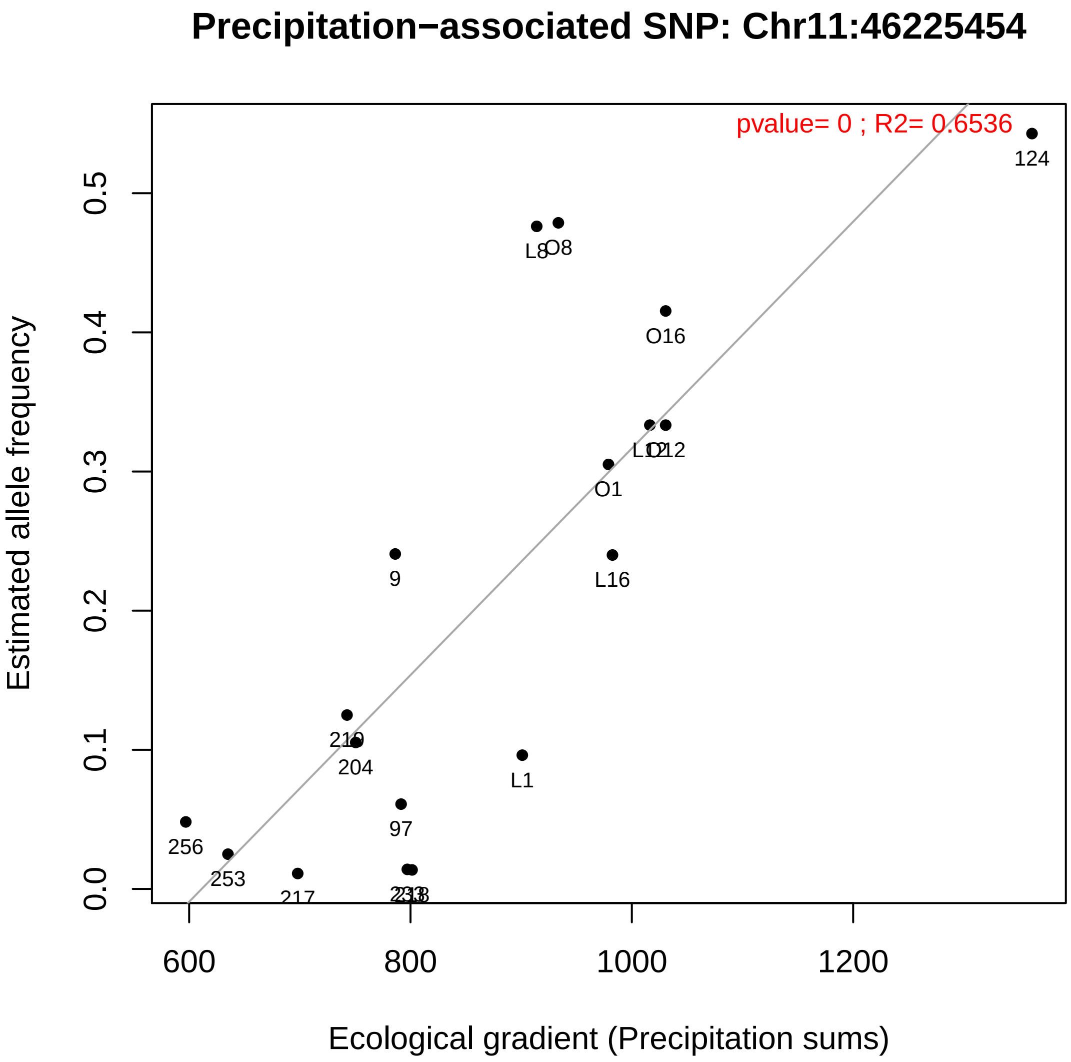
Precipitation−associated SNP: Chr11:46225454
pvalue= 0 ; R2= 0.6536
Estimated allele frequency
0.0
0.1
0.2
0.3
0.4
0.5
Ecological gradient (Precipitation sums)
600
800
1000
1200
124
L8
O8
O16
O12
O1
9
L16
204
L1
97
256
253
217

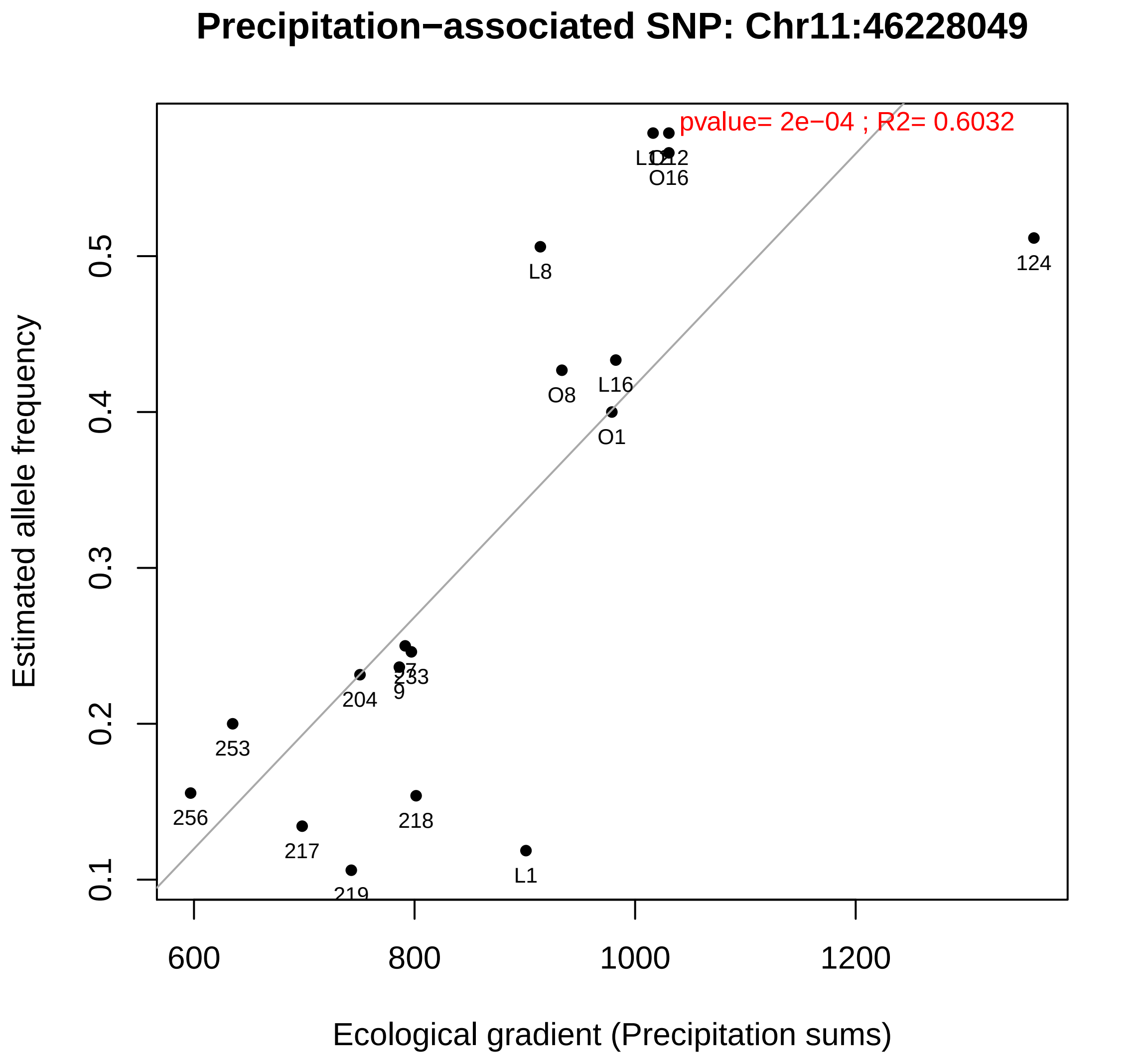

Precipitation−associated SNP: Chr11:46228049
pvalue= 2e−04 ; R2= 0.6032
Estimated allele frequency
0.1
0.2
0.3
0.4
0.5
600
800
1000
1200
Ecological gradient (Precipitation sums)
O16
L8
124
L16
O8
O1
233
9
204
253
256
218
217
L1

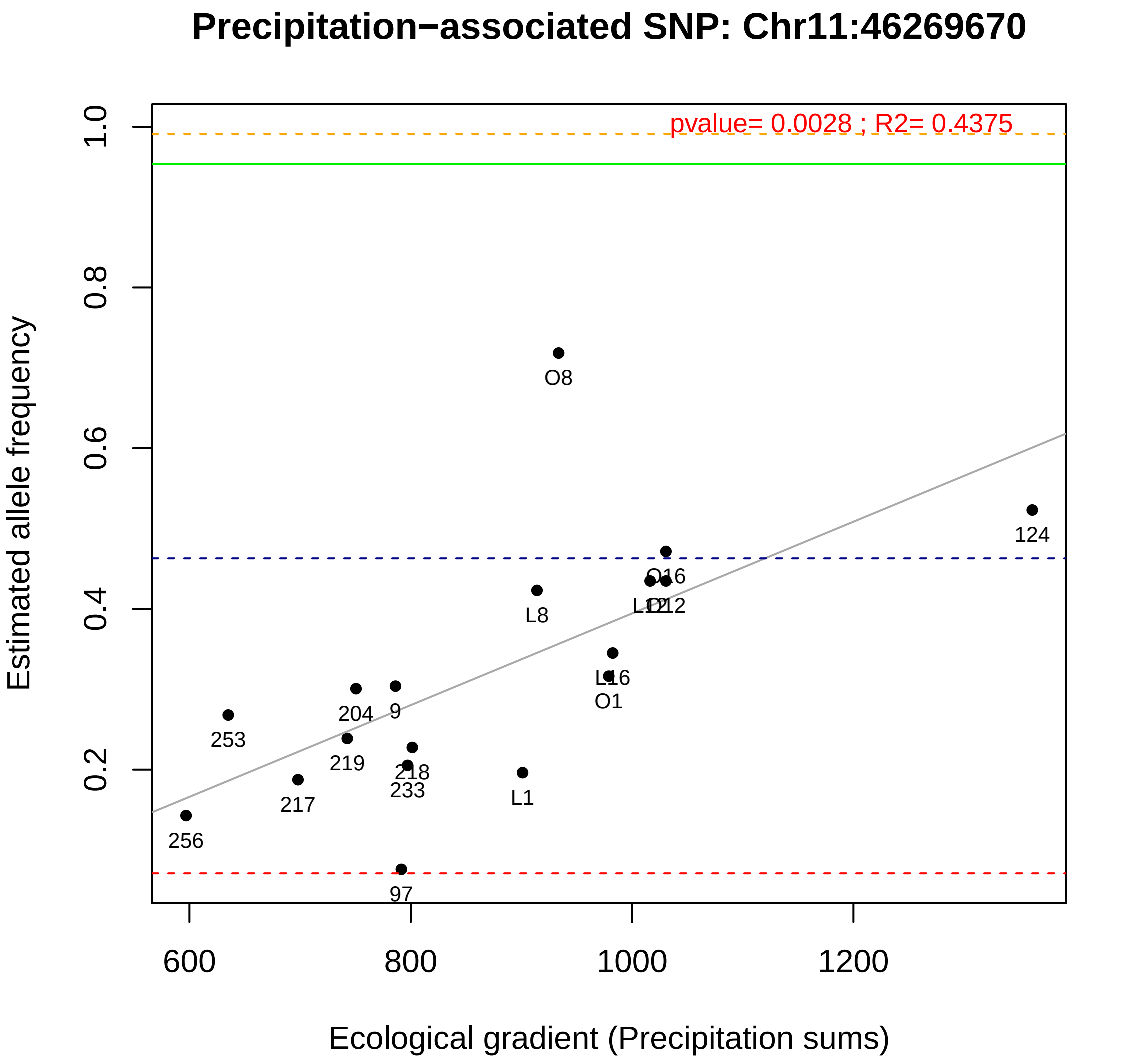
Precipitation−associated SNP: Chr11:46269670
pvalue= 0.0028 ; R2= 0.4375
Estimated allele frequency
Ecological gradient (Precipitation sums)
0.2
0.4
0.6
0.8
1.0
600
800
1000
1200
O8
124
O16
L12
O12
L8
L16
O1
204
9
253
219
218
233
217
L1
256
97

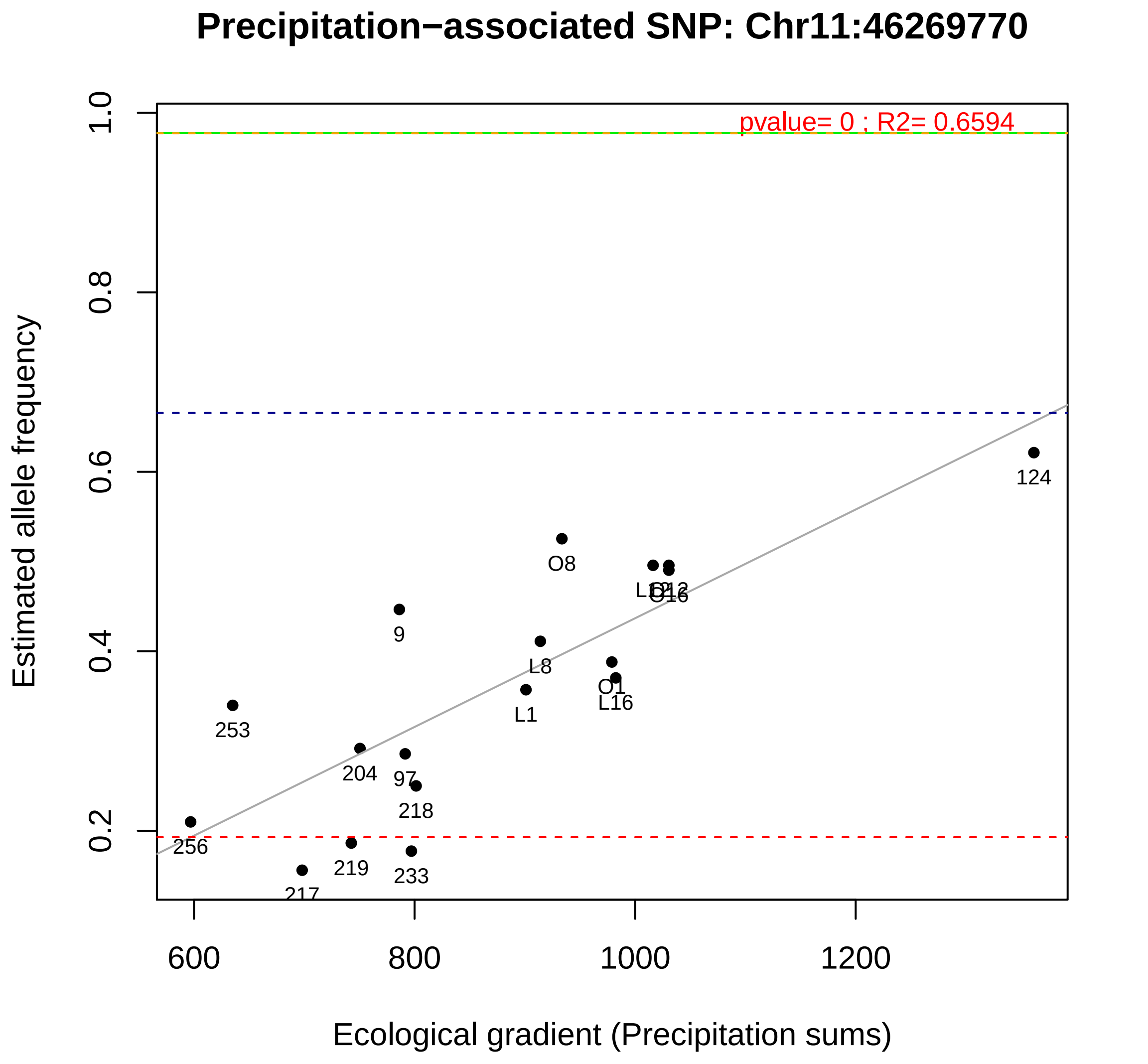
Precipitation−associated SNP: Chr11:46269770
pvalue= 0 ; R2= 0.6594
Estimated allele frequency
Ecological gradient (Precipitation sums)
0.2
0.4
0.6
0.8
1.0
600
800
1000
1200
124
O8
9
L8
L1
O1
L16
253
204
97
218
256
219
233
217

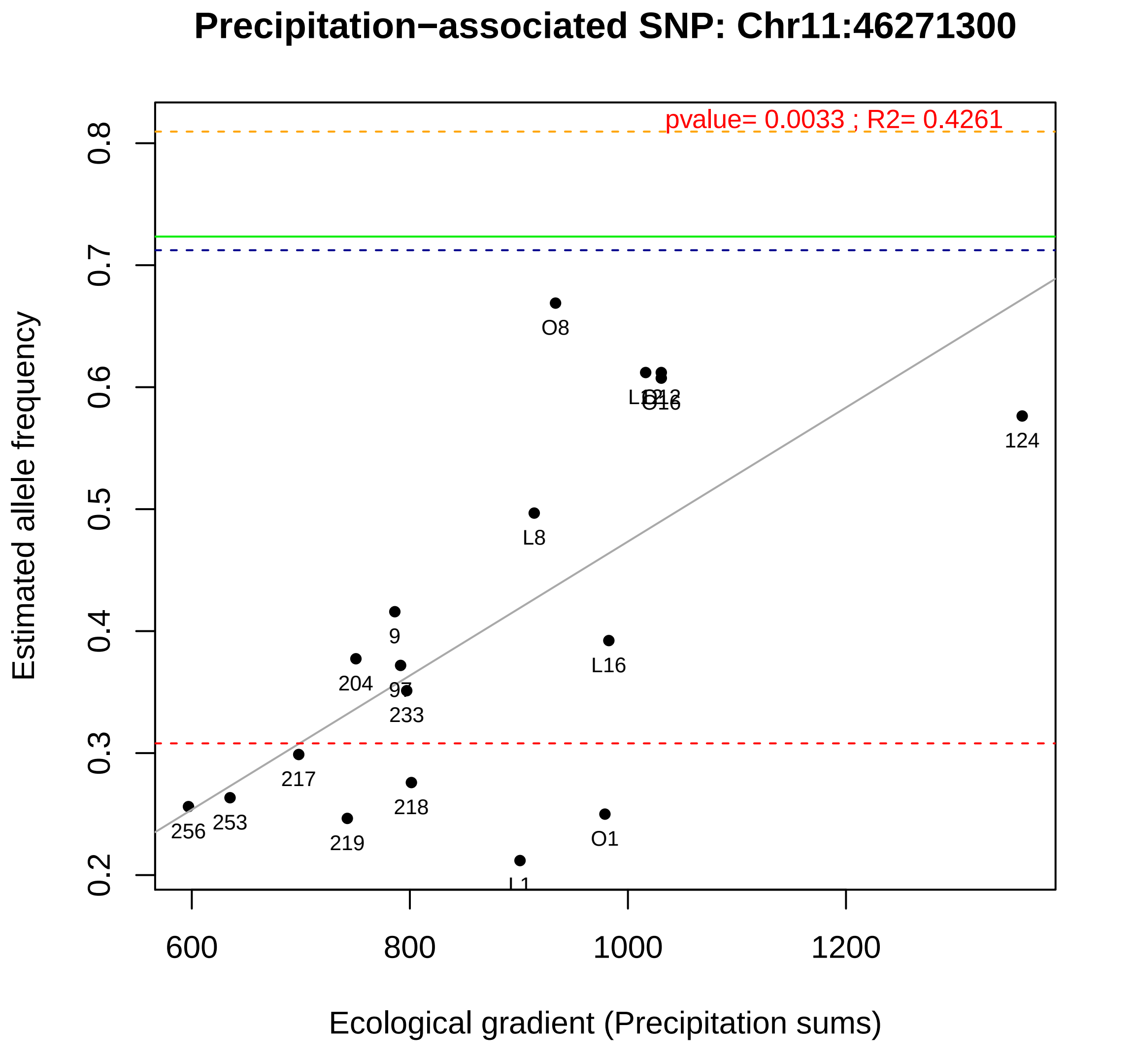

Precipitation−associated SNP: Chr11:46271300
pvalue= 0.0033 ; R2= 0.4261
Estimated allele frequency
0.2
0.3
0.4
0.5
0.6
0.7
0.8
600
800
1000
1200
Ecological gradient (Precipitation sums)
O8
L8
124
9
204
L16
233
217
218
256
253
219
O1
O16

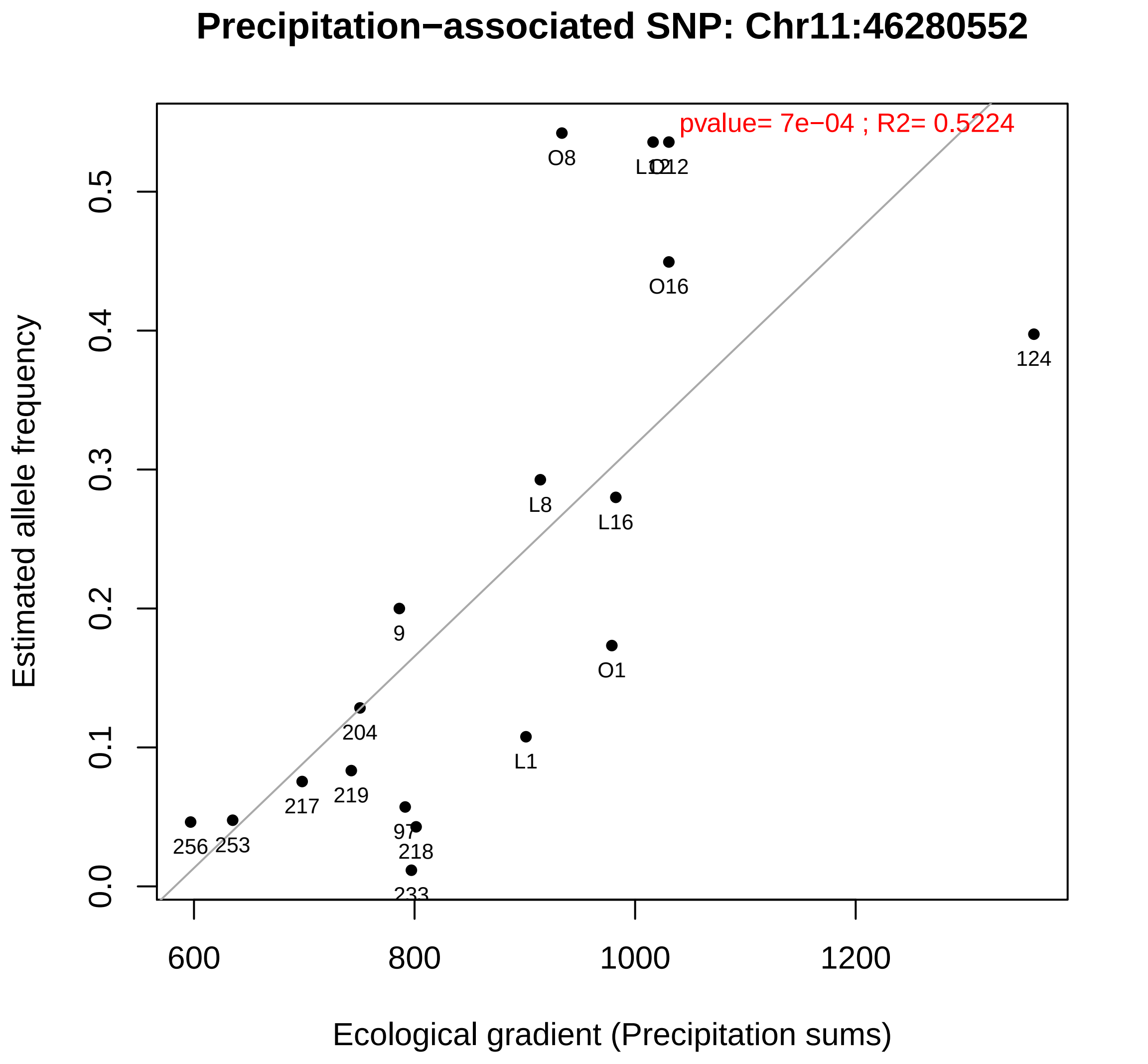

Precipitation−associated SNP: Chr11:46280552
pvalue= 7e−04 ; R2= 0.5224
O8
L10
O12
O16
124
L8
L16
9
O1
204
L1
217
219
256
253
97
218
233
0.0
0.1
0.2
0.3
0.4
0.5
600
800
1000
1200
Estimated allele frequency
Ecological gradient (Precipitation sums)

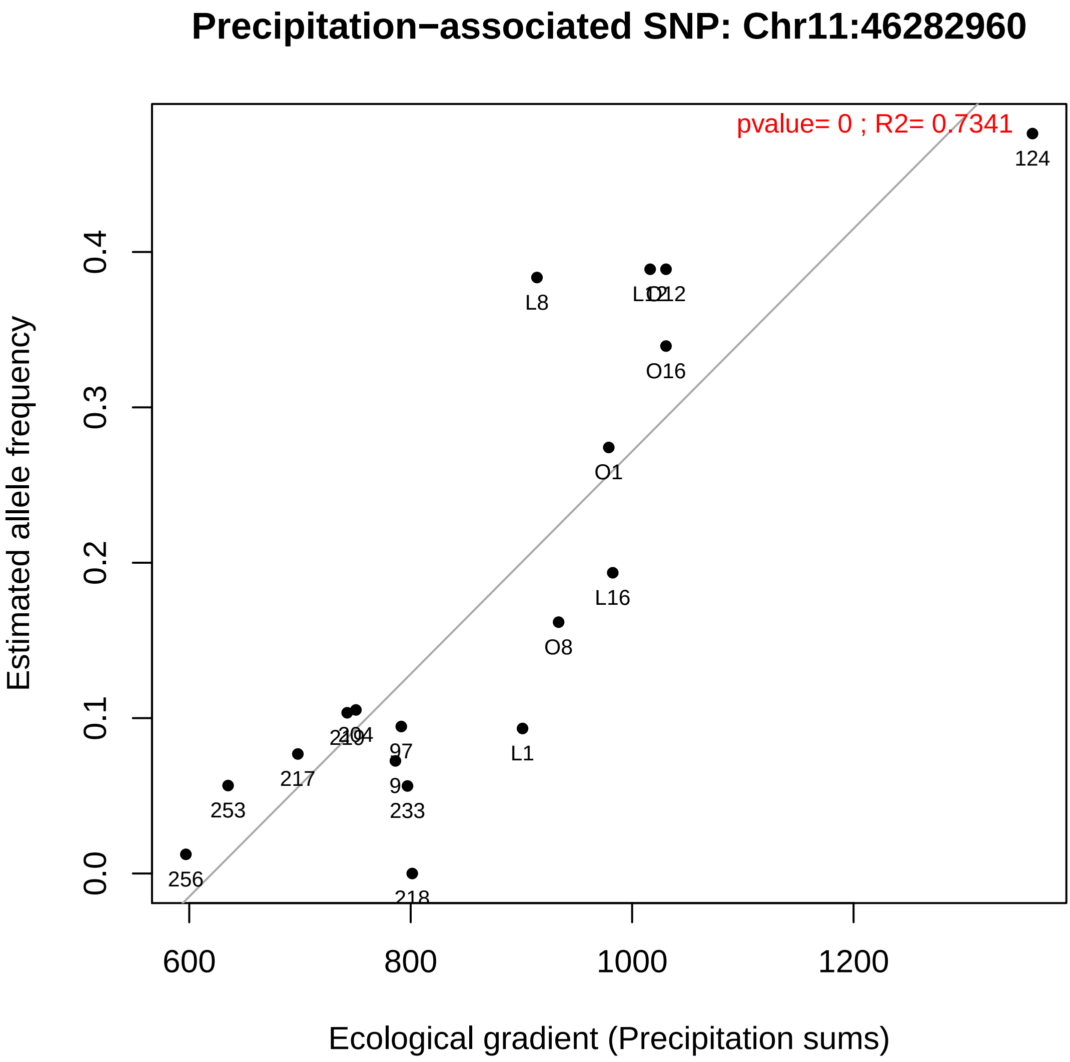

Precipitation−associated SNP: Chr11:46282960
pvalue= 0 ; R2= 0.7341
124
L8
L12
O12
O16
O1
L16
O8
210
204
97
L1
217
9
233
253
256
218
0.0
0.1
0.2
0.3
0.4
600
800
1000
1200
Estimated allele frequency
Ecological gradient (Precipitation sums)

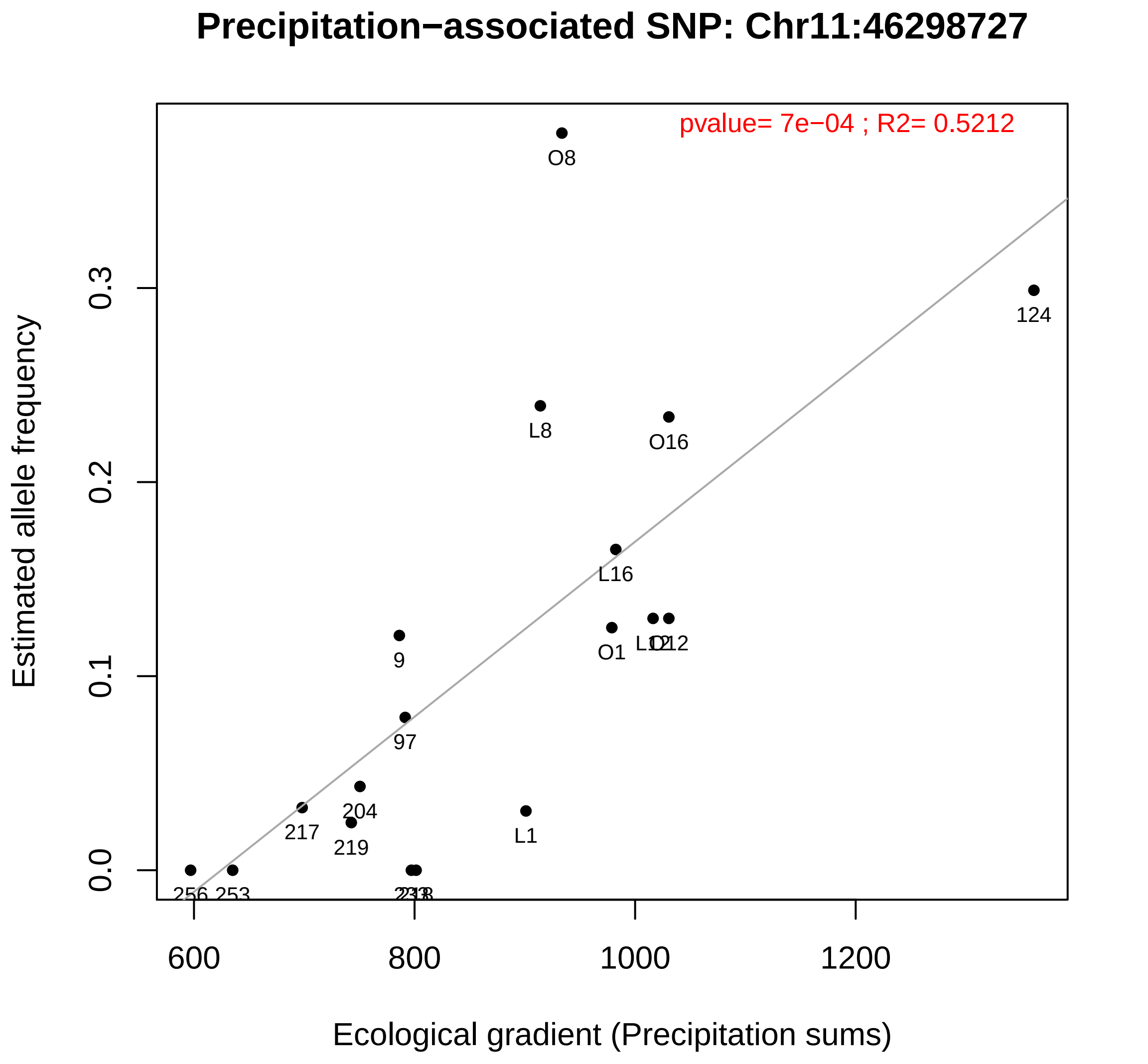

Precipitation−associated SNP: Chr11:46298727
pvalue= 7e−04 ; R2= 0.5212
O8
124
L8
O16
L16
9
O1
L12
O12
97
204
217
219
L1
256
253
0.0
0.1
0.2
0.3
600
800
1000
1200
Estimated allele frequency
Ecological gradient (Precipitation sums)

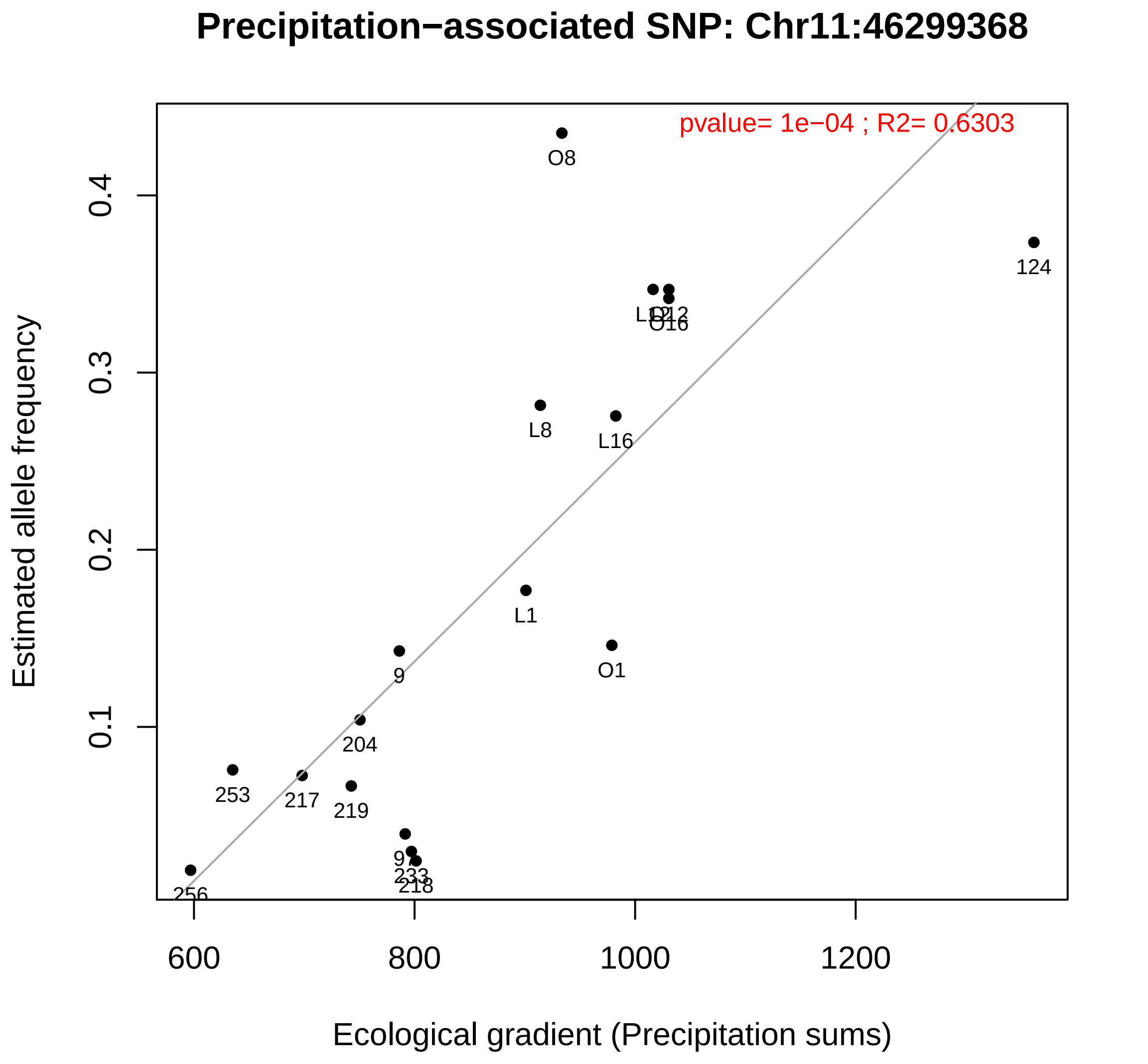

Precipitation−associated SNP: Chr11:46299368
pvalue= 1e−04 ; R2= 0.6303
Estimated allele frequency
0.1
0.2
0.3
0.4
600
800
1000
1200
Ecological gradient (Precipitation sums)
O8
124
L8
L16
L1
O1
9
204
253
217
219
256
O16

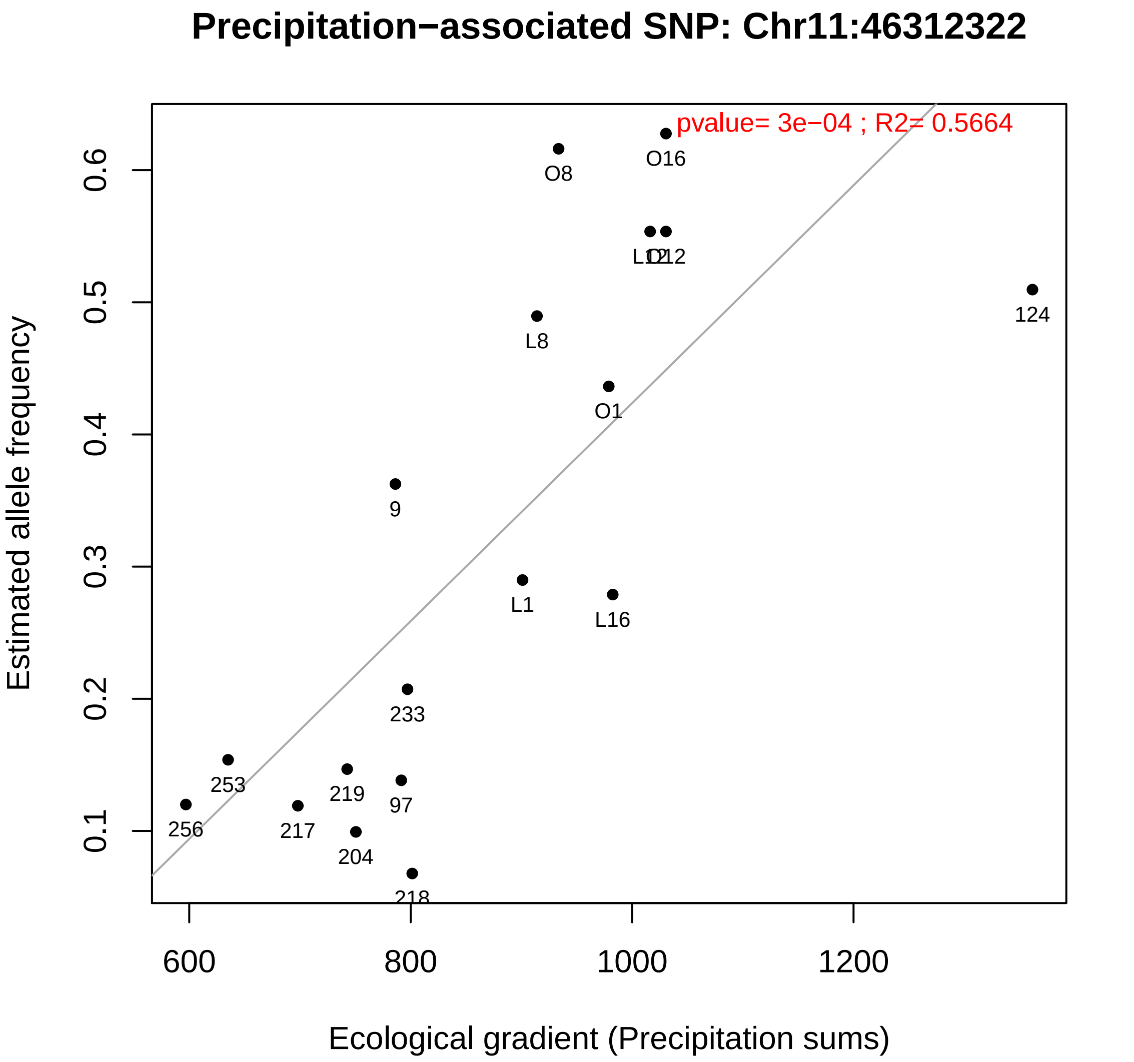
Precipitation−associated SNP: Chr11:46312322
pvalue= 3e−04 ; R2= 0.5664
Estimated allele frequency
Ecological gradient (Precipitation sums)
0.1
0.2
0.3
0.4
0.5
0.6
600
800
1000
1200
O16
O8
L12
O12
124
L8
O1
9
L1
L16
233
253
219
97
256
217
204
218

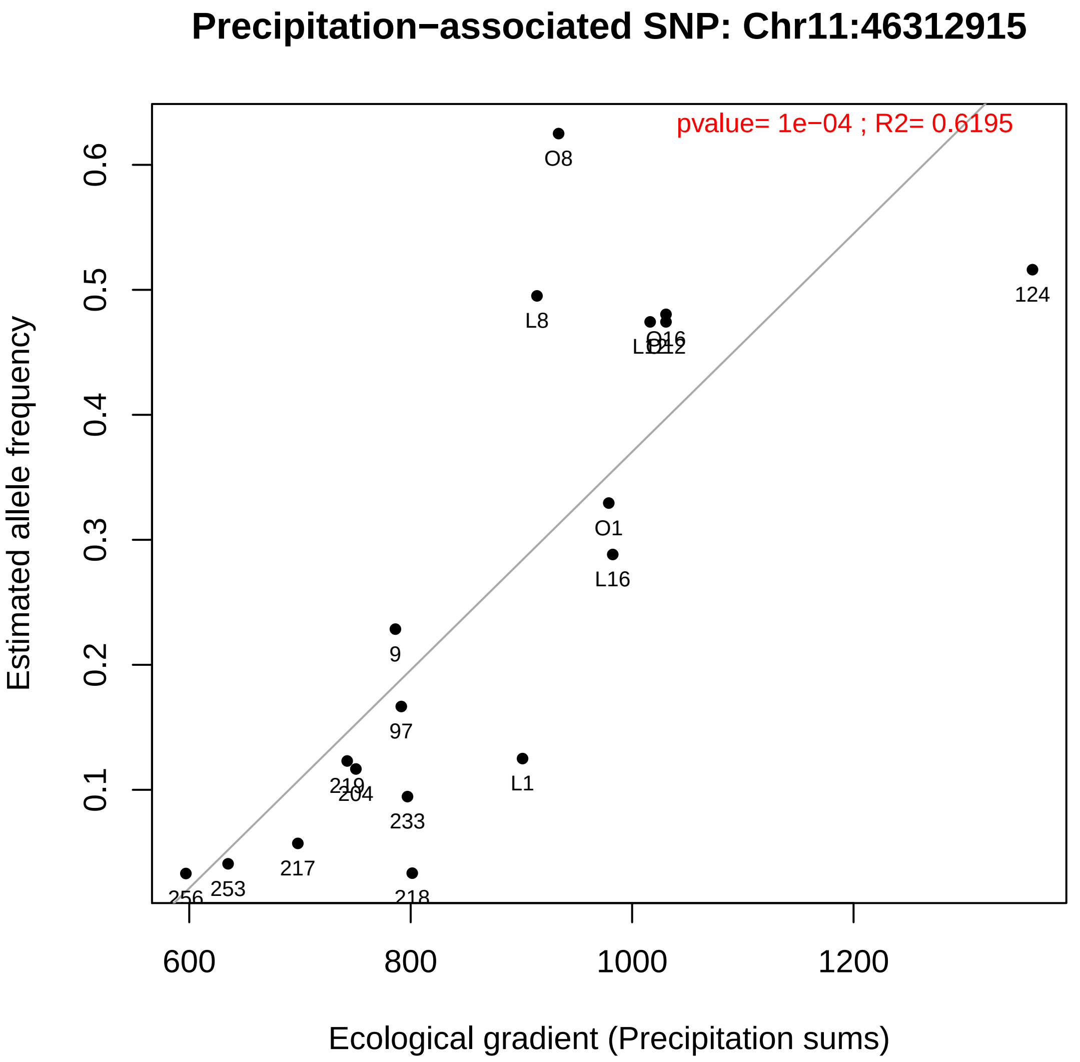

Precipitation−associated SNP: Chr11:46312915
pvalue= 1e−04 ; R2= 0.6195
O8
L8
124
O16
L10
O12
O1
L16
9
97
219
204
L1
233
217
256
253
218
Estimated allele frequency
0.1
0.2
0.3
0.4
0.5
0.6
600
800
1000
1200
Ecological gradient (Precipitation sums)

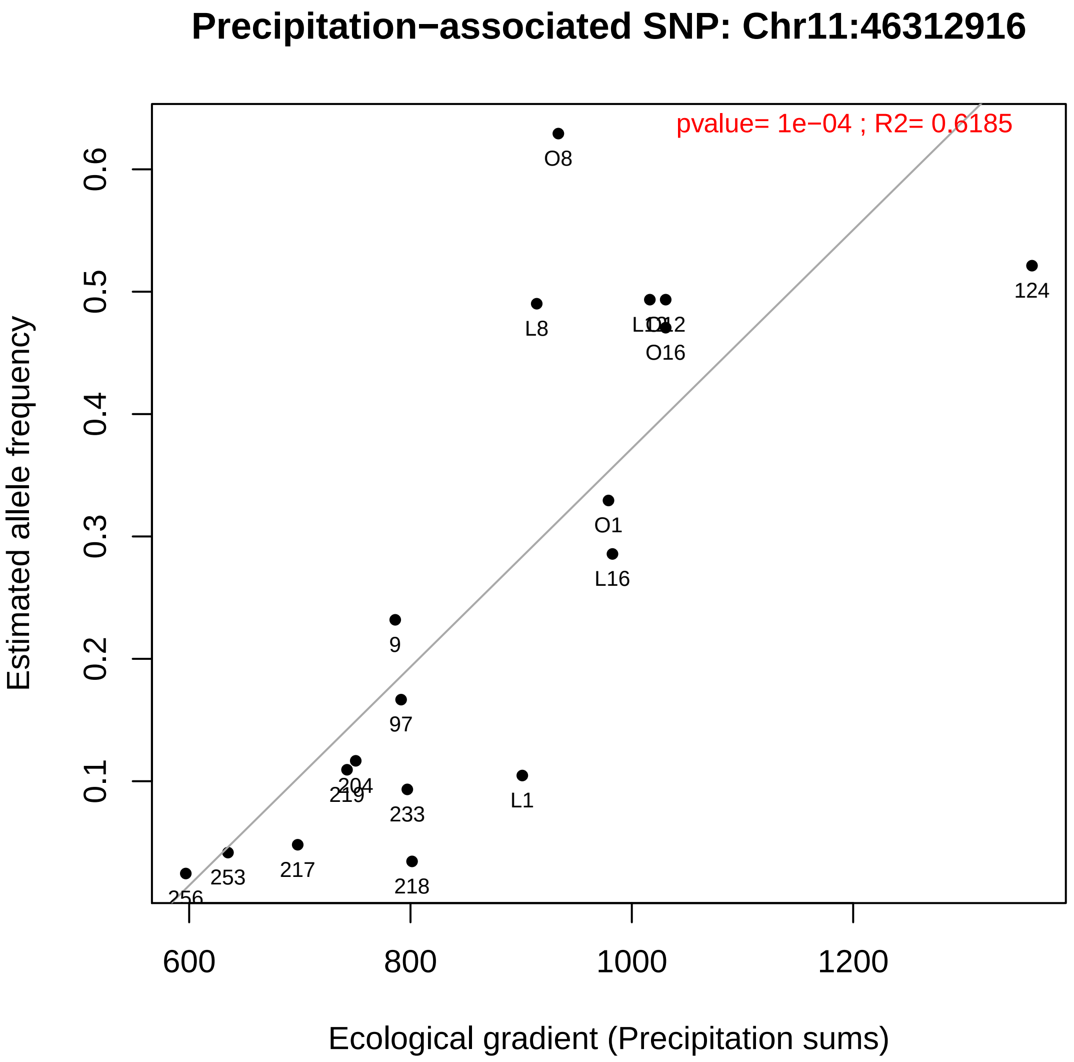

Precipitation−associated SNP: Chr11:46312916
pvalue= 1e−04 ; R2= 0.6185
Estimated allele frequency
0.1
0.2
0.3
0.4
0.5
0.6
600
800
1000
1200
Ecological gradient (Precipitation sums)
O8
124
L8
O16
O1
L16
9
97
204
219
233
L1
253
217
218
256

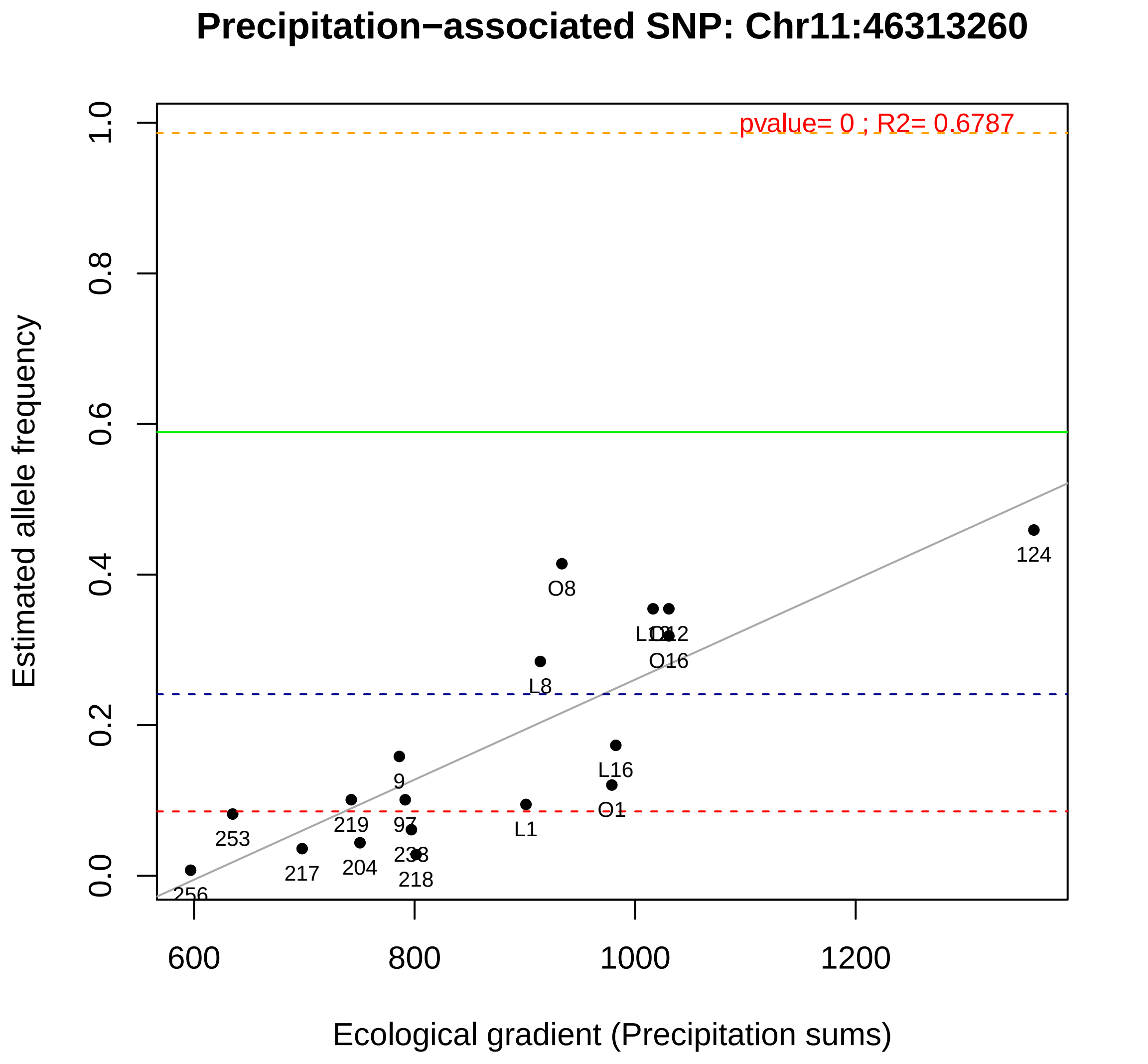

Precipitation−associated SNP: Chr11:46313260
pvalue= 0 ; R2= 0.6787
Estimated allele frequency
0.0
0.2
0.4
0.6
0.8
1.0
600
800
1000
1200
Ecological gradient (Precipitation sums)
124
O8
L12
O12
O16
L8
L16
9
O1
219
97
L1
253
204
217
218
256

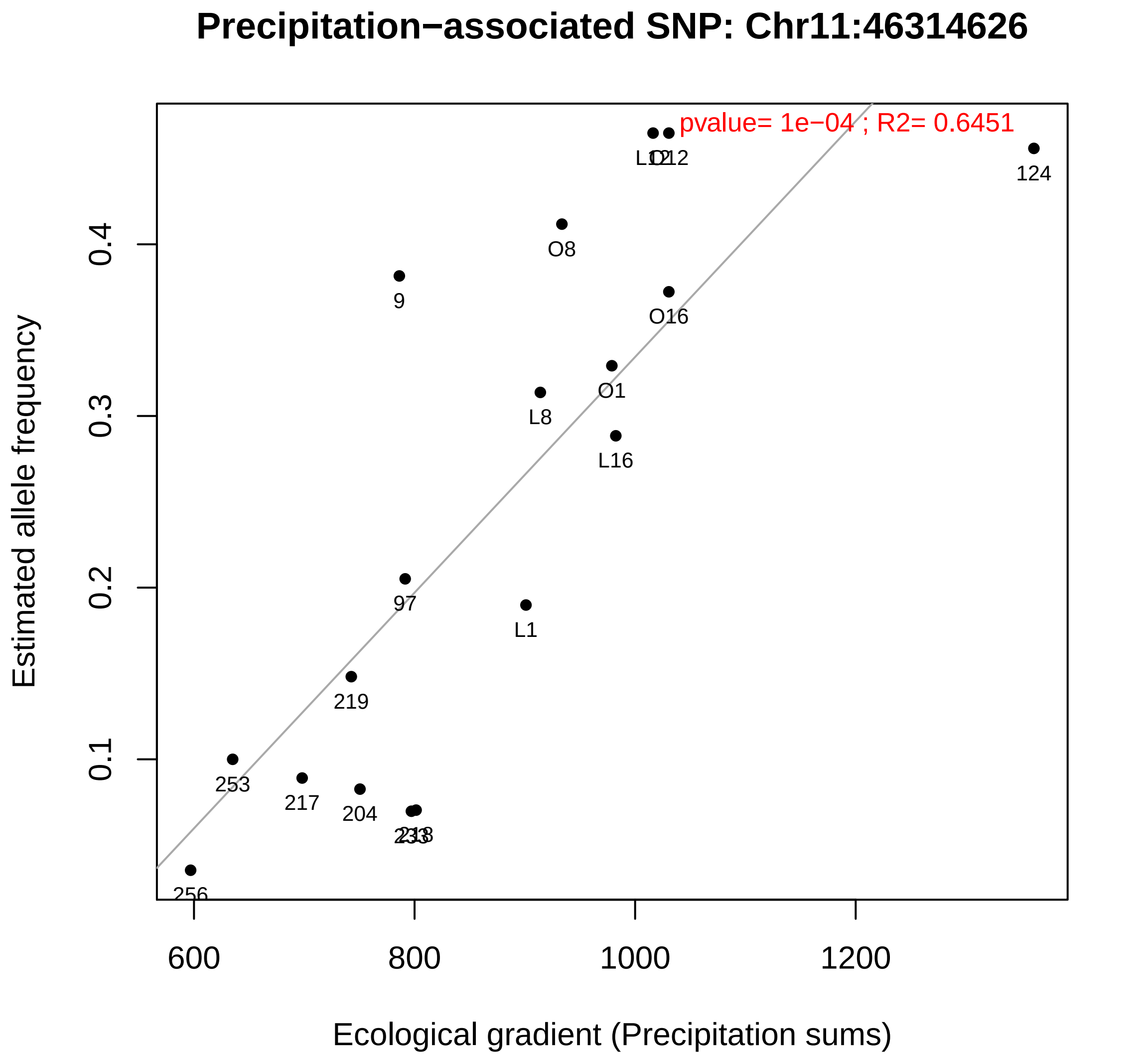

Precipitation−associated SNP: Chr11:46314626
pvalue= 1e−04 ; R2= 0.6451
Estimated allele frequency
Ecological gradient (Precipitation sums)
0.1
0.2
0.3
0.4
600
800
1000
1200
124
O8
9
O16
O1
L8
L16
97
L1
219
253
217
204
256

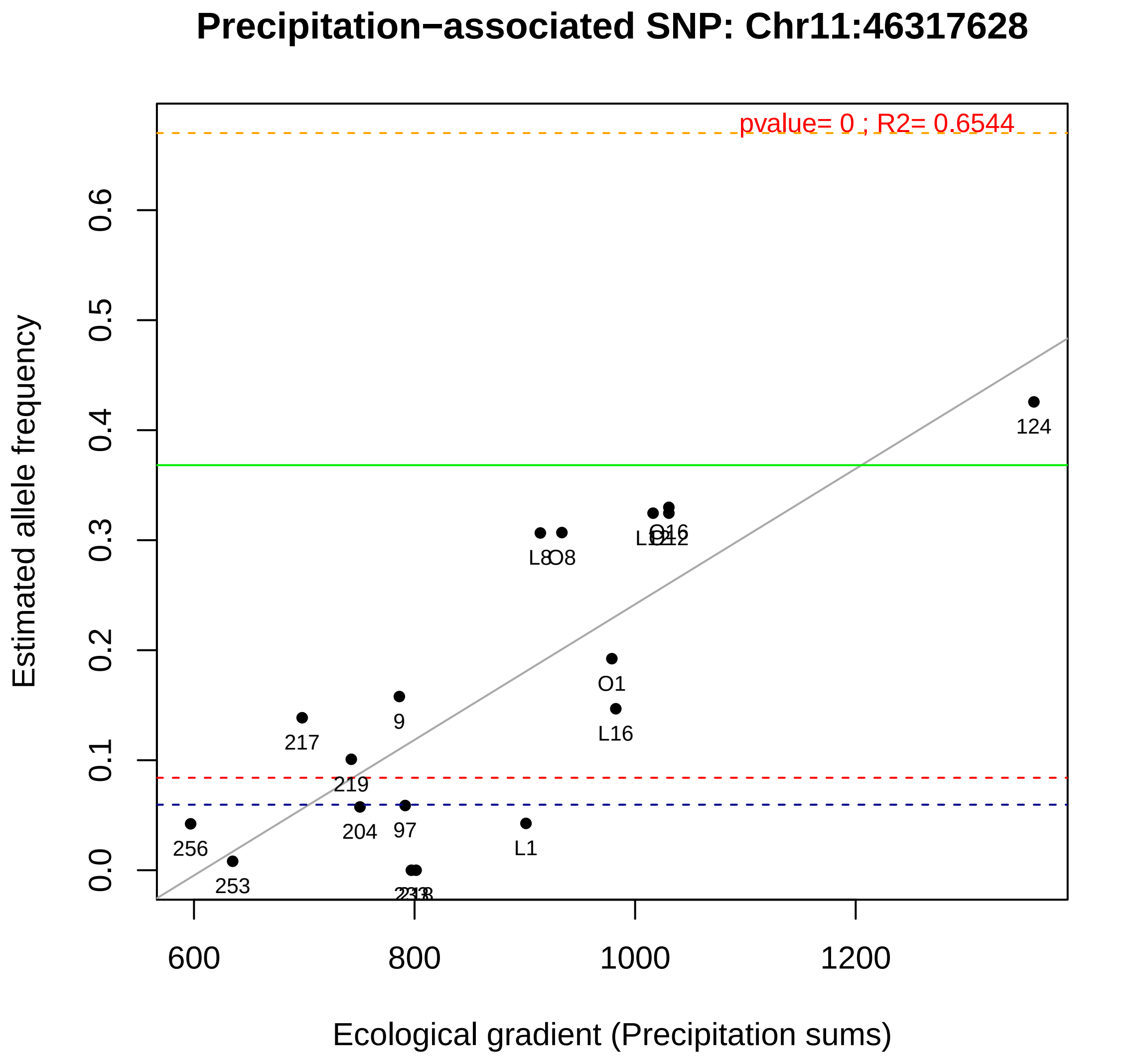
Precipitation−associated SNP: Chr11:46317628
pvalue= 0 ; R2= 0.6544
Estimated allele frequency
0.0
0.1
0.2
0.3
0.4
0.5
0.6
600
800
1000
1200
Ecological gradient (Precipitation sums)
124
L8
O8
O1
L16
9
217
219
204
97
256
253
L1

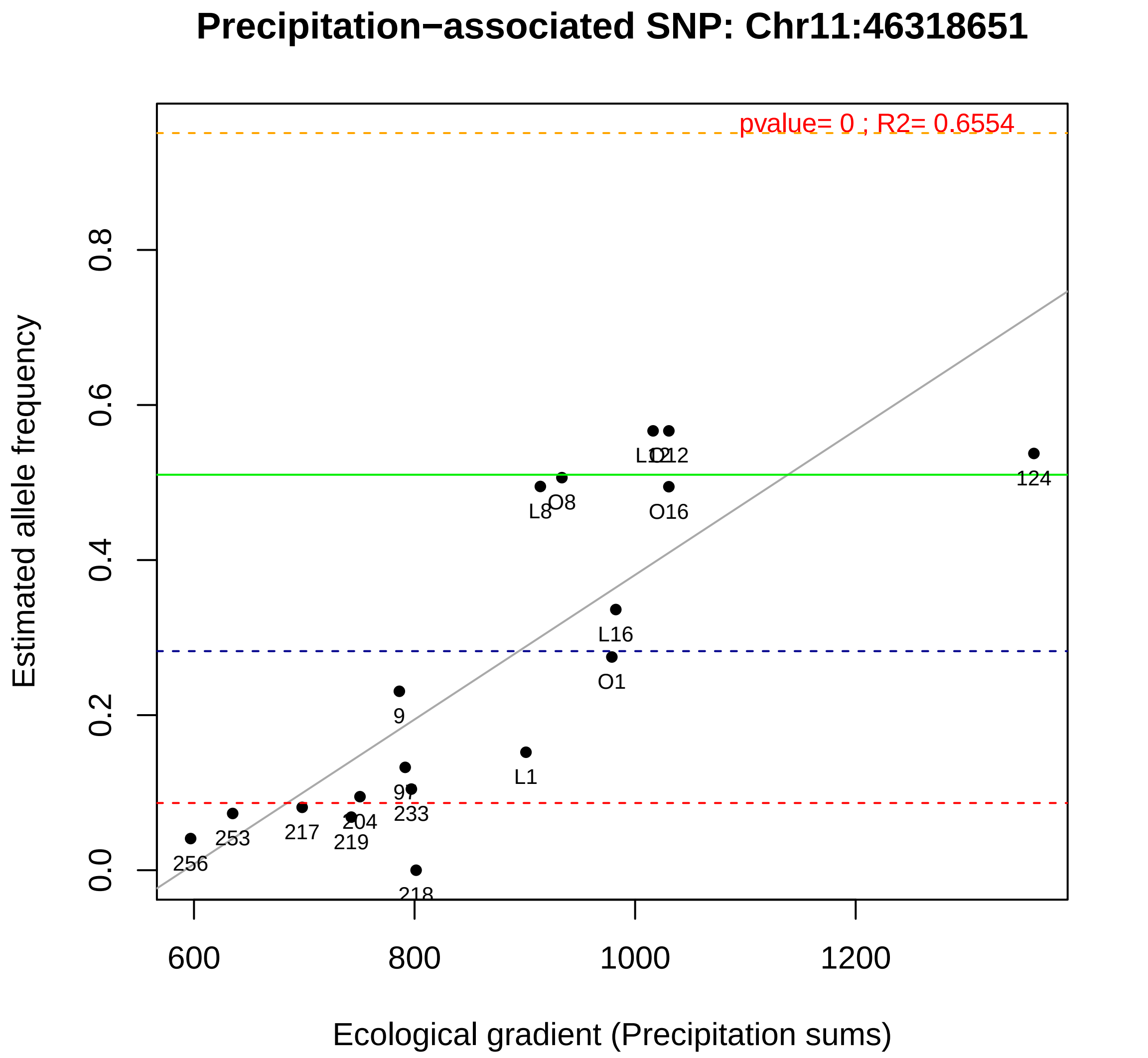

Precipitation−associated SNP: Chr11:46318651
pvalue= 0 ; R2= 0.6554
Estimated allele frequency
0.0
0.2
0.4
0.6
0.8
600
800
1000
1200
Ecological gradient (Precipitation sums)
L12
O12
124
L8
O8
O16
L16
O1
9
L1
97
233
204
219
217
253
256
218

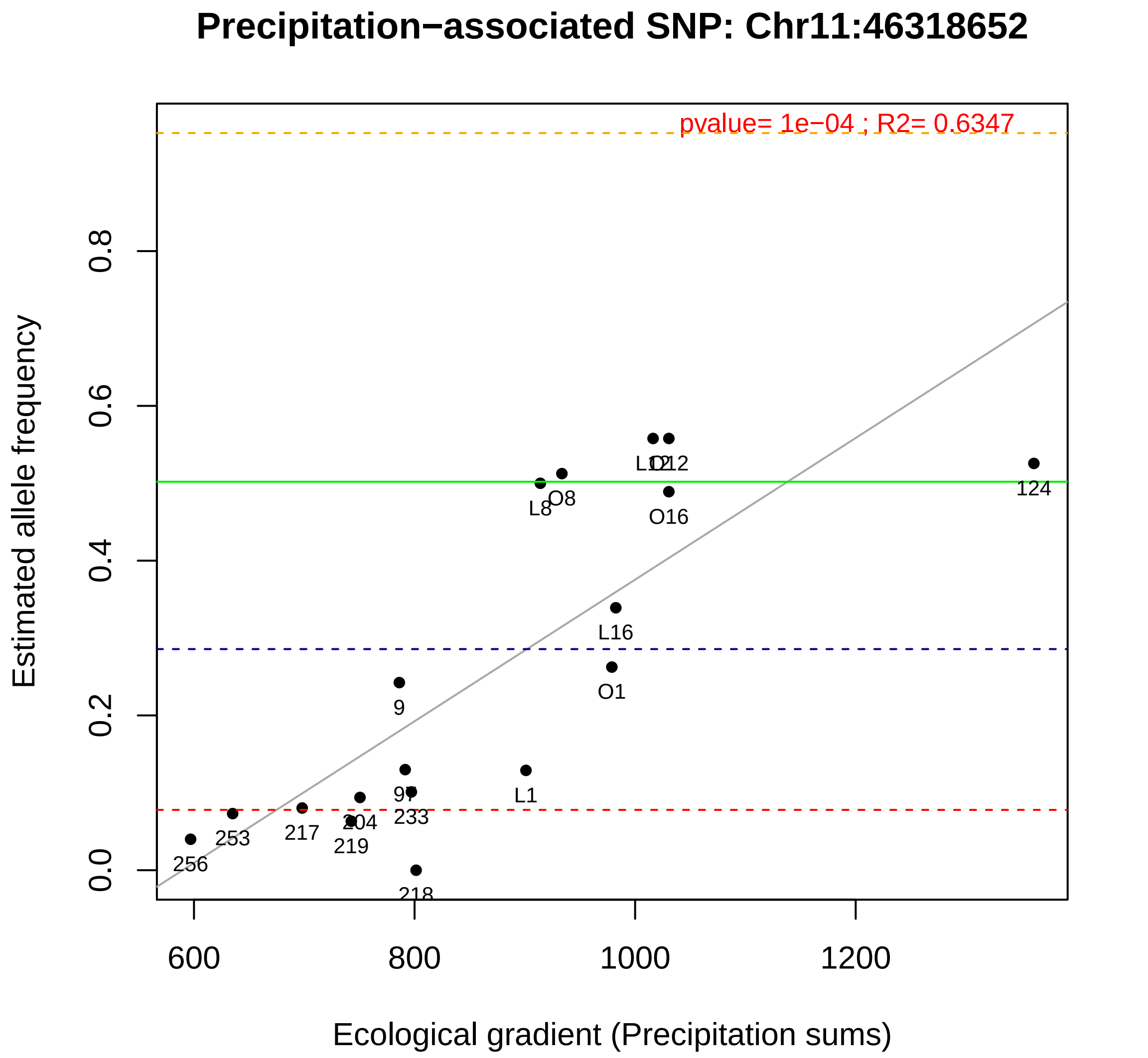
Precipitation−associated SNP: Chr11:46318652
pvalue= 1e−04 ; R2= 0.6347
Estimated allele frequency
0.0
0.2
0.4
0.6
0.8
Ecological gradient (Precipitation sums)
600
800
1000
1200
256
253
217
219
204
9
97
233
218
L1
L8
O8
L16
O1
O16
124

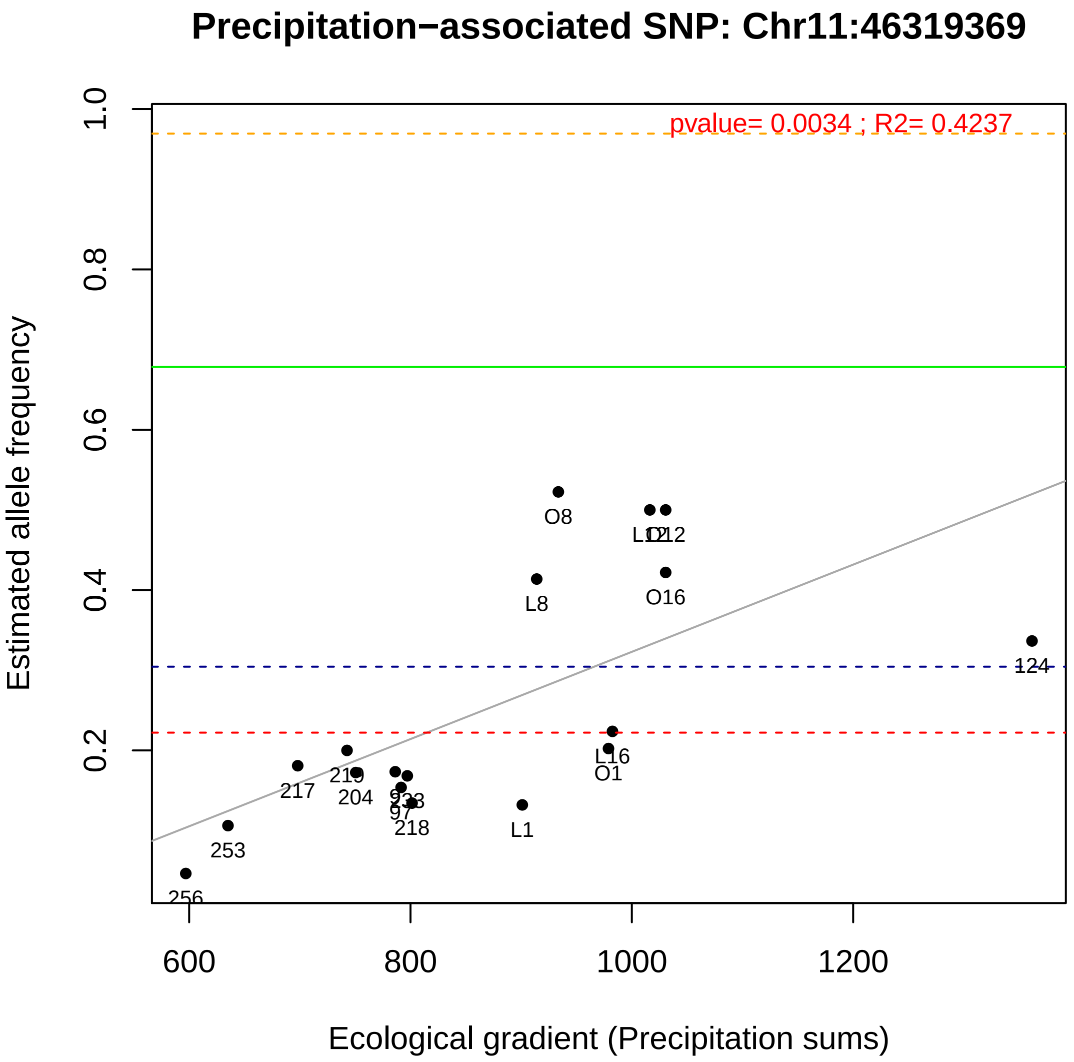

Precipitation−associated SNP: Chr11:46319369
pvalue= 0.0034 ; R2= 0.4237
Estimated allele frequency
Ecological gradient (Precipitation sums)
0.2
0.4
0.6
0.8
1.0
600
800
1000
1200
O8
O16
L8
124
L16
O1
217
204
218
L1
253
256

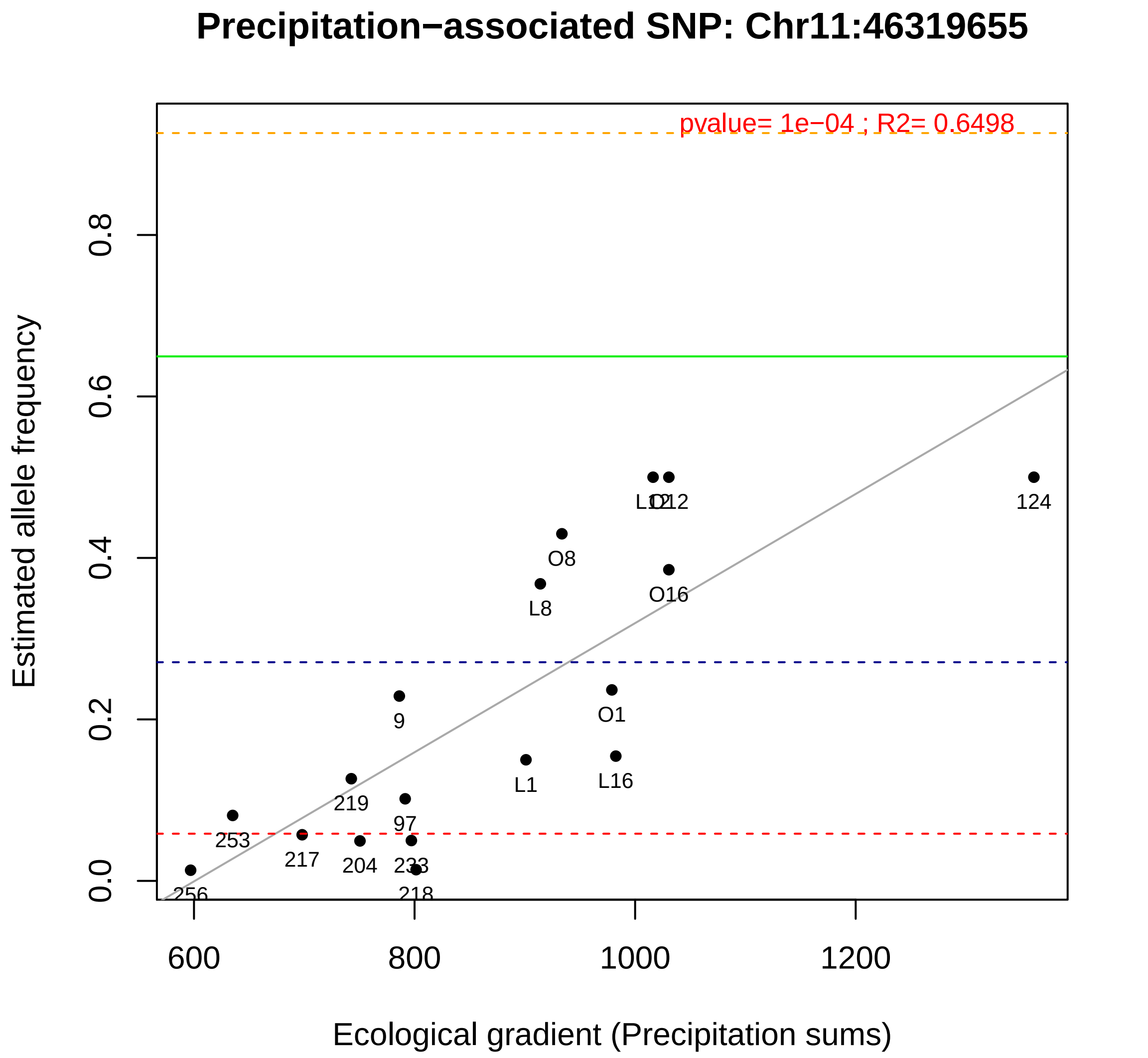
Precipitation−associated SNP: Chr11:46319655
pvalue= 1e−04 ; R2= 0.6498
Estimated allele frequency
0.0
0.2
0.4
0.6
0.8
600
800
1000
1200
Ecological gradient (Precipitation sums)
124
O8
O16
L8
9
O1
L1
L16
219
97
253
217
204
233
218
256

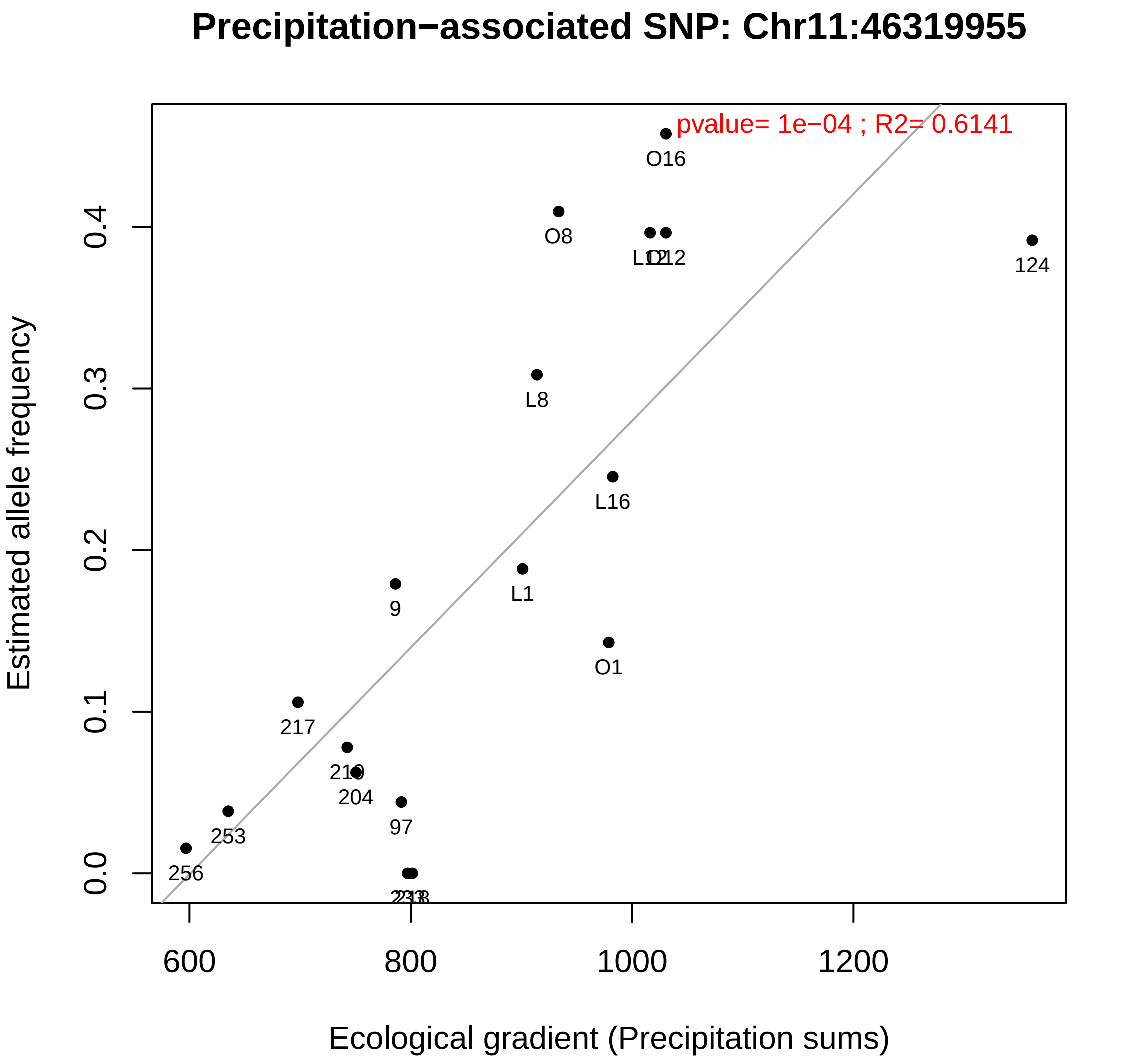
Precipitation−associated SNP: Chr11:46319955
pvalue= 1e−04 ; R2= 0.6141
O16
O8
L12
O12
124
L8
L16
L1
9
O1
217
210
204
97
253
256
Estimated allele frequency
0.0
0.1
0.2
0.3
0.4
600
800
1000
1200
Ecological gradient (Precipitation sums)

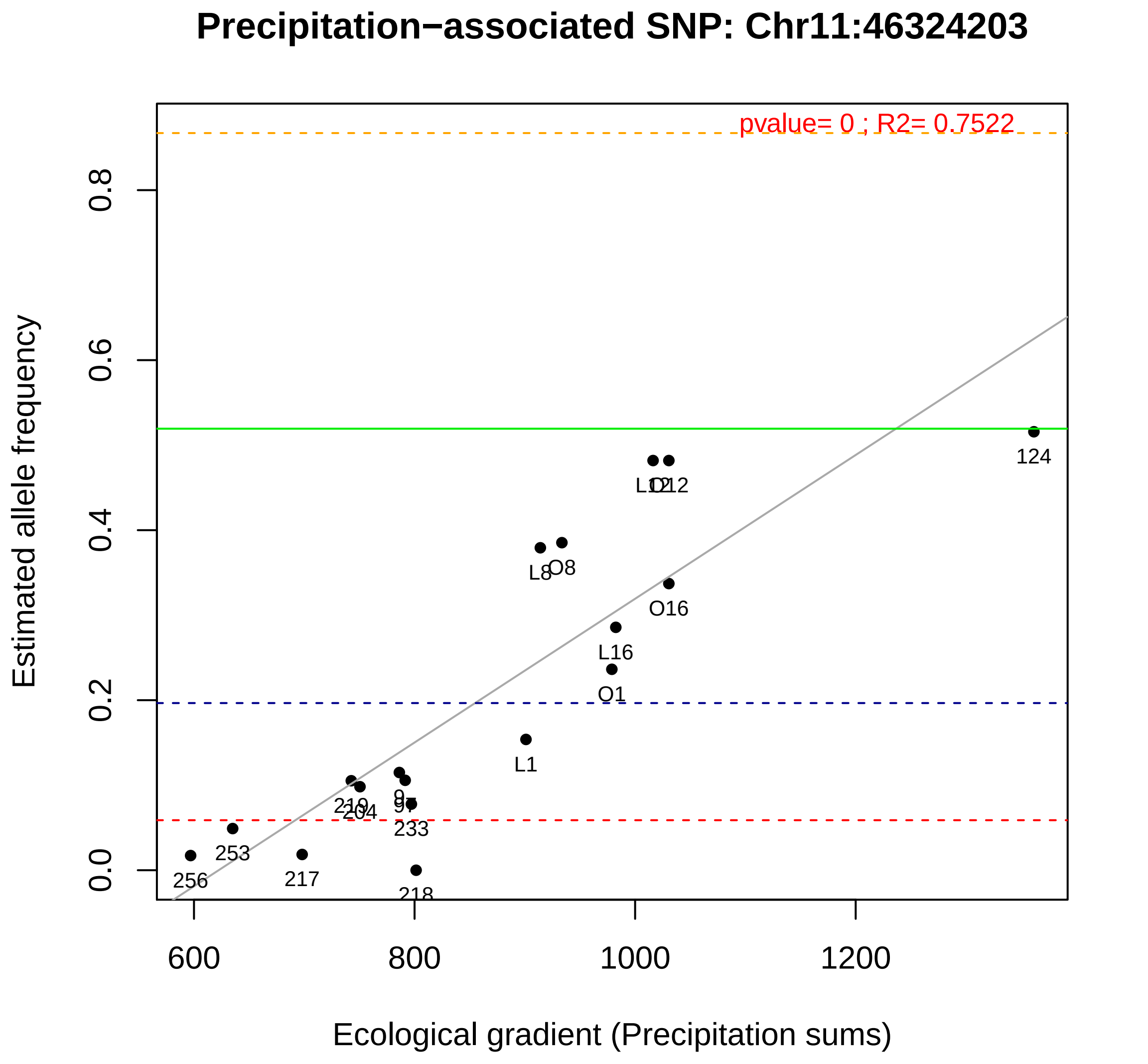
Precipitation−associated SNP: Chr11:46324203
pvalue= 0 ; R2= 0.7522
Estimated allele frequency
0.0
0.2
0.4
0.6
0.8
600
800
1000
1200
Ecological gradient (Precipitation sums)
124
L12
O12
L8
O8
O16
L16
O1
L1
219
204
9
97
233
253
256
217
218

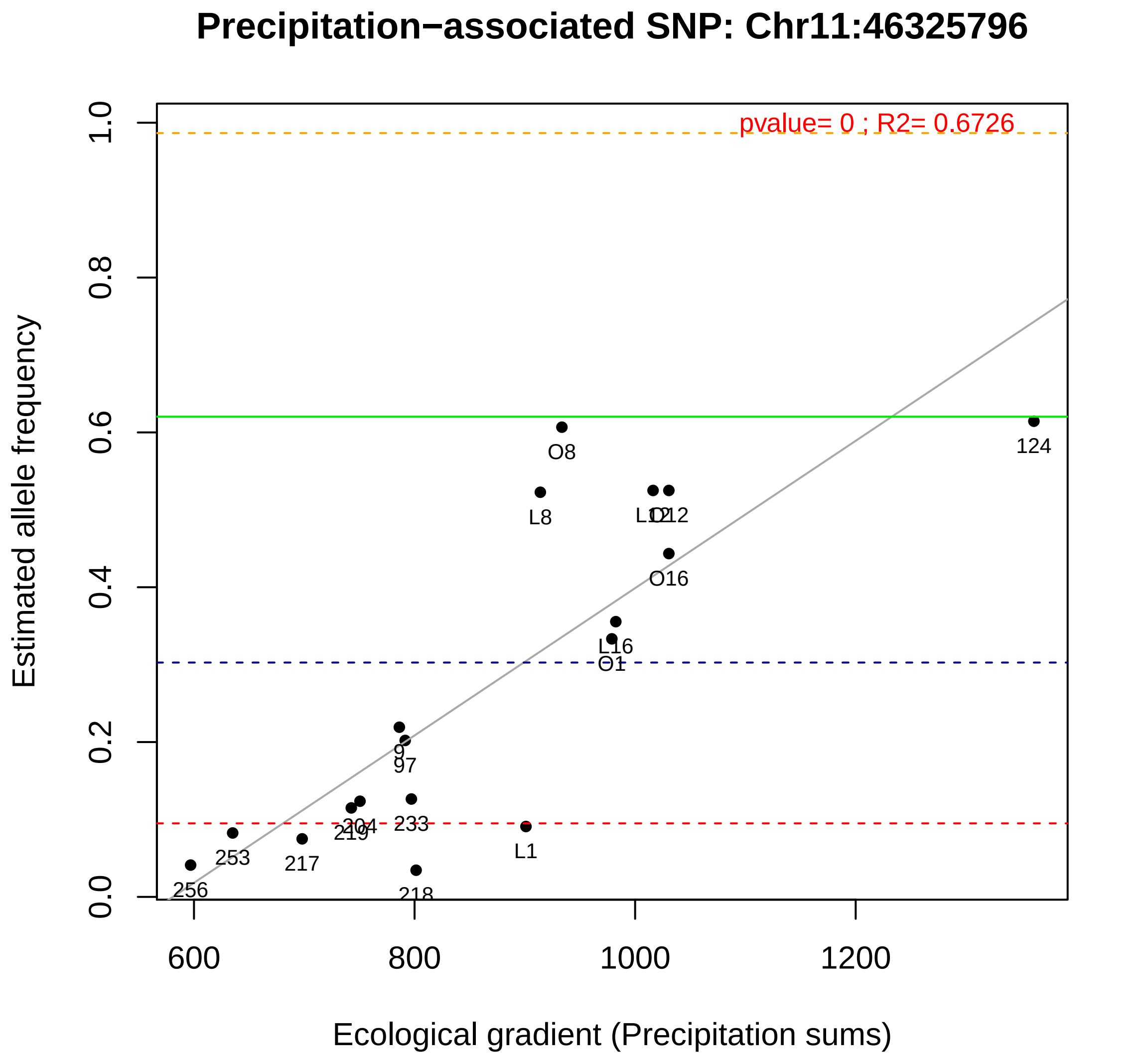

Precipitation−associated SNP: Chr11:46325796
pvalue= 0 ; R2= 0.6726
Estimated allele frequency
Ecological gradient (Precipitation sums)
0.0
0.2
0.4
0.6
0.8
1.0
600
800
1000
1200
O8
124
L8
L12
O12
O16
L16
O1
9
97
204
219
233
L1
253
217
256
218

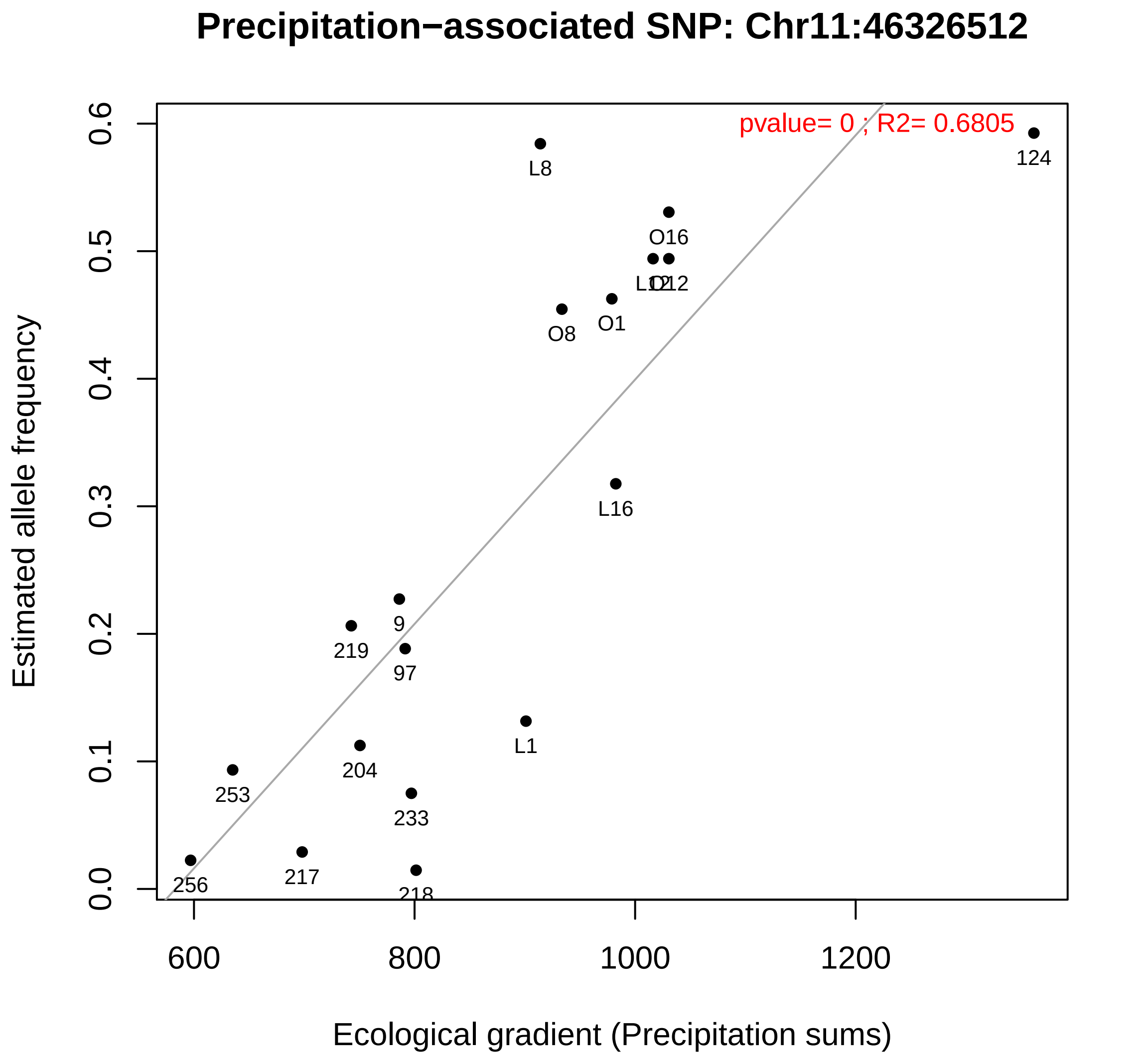

Precipitation−associated SNP: Chr11:46326512
pvalue= 0 ; R2= 0.6805
Estimated allele frequency
Ecological gradient (Precipitation sums)
0.0
0.1
0.2
0.3
0.4
0.5
0.6
600
800
1000
1200
L8
124
O16
L12
O12
O1
O8
L16
9
219
97
L1
204
253
233
256
217
218

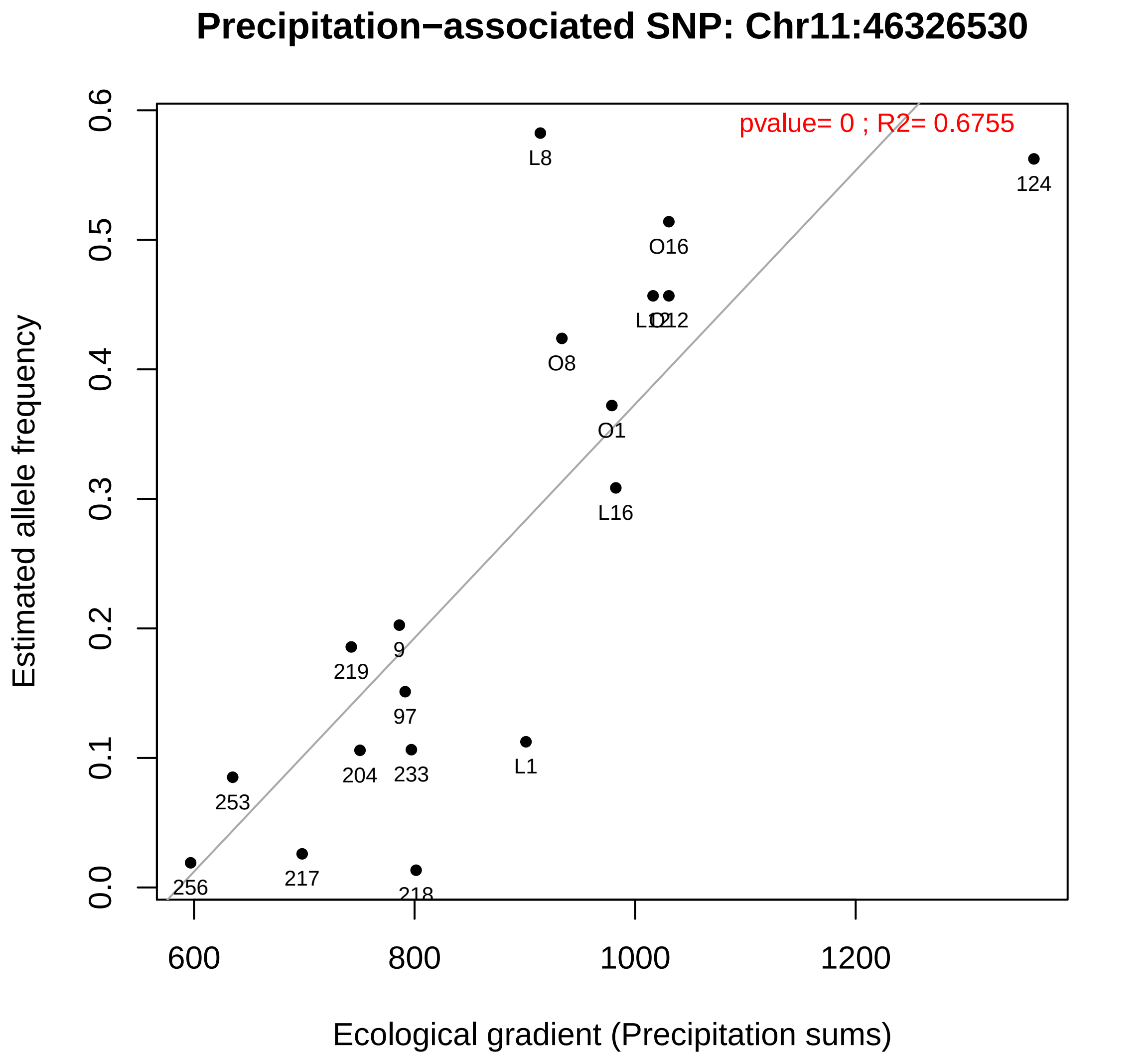

Precipitation−associated SNP: Chr11:46326530
pvalue= 0 ; R2= 0.6755
Estimated allele frequency
Ecological gradient (Precipitation sums)
0.0
0.1
0.2
0.3
0.4
0.5
0.6
600
800
1000
1200
L8
124
O16
O12
O8
O1
L16
9
219
97
204
233
L1
253
256
217
218

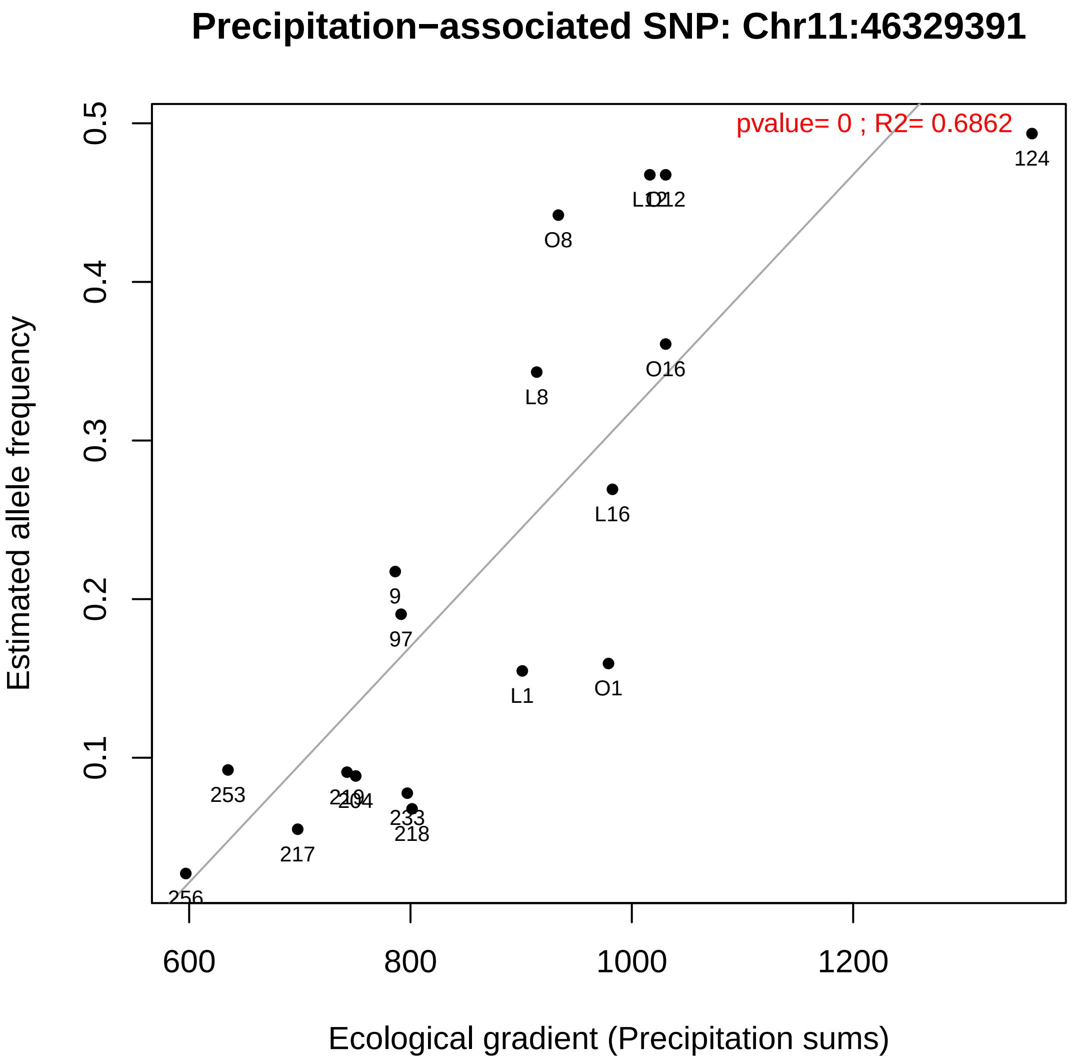

Precipitation−associated SNP: Chr11:46329391
pvalue= 0 ; R2= 0.6862
Estimated allele frequency
0.1
0.2
0.3
0.4
0.5
Ecological gradient (Precipitation sums)
600
800
1000
1200
124
O8
O16
L8
L16
9
97
L1
O1
253
217
256
233
218

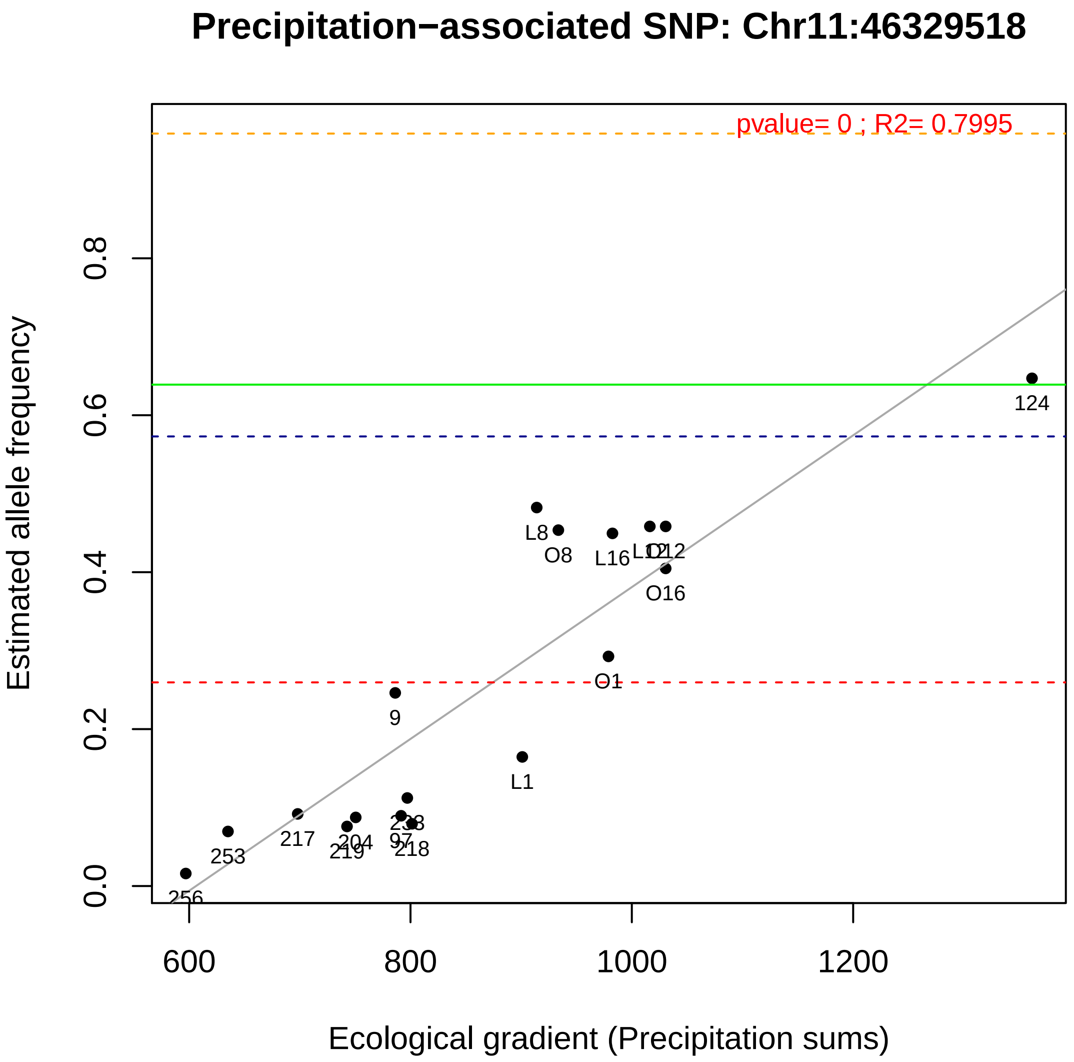
Precipitation−associated SNP: Chr11:46329518
pvalue= 0 ; R2= 0.7995
Estimated allele frequency
0.0
0.2
0.4
0.6
0.8
Ecological gradient (Precipitation sums)
600
800
1000
1200
124
L8
O8
L16
O16
O1
9
L1
217
253
204
219
97
218
256

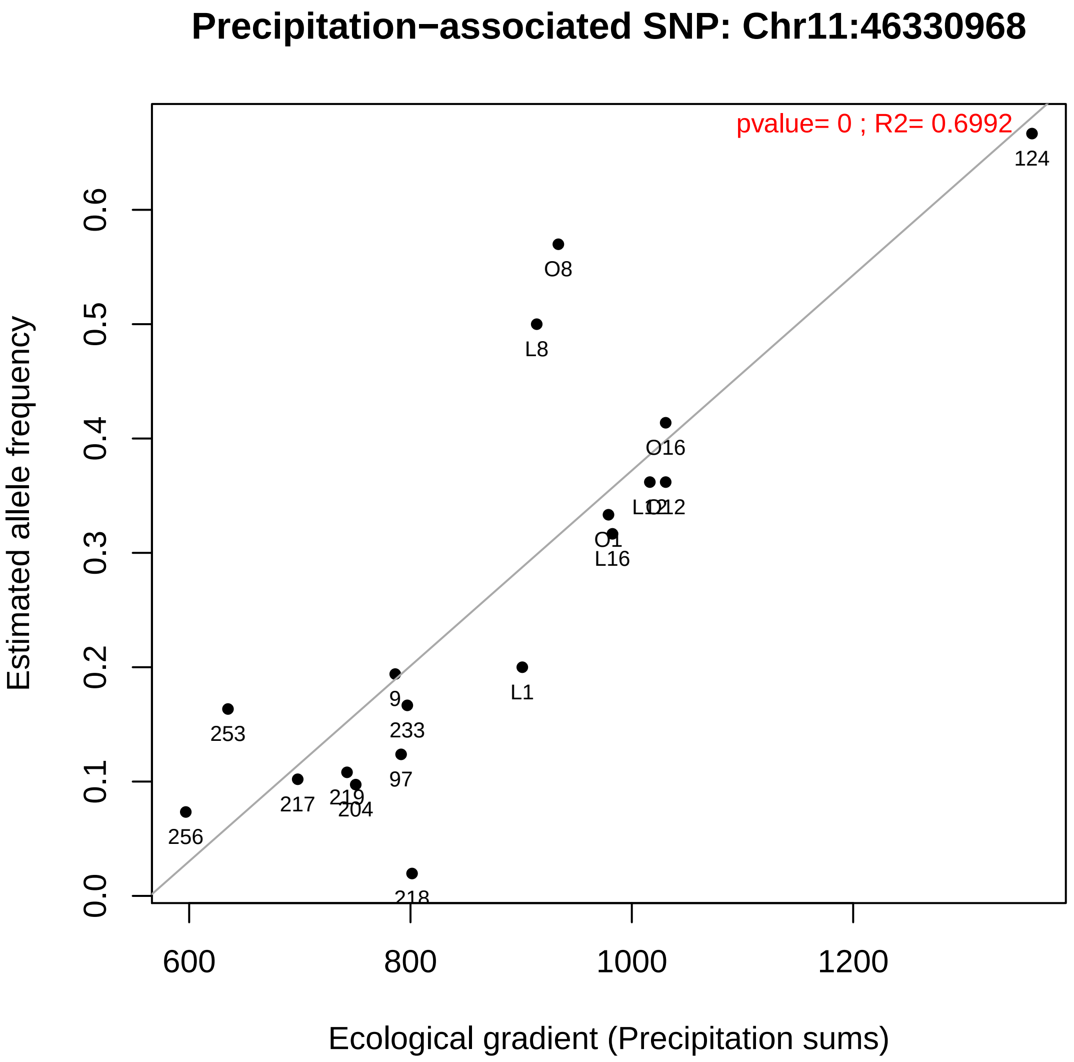
Precipitation−associated SNP: Chr11:46330968
pvalue= 0 ; R2= 0.6992
Estimated allele frequency
Ecological gradient (Precipitation sums)
0.0
0.1
0.2
0.3
0.4
0.5
0.6
600
800
1000
1200
124
O8
L8
O16
L12
O12
O1
L16
L1
9
233
253
97
217
219
204
256
218

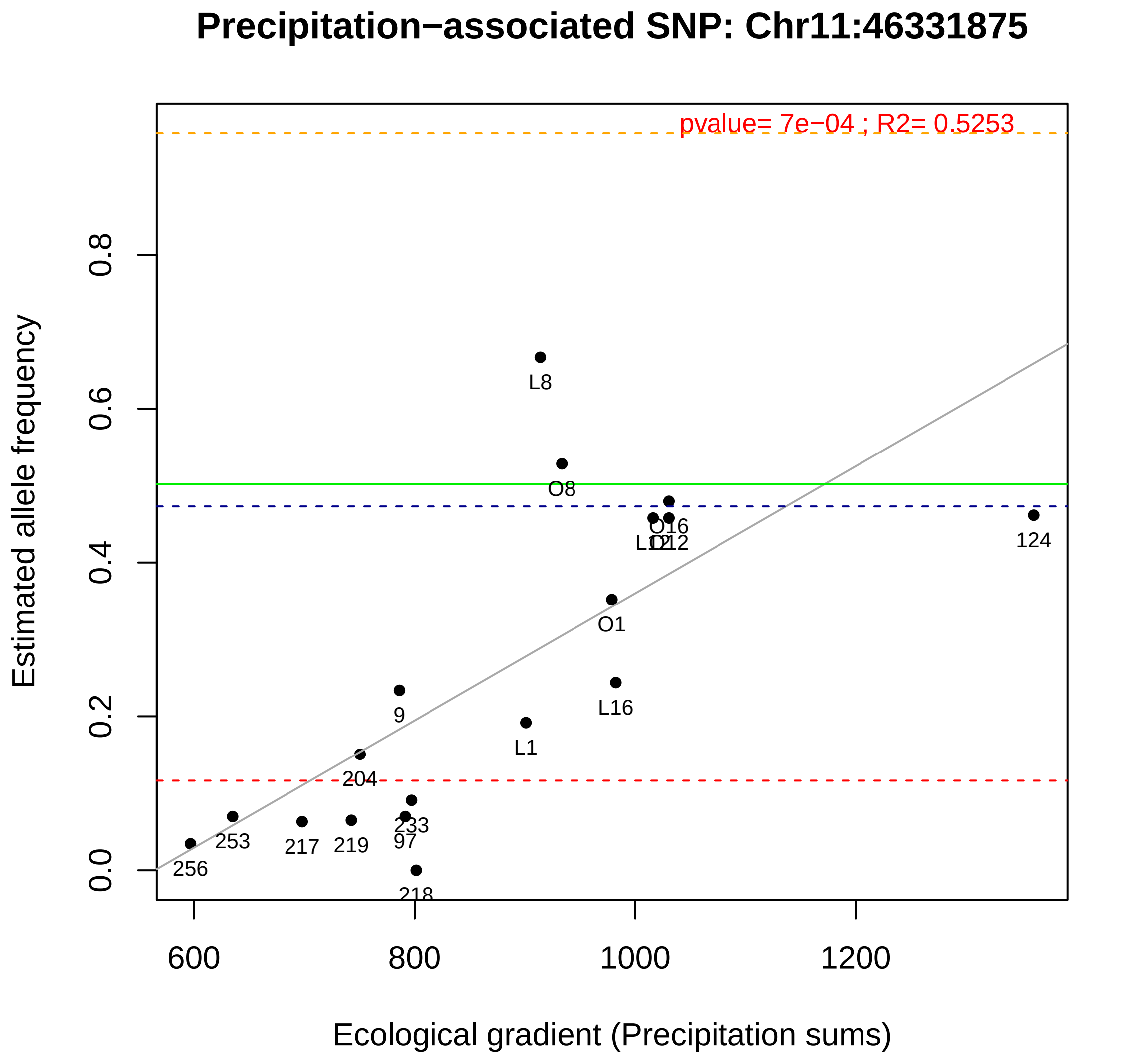
Precipitation−associated SNP: Chr11:46331875
pvalue= 7e−04 ; R2= 0.5253
Estimated allele frequency
0.0
0.2
0.4
0.6
0.8
600
800
1000
1200
Ecological gradient (Precipitation sums)
L8
O8
O16
L12
O12
124
O1
L16
9
L1
204
253
217
219
233
97
256
218

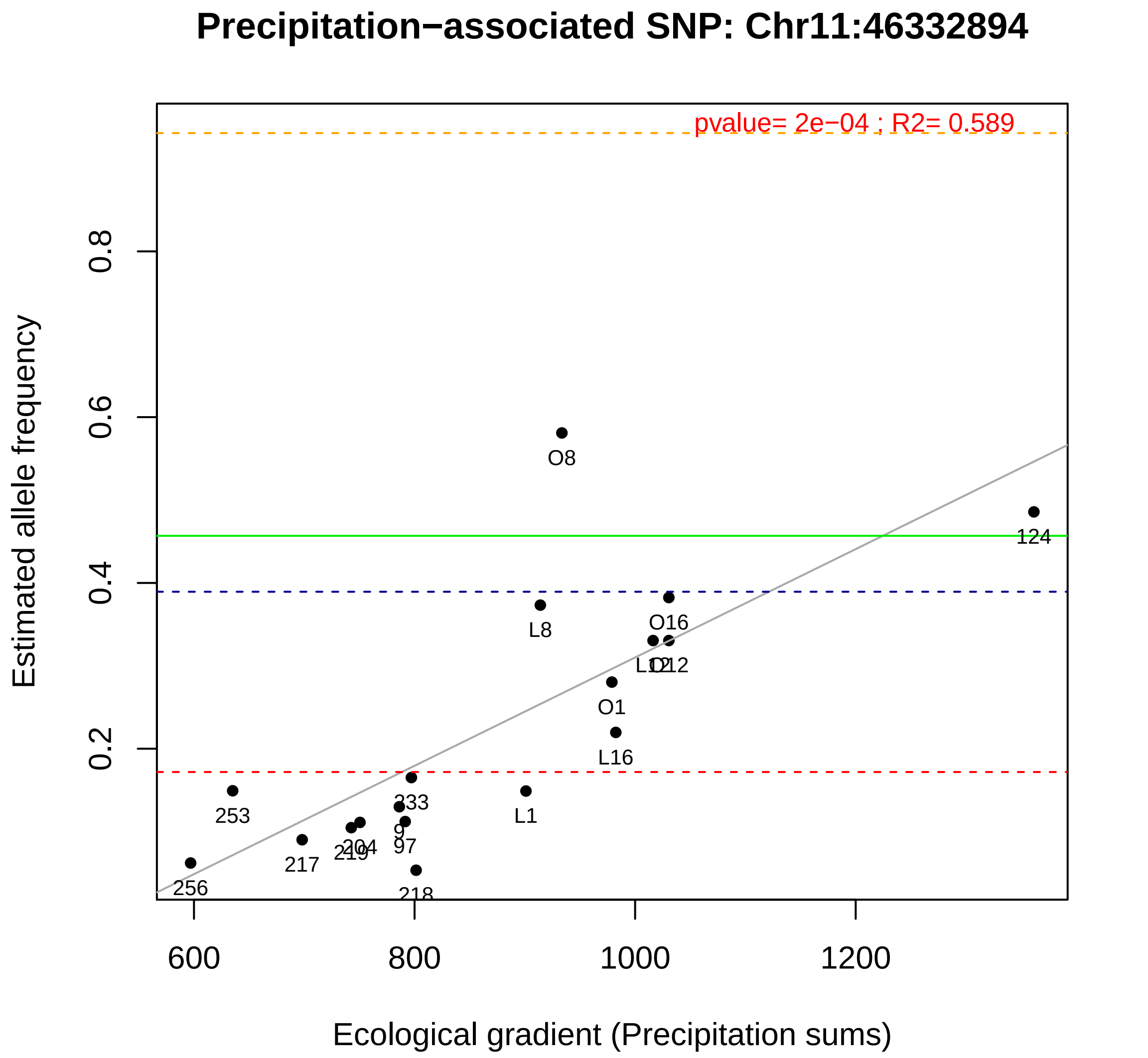

Precipitation−associated SNP: Chr11:46332894
pvalue= 2e−04 ; R2= 0.589
Estimated allele frequency
0.2
0.4
0.6
0.8
600
800
1000
1200
Ecological gradient (Precipitation sums)
O8
124
L8
O16
O12
O1
L16
253
233
L1
9
97
217
204
219
256
218

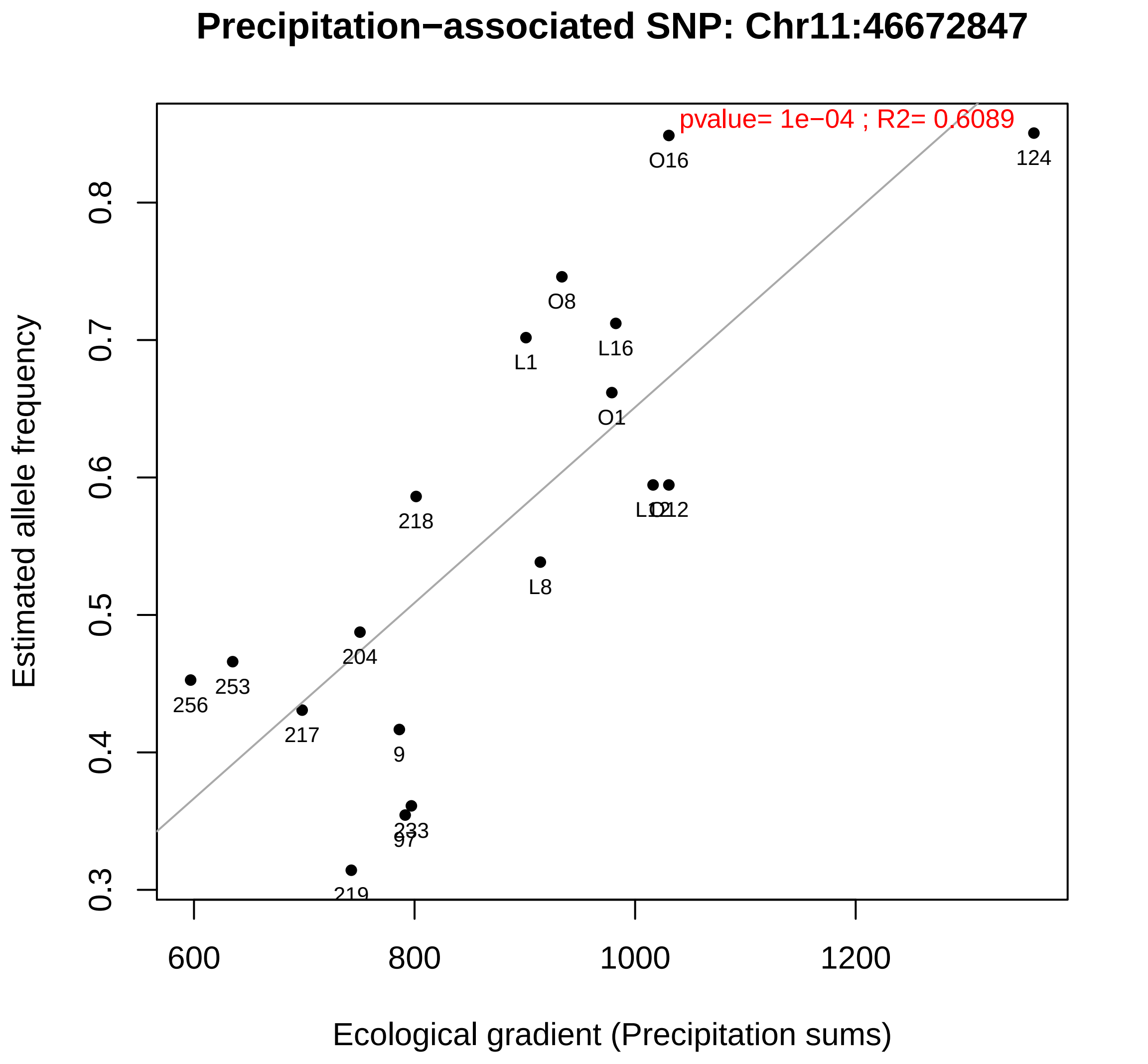

Precipitation−associated SNP: Chr11:46672847
pvalue= 1e−04 ; R2= 0.6089
O16
124
O8
L16
L1
O1
218
L12
O12
L8
204
253
256
217
9
233
97
219
0.3
0.4
0.5
0.6
0.7
0.8
600
800
1000
1200
Estimated allele frequency
Ecological gradient (Precipitation sums)

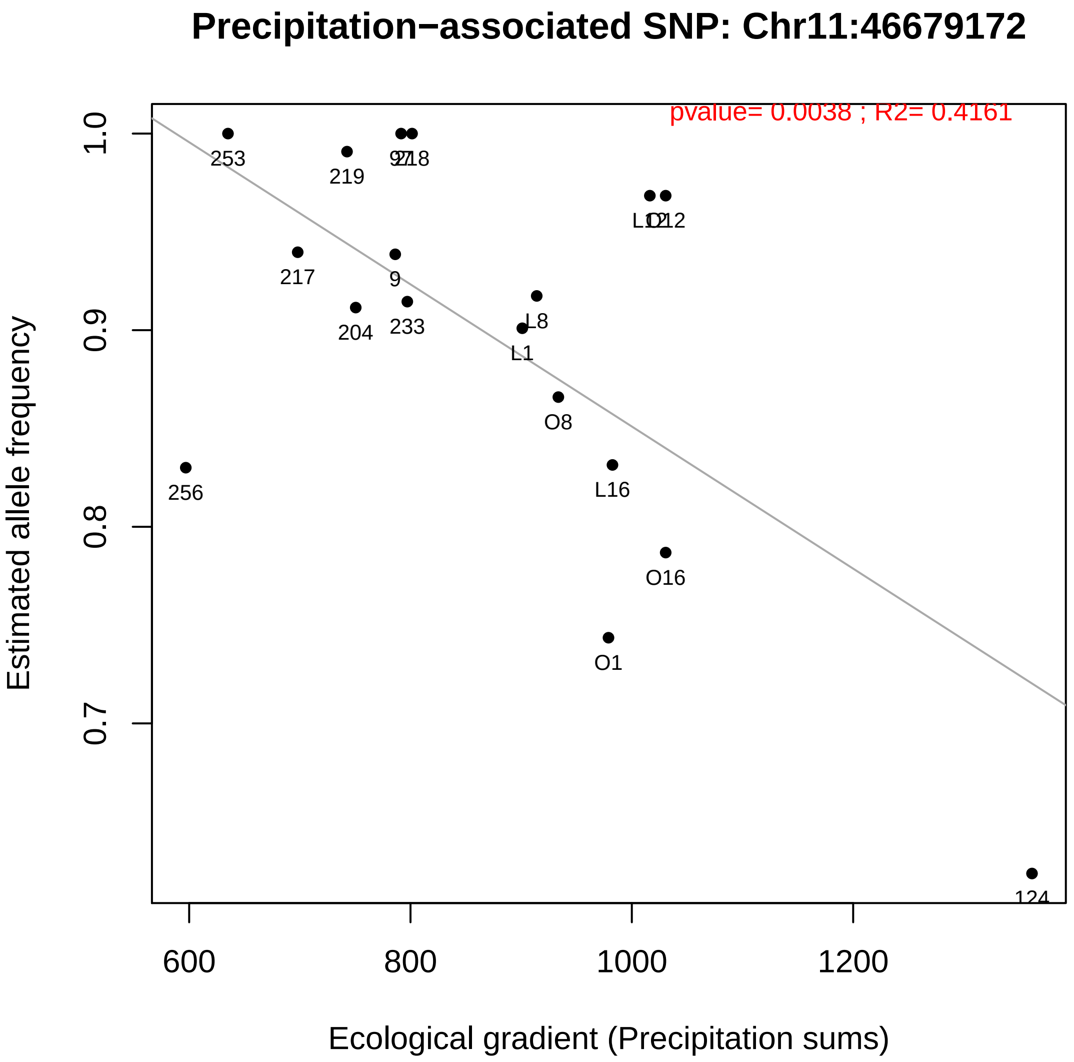

Precipitation−associated SNP: Chr11:46679172
pvalue= 0.0038 ; R2= 0.4161
Estimated allele frequency
Ecological gradient (Precipitation sums)
0.7
0.8
0.9
1.0
600
800
1000
1200
253
219
97
218
L12
O12
217
9
204
233
L8
L1
O8
256
L16
O16
O1
124

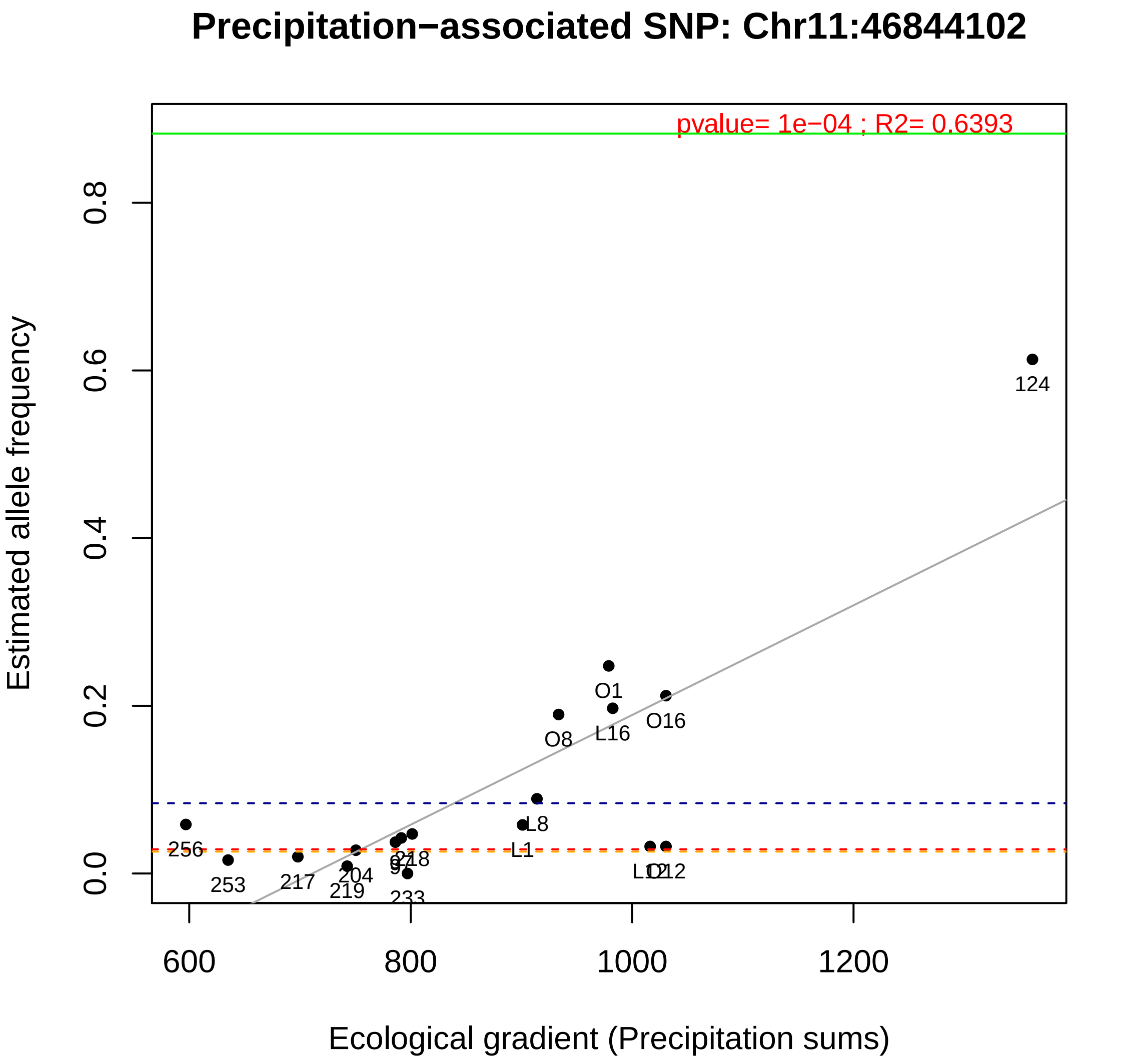
Precipitation−associated SNP: Chr11:46844102
pvalue= 1e−04 ; R2= 0.6393
Estimated allele frequency
0.0
0.2
0.4
0.6
0.8
Ecological gradient (Precipitation sums)
600
800
1000
1200
124
O1
O8
L16
O16
L8
L1
256
253
217
204
219
218
233
O12

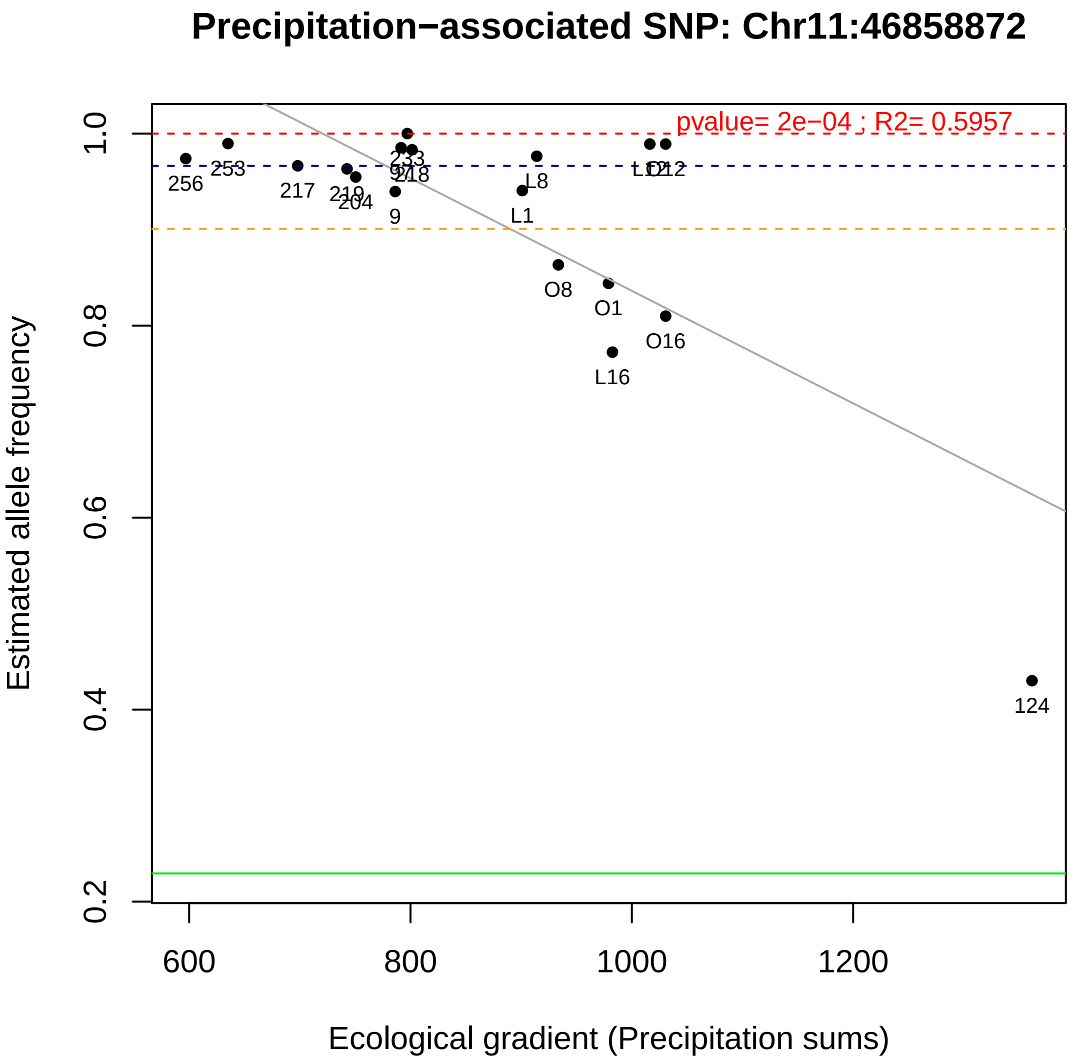

Precipitation−associated SNP: Chr11:46858872
pvalue= 2e−04 ; R2= 0.5957
Estimated allele frequency
Ecological gradient (Precipitation sums)
0.2
0.4
0.6
0.8
1.0
600
800
1000
1200
256
253
217
219
204
233
97
218
9
L8
L1
L12
O12
O8
O1
O16
L16
124

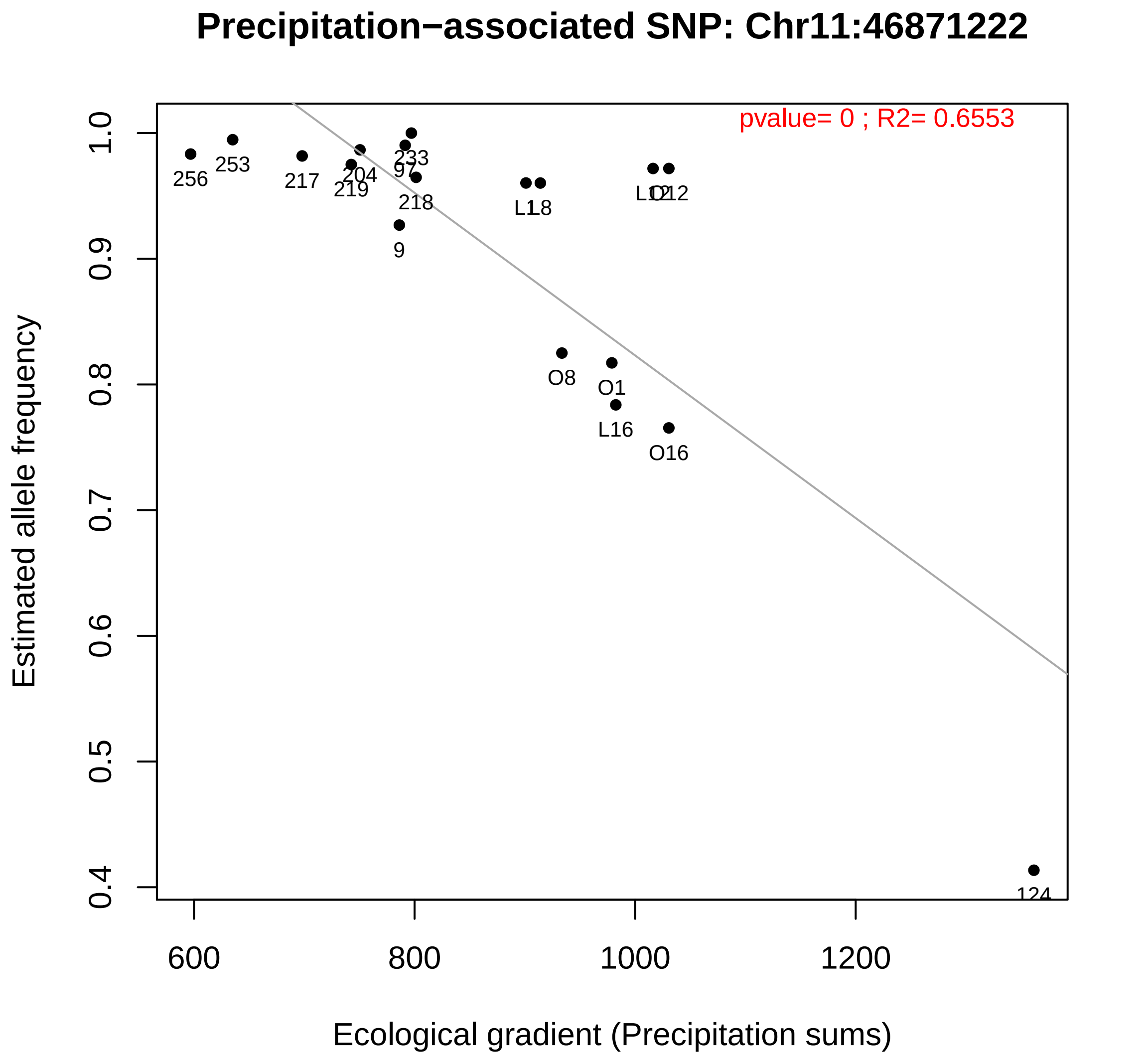
Precipitation−associated SNP: Chr11:46871222
pvalue= 0 ; R2= 0.6553
Estimated allele frequency
Ecological gradient (Precipitation sums)
0.4
0.5
0.6
0.7
0.8
0.9
1.0
600
800
1000
1200
256
253
217
204
219
233
97
218
9
L1
L8
L12
O12
O8
O1
L16
O16
124

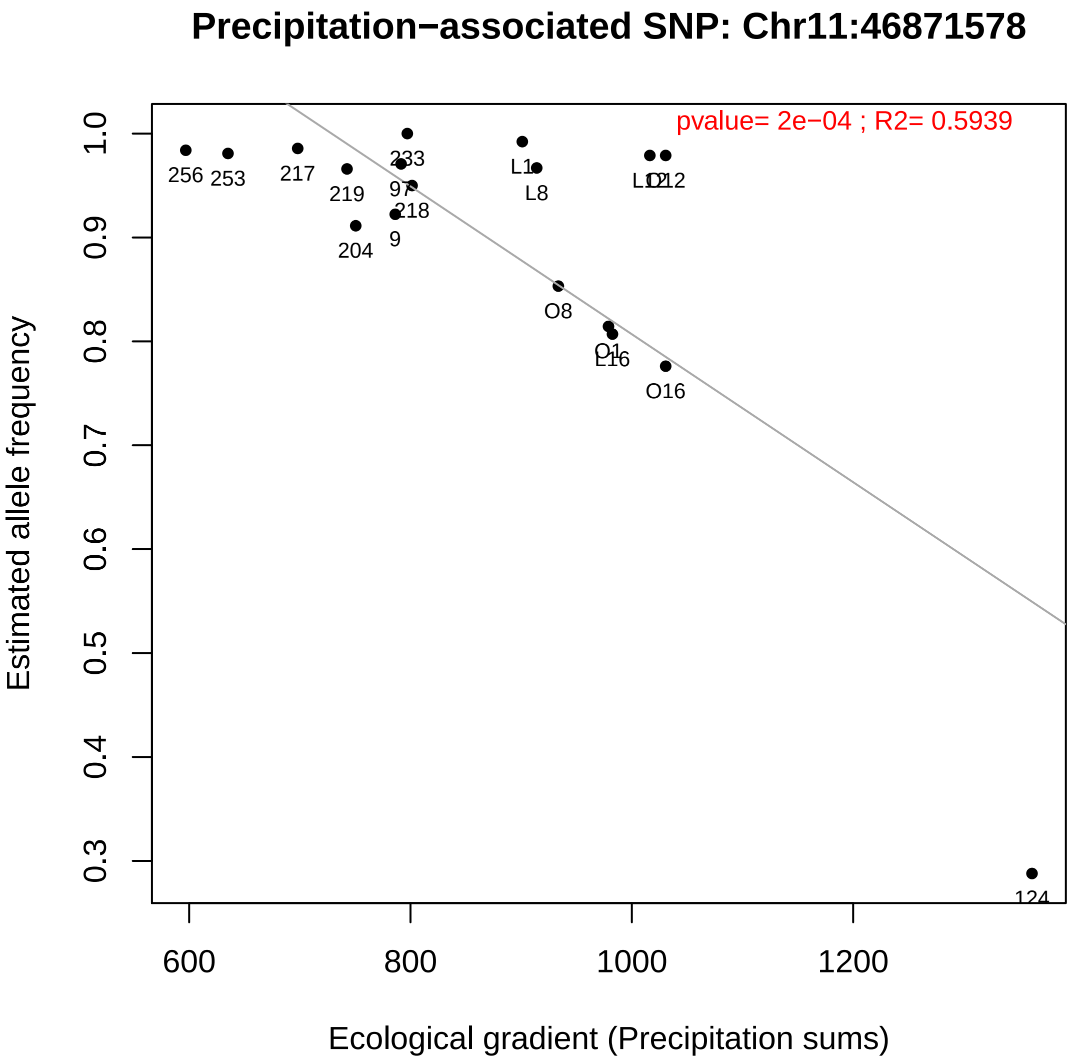
Precipitation−associated SNP: Chr11:46871578
pvalue= 2e−04 ; R2= 0.5939
Estimated allele frequency
Ecological gradient (Precipitation sums)
1.0
0.9
0.8
0.7
0.6
0.5
0.4
0.3
600
800
1000
1200
256
253
217
219
233
97
218
9
204
L1
L8
L12
O12
O8
O1
L16
O16
124

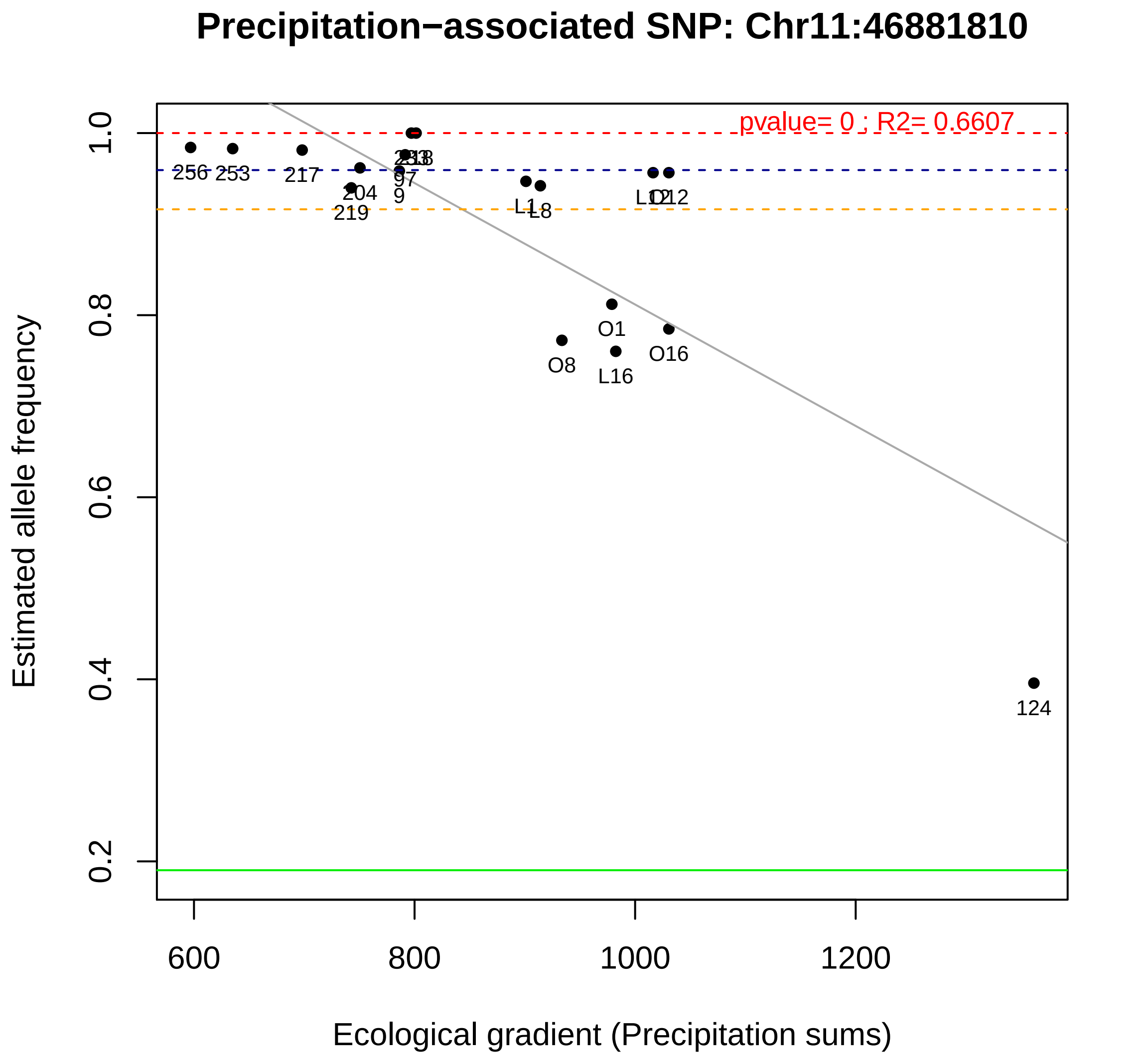

Precipitation−associated SNP: Chr11:46881810
pvalue= 0 ; R2= 0.6607
Estimated allele frequency
Ecological gradient (Precipitation sums)
0.2
0.4
0.6
0.8
1.0
600
800
1000
1200
256
253
217
204
219
97
9
L1
L8
O1
O8
L16
O16
124

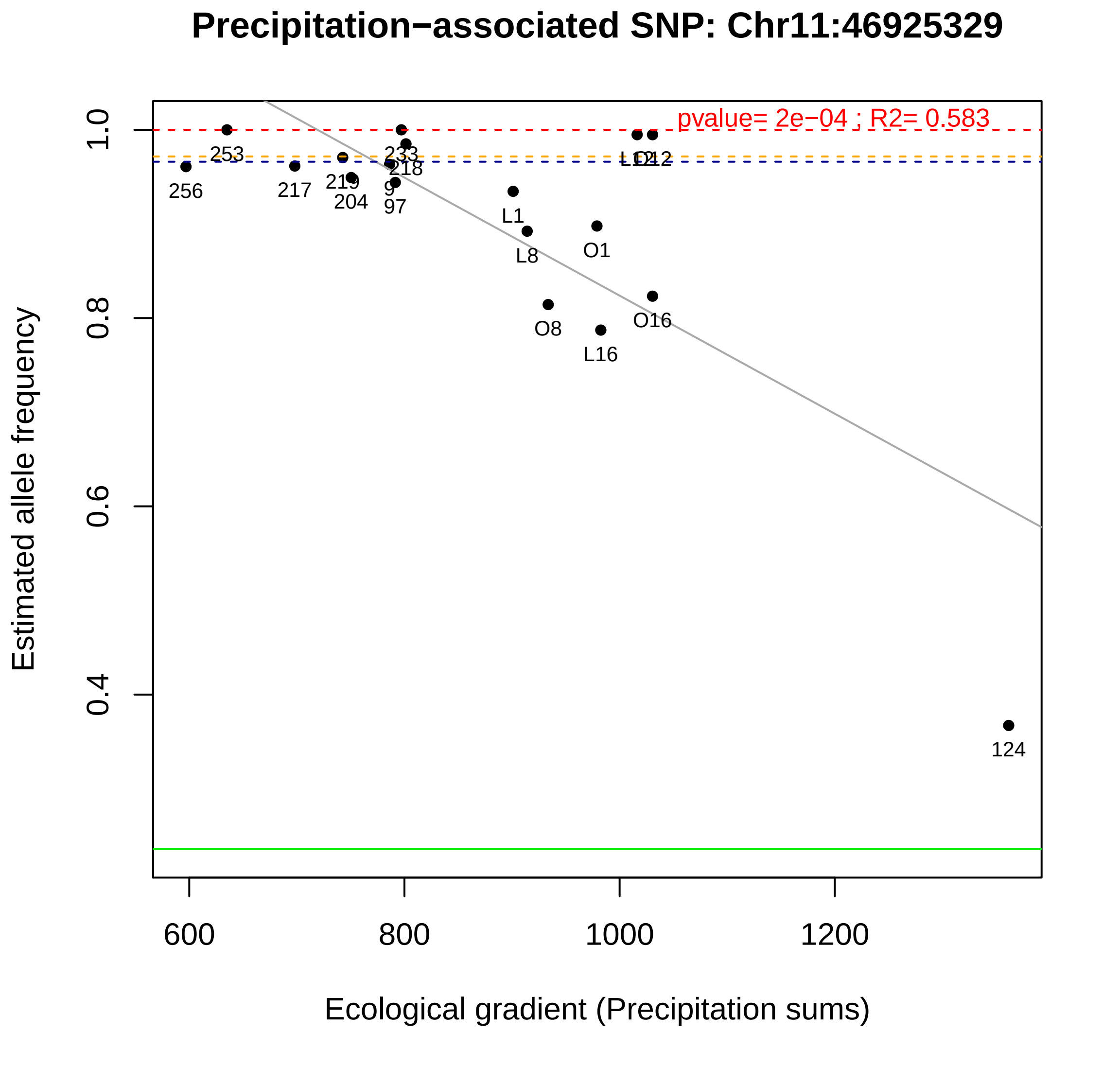

Precipitation−associated SNP: Chr11:46925329
pvalue= 2e−04 ; R2= 0.583
Estimated allele frequency
Ecological gradient (Precipitation sums)
1.0
0.8
0.6
0.4
600
800
1000
1200
256
253
217
219
204
233
218
9
97
L1
L8
O1
L12
O12
O8
L16
O16
124

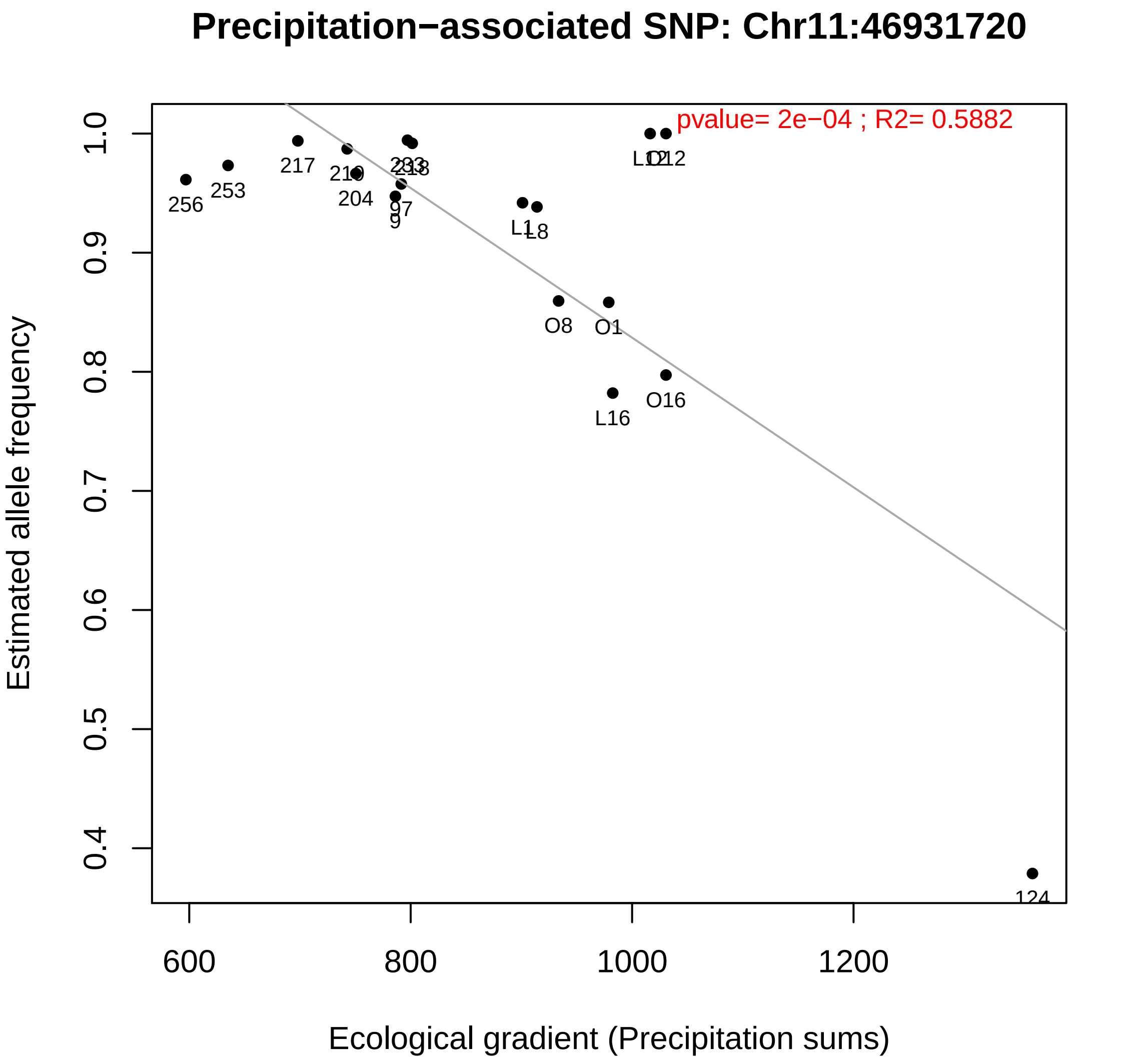

Precipitation−associated SNP: Chr11:46931720
pvalue= 2e−04 ; R2= 0.5882
Estimated allele frequency
Ecological gradient (Precipitation sums)
0.4
0.5
0.6
0.7
0.8
0.9
1.0
600
800
1000
1200
256
253
217
210
204
233
218
97
9
L1
L8
O8
O1
L12
O12
L16
O16
124

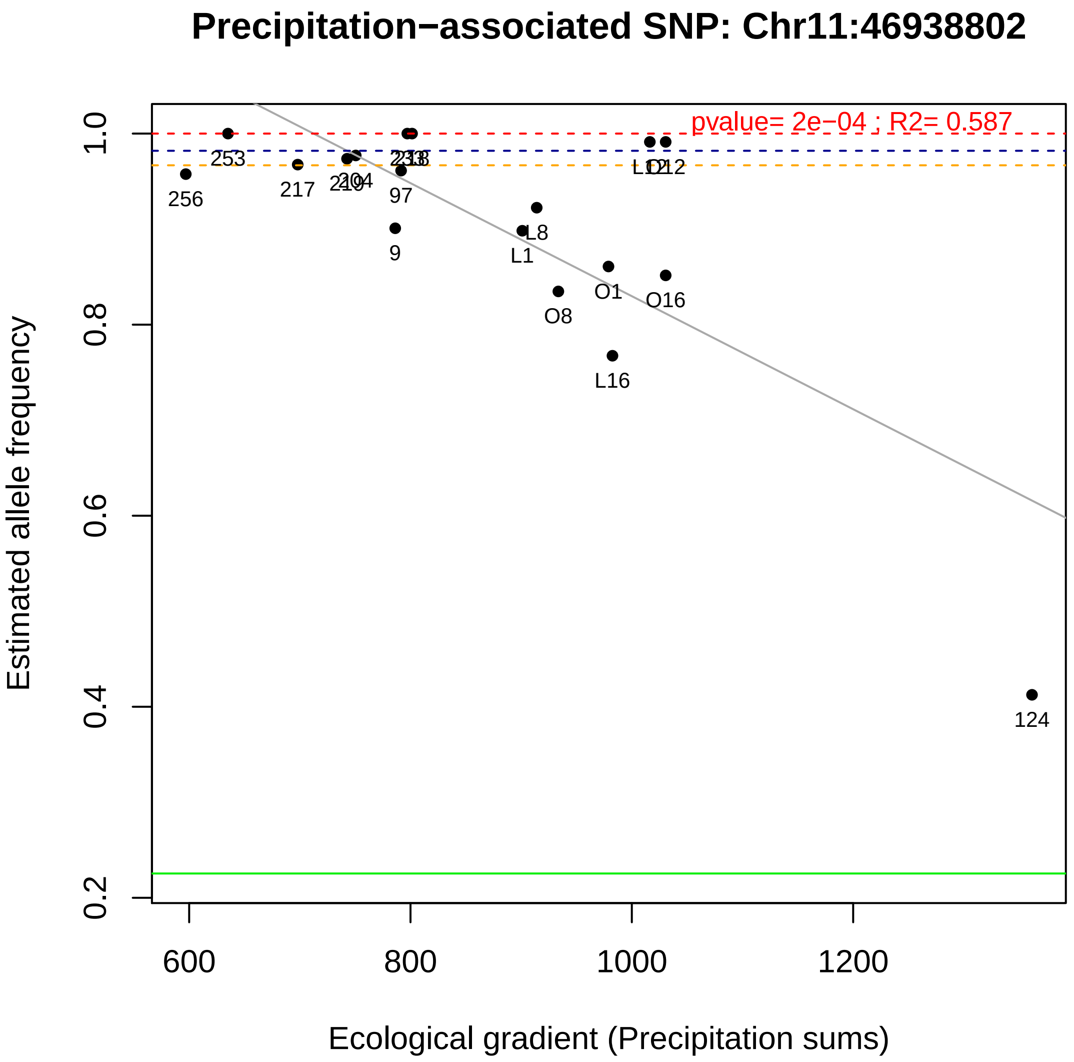
Precipitation−associated SNP: Chr11:46938802
pvalue= 2e−04 ; R2= 0.587
Estimated allele frequency
1.0
0.8
0.6
0.4
0.2
600
800
1000
1200
Ecological gradient (Precipitation sums)
253
256
217
97
9
L8
L1
O1
O16
O8
L16
124

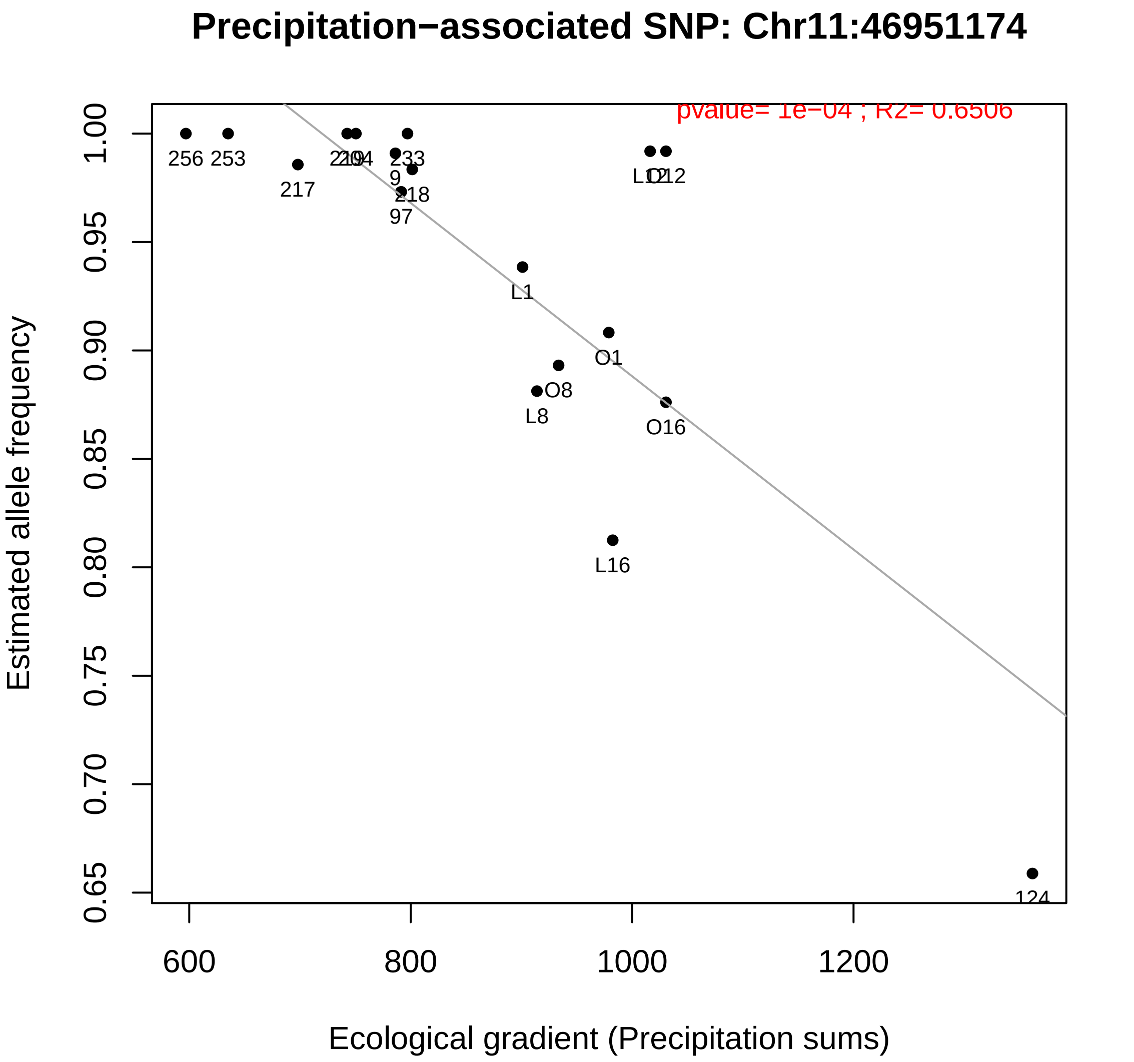

Precipitation−associated SNP: Chr11:46951174
pvalue= 1e−04 ; R2= 0.6506
Estimated allele frequency
1.00
0.95
0.90
0.85
0.80
0.75
0.70
0.65
600
800
1000
1200
Ecological gradient (Precipitation sums)
256
253
217
233
9
218
97
L1
O1
O8
L8
O16
L16
124

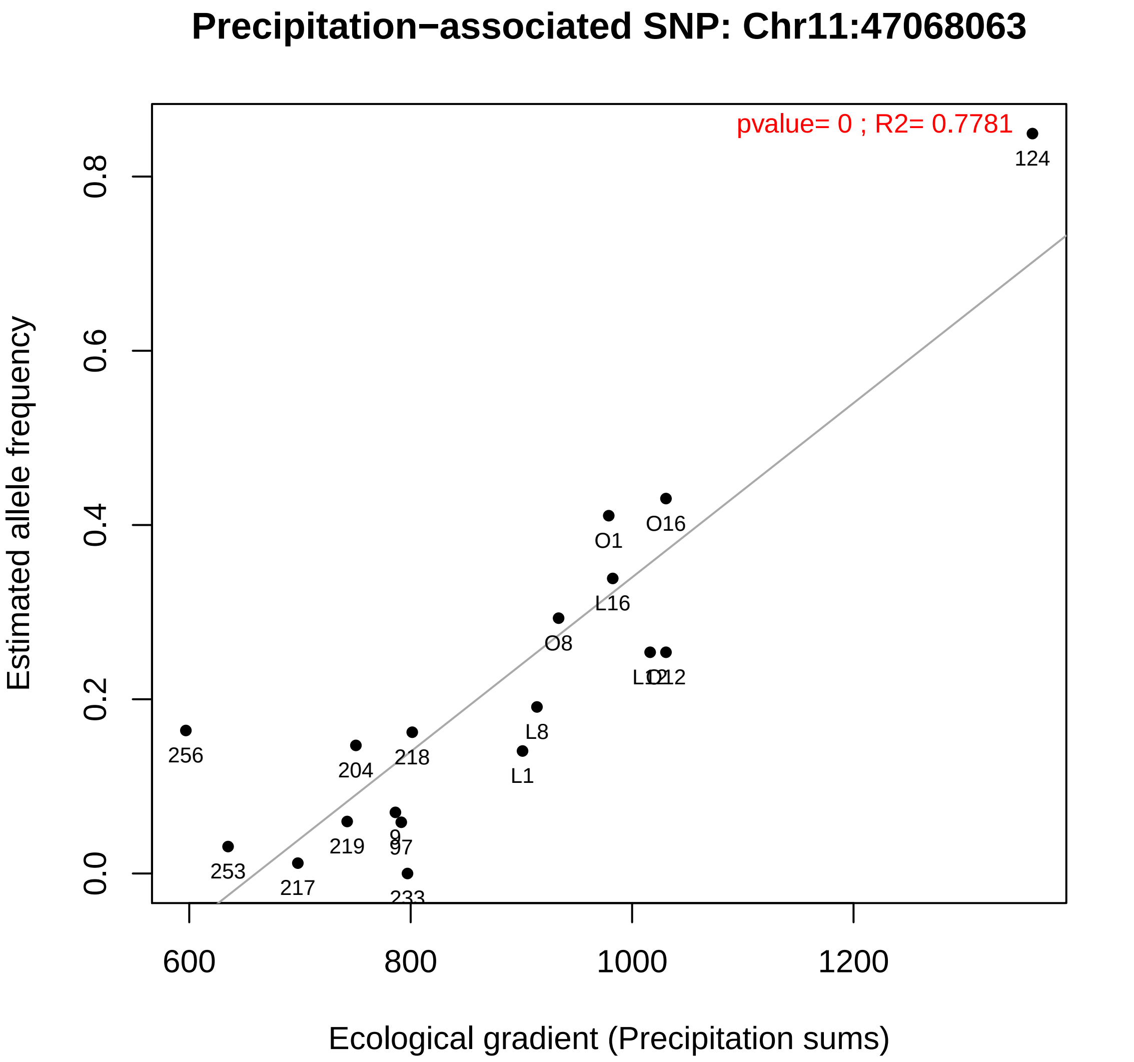
Precipitation−associated SNP: Chr11:47068063
pvalue= 0 ; R2= 0.7781
124
O16
O1
L16
O8
L10
O12
L8
L1
256
204
218
219
9
97
253
217
233
0.0
0.2
0.4
0.6
0.8
600
800
1000
1200
Estimated allele frequency
Ecological gradient (Precipitation sums)

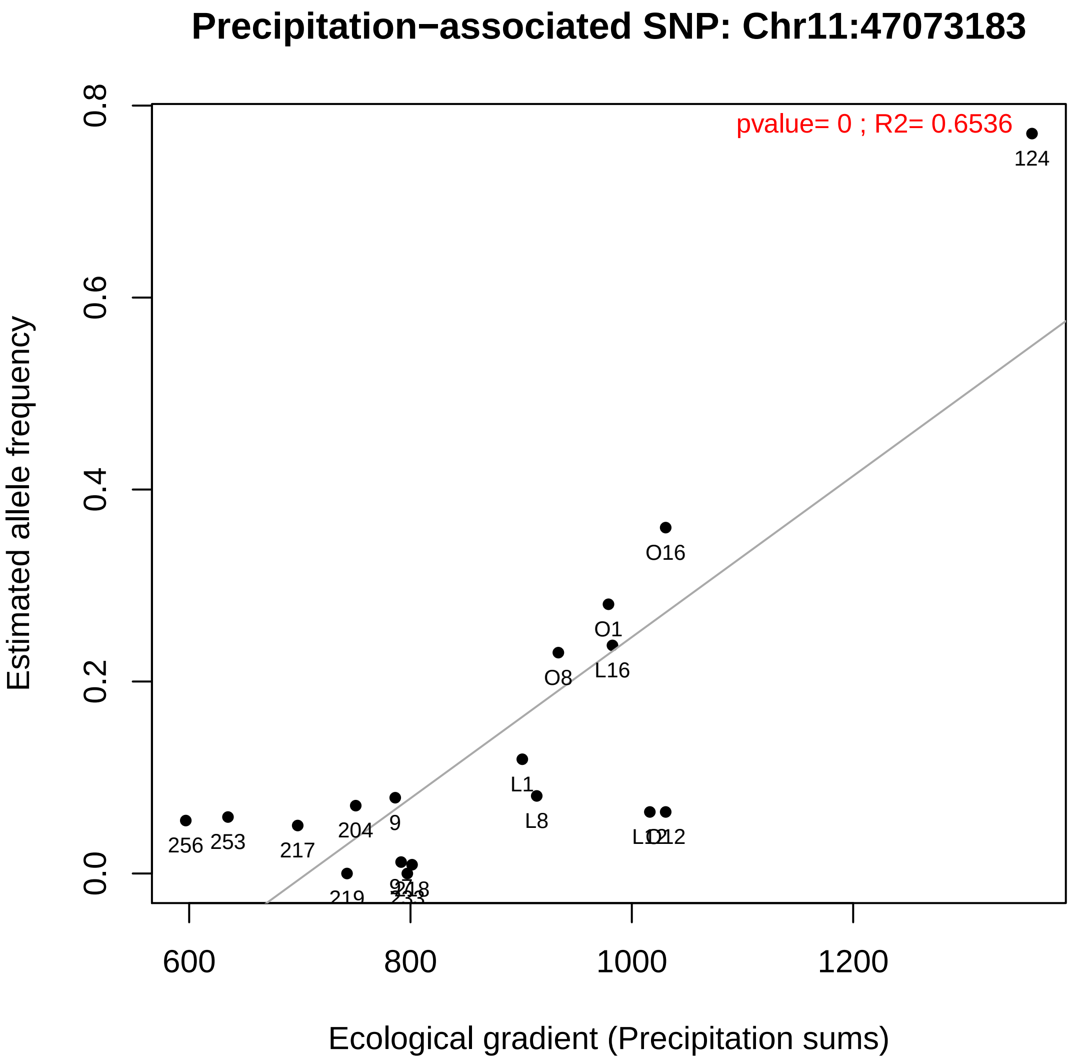

Precipitation−associated SNP: Chr11:47073183
pvalue= 0 ; R2= 0.6536
Estimated allele frequency
0.0
0.2
0.4
0.6
0.8
Ecological gradient (Precipitation sums)
600
800
1000
1200
124
O16
O1
L16
O8
L1
L8
256
253
217
204
9
219

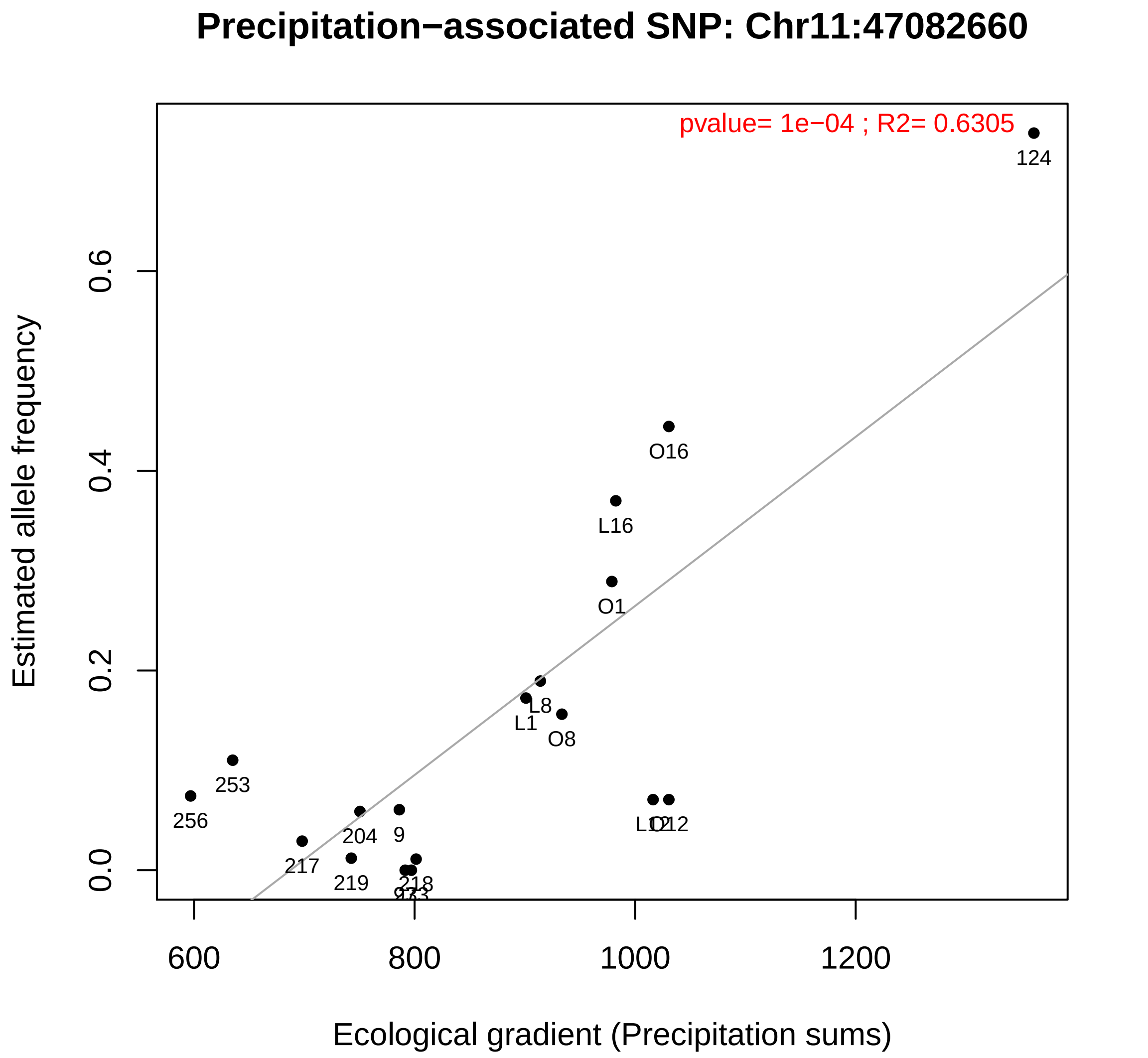

Precipitation−associated SNP: Chr11:47082660
pvalue= 1e−04 ; R2= 0.6305
Estimated allele frequency
0.0
0.2
0.4
0.6
600
800
1000
1200
Ecological gradient (Precipitation sums)
124
O16
L16
O1
L8
L1
O8
253
256
204
9
217
219
218

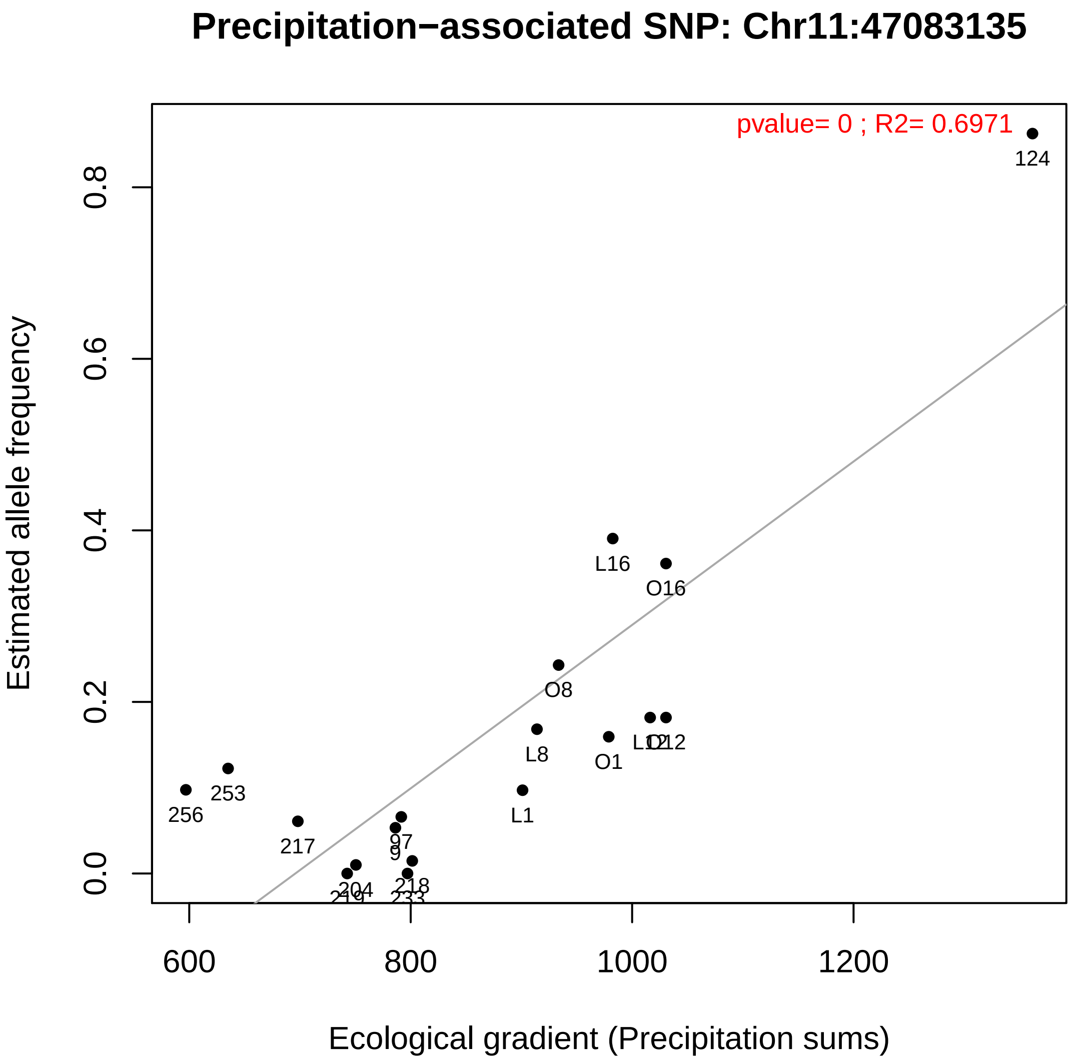
Precipitation−associated SNP: Chr11:47083135
pvalue= 0 ; R2= 0.6971
124
L16
O16
O8
L8
O1
L1
253
256
217
97
9
204
218
233
0.0
0.2
0.4
0.6
0.8
600
800
1000
1200
Estimated allele frequency
Ecological gradient (Precipitation sums)

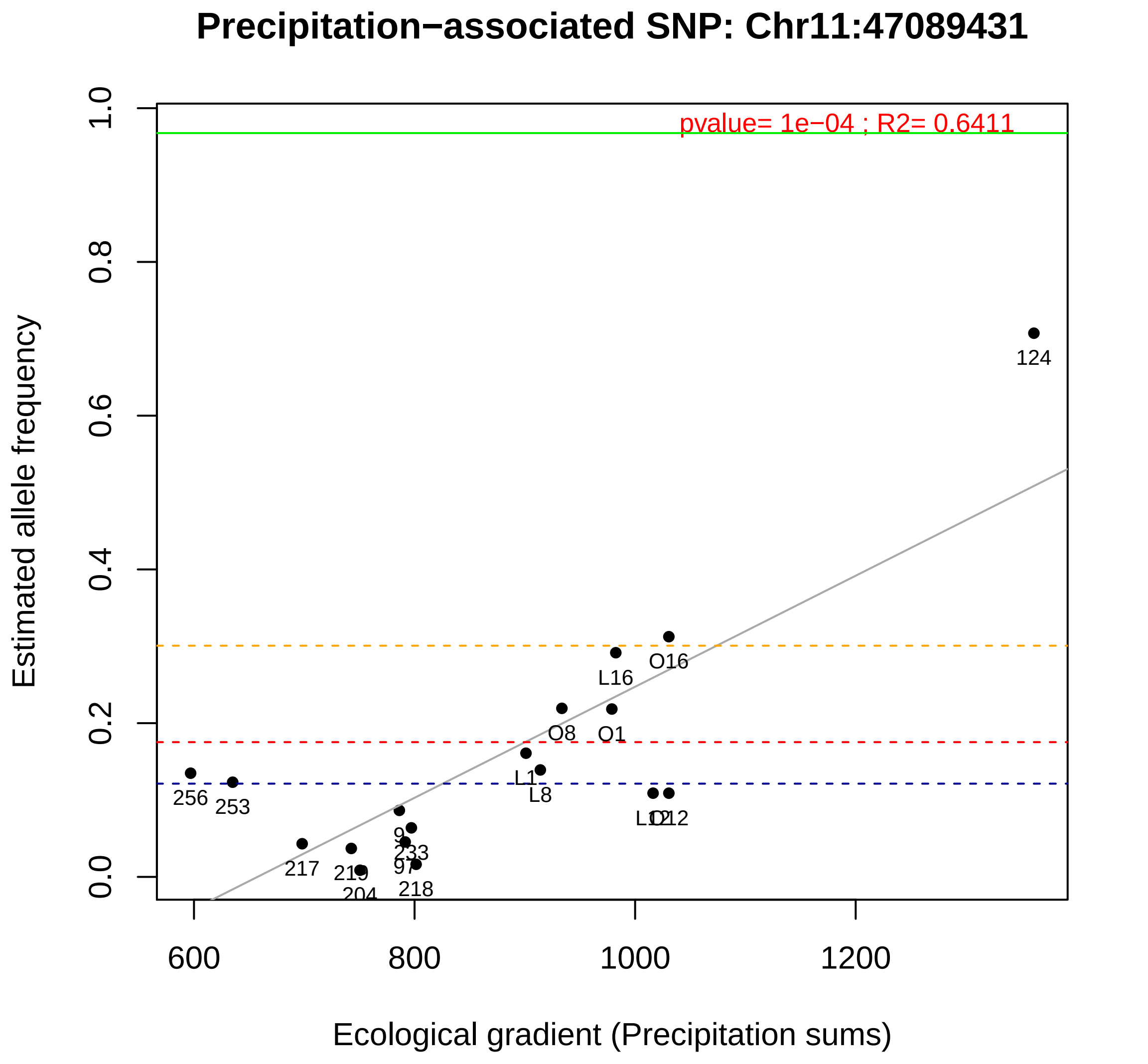

Precipitation−associated SNP: Chr11:47089431
pvalue= 1e−04 ; R2= 0.6411
Estimated allele frequency
Ecological gradient (Precipitation sums)
0.0
0.2
0.4
0.6
0.8
1.0
600
800
1000
1200
124
O16
L16
O8
O1
L1
L8
256
253
217
219
204
233
97
218
L12
O12

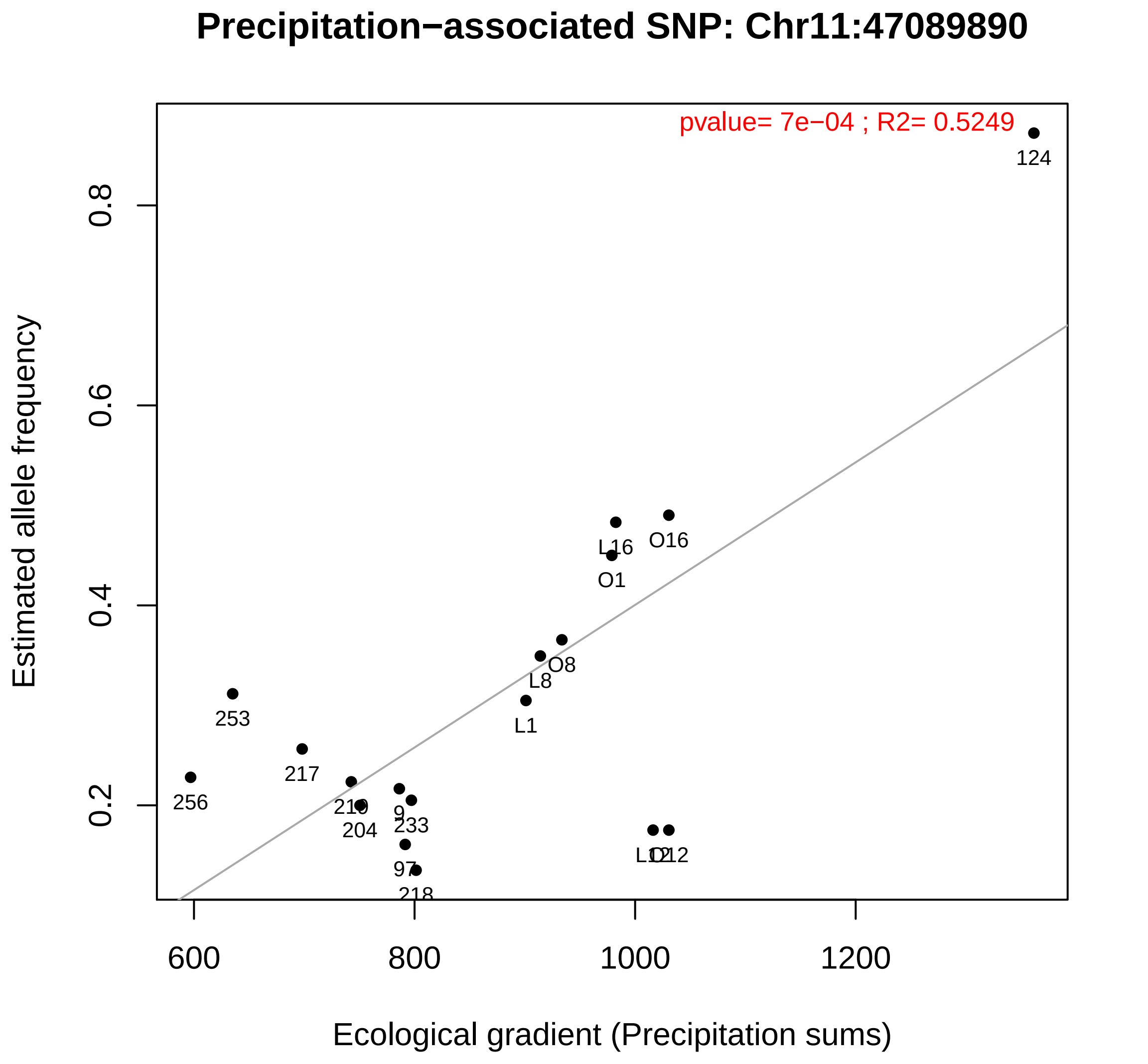

Precipitation−associated SNP: Chr11:47089890
pvalue= 7e−04 ; R2= 0.5249
124
L16
O16
O1
O8
L8
L1
253
217
256
210
204
9
233
97
218
L12
O12
0.2
0.4
0.6
0.8
600
800
1000
1200
Estimated allele frequency
Ecological gradient (Precipitation sums)

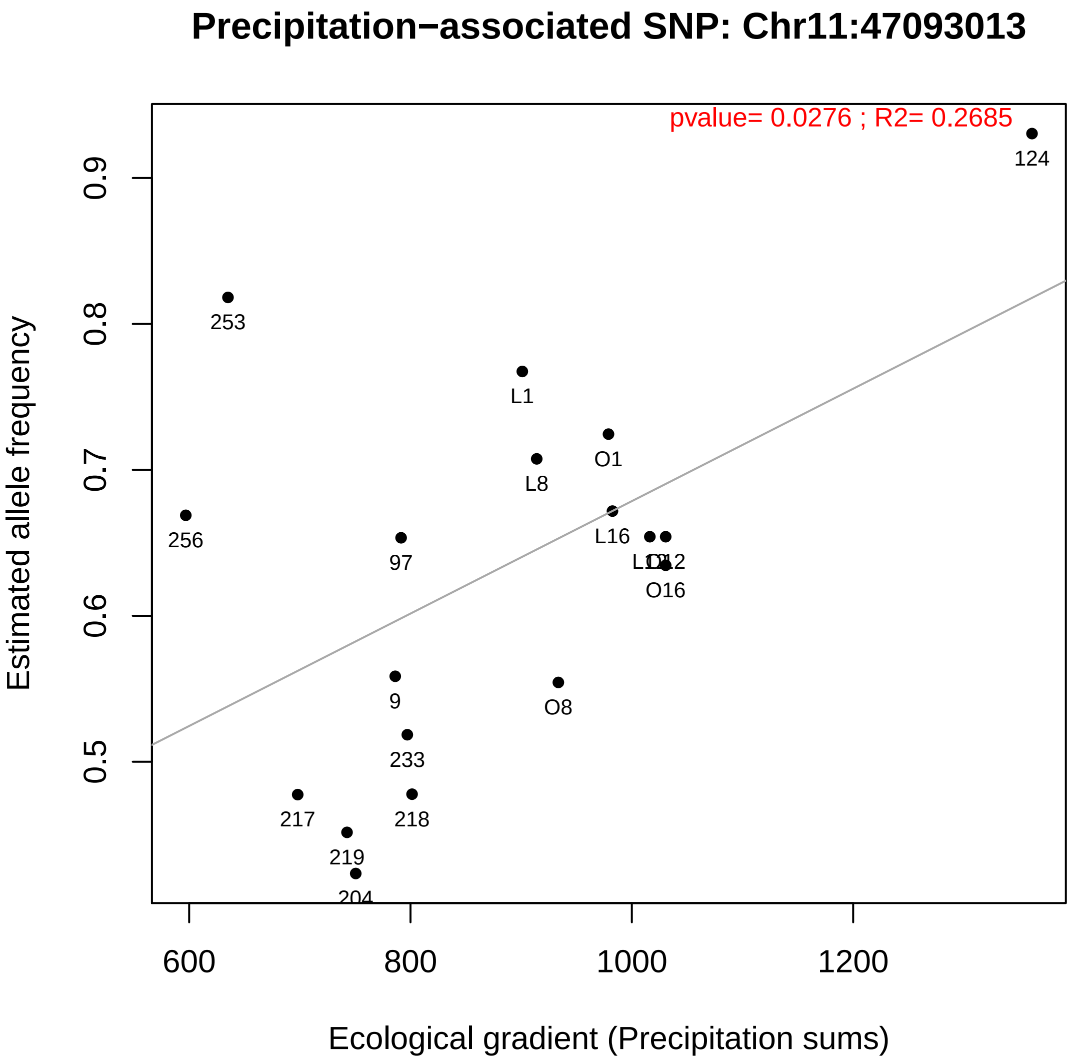
Precipitation−associated SNP: Chr11:47093013
pvalue= 0.0276 ; R2= 0.2685
124
253
L1
O1
L8
L16
256
97
O12
O16
9
O8
233
217
218
219
204
0.9
0.8
0.7
0.6
0.5
600
800
1000
1200
Estimated allele frequency
Ecological gradient (Precipitation sums)

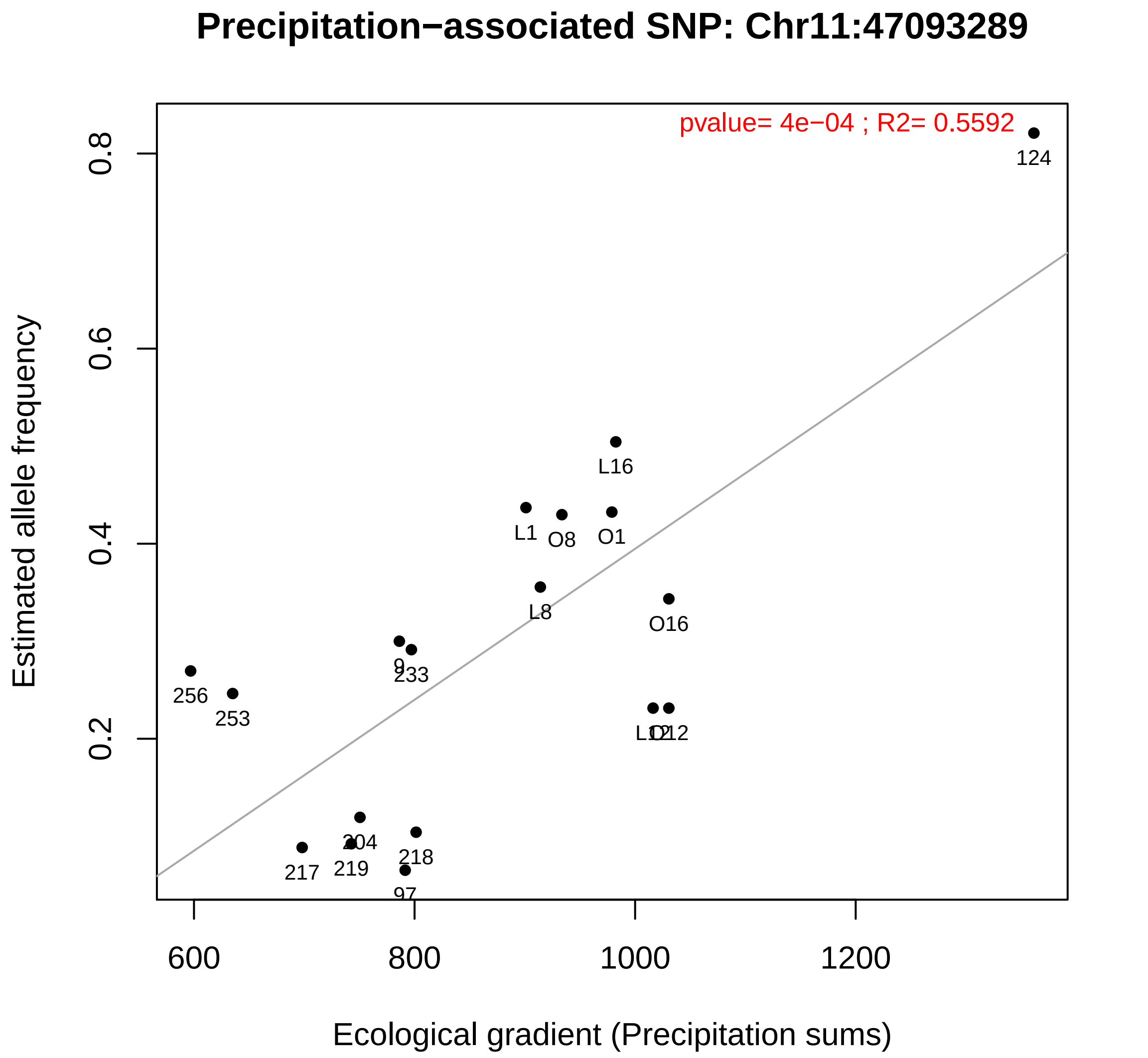

Precipitation−associated SNP: Chr11:47093289
pvalue= 4e−04 ; R2= 0.5592
124
L16
L1
O8
O1
L8
O16
9
233
256
253
L12
O12
204
218
217
219
97
0.8
0.6
0.4
0.2
600
800
1000
1200
Estimated allele frequency
Ecological gradient (Precipitation sums)

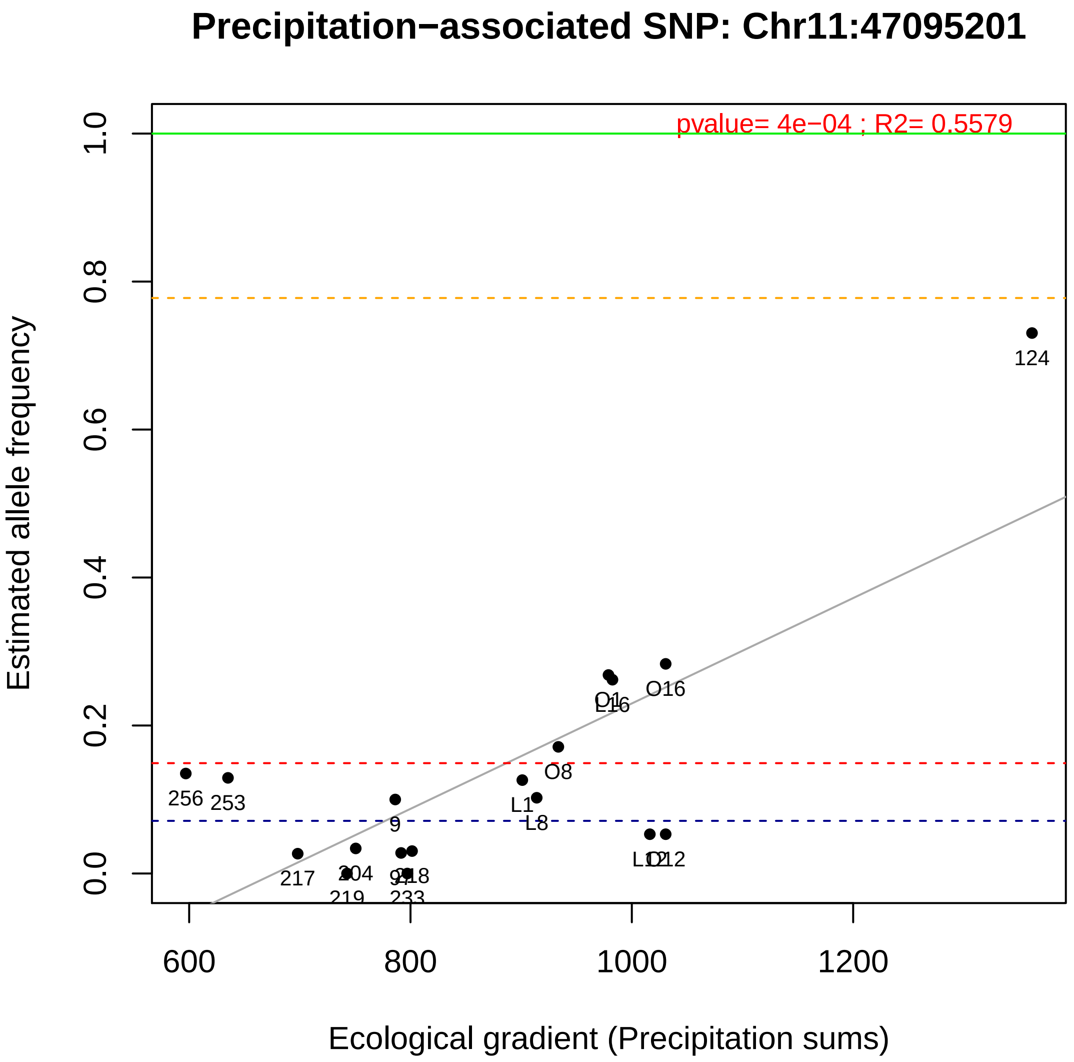

Precipitation−associated SNP: Chr11:47095201
pvalue= 4e−04 ; R2= 0.5579
Estimated allele frequency
0.0
0.2
0.4
0.6
0.8
1.0
600
800
1000
1200
Ecological gradient (Precipitation sums)
124
O16
O8
L1
L8
9
256
253
217
204
219
218
233

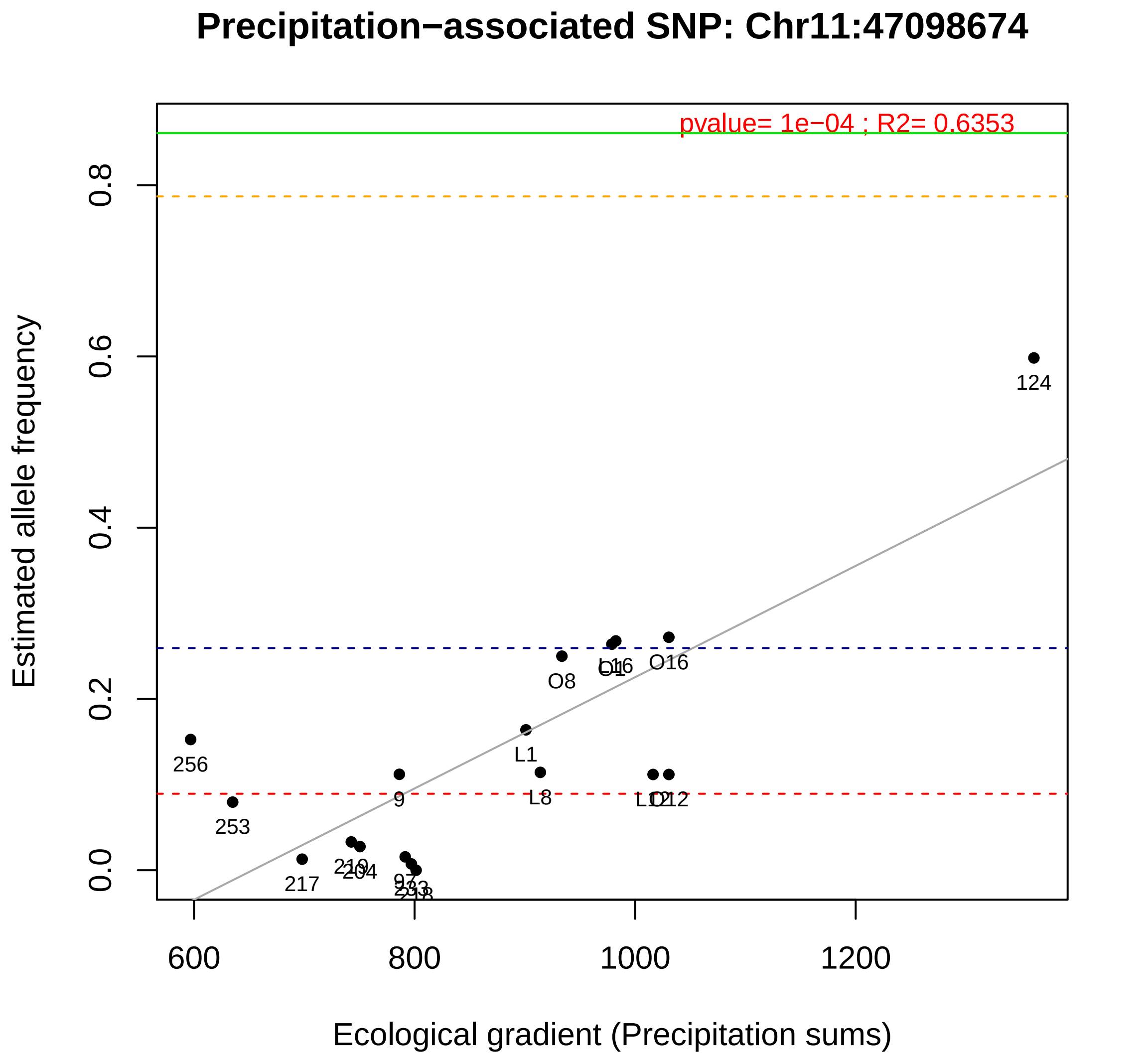
Precipitation−associated SNP: Chr11:47098674
pvalue= 1e−04 ; R2= 0.6353
Estimated allele frequency
0.0
0.2
0.4
0.6
0.8
600
800
1000
1200
Ecological gradient (Precipitation sums)
124
O16
O8
L1
256
L8
9
253
217
204
O12

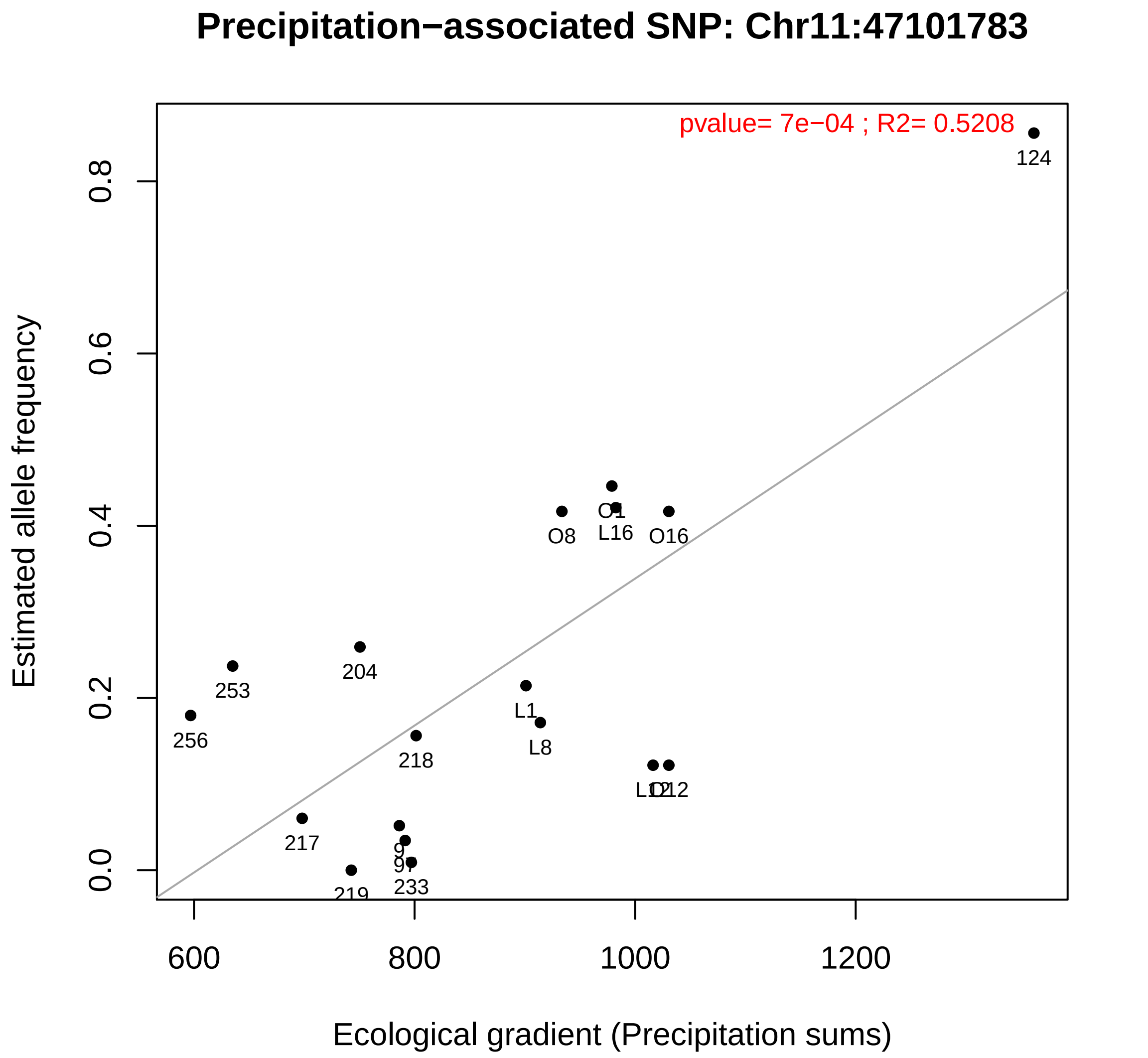
Precipitation−associated SNP: Chr11:47101783
pvalue= 7e−04 ; R2= 0.5208
Estimated allele frequency
0.0
0.2
0.4
0.6
0.8
600
800
1000
1200
Ecological gradient (Precipitation sums)
124
O8
O1
L16
O16
204
253
256
L1
L8
218
O12
217
233
219

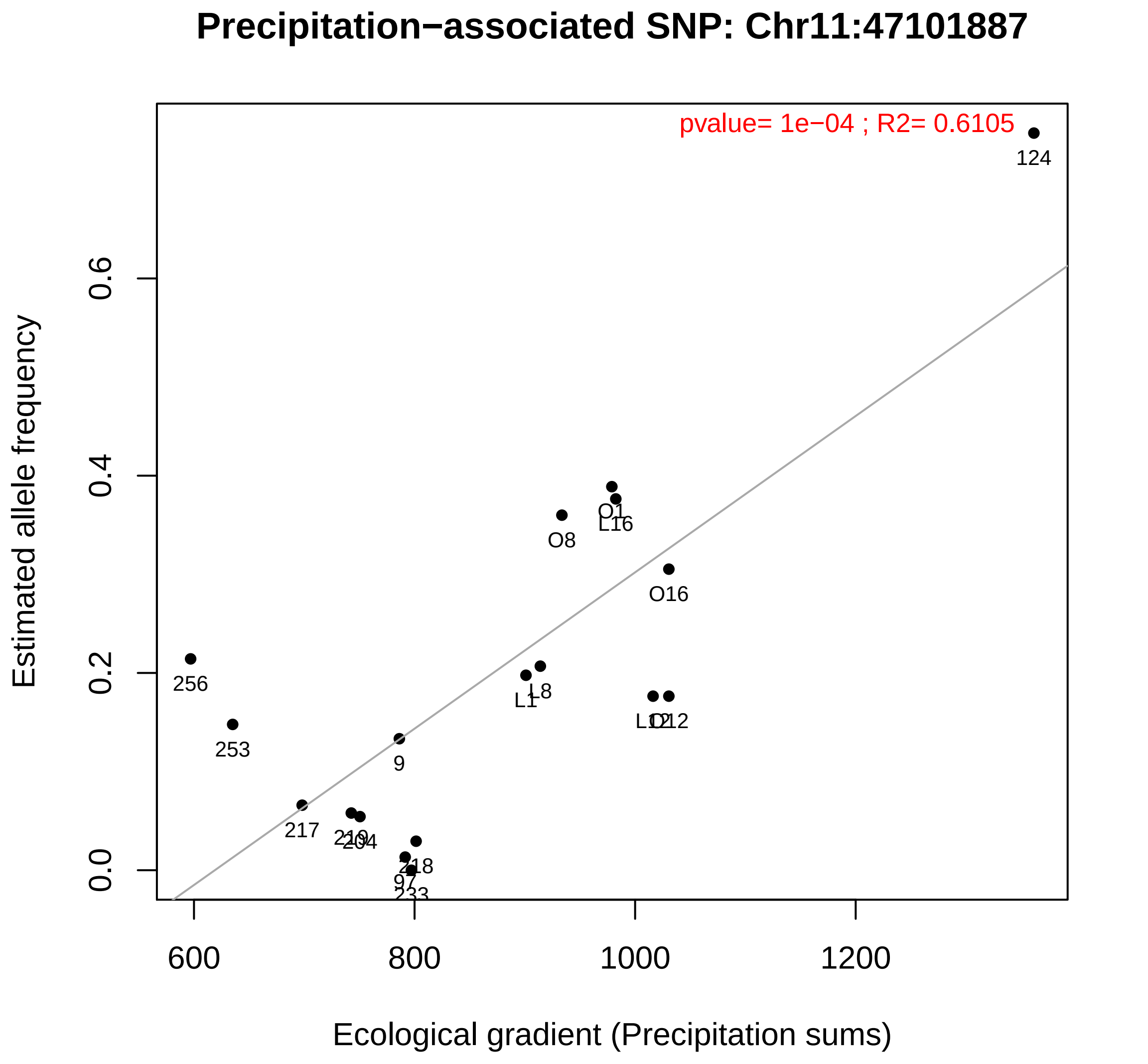
Precipitation−associated SNP: Chr11:47101887
pvalue= 1e−04 ; R2= 0.6105
124
O1
L16
O8
O16
256
L1
L8
L12
O12
253
9
217
210
204
218
97
233
0.0
0.2
0.4
0.6
600
800
1000
1200
Estimated allele frequency
Ecological gradient (Precipitation sums)

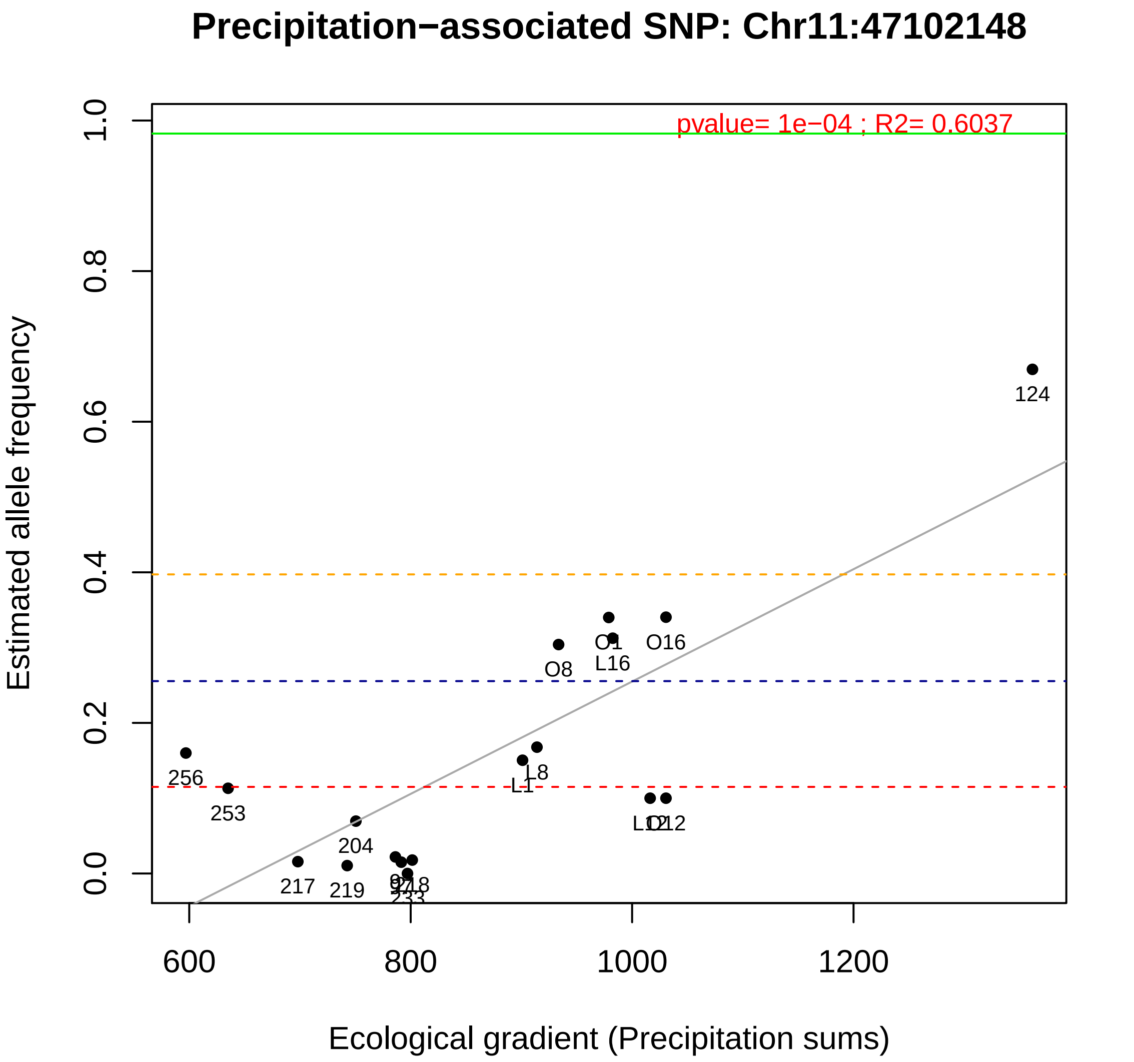

Precipitation−associated SNP: Chr11:47102148
pvalue= 1e−04 ; R2= 0.6037
Estimated allele frequency
0.0
0.2
0.4
0.6
0.8
1.0
600
800
1000
1200
Ecological gradient (Precipitation sums)
124
O1
L16
O16
O8
L8
L1
256
253
L12
O12
204
217
219
218
233

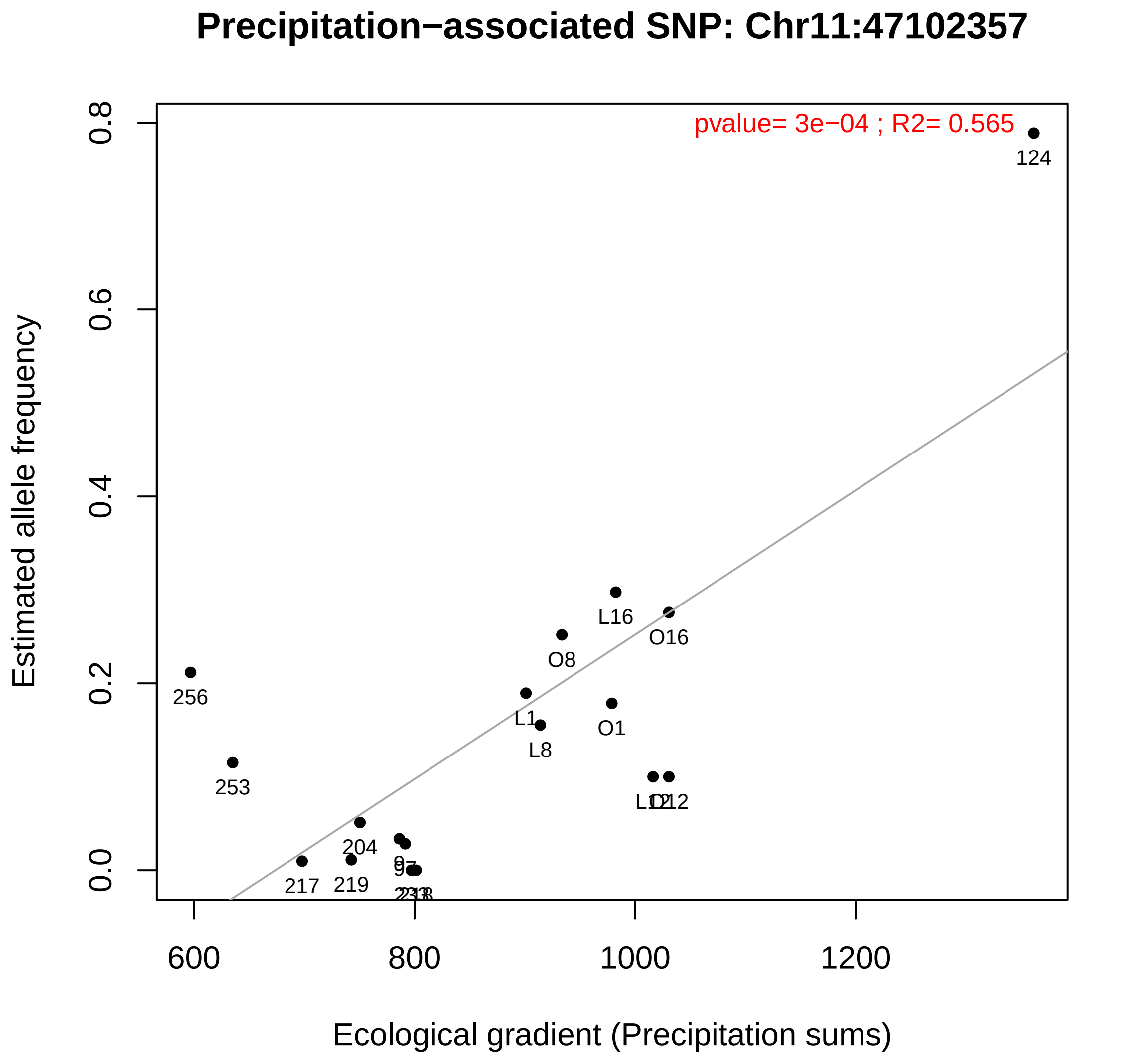
Precipitation−associated SNP: Chr11:47102357
pvalue= 3e−04 ; R2= 0.565
Estimated allele frequency
Ecological gradient (Precipitation sums)
0.0
0.2
0.4
0.6
0.8
600
800
1000
1200
124
L16
O16
O8
256
L1
L8
O1
253
204
217
219

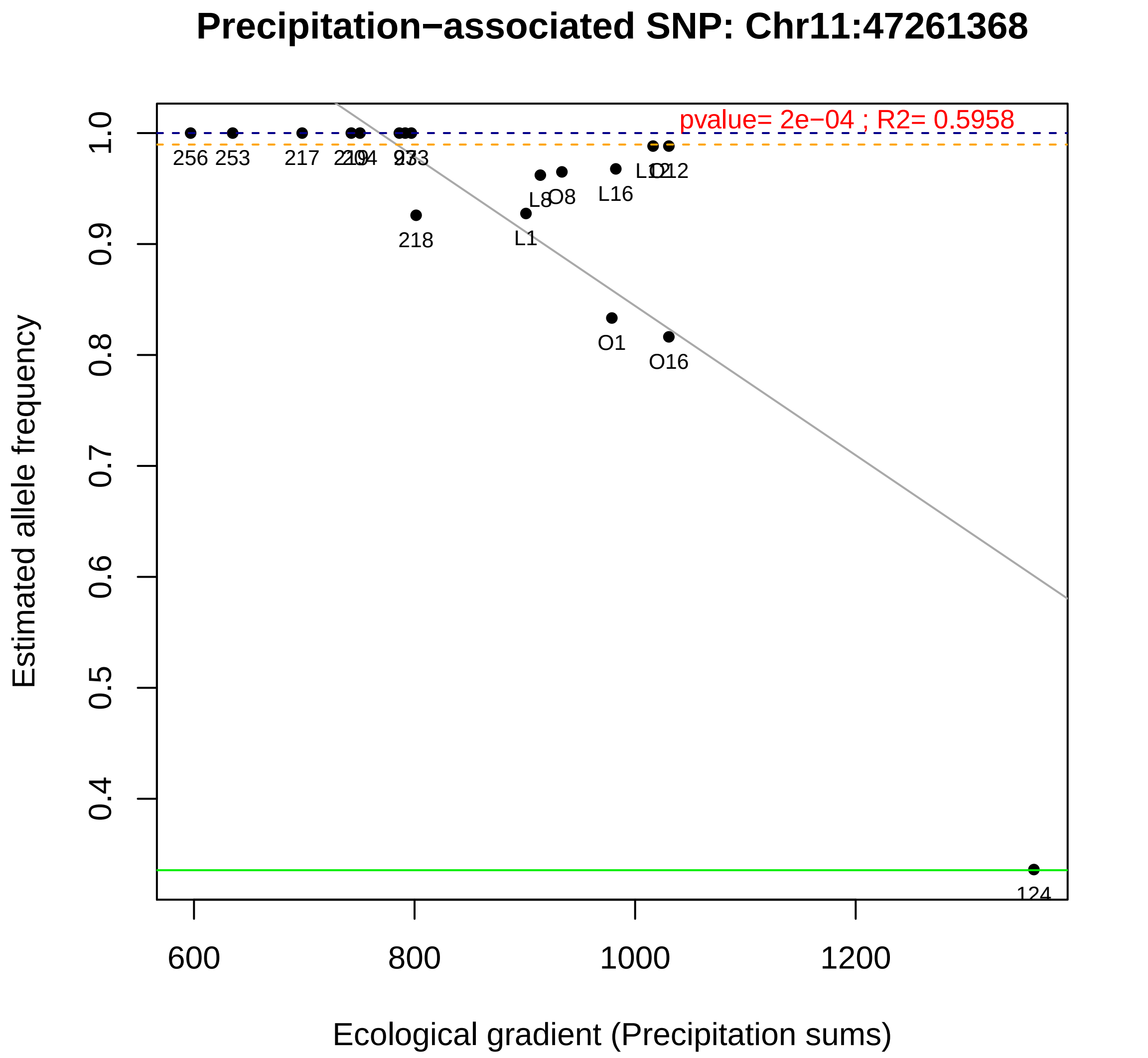
Precipitation−associated SNP: Chr11:47261368
pvalue= 2e−04 ; R2= 0.5958
Estimated allele frequency
Ecological gradient (Precipitation sums)
256
253
217
218
L1
L8
O8
L16
O1
O16
124
0.4
0.5
0.6
0.7
0.8
0.9
1.0
600
800
1000
1200

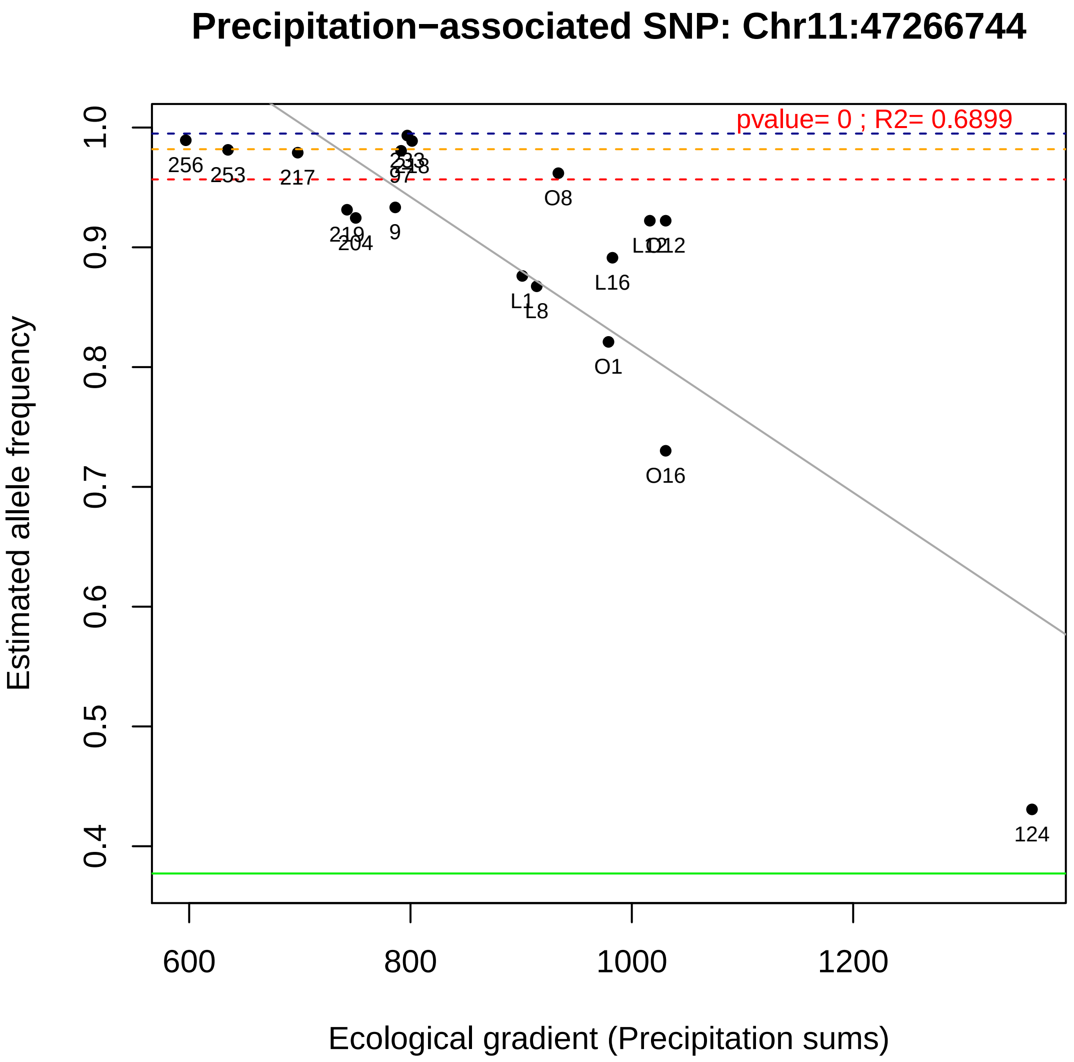

Precipitation−associated SNP: Chr11:47266744
pvalue= 0 ; R2= 0.6899
Estimated allele frequency
Ecological gradient (Precipitation sums)
1.0
0.9
0.8
0.7
0.6
0.5
0.4
600
800
1000
1200
256
253
217
219
204
9
O8
L1
L8
L16
O1
L12
O12
O16
124

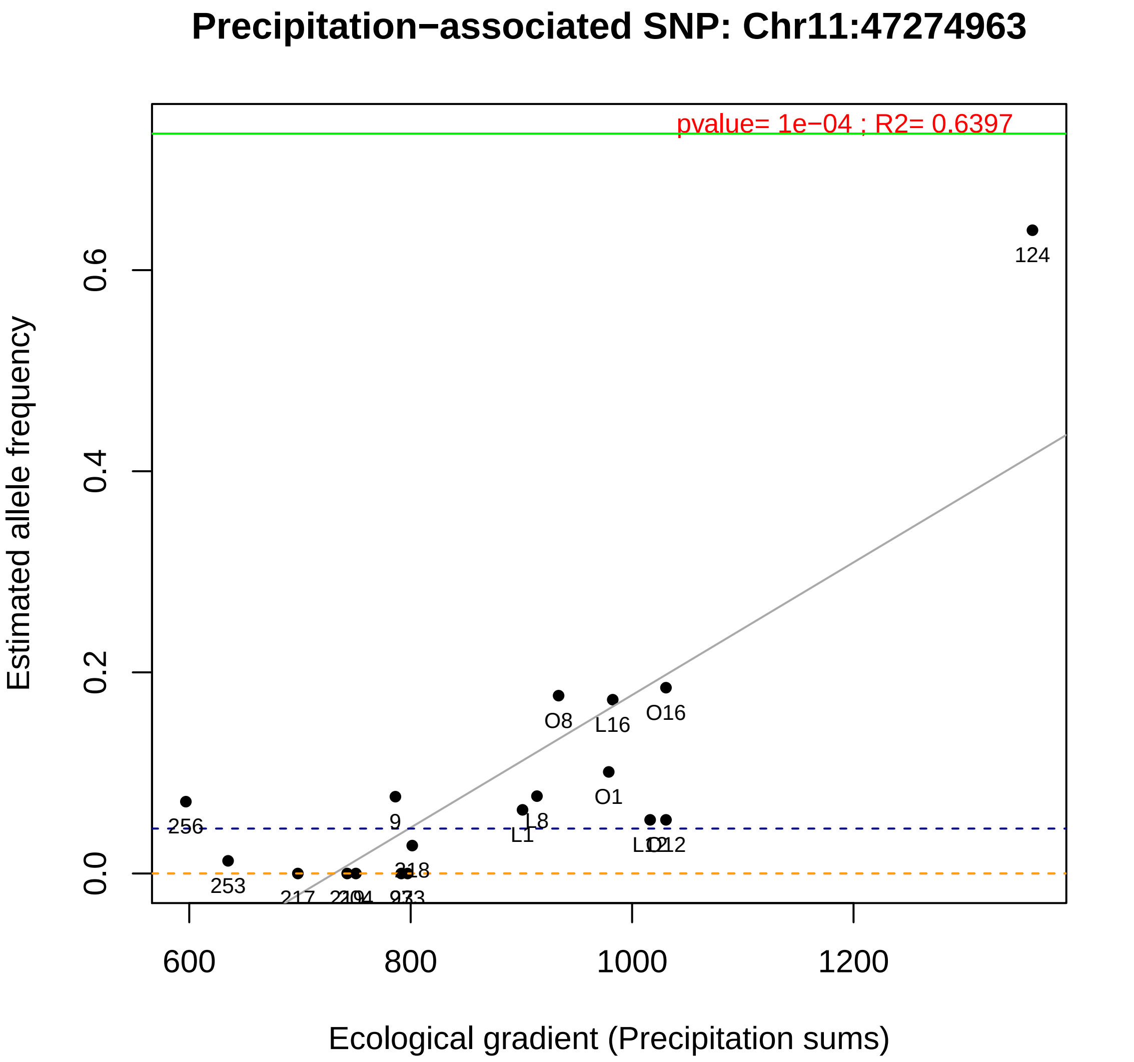
Precipitation−associated SNP: Chr11:47274963
pvalue= 1e−04 ; R2= 0.6397
Estimated allele frequency
0.0
0.2
0.4
0.6
600
800
1000
1200
Ecological gradient (Precipitation sums)
124
O8
L16
O16
O1
256
9
L8
L1
L12
O12
253
218
217
97
273

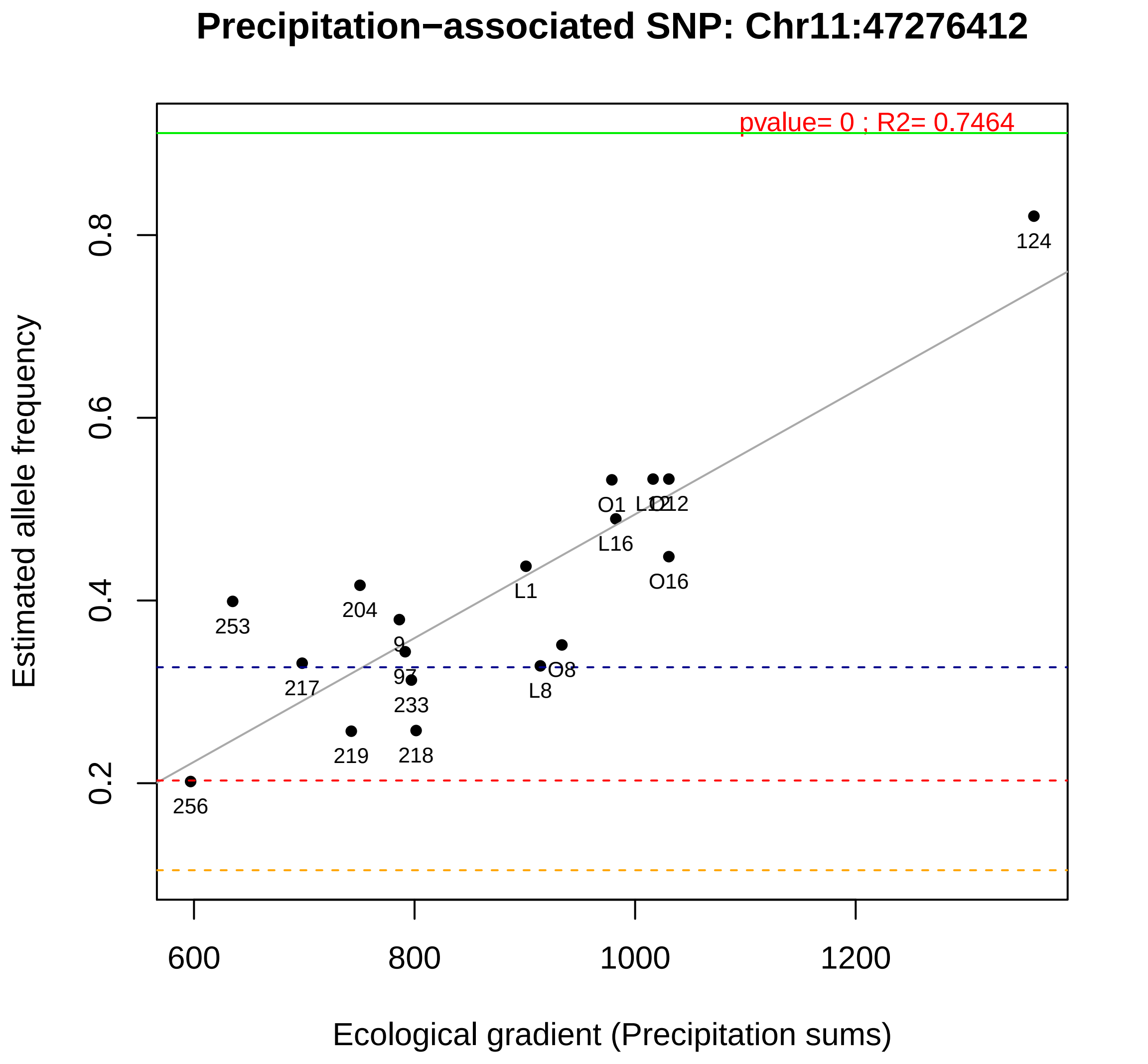

Precipitation−associated SNP: Chr11:47276412
pvalue= 0 ; R2= 0.7464
124
O1
L12
O12
L16
O16
L1
204
253
9
97
233
O8
L8
217
219
218
256
0.2
0.4
0.6
0.8
600
800
1000
1200
Estimated allele frequency
Ecological gradient (Precipitation sums)

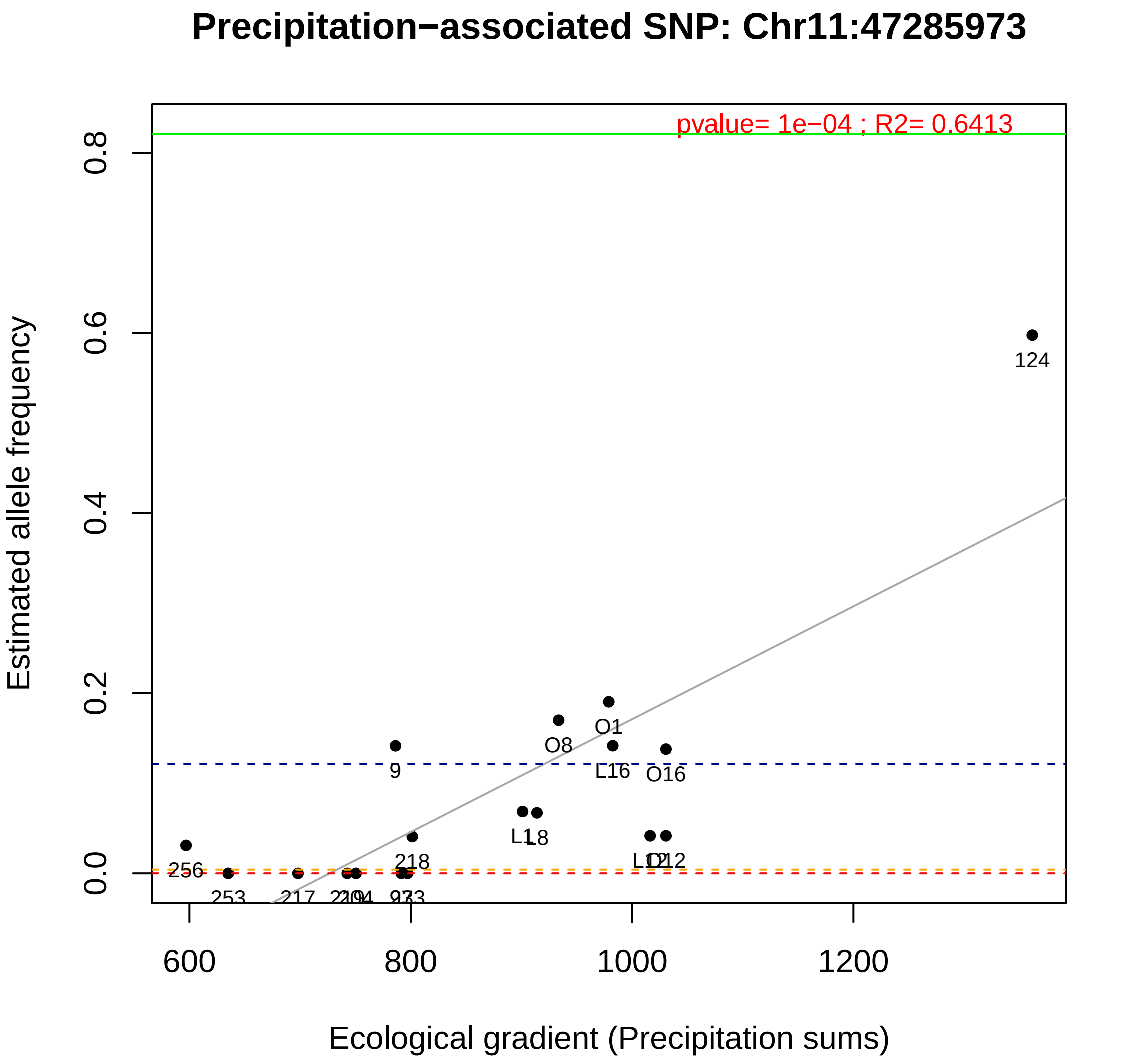

Precipitation−associated SNP: Chr11:47285973
pvalue= 1e−04 ; R2= 0.6413
Estimated allele frequency
0.0
0.2
0.4
0.6
0.8
600
800
1000
1200
Ecological gradient (Precipitation sums)
124
O1
O8
9
L16
O16
218
256
253
217

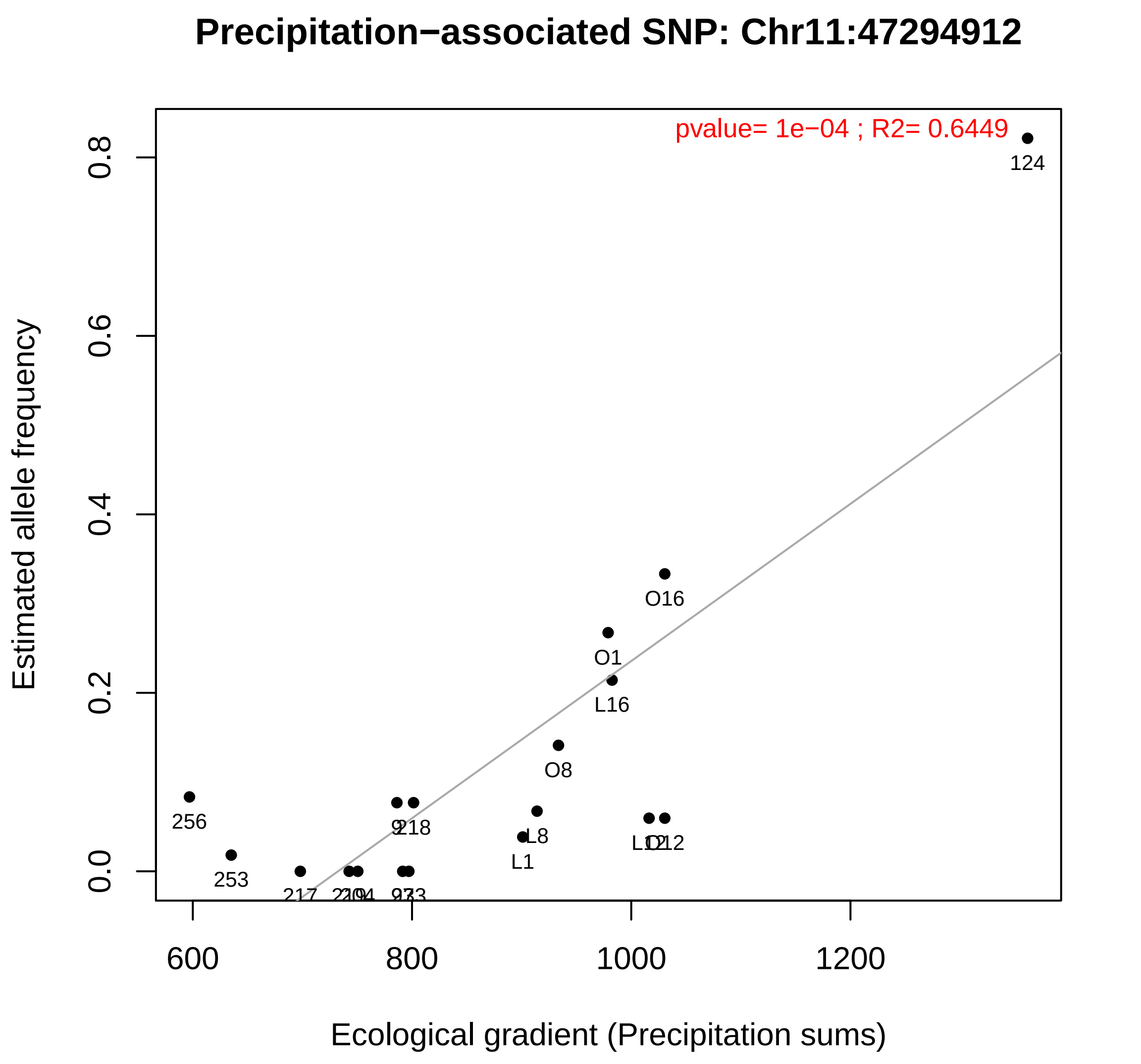

Precipitation−associated SNP: Chr11:47294912
pvalue= 1e−04 ; R2= 0.6449
Estimated allele frequency
0.0
0.2
0.4
0.6
0.8
600
800
1000
1200
Ecological gradient (Precipitation sums)
124
O16
O1
L16
O8
256
253
217
218
L8
L1
O12

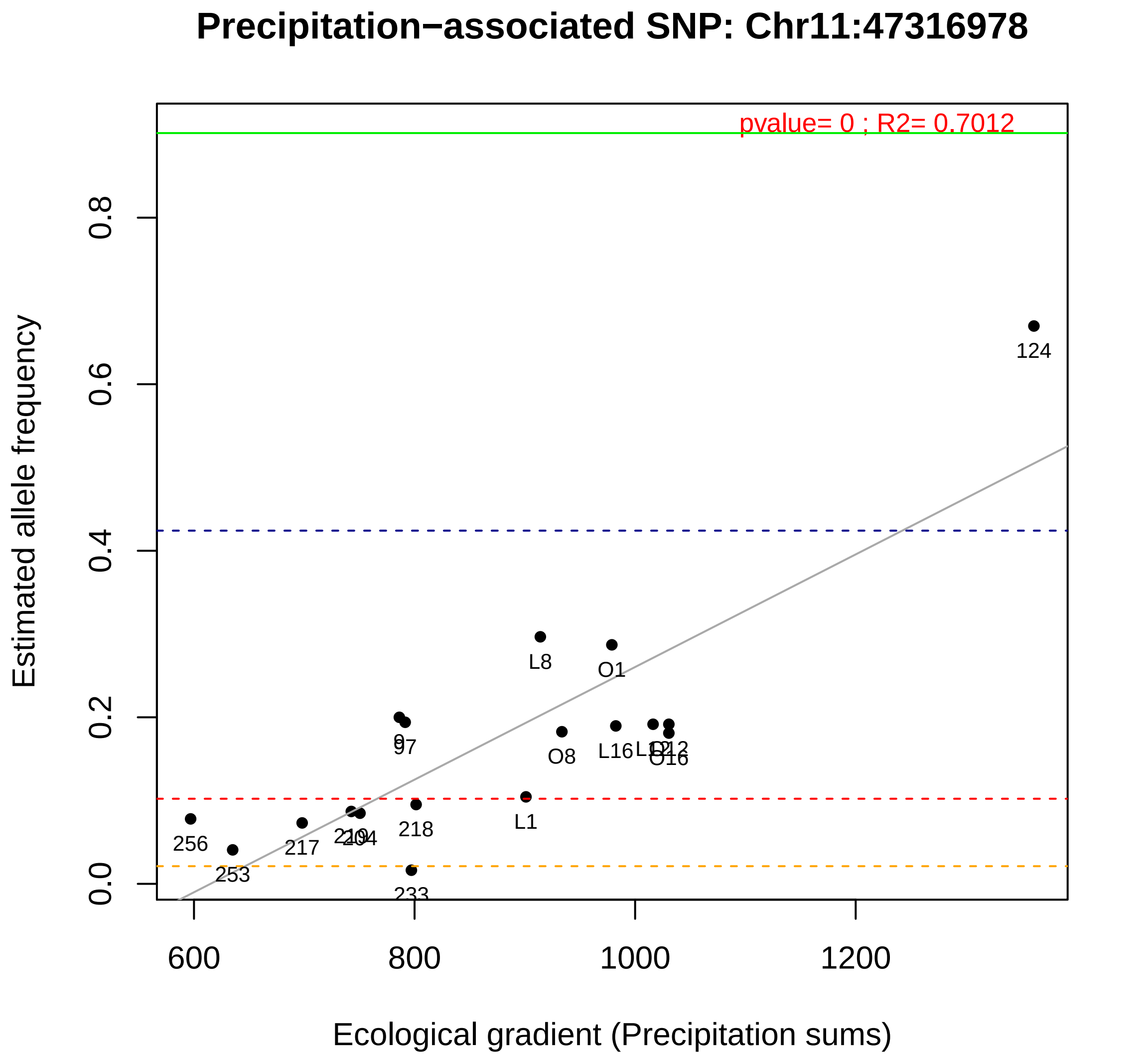

Precipitation−associated SNP: Chr11:47316978
pvalue= 0 ; R2= 0.7012
Estimated allele frequency
0.0
0.2
0.4
0.6
0.8
600
800
1000
1200
Ecological gradient (Precipitation sums)
124
L8
O1
O8
L16
97
L1
218
256
253
217
233

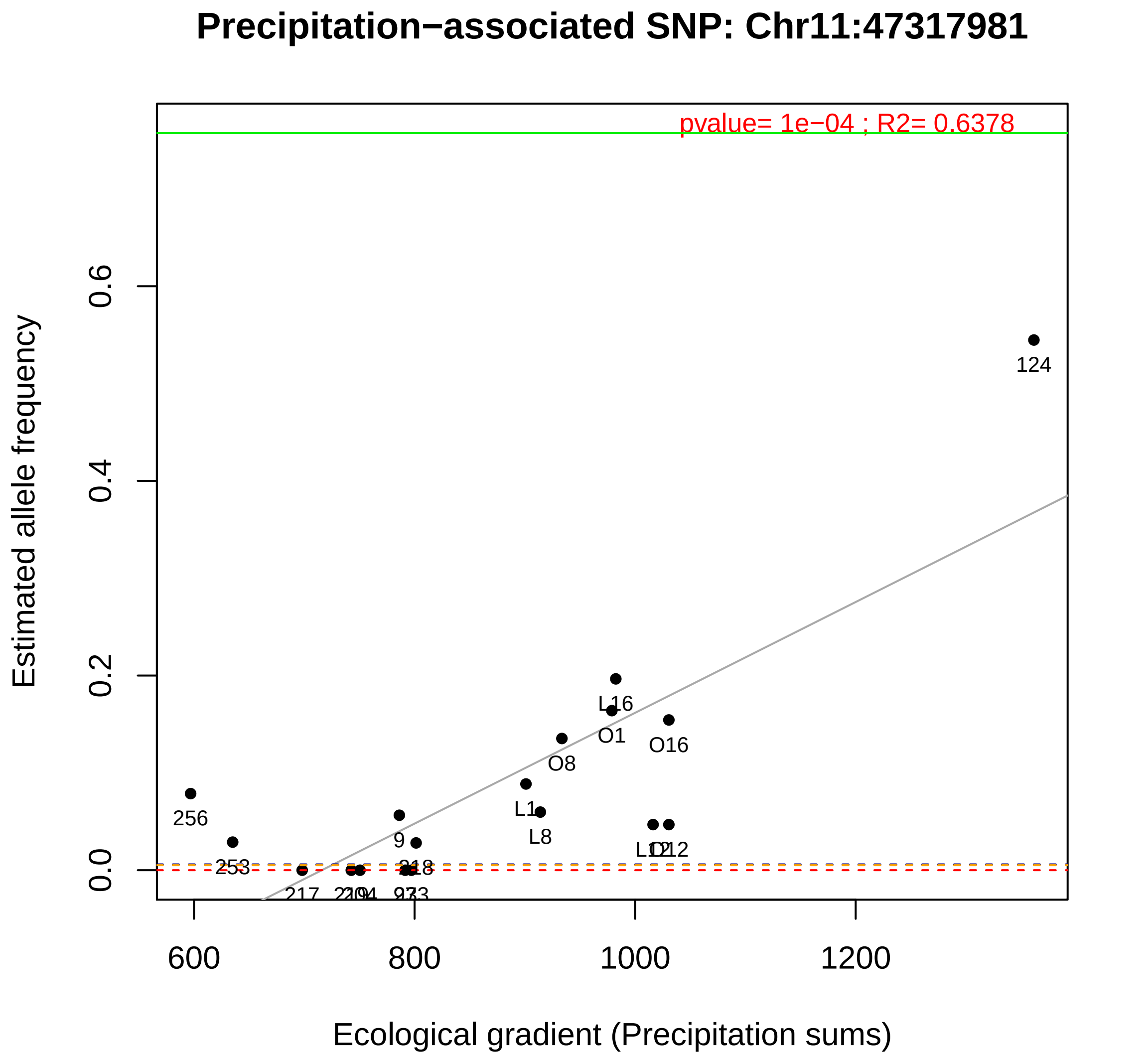

Precipitation−associated SNP: Chr11:47317981
pvalue= 1e−04 ; R2= 0.6378
Estimated allele frequency
0.0
0.2
0.4
0.6
600
800
1000
1200
Ecological gradient (Precipitation sums)
124
L16
O1
O16
O8
256
L1
L8
9
253
L12
O12

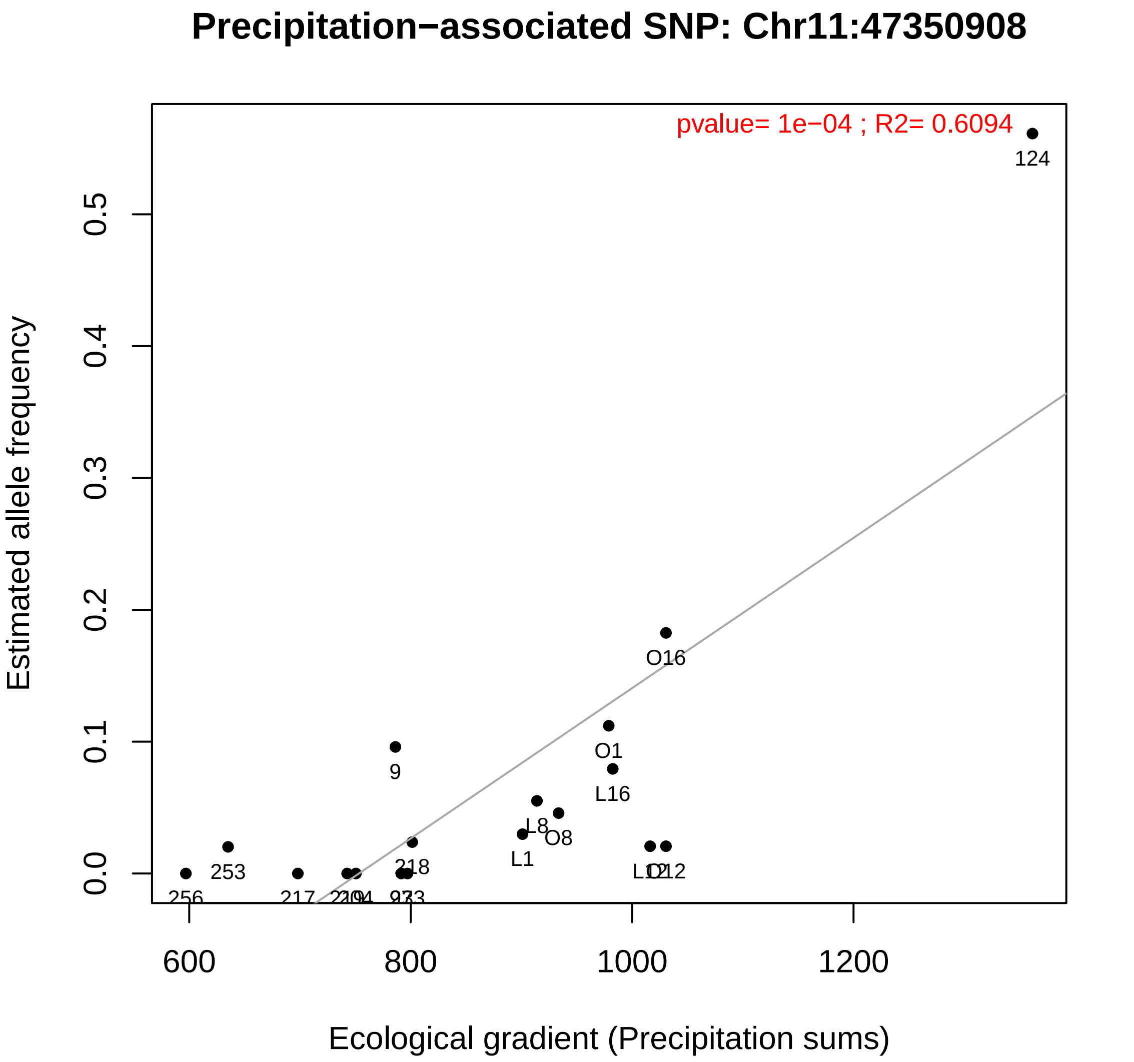
Precipitation−associated SNP: Chr11:47350908
pvalue= 1e−04 ; R2= 0.6094
124
O16
O1
L16
9
L8
O8
L1
218
253
256
217
Estimated allele frequency
0.0
0.1
0.2
0.3
0.4
0.5
600
800
1000
1200
Ecological gradient (Precipitation sums)

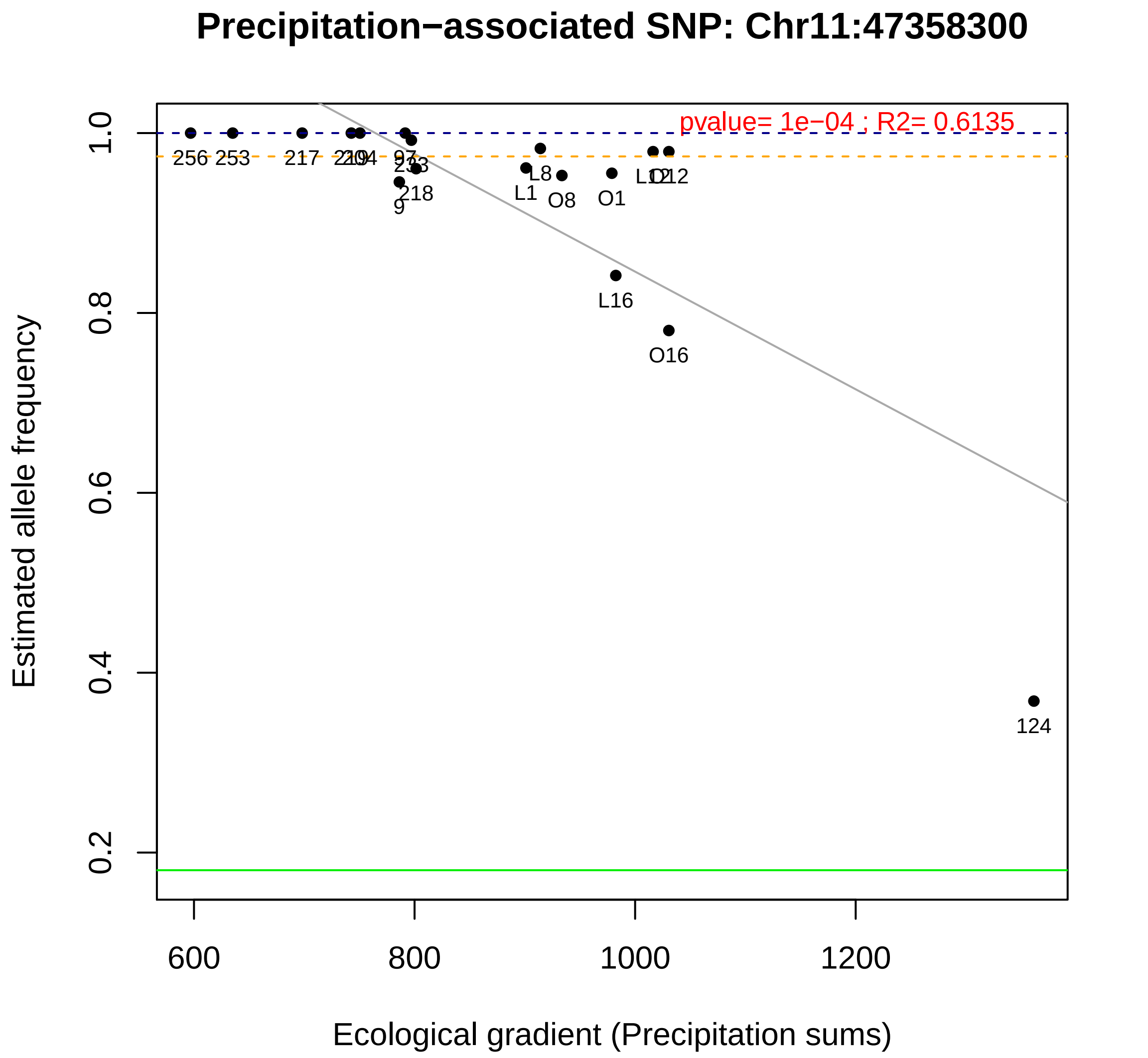

Precipitation−associated SNP: Chr11:47358300
pvalue= 1e−04 ; R2= 0.6135
Estimated allele frequency
Ecological gradient (Precipitation sums)
1.0
0.8
0.6
0.4
0.2
600
800
1000
1200
256
253
217
218
L16
O16
124
L8
L1
O8
O1
9

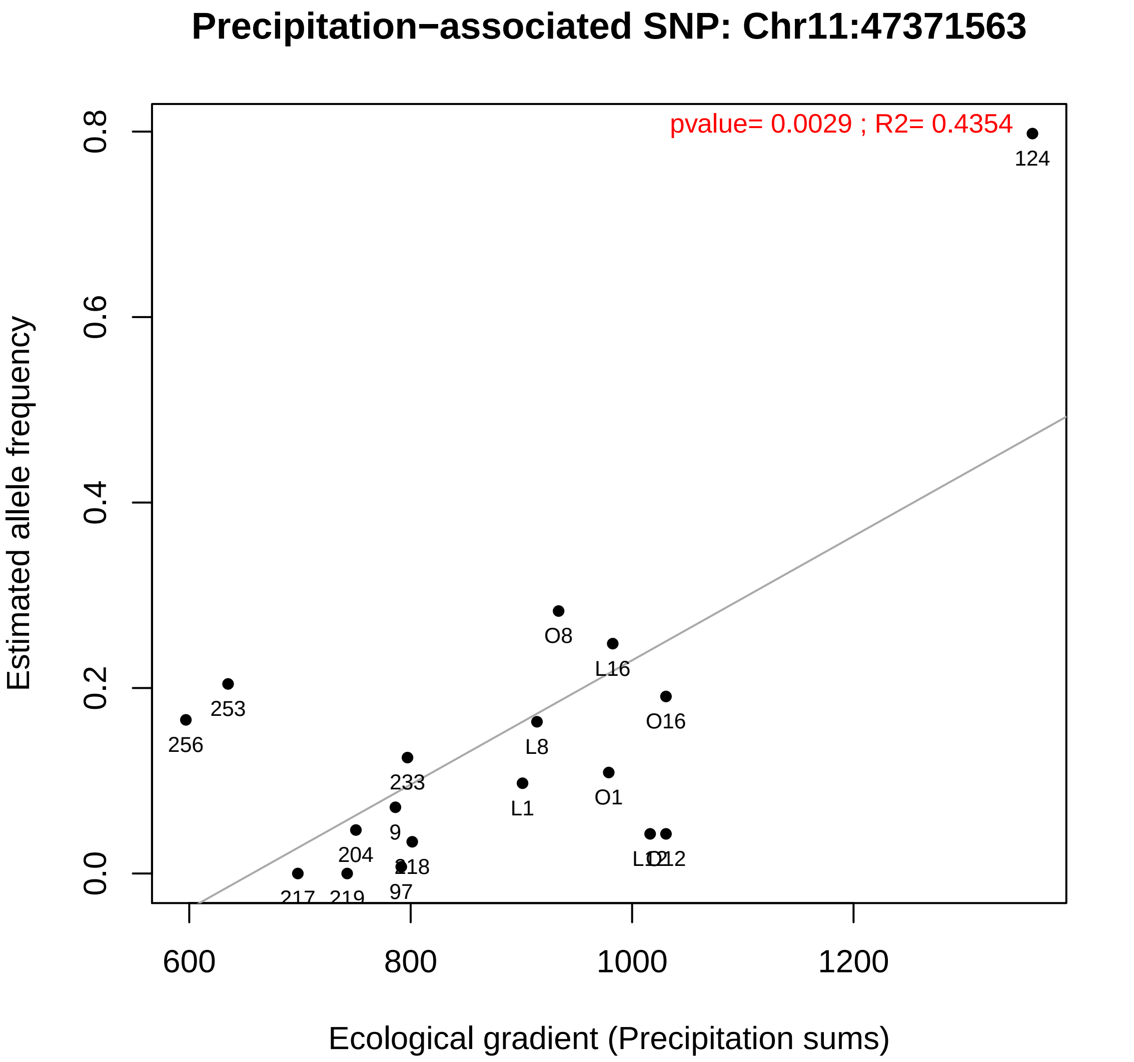

Precipitation−associated SNP: Chr11:47371563
pvalue= 0.0029 ; R2= 0.4354
Estimated allele frequency
Ecological gradient (Precipitation sums)
0.0
0.2
0.4
0.6
0.8
600
800
1000
1200
124
O8
L16
253
256
O16
L8
233
L1
O1
9
204
218
97
217
219
L12
O12

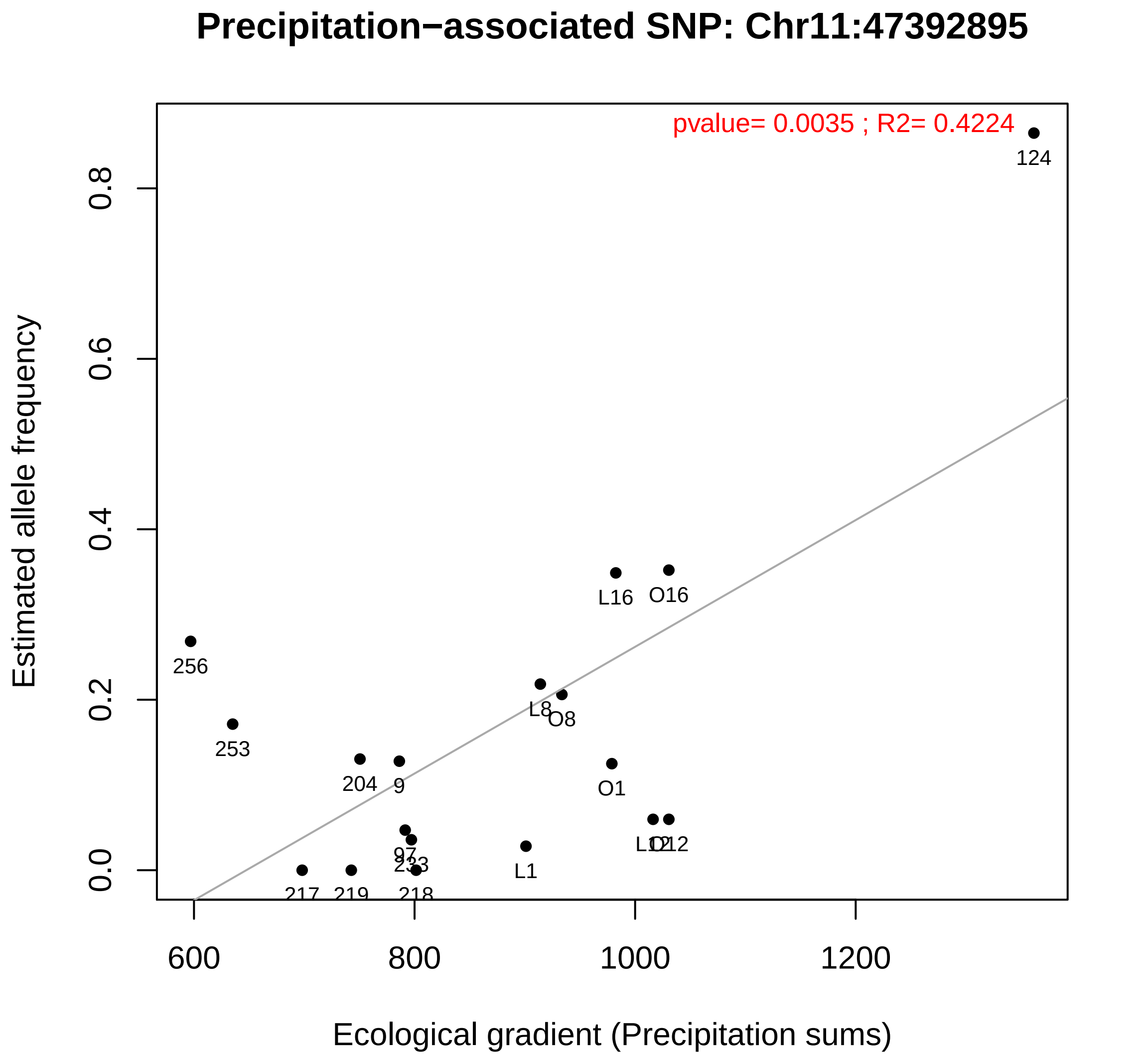
Precipitation−associated SNP: Chr11:47392895
pvalue= 0.0035 ; R2= 0.4224
124
L16
O16
256
L8
O8
253
204
9
O1
L12
O12
97
233
L1
217
219
218
0.0
0.2
0.4
0.6
0.8
600
800
1000
1200
Estimated allele frequency
Ecological gradient (Precipitation sums)

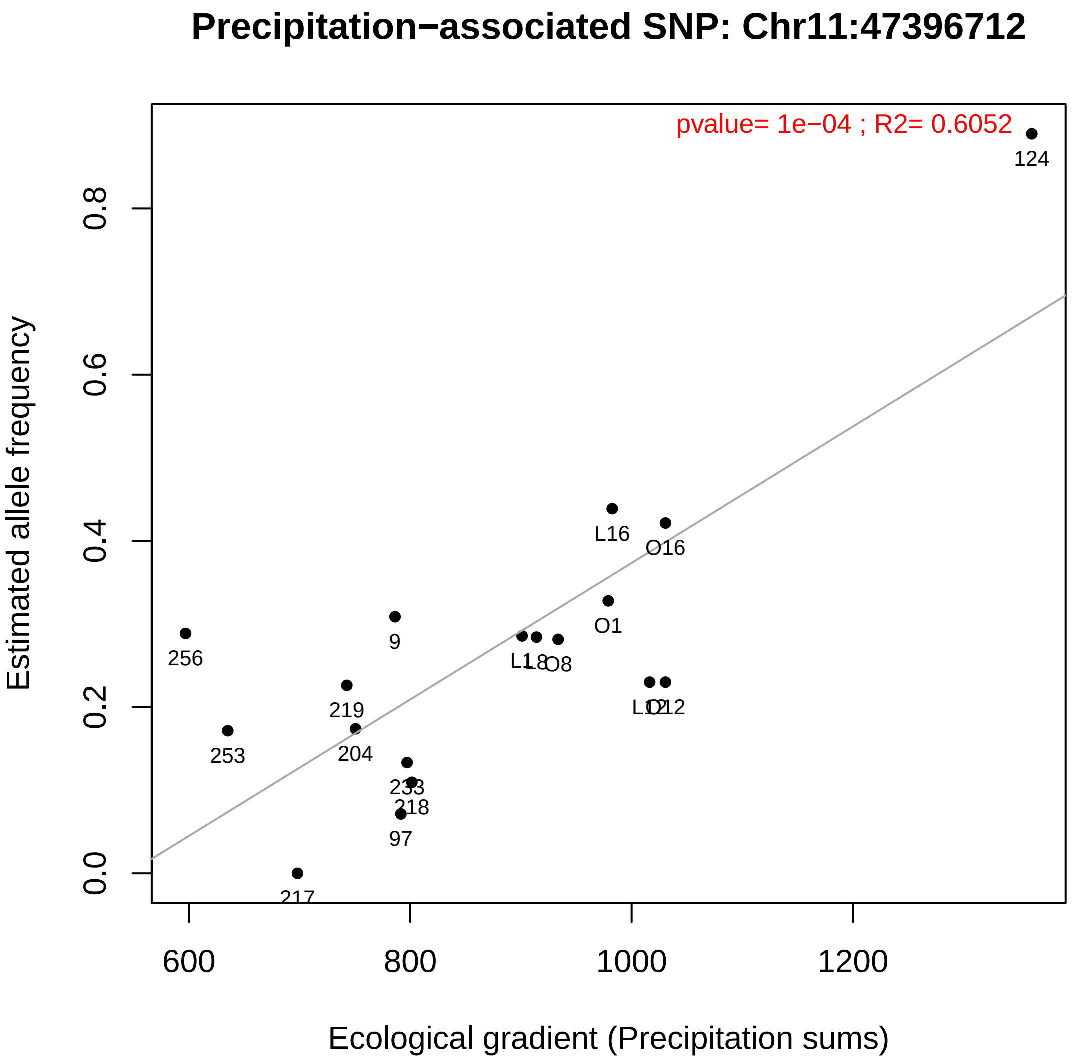

Precipitation−associated SNP: Chr11:47396712
pvalue= 1e−04 ; R2= 0.6052
Estimated allele frequency
Ecological gradient (Precipitation sums)
0.0
0.2
0.4
0.6
0.8
600
800
1000
1200
124
L16
O16
O1
9
256
L1
L8
O8
L12
O12
219
253
204
233
218
97
217

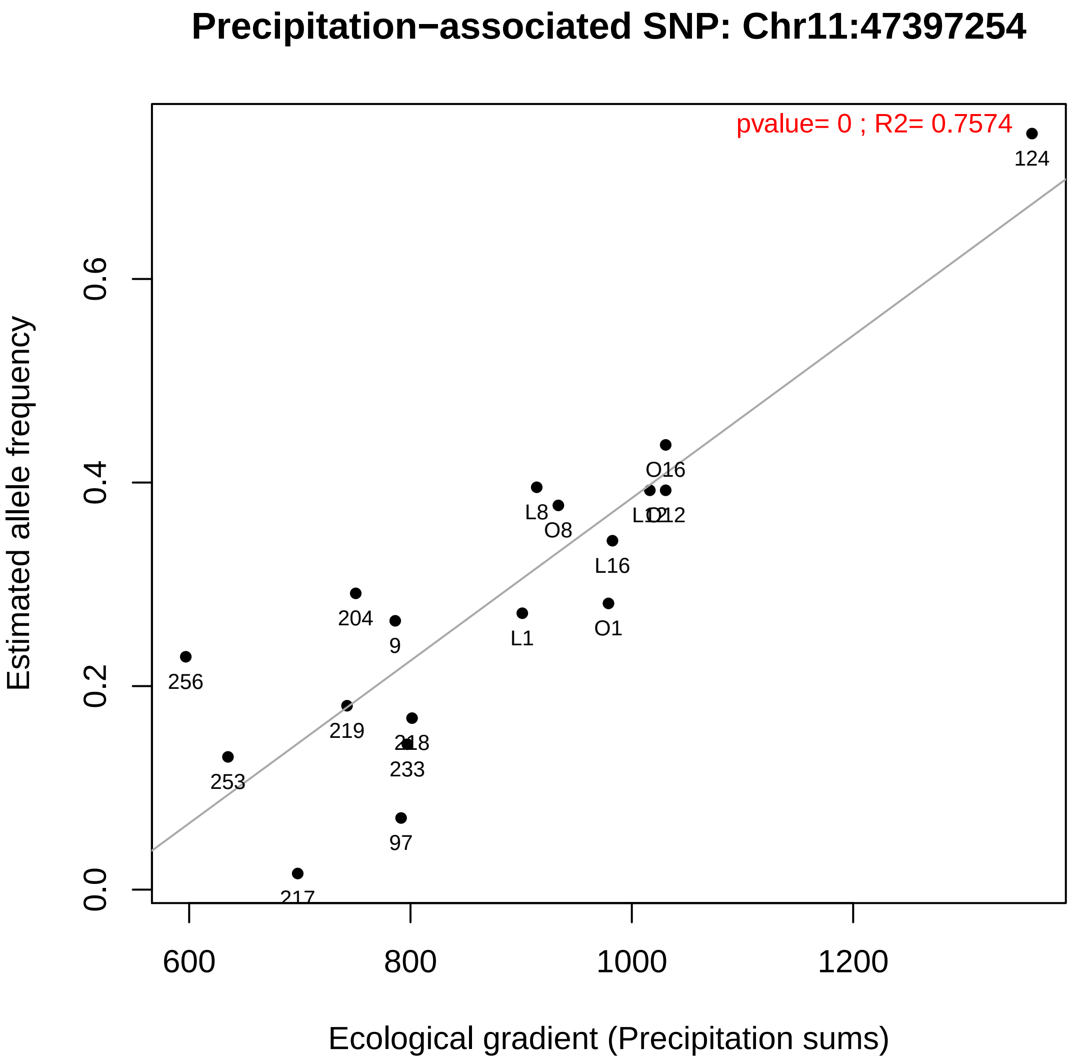
Precipitation−associated SNP: Chr11:47397254
pvalue= 0 ; R2= 0.7574
124
O16
L8
O8
L12
O12
L16
204
9
L1
O1
256
219
218
233
253
97
217
0.0
0.2
0.4
0.6
600
800
1000
1200
Estimated allele frequency
Ecological gradient (Precipitation sums)

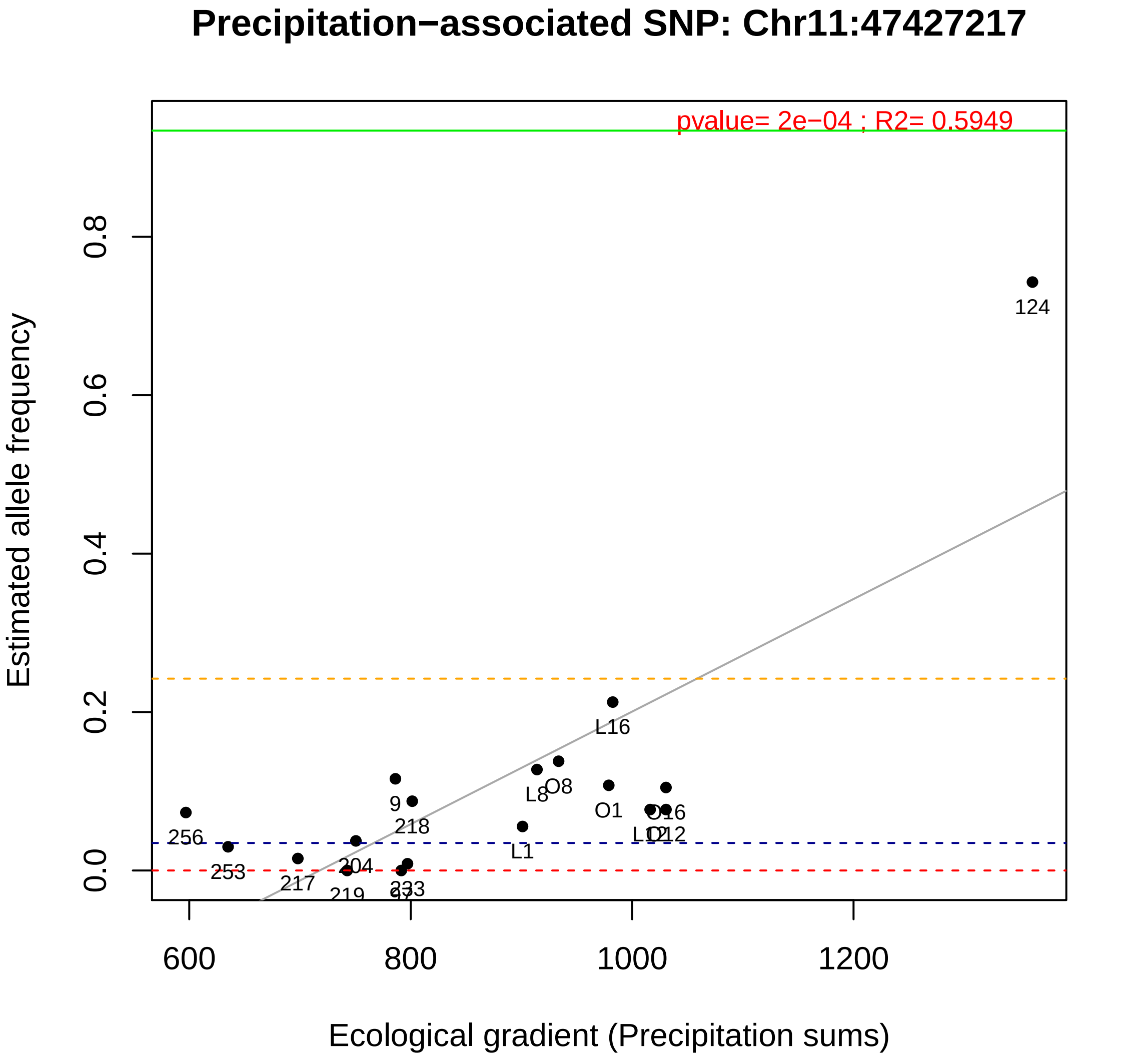

Precipitation−associated SNP: Chr11:47427217
pvalue= 2e−04 ; R2= 0.5949
Estimated allele frequency
0.0
0.2
0.4
0.6
0.8
600
800
1000
1200
Ecological gradient (Precipitation sums)
124
L16
O8
L8
O1
O16
O12
9
218
256
253
217
204
219
233
L1

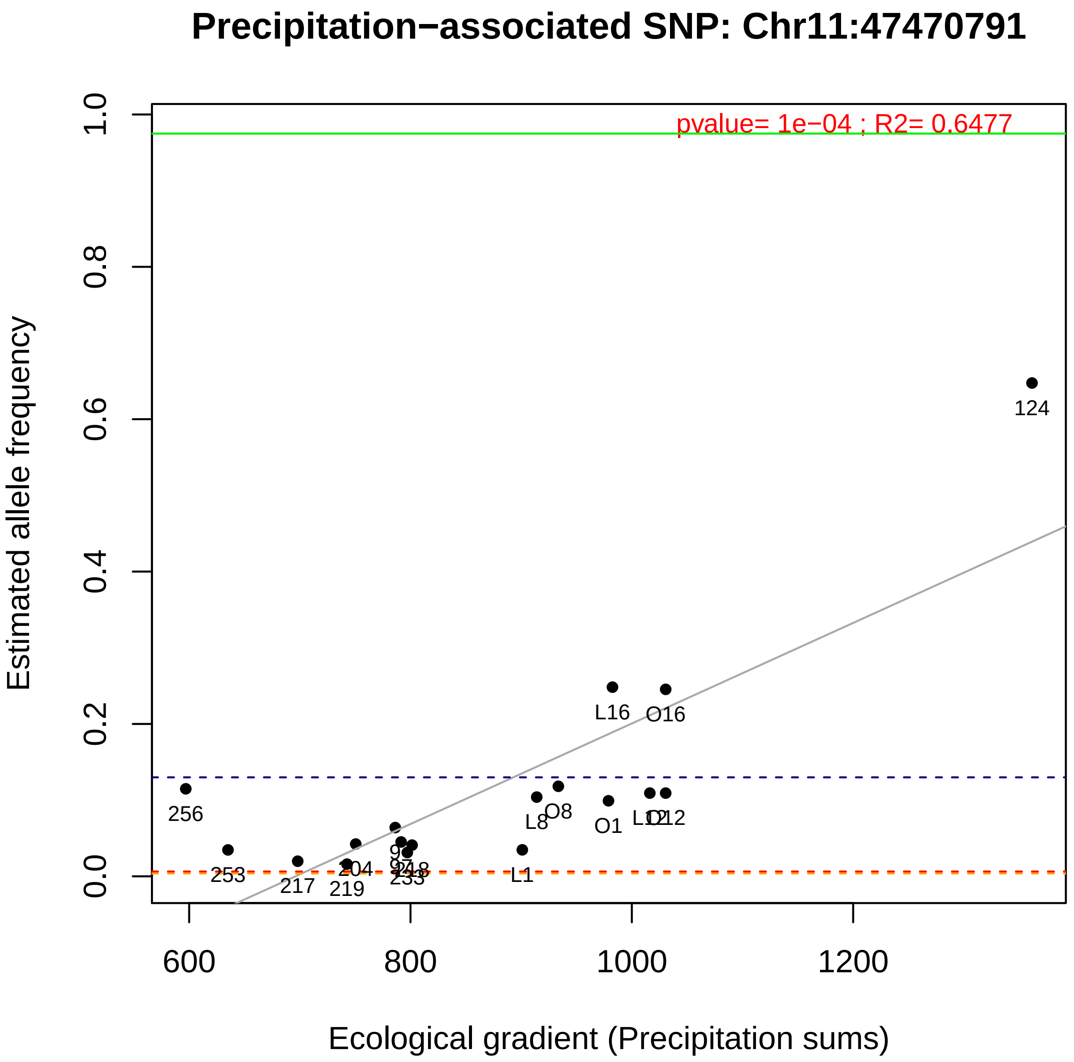

Precipitation−associated SNP: Chr11:47470791
pvalue= 1e−04 ; R2= 0.6477
Estimated allele frequency
0.0
0.2
0.4
0.6
0.8
1.0
Ecological gradient (Precipitation sums)
600
800
1000
1200
124
L16
O16
256
L8
O8
O1
253
217
219
204
L1
233

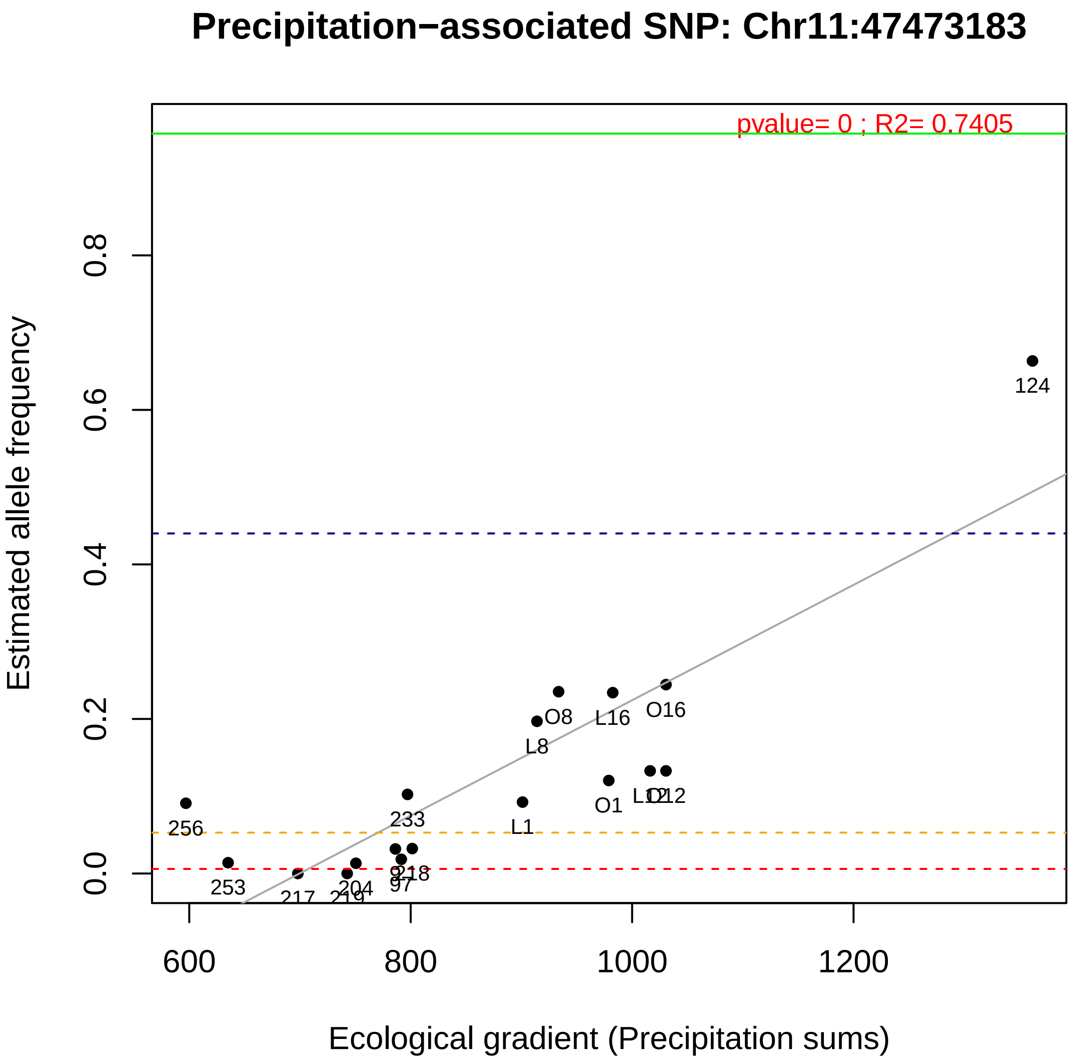
Precipitation−associated SNP: Chr11:47473183
pvalue= 0 ; R2= 0.7405
Estimated allele frequency
0.0
0.2
0.4
0.6
0.8
600
800
1000
1200
Ecological gradient (Precipitation sums)
124
O8
L16
O16
L8
O1
L12
O12
233
L1
256
253
217
219
204
218
97

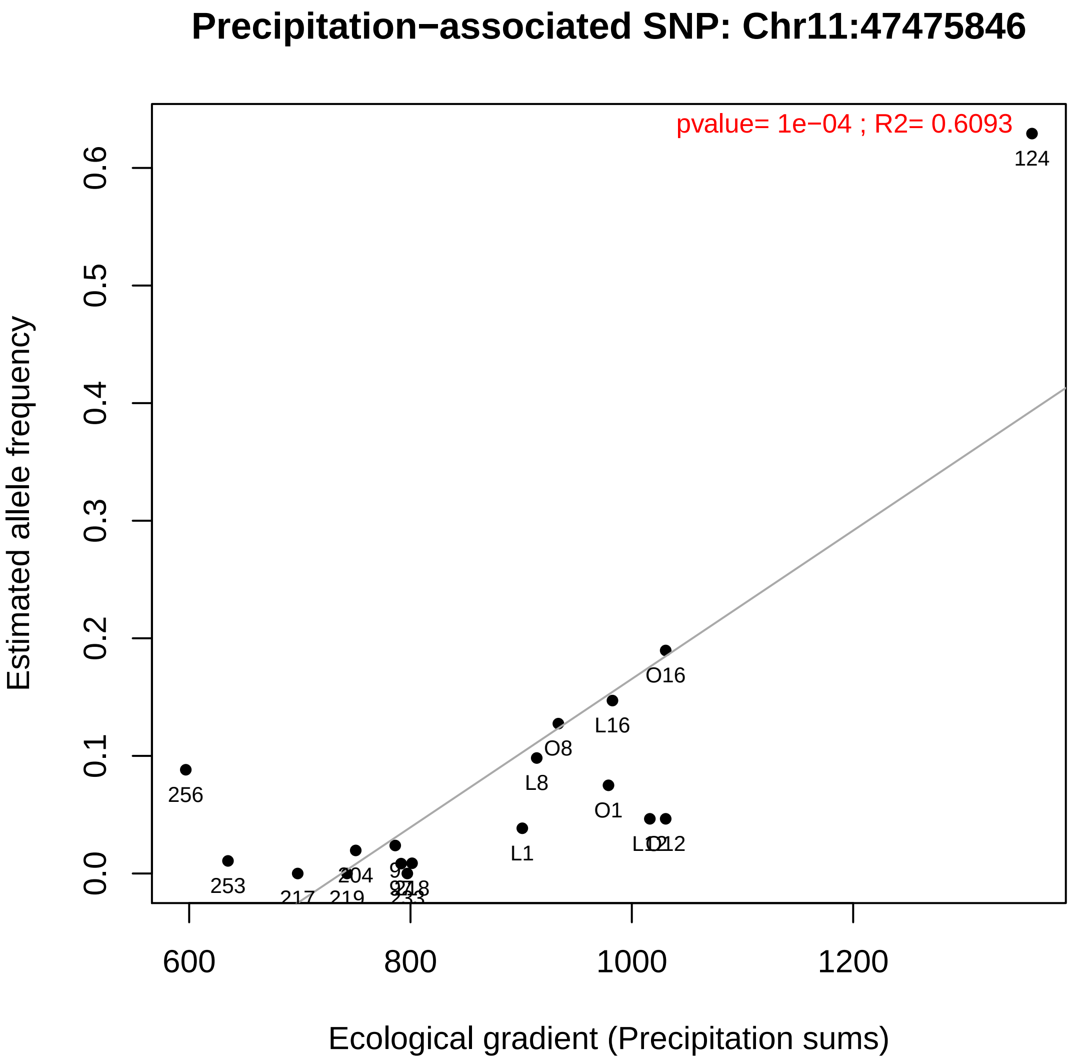

Precipitation−associated SNP: Chr11:47475846
pvalue= 1e−04 ; R2= 0.6093
Estimated allele frequency
Ecological gradient (Precipitation sums)
0.0
0.1
0.2
0.3
0.4
0.5
0.6
600
800
1000
1200
124
O16
L16
O8
L8
O1
256
L1
253
204
217
219

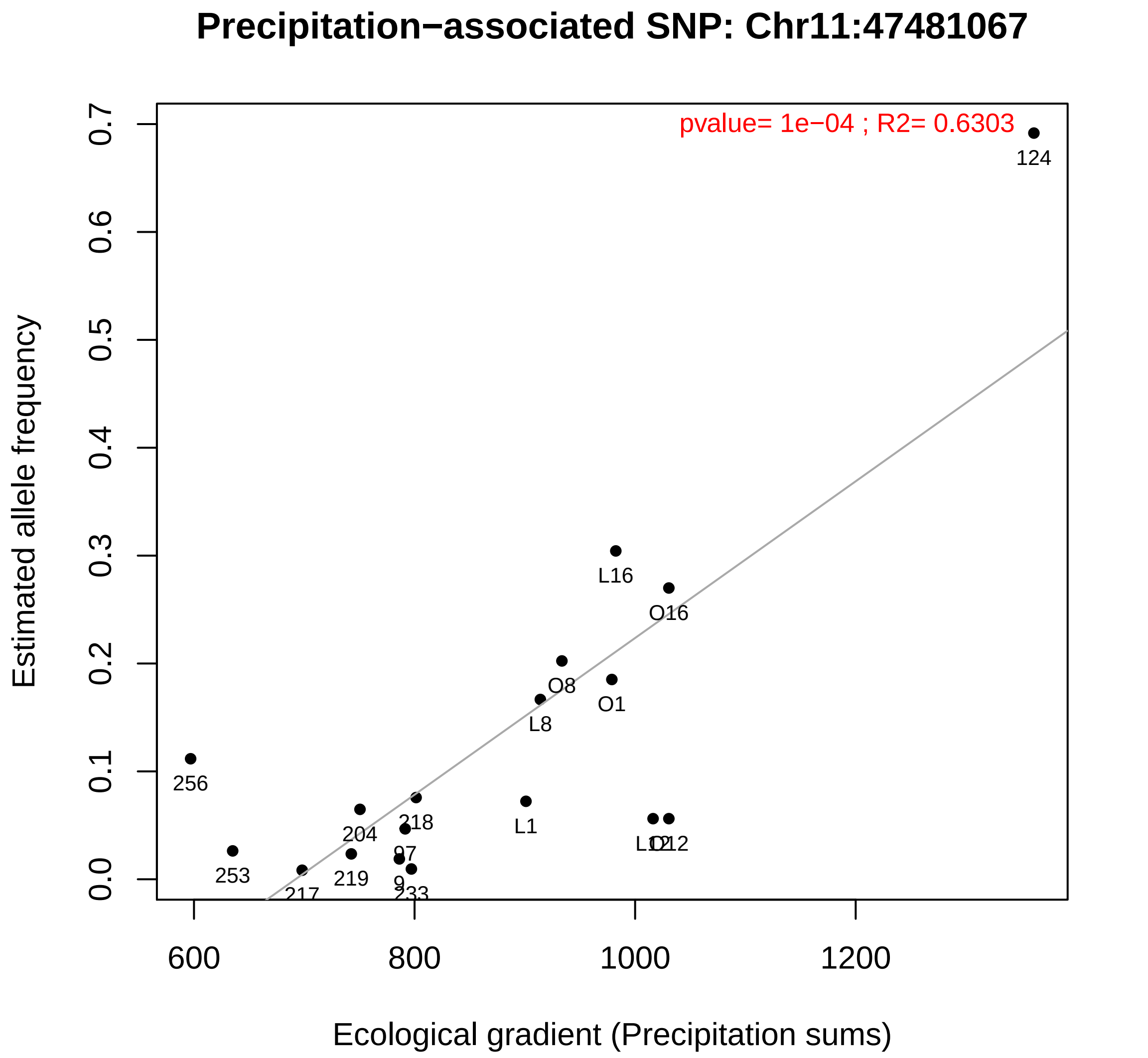
Precipitation−associated SNP: Chr11:47481067
pvalue= 1e−04 ; R2= 0.6303
Estimated allele frequency
Ecological gradient (Precipitation sums)
0.0
0.1
0.2
0.3
0.4
0.5
0.6
0.7
600
800
1000
1200
124
L16
O16
O8
O1
L8
256
L1
218
204
97
253
219
9
217
233

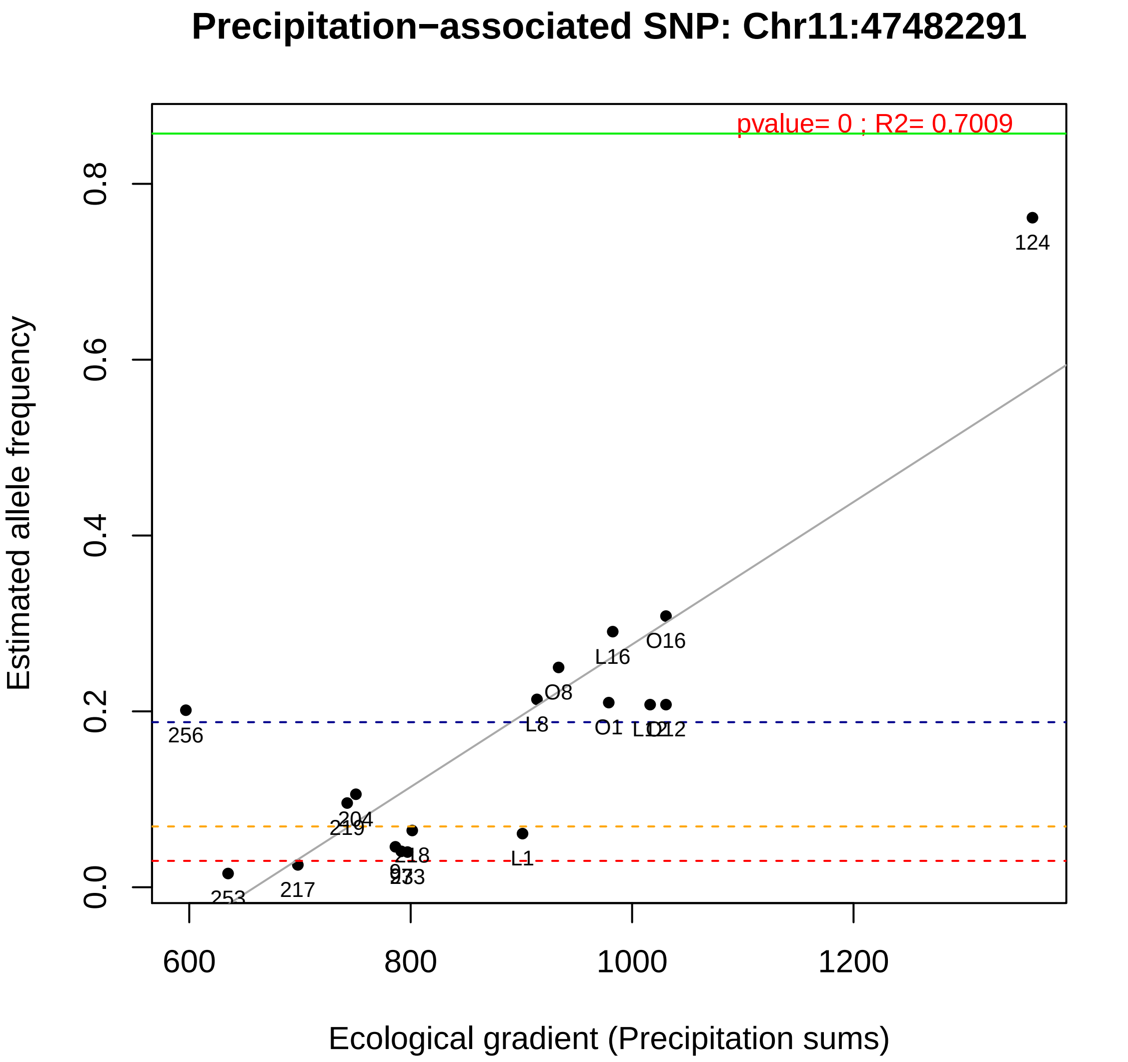

Precipitation−associated SNP: Chr11:47482291
pvalue= 0 ; R2= 0.7009
Estimated allele frequency
0.0
0.2
0.4
0.6
0.8
600
800
1000
1200
Ecological gradient (Precipitation sums)
124
O16
L16
O8
L8
O1
L12
O12
256
204
219
218
L1
217
253

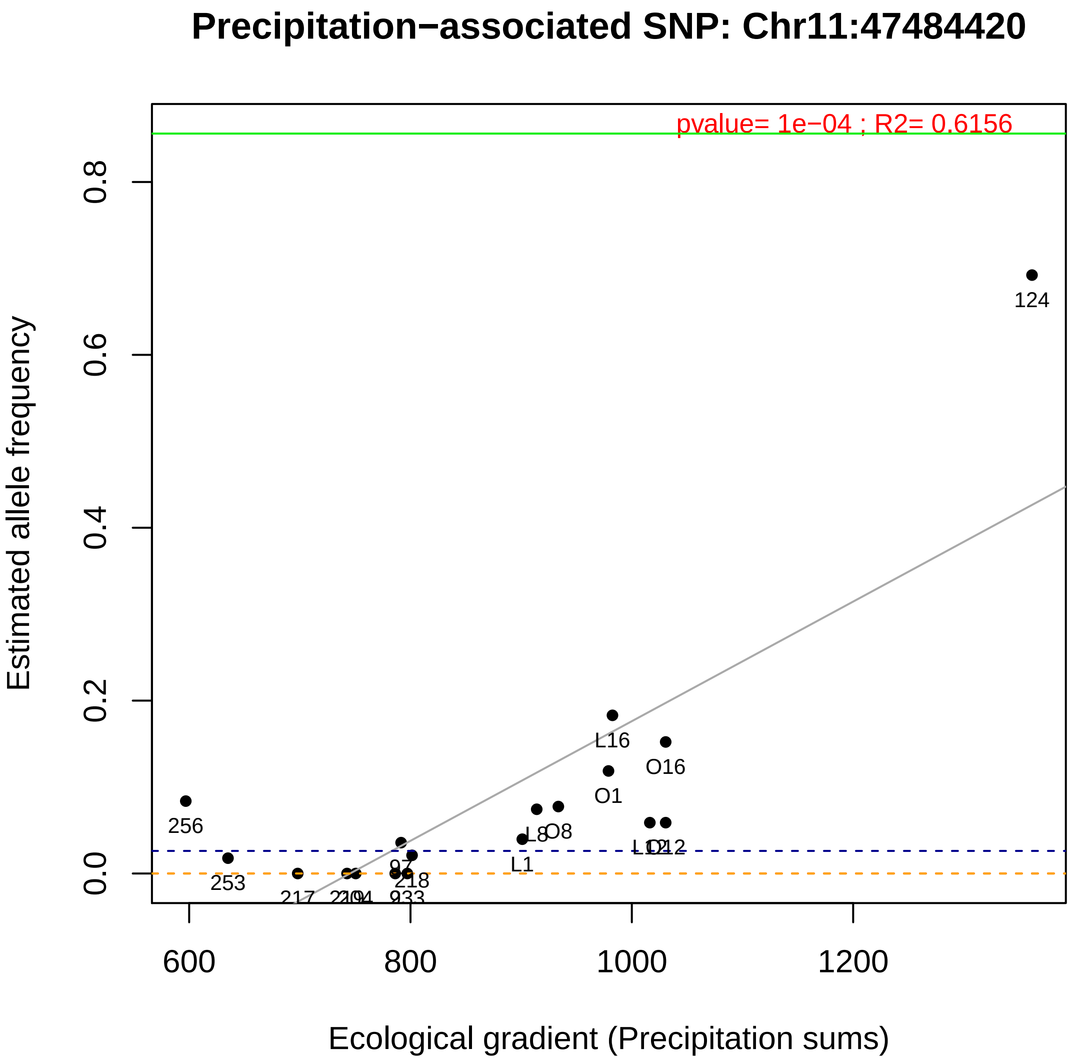

Precipitation−associated SNP: Chr11:47484420
pvalue= 1e−04 ; R2= 0.6156
Estimated allele frequency
0.0
0.2
0.4
0.6
0.8
600
800
1000
1200
Ecological gradient (Precipitation sums)
124
L16
O16
O1
256
253
L8
O8
L1
218

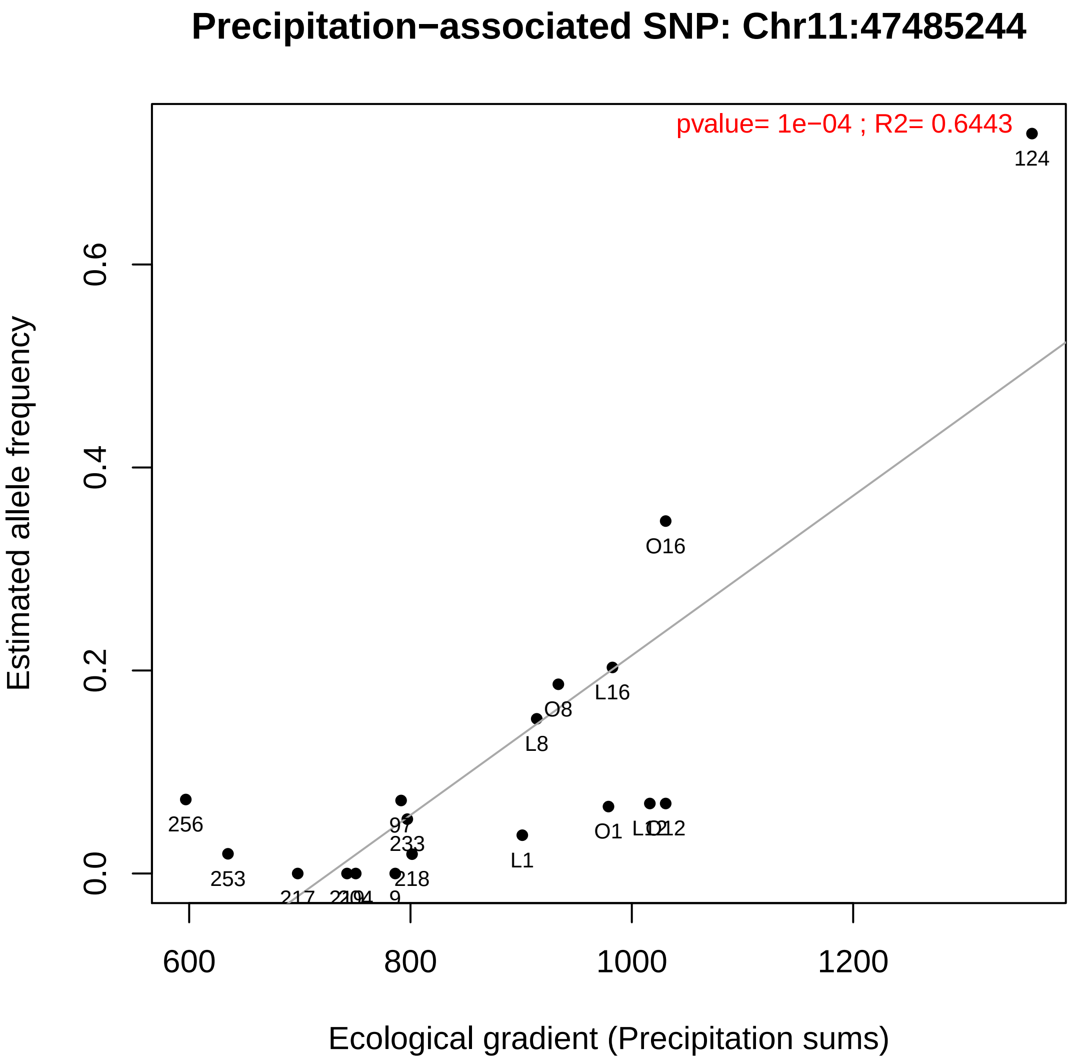
Precipitation−associated SNP: Chr11:47485244
pvalue= 1e−04 ; R2= 0.6443
Estimated allele frequency
0.0
0.2
0.4
0.6
600
800
1000
1200
Ecological gradient (Precipitation sums)
124
O16
L16
O8
L8
256
253
217
97
233
218
9
L1
O1
O12

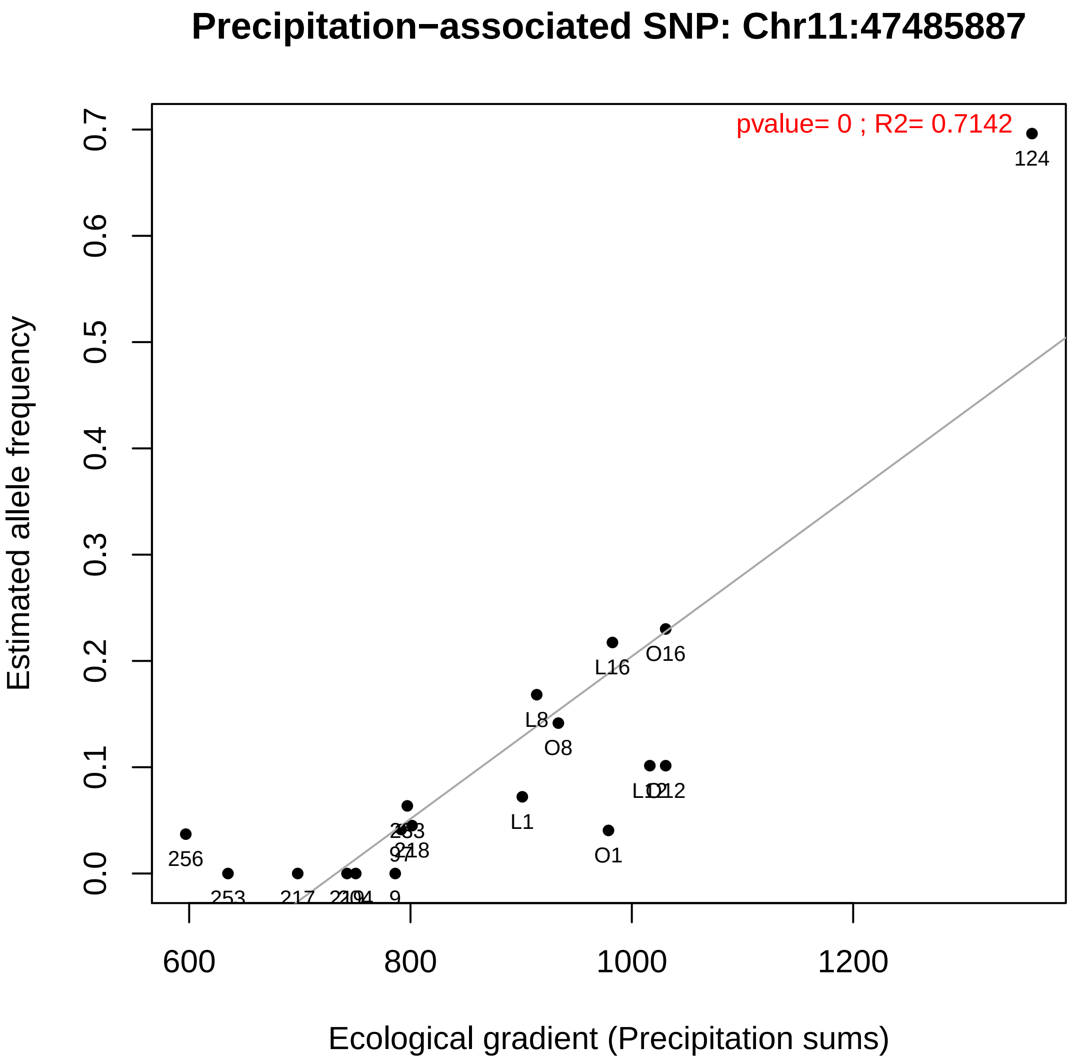

Precipitation−associated SNP: Chr11:47485887
pvalue= 0 ; R2= 0.7142
Estimated allele frequency
0.0
0.1
0.2
0.3
0.4
0.5
0.6
0.7
600
800
1000
1200
Ecological gradient (Precipitation sums)
124
O16
L16
L8
O8
L1
O1
256
253
217
9

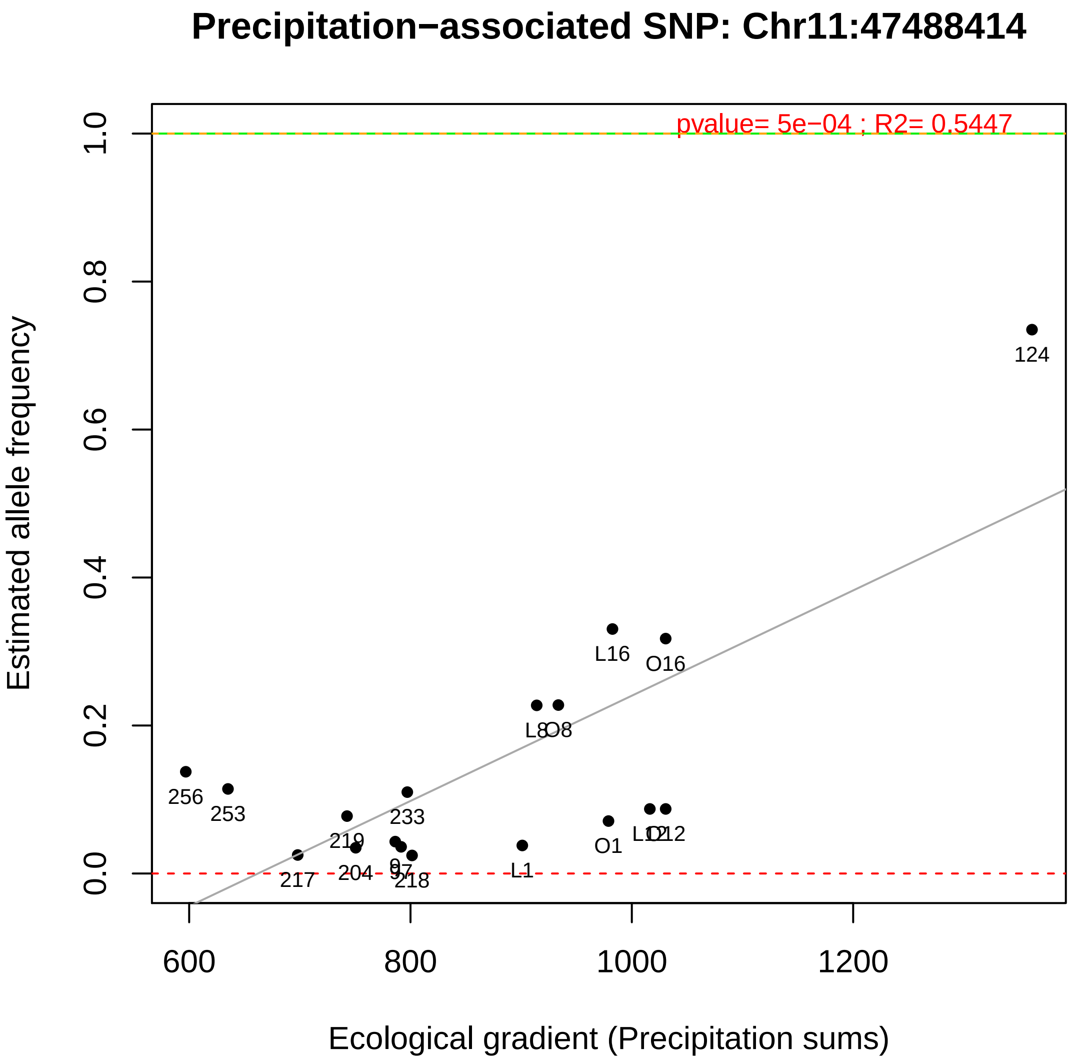

Precipitation−associated SNP: Chr11:47488414
pvalue= 5e−04 ; R2= 0.5447
Estimated allele frequency
0.0
0.2
0.4
0.6
0.8
1.0
600
800
1000
1200
Ecological gradient (Precipitation sums)
124
L16
O16
L8
O8
256
253
233
219
204
217
218
L1
O1
O12

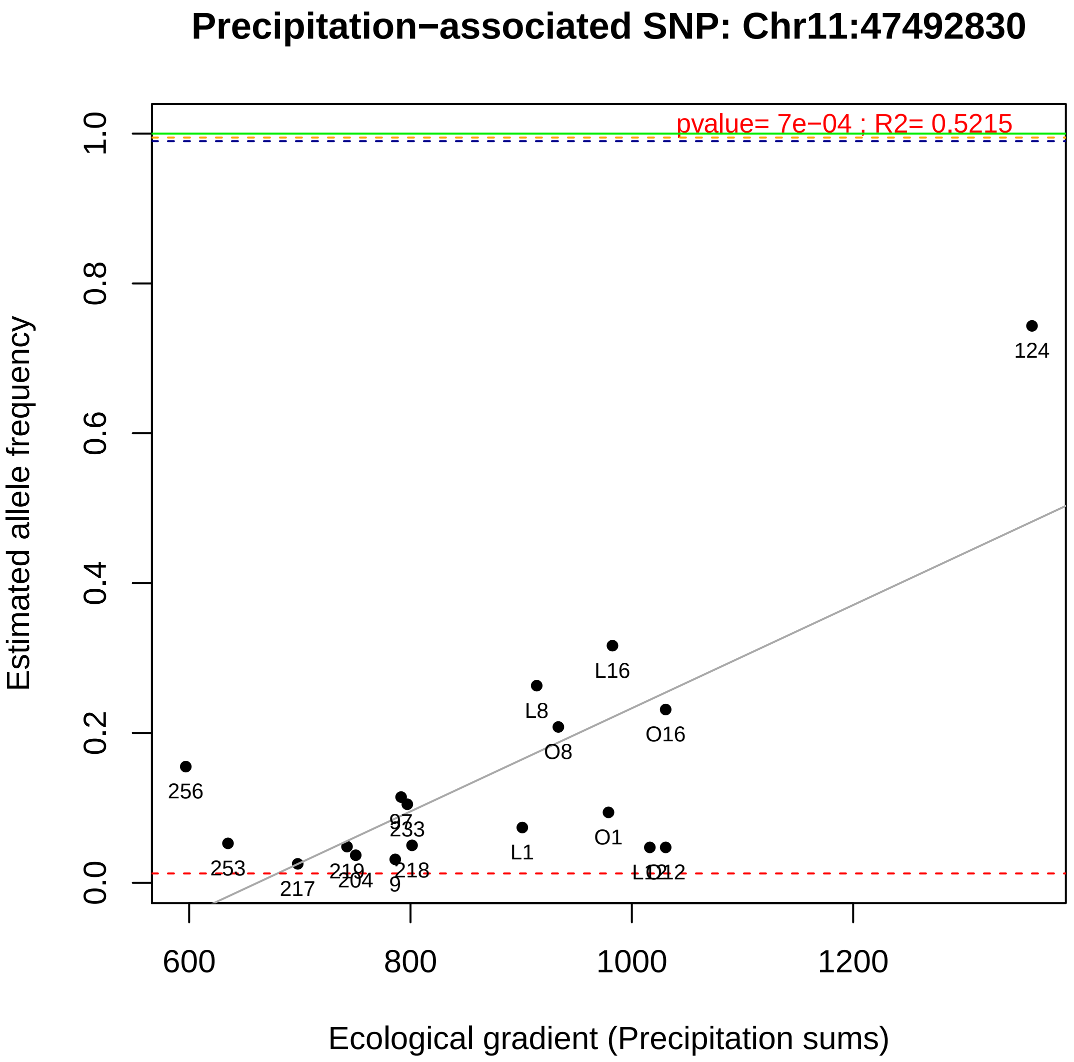

Precipitation−associated SNP: Chr11:47492830
pvalue= 7e−04 ; R2= 0.5215
Estimated allele frequency
0.0
0.2
0.4
0.6
0.8
1.0
600
800
1000
1200
Ecological gradient (Precipitation sums)
124
L16
L8
O8
O16
256
97
233
L1
O1
253
219
204
218
217
9

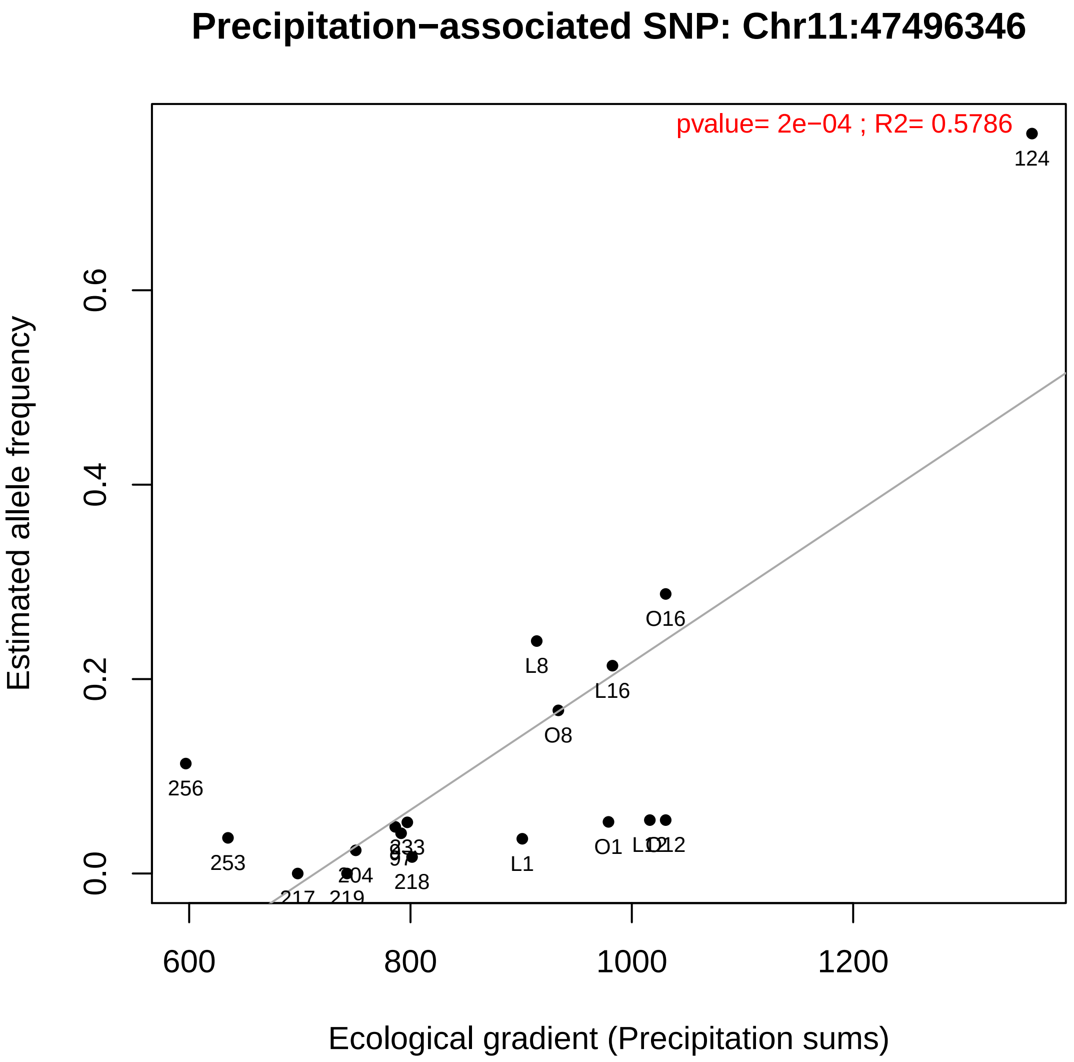
Precipitation−associated SNP: Chr11:47496346
pvalue= 2e−04 ; R2= 0.5786
124
O16
L8
L16
O8
256
253
217
219
204
233
97
218
L1
O1
L12
O12
0.0
0.2
0.4
0.6
600
800
1000
1200
Estimated allele frequency
Ecological gradient (Precipitation sums)

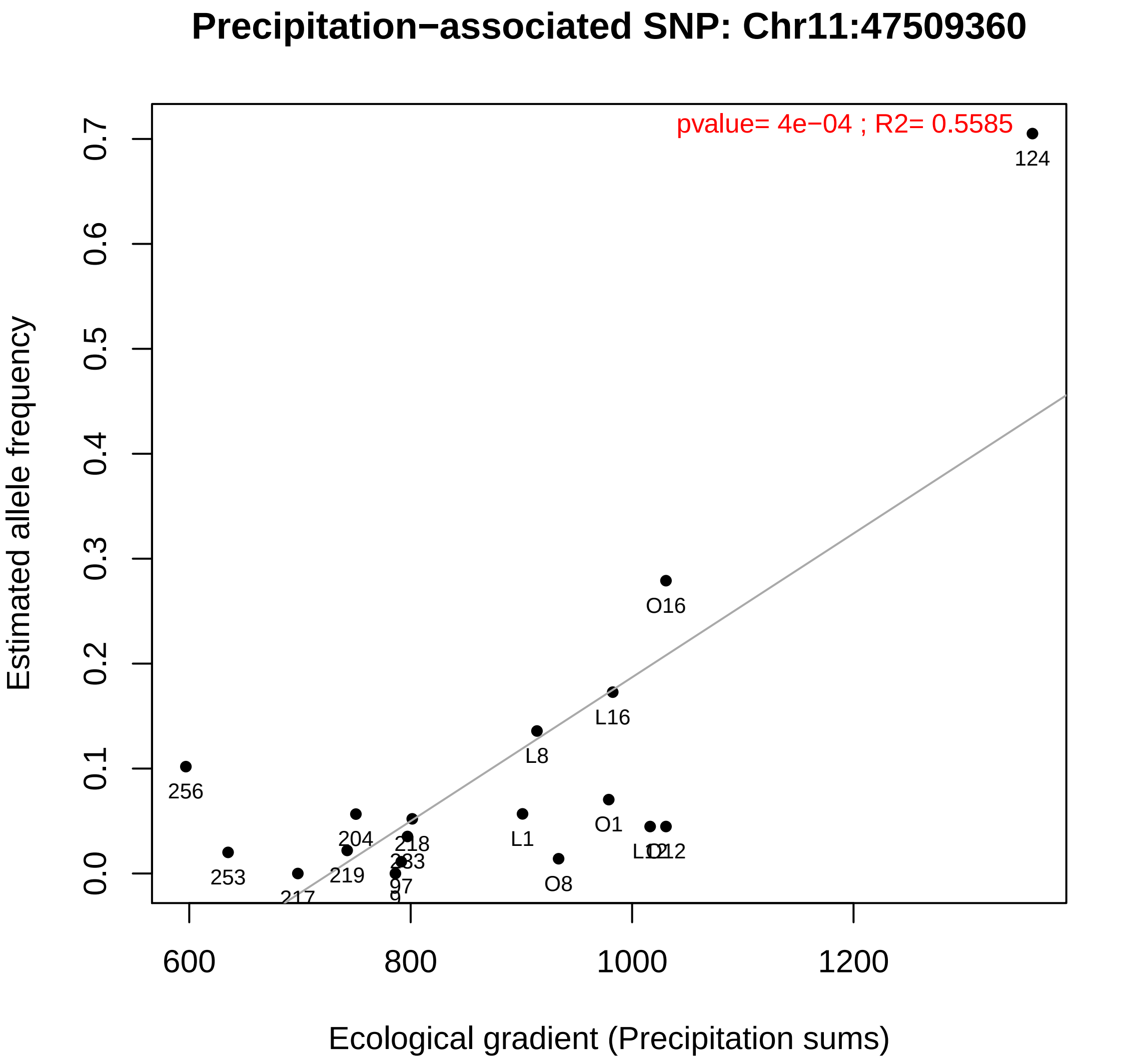

Precipitation−associated SNP: Chr11:47509360
pvalue= 4e−04 ; R2= 0.5585
Estimated allele frequency
Ecological gradient (Precipitation sums)
0.0
0.1
0.2
0.3
0.4
0.5
0.6
0.7
600
800
1000
1200
124
O16
L16
L8
256
O1
L1
204
218
233
219
253
97
217
O8

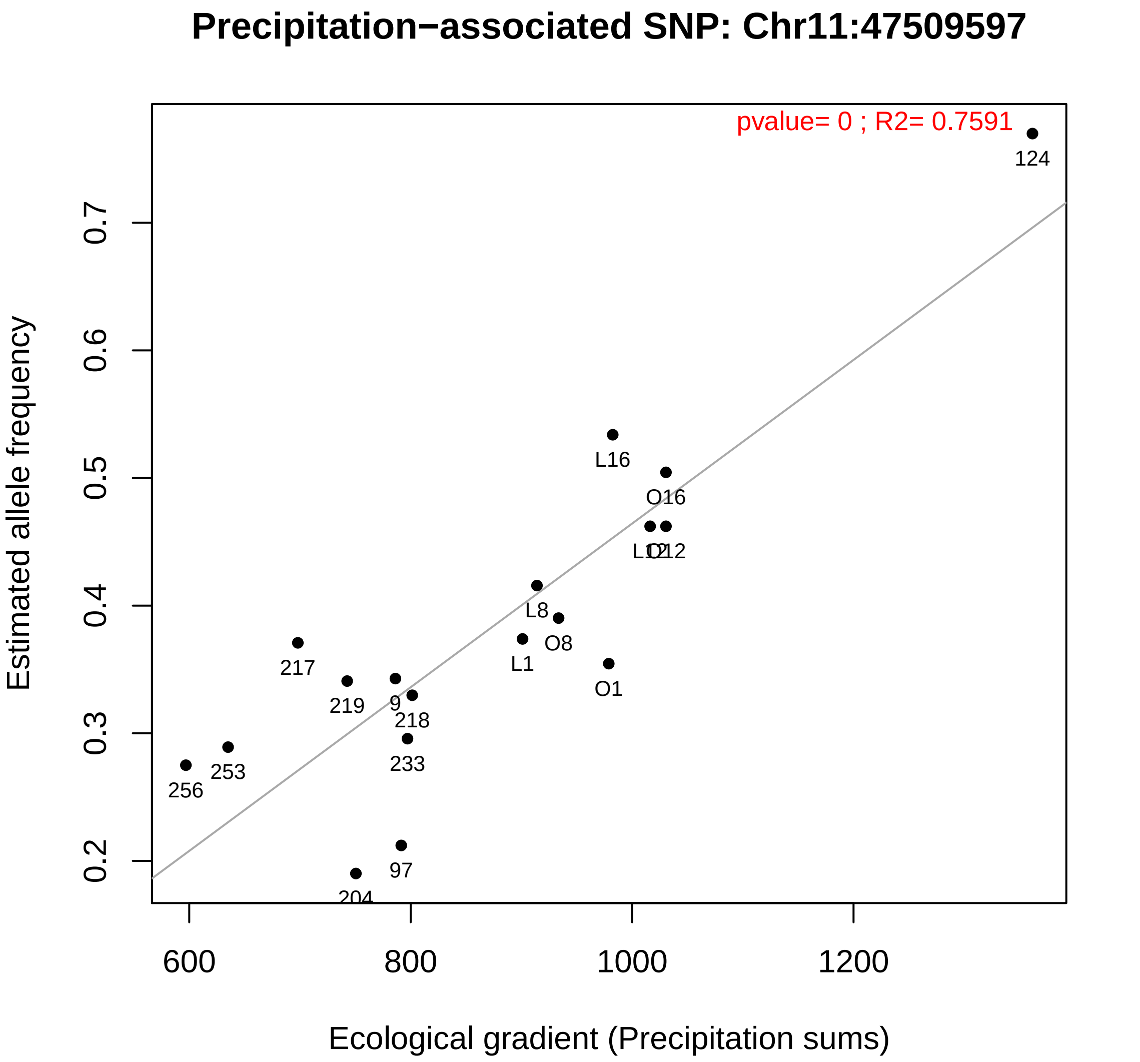

Precipitation−associated SNP: Chr11:47509597
pvalue= 0 ; R2= 0.7591
124
L16
O16
L12
O12
L8
O8
L1
O1
217
219
9
218
233
253
256
97
204
0.2
0.3
0.4
0.5
0.6
0.7
600
800
1000
1200
Estimated allele frequency
Ecological gradient (Precipitation sums)

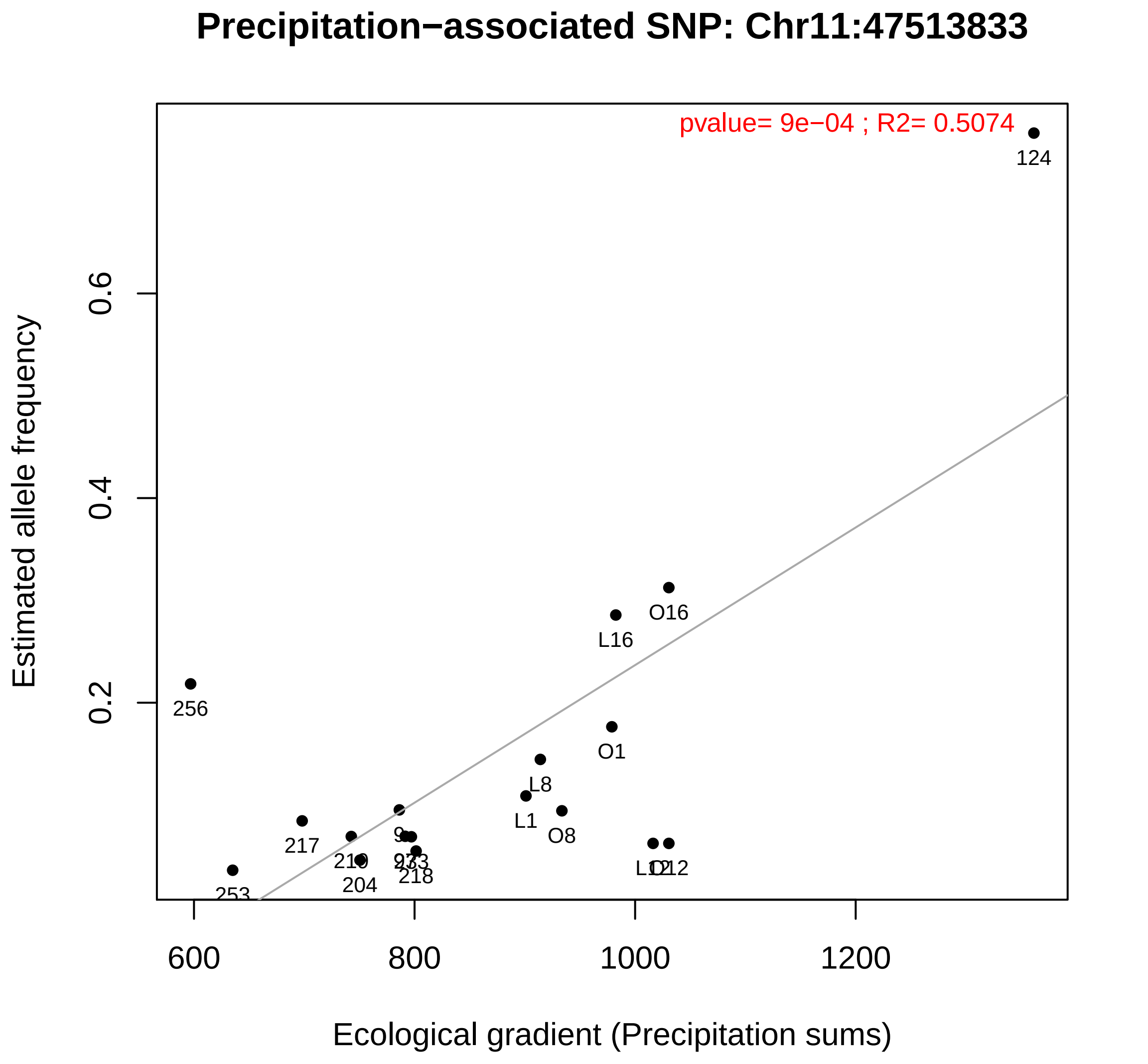

Precipitation−associated SNP: Chr11:47513833
pvalue= 9e−04 ; R2= 0.5074
124
O16
L16
256
O1
L8
L1
O8
217
210
204
253
218
L12
O12
0.2
0.4
0.6
600
800
1000
1200
Estimated allele frequency
Ecological gradient (Precipitation sums)

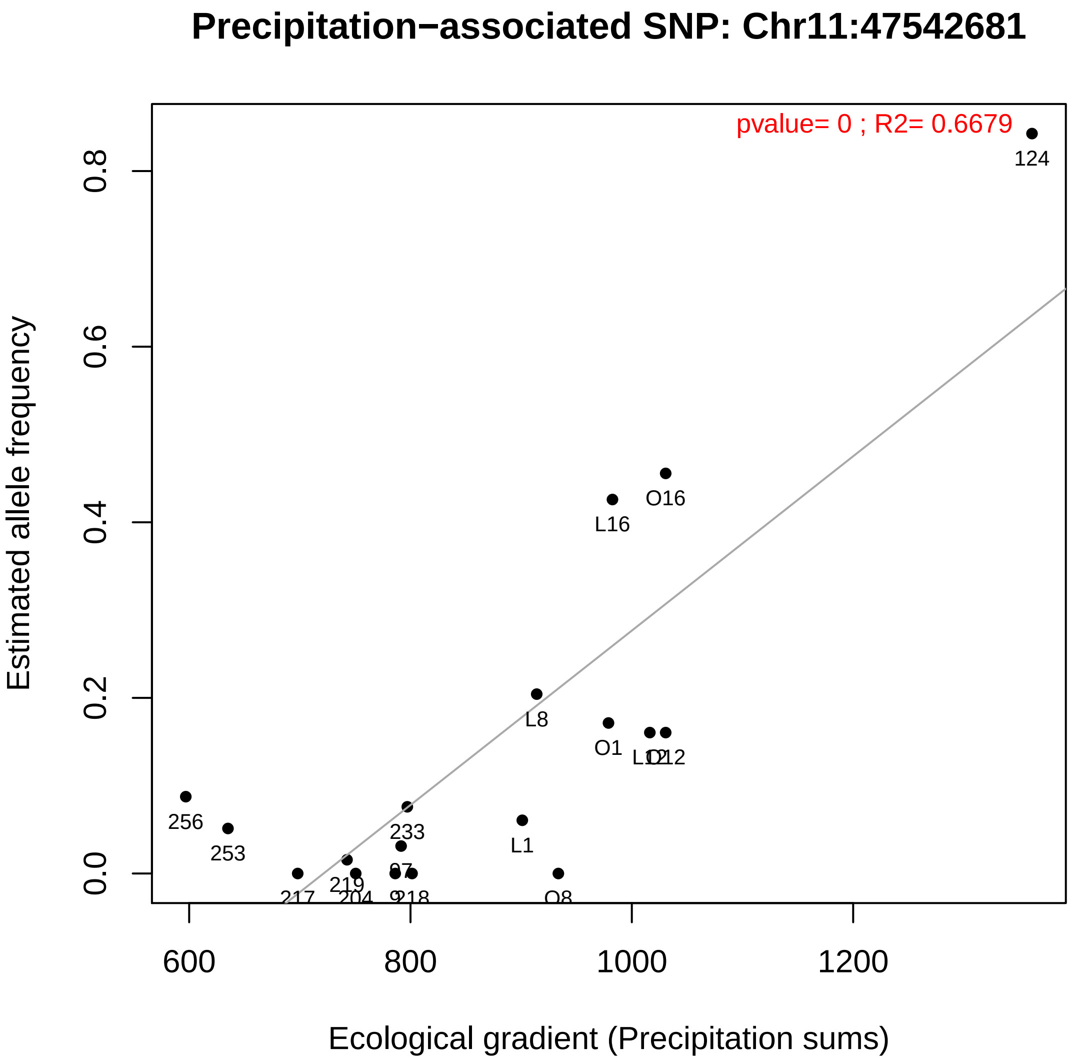

Precipitation−associated SNP: Chr11:47542681
pvalue= 0 ; R2= 0.6679
124
O16
L16
L8
O1
L12
O12
256
253
233
L1
219
217
204
218
O8
Estimated allele frequency
0.0
0.2
0.4
0.6
0.8
600
800
1000
1200
Ecological gradient (Precipitation sums)

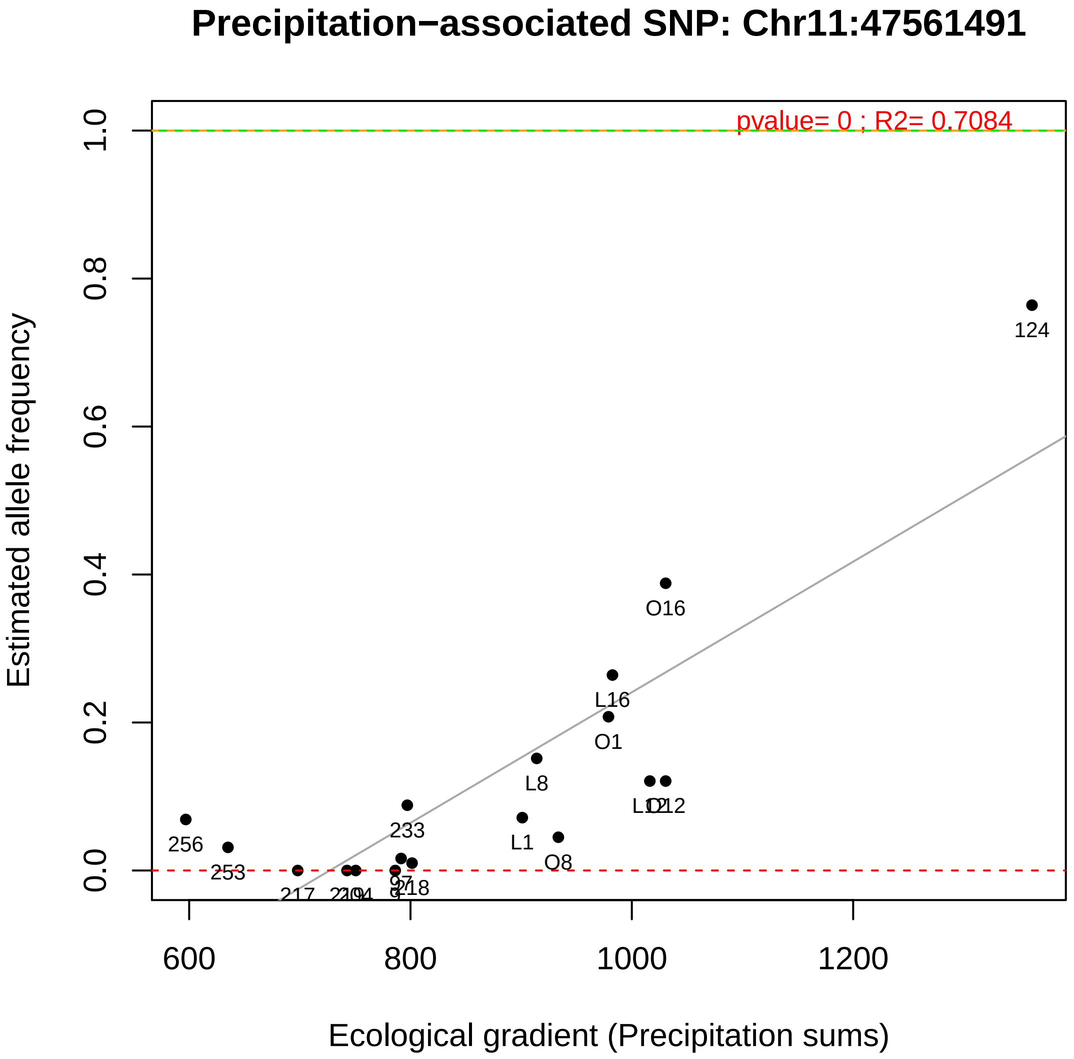

Precipitation−associated SNP: Chr11:47561491
pvalue= 0 ; R2= 0.7084
Estimated allele frequency
Ecological gradient (Precipitation sums)
0.0
0.2
0.4
0.6
0.8
1.0
600
800
1000
1200
124
O16
L16
O1
L8
233
L1
O8
256
253
217
218

**Precipitation−associated SNP: Chr11:47561494**

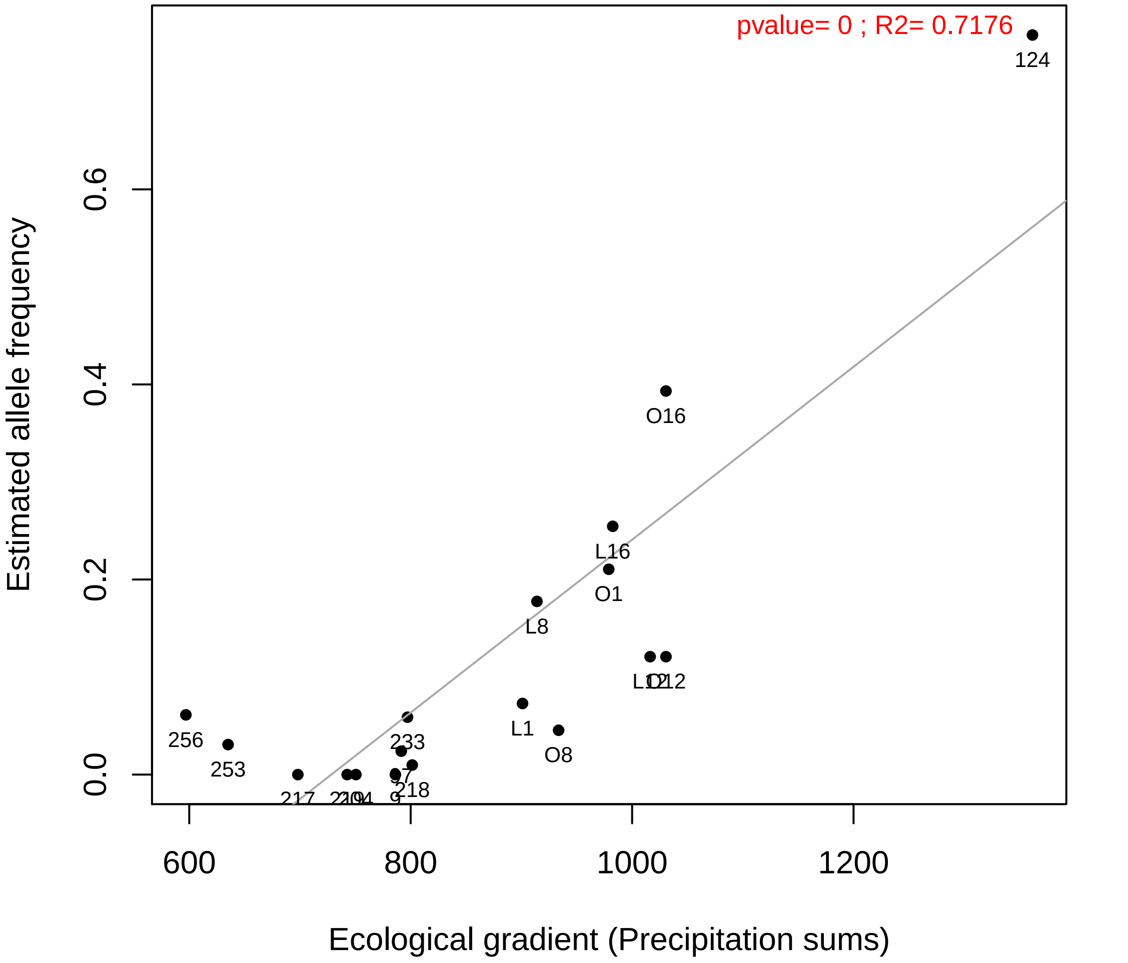

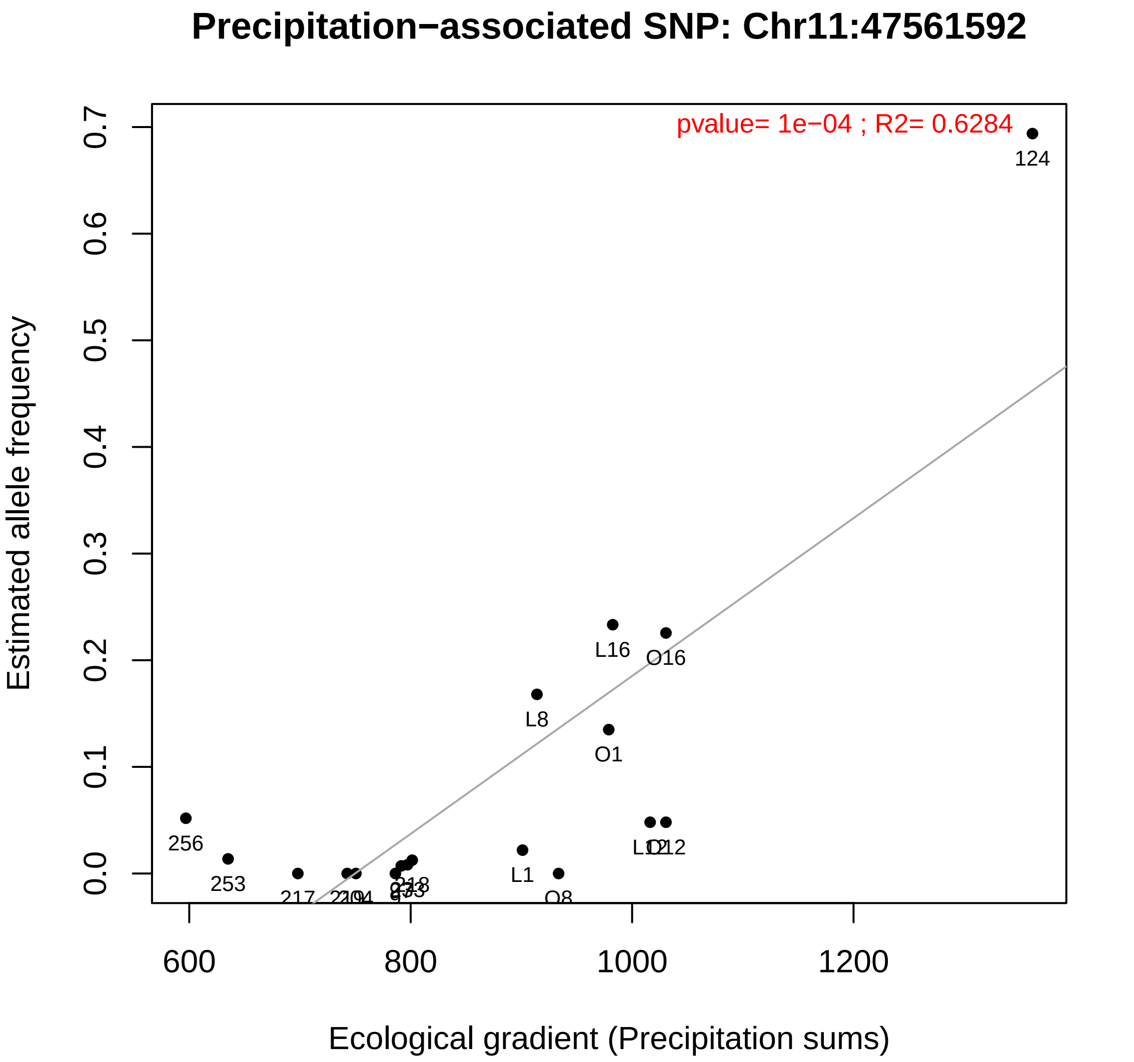

Precipitation−associated SNP: Chr11:47561592
pvalue= 1e−04 ; R2= 0.6284
Estimated allele frequency
Ecological gradient (Precipitation sums)
0.0
0.1
0.2
0.3
0.4
0.5
0.6
0.7
600
800
1000
1200
124
L16
O16
L8
O1
256
253
217
L1
O8

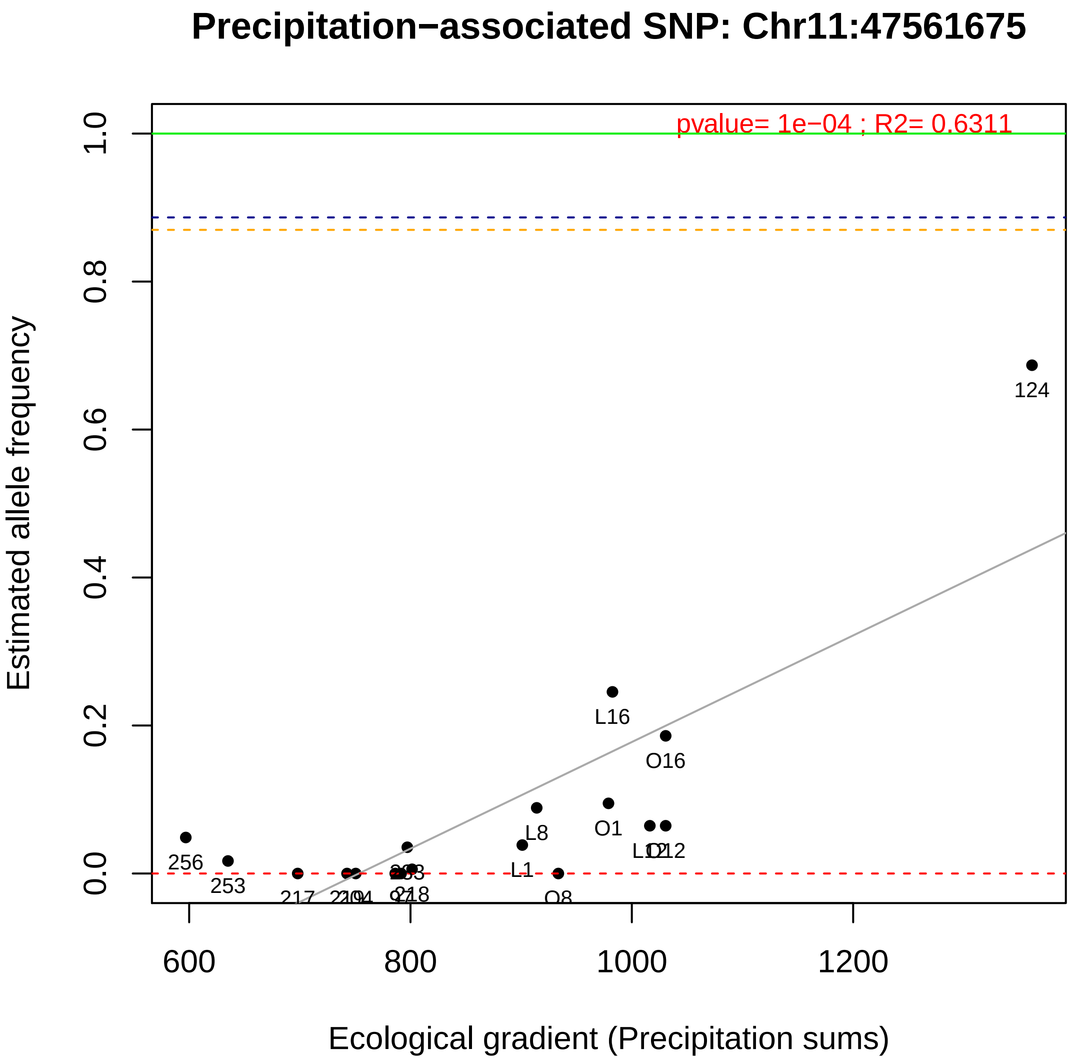

Precipitation−associated SNP: Chr11:47561675
pvalue= 1e−04 ; R2= 0.6311
Estimated allele frequency
0.0
0.2
0.4
0.6
0.8
1.0
600
800
1000
1200
Ecological gradient (Precipitation sums)
124
L16
O16
L8
O1
L1
O8
256
253
217

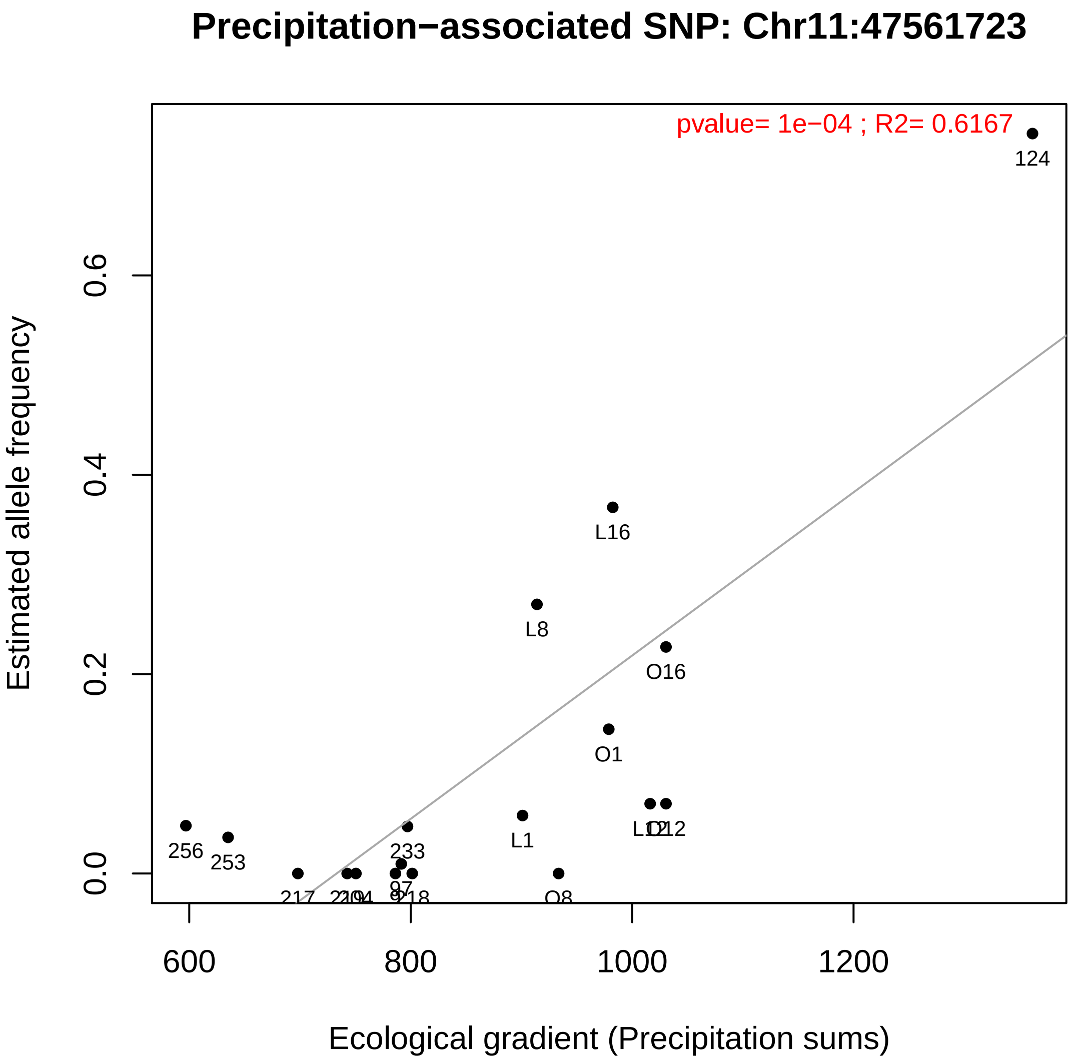
Precipitation−associated SNP: Chr11:47561723
pvalue= 1e−04 ; R2= 0.6167
124
L16
L8
O16
O1
L1
256
253
233
97
O8
217
218
Estimated allele frequency
0.0
0.2
0.4
0.6
600
800
1000
1200
Ecological gradient (Precipitation sums)

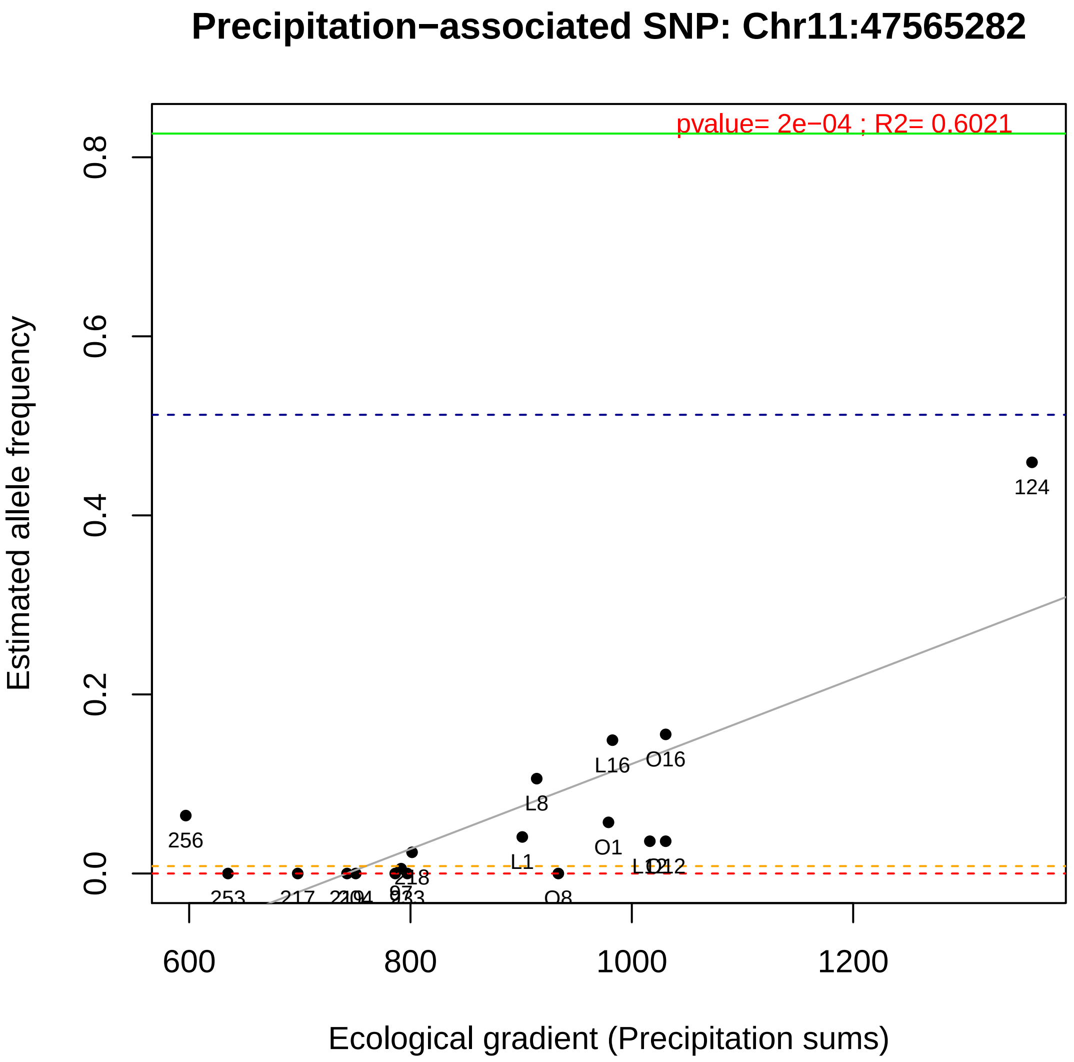
Precipitation−associated SNP: Chr11:47565282
pvalue= 2e−04 ; R2= 0.6021
Estimated allele frequency
0.0
0.2
0.4
0.6
0.8
Ecological gradient (Precipitation sums)
600
800
1000
1200
124
O16
L16
L8
256
O1
L1
218
253
217
O8

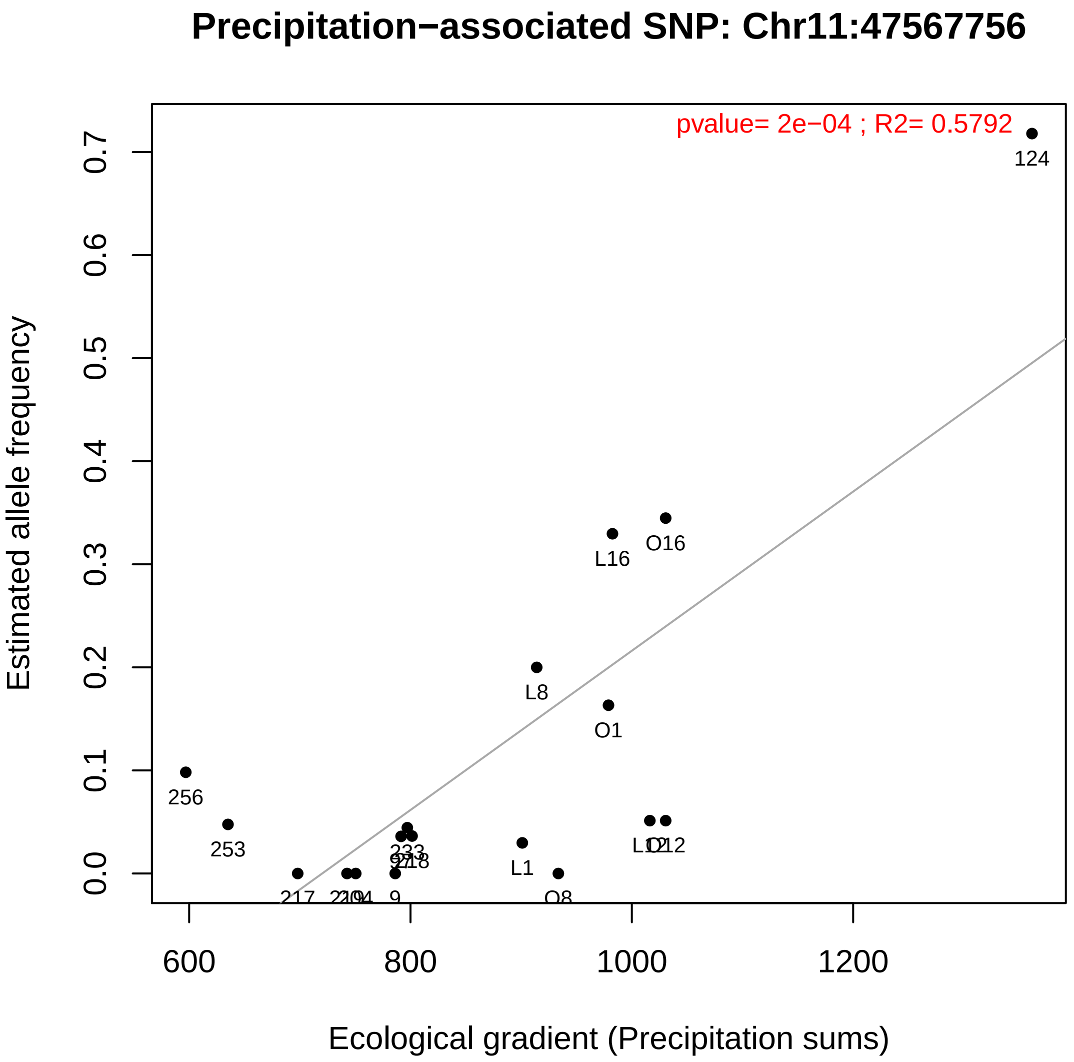

Precipitation−associated SNP: Chr11:47567756
pvalue= 2e−04 ; R2= 0.5792
Estimated allele frequency
0.0
0.1
0.2
0.3
0.4
0.5
0.6
0.7
600
800
1000
1200
Ecological gradient (Precipitation sums)
124
O16
L16
L8
O1
256
253
L1
217
O8

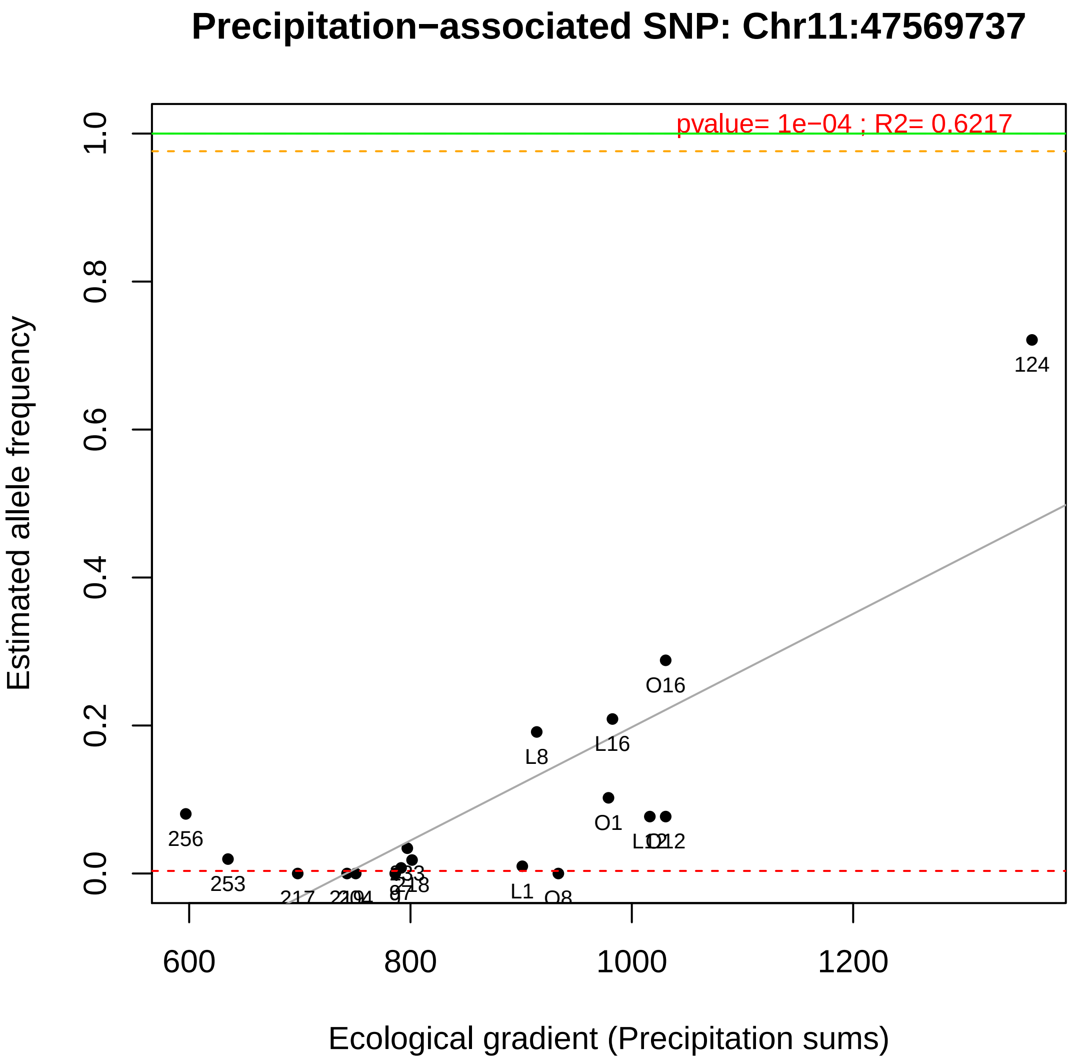
Precipitation−associated SNP: Chr11:47569737
pvalue= 1e−04 ; R2= 0.6217
Estimated allele frequency
0.0
0.2
0.4
0.6
0.8
1.0
600
800
1000
1200
Ecological gradient (Precipitation sums)
124
O16
L16
L8
O1
256
253
L1
218

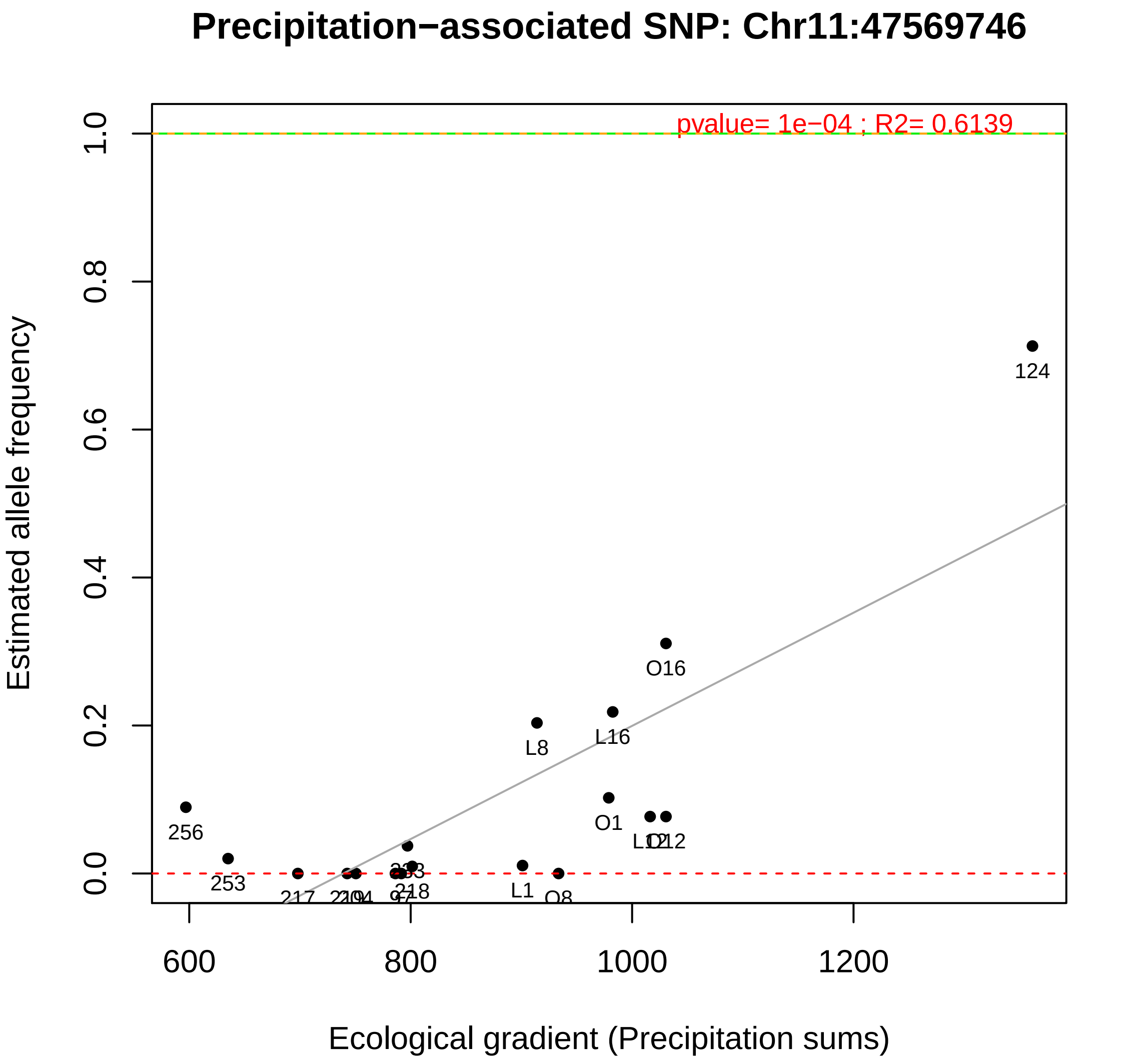

Precipitation−associated SNP: Chr11:47569746
pvalue= 1e−04 ; R2= 0.6139
Estimated allele frequency
0.0
0.2
0.4
0.6
0.8
1.0
600
800
1000
1200
Ecological gradient (Precipitation sums)
124
O16
L8
L16
O1
256
253
217
218
L1
O8

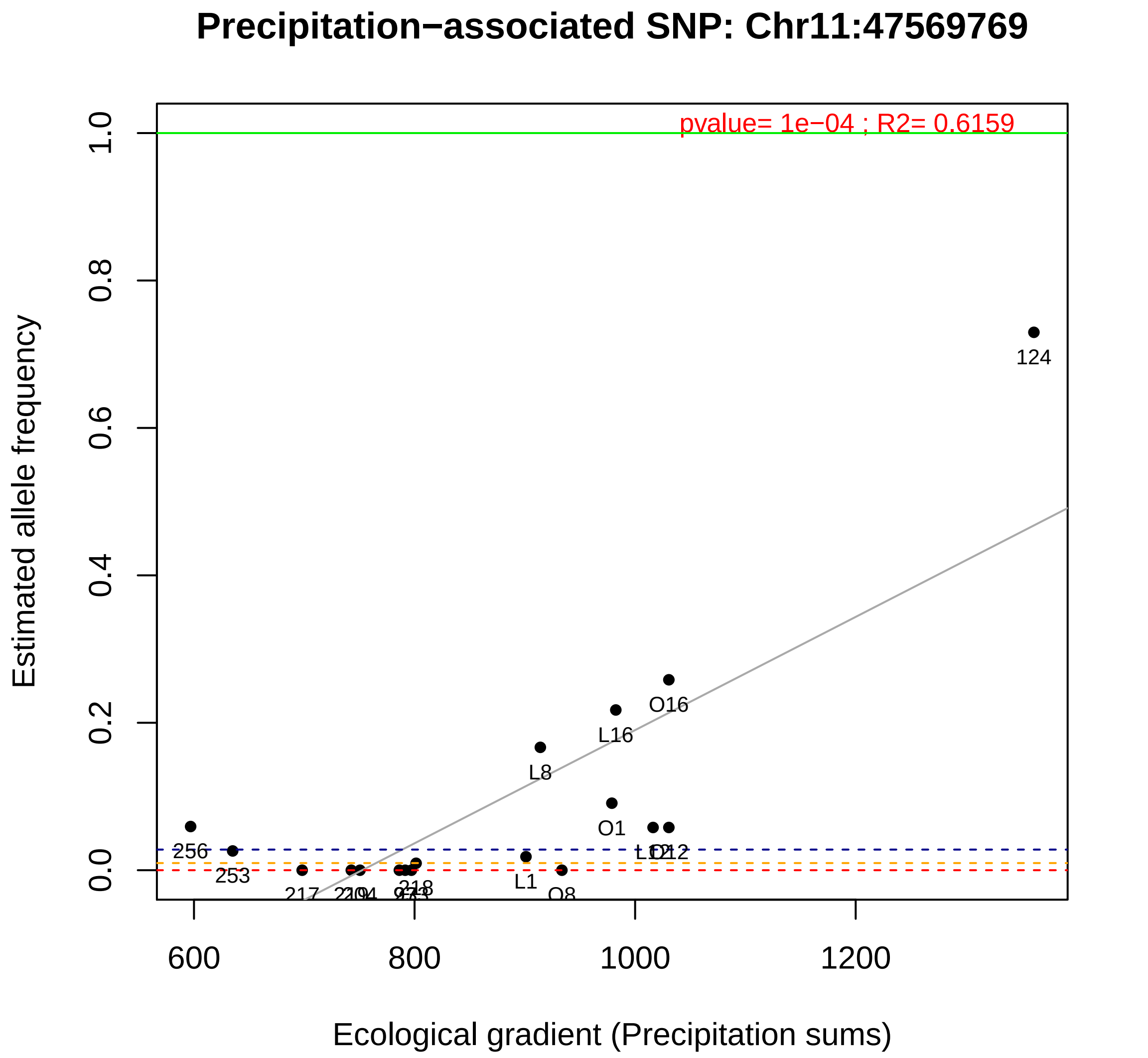

Precipitation−associated SNP: Chr11:47569769
pvalue= 1e−04 ; R2= 0.6159
Estimated allele frequency
Ecological gradient (Precipitation sums)
0.0
0.2
0.4
0.6
0.8
1.0
600
800
1000
1200
124
O16
L16
L8
O1
256
253
217
218
L1
O8

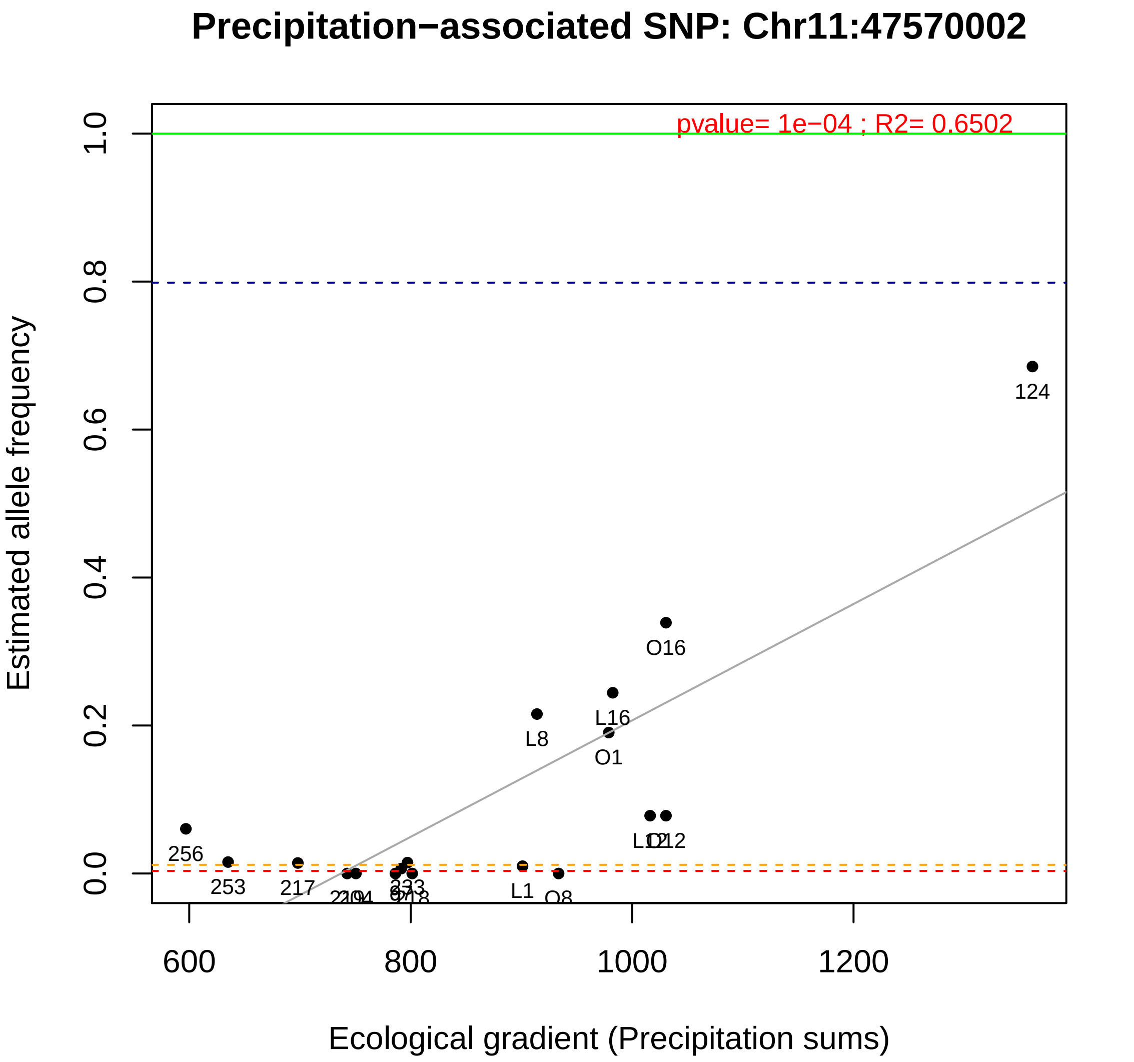
Precipitation−associated SNP: Chr11:47570002
pvalue= 1e−04 ; R2= 0.6502
Estimated allele frequency
0.0
0.2
0.4
0.6
0.8
1.0
600
800
1000
1200
Ecological gradient (Precipitation sums)
124
O16
L16
L8
O1
256
253
217
L1
O8

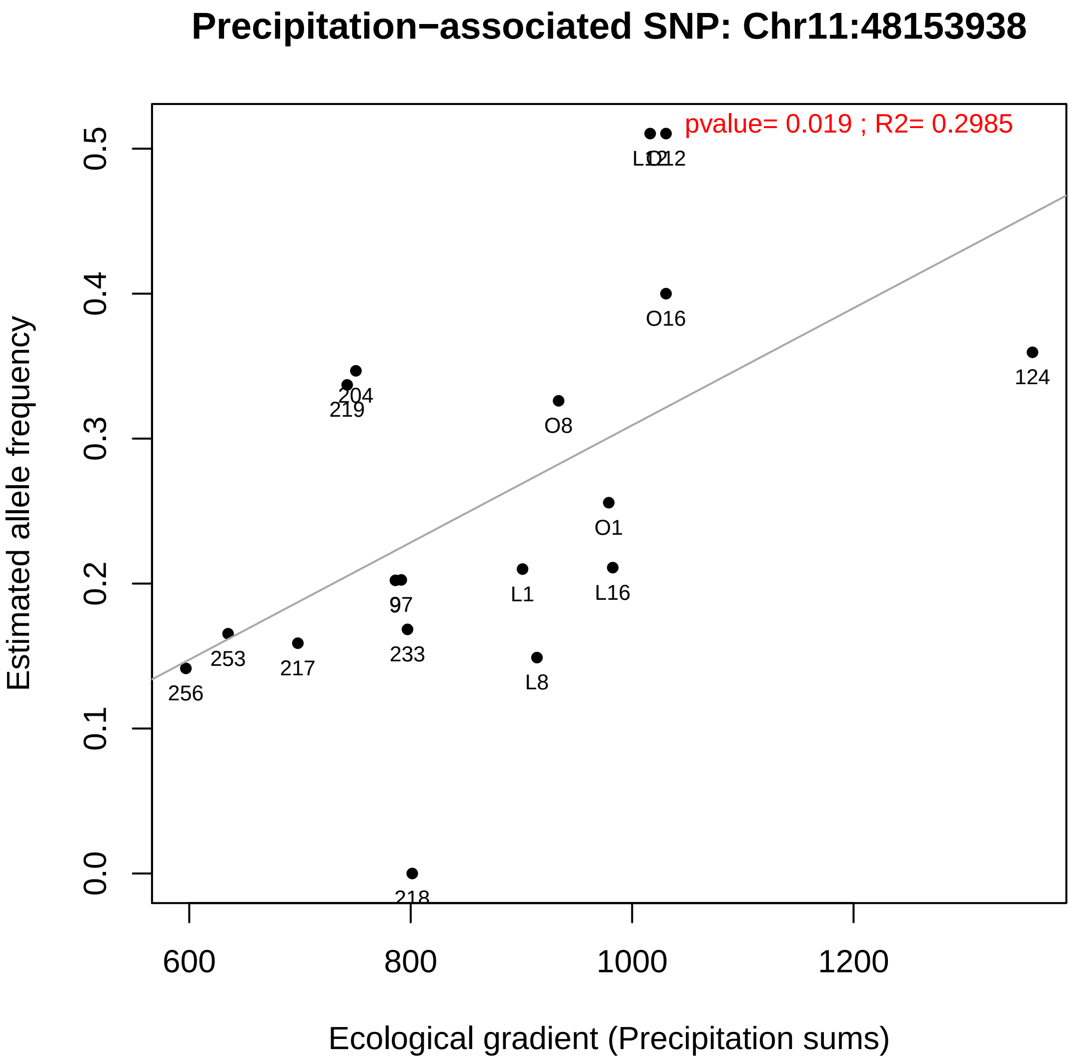

Precipitation−associated SNP: Chr11:48153938
pvalue= 0.019 ; R2= 0.2985
Estimated allele frequency
Ecological gradient (Precipitation sums)
0.0
0.1
0.2
0.3
0.4
0.5
600
800
1000
1200
L12
O12
O16
124
204
219
O8
O1
L1
L16
97
233
L8
253
217
256
218

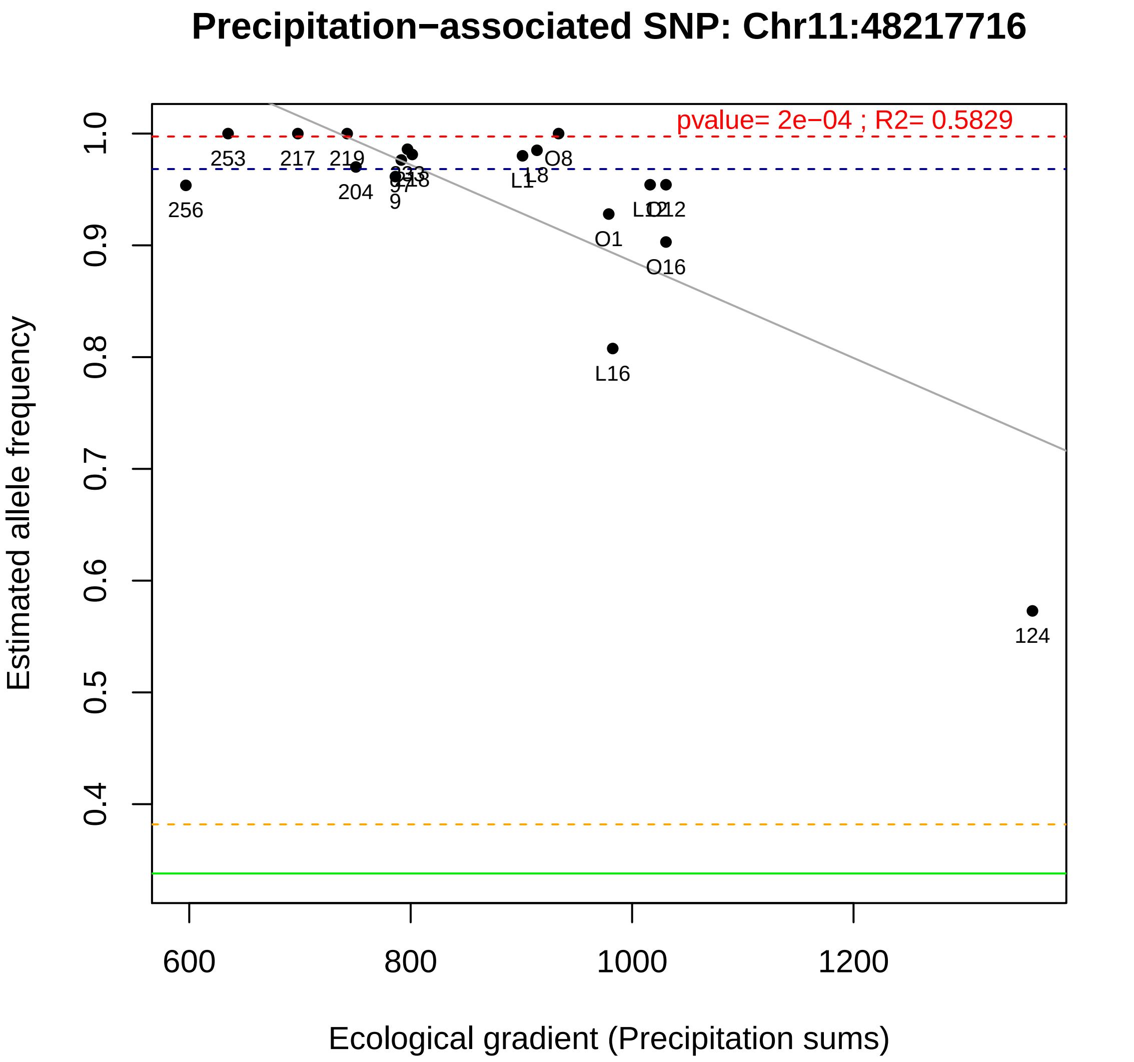

Precipitation−associated SNP: Chr11:48217716
pvalue= 2e−04 ; R2= 0.5829
Estimated allele frequency
Ecological gradient (Precipitation sums)
0.4
0.5
0.6
0.7
0.8
0.9
1.0
600
800
1000
1200
253
217
219
204
256
O8
L1
L8
O1
O12
O16
L16
124

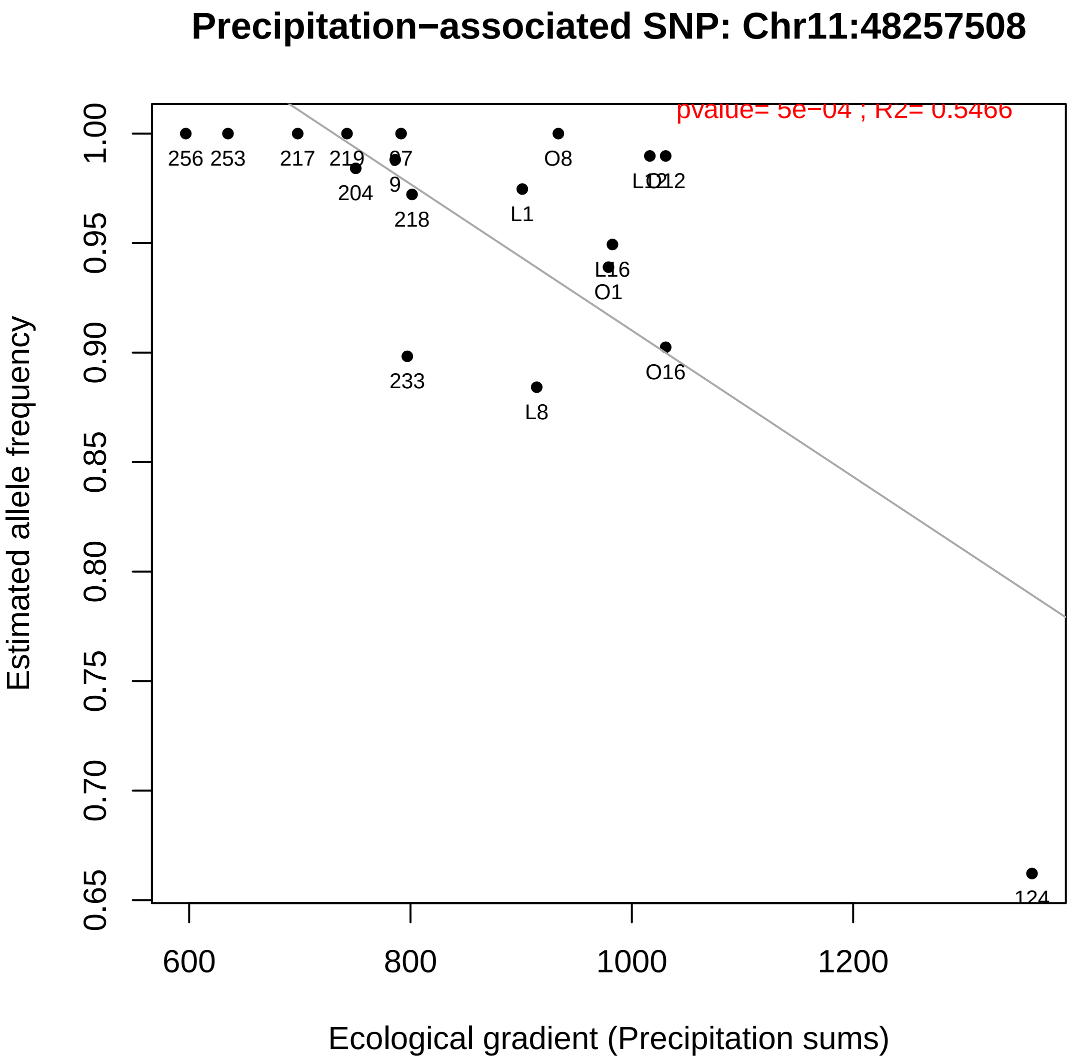
Precipitation−associated SNP: Chr11:48257508
pvalue= 5e−04 ; R2= 0.5466
Estimated allele frequency
Ecological gradient (Precipitation sums)
1.00
0.95
0.90
0.85
0.80
0.75
0.70
0.65
600
800
1000
1200
256
253
217
219
204
9
218
233
L1
O8
L8
L16
O1
O16
124

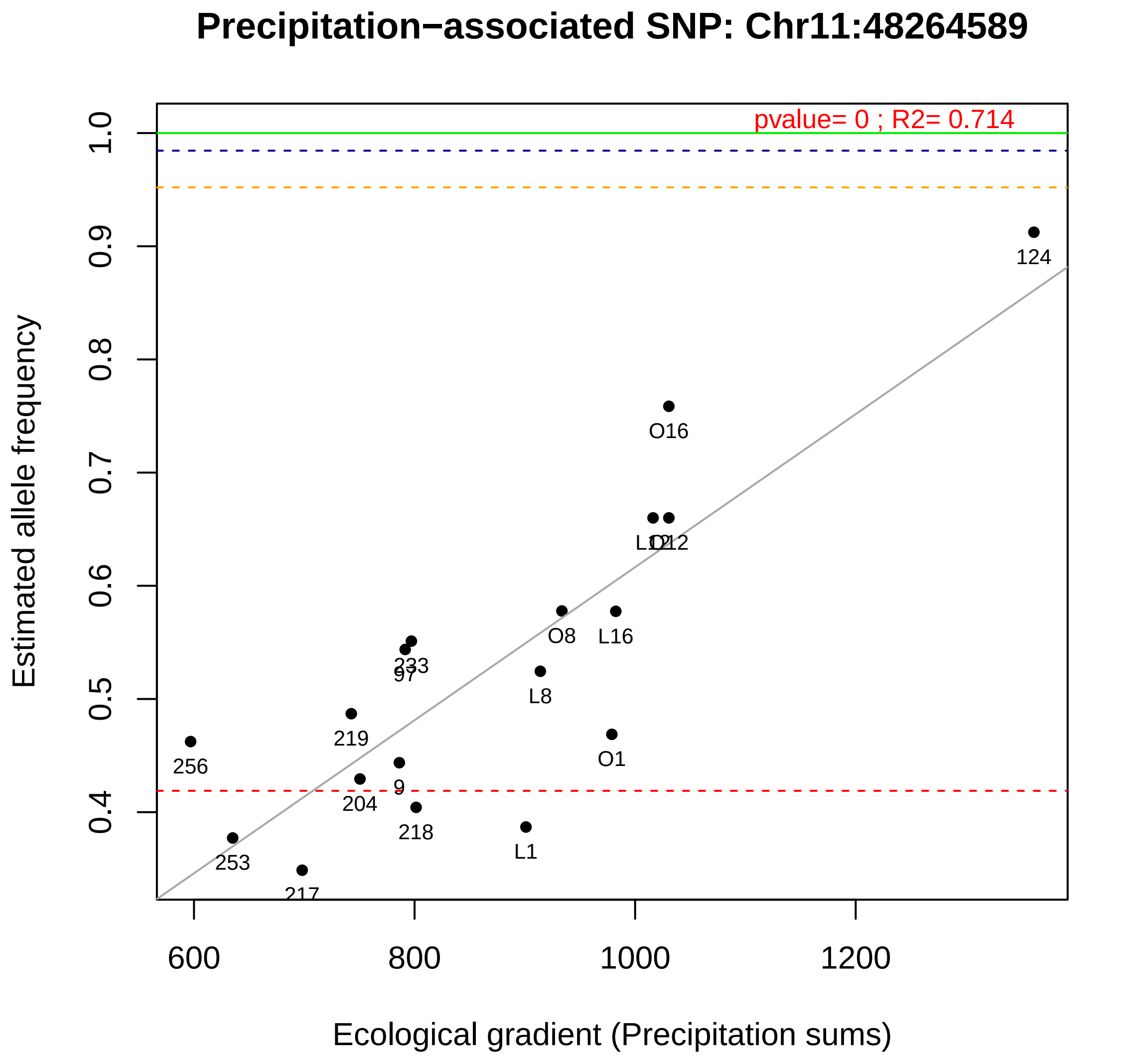
Precipitation−associated SNP: Chr11:48264589
pvalue= 0 ; R2= 0.714
Estimated allele frequency
0.4
0.5
0.6
0.7
0.8
0.9
1.0
Ecological gradient (Precipitation sums)
600
800
1000
1200
124
O16
L12
O12
O8
L16
233
97
L8
219
256
O1
9
204
218
L1
253
217

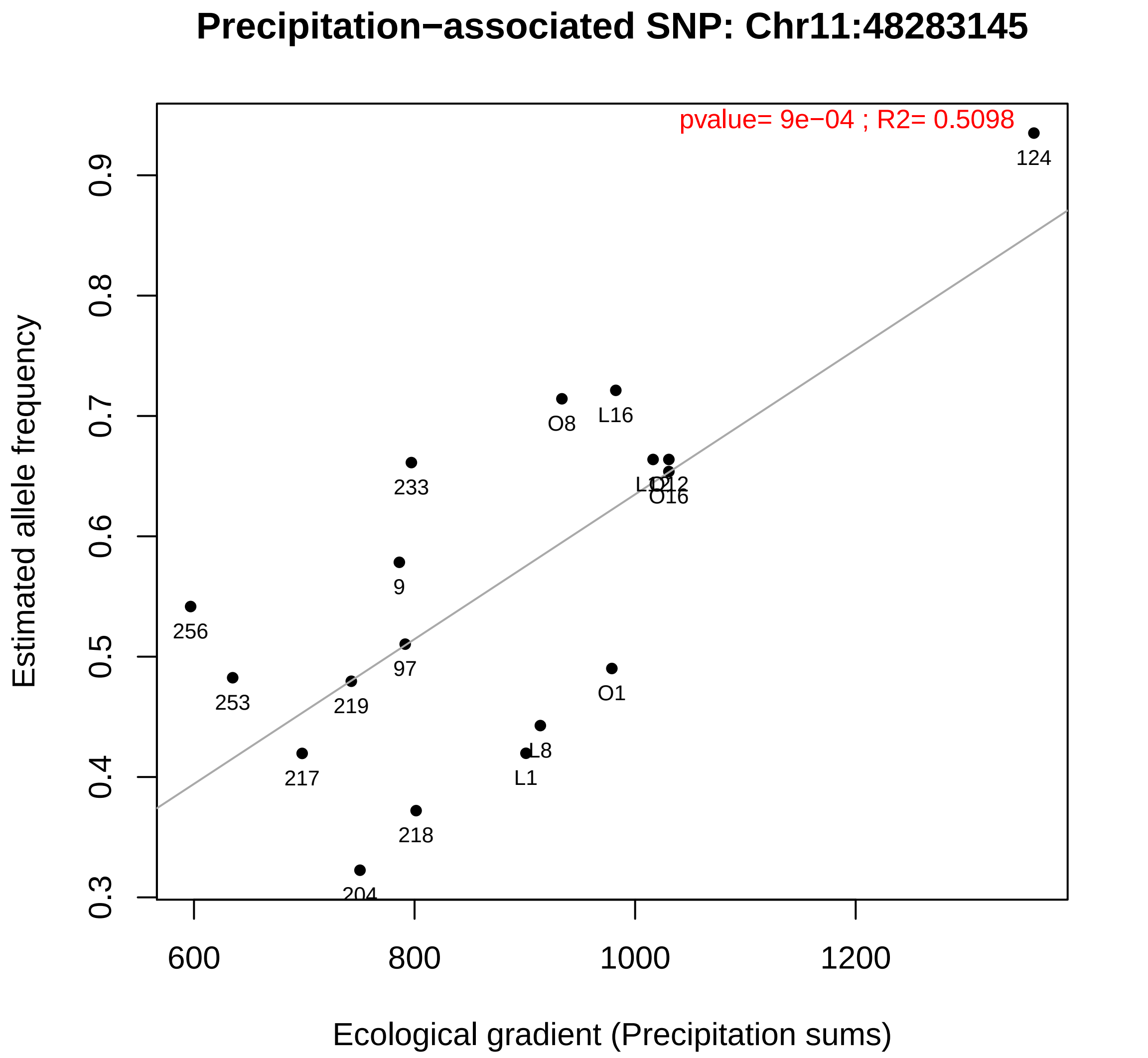

Precipitation−associated SNP: Chr11:48283145
pvalue= 9e−04 ; R2= 0.5098
Estimated allele frequency
Ecological gradient (Precipitation sums)
0.3
0.4
0.5
0.6
0.7
0.8
0.9
600
800
1000
1200
124
O8
L16
233
O16
9
256
97
253
219
O1
217
L8
L1
218
204

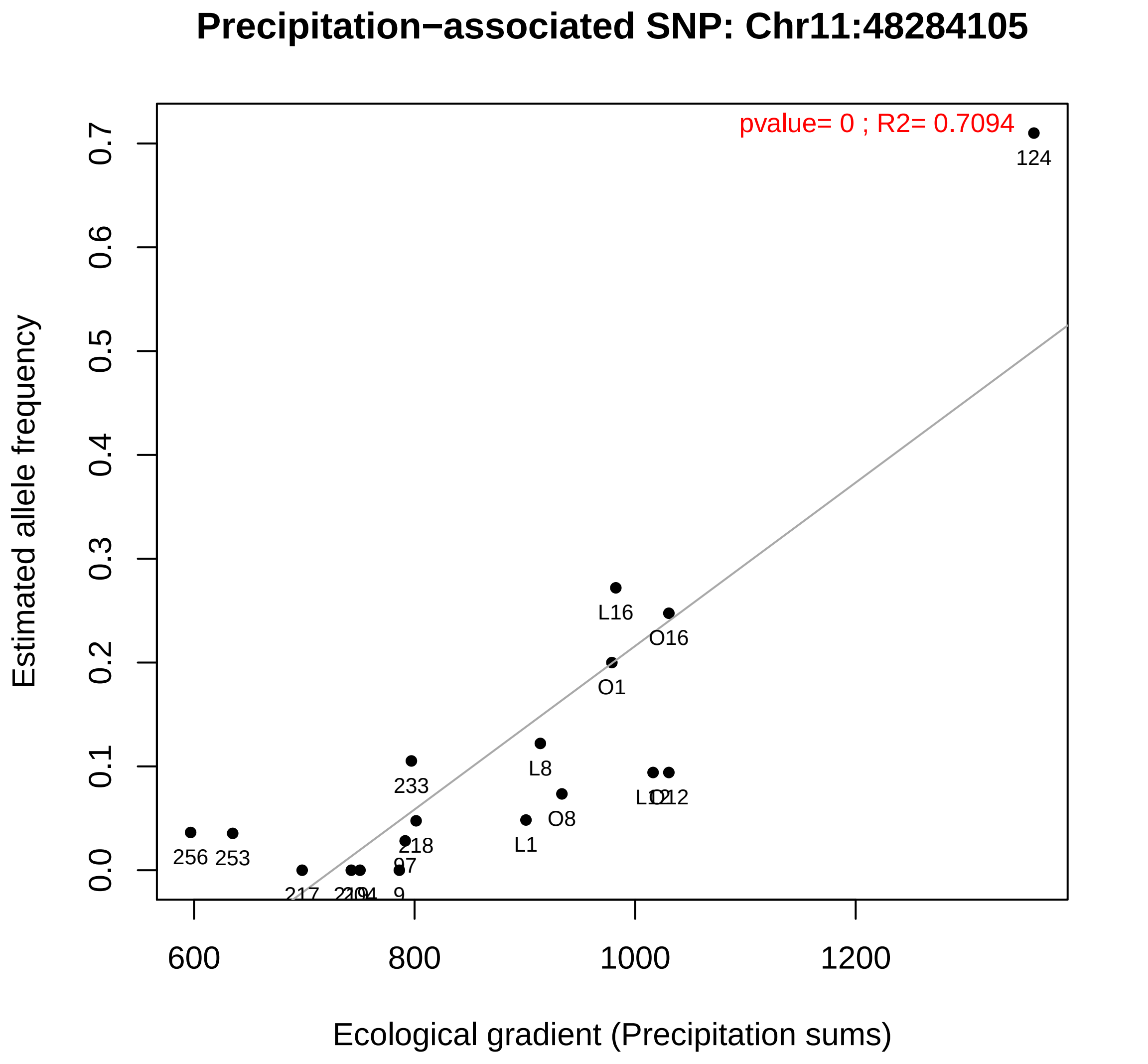
Precipitation−associated SNP: Chr11:48284105
pvalue= 0 ; R2= 0.7094
Estimated allele frequency
0.0
0.1
0.2
0.3
0.4
0.5
0.6
0.7
600
800
1000
1200
Ecological gradient (Precipitation sums)
124
L16
O16
O1
L8
233
O8
L1
218
256
253
217
9

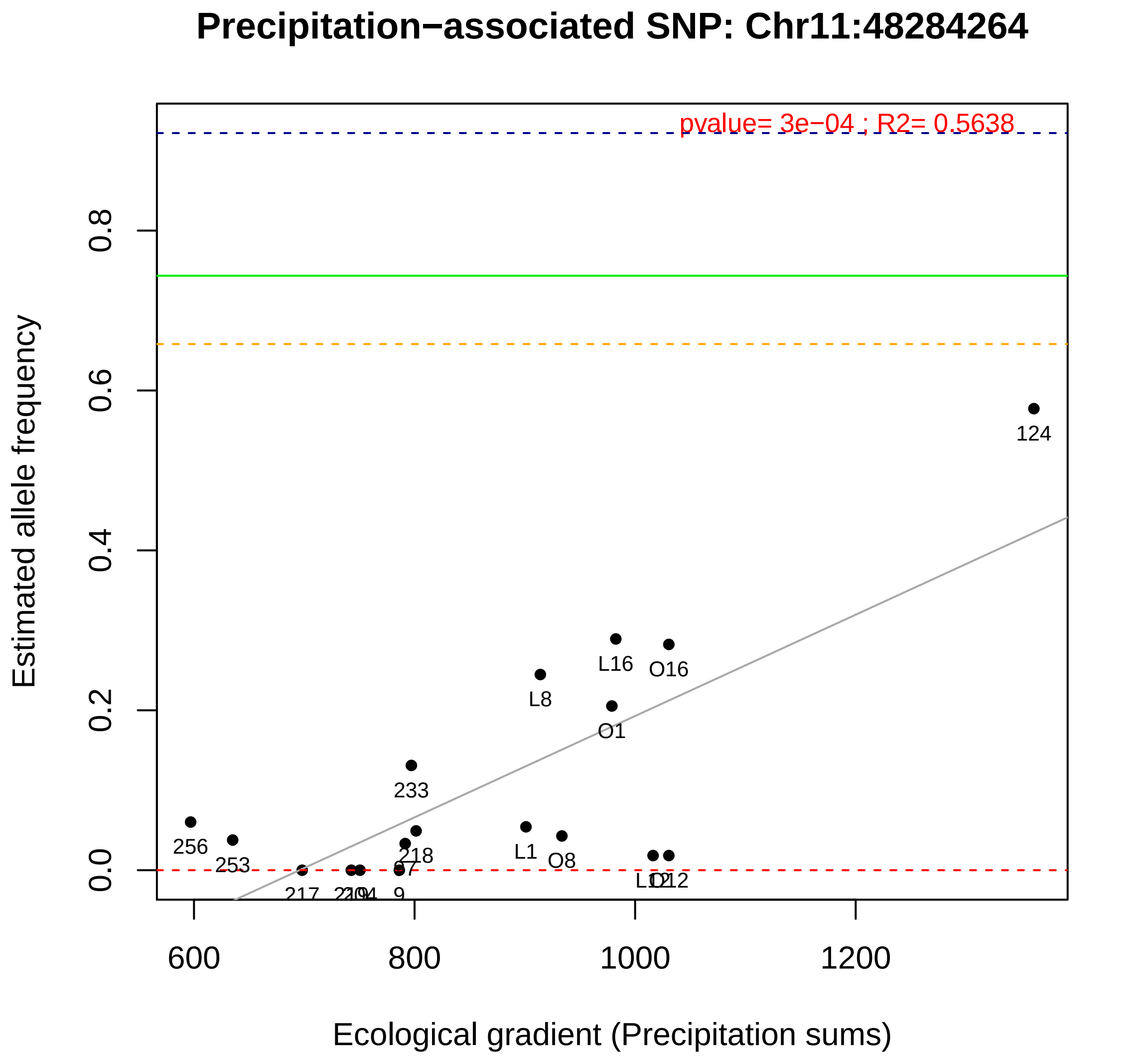

Precipitation−associated SNP: Chr11:48284264
pvalue= 3e−04 ; R2= 0.5638
Estimated allele frequency
0.0
0.2
0.4
0.6
0.8
600
800
1000
1200
Ecological gradient (Precipitation sums)
124
L16
O16
L8
O1
233
256
253
218
L1
O8
217
9

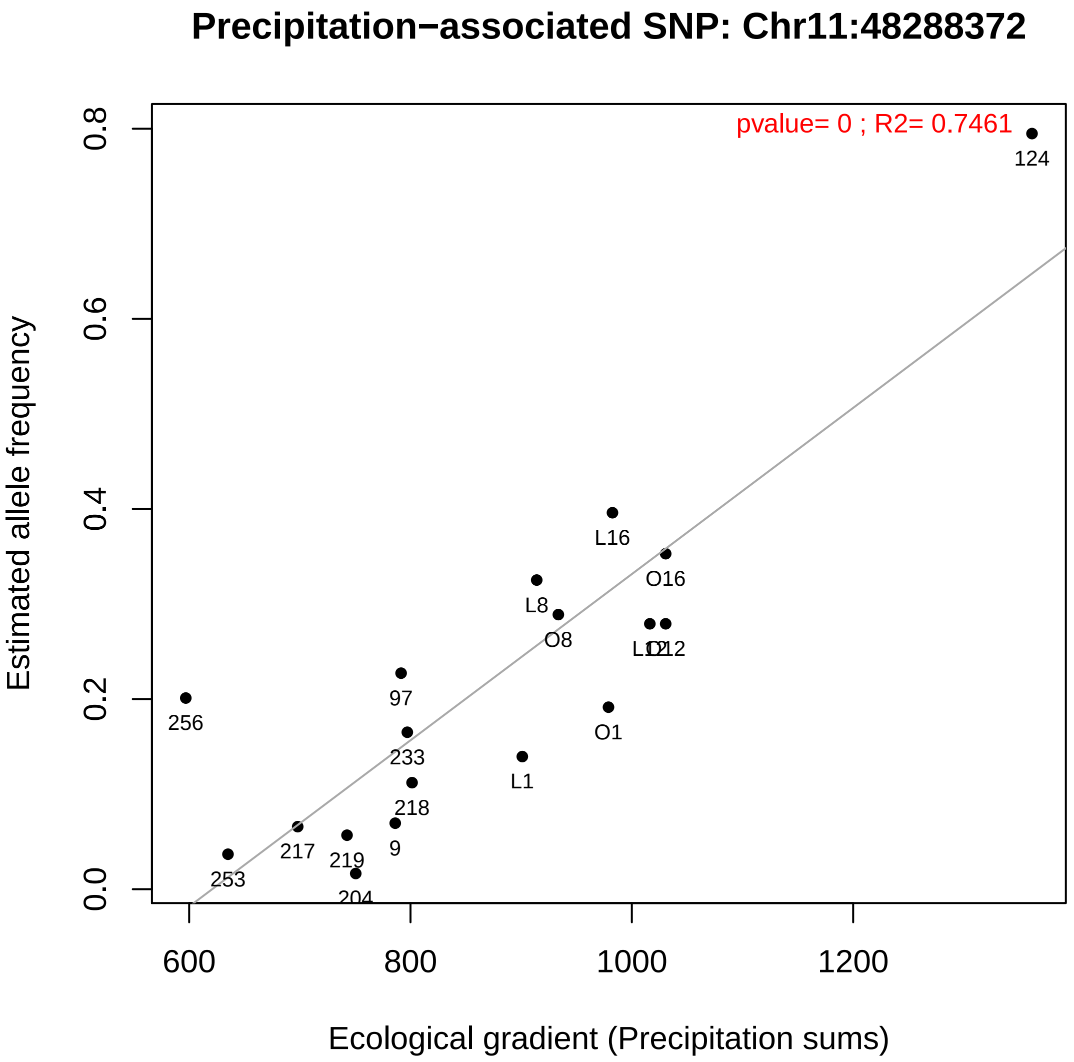
Precipitation−associated SNP: Chr11:48288372
pvalue= 0 ; R2= 0.7461
124
L16
O16
L8
O8
L12
O12
97
256
O1
233
L1
218
217
9
219
253
204
0.0
0.2
0.4
0.6
0.8
600
800
1000
1200
Estimated allele frequency
Ecological gradient (Precipitation sums)

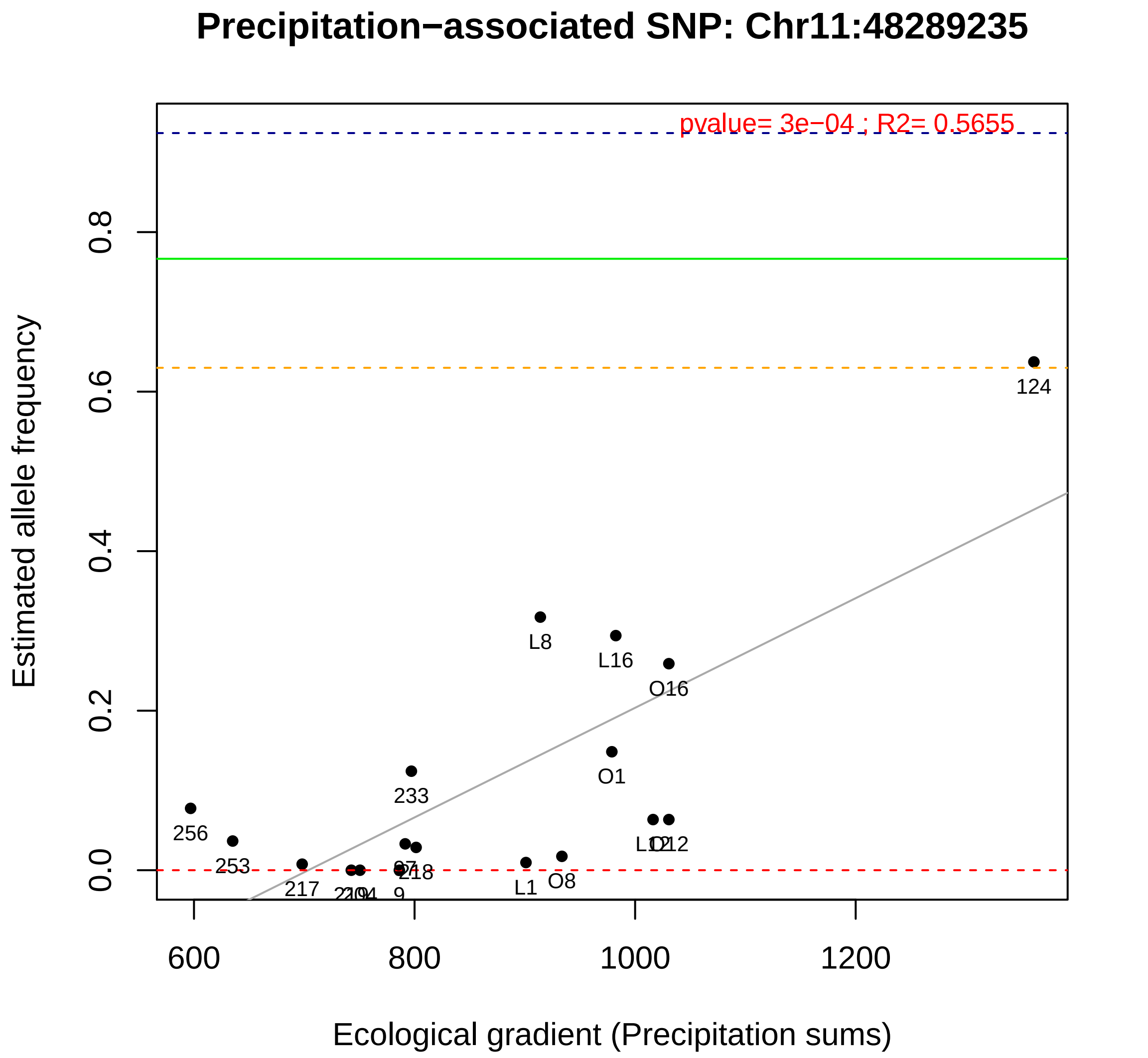

Precipitation−associated SNP: Chr11:48289235
pvalue= 3e−04 ; R2= 0.5655
Estimated allele frequency
0.0
0.2
0.4
0.6
0.8
600
800
1000
1200
Ecological gradient (Precipitation sums)
124
L8
L16
O16
O1
233
256
253
217
L1
O8

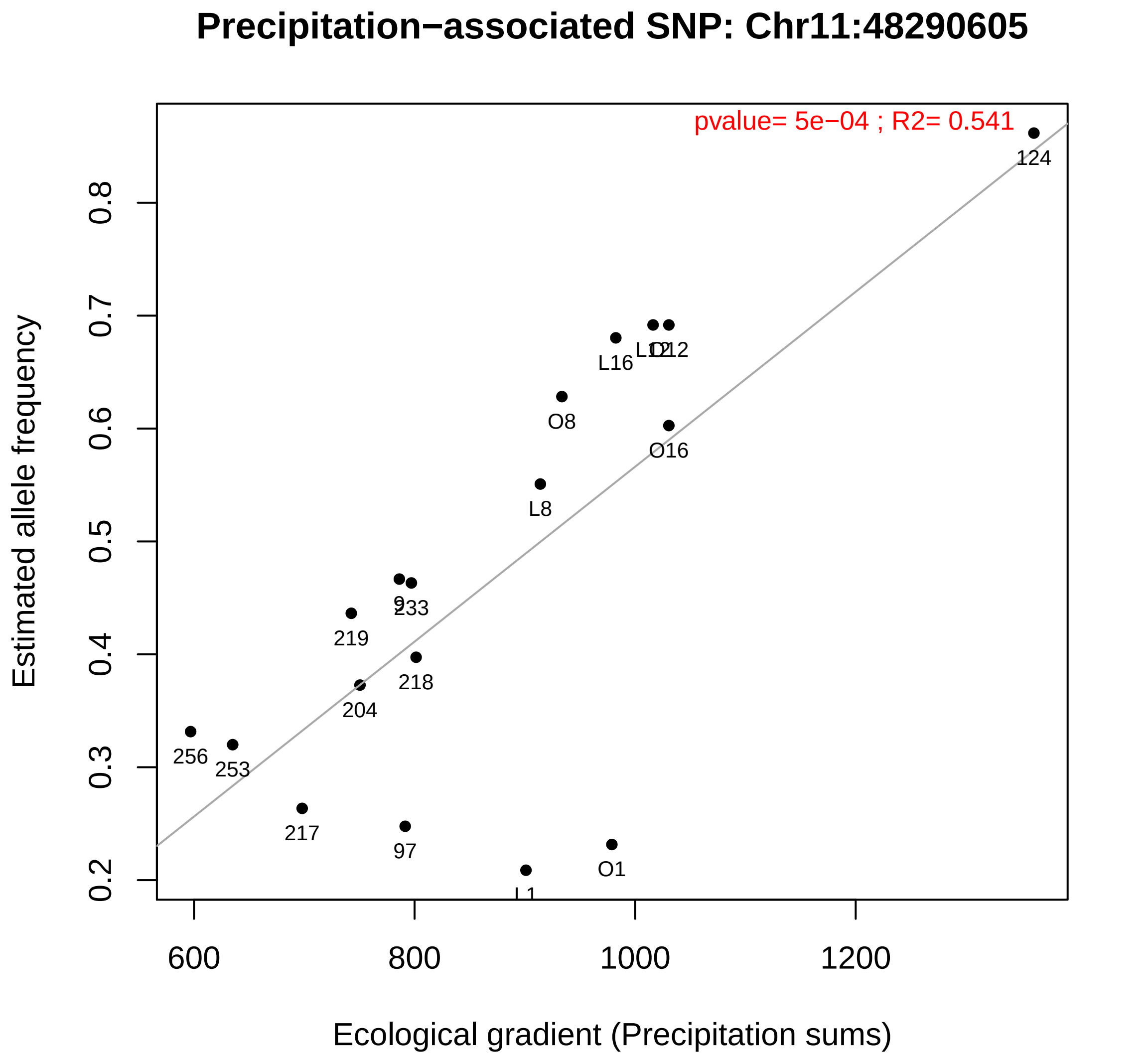
Precipitation−associated SNP: Chr11:48290605
pvalue= 5e−04 ; R2= 0.541
Estimated allele frequency
Ecological gradient (Precipitation sums)
0.2
0.3
0.4
0.5
0.6
0.7
0.8
600
800
1000
1200
124
L16
O12
O8
O16
L8
233
219
218
204
256
253
217
97
O1

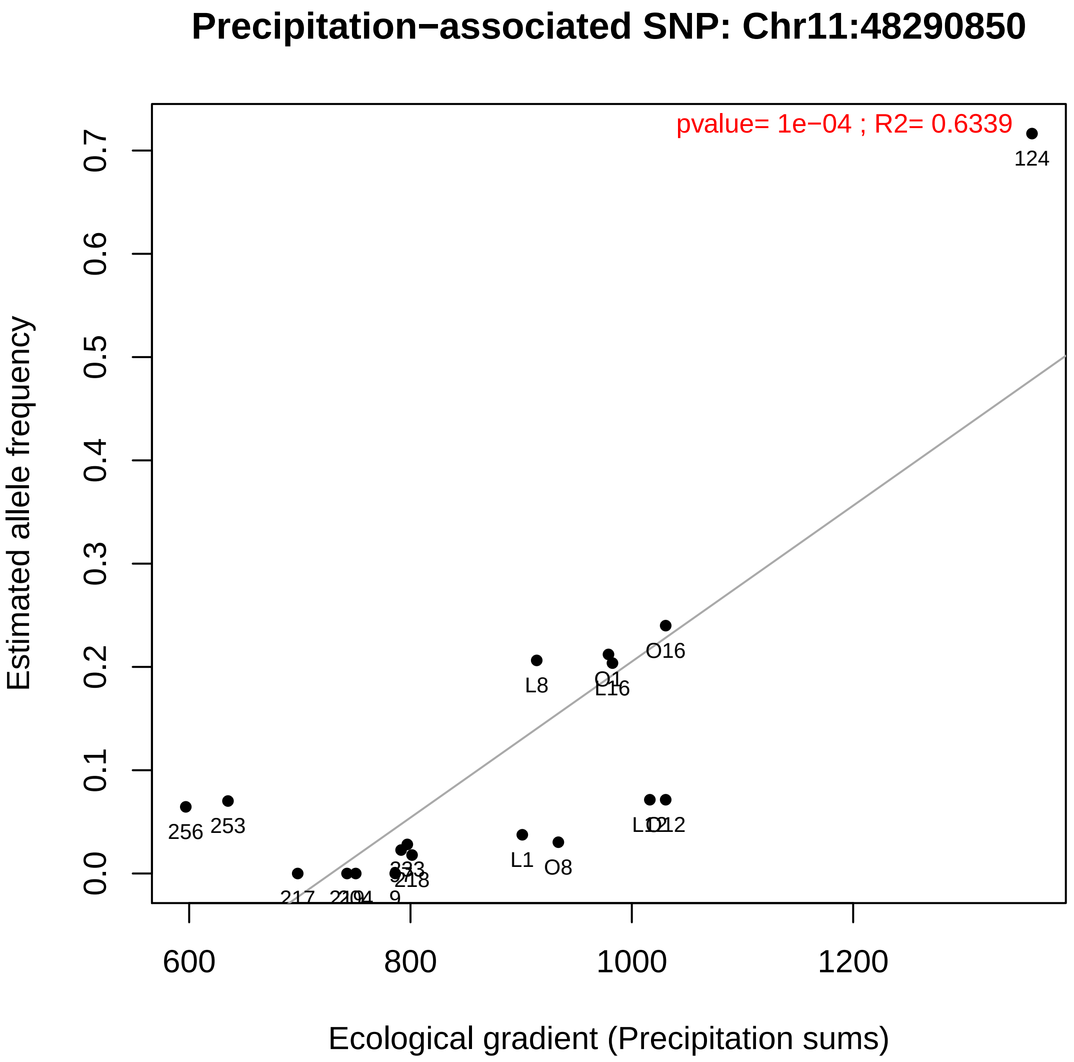
Precipitation−associated SNP: Chr11:48290850
pvalue= 1e−04 ; R2= 0.6339
Estimated allele frequency
Ecological gradient (Precipitation sums)
0.0
0.1
0.2
0.3
0.4
0.5
0.6
0.7
600
800
1000
1200
124
O16
L8
O1
L16
256
253
217
L1
O8
L12
O12
233
218
9

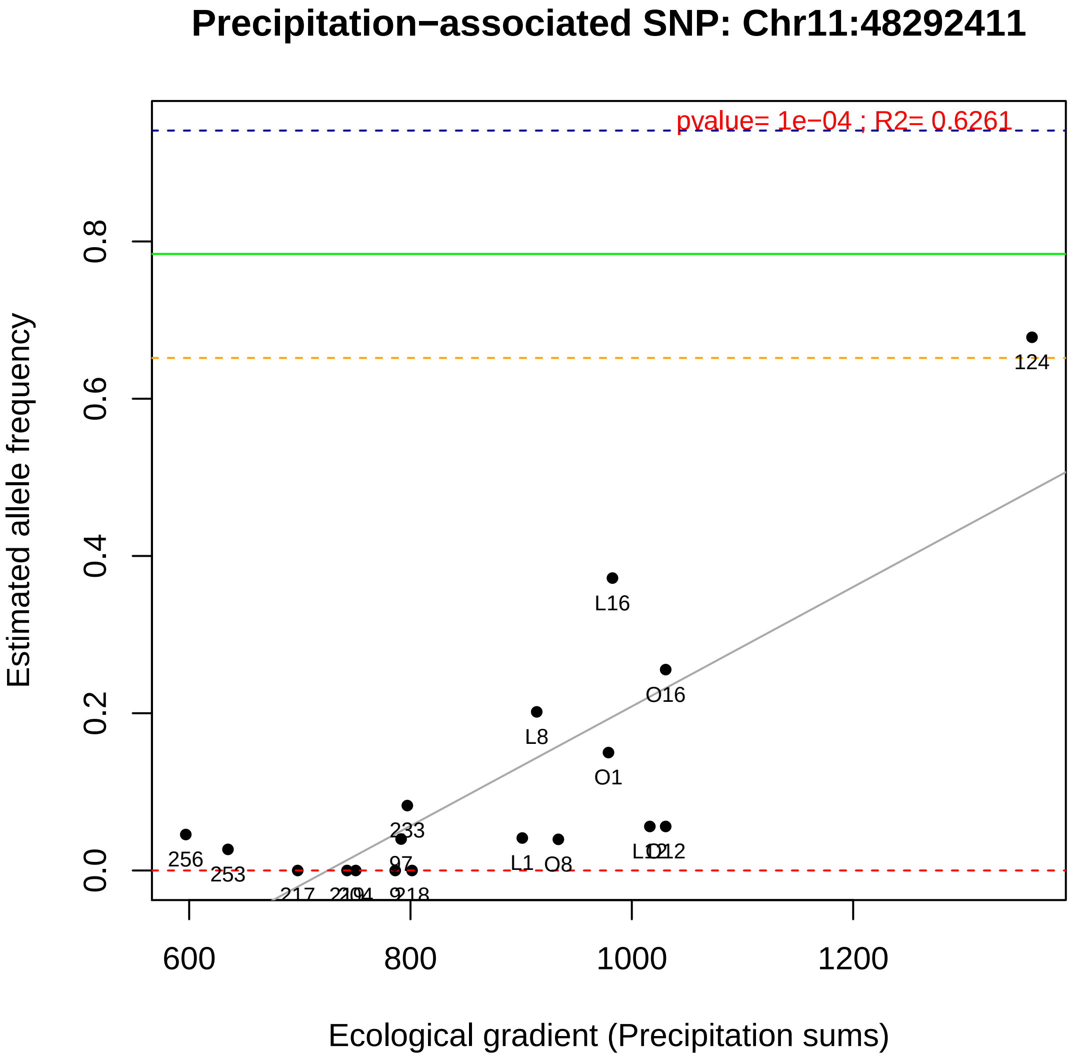
Precipitation−associated SNP: Chr11:48292411
pvalue= 1e−04 ; R2= 0.6261
Estimated allele frequency
0.0
0.2
0.4
0.6
0.8
600
800
1000
1200
Ecological gradient (Precipitation sums)
124
L16
O16
L8
O1
233
97
256
253
217
218
L1
O8

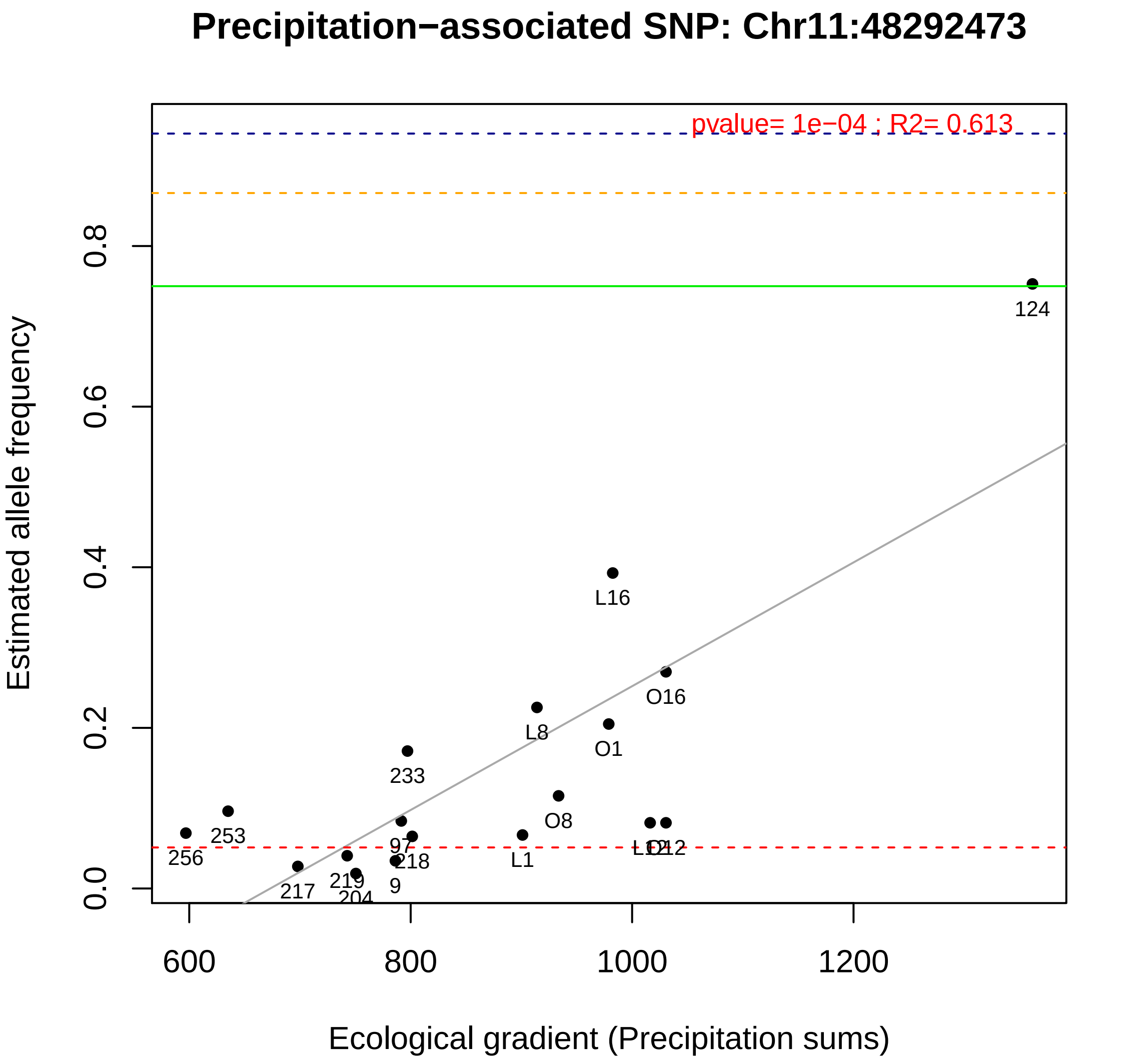

Precipitation−associated SNP: Chr11:48292473
pvalue= 1e−04 ; R2= 0.613
Estimated allele frequency
0.0
0.2
0.4
0.6
0.8
600
800
1000
1200
Ecological gradient (Precipitation sums)
124
L16
O16
L8
O1
233
O8
253
256
217
219
204
97
218
9
L1

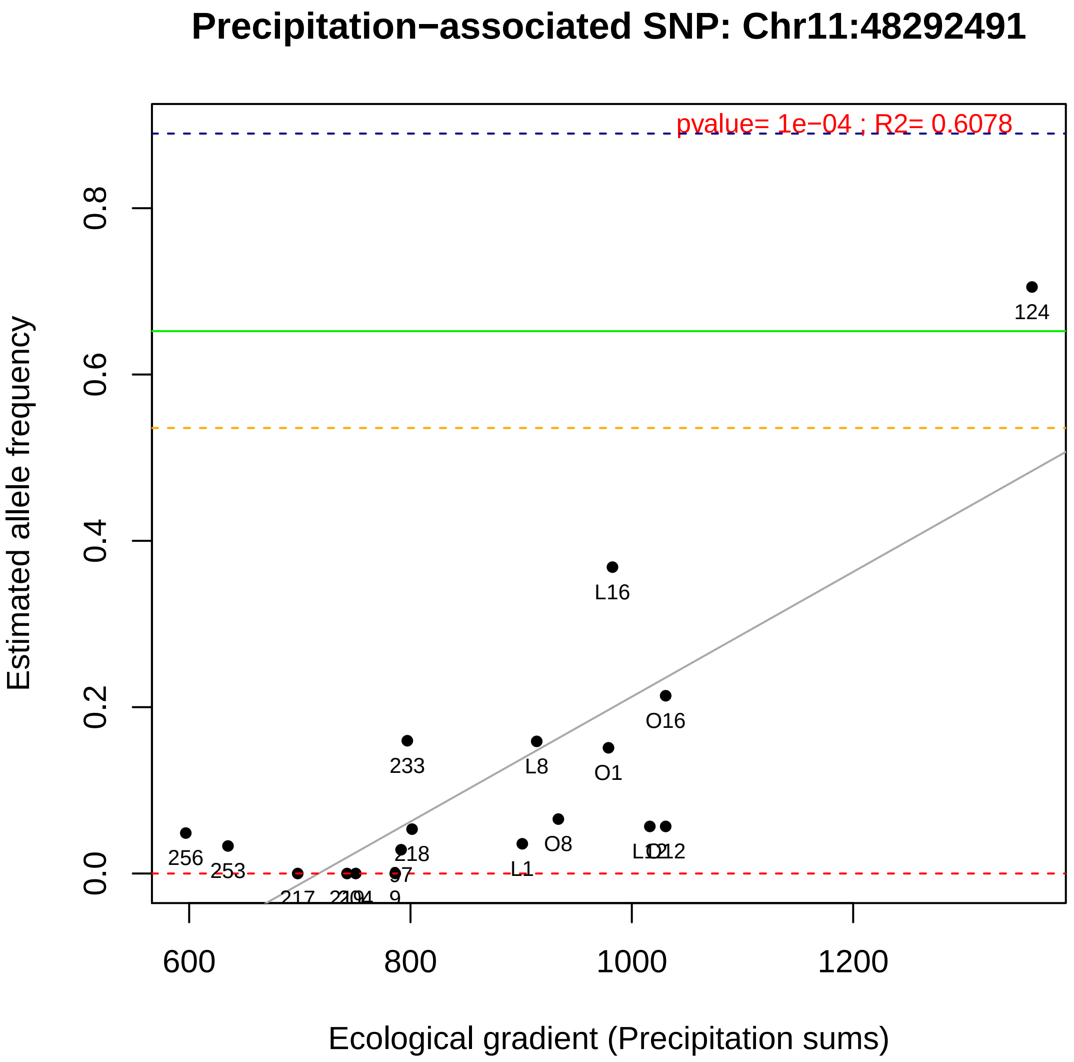
Precipitation−associated SNP: Chr11:48292491
pvalue= 1e−04 ; R2= 0.6078
Estimated allele frequency
Ecological gradient (Precipitation sums)
0.0
0.2
0.4
0.6
0.8
600
800
1000
1200
124
L16
O16
233
L8
O1
O8
256
253
218
L1
217
97
9

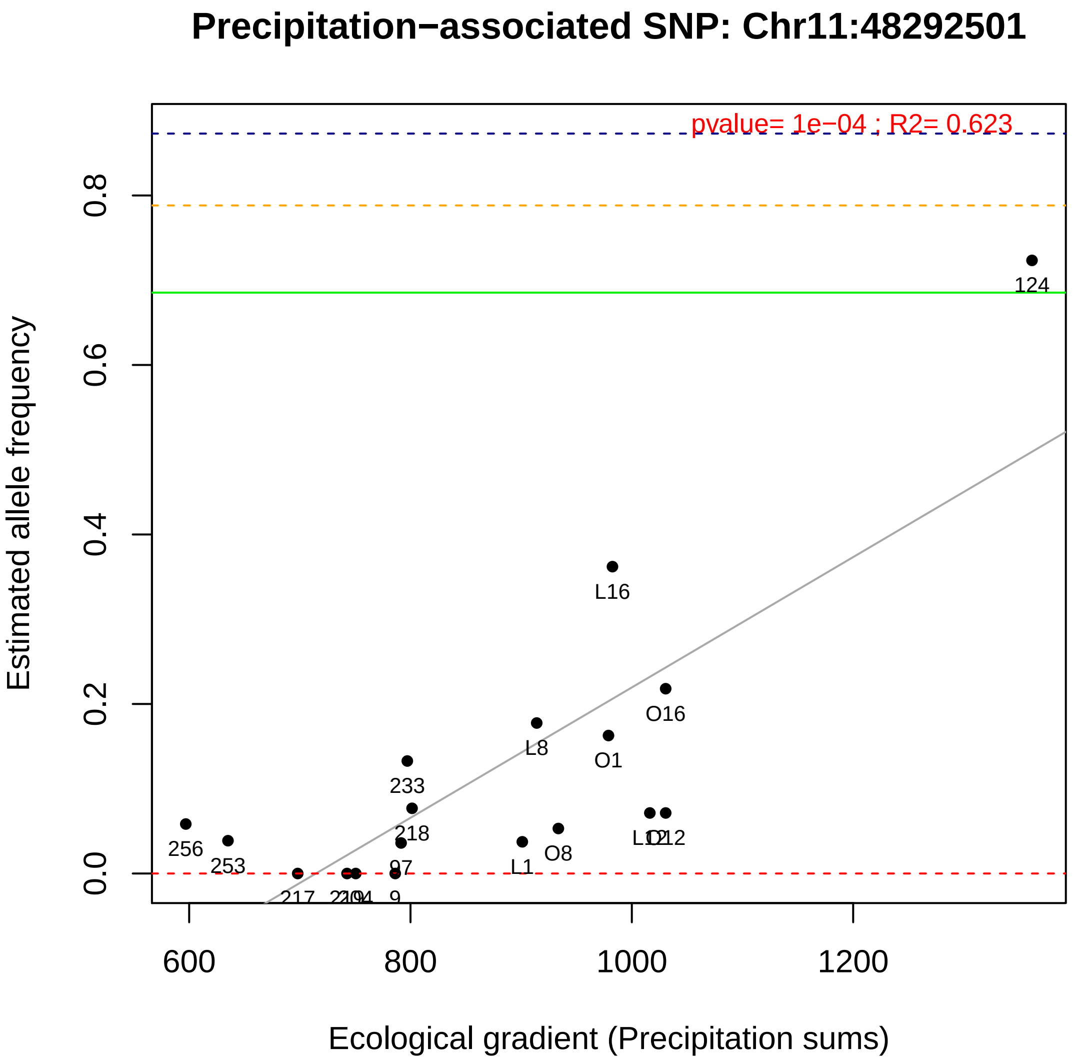

Precipitation−associated SNP: Chr11:48292501
pvalue= 1e−04 ; R2= 0.623
Estimated allele frequency
0.0
0.2
0.4
0.6
0.8
600
800
1000
1200
Ecological gradient (Precipitation sums)
124
L16
O16
L8
O1
233
218
256
253
O8
L1
97
217
9

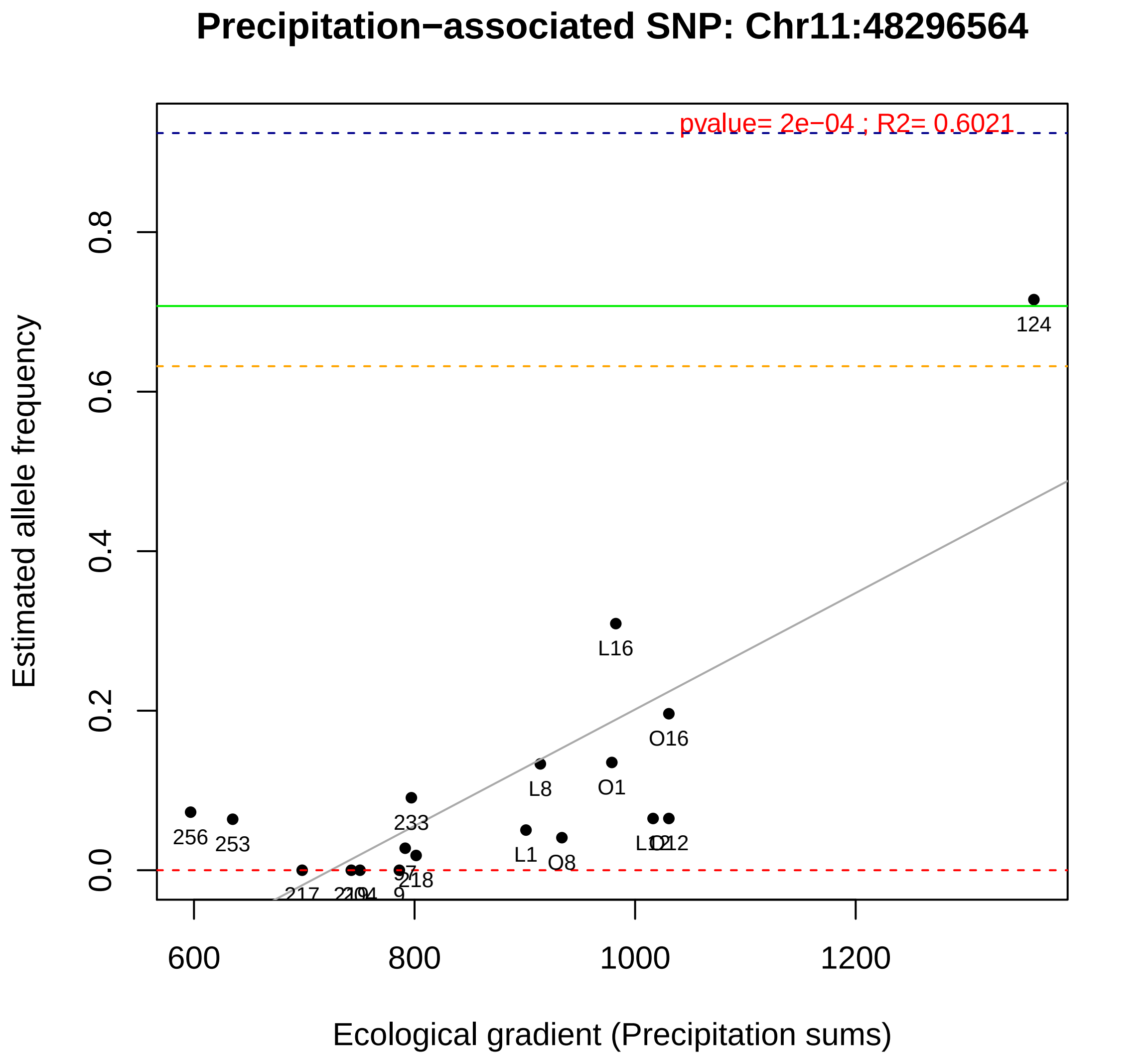
Precipitation−associated SNP: Chr11:48296564
pvalue= 2e−04 ; R2= 0.6021
Estimated allele frequency
Ecological gradient (Precipitation sums)
0.0
0.2
0.4
0.6
0.8
600
800
1000
1200
124
L16
O16
L8
O1
233
256
253
L1
O8
218
217
9

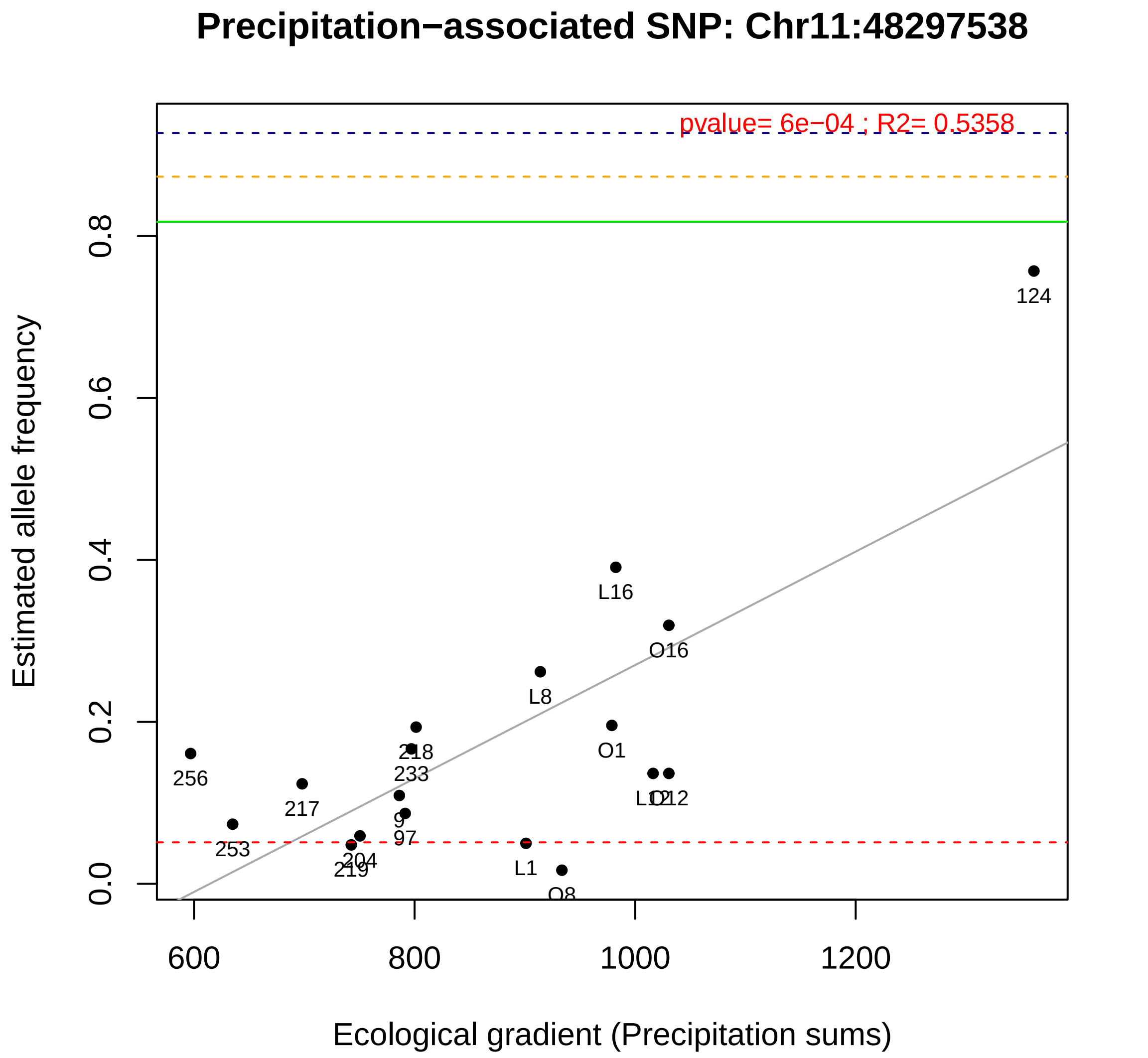

Precipitation−associated SNP: Chr11:48297538
pvalue= 6e−04 ; R2= 0.5358
Estimated allele frequency
Ecological gradient (Precipitation sums)
0.0
0.2
0.4
0.6
0.8
600
800
1000
1200
124
L16
O16
L8
O1
218
233
256
217
253
9
97
204
219
L1
O8
L12
O12

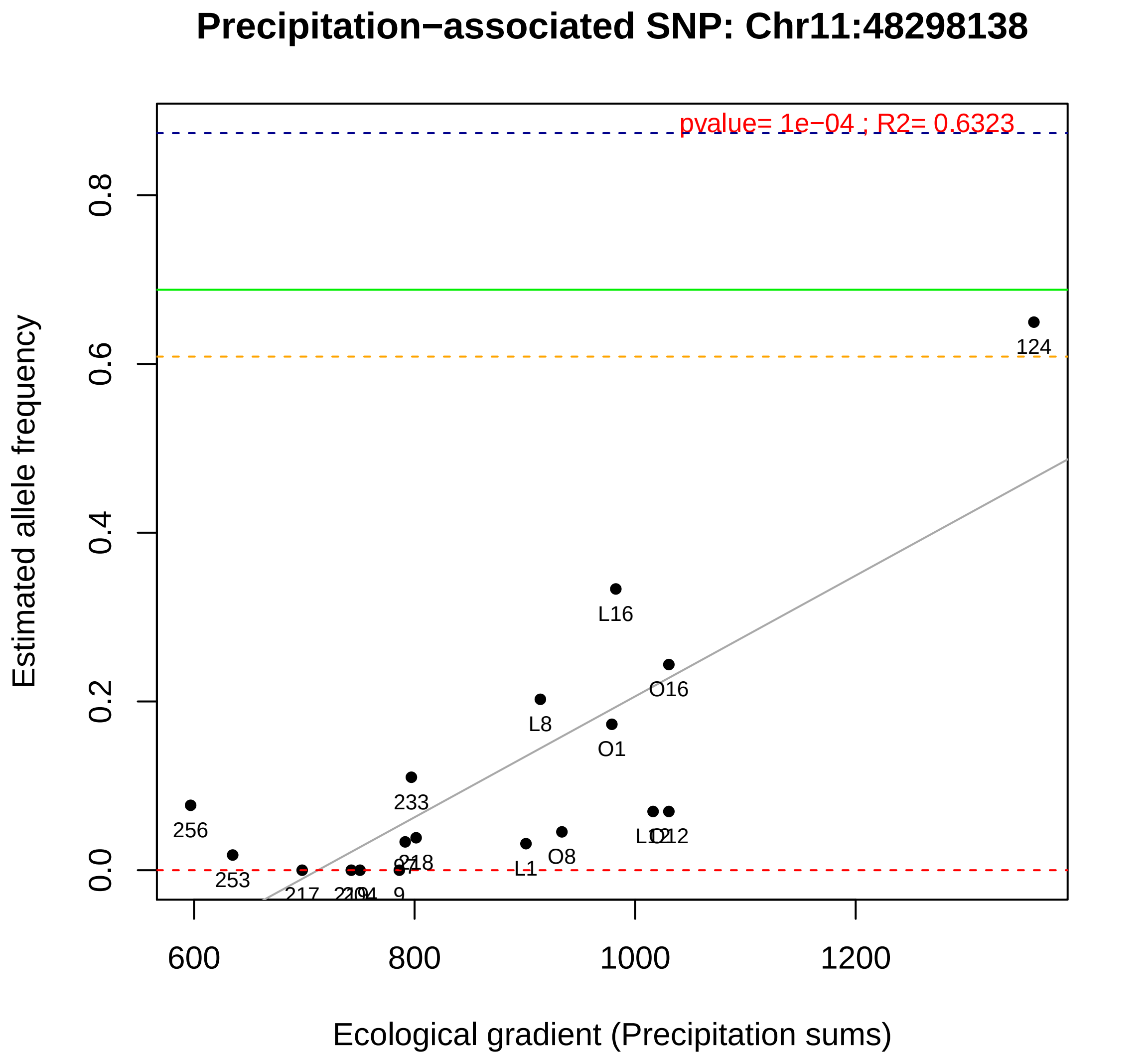
Precipitation−associated SNP: Chr11:48298138
pvalue= 1e−04 ; R2= 0.6323
Estimated allele frequency
Ecological gradient (Precipitation sums)
0.0
0.2
0.4
0.6
0.8
600
800
1000
1200
124
L16
O16
L8
O1
233
256
253
217
218
9
L1
O8

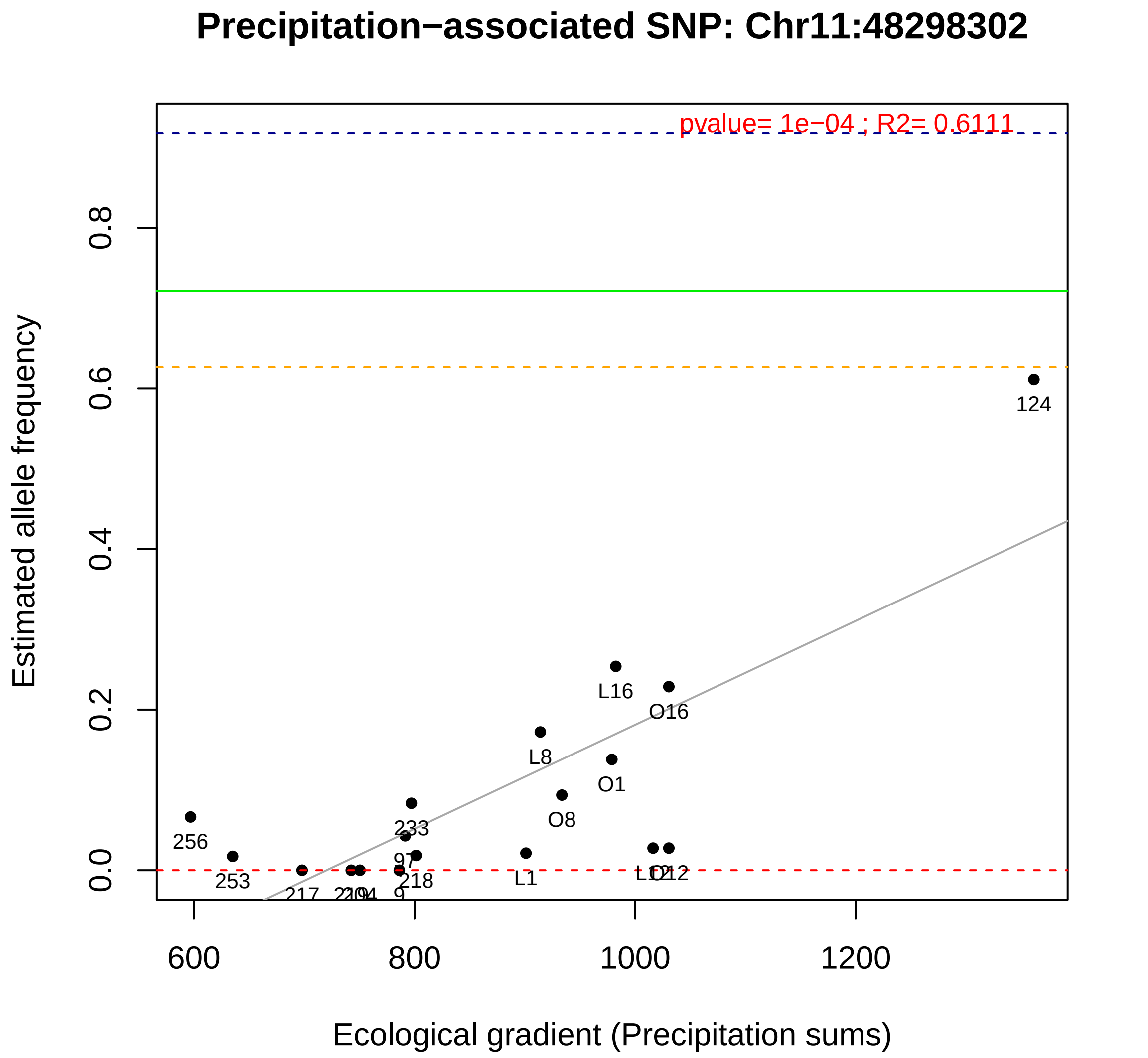

Precipitation−associated SNP: Chr11:48298302
pvalue= 1e−04 ; R2= 0.6111
Estimated allele frequency
0.0
0.2
0.4
0.6
0.8
600
800
1000
1200
Ecological gradient (Precipitation sums)
124
L16
O16
L8
O1
O8
233
256
253
97
218
L1
217
9

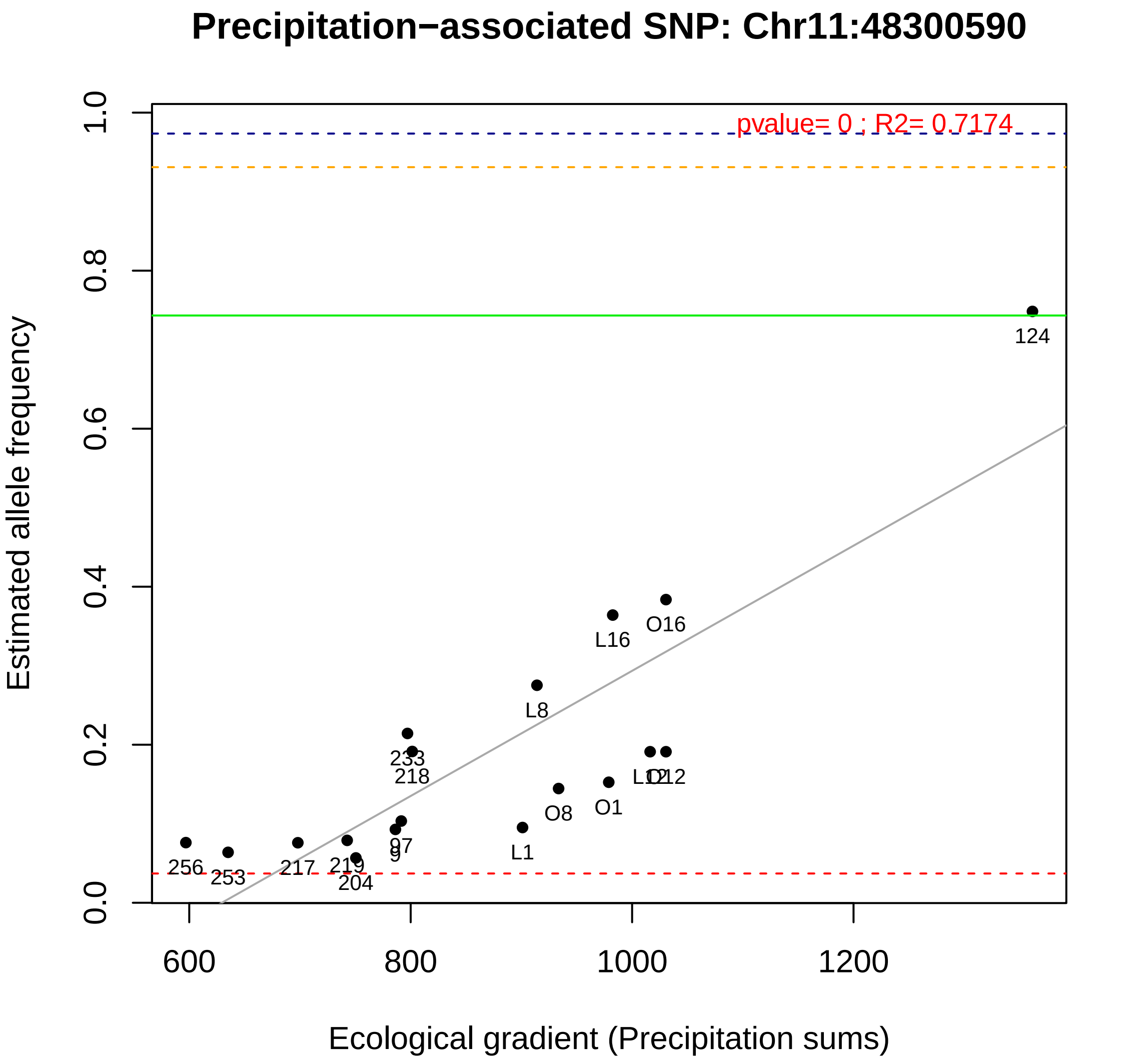
Precipitation−associated SNP: Chr11:48300590
pvalue= 0 ; R2= 0.7174
Estimated allele frequency
Ecological gradient (Precipitation sums)
0.0
0.2
0.4
0.6
0.8
1.0
600
800
1000
1200
124
O16
L16
L8
233
218
L12
O12
O1
O8
97
9
L1
256
253
217
219
204

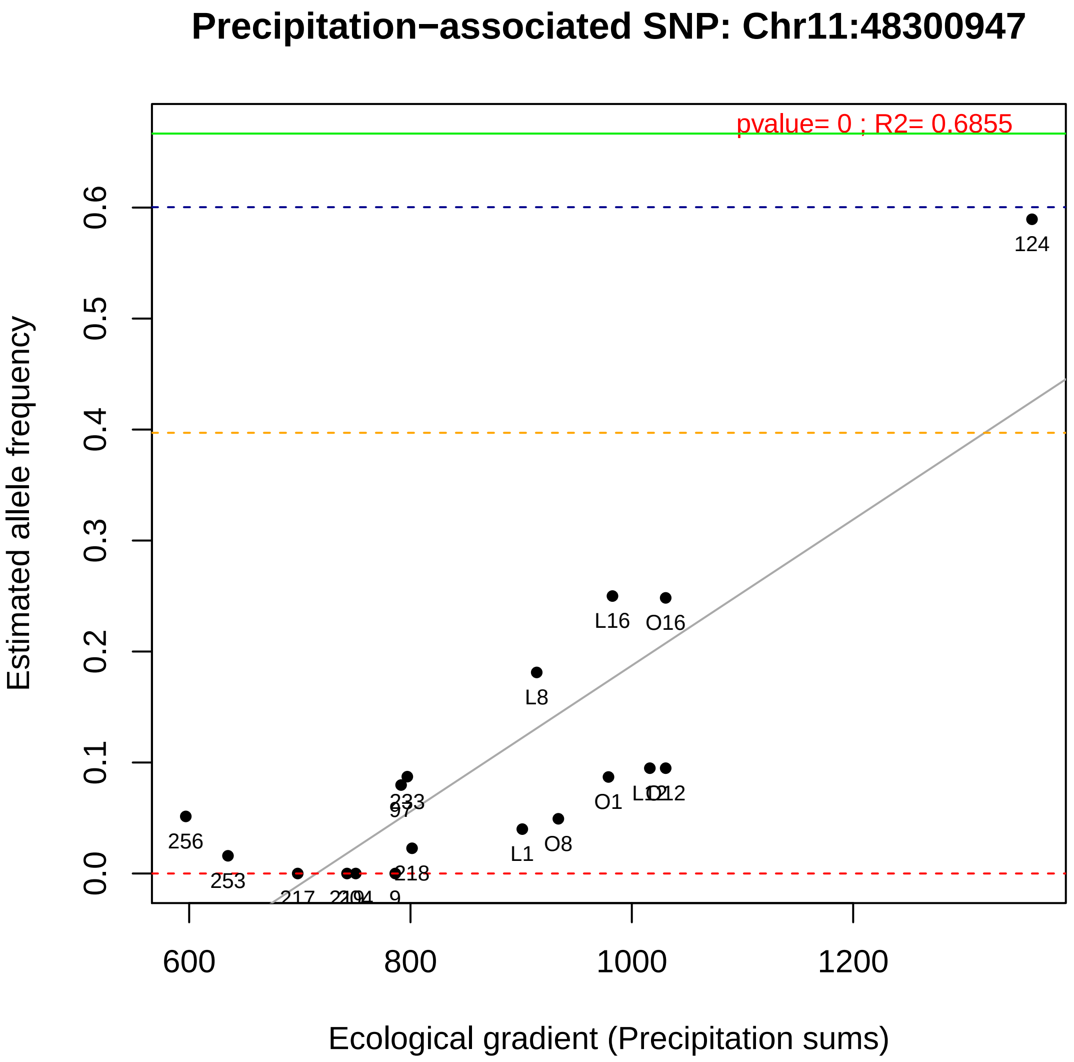
Precipitation−associated SNP: Chr11:48300947
pvalue= 0 ; R2= 0.6855
Estimated allele frequency
Ecological gradient (Precipitation sums)
0.0
0.1
0.2
0.3
0.4
0.5
0.6
600
800
1000
1200
124
L16
O16
L8
233
97
O1
O12
256
253
217
218
9
L1
O8

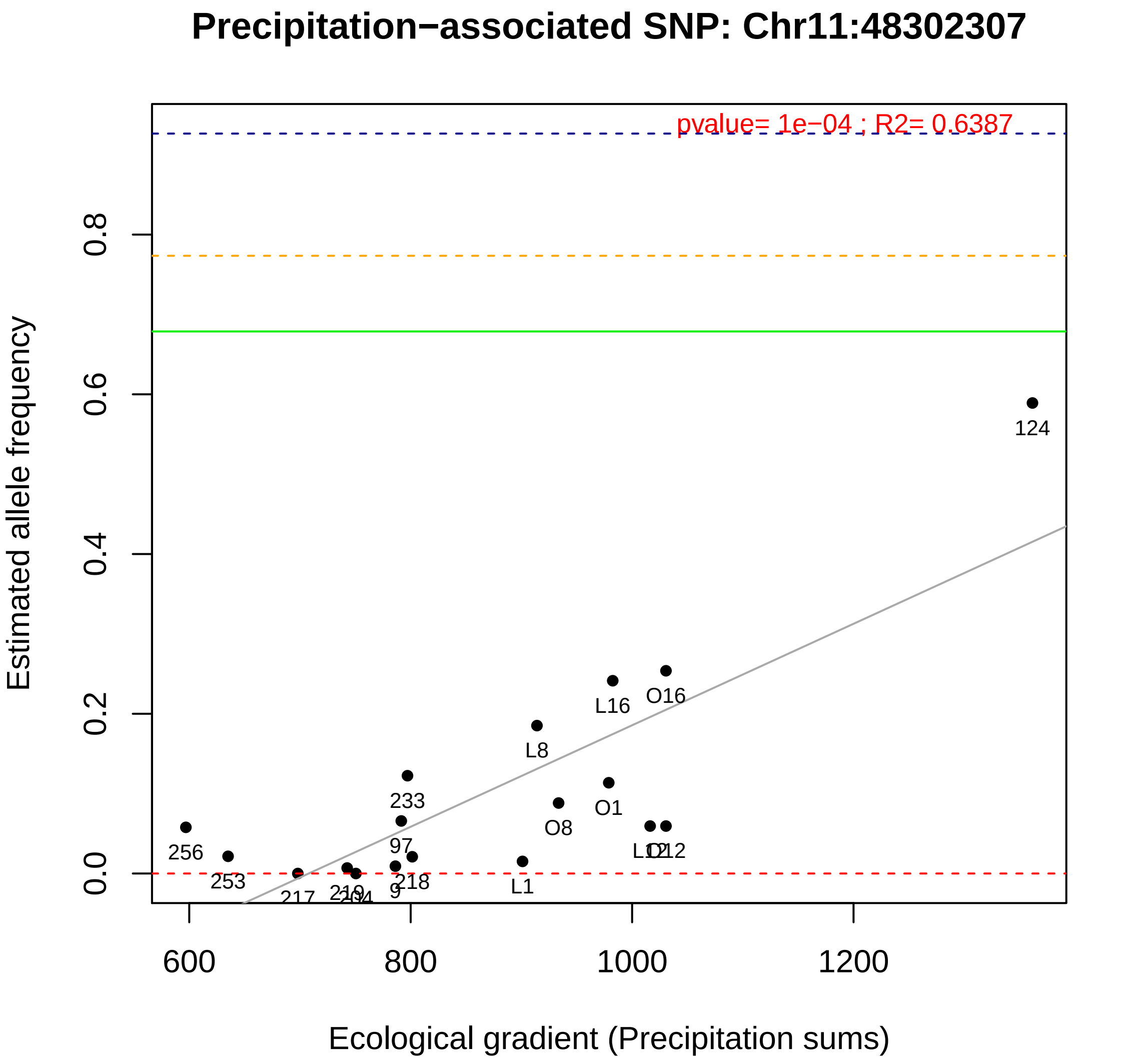
Precipitation−associated SNP: Chr11:48302307
pvalue= 1e−04 ; R2= 0.6387
Estimated allele frequency
0.0
0.2
0.4
0.6
0.8
600
800
1000
1200
Ecological gradient (Precipitation sums)
124
O16
L16
L8
233
O1
O8
97
256
253
218
217
9
L1

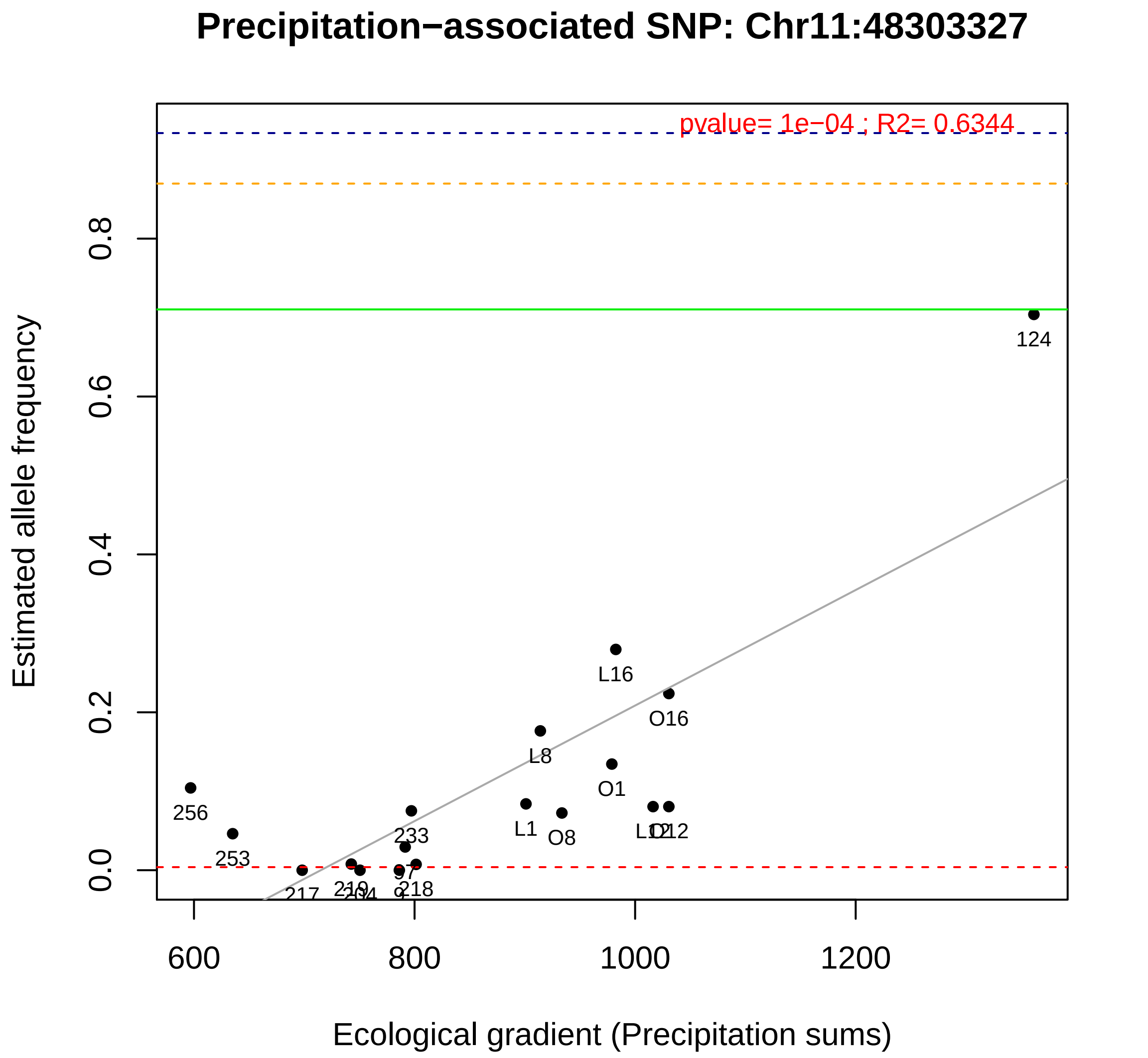
Precipitation−associated SNP: Chr11:48303327
pvalue= 1e−04 ; R2= 0.6344
Estimated allele frequency
Ecological gradient (Precipitation sums)
0.0
0.2
0.4
0.6
0.8
600
800
1000
1200
124
L16
O16
L8
O1
256
L1
O8
233
253
97
218
217

**Precipitation−associated SNP: Chr11:48304187**

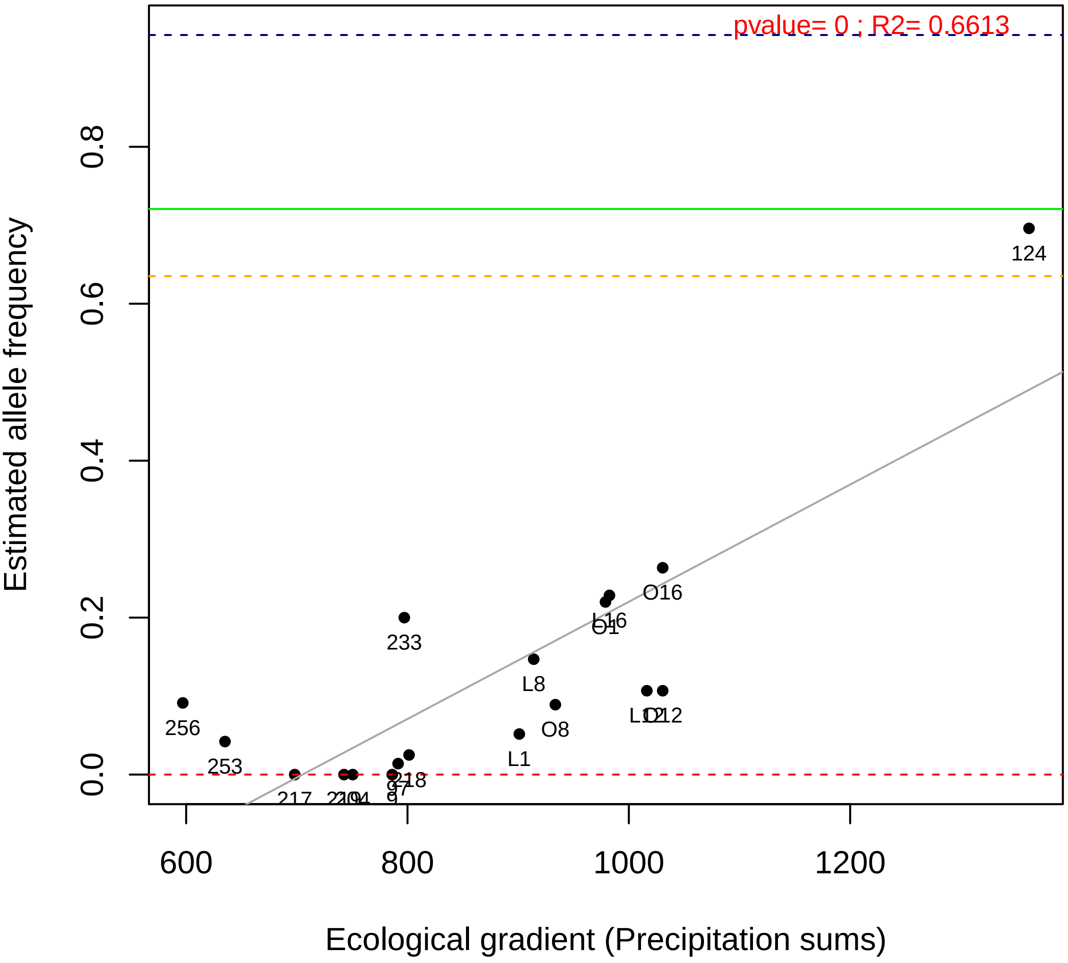

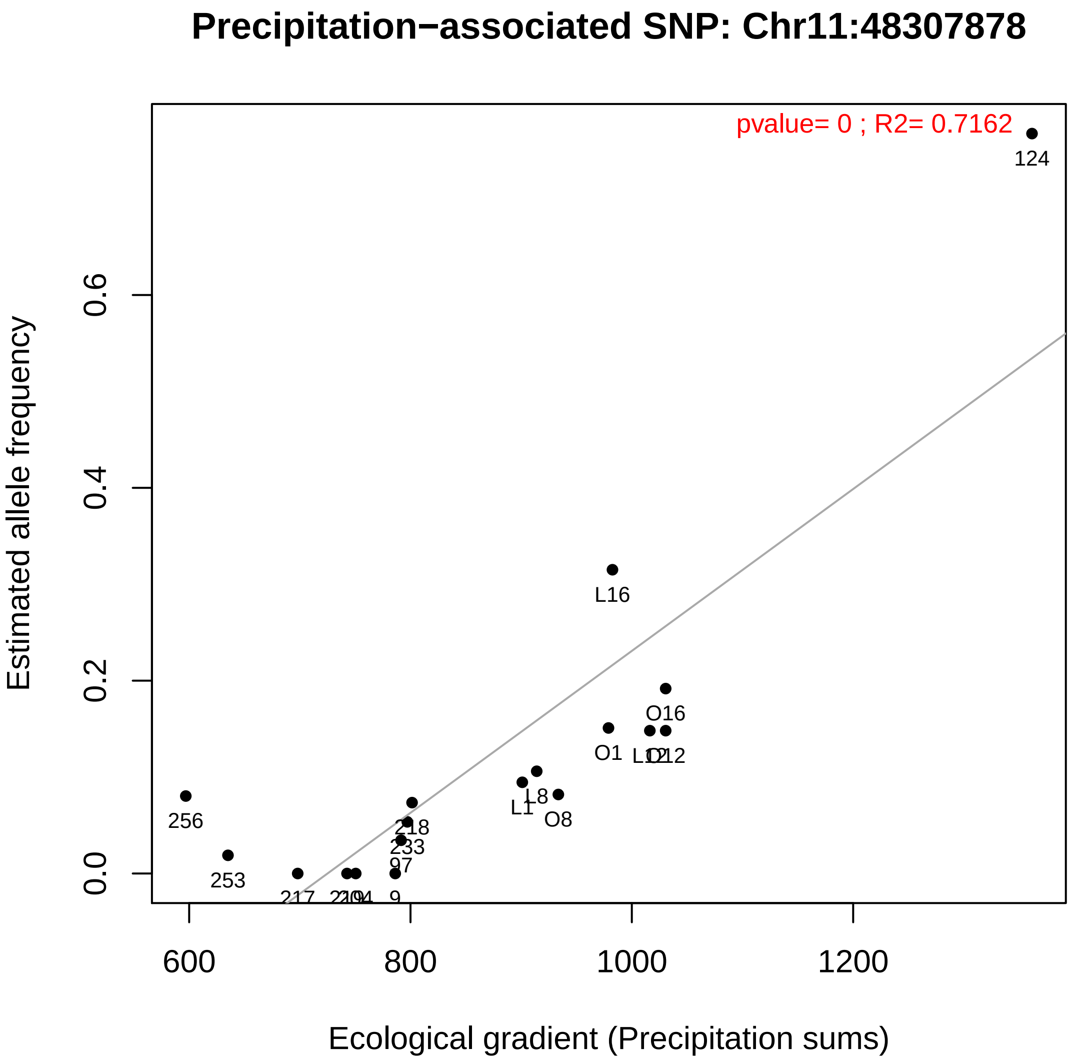

Precipitation−associated SNP: Chr11:48307878
pvalue= 0 ; R2= 0.7162
Estimated allele frequency
Ecological gradient (Precipitation sums)
0.0
0.2
0.4
0.6
600
800
1000
1200
124
L16
O16
O1
256
253
217
218
233
97
9
L8
L1
O8

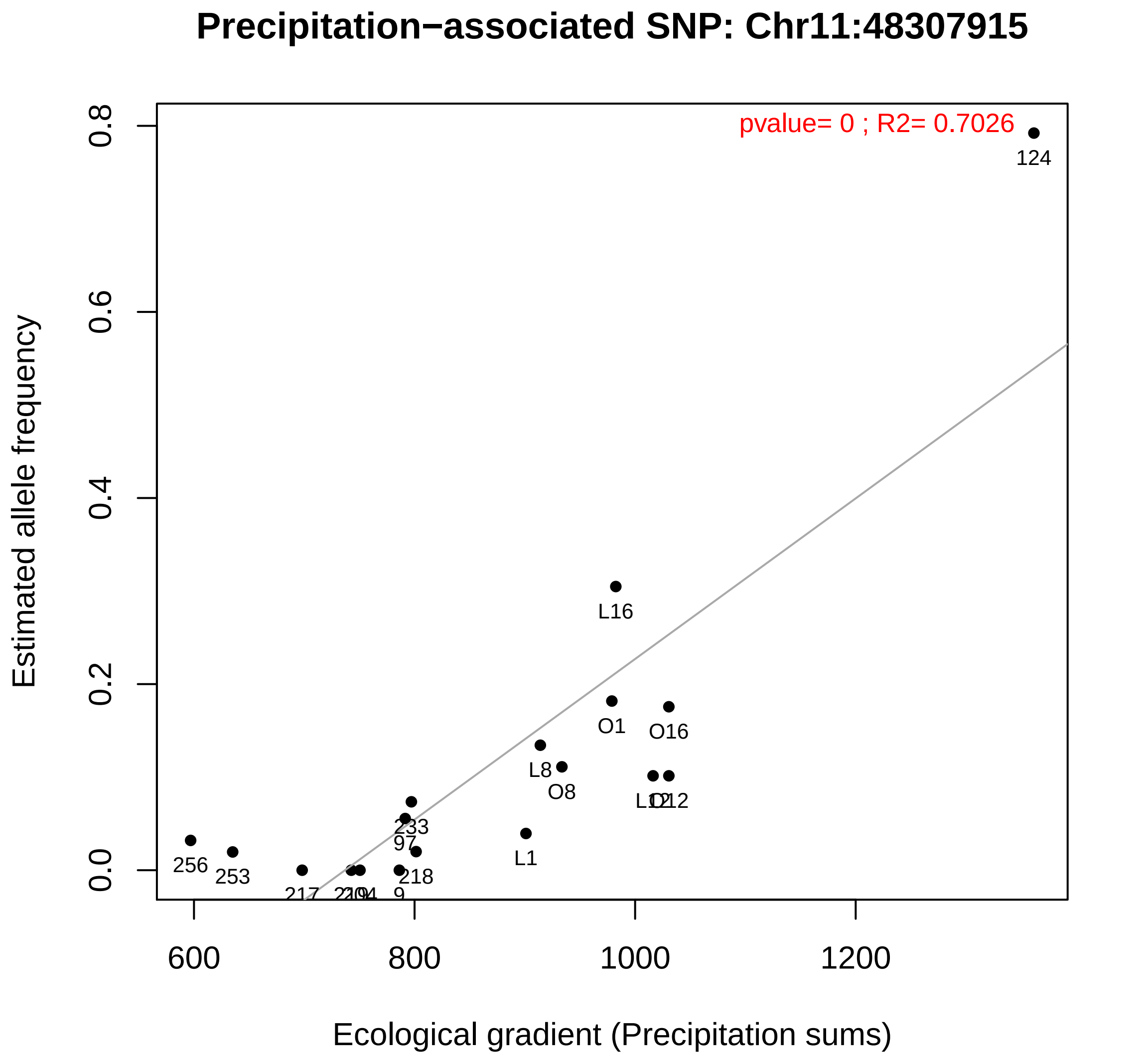

Precipitation−associated SNP: Chr11:48307915
pvalue= 0 ; R2= 0.7026
124
L16
O1
O16
L8
O8
233
97
218
L1
256
253
217
9
0.0
0.2
0.4
0.6
0.8
600
800
1000
1200
Estimated allele frequency
Ecological gradient (Precipitation sums)

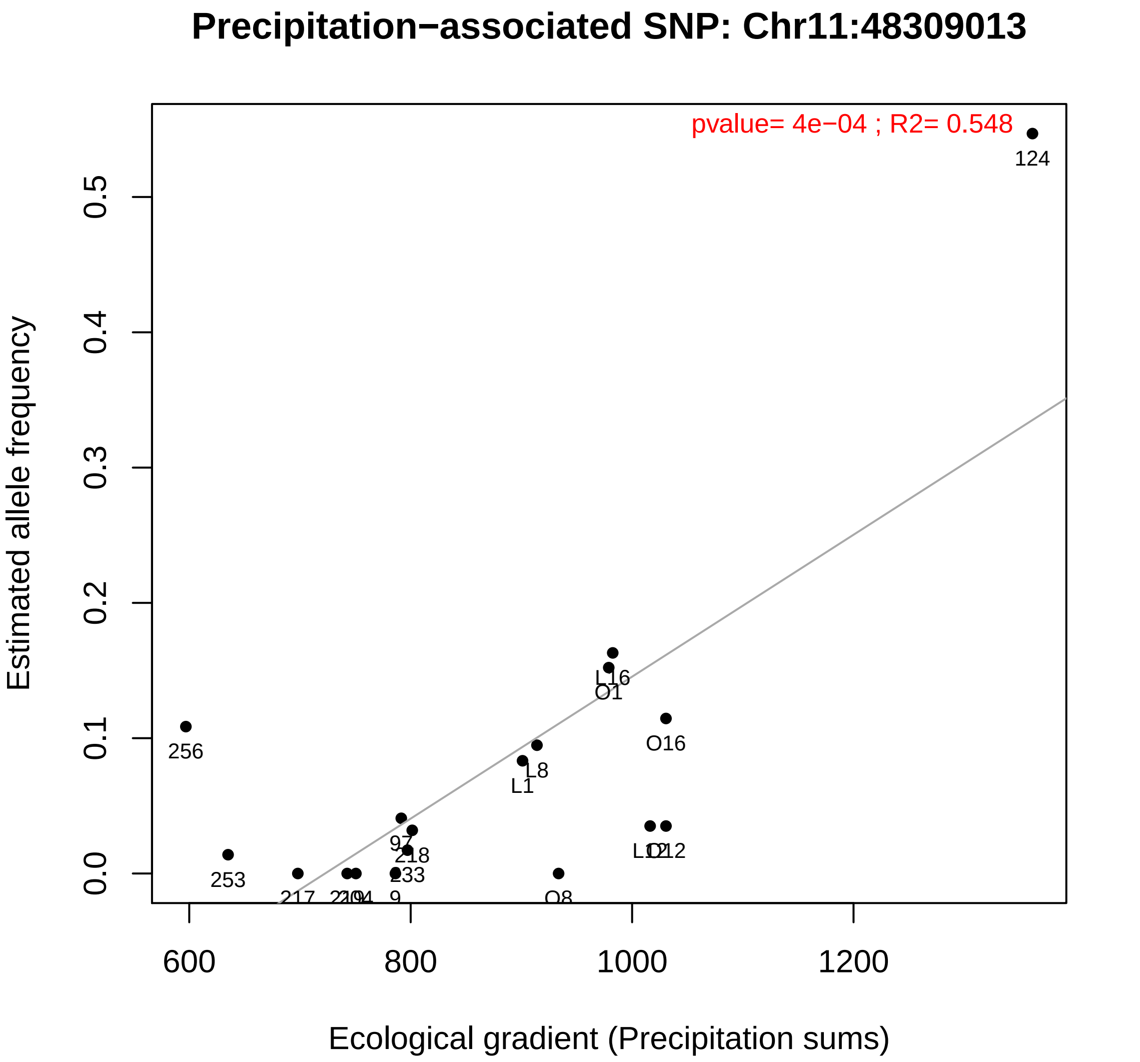
Precipitation−associated SNP: Chr11:48309013
pvalue= 4e−04 ; R2= 0.548
124
L16
O1
O16
256
L8
L1
97
218
233
253
217
9
O8
O12
0.0
0.1
0.2
0.3
0.4
0.5
600
800
1000
1200
Estimated allele frequency
Ecological gradient (Precipitation sums)

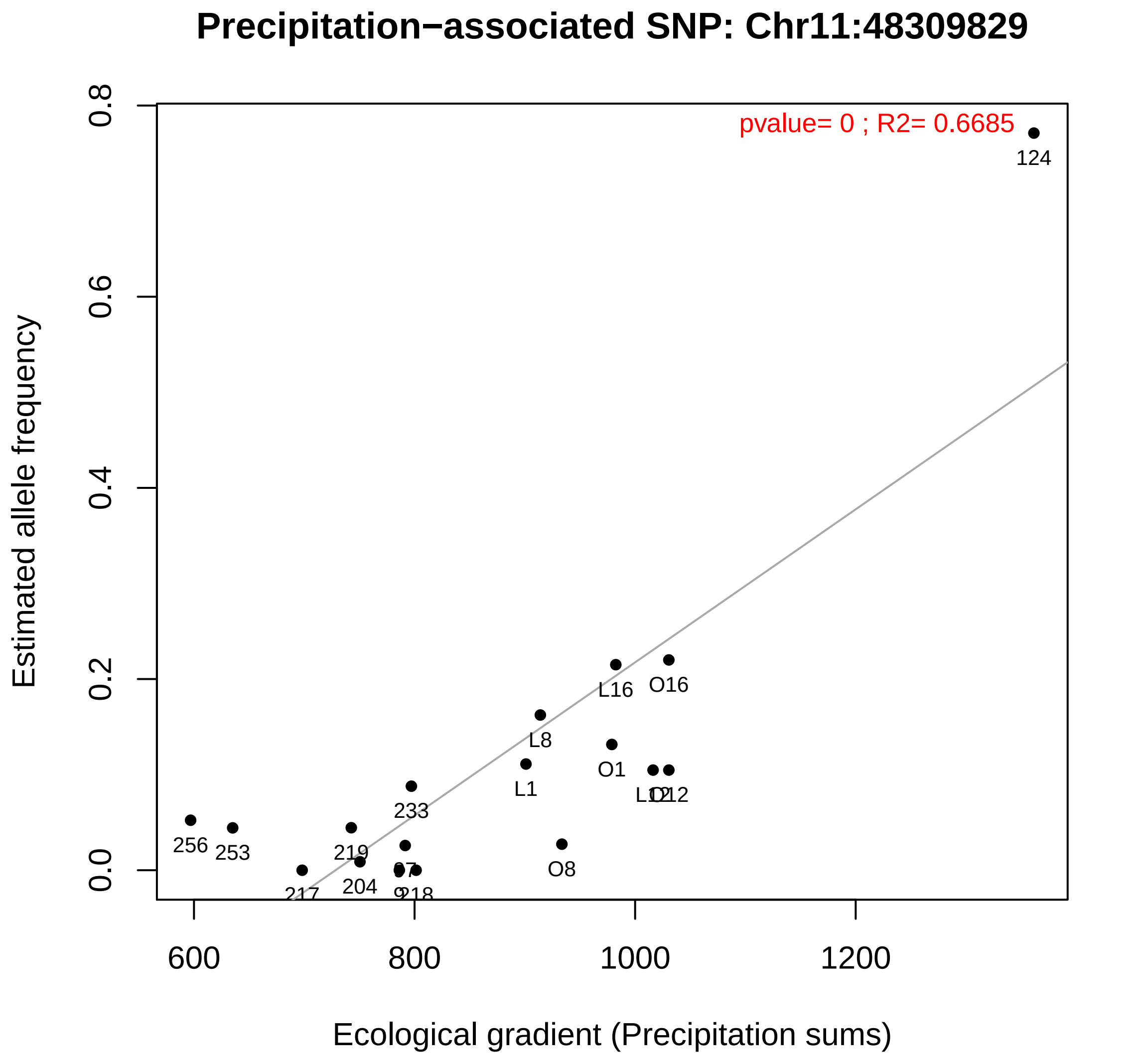

Precipitation−associated SNP: Chr11:48309829
pvalue= 0 ; R2= 0.6685
Estimated allele frequency
Ecological gradient (Precipitation sums)
0.0
0.2
0.4
0.6
0.8
600
800
1000
1200
124
L16
O16
L8
O1
L1
233
256
253
219
204
217
O8
218

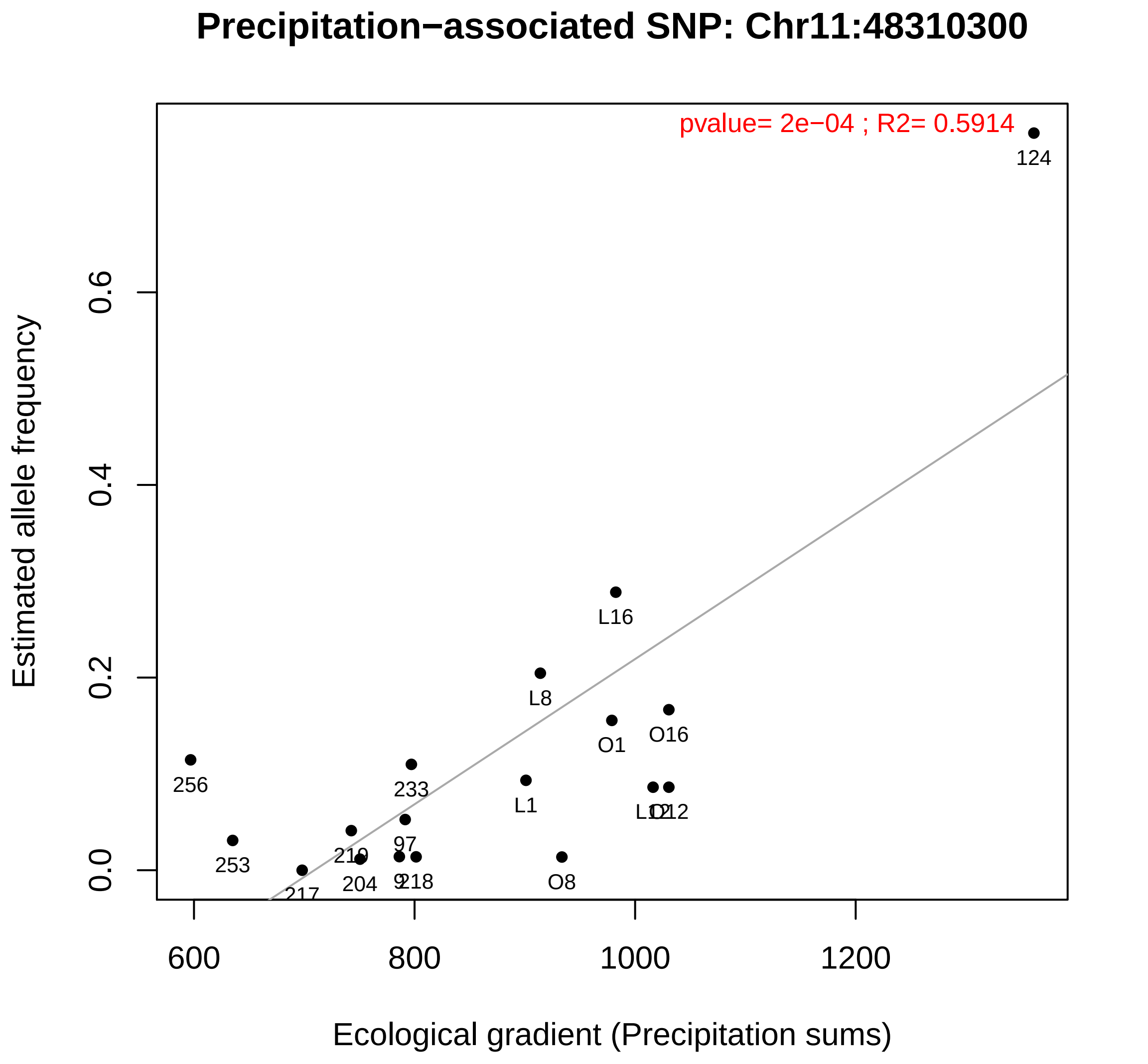

Precipitation−associated SNP: Chr11:48310300
pvalue= 2e−04 ; R2= 0.5914
Estimated allele frequency
0.0
0.2
0.4
0.6
600
800
1000
1200
Ecological gradient (Precipitation sums)
124
L16
L8
O1
O16
256
233
L1
L12
O12
97
253
210
204
217
9
218
O8

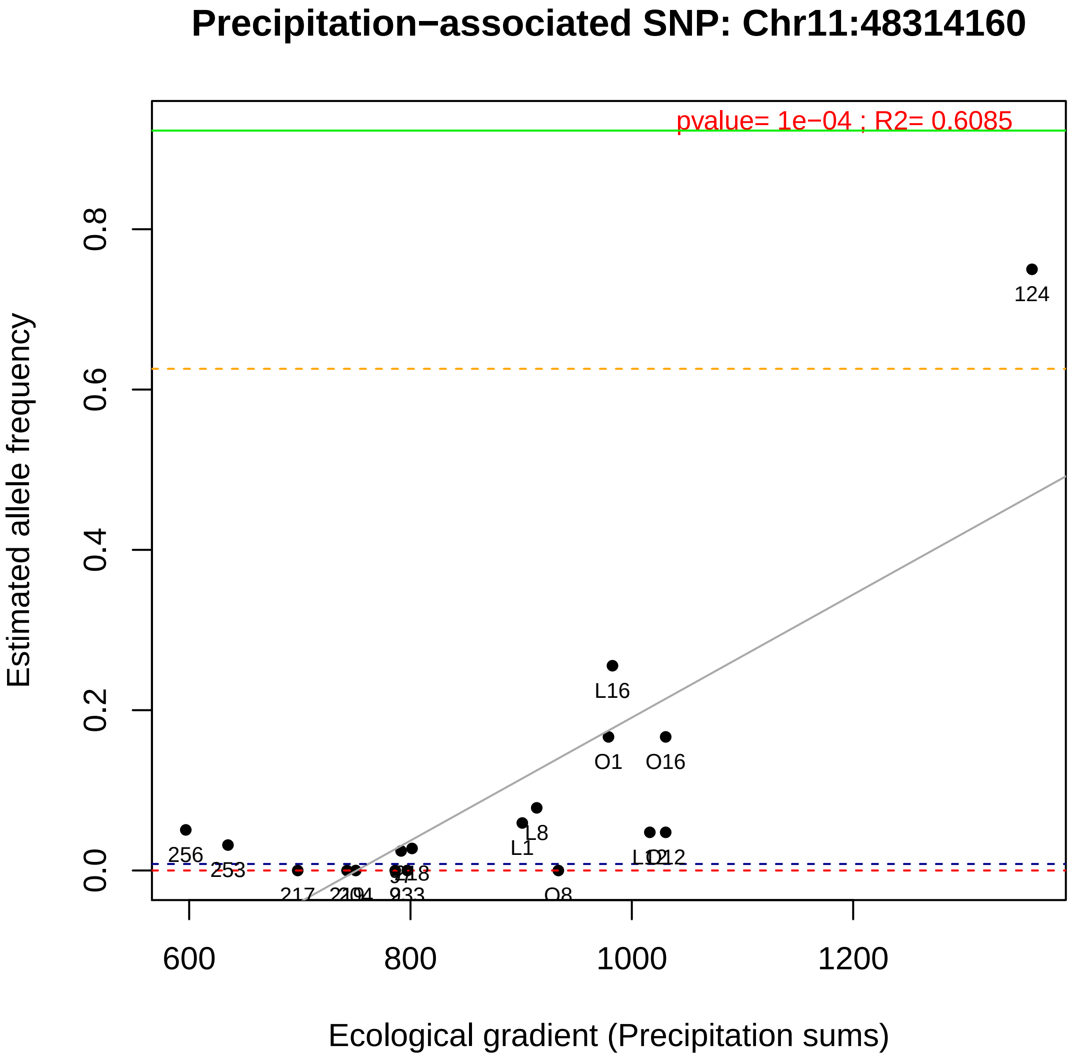
Precipitation−associated SNP: Chr11:48314160
pvalue= 1e−04 ; R2= 0.6085
Estimated allele frequency
0.0
0.2
0.4
0.6
0.8
124
L16
O1
O16
L8
L1
256
253
217
O8
600
800
1000
1200
Ecological gradient (Precipitation sums)

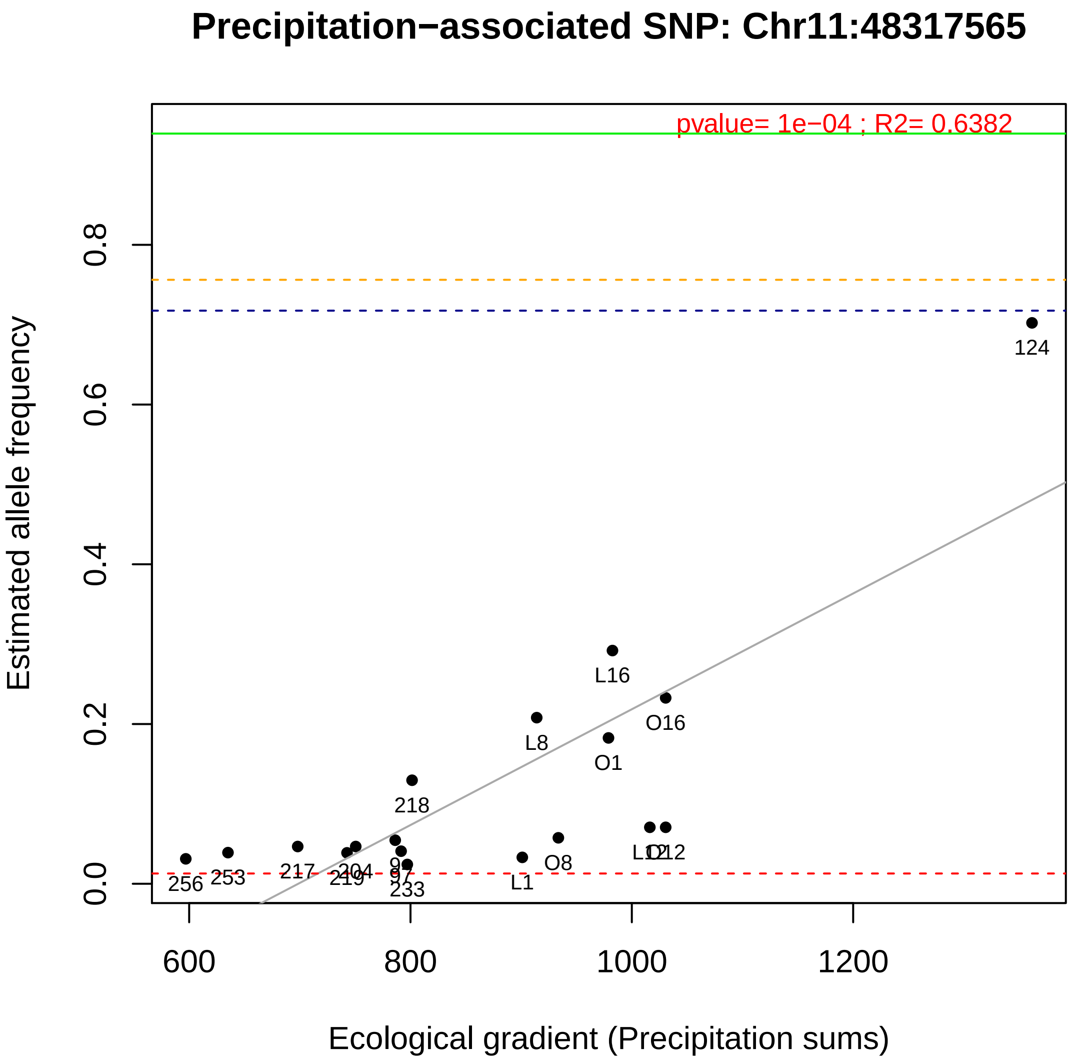
Precipitation−associated SNP: Chr11:48317565
pvalue= 1e−04 ; R2= 0.6382
Estimated allele frequency
0.0
0.2
0.4
0.6
0.8
600
800
1000
1200
Ecological gradient (Precipitation sums)
124
L16
O16
L8
O1
218
O8
L1
256
253
217
204
219
233

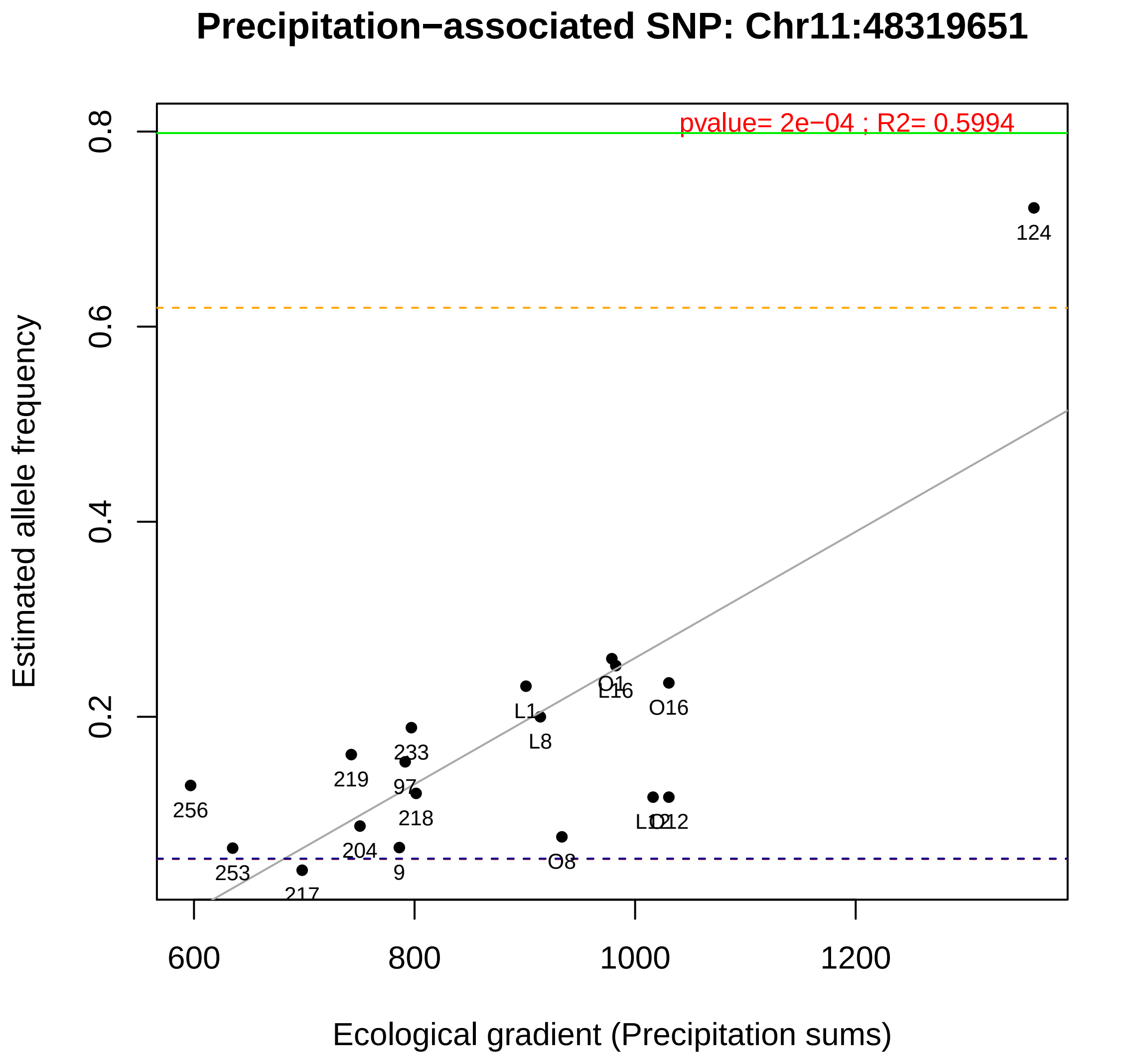

Precipitation−associated SNP: Chr11:48319651
pvalue= 2e−04 ; R2= 0.5994
Estimated allele frequency
Ecological gradient (Precipitation sums)
0.2
0.4
0.6
0.8
600
800
1000
1200
124
O1
L16
O16
L1
L8
233
219
97
218
256
204
253
9
217
O8
L12
O12

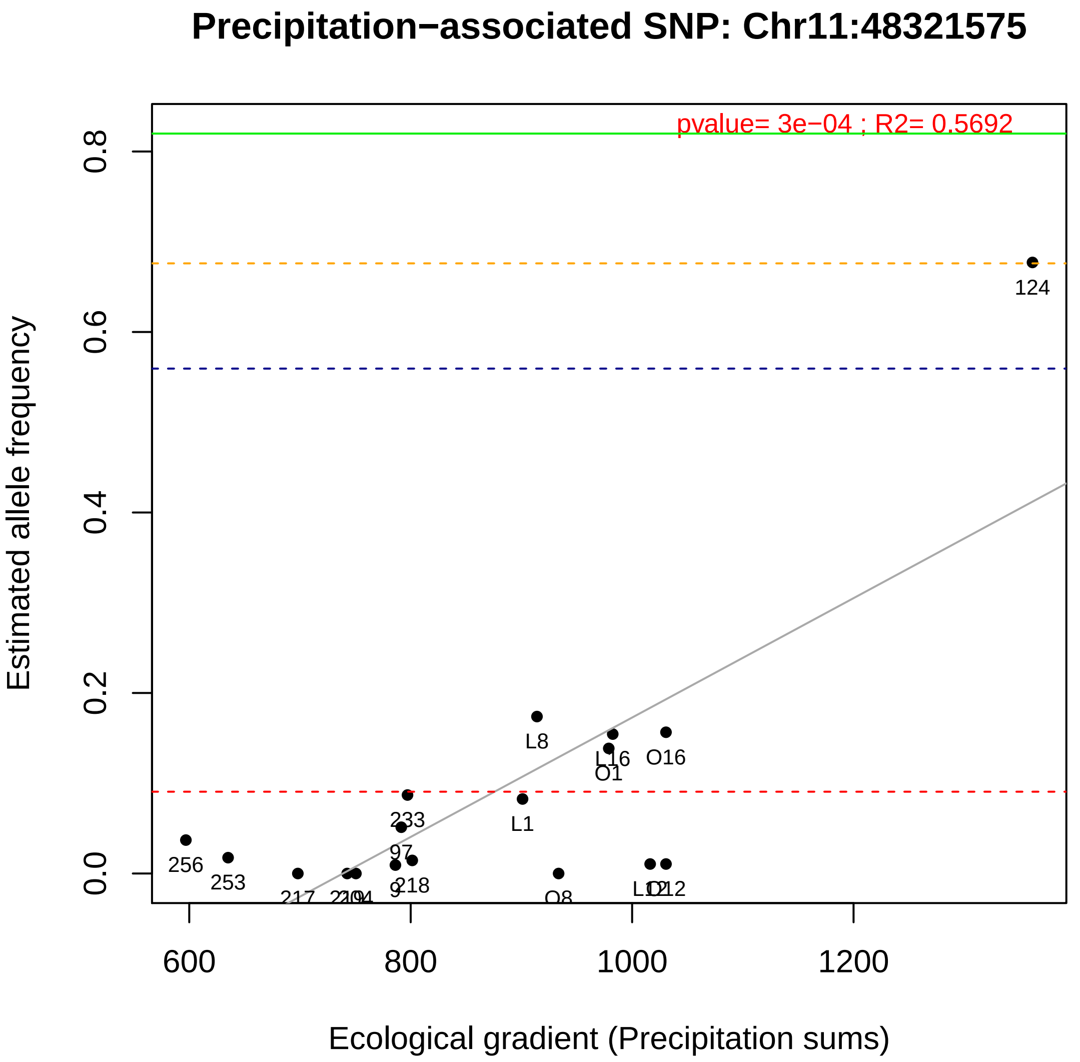

Precipitation−associated SNP: Chr11:48321575
pvalue= 3e−04 ; R2= 0.5692
Estimated allele frequency
0.0
0.2
0.4
0.6
0.8
600
800
1000
1200
Ecological gradient (Precipitation sums)
124
L8
L16
O1
O16
233
L1
97
256
253
218

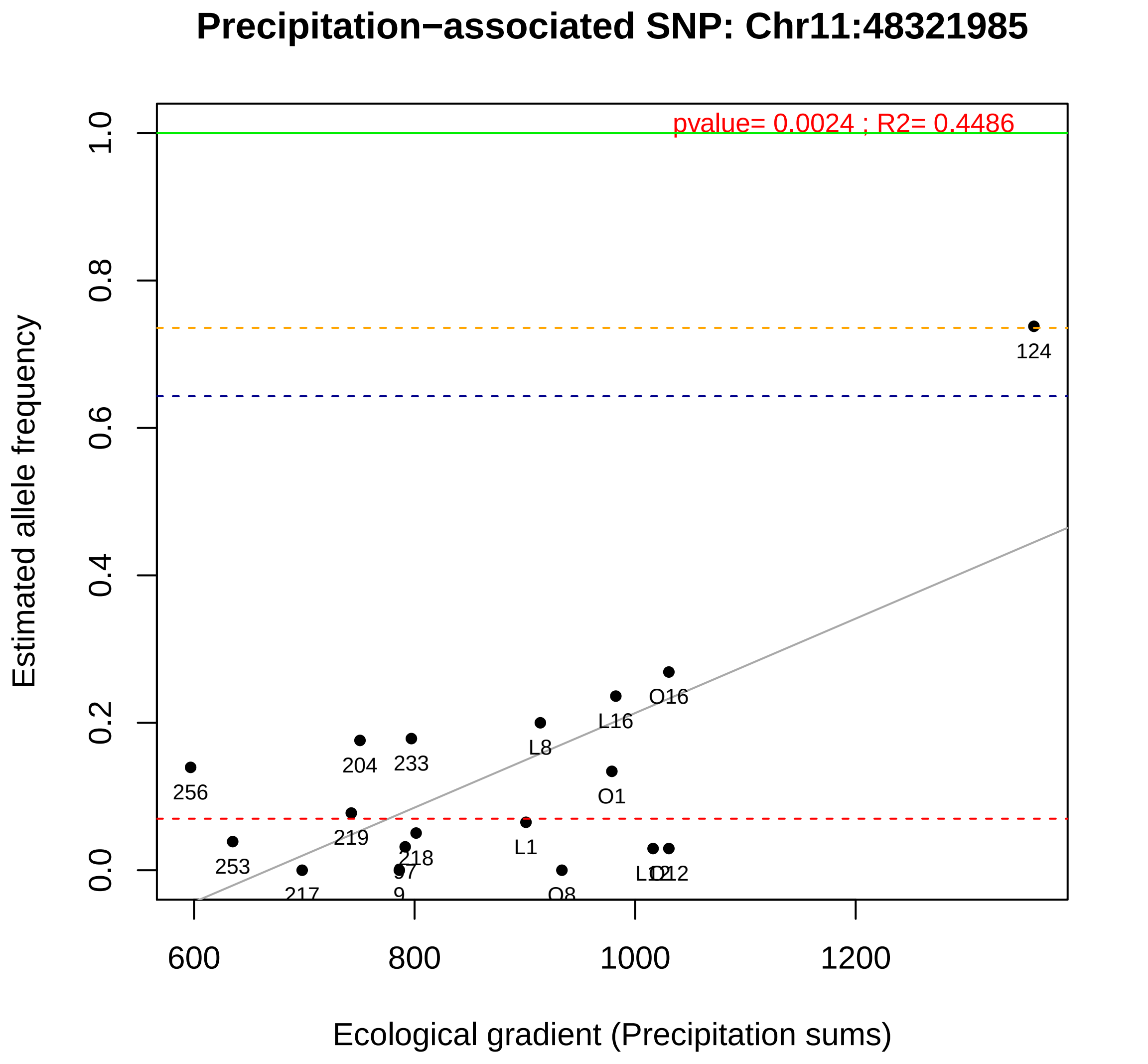

Precipitation−associated SNP: Chr11:48321985
pvalue= 0.0024 ; R2= 0.4486
Estimated allele frequency
Ecological gradient (Precipitation sums)
0.0
0.2
0.4
0.6
0.8
1.0
600
800
1000
1200
124
O16
L16
L8
233
204
256
O1
219
253
218
L1
97
217
9
O8
L10
O12

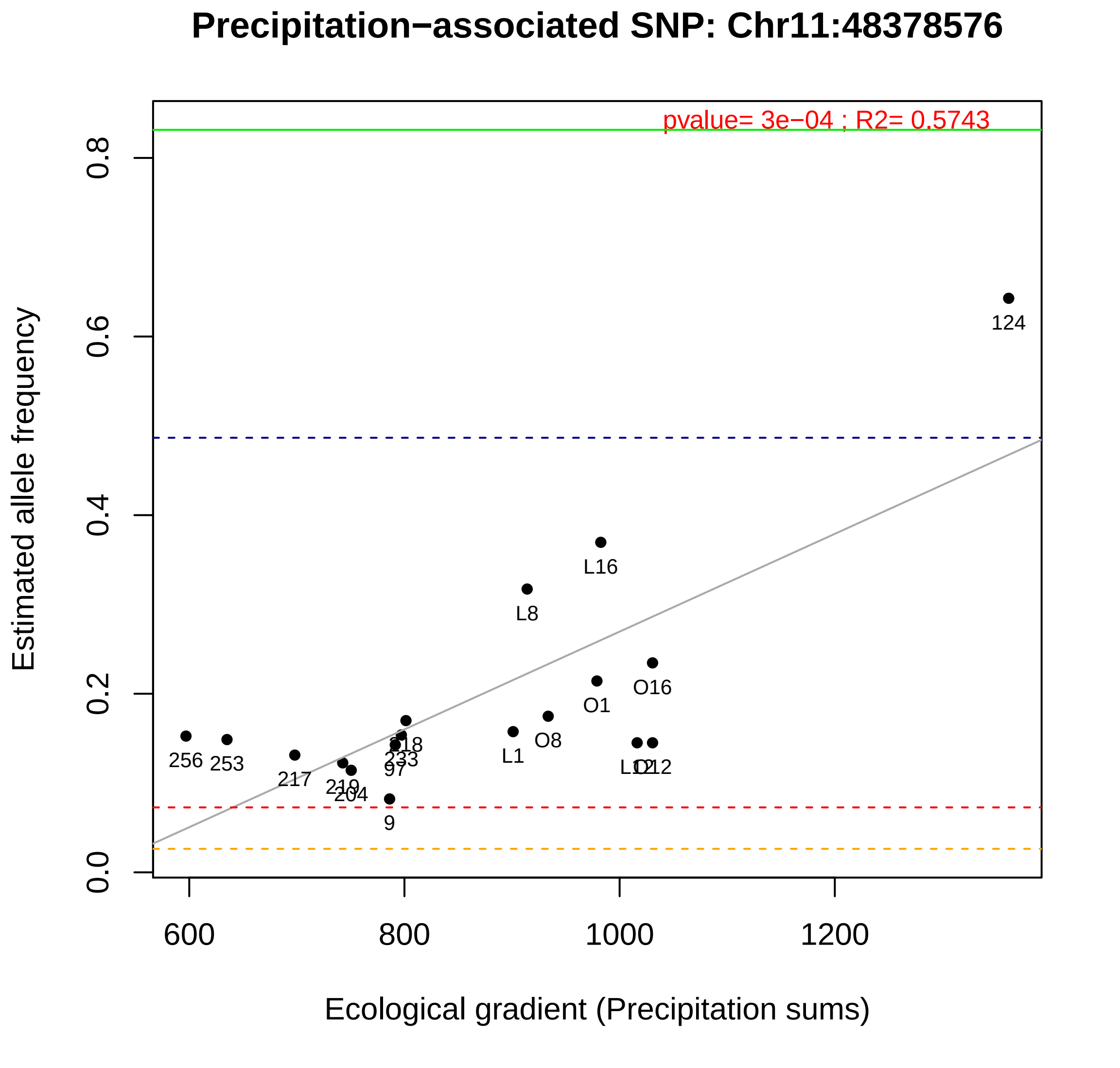
Precipitation−associated SNP: Chr11:48378576
pvalue= 3e−04 ; R2= 0.5743
Estimated allele frequency
0.0
0.2
0.4
0.6
0.8
600
800
1000
1200
Ecological gradient (Precipitation sums)
124
L16
L8
O16
O1
O8
L1
256
253
217
219
204
233
97
9
L12
O12

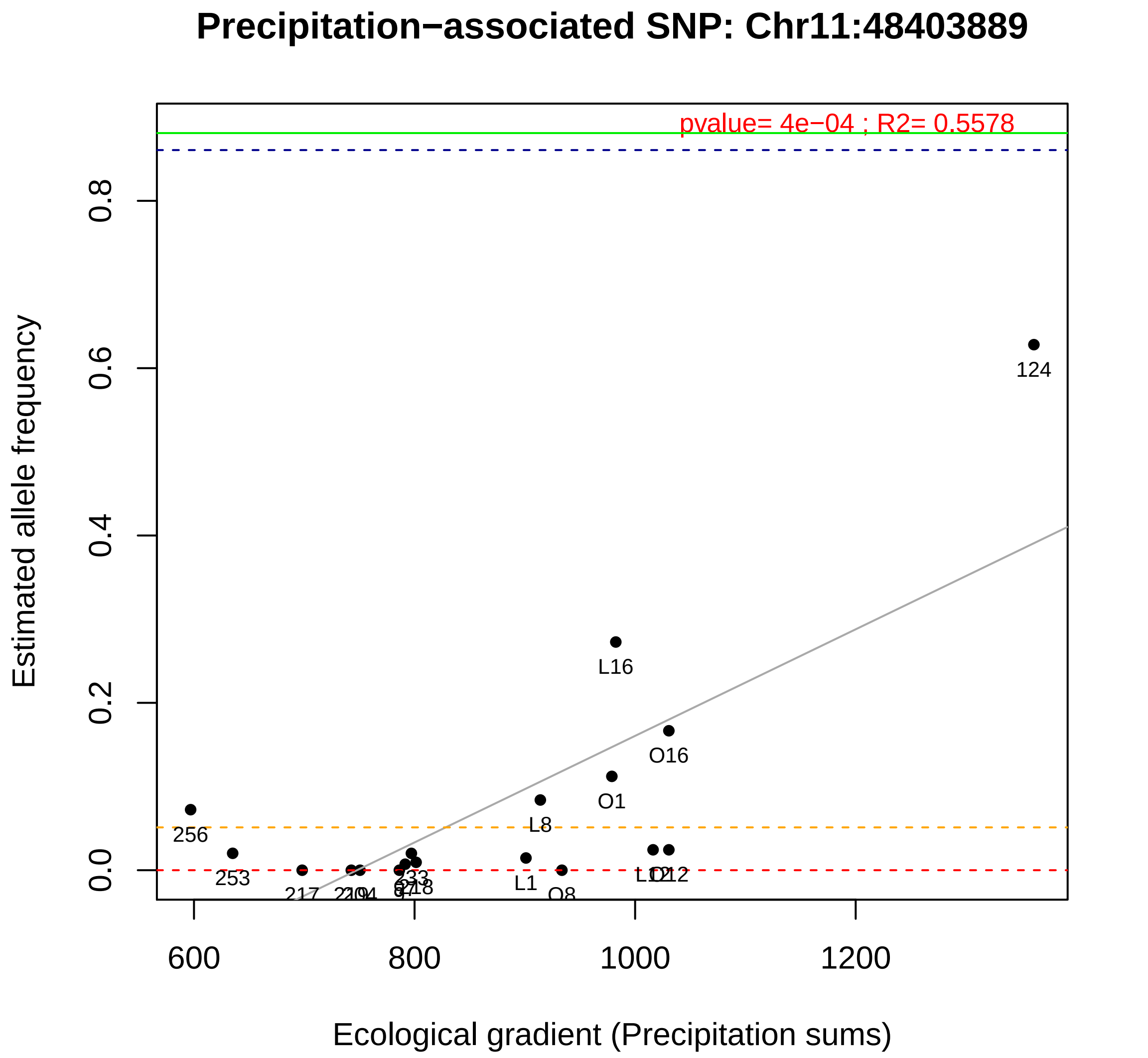

Precipitation−associated SNP: Chr11:48403889
pvalue= 4e−04 ; R2= 0.5578
Estimated allele frequency
Ecological gradient (Precipitation sums)
0.0
0.2
0.4
0.6
0.8
600
800
1000
1200
124
L16
O16
O1
L8
256
253
217
L1
O8

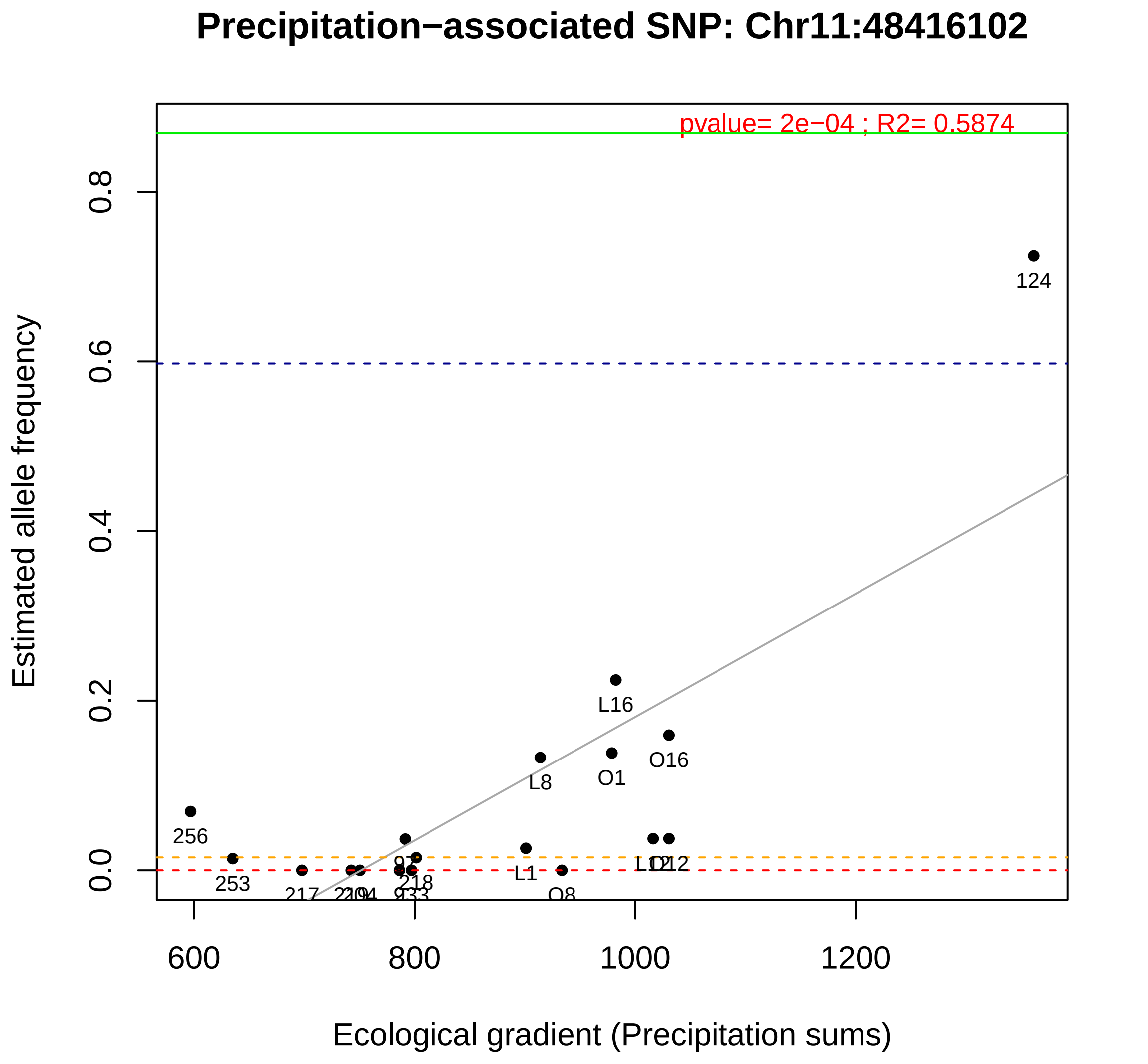

Precipitation−associated SNP: Chr11:48416102
pvalue= 2e−04 ; R2= 0.5874
Estimated allele frequency
Ecological gradient (Precipitation sums)
0.0
0.2
0.4
0.6
0.8
600
800
1000
1200
124
L16
O16
O1
L8
256
253
217
218
L1
O8

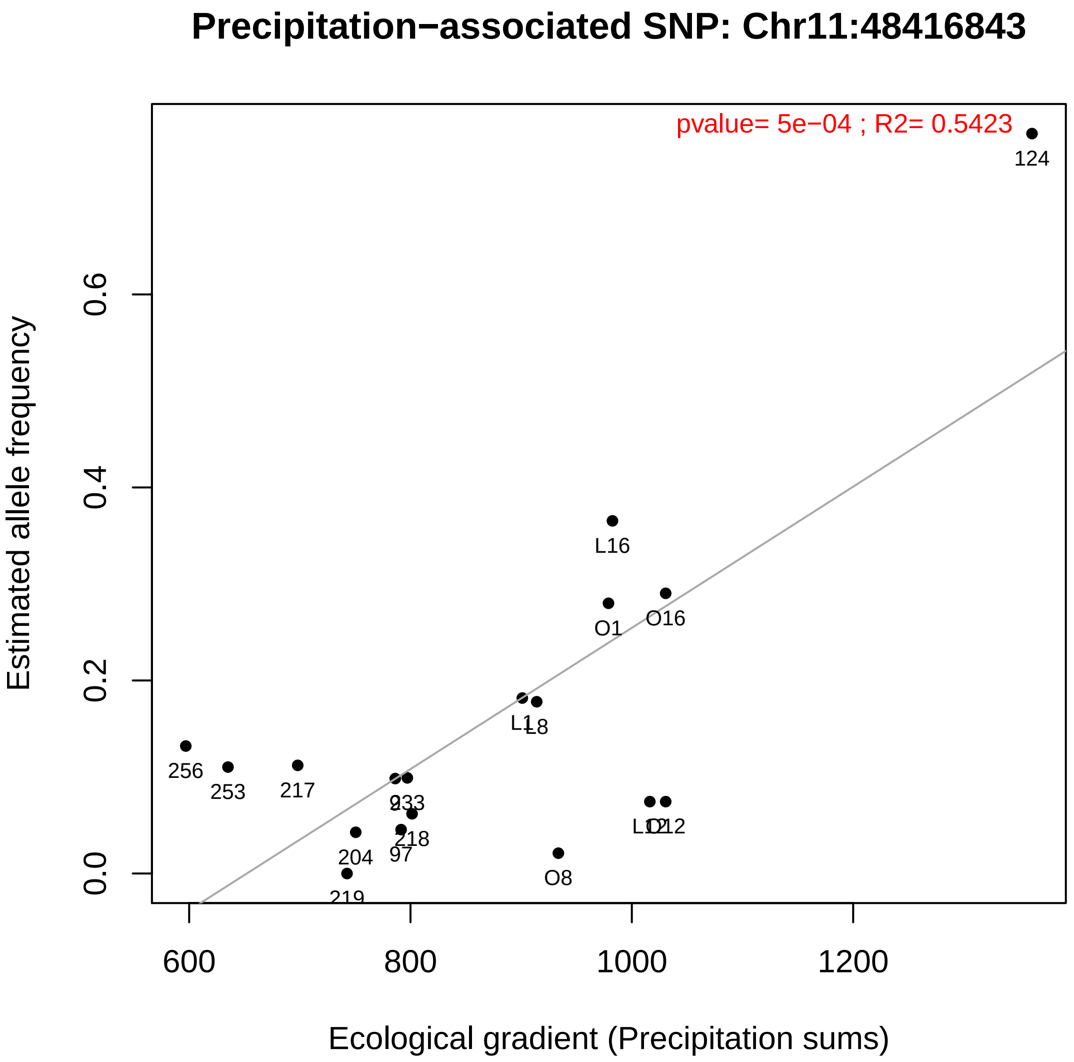

Precipitation−associated SNP: Chr11:48416843
pvalue= 5e−04 ; R2= 0.5423
124
L16
O1
O16
L1
L8
256
253
217
233
218
204
97
219
O8
O12
0.0
0.2
0.4
0.6
600
800
1000
1200
Estimated allele frequency
Ecological gradient (Precipitation sums)

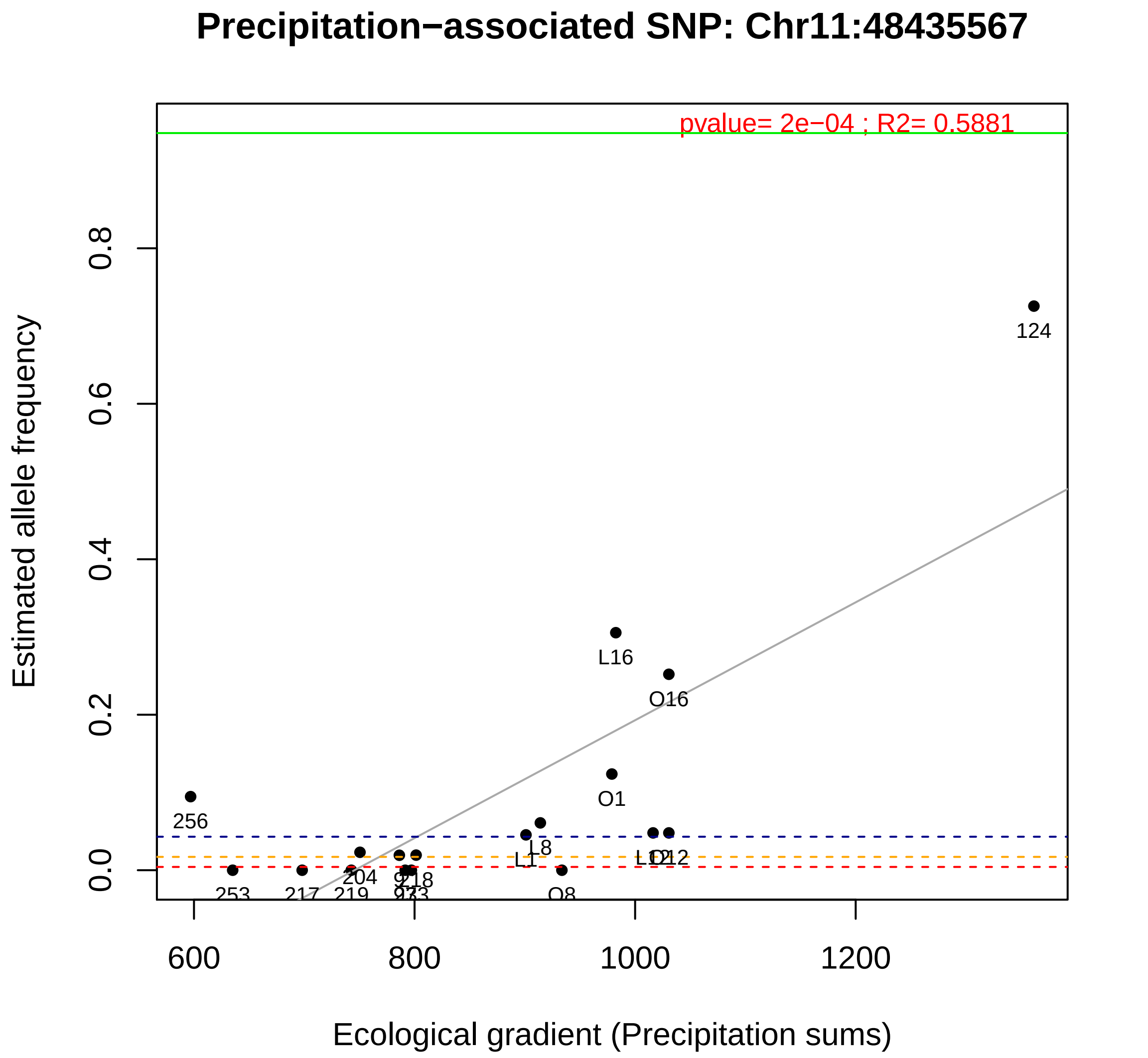

Precipitation−associated SNP: Chr11:48435567
pvalue= 2e−04 ; R2= 0.5881
Estimated allele frequency
0.0
0.2
0.4
0.6
0.8
600
800
1000
1200
Ecological gradient (Precipitation sums)
124
L16
O16
O1
256
L8
L1
O12
204
218
253
217
219
O8

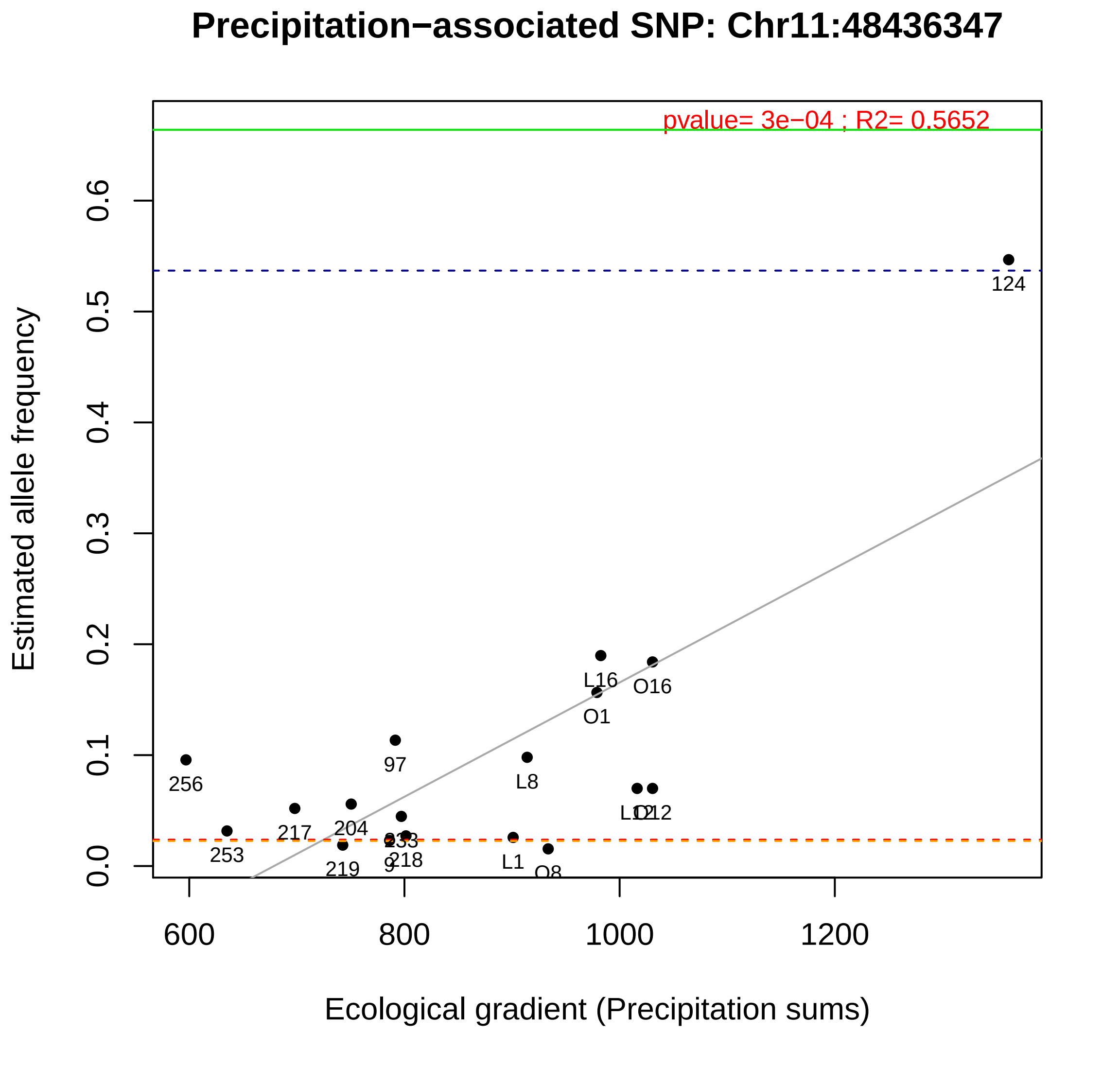
Precipitation−associated SNP: Chr11:48436347
pvalue= 3e−04 ; R2= 0.5652
Estimated allele frequency
Ecological gradient (Precipitation sums)
0.0
0.1
0.2
0.3
0.4
0.5
0.6
600
800
1000
1200
124
L16
O16
O1
97
256
L8
217
204
253
219
218
L1

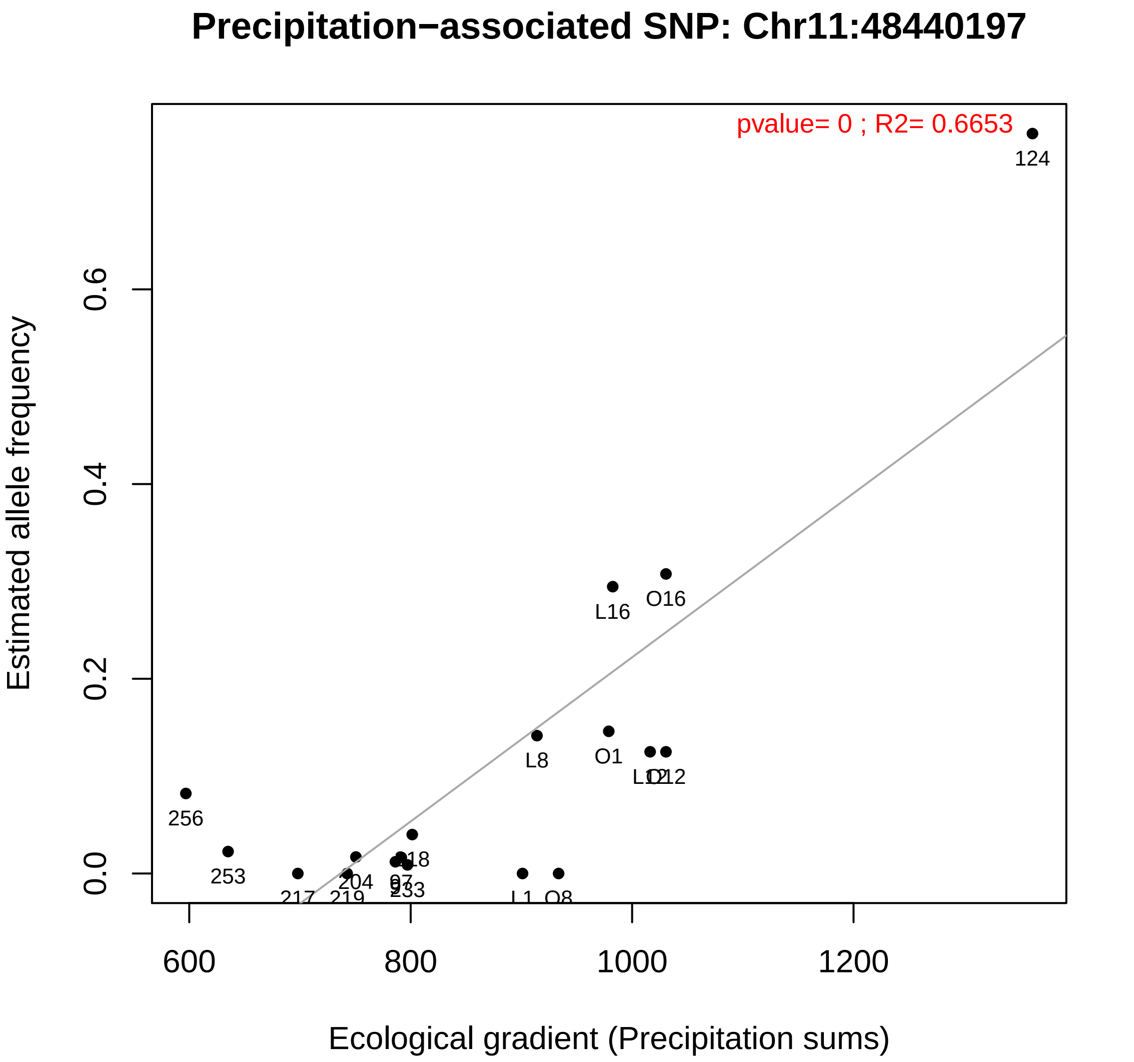

Precipitation−associated SNP: Chr11:48440197
pvalue= 0 ; R2= 0.6653
124
O16
L16
L8
O1
O12
256
253
218
204
233
Estimated allele frequency
0.0
0.2
0.4
0.6
600
800
1000
1200
Ecological gradient (Precipitation sums)

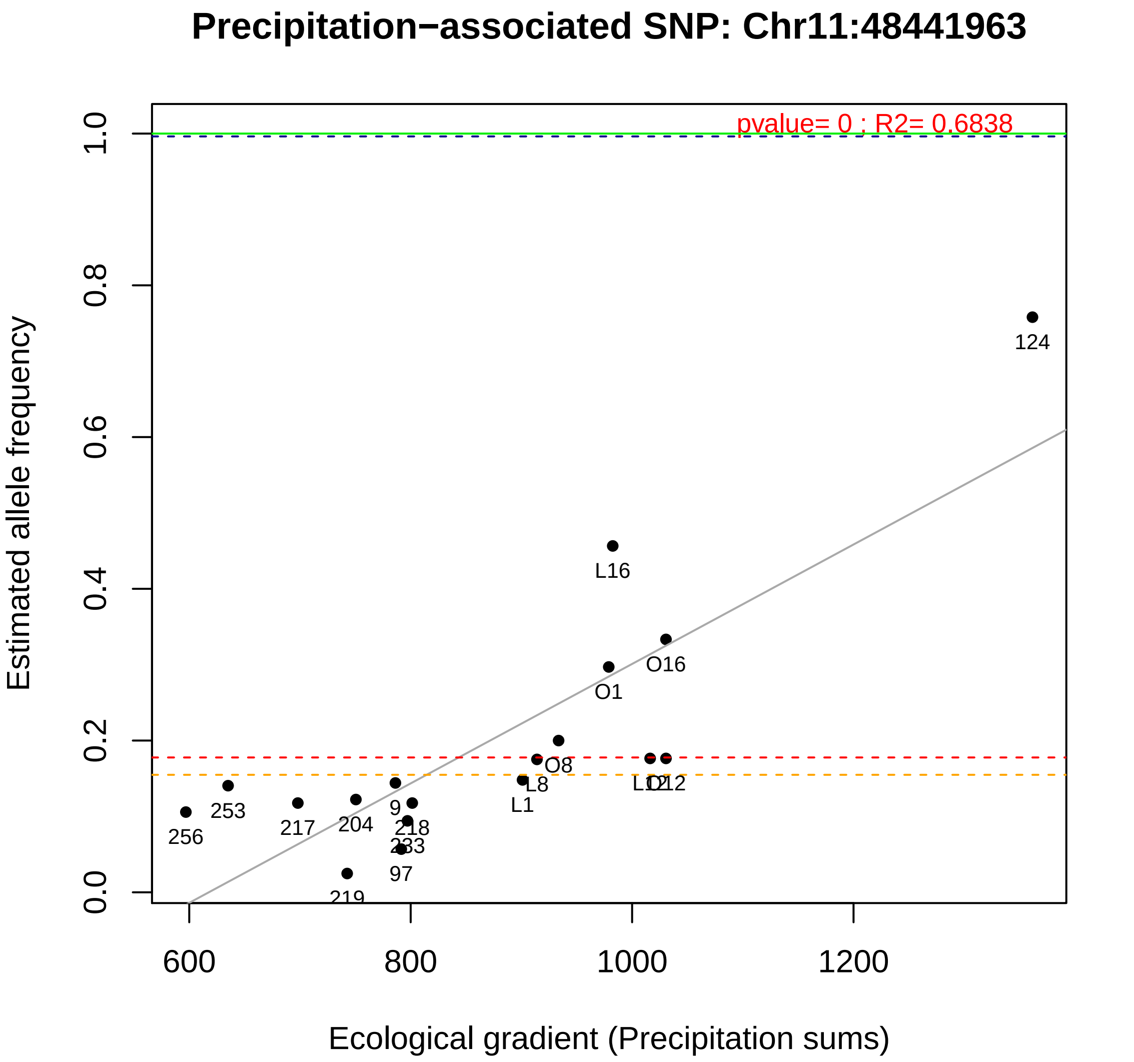

Precipitation−associated SNP: Chr11:48441963
pvalue= 0 ; R2= 0.6838
Estimated allele frequency
Ecological gradient (Precipitation sums)
0.0
0.2
0.4
0.6
0.8
1.0
600
800
1000
1200
124
L16
O16
O1
O8
L8
L1
253
256
217
204
9
218
233
97
219

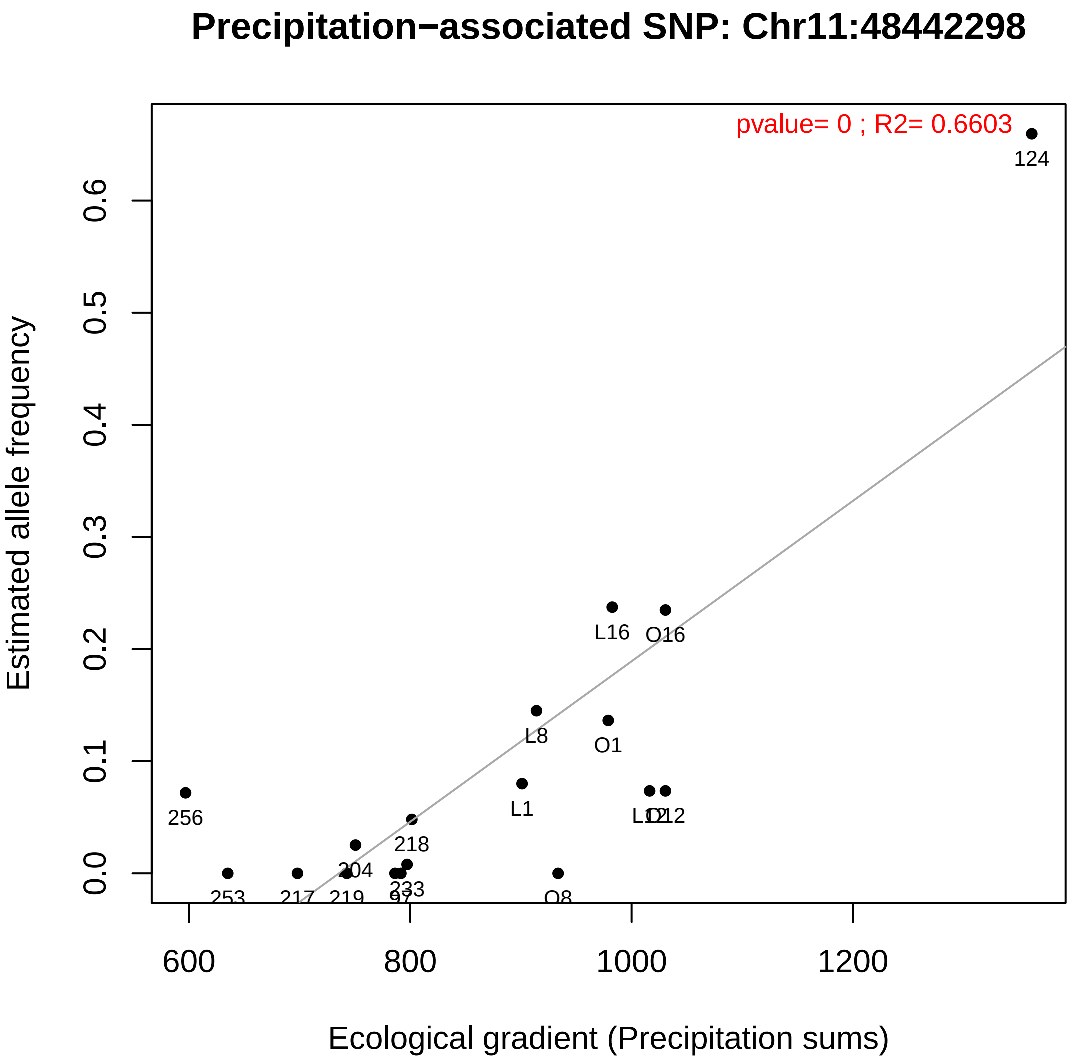
Precipitation−associated SNP: Chr11:48442298
pvalue= 0 ; R2= 0.6603
Estimated allele frequency
Ecological gradient (Precipitation sums)
0.0
0.1
0.2
0.3
0.4
0.5
0.6
600
800
1000
1200
124
L16
O16
L8
O1
256
L1
218
204
233
253
217
219
O8

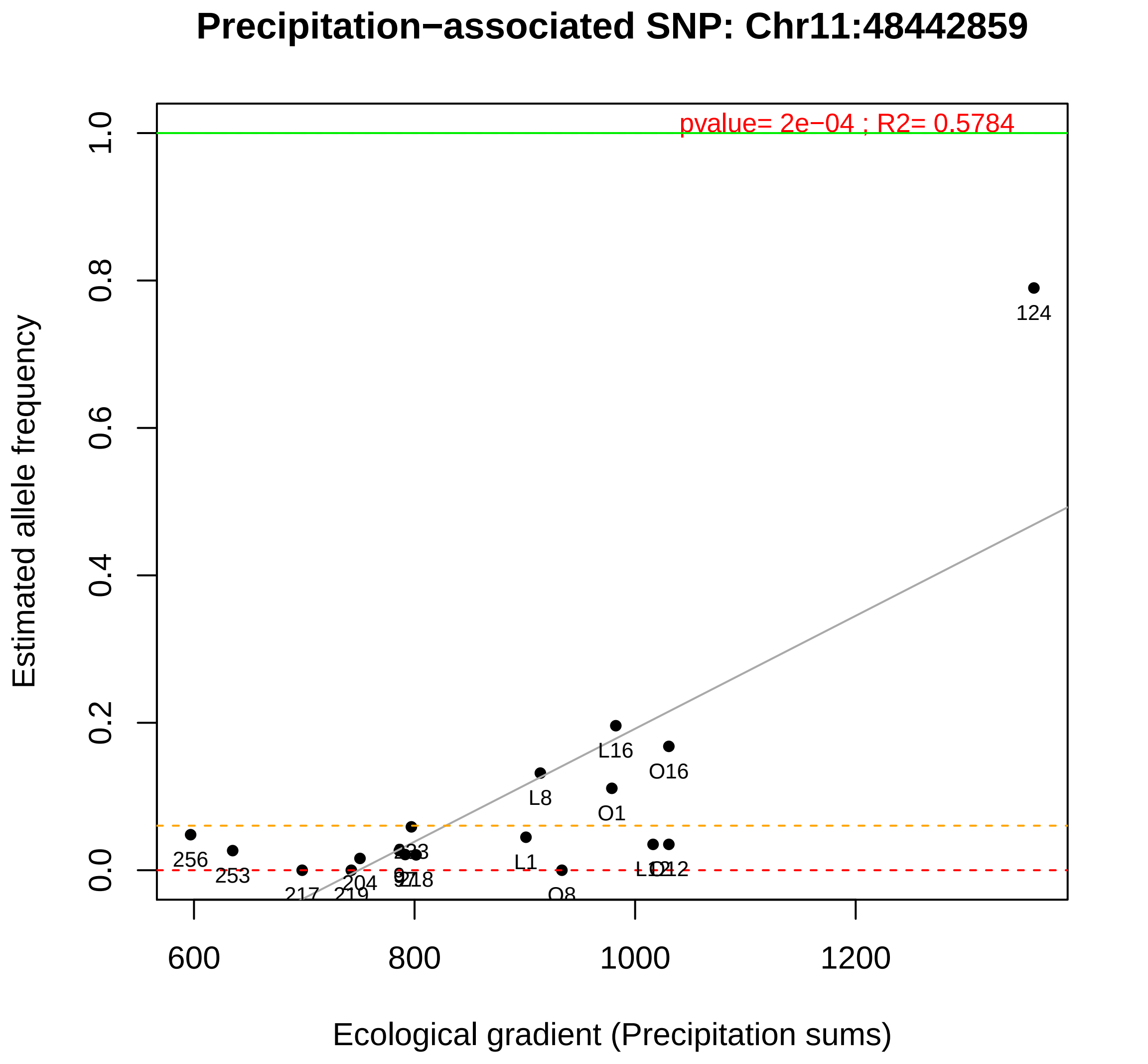
Precipitation−associated SNP: Chr11:48442859
pvalue= 2e−04 ; R2= 0.5784
Estimated allele frequency
0.0
0.2
0.4
0.6
0.8
1.0
Ecological gradient (Precipitation sums)
600
800
1000
1200
124
L16
O16
L8
O1
256
253
L1
217
219
204
218
O8
O12

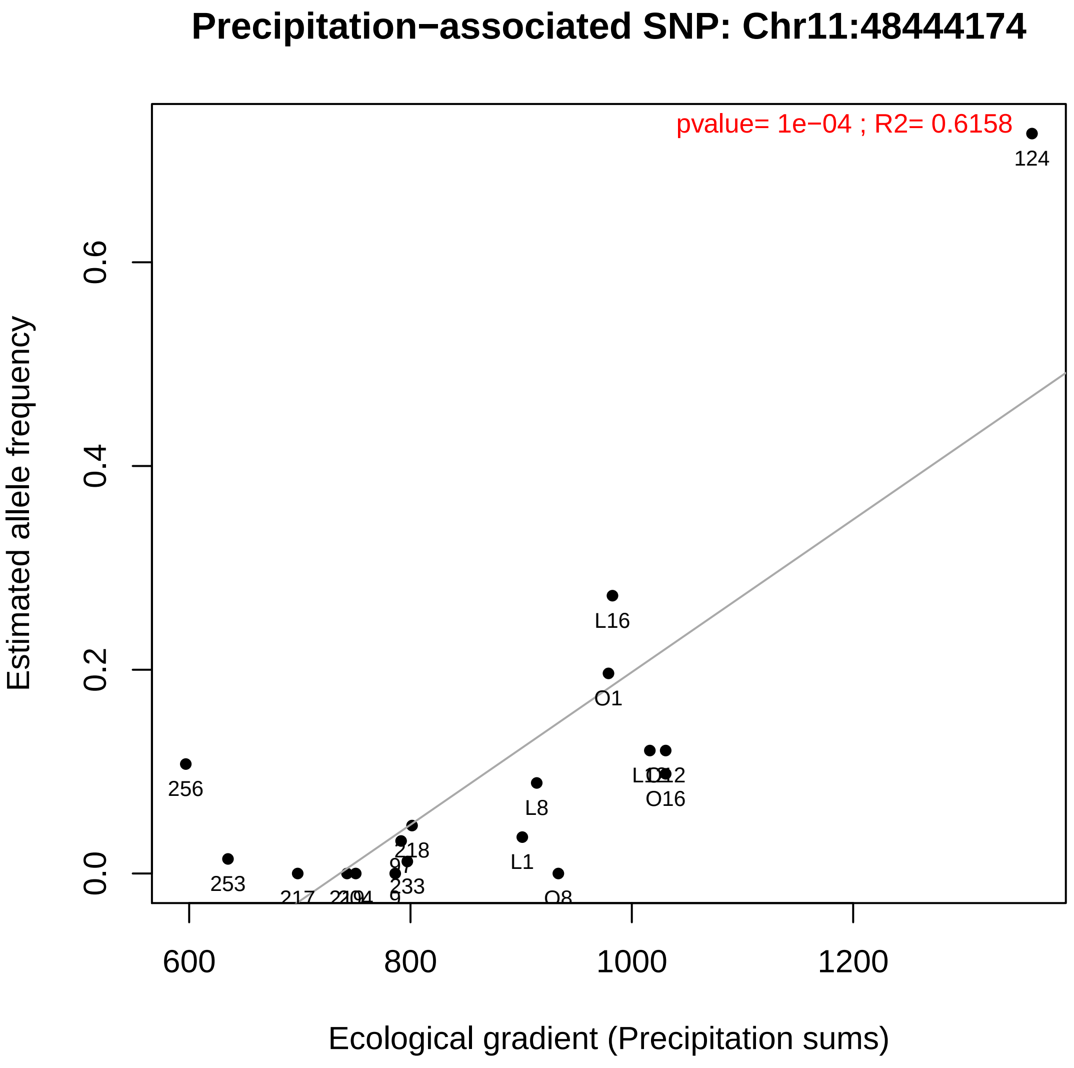

Precipitation−associated SNP: Chr11:48444174
pvalue= 1e−04 ; R2= 0.6158
Estimated allele frequency
Ecological gradient (Precipitation sums)
0.0
0.2
0.4
0.6
600
800
1000
1200
124
L16
O1
256
L8
O16
218
L1
253
233
O8
217

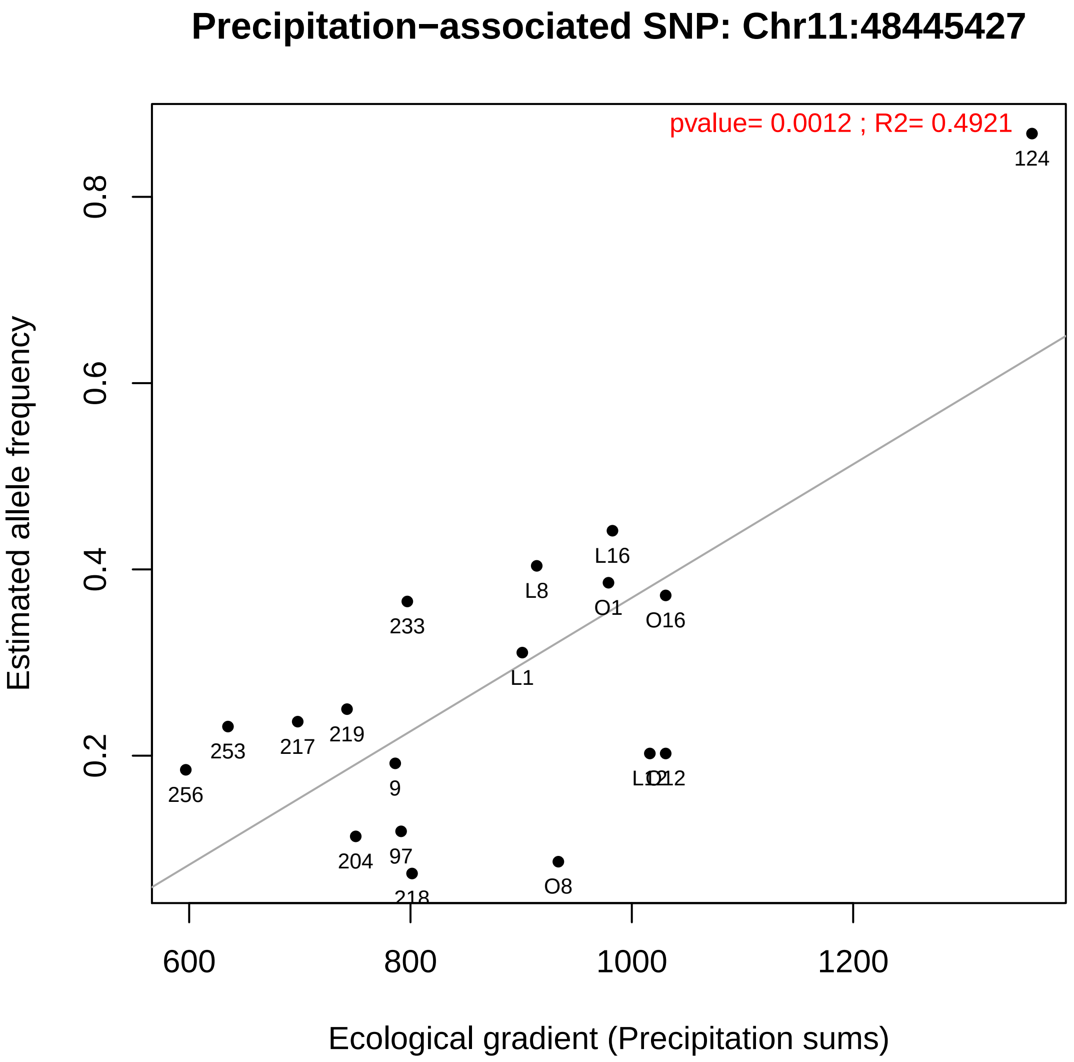

Precipitation−associated SNP: Chr11:48445427
pvalue= 0.0012 ; R2= 0.4921
124
L16
L8
O1
O16
233
L1
219
253
217
256
9
L12
O12
204
97
218
O8
0.2
0.4
0.6
0.8
600
800
1000
1200
Estimated allele frequency
Ecological gradient (Precipitation sums)

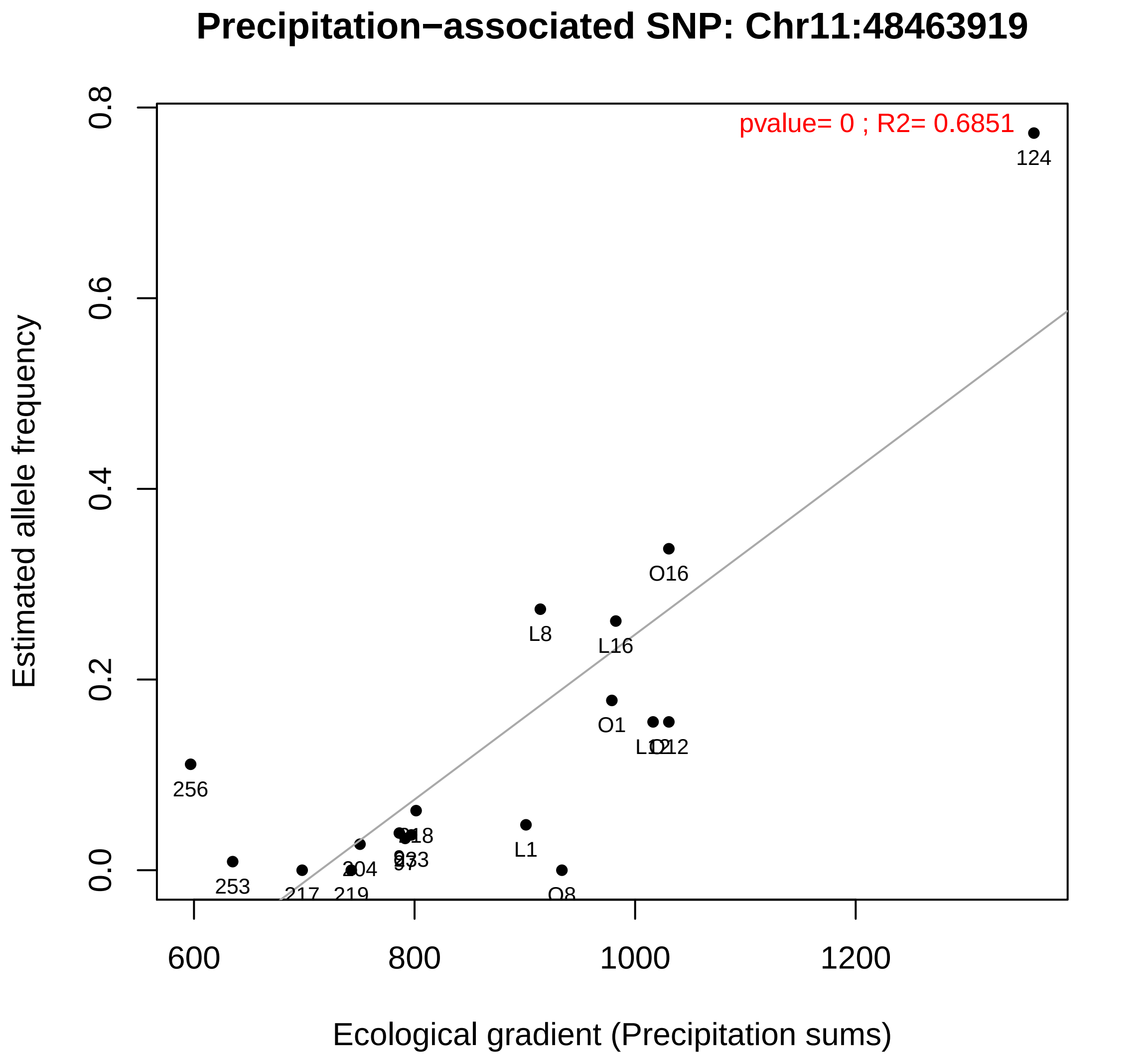

Precipitation−associated SNP: Chr11:48463919
pvalue= 0 ; R2= 0.6851
Estimated allele frequency
Ecological gradient (Precipitation sums)
0.0
0.2
0.4
0.6
0.8
600
800
1000
1200
124
O16
L8
L16
O1
256
L1
253
204
217
219
O8

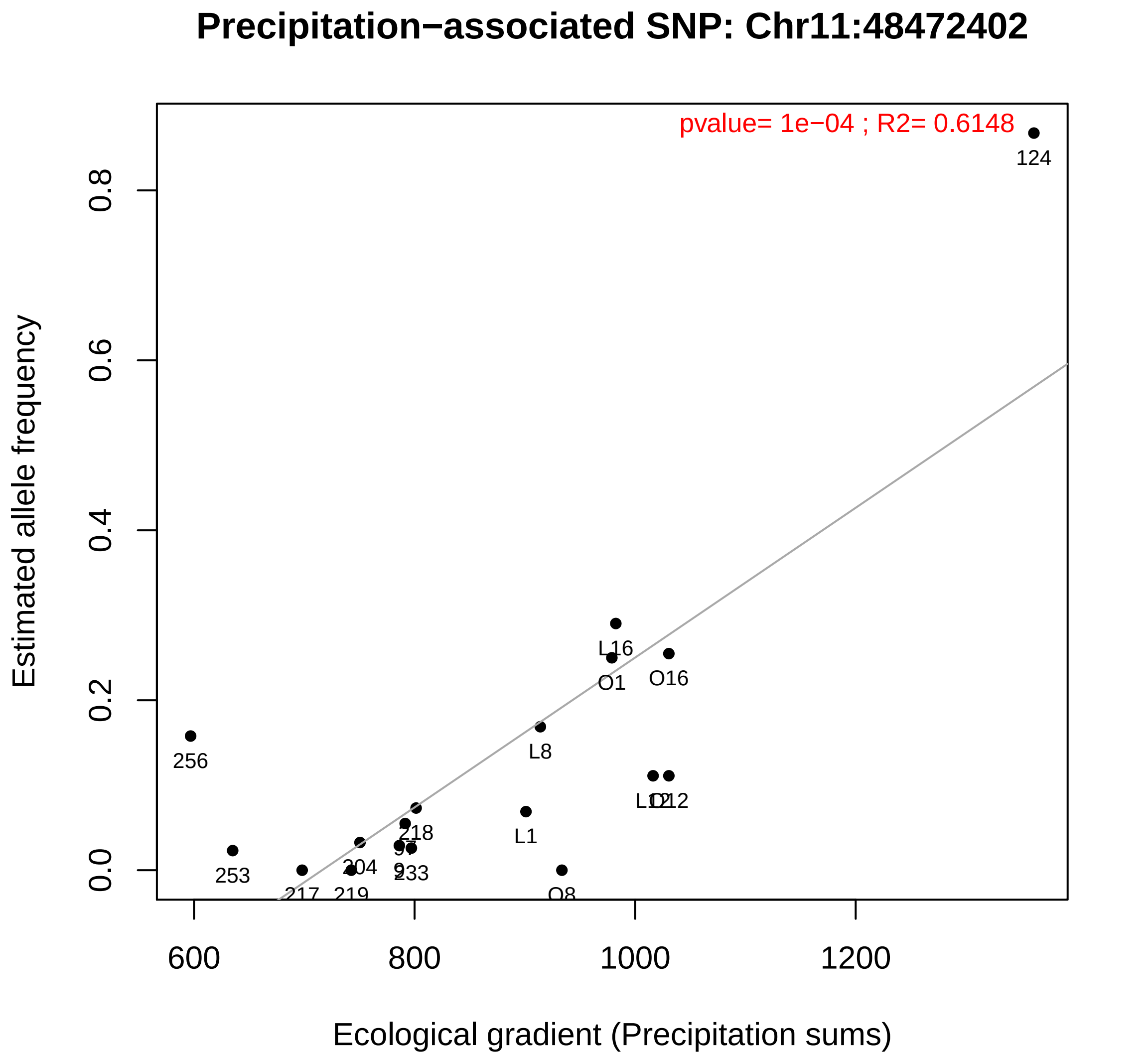

Precipitation−associated SNP: Chr11:48472402
pvalue= 1e−04 ; R2= 0.6148
124
L16
O1
O16
L8
256
L10
O12
218
L1
97
204
233
253
217
219
9
O8
0.0
0.2
0.4
0.6
0.8
600
800
1000
1200
Estimated allele frequency
Ecological gradient (Precipitation sums)

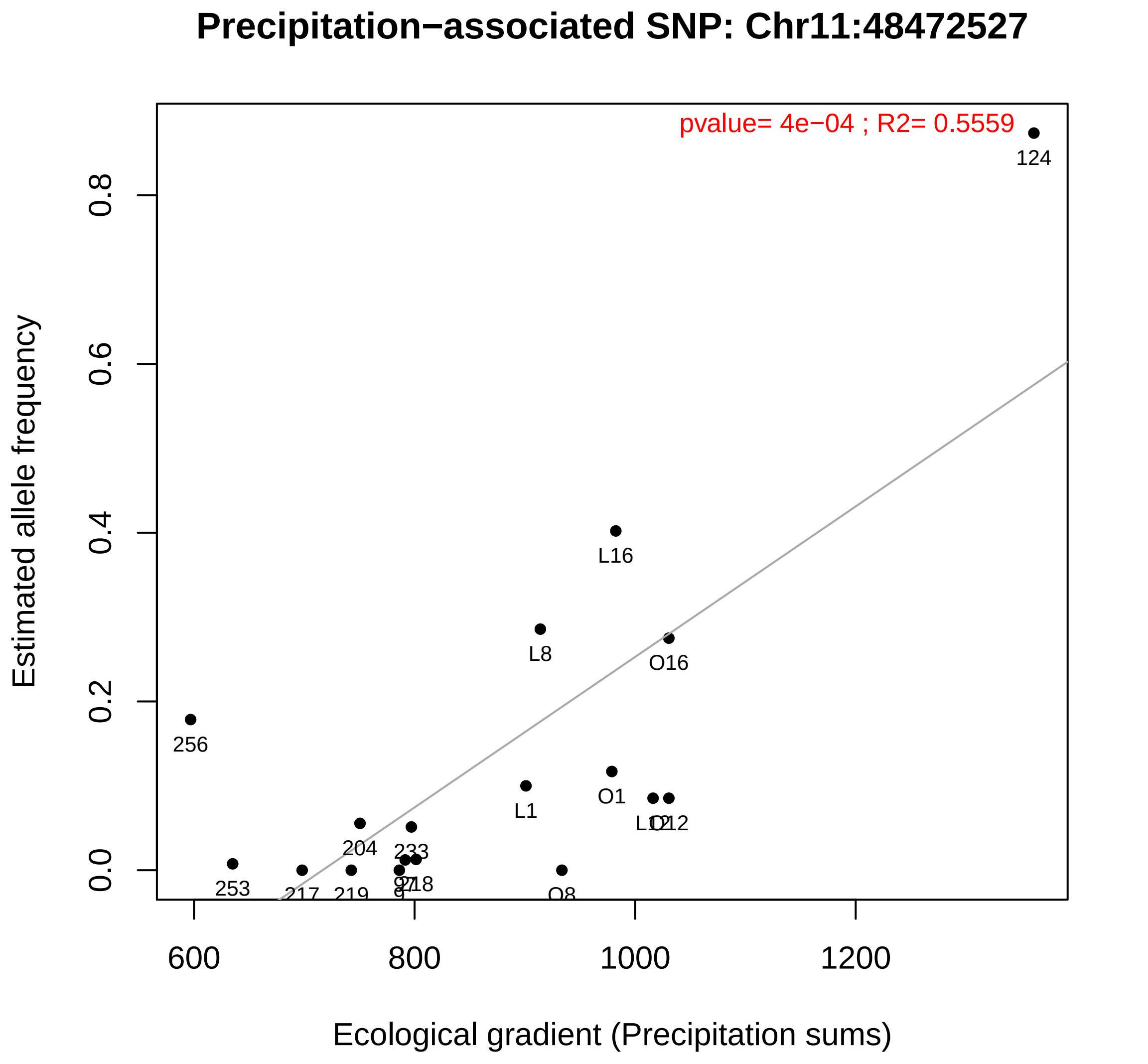
Precipitation−associated SNP: Chr11:48472527
pvalue= 4e−04 ; R2= 0.5559
Estimated allele frequency
0.0
0.2
0.4
0.6
0.8
600
800
1000
1200
Ecological gradient (Precipitation sums)
124
L16
L8
O16
256
L1
O1
204
233
253
217
219
218
O8

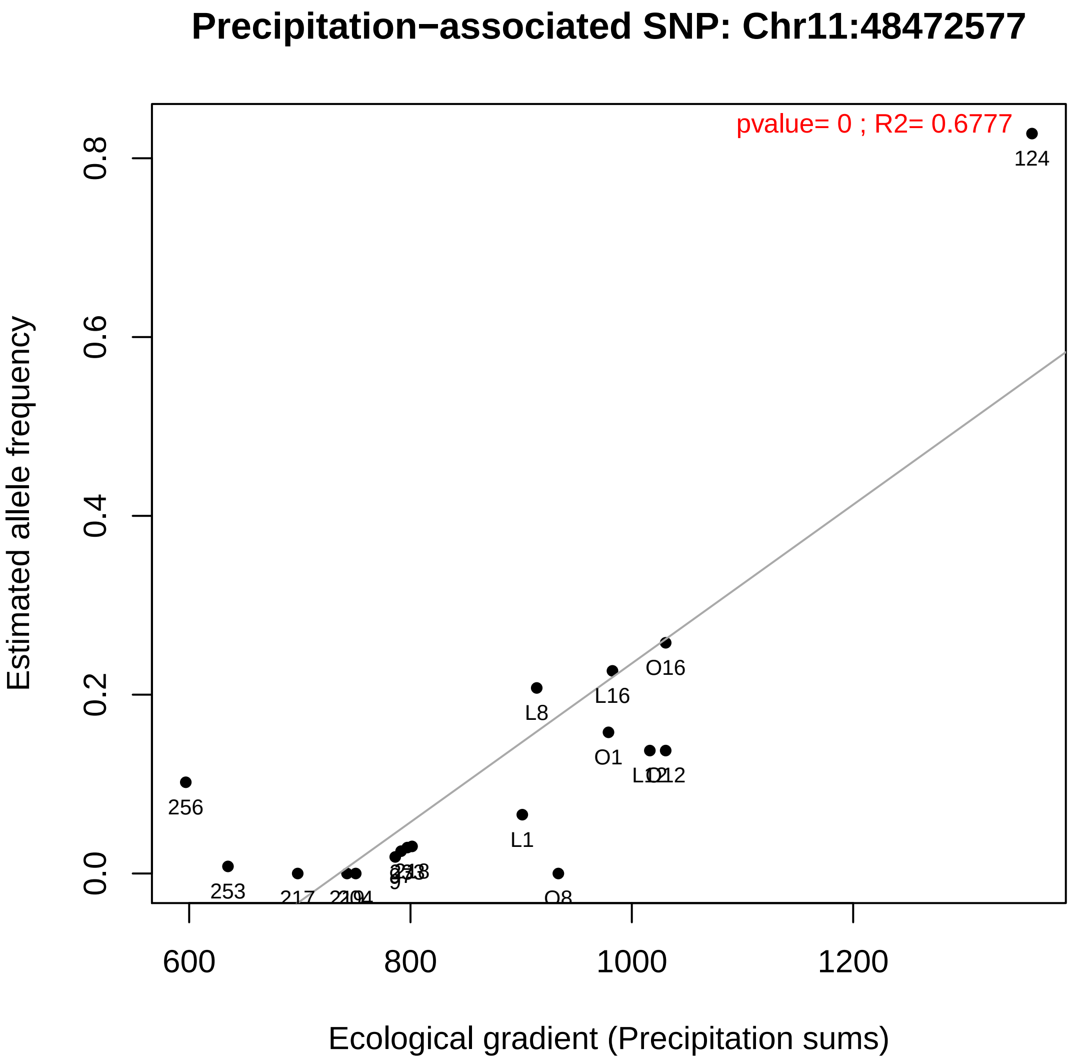
Precipitation−associated SNP: Chr11:48472577
pvalue= 0 ; R2= 0.6777
Estimated allele frequency
Ecological gradient (Precipitation sums)
0.0
0.2
0.4
0.6
0.8
600
800
1000
1200
124
O16
L16
L8
O1
O12
L1
256
253
217
O8

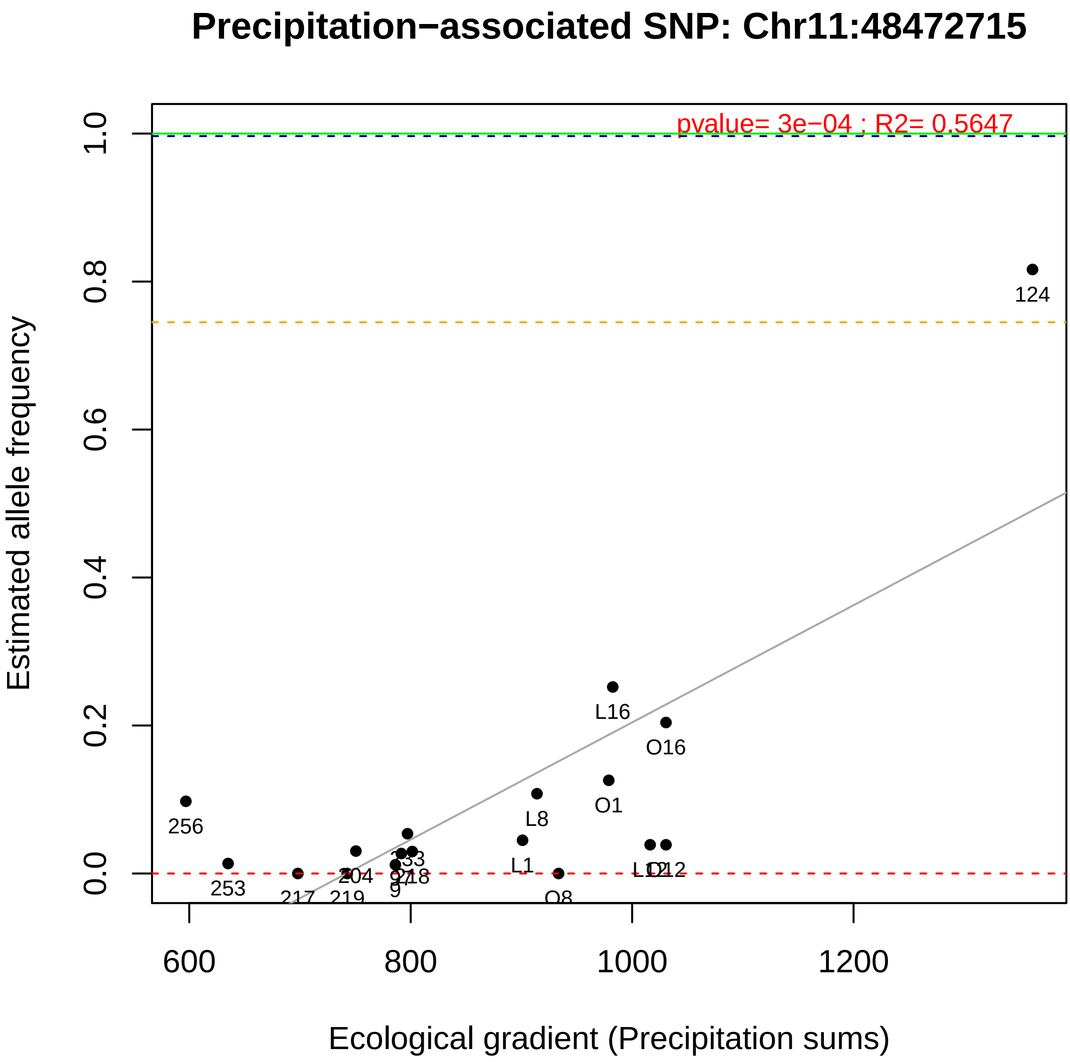

Precipitation−associated SNP: Chr11:48472715
pvalue= 3e−04 ; R2= 0.5647
Estimated allele frequency
Ecological gradient (Precipitation sums)
0.0
0.2
0.4
0.6
0.8
1.0
600
800
1000
1200
124
L16
O16
O1
L8
256
253
217
219
204
L1
O8
O12

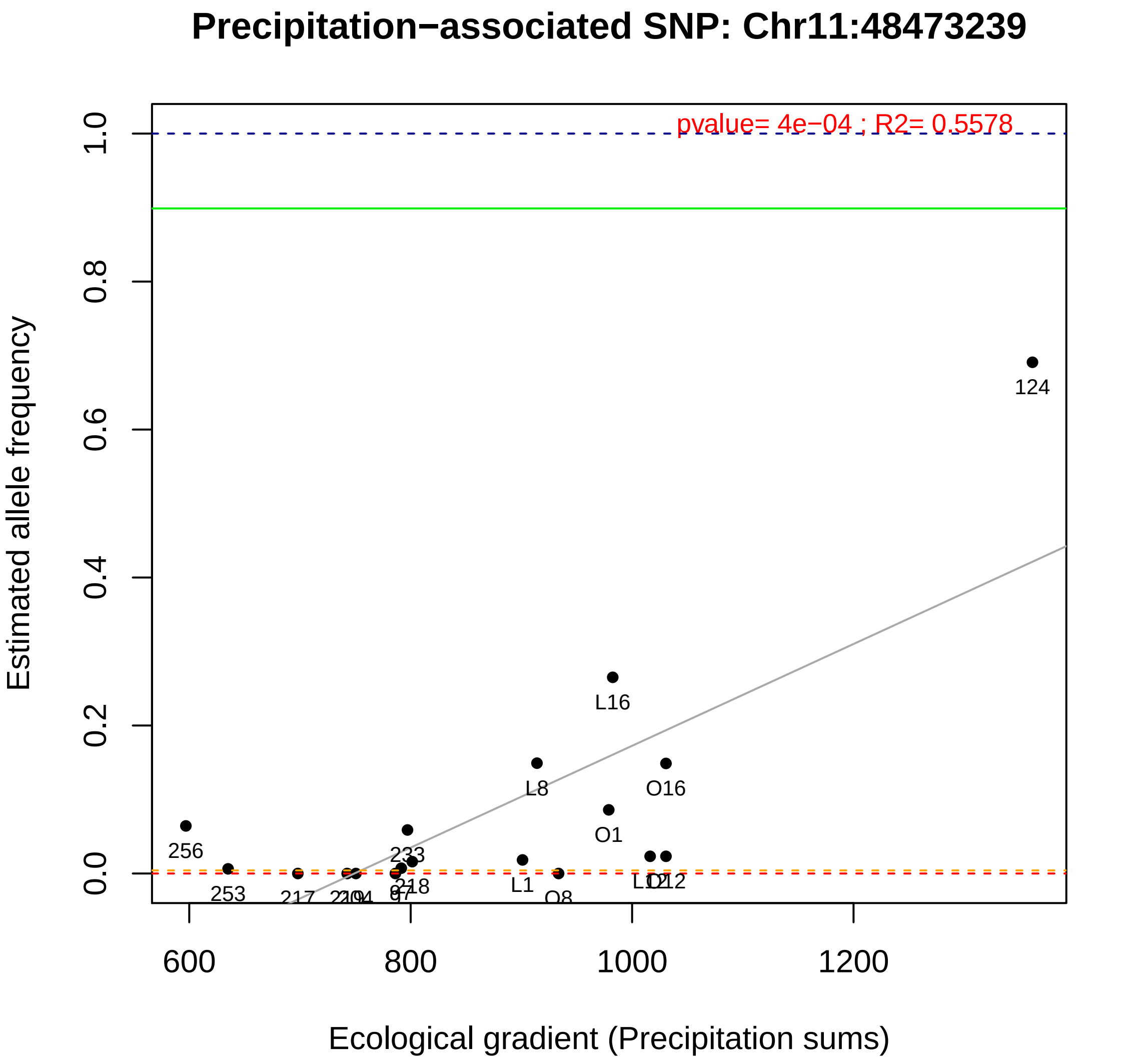

Precipitation−associated SNP: Chr11:48473239
pvalue= 4e−04 ; R2= 0.5578
Estimated allele frequency
Ecological gradient (Precipitation sums)
0.0
0.2
0.4
0.6
0.8
1.0
600
800
1000
1200
124
L16
L8
O16
O1
256
233
253
218
L1
O12

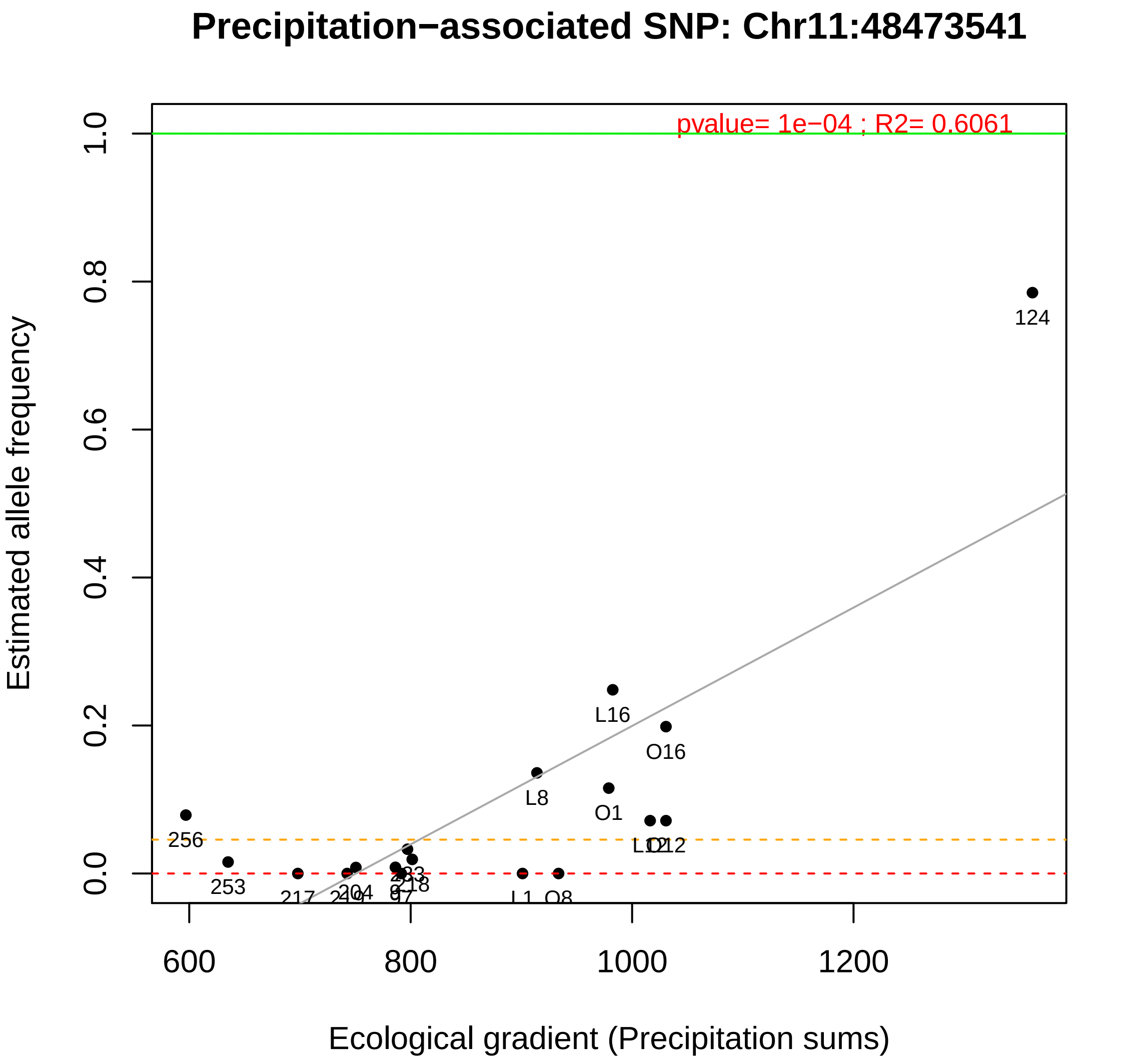
Precipitation−associated SNP: Chr11:48473541
pvalue= 1e−04 ; R2= 0.6061
Estimated allele frequency
0.0
0.2
0.4
0.6
0.8
1.0
600
800
1000
1200
Ecological gradient (Precipitation sums)
124
L16
O16
L8
O1
256
253
217
204
218
L1
O8

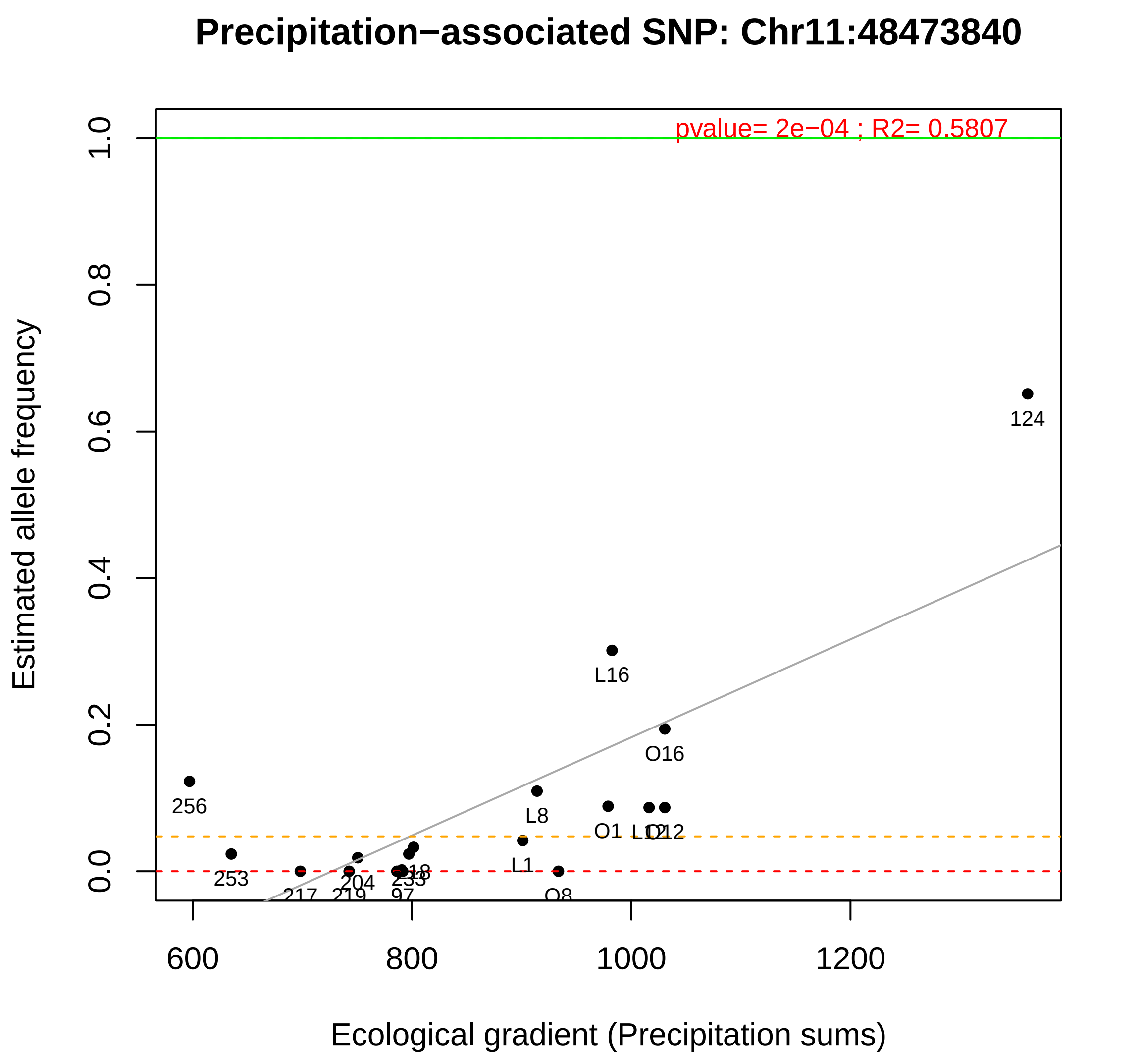
Precipitation−associated SNP: Chr11:48473840
pvalue= 2e−04 ; R2= 0.5807
Estimated allele frequency
Ecological gradient (Precipitation sums)
0.0
0.2
0.4
0.6
0.8
1.0
600
800
1000
1200
124
L16
O16
256
L8
O1
253
L1
204
O8
217
219
97

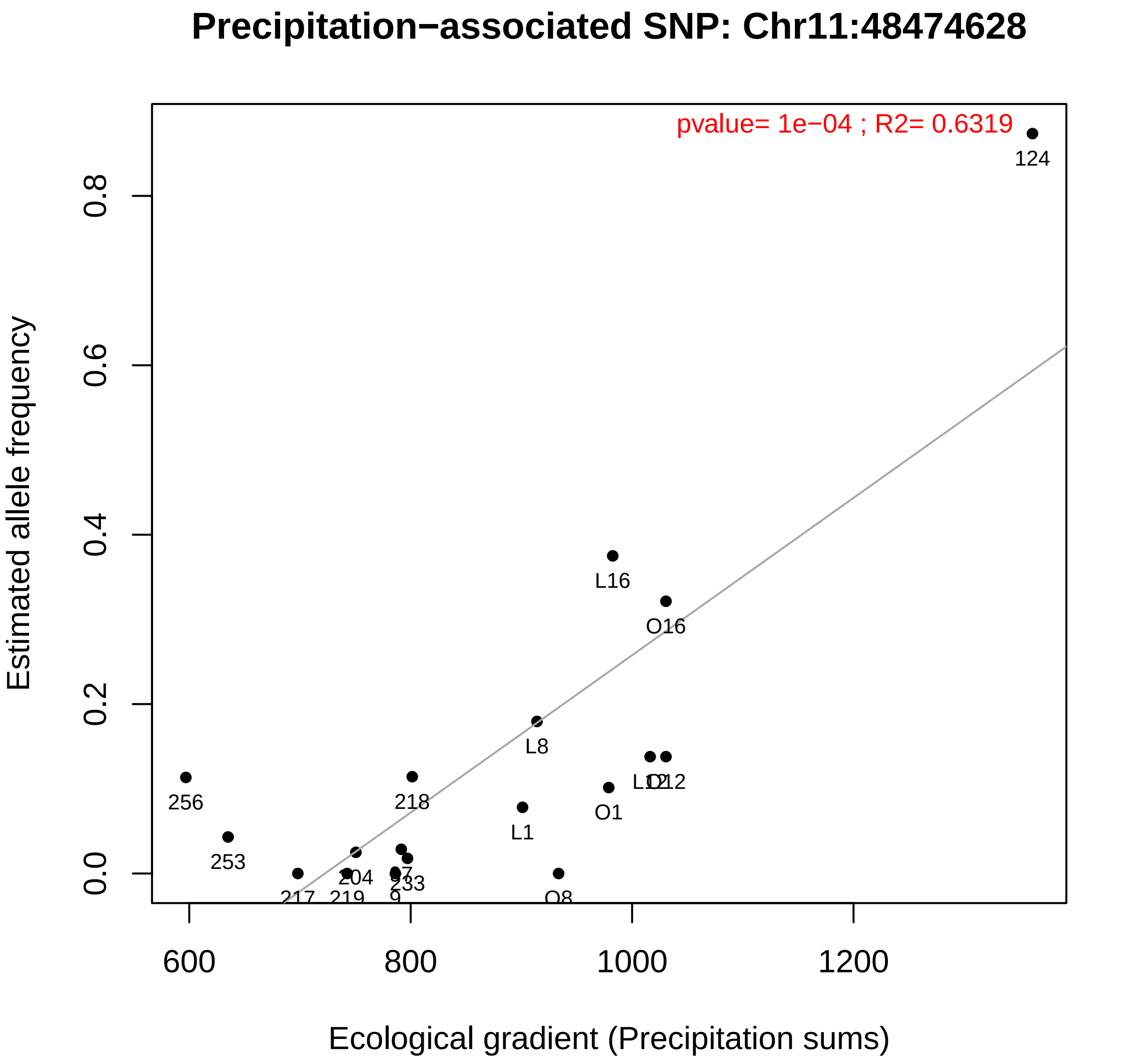

Precipitation−associated SNP: Chr11:48474628
pvalue= 1e−04 ; R2= 0.6319
Estimated allele frequency
0.0
0.2
0.4
0.6
0.8
600
800
1000
1200
Ecological gradient (Precipitation sums)
124
L16
O16
L8
256
218
L1
O1
253
204
233
O8
217
219
9

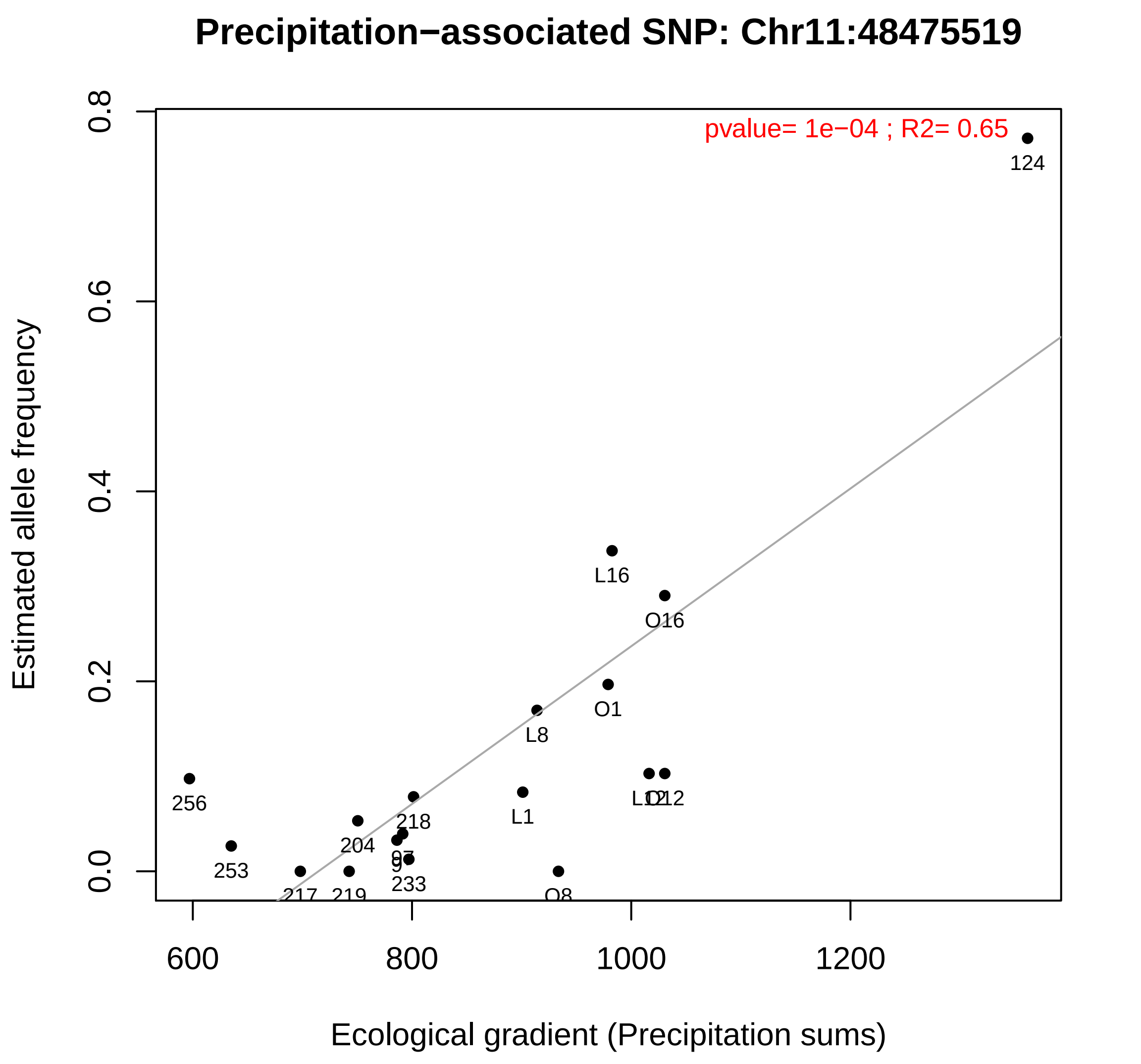
Precipitation−associated SNP: Chr11:48475519
pvalue= 1e−04 ; R2= 0.65
Estimated allele frequency
Ecological gradient (Precipitation sums)
0.0
0.2
0.4
0.6
0.8
600
800
1000
1200
124
L16
O16
O1
L8
256
253
217
219
204
218
97
9
233
L1
O8
L12
O12

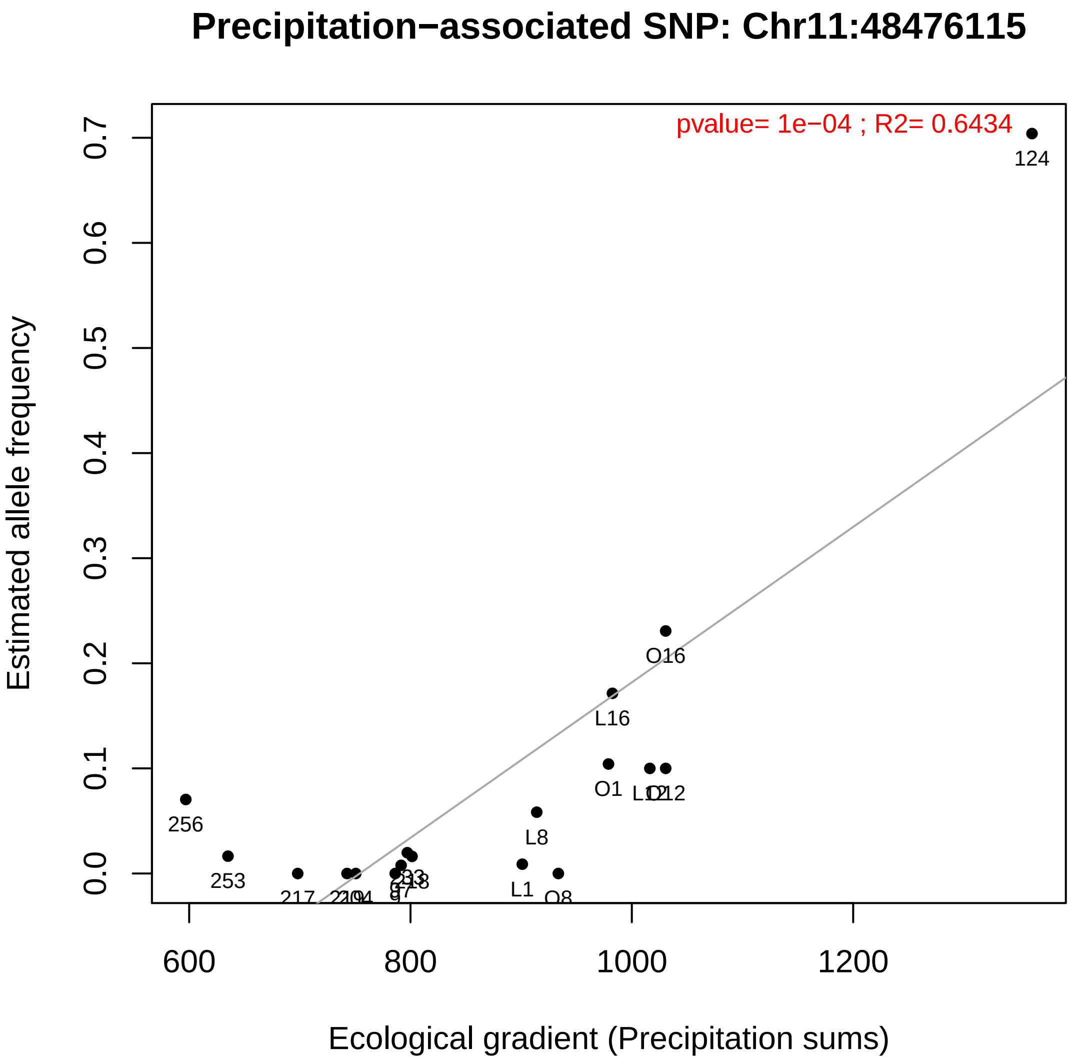

Precipitation−associated SNP: Chr11:48476115
pvalue= 1e−04 ; R2= 0.6434
Estimated allele frequency
Ecological gradient (Precipitation sums)
0.0
0.1
0.2
0.3
0.4
0.5
0.6
0.7
600
800
1000
1200
124
O16
L16
O1
O12
L8
L1
O8
256
253
217

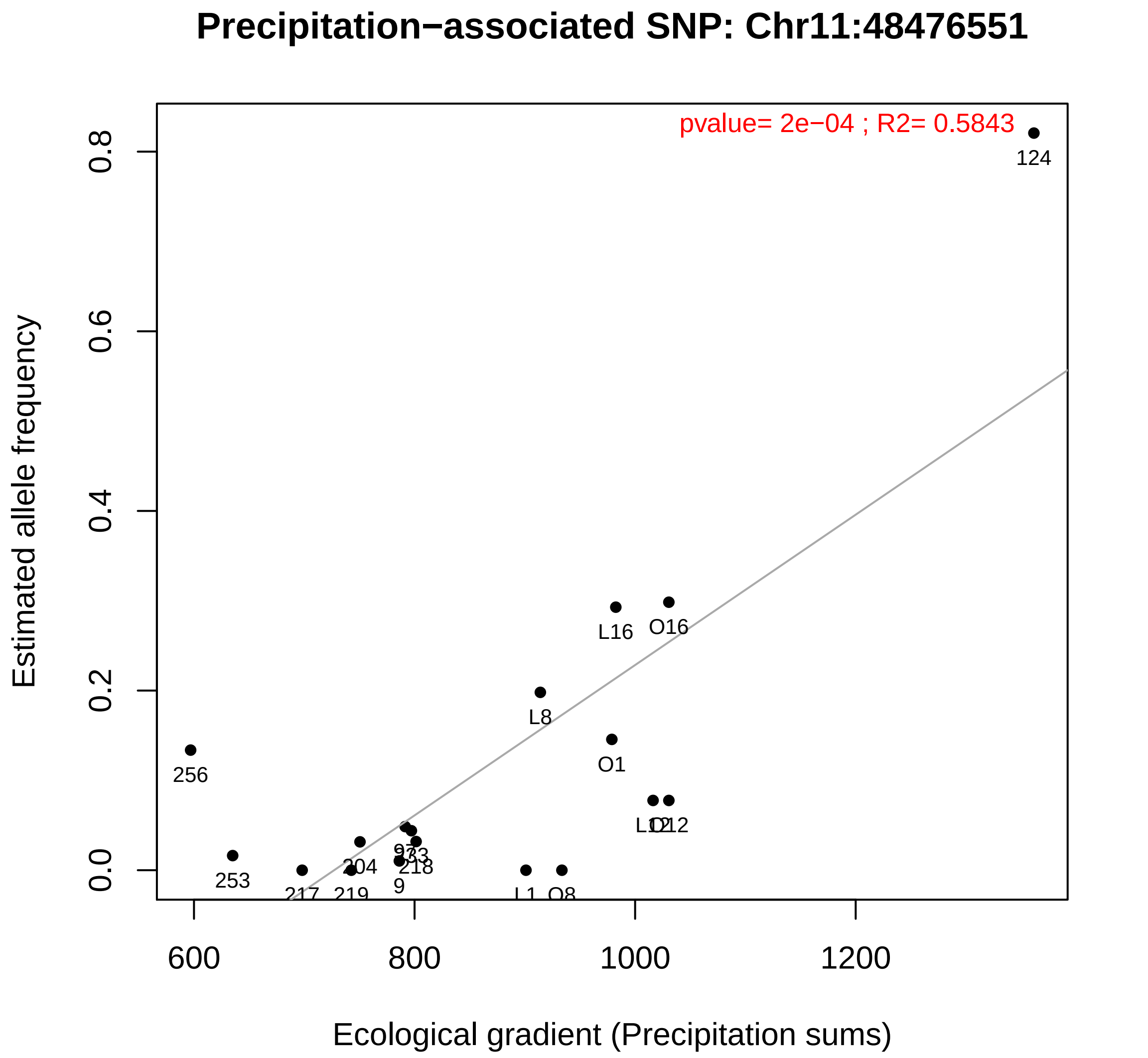
Precipitation−associated SNP: Chr11:48476551
pvalue= 2e−04 ; R2= 0.5843
Estimated allele frequency
Ecological gradient (Precipitation sums)
0.0
0.2
0.4
0.6
0.8
600
800
1000
1200
124
L16
O16
L8
O1
256
253
204
217
219
218
9
L1
O8

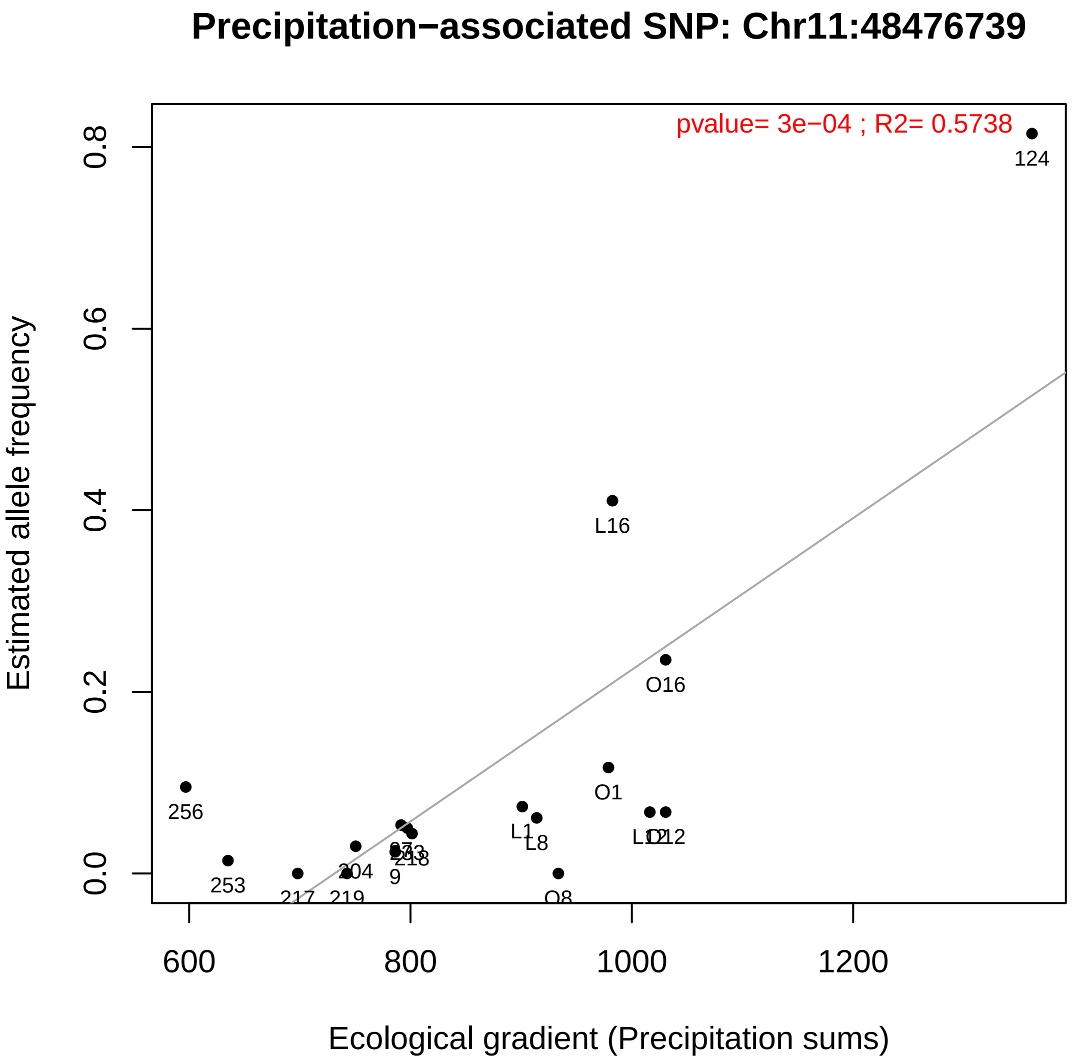

Precipitation−associated SNP: Chr11:48476739
pvalue= 3e−04 ; R2= 0.5738
Estimated allele frequency
Ecological gradient (Precipitation sums)
0.0
0.2
0.4
0.6
0.8
600
800
1000
1200
124
L16
O16
O1
256
253
217
219
204
218
9
L1
L8
O8
O12

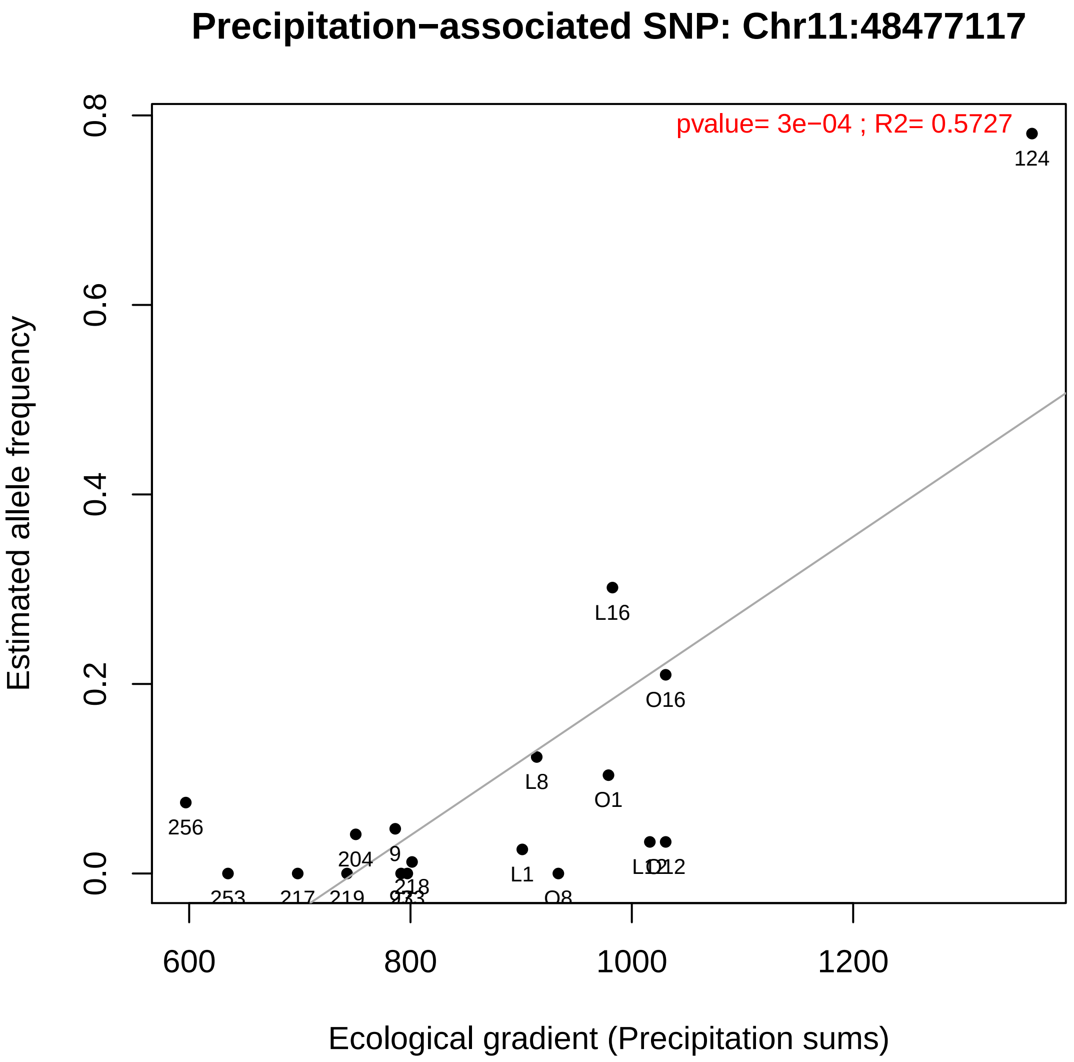

Precipitation−associated SNP: Chr11:48477117
pvalue= 3e−04 ; R2= 0.5727
Estimated allele frequency
Ecological gradient (Precipitation sums)
0.0
0.2
0.4
0.6
0.8
600
800
1000
1200
124
L16
O16
L8
O1
256
204
9
L1
218
253
217
219
O8

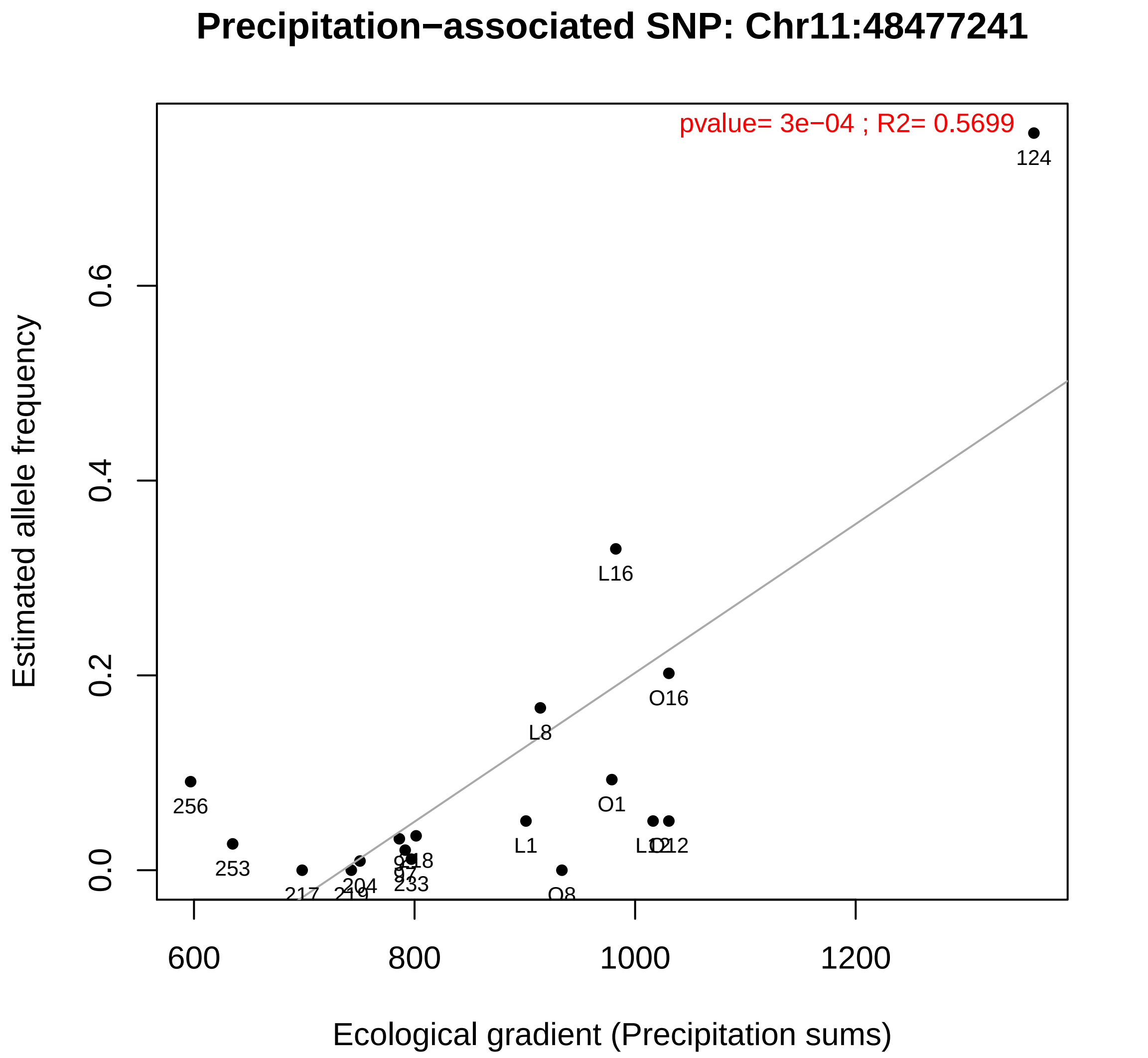

Precipitation−associated SNP: Chr11:48477241
pvalue= 3e−04 ; R2= 0.5699
Estimated allele frequency
0.0
0.2
0.4
0.6
600
800
1000
1200
Ecological gradient (Precipitation sums)
124
L16
O16
L8
256
O1
L1
253

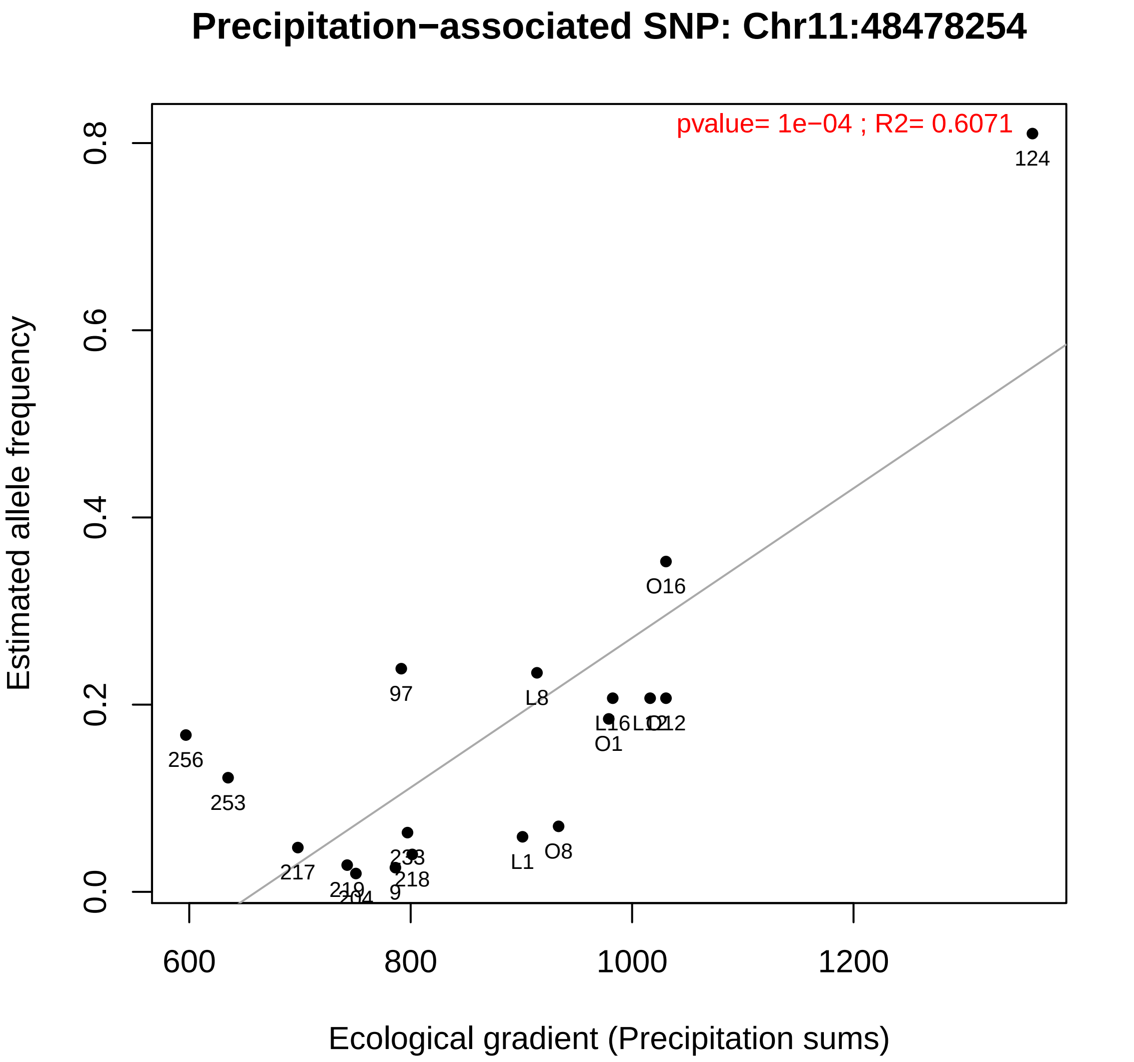
Precipitation−associated SNP: Chr11:48478254
pvalue= 1e−04 ; R2= 0.6071
Estimated allele frequency
Ecological gradient (Precipitation sums)
0.0
0.2
0.4
0.6
0.8
600
800
1000
1200
124
O16
97
L8
L16
O1
256
253
217
L1
O8
218
9

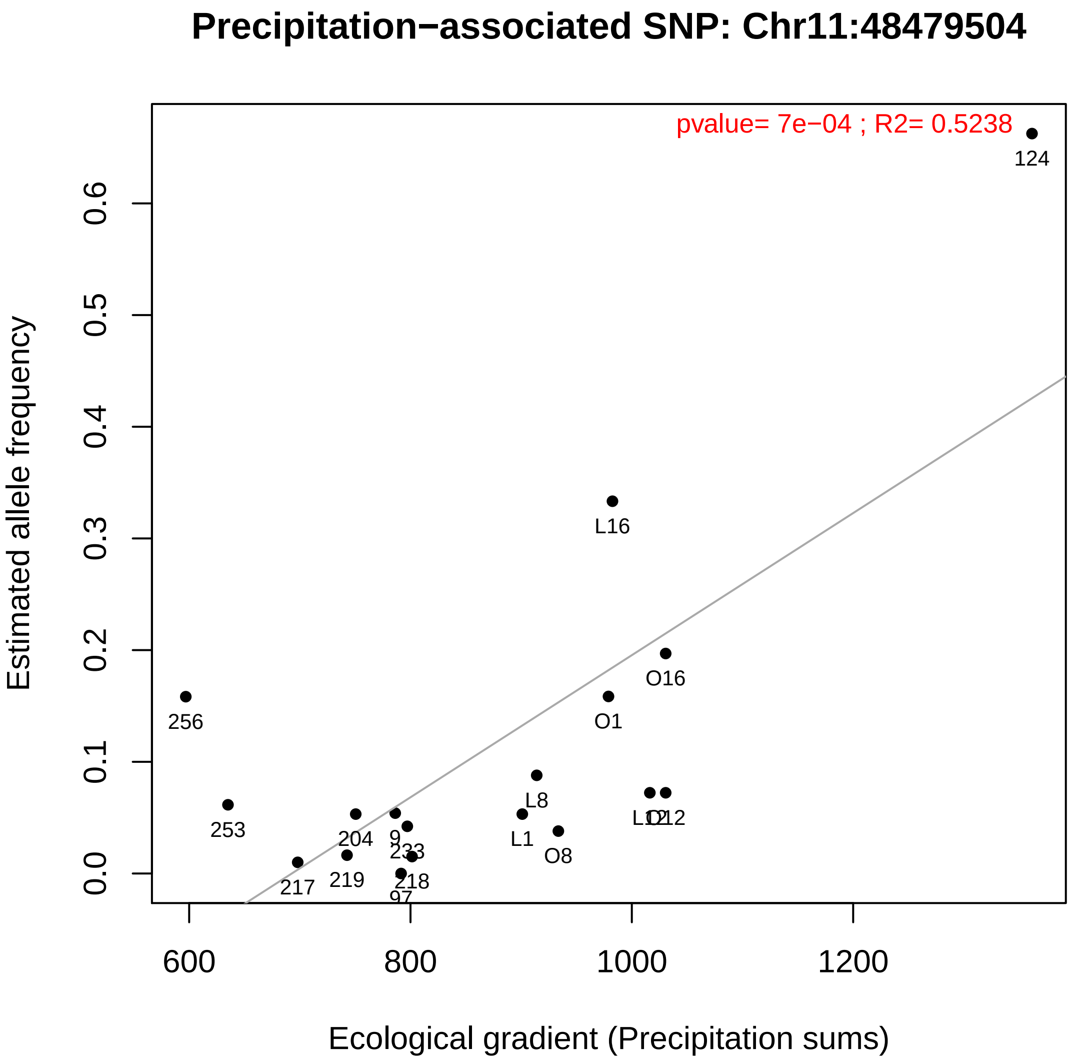
Precipitation−associated SNP: Chr11:48479504
pvalue= 7e−04 ; R2= 0.5238
124
L16
O16
O1
256
L8
253
204
9
233
L1
O8
L12
O12
217
219
218
97
0.0
0.1
0.2
0.3
0.4
0.5
0.6
600
800
1000
1200
Estimated allele frequency
Ecological gradient (Precipitation sums)

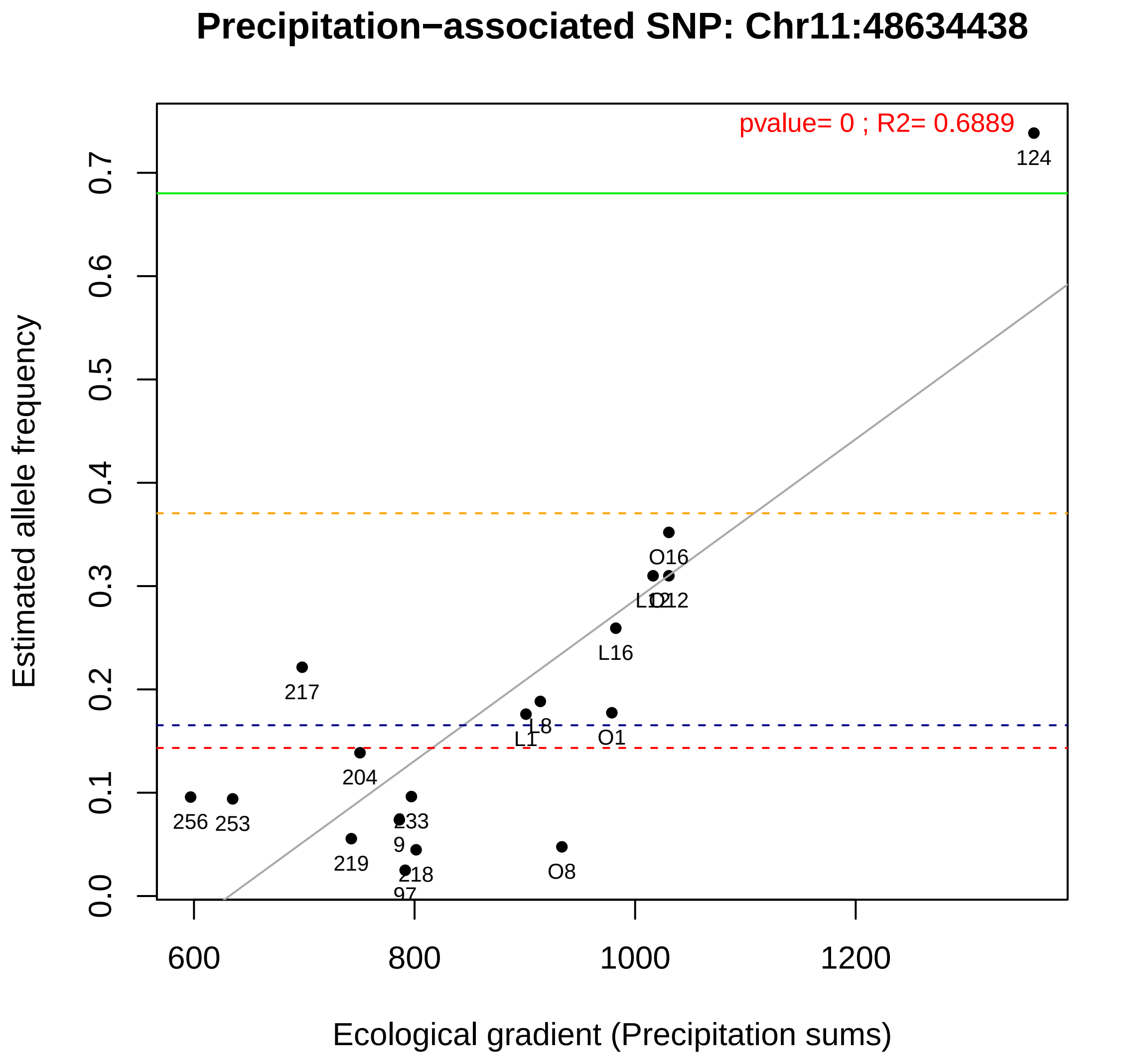

Precipitation−associated SNP: Chr11:48634438
pvalue= 0 ; R2= 0.6889
Estimated allele frequency
Ecological gradient (Precipitation sums)
0.0
0.1
0.2
0.3
0.4
0.5
0.6
0.7
600
800
1000
1200
124
O16
L12
O12
L16
217
L1
18
O1
204
256
253
233
9
219
218
97
O8

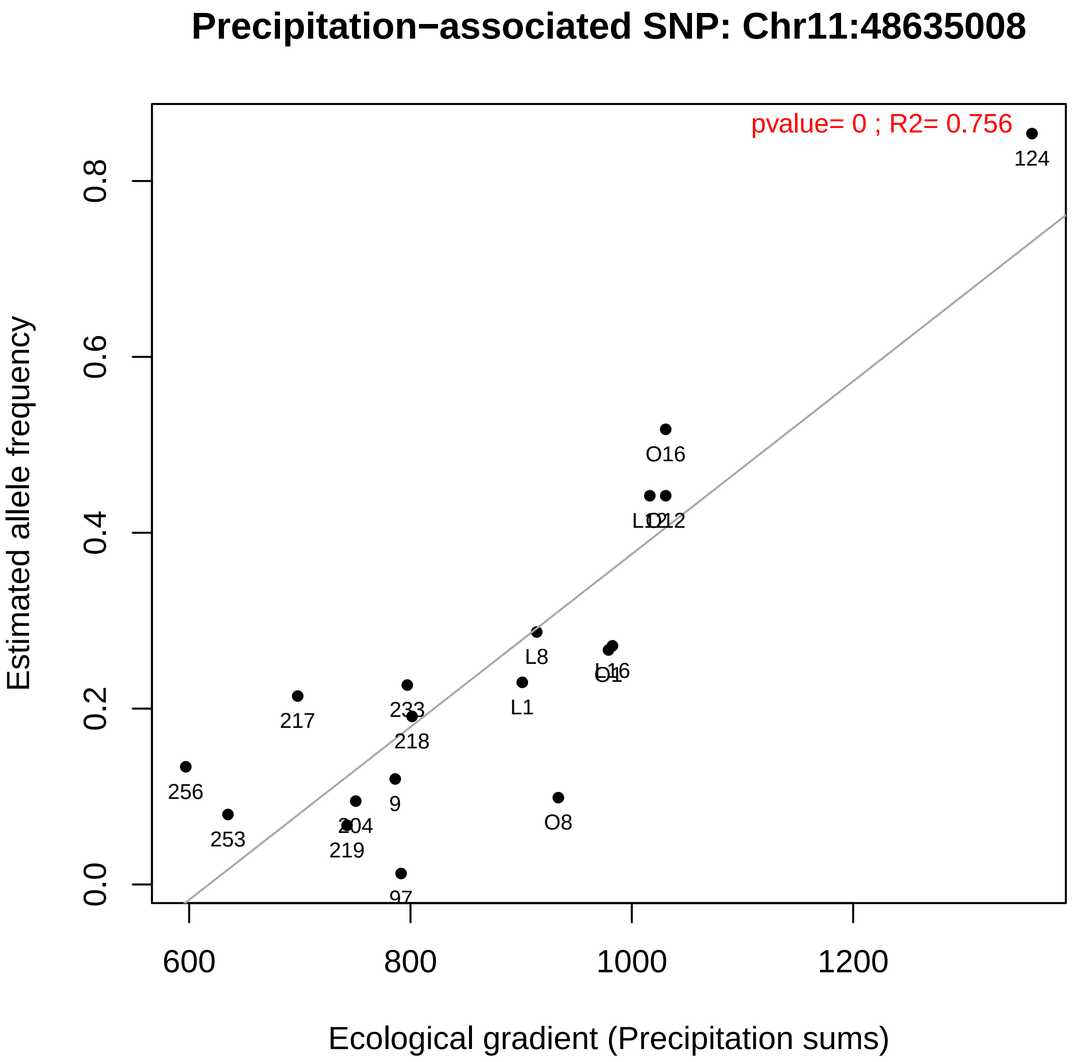
Precipitation−associated SNP: Chr11:48635008
pvalue= 0 ; R2= 0.756
124
O16
L10
O12
L8
L16
O1
233
217
L1
218
256
9
O8
204
253
219
97
0.0
0.2
0.4
0.6
0.8
600
800
1000
1200
Estimated allele frequency
Ecological gradient (Precipitation sums)

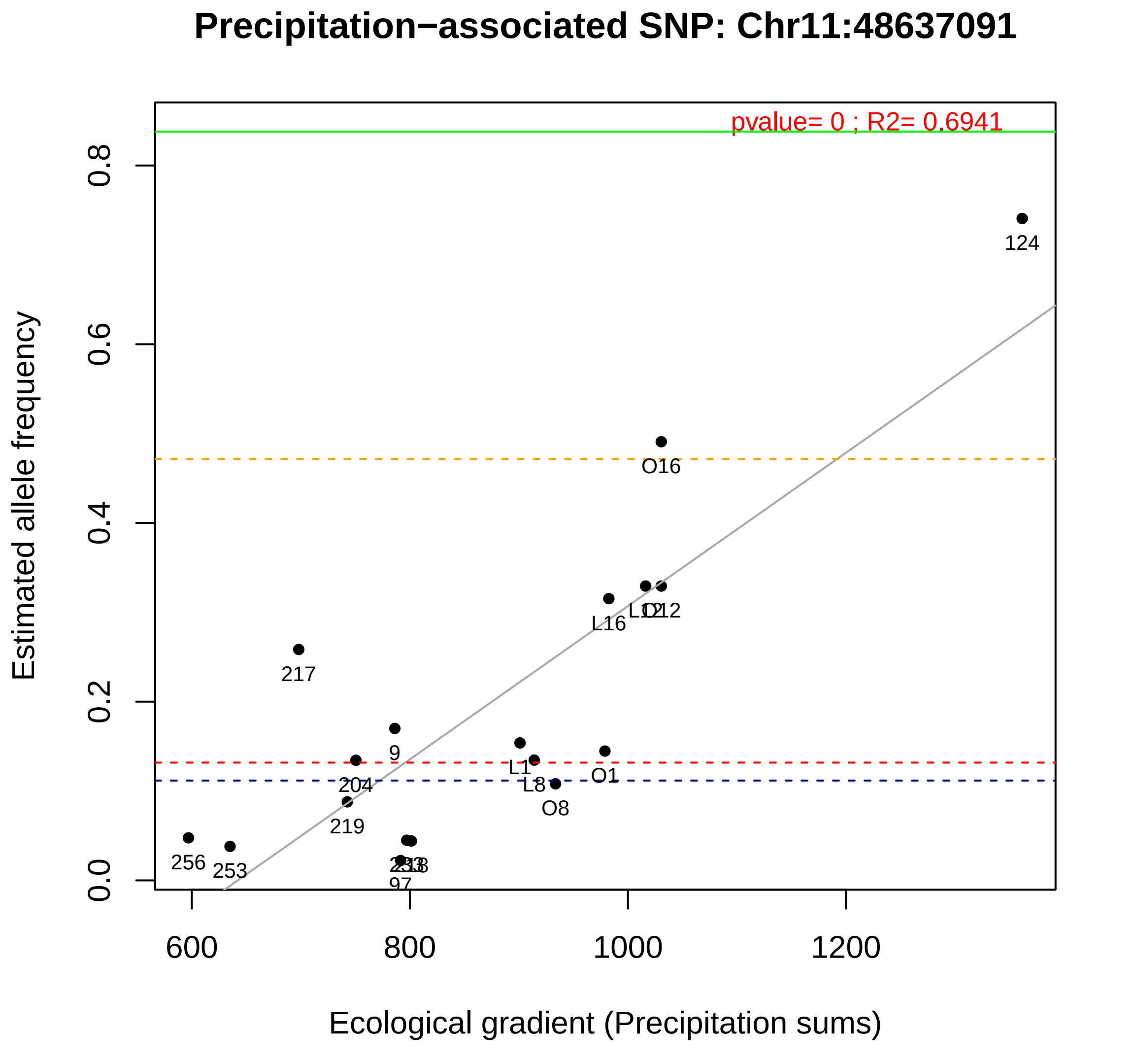

Precipitation−associated SNP: Chr11:48637091
pvalue= 0 ; R2= 0.6941
Estimated allele frequency
0.0
0.2
0.4
0.6
0.8
600
800
1000
1200
Ecological gradient (Precipitation sums)
124
O16
L16
217
9
204
219
256
253
97
L1
L8
O8
O1

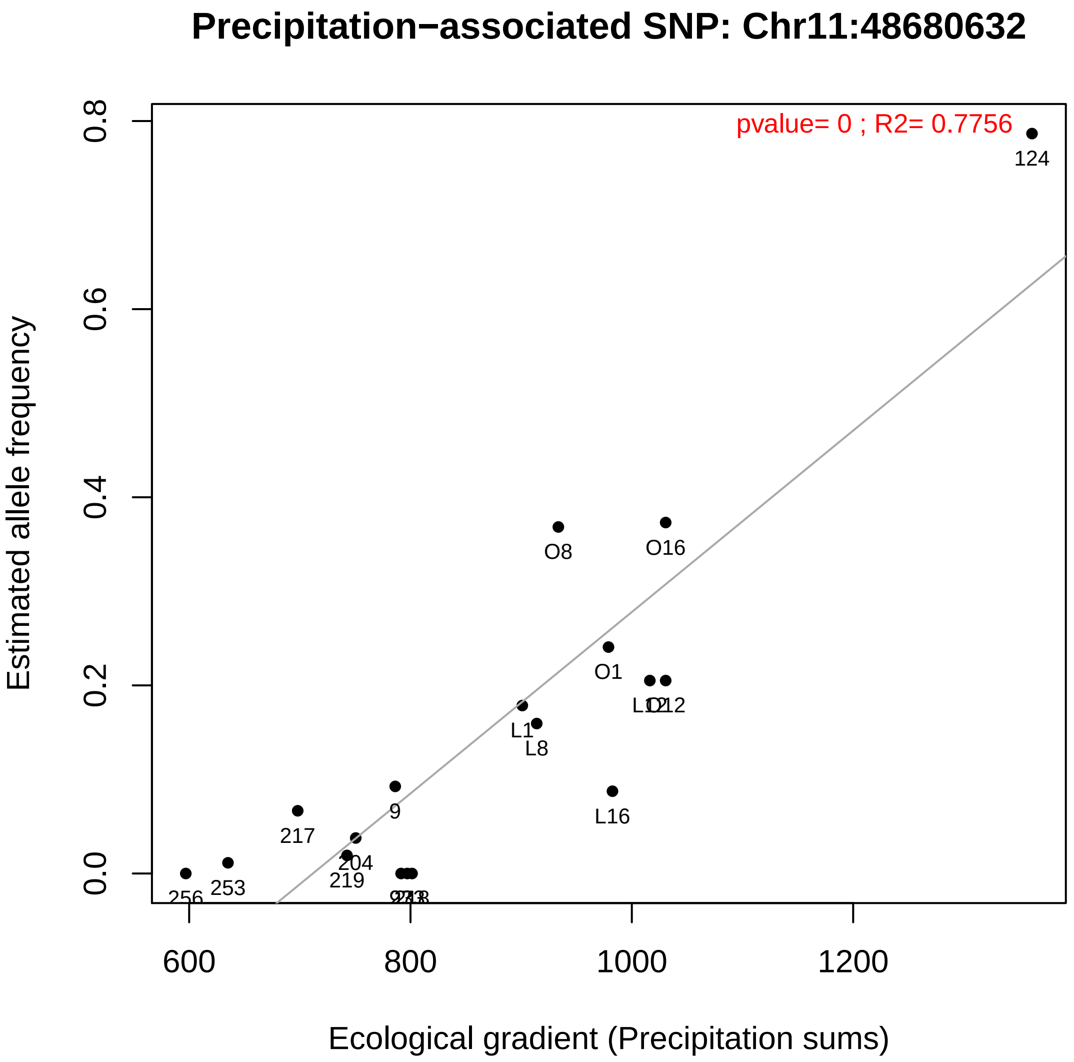

Precipitation−associated SNP: Chr11:48680632
pvalue= 0 ; R2= 0.7756
Estimated allele frequency
0.0
0.2
0.4
0.6
0.8
600
800
1000
1200
Ecological gradient (Precipitation sums)
124
O8
O16
O1
L12
O12
L1
L8
L16
9
217
204
219
253
256

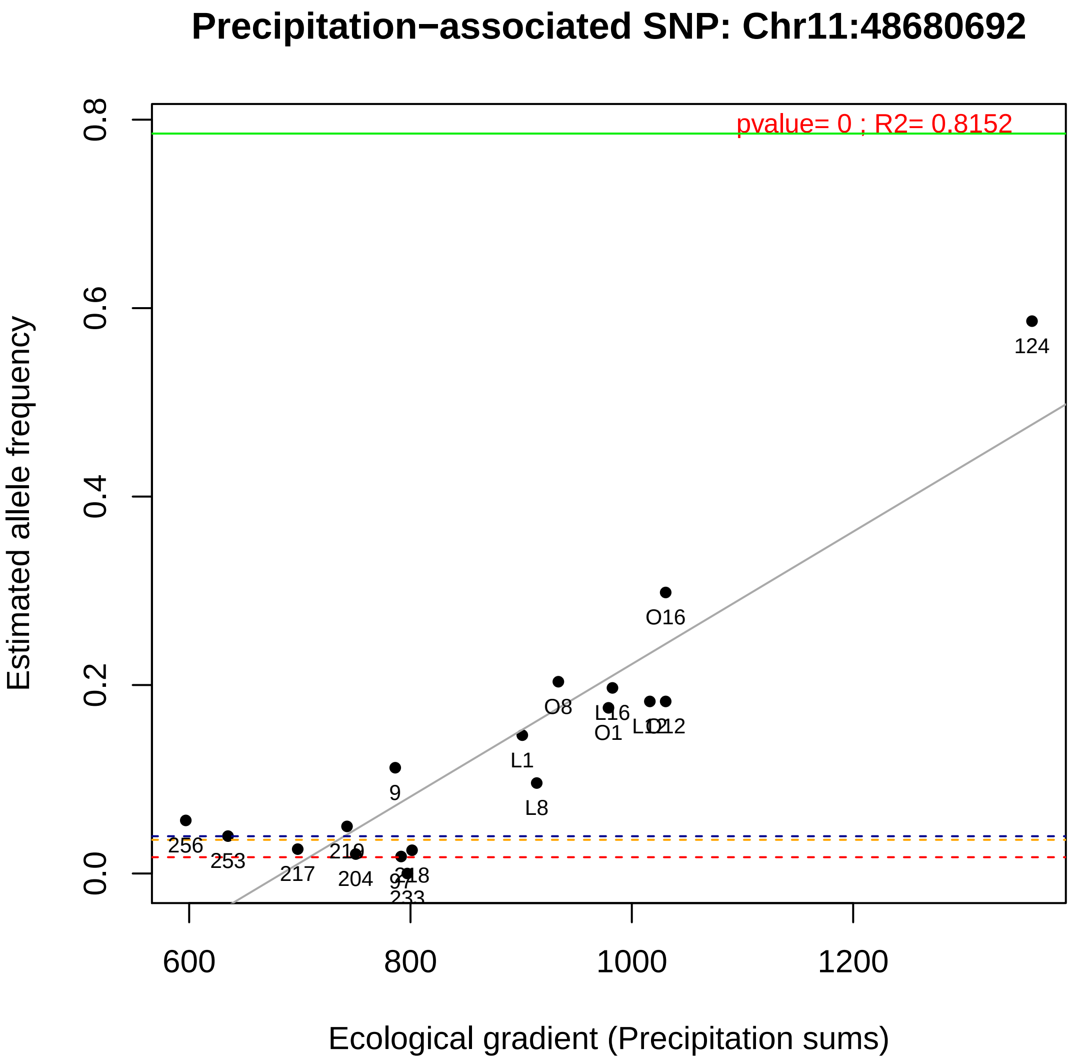

Precipitation−associated SNP: Chr11:48680692
pvalue= 0 ; R2= 0.8152
Estimated allele frequency
0.0
0.2
0.4
0.6
0.8
600
800
1000
1200
Ecological gradient (Precipitation sums)
124
O16
O8
L16
O1
L1
L8
9
256
253
217
210
204
218
97
233

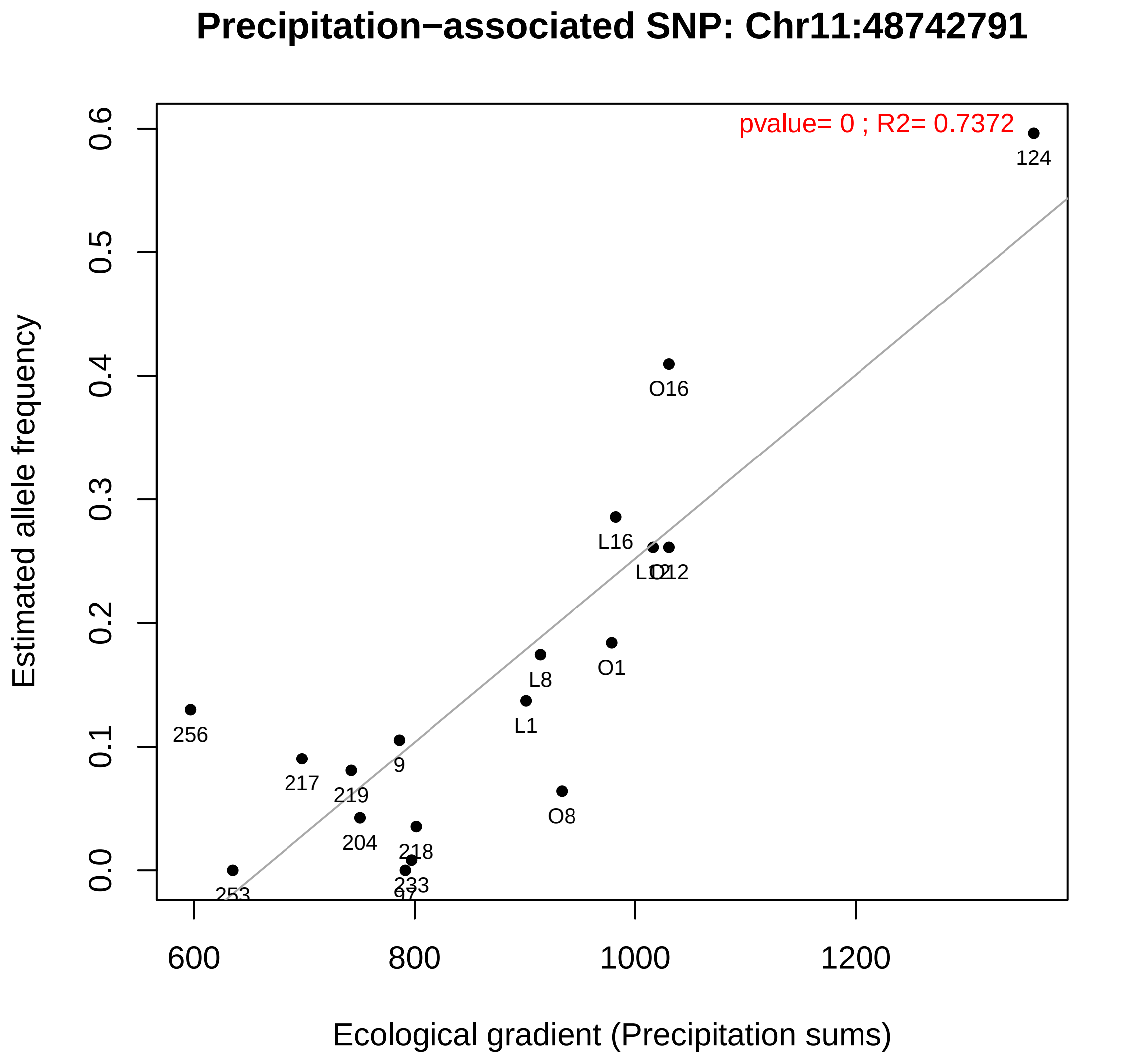

Precipitation−associated SNP: Chr11:48742791
pvalue= 0 ; R2= 0.7372
124
O16
L16
L12
O12
O1
L8
L1
256
217
219
9
204
O8
218
233
253
0.0
0.1
0.2
0.3
0.4
0.5
0.6
600
800
1000
1200
Estimated allele frequency
Ecological gradient (Precipitation sums)

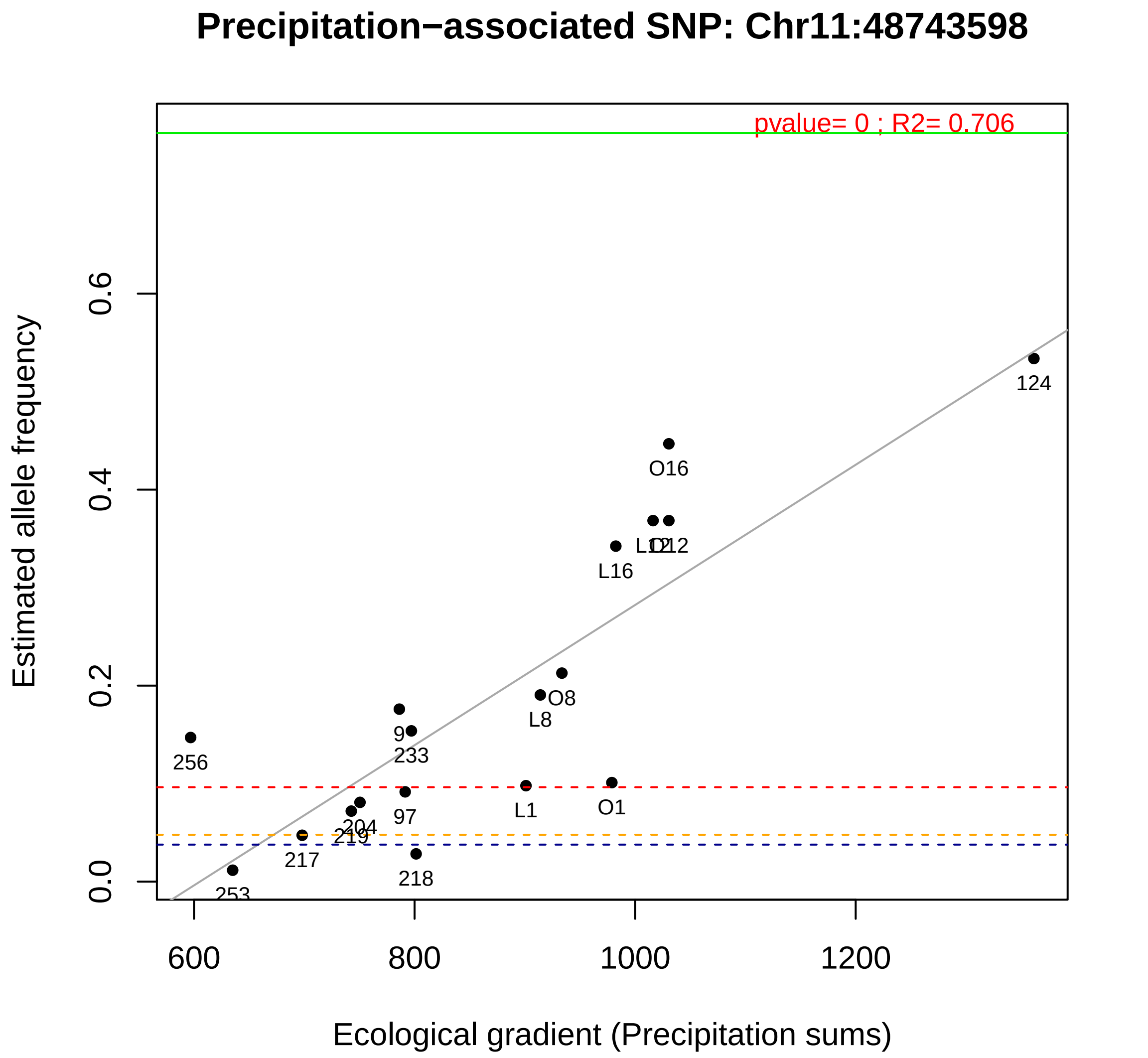
Precipitation−associated SNP: Chr11:48743598
pvalue= 0 ; R2= 0.706
Estimated allele frequency
0.0
0.2
0.4
0.6
600
800
1000
1200
Ecological gradient (Precipitation sums)
124
O16
L12
O12
L16
O8
L8
256
233
L1
O1
97
204
219
217
218
253

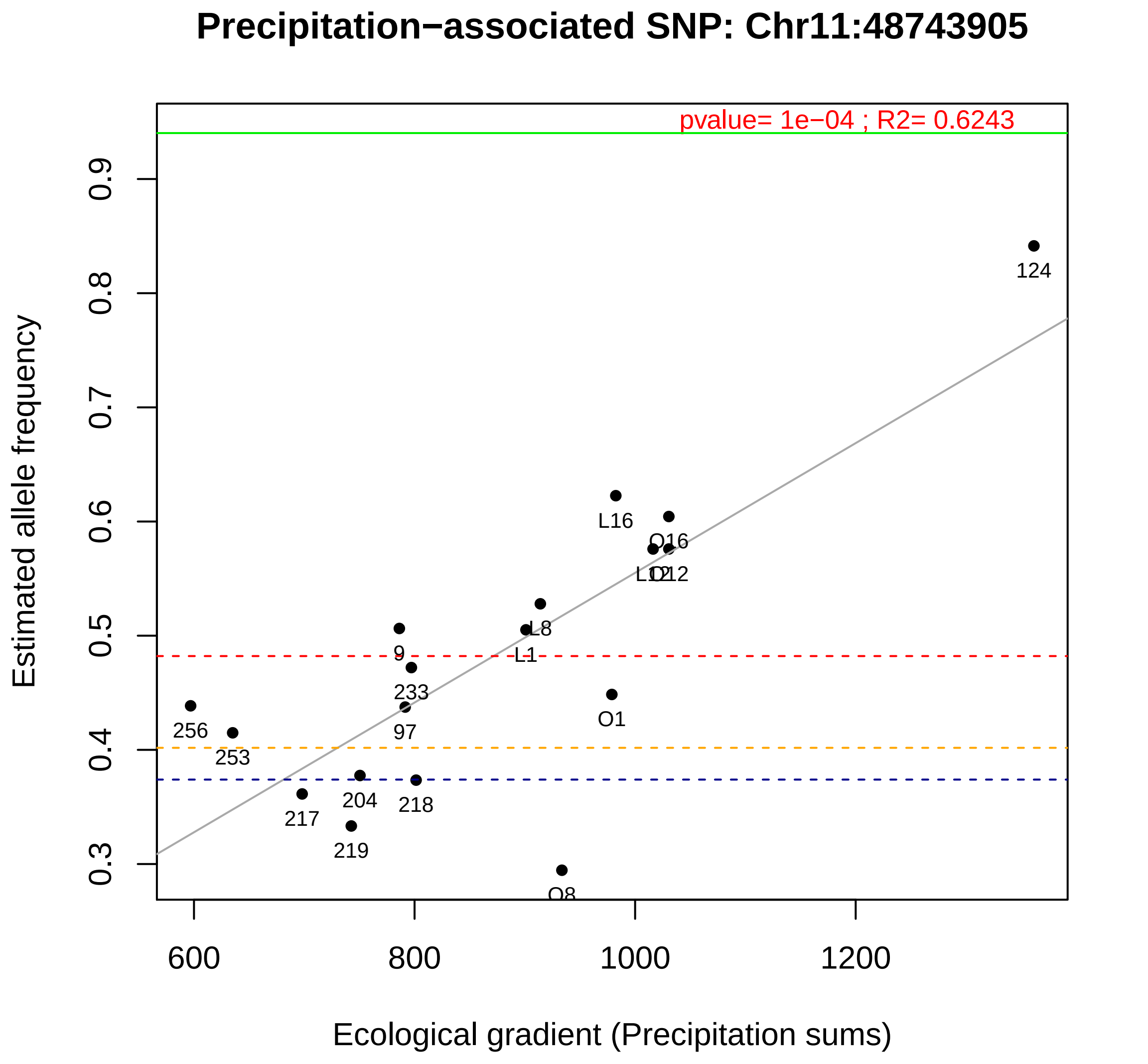
Precipitation−associated SNP: Chr11:48743905
pvalue= 1e−04 ; R2= 0.6243
Estimated allele frequency
0.3
0.4
0.5
0.6
0.7
0.8
0.9
600
800
1000
1200
Ecological gradient (Precipitation sums)
124
L16
O16
L10
O12
L8
L1
9
233
97
O1
256
253
204
218
217
219
O8

Precipitation−associated SNP: Chr11:48761199

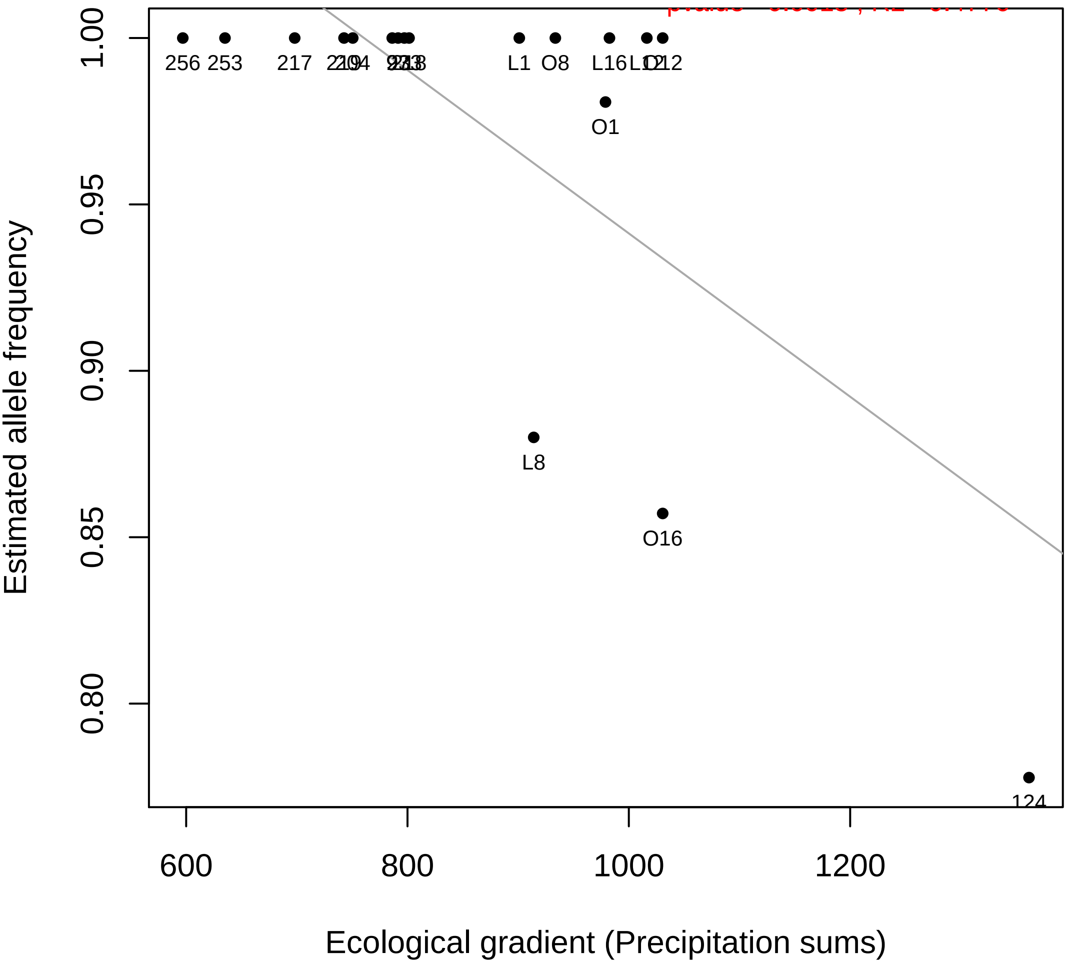

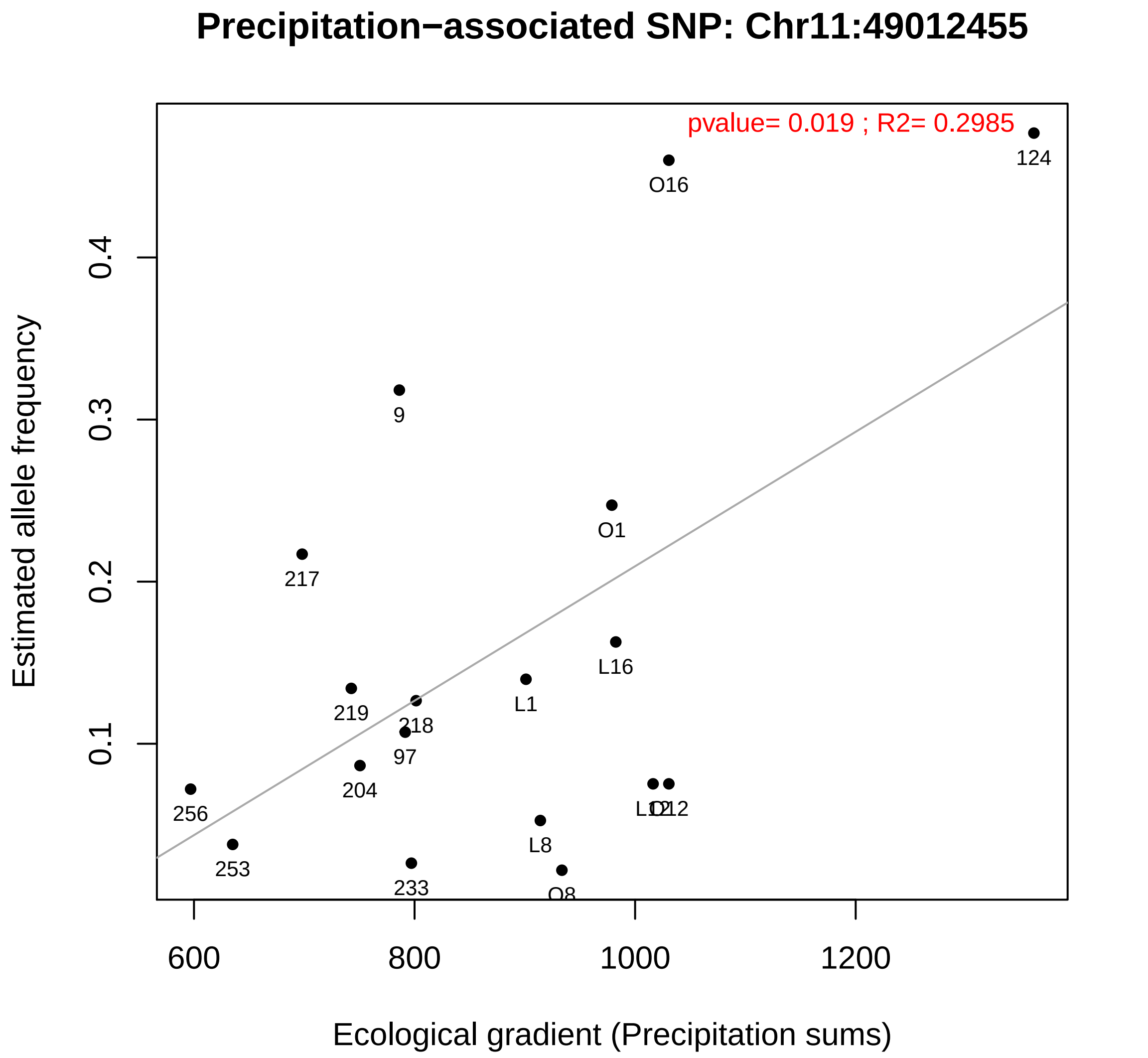
Precipitation−associated SNP: Chr11:49012455
pvalue= 0.019 ; R2= 0.2985
Estimated allele frequency
0.1
0.2
0.3
0.4
600
800
1000
1200
Ecological gradient (Precipitation sums)
124
O16
9
O1
217
L16
L1
219
218
97
204
256
L8
253
233
O8

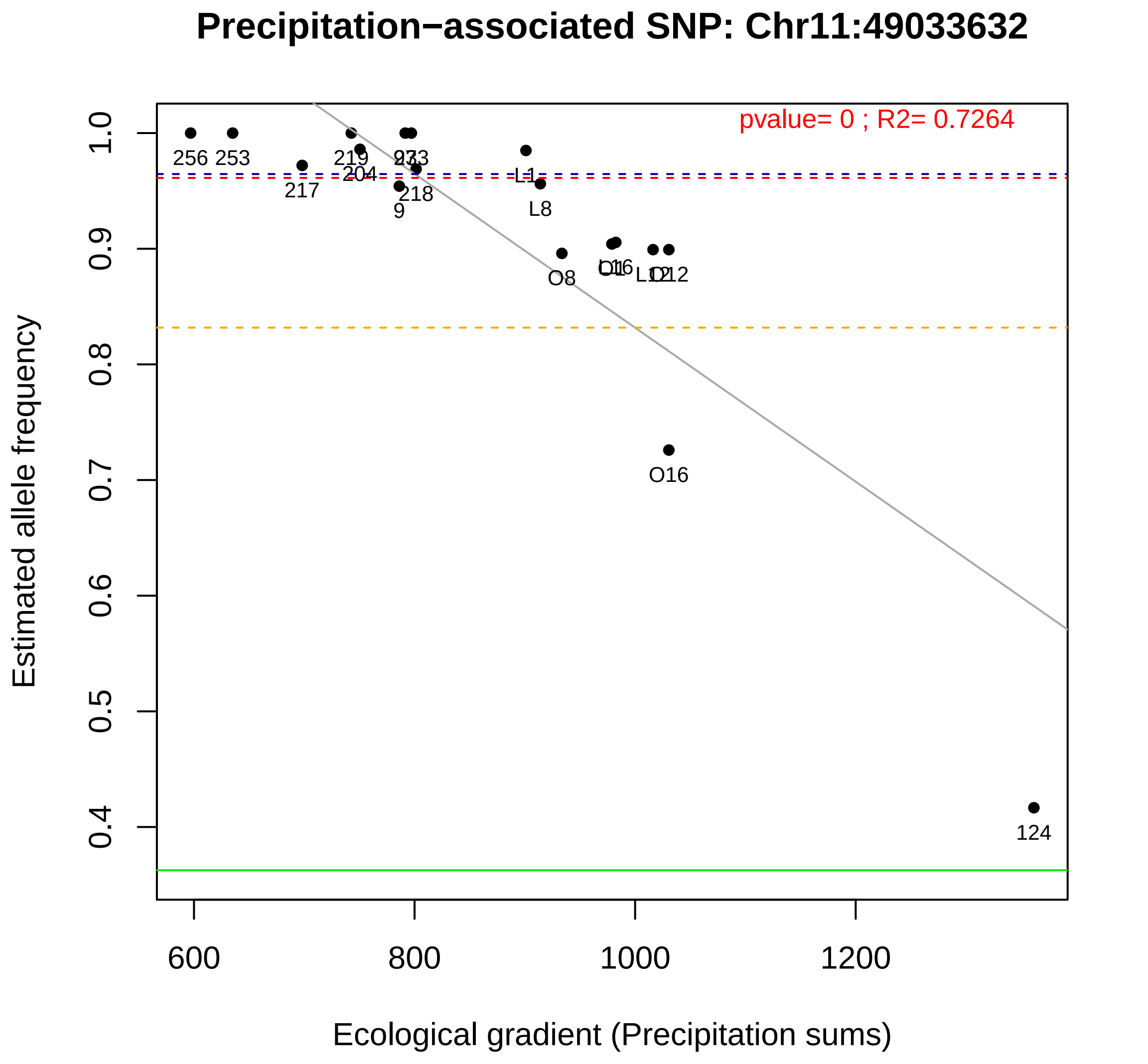

Precipitation−associated SNP: Chr11:49033632
pvalue= 0 ; R2= 0.7264
Estimated allele frequency
Ecological gradient (Precipitation sums)
1.0
0.9
0.8
0.7
0.6
0.5
0.4
600
800
1000
1200
256
253
219
204
217
218
9
L1
L8
O8
O16
124

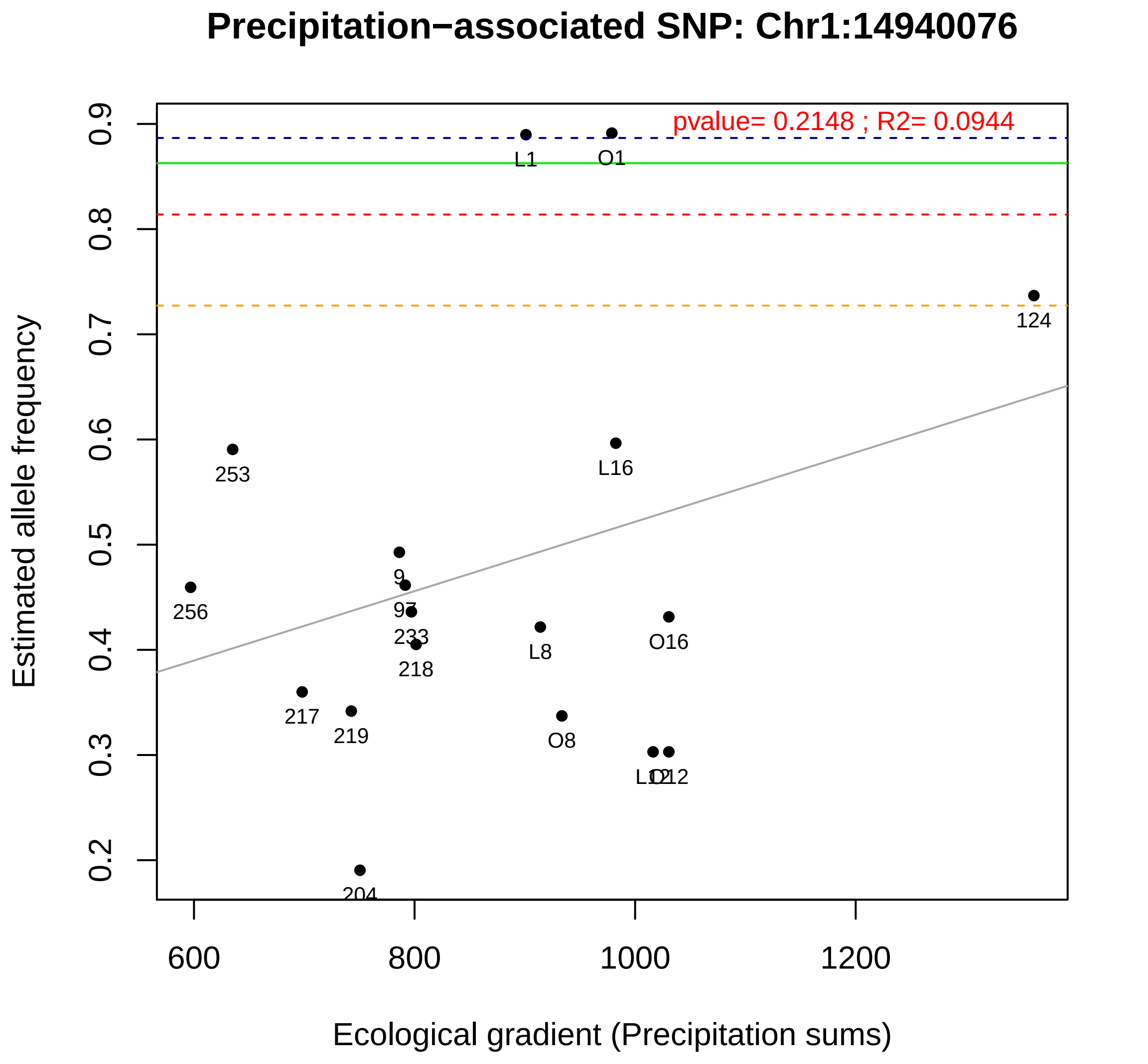
Precipitation−associated SNP: Chr1:14940076
pvalue= 0.2148 ; R2= 0.0944
Estimated allele frequency
Ecological gradient (Precipitation sums)
0.2
0.3
0.4
0.5
0.6
0.7
0.8
0.9
600
800
1000
1200
L1
O1
124
253
L16
9
97
256
233
218
L8
O16
217
219
O8
L12
O12
204

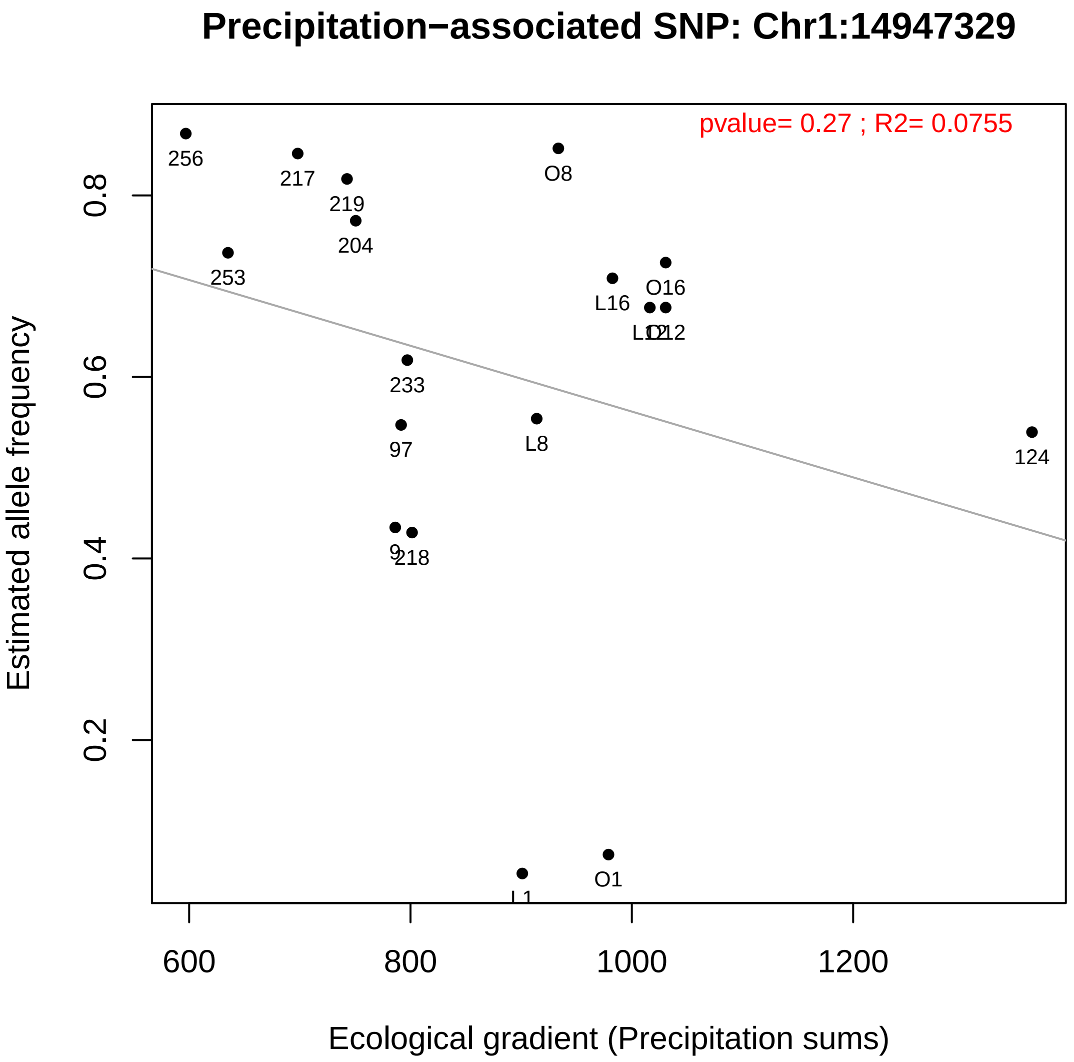
Precipitation−associated SNP: Chr1:14947329
pvalue= 0.27 ; R2= 0.0755
Estimated allele frequency
0.2
0.4
0.6
0.8
600
800
1000
1200
Ecological gradient (Precipitation sums)
256
217
219
204
253
O8
O16
L16
L12
O12
233
97
L8
124
9
218
O1
L1

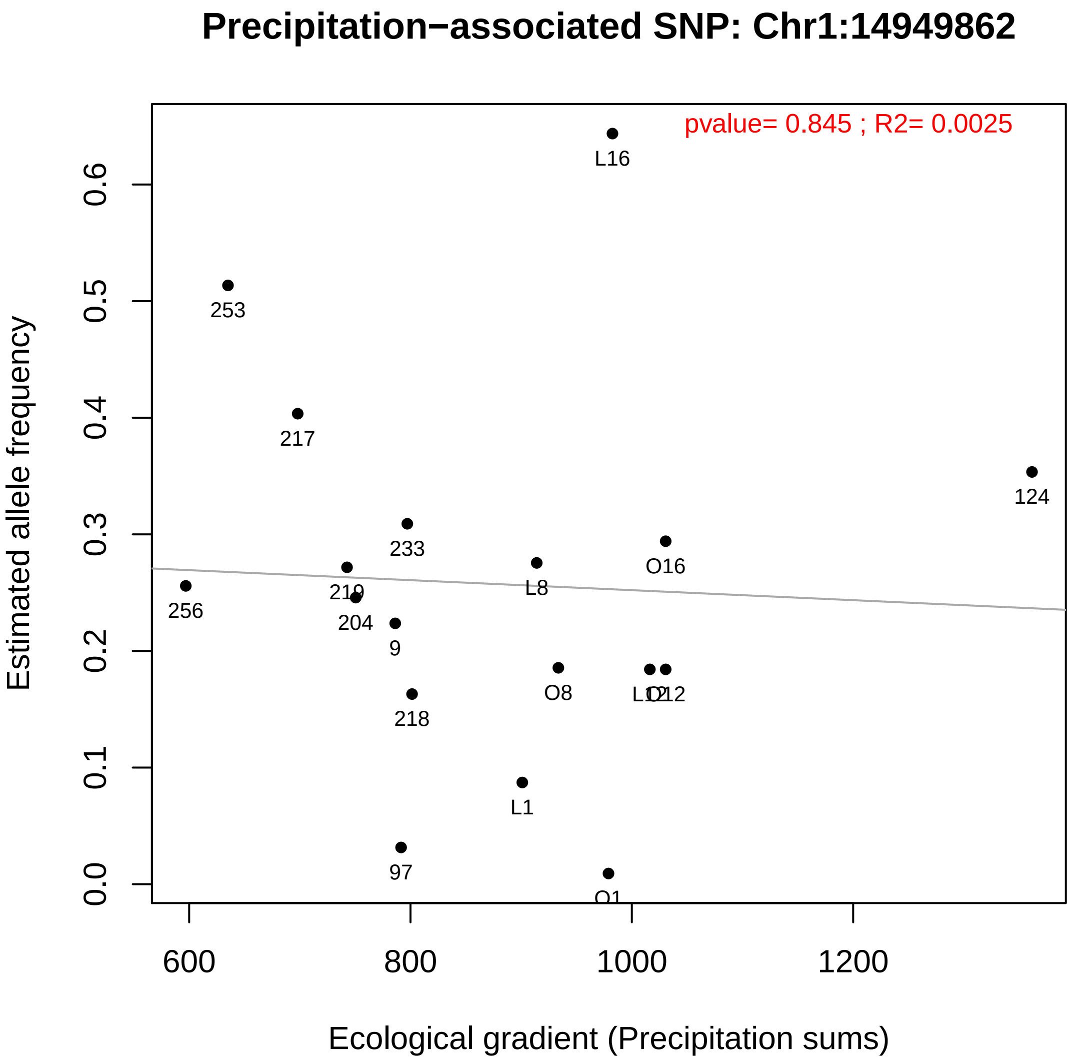
Precipitation−associated SNP: Chr1:14949862
pvalue= 0.845 ; R2= 0.0025
L16
253
217
124
233
O16
L8
219
256
204
9
O8
L10
O12
218
L1
97
O1
0.0
0.1
0.2
0.3
0.4
0.5
0.6
Estimated allele frequency
600
800
1000
1200
Ecological gradient (Precipitation sums)

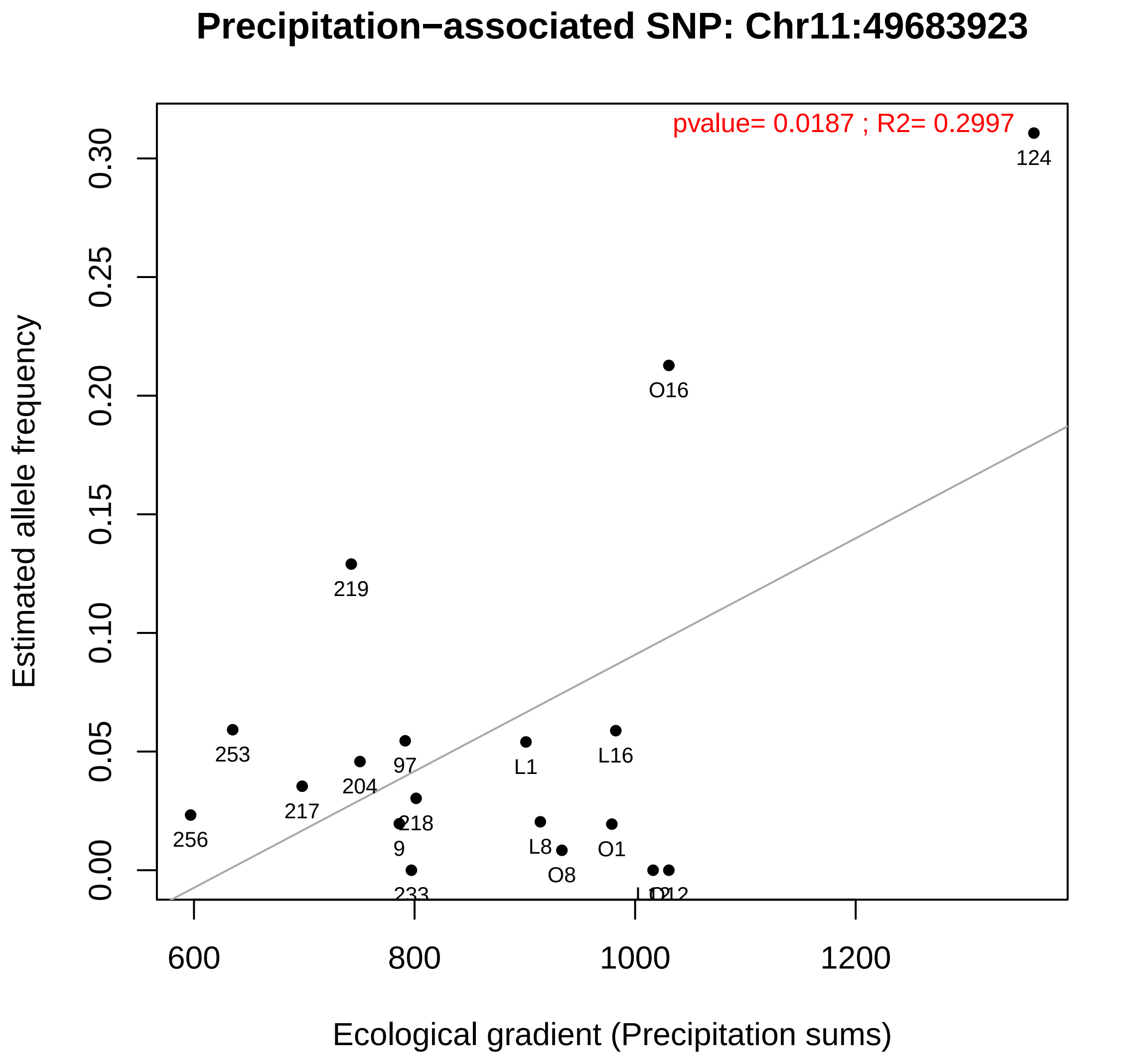

Precipitation−associated SNP: Chr11:49683923
pvalue= 0.0187 ; R2= 0.2997
Estimated allele frequency
Ecological gradient (Precipitation sums)
0.00
0.05
0.10
0.15
0.20
0.25
0.30
600
800
1000
1200
124
O16
219
253
97
L1
L16
204
217
218
256
9
L8
O1
O8
233
O12

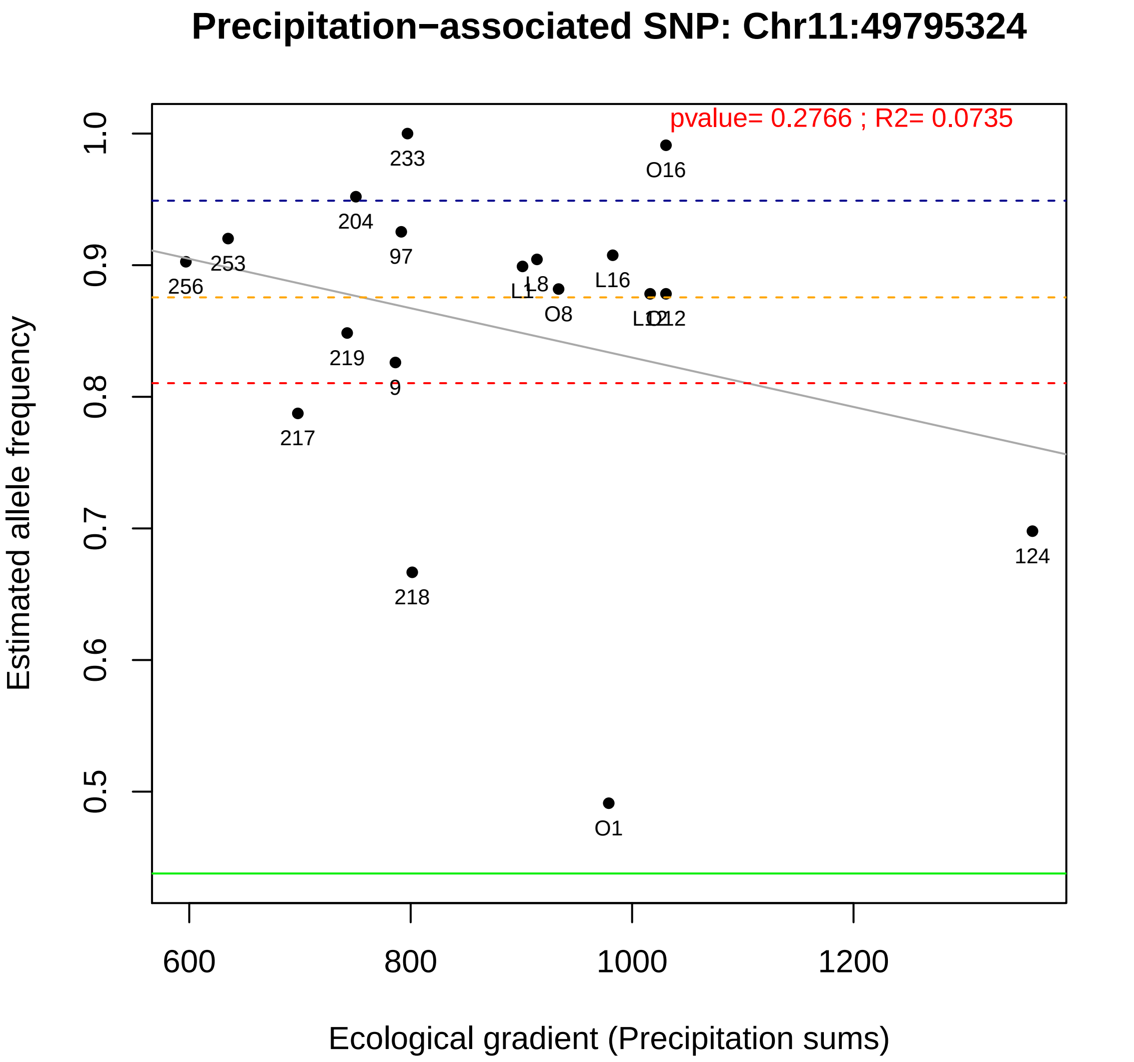
Precipitation−associated SNP: Chr11:49795324
pvalue= 0.2766 ; R2= 0.0735
Estimated allele frequency
Ecological gradient (Precipitation sums)
1.0
0.9
0.8
0.7
0.6
0.5
600
800
1000
1200
233
O16
204
97
253
256
L1
L8
O8
L16
L12
O12
219
9
217
124
218
O1

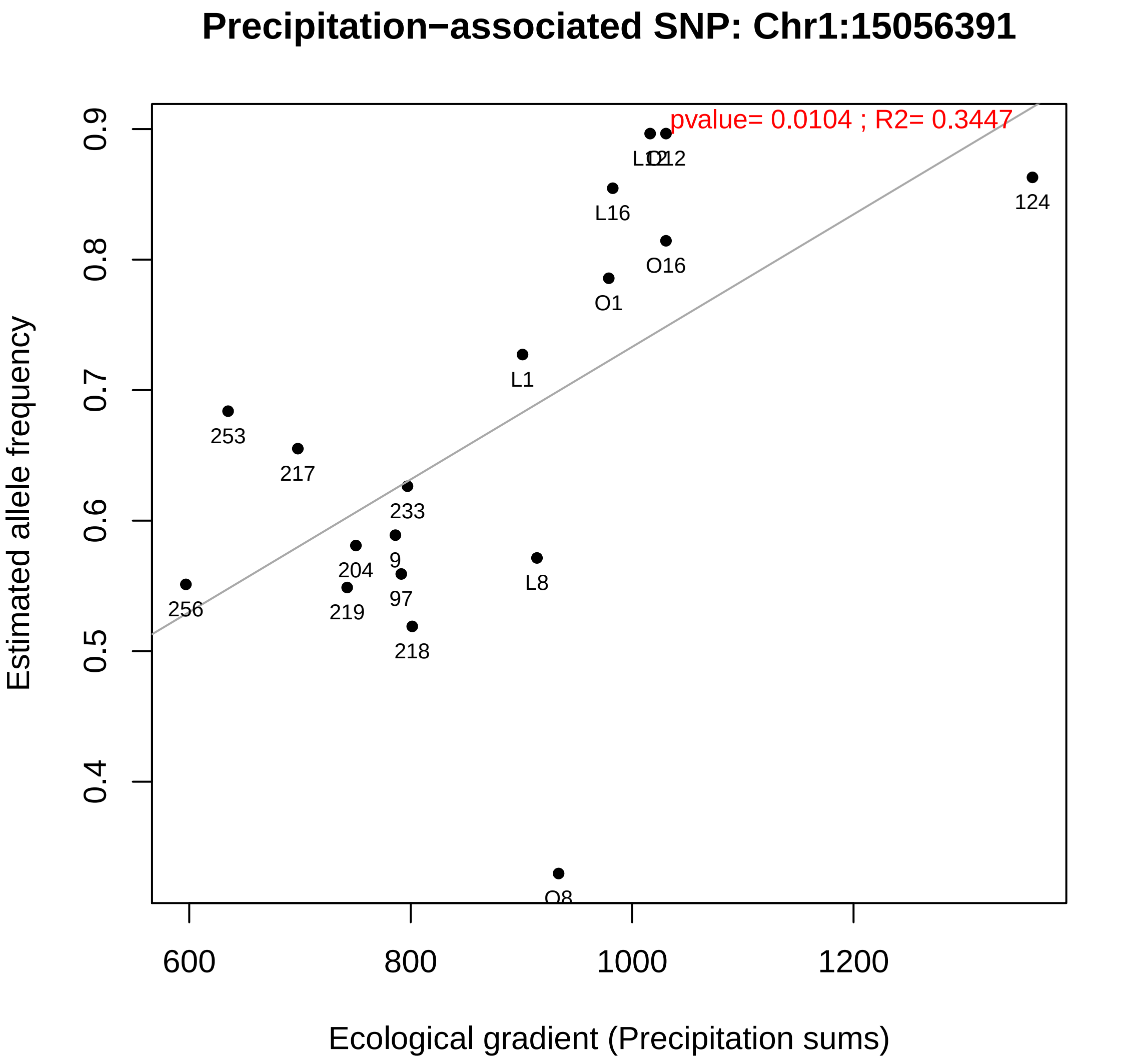

Precipitation−associated SNP: Chr1:15056391
pvalue= 0.0104 ; R2= 0.3447
Estimated allele frequency
0.4
0.5
0.6
0.7
0.8
0.9
600
800
1000
1200
Ecological gradient (Precipitation sums)
124
L16
O16
O1
L1
253
217
233
204
9
97
256
219
218
L8
O8

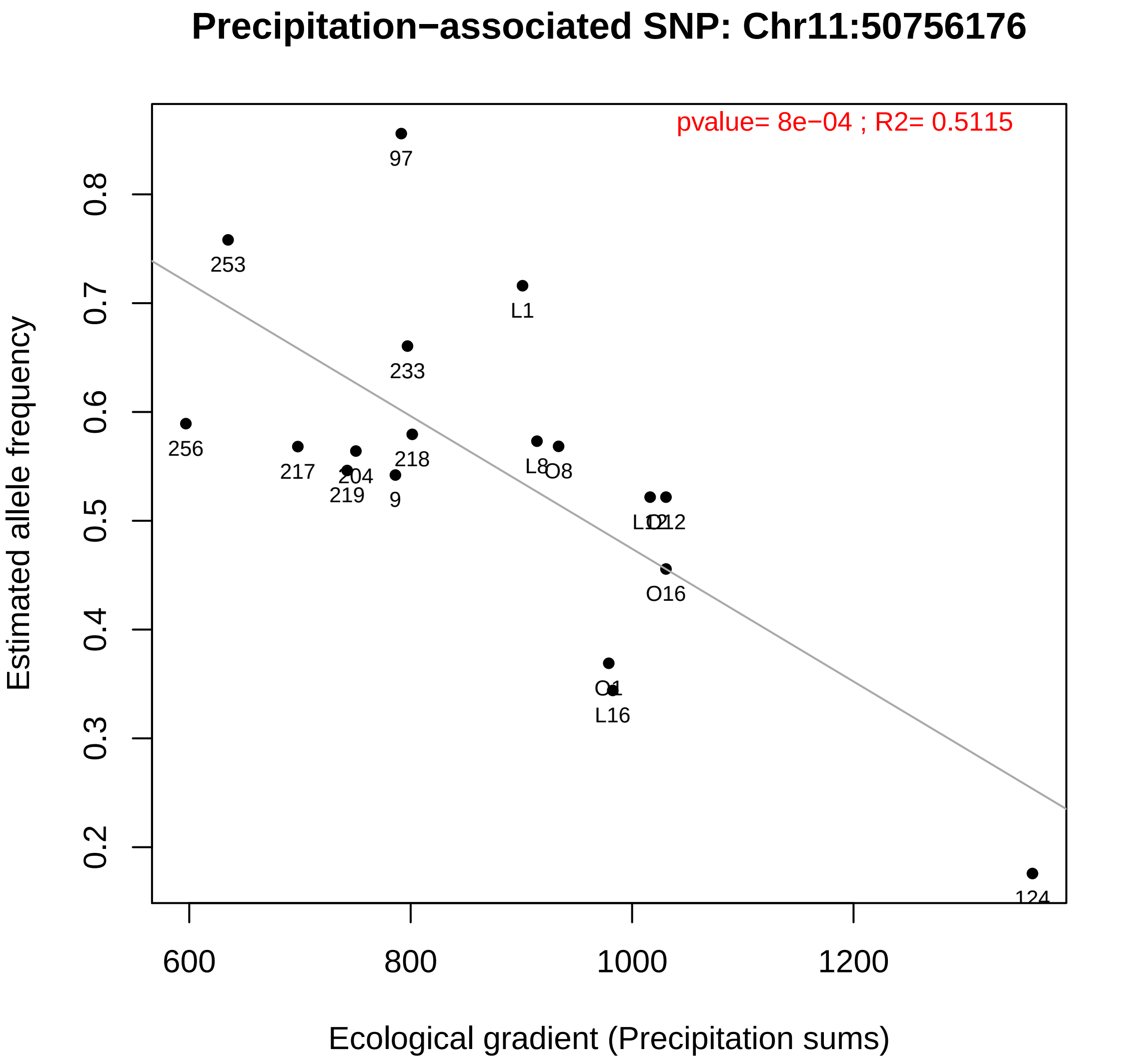
Precipitation−associated SNP: Chr11:50756176
pvalue= 8e−04 ; R2= 0.5115
97
253
L1
233
256
217
204
219
9
218
L8
O8
L12
O12
O16
O1
L16
124
Estimated allele frequency
0.2
0.3
0.4
0.5
0.6
0.7
0.8
600
800
1000
1200
Ecological gradient (Precipitation sums)

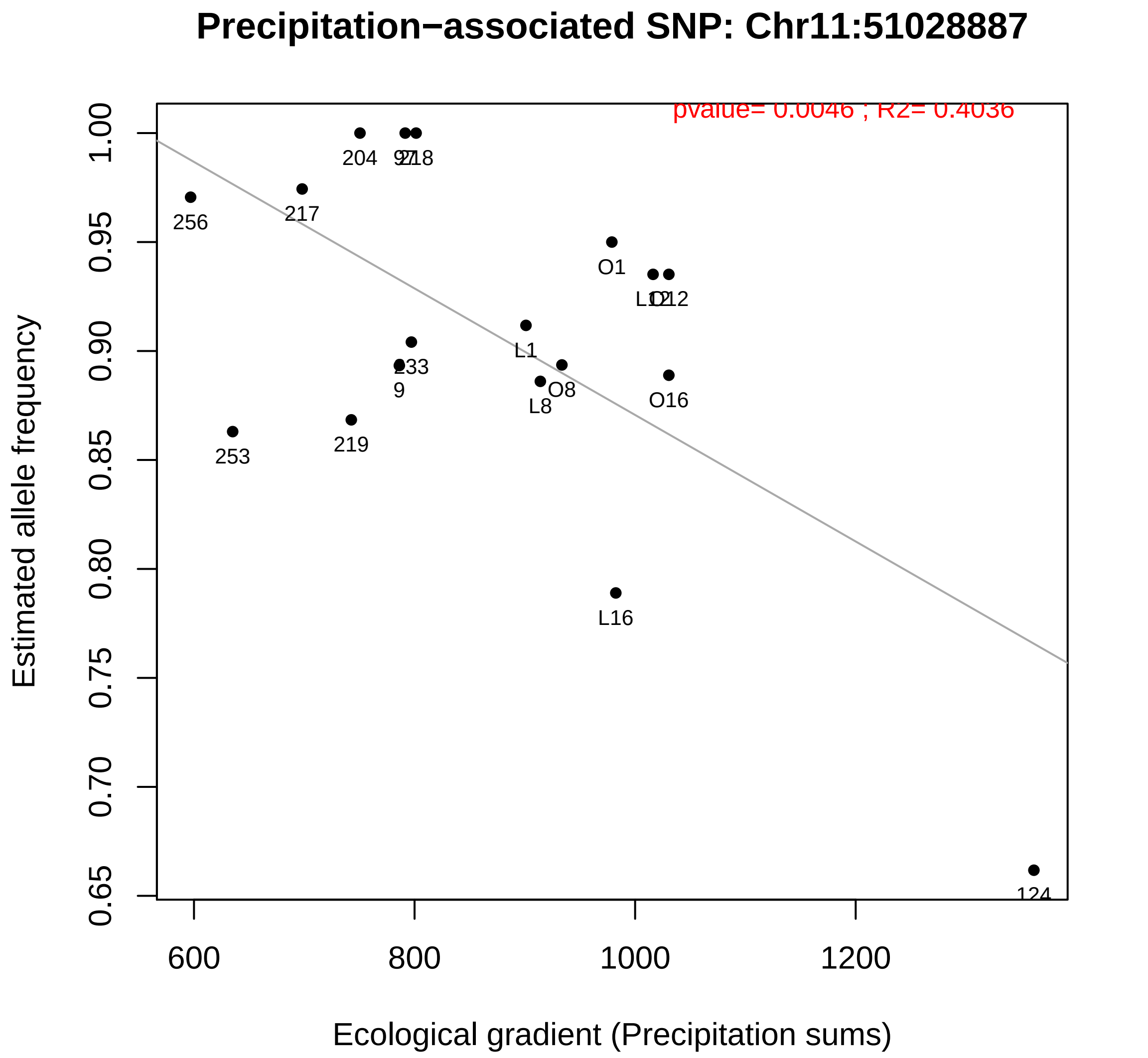
Precipitation−associated SNP: Chr11:51028887
pvalue= 0.0046 ; R2= 0.4036
Estimated allele frequency
Ecological gradient (Precipitation sums)
0.65
0.70
0.75
0.80
0.85
0.90
0.95
1.00
600
800
1000
1200
204
218
256
217
O1
O12
L1
233
9
O8
L8
O16
219
253
L16
124

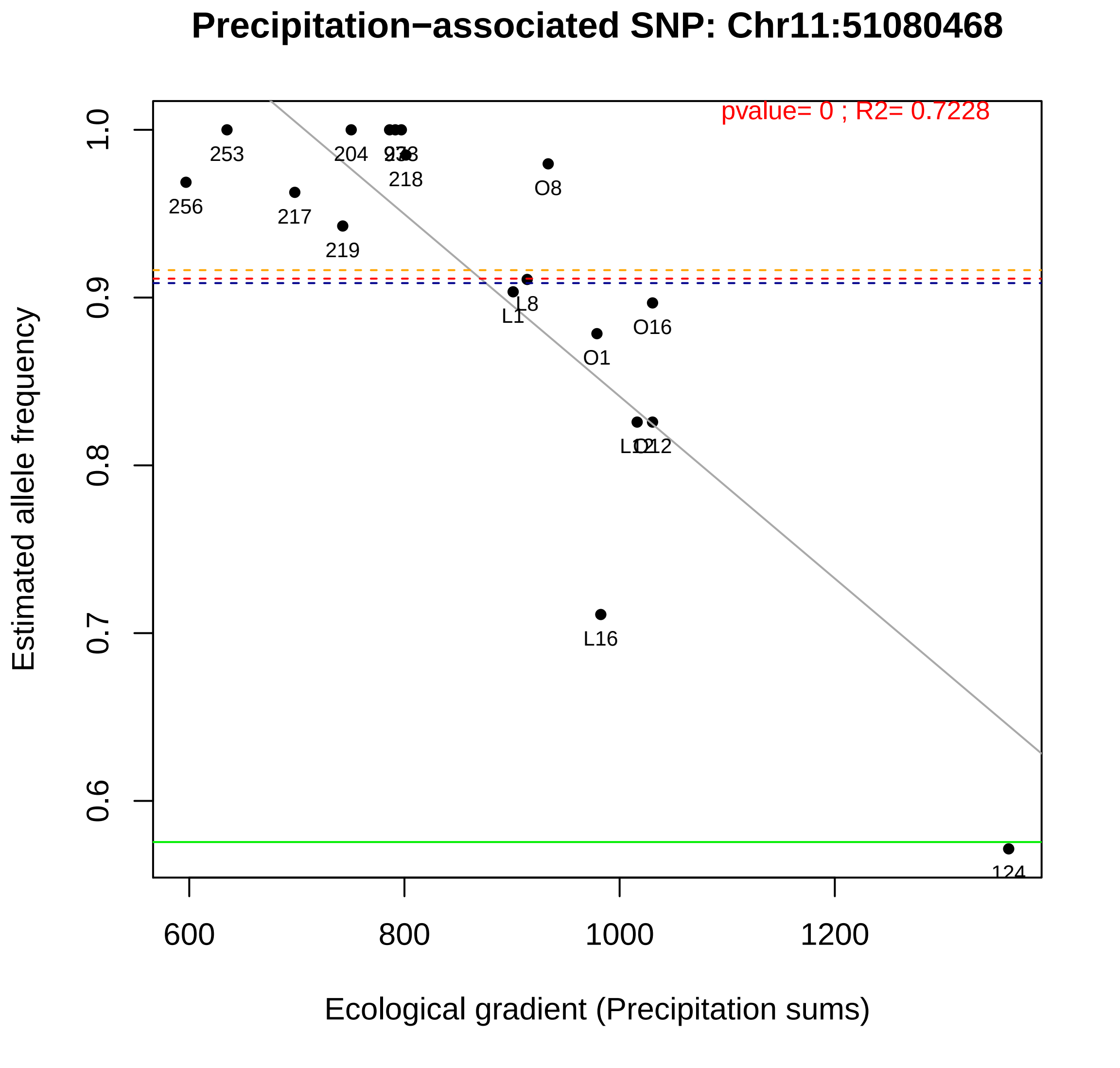

Precipitation−associated SNP: Chr11:51080468
pvalue= 0 ; R2= 0.7228
Estimated allele frequency
Ecological gradient (Precipitation sums)
0.6
0.7
0.8
0.9
1.0
600
800
1000
1200
253
204
218
O8
256
217
219
L8
L1
O16
O1
O12
L16
124

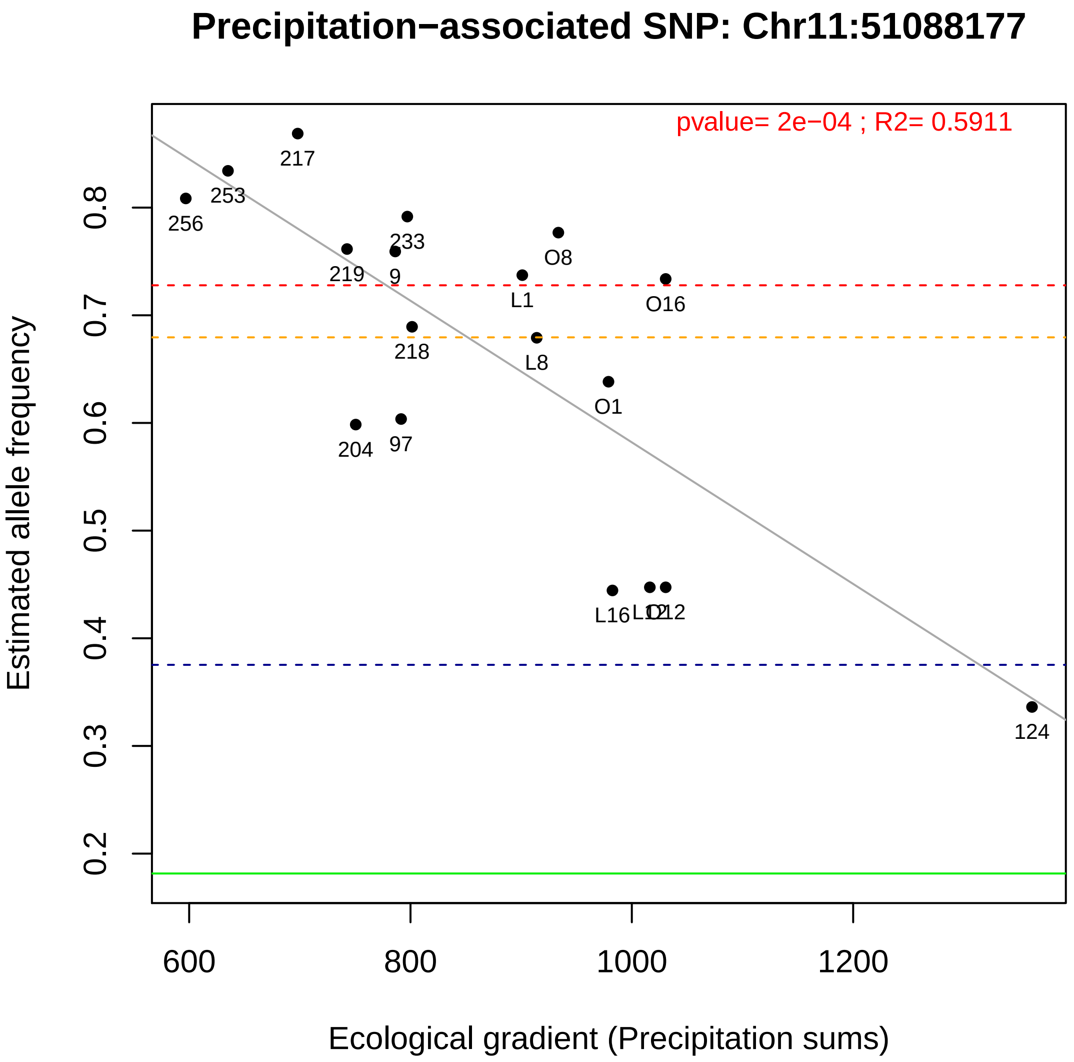

Precipitation−associated SNP: Chr11:51088177
pvalue= 2e−04 ; R2= 0.5911
Estimated allele frequency
Ecological gradient (Precipitation sums)
0.2
0.3
0.4
0.5
0.6
0.7
0.8
600
800
1000
1200
217
253
256
233
219
9
O8
L1
O16
218
L8
O1
204
97
L16
L12
O12
124

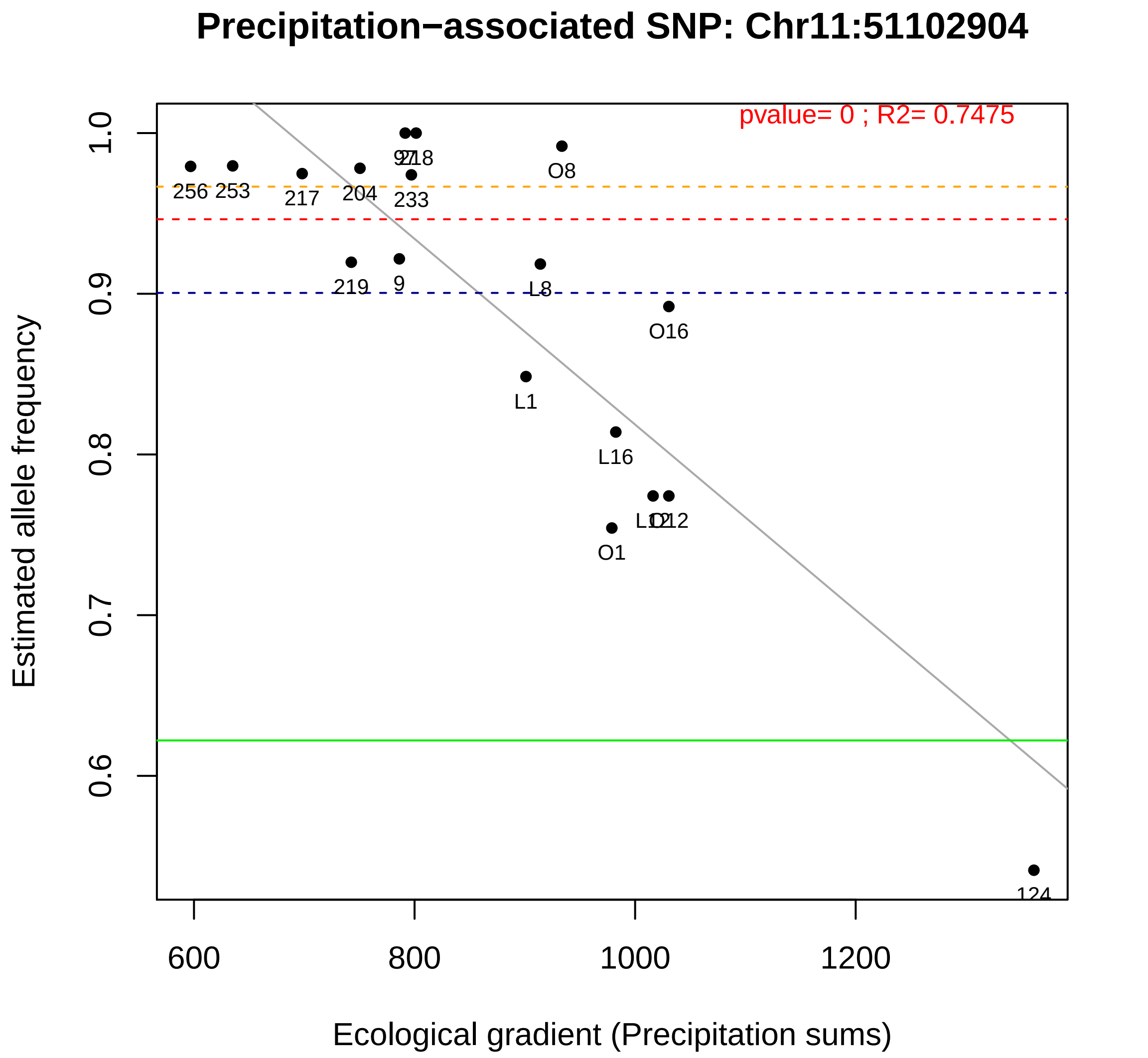

Precipitation−associated SNP: Chr11:51102904
pvalue= 0 ; R2= 0.7475
Estimated allele frequency
Ecological gradient (Precipitation sums)
1.0
0.9
0.8
0.7
0.6
600
800
1000
1200
256
253
217
204
233
O8
219
9
L8
O16
L1
L16
O1
124

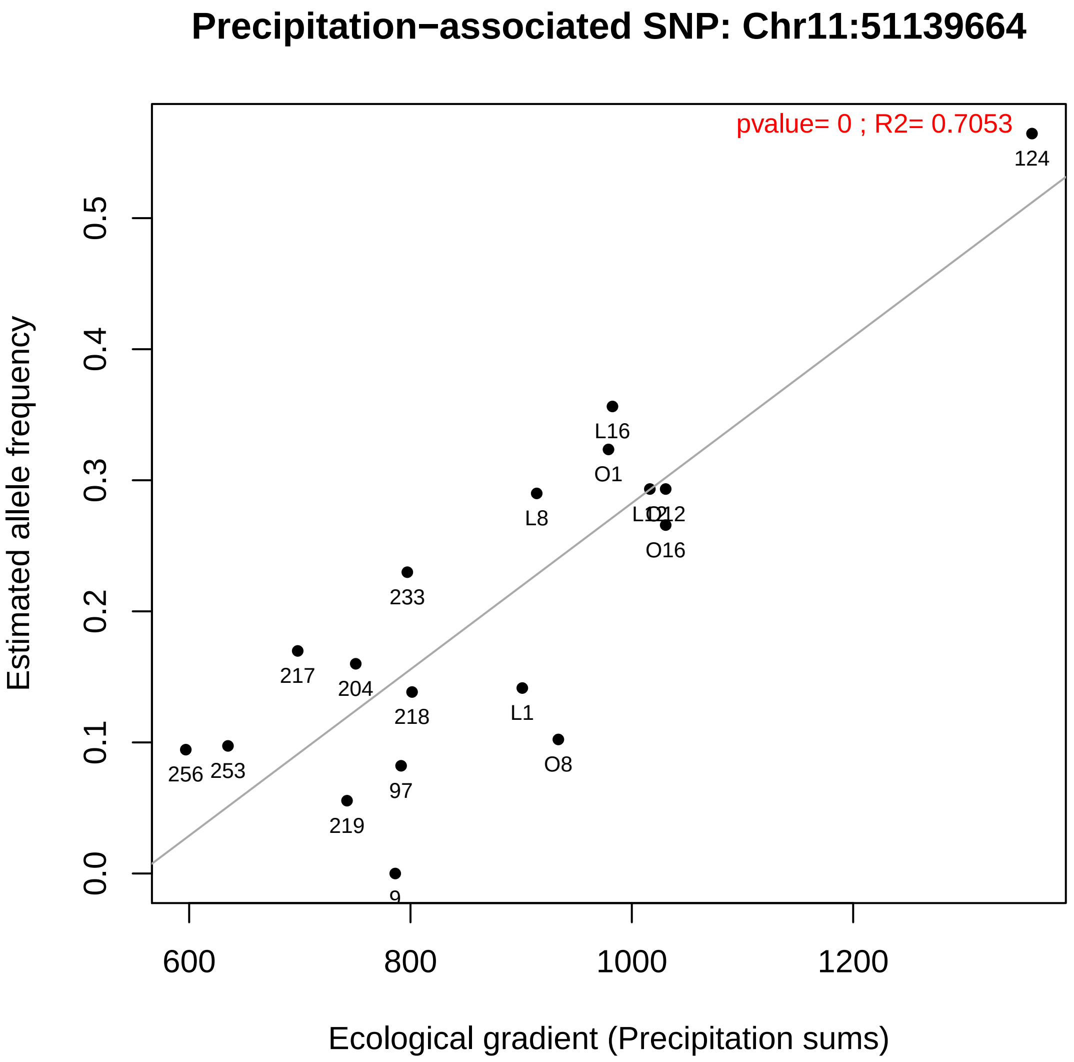

Precipitation−associated SNP: Chr11:51139664
pvalue= 0 ; R2= 0.7053
124
L16
O1
L8
L12
O12
O16
233
217
204
218
L1
256
253
O8
97
219
9
0.0
0.1
0.2
0.3
0.4
0.5
600
800
1000
1200
Estimated allele frequency
Ecological gradient (Precipitation sums)

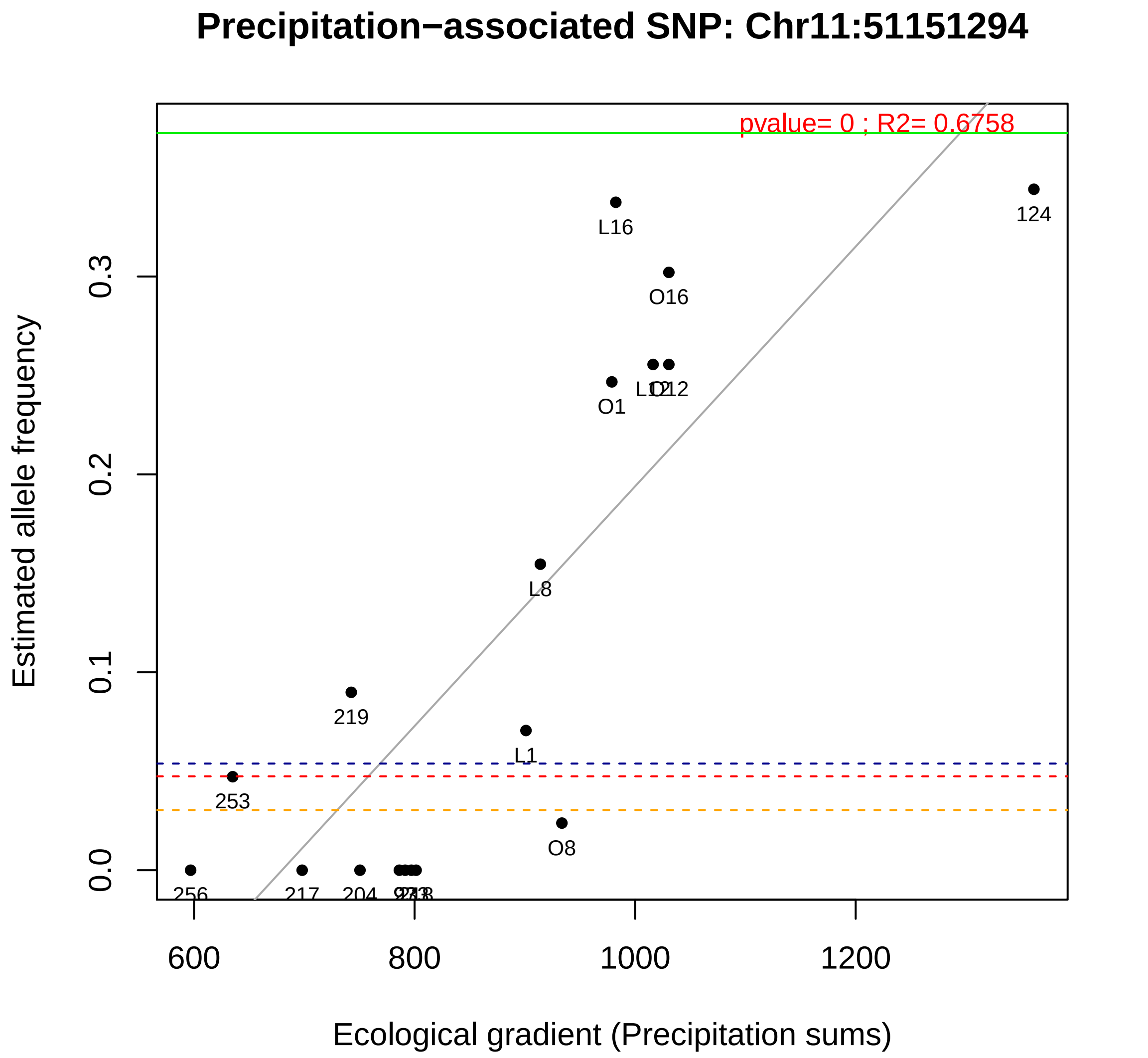
Precipitation−associated SNP: Chr11:51151294
pvalue= 0 ; R2= 0.6758
Estimated allele frequency
0.0
0.1
0.2
0.3
600
800
1000
1200
Ecological gradient (Precipitation sums)
L16
124
O16
O1
L8
219
L1
253
O8
256
217
204

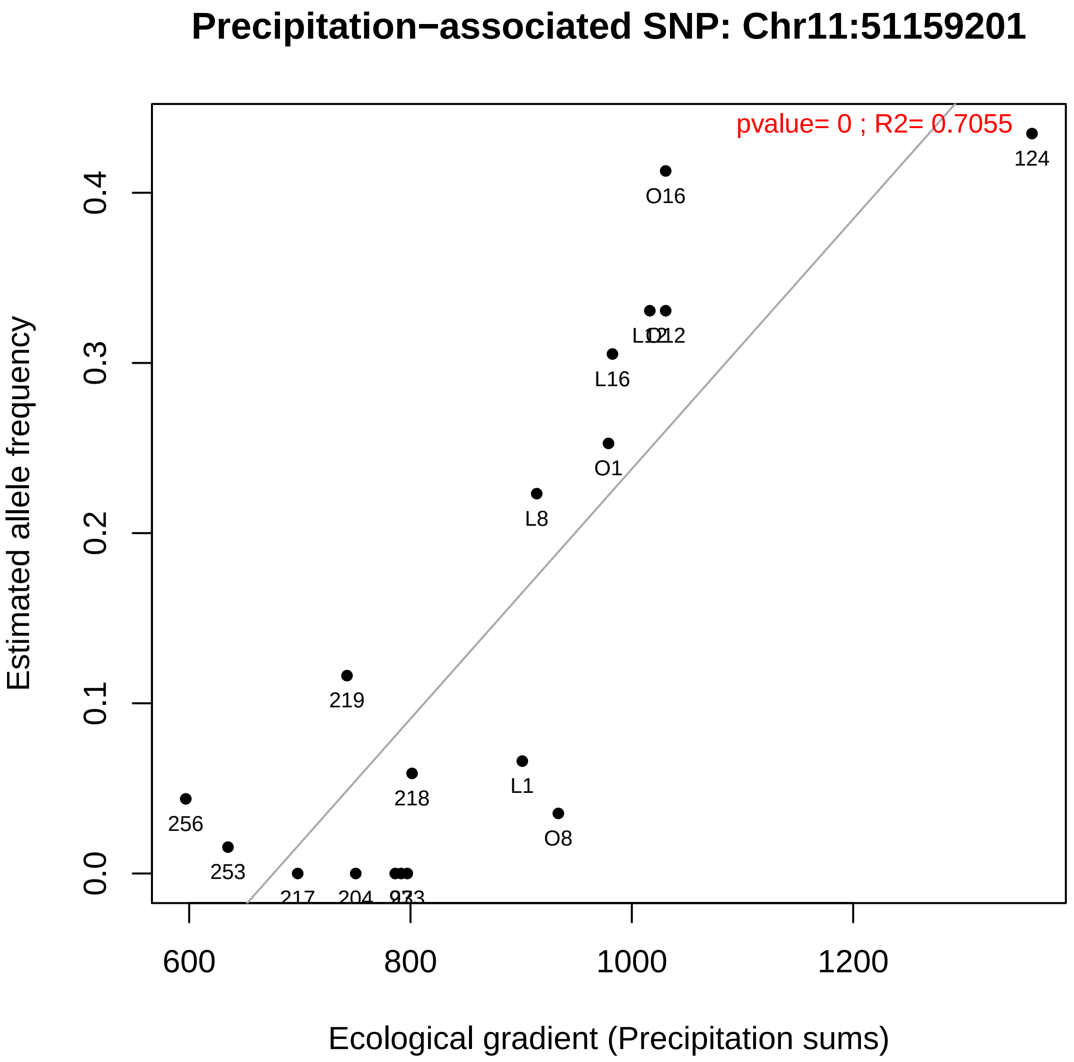
Precipitation−associated SNP: Chr11:51159201
pvalue= 0 ; R2= 0.7055
124
O16
L10
O12
L16
O1
L8
219
218
L1
256
O8
253
217
204
0.0
0.1
0.2
0.3
0.4
Estimated allele frequency
600
800
1000
1200
Ecological gradient (Precipitation sums)

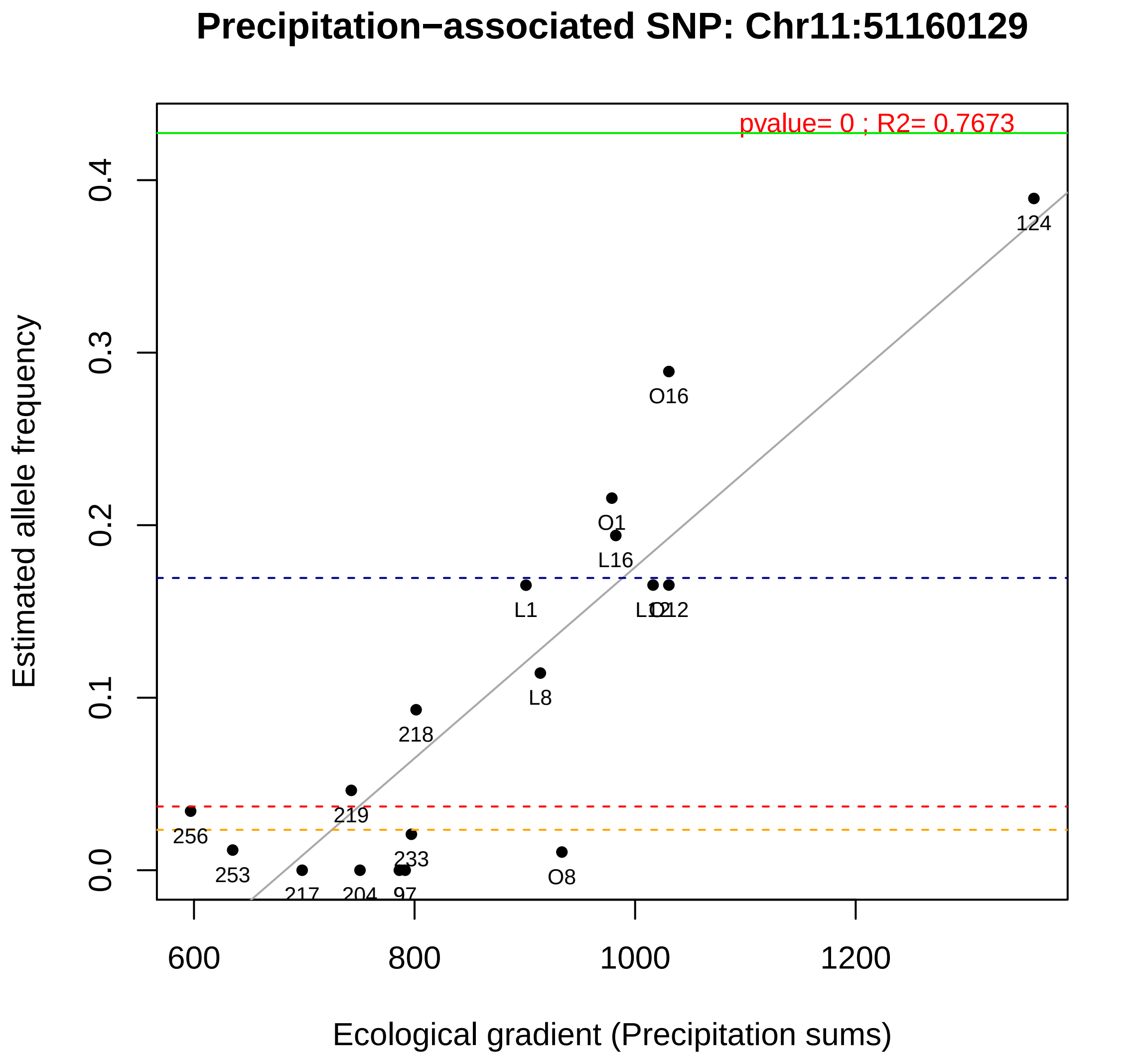

Precipitation−associated SNP: Chr11:51160129
pvalue= 0 ; R2= 0.7673
Estimated allele frequency
0.0
0.1
0.2
0.3
0.4
600
800
1000
1200
Ecological gradient (Precipitation sums)
124
O16
O1
L16
L1
L8
218
219
256
253
233
217
204
97
O8

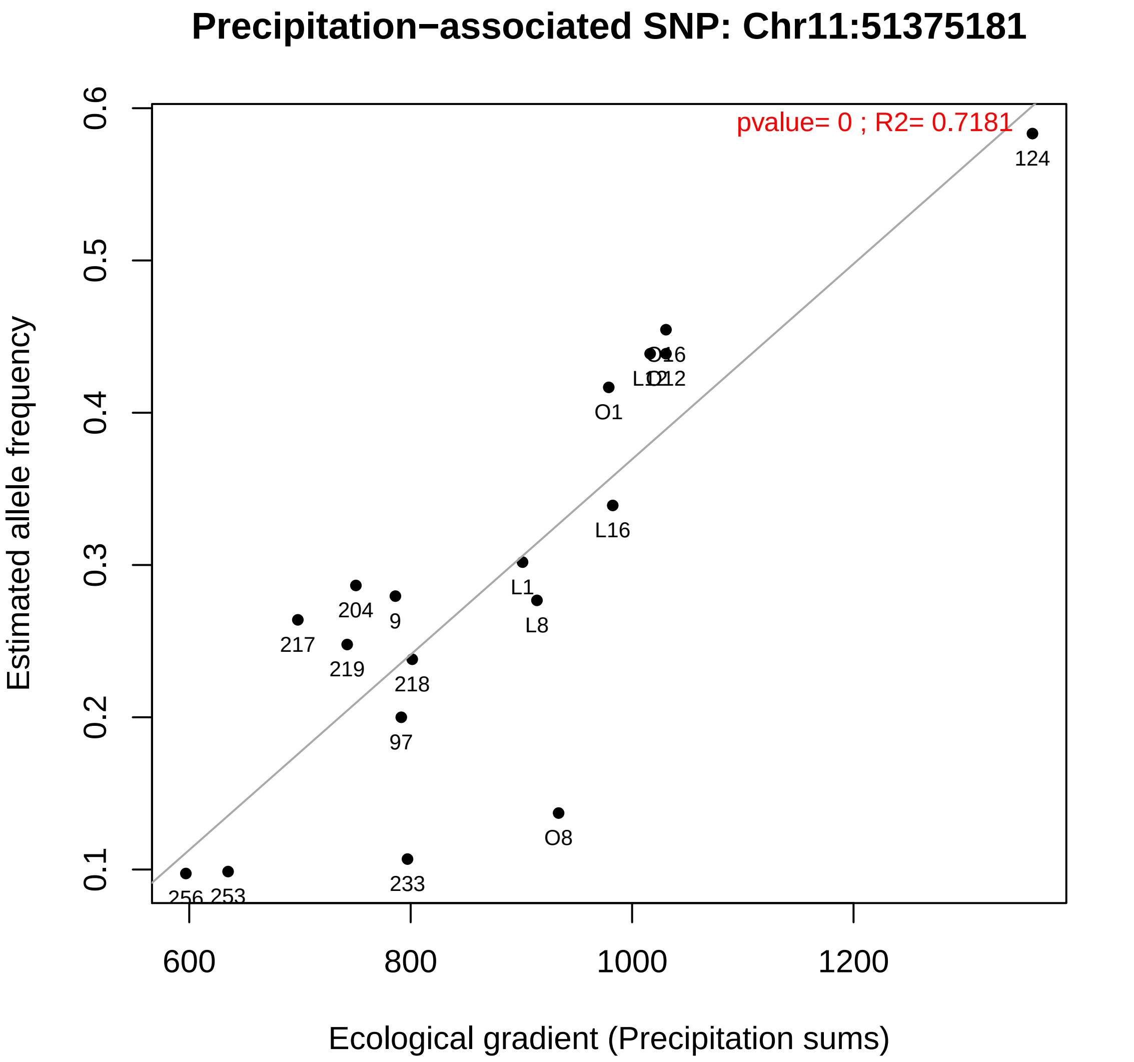

Precipitation−associated SNP: Chr11:51375181
pvalue= 0 ; R2= 0.7181
Estimated allele frequency
Ecological gradient (Precipitation sums)
0.1
0.2
0.3
0.4
0.5
0.6
600
800
1000
1200
124
O16
L12
O12
O1
L16
L1
L8
204
9
217
219
218
97
O8
233
256
253

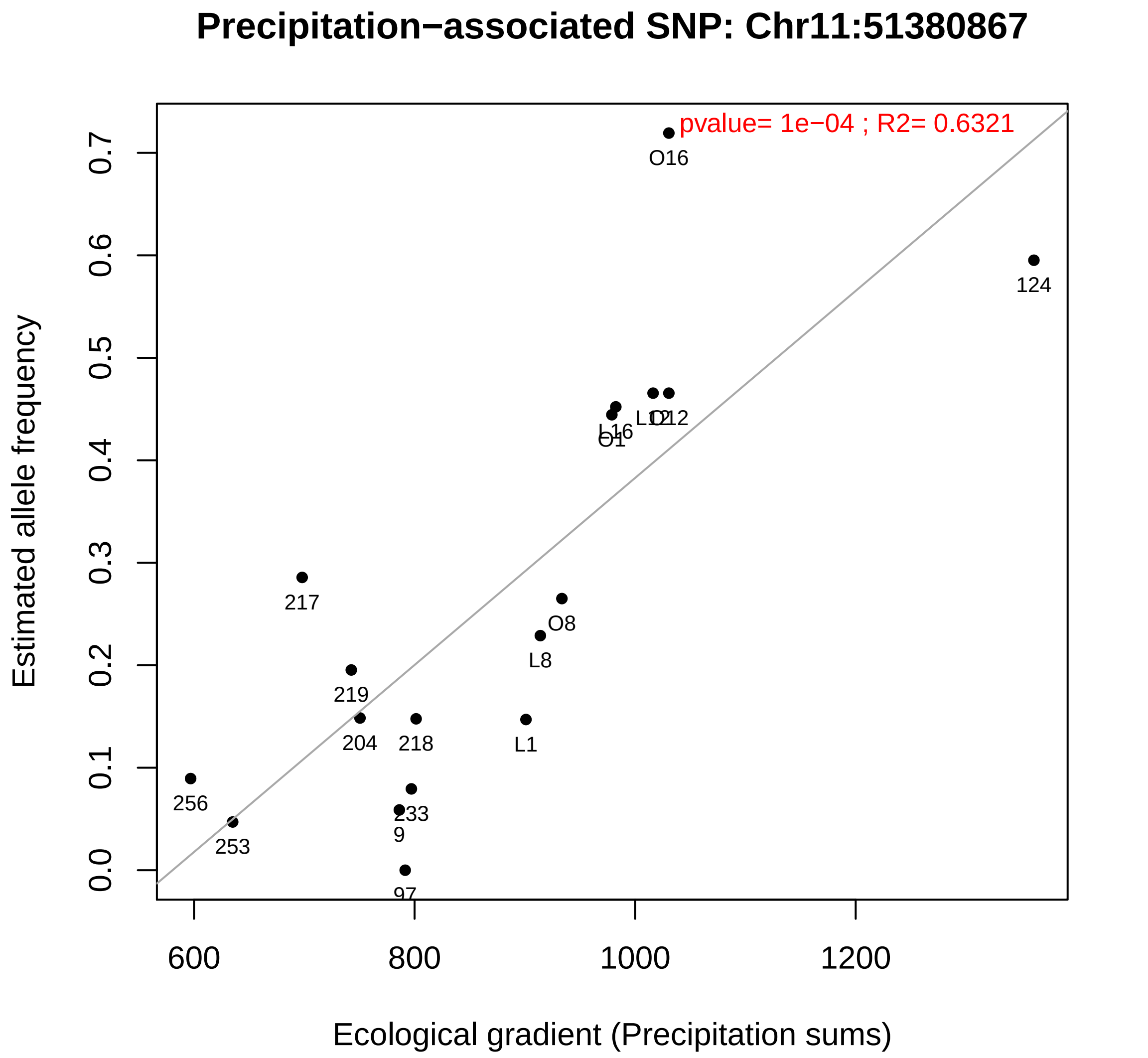

Precipitation−associated SNP: Chr11:51380867
pvalue= 1e−04 ; R2= 0.6321
O16
124
L16
O1
L12
O12
217
O8
L8
219
204
218
L1
256
233
9
253
97
0.0
0.1
0.2
0.3
0.4
0.5
0.6
0.7
600
800
1000
1200
Estimated allele frequency
Ecological gradient (Precipitation sums)

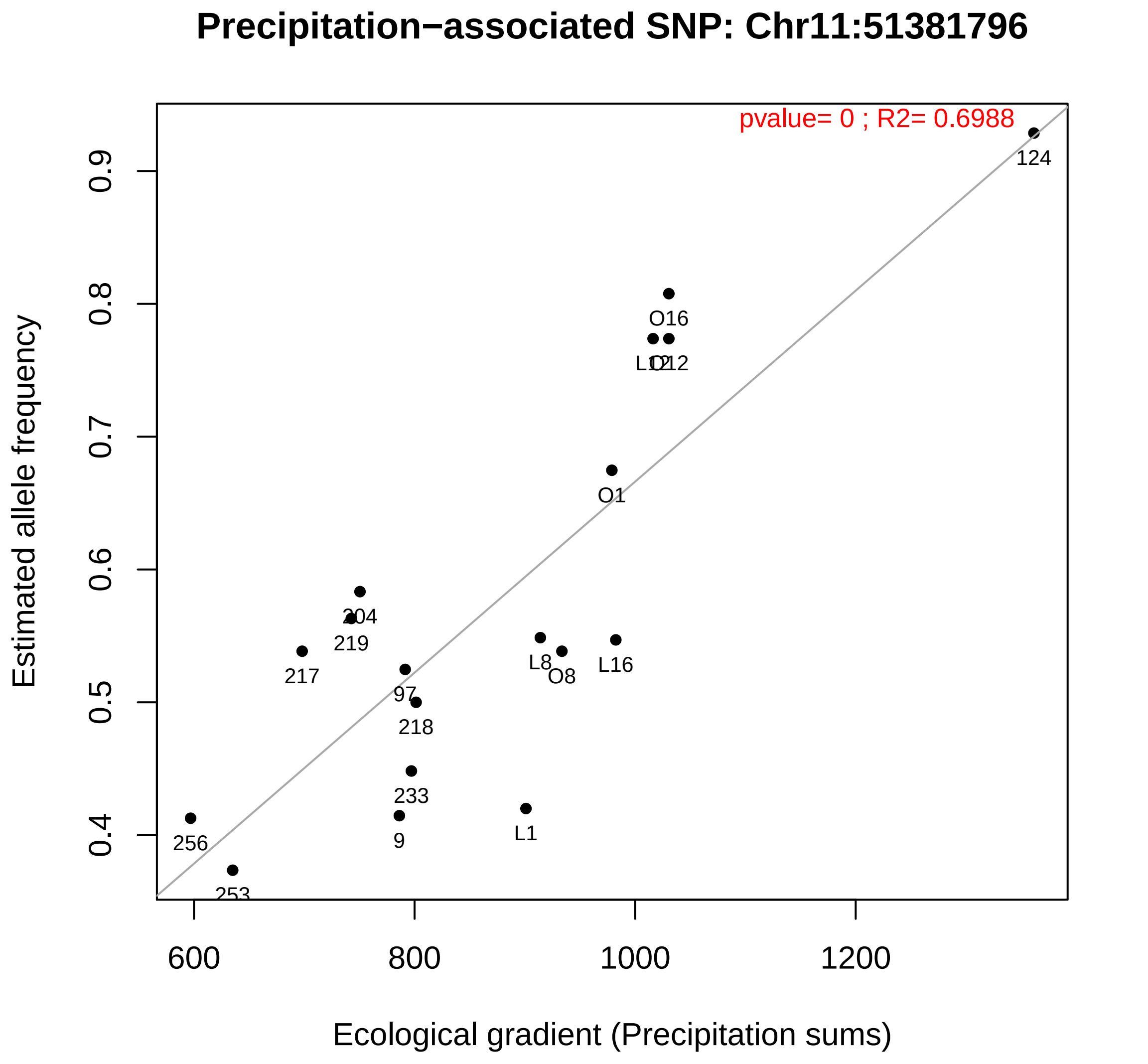

Precipitation−associated SNP: Chr11:51381796
pvalue= 0 ; R2= 0.6988
Estimated allele frequency
Ecological gradient (Precipitation sums)
0.4
0.5
0.6
0.7
0.8
0.9
600
800
1000
1200
124
O16
L12
O12
O1
204
219
217
97
218
L8
O8
L16
233
9
L1
256
253

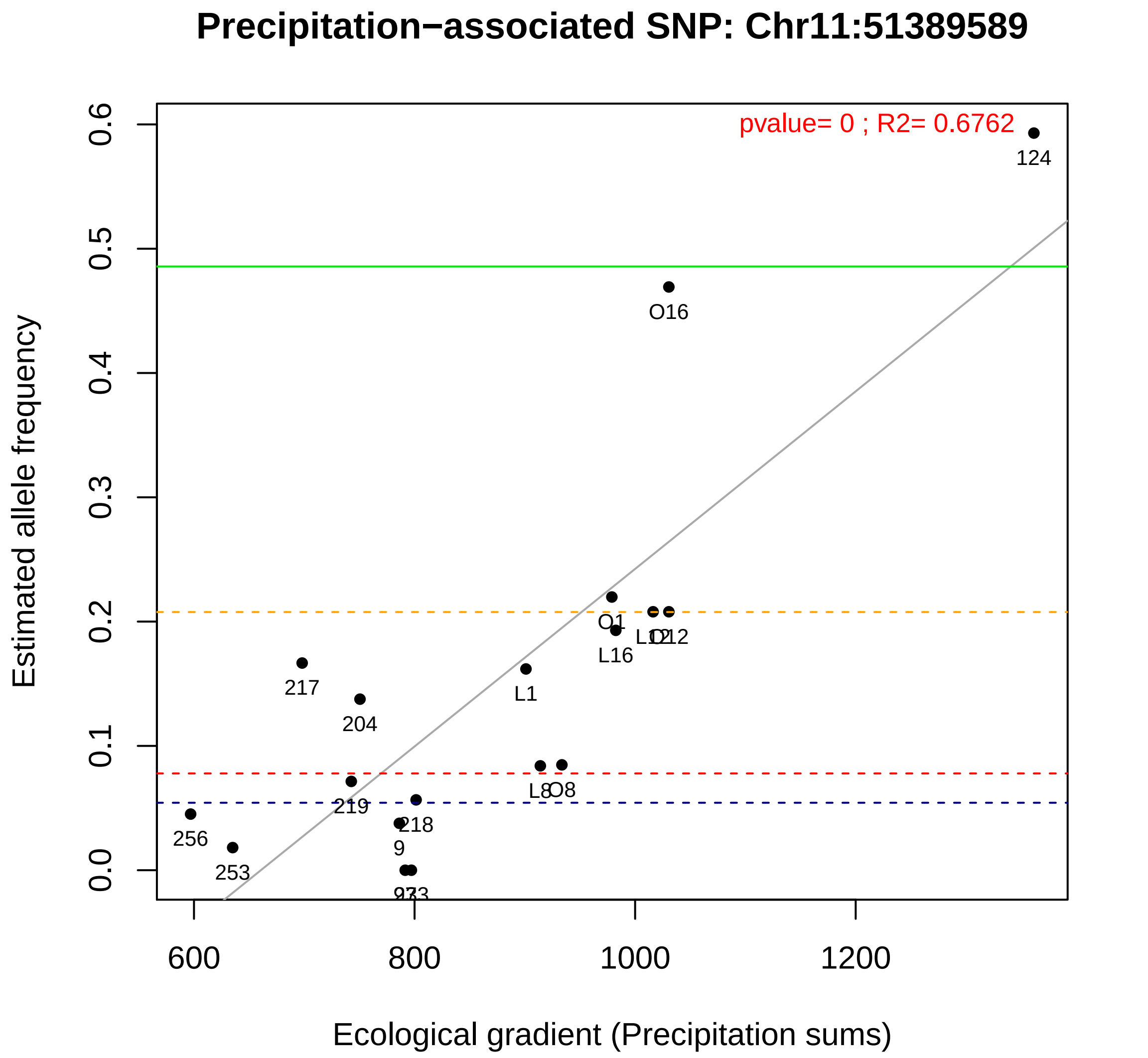
Precipitation−associated SNP: Chr11:51389589
pvalue= 0 ; R2= 0.6762
124
O16
O1
L12
O12
L16
217
L1
204
L8
O8
219
218
256
9
253
Estimated allele frequency
0.0
0.1
0.2
0.3
0.4
0.5
0.6
600
800
1000
1200
Ecological gradient (Precipitation sums)

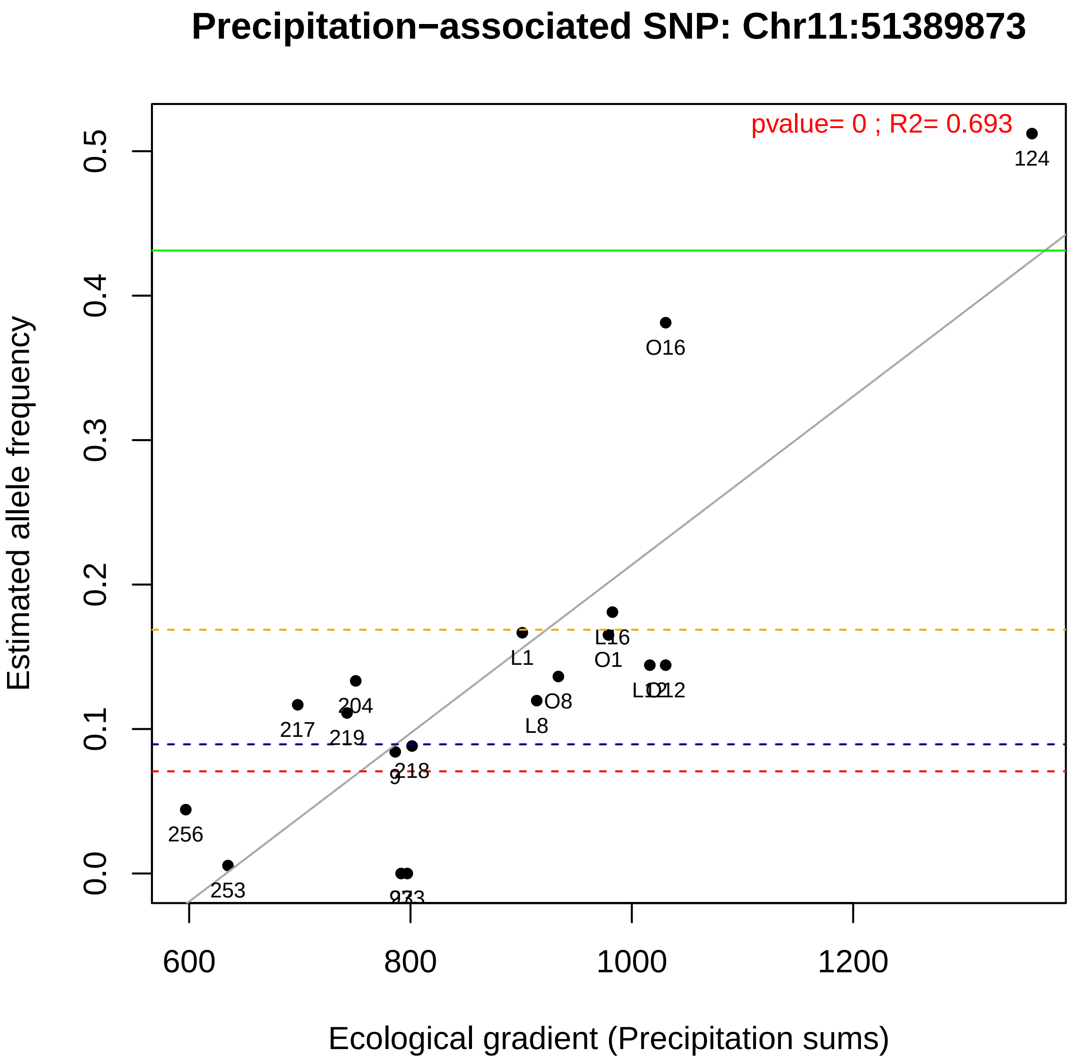

Precipitation−associated SNP: Chr11:51389873
pvalue= 0 ; R2= 0.693
Estimated allele frequency
Ecological gradient (Precipitation sums)
0.0
0.1
0.2
0.3
0.4
0.5
600
800
1000
1200
124
O16
L16
O1
L1
L12
O12
O8
L8
204
217
219
218
9
256
253

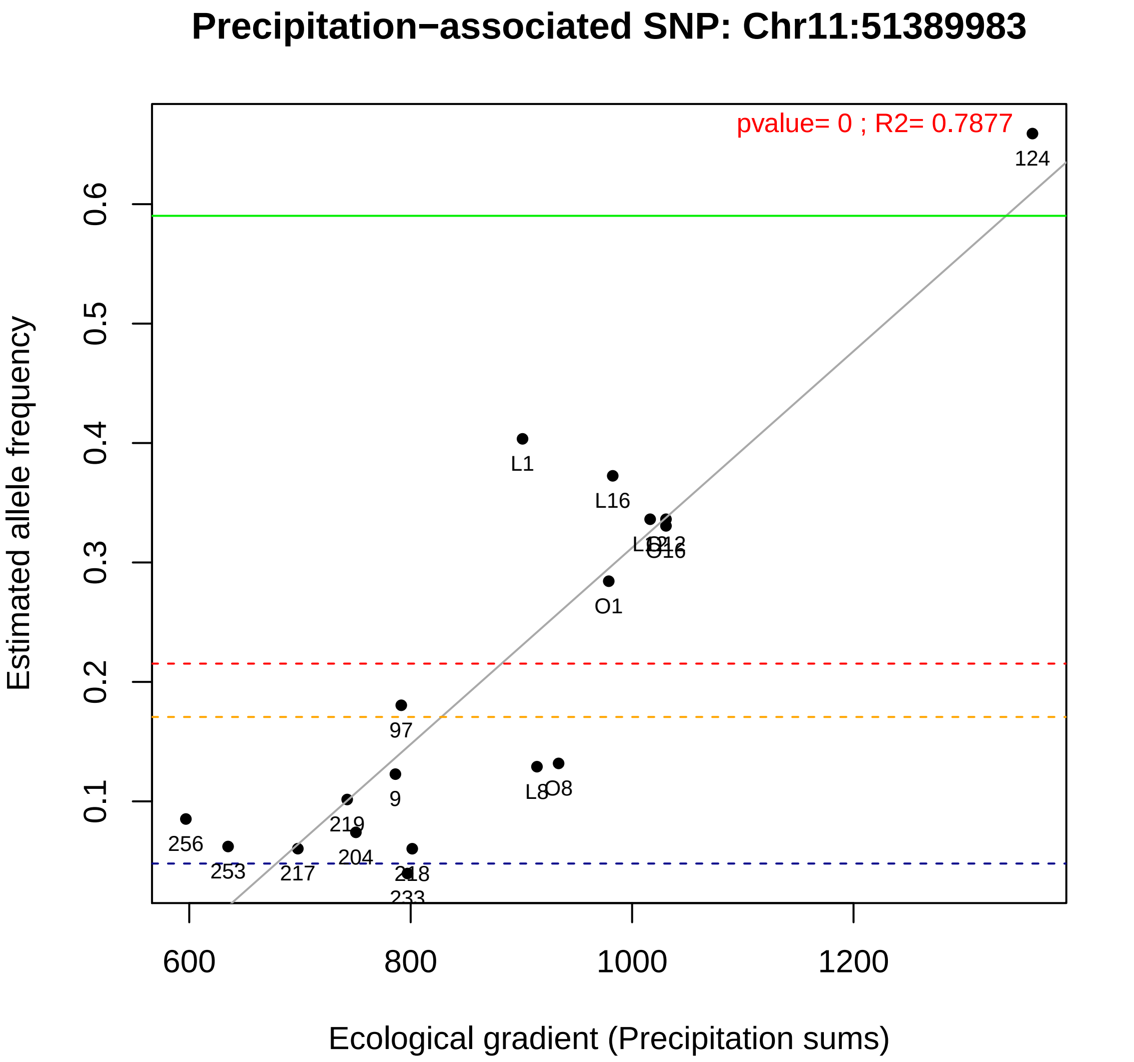

Precipitation−associated SNP: Chr11:51389983
pvalue= 0 ; R2= 0.7877
Estimated allele frequency
Ecological gradient (Precipitation sums)
0.1
0.2
0.3
0.4
0.5
0.6
600
800
1000
1200
124
L1
L16
O1
97
9
219
204
256
253
217
218
233
L8
O8

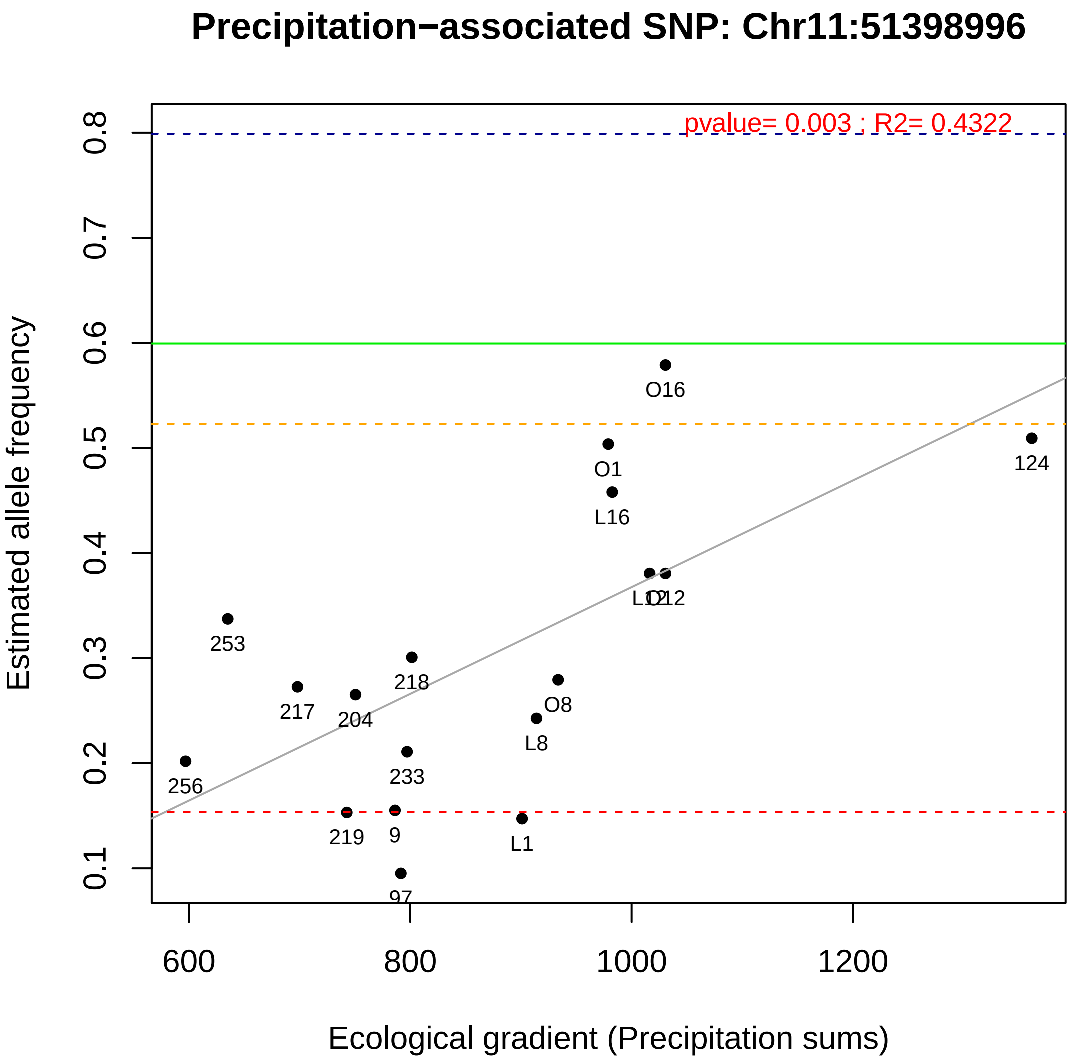

Precipitation−associated SNP: Chr11:51398996
pvalue= 0.003 ; R2= 0.4322
Estimated allele frequency
0.1
0.2
0.3
0.4
0.5
0.6
0.7
0.8
600
800
1000
1200
Ecological gradient (Precipitation sums)
O16
O1
124
L16
L12
O12
253
218
217
204
O8
L8
233
256
219
9
L1
97

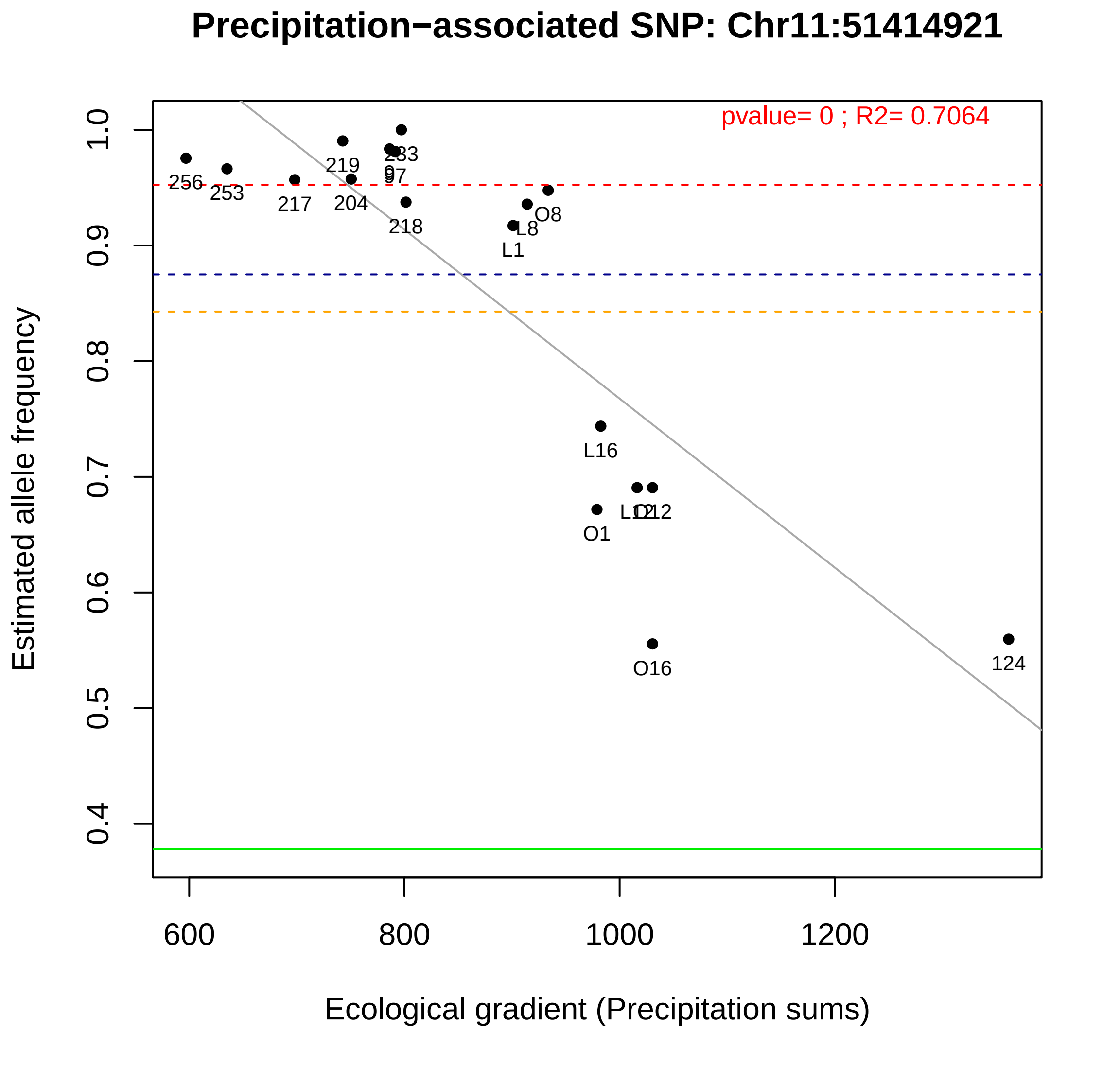

Precipitation−associated SNP: Chr11:51414921
pvalue= 0 ; R2= 0.7064
Estimated allele frequency
Ecological gradient (Precipitation sums)
1.0
0.9
0.8
0.7
0.6
0.5
0.4
600
800
1000
1200
256
253
217
219
204
233
97
218
L1
L8
O8
L16
O1
O12
O16
124

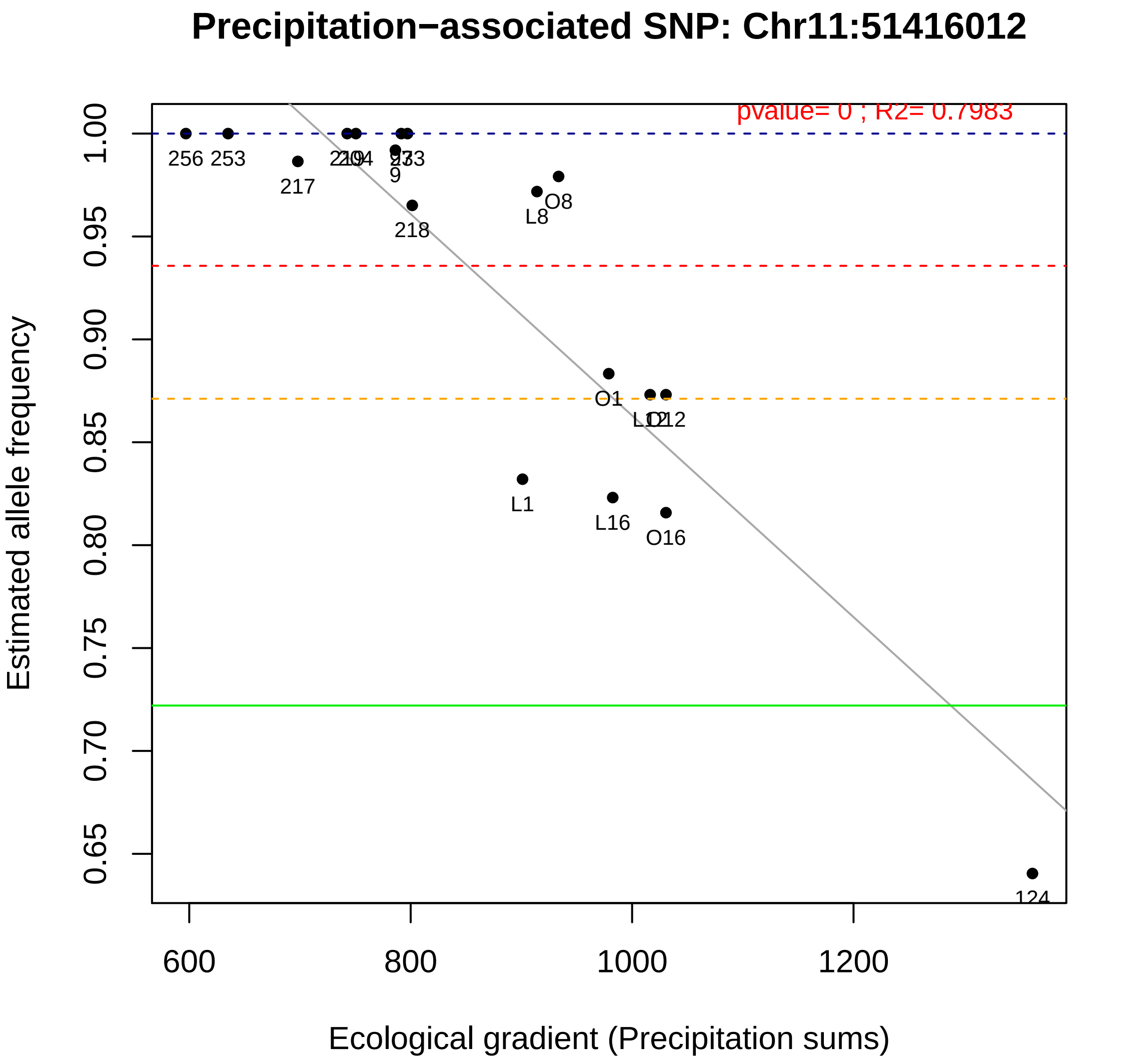

Precipitation−associated SNP: Chr11:51416012
pvalue= 0 ; R2= 0.7983
Estimated allele frequency
Ecological gradient (Precipitation sums)
0.65
0.70
0.75
0.80
0.85
0.90
0.95
1.00
600
800
1000
1200
256
253
217
218
L8
O8
O1
L1
L16
O16
124

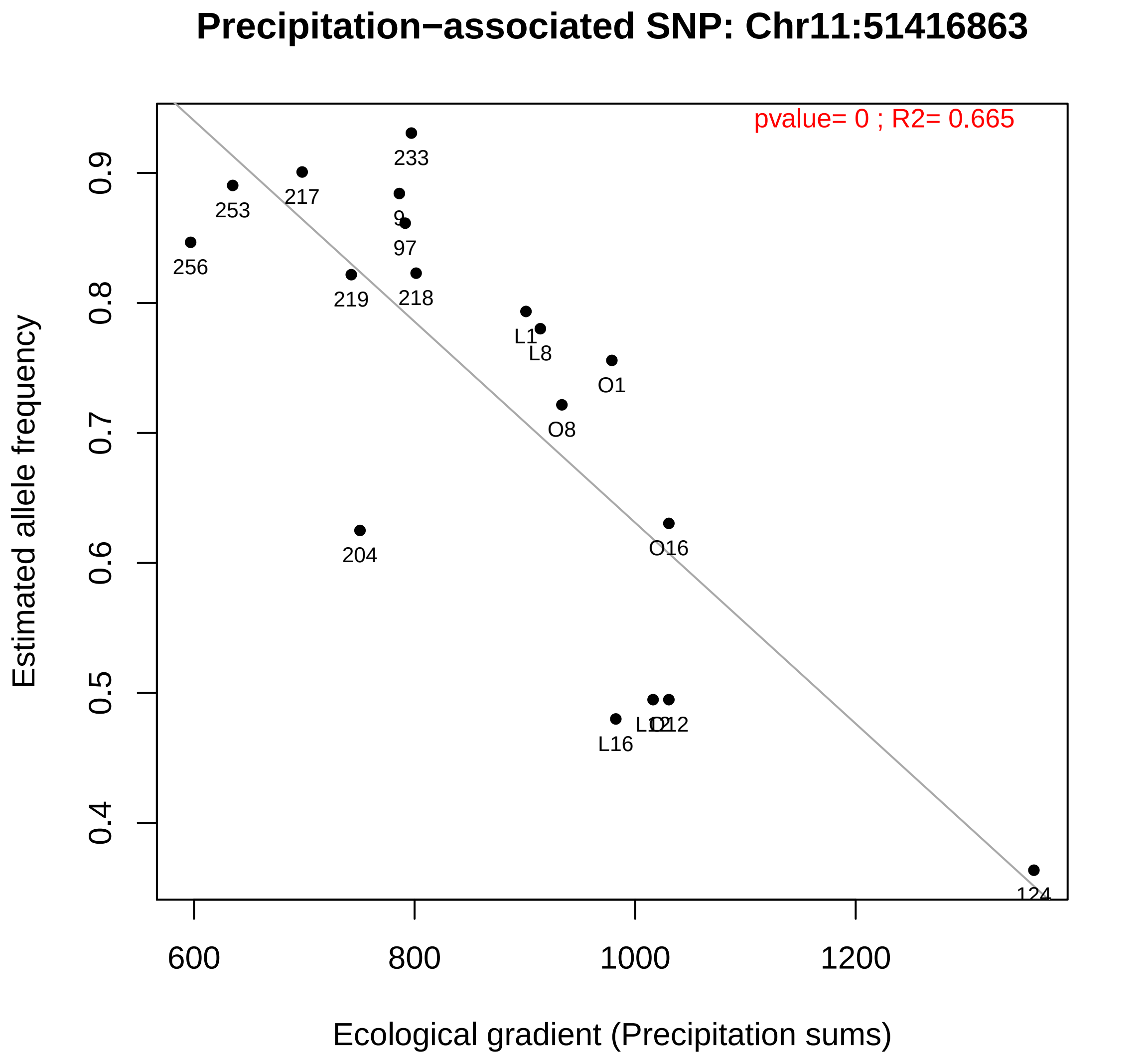

Precipitation−associated SNP: Chr11:51416863
pvalue= 0 ; R2= 0.665
Estimated allele frequency
Ecological gradient (Precipitation sums)
0.4
0.5
0.6
0.7
0.8
0.9
600
800
1000
1200
233
217
253
9
97
256
219
218
L1
L8
O1
O8
204
O16
L16
L12
O12
124

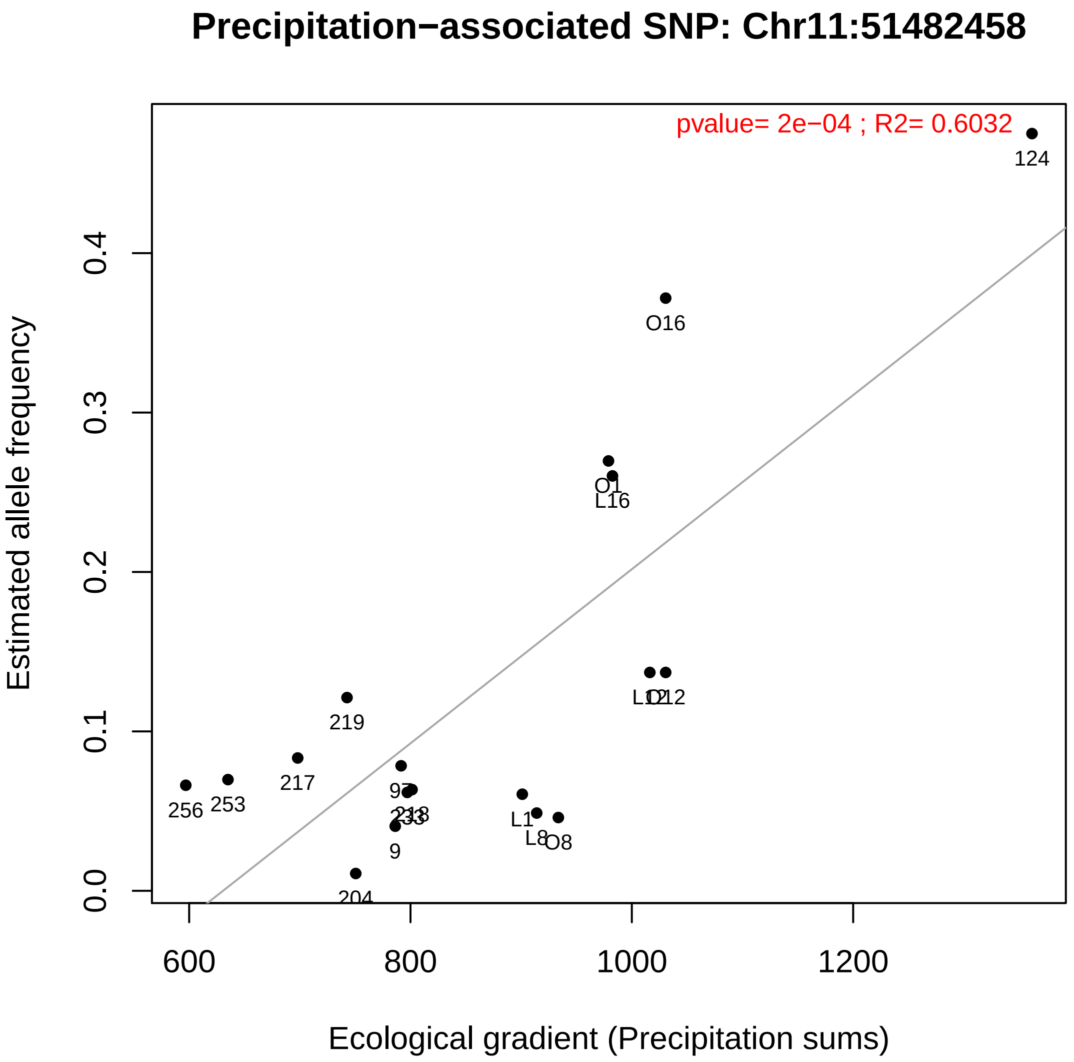

Precipitation−associated SNP: Chr11:51482458
pvalue= 2e−04 ; R2= 0.6032
124
O16
O1
L16
L12
O12
219
217
256
253
97
233
218
9
204
L1
L8
O8
0.0
0.1
0.2
0.3
0.4
600
800
1000
1200
Estimated allele frequency
Ecological gradient (Precipitation sums)

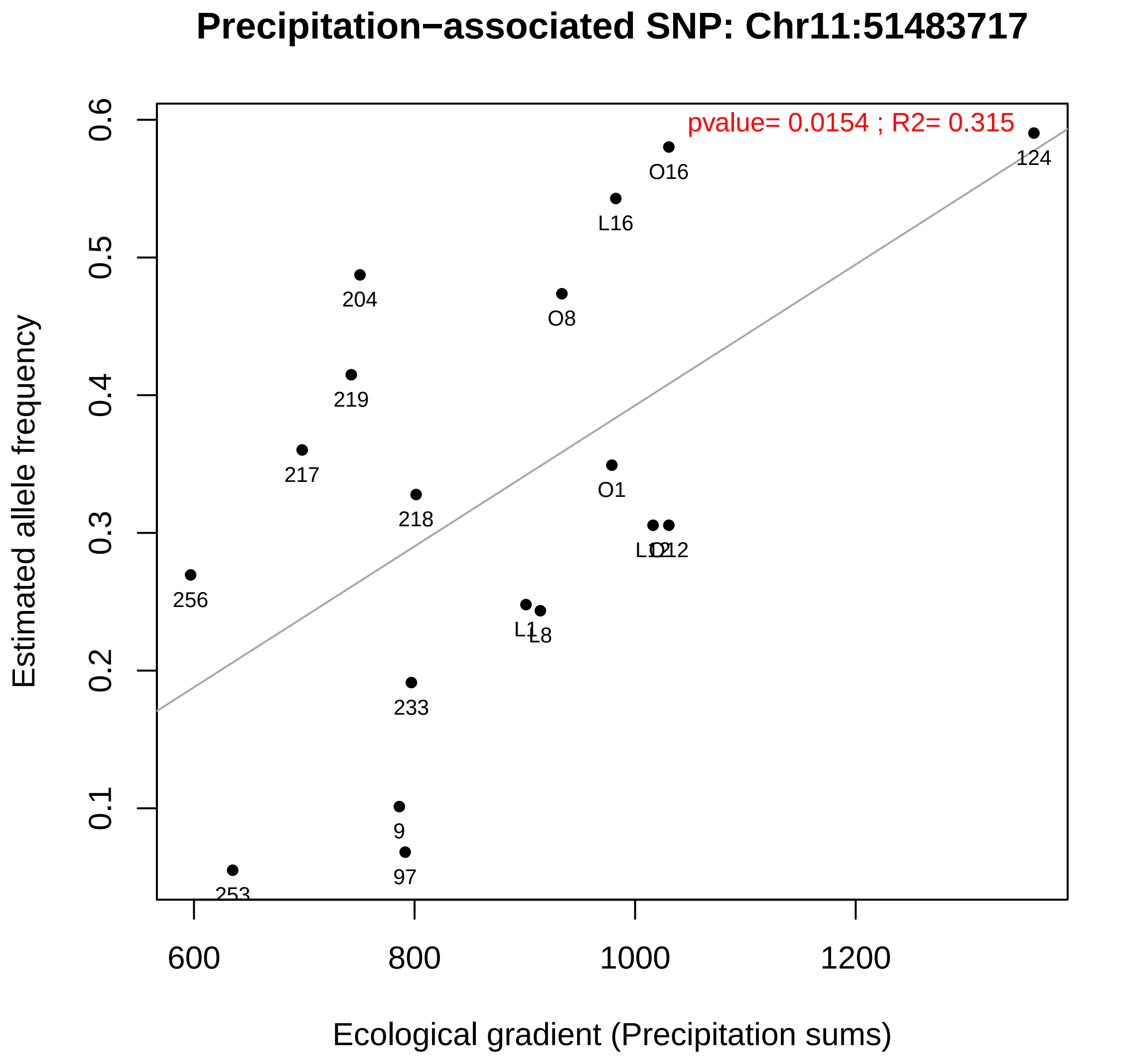

Precipitation−associated SNP: Chr11:51483717
pvalue= 0.0154 ; R2= 0.315
Estimated allele frequency
0.1
0.2
0.3
0.4
0.5
0.6
600
800
1000
1200
Ecological gradient (Precipitation sums)
124
O16
L16
204
O8
219
217
218
O1
L10
O12
256
L1
L8
233
9
97
253

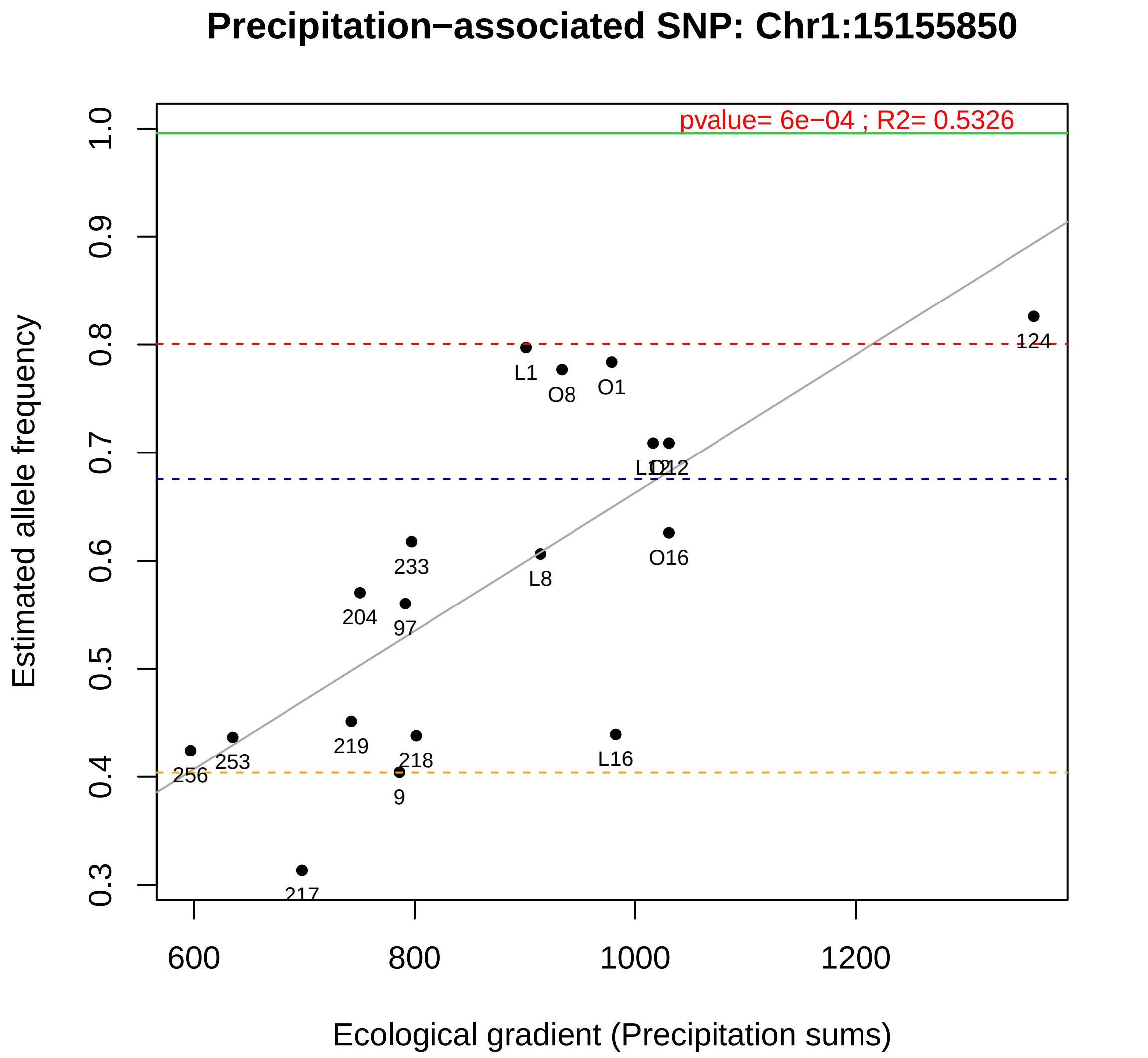
Precipitation−associated SNP: Chr1:15155850
pvalue= 6e−04 ; R2= 0.5326
Estimated allele frequency
Ecological gradient (Precipitation sums)
0.3
0.4
0.5
0.6
0.7
0.8
0.9
1.0
600
800
1000
1200
124
L1
O8
O1
O12
O16
233
L8
204
97
219
218
253
256
L16
9
217

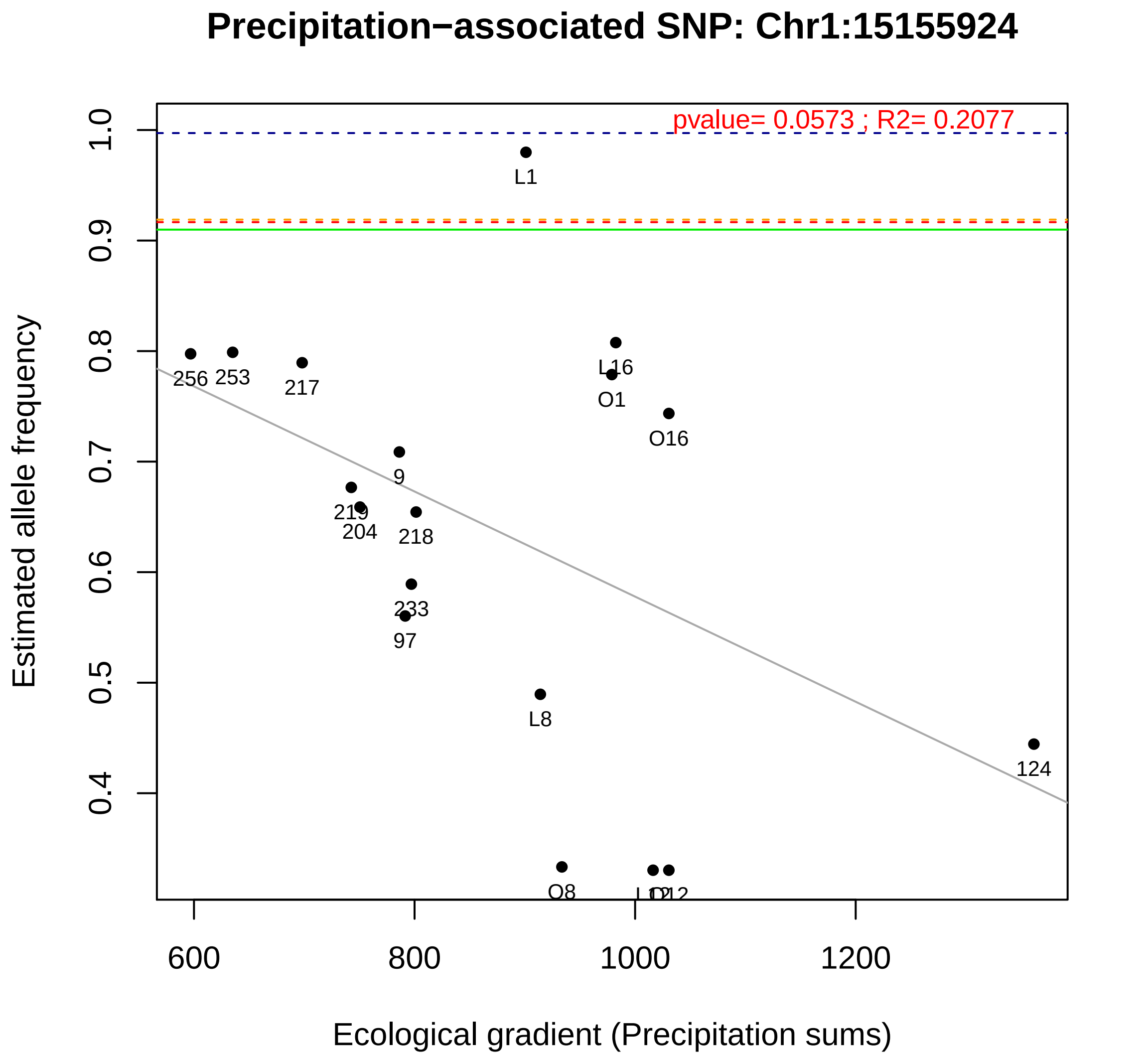

Precipitation−associated SNP: Chr1:15155924
pvalue= 0.0573 ; R2= 0.2077
Estimated allele frequency
0.4
0.5
0.6
0.7
0.8
0.9
1.0
600
800
1000
1200
Ecological gradient (Precipitation sums)
L1
256
253
217
L16
O1
O16
9
219
204
218
233
97
L8
124
O8

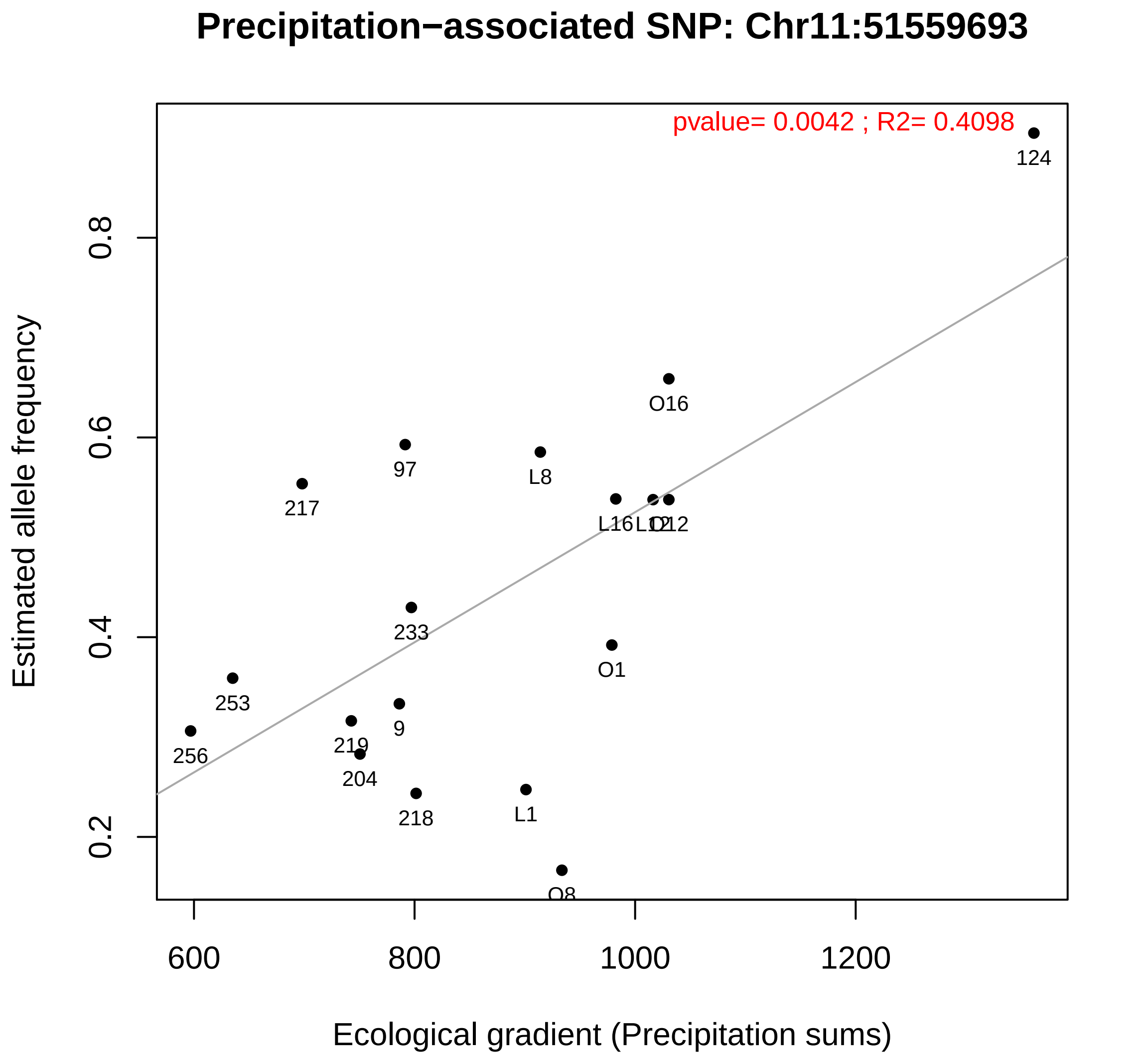

Precipitation−associated SNP: Chr11:51559693
pvalue= 0.0042 ; R2= 0.4098
124
O16
97
L8
217
L16
L12
O12
233
O1
253
9
256
219
204
218
L1
O8
0.2
0.4
0.6
0.8
600
800
1000
1200
Estimated allele frequency
Ecological gradient (Precipitation sums)

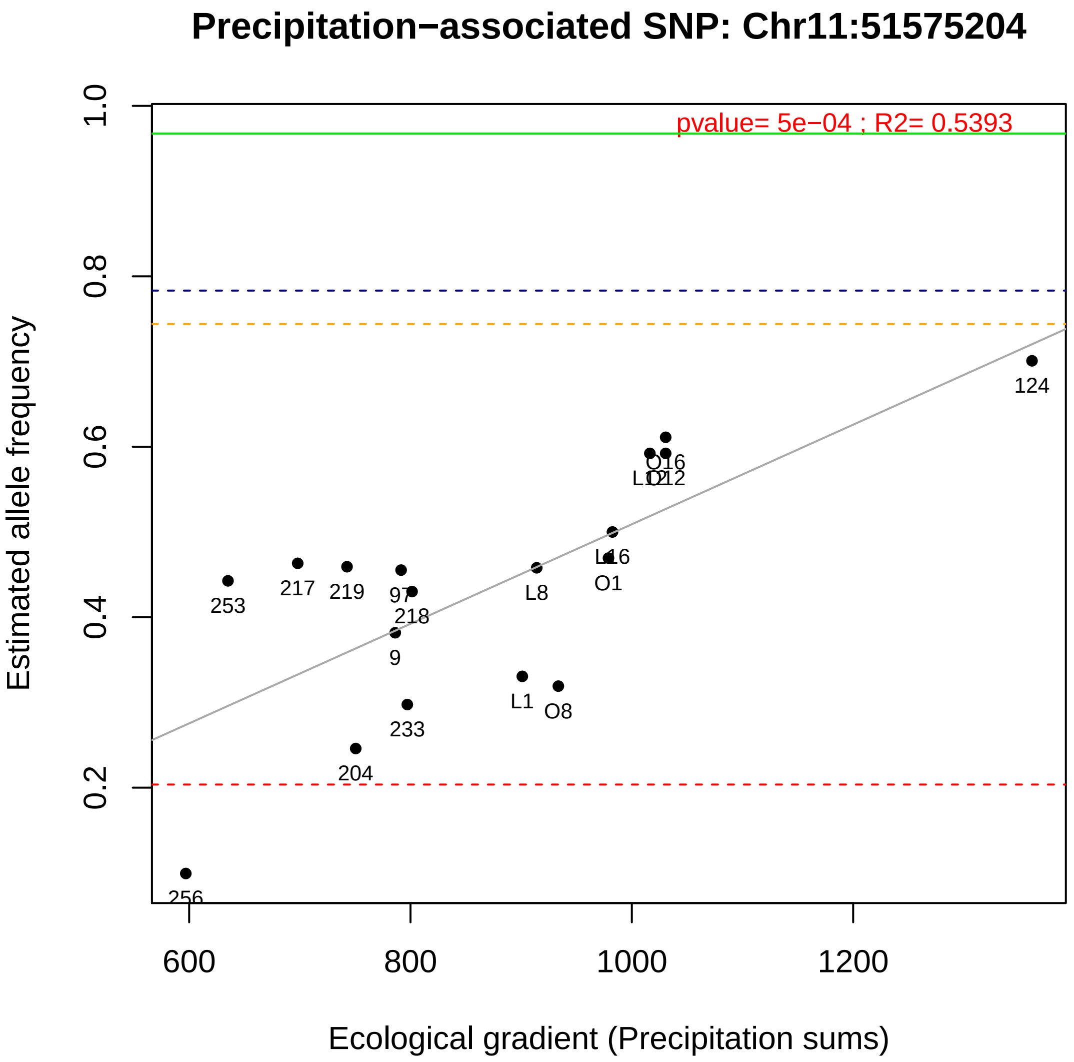
Precipitation−associated SNP: Chr11:51575204
pvalue= 5e−04 ; R2= 0.5393
Estimated allele frequency
Ecological gradient (Precipitation sums)
0.2
0.4
0.6
0.8
1.0
600
800
1000
1200
124
O16
L12
O12
L16
O1
L8
253
217
219
97
218
9
L1
O8
233
204
256

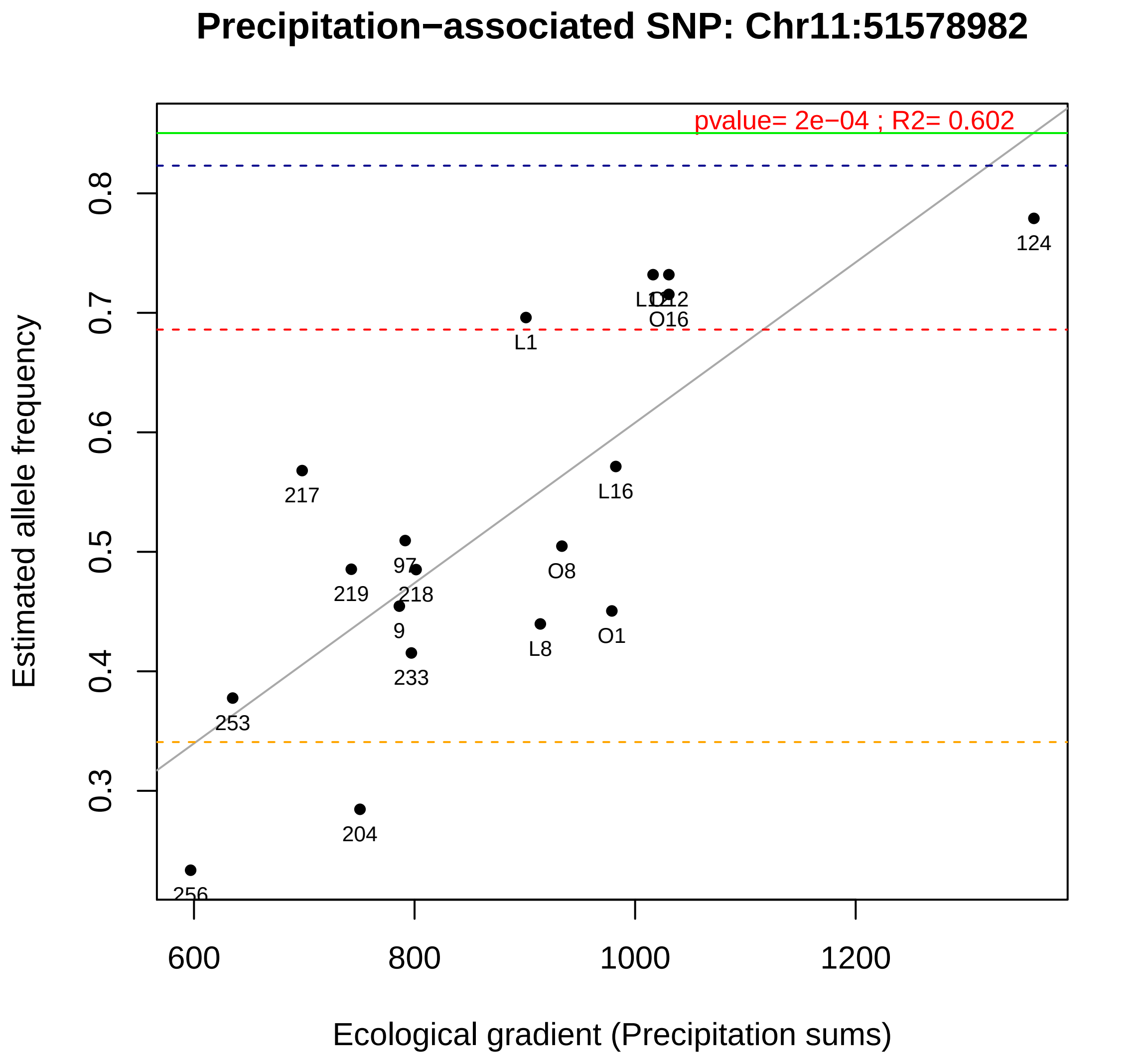

Precipitation−associated SNP: Chr11:51578982
pvalue= 2e−04 ; R2= 0.602
Estimated allele frequency
0.3
0.4
0.5
0.6
0.7
0.8
600
800
1000
1200
Ecological gradient (Precipitation sums)
124
O16
L1
217
L16
97
O8
219
218
9
L8
O1
233
253
204
256

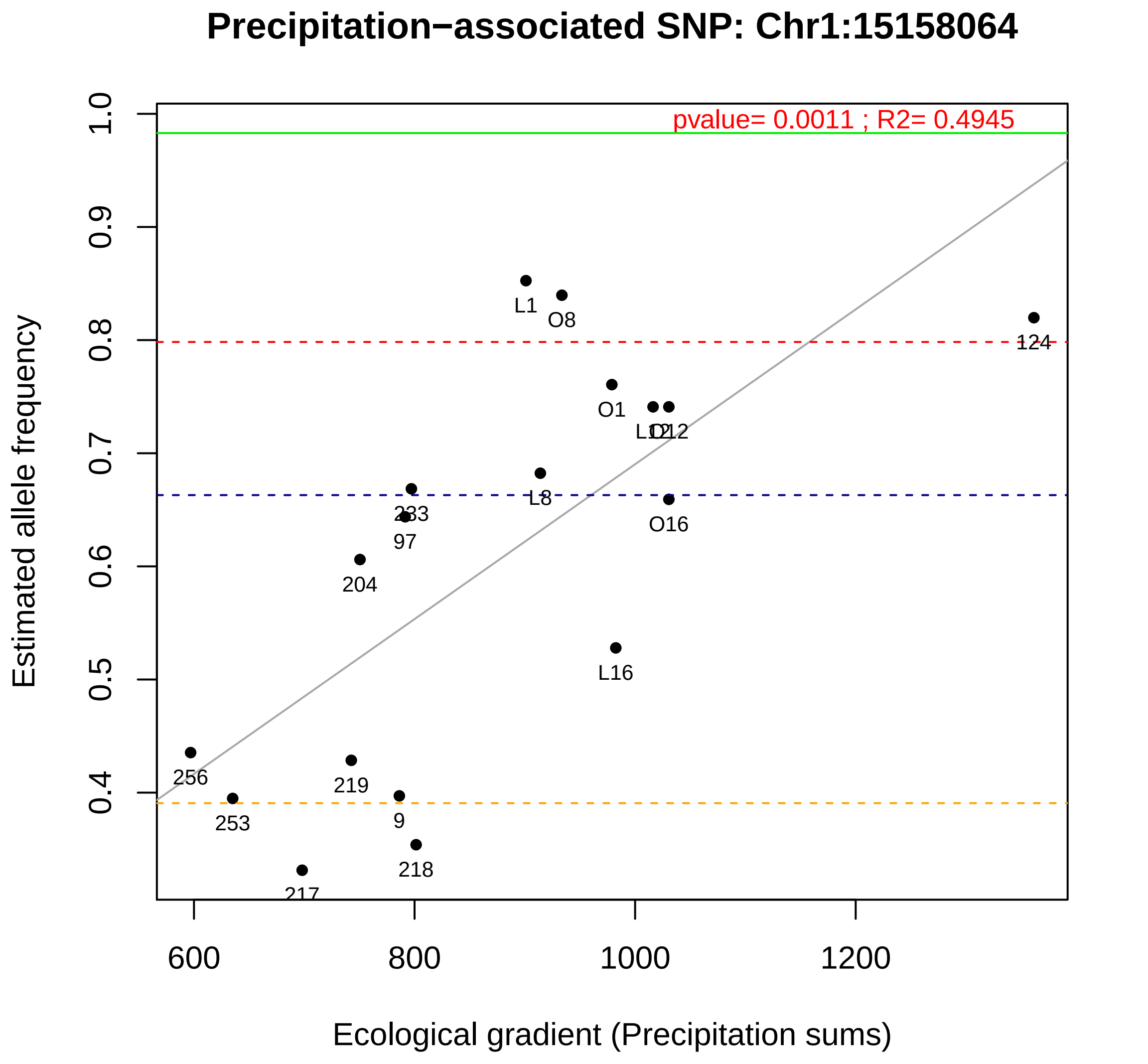

Precipitation−associated SNP: Chr1:15158064
pvalue= 0.0011 ; R2= 0.4945
Estimated allele frequency
Ecological gradient (Precipitation sums)
0.4
0.5
0.6
0.7
0.8
0.9
1.0
600
800
1000
1200
L1
O8
124
O1
L12
O12
L8
233
97
O16
204
L16
256
219
253
9
218
217

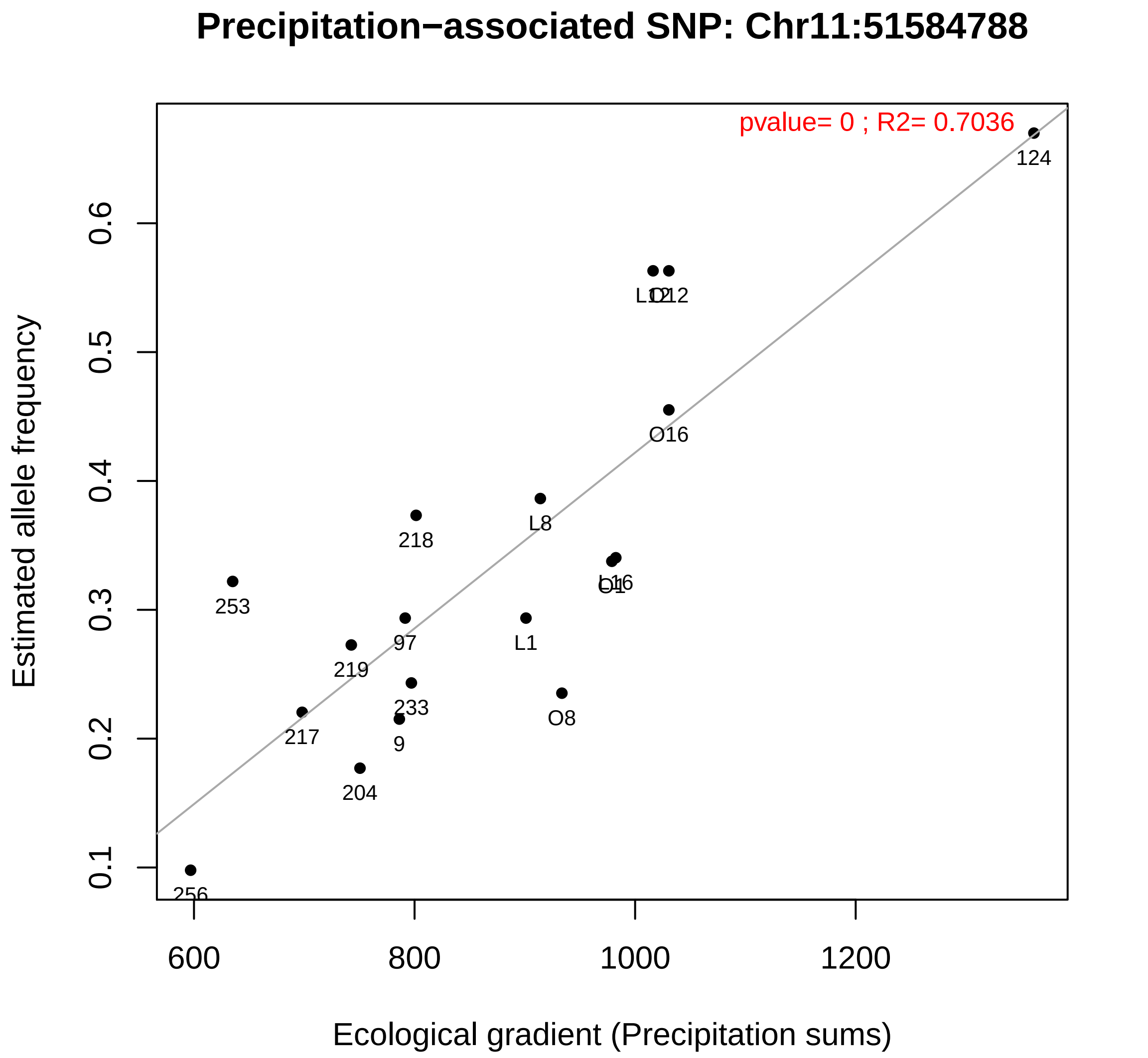

Precipitation−associated SNP: Chr11:51584788
pvalue= 0 ; R2= 0.7036
Estimated allele frequency
Ecological gradient (Precipitation sums)
0.1
0.2
0.3
0.4
0.5
0.6
600
800
1000
1200
124
L12
O12
O16
L8
218
L16
O1
253
97
L1
219
233
O8
217
9
204
256

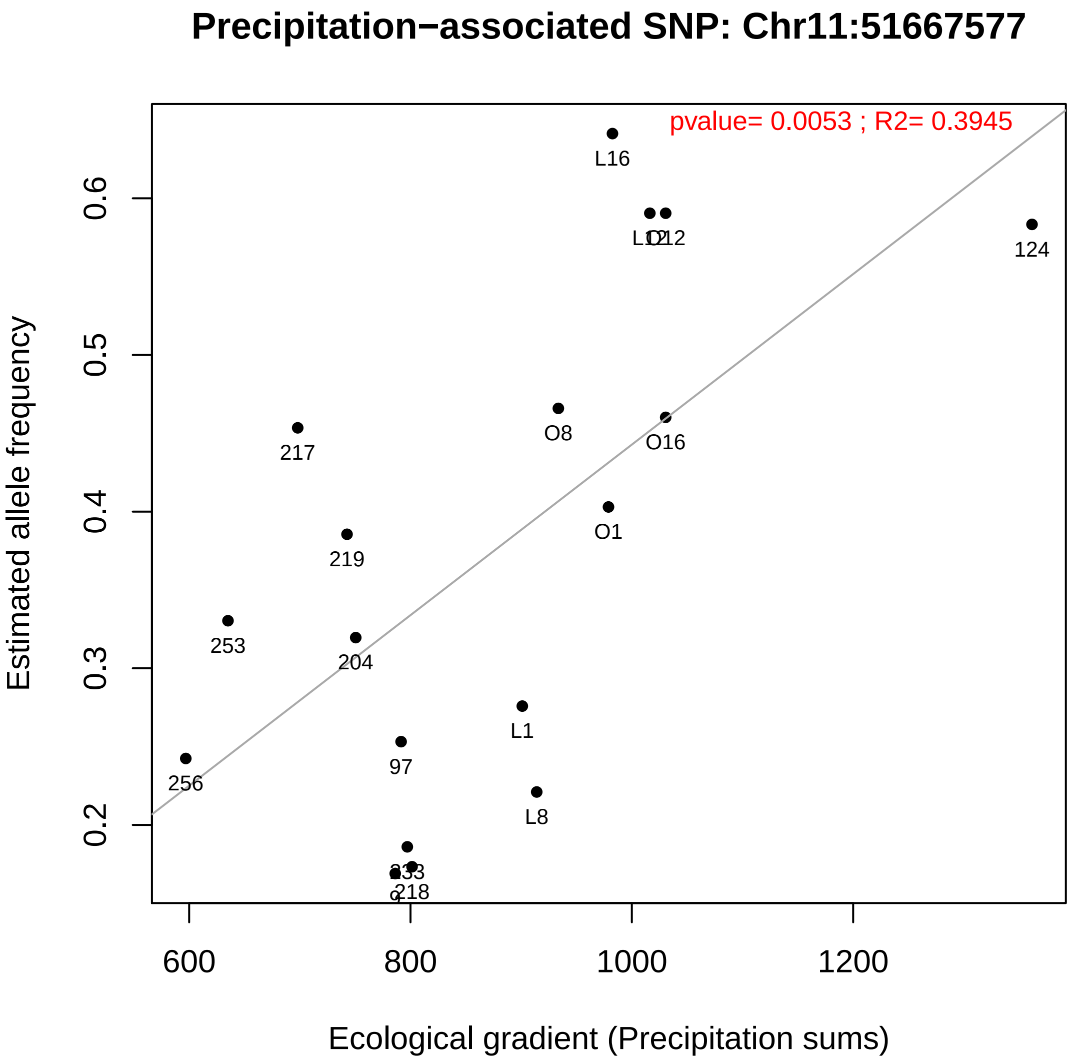
Precipitation−associated SNP: Chr11:51667577
pvalue= 0.0053 ; R2= 0.3945
Estimated allele frequency
Ecological gradient (Precipitation sums)
0.2
0.3
0.4
0.5
0.6
600
800
1000
1200
L16
124
O8
O16
217
O1
219
253
204
L1
97
256
L8
233
218

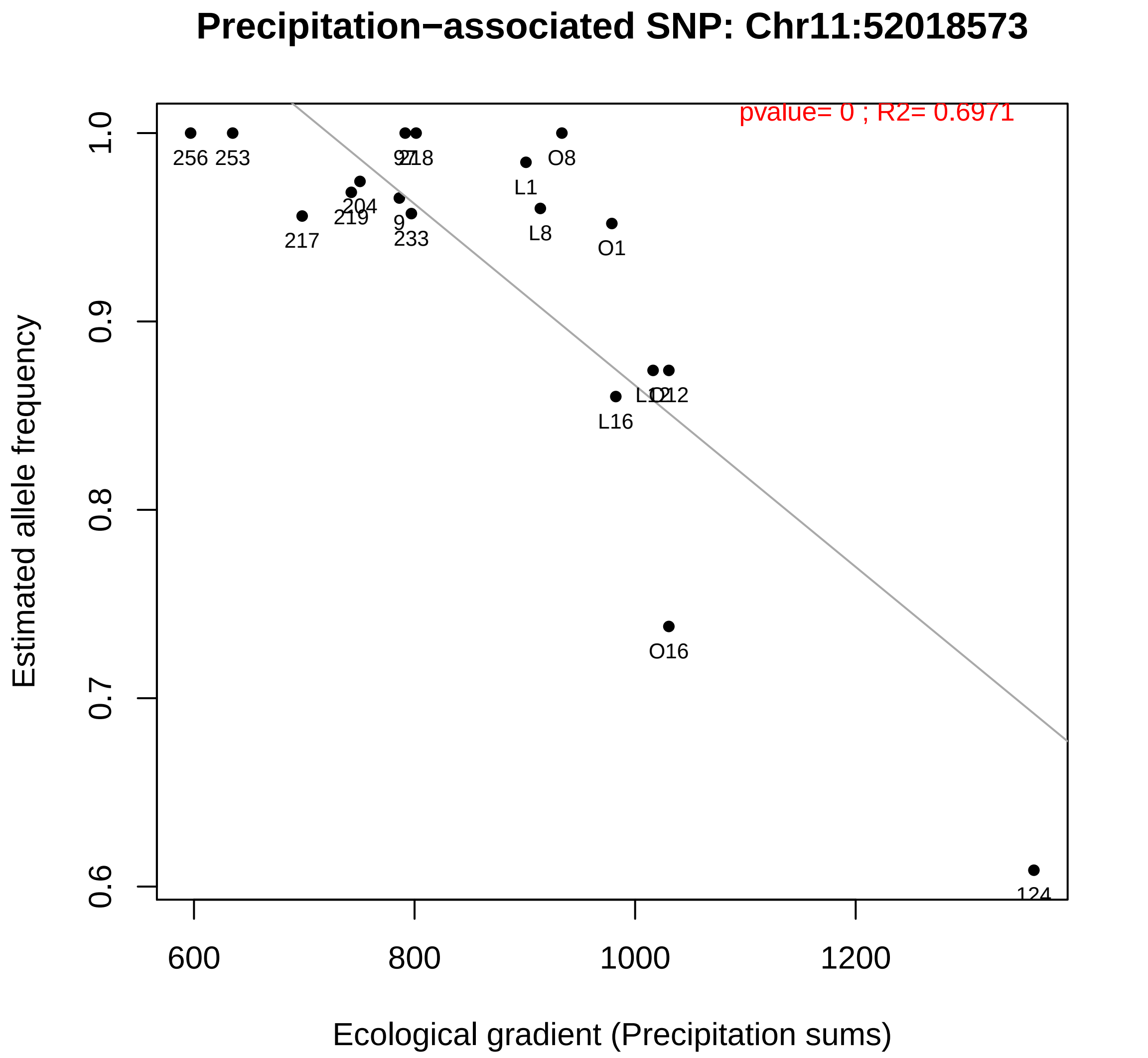

Precipitation−associated SNP: Chr11:52018573
pvalue= 0 ; R2= 0.6971
Estimated allele frequency
Ecological gradient (Precipitation sums)
0.6
0.7
0.8
0.9
1.0
600
800
1000
1200
256
253
97
218
O8
L1
204
219
9
233
217
L8
O1
L12
O12
L16
O16
124

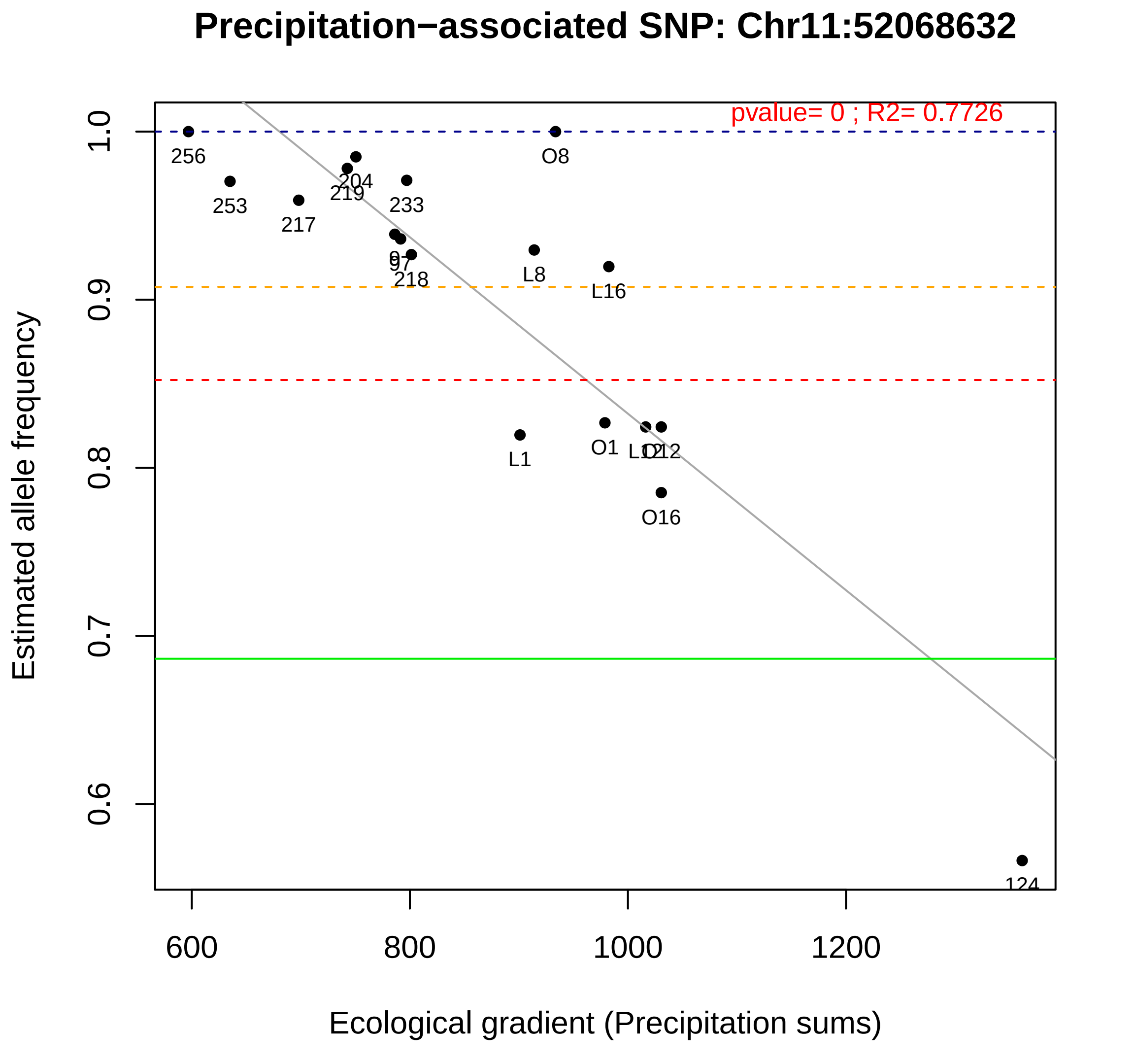
Precipitation−associated SNP: Chr11:52068632
pvalue= 0 ; R2= 0.7726
Estimated allele frequency
1.0
0.9
0.8
0.7
0.6
600
800
1000
1200
Ecological gradient (Precipitation sums)
256
253
217
204
219
233
97
218
L8
L16
O8
L1
O1
O12
O16
124

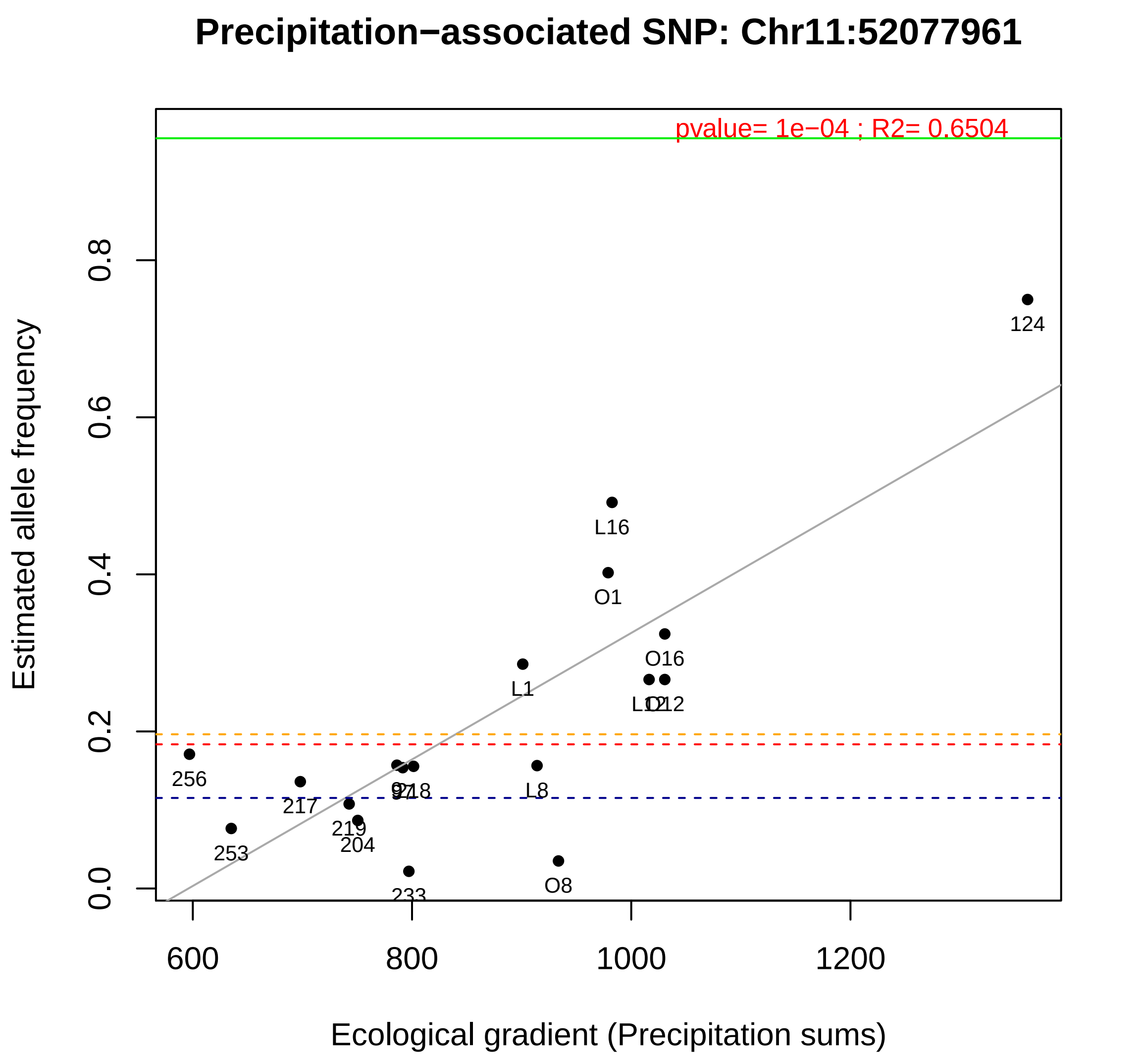

Precipitation−associated SNP: Chr11:52077961
pvalue= 1e−04 ; R2= 0.6504
Estimated allele frequency
0.0
0.2
0.4
0.6
0.8
600
800
1000
1200
Ecological gradient (Precipitation sums)
124
L16
O1
O16
L1
L8
O8
256
217
219
204
253
233

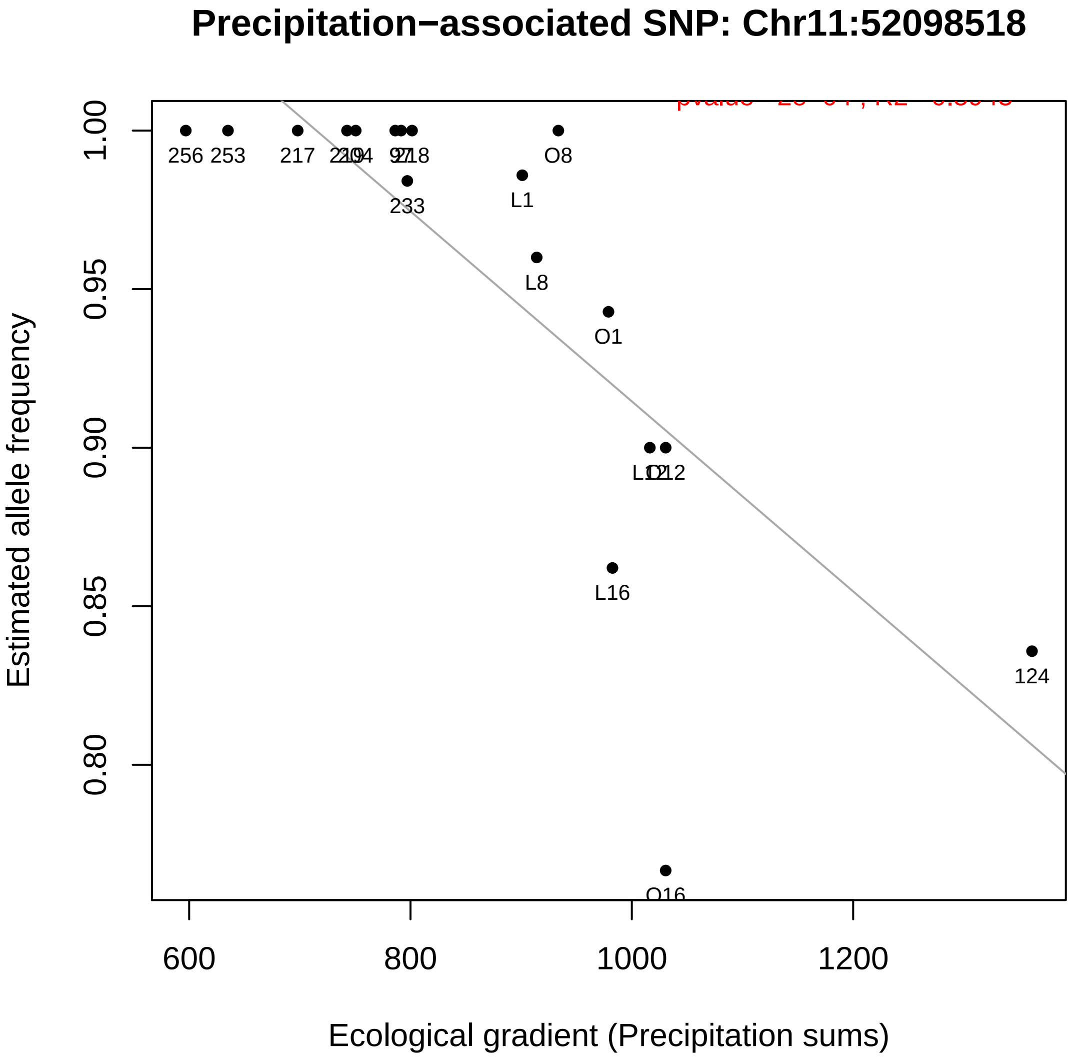

Precipitation−associated SNP: Chr11:52098518
Estimated allele frequency
Ecological gradient (Precipitation sums)
1.00
0.95
0.90
0.85
0.80
600
800
1000
1200
256
253
217
210
194
97
218
O8
233
L1
L8
O1
L12
O12
L16
124
O16

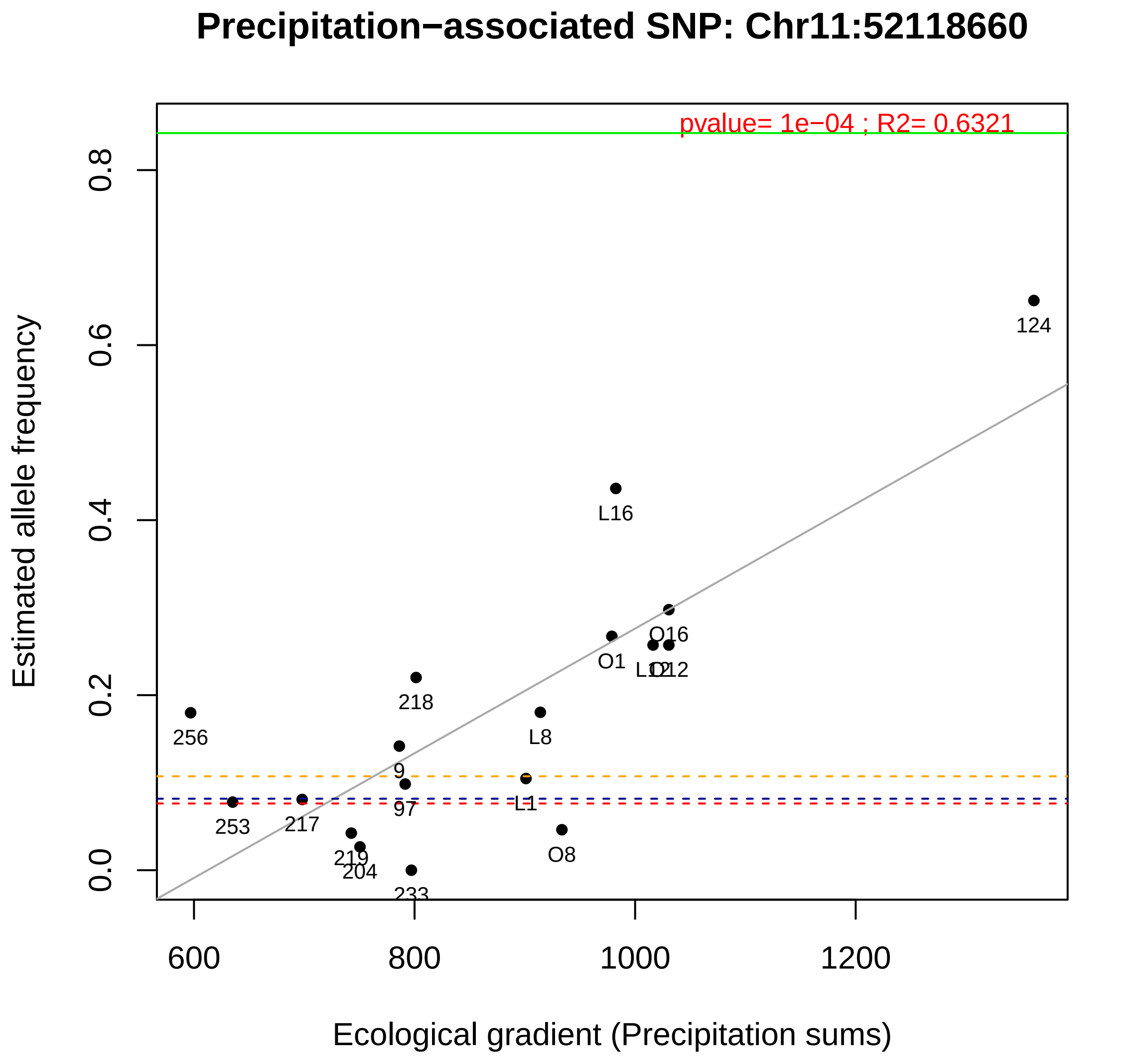

Precipitation−associated SNP: Chr11:52118660
pvalue= 1e−04 ; R2= 0.6321
Estimated allele frequency
0.0
0.2
0.4
0.6
0.8
600
800
1000
1200
Ecological gradient (Precipitation sums)
124
L16
O16
O1
L10
O12
218
256
L8
9
97
L1
253
217
219
204
O8
233

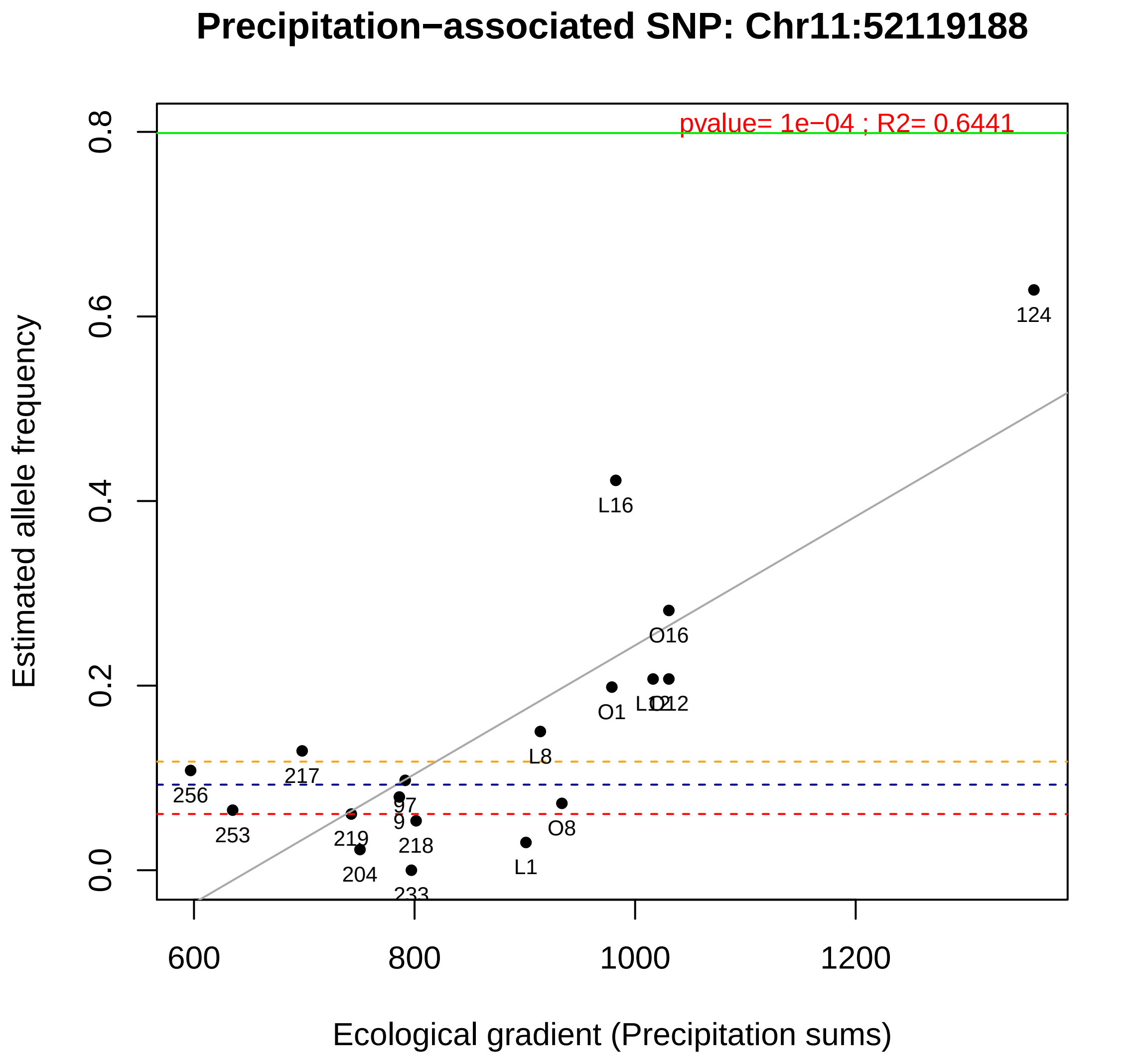

Precipitation−associated SNP: Chr11:52119188
pvalue= 1e−04 ; R2= 0.6441
Estimated allele frequency
0.0
0.2
0.4
0.6
0.8
Ecological gradient (Precipitation sums)
600
800
1000
1200
124
L16
O16
O1
O12
L8
217
256
253
219
204
97
9
218
233
O8
L1

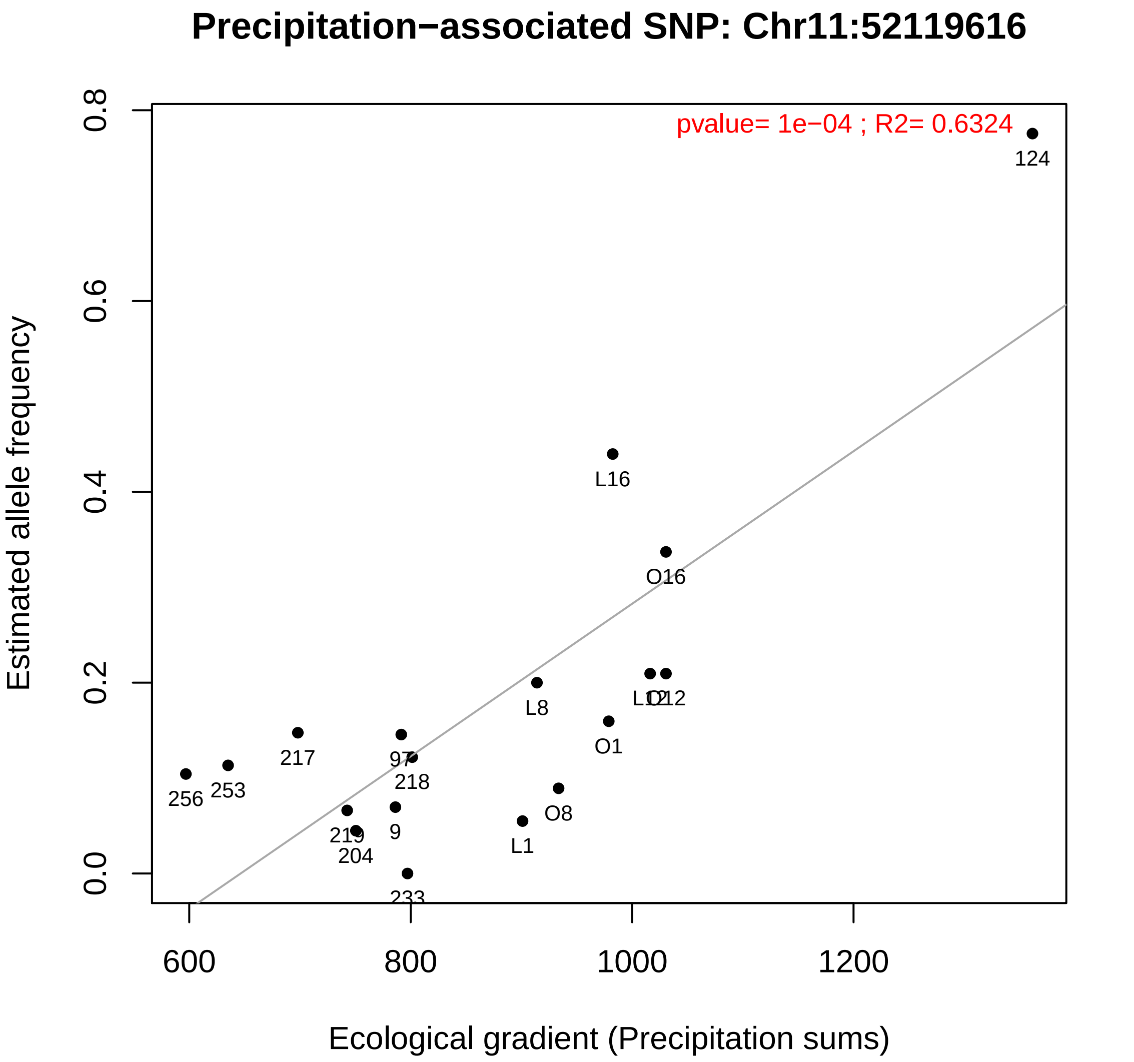
Precipitation−associated SNP: Chr11:52119616
pvalue= 1e−04 ; R2= 0.6324
Estimated allele frequency
0.0
0.2
0.4
0.6
0.8
600
800
1000
1200
Ecological gradient (Precipitation sums)
124
L16
O16
L8
O1
217
256
253
218
O8
9
L1
204
233

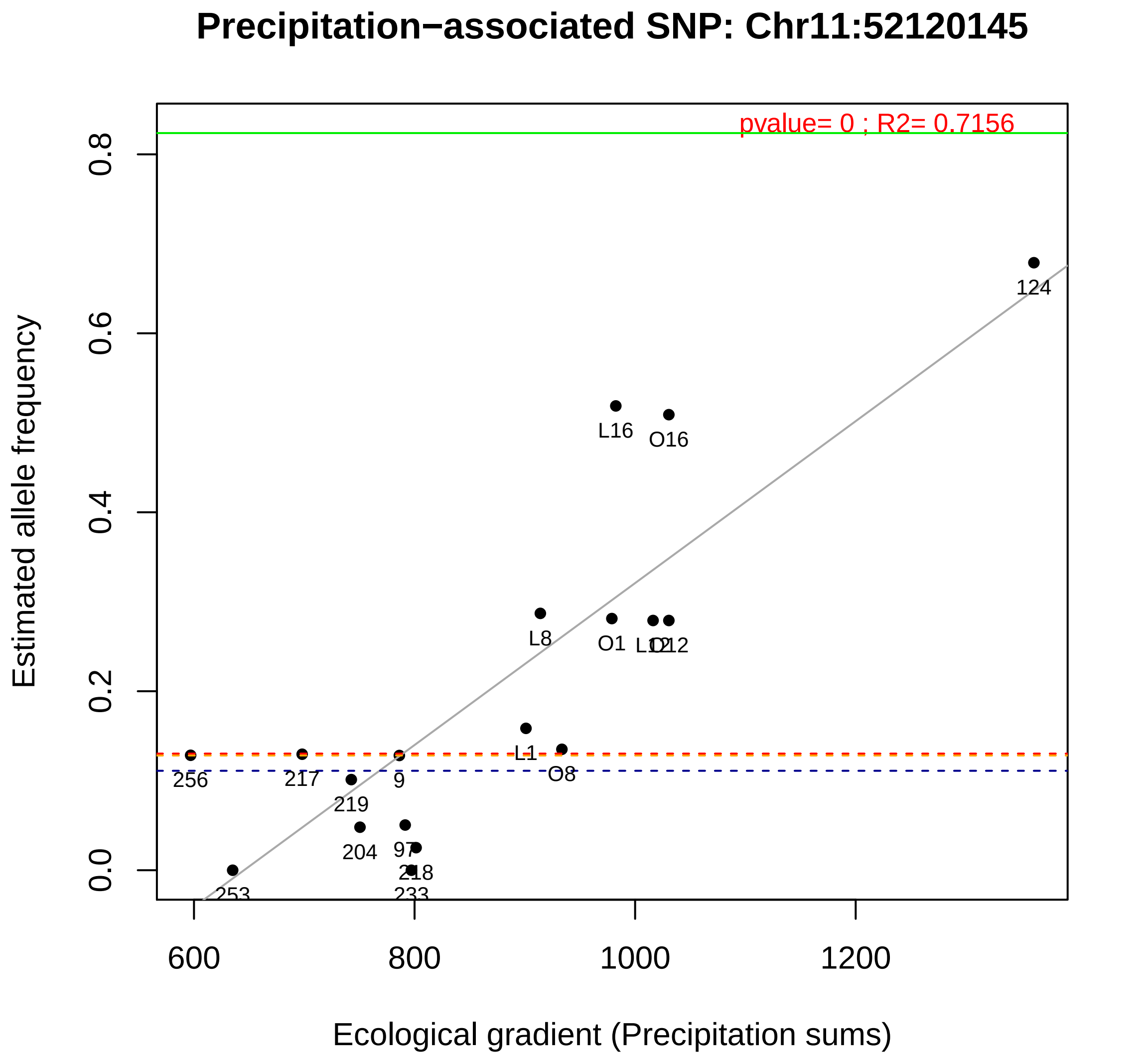

Precipitation−associated SNP: Chr11:52120145
pvalue= 0 ; R2= 0.7156
Estimated allele frequency
0.0
0.2
0.4
0.6
0.8
600
800
1000
1200
Ecological gradient (Precipitation sums)
124
L16
O16
L8
O1
O12
L1
O8
256
217
9
219
204
97
218
233
253

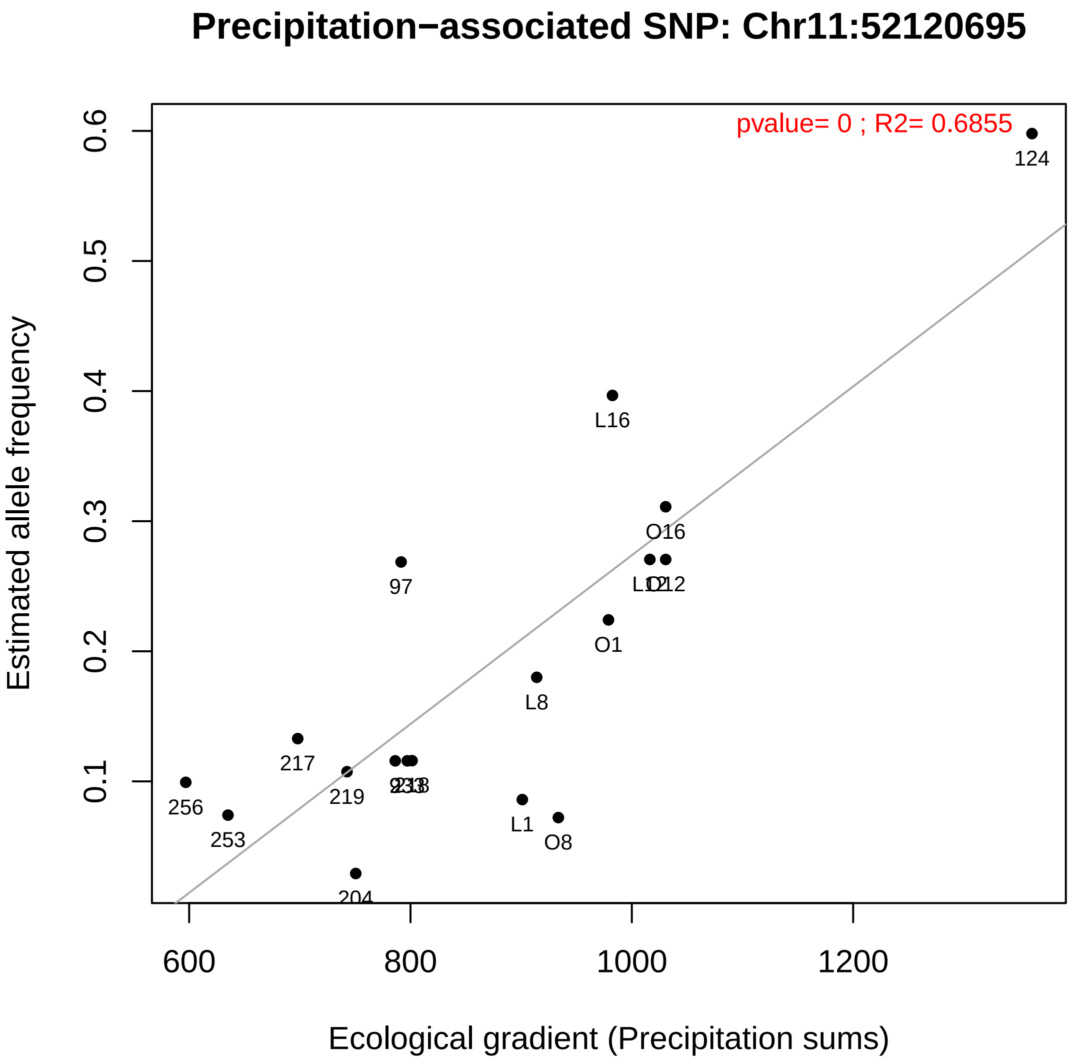

Precipitation−associated SNP: Chr11:52120695
pvalue= 0 ; R2= 0.6855
Estimated allele frequency
0.1
0.2
0.3
0.4
0.5
0.6
600
800
1000
1200
Ecological gradient (Precipitation sums)
124
L16
O16
97
O1
L8
217
219
256
253
204
L1
O8

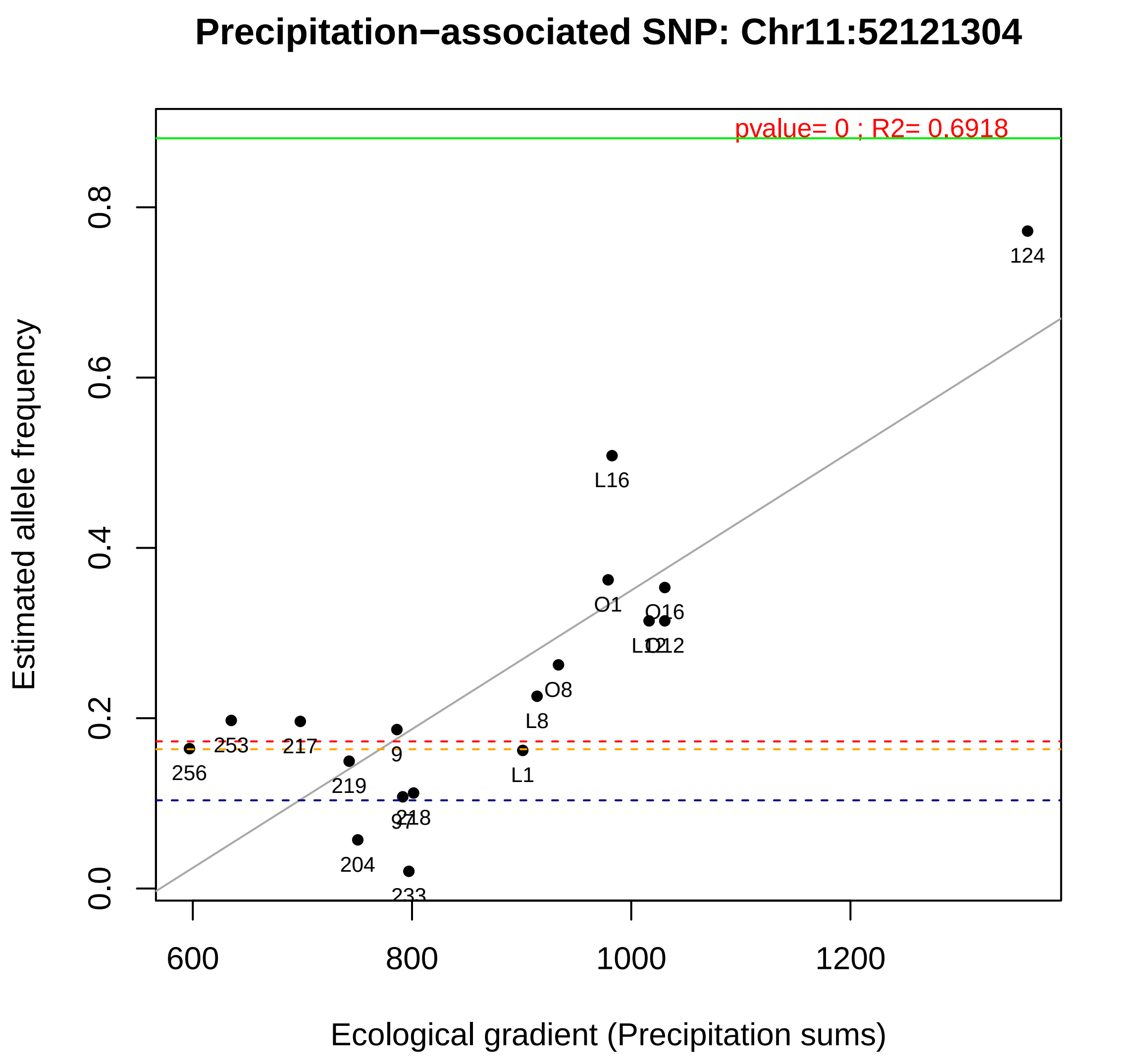

Precipitation−associated SNP: Chr11:52121304
pvalue= 0 ; R2= 0.6918
Estimated allele frequency
Ecological gradient (Precipitation sums)
0.0
0.2
0.4
0.6
0.8
600
800
1000
1200
124
L16
O1
O16
L10
O12
O8
L8
253
217
256
9
219
L1
218
97
204
233

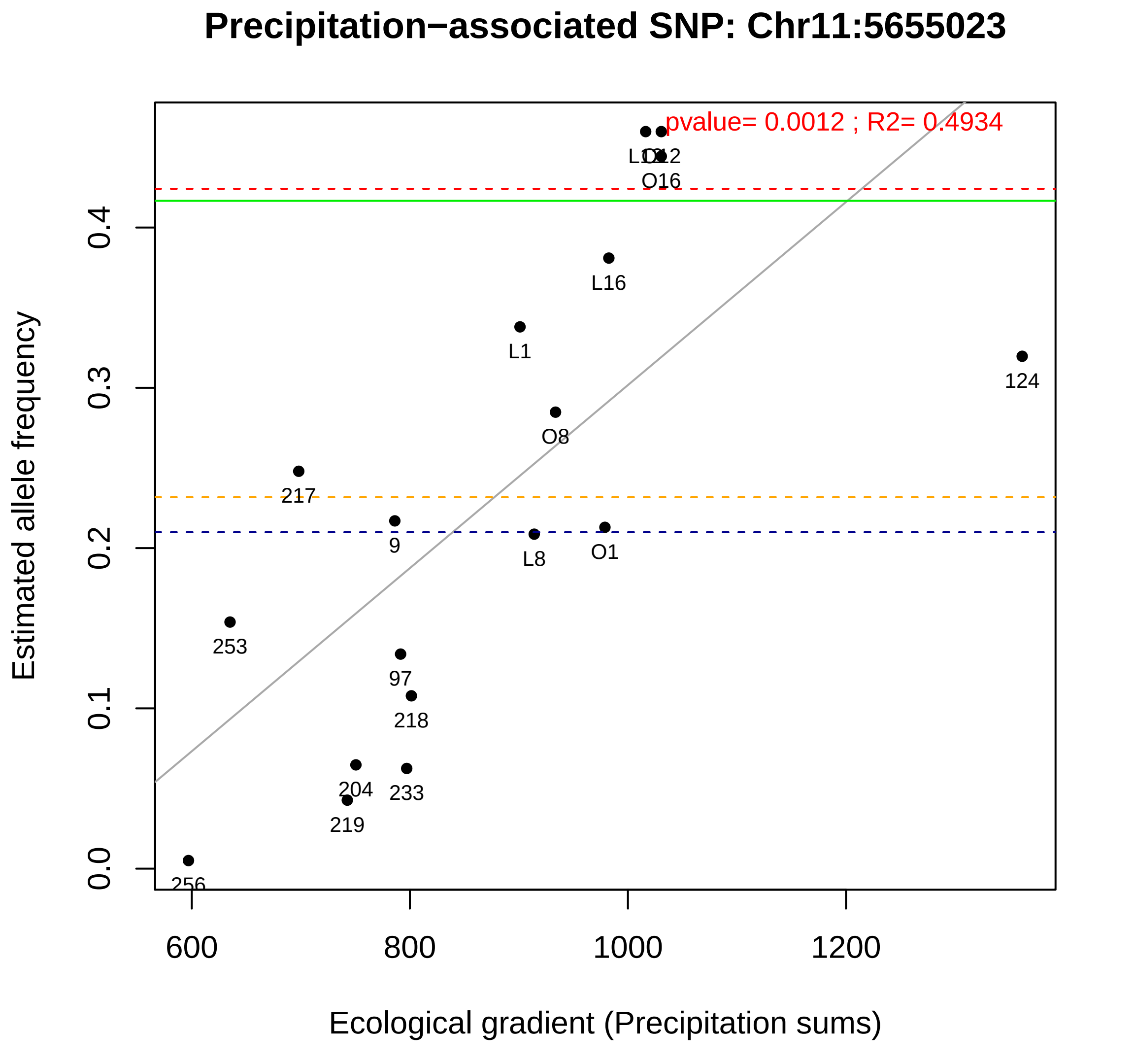

Precipitation−associated SNP: Chr11:5655023
pvalue= 0.0012 ; R2= 0.4934
Estimated allele frequency
Ecological gradient (Precipitation sums)
0.0
0.1
0.2
0.3
0.4
600
800
1000
1200
O16
L16
L1
124
O8
217
9
L8
O1
253
97
218
204
233
219
256

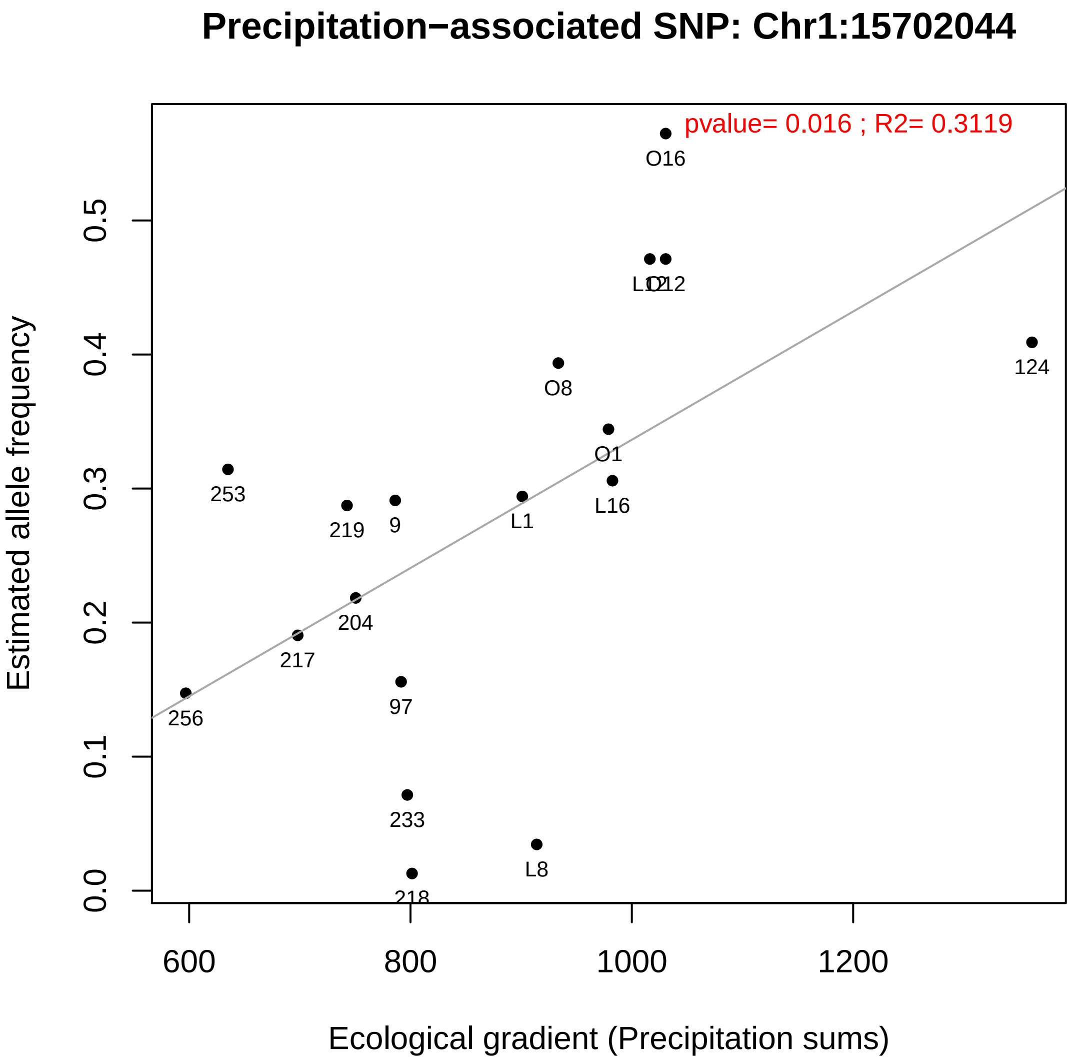

Precipitation−associated SNP: Chr1:15702044
pvalue= 0.016 ; R2= 0.3119
O16
L12
O12
124
O8
O1
253
L16
219
9
L1
204
217
97
256
233
L8
218
0.0
0.1
0.2
0.3
0.4
0.5
600
800
1000
1200
Estimated allele frequency
Ecological gradient (Precipitation sums)

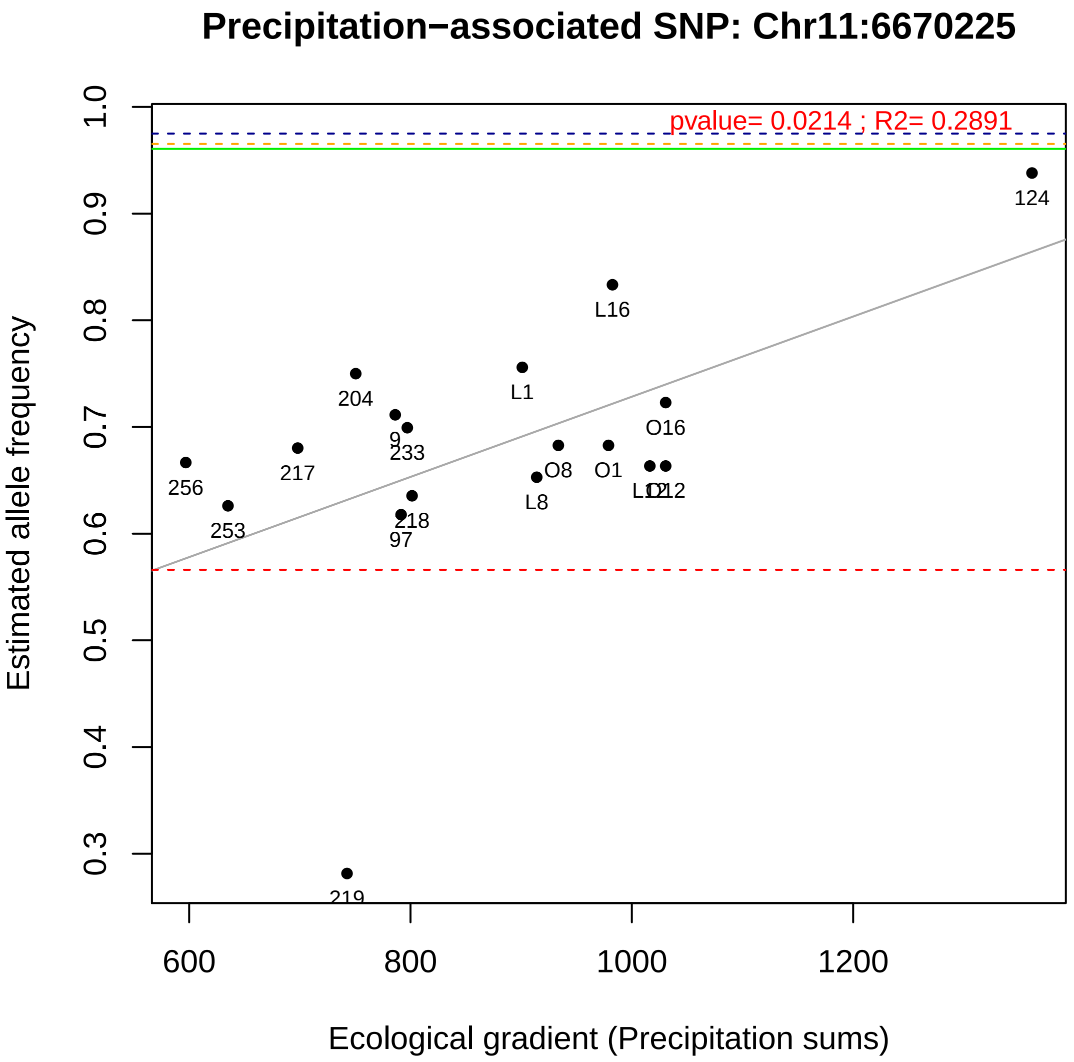

Precipitation−associated SNP: Chr11:6670225
pvalue= 0.0214 ; R2= 0.2891
Estimated allele frequency
1.0
0.9
0.8
0.7
0.6
0.5
0.4
0.3
600
800
1000
1200
Ecological gradient (Precipitation sums)
124
L16
204
L1
O16
9
233
256
217
O8
O1
L8
253
218
97
219

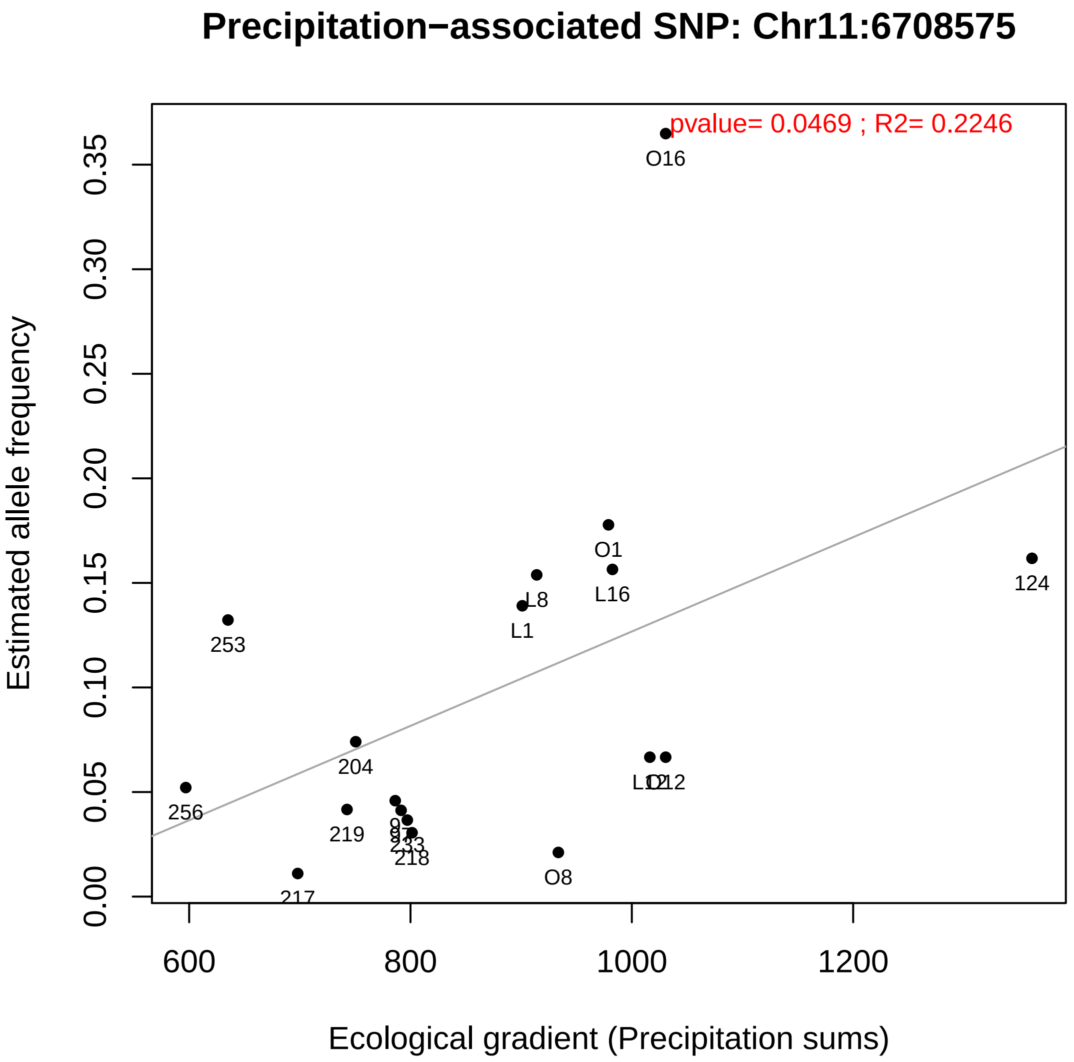

Precipitation−associated SNP: Chr11:6708575
pvalue= 0.0469 ; R2= 0.2246
Estimated allele frequency
Ecological gradient (Precipitation sums)
0.00
0.05
0.10
0.15
0.20
0.25
0.30
0.35
600
800
1000
1200
O16
O1
L16
124
L8
L1
253
204
256
219
233
218
217
O8

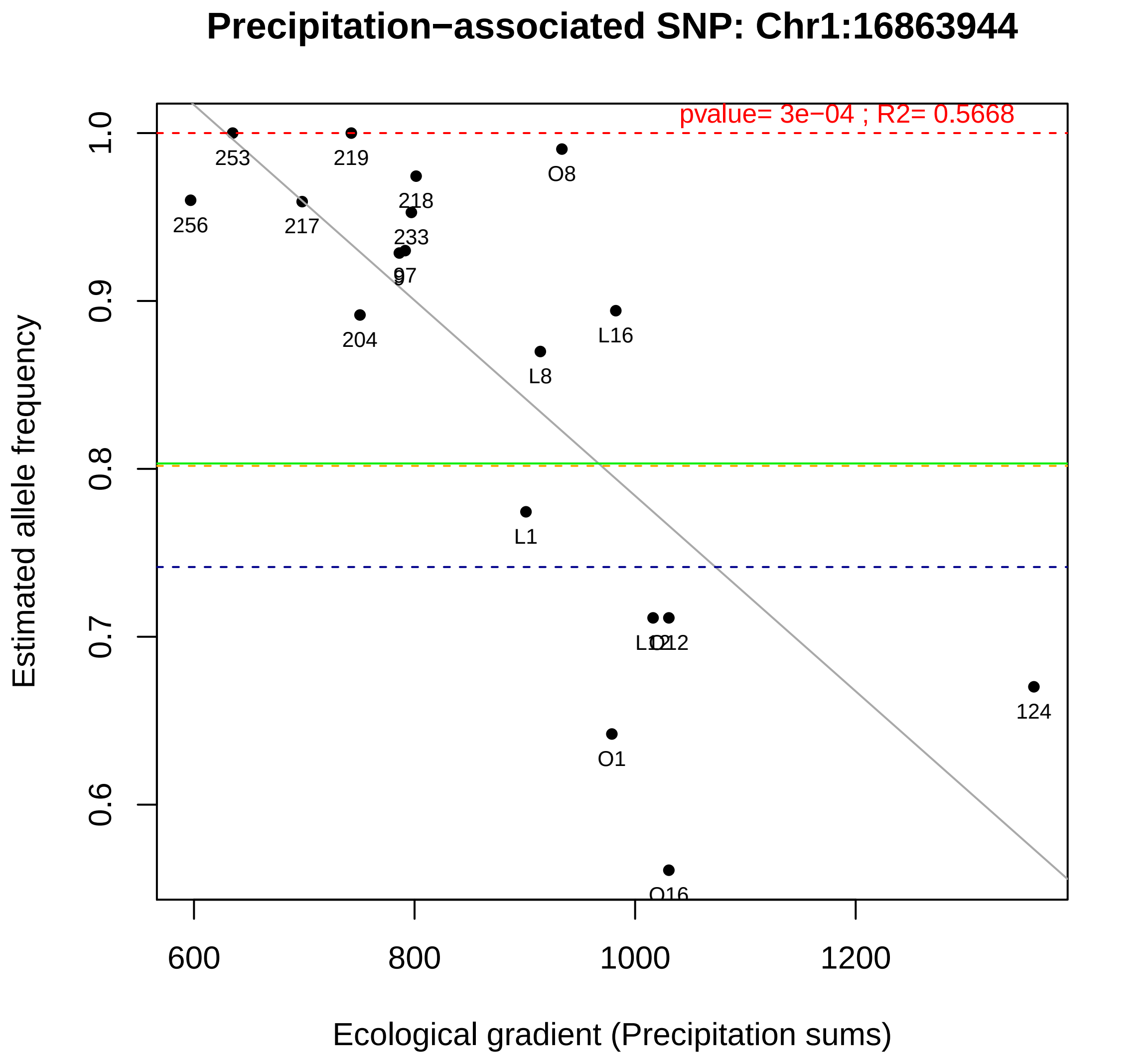

Precipitation−associated SNP: Chr1:16863944
pvalue= 3e−04 ; R2= 0.5668
Estimated allele frequency
Ecological gradient (Precipitation sums)
0.6
0.7
0.8
0.9
1.0
600
800
1000
1200
253
219
O8
256
217
218
233
97
204
L16
L8
L1
L12
O12
124
O1
O16

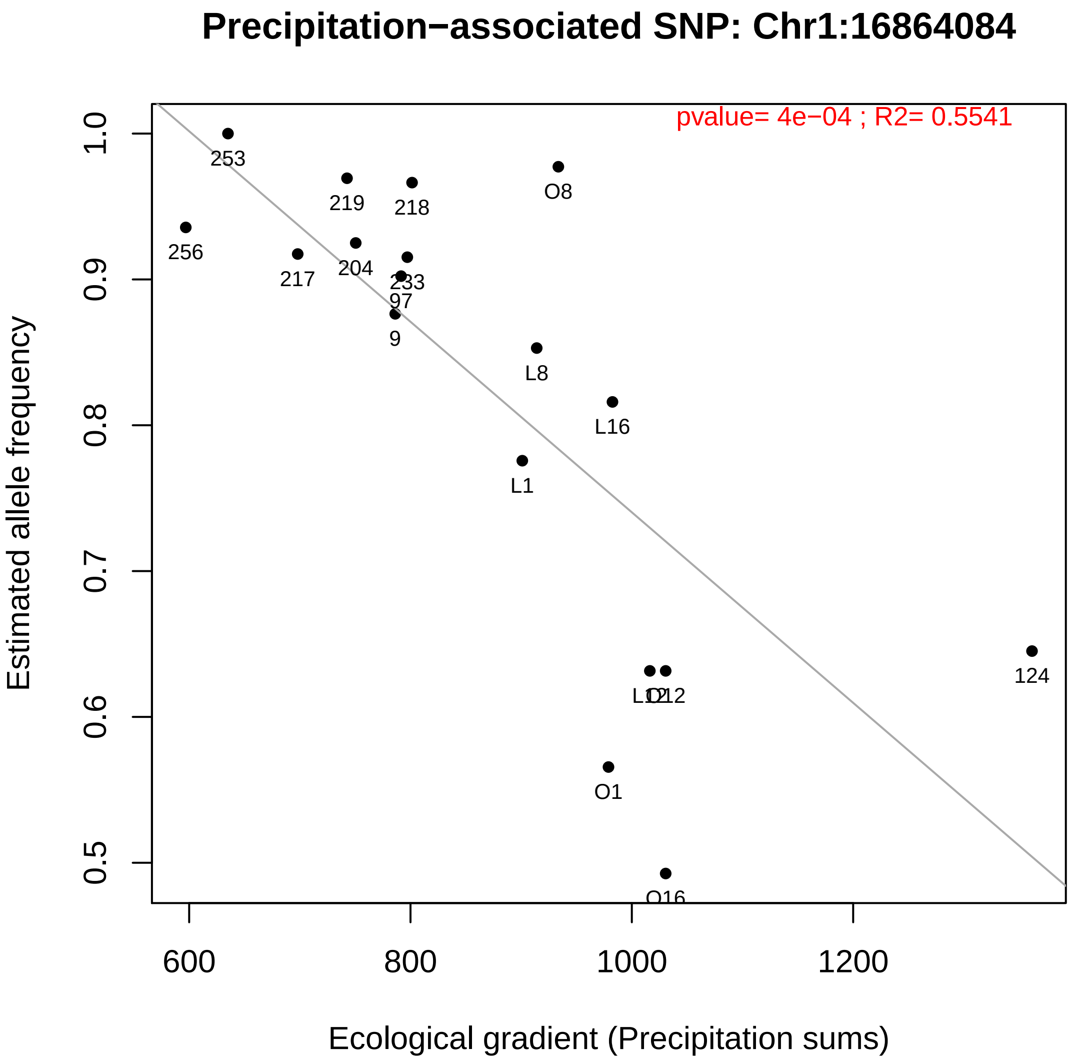

Precipitation−associated SNP: Chr1:16864084
pvalue= 4e−04 ; R2= 0.5541
Estimated allele frequency
Ecological gradient (Precipitation sums)
0.5
0.6
0.7
0.8
0.9
1.0
600
800
1000
1200
253
219
218
O8
256
217
204
233
97
9
L8
L16
L1
124
O12
O1
O16

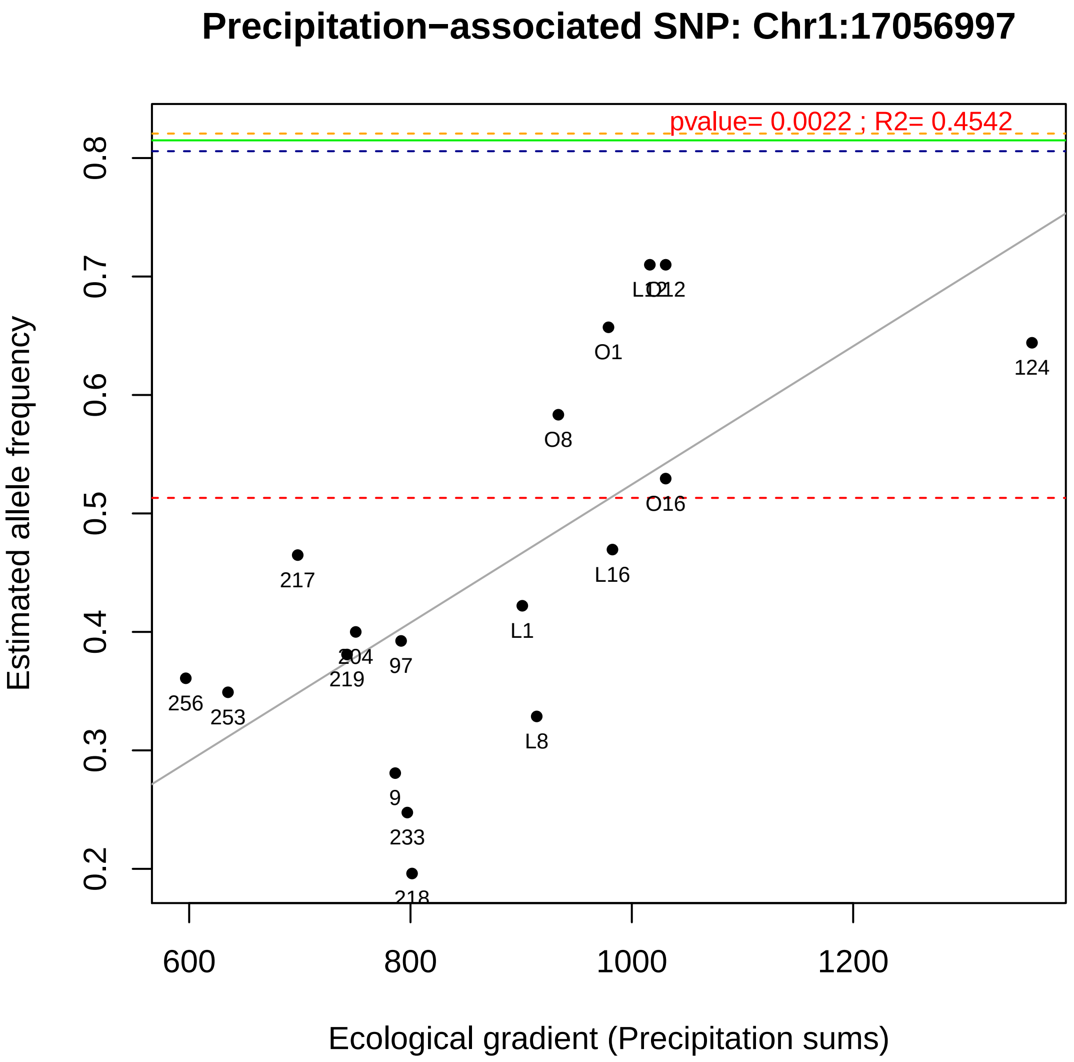
Precipitation−associated SNP: Chr1:17056997
pvalue= 0.0022 ; R2= 0.4542
Estimated allele frequency
0.2
0.3
0.4
0.5
0.6
0.7
0.8
600
800
1000
1200
Ecological gradient (Precipitation sums)
L12
O12
O1
124
O8
O16
L16
217
L1
204
219
97
256
253
L8
9
233
218

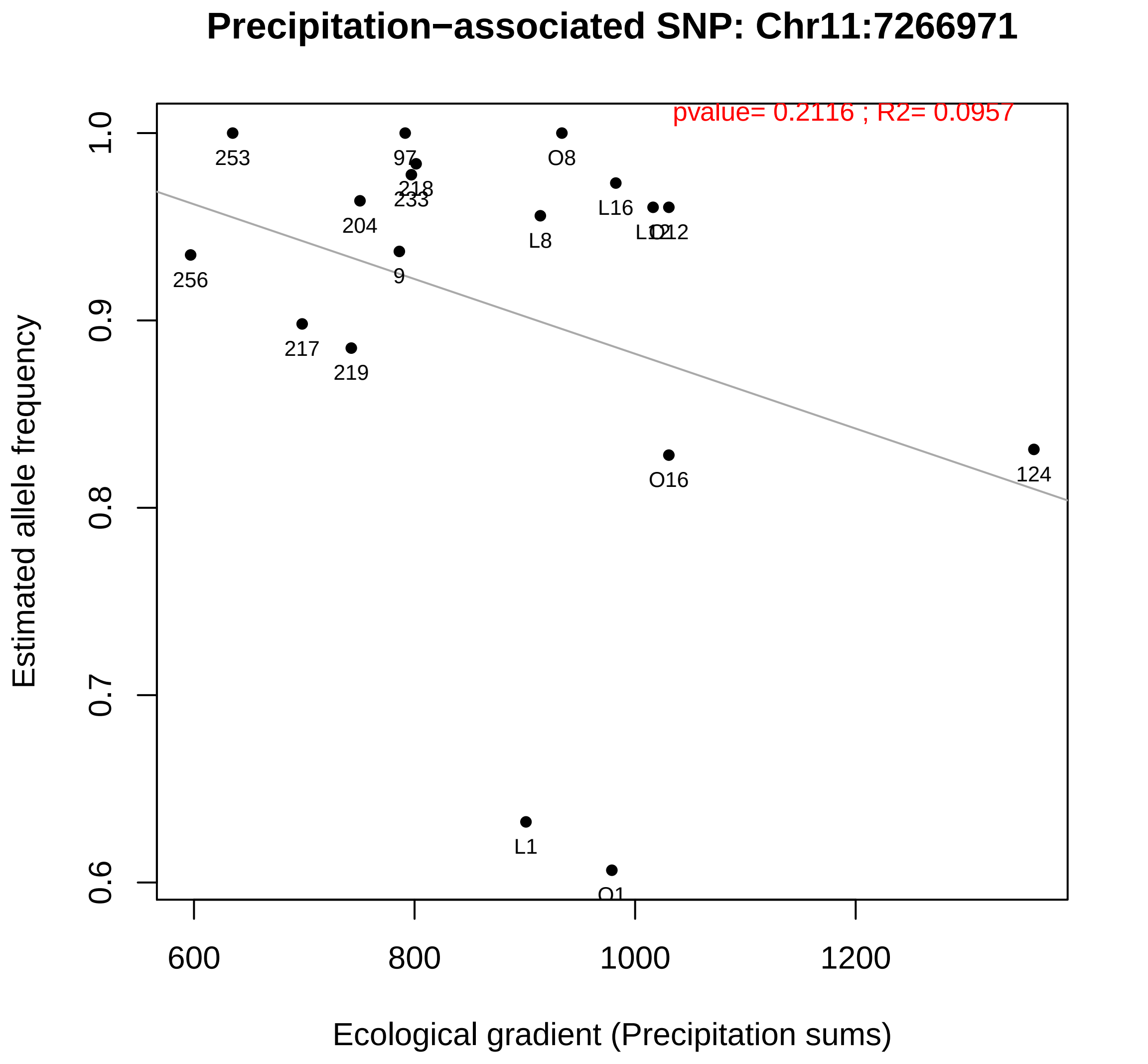
Precipitation−associated SNP: Chr11:7266971
pvalue= 0.2116 ; R2= 0.0957
Estimated allele frequency
1.0
0.9
0.8
0.7
0.6
600
800
1000
1200
Ecological gradient (Precipitation sums)
253
97
218
233
204
256
9
217
219
O8
L16
L8
L12
O12
O16
124
L1
O1

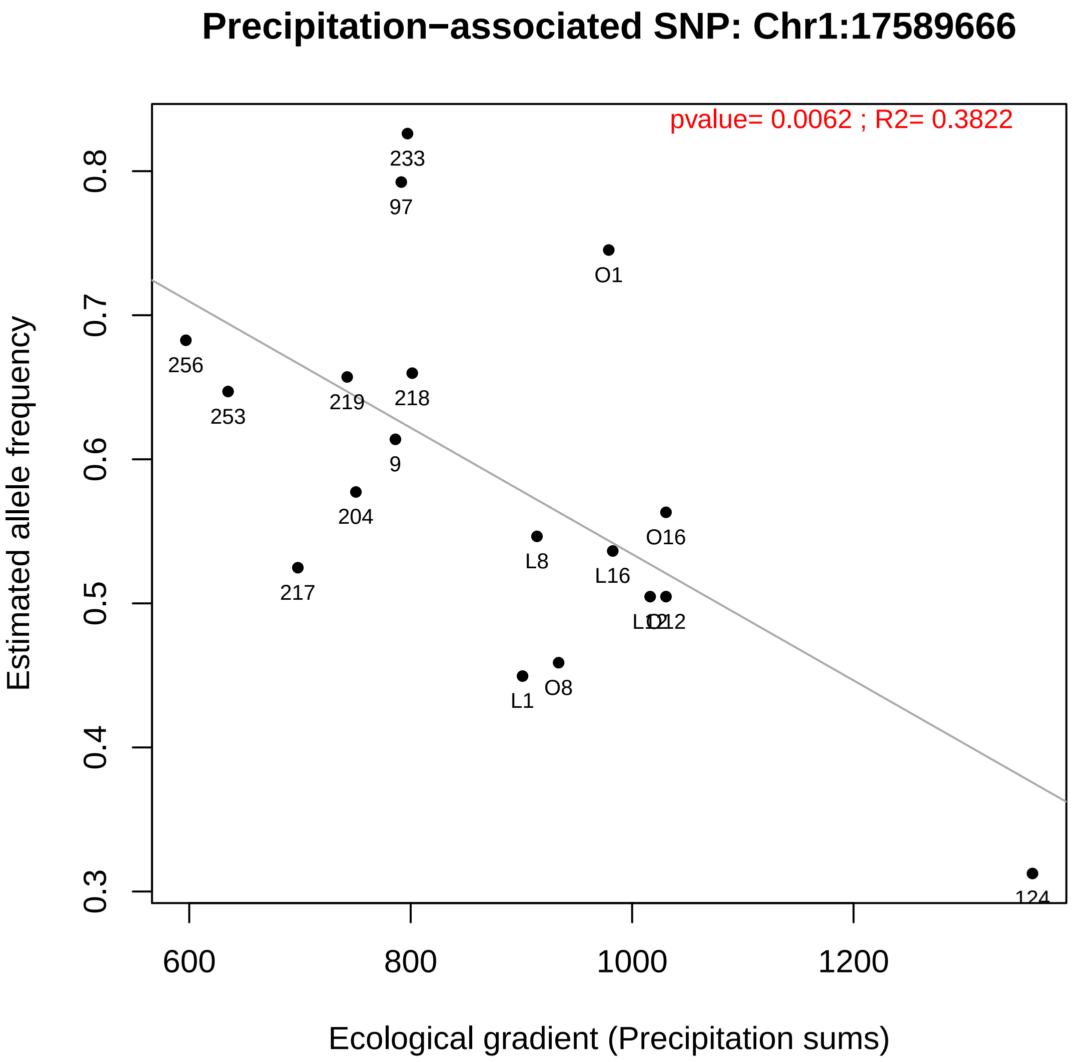
Precipitation−associated SNP: Chr1:17589666
pvalue= 0.0062 ; R2= 0.3822
Estimated allele frequency
0.3
0.4
0.5
0.6
0.7
0.8
600
800
1000
1200
Ecological gradient (Precipitation sums)
233
97
O1
256
253
219
218
9
204
217
L8
L16
O16
L1
O8
124

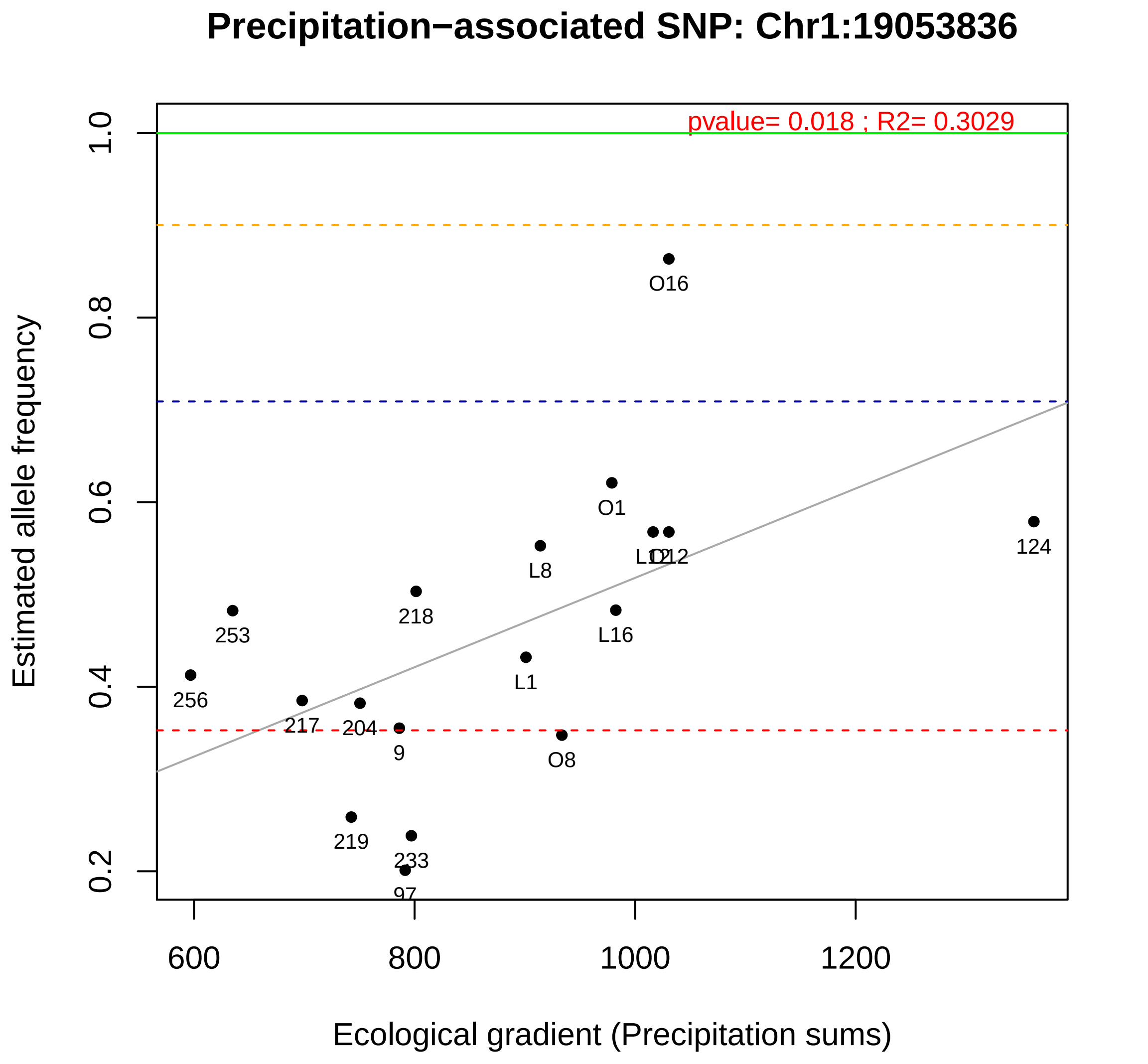
Precipitation−associated SNP: Chr1:19053836
pvalue= 0.018 ; R2= 0.3029
Estimated allele frequency
Ecological gradient (Precipitation sums)
0.2
0.4
0.6
0.8
1.0
600
800
1000
1200
O16
O1
L10
O12
124
L8
218
253
L16
L1
256
217
204
9
O8
219
233
97

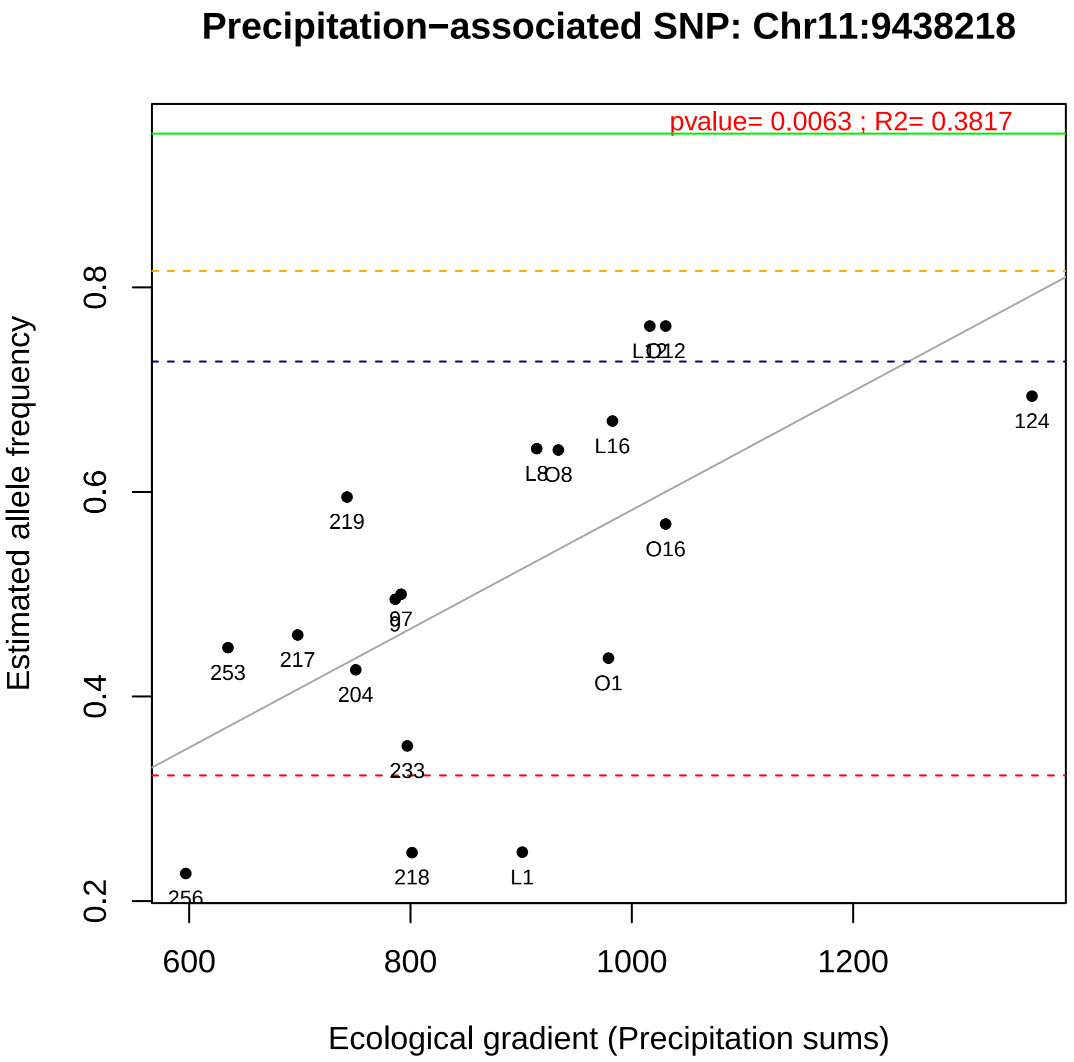
Precipitation−associated SNP: Chr11:9438218
pvalue= 0.0063 ; R2= 0.3817
Estimated allele frequency
0.2
0.4
0.6
0.8
600
800
1000
1200
Ecological gradient (Precipitation sums)
256
253
217
219
204
233
218
L1
L8
O8
L16
O16
O1
124
97

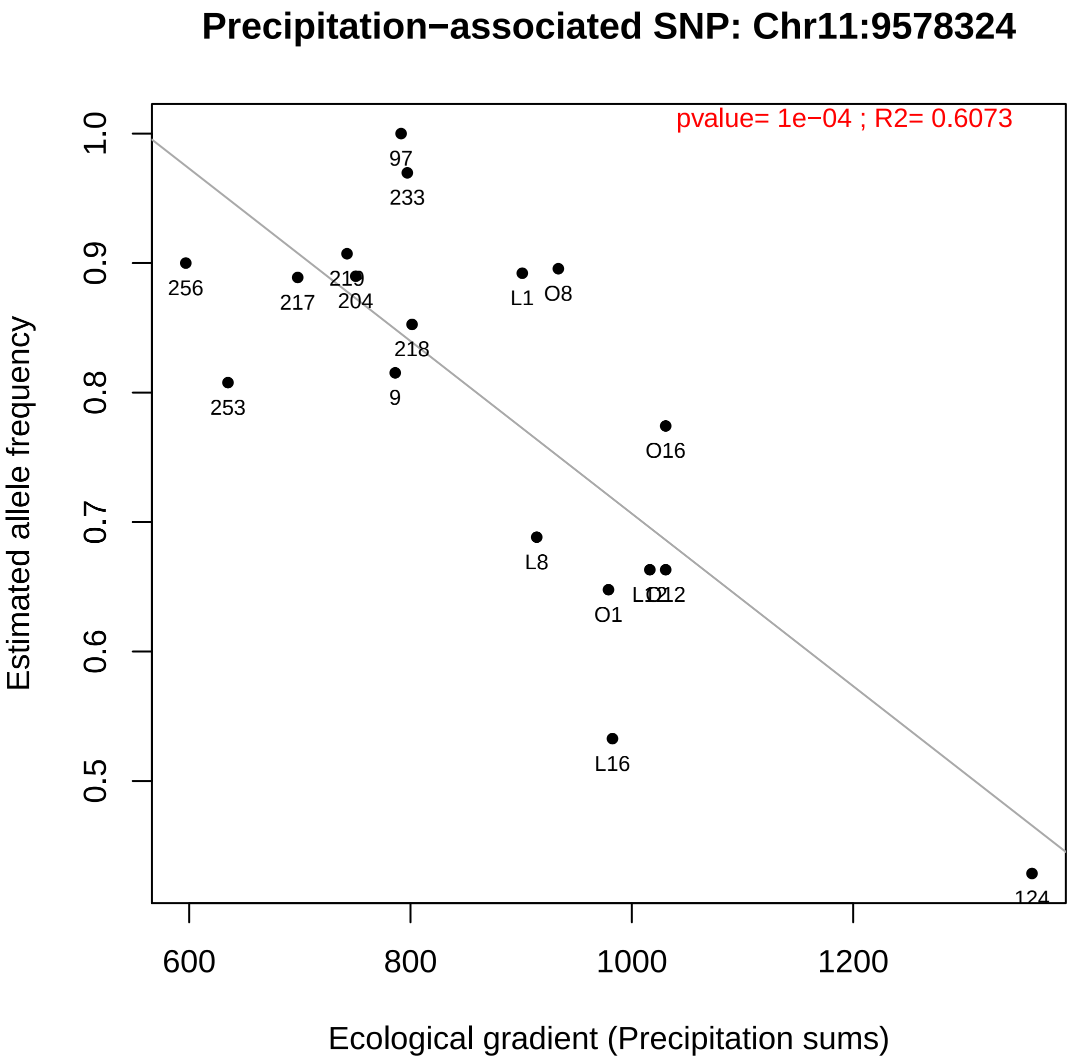

Precipitation−associated SNP: Chr11:9578324
pvalue= 1e−04 ; R2= 0.6073
Estimated allele frequency
Ecological gradient (Precipitation sums)
1.0
0.9
0.8
0.7
0.6
0.5
600
800
1000
1200
97
233
256
217
219
204
218
253
9
L1
O8
O16
L8
L12
O12
O1
L16
124

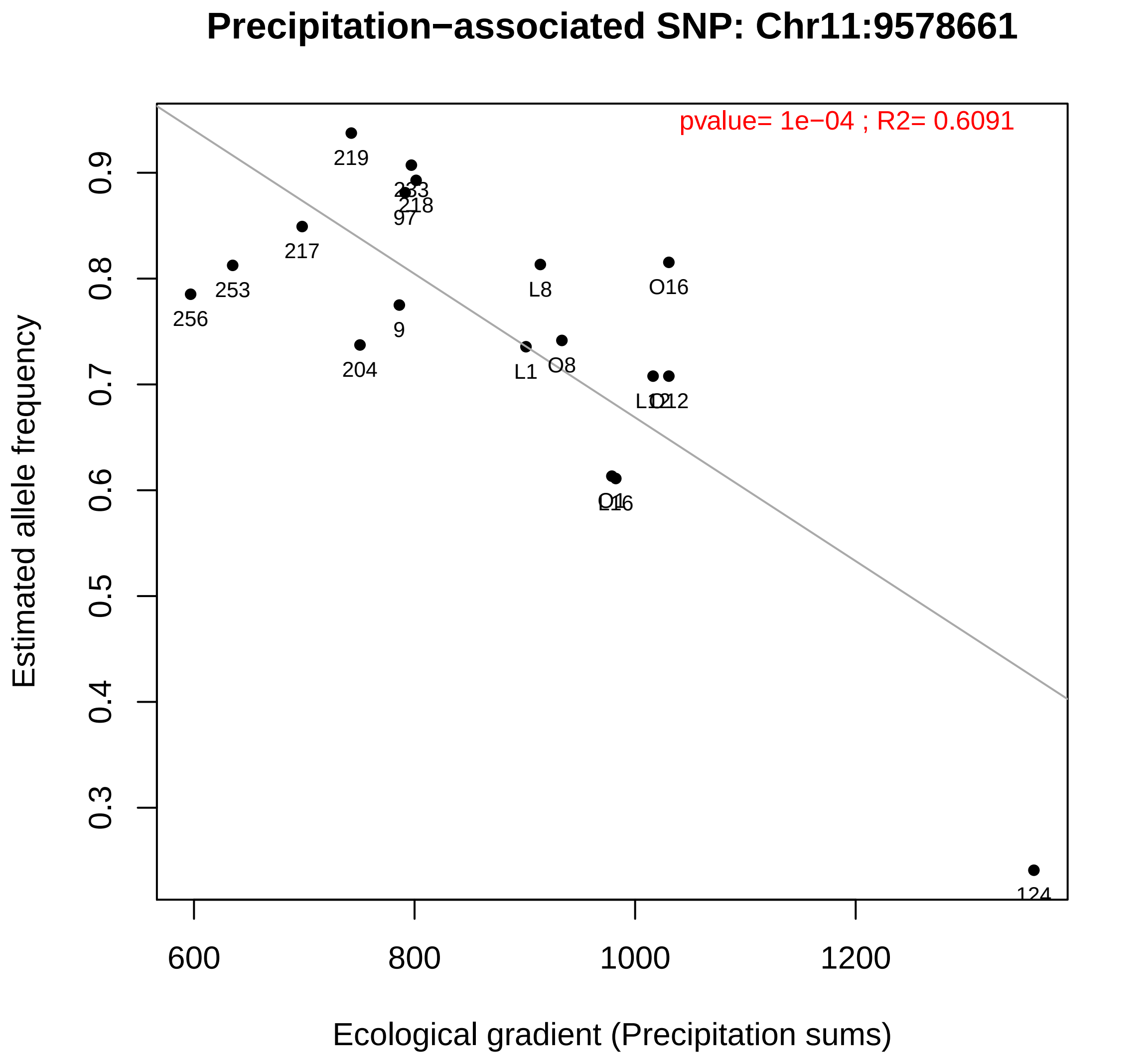

Precipitation−associated SNP: Chr11:9578661
pvalue= 1e−04 ; R2= 0.6091
Estimated allele frequency
Ecological gradient (Precipitation sums)
0.3
0.4
0.5
0.6
0.7
0.8
0.9
600
800
1000
1200
219
233
218
97
217
253
256
204
9
L8
O16
L1
O8
L12
O12
O1
L16
124

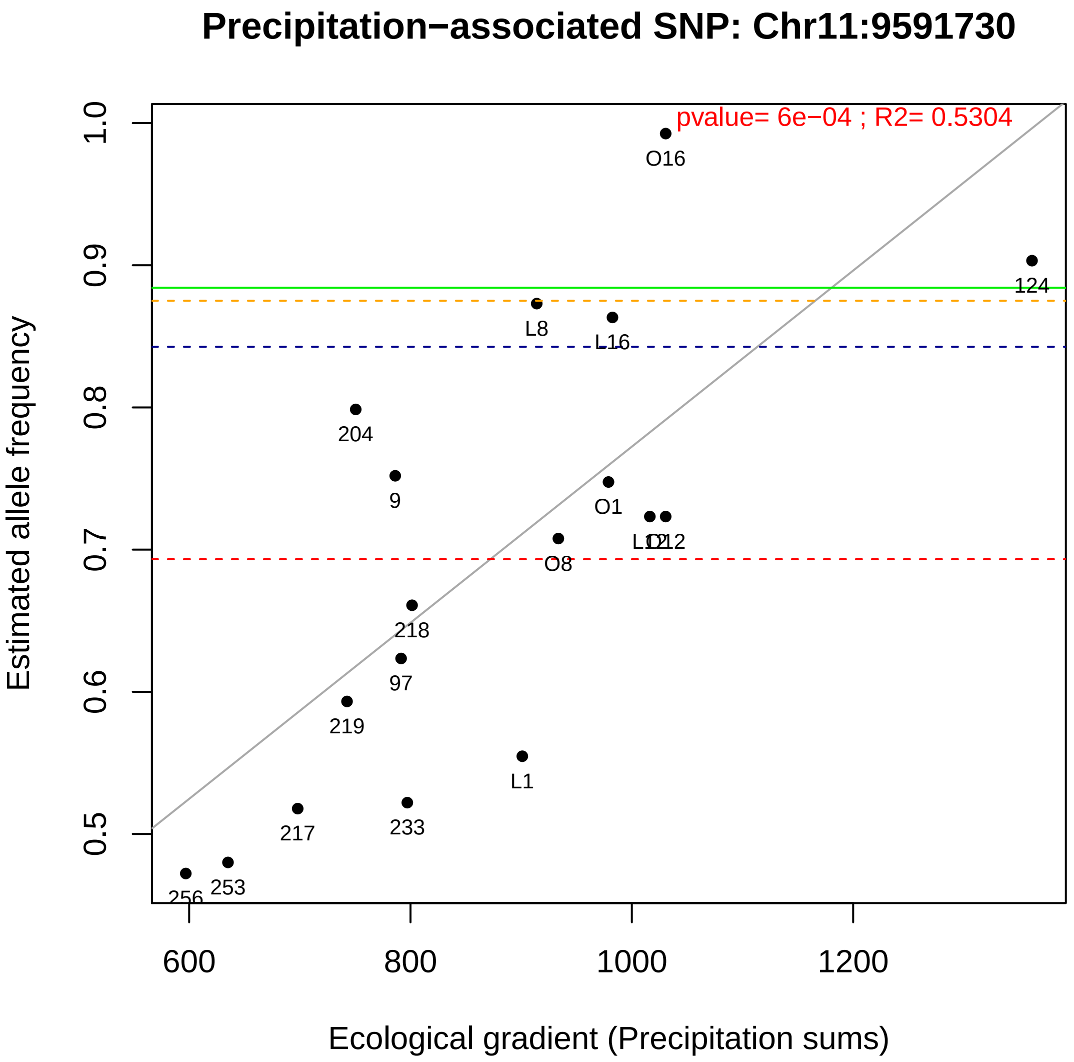

Precipitation−associated SNP: Chr11:9591730
pvalue= 6e−04 ; R2= 0.5304
Estimated allele frequency
Ecological gradient (Precipitation sums)
1.0
0.9
0.8
0.7
0.6
0.5
600
800
1000
1200
O16
124
L8
L16
204
9
O1
L12
O12
O8
218
97
219
L1
217
233
253
256

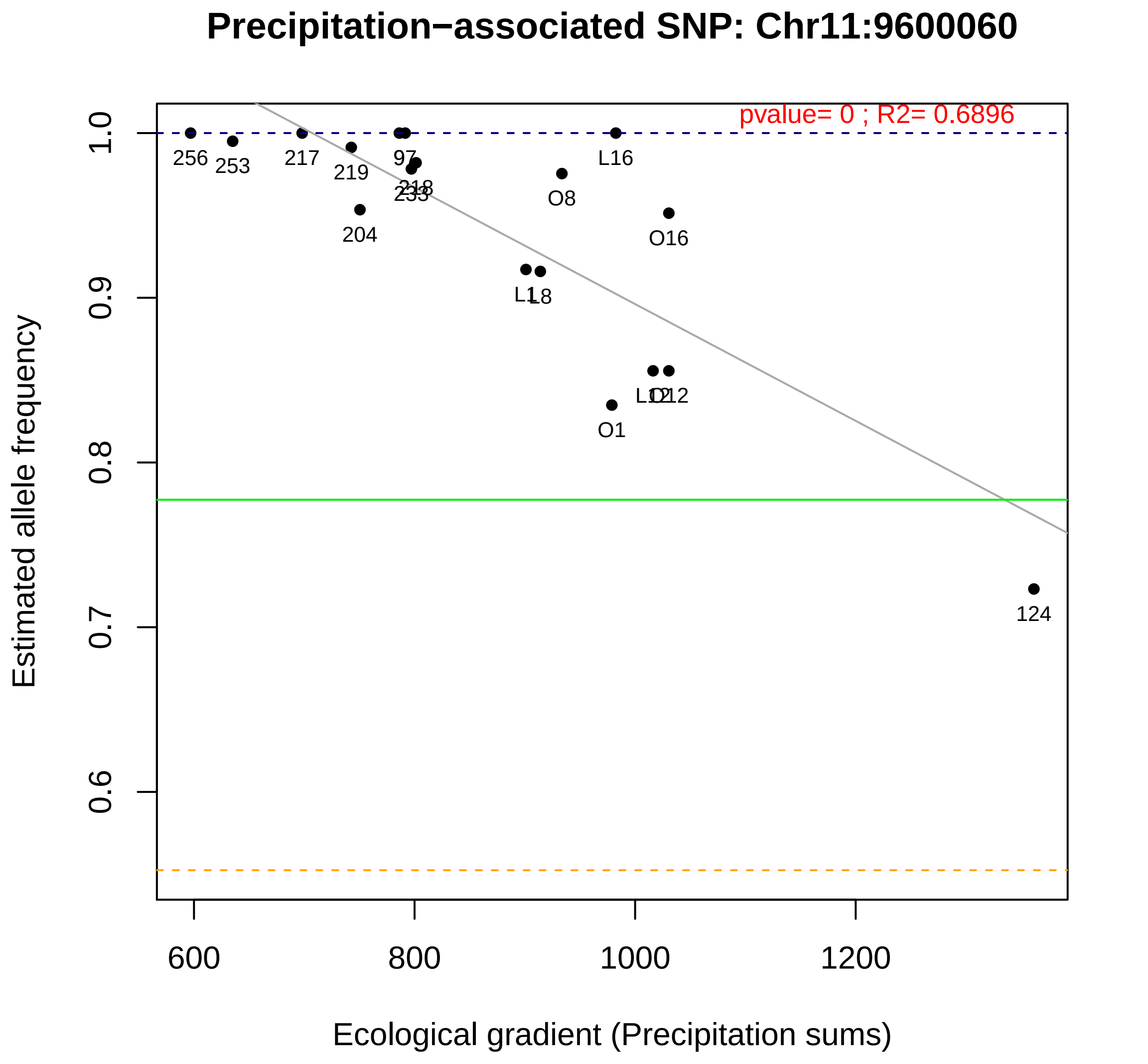
Precipitation−associated SNP: Chr11:9600060
pvalue= 0 ; R2= 0.6896
Estimated allele frequency
Ecological gradient (Precipitation sums)
0.6
0.7
0.8
0.9
1.0
600
800
1000
1200
256
253
217
219
97
218
233
204
L16
O8
O16
L1
L8
L12
O12
O1
124

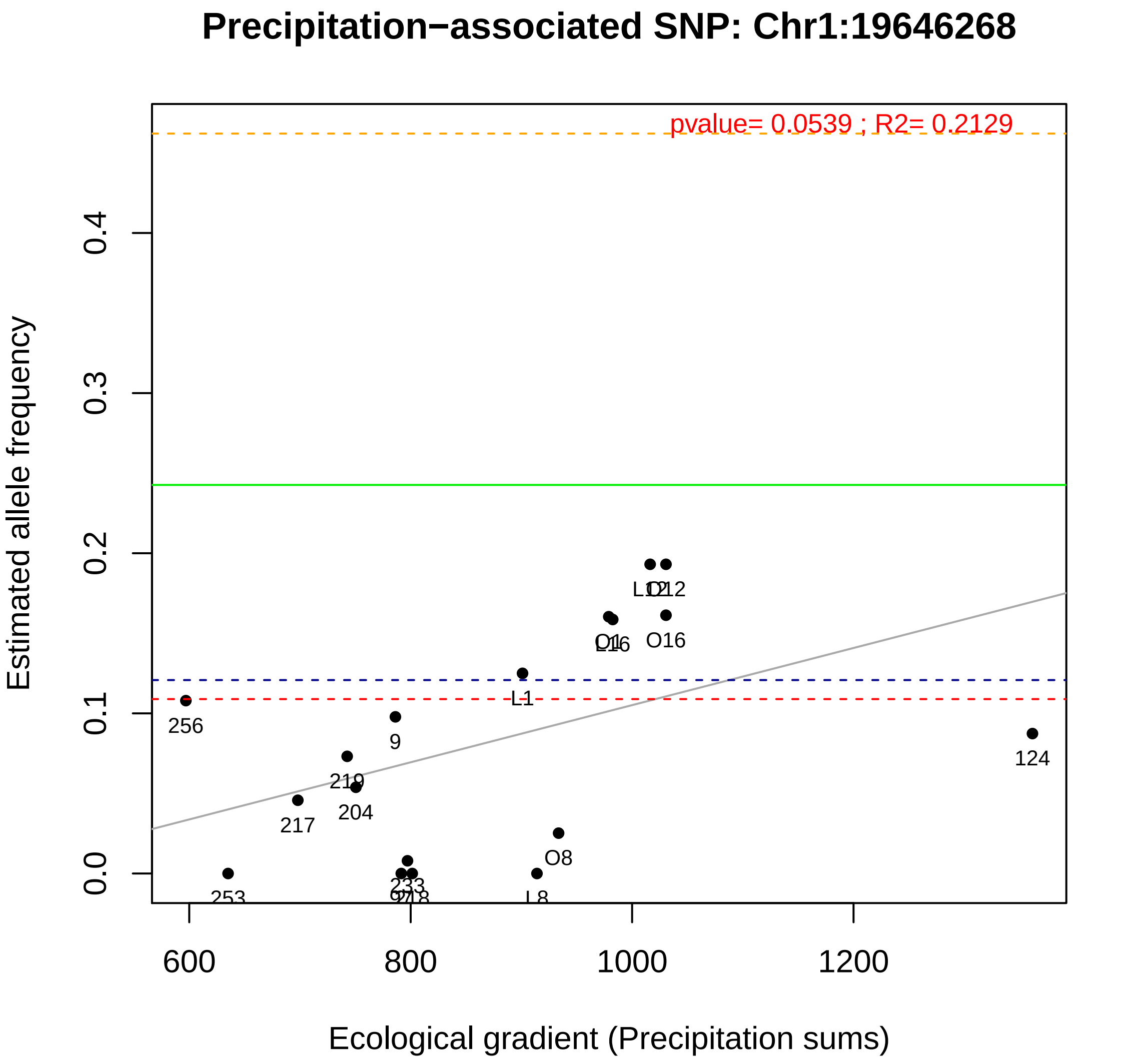

Precipitation−associated SNP: Chr1:19646268
pvalue= 0.0539 ; R2= 0.2129
Estimated allele frequency
0.0
0.1
0.2
0.3
0.4
600
800
1000
1200
Ecological gradient (Precipitation sums)
256
9
219
204
217
253
233
L1
O8
O16
124

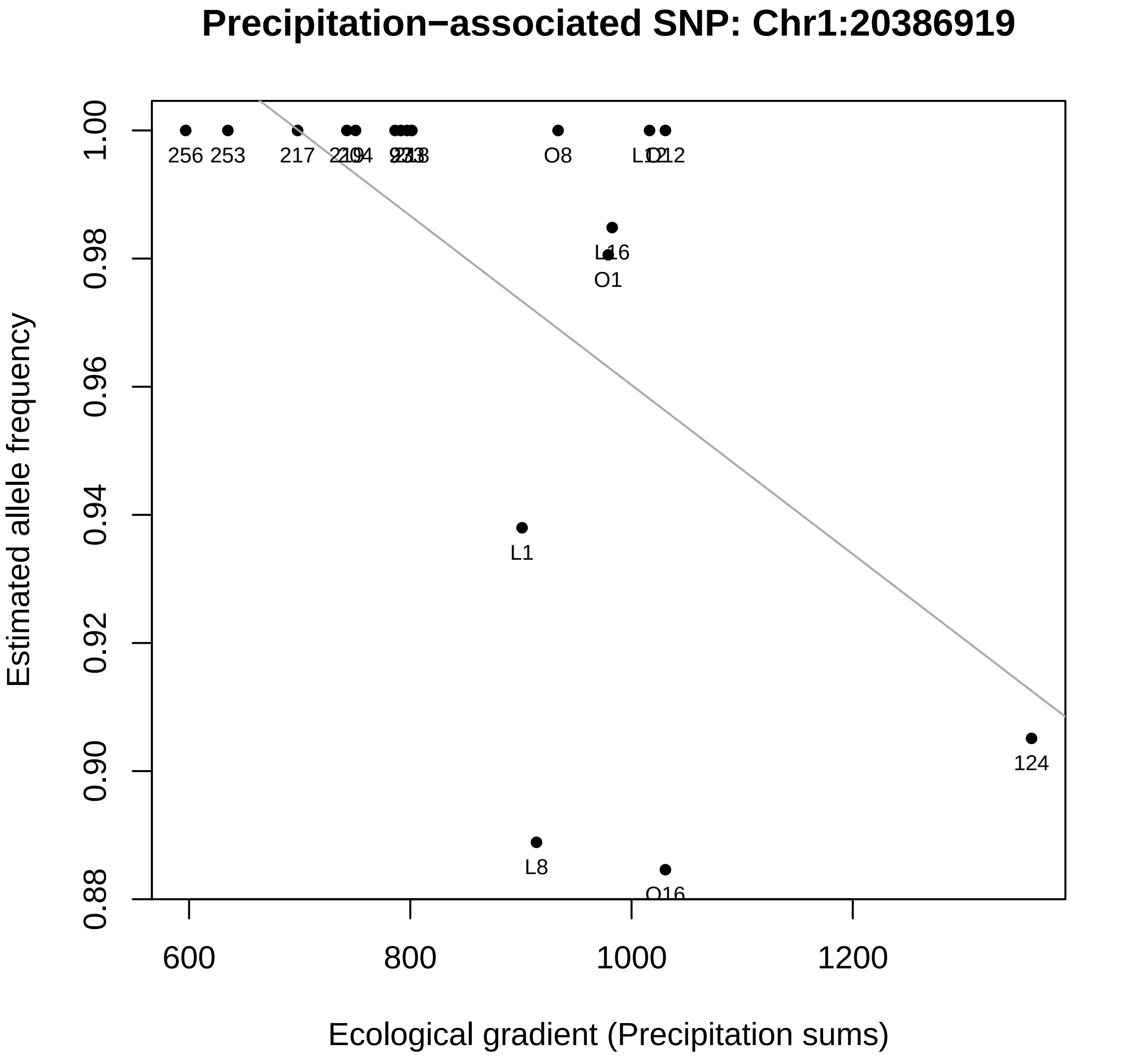
Precipitation−associated SNP: Chr1:20386919
Estimated allele frequency
Ecological gradient (Precipitation sums)
1.00
0.98
0.96
0.94
0.92
0.90
0.88
600
800
1000
1200
256
253
217
O8
L16
O1
L1
124
L8
O16

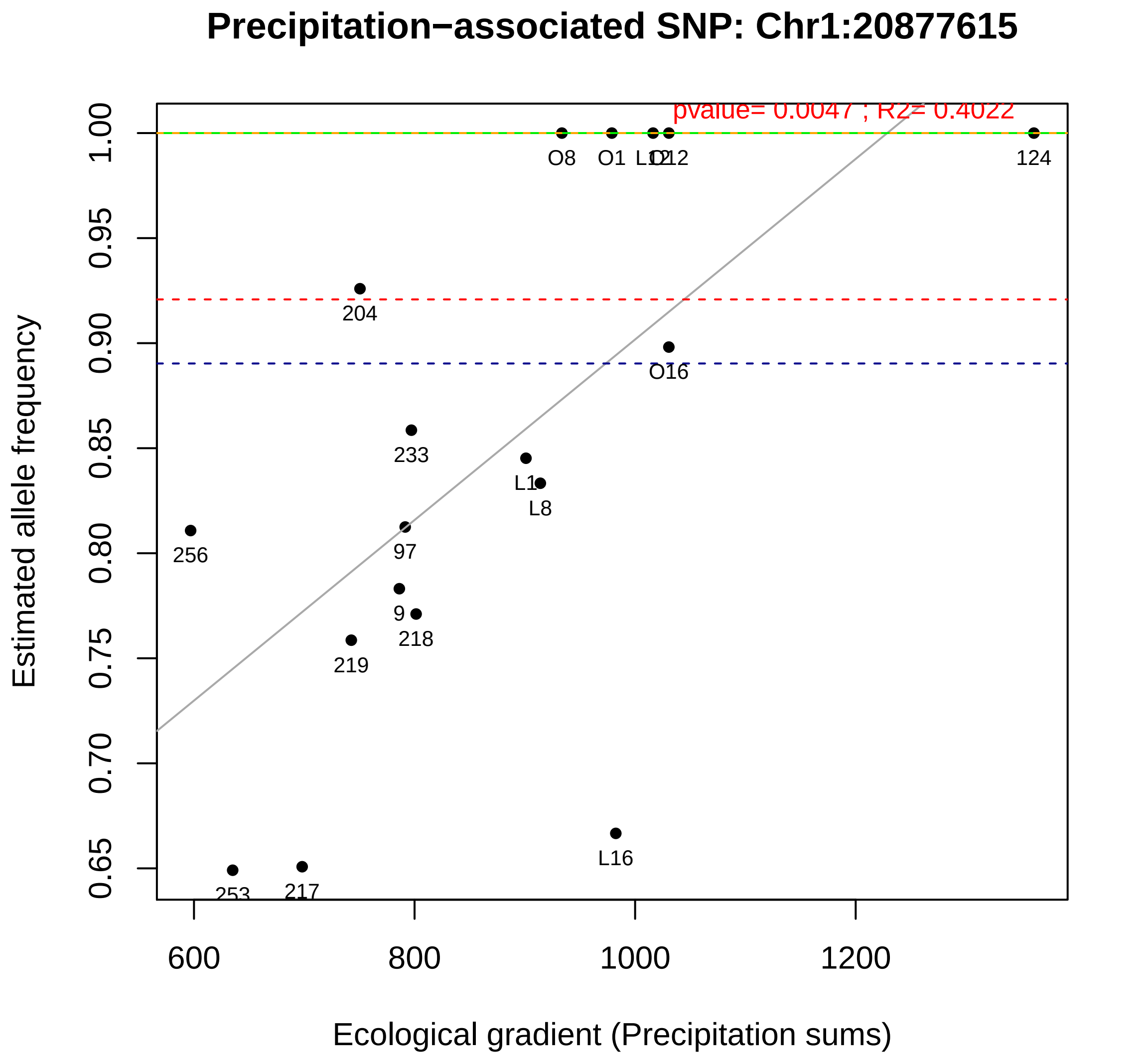

Precipitation−associated SNP: Chr1:20877615
pvalue= 0.0047 ; R2= 0.4022
Estimated allele frequency
Ecological gradient (Precipitation sums)
1.00
0.95
0.90
0.85
0.80
0.75
0.70
0.65
600
800
1000
1200
O8
O1
O12
124
204
O16
233
L1
L8
256
97
9
218
219
L16
253
217

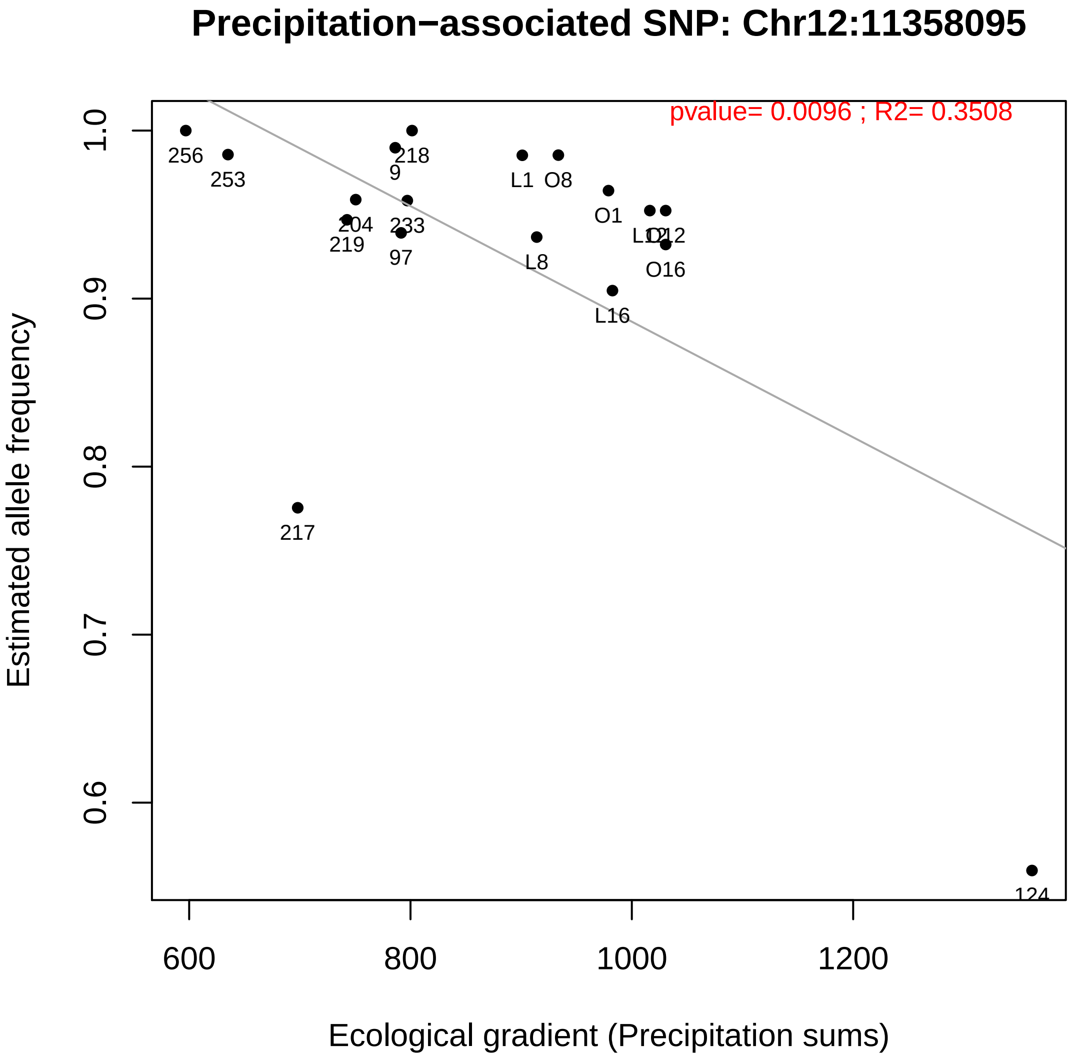
Precipitation−associated SNP: Chr12:11358095
pvalue= 0.0096 ; R2= 0.3508
Estimated allele frequency
Ecological gradient (Precipitation sums)
0.6
0.7
0.8
0.9
1.0
600
800
1000
1200
256
253
218
9
L1
O8
O1
204
219
233
97
L8
O16
L16
217
124

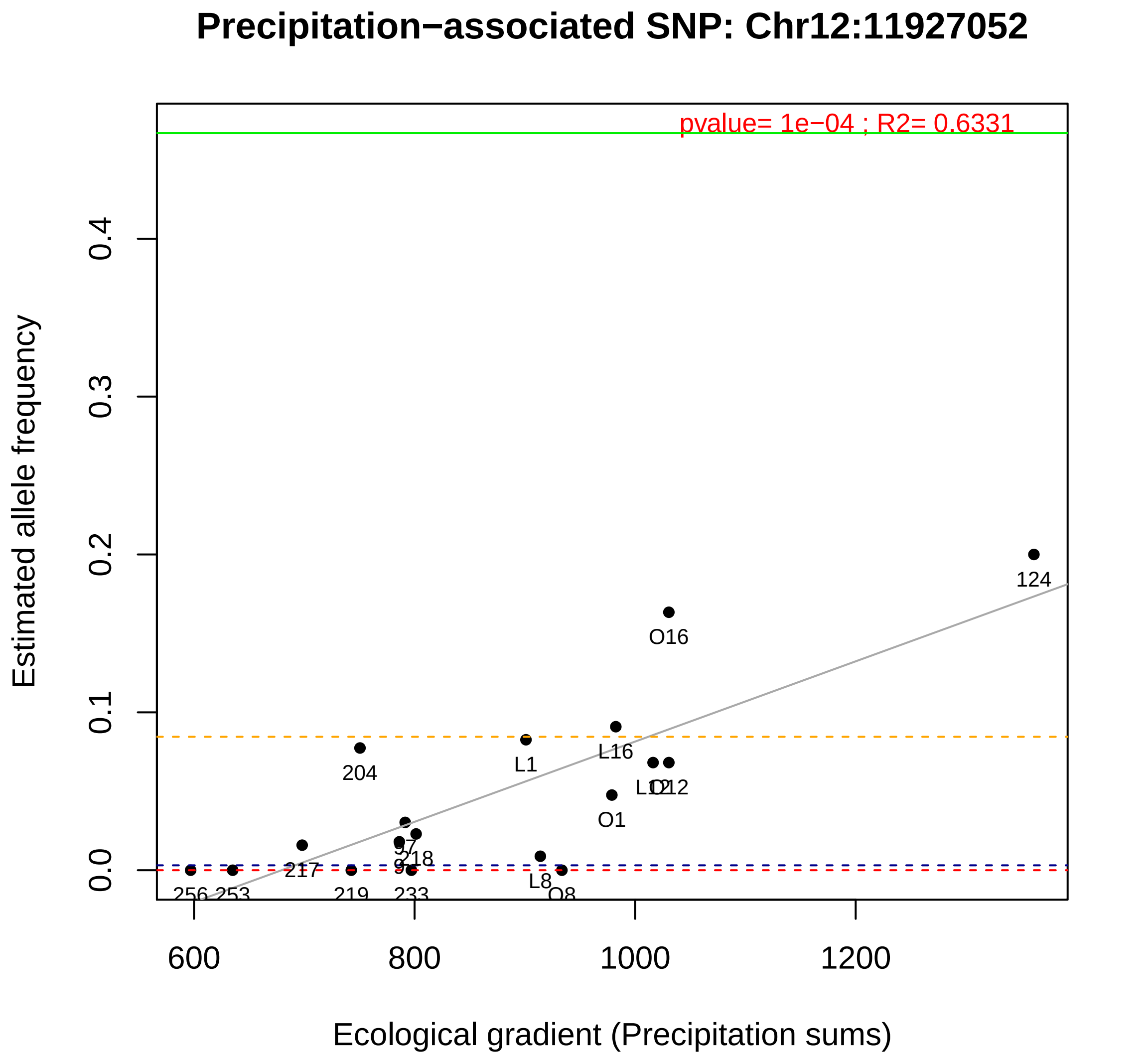

Precipitation−associated SNP: Chr12:11927052
pvalue= 1e−04 ; R2= 0.6331
Estimated allele frequency
0.0
0.1
0.2
0.3
0.4
600
800
1000
1200
Ecological gradient (Precipitation sums)
124
O16
L16
L1
204
L12
O12
O1
97
218
217
L8
O8
256
253
219
233

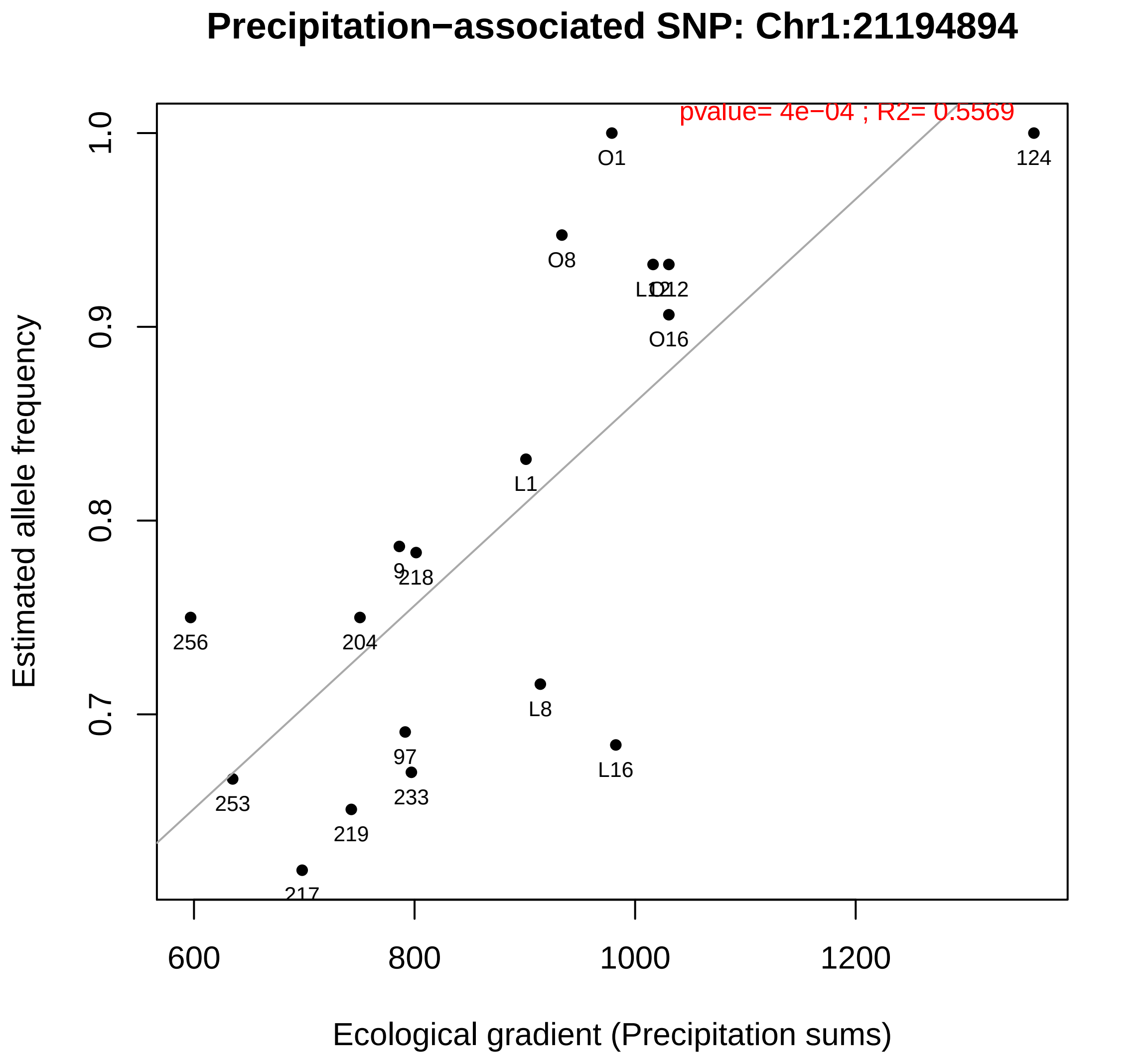
Precipitation−associated SNP: Chr1:21194894
pvalue= 4e−04 ; R2= 0.5569
Estimated allele frequency
Ecological gradient (Precipitation sums)
1.0
0.9
0.8
0.7
600
800
1000
1200
O1
124
O8
L12
O12
O16
L1
9
218
256
204
L8
97
L16
233
253
219
217

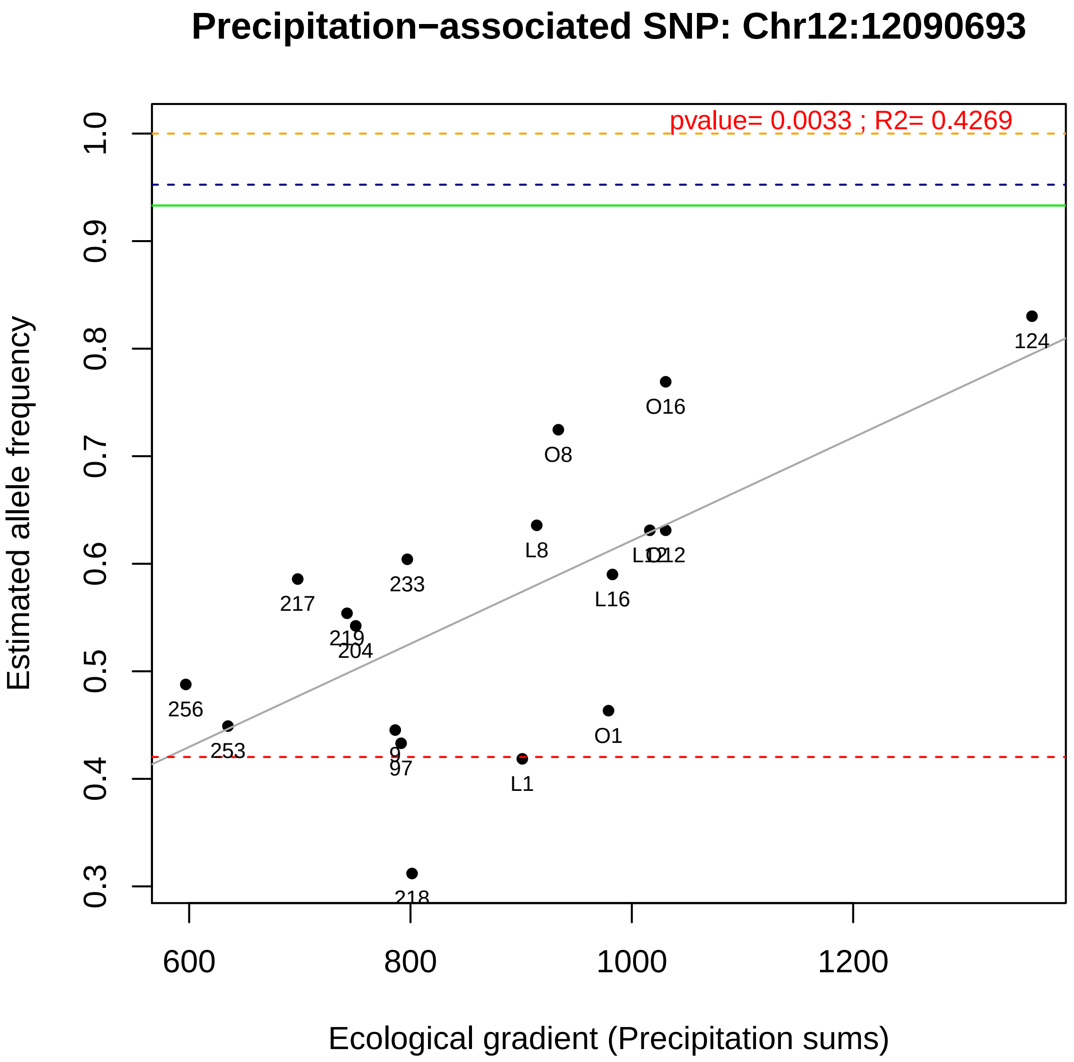

Precipitation−associated SNP: Chr12:12090693
pvalue= 0.0033 ; R2= 0.4269
Estimated allele frequency
0.3
0.4
0.5
0.6
0.7
0.8
0.9
1.0
600
800
1000
1200
Ecological gradient (Precipitation sums)
124
O16
O8
L8
L12
O12
L16
233
217
219
204
256
253
O1
9
97
L1
218

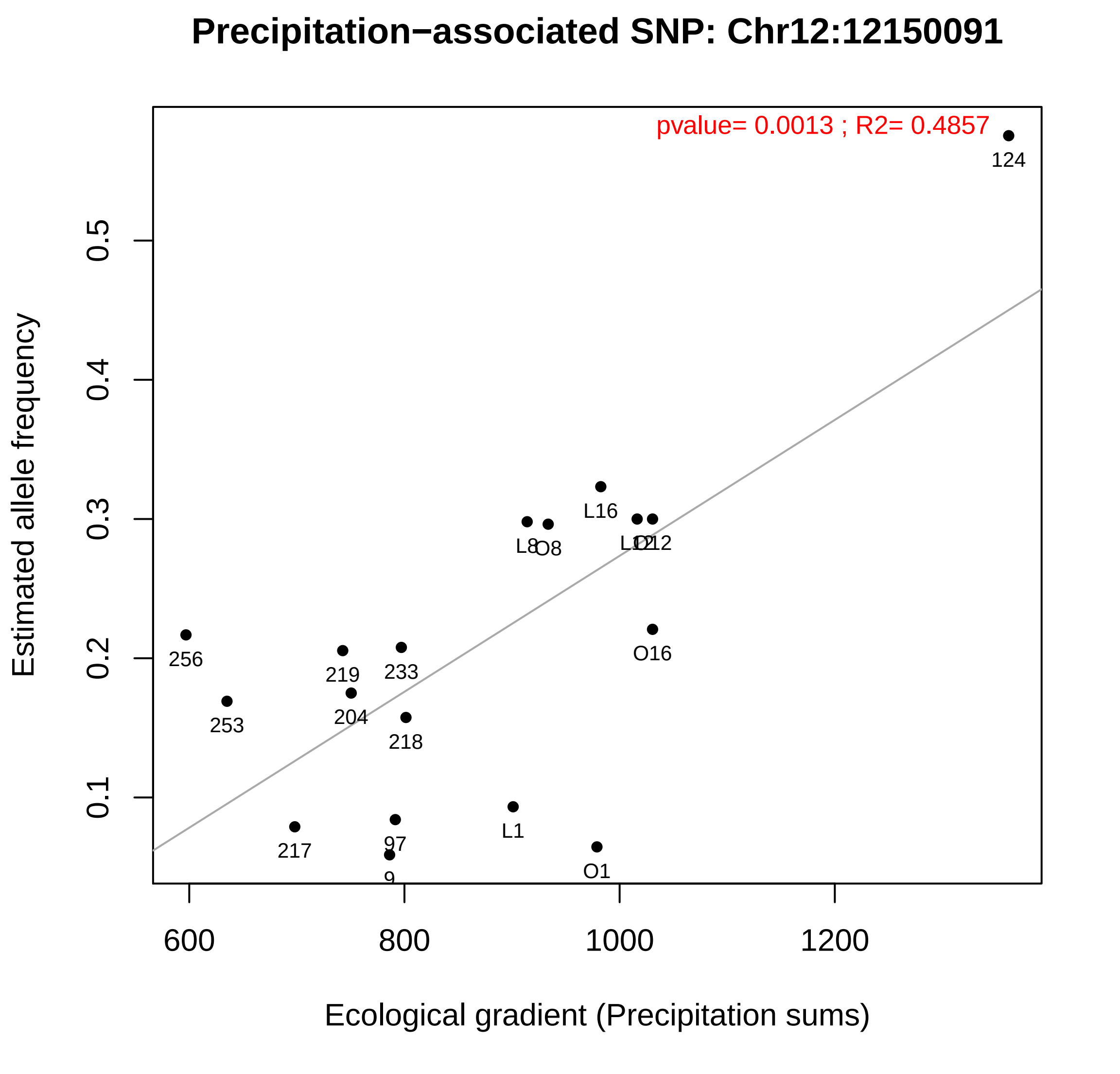
Precipitation−associated SNP: Chr12:12150091
pvalue= 0.0013 ; R2= 0.4857
Estimated allele frequency
Ecological gradient (Precipitation sums)
0.1
0.2
0.3
0.4
0.5
600
800
1000
1200
124
L16
L8
O8
L12
O12
O16
256
219
233
253
204
218
217
97
9
L1
O1

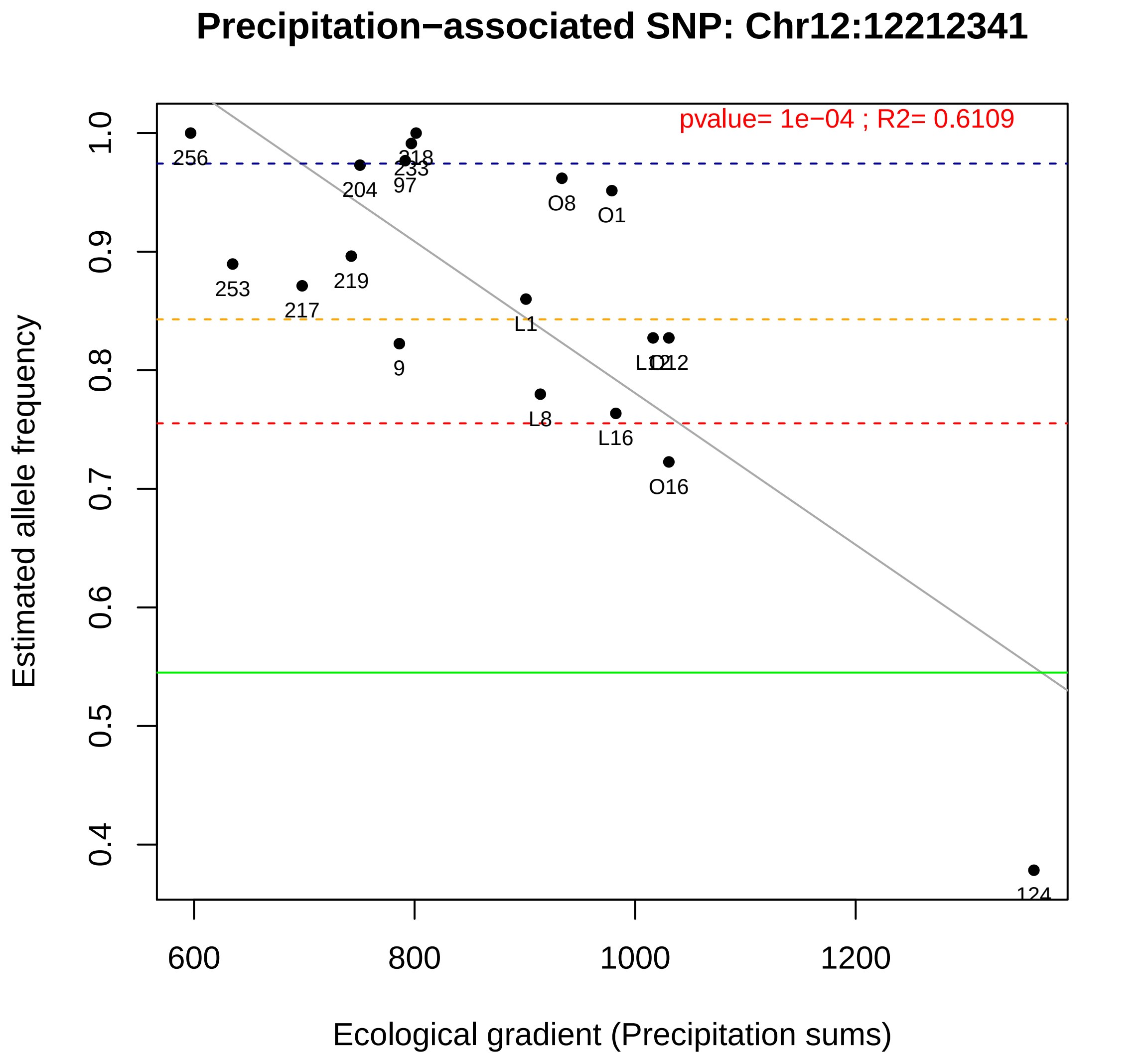

Precipitation−associated SNP: Chr12:12212341
pvalue= 1e−04 ; R2= 0.6109
Estimated allele frequency
Ecological gradient (Precipitation sums)
1.0
0.9
0.8
0.7
0.6
0.5
0.4
600
800
1000
1200
256
218
233
97
204
O8
O1
253
219
217
L1
9
L12
O12
L8
L16
O16
124

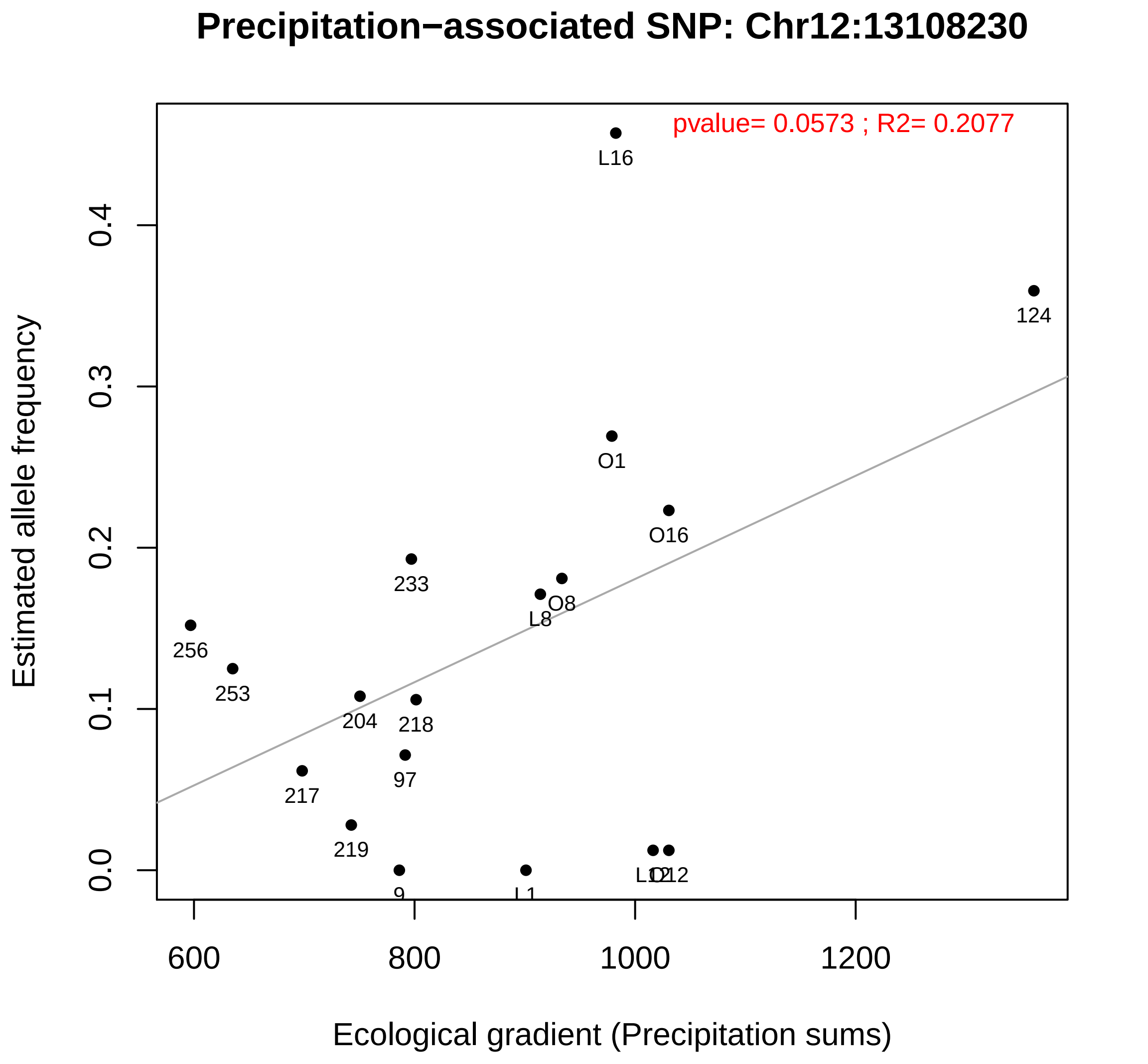
Precipitation−associated SNP: Chr12:13108230
pvalue= 0.0573 ; R2= 0.2077
L16
124
O1
O16
233
O8
L8
256
253
204
218
97
217
219
9
L1
L12
O12
0.0
0.1
0.2
0.3
0.4
600
800
1000
1200
Estimated allele frequency
Ecological gradient (Precipitation sums)

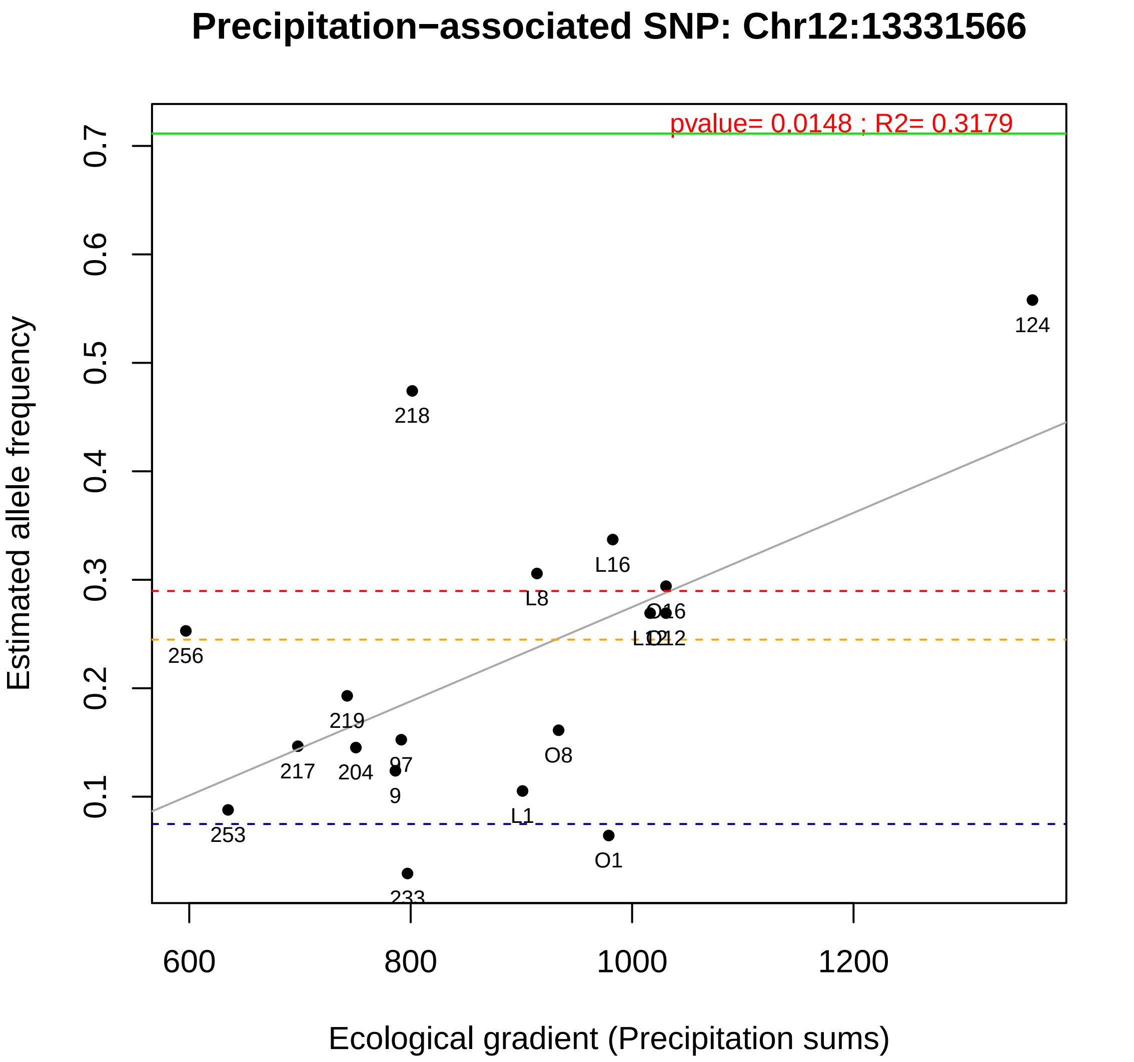
Precipitation−associated SNP: Chr12:13331566
pvalue= 0.0148 ; R2= 0.3179
Estimated allele frequency
Ecological gradient (Precipitation sums)
0.1
0.2
0.3
0.4
0.5
0.6
0.7
600
800
1000
1200
124
218
L16
L8
256
219
217
204
97
9
O8
L1
253
O1
233

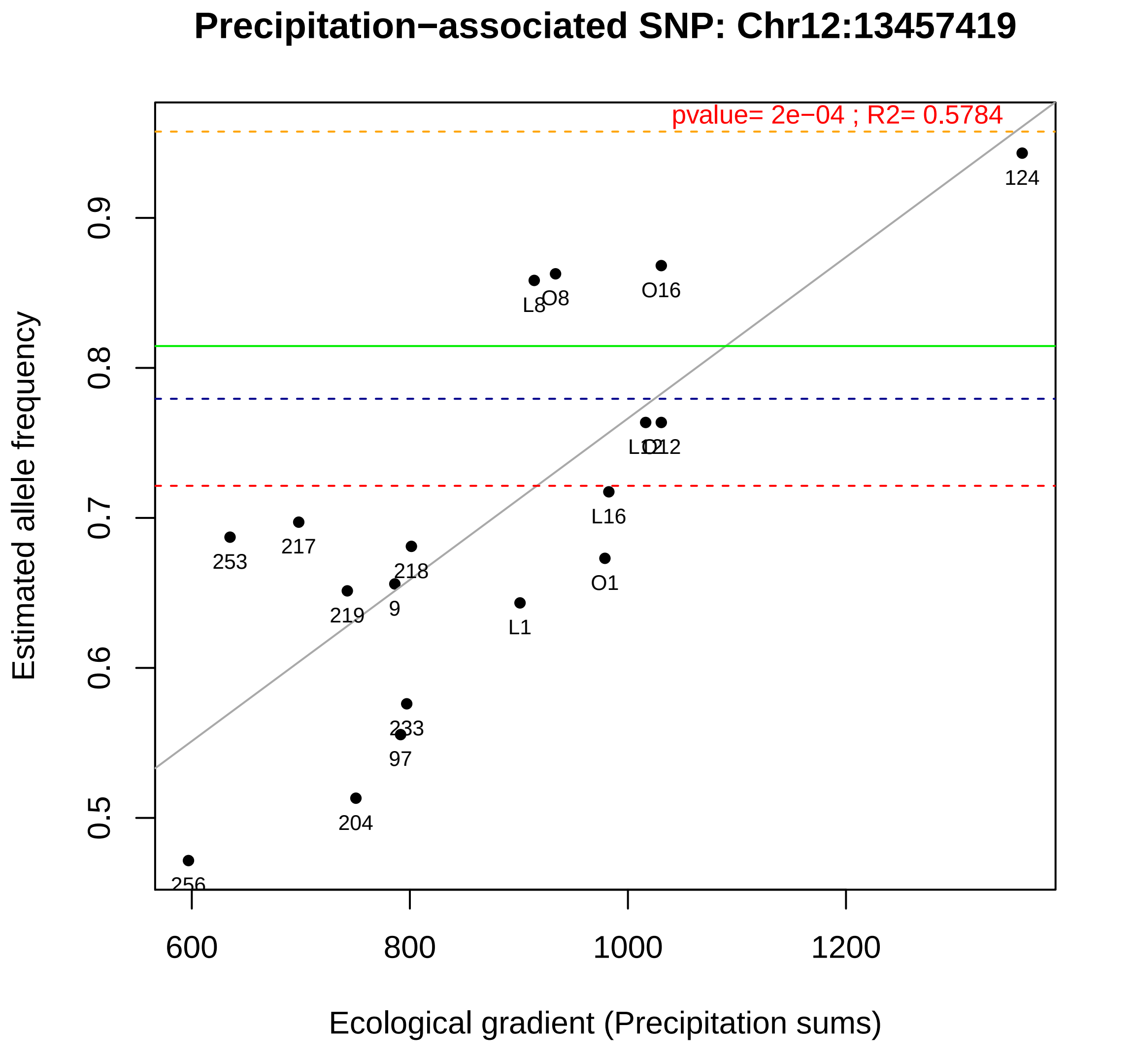
Precipitation−associated SNP: Chr12:13457419
pvalue= 2e−04 ; R2= 0.5784
Estimated allele frequency
Ecological gradient (Precipitation sums)
0.5
0.6
0.7
0.8
0.9
600
800
1000
1200
124
L8
O8
O16
L12
O12
L16
253
217
218
219
9
L1
O1
233
97
204
256

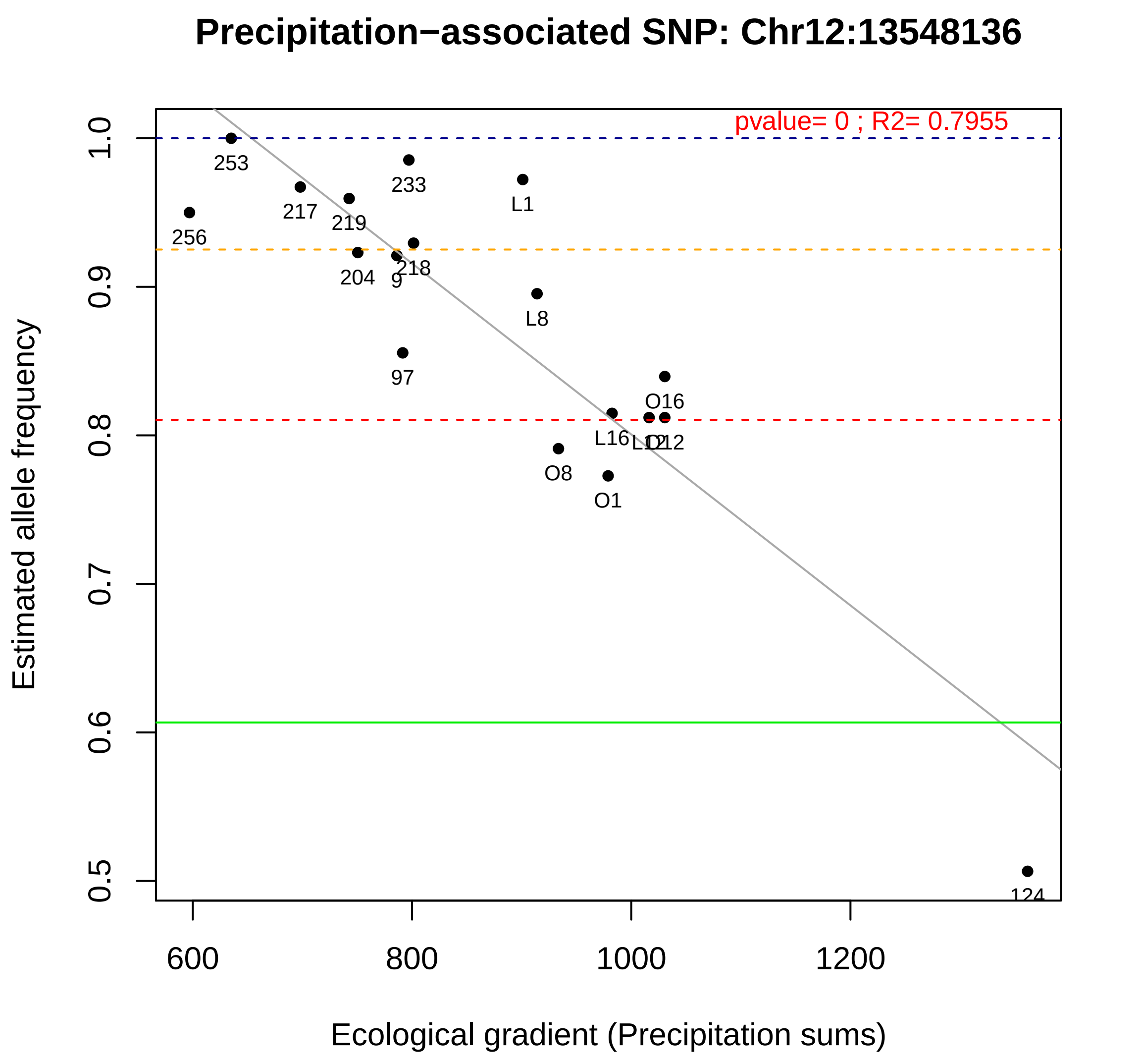

Precipitation−associated SNP: Chr12:13548136
pvalue= 0 ; R2= 0.7955
Estimated allele frequency
1.0
0.9
0.8
0.7
0.6
0.5
600
800
1000
1200
Ecological gradient (Precipitation sums)
253
233
L1
217
219
256
204
9
218
L8
97
O16
L16
L12
O12
O8
O1
124

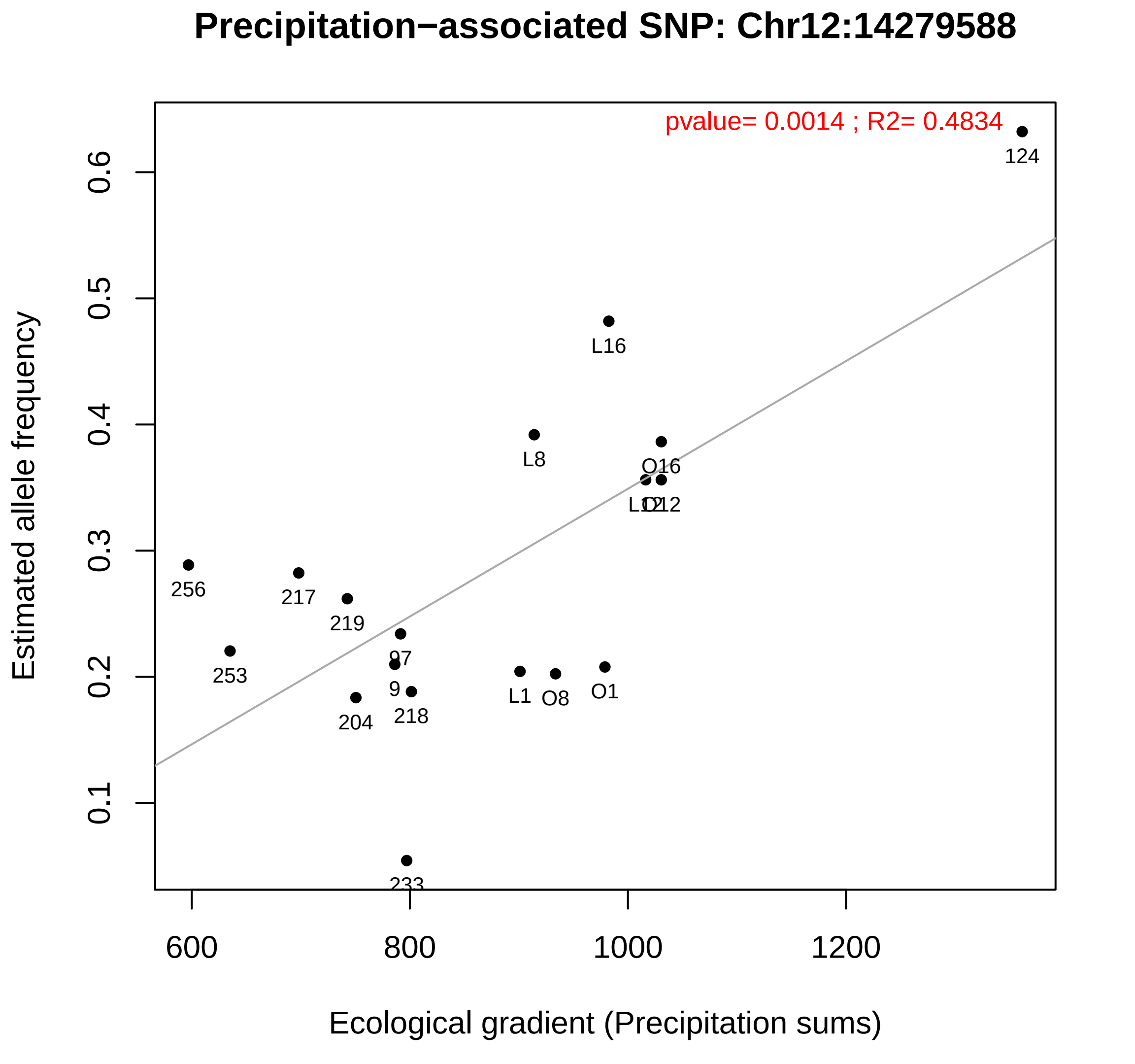
Precipitation−associated SNP: Chr12:14279588
pvalue= 0.0014 ; R2= 0.4834
124
L16
L8
O16
L12
O12
256
217
219
97
9
253
204
218
L1
O8
O1
233
0.1
0.2
0.3
0.4
0.5
0.6
600
800
1000
1200
Estimated allele frequency
Ecological gradient (Precipitation sums)

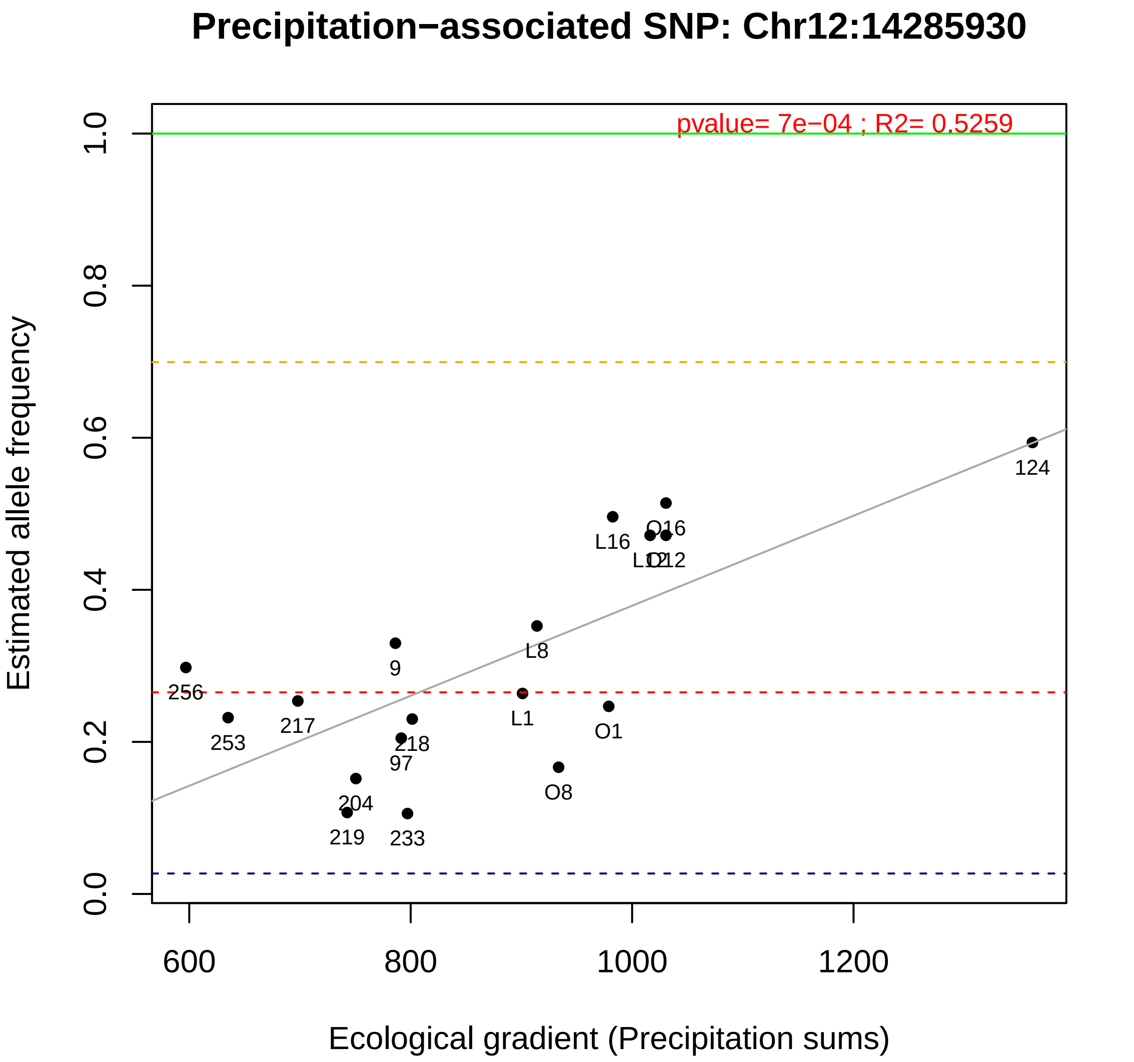

Precipitation−associated SNP: Chr12:14285930
pvalue= 7e−04 ; R2= 0.5259
Estimated allele frequency
0.0
0.2
0.4
0.6
0.8
1.0
Ecological gradient (Precipitation sums)
600
800
1000
1200
124
L16
O16
L12
O12
L8
9
256
253
217
218
97
L1
O1
O8
204
219
233

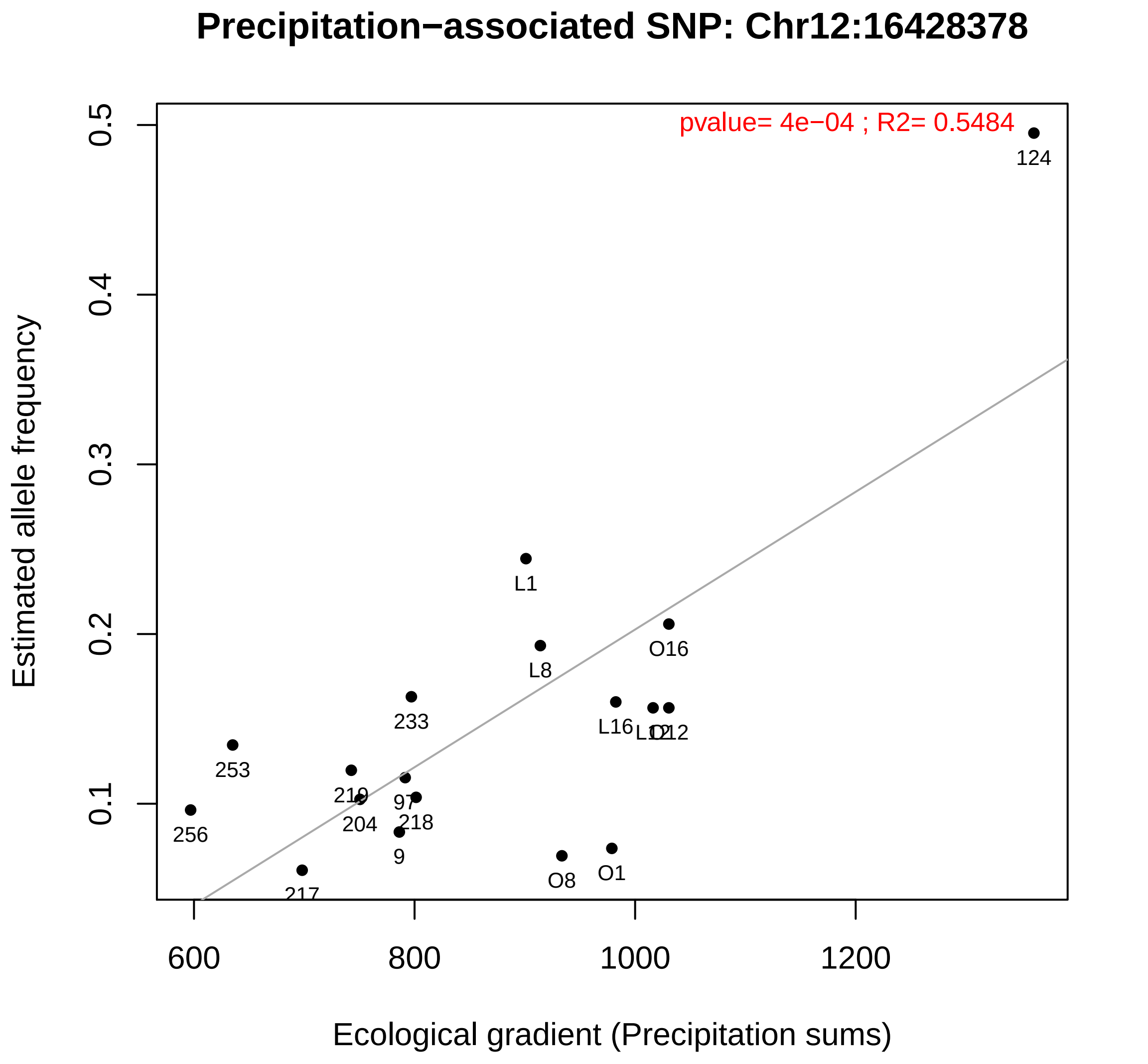
Precipitation−associated SNP: Chr12:16428378
pvalue= 4e−04 ; R2= 0.5484
Estimated allele frequency
Ecological gradient (Precipitation sums)
0.1
0.2
0.3
0.4
0.5
600
800
1000
1200
124
L1
O16
L8
233
L16
O12
253
256
204
218
9
O8
O1
217

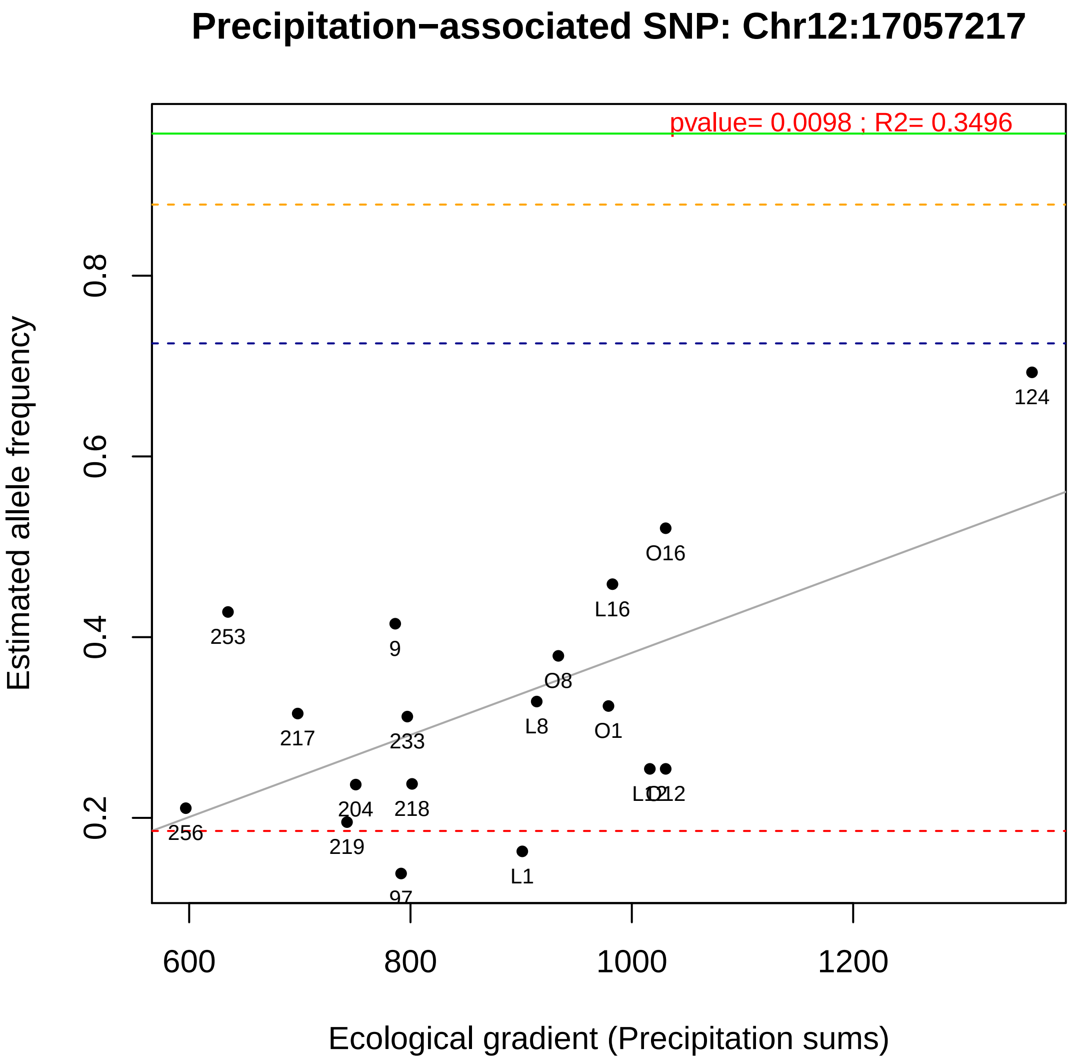

Precipitation−associated SNP: Chr12:17057217
pvalue= 0.0098 ; R2= 0.3496
Estimated allele frequency
0.2
0.4
0.6
0.8
600
800
1000
1200
Ecological gradient (Precipitation sums)
124
O16
L16
253
9
O8
L8
O1
217
233
L10
O12
204
218
256
219
L1
97

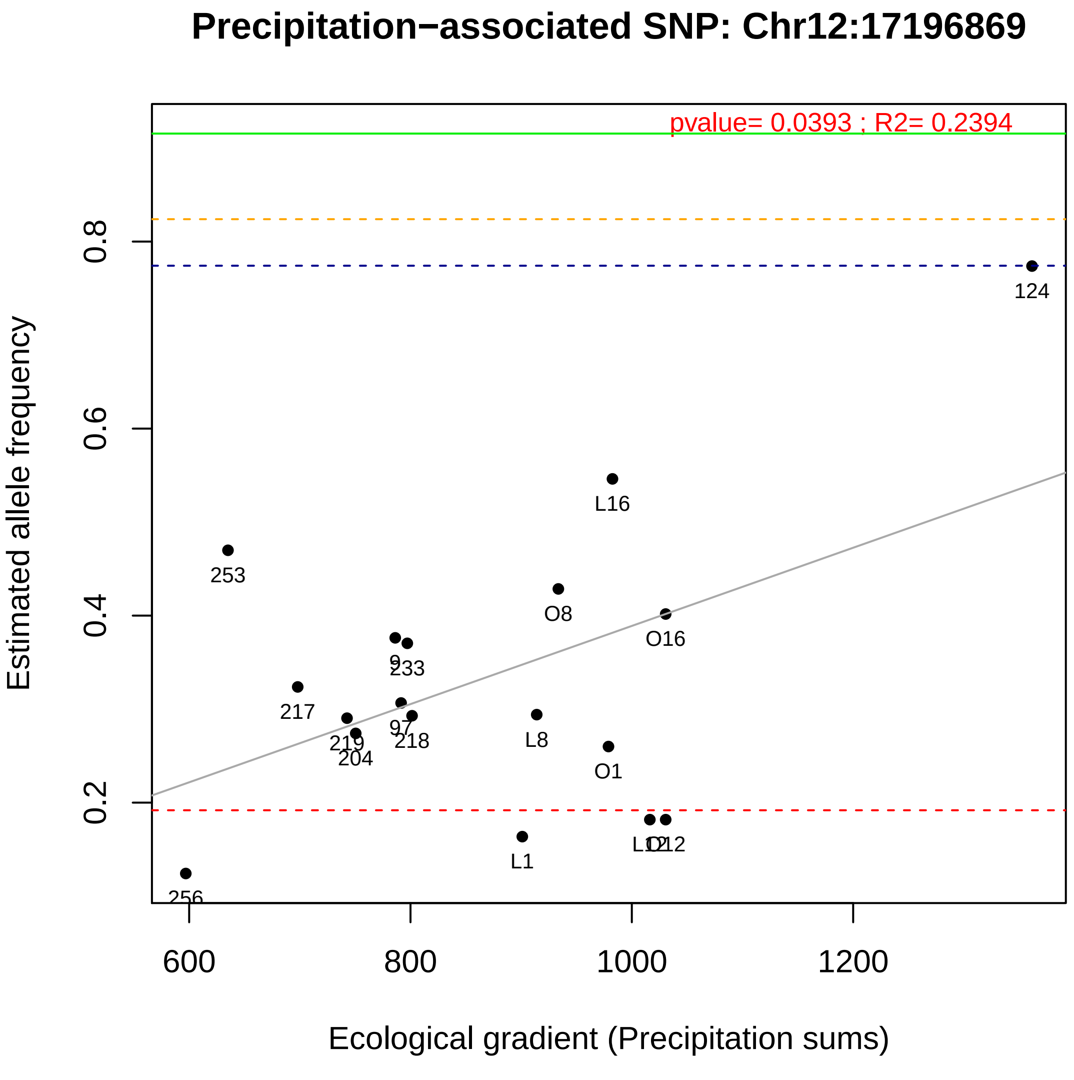

Precipitation−associated SNP: Chr12:17196869
pvalue= 0.0393 ; R2= 0.2394
Estimated allele frequency
0.2
0.4
0.6
0.8
600
800
1000
1200
Ecological gradient (Precipitation sums)
124
L16
253
O8
O16
9
233
217
97
218
219
204
L8
O1
L12
O12
L1
256

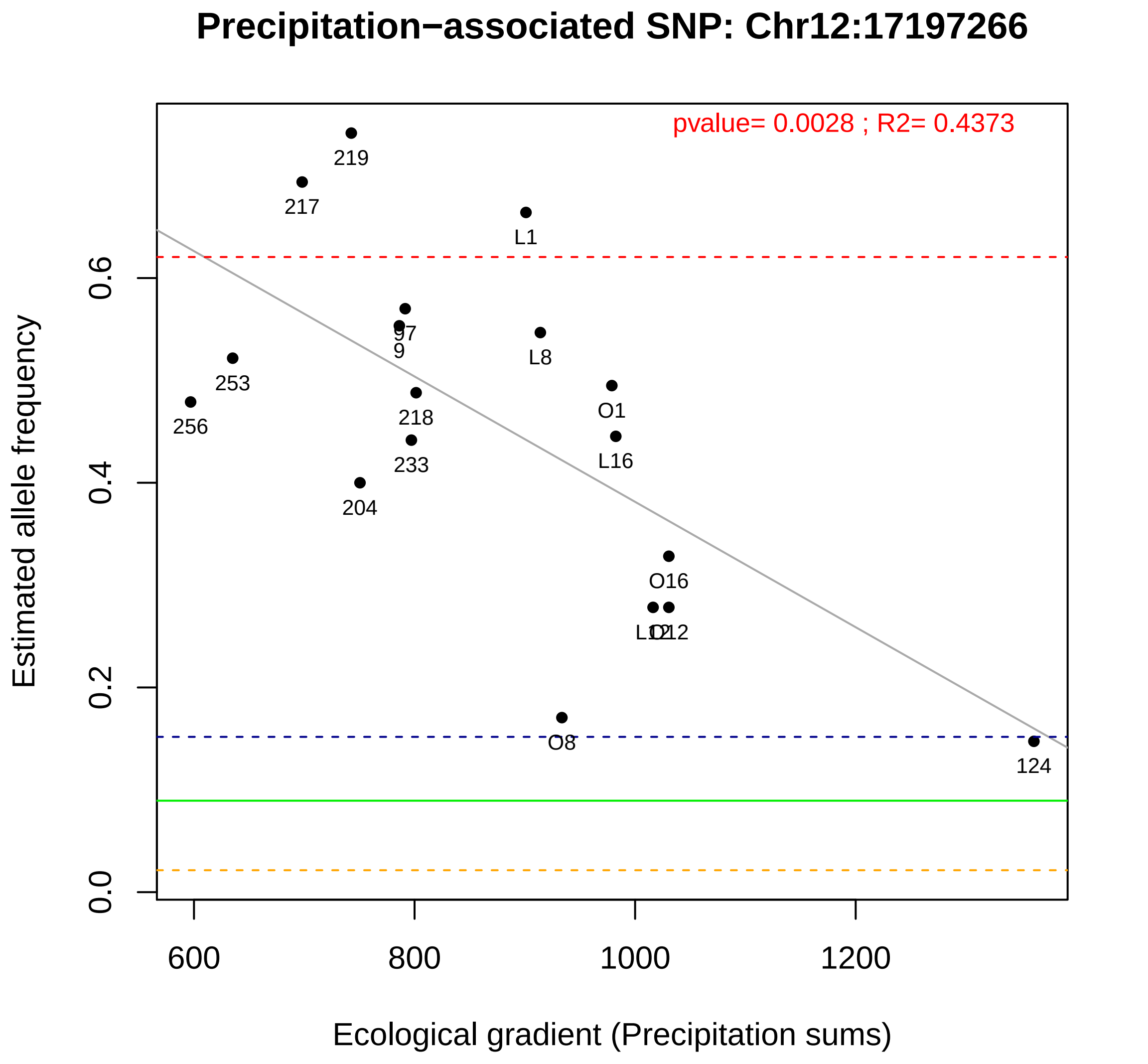

Precipitation−associated SNP: Chr12:17197266
pvalue= 0.0028 ; R2= 0.4373
219
217
L1
97
9
L8
253
256
218
O1
233
L16
204
O16
L12
O12
O8
124
0.0
0.2
0.4
0.6
600
800
1000
1200
Estimated allele frequency
Ecological gradient (Precipitation sums)

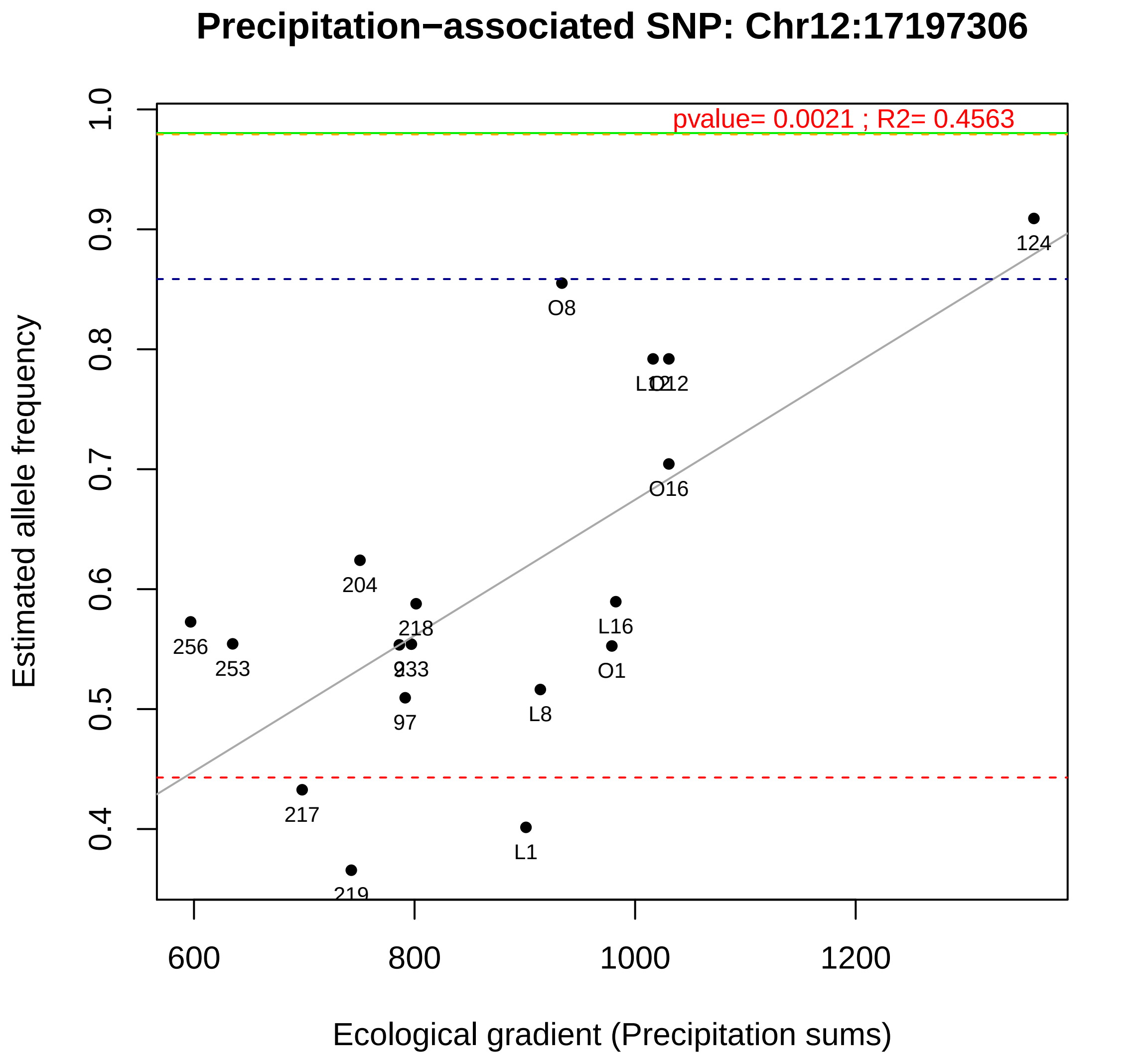

Precipitation−associated SNP: Chr12:17197306
pvalue= 0.0021 ; R2= 0.4563
Estimated allele frequency
0.4
0.5
0.6
0.7
0.8
0.9
1.0
Ecological gradient (Precipitation sums)
600
800
1000
1200
124
O8
L12
O12
O16
204
218
L16
256
253
233
O1
L8
97
217
L1
219

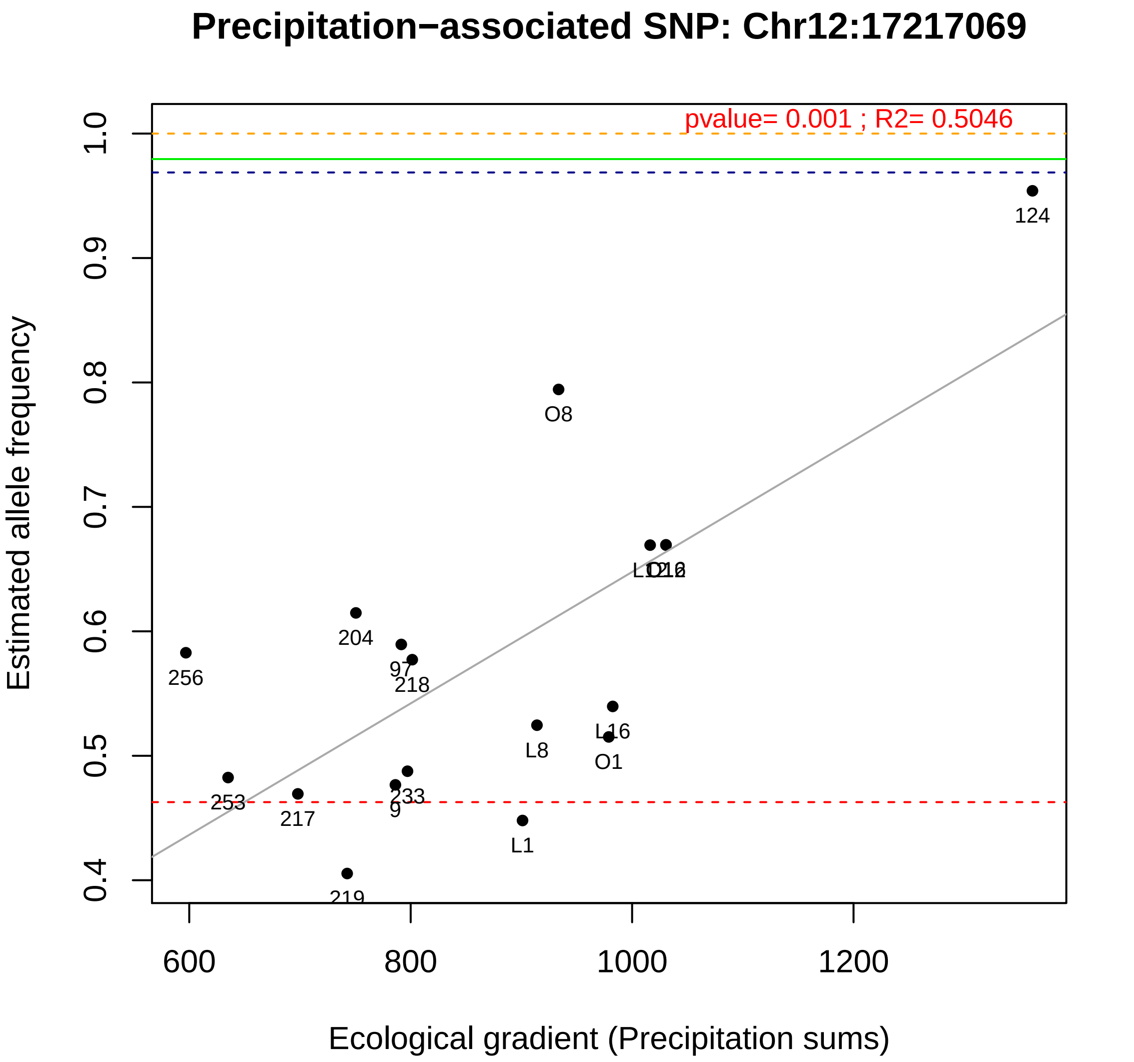

Precipitation−associated SNP: Chr12:17217069
pvalue= 0.001 ; R2= 0.5046
Estimated allele frequency
Ecological gradient (Precipitation sums)
0.4
0.5
0.6
0.7
0.8
0.9
1.0
600
800
1000
1200
124
O8
204
256
97
218
L16
L8
O1
253
233
9
217
L1
219

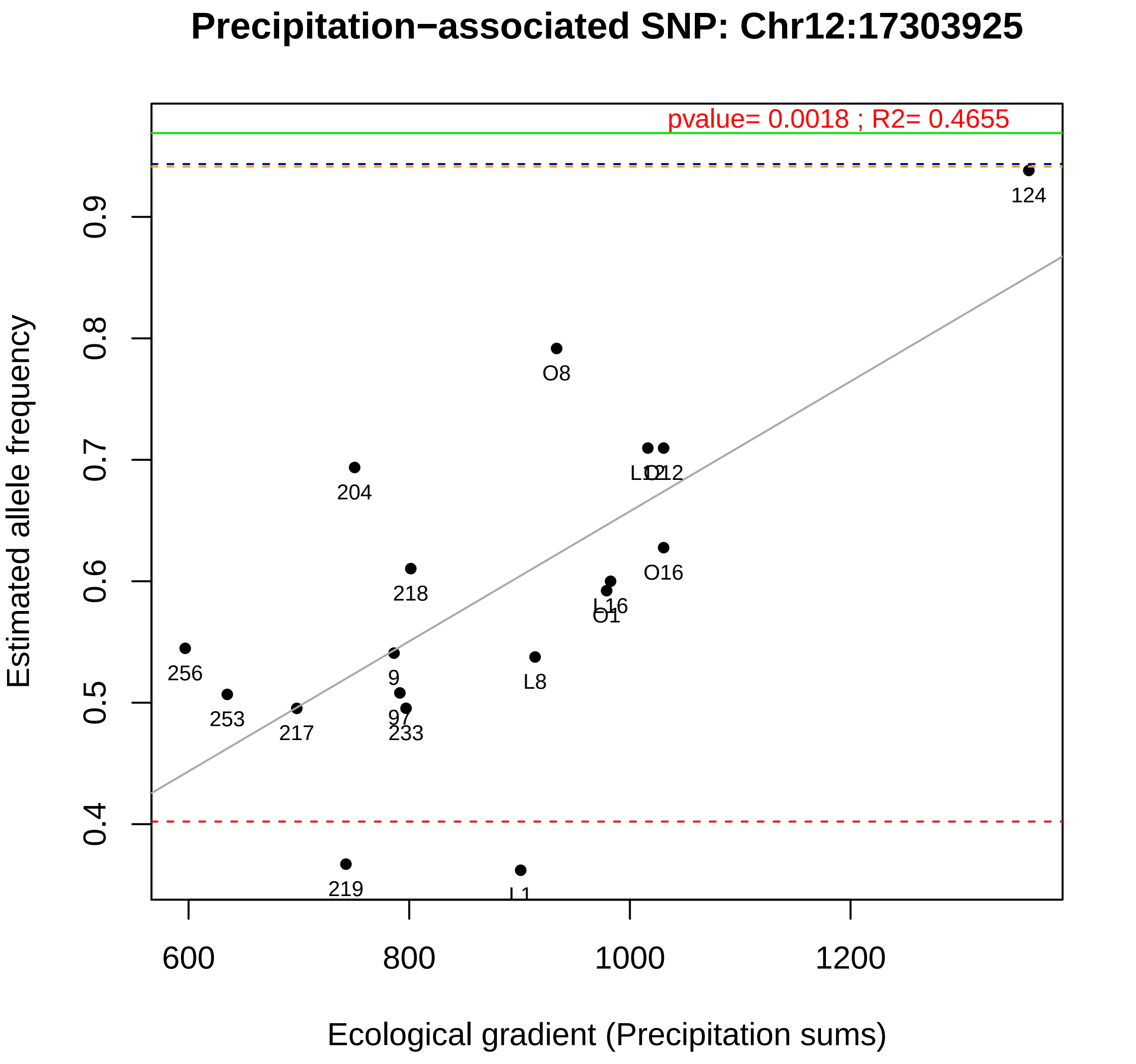

Precipitation−associated SNP: Chr12:17303925
pvalue= 0.0018 ; R2= 0.4655
Estimated allele frequency
0.4
0.5
0.6
0.7
0.8
0.9
600
800
1000
1200
Ecological gradient (Precipitation sums)
124
O8
204
L12
O12
O16
218
L16
O1
256
9
L8
253
217
97
233
219
L1

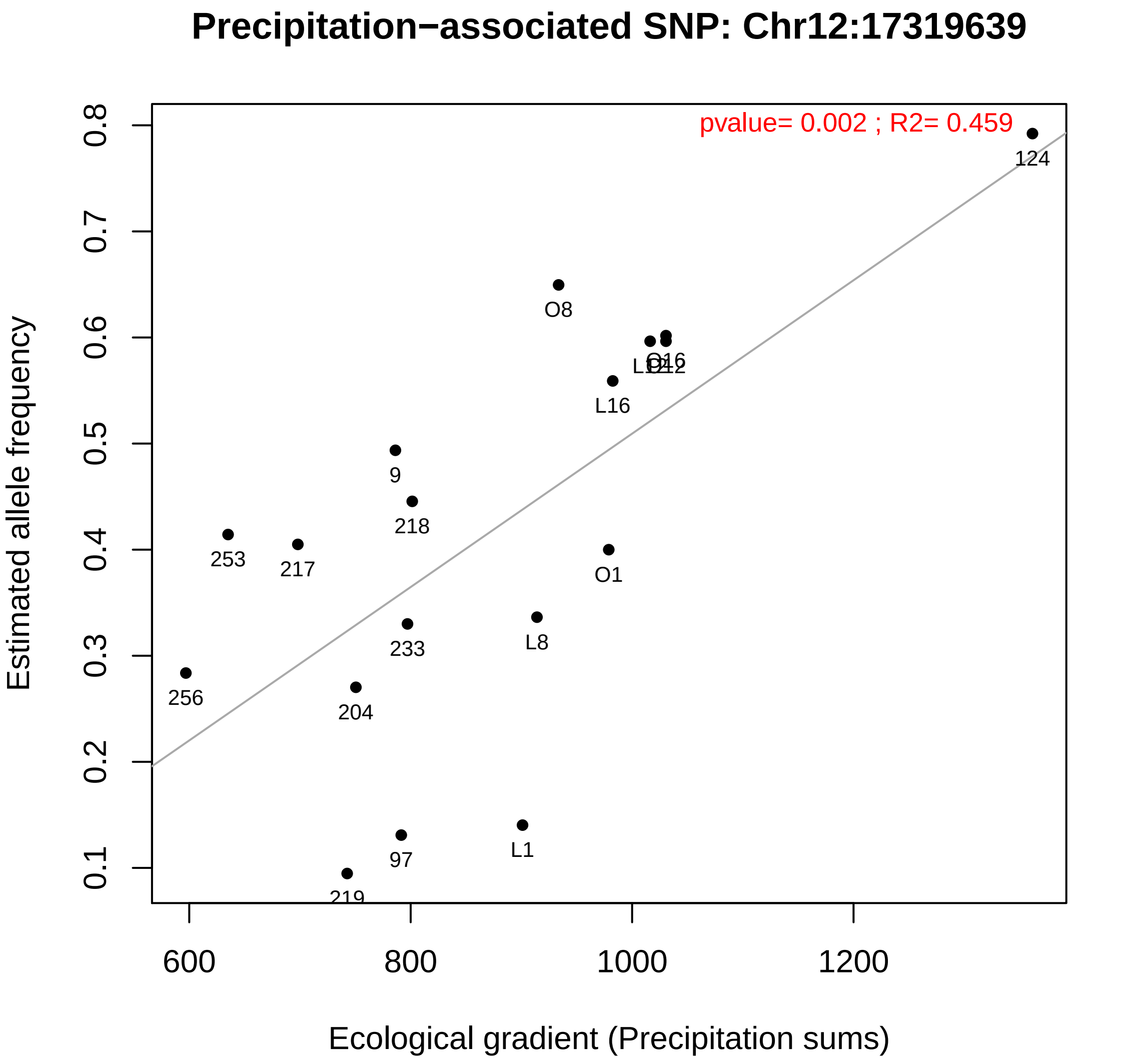
Precipitation−associated SNP: Chr12:17319639
pvalue= 0.002 ; R2= 0.459
Estimated allele frequency
0.1
0.2
0.3
0.4
0.5
0.6
0.7
0.8
600
800
1000
1200
Ecological gradient (Precipitation sums)
124
O8
O16
L16
9
218
253
217
O1
233
L8
256
204
97
L1
219

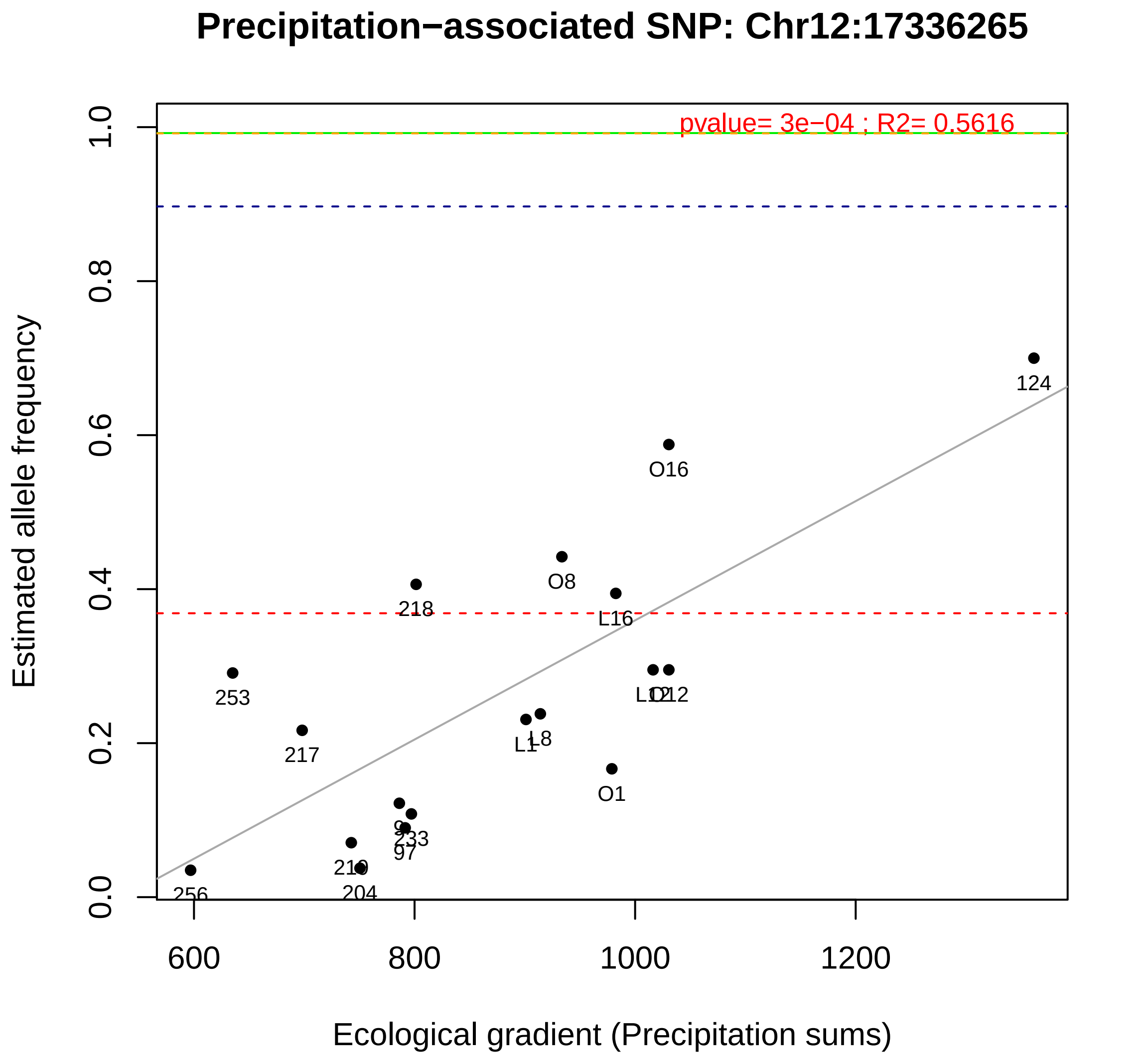
Precipitation−associated SNP: Chr12:17336265
pvalue= 3e−04 ; R2= 0.5616
Estimated allele frequency
0.0
0.2
0.4
0.6
0.8
1.0
600
800
1000
1200
Ecological gradient (Precipitation sums)
124
O16
O8
218
L16
253
217
O1
256
204
233
97

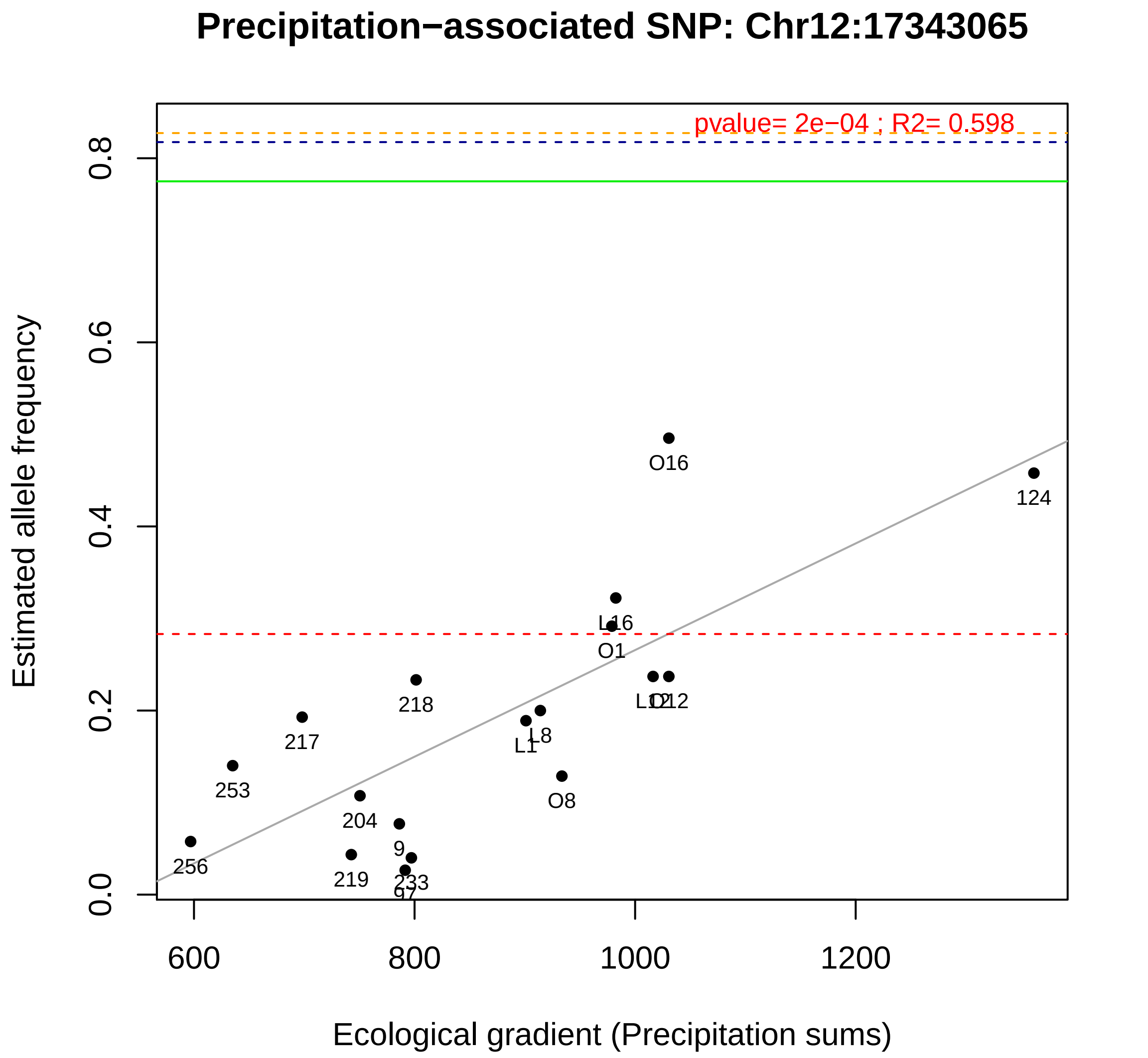

Precipitation−associated SNP: Chr12:17343065
pvalue= 2e−04 ; R2= 0.598
Estimated allele frequency
0.0
0.2
0.4
0.6
0.8
600
800
1000
1200
Ecological gradient (Precipitation sums)
O16
124
L16
O1
L12
O12
218
217
253
204
256
219
9
233
97
L1
L8
O8

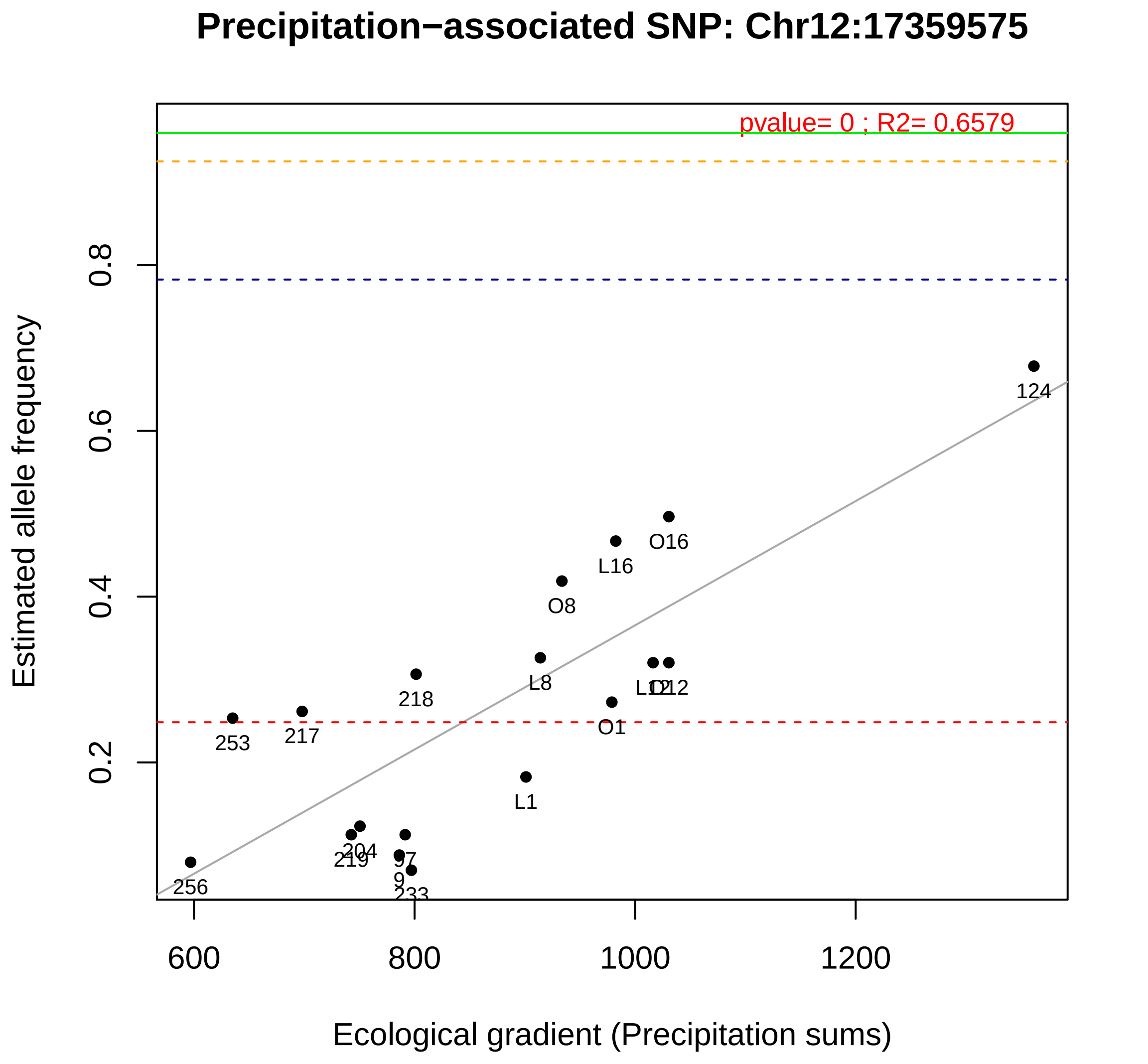

Precipitation−associated SNP: Chr12:17359575
pvalue= 0 ; R2= 0.6579
Estimated allele frequency
0.2
0.4
0.6
0.8
600
800
1000
1200
Ecological gradient (Precipitation sums)
124
O16
L16
O8
L8
218
253
217
O1
L1
256
219
204
97
9
233

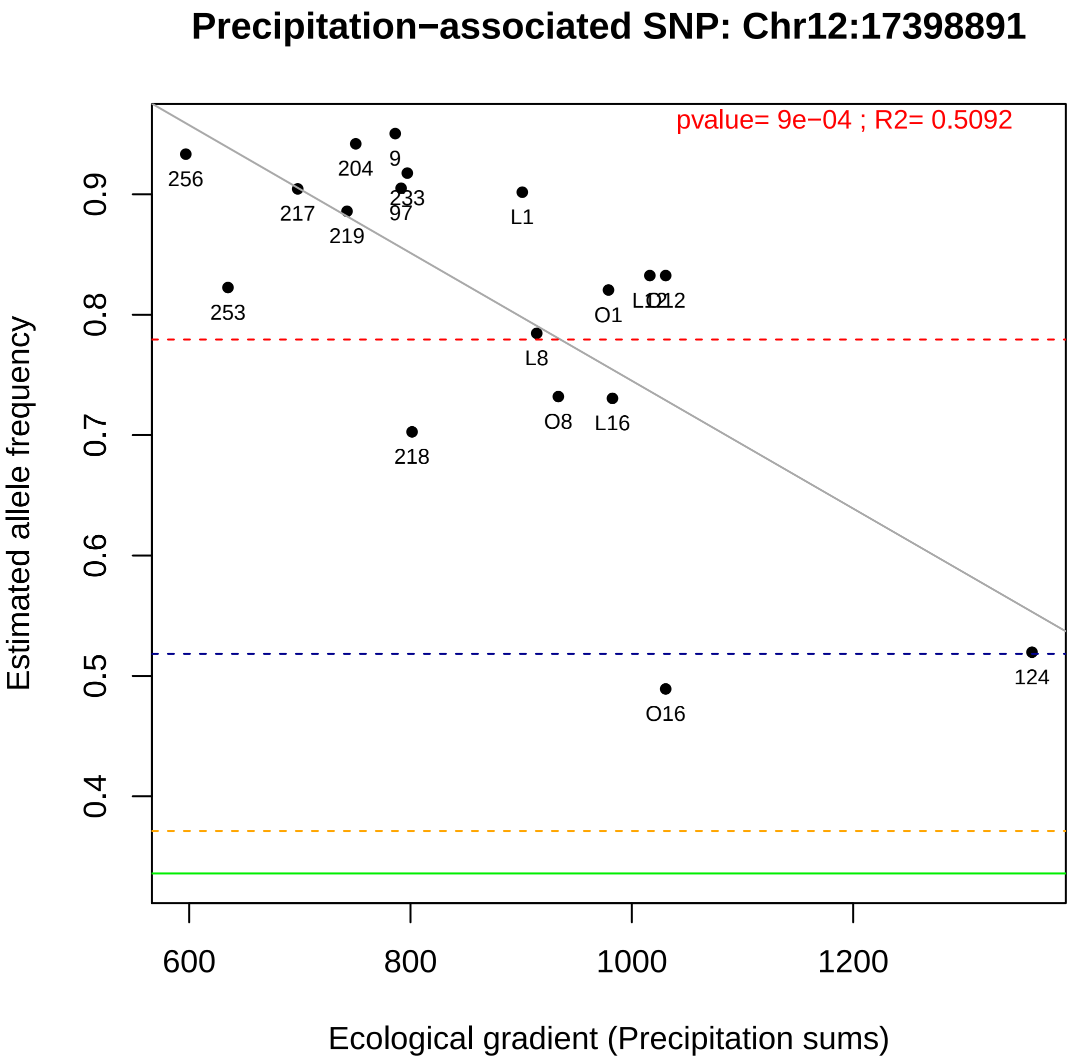

Precipitation−associated SNP: Chr12:17398891
pvalue= 9e−04 ; R2= 0.5092
Estimated allele frequency
Ecological gradient (Precipitation sums)
0.4
0.5
0.6
0.7
0.8
0.9
600
800
1000
1200
256
204
9
217
233
97
219
L1
253
O1
L8
O8
L16
218
124
O16

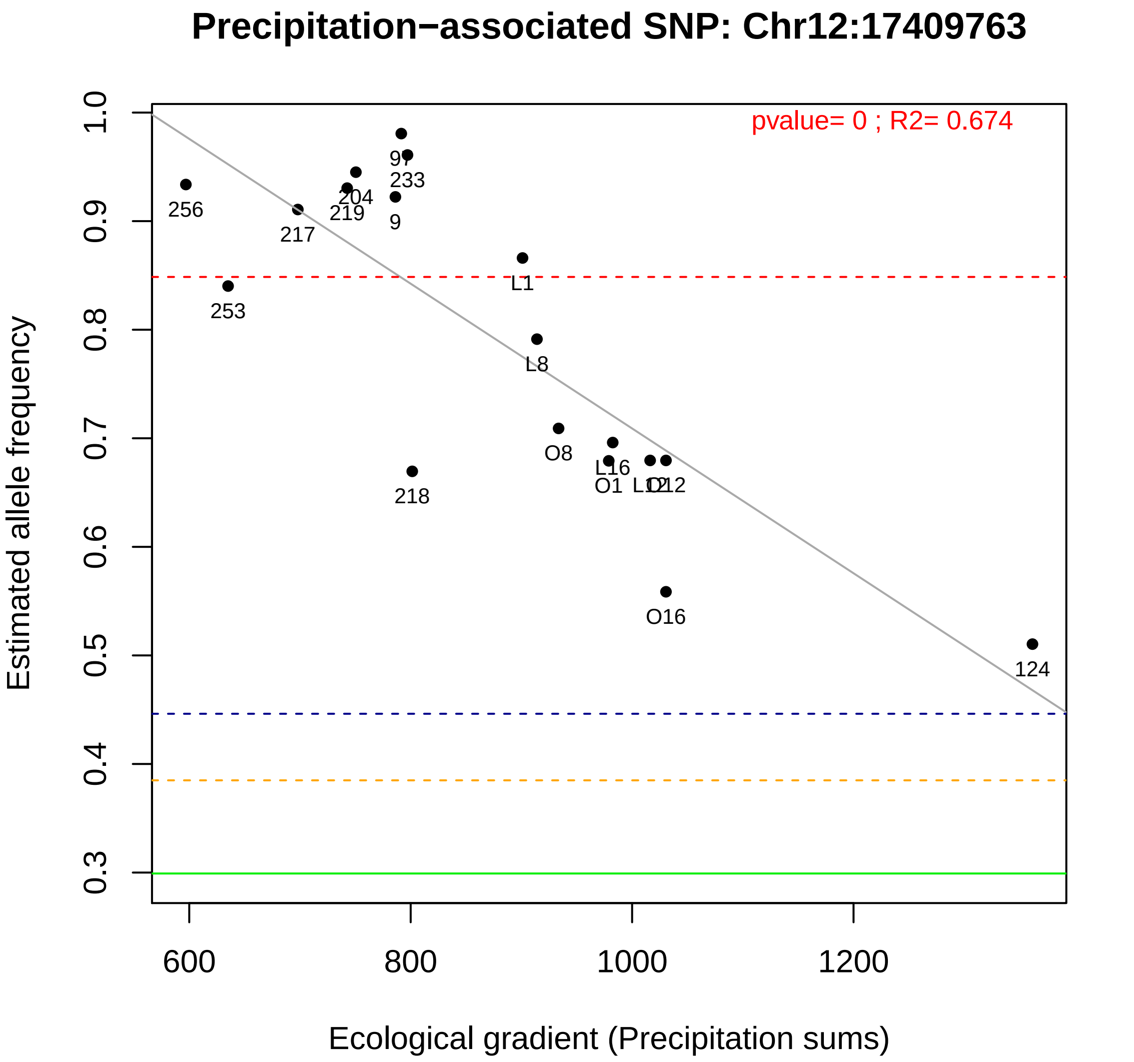

Precipitation−associated SNP: Chr12:17409763
pvalue= 0 ; R2= 0.674
Estimated allele frequency
Ecological gradient (Precipitation sums)
0.3
0.4
0.5
0.6
0.7
0.8
0.9
1.0
600
800
1000
1200
256
253
217
219
204
97
233
9
218
L1
L8
O8
L16
O1
L12
O12
O16
124

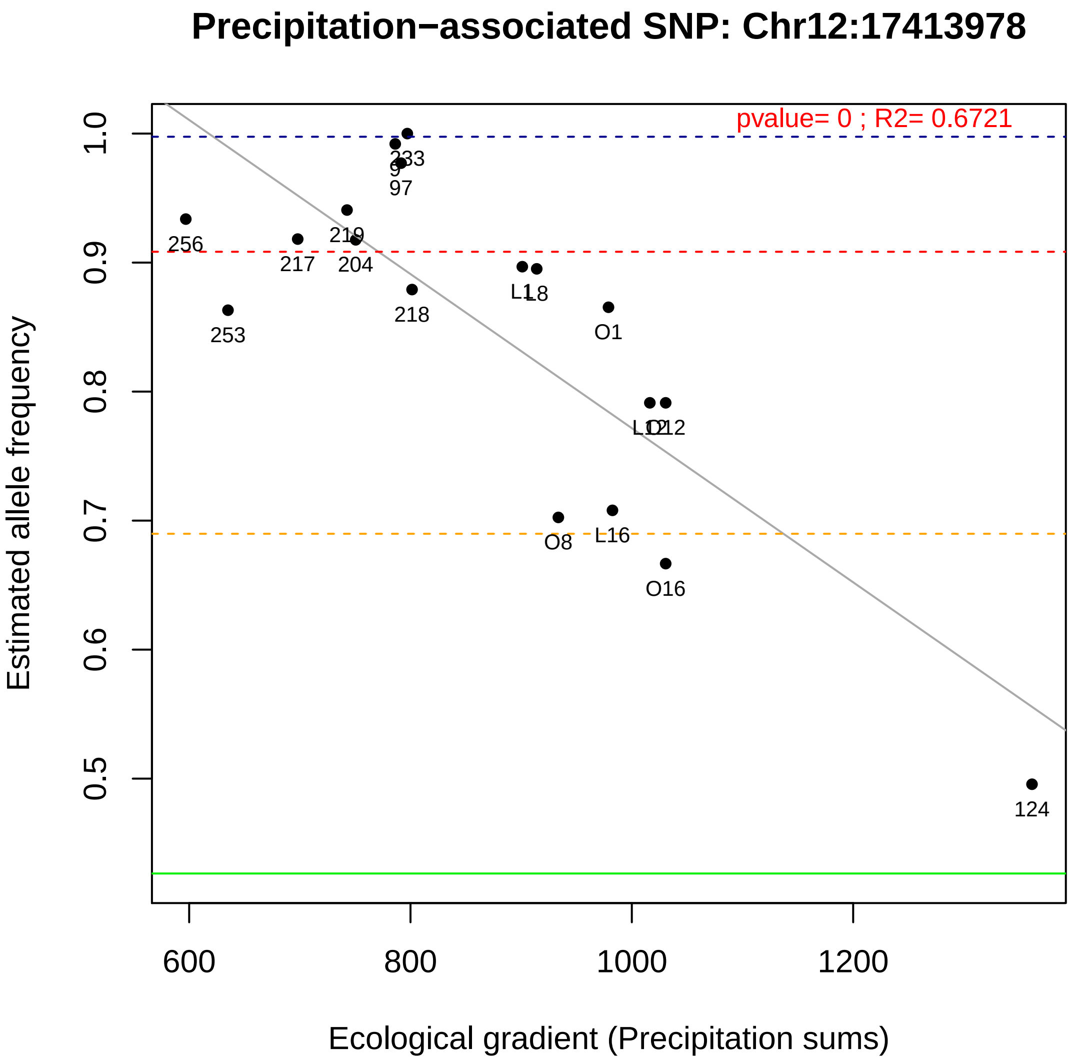

Precipitation−associated SNP: Chr12:17413978
pvalue= 0 ; R2= 0.6721
Estimated allele frequency
Ecological gradient (Precipitation sums)
1.0
0.9
0.8
0.7
0.6
0.5
600
800
1000
1200
233
97
256
219
217
204
218
L18
O1
O12
O8
L16
O16
124
253

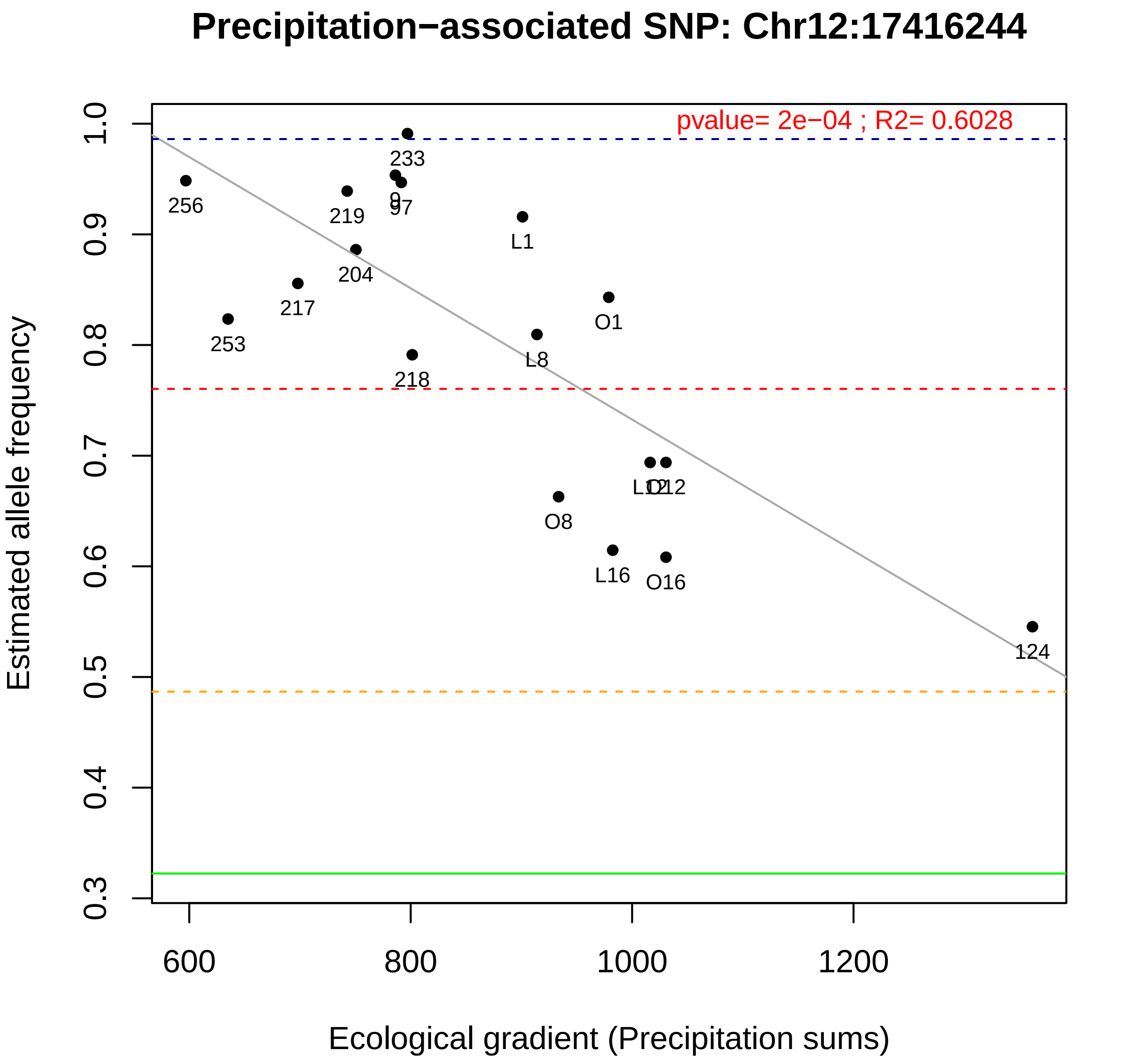

Precipitation−associated SNP: Chr12:17416244
pvalue= 2e−04 ; R2= 0.6028
Estimated allele frequency
0.3
0.4
0.5
0.6
0.7
0.8
0.9
1.0
600
800
1000
1200
Ecological gradient (Precipitation sums)
233
256
219
9
97
L1
204
217
253
O1
L8
218
L12
O12
O8
L16
O16
124

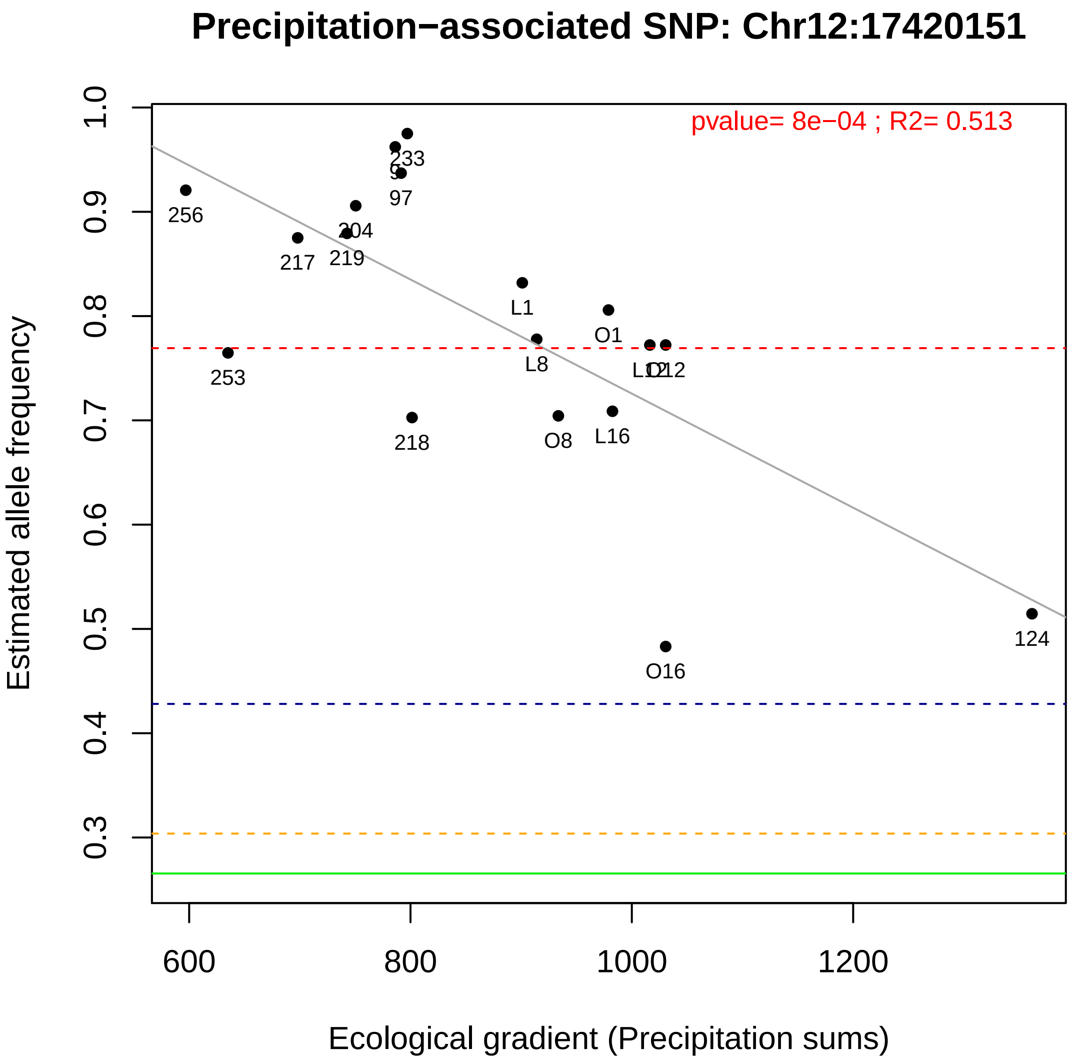

Precipitation−associated SNP: Chr12:17420151
pvalue= 8e−04 ; R2= 0.513
Estimated allele frequency
Ecological gradient (Precipitation sums)
0.3
0.4
0.5
0.6
0.7
0.8
0.9
1.0
600
800
1000
1200
256
253
217
219
204
233
97
218
L1
L8
O1
O8
L16
O16
124

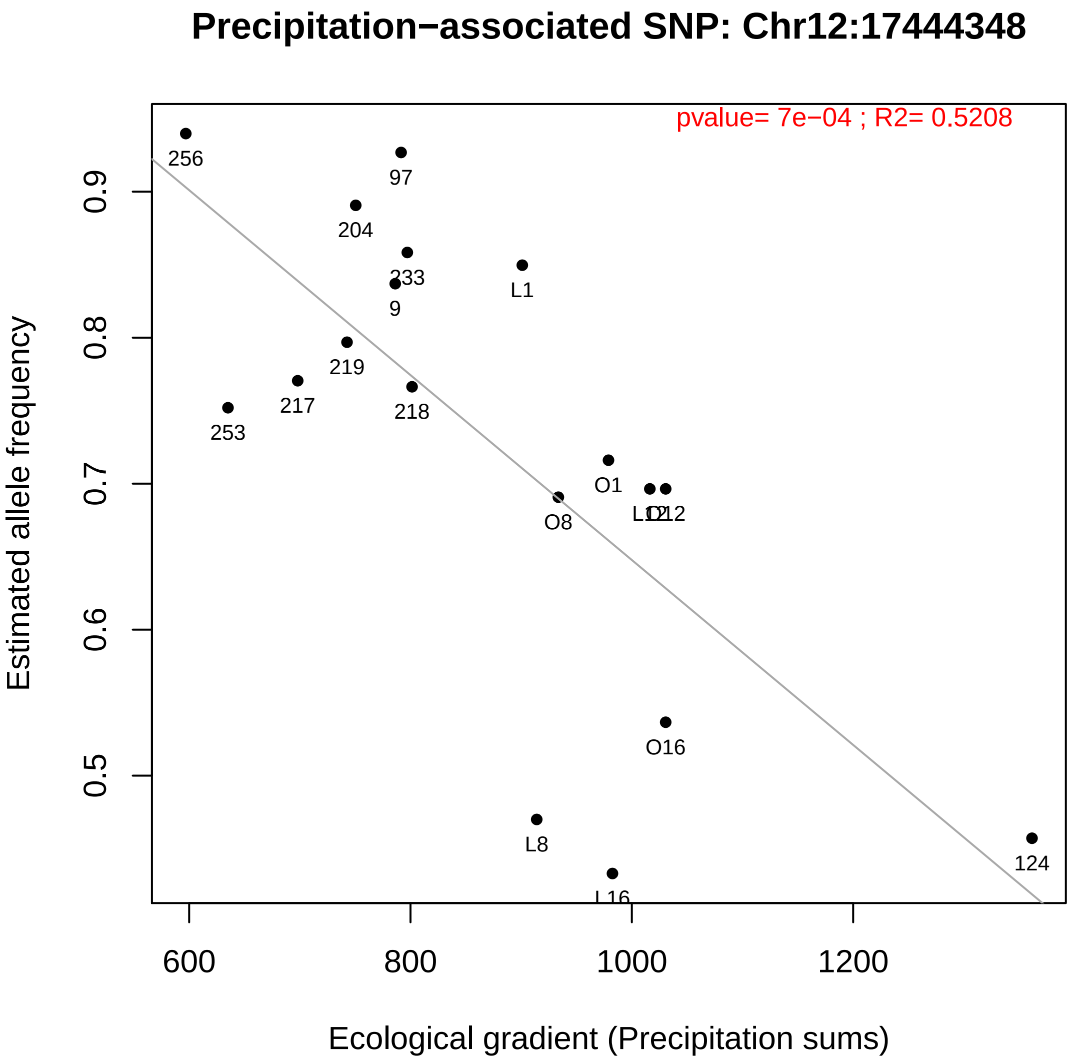

Precipitation−associated SNP: Chr12:17444348
pvalue= 7e−04 ; R2= 0.5208
Estimated allele frequency
0.5
0.6
0.7
0.8
0.9
600
800
1000
1200
Ecological gradient (Precipitation sums)
256
97
204
233
9
L1
219
217
218
253
O1
O8
L12
O12
O16
L8
L16
124

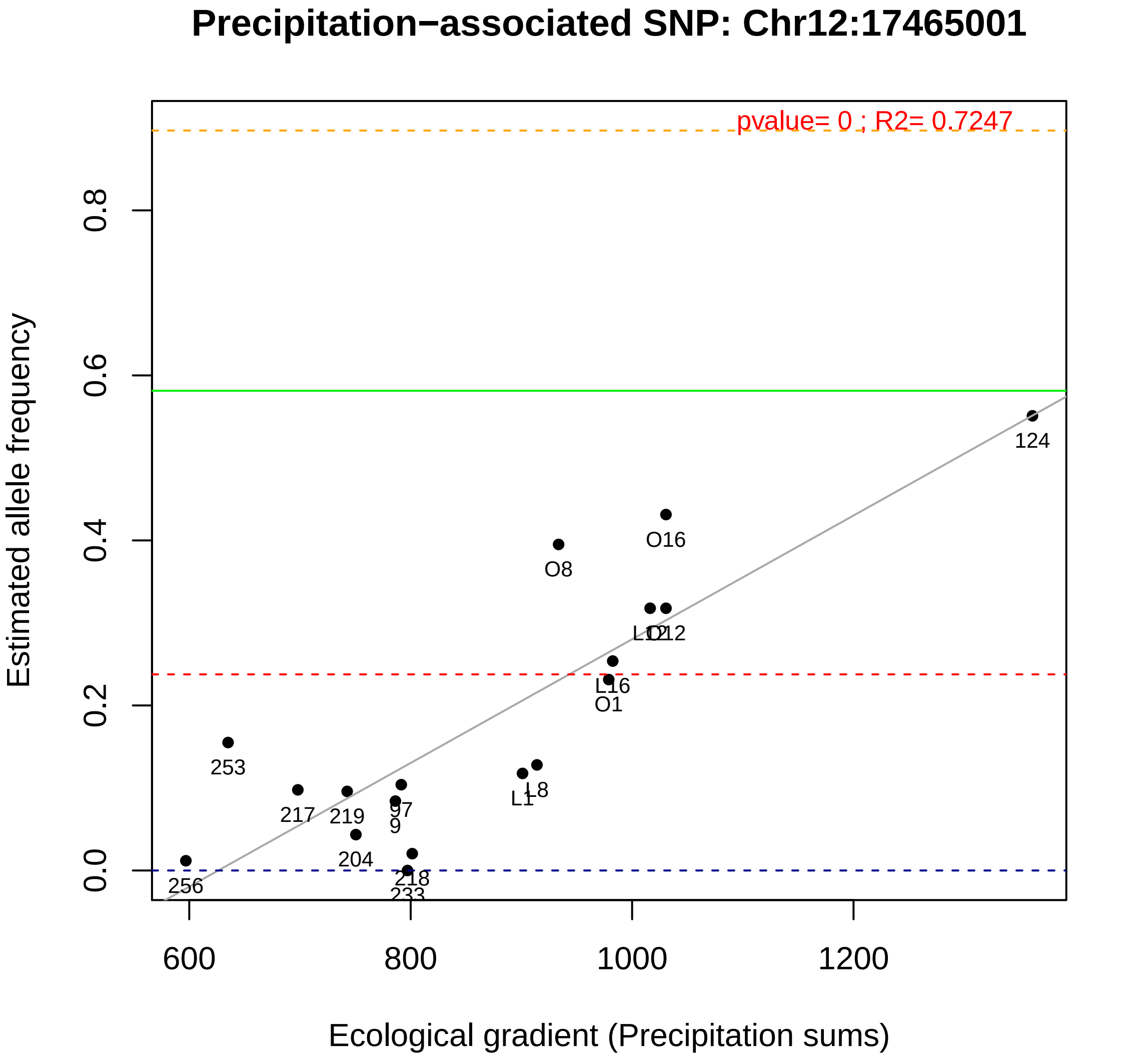
Precipitation−associated SNP: Chr12:17465001
pvalue= 0 ; R2= 0.7247
Estimated allele frequency
Ecological gradient (Precipitation sums)
0.0
0.2
0.4
0.6
0.8
600
800
1000
1200
124
O16
O8
O12
L16
O1
253
L8
L1
217
219
97
9
204
218
233
256

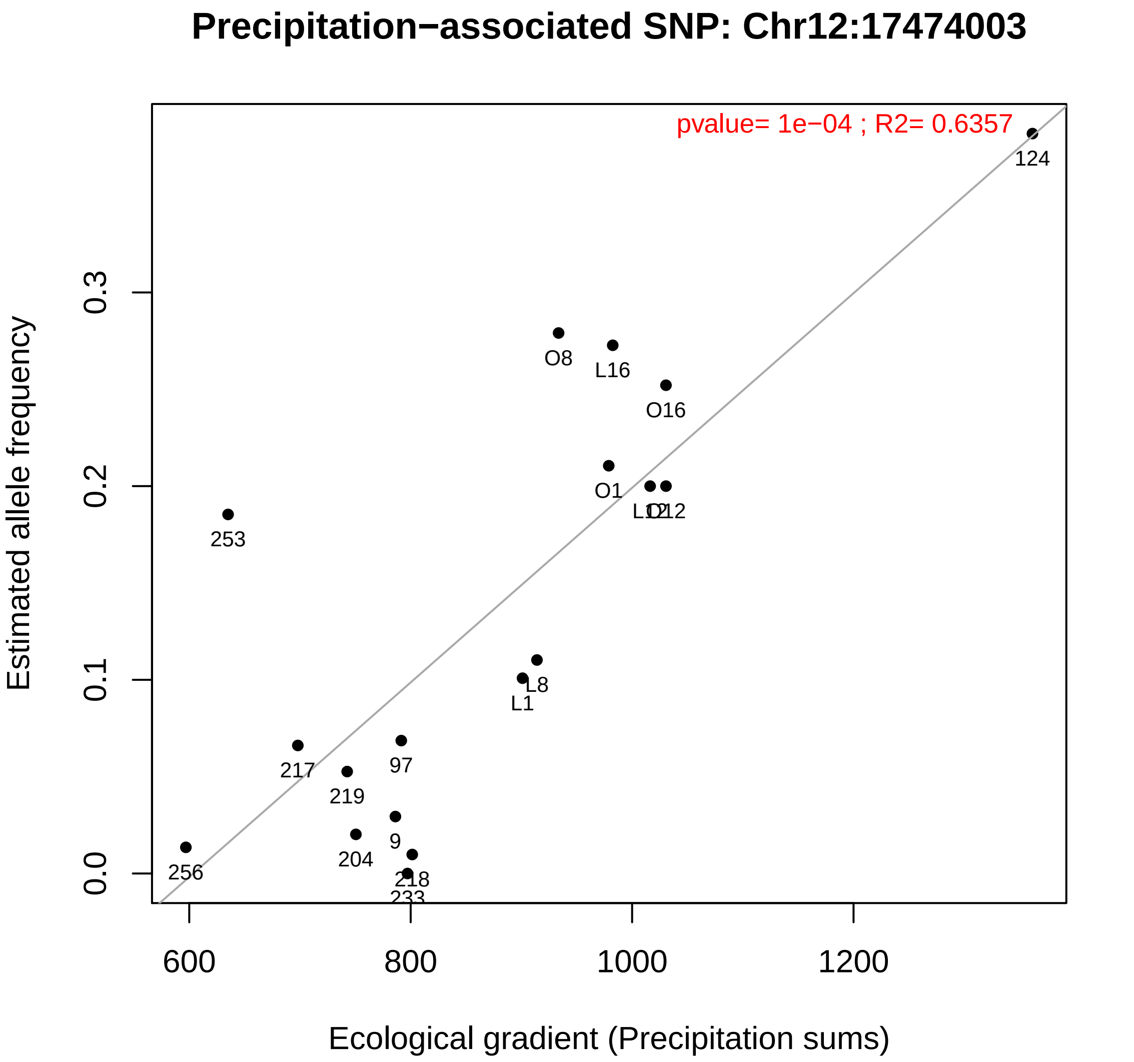
Precipitation−associated SNP: Chr12:17474003
pvalue= 1e−04 ; R2= 0.6357
124
O8
L16
O16
O1
L10
O12
253
L8
L1
217
97
219
9
204
256
218
233
0.0
0.1
0.2
0.3
600
800
1000
1200
Estimated allele frequency
Ecological gradient (Precipitation sums)

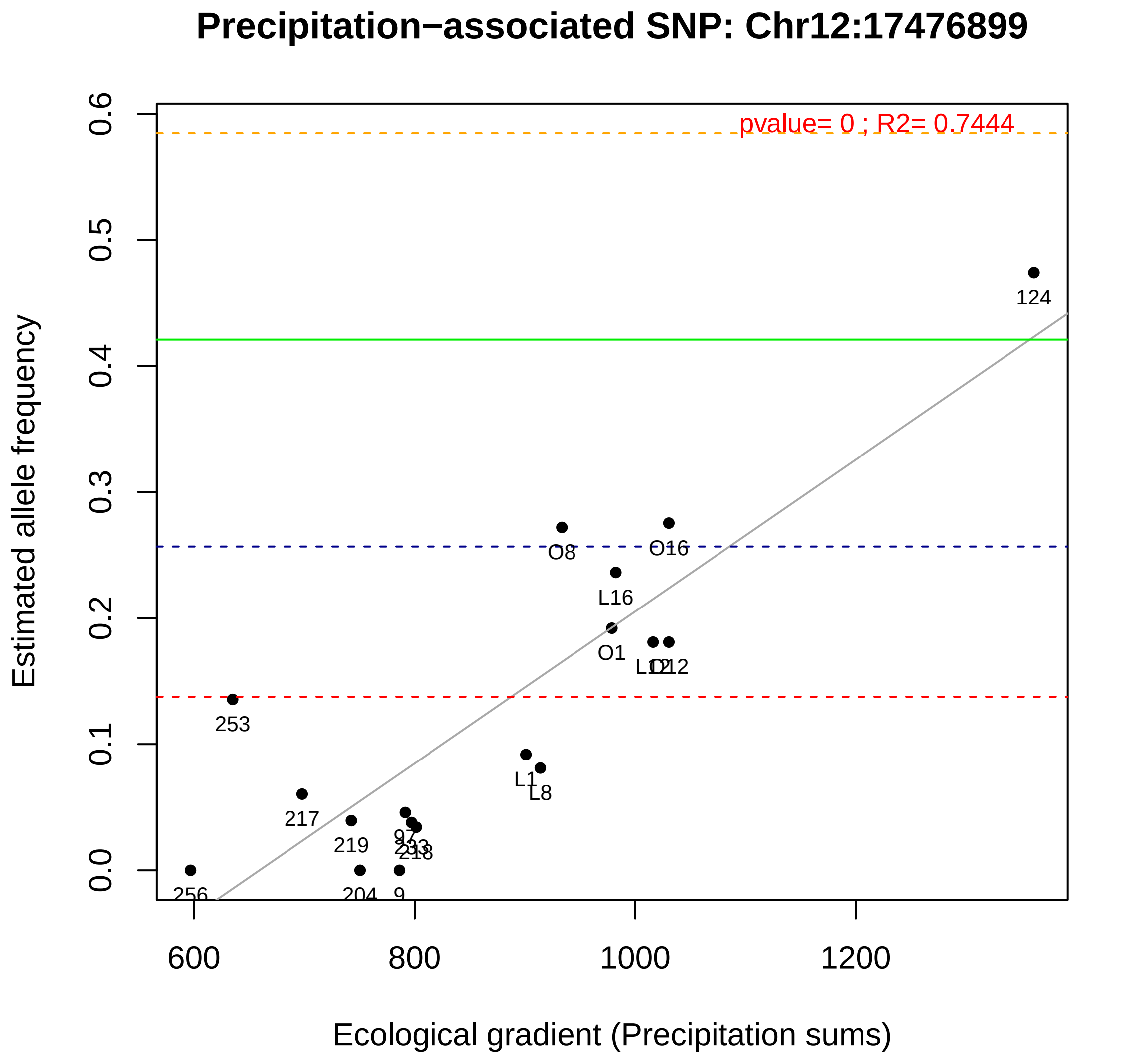

Precipitation−associated SNP: Chr12:17476899
pvalue= 0 ; R2= 0.7444
Estimated allele frequency
Ecological gradient (Precipitation sums)
0.0
0.1
0.2
0.3
0.4
0.5
0.6
600
800
1000
1200
124
O8
O16
L16
O1
L12
O12
253
L1
L8
217
219
97
233
218
256
204
9

**Precipitation−associated SNP: Chr12:17487856**

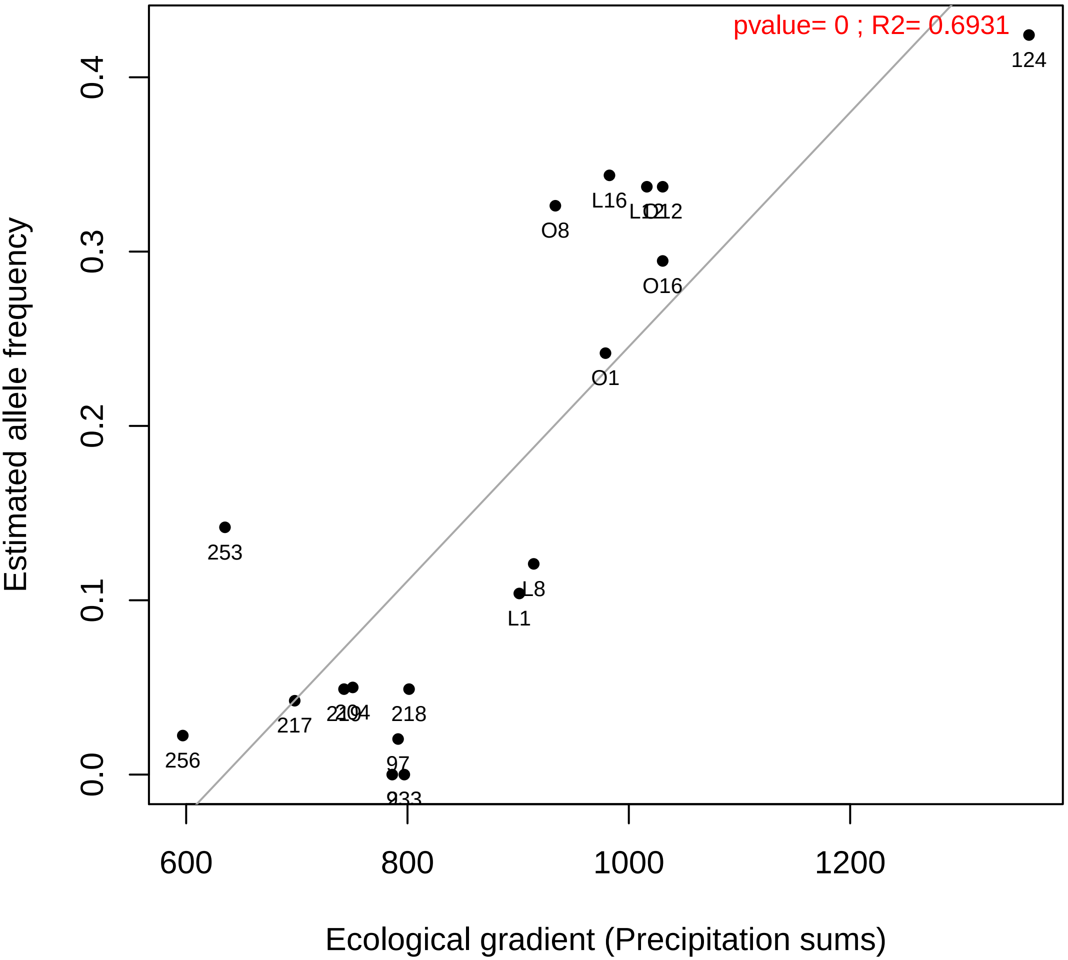

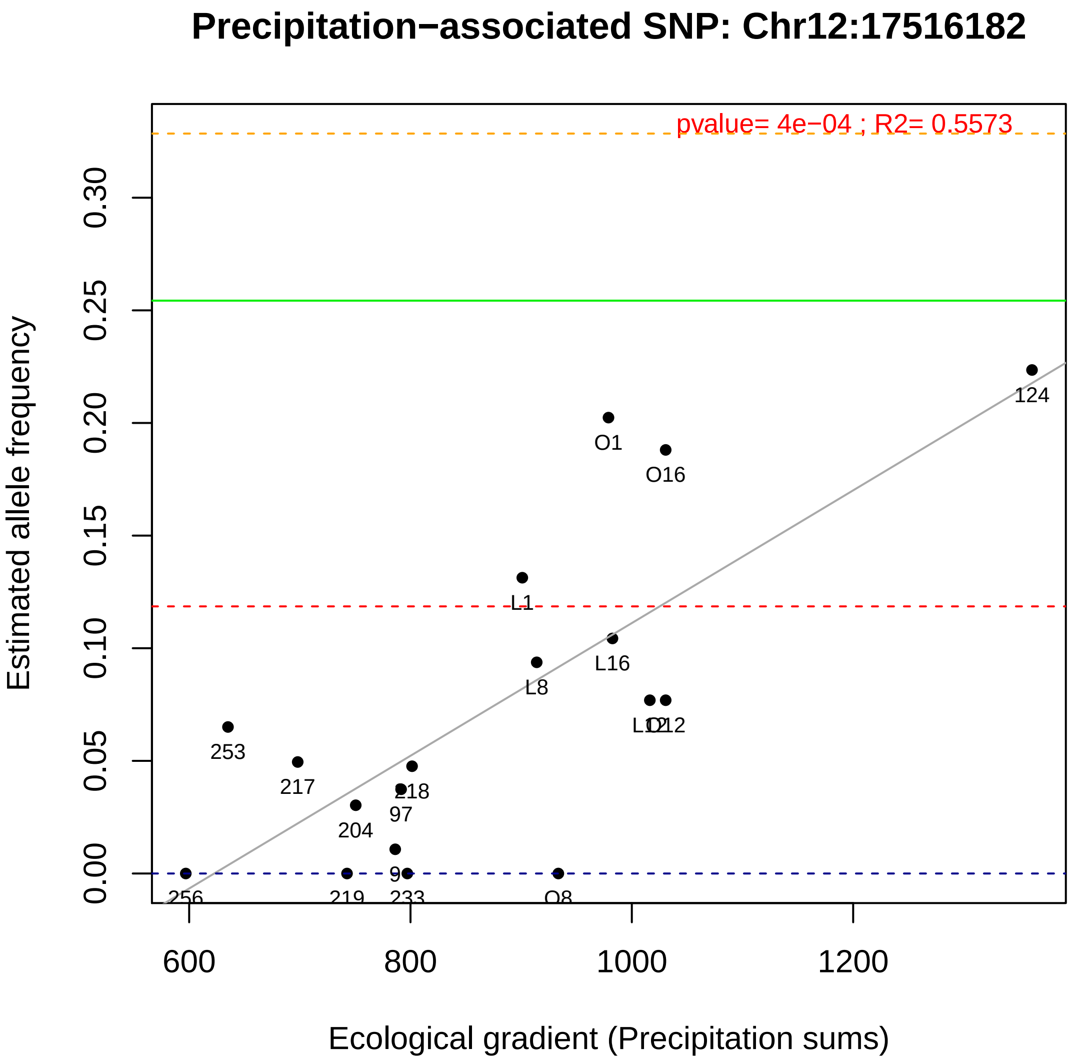

Precipitation−associated SNP: Chr12:17516182
pvalue= 4e−04 ; R2= 0.5573
Estimated allele frequency
0.00
0.05
0.10
0.15
0.20
0.25
0.30
600
800
1000
1200
Ecological gradient (Precipitation sums)
124
O1
O16
L1
L16
L8
253
217
218
97
204
219
233
O8
256

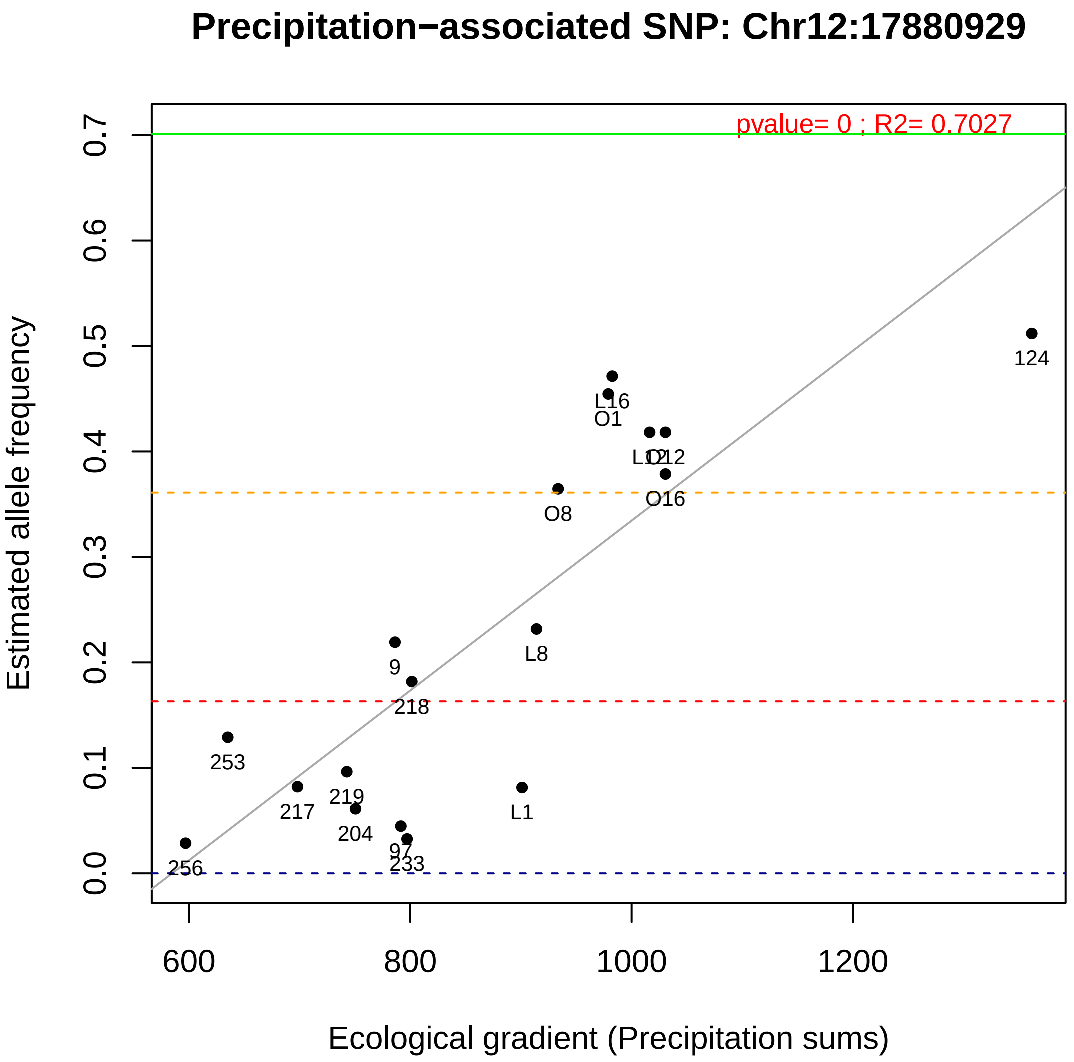

Precipitation−associated SNP: Chr12:17880929
pvalue= 0 ; R2= 0.7027
Estimated allele frequency
Ecological gradient (Precipitation sums)
0.0
0.1
0.2
0.3
0.4
0.5
0.6
0.7
600
800
1000
1200
124
L16
O1
L12
O12
O16
O8
L8
9
218
253
219
217
204
L1
97
233
256

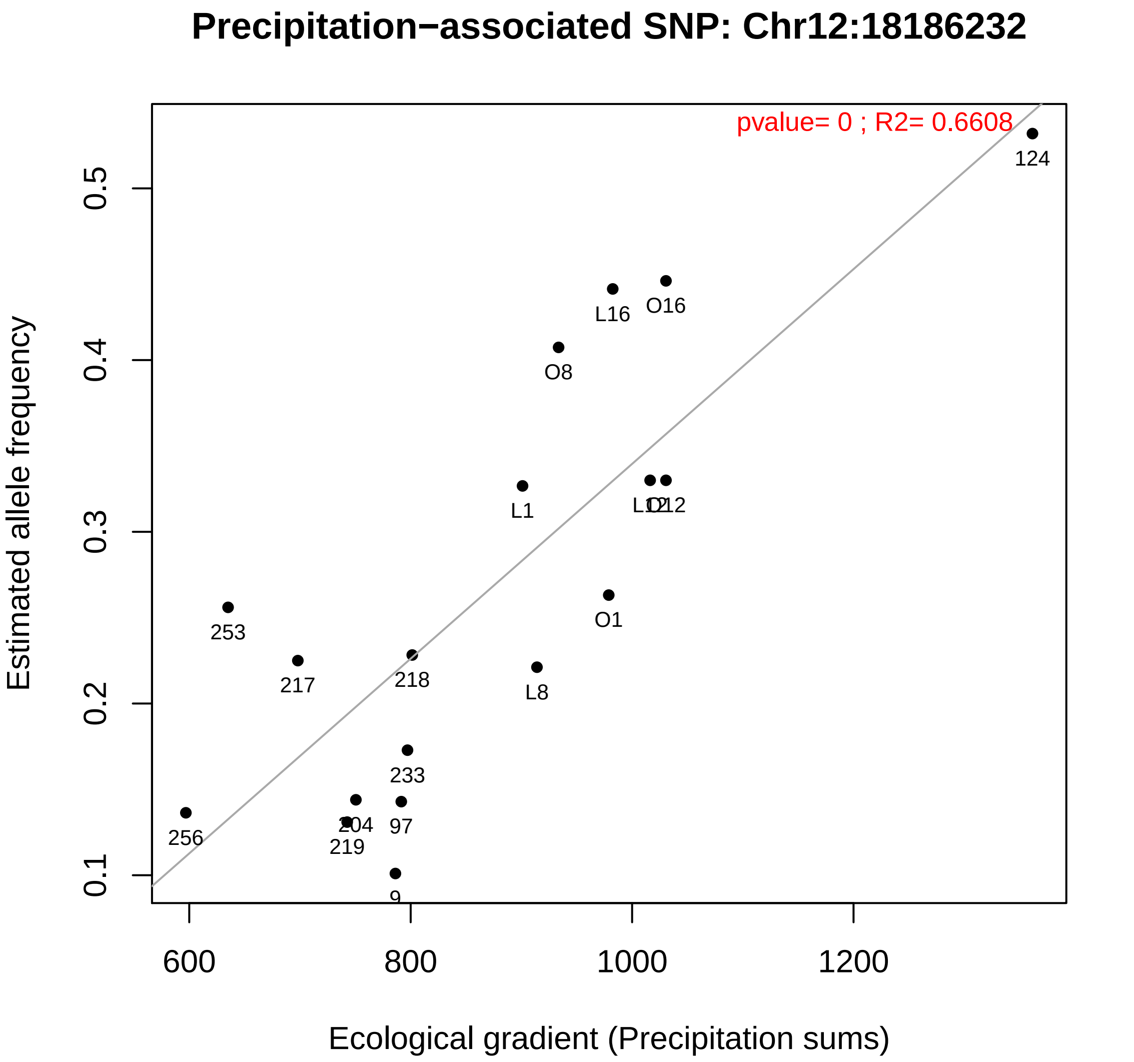

Precipitation−associated SNP: Chr12:18186232
pvalue= 0 ; R2= 0.6608
124
O16
L16
O8
L1
L12
O12
O1
253
217
218
L8
233
204
219
97
256
9
0.5
0.4
0.3
0.2
0.1
600
800
1000
1200
Estimated allele frequency
Ecological gradient (Precipitation sums)

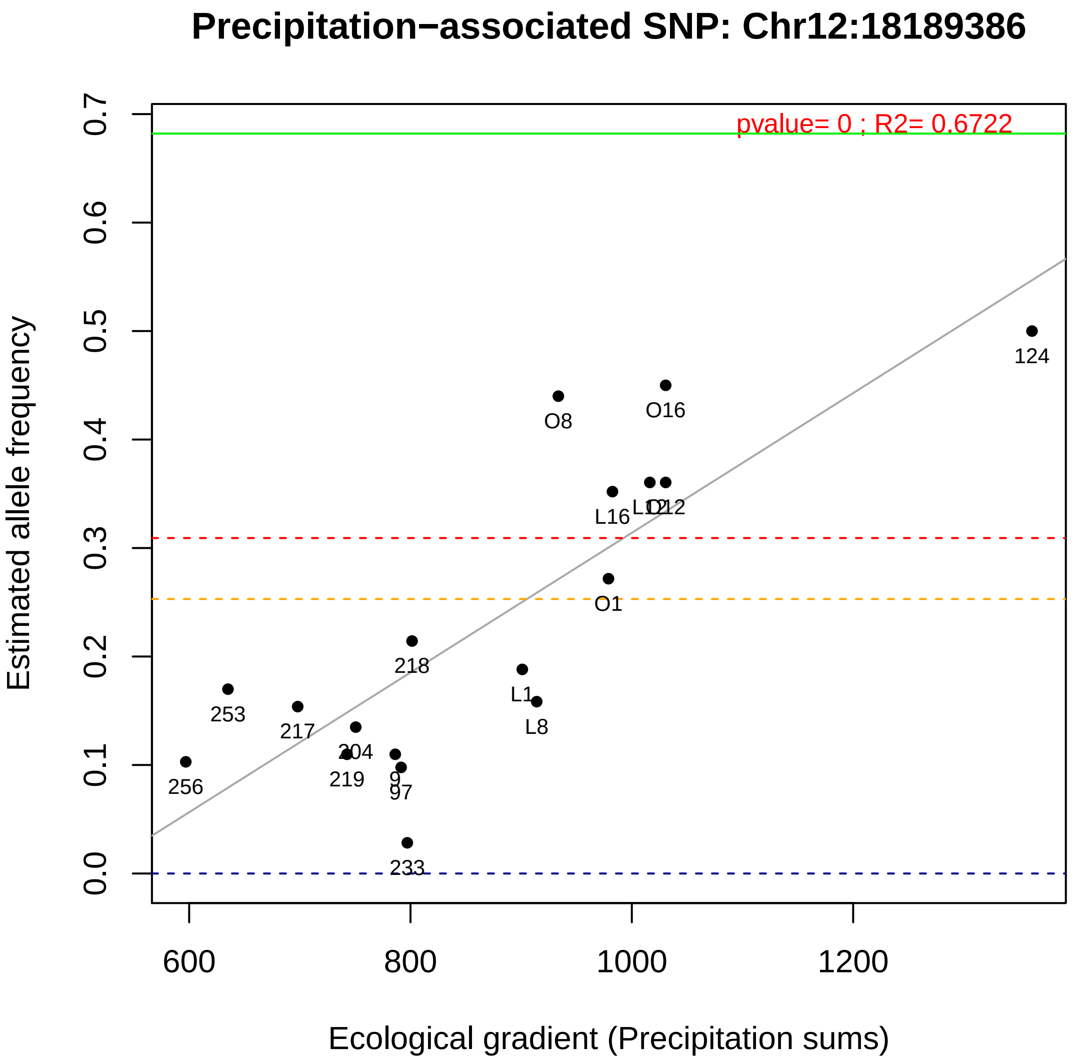
Precipitation−associated SNP: Chr12:18189386
pvalue= 0 ; R2= 0.6722
Estimated allele frequency
0.0
0.1
0.2
0.3
0.4
0.5
0.6
0.7
600
800
1000
1200
Ecological gradient (Precipitation sums)
124
O8
O16
L16
O1
218
L1
L8
253
217
204
219
256
97
233

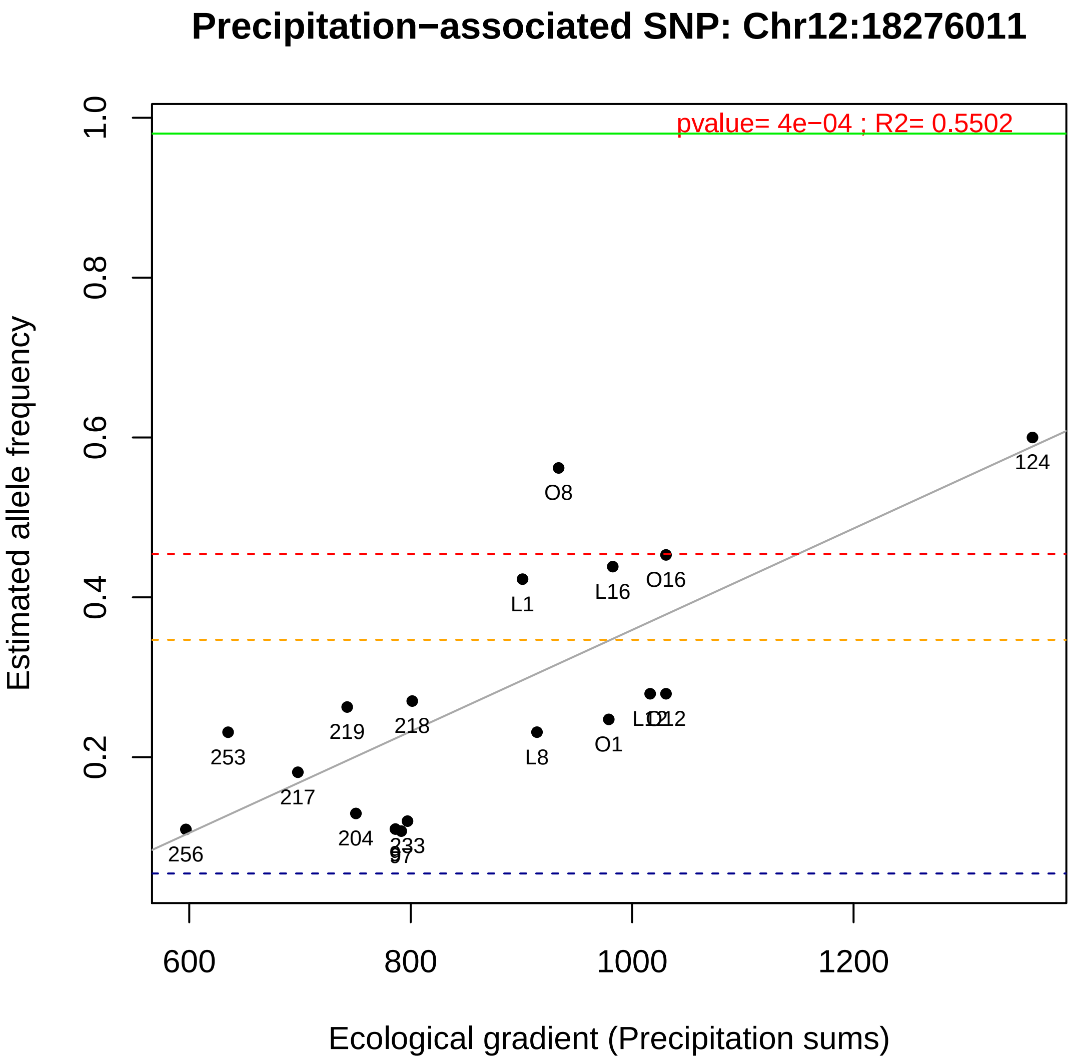

Precipitation−associated SNP: Chr12:18276011
pvalue= 4e−04 ; R2= 0.5502
Estimated allele frequency
0.2
0.4
0.6
0.8
1.0
600
800
1000
1200
Ecological gradient (Precipitation sums)
124
O8
O16
L16
L1
L12
O12
O1
L8
219
218
253
217
204
233
97
256

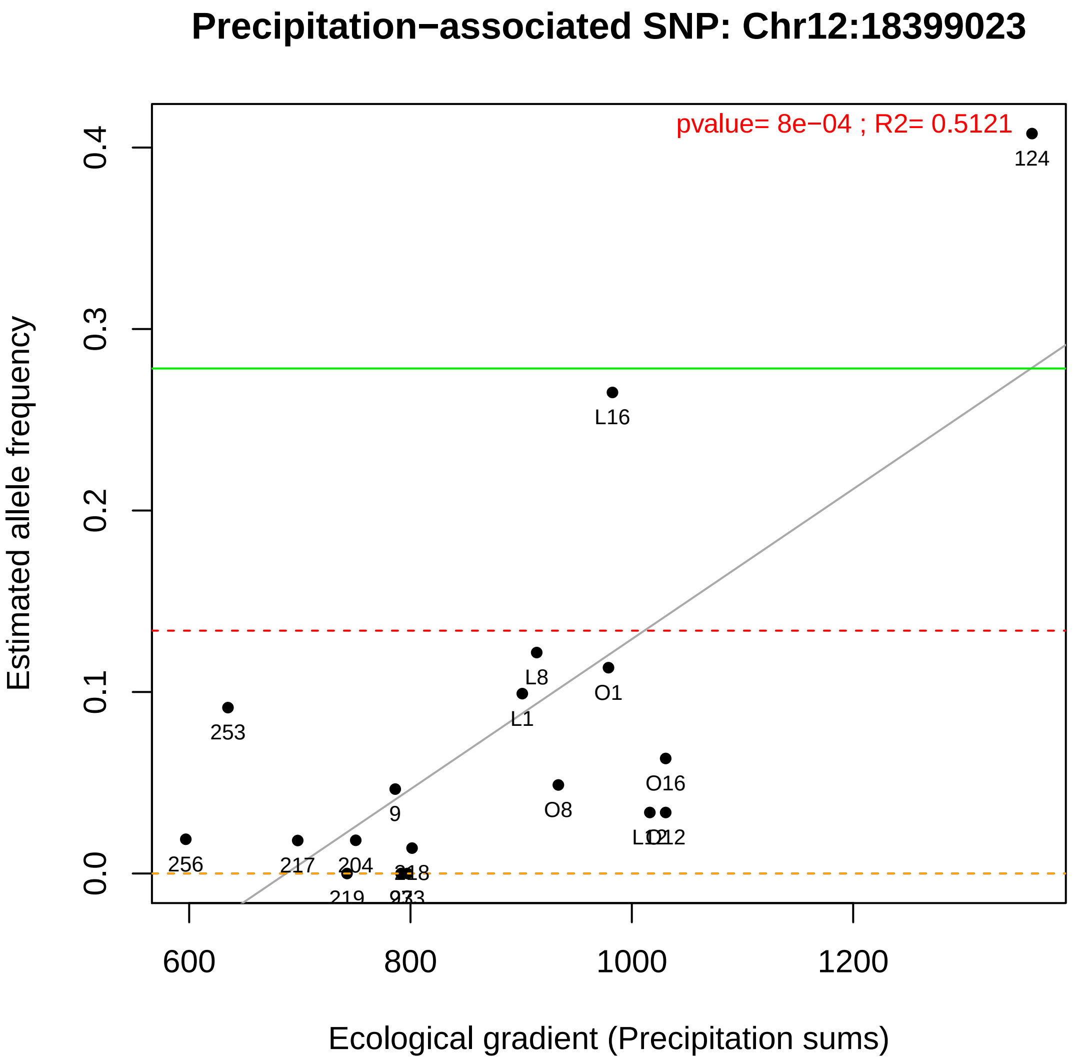
Precipitation−associated SNP: Chr12:18399023
pvalue= 8e−04 ; R2= 0.5121
Estimated allele frequency
0.0
0.1
0.2
0.3
0.4
Ecological gradient (Precipitation sums)
600
800
1000
1200
124
L16
L8
O1
L1
253
O16
9
O8
256
217
204
219

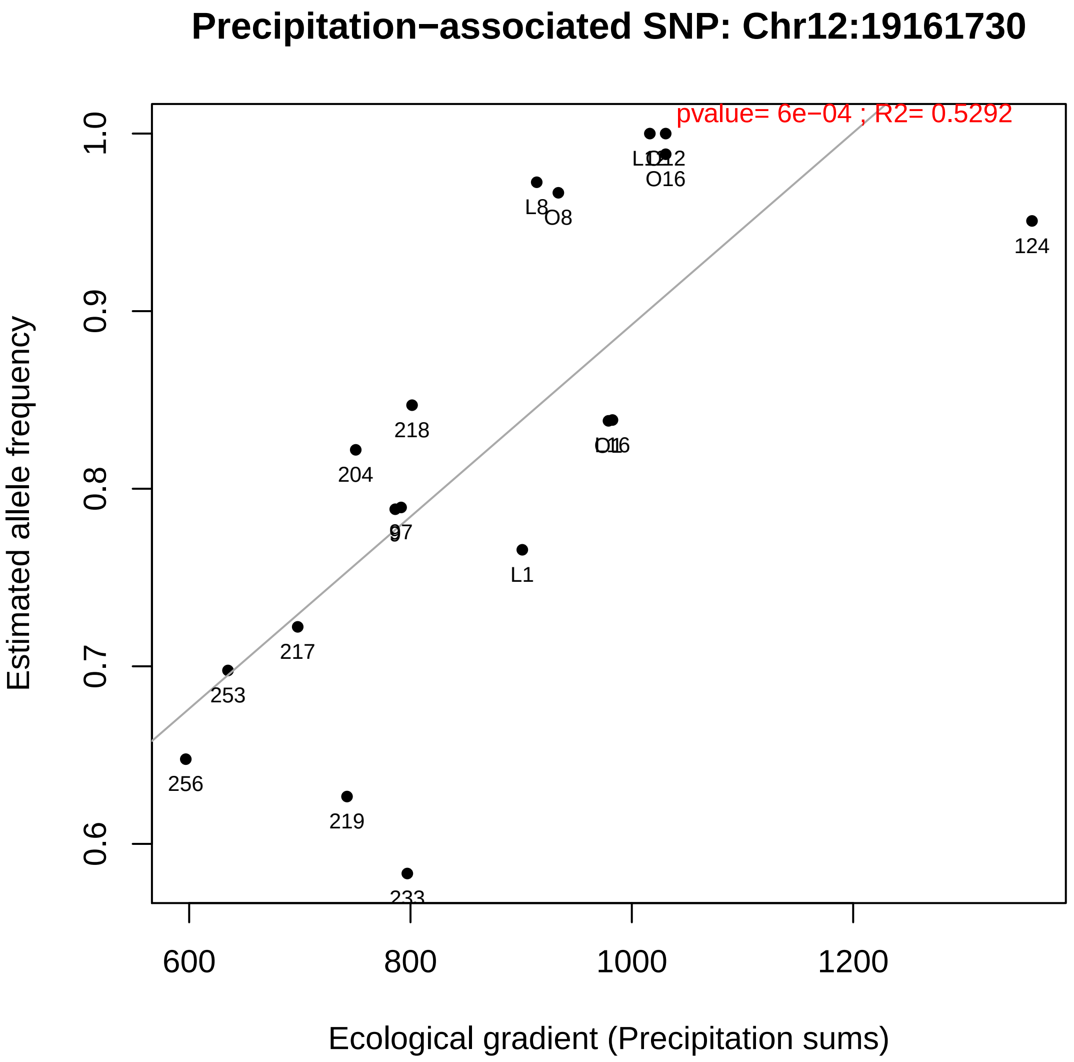
Precipitation−associated SNP: Chr12:19161730
pvalue= 6e−04 ; R2= 0.5292
Estimated allele frequency
Ecological gradient (Precipitation sums)
0.6
0.7
0.8
0.9
1.0
600
800
1000
1200
O16
L8
O8
124
218
204
217
253
256
219
233
L1

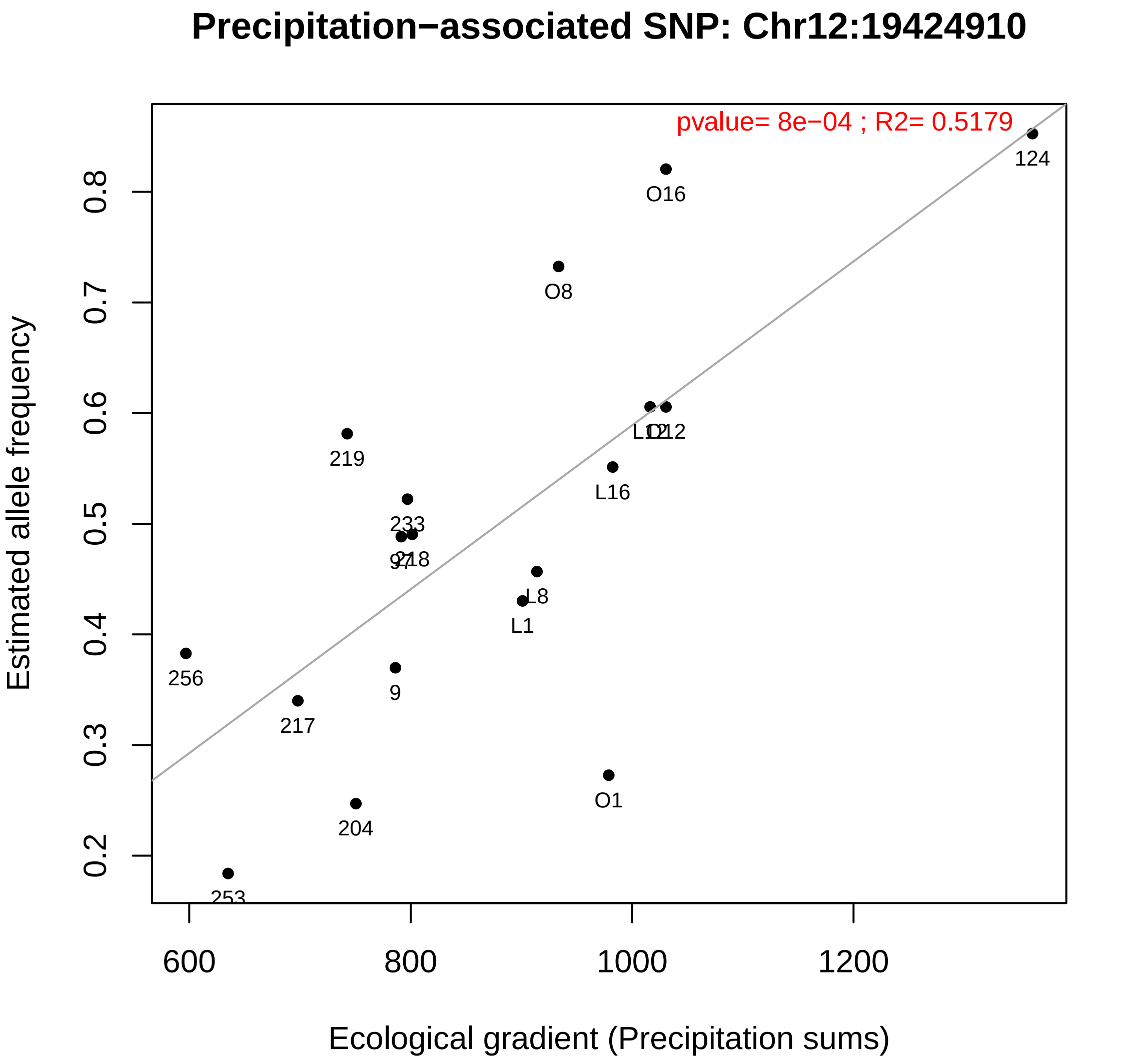
Precipitation−associated SNP: Chr12:19424910
pvalue= 8e−04 ; R2= 0.5179
Estimated allele frequency
0.2
0.3
0.4
0.5
0.6
0.7
0.8
600
800
1000
1200
Ecological gradient (Precipitation sums)
124
O16
O8
219
233
218
L16
L8
L1
256
9
217
O1
204
253

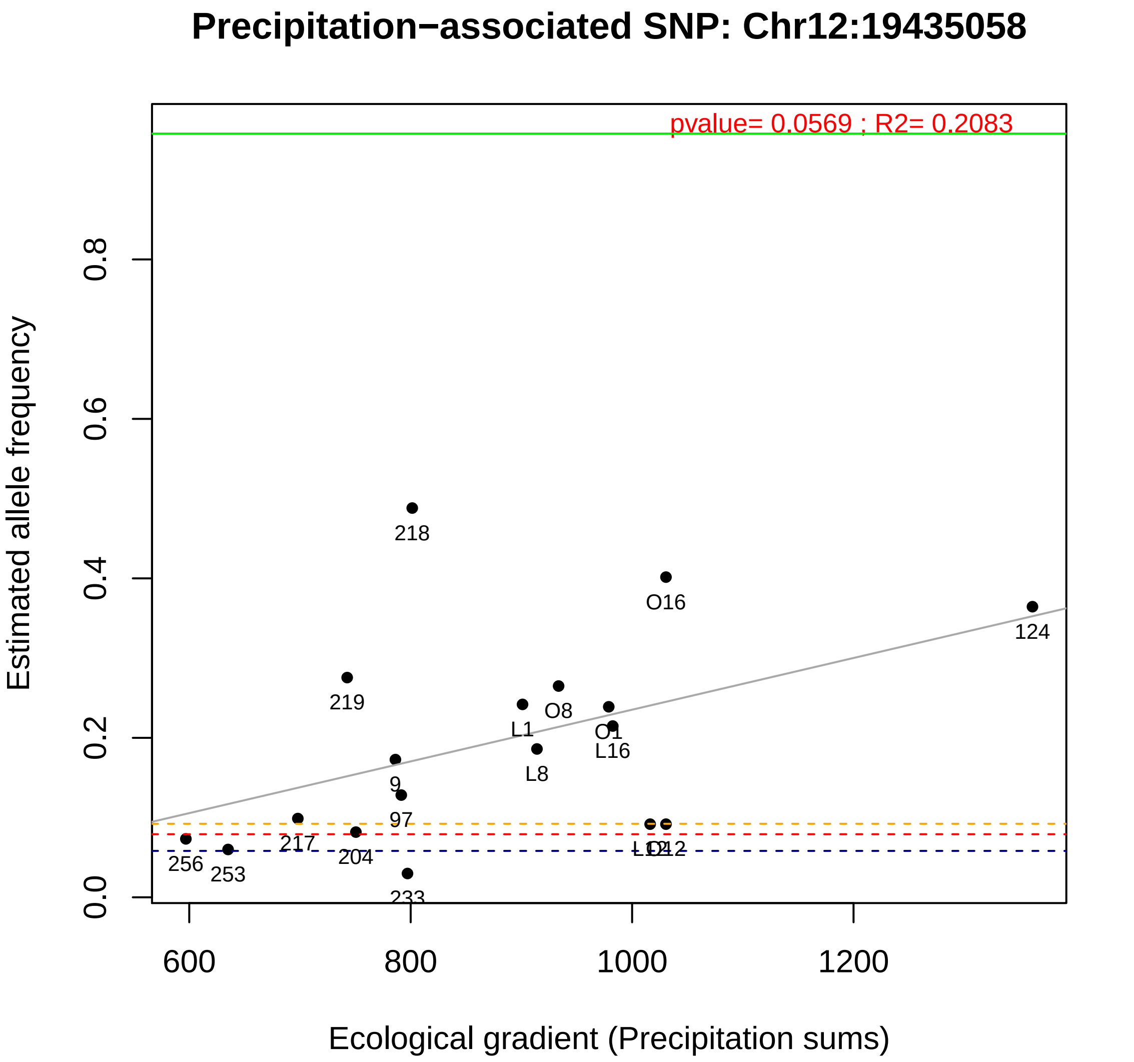

Precipitation−associated SNP: Chr12:19435058
pvalue= 0.0569 ; R2= 0.2083
Estimated allele frequency
0.0
0.2
0.4
0.6
0.8
600
800
1000
1200
Ecological gradient (Precipitation sums)
218
O16
124
219
O8
L1
O1
L16
L8
9
97
217
204
256
253
233

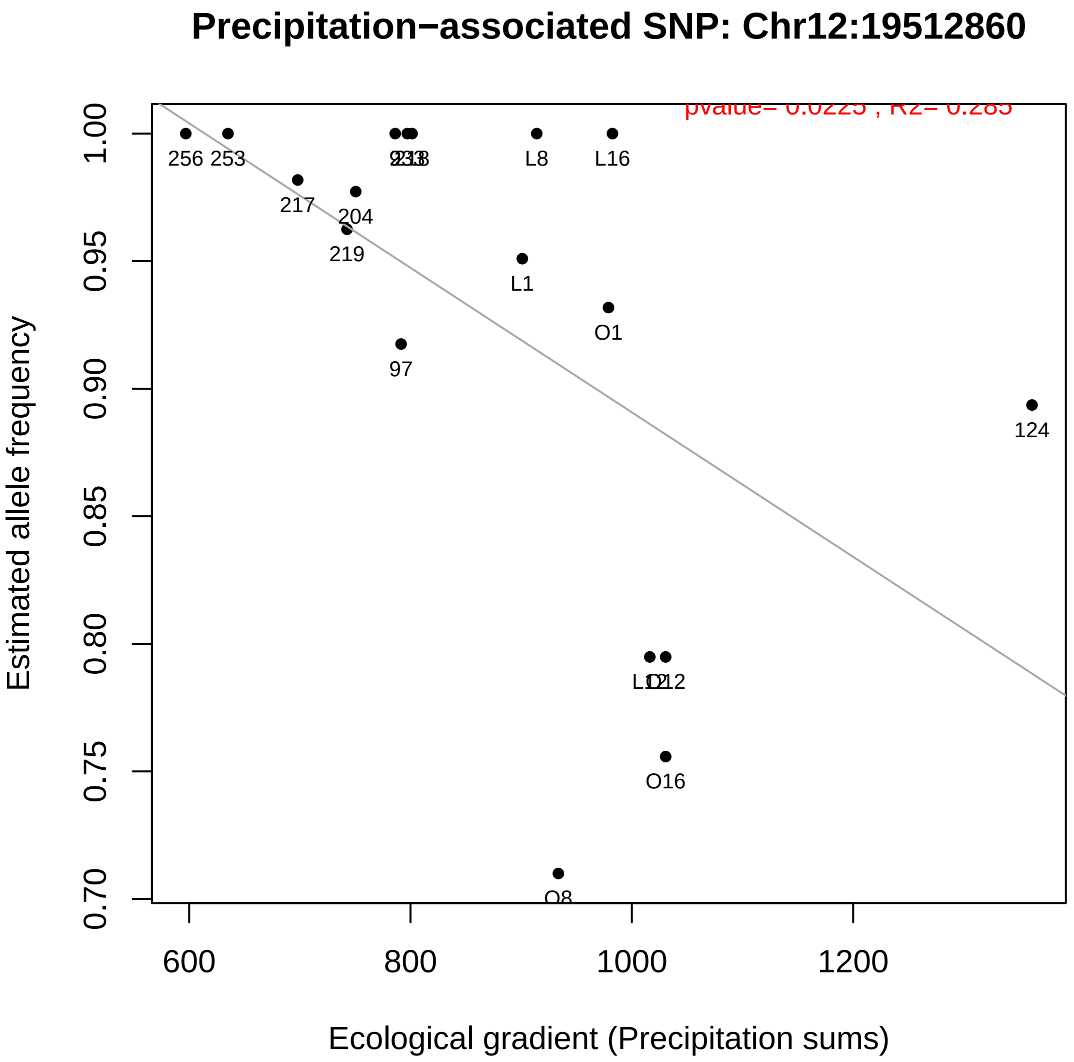

Precipitation−associated SNP: Chr12:19512860
pvalue= 0.0225 , R2= 0.285
Estimated allele frequency
Ecological gradient (Precipitation sums)
1.00
0.95
0.90
0.85
0.80
0.75
0.70
600
800
1000
1200
256
253
217
204
219
L8
L16
L1
O1
97
124
O16
O8

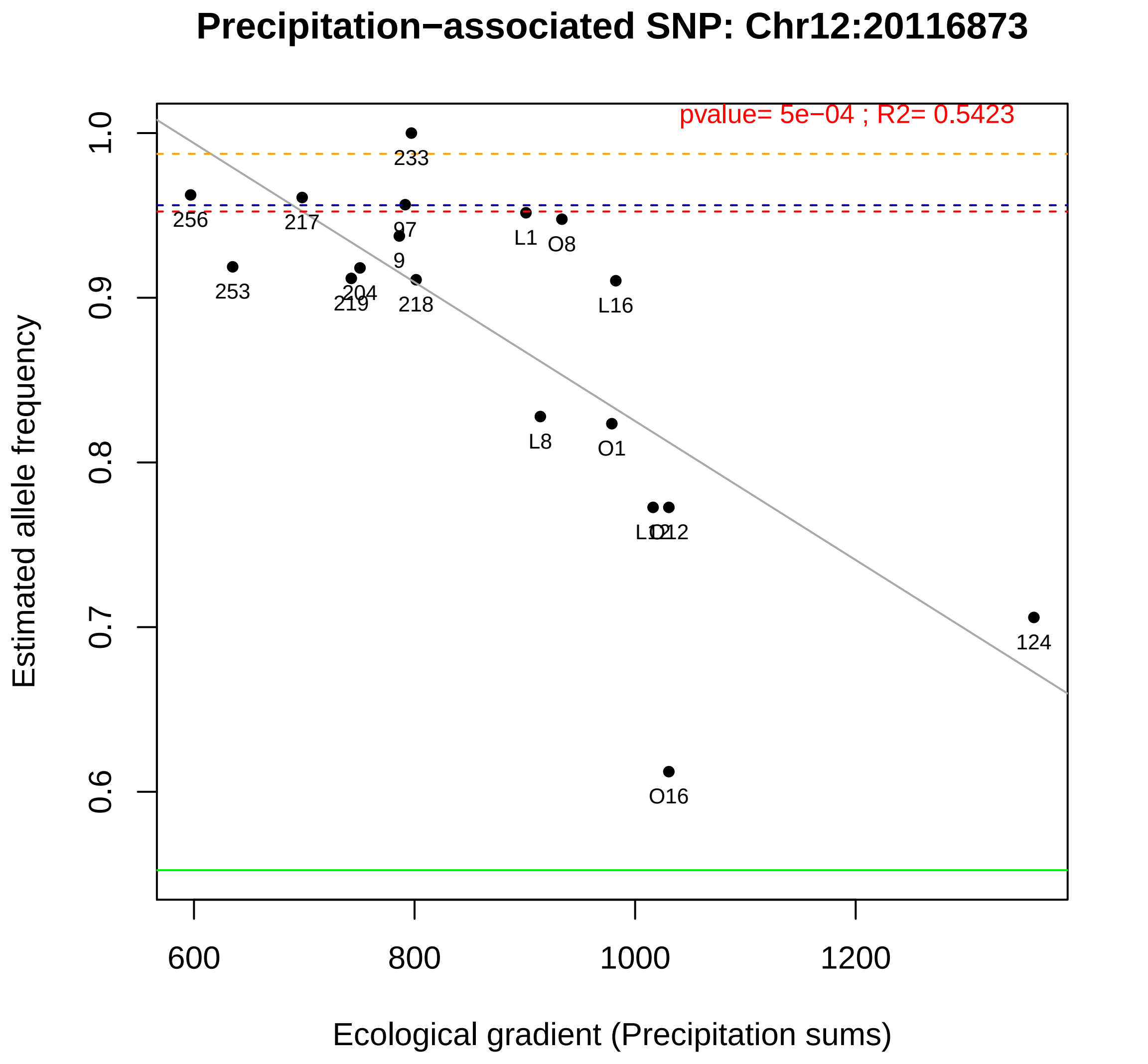

Precipitation−associated SNP: Chr12:20116873
pvalue= 5e−04 ; R2= 0.5423
Estimated allele frequency
Ecological gradient (Precipitation sums)
0.6
0.7
0.8
0.9
1.0
600
800
1000
1200
233
256
217
97
L1
O8
9
253
204
219
218
L16
L8
O1
L12
O12
124
O16

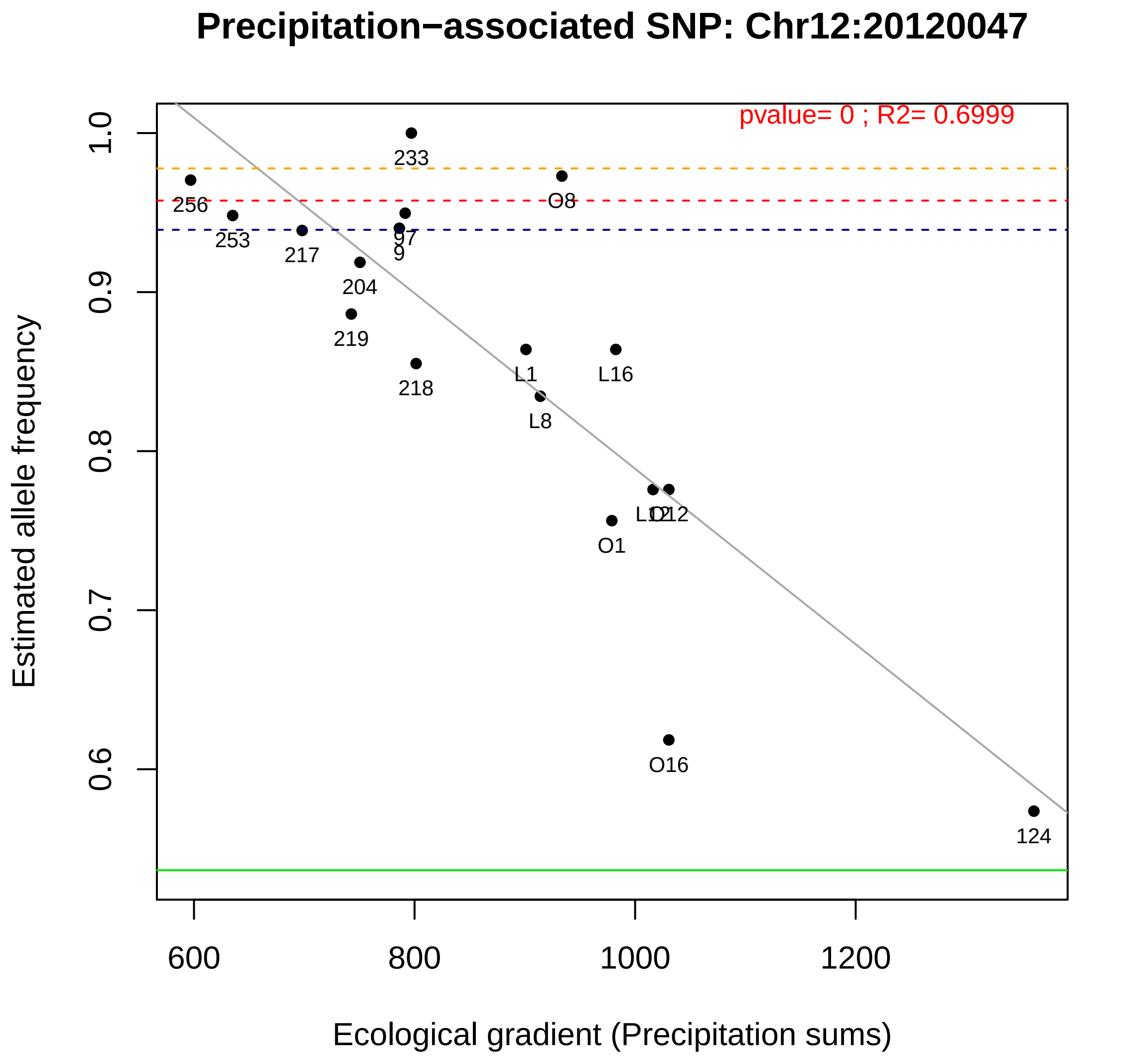
Precipitation−associated SNP: Chr12:20120047
pvalue= 0 ; R2= 0.6999
Estimated allele frequency
Ecological gradient (Precipitation sums)
0.6
0.7
0.8
0.9
1.0
600
800
1000
1200
233
256
O8
253
97
217
9
204
219
218
L1
L16
L8
L12
O12
O1
O16
124

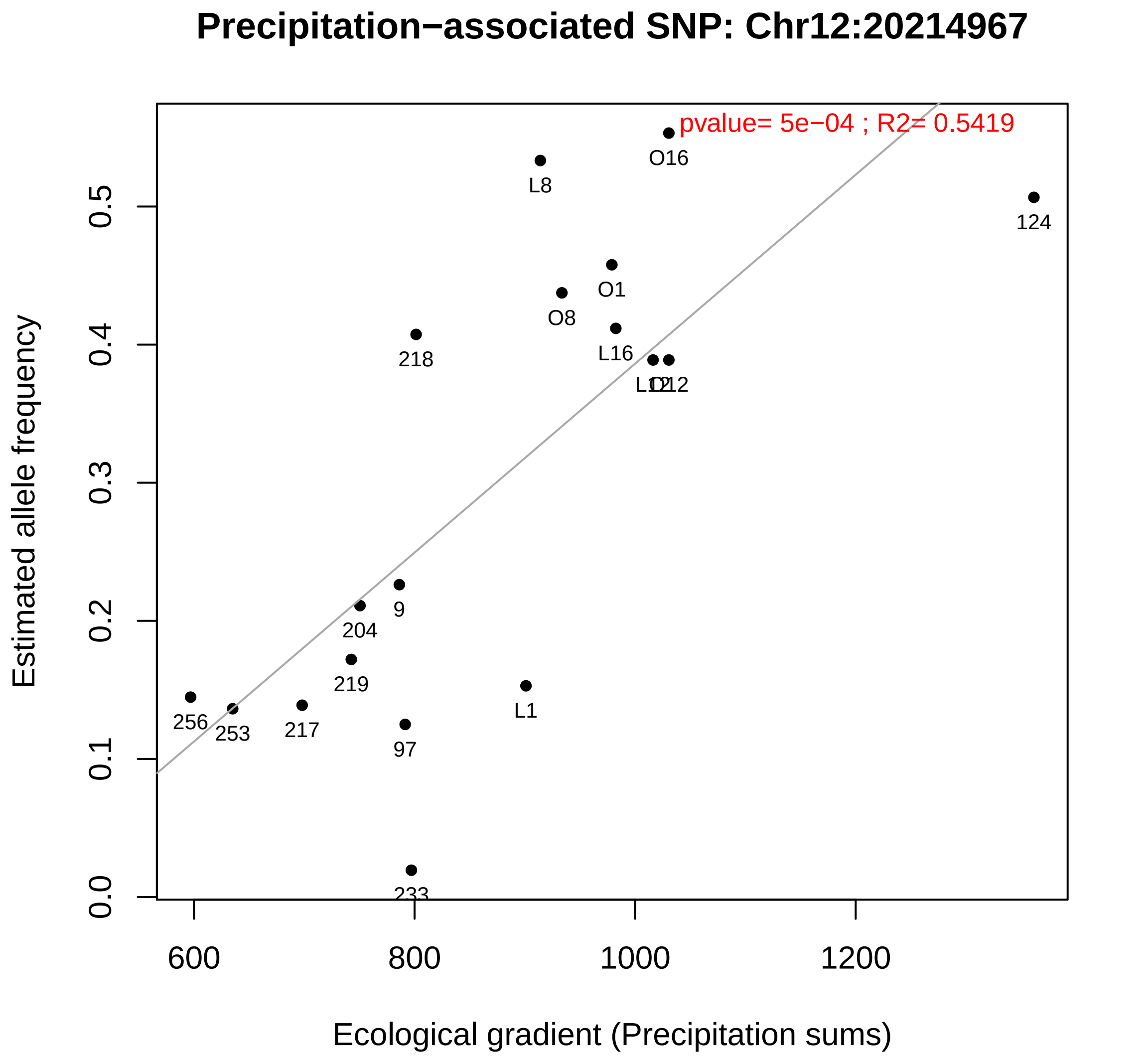
Precipitation−associated SNP: Chr12:20214967
pvalue= 5e−04 ; R2= 0.5419
O16
L8
124
O1
O8
L16
218
L12
O12
9
204
219
L1
256
253
217
97
233
0.0
0.1
0.2
0.3
0.4
0.5
600
800
1000
1200
Estimated allele frequency
Ecological gradient (Precipitation sums)

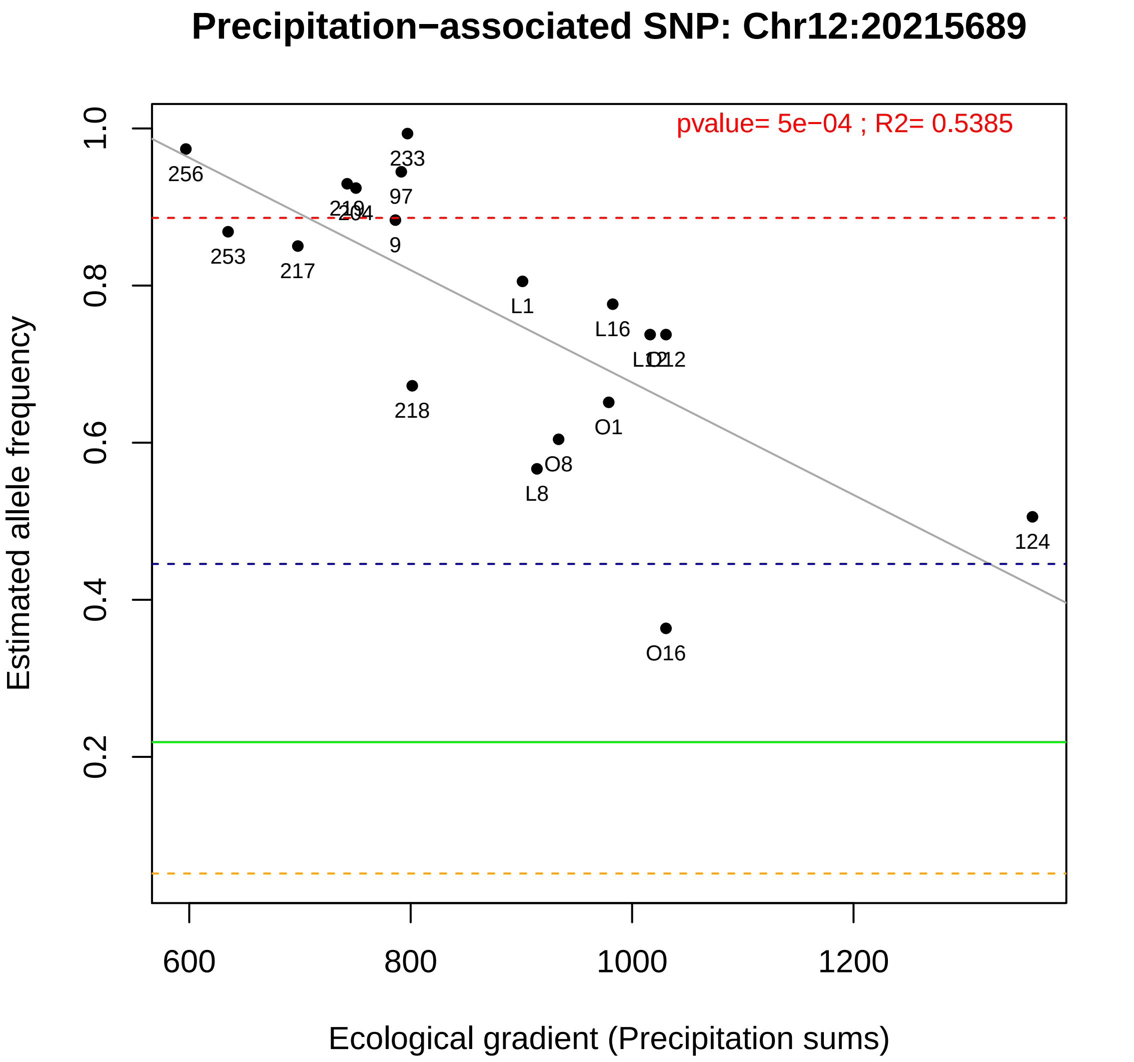

Precipitation−associated SNP: Chr12:20215689
pvalue= 5e−04 ; R2= 0.5385
Estimated allele frequency
0.2
0.4
0.6
0.8
1.0
Ecological gradient (Precipitation sums)
600
800
1000
1200
256
233
97
210
204
9
253
217
L1
L16
L12
O12
218
O1
O8
L8
124
O16

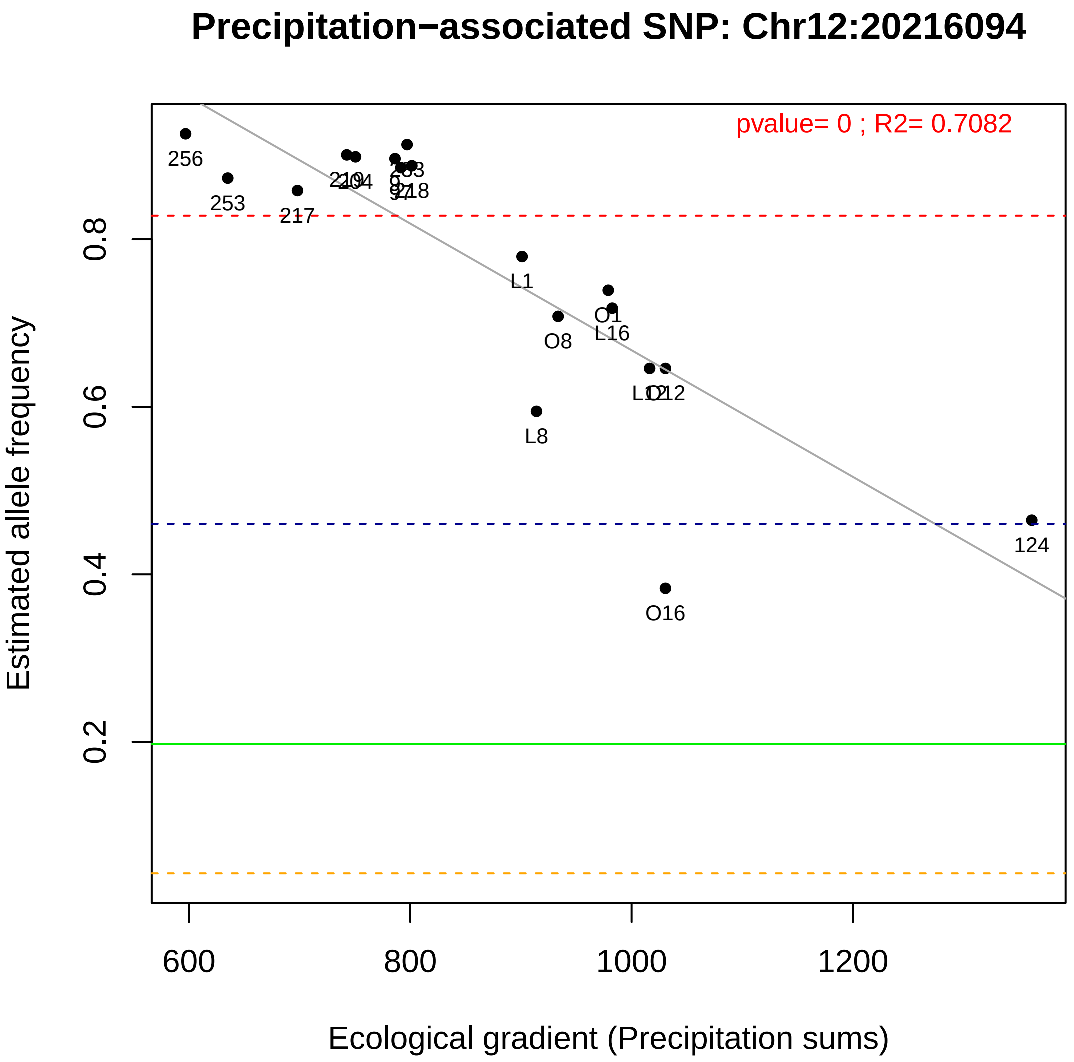
Precipitation−associated SNP: Chr12:20216094
pvalue= 0 ; R2= 0.7082
256
253
217
204
283
218
L1
O1
L16
O8
O12
L8
124
O16
0.8
0.6
0.4
0.2
600
800
1000
1200
Estimated allele frequency
Ecological gradient (Precipitation sums)

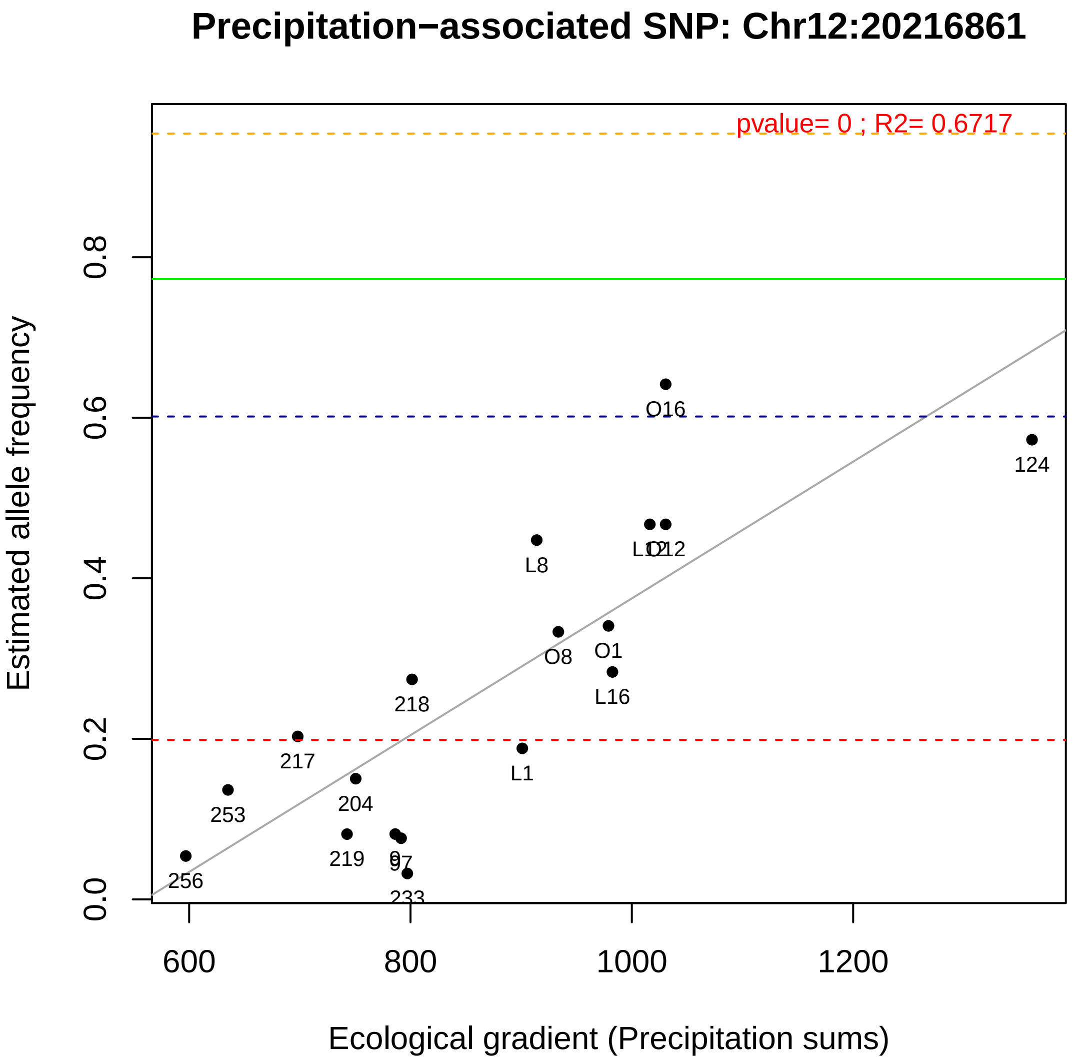
Precipitation−associated SNP: Chr12:20216861
pvalue= 0 ; R2= 0.6717
Estimated allele frequency
0.0
0.2
0.4
0.6
0.8
600
800
1000
1200
Ecological gradient (Precipitation sums)
O16
124
L8
L12
O12
O8
O1
L16
218
217
L1
204
253
219
97
256
233

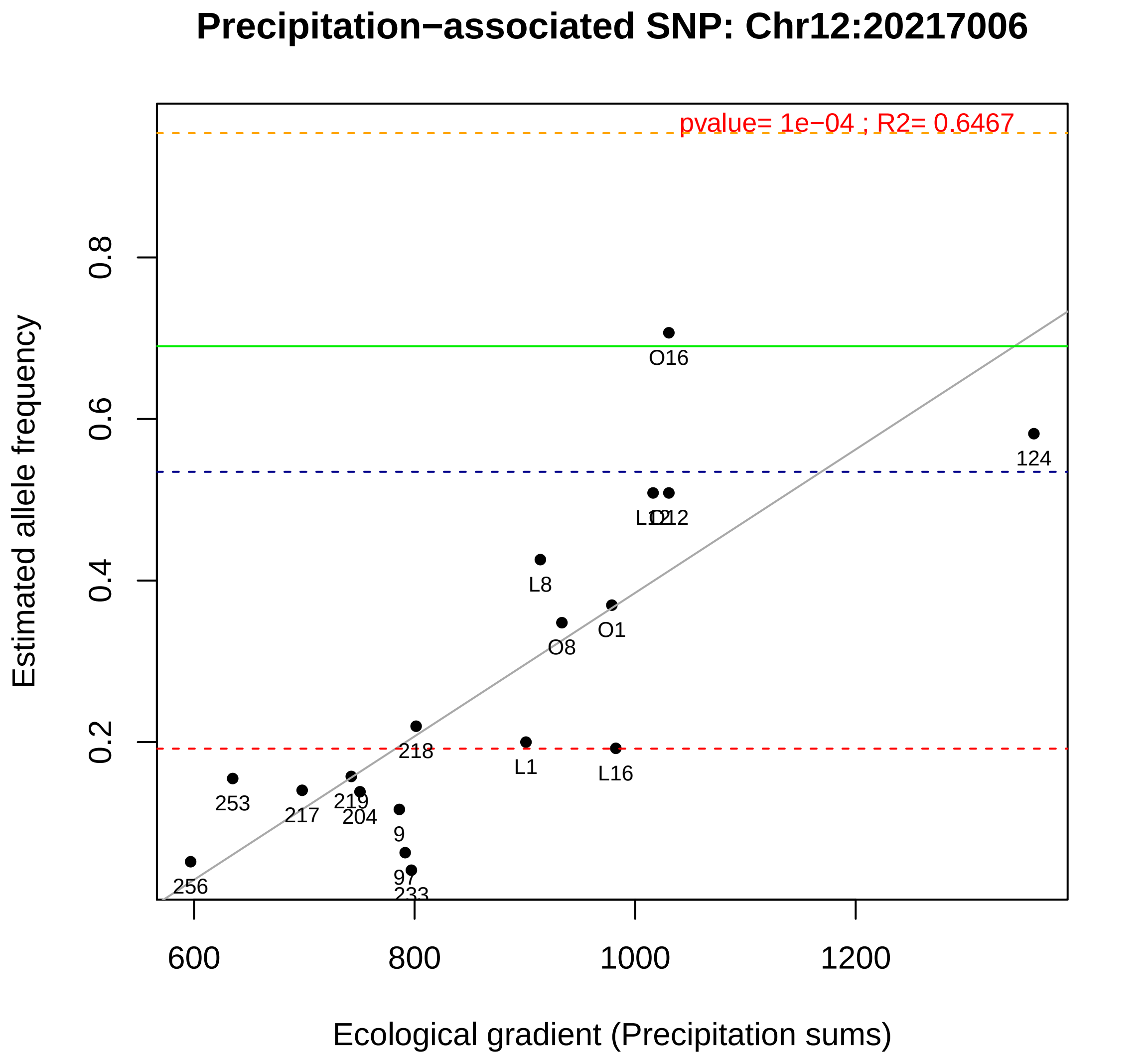

Precipitation−associated SNP: Chr12:20217006
pvalue= 1e−04 ; R2= 0.6467
Estimated allele frequency
0.2
0.4
0.6
0.8
600
800
1000
1200
Ecological gradient (Precipitation sums)
O16
124
L12
O12
L8
O1
O8
218
L1
L16
253
217
219
204
9
97
233
256

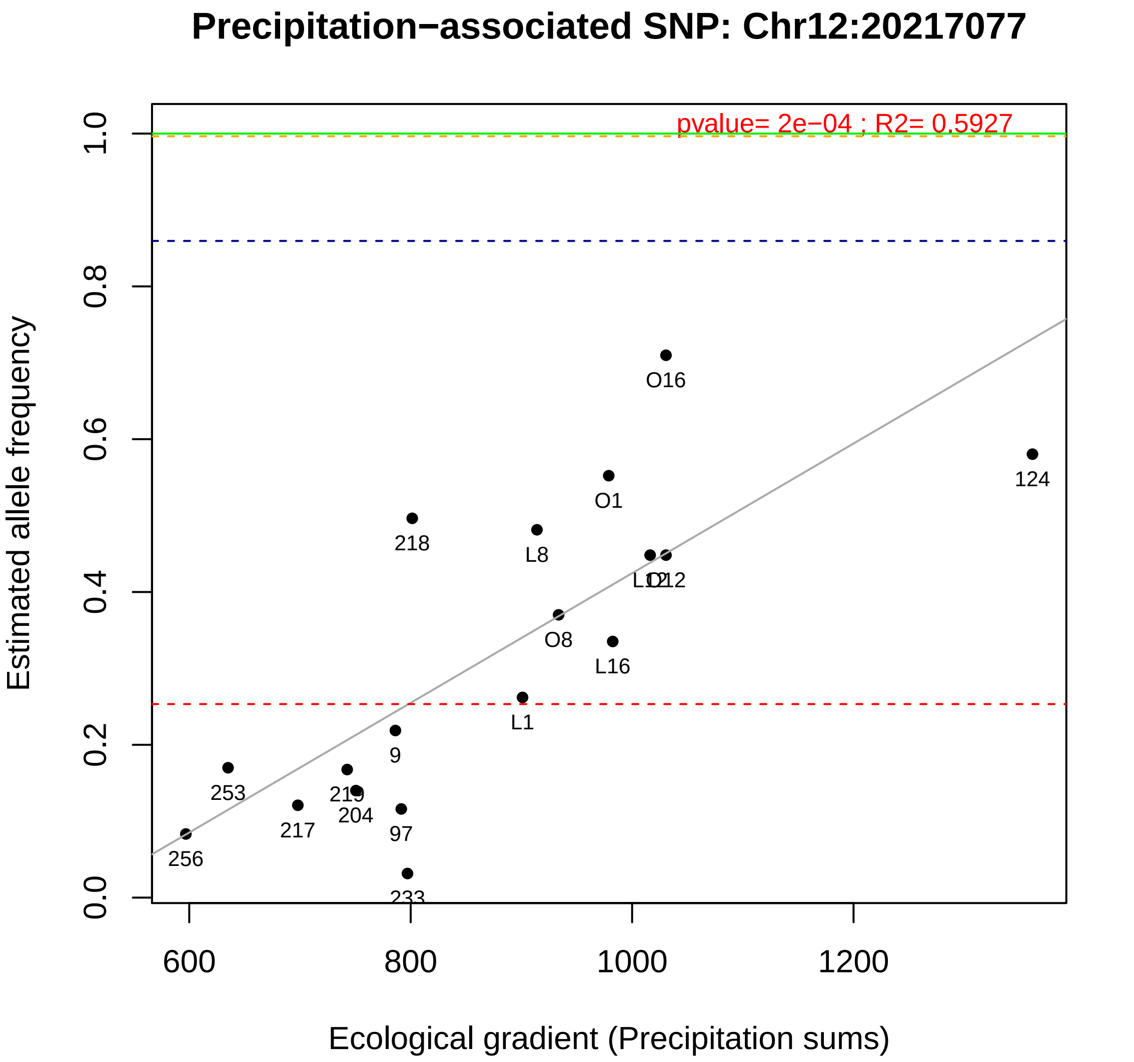

Precipitation−associated SNP: Chr12:20217077
pvalue= 2e−04 ; R2= 0.5927
Estimated allele frequency
0.0
0.2
0.4
0.6
0.8
1.0
600
800
1000
1200
Ecological gradient (Precipitation sums)
O16
124
O1
218
L8
L12
O12
O8
L16
L1
9
253
219
204
217
97
256
233

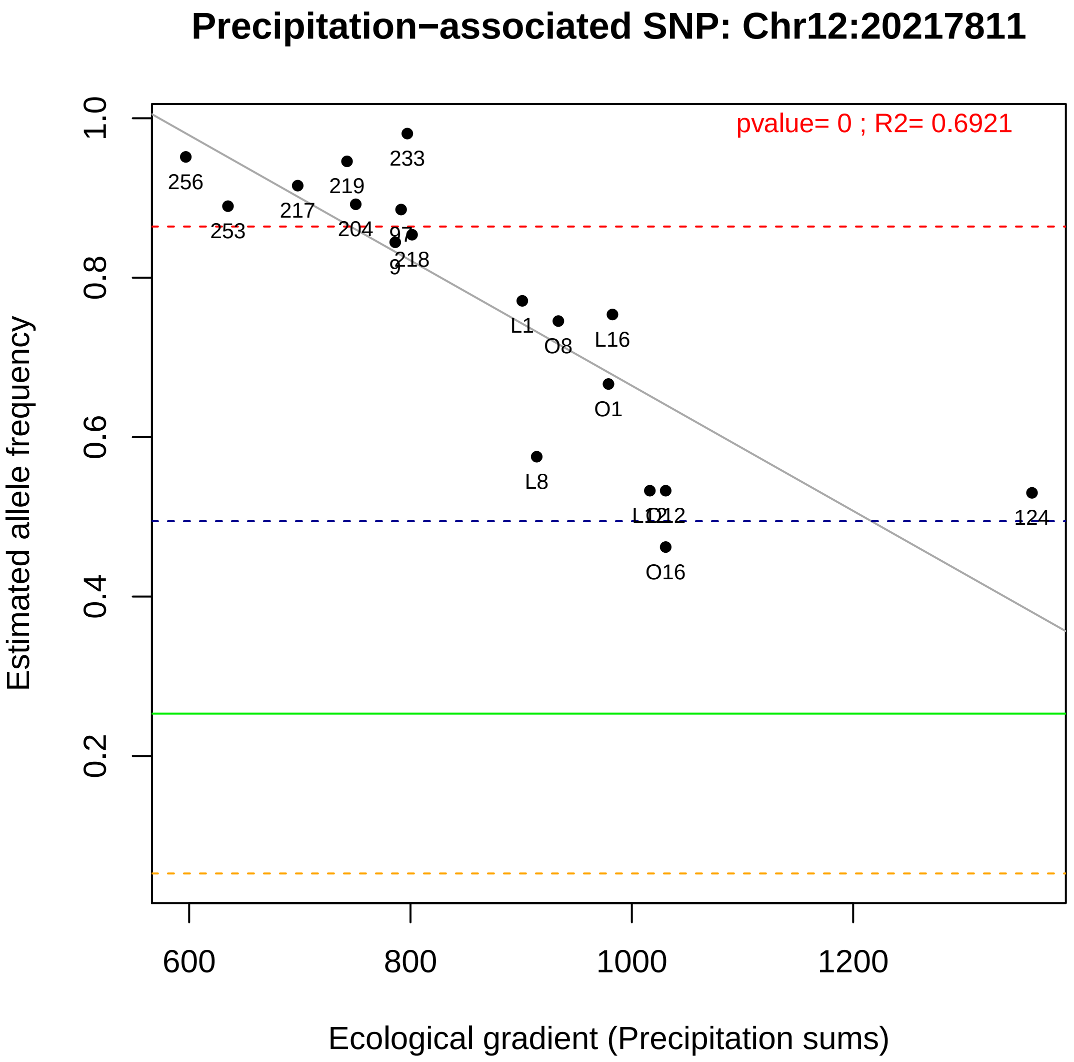

Precipitation−associated SNP: Chr12:20217811
pvalue= 0 ; R2= 0.6921
Estimated allele frequency
Ecological gradient (Precipitation sums)
1.0
0.8
0.6
0.4
0.2
600
800
1000
1200
256
253
217
219
204
233
97
9
218
L1
O8
L16
O1
L8
L12
O12
O16
124

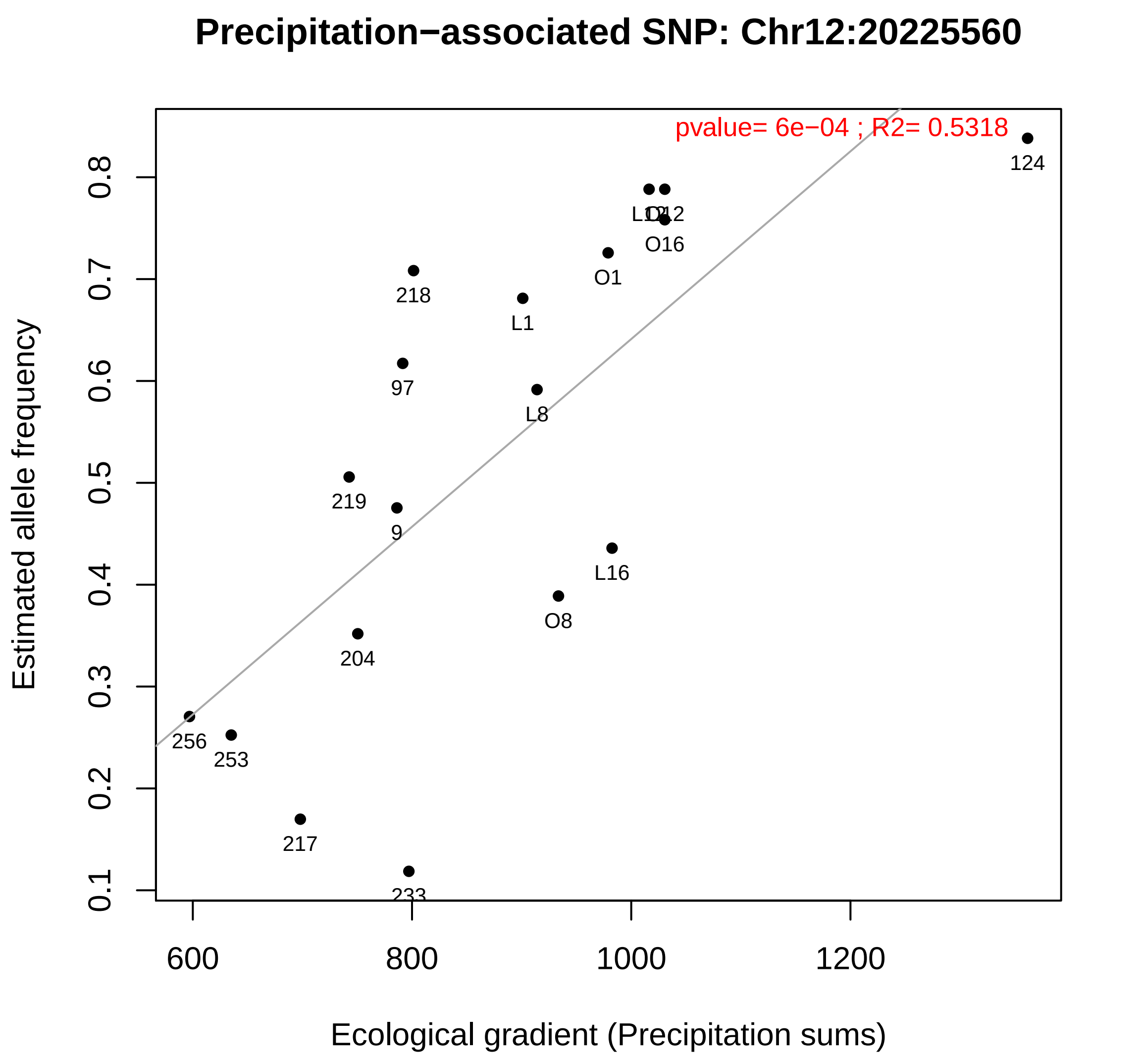
Precipitation−associated SNP: Chr12:20225560
pvalue= 6e−04 ; R2= 0.5318
Estimated allele frequency
Ecological gradient (Precipitation sums)
0.1
0.2
0.3
0.4
0.5
0.6
0.7
0.8
600
800
1000
1200
124
O12
O16
O1
218
L1
97
L8
219
9
L16
O8
204
256
253
217
233

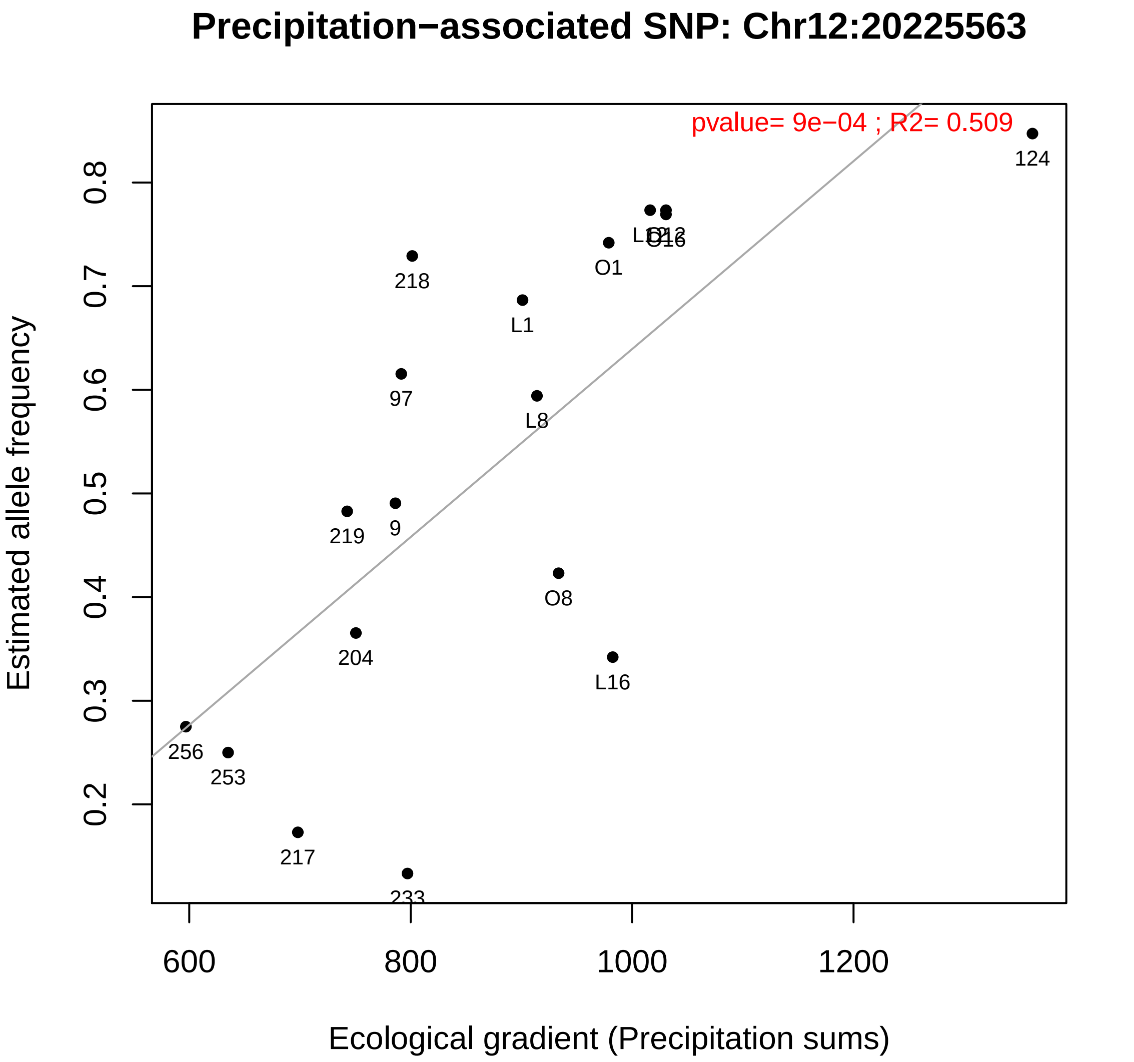
Precipitation−associated SNP: Chr12:20225563
pvalue= 9e−04 ; R2= 0.509
124
L12
O12
O16
O1
218
L1
97
L8
219
9
O8
204
L16
256
253
217
233
0.2
0.3
0.4
0.5
0.6
0.7
0.8
600
800
1000
1200
Estimated allele frequency
Ecological gradient (Precipitation sums)

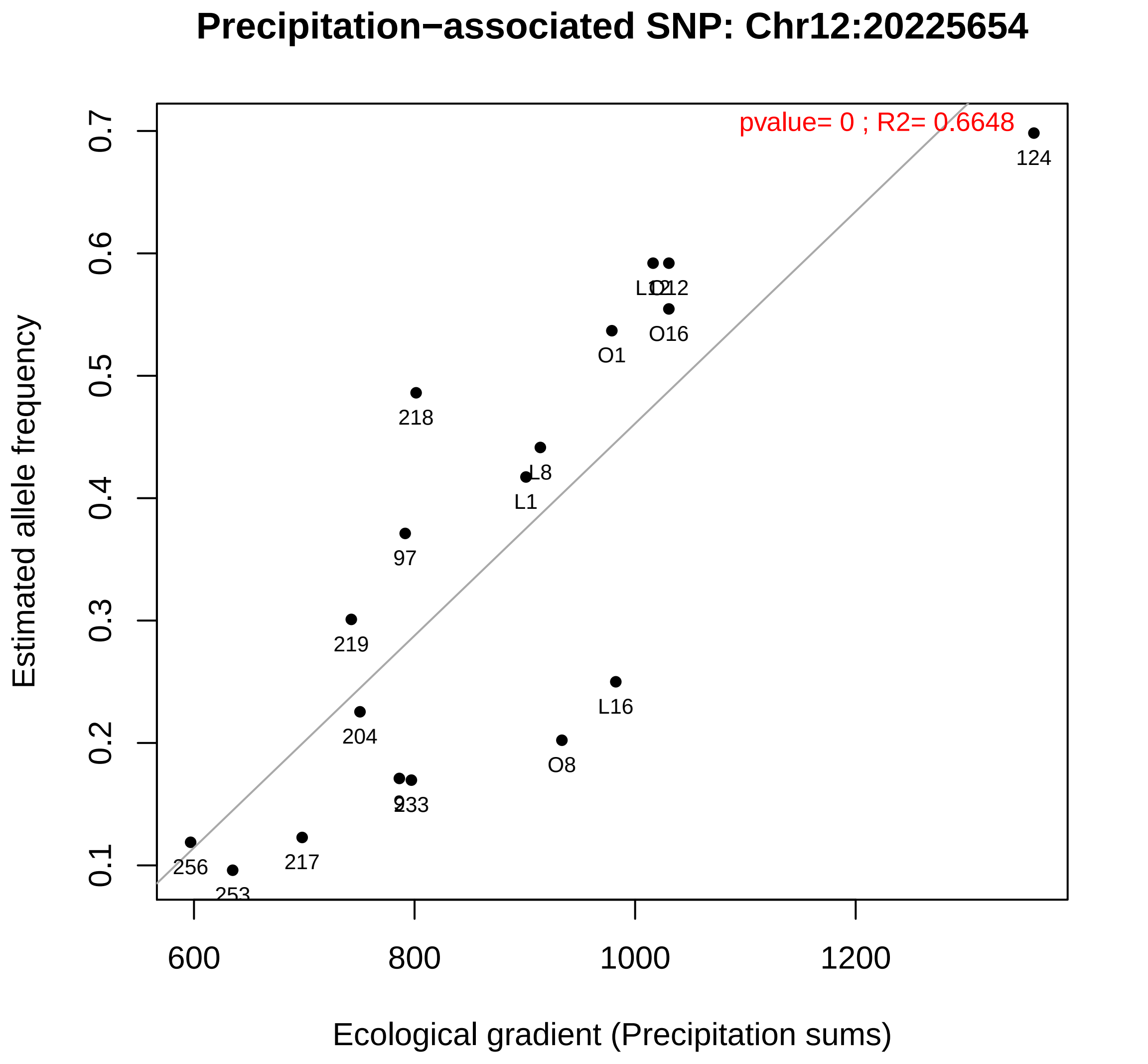
Precipitation−associated SNP: Chr12:20225654
pvalue= 0 ; R2= 0.6648
124
L12
O12
O16
O1
218
L8
L1
97
219
L16
204
O8
2
233
256
217
253
0.1
0.2
0.3
0.4
0.5
0.6
0.7
600
800
1000
1200
Estimated allele frequency
Ecological gradient (Precipitation sums)

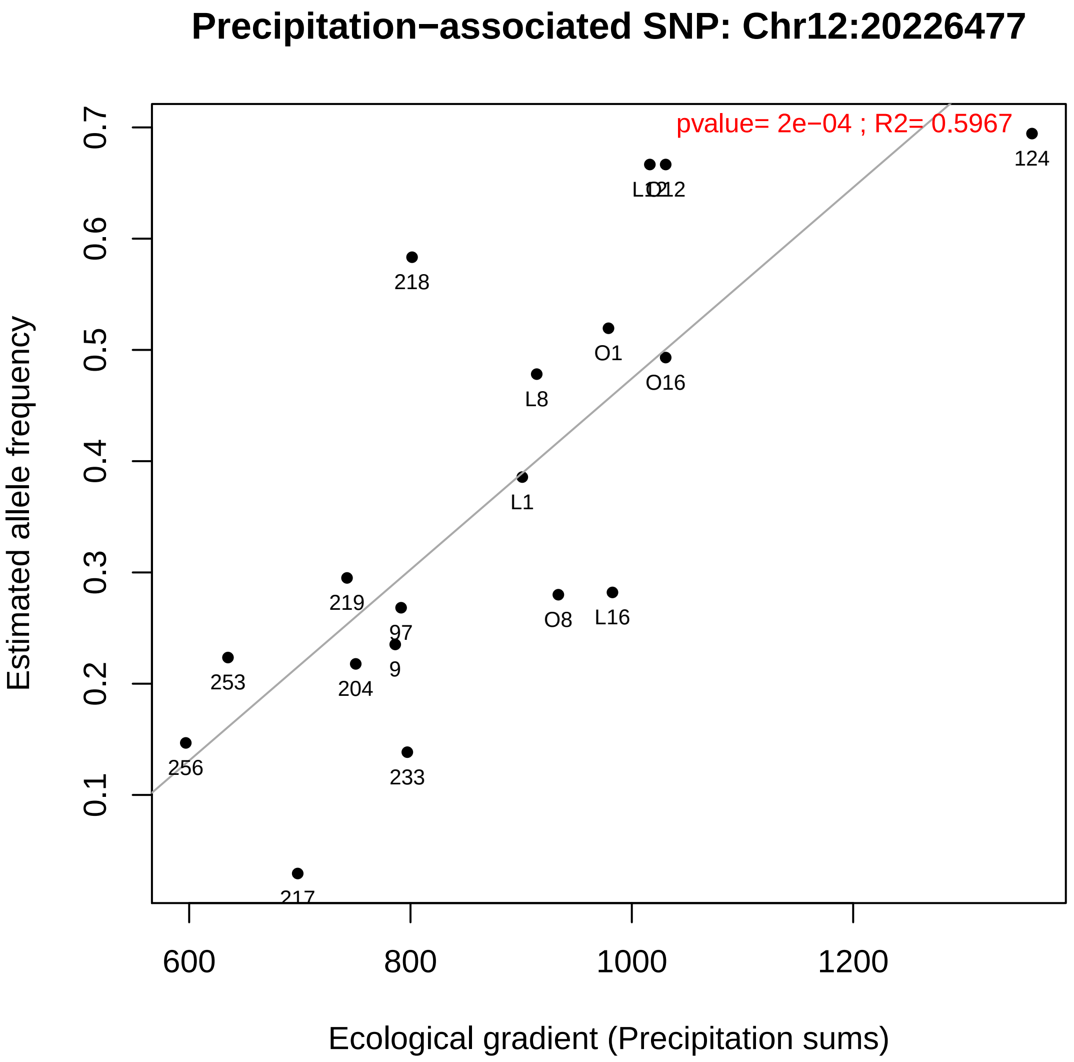

Precipitation−associated SNP: Chr12:20226477
pvalue= 2e−04 ; R2= 0.5967
124
L12
O12
218
O1
O16
L8
L1
219
97
O8
L16
253
204
9
256
233
217
0.1
0.2
0.3
0.4
0.5
0.6
0.7
600
800
1000
1200
Estimated allele frequency
Ecological gradient (Precipitation sums)

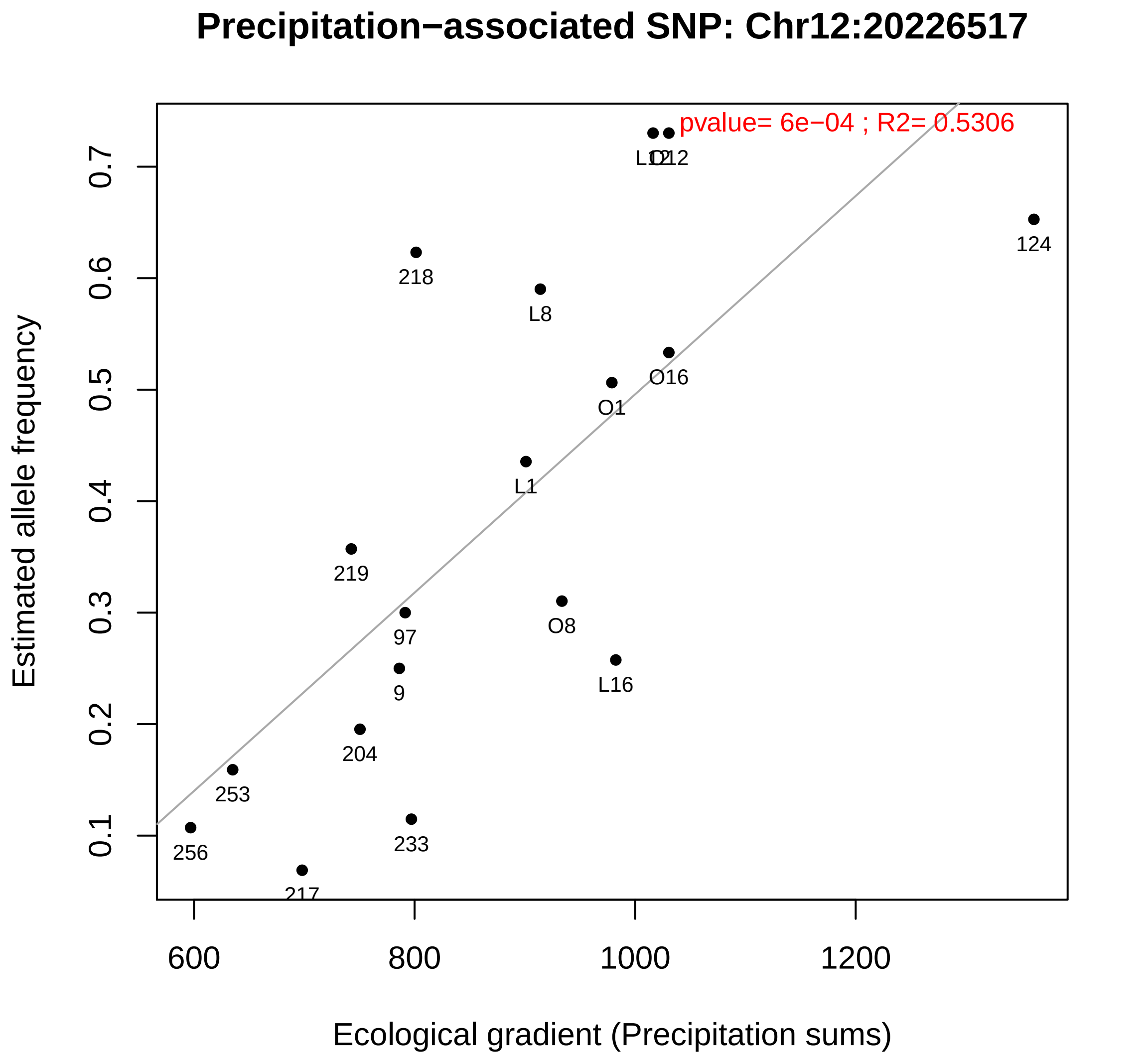
Precipitation−associated SNP: Chr12:20226517
pvalue= 6e−04 ; R2= 0.5306
Estimated allele frequency
0.1
0.2
0.3
0.4
0.5
0.6
0.7
600
800
1000
1200
Ecological gradient (Precipitation sums)
L12
O12
124
218
L8
O16
O1
L1
219
97
O8
9
L16
204
253
256
233
217

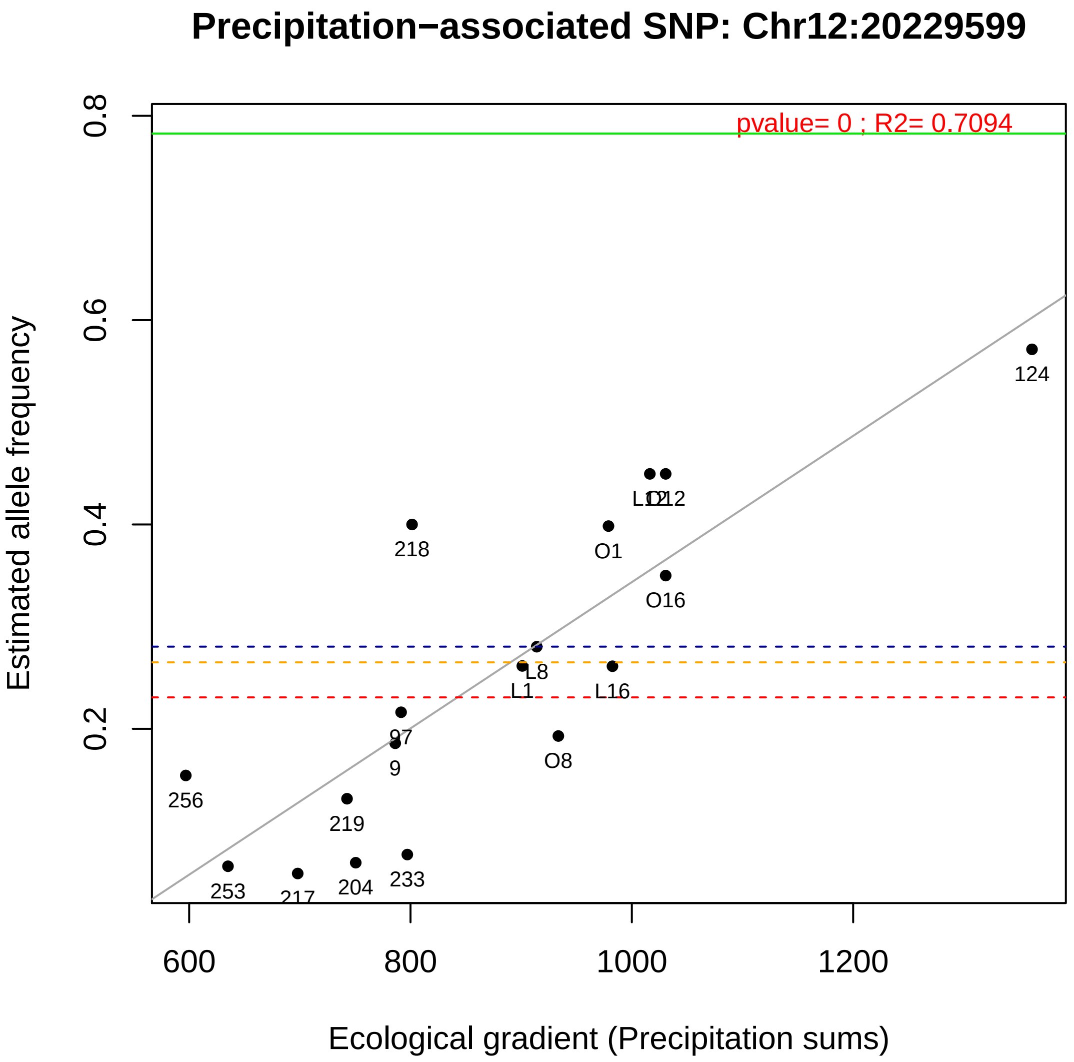
Precipitation−associated SNP: Chr12:20229599
pvalue= 0 ; R2= 0.7094
Estimated allele frequency
0.2
0.4
0.6
0.8
Ecological gradient (Precipitation sums)
600
800
1000
1200
124
L12
O12
218
O1
O16
L8
L1
L16
97
9
O8
256
219
253
217
204
233

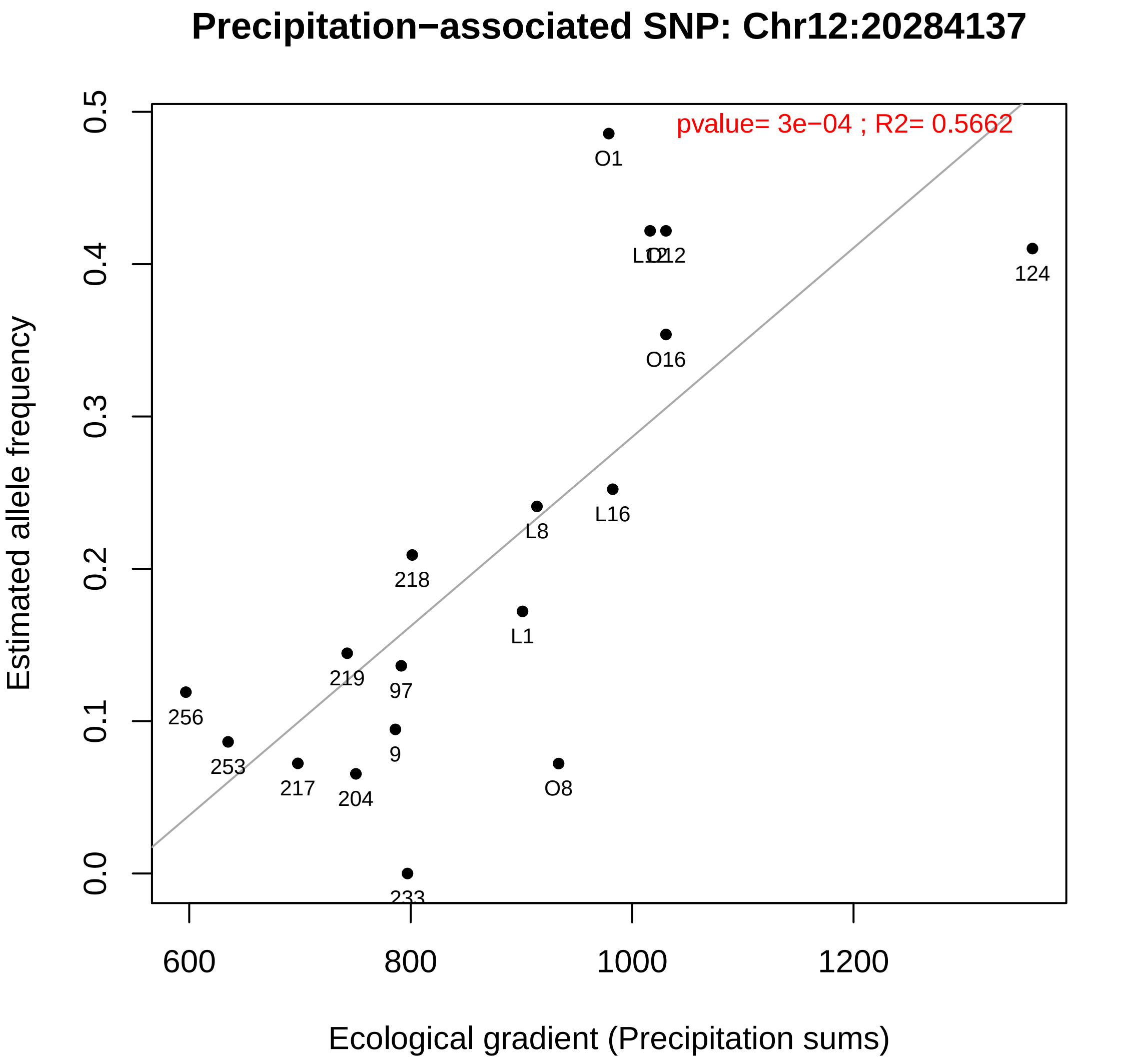

Precipitation−associated SNP: Chr12:20284137
pvalue= 3e−04 ; R2= 0.5662
Estimated allele frequency
0.0
0.1
0.2
0.3
0.4
0.5
Ecological gradient (Precipitation sums)
600
800
1000
1200
O1
L12
O12
124
O16
L16
L8
218
L1
219
97
256
253
9
217
204
O8
233

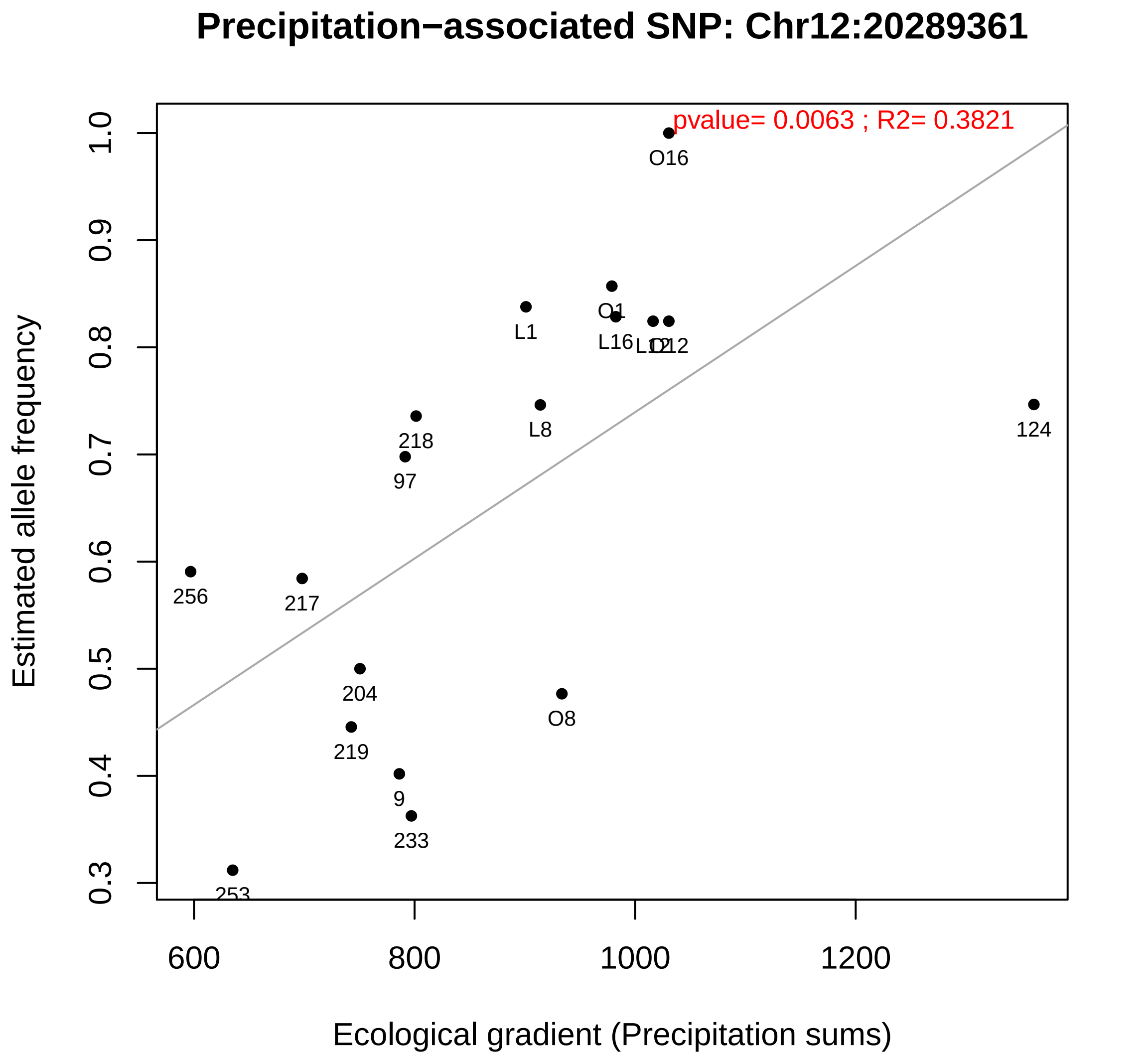

Precipitation−associated SNP: Chr12:20289361
pvalue= 0.0063 ; R2= 0.3821
Estimated allele frequency
Ecological gradient (Precipitation sums)
1.0
0.9
0.8
0.7
0.6
0.5
0.4
0.3
600
800
1000
1200
O16
O1
L1
L16
O12
L8
124
218
97
256
217
204
O8
219
9
233

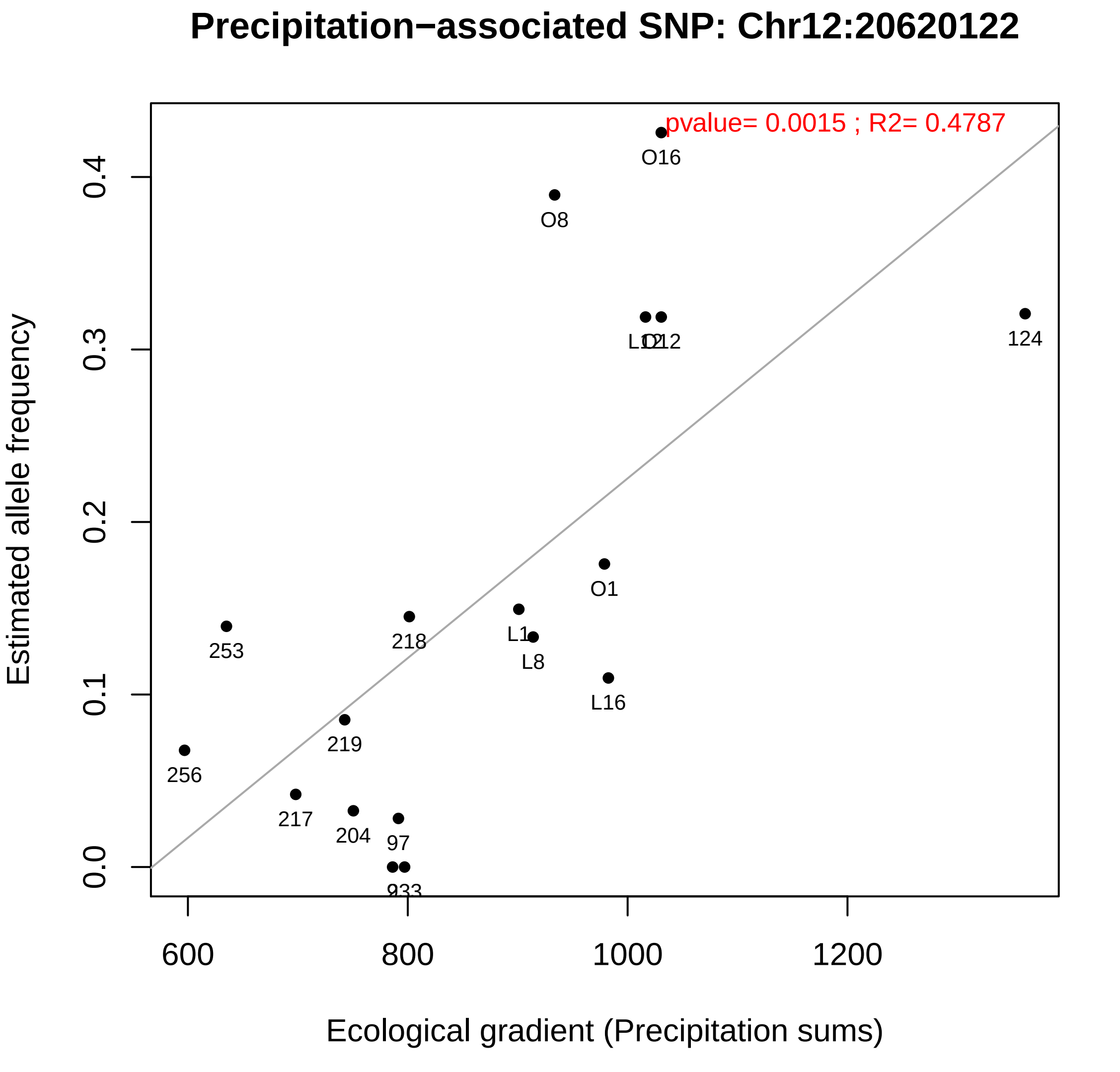
Precipitation−associated SNP: Chr12:20620122
pvalue= 0.0015 ; R2= 0.4787
Estimated allele frequency
Ecological gradient (Precipitation sums)
0.0
0.1
0.2
0.3
0.4
600
800
1000
1200
O16
O8
O12
124
O1
218
253
L1
L8
L16
219
256
217
204
97

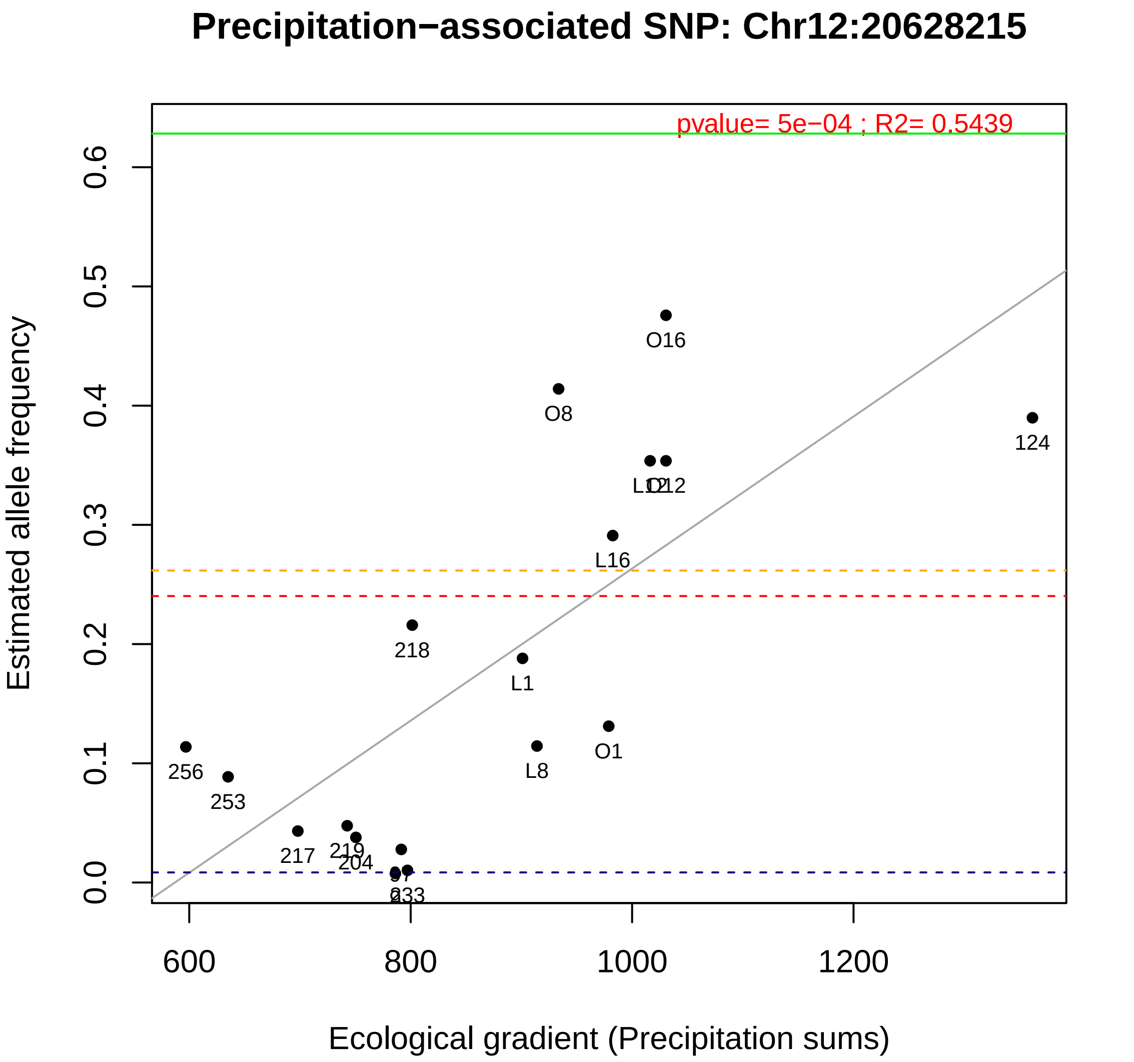
Precipitation−associated SNP: Chr12:20628215
pvalue= 5e−04 ; R2= 0.5439
Estimated allele frequency
0.0
0.1
0.2
0.3
0.4
0.5
0.6
600
800
1000
1200
Ecological gradient (Precipitation sums)
O16
O8
124
O12
L16
218
L1
O1
L8
256
253
217
219
204
233

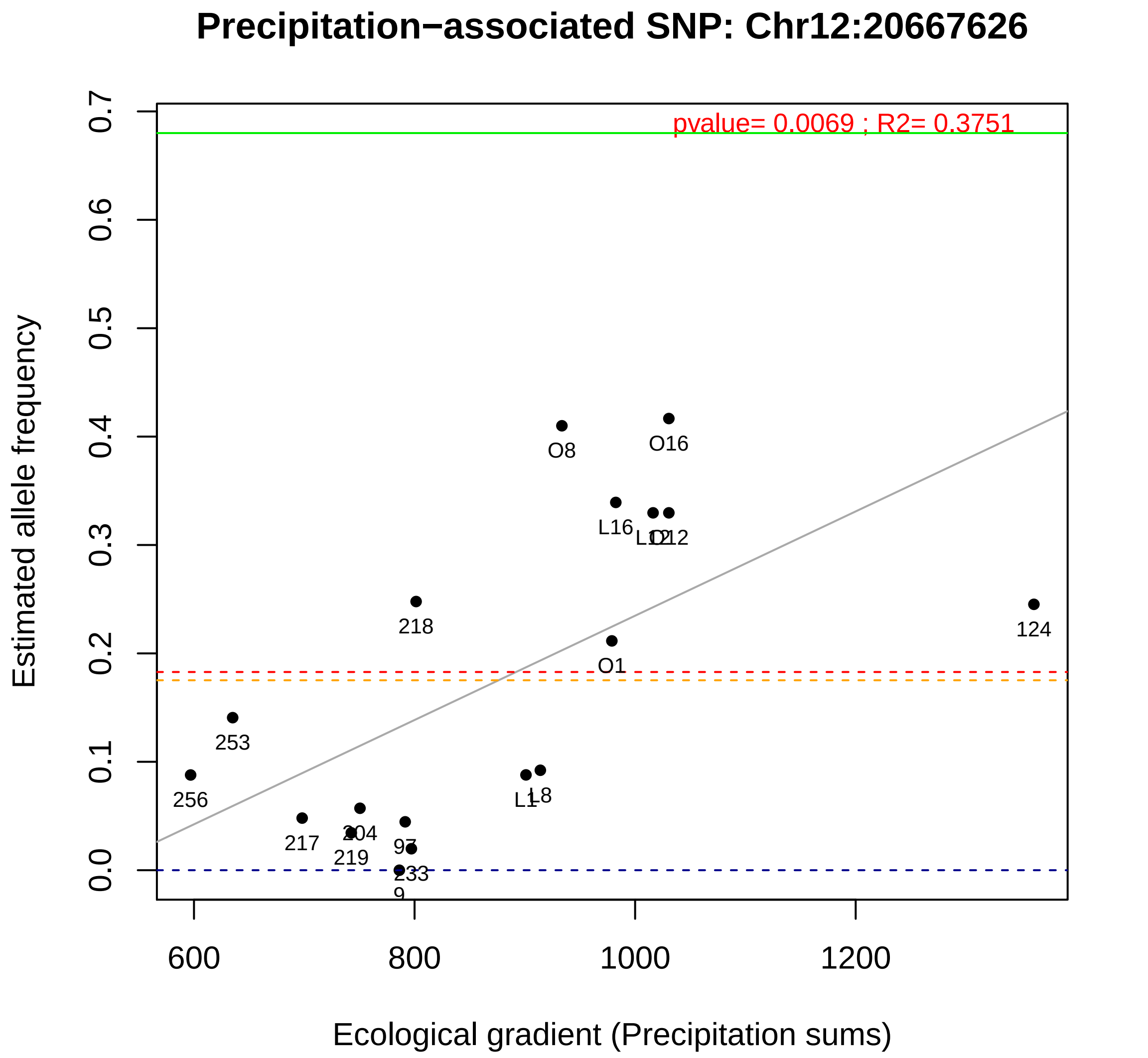

Precipitation−associated SNP: Chr12:20667626
pvalue= 0.0069 ; R2= 0.3751
Estimated allele frequency
Ecological gradient (Precipitation sums)
0.0
0.1
0.2
0.3
0.4
0.5
0.6
0.7
600
800
1000
1200
O8
O16
L16
L12
O12
218
124
O1
253
256
L1
L8
217
204
219
97
233
9

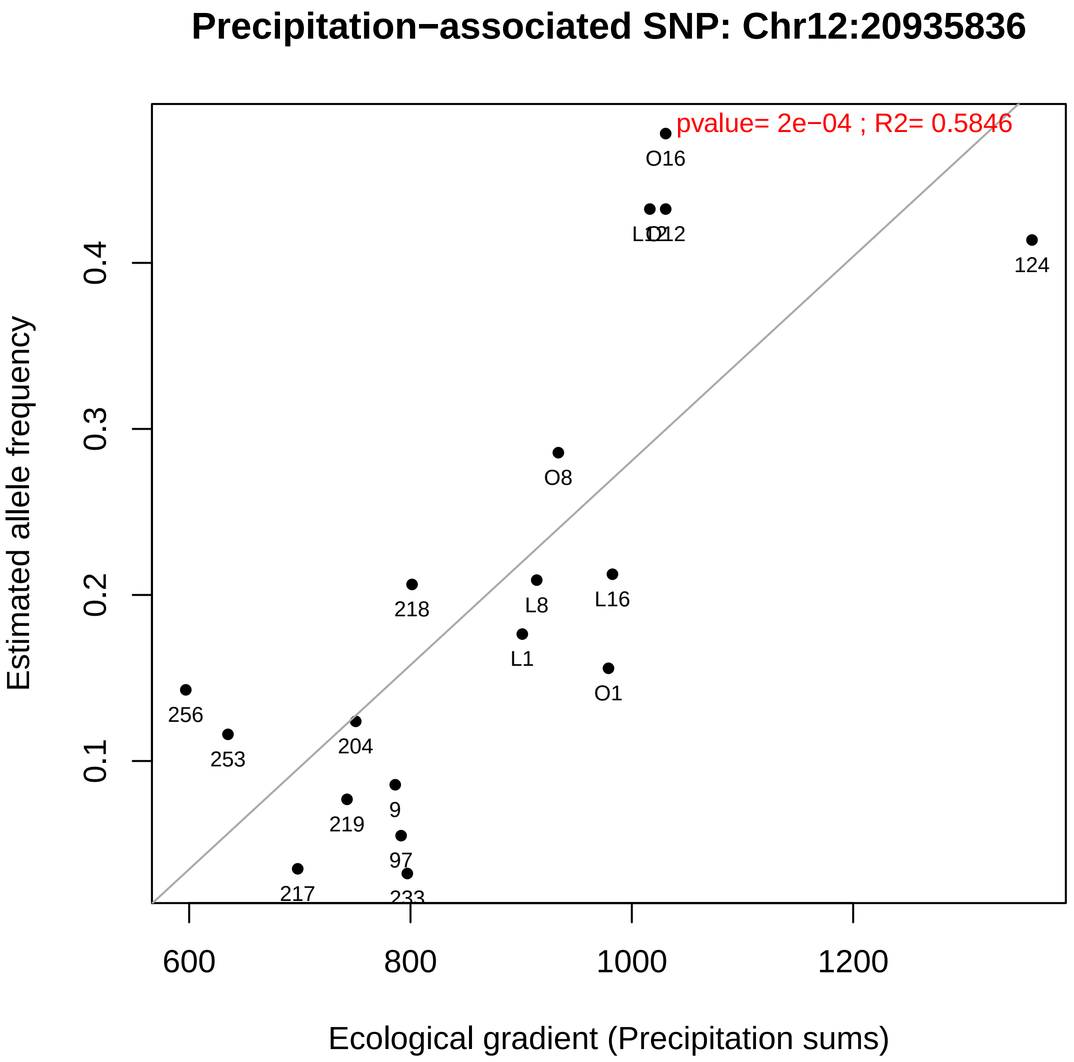

Precipitation−associated SNP: Chr12:20935836
pvalue= 2e−04 ; R2= 0.5846
O16
L12
O12
124
O8
218
L8
L16
L1
O1
256
253
204
219
9
97
217
233
0.1
0.2
0.3
0.4
Estimated allele frequency
600
800
1000
1200
Ecological gradient (Precipitation sums)

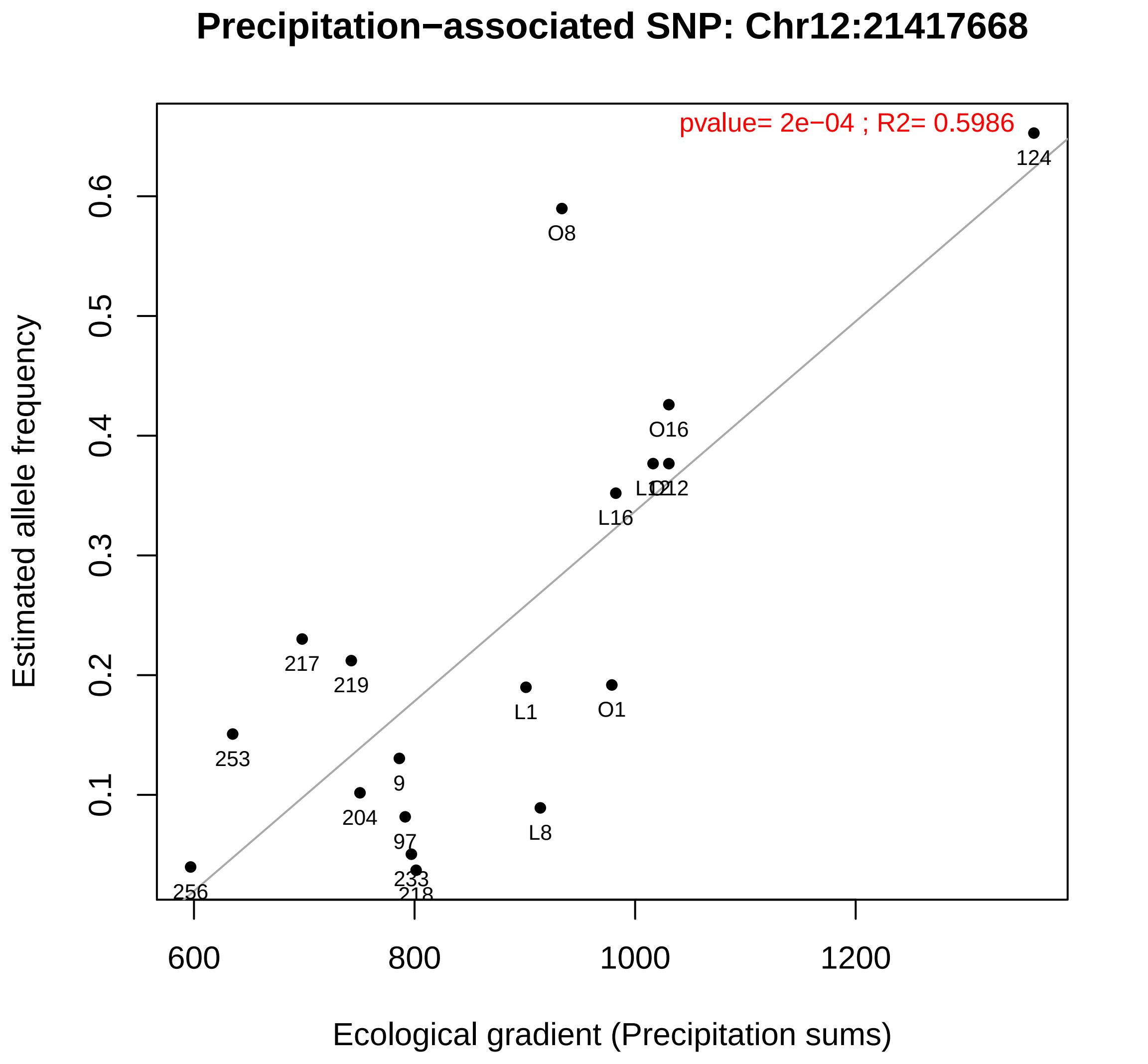

Precipitation−associated SNP: Chr12:21417668
pvalue= 2e−04 ; R2= 0.5986
Estimated allele frequency
0.1
0.2
0.3
0.4
0.5
0.6
600
800
1000
1200
Ecological gradient (Precipitation sums)
124
O8
O16
L12
O12
L16
217
219
L1
O1
253
9
204
97
L8
233
218
256

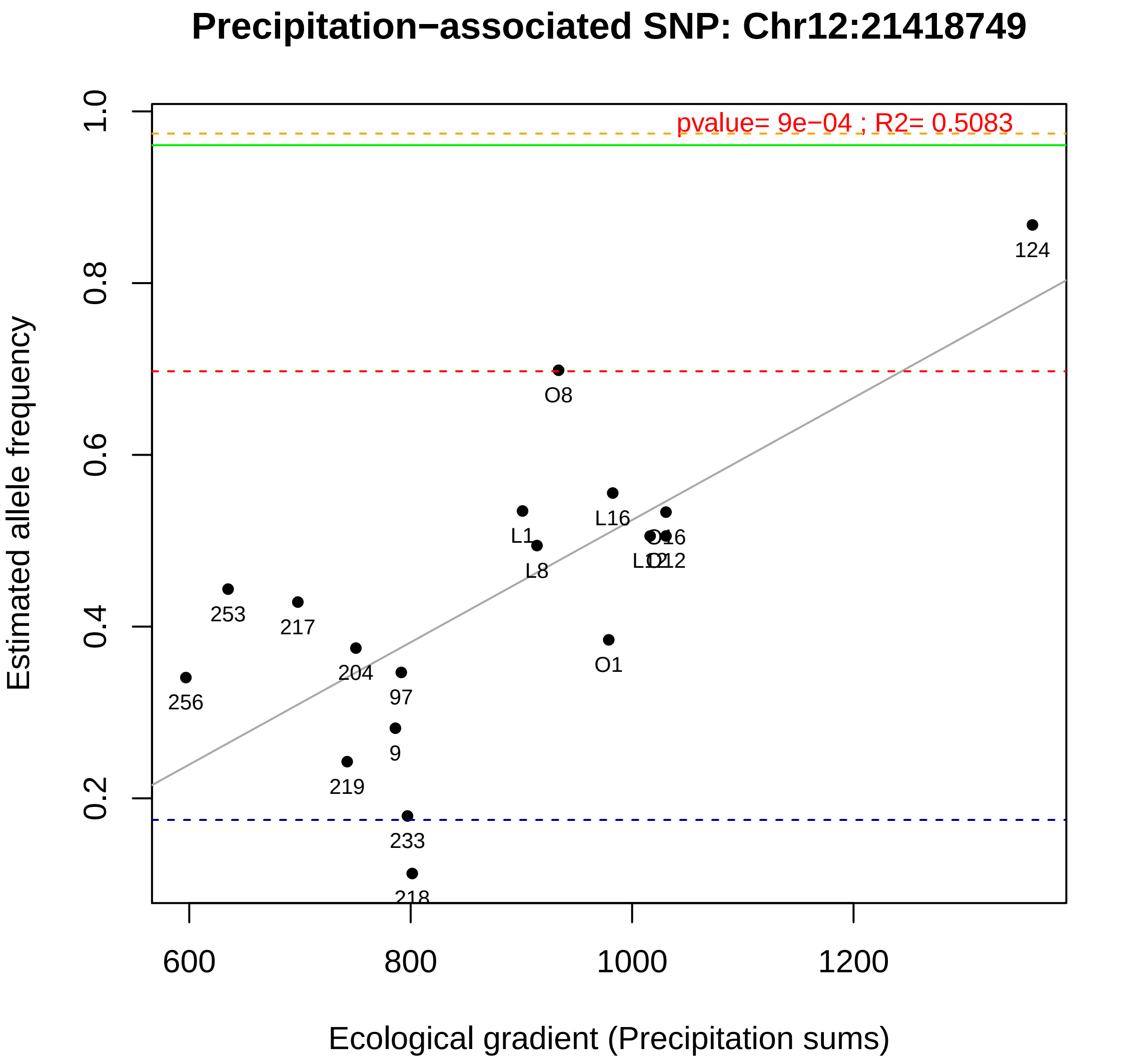

Precipitation−associated SNP: Chr12:21418749
pvalue= 9e−04 ; R2= 0.5083
Estimated allele frequency
Ecological gradient (Precipitation sums)
0.2
0.4
0.6
0.8
1.0
600
800
1000
1200
124
O8
L16
L1
L8
O16
L12
O12
O1
253
217
204
256
97
9
219
233
218

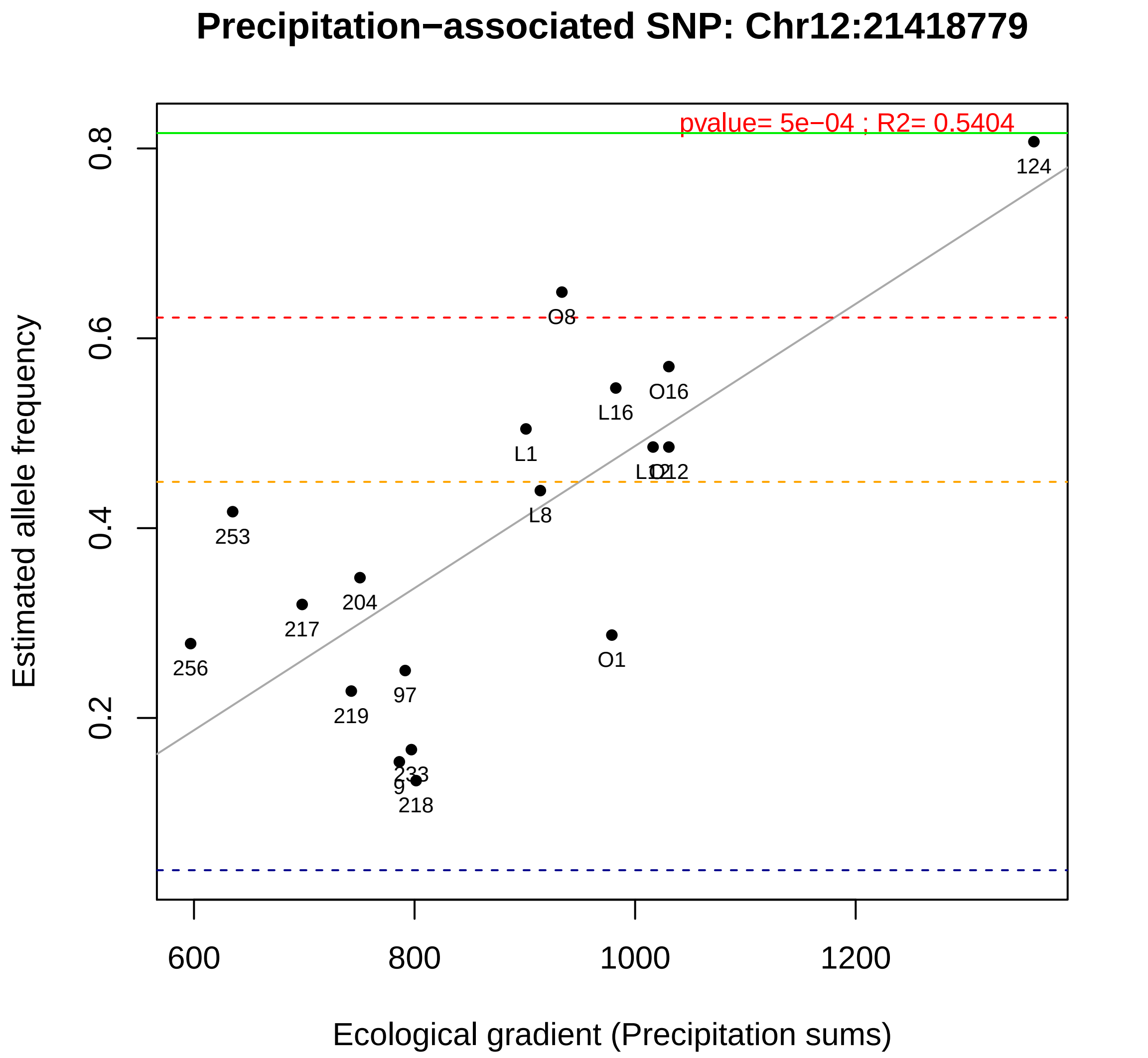
Precipitation−associated SNP: Chr12:21418779
pvalue= 5e−04 ; R2= 0.5404
Estimated allele frequency
0.2
0.4
0.6
0.8
600
800
1000
1200
Ecological gradient (Precipitation sums)
124
O8
O16
L16
L1
L12
O12
L8
253
204
217
256
O1
97
219
233
9
218

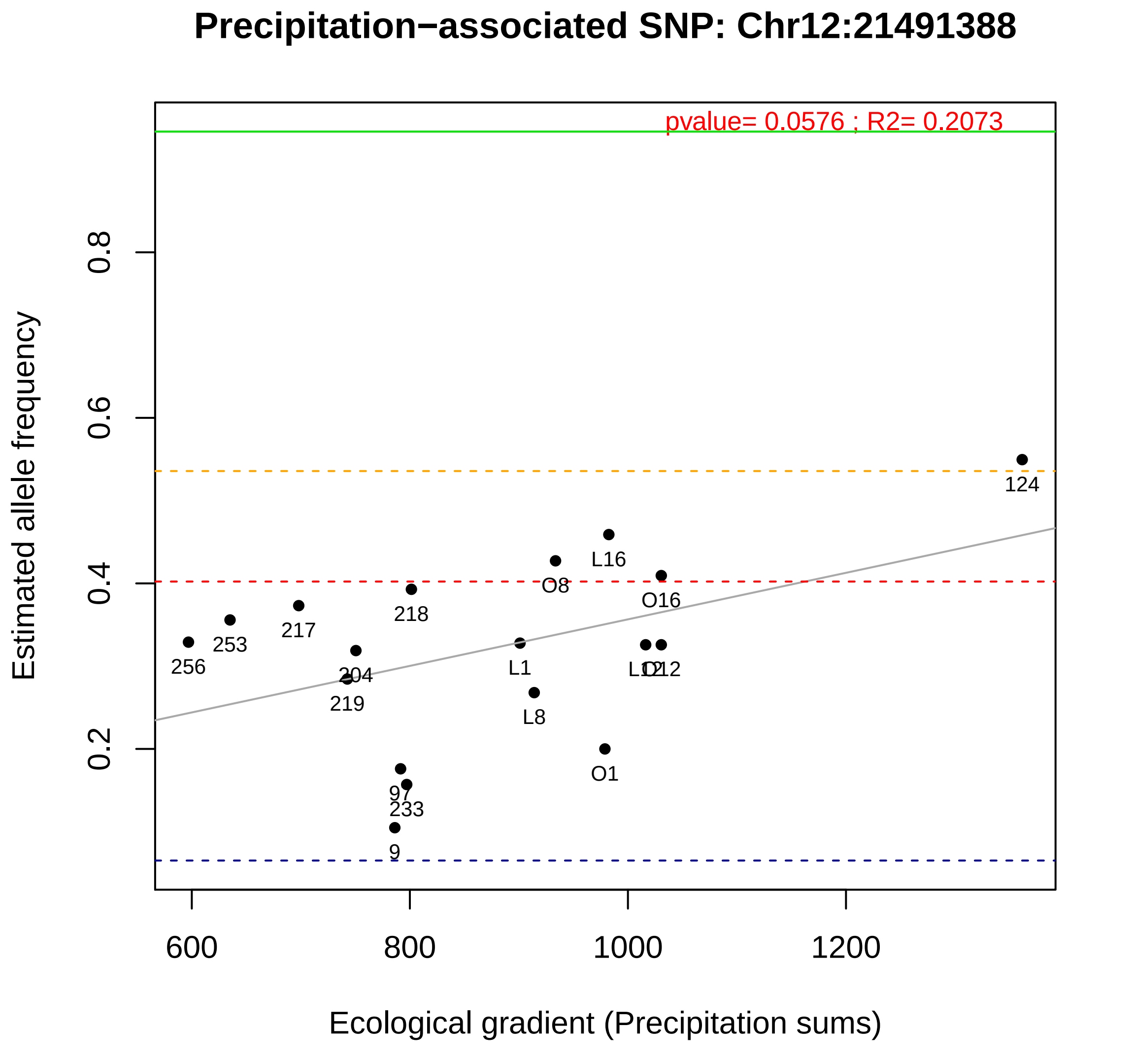
Precipitation−associated SNP: Chr12:21491388
pvalue= 0.0576 ; R2= 0.2073
Estimated allele frequency
0.2
0.4
0.6
0.8
600
800
1000
1200
Ecological gradient (Precipitation sums)
124
L16
O8
O16
218
217
253
256
204
219
L1
L8
L12
O12
O1
97
233
9

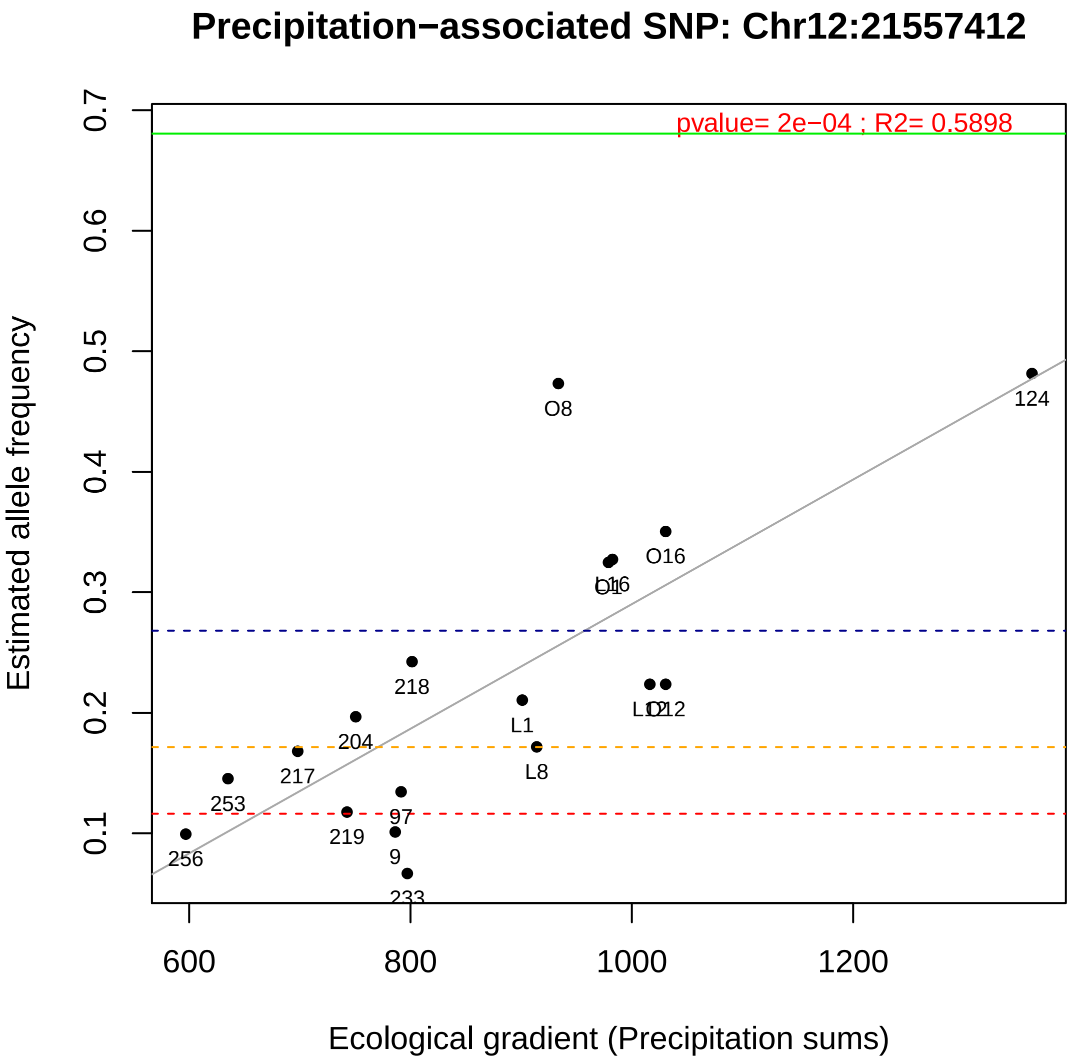
Precipitation−associated SNP: Chr12:21557412
pvalue= 2e−04 ; R2= 0.5898
Estimated allele frequency
0.1
0.2
0.3
0.4
0.5
0.6
0.7
600
800
1000
1200
Ecological gradient (Precipitation sums)
O8
124
O16
L16
O1
218
L1
L10
O12
204
217
L8
253
97
219
9
256
233

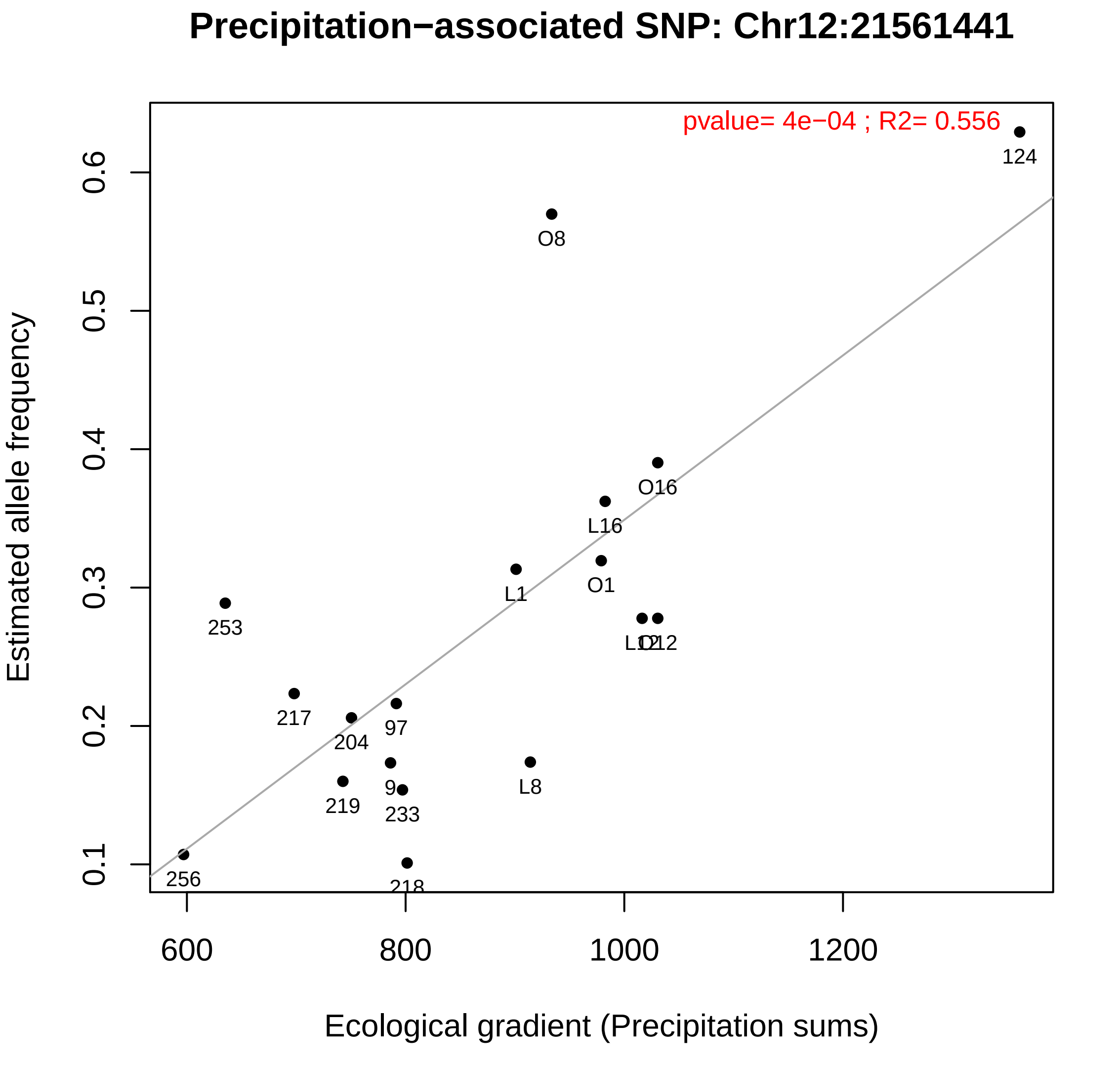
Precipitation−associated SNP: Chr12:21561441
pvalue= 4e−04 ; R2= 0.556
Estimated allele frequency
Ecological gradient (Precipitation sums)
0.1
0.2
0.3
0.4
0.5
0.6
600
800
1000
1200
124
O8
O16
L16
O1
L1
253
217
204
97
9
219
233
L8
256
218

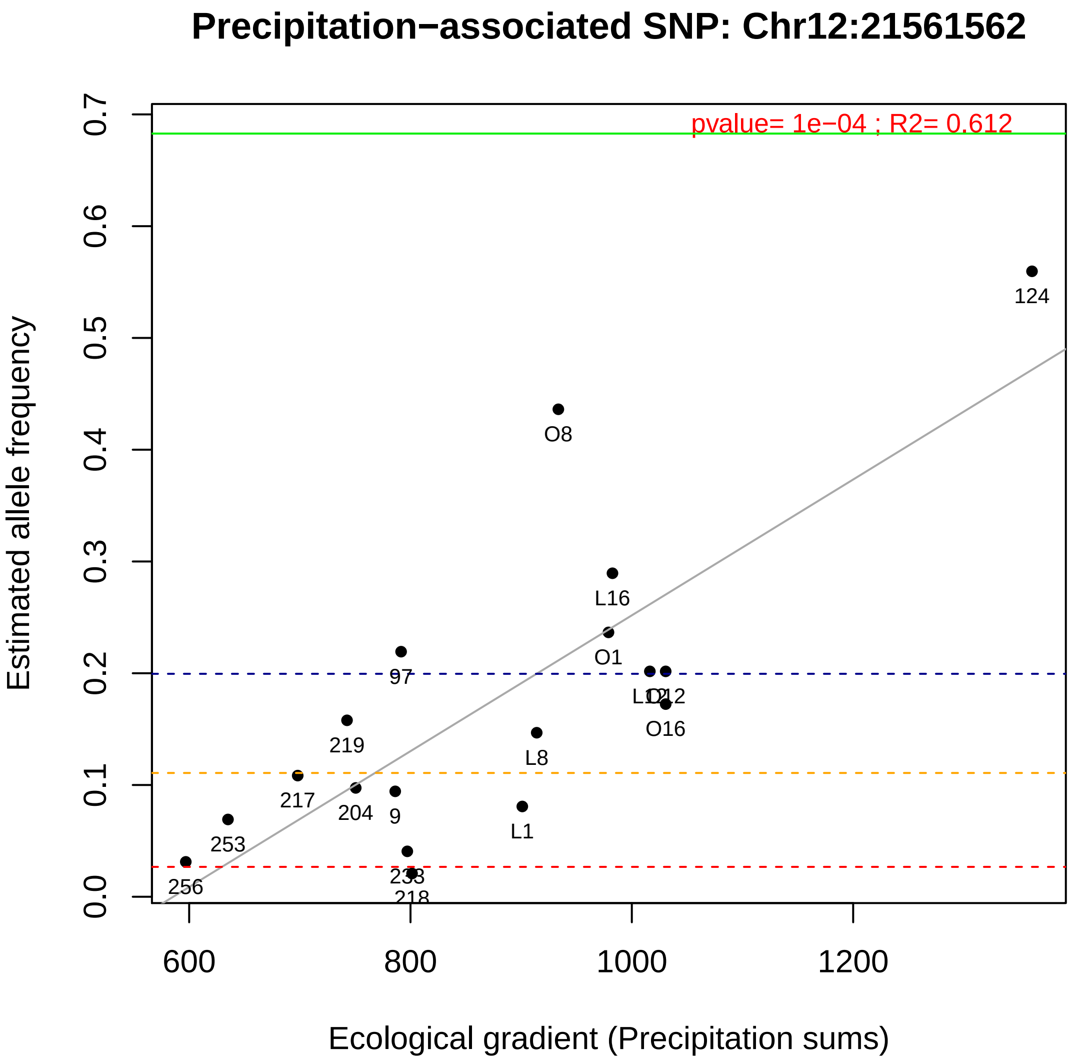

Precipitation−associated SNP: Chr12:21561562
pvalue= 1e−04 ; R2= 0.612
Estimated allele frequency
0.0
0.1
0.2
0.3
0.4
0.5
0.6
0.7
124
O8
L16
O1
97
L12
O12
O16
219
L8
217
204
9
L1
253
256
233
218
600
800
1000
1200
Ecological gradient (Precipitation sums)

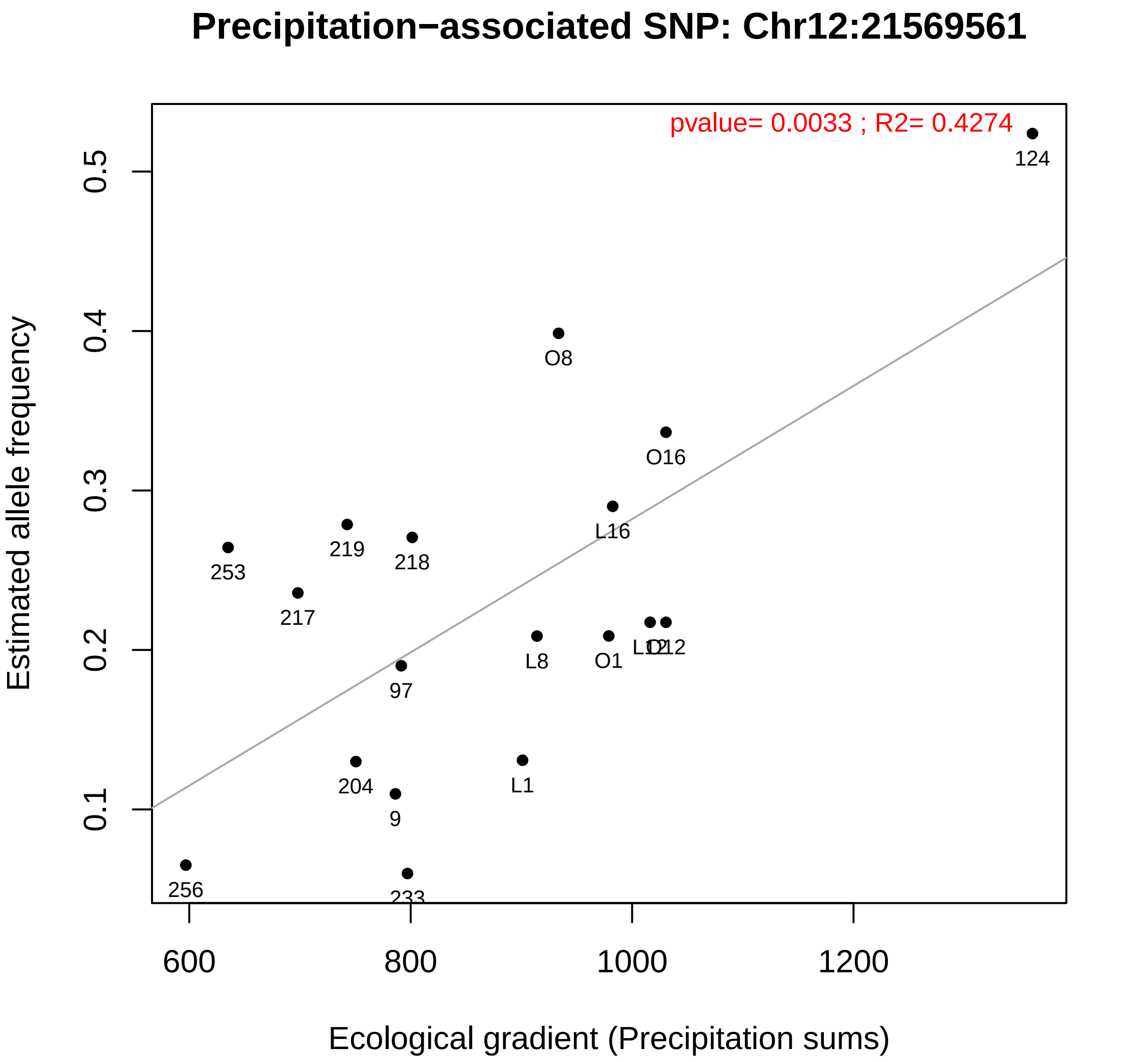
Precipitation−associated SNP: Chr12:21569561
pvalue= 0.0033 ; R2= 0.4274
Estimated allele frequency
0.1
0.2
0.3
0.4
0.5
600
800
1000
1200
Ecological gradient (Precipitation sums)
124
O8
O16
L16
219
218
253
217
L8
O1
O12
97
204
L1
9
256
233

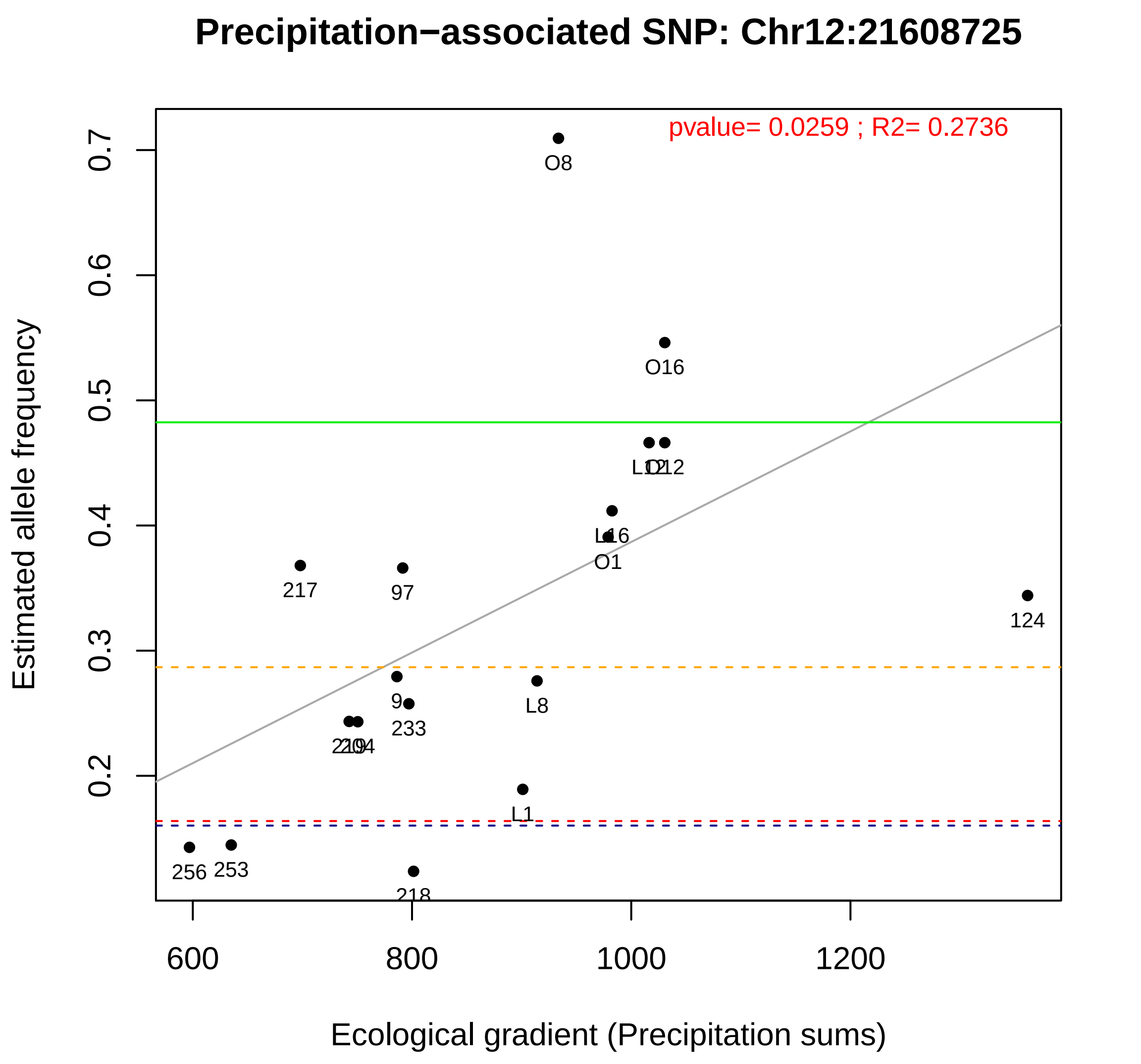

Precipitation−associated SNP: Chr12:21608725
pvalue= 0.0259 ; R2= 0.2736
Estimated allele frequency
Ecological gradient (Precipitation sums)
0.2
0.3
0.4
0.5
0.6
0.7
600
800
1000
1200
O8
O16
L12
O12
L16
O1
217
97
124
9
233
L8
210
194
L1
256
253
218

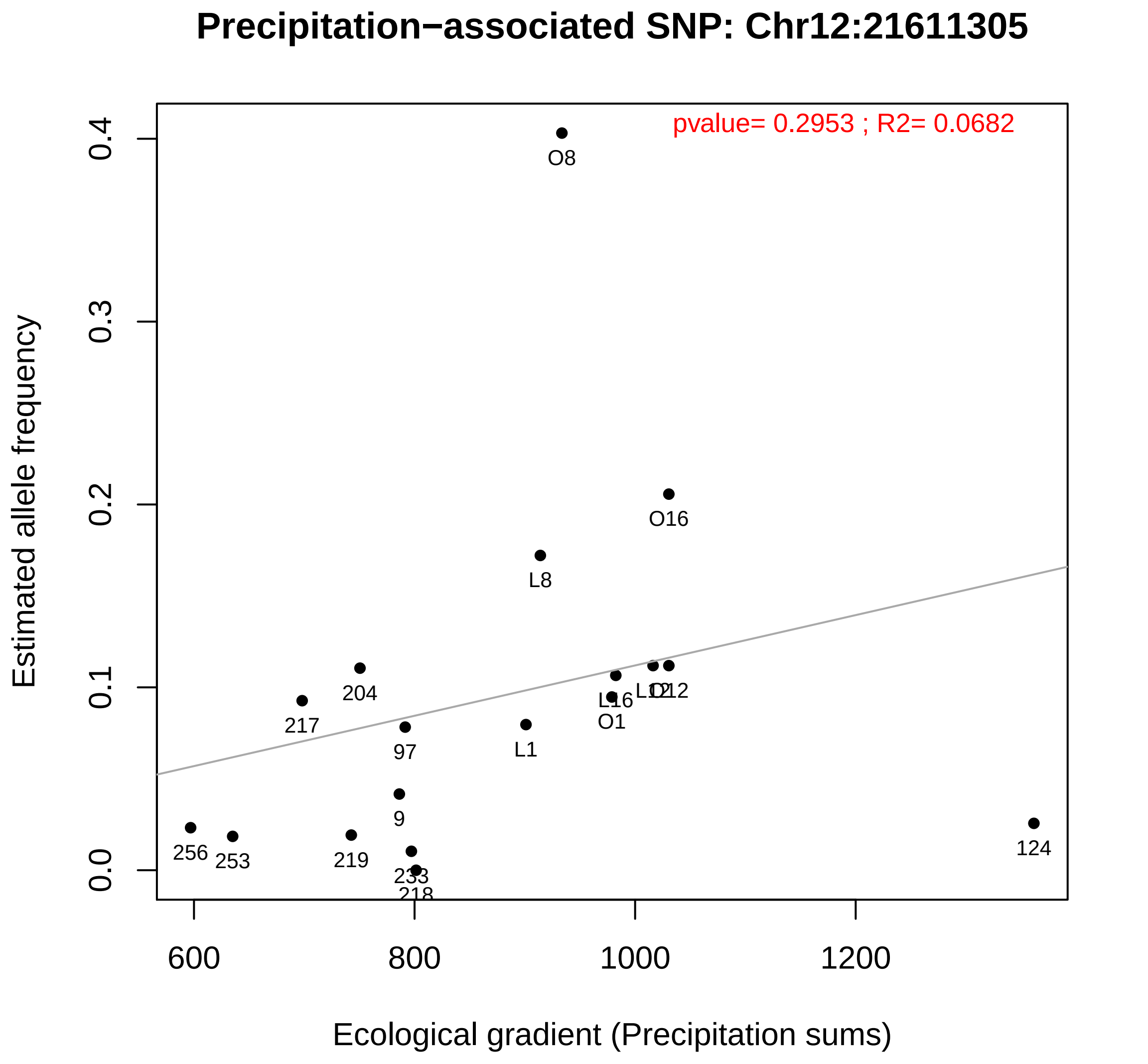
Precipitation−associated SNP: Chr12:21611305
pvalue= 0.2953 ; R2= 0.0682
Estimated allele frequency
0.0
0.1
0.2
0.3
0.4
600
800
1000
1200
Ecological gradient (Precipitation sums)
O8
O16
L8
204
217
97
L1
L16
O1
L12
O12
9
256
253
219
233
218
124

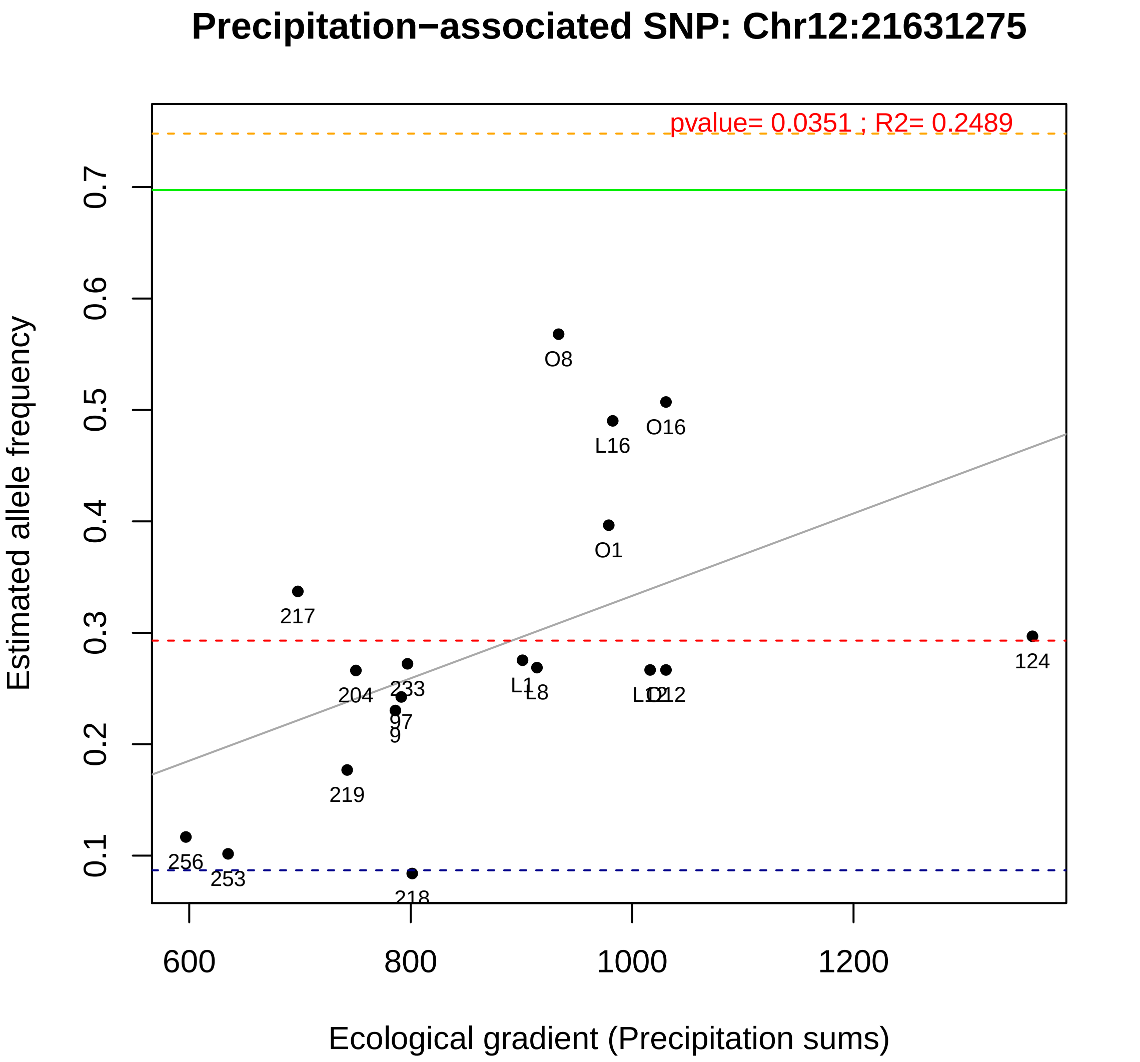

Precipitation−associated SNP: Chr12:21631275
pvalue= 0.0351 ; R2= 0.2489
Estimated allele frequency
Ecological gradient (Precipitation sums)
0.1
0.2
0.3
0.4
0.5
0.6
0.7
600
800
1000
1200
O8
O16
L16
O1
217
124
204
233
L1
L8
L12
O12
97
9
219
256
253
218

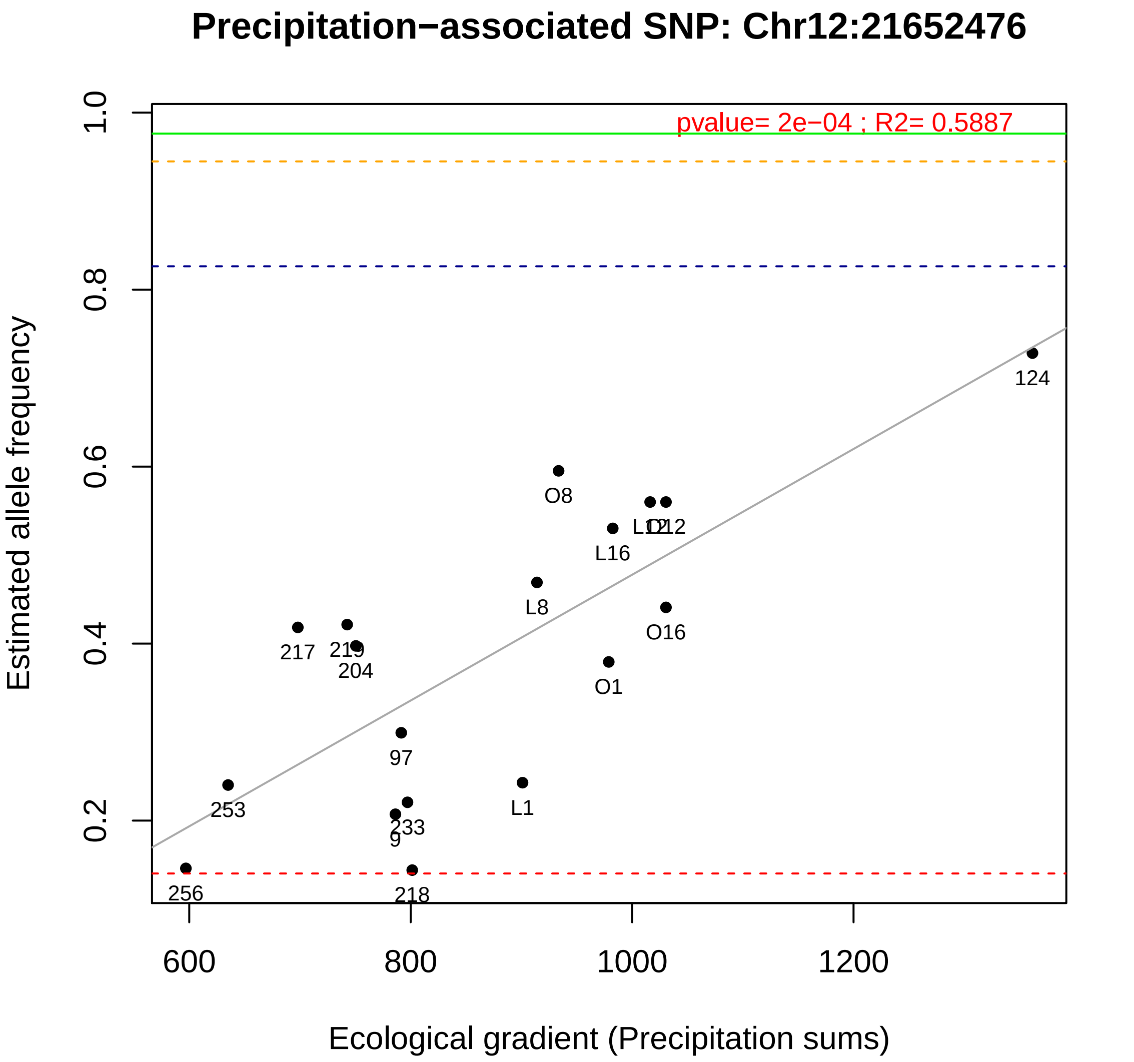

Precipitation−associated SNP: Chr12:21652476
pvalue= 2e−04 ; R2= 0.5887
Estimated allele frequency
Ecological gradient (Precipitation sums)
0.2
0.4
0.6
0.8
1.0
600
800
1000
1200
124
O8
L12
O12
L16
L8
O16
O1
217
219
204
97
253
L1
233
9
256
218

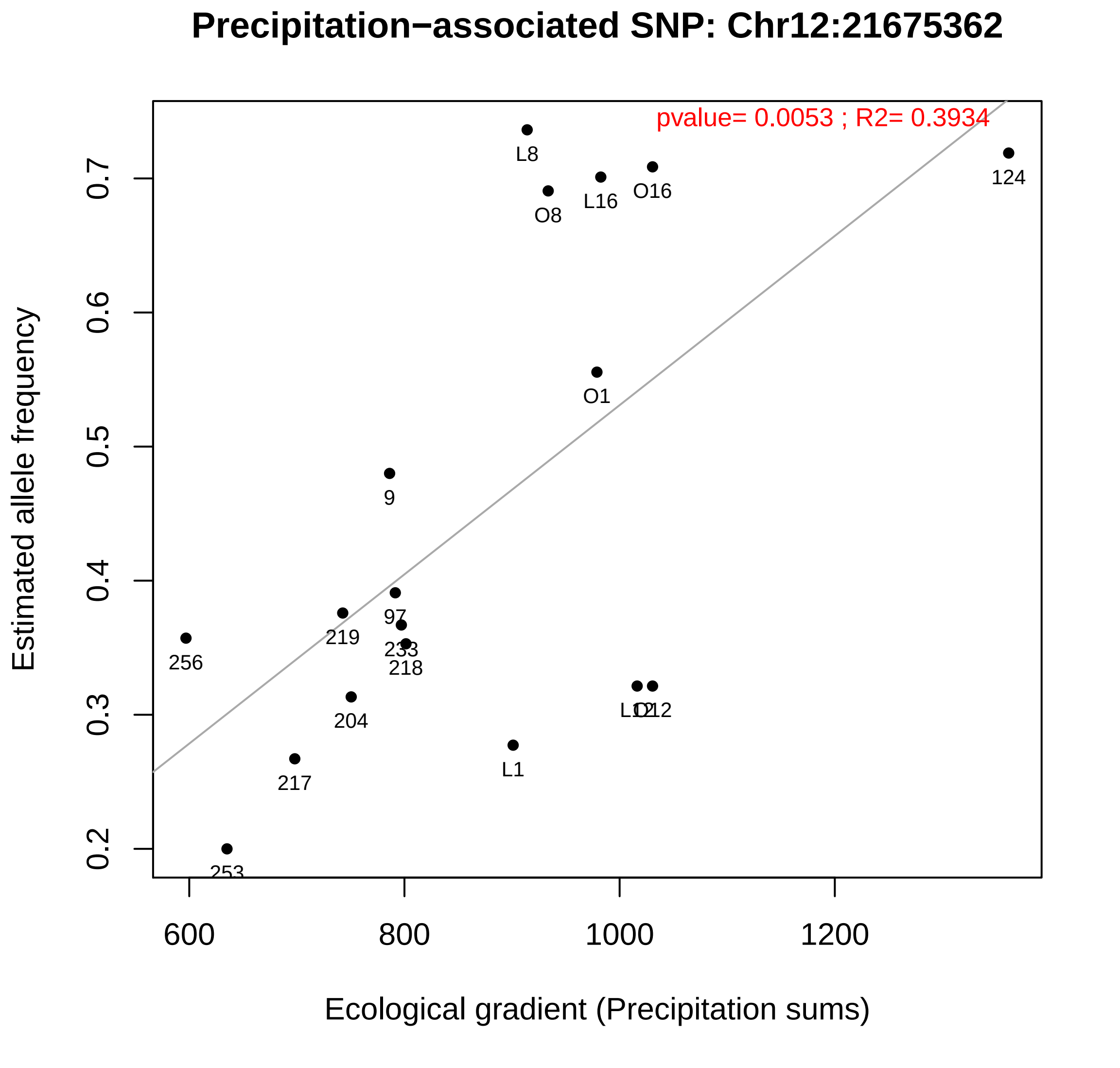
Precipitation−associated SNP: Chr12:21675362
pvalue= 0.0053 ; R2= 0.3934
Estimated allele frequency
0.2
0.3
0.4
0.5
0.6
0.7
600
800
1000
1200
Ecological gradient (Precipitation sums)
L8
O8
L16
O16
124
O1
9
97
219
233
218
256
204
L12
O12
217
L1
253

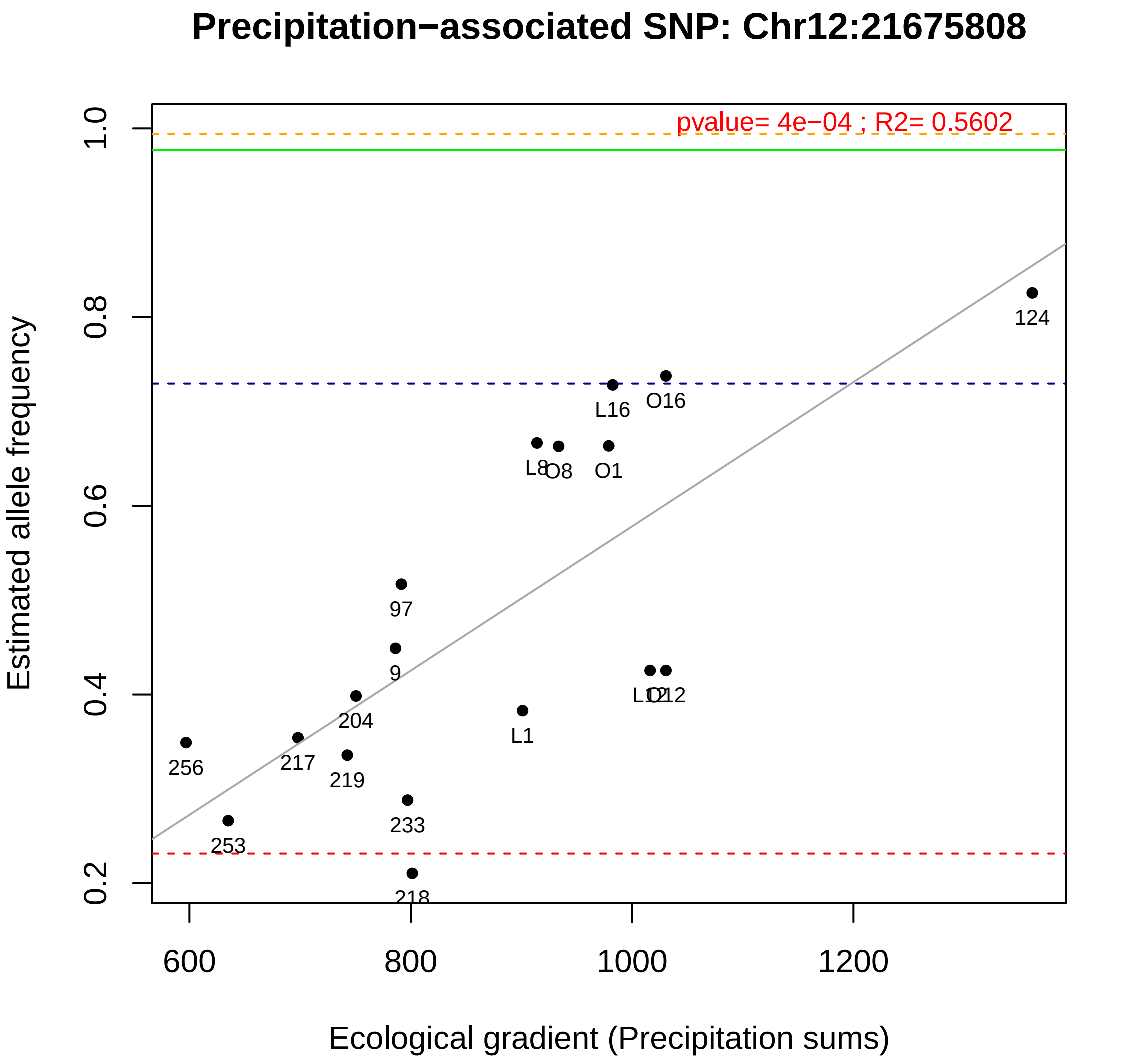

Precipitation−associated SNP: Chr12:21675808
pvalue= 4e−04 ; R2= 0.5602
Estimated allele frequency
Ecological gradient (Precipitation sums)
0.2
0.4
0.6
0.8
1.0
600
800
1000
1200
124
O16
L16
L8
O8
O1
97
9
204
L1
L12
O12
256
217
219
233
253
218

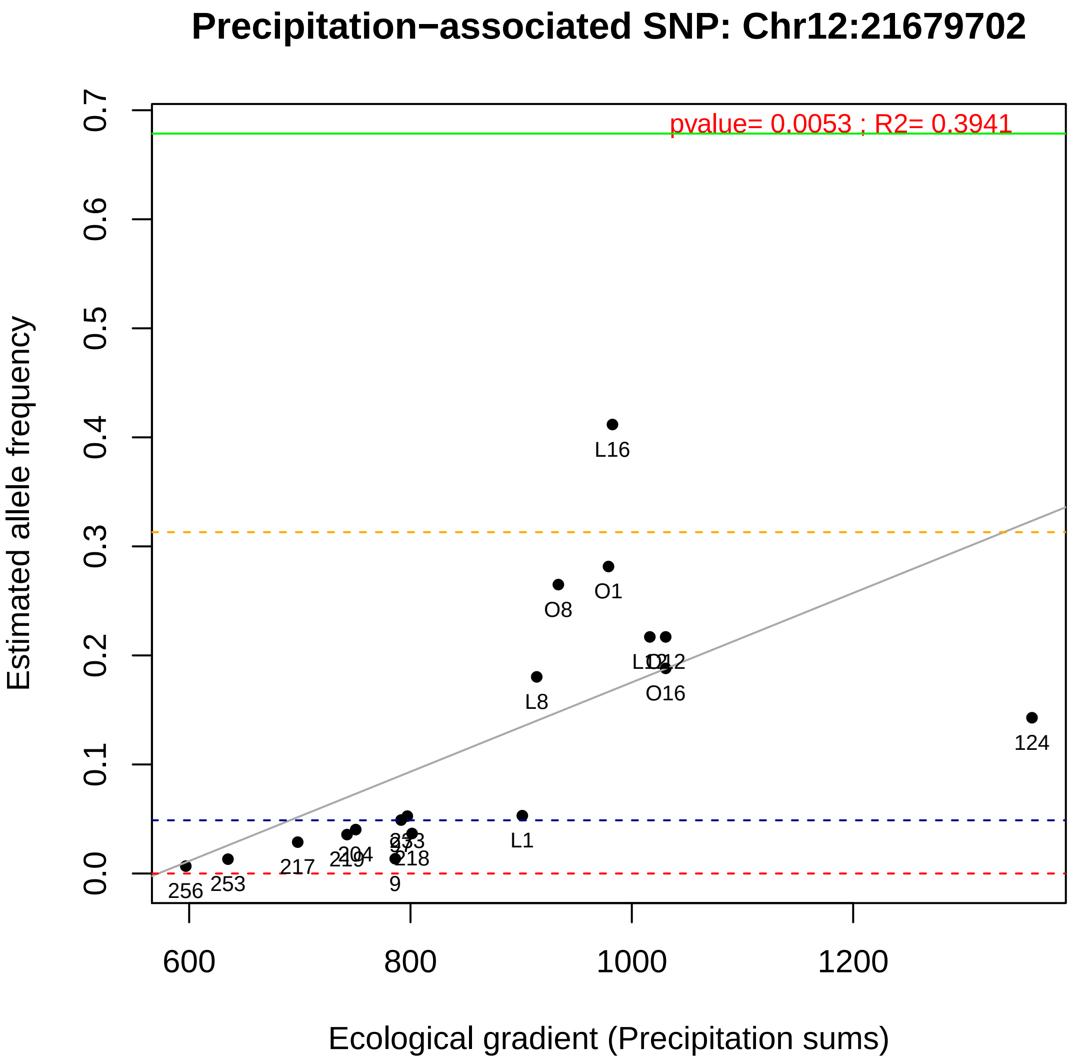
Precipitation−associated SNP: Chr12:21679702
pvalue= 0.0053 ; R2= 0.3941
Estimated allele frequency
Ecological gradient (Precipitation sums)
0.0
0.1
0.2
0.3
0.4
0.5
0.6
0.7
600
800
1000
1200
L16
O1
O8
L8
O16
124
L1
256
253
217
219
204
218
9

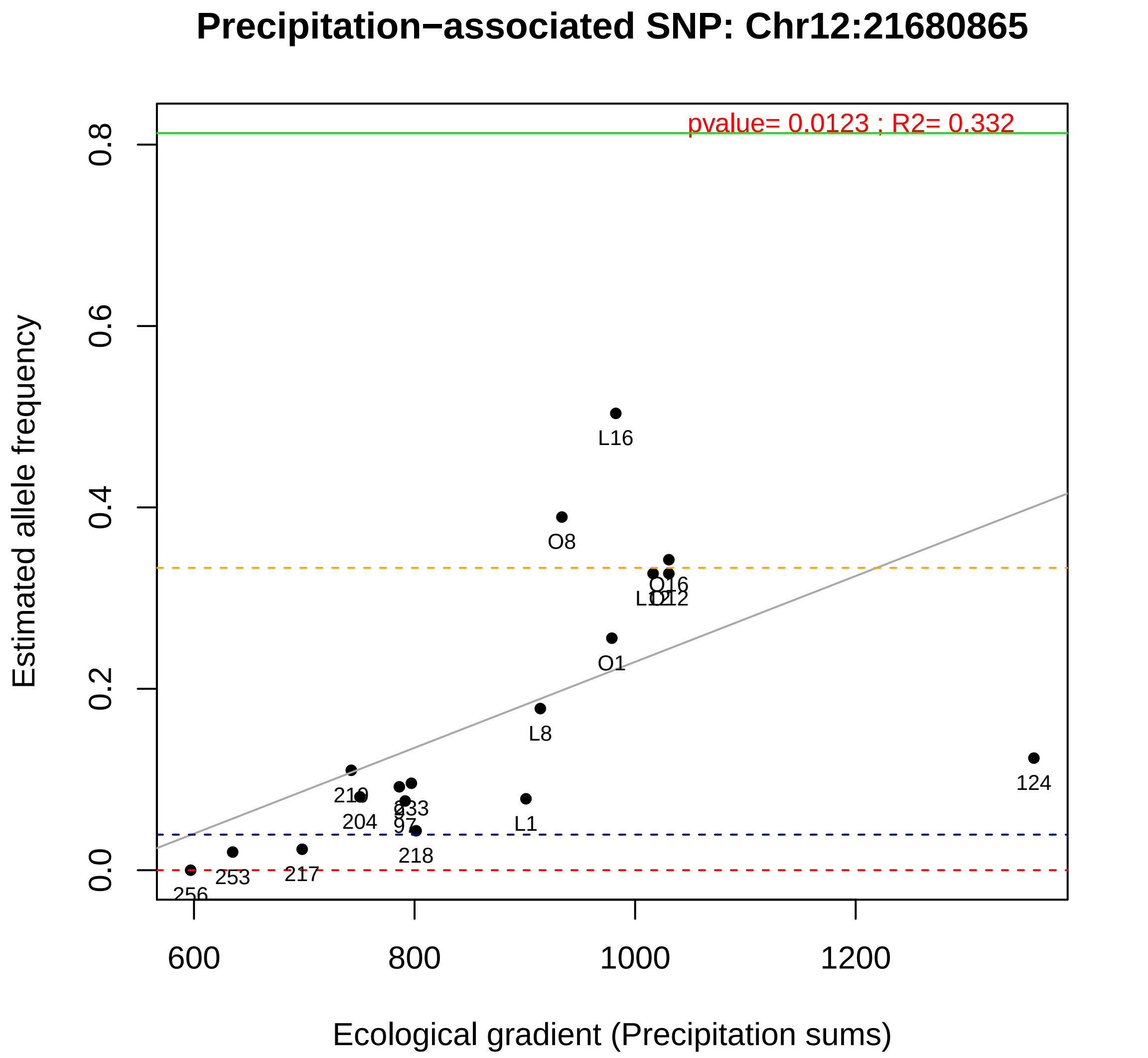
Precipitation−associated SNP: Chr12:21680865
pvalue= 0.0123 ; R2= 0.332
Estimated allele frequency
0.0
0.2
0.4
0.6
0.8
600
800
1000
1200
Ecological gradient (Precipitation sums)
L16
O8
O16
O12
O1
L8
124
204
233
97
218
L1
253
217
256

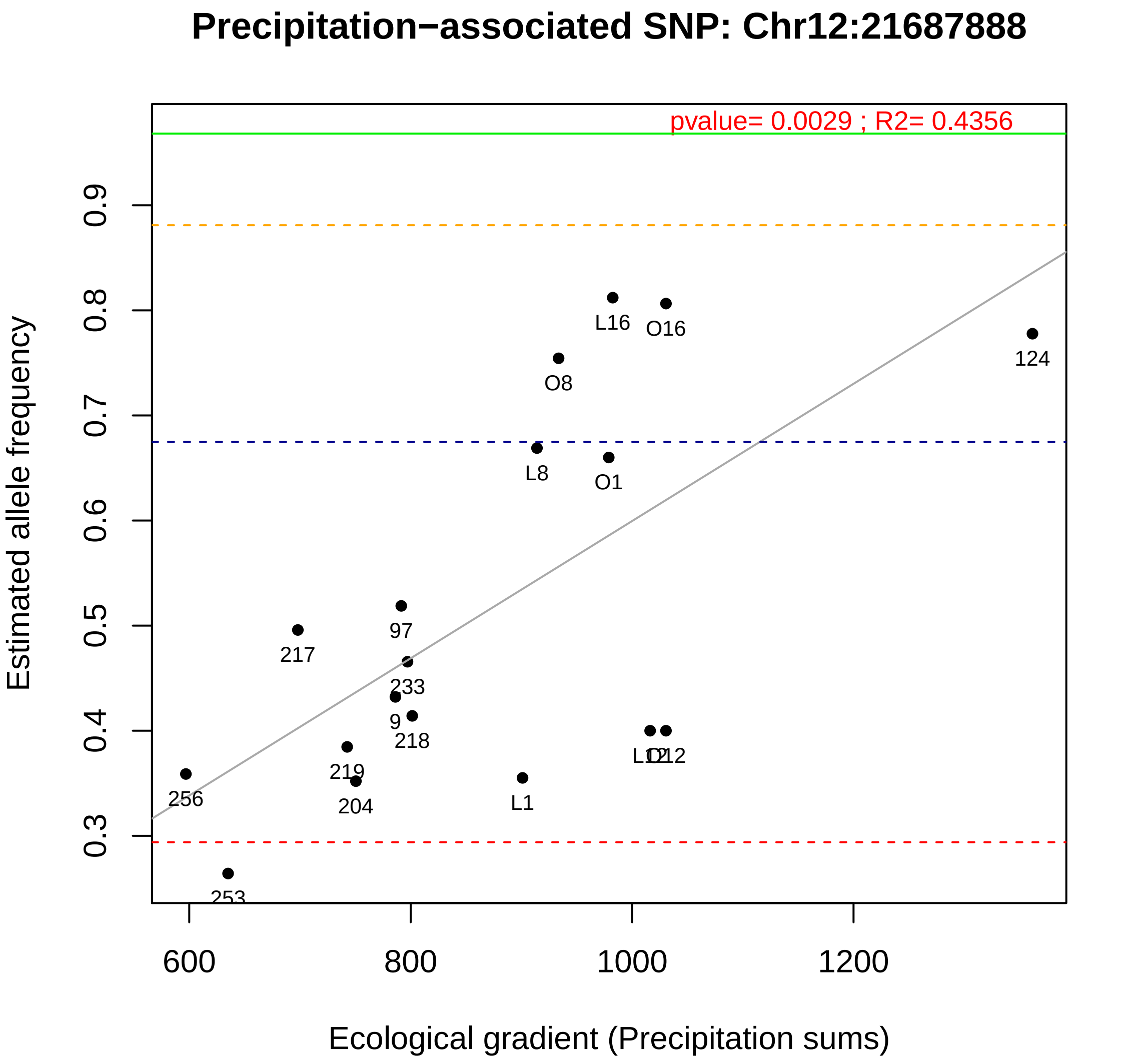
Precipitation−associated SNP: Chr12:21687888
pvalue= 0.0029 ; R2= 0.4356
Estimated allele frequency
0.3
0.4
0.5
0.6
0.7
0.8
0.9
600
800
1000
1200
Ecological gradient (Precipitation sums)
L16
O16
124
O8
L8
O1
97
217
233
9
218
L12
O12
219
256
204
L1
253

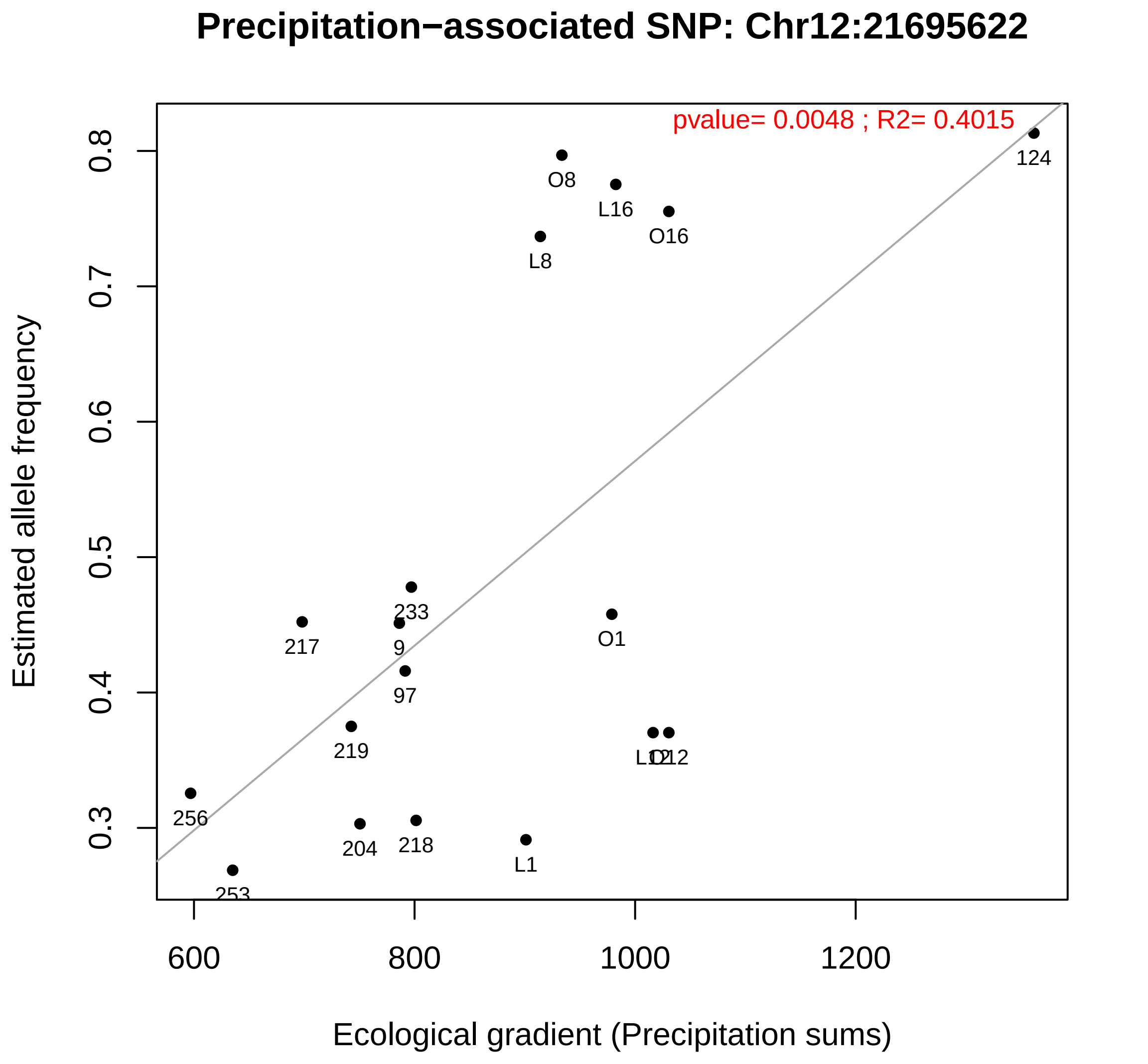
Precipitation−associated SNP: Chr12:21695622
pvalue= 0.0048 ; R2= 0.4015
Estimated allele frequency
0.3
0.4
0.5
0.6
0.7
0.8
600
800
1000
1200
Ecological gradient (Precipitation sums)
124
O8
L16
O16
L8
233
217
9
O1
97
219
O12
256
204
218
L1
253

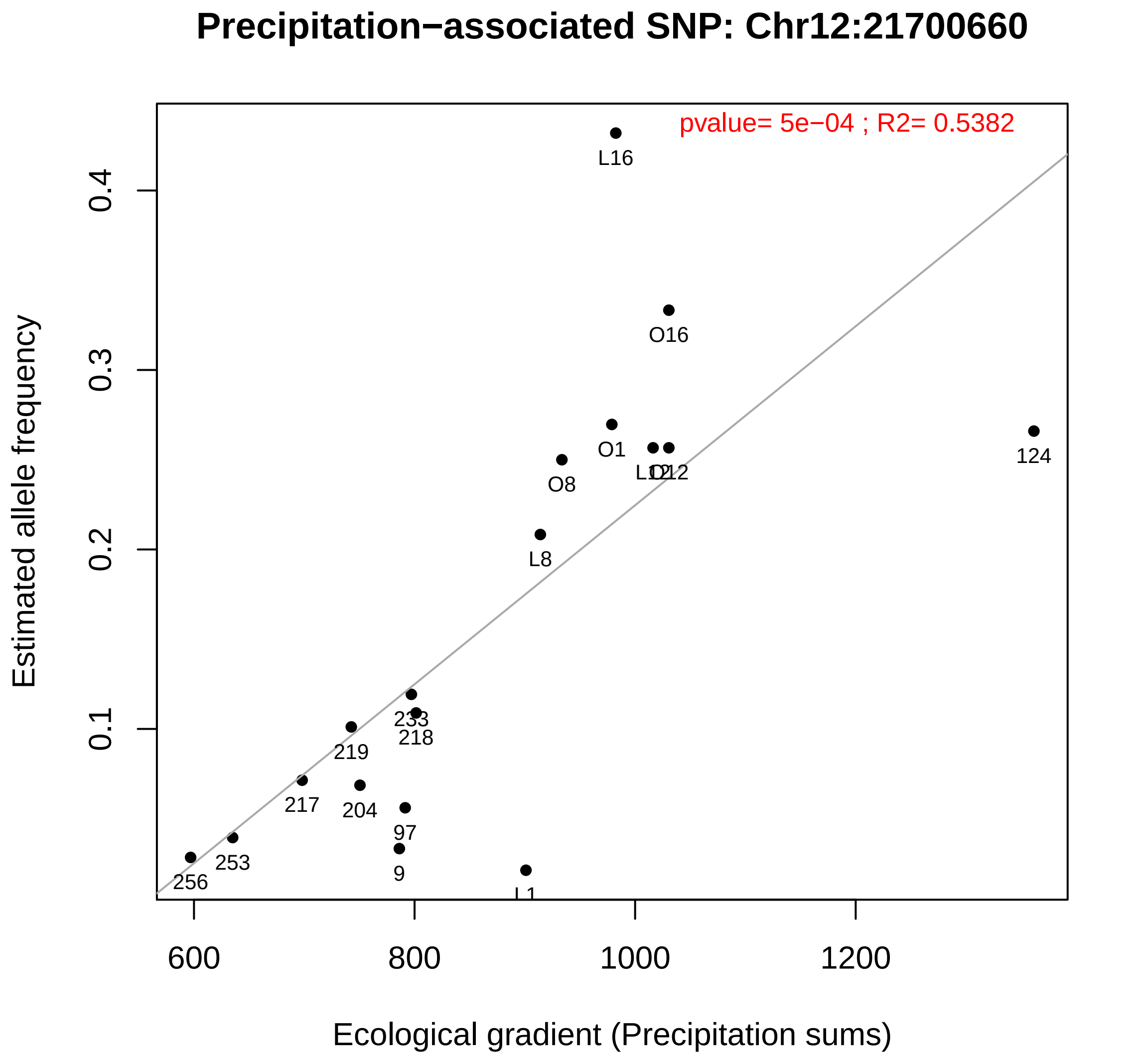

Precipitation−associated SNP: Chr12:21700660
pvalue= 5e−04 ; R2= 0.5382
Estimated allele frequency
0.1
0.2
0.3
0.4
600
800
1000
1200
Ecological gradient (Precipitation sums)
L16
O16
O1
124
O8
L8
233
218
219
217
204
97
253
256
9

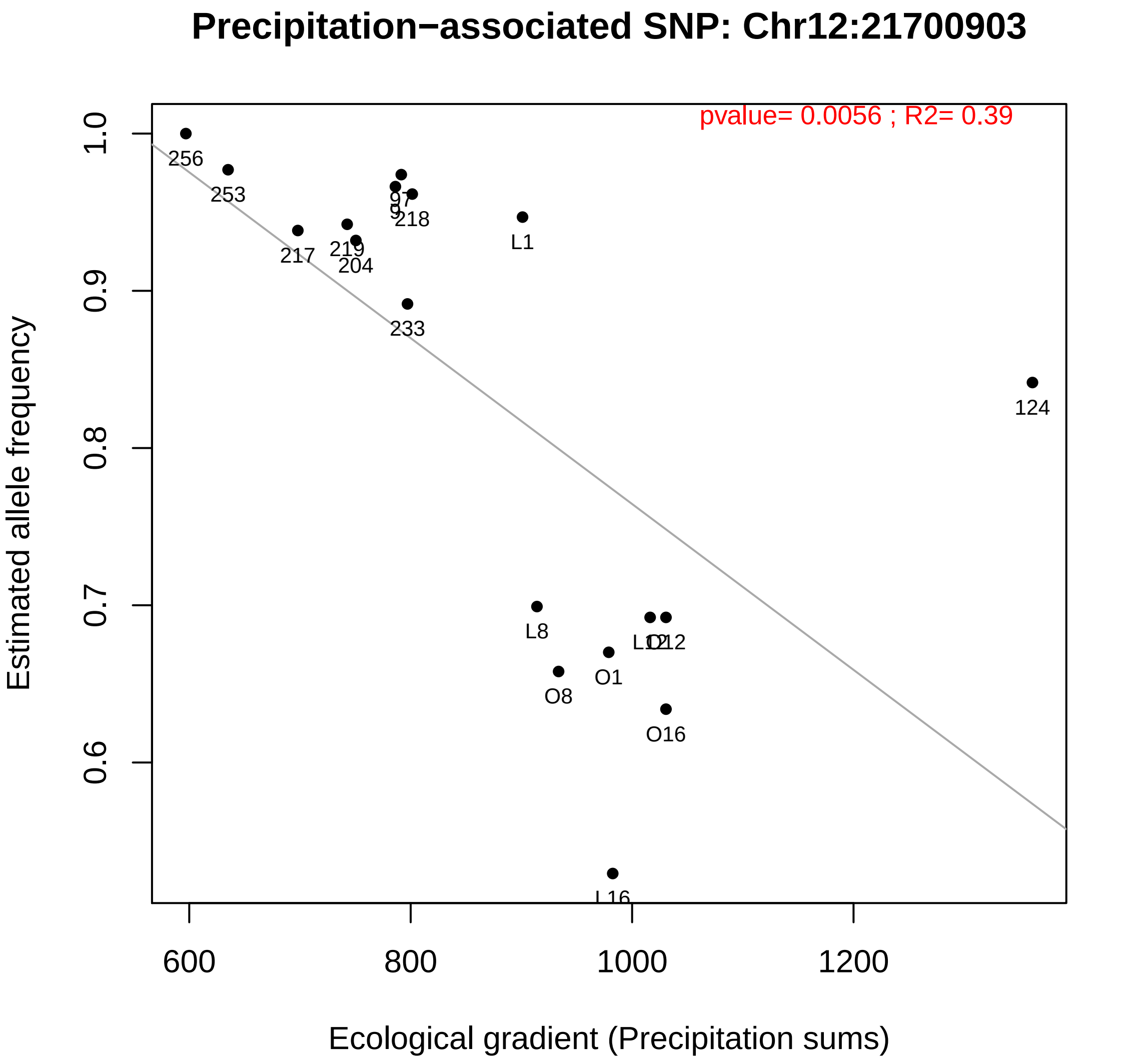
Precipitation−associated SNP: Chr12:21700903
pvalue= 0.0056 ; R2= 0.39
Estimated allele frequency
Ecological gradient (Precipitation sums)
1.0
0.9
0.8
0.7
0.6
600
800
1000
1200
256
253
217
219
204
97
218
233
L1
124
L8
O1
O8
O12
O16
L16

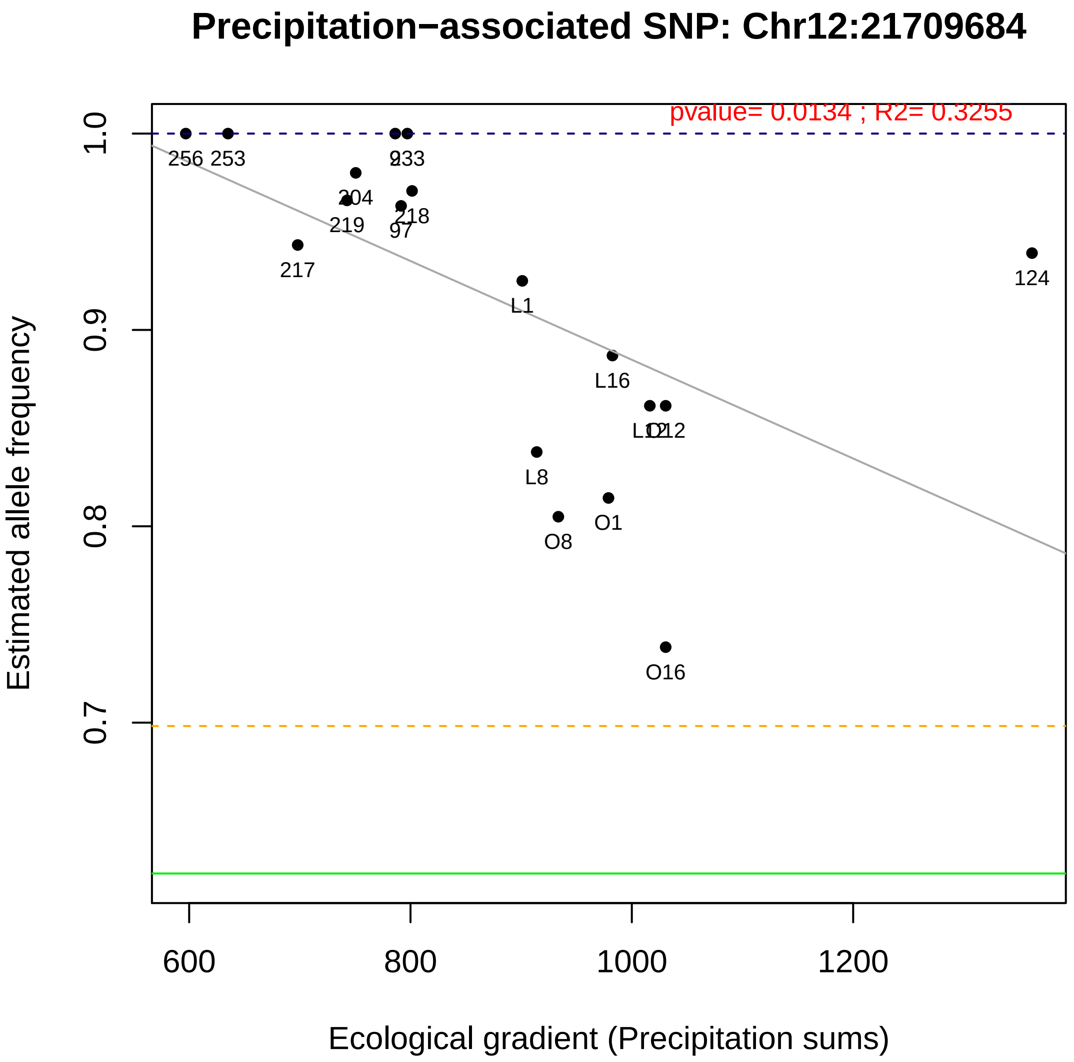
Precipitation−associated SNP: Chr12:21709684
pvalue= 0.0134 ; R2= 0.3255
Estimated allele frequency
Ecological gradient (Precipitation sums)
1.0
0.9
0.8
0.7
600
800
1000
1200
256
253
233
204
219
218
97
217
124
L1
L16
L12
O12
L8
O1
O8
O16

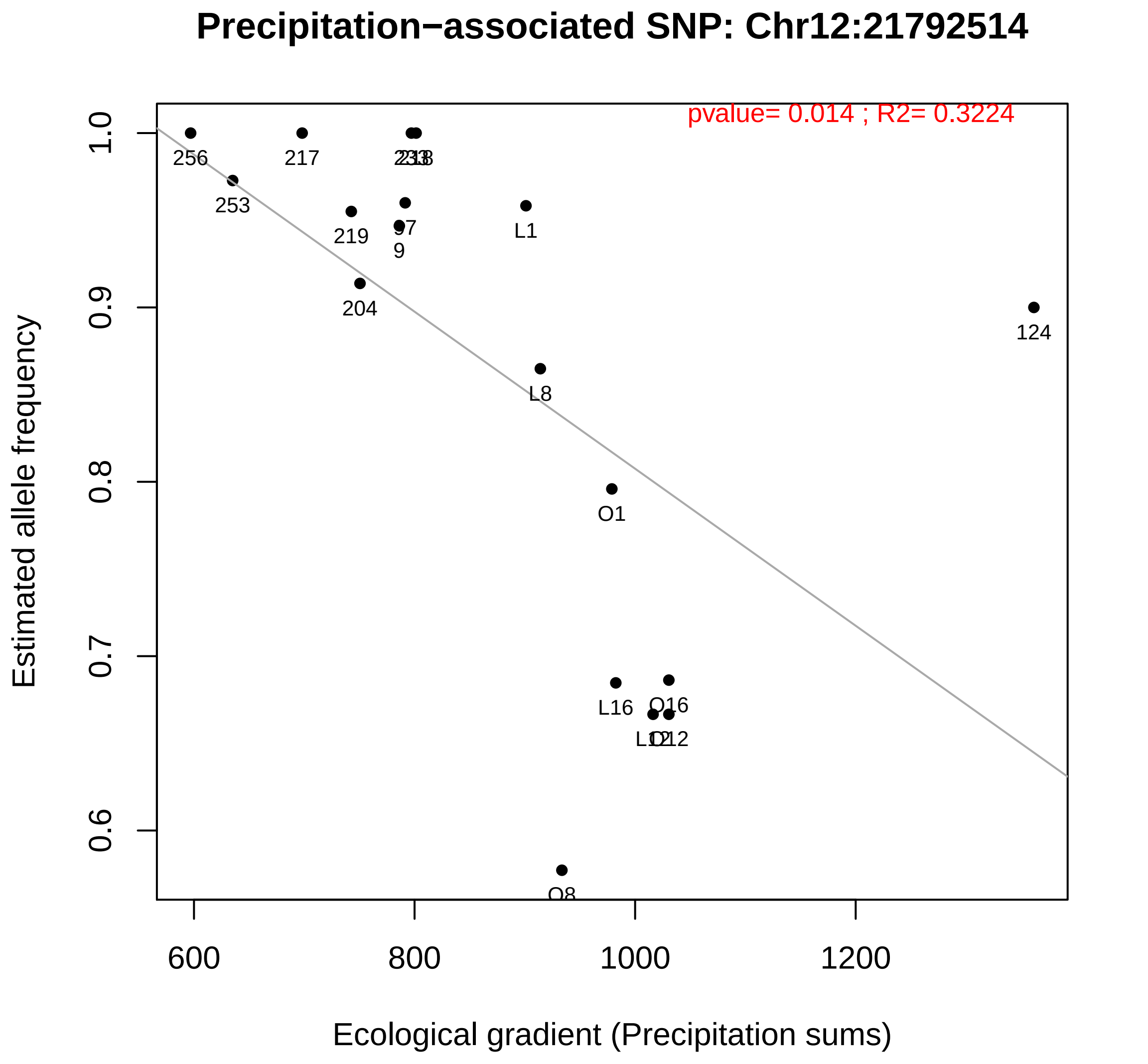
Precipitation−associated SNP: Chr12:21792514
pvalue= 0.014 ; R2= 0.3224
Estimated allele frequency
Ecological gradient (Precipitation sums)
1.0
0.9
0.8
0.7
0.6
600
800
1000
1200
256
217
253
219
204
L1
124
L8
O1
L16
O16
O8

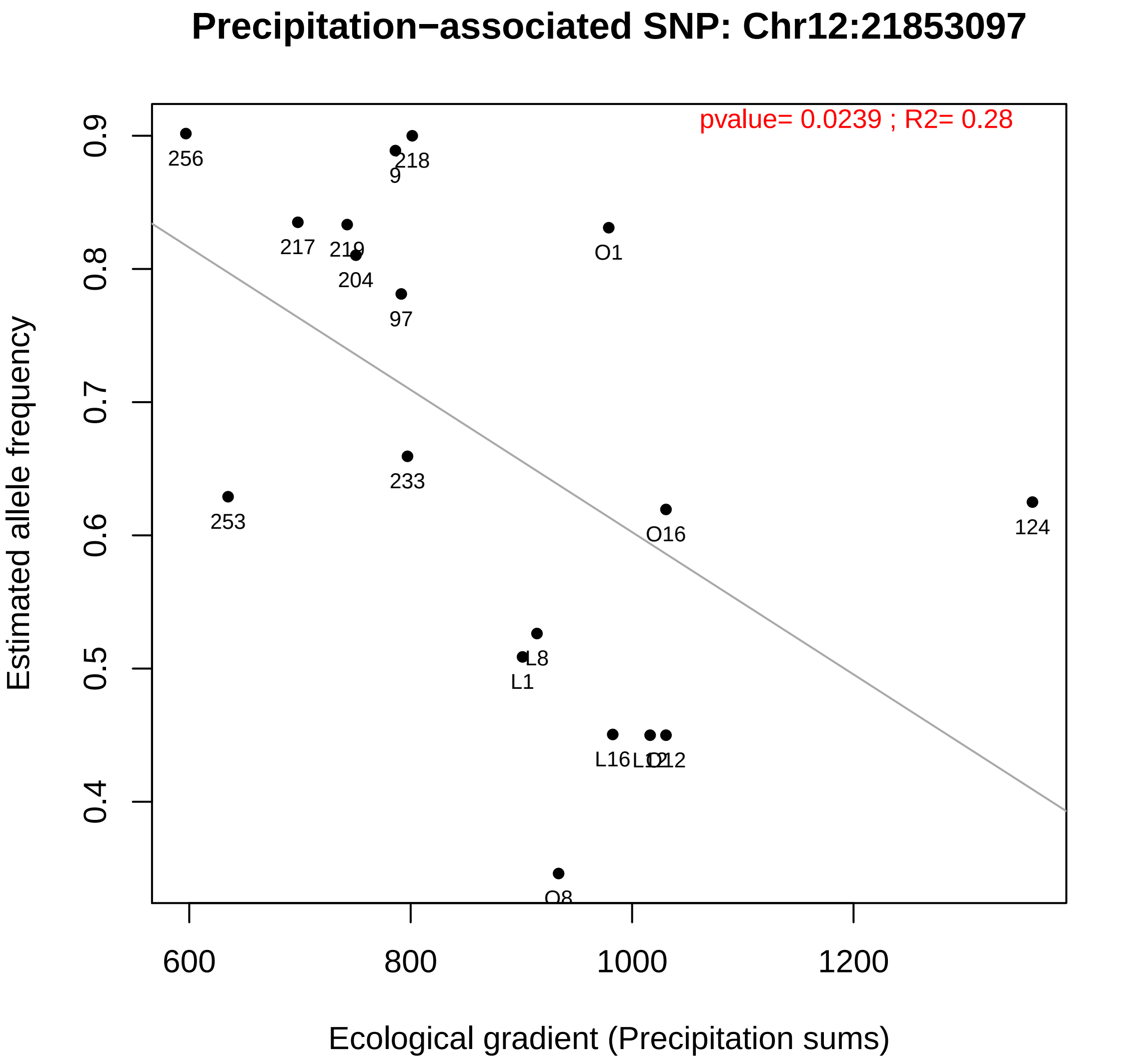

Precipitation−associated SNP: Chr12:21853097
pvalue= 0.0239 ; R2= 0.28
Estimated allele frequency
0.4
0.5
0.6
0.7
0.8
0.9
600
800
1000
1200
Ecological gradient (Precipitation sums)
256
218
9
217
219
204
O1
97
233
253
O16
124
L8
L1
L16
L12
O12
O8

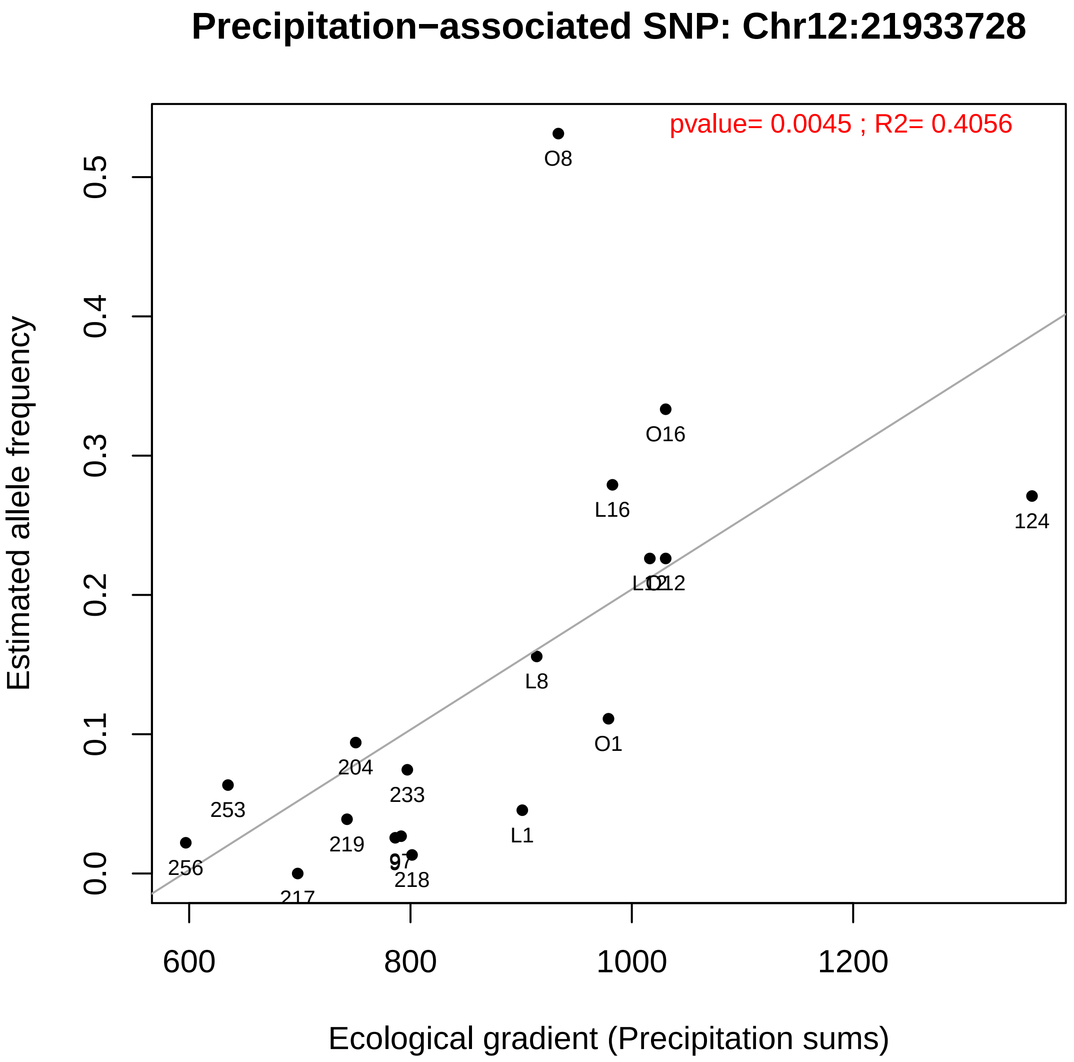

Precipitation−associated SNP: Chr12:21933728
pvalue= 0.0045 ; R2= 0.4056
Estimated allele frequency
Ecological gradient (Precipitation sums)
0.0
0.1
0.2
0.3
0.4
0.5
600
800
1000
1200
O8
O16
L16
124
L12
O12
L8
O1
204
233
253
L1
219
256
97
218
217

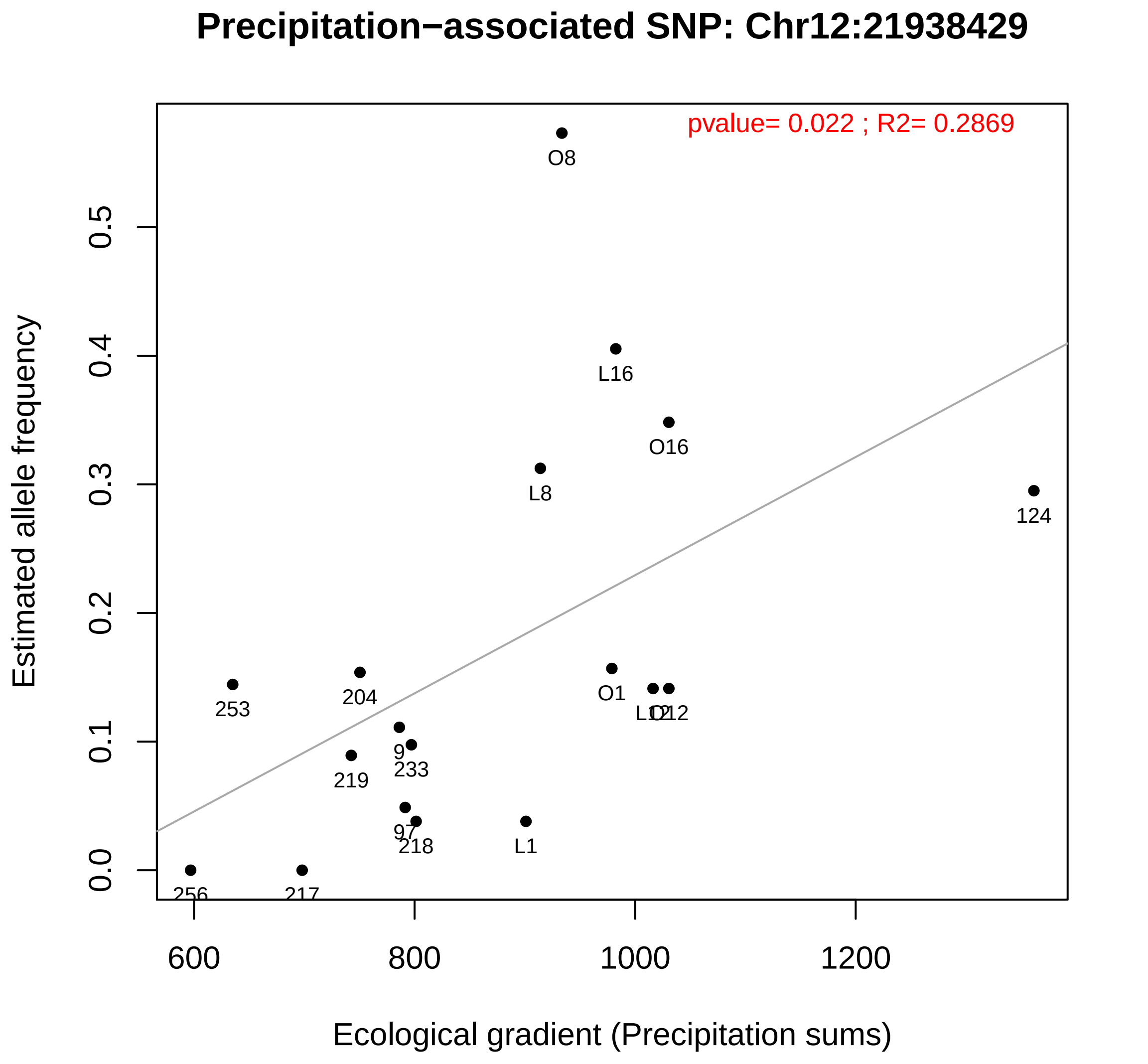
Precipitation−associated SNP: Chr12:21938429
pvalue= 0.022 ; R2= 0.2869
Estimated allele frequency
Ecological gradient (Precipitation sums)
0.0
0.1
0.2
0.3
0.4
0.5
600
800
1000
1200
O8
L16
O16
L8
124
253
204
O1
9
233
219
97
218
L1
256
217

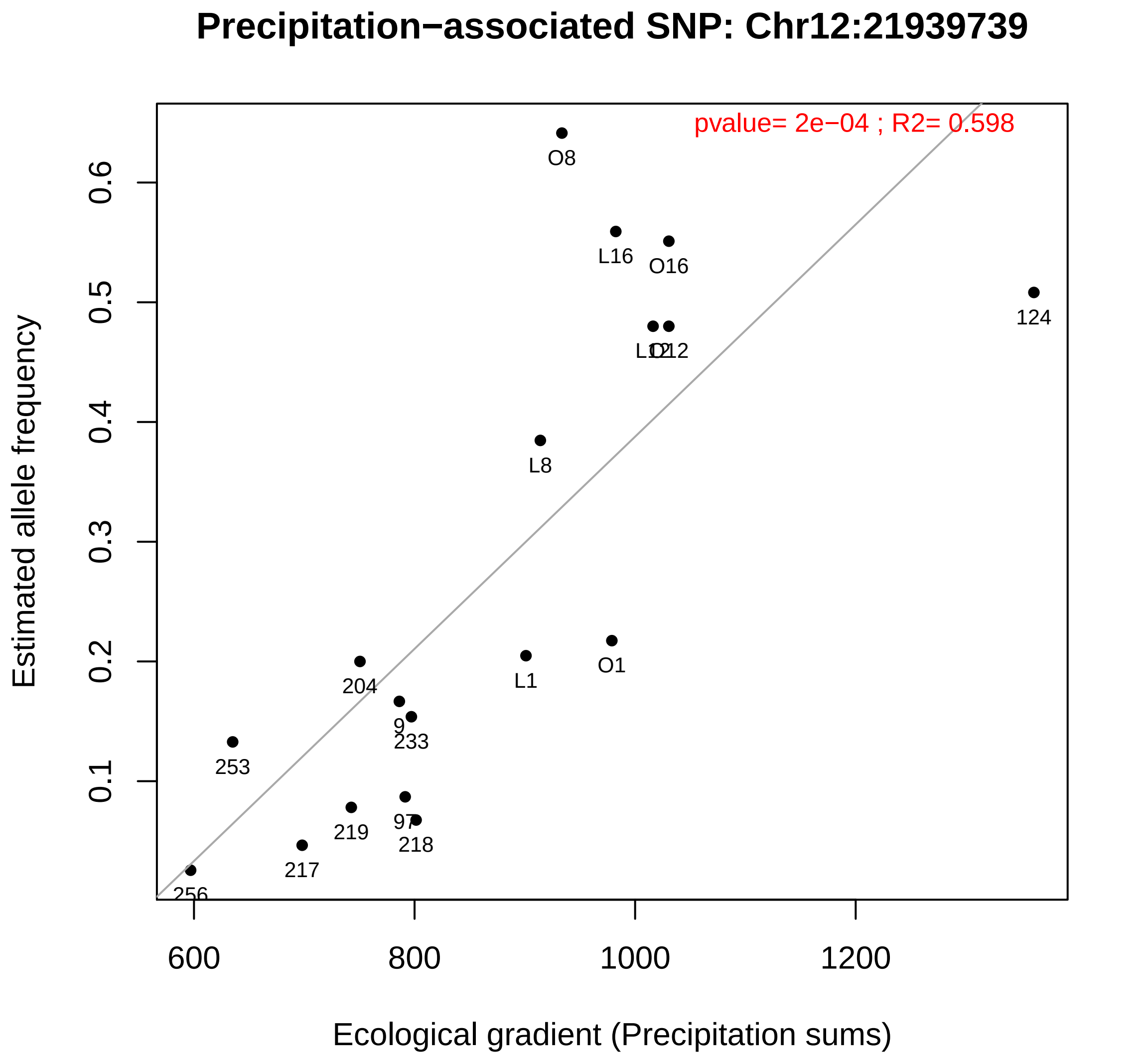

Precipitation−associated SNP: Chr12:21939739
pvalue= 2e−04 ; R2= 0.598
Estimated allele frequency
Ecological gradient (Precipitation sums)
0.1
0.2
0.3
0.4
0.5
0.6
600
800
1000
1200
O8
L16
O16
124
L12
O12
L8
L1
O1
204
9
233
253
219
97
218
217
256

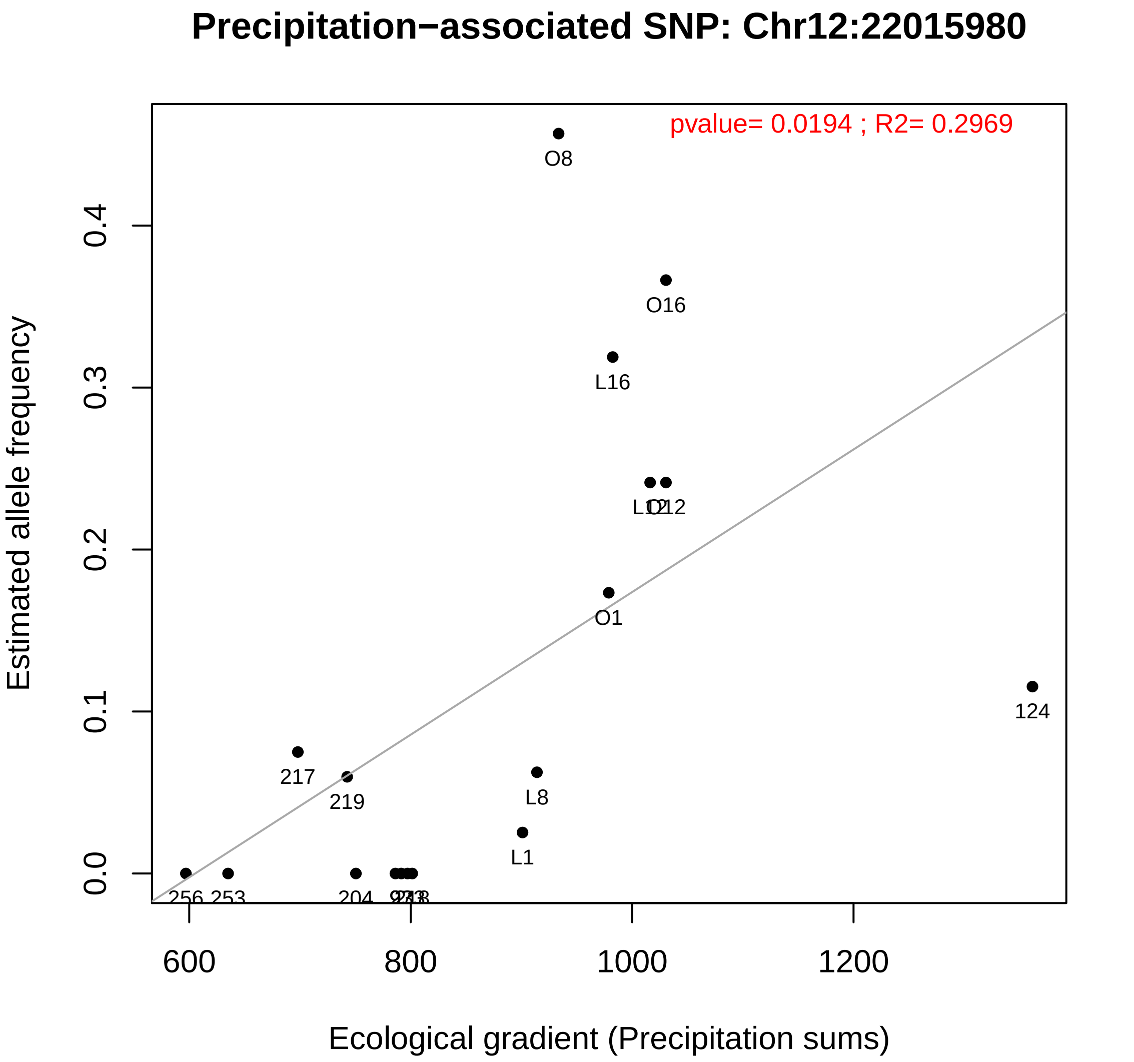

Precipitation−associated SNP: Chr12:22015980
pvalue= 0.0194 ; R2= 0.2969
O8
O16
L16
L12
O12
O1
124
217
219
L8
L1
256
253
204
0.0
0.1
0.2
0.3
0.4
600
800
1000
1200
Estimated allele frequency
Ecological gradient (Precipitation sums)

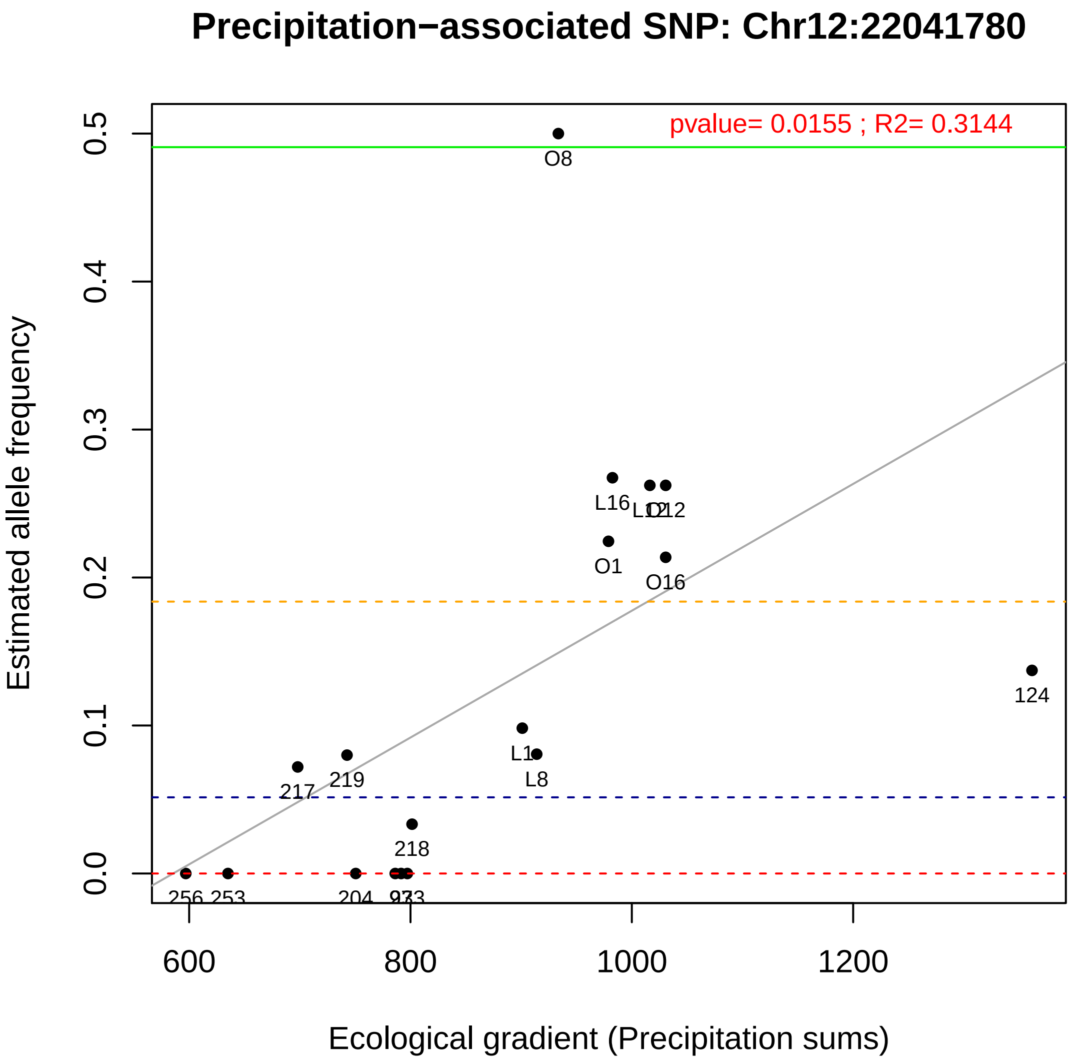

Precipitation−associated SNP: Chr12:22041780
pvalue= 0.0155 ; R2= 0.3144
Estimated allele frequency
Ecological gradient (Precipitation sums)
0.0
0.1
0.2
0.3
0.4
0.5
600
800
1000
1200
O8
L16
O12
O1
O16
124
L1
L8
217
219
218
256
253
204

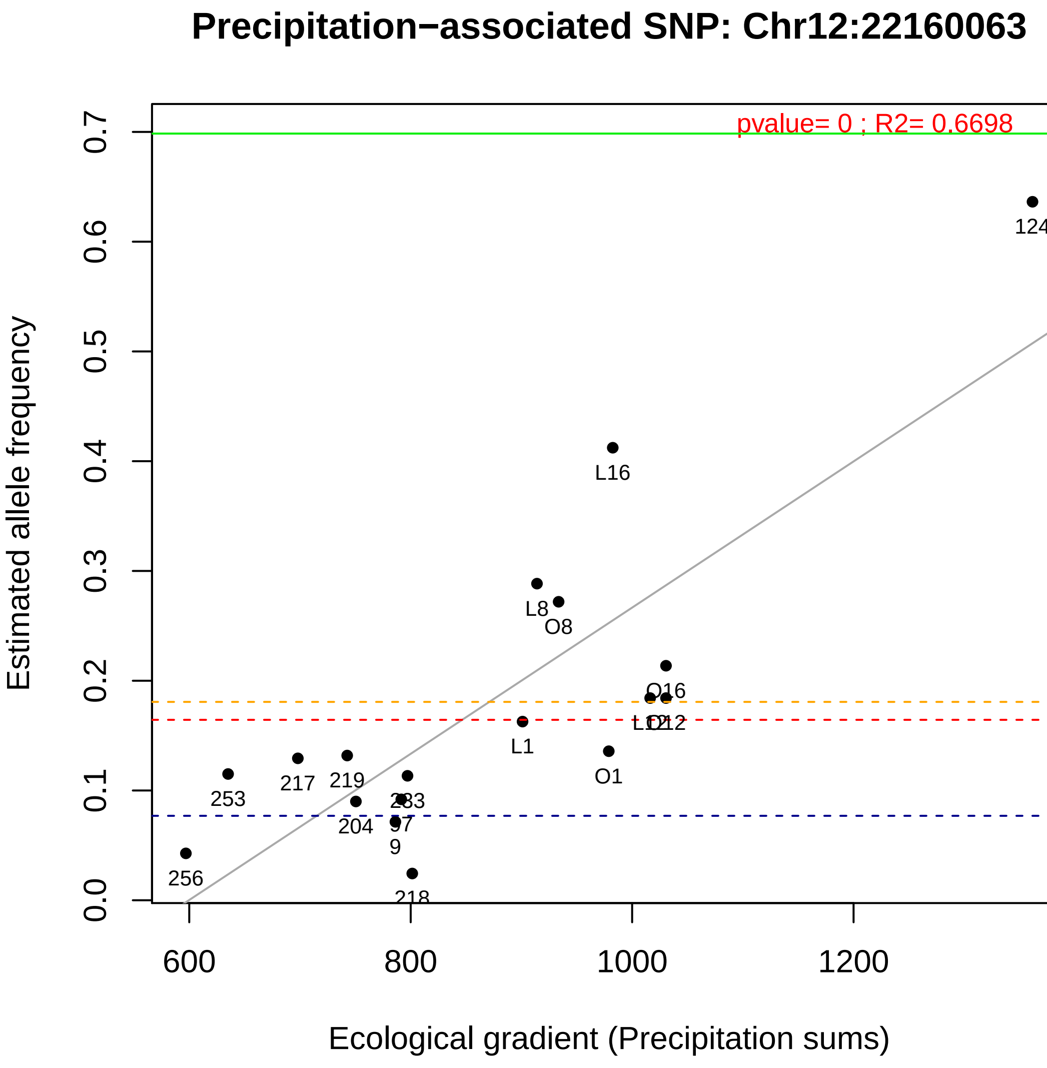
Precipitation−associated SNP: Chr12:22160063
pvalue= 0 ; R2= 0.6698
Estimated allele frequency
Ecological gradient (Precipitation sums)
0.0
0.1
0.2
0.3
0.4
0.5
0.6
0.7
600
800
1000
1200
124
L16
L8
O8
O16
L10
O12
L1
O1
217
219
253
204
233
97
9
256
218

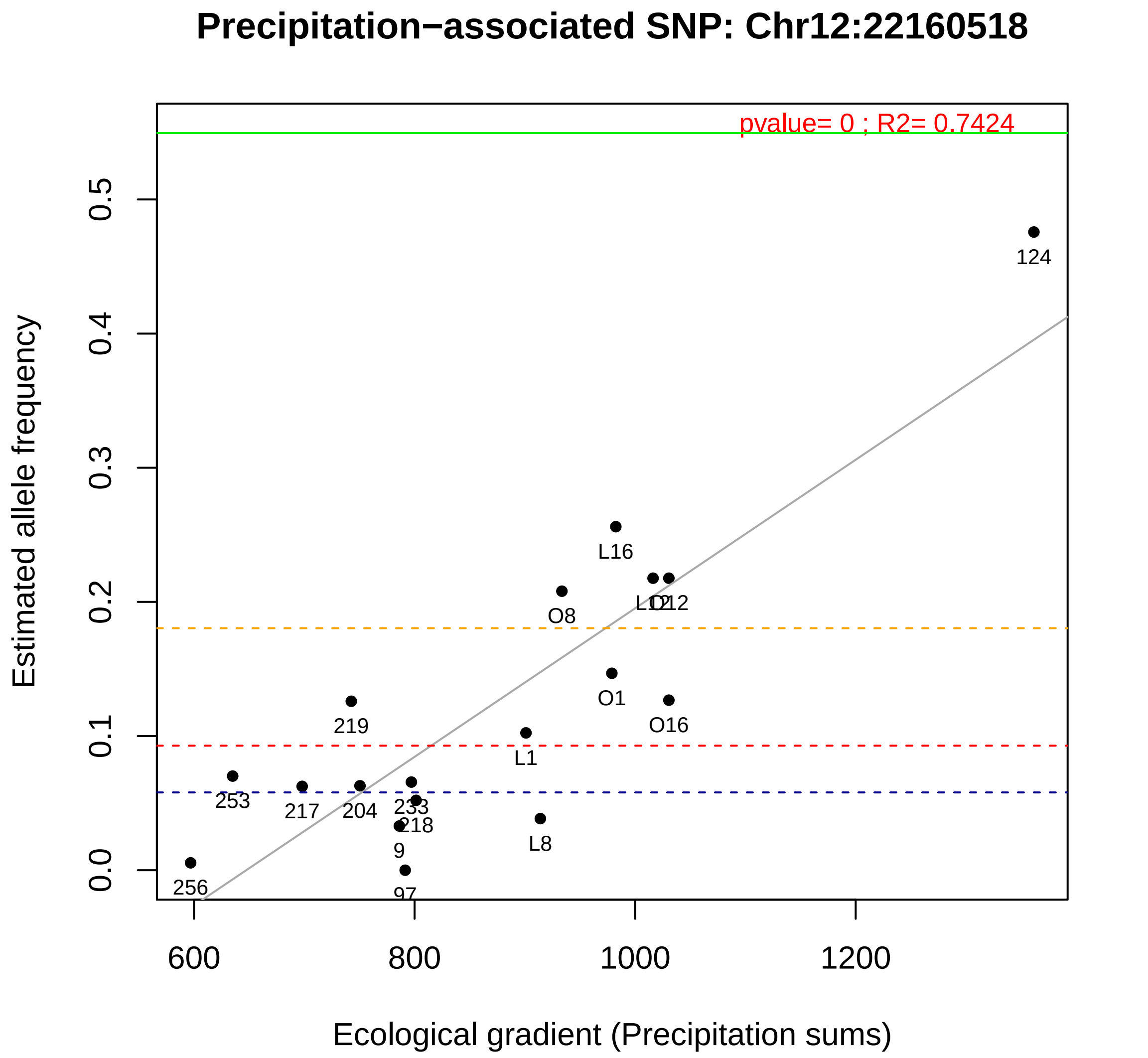
Precipitation−associated SNP: Chr12:22160518
pvalue= 0 ; R2= 0.7424
Estimated allele frequency
Ecological gradient (Precipitation sums)
0.0
0.1
0.2
0.3
0.4
0.5
600
800
1000
1200
124
L16
O8
O12
O1
O16
219
L1
253
217
204
233
218
9
L8
256
97

#### Precipitation−associated SNP: Chr12:22395766

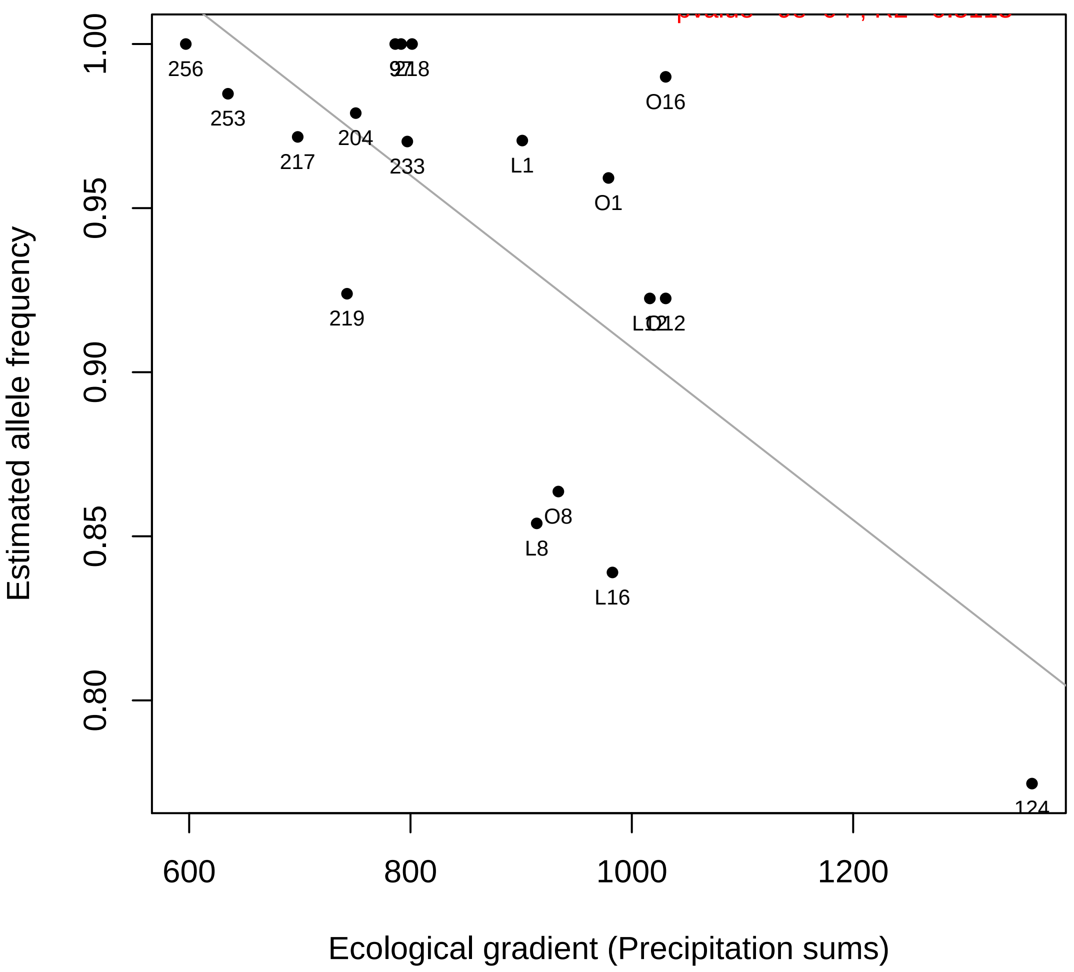

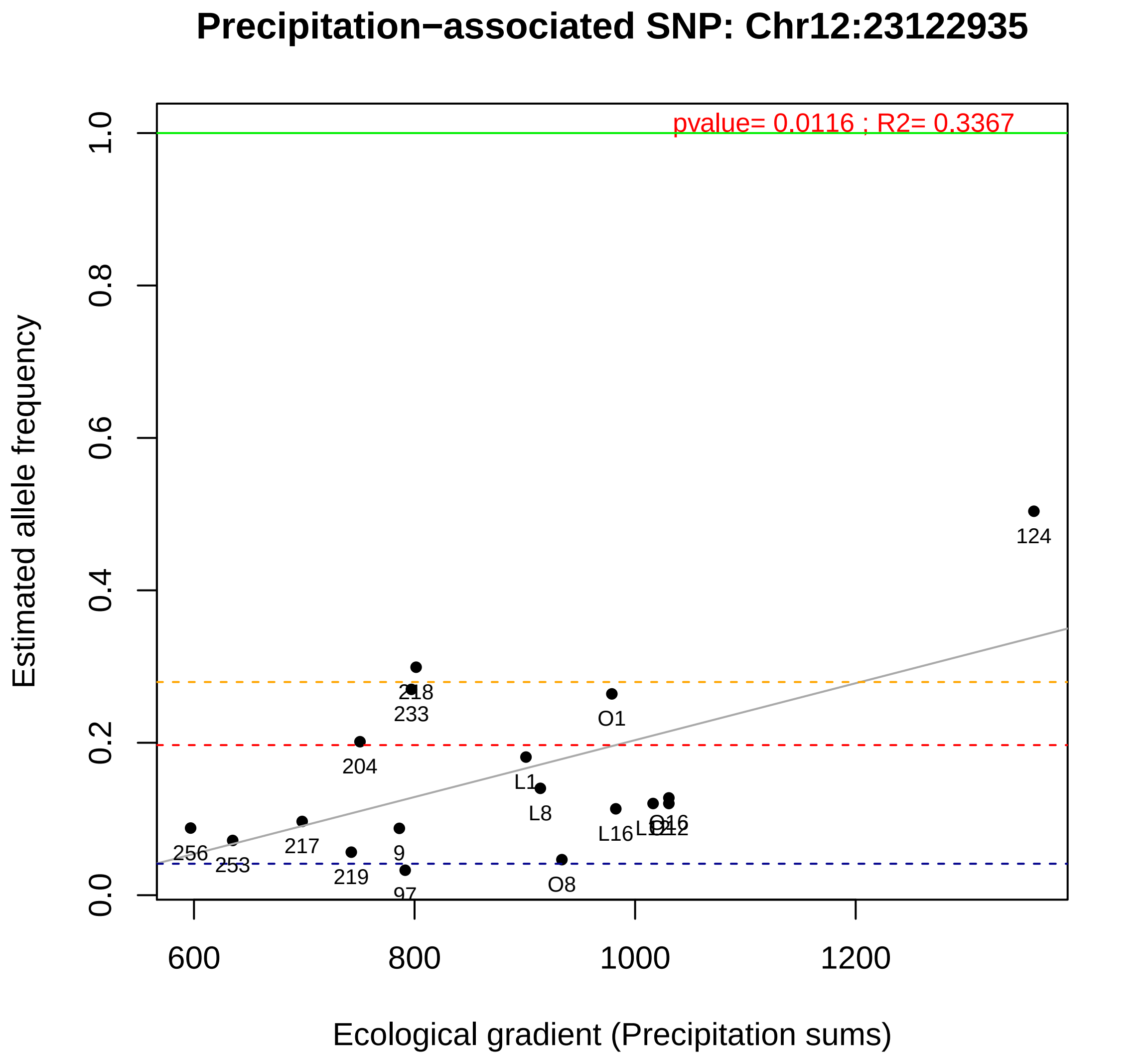

Precipitation−associated SNP: Chr12:23122935
pvalue= 0.0116 ; R2= 0.3367
Estimated allele frequency
0.0
0.2
0.4
0.6
0.8
1.0
600
800
1000
1200
Ecological gradient (Precipitation sums)
124
218
233
O1
204
L1
L8
L16
O16
256
253
217
219
9
97
O8

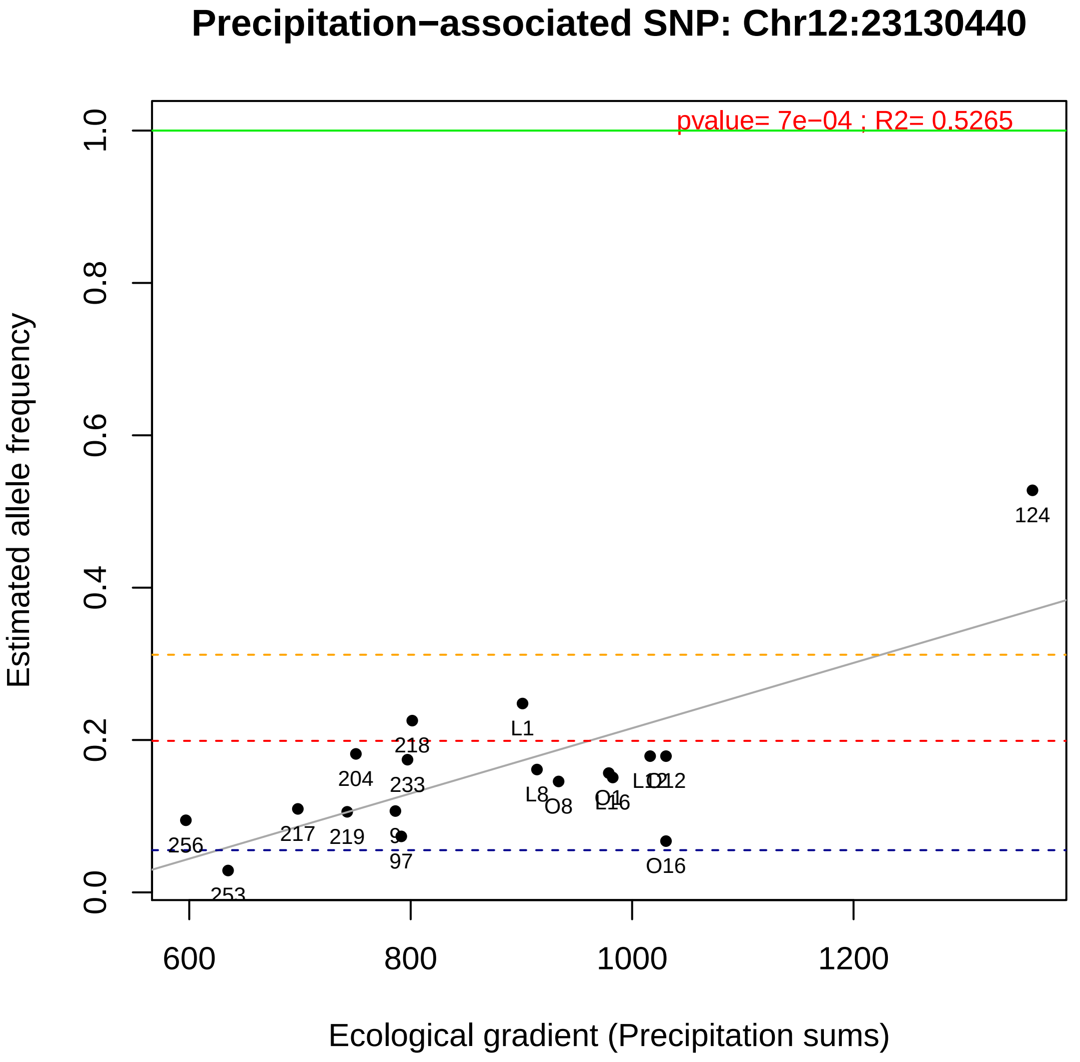
Precipitation−associated SNP: Chr12:23130440
pvalue= 7e−04 ; R2= 0.5265
Estimated allele frequency
0.0
0.2
0.4
0.6
0.8
1.0
600
800
1000
1200
Ecological gradient (Precipitation sums)
124
L1
218
204
233
L8
O8
217
219
256
253
97
O16

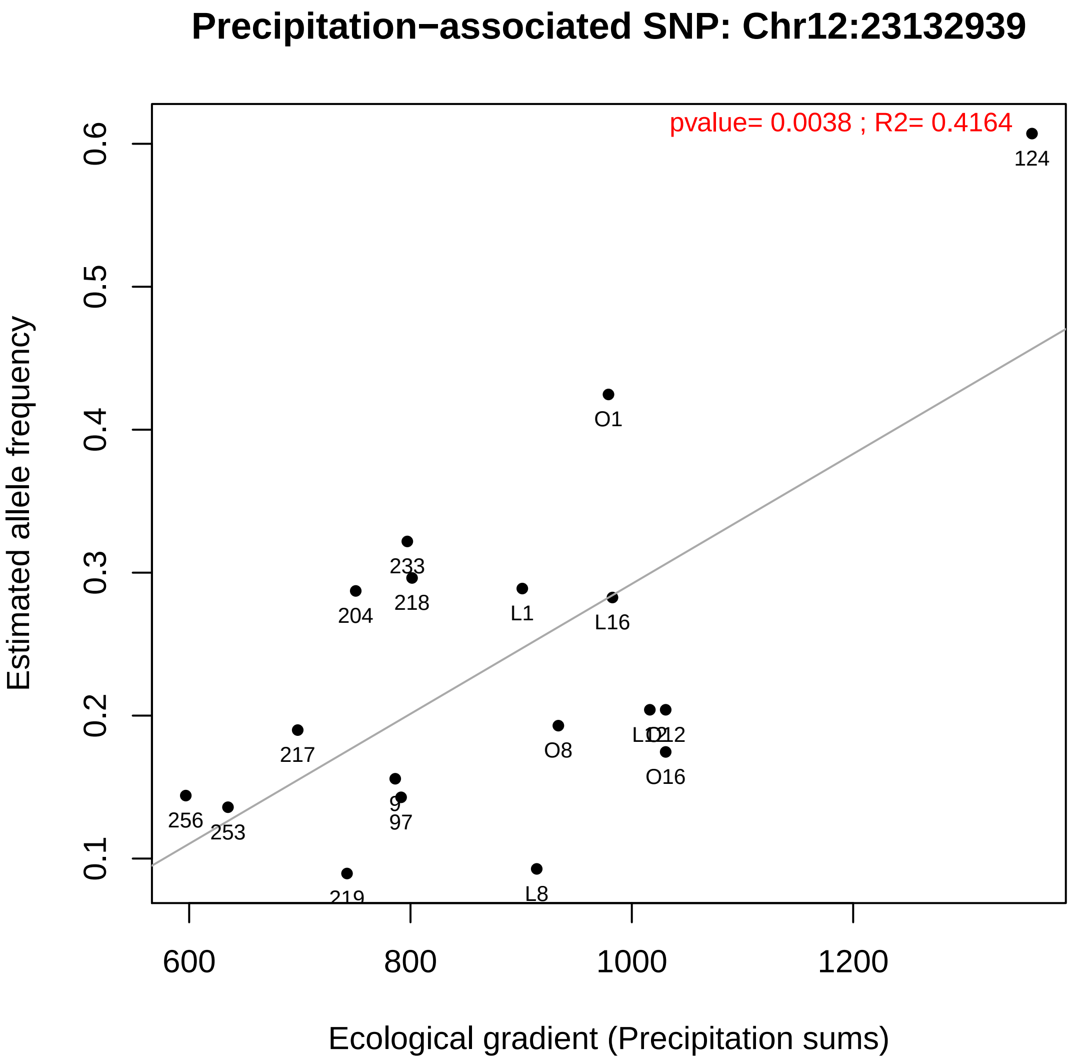

Precipitation−associated SNP: Chr12:23132939
pvalue= 0.0038 ; R2= 0.4164
Estimated allele frequency
Ecological gradient (Precipitation sums)
0.1
0.2
0.3
0.4
0.5
0.6
600
800
1000
1200
124
O1
233
218
204
L1
L16
217
O8
O12
O16
256
253
9
97
219
L8

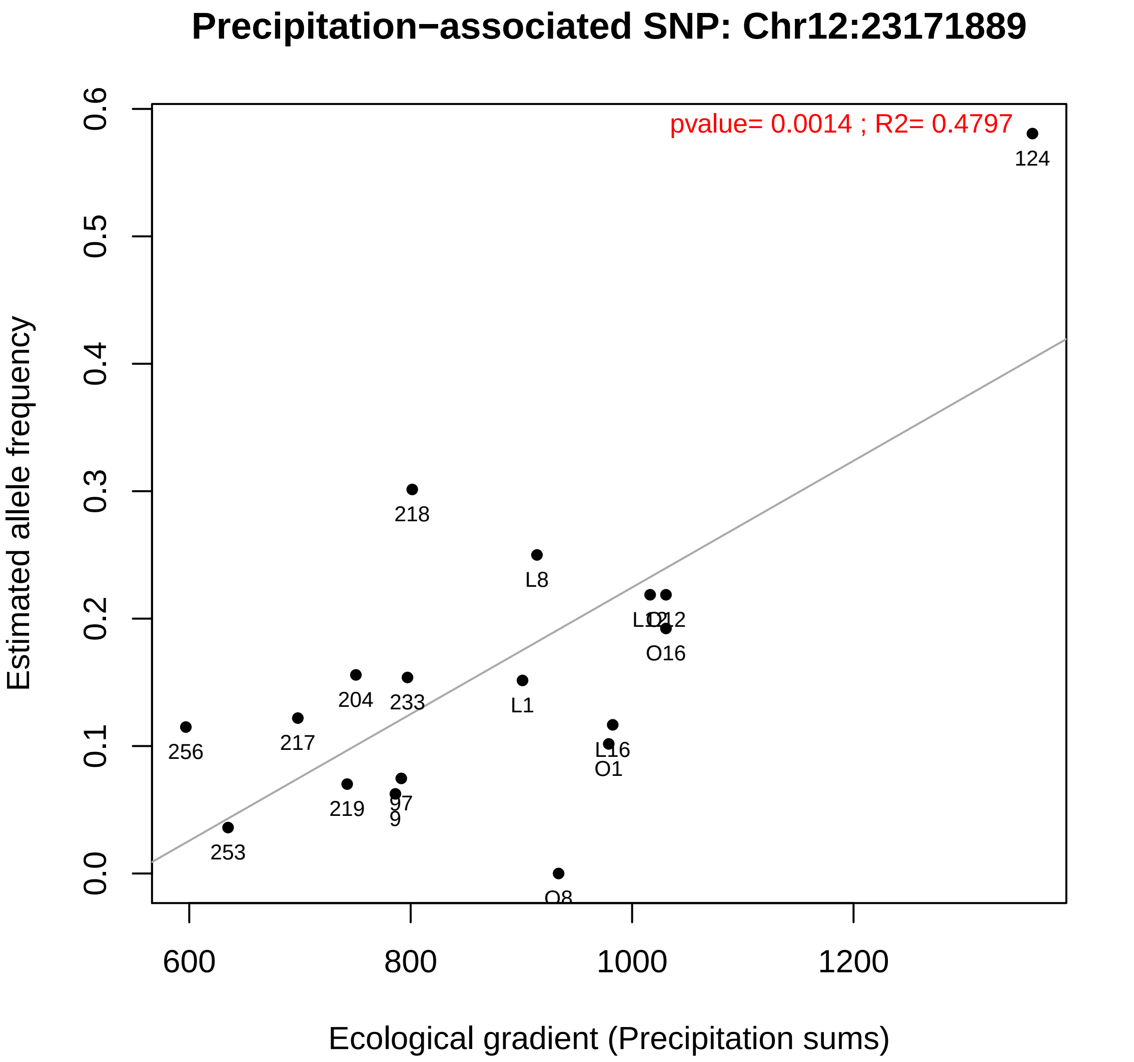
Precipitation−associated SNP: Chr12:23171889
pvalue= 0.0014 ; R2= 0.4797
Estimated allele frequency
Ecological gradient (Precipitation sums)
0.0
0.1
0.2
0.3
0.4
0.5
0.6
600
800
1000
1200
124
218
L8
L12
O12
O16
204
233
L1
256
217
L16
O1
219
97
9
253
O8

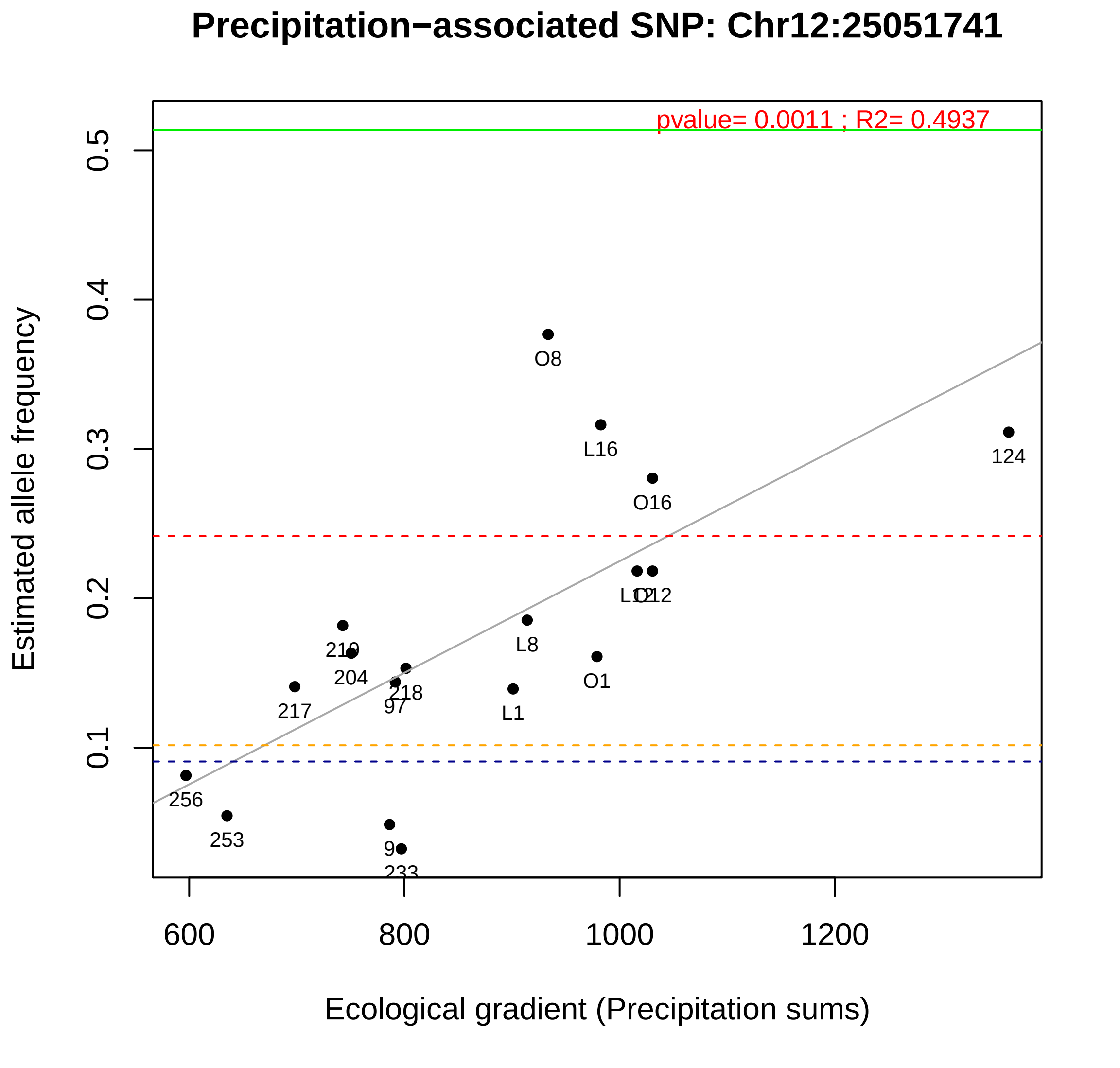

Precipitation−associated SNP: Chr12:25051741
pvalue= 0.0011 ; R2= 0.4937
Estimated allele frequency
0.1
0.2
0.3
0.4
0.5
Ecological gradient (Precipitation sums)
600
800
1000
1200
O8
L16
124
O16
L12
O12
L8
O1
219
204
218
97
217
L1
256
253
9
233

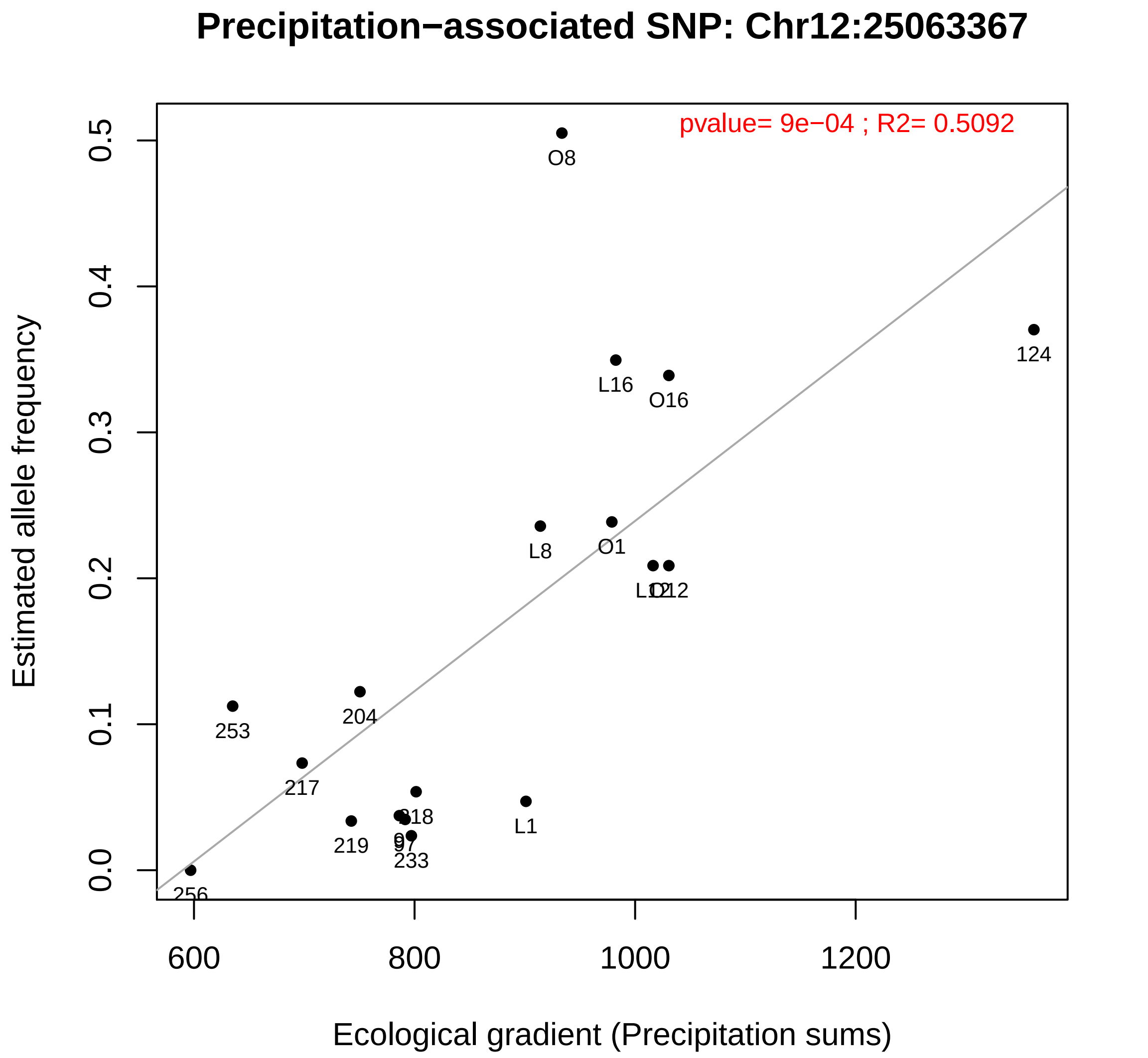
Precipitation−associated SNP: Chr12:25063367
pvalue= 9e−04 ; R2= 0.5092
Estimated allele frequency
0.0
0.1
0.2
0.3
0.4
0.5
600
800
1000
1200
Ecological gradient (Precipitation sums)
O8
124
L16
O16
L8
O1
L12
O12
204
253
217
219
218
97
233
L1
256

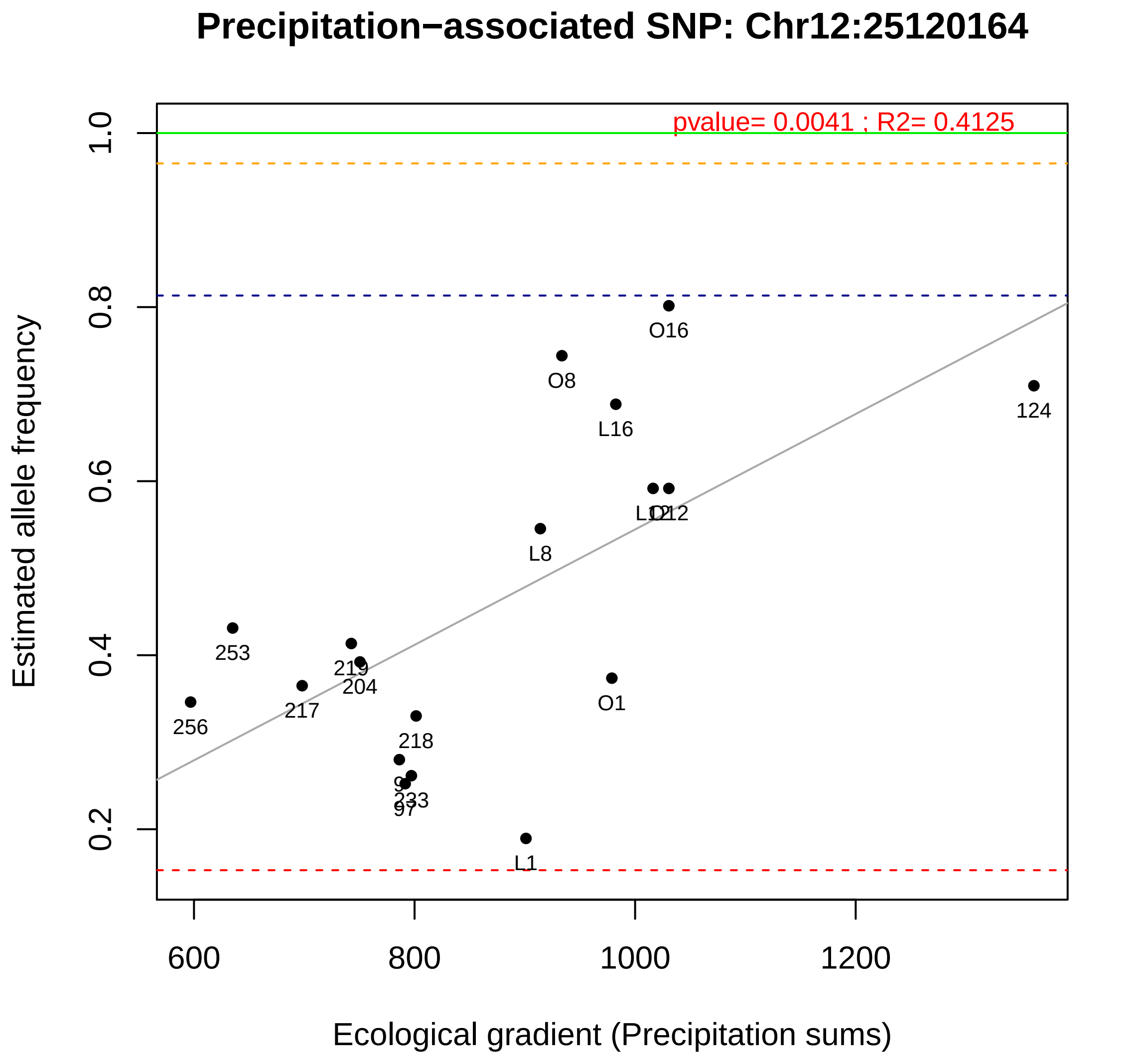
Precipitation−associated SNP: Chr12:25120164
pvalue= 0.0041 ; R2= 0.4125
Estimated allele frequency
Ecological gradient (Precipitation sums)
0.2
0.4
0.6
0.8
1.0
600
800
1000
1200
O16
O8
124
L16
O12
L8
253
219
204
217
256
O1
218
233
97
L1

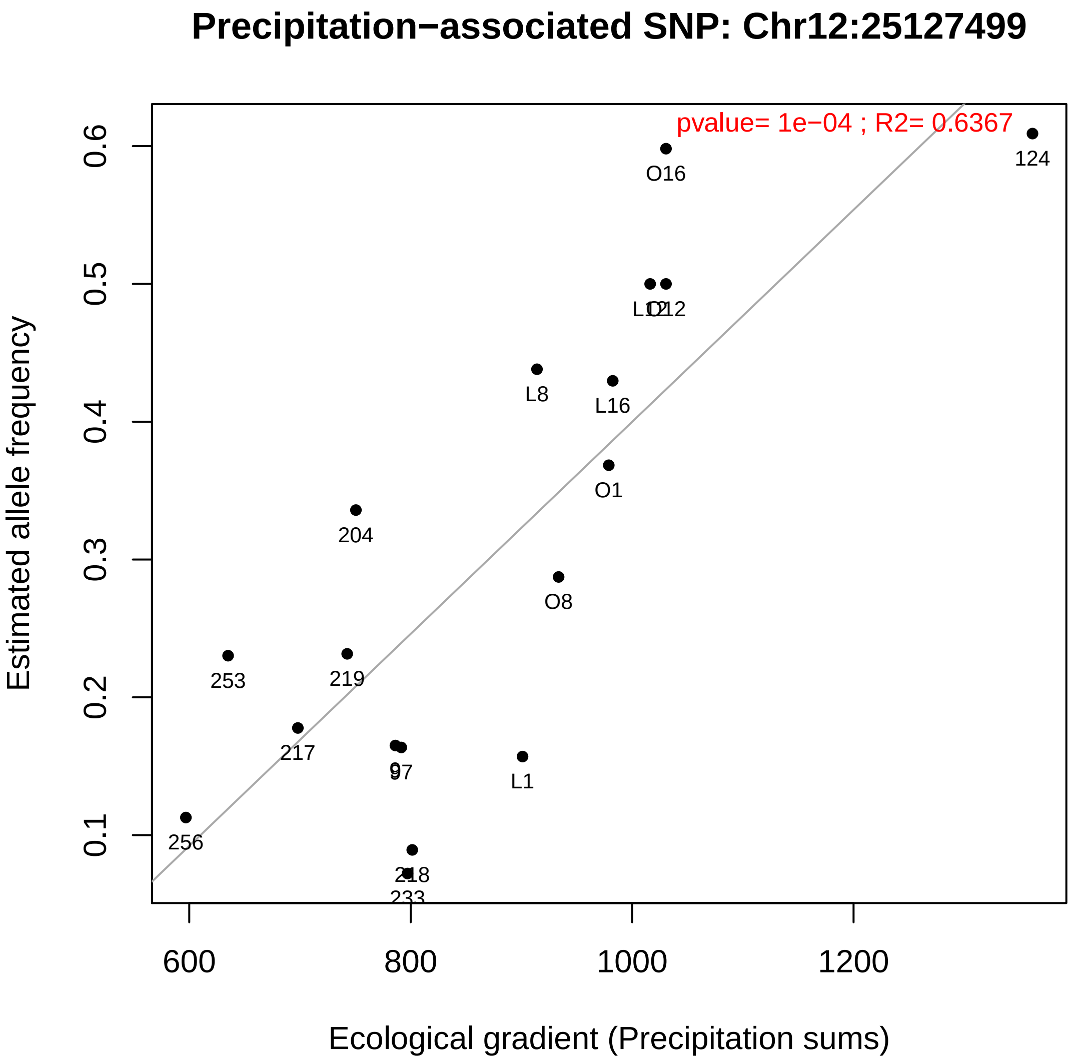

Precipitation−associated SNP: Chr12:25127499
pvalue= 1e−04 ; R2= 0.6367
Estimated allele frequency
Ecological gradient (Precipitation sums)
0.1
0.2
0.3
0.4
0.5
0.6
600
800
1000
1200
124
O16
L12
O12
L8
L16
O1
204
O8
253
219
217
9
97
L1
256
218
233

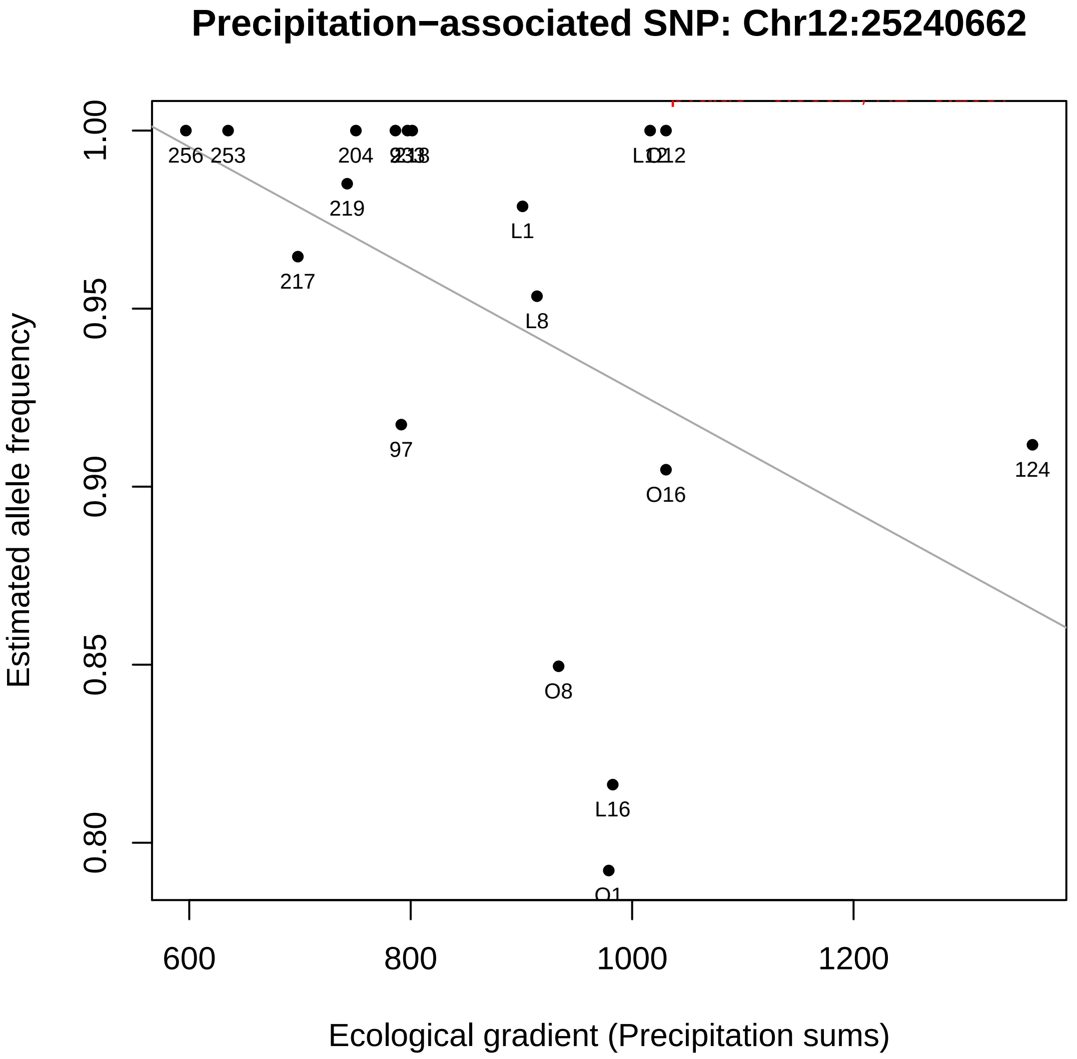

Precipitation−associated SNP: Chr12:25240662
Estimated allele frequency
1.00
0.95
0.90
0.85
0.80
600
800
1000
1200
Ecological gradient (Precipitation sums)
256
253
204
219
217
L1
L8
97
O16
124
O8
L16

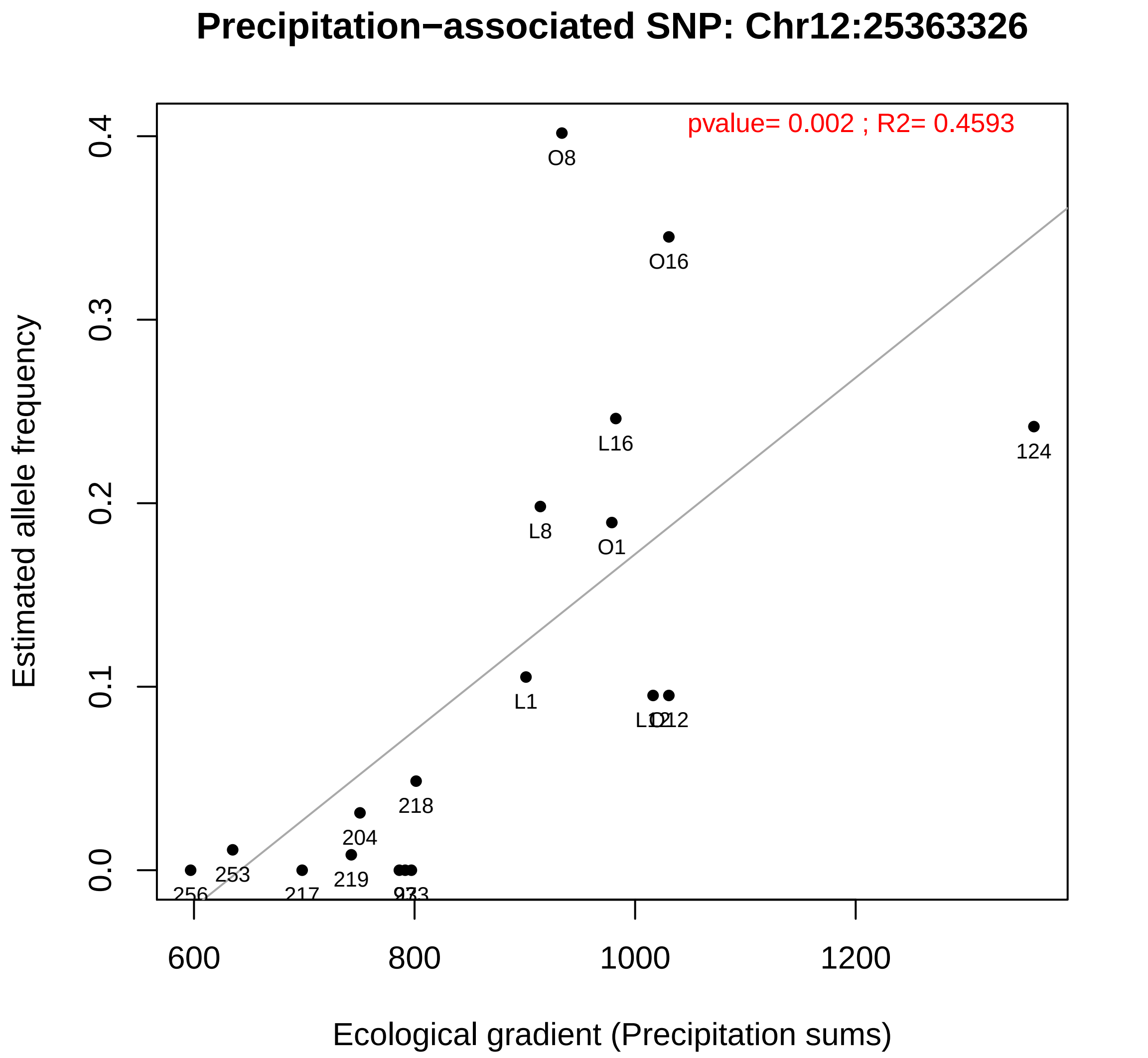

Precipitation−associated SNP: Chr12:25363326
pvalue= 0.002 ; R2= 0.4593
Estimated allele frequency
Ecological gradient (Precipitation sums)
0.0
0.1
0.2
0.3
0.4
600
800
1000
1200
O8
O16
L16
124
L8
O1
L1
L12
O12
218
204
253
219
256
217

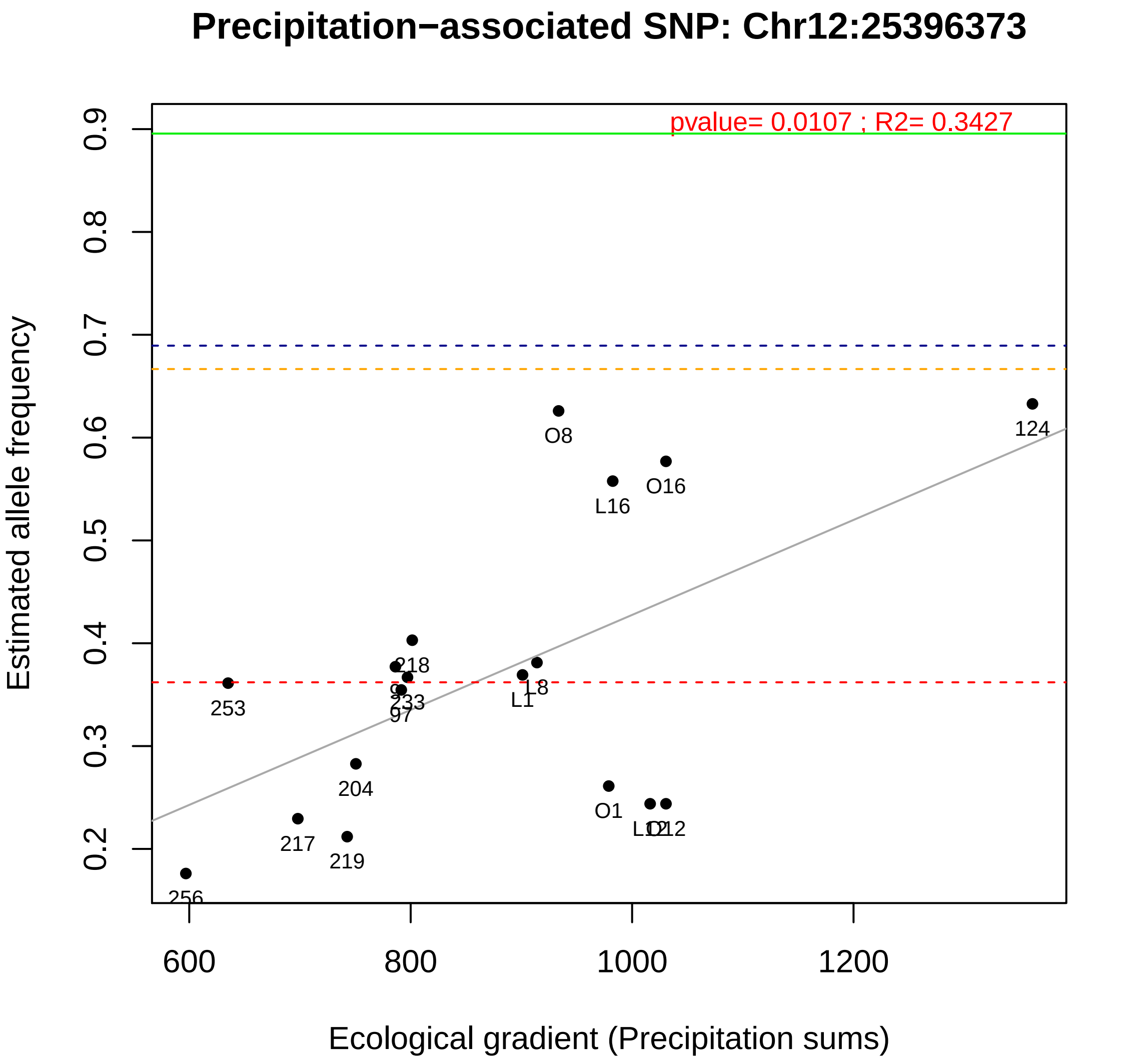

Precipitation−associated SNP: Chr12:25396373
pvalue= 0.0107 ; R2= 0.3427
Estimated allele frequency
Ecological gradient (Precipitation sums)
0.2
0.3
0.4
0.5
0.6
0.7
0.8
0.9
600
800
1000
1200
O8
124
L16
O16
218
253
233
97
L1
18
204
O1
217
219
256

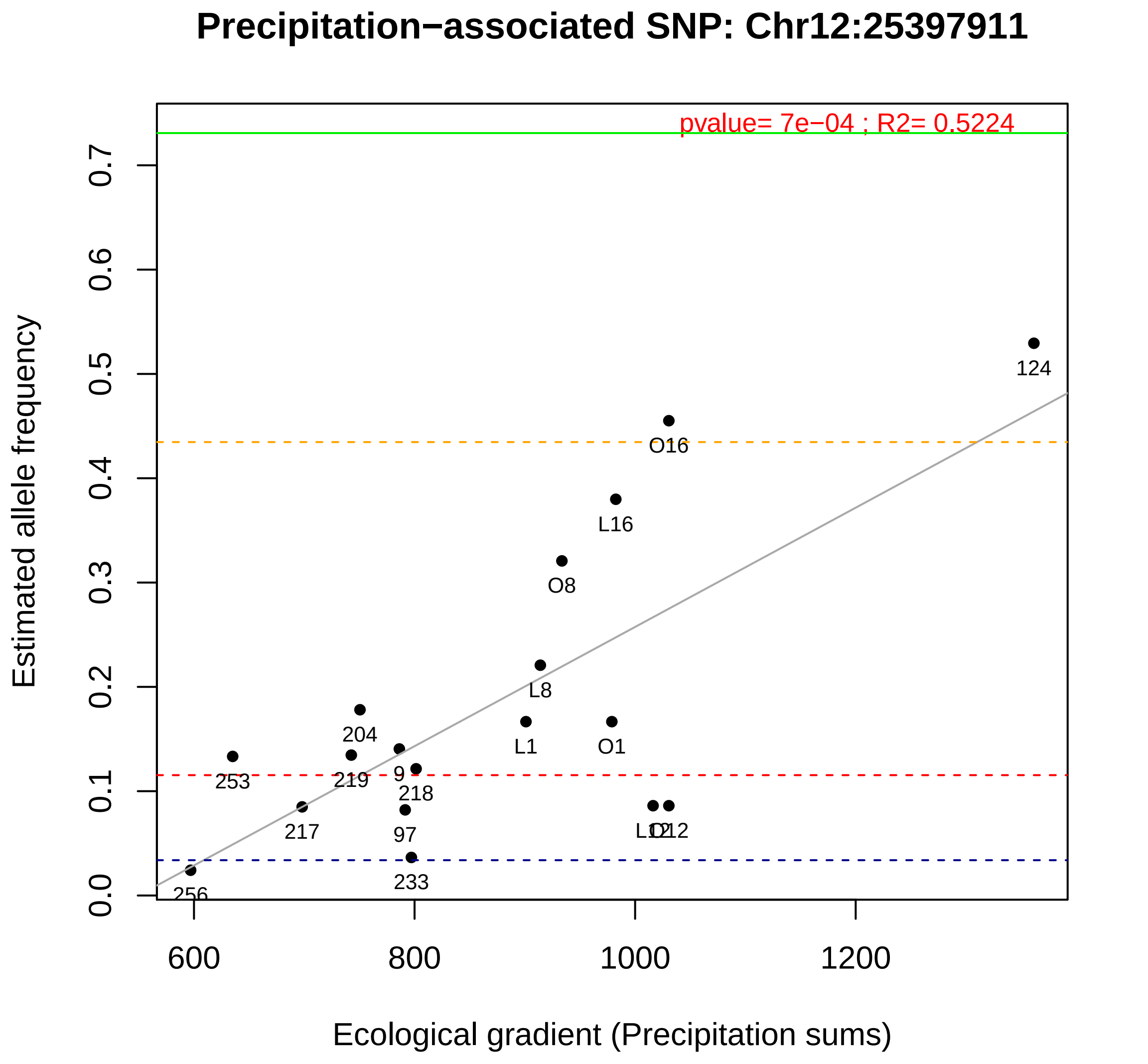
Precipitation−associated SNP: Chr12:25397911
pvalue= 7e−04 ; R2= 0.5224
Estimated allele frequency
0.0
0.1
0.2
0.3
0.4
0.5
0.6
0.7
600
800
1000
1200
Ecological gradient (Precipitation sums)
124
O16
L16
O8
L8
204
L1
O1
253
219
9
218
217
97
233
256

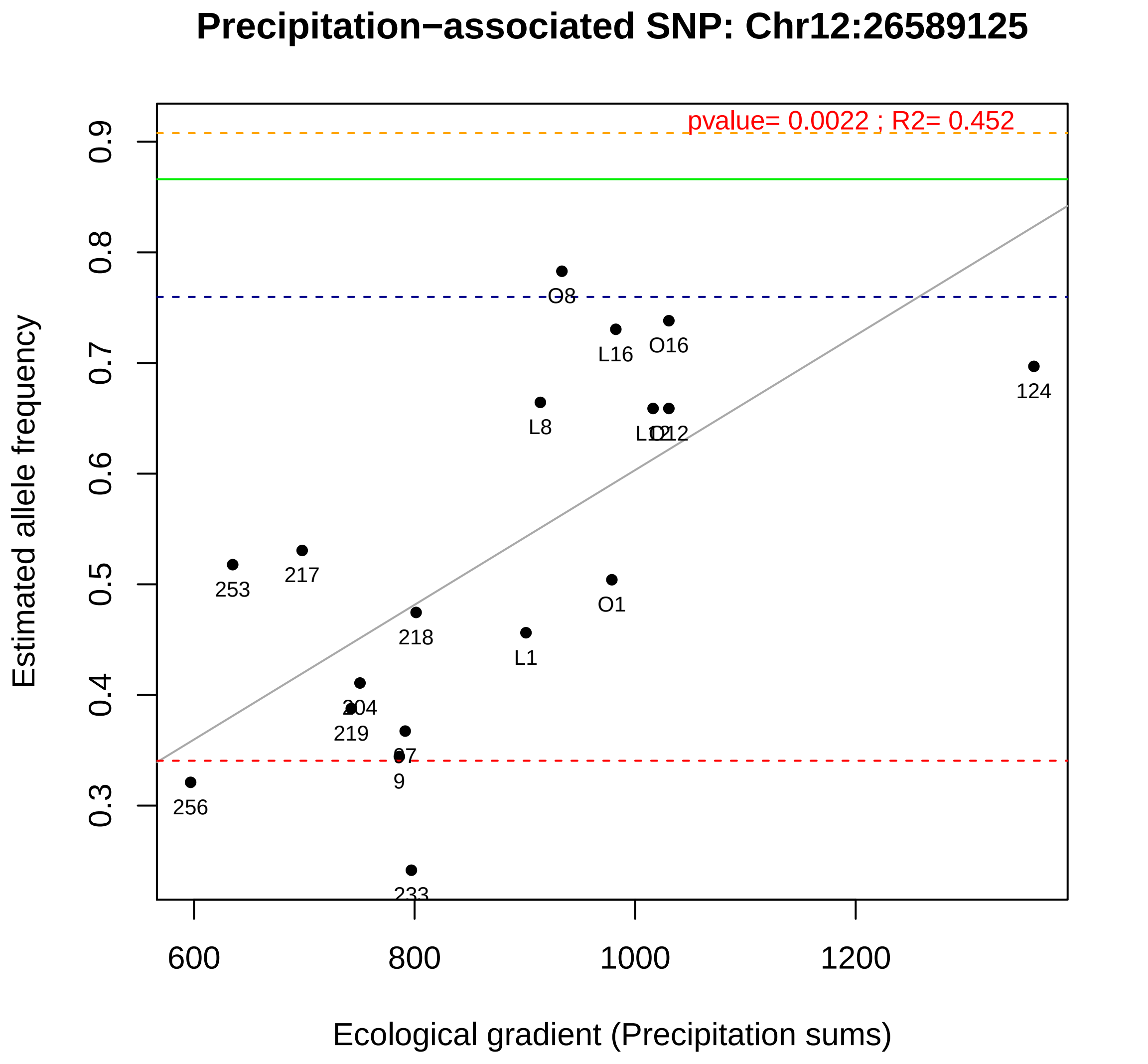

Precipitation−associated SNP: Chr12:26589125
pvalue= 0.0022 ; R2= 0.452
Estimated allele frequency
0.3
0.4
0.5
0.6
0.7
0.8
0.9
600
800
1000
1200
Ecological gradient (Precipitation sums)
O8
L16
O16
124
L8
253
217
O1
218
L1
204
219
256
9
233

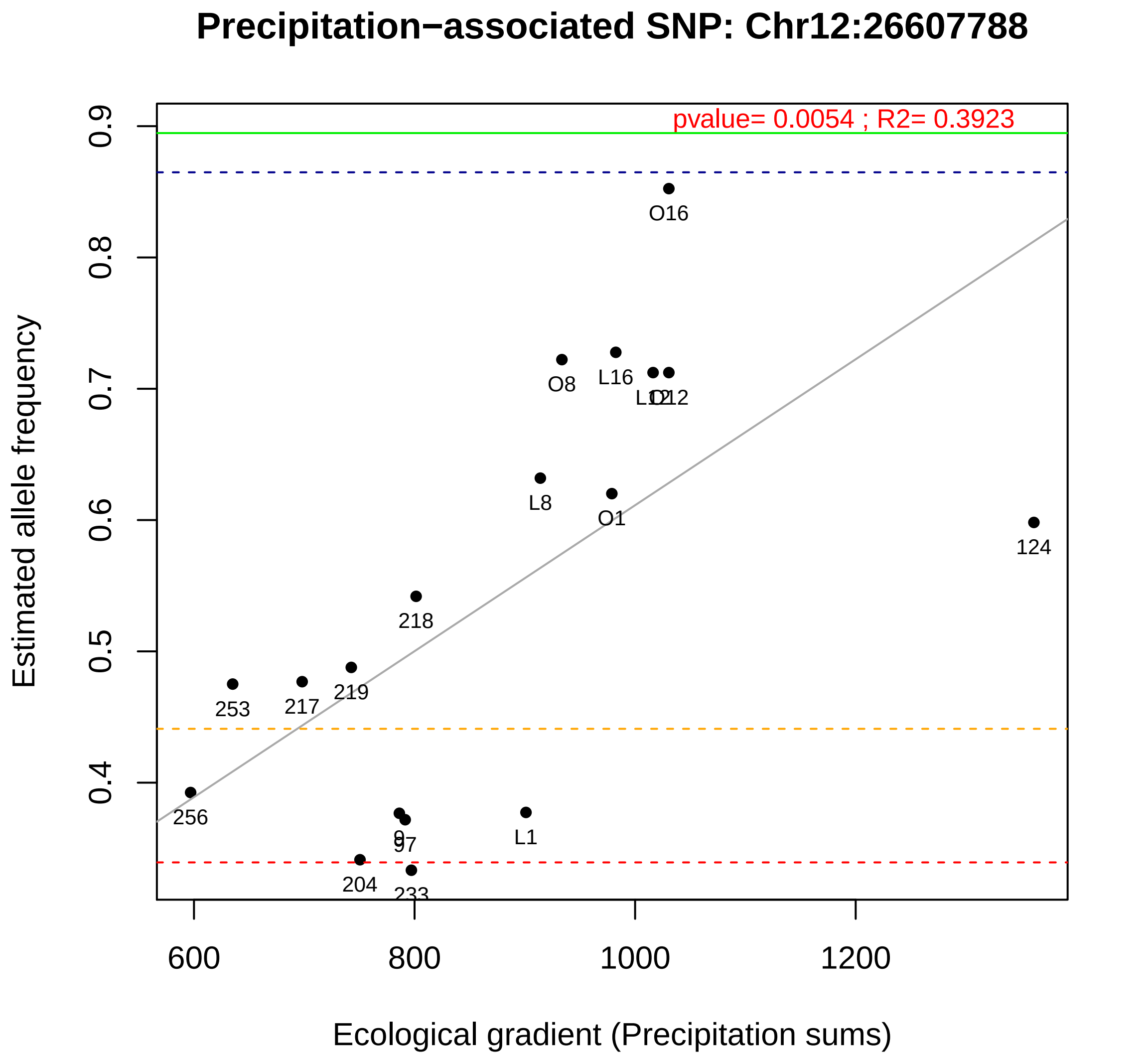
Precipitation−associated SNP: Chr12:26607788
pvalue= 0.0054 ; R2= 0.3923
Estimated allele frequency
0.4
0.5
0.6
0.7
0.8
0.9
600
800
1000
1200
Ecological gradient (Precipitation sums)
O16
O8
L16
L12
O12
L8
O1
124
218
253
217
219
256
97
L1
204
233

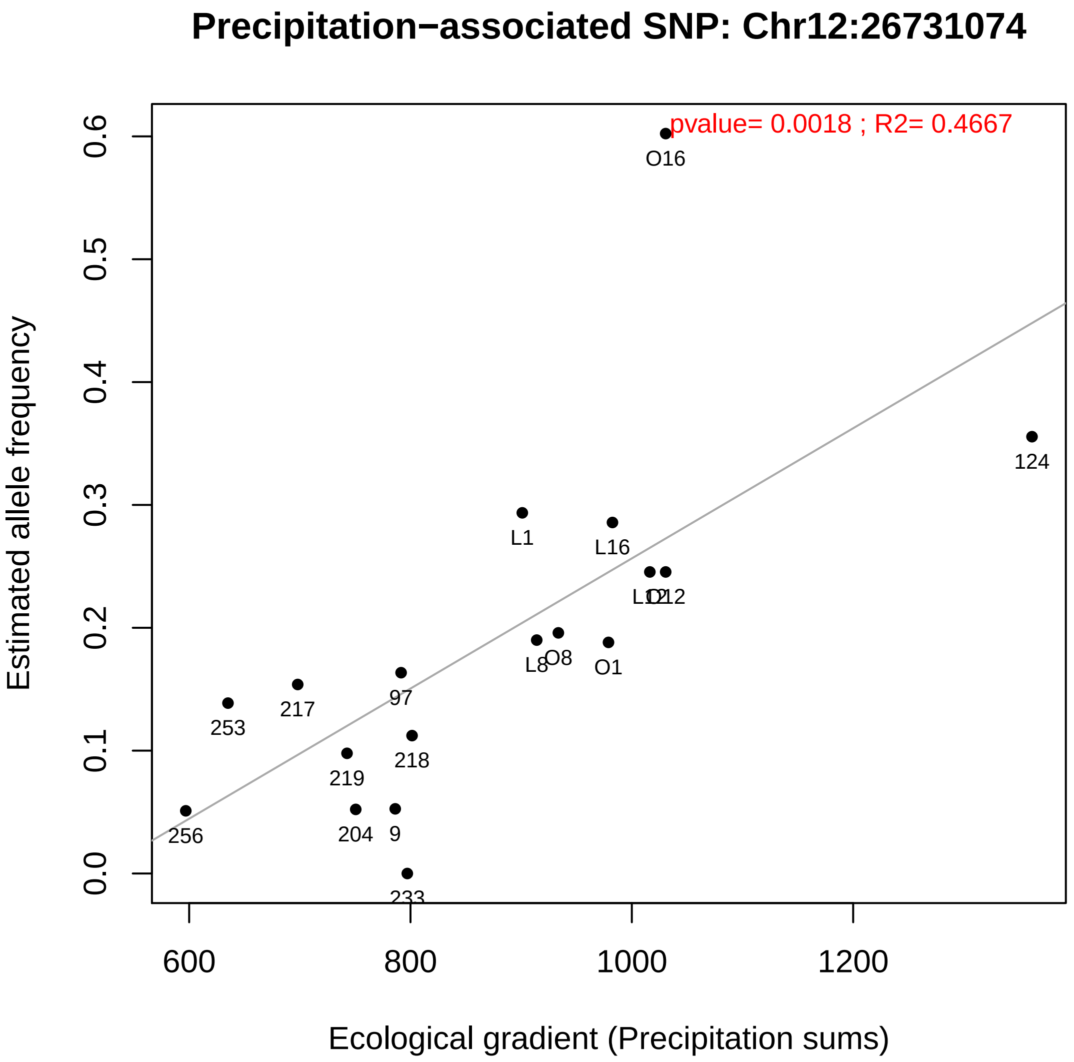

Precipitation−associated SNP: Chr12:26731074
pvalue= 0.0018 ; R2= 0.4667
Estimated allele frequency
Ecological gradient (Precipitation sums)
0.0
0.1
0.2
0.3
0.4
0.5
0.6
600
800
1000
1200
O16
124
L1
L16
O12
O8
L8
O1
97
217
253
218
219
256
204
9
233

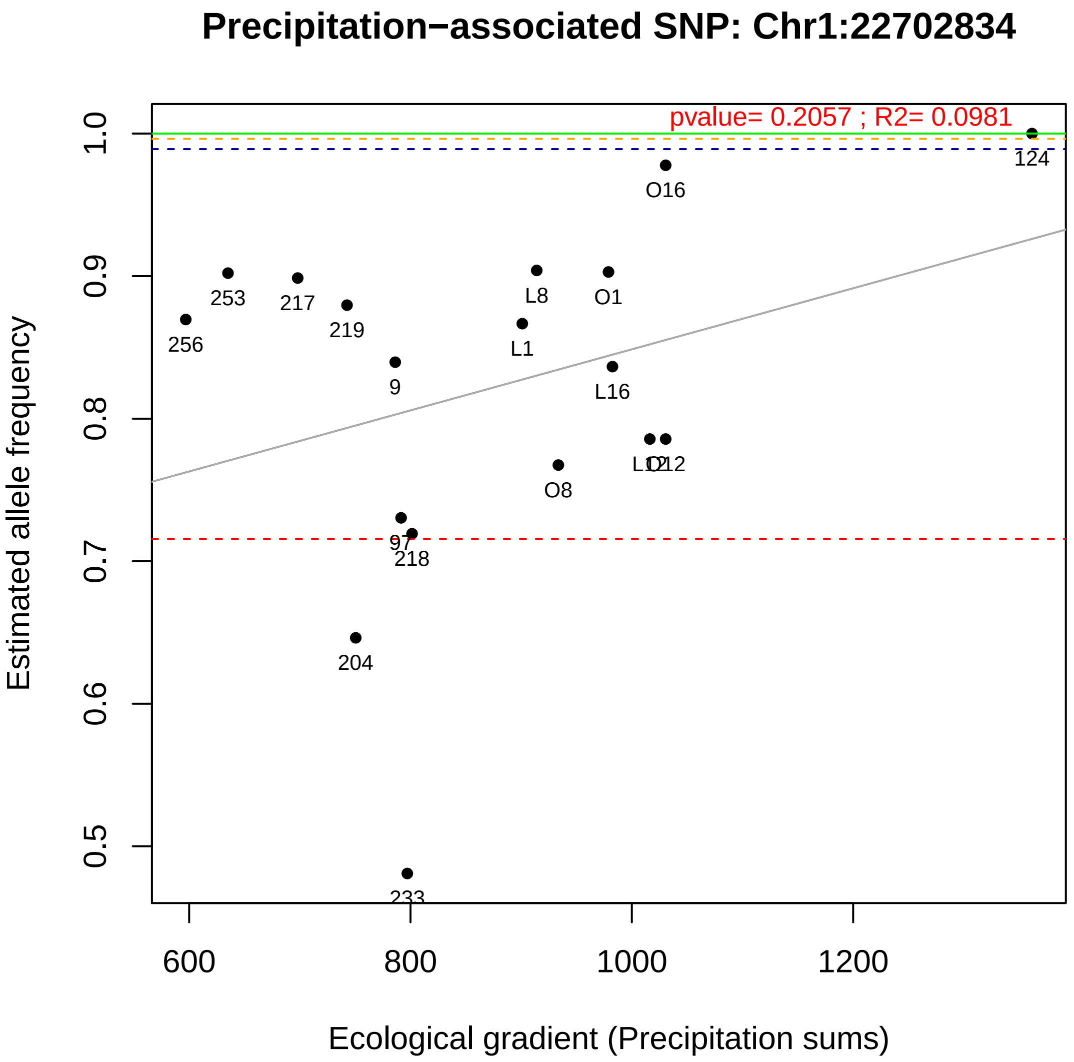
Precipitation−associated SNP: Chr1:22702834
pvalue= 0.2057 ; R2= 0.0981
Estimated allele frequency
Ecological gradient (Precipitation sums)
0.5
0.6
0.7
0.8
0.9
1.0
600
800
1000
1200
124
O16
253
217
219
256
L8
O1
L1
9
L16
L12
O12
O8
97
218
204
233

**Precipitation−associated SNP: Chr12:27066866**

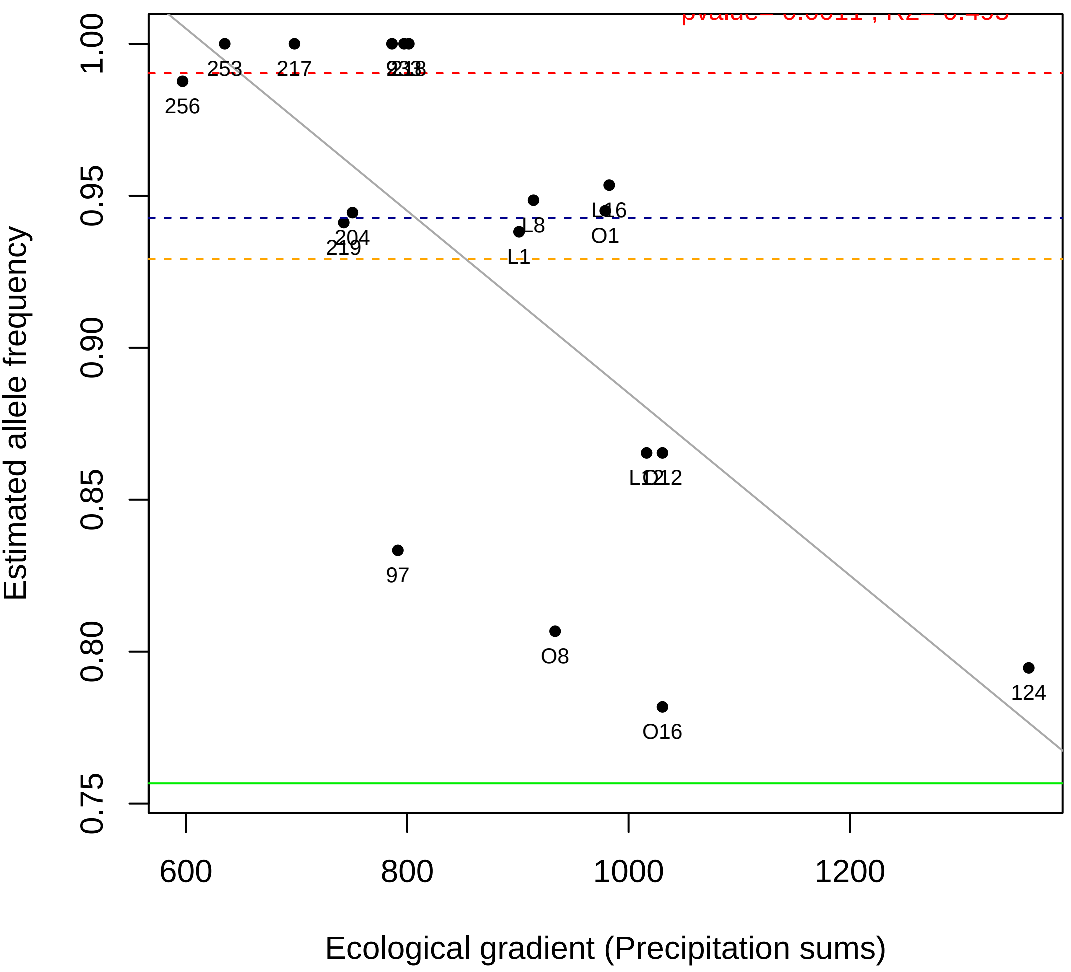

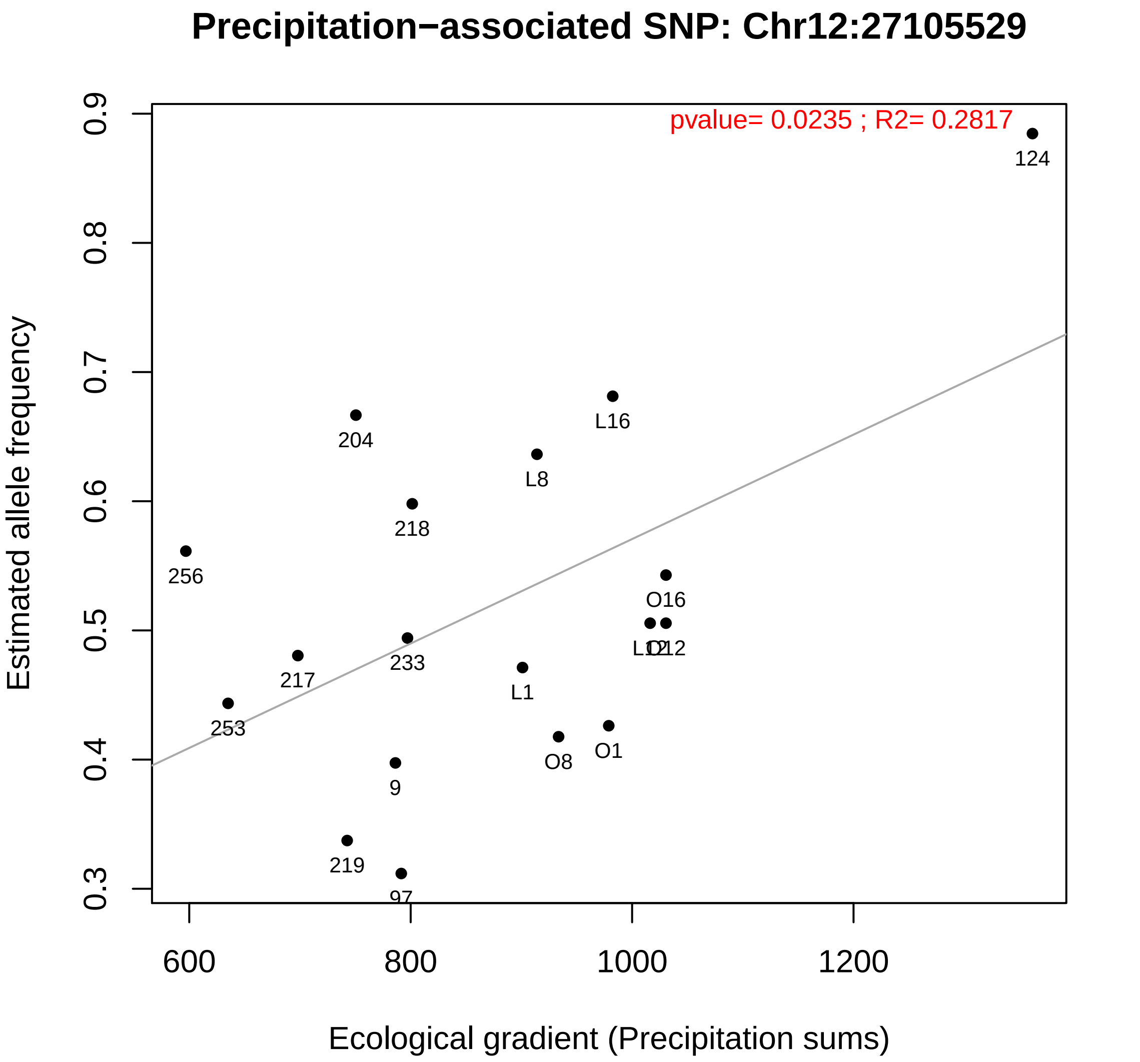

Precipitation−associated SNP: Chr12:27105529
pvalue= 0.0235 ; R2= 0.2817
Estimated allele frequency
0.3
0.4
0.5
0.6
0.7
0.8
0.9
600
800
1000
1200
Ecological gradient (Precipitation sums)
124
L16
204
L8
218
256
O16
L12
O12
233
217
L1
253
O1
O8
9
219
97

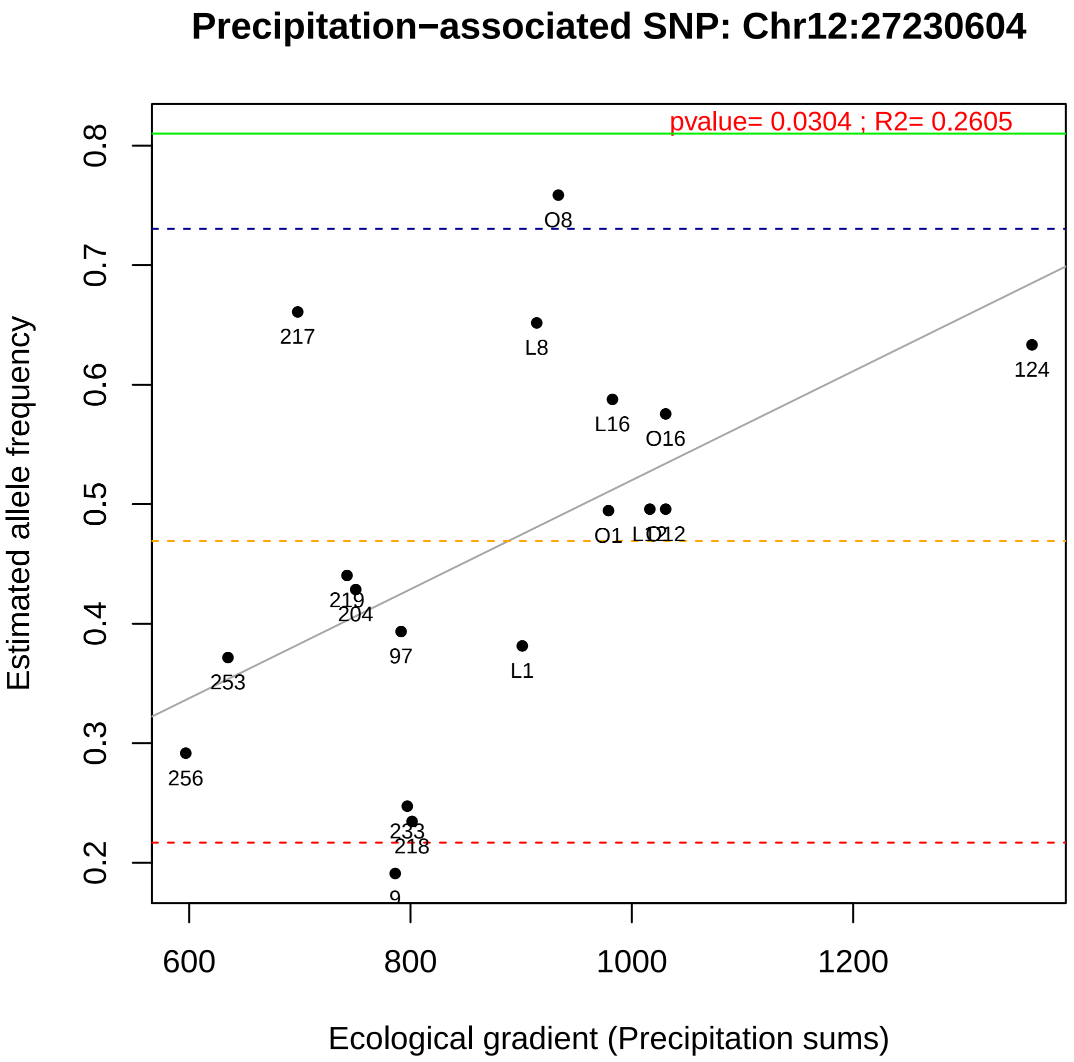

Precipitation−associated SNP: Chr12:27230604
pvalue= 0.0304 ; R2= 0.2605
Estimated allele frequency
0.2
0.3
0.4
0.5
0.6
0.7
0.8
600
800
1000
1200
Ecological gradient (Precipitation sums)
O8
217
L8
124
L16
O16
O1
O12
219
204
97
L1
253
256
233
218
9

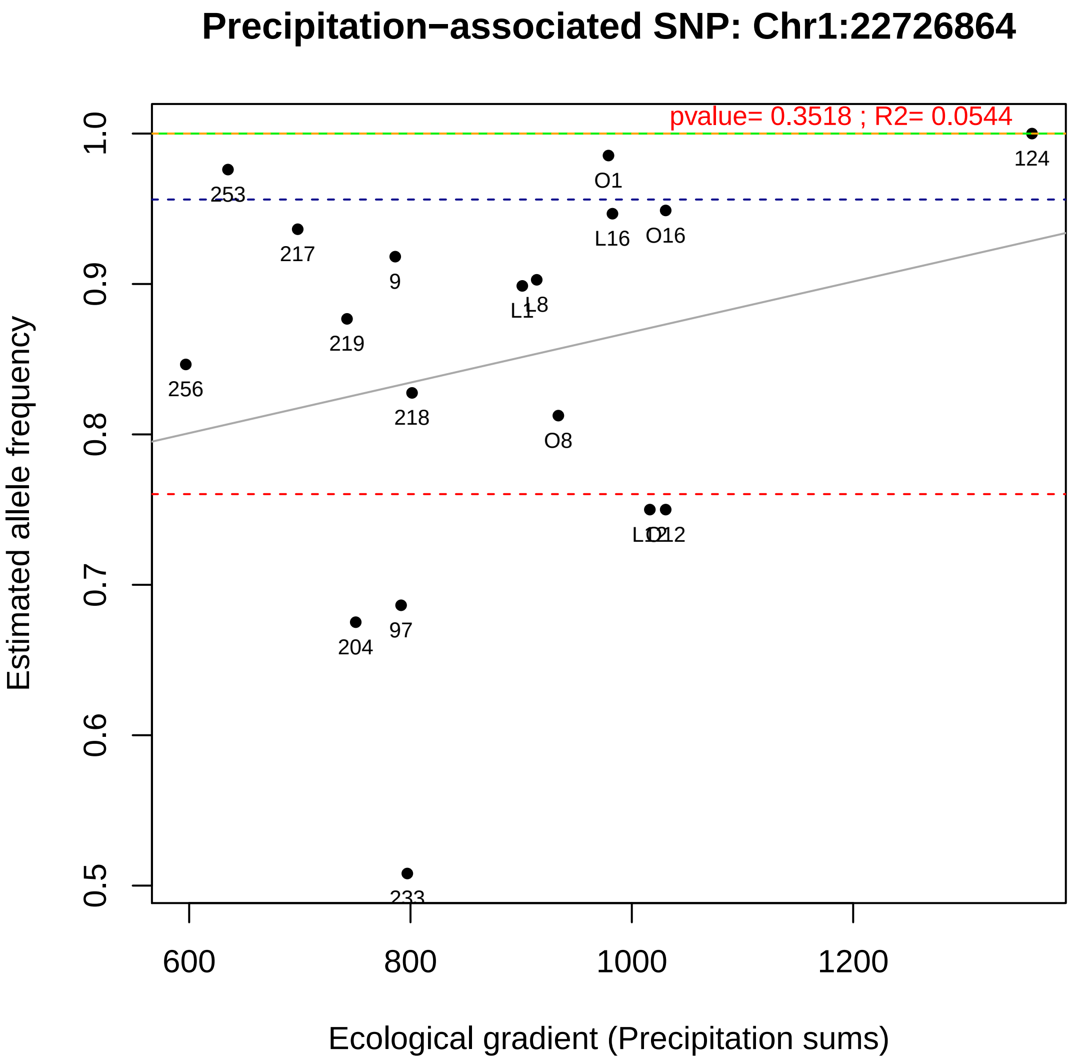
Precipitation−associated SNP: Chr1:22726864
pvalue= 0.3518 ; R2= 0.0544
Estimated allele frequency
Ecological gradient (Precipitation sums)
1.0
0.9
0.8
0.7
0.6
0.5
600
800
1000
1200
124
O1
253
L16
O16
217
9
L1
L8
219
256
218
O8
L12
O12
97
204
233

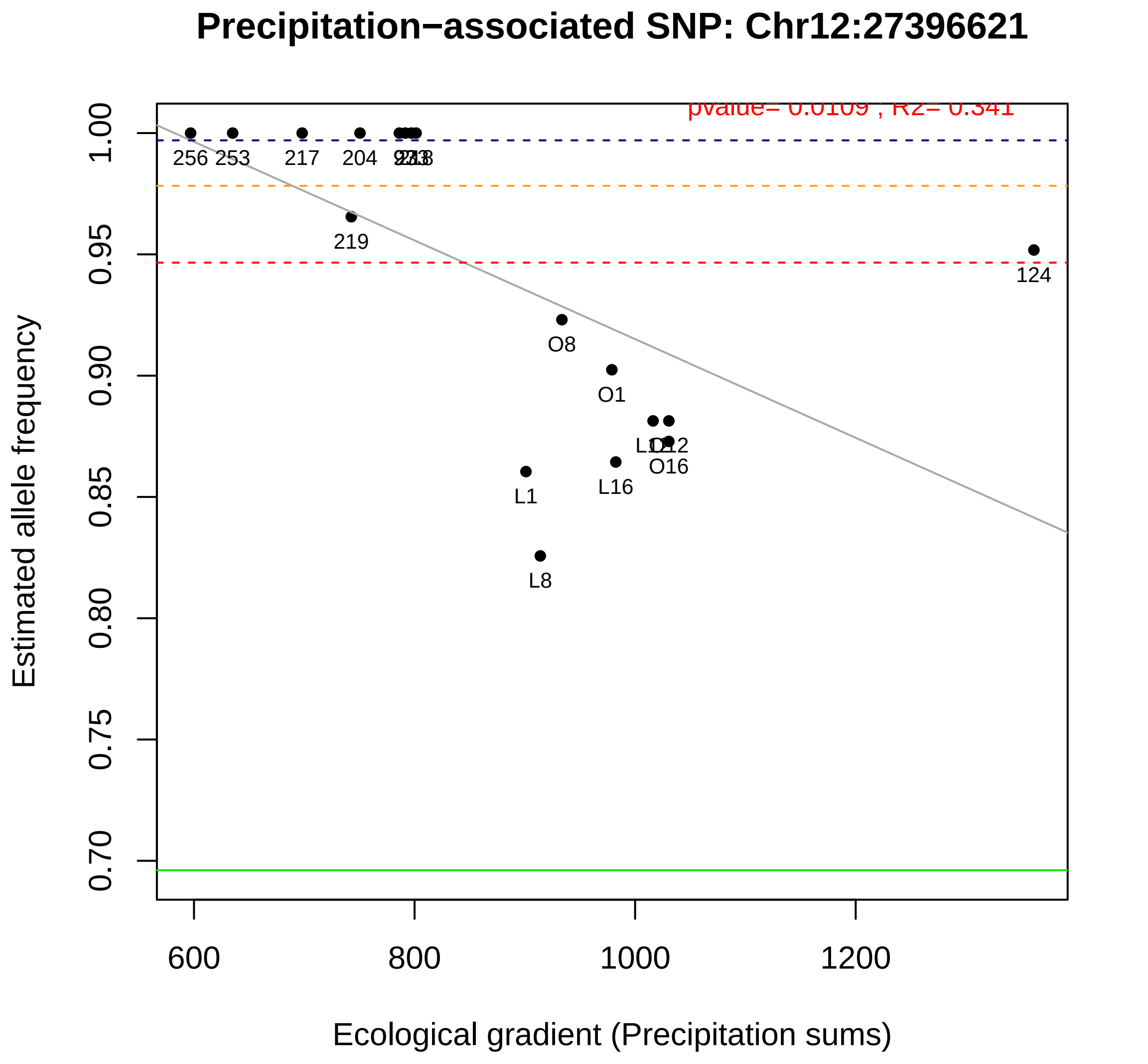

Precipitation−associated SNP: Chr12:27396621
pvalue= 0.0109 , R2= 0.341
Estimated allele frequency
Ecological gradient (Precipitation sums)
1.00
0.95
0.90
0.85
0.80
0.75
0.70
600
800
1000
1200
256
253
217
204
219
124
O8
O1
L1
L16
O16
L8

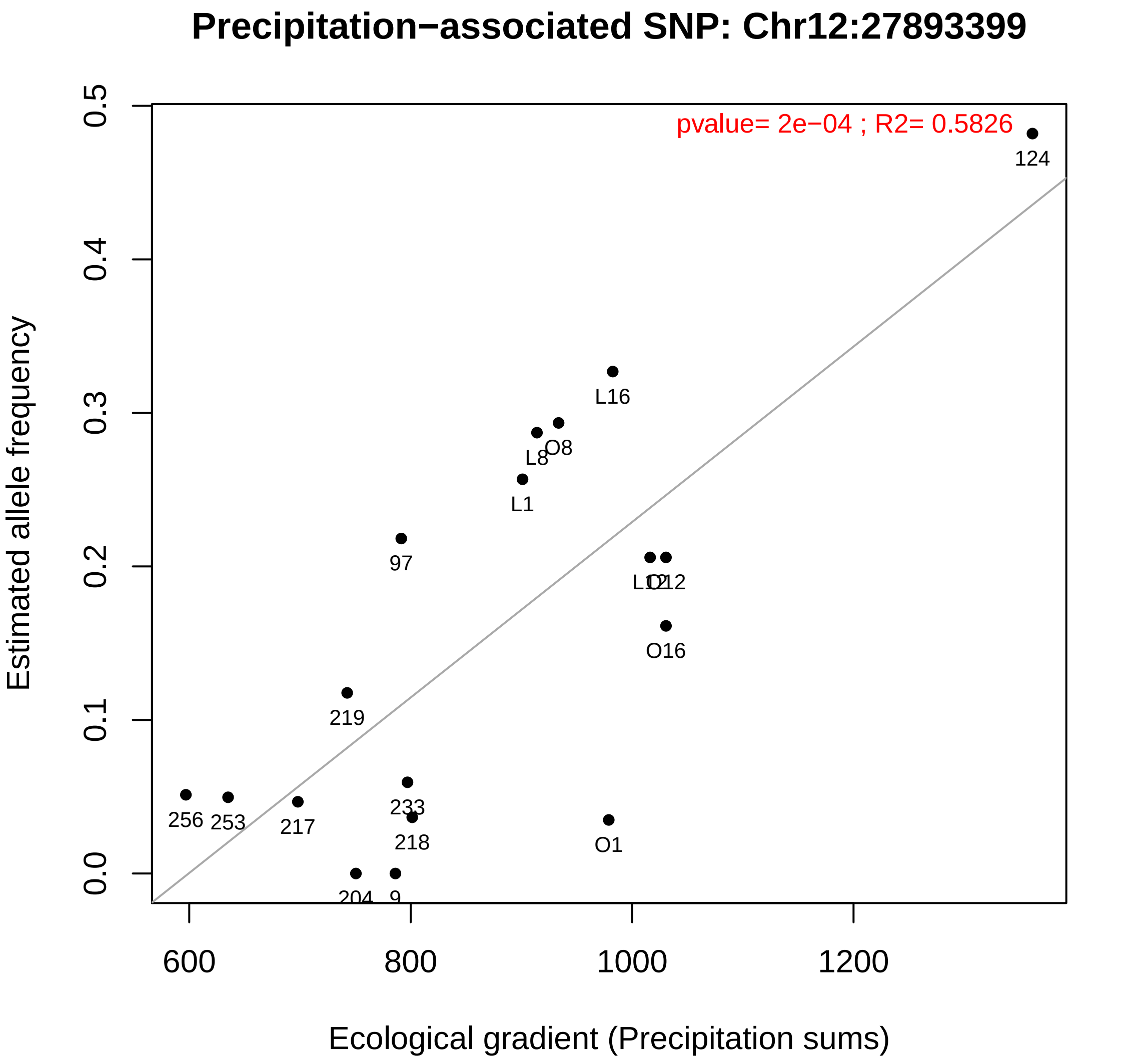

Precipitation−associated SNP: Chr12:27893399
pvalue= 2e−04 ; R2= 0.5826
Estimated allele frequency
Ecological gradient (Precipitation sums)
0.0
0.1
0.2
0.3
0.4
0.5
600
800
1000
1200
124
L16
O8
L8
L1
97
O16
219
233
218
256
253
217
O1
204
9

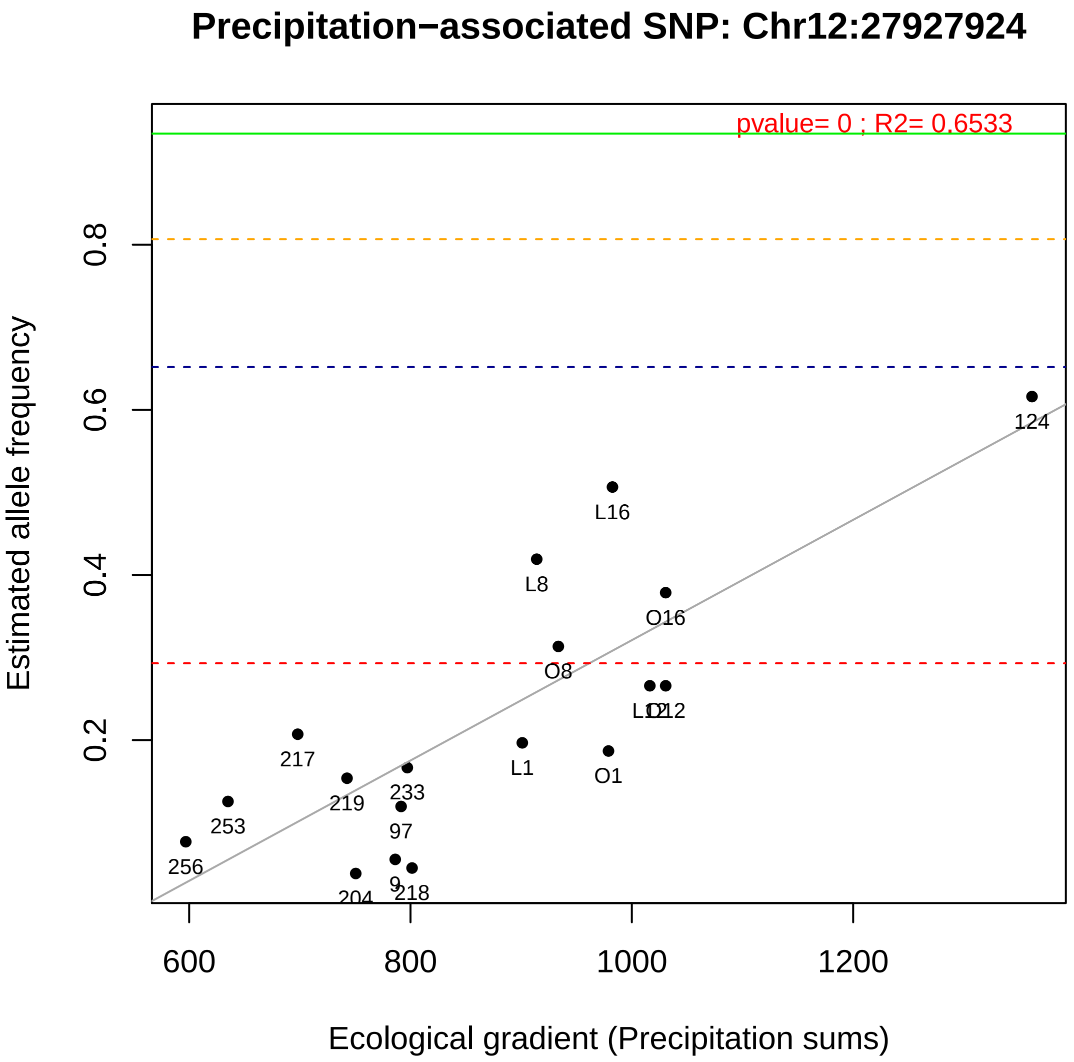

Precipitation−associated SNP: Chr12:27927924
pvalue= 0 ; R2= 0.6533
Estimated allele frequency
0.2
0.4
0.6
0.8
600
800
1000
1200
Ecological gradient (Precipitation sums)
124
L16
L8
O16
O8
L12
O12
217
L1
O1
219
233
253
97
256
9
218
204

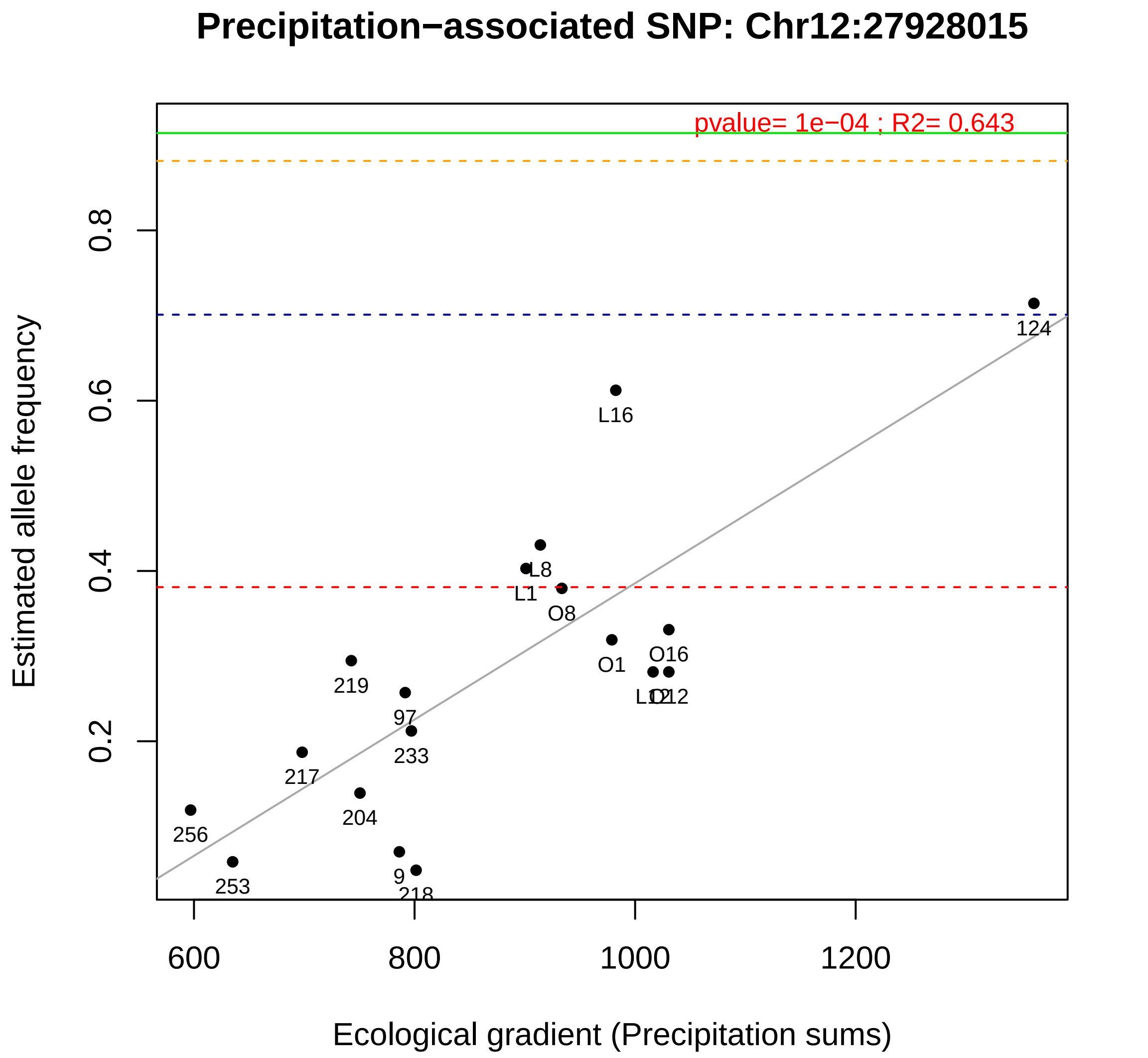

Precipitation−associated SNP: Chr12:27928015
pvalue= 1e−04 ; R2= 0.643
Estimated allele frequency
0.2
0.4
0.6
0.8
Ecological gradient (Precipitation sums)
600
800
1000
1200
124
L16
L8
L1
O8
O16
O1
L12
O12
219
97
233
217
204
256
253
9
218

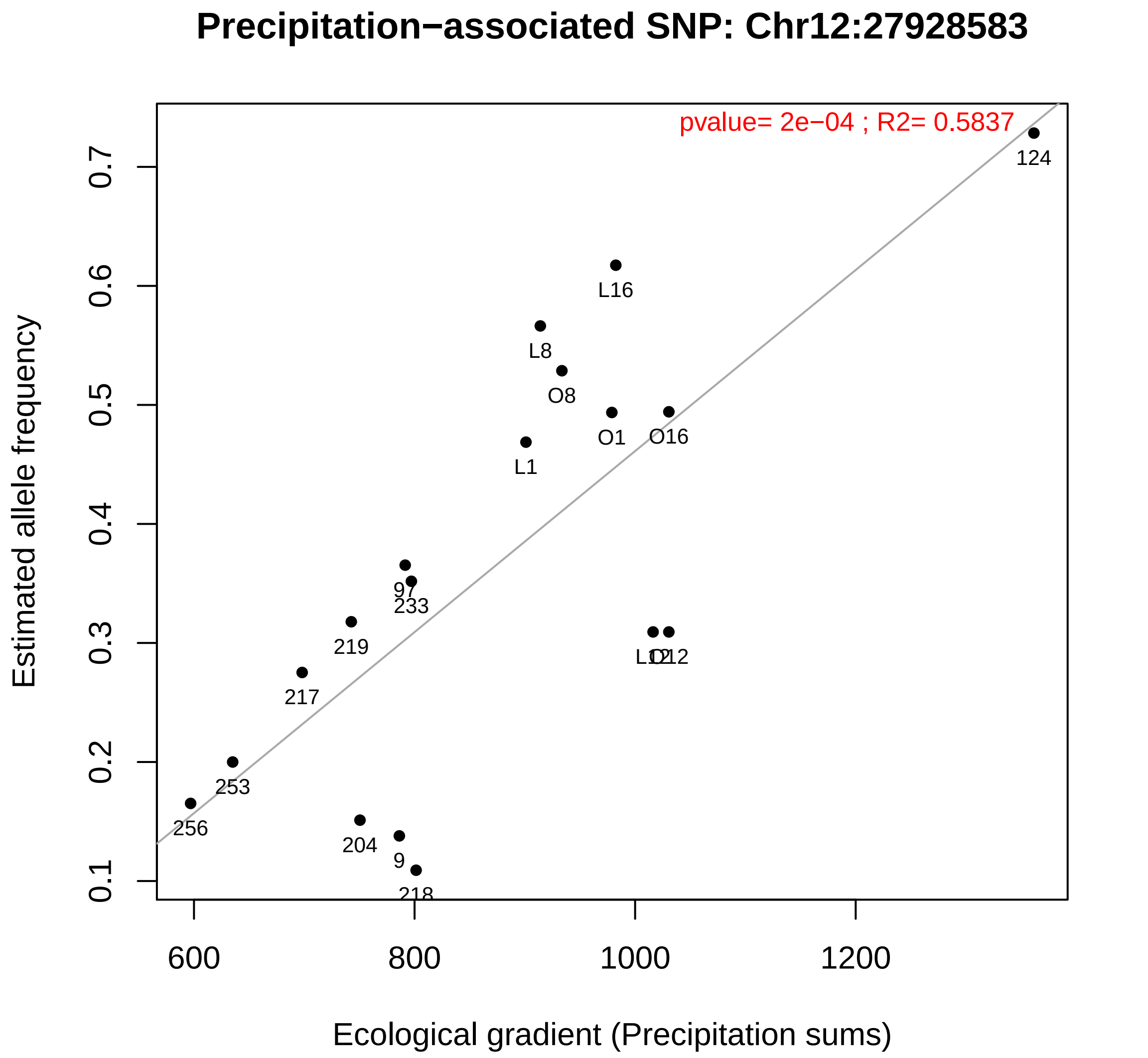

Precipitation−associated SNP: Chr12:27928583
pvalue= 2e−04 ; R2= 0.5837
Estimated allele frequency
Ecological gradient (Precipitation sums)
0.1
0.2
0.3
0.4
0.5
0.6
0.7
600
800
1000
1200
124
L16
L8
O8
O1
O16
L1
97
233
219
217
253
256
204
9
218

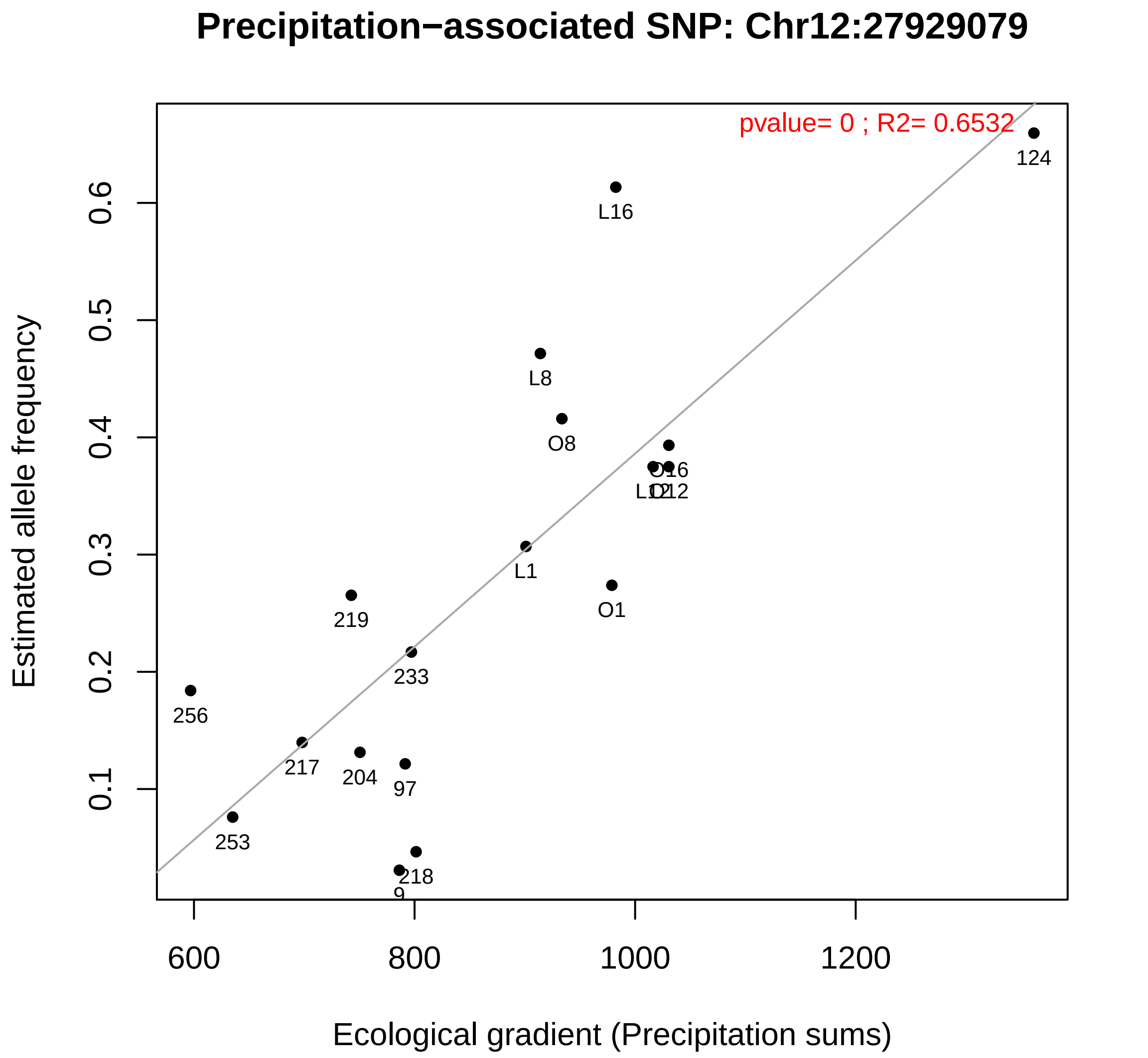

Precipitation−associated SNP: Chr12:27929079
pvalue= 0 ; R2= 0.6532
124
L16
L8
O8
O16
L12
O12
L1
O1
219
233
256
217
204
97
253
218
9
0.1
0.2
0.3
0.4
0.5
0.6
600
800
1000
1200
Estimated allele frequency
Ecological gradient (Precipitation sums)

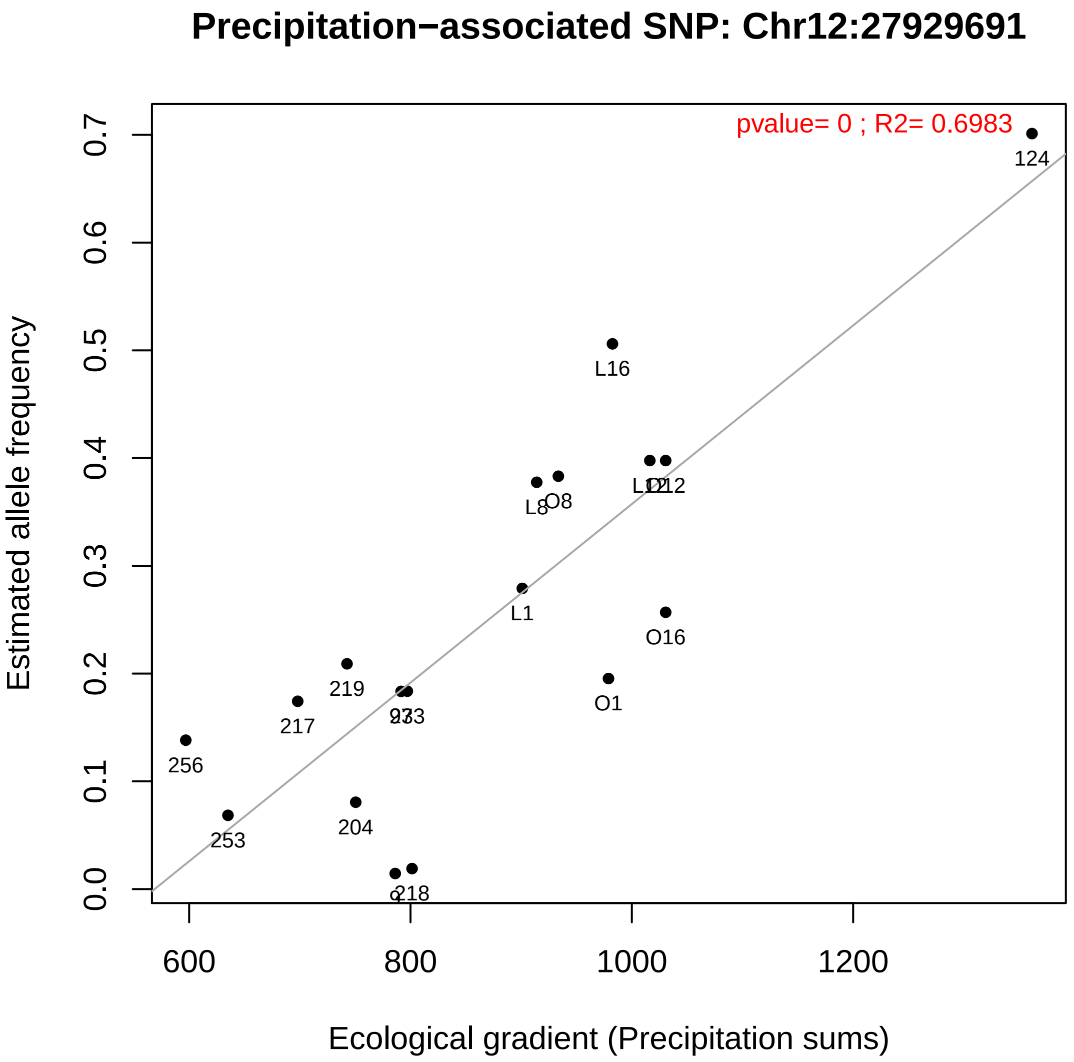

Precipitation−associated SNP: Chr12:27929691
pvalue= 0 ; R2= 0.6983
124
L16
L12
O12
L8
O8
L1
O16
O1
219
217
256
253
204
218
Estimated allele frequency
0.0
0.1
0.2
0.3
0.4
0.5
0.6
0.7
600
800
1000
1200
Ecological gradient (Precipitation sums)

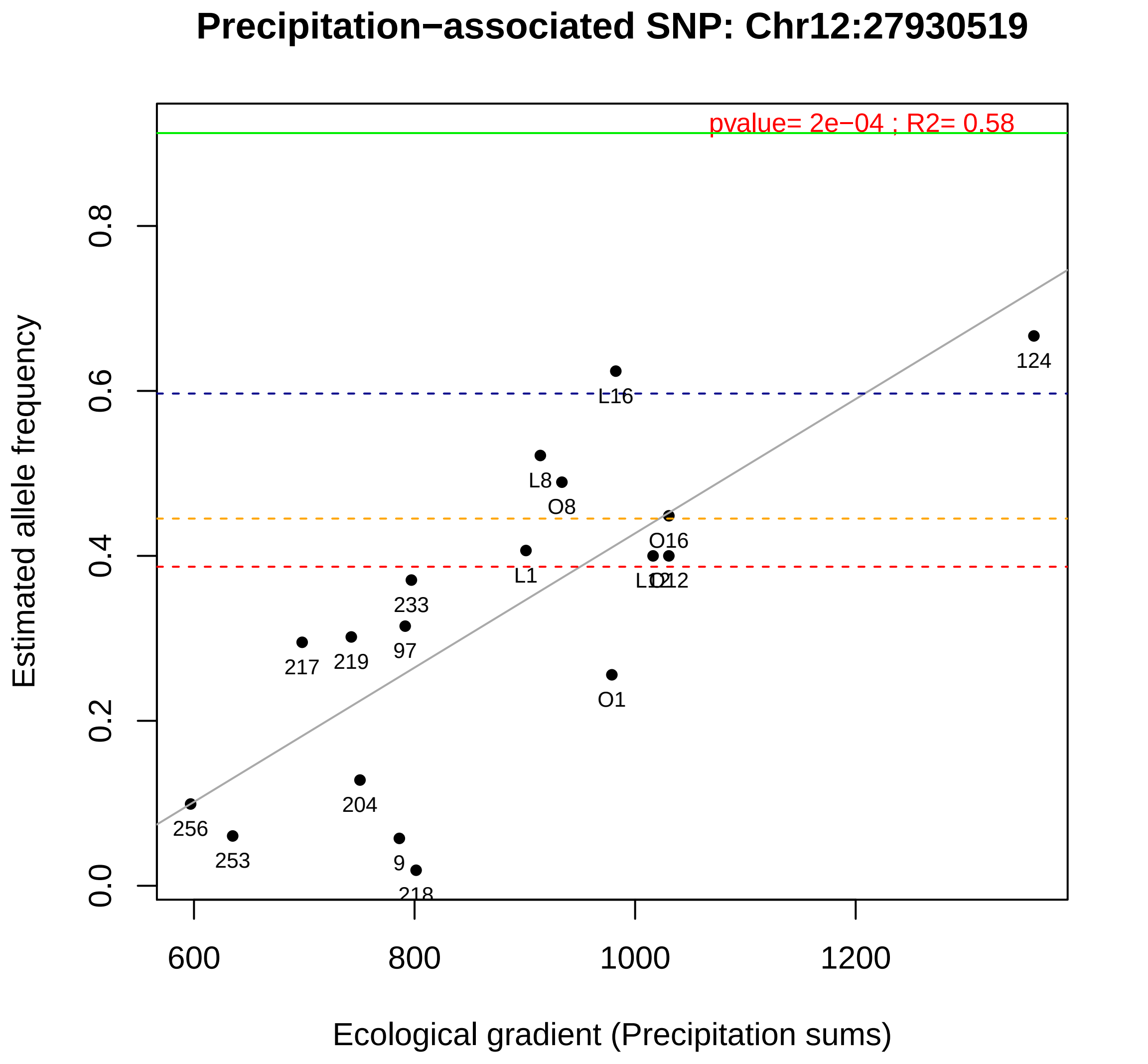

Precipitation−associated SNP: Chr12:27930519
pvalue= 2e−04 ; R2= 0.58
Estimated allele frequency
0.0
0.2
0.4
0.6
0.8
600
800
1000
1200
Ecological gradient (Precipitation sums)
124
L16
L8
O8
O16
L1
L12
O12
233
97
217
219
O1
204
256
253
9
218

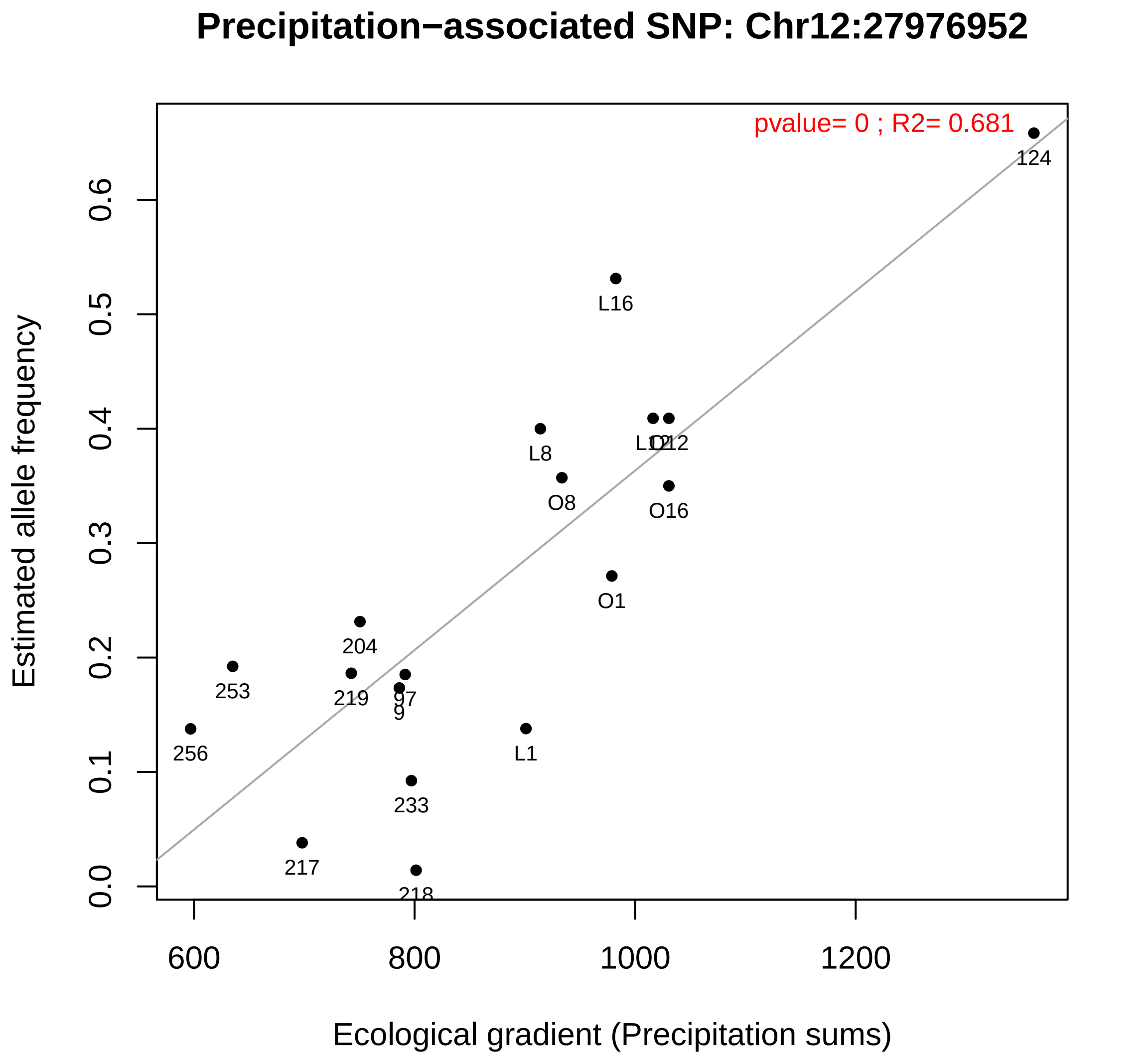
Precipitation−associated SNP: Chr12:27976952
pvalue= 0 ; R2= 0.681
Estimated allele frequency
Ecological gradient (Precipitation sums)
0.0
0.1
0.2
0.3
0.4
0.5
0.6
600
800
1000
1200
124
L16
L8
O8
O16
O1
204
253
219
97
9
256
L1
233
217
218

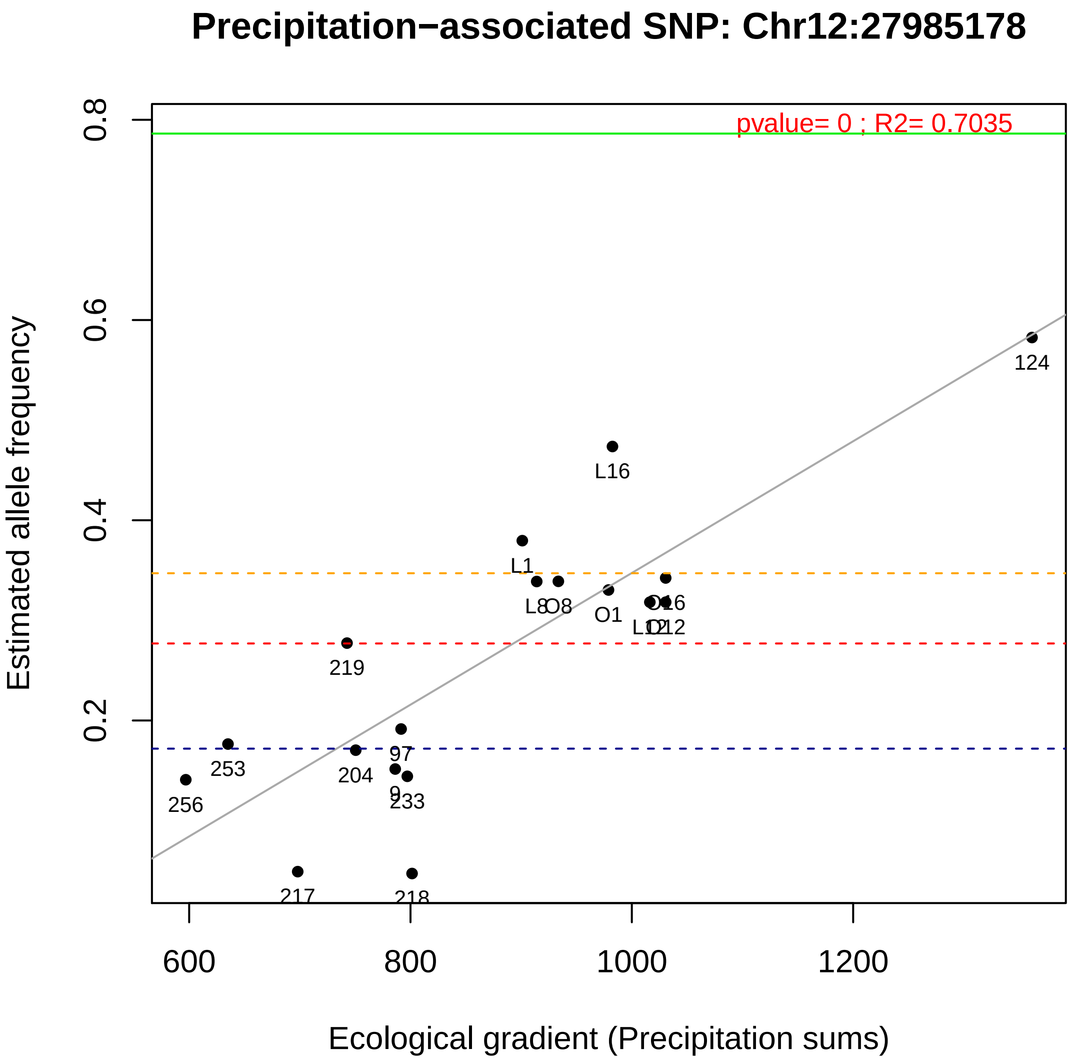
Precipitation−associated SNP: Chr12:27985178
pvalue= 0 ; R2= 0.7035
Estimated allele frequency
0.2
0.4
0.6
0.8
600
800
1000
1200
Ecological gradient (Precipitation sums)
124
L16
L1
L8
O8
O1
O16
L12
O12
219
97
253
204
256
9
233
217
218

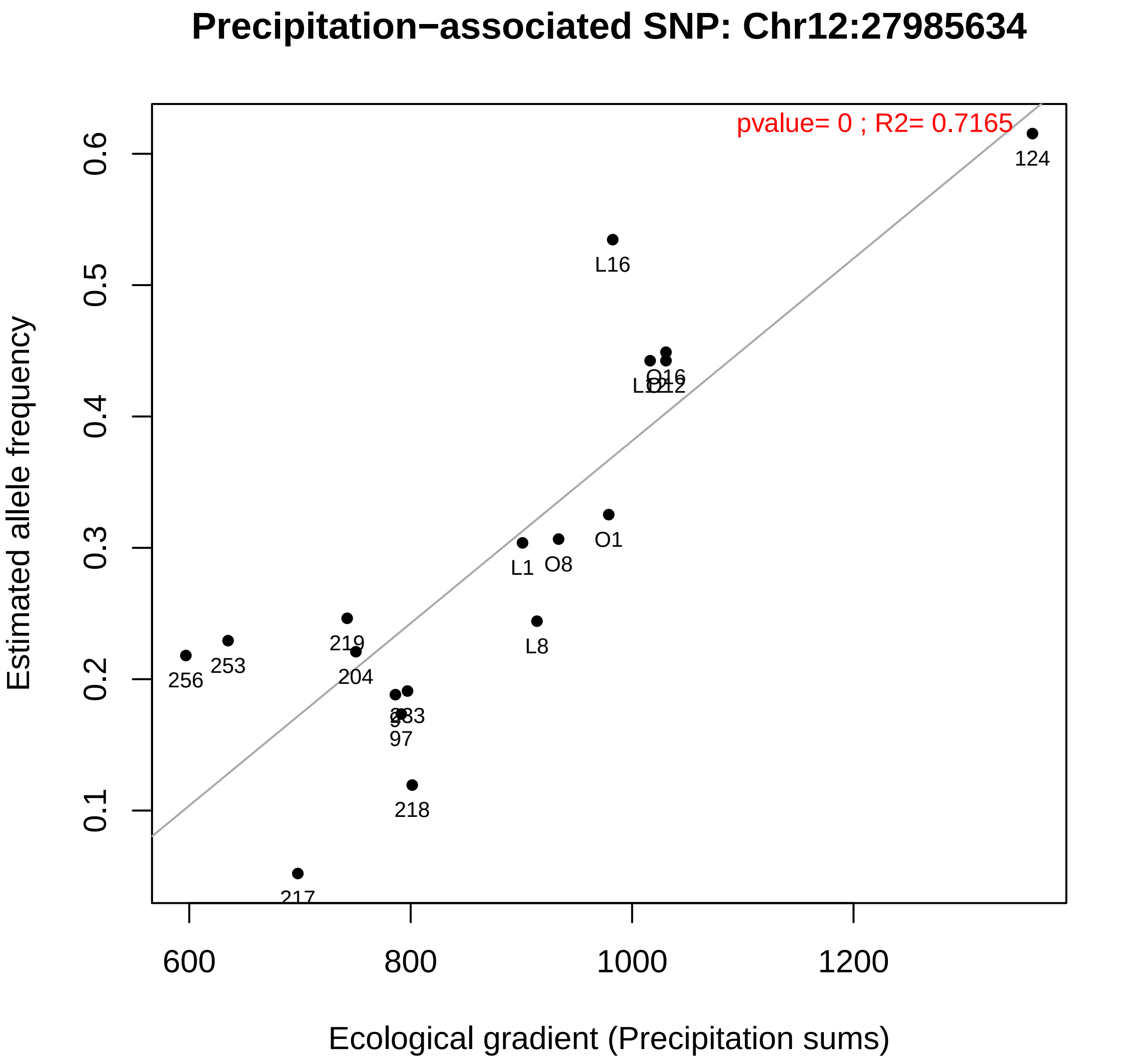
Precipitation−associated SNP: Chr12:27985634
pvalue= 0 ; R2= 0.7165
Estimated allele frequency
Ecological gradient (Precipitation sums)
0.1
0.2
0.3
0.4
0.5
0.6
600
800
1000
1200
124
L16
O16
O12
L1
O8
O1
L8
219
204
253
256
233
97
218
217

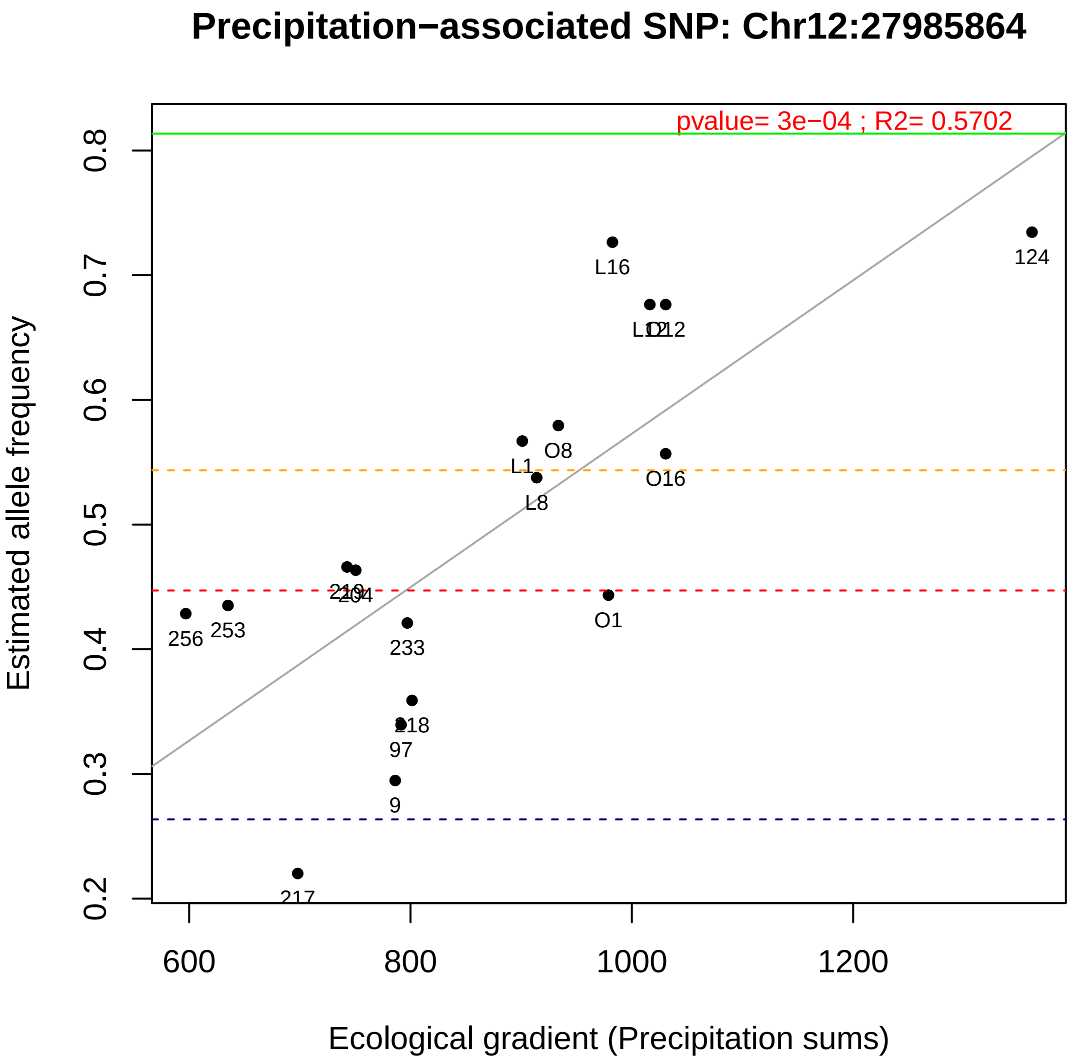
Precipitation−associated SNP: Chr12:27985864
pvalue= 3e−04 ; R2= 0.5702
Estimated allele frequency
0.2
0.3
0.4
0.5
0.6
0.7
0.8
Ecological gradient (Precipitation sums)
600
800
1000
1200
L16
124
L12
O12
O8
L1
L8
O16
210
204
256
253
233
O1
218
97
9
217

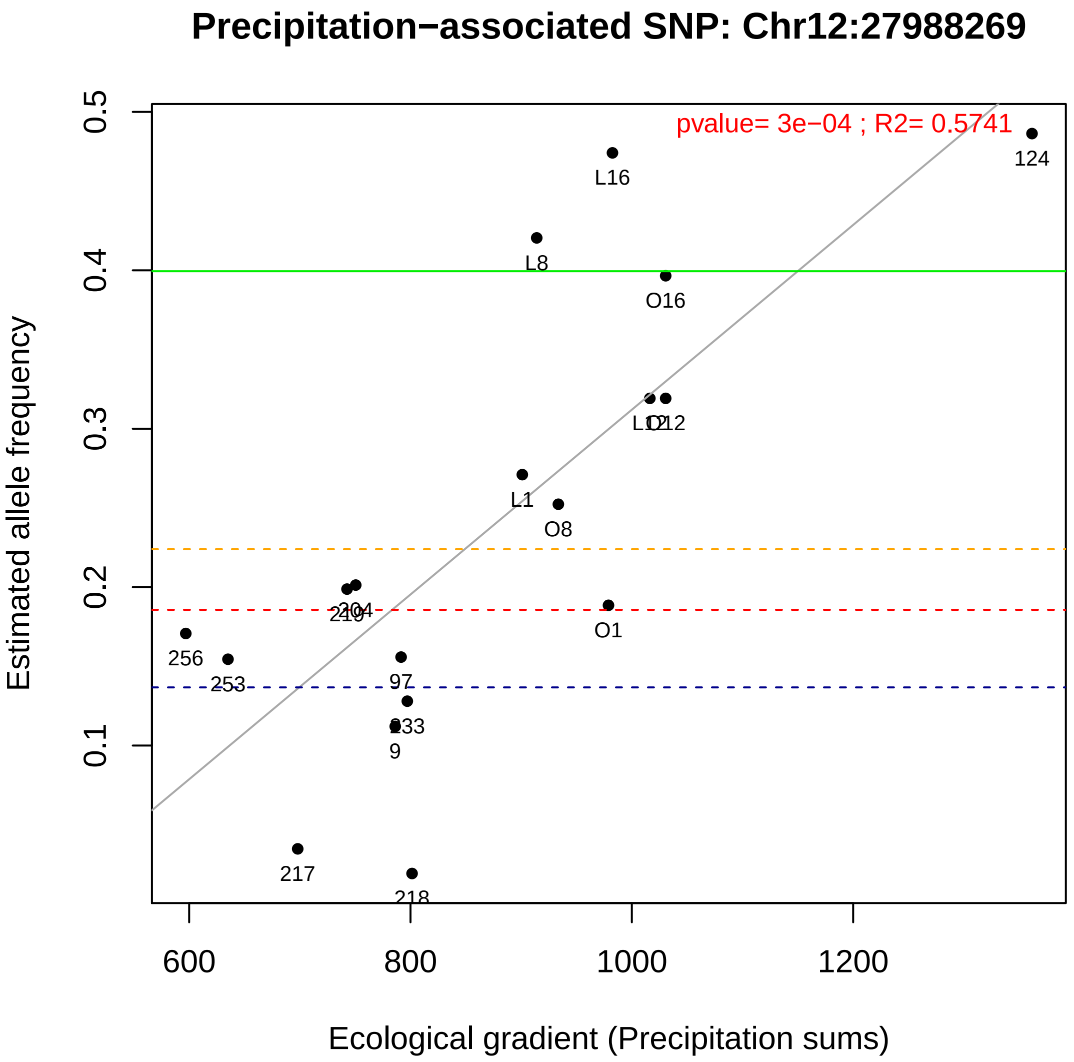
Precipitation−associated SNP: Chr12:27988269
pvalue= 3e−04 ; R2= 0.5741
Estimated allele frequency
0.1
0.2
0.3
0.4
0.5
Ecological gradient (Precipitation sums)
600
800
1000
1200
124
L16
L8
O16
L12
O12
L1
O8
204
219
O1
256
253
97
233
9
217
218

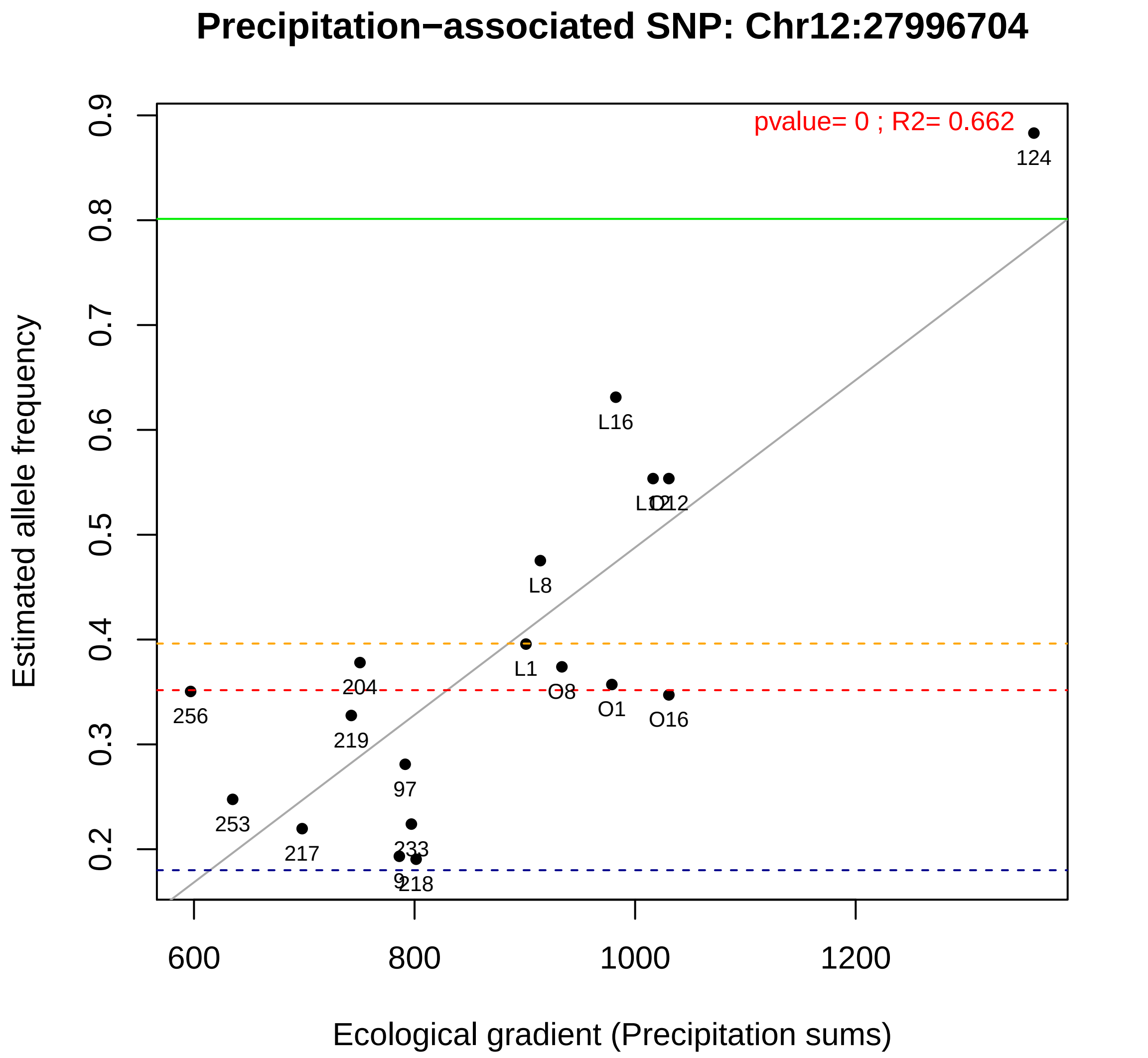
Precipitation−associated SNP: Chr12:27996704
pvalue= 0 ; R2= 0.662
Estimated allele frequency
Ecological gradient (Precipitation sums)
0.2
0.3
0.4
0.5
0.6
0.7
0.8
0.9
600
800
1000
1200
124
L16
L12
O12
L8
L1
O8
O1
O16
204
256
219
97
253
217
233
9
218

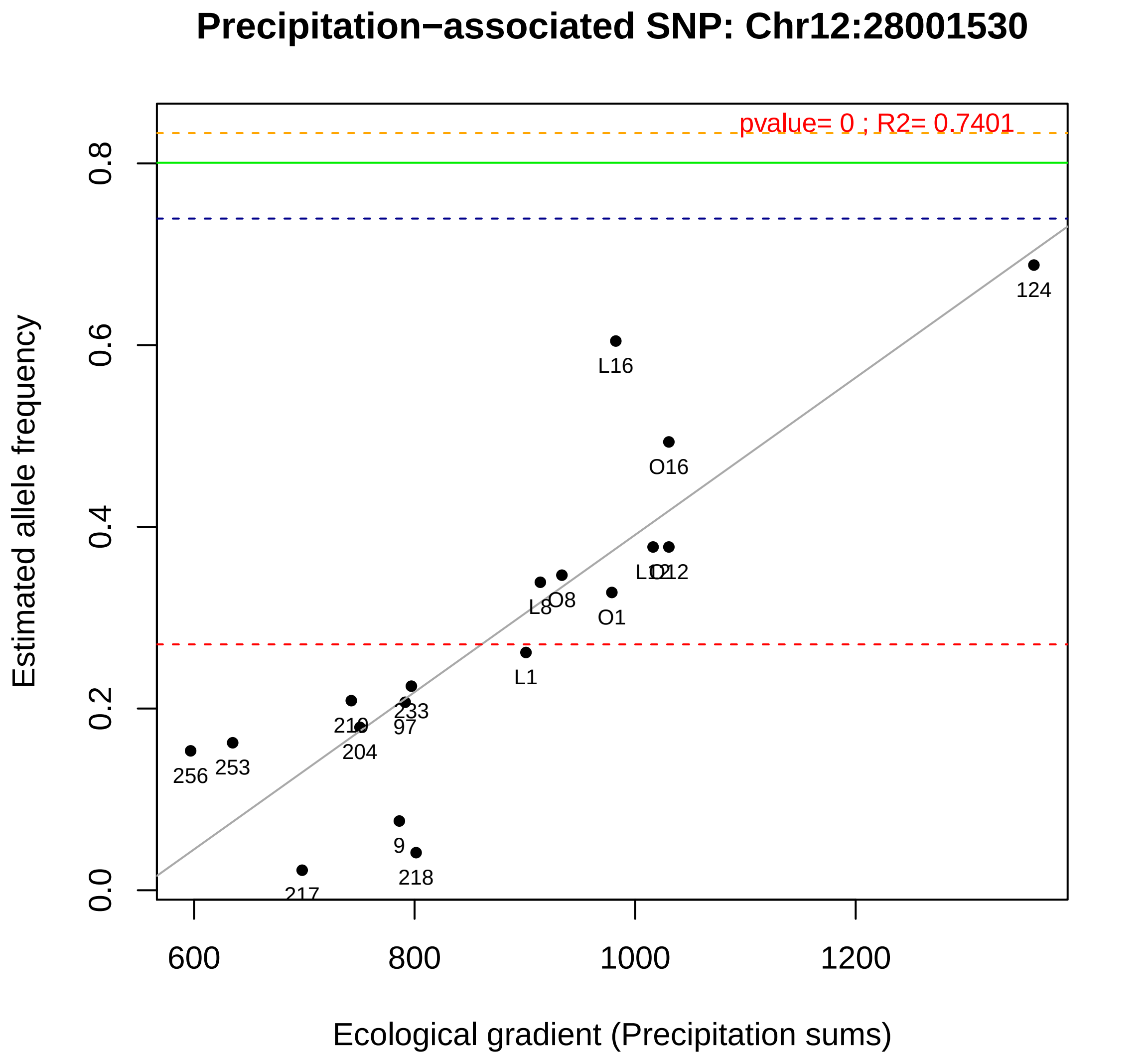
Precipitation−associated SNP: Chr12:28001530
pvalue= 0 ; R2= 0.7401
Estimated allele frequency
Ecological gradient (Precipitation sums)
0.0
0.2
0.4
0.6
0.8
600
800
1000
1200
124
L16
O16
L12
O12
L8
O8
O1
L1
233
97
210
204
256
253
9
218
217

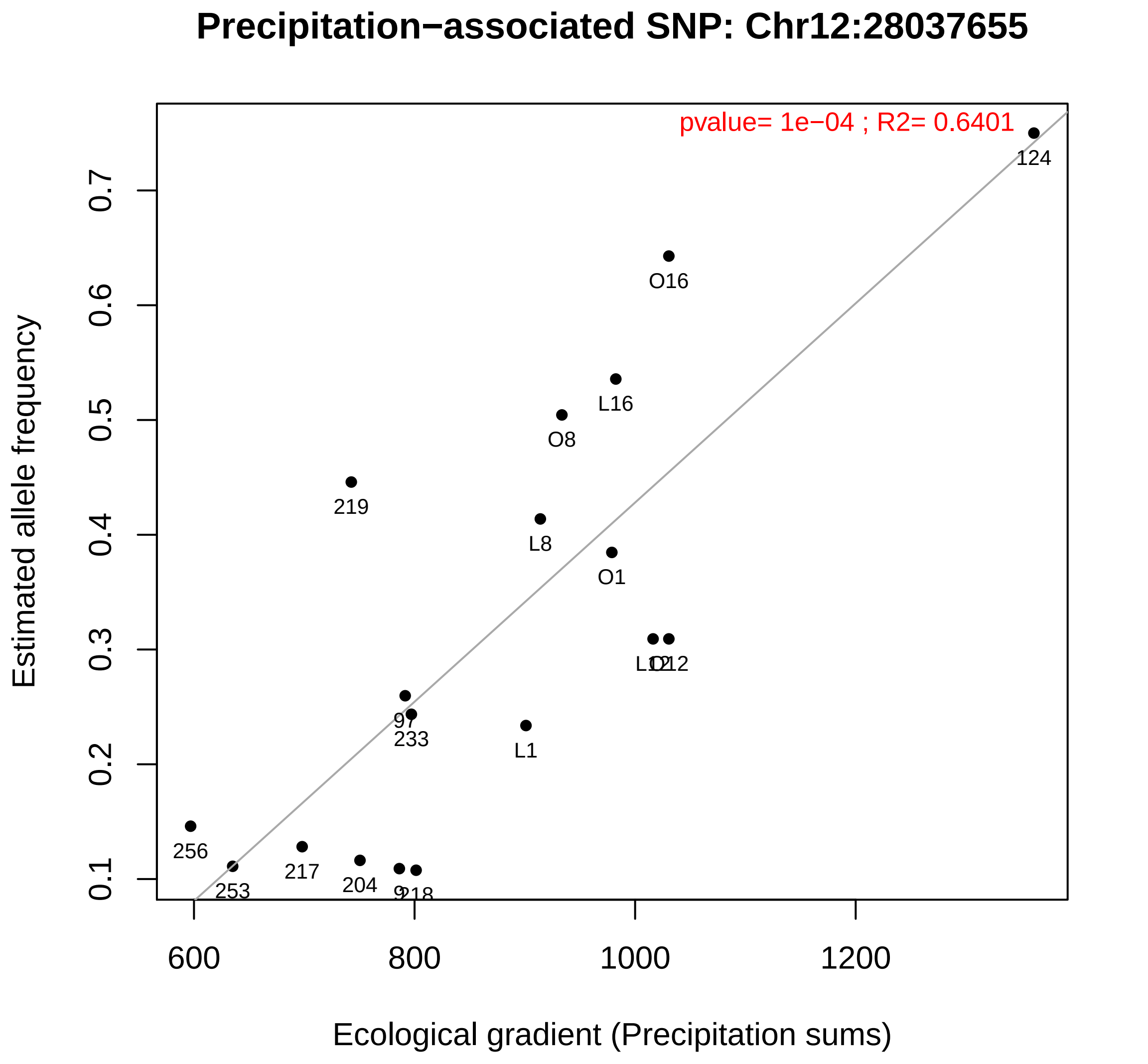
Precipitation−associated SNP: Chr12:28037655
pvalue= 1e−04 ; R2= 0.6401
124
O16
L16
O8
219
L8
O1
L12
O12
97
233
L1
256
253
217
204
9
218
Estimated allele frequency
0.1
0.2
0.3
0.4
0.5
0.6
0.7
600
800
1000
1200
Ecological gradient (Precipitation sums)

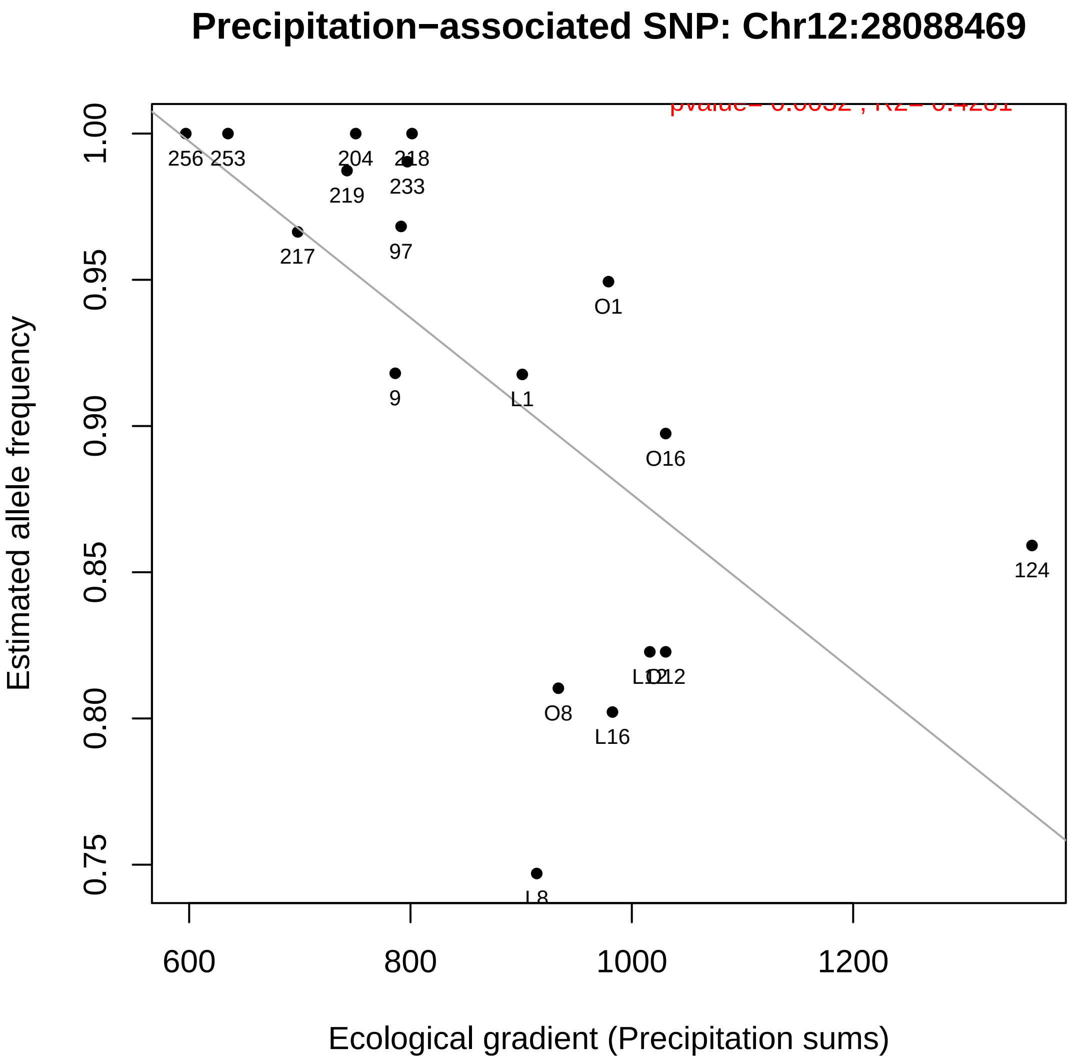
Precipitation−associated SNP: Chr12:28088469
Estimated allele frequency
Ecological gradient (Precipitation sums)
0.75
0.80
0.85
0.90
0.95
1.00
600
800
1000
1200
256
253
204
218
219
233
217
97
O1
9
L1
O16
124
O8
L16

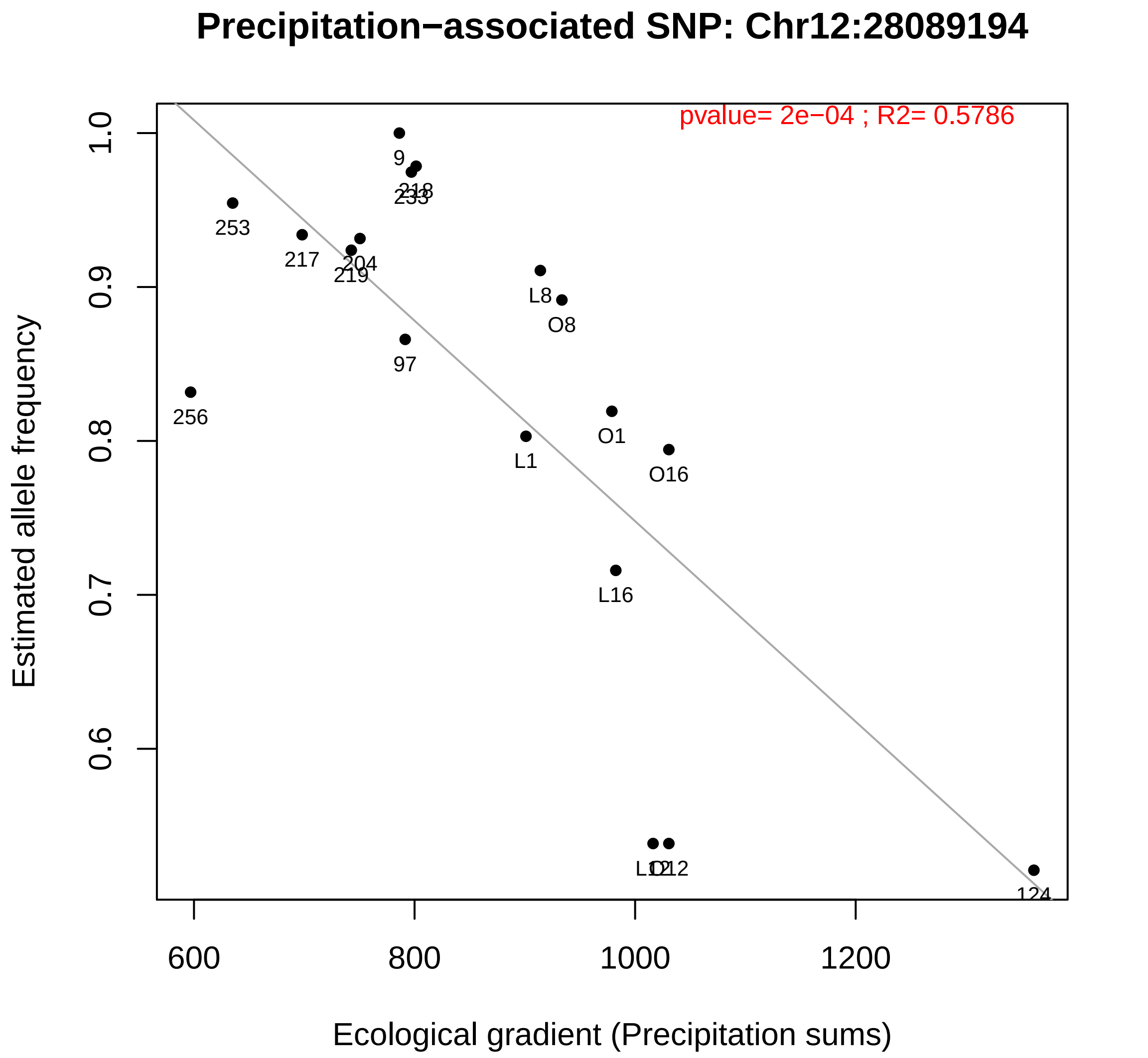
Precipitation−associated SNP: Chr12:28089194
pvalue= 2e−04 ; R2= 0.5786
Estimated allele frequency
Ecological gradient (Precipitation sums)
1.0
0.9
0.8
0.7
0.6
600
800
1000
1200
9
218
233
253
217
204
219
L8
O8
97
256
O1
L1
O16
L16
L12
O12
124

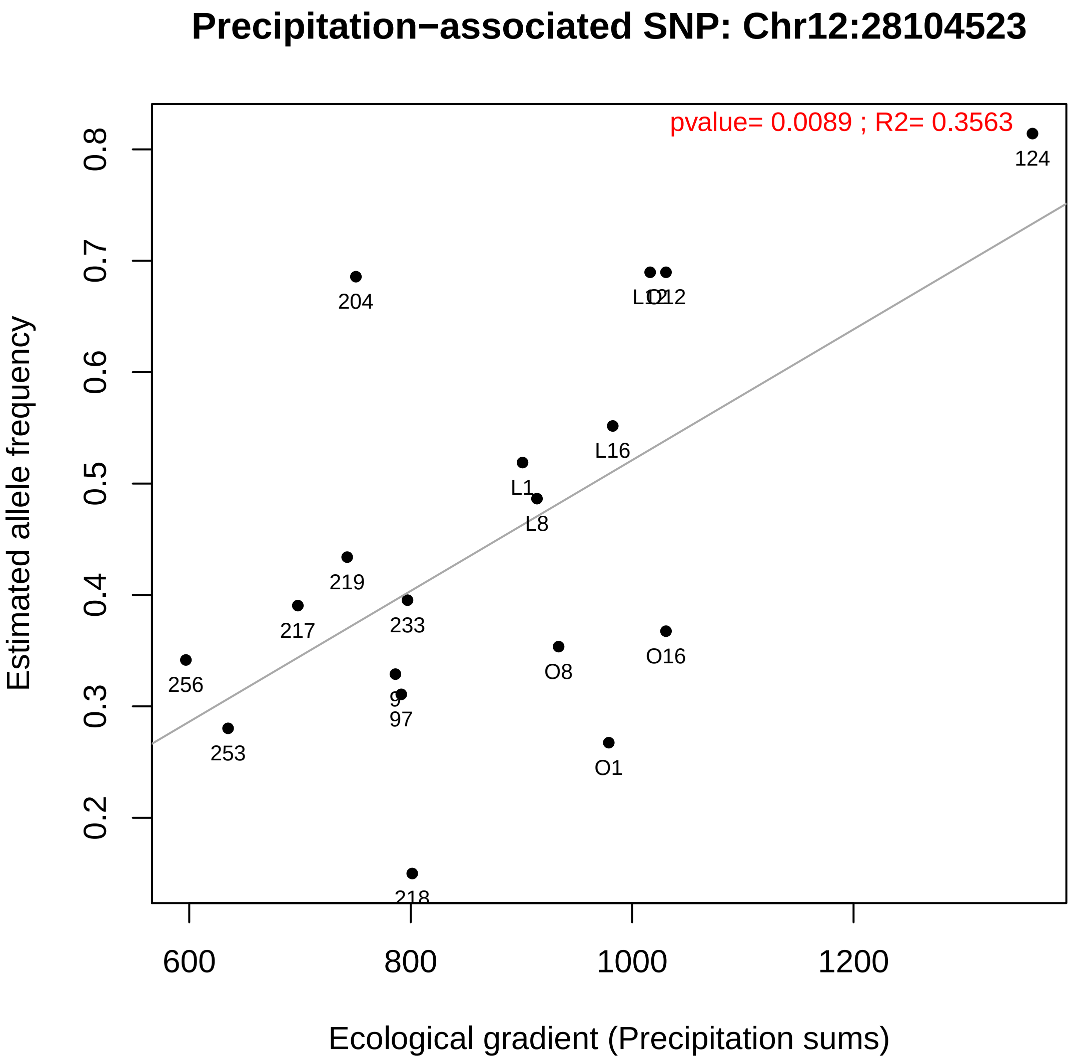

Precipitation−associated SNP: Chr12:28104523
pvalue= 0.0089 ; R2= 0.3563
Estimated allele frequency
Ecological gradient (Precipitation sums)
0.2
0.3
0.4
0.5
0.6
0.7
0.8
600
800
1000
1200
124
204
L16
L1
L8
219
217
233
256
253
O8
O16
O1
97
218

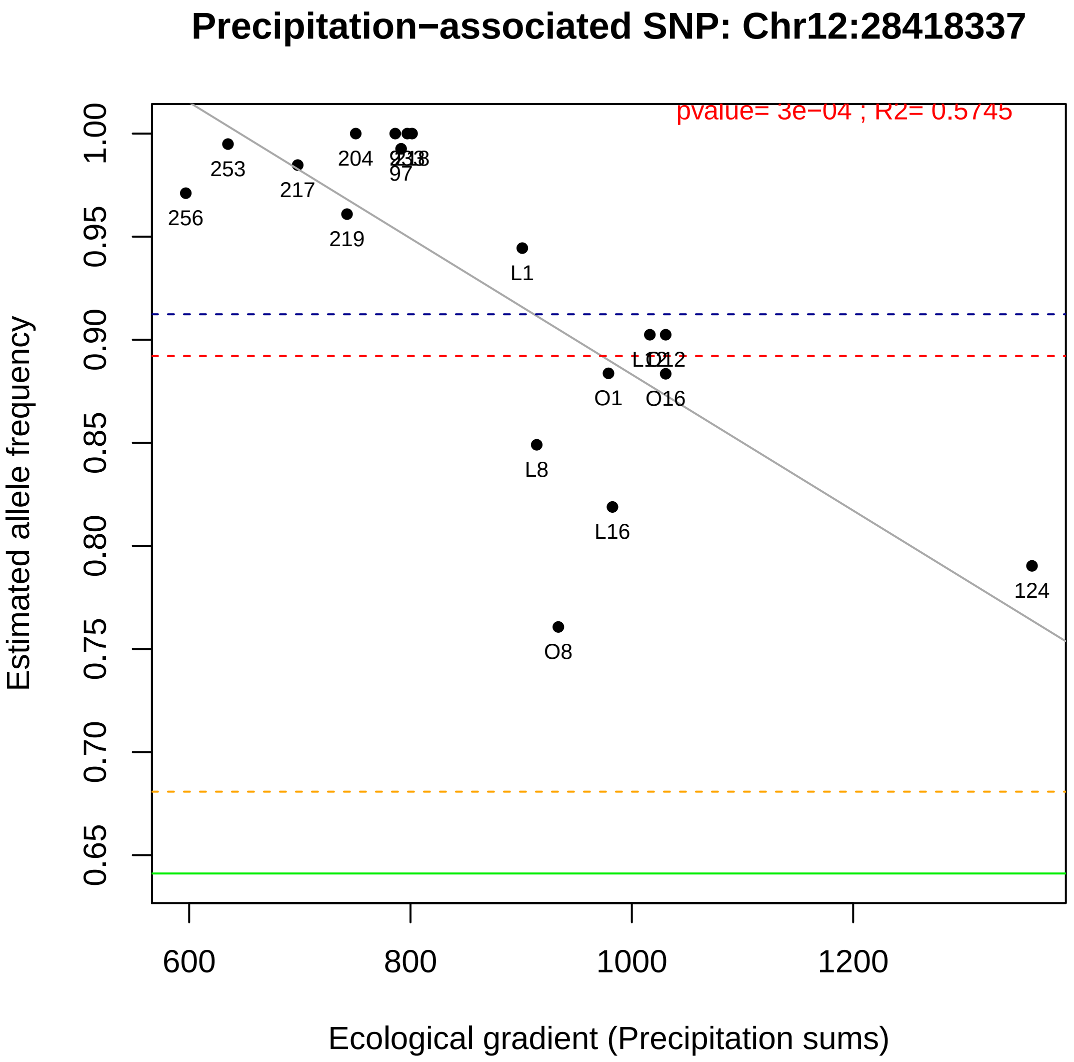
Precipitation−associated SNP: Chr12:28418337
pvalue= 3e−04 ; R2= 0.5745
Estimated allele frequency
Ecological gradient (Precipitation sums)
0.65
0.70
0.75
0.80
0.85
0.90
0.95
1.00
600
800
1000
1200
256
253
217
219
204
97
L1
L8
O1
O16
L16
O8
124

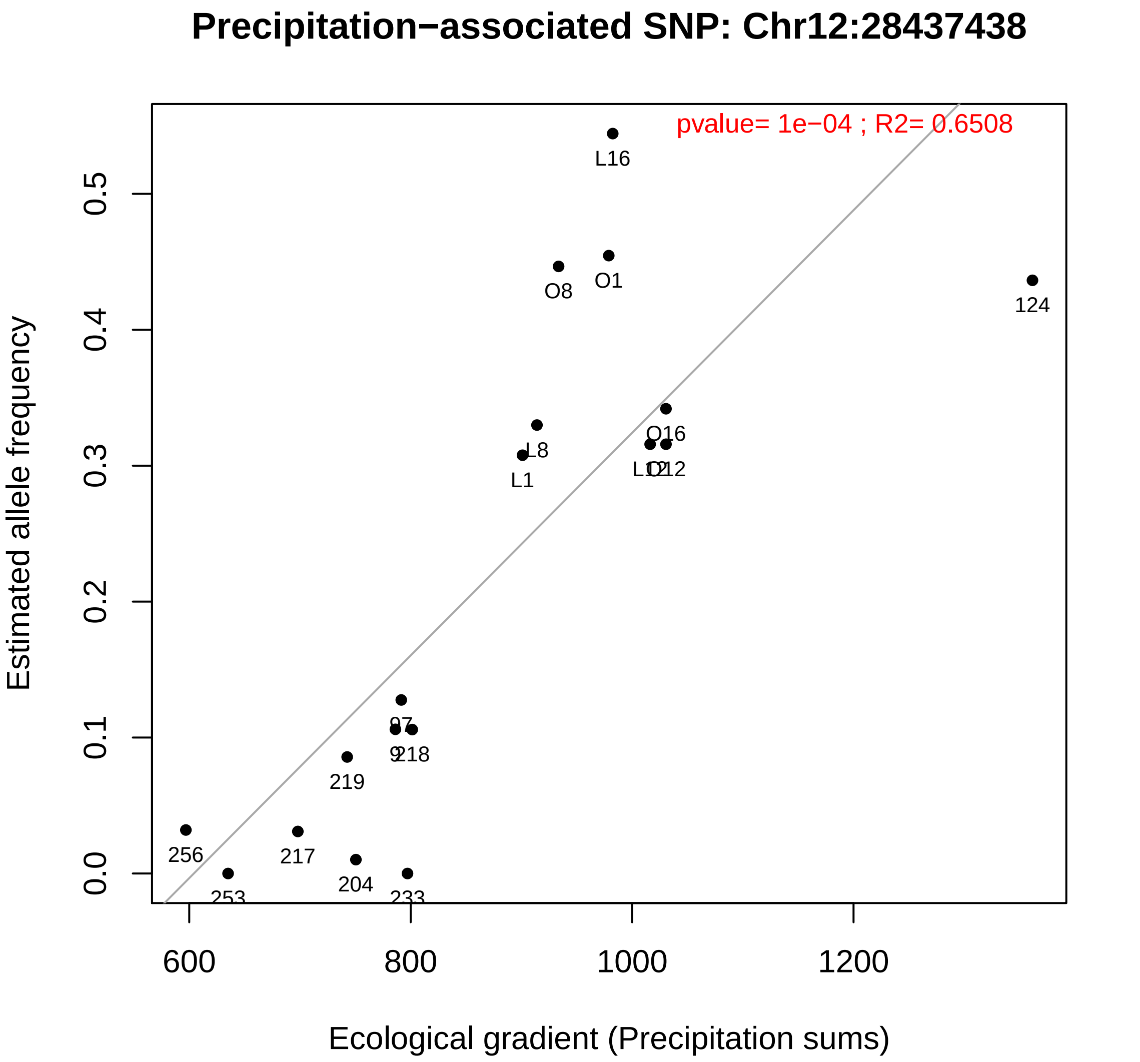

Precipitation−associated SNP: Chr12:28437438
pvalue= 1e−04 ; R2= 0.6508
L16
O8
O1
124
L8
L1
O16
L12
O12
97
9
218
219
256
253
217
204
233
0.0
0.1
0.2
0.3
0.4
0.5
600
800
1000
1200
Estimated allele frequency
Ecological gradient (Precipitation sums)

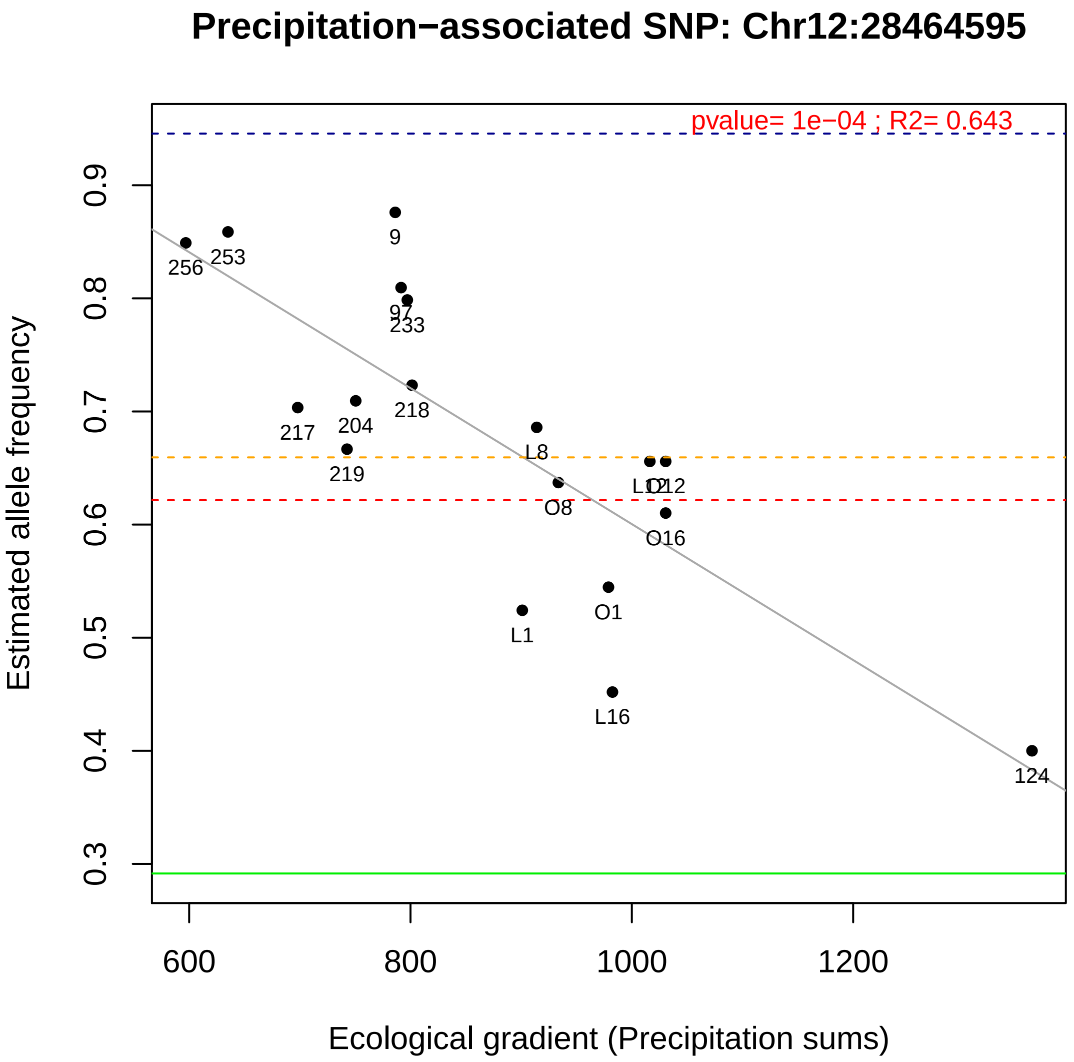

Precipitation−associated SNP: Chr12:28464595
pvalue= 1e−04 ; R2= 0.643
Estimated allele frequency
0.3
0.4
0.5
0.6
0.7
0.8
0.9
600
800
1000
1200
Ecological gradient (Precipitation sums)
256
253
9
97
233
218
217
204
219
L8
O8
L12
O12
O16
O1
L1
L16
124

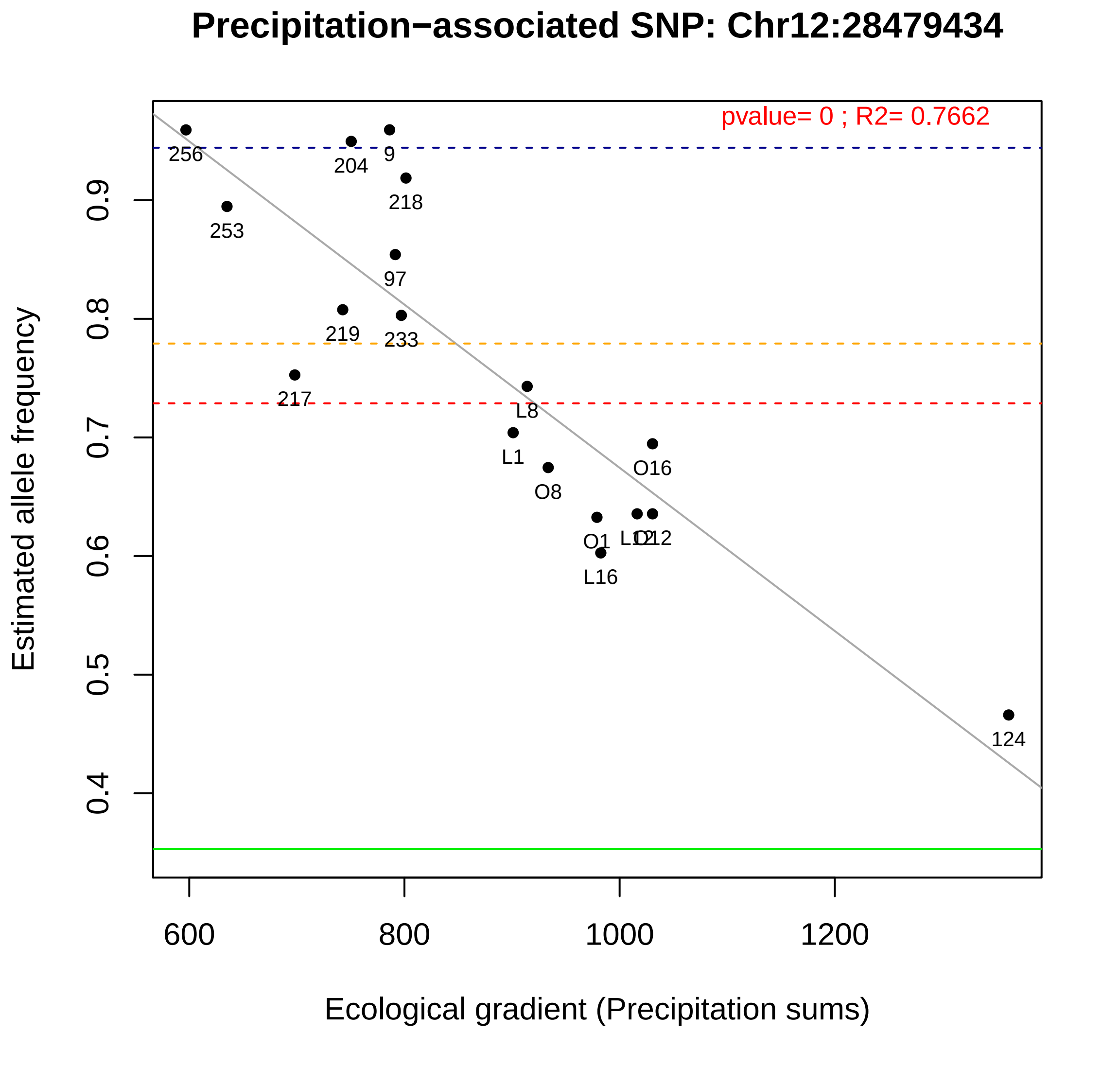

Precipitation−associated SNP: Chr12:28479434
pvalue= 0 ; R2= 0.7662
256
204
9
218
253
97
219
233
217
L8
L1
O16
O8
O1
L12
O12
L16
124
0.4
0.5
0.6
0.7
0.8
0.9
600
800
1000
1200
Estimated allele frequency
Ecological gradient (Precipitation sums)

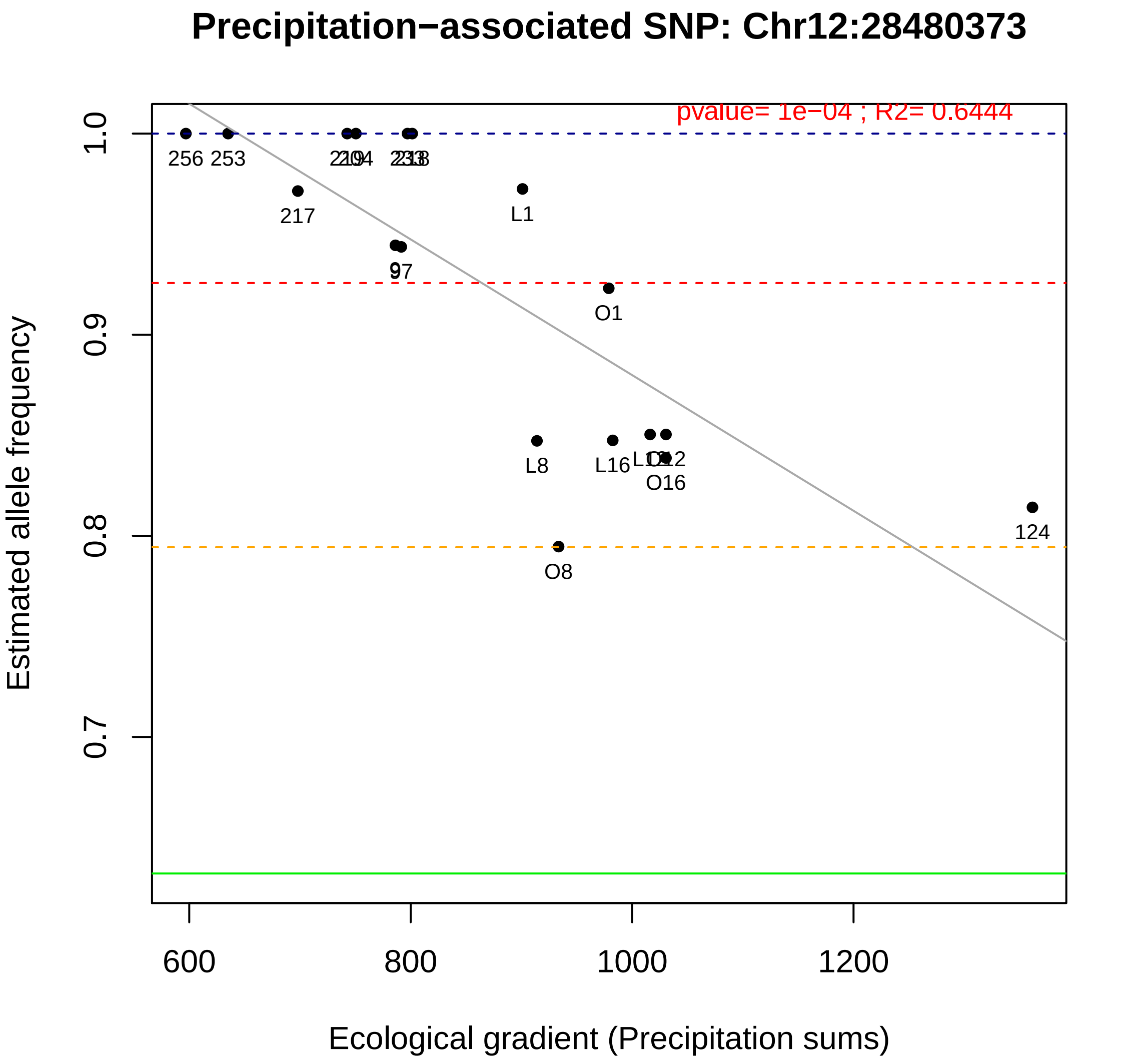

Precipitation−associated SNP: Chr12:28480373
pvalue= 1e−04 ; R2= 0.6444
Estimated allele frequency
1.0
0.9
0.8
0.7
256
253
217
L1
O1
L8
L16
O16
O8
124
600
800
1000
1200
Ecological gradient (Precipitation sums)

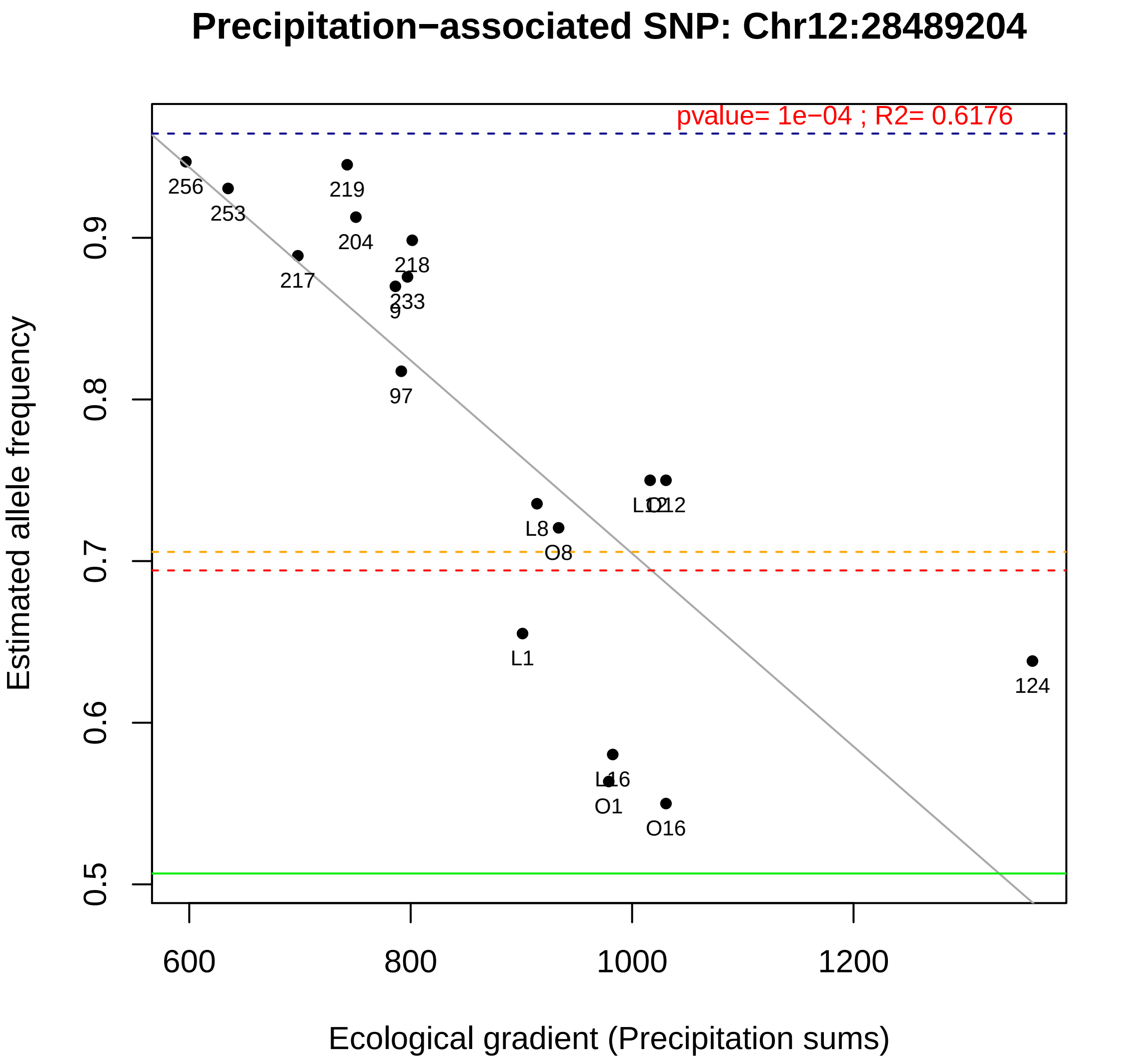

Precipitation−associated SNP: Chr12:28489204
pvalue= 1e−04 ; R2= 0.6176
Estimated allele frequency
0.5
0.6
0.7
0.8
0.9
600
800
1000
1200
Ecological gradient (Precipitation sums)
256
253
219
204
217
218
233
9
97
L8
O8
L12
O12
L1
124
L16
O1
O16

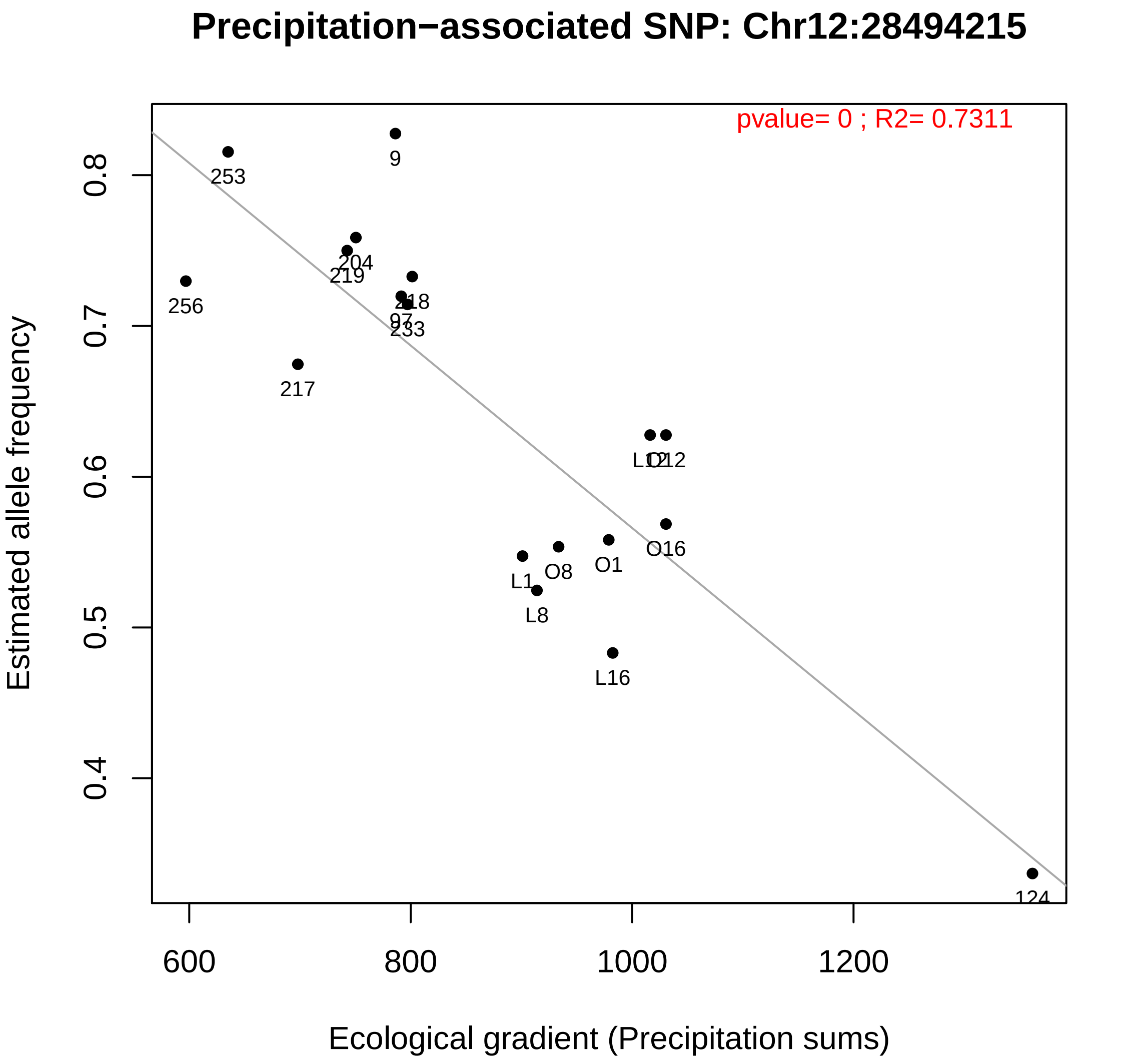
Precipitation−associated SNP: Chr12:28494215
pvalue= 0 ; R2= 0.7311
Estimated allele frequency
Ecological gradient (Precipitation sums)
0.4
0.5
0.6
0.7
0.8
600
800
1000
1200
9
253
204
219
256
218
97
233
217
L12
O12
O16
O1
O8
L1
L8
L16
124

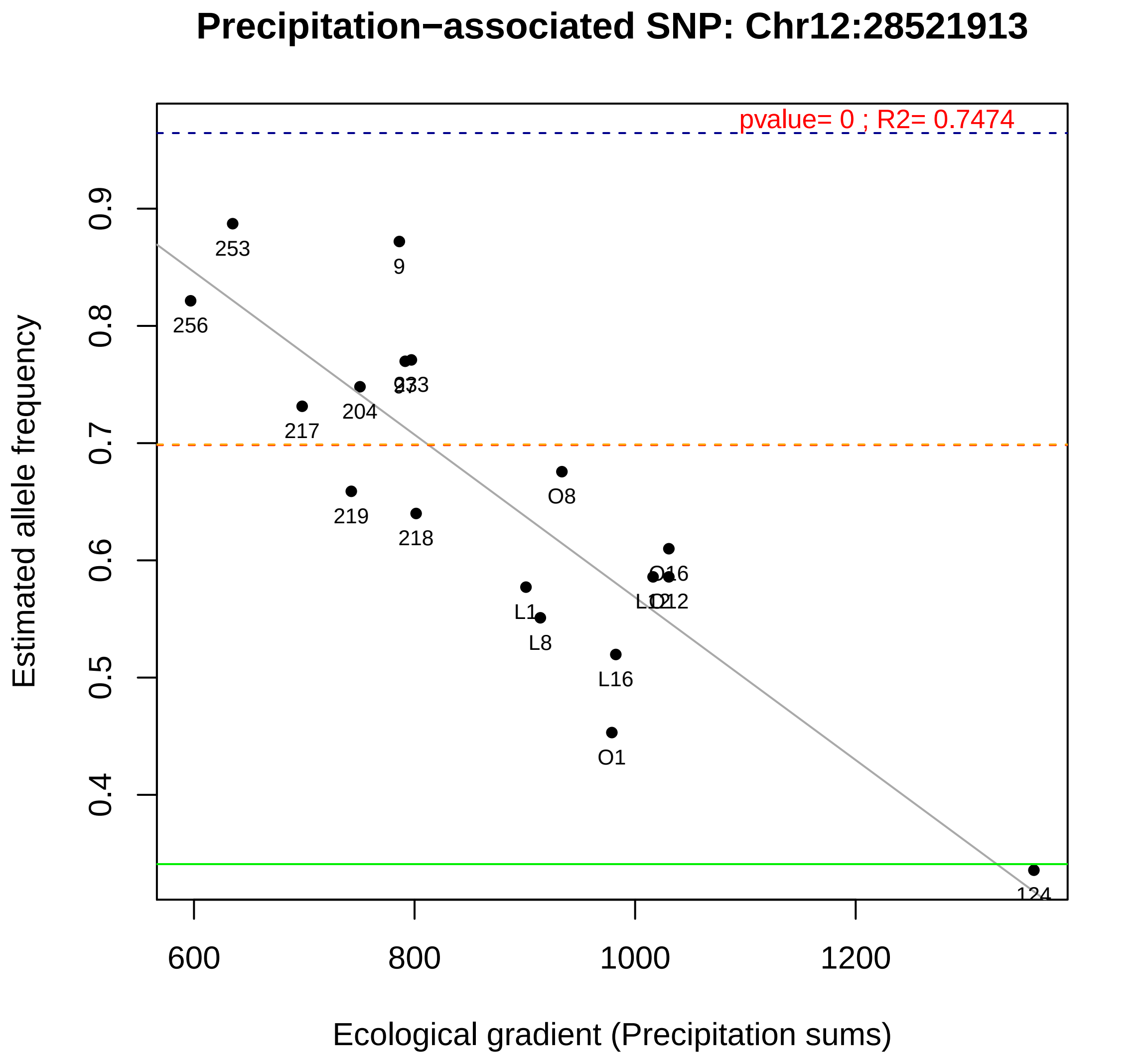
Precipitation−associated SNP: Chr12:28521913
pvalue= 0 ; R2= 0.7474
Estimated allele frequency
Ecological gradient (Precipitation sums)
0.4
0.5
0.6
0.7
0.8
0.9
600
800
1000
1200
253
9
256
204
217
219
218
O8
L1
L8
L16
O1
O16
L12
O12
124

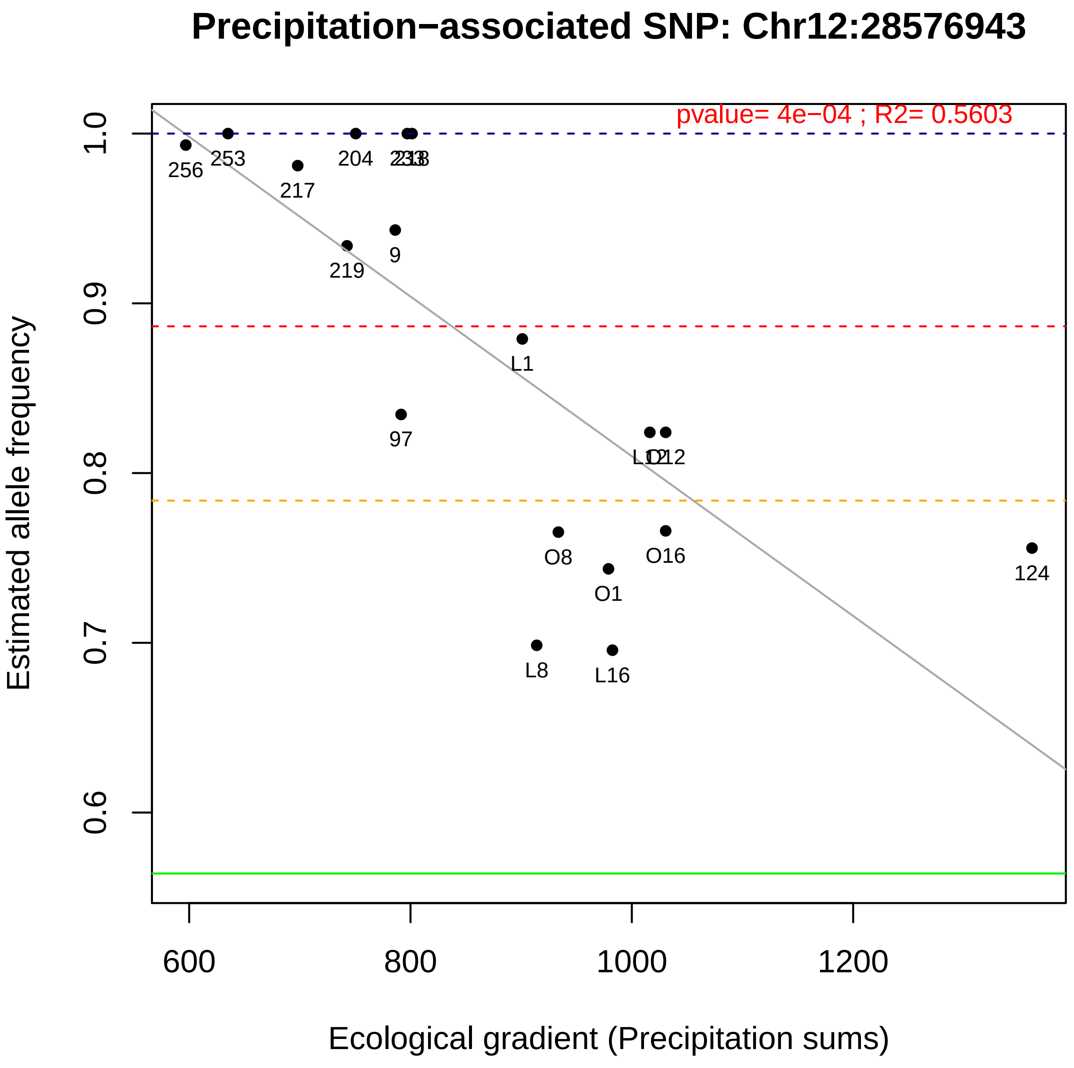
Precipitation−associated SNP: Chr12:28576943
pvalue= 4e−04 ; R2= 0.5603
Estimated allele frequency
Ecological gradient (Precipitation sums)
0.6
0.7
0.8
0.9
1.0
600
800
1000
1200
256
253
204
217
219
9
L1
97
O8
O1
O16
124
L8
L16

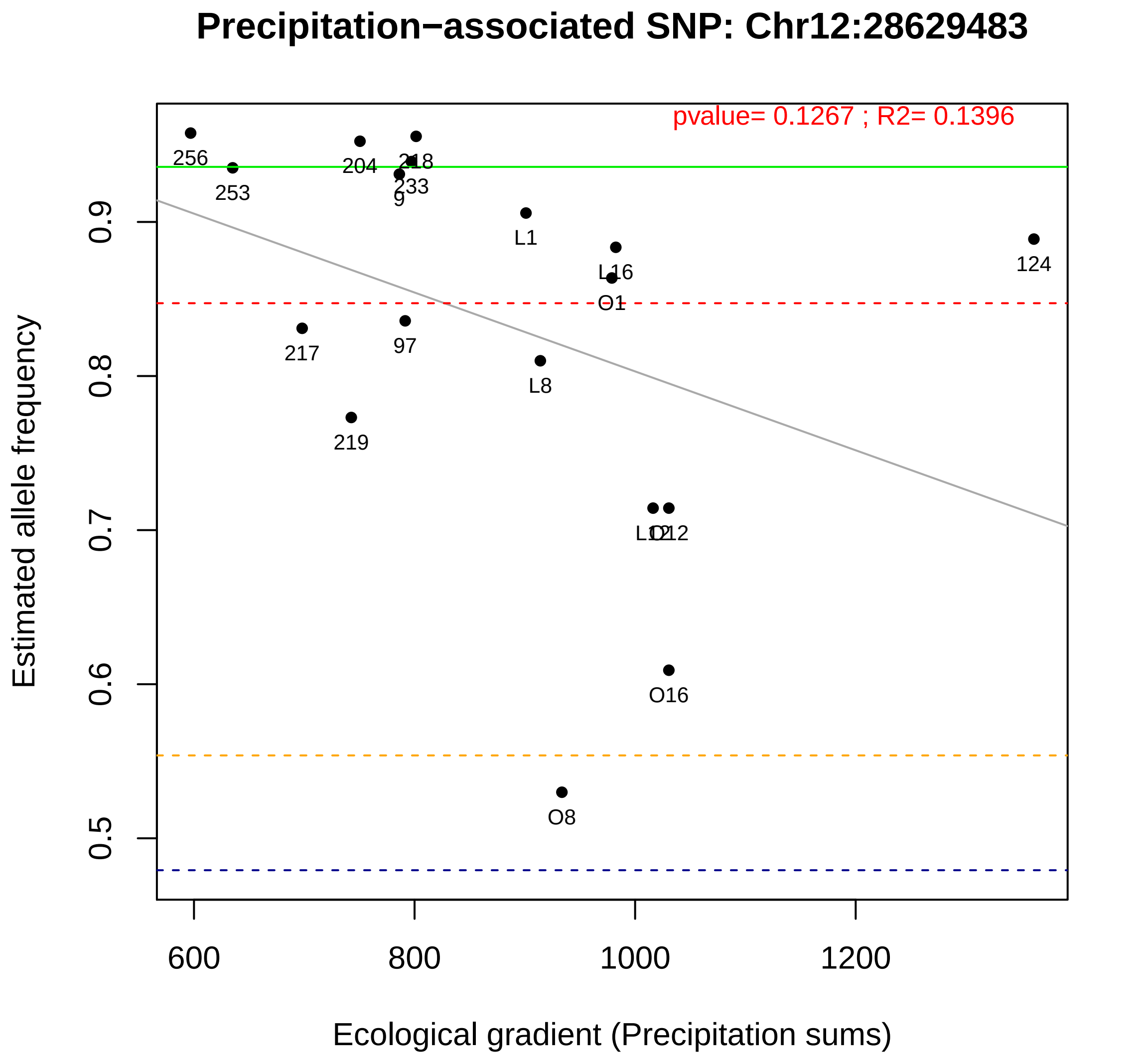

Precipitation−associated SNP: Chr12:28629483
pvalue= 0.1267 ; R2= 0.1396
256
253
204
218
233
9
L1
L16
O1
124
217
97
L8
219
L12
O12
O16
O8
Estimated allele frequency
0.5
0.6
0.7
0.8
0.9
600
800
1000
1200
Ecological gradient (Precipitation sums)

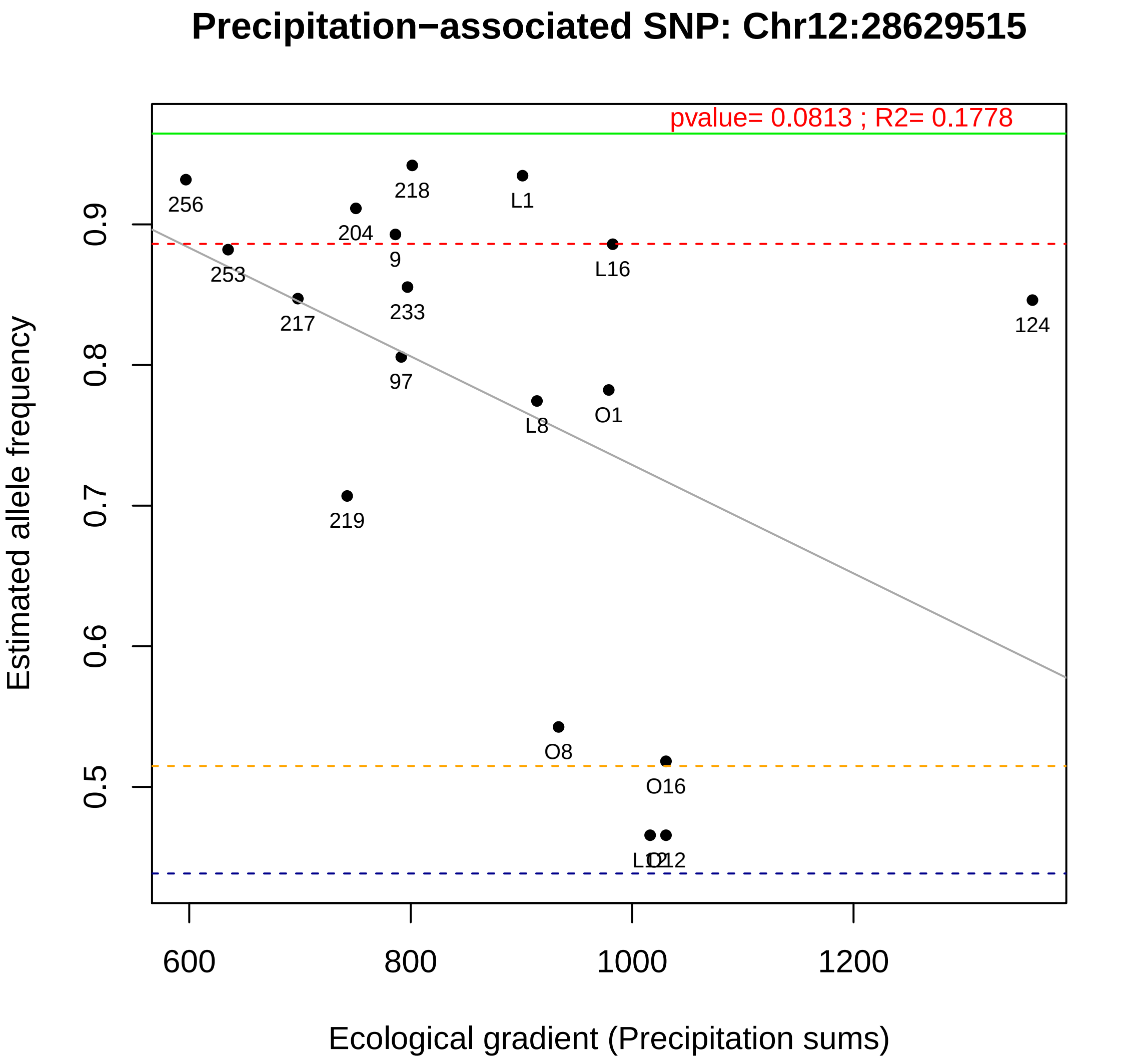

Precipitation−associated SNP: Chr12:28629515
pvalue= 0.0813 ; R2= 0.1778
Estimated allele frequency
0.5
0.6
0.7
0.8
0.9
600
800
1000
1200
Ecological gradient (Precipitation sums)
256
218
L1
204
9
L16
253
217
233
124
97
L8
O1
219
O8
O16

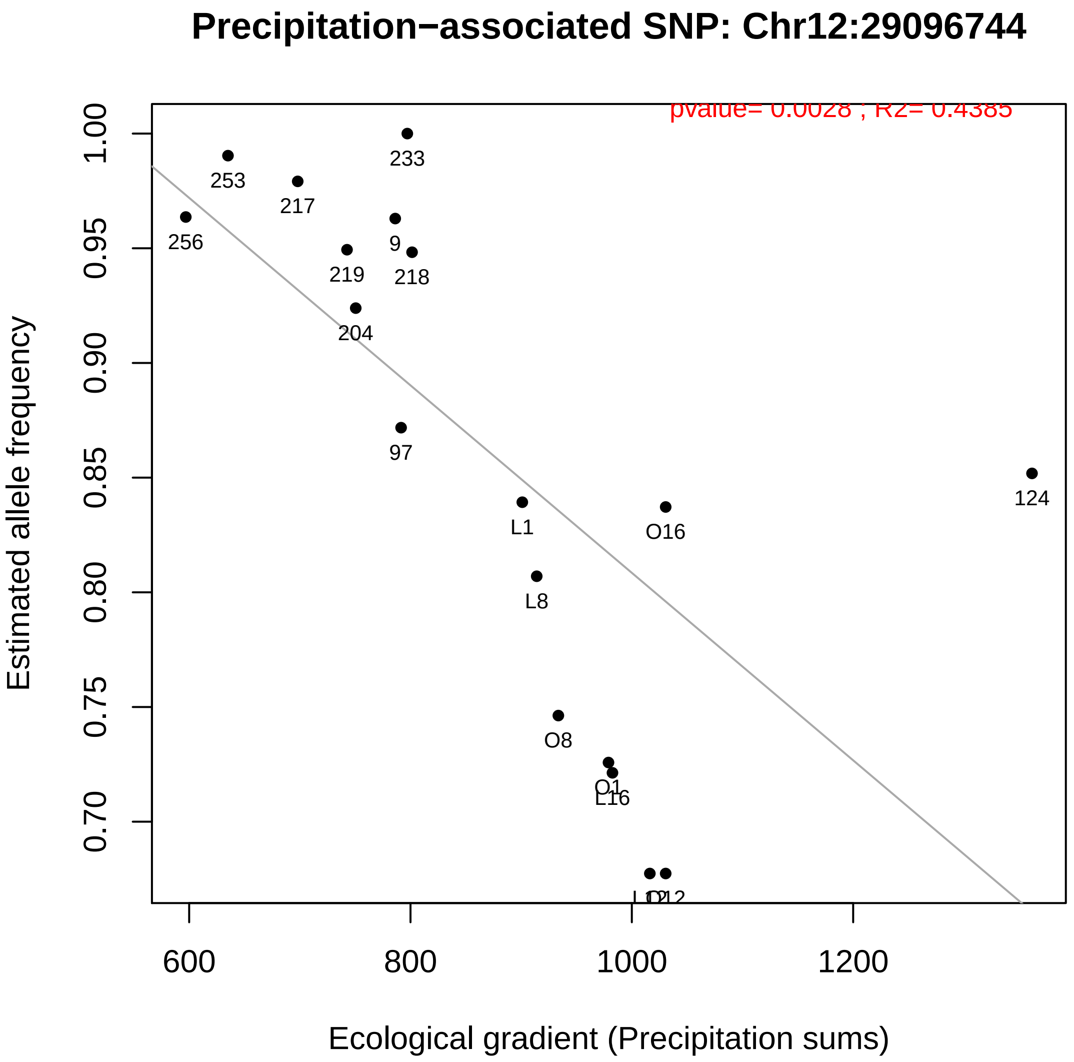

Precipitation−associated SNP: Chr12:29096744
pvalue= 0.0028 ; R2= 0.4385
Estimated allele frequency
1.00
0.95
0.90
0.85
0.80
0.75
0.70
600
800
1000
1200
Ecological gradient (Precipitation sums)
233
253
217
256
9
219
218
204
97
L1
O16
124
L8
O8
O1
L16

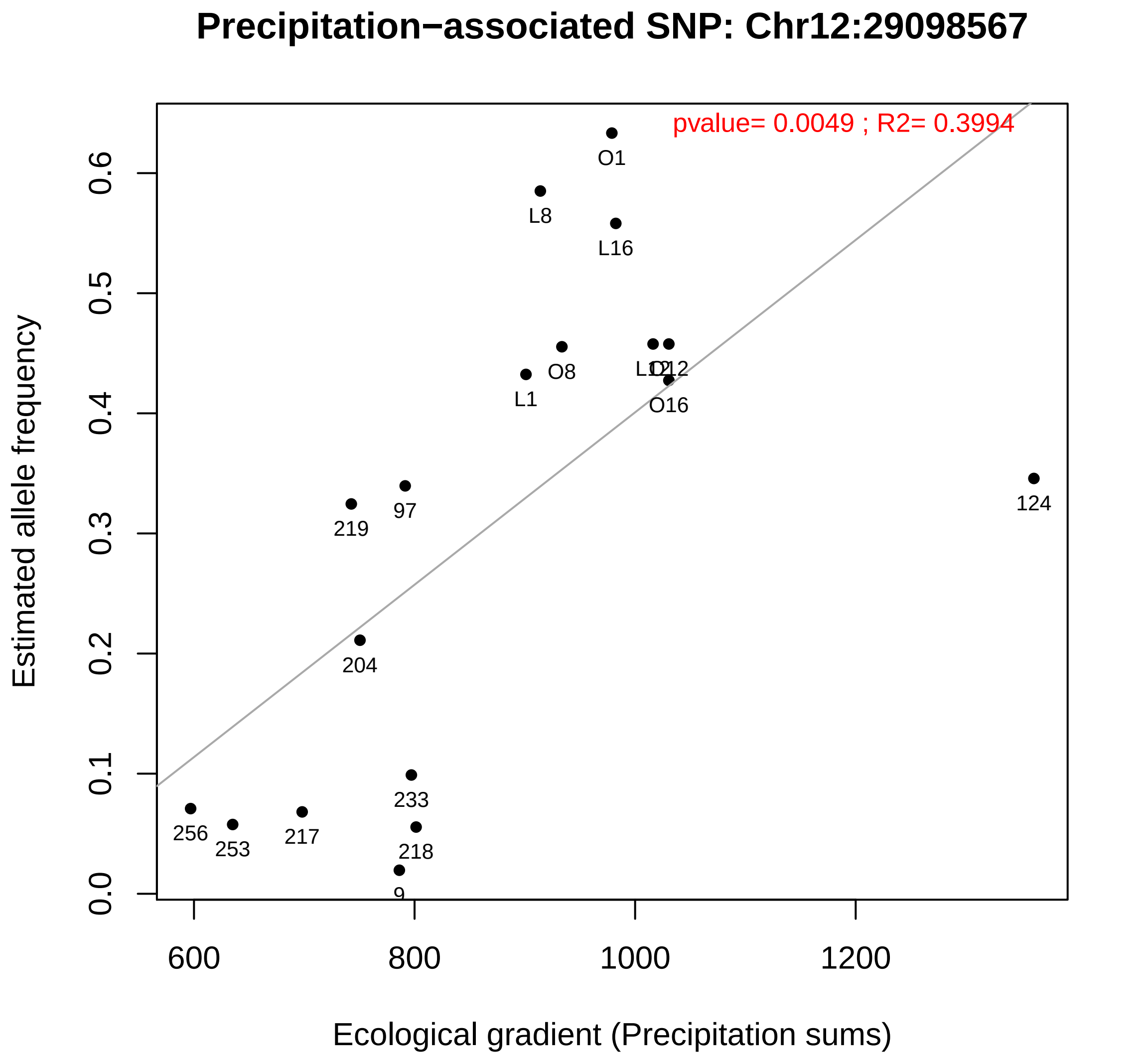

Precipitation−associated SNP: Chr12:29098567
pvalue= 0.0049 ; R2= 0.3994
O1
L8
L16
O8
L12
O12
L1
O16
97
219
124
204
233
256
253
217
218
9
0.0
0.1
0.2
0.3
0.4
0.5
0.6
600
800
1000
1200
Estimated allele frequency
Ecological gradient (Precipitation sums)

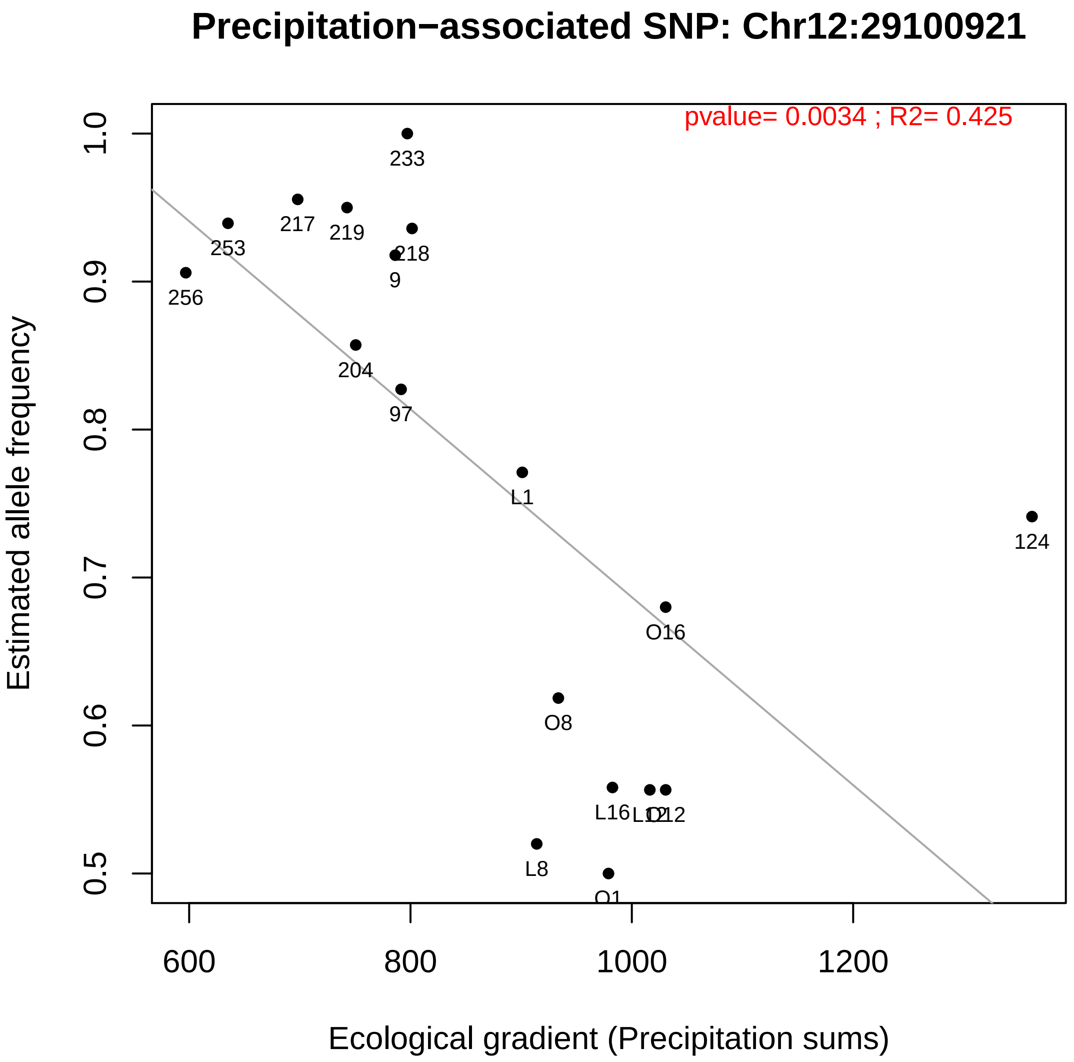

Precipitation−associated SNP: Chr12:29100921
pvalue= 0.0034 ; R2= 0.425
Estimated allele frequency
Ecological gradient (Precipitation sums)
1.0
0.9
0.8
0.7
0.6
0.5
600
800
1000
1200
233
217
219
253
218
9
256
204
97
L1
124
O16
O8
L16
O12
L8

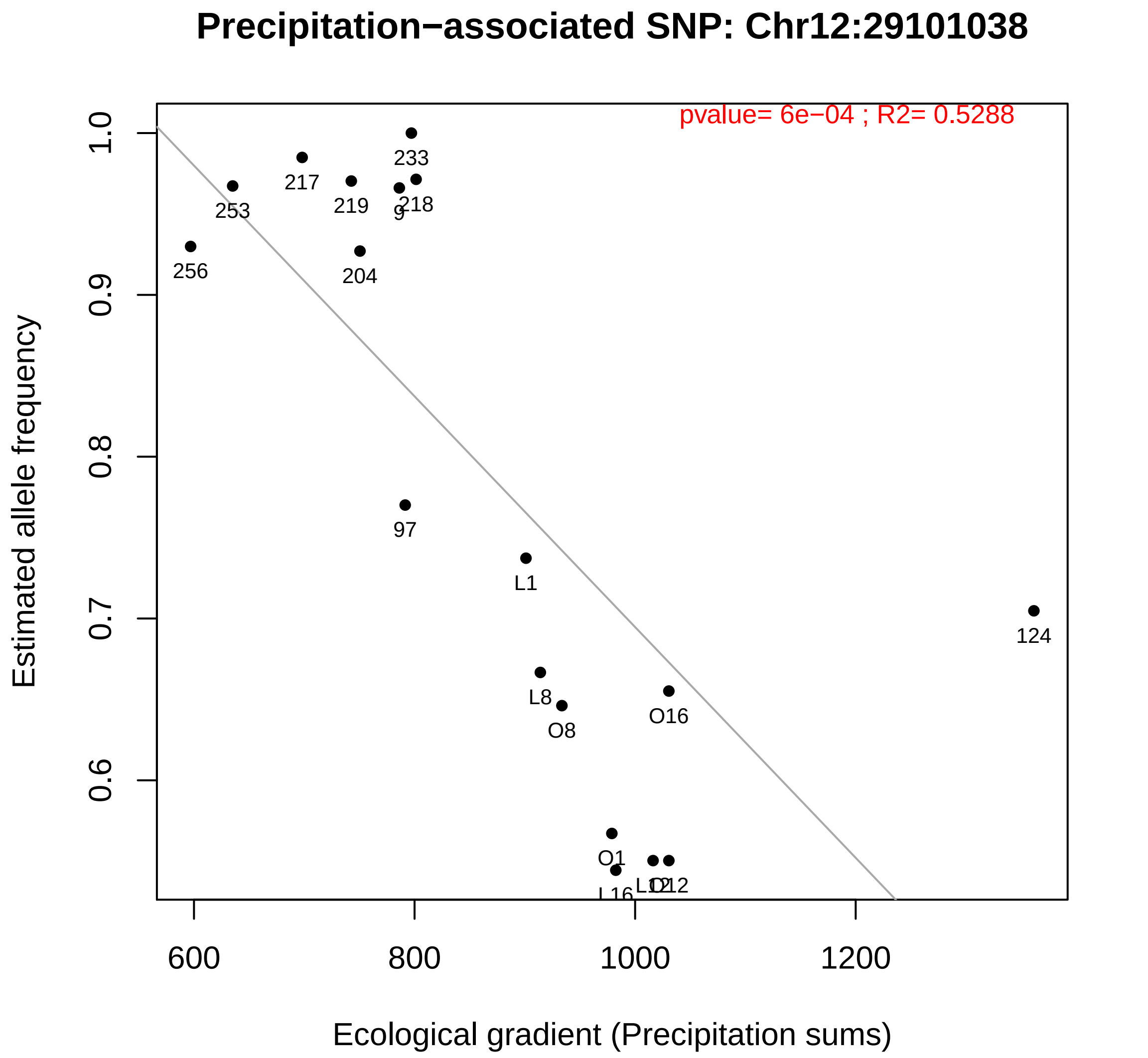

Precipitation−associated SNP: Chr12:29101038
pvalue= 6e−04 ; R2= 0.5288
Estimated allele frequency
Ecological gradient (Precipitation sums)
0.6
0.7
0.8
0.9
1.0
600
800
1000
1200
233
217
253
219
9
218
256
204
97
L1
124
L8
O8
O16
O1
L16
L12
O12

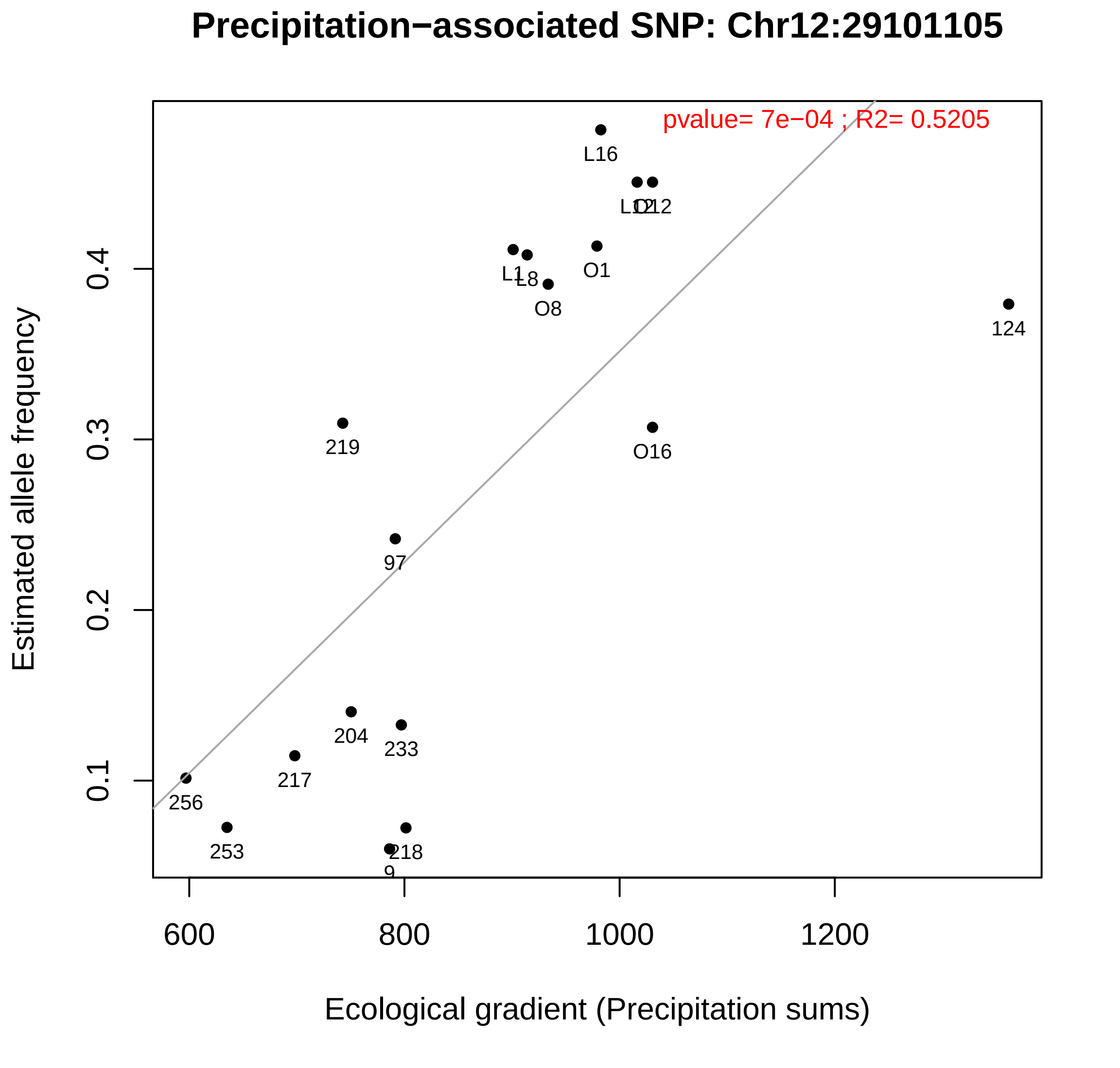

Precipitation−associated SNP: Chr12:29101105
pvalue= 7e−04 ; R2= 0.5205
Estimated allele frequency
0.1
0.2
0.3
0.4
Ecological gradient (Precipitation sums)
600
800
1000
1200
L16
L12
O12
O1
L1
L8
O8
124
219
O16
97
204
233
217
256
253
218
9

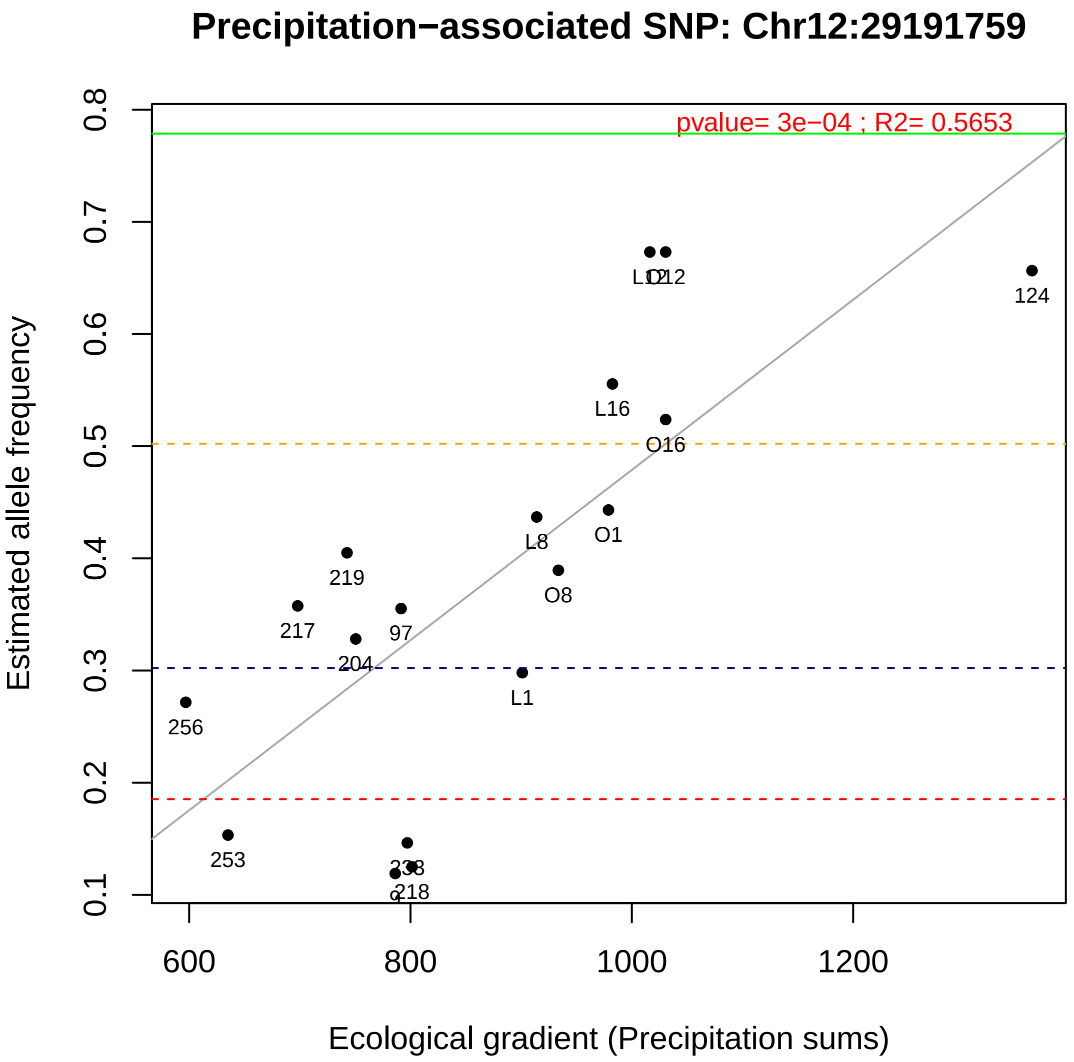
Precipitation−associated SNP: Chr12:29191759
pvalue= 3e−04 ; R2= 0.5653
Estimated allele frequency
0.1
0.2
0.3
0.4
0.5
0.6
0.7
0.8
Ecological gradient (Precipitation sums)
600
800
1000
1200
124
L16
O16
L8
O1
O8
219
217
97
204
L1
256
253
218

**Precipitation−associated SNP: Chr12:29204058**

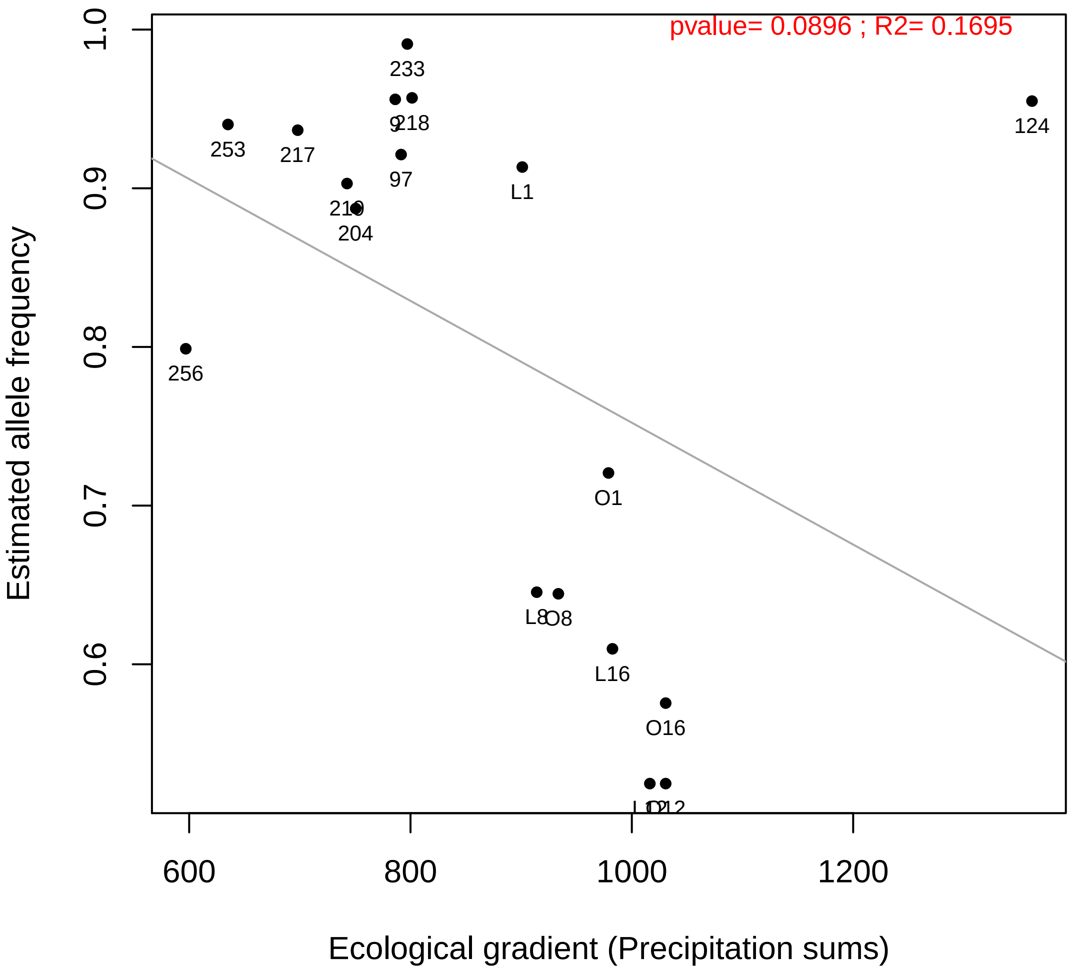

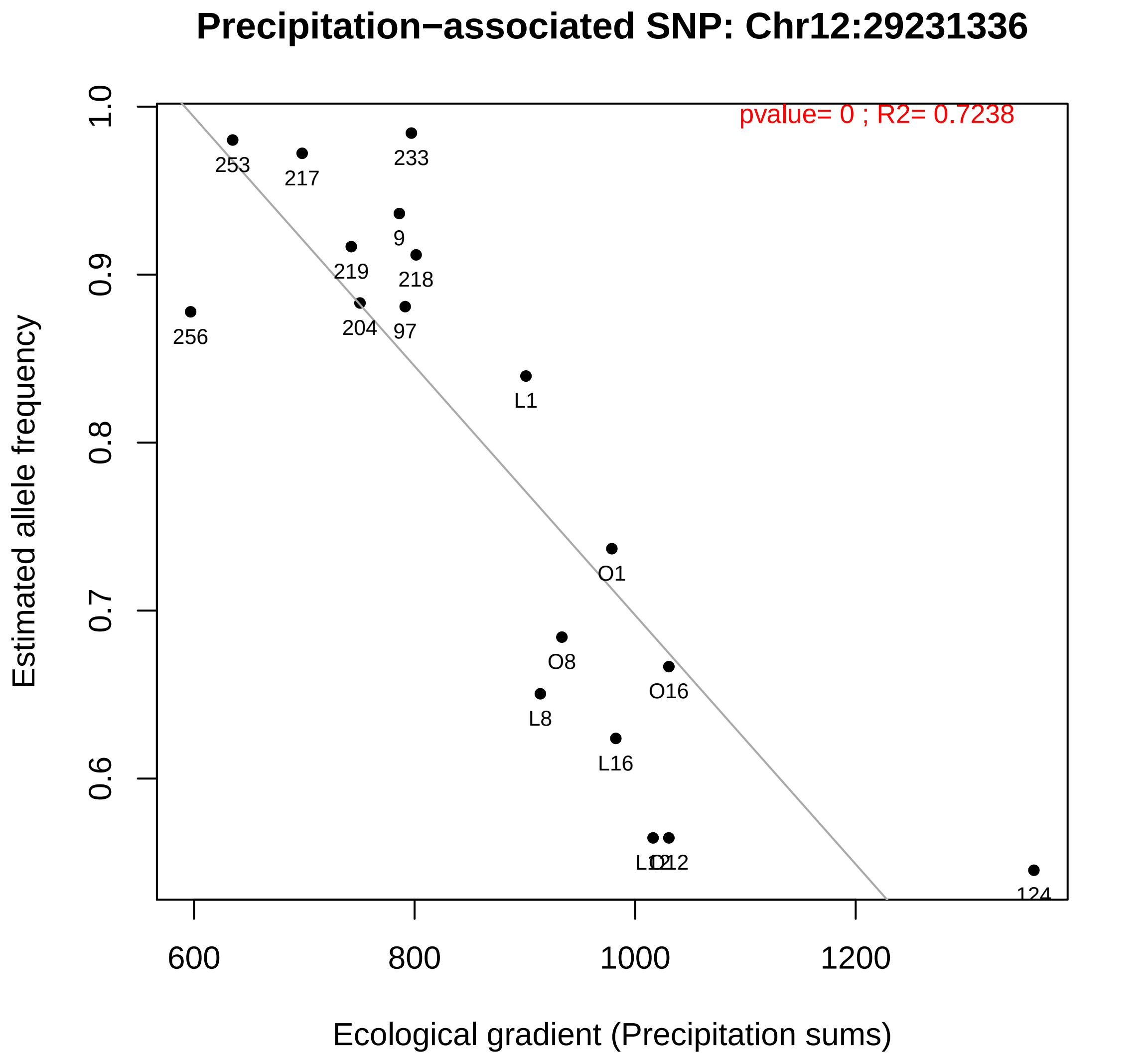

Precipitation−associated SNP: Chr12:29231336
pvalue= 0 ; R2= 0.7238
Estimated allele frequency
Ecological gradient (Precipitation sums)
1.0
0.9
0.8
0.7
0.6
600
800
1000
1200
253
217
233
9
219
218
256
204
97
L1
O1
O8
O16
L8
L16
L12
O12
124

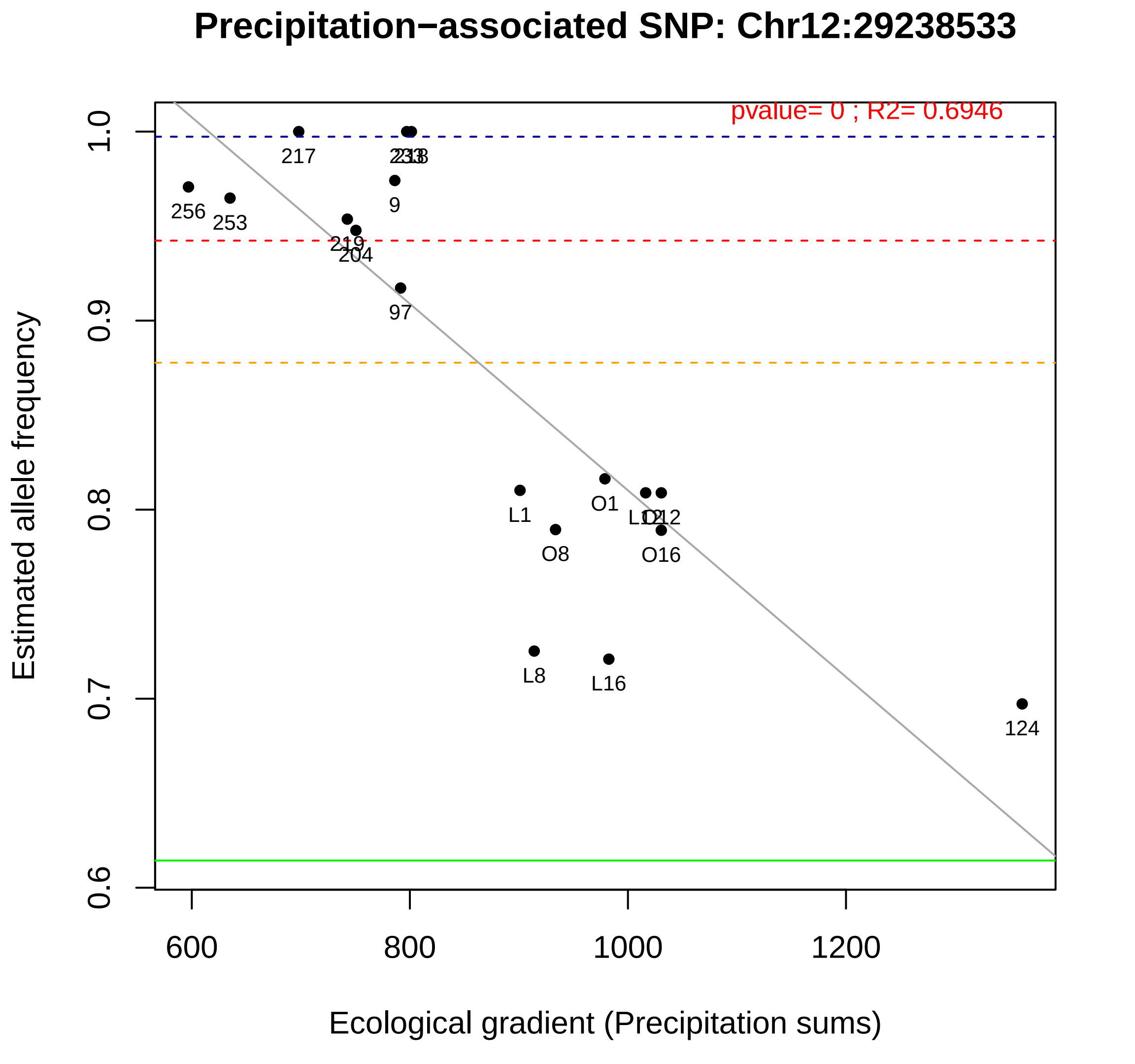
Precipitation−associated SNP: Chr12:29238533
pvalue= 0 ; R2= 0.6946
Estimated allele frequency
0.6
0.7
0.8
0.9
1.0
Ecological gradient (Precipitation sums)
600
800
1000
1200
217
256
253
9
219
204
97
L1
O1
O8
O16
L8
L16
124

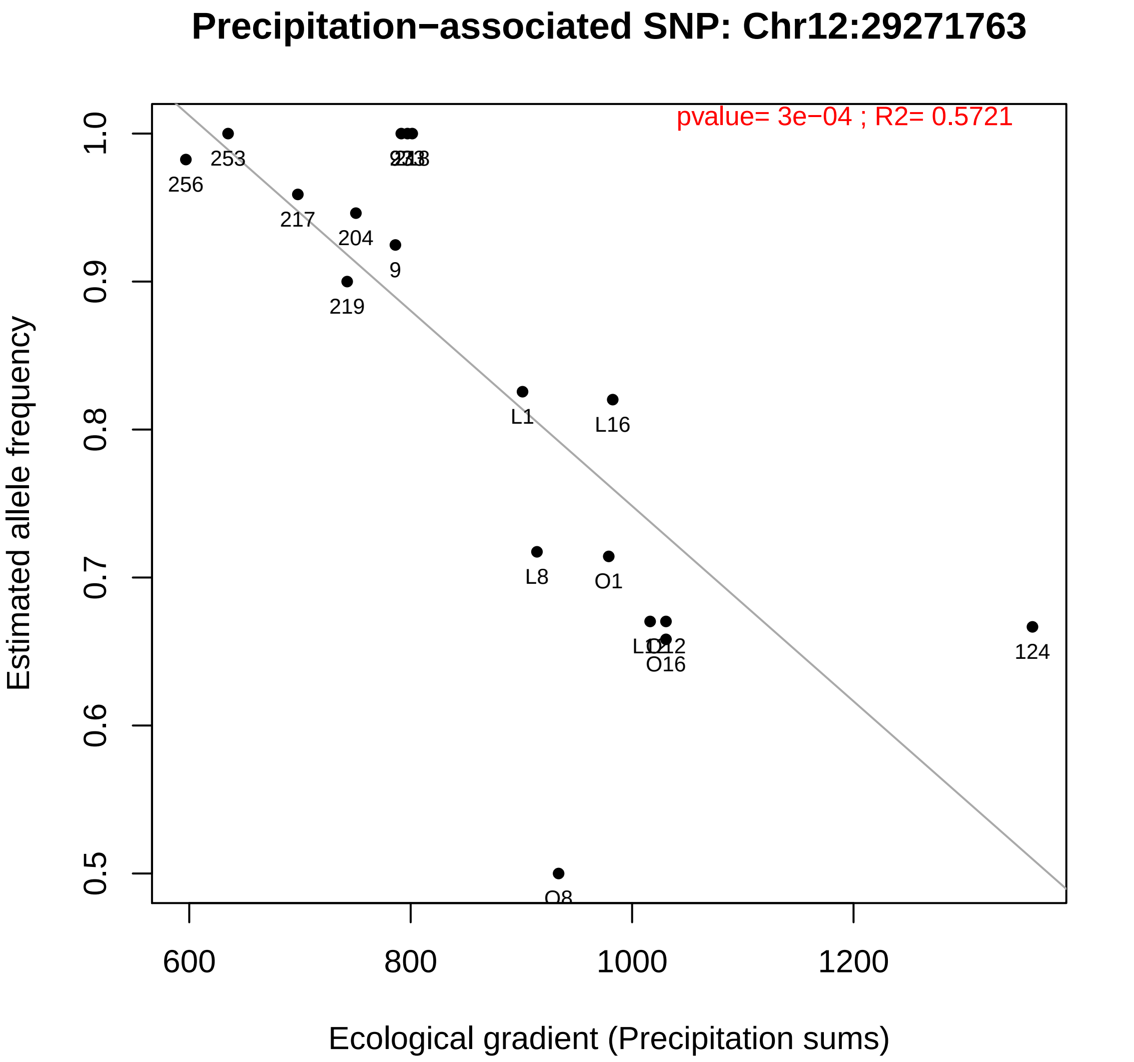
Precipitation−associated SNP: Chr12:29271763
pvalue= 3e−04 ; R2= 0.5721
Estimated allele frequency
Ecological gradient (Precipitation sums)
0.5
0.6
0.7
0.8
0.9
1.0
600
800
1000
1200
253
256
217
204
9
219
L1
L16
L8
O1
O16
124
O8

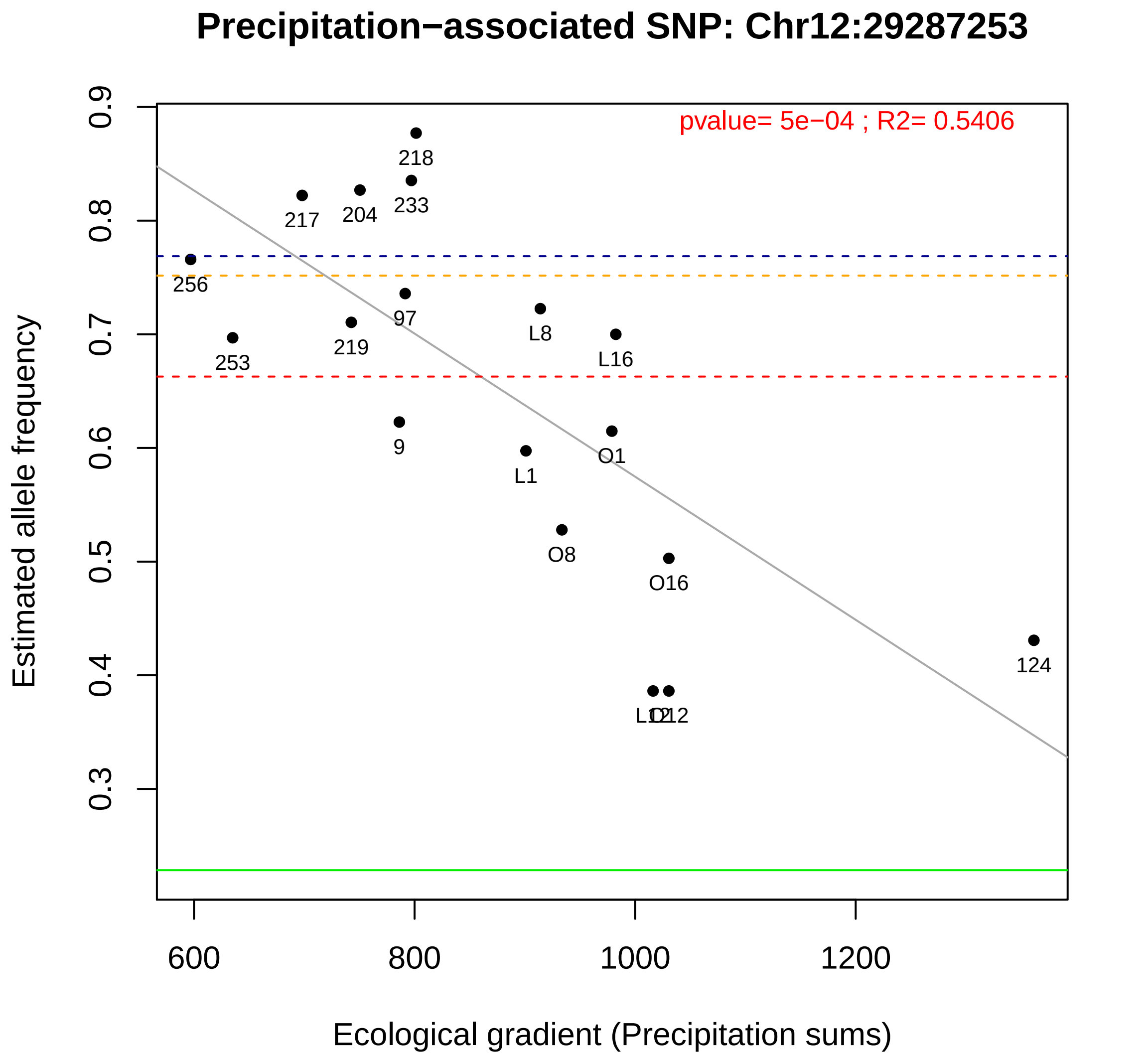
Precipitation−associated SNP: Chr12:29287253
pvalue= 5e−04 ; R2= 0.5406
Estimated allele frequency
0.3
0.4
0.5
0.6
0.7
0.8
0.9
600
800
1000
1200
Ecological gradient (Precipitation sums)
218
217
204
233
256
97
L8
219
L16
253
9
O1
L1
O8
O16
124
L12
O12

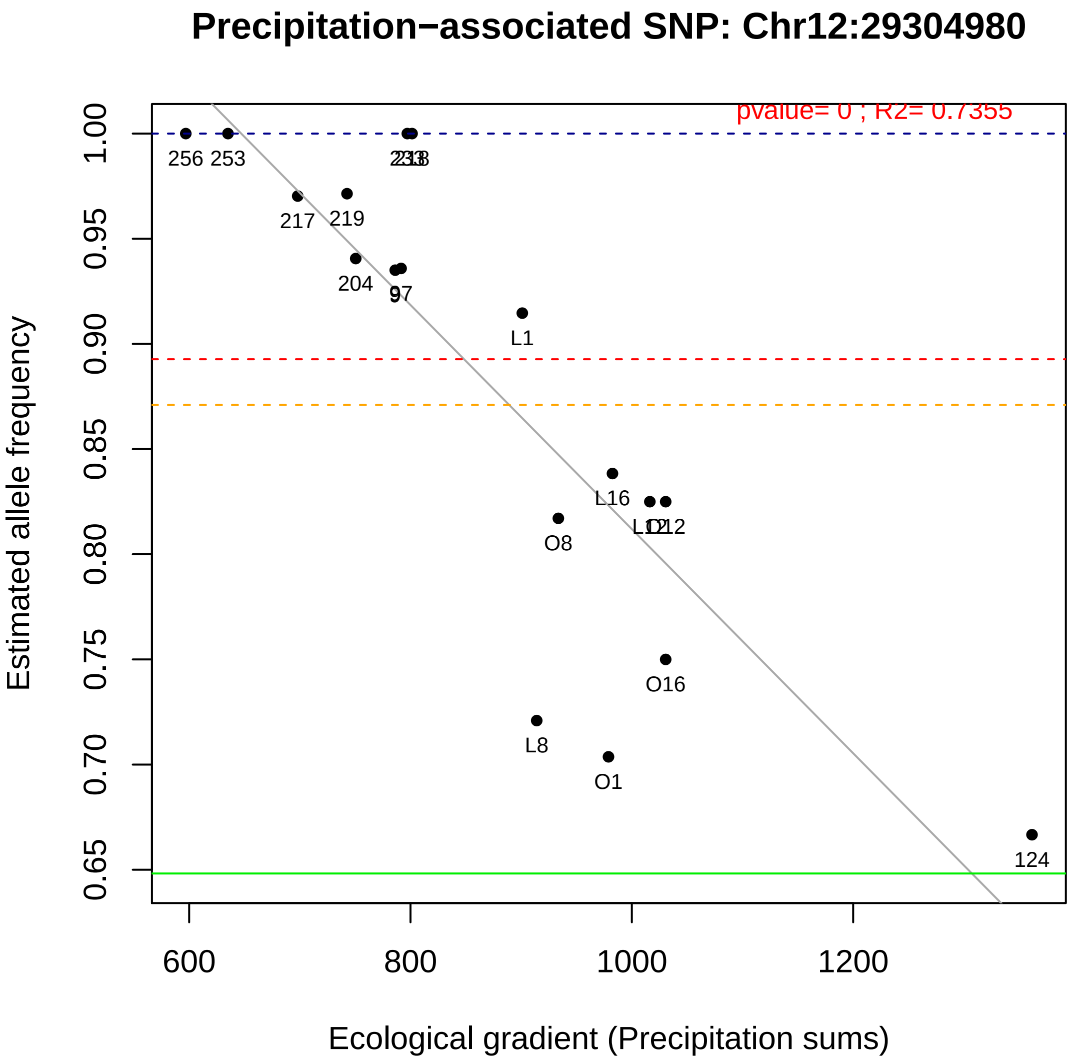

Precipitation−associated SNP: Chr12:29304980
pvalue= 0 ; R2= 0.7355
Estimated allele frequency
Ecological gradient (Precipitation sums)
1.00
0.95
0.90
0.85
0.80
0.75
0.70
0.65
600
800
1000
1200
256
253
217
219
204
97
L1
L16
L12
O12
O8
O16
L8
O1
124

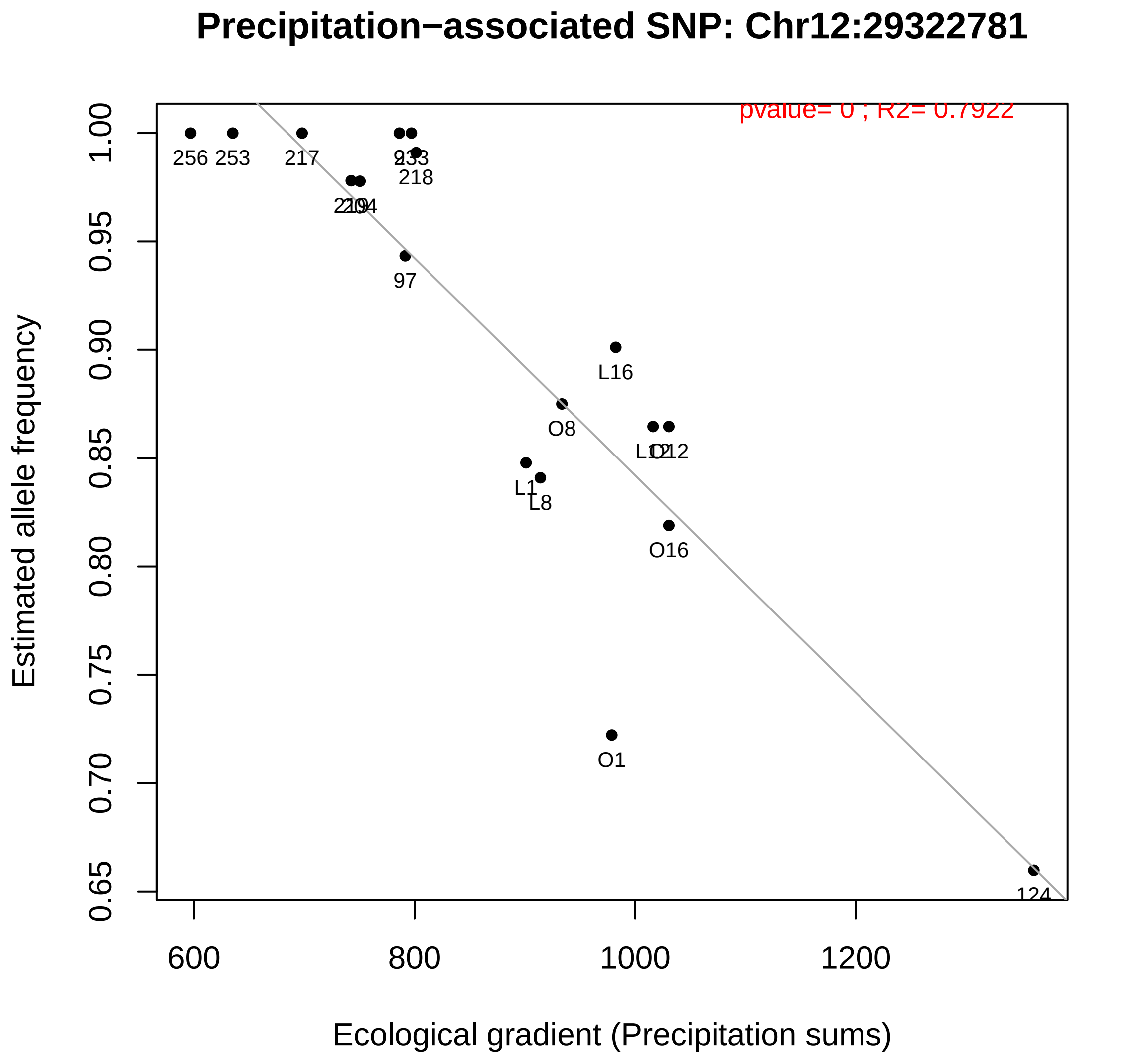

Precipitation−associated SNP: Chr12:29322781
pvalue= 0 ; R2= 0.7922
Estimated allele frequency
Ecological gradient (Precipitation sums)
0.65
0.70
0.75
0.80
0.85
0.90
0.95
1.00
600
800
1000
1200
256
253
217
233
218
97
L16
O8
L1
L8
O12
O16
O1
124

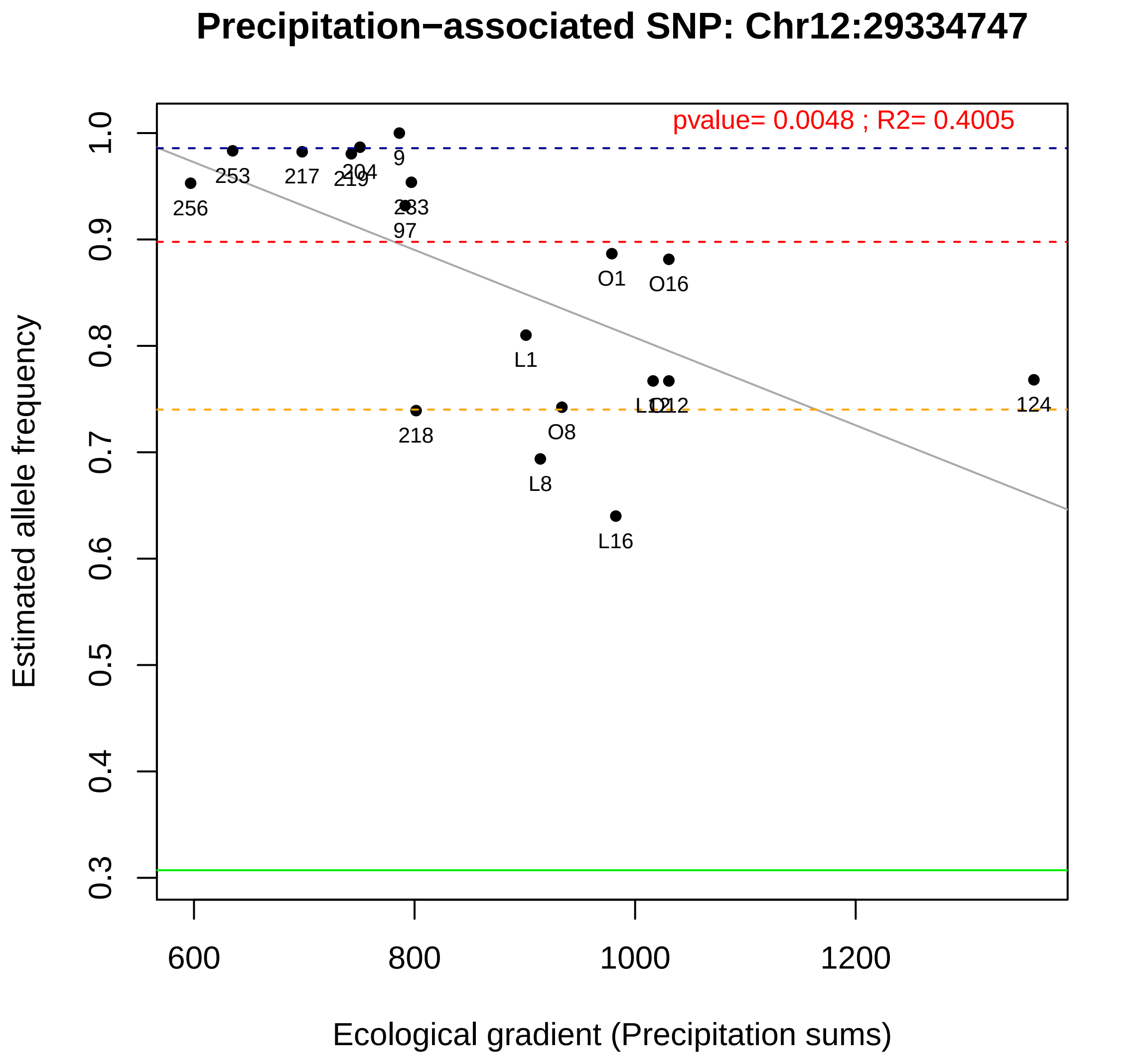

Precipitation−associated SNP: Chr12:29334747
pvalue= 0.0048 ; R2= 0.4005
Estimated allele frequency
Ecological gradient (Precipitation sums)
1.0
0.9
0.8
0.7
0.6
0.5
0.4
0.3
600
800
1000
1200
256
253
217
219
204
9
283
97
218
L1
O1
O16
O8
L8
L16
L12
O12
124

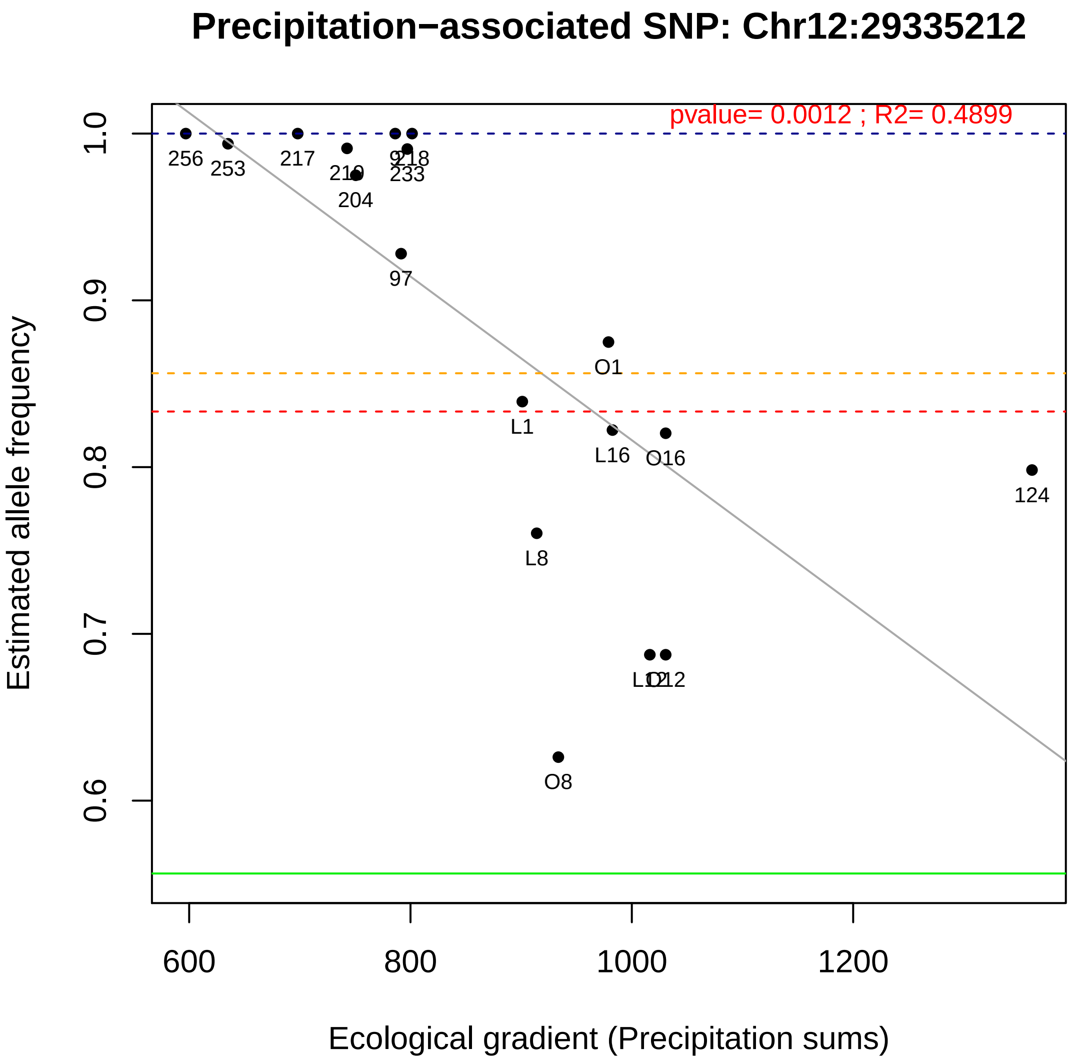
Precipitation−associated SNP: Chr12:29335212
pvalue= 0.0012 ; R2= 0.4899
Estimated allele frequency
Ecological gradient (Precipitation sums)
1.0
0.9
0.8
0.7
0.6
600
800
1000
1200
256
253
217
210
204
218
233
97
O1
L1
L16
O16
124
L8
O12
O8

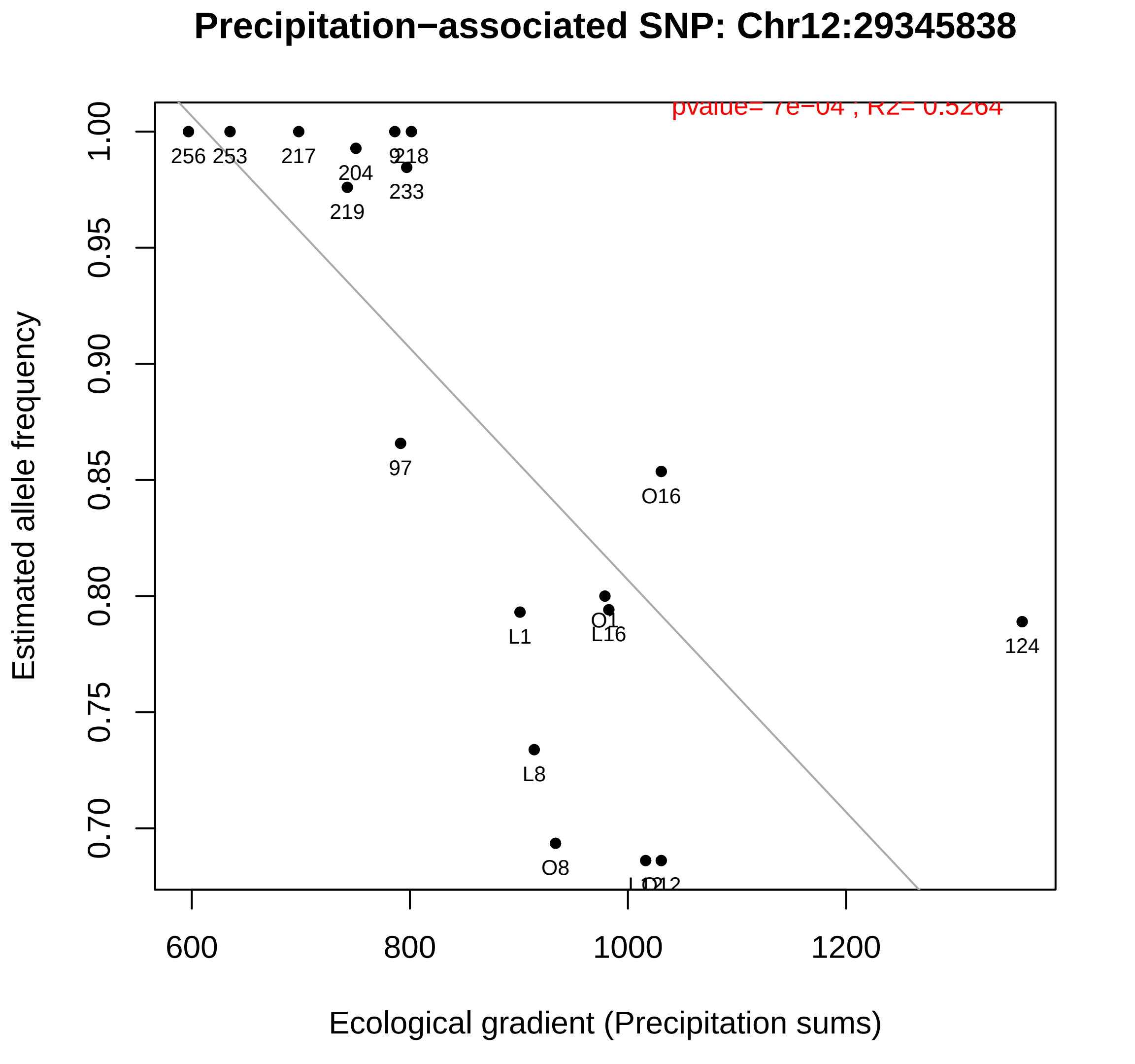

Precipitation−associated SNP: Chr12:29345838
pvalue= 7e−04 ; R2= 0.5264
Estimated allele frequency
Ecological gradient (Precipitation sums)
1.00
0.95
0.90
0.85
0.80
0.75
0.70
600
800
1000
1200
256
253
217
204
218
233
219
97
O16
L1
O1
L16
124
L8
O8
O12

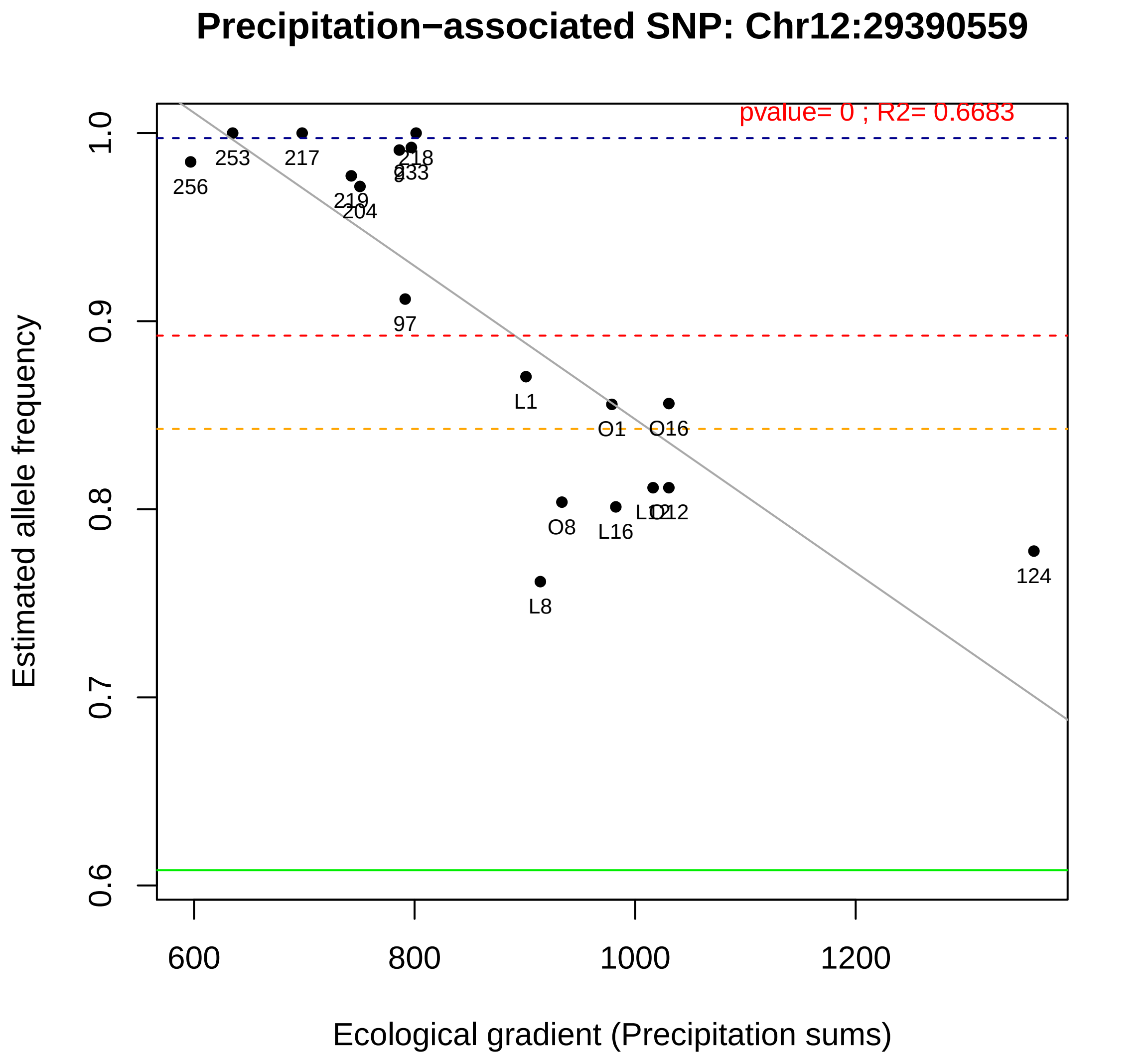

Precipitation−associated SNP: Chr12:29390559
pvalue= 0 ; R2= 0.6683
Estimated allele frequency
Ecological gradient (Precipitation sums)
0.6
0.7
0.8
0.9
1.0
600
800
1000
1200
256
253
217
218
233
219
204
97
L1
O1
O16
O8
L16
O12
L8
124

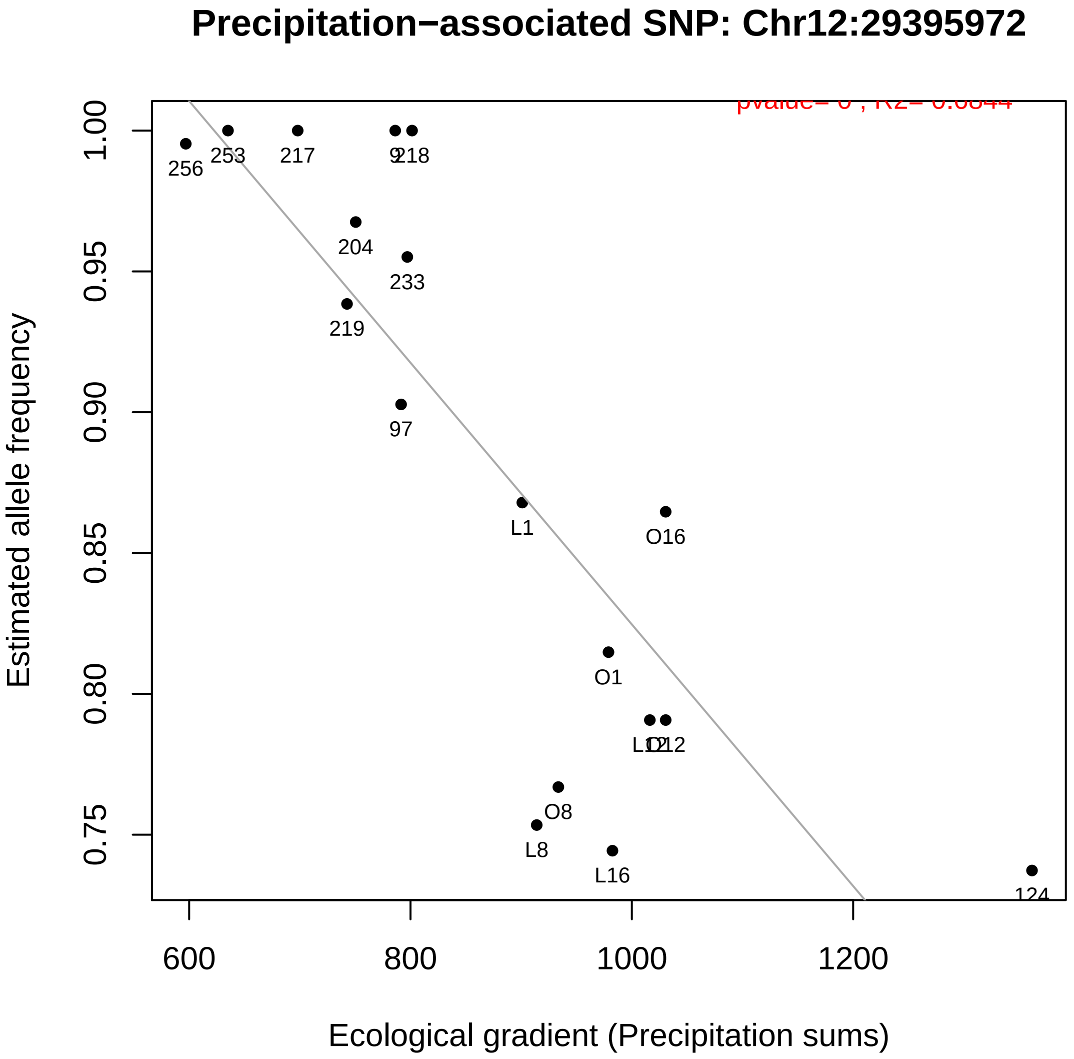

Precipitation−associated SNP: Chr12:29395972
Estimated allele frequency
Ecological gradient (Precipitation sums)
1.00
0.95
0.90
0.85
0.80
0.75
600
800
1000
1200
256
253
217
218
204
233
219
97
L1
O16
O1
O8
L8
L16
124

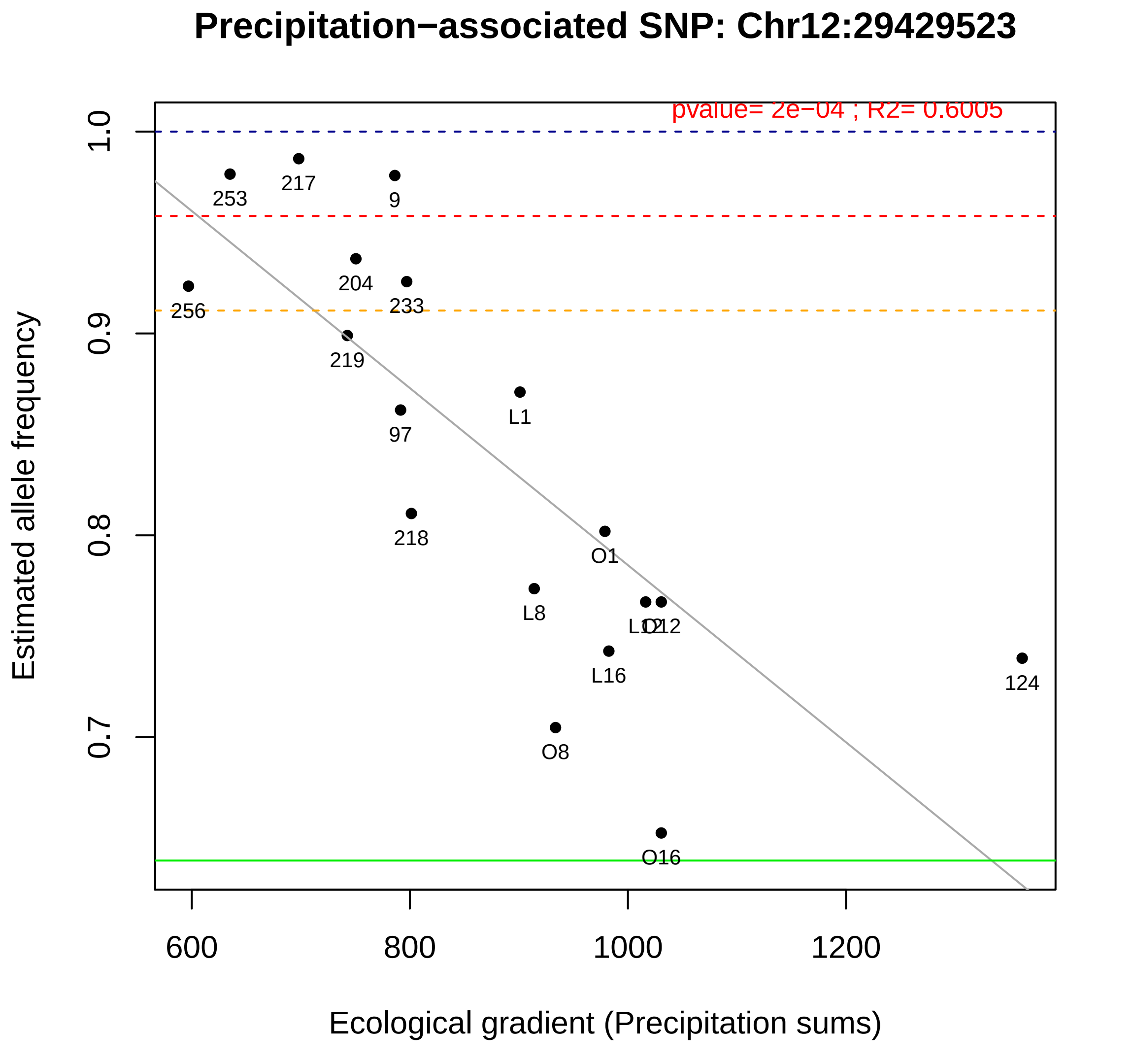
Precipitation−associated SNP: Chr12:29429523
pvalue= 2e−04 ; R2= 0.6005
Estimated allele frequency
Ecological gradient (Precipitation sums)
1.0
0.9
0.8
0.7
600
800
1000
1200
253
217
9
204
256
233
219
97
L1
218
O1
L8
L16
124
O8
O16

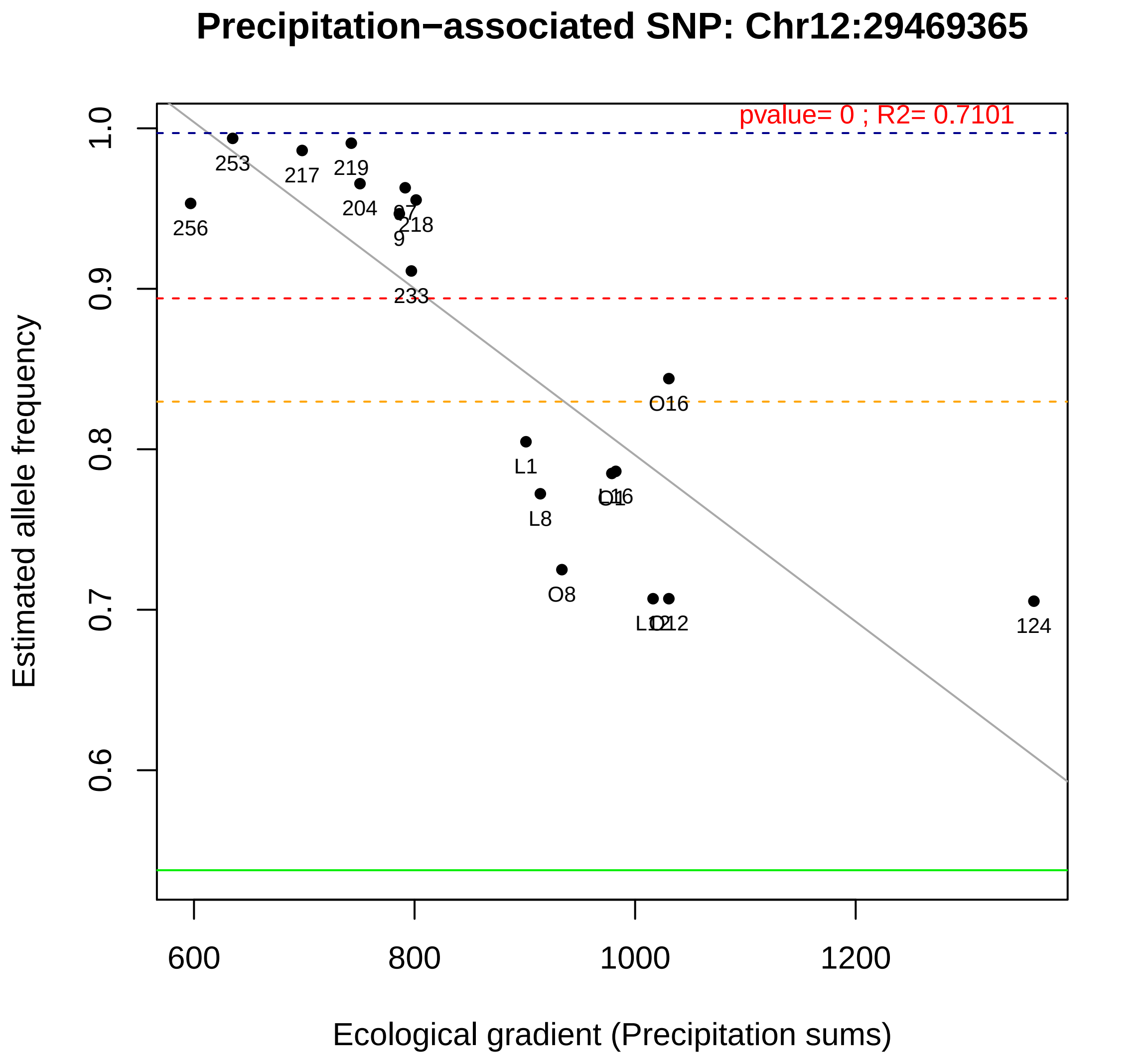

Precipitation−associated SNP: Chr12:29469365
pvalue= 0 ; R2= 0.7101
Estimated allele frequency
Ecological gradient (Precipitation sums)
1.0
0.9
0.8
0.7
0.6
600
800
1000
1200
253
217
219
204
256
97
218
9
233
O16
L1
L16
O1
L8
O8
L12
O12
124

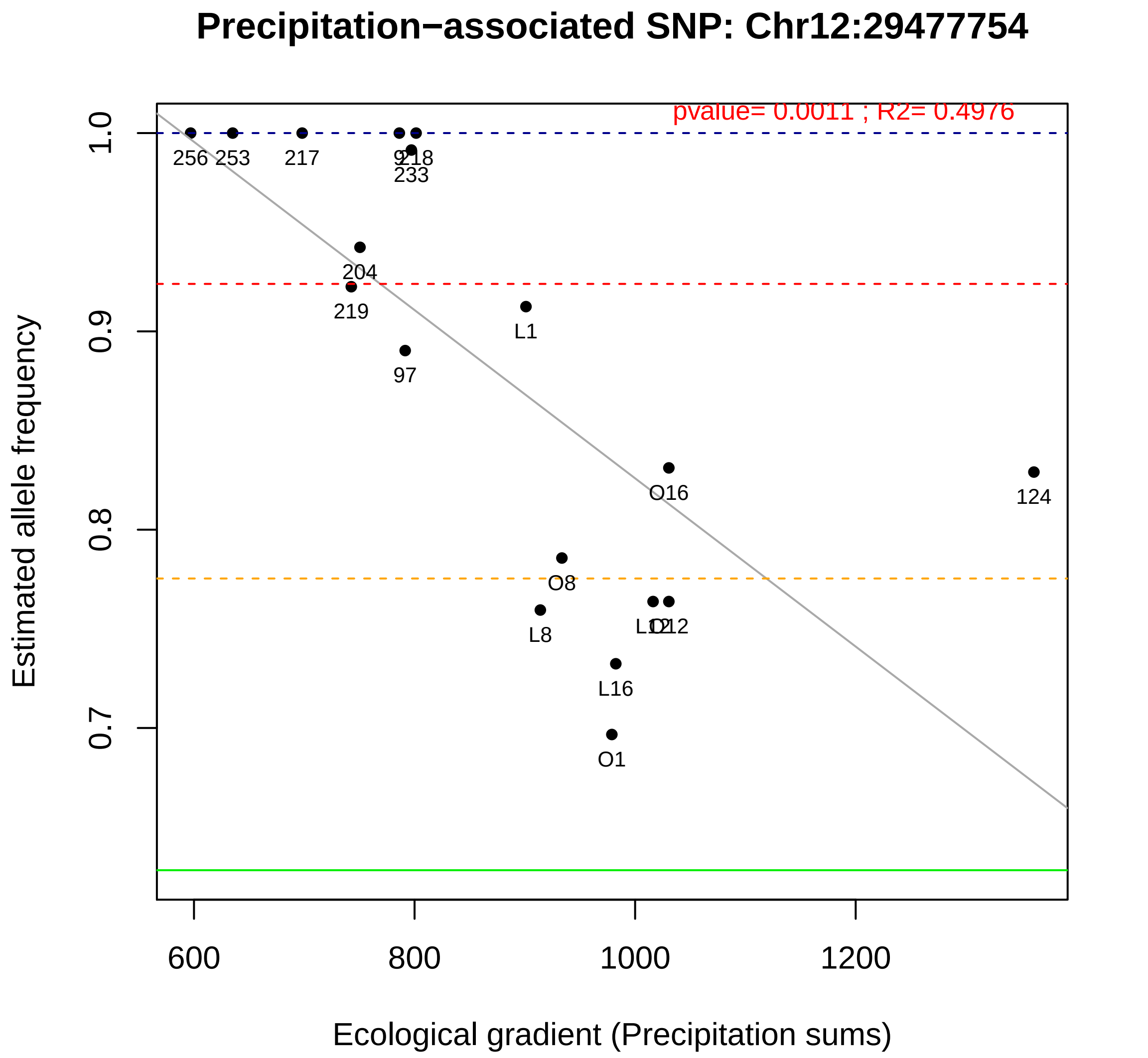
Precipitation−associated SNP: Chr12:29477754
pvalue= 0.0011 ; R2= 0.4976
Estimated allele frequency
Ecological gradient (Precipitation sums)
1.0
0.9
0.8
0.7
600
800
1000
1200
256
253
217
218
233
204
219
97
L1
O16
124
O8
L8
O12
L16
O1

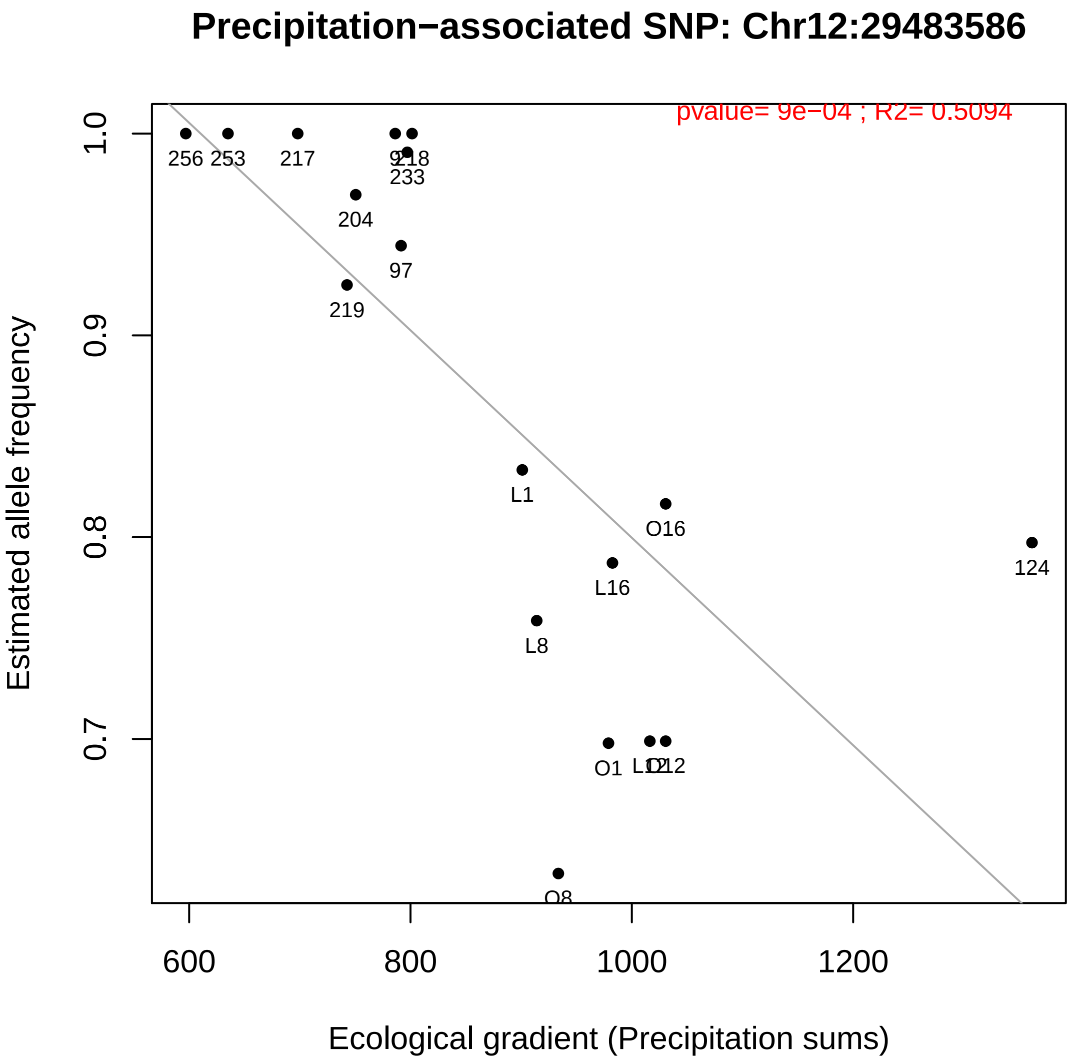

Precipitation−associated SNP: Chr12:29483586
pvalue= 9e−04 ; R2= 0.5094
Estimated allele frequency
1.0
0.9
0.8
0.7
600
800
1000
1200
Ecological gradient (Precipitation sums)
256
253
217
218
233
204
97
219
L1
O16
124
L16
L8
O1
O12
O8

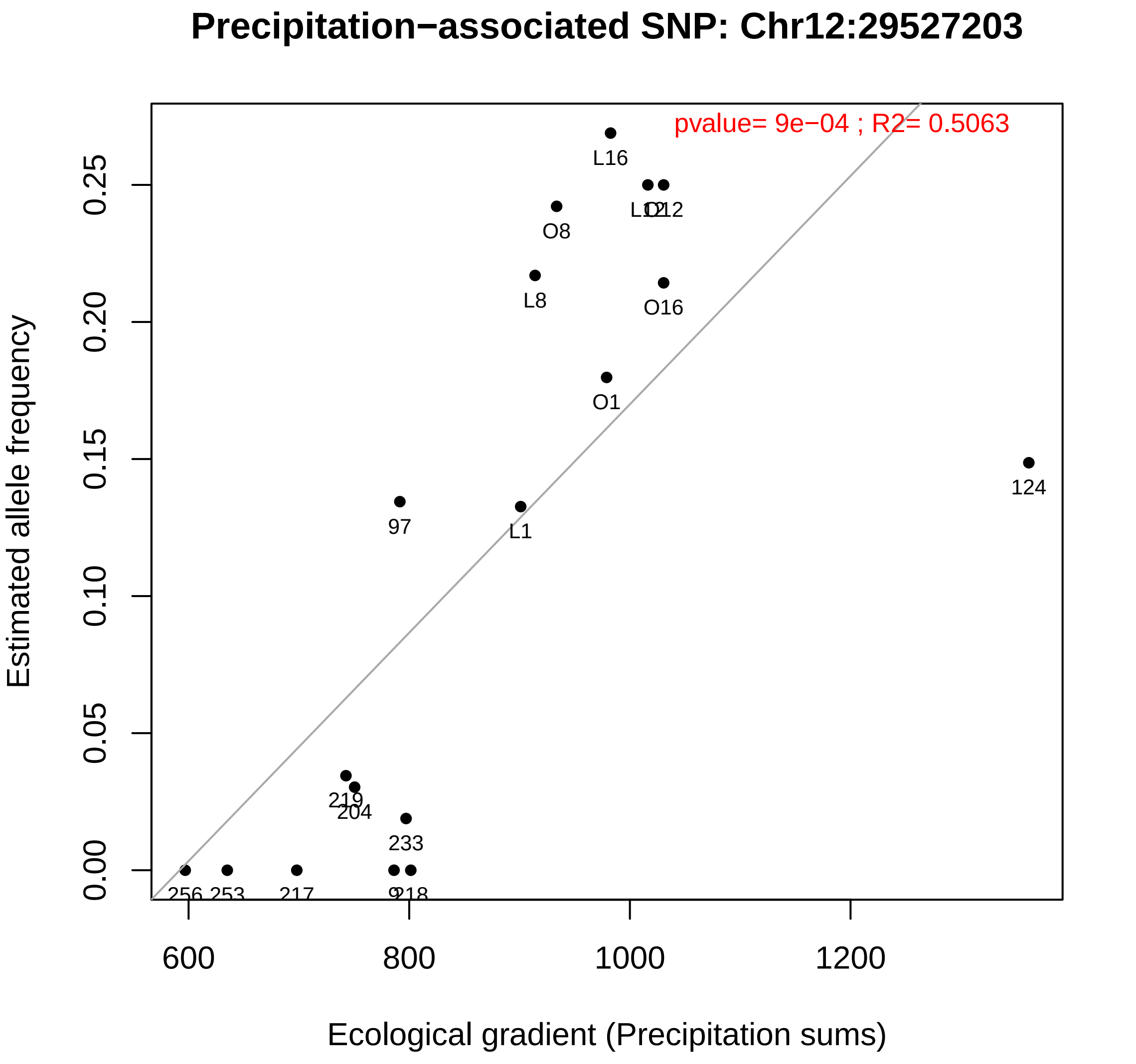

Precipitation−associated SNP: Chr12:29527203
pvalue= 9e−04 ; R2= 0.5063
Estimated allele frequency
0.00
0.05
0.10
0.15
0.20
0.25
600
800
1000
1200
Ecological gradient (Precipitation sums)
L16
O8
L8
O16
O1
97
L1
124
219
204
233
256
253
217
218

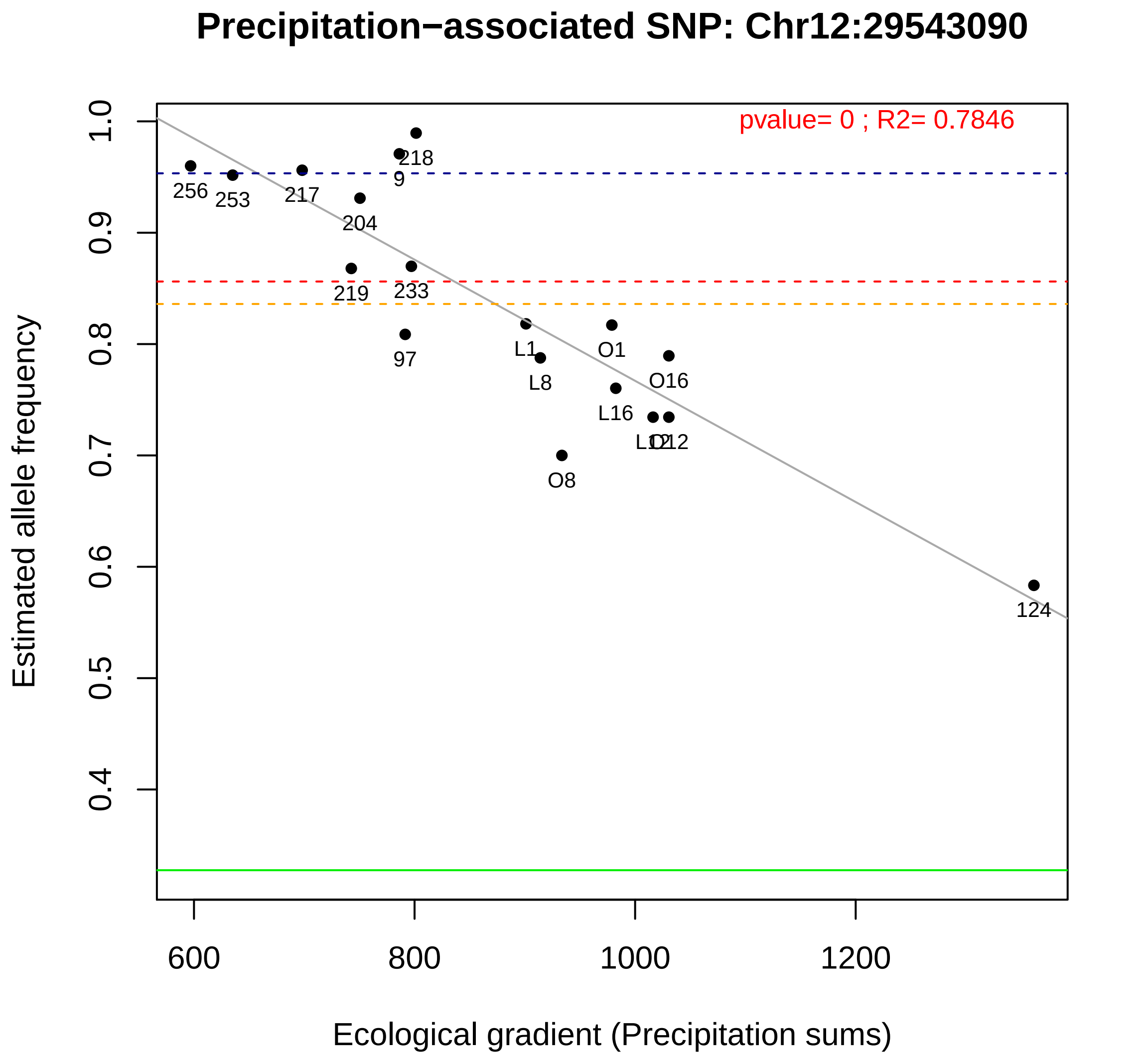

Precipitation−associated SNP: Chr12:29543090
pvalue= 0 ; R2= 0.7846
Estimated allele frequency
Ecological gradient (Precipitation sums)
0.4
0.5
0.6
0.7
0.8
0.9
1.0
600
800
1000
1200
256
253
217
218
9
204
219
233
97
L1
L8
O1
O16
L16
L12
O12
O8
124

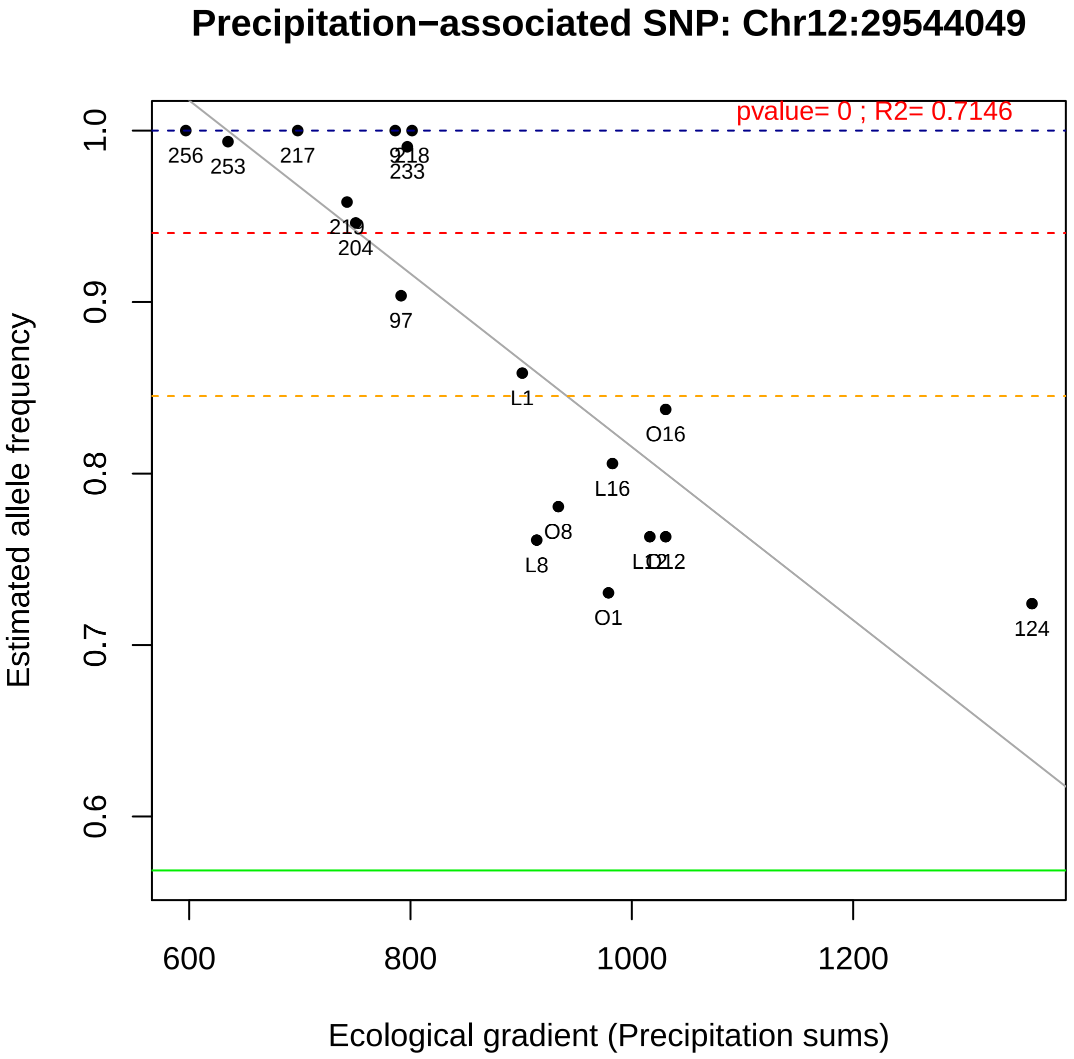
Precipitation−associated SNP: Chr12:29544049
pvalue= 0 ; R2= 0.7146
Estimated allele frequency
Ecological gradient (Precipitation sums)
0.6
0.7
0.8
0.9
1.0
600
800
1000
1200
256
253
217
218
233
219
204
97
L1
O16
L16
O8
L8
O12
O1
124

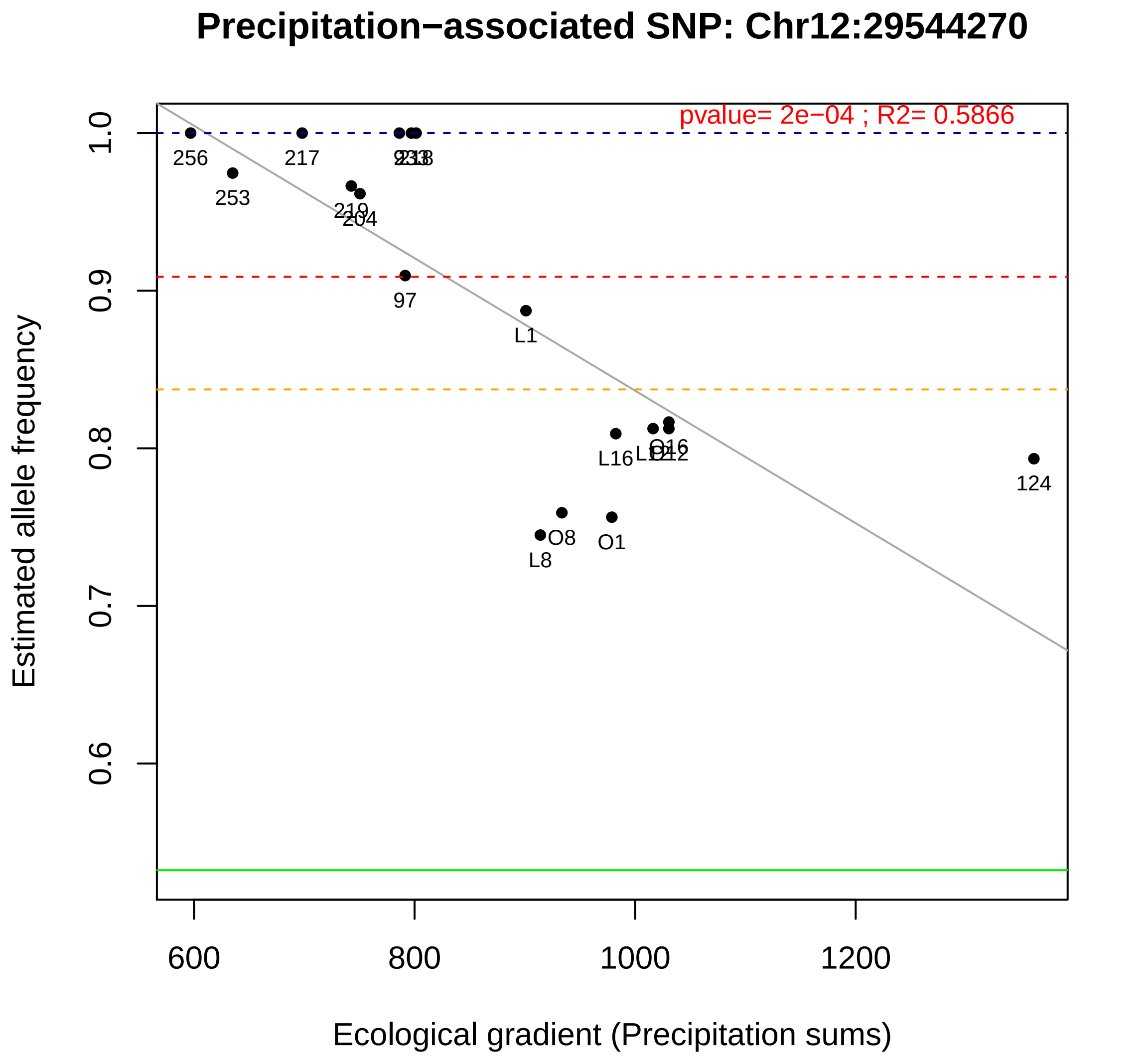

Precipitation−associated SNP: Chr12:29544270
pvalue= 2e−04 ; R2= 0.5866
Estimated allele frequency
Ecological gradient (Precipitation sums)
1.0
0.9
0.8
0.7
0.6
600
800
1000
1200
256
217
253
219
204
97
L1
L16
O16
124
O8
O1
L8

**Precipitation−associated SNP: Chr12:29556323**

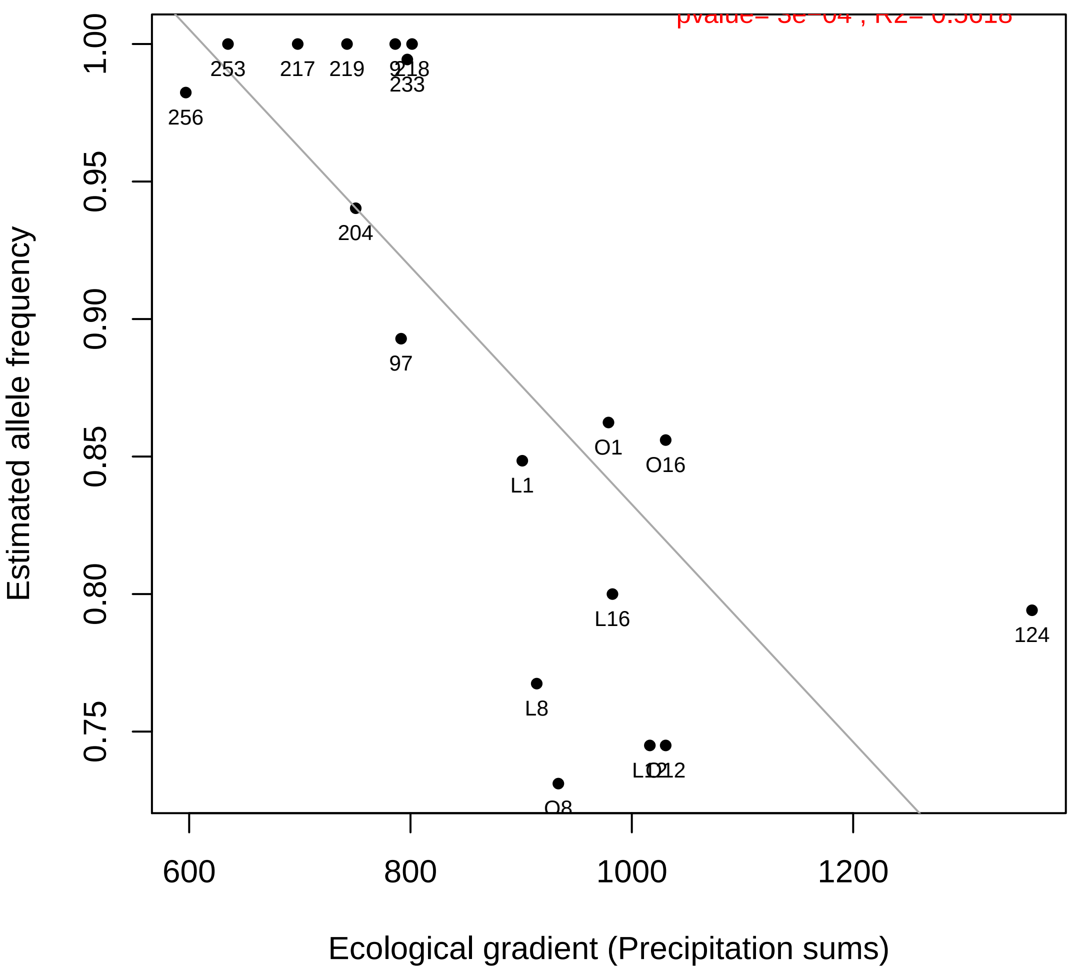

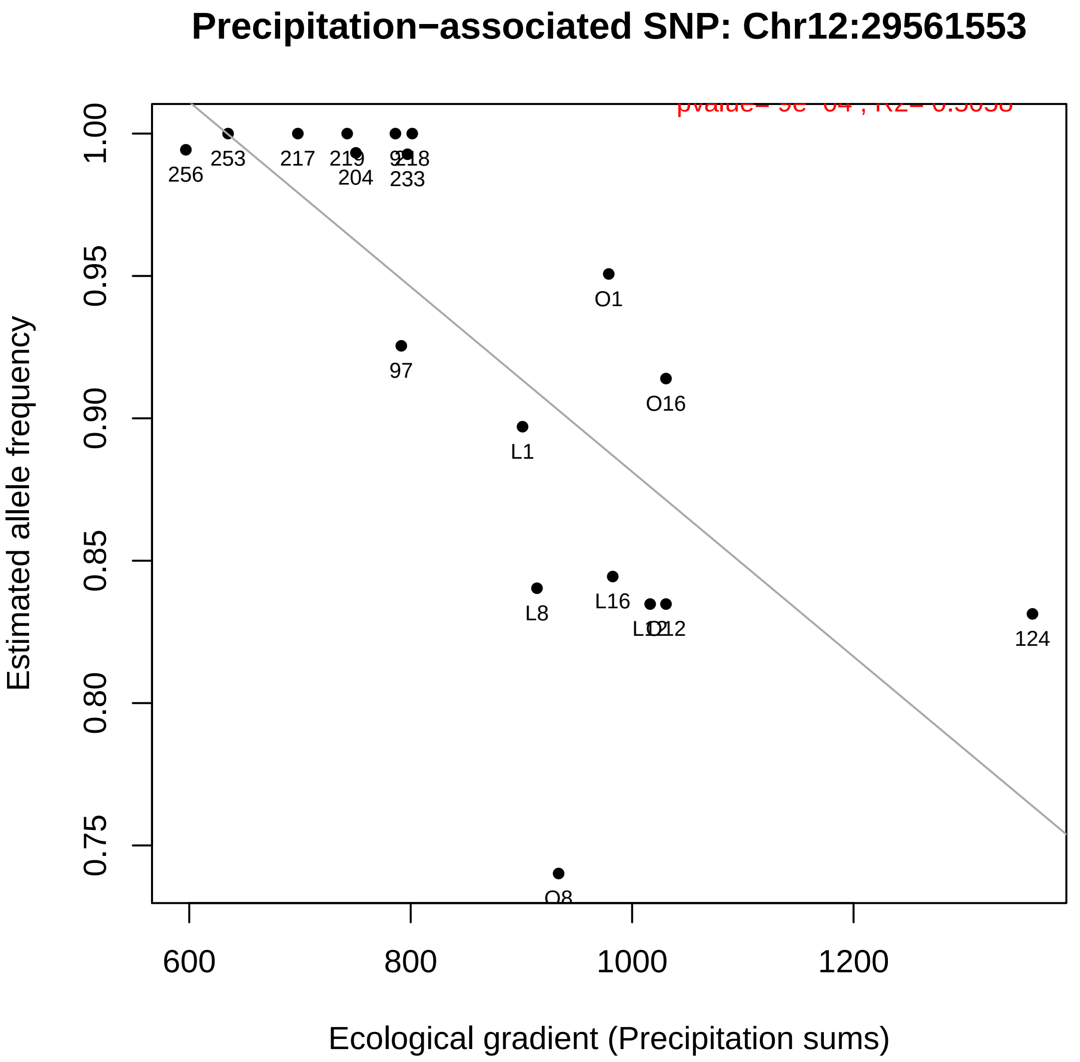
Precipitation−associated SNP: Chr12:29561553
Estimated allele frequency
Ecological gradient (Precipitation sums)
1.00
0.95
0.90
0.85
0.80
0.75
600
800
1000
1200
256
253
217
219
204
218
233
97
O1
O16
L1
L8
L16
O12
124

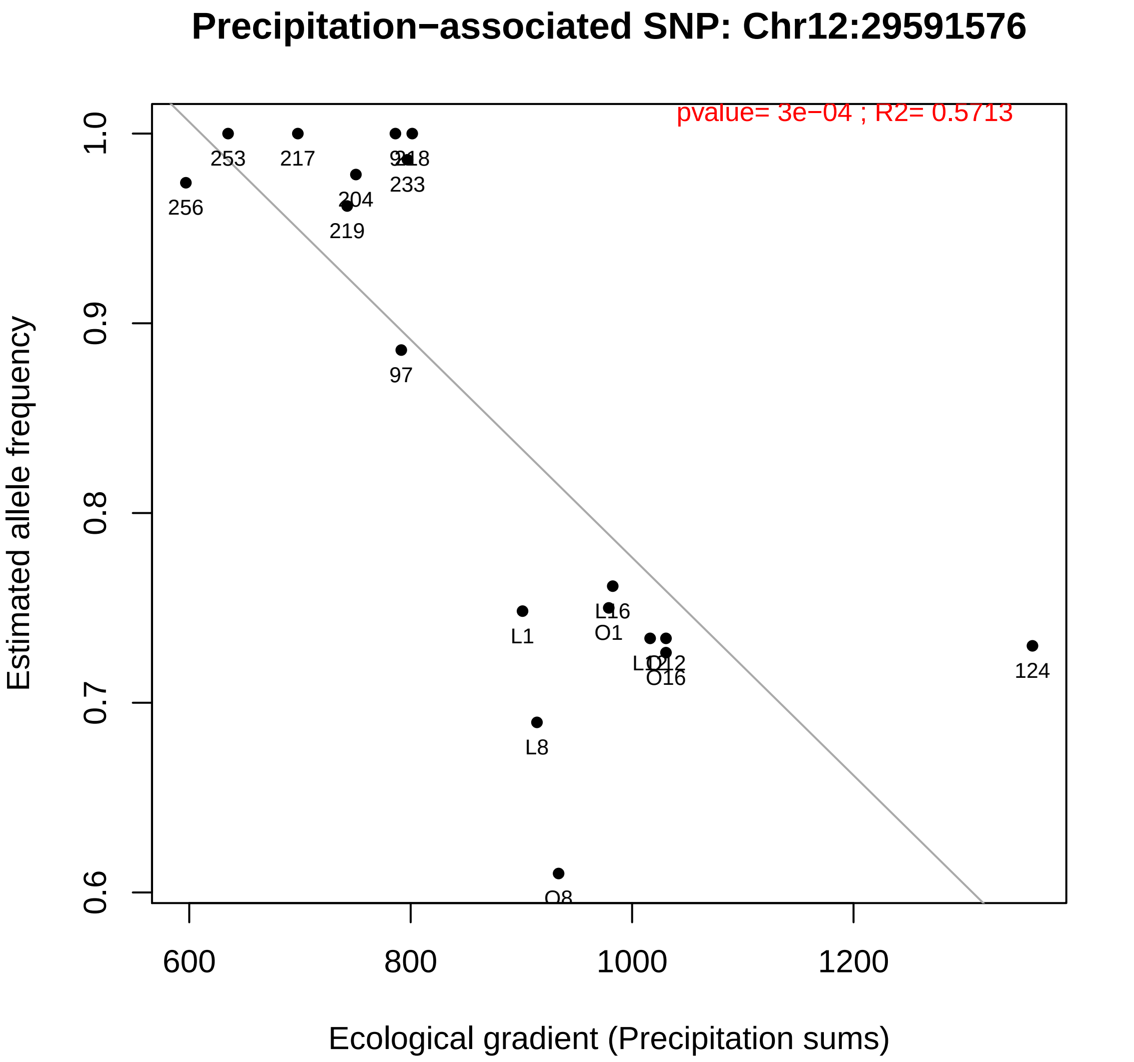
Precipitation−associated SNP: Chr12:29591576
pvalue= 3e−04 ; R2= 0.5713
Estimated allele frequency
Ecological gradient (Precipitation sums)
0.6
0.7
0.8
0.9
1.0
600
800
1000
1200
253
217
218
233
256
204
219
97
L16
O1
L1
O12
O16
124
L8
O8

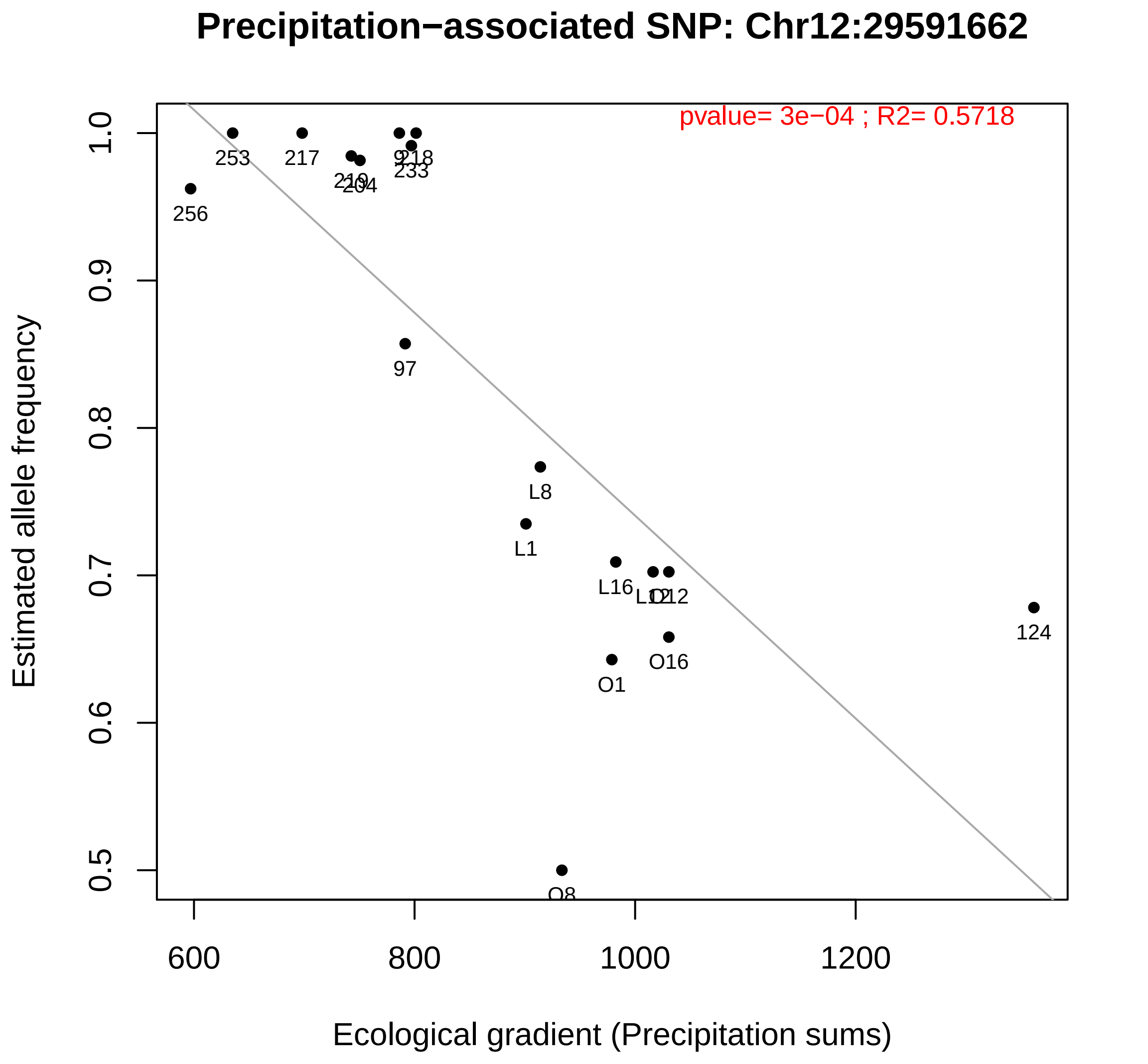

Precipitation−associated SNP: Chr12:29591662
pvalue= 3e−04 ; R2= 0.5718
Estimated allele frequency
Ecological gradient (Precipitation sums)
1.0
0.9
0.8
0.7
0.6
0.5
600
800
1000
1200
253
217
233
256
97
L8
L1
L16
124
O16
O1

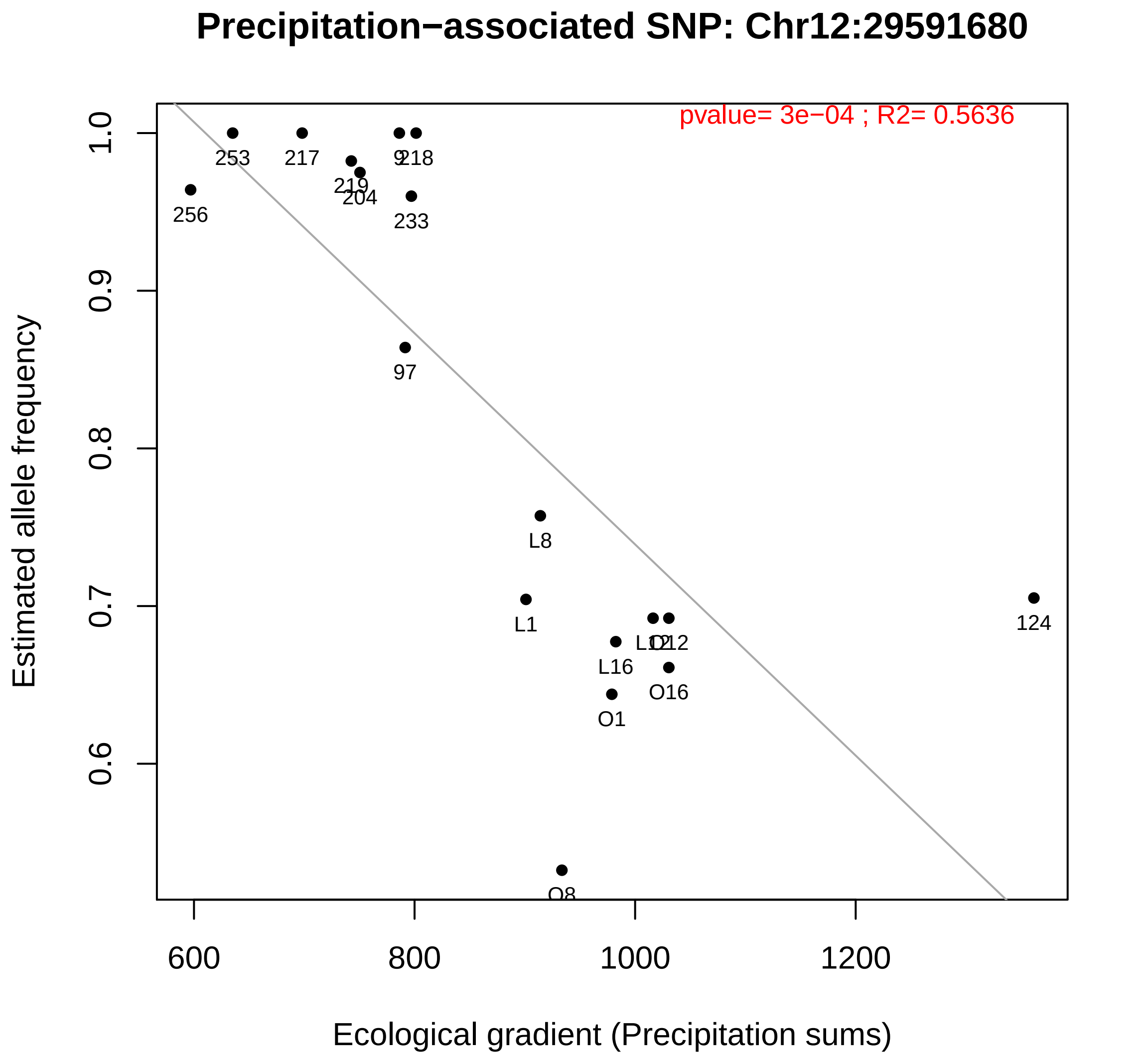

Precipitation−associated SNP: Chr12:29591680
pvalue= 3e−04 ; R2= 0.5636
Estimated allele frequency
Ecological gradient (Precipitation sums)
1.0
0.9
0.8
0.7
0.6
600
800
1000
1200
253
217
9
218
219
204
256
233
97
L8
L1
124
L16
O16
O1
O8

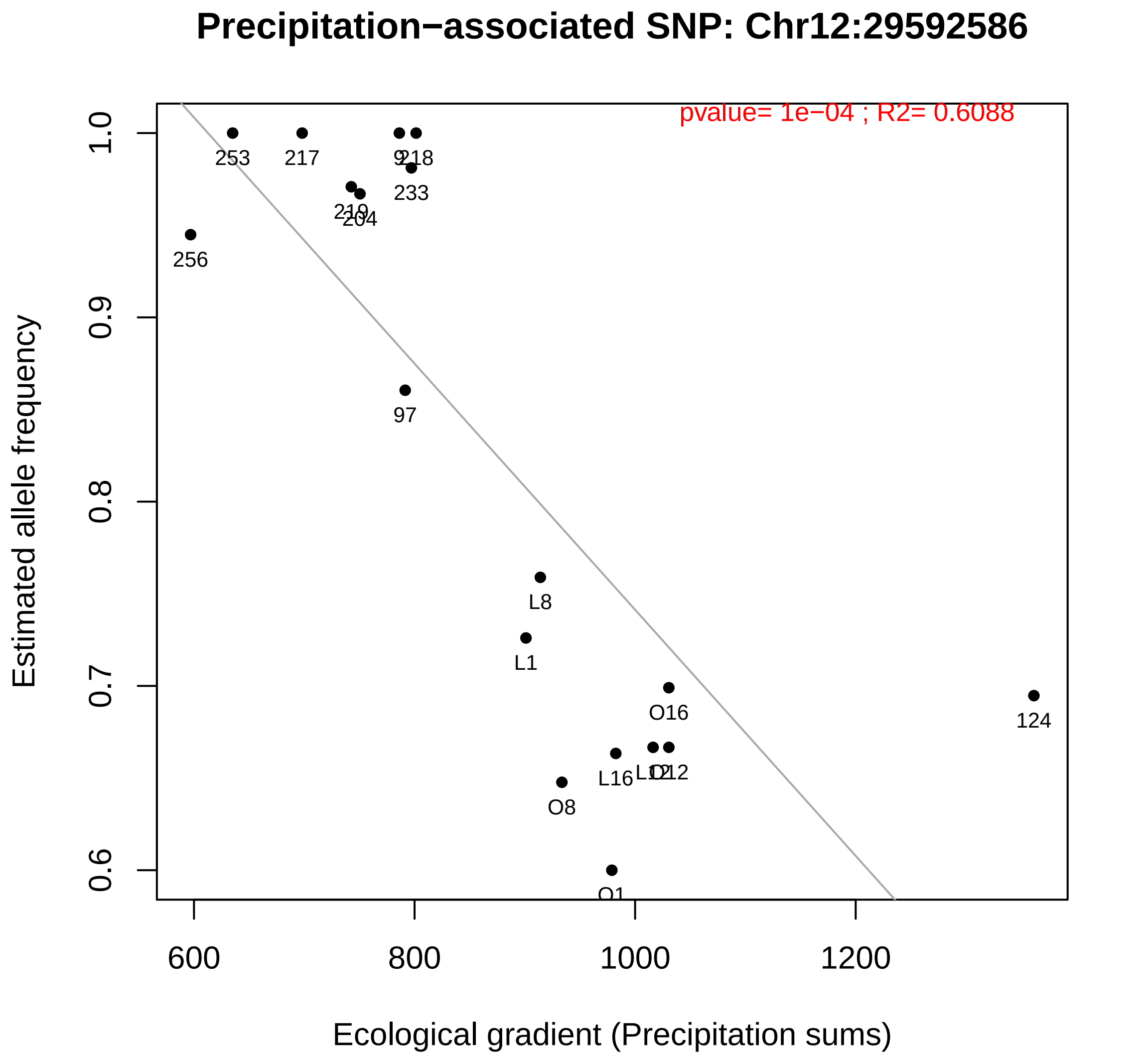

Precipitation−associated SNP: Chr12:29592586
pvalue= 1e−04 ; R2= 0.6088
Estimated allele frequency
Ecological gradient (Precipitation sums)
1.0
0.9
0.8
0.7
0.6
600
800
1000
1200
253
217
218
233
219
204
256
97
L8
L1
O16
124
L16
O12
O8

**Precipitation−associated SNP: Chr12:29594169**

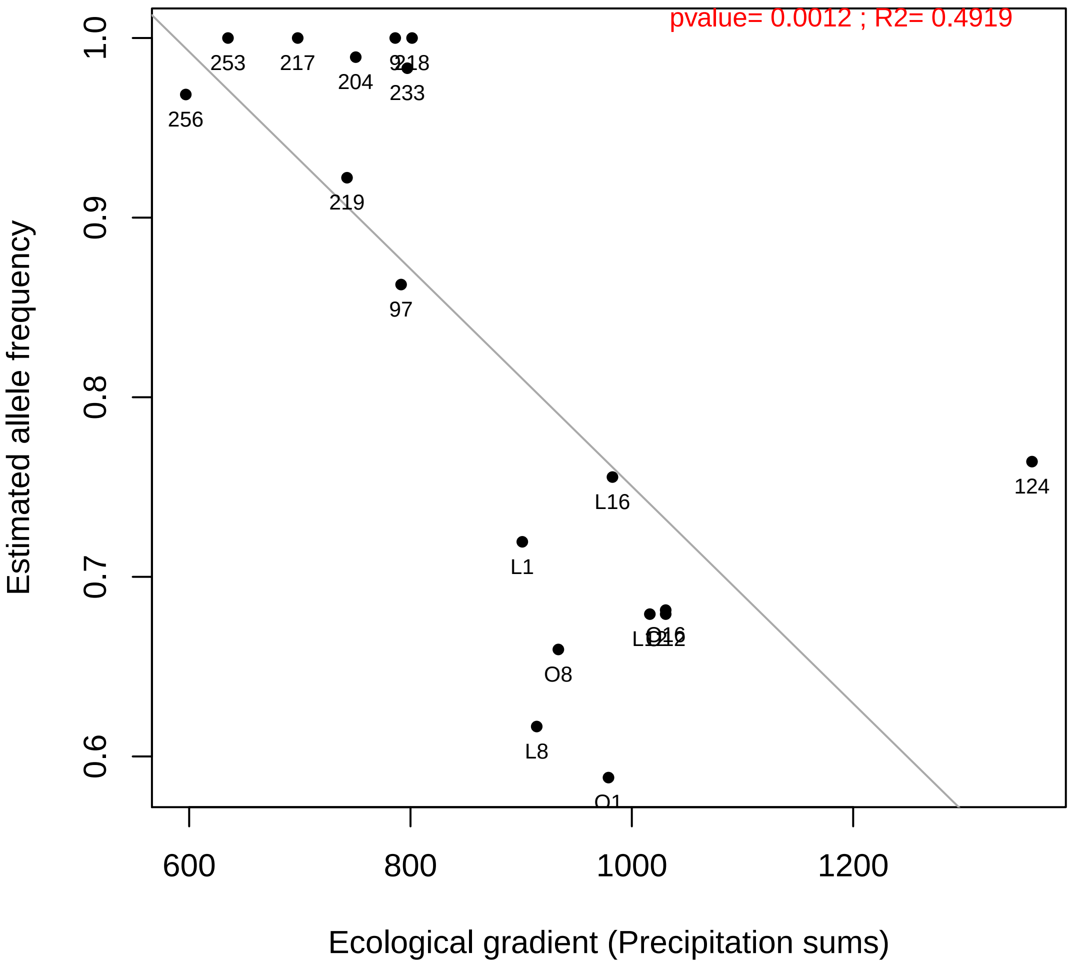

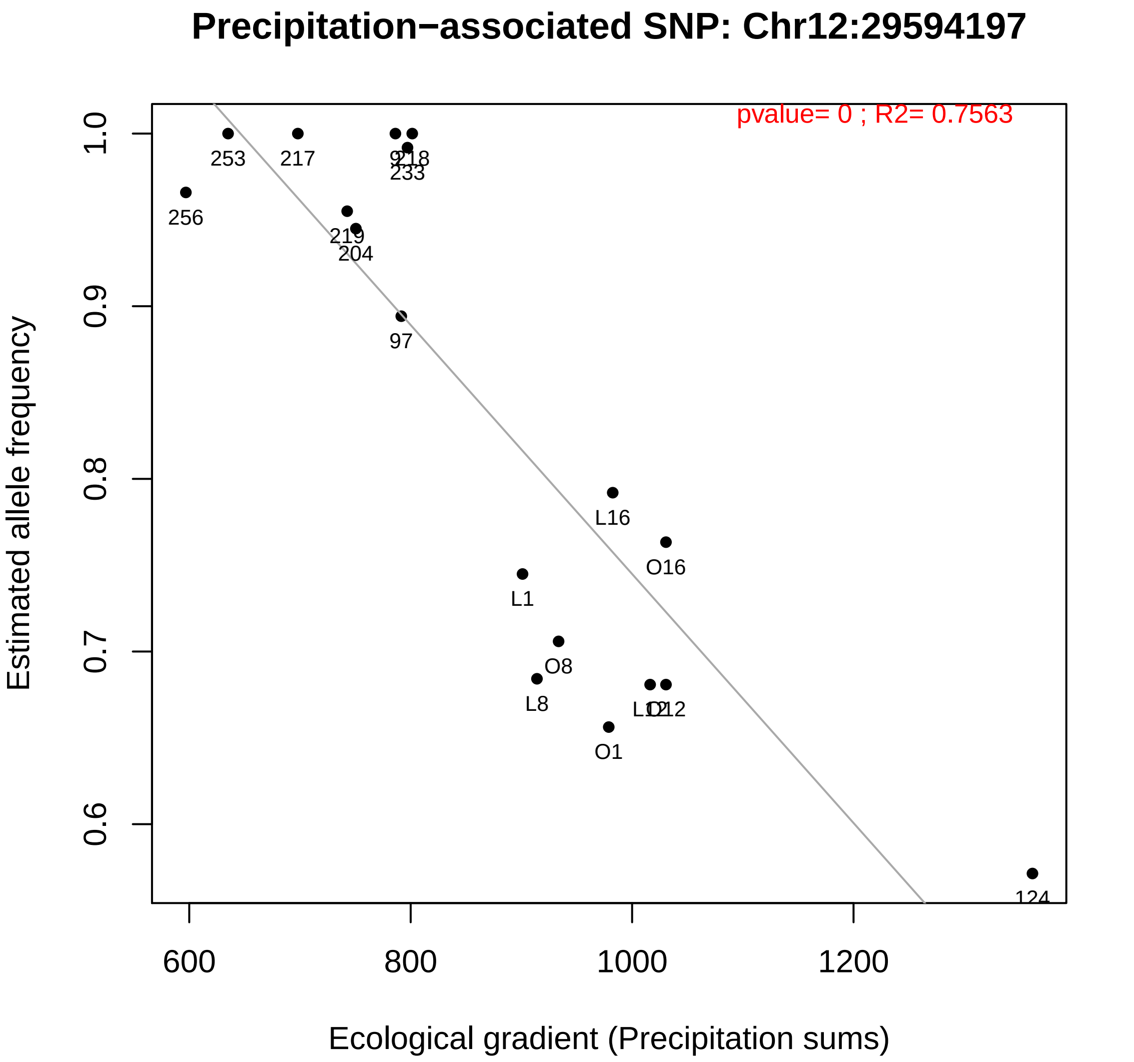

Precipitation−associated SNP: Chr12:29594197
pvalue= 0 ; R2= 0.7563
Estimated allele frequency
Ecological gradient (Precipitation sums)
1.0
0.9
0.8
0.7
0.6
600
800
1000
1200
253
217
218
233
256
219
204
97
L16
O16
L1
O8
L8
O12
O1
124

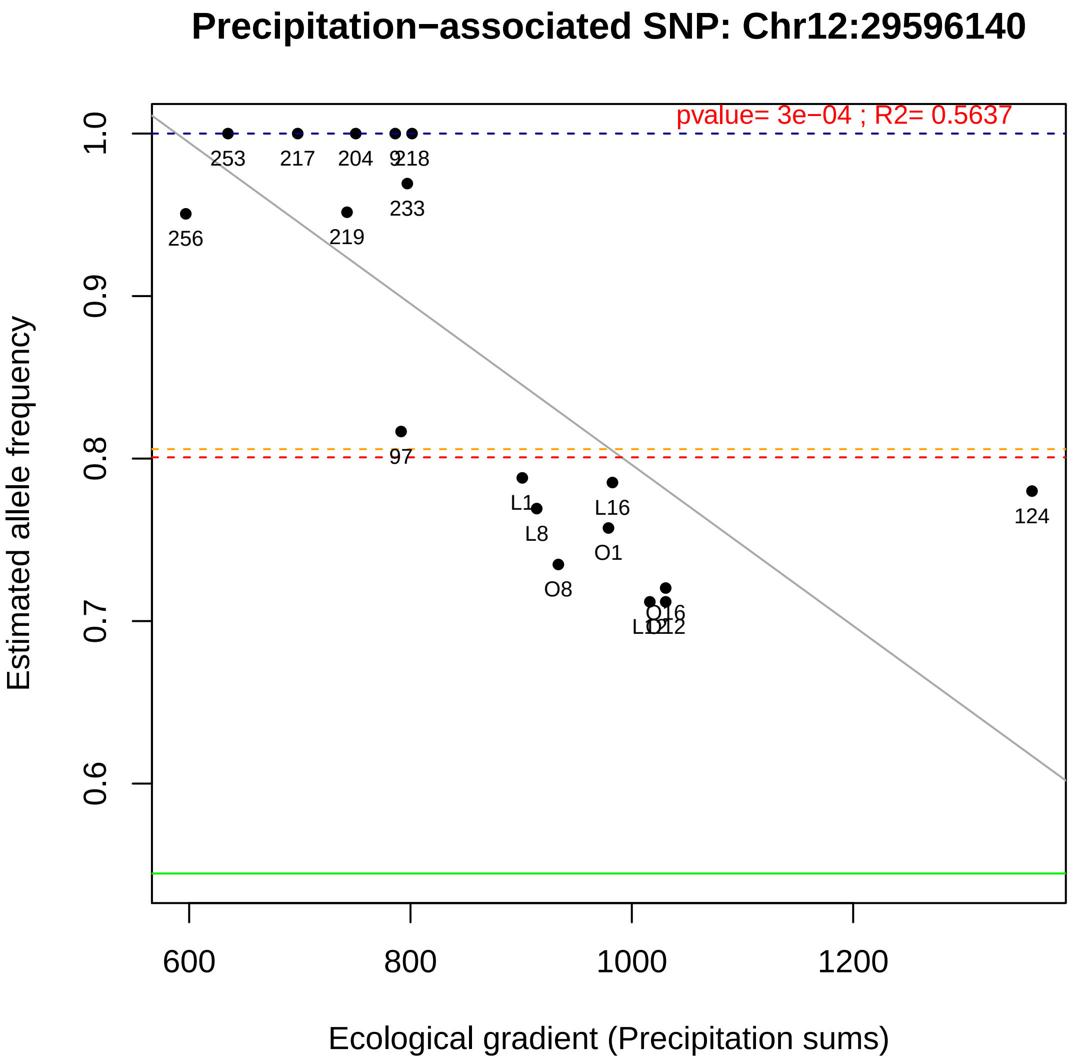
Precipitation−associated SNP: Chr12:29596140
pvalue= 3e−04 ; R2= 0.5637
Estimated allele frequency
Ecological gradient (Precipitation sums)
1.0
0.9
0.8
0.7
0.6
600
800
1000
1200
253
217
204
218
233
256
219
97
L1
L8
L16
O1
O8
O16
O12
124

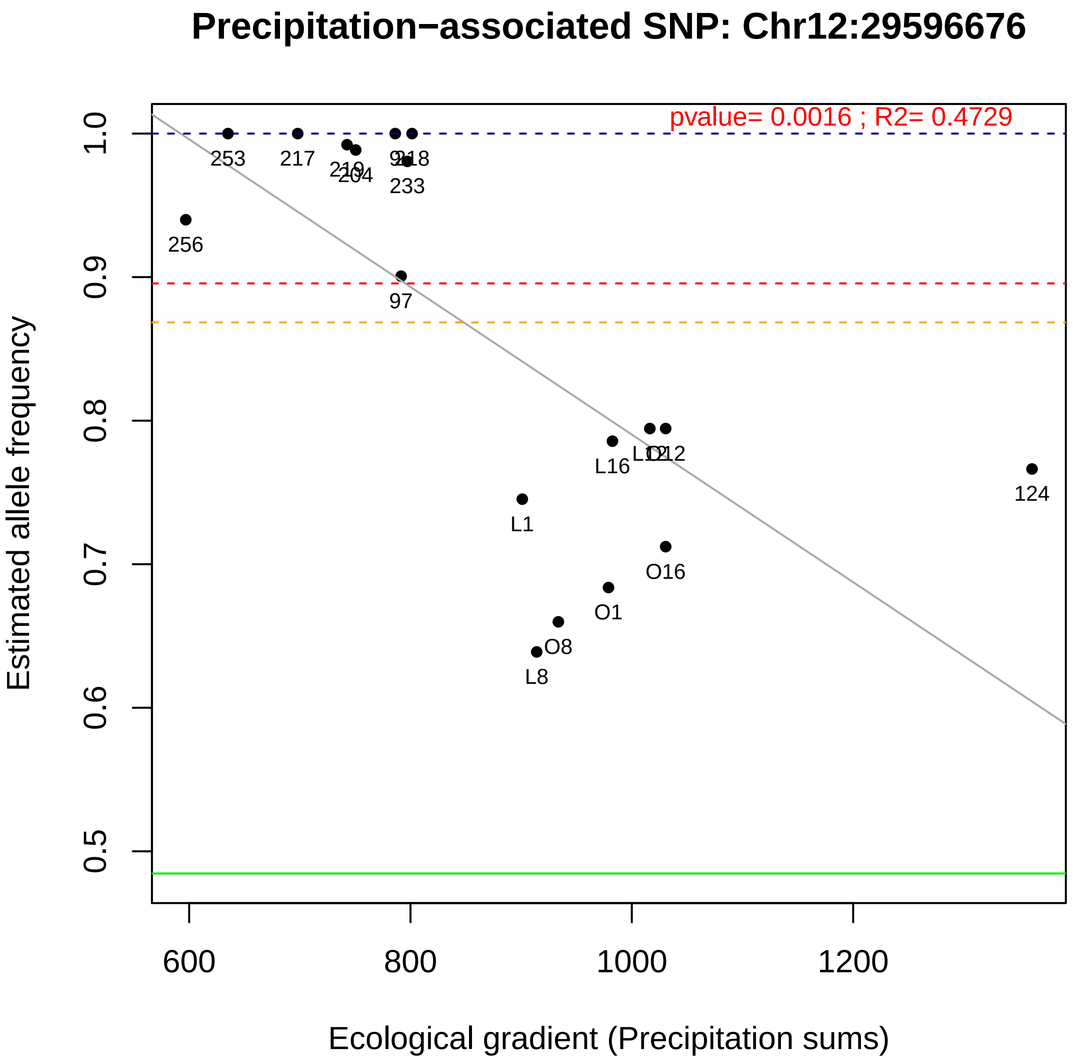

Precipitation−associated SNP: Chr12:29596676
pvalue= 0.0016 ; R2= 0.4729
Estimated allele frequency
1.0
0.9
0.8
0.7
0.6
0.5
253
217
219
204
92
218
233
256
97
L16
L12
O12
124
L1
O16
O1
O8
L8
600
800
1000
1200
Ecological gradient (Precipitation sums)

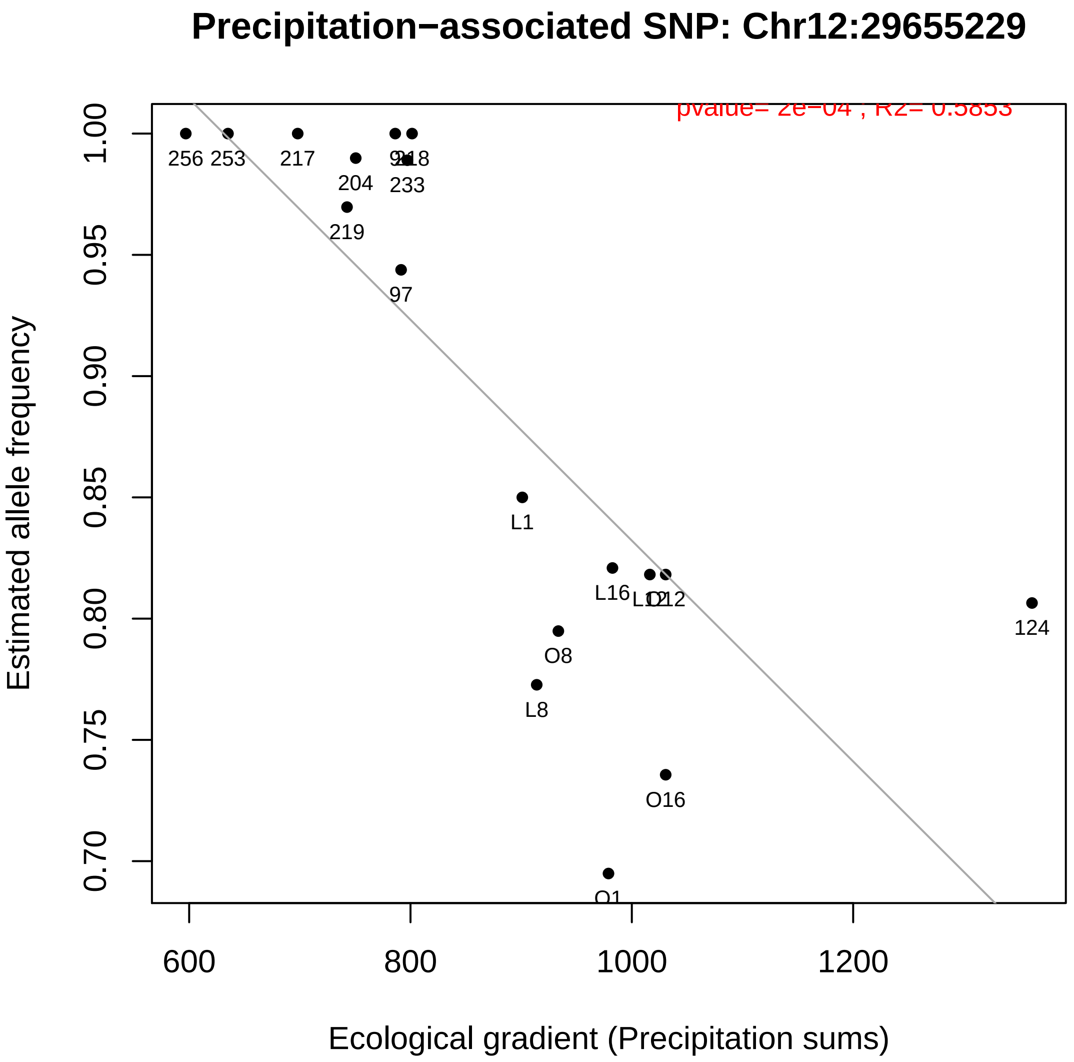
Precipitation−associated SNP: Chr12:29655229
pvalue= 2e−04 ; R2= 0.5853
Estimated allele frequency
Ecological gradient (Precipitation sums)
1.00
0.95
0.90
0.85
0.80
0.75
0.70
600
800
1000
1200
256
253
217
204
233
219
97
L1
L16
124
O8
L8
O16

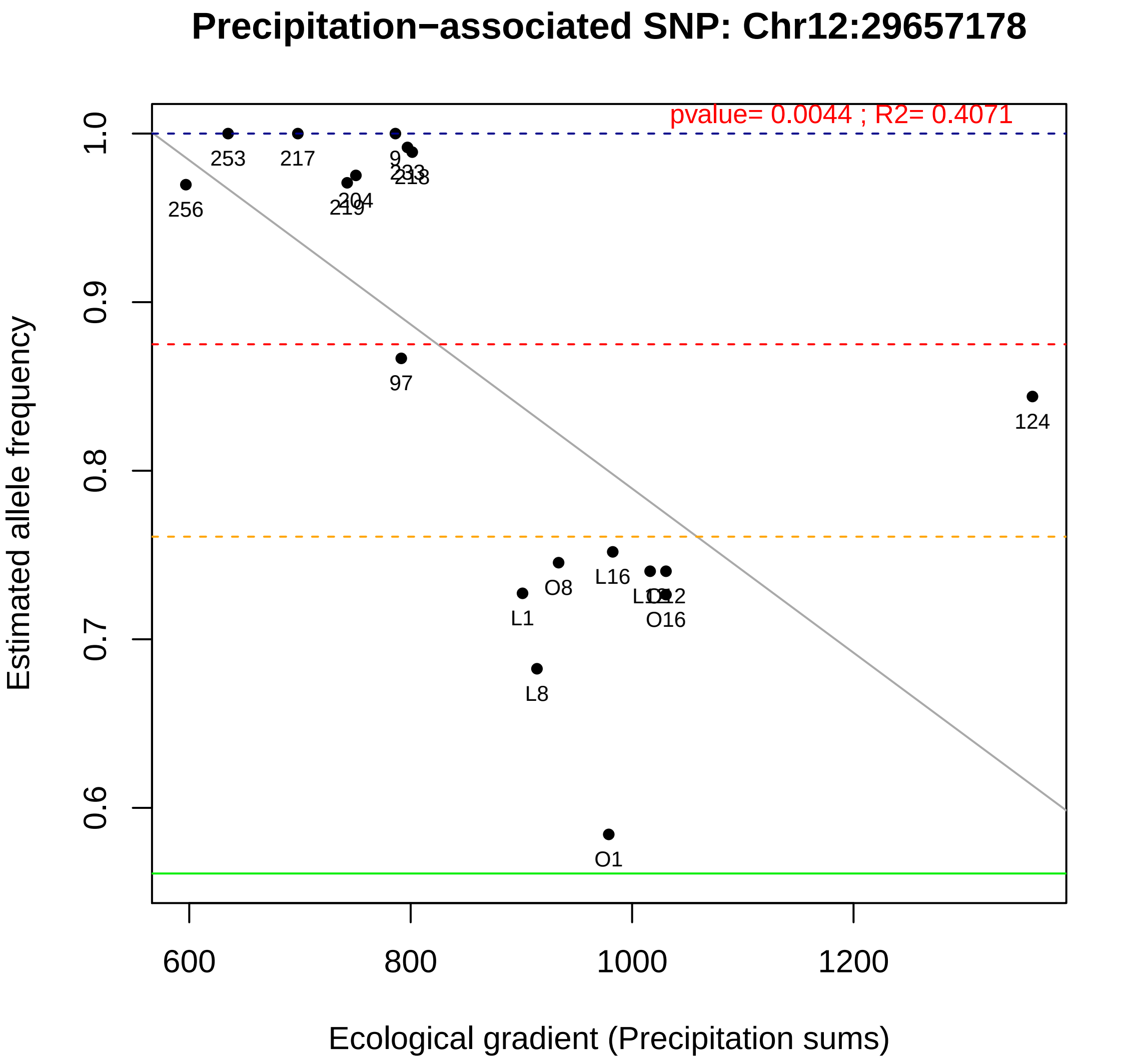
Precipitation−associated SNP: Chr12:29657178
pvalue= 0.0044 ; R2= 0.4071
Estimated allele frequency
Ecological gradient (Precipitation sums)
0.6
0.7
0.8
0.9
1.0
600
800
1000
1200
256
253
217
219
204
9
233
218
97
124
O8
L16
L1
O12
O16
L8
O1

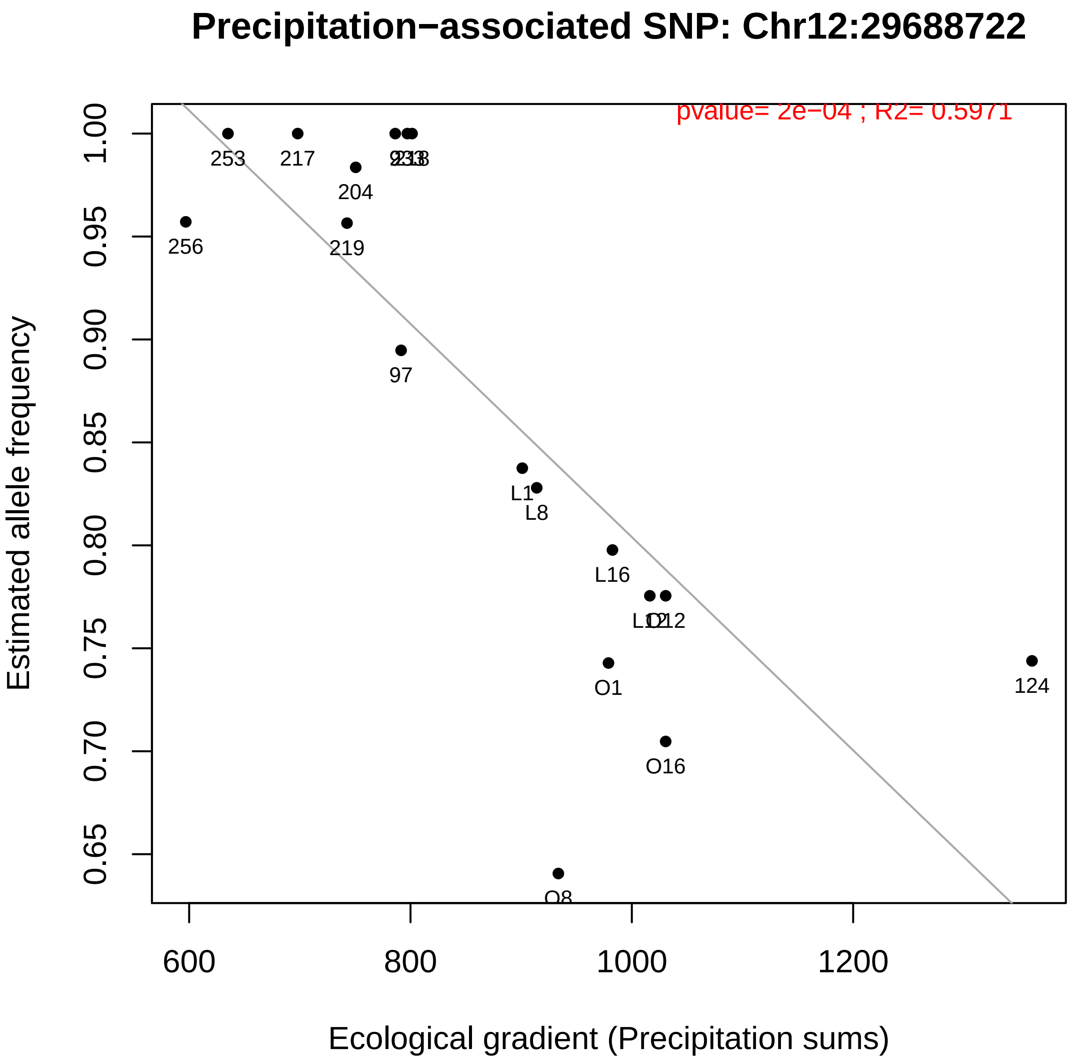

Precipitation−associated SNP: Chr12:29688722
pvalue= 2e−04 ; R2= 0.5971
Estimated allele frequency
Ecological gradient (Precipitation sums)
0.65
0.70
0.75
0.80
0.85
0.90
0.95
1.00
600
800
1000
1200
253
217
204
256
219
97
L1
L8
L16
O12
O1
124
O16
O8

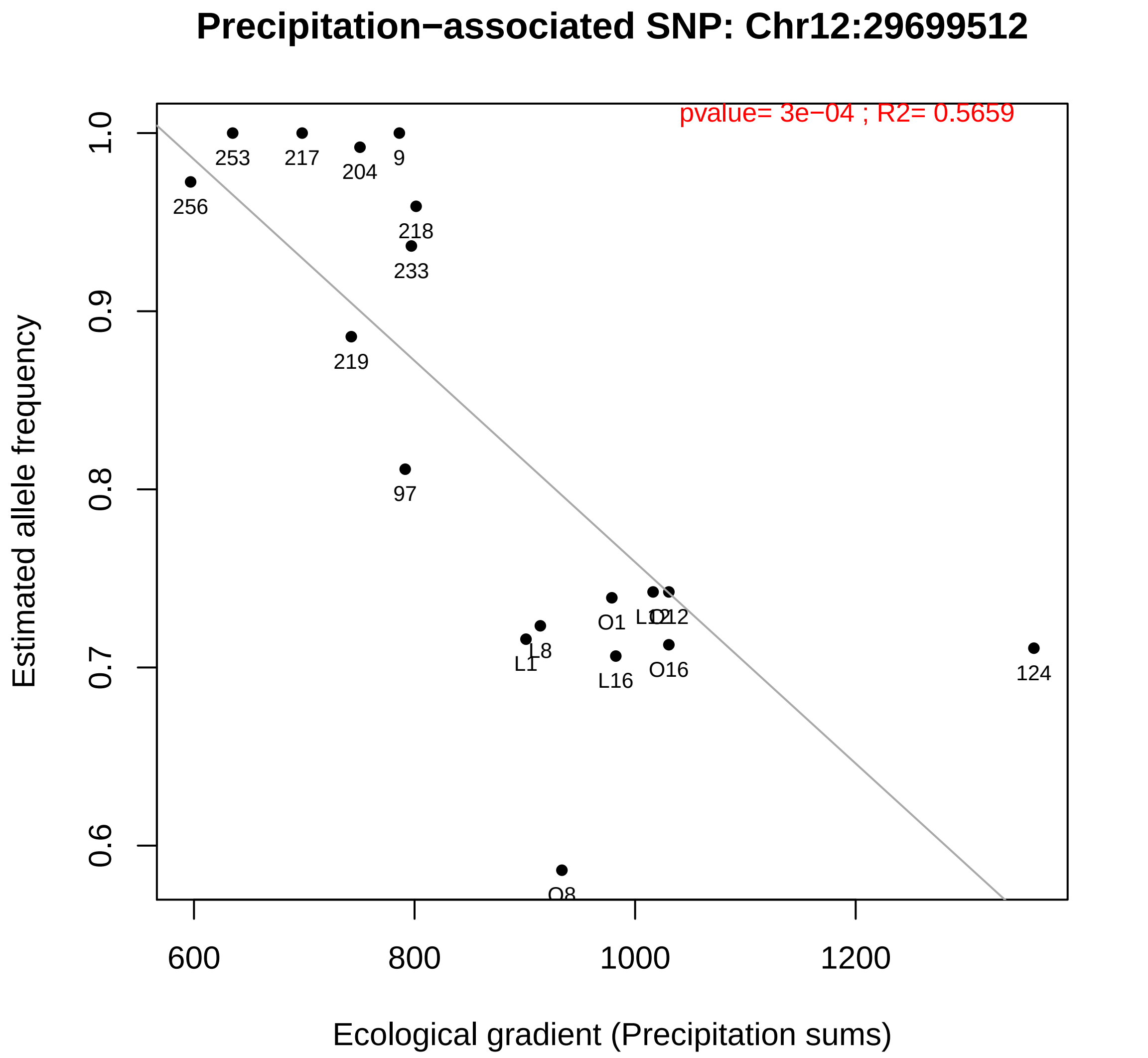
Precipitation−associated SNP: Chr12:29699512
pvalue= 3e−04 ; R2= 0.5659
Estimated allele frequency
Ecological gradient (Precipitation sums)
1.0
0.9
0.8
0.7
0.6
600
800
1000
1200
253
217
204
9
256
218
233
219
97
O1
L12
O12
L8
L1
L16
O16
124
O8

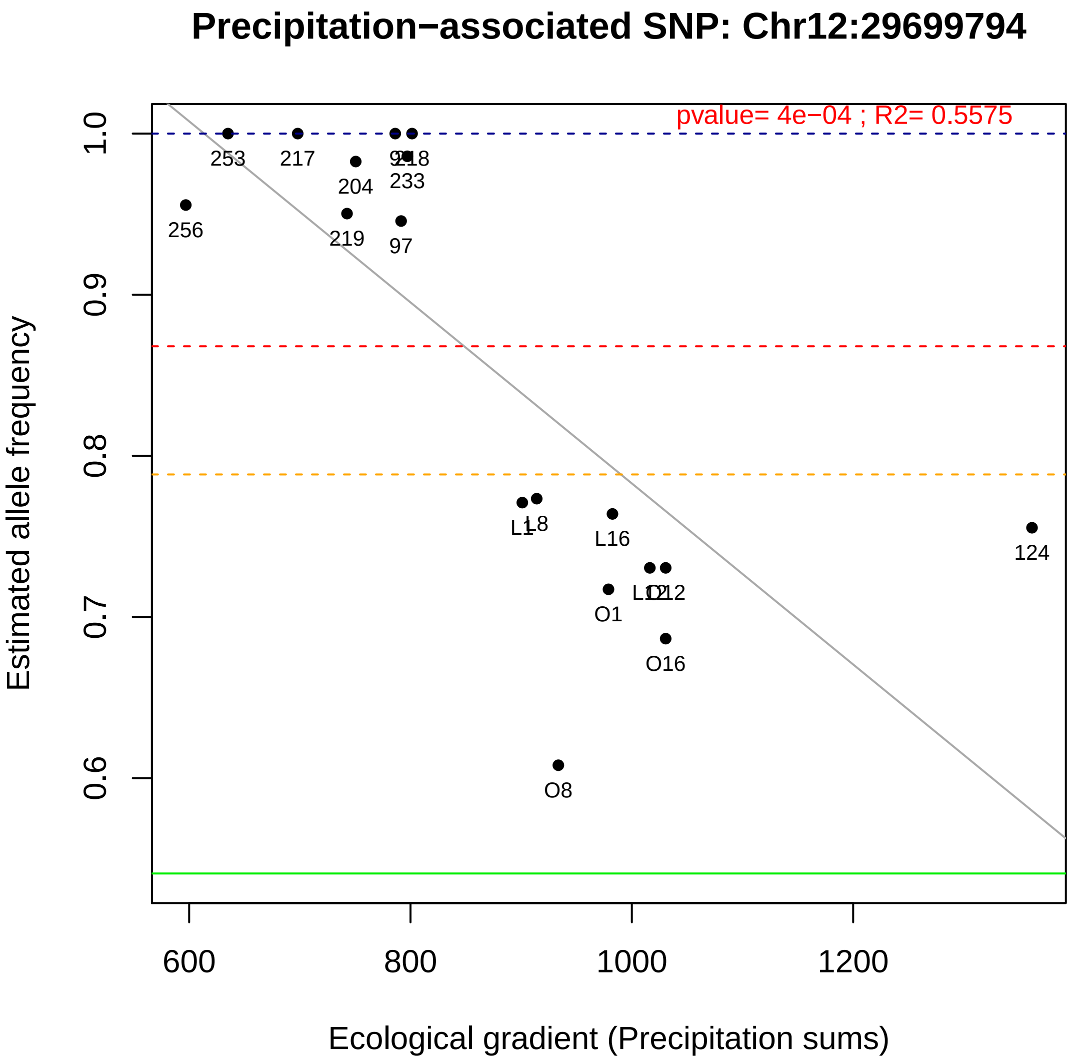

Precipitation−associated SNP: Chr12:29699794
pvalue= 4e−04 ; R2= 0.5575
Estimated allele frequency
1.0
0.9
0.8
0.7
0.6
600
800
1000
1200
Ecological gradient (Precipitation sums)
253
217
204
233
256
219
97
L1
L8
L16
124
O1
O12
O16
O8

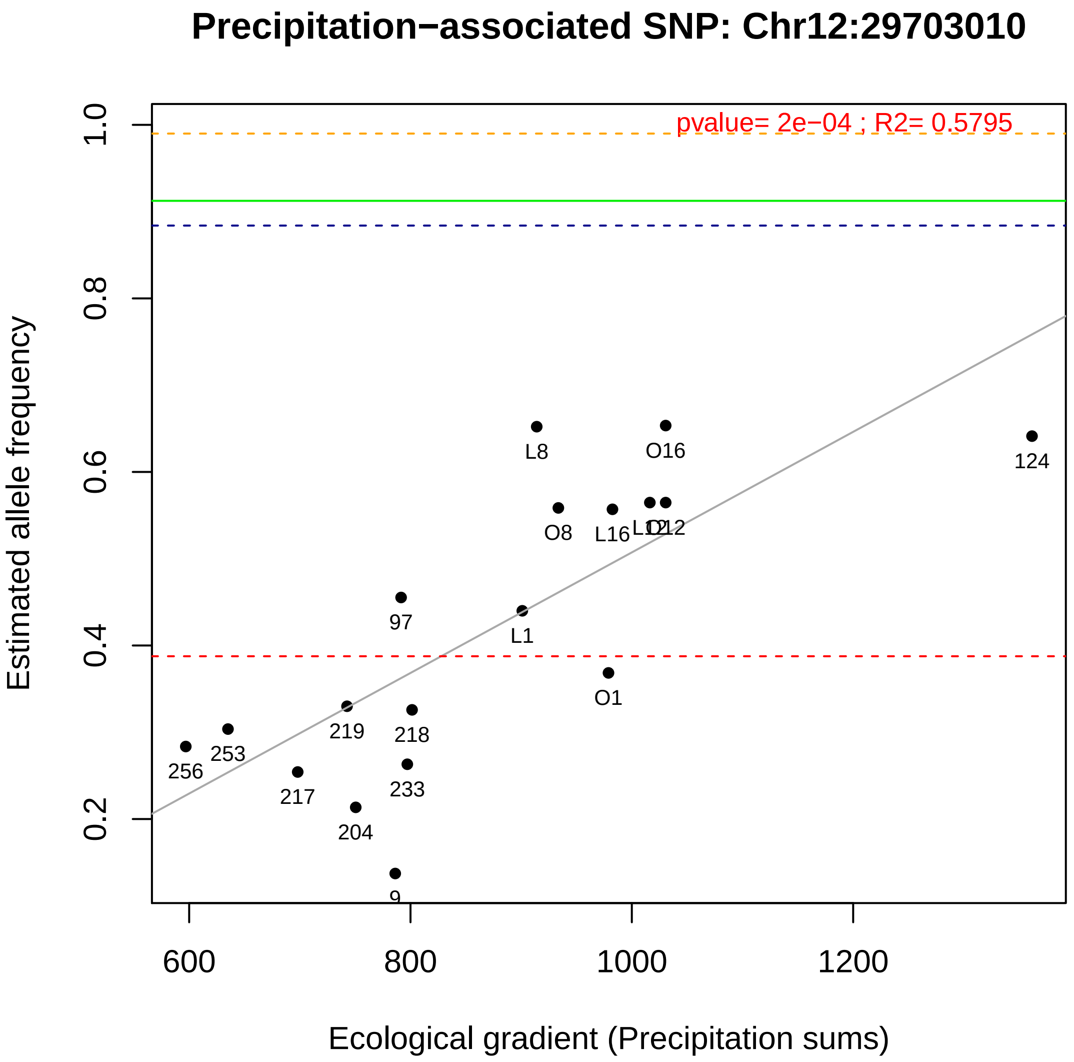

Precipitation−associated SNP: Chr12:29703010
pvalue= 2e−04 ; R2= 0.5795
Estimated allele frequency
Ecological gradient (Precipitation sums)
0.2
0.4
0.6
0.8
1.0
600
800
1000
1200
L8
O16
124
O8
L16
L1
97
O1
219
218
253
256
217
233
204
9

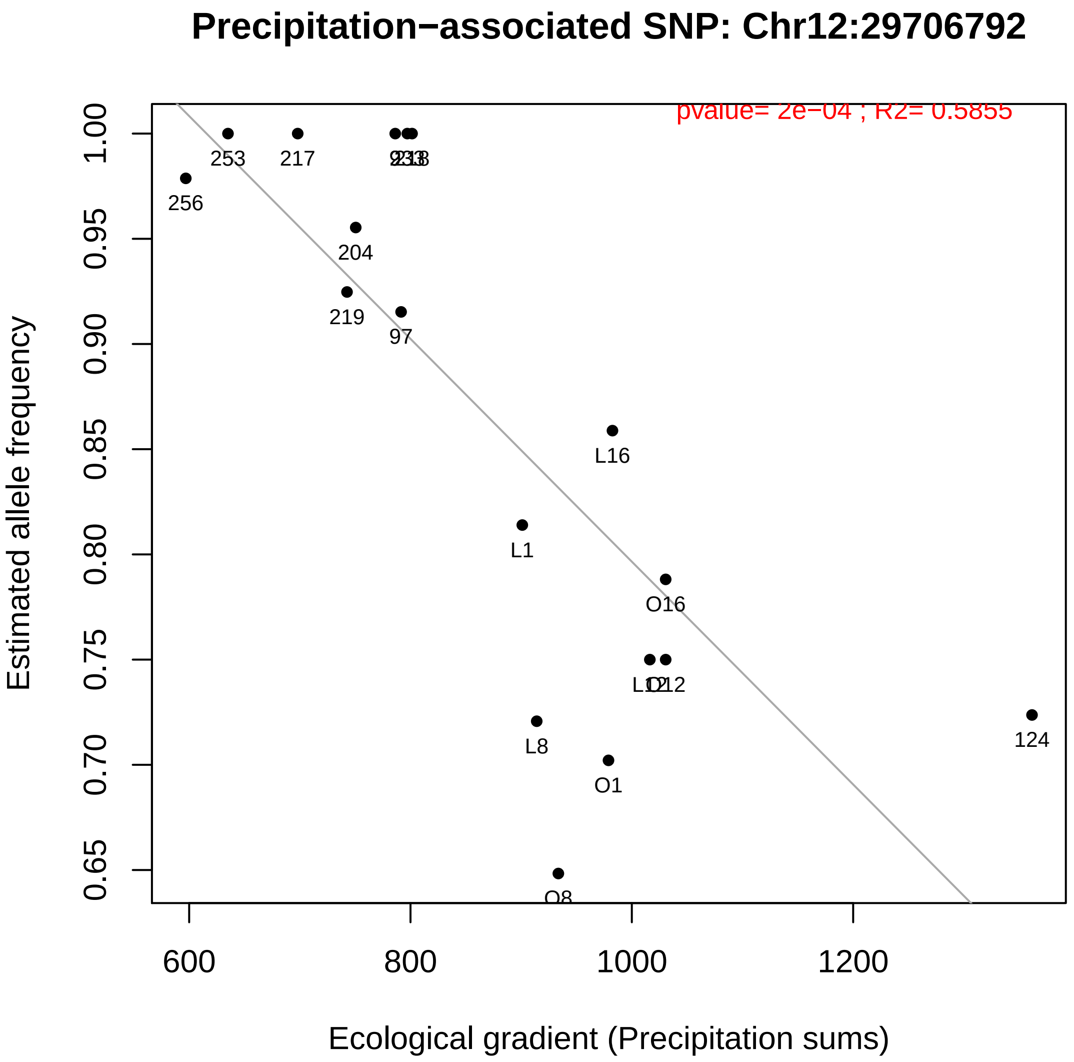

Precipitation−associated SNP: Chr12:29706792
pvalue= 2e−04 ; R2= 0.5855
Estimated allele frequency
Ecological gradient (Precipitation sums)
0.65
0.70
0.75
0.80
0.85
0.90
0.95
1.00
600
800
1000
1200
253
217
256
204
219
97
L16
L1
O16
L8
O1
O8
124

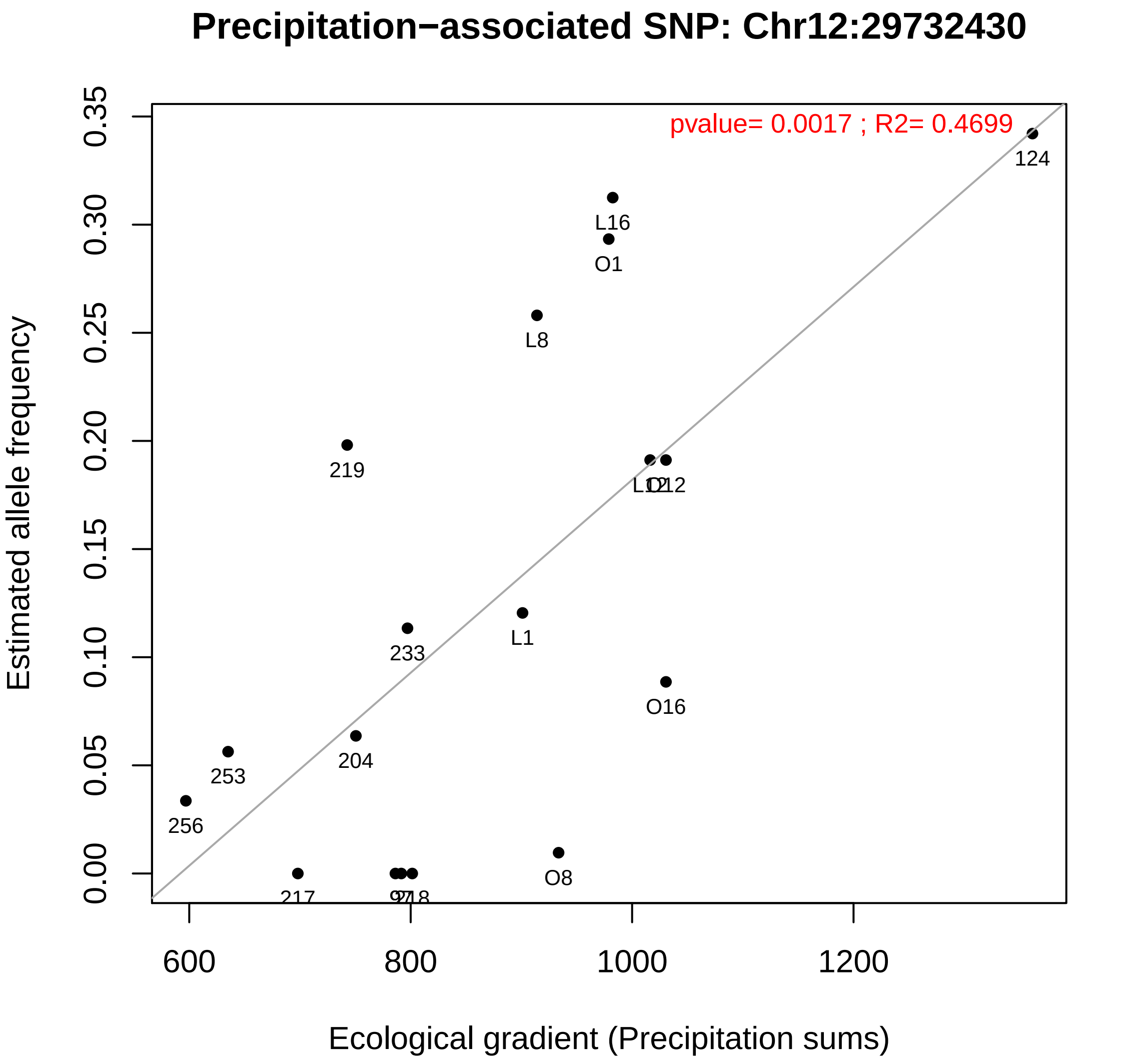
Precipitation−associated SNP: Chr12:29732430
pvalue= 0.0017 ; R2= 0.4699
Estimated allele frequency
Ecological gradient (Precipitation sums)
0.00
0.05
0.10
0.15
0.20
0.25
0.30
0.35
600
800
1000
1200
124
L16
O1
L8
219
L12
O12
L1
233
O16
204
253
256
217
O8

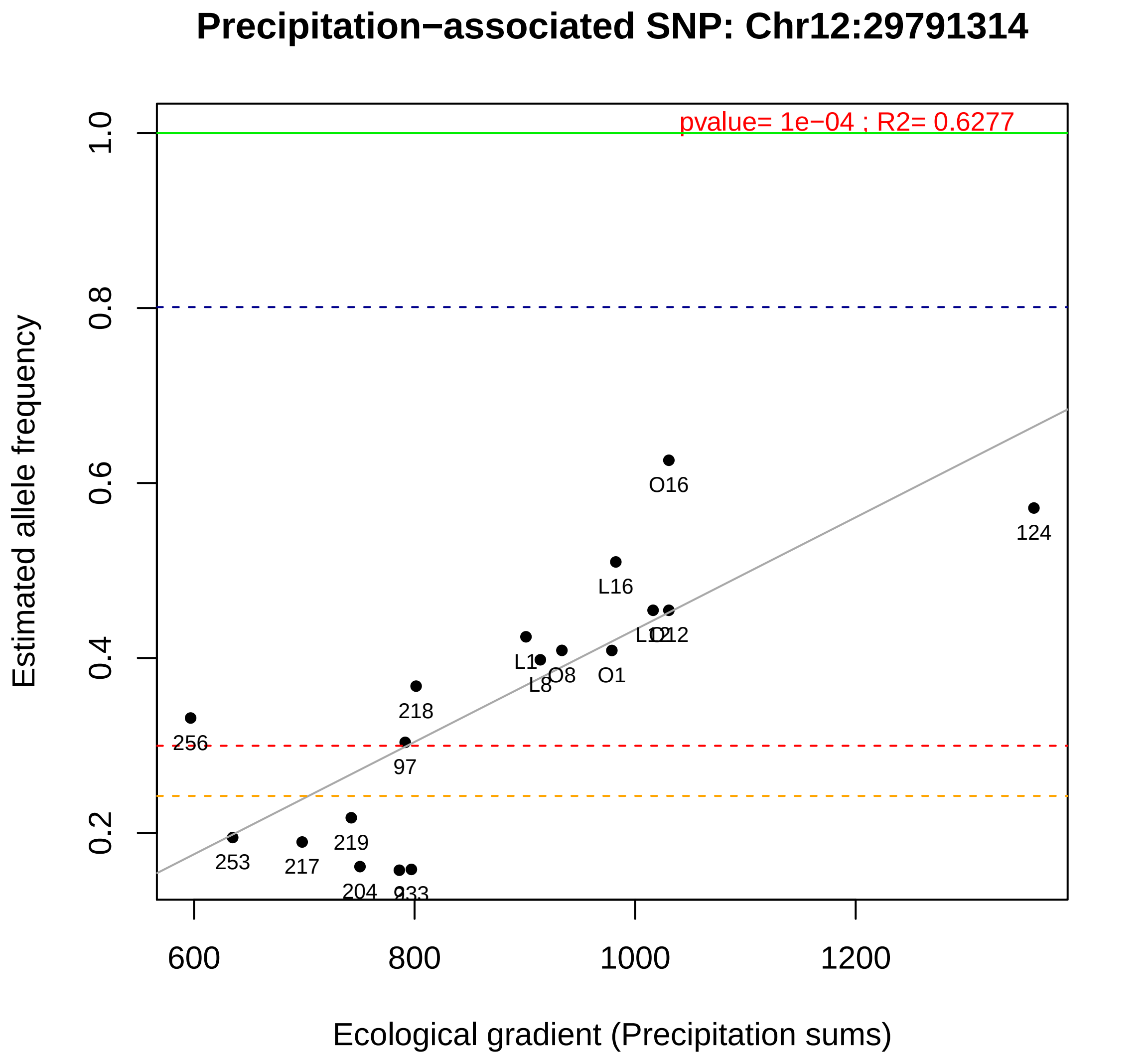

Precipitation−associated SNP: Chr12:29791314
pvalue= 1e−04 ; R2= 0.6277
Estimated allele frequency
0.2
0.4
0.6
0.8
1.0
600
800
1000
1200
Ecological gradient (Precipitation sums)
O16
124
L16
L12
O12
L1
L8
O8
O1
218
256
97
219
253
217
204
233

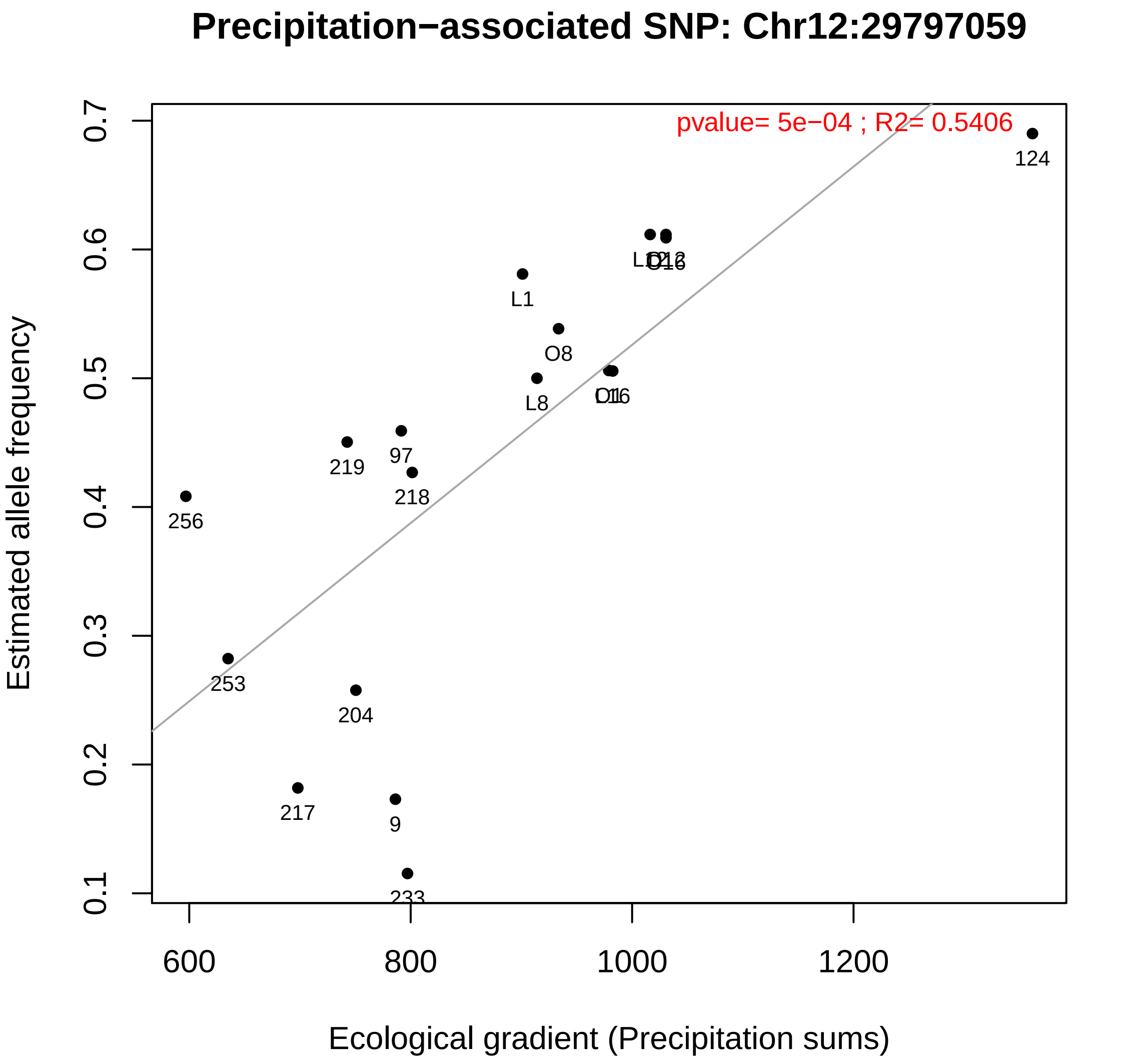

Precipitation−associated SNP: Chr12:29797059
pvalue= 5e−04 ; R2= 0.5406
Estimated allele frequency
Ecological gradient (Precipitation sums)
0.1
0.2
0.3
0.4
0.5
0.6
0.7
600
800
1000
1200
124
L1
O8
L8
97
219
218
256
253
204
217
9
233

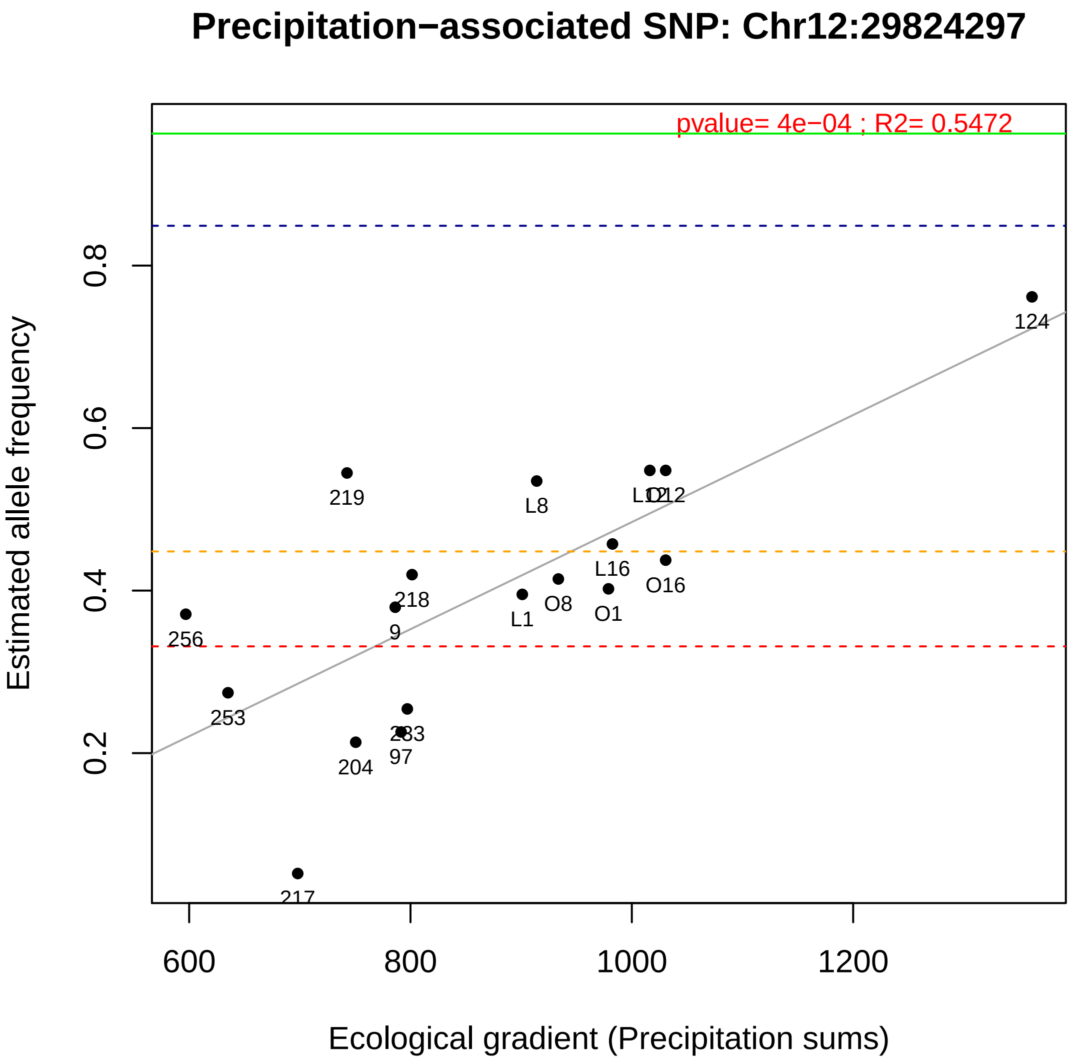

Precipitation−associated SNP: Chr12:29824297
pvalue= 4e−04 ; R2= 0.5472
Estimated allele frequency
0.2
0.4
0.6
0.8
600
800
1000
1200
Ecological gradient (Precipitation sums)
219
L8
L12
O12
124
L16
O16
218
O8
O1
L1
9
256
253
283
97
204
217

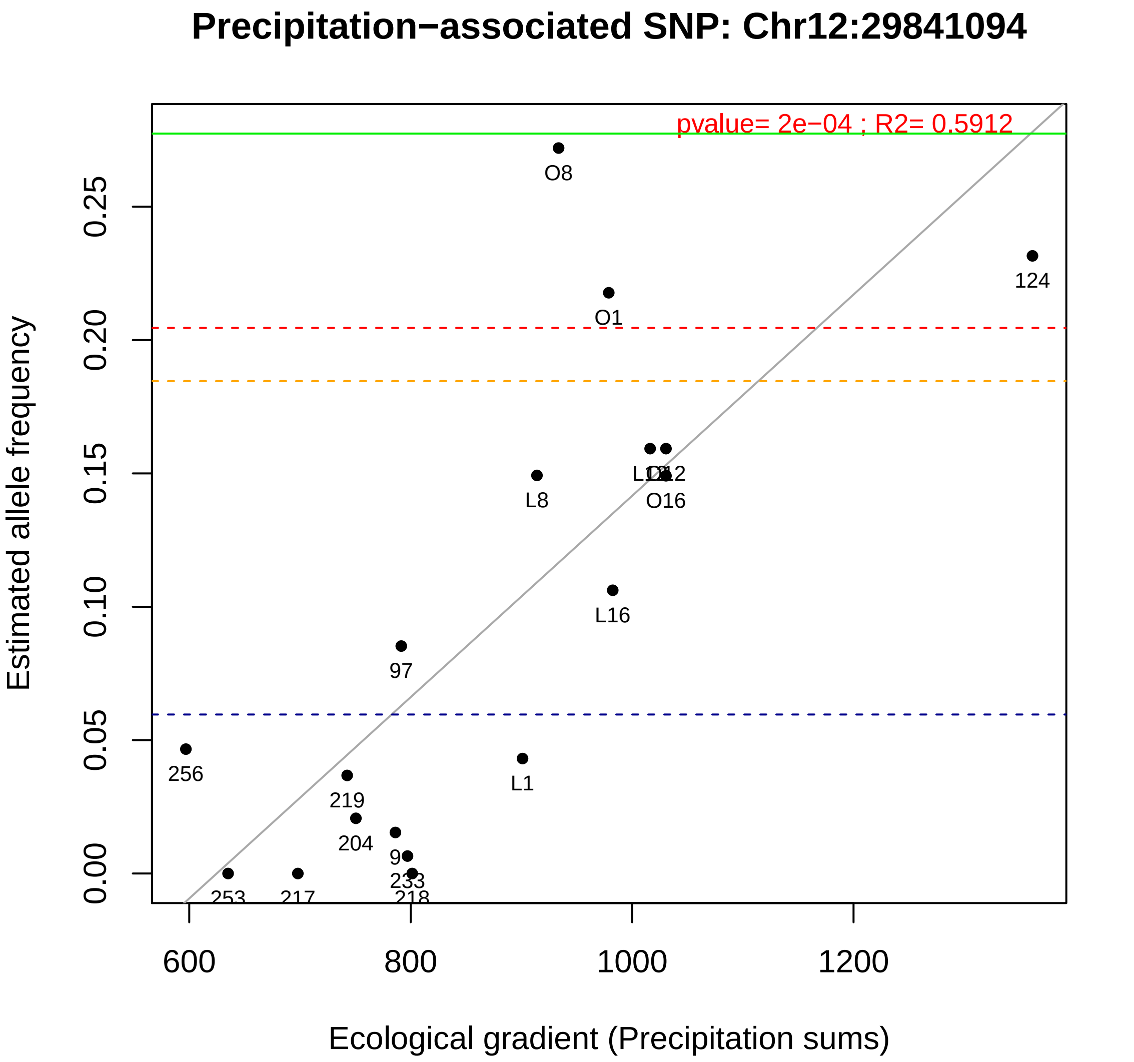

Precipitation−associated SNP: Chr12:29841094
pvalue= 2e−04 ; R2= 0.5912
Estimated allele frequency
0.00
0.05
0.10
0.15
0.20
0.25
600
800
1000
1200
Ecological gradient (Precipitation sums)
O8
124
O1
L8
O16
L16
97
256
219
204
L1
233
218
253
217

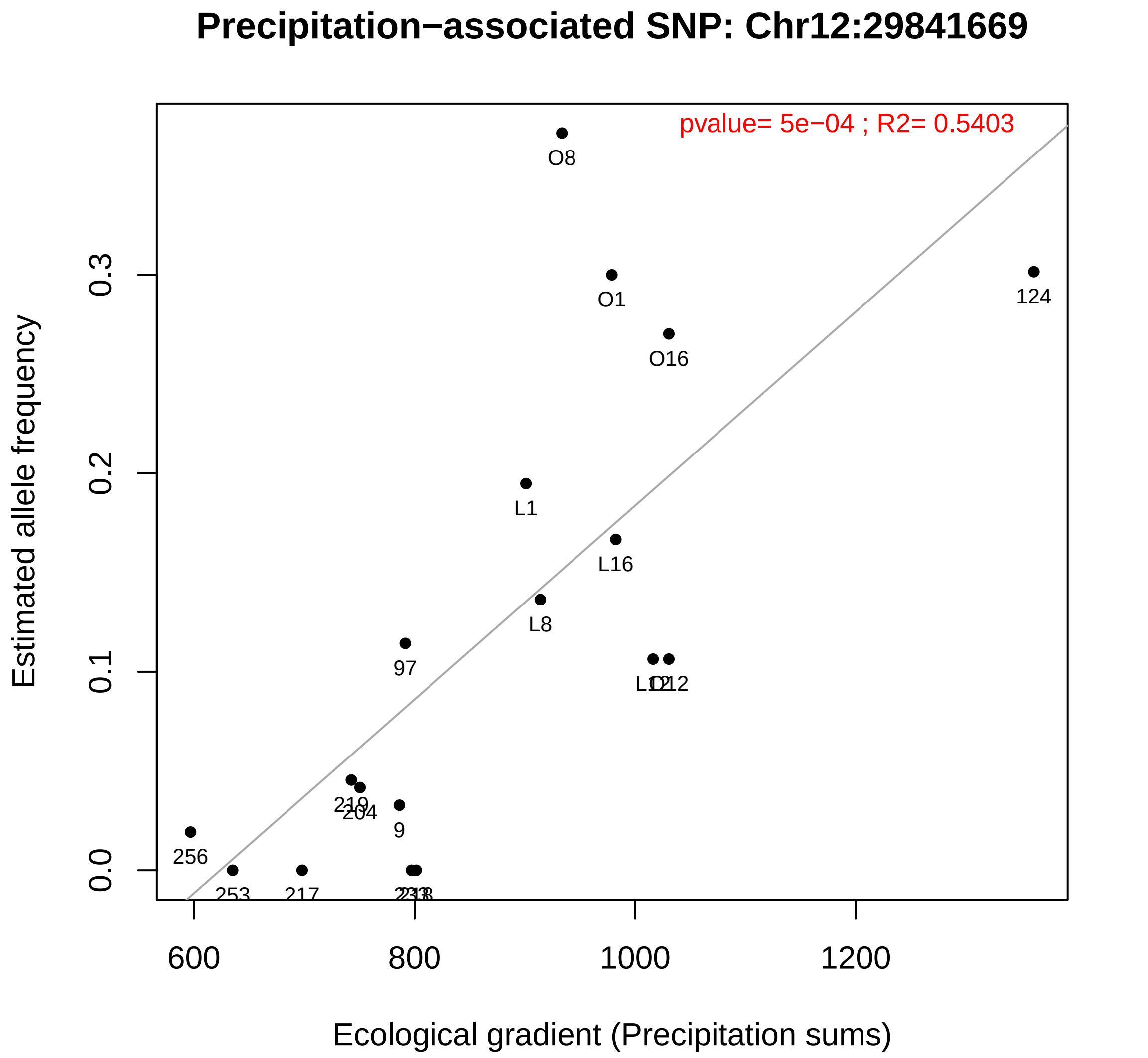

Precipitation−associated SNP: Chr12:29841669
pvalue= 5e−04 ; R2= 0.5403
O8
O1
124
O16
L1
L16
L8
97
O12
219
204
9
256
253
217
0.0
0.1
0.2
0.3
600
800
1000
1200
Estimated allele frequency
Ecological gradient (Precipitation sums)

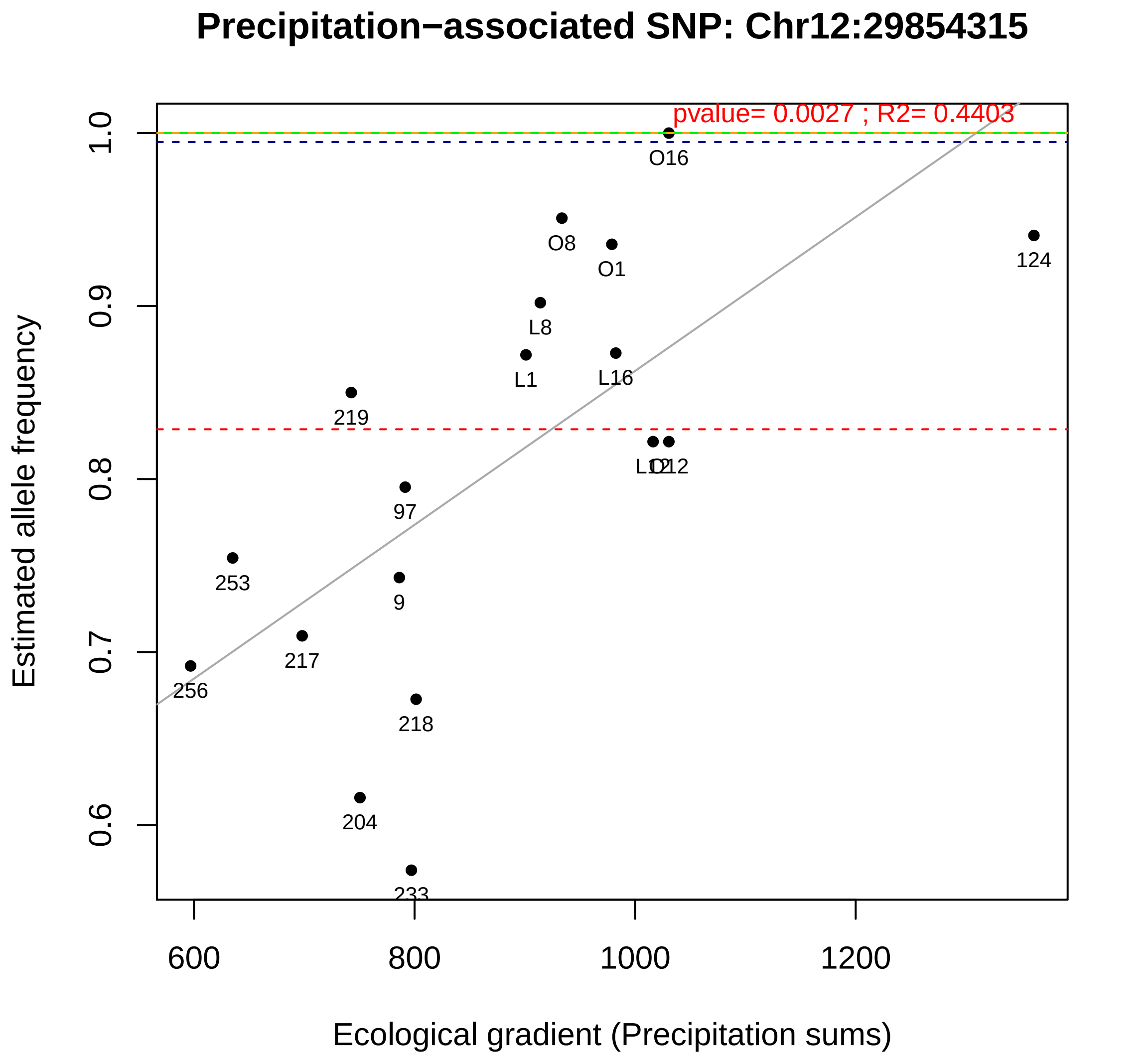

Precipitation−associated SNP: Chr12:29854315
pvalue= 0.0027 ; R2= 0.4403
Estimated allele frequency
Ecological gradient (Precipitation sums)
0.6
0.7
0.8
0.9
1.0
600
800
1000
1200
O16
O8
O1
124
L8
L1
L16
219
L12
O12
97
253
9
217
256
218
204
233

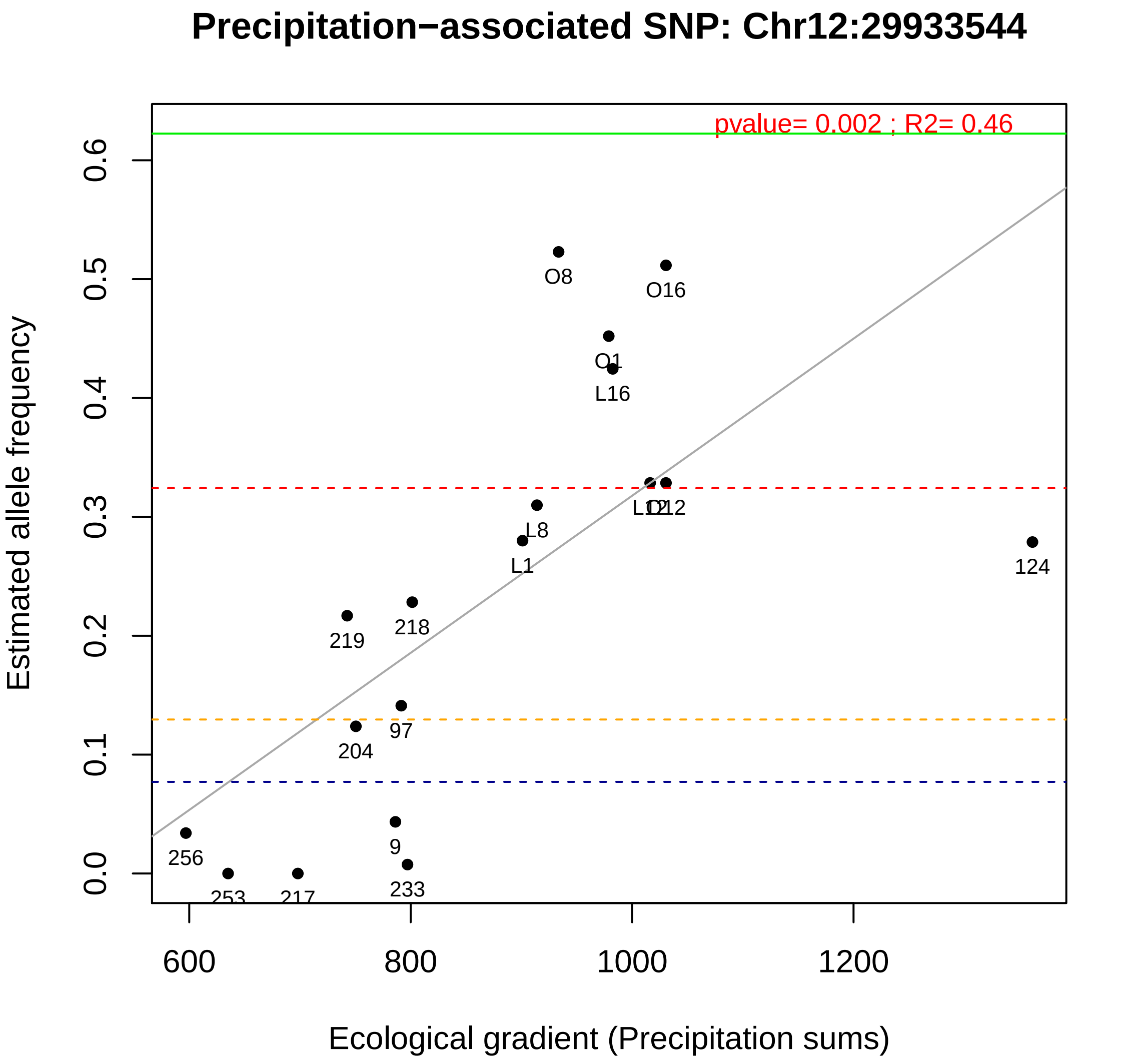

Precipitation−associated SNP: Chr12:29933544
pvalue= 0.002 ; R2= 0.46
Estimated allele frequency
Ecological gradient (Precipitation sums)
0.0
0.1
0.2
0.3
0.4
0.5
0.6
600
800
1000
1200
O8
O16
O1
L16
L12
O12
L8
L1
124
219
218
97
204
256
9
253
217
233

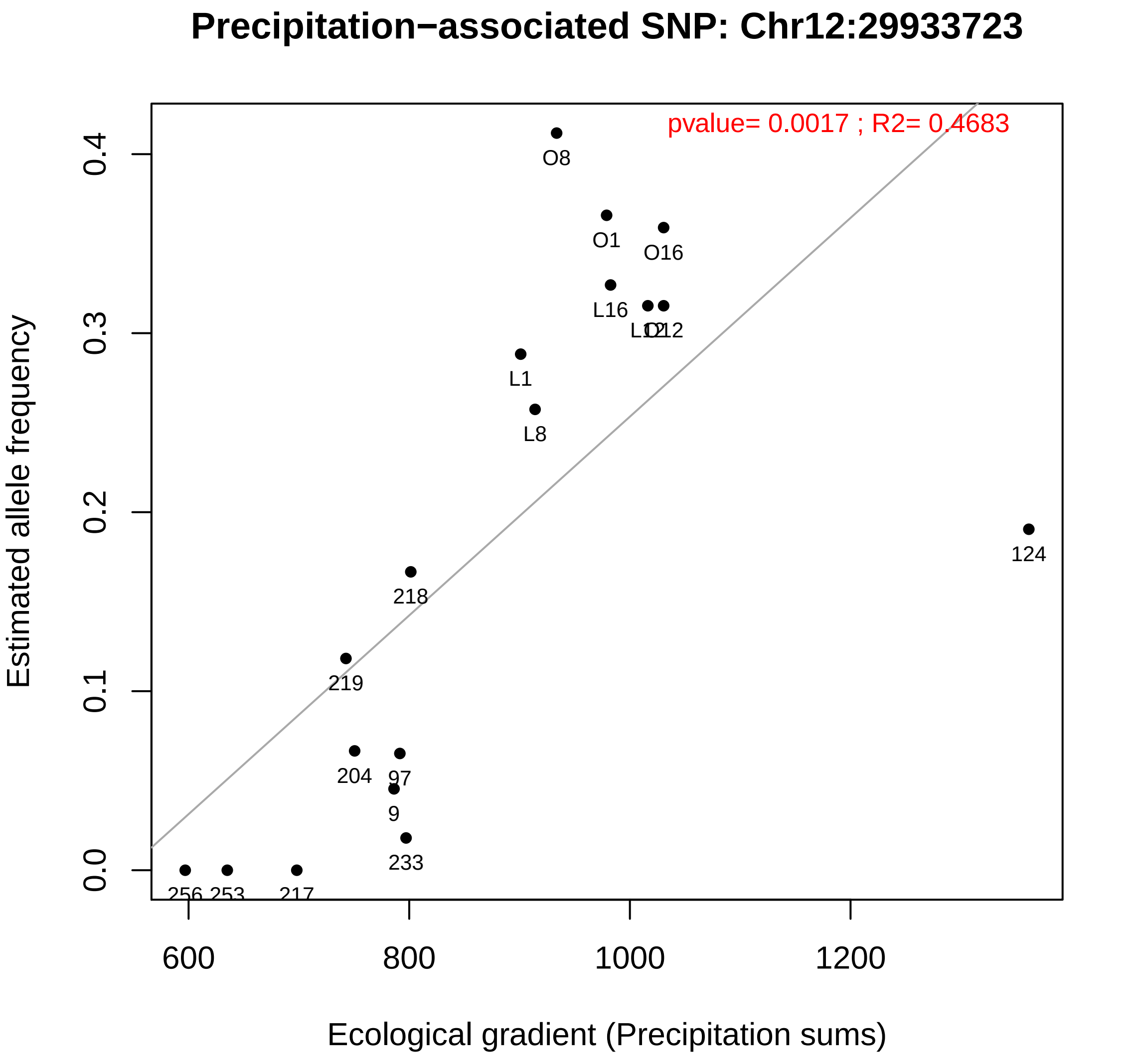
Precipitation−associated SNP: Chr12:29933723
pvalue= 0.0017 ; R2= 0.4683
O8
O1
O16
L16
L1
L8
124
218
219
204
97
9
233
256
253
217
0.0
0.1
0.2
0.3
0.4
600
800
1000
1200
Estimated allele frequency
Ecological gradient (Precipitation sums)

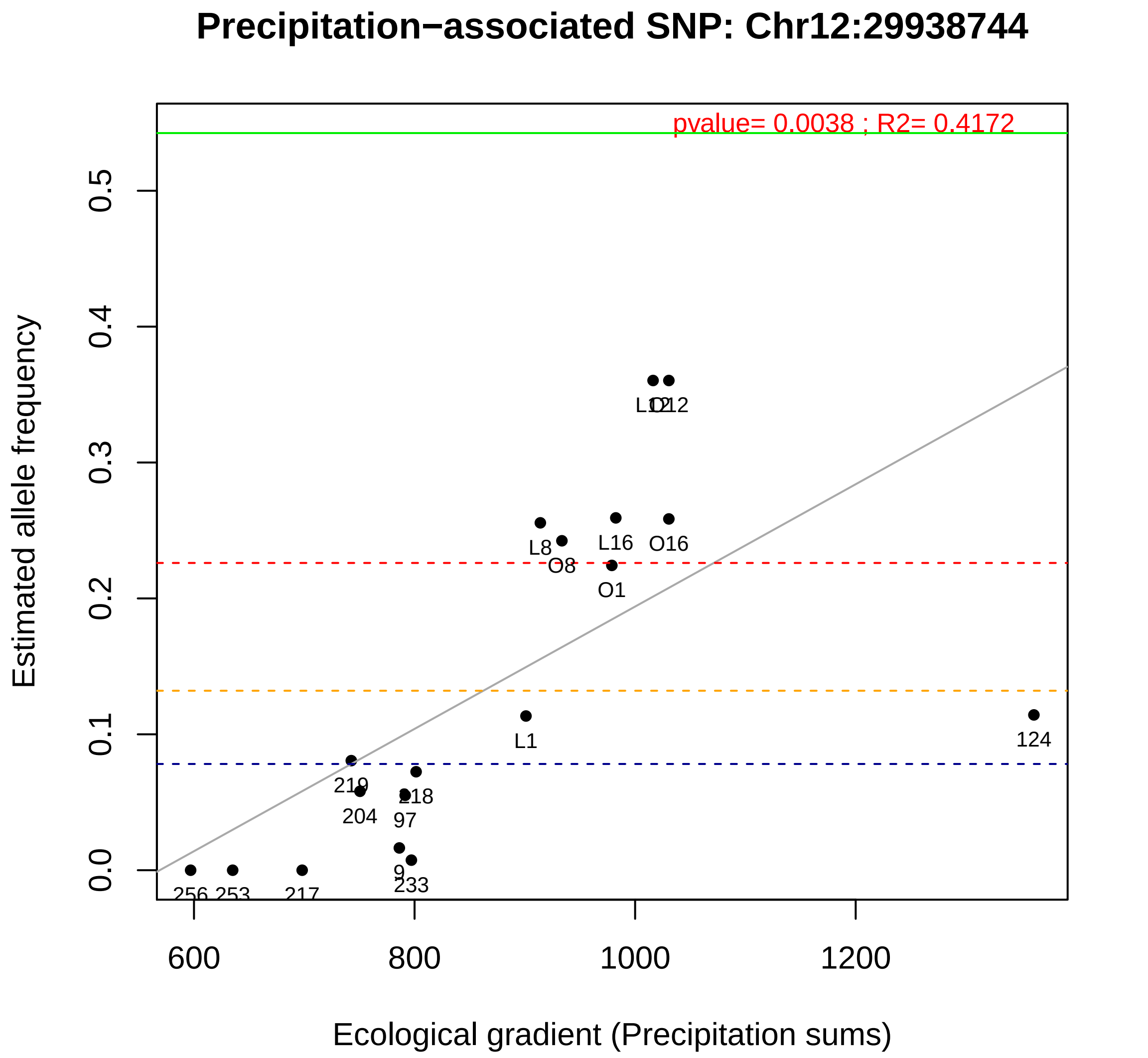

Precipitation−associated SNP: Chr12:29938744
pvalue= 0.0038 ; R2= 0.4172
Estimated allele frequency
0.0
0.1
0.2
0.3
0.4
0.5
600
800
1000
1200
Ecological gradient (Precipitation sums)
L12
O12
L8
O8
L16
O16
O1
L1
124
219
204
218
97
9
233
256
253
217

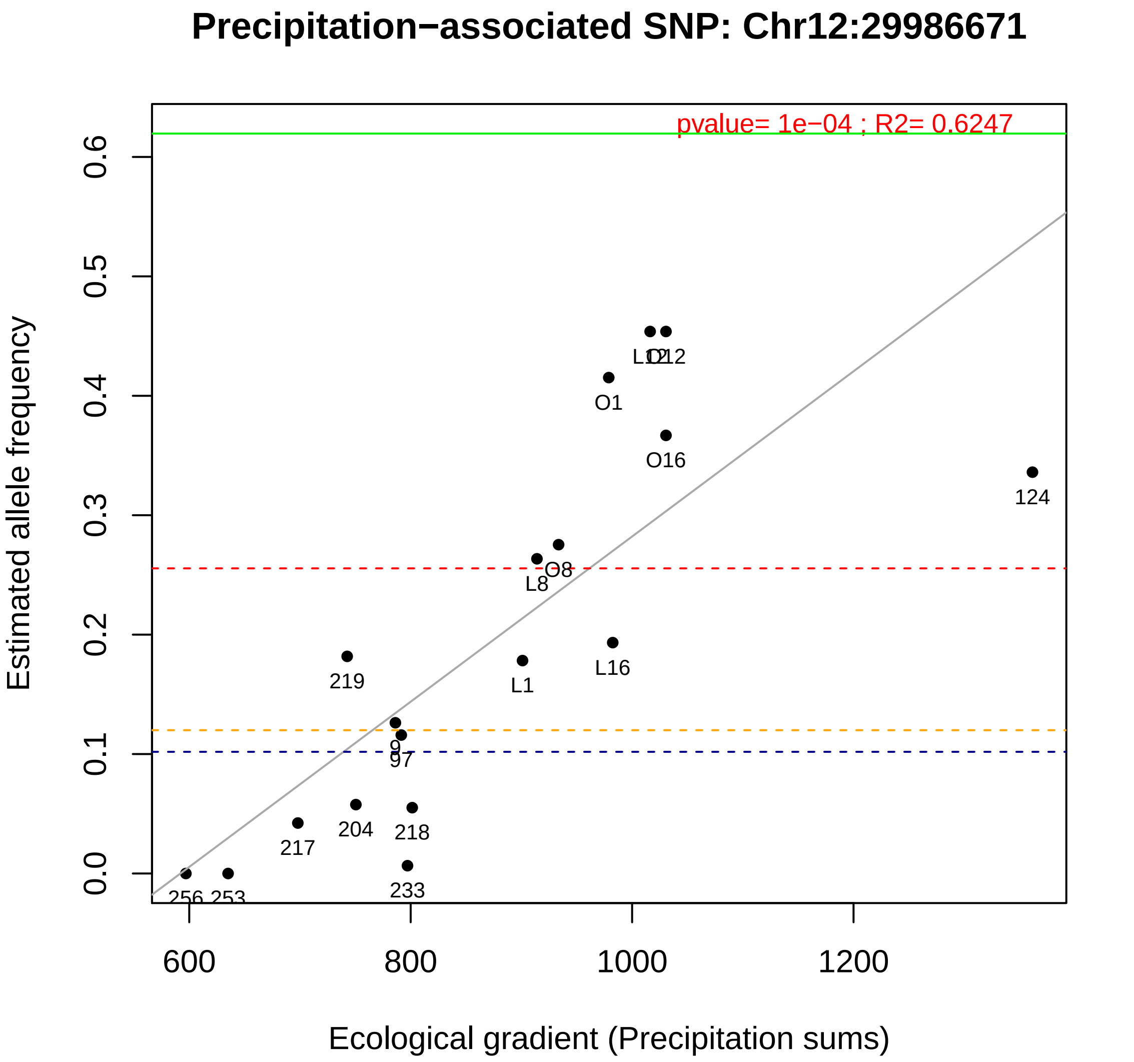

Precipitation−associated SNP: Chr12:29986671
pvalue= 1e−04 ; R2= 0.6247
Estimated allele frequency
0.0
0.1
0.2
0.3
0.4
0.5
0.6
600
800
1000
1200
Ecological gradient (Precipitation sums)
L12
O12
O1
O16
124
O8
L8
L16
219
L1
9
97
204
218
217
233
256
253

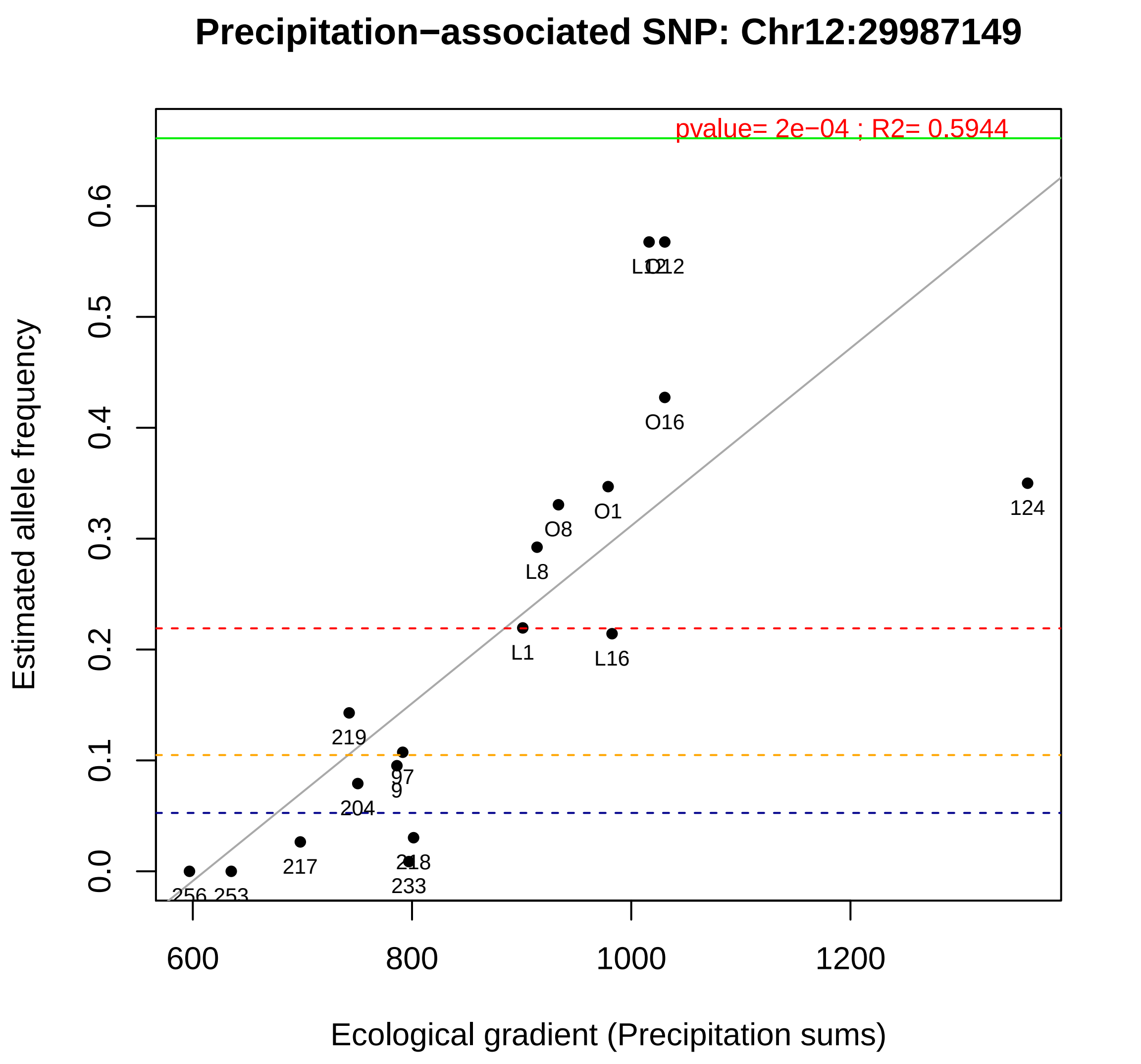

Precipitation−associated SNP: Chr12:29987149
pvalue= 2e−04 ; R2= 0.5944
Estimated allele frequency
0.0
0.1
0.2
0.3
0.4
0.5
0.6
600
800
1000
1200
Ecological gradient (Precipitation sums)
O12
O16
124
O1
O8
L8
L1
L16
219
97
9
204
217
233

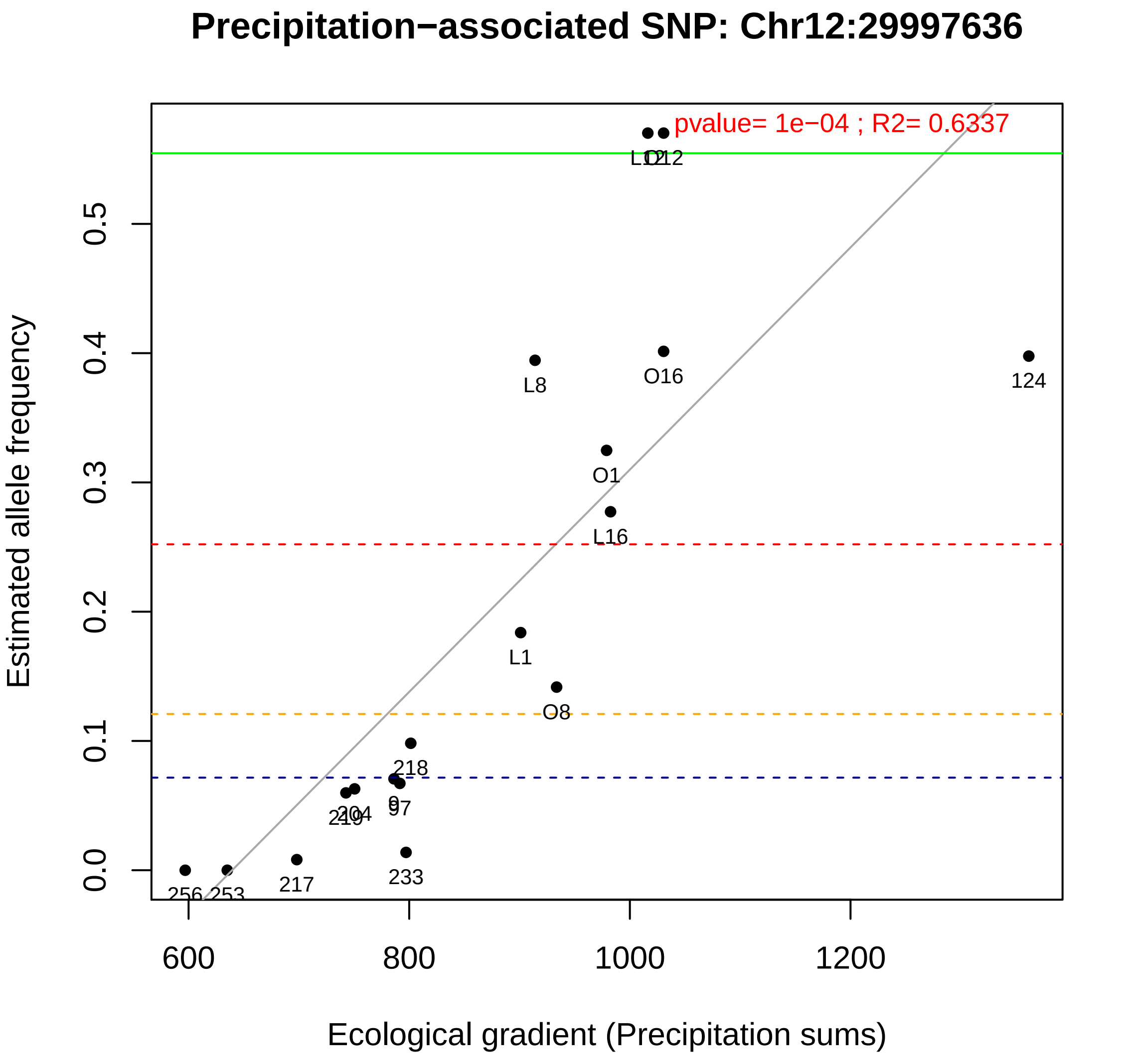
Precipitation−associated SNP: Chr12:29997636
pvalue= 1e−04 ; R2= 0.6337
Estimated allele frequency
Ecological gradient (Precipitation sums)
0.0
0.1
0.2
0.3
0.4
0.5
600
800
1000
1200
L8
O16
124
O1
L16
L1
O8
218
217
233
97
256
253

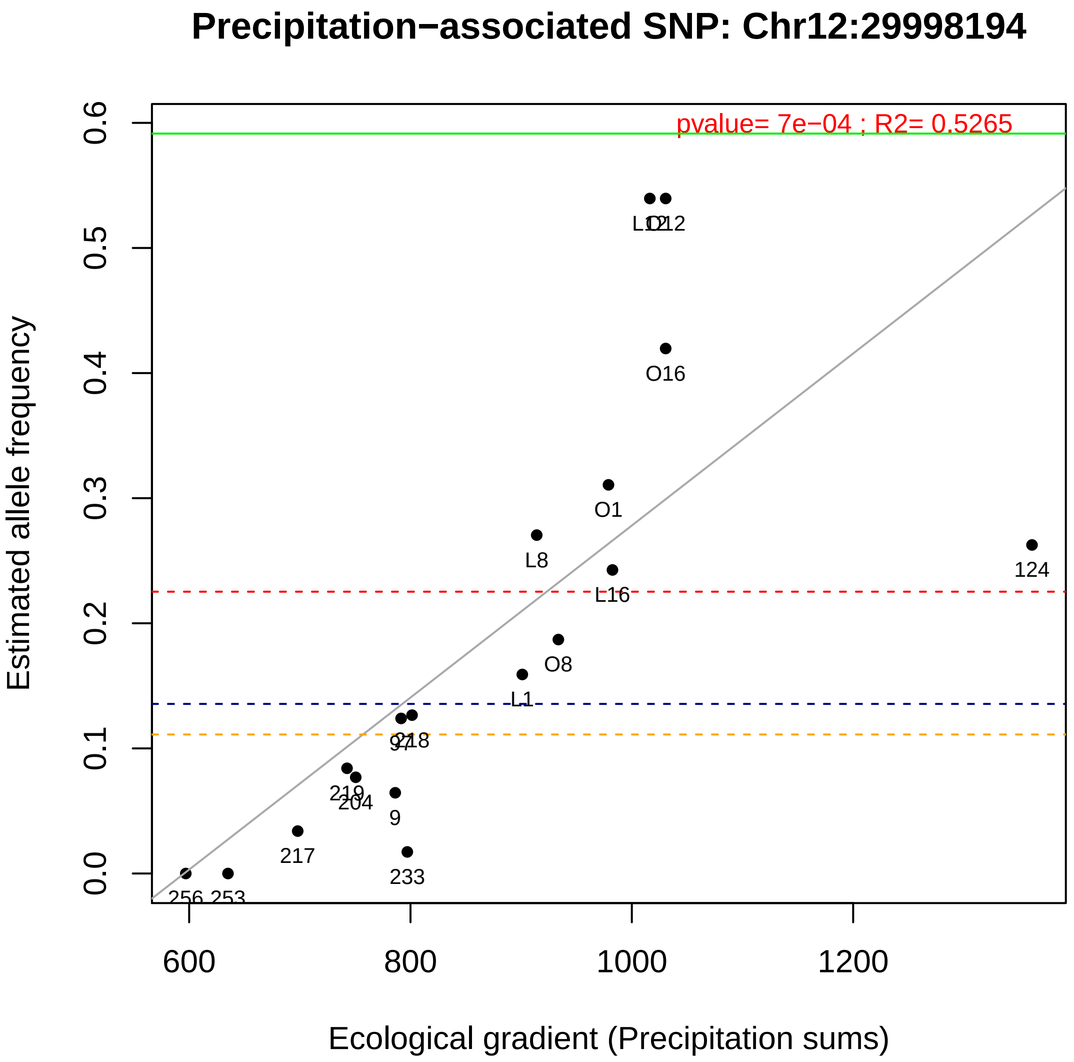
Precipitation−associated SNP: Chr12:29998194
pvalue= 7e−04 ; R2= 0.5265
Estimated allele frequency
Ecological gradient (Precipitation sums)
0.0
0.1
0.2
0.3
0.4
0.5
0.6
600
800
1000
1200
L12
O12
O16
O1
L8
L16
124
O8
L1
97
218
219
204
9
217
233
256
253

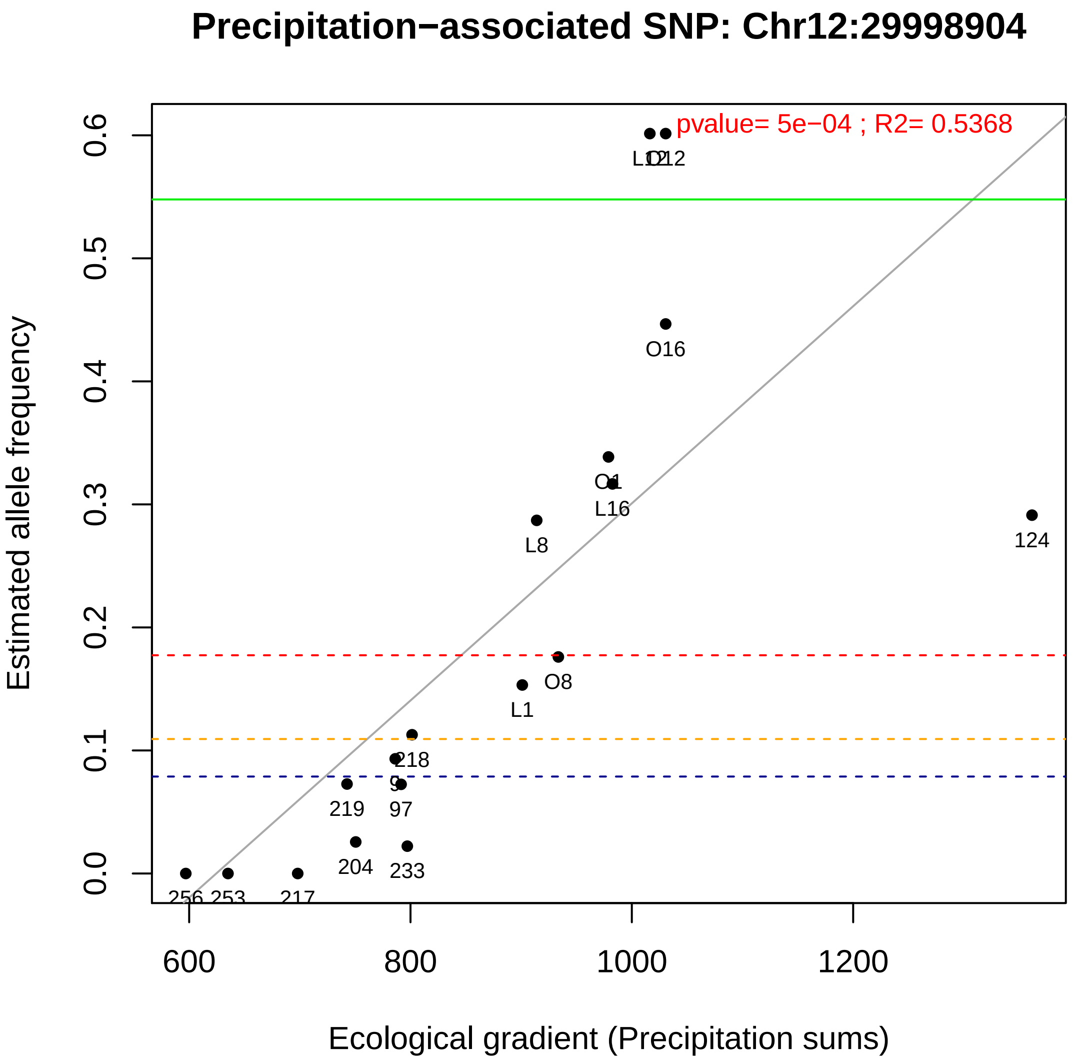
Precipitation−associated SNP: Chr12:29998904
pvalue= 5e−04 ; R2= 0.5368
Estimated allele frequency
0.0
0.1
0.2
0.3
0.4
0.5
0.6
Ecological gradient (Precipitation sums)
600
800
1000
1200
L10
O12
O16
O1
L16
L8
124
O8
L1
218
9
97
219
204
233
256
253
217

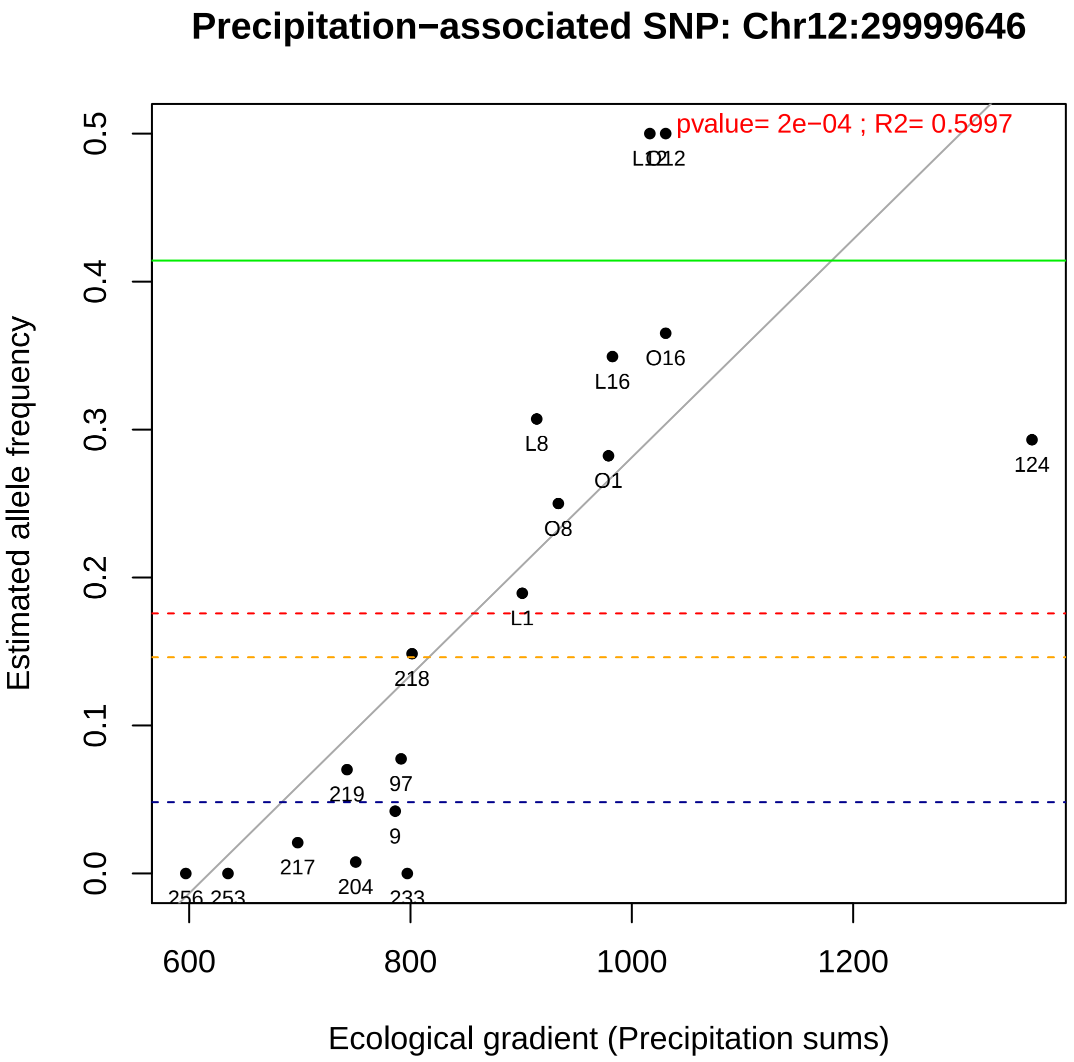
Precipitation−associated SNP: Chr12:29999646
pvalue= 2e−04 ; R2= 0.5997
Estimated allele frequency
Ecological gradient (Precipitation sums)
0.0
0.1
0.2
0.3
0.4
0.5
600
800
1000
1200
L12
O12
O16
L16
L8
O1
124
O8
L1
218
219
97
9
217
204
256
253
233

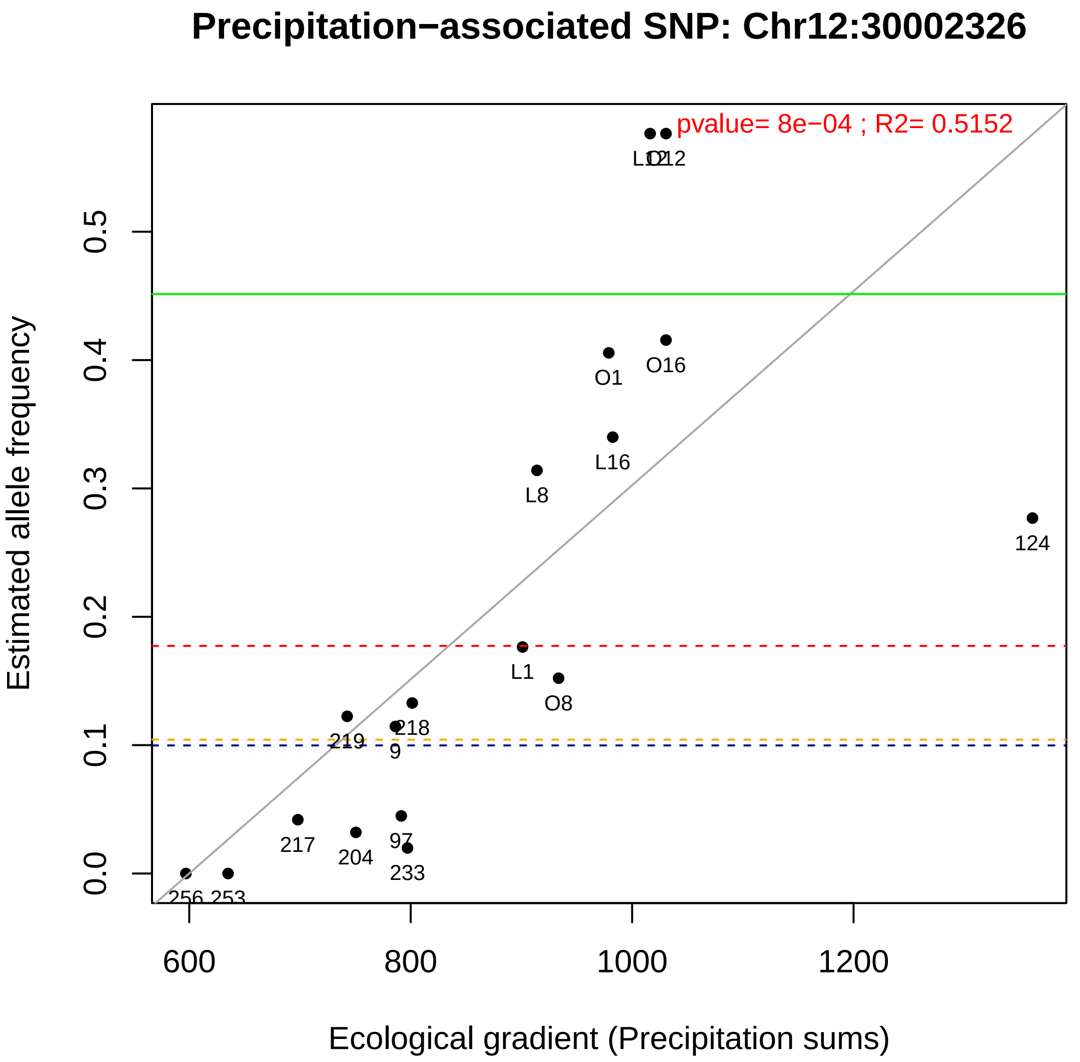

Precipitation−associated SNP: Chr12:30002326
pvalue= 8e−04 ; R2= 0.5152
Estimated allele frequency
Ecological gradient (Precipitation sums)
0.0
0.1
0.2
0.3
0.4
0.5
600
800
1000
1200
L12
O12
O16
O1
L16
L8
124
L1
O8
218
219
9
217
204
97
233
256
253

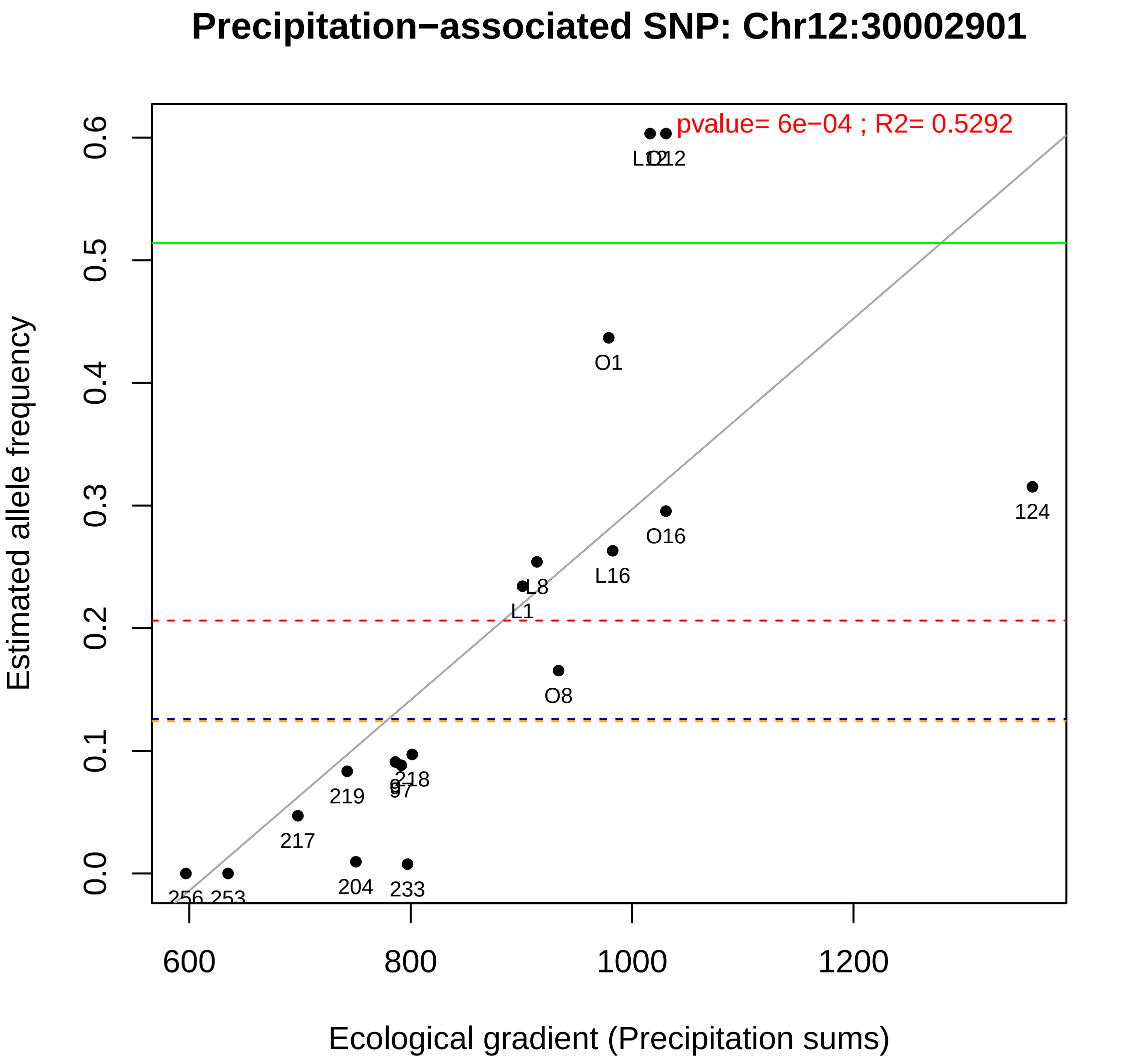
Precipitation−associated SNP: Chr12:30002901
pvalue= 6e−04 ; R2= 0.5292
L12
O12
O1
124
O16
L16
L8
L1
O8
218
219
97
217
204
233
256
253
Estimated allele frequency
0.0
0.1
0.2
0.3
0.4
0.5
0.6
600
800
1000
1200
Ecological gradient (Precipitation sums)

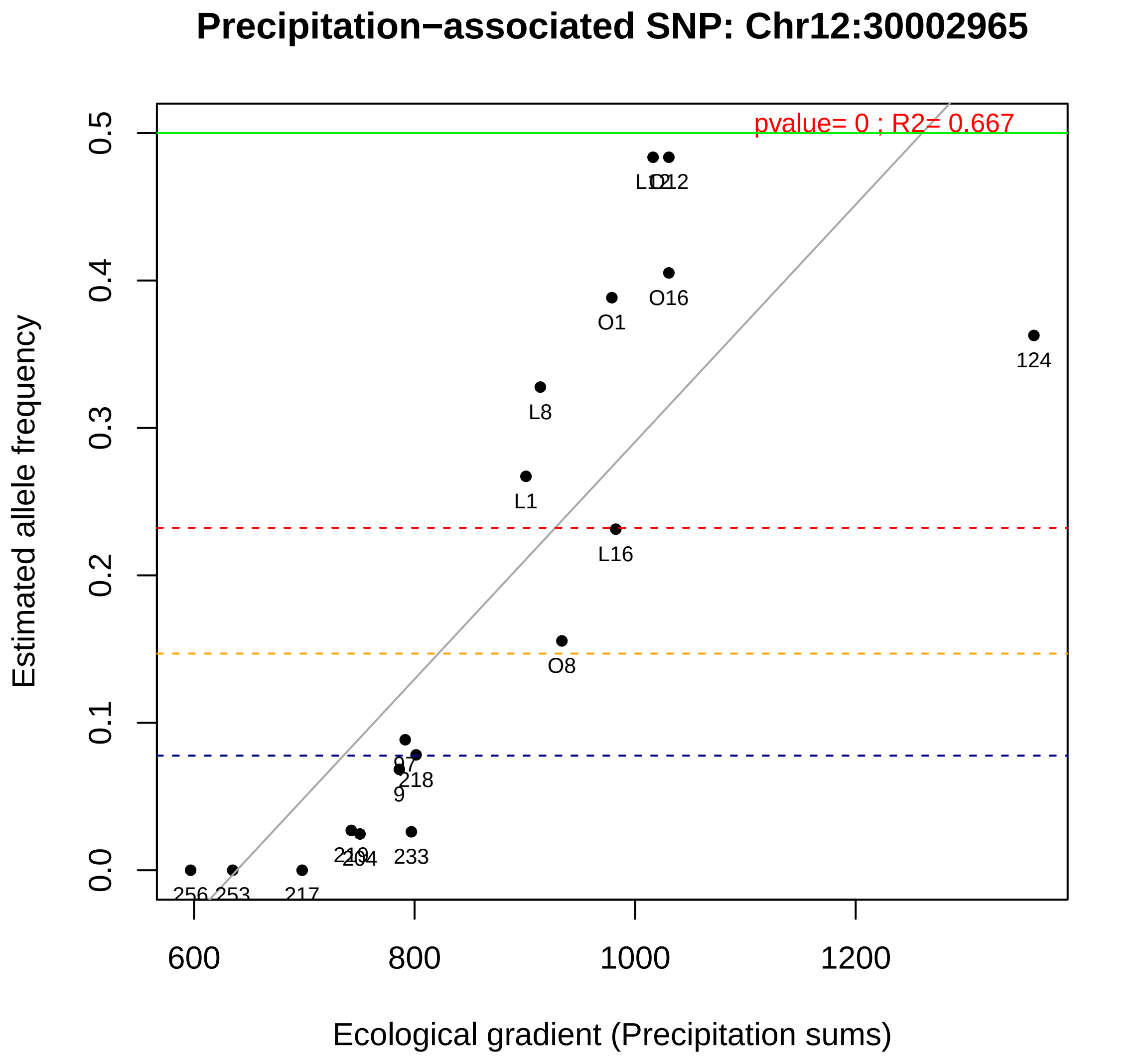
Precipitation−associated SNP: Chr12:30002965
pvalue= 0 ; R2= 0.667
Estimated allele frequency
Ecological gradient (Precipitation sums)
0.0
0.1
0.2
0.3
0.4
0.5
600
800
1000
1200
L12
O12
O16
O1
124
L8
L1
L16
O8
O7
218
9
210
204
233
256
253
217

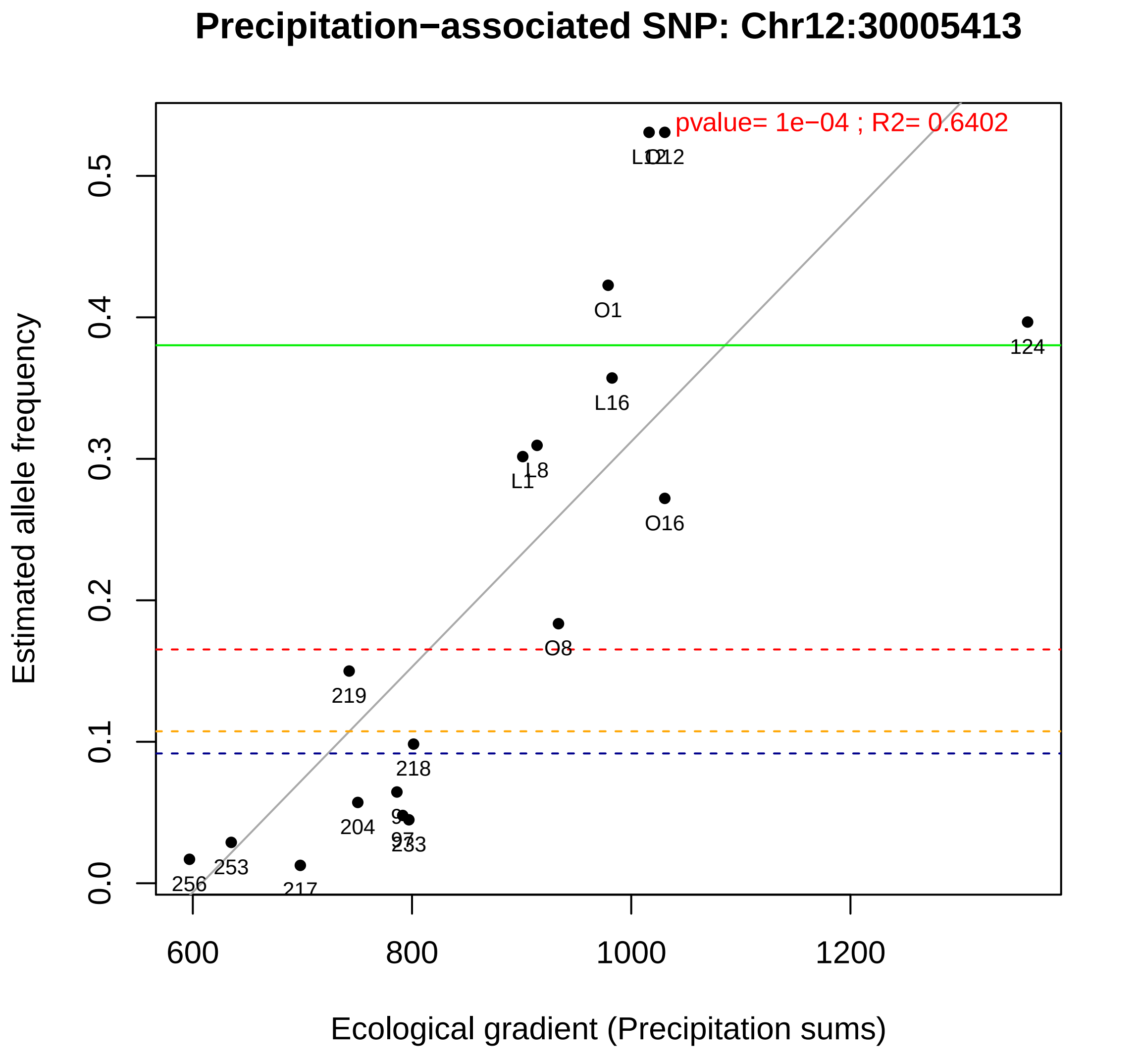
Precipitation−associated SNP: Chr12:30005413
pvalue= 1e−04 ; R2= 0.6402
Estimated allele frequency
Ecological gradient (Precipitation sums)
0.0
0.1
0.2
0.3
0.4
0.5
600
800
1000
1200
L12
O12
O1
124
L16
L8
L1
O16
O8
219
218
204
233
253
256
217

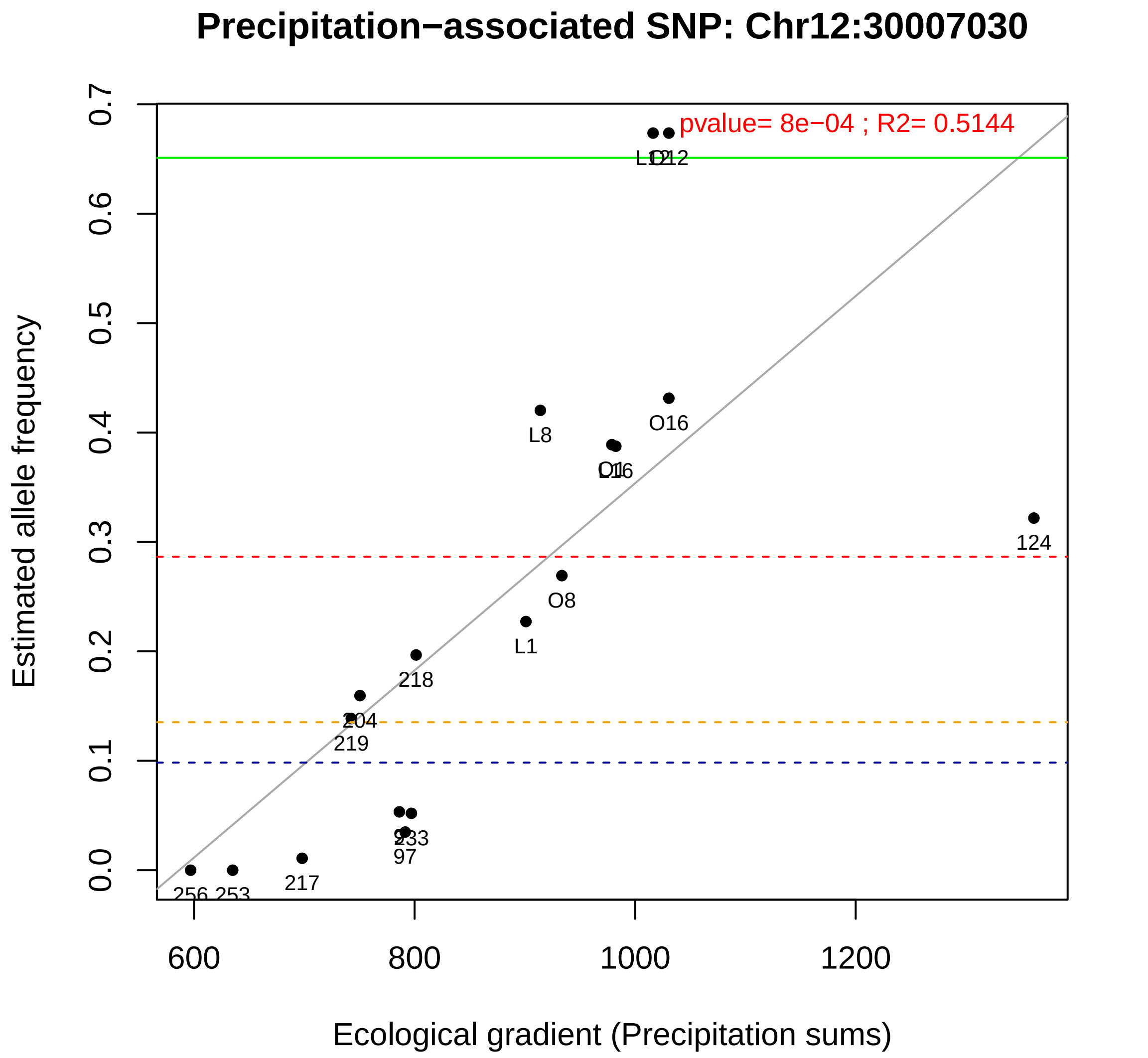

Precipitation−associated SNP: Chr12:30007030
pvalue= 8e−04 ; R2= 0.5144
Estimated allele frequency
Ecological gradient (Precipitation sums)
0.0
0.1
0.2
0.3
0.4
0.5
0.6
0.7
600
800
1000
1200
L8
O16
O8
L1
218
204
219
124
233
97
217
256
253

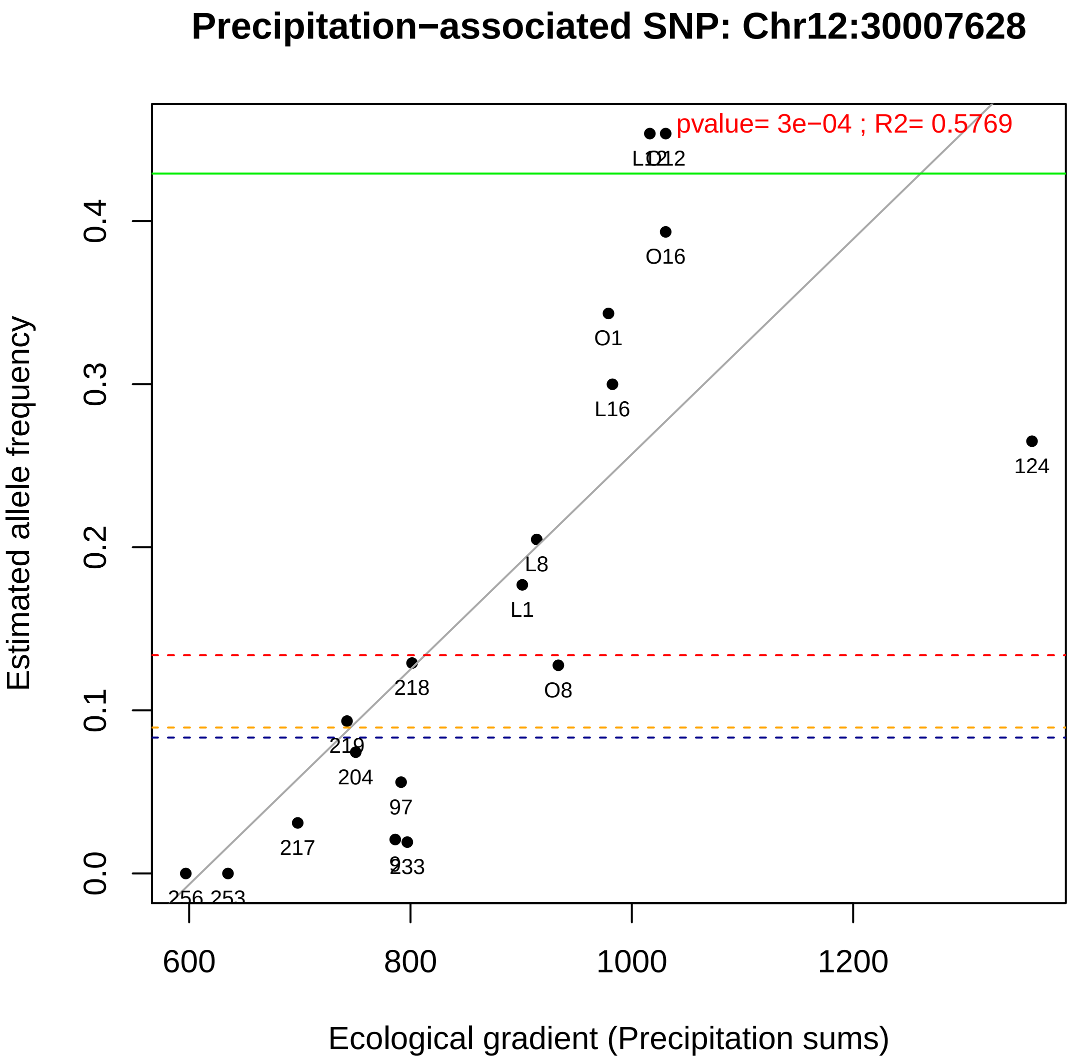
Precipitation−associated SNP: Chr12:30007628
pvalue= 3e−04 ; R2= 0.5769
Estimated allele frequency
Ecological gradient (Precipitation sums)
0.0
0.1
0.2
0.3
0.4
600
800
1000
1200
O16
O1
L16
124
L8
L1
218
O8
204
97
217
256
253

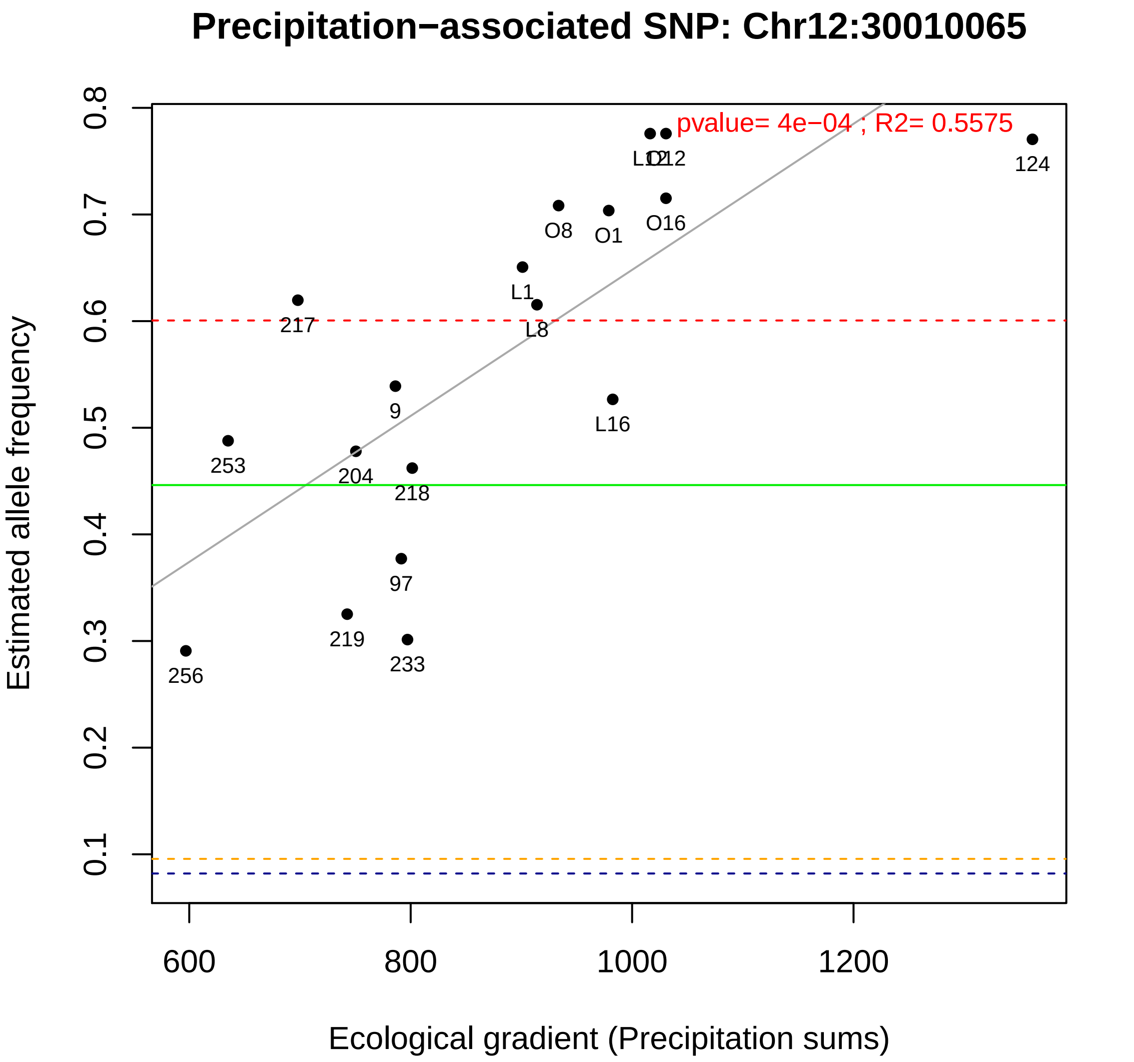
Precipitation−associated SNP: Chr12:30010065
pvalue= 4e−04 ; R2= 0.5575
Estimated allele frequency
0.1
0.2
0.3
0.4
0.5
0.6
0.7
0.8
Ecological gradient (Precipitation sums)
600
800
1000
1200
124
O16
O8
O1
L1
L8
217
9
L16
253
204
218
97
219
233
256

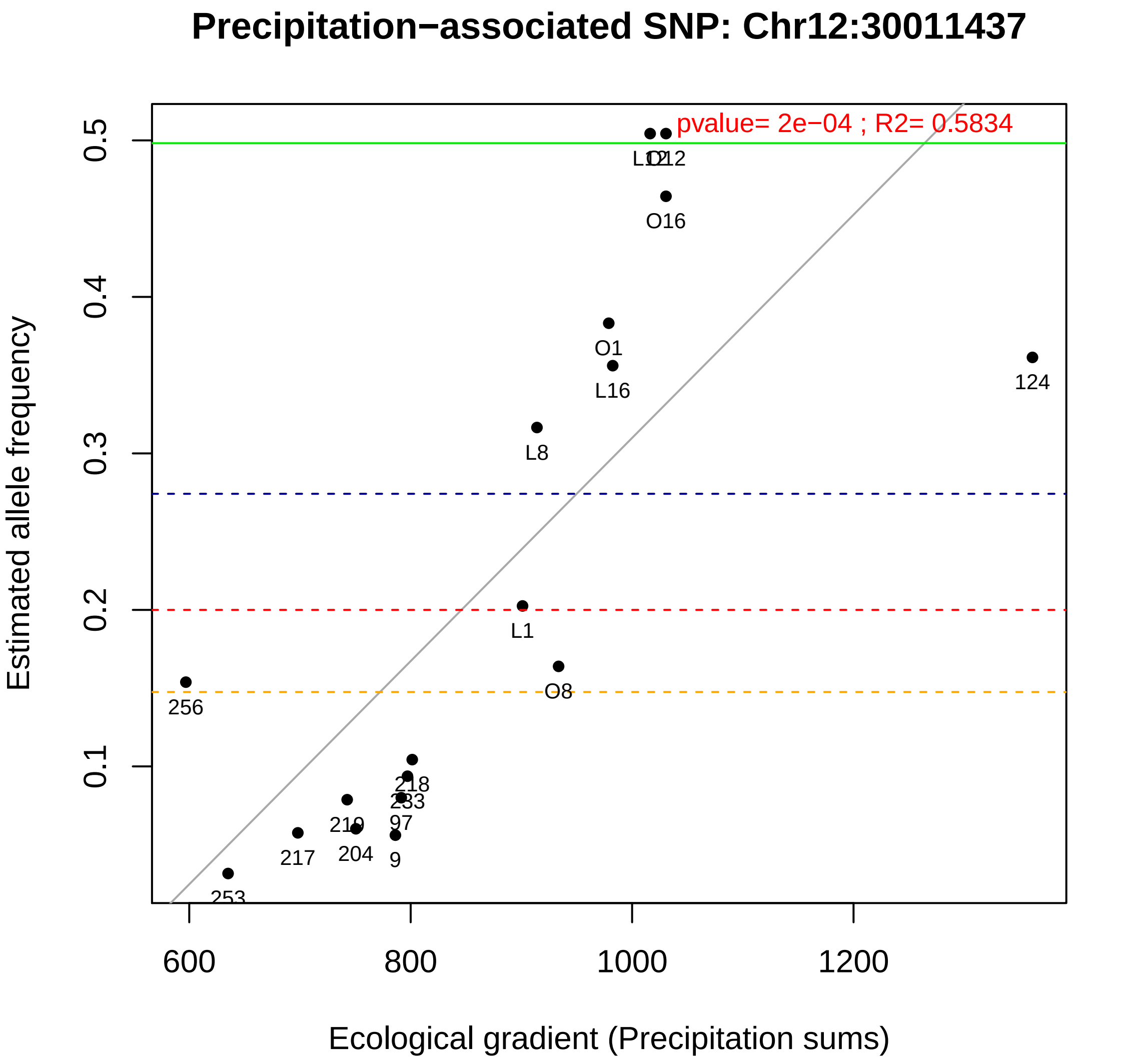

Precipitation−associated SNP: Chr12:30011437
pvalue= 2e−04 ; R2= 0.5834
Estimated allele frequency
Ecological gradient (Precipitation sums)
0.1
0.2
0.3
0.4
0.5
600
800
1000
1200
O12
O16
O1
L16
124
L8
L1
O8
256
218
233
210
97
204
217
9
253

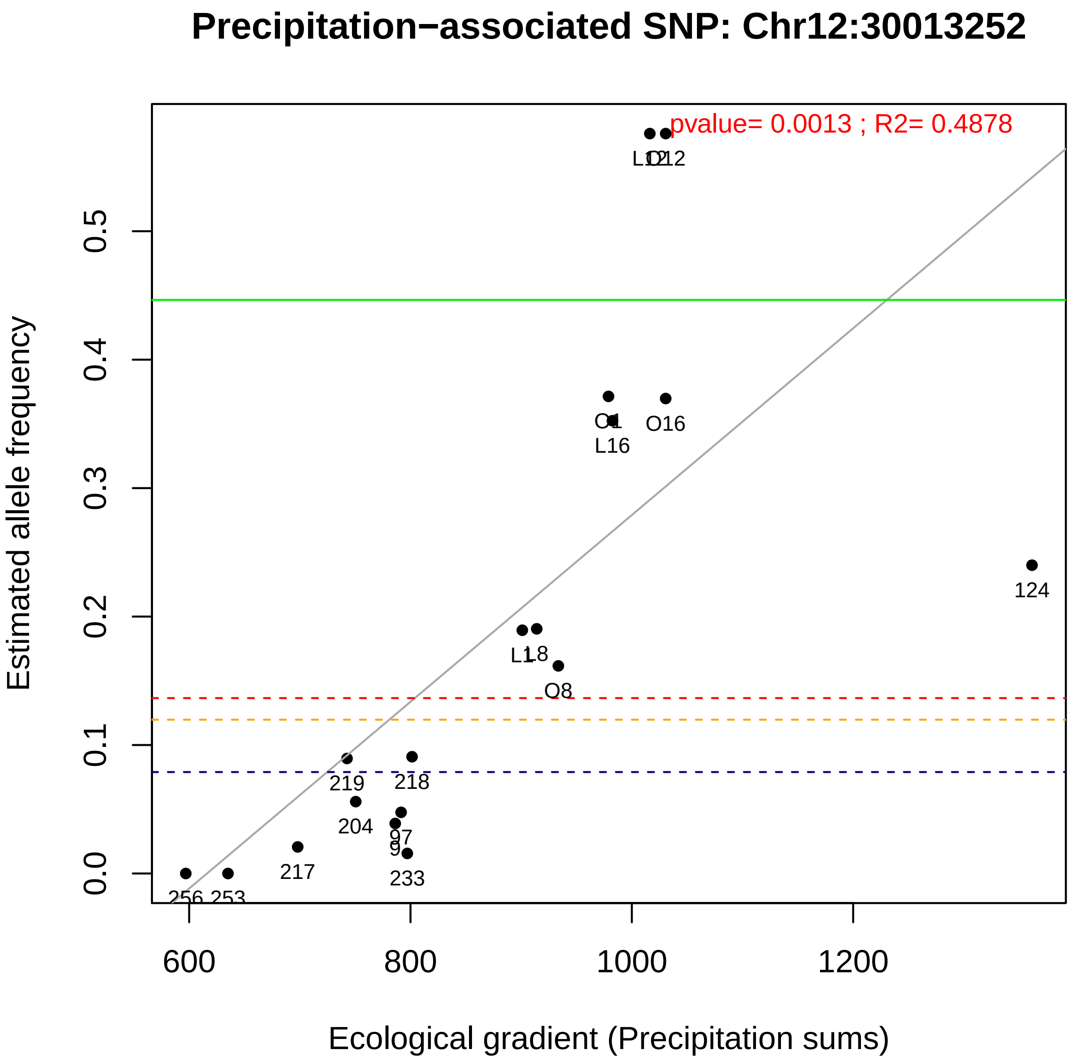
Precipitation−associated SNP: Chr12:30013252
pvalue= 0.0013 ; R2= 0.4878
Estimated allele frequency
0.0
0.1
0.2
0.3
0.4
0.5
600
800
1000
1200
Ecological gradient (Precipitation sums)
L16
O16
124
O8
219
218
204
217
233
256
253

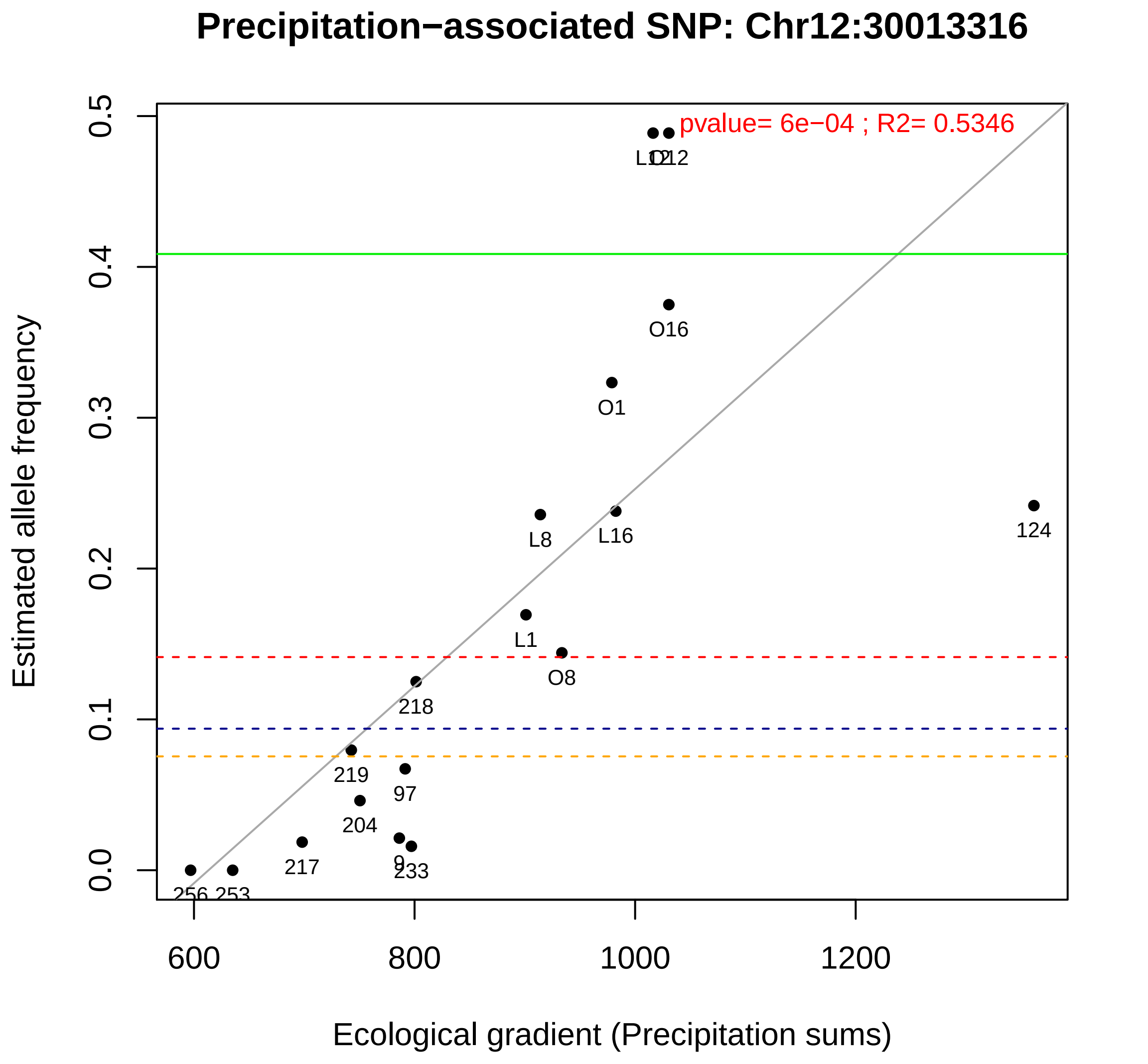
Precipitation−associated SNP: Chr12:30013316
pvalue= 6e−04 ; R2= 0.5346
Estimated allele frequency
Ecological gradient (Precipitation sums)
0.0
0.1
0.2
0.3
0.4
0.5
600
800
1000
1200
O16
O1
L8
L16
124
L1
O8
218
219
97
204
217
233
256
253

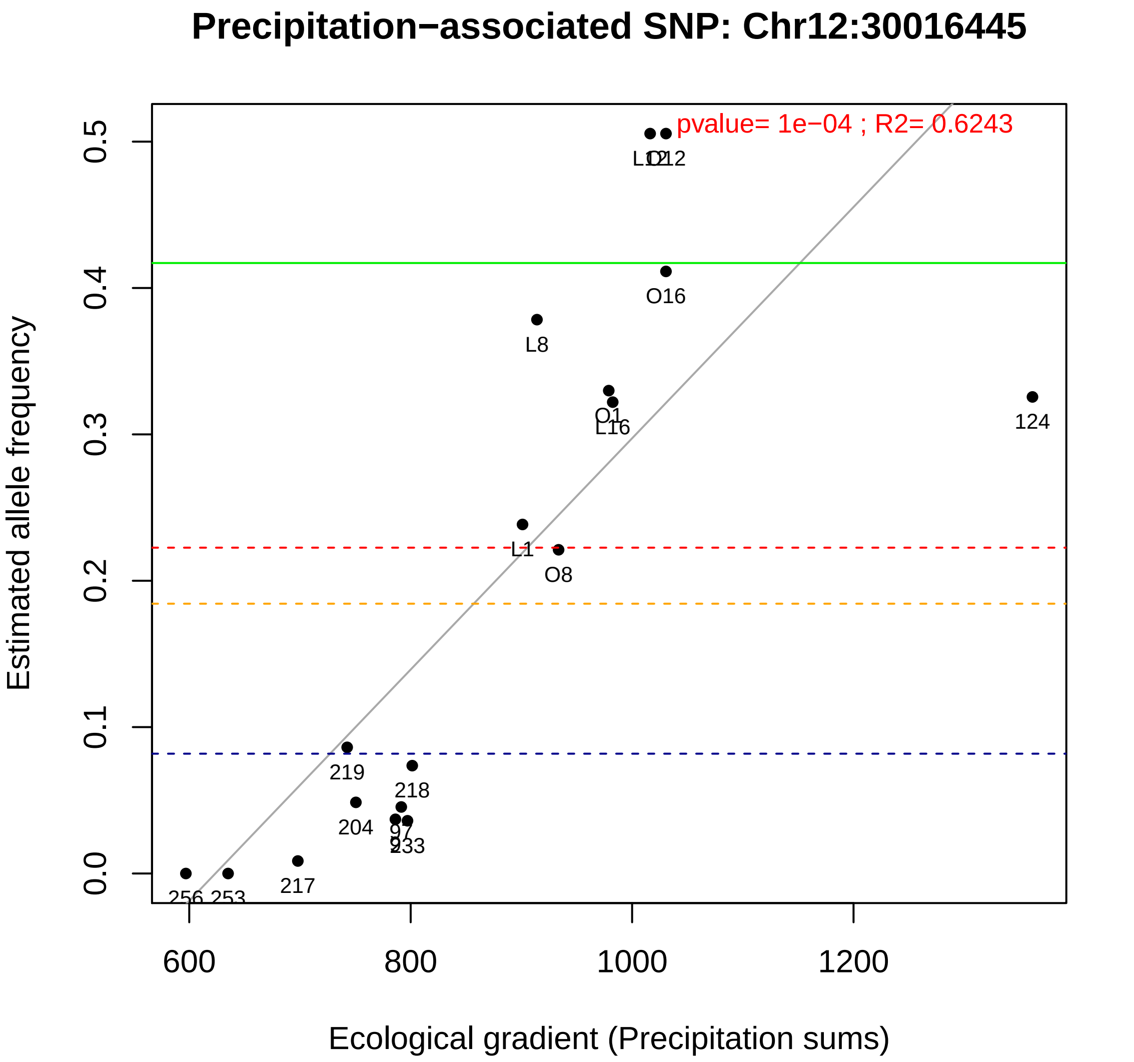

Precipitation−associated SNP: Chr12:30016445
pvalue= 1e−04 ; R2= 0.6243
Estimated allele frequency
Ecological gradient (Precipitation sums)
0.0
0.1
0.2
0.3
0.4
0.5
600
800
1000
1200
L10
O12
O16
L8
O1
L16
124
L1
O8
219
218
204
97
233
217
256
253

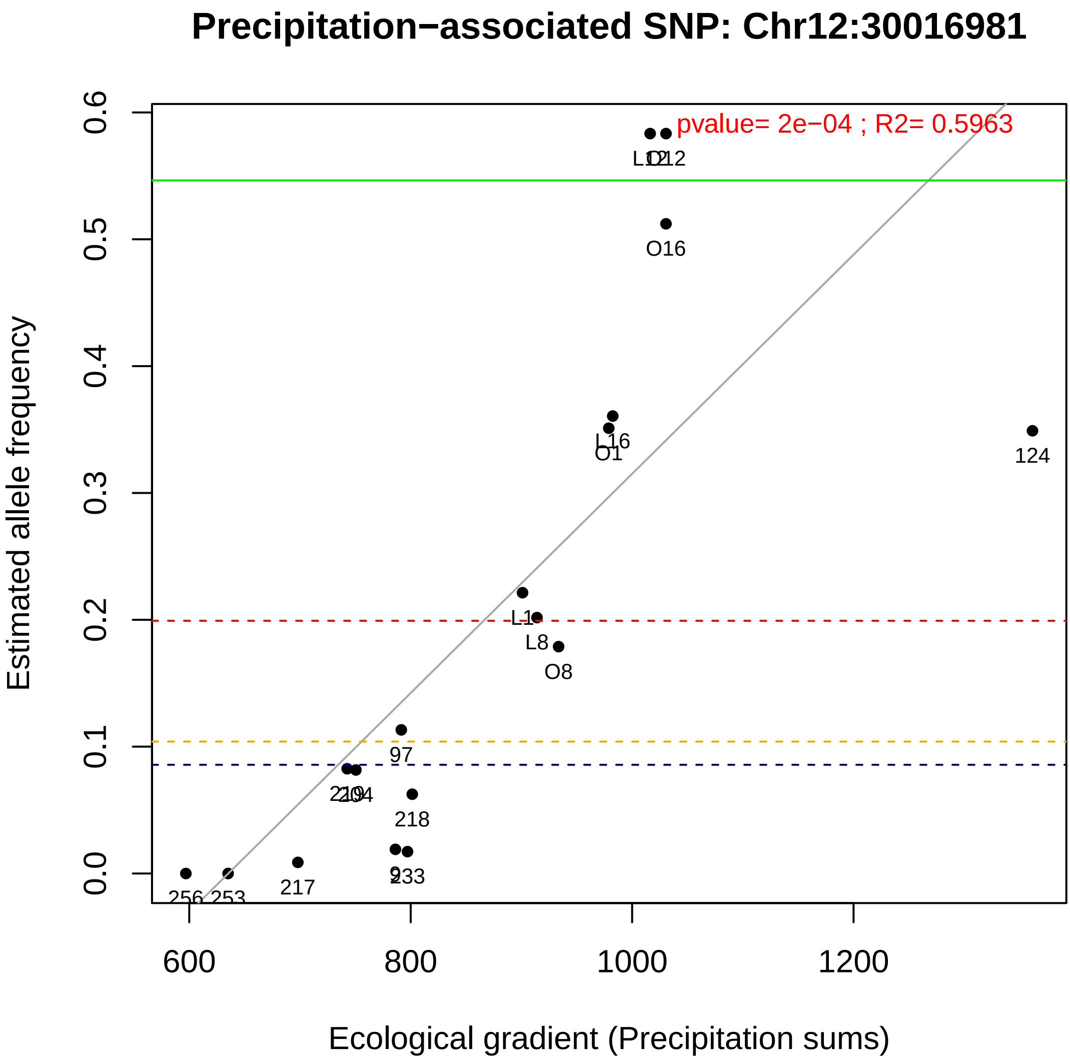
Precipitation−associated SNP: Chr12:30016981
pvalue= 2e−04 ; R2= 0.5963
Estimated allele frequency
Ecological gradient (Precipitation sums)
0.0
0.1
0.2
0.3
0.4
0.5
0.6
600
800
1000
1200
L12
O12
O16
L16
O1
124
L1
L8
O8
97
210
204
218
217
233
256
253

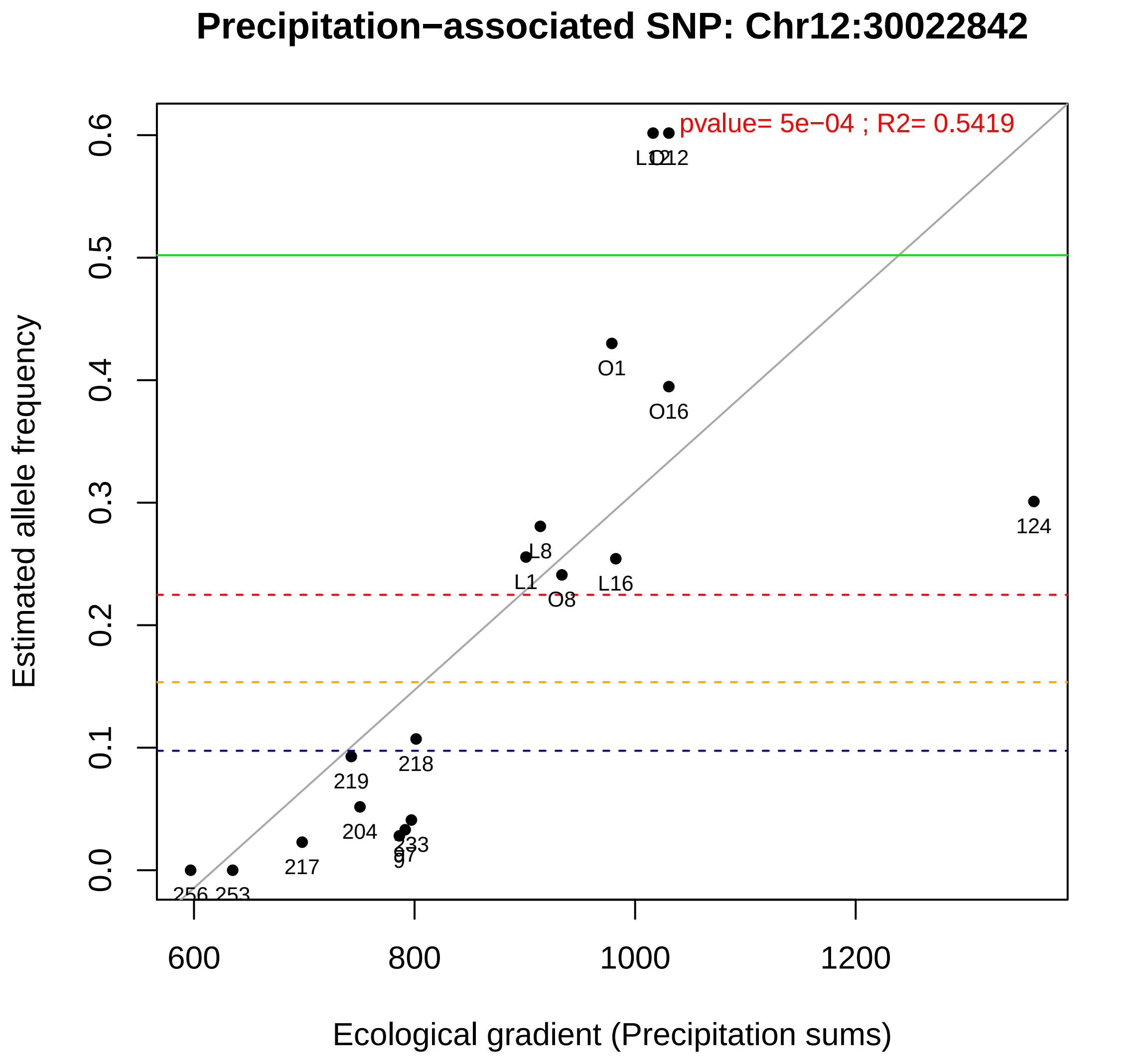

Precipitation−associated SNP: Chr12:30022842
pvalue= 5e−04 ; R2= 0.5419
Estimated allele frequency
Ecological gradient (Precipitation sums)
0.0
0.1
0.2
0.3
0.4
0.5
0.6
600
800
1000
1200
O12
O1
O16
124
L8
L1
L16
O8
218
219
204
233
217
256
253

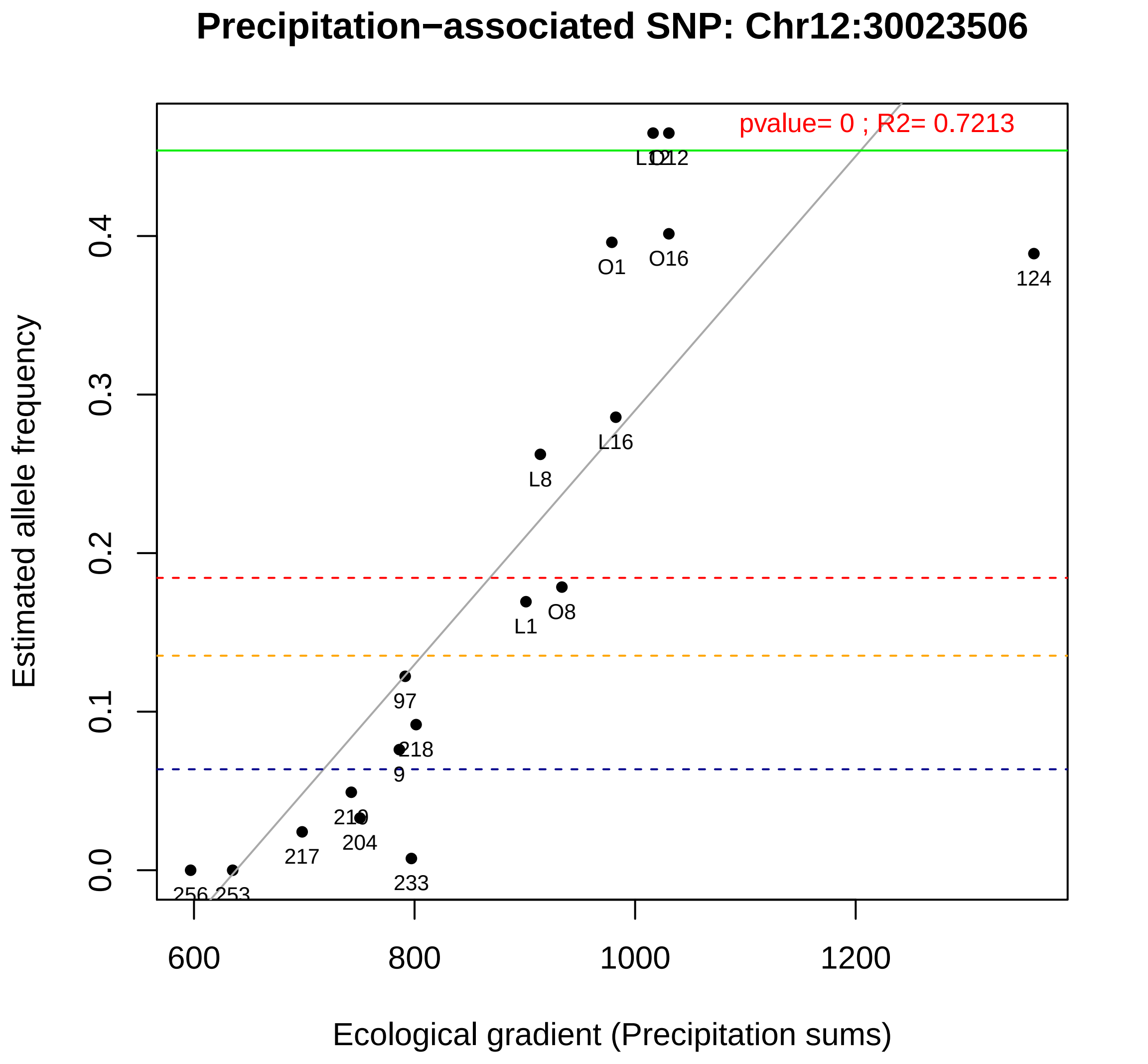

Precipitation−associated SNP: Chr12:30023506
pvalue= 0 ; R2= 0.7213
L12
O12
O1
O16
124
L16
L8
L1
O8
97
218
9
210
204
217
233
256
253
0.0
0.1
0.2
0.3
0.4
600
800
1000
1200
Estimated allele frequency
Ecological gradient (Precipitation sums)

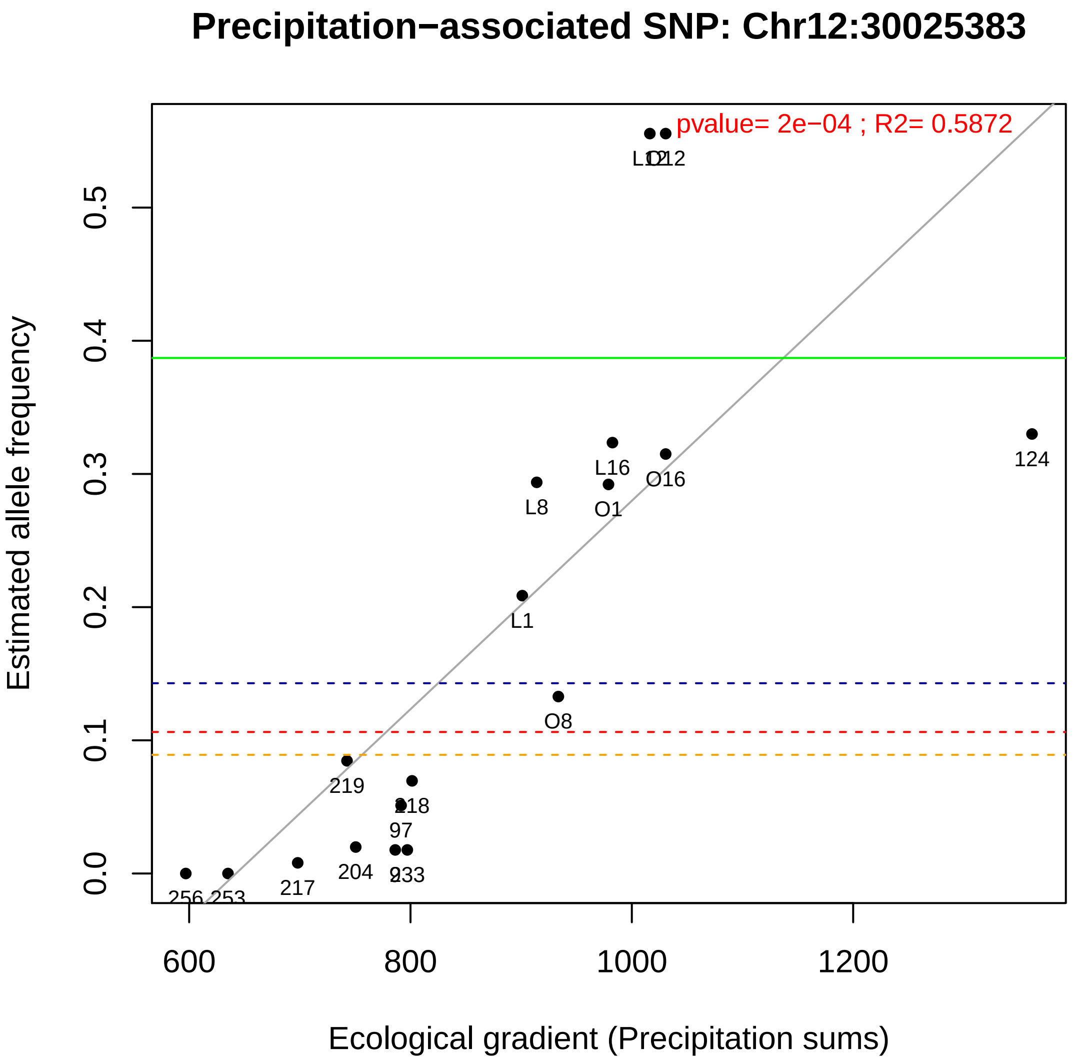

Precipitation−associated SNP: Chr12:30025383
pvalue= 2e−04 ; R2= 0.5872
Estimated allele frequency
Ecological gradient (Precipitation sums)
0.0
0.1
0.2
0.3
0.4
0.5
600
800
1000
1200
L12
O12
L16
O16
124
L8
O1
L1
O8
219
218
97
204
233
217
256
253

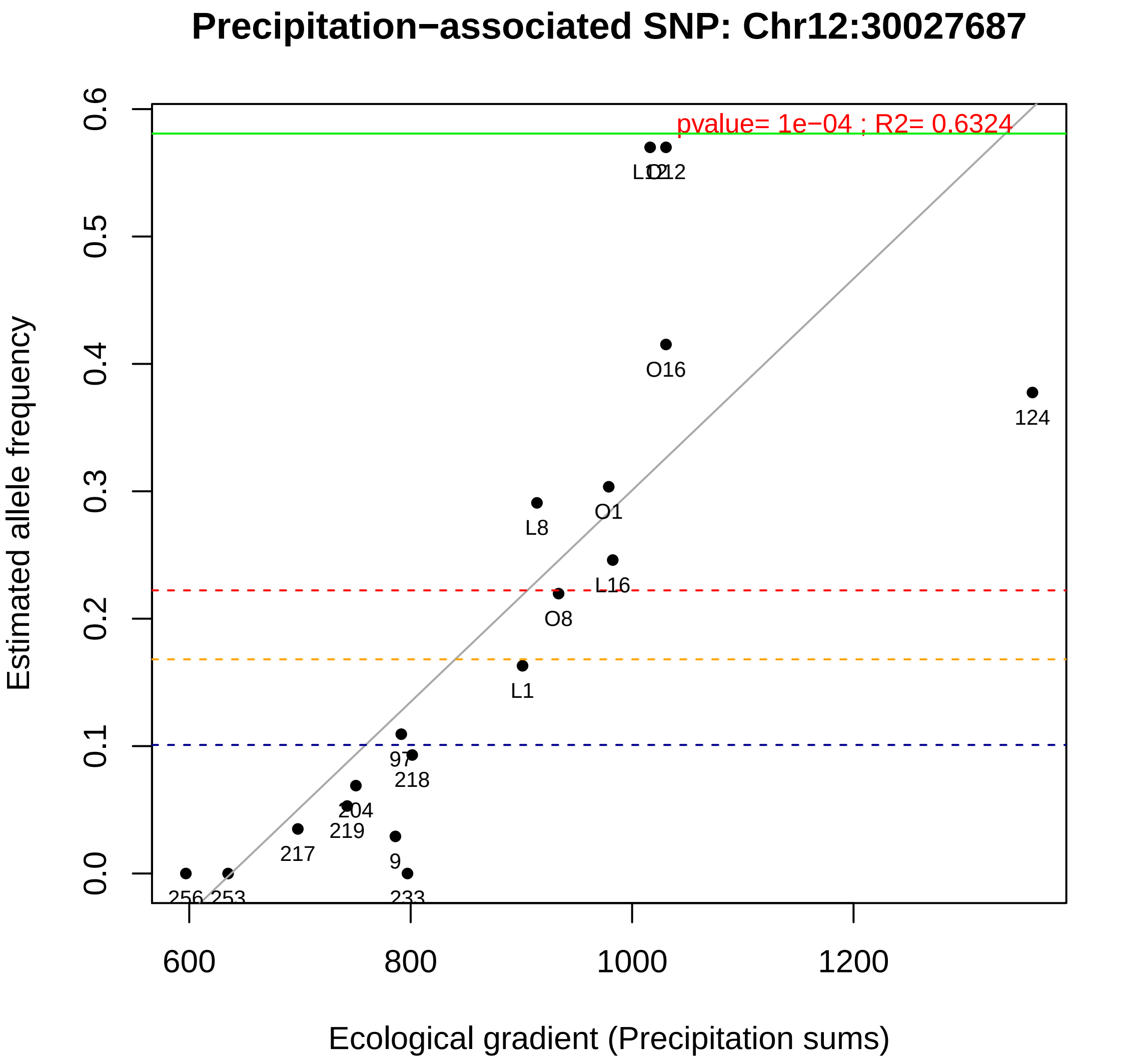

Precipitation−associated SNP: Chr12:30027687
pvalue= 1e−04 ; R2= 0.6324
Estimated allele frequency
Ecological gradient (Precipitation sums)
0.0
0.1
0.2
0.3
0.4
0.5
0.6
600
800
1000
1200
L12
O12
O16
124
O1
L8
L16
O8
L1
97
218
204
219
217
9
233
256
253

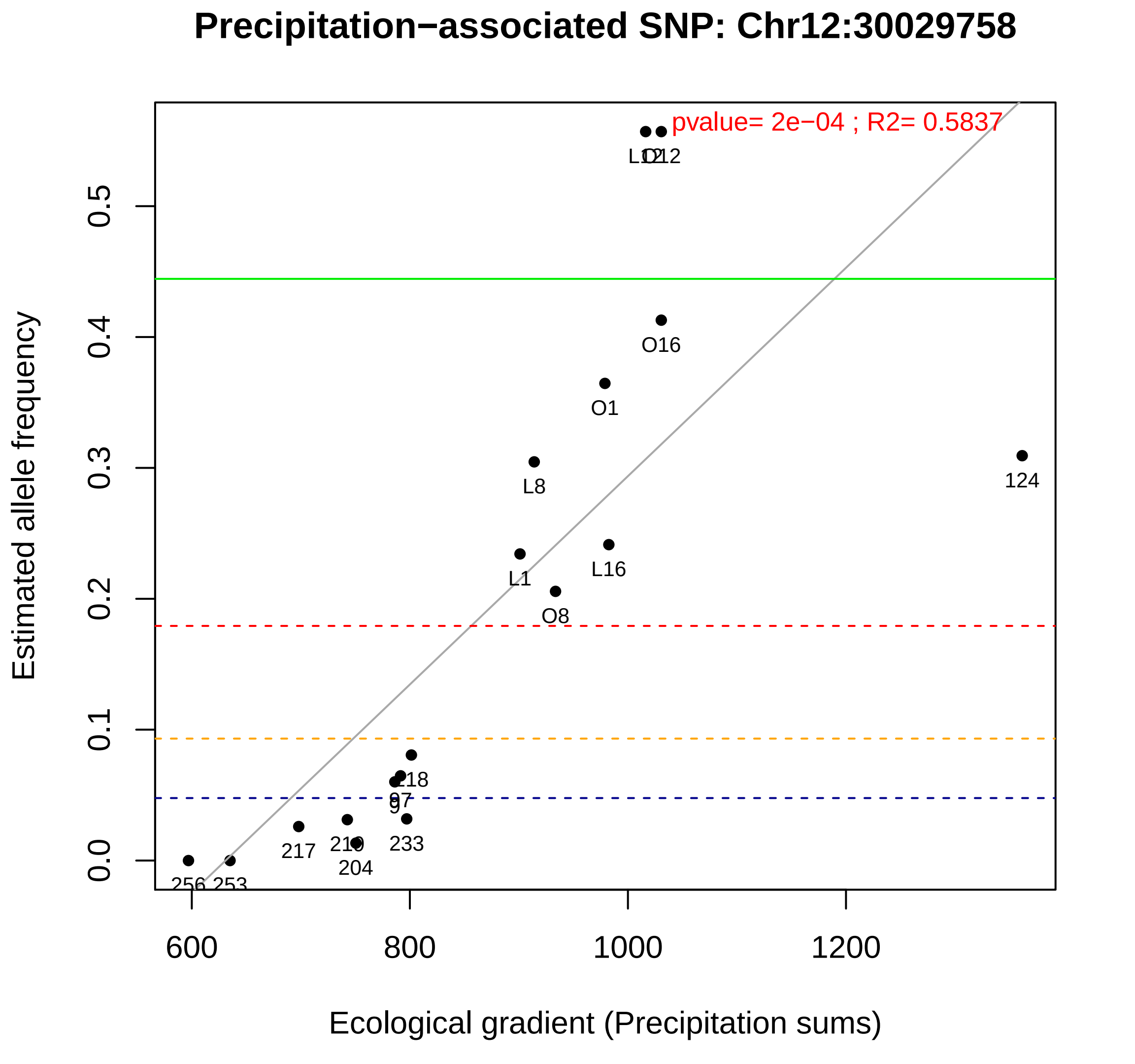
Precipitation−associated SNP: Chr12:30029758
pvalue= 2e−04 ; R2= 0.5837
Estimated allele frequency
Ecological gradient (Precipitation sums)
0.0
0.1
0.2
0.3
0.4
0.5
600
800
1000
1200
O16
O1
L8
124
L1
L16
O8
218
97
233
217
204
256
253

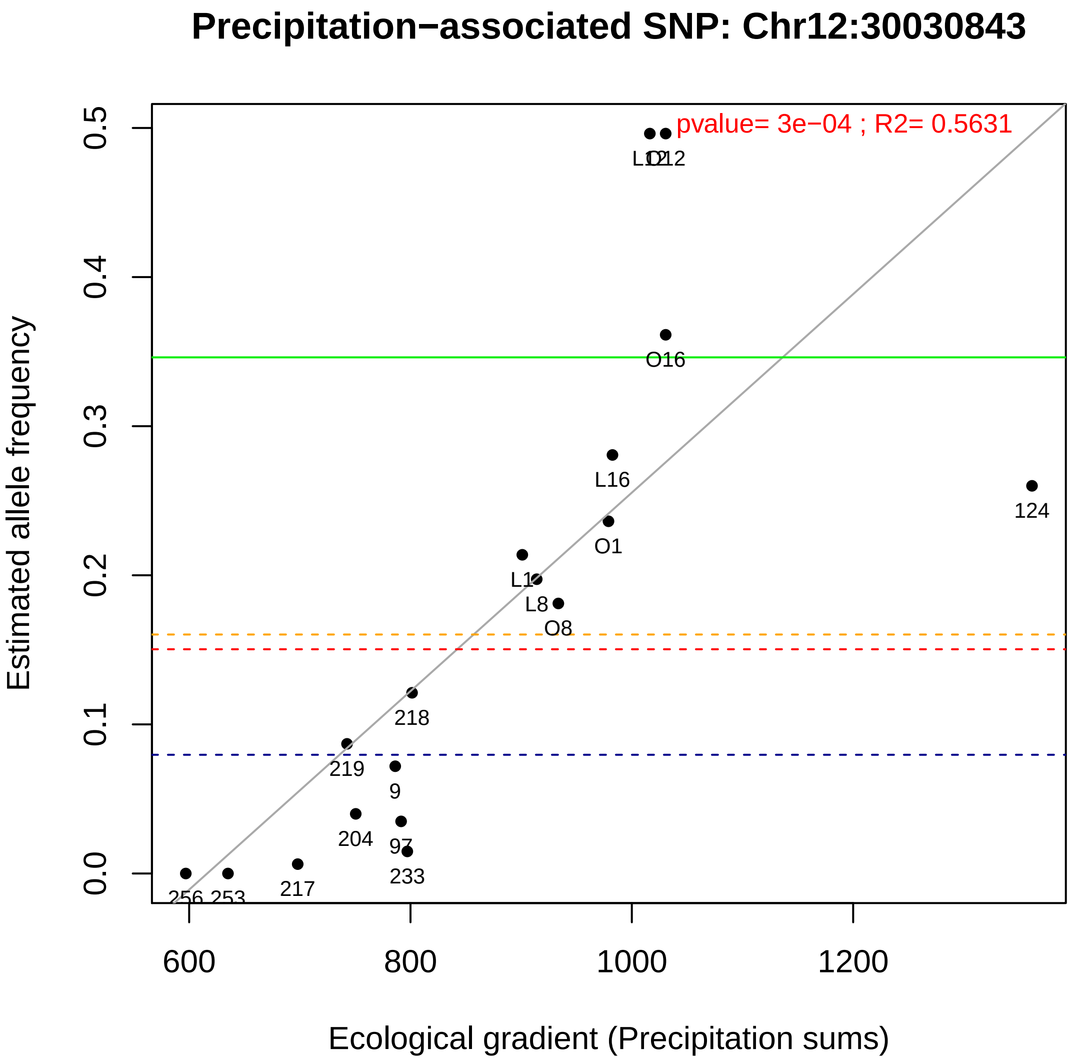

Precipitation−associated SNP: Chr12:30030843
pvalue= 3e−04 ; R2= 0.5631
Estimated allele frequency
0.0
0.1
0.2
0.3
0.4
0.5
600
800
1000
1200
Ecological gradient (Precipitation sums)
L12
O12
O16
L16
124
O1
L1
L8
O8
218
219
9
204
97
233
217
256
253

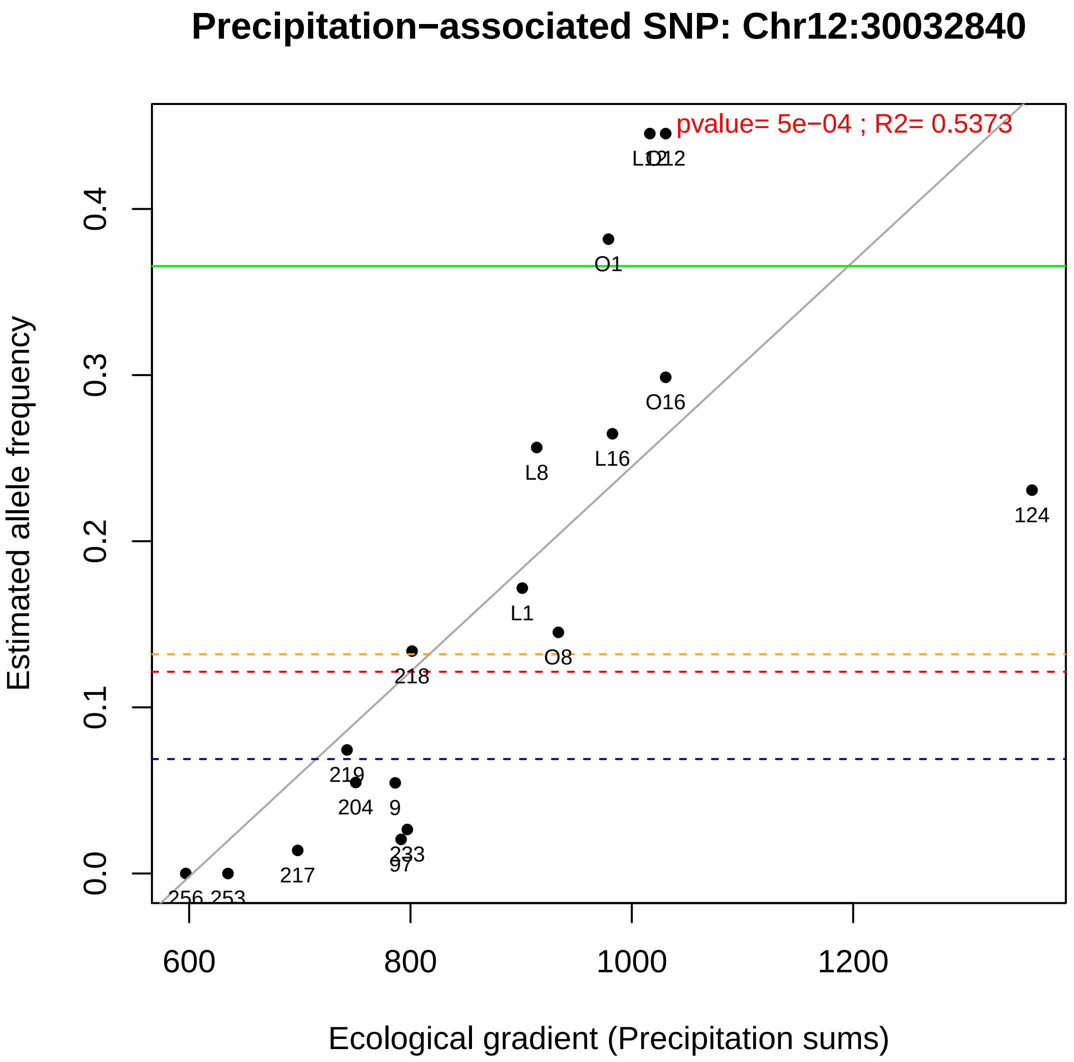
Precipitation−associated SNP: Chr12:30032840
pvalue= 5e−04 ; R2= 0.5373
Estimated allele frequency
Ecological gradient (Precipitation sums)
0.0
0.1
0.2
0.3
0.4
600
800
1000
1200
L12
O12
O1
O16
L16
L8
124
L1
O8
218
219
204
9
233
97
217
256
253

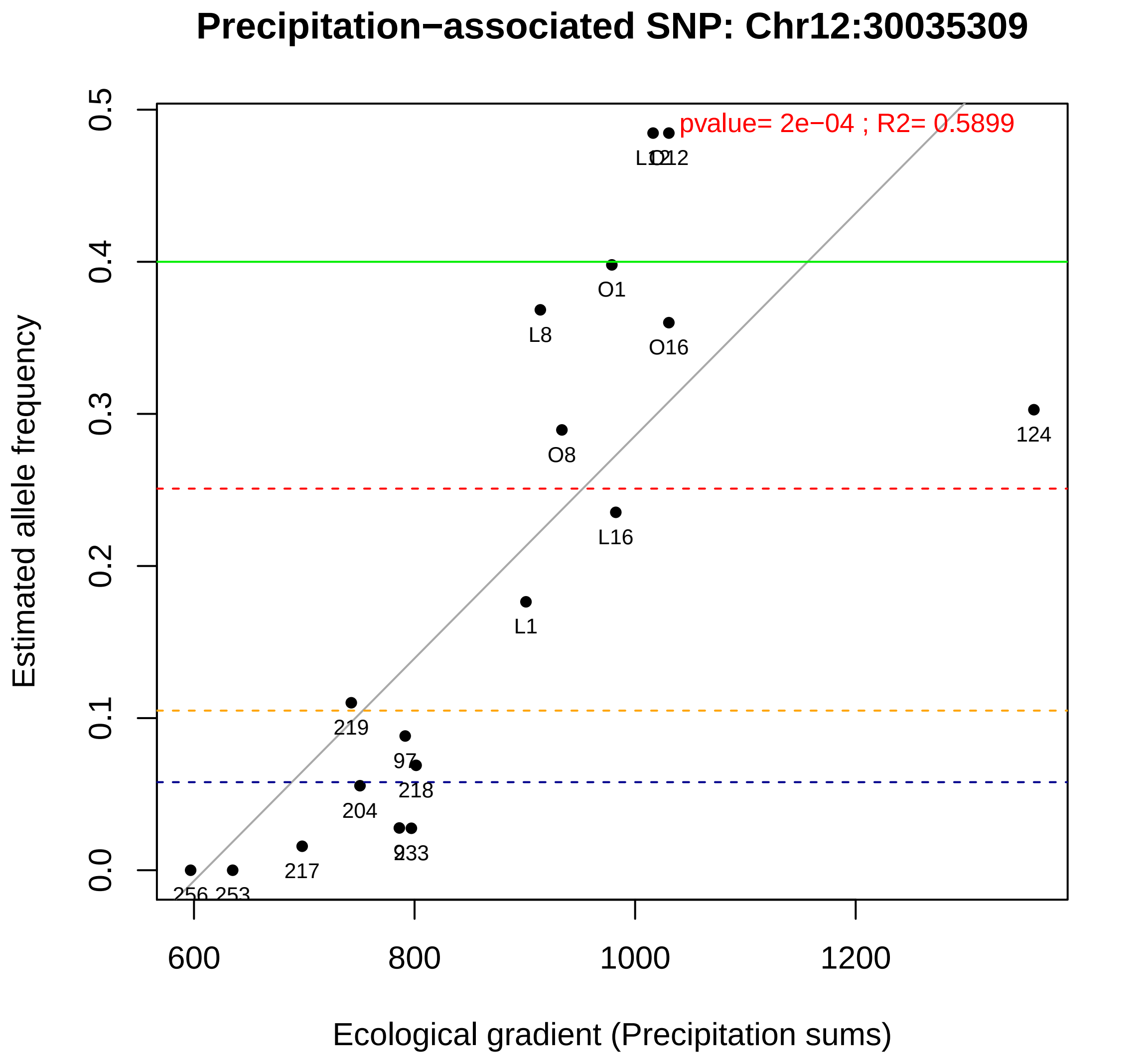
Precipitation−associated SNP: Chr12:30035309
pvalue= 2e−04 ; R2= 0.5899
Estimated allele frequency
0.0
0.1
0.2
0.3
0.4
0.5
600
800
1000
1200
Ecological gradient (Precipitation sums)
O1
L8
O16
124
O8
L16
L1
219
97
218
204
217
256
253

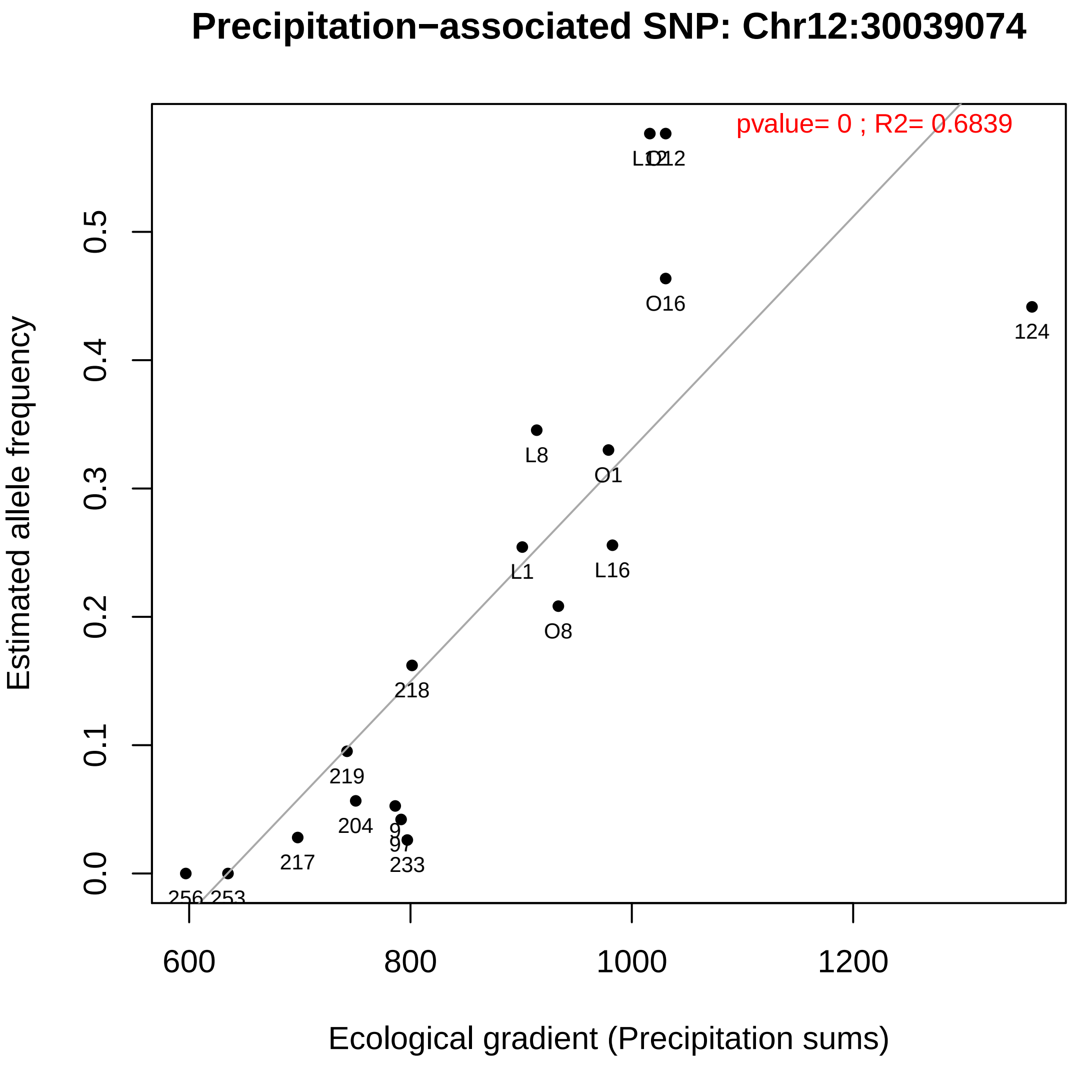

Precipitation−associated SNP: Chr12:30039074
pvalue= 0 ; R2= 0.6839
Estimated allele frequency
Ecological gradient (Precipitation sums)
0.0
0.1
0.2
0.3
0.4
0.5
600
800
1000
1200
L12
O12
O16
124
L8
O1
L1
L16
O8
218
219
204
9
97
233
217
256
253

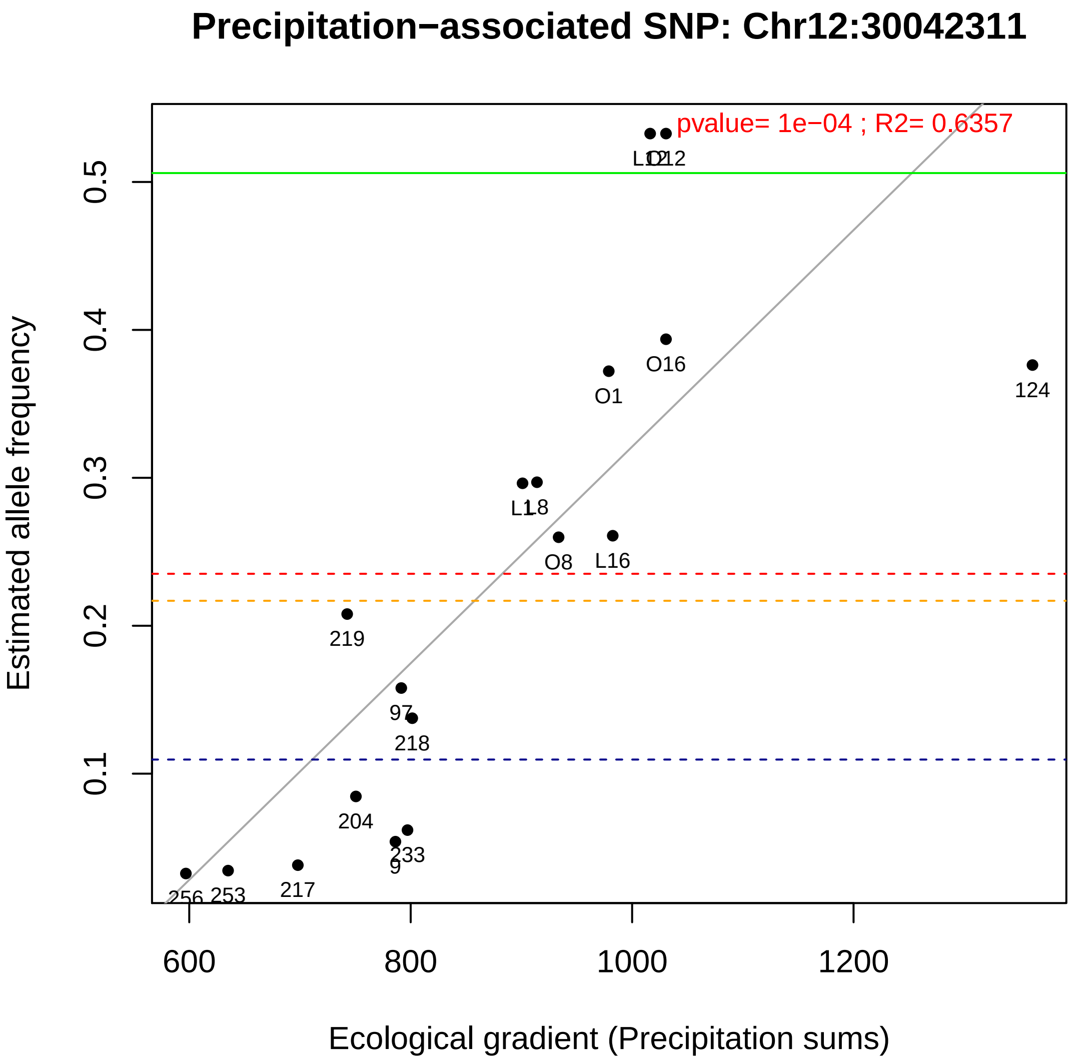
Precipitation−associated SNP: Chr12:30042311
pvalue= 1e−04 ; R2= 0.6357
Estimated allele frequency
0.1
0.2
0.3
0.4
0.5
600
800
1000
1200
Ecological gradient (Precipitation sums)
O16
O1
124
O8
L16
219
97
218
204
233
9
256
253
217

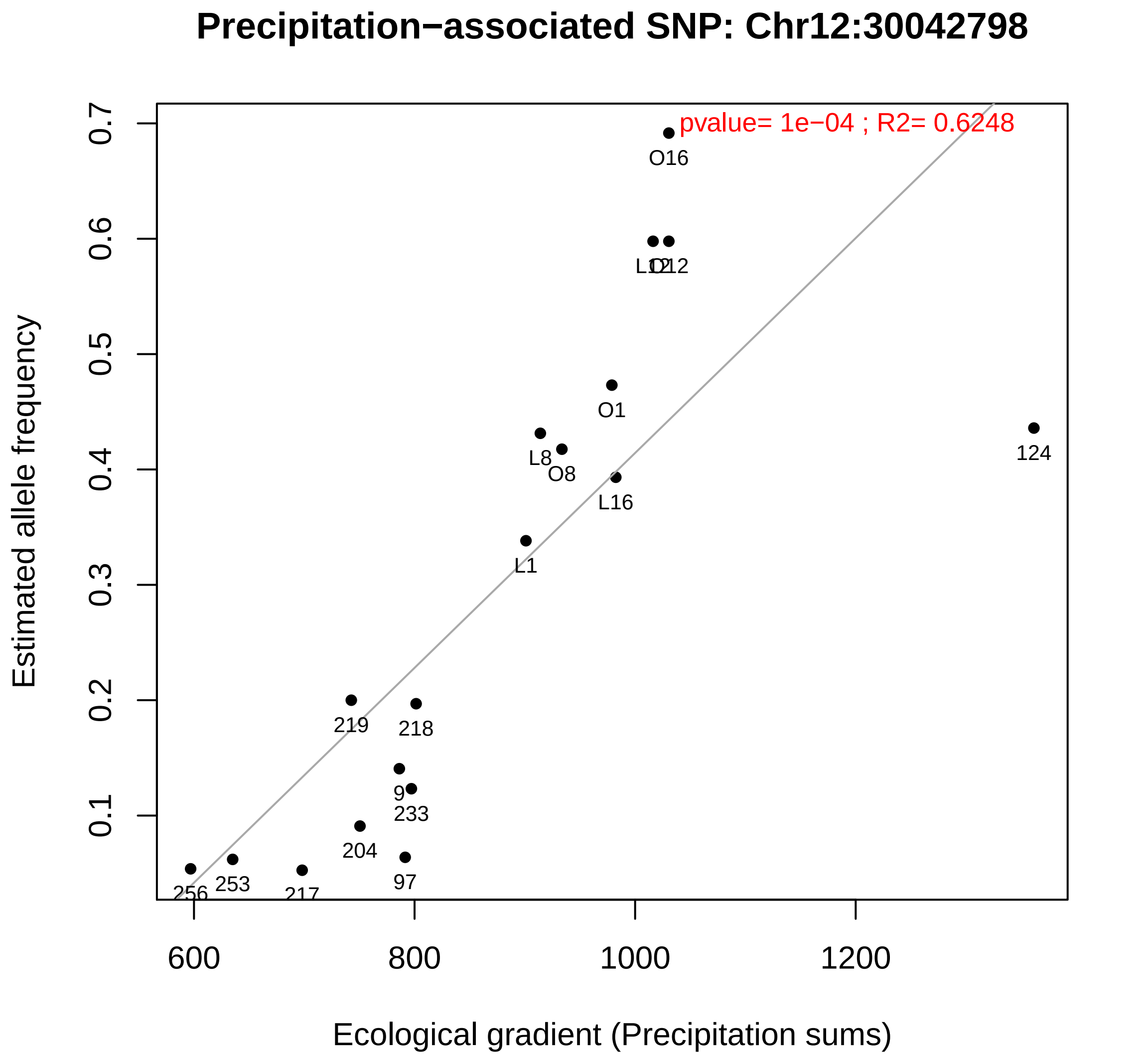
Precipitation−associated SNP: Chr12:30042798
pvalue= 1e−04 ; R2= 0.6248
O16
L12
O12
O1
124
L8
O8
L16
L1
219
218
9
233
204
97
256
253
217
0.1
0.2
0.3
0.4
0.5
0.6
0.7
600
800
1000
1200
Estimated allele frequency
Ecological gradient (Precipitation sums)

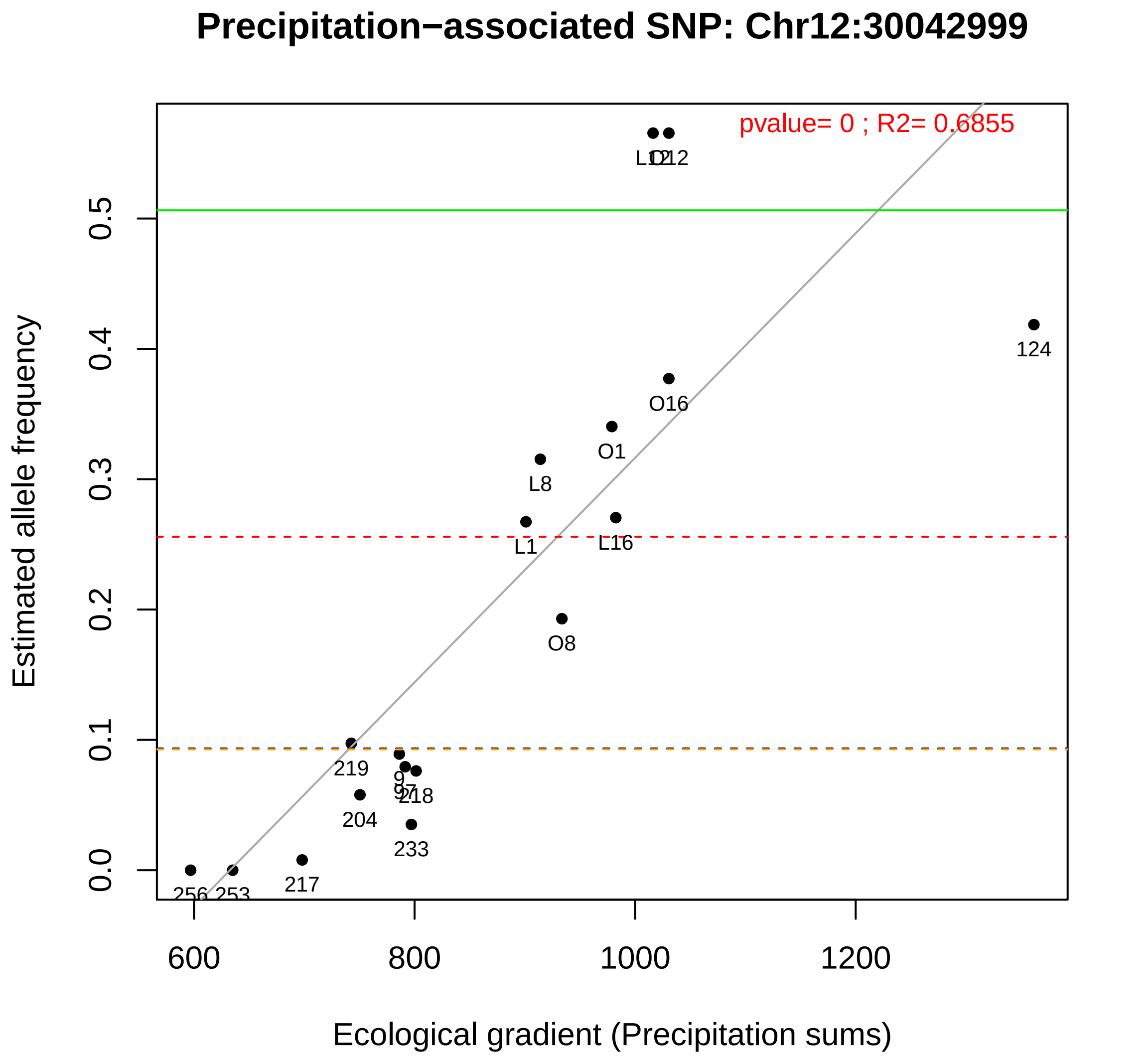
Precipitation−associated SNP: Chr12:30042999
pvalue= 0 ; R2= 0.6855
Estimated allele frequency
Ecological gradient (Precipitation sums)
0.0
0.1
0.2
0.3
0.4
0.5
600
800
1000
1200
L12
O12
124
O16
O1
L8
L1
L16
O8
219
9
97
218
204
233
217
256
253

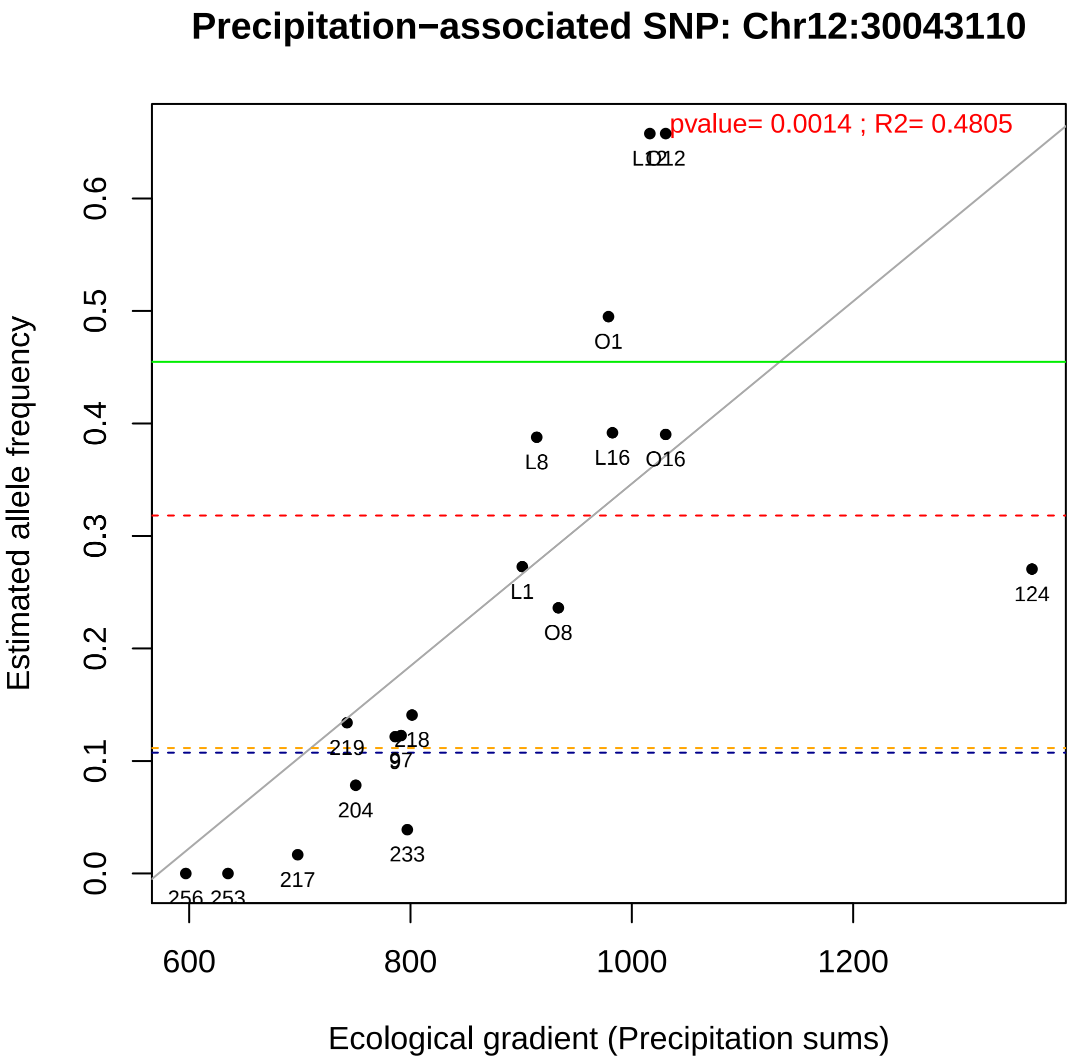

Precipitation−associated SNP: Chr12:30043110
pvalue= 0.0014 ; R2= 0.4805
Estimated allele frequency
Ecological gradient (Precipitation sums)
0.0
0.1
0.2
0.3
0.4
0.5
0.6
600
800
1000
1200
O1
L8
L16
O16
L1
O8
124
219
218
204
233
217
256
253

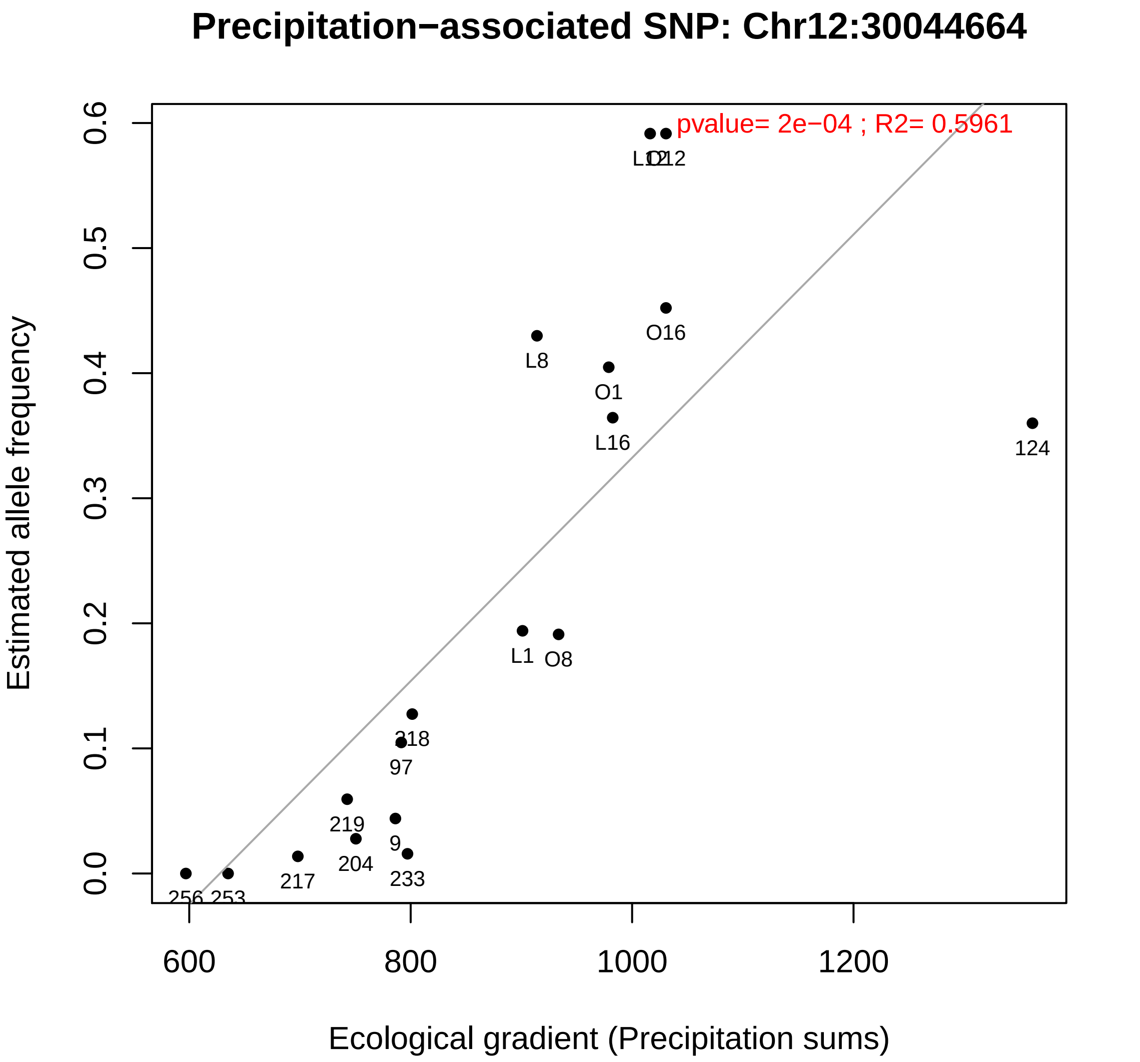

Precipitation−associated SNP: Chr12:30044664
pvalue= 2e−04 ; R2= 0.5961
Estimated allele frequency
Ecological gradient (Precipitation sums)
0.0
0.1
0.2
0.3
0.4
0.5
0.6
600
800
1000
1200
L12
O12
O16
L8
O1
L16
124
L1
O8
218
97
219
9
204
233
217
256
253

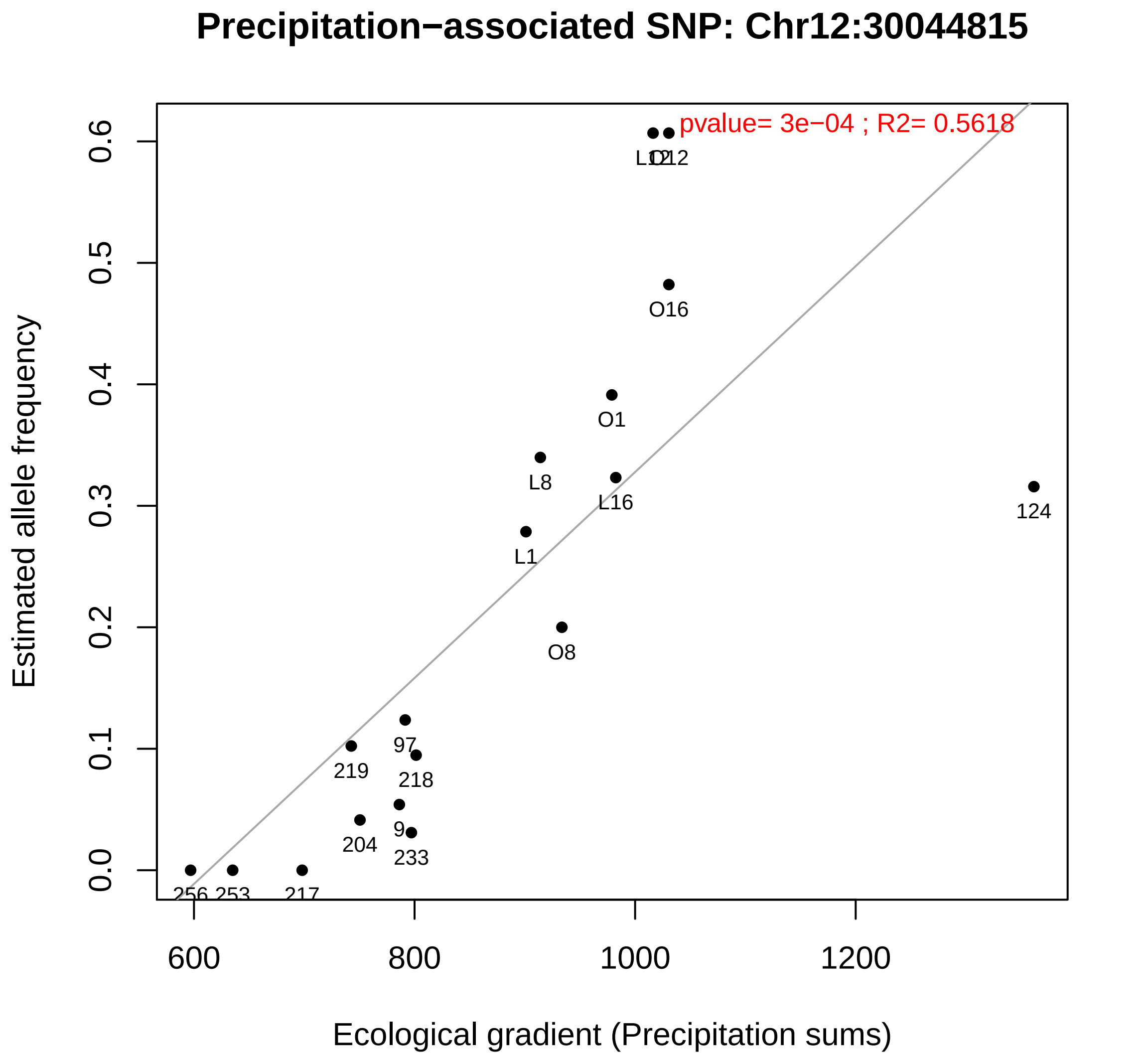
Precipitation−associated SNP: Chr12:30044815
pvalue= 3e−04 ; R2= 0.5618
Estimated allele frequency
Ecological gradient (Precipitation sums)
0.0
0.1
0.2
0.3
0.4
0.5
0.6
600
800
1000
1200
O12
O16
O1
L8
L16
124
L1
O8
97
219
218
204
9
233
256
253
217

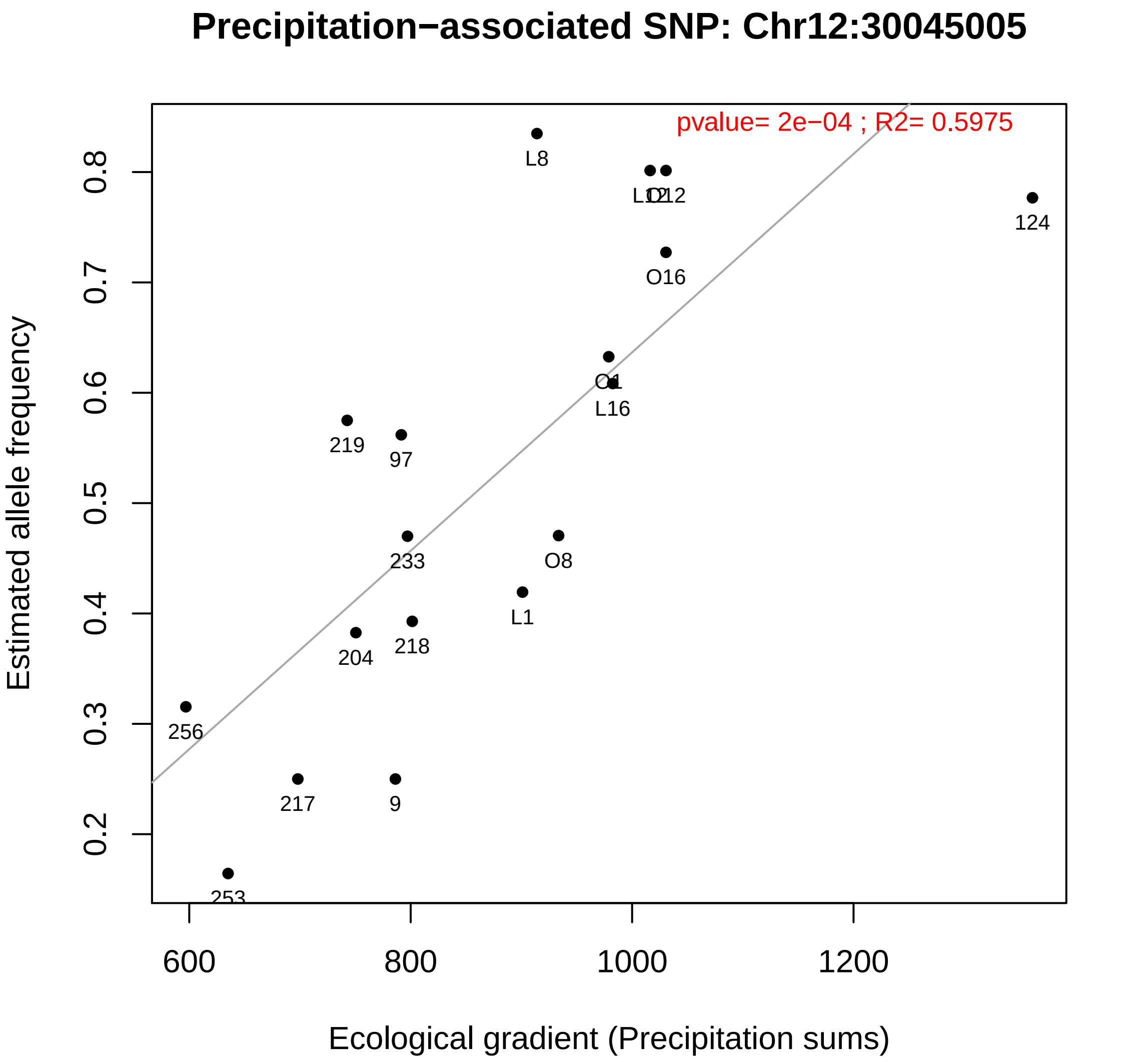
Precipitation−associated SNP: Chr12:30045005
pvalue= 2e−04 ; R2= 0.5975
Estimated allele frequency
0.2
0.3
0.4
0.5
0.6
0.7
0.8
600
800
1000
1200
Ecological gradient (Precipitation sums)
L8
L12
O12
124
O16
O1
L16
219
97
233
O8
L1
204
218
256
217
9
253

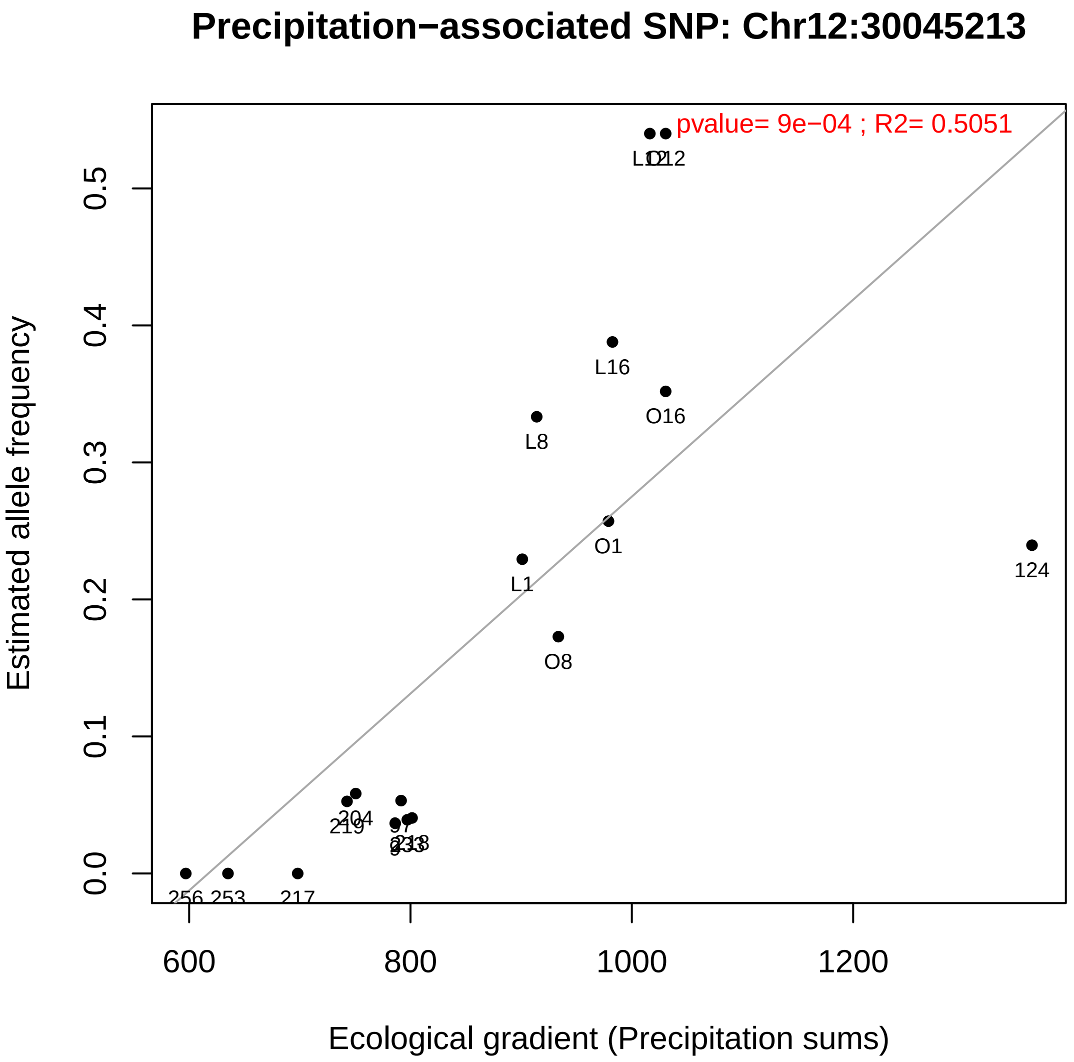
Precipitation−associated SNP: Chr12:30045213
pvalue= 9e−04 ; R2= 0.5051
Estimated allele frequency
0.0
0.1
0.2
0.3
0.4
0.5
Ecological gradient (Precipitation sums)
600
800
1000
1200
L16
O16
L8
O1
L1
O8
124
204
219
97
256
253
217

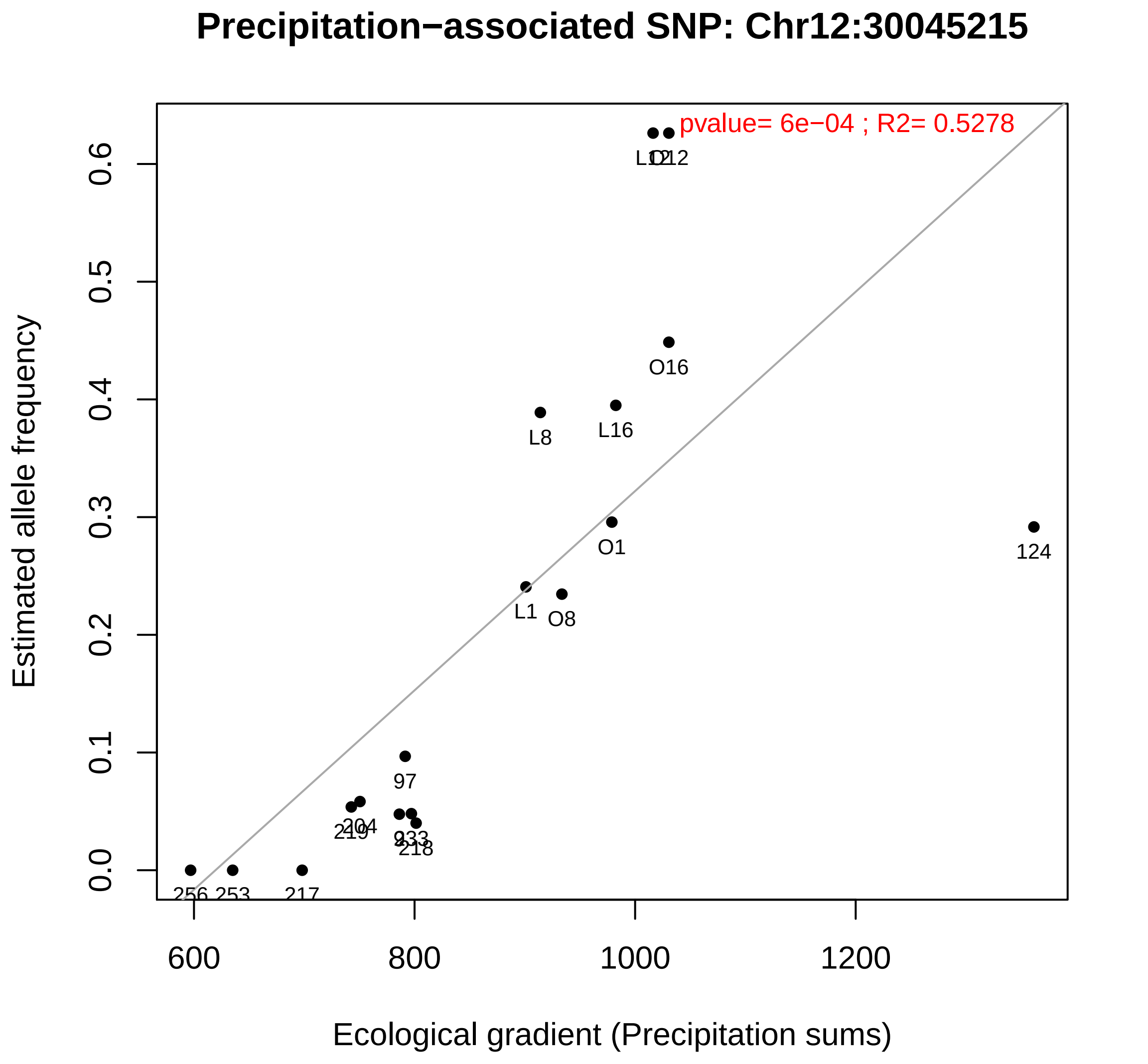

Precipitation−associated SNP: Chr12:30045215
pvalue= 6e−04 ; R2= 0.5278
Estimated allele frequency
Ecological gradient (Precipitation sums)
0.0
0.1
0.2
0.3
0.4
0.5
0.6
600
800
1000
1200
O16
L8
L16
O1
124
L1
O8
97

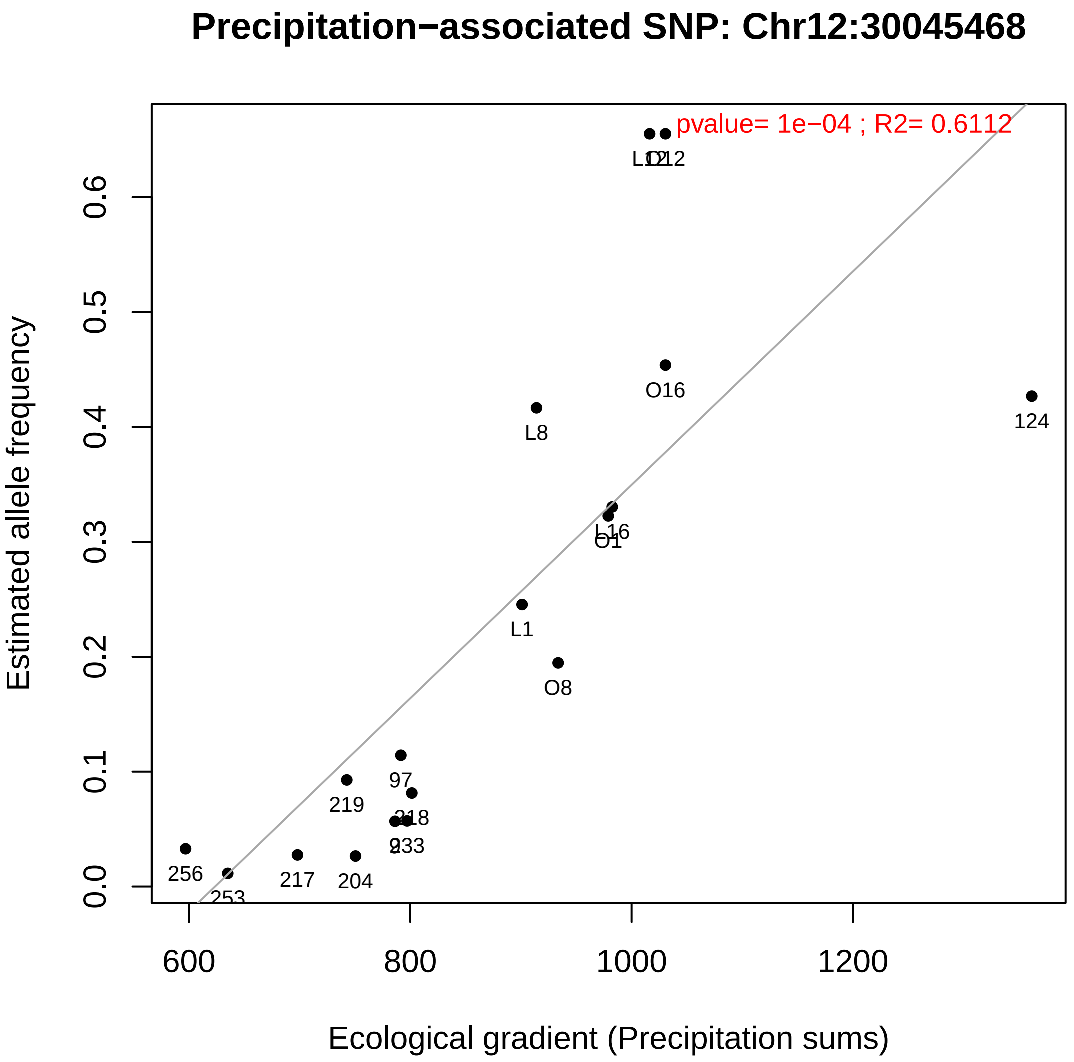
Precipitation−associated SNP: Chr12:30045468
pvalue= 1e−04 ; R2= 0.6112
Estimated allele frequency
Ecological gradient (Precipitation sums)
0.0
0.1
0.2
0.3
0.4
0.5
0.6
600
800
1000
1200
O16
124
L8
L16
O1
L1
O8
97
219
218
233
256
217
204
253

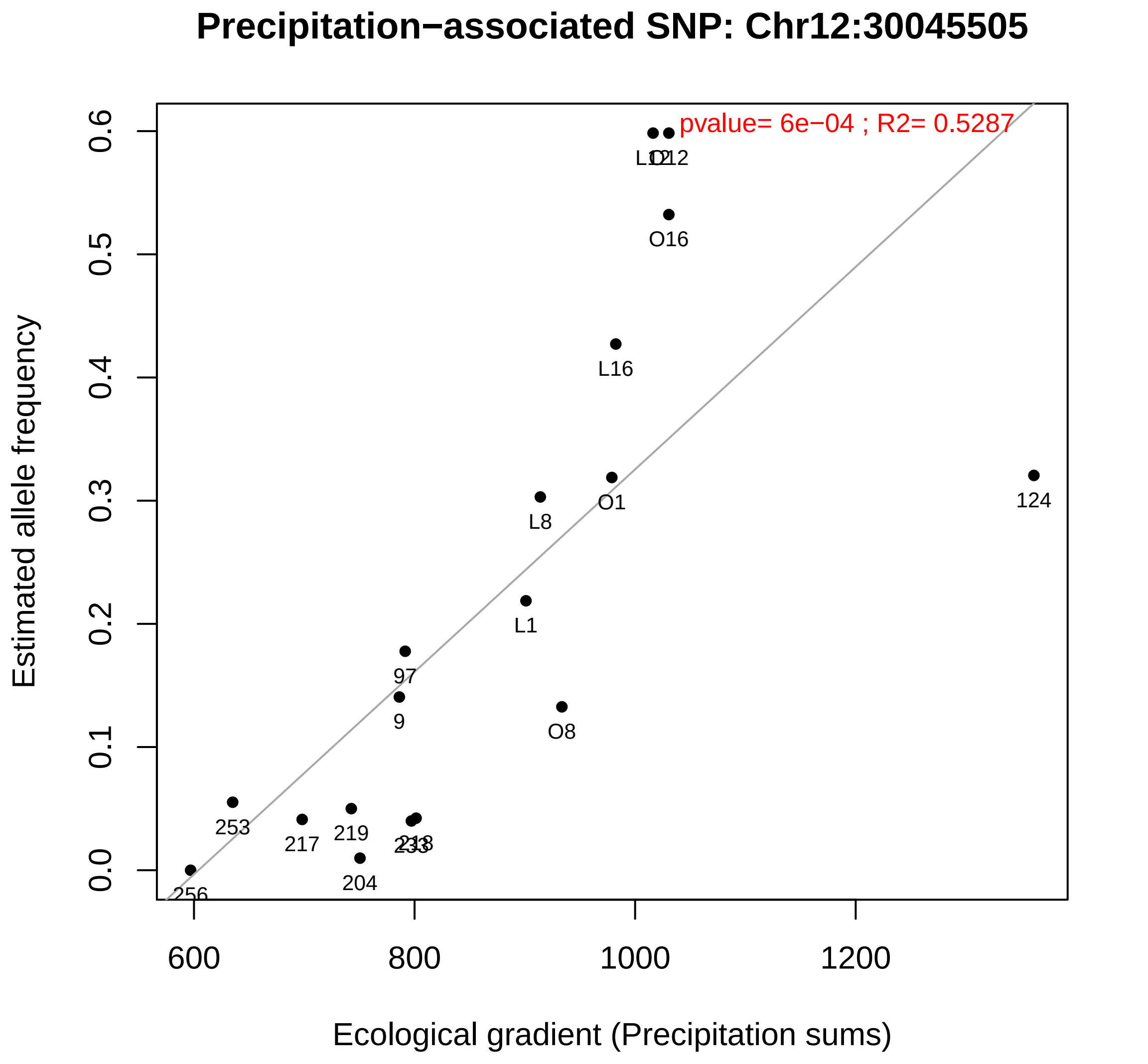

Precipitation−associated SNP: Chr12:30045505
pvalue= 6e−04 ; R2= 0.5287
Estimated allele frequency
Ecological gradient (Precipitation sums)
0.0
0.1
0.2
0.3
0.4
0.5
0.6
600
800
1000
1200
O16
L16
O1
124
L8
L1
97
9
O8
253
217
219
204
256

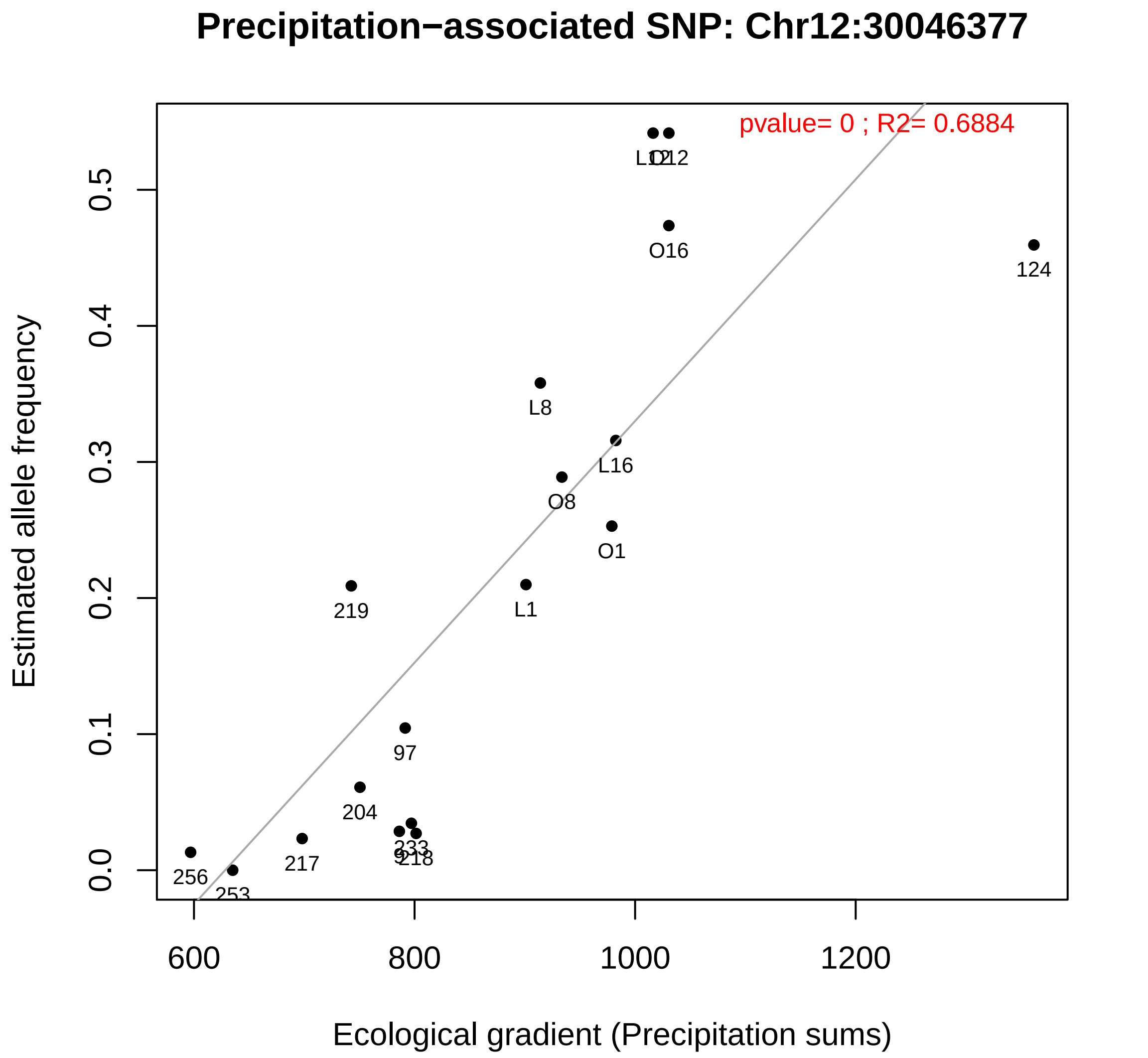

Precipitation−associated SNP: Chr12:30046377
pvalue= 0 ; R2= 0.6884
Estimated allele frequency
Ecological gradient (Precipitation sums)
0.0
0.1
0.2
0.3
0.4
0.5
600
800
1000
1200
O16
124
L8
L16
O8
O1
219
L1
97
204
217
256
233
218

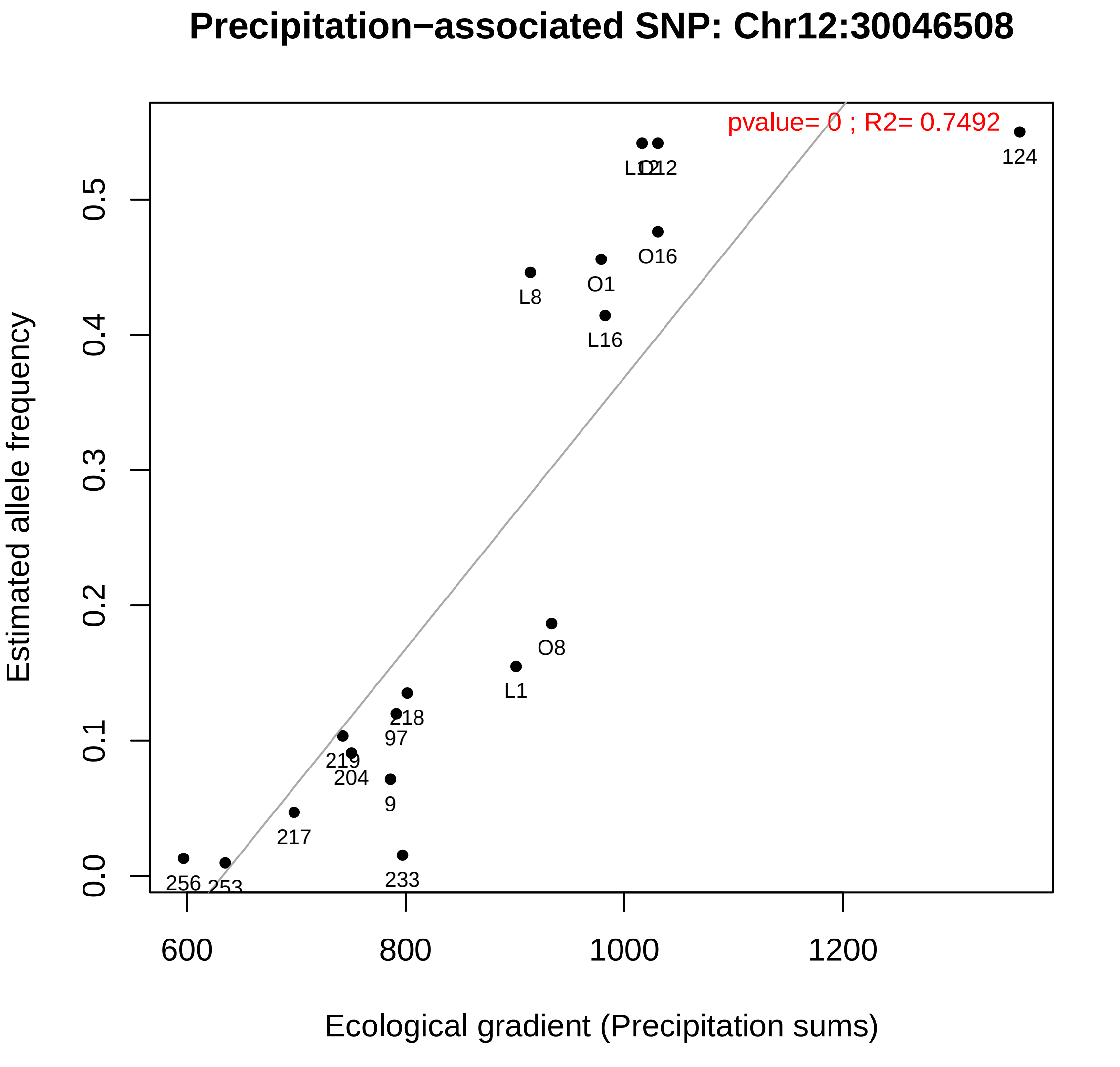

Precipitation−associated SNP: Chr12:30046508
pvalue= 0 ; R2= 0.7492
Estimated allele frequency
Ecological gradient (Precipitation sums)
0.0
0.1
0.2
0.3
0.4
0.5
600
800
1000
1200
124
O16
O1
L8
L16
O8
L1
218
97
219
204
9
217
233
256
253

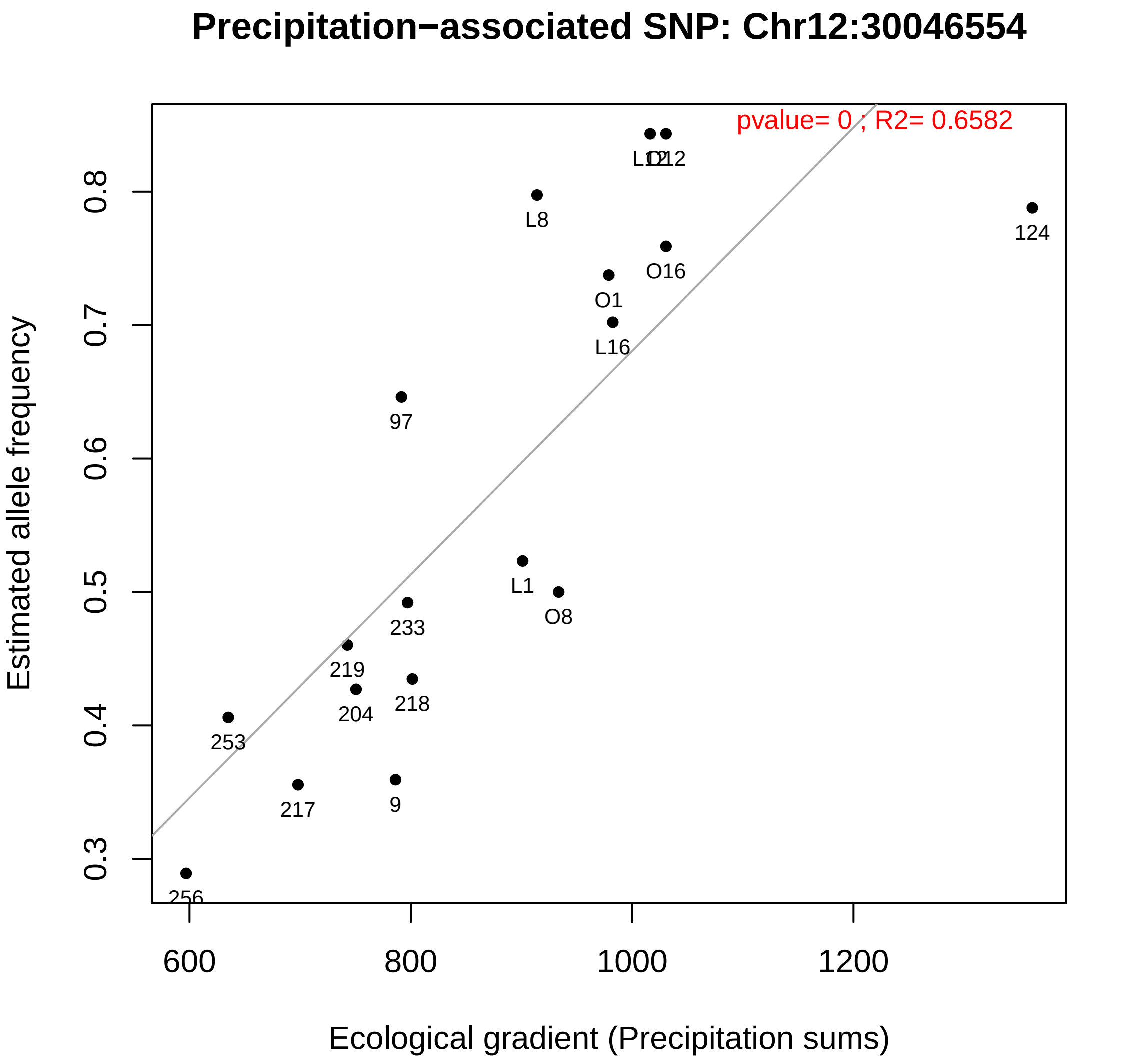
Precipitation−associated SNP: Chr12:30046554
pvalue= 0 ; R2= 0.6582
Estimated allele frequency
0.3
0.4
0.5
0.6
0.7
0.8
600
800
1000
1200
Ecological gradient (Precipitation sums)
O12
L8
124
O16
O1
L16
97
L1
O8
233
219
204
218
253
217
9
256

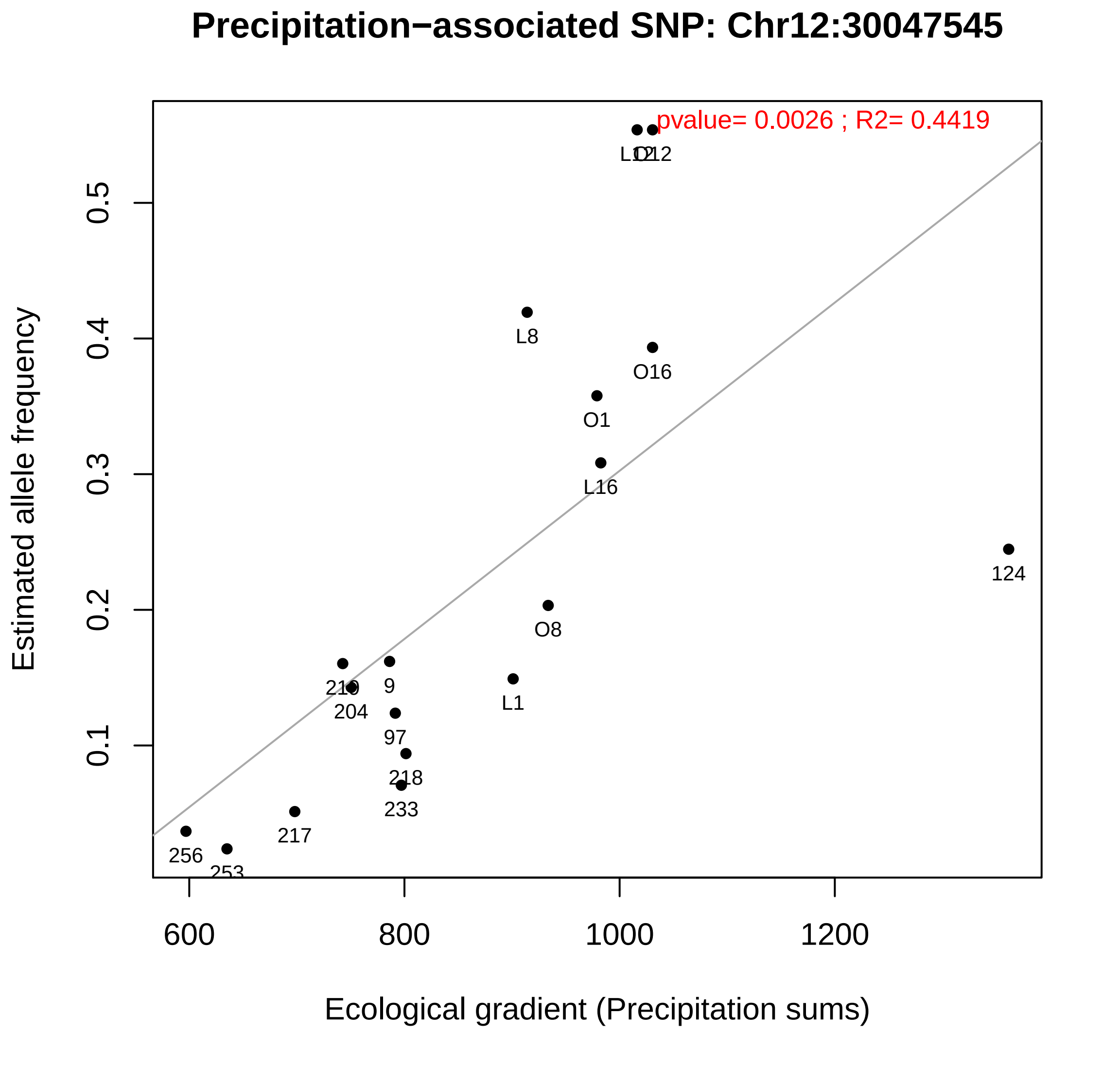

Precipitation−associated SNP: Chr12:30047545
pvalue= 0.0026 ; R2= 0.4419
Estimated allele frequency
Ecological gradient (Precipitation sums)
0.1
0.2
0.3
0.4
0.5
600
800
1000
1200
L8
O16
O1
L16
124
O8
9
204
97
L1
218
233
217
256
253

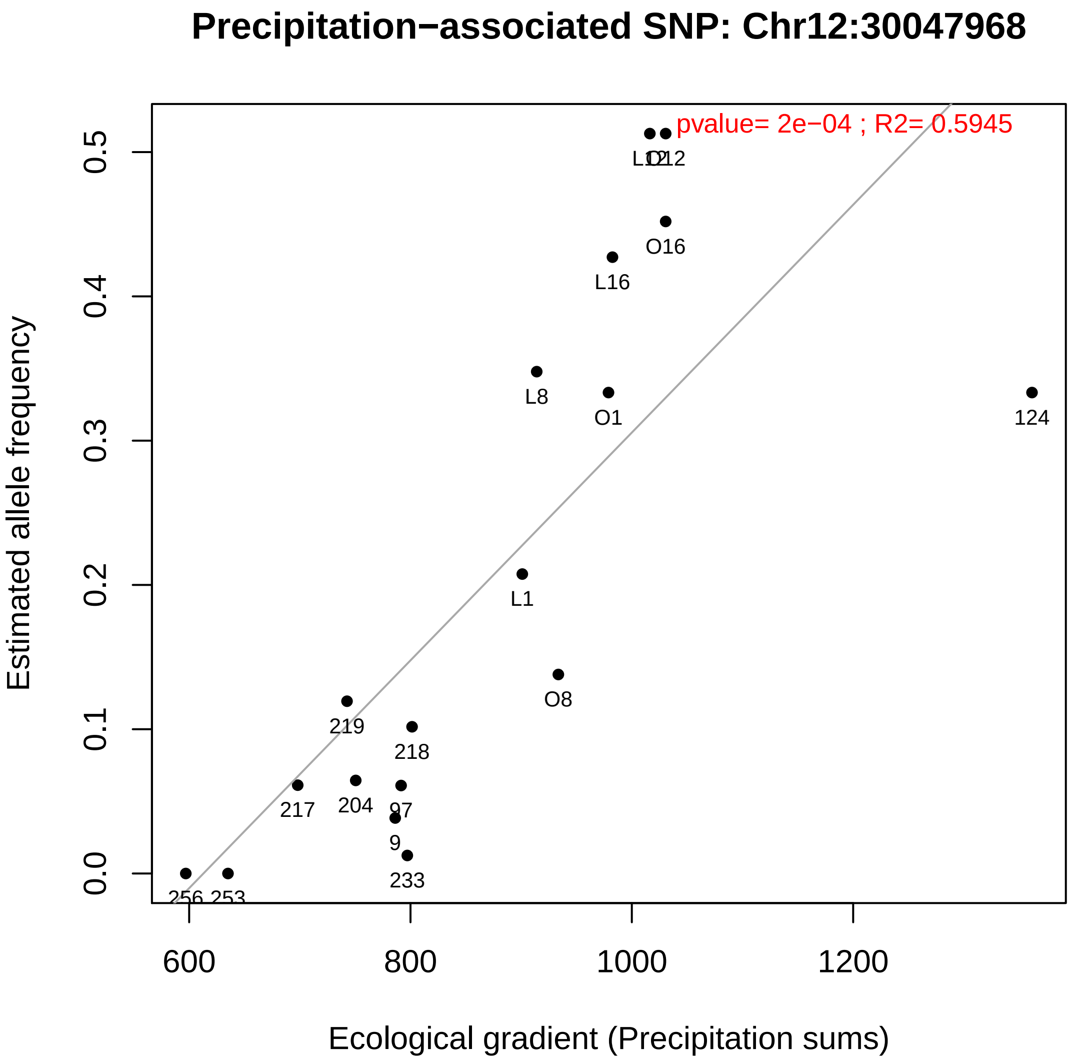
Precipitation−associated SNP: Chr12:30047968
pvalue= 2e−04 ; R2= 0.5945
Estimated allele frequency
Ecological gradient (Precipitation sums)
0.0
0.1
0.2
0.3
0.4
0.5
600
800
1000
1200
O12
O16
L16
L8
O1
124
L1
O8
219
218
217
204
97
9
233
253

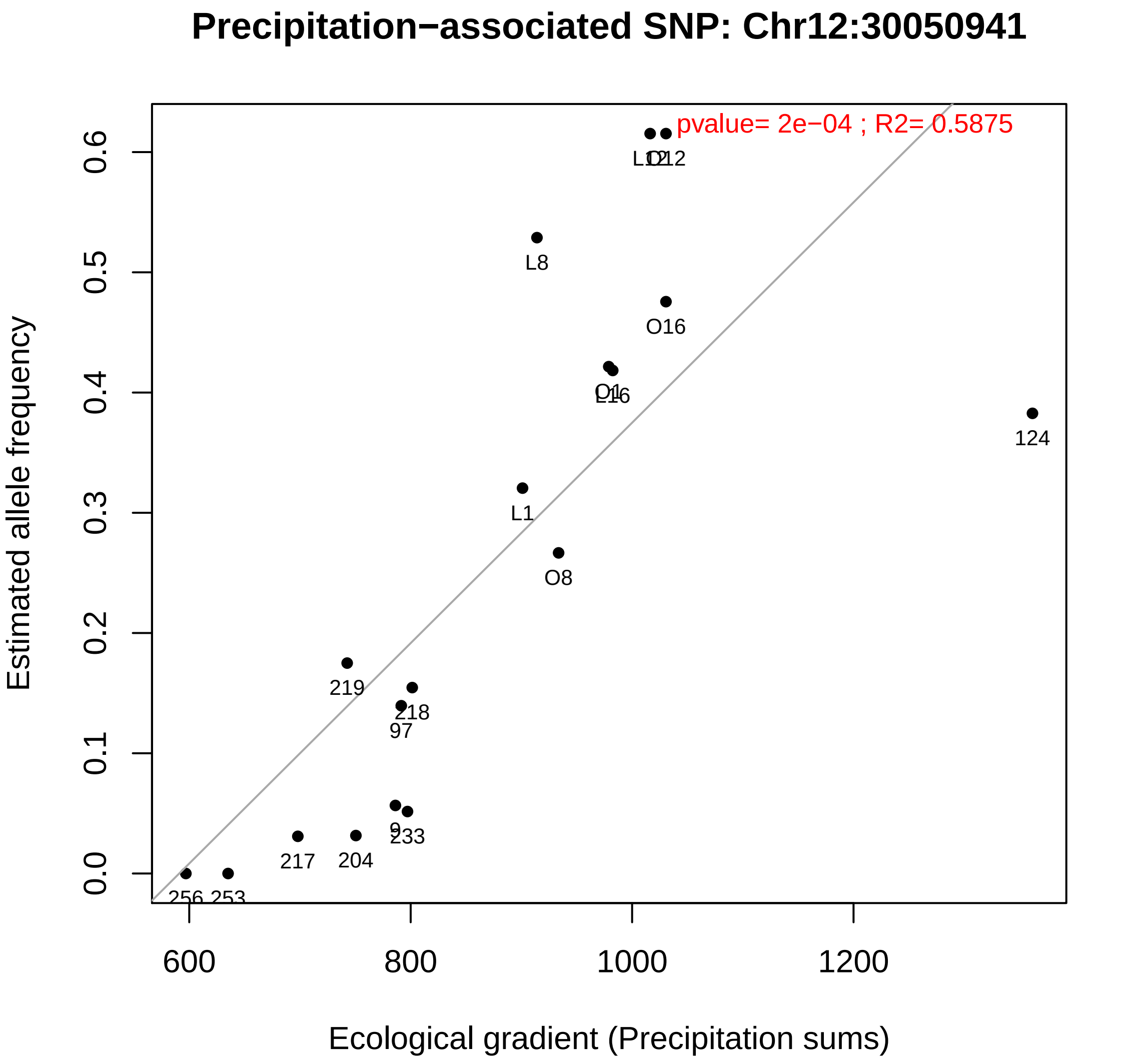

Precipitation−associated SNP: Chr12:30050941
pvalue= 2e−04 ; R2= 0.5875
L10
O12
L8
O16
O1
L16
124
L1
O8
219
218
97
9
233
217
204
256
253
0.0
0.1
0.2
0.3
0.4
0.5
0.6
600
800
1000
1200
Estimated allele frequency
Ecological gradient (Precipitation sums)

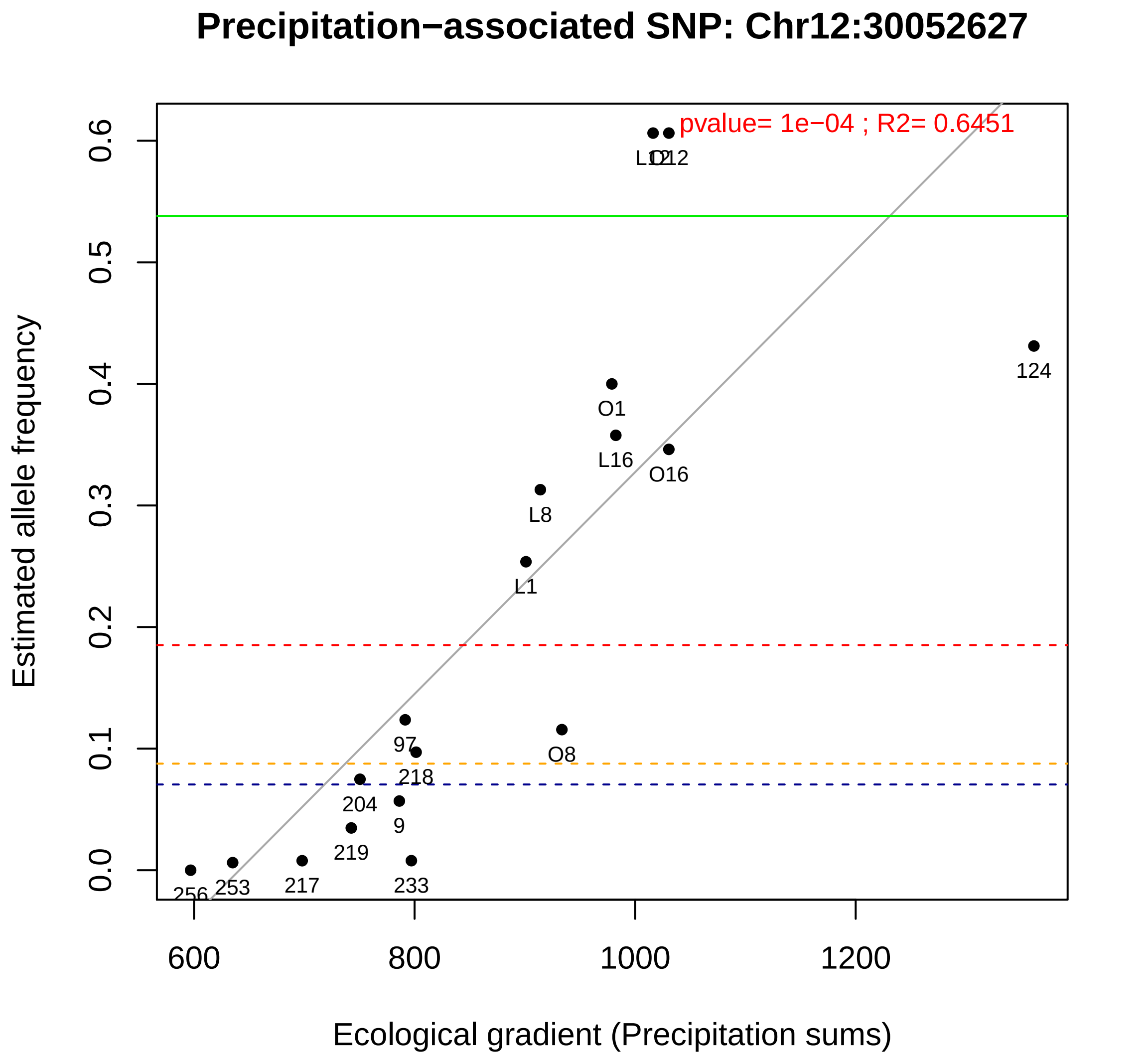

Precipitation−associated SNP: Chr12:30052627
pvalue= 1e−04 ; R2= 0.6451
Estimated allele frequency
Ecological gradient (Precipitation sums)
0.0
0.1
0.2
0.3
0.4
0.5
0.6
600
800
1000
1200
L12
O12
124
O1
L16
O16
L8
L1
97
O8
218
204
9
219
233
217
253
256

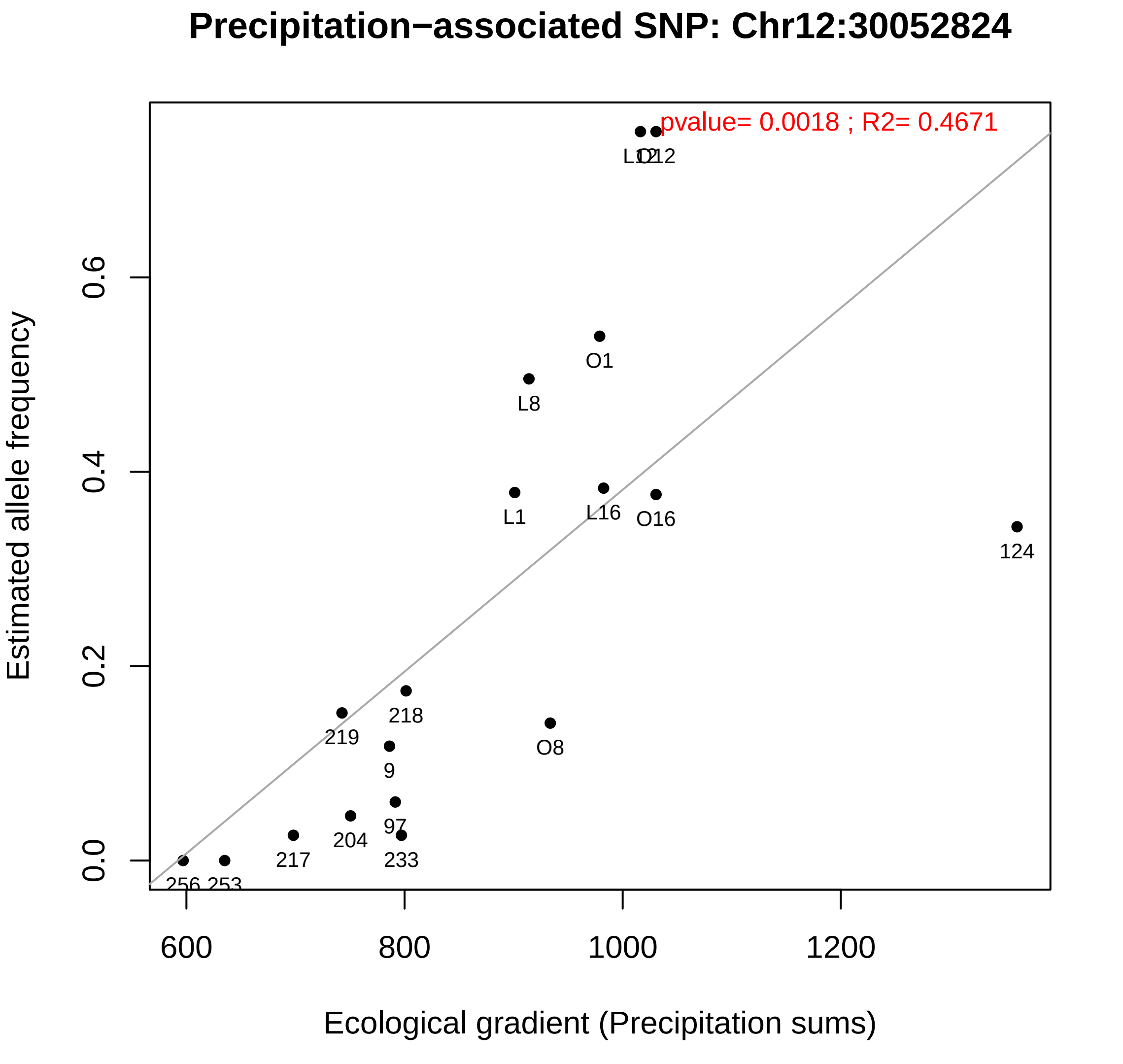

Precipitation−associated SNP: Chr12:30052824
pvalue= 0.0018 ; R2= 0.4671
Estimated allele frequency
Ecological gradient (Precipitation sums)
0.0
0.2
0.4
0.6
600
800
1000
1200
O1
L8
L1
L16
O16
124
218
219
O8
9
97
204
233
217

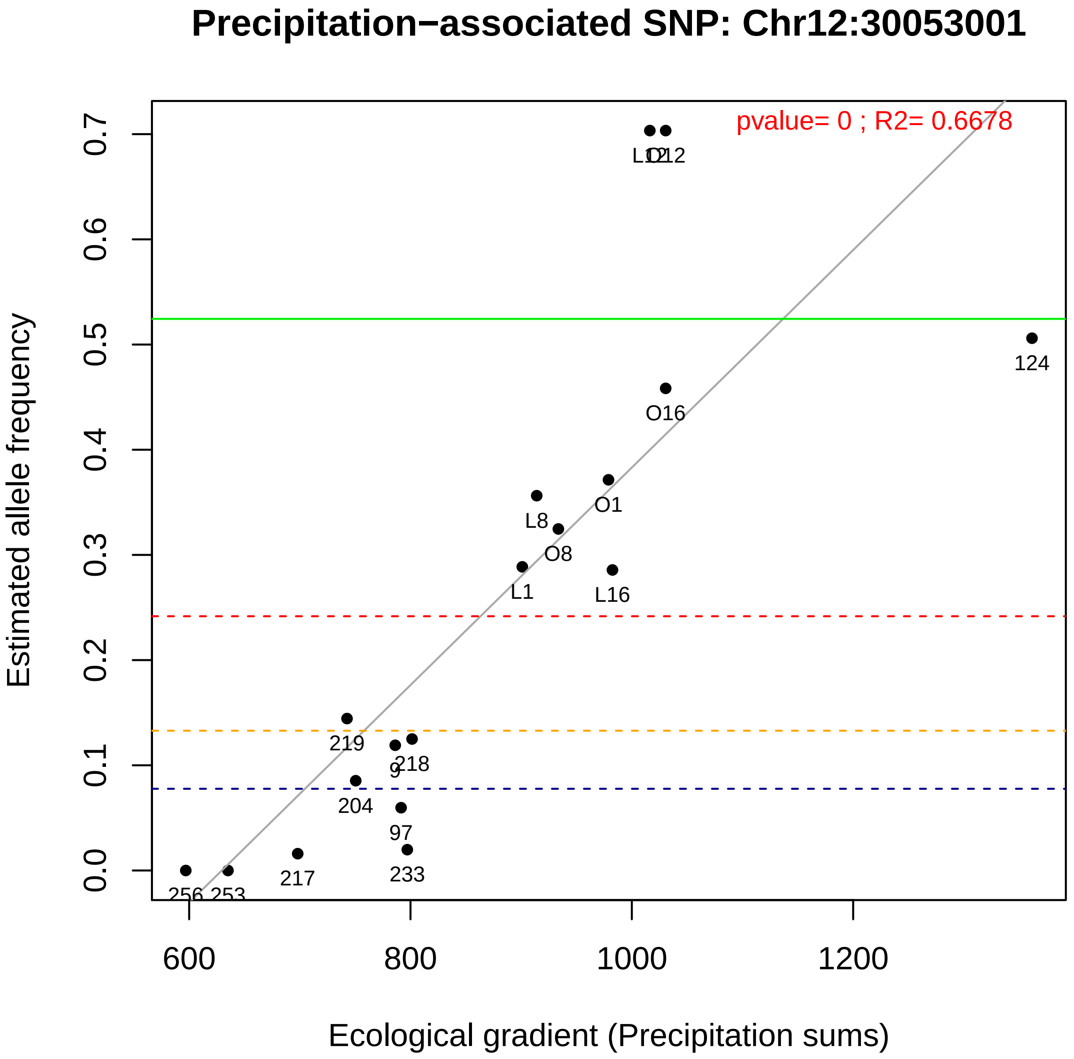
Precipitation−associated SNP: Chr12:30053001
pvalue= 0 ; R2= 0.6678
Estimated allele frequency
0.0
0.1
0.2
0.3
0.4
0.5
0.6
0.7
Ecological gradient (Precipitation sums)
600
800
1000
1200
124
O16
O1
L8
O8
L1
L16
219
218
9
204
97
233
217
256
253

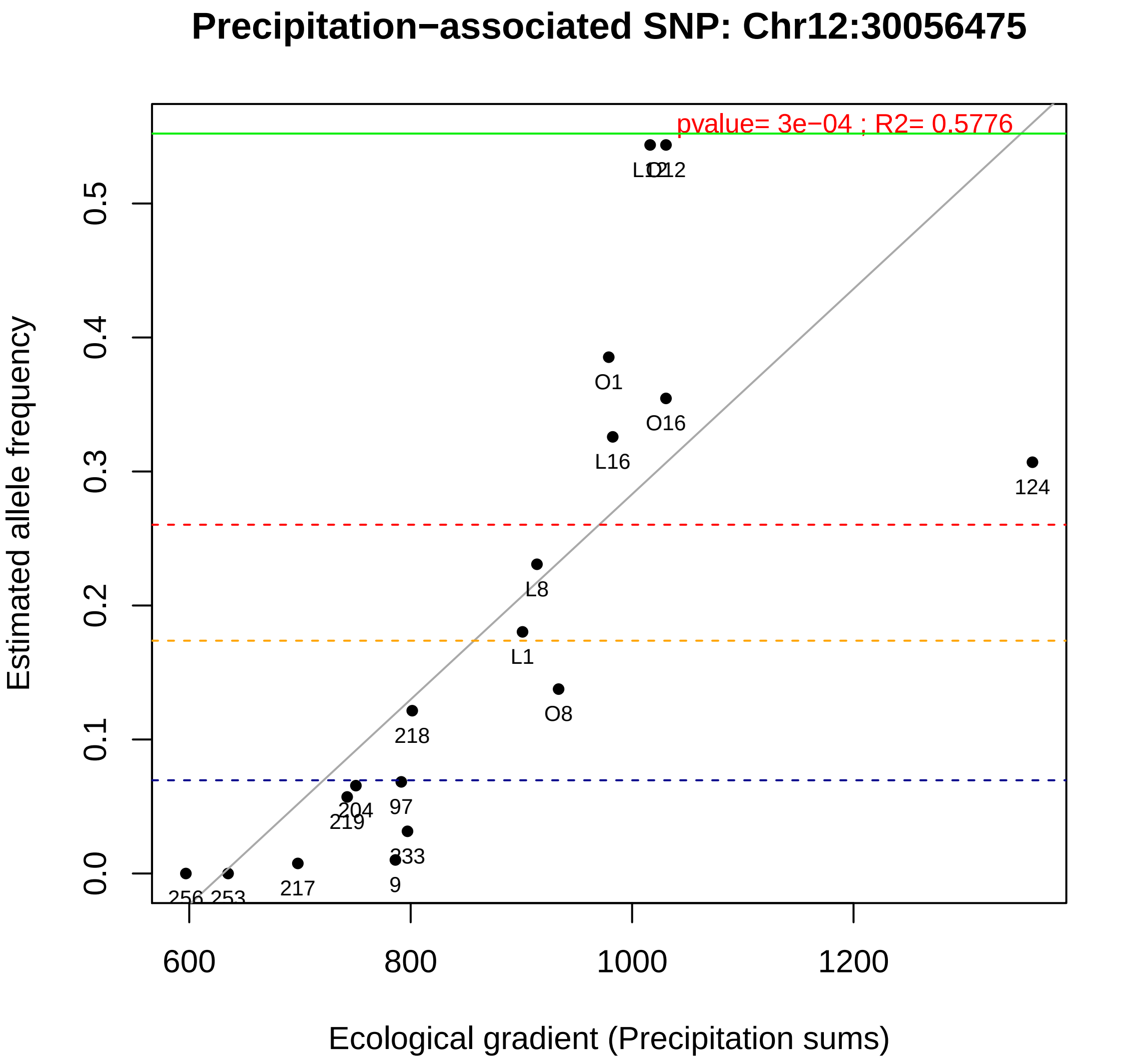
Precipitation−associated SNP: Chr12:30056475
pvalue= 3e−04 ; R2= 0.5776
Estimated allele frequency
Ecological gradient (Precipitation sums)
0.0
0.1
0.2
0.3
0.4
0.5
600
800
1000
1200
L12
O12
O1
O16
L16
124
L8
L1
O8
218
204
219
97
333
9
217
256
253

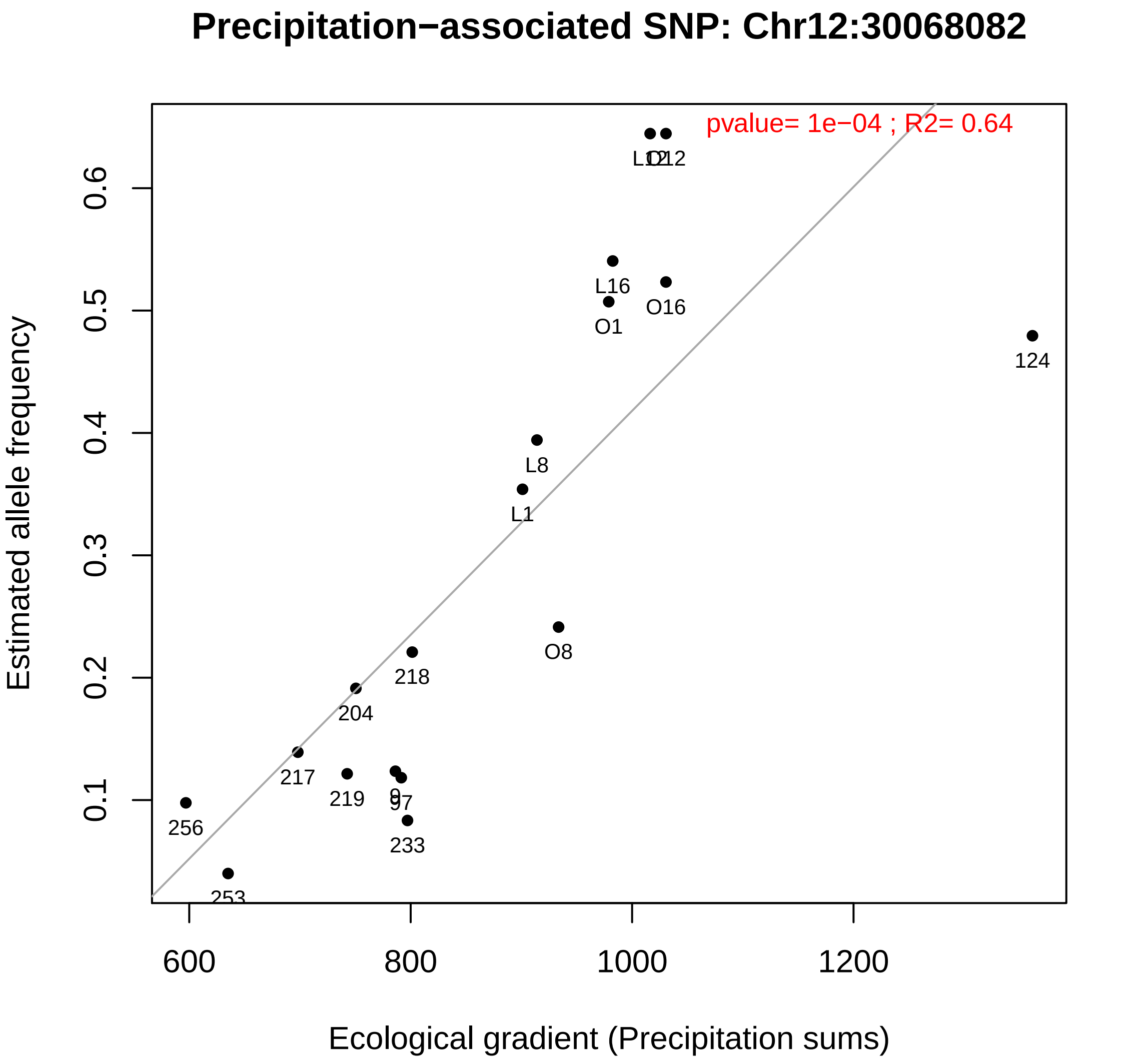

Precipitation−associated SNP: Chr12:30068082
pvalue= 1e−04 ; R2= 0.64
Estimated allele frequency
Ecological gradient (Precipitation sums)
0.1
0.2
0.3
0.4
0.5
0.6
600
800
1000
1200
L12
O12
L16
O16
O1
124
L8
L1
O8
218
204
217
219
97
233
256
253

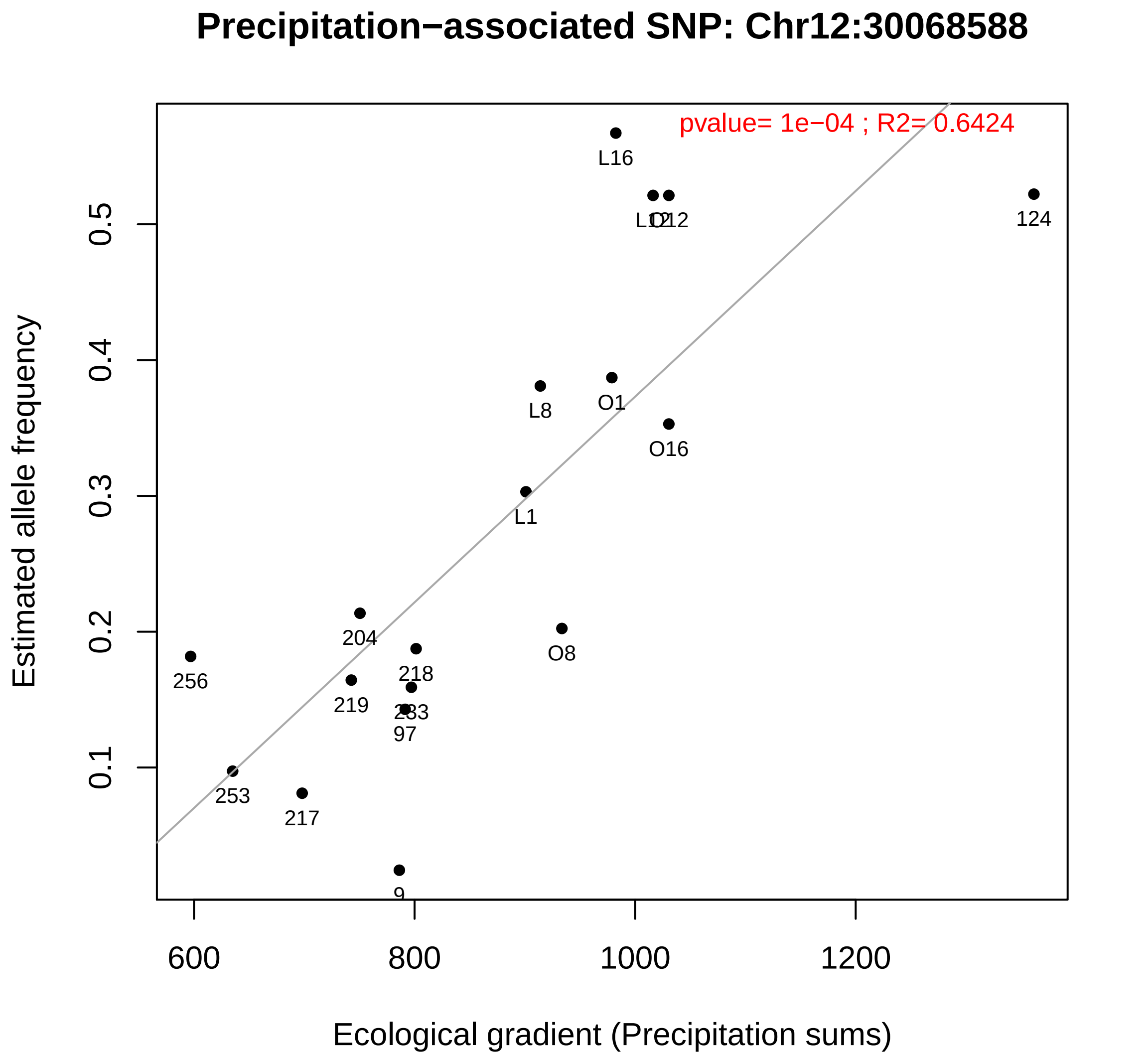

Precipitation−associated SNP: Chr12:30068588
pvalue= 1e−04 ; R2= 0.6424
Estimated allele frequency
Ecological gradient (Precipitation sums)
0.1
0.2
0.3
0.4
0.5
600
800
1000
1200
L16
L12
O12
124
L8
O1
O16
L1
204
O8
256
218
219
233
97
253
217
9

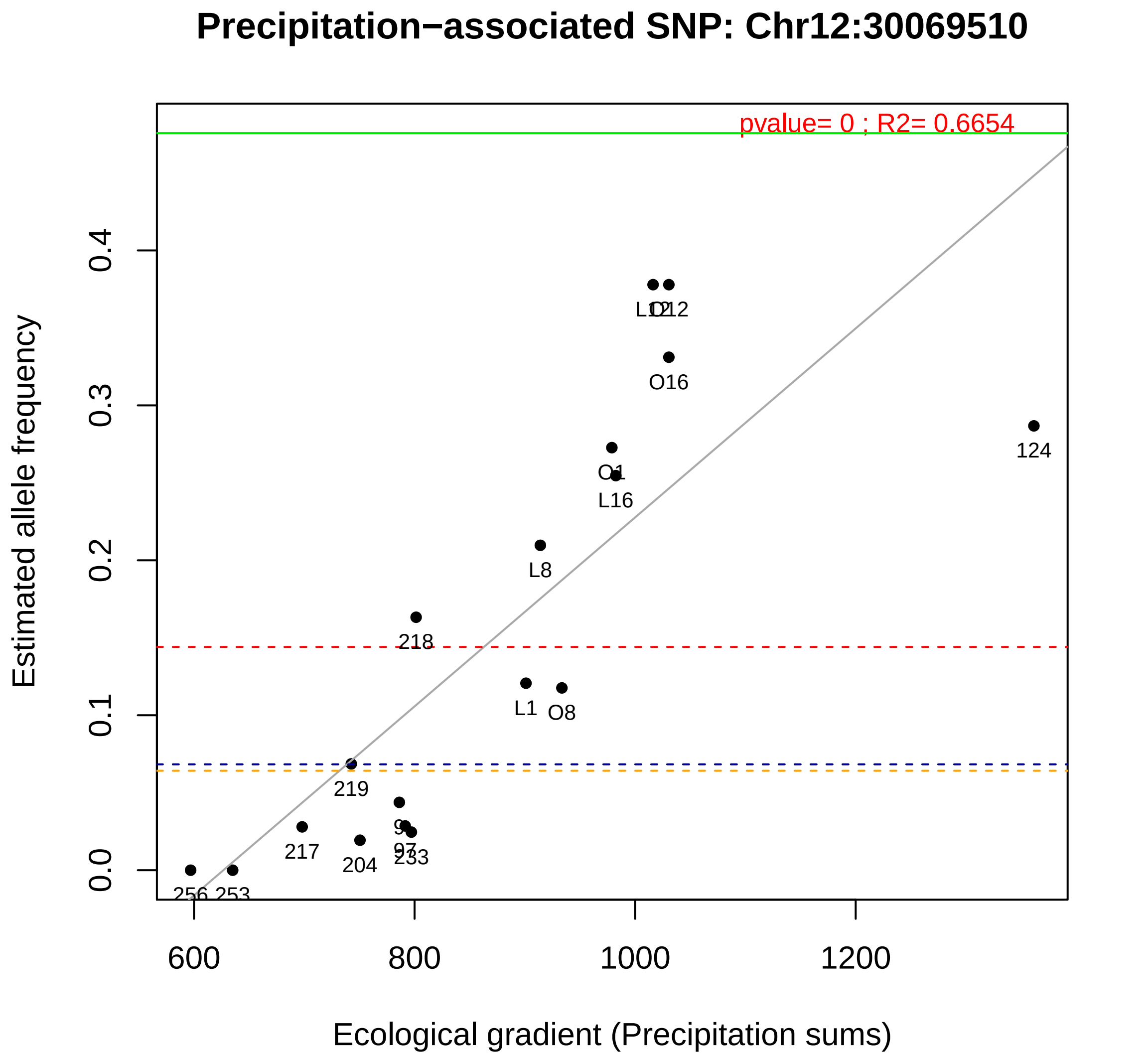

Precipitation−associated SNP: Chr12:30069510
pvalue= 0 ; R2= 0.6654
Estimated allele frequency
0.0
0.1
0.2
0.3
0.4
Ecological gradient (Precipitation sums)
600
800
1000
1200
L12
O12
O16
124
O1
L16
L8
218
L1
O8
219
217
204
233
256
253

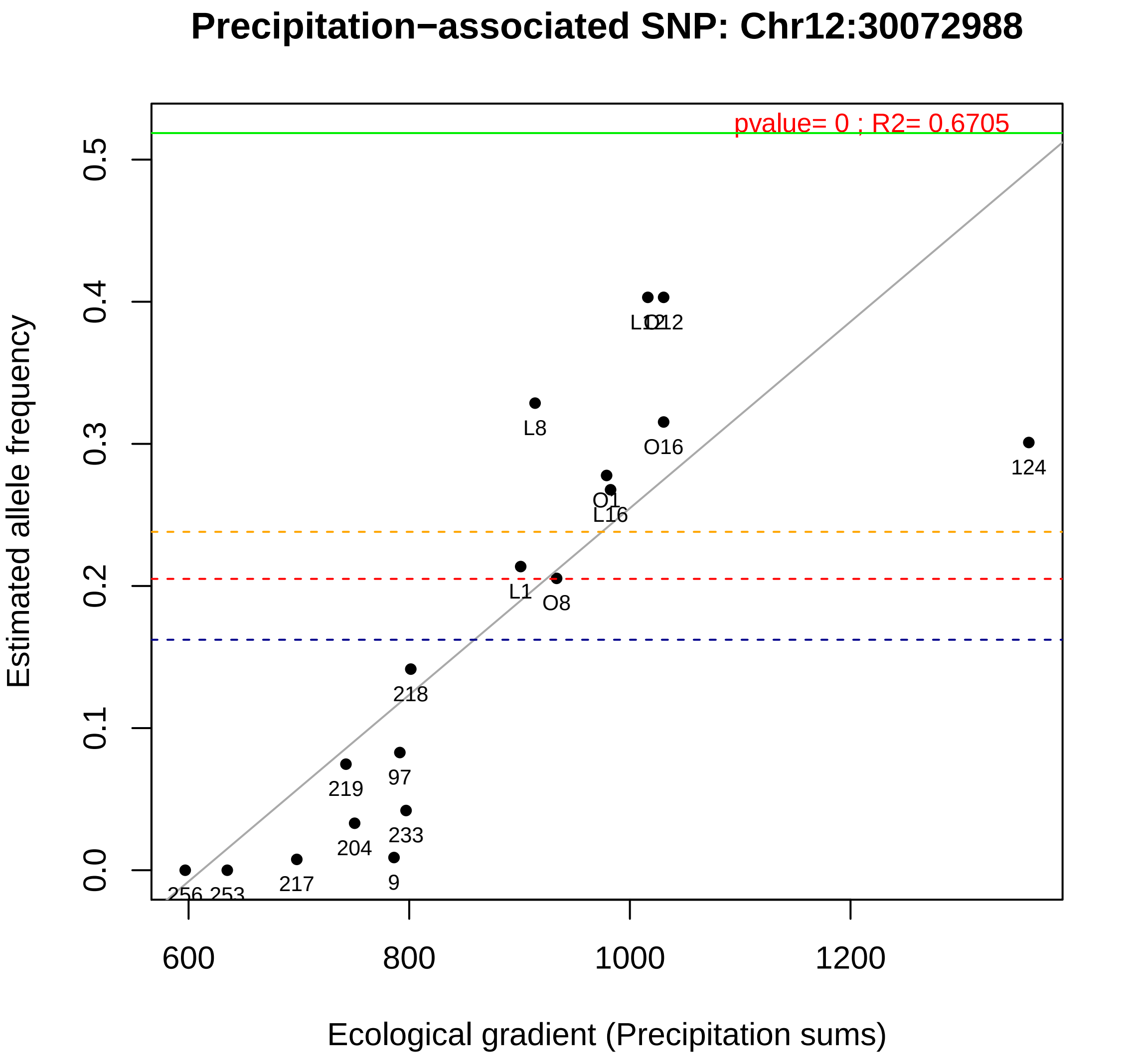

Precipitation−associated SNP: Chr12:30072988
pvalue= 0 ; R2= 0.6705
Estimated allele frequency
0.0
0.1
0.2
0.3
0.4
0.5
Ecological gradient (Precipitation sums)
600
800
1000
1200
L8
O16
124
O1
L16
L1
O8
218
219
97
204
233
217
9
256
253

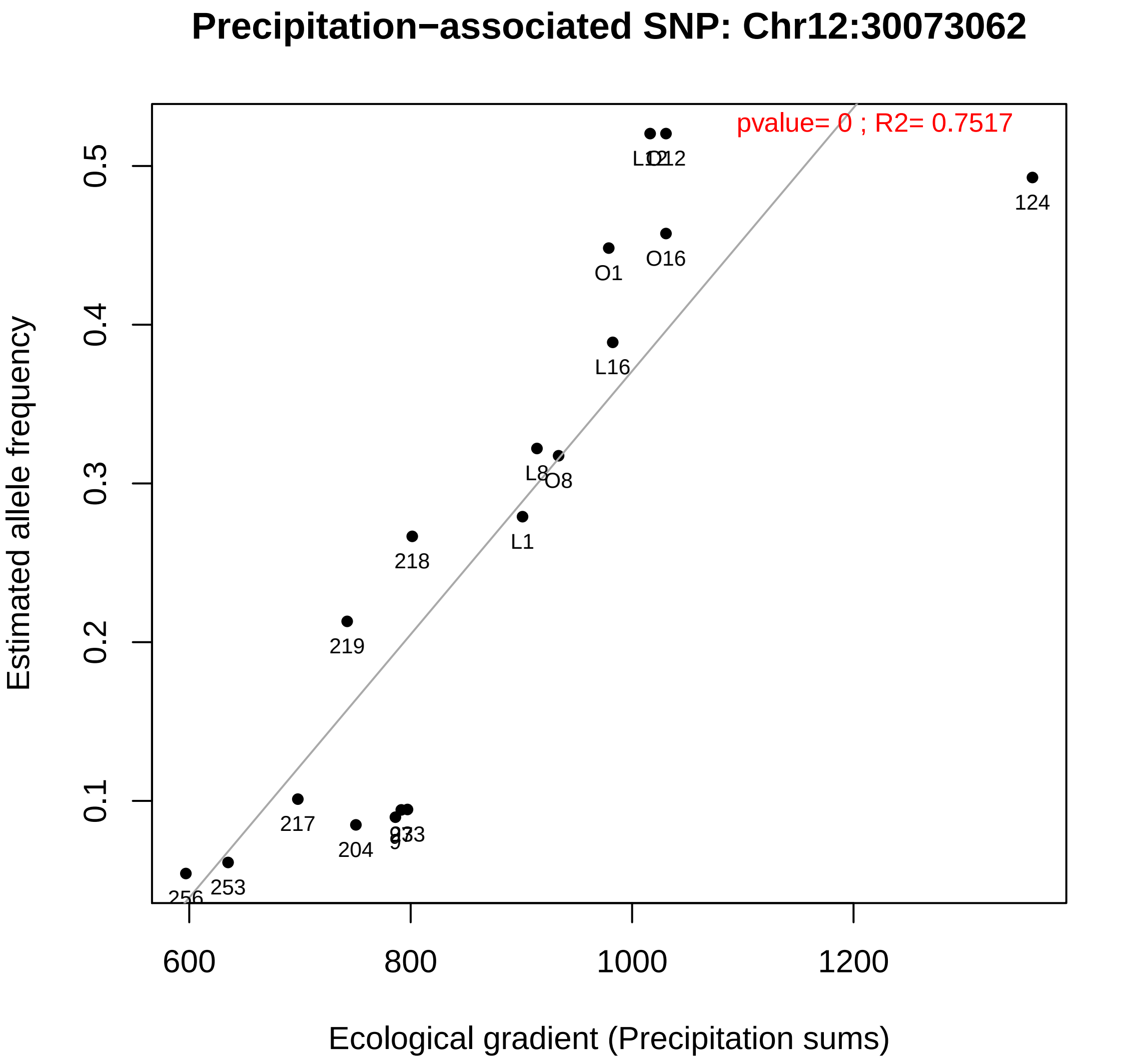

Precipitation−associated SNP: Chr12:30073062
pvalue= 0 ; R2= 0.7517
Estimated allele frequency
0.1
0.2
0.3
0.4
0.5
600
800
1000
1200
Ecological gradient (Precipitation sums)
L12
O12
124
O16
O1
L16
L8
O8
L1
218
219
217
204
253
256

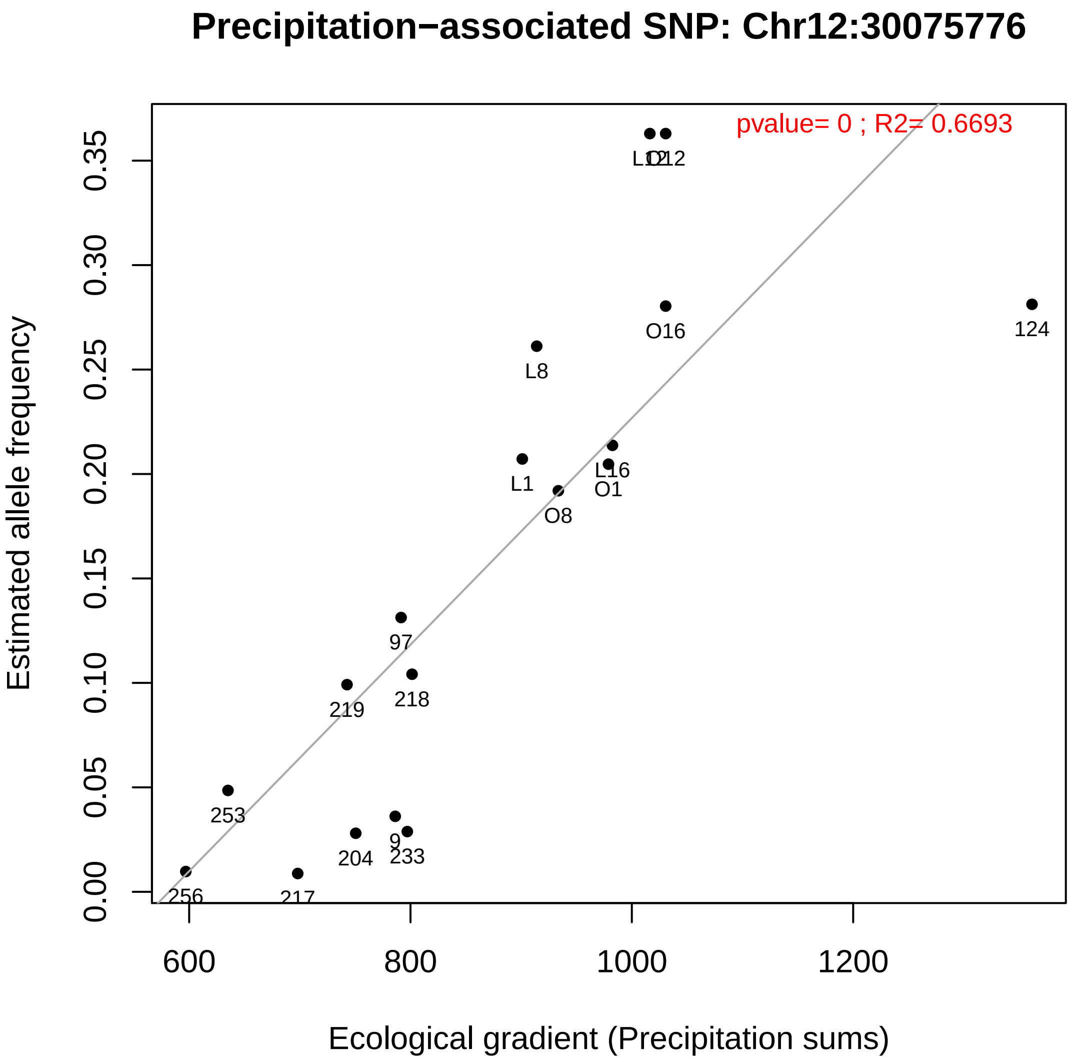
Precipitation−associated SNP: Chr12:30075776
pvalue= 0 ; R2= 0.6693
Estimated allele frequency
Ecological gradient (Precipitation sums)
0.00
0.05
0.10
0.15
0.20
0.25
0.30
0.35
600
800
1000
1200
L12
O12
O16
124
L8
L1
L16
O1
O8
97
219
218
253
204
9
233
256
217

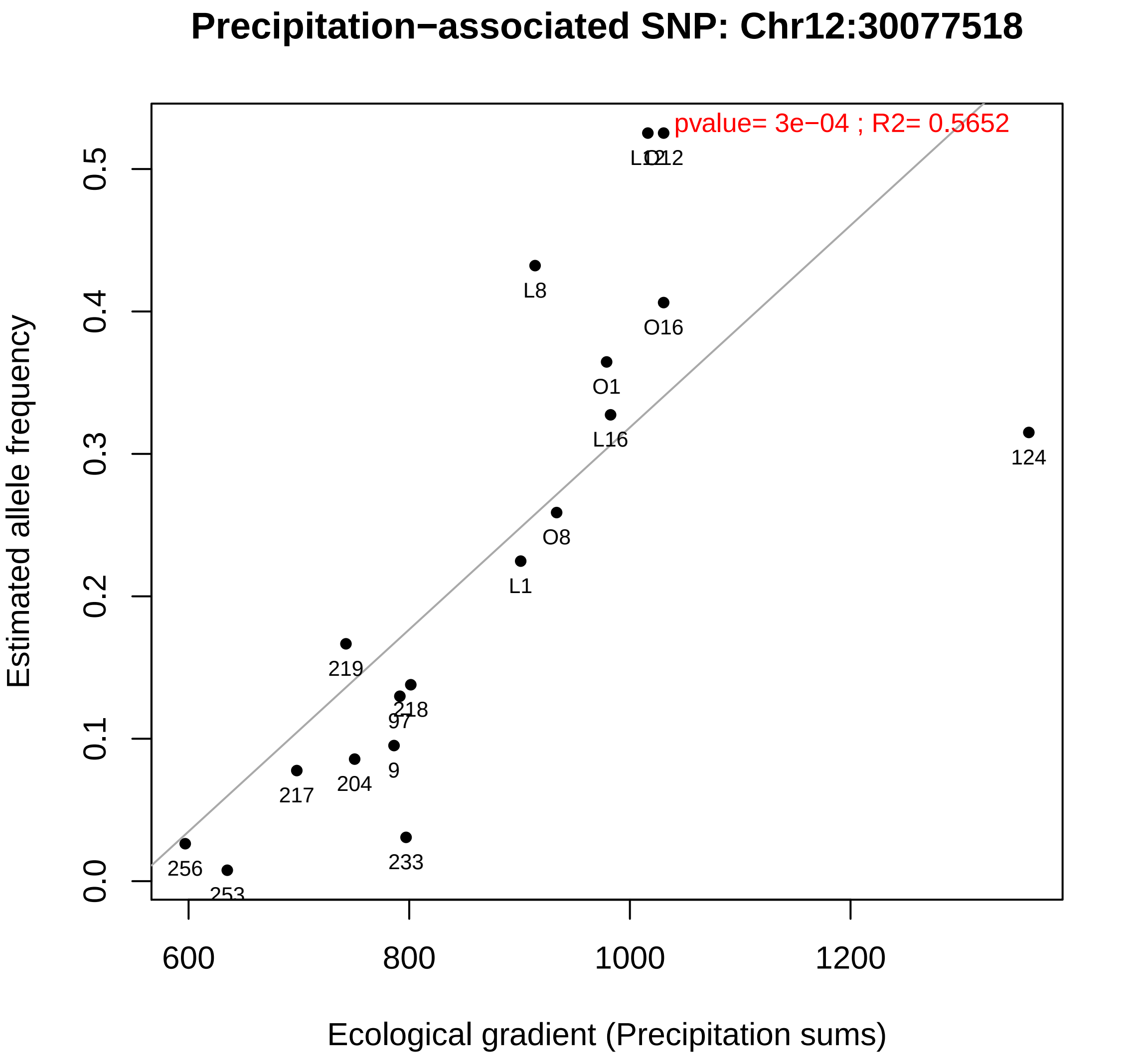

Precipitation−associated SNP: Chr12:30077518
pvalue= 3e−04 ; R2= 0.5652
Estimated allele frequency
Ecological gradient (Precipitation sums)
0.0
0.1
0.2
0.3
0.4
0.5
600
800
1000
1200
L8
O16
O1
L16
124
O8
L1
219
218
97
9
204
217
233
256

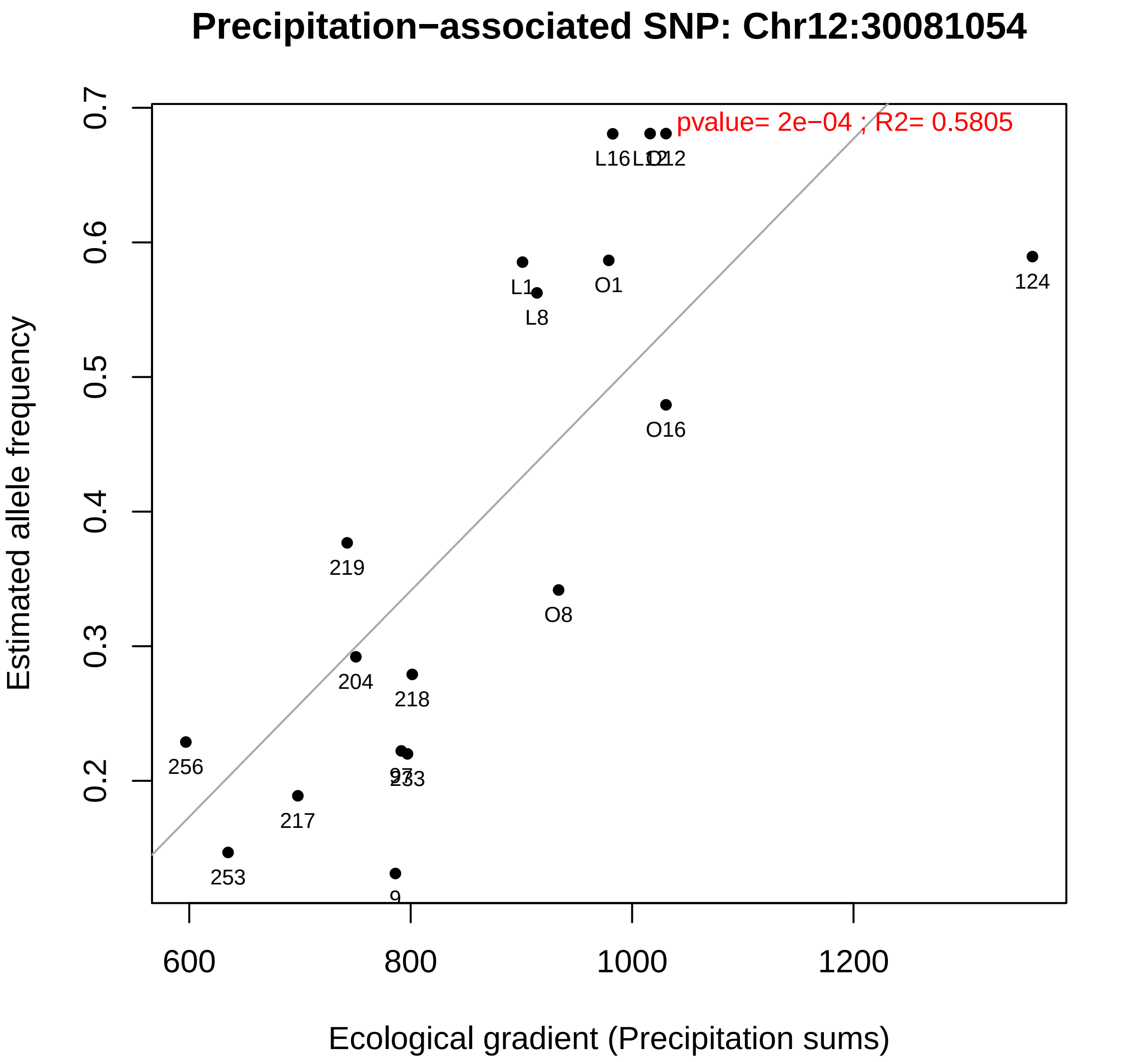

Precipitation−associated SNP: Chr12:30081054
pvalue= 2e−04 ; R2= 0.5805
Estimated allele frequency
Ecological gradient (Precipitation sums)
0.2
0.3
0.4
0.5
0.6
0.7
600
800
1000
1200
L16
O12
L1
L8
O1
124
O16
219
O8
204
218
256
217
253
9

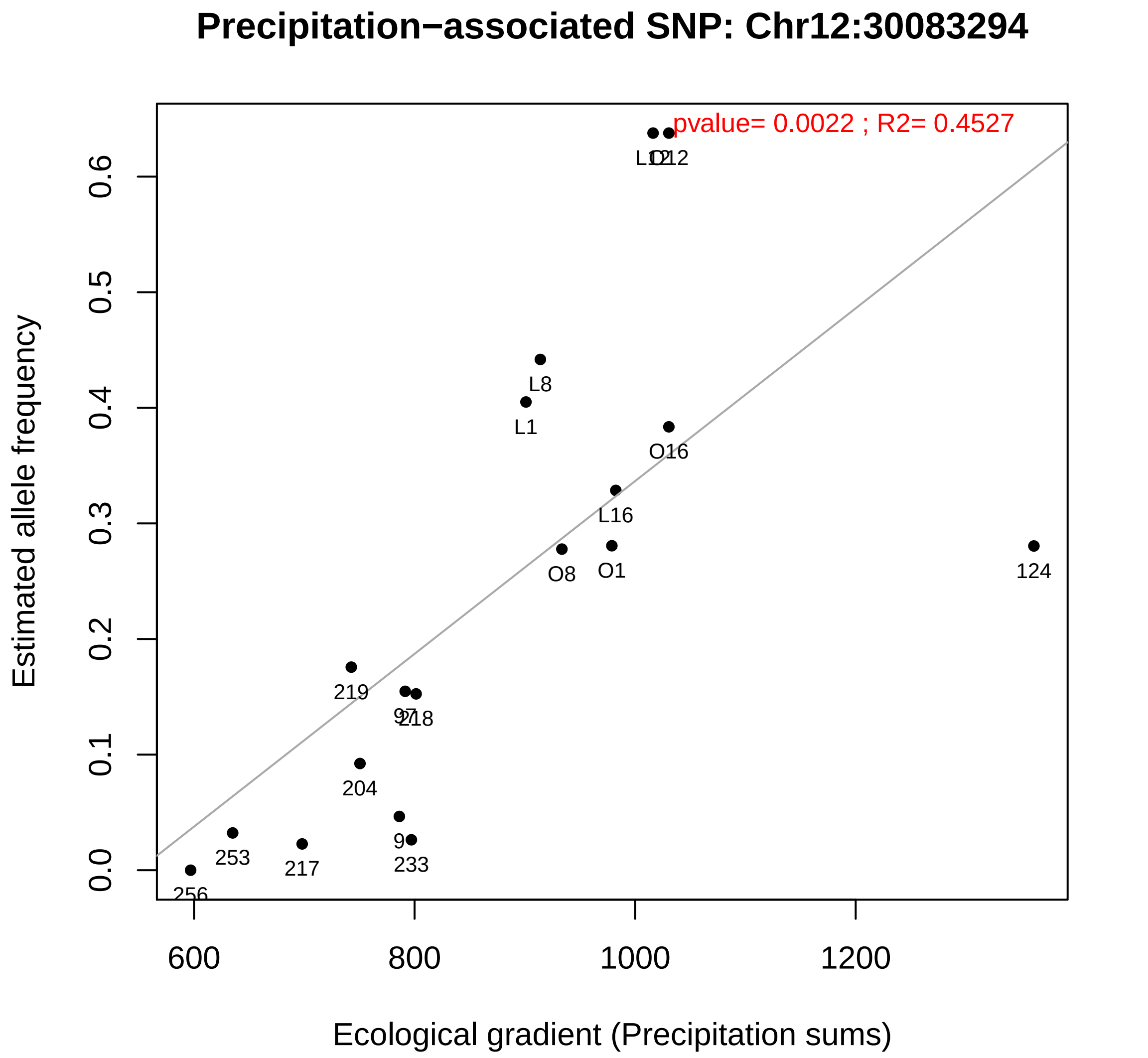

Precipitation−associated SNP: Chr12:30083294
pvalue= 0.0022 ; R2= 0.4527
Estimated allele frequency
Ecological gradient (Precipitation sums)
0.0
0.1
0.2
0.3
0.4
0.5
0.6
600
800
1000
1200
L12
O2
L8
L1
O16
L16
O8
O1
124
219
97
218
204
253
217
9
233
256

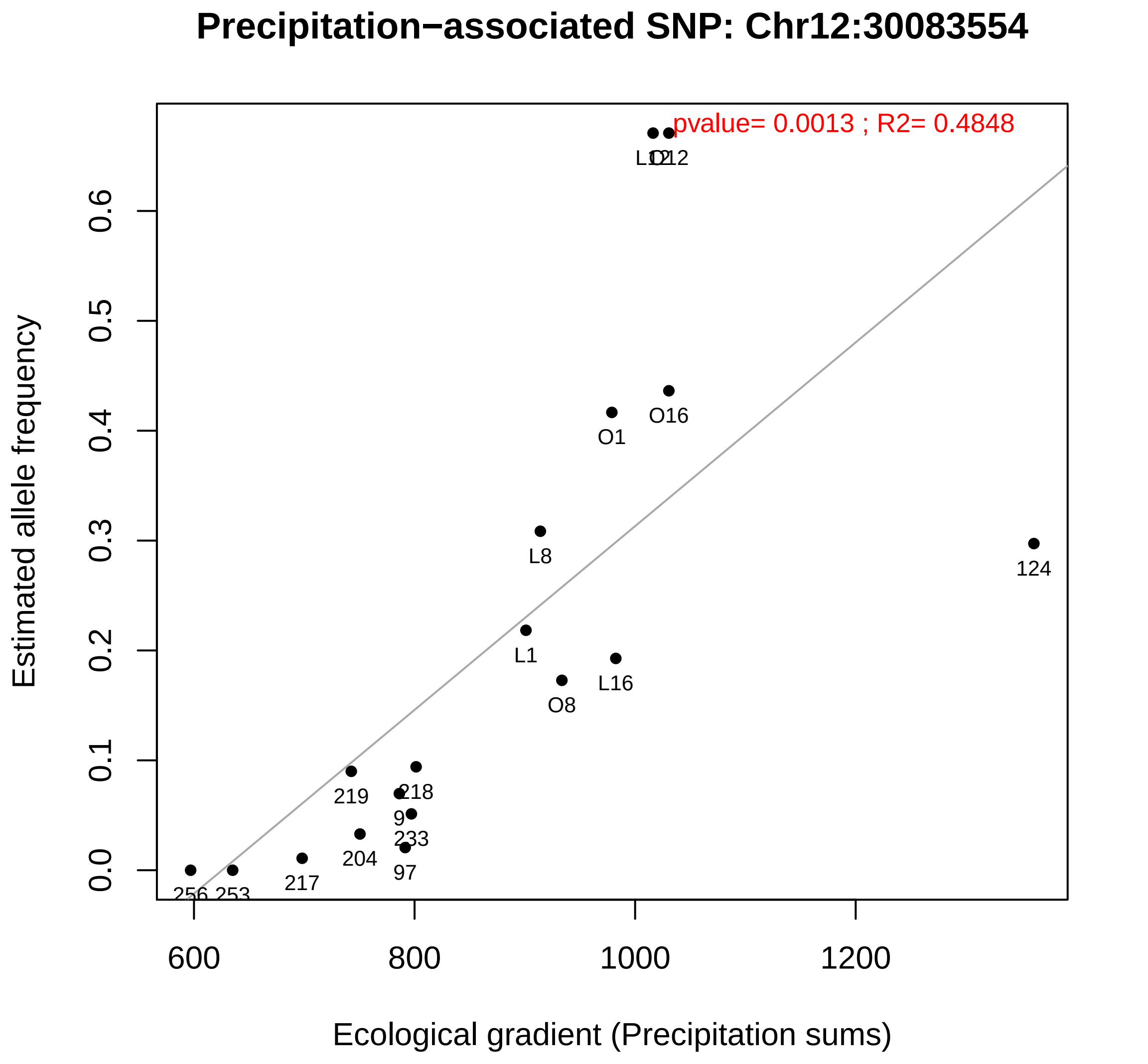

Precipitation−associated SNP: Chr12:30083554
pvalue= 0.0013 ; R2= 0.4848
Estimated allele frequency
Ecological gradient (Precipitation sums)
0.0
0.1
0.2
0.3
0.4
0.5
0.6
600
800
1000
1200
L12
O12
O16
O1
L8
124
L1
L16
O8
219
218
9
233
204
97
217
256
253

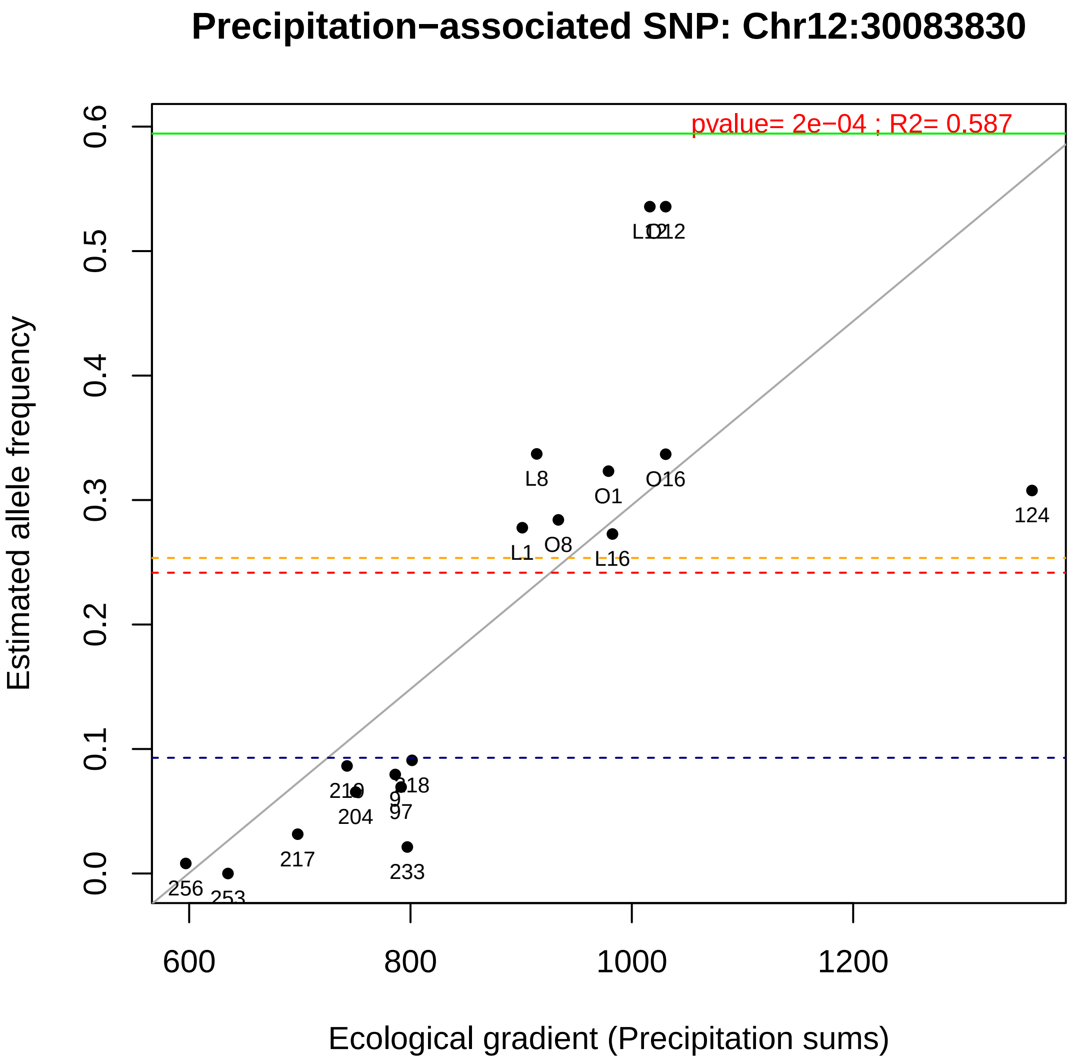

Precipitation−associated SNP: Chr12:30083830
pvalue= 2e−04 ; R2= 0.587
Estimated allele frequency
Ecological gradient (Precipitation sums)
0.0
0.1
0.2
0.3
0.4
0.5
0.6
600
800
1000
1200
L8
O1
O16
124
L1
O8
L16
204
97
217
233
256

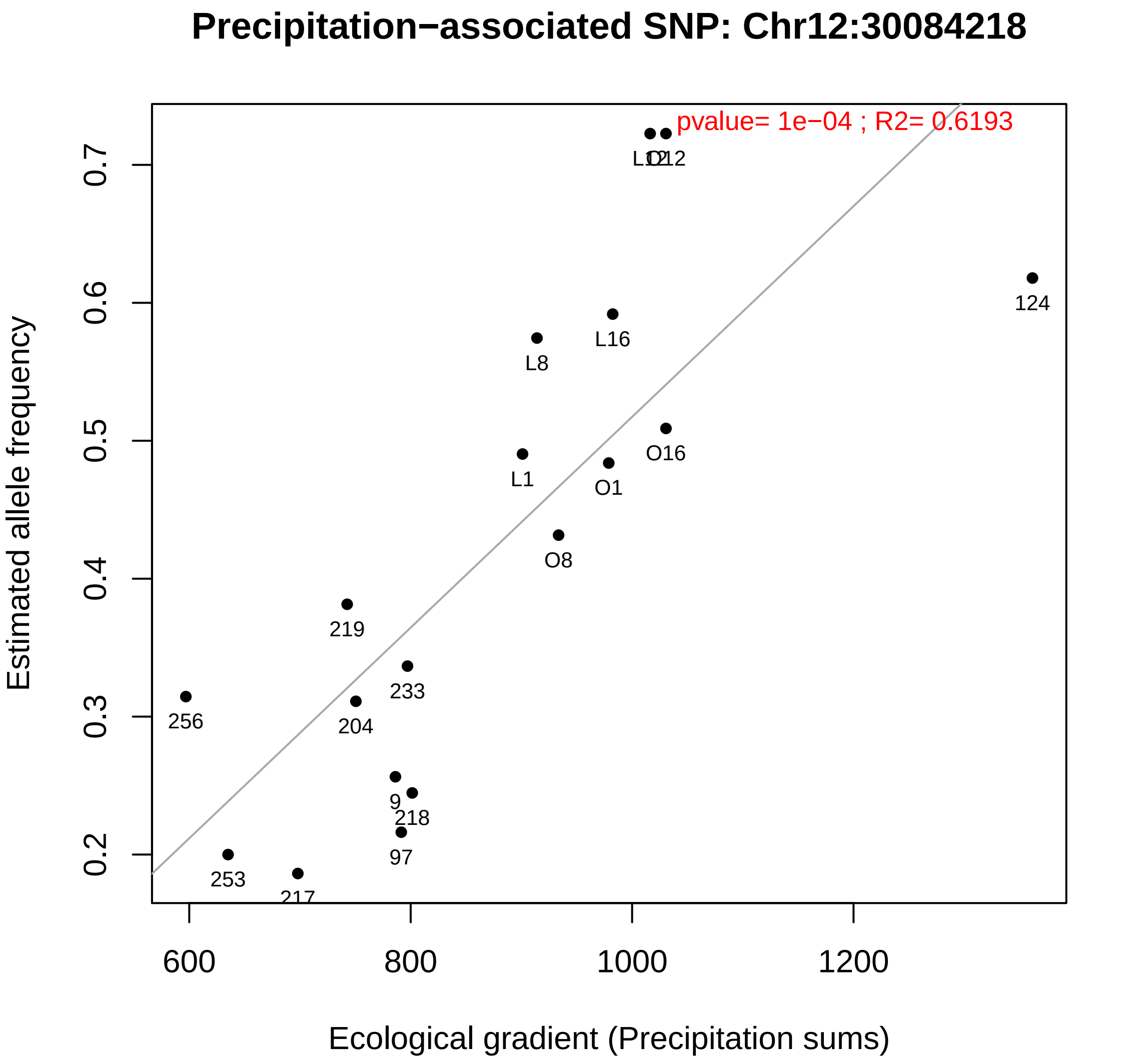
Precipitation−associated SNP: Chr12:30084218
pvalue= 1e−04 ; R2= 0.6193
Estimated allele frequency
Ecological gradient (Precipitation sums)
0.2
0.3
0.4
0.5
0.6
0.7
600
800
1000
1200
124
L16
L8
O16
L1
O1
O8
219
233
256
204
9
218
97
253
217

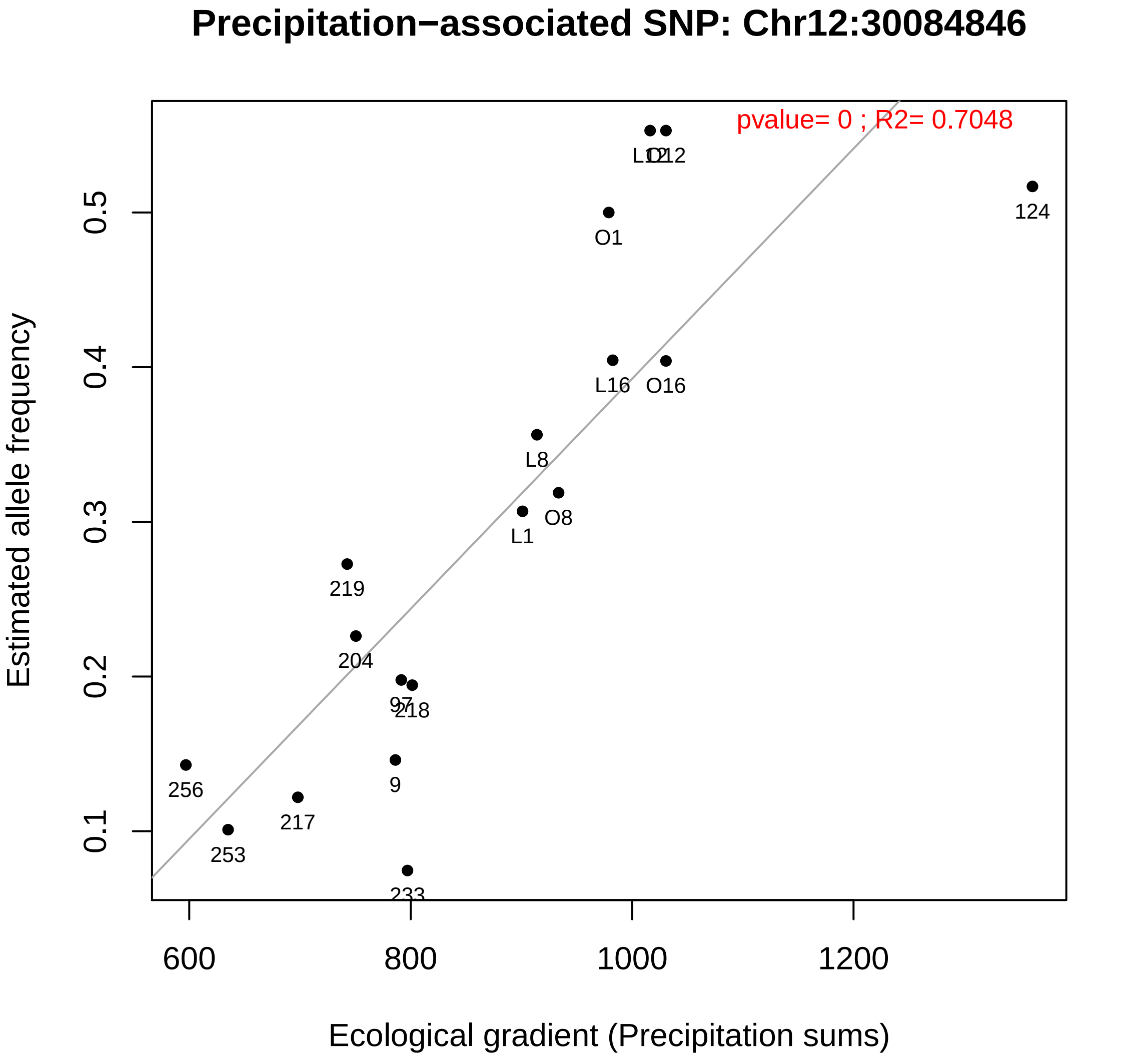

Precipitation−associated SNP: Chr12:30084846
pvalue= 0 ; R2= 0.7048
Estimated allele frequency
Ecological gradient (Precipitation sums)
0.1
0.2
0.3
0.4
0.5
600
800
1000
1200
L12
O12
124
O1
L16
O16
L8
O8
L1
219
204
97
218
256
9
217
253
233

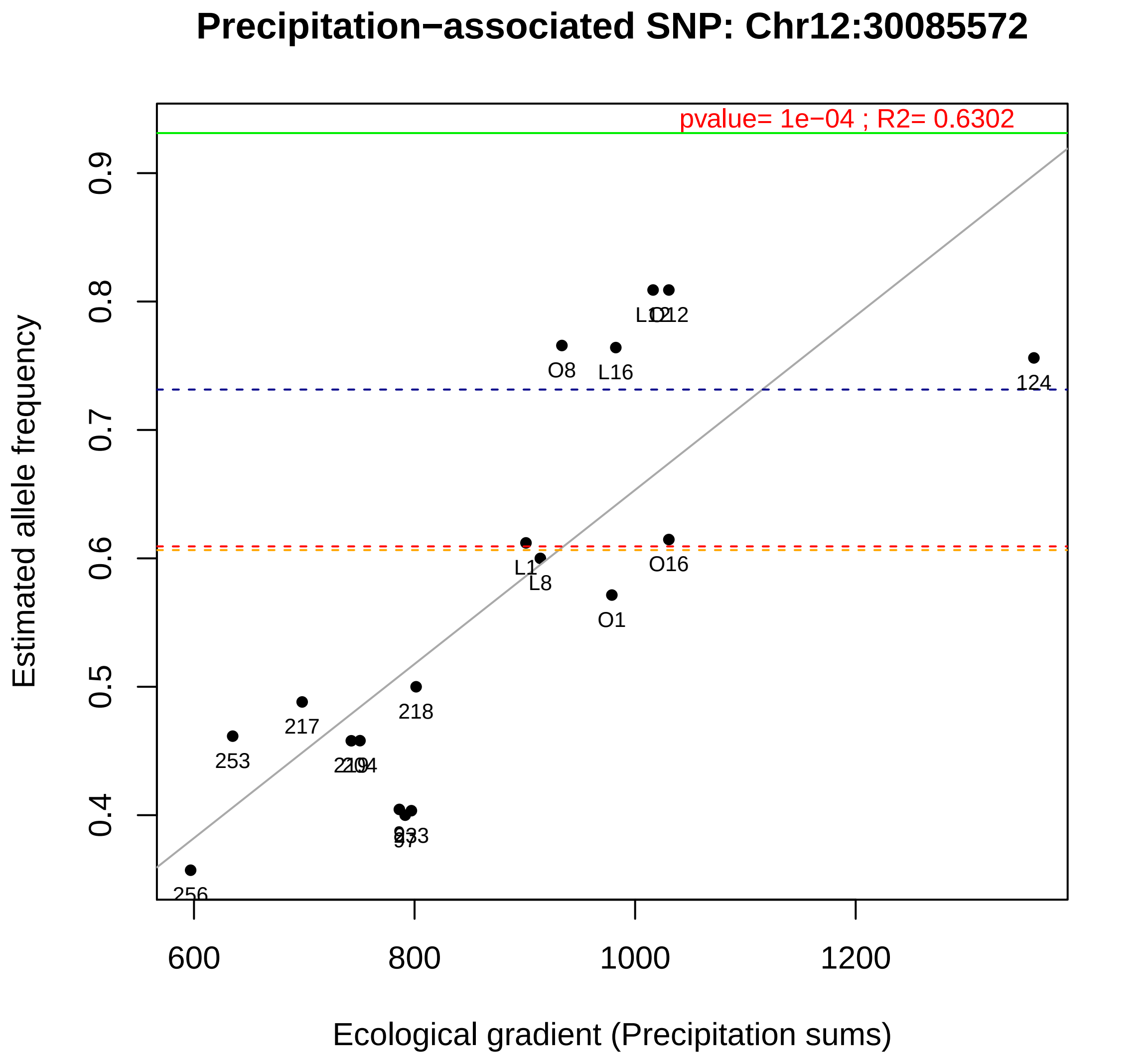

Precipitation−associated SNP: Chr12:30085572
pvalue= 1e−04 ; R2= 0.6302
Estimated allele frequency
Ecological gradient (Precipitation sums)
0.4
0.5
0.6
0.7
0.8
0.9
600
800
1000
1200
L12
O12
O8
L16
124
L1
L8
O16
O1
218
217
253
210
214
256

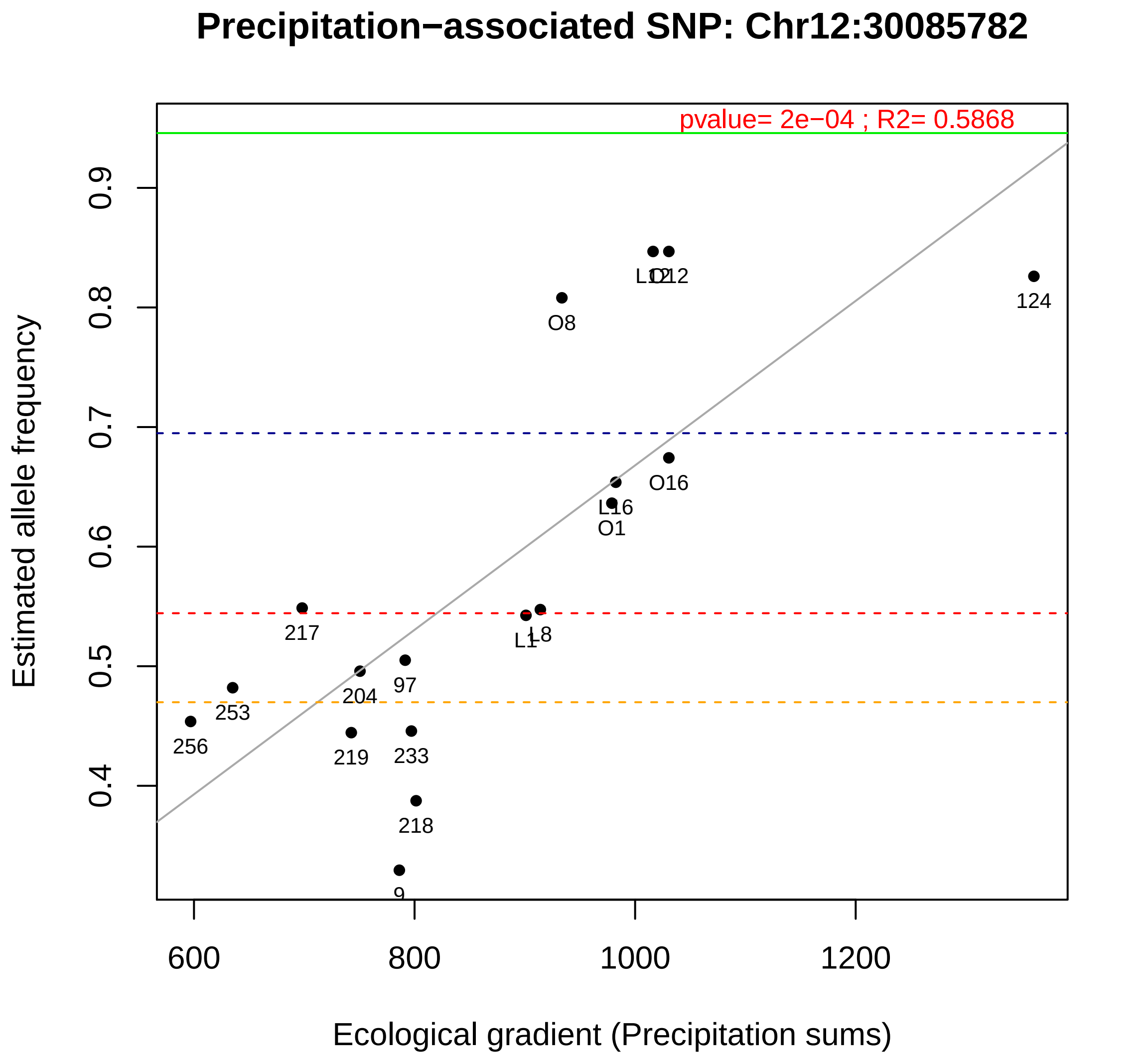

Precipitation−associated SNP: Chr12:30085782
pvalue= 2e−04 ; R2= 0.5868
Estimated allele frequency
Ecological gradient (Precipitation sums)
0.4
0.5
0.6
0.7
0.8
0.9
600
800
1000
1200
L12
O12
124
O8
O16
L16
O1
217
L1
L8
97
204
253
256
219
233
218
9

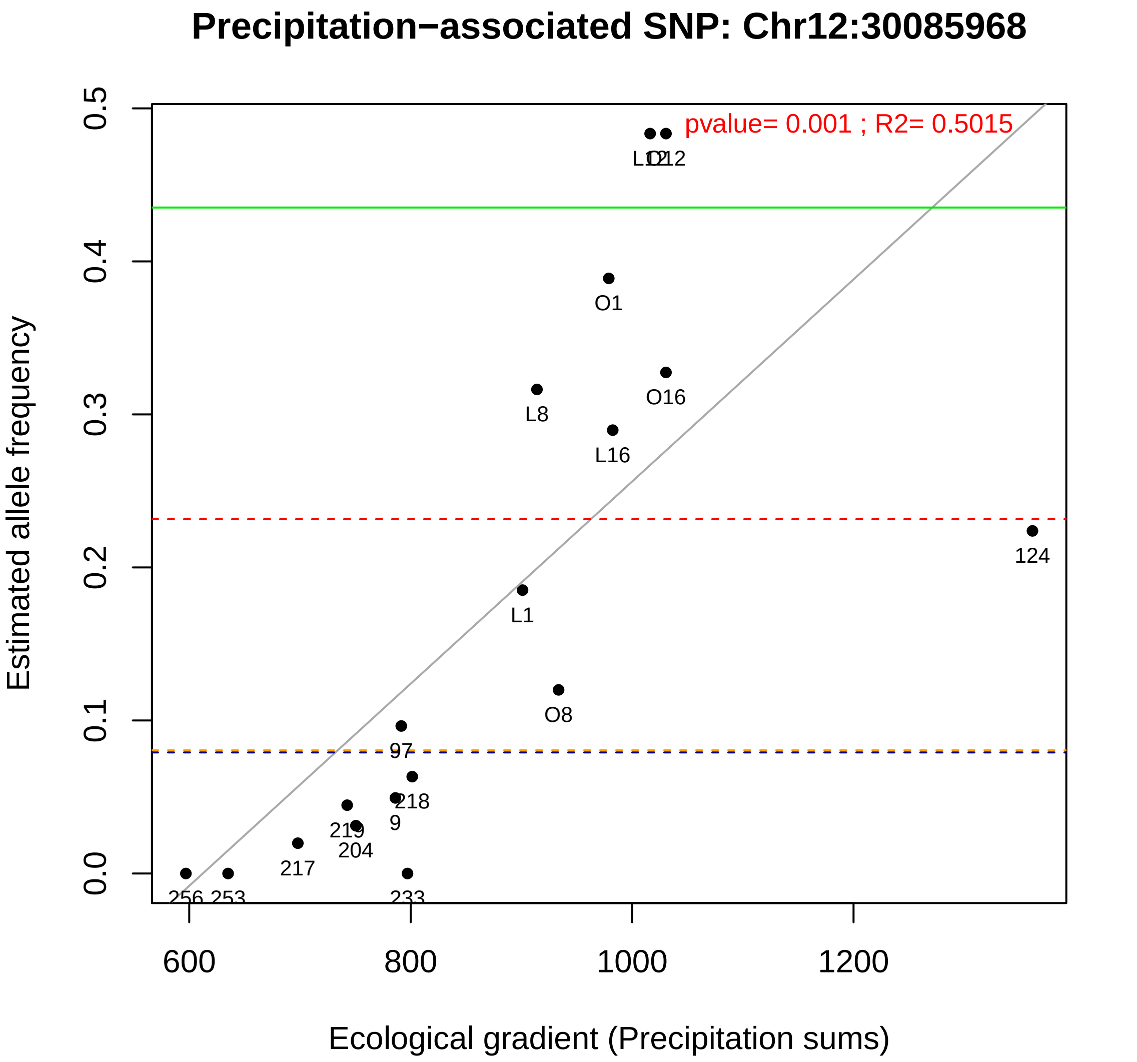

Precipitation−associated SNP: Chr12:30085968
pvalue= 0.001 ; R2= 0.5015
Estimated allele frequency
Ecological gradient (Precipitation sums)
0.0
0.1
0.2
0.3
0.4
0.5
600
800
1000
1200
O1
O16
L8
L16
124
L1
O8
97
218
9
219
204
217
256
253
233

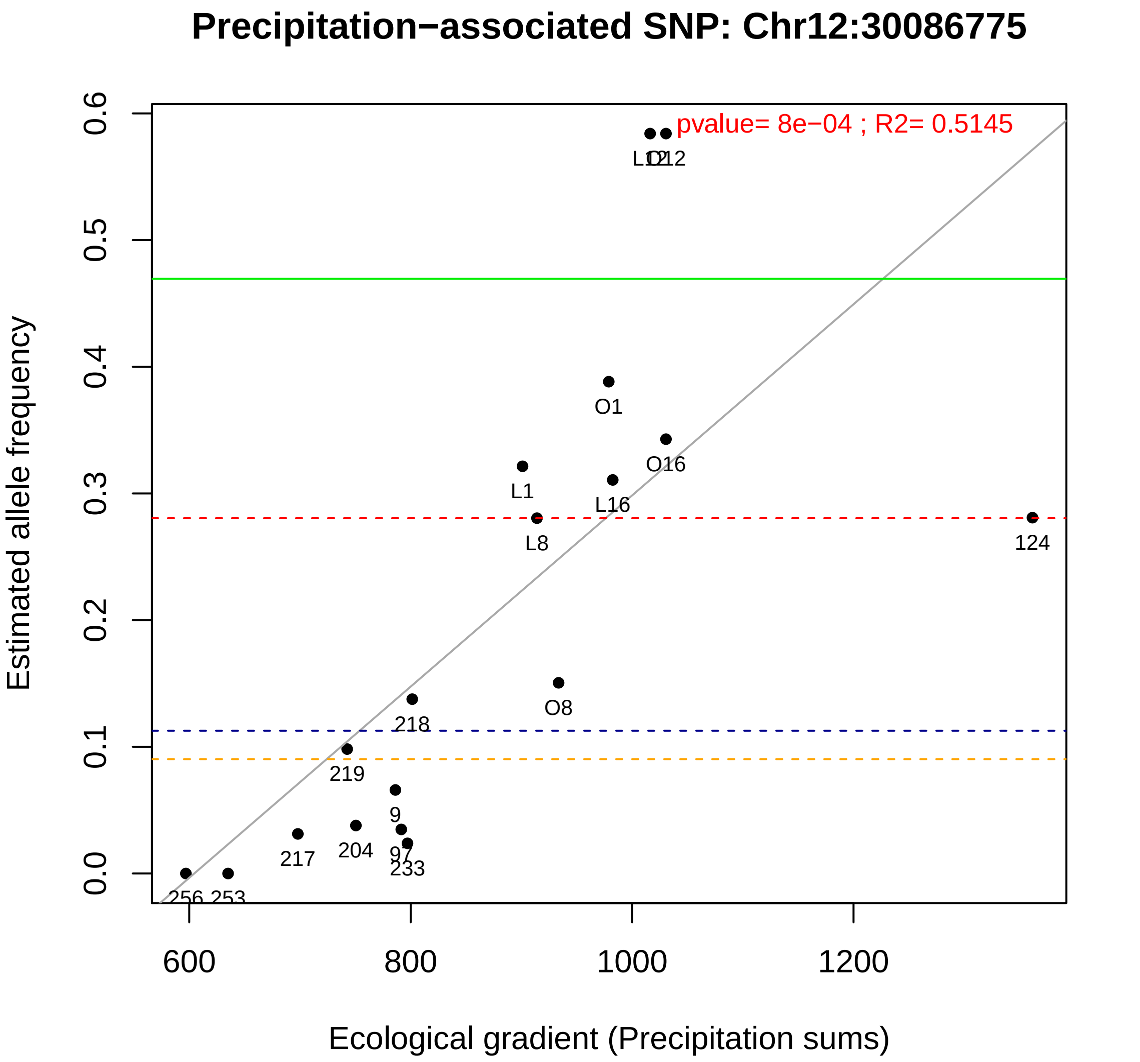
Precipitation−associated SNP: Chr12:30086775
pvalue= 8e−04 ; R2= 0.5145
Estimated allele frequency
Ecological gradient (Precipitation sums)
0.0
0.1
0.2
0.3
0.4
0.5
0.6
600
800
1000
1200
L12
O12
O1
O16
L1
L16
L8
124
O8
218
219
9
217
204
97
233
256
253

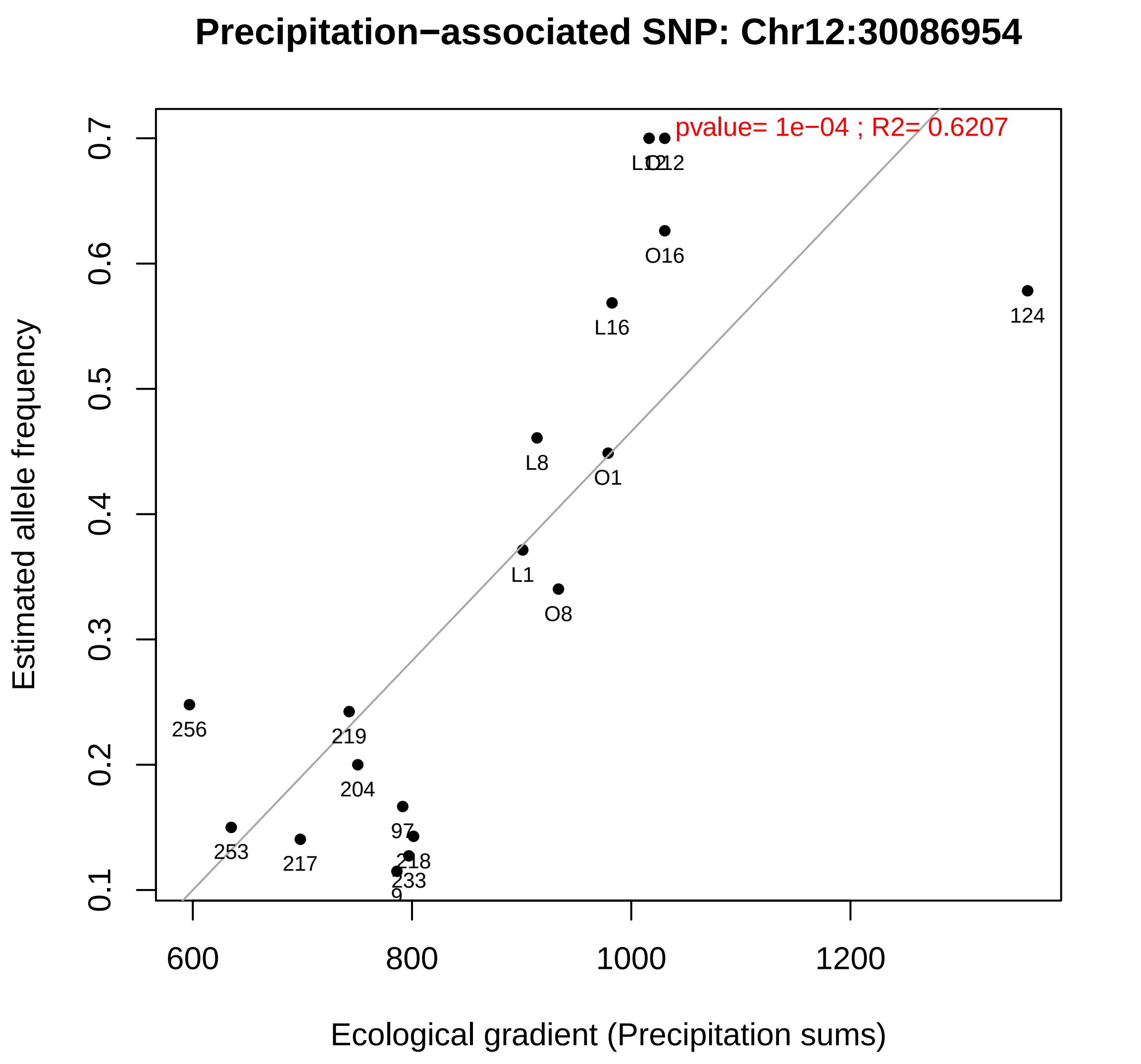
Precipitation−associated SNP: Chr12:30086954
pvalue= 1e−04 ; R2= 0.6207
Estimated allele frequency
Ecological gradient (Precipitation sums)
0.1
0.2
0.3
0.4
0.5
0.6
0.7
600
800
1000
1200
L12
O12
O16
L16
124
L8
O1
L1
O8
256
219
204
253
217
97
218
233
9

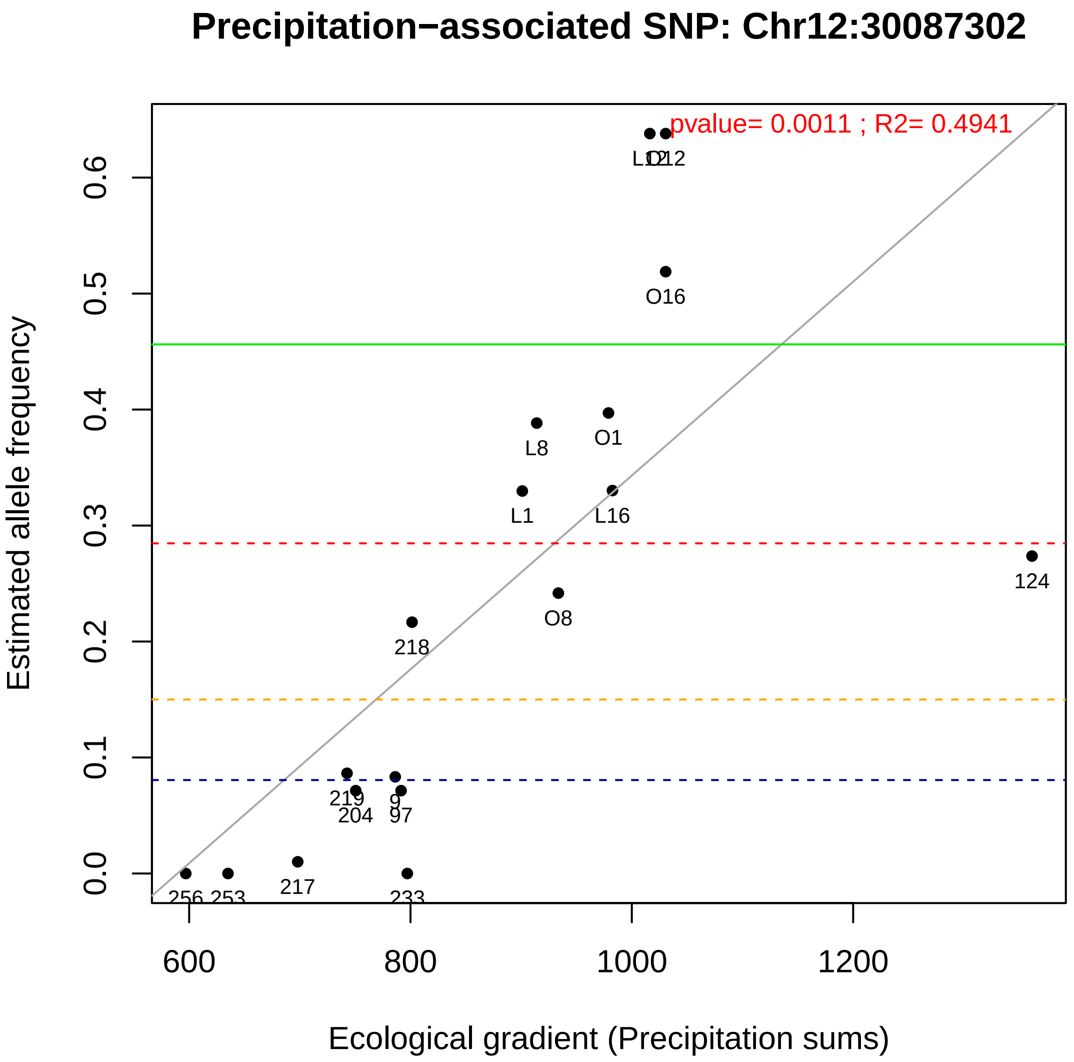
Precipitation−associated SNP: Chr12:30087302
pvalue= 0.0011 ; R2= 0.4941
Estimated allele frequency
Ecological gradient (Precipitation sums)
0.0
0.1
0.2
0.3
0.4
0.5
0.6
600
800
1000
1200
O16
L8
O1
L1
L16
124
O8
218
219
204
9
97
217
233

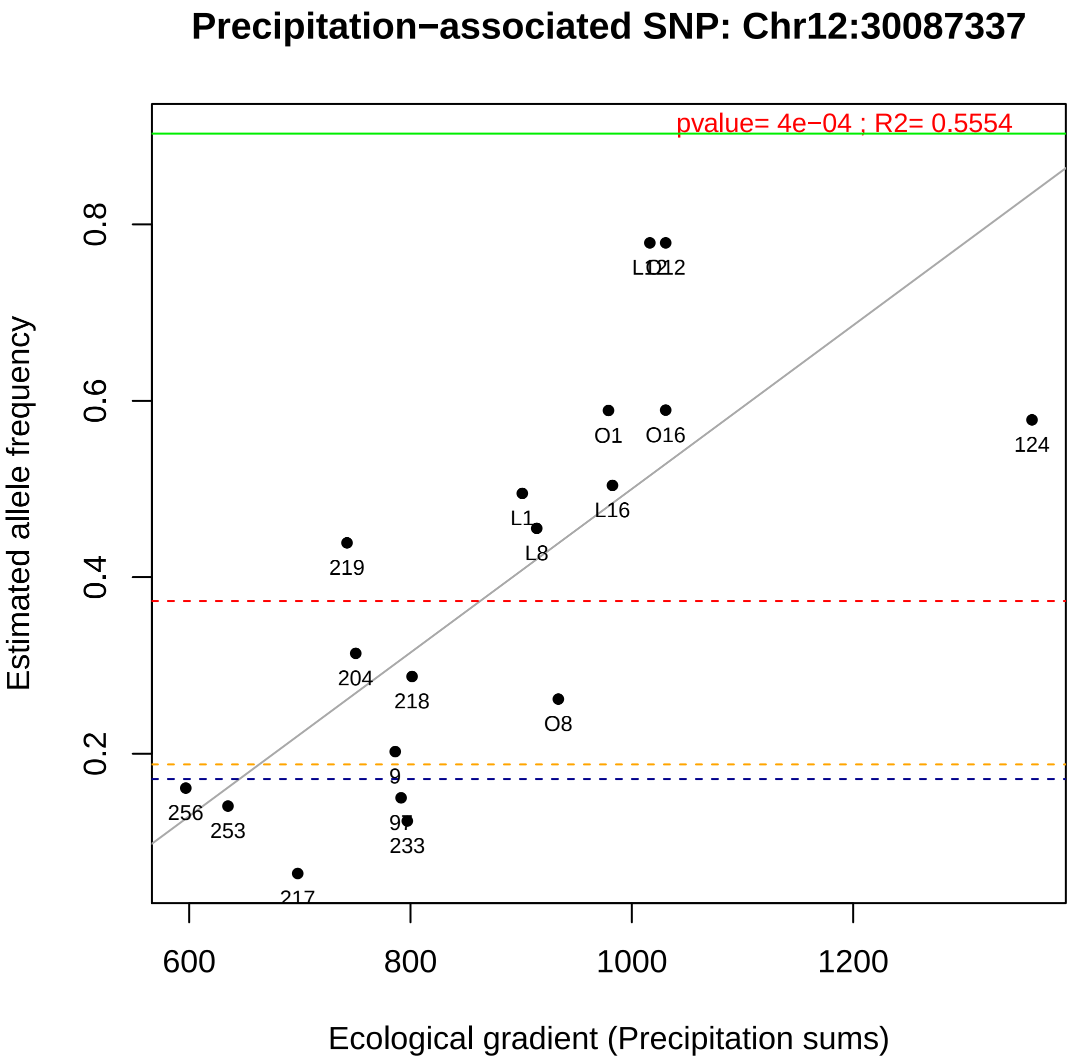

Precipitation−associated SNP: Chr12:30087337
pvalue= 4e−04 ; R2= 0.5554
Estimated allele frequency
Ecological gradient (Precipitation sums)
0.2
0.4
0.6
0.8
600
800
1000
1200
L12
O12
O1
O16
124
L1
L16
L8
219
204
218
O8
9
256
253
97
233
217

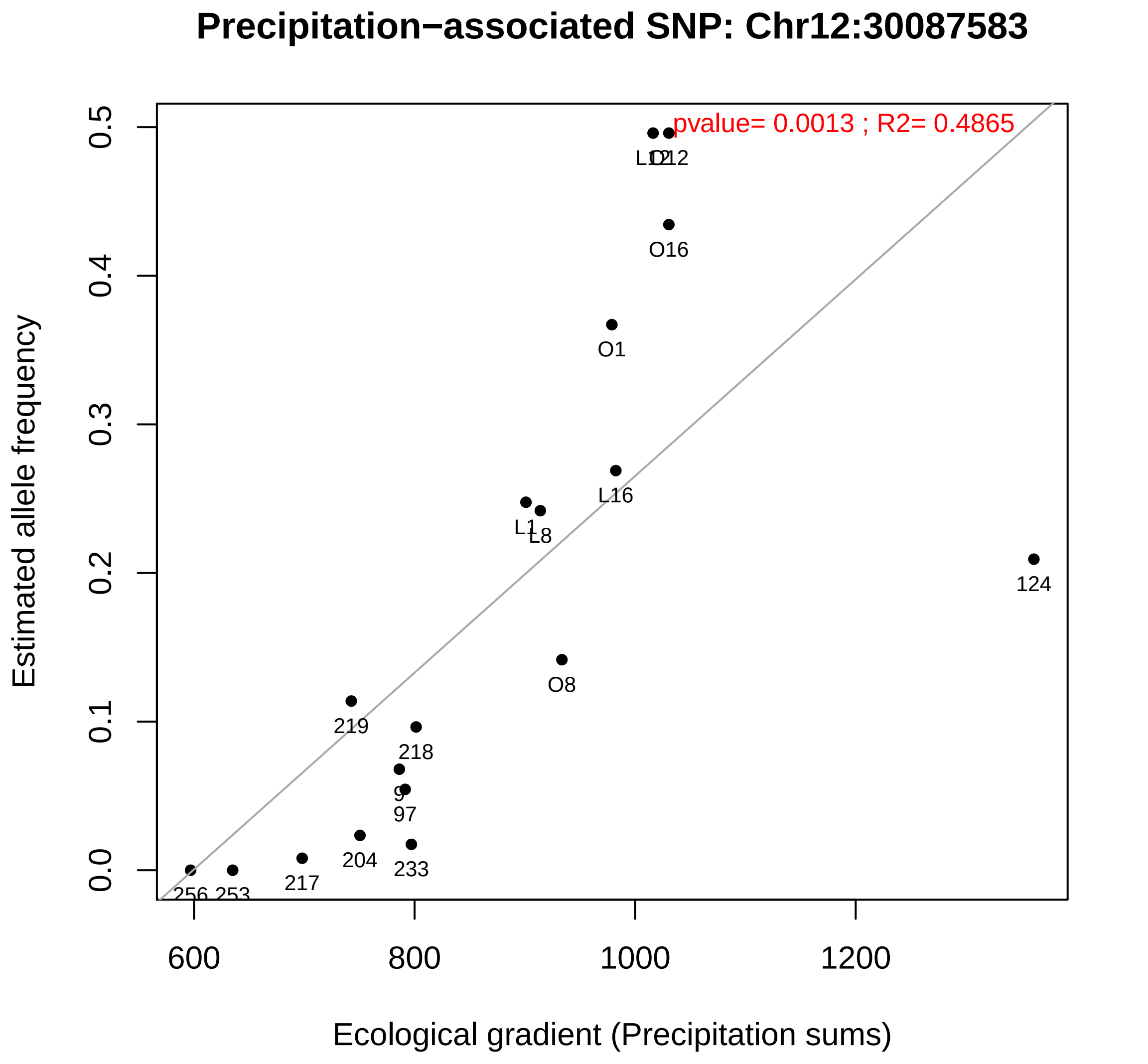
Precipitation−associated SNP: Chr12:30087583
pvalue= 0.0013 ; R2= 0.4865
Estimated allele frequency
Ecological gradient (Precipitation sums)
0.0
0.1
0.2
0.3
0.4
0.5
600
800
1000
1200
O16
O1
L16
L1
L8
124
O8
219
218
9
97
204
233
217
256
253

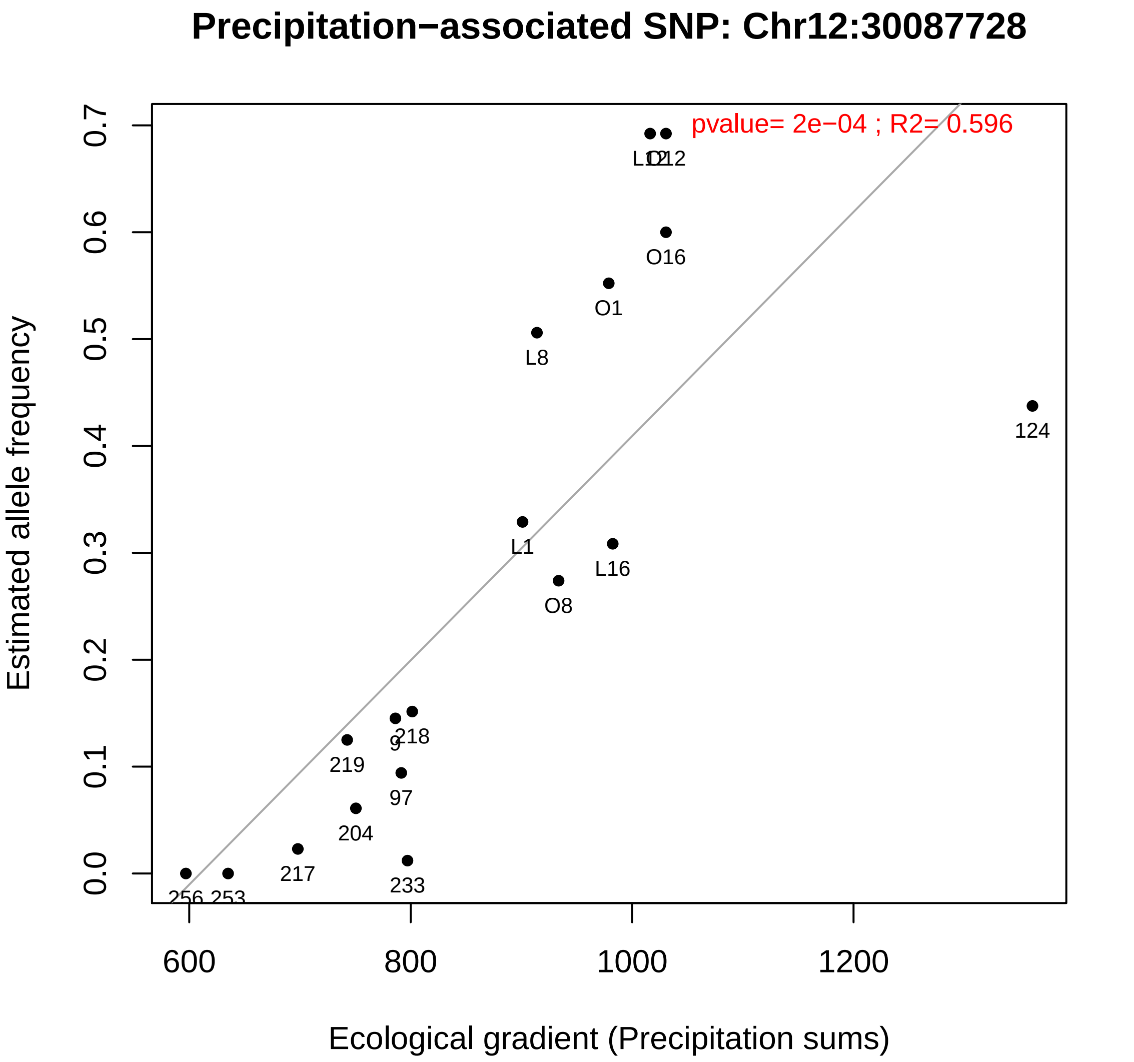
Precipitation−associated SNP: Chr12:30087728
pvalue= 2e−04 ; R2= 0.596
Estimated allele frequency
0.0
0.1
0.2
0.3
0.4
0.5
0.6
0.7
Ecological gradient (Precipitation sums)
600
800
1000
1200
O16
O1
L8
124
L1
L16
O8
218
219
97
204
217
233
256
253

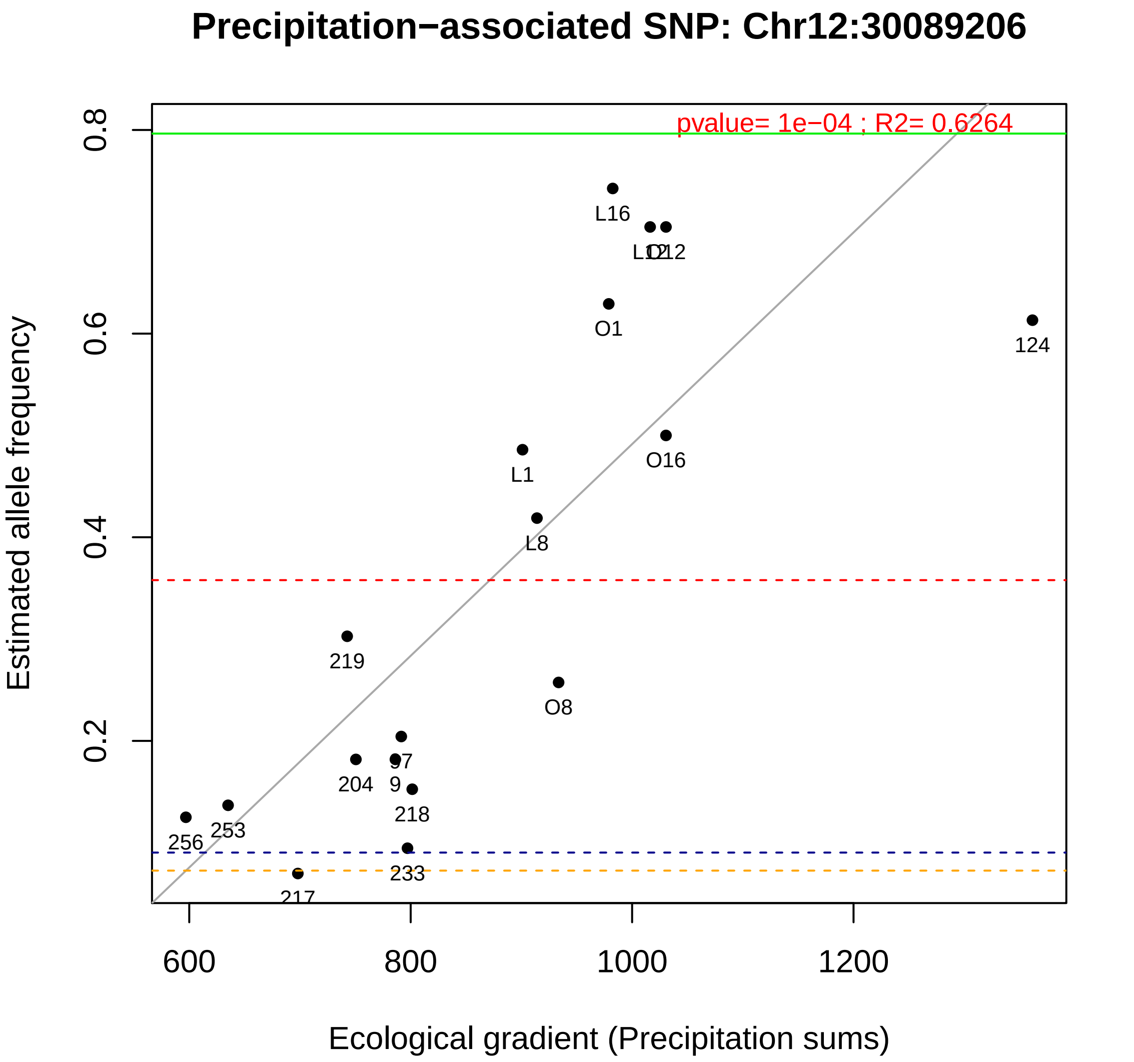
Precipitation−associated SNP: Chr12:30089206
pvalue= 1e−04 ; R2= 0.6264
Estimated allele frequency
Ecological gradient (Precipitation sums)
0.2
0.4
0.6
0.8
600
800
1000
1200
L16
O1
124
O16
L1
L8
219
O8
204
218
256
253
233

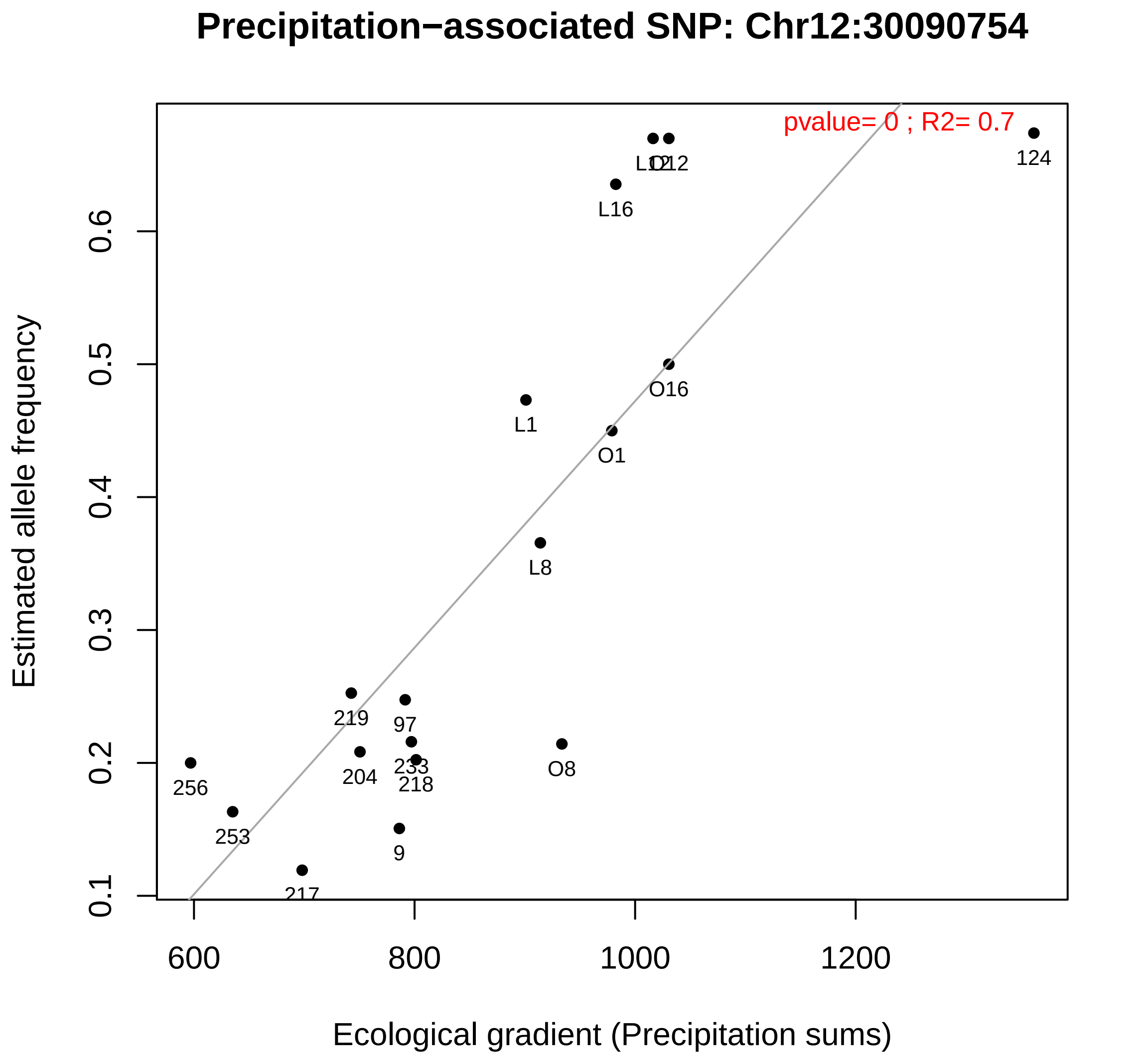
Precipitation−associated SNP: Chr12:30090754
pvalue= 0 ; R2= 0.7
Estimated allele frequency
Ecological gradient (Precipitation sums)
0.1
0.2
0.3
0.4
0.5
0.6
600
800
1000
1200
124
L16
O16
L1
O1
L8
219
97
204
233
218
O8
256
253
9
217

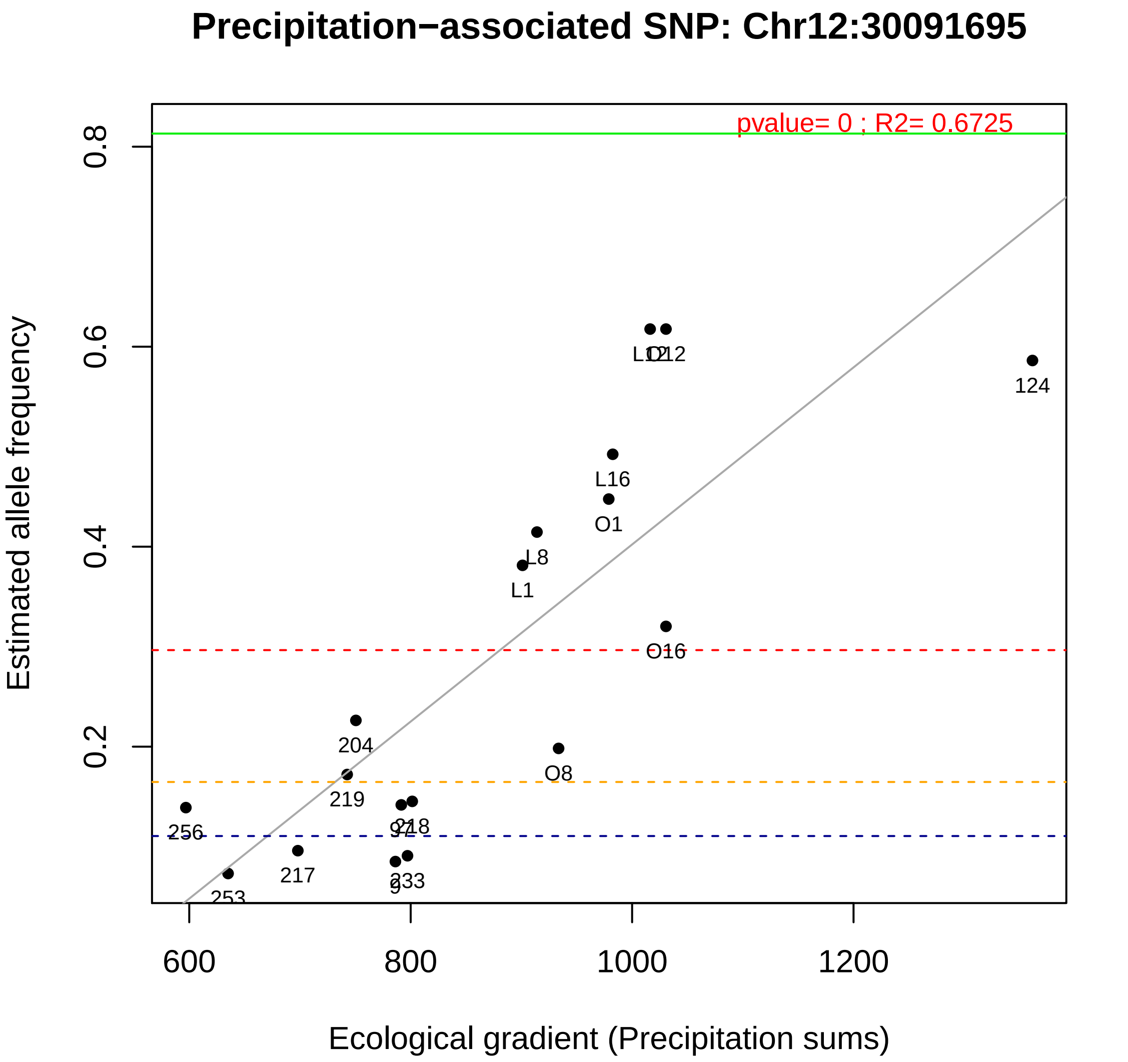
Precipitation−associated SNP: Chr12:30091695
pvalue= 0 ; R2= 0.6725
Estimated allele frequency
0.2
0.4
0.6
0.8
600
800
1000
1200
Ecological gradient (Precipitation sums)
L12
O12
124
L16
O1
L8
L1
O16
204
O8
219
256
218
217
233
253

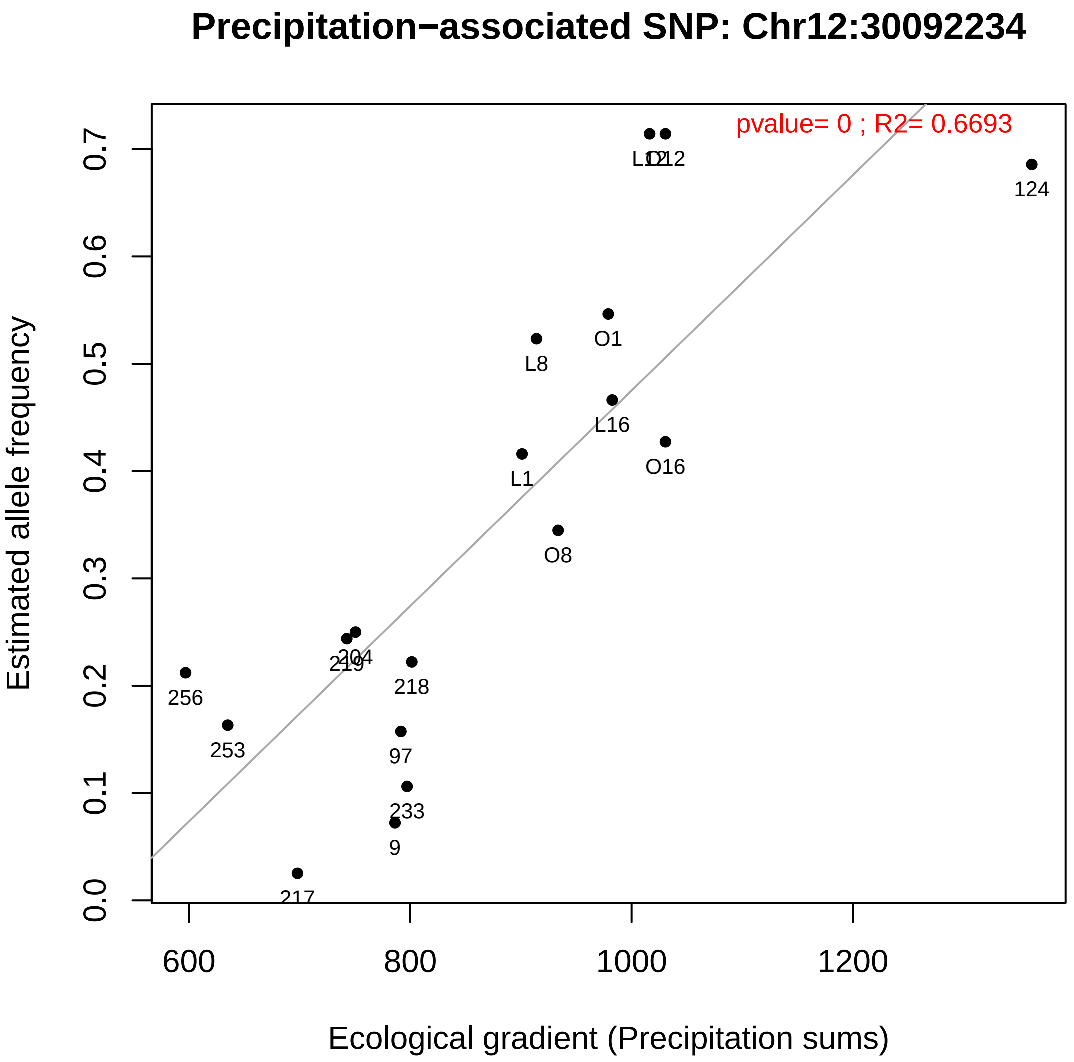
Precipitation−associated SNP: Chr12:30092234
pvalue= 0 ; R2= 0.6693
Estimated allele frequency
Ecological gradient (Precipitation sums)
0.0
0.1
0.2
0.3
0.4
0.5
0.6
0.7
600
800
1000
1200
124
O1
L8
L16
O16
L1
O8
204
219
218
256
253
97
233
9
217

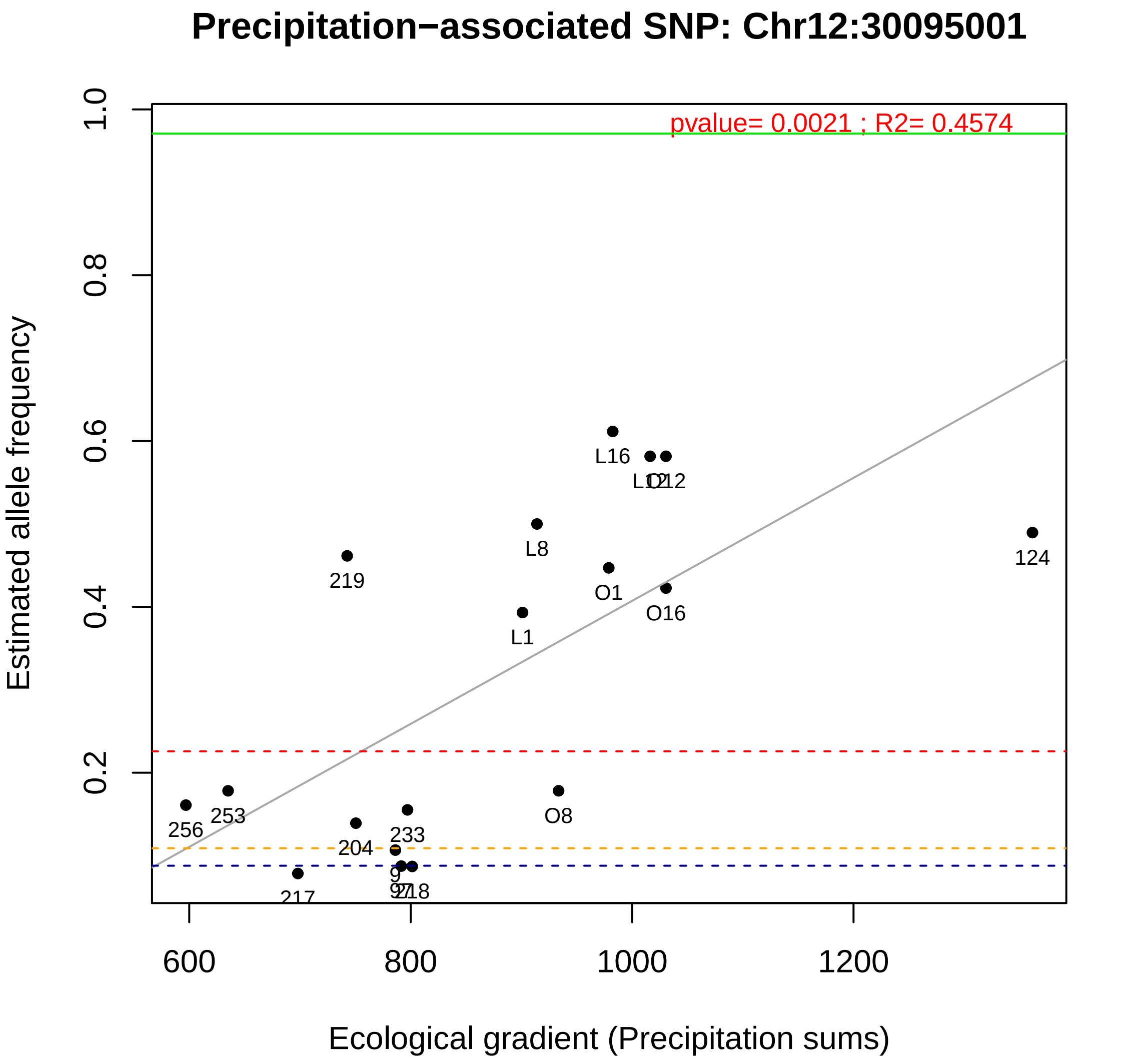

Precipitation−associated SNP: Chr12:30095001
pvalue= 0.0021 ; R2= 0.4574
Estimated allele frequency
0.2
0.4
0.6
0.8
1.0
600
800
1000
1200
Ecological gradient (Precipitation sums)
L16
L12
O12
L8
219
124
O1
O16
L1
256
253
204
233
O8
217
9
97
218

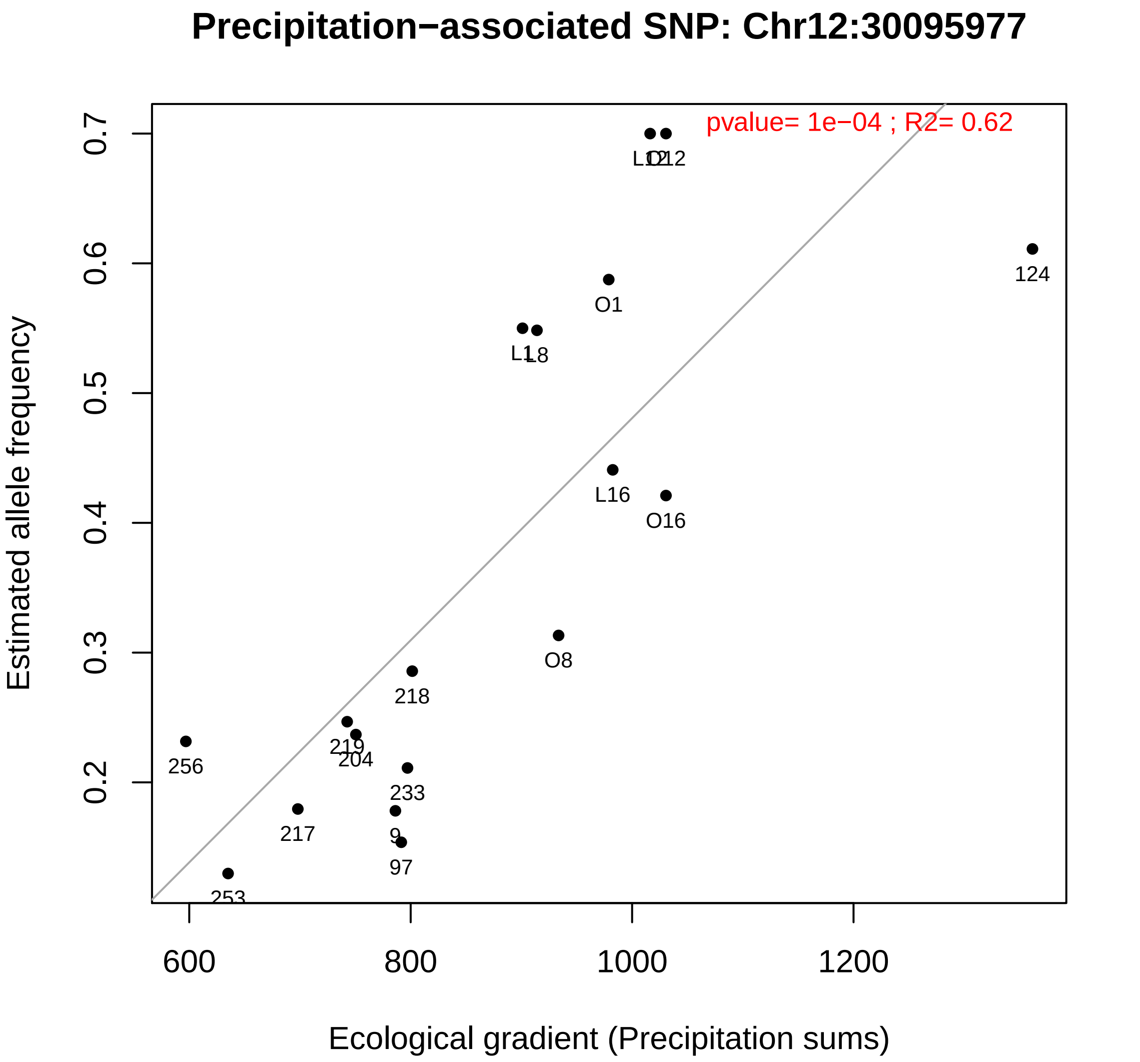
Precipitation−associated SNP: Chr12:30095977
pvalue= 1e−04 ; R2= 0.62
Estimated allele frequency
Ecological gradient (Precipitation sums)
0.2
0.3
0.4
0.5
0.6
0.7
600
800
1000
1200
124
O1
L18
L16
O16
O8
218
219
204
256
233
217
97
253

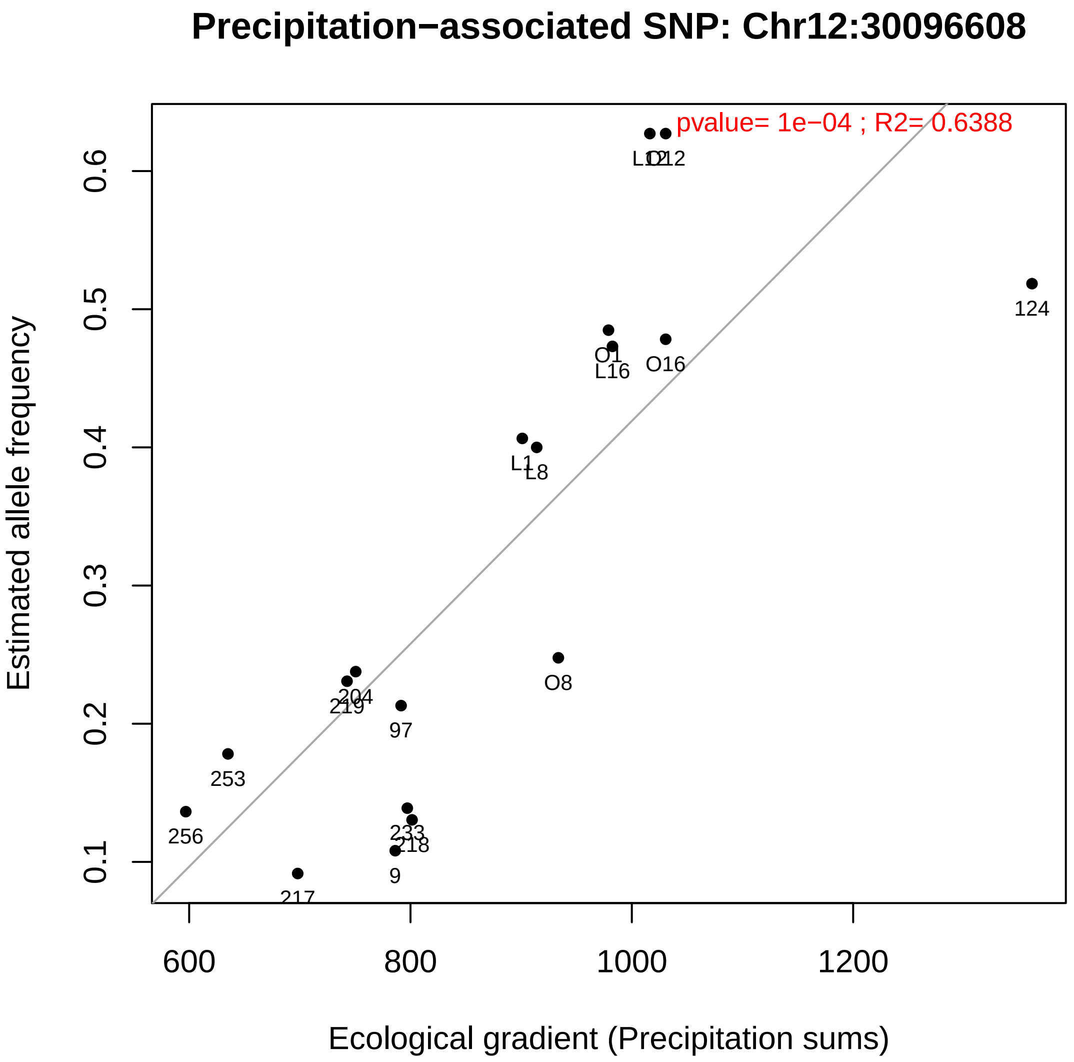
Precipitation−associated SNP: Chr12:30096608
pvalue= 1e−04 ; R2= 0.6388
Estimated allele frequency
Ecological gradient (Precipitation sums)
0.1
0.2
0.3
0.4
0.5
0.6
600
800
1000
1200
124
O16
L16
O8
L8
97
253
256
233
218
9
217
204
219

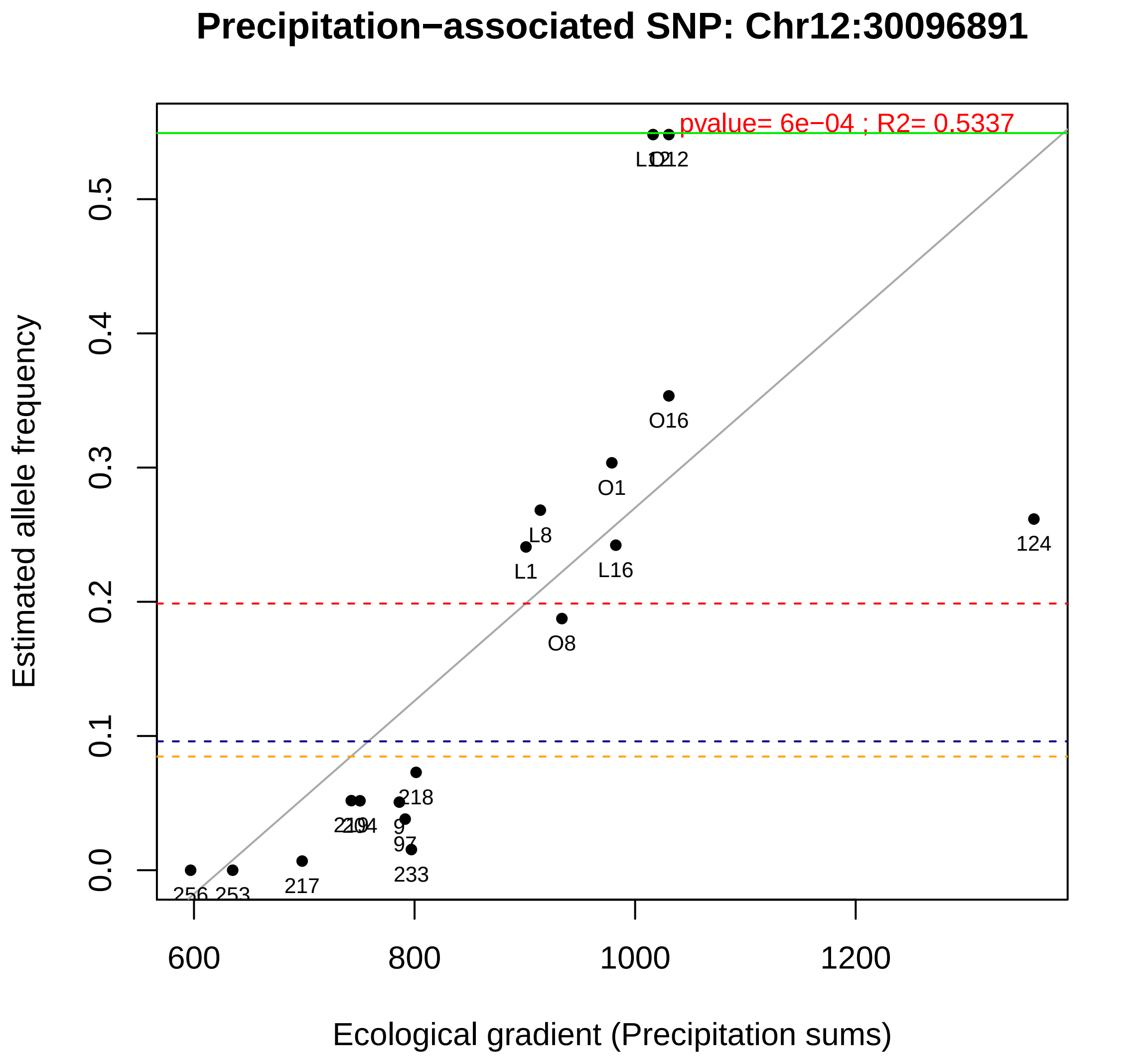

Precipitation−associated SNP: Chr12:30096891
pvalue= 6e−04 ; R2= 0.5337
Estimated allele frequency
0.0
0.1
0.2
0.3
0.4
0.5
600
800
1000
1200
Ecological gradient (Precipitation sums)
O16
O1
L8
L1
L16
124
O8
218
9
97
233
217

**Precipitation−associated SNP: Chr12:30098102**

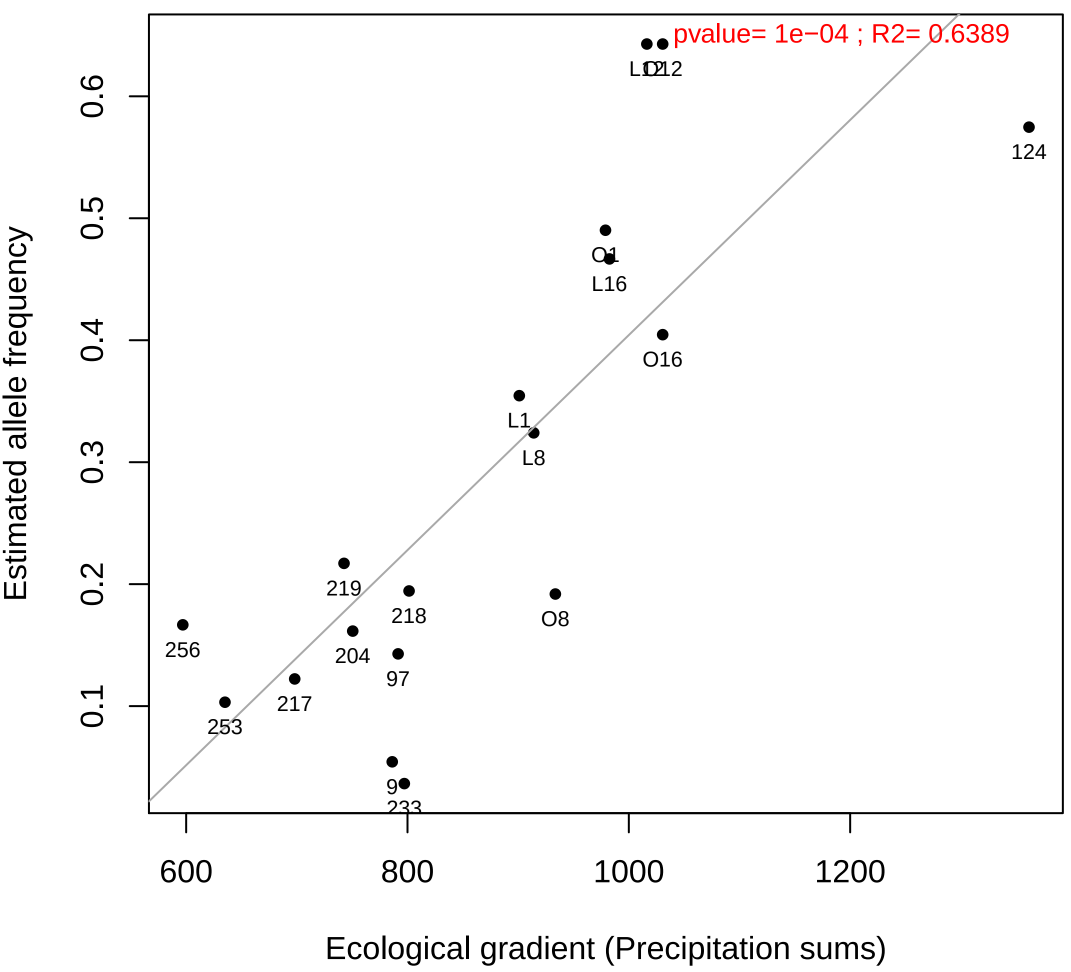

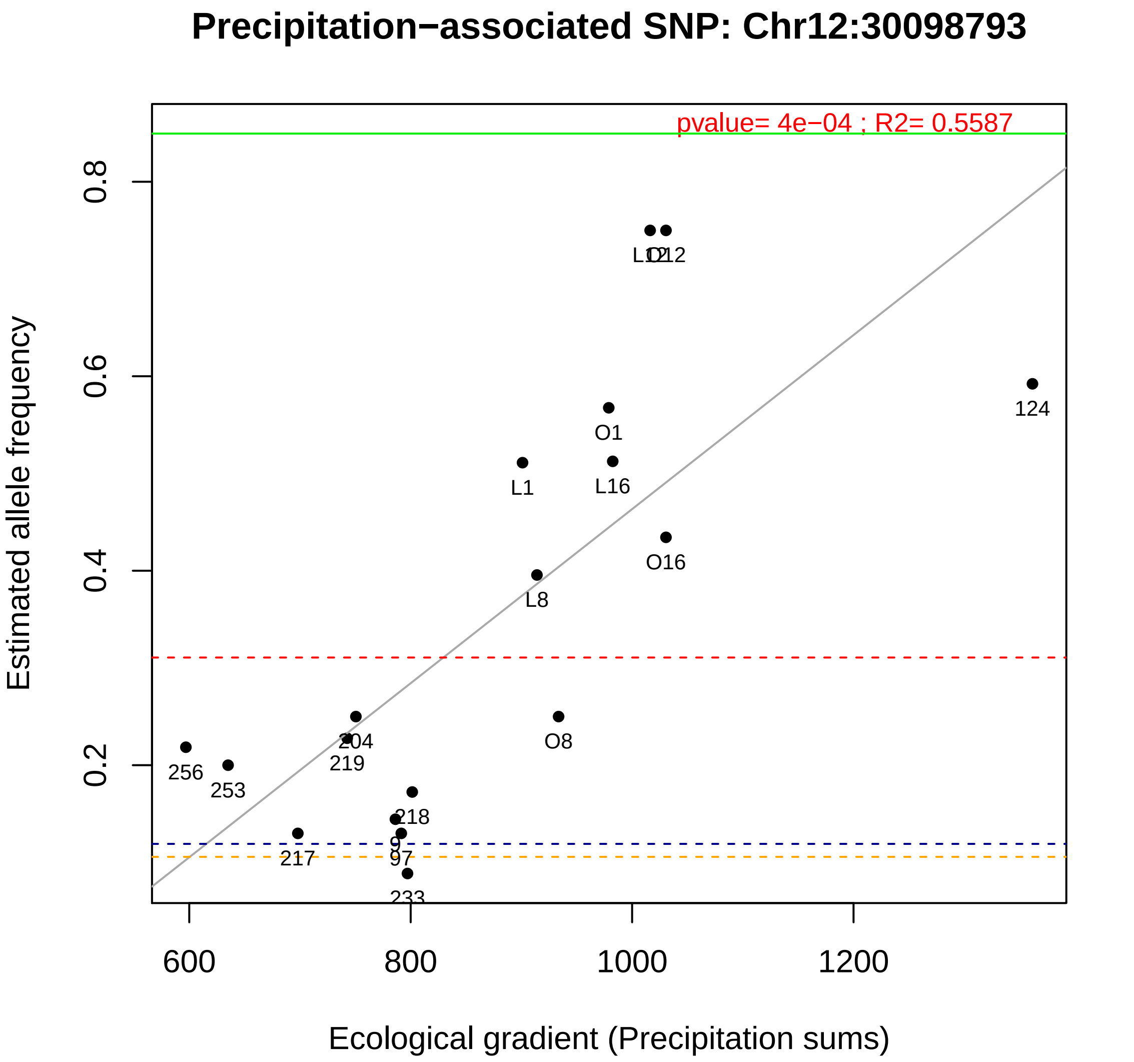

Precipitation−associated SNP: Chr12:30098793
pvalue= 4e−04 ; R2= 0.5587
Estimated allele frequency
0.2
0.4
0.6
0.8
600
800
1000
1200
Ecological gradient (Precipitation sums)
124
O1
L1
L16
O16
L8
O8
204
219
256
253
218
217
9
97
233

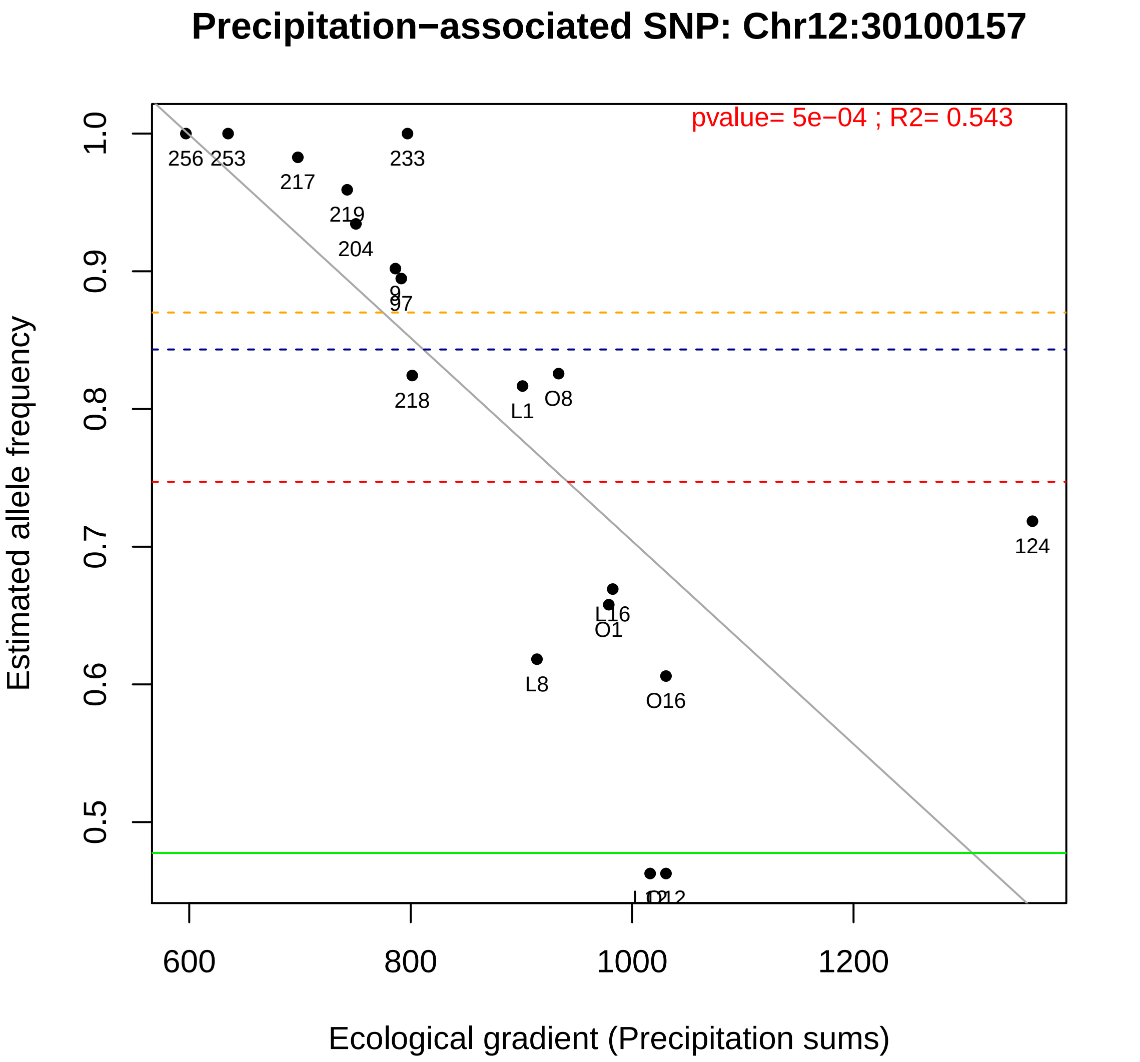
Precipitation−associated SNP: Chr12:30100157
pvalue= 5e−04 ; R2= 0.543
Estimated allele frequency
Ecological gradient (Precipitation sums)
1.0
0.9
0.8
0.7
0.6
0.5
600
800
1000
1200
256
253
217
233
219
204
9
97
218
L1
O8
124
L16
O1
L8
O16

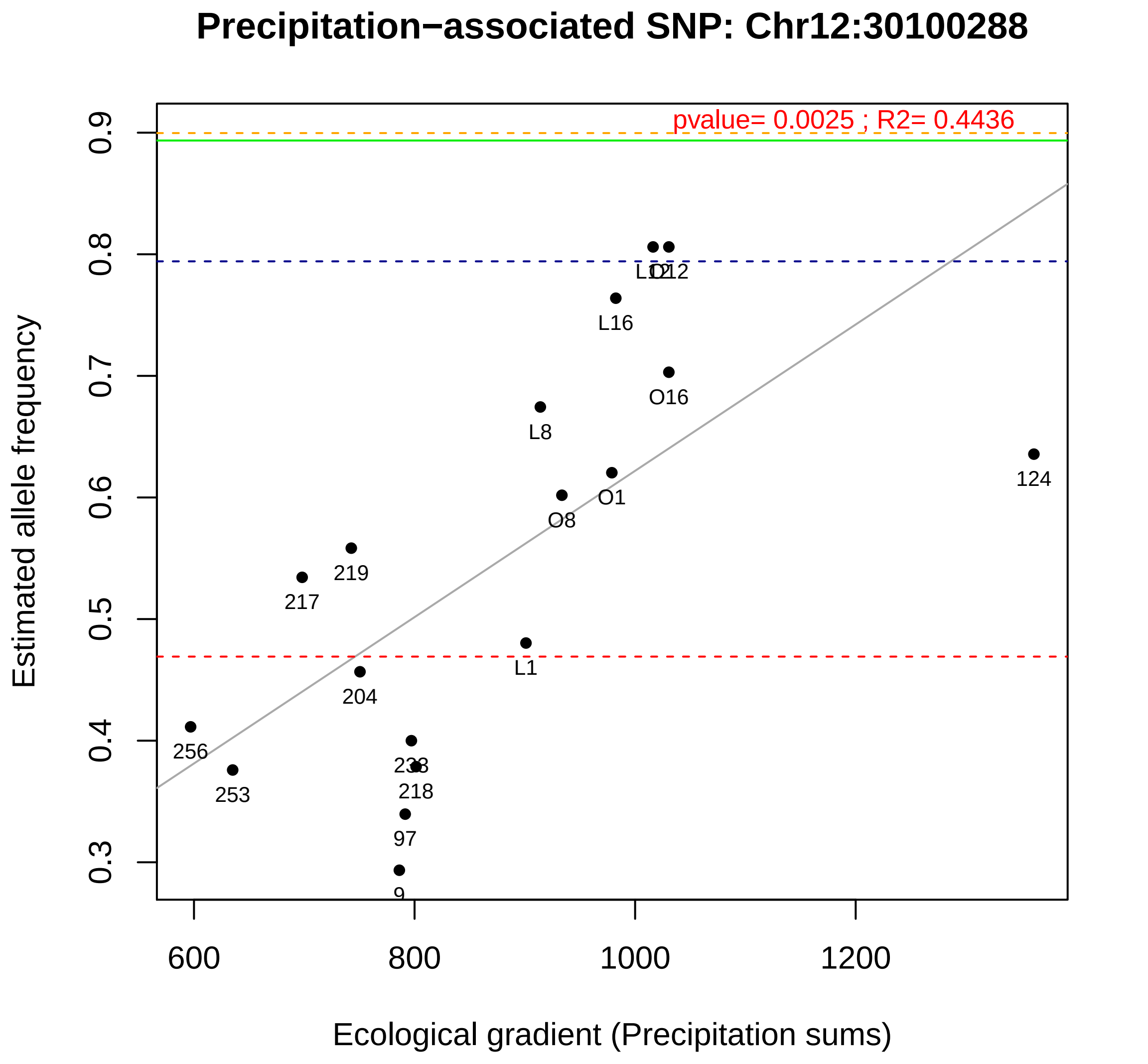
Precipitation−associated SNP: Chr12:30100288
pvalue= 0.0025 ; R2= 0.4436
Estimated allele frequency
0.3
0.4
0.5
0.6
0.7
0.8
0.9
600
800
1000
1200
Ecological gradient (Precipitation sums)
256
253
217
219
204
97
9
218
L1
L8
O8
O1
L16
O16
124

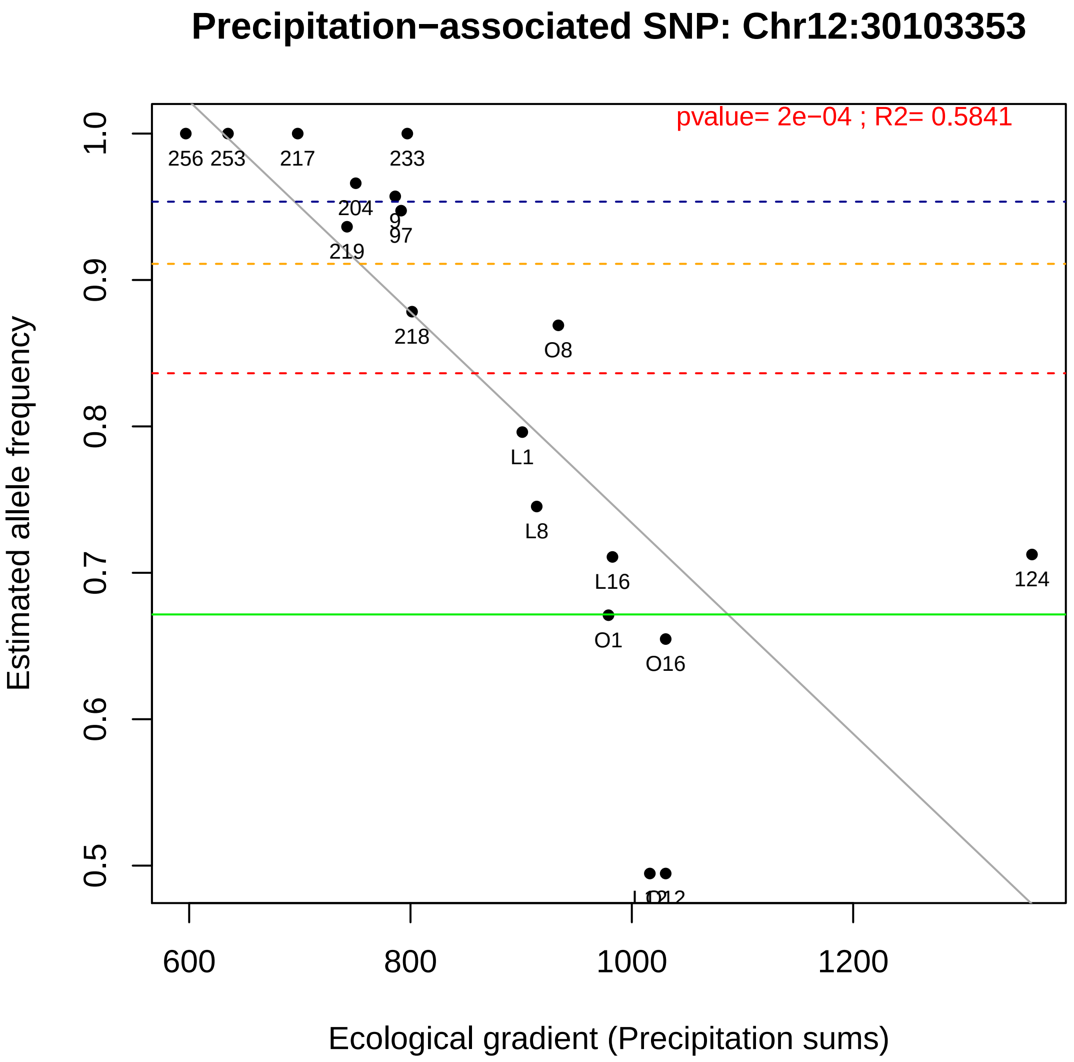
Precipitation−associated SNP: Chr12:30103353
pvalue= 2e−04 ; R2= 0.5841
Estimated allele frequency
Ecological gradient (Precipitation sums)
1.0
0.9
0.8
0.7
0.6
0.5
600
800
1000
1200
256
253
217
233
204
9
97
219
218
O8
L1
L8
L16
124
O1
O16

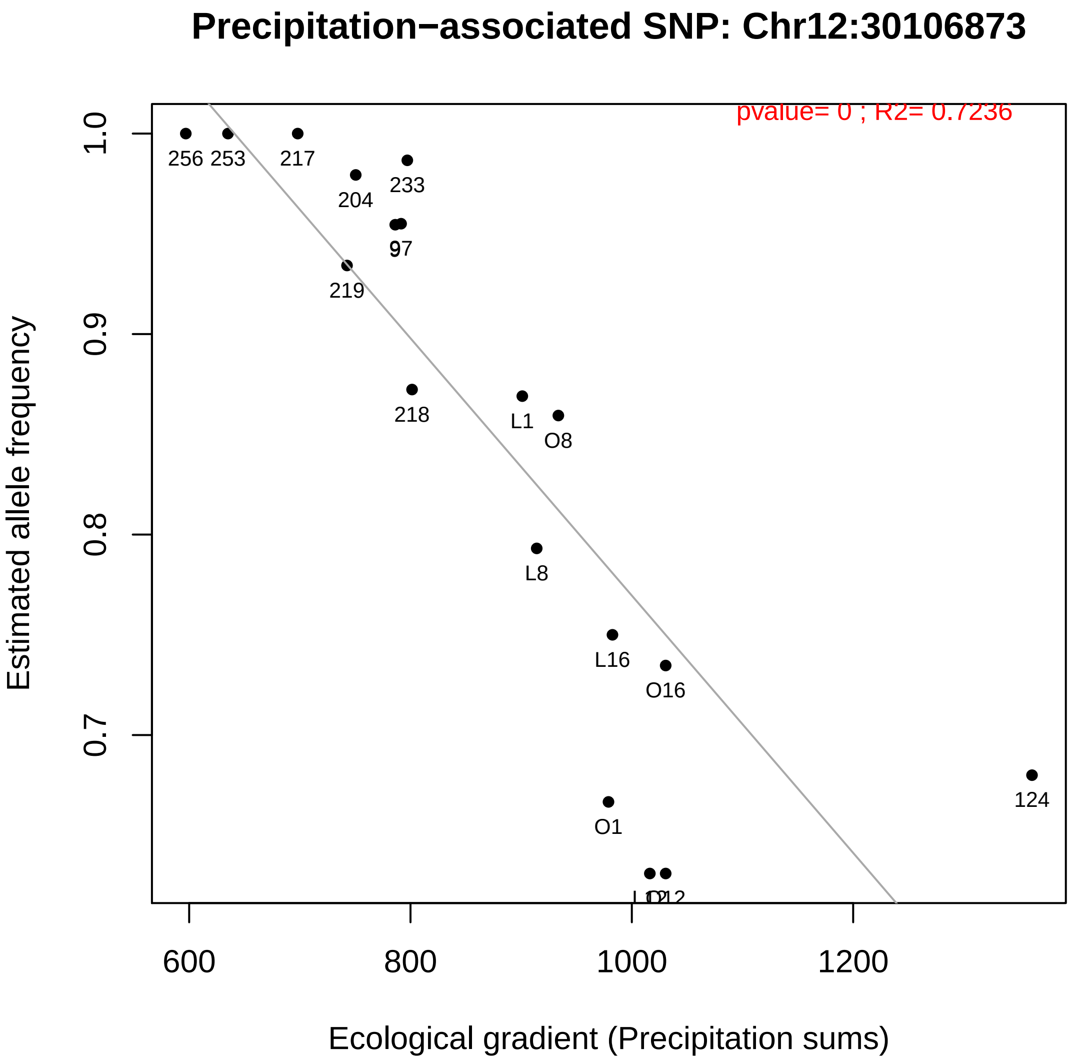
Precipitation−associated SNP: Chr12:30106873
pvalue= 0 ; R2= 0.7236
Estimated allele frequency
Ecological gradient (Precipitation sums)
1.0
0.9
0.8
0.7
600
800
1000
1200
256
253
217
233
204
97
219
218
L1
O8
L8
L16
O16
O1
124

#### Precipitation−associated SNP: Chr12:30107530

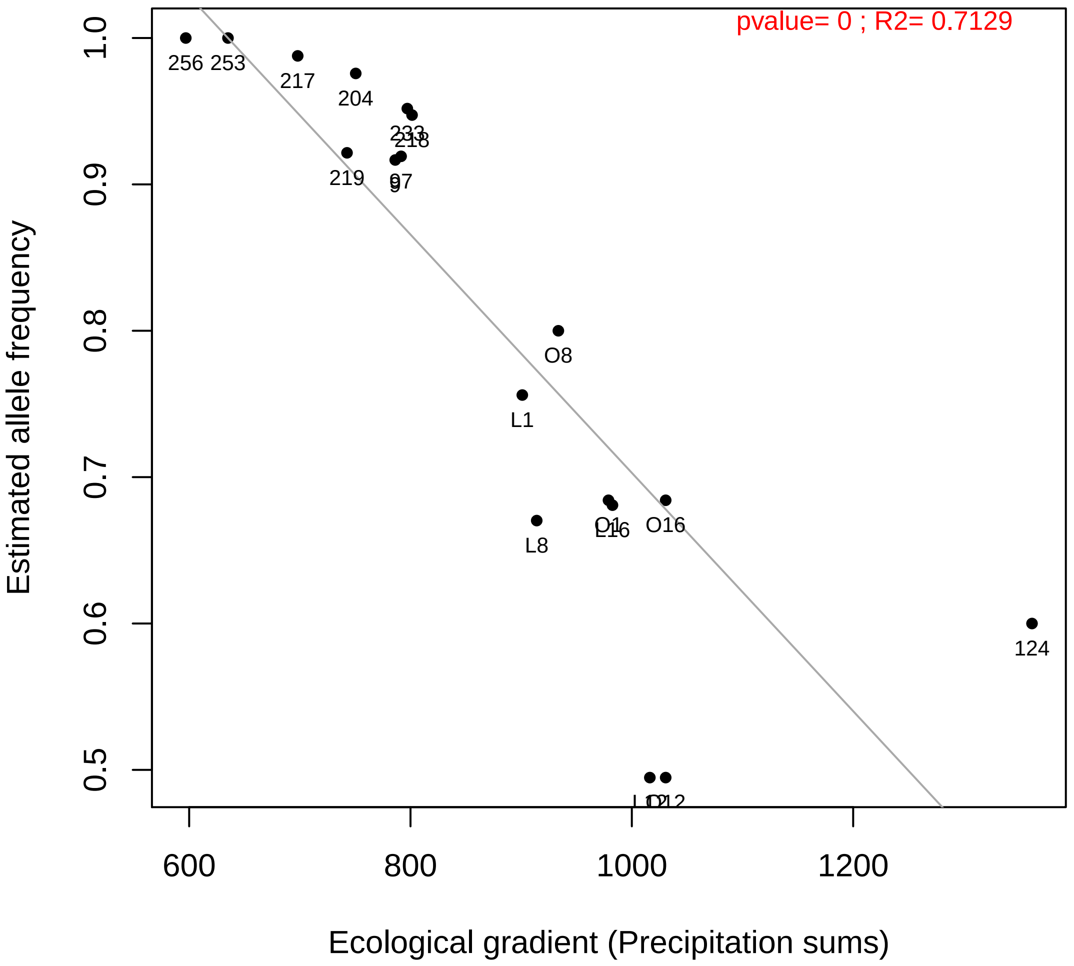

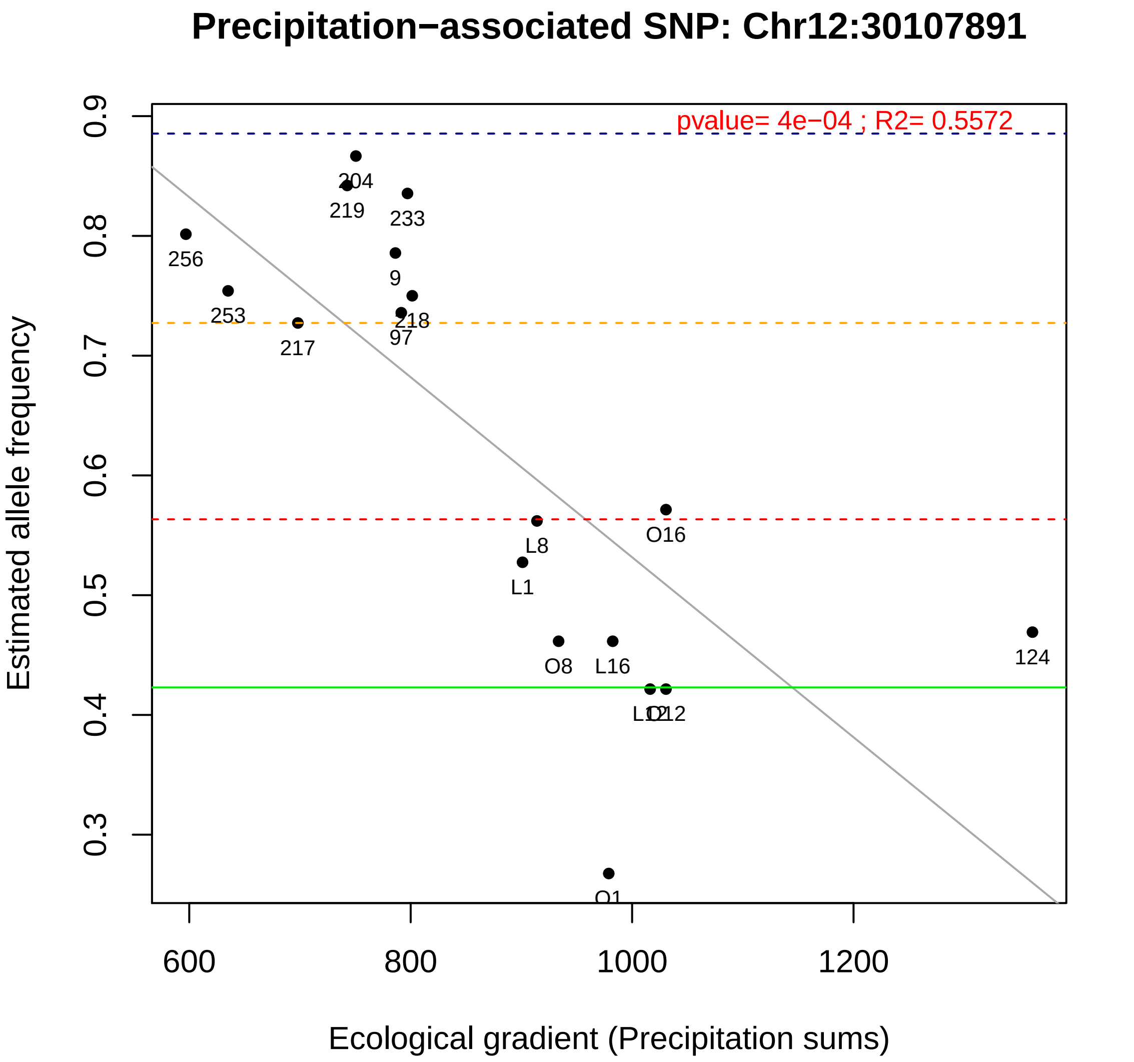
Precipitation−associated SNP: Chr12:30107891
pvalue= 4e−04 ; R2= 0.5572
Estimated allele frequency
0.3
0.4
0.5
0.6
0.7
0.8
0.9
600
800
1000
1200
Ecological gradient (Precipitation sums)
204
219
233
256
9
253
218
217
97
L8
O16
L1
O8
L16
124
O12
O1

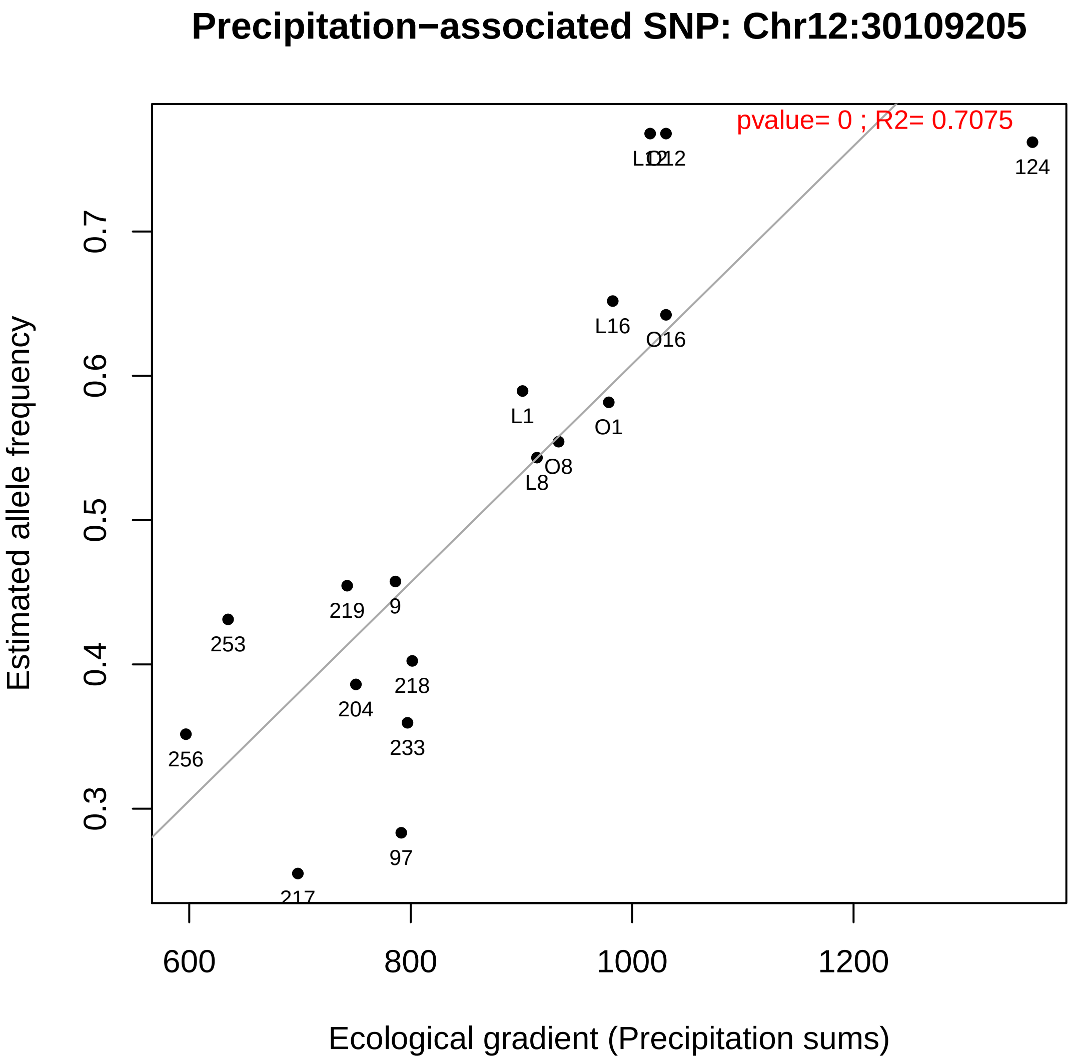
Precipitation−associated SNP: Chr12:30109205
pvalue= 0 ; R2= 0.7075
Estimated allele frequency
0.3
0.4
0.5
0.6
0.7
600
800
1000
1200
Ecological gradient (Precipitation sums)
124
L16
O16
L1
O1
O8
L8
219
9
253
218
204
233
256
97
217

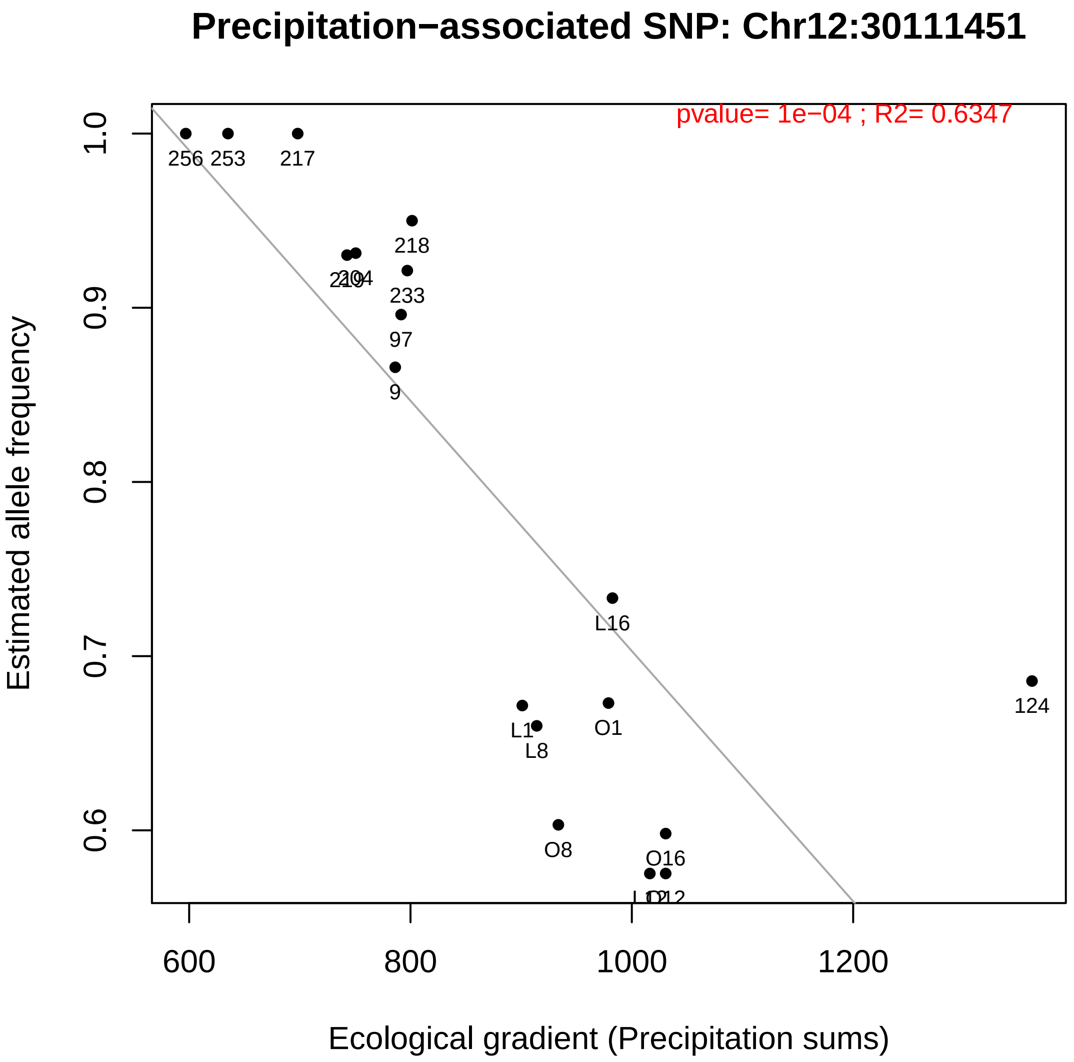
Precipitation−associated SNP: Chr12:30111451
pvalue= 1e−04 ; R2= 0.6347
Estimated allele frequency
Ecological gradient (Precipitation sums)
0.6
0.7
0.8
0.9
1.0
600
800
1000
1200
256
253
217
218
233
97
9
L16
124
L1
L8
O1
O8
O16

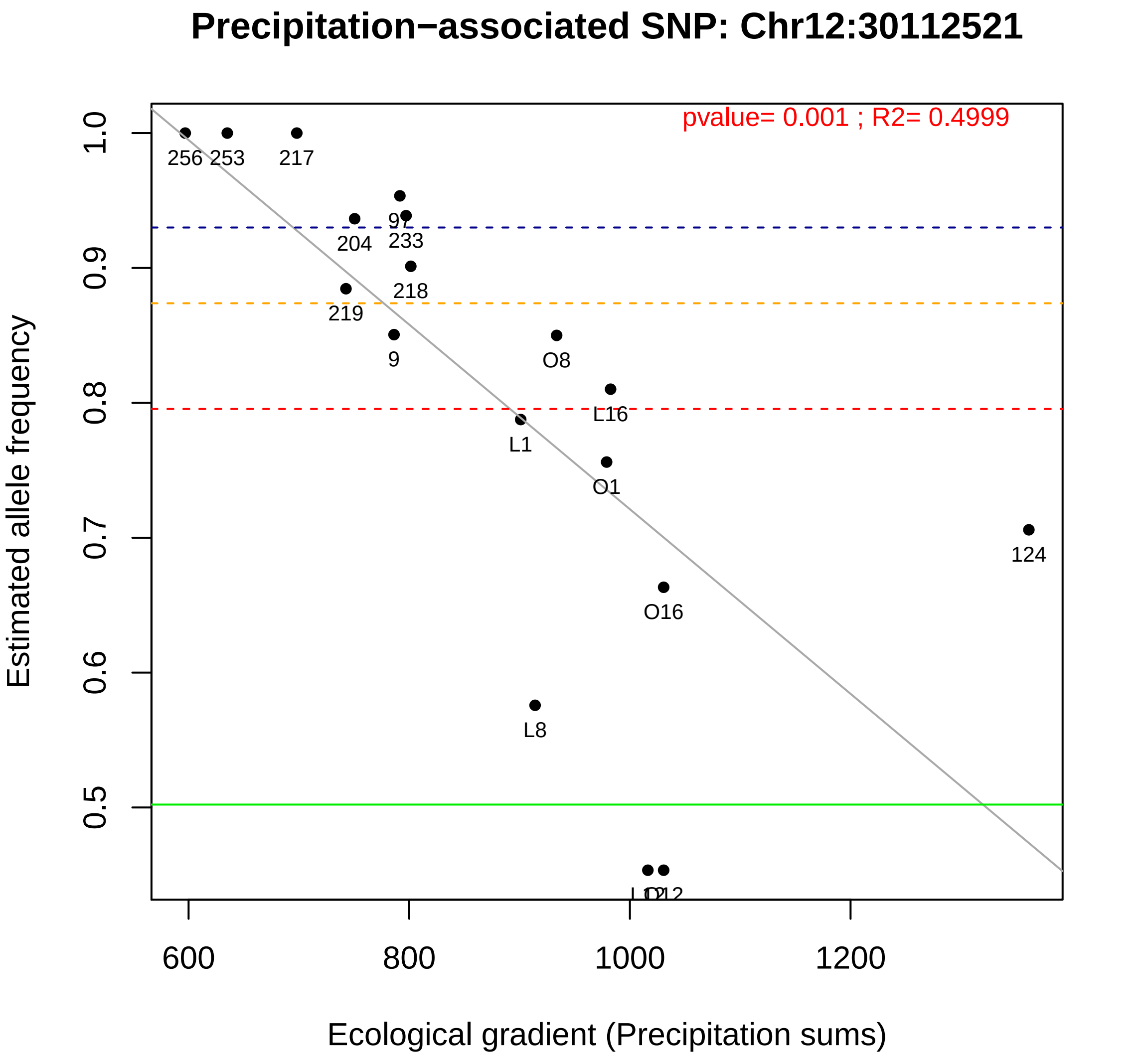
Precipitation−associated SNP: Chr12:30112521
pvalue= 0.001 ; R2= 0.4999
Estimated allele frequency
1.0
0.9
0.8
0.7
0.6
0.5
600
800
1000
1200
Ecological gradient (Precipitation sums)
256
253
217
97
233
204
218
219
9
O8
L16
L1
O1
124
O16
L8

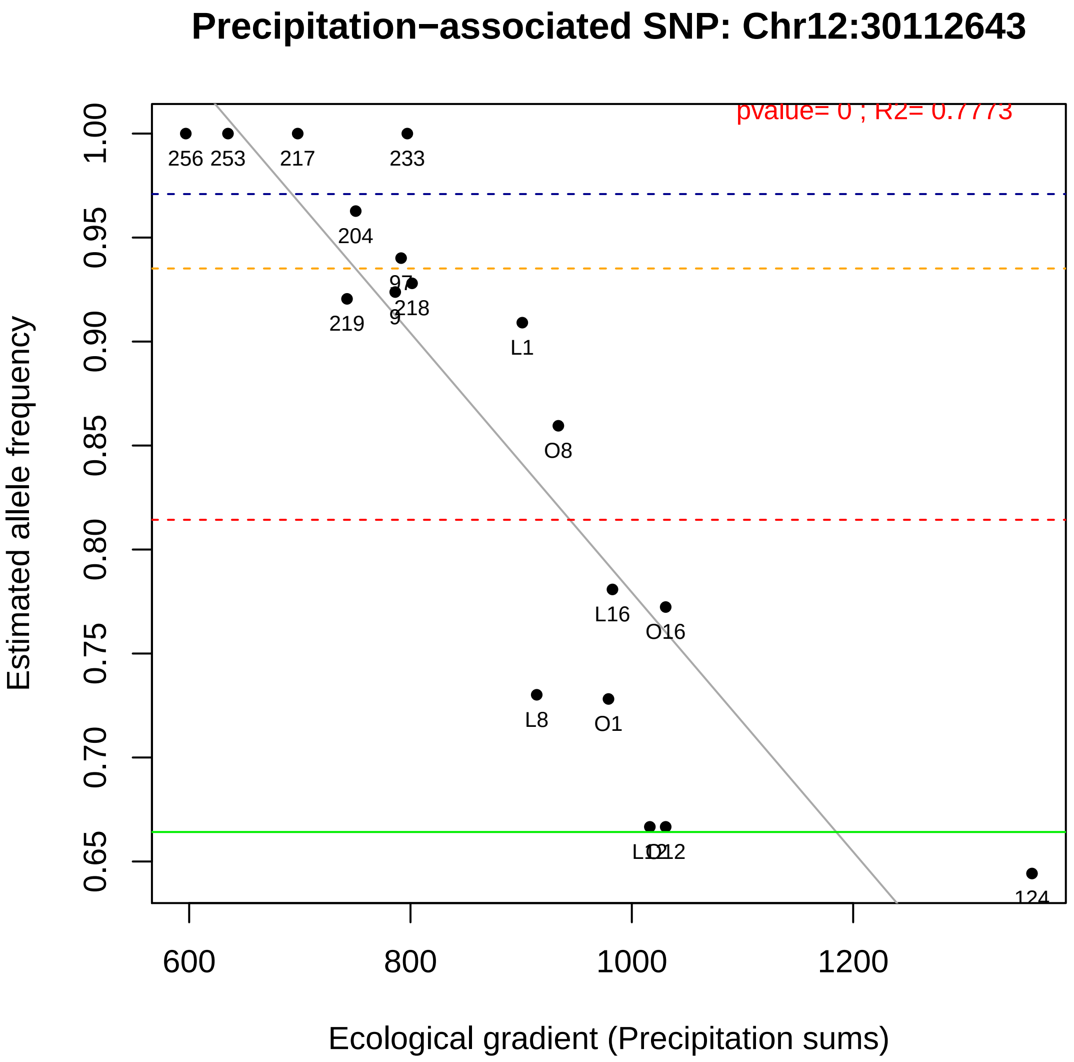

Precipitation−associated SNP: Chr12:30112643
pvalue= 0 ; R2= 0.7773
Estimated allele frequency
1.00
0.95
0.90
0.85
0.80
0.75
0.70
0.65
600
800
1000
1200
Ecological gradient (Precipitation sums)
256
253
217
233
204
97
218
219
9
L1
O8
L16
O16
L8
O1
L12
O12
124

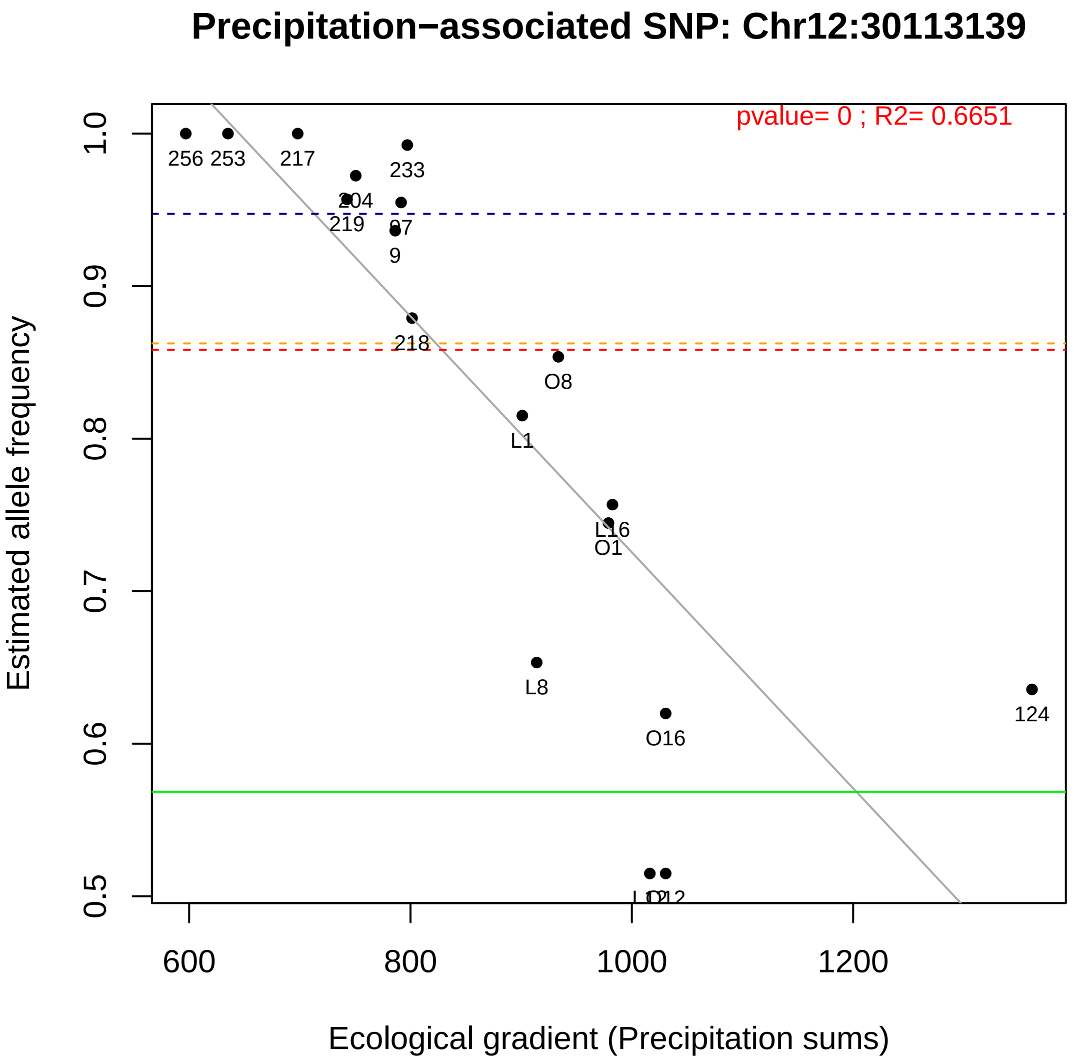

Precipitation−associated SNP: Chr12:30113139
pvalue= 0 ; R2= 0.6651
Estimated allele frequency
Ecological gradient (Precipitation sums)
1.0
0.9
0.8
0.7
0.6
0.5
600
800
1000
1200
256
253
217
233
204
219
9
218
O8
L1
L16
O1
L8
O16
124

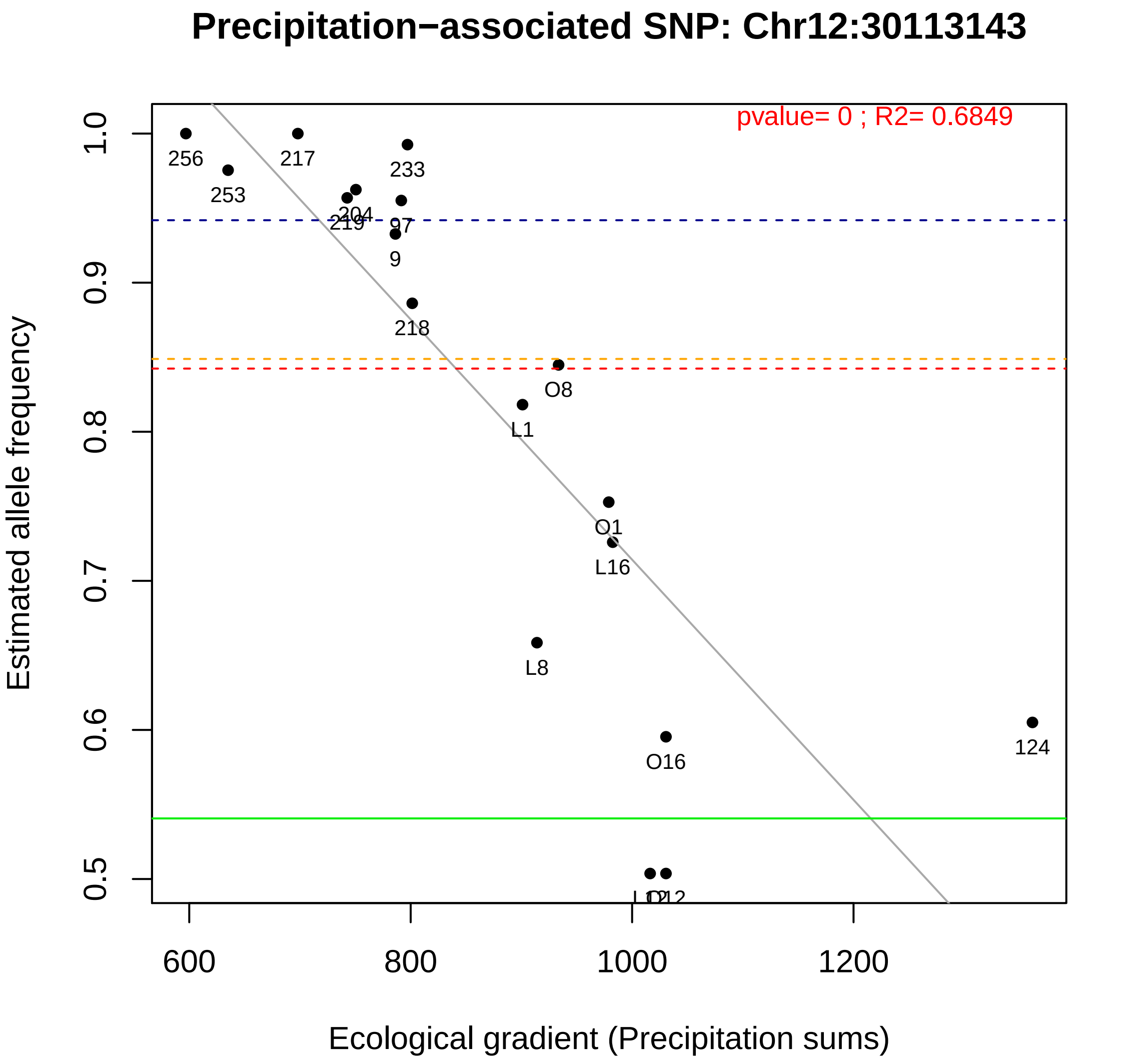

Precipitation−associated SNP: Chr12:30113143
pvalue= 0 ; R2= 0.6849
Estimated allele frequency
Ecological gradient (Precipitation sums)
1.0
0.9
0.8
0.7
0.6
0.5
600
800
1000
1200
256
253
217
233
204
219
97
9
218
O8
L1
O1
L16
L8
O16
124

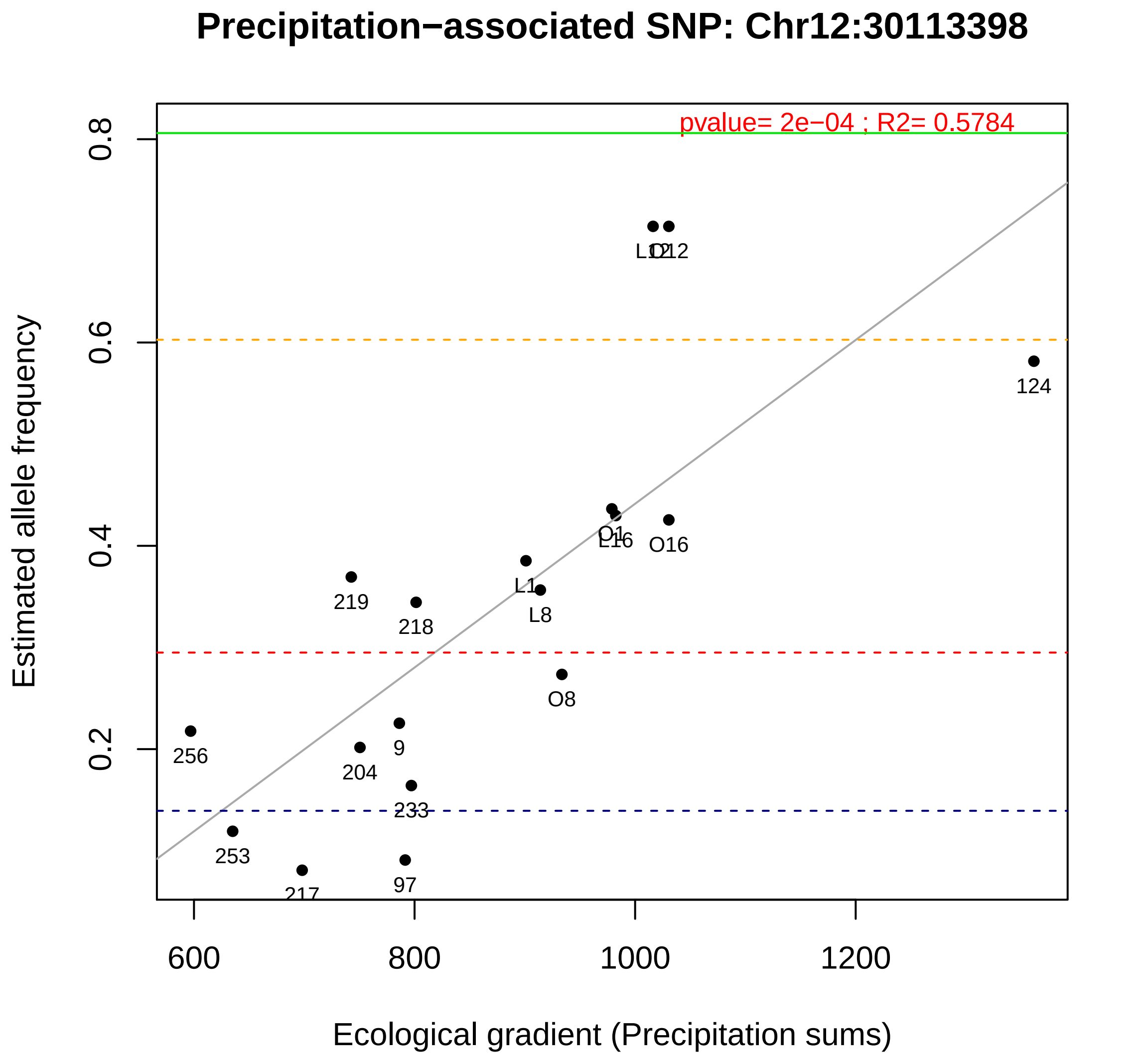

Precipitation−associated SNP: Chr12:30113398
pvalue= 2e−04 ; R2= 0.5784
Estimated allele frequency
0.2
0.4
0.6
0.8
Ecological gradient (Precipitation sums)
600
800
1000
1200
124
O16
O8
L8
L1
219
218
256
204
9
233
253
217
97

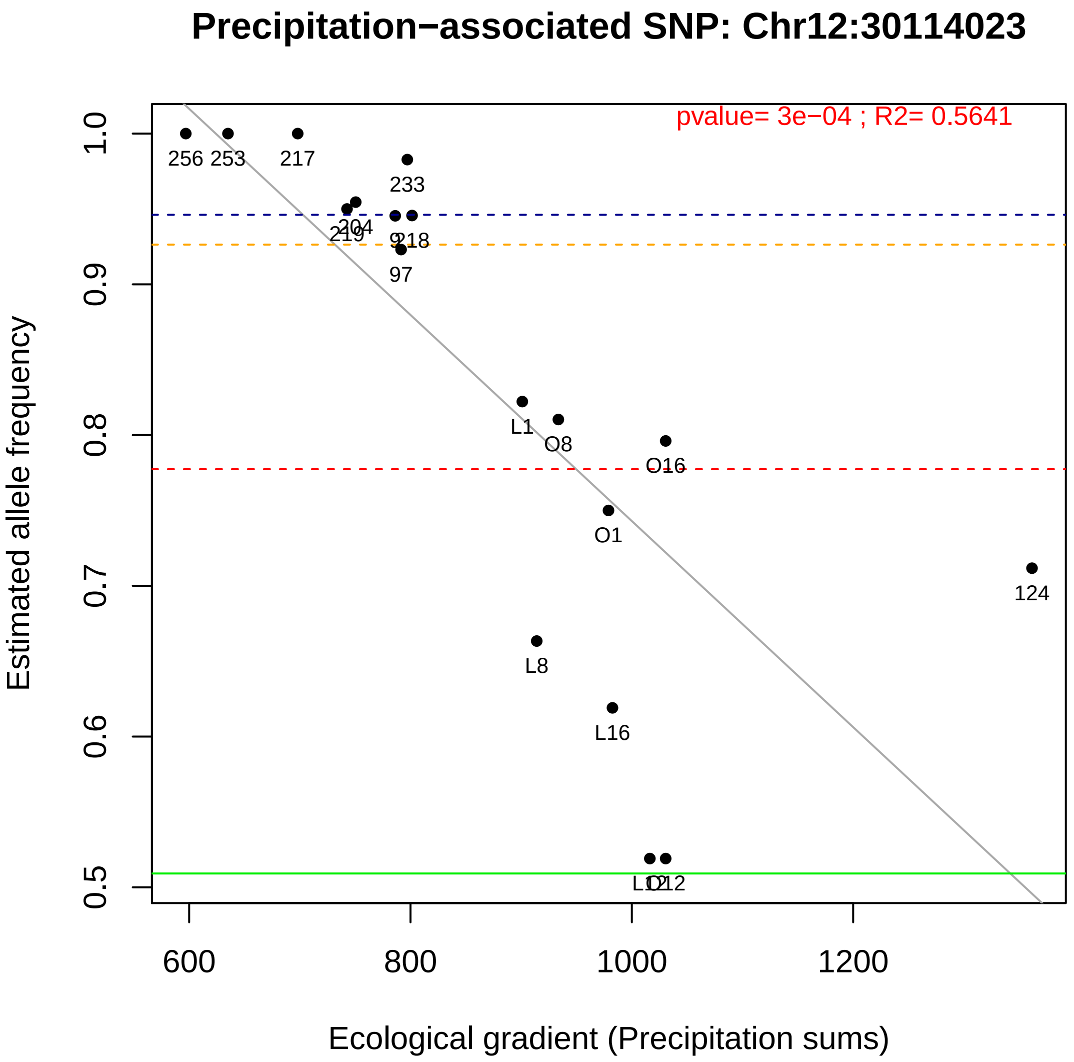

Precipitation−associated SNP: Chr12:30114023
pvalue= 3e−04 ; R2= 0.5641
Estimated allele frequency
Ecological gradient (Precipitation sums)
1.0
0.9
0.8
0.7
0.6
0.5
600
800
1000
1200
256
253
217
233
204
219
218
97
L1
O8
O16
O1
124
L8
L16
O12

**Precipitation−associated SNP: Chr12:30114593**

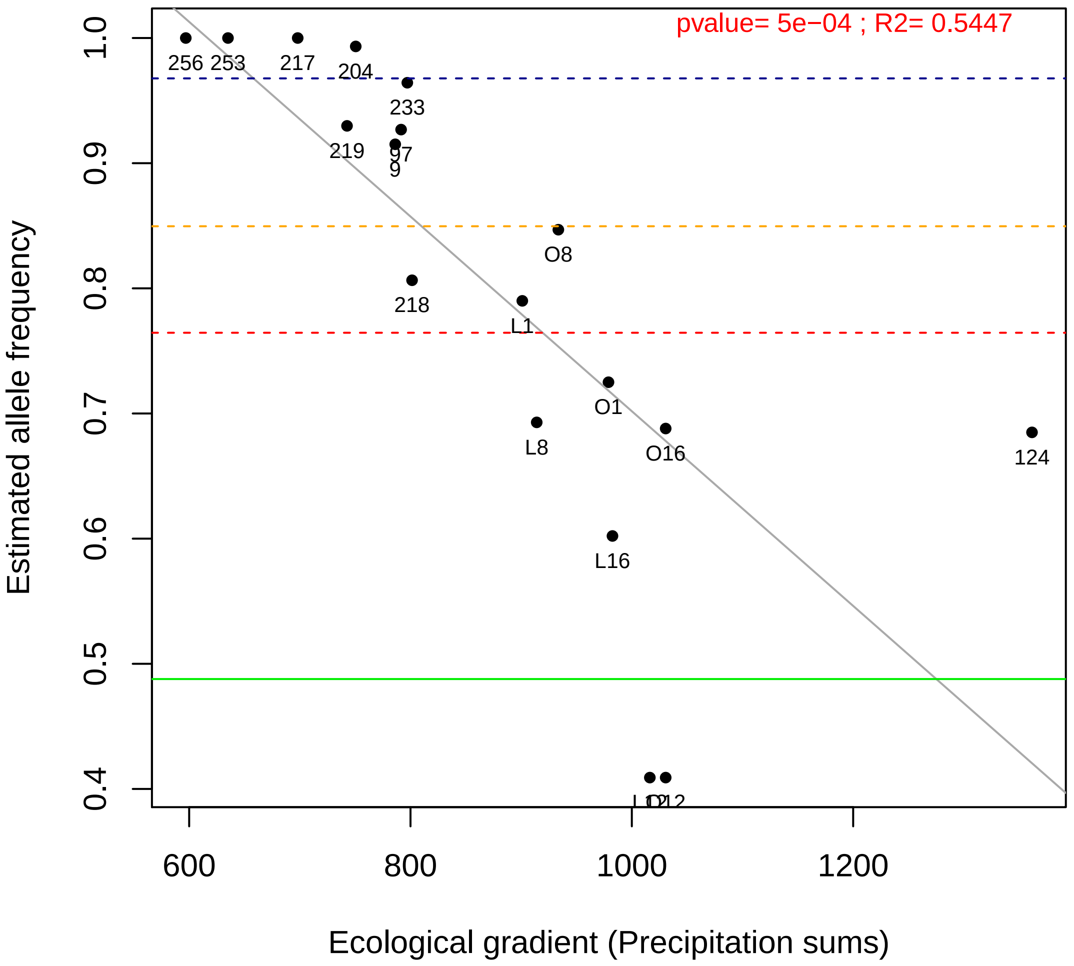

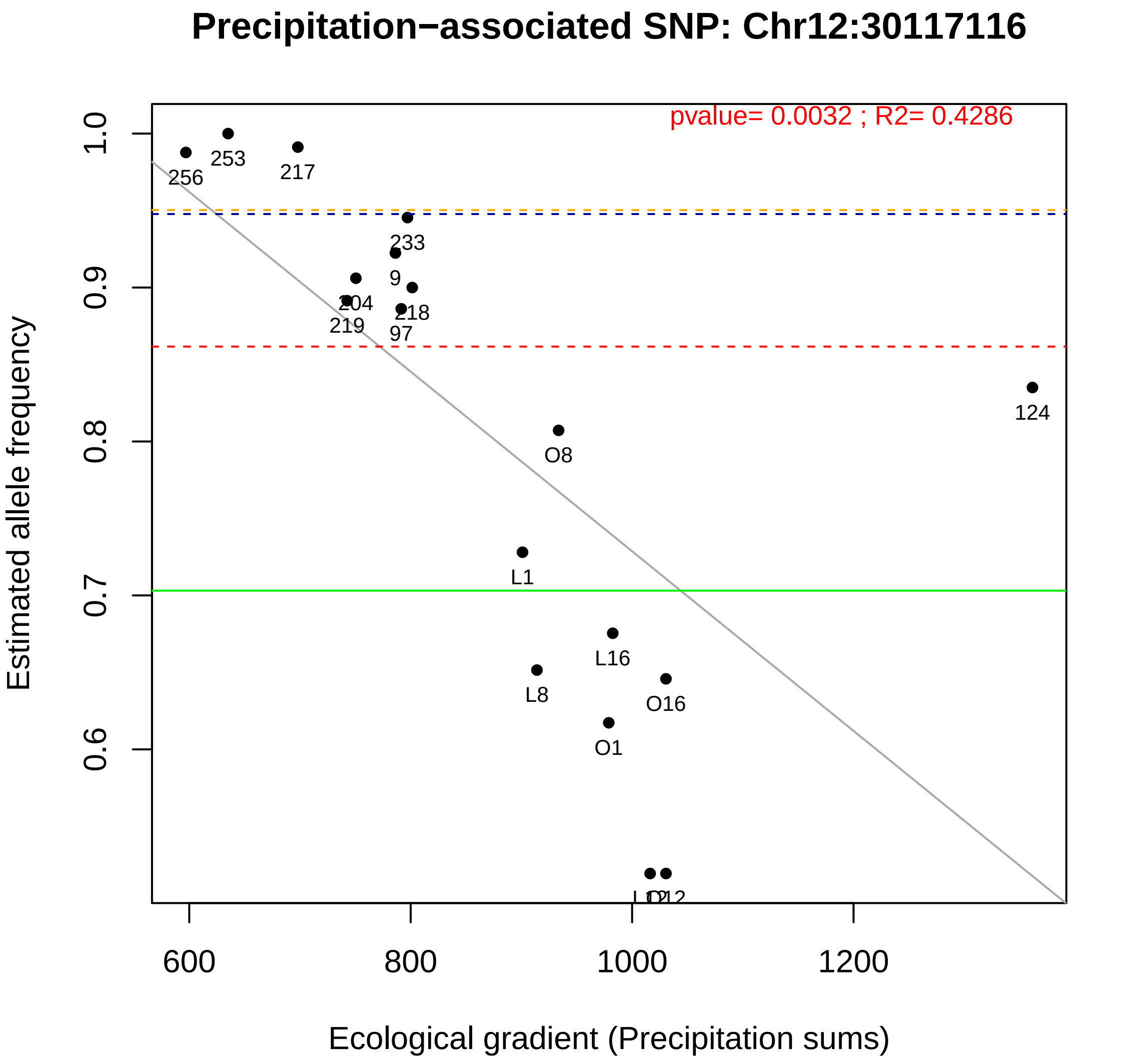

Precipitation−associated SNP: Chr12:30117116
pvalue= 0.0032 ; R2= 0.4286
Estimated allele frequency
Ecological gradient (Precipitation sums)
0.6
0.7
0.8
0.9
1.0
600
800
1000
1200
256
253
217
233
9
204
218
219
97
124
O8
L1
L16
L8
O16
O1

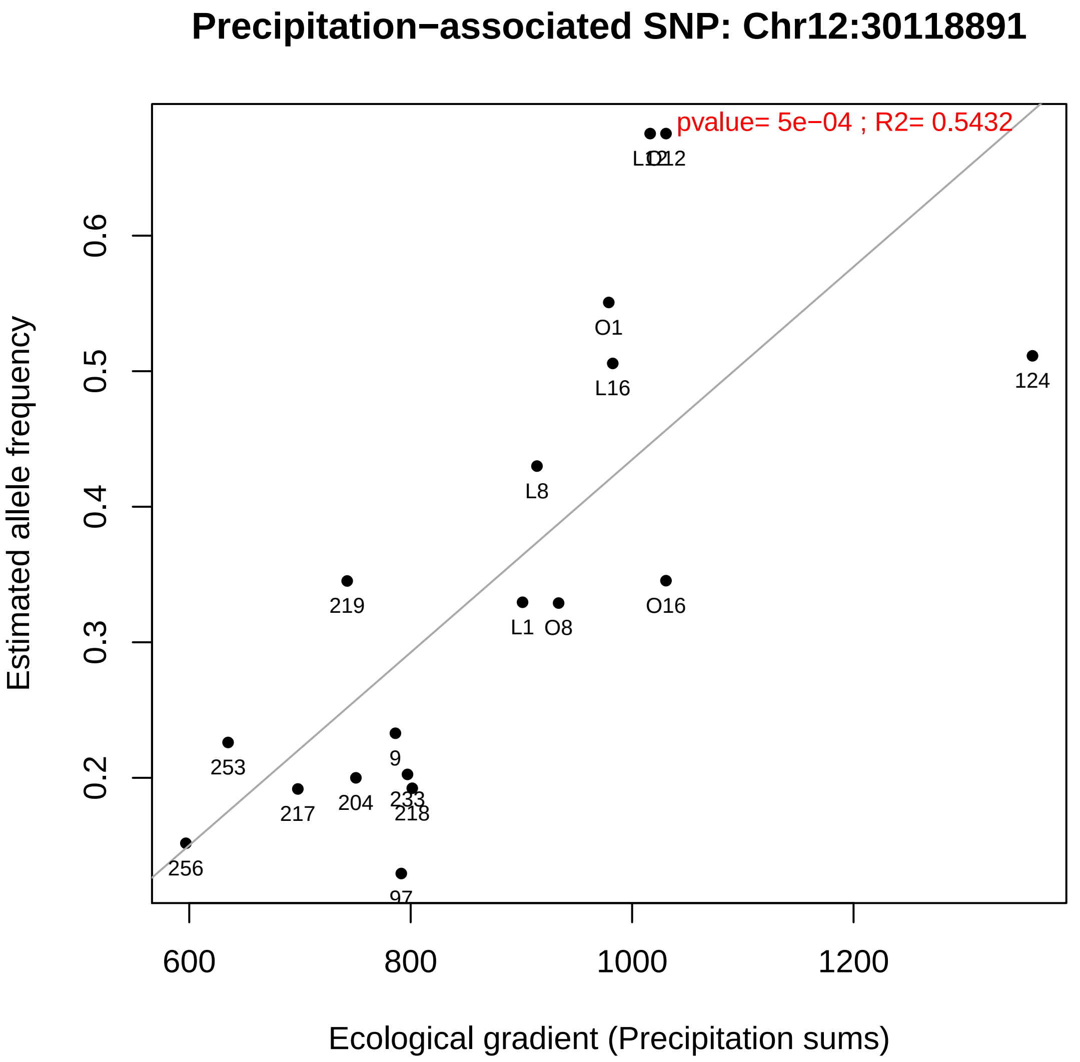

Precipitation−associated SNP: Chr12:30118891
pvalue= 5e−04 ; R2= 0.5432
Estimated allele frequency
Ecological gradient (Precipitation sums)
0.2
0.3
0.4
0.5
0.6
600
800
1000
1200
O12
O1
L16
124
L8
219
L1
O8
O16
253
9
204
217
233
218
256
97

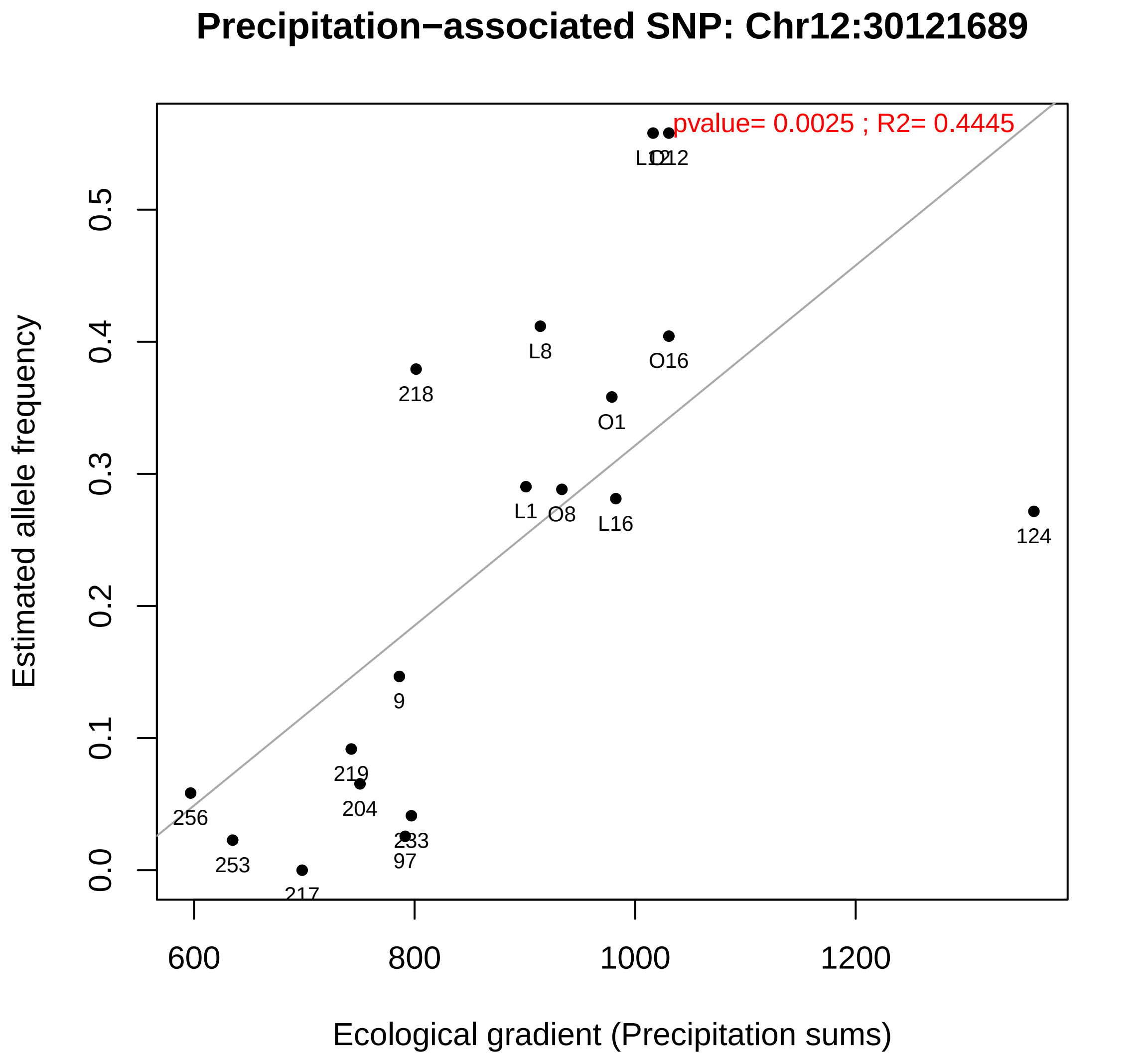
Precipitation−associated SNP: Chr12:30121689
pvalue= 0.0025 ; R2= 0.4445
Estimated allele frequency
Ecological gradient (Precipitation sums)
0.0
0.1
0.2
0.3
0.4
0.5
600
800
1000
1200
L8
O16
218
O1
L1
O8
L16
124
9
219
204
256
253
217
233
97

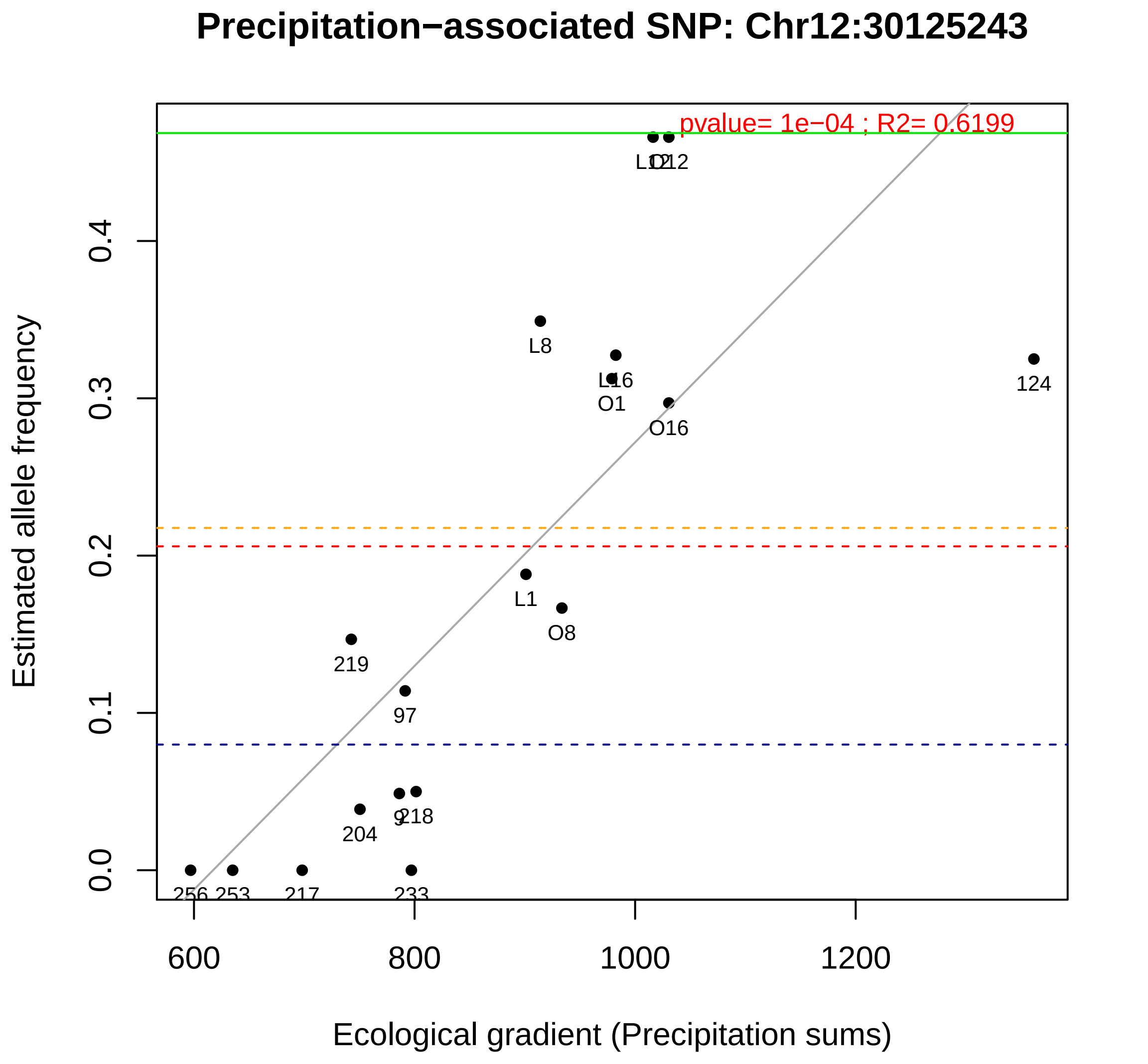

Precipitation−associated SNP: Chr12:30125243
pvalue= 1e−04 ; R2= 0.6199
Estimated allele frequency
Ecological gradient (Precipitation sums)
0.0
0.1
0.2
0.3
0.4
600
800
1000
1200
L8
L16
O1
O16
124
L1
O8
219
97
204
218
256
253
217
233

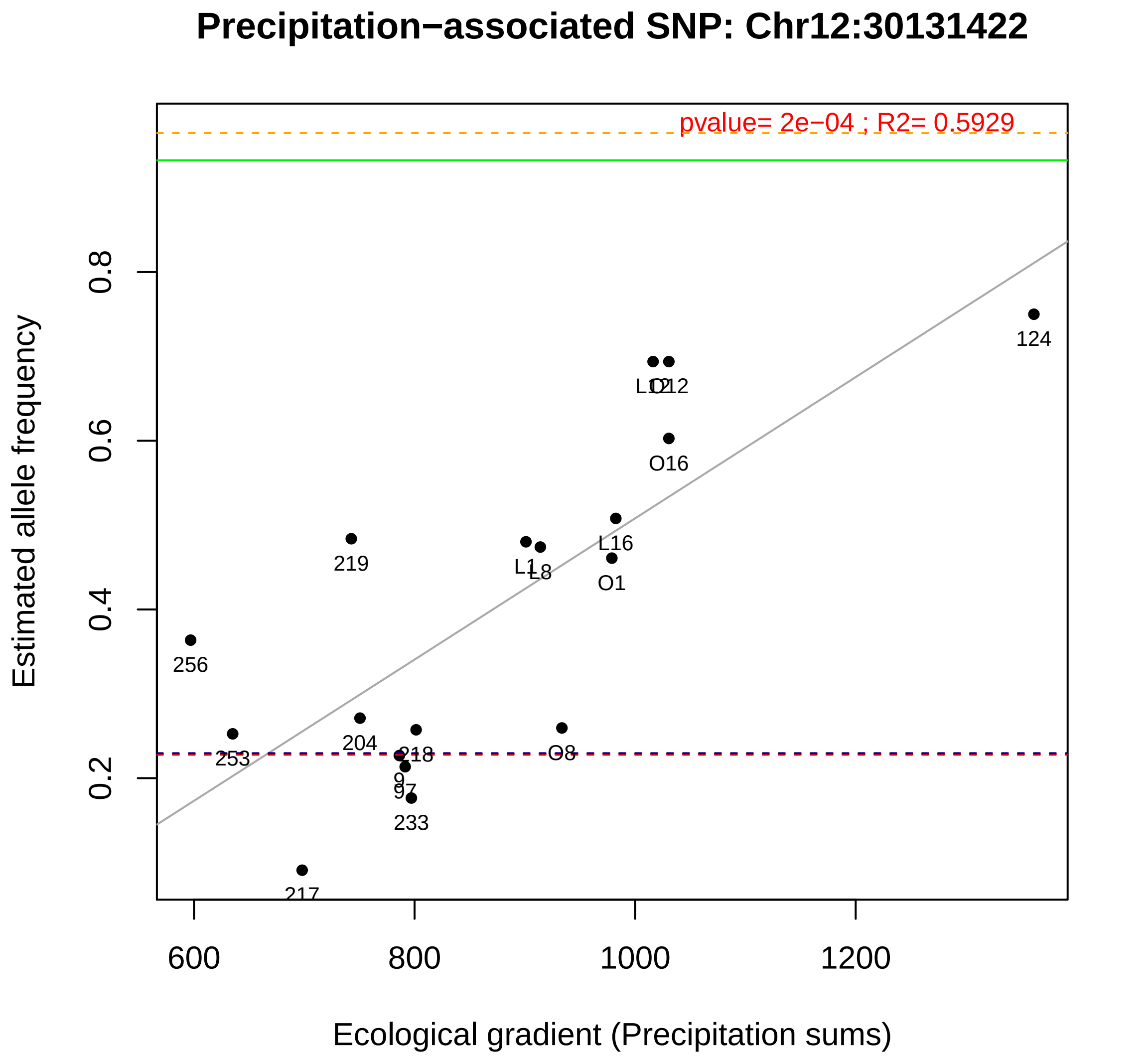
Precipitation−associated SNP: Chr12:30131422
pvalue= 2e−04 ; R2= 0.5929
Estimated allele frequency
Ecological gradient (Precipitation sums)
0.2
0.4
0.6
0.8
600
800
1000
1200
124
L12
O12
O16
L16
O1
L1
L8
219
256
204
253
218
O8
9
97
233
217

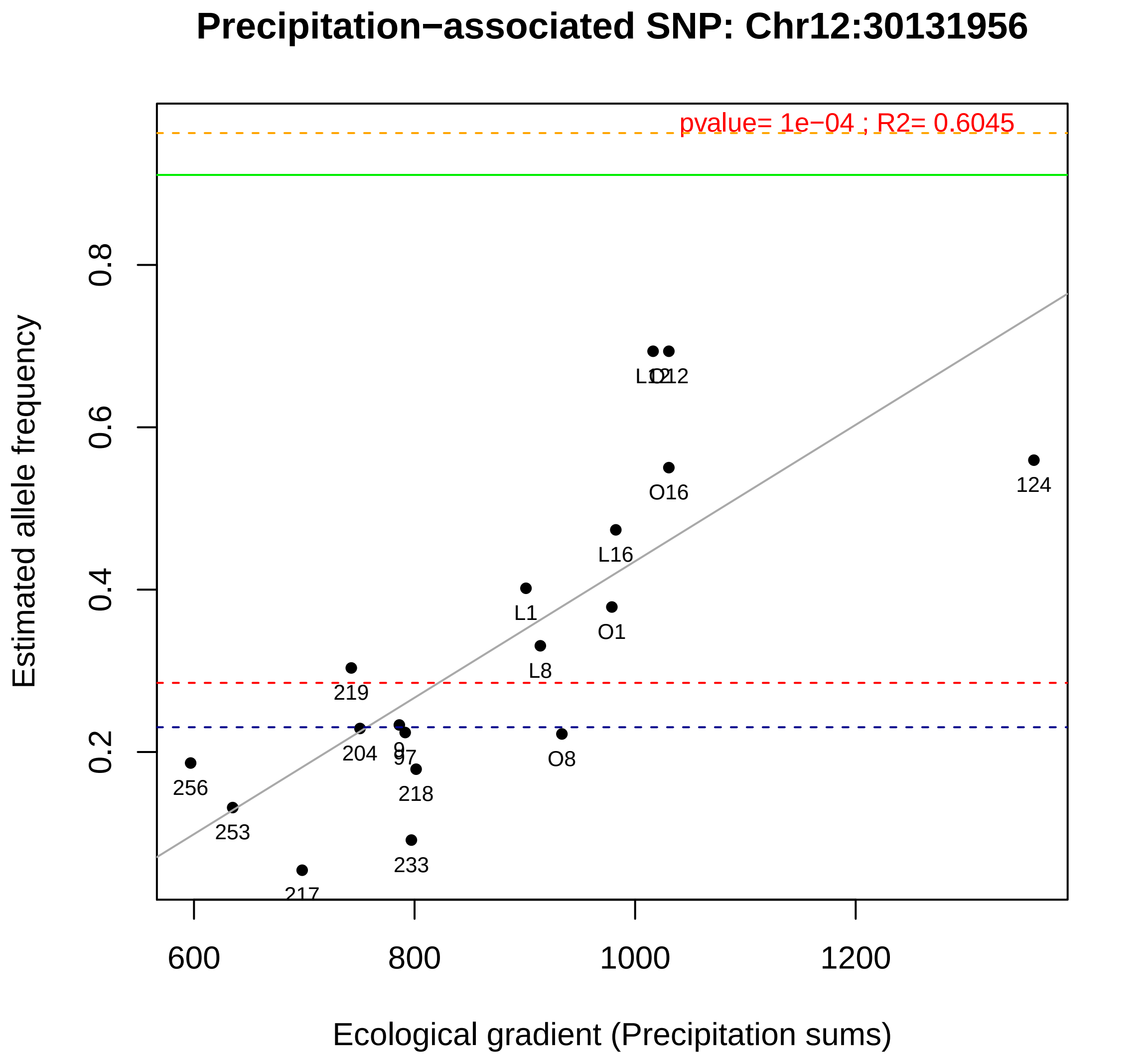

Precipitation−associated SNP: Chr12:30131956
pvalue= 1e−04 ; R2= 0.6045
Estimated allele frequency
0.2
0.4
0.6
0.8
Ecological gradient (Precipitation sums)
600
800
1000
1200
L12
O12
124
O16
L16
L1
O1
L8
219
204
9
97
O8
218
256
253
233
217

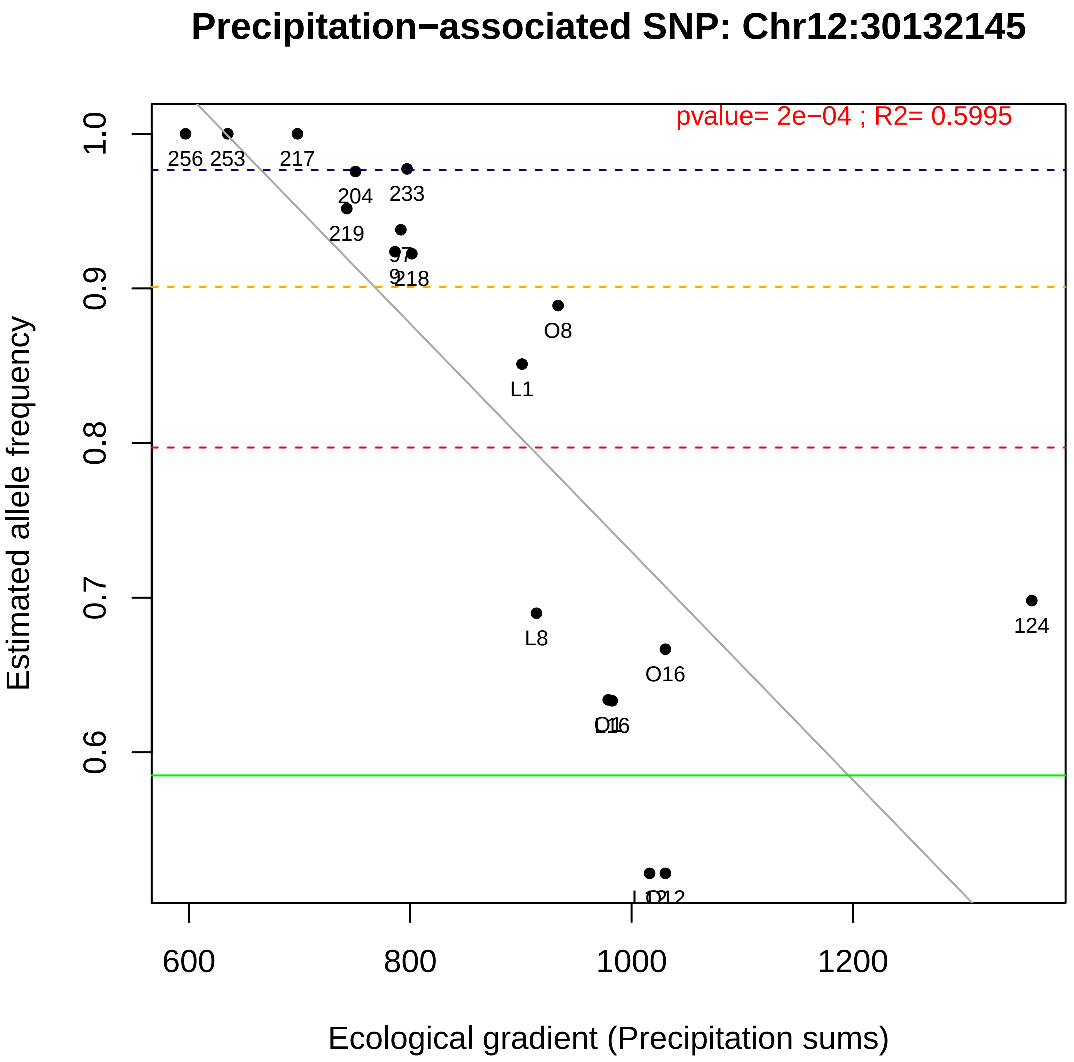

Precipitation−associated SNP: Chr12:30132145
pvalue= 2e−04 ; R2= 0.5995
Estimated allele frequency
Ecological gradient (Precipitation sums)
1.0
0.9
0.8
0.7
0.6
600
800
1000
1200
256
253
217
204
233
219
218
O8
L1
L8
124
O16

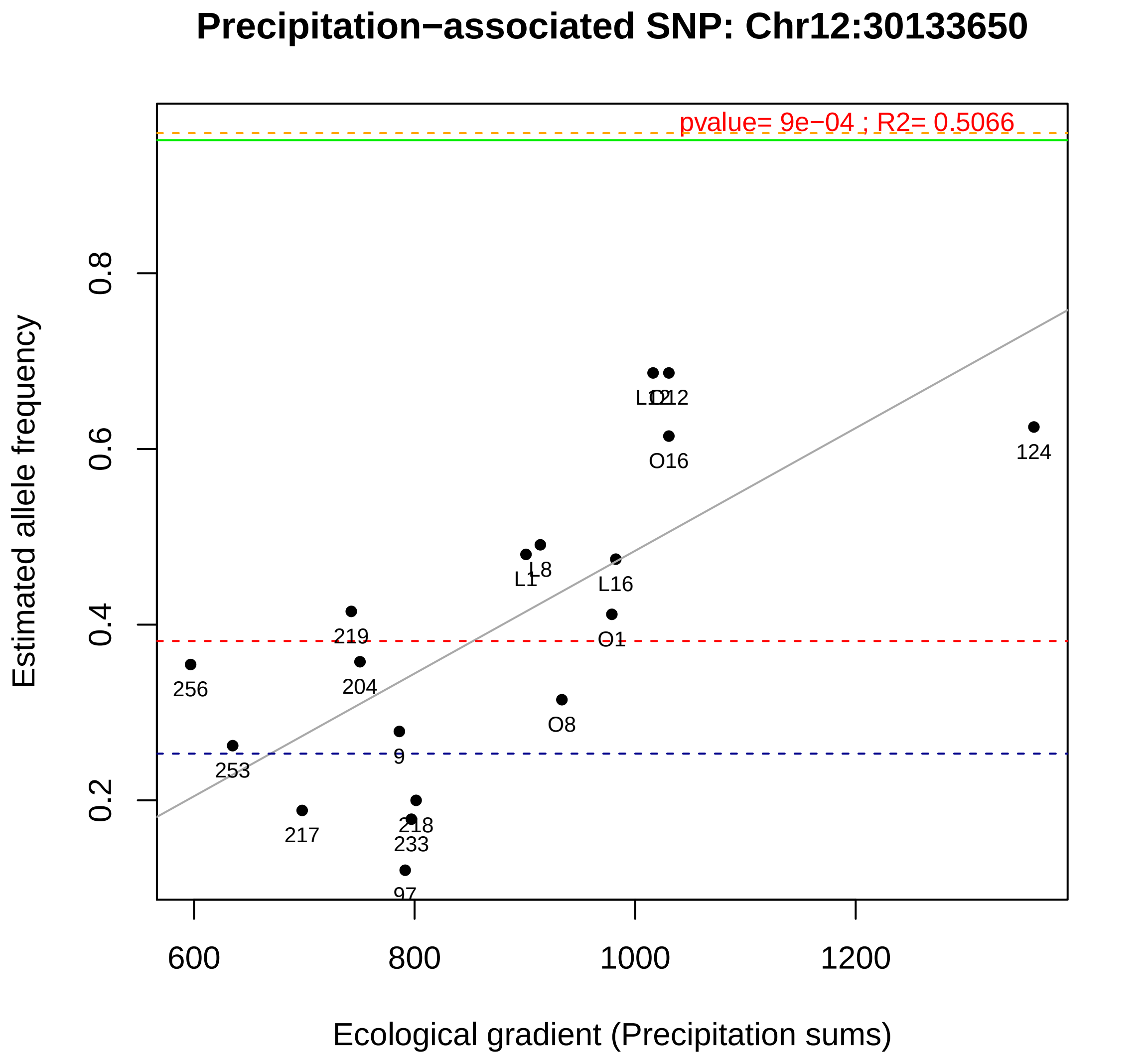
Precipitation−associated SNP: Chr12:30133650
pvalue= 9e−04 ; R2= 0.5066
Estimated allele frequency
0.2
0.4
0.6
0.8
600
800
1000
1200
Ecological gradient (Precipitation sums)
256
253
217
219
204
9
218
233
97
L1
L8
L16
O1
O8
O16
124

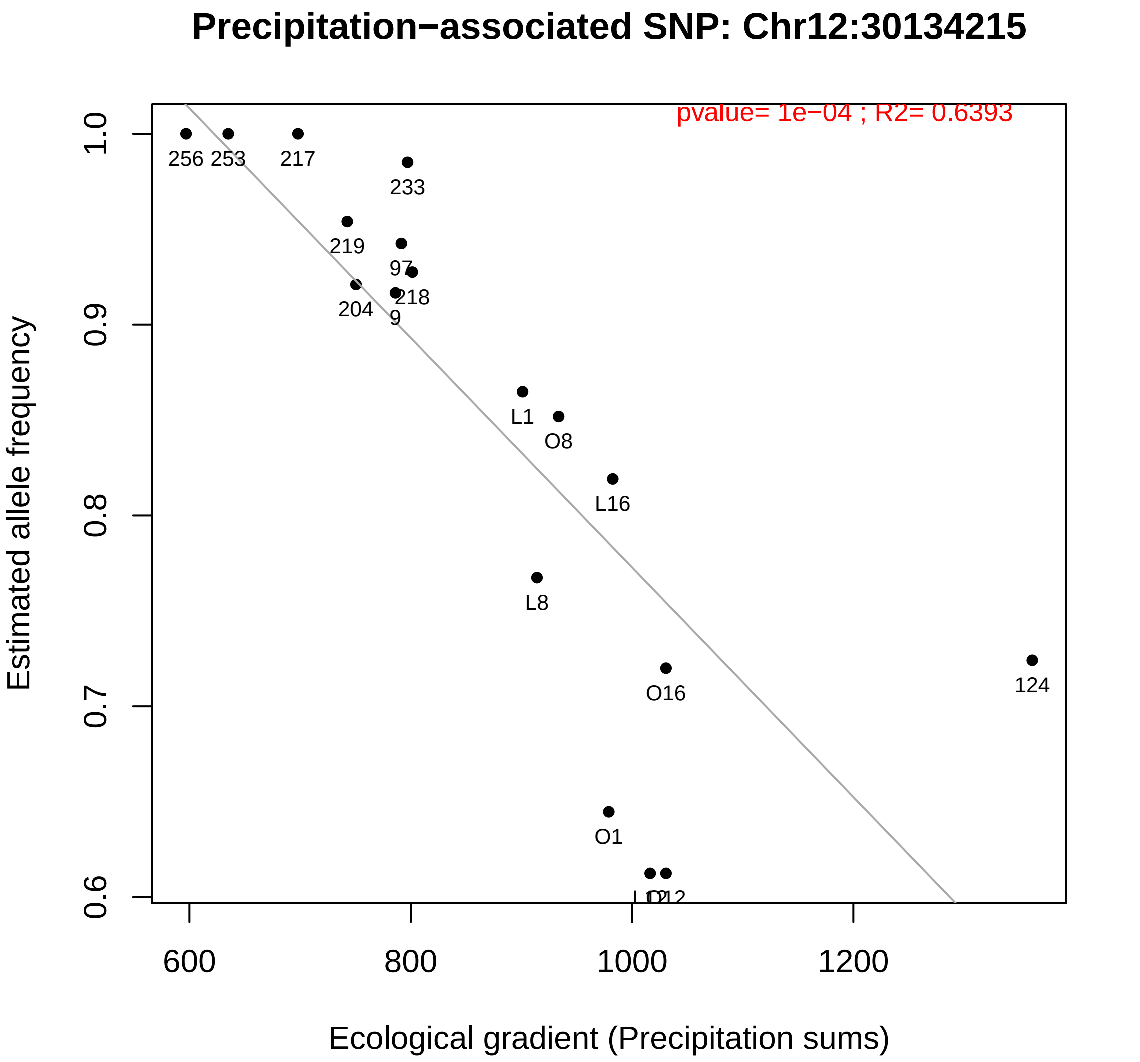
Precipitation−associated SNP: Chr12:30134215
pvalue= 1e−04 ; R2= 0.6393
Estimated allele frequency
Ecological gradient (Precipitation sums)
1.0
0.9
0.8
0.7
0.6
600
800
1000
1200
256
253
217
233
219
97
218
204
9
L1
O8
L16
L8
O16
124
O1

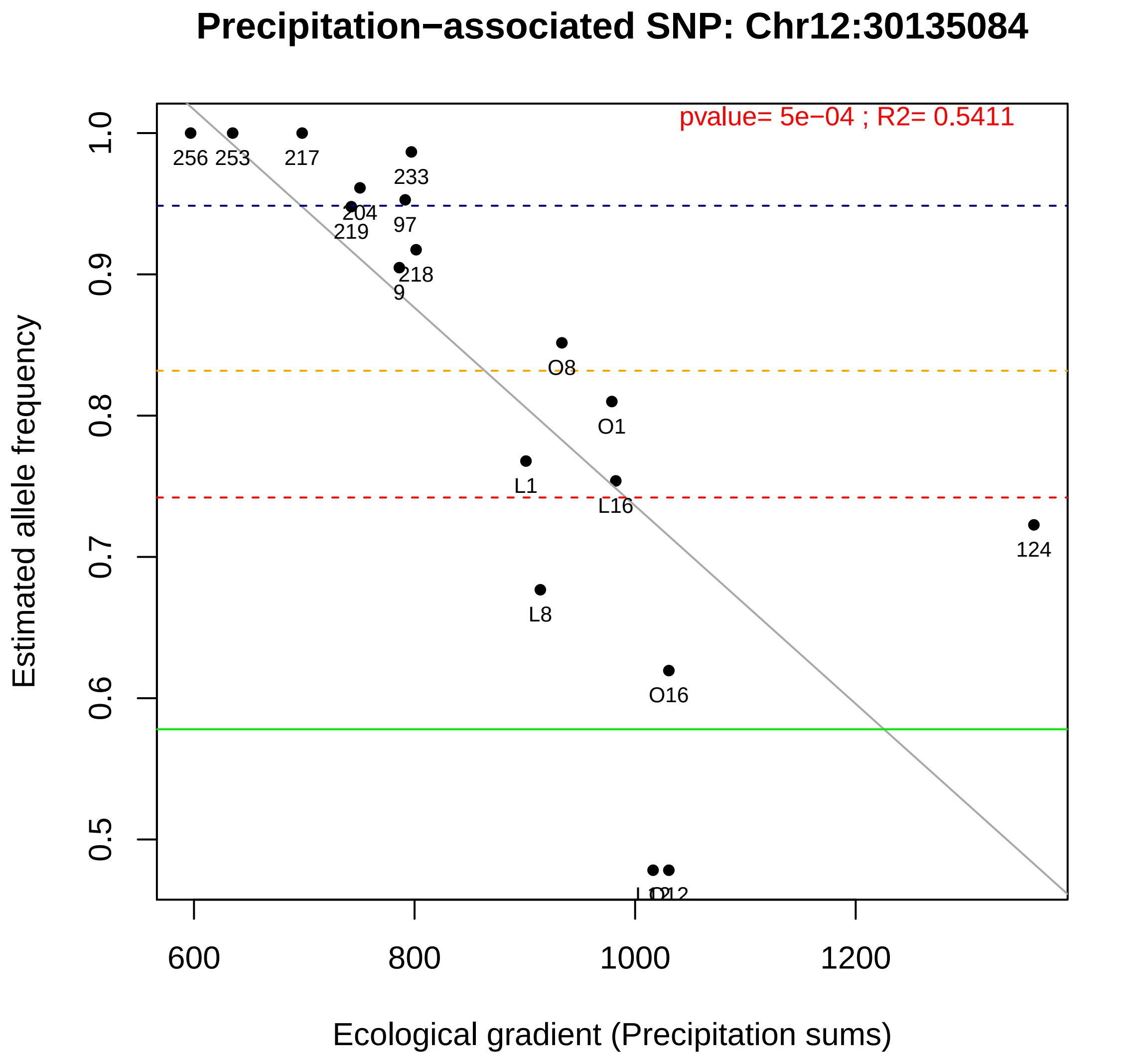

Precipitation−associated SNP: Chr12:30135084
pvalue= 5e−04 ; R2= 0.5411
Estimated allele frequency
Ecological gradient (Precipitation sums)
1.0
0.9
0.8
0.7
0.6
0.5
600
800
1000
1200
256
253
217
233
204
219
97
218
9
O8
O1
L1
L16
124
L8
O16

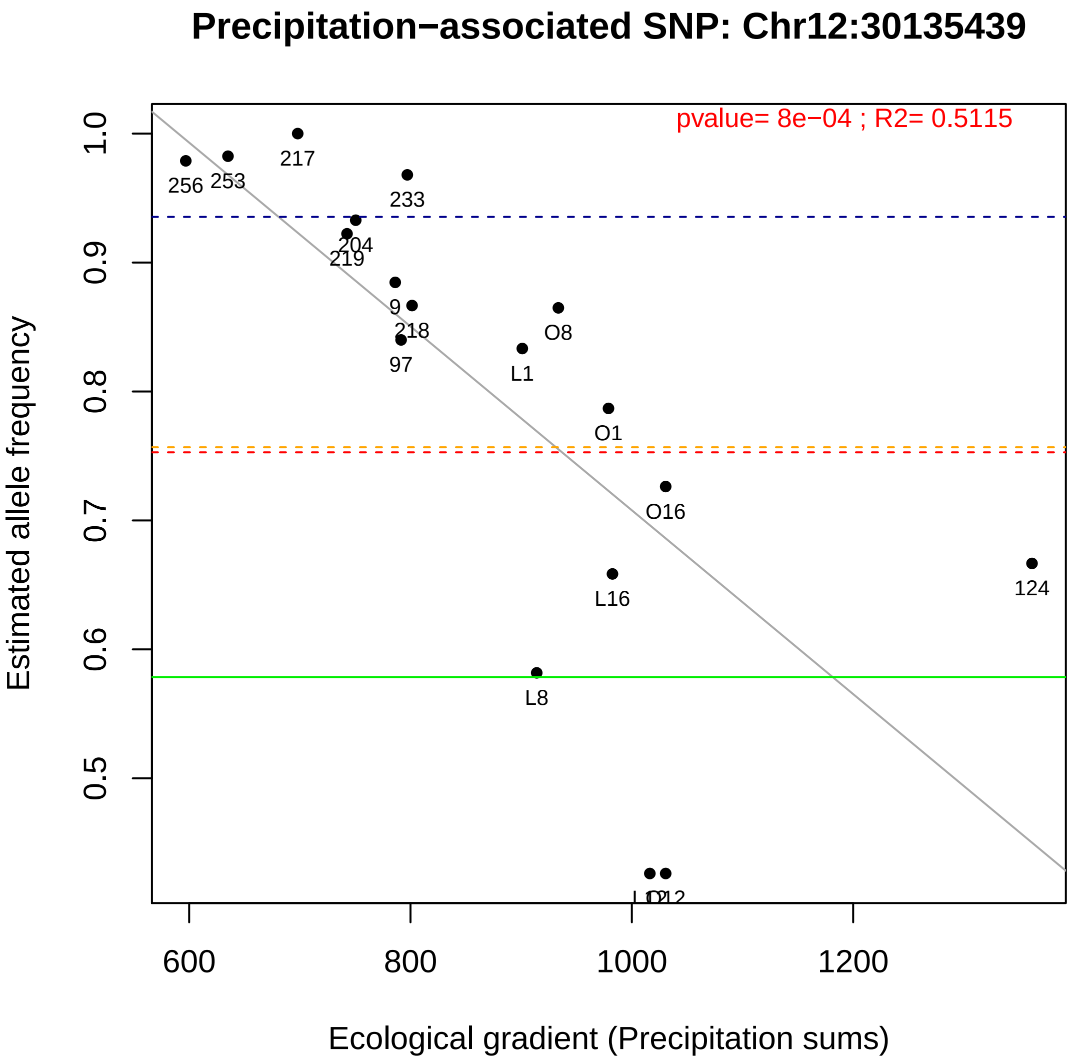

Precipitation−associated SNP: Chr12:30135439
pvalue= 8e−04 ; R2= 0.5115
Estimated allele frequency
Ecological gradient (Precipitation sums)
1.0
0.9
0.8
0.7
0.6
0.5
600
800
1000
1200
256
253
217
233
204
219
9
218
97
L1
O8
O1
O16
L16
124
L8

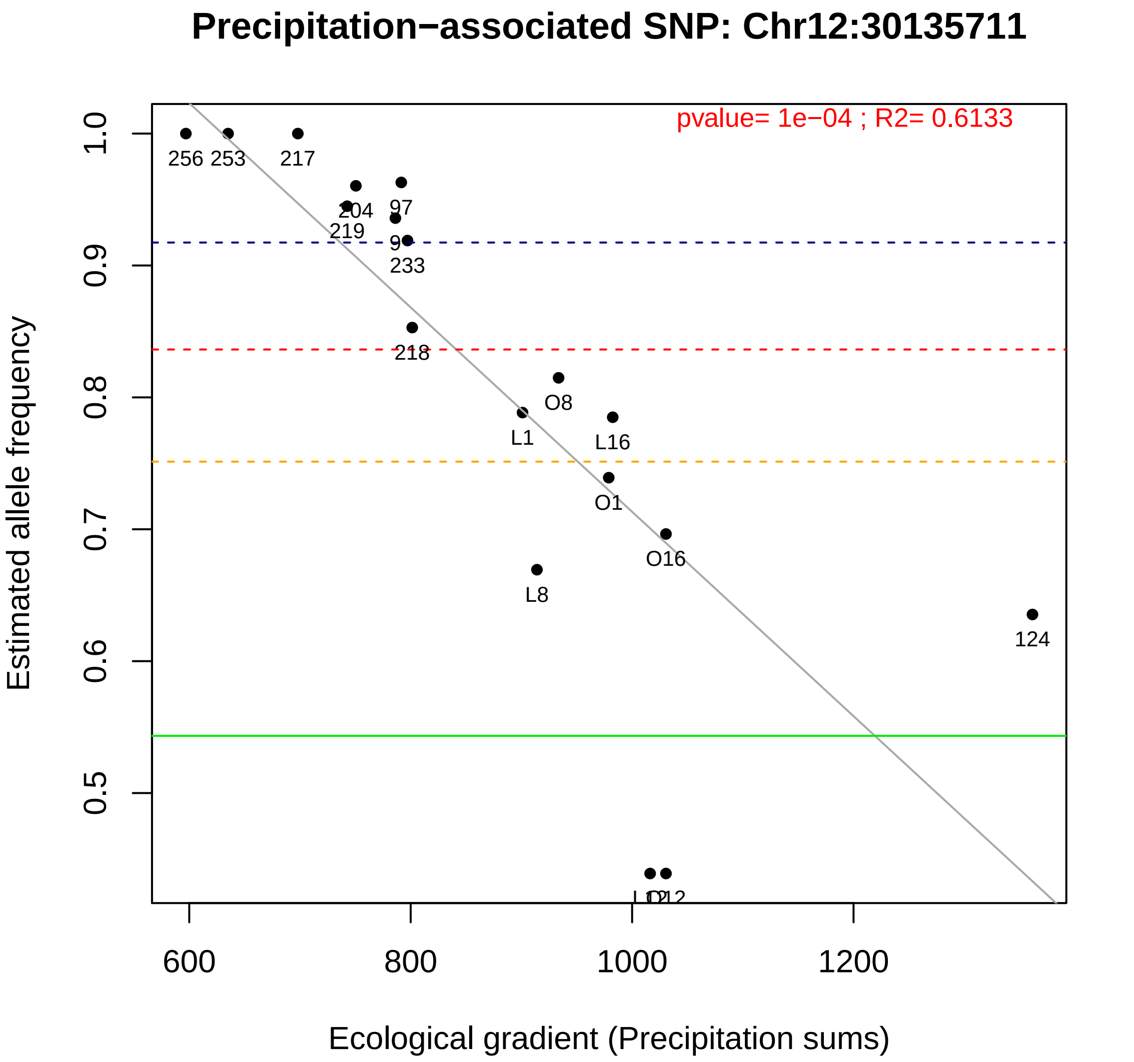

Precipitation−associated SNP: Chr12:30135711
pvalue= 1e−04 ; R2= 0.6133
Estimated allele frequency
Ecological gradient (Precipitation sums)
1.0
0.9
0.8
0.7
0.6
0.5
600
800
1000
1200
256
253
217
204
97
219
233
218
O8
L1
L16
O1
O16
L8
124

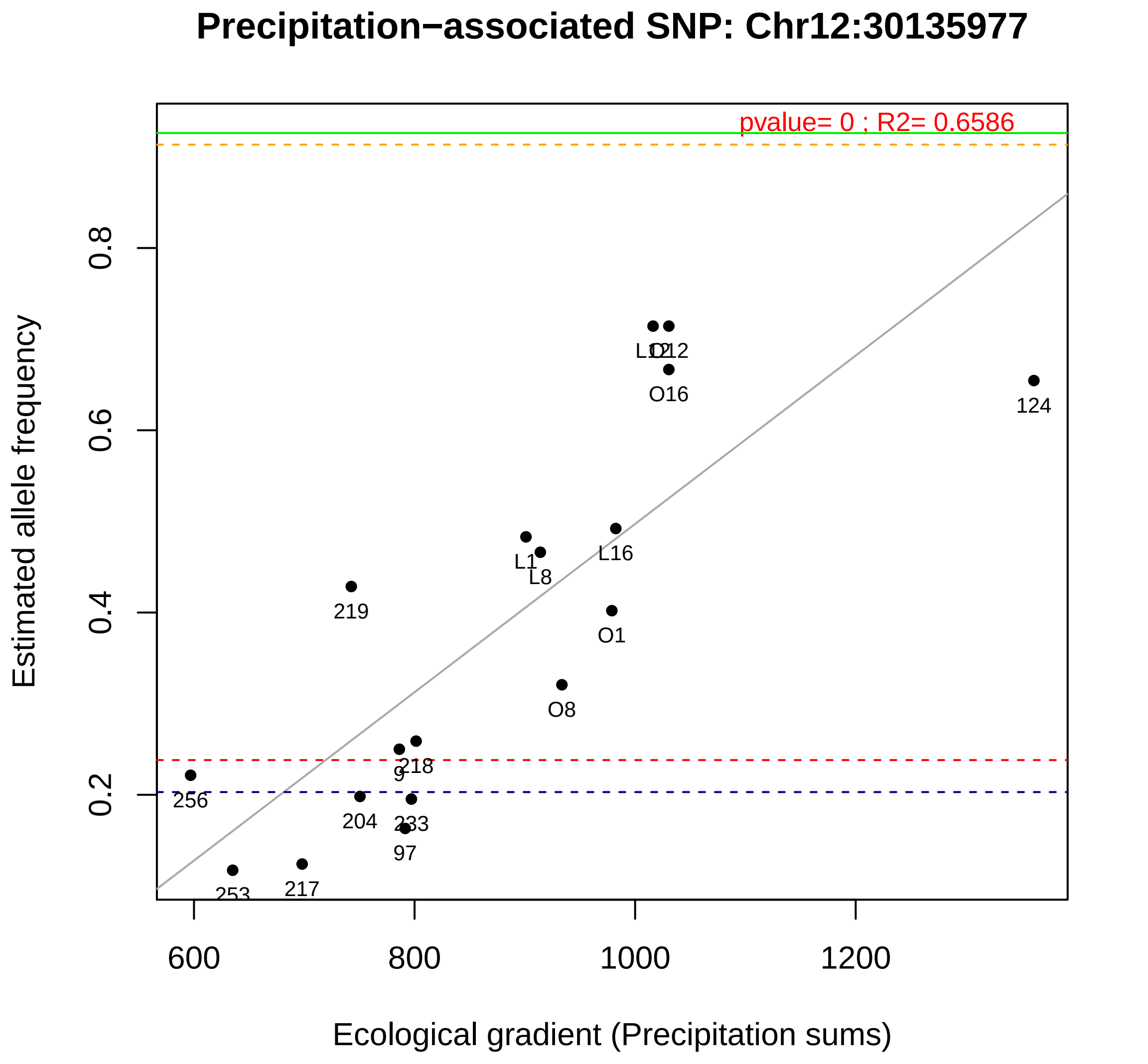

Precipitation−associated SNP: Chr12:30135977
pvalue= 0 ; R2= 0.6586
Estimated allele frequency
0.2
0.4
0.6
0.8
600
800
1000
1200
Ecological gradient (Precipitation sums)
L12
O12
O16
124
L16
L1
L8
219
O1
O8
218
9
256
204
233
97
253
217

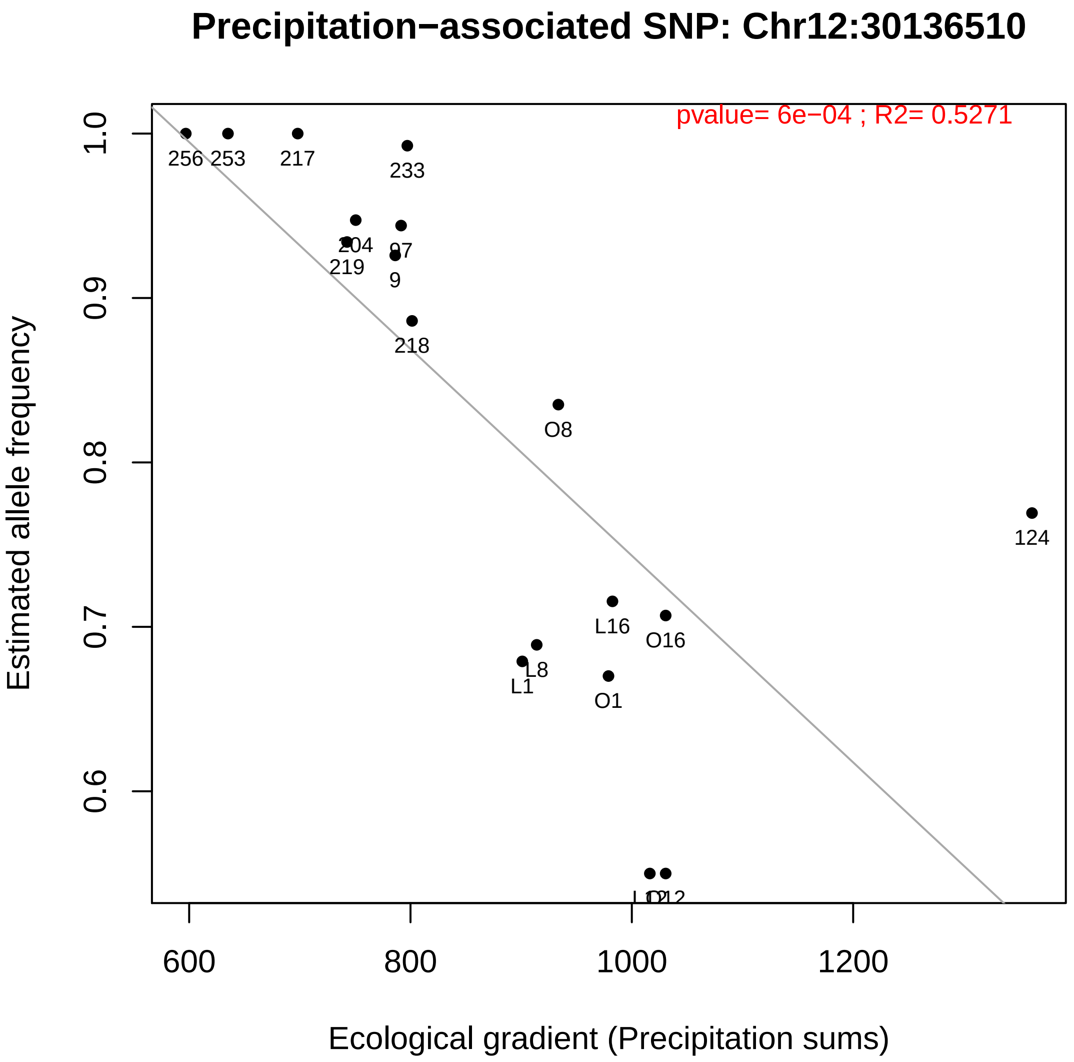

Precipitation−associated SNP: Chr12:30136510
pvalue= 6e−04 ; R2= 0.5271
256
253
217
233
204
97
219
9
218
O8
124
L16
O16
L8
L1
O1
O12
1.0
0.9
0.8
0.7
0.6
600
800
1000
1200
Estimated allele frequency
Ecological gradient (Precipitation sums)

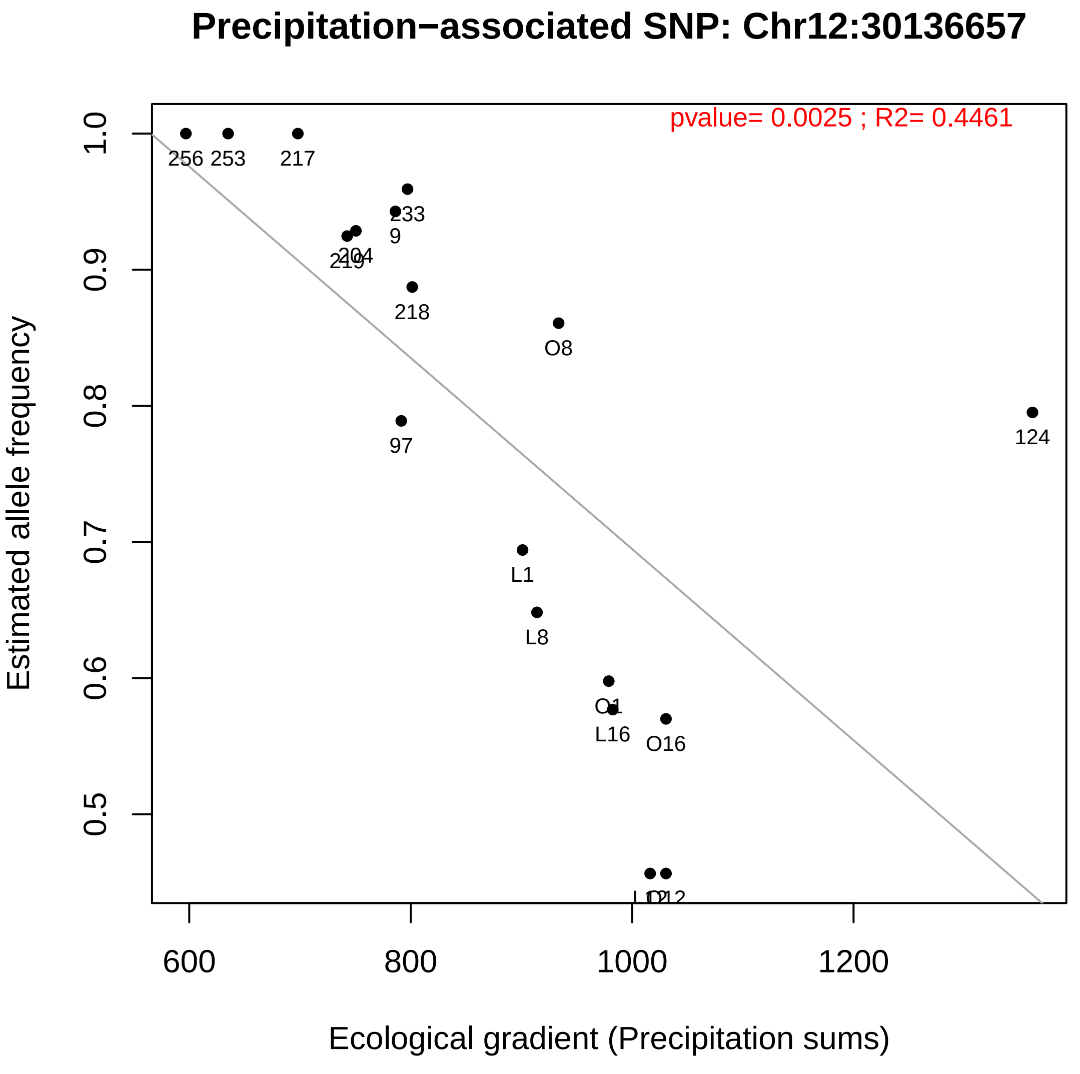
Precipitation−associated SNP: Chr12:30136657
pvalue= 0.0025 ; R2= 0.4461
Estimated allele frequency
Ecological gradient (Precipitation sums)
1.0
0.9
0.8
0.7
0.6
0.5
600
800
1000
1200
256
253
217
233
9
219
204
218
O8
97
124
L1
L8
O1
L16
O16
O12

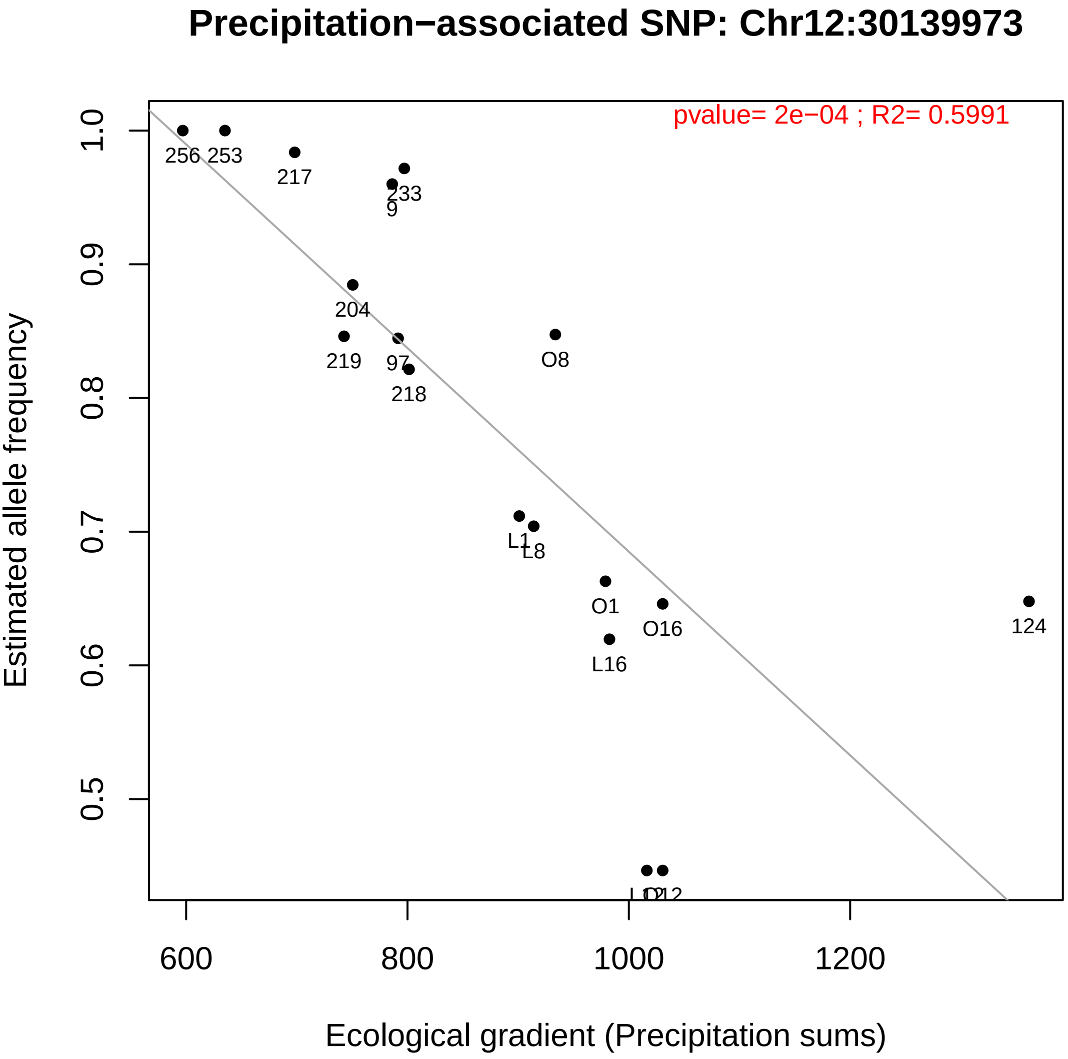
Precipitation−associated SNP: Chr12:30139973
pvalue= 2e−04 ; R2= 0.5991
Estimated allele frequency
Ecological gradient (Precipitation sums)
1.0
0.9
0.8
0.7
0.6
0.5
600
800
1000
1200
256
253
217
233
9
204
219
97
218
O8
L1
L8
O1
O16
L16
124

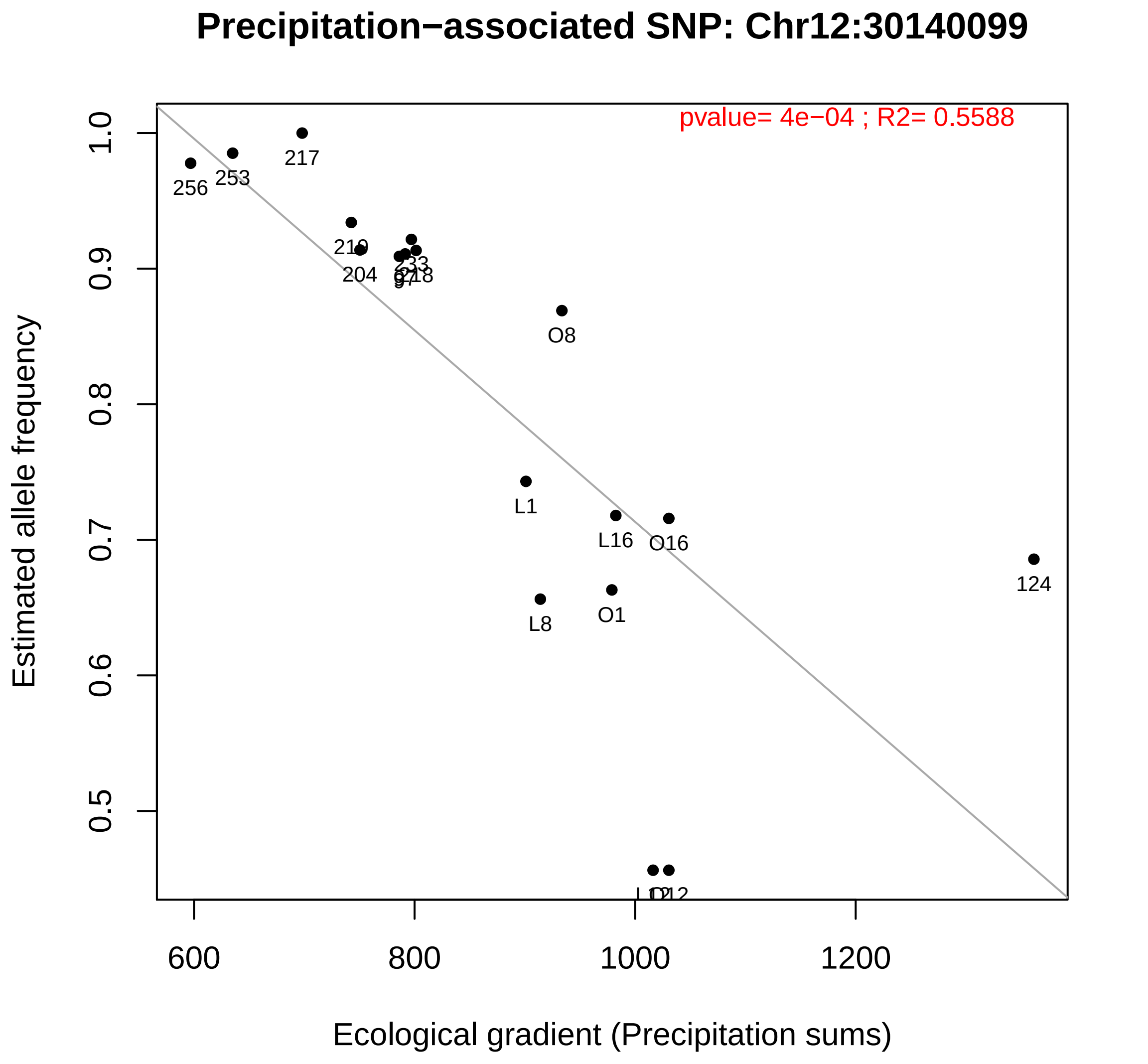
Precipitation−associated SNP: Chr12:30140099
pvalue= 4e−04 ; R2= 0.5588
Estimated allele frequency
Ecological gradient (Precipitation sums)
1.0
0.9
0.8
0.7
0.6
0.5
600
800
1000
1200
217
253
256
210
204
218
O8
L1
L16
O16
124
O1
L8

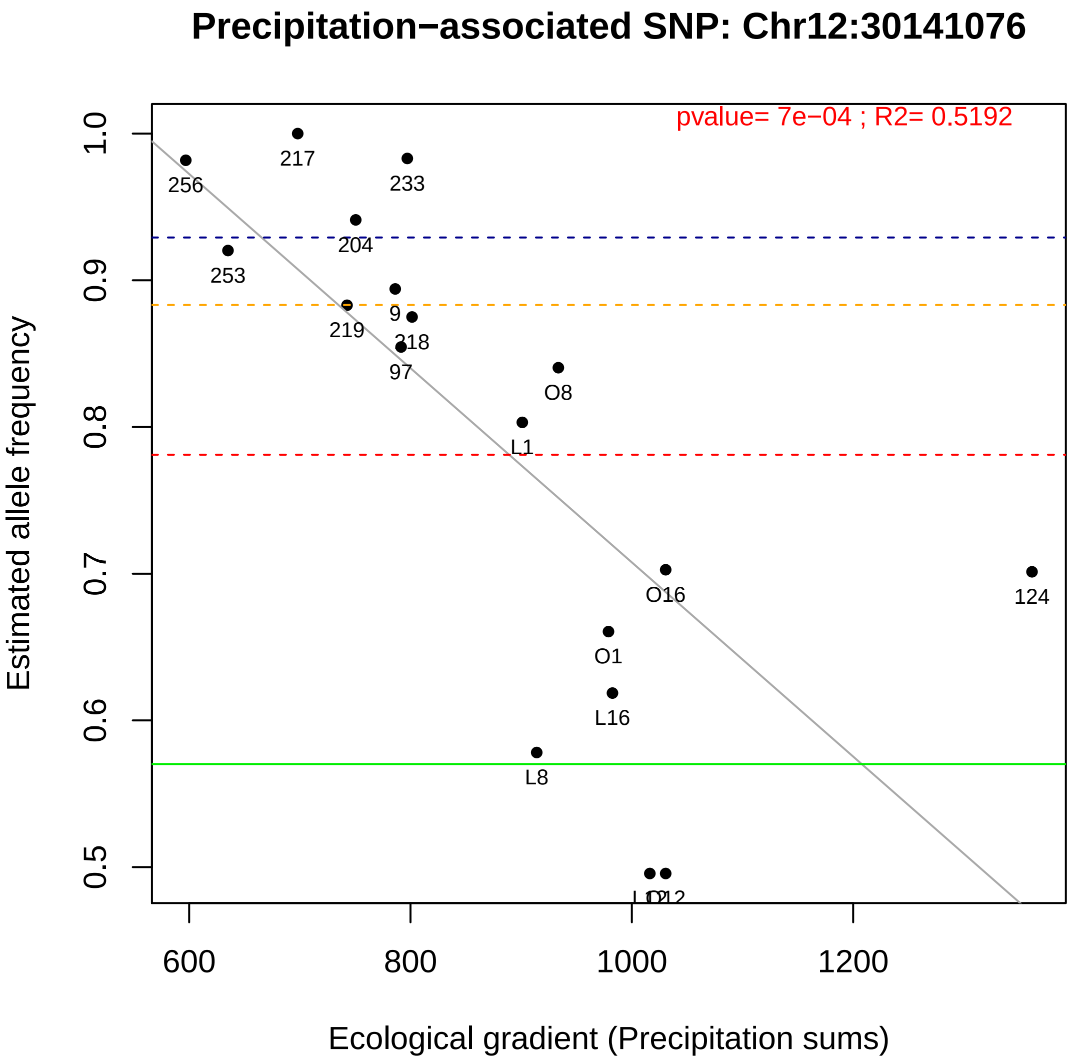
Precipitation−associated SNP: Chr12:30141076
pvalue= 7e−04 ; R2= 0.5192
Estimated allele frequency
Ecological gradient (Precipitation sums)
0.5
0.6
0.7
0.8
0.9
1.0
600
800
1000
1200
256
217
233
204
253
219
9
218
97
O8
L1
O16
124
O1
L16
L8

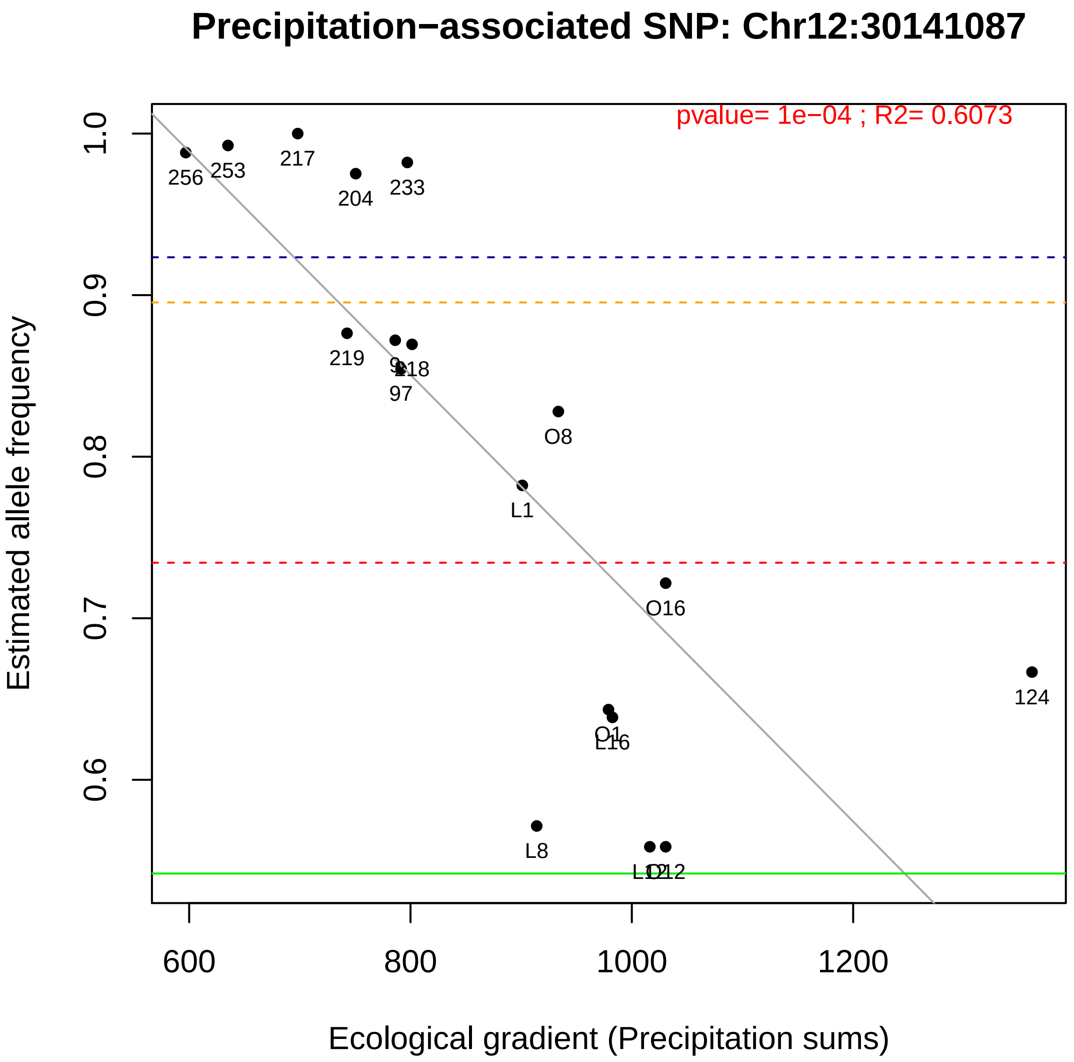
Precipitation−associated SNP: Chr12:30141087
pvalue= 1e−04 ; R2= 0.6073
Estimated allele frequency
Ecological gradient (Precipitation sums)
1.0
0.9
0.8
0.7
0.6
600
800
1000
1200
256
253
217
204
233
219
218
97
O8
L1
O16
124
O1
L16
L8
O12

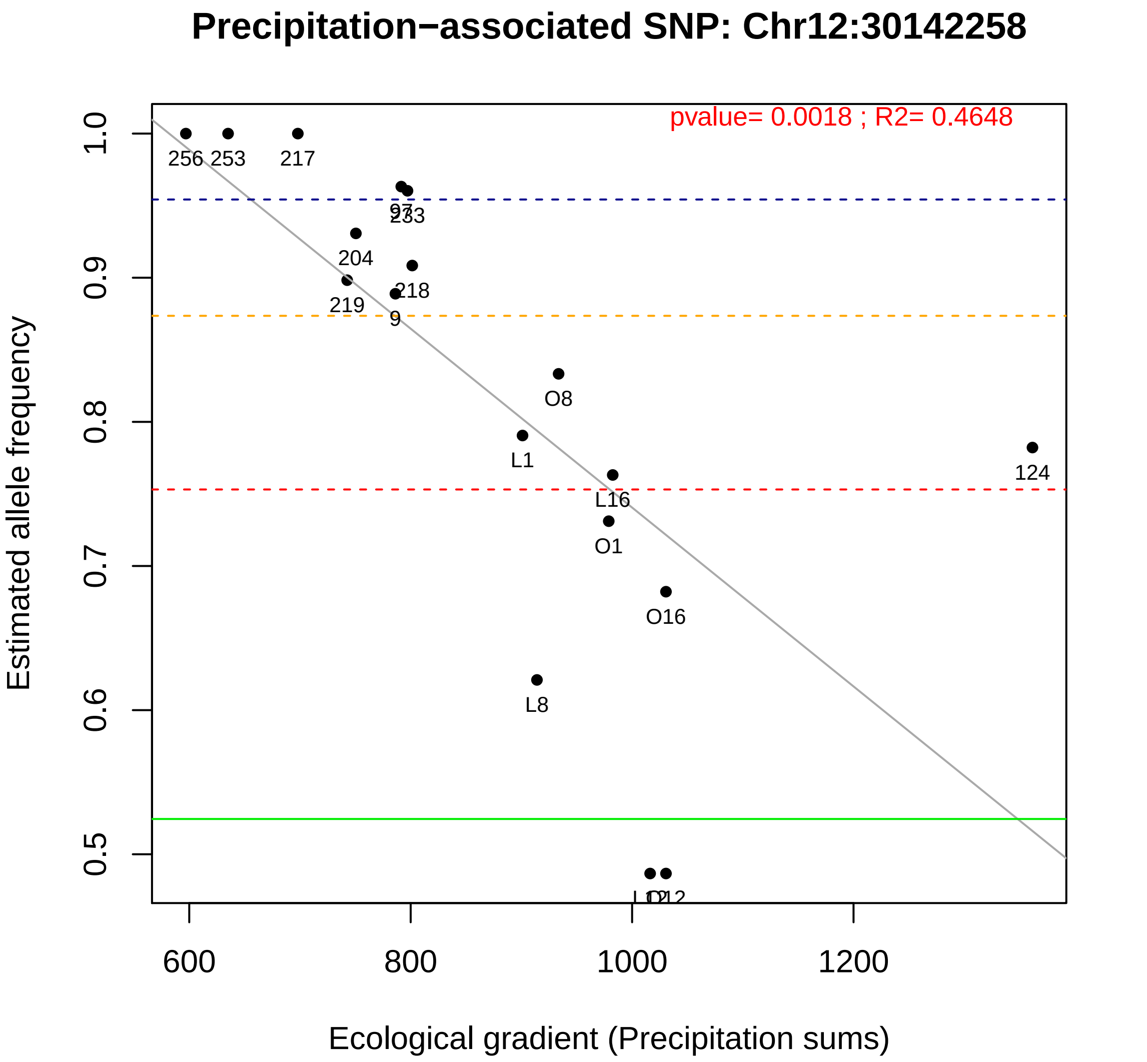

Precipitation−associated SNP: Chr12:30142258
pvalue= 0.0018 ; R2= 0.4648
Estimated allele frequency
Ecological gradient (Precipitation sums)
0.5
0.6
0.7
0.8
0.9
1.0
600
800
1000
1200
256
253
217
97
233
204
219
218
9
O8
L1
L16
124
O1
O16
L8

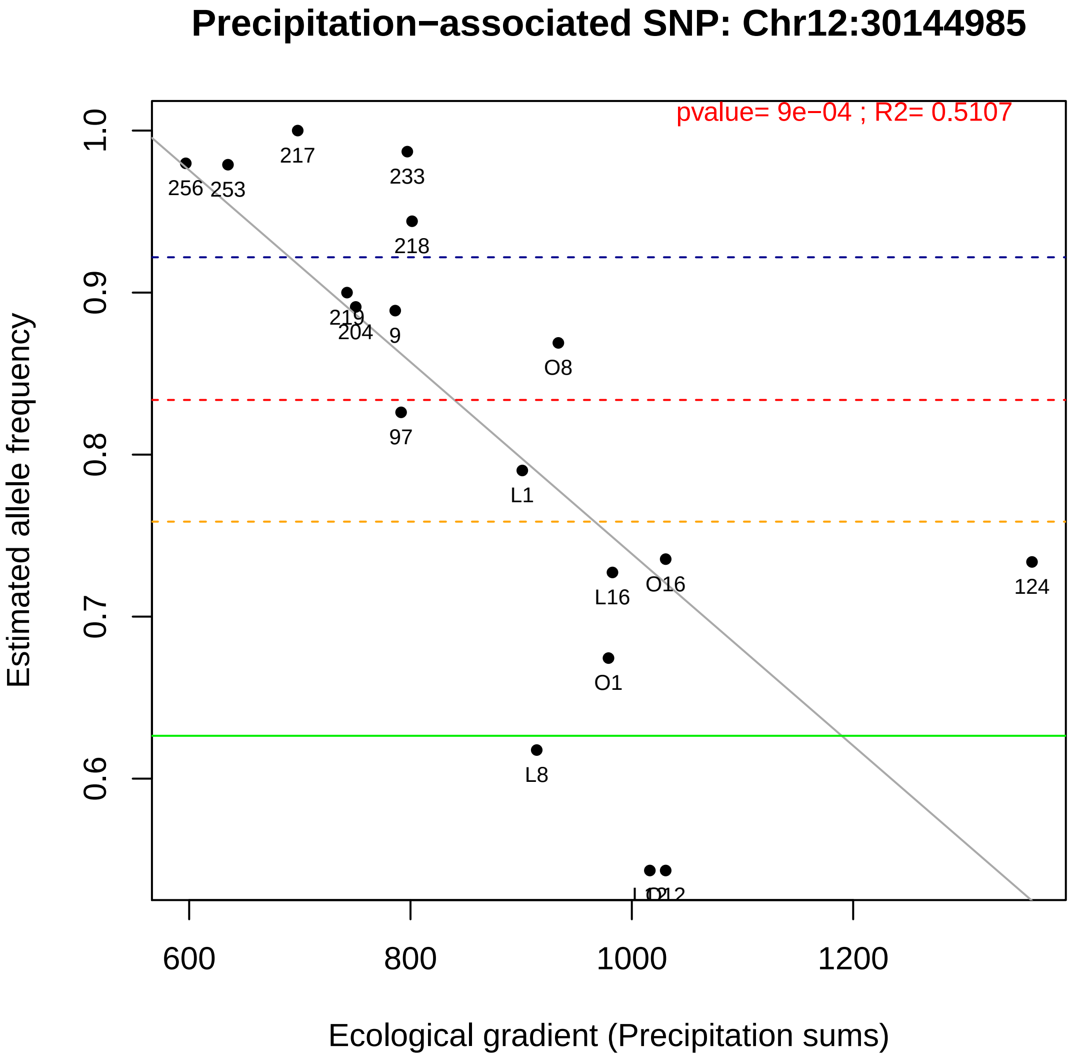

Precipitation−associated SNP: Chr12:30144985
pvalue= 9e−04 ; R2= 0.5107
Estimated allele frequency
Ecological gradient (Precipitation sums)
0.6
0.7
0.8
0.9
1.0
600
800
1000
1200
256
253
217
233
218
219
204
9
O8
97
L1
L16
O16
124
O1
L8

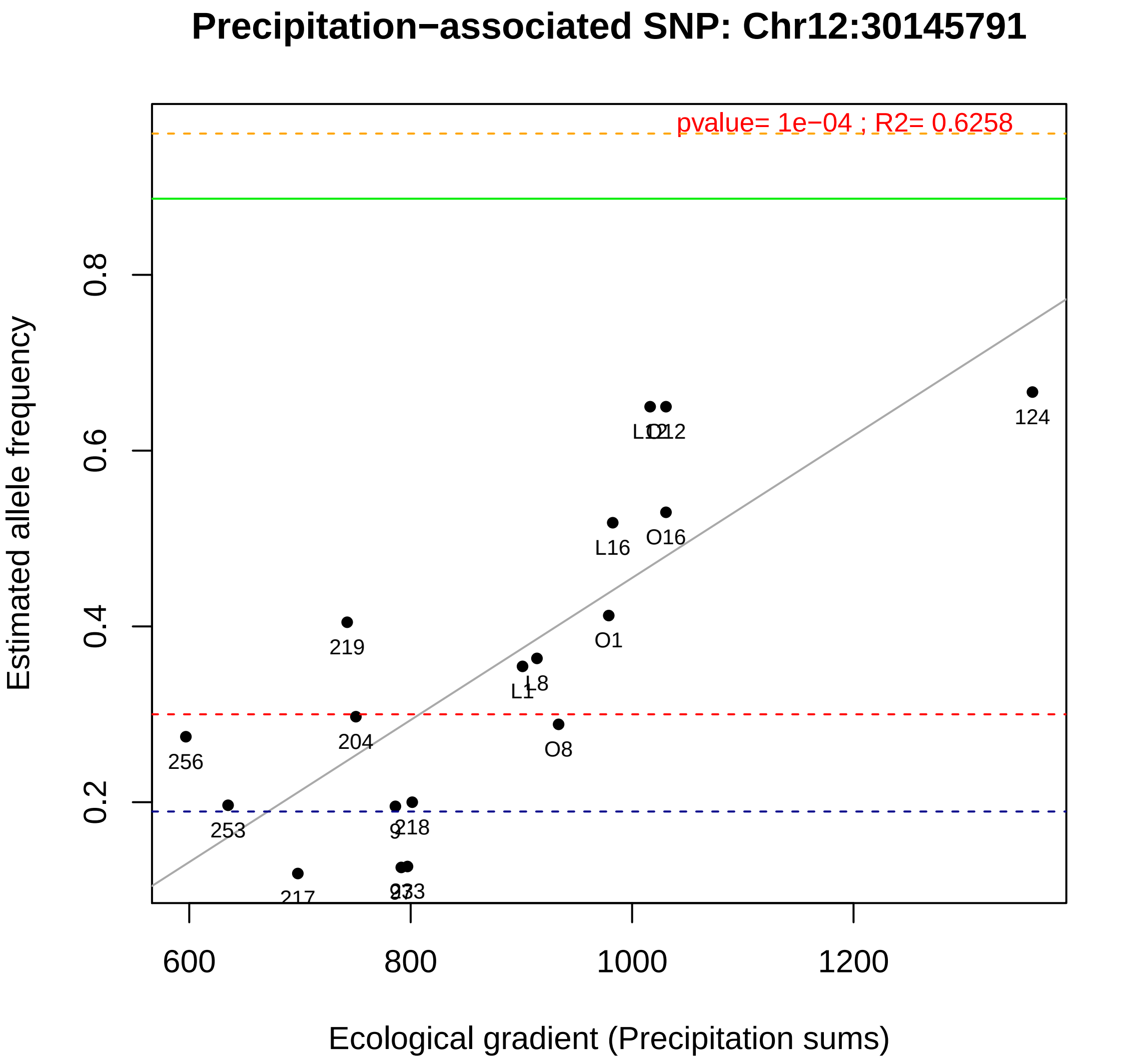
Precipitation−associated SNP: Chr12:30145791
pvalue= 1e−04 ; R2= 0.6258
Estimated allele frequency
0.2
0.4
0.6
0.8
600
800
1000
1200
Ecological gradient (Precipitation sums)
124
L12
O12
O16
L16
O1
219
L1
L8
204
O8
256
253
218
9
217
233

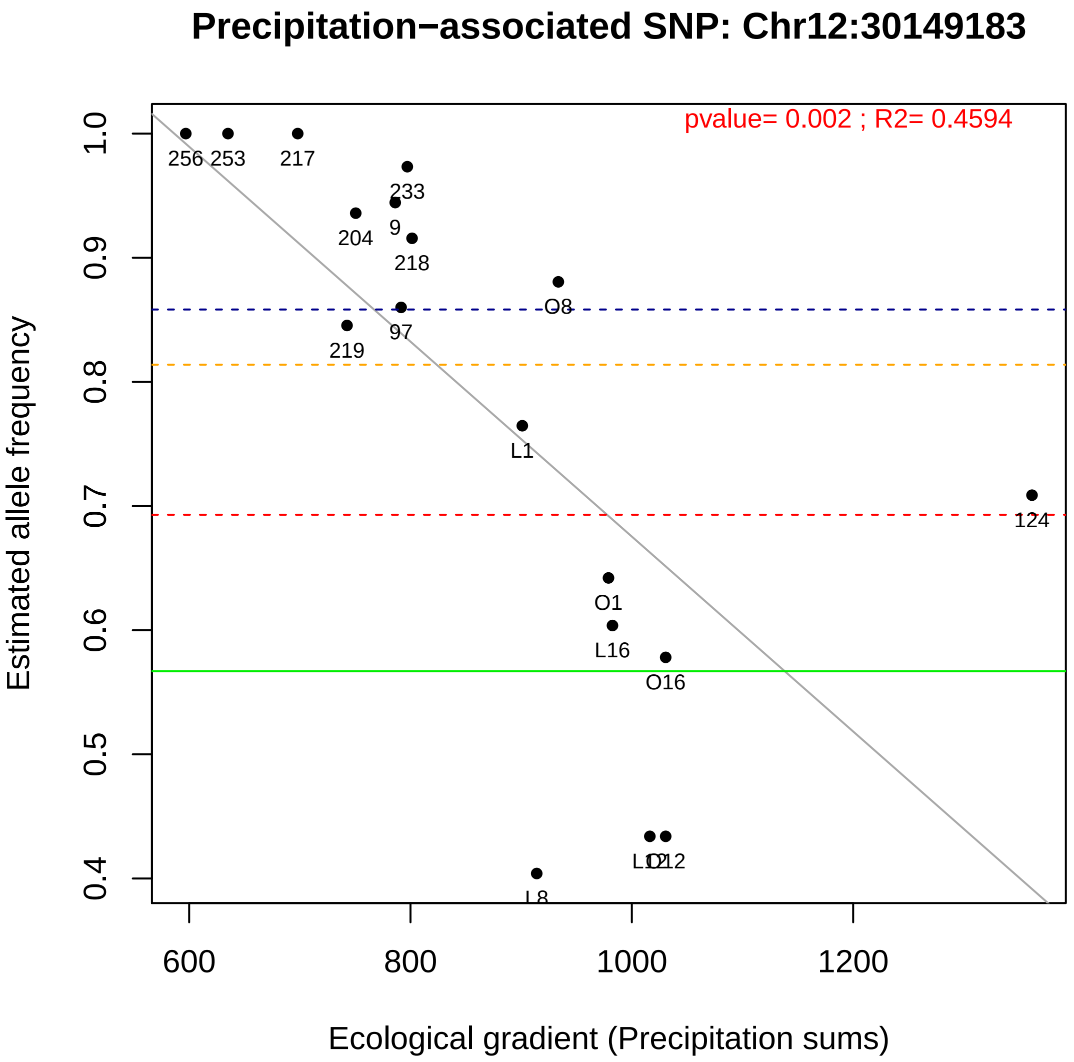
Precipitation−associated SNP: Chr12:30149183
pvalue= 0.002 ; R2= 0.4594
Estimated allele frequency
Ecological gradient (Precipitation sums)
0.4
0.5
0.6
0.7
0.8
0.9
1.0
600
800
1000
1200
256
253
217
233
9
204
218
O8
97
219
L1
124
O1
L16
O16
L8

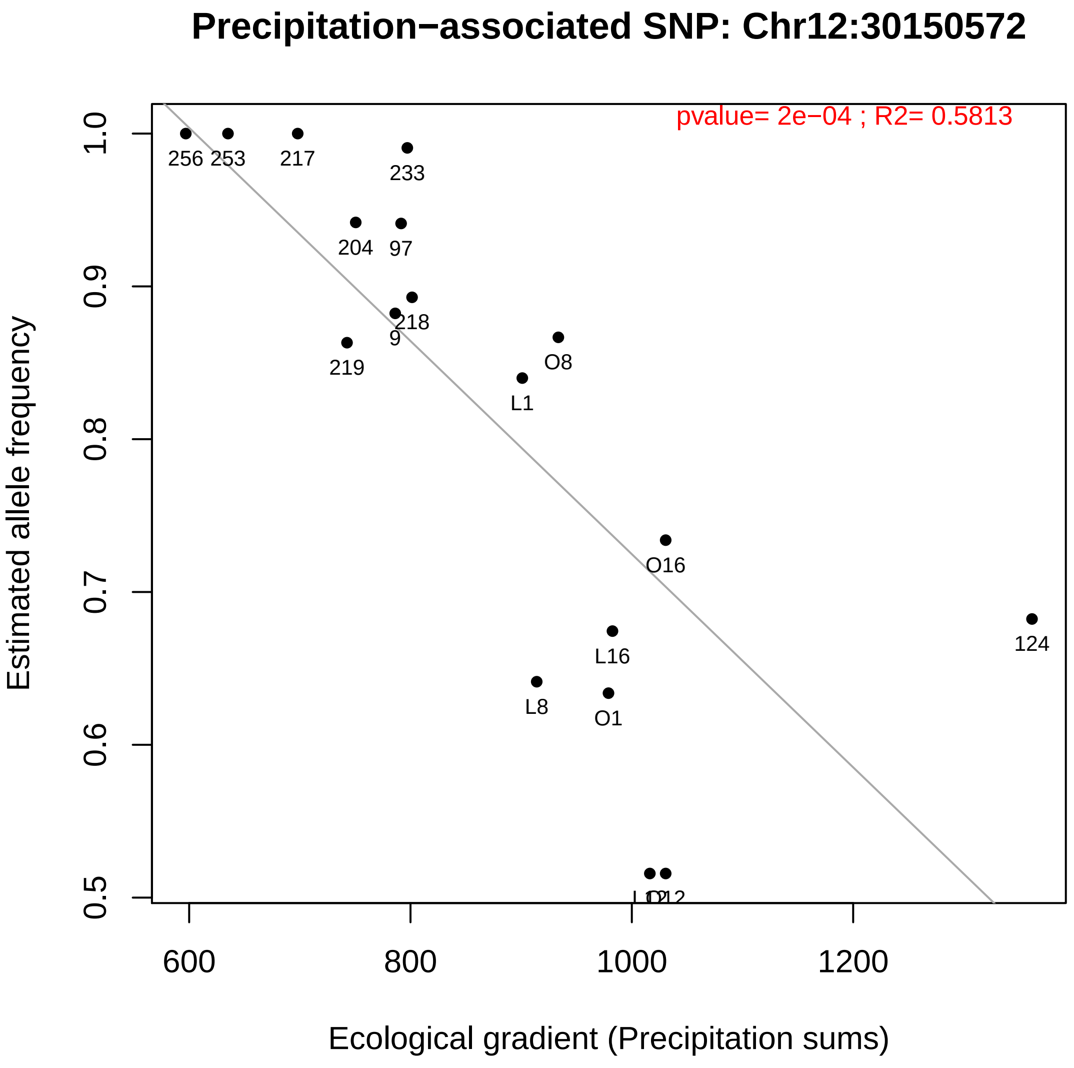
Precipitation−associated SNP: Chr12:30150572
pvalue= 2e−04 ; R2= 0.5813
Estimated allele frequency
Ecological gradient (Precipitation sums)
1.0
0.9
0.8
0.7
0.6
0.5
600
800
1000
1200
256
253
217
233
204
97
218
9
219
O8
L1
O16
124
L16
L8
O1

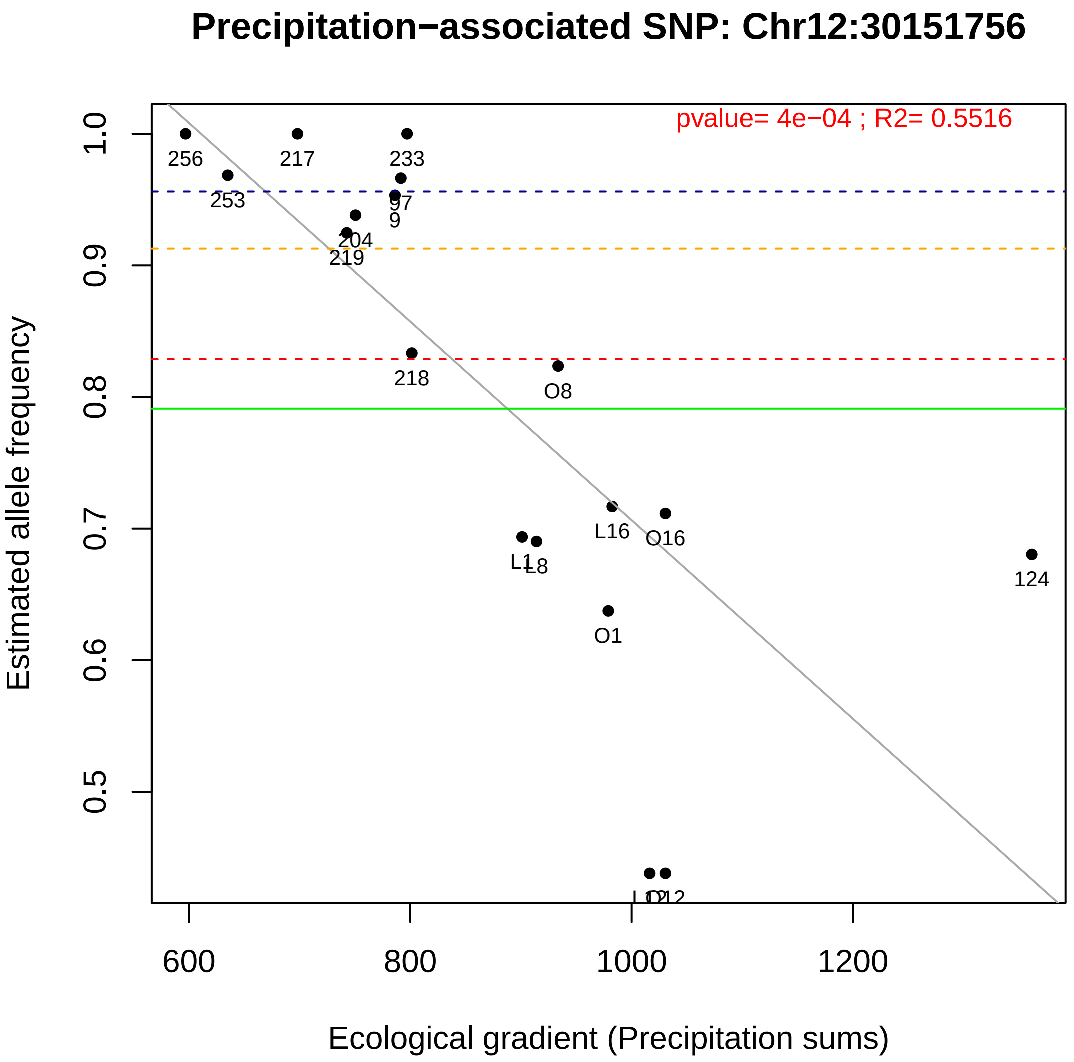

Precipitation−associated SNP: Chr12:30151756
pvalue= 4e−04 ; R2= 0.5516
256
217
233
253
97
9
204
219
218
O8
L16
O16
L1
L8
124
O1
L12
O12
Estimated allele frequency
1.0
0.9
0.8
0.7
0.6
0.5
600
800
1000
1200
Ecological gradient (Precipitation sums)

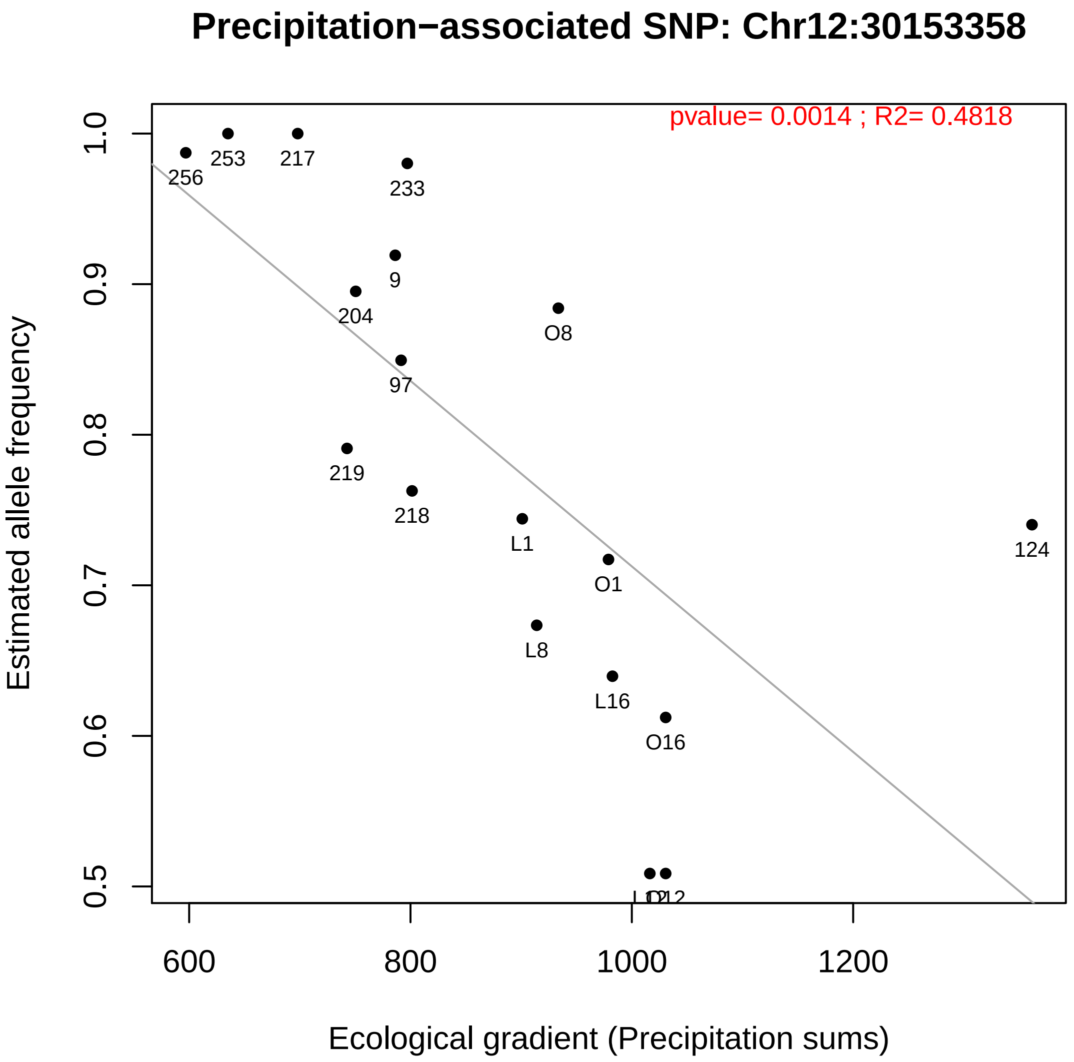
Precipitation−associated SNP: Chr12:30153358
pvalue= 0.0014 ; R2= 0.4818
Estimated allele frequency
Ecological gradient (Precipitation sums)
1.0
0.9
0.8
0.7
0.6
0.5
600
800
1000
1200
256
253
217
233
9
204
O8
97
219
218
L1
124
O1
L8
L16
O16
O12

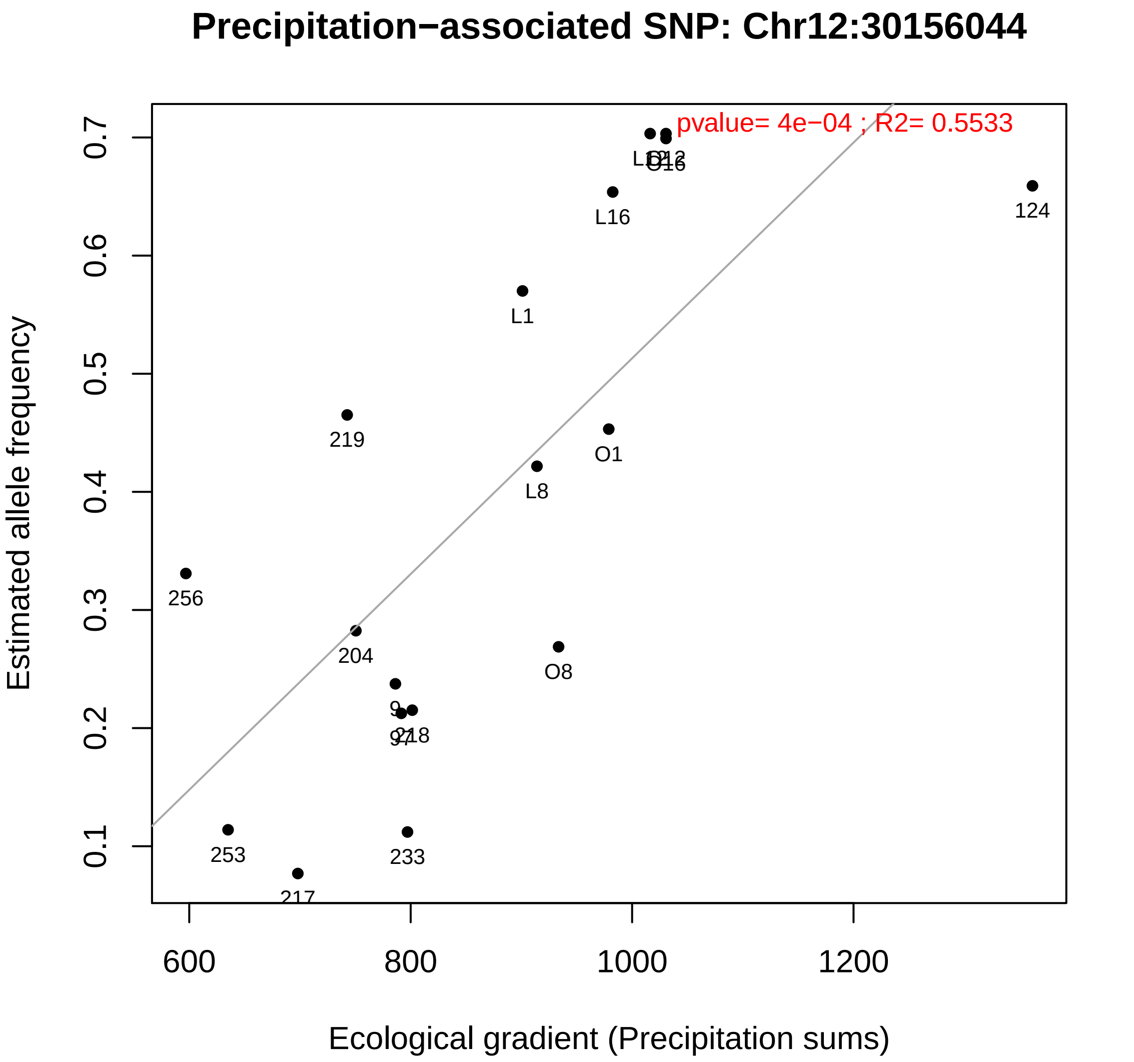
Precipitation−associated SNP: Chr12:30156044
pvalue= 4e−04 ; R2= 0.5533
Estimated allele frequency
Ecological gradient (Precipitation sums)
0.1
0.2
0.3
0.4
0.5
0.6
0.7
600
800
1000
1200
L16
124
L1
219
O1
L8
256
204
O8
218
253
233
217
O16

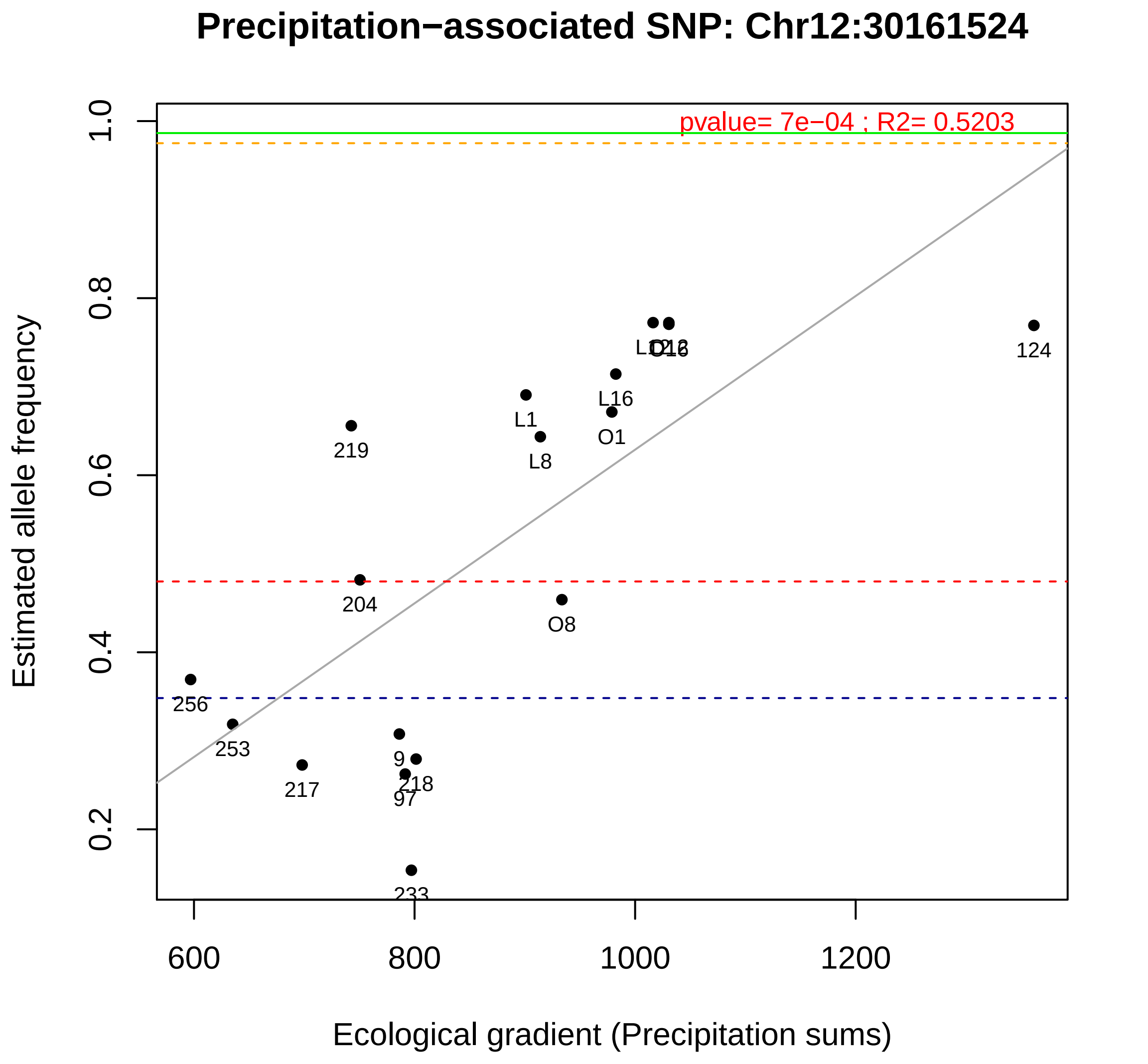

Precipitation−associated SNP: Chr12:30161524
pvalue= 7e−04 ; R2= 0.5203
Estimated allele frequency
0.2
0.4
0.6
0.8
1.0
600
800
1000
1200
Ecological gradient (Precipitation sums)
124
L16
L1
O1
L8
219
204
O8
256
253
217
9
218
97
233

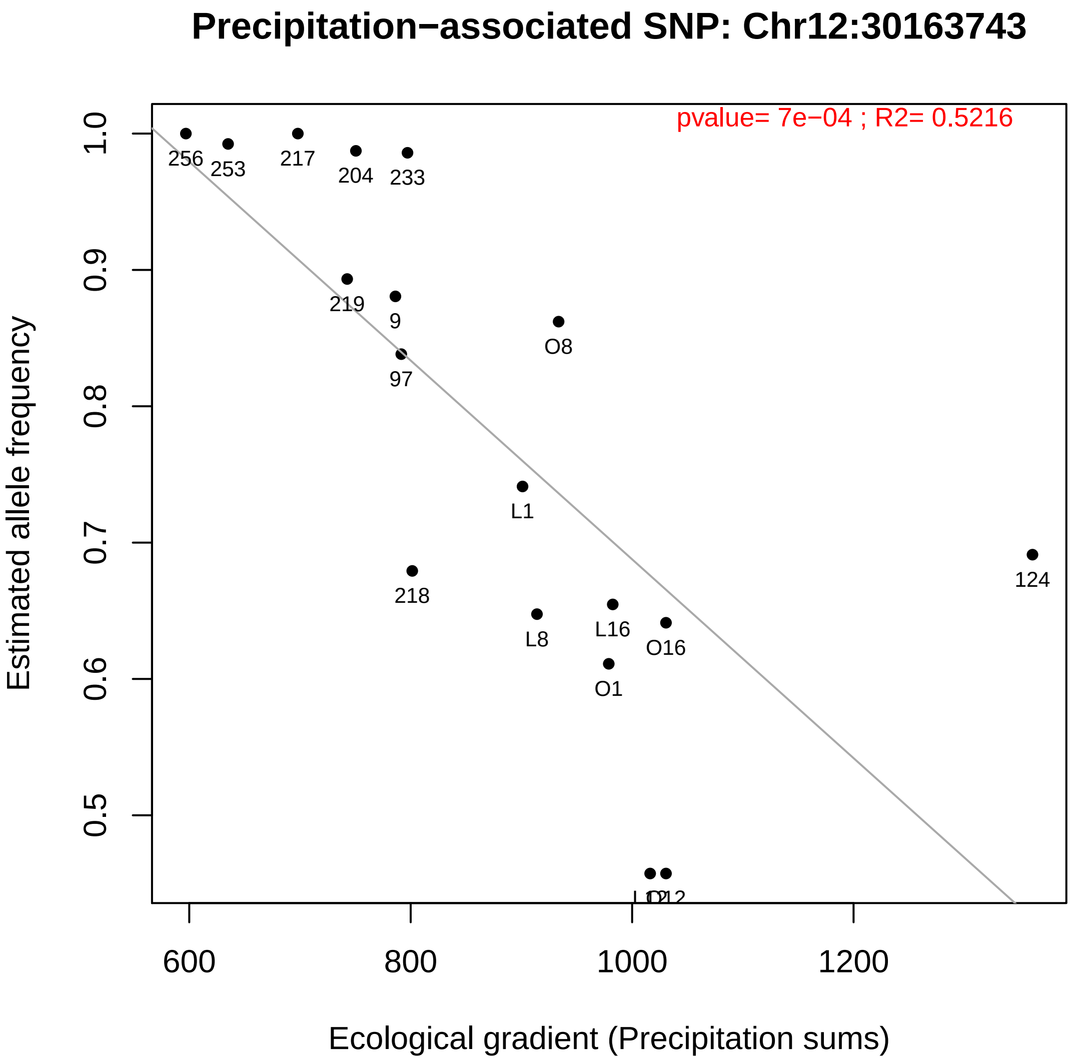
Precipitation−associated SNP: Chr12:30163743
pvalue= 7e−04 ; R2= 0.5216
Estimated allele frequency
Ecological gradient (Precipitation sums)
1.0
0.9
0.8
0.7
0.6
0.5
600
800
1000
1200
256
253
217
204
233
219
9
97
O8
L1
218
L8
L16
O16
O1
124

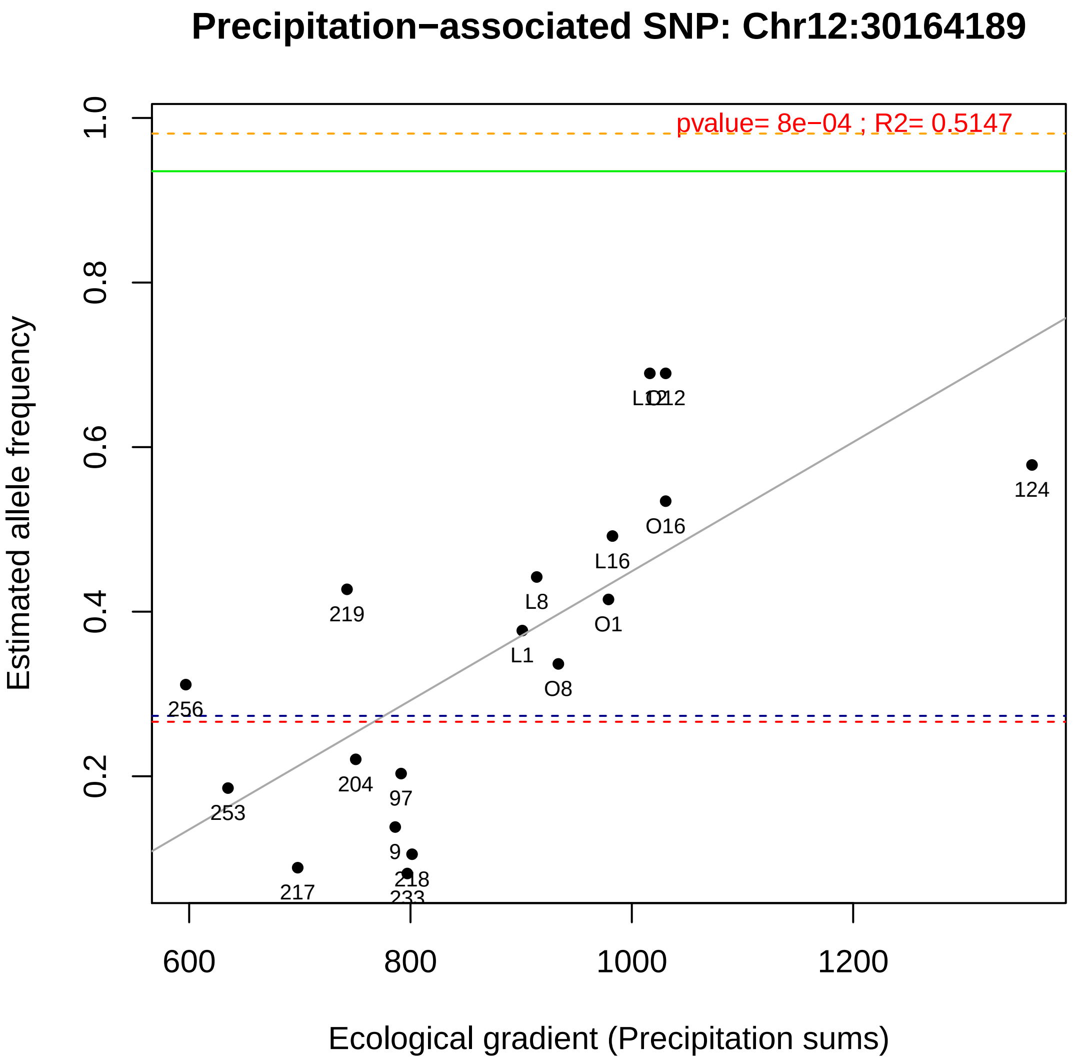

Precipitation−associated SNP: Chr12:30164189
pvalue= 8e−04 ; R2= 0.5147
Estimated allele frequency
0.2
0.4
0.6
0.8
1.0
600
800
1000
1200
Ecological gradient (Precipitation sums)
L12
O12
124
O16
L16
L8
O1
219
L1
O8
256
204
97
253
9
218
217
233

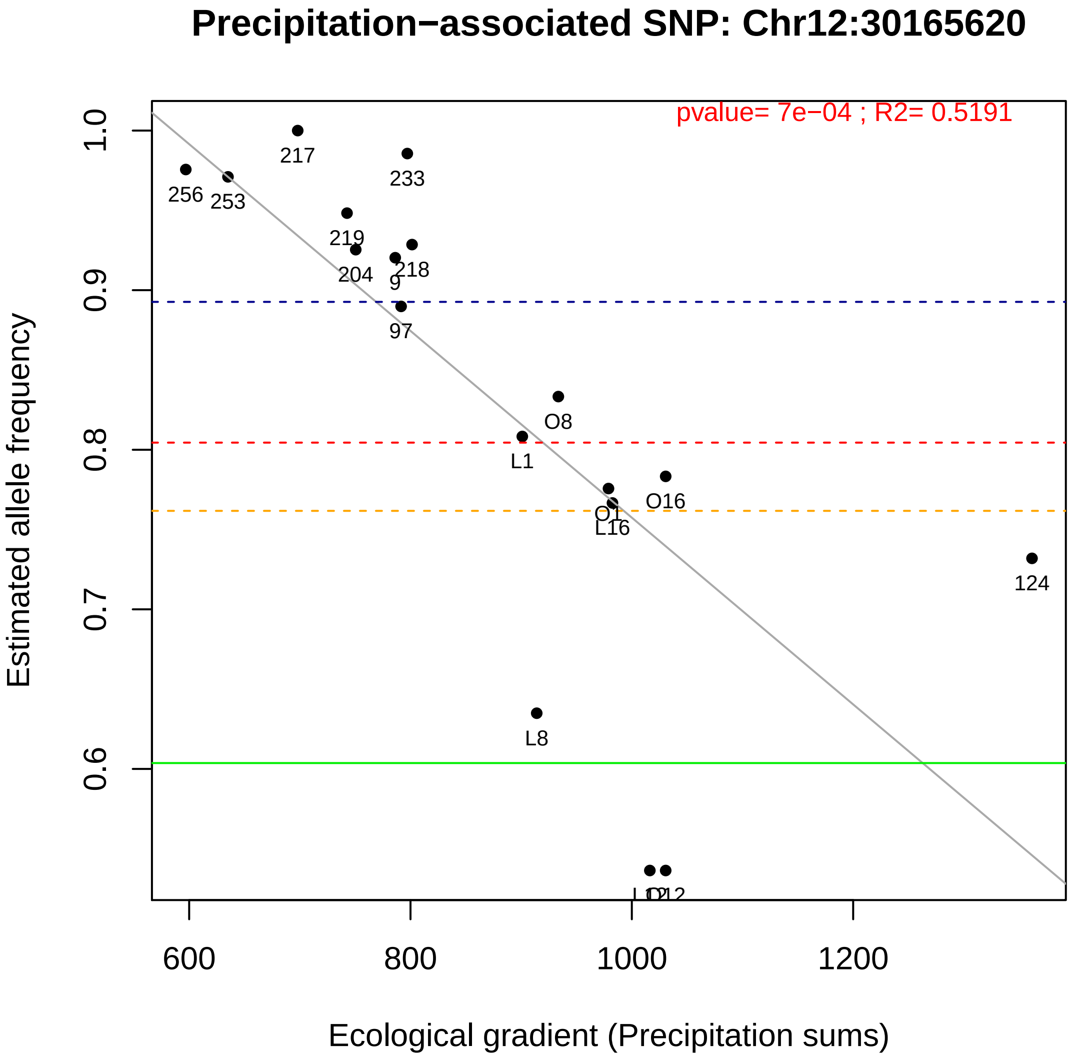
Precipitation−associated SNP: Chr12:30165620
pvalue= 7e−04 ; R2= 0.5191
Estimated allele frequency
1.0
0.9
0.8
0.7
0.6
600
800
1000
1200
Ecological gradient (Precipitation sums)
217
233
256
253
219
204
218
9
97
O8
L1
O16
O1
L16
124
L8

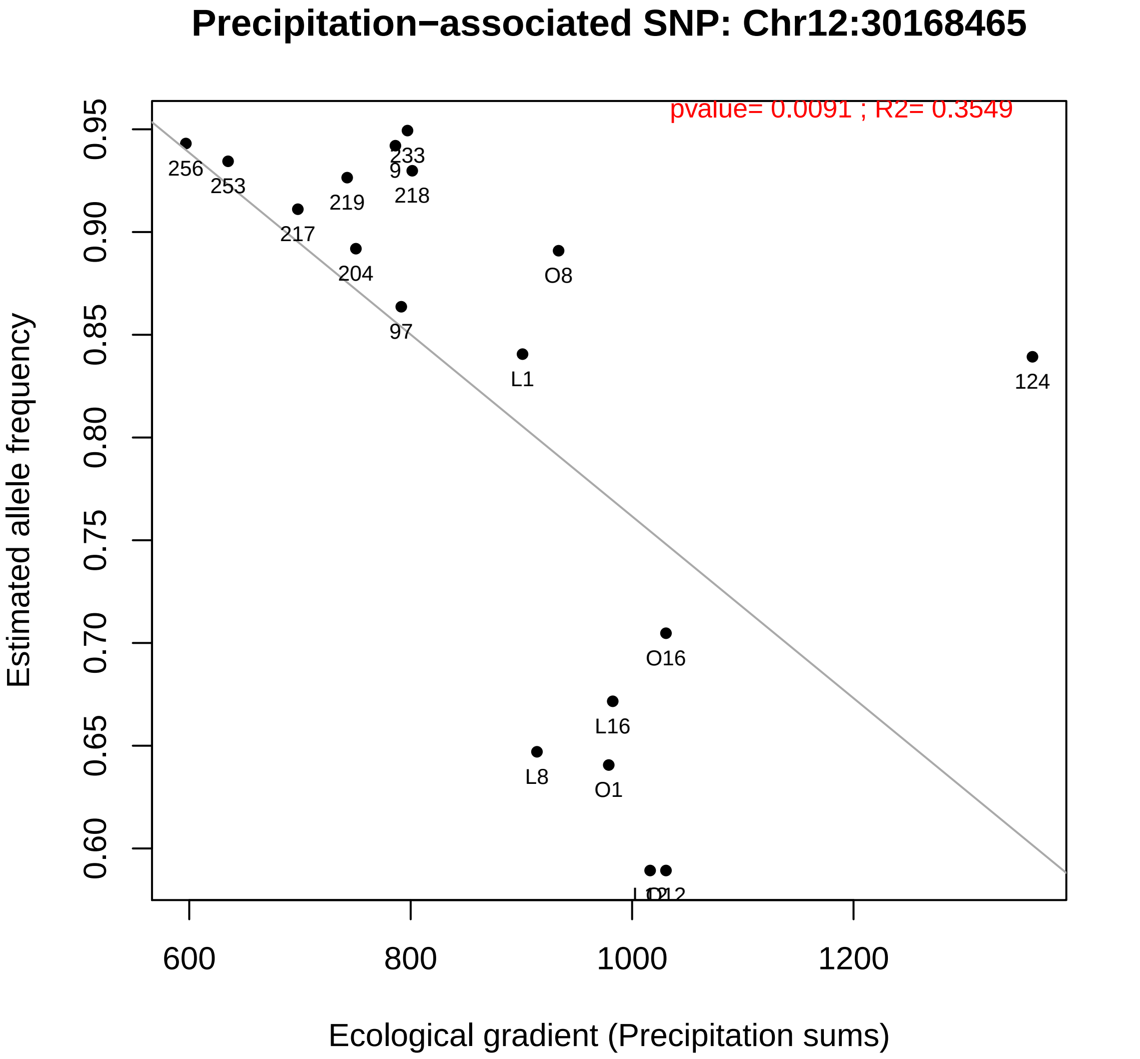

Precipitation−associated SNP: Chr12:30168465
pvalue= 0.0091 ; R2= 0.3549
Estimated allele frequency
0.60
0.65
0.70
0.75
0.80
0.85
0.90
0.95
600
800
1000
1200
Ecological gradient (Precipitation sums)
256
253
233
9
218
219
217
204
O8
97
L1
124
O16
L16
L8
O1
O12

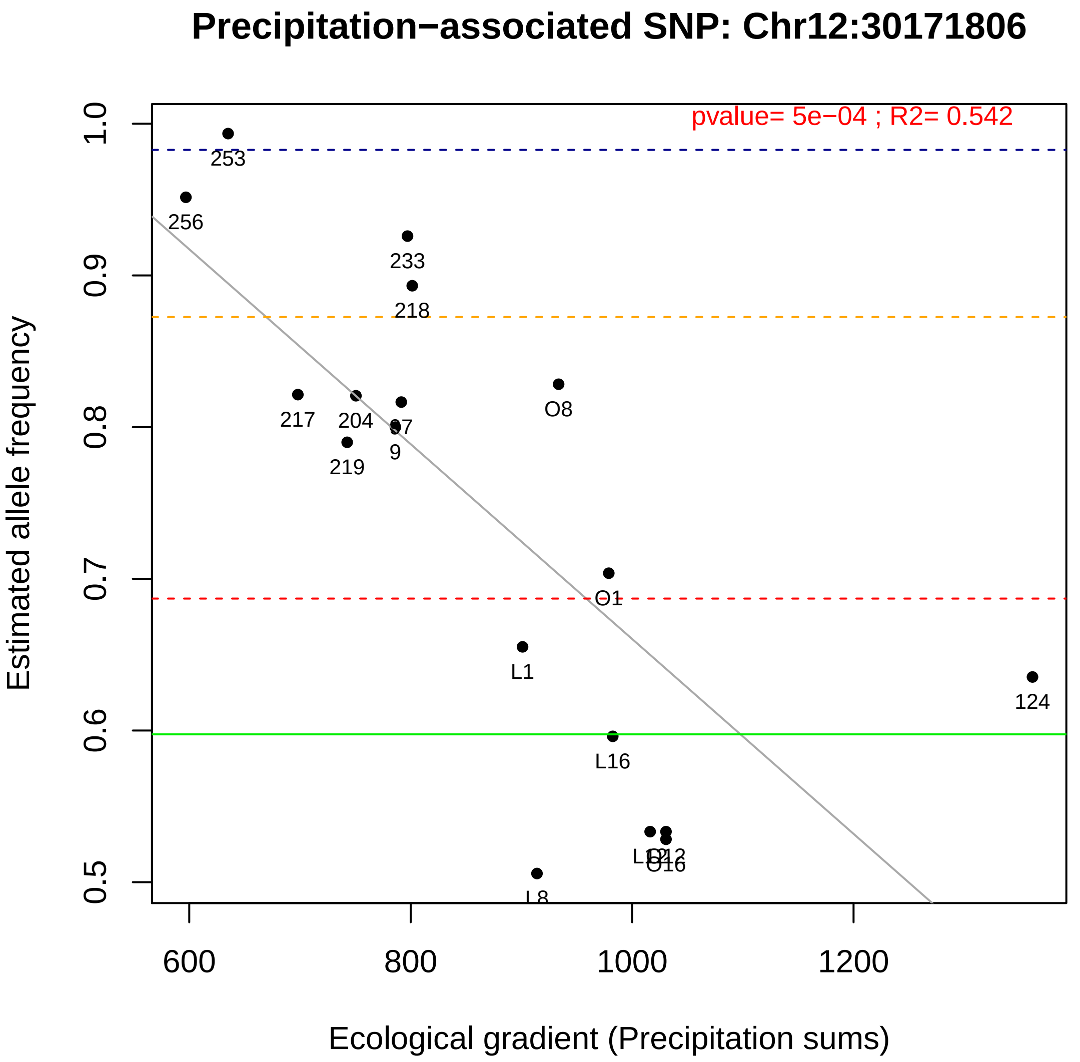

Precipitation−associated SNP: Chr12:30171806
pvalue= 5e−04 ; R2= 0.542
Estimated allele frequency
Ecological gradient (Precipitation sums)
1.0
0.9
0.8
0.7
0.6
0.5
600
800
1000
1200
253
256
233
218
217
204
219
O8
O1
L1
124
L16
L8

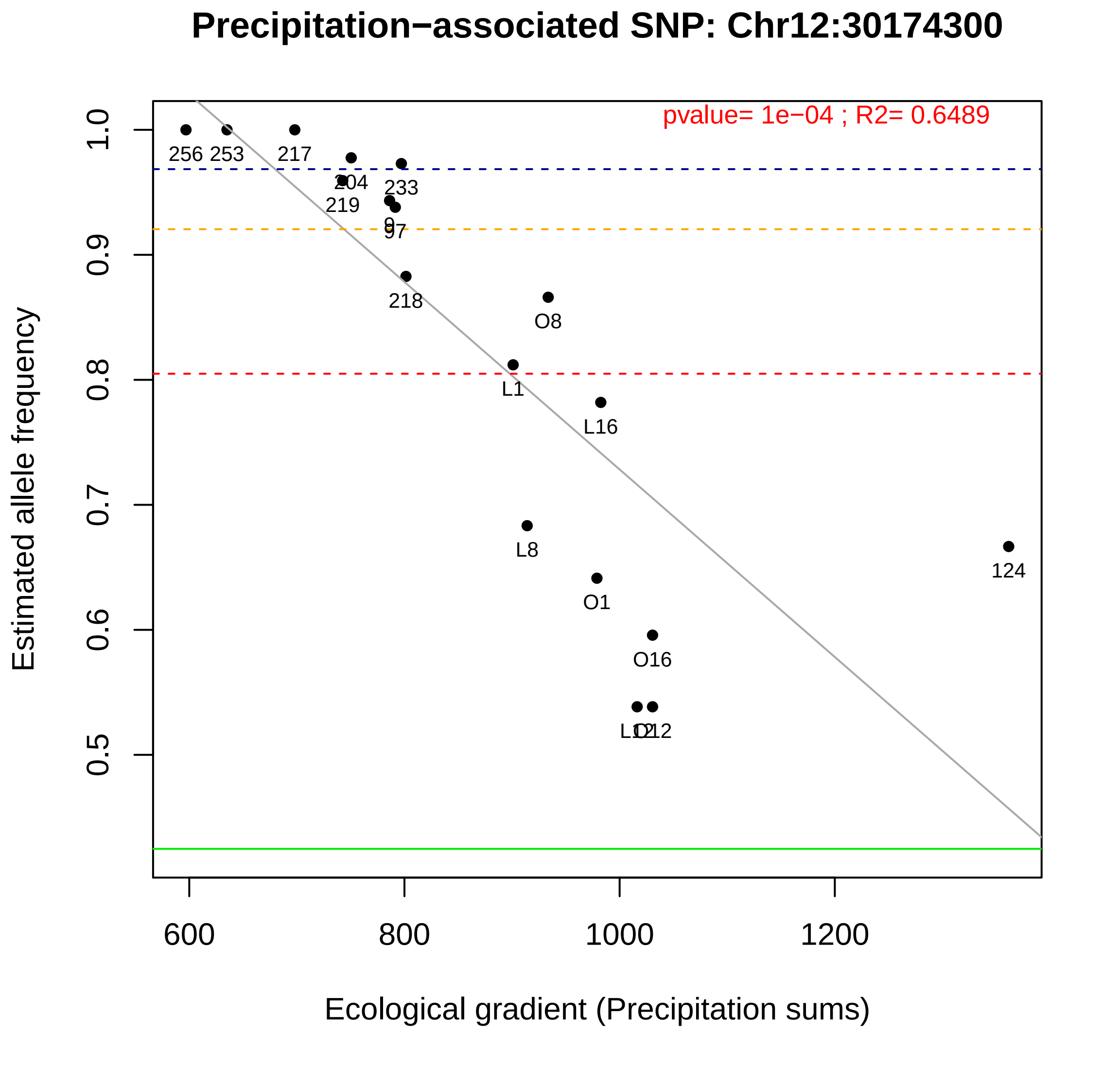

Precipitation−associated SNP: Chr12:30174300
pvalue= 1e−04 ; R2= 0.6489
Estimated allele frequency
Ecological gradient (Precipitation sums)
0.5
0.6
0.7
0.8
0.9
1.0
600
800
1000
1200
256
253
217
204
233
219
9
97
218
O8
L1
L16
L8
124
O1
O16
L12
O12

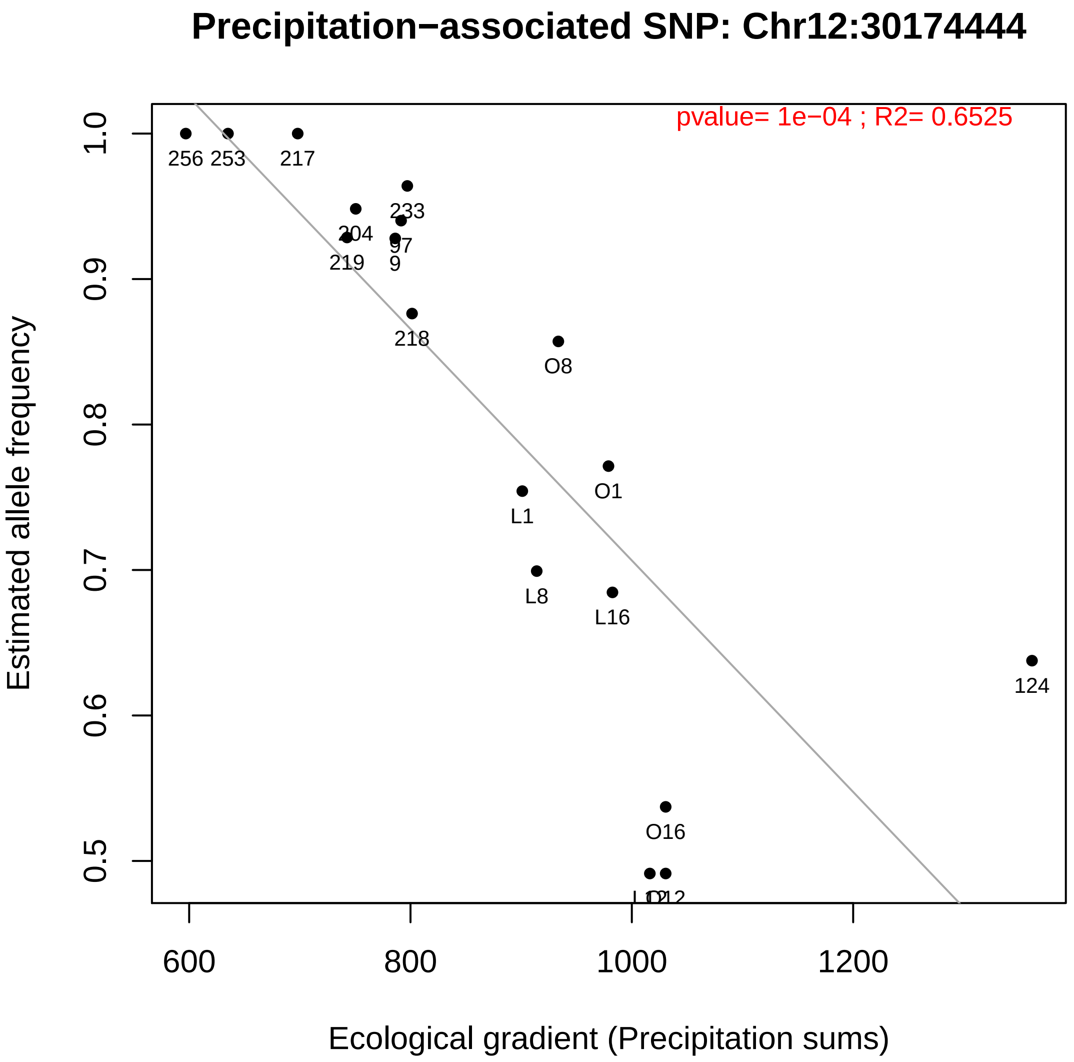
Precipitation−associated SNP: Chr12:30174444
pvalue= 1e−04 ; R2= 0.6525
Estimated allele frequency
0.5
0.6
0.7
0.8
0.9
1.0
Ecological gradient (Precipitation sums)
600
800
1000
1200
256
253
217
233
204
97
219
9
218
O8
O1
L1
L8
L16
124
O16

**Precipitation−associated SNP: Chr12:30176072**

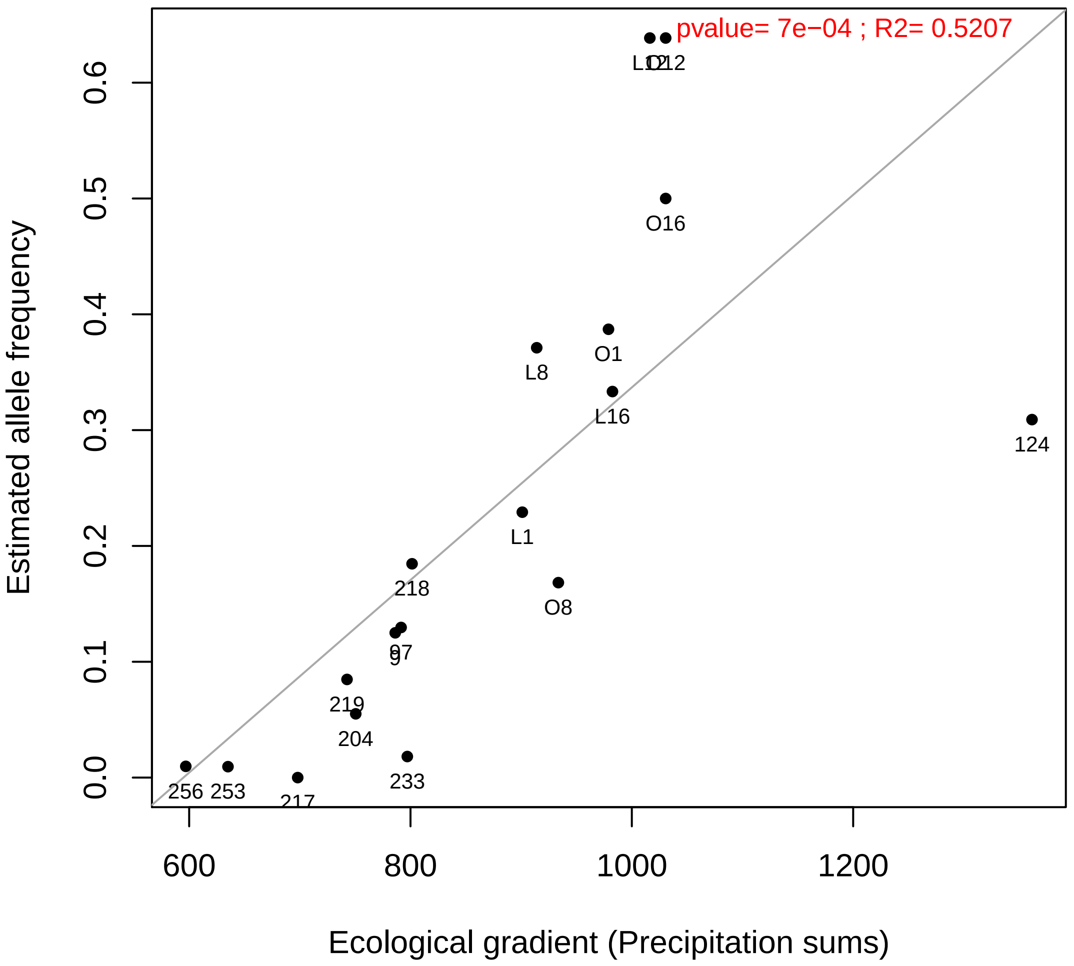

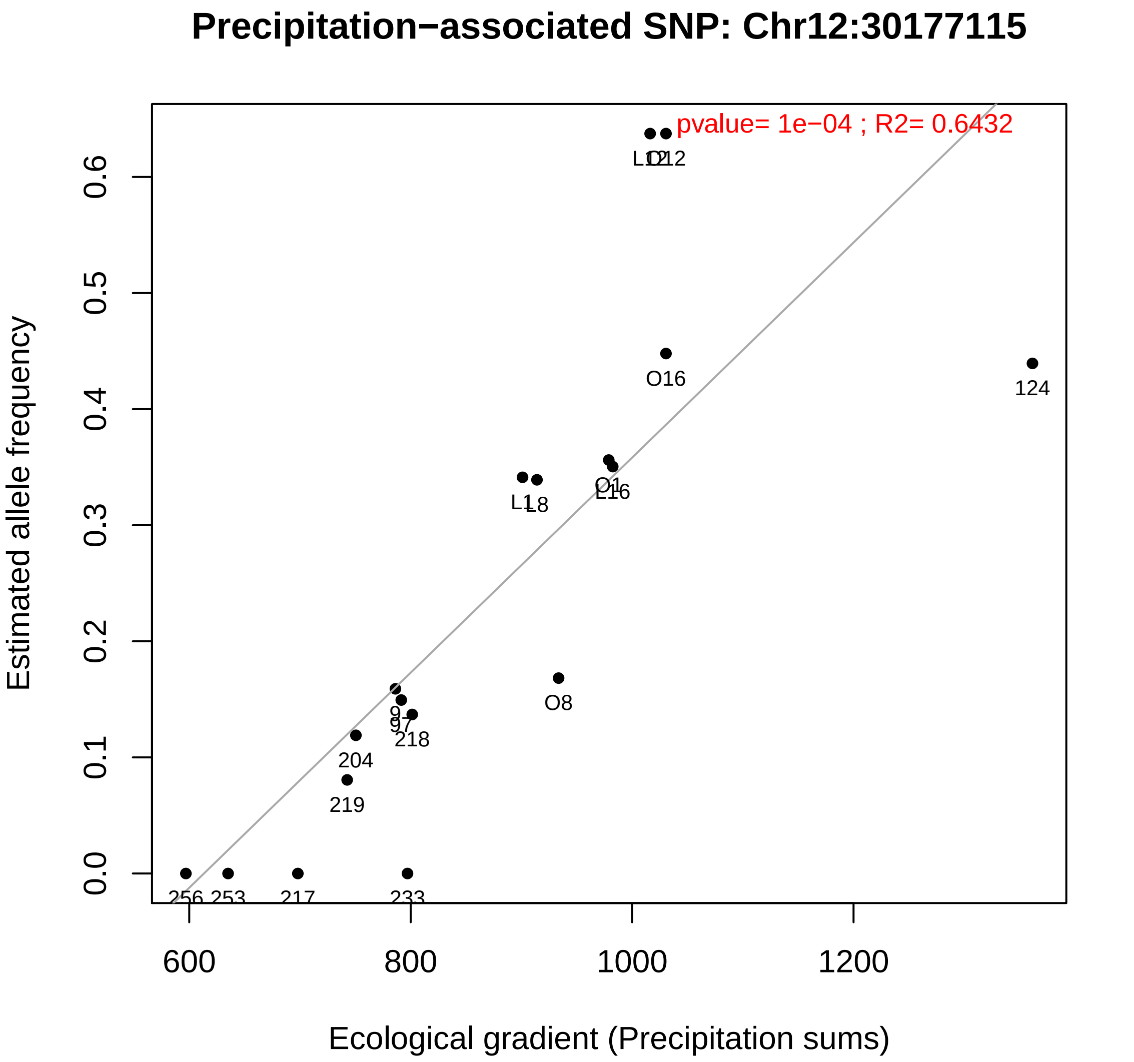
Precipitation−associated SNP: Chr12:30177115
pvalue= 1e−04 ; R2= 0.6432
Estimated allele frequency
0.0
0.1
0.2
0.3
0.4
0.5
0.6
Ecological gradient (Precipitation sums)
600
800
1000
1200
O12
O16
124
L16
L18
O8
97
218
204
219
256
253
217
233

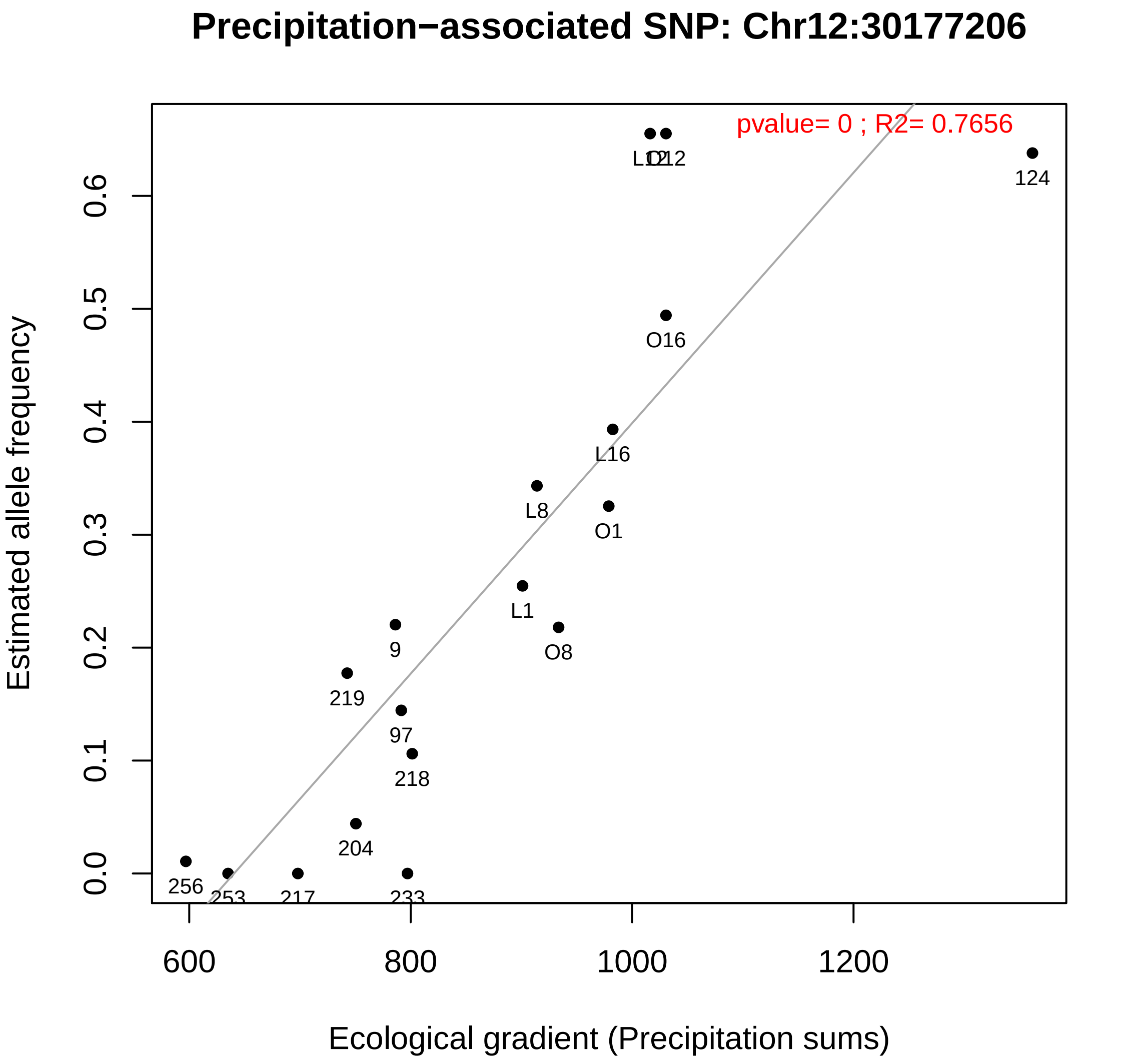
Precipitation−associated SNP: Chr12:30177206
pvalue= 0 ; R2= 0.7656
Estimated allele frequency
Ecological gradient (Precipitation sums)
0.0
0.1
0.2
0.3
0.4
0.5
0.6
600
800
1000
1200
124
O16
L16
L8
O1
L1
O8
9
219
97
218
204
256
253
217
233

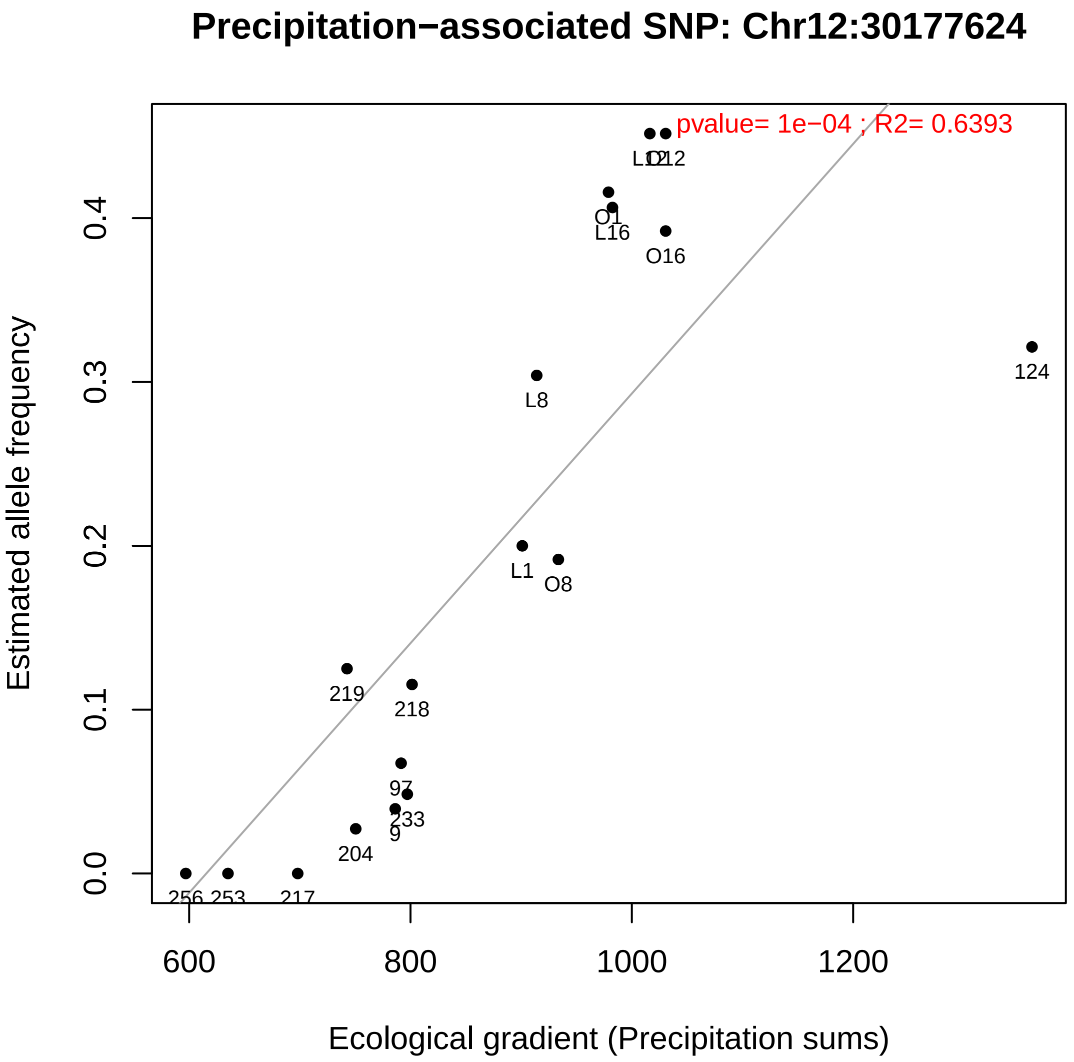
Precipitation−associated SNP: Chr12:30177624
pvalue= 1e−04 ; R2= 0.6393
Estimated allele frequency
0.0
0.1
0.2
0.3
0.4
600
800
1000
1200
Ecological gradient (Precipitation sums)
L12
O12
O1
L16
O16
124
L8
L1
O8
219
218
97
233
9
204
256
253
217

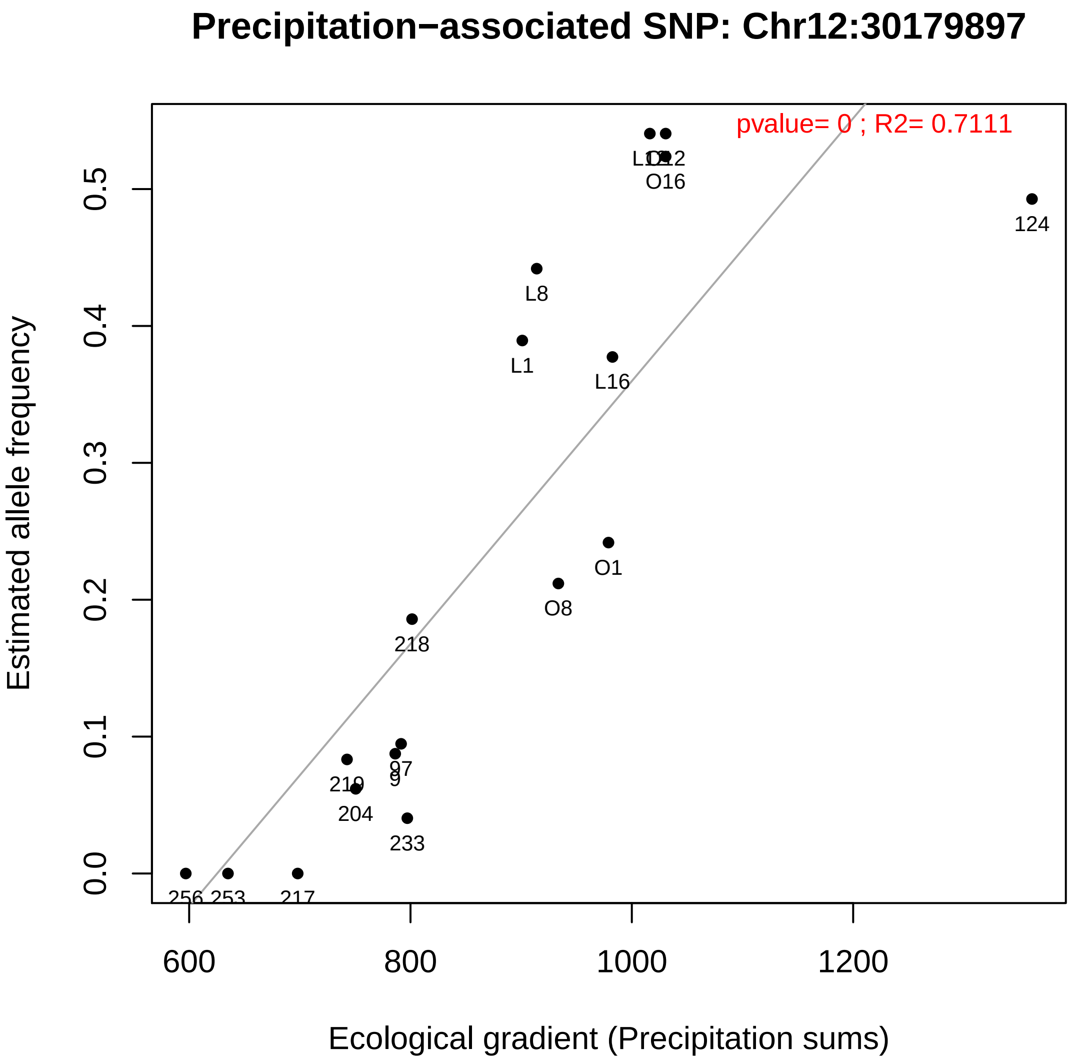

Precipitation−associated SNP: Chr12:30179897
pvalue= 0 ; R2= 0.7111
Estimated allele frequency
Ecological gradient (Precipitation sums)
0.0
0.1
0.2
0.3
0.4
0.5
600
800
1000
1200
O16
124
L8
L1
L16
O1
O8
218
219
97
9
204
233
256
253
217

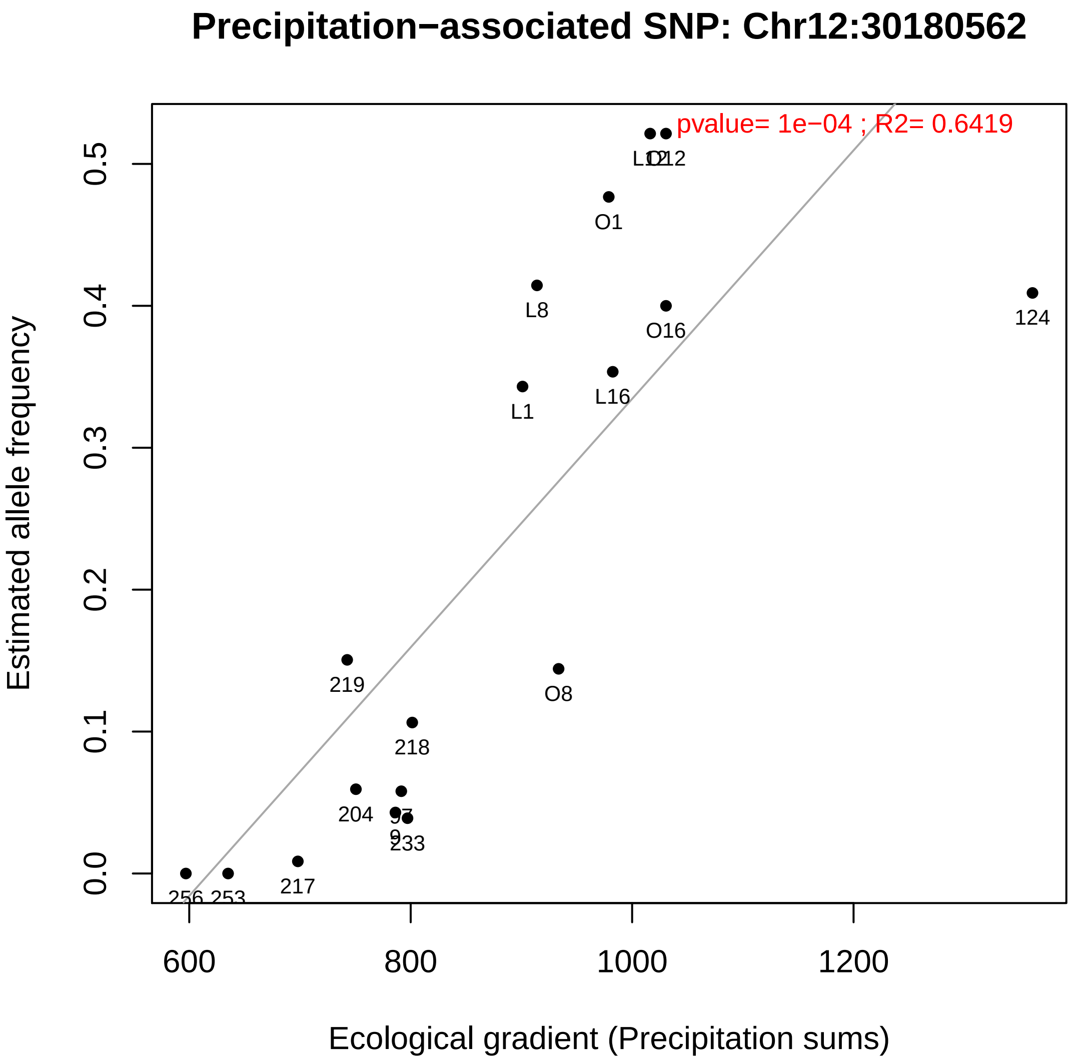

Precipitation−associated SNP: Chr12:30180562
pvalue= 1e−04 ; R2= 0.6419
Estimated allele frequency
0.0
0.1
0.2
0.3
0.4
0.5
600
800
1000
1200
Ecological gradient (Precipitation sums)
L12
O12
O1
L8
O16
124
L16
L1
219
O8
218
204
233
217
256
253

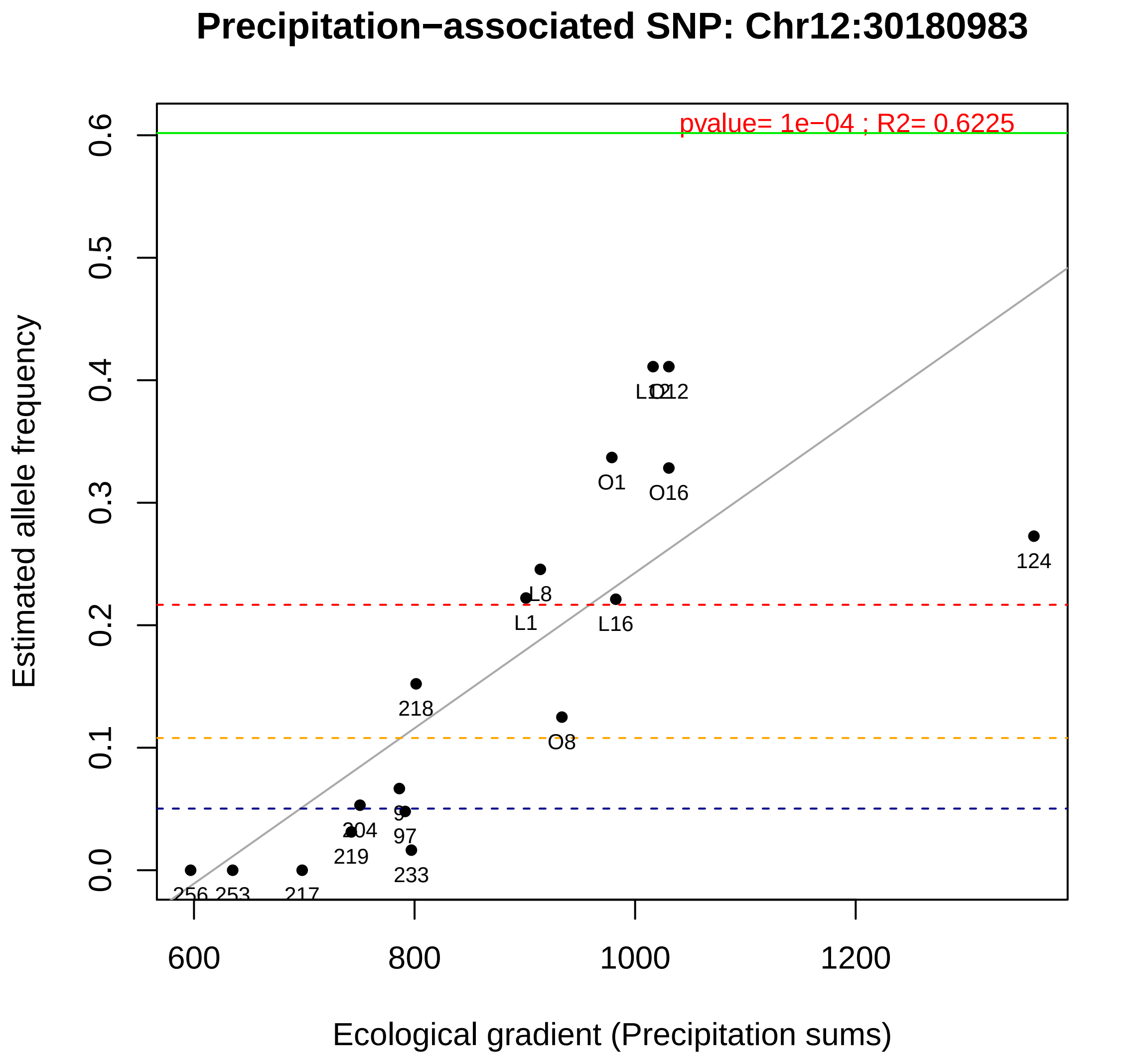

Precipitation−associated SNP: Chr12:30180983
pvalue= 1e−04 ; R2= 0.6225
Estimated allele frequency
Ecological gradient (Precipitation sums)
0.0
0.1
0.2
0.3
0.4
0.5
0.6
600
800
1000
1200
L12
O12
O1
O16
124
L8
L1
L16
218
O8
9
204
97
219
233
256
253
217

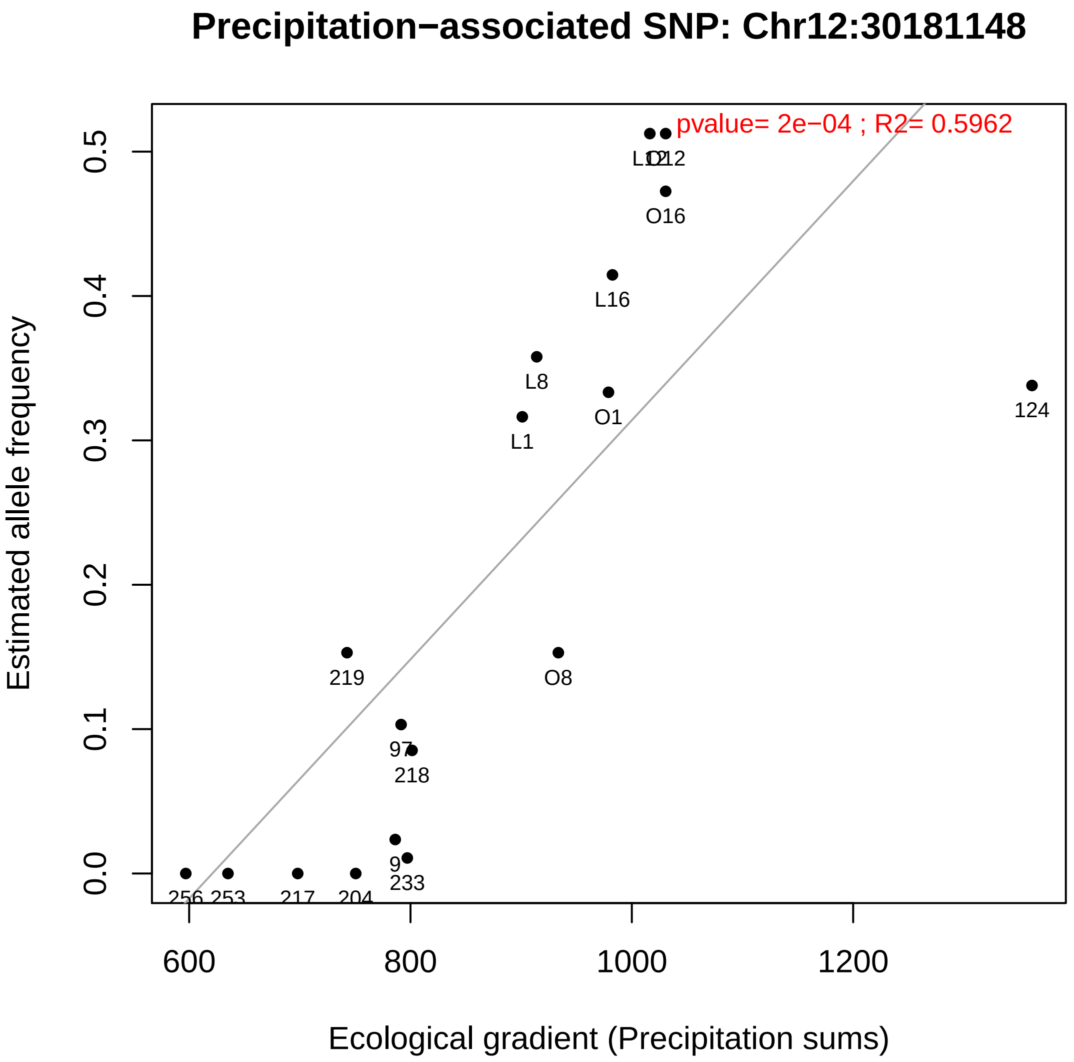

Precipitation−associated SNP: Chr12:30181148
pvalue= 2e−04 ; R2= 0.5962
Estimated allele frequency
0.0
0.1
0.2
0.3
0.4
0.5
600
800
1000
1200
Ecological gradient (Precipitation sums)
L12
O12
O16
L16
L8
O1
L1
124
219
O8
97
218
9
233
256
253
217
204

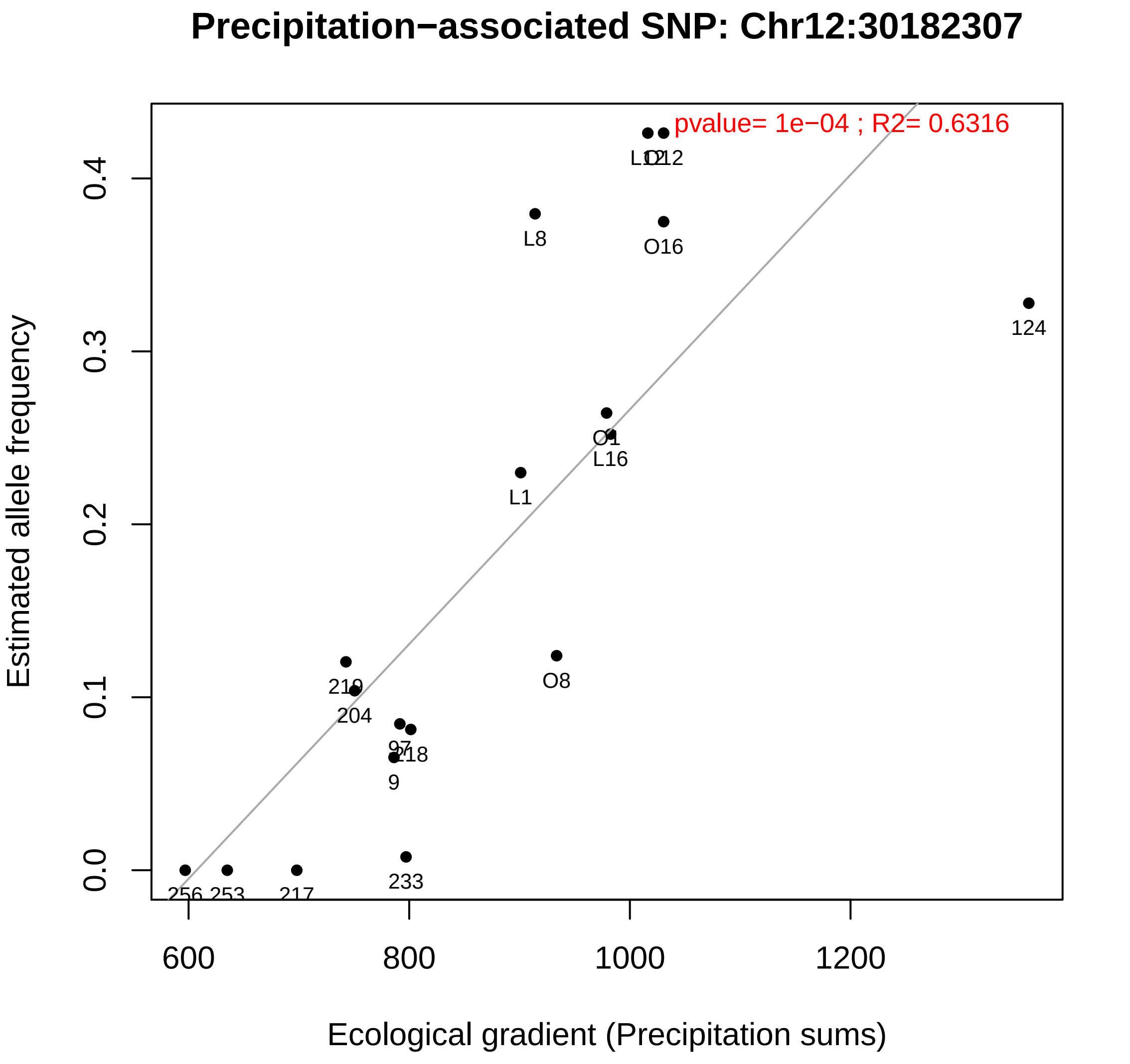

Precipitation−associated SNP: Chr12:30182307
pvalue= 1e−04 ; R2= 0.6316
Estimated allele frequency
Ecological gradient (Precipitation sums)
0.0
0.1
0.2
0.3
0.4
600
800
1000
1200
L8
O16
124
O1
L16
L1
O8
204
9
233

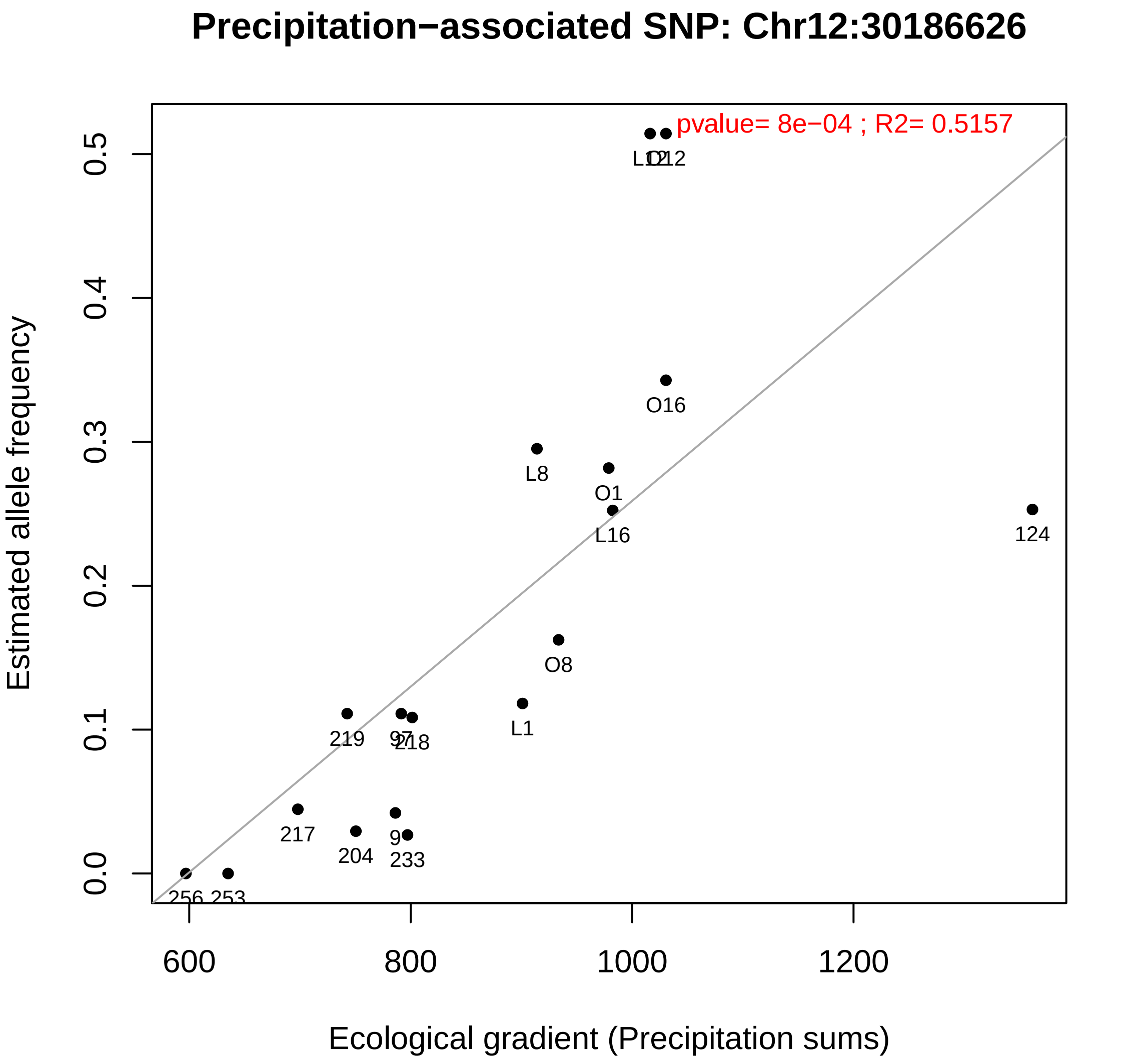

Precipitation−associated SNP: Chr12:30186626
pvalue= 8e−04 ; R2= 0.5157
Estimated allele frequency
Ecological gradient (Precipitation sums)
0.0
0.1
0.2
0.3
0.4
0.5
600
800
1000
1200
L10
O12
O16
L8
O1
L16
124
O8
L1
219
97
218
217
204
9
233
256
253

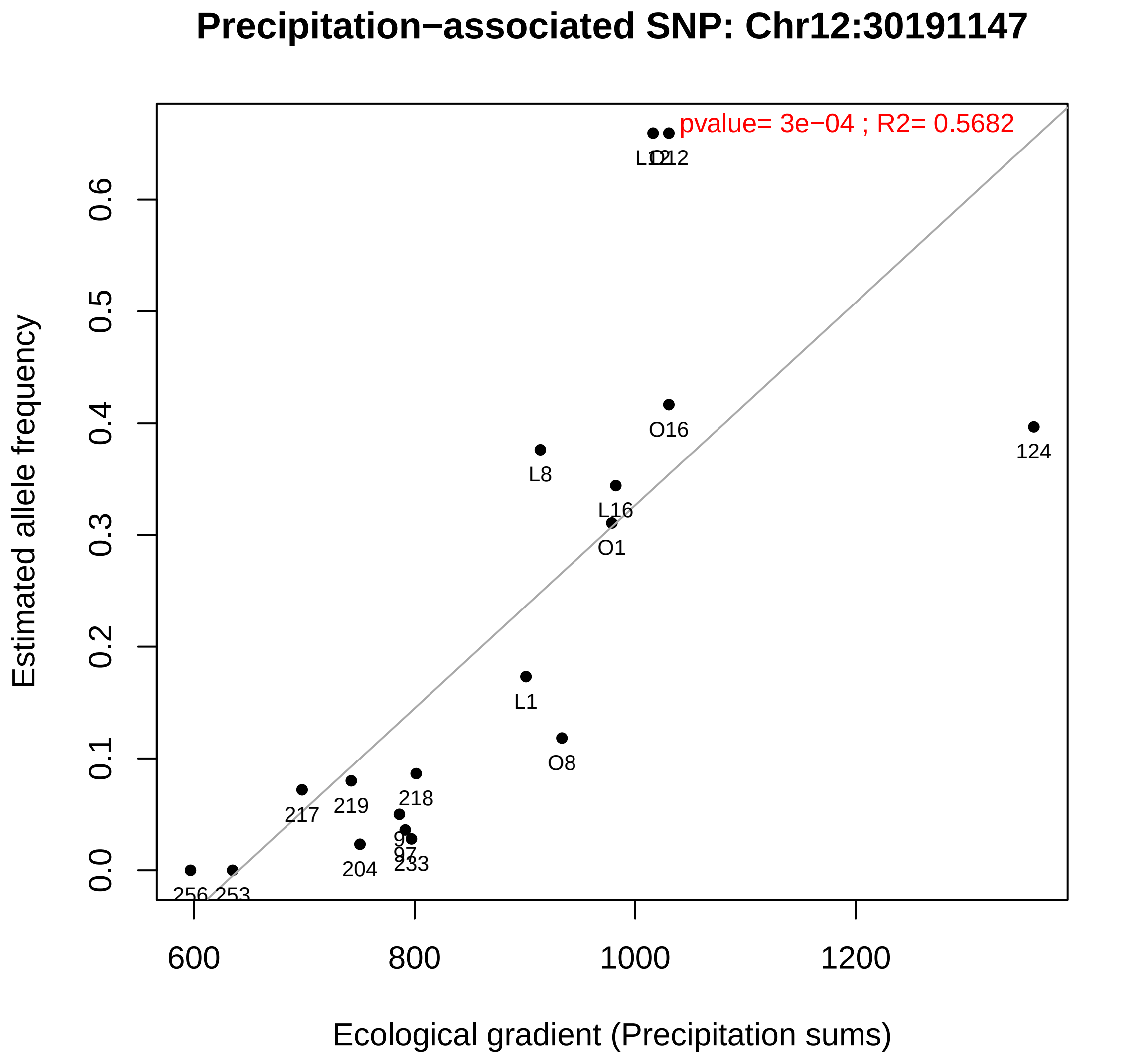
Precipitation−associated SNP: Chr12:30191147
pvalue= 3e−04 ; R2= 0.5682
Estimated allele frequency
Ecological gradient (Precipitation sums)
0.0
0.1
0.2
0.3
0.4
0.5
0.6
600
800
1000
1200
O16
124
L8
L16
O1
L1
O8
218
217
219
204
233
256
253

### Precipitation−associated SNP: Chr12:30192174

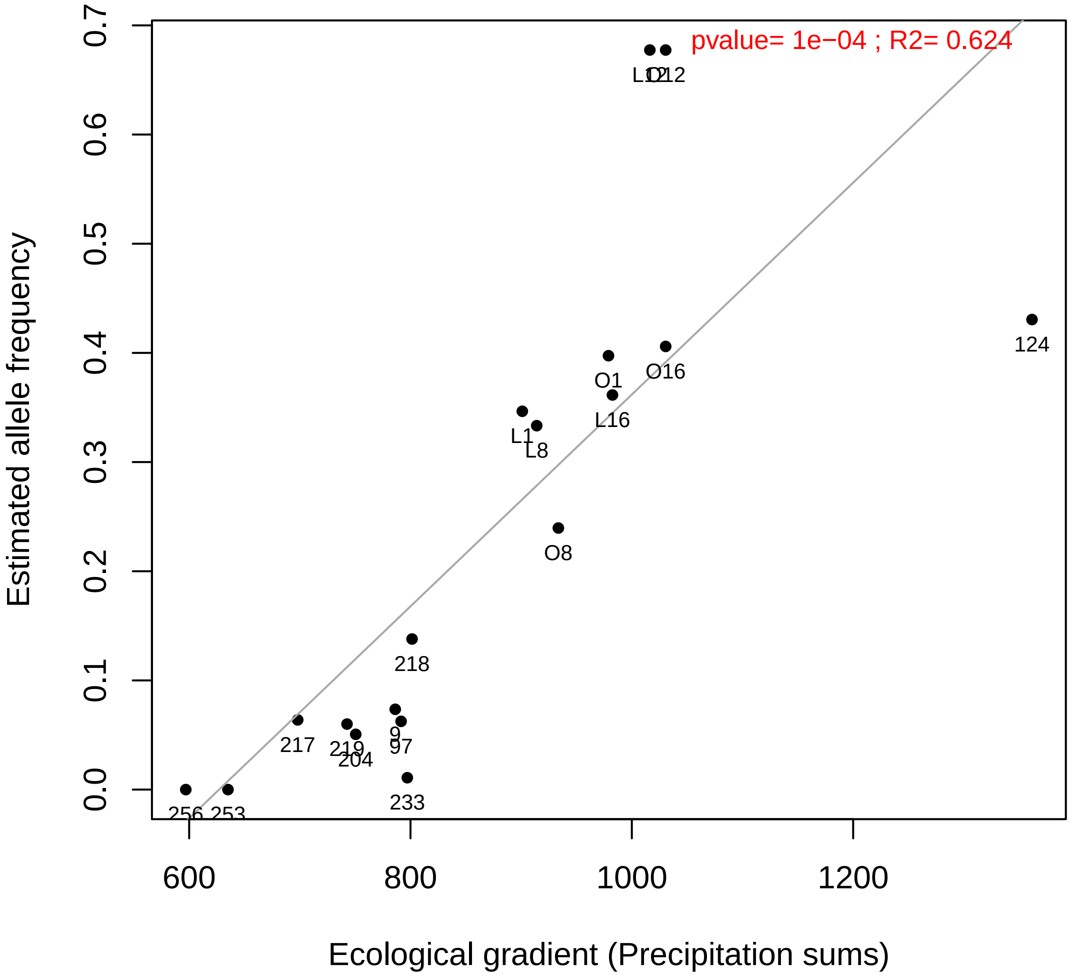

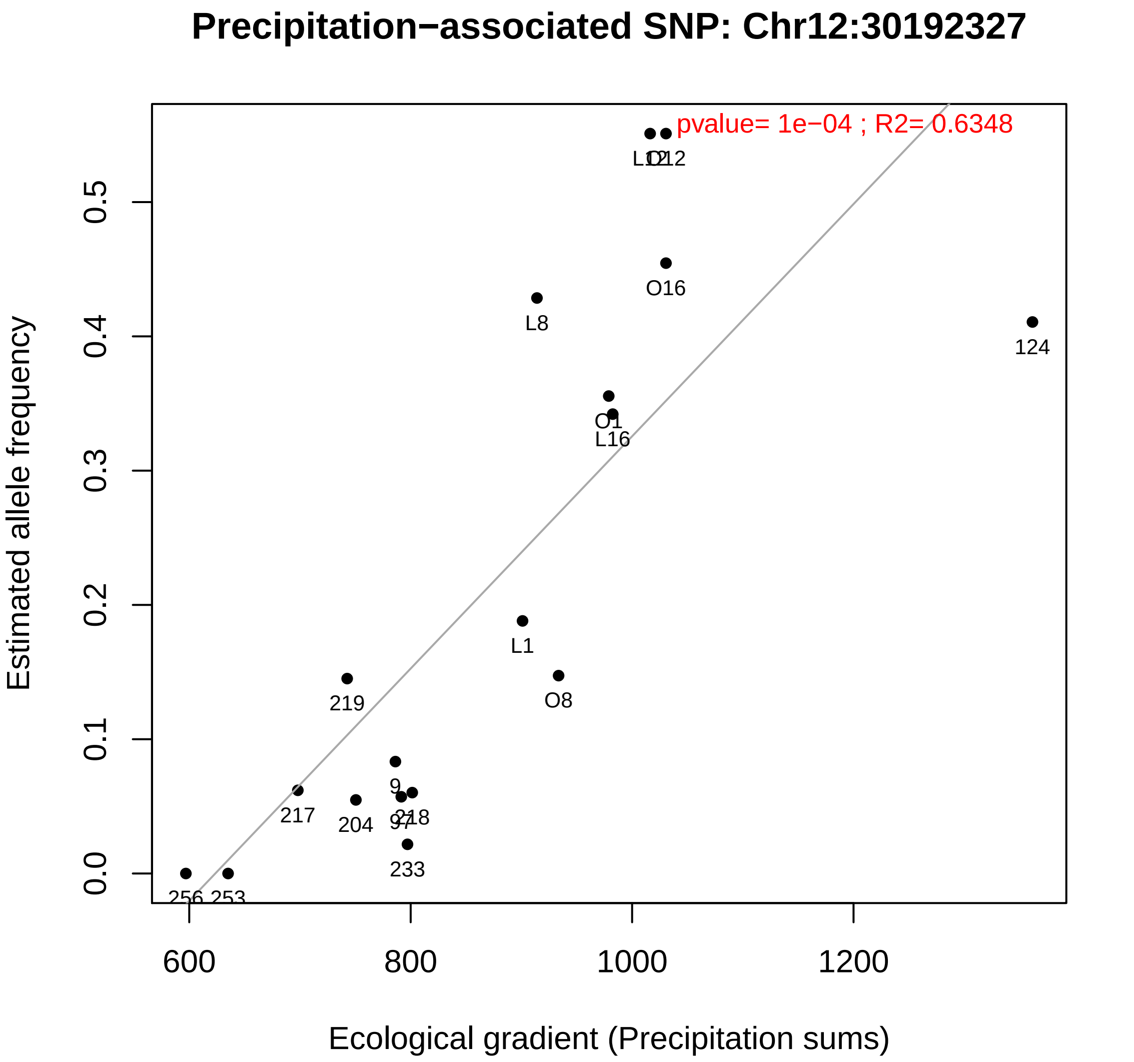

Precipitation−associated SNP: Chr12:30192327
pvalue= 1e−04 ; R2= 0.6348
Estimated allele frequency
Ecological gradient (Precipitation sums)
0.0
0.1
0.2
0.3
0.4
0.5
600
800
1000
1200
L12
O12
O16
L8
124
O1
L16
L1
O8
219
9
217
204
97
218
233
256
253

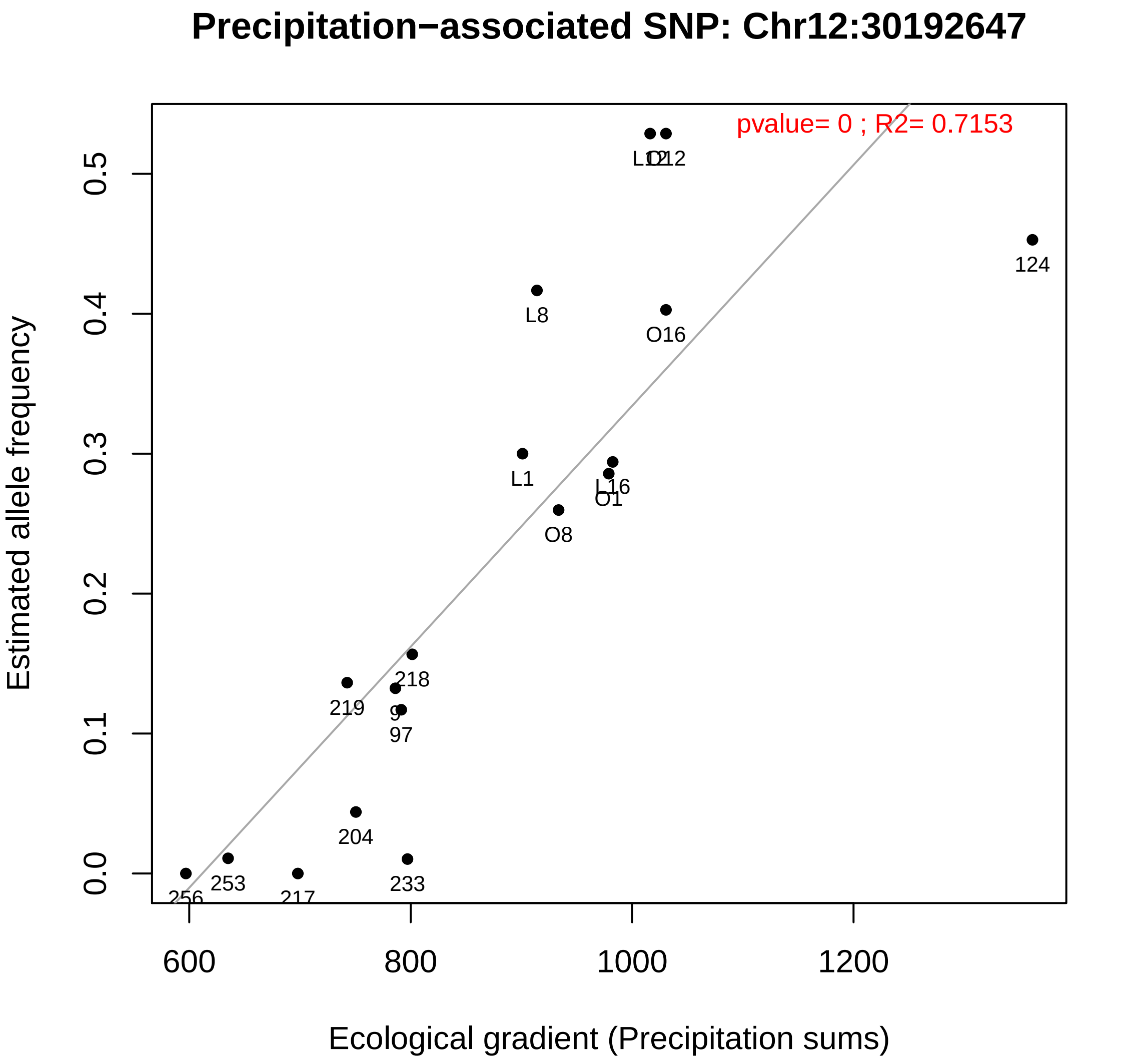

Precipitation−associated SNP: Chr12:30192647
pvalue= 0 ; R2= 0.7153
Estimated allele frequency
Ecological gradient (Precipitation sums)
0.0
0.1
0.2
0.3
0.4
0.5
600
800
1000
1200
L12
O12
124
L8
O16
L1
L16
O1
O8
218
219
9
97
204
253
233
256
217

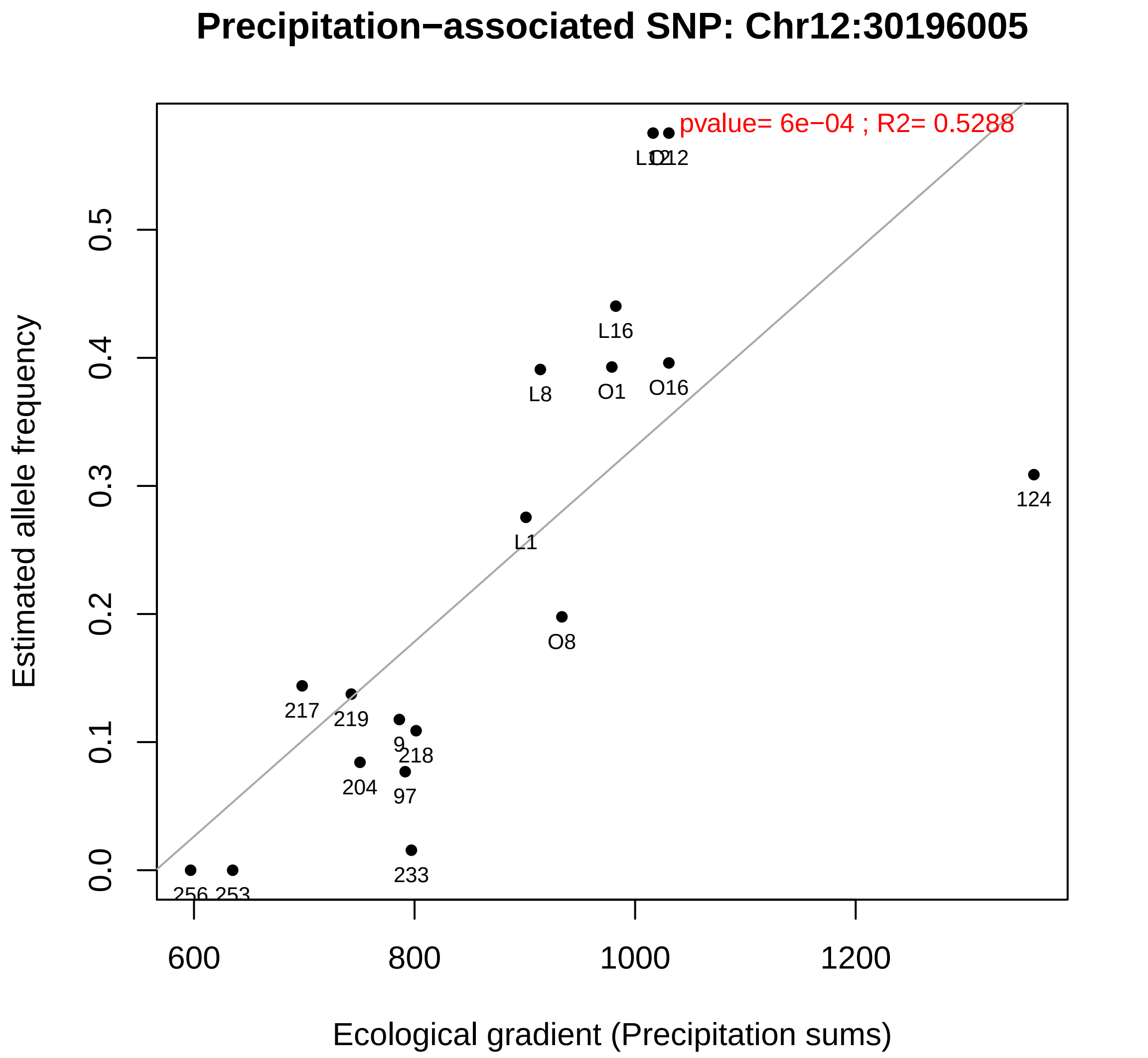
Precipitation−associated SNP: Chr12:30196005
pvalue= 6e−04 ; R2= 0.5288
Estimated allele frequency
Ecological gradient (Precipitation sums)
0.0
0.1
0.2
0.3
0.4
0.5
600
800
1000
1200
L12
O12
L16
L8
O1
O16
124
L1
O8
217
219
9
218
204
97
233
256
253

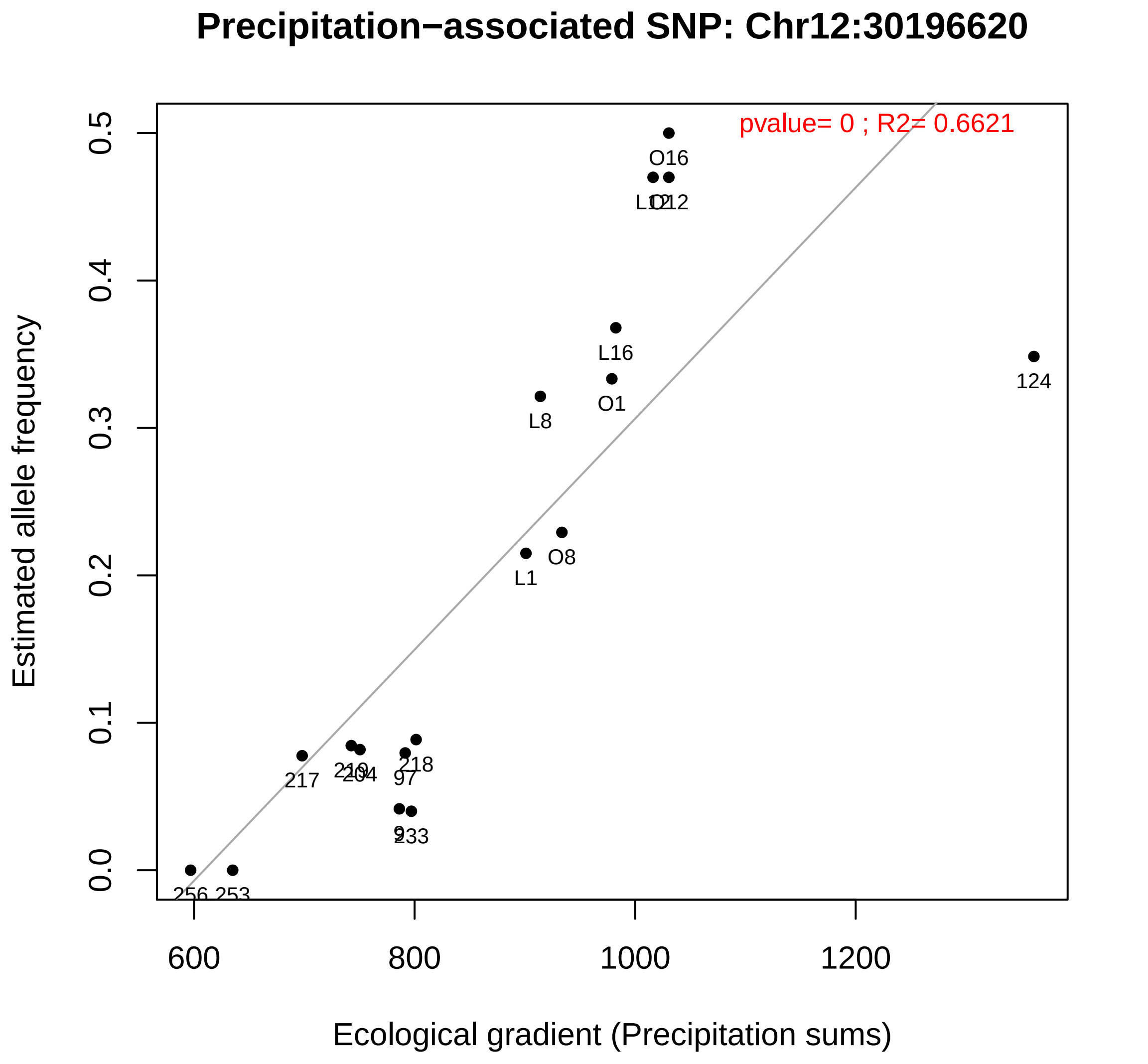

Precipitation−associated SNP: Chr12:30196620
pvalue= 0 ; R2= 0.6621
O16
L10
O12
L16
124
O1
L8
O8
L1
217
219
204
218
97
9
233
256
253
0.0
0.1
0.2
0.3
0.4
0.5
600
800
1000
1200
Estimated allele frequency
Ecological gradient (Precipitation sums)

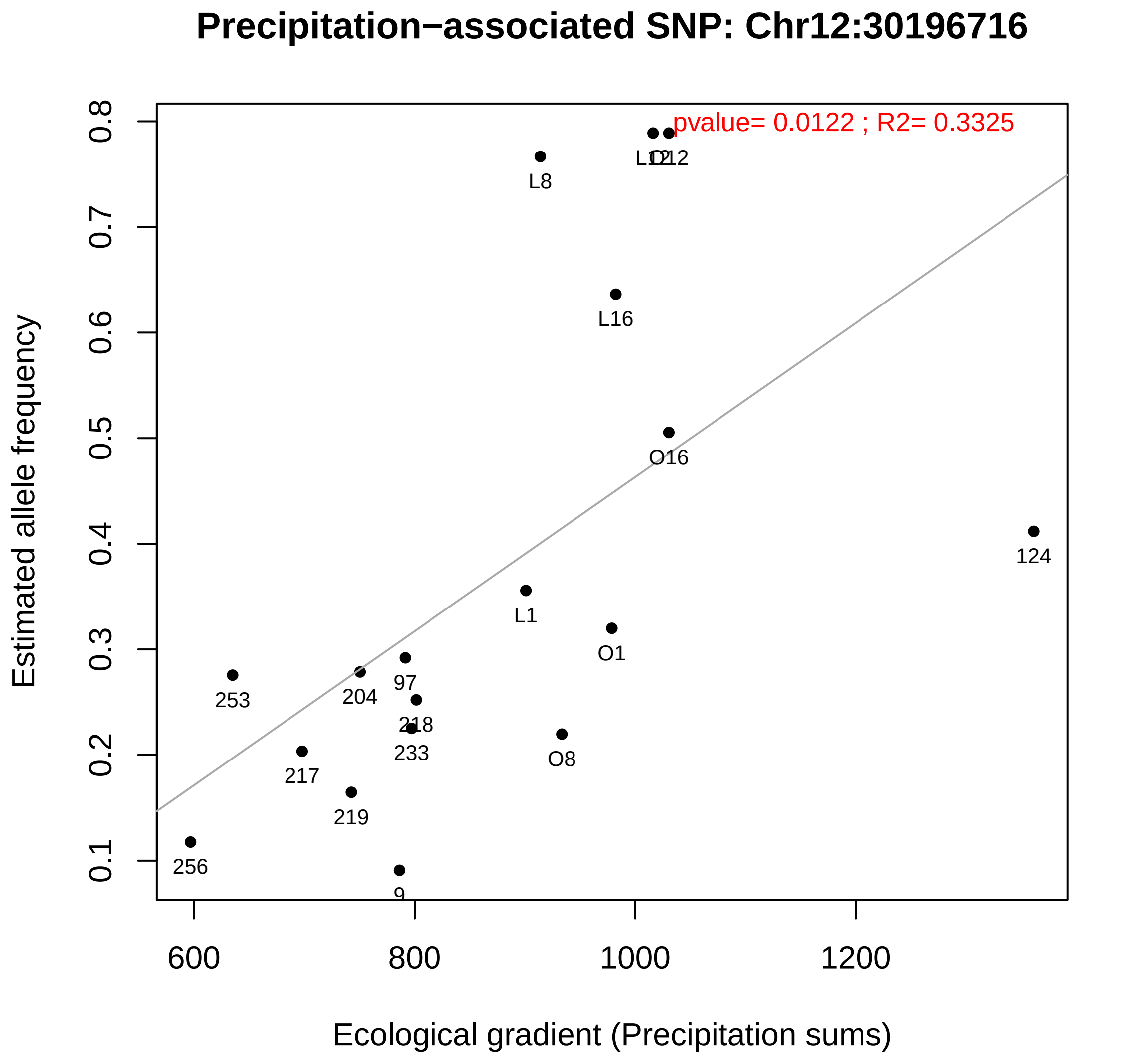
Precipitation−associated SNP: Chr12:30196716
pvalue= 0.0122 ; R2= 0.3325
Estimated allele frequency
0.1
0.2
0.3
0.4
0.5
0.6
0.7
0.8
600
800
1000
1200
Ecological gradient (Precipitation sums)
L8
L12
O12
L16
O16
124
L1
O1
253
204
97
218
233
O8
217
219
256
9

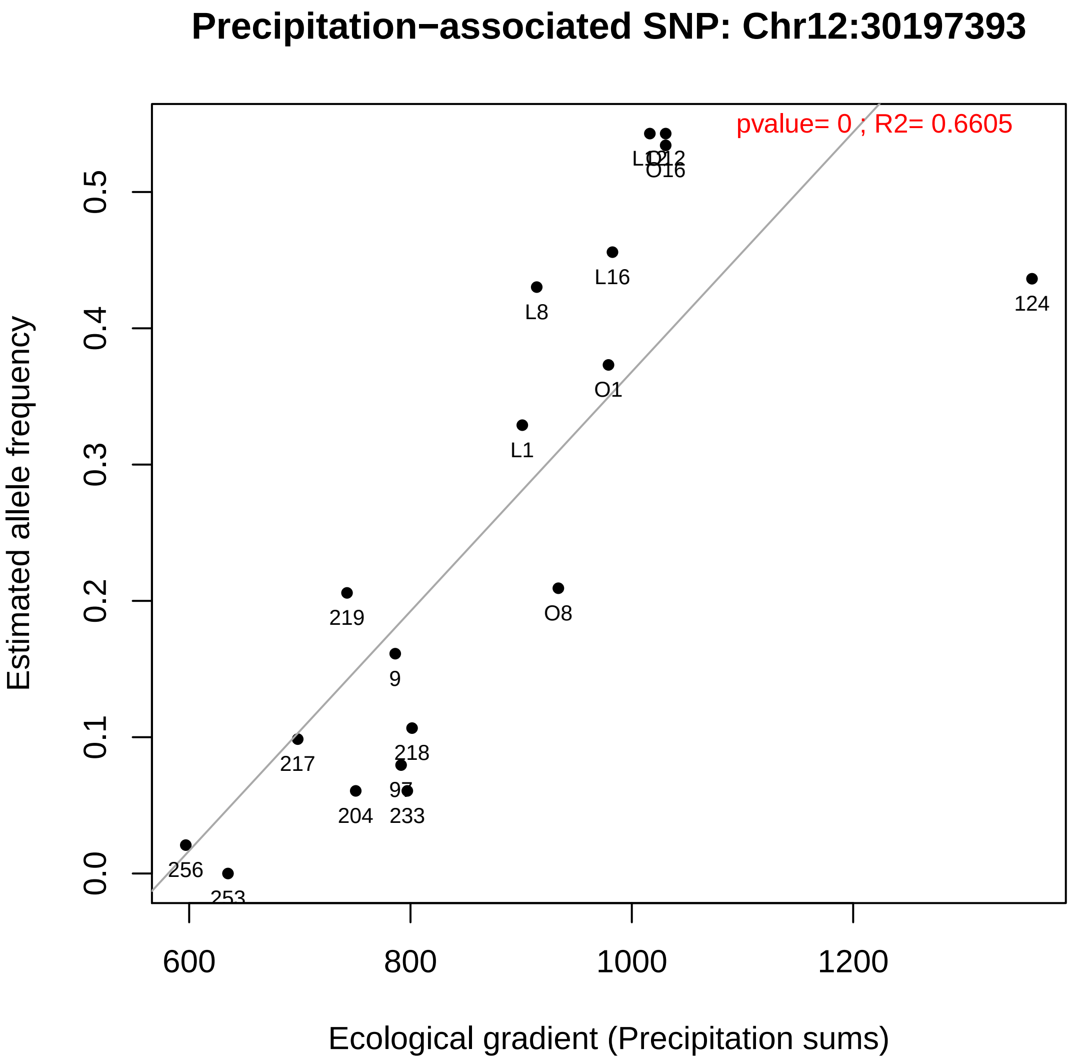
Precipitation−associated SNP: Chr12:30197393
pvalue= 0 ; R2= 0.6605
L12
O12
O16
L16
124
L8
O1
L1
219
O8
9
218
217
97
204
233
256
253
0.0
0.1
0.2
0.3
0.4
0.5
600
800
1000
1200
Estimated allele frequency
Ecological gradient (Precipitation sums)

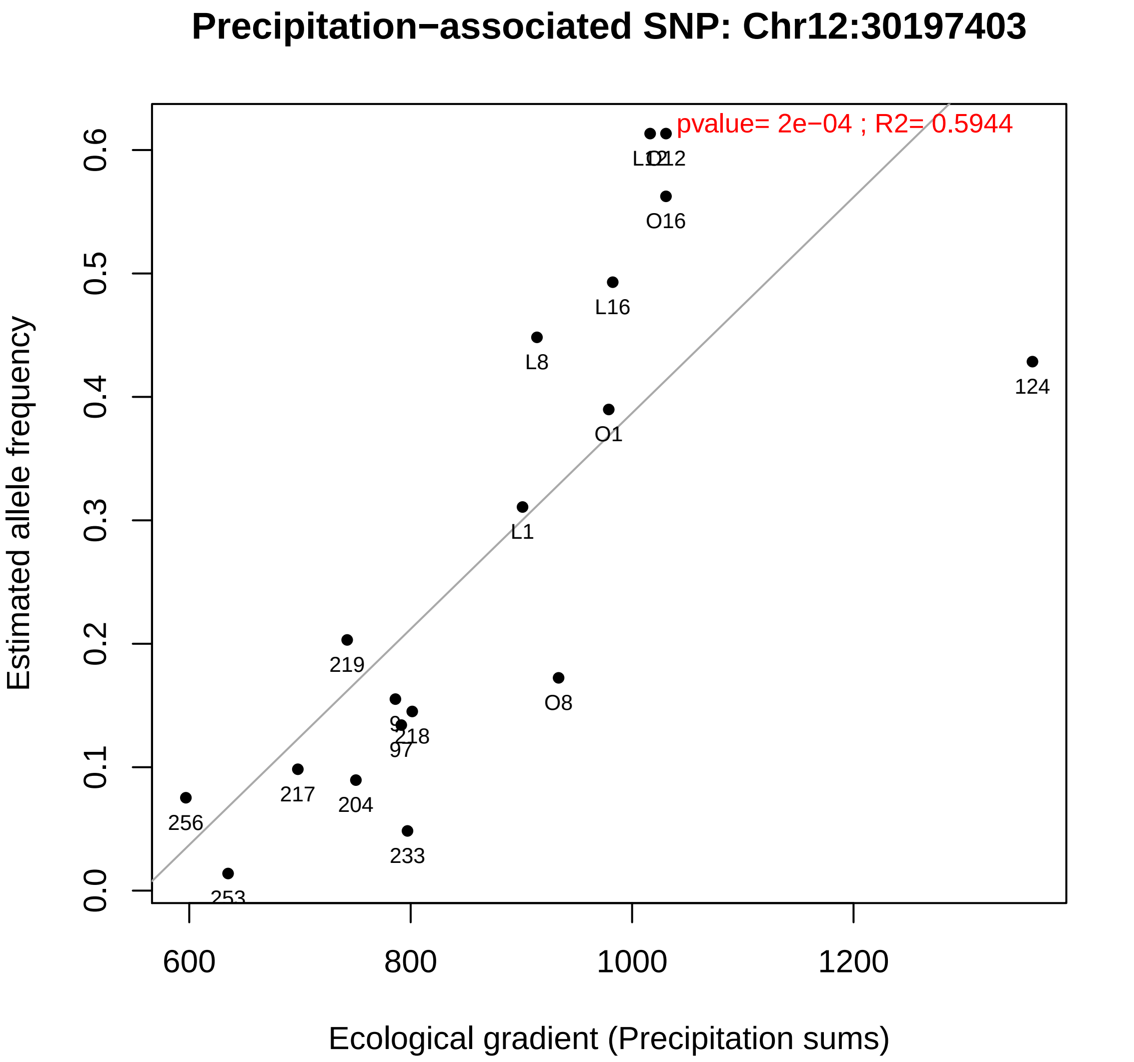

Precipitation−associated SNP: Chr12:30197403
pvalue= 2e−04 ; R2= 0.5944
Estimated allele frequency
Ecological gradient (Precipitation sums)
0.0
0.1
0.2
0.3
0.4
0.5
0.6
600
800
1000
1200
L12
O12
O16
L16
L8
124
O1
L1
219
O8
9
218
97
217
204
256
233
253

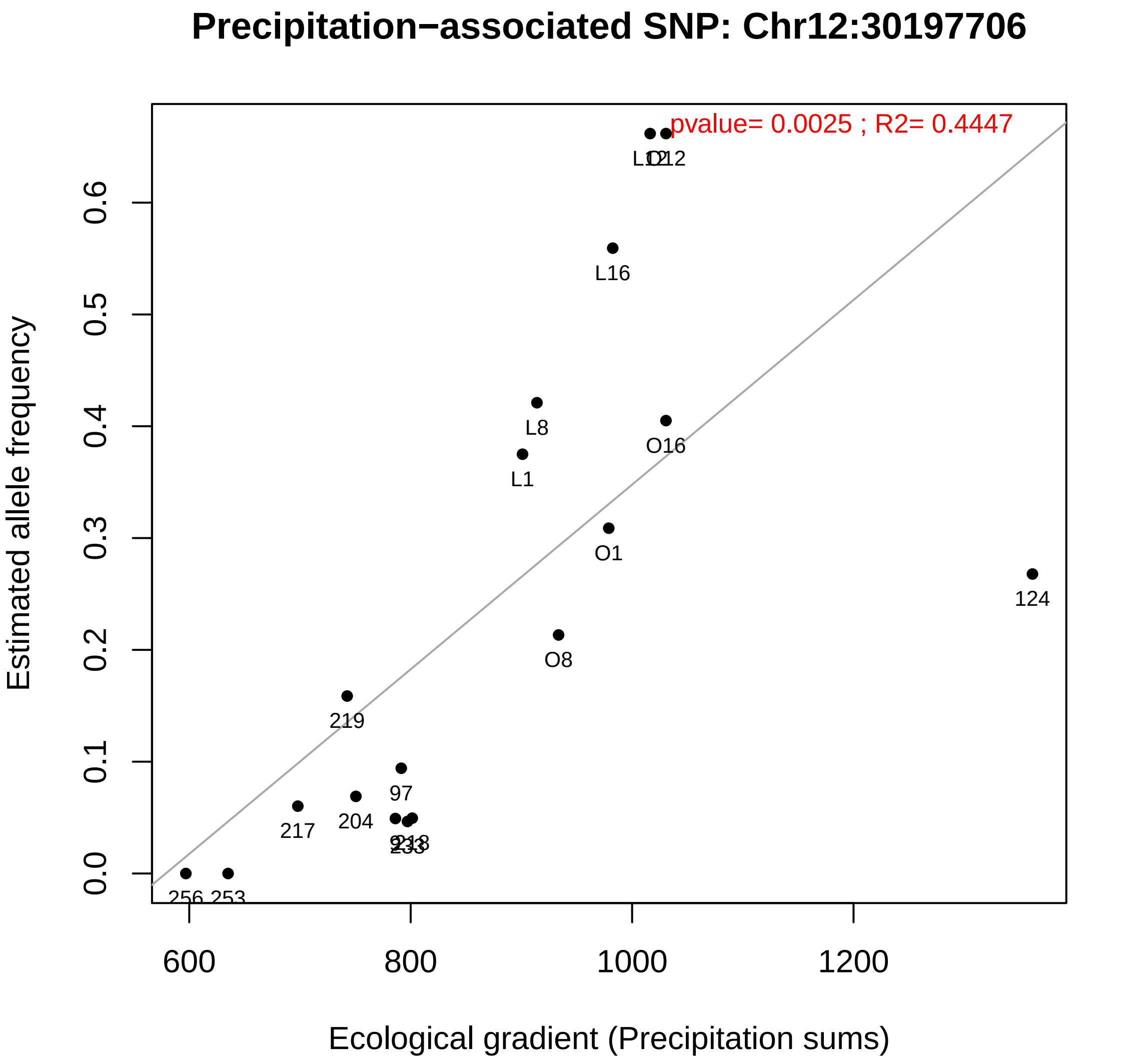

Precipitation−associated SNP: Chr12:30197706
pvalue= 0.0025 ; R2= 0.4447
Estimated allele frequency
Ecological gradient (Precipitation sums)
0.0
0.1
0.2
0.3
0.4
0.5
0.6
600
800
1000
1200
L16
L8
O16
L1
O1
124
O8
219
97
204
217

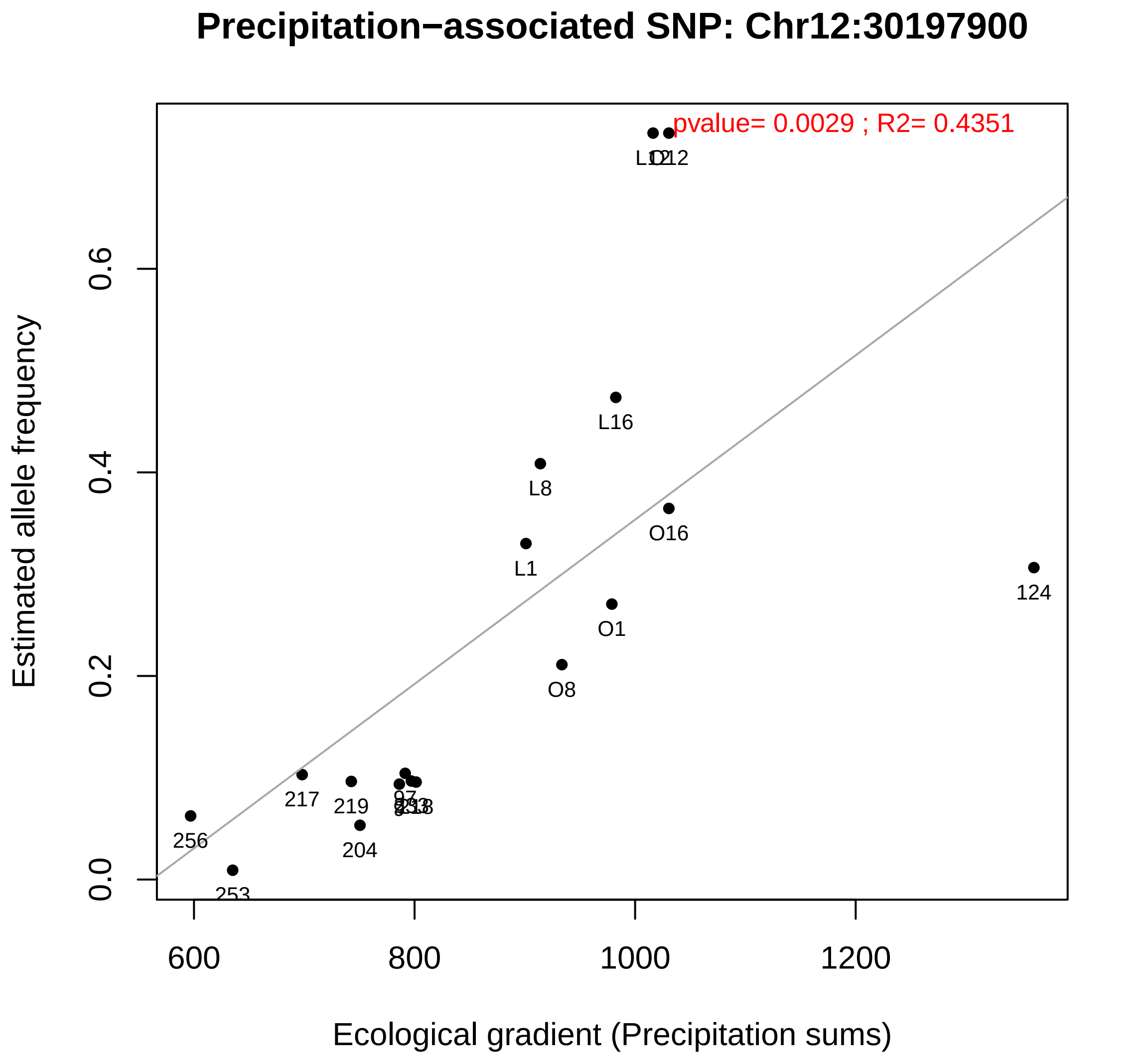
Precipitation−associated SNP: Chr12:30197900
pvalue= 0.0029 ; R2= 0.4351
Estimated allele frequency
Ecological gradient (Precipitation sums)
0.0
0.2
0.4
0.6
600
800
1000
1200
L16
L8
O16
L1
124
O1
O8
217
219
204
256

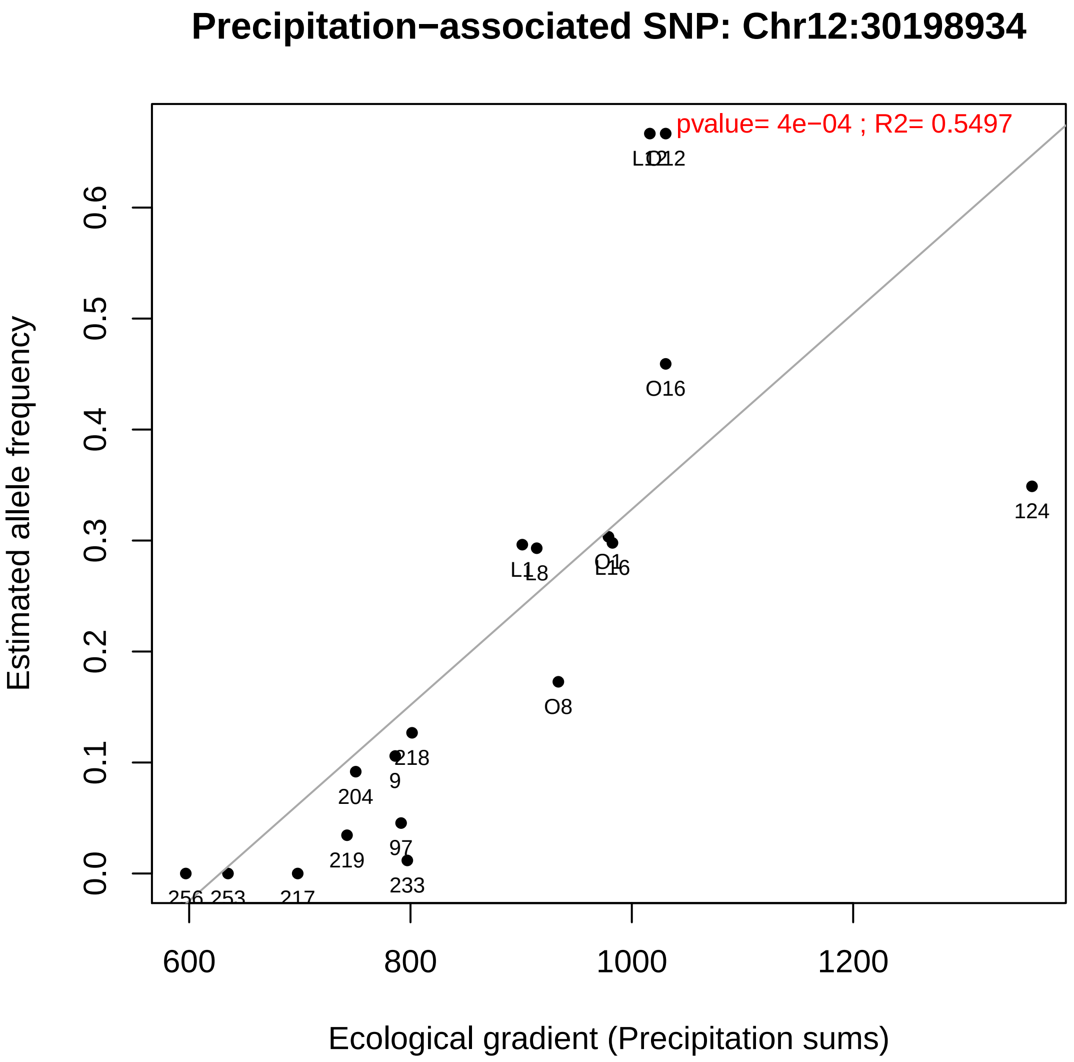

Precipitation−associated SNP: Chr12:30198934
pvalue= 4e−04 ; R2= 0.5497
Estimated allele frequency
Ecological gradient (Precipitation sums)
0.0
0.1
0.2
0.3
0.4
0.5
0.6
600
800
1000
1200
O16
124
O8
218
9
204
97
219
233
256
253
217

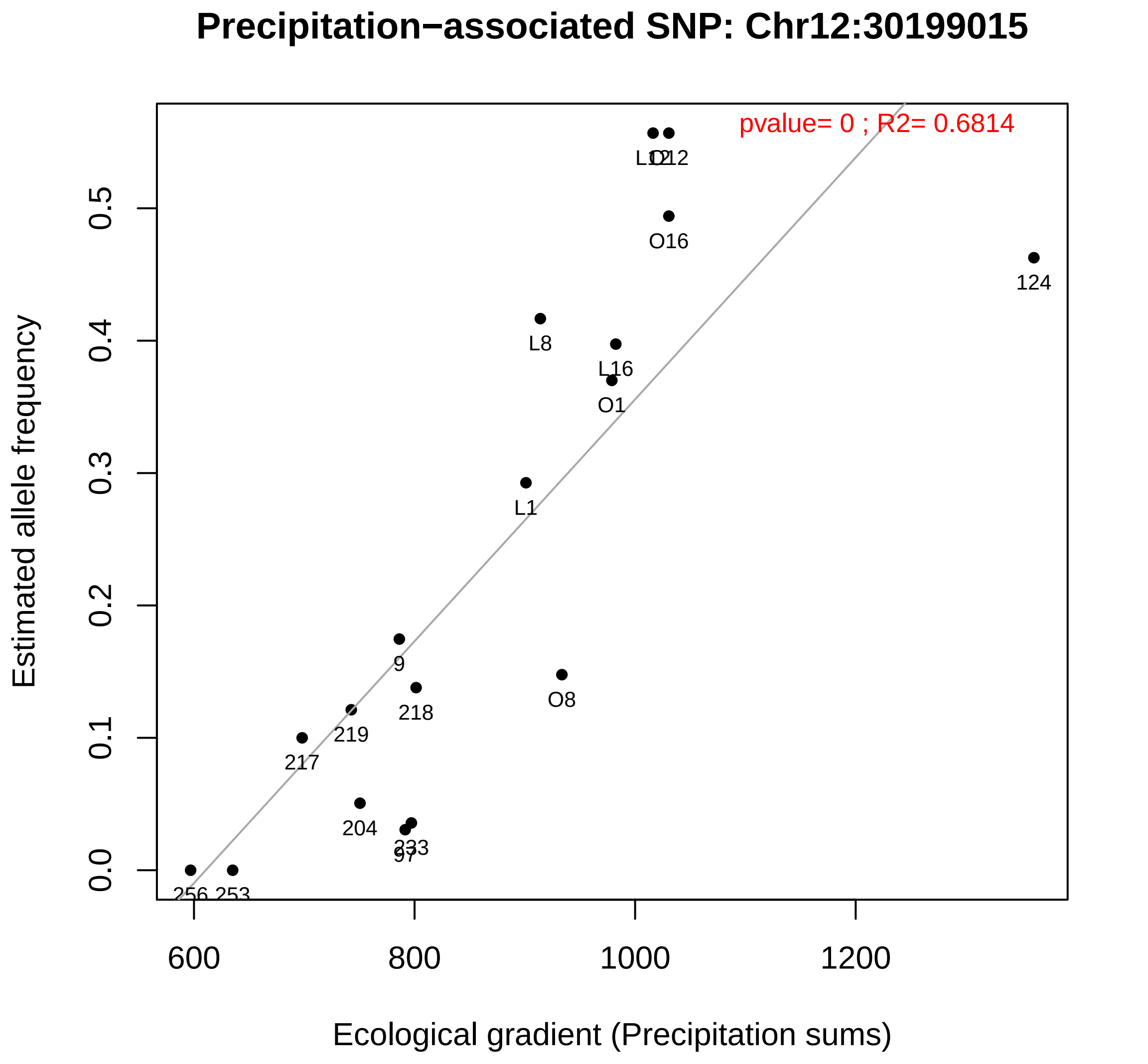

Precipitation−associated SNP: Chr12:30199015
pvalue= 0 ; R2= 0.6814
Estimated allele frequency
Ecological gradient (Precipitation sums)
0.0
0.1
0.2
0.3
0.4
0.5
600
800
1000
1200
L12
O12
O16
124
L8
L16
O1
L1
9
218
O8
219
217
204
233
97
256
253

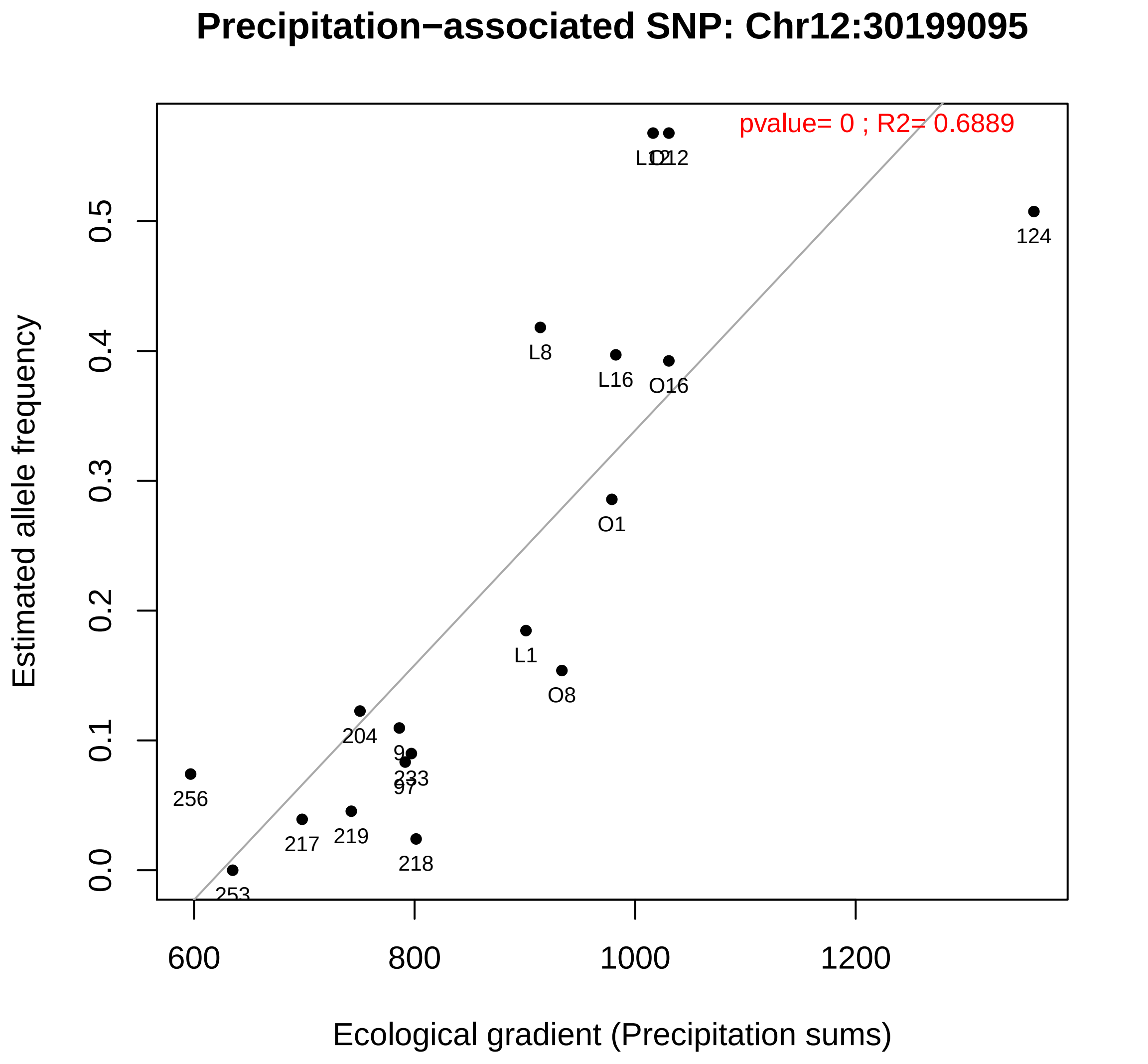
Precipitation−associated SNP: Chr12:30199095
pvalue= 0 ; R2= 0.6889
Estimated allele frequency
Ecological gradient (Precipitation sums)
0.0
0.1
0.2
0.3
0.4
0.5
600
800
1000
1200
L12
O12
124
L8
L16
O16
O1
L1
O8
204
9
233
97
256
217
219
218
253

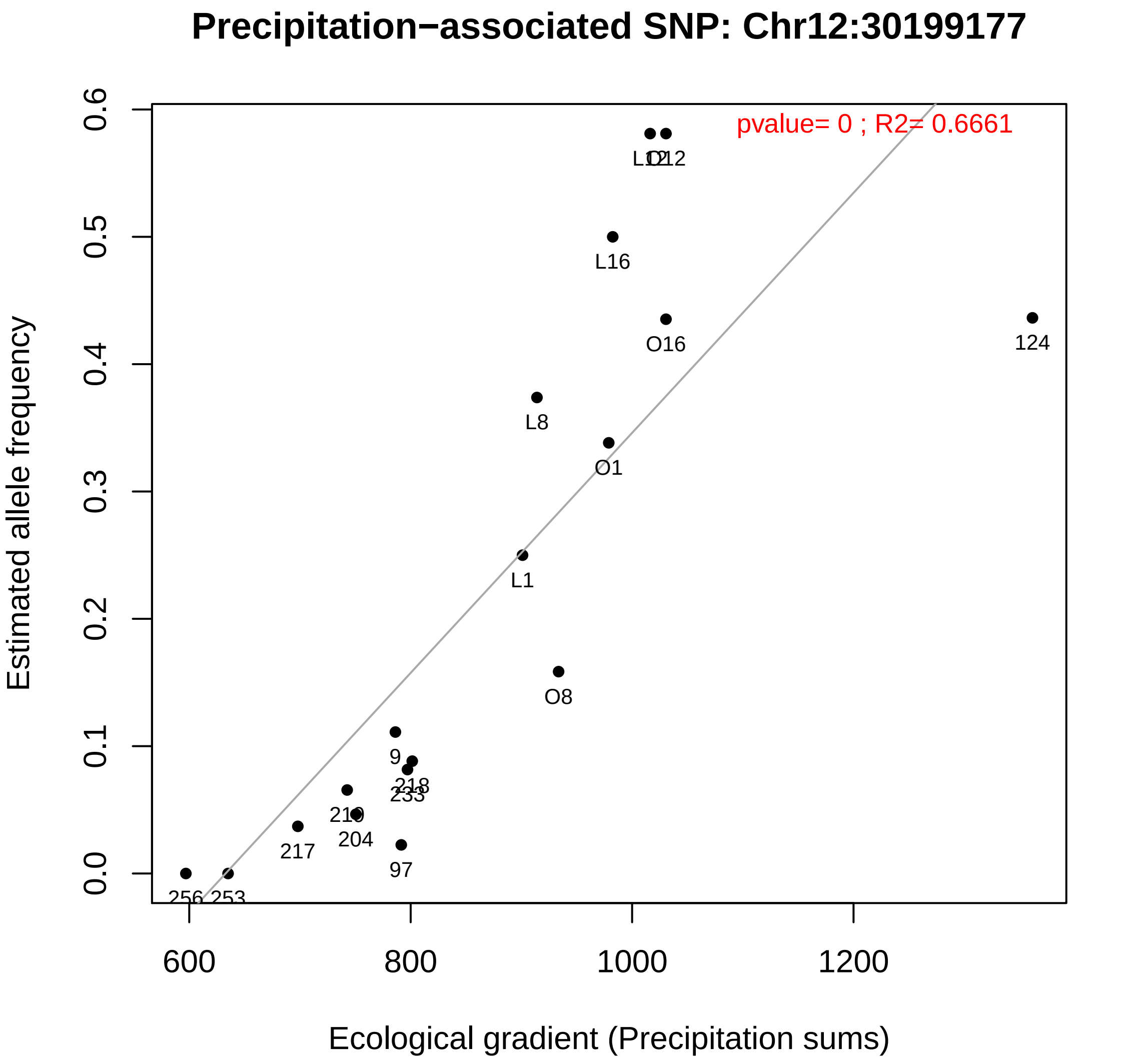

Precipitation−associated SNP: Chr12:30199177
pvalue= 0 ; R2= 0.6661
Estimated allele frequency
Ecological gradient (Precipitation sums)
0.0
0.1
0.2
0.3
0.4
0.5
0.6
600
800
1000
1200
O12
L16
O16
124
L8
O1
L1
O8
9
218
233
204
217
97
256
253

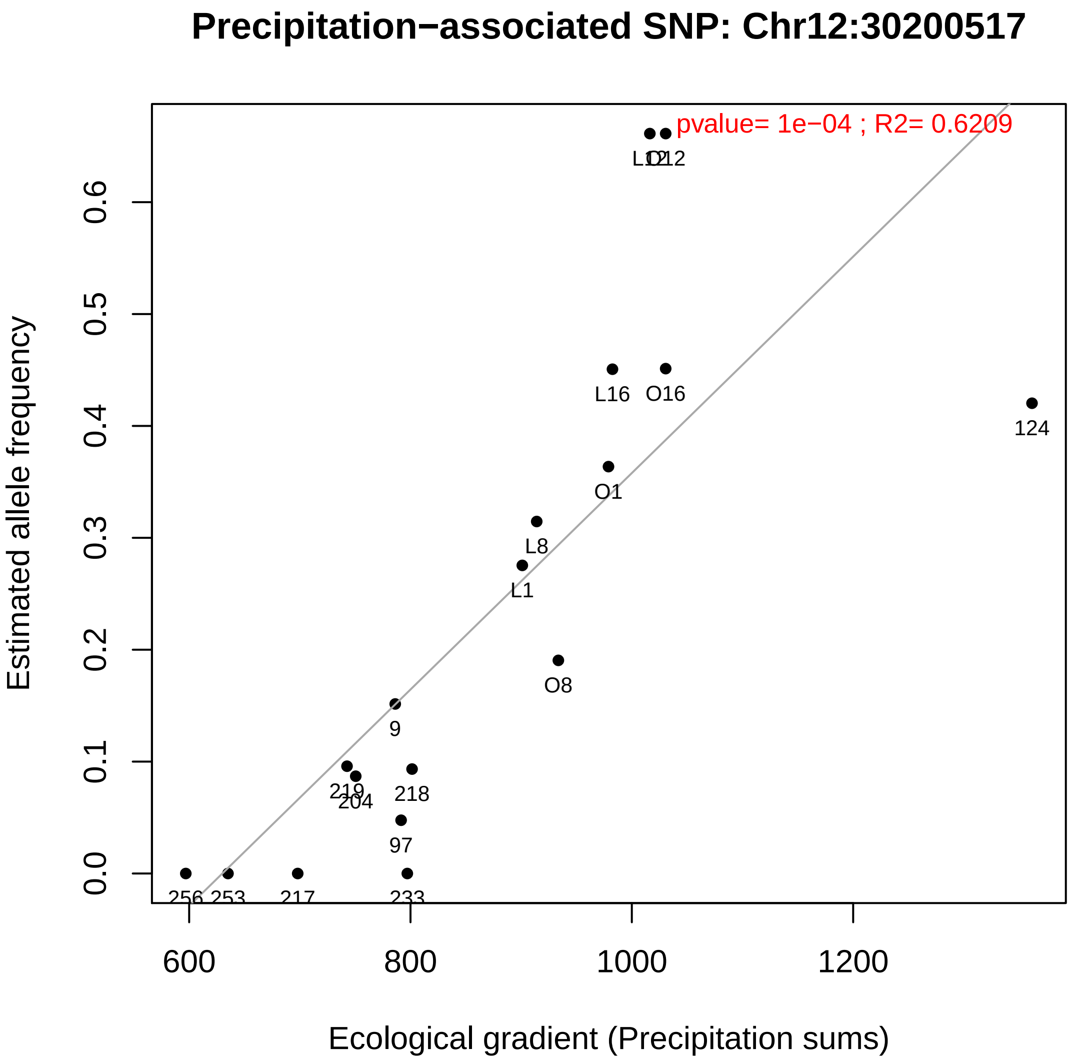

Precipitation−associated SNP: Chr12:30200517
pvalue= 1e−04 ; R2= 0.6209
L12
O12
L16
O16
124
O1
L8
L1
O8
9
219
204
218
97
256
253
217
233
0.0
0.1
0.2
0.3
0.4
0.5
0.6
600
800
1000
1200
Estimated allele frequency
Ecological gradient (Precipitation sums)

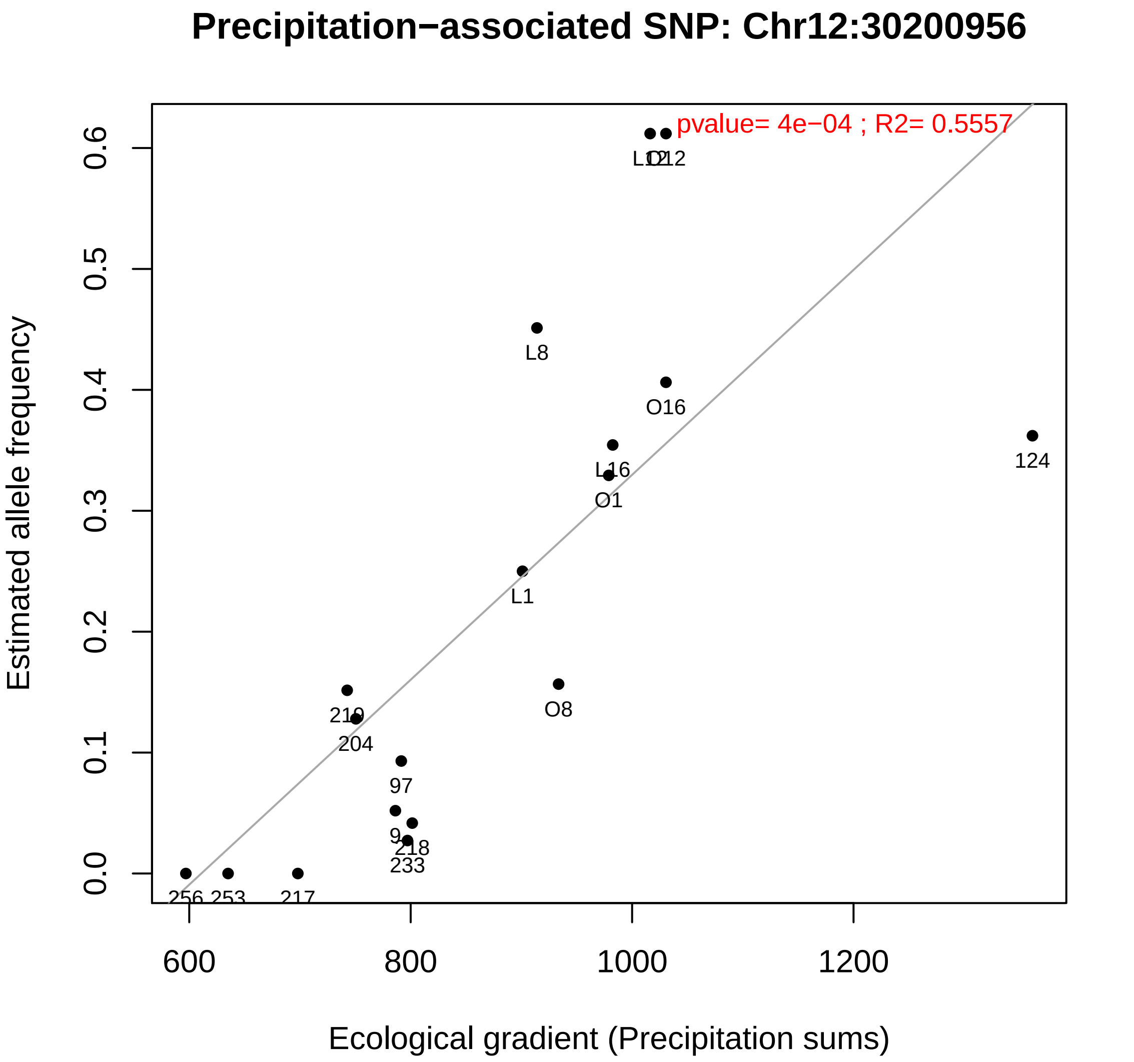
Precipitation−associated SNP: Chr12:30200956
pvalue= 4e−04 ; R2= 0.5557
Estimated allele frequency
Ecological gradient (Precipitation sums)
0.0
0.1
0.2
0.3
0.4
0.5
0.6
600
800
1000
1200
L12
O12
L8
O16
124
L16
O1
L1
O8
210
204
97
9
218
233
256
253
217

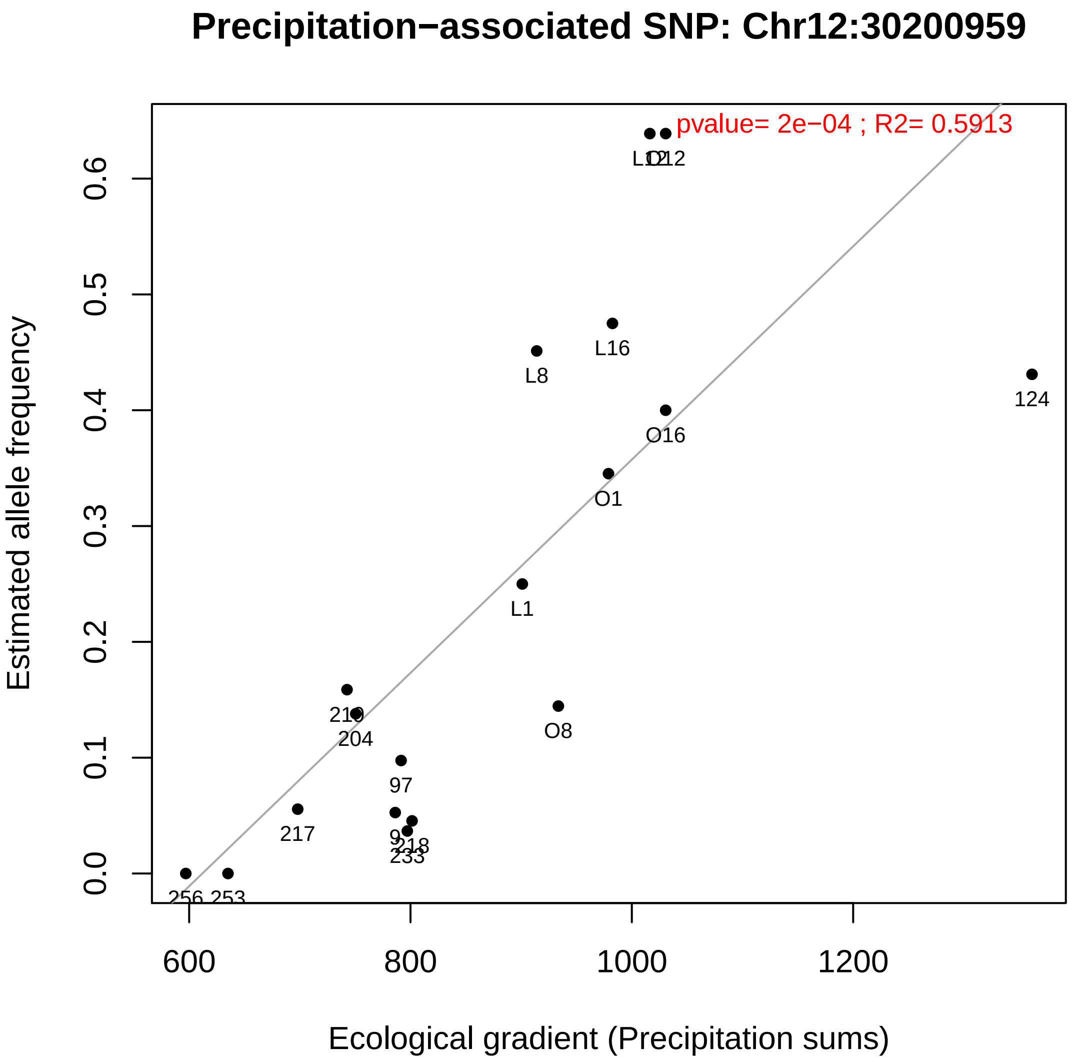

Precipitation−associated SNP: Chr12:30200959
pvalue= 2e−04 ; R2= 0.5913
Estimated allele frequency
Ecological gradient (Precipitation sums)
0.0
0.1
0.2
0.3
0.4
0.5
0.6
600
800
1000
1200
O12
L16
L8
124
O16
O1
L1
204
O8
97
217
233
256
253

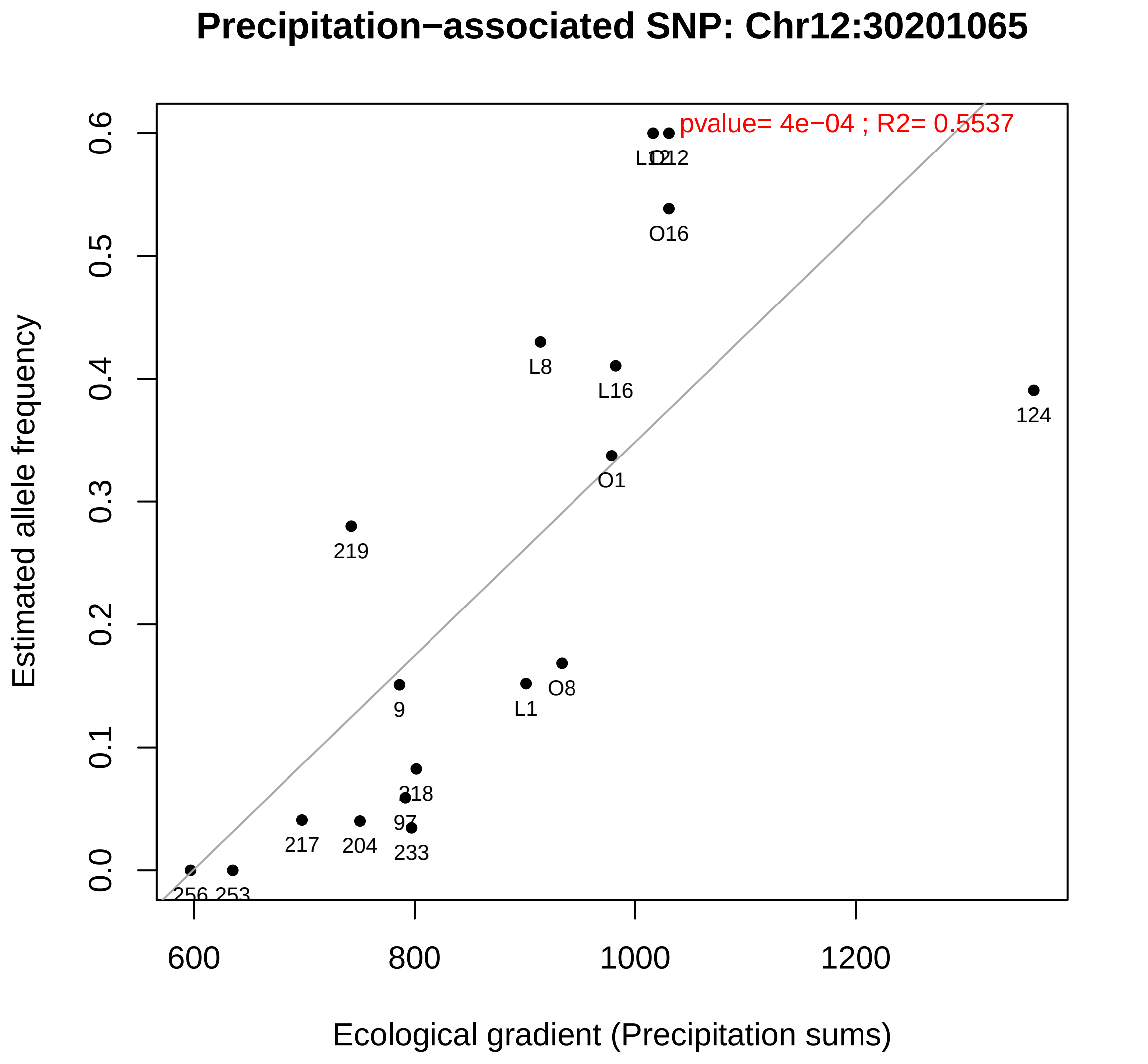

Precipitation−associated SNP: Chr12:30201065
pvalue= 4e−04 ; R2= 0.5537
L10
O12
O16
L8
L16
124
O1
219
O8
9
L1
218
97
217
204
233
256
253
0.0
0.1
0.2
0.3
0.4
0.5
0.6
600
800
1000
1200
Estimated allele frequency
Ecological gradient (Precipitation sums)

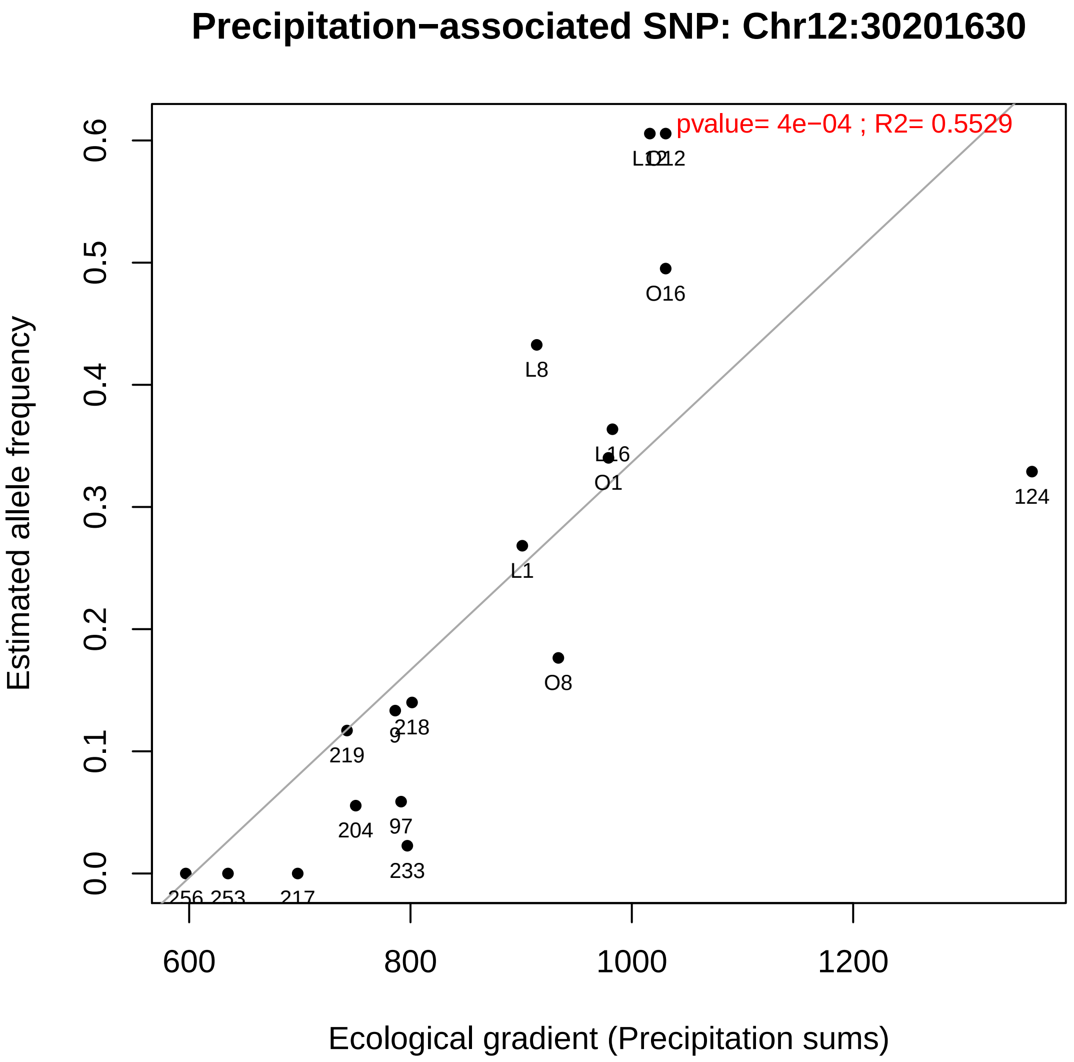

Precipitation−associated SNP: Chr12:30201630
pvalue= 4e−04 ; R2= 0.5529
Estimated allele frequency
Ecological gradient (Precipitation sums)
0.0
0.1
0.2
0.3
0.4
0.5
0.6
600
800
1000
1200
L12
O12
O16
L8
L16
O1
124
L1
O8
218
9
219
204
97
233
256
253
217

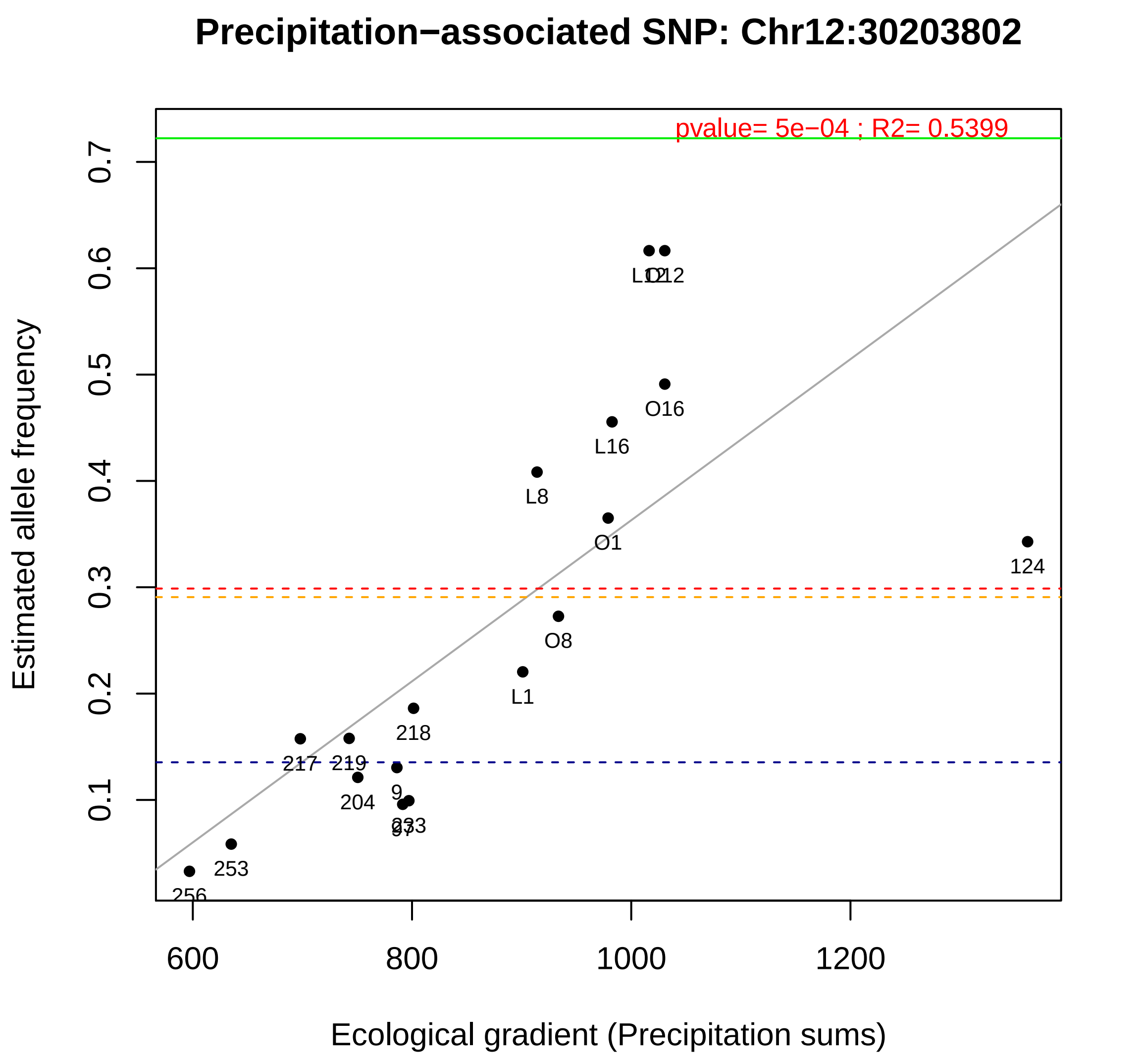

Precipitation−associated SNP: Chr12:30203802
pvalue= 5e−04 ; R2= 0.5399
Estimated allele frequency
0.1
0.2
0.3
0.4
0.5
0.6
0.7
600
800
1000
1200
Ecological gradient (Precipitation sums)
L12
O12
O16
L16
L8
O1
124
O8
L1
218
217
219
204
9
233
253
256

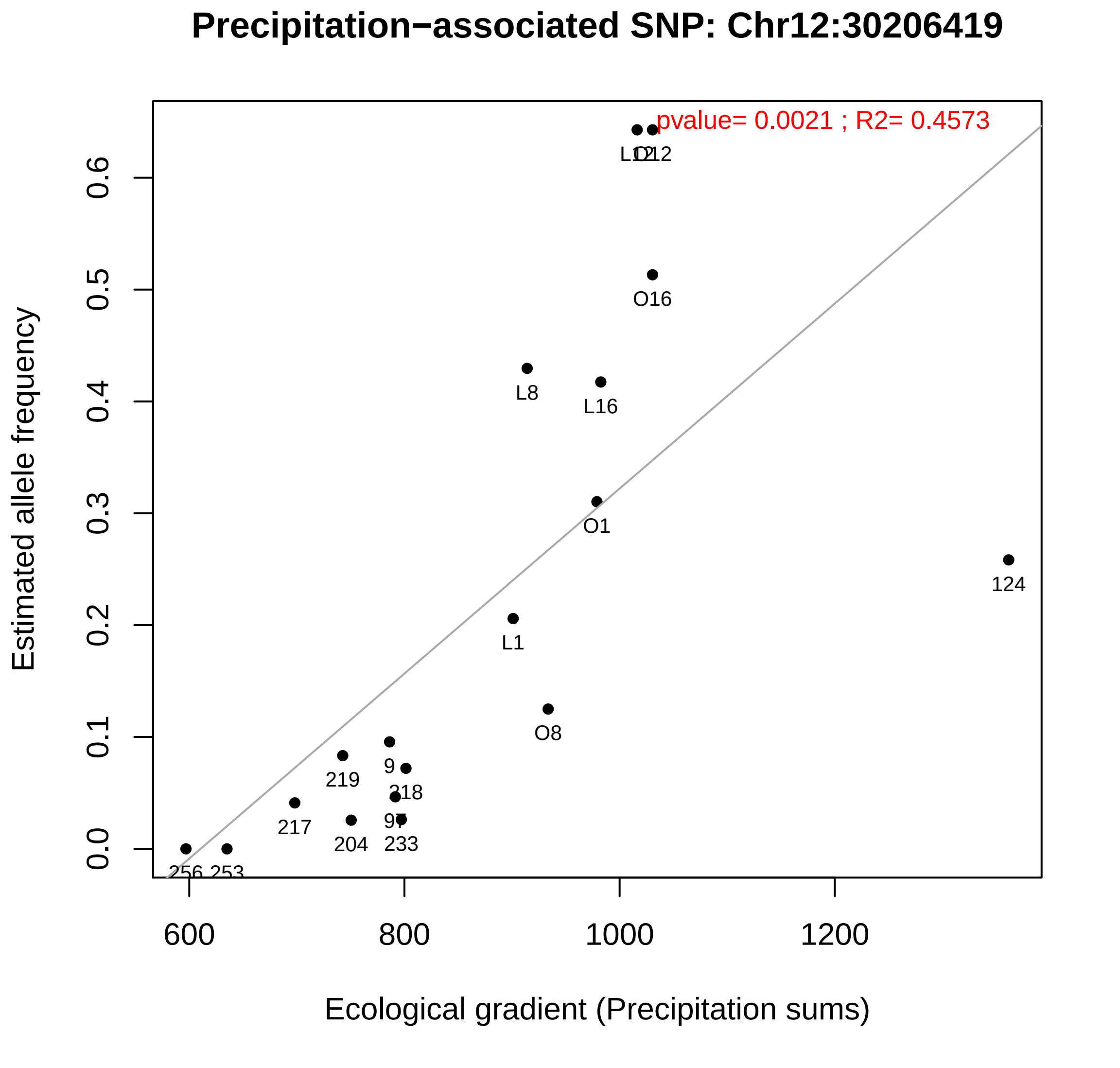

Precipitation−associated SNP: Chr12:30206419
pvalue= 0.0021 ; R2= 0.4573
Estimated allele frequency
Ecological gradient (Precipitation sums)
0.0
0.1
0.2
0.3
0.4
0.5
0.6
600
800
1000
1200
L12
O12
O16
L8
L16
O1
124
L1
O8
9
219
218
217
97
204
233
256
253

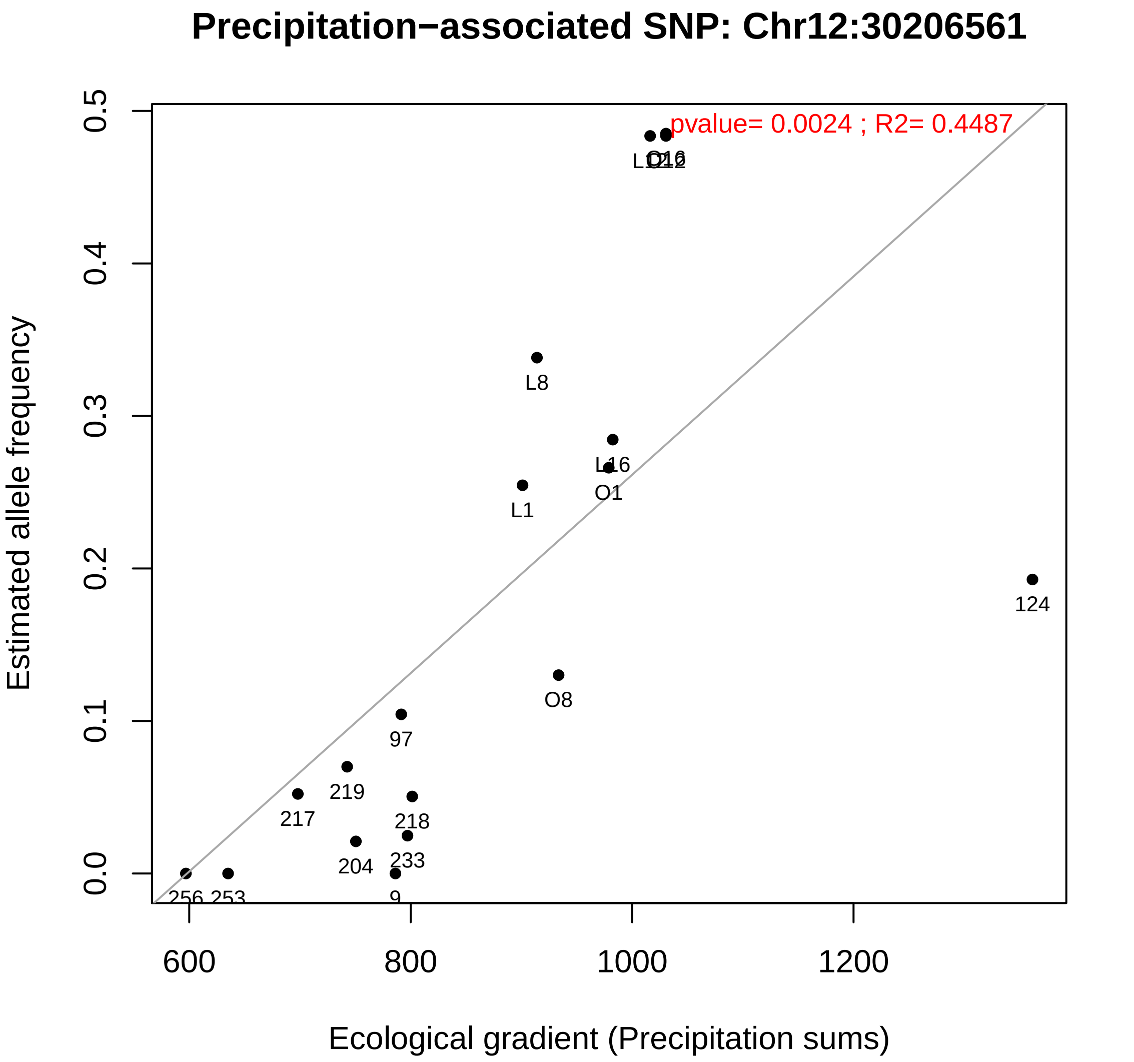

Precipitation−associated SNP: Chr12:30206561
pvalue= 0.0024 ; R2= 0.4487
Estimated allele frequency
Ecological gradient (Precipitation sums)
0.0
0.1
0.2
0.3
0.4
0.5
600
800
1000
1200
L8
L16
O1
L1
124
O8
97
219
217
218
204
233
9

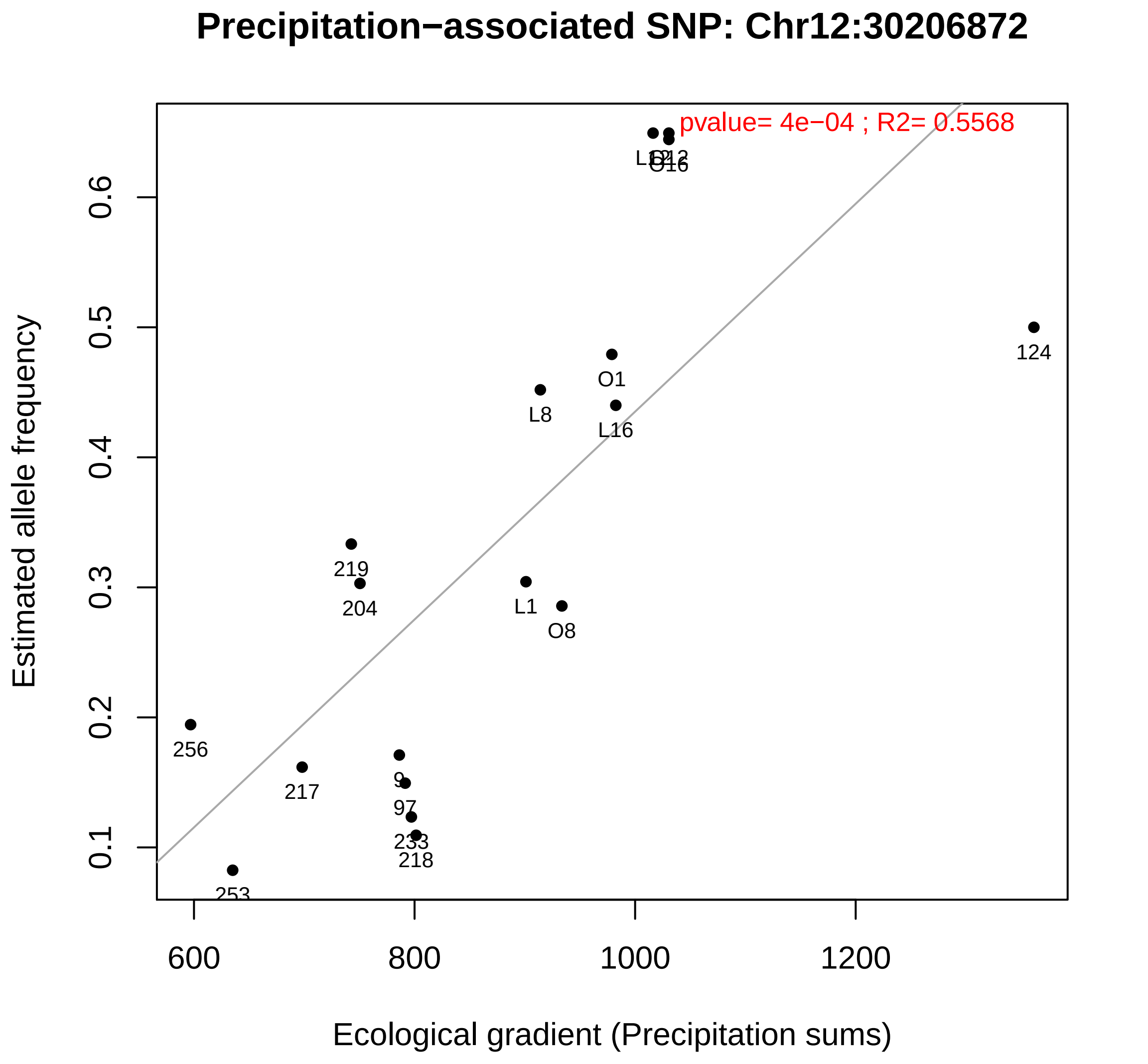

Precipitation−associated SNP: Chr12:30206872
pvalue= 4e−04 ; R2= 0.5568
Estimated allele frequency
Ecological gradient (Precipitation sums)
0.1
0.2
0.3
0.4
0.5
0.6
600
800
1000
1200
124
O1
L8
L16
219
204
L1
O8
256
217
97
233
218
253
O16

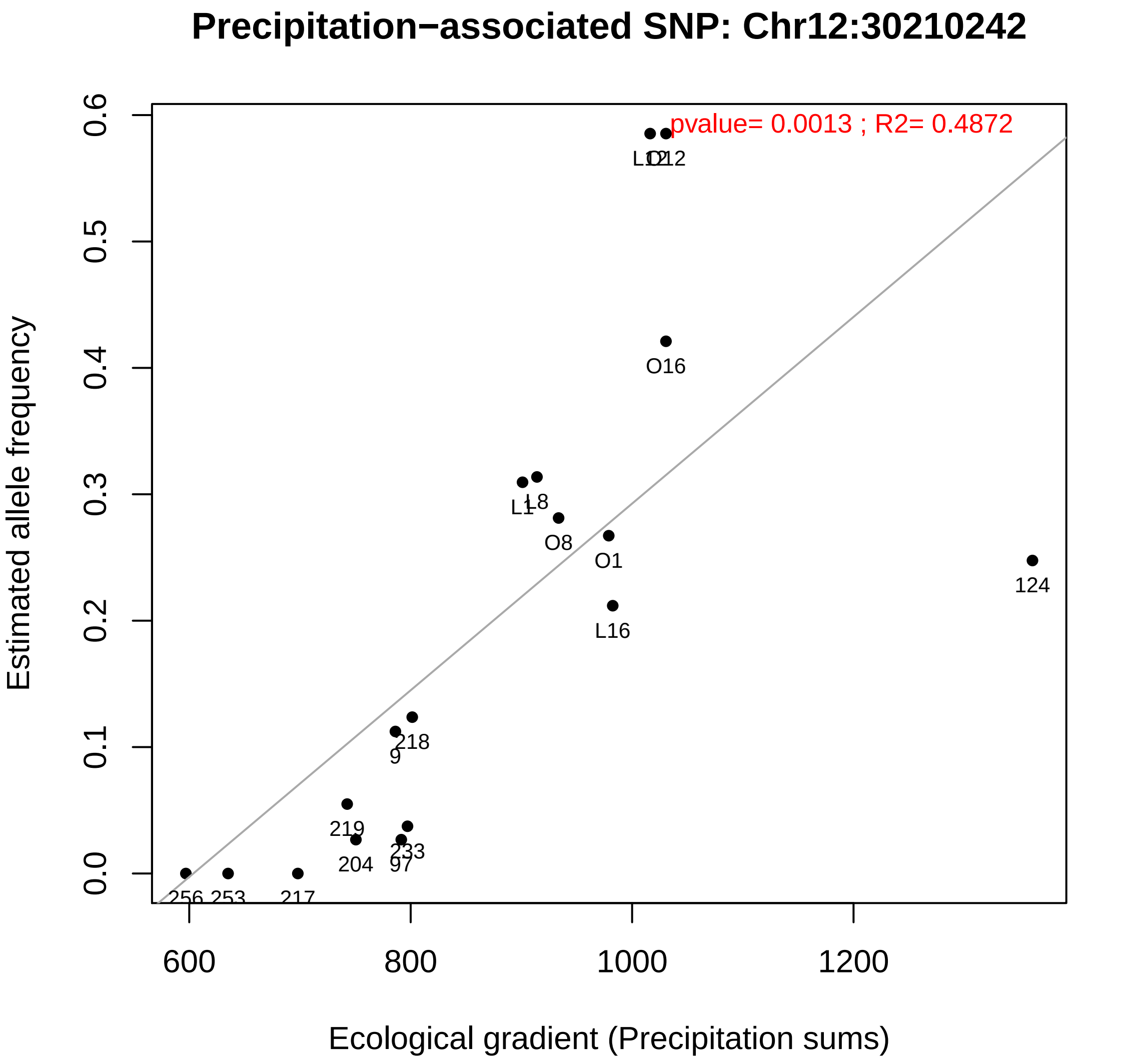
Precipitation−associated SNP: Chr12:30210242
pvalue= 0.0013 ; R2= 0.4872
Estimated allele frequency
Ecological gradient (Precipitation sums)
0.0
0.1
0.2
0.3
0.4
0.5
0.6
600
800
1000
1200
O16
L1
L8
O8
O1
124
L16
218
9
219
233
204
97
256
253
217

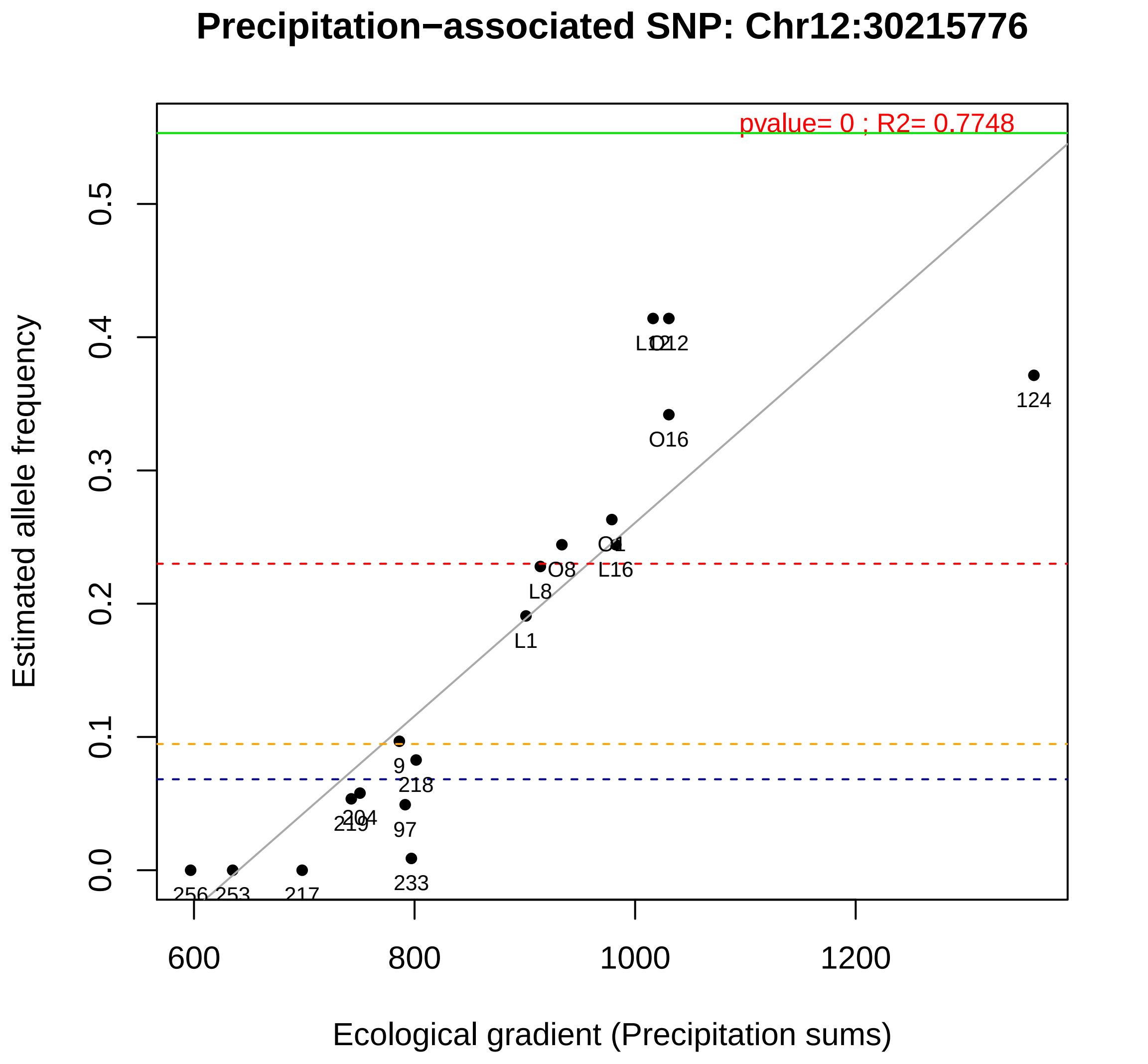

Precipitation−associated SNP: Chr12:30215776
pvalue= 0 ; R2= 0.7748
Estimated allele frequency
Ecological gradient (Precipitation sums)
0.0
0.1
0.2
0.3
0.4
0.5
600
800
1000
1200
L12
O12
124
O16
O1
L16
O8
L8
L1
9
218
204
219
97
233
256
253
217

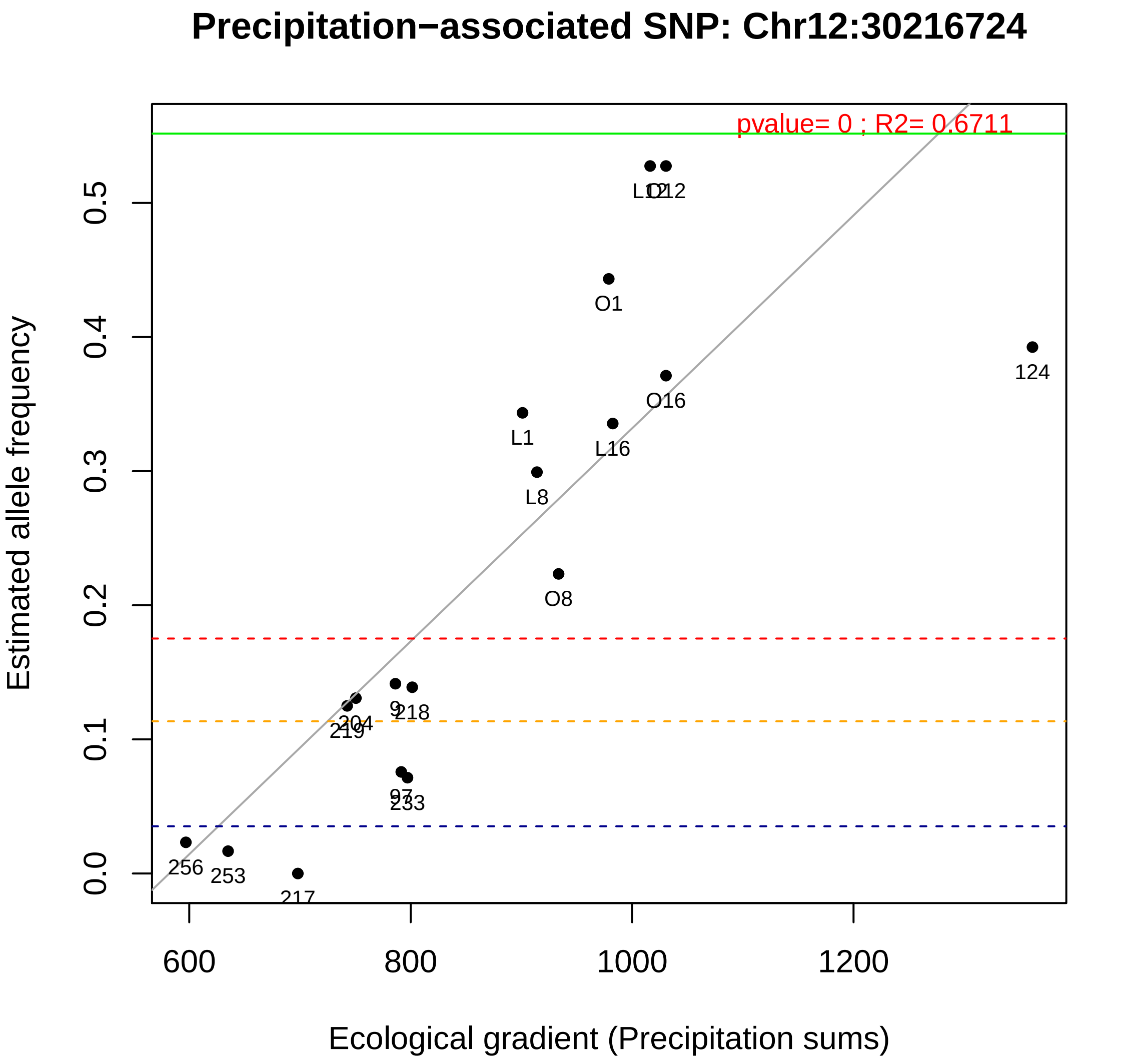
Precipitation−associated SNP: Chr12:30216724
pvalue= 0 ; R2= 0.6711
Estimated allele frequency
Ecological gradient (Precipitation sums)
0.0
0.1
0.2
0.3
0.4
0.5
600
800
1000
1200
L12
O12
O1
124
O16
L1
L16
L8
O8
9
218
204
219
97
233
256
253
217

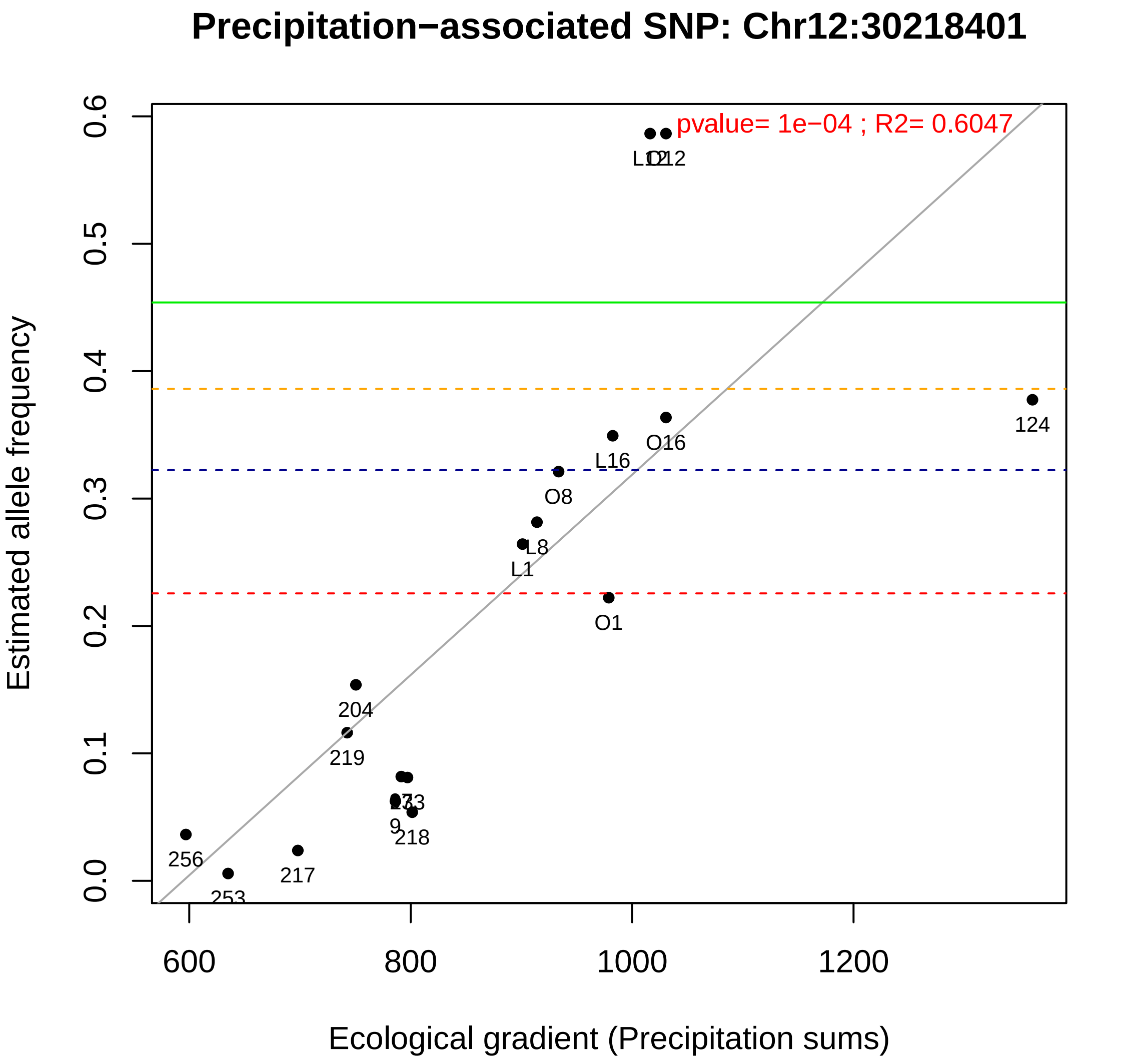

Precipitation−associated SNP: Chr12:30218401
pvalue= 1e−04 ; R2= 0.6047
Estimated allele frequency
Ecological gradient (Precipitation sums)
0.0
0.1
0.2
0.3
0.4
0.5
0.6
600
800
1000
1200
L12
O12
124
O16
L16
O8
L8
L1
O1
204
219
218
256
253
217

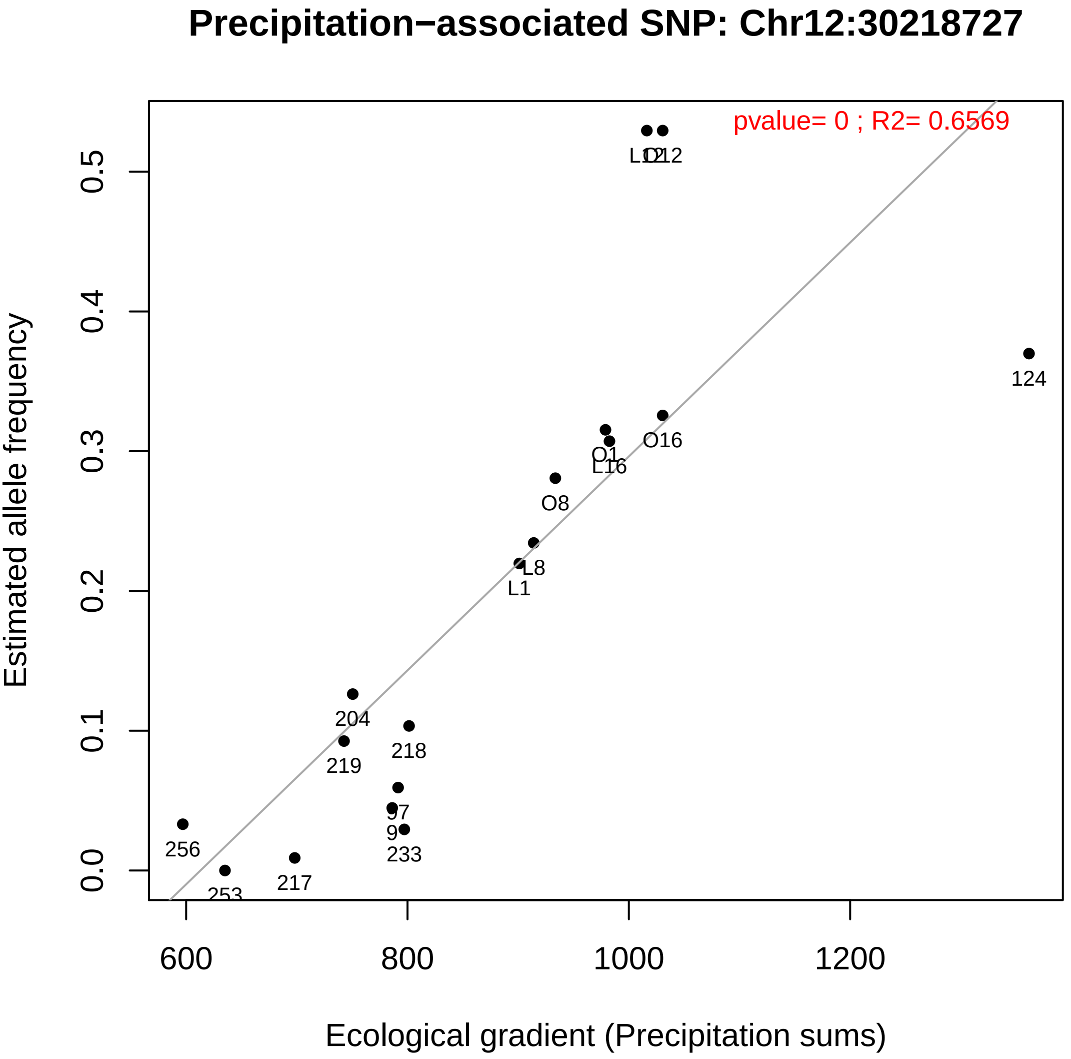
Precipitation−associated SNP: Chr12:30218727
pvalue= 0 ; R2= 0.6569
Estimated allele frequency
0.0
0.1
0.2
0.3
0.4
0.5
600
800
1000
1200
Ecological gradient (Precipitation sums)
O12
124
O16
O1
L16
O8
L8
L1
204
218
219
97
9
233
256
253
217

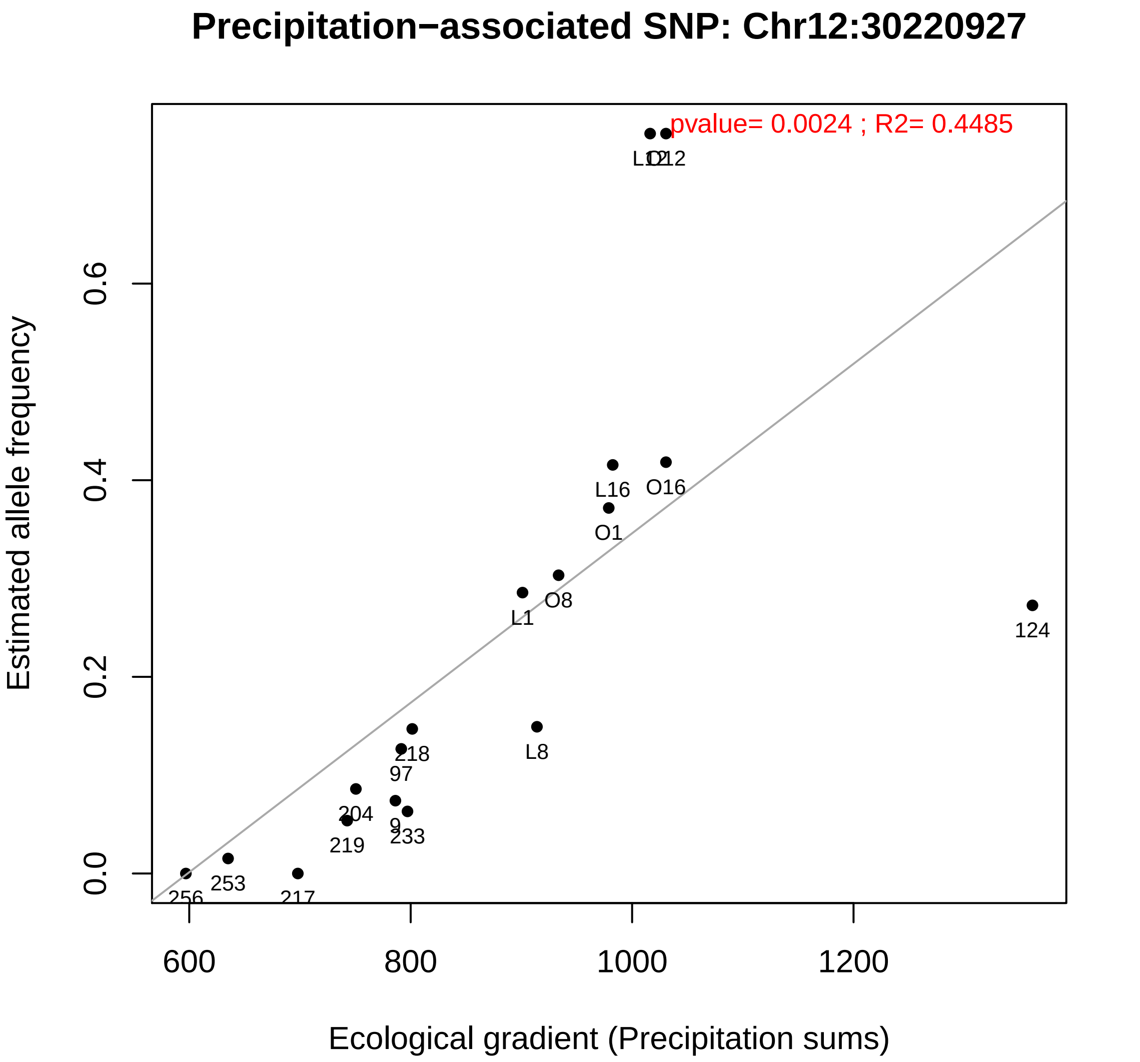

Precipitation−associated SNP: Chr12:30220927
pvalue= 0.0024 ; R2= 0.4485
Estimated allele frequency
0.0
0.2
0.4
0.6
600
800
1000
1200
Ecological gradient (Precipitation sums)
L1
O8
O1
L16
O16
L8
124
218
97
204
9
233
219
253
256
217

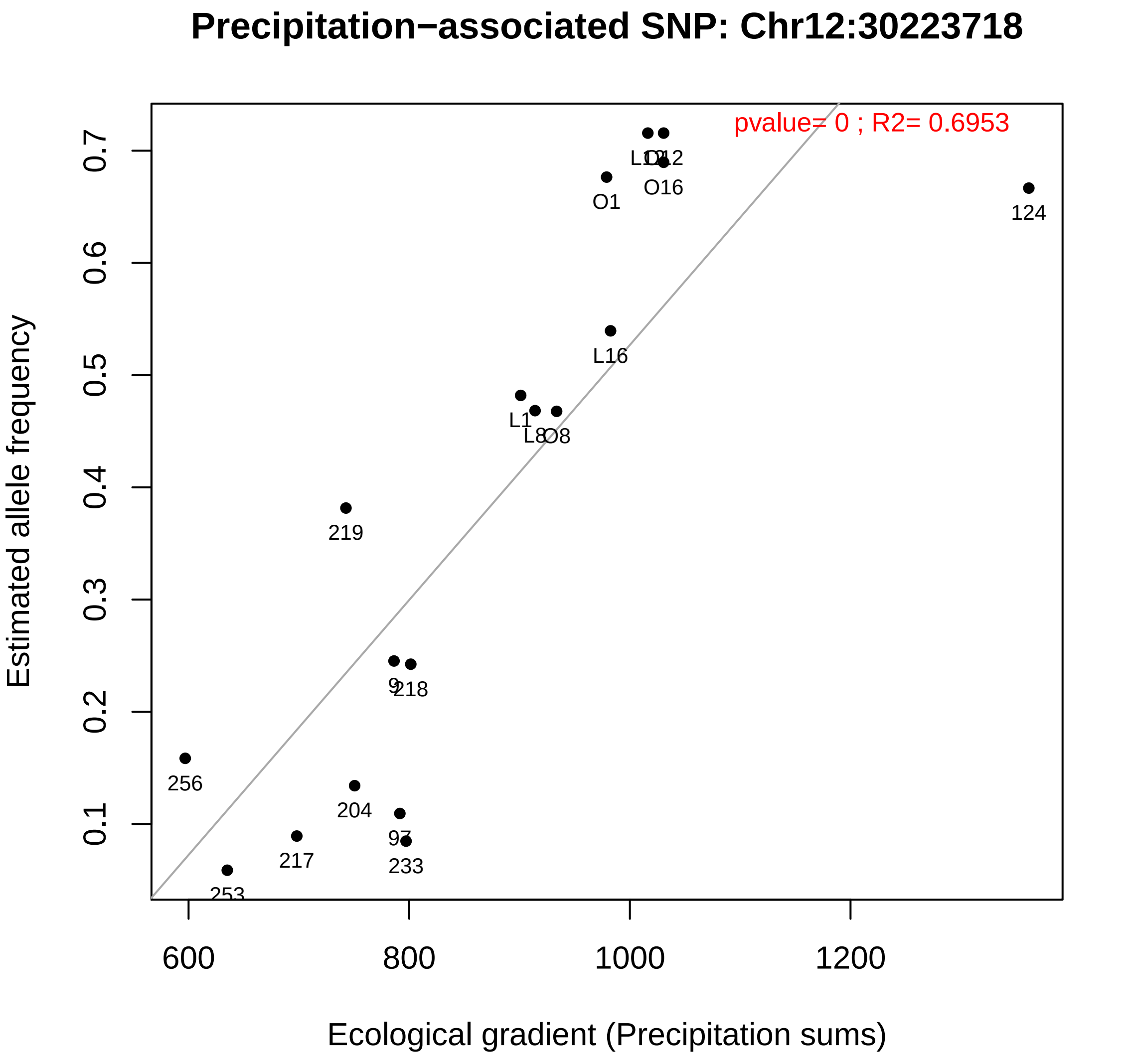
Precipitation−associated SNP: Chr12:30223718
pvalue= 0 ; R2= 0.6953
Estimated allele frequency
0.1
0.2
0.3
0.4
0.5
0.6
0.7
600
800
1000
1200
Ecological gradient (Precipitation sums)
O12
O16
O1
124
L16
L1
L8
O8
219
218
256
204
217
233
253

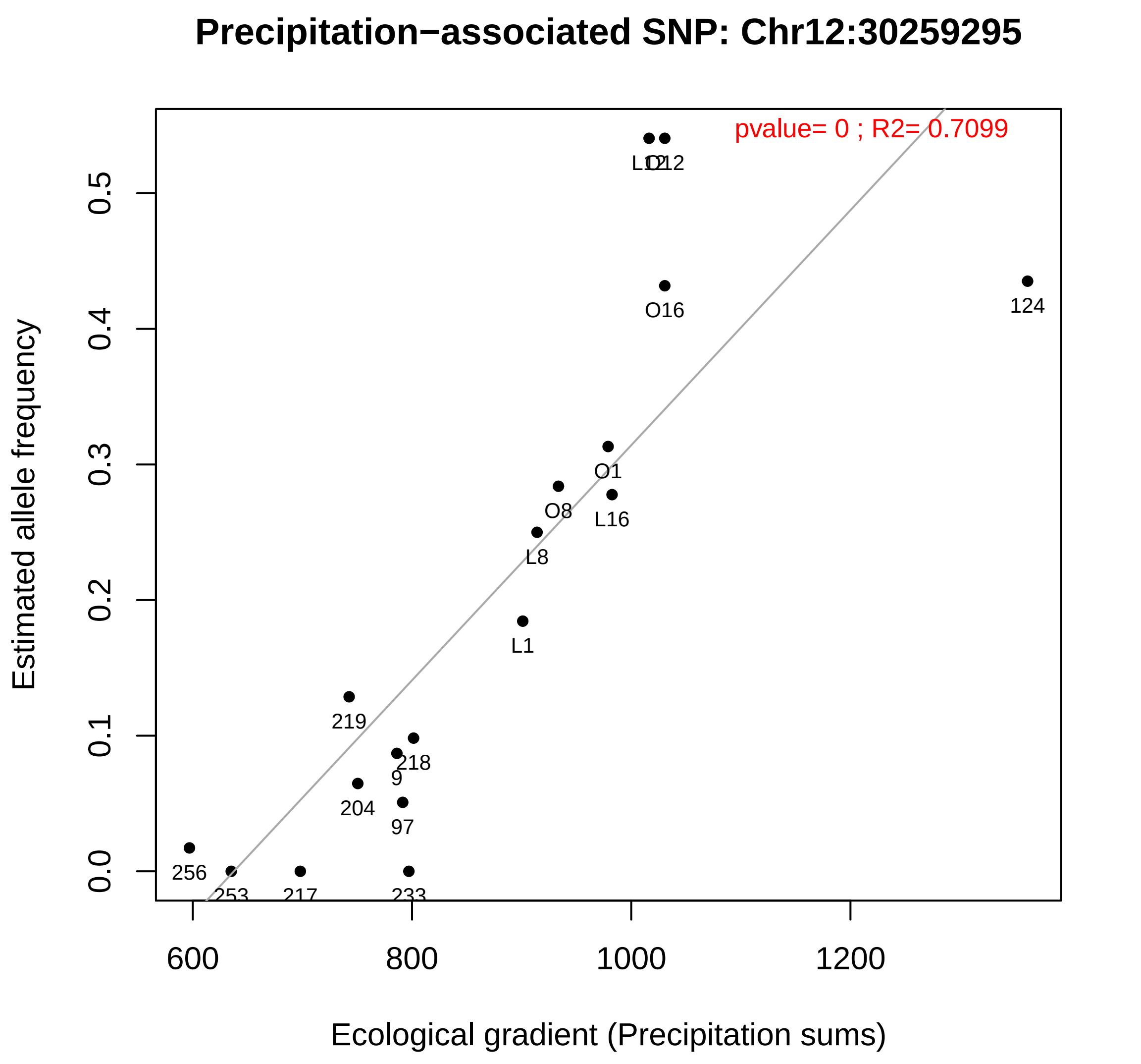

Precipitation−associated SNP: Chr12:30259295
pvalue= 0 ; R2= 0.7099
Estimated allele frequency
0.0
0.1
0.2
0.3
0.4
0.5
600
800
1000
1200
Ecological gradient (Precipitation sums)
L12
O12
O16
124
O1
O8
L16
L8
L1
219
218
9
204
97
256
253
217
233

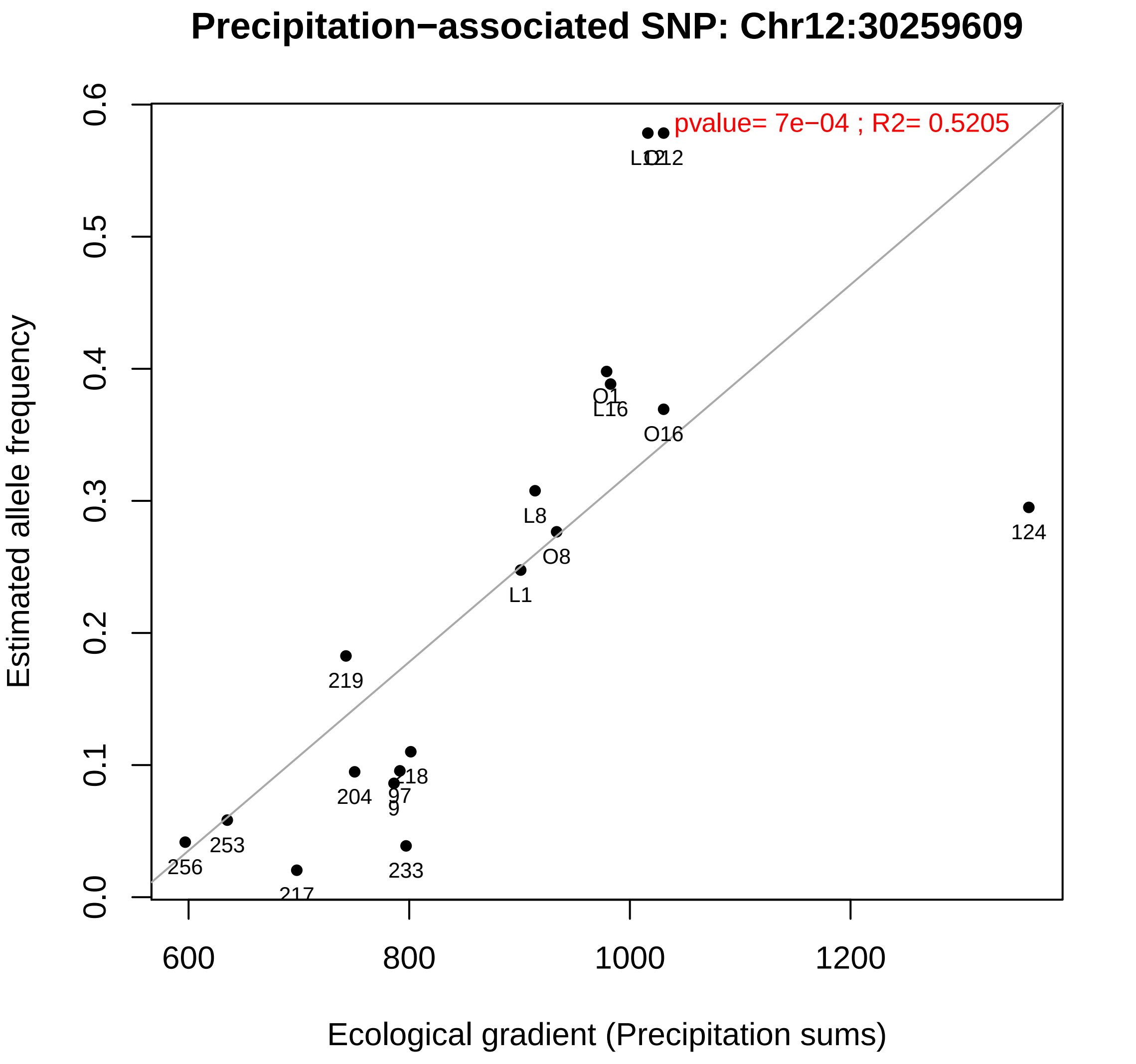
Precipitation−associated SNP: Chr12:30259609
pvalue= 7e−04 ; R2= 0.5205
Estimated allele frequency
Ecological gradient (Precipitation sums)
0.0
0.1
0.2
0.3
0.4
0.5
0.6
600
800
1000
1200
L12
O12
O1
L16
O16
L8
O8
L1
124
219
204
118
97
9
253
256
217
233

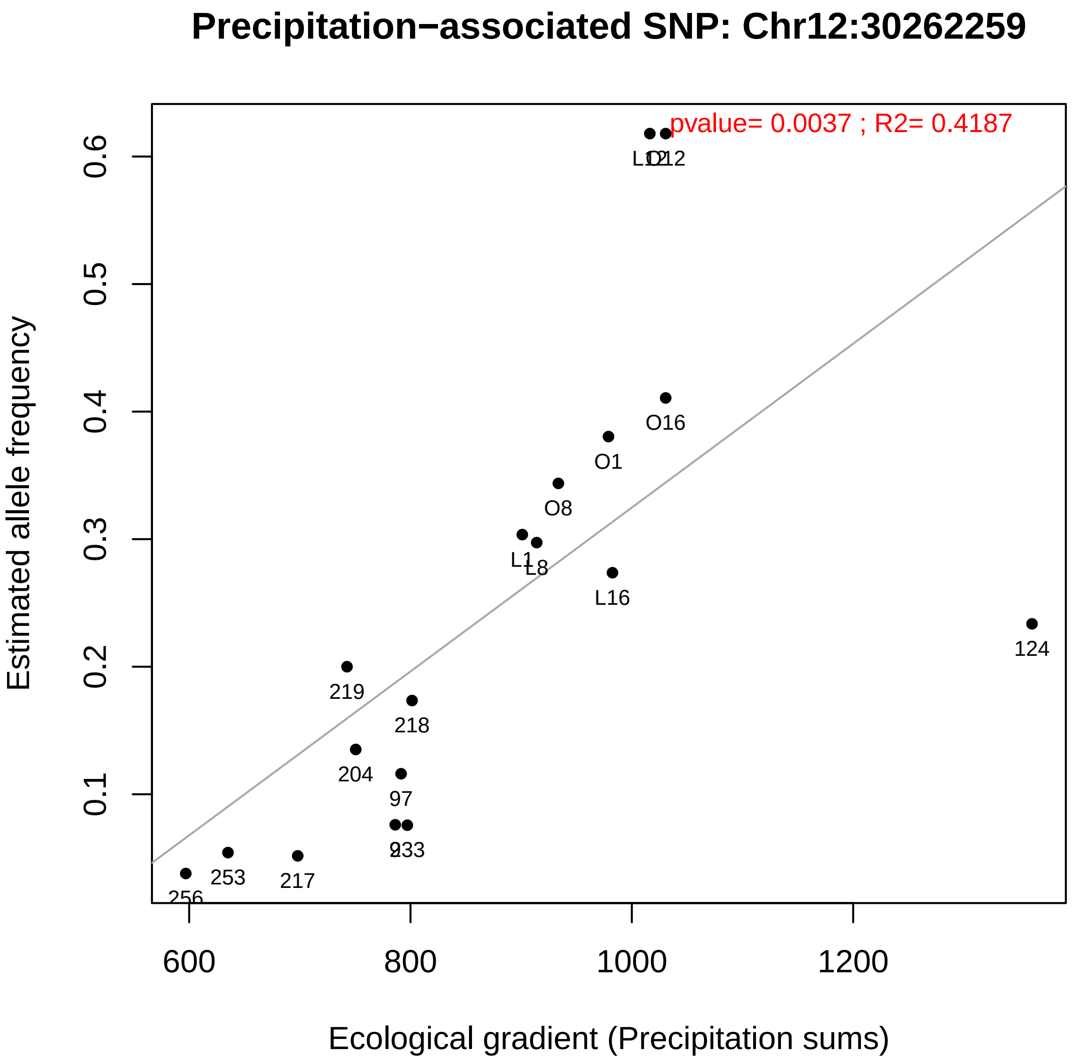

Precipitation−associated SNP: Chr12:30262259
pvalue= 0.0037 ; R2= 0.4187
Estimated allele frequency
Ecological gradient (Precipitation sums)
0.1
0.2
0.3
0.4
0.5
0.6
600
800
1000
1200
O12
O16
O1
O8
L1
L8
L16
124
219
218
204
97
233
253
217
256

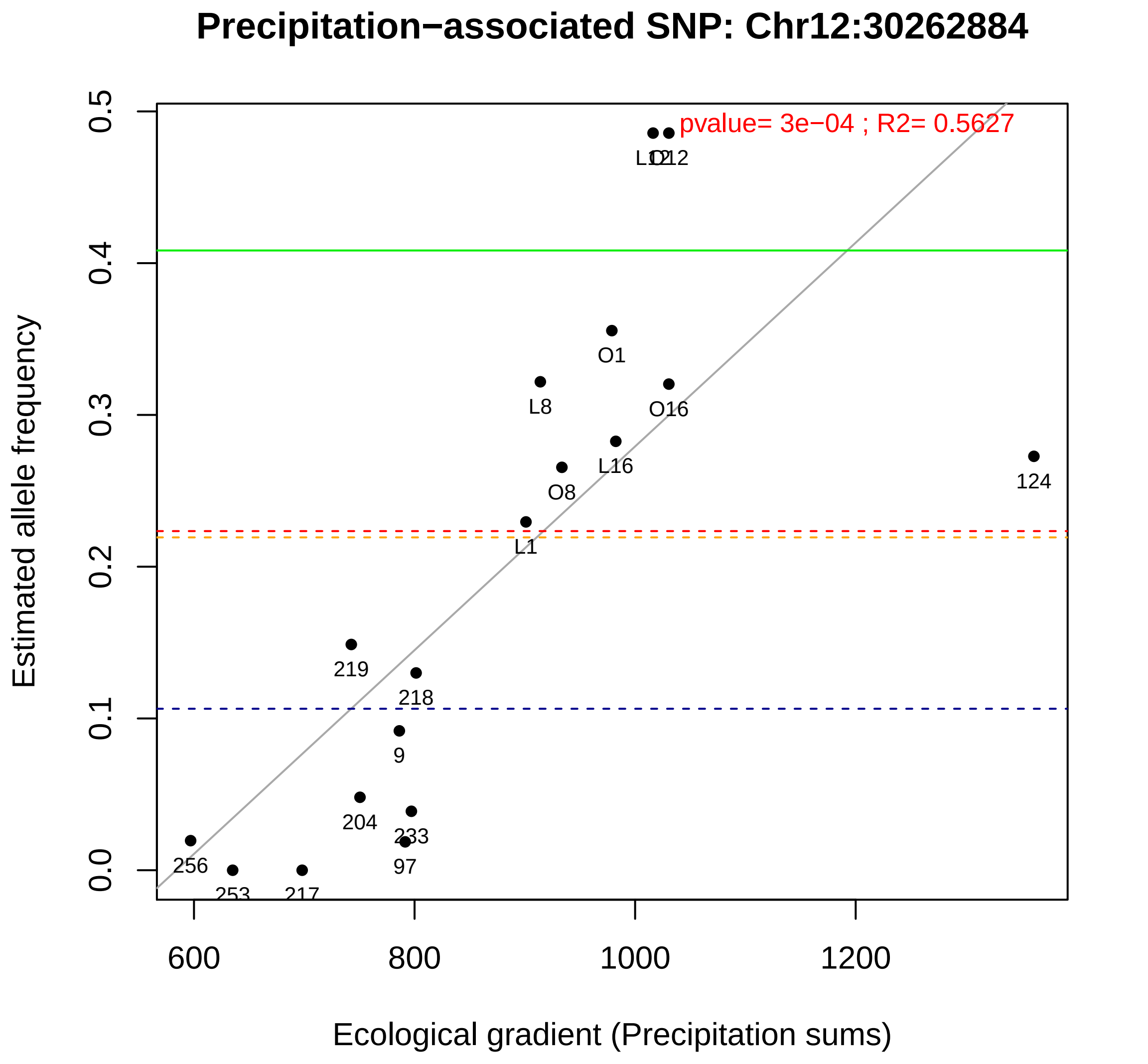
Precipitation−associated SNP: Chr12:30262884
pvalue= 3e−04 ; R2= 0.5627
Estimated allele frequency
Ecological gradient (Precipitation sums)
0.0
0.1
0.2
0.3
0.4
0.5
600
800
1000
1200
O1
L8
O16
L16
O8
124
L1
219
218
9
204
233
97
256
253
217

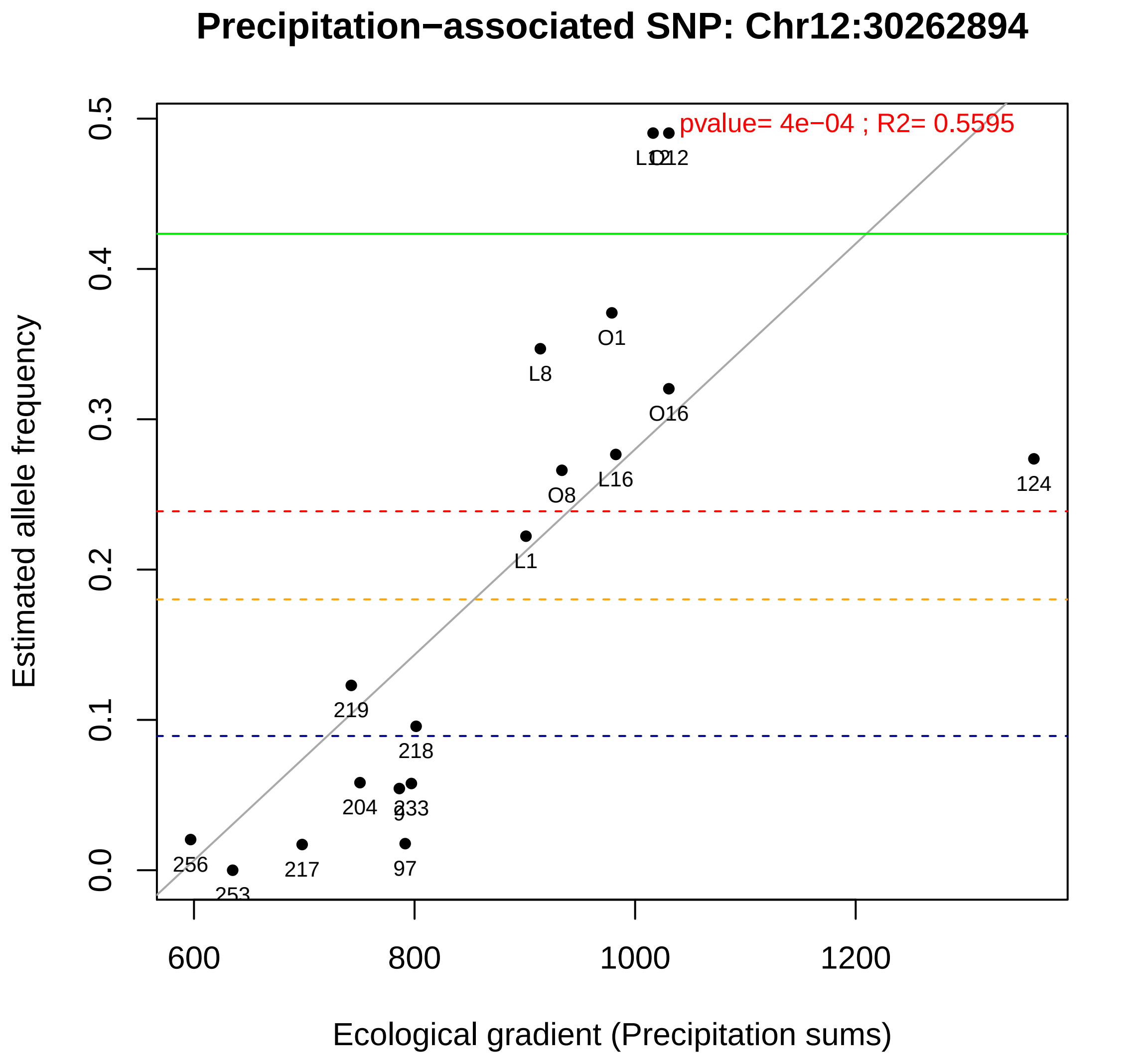
Precipitation−associated SNP: Chr12:30262894
pvalue= 4e−04 ; R2= 0.5595
Estimated allele frequency
Ecological gradient (Precipitation sums)
0.0
0.1
0.2
0.3
0.4
0.5
600
800
1000
1200
O1
L8
O16
L16
O8
124
L1
219
218
204
233
97
256
253
217

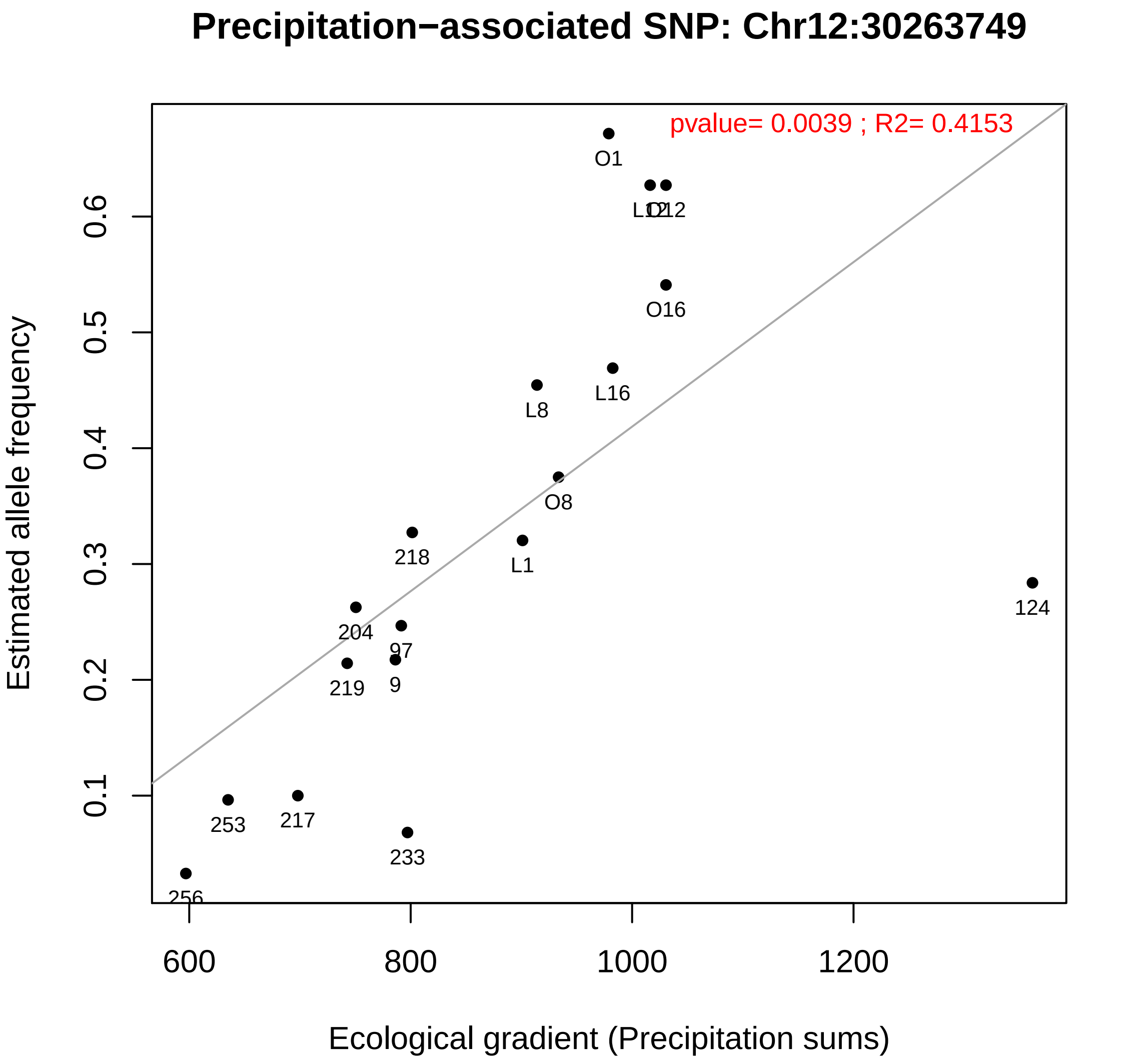
Precipitation−associated SNP: Chr12:30263749
pvalue= 0.0039 ; R2= 0.4153
Estimated allele frequency
0.1
0.2
0.3
0.4
0.5
0.6
600
800
1000
1200
Ecological gradient (Precipitation sums)
O1
O16
L16
L8
O8
218
L1
124
204
97
219
9
253
217
233
256

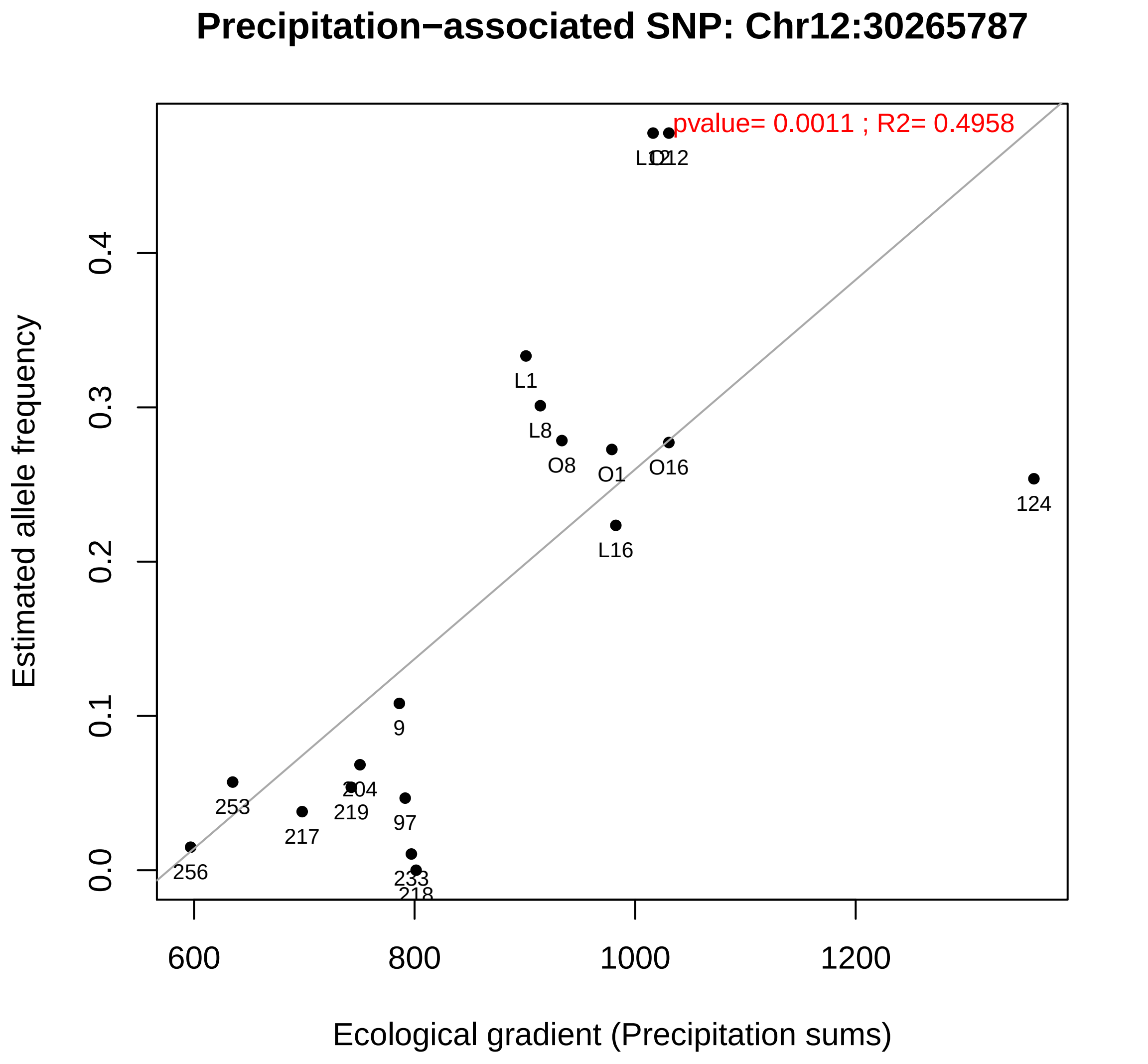

Precipitation−associated SNP: Chr12:30265787
pvalue= 0.0011 ; R2= 0.4958
L1
L8
O8
O1
O16
L16
124
9
253
204
219
217
97
256
233
218
0.0
0.1
0.2
0.3
0.4
600
800
1000
1200
Estimated allele frequency
Ecological gradient (Precipitation sums)

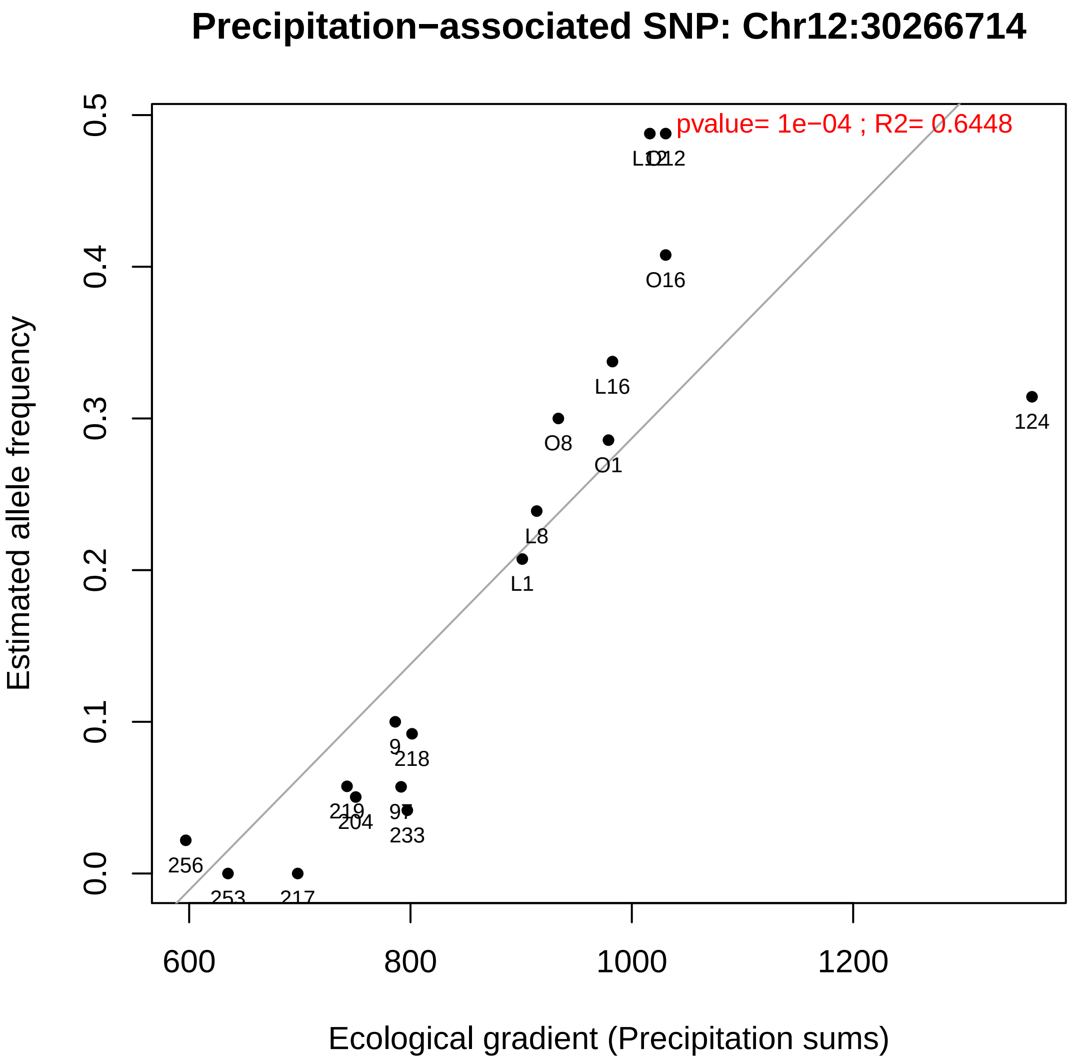

Precipitation−associated SNP: Chr12:30266714
pvalue= 1e−04 ; R2= 0.6448
Estimated allele frequency
Ecological gradient (Precipitation sums)
0.0
0.1
0.2
0.3
0.4
0.5
600
800
1000
1200
O16
L16
124
O8
O1
L8
L1
9
218
219
204
233
256
253
217

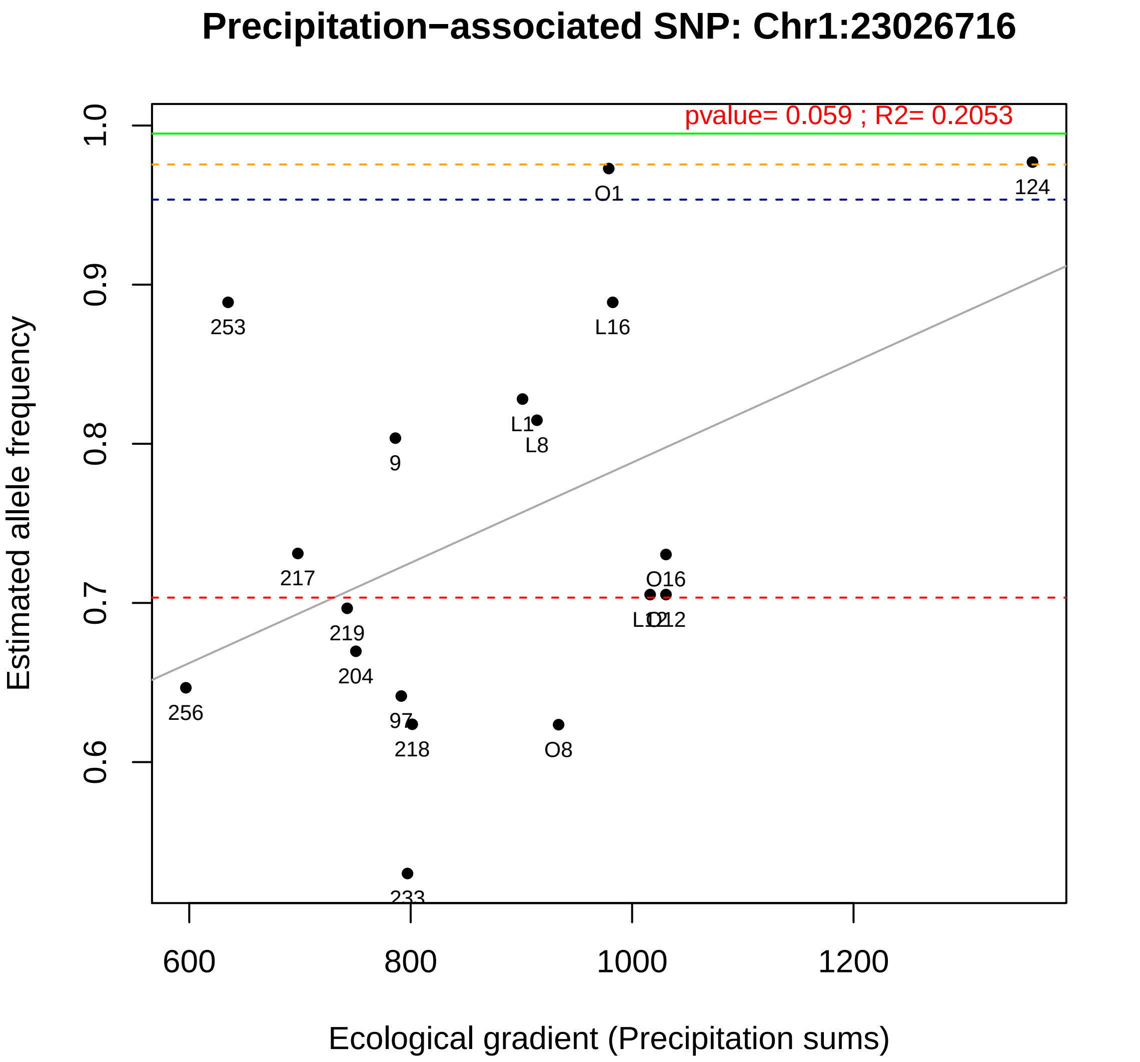
Precipitation−associated SNP: Chr1:23026716
pvalue= 0.059 ; R2= 0.2053
Estimated allele frequency
Ecological gradient (Precipitation sums)
0.6
0.7
0.8
0.9
1.0
600
800
1000
1200
O1
124
253
L16
L1
L8
9
217
O16
219
L10
O12
204
256
97
218
O8
233

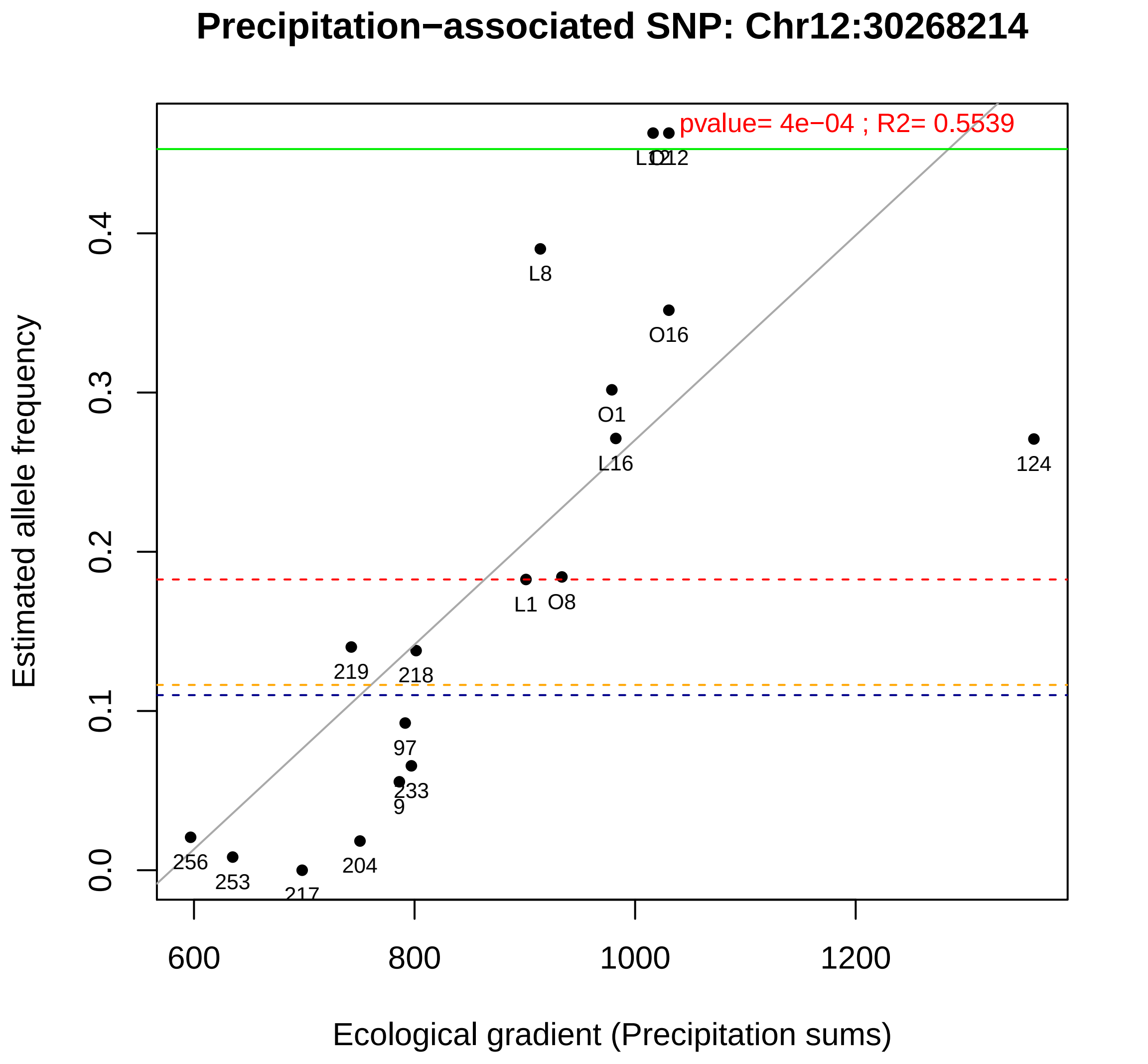

Precipitation−associated SNP: Chr12:30268214
pvalue= 4e−04 ; R2= 0.5539
Estimated allele frequency
Ecological gradient (Precipitation sums)
0.0
0.1
0.2
0.3
0.4
600
800
1000
1200
L8
O16
O1
L16
124
L1
O8
219
218
97
233
9
256
253
217
204

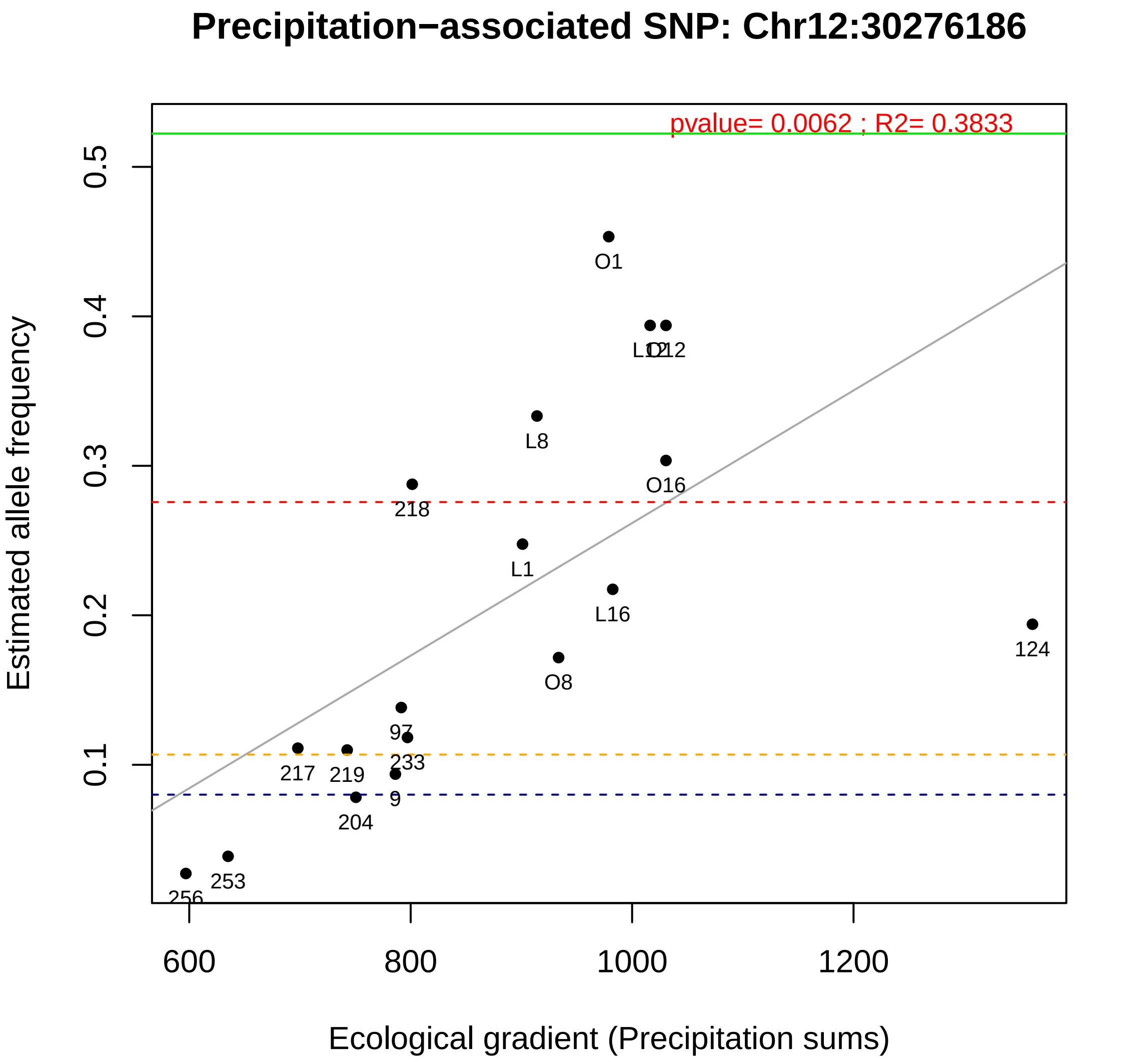

Precipitation−associated SNP: Chr12:30276186
pvalue= 0.0062 ; R2= 0.3833
Estimated allele frequency
0.1
0.2
0.3
0.4
0.5
600
800
1000
1200
Ecological gradient (Precipitation sums)
O1
L8
O16
218
L1
L16
124
O8
97
233
217
219
9
204
253
256

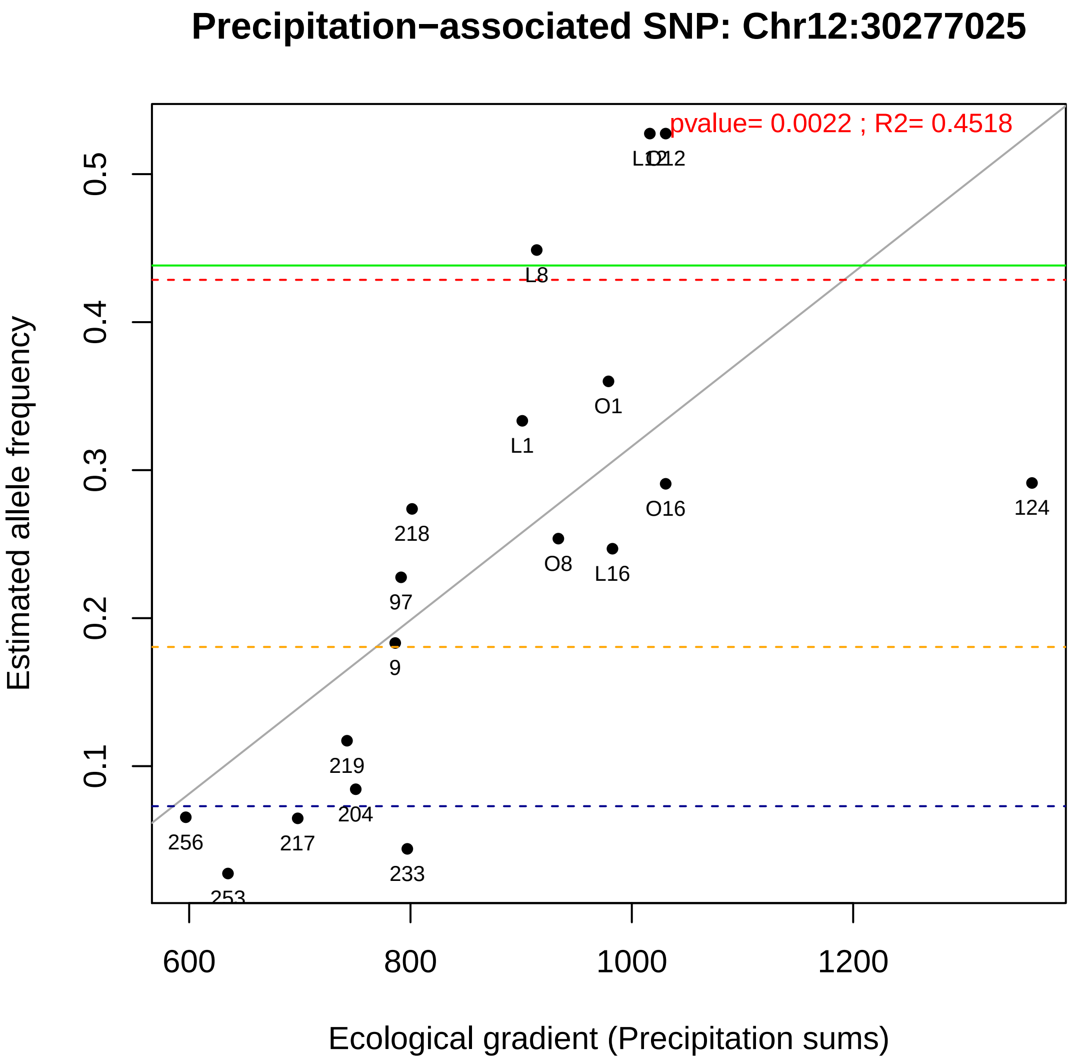

Precipitation−associated SNP: Chr12:30277025
pvalue= 0.0022 ; R2= 0.4518
Estimated allele frequency
Ecological gradient (Precipitation sums)
0.1
0.2
0.3
0.4
0.5
600
800
1000
1200
L8
O1
L1
O16
124
218
O8
L16
97
9
219
204
256
217
233

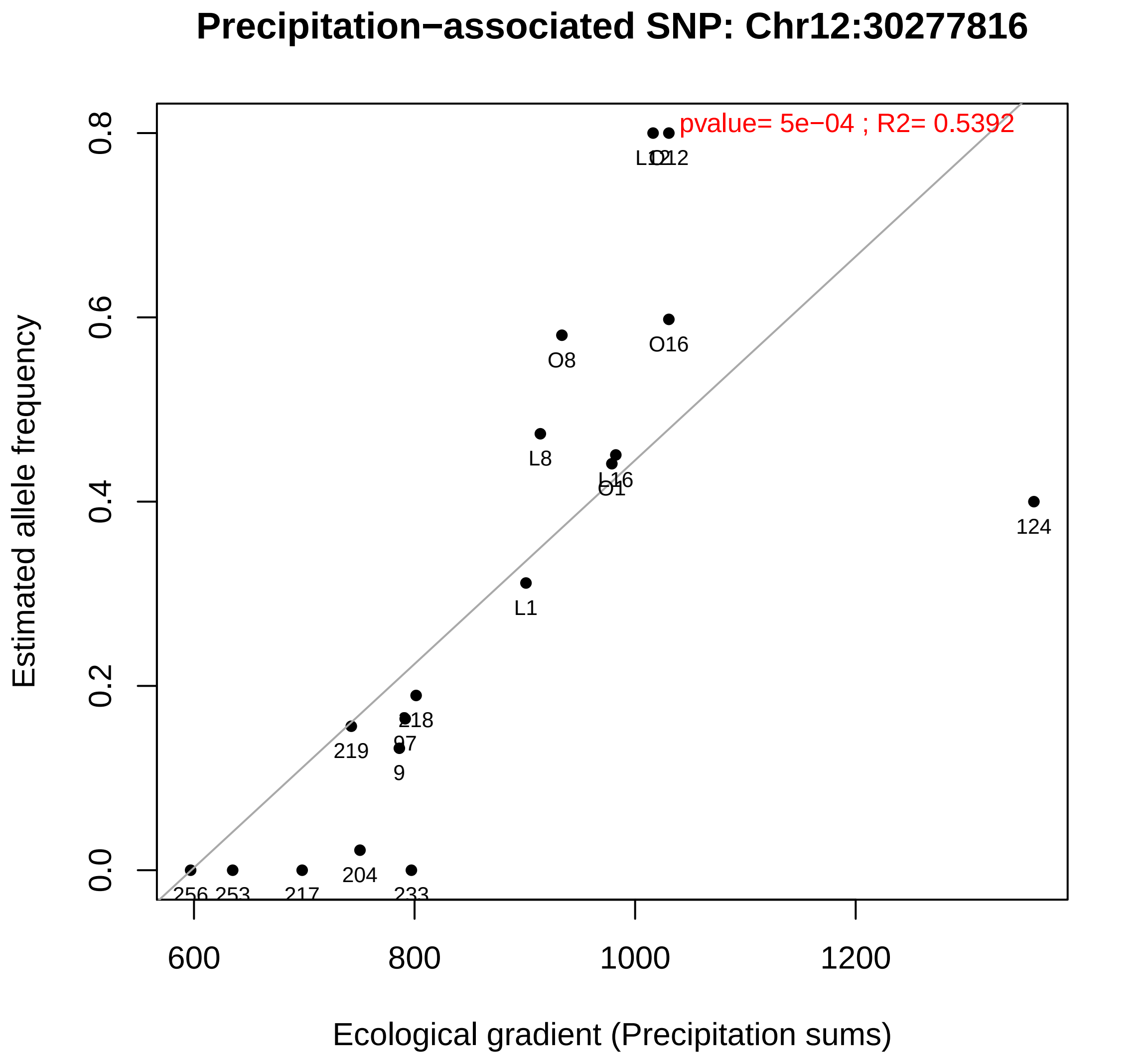

Precipitation−associated SNP: Chr12:30277816
pvalue= 5e−04 ; R2= 0.5392
Estimated allele frequency
Ecological gradient (Precipitation sums)
0.0
0.2
0.4
0.6
0.8
600
800
1000
1200
L12
O12
O16
O8
L8
L16
O1
124
L1
218
97
219
9
204
256
253
217
233

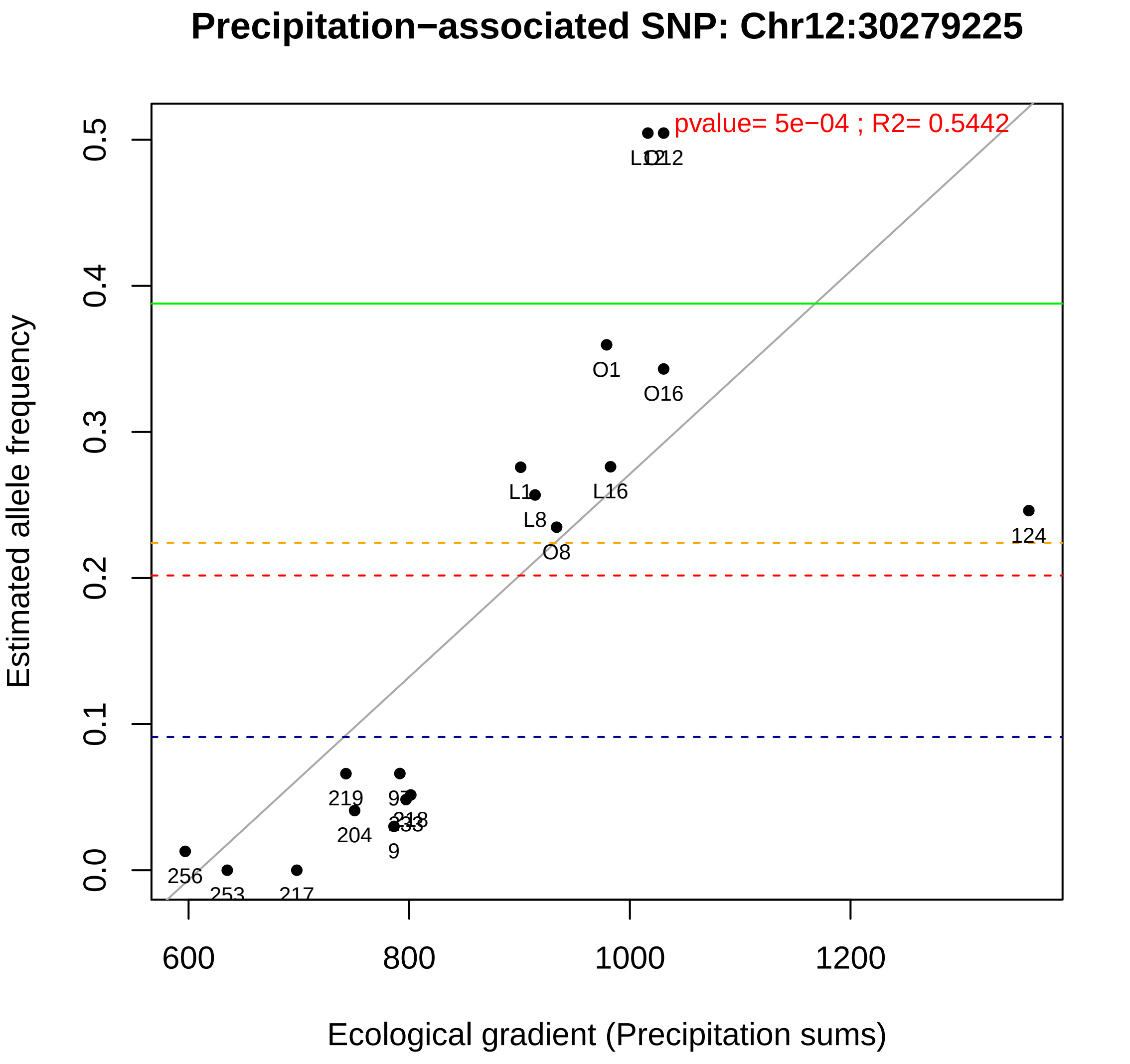

Precipitation−associated SNP: Chr12:30279225
pvalue= 5e−04 ; R2= 0.5442
Estimated allele frequency
Ecological gradient (Precipitation sums)
0.0
0.1
0.2
0.3
0.4
0.5
600
800
1000
1200
O1
O16
L1
L16
L8
O8
124
219
204
9
256
253
217

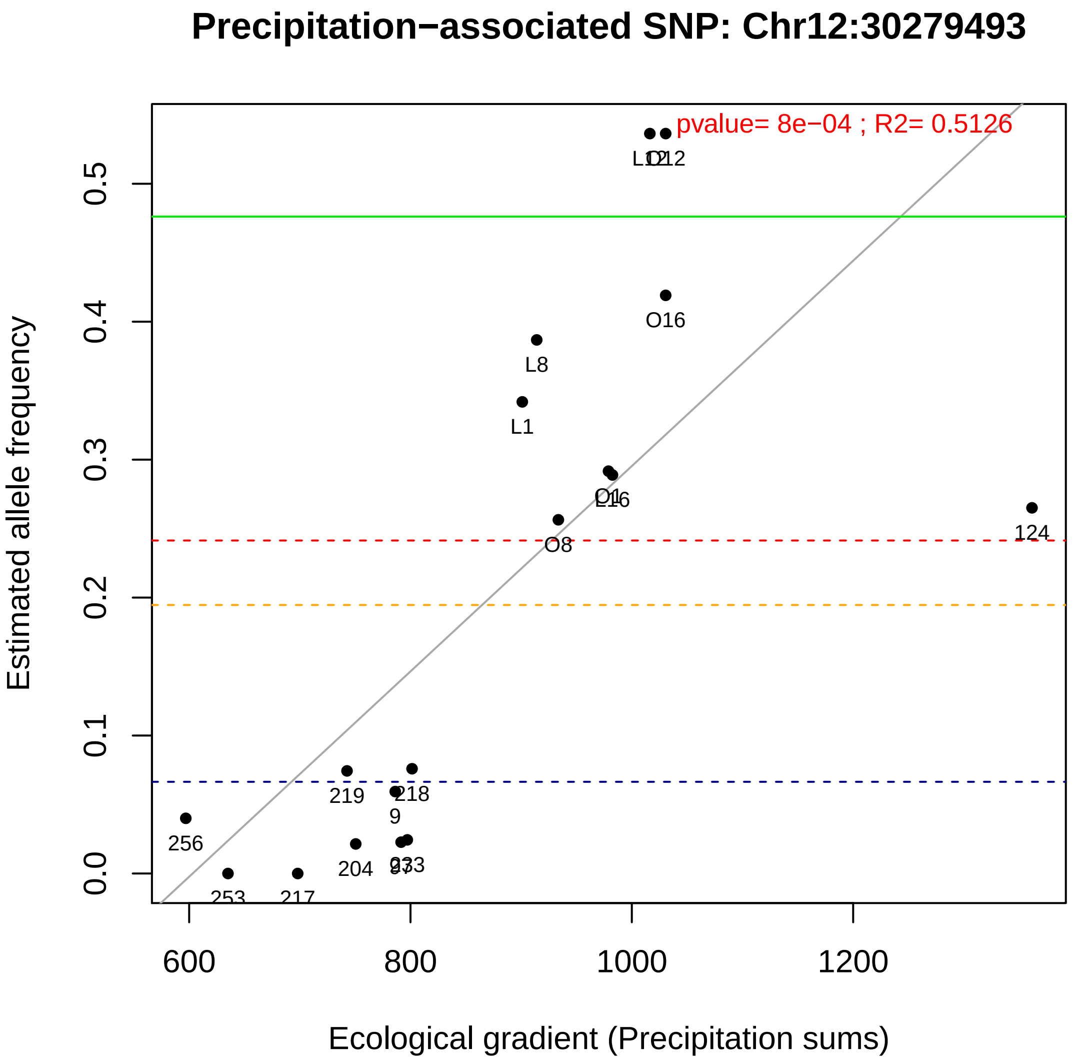

Precipitation−associated SNP: Chr12:30279493
pvalue= 8e−04 ; R2= 0.5126
Estimated allele frequency
Ecological gradient (Precipitation sums)
0.0
0.1
0.2
0.3
0.4
0.5
600
800
1000
1200
L12
O12
O16
L8
L1
O1
L16
O8
124
219
218
9
256
204
253
217
233

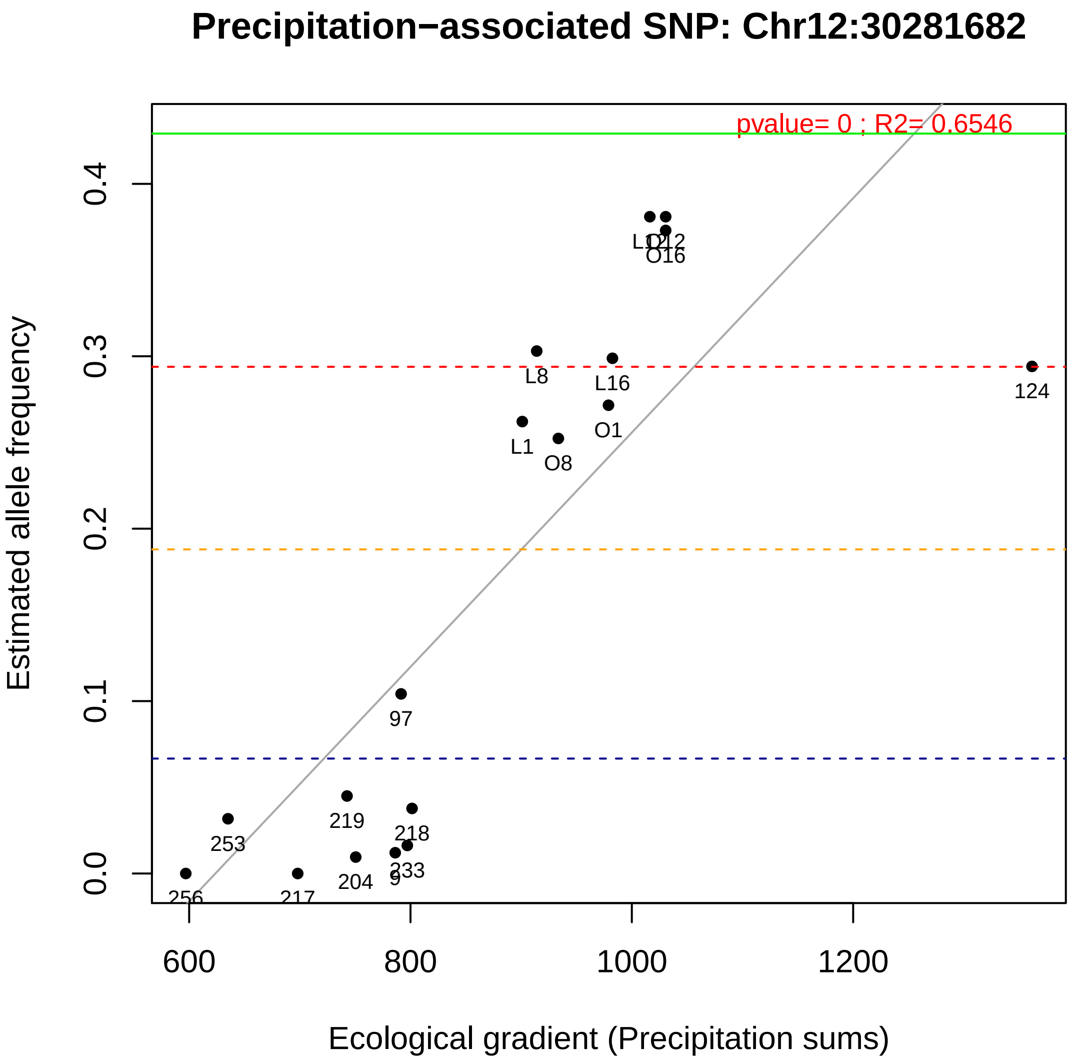

Precipitation−associated SNP: Chr12:30281682
pvalue= 0 ; R2= 0.6546
Estimated allele frequency
Ecological gradient (Precipitation sums)
0.0
0.1
0.2
0.3
0.4
600
800
1000
1200
L8
L16
124
O1
L1
O8
O16
97
219
253
218
233
204
9
256
217

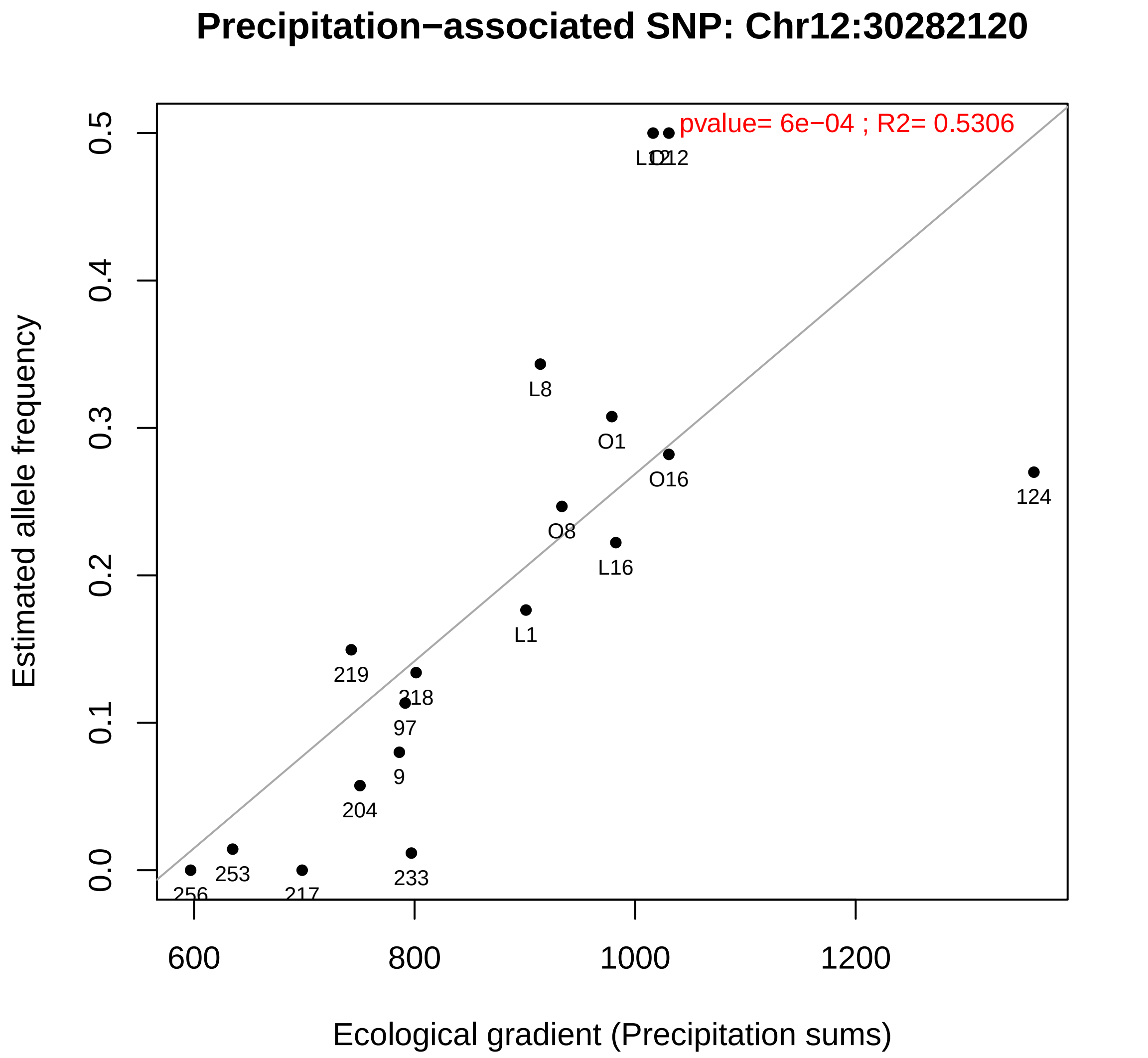

Precipitation−associated SNP: Chr12:30282120
pvalue= 6e−04 ; R2= 0.5306
Estimated allele frequency
Ecological gradient (Precipitation sums)
0.0
0.1
0.2
0.3
0.4
0.5
600
800
1000
1200
L12
O12
L8
O1
O16
124
O8
L16
L1
219
218
97
9
204
253
233
256
217

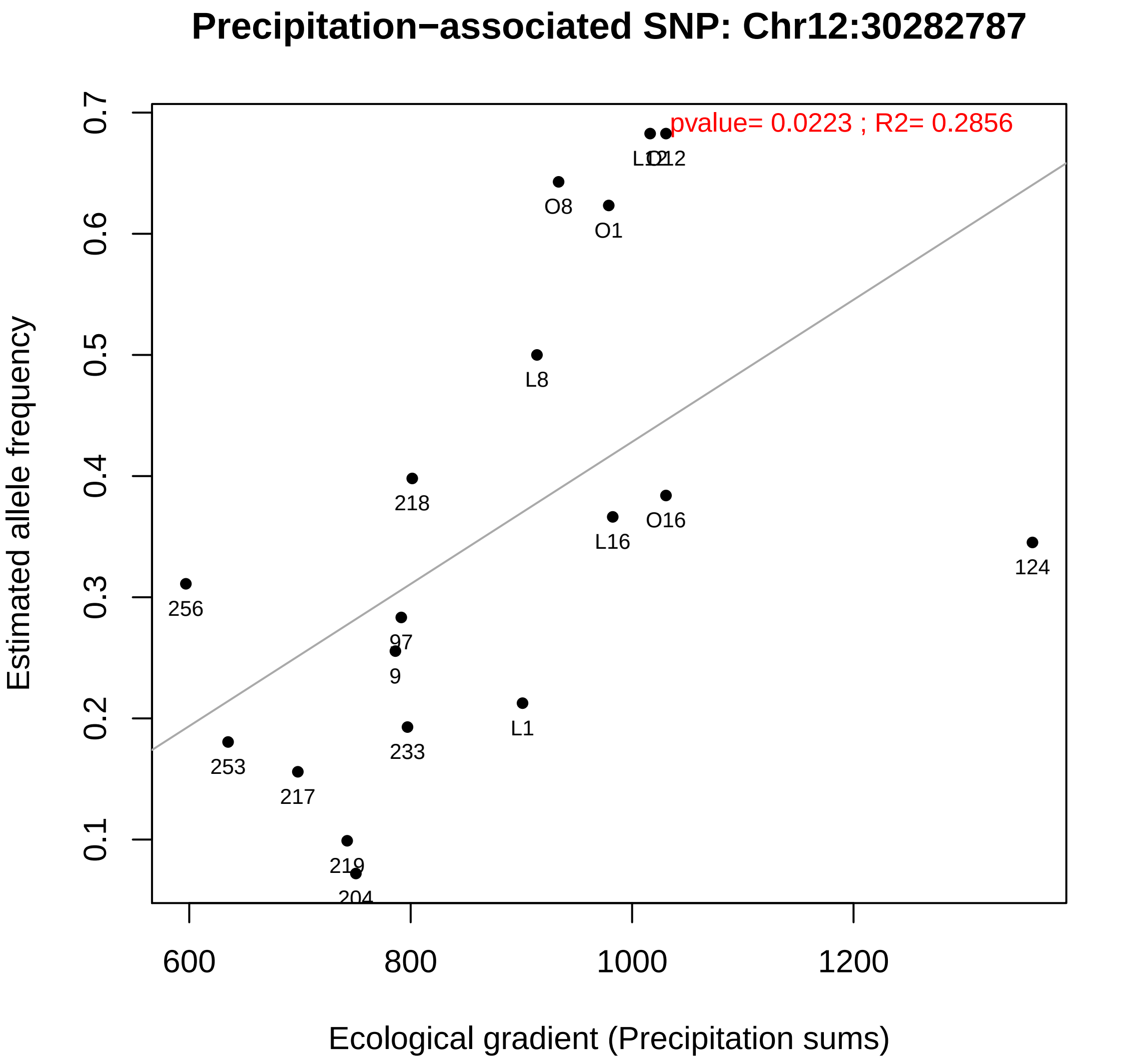

Precipitation−associated SNP: Chr12:30282787
pvalue= 0.0223 ; R2= 0.2856
Estimated allele frequency
Ecological gradient (Precipitation sums)
0.1
0.2
0.3
0.4
0.5
0.6
0.7
600
800
1000
1200
L12
O12
O8
O1
L8
218
O16
L16
124
256
97
9
L1
233
253
217
219
204

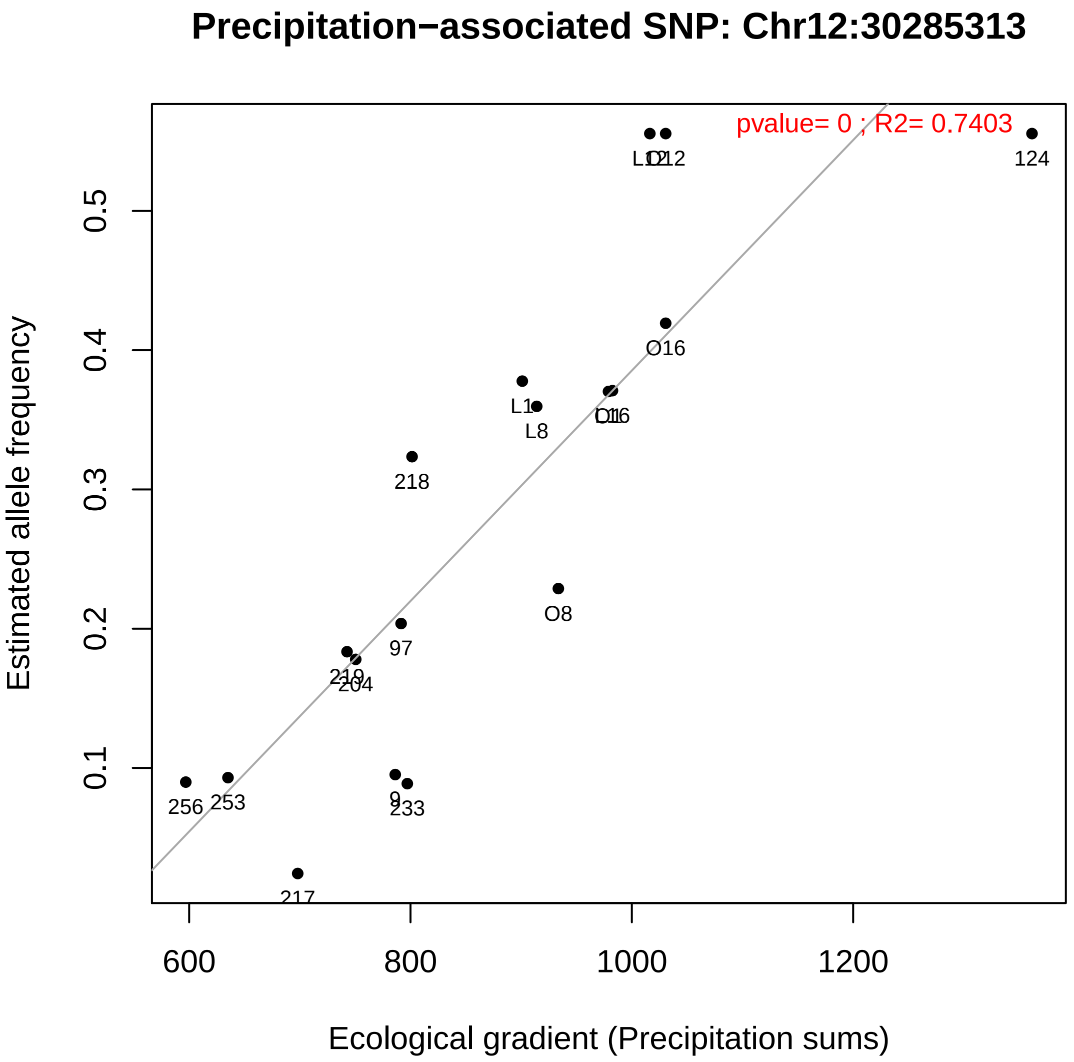
Precipitation−associated SNP: Chr12:30285313
pvalue= 0 ; R2= 0.7403
Estimated allele frequency
0.1
0.2
0.3
0.4
0.5
600
800
1000
1200
Ecological gradient (Precipitation sums)
124
O16
218
L8
O8
97
204
256
253
233
217

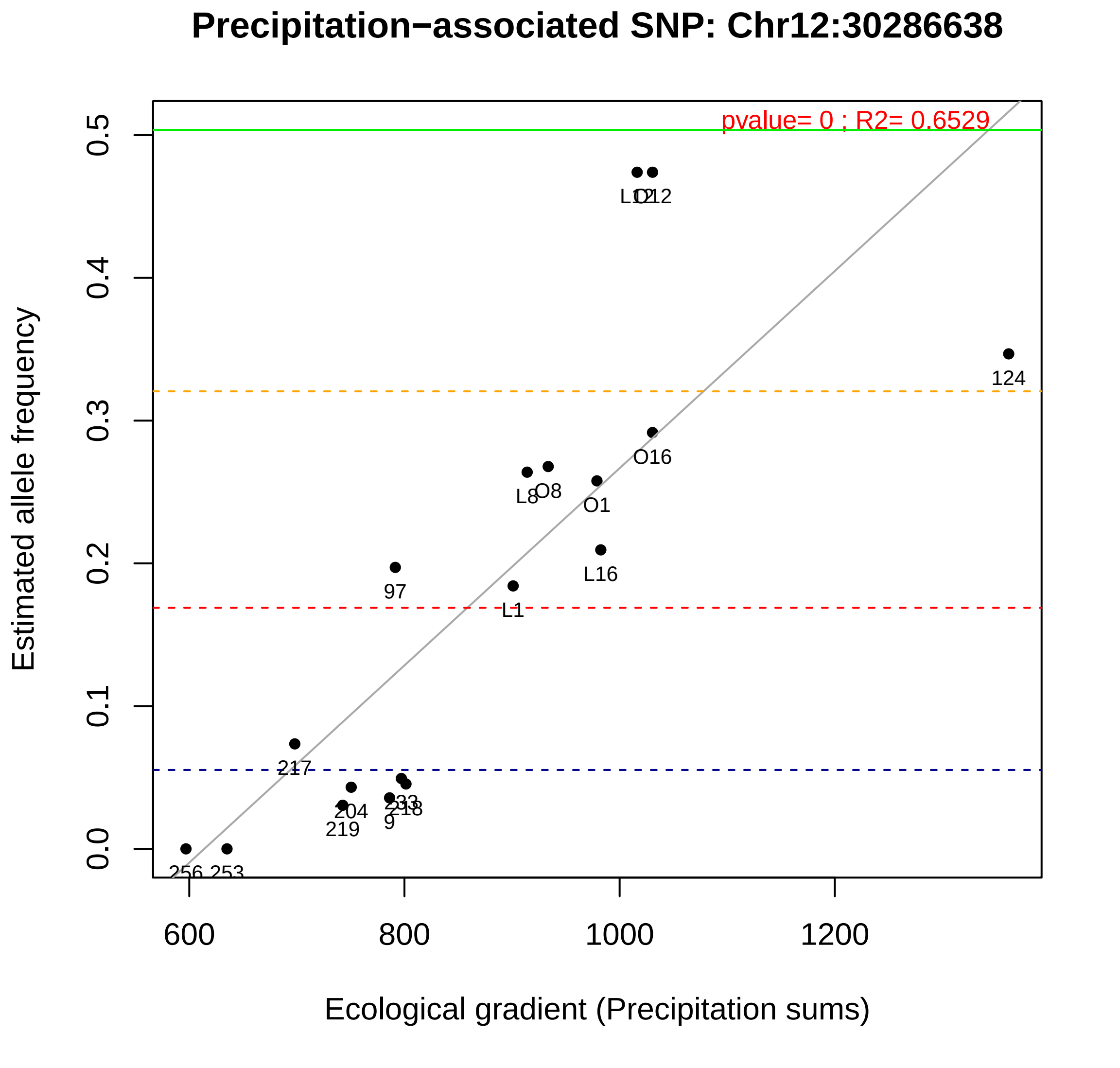

Precipitation−associated SNP: Chr12:30286638
pvalue= 0 ; R2= 0.6529
Estimated allele frequency
Ecological gradient (Precipitation sums)
0.0
0.1
0.2
0.3
0.4
0.5
600
800
1000
1200
O16
L8
O8
O1
L16
97
L1
124
217
204
219
218
9

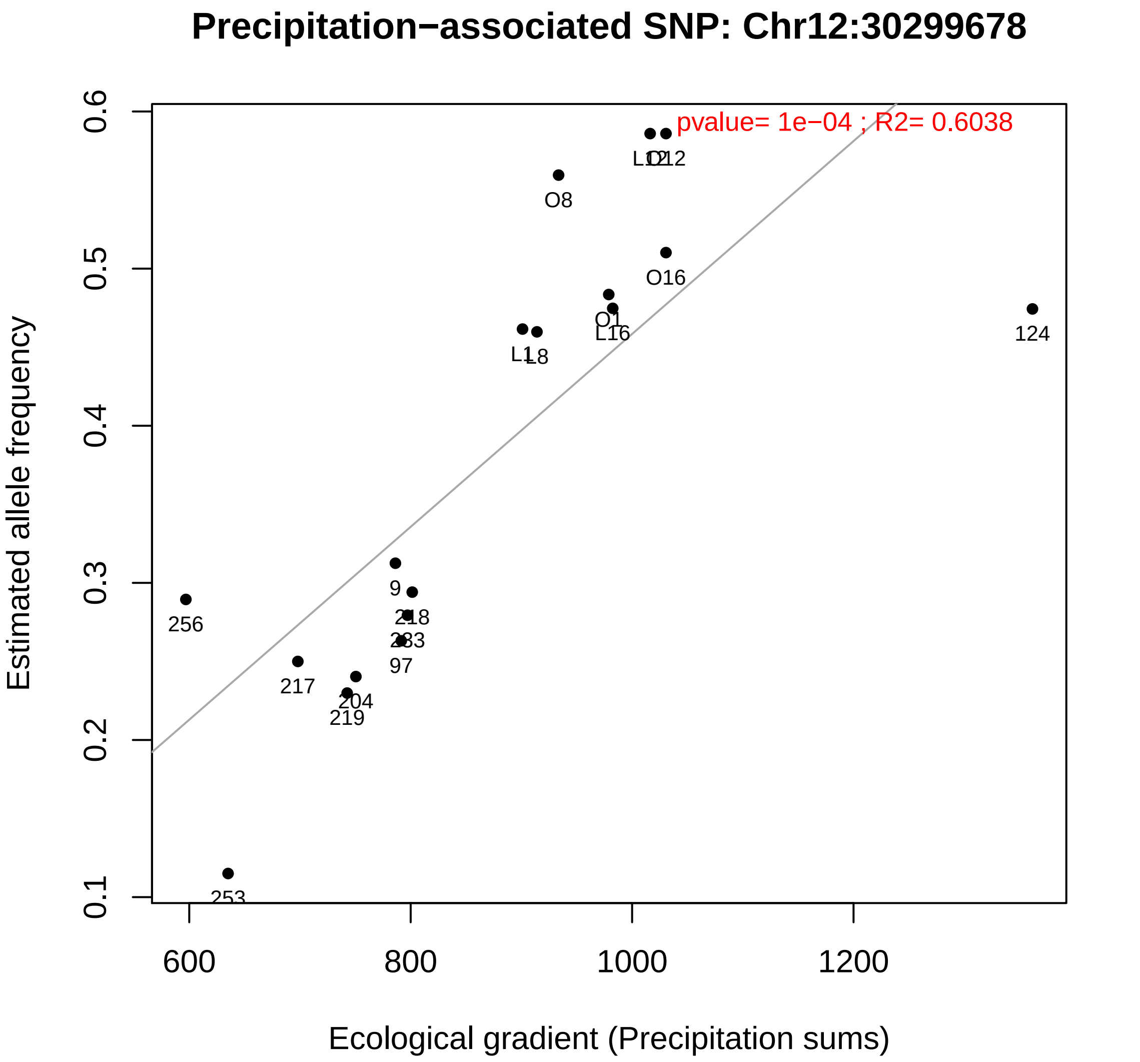

Precipitation−associated SNP: Chr12:30299678
pvalue= 1e−04 ; R2= 0.6038
Estimated allele frequency
0.1
0.2
0.3
0.4
0.5
0.6
600
800
1000
1200
Ecological gradient (Precipitation sums)
L12
O2
O8
O16
O1
L16
124
L1
L8
9
218
233
97
256
217
204
219
253

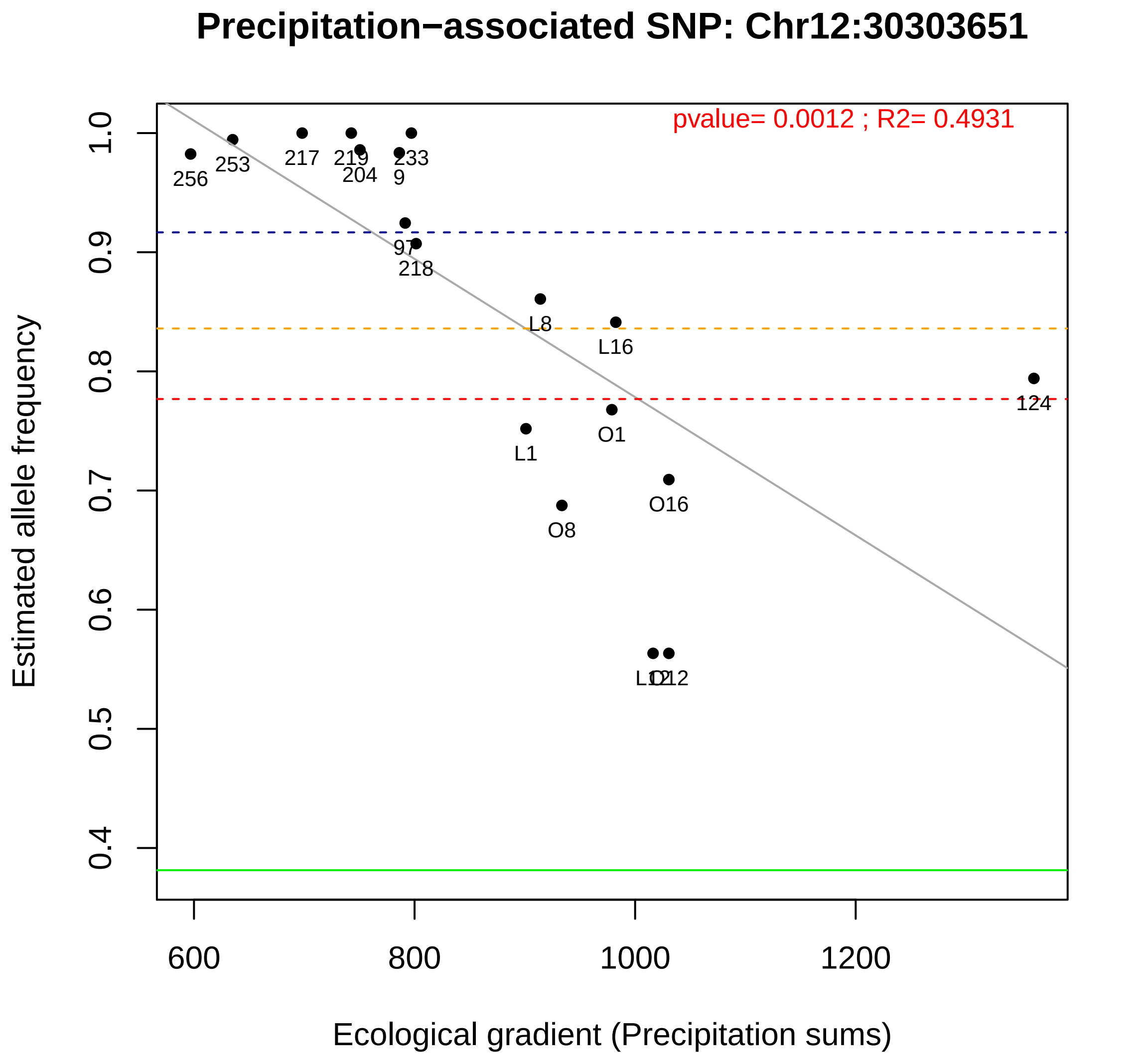

Precipitation−associated SNP: Chr12:30303651
pvalue= 0.0012 ; R2= 0.4931
Estimated allele frequency
Ecological gradient (Precipitation sums)
0.4
0.5
0.6
0.7
0.8
0.9
1.0
600
800
1000
1200
256
253
217
219
204
233
9
97
218
L8
L16
124
O1
L1
O16
O8
L12
O12

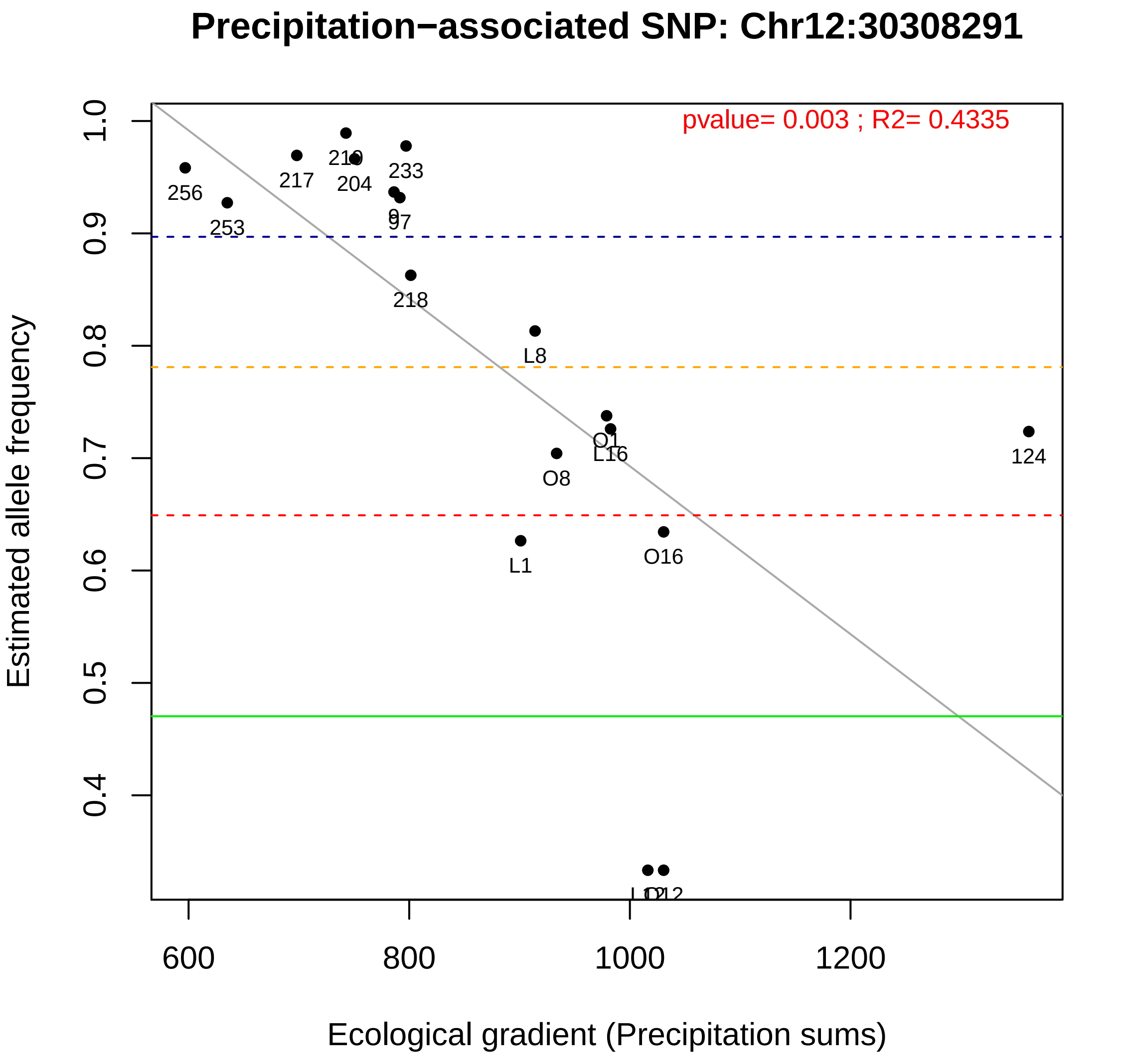

Precipitation−associated SNP: Chr12:30308291
pvalue= 0.003 ; R2= 0.4335
Estimated allele frequency
Ecological gradient (Precipitation sums)
1.0
0.9
0.8
0.7
0.6
0.5
0.4
600
800
1000
1200
256
253
217
210
204
233
9
97
218
L8
O1
L16
O8
124
L1
O16
O12

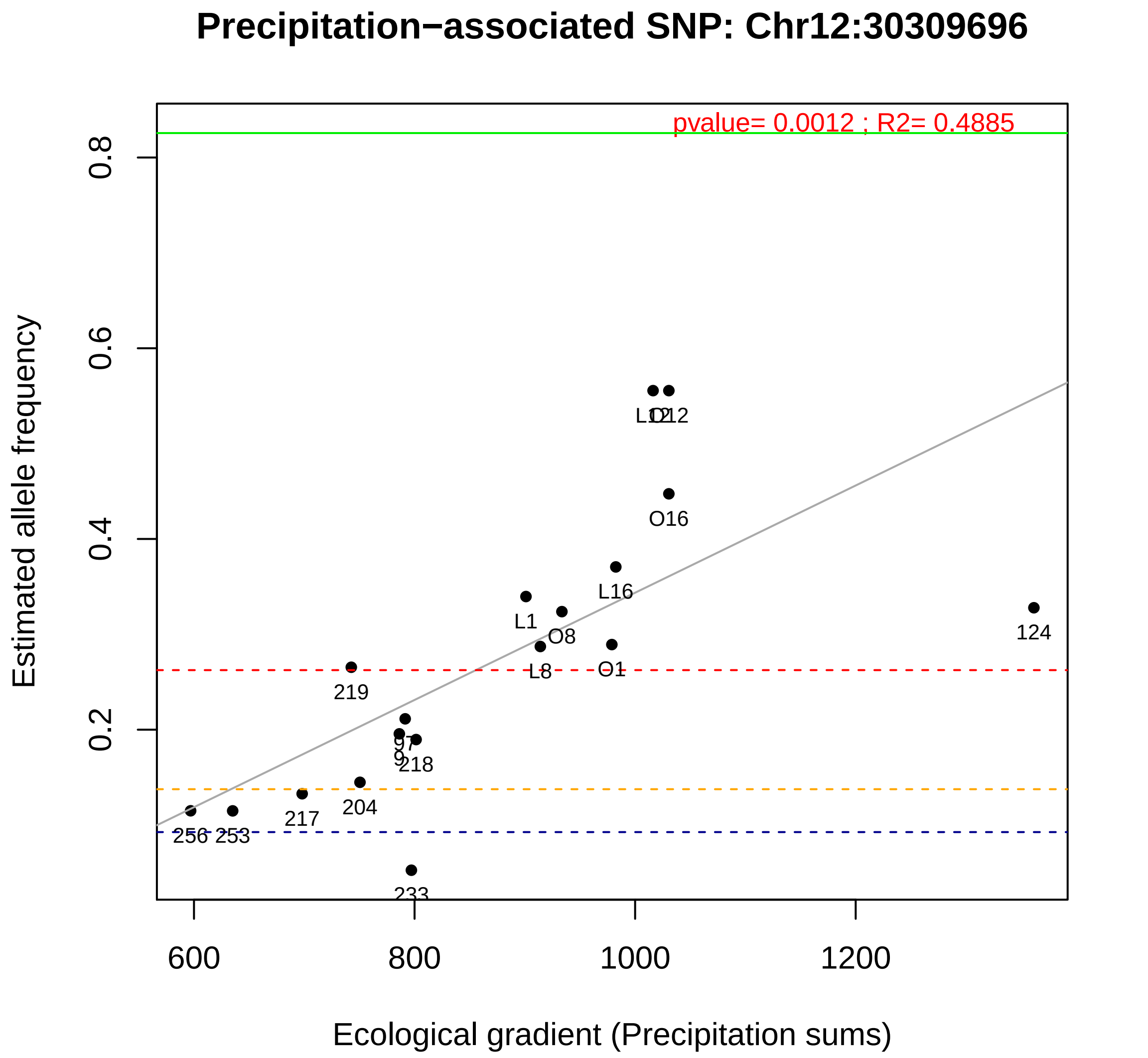
Precipitation−associated SNP: Chr12:30309696
pvalue= 0.0012 ; R2= 0.4885
Estimated allele frequency
Ecological gradient (Precipitation sums)
0.2
0.4
0.6
0.8
600
800
1000
1200
L12
O12
O16
L16
L1
O8
L8
O1
124
219
97
9
218
204
217
256
253
233

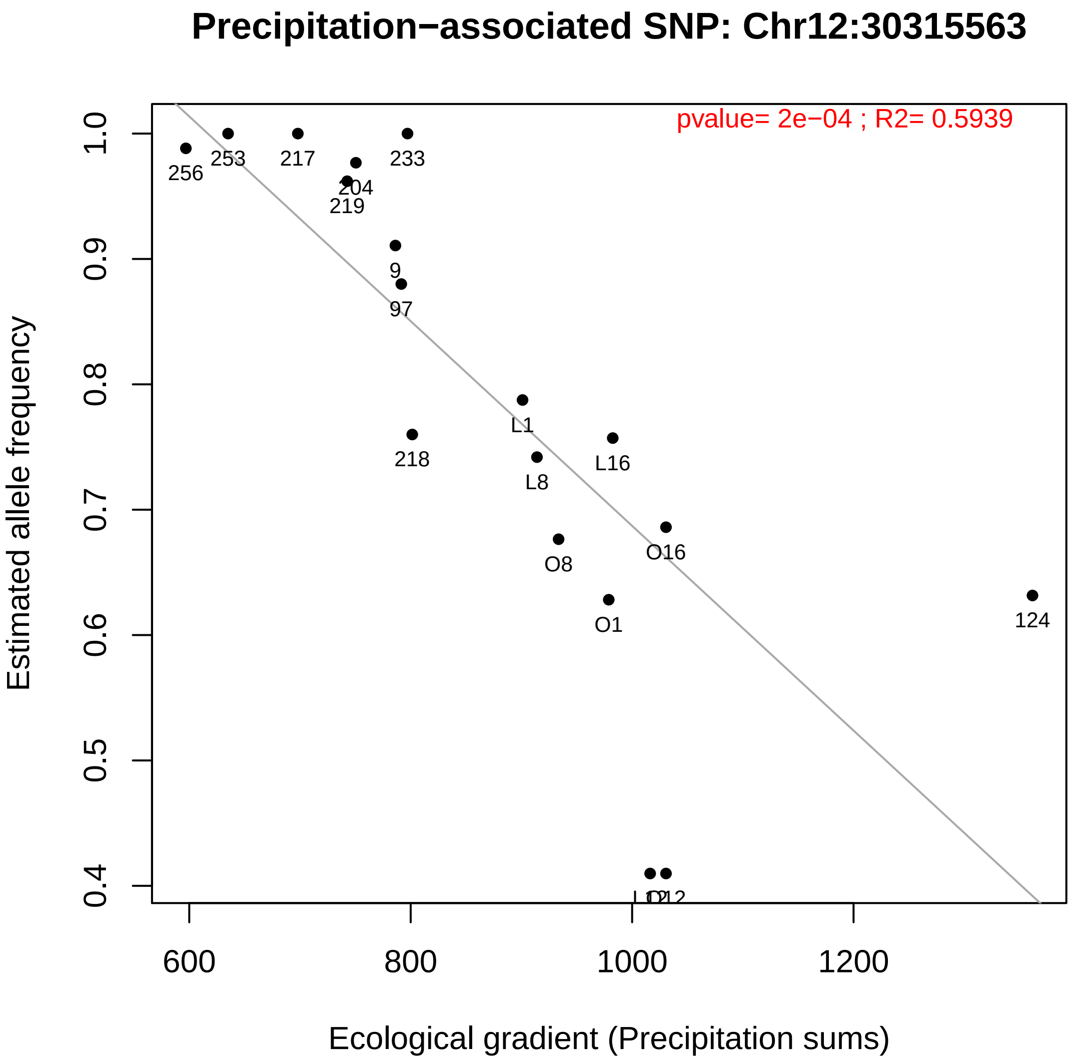
Precipitation−associated SNP: Chr12:30315563
pvalue= 2e−04 ; R2= 0.5939
Estimated allele frequency
Ecological gradient (Precipitation sums)
0.4
0.5
0.6
0.7
0.8
0.9
1.0
600
800
1000
1200
256
253
217
233
204
219
9
97
218
L1
L8
L16
O16
O8
O1
124

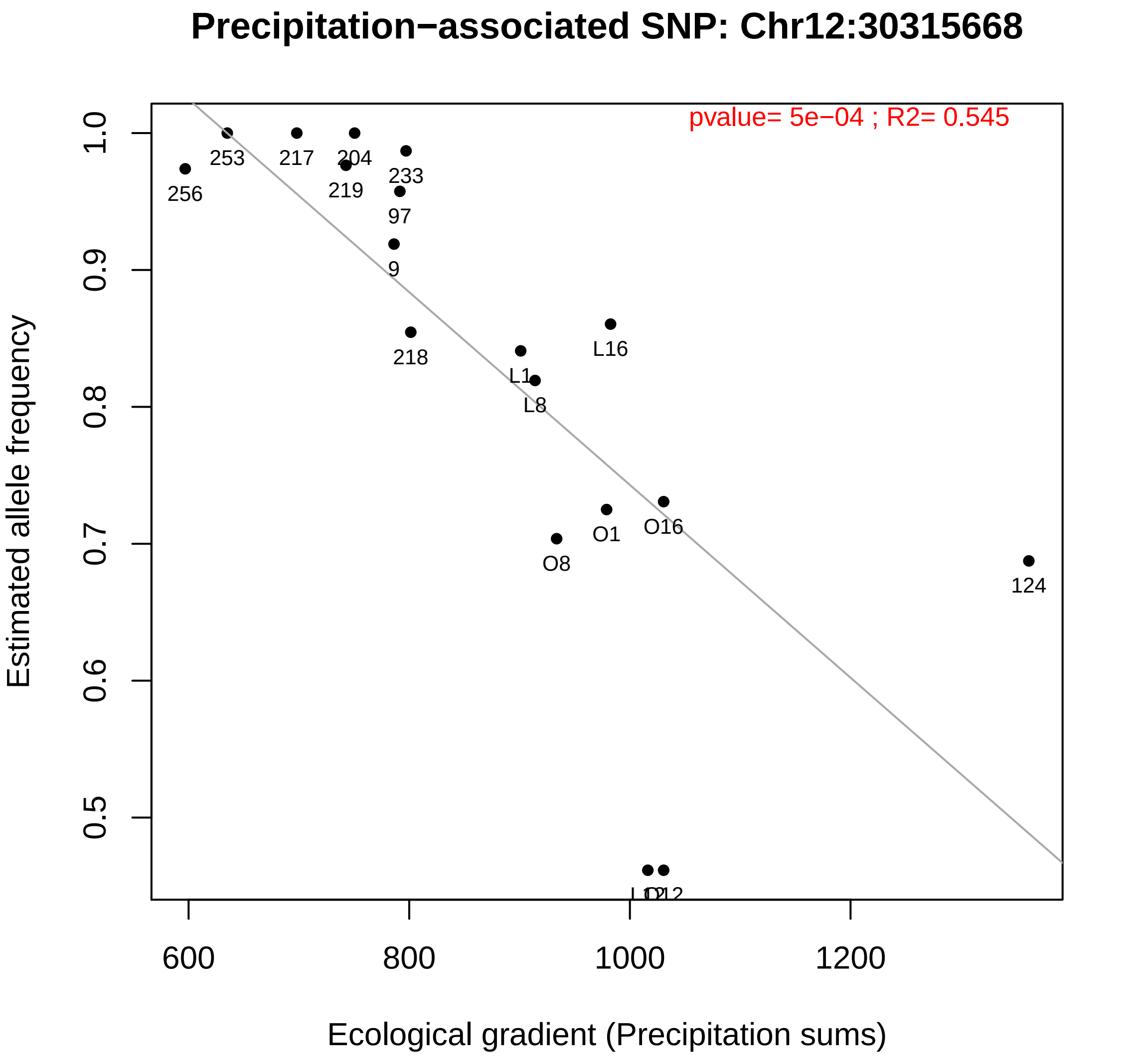
Precipitation−associated SNP: Chr12:30315668
pvalue= 5e−04 ; R2= 0.545
Estimated allele frequency
Ecological gradient (Precipitation sums)
0.5
0.6
0.7
0.8
0.9
1.0
600
800
1000
1200
256
253
217
204
219
233
97
9
218
L1
L8
L16
O8
O1
O16
124

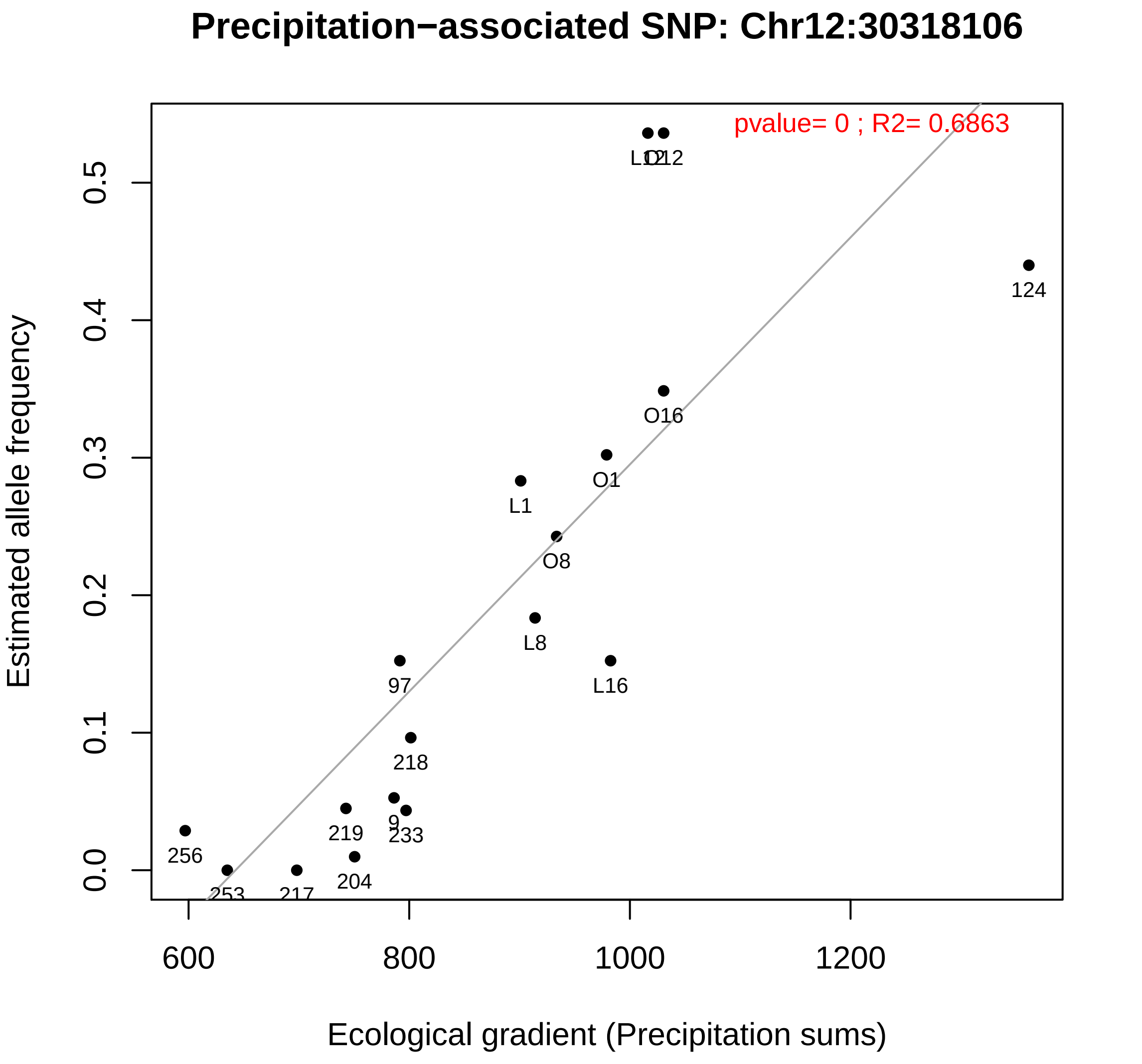
Precipitation−associated SNP: Chr12:30318106
pvalue= 0 ; R2= 0.6863
Estimated allele frequency
Ecological gradient (Precipitation sums)
0.0
0.1
0.2
0.3
0.4
0.5
600
800
1000
1200
L12
O12
124
O16
O1
L1
O8
L8
L16
97
218
219
9
233
256
204
253
217

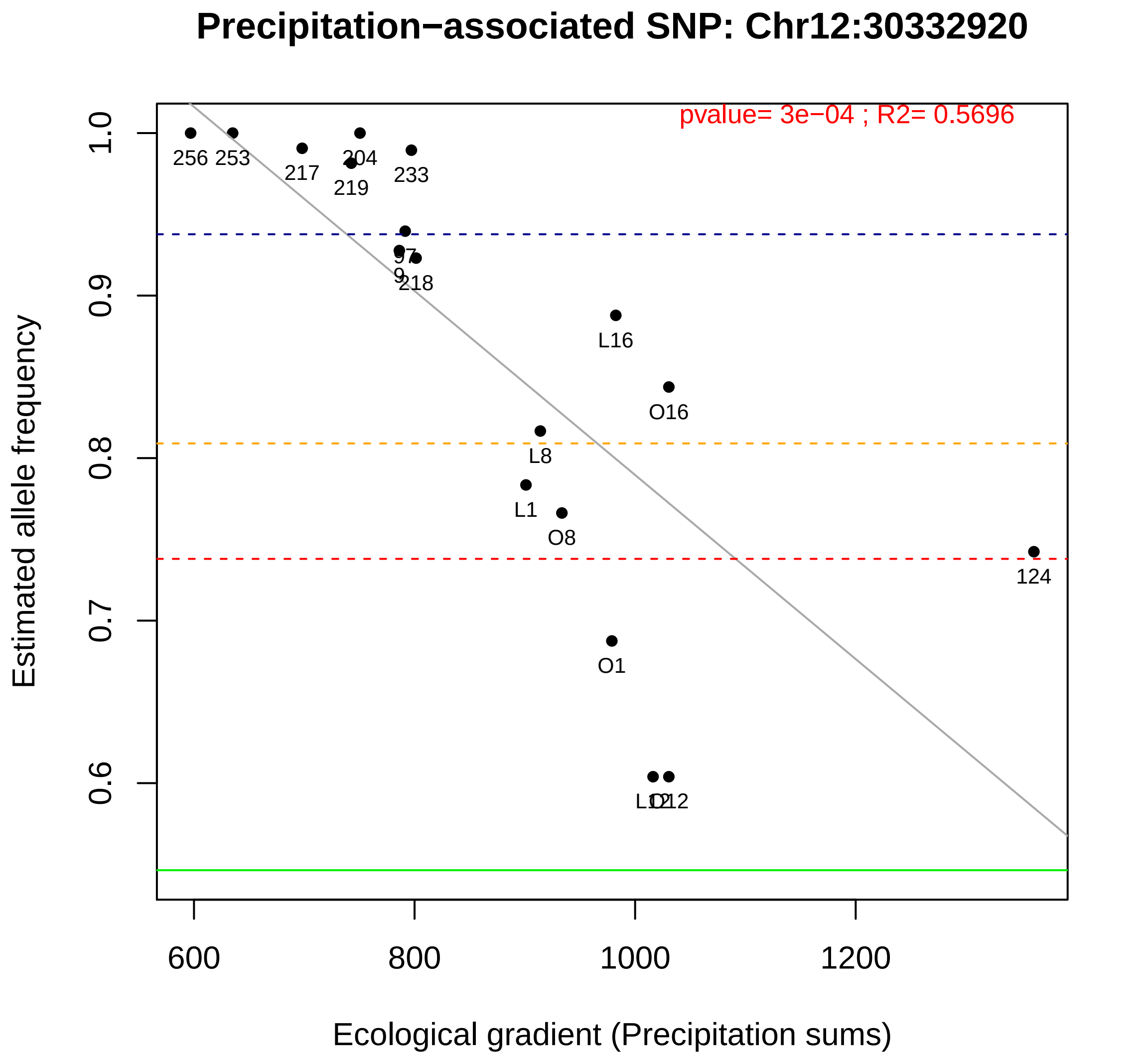

Precipitation−associated SNP: Chr12:30332920
pvalue= 3e−04 ; R2= 0.5696
Estimated allele frequency
Ecological gradient (Precipitation sums)
1.0
0.9
0.8
0.7
0.6
600
800
1000
1200
256
253
217
204
219
233
97
9
218
L16
O16
L8
L1
O8
124
O1
L12
O12

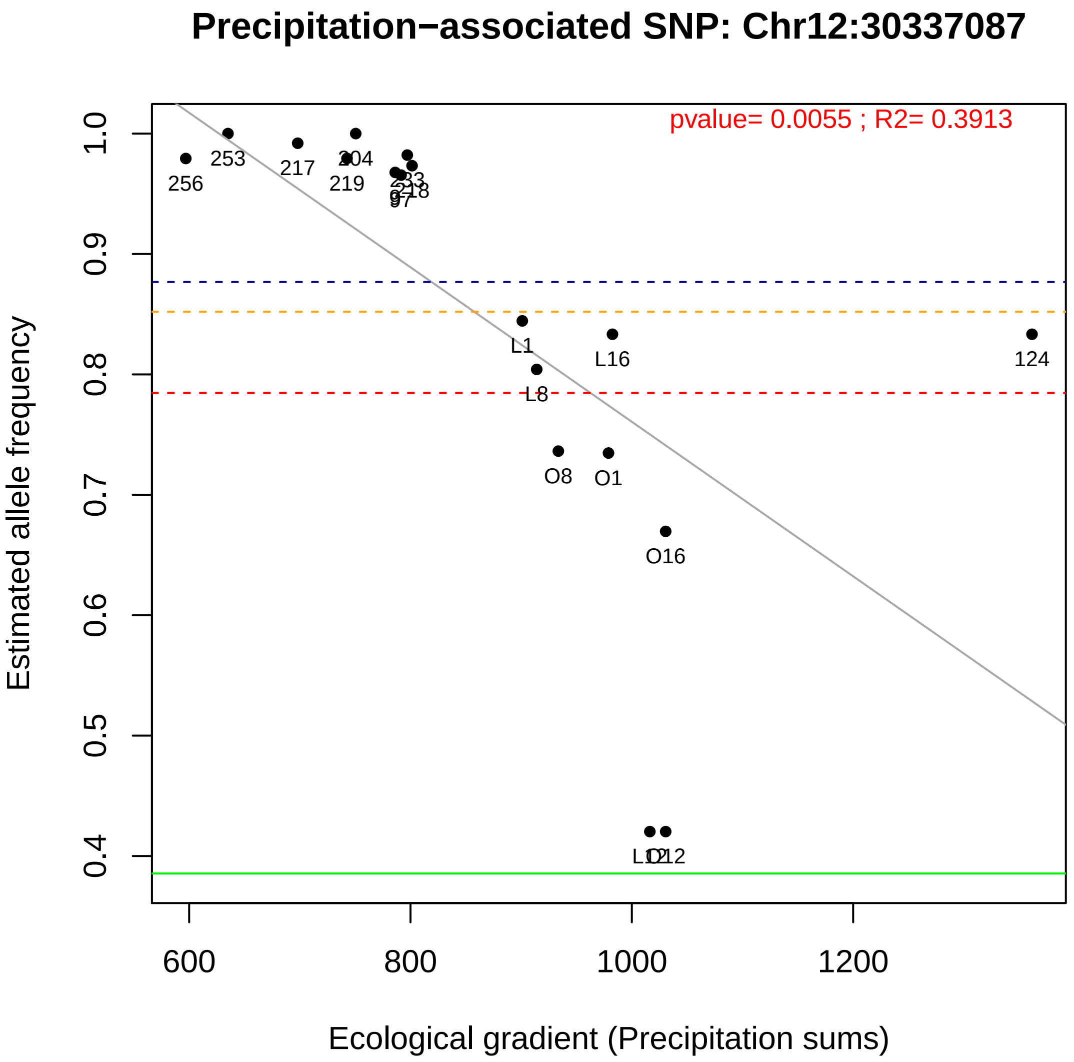
Precipitation−associated SNP: Chr12:30337087
pvalue= 0.0055 ; R2= 0.3913
Estimated allele frequency
Ecological gradient (Precipitation sums)
0.4
0.5
0.6
0.7
0.8
0.9
1.0
600
800
1000
1200
256
253
217
204
219
233
218
97
L1
L16
124
L8
O8
O1
O16
O12

**Precipitation−associated SNP: Chr12:30344934**

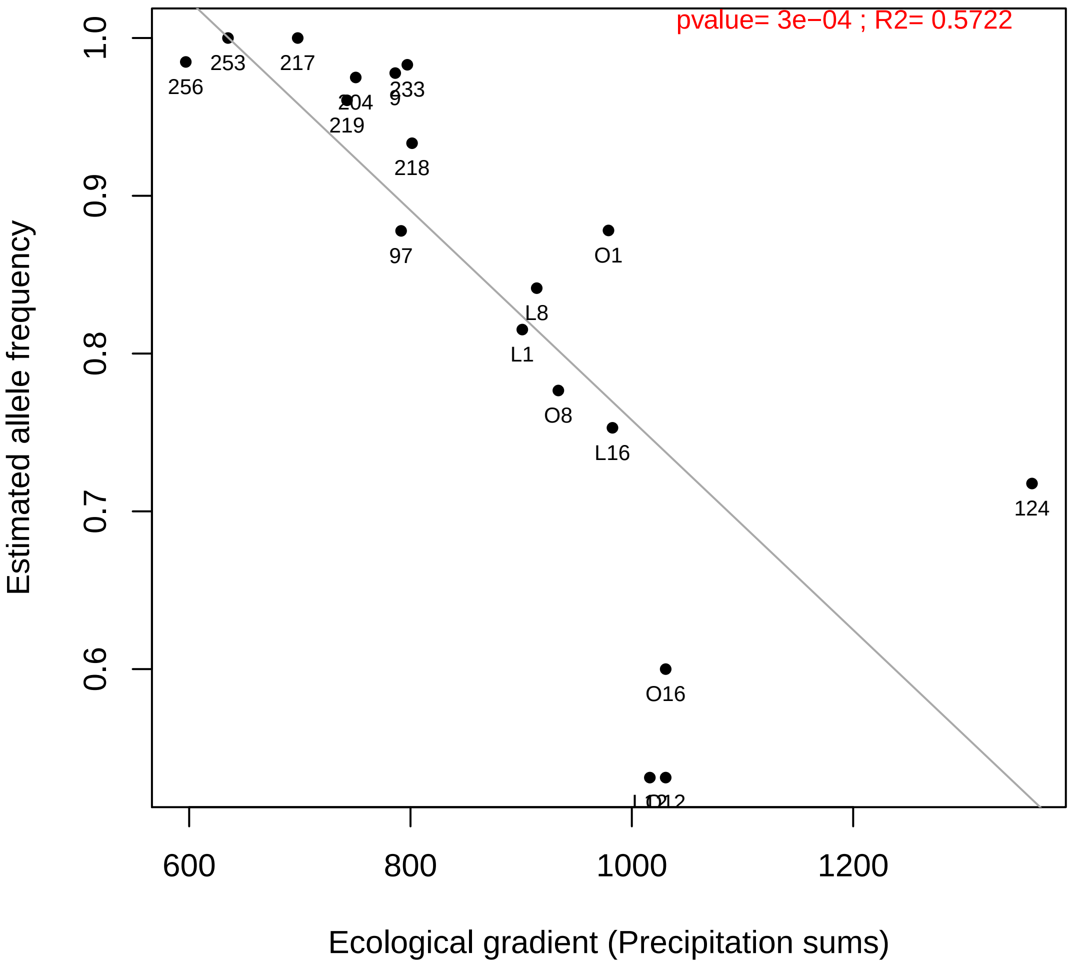

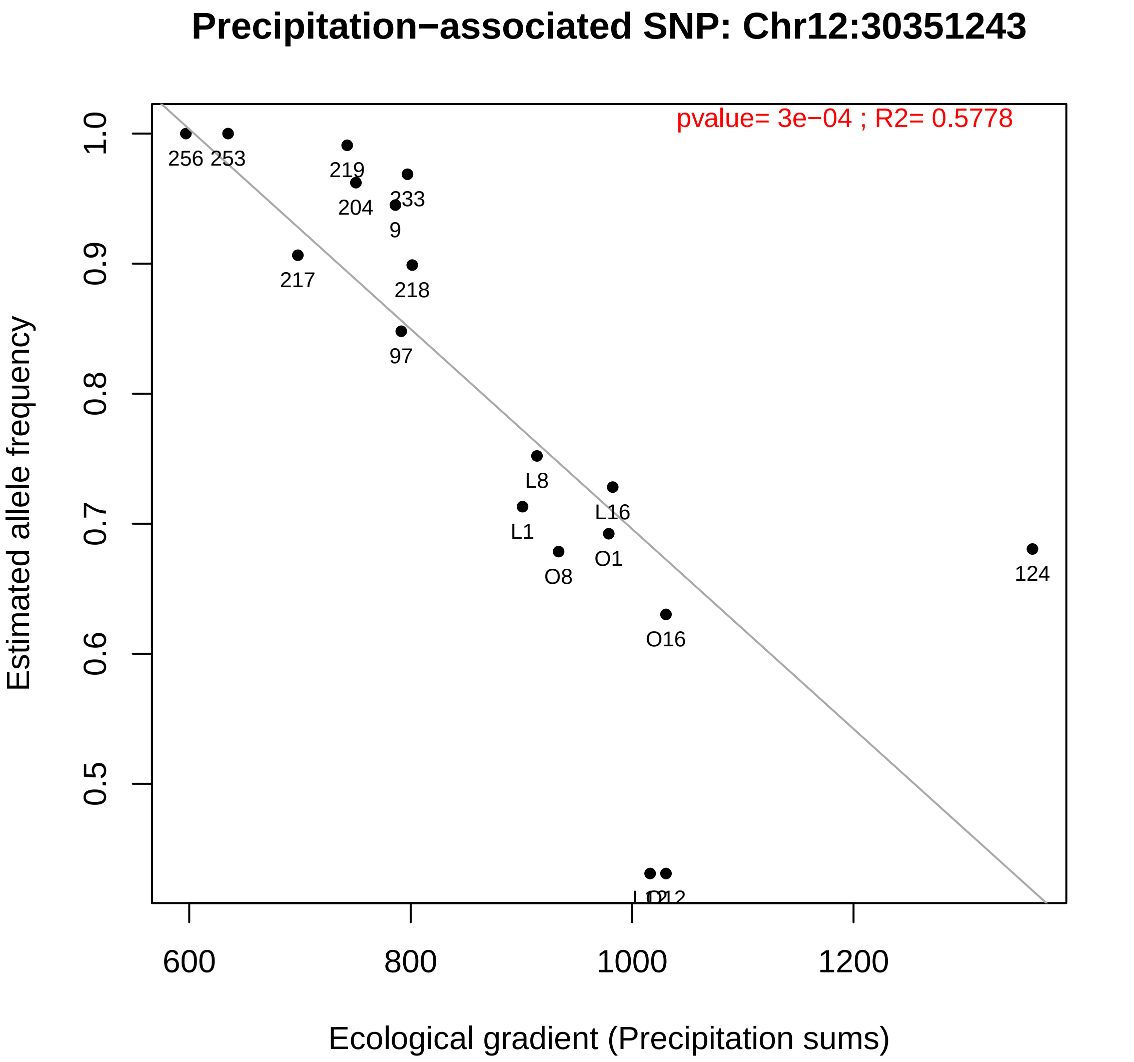
Precipitation−associated SNP: Chr12:30351243
pvalue= 3e−04 ; R2= 0.5778
Estimated allele frequency
Ecological gradient (Precipitation sums)
1.0
0.9
0.8
0.7
0.6
0.5
600
800
1000
1200
256
253
219
204
233
9
217
218
97
L8
L16
L1
O1
O8
124
O16

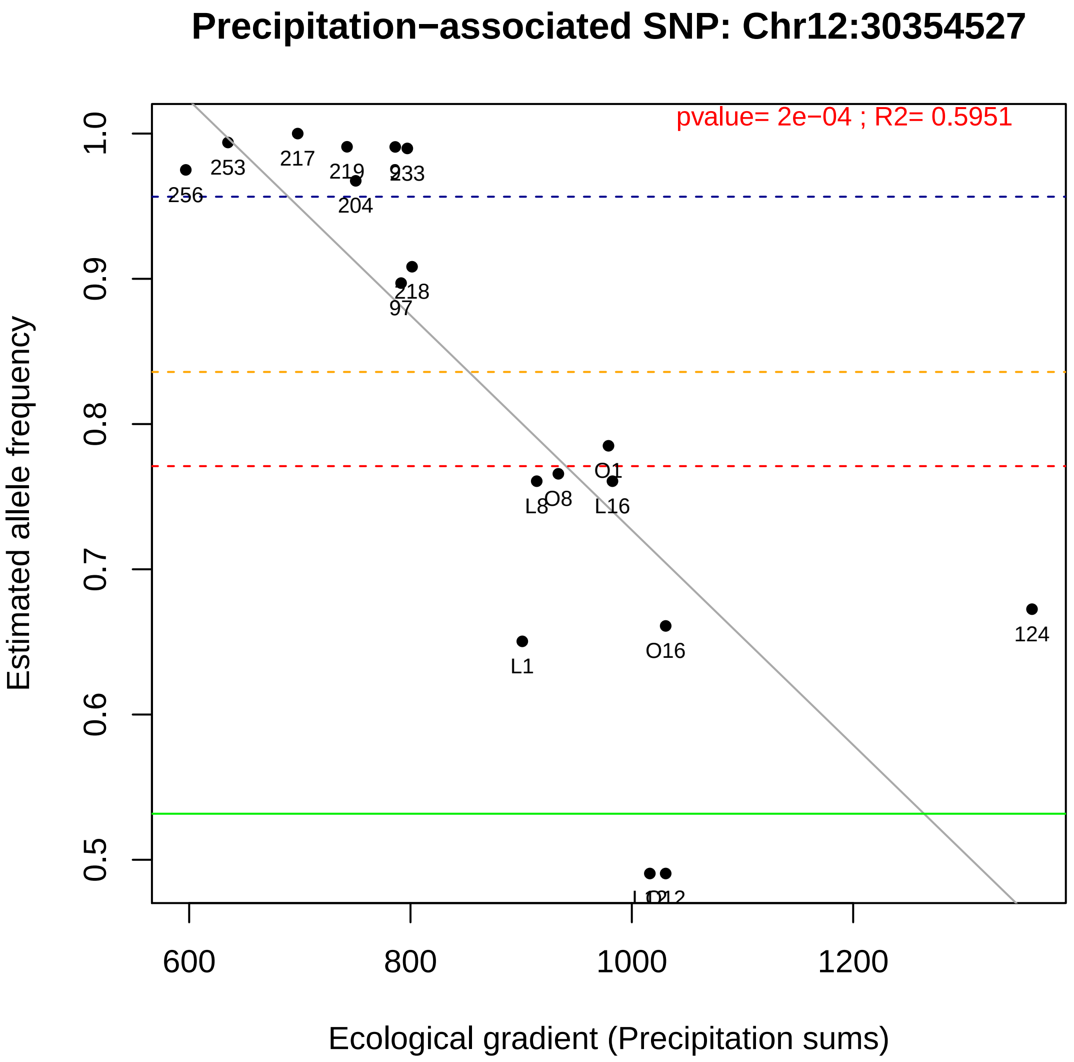

Precipitation−associated SNP: Chr12:30354527
pvalue= 2e−04 ; R2= 0.5951
Estimated allele frequency
Ecological gradient (Precipitation sums)
1.0
0.9
0.8
0.7
0.6
0.5
600
800
1000
1200
256
253
217
219
204
233
218
97
L8
O8
O1
L16
L1
O16
124
O12

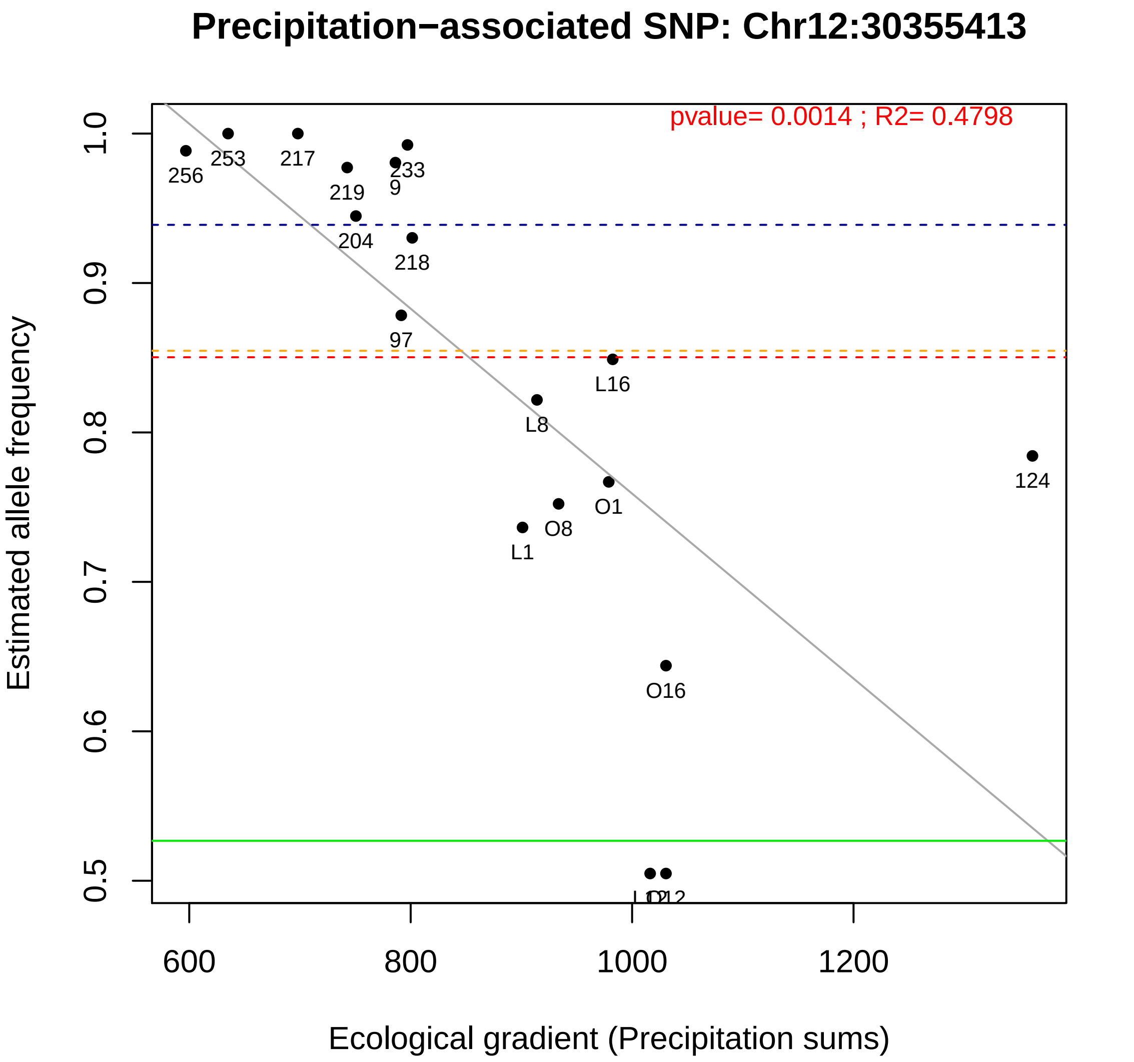

Precipitation−associated SNP: Chr12:30355413
pvalue= 0.0014 ; R2= 0.4798
Estimated allele frequency
Ecological gradient (Precipitation sums)
1.0
0.9
0.8
0.7
0.6
0.5
600
800
1000
1200
256
253
217
219
233
9
204
218
97
L16
L8
124
O1
O8
L1
O16

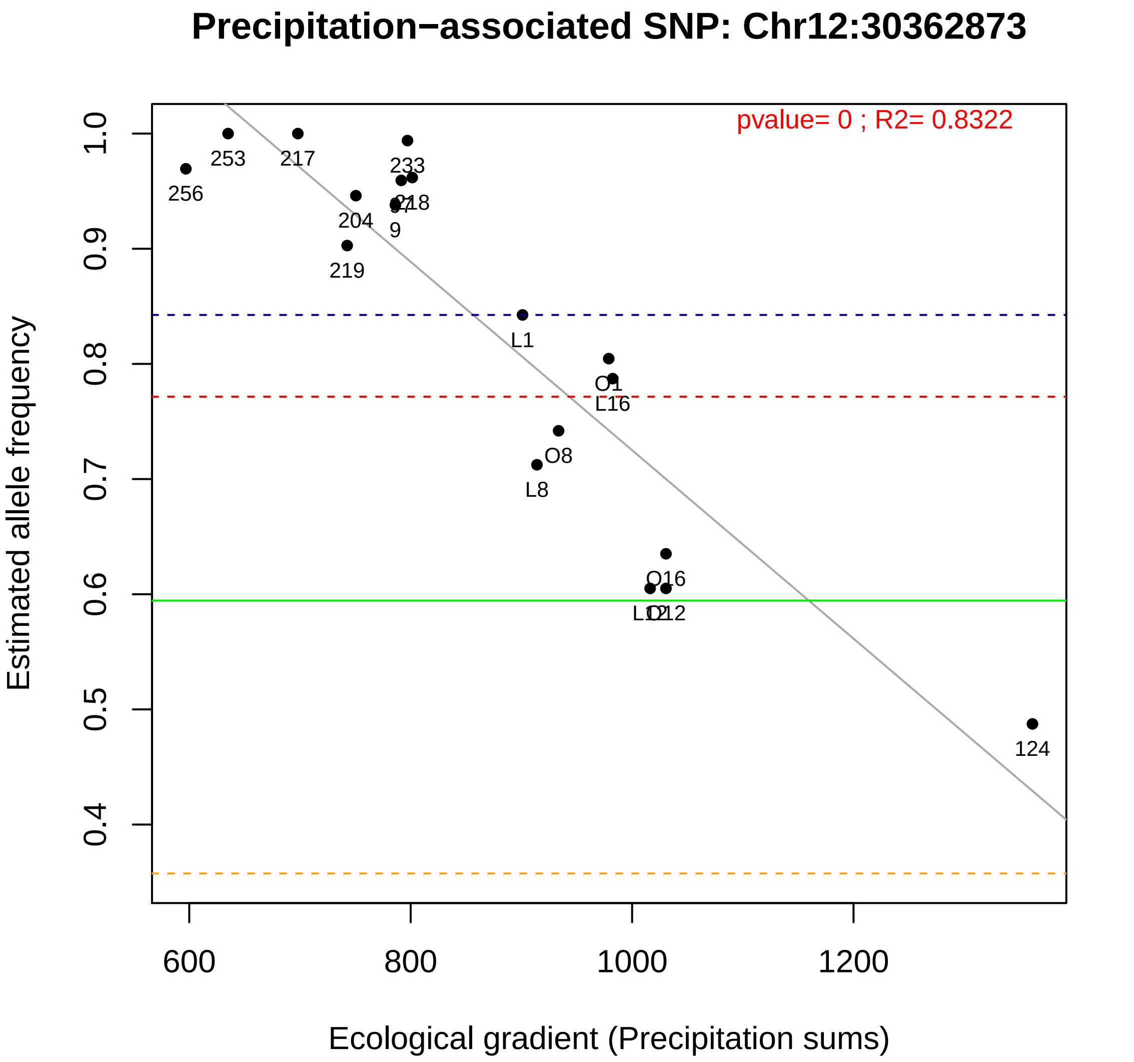

Precipitation−associated SNP: Chr12:30362873
pvalue= 0 ; R2= 0.8322
Estimated allele frequency
Ecological gradient (Precipitation sums)
0.4
0.5
0.6
0.7
0.8
0.9
1.0
600
800
1000
1200
256
253
217
233
218
204
9
219
L1
O1
L16
O8
L8
O16
O12
124

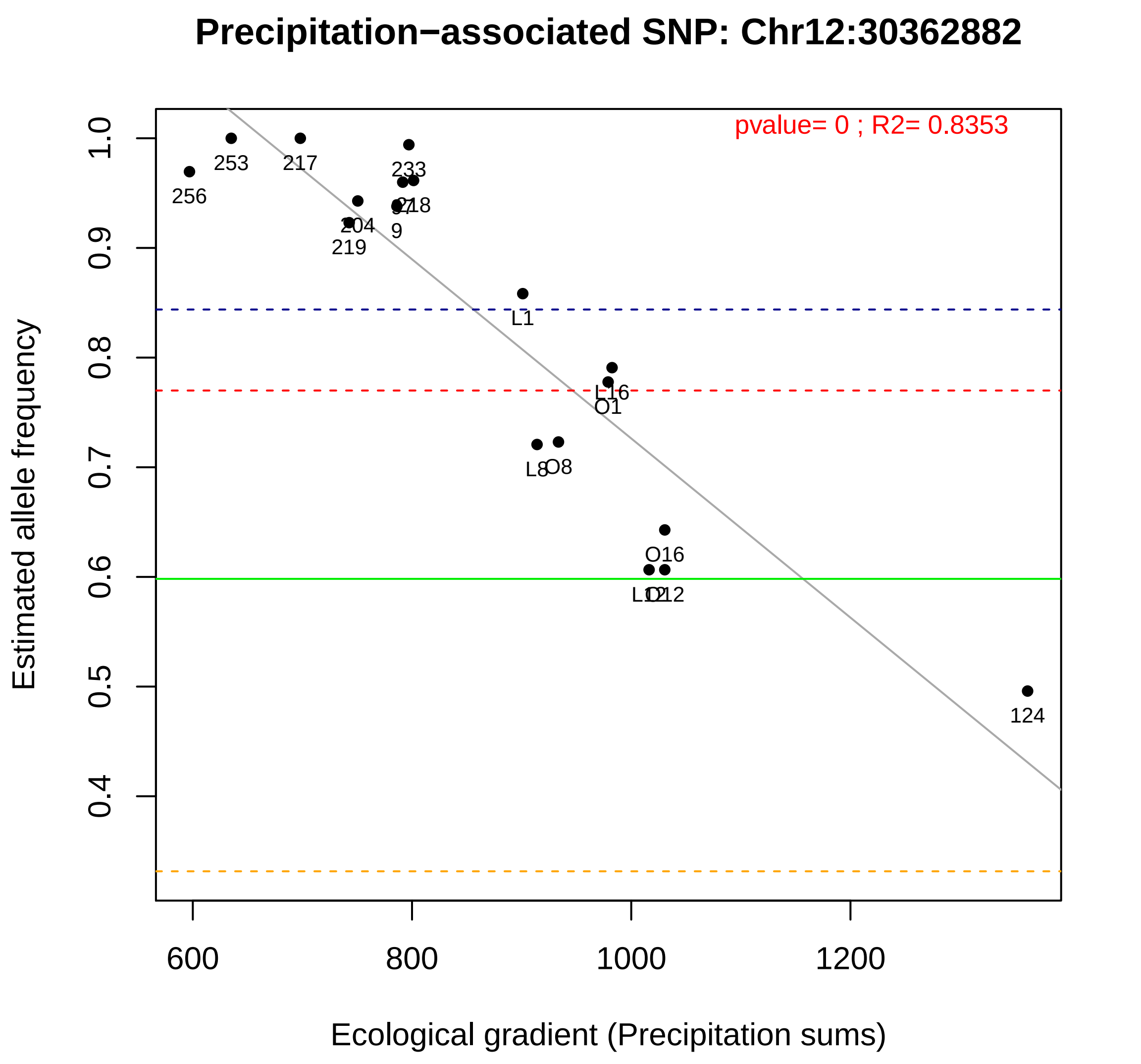
Precipitation−associated SNP: Chr12:30362882
pvalue= 0 ; R2= 0.8353
Estimated allele frequency
Ecological gradient (Precipitation sums)
0.4
0.5
0.6
0.7
0.8
0.9
1.0
600
800
1000
1200
256
253
217
233
218
204
219
9
L1
L16
O1
L8
O8
O16
124

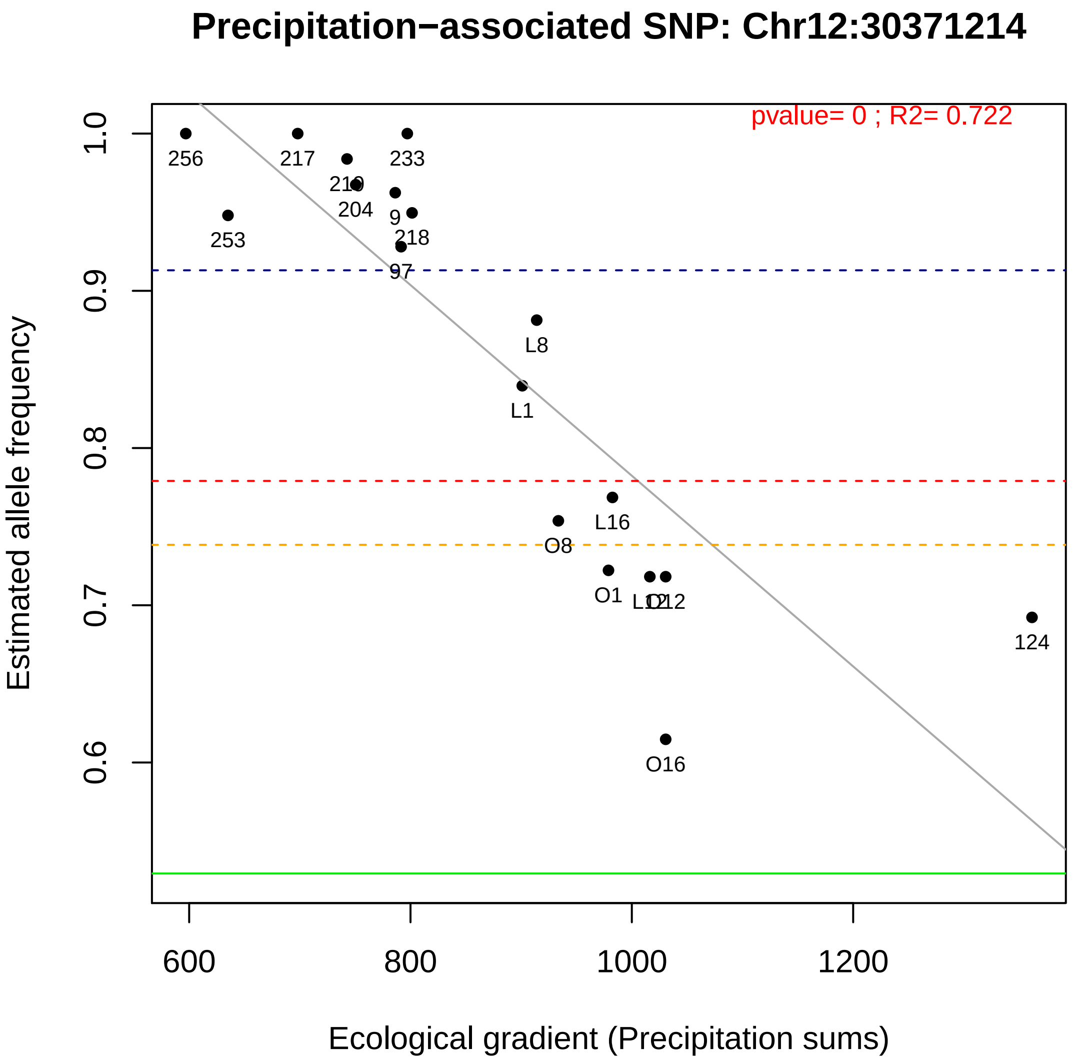

Precipitation−associated SNP: Chr12:30371214
pvalue= 0 ; R2= 0.722
Estimated allele frequency
Ecological gradient (Precipitation sums)
1.0
0.9
0.8
0.7
0.6
600
800
1000
1200
256
217
233
210
204
9
218
253
97
L8
L1
L16
O8
O1
L12
O12
124
O16

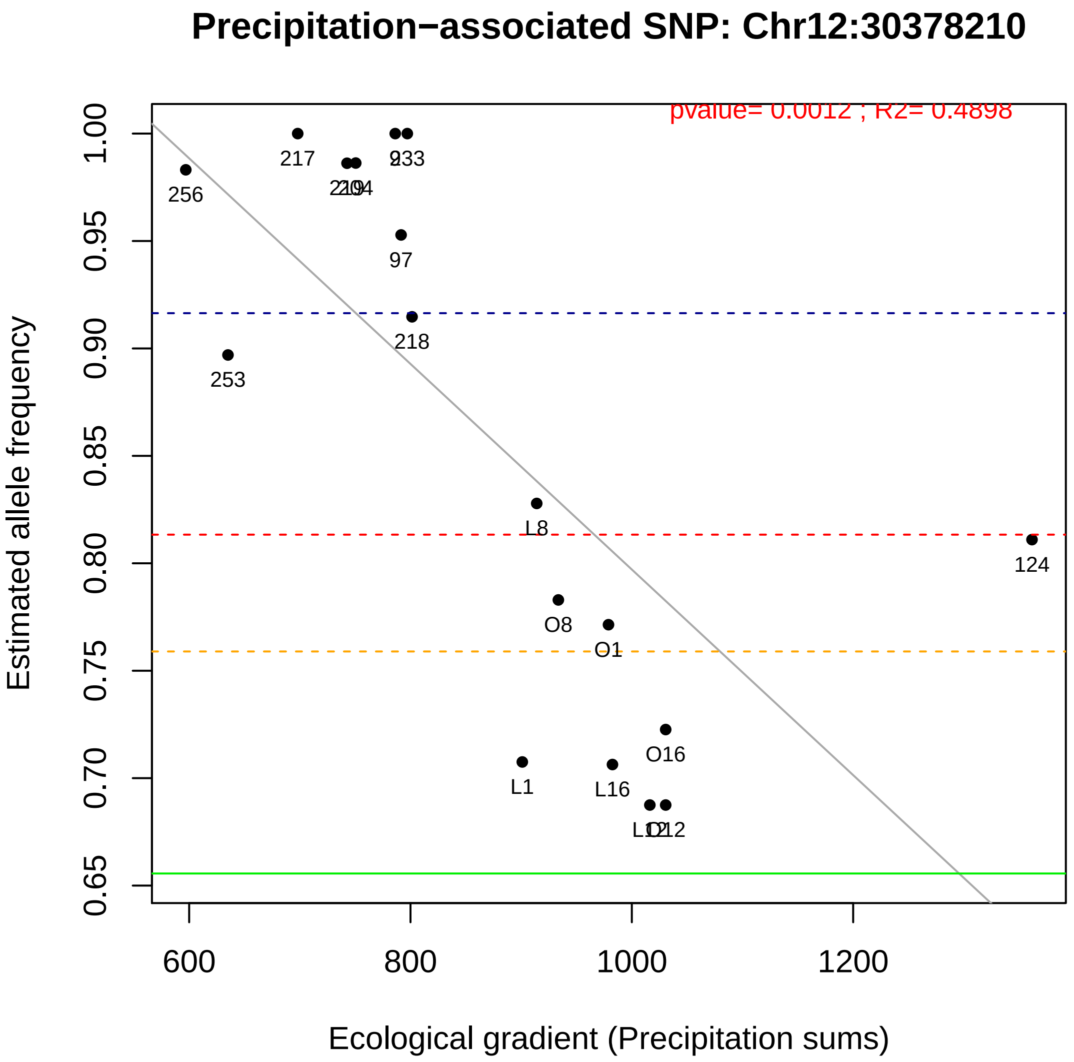
Precipitation−associated SNP: Chr12:30378210
pvalue= 0.0012 ; R2= 0.4898
Estimated allele frequency
Ecological gradient (Precipitation sums)
1.00
0.95
0.90
0.85
0.80
0.75
0.70
0.65
600
800
1000
1200
217
256
97
218
253
L8
124
O8
O1
O16
L1
L16

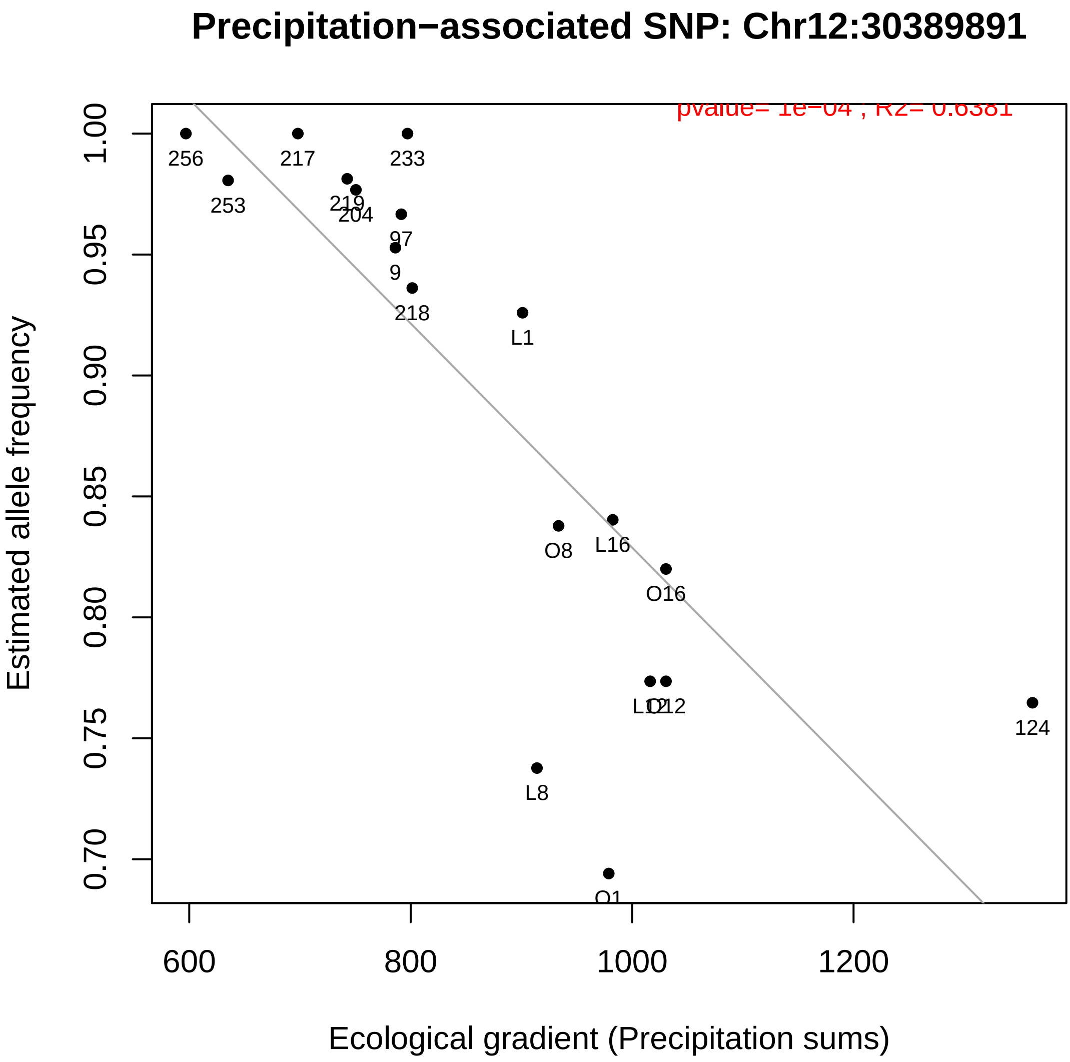
Precipitation−associated SNP: Chr12:30389891
pvalue= 1e−04 ; R2= 0.6381
Estimated allele frequency
Ecological gradient (Precipitation sums)
0.70
0.75
0.80
0.85
0.90
0.95
1.00
600
800
1000
1200
256
253
217
219
204
233
97
9
218
L1
O8
L16
O16
O12
124
L8
O1

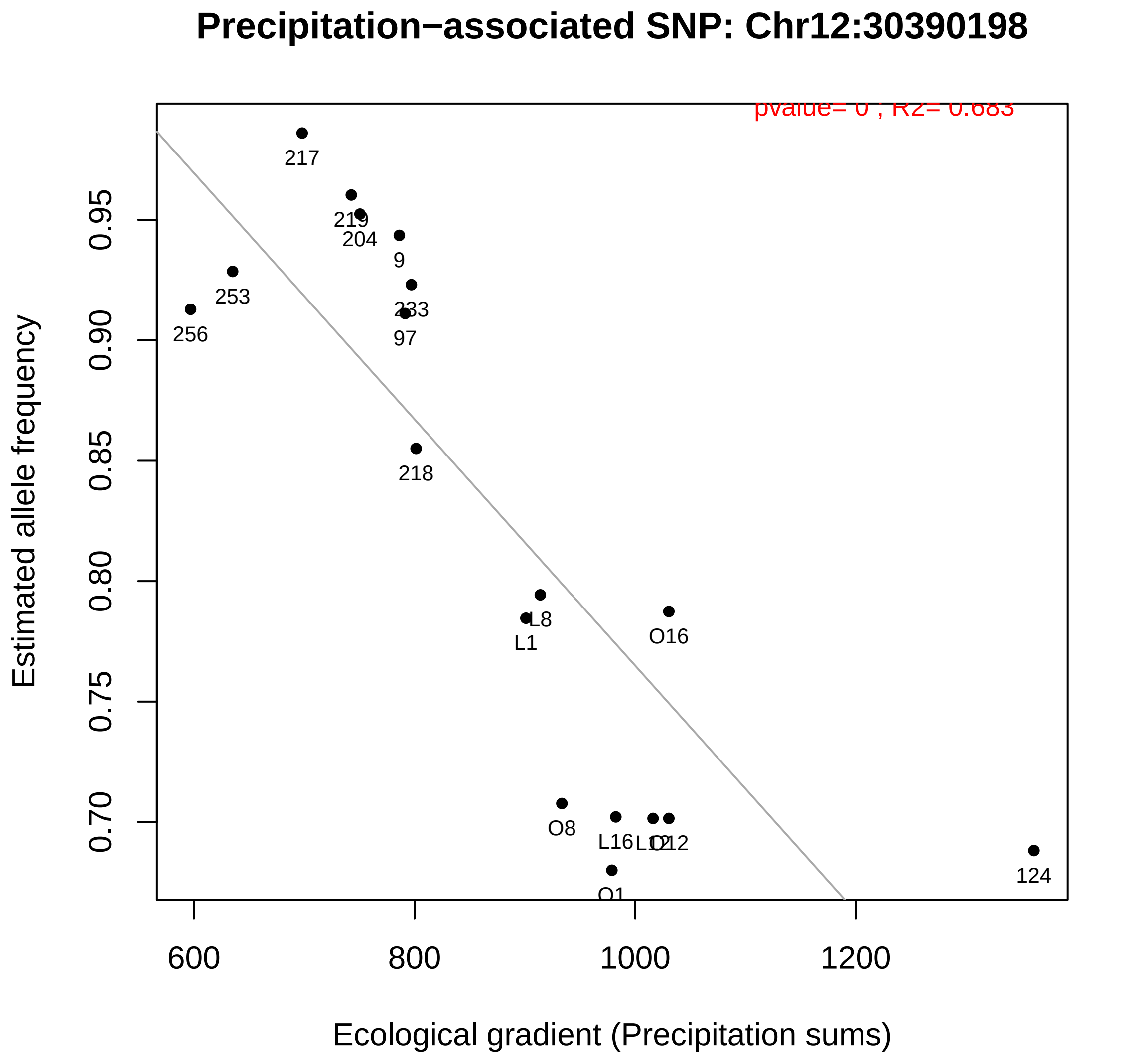

Precipitation−associated SNP: Chr12:30390198
pvalue= 0 ; R2= 0.683
Estimated allele frequency
0.70
0.75
0.80
0.85
0.90
0.95
600
800
1000
1200
Ecological gradient (Precipitation sums)
217
219
204
9
253
256
233
97
218
L8
L1
O16
O8
L16
O12
124

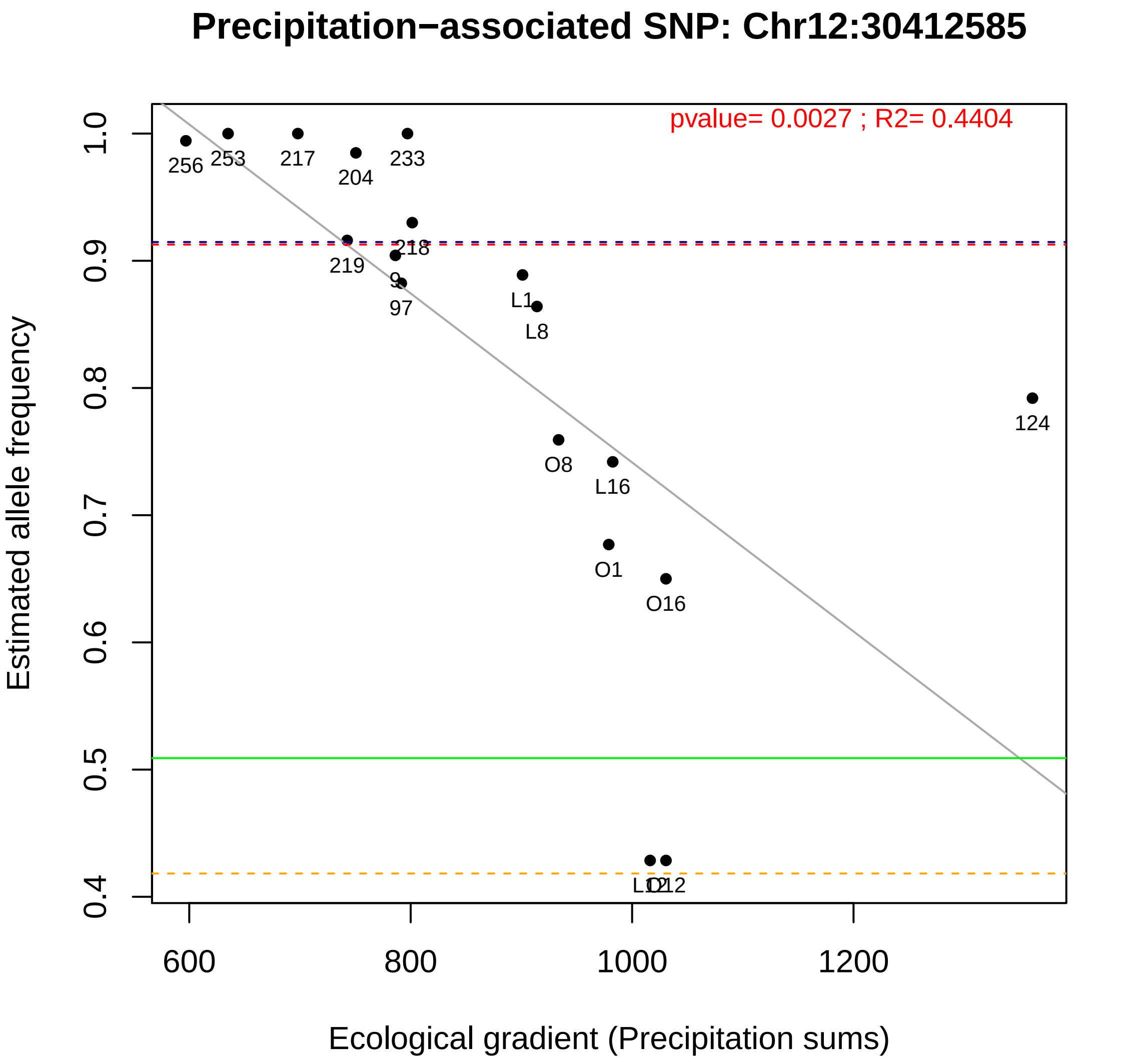

Precipitation−associated SNP: Chr12:30412585
pvalue= 0.0027 ; R2= 0.4404
Estimated allele frequency
Ecological gradient (Precipitation sums)
1.0
0.9
0.8
0.7
0.6
0.5
0.4
600
800
1000
1200
256
253
217
204
233
218
219
9
97
L1
L8
124
O8
L16
O1
O16
L12
O12

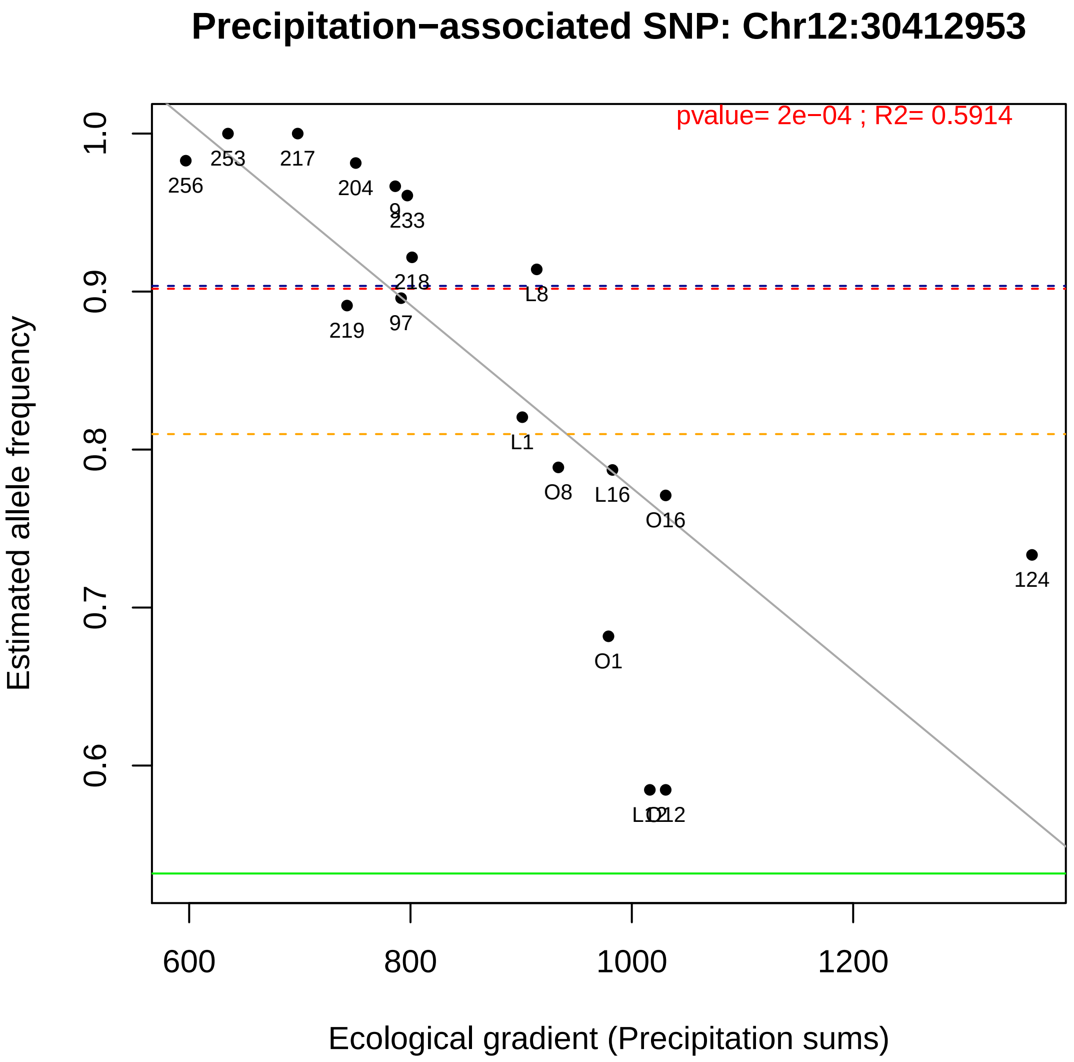
Precipitation−associated SNP: Chr12:30412953
pvalue= 2e−04 ; R2= 0.5914
Estimated allele frequency
1.0
0.9
0.8
0.7
0.6
600
800
1000
1200
Ecological gradient (Precipitation sums)
256
253
217
204
9
233
218
219
97
L8
L1
O8
L16
O16
124
O1
L12
O12

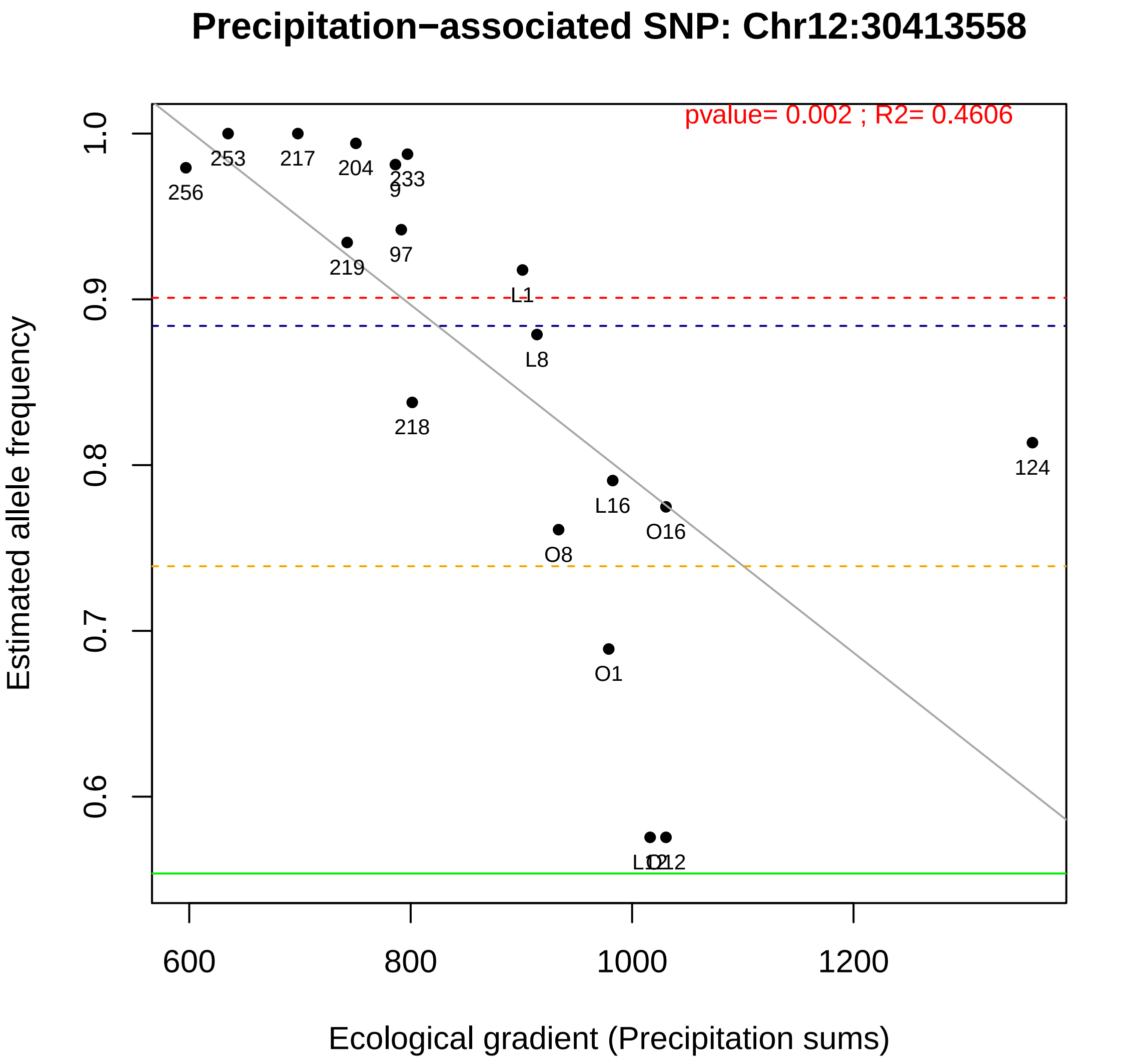
Precipitation−associated SNP: Chr12:30413558
pvalue= 0.002 ; R2= 0.4606
Estimated allele frequency
Ecological gradient (Precipitation sums)
1.0
0.9
0.8
0.7
0.6
600
800
1000
1200
256
253
217
204
233
9
219
97
L1
L8
218
124
L16
O16
O8
O1
L10
O12

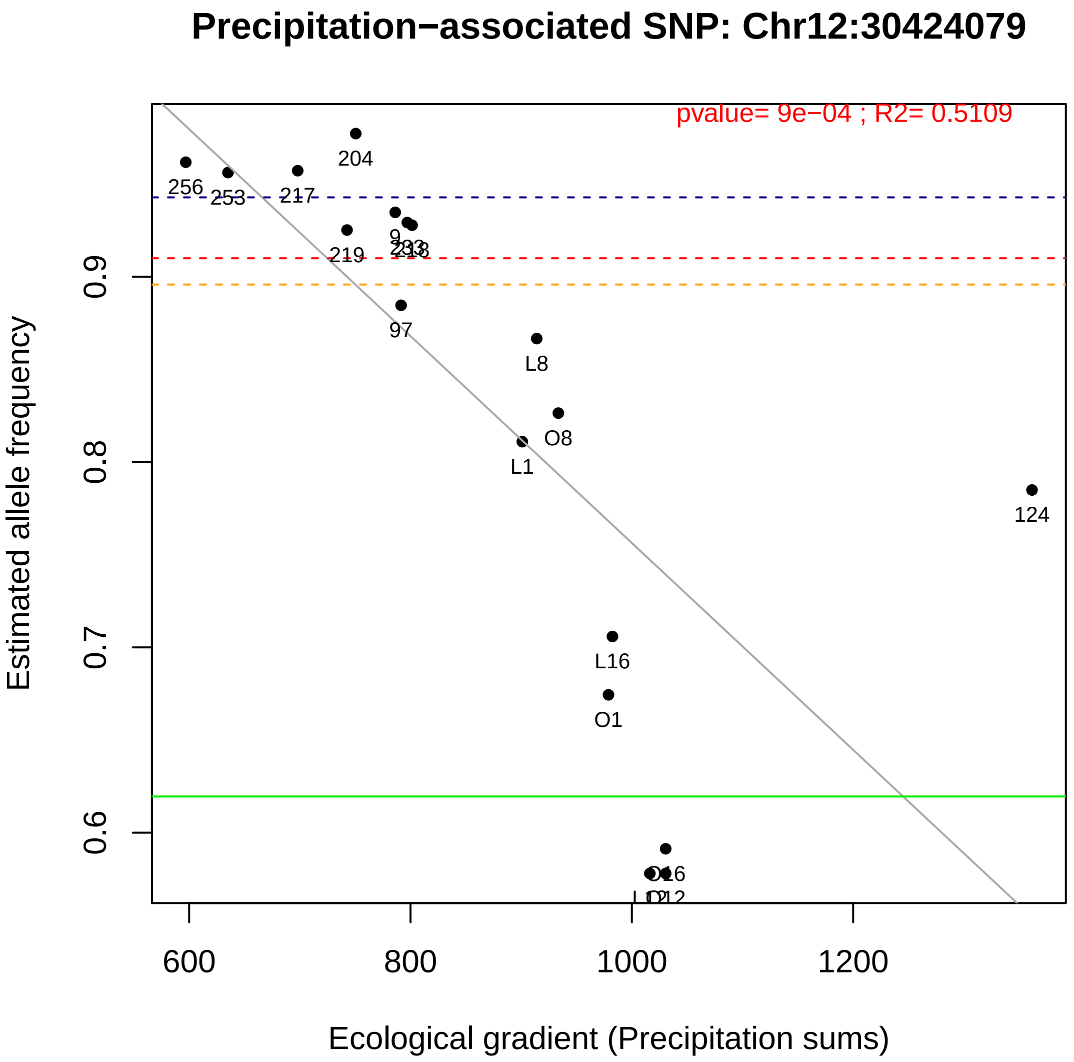

Precipitation−associated SNP: Chr12:30424079
pvalue= 9e−04 ; R2= 0.5109
204
256
253
217
9
233
218
219
97
L8
O8
L1
124
L16
O1
O16
L12
O12
0.6
0.7
0.8
0.9
600
800
1000
1200
Estimated allele frequency
Ecological gradient (Precipitation sums)

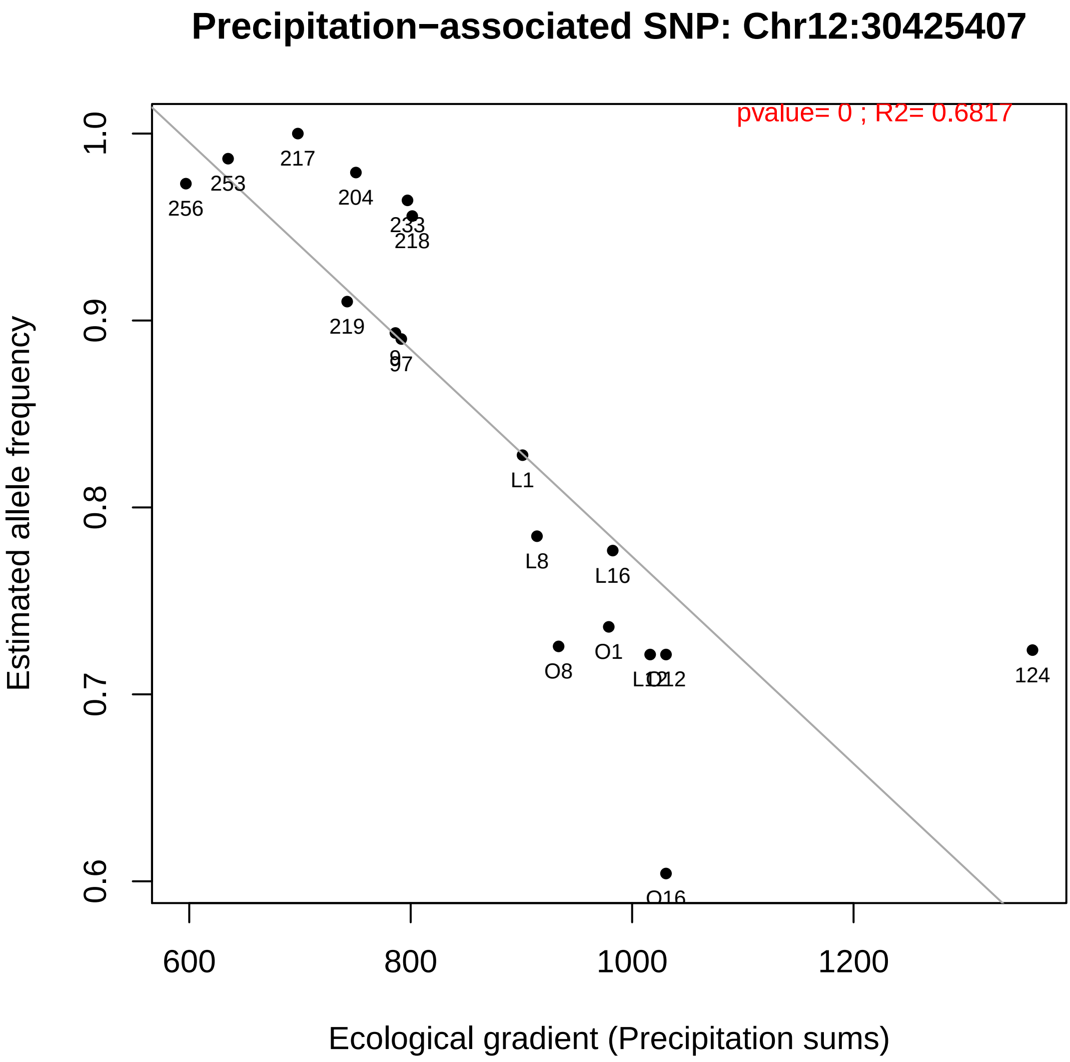

Precipitation−associated SNP: Chr12:30425407
pvalue= 0 ; R2= 0.6817
Estimated allele frequency
Ecological gradient (Precipitation sums)
1.0
0.9
0.8
0.7
0.6
600
800
1000
1200
217
253
256
204
233
218
219
97
L1
L8
L16
O1
O8
O12
124
O16

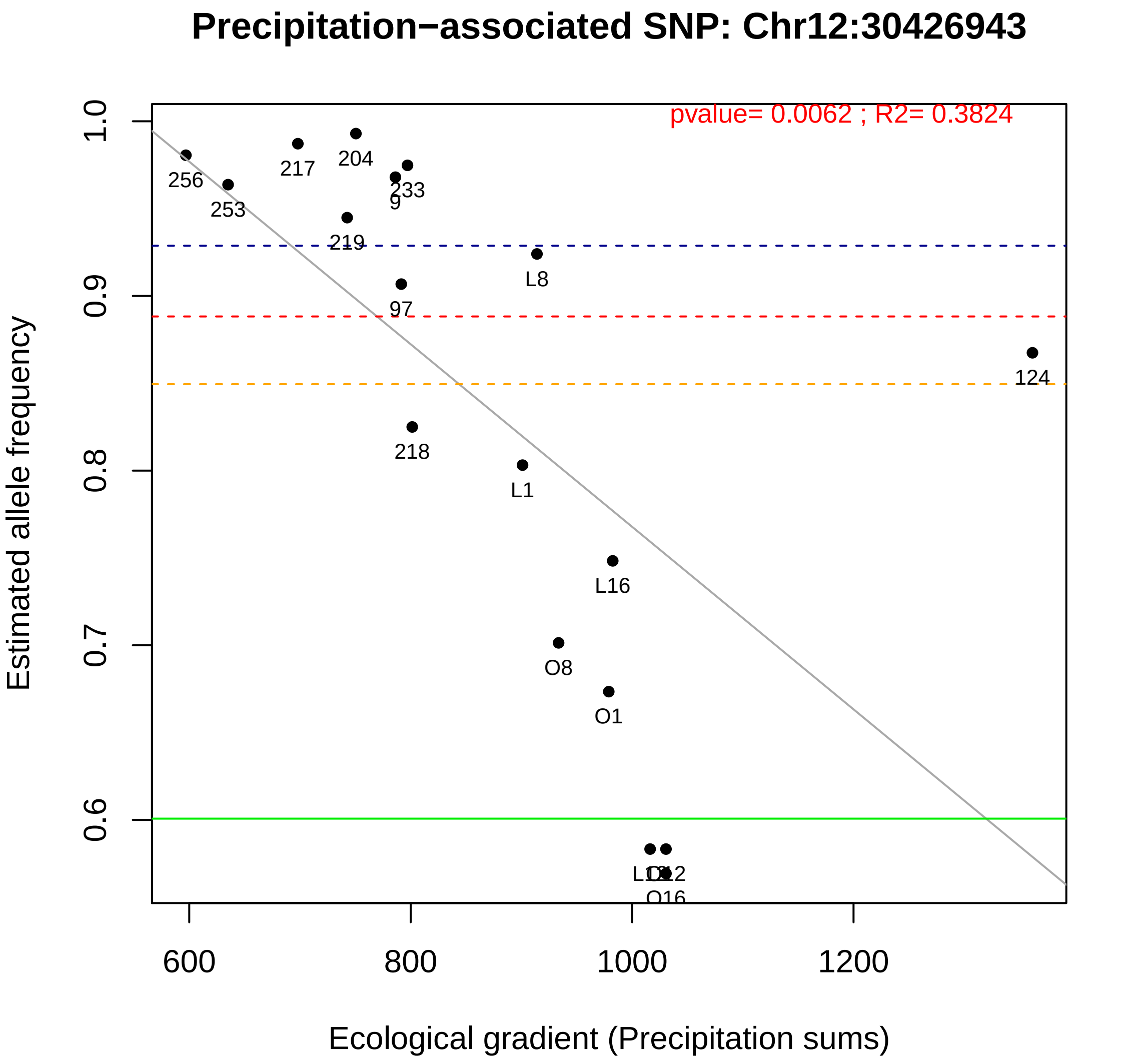
Precipitation−associated SNP: Chr12:30426943
pvalue= 0.0062 ; R2= 0.3824
Estimated allele frequency
Ecological gradient (Precipitation sums)
1.0
0.9
0.8
0.7
0.6
600
800
1000
1200
256
253
217
204
233
9
219
L8
97
124
218
L1
L16
O8
O1
O16

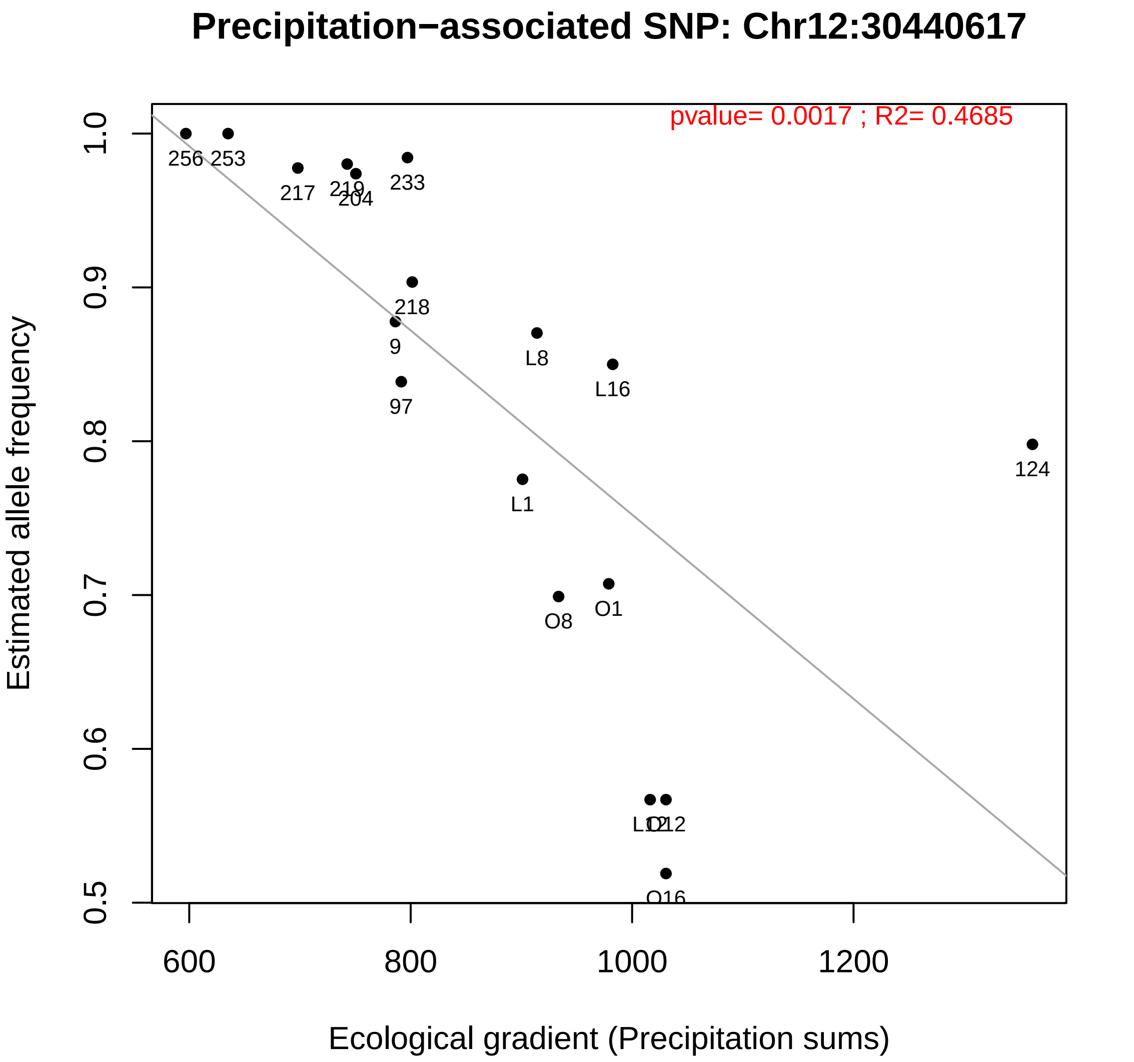

Precipitation−associated SNP: Chr12:30440617
pvalue= 0.0017 ; R2= 0.4685
Estimated allele frequency
Ecological gradient (Precipitation sums)
1.0
0.9
0.8
0.7
0.6
0.5
600
800
1000
1200
256
253
217
219
204
233
218
9
97
L8
L16
L1
124
O8
O1
L12
O12
O16

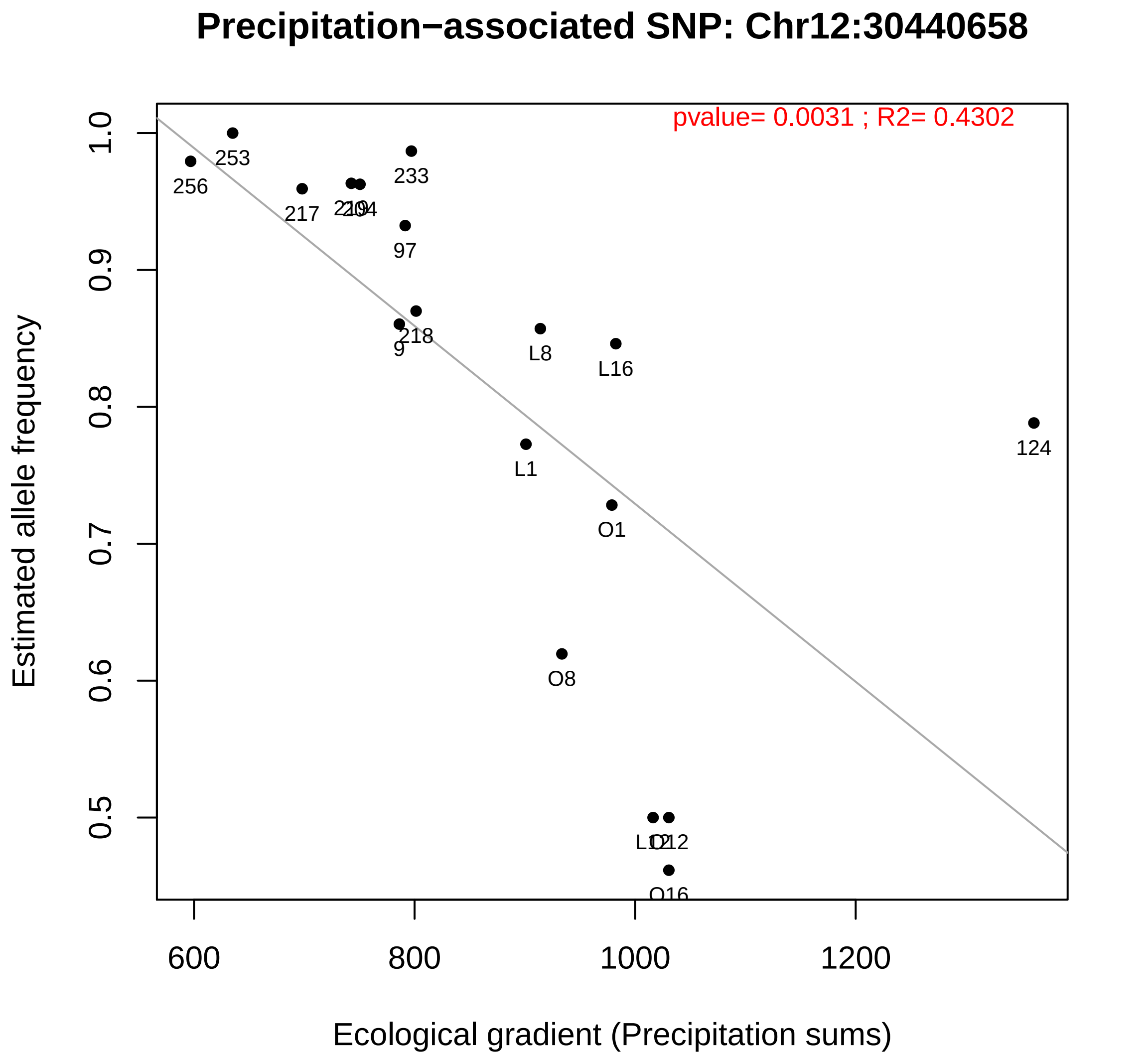

Precipitation−associated SNP: Chr12:30440658
pvalue= 0.0031 ; R2= 0.4302
Estimated allele frequency
Ecological gradient (Precipitation sums)
0.5
0.6
0.7
0.8
0.9
1.0
600
800
1000
1200
253
256
233
217
97
218
9
L8
L16
124
L1
O1
O8
O16

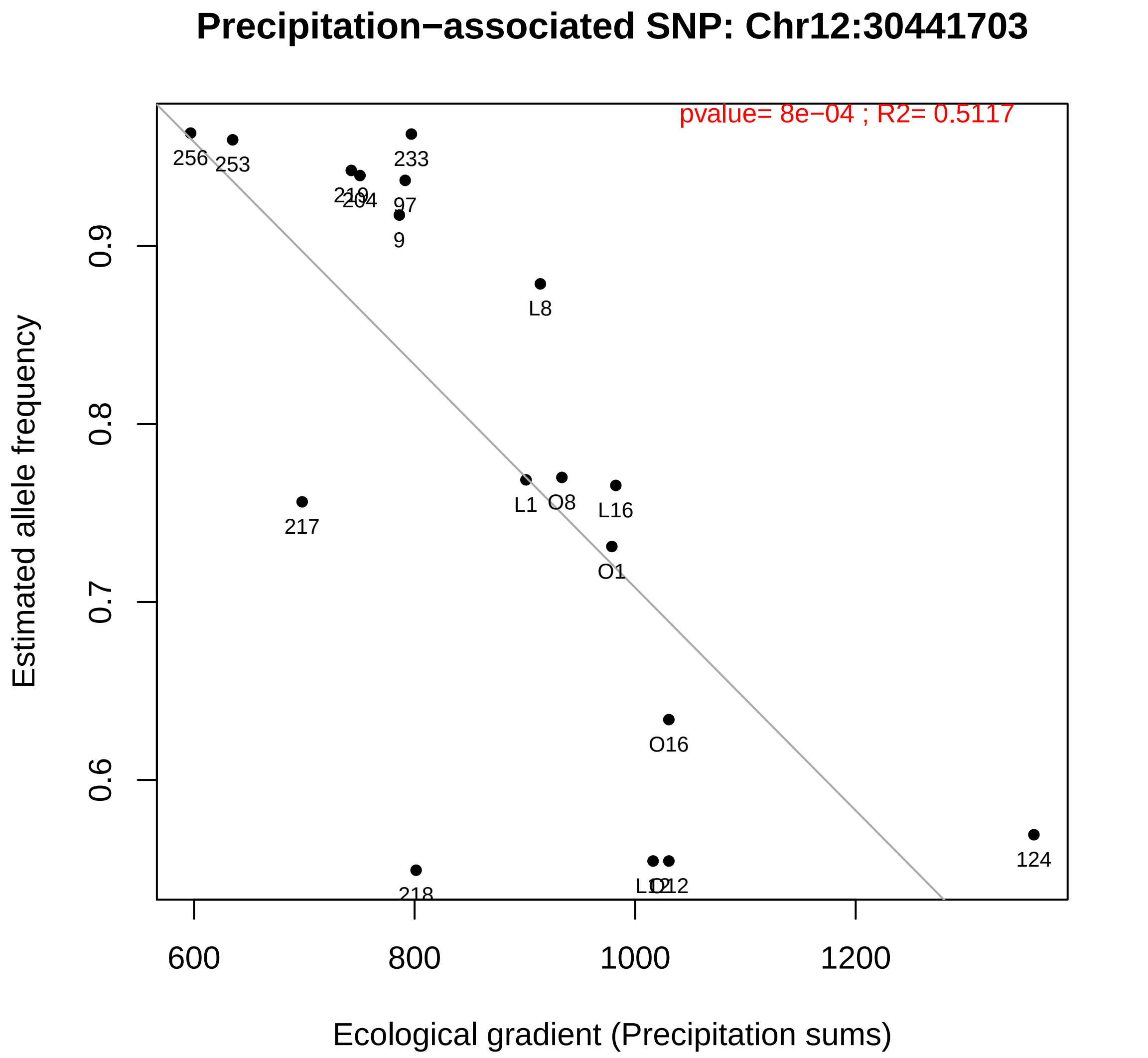
Precipitation−associated SNP: Chr12:30441703
pvalue= 8e−04 ; R2= 0.5117
Estimated allele frequency
Ecological gradient (Precipitation sums)
0.6
0.7
0.8
0.9
600
800
1000
1200
256
253
233
219
204
97
9
L8
217
L1
O8
L16
O1
O16
218
124
O12

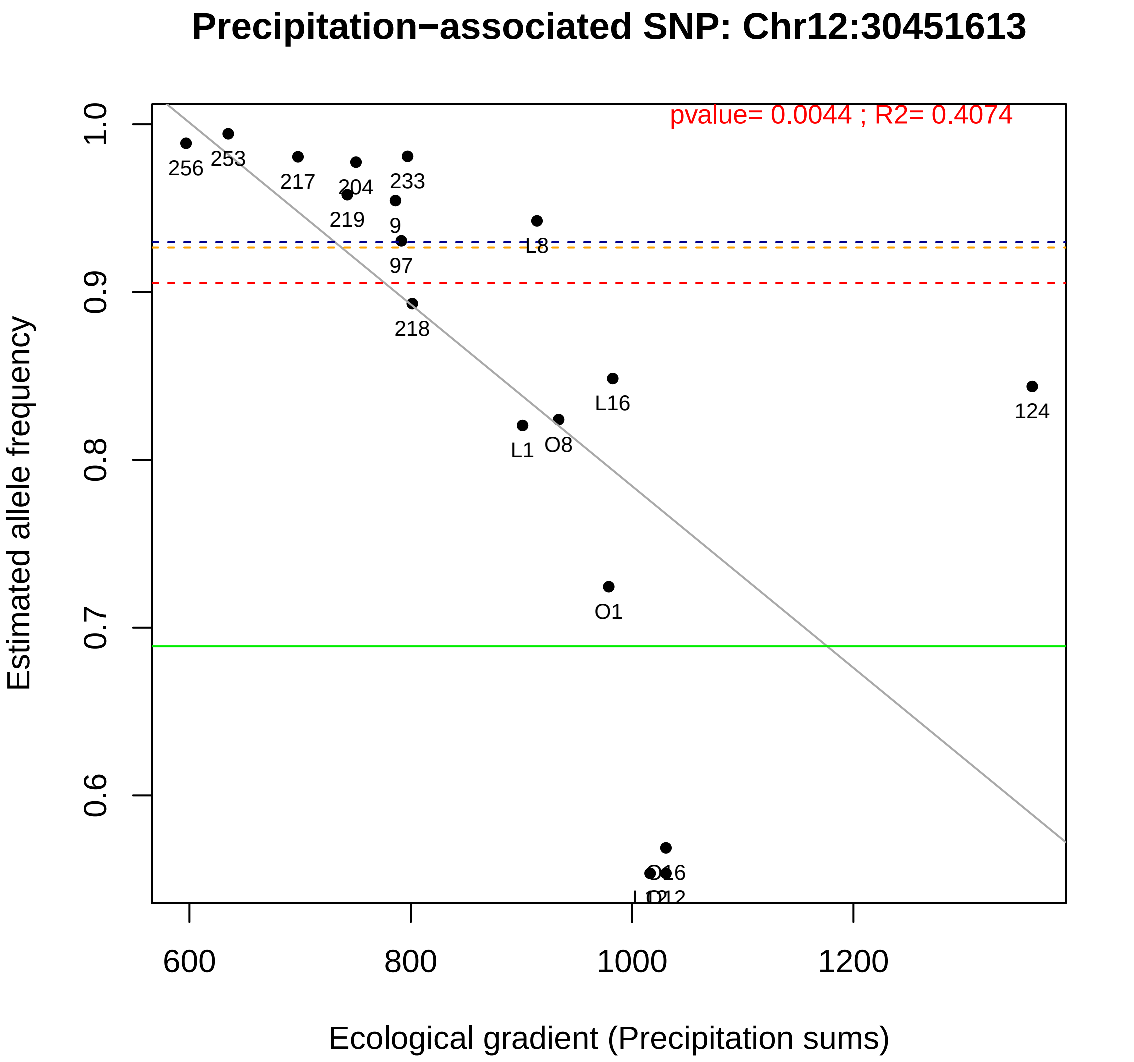
Precipitation−associated SNP: Chr12:30451613
pvalue= 0.0044 ; R2= 0.4074
Estimated allele frequency
Ecological gradient (Precipitation sums)
1.0
0.9
0.8
0.7
0.6
600
800
1000
1200
256
253
217
204
233
219
9
97
L8
218
L16
124
L1
O8
O1
O16
O12

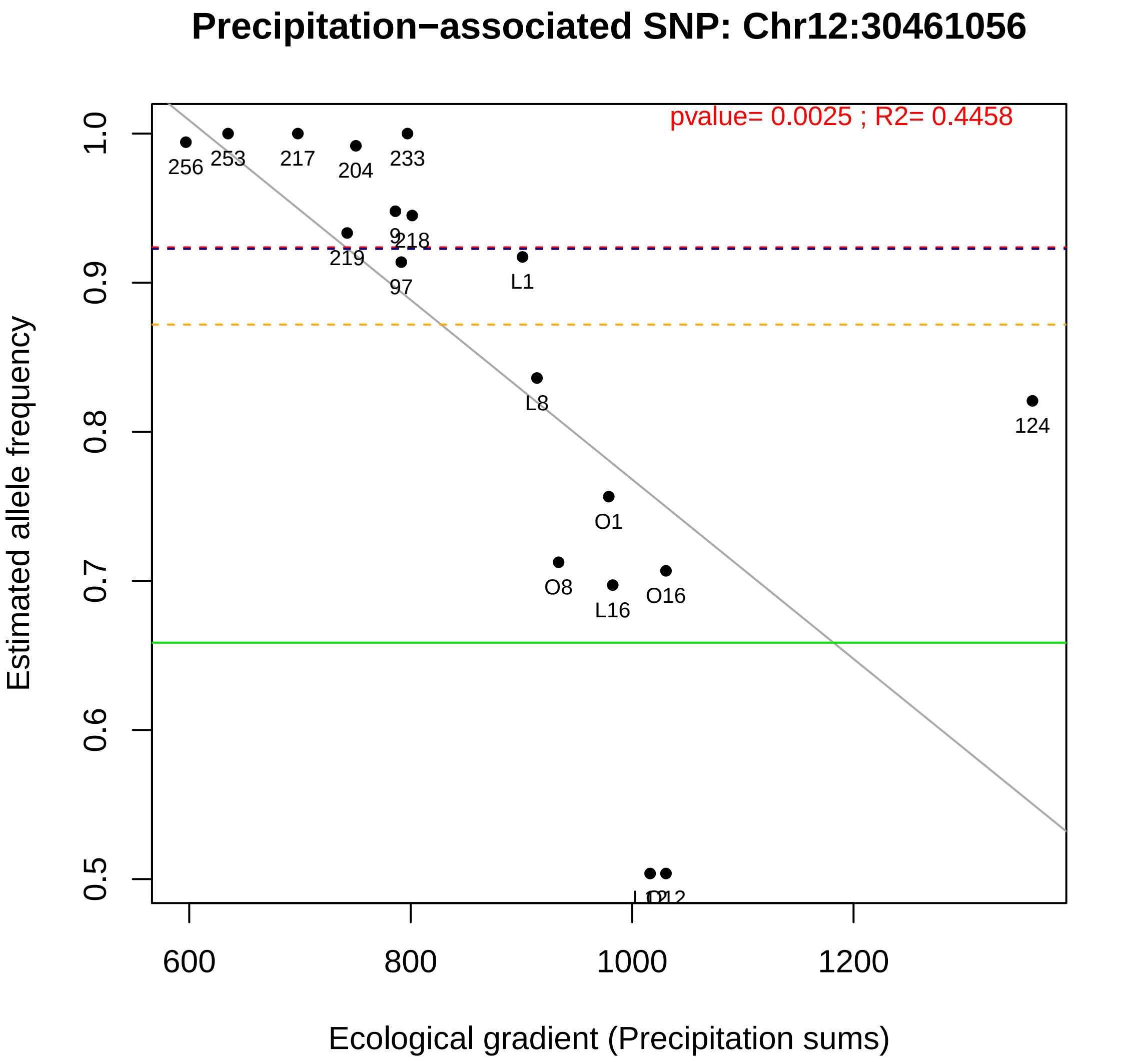
Precipitation−associated SNP: Chr12:30461056
pvalue= 0.0025 ; R2= 0.4458
Estimated allele frequency
Ecological gradient (Precipitation sums)
0.5
0.6
0.7
0.8
0.9
1.0
600
800
1000
1200
256
253
217
204
233
219
218
97
L1
L8
124
O1
O8
L16
O16

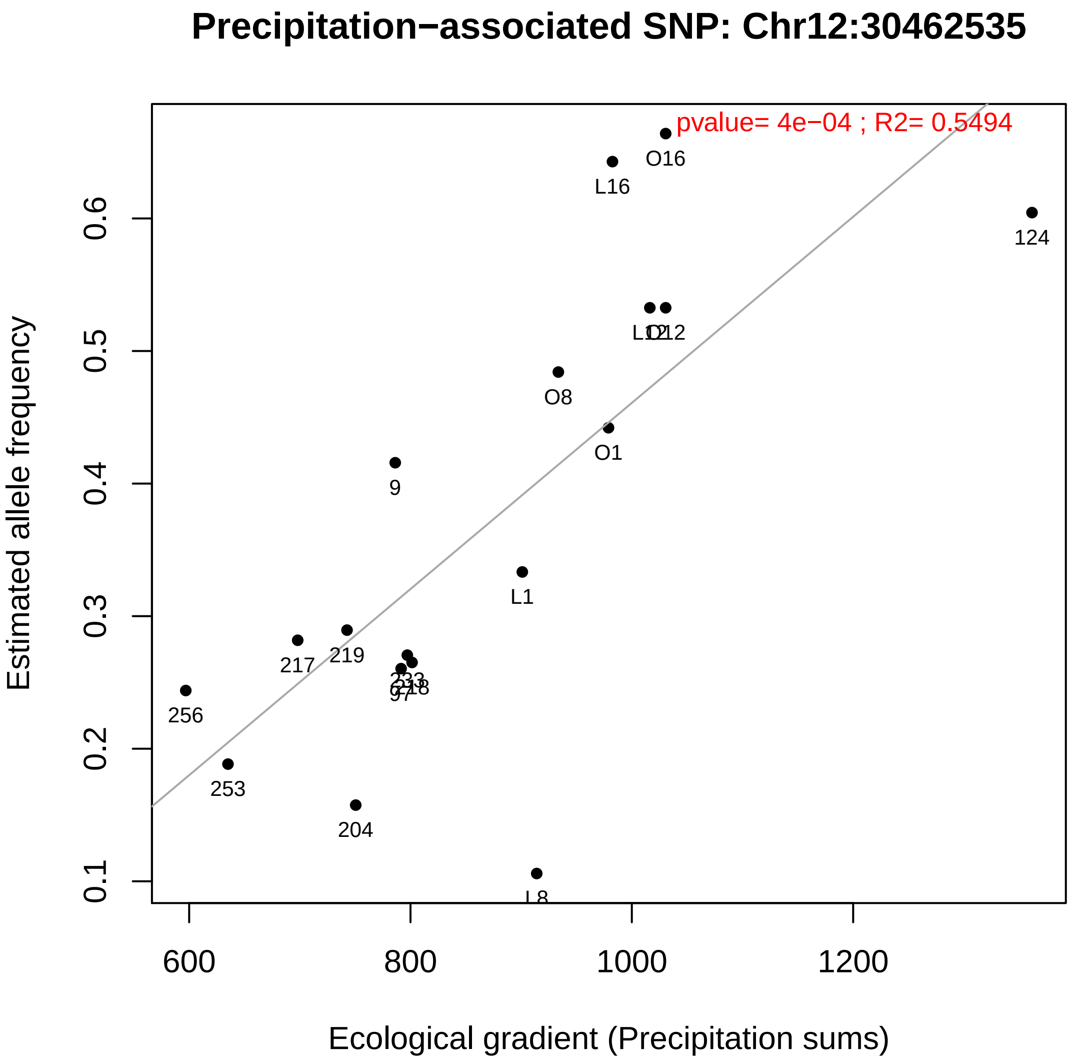
Precipitation−associated SNP: Chr12:30462535
pvalue= 4e−04 ; R2= 0.5494
O16
L16
124
L12
O12
O8
O1
9
L1
219
217
233
218
97
256
253
204
L8
0.1
0.2
0.3
0.4
0.5
0.6
600
800
1000
1200
Estimated allele frequency
Ecological gradient (Precipitation sums)

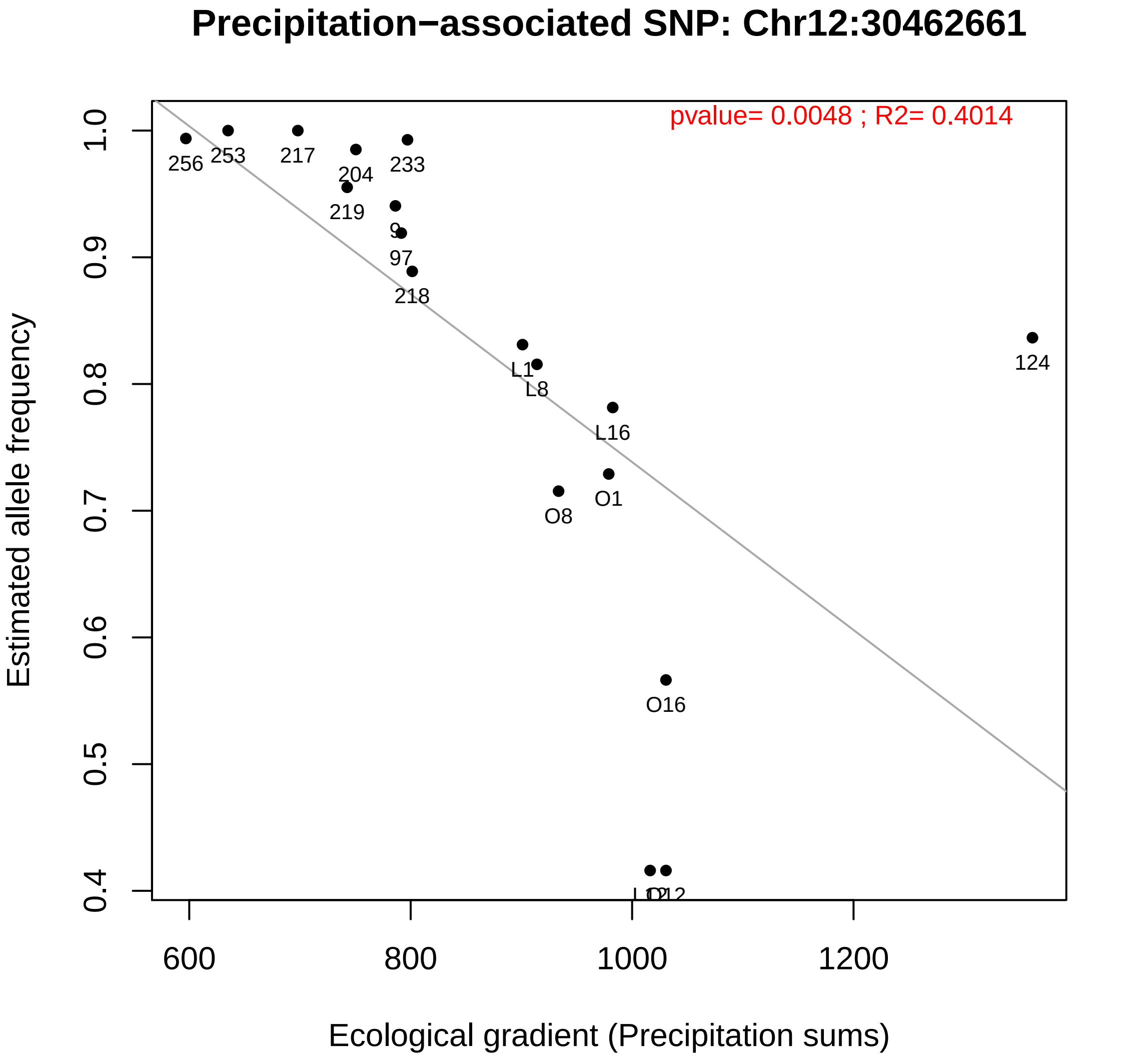
Precipitation−associated SNP: Chr12:30462661
pvalue= 0.0048 ; R2= 0.4014
Estimated allele frequency
Ecological gradient (Precipitation sums)
0.4
0.5
0.6
0.7
0.8
0.9
1.0
600
800
1000
1200
256
253
217
204
233
219
97
218
L1
L8
L16
O1
O8
124
O16

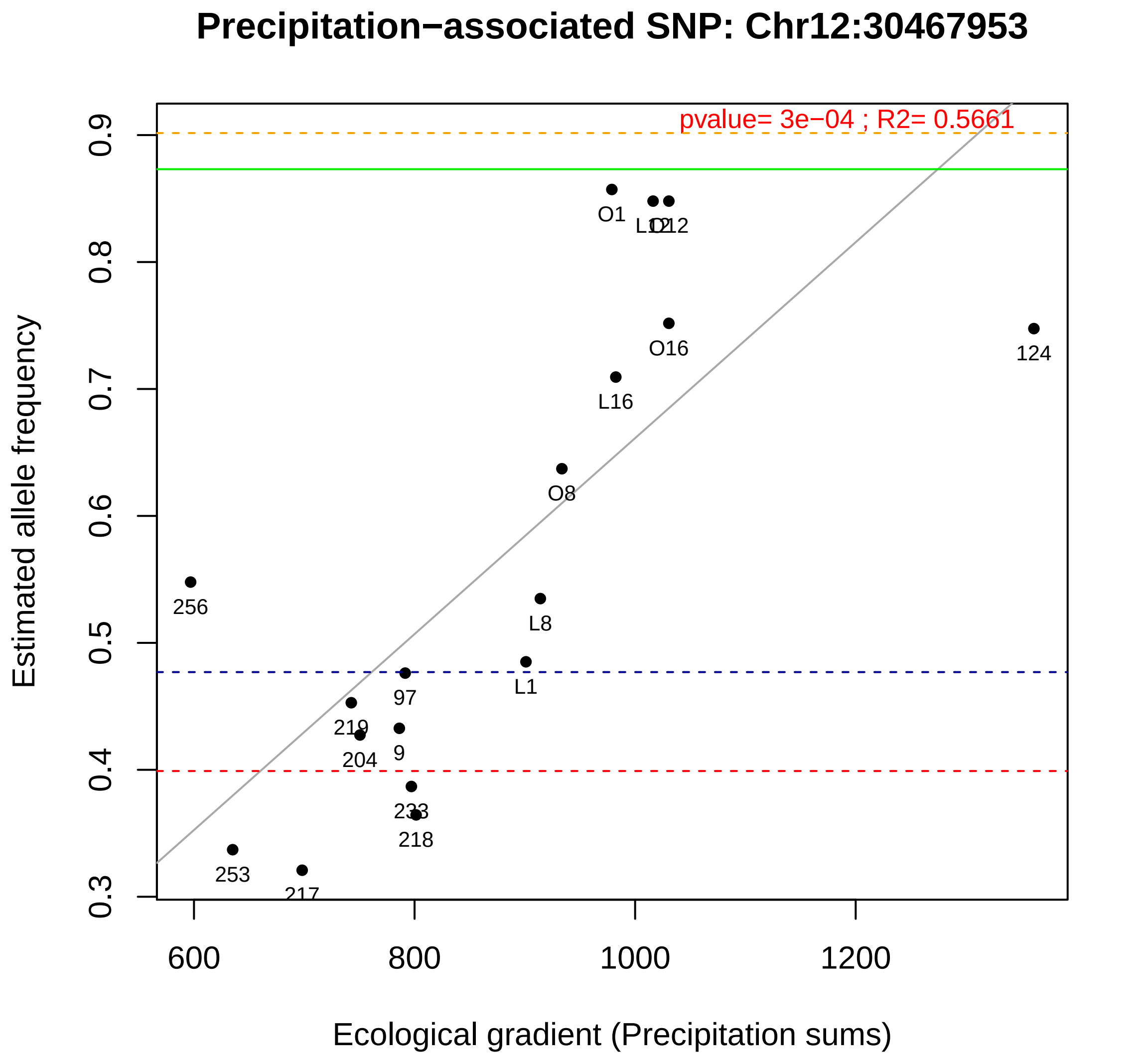

Precipitation−associated SNP: Chr12:30467953
pvalue= 3e−04 ; R2= 0.5661
Estimated allele frequency
0.3
0.4
0.5
0.6
0.7
0.8
0.9
600
800
1000
1200
Ecological gradient (Precipitation sums)
O1
L12
O12
O16
124
L16
O8
256
L8
L1
97
219
204
9
233
218
253
217

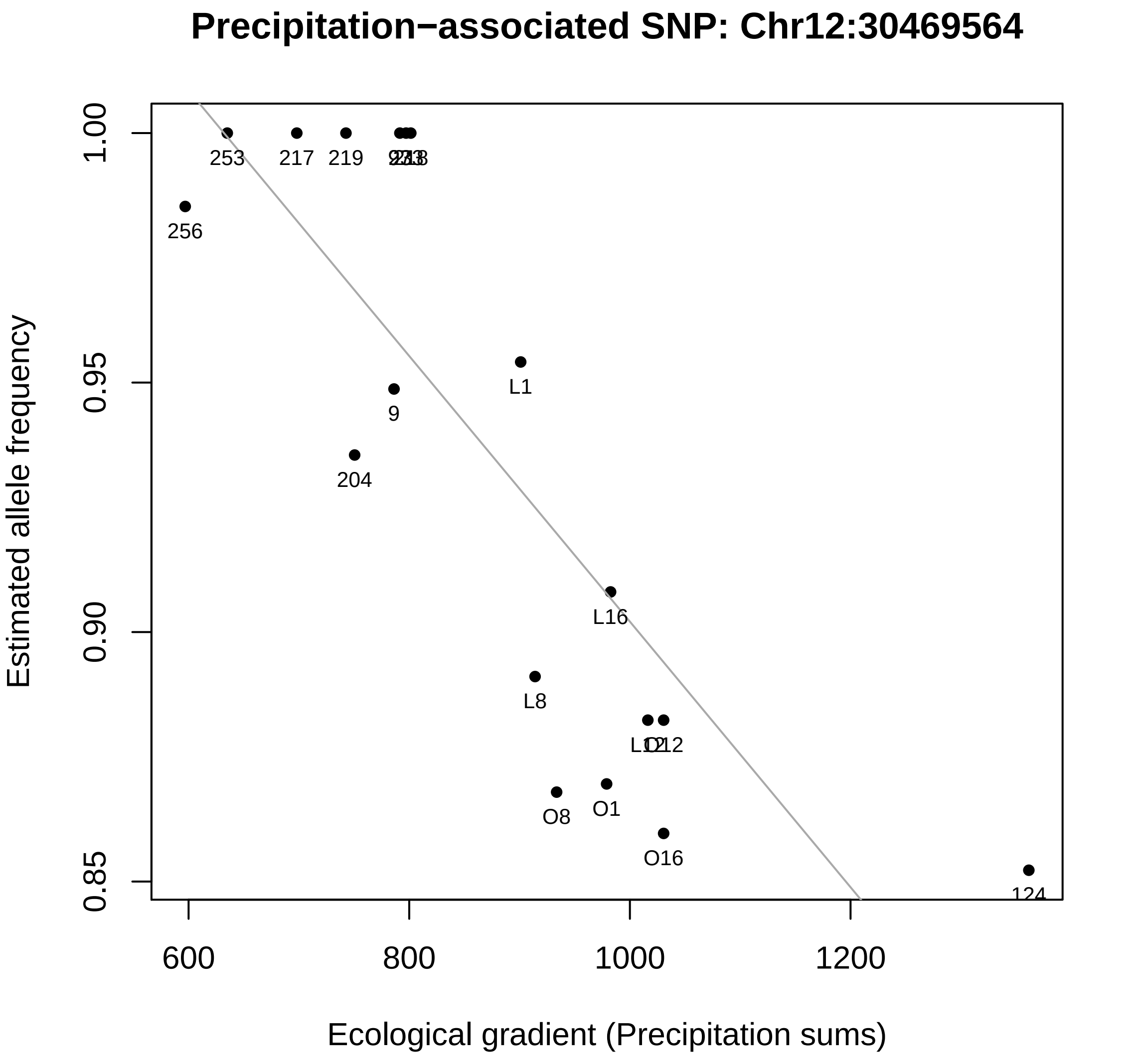
Precipitation−associated SNP: Chr12:30469564
Estimated allele frequency
Ecological gradient (Precipitation sums)
1.00
0.95
0.90
0.85
600
800
1000
1200
253
217
219
256
L1
9
204
L16
L8
O8
O1
O16
124

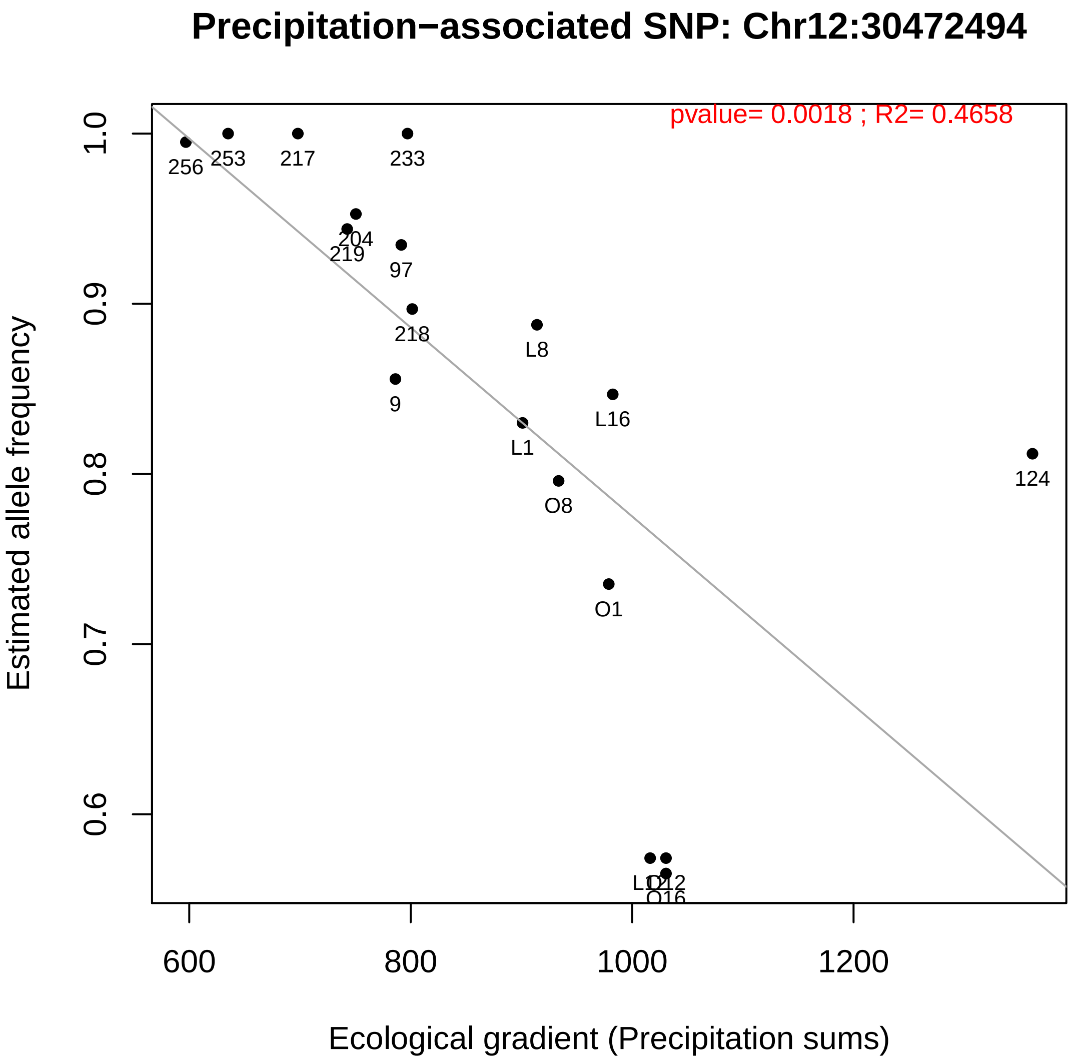
Precipitation−associated SNP: Chr12:30472494
pvalue= 0.0018 ; R2= 0.4658
Estimated allele frequency
1.0
0.9
0.8
0.7
0.6
600
800
1000
1200
Ecological gradient (Precipitation sums)
256
253
217
233
204
219
97
218
L8
9
L16
L1
124
O8
O1

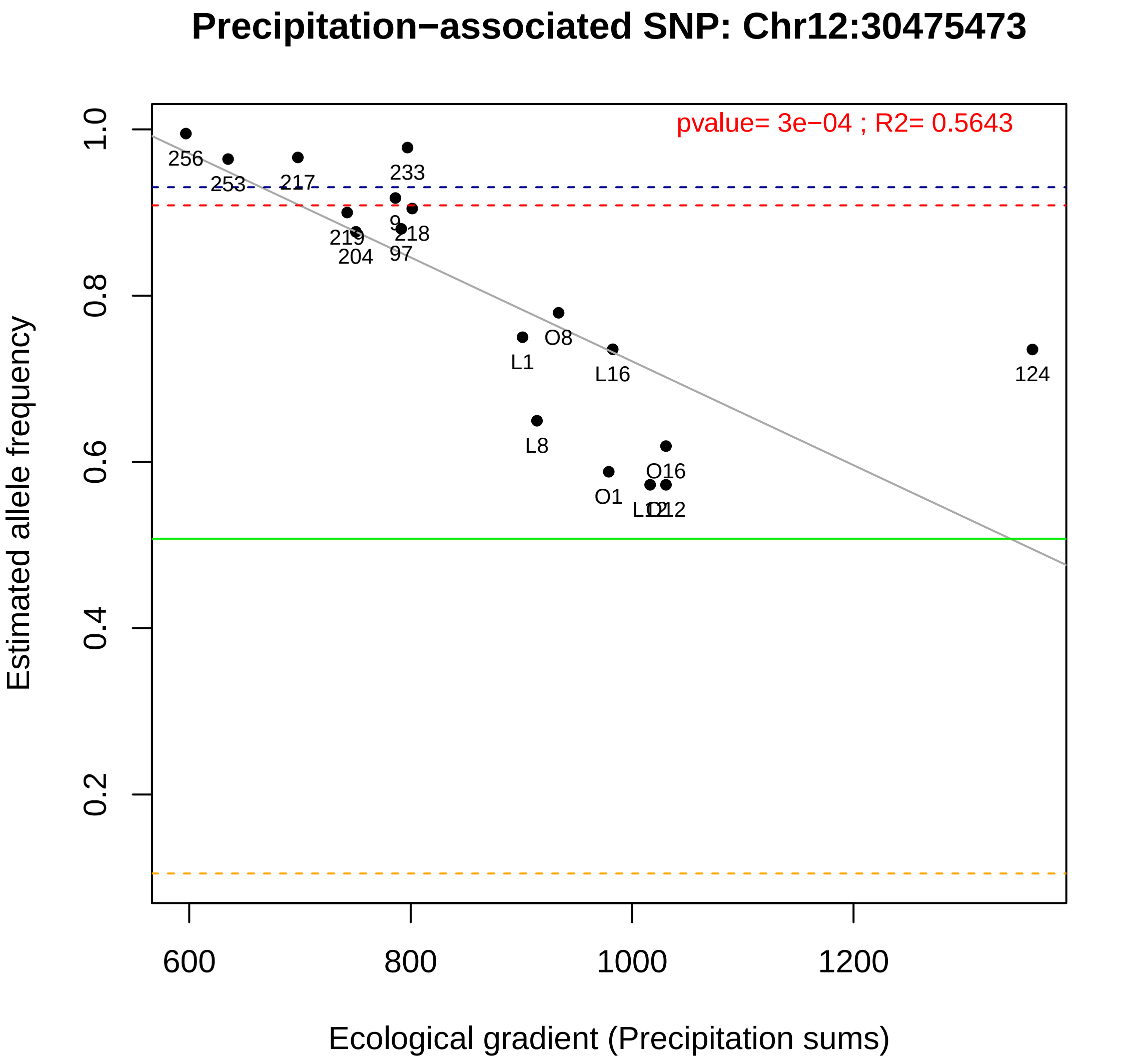
Precipitation−associated SNP: Chr12:30475473
pvalue= 3e−04 ; R2= 0.5643
Estimated allele frequency
Ecological gradient (Precipitation sums)
0.2
0.4
0.6
0.8
1.0
600
800
1000
1200
256
253
217
233
9
218
219
204
97
L1
O8
L16
124
L8
O16
O1

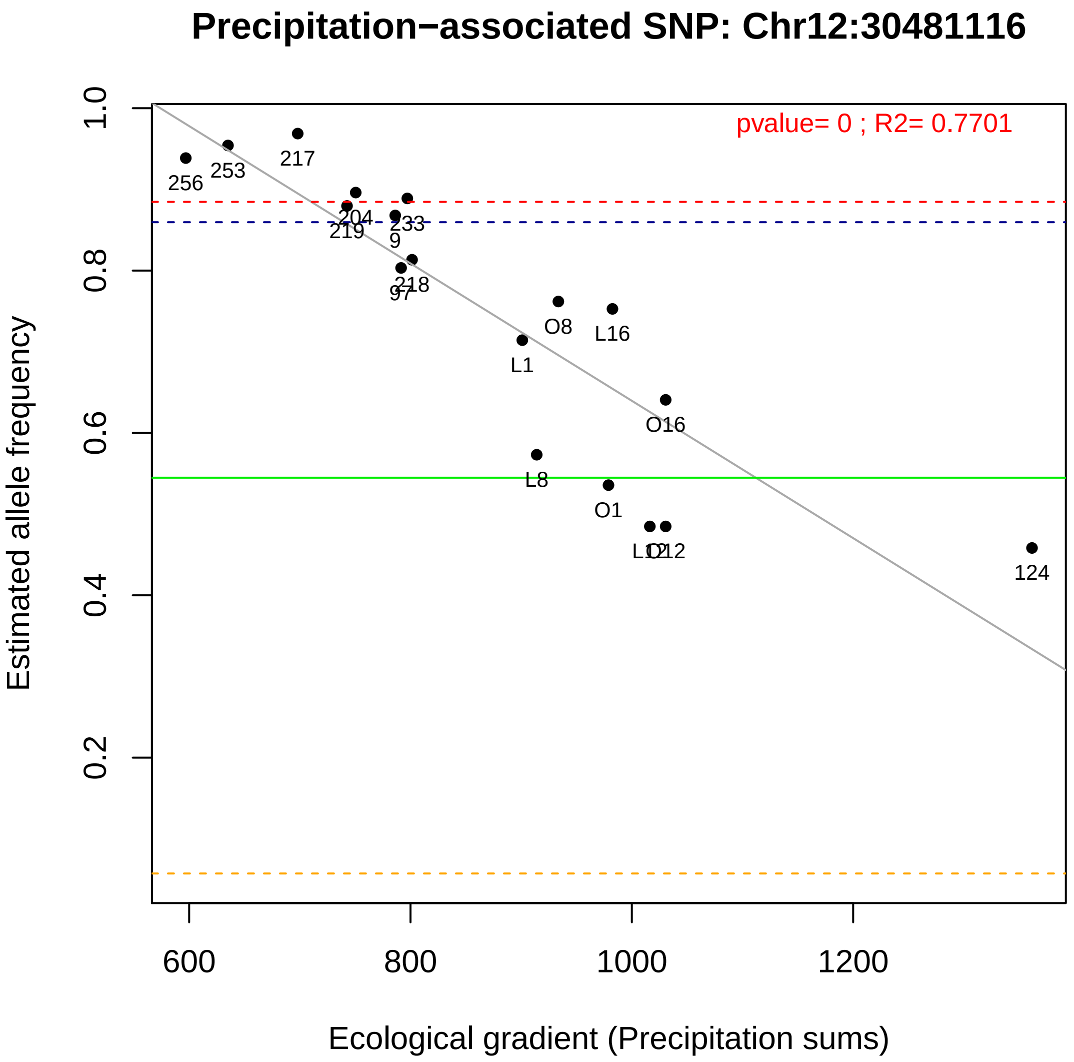
Precipitation−associated SNP: Chr12:30481116
pvalue= 0 ; R2= 0.7701
Estimated allele frequency
Ecological gradient (Precipitation sums)
0.2
0.4
0.6
0.8
1.0
600
800
1000
1200
256
253
217
204
219
233
9
218
97
O8
L16
L1
O16
L8
O1
L12
O12
124

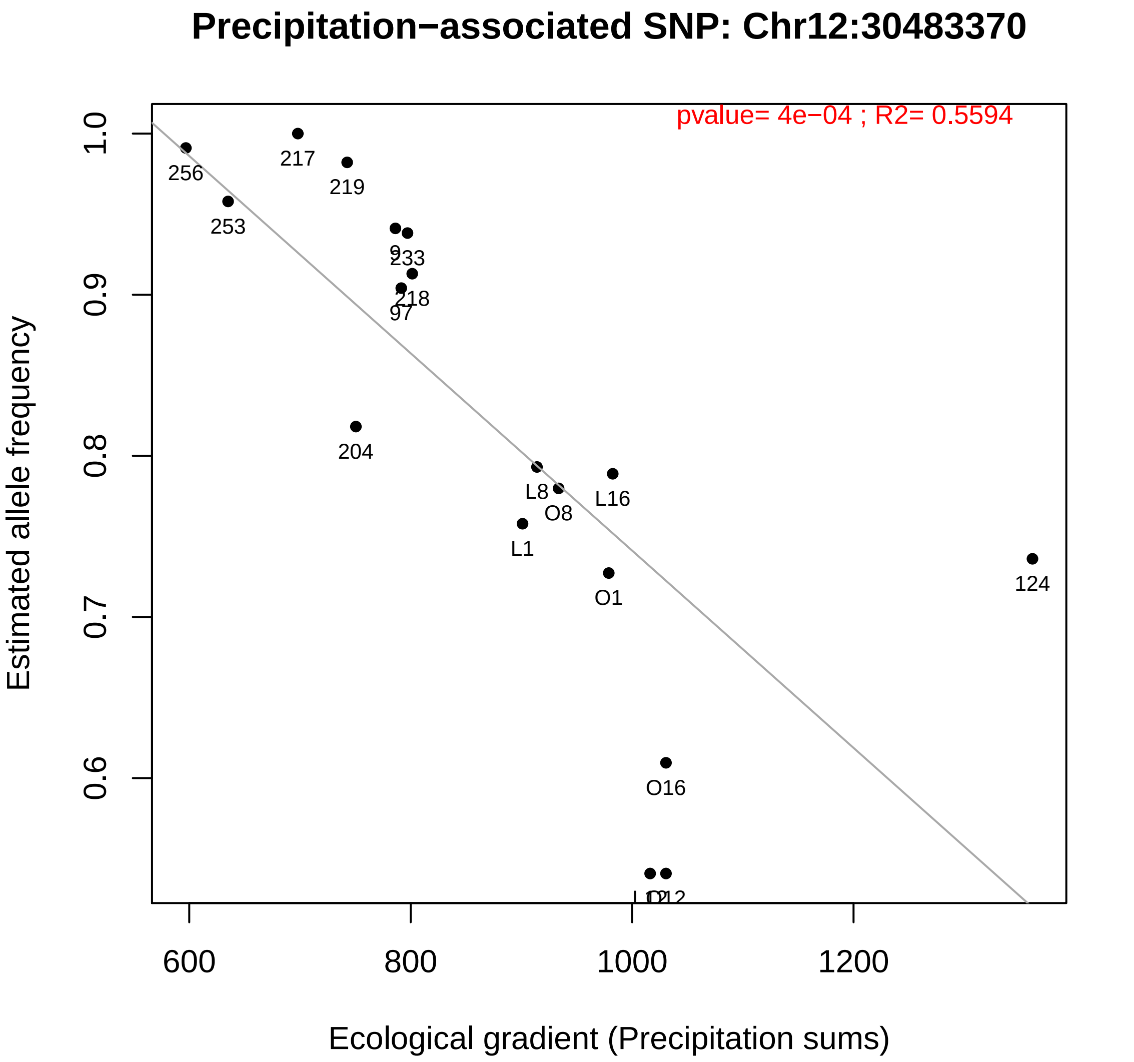

Precipitation−associated SNP: Chr12:30483370
pvalue= 4e−04 ; R2= 0.5594
Estimated allele frequency
1.0
0.9
0.8
0.7
0.6
600
800
1000
1200
Ecological gradient (Precipitation sums)
256
217
219
253
9
233
218
97
204
L8
O8
L16
L1
O1
124
O16
O12

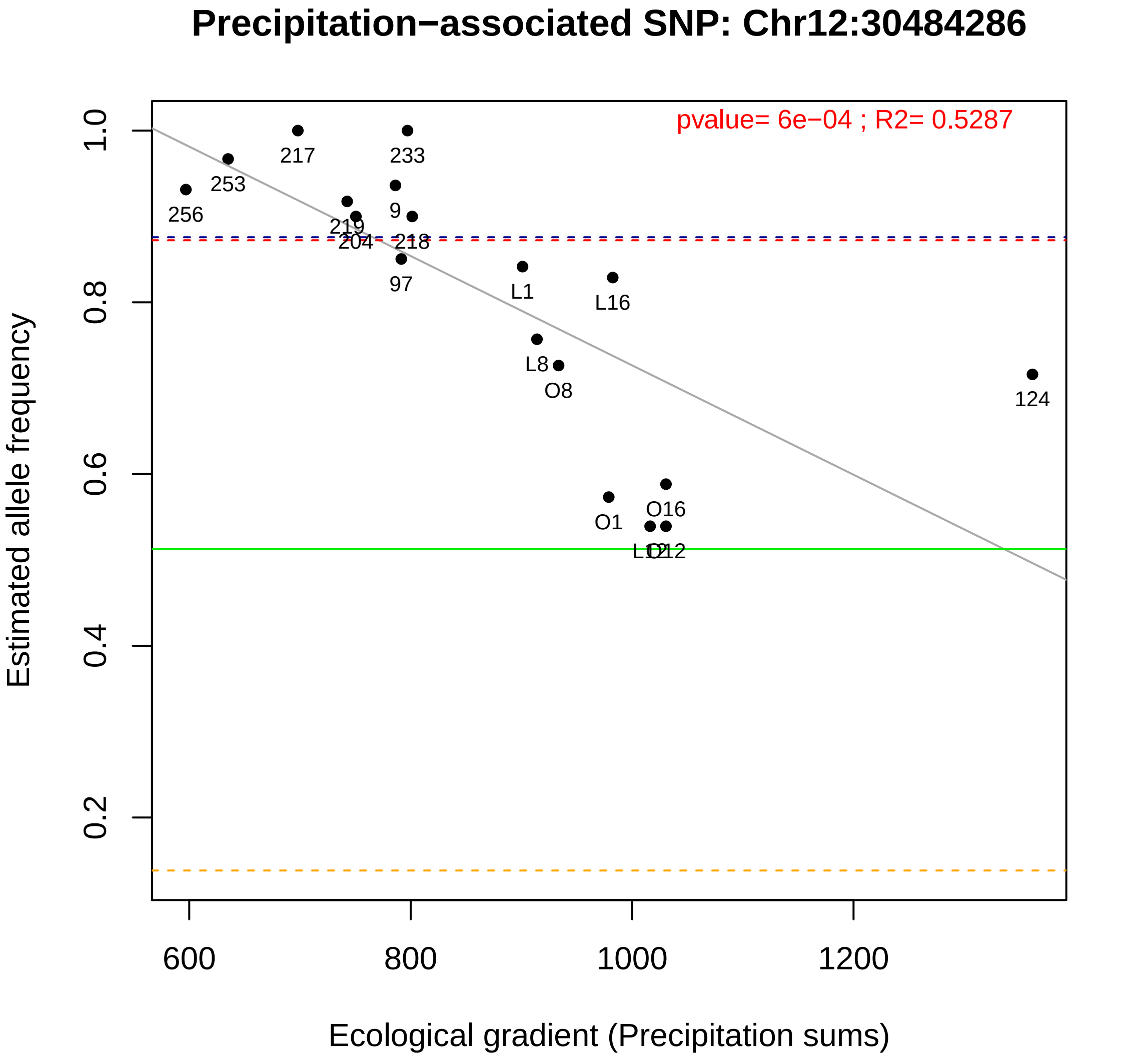
Precipitation−associated SNP: Chr12:30484286
pvalue= 6e−04 ; R2= 0.5287
Estimated allele frequency
0.2
0.4
0.6
0.8
1.0
600
800
1000
1200
Ecological gradient (Precipitation sums)
217
233
253
256
9
219
204
218
97
L1
L16
L8
O8
124
O16
O1

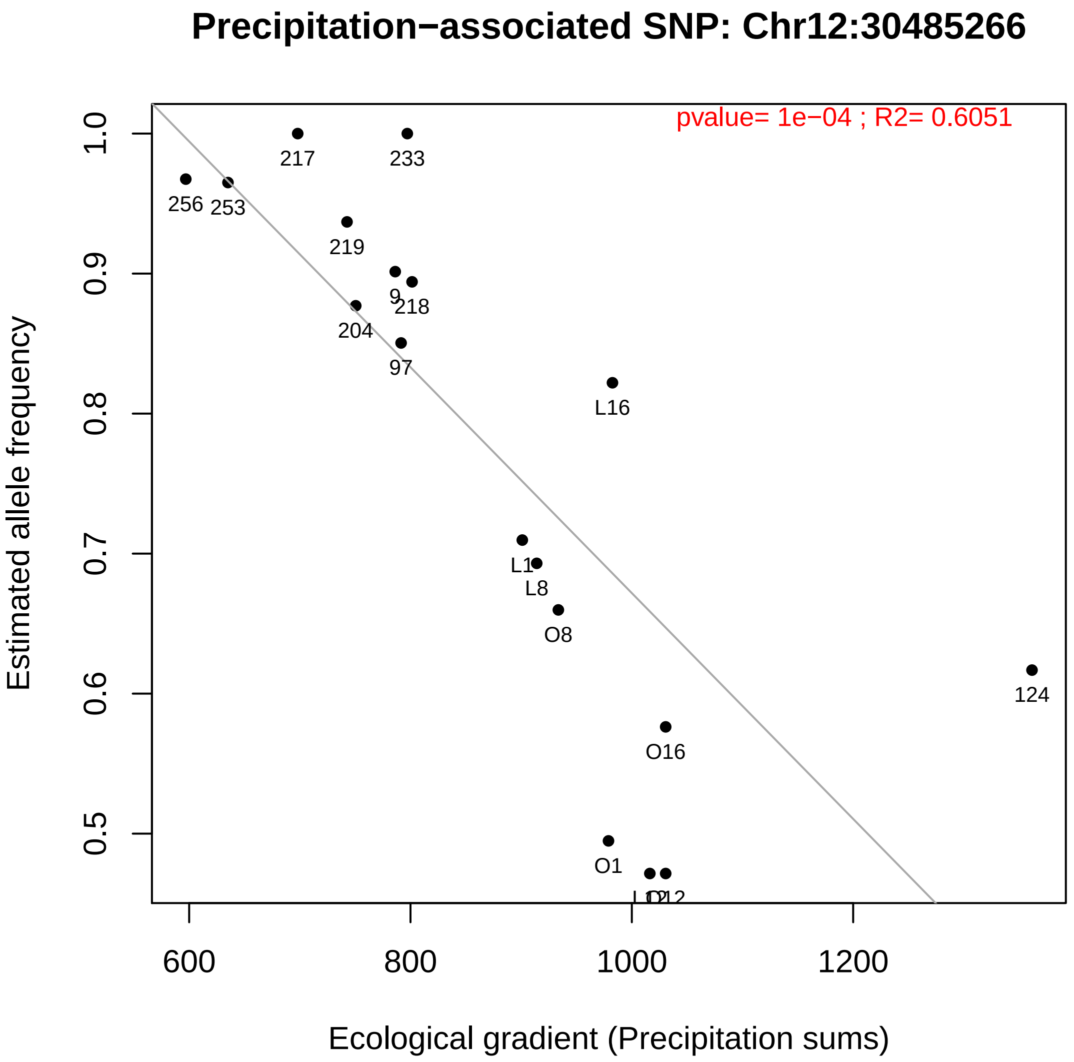

Precipitation−associated SNP: Chr12:30485266
pvalue= 1e−04 ; R2= 0.6051
Estimated allele frequency
Ecological gradient (Precipitation sums)
0.5
0.6
0.7
0.8
0.9
1.0
600
800
1000
1200
217
233
256
253
219
9
218
204
97
L16
L1
L8
O8
124
O16
O1
O12

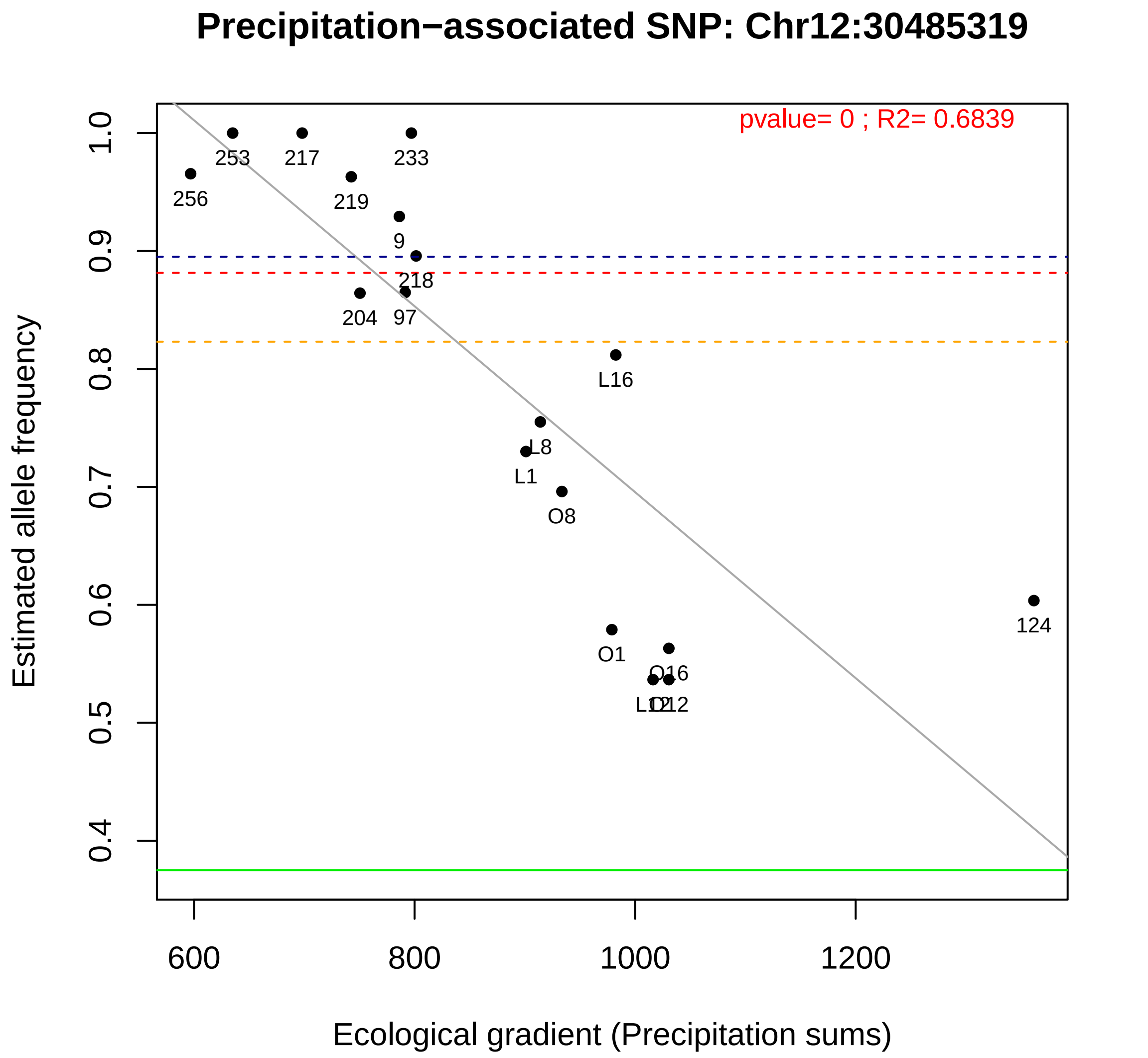

Precipitation−associated SNP: Chr12:30485319
pvalue= 0 ; R2= 0.6839
Estimated allele frequency
Ecological gradient (Precipitation sums)
1.0
0.9
0.8
0.7
0.6
0.5
0.4
600
800
1000
1200
253
217
233
256
219
9
218
204
97
L16
L8
L1
O8
O1
O16
L12
O12
124

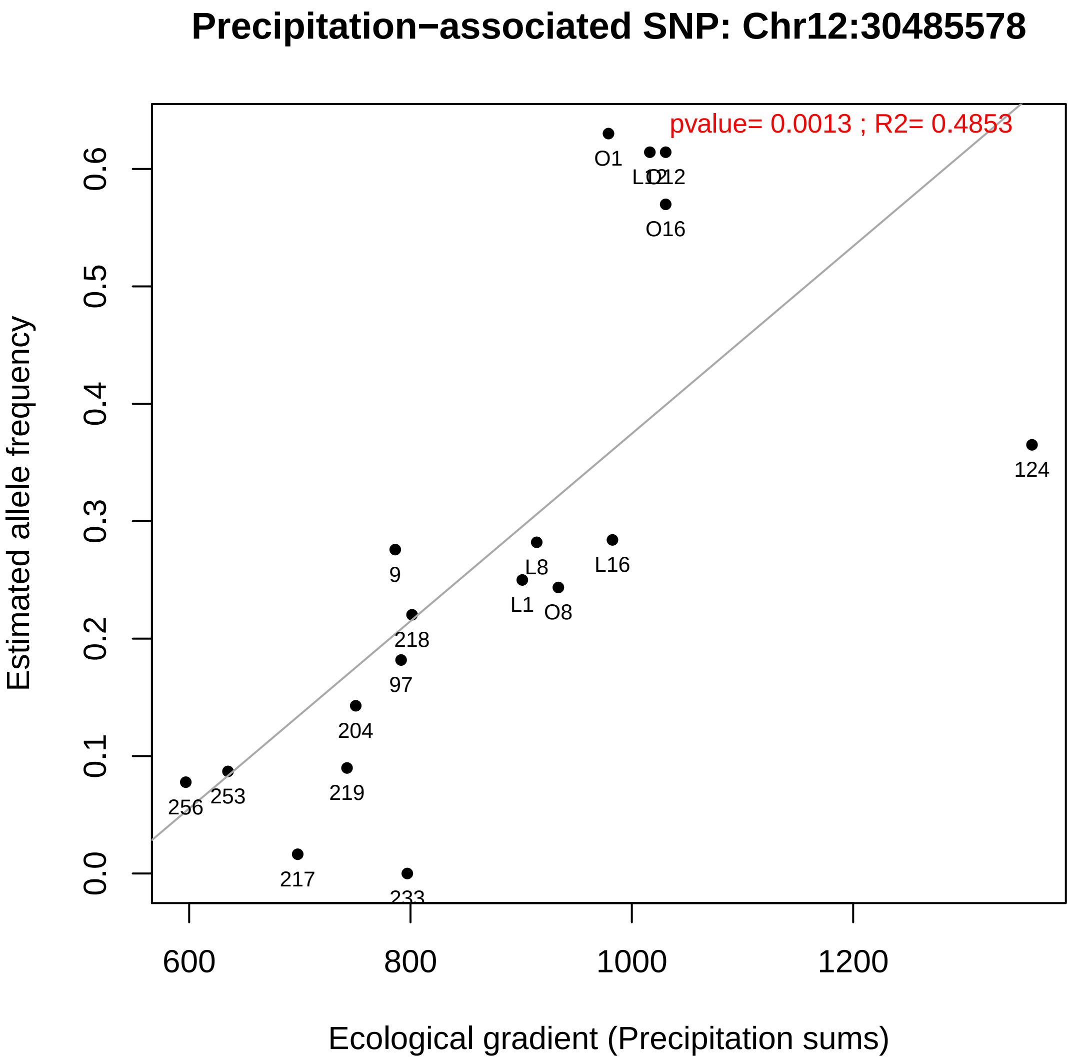

Precipitation−associated SNP: Chr12:30485578
pvalue= 0.0013 ; R2= 0.4853
Estimated allele frequency
Ecological gradient (Precipitation sums)
0.0
0.1
0.2
0.3
0.4
0.5
0.6
600
800
1000
1200
O1
L12
O12
O16
124
9
L8
L16
L1
O8
218
97
204
219
256
253
217
233

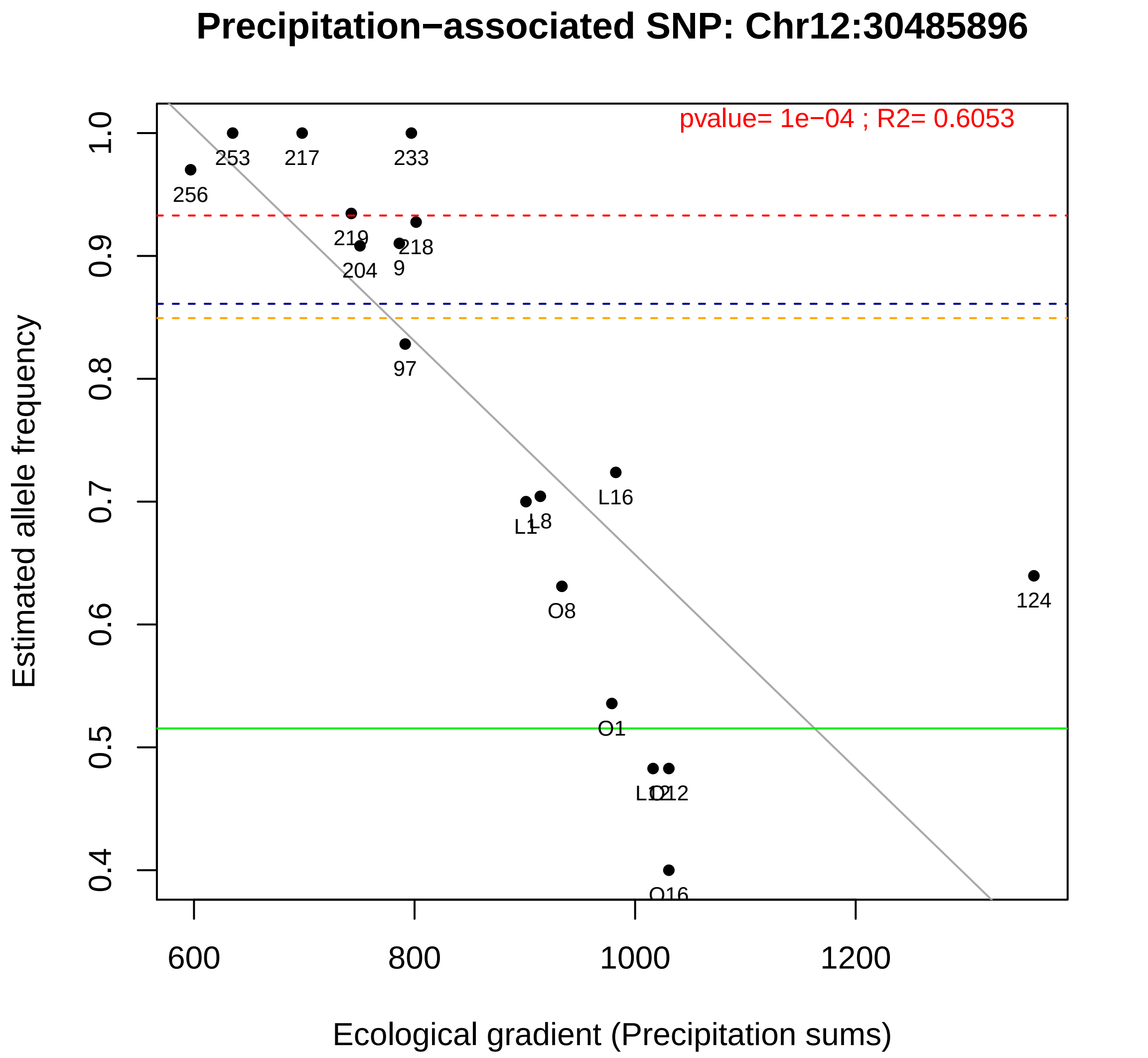

Precipitation−associated SNP: Chr12:30485896
pvalue= 1e−04 ; R2= 0.6053
Estimated allele frequency
Ecological gradient (Precipitation sums)
0.4
0.5
0.6
0.7
0.8
0.9
1.0
600
800
1000
1200
256
253
217
233
219
218
204
9
97
L16
L1
L8
O8
124
O1
L12
O12
O16

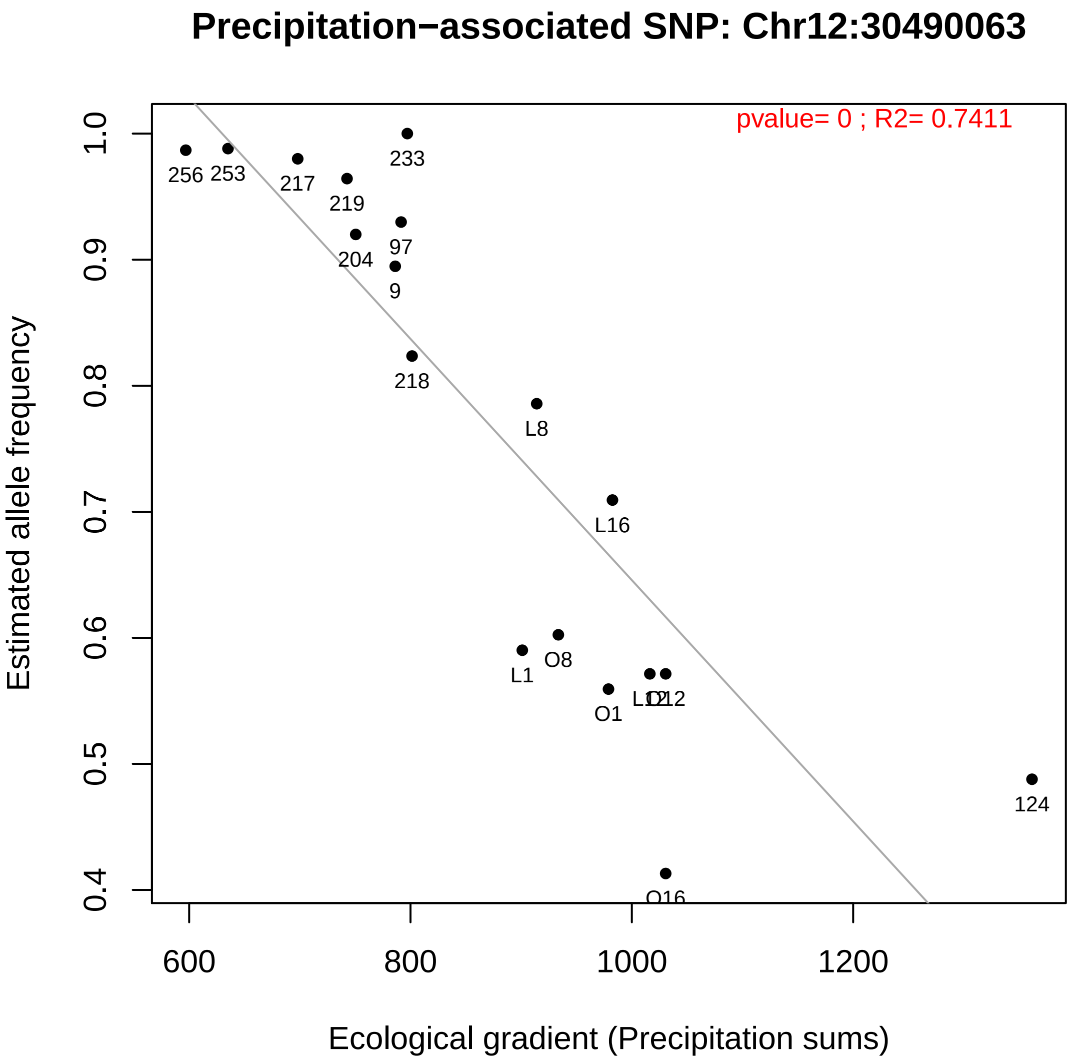
Precipitation−associated SNP: Chr12:30490063
pvalue= 0 ; R2= 0.7411
256
253
217
219
233
97
204
9
218
L8
L16
O8
L1
O1
O12
124
O16
0.4
0.5
0.6
0.7
0.8
0.9
1.0
600
800
1000
1200
Estimated allele frequency
Ecological gradient (Precipitation sums)

**Precipitation−associated SNP: Chr12:30491226**

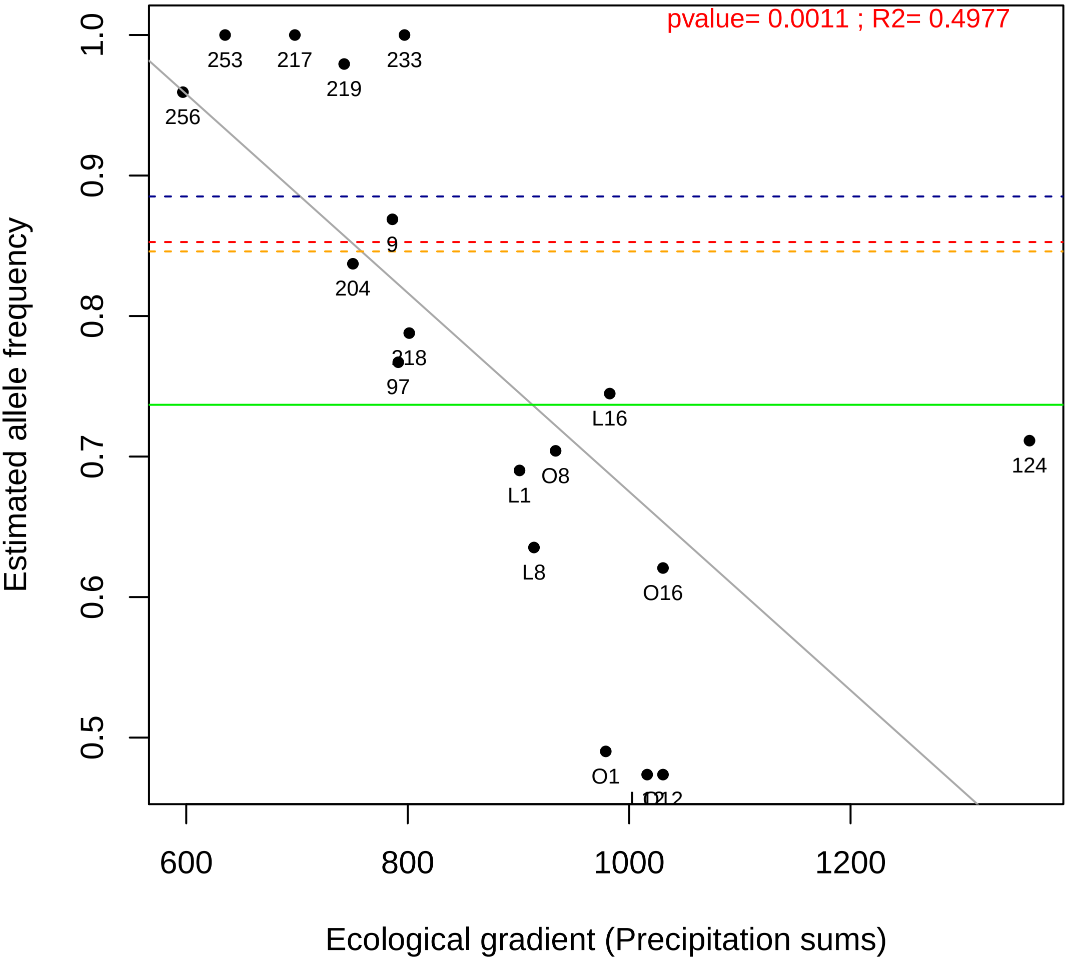

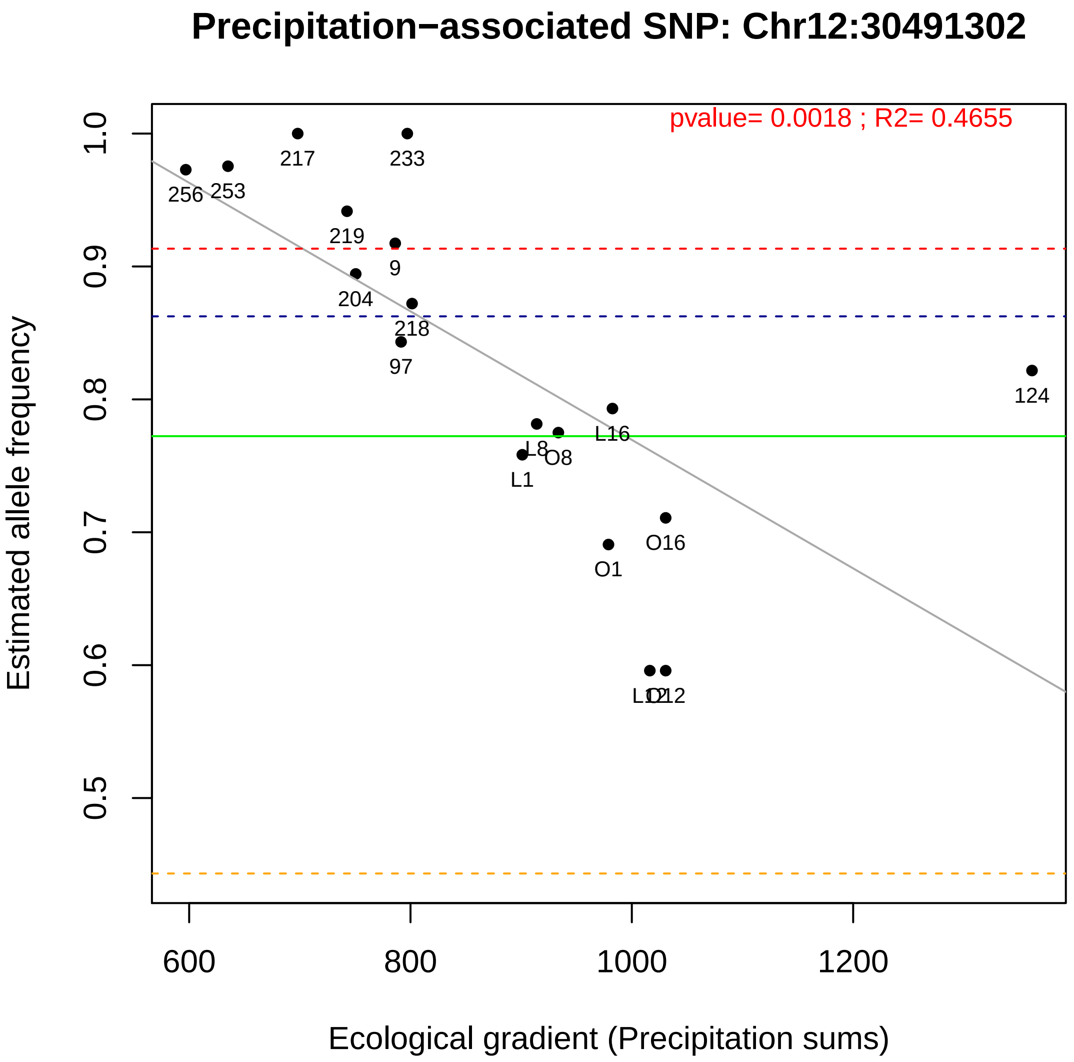
Precipitation−associated SNP: Chr12:30491302
pvalue= 0.0018 ; R2= 0.4655
Estimated allele frequency
Ecological gradient (Precipitation sums)
1.0
0.9
0.8
0.7
0.6
0.5
600
800
1000
1200
217
233
256
253
219
9
204
218
97
124
L16
L8
O8
L1
O16
O1
L12
O12

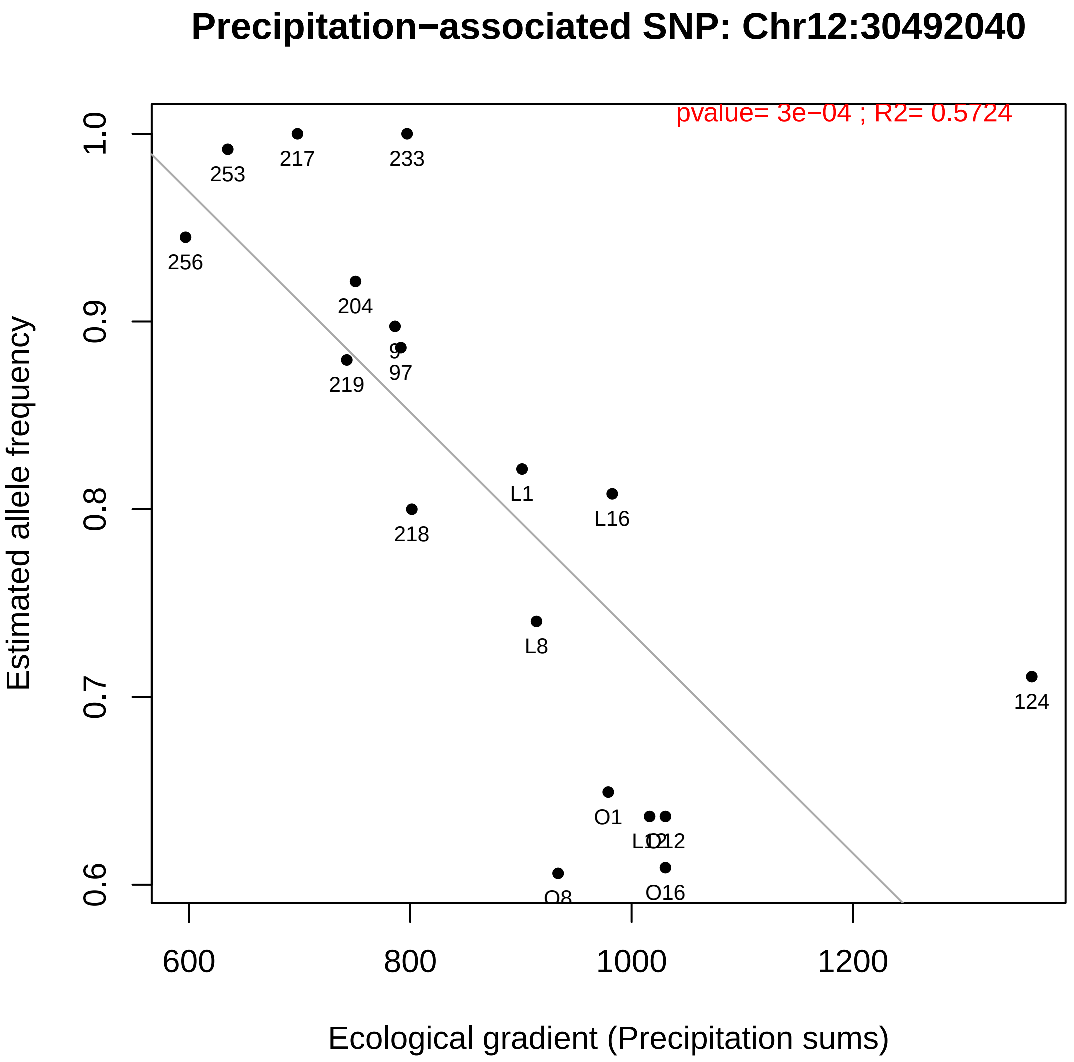

Precipitation−associated SNP: Chr12:30492040
pvalue= 3e−04 ; R2= 0.5724
253
217
233
256
204
9
97
219
L1
L16
218
L8
124
O1
L12
O12
O8
O16
Estimated allele frequency
0.6
0.7
0.8
0.9
1.0
600
800
1000
1200
Ecological gradient (Precipitation sums)

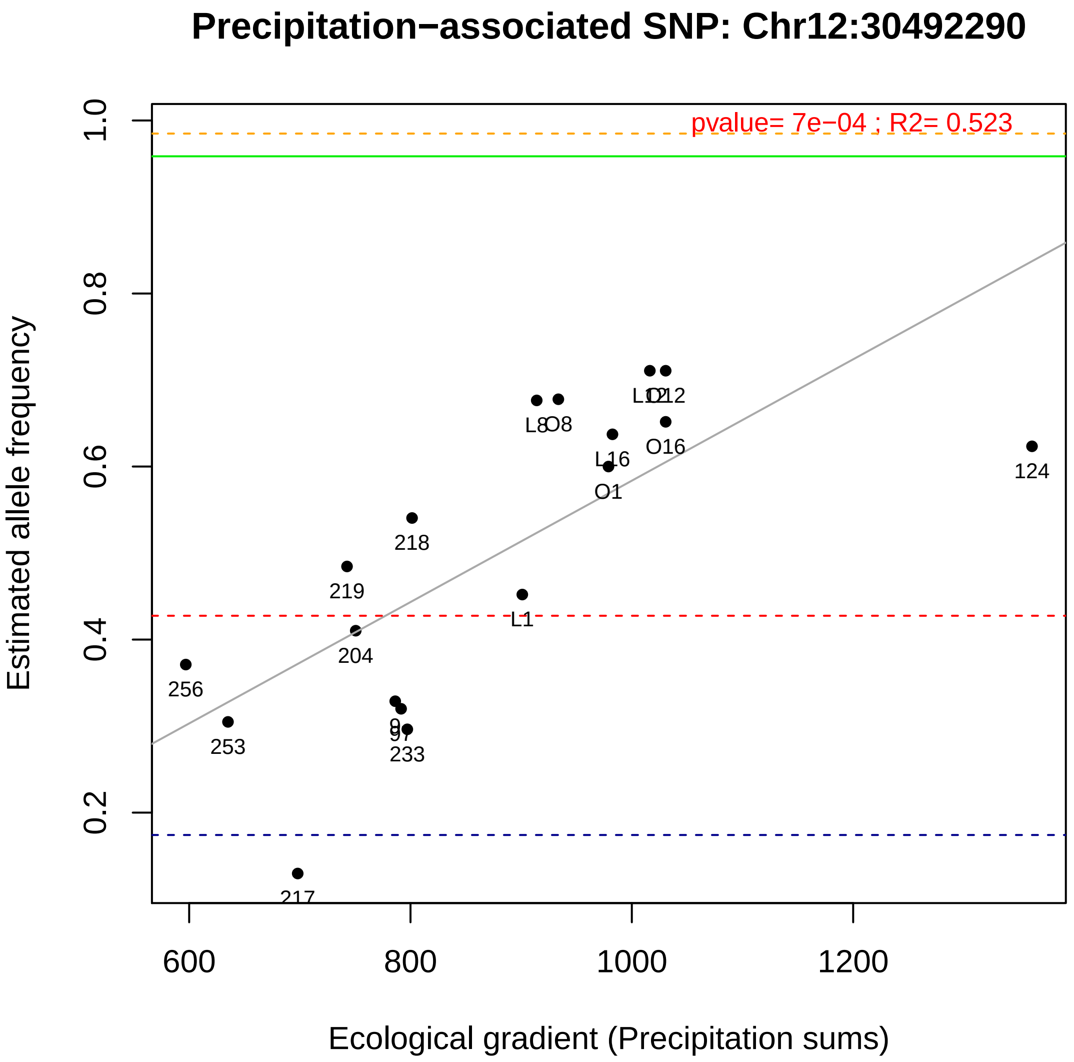

Precipitation−associated SNP: Chr12:30492290
pvalue= 7e−04 ; R2= 0.523
Estimated allele frequency
Ecological gradient (Precipitation sums)
0.2
0.4
0.6
0.8
1.0
600
800
1000
1200
L12
O12
L8
O8
O16
L16
124
O1
218
219
L1
204
256
253
233
217

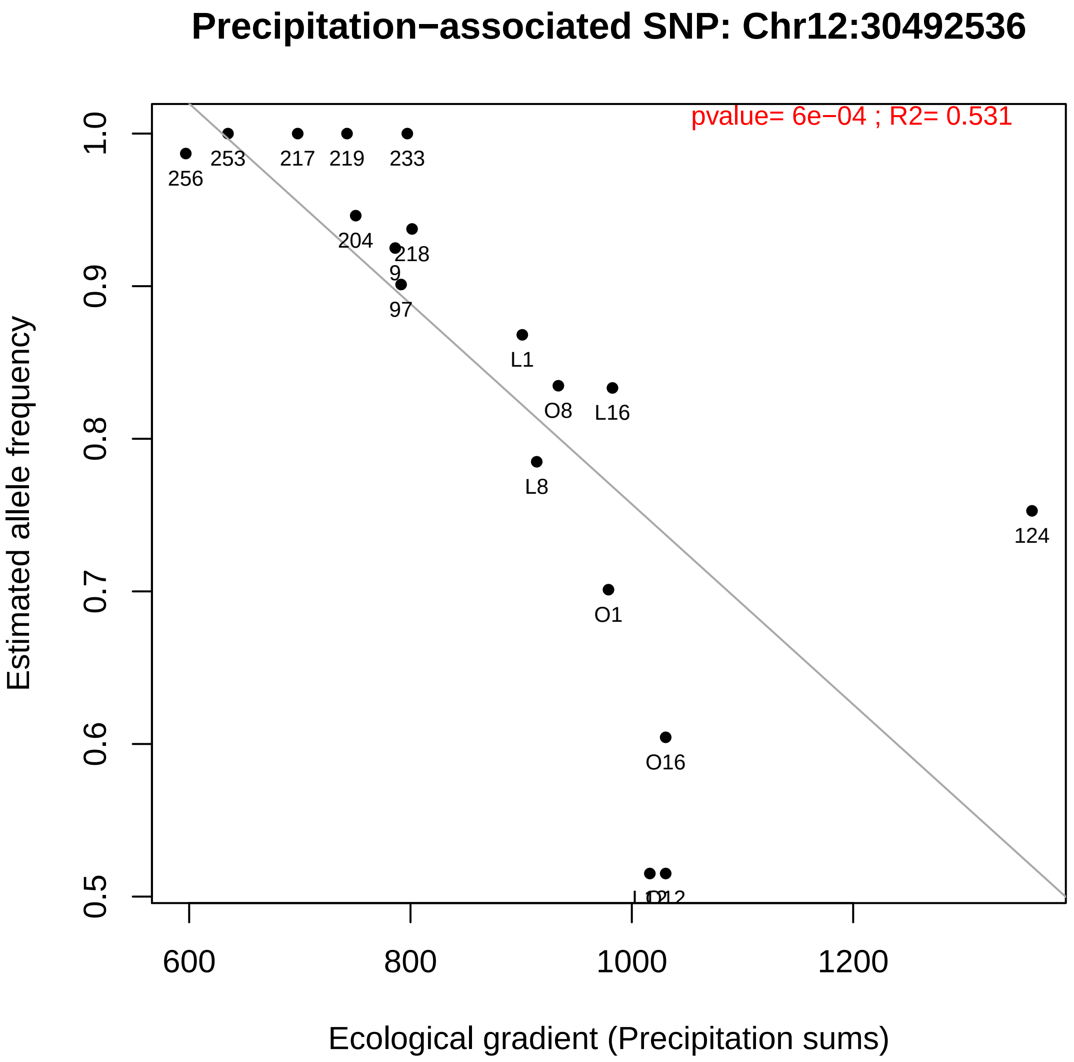

Precipitation−associated SNP: Chr12:30492536
pvalue= 6e−04 ; R2= 0.531
Estimated allele frequency
Ecological gradient (Precipitation sums)
1.0
0.9
0.8
0.7
0.6
0.5
600
800
1000
1200
256
253
217
219
233
204
218
9
97
L1
O8
L16
L8
124
O1
O16

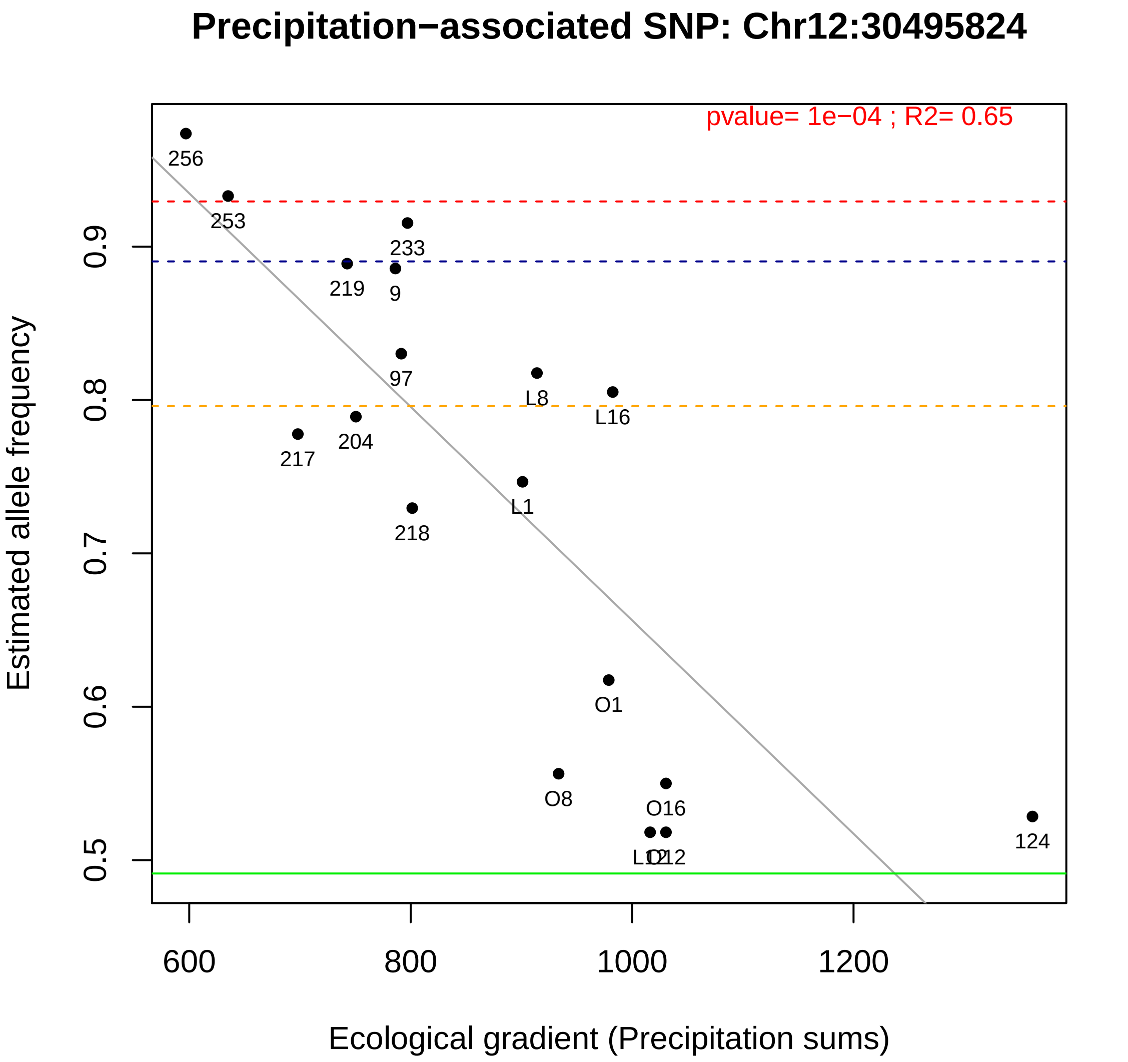

Precipitation−associated SNP: Chr12:30495824
pvalue= 1e−04 ; R2= 0.65
256
253
233
219
9
97
L8
L16
204
217
L1
218
O1
O8
O16
L12
O12
124
0.5
0.6
0.7
0.8
0.9
Estimated allele frequency
600
800
1000
1200
Ecological gradient (Precipitation sums)

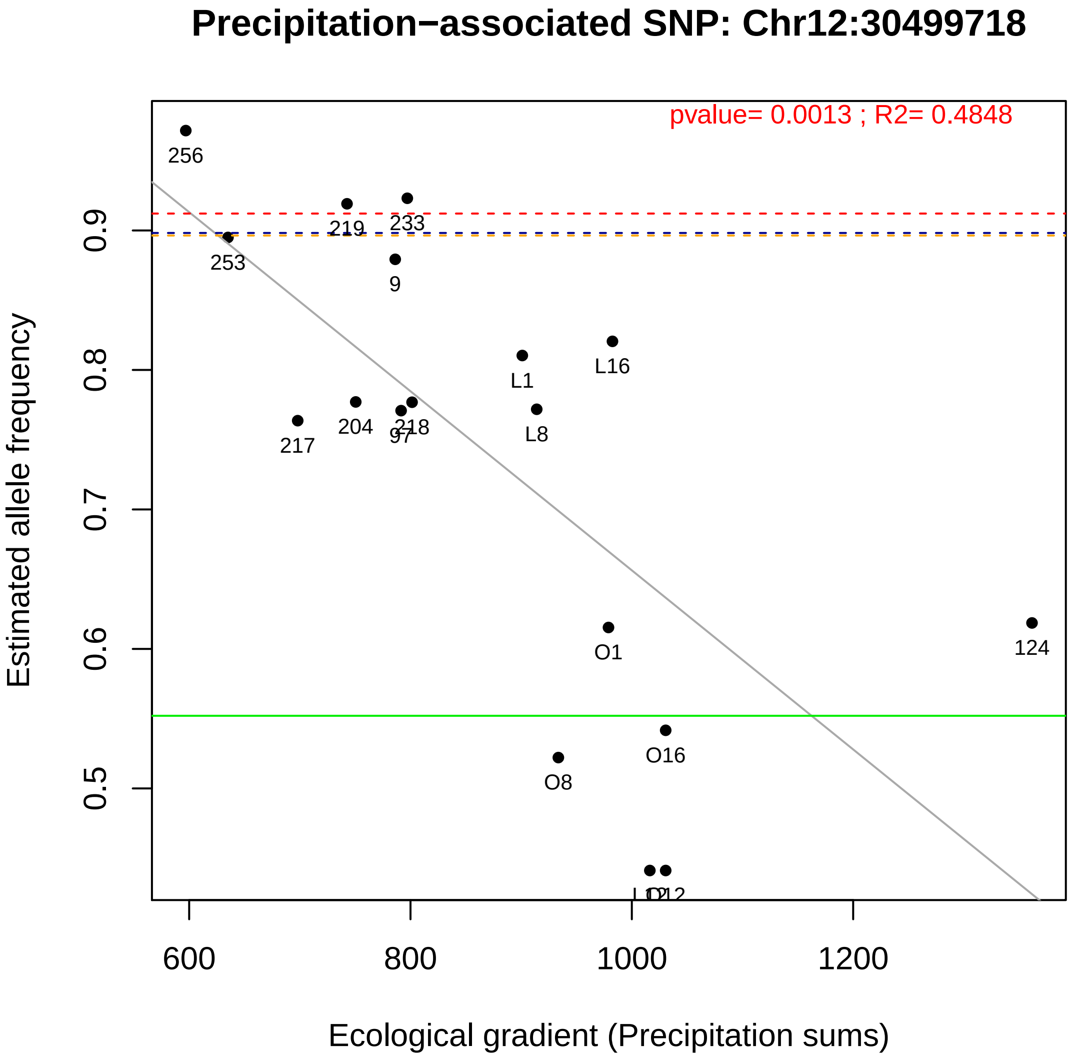
Precipitation−associated SNP: Chr12:30499718
pvalue= 0.0013 ; R2= 0.4848
Estimated allele frequency
0.5
0.6
0.7
0.8
0.9
600
800
1000
1200
Ecological gradient (Precipitation sums)
256
219
233
253
9
L16
L1
204
218
97
217
L8
O1
124
O16
O8

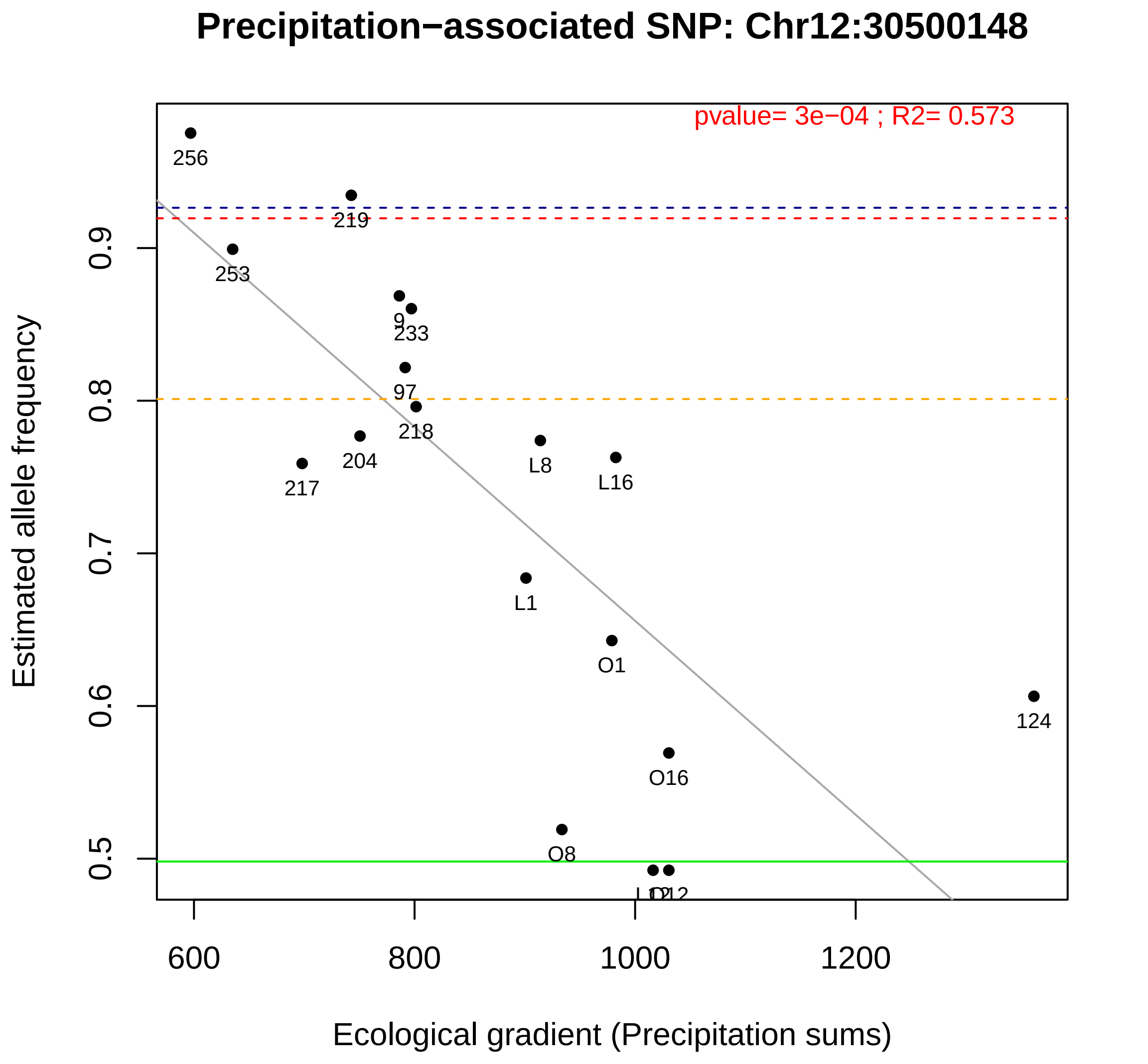

Precipitation−associated SNP: Chr12:30500148
pvalue= 3e−04 ; R2= 0.573
256
219
253
9
233
97
218
204
217
L8
L16
L1
O1
124
O16
O8
L12
O12
0.5
0.6
0.7
0.8
0.9
600
800
1000
1200
Estimated allele frequency
Ecological gradient (Precipitation sums)

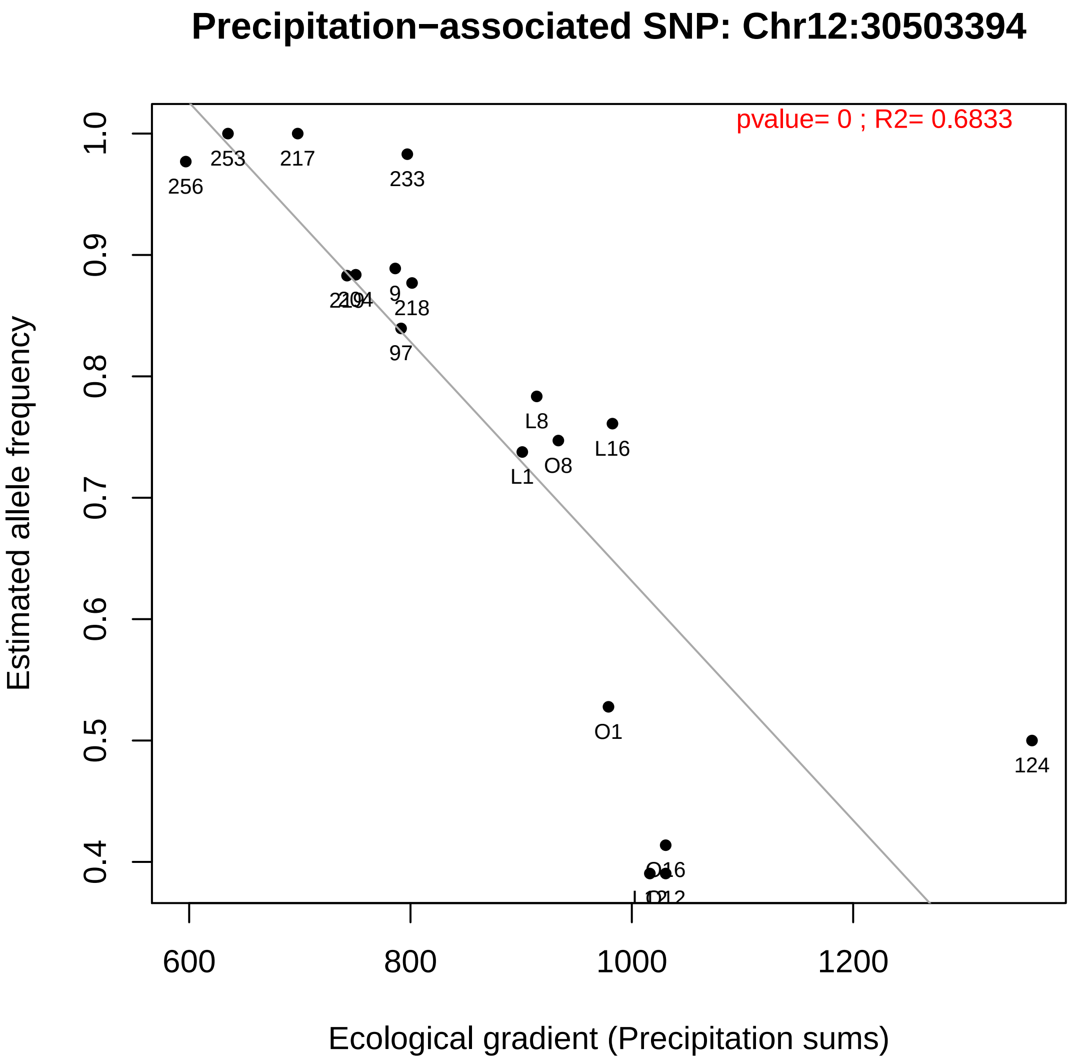

Precipitation−associated SNP: Chr12:30503394
pvalue= 0 ; R2= 0.6833
Estimated allele frequency
Ecological gradient (Precipitation sums)
0.4
0.5
0.6
0.7
0.8
0.9
1.0
600
800
1000
1200
256
253
217
233
9
218
97
L8
L16
L1
O8
O1
124
O16

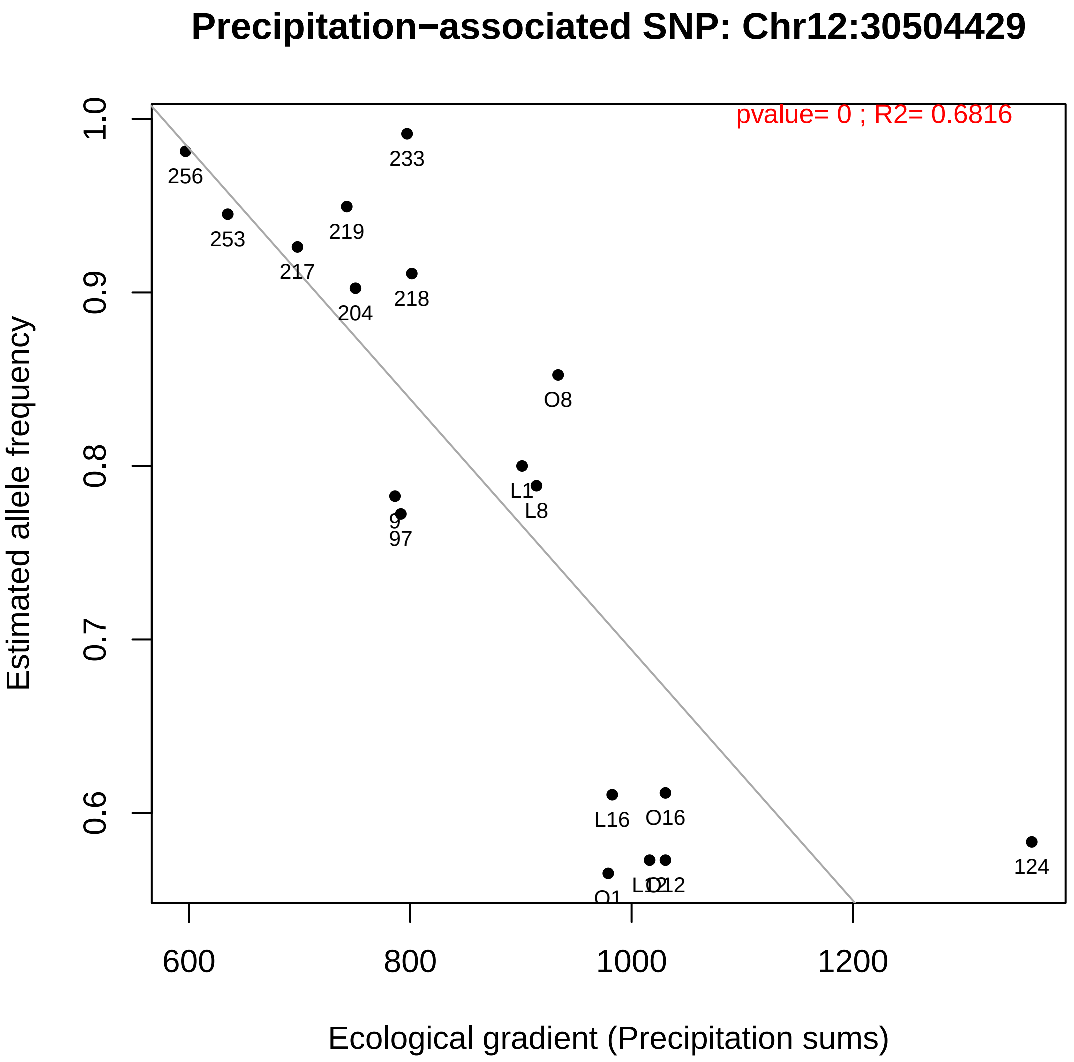

Precipitation−associated SNP: Chr12:30504429
pvalue= 0 ; R2= 0.6816
Estimated allele frequency
Ecological gradient (Precipitation sums)
1.0
0.9
0.8
0.7
0.6
600
800
1000
1200
256
233
253
219
217
204
218
O8
L1
L8
9
97
L16
O16
124
O1
L12
O12

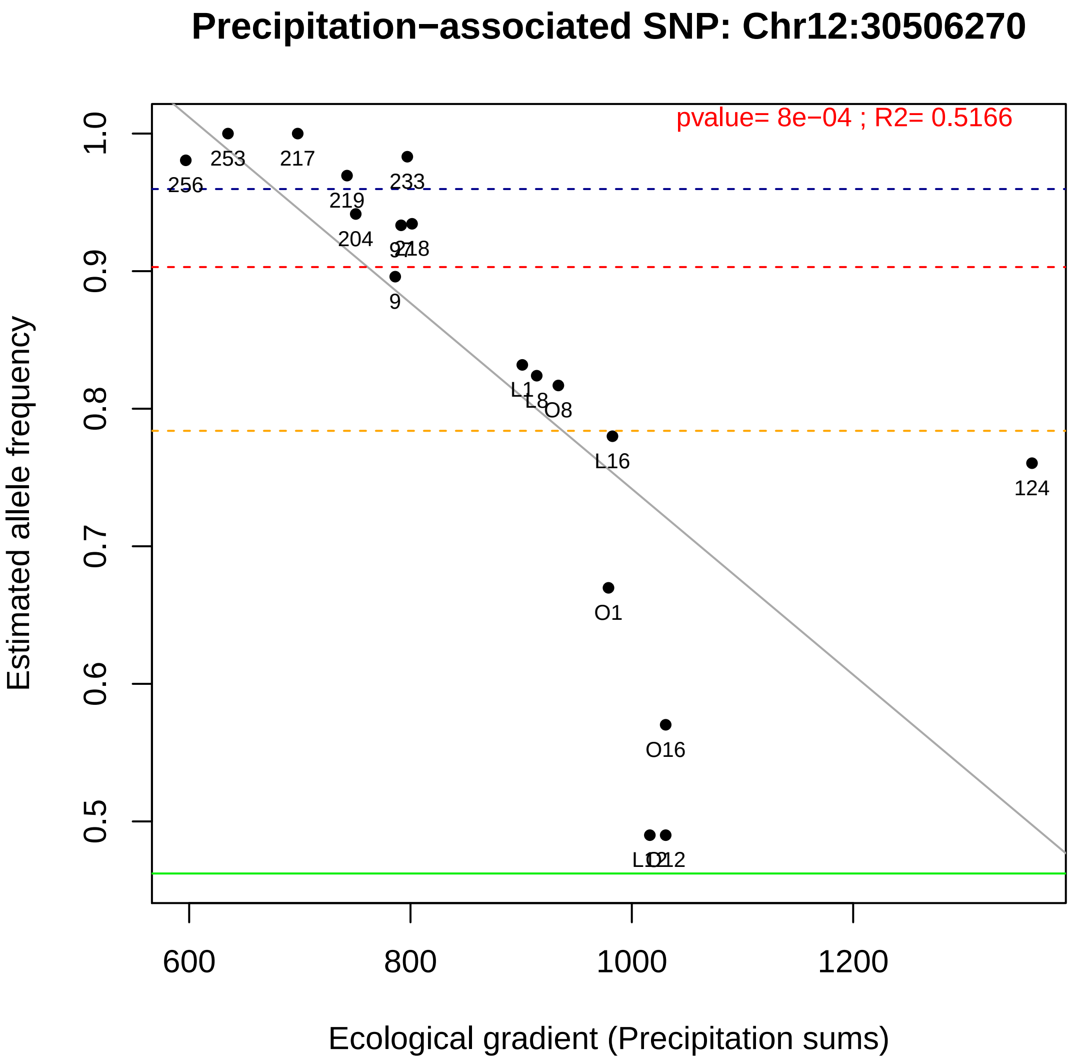
Precipitation−associated SNP: Chr12:30506270
pvalue= 8e−04 ; R2= 0.5166
Estimated allele frequency
Ecological gradient (Precipitation sums)
1.0
0.9
0.8
0.7
0.6
0.5
600
800
1000
1200
256
253
217
219
233
204
218
9
L1
L8
O8
L16
124
O1
O16
O12

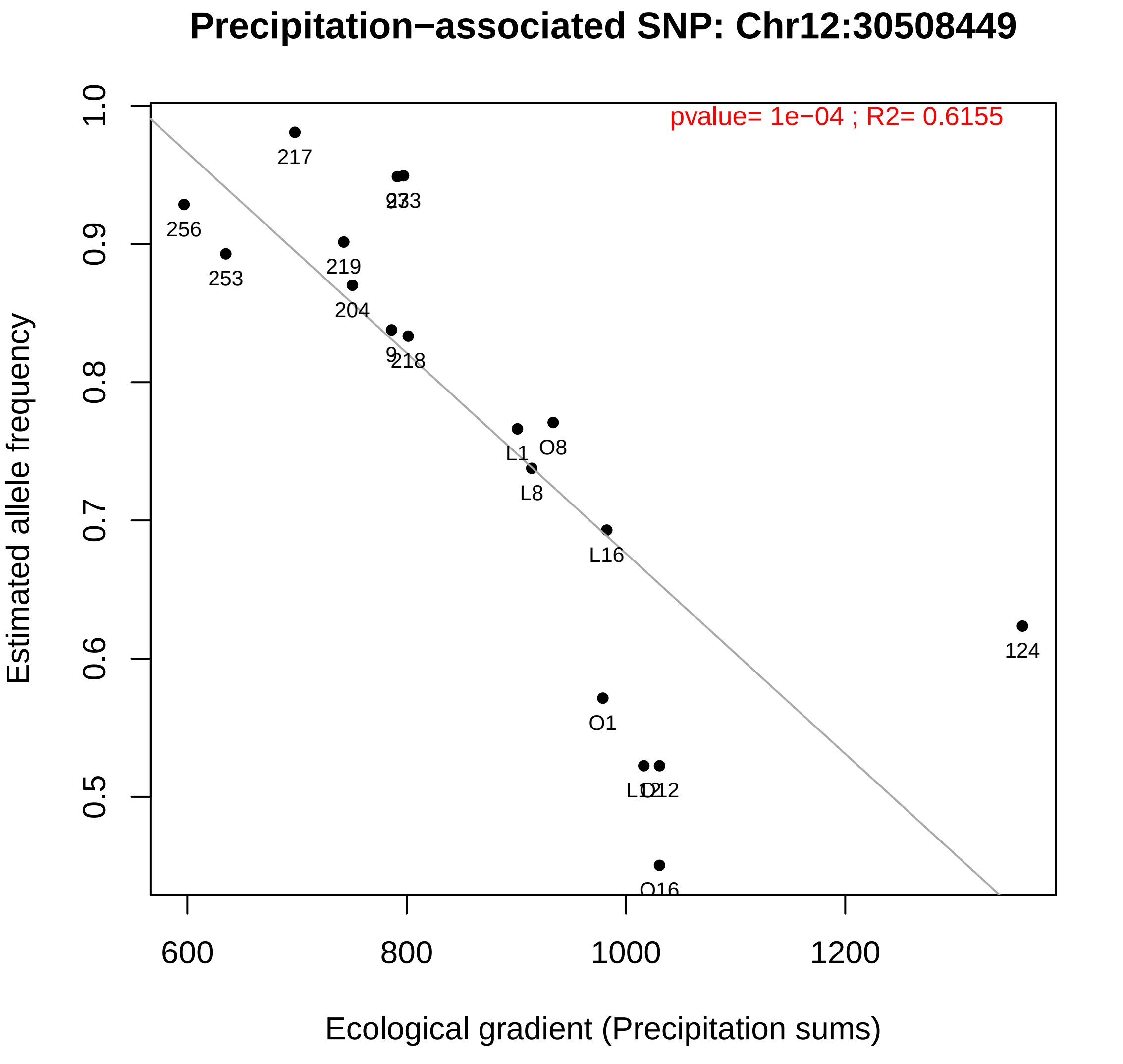
Precipitation−associated SNP: Chr12:30508449
pvalue= 1e−04 ; R2= 0.6155
Estimated allele frequency
Ecological gradient (Precipitation sums)
1.0
0.9
0.8
0.7
0.6
0.5
600
800
1000
1200
217
256
253
219
204
218
L1
O8
L8
L16
124
O1
O16

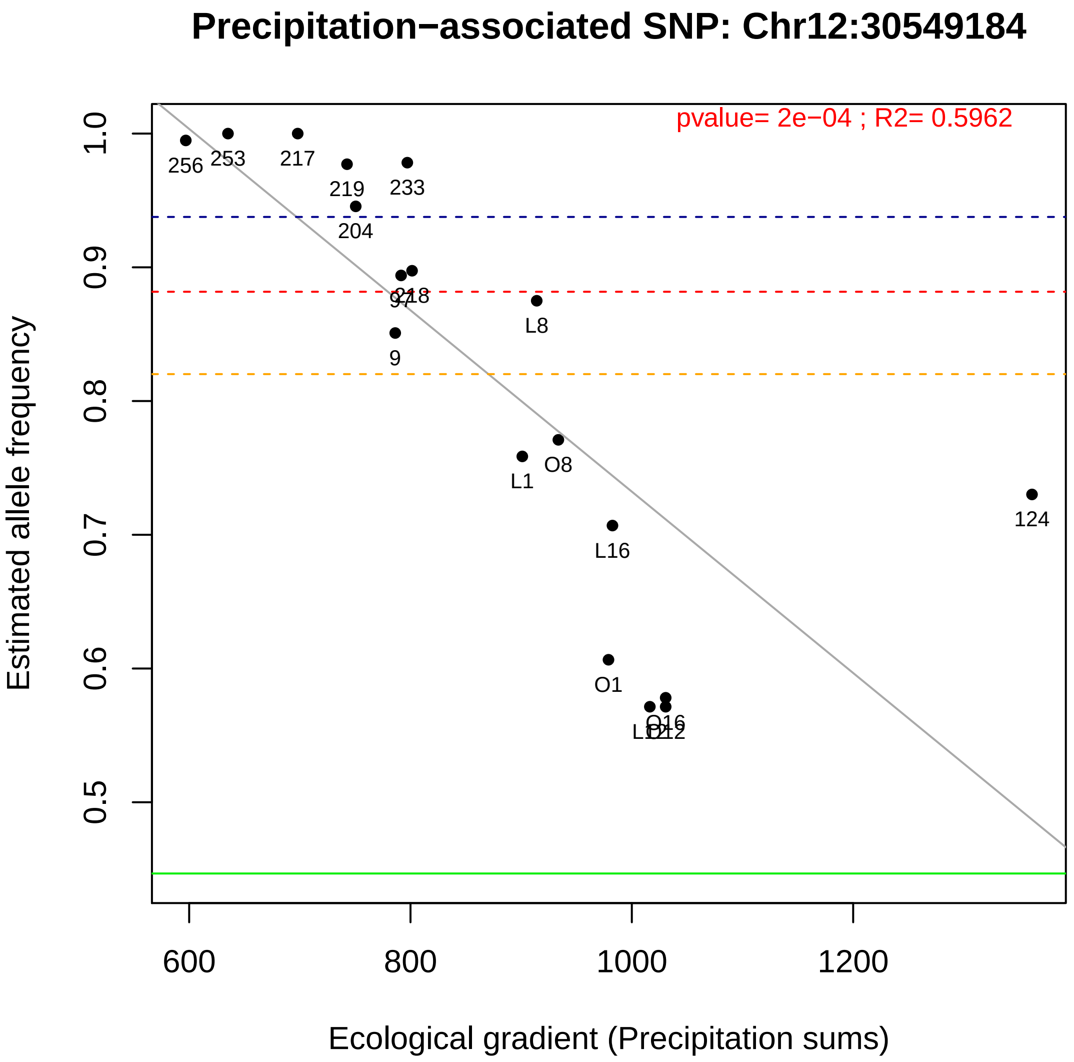

Precipitation−associated SNP: Chr12:30549184
pvalue= 2e−04 ; R2= 0.5962
256
253
217
219
233
204
97
218
L8
9
L1
O8
124
L16
O1
O16
L10
O12
1.0
0.9
0.8
0.7
0.6
0.5
600
800
1000
1200
Estimated allele frequency
Ecological gradient (Precipitation sums)

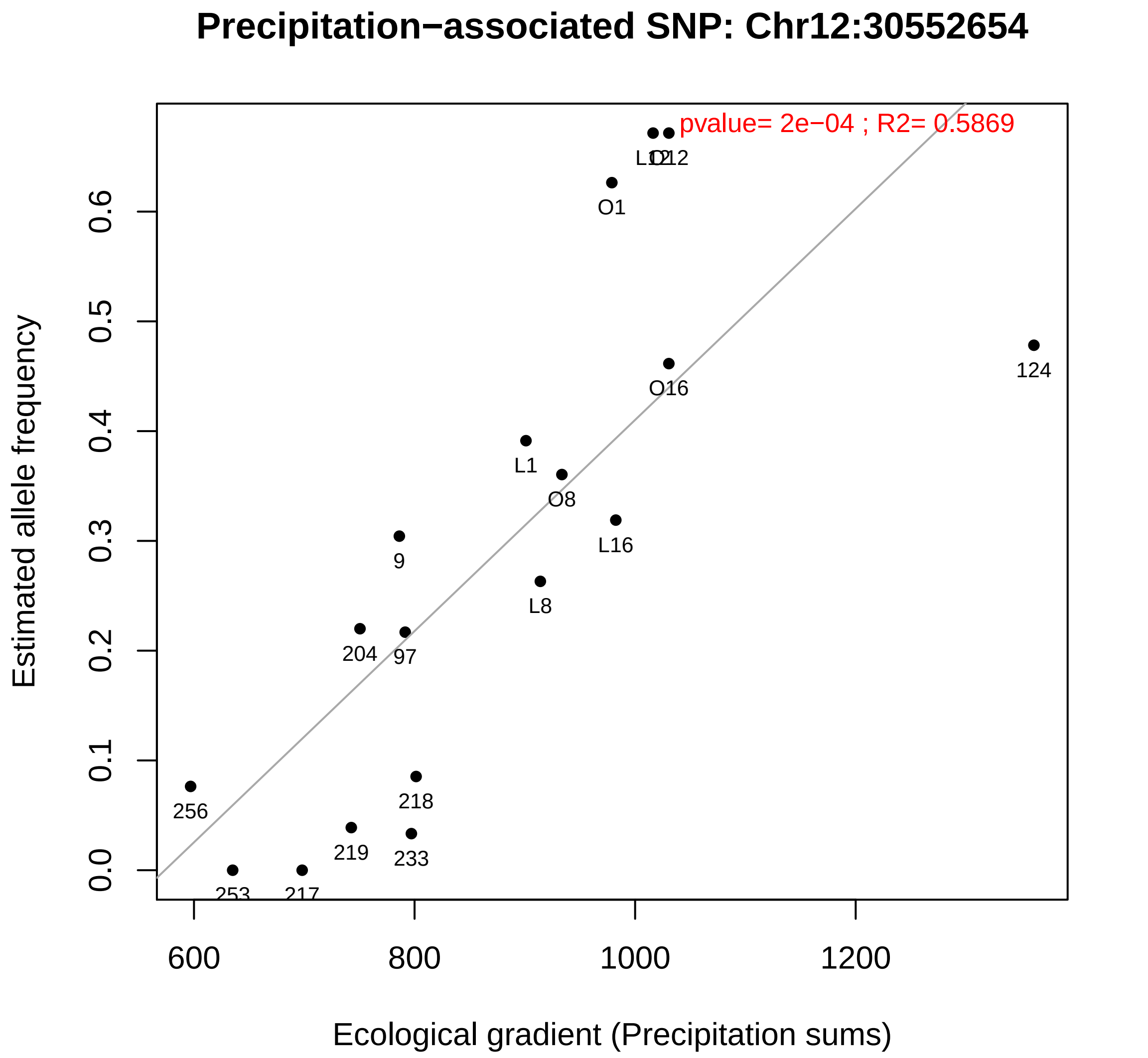
Precipitation−associated SNP: Chr12:30552654
pvalue= 2e−04 ; R2= 0.5869
Estimated allele frequency
Ecological gradient (Precipitation sums)
0.0
0.1
0.2
0.3
0.4
0.5
0.6
600
800
1000
1200
L12
O12
O1
124
O16
L1
O8
L16
9
L8
204
97
256
218
219
233
253
217

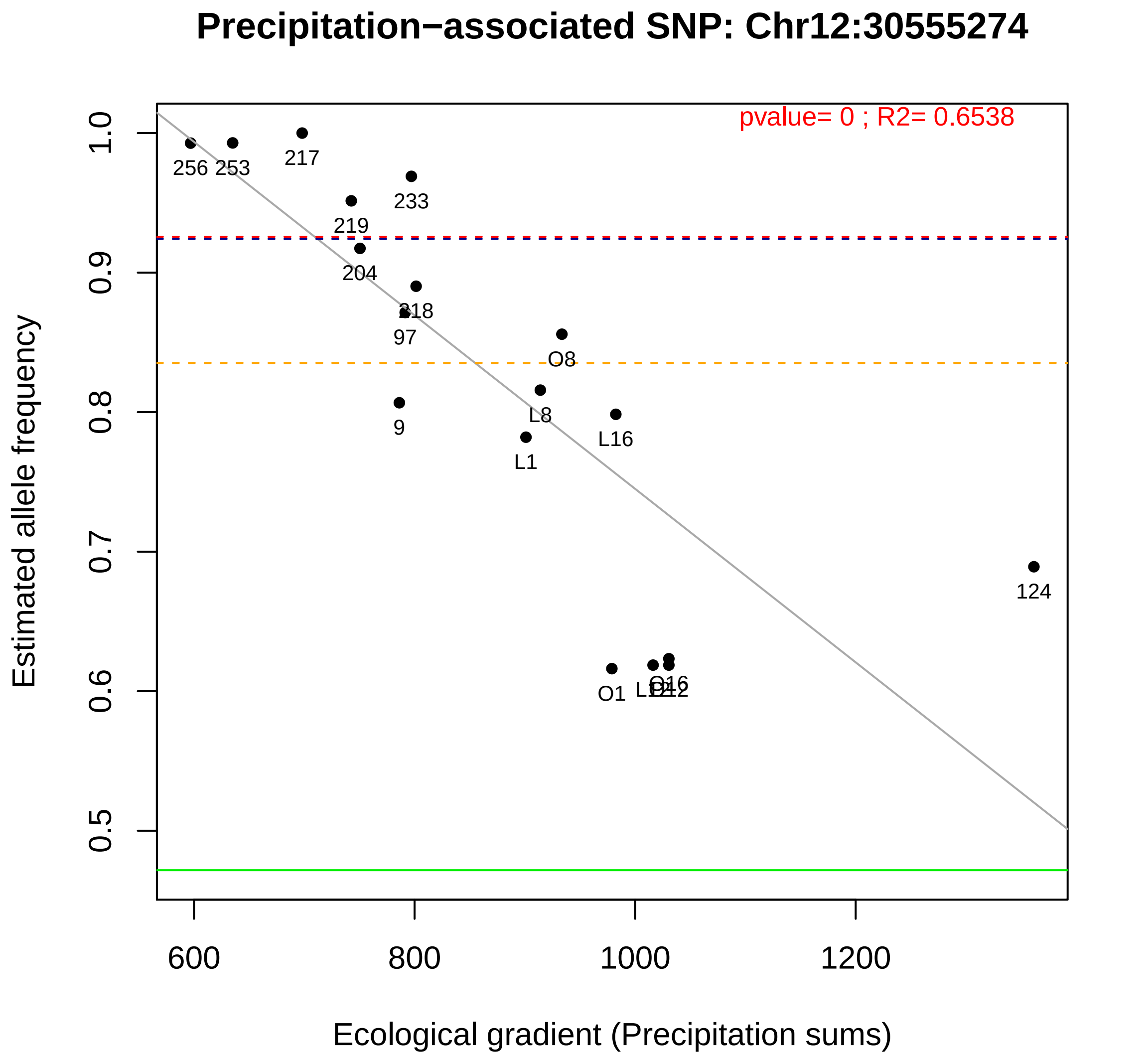
Precipitation−associated SNP: Chr12:30555274
pvalue= 0 ; R2= 0.6538
Estimated allele frequency
Ecological gradient (Precipitation sums)
1.0
0.9
0.8
0.7
0.6
0.5
600
800
1000
1200
256
253
217
233
219
204
218
97
O8
9
L8
L16
L1
124
O1
O16
O12

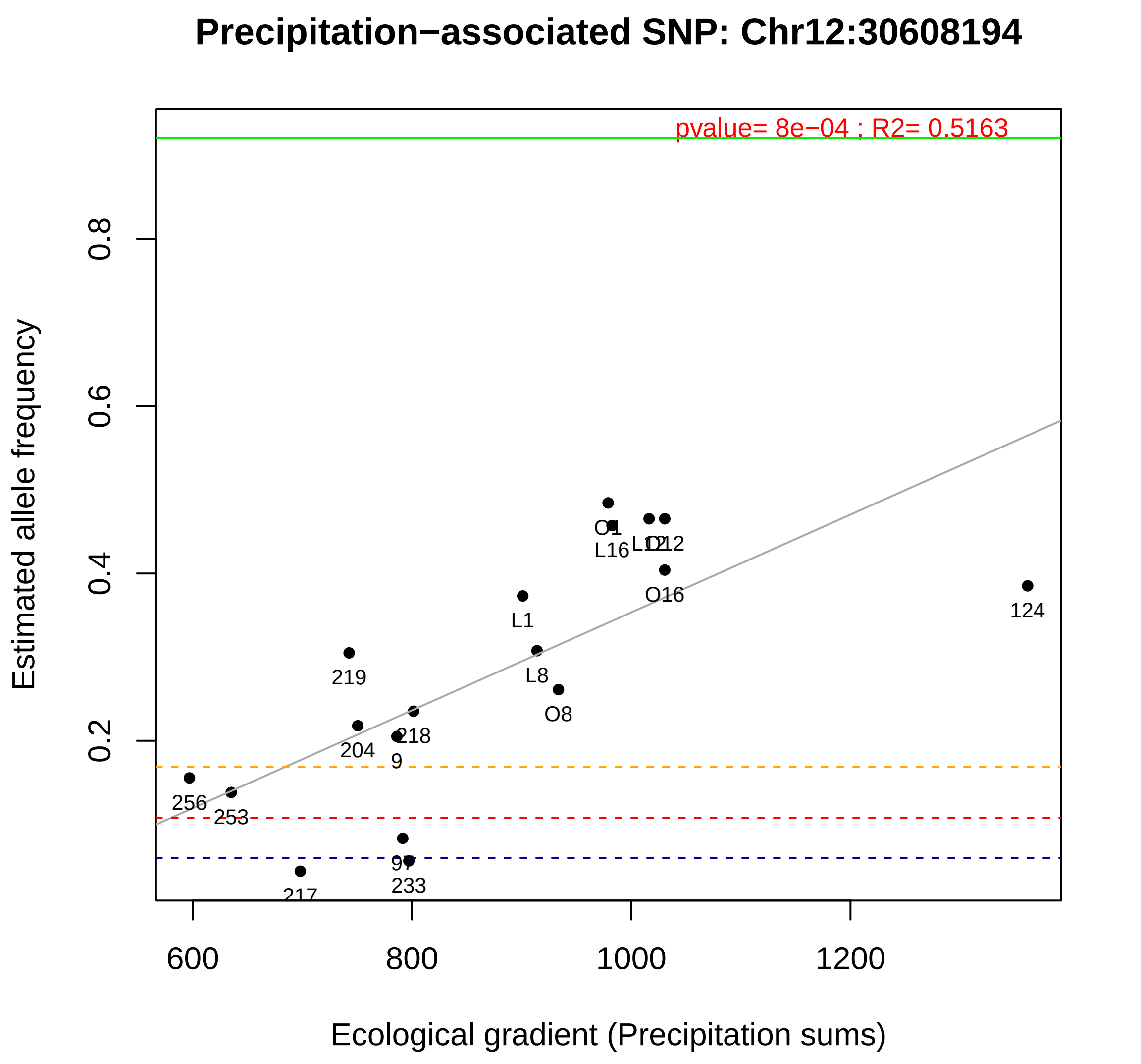

Precipitation−associated SNP: Chr12:30608194
pvalue= 8e−04 ; R2= 0.5163
Estimated allele frequency
0.2
0.4
0.6
0.8
600
800
1000
1200
Ecological gradient (Precipitation sums)
O1
L16
L10
O12
O16
124
L1
219
L8
O8
218
204
9
256
253
97
233
217

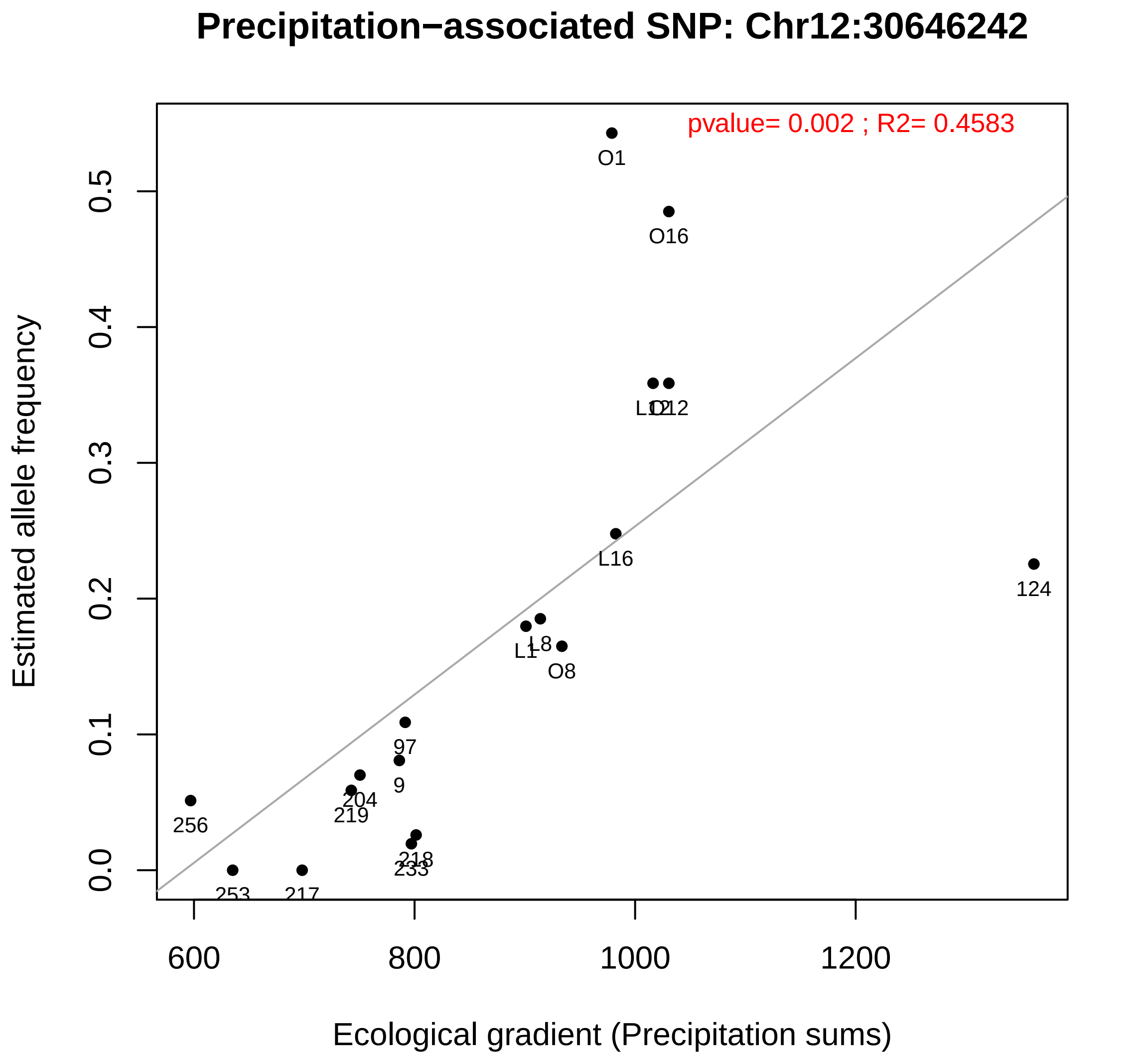

Precipitation−associated SNP: Chr12:30646242
pvalue= 0.002 ; R2= 0.4583
O1
O16
L10
O12
L16
124
L1
L8
O8
97
9
204
219
256
218
233
253
217
0.0
0.1
0.2
0.3
0.4
0.5
600
800
1000
1200
Estimated allele frequency
Ecological gradient (Precipitation sums)

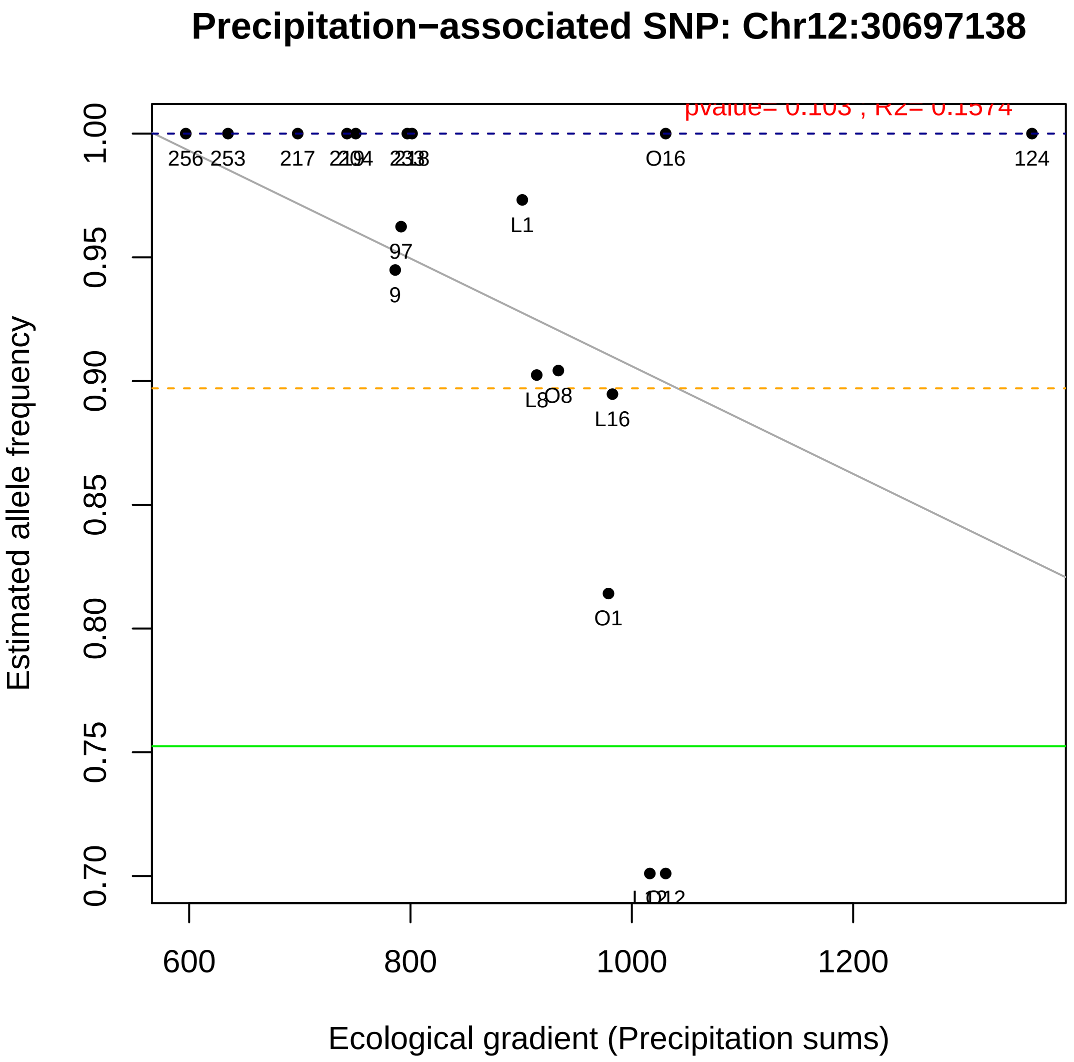

Precipitation−associated SNP: Chr12:30697138
pvalue= 0.103 , R2= 0.1574
Estimated allele frequency
Ecological gradient (Precipitation sums)
0.70
0.75
0.80
0.85
0.90
0.95
1.00
600
800
1000
1200
256
253
217
O16
124
L1
97
9
L8
O8
L16
O1

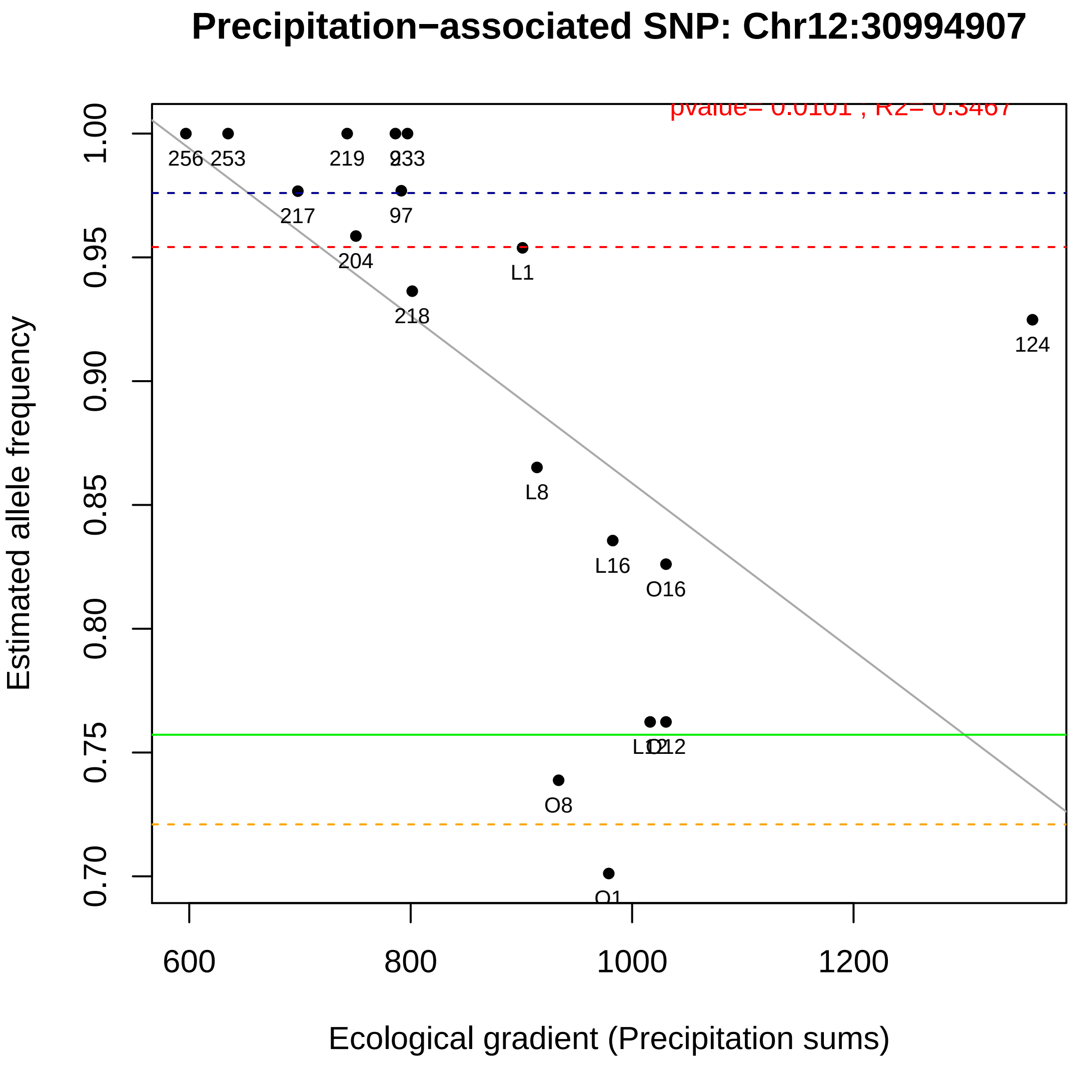

Precipitation−associated SNP: Chr12:30994907
pvalue= 0.0101 , R2= 0.3467
Estimated allele frequency
1.00
0.95
0.90
0.85
0.80
0.75
0.70
600
800
1000
1200
Ecological gradient (Precipitation sums)
256
253
219
217
97
204
L1
218
124
L8
L16
O16
O8
O1

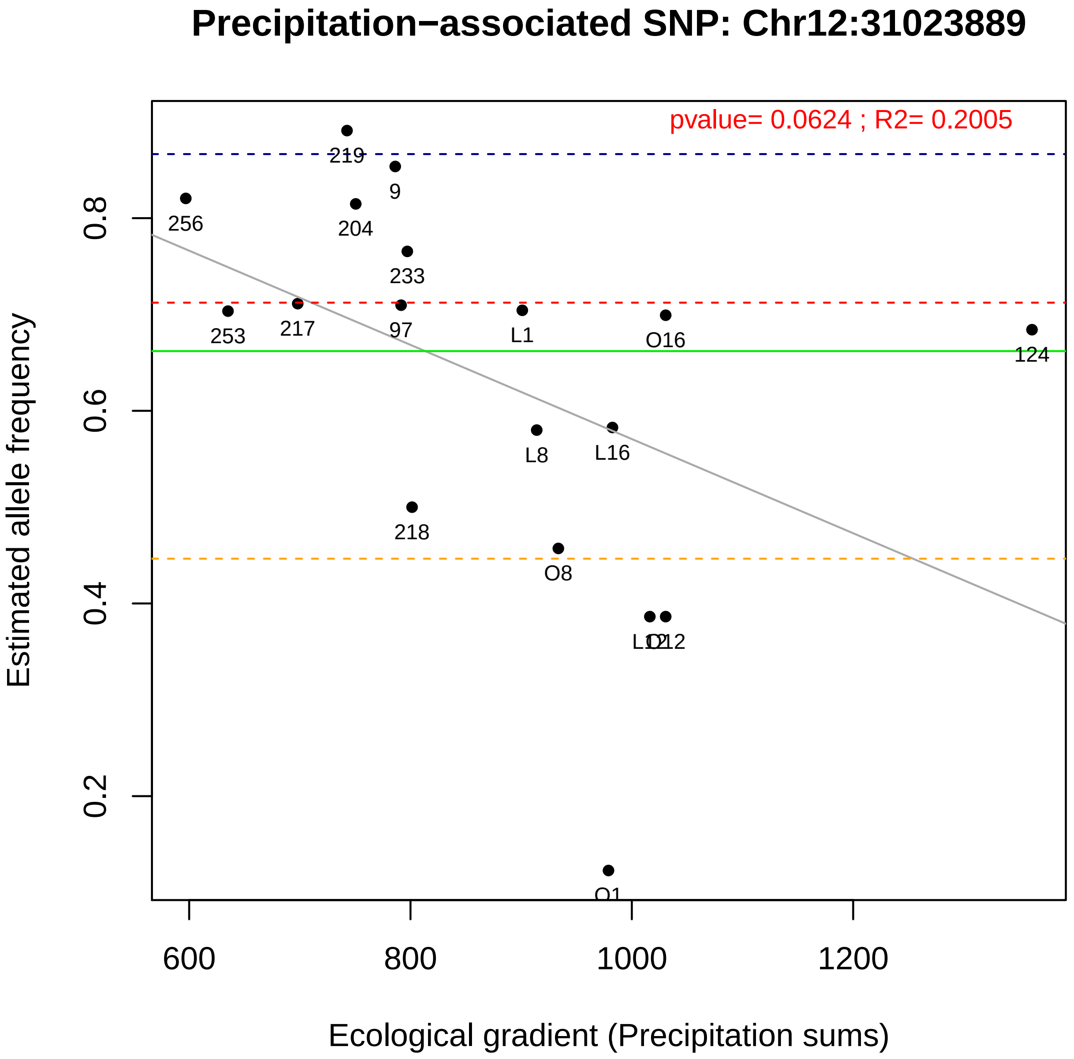
Precipitation−associated SNP: Chr12:31023889
pvalue= 0.0624 ; R2= 0.2005
Estimated allele frequency
0.2
0.4
0.6
0.8
600
800
1000
1200
Ecological gradient (Precipitation sums)
219
9
256
204
233
253
217
97
L1
O16
124
L8
L16
218
O8
O12

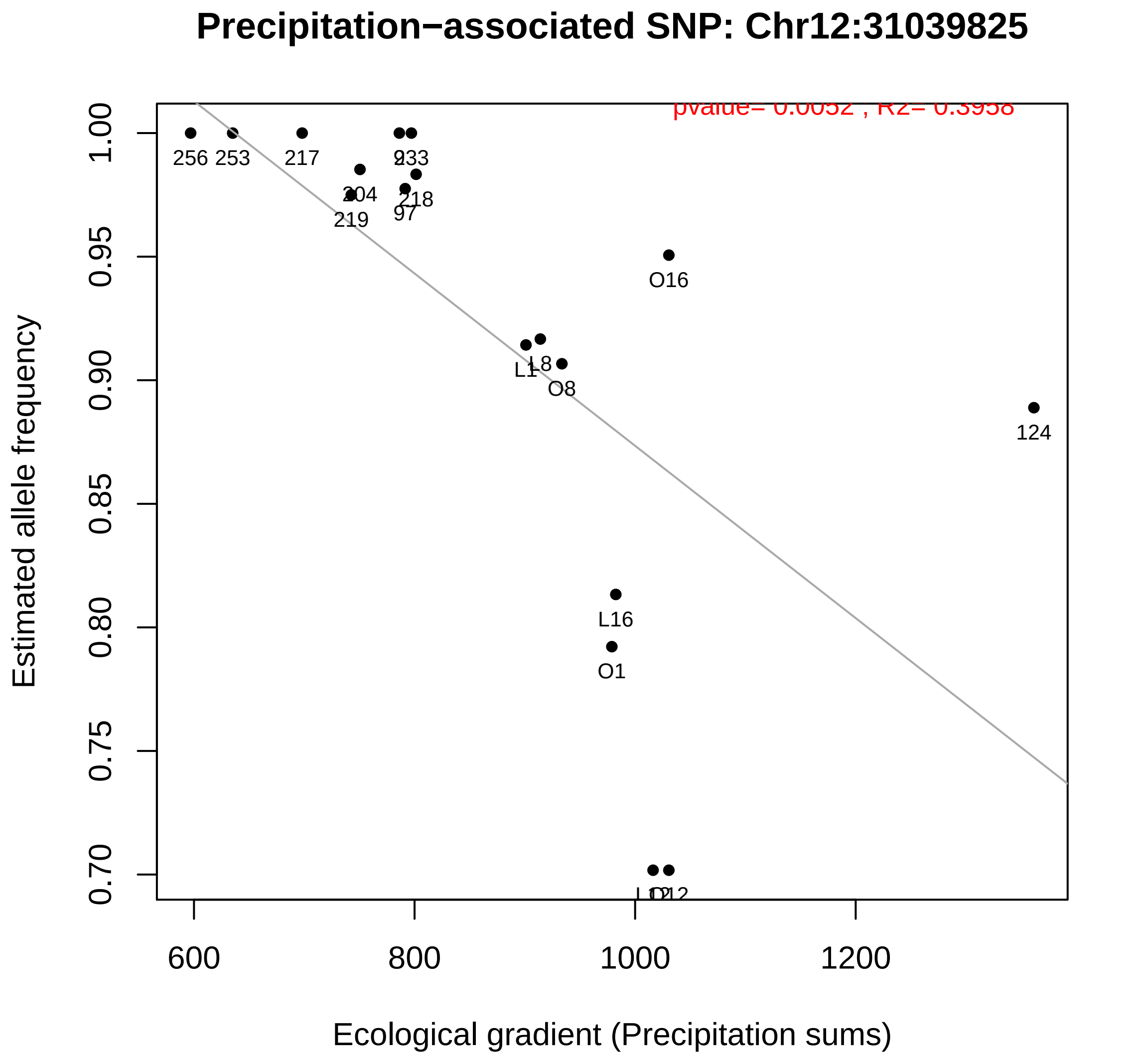
Precipitation−associated SNP: Chr12:31039825
pvalue= 0.0052 , R2= 0.3958
Estimated allele frequency
Ecological gradient (Precipitation sums)
1.00
0.95
0.90
0.85
0.80
0.75
0.70
600
800
1000
1200
256
253
217
233
204
218
219
97
O16
L8
L1
O8
124
L16
O1
O12

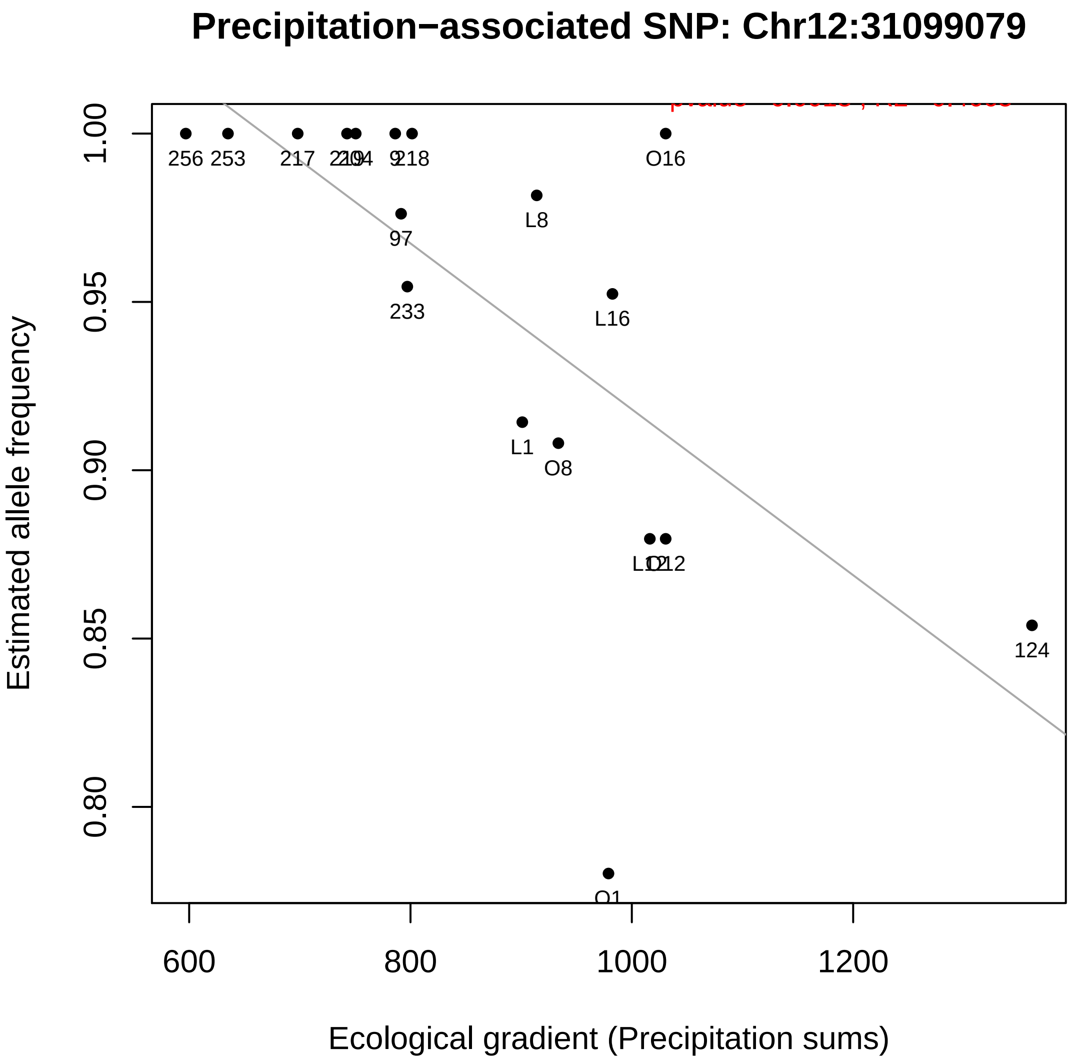

Precipitation−associated SNP: Chr12:31099079
Estimated allele frequency
Ecological gradient (Precipitation sums)
1.00
0.95
0.90
0.85
0.80
600
800
1000
1200
256
253
217
218
O16
L8
97
233
L16
L1
O8
124

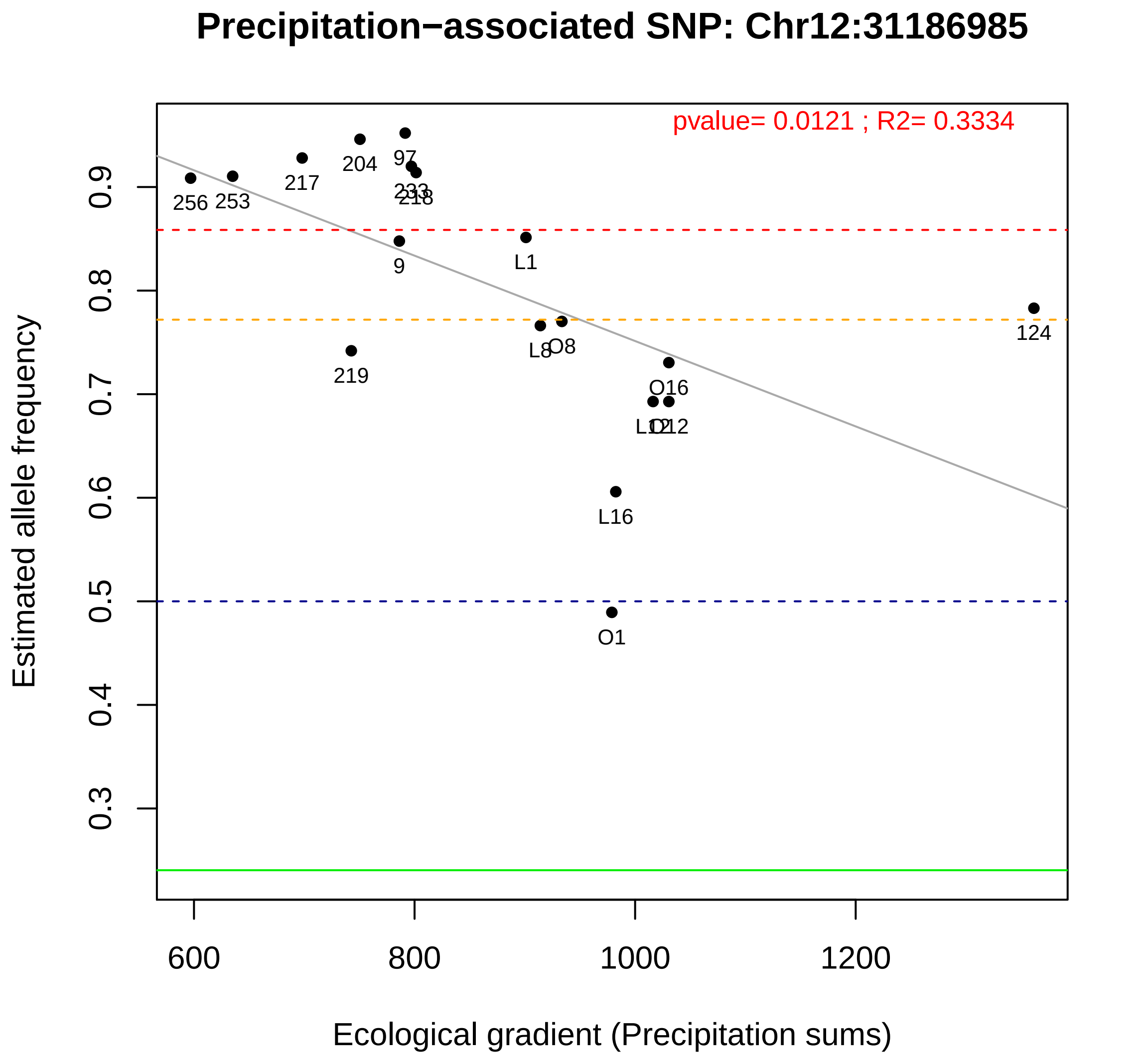
Precipitation−associated SNP: Chr12:31186985
pvalue= 0.0121 ; R2= 0.3334
Estimated allele frequency
Ecological gradient (Precipitation sums)
0.3
0.4
0.5
0.6
0.7
0.8
0.9
600
800
1000
1200
256
253
217
204
97
233
218
9
L1
219
L8
O8
124
O16
L12
O12
L16
O1

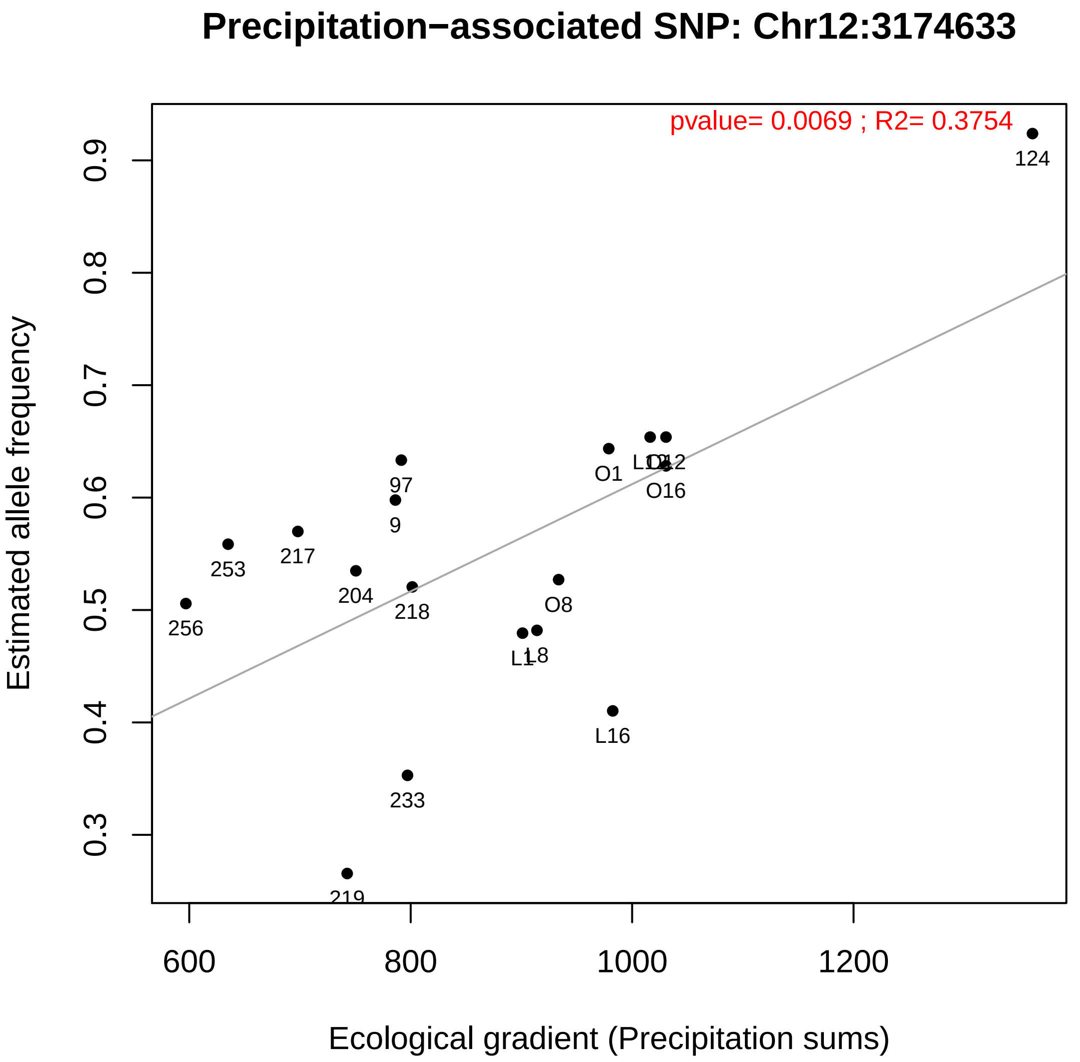
Precipitation−associated SNP: Chr12:3174633
pvalue= 0.0069 ; R2= 0.3754
Estimated allele frequency
Ecological gradient (Precipitation sums)
0.3
0.4
0.5
0.6
0.7
0.8
0.9
600
800
1000
1200
124
O1
L12
O12
O16
97
9
253
217
204
218
256
O8
L1
L8
L16
233
219

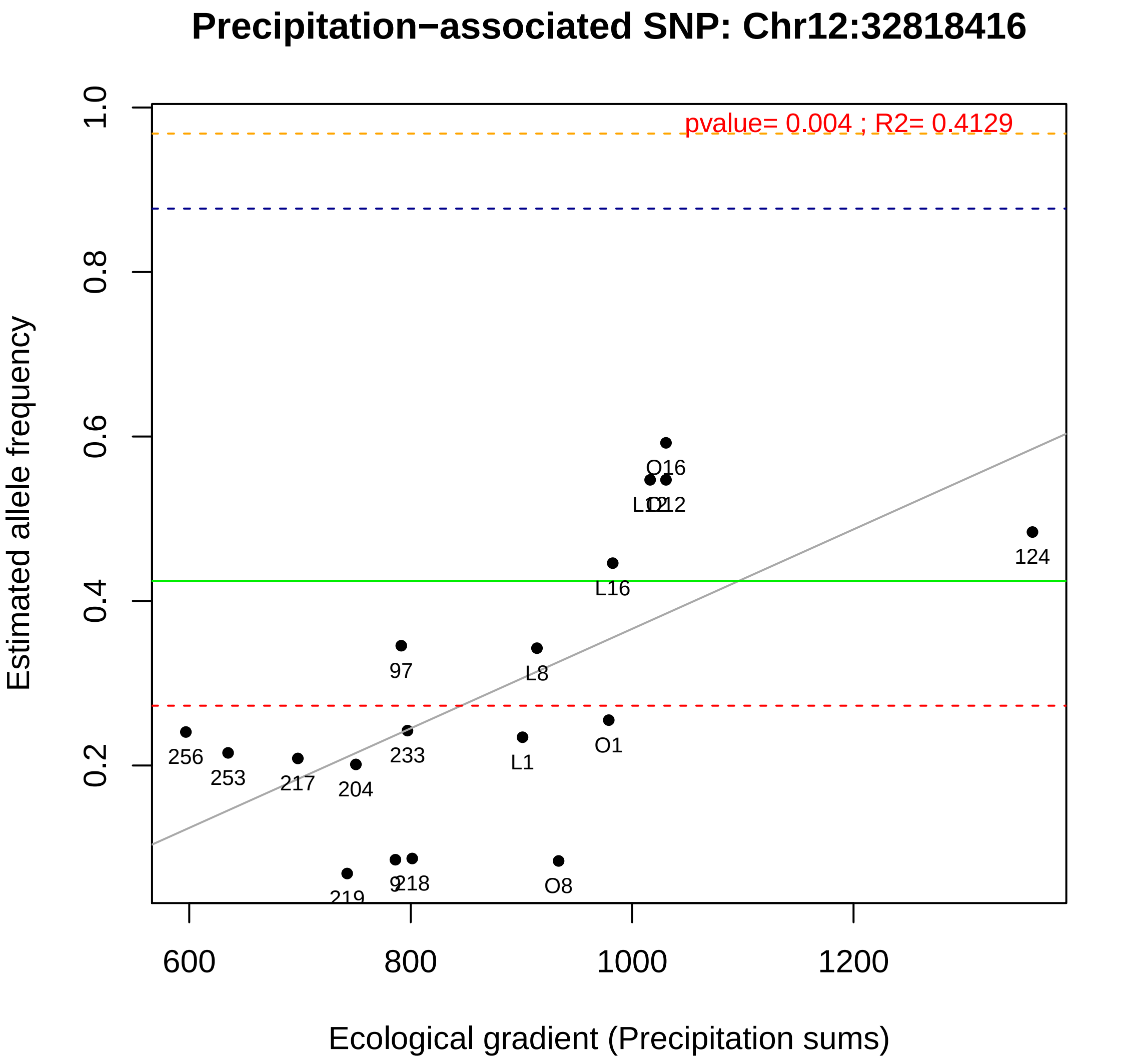
Precipitation−associated SNP: Chr12:32818416
pvalue= 0.004 ; R2= 0.4129
Estimated allele frequency
0.2
0.4
0.6
0.8
1.0
600
800
1000
1200
Ecological gradient (Precipitation sums)
O16
L12
O12
124
L16
97
L8
256
253
217
204
233
L1
O1
219
9
218
O8

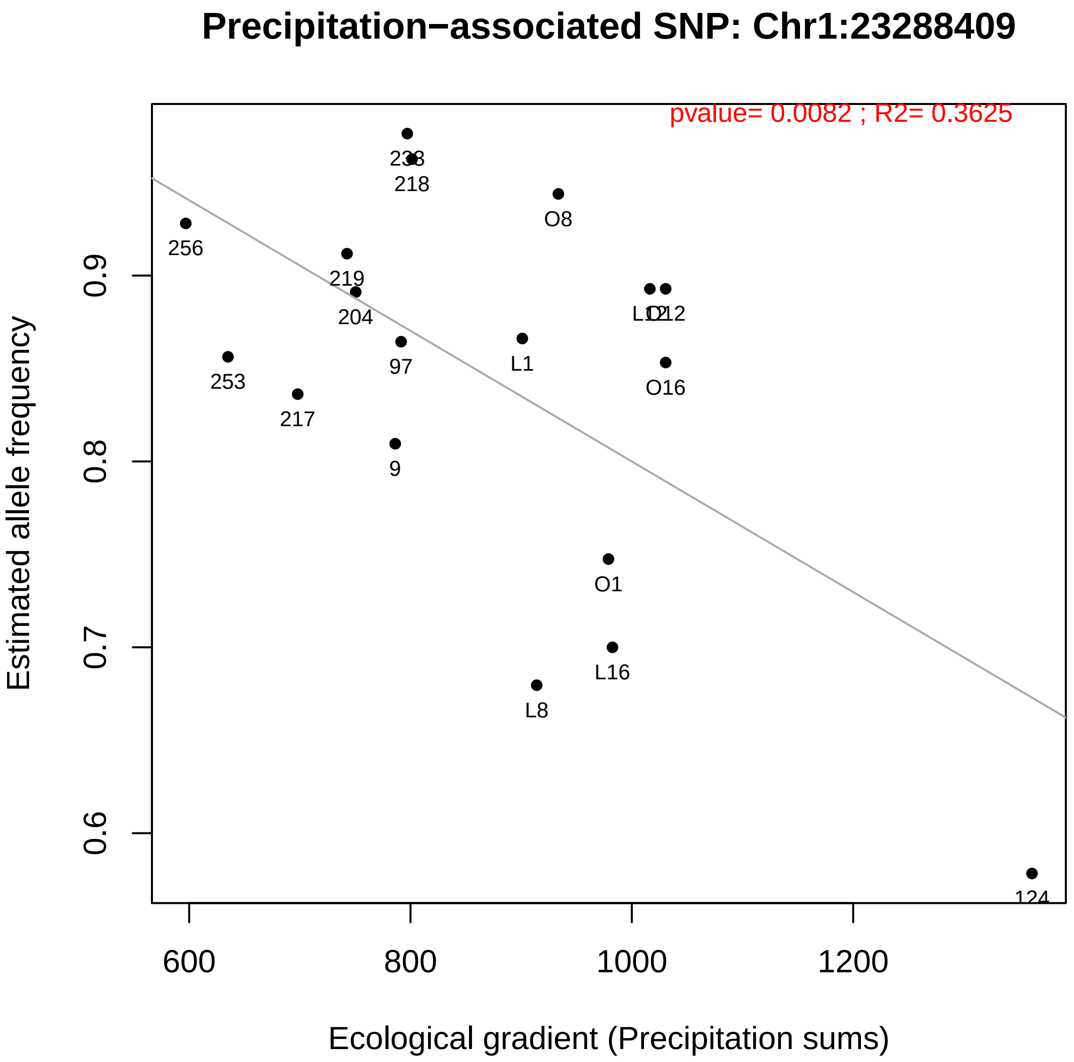

Precipitation−associated SNP: Chr1:23288409
pvalue= 0.0082 ; R2= 0.3625
Estimated allele frequency
0.6
0.7
0.8
0.9
600
800
1000
1200
Ecological gradient (Precipitation sums)
256
219
204
218
97
253
217
9
O8
L1
O16
O1
L16
L8
124

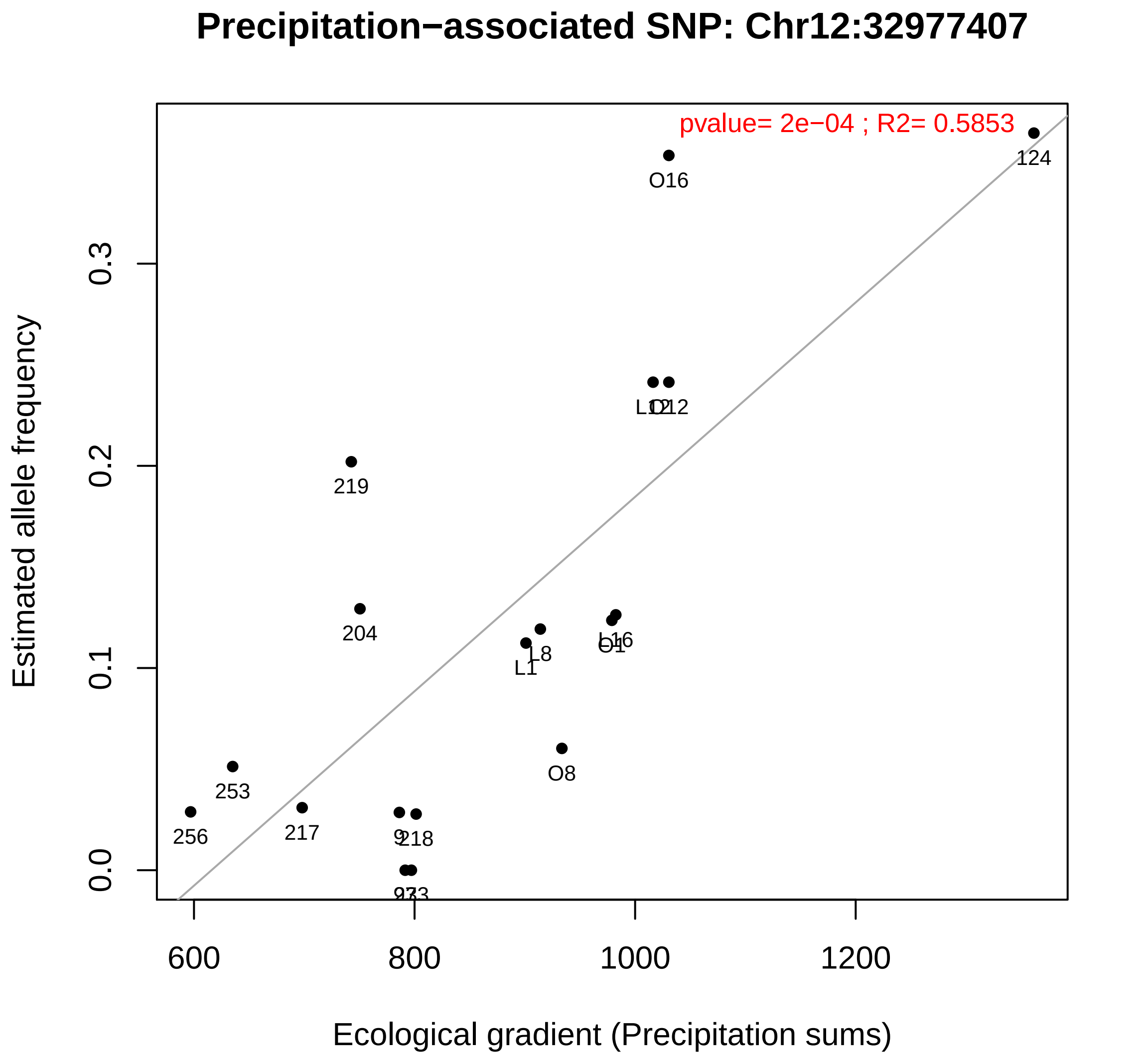
Precipitation−associated SNP: Chr12:32977407
pvalue= 2e−04 ; R2= 0.5853
124
O16
L10
O12
219
204
L16
O1
L8
L1
O8
253
256
217
9
218
0.0
0.1
0.2
0.3
600
800
1000
1200
Estimated allele frequency
Ecological gradient (Precipitation sums)

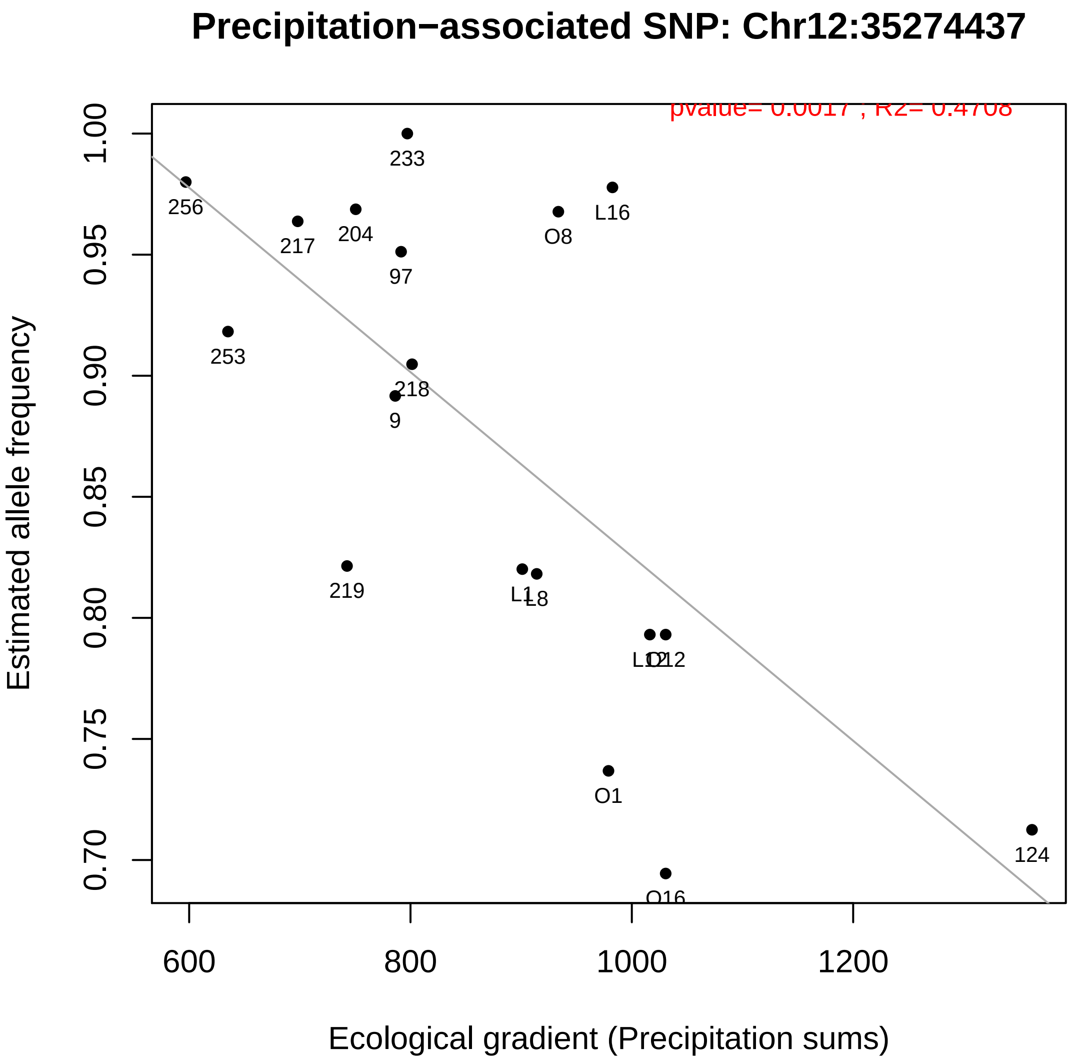
Precipitation−associated SNP: Chr12:35274437
pvalue= 0.0017 ; R2= 0.4708
Estimated allele frequency
1.00
0.95
0.90
0.85
0.80
0.75
0.70
600
800
1000
1200
Ecological gradient (Precipitation sums)
233
256
217
204
97
O8
L16
253
218
9
219
L1
L8
L12
O12
O1
124
O16

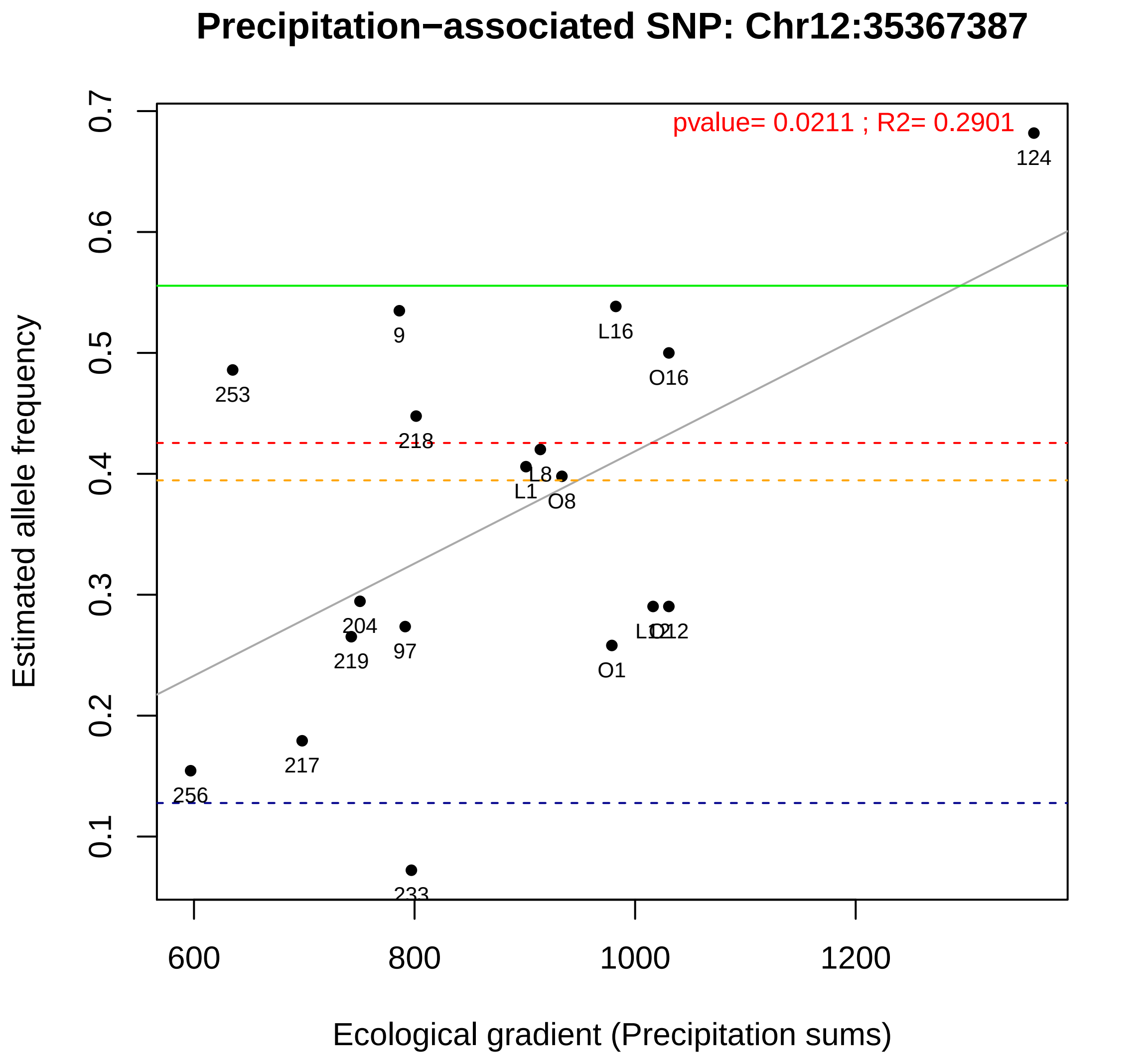

Precipitation−associated SNP: Chr12:35367387
pvalue= 0.0211 ; R2= 0.2901
Estimated allele frequency
Ecological gradient (Precipitation sums)
0.1
0.2
0.3
0.4
0.5
0.6
0.7
600
800
1000
1200
124
9
L16
O16
253
218
L8
L1
O8
204
97
219
L12
O12
O1
217
256
233

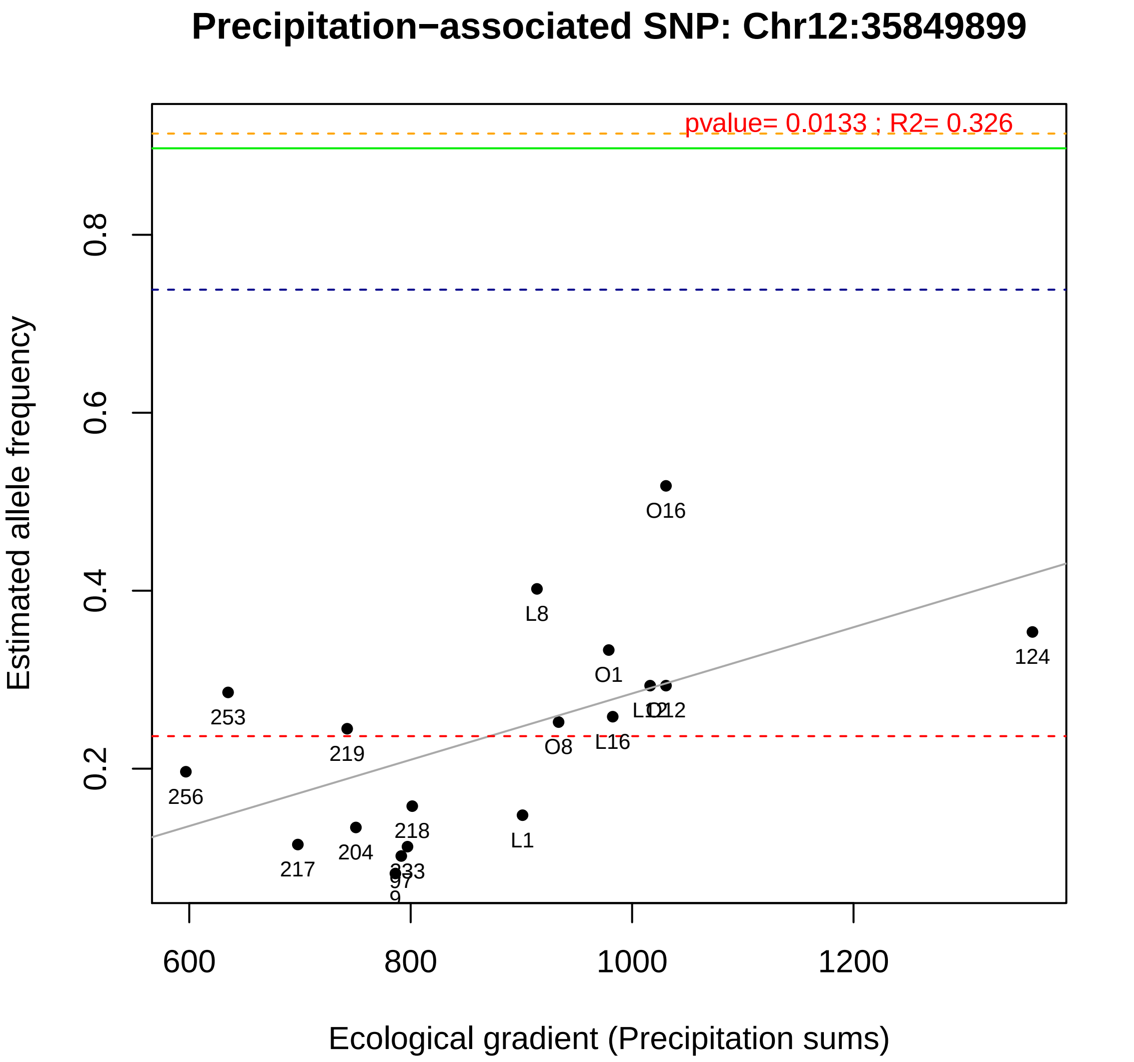
Precipitation−associated SNP: Chr12:35849899
pvalue= 0.0133 ; R2= 0.326
Estimated allele frequency
0.2
0.4
0.6
0.8
600
800
1000
1200
Ecological gradient (Precipitation sums)
O16
L8
124
O1
L12
O12
253
219
O8
L16
256
218
L1
204
217
333
97
9

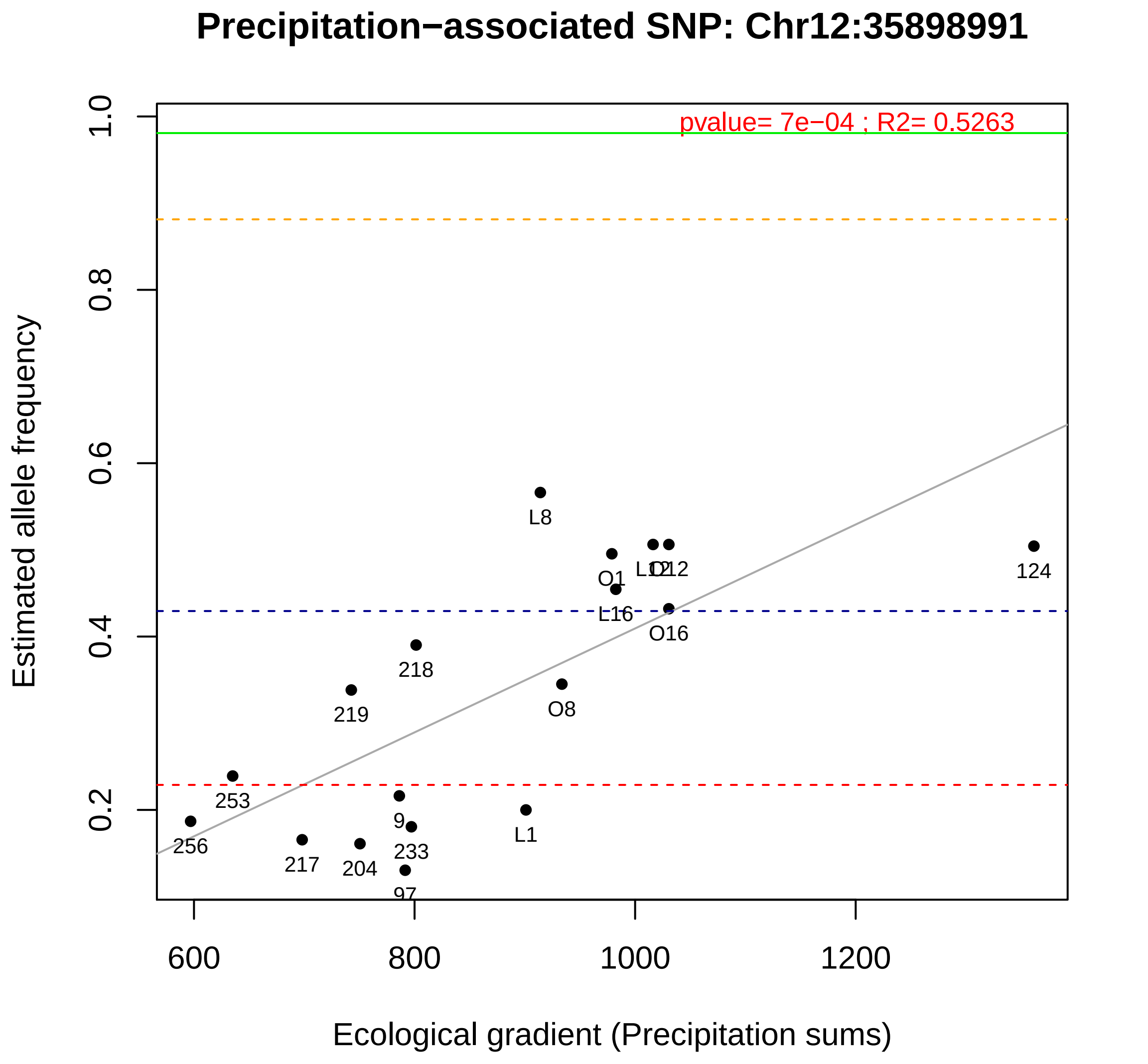
Precipitation−associated SNP: Chr12:35898991
pvalue= 7e−04 ; R2= 0.5263
Estimated allele frequency
0.2
0.4
0.6
0.8
1.0
600
800
1000
1200
Ecological gradient (Precipitation sums)
L8
O1
L12
O12
124
L16
O16
218
219
O8
253
256
9
233
L1
217
204
97

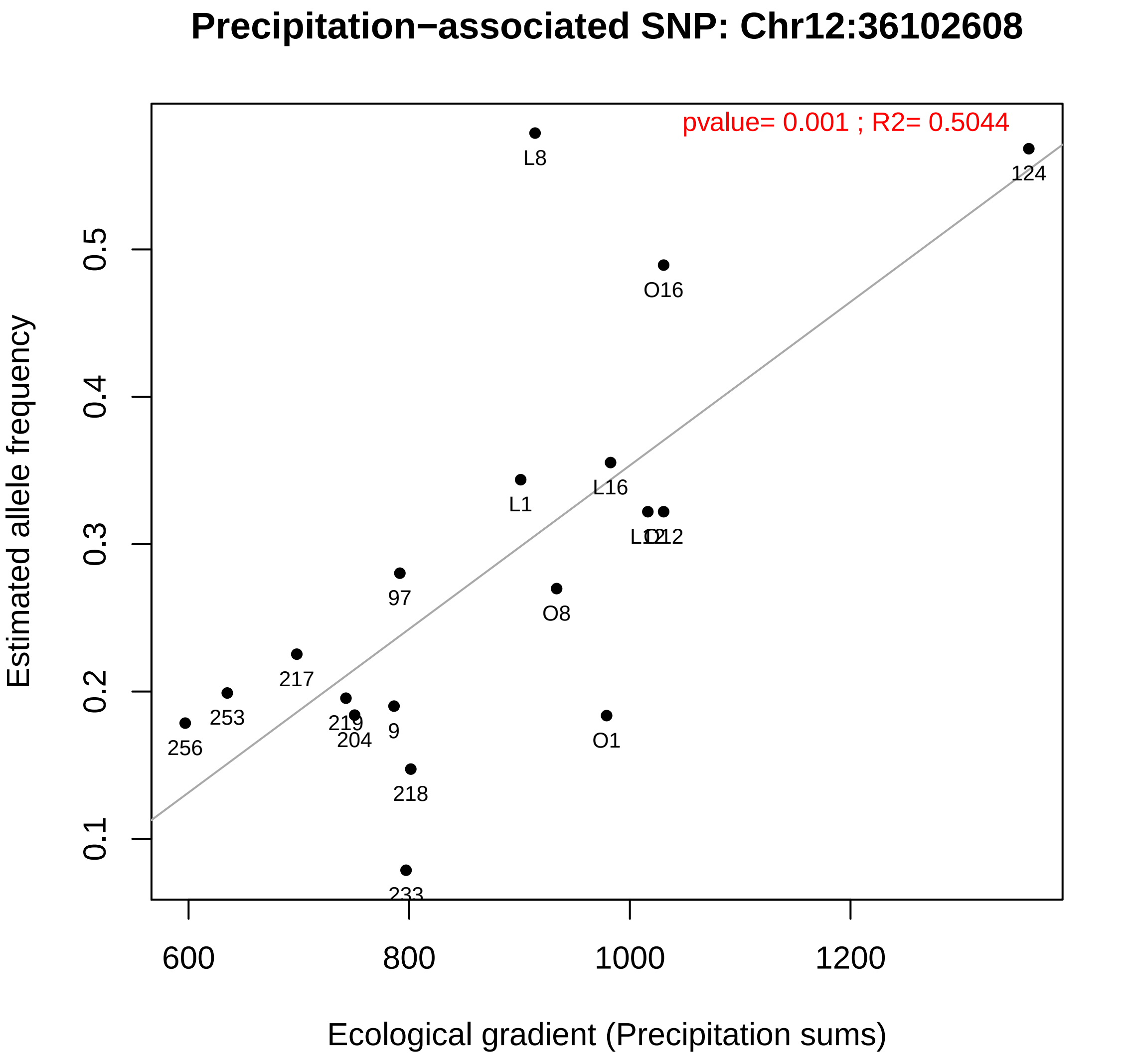
Precipitation−associated SNP: Chr12:36102608
pvalue= 0.001 ; R2= 0.5044
Estimated allele frequency
0.1
0.2
0.3
0.4
0.5
600
800
1000
1200
Ecological gradient (Precipitation sums)
L8
124
O16
L16
L1
L12
O12
97
O8
217
253
256
219
204
9
O1
218
233

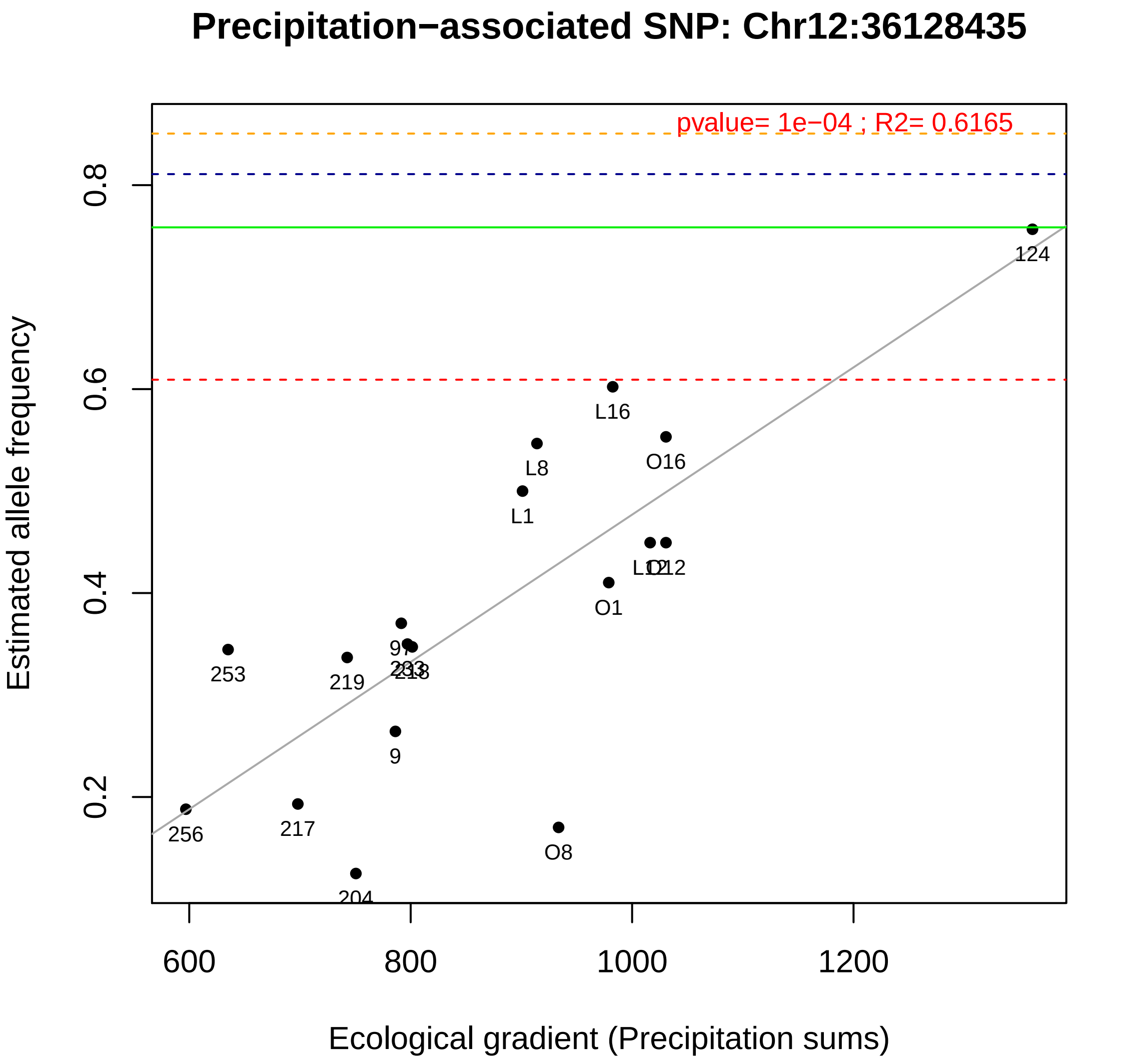
Precipitation−associated SNP: Chr12:36128435
pvalue= 1e−04 ; R2= 0.6165
Estimated allele frequency
0.2
0.4
0.6
0.8
600
800
1000
1200
Ecological gradient (Precipitation sums)
124
L16
O16
L8
L1
O12
O1
97
253
219
9
256
217
O8
204

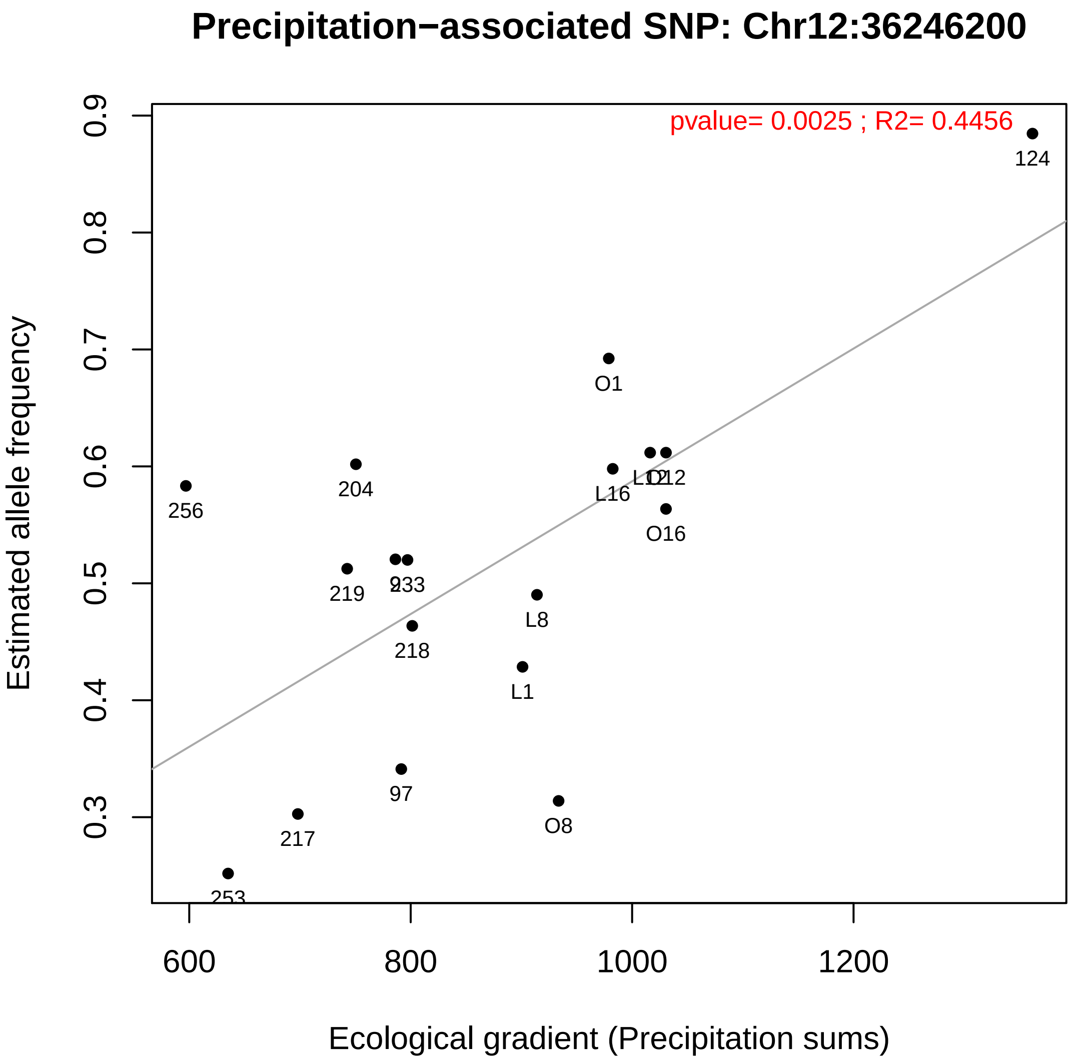
Precipitation−associated SNP: Chr12:36246200
pvalue= 0.0025 ; R2= 0.4456
124
O1
204
256
L16
O16
219
L8
218
L1
97
217
O8
253
0.9
0.8
0.7
0.6
0.5
0.4
0.3
Estimated allele frequency
600
800
1000
1200
Ecological gradient (Precipitation sums)

**Precipitation−associated SNP: Chr12:36253066**

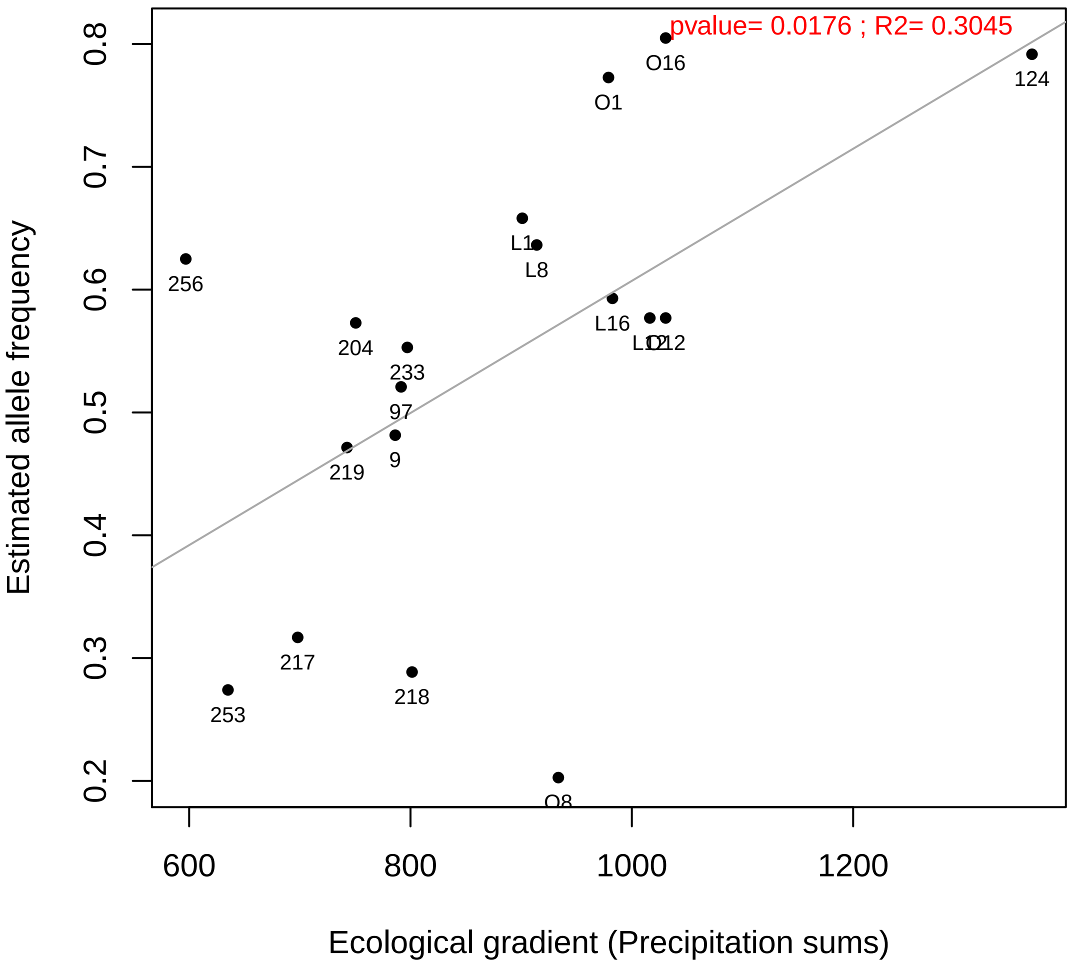

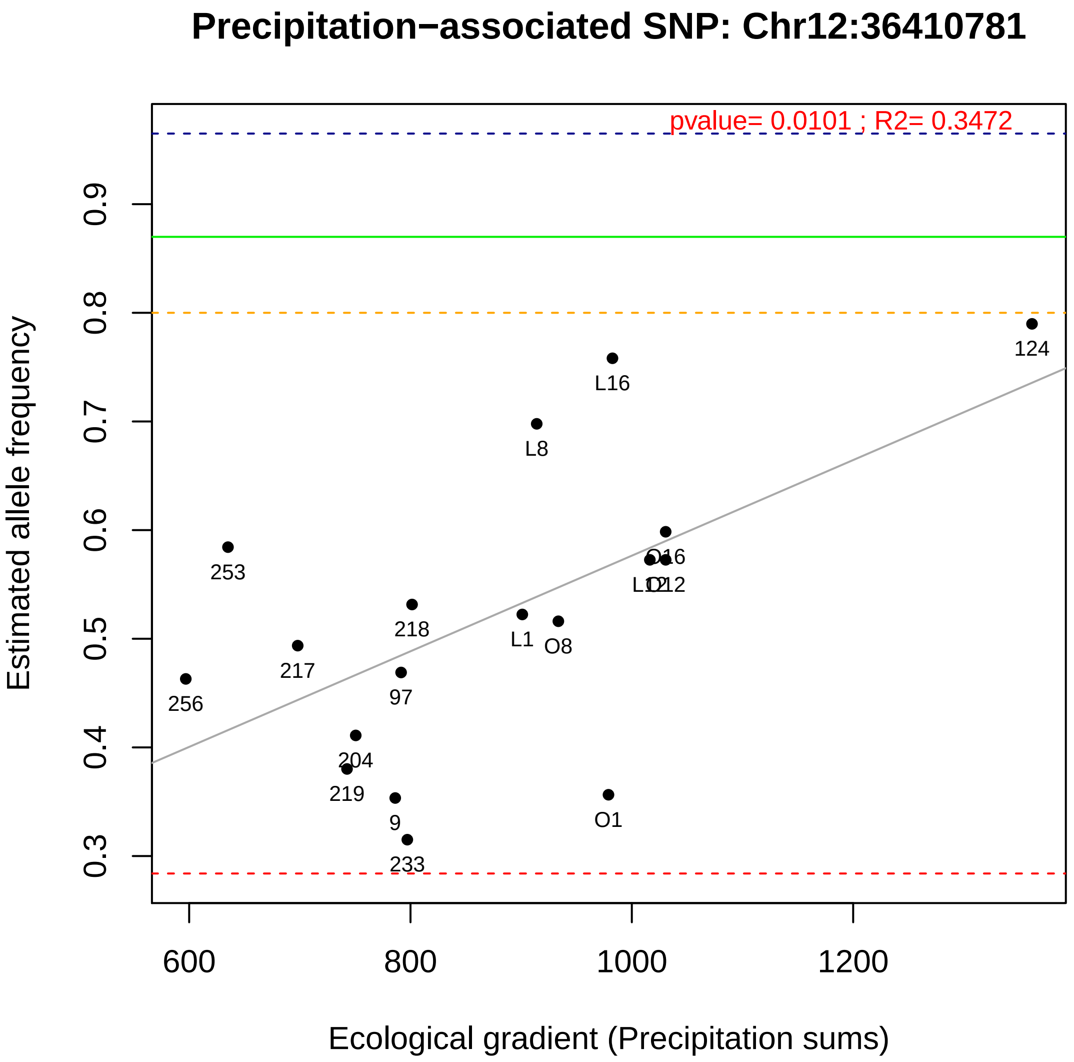

Precipitation−associated SNP: Chr12:36410781
pvalue= 0.0101 ; R2= 0.3472
Estimated allele frequency
0.3
0.4
0.5
0.6
0.7
0.8
0.9
600
800
1000
1200
Ecological gradient (Precipitation sums)
124
L16
L8
253
O16
L12
O12
218
L1
O8
217
97
256
204
219
9
O1
233

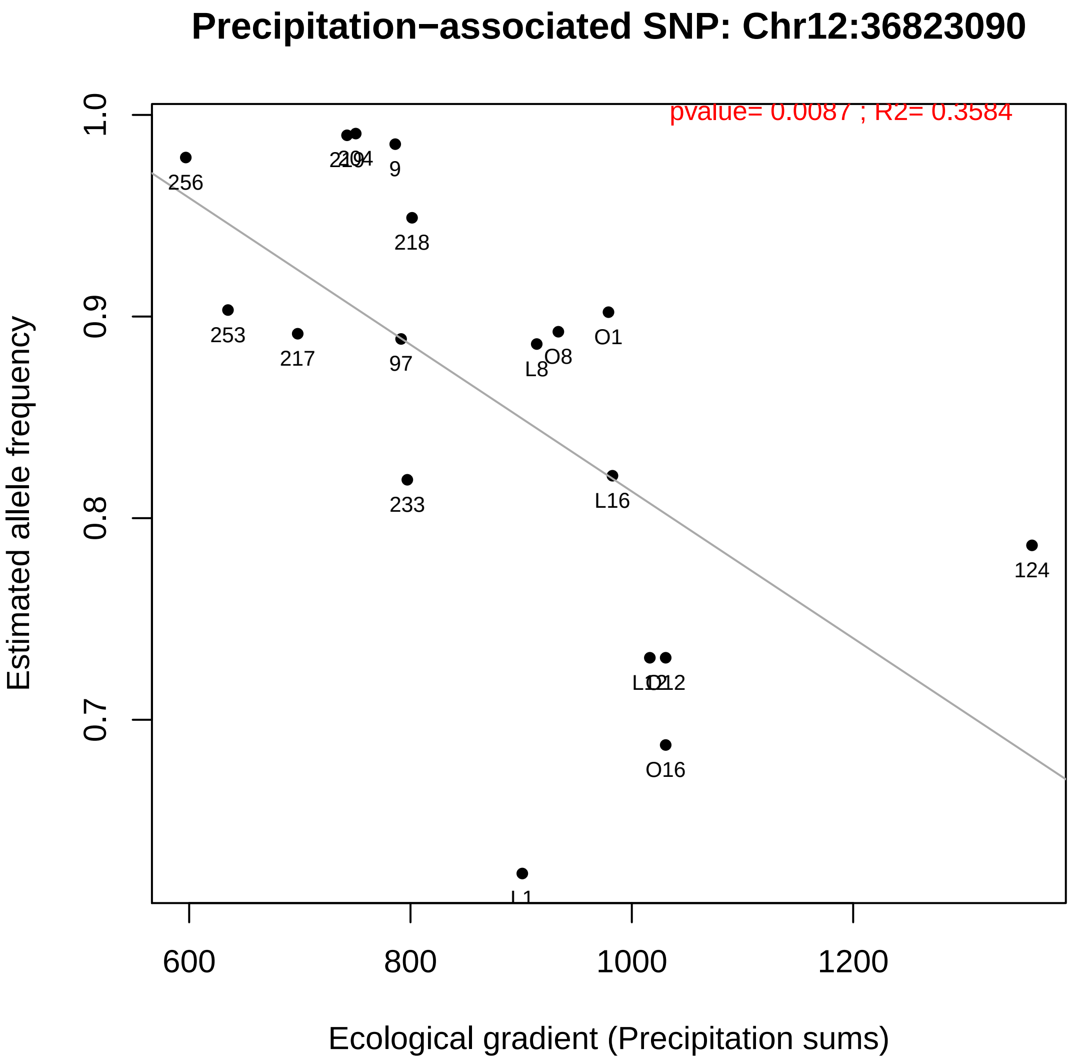

Precipitation−associated SNP: Chr12:36823090
pvalue= 0.0087 ; R2= 0.3584
Estimated allele frequency
Ecological gradient (Precipitation sums)
1.0
0.9
0.8
0.7
600
800
1000
1200
256
219
204
9
218
253
217
97
L8
O8
O1
233
L16
124
L12
O12
O16
L1

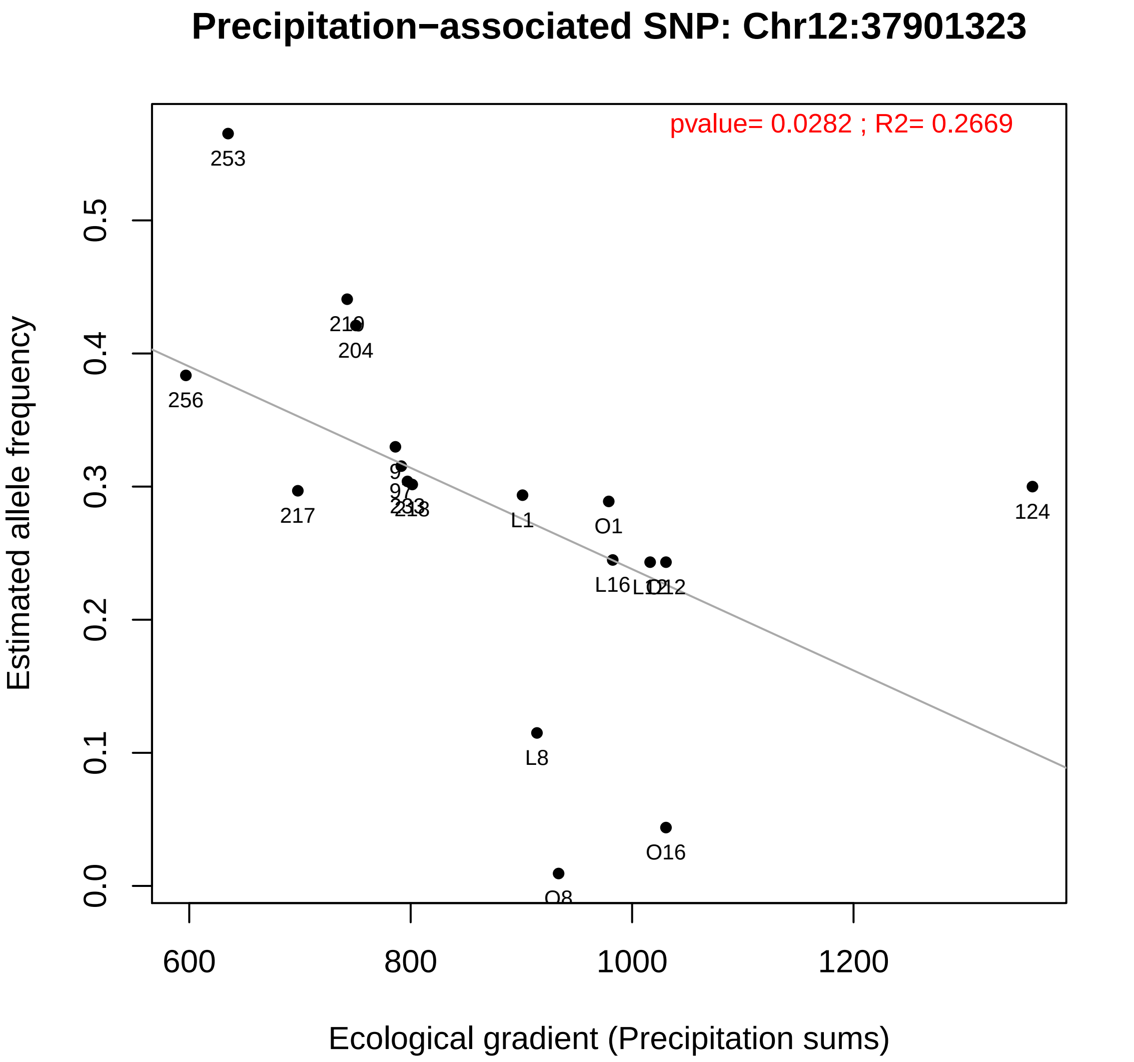

Precipitation−associated SNP: Chr12:37901323
pvalue= 0.0282 ; R2= 0.2669
253
210
204
256
9
97
233
218
217
L1
O1
124
L16
L12
O12
L8
O16
O8
0.0
0.1
0.2
0.3
0.4
0.5
Estimated allele frequency
600
800
1000
1200
Ecological gradient (Precipitation sums)

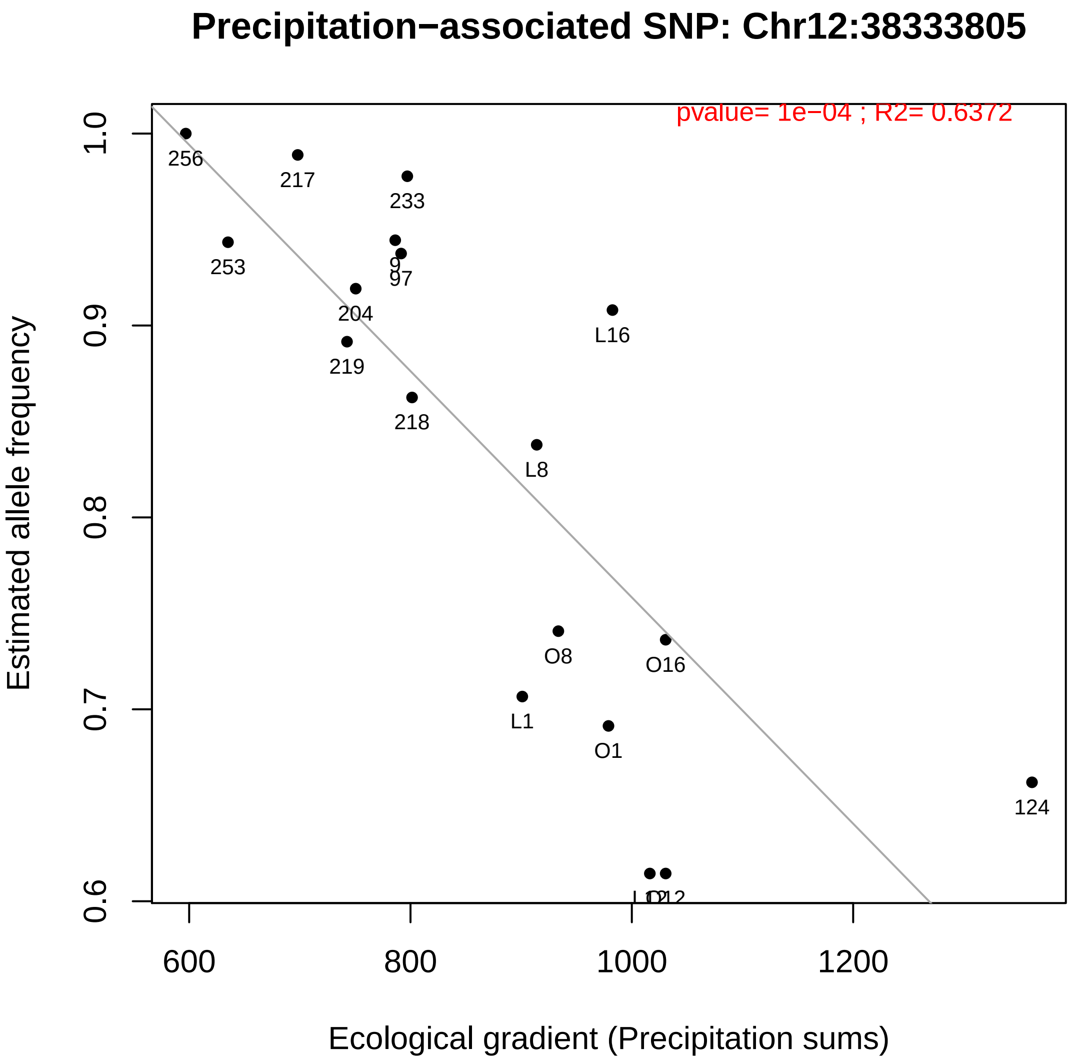

Precipitation−associated SNP: Chr12:38333805
pvalue= 1e−04 ; R2= 0.6372
Estimated allele frequency
Ecological gradient (Precipitation sums)
1.0
0.9
0.8
0.7
0.6
600
800
1000
1200
256
217
233
253
9
97
204
219
218
L16
L8
O8
O16
L1
O1
124

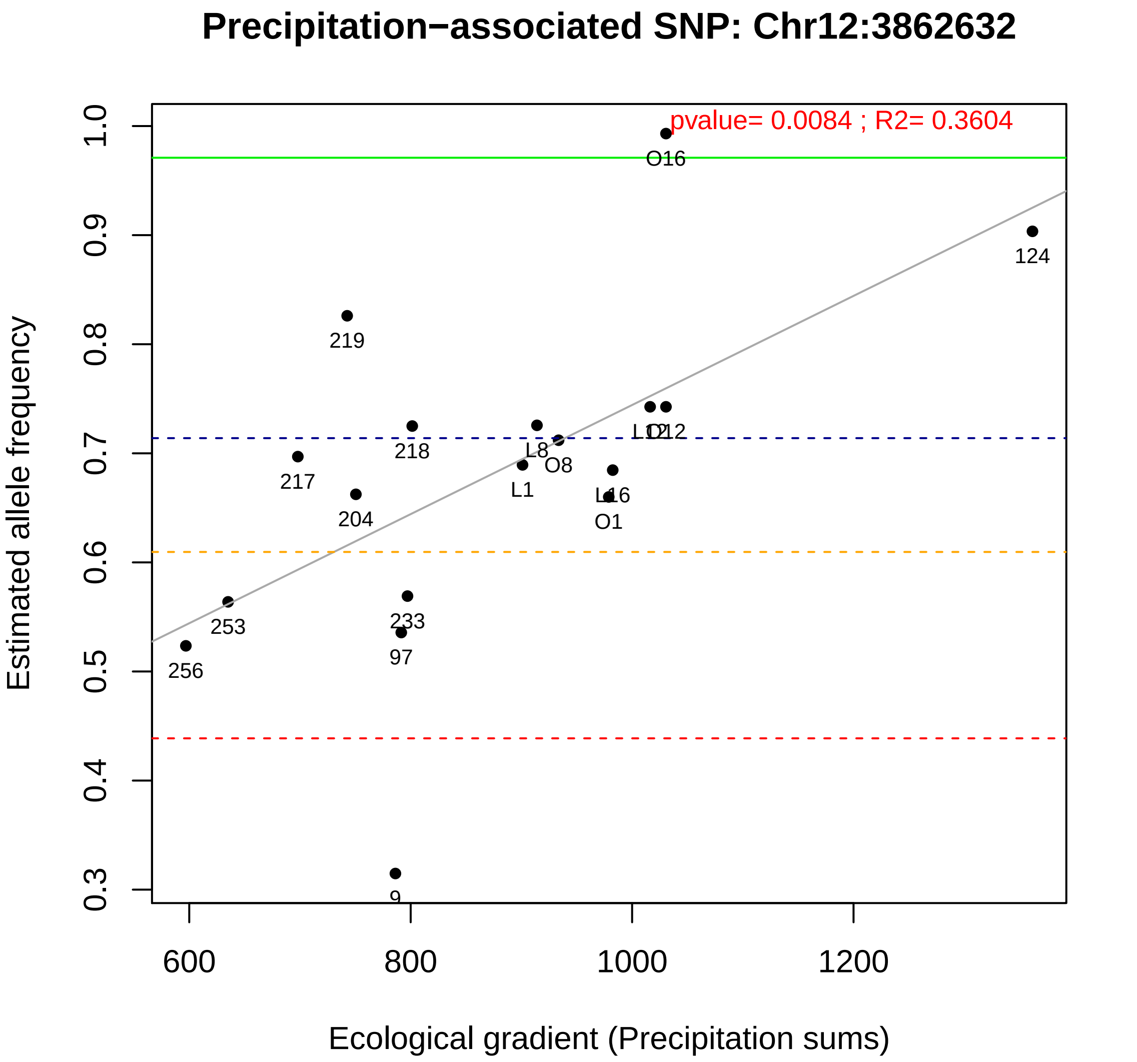

Precipitation−associated SNP: Chr12:3862632
pvalue= 0.0084 ; R2= 0.3604
Estimated allele frequency
Ecological gradient (Precipitation sums)
0.3
0.4
0.5
0.6
0.7
0.8
0.9
1.0
600
800
1000
1200
O16
124
219
218
217
204
L8
O8
L1
L16
O1
L12
O12
253
256
233
97
9

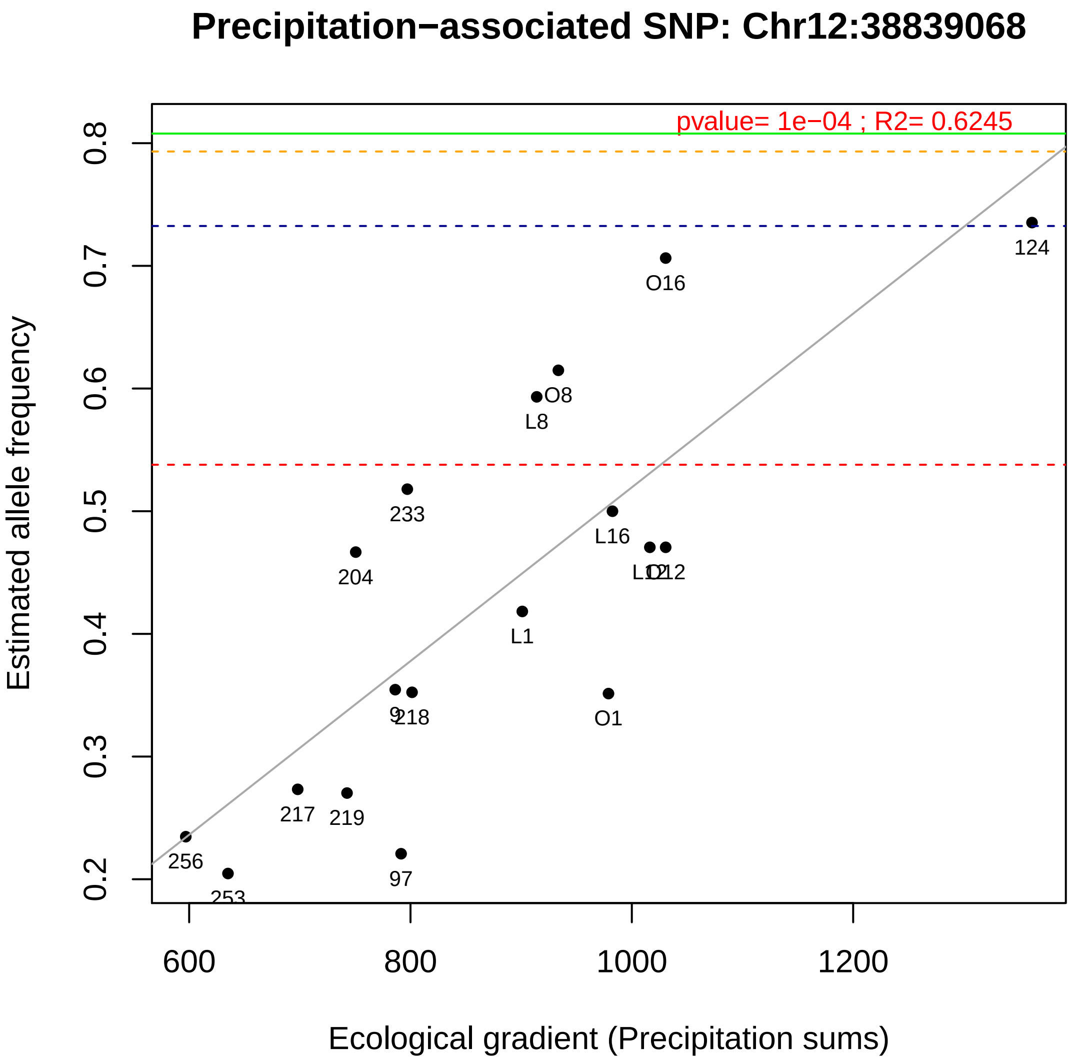
Precipitation−associated SNP: Chr12:38839068
pvalue= 1e−04 ; R2= 0.6245
Estimated allele frequency
Ecological gradient (Precipitation sums)
0.2
0.3
0.4
0.5
0.6
0.7
0.8
600
800
1000
1200
124
O16
O8
L8
233
204
L16
O12
L1
218
O1
217
219
256
97

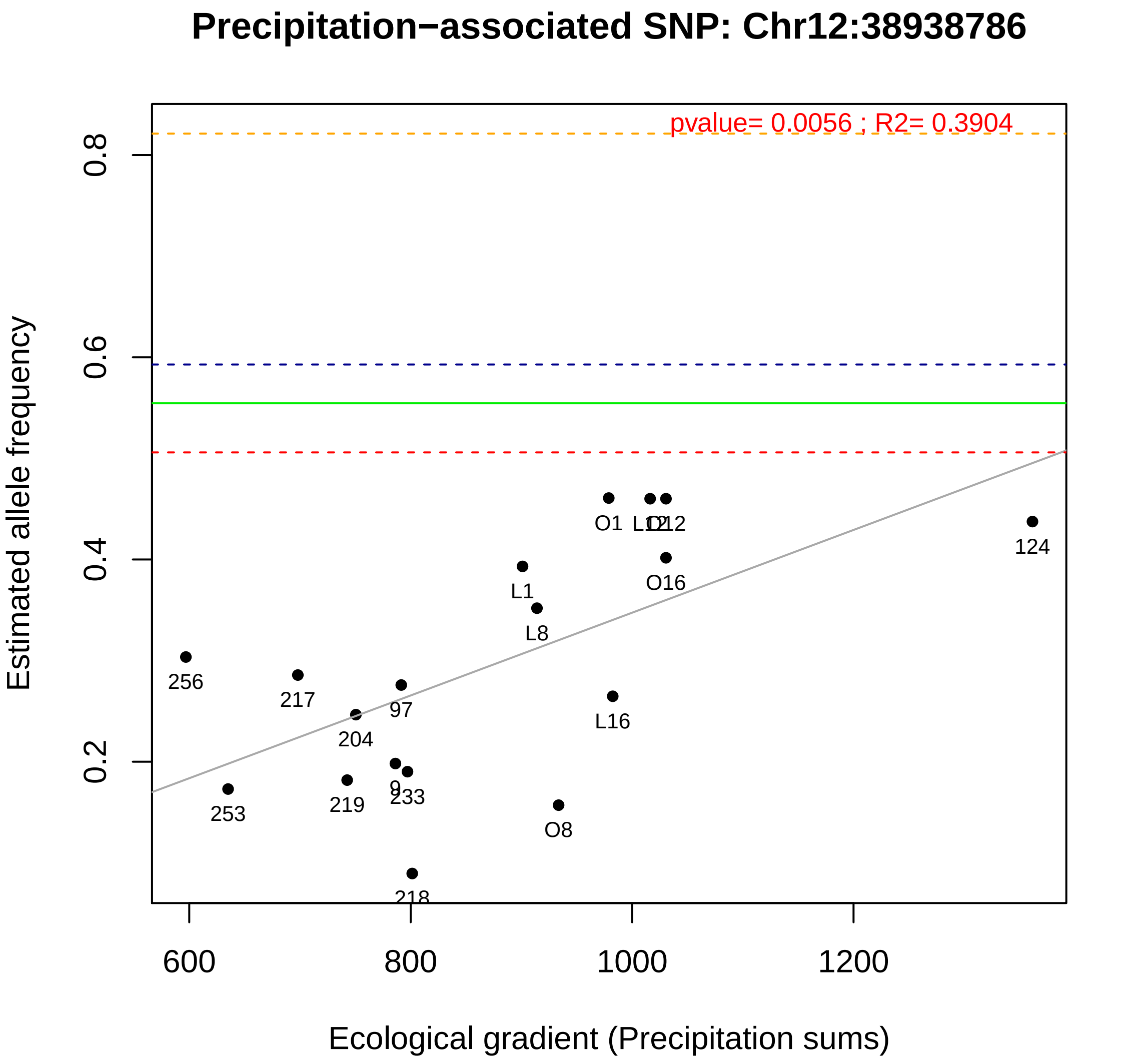
Precipitation−associated SNP: Chr12:38938786
pvalue= 0.0056 ; R2= 0.3904
Estimated allele frequency
0.2
0.4
0.6
0.8
600
800
1000
1200
Ecological gradient (Precipitation sums)
256
253
217
204
219
97
233
218
L1
L8
O8
O1
L16
O16
124

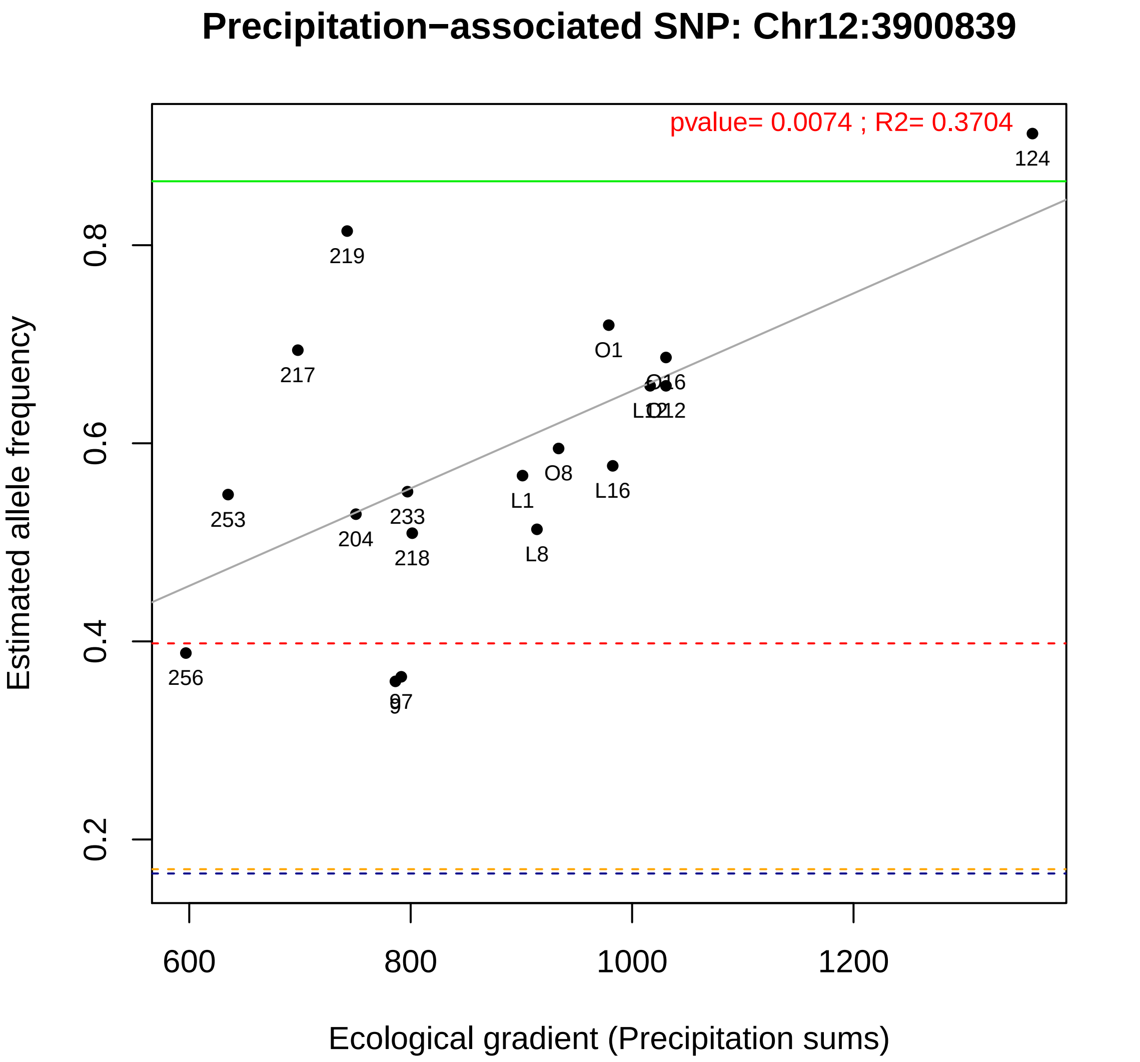
Precipitation−associated SNP: Chr12:3900839
pvalue= 0.0074 ; R2= 0.3704
Estimated allele frequency
0.2
0.4
0.6
0.8
600
800
1000
1200
Ecological gradient (Precipitation sums)
124
219
217
O1
O16
L10
O12
O8
L16
L1
253
204
233
218
L8
256
97

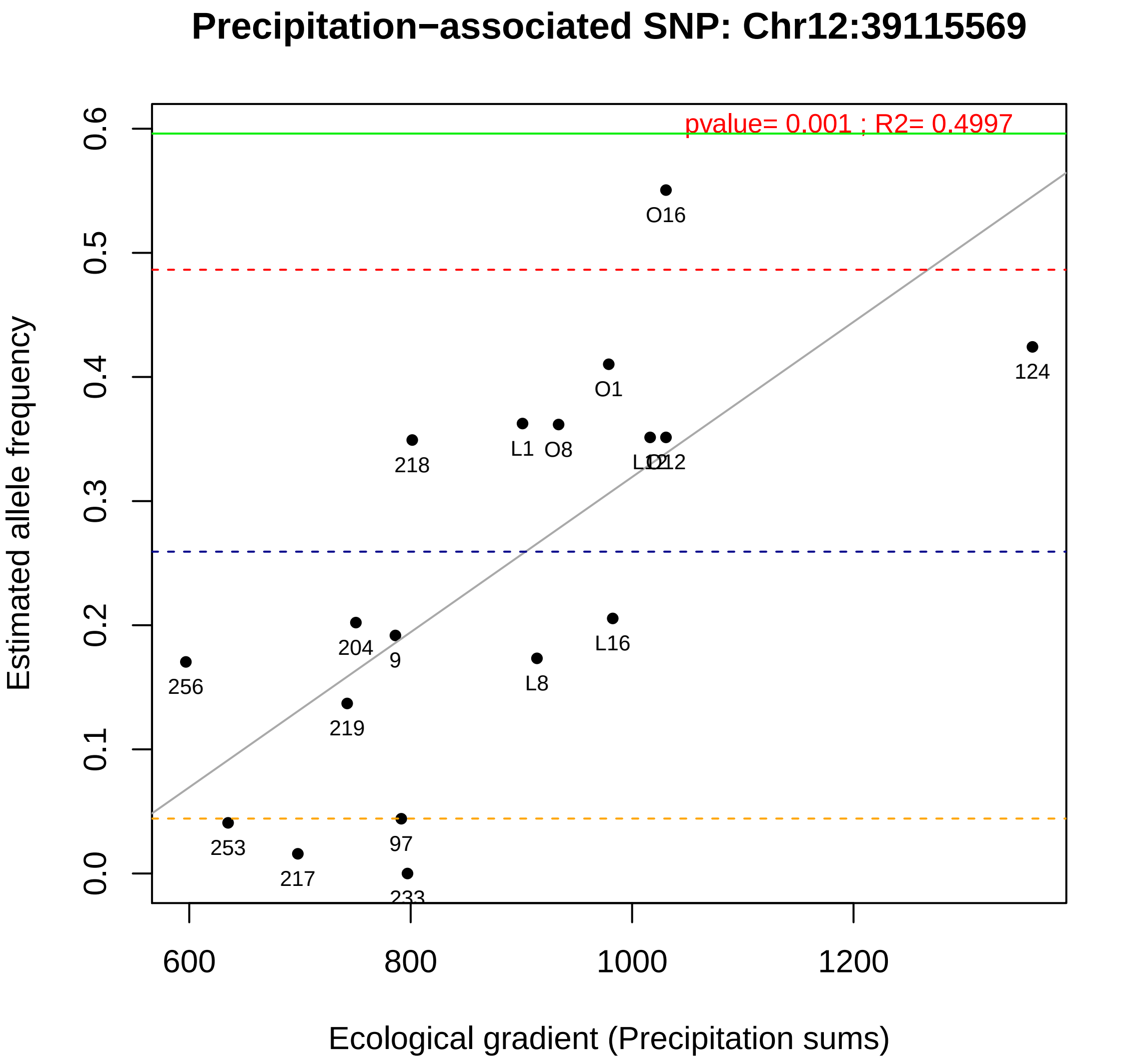
Precipitation−associated SNP: Chr12:39115569
pvalue= 0.001 ; R2= 0.4997
Estimated allele frequency
0.0
0.1
0.2
0.3
0.4
0.5
0.6
600
800
1000
1200
Ecological gradient (Precipitation sums)
O16
124
O1
218
L1
O8
256
204
9
219
L8
L16
253
217
97
233

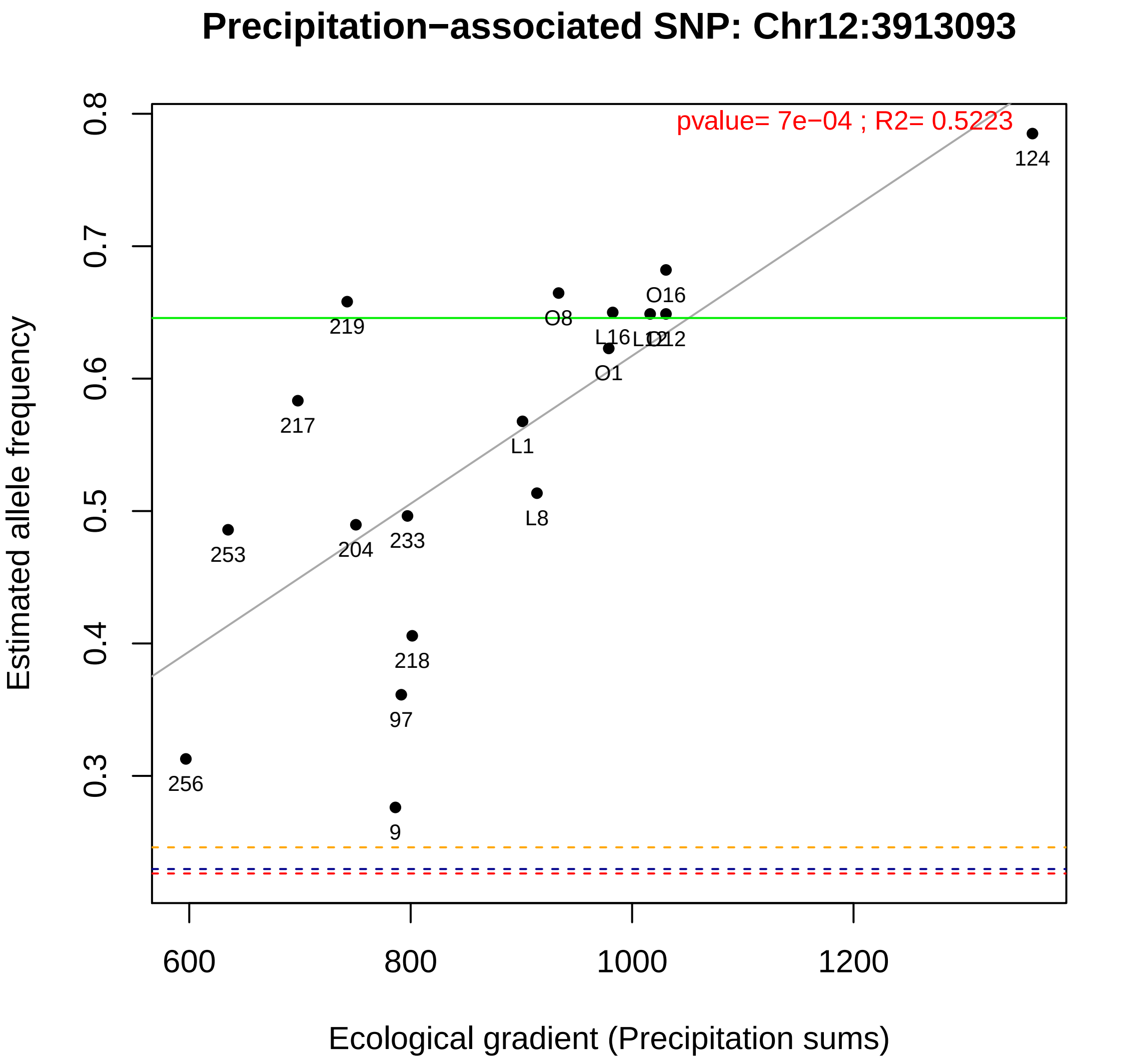

Precipitation−associated SNP: Chr12:3913093
pvalue= 7e−04 ; R2= 0.5223
Estimated allele frequency
Ecological gradient (Precipitation sums)
0.3
0.4
0.5
0.6
0.7
0.8
600
800
1000
1200
124
O16
219
O8
L16
L12
O12
O1
217
L1
L8
253
204
233
218
97
256
9

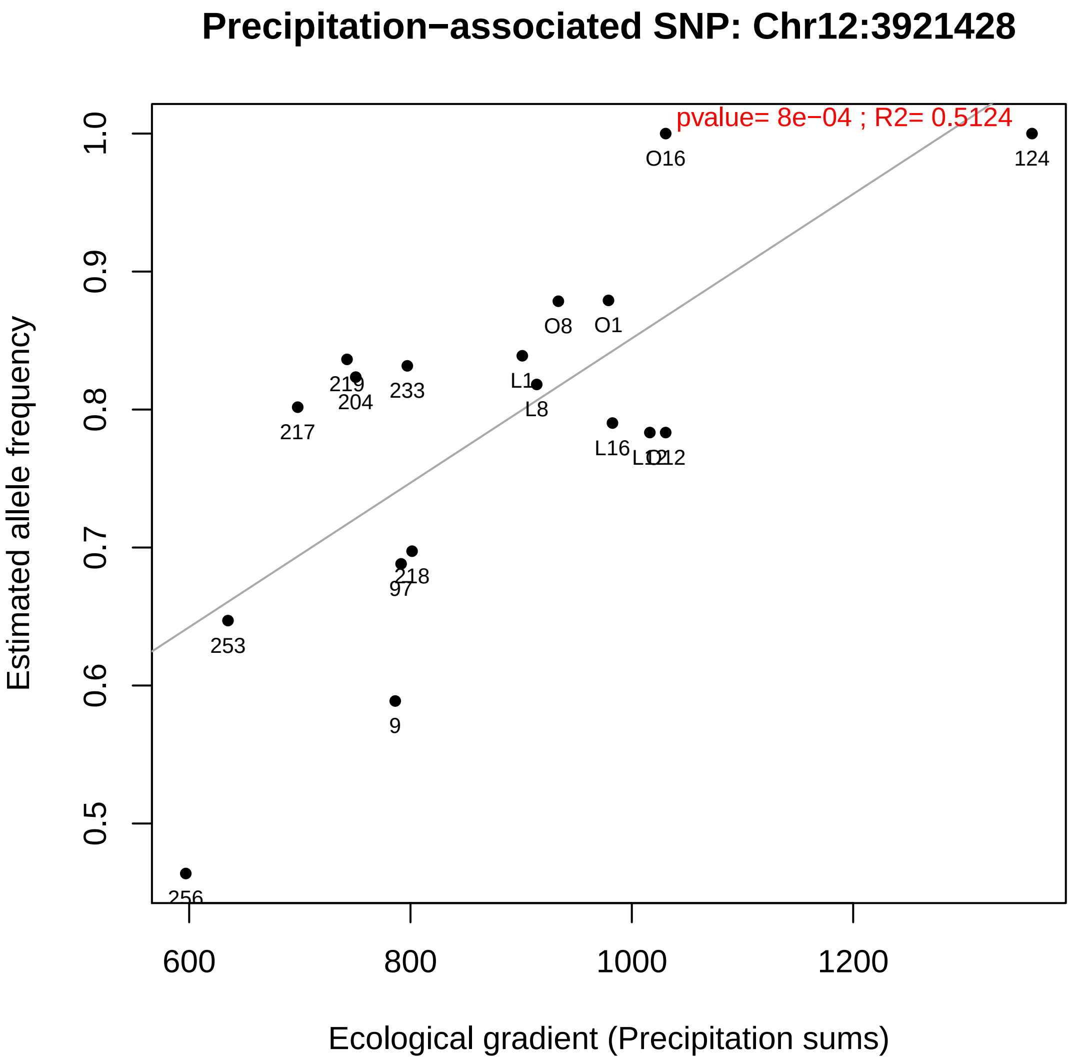

Precipitation−associated SNP: Chr12:3921428
pvalue= 8e−04 ; R2= 0.5124
Estimated allele frequency
Ecological gradient (Precipitation sums)
0.5
0.6
0.7
0.8
0.9
1.0
600
800
1000
1200
O16
124
O8
O1
219
204
233
L1
L8
217
L16
L12
O12
218
97
253
9
256

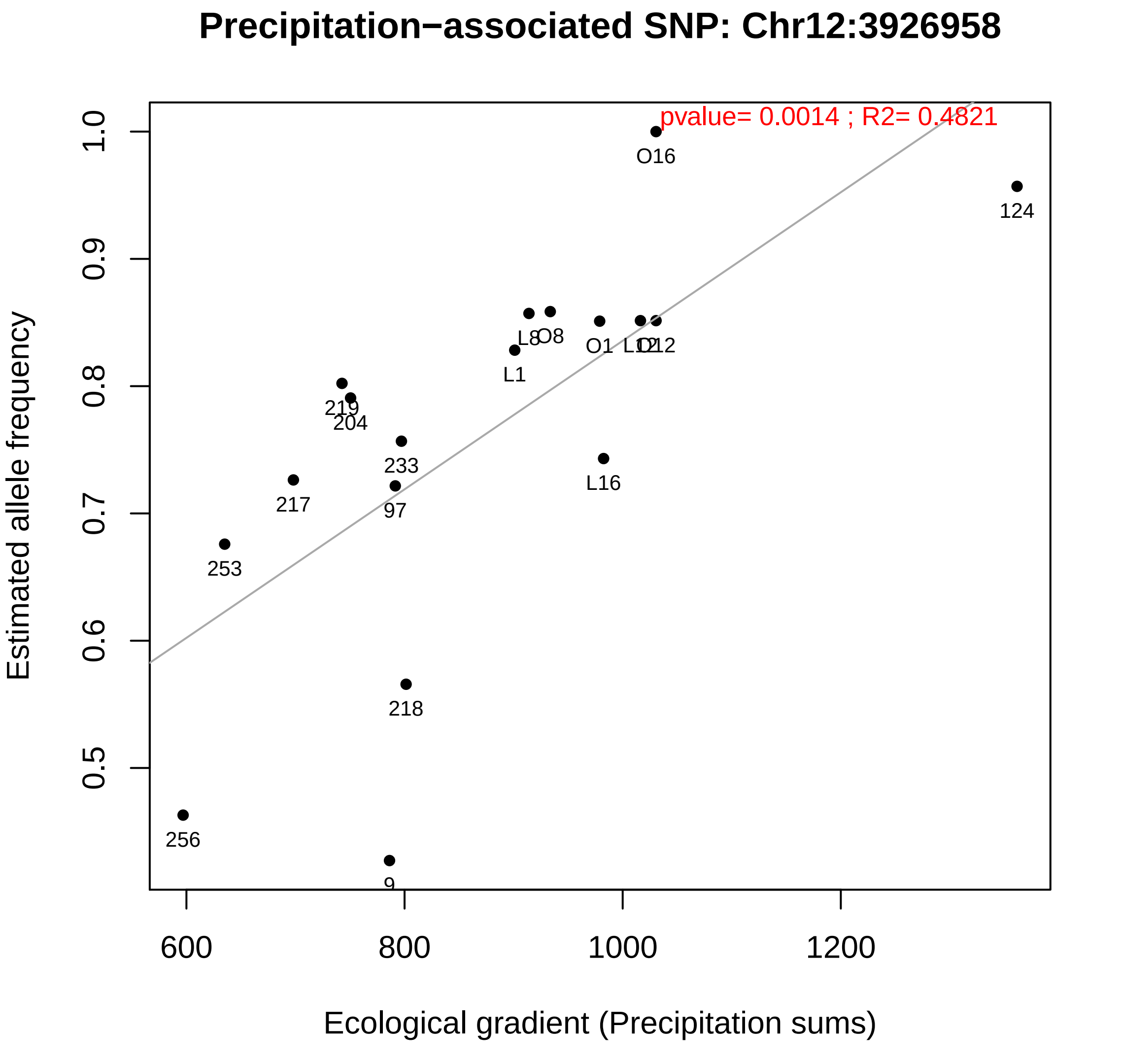

Precipitation−associated SNP: Chr12:3926958
pvalue= 0.0014 ; R2= 0.4821
Estimated allele frequency
Ecological gradient (Precipitation sums)
0.5
0.6
0.7
0.8
0.9
1.0
600
800
1000
1200
O16
124
L8
O8
O1
L1
219
204
233
217
97
L16
253
218
256
9

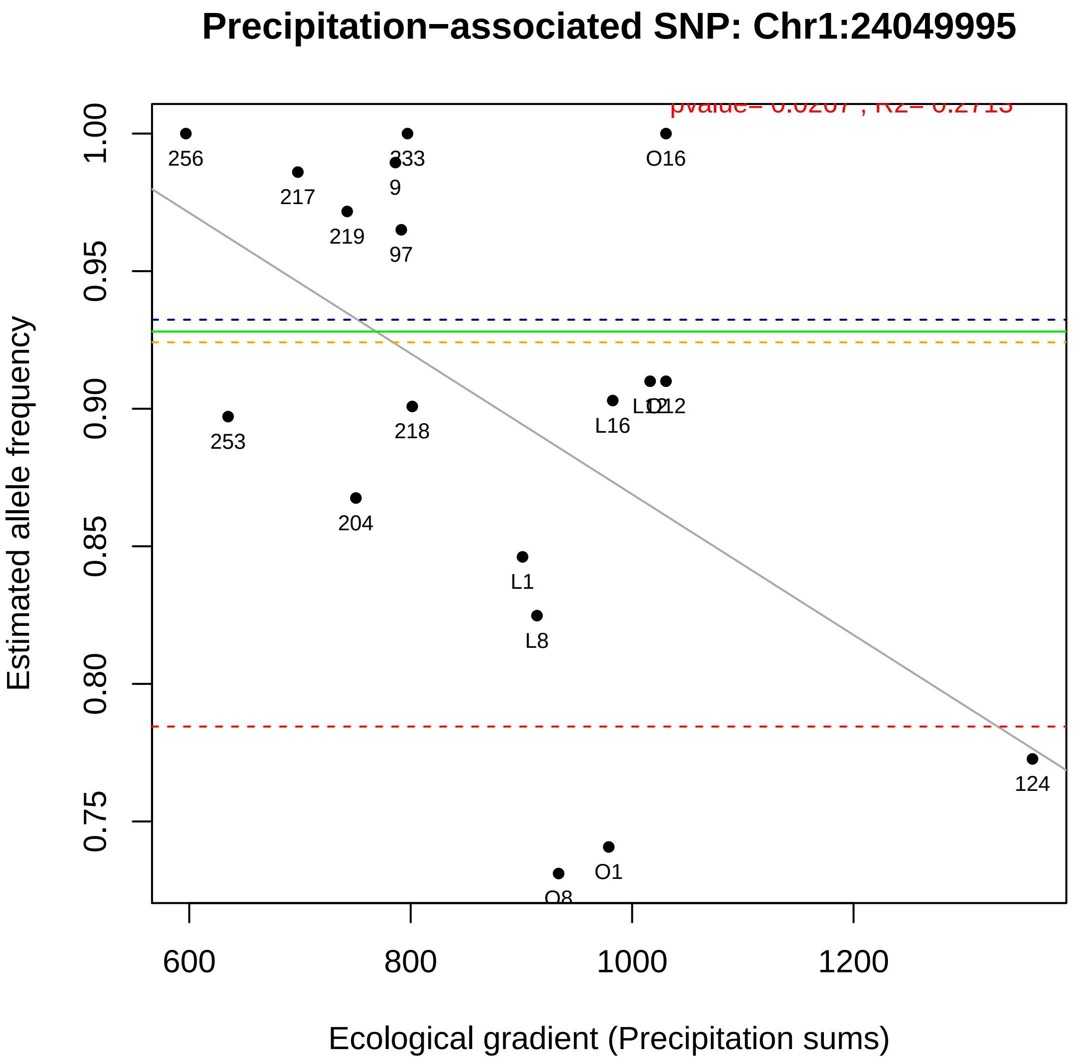

Precipitation−associated SNP: Chr1:24049995
pvalue= 0.0267 , R2= 0.2713
Estimated allele frequency
1.00
0.95
0.90
0.85
0.80
0.75
600
800
1000
1200
Ecological gradient (Precipitation sums)
256
217
219
233
9
97
O16
253
218
204
L16
L12
O12
L1
L8
124
O1
O8

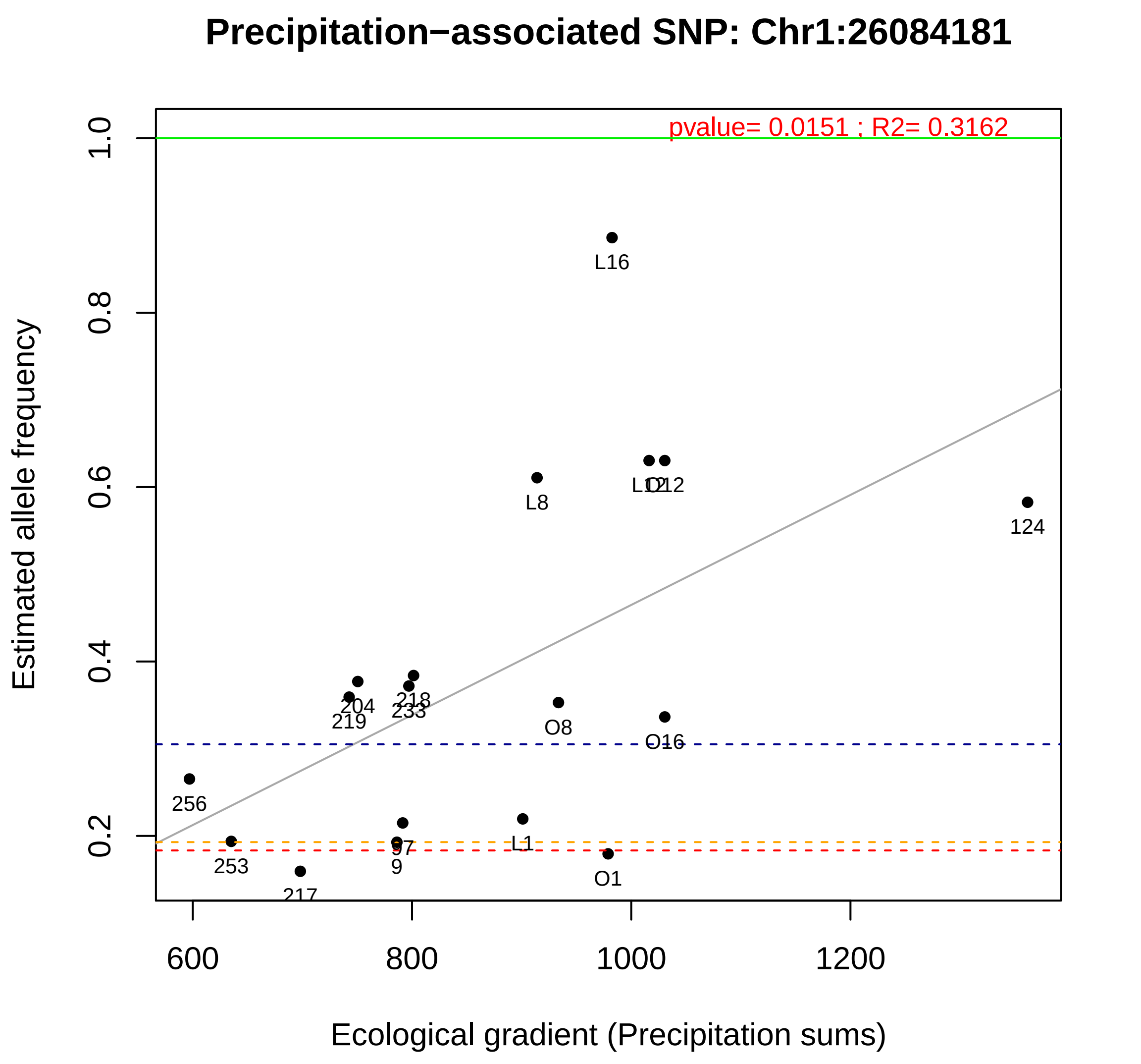

Precipitation−associated SNP: Chr1:26084181
pvalue= 0.0151 ; R2= 0.3162
Estimated allele frequency
0.2
0.4
0.6
0.8
1.0
600
800
1000
1200
Ecological gradient (Precipitation sums)
L16
L8
O12
124
204
219
218
233
O8
O16
256
253
217
9
L1
O1

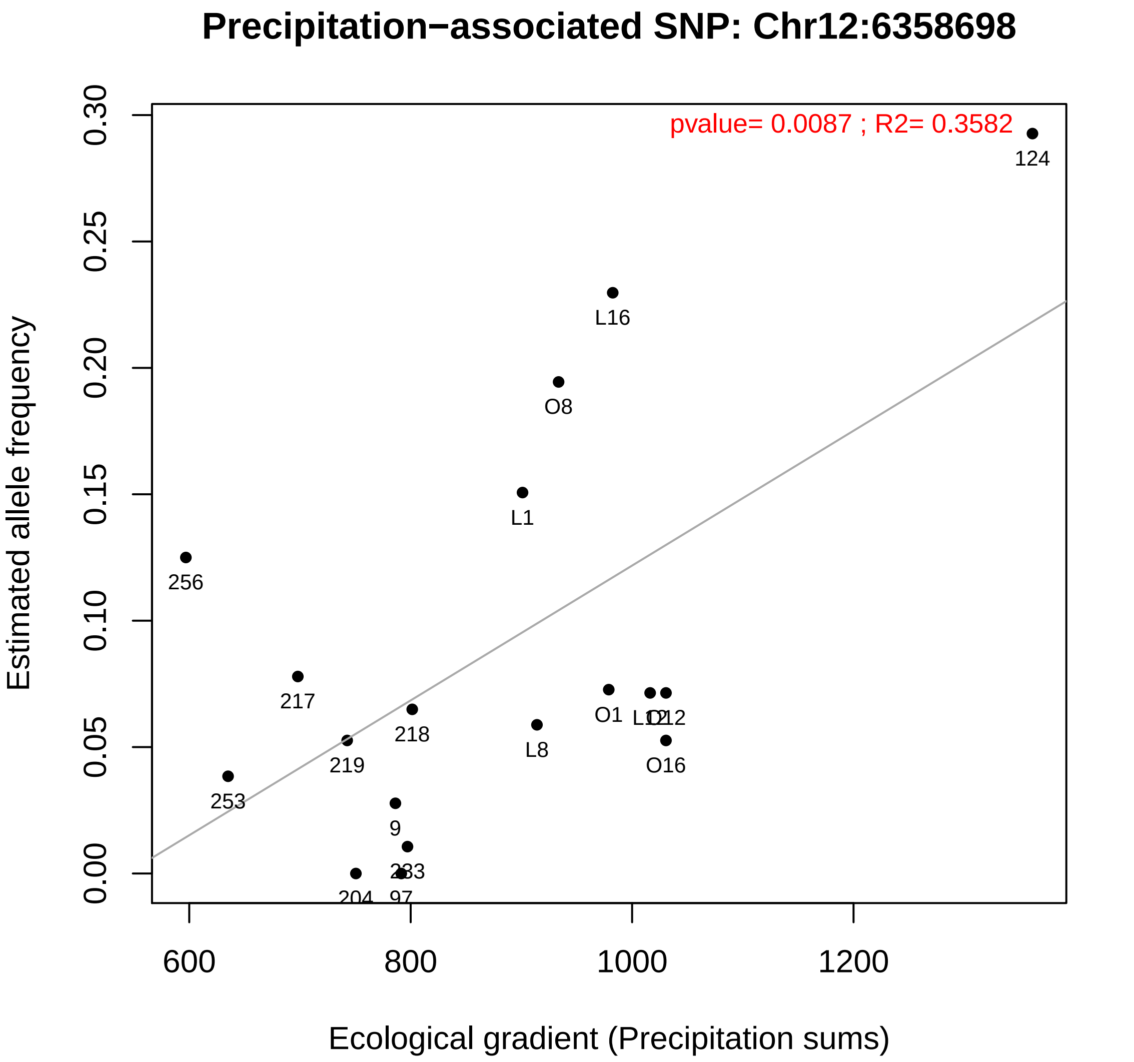
Precipitation−associated SNP: Chr12:6358698
pvalue= 0.0087 ; R2= 0.3582
Estimated allele frequency
Ecological gradient (Precipitation sums)
0.00
0.05
0.10
0.15
0.20
0.25
0.30
600
800
1000
1200
124
L16
O8
L1
256
217
218
219
253
9
233
204
97
L8
O1
O12
O16

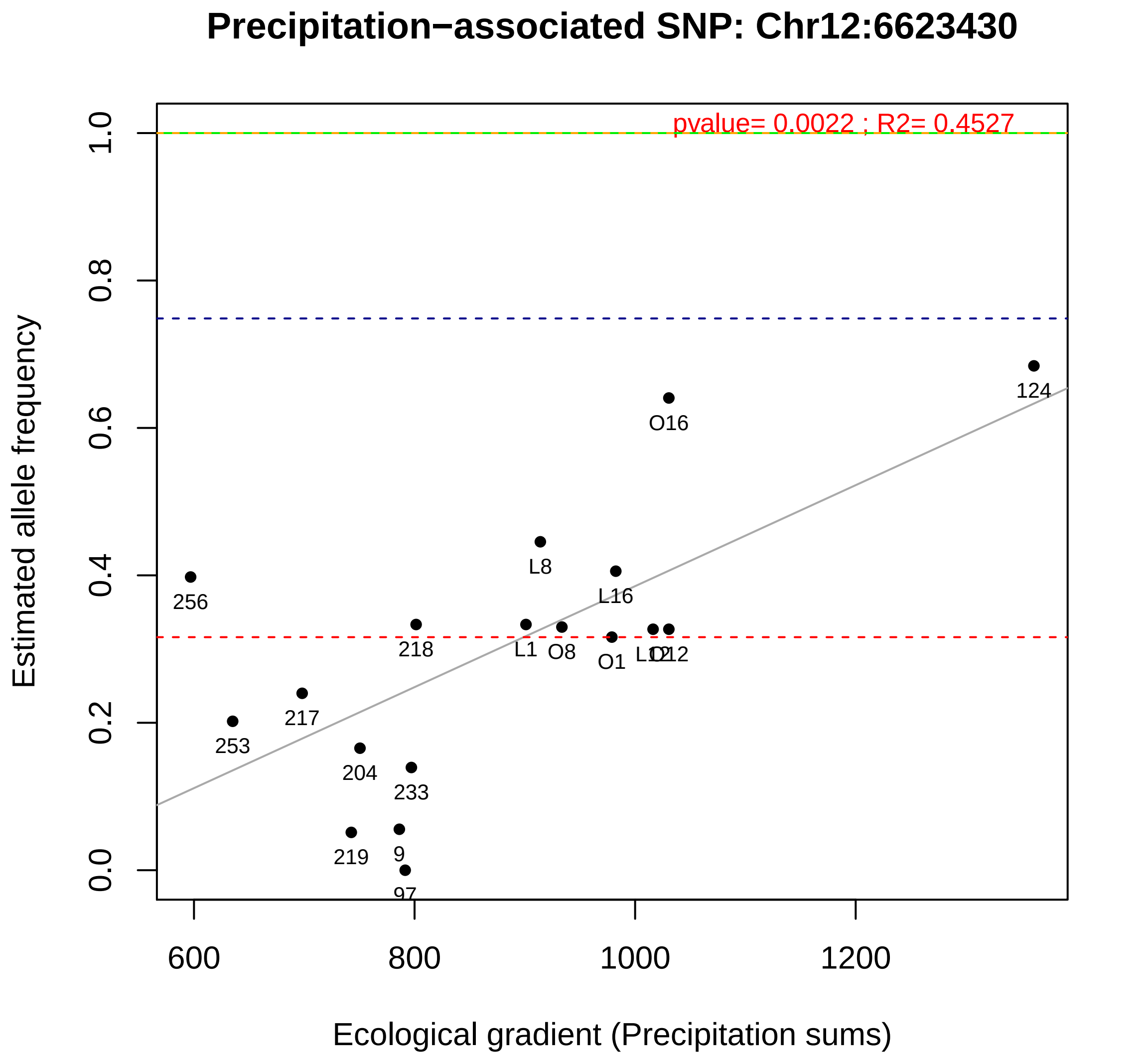
Precipitation−associated SNP: Chr12:6623430
pvalue= 0.0022 ; R2= 0.4527
Estimated allele frequency
0.0
0.2
0.4
0.6
0.8
1.0
124
O16
L8
L16
256
218
L1
O8
O1
L12
O12
217
253
204
233
219
9
97
600
800
1000
1200
Ecological gradient (Precipitation sums)

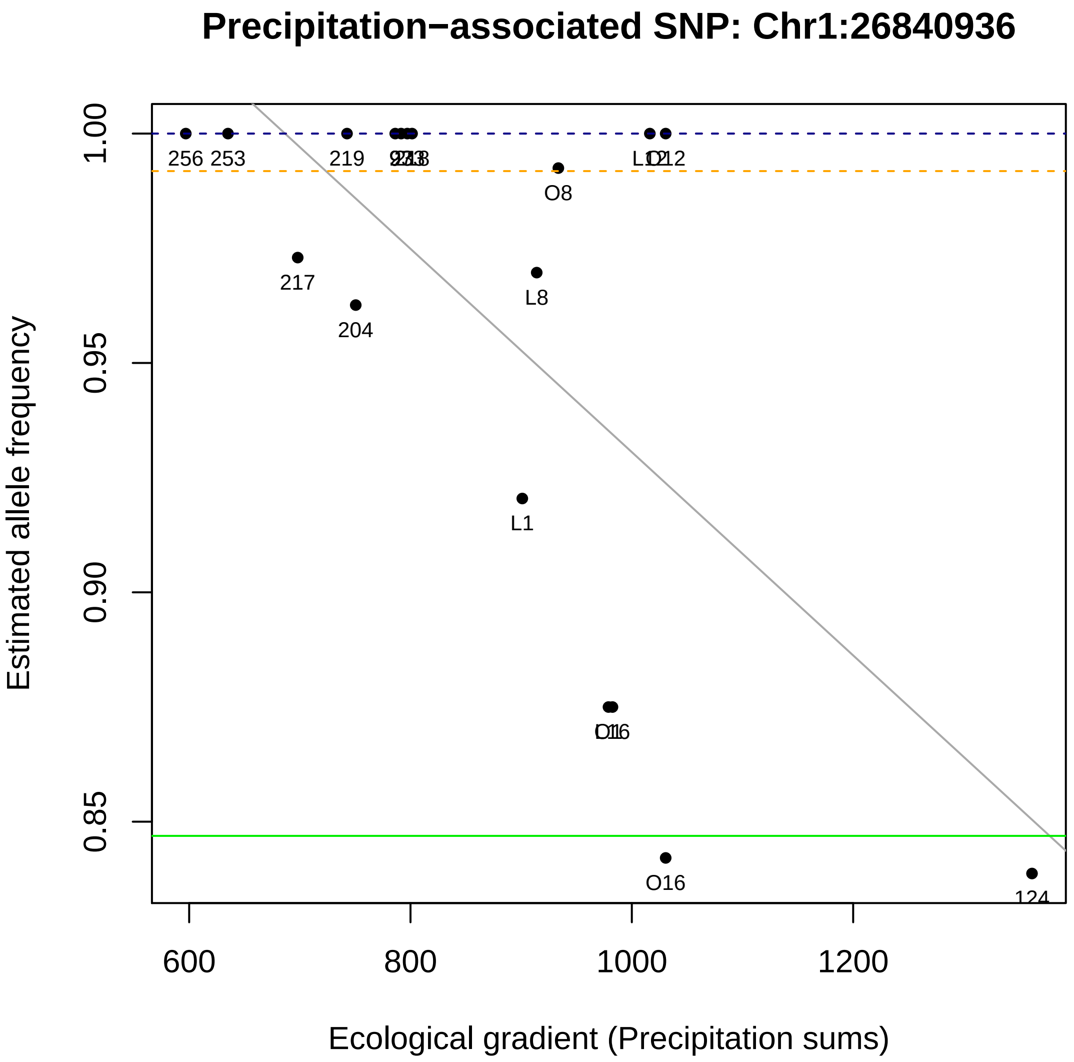
Precipitation−associated SNP: Chr1:26840936
Estimated allele frequency
Ecological gradient (Precipitation sums)
1.00
0.95
0.90
0.85
600
800
1000
1200
256
253
219
217
204
O8
L8
L1
O16
124

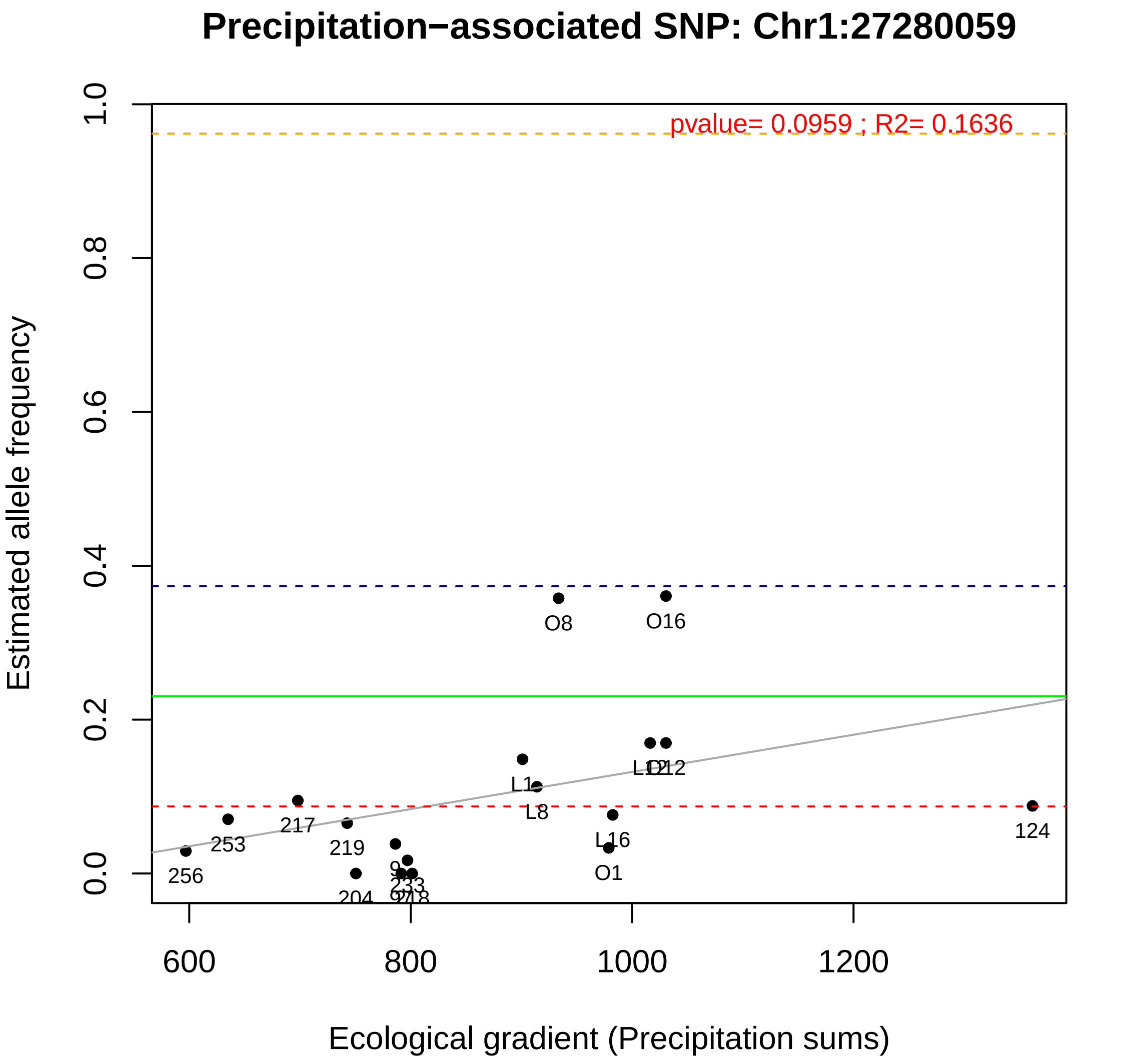

Precipitation−associated SNP: Chr1:27280059
pvalue= 0.0959 ; R2= 0.1636
Estimated allele frequency
Ecological gradient (Precipitation sums)
0.0
0.2
0.4
0.6
0.8
1.0
600
800
1000
1200
O8
O16
L1
L8
L16
O1
O12
124
217
253
256
219
233
204

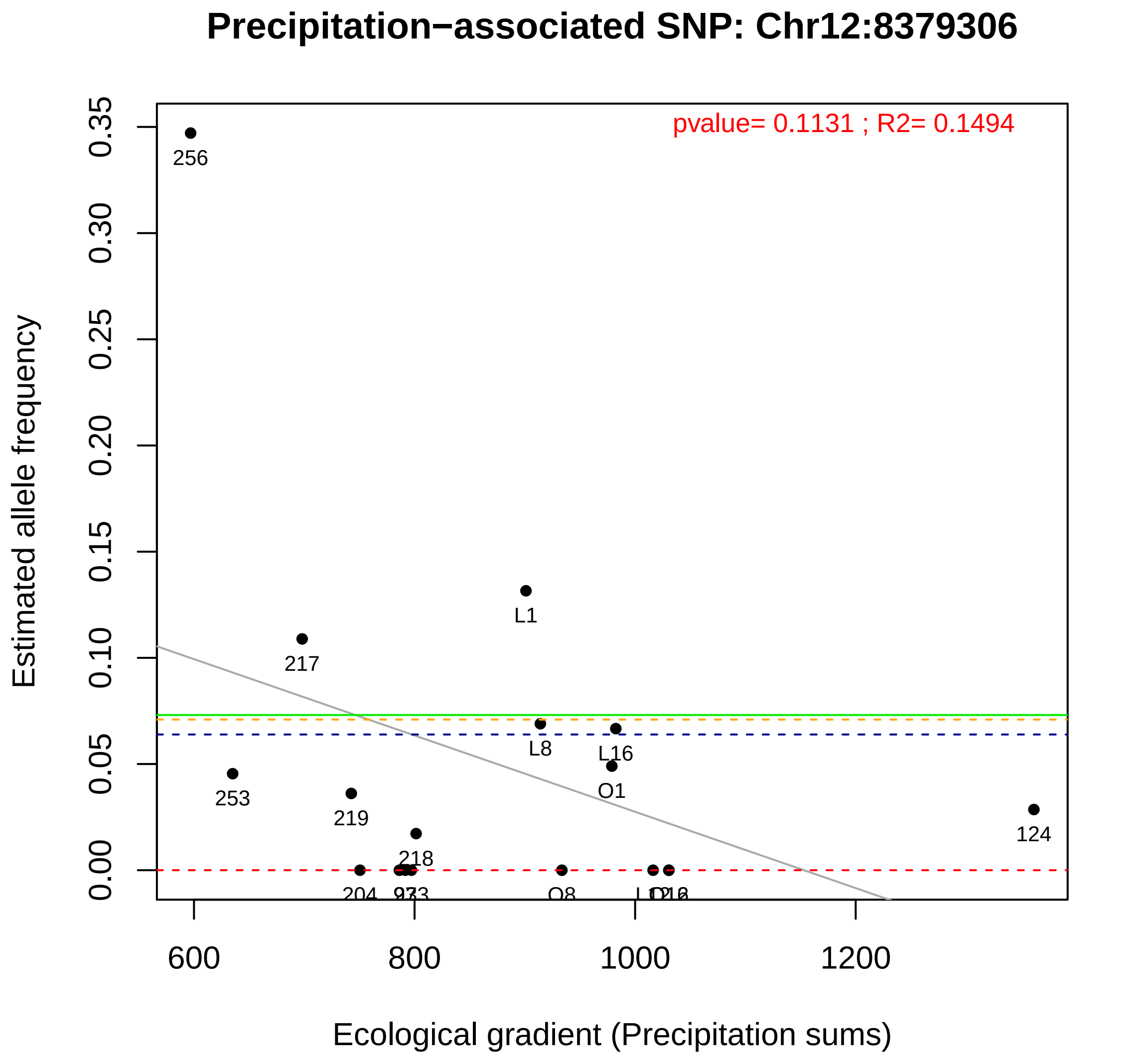
Precipitation−associated SNP: Chr12:8379306
pvalue= 0.1131 ; R2= 0.1494
Estimated allele frequency
Ecological gradient (Precipitation sums)
0.00
0.05
0.10
0.15
0.20
0.25
0.30
0.35
600
800
1000
1200
256
217
L1
L8
L16
O1
253
219
218
124
204
O8

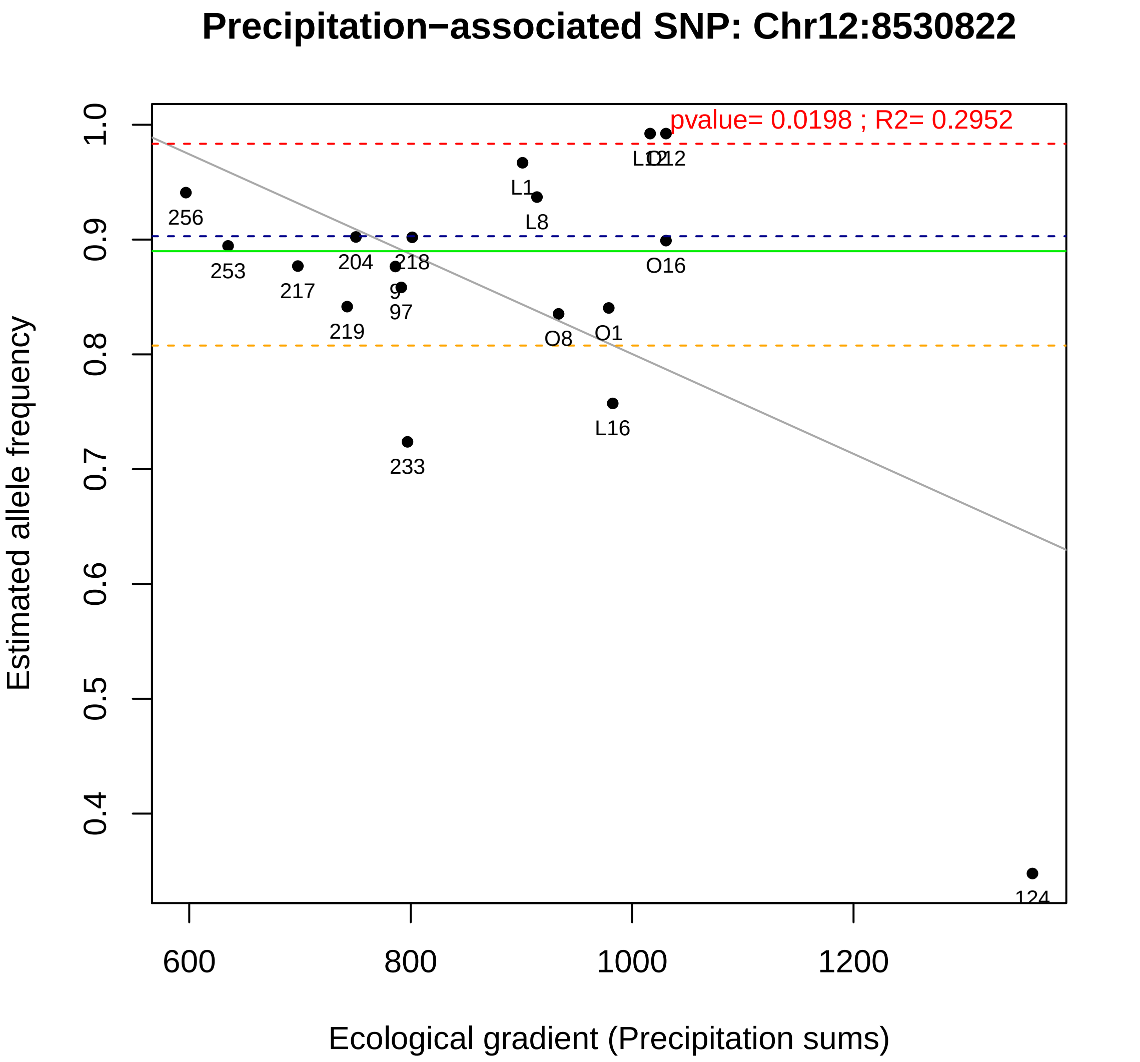

Precipitation−associated SNP: Chr12:8530822
pvalue= 0.0198 ; R2= 0.2952
Estimated allele frequency
Ecological gradient (Precipitation sums)
0.4
0.5
0.6
0.7
0.8
0.9
1.0
600
800
1000
1200
256
253
217
204
219
218
9
97
233
L1
L8
O8
O1
L16
O16
L12
O12
124

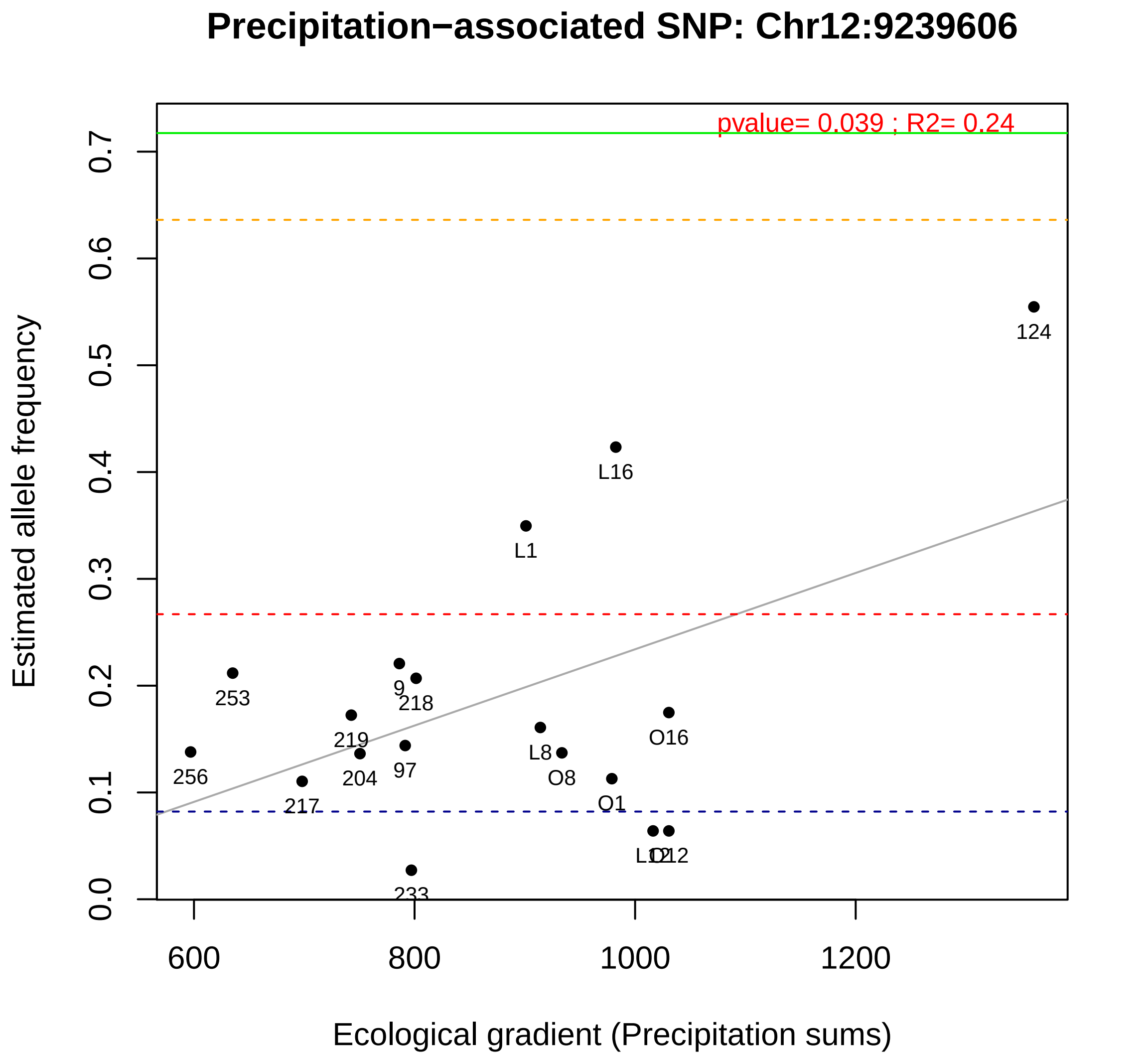
Precipitation−associated SNP: Chr12:9239606
pvalue= 0.039 ; R2= 0.24
Estimated allele frequency
0.0
0.1
0.2
0.3
0.4
0.5
0.6
0.7
600
800
1000
1200
Ecological gradient (Precipitation sums)
124
L16
L1
253
9
218
219
256
204
97
217
233
L8
O8
O16
O1
L12
O12

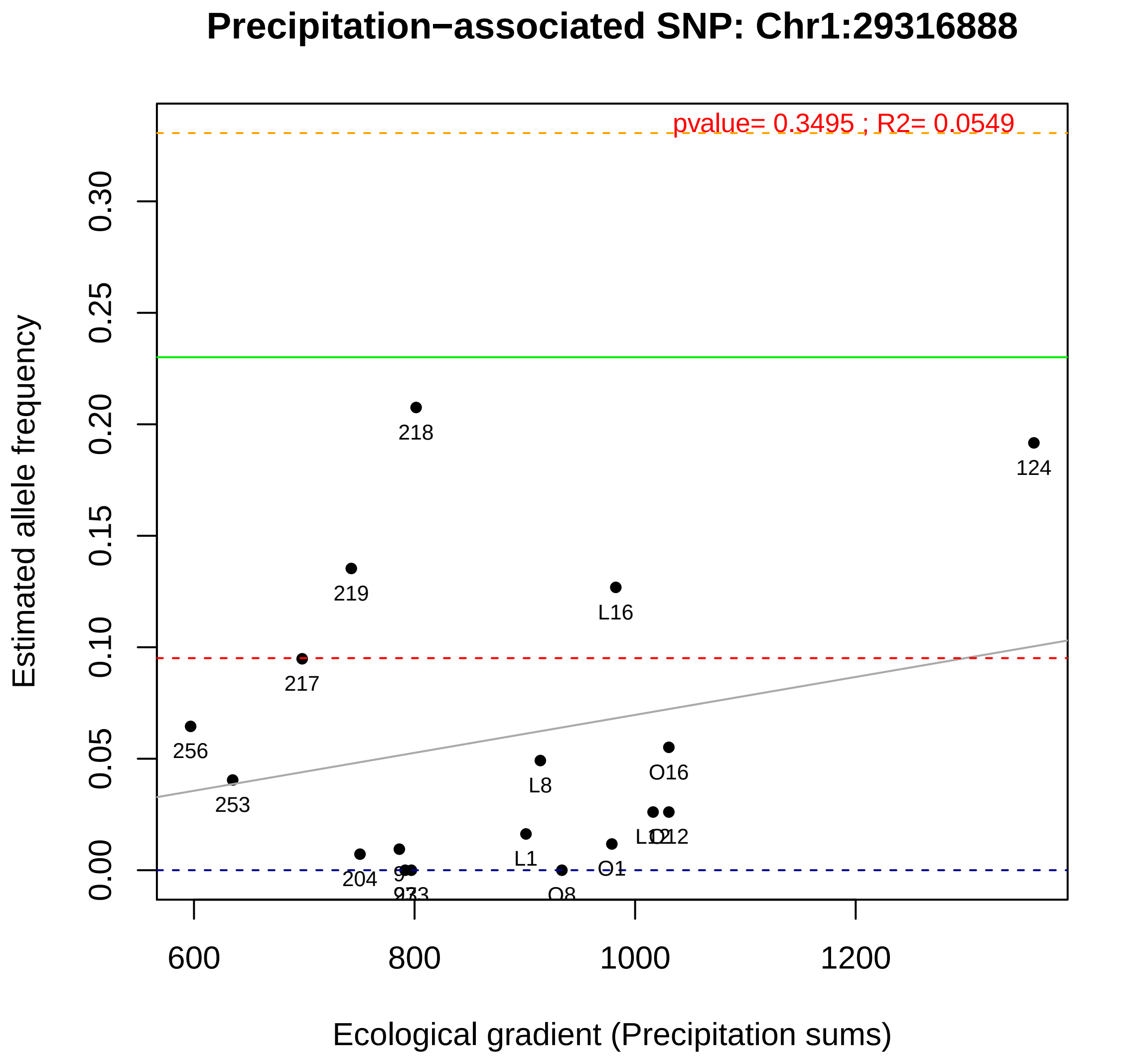
Precipitation−associated SNP: Chr1:29316888
pvalue= 0.3495 ; R2= 0.0549
Estimated allele frequency
0.00
0.05
0.10
0.15
0.20
0.25
0.30
600
800
1000
1200
Ecological gradient (Precipitation sums)
218
124
219
L16
217
256
253
L8
O16
204
L1
O1
O8
L12
O12

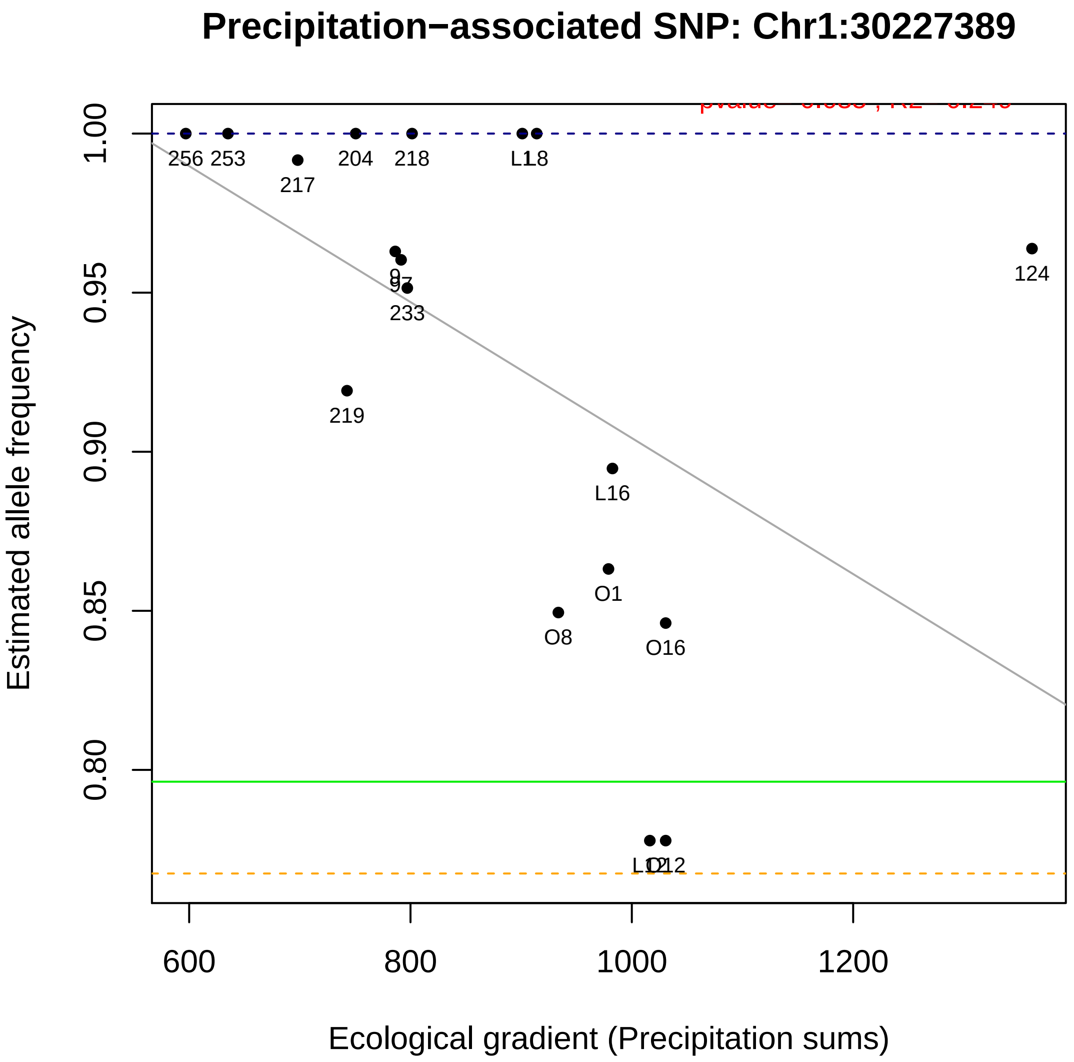

Precipitation−associated SNP: Chr1:30227389
Estimated allele frequency
Ecological gradient (Precipitation sums)
1.00
0.95
0.90
0.85
0.80
600
800
1000
1200
256
253
217
204
218
219
233
124
L16
O1
O8
O16

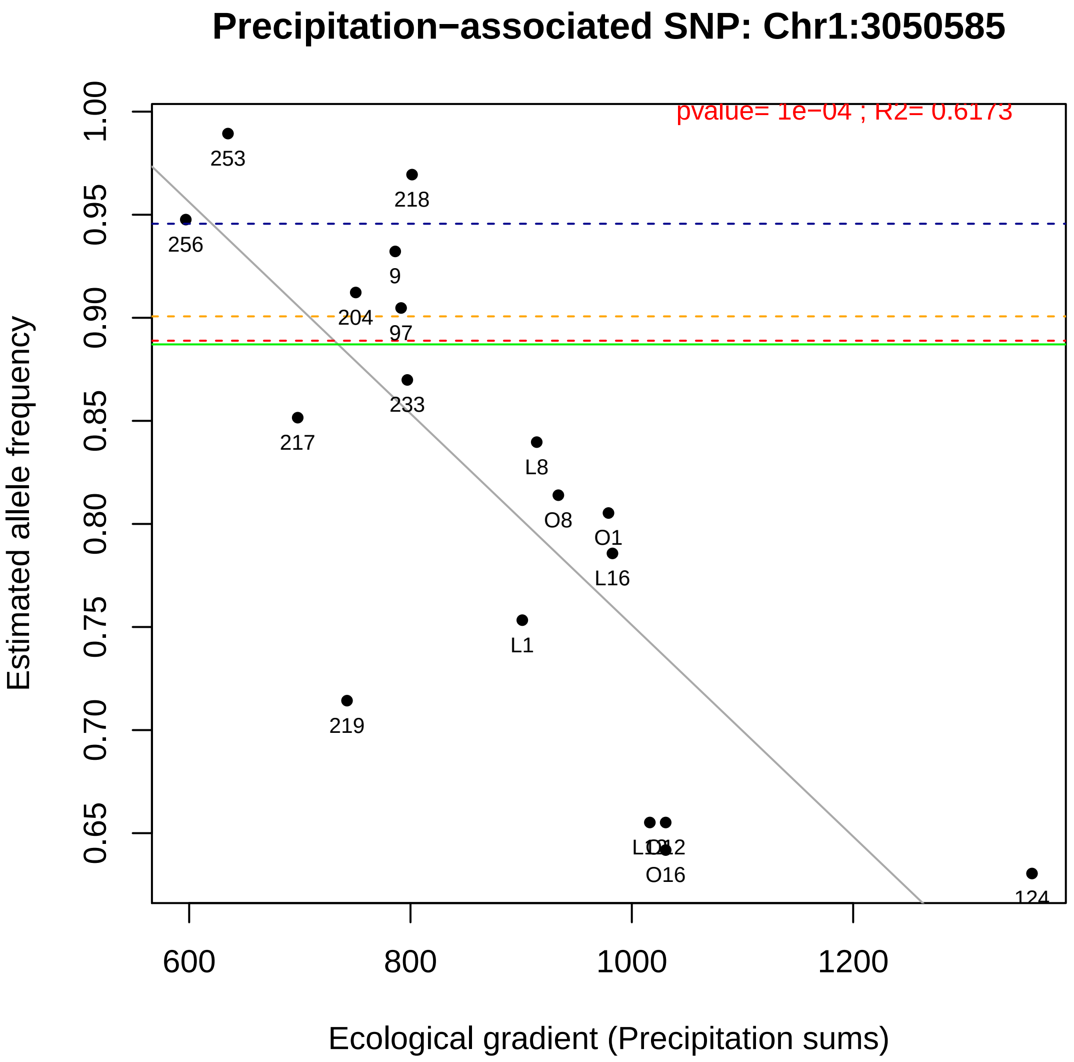

Precipitation−associated SNP: Chr1:3050585
pvalue= 1e−04 ; R2= 0.6173
Estimated allele frequency
Ecological gradient (Precipitation sums)
0.65
0.70
0.75
0.80
0.85
0.90
0.95
1.00
600
800
1000
1200
253
218
256
9
204
97
233
217
L8
O8
O1
L16
L1
219
O16
124

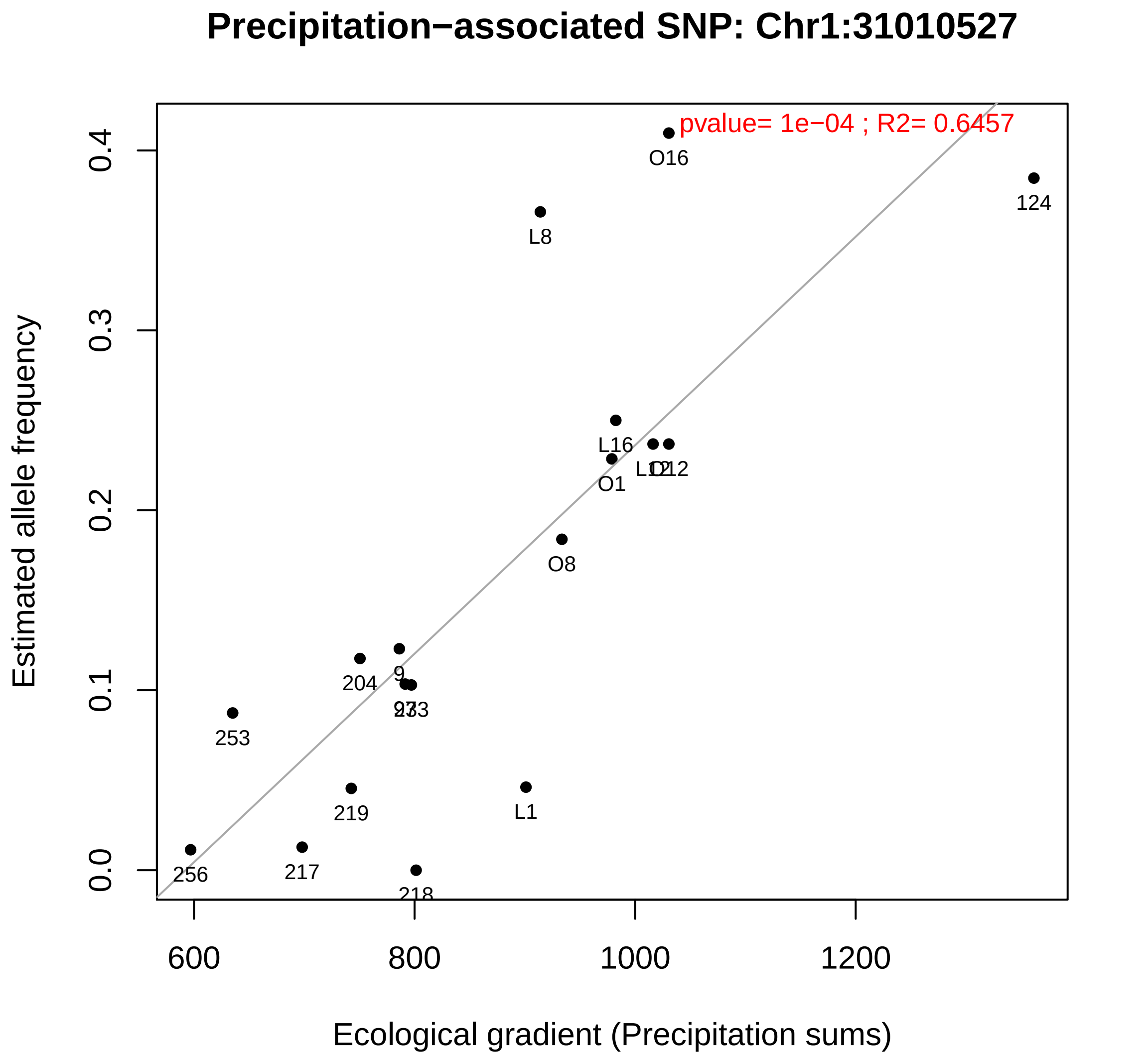

Precipitation−associated SNP: Chr1:31010527
pvalue= 1e−04 ; R2= 0.6457
Estimated allele frequency
Ecological gradient (Precipitation sums)
0.0
0.1
0.2
0.3
0.4
600
800
1000
1200
O16
124
L8
L16
L12
O12
O1
O8
204
9
27
233
253
219
L1
256
217
218

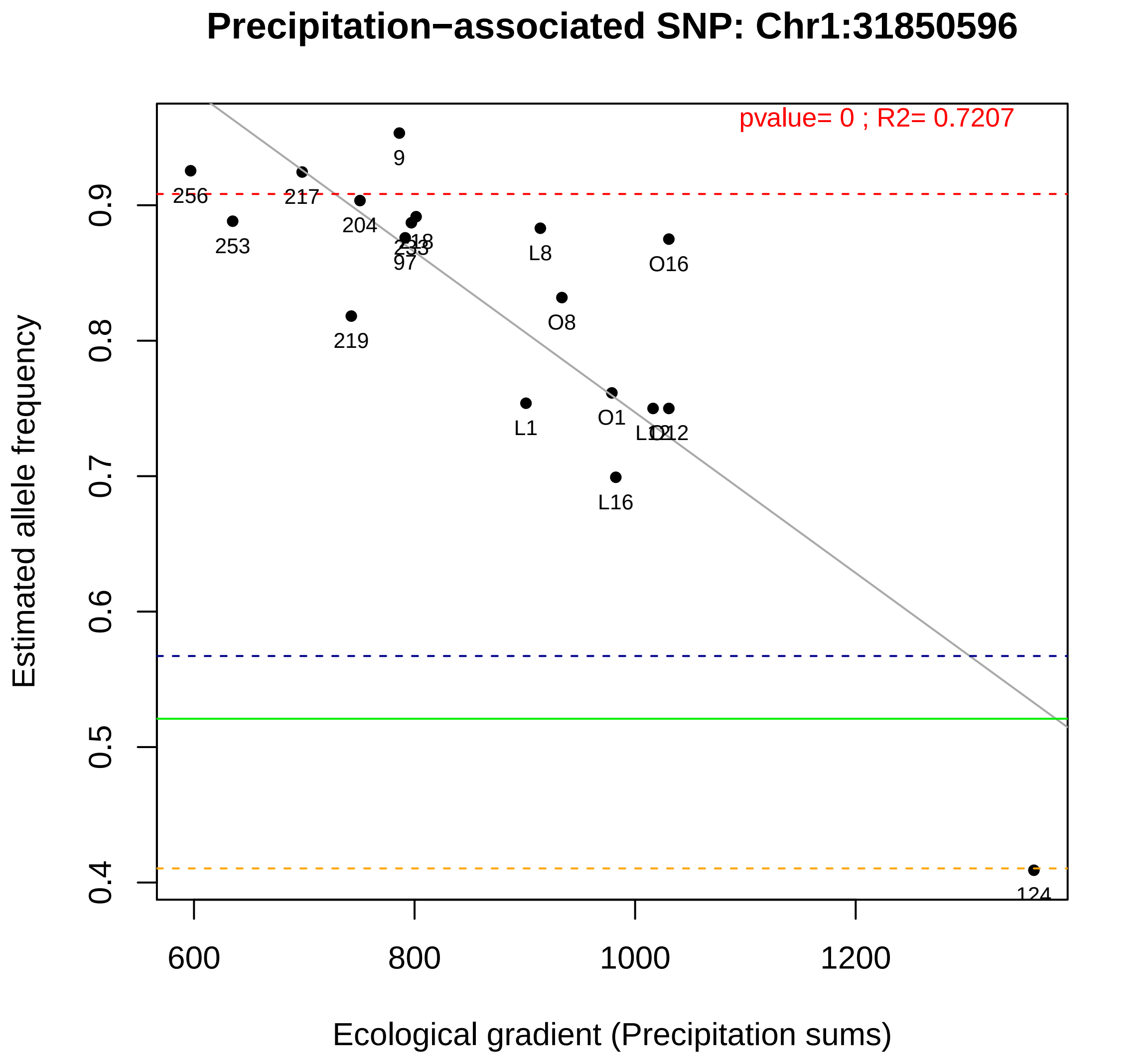

Precipitation−associated SNP: Chr1:31850596
pvalue= 0 ; R2= 0.7207
Estimated allele frequency
Ecological gradient (Precipitation sums)
0.4
0.5
0.6
0.7
0.8
0.9
600
800
1000
1200
9
256
217
204
253
97
L8
O16
O8
219
L1
O1
L16
124

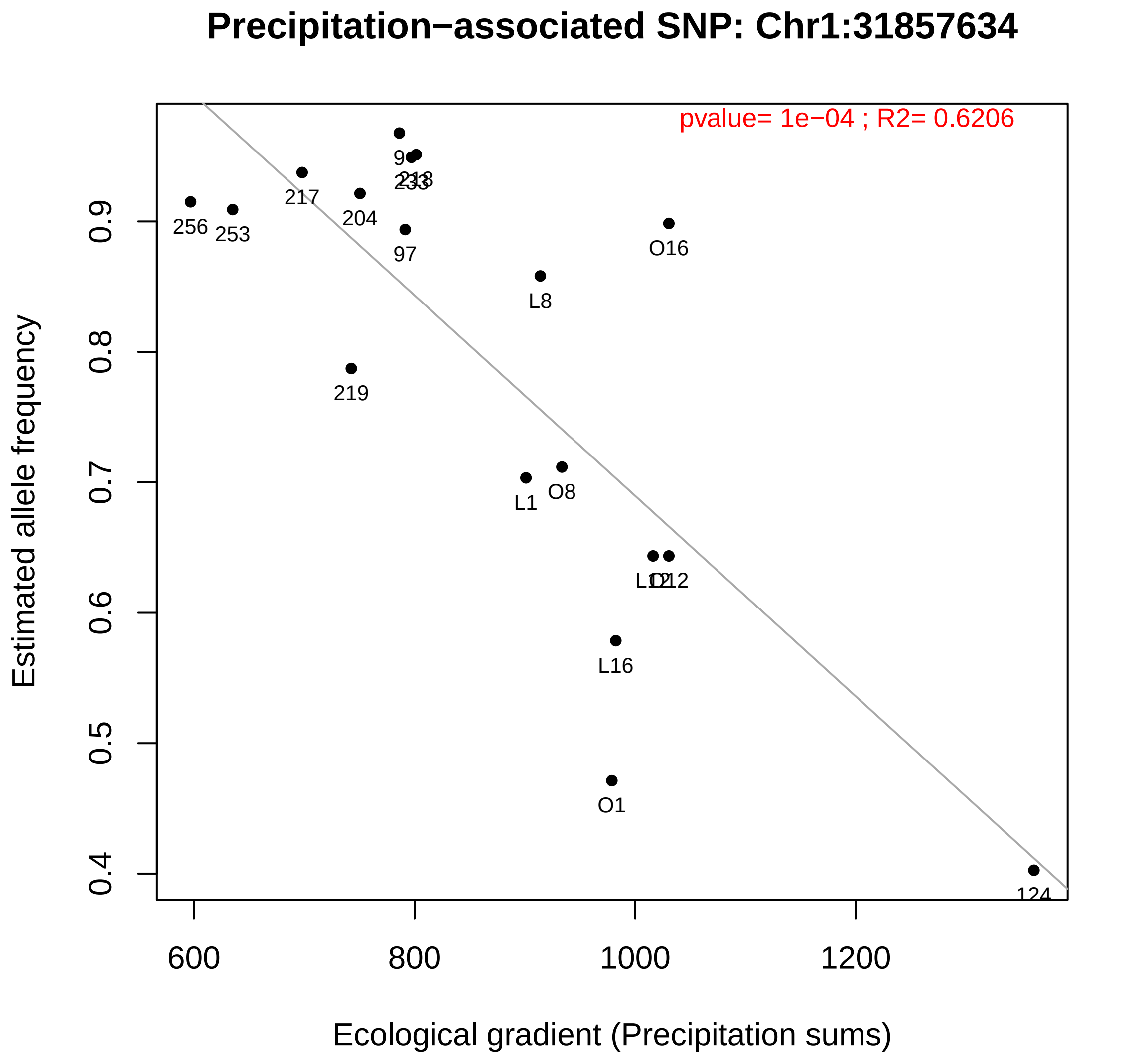

Precipitation−associated SNP: Chr1:31857634
pvalue= 1e−04 ; R2= 0.6206
Estimated allele frequency
0.4
0.5
0.6
0.7
0.8
0.9
600
800
1000
1200
Ecological gradient (Precipitation sums)
256
253
217
204
219
97
233
218
L8
O16
L1
O8
L12
O12
L16
O1
124

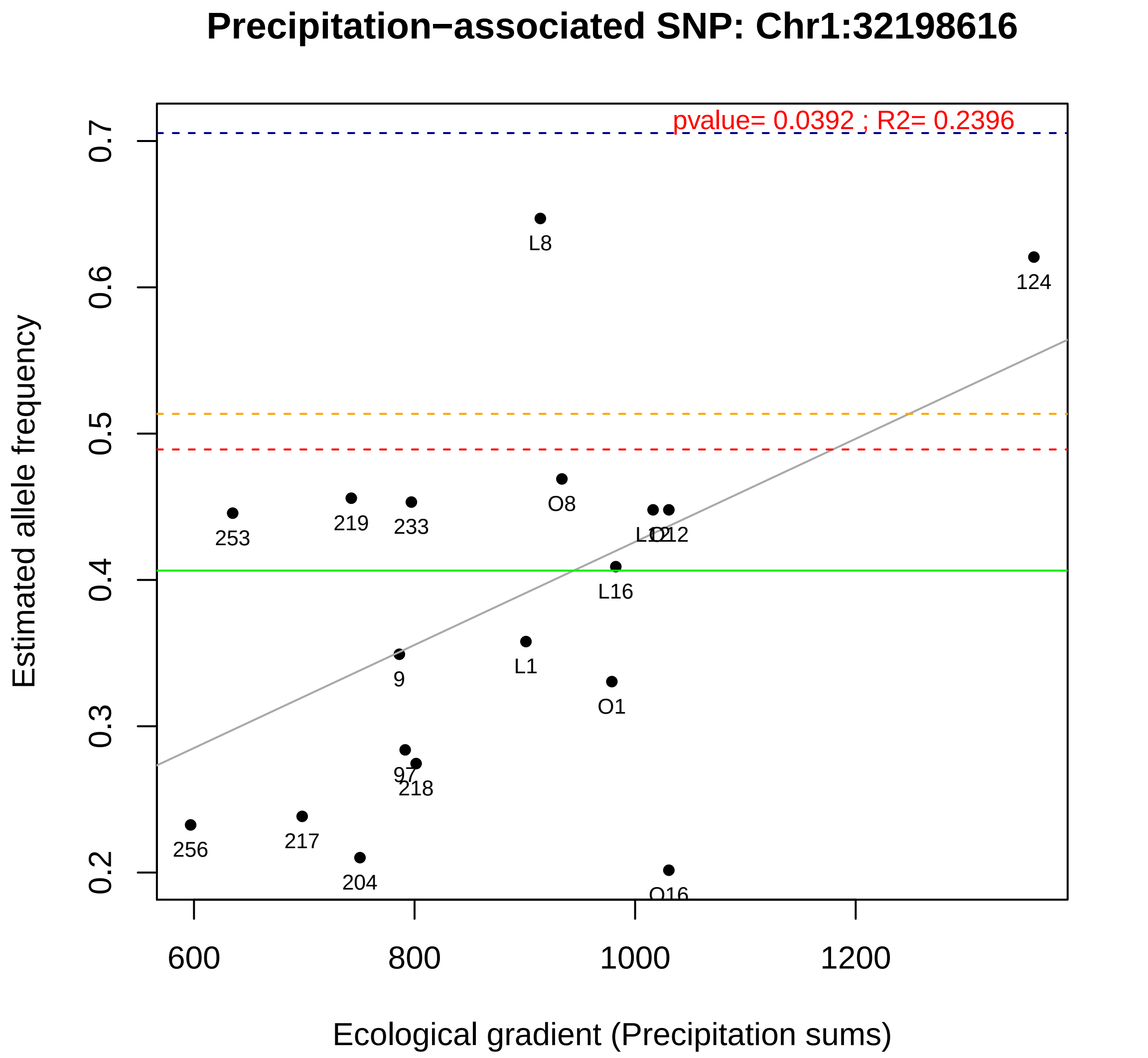

Precipitation−associated SNP: Chr1:32198616
pvalue= 0.0392 ; R2= 0.2396
Estimated allele frequency
Ecological gradient (Precipitation sums)
0.2
0.3
0.4
0.5
0.6
0.7
600
800
1000
1200
L8
124
O8
253
219
233
L12
O12
L16
L1
9
O1
97
218
256
217
204
O16

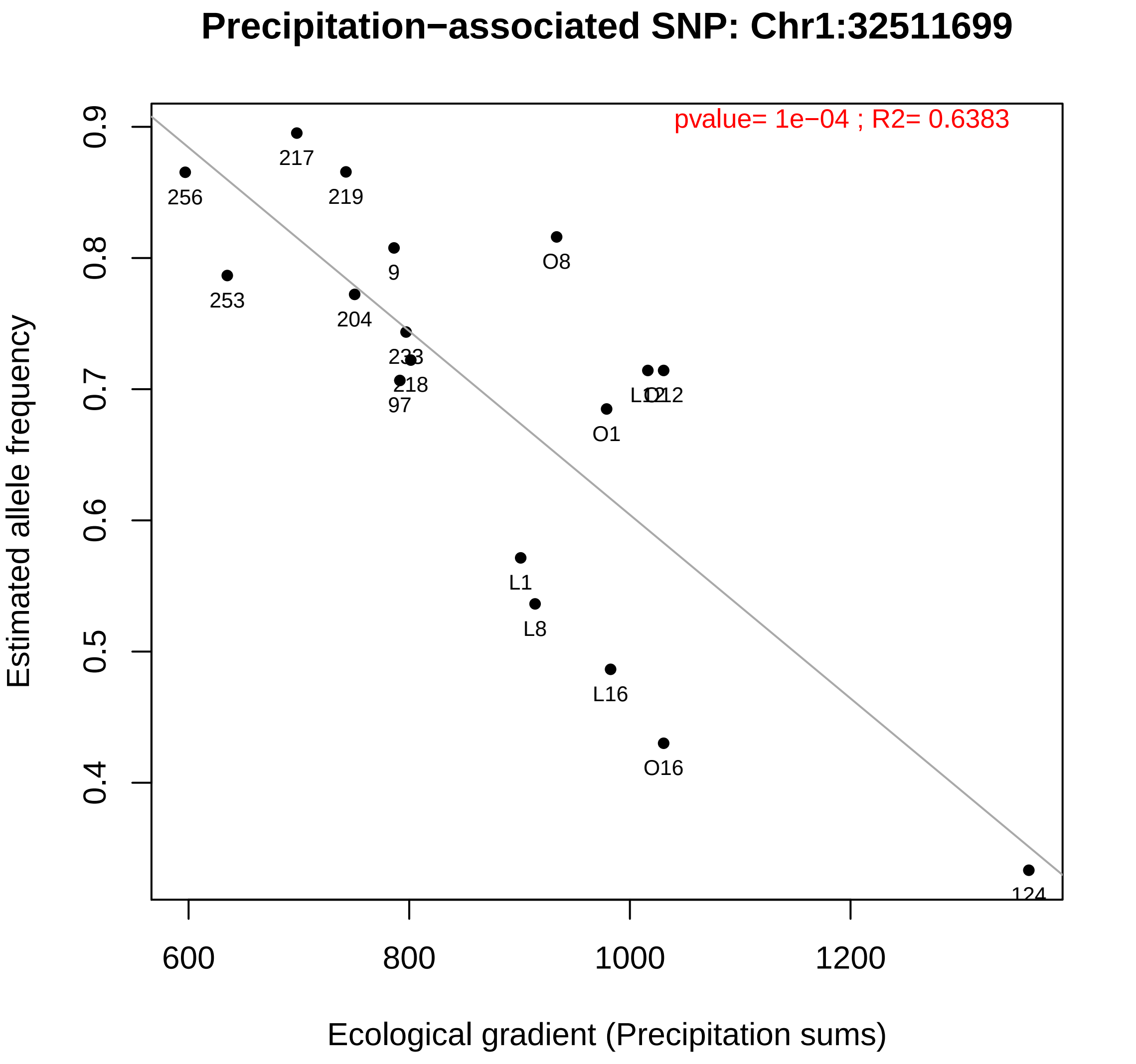

Precipitation−associated SNP: Chr1:32511699
pvalue= 1e−04 ; R2= 0.6383
Estimated allele frequency
0.4
0.5
0.6
0.7
0.8
0.9
600
800
1000
1200
Ecological gradient (Precipitation sums)
256
217
219
253
9
O8
204
233
218
97
L12
O12
O1
L1
L8
L16
O16
124

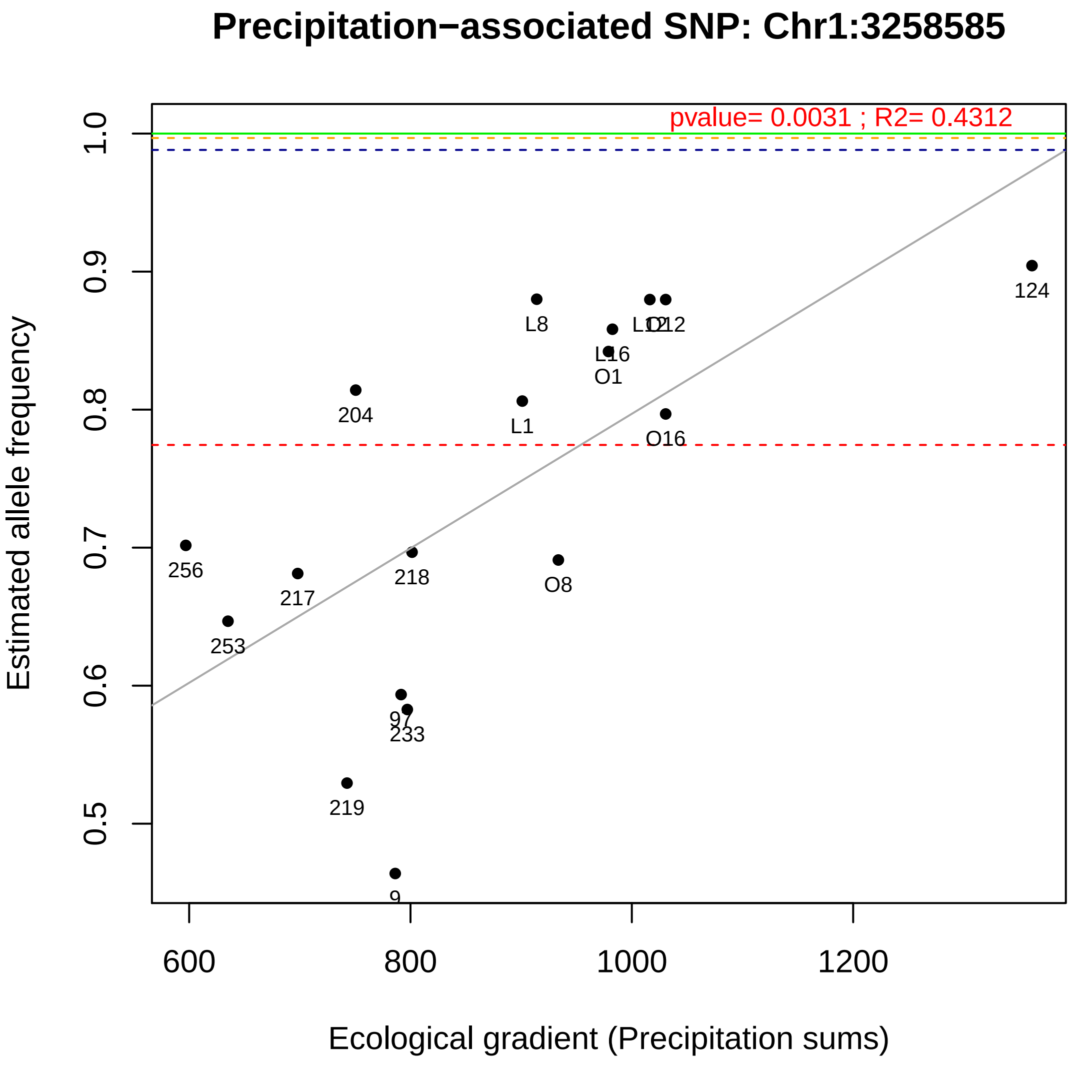

Precipitation−associated SNP: Chr1:3258585
pvalue= 0.0031 ; R2= 0.4312
Estimated allele frequency
Ecological gradient (Precipitation sums)
0.5
0.6
0.7
0.8
0.9
1.0
600
800
1000
1200
124
L8
L16
O1
204
L1
O16
256
217
218
O8
253
97
233
219
9

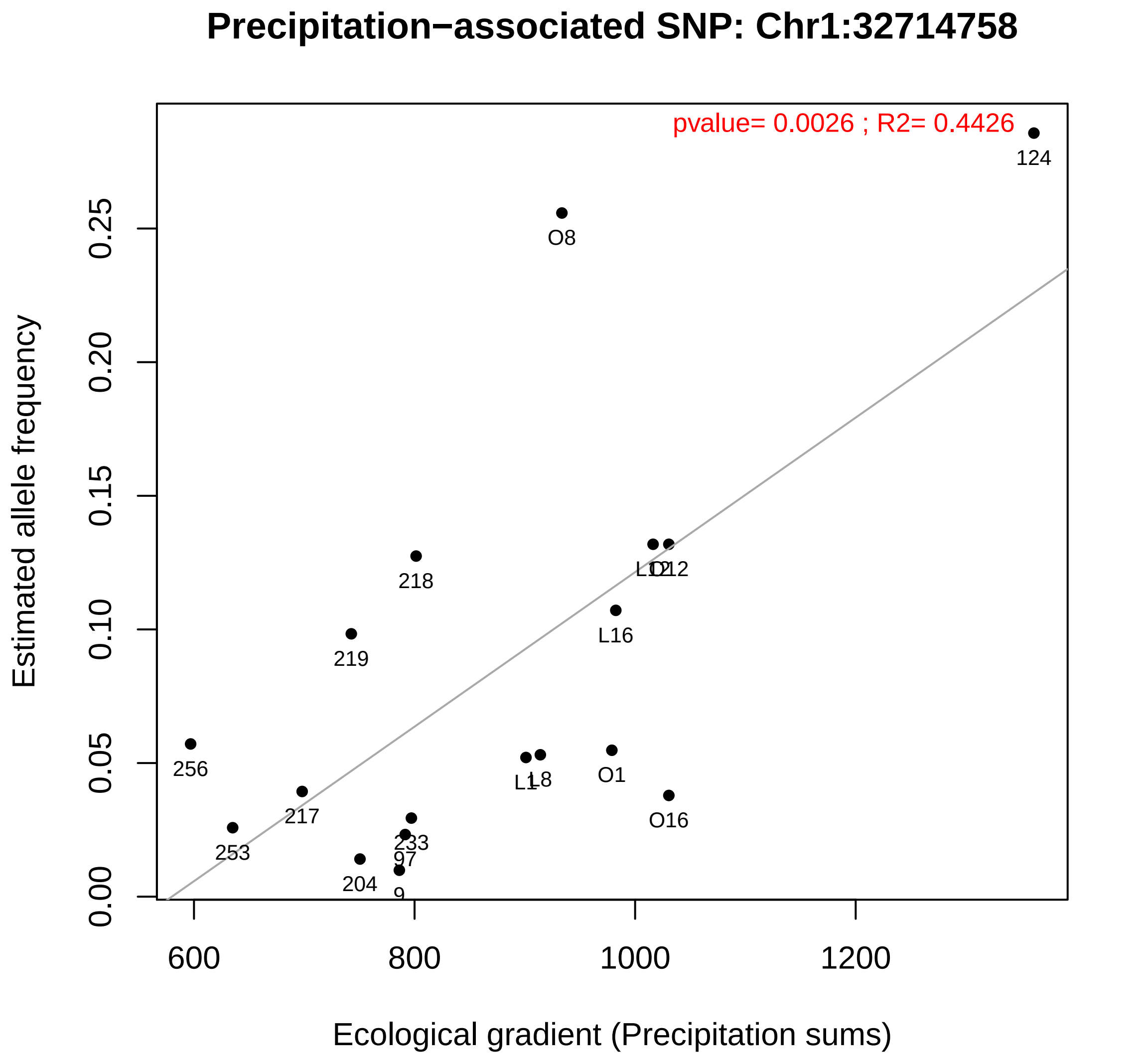

Precipitation−associated SNP: Chr1:32714758
pvalue= 0.0026 ; R2= 0.4426
124
O8
218
219
L16
256
217
253
204
233
97
9
O1
O16
0.00
0.05
0.10
0.15
0.20
0.25
600
800
1000
1200
Estimated allele frequency
Ecological gradient (Precipitation sums)

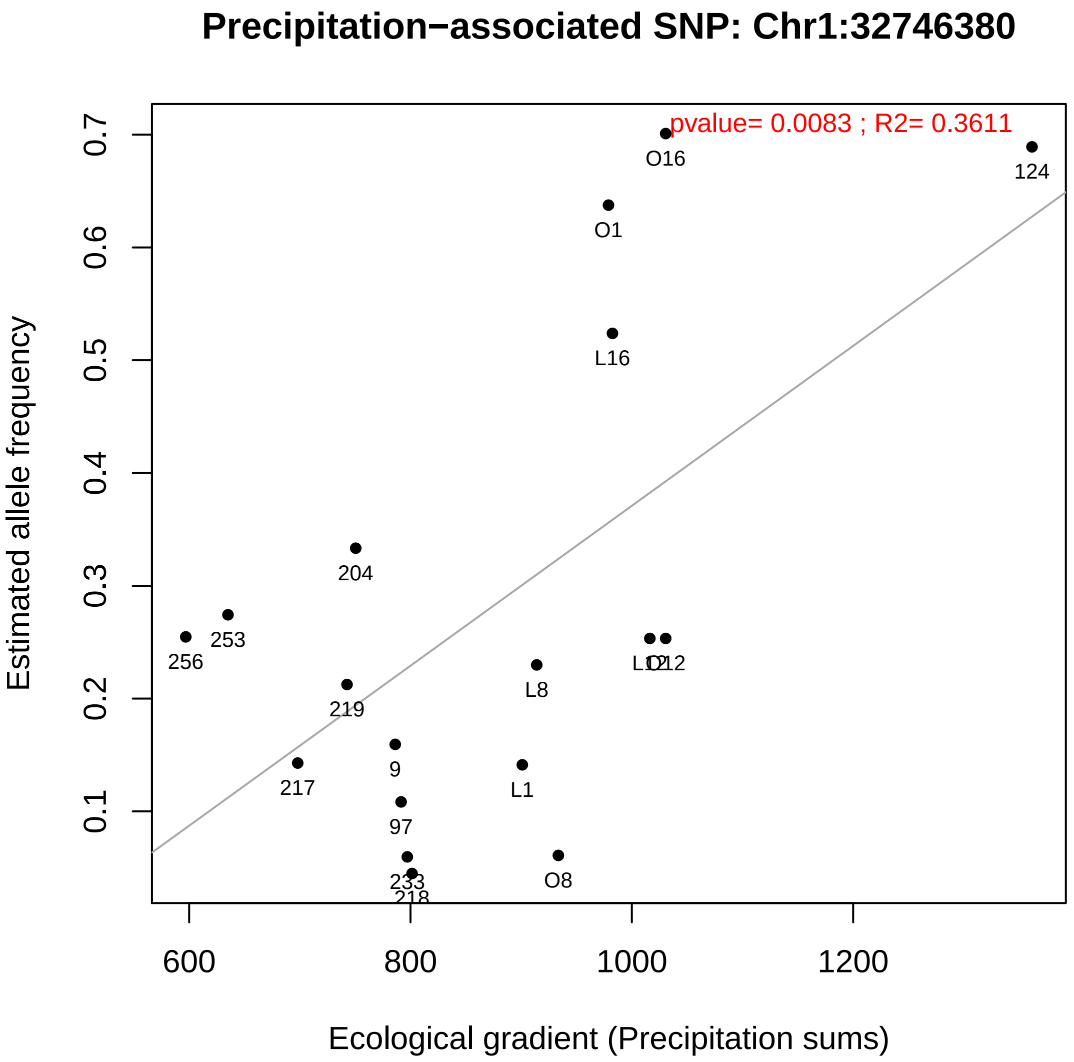
Precipitation−associated SNP: Chr1:32746380
pvalue= 0.0083 ; R2= 0.3611
Estimated allele frequency
Ecological gradient (Precipitation sums)
0.1
0.2
0.3
0.4
0.5
0.6
0.7
600
800
1000
1200
O16
124
O1
L16
204
253
256
219
L8
217
9
L1
97
233
218
O8

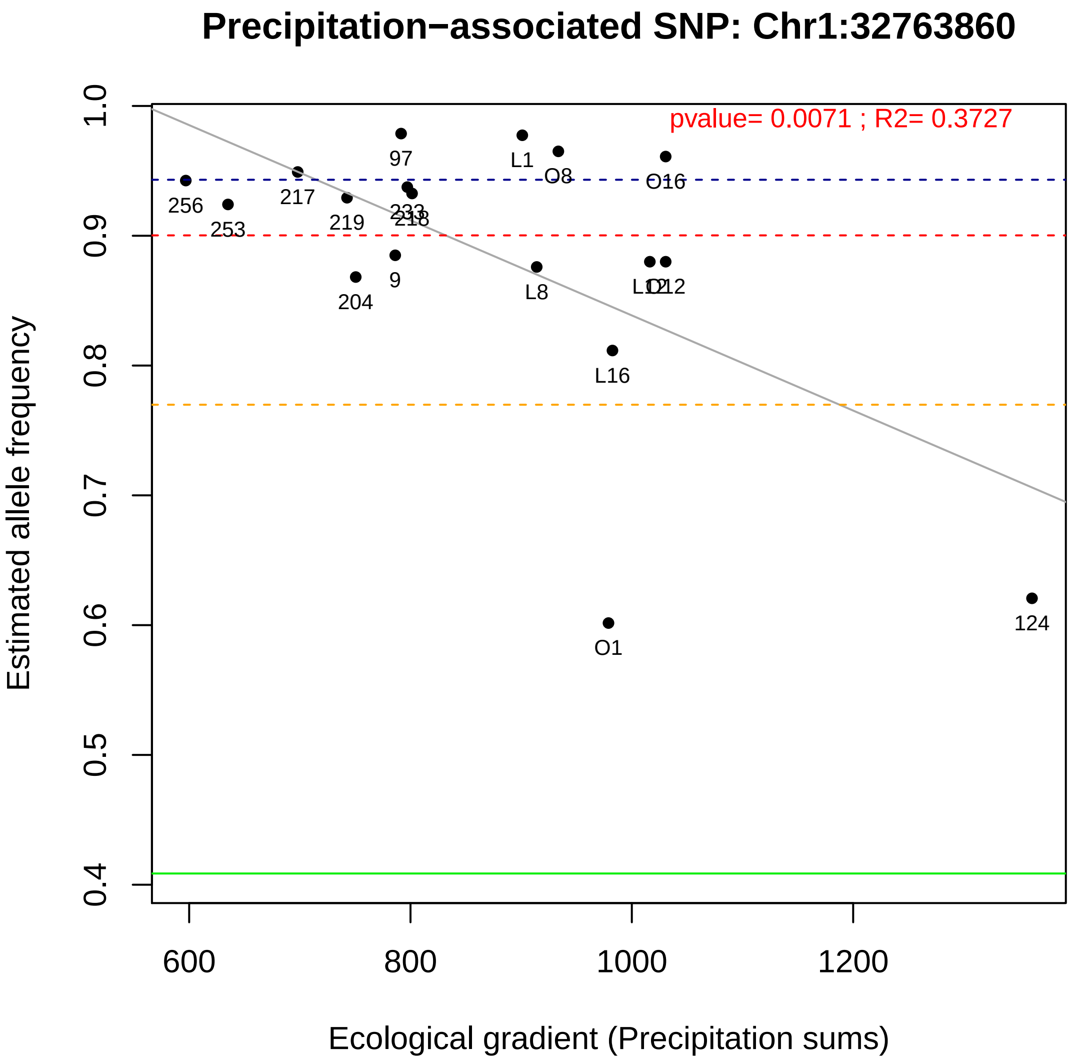

Precipitation−associated SNP: Chr1:32763860
pvalue= 0.0071 ; R2= 0.3727
Estimated allele frequency
Ecological gradient (Precipitation sums)
1.0
0.9
0.8
0.7
0.6
0.5
0.4
600
800
1000
1200
97
L1
O8
O16
256
217
253
219
233
218
9
204
L8
L12
O12
L16
O1
124

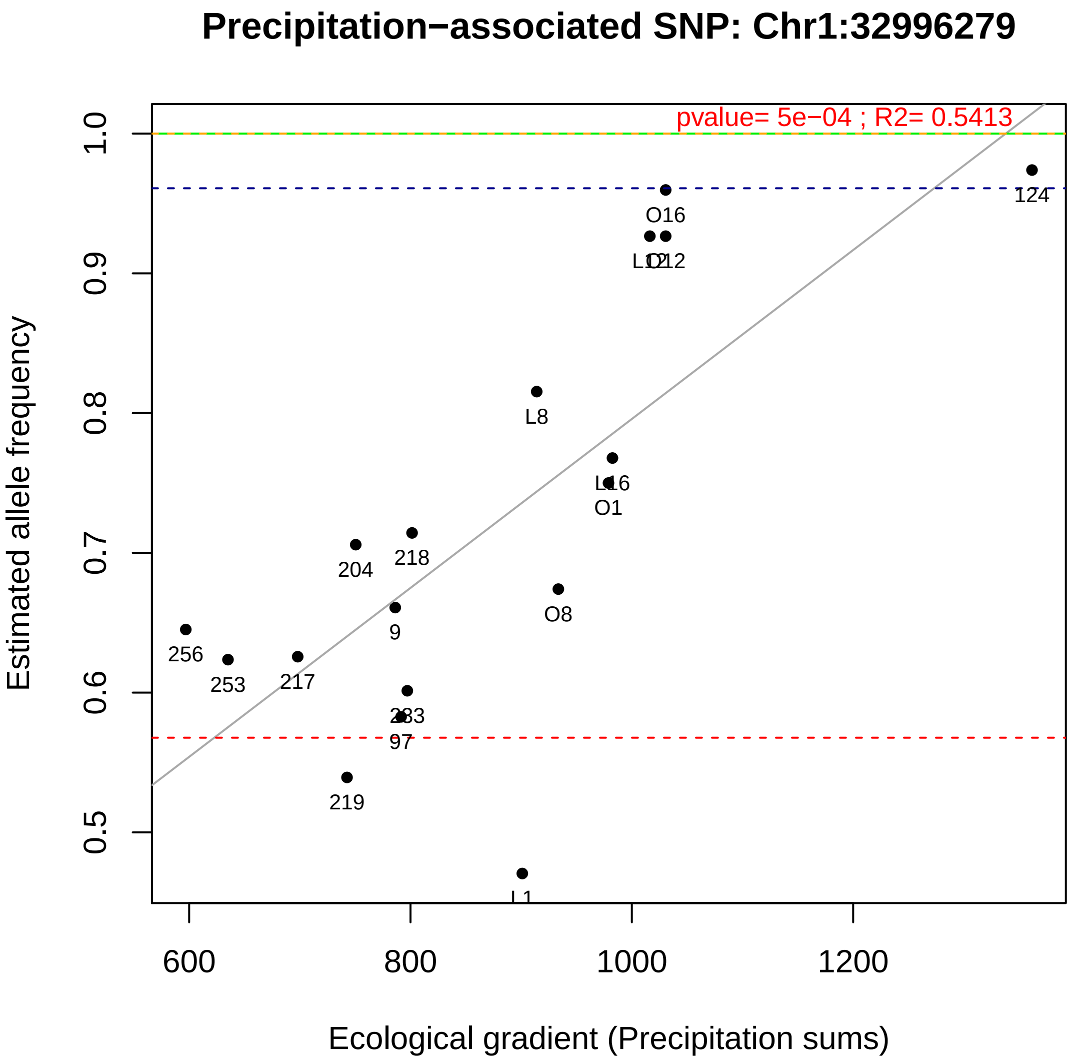
Precipitation−associated SNP: Chr1:32996279
pvalue= 5e−04 ; R2= 0.5413
Estimated allele frequency
Ecological gradient (Precipitation sums)
0.5
0.6
0.7
0.8
0.9
1.0
600
800
1000
1200
124
O16
L10
O12
L8
L16
O1
218
204
O8
9
256
253
217
233
97
219
L1

#### Precipitation−associated SNP: Chr1:33064558

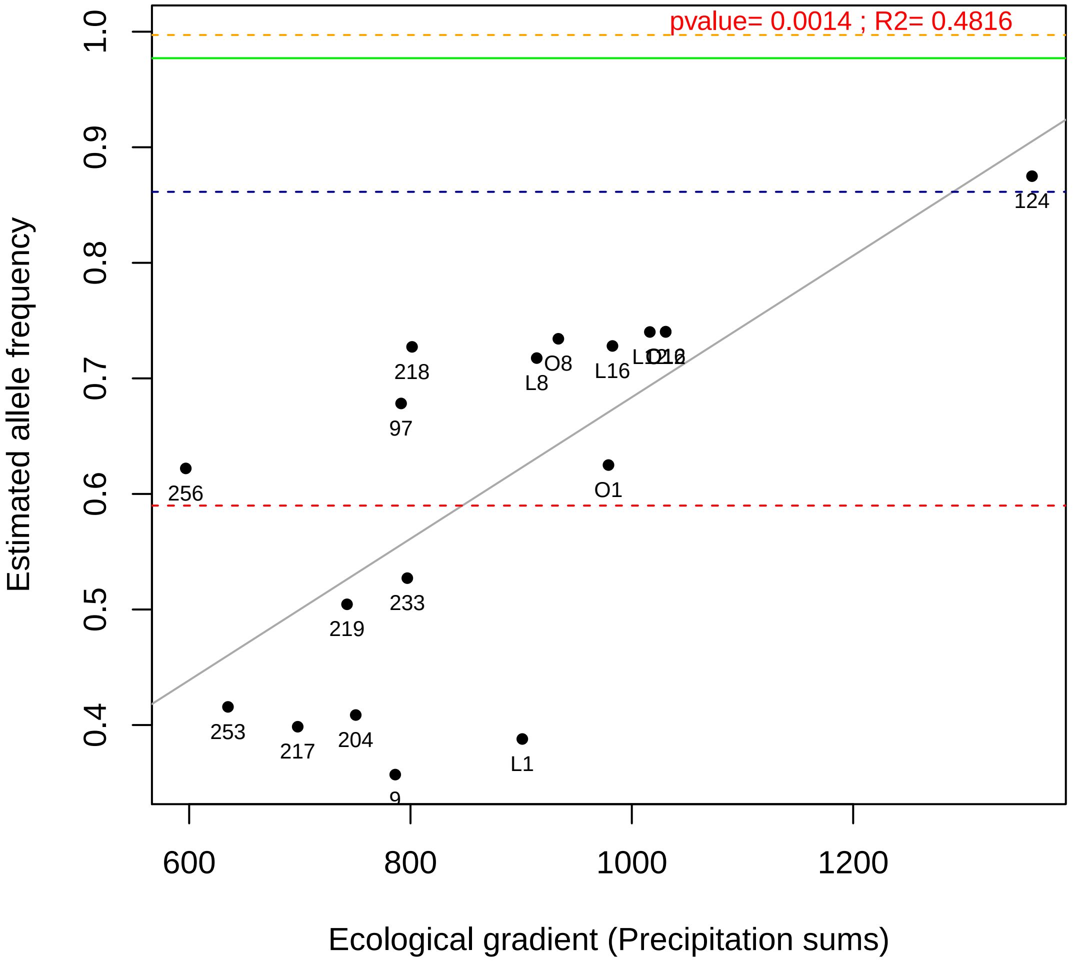

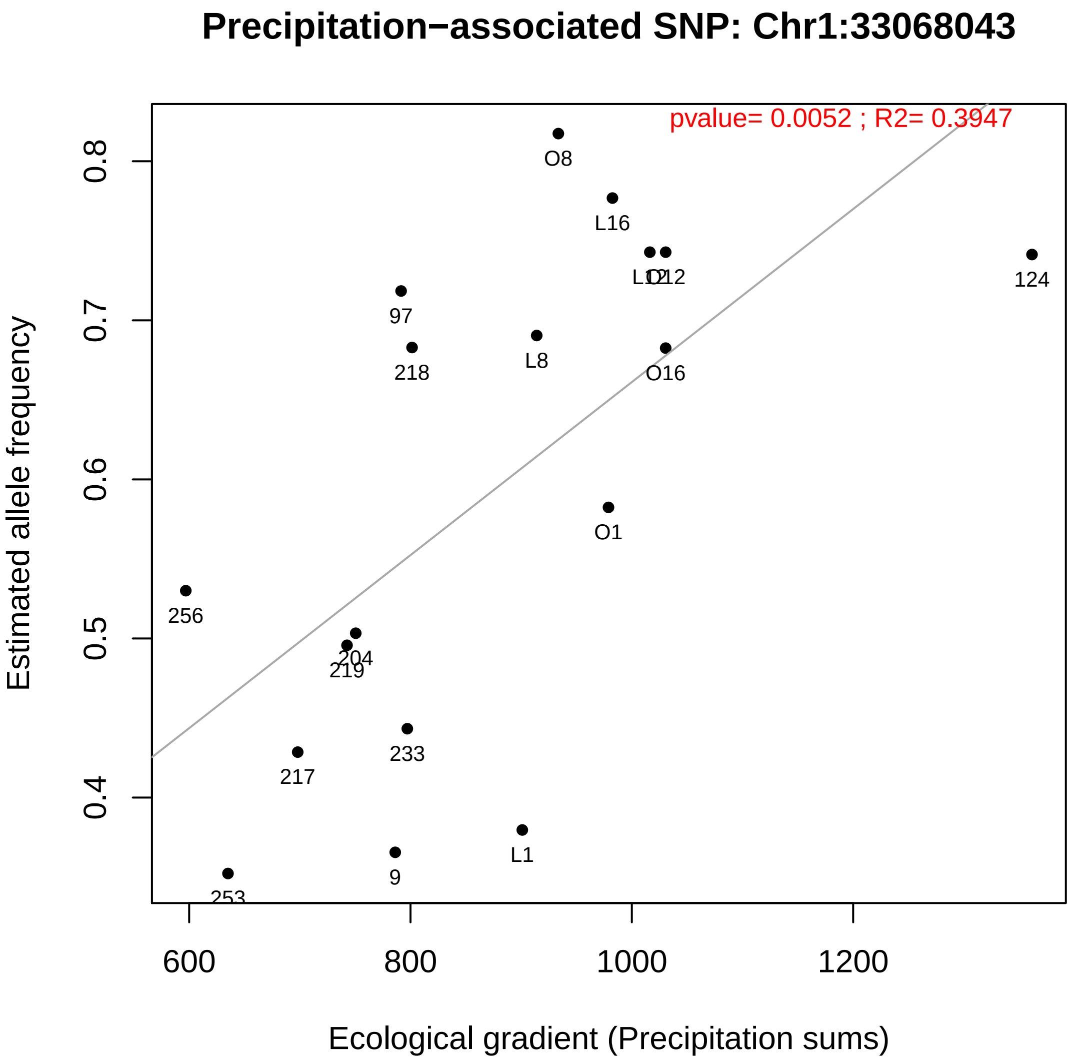

Precipitation−associated SNP: Chr1:33068043
pvalue= 0.0052 ; R2= 0.3947
Estimated allele frequency
0.4
0.5
0.6
0.7
0.8
600
800
1000
1200
Ecological gradient (Precipitation sums)
O8
L16
L12
O12
124
97
218
L8
O16
O1
256
204
219
233
217
L1
9
253

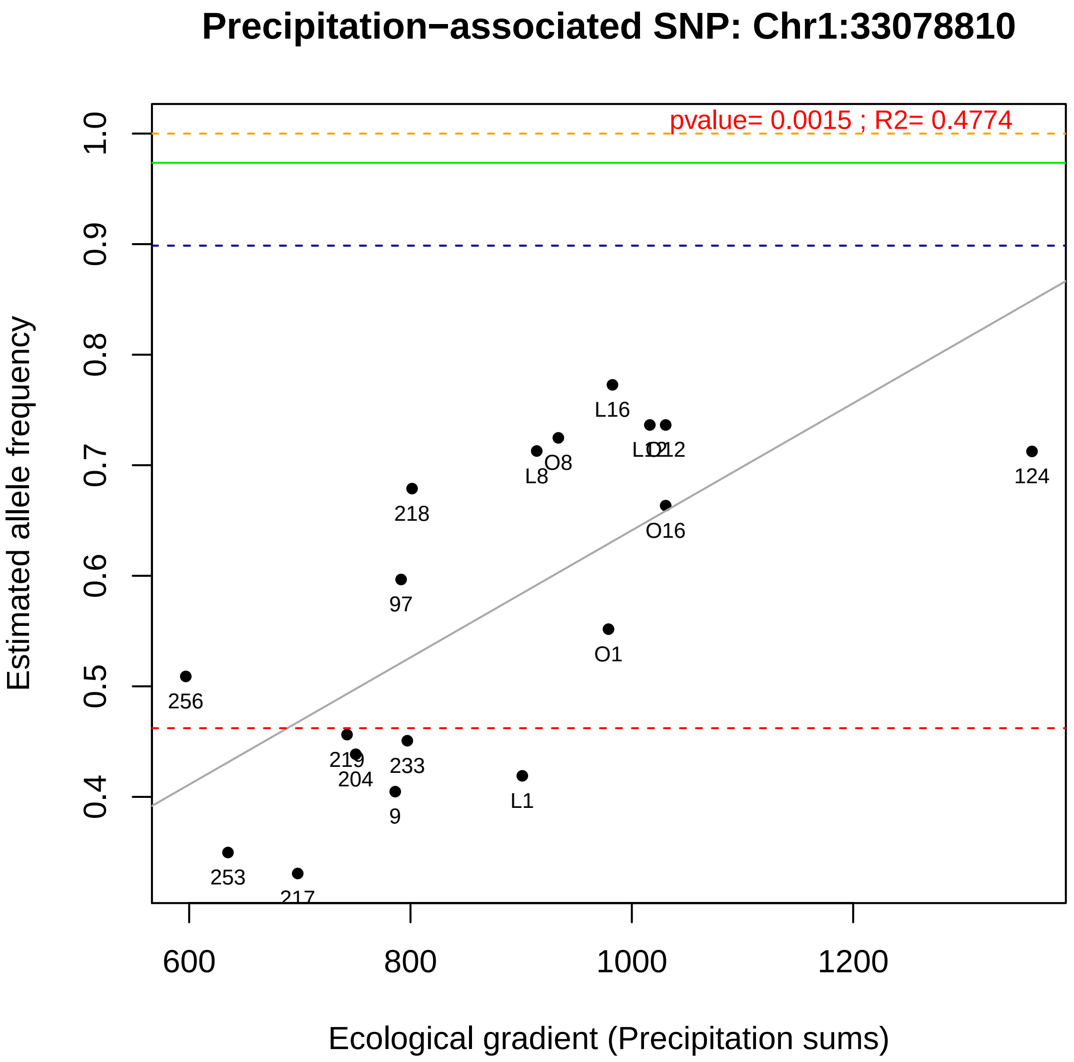
Precipitation−associated SNP: Chr1:33078810
pvalue= 0.0015 ; R2= 0.4774
Estimated allele frequency
Ecological gradient (Precipitation sums)
0.4
0.5
0.6
0.7
0.8
0.9
1.0
600
800
1000
1200
L16
L12
O12
O8
L8
124
218
O16
97
O1
256
219
204
233
L1
9
253
217

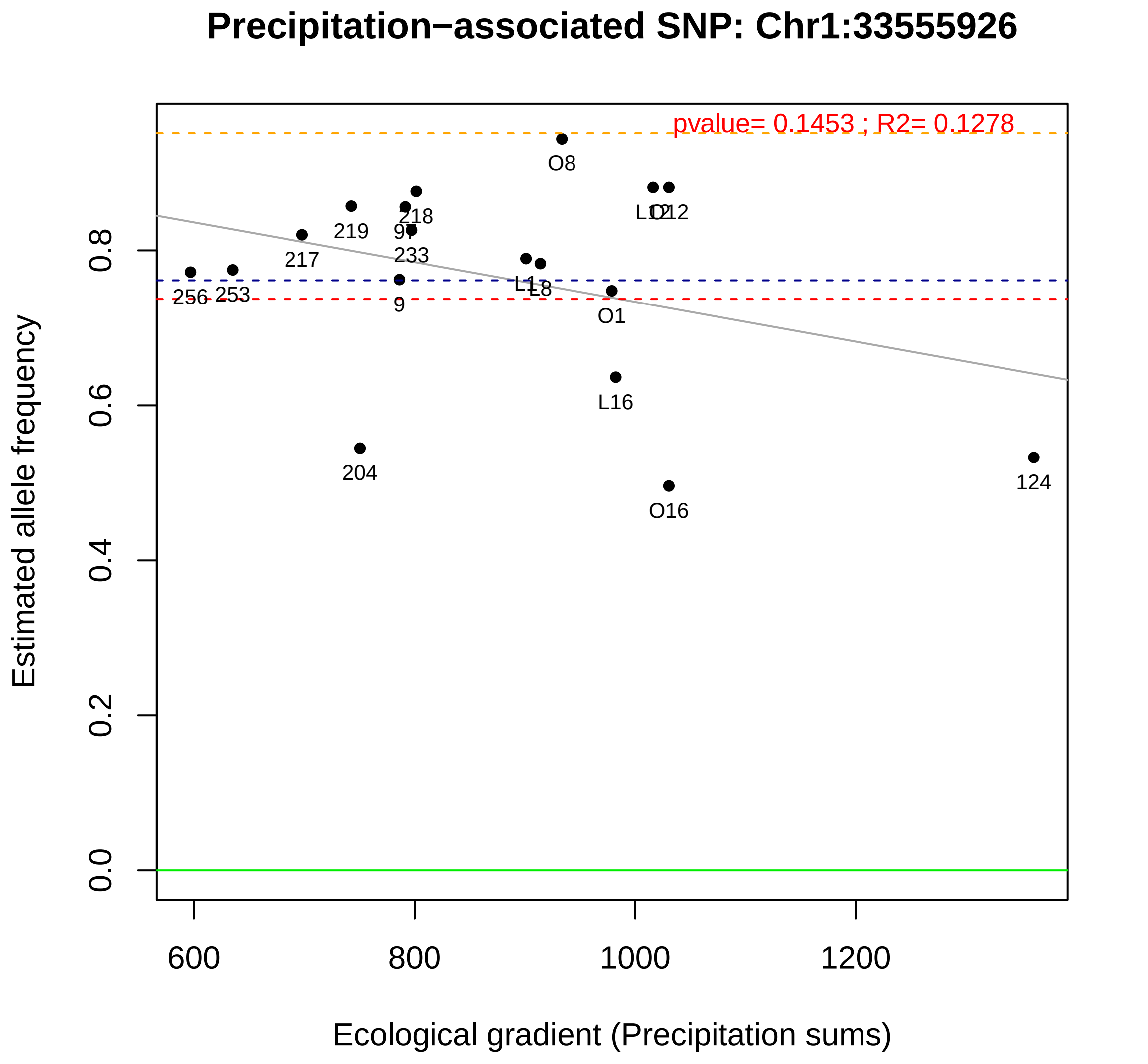

Precipitation−associated SNP: Chr1:33555926
pvalue= 0.1453 ; R2= 0.1278
Estimated allele frequency
Ecological gradient (Precipitation sums)
0.0
0.2
0.4
0.6
0.8
600
800
1000
1200
O8
L12
O12
218
219
97
233
217
256
253
9
L1
L8
O1
L16
204
O16
124

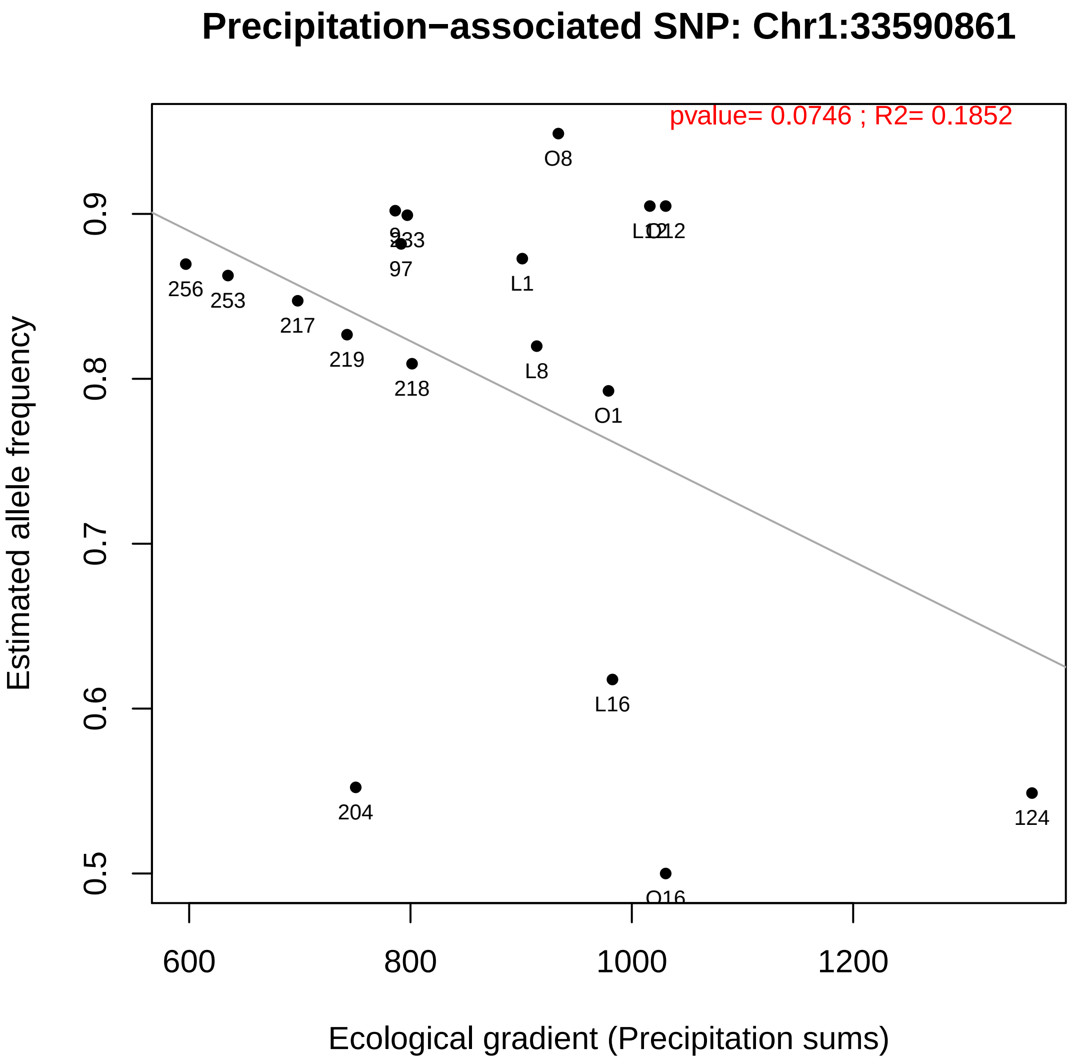
Precipitation−associated SNP: Chr1:33590861
pvalue= 0.0746 ; R2= 0.1852
Estimated allele frequency
0.5
0.6
0.7
0.8
0.9
600
800
1000
1200
Ecological gradient (Precipitation sums)
O8
256
253
217
219
218
97
L1
L8
O1
L16
204
124
O16

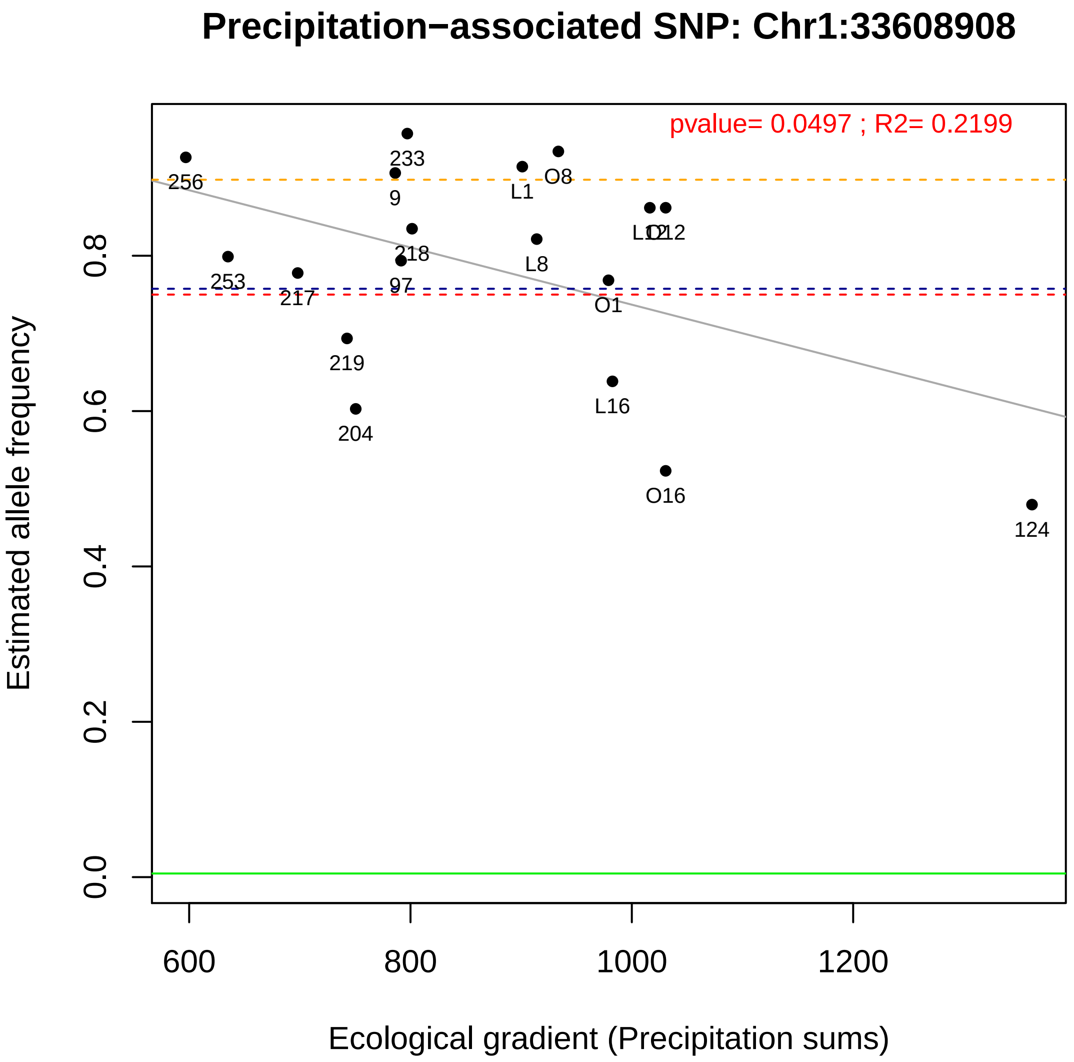
Precipitation−associated SNP: Chr1:33608908
pvalue= 0.0497 ; R2= 0.2199
Estimated allele frequency
Ecological gradient (Precipitation sums)
0.0
0.2
0.4
0.6
0.8
600
800
1000
1200
256
253
217
219
204
233
9
218
97
L1
O8
L8
L10
O12
O1
L16
O16
124

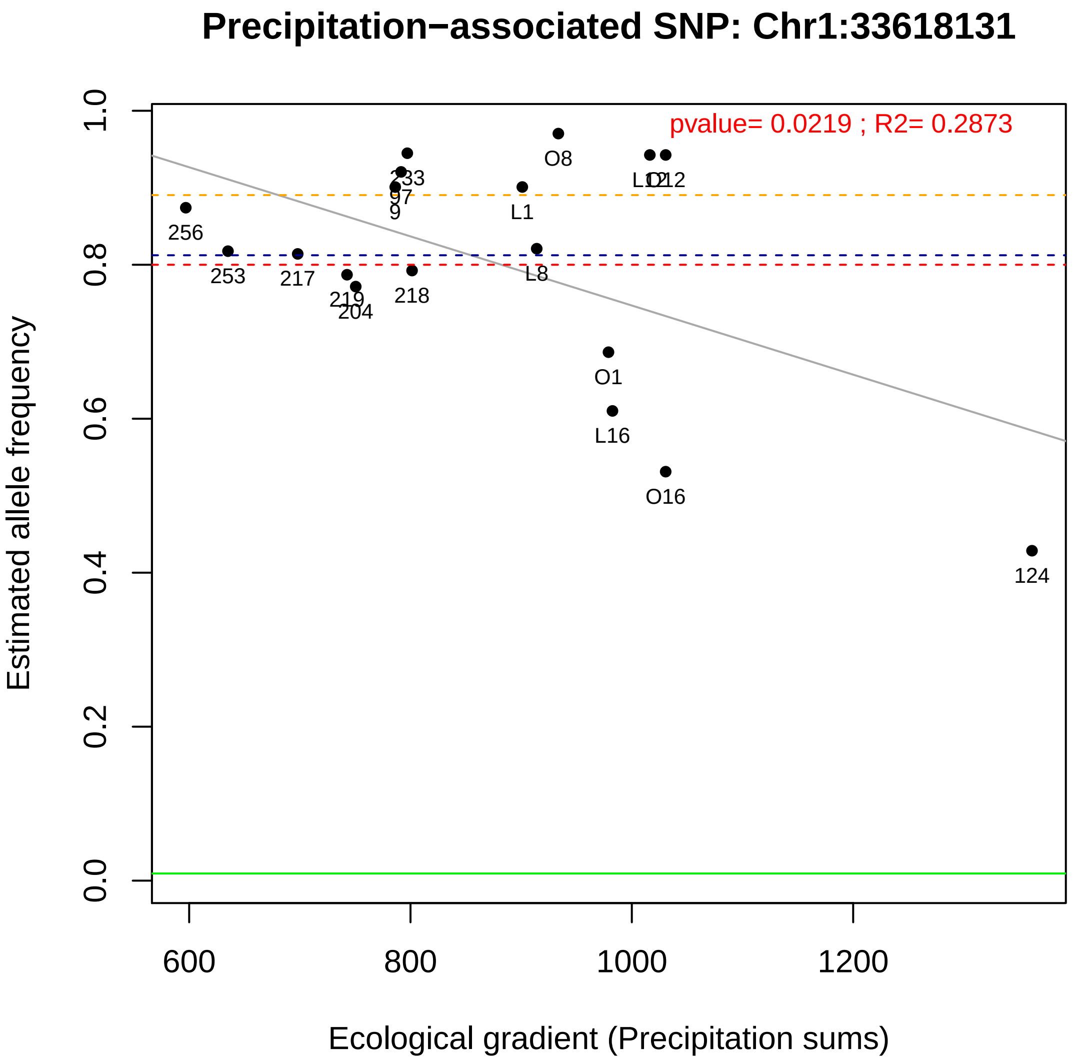

Precipitation−associated SNP: Chr1:33618131
pvalue= 0.0219 ; R2= 0.2873
Estimated allele frequency
Ecological gradient (Precipitation sums)
0.0
0.2
0.4
0.6
0.8
1.0
600
800
1000
1200
256
253
217
219
204
233
97
9
218
L1
L8
O8
L12
O12
O1
L16
O16
124

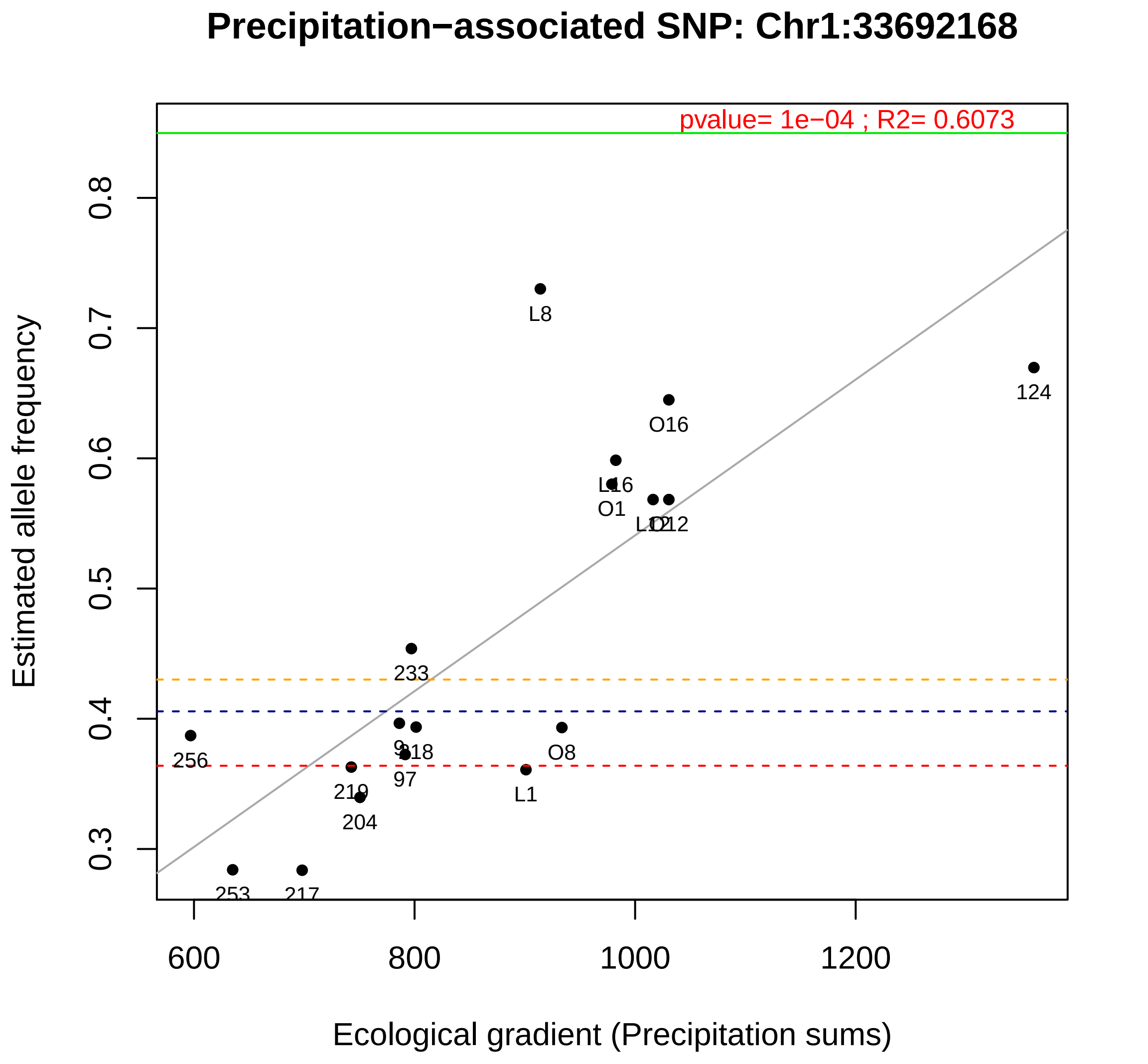

Precipitation−associated SNP: Chr1:33692168
pvalue= 1e−04 ; R2= 0.6073
Estimated allele frequency
0.3
0.4
0.5
0.6
0.7
0.8
600
800
1000
1200
Ecological gradient (Precipitation sums)
L8
124
O16
L16
O1
O12
233
256
O8
97
219
204
L1
253
217

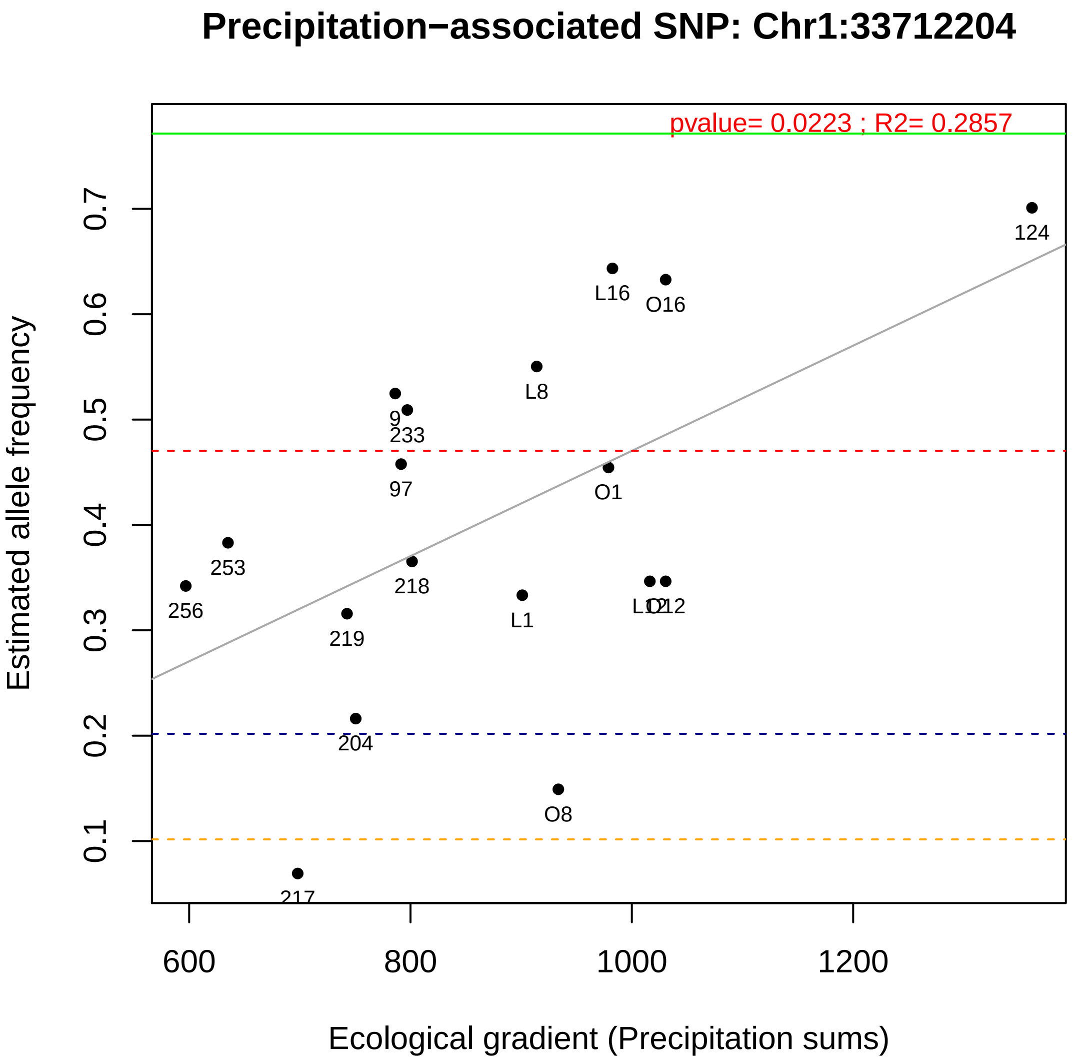
Precipitation−associated SNP: Chr1:33712204
pvalue= 0.0223 ; R2= 0.2857
Estimated allele frequency
Ecological gradient (Precipitation sums)
0.1
0.2
0.3
0.4
0.5
0.6
0.7
600
800
1000
1200
124
L16
O16
L8
9
233
97
O1
253
218
256
L1
219
204
O8
217

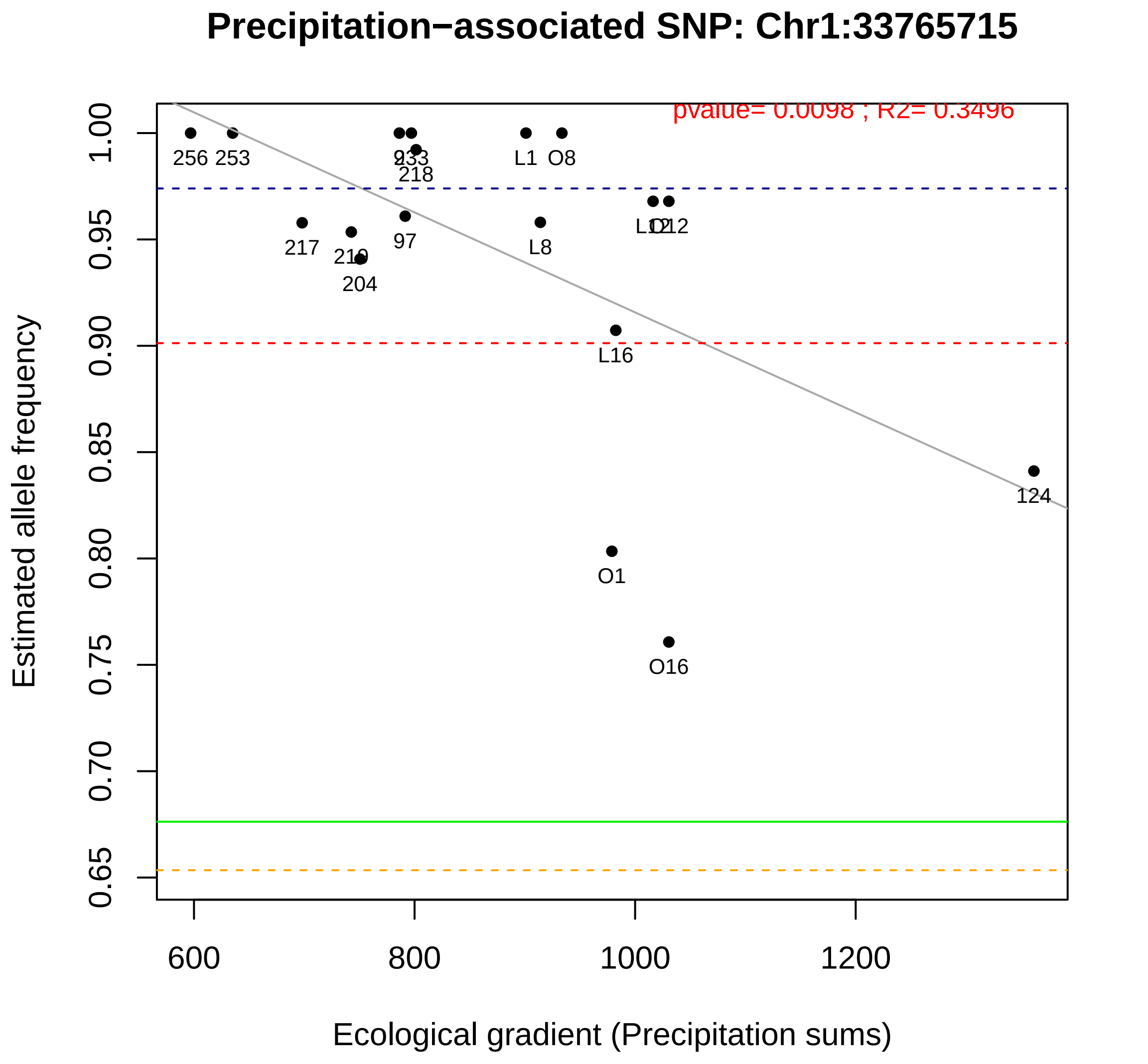

Precipitation−associated SNP: Chr1:33765715
pvalue= 0.0098 ; R2= 0.3496
Estimated allele frequency
Ecological gradient (Precipitation sums)
0.65
0.70
0.75
0.80
0.85
0.90
0.95
1.00
600
800
1000
1200
256
253
233
218
L1
O8
217
210
204
97
L8
L12
O12
L16
124
O1
O16

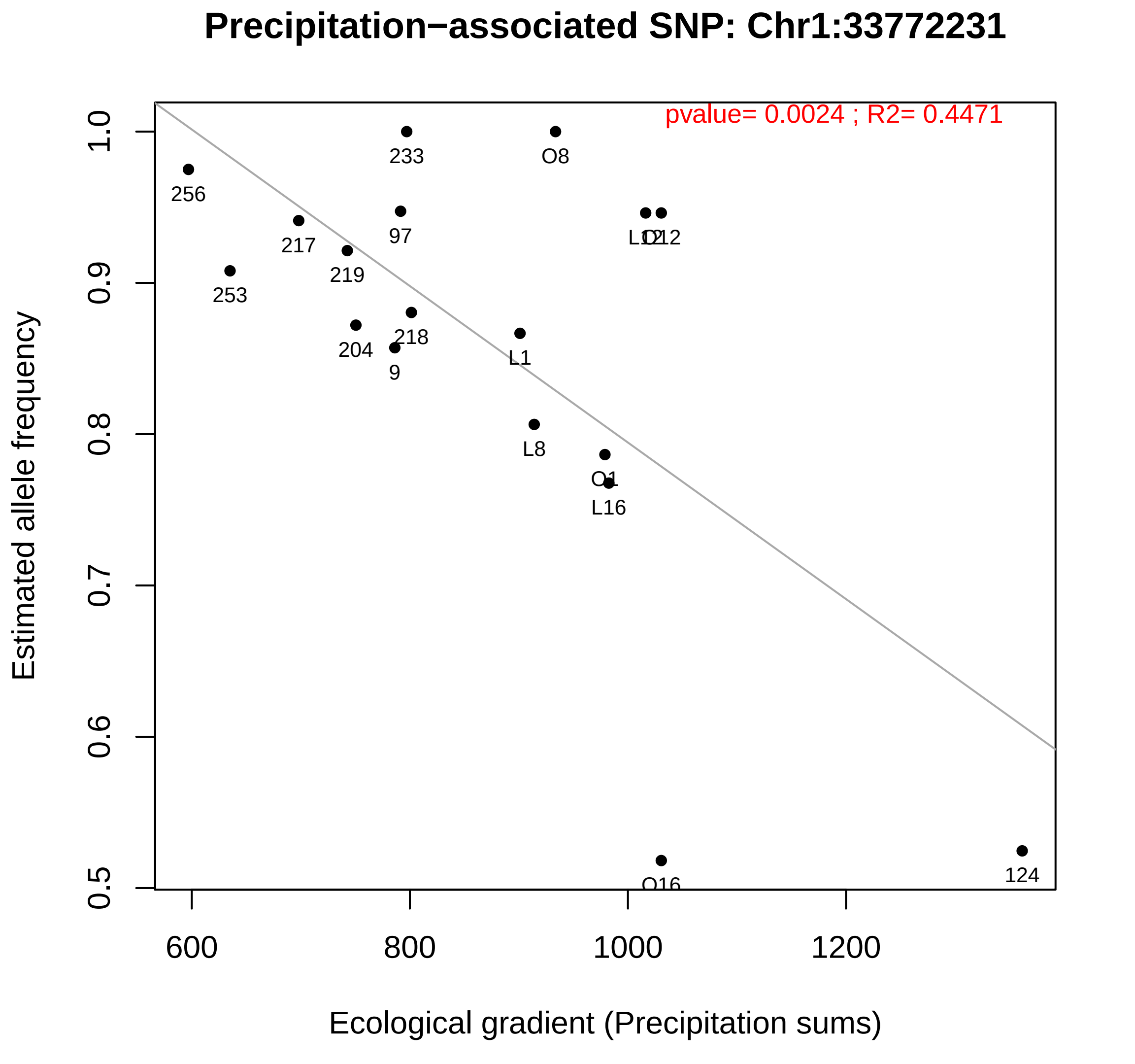

Precipitation−associated SNP: Chr1:33772231
pvalue= 0.0024 ; R2= 0.4471
Estimated allele frequency
Ecological gradient (Precipitation sums)
0.5
0.6
0.7
0.8
0.9
1.0
600
800
1000
1200
233
O8
256
97
217
L12
O12
219
253
218
204
L1
9
L8
O1
L16
O16
124

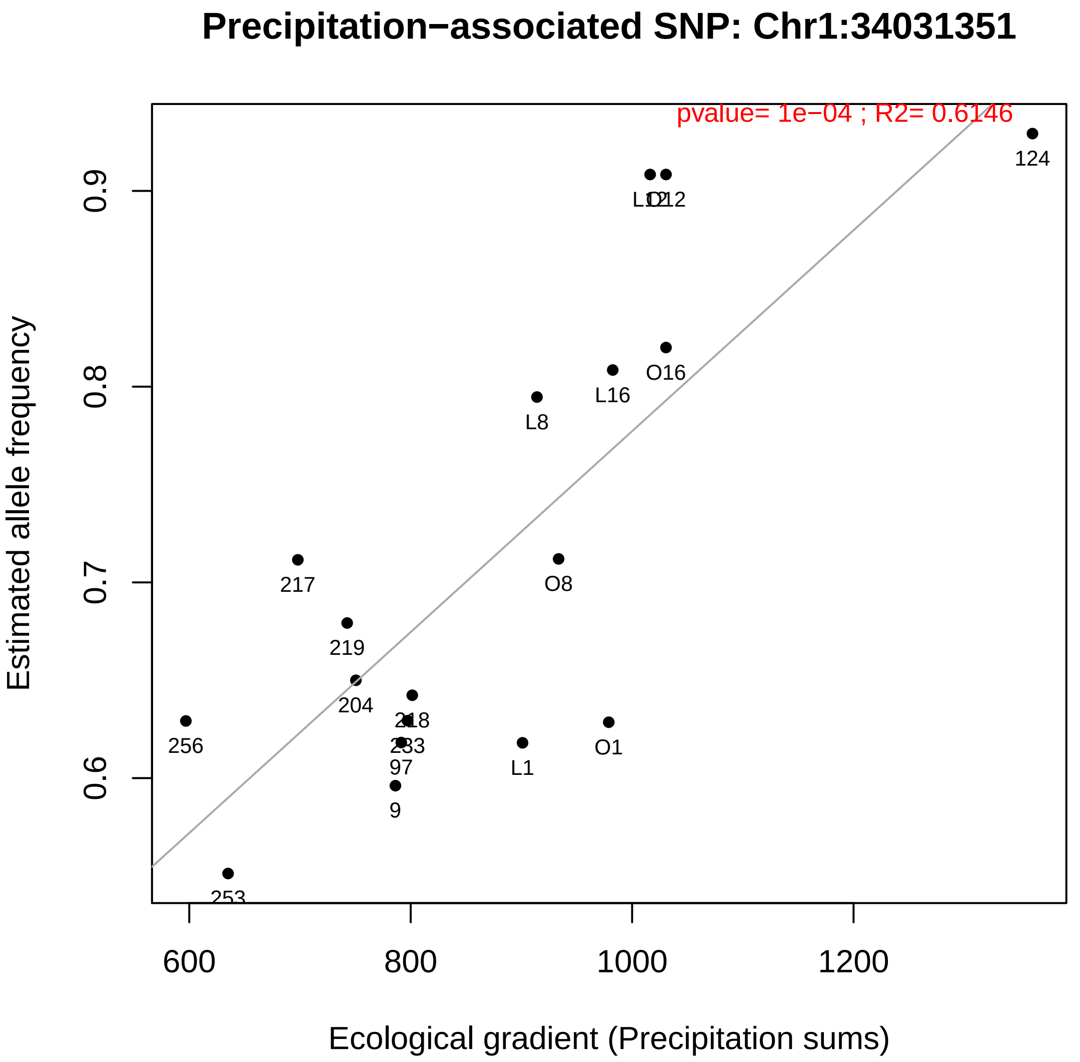
Precipitation−associated SNP: Chr1:34031351
pvalue= 1e−04 ; R2= 0.6146
Estimated allele frequency
0.6
0.7
0.8
0.9
600
800
1000
1200
Ecological gradient (Precipitation sums)
124
L12
O12
O16
L16
L8
217
O8
219
204
218
256
233
97
L1
O1
9
253

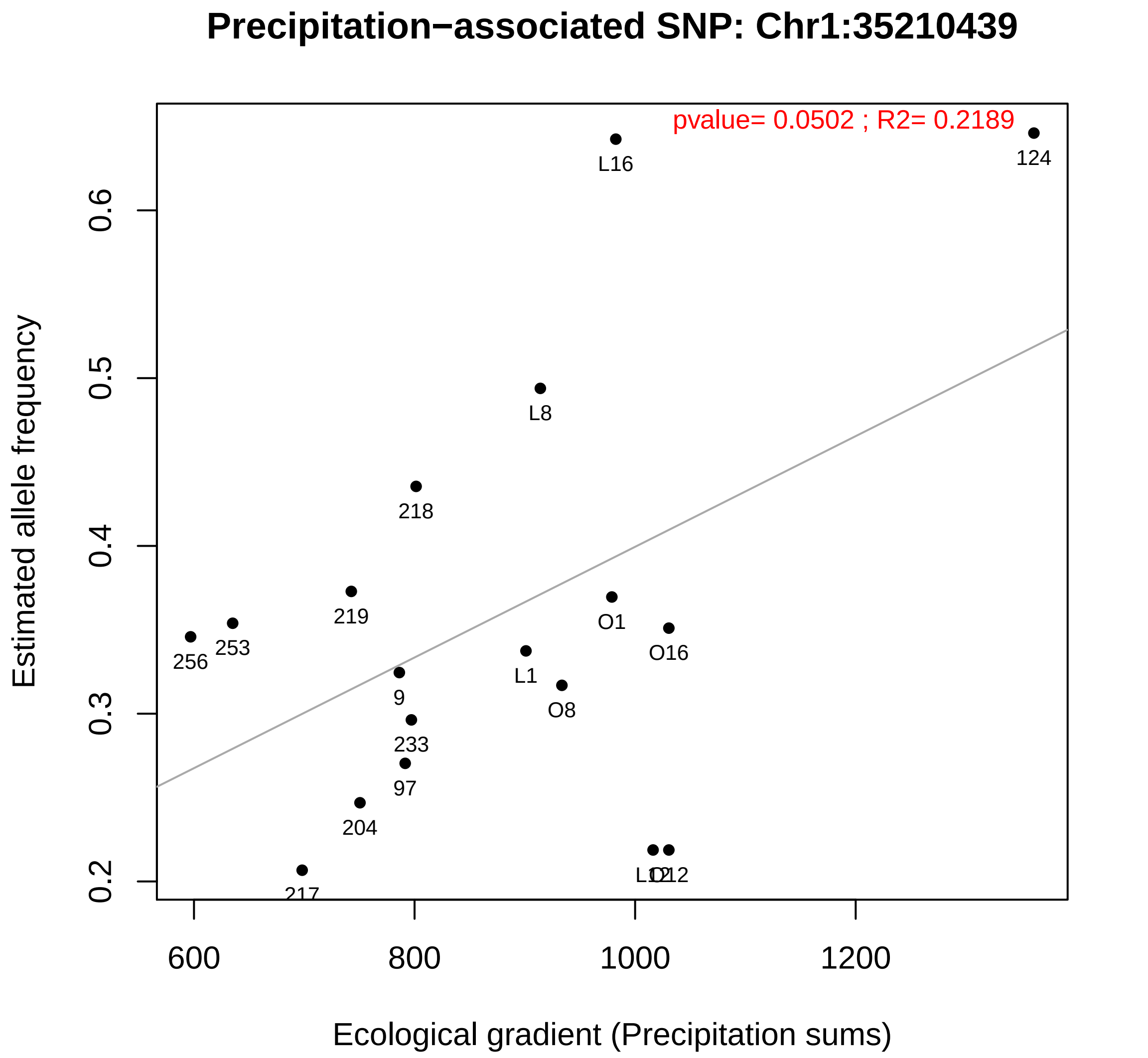

Precipitation−associated SNP: Chr1:35210439
pvalue= 0.0502 ; R2= 0.2189
Estimated allele frequency
0.2
0.3
0.4
0.5
0.6
600
800
1000
1200
Ecological gradient (Precipitation sums)
L16
124
L8
218
219
253
256
9
L1
O1
O16
O8
233
97
204
217
L10
O12

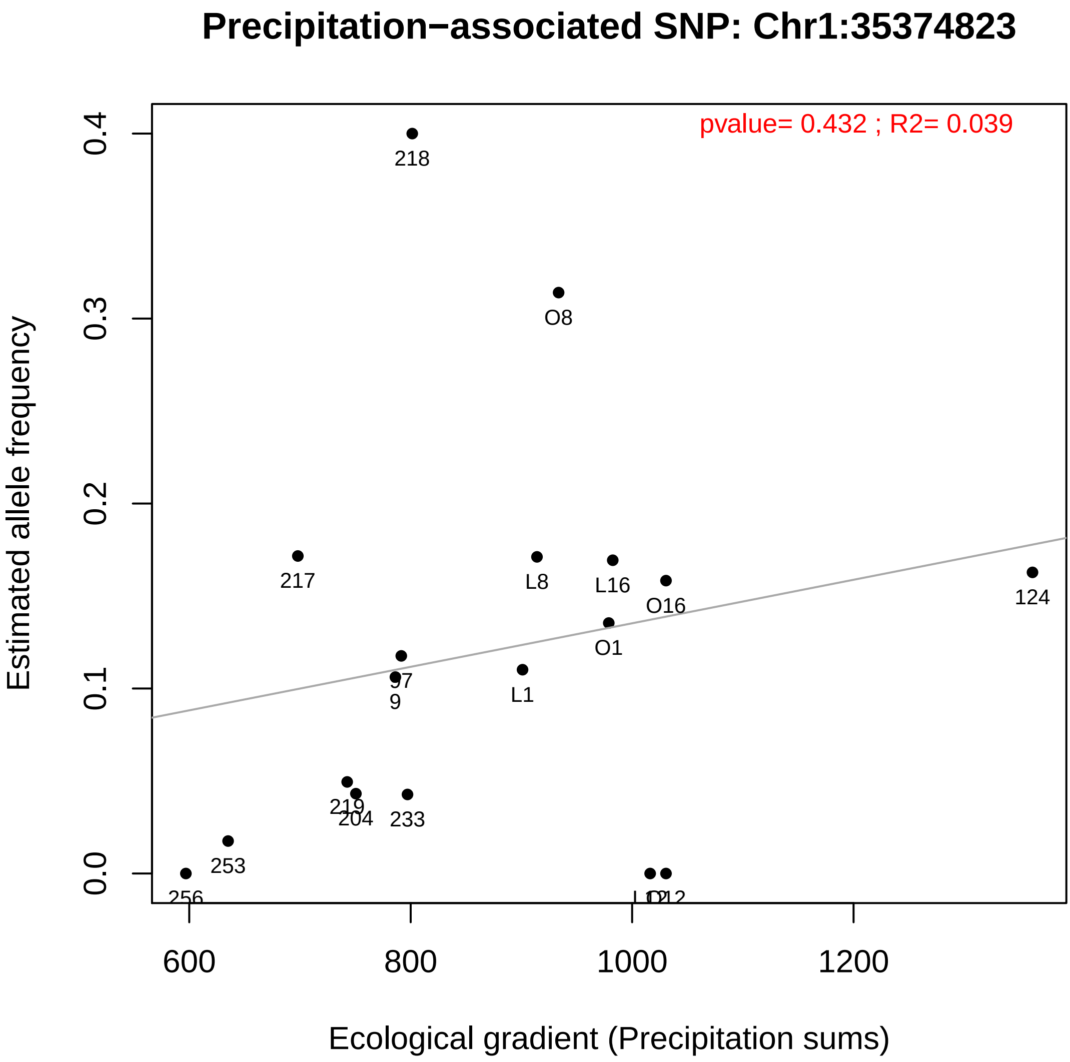

Precipitation−associated SNP: Chr1:35374823
pvalue= 0.432 ; R2= 0.039
Estimated allele frequency
0.0
0.1
0.2
0.3
0.4
600
800
1000
1200
Ecological gradient (Precipitation sums)
218
O8
217
L8
L16
O16
124
O1
97
9
L1
219
204
233
253
256

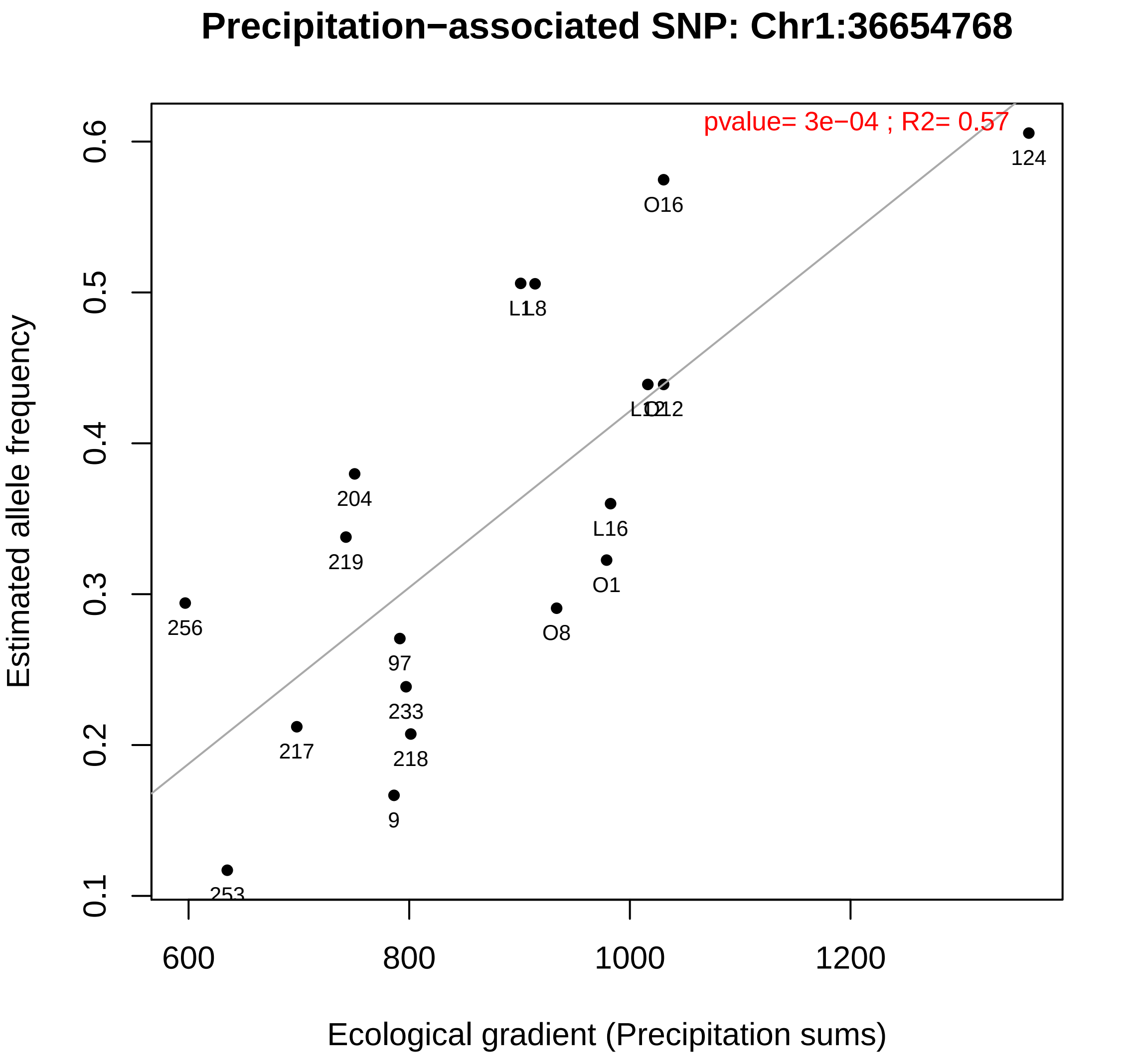

Precipitation−associated SNP: Chr1:36654768
pvalue= 3e−04 ; R2= 0.57
124
O16
L18
L10
O12
204
L16
219
O1
256
O8
97
233
217
218
9
253
0.6
0.5
0.4
0.3
0.2
0.1
600
800
1000
1200
Estimated allele frequency
Ecological gradient (Precipitation sums)

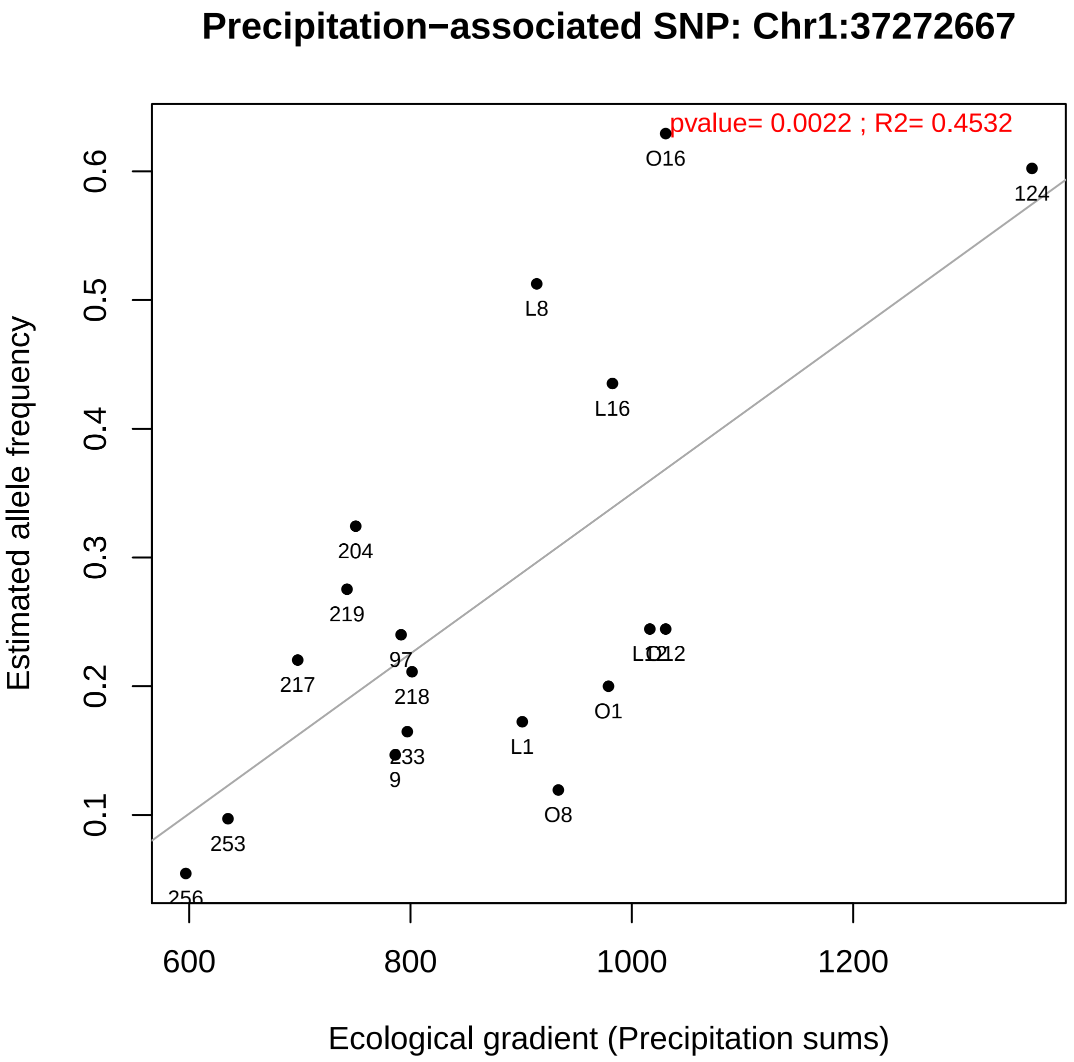

Precipitation−associated SNP: Chr1:37272667
pvalue= 0.0022 ; R2= 0.4532
Estimated allele frequency
0.1
0.2
0.3
0.4
0.5
0.6
600
800
1000
1200
Ecological gradient (Precipitation sums)
O16
124
L8
L16
204
219
97
217
218
O1
L1
233
9
O8
253

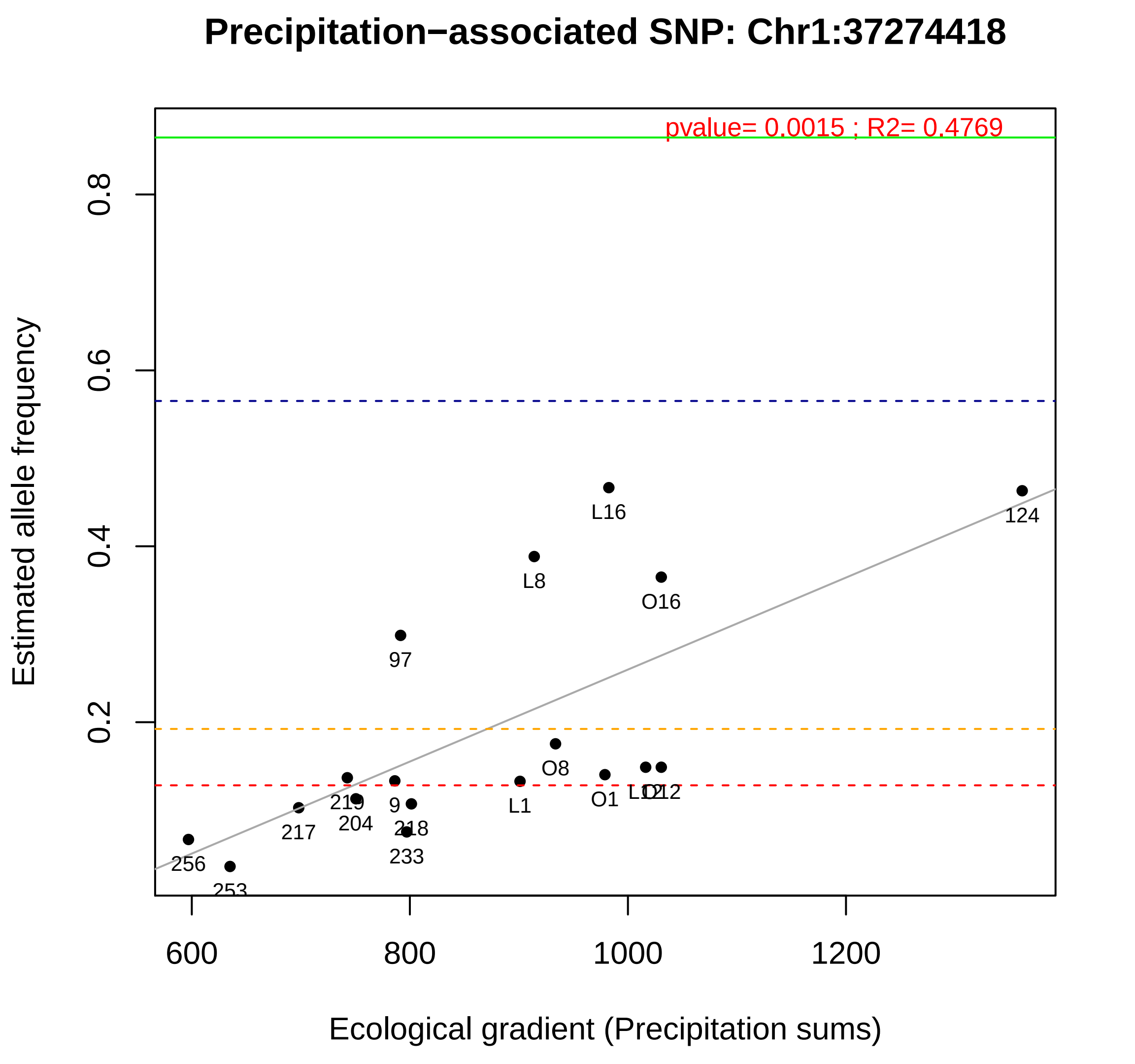
Precipitation−associated SNP: Chr1:37274418
pvalue= 0.0015 ; R2= 0.4769
Estimated allele frequency
0.2
0.4
0.6
0.8
600
800
1000
1200
Ecological gradient (Precipitation sums)
L16
124
L8
O16
97
O8
O1
L1
217
204
233
256
253
9

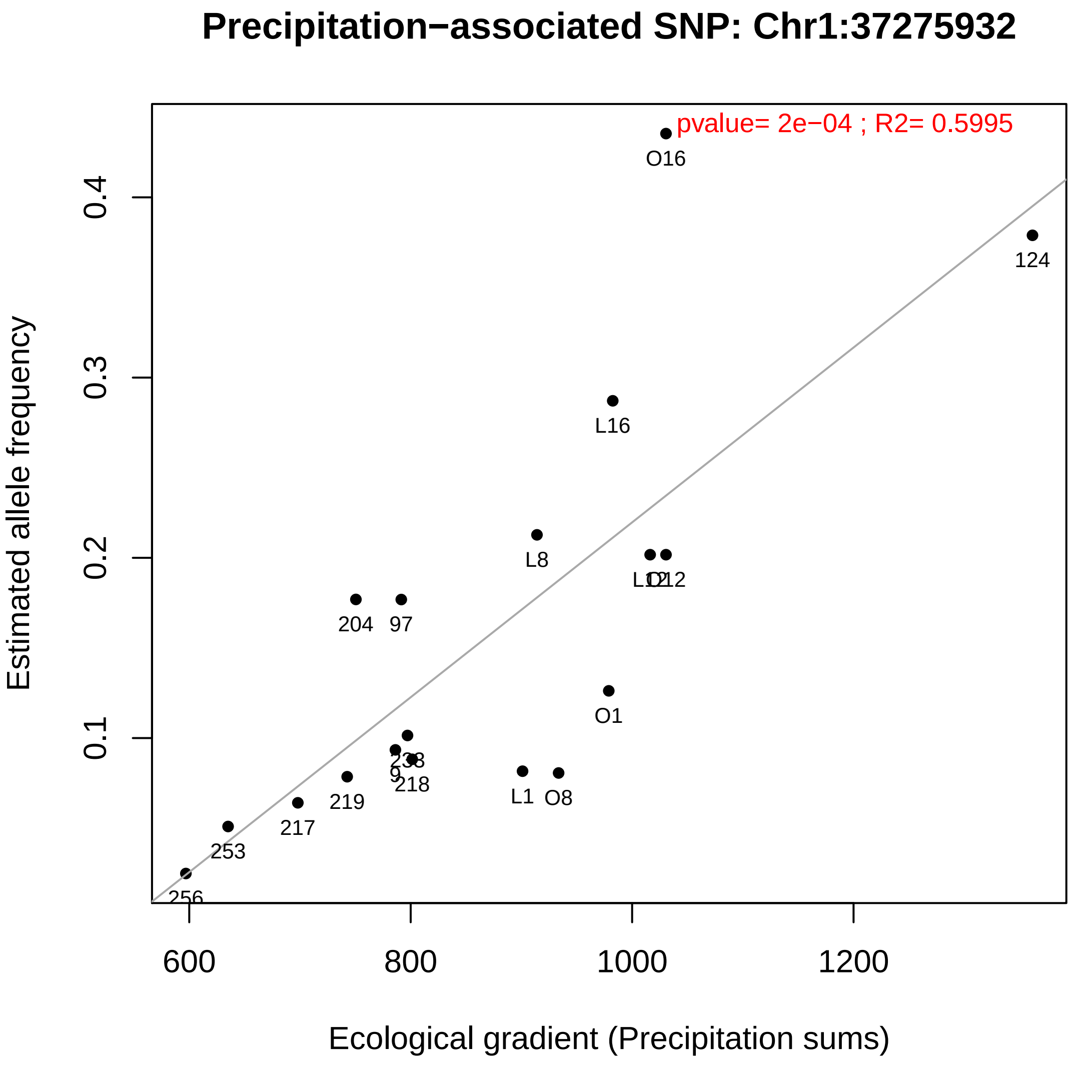

Precipitation−associated SNP: Chr1:37275932
pvalue= 2e−04 ; R2= 0.5995
Estimated allele frequency
0.1
0.2
0.3
0.4
Ecological gradient (Precipitation sums)
600
800
1000
1200
O16
124
L16
L8
204
97
O1
219
217
253
256
218
L1
O8

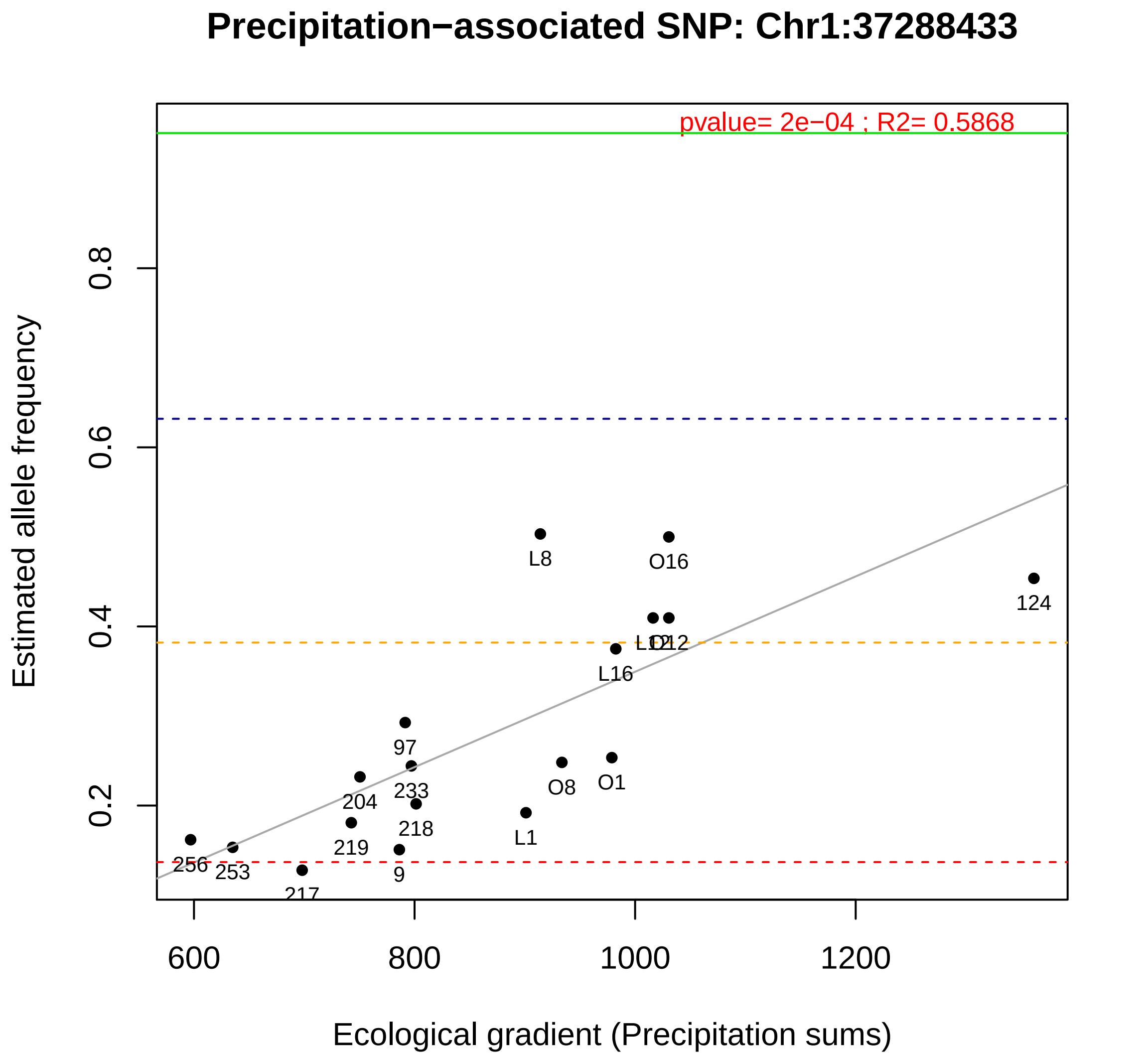

Precipitation−associated SNP: Chr1:37288433
pvalue= 2e−04 ; R2= 0.5868
Estimated allele frequency
0.2
0.4
0.6
0.8
600
800
1000
1200
Ecological gradient (Precipitation sums)
L8
O16
124
L10
O12
L16
97
233
204
218
219
9
256
253
217
L1
O8
O1

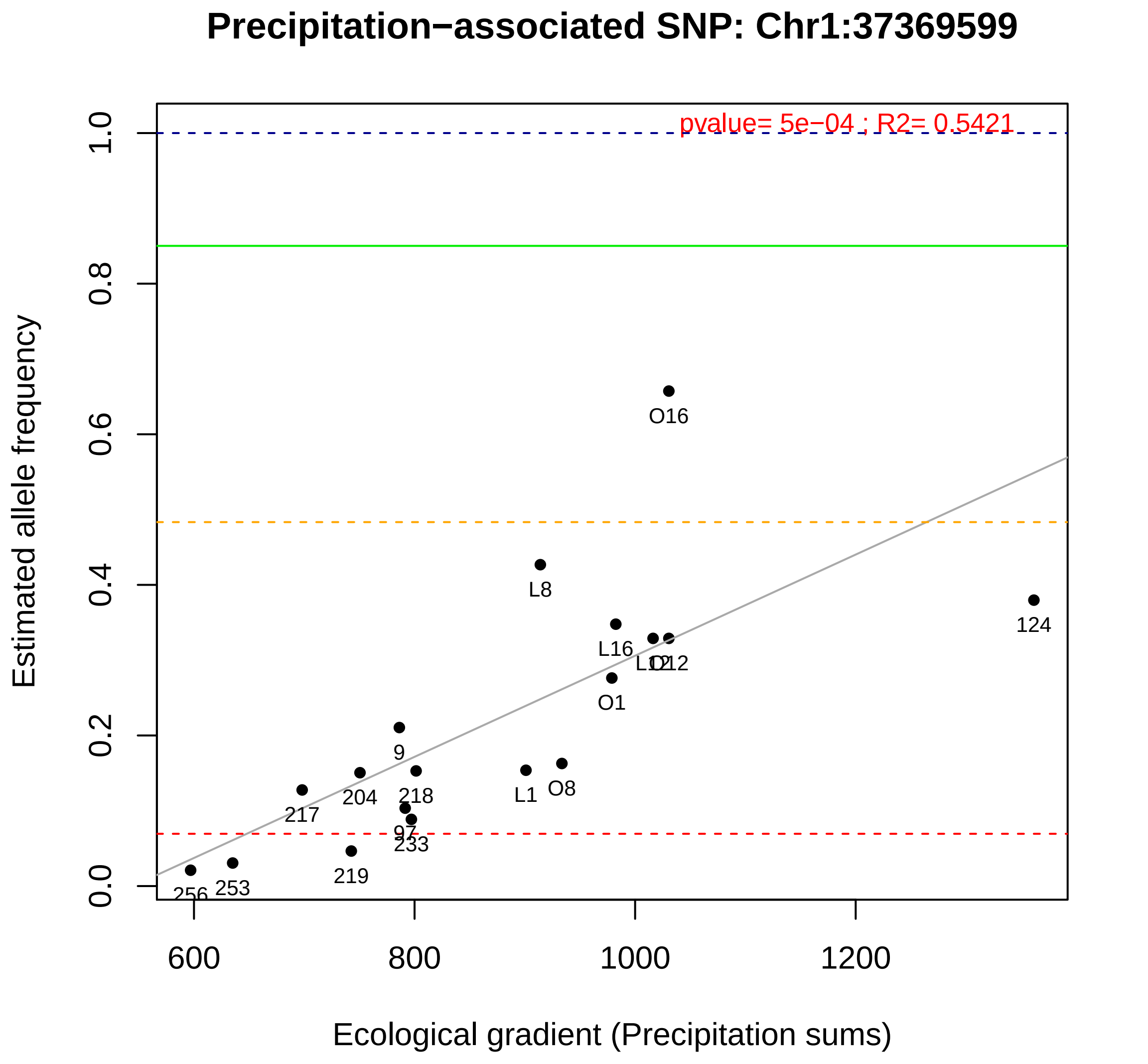
Precipitation−associated SNP: Chr1:37369599
pvalue= 5e−04 ; R2= 0.5421
Estimated allele frequency
Ecological gradient (Precipitation sums)
0.0
0.2
0.4
0.6
0.8
1.0
600
800
1000
1200
O16
L8
124
L16
L12
O12
O1
9
204
218
217
L1
O8
97
233
219
253
256

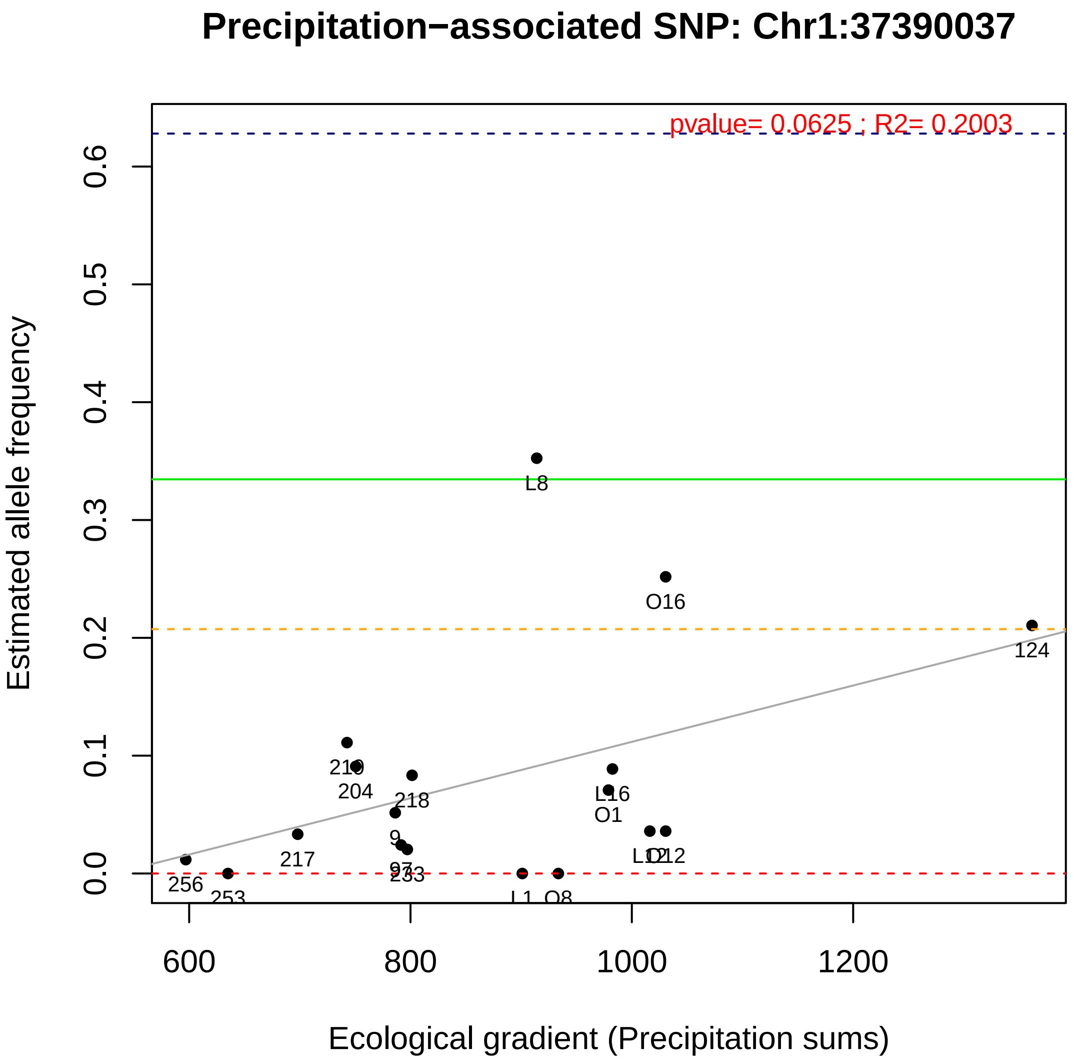

Precipitation−associated SNP: Chr1:37390037
pvalue= 0.0625 ; R2= 0.2003
Estimated allele frequency
0.0
0.1
0.2
0.3
0.4
0.5
0.6
600
800
1000
1200
Ecological gradient (Precipitation sums)
L8
O16
124
210
204
218
L16
O1
9
217
256
97
233
L10
O12

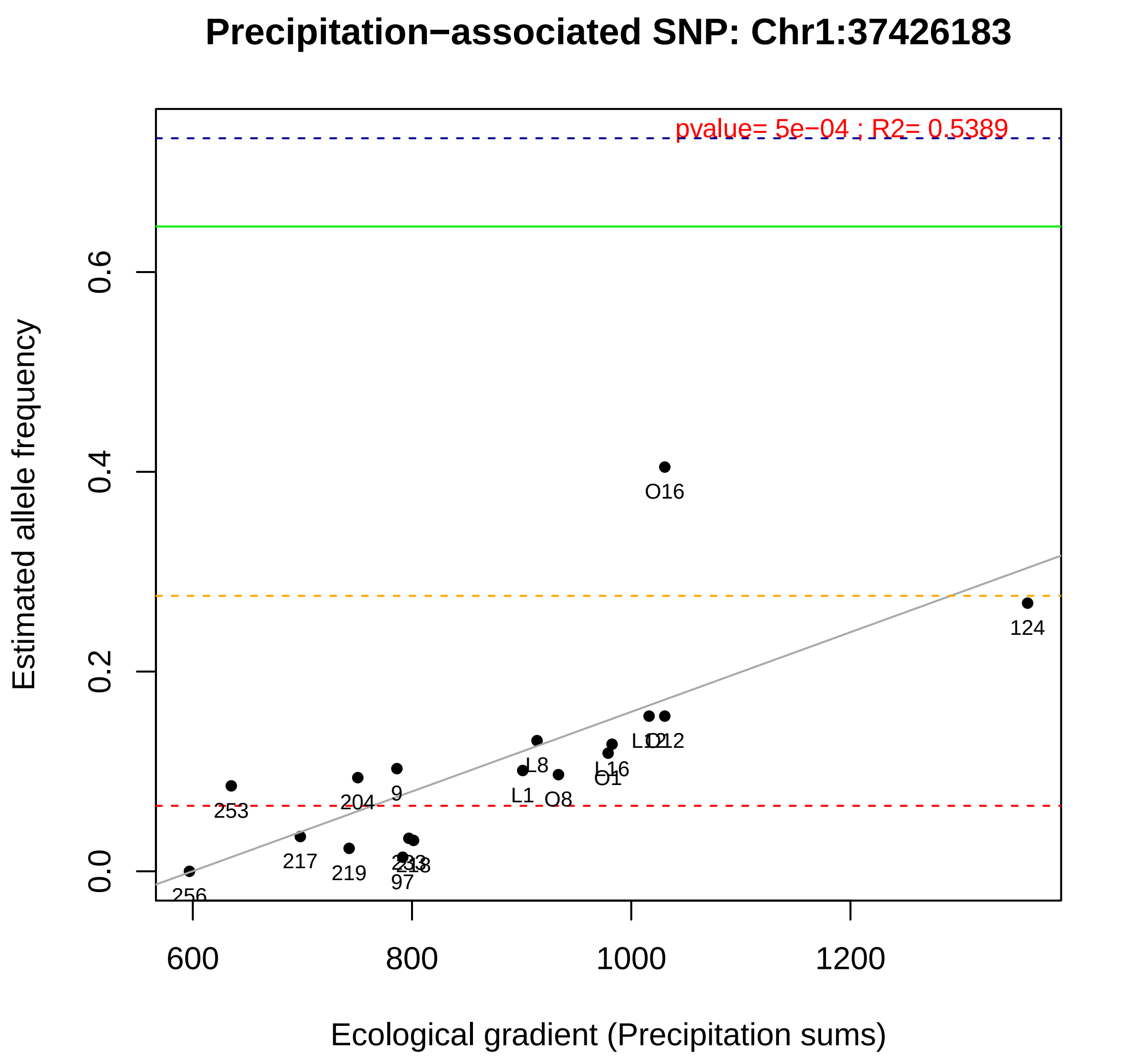
Precipitation−associated SNP: Chr1:37426183
pvalue= 5e−04 ; R2= 0.5389
Estimated allele frequency
0.0
0.2
0.4
0.6
600
800
1000
1200
Ecological gradient (Precipitation sums)
O16
124
L12
O12
L8
L16
O1
L1
O8
9
204
253
217
219
97
256

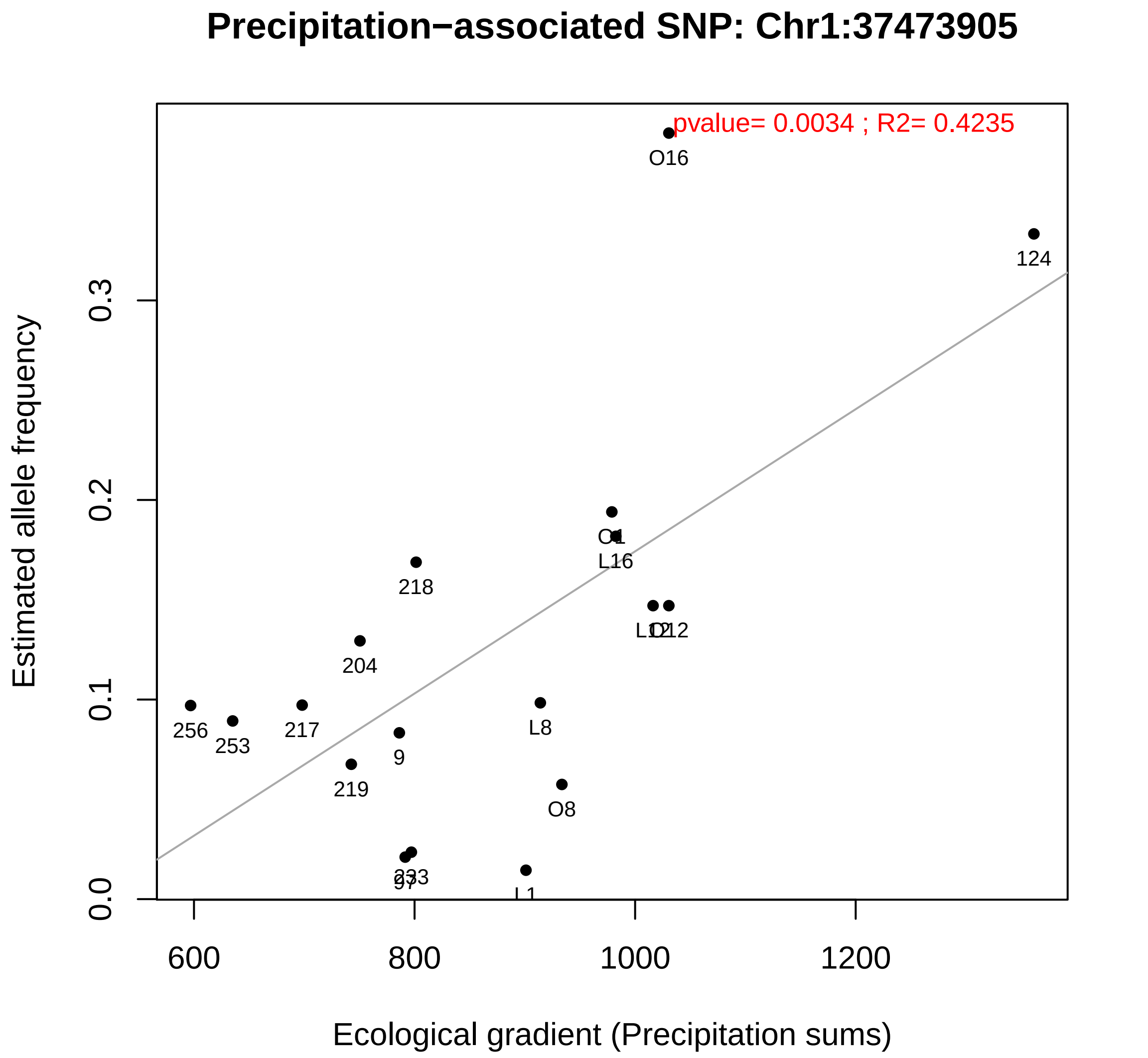
Precipitation−associated SNP: Chr1:37473905
pvalue= 0.0034 ; R2= 0.4235
Estimated allele frequency
Ecological gradient (Precipitation sums)
0.0
0.1
0.2
0.3
600
800
1000
1200
O16
124
218
204
256
253
217
219
9
L8
O8
L16
L1

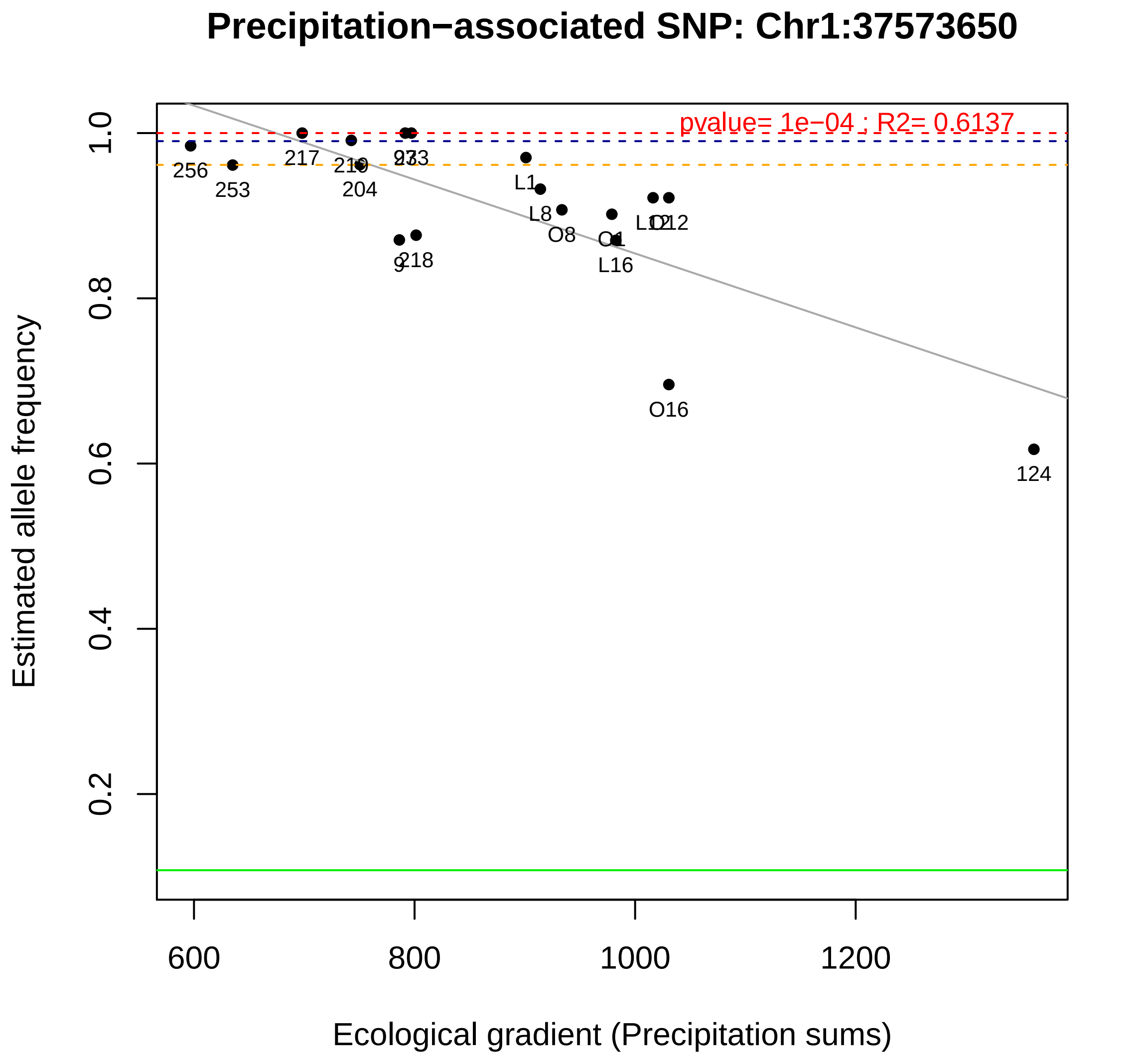

Precipitation−associated SNP: Chr1:37573650
pvalue= 1e−04 ; R2= 0.6137
Estimated allele frequency
Ecological gradient (Precipitation sums)
0.2
0.4
0.6
0.8
1.0
600
800
1000
1200
256
253
217
210
204
97
233
9
218
L1
L8
O8
O1
L16
L12
O12
O16
124

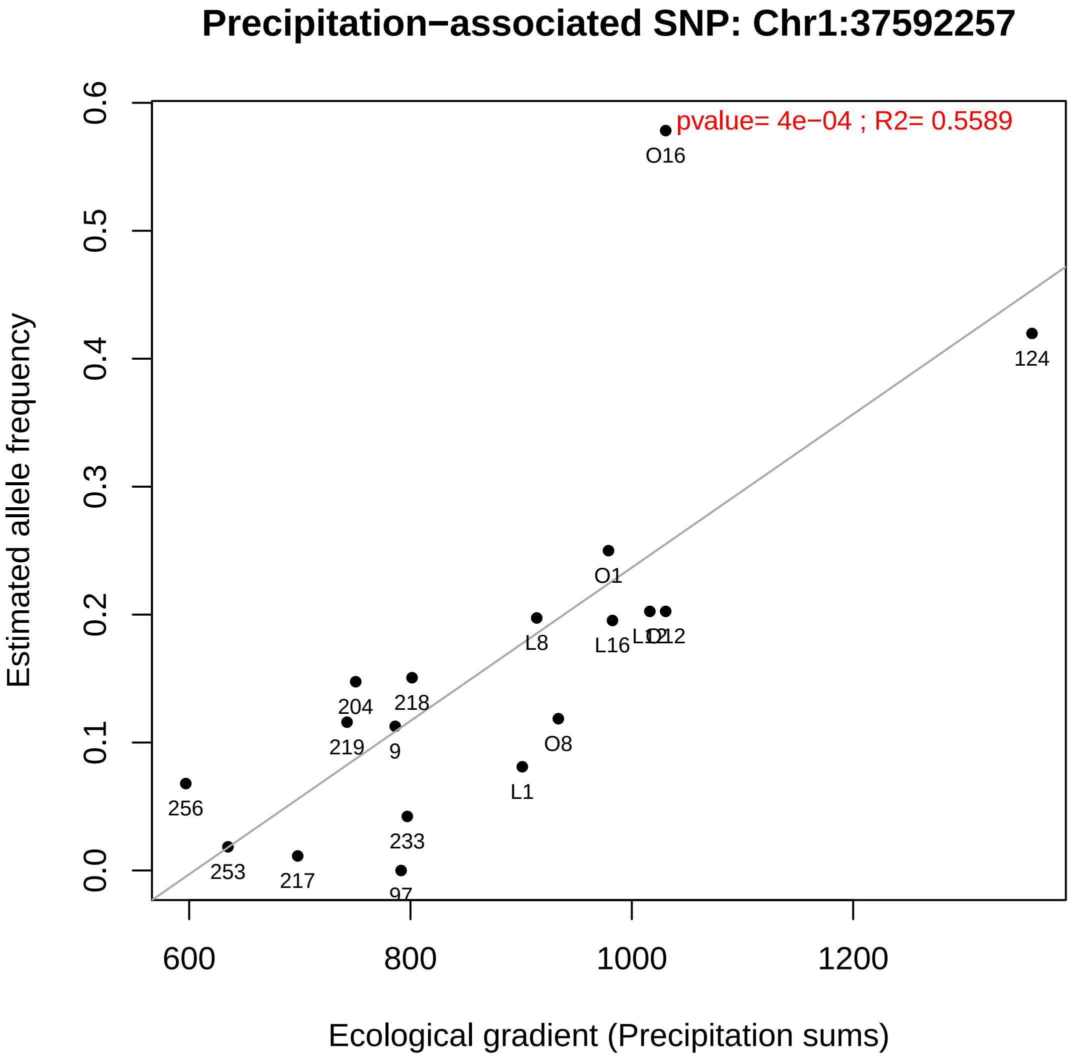
Precipitation−associated SNP: Chr1:37592257
pvalue= 4e−04 ; R2= 0.5589
Estimated allele frequency
Ecological gradient (Precipitation sums)
0.0
0.1
0.2
0.3
0.4
0.5
0.6
600
800
1000
1200
O16
124
O1
L8
L16
L12
O12
204
218
219
9
O8
L1
256
233
253
217
97

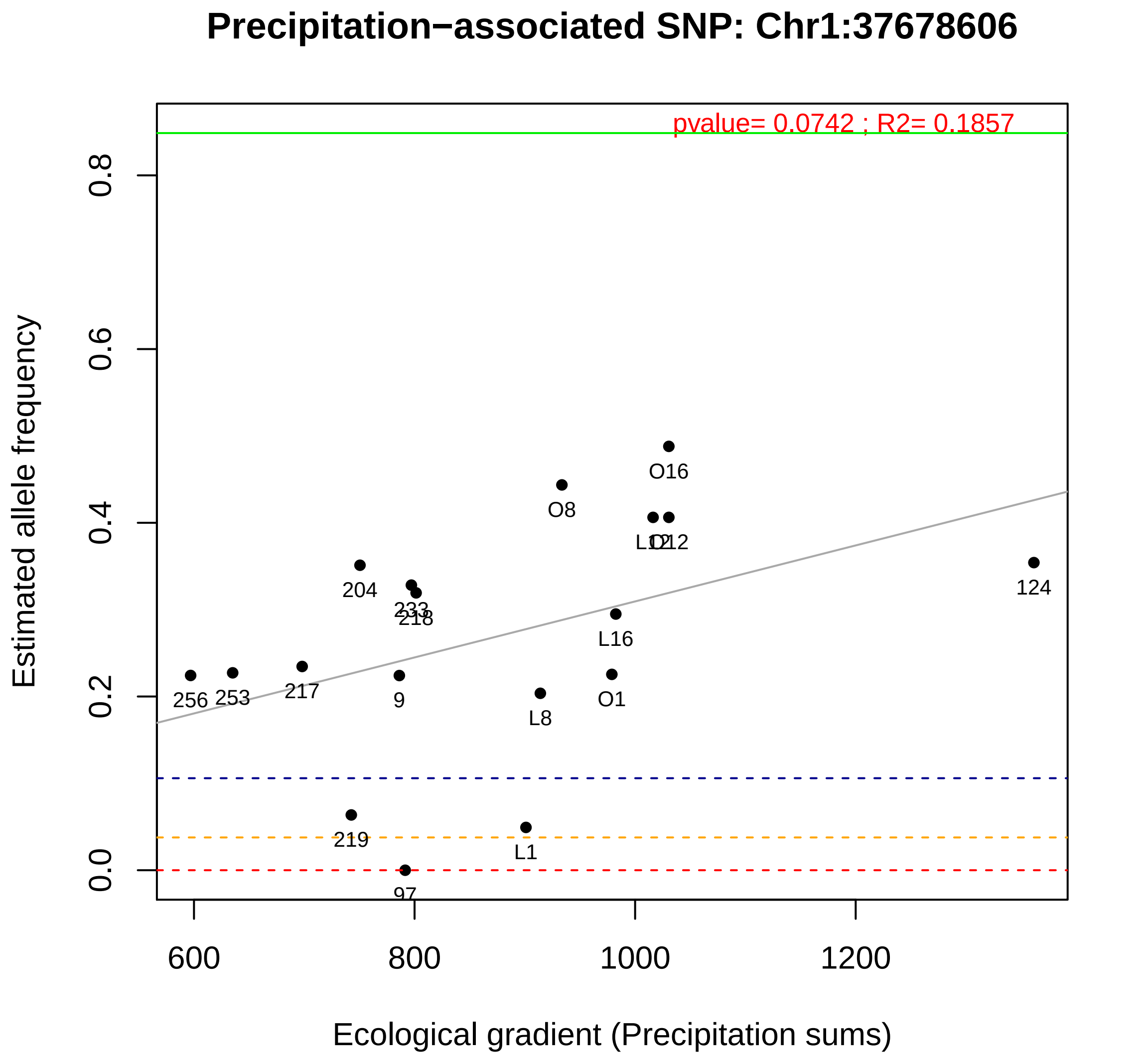

Precipitation−associated SNP: Chr1:37678606
pvalue= 0.0742 ; R2= 0.1857
Estimated allele frequency
0.0
0.2
0.4
0.6
0.8
600
800
1000
1200
Ecological gradient (Precipitation sums)
O16
O8
L12
O12
204
233
218
124
L16
256
253
217
9
L8
O1
219
L1
97

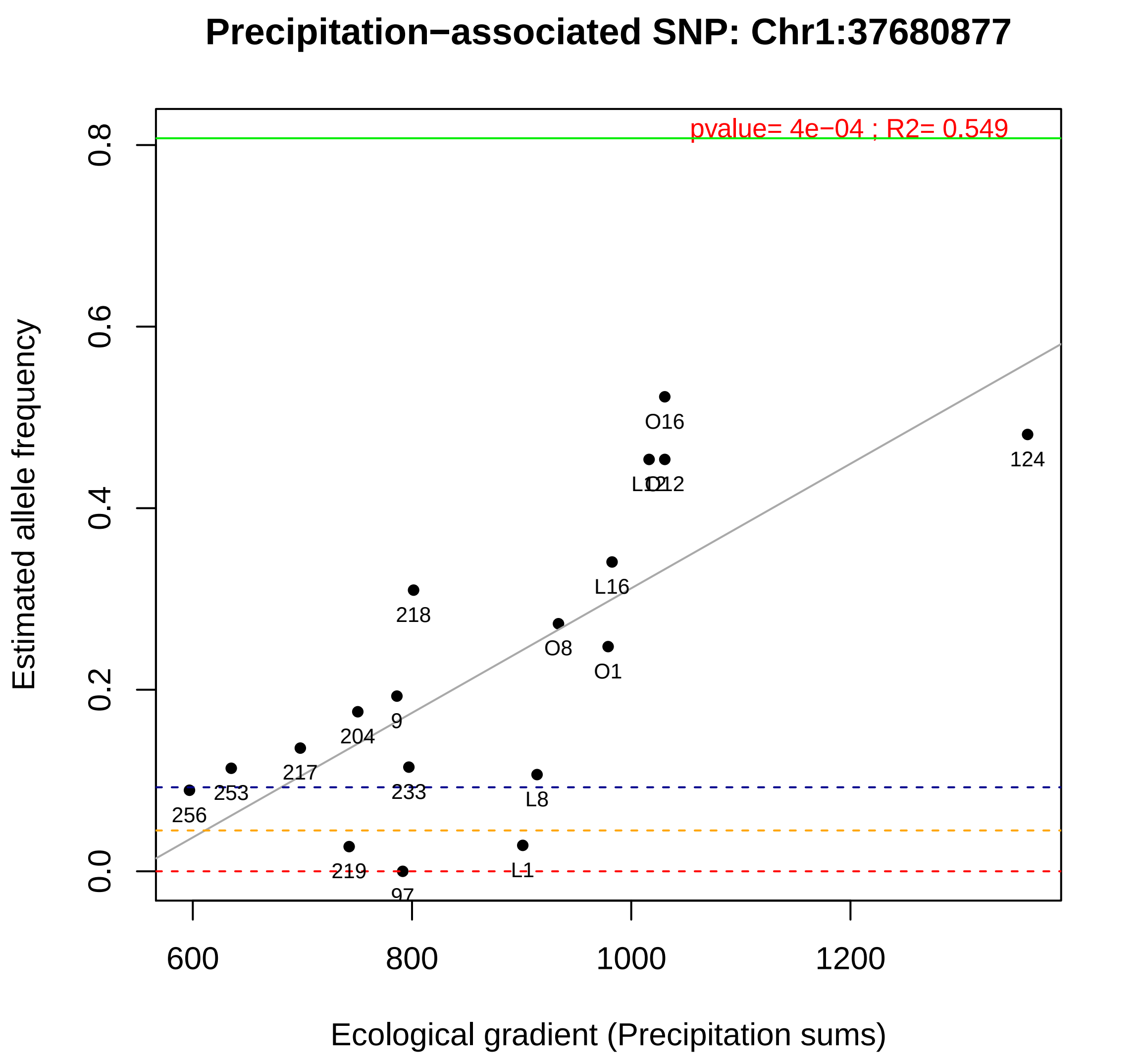

Precipitation−associated SNP: Chr1:37680877
pvalue= 4e−04 ; R2= 0.549
Estimated allele frequency
0.0
0.2
0.4
0.6
0.8
600
800
1000
1200
Ecological gradient (Precipitation sums)
O16
124
L12
O12
L16
218
O8
O1
9
204
217
253
233
L8
256
219
L1
97

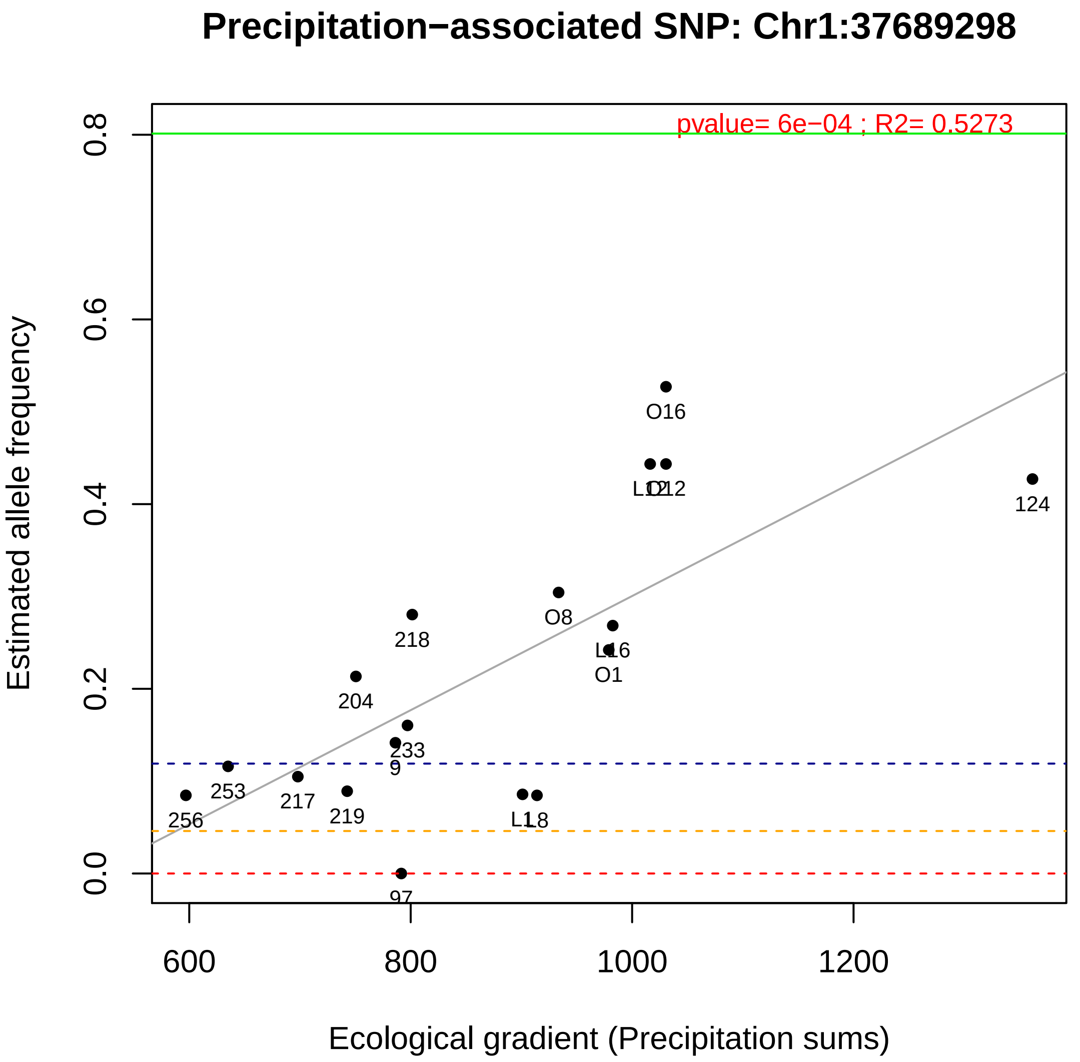
Precipitation−associated SNP: Chr1:37689298
pvalue= 6e−04 ; R2= 0.5273
Estimated allele frequency
Ecological gradient (Precipitation sums)
0.0
0.2
0.4
0.6
0.8
600
800
1000
1200
O16
124
O8
218
L16
O1
204
233
9
253
217
219
256
97

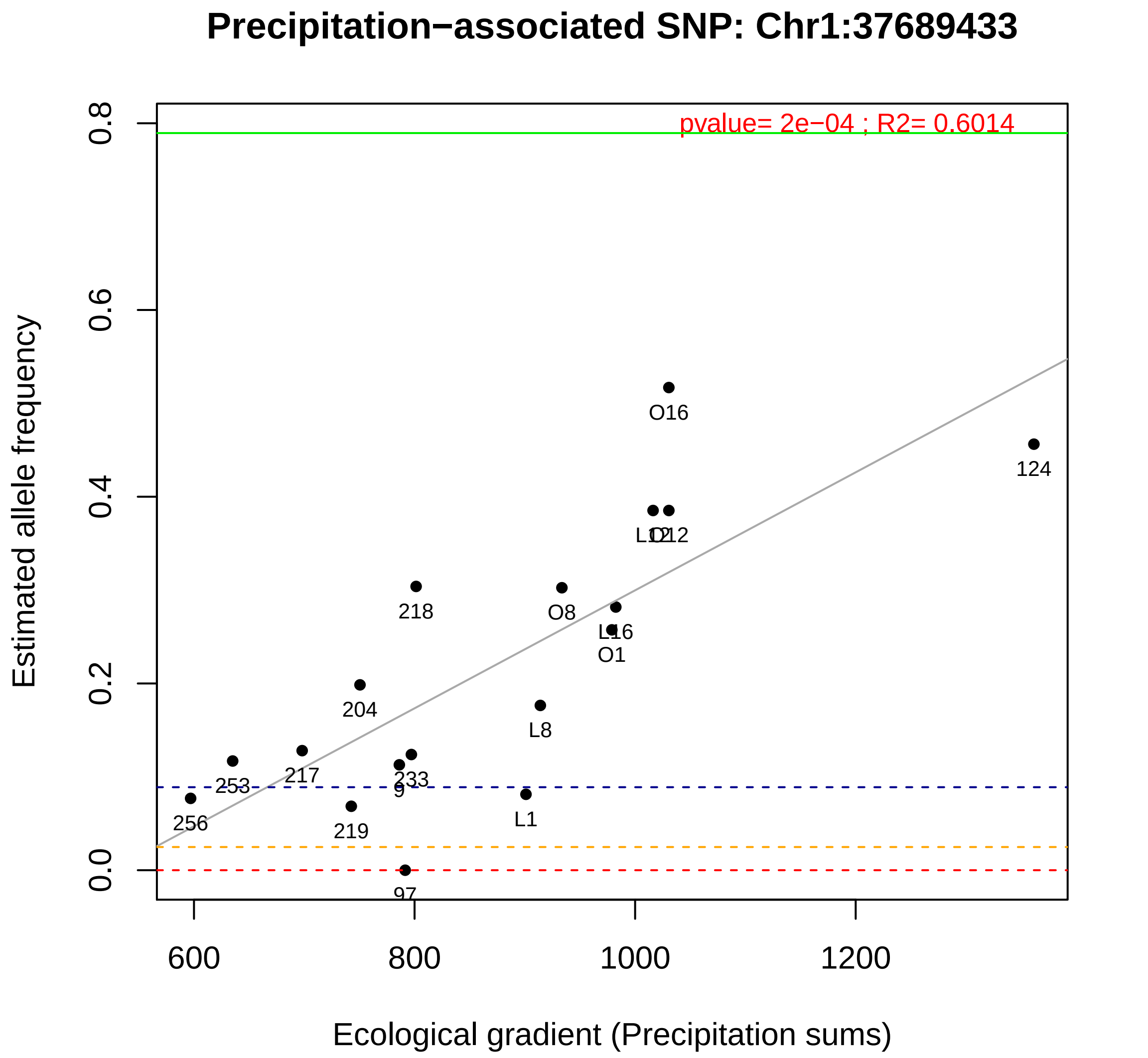
Precipitation−associated SNP: Chr1:37689433
pvalue= 2e−04 ; R2= 0.6014
Estimated allele frequency
0.0
0.2
0.4
0.6
0.8
600
800
1000
1200
Ecological gradient (Precipitation sums)
O16
124
L12
O12
218
O8
L16
O1
204
L8
217
253
233
9
256
219
L1
97

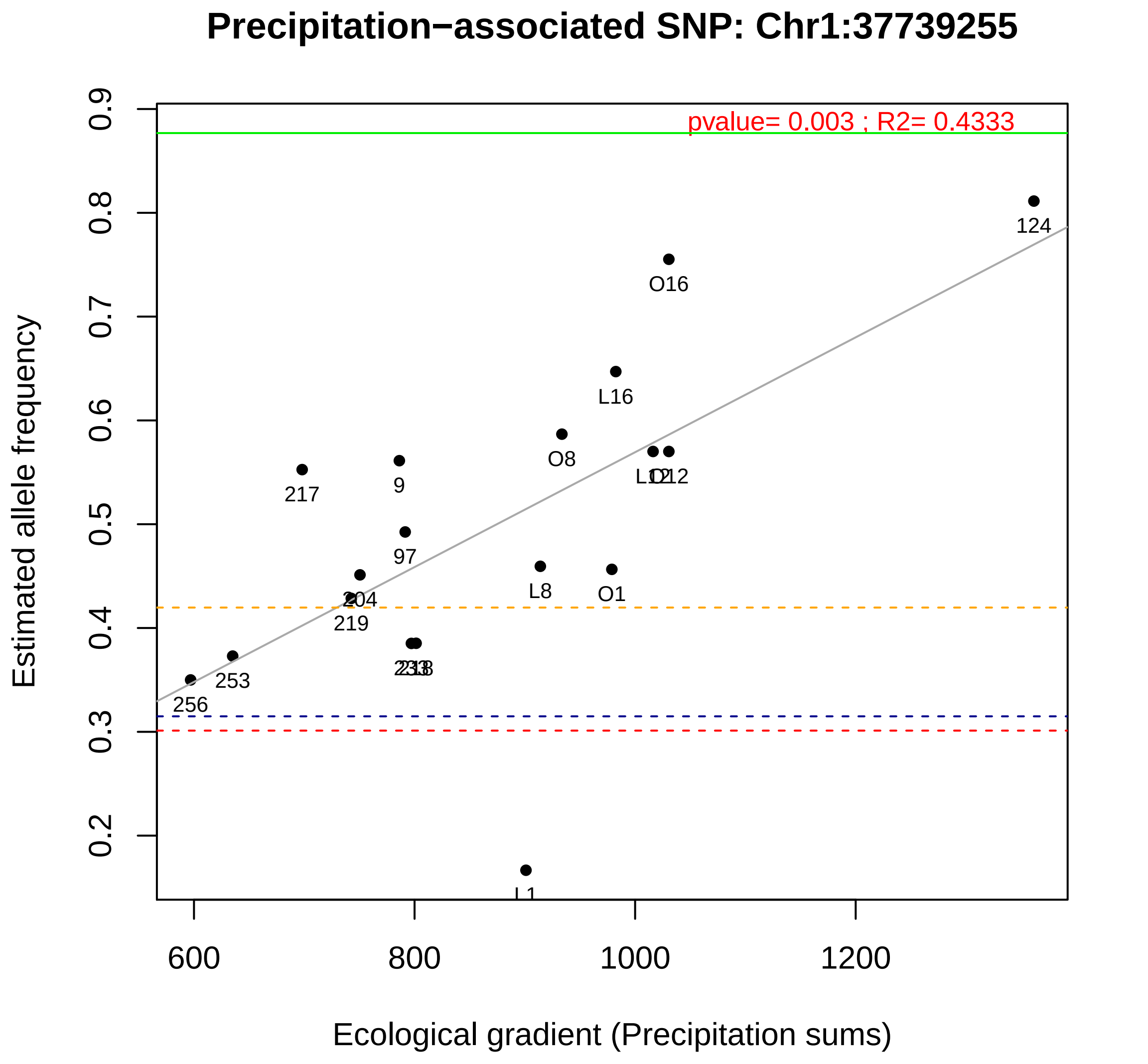

Precipitation−associated SNP: Chr1:37739255
pvalue= 0.003 ; R2= 0.4333
Estimated allele frequency
Ecological gradient (Precipitation sums)
0.2
0.3
0.4
0.5
0.6
0.7
0.8
0.9
600
800
1000
1200
124
O16
L16
O8
217
9
97
L8
O1
219
253
256
L1

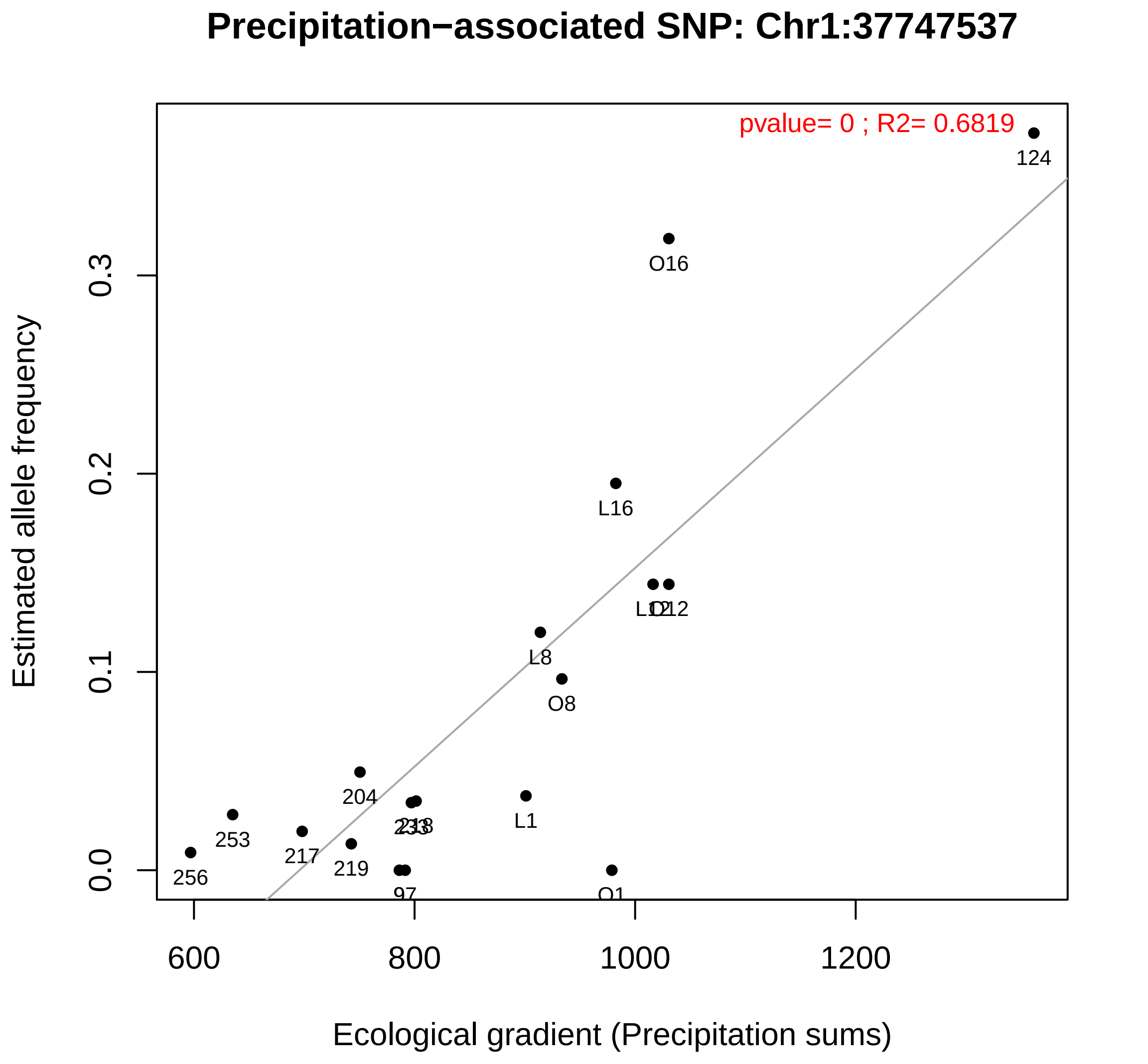
Precipitation−associated SNP: Chr1:37747537
pvalue= 0 ; R2= 0.6819
Estimated allele frequency
Ecological gradient (Precipitation sums)
0.0
0.1
0.2
0.3
600
800
1000
1200
124
O16
L16
O12
L8
O8
204
L1
253
217
219
256

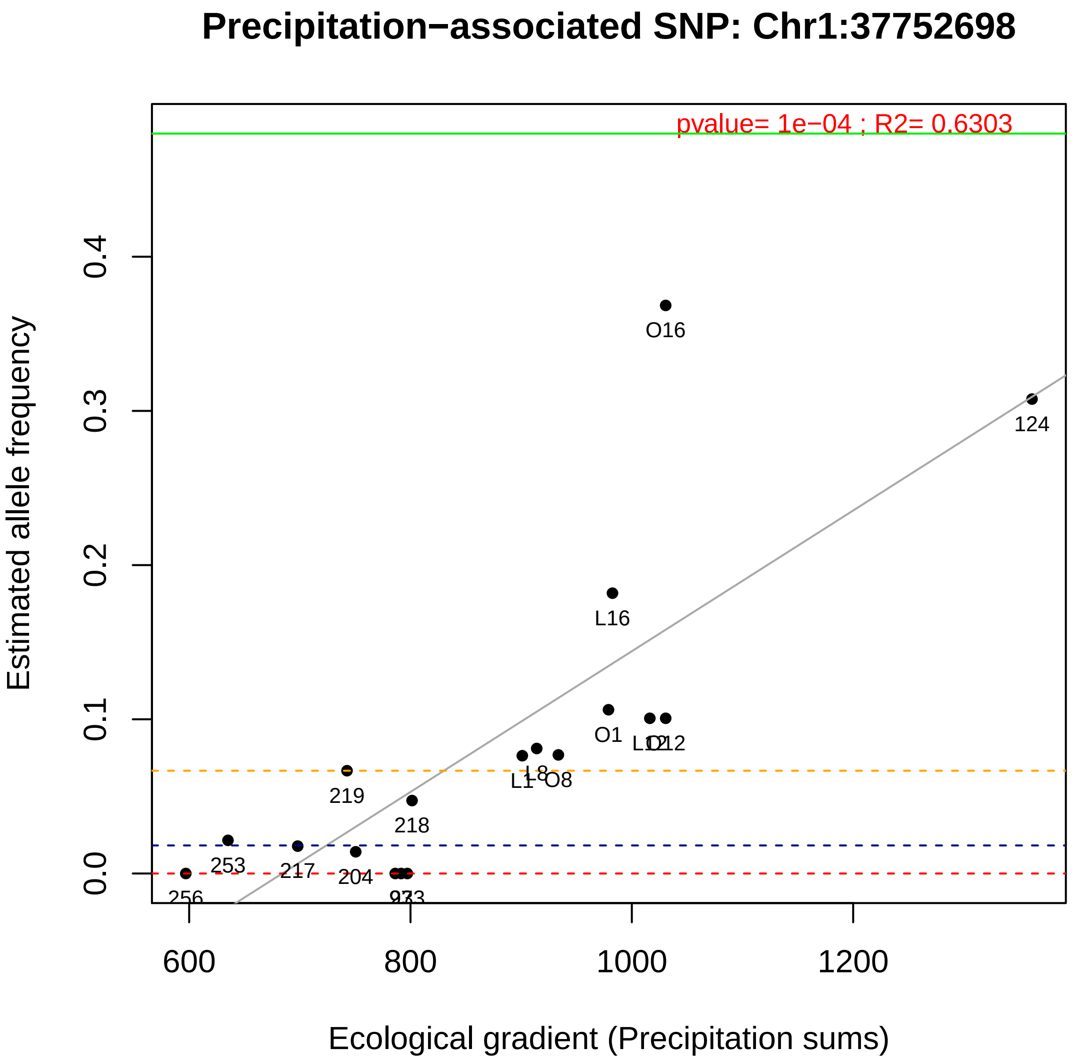

Precipitation−associated SNP: Chr1:37752698
pvalue= 1e−04 ; R2= 0.6303
Estimated allele frequency
0.0
0.1
0.2
0.3
0.4
O16
124
L16
O1
L12
O12
L1
L8
O8
219
218
253
217
204
256
600
800
1000
1200
Ecological gradient (Precipitation sums)

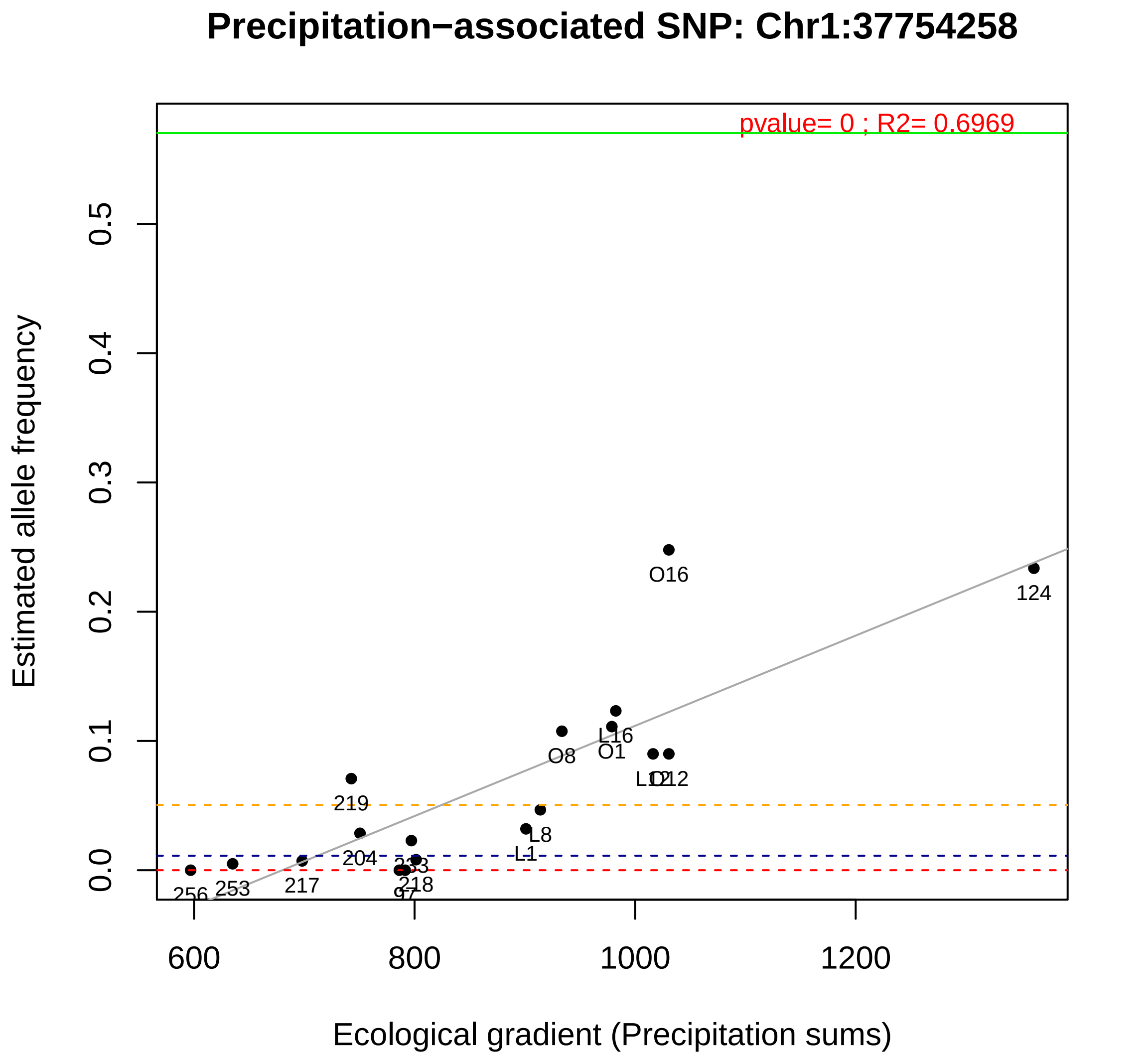
Precipitation−associated SNP: Chr1:37754258
pvalue= 0 ; R2= 0.6969
Estimated allele frequency
0.0
0.1
0.2
0.3
0.4
0.5
600
800
1000
1200
Ecological gradient (Precipitation sums)
O16
124
L16
O1
O8
O12
219
L8
L1
204
218
217
253

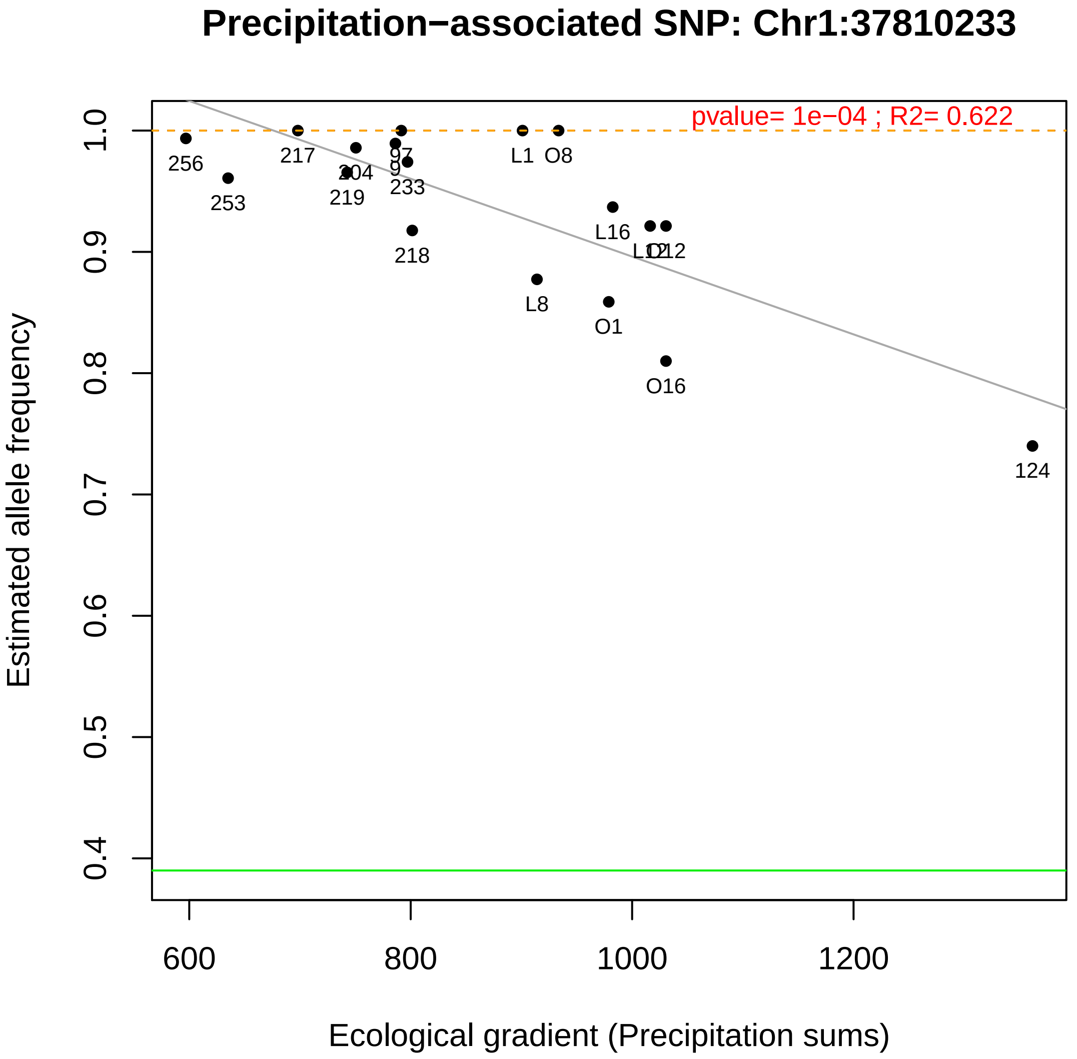

Precipitation−associated SNP: Chr1:37810233
pvalue= 1e−04 ; R2= 0.622
Estimated allele frequency
Ecological gradient (Precipitation sums)
1.0
0.9
0.8
0.7
0.6
0.5
0.4
600
800
1000
1200
256
253
217
204
219
97
9
233
218
L1
O8
L8
L16
L12
O12
O1
O16
124

**Precipitation−associated SNP: Chr1:38027641**

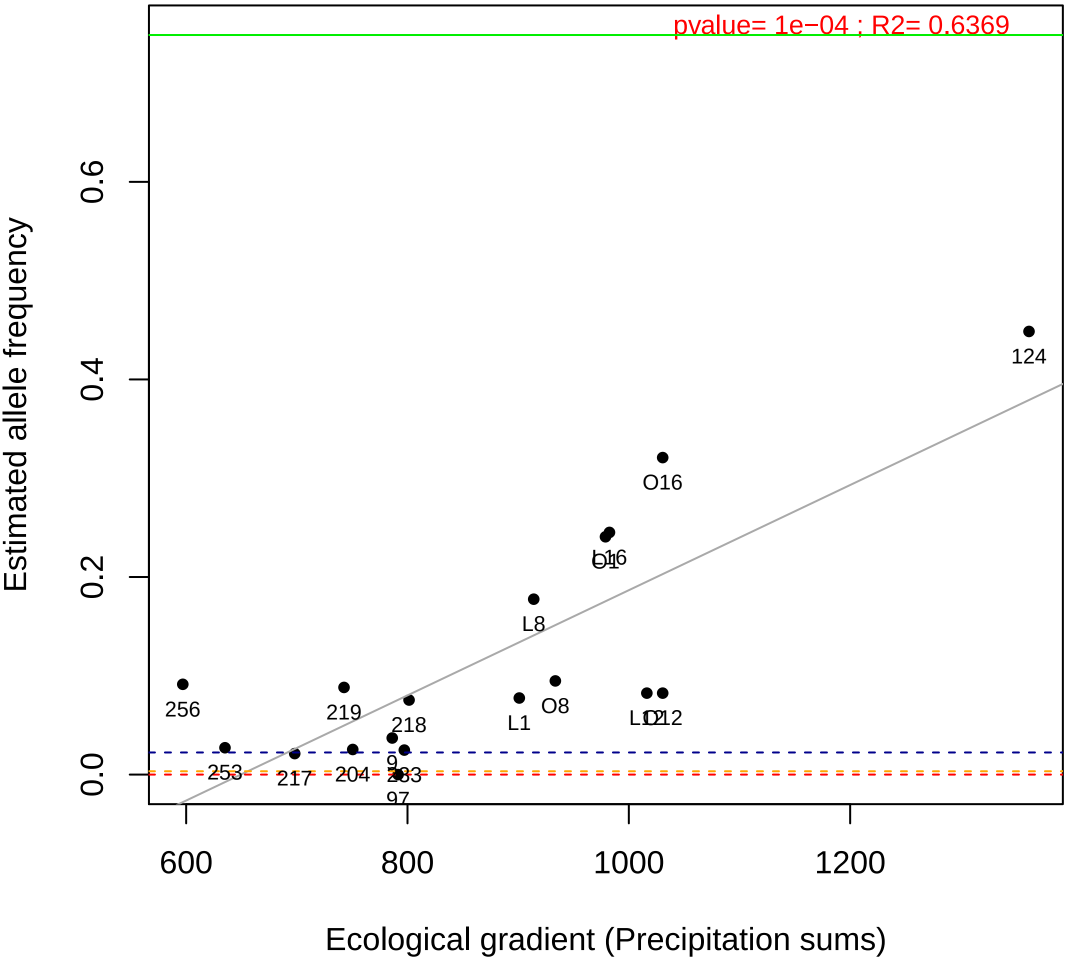

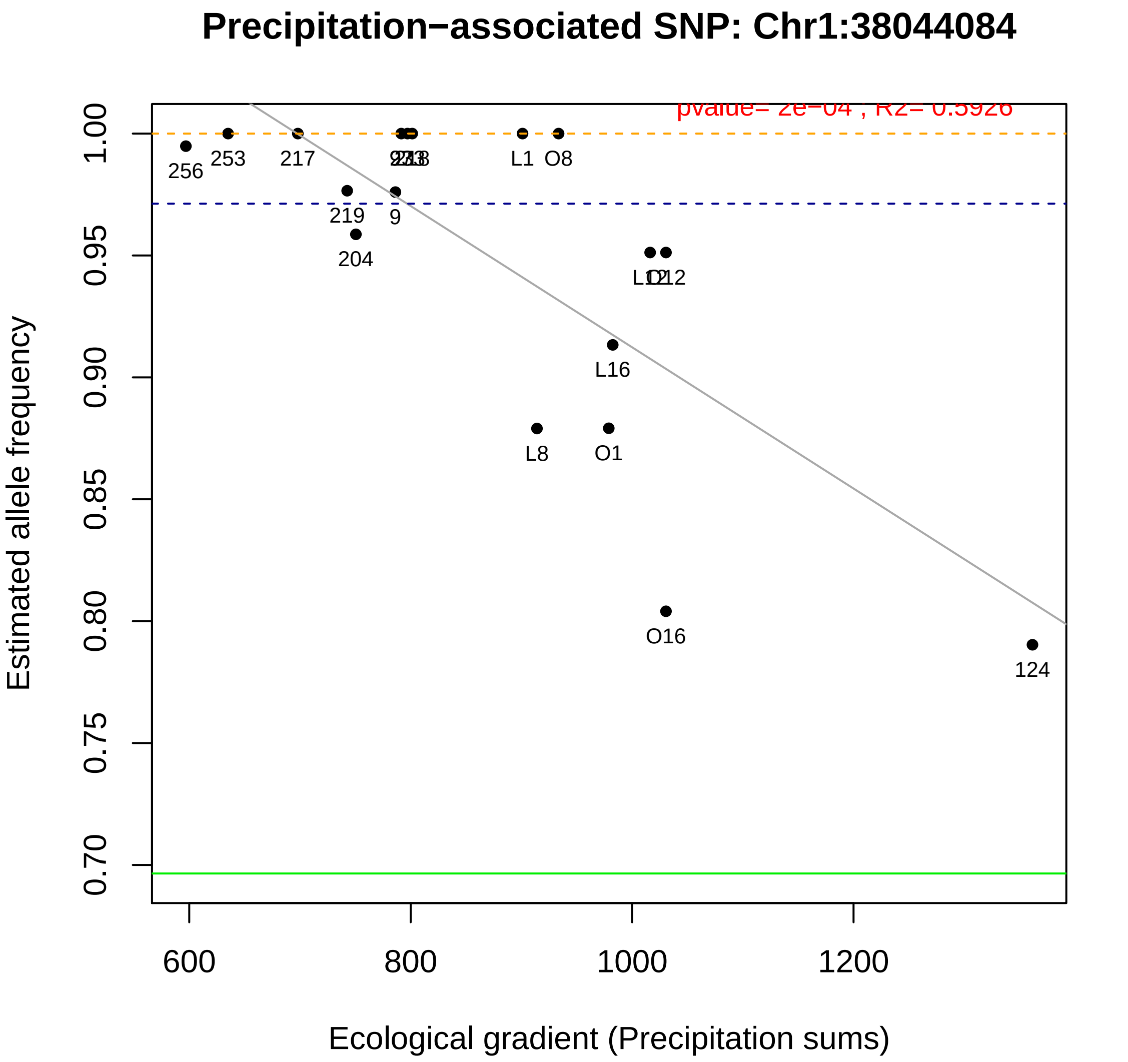

Precipitation−associated SNP: Chr1:38044084
pvalue= 2e−04 ; R2= 0.5926
Estimated allele frequency
Ecological gradient (Precipitation sums)
0.70
0.75
0.80
0.85
0.90
0.95
1.00
600
800
1000
1200
256
253
217
L1
O8
219
9
204
L16
L8
O1
O16
124

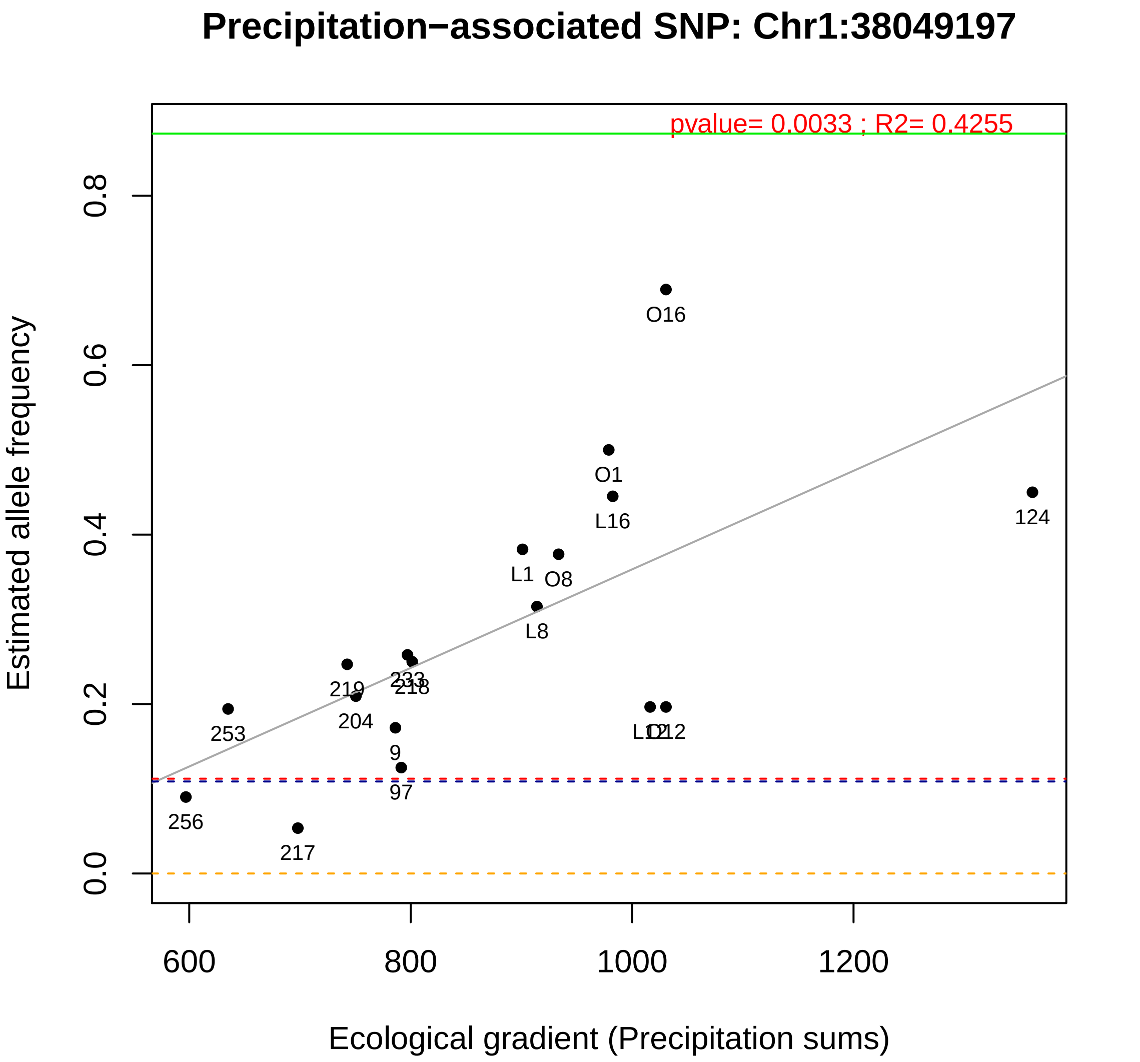

Precipitation−associated SNP: Chr1:38049197
pvalue= 0.0033 ; R2= 0.4255
Estimated allele frequency
0.0
0.2
0.4
0.6
0.8
600
800
1000
1200
Ecological gradient (Precipitation sums)
O16
O1
L16
124
L1
O8
L8
233
218
219
204
253
9
97
256
217
L12
O12

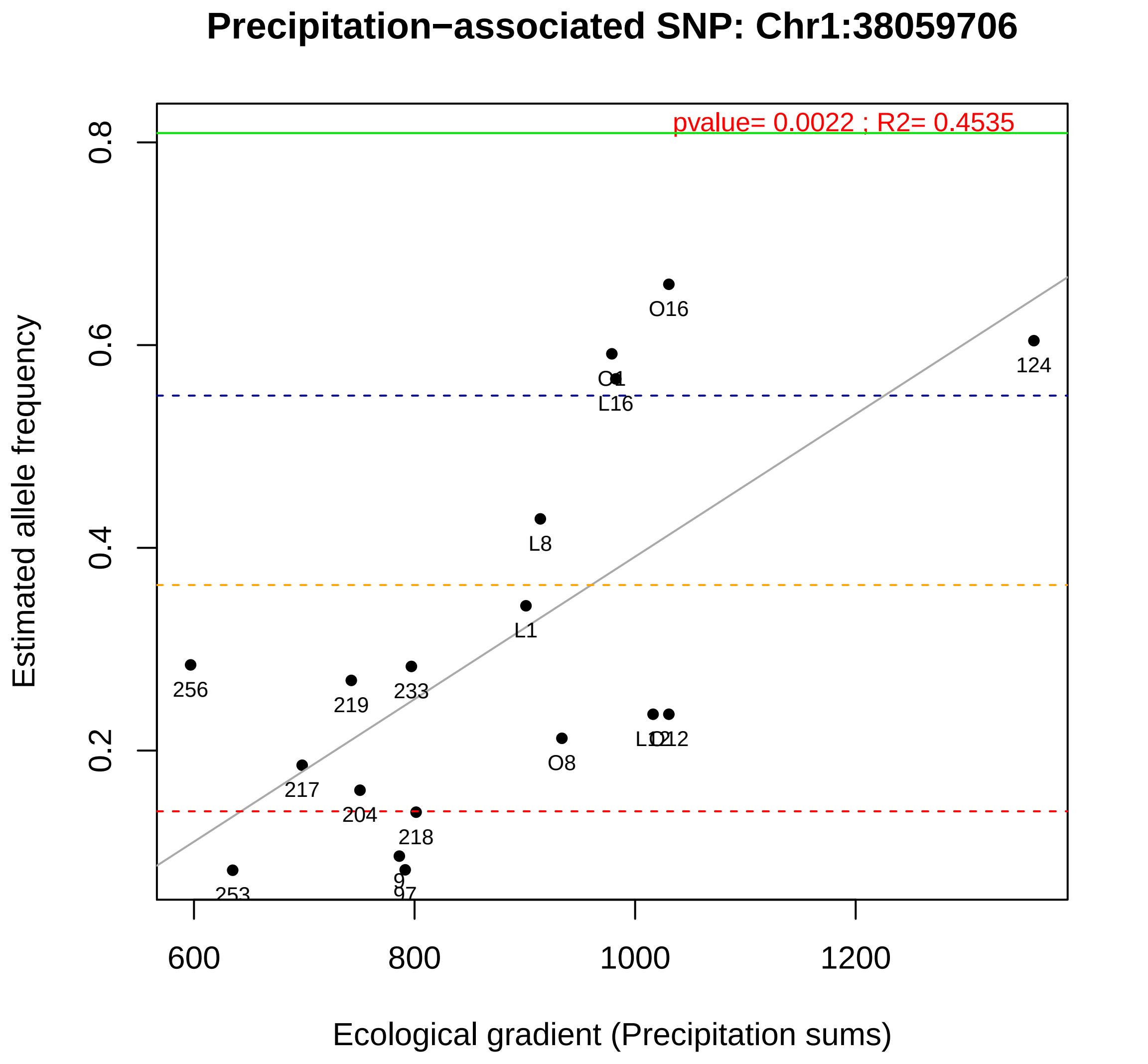

Precipitation−associated SNP: Chr1:38059706
pvalue= 0.0022 ; R2= 0.4535
Estimated allele frequency
0.2
0.4
0.6
0.8
600
800
1000
1200
Ecological gradient (Precipitation sums)
O16
124
O1
L16
L8
L1
256
219
233
L12
O12
O8
217
204
218
9
97
253

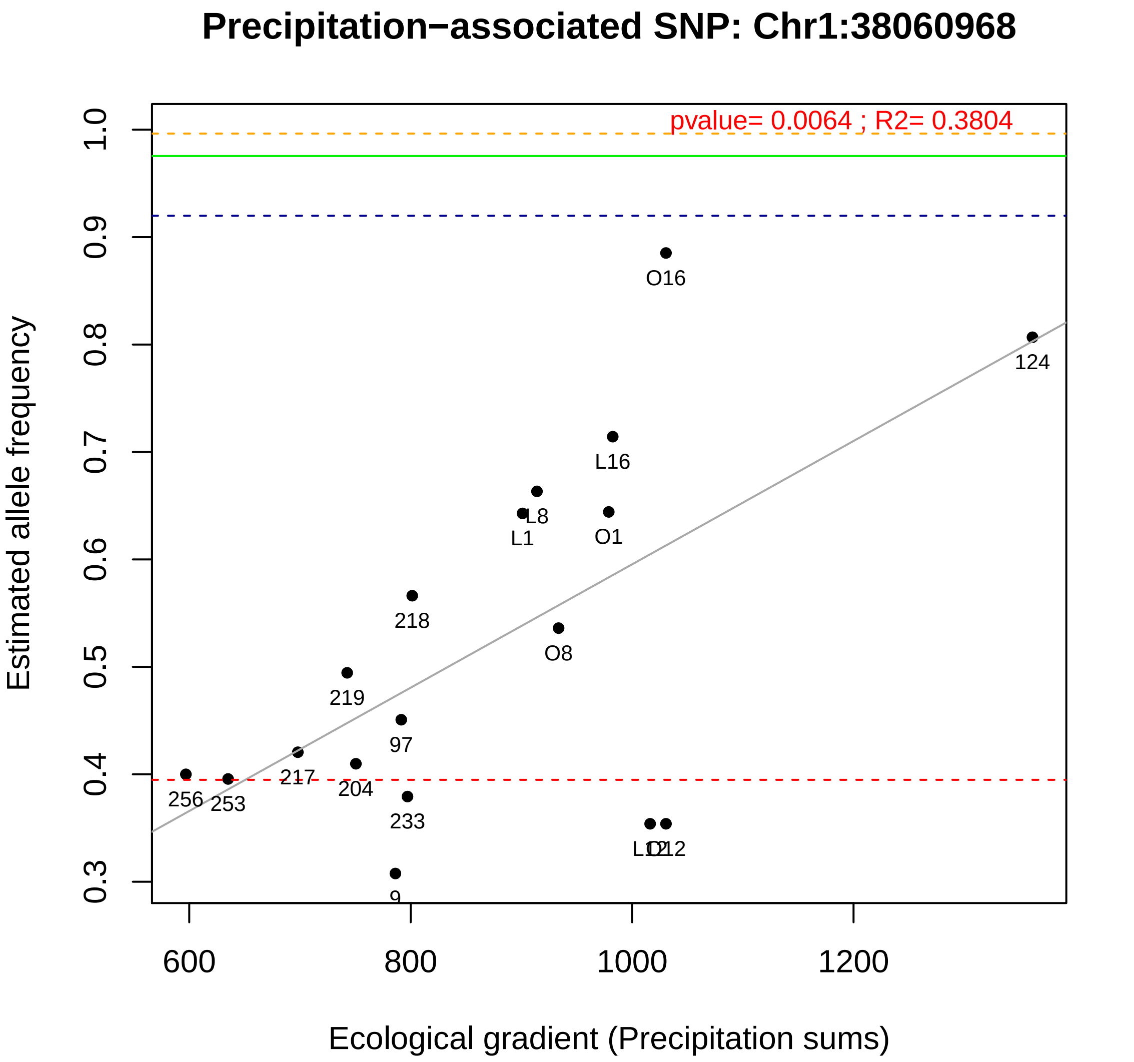
Precipitation−associated SNP: Chr1:38060968
pvalue= 0.0064 ; R2= 0.3804
Estimated allele frequency
0.3
0.4
0.5
0.6
0.7
0.8
0.9
1.0
O16
124
L16
L8
L1
O1
218
O8
219
97
217
204
256
253
233
L12
O12
9
600
800
1000
1200
Ecological gradient (Precipitation sums)

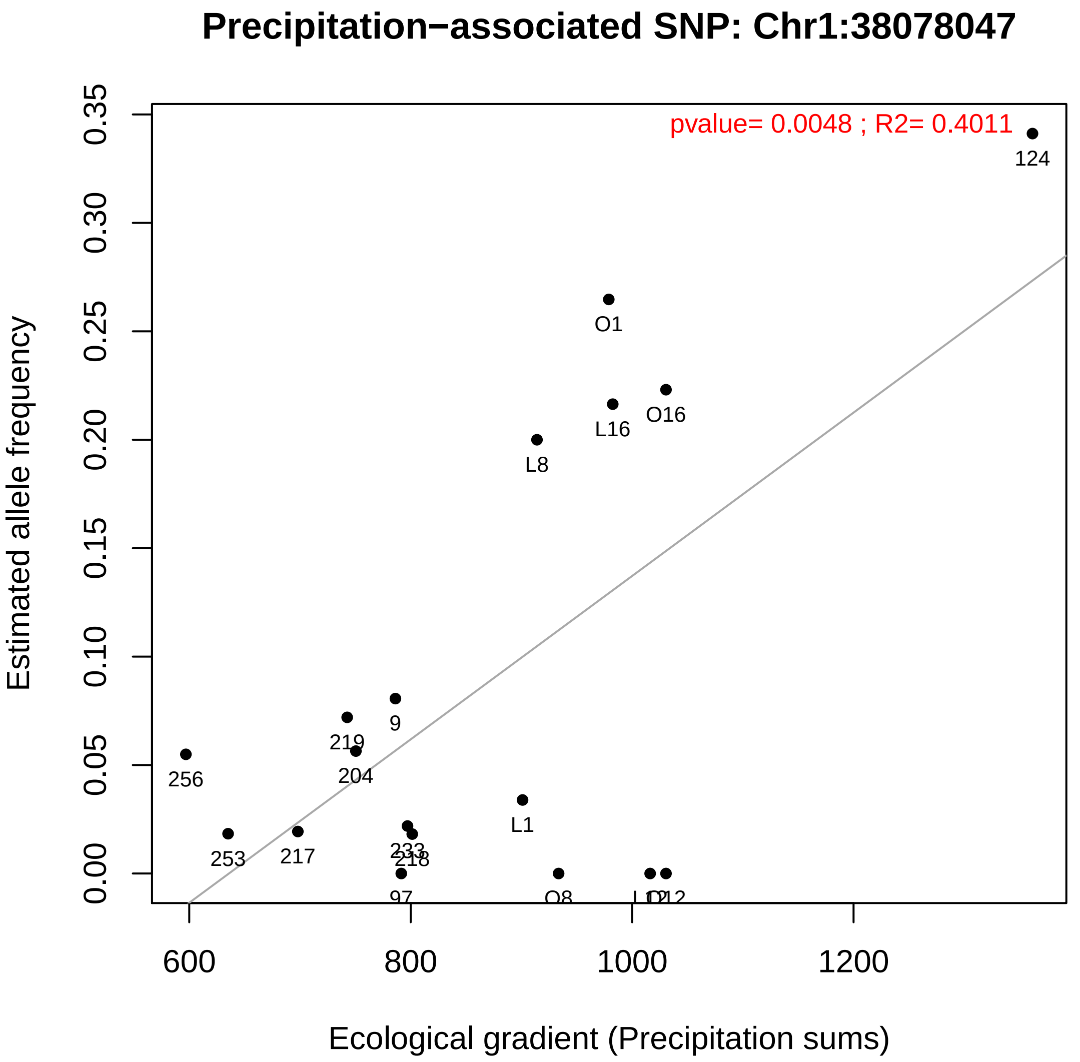
Precipitation−associated SNP: Chr1:38078047
pvalue= 0.0048 ; R2= 0.4011
Estimated allele frequency
0.00
0.05
0.10
0.15
0.20
0.25
0.30
0.35
600
800
1000
1200
Ecological gradient (Precipitation sums)
124
O1
O16
L16
L8
9
219
204
256
253
217
233
218
97
L1
O8

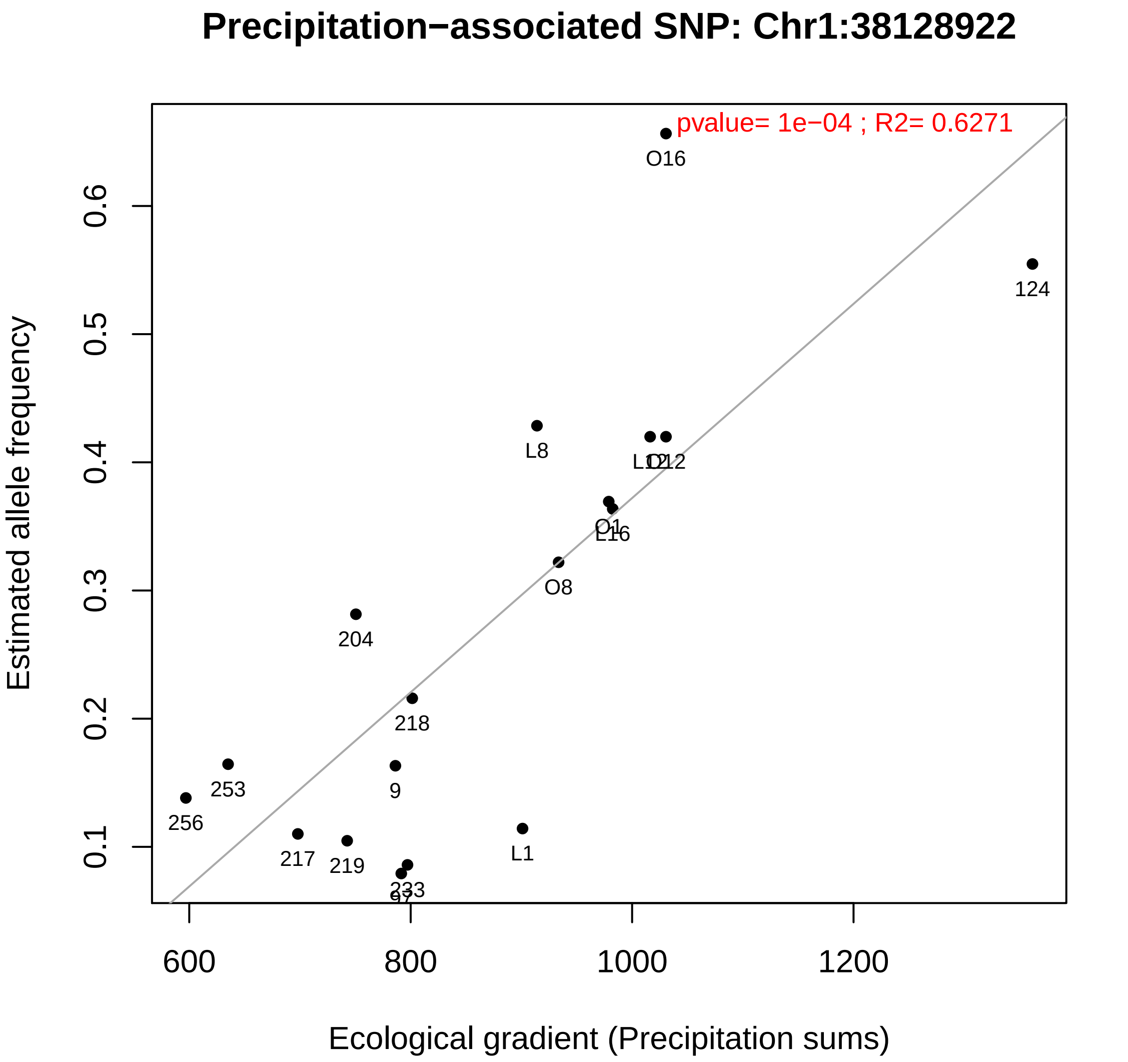

Precipitation−associated SNP: Chr1:38128922
pvalue= 1e−04 ; R2= 0.6271
O16
124
L8
L12
O12
O1
L16
O8
204
218
253
9
256
217
219
L1
233
97
0.1
0.2
0.3
0.4
0.5
0.6
600
800
1000
1200
Estimated allele frequency
Ecological gradient (Precipitation sums)

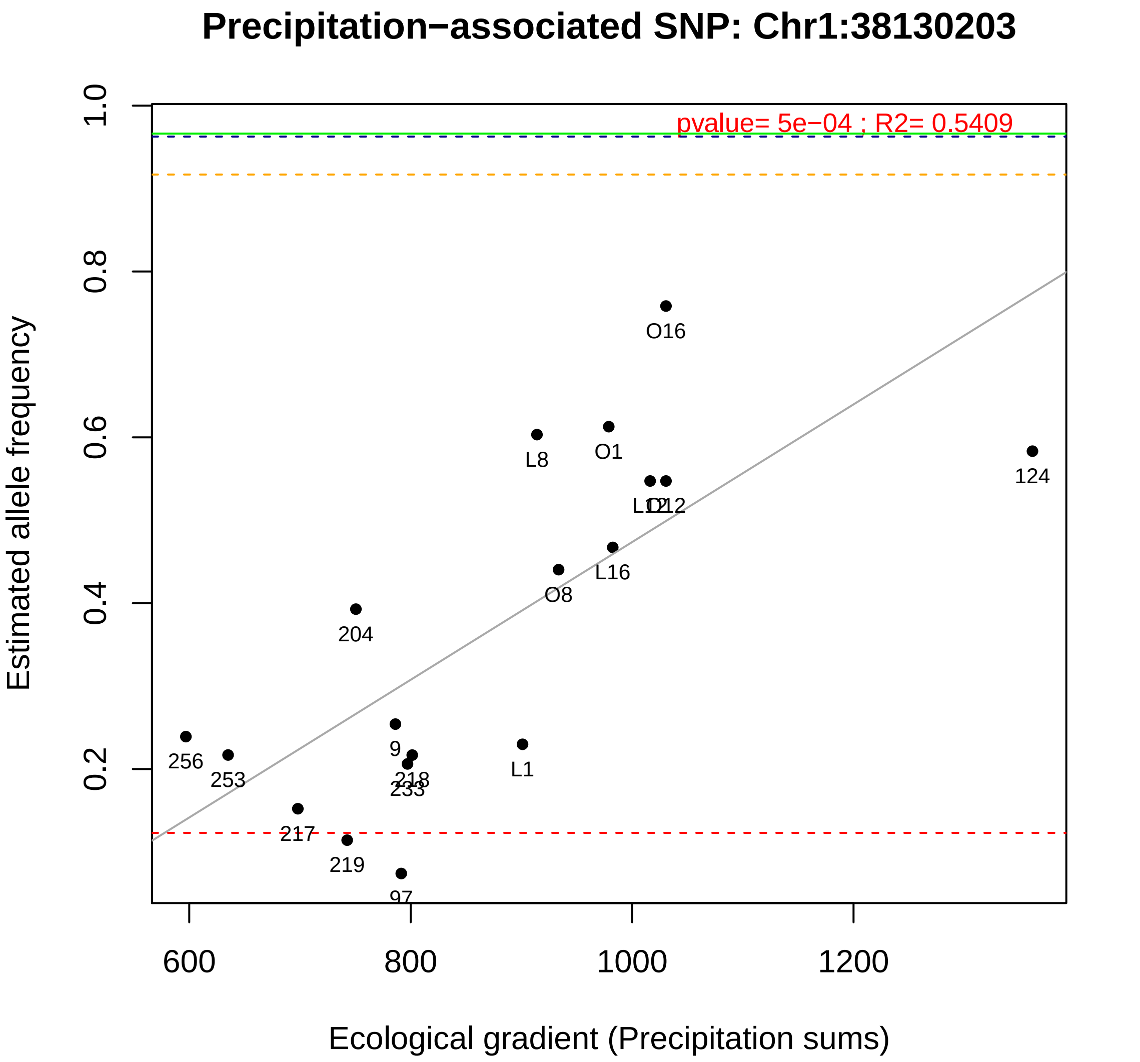
Precipitation−associated SNP: Chr1:38130203
pvalue= 5e−04 ; R2= 0.5409
Estimated allele frequency
0.2
0.4
0.6
0.8
1.0
600
800
1000
1200
Ecological gradient (Precipitation sums)
O16
L8
O1
124
L12
O12
L16
O8
204
9
218
233
L1
256
253
217
219
97

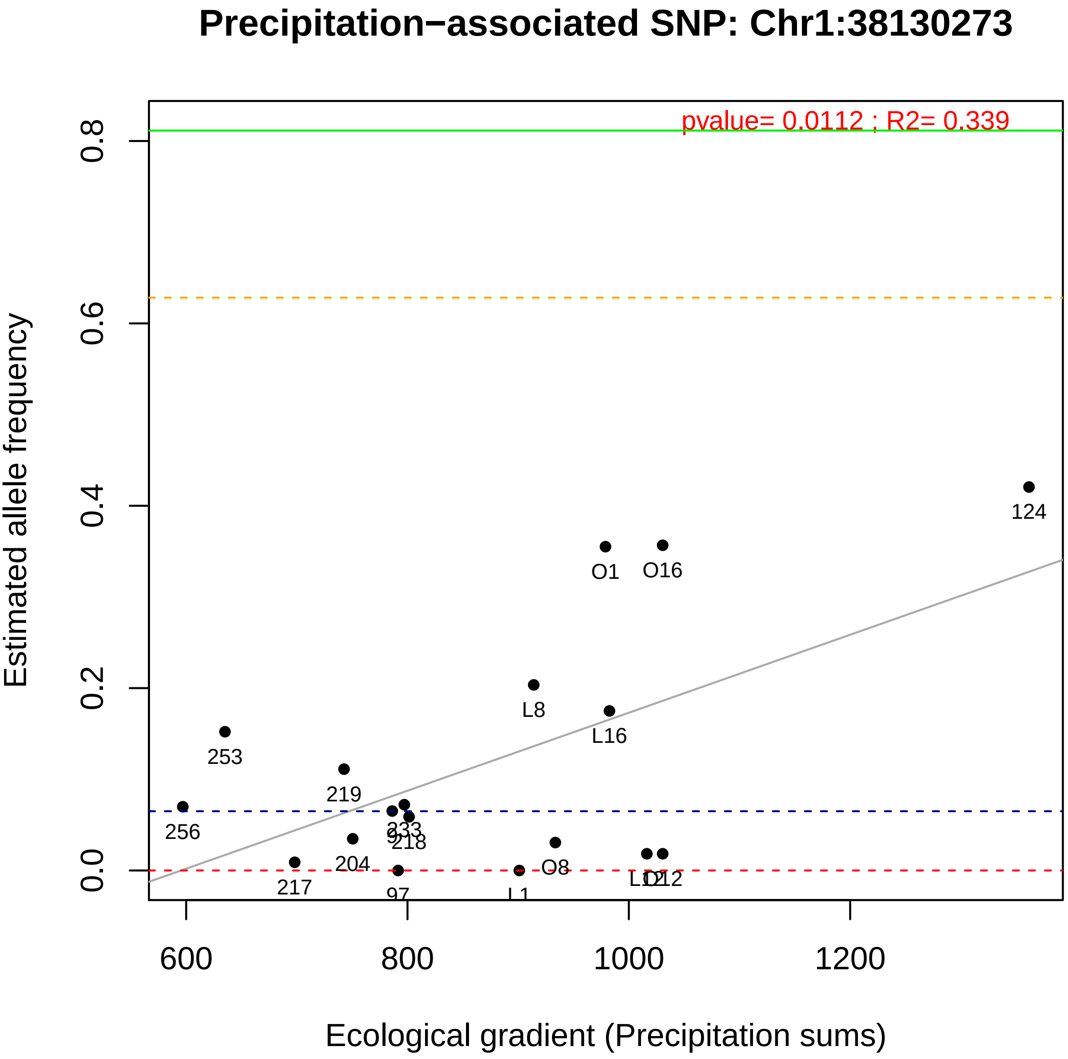
Precipitation−associated SNP: Chr1:38130273
pvalue= 0.0112 ; R2= 0.339
Estimated allele frequency
0.0
0.2
0.4
0.6
0.8
600
800
1000
1200
Ecological gradient (Precipitation sums)
124
O1
O16
L8
L16
253
219
256
233
218
204
217
O8

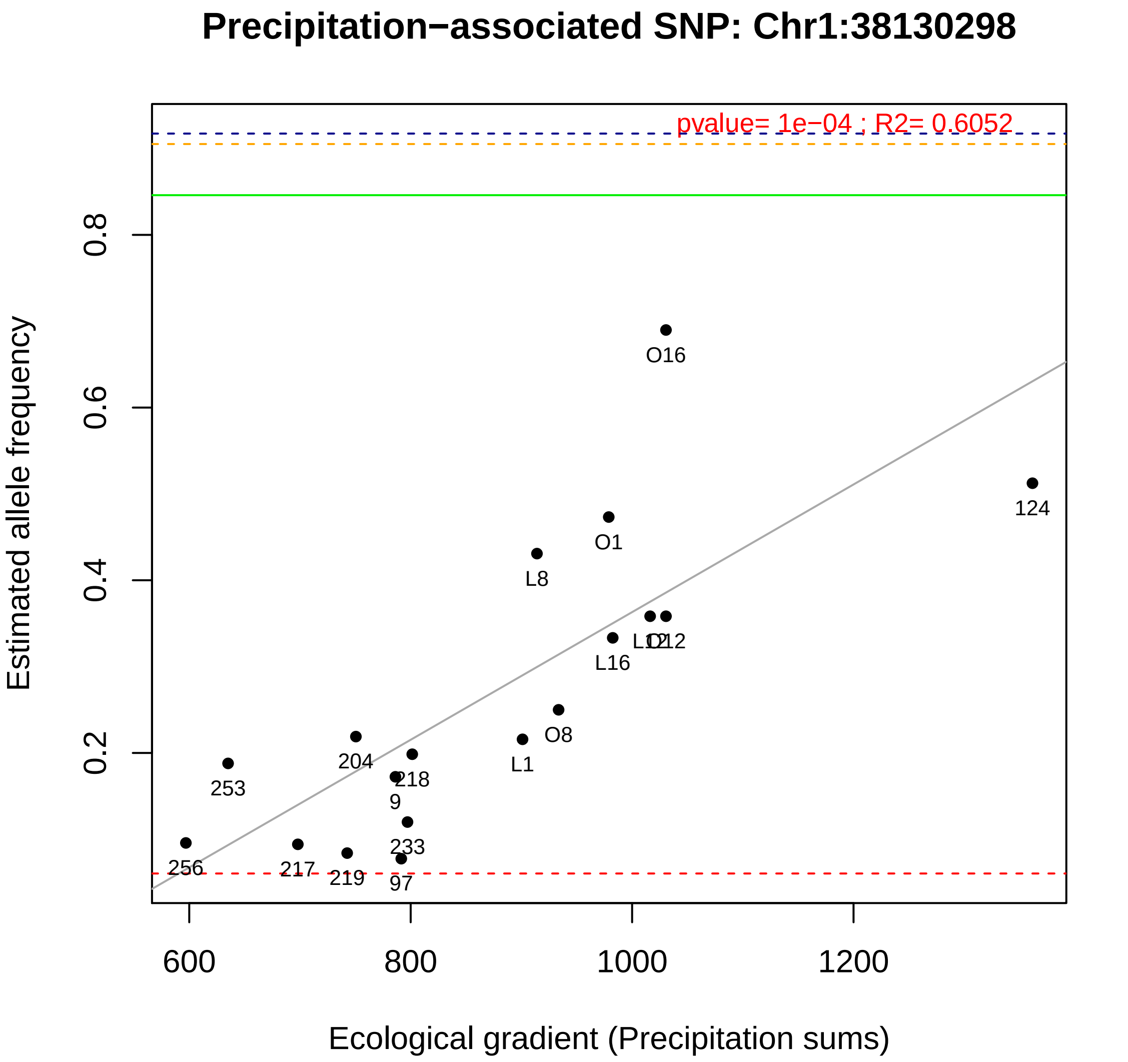
Precipitation−associated SNP: Chr1:38130298
pvalue= 1e−04 ; R2= 0.6052
Estimated allele frequency
0.2
0.4
0.6
0.8
600
800
1000
1200
Ecological gradient (Precipitation sums)
O16
124
O1
L8
L16
L12
O12
O8
L1
204
253
218
9
233
256
217
219
97

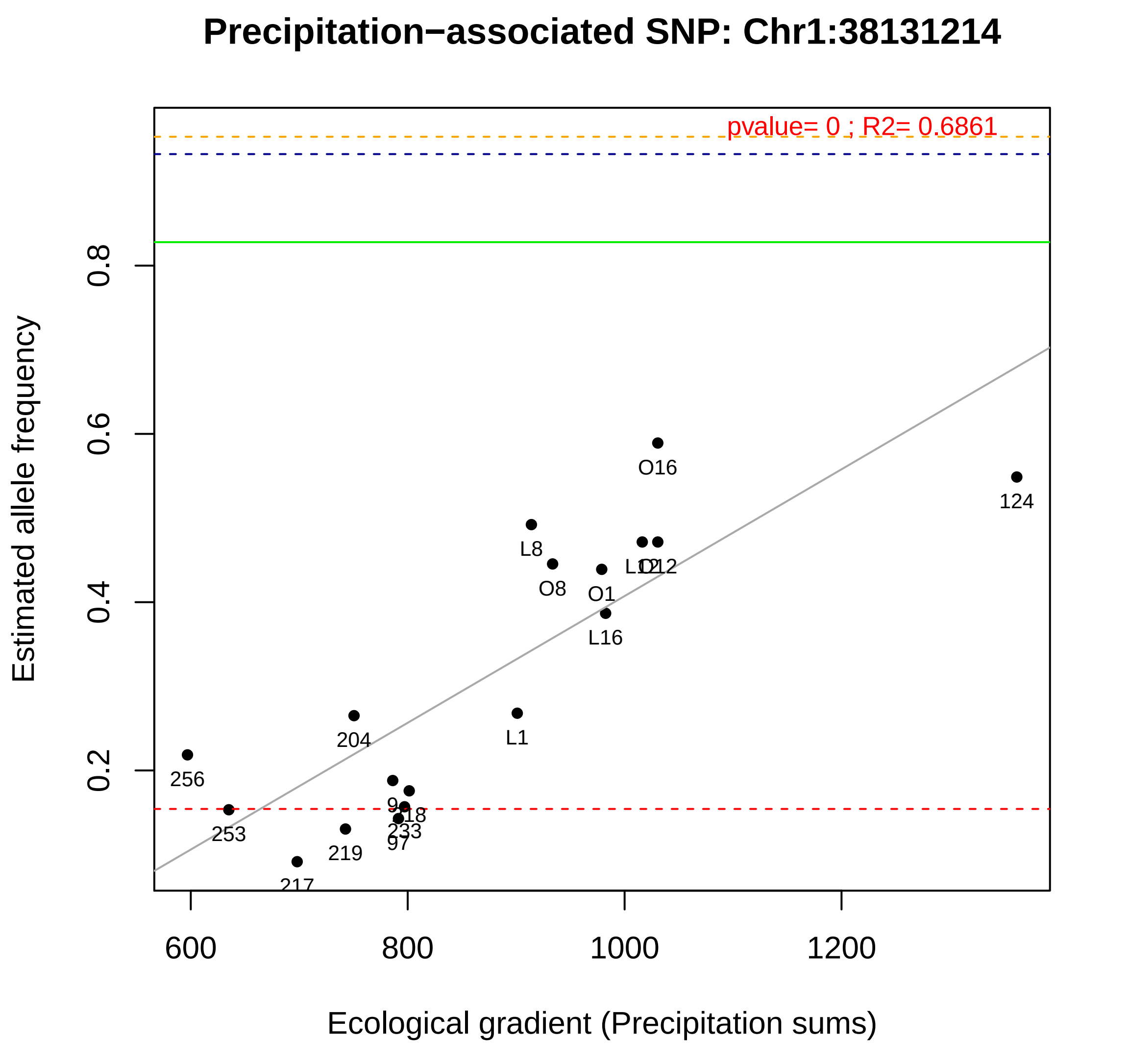

Precipitation−associated SNP: Chr1:38131214
pvalue= 0 ; R2= 0.6861
Estimated allele frequency
0.2
0.4
0.6
0.8
600
800
1000
1200
Ecological gradient (Precipitation sums)
O16
124
L8
O8
O1
L16
204
L1
256
253
219
217
233
97

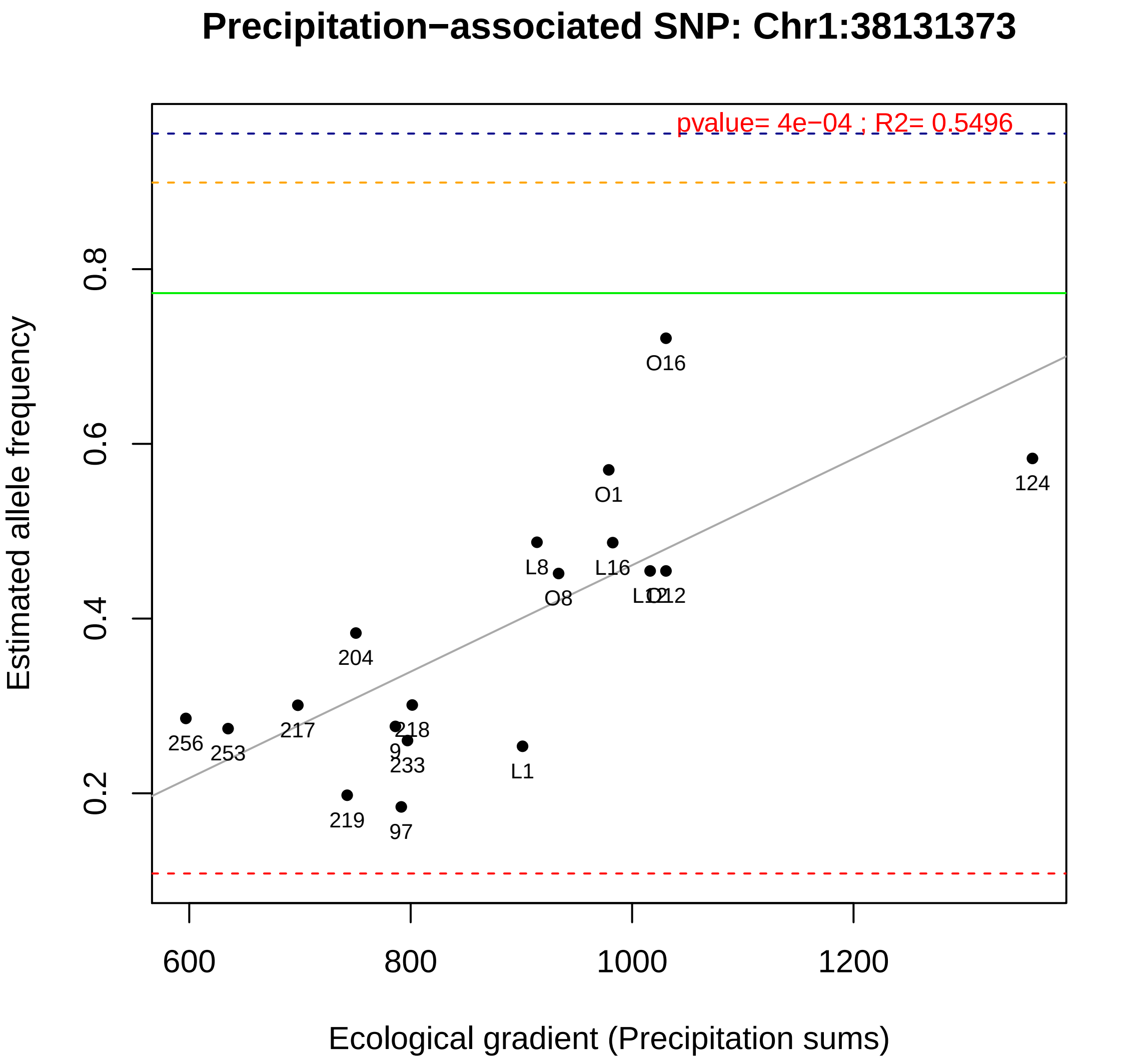

Precipitation−associated SNP: Chr1:38131373
pvalue= 4e−04 ; R2= 0.5496
Estimated allele frequency
0.2
0.4
0.6
0.8
600
800
1000
1200
Ecological gradient (Precipitation sums)
O16
124
O1
L8
L16
O8
L12
O12
204
217
218
9
233
256
253
L1
219
97

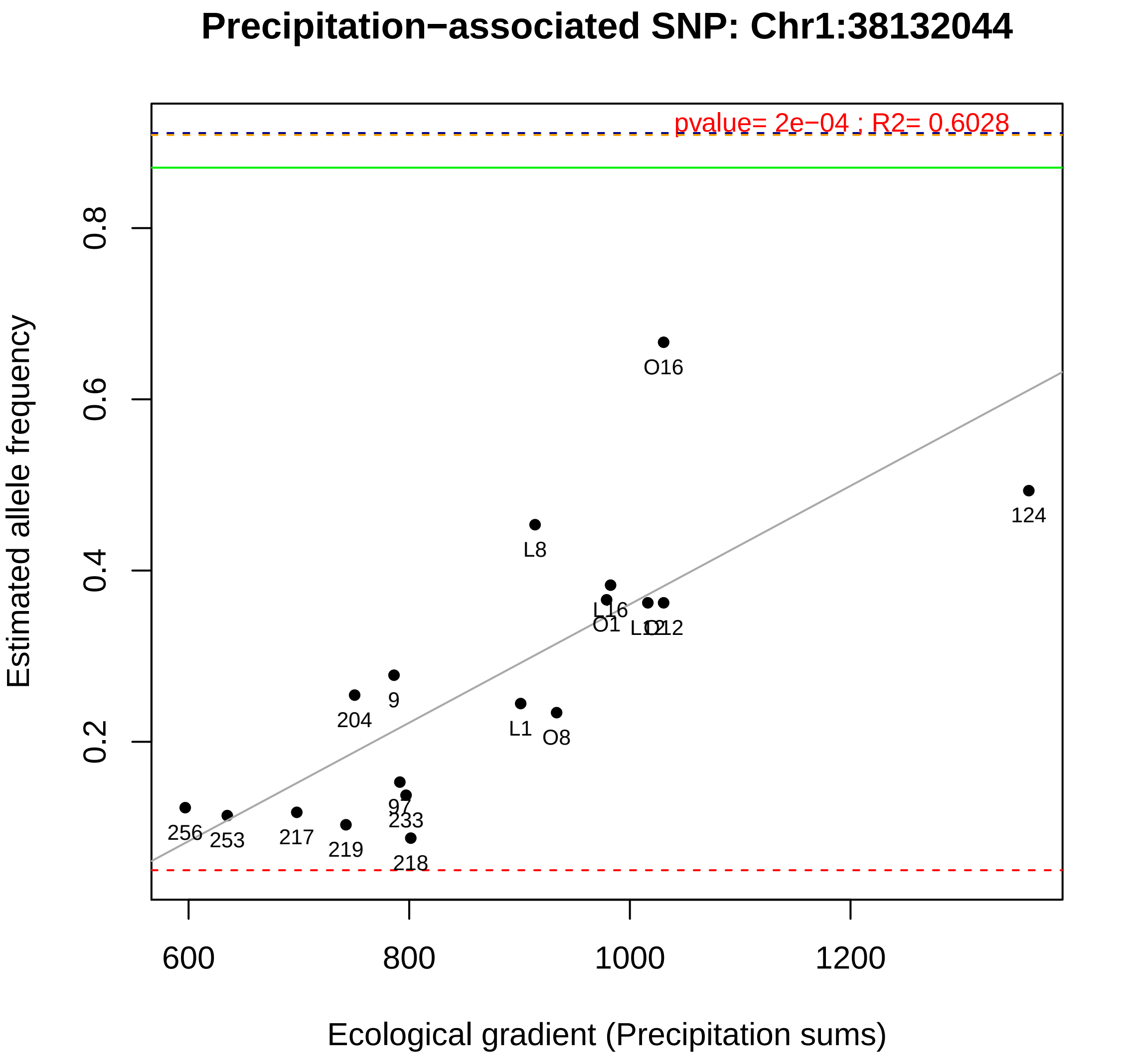

Precipitation−associated SNP: Chr1:38132044
pvalue= 2e−04 ; R2= 0.6028
Estimated allele frequency
0.2
0.4
0.6
0.8
600
800
1000
1200
Ecological gradient (Precipitation sums)
O16
124
L8
L16
O1
L12
O12
9
204
L1
O8
97
233
256
253
217
219
218

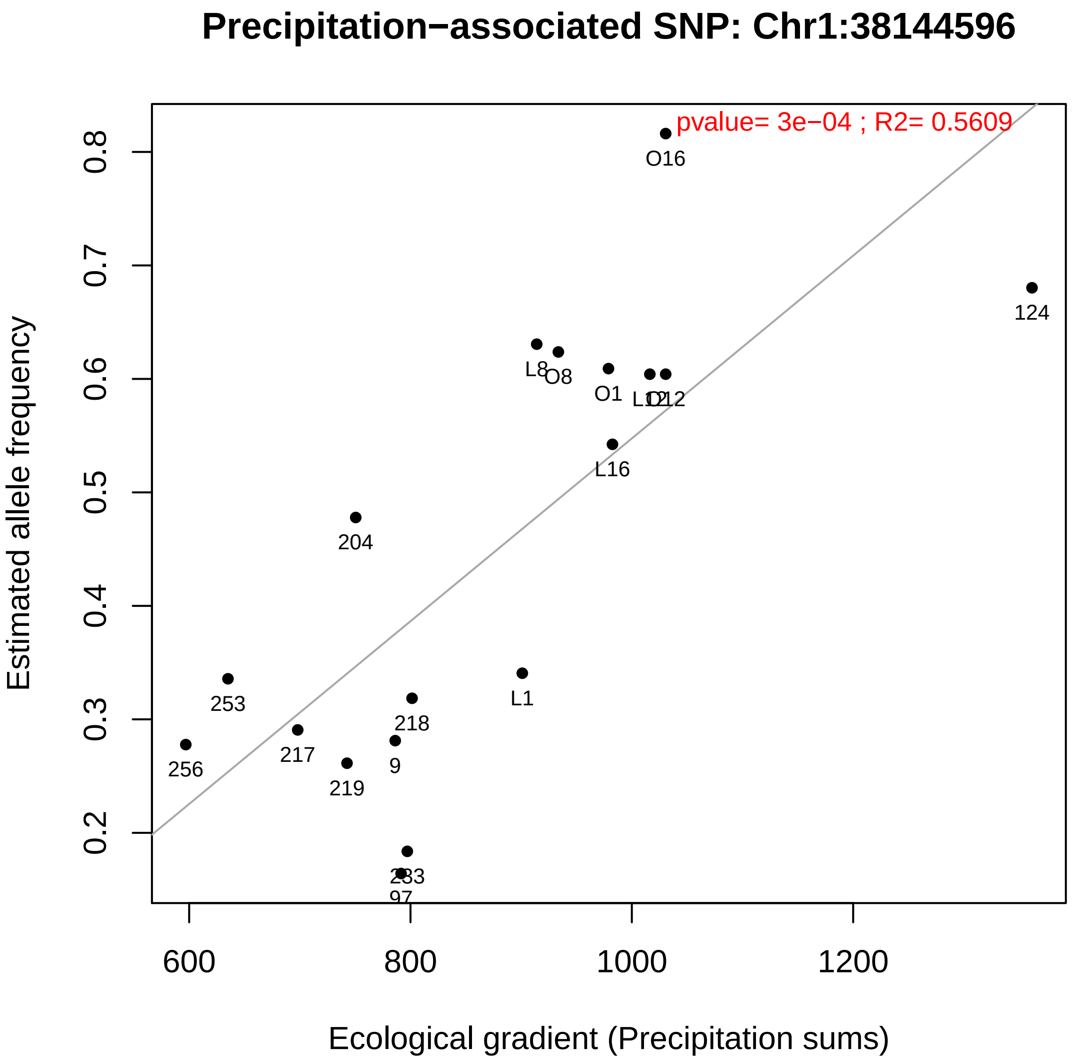
Precipitation−associated SNP: Chr1:38144596
pvalue= 3e−04 ; R2= 0.5609
O16
124
L8
O8
O1
L12
O12
L16
204
253
L1
218
217
256
9
219
233
97
0.2
0.3
0.4
0.5
0.6
0.7
0.8
600
800
1000
1200
Estimated allele frequency
Ecological gradient (Precipitation sums)

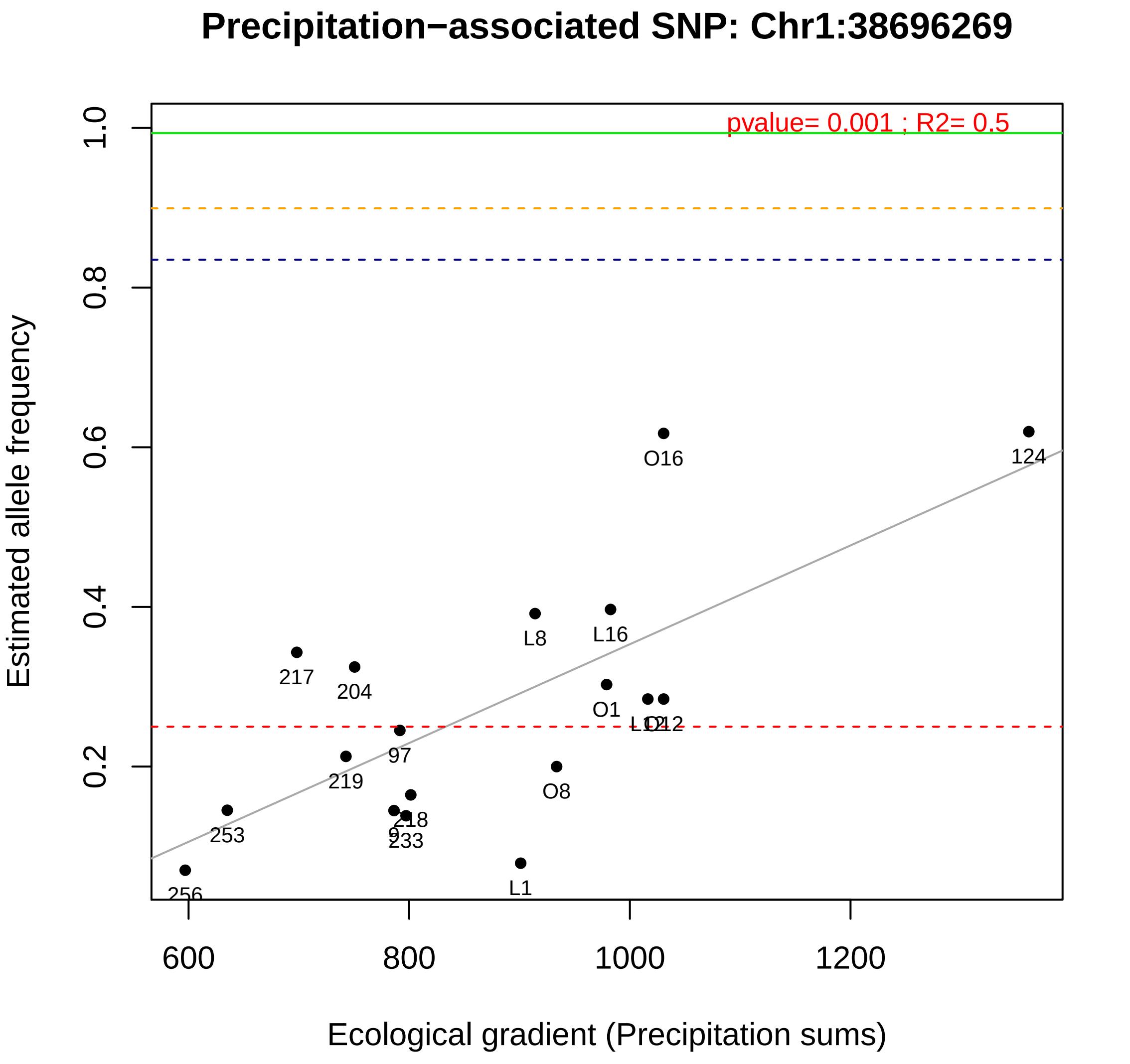

Precipitation−associated SNP: Chr1:38696269
pvalue= 0.001 ; R2= 0.5
Estimated allele frequency
Ecological gradient (Precipitation sums)
0.2
0.4
0.6
0.8
1.0
600
800
1000
1200
O16
124
L8
L16
217
204
O1
97
219
O8
253
218
233
L1

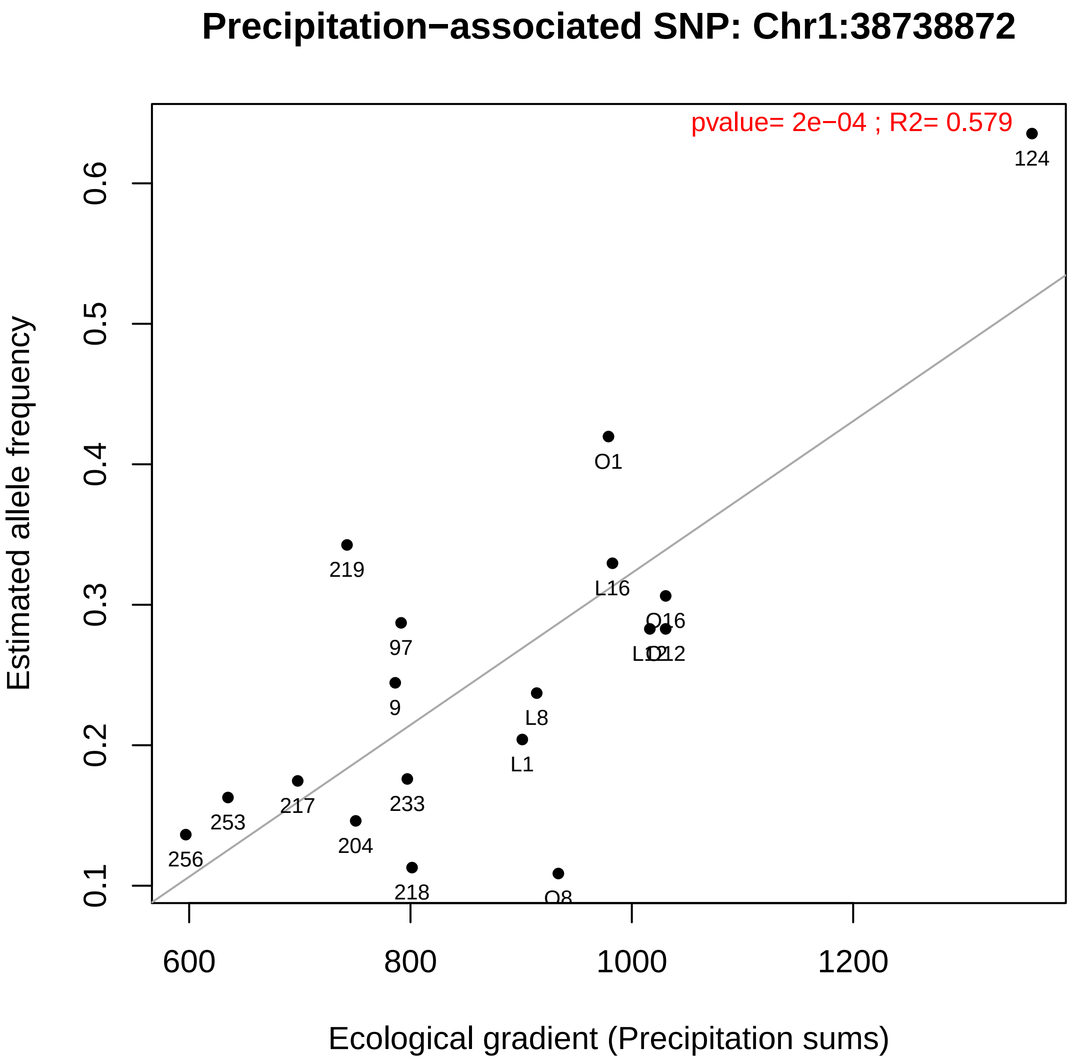

Precipitation−associated SNP: Chr1:38738872
pvalue= 2e−04 ; R2= 0.579
Estimated allele frequency
0.1
0.2
0.3
0.4
0.5
0.6
600
800
1000
1200
Ecological gradient (Precipitation sums)
124
O1
219
L16
O16
97
9
L8
L1
217
233
253
204
256
218
O8

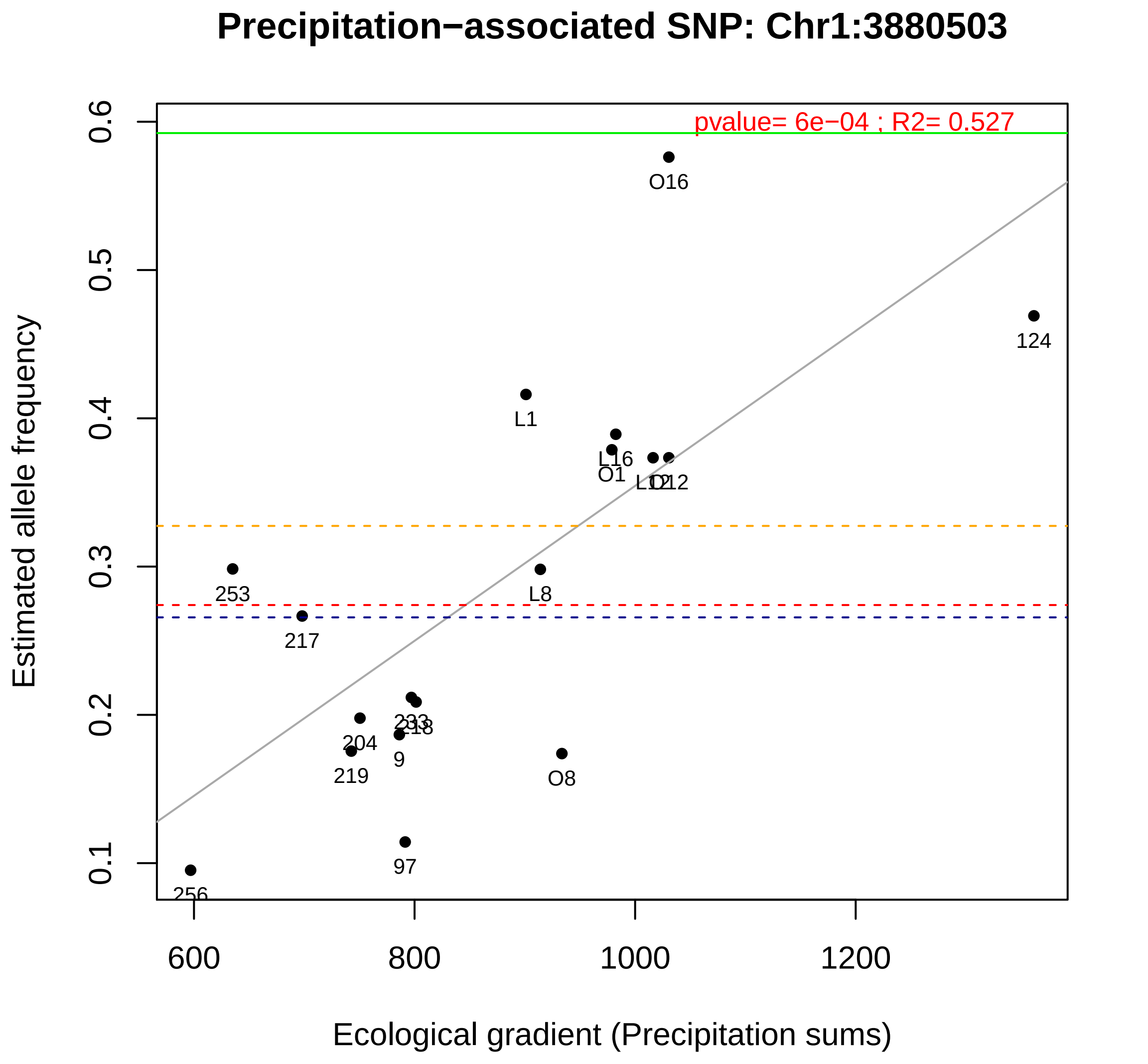
Precipitation−associated SNP: Chr1:3880503
pvalue= 6e−04 ; R2= 0.527
Estimated allele frequency
Ecological gradient (Precipitation sums)
0.1
0.2
0.3
0.4
0.5
0.6
600
800
1000
1200
O16
124
L1
L16
O1
L8
253
217
233
218
204
9
219
O8
97
256

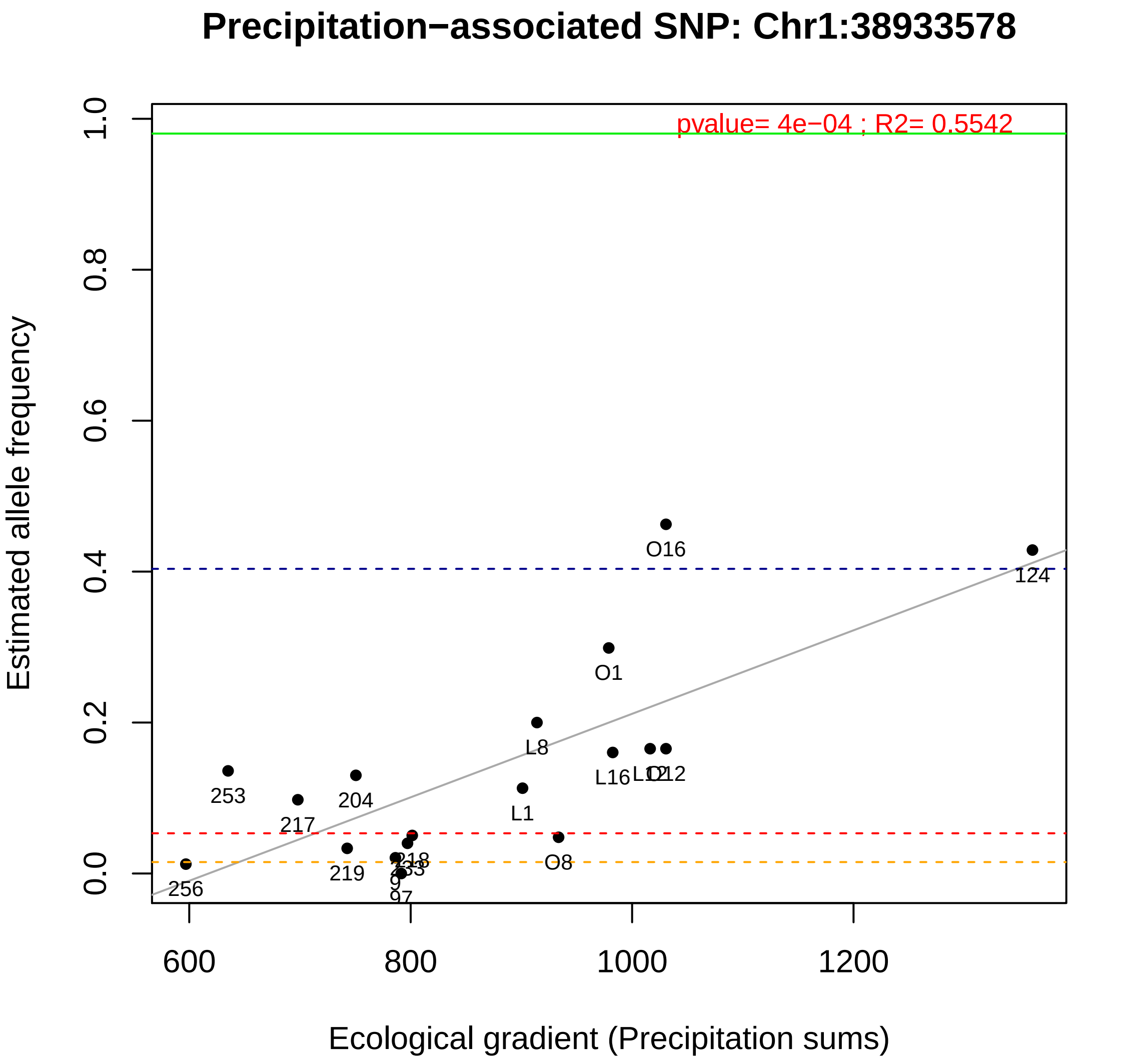
Precipitation−associated SNP: Chr1:38933578
pvalue= 4e−04 ; R2= 0.5542
Estimated allele frequency
0.0
0.2
0.4
0.6
0.8
1.0
600
800
1000
1200
Ecological gradient (Precipitation sums)
O16
124
O1
L8
L16
L1
253
204
217
219
256
O8
218
233
9
97

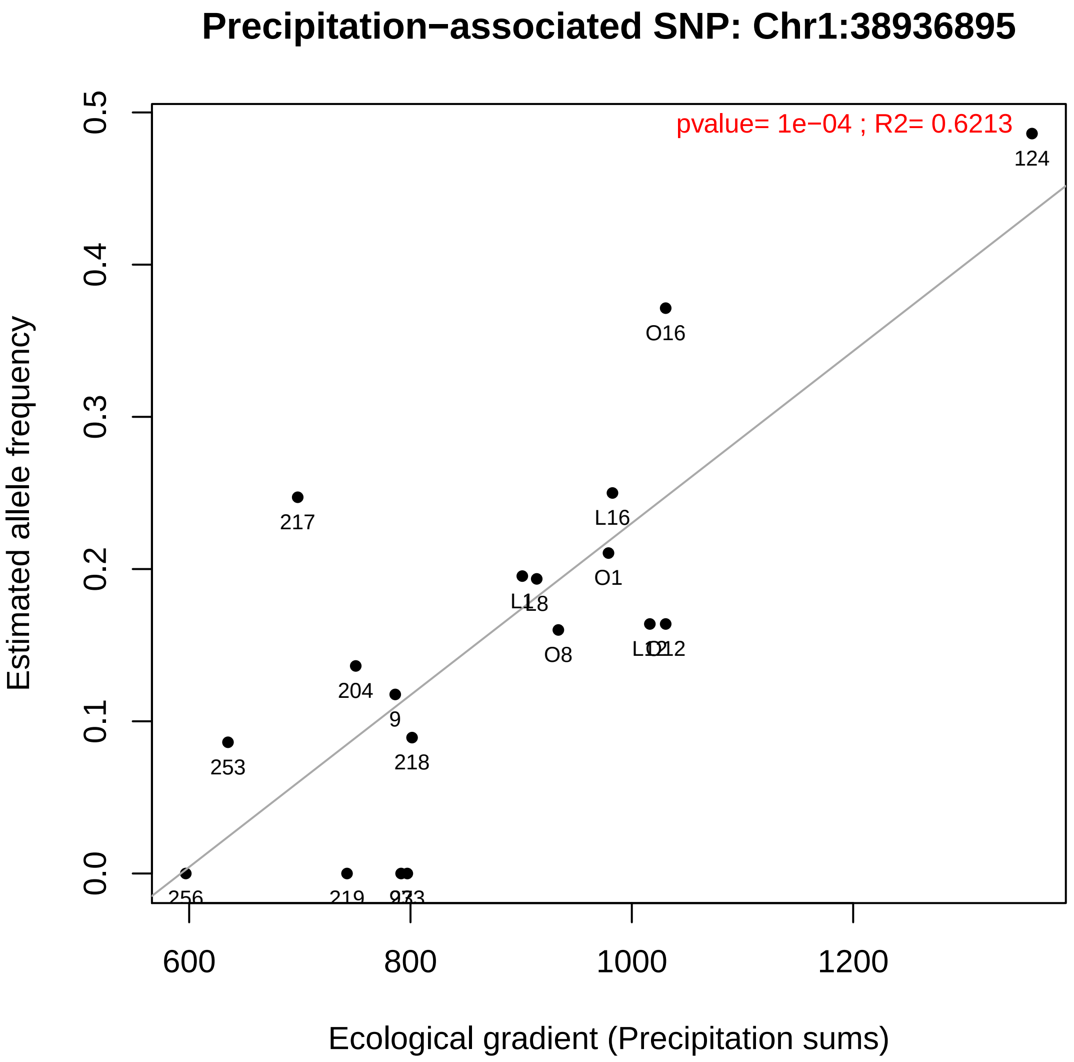
Precipitation−associated SNP: Chr1:38936895
pvalue= 1e−04 ; R2= 0.6213
Estimated allele frequency
Ecological gradient (Precipitation sums)
0.0
0.1
0.2
0.3
0.4
0.5
600
800
1000
1200
124
O16
217
L16
O1
L1
L8
O8
L12
O12
204
9
218
253
256
219
97
73

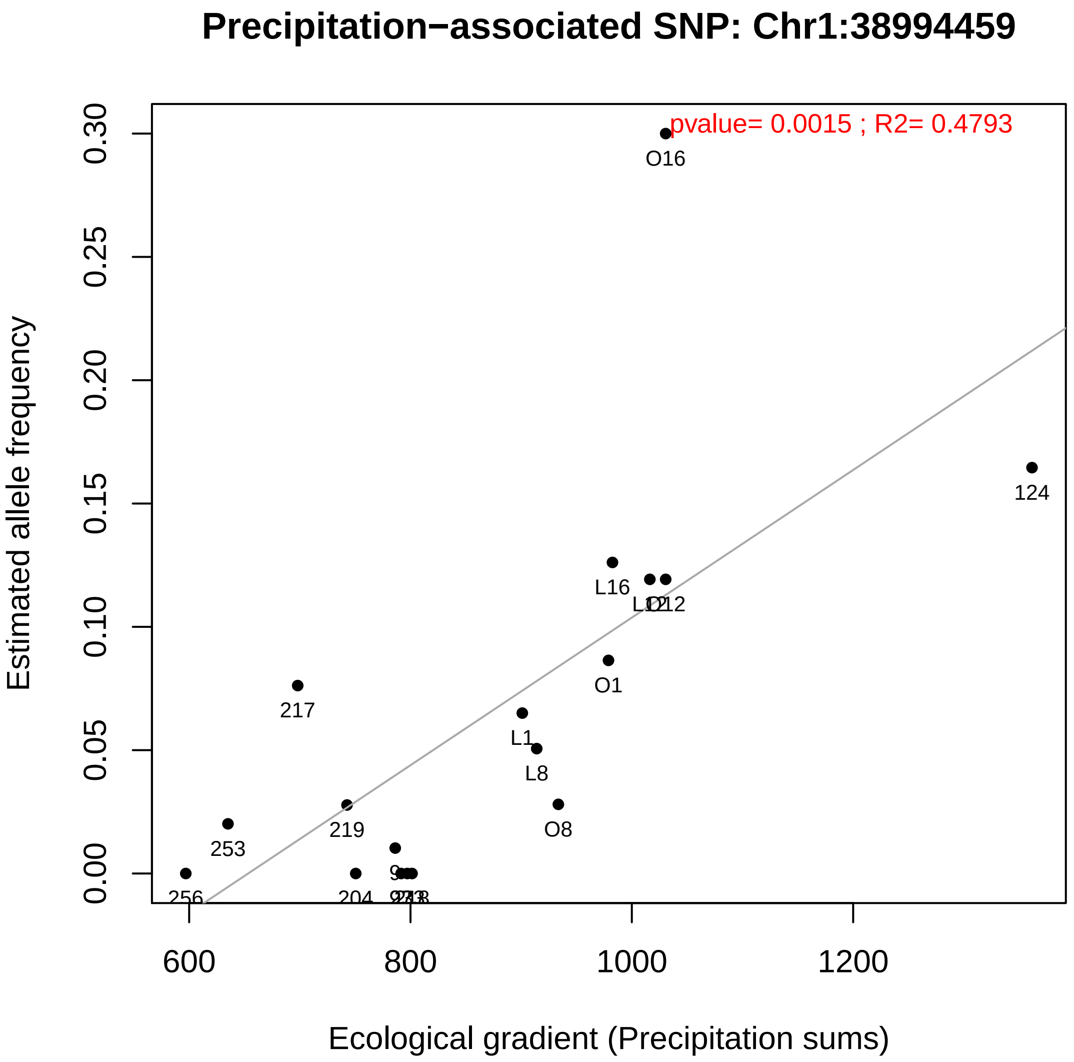

Precipitation−associated SNP: Chr1:38994459
pvalue= 0.0015 ; R2= 0.4793
O16
124
L16
O12
O1
217
L1
L8
219
O8
253
256
204
0.00
0.05
0.10
0.15
0.20
0.25
0.30
600
800
1000
1200
Estimated allele frequency
Ecological gradient (Precipitation sums)

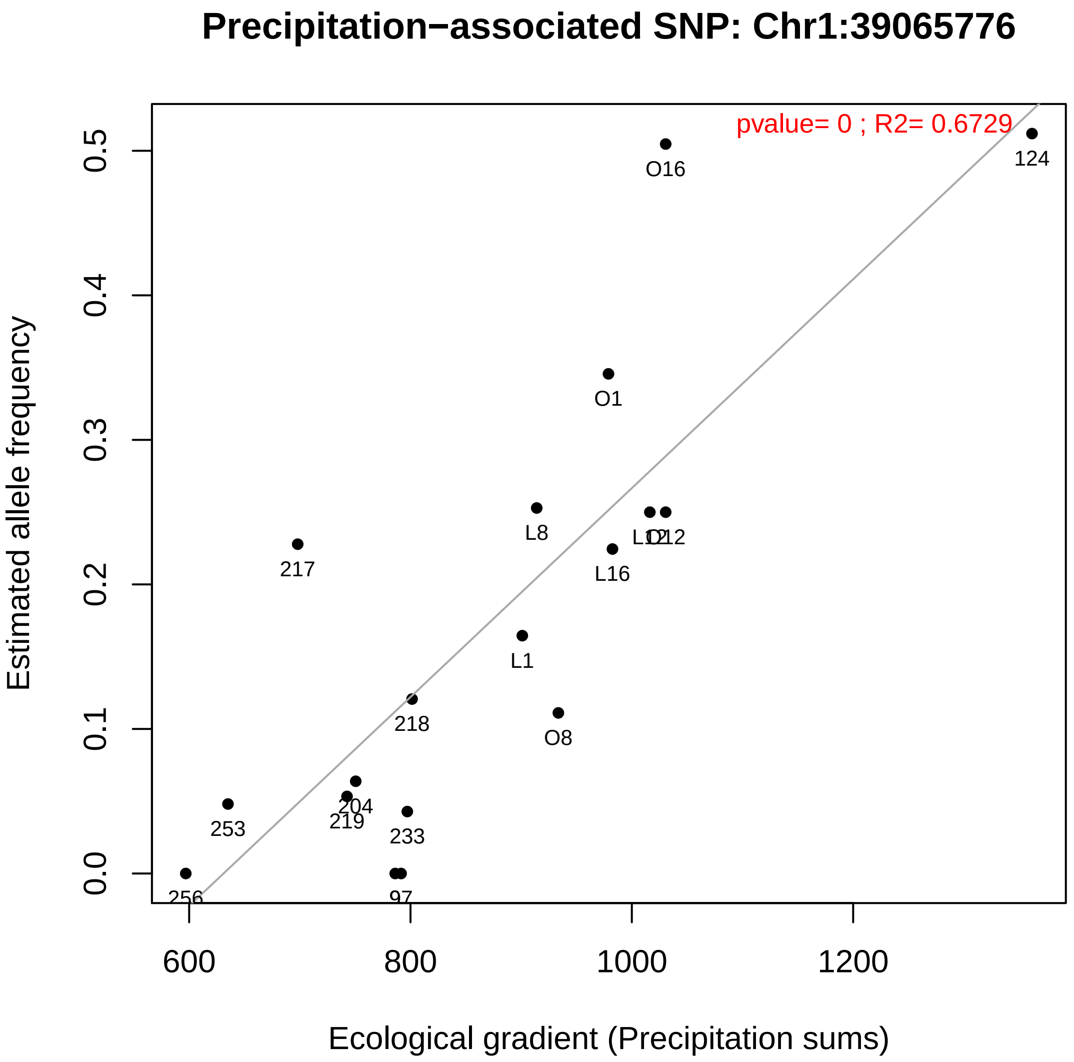
Precipitation−associated SNP: Chr1:39065776
pvalue= 0 ; R2= 0.6729
Estimated allele frequency
0.0
0.1
0.2
0.3
0.4
0.5
600
800
1000
1200
Ecological gradient (Precipitation sums)
O16
124
O1
L8
L12
O12
L16
217
L1
218
O8
204
219
253
233
256
97

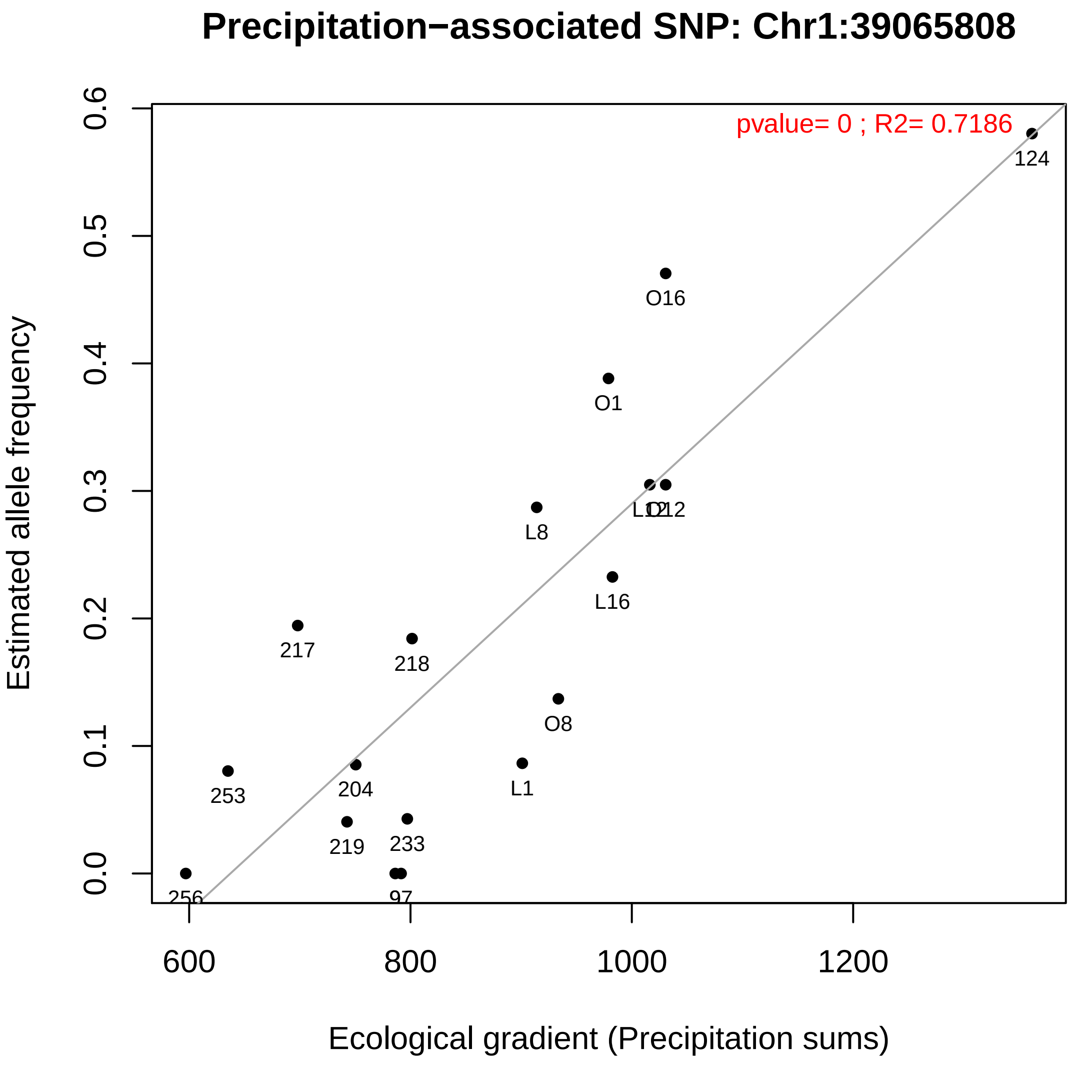
Precipitation−associated SNP: Chr1:39065808
pvalue= 0 ; R2= 0.7186
Estimated allele frequency
Ecological gradient (Precipitation sums)
0.0
0.1
0.2
0.3
0.4
0.5
0.6
600
800
1000
1200
124
O16
O1
L8
L16
217
218
O8
253
204
L1
219
233
256
97

**Precipitation−associated SNP: Chr1:39065847**

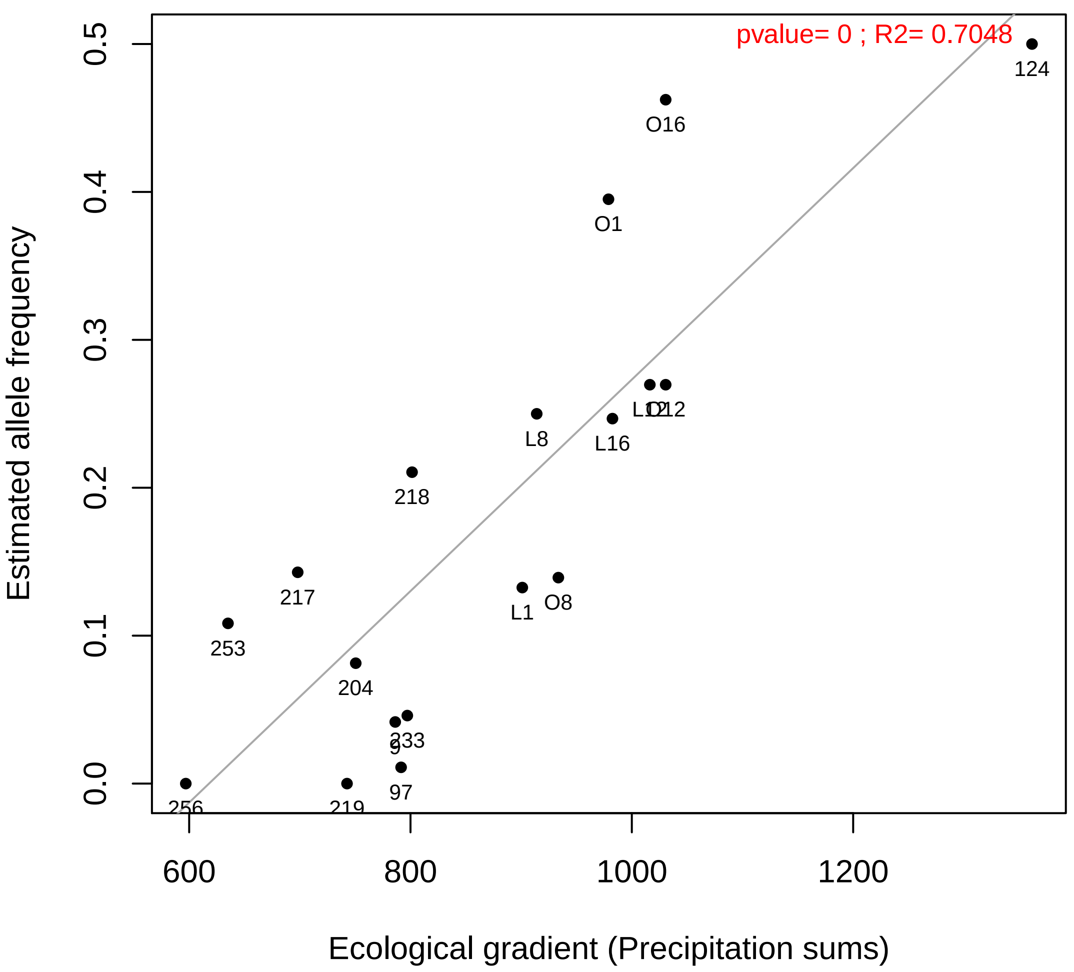

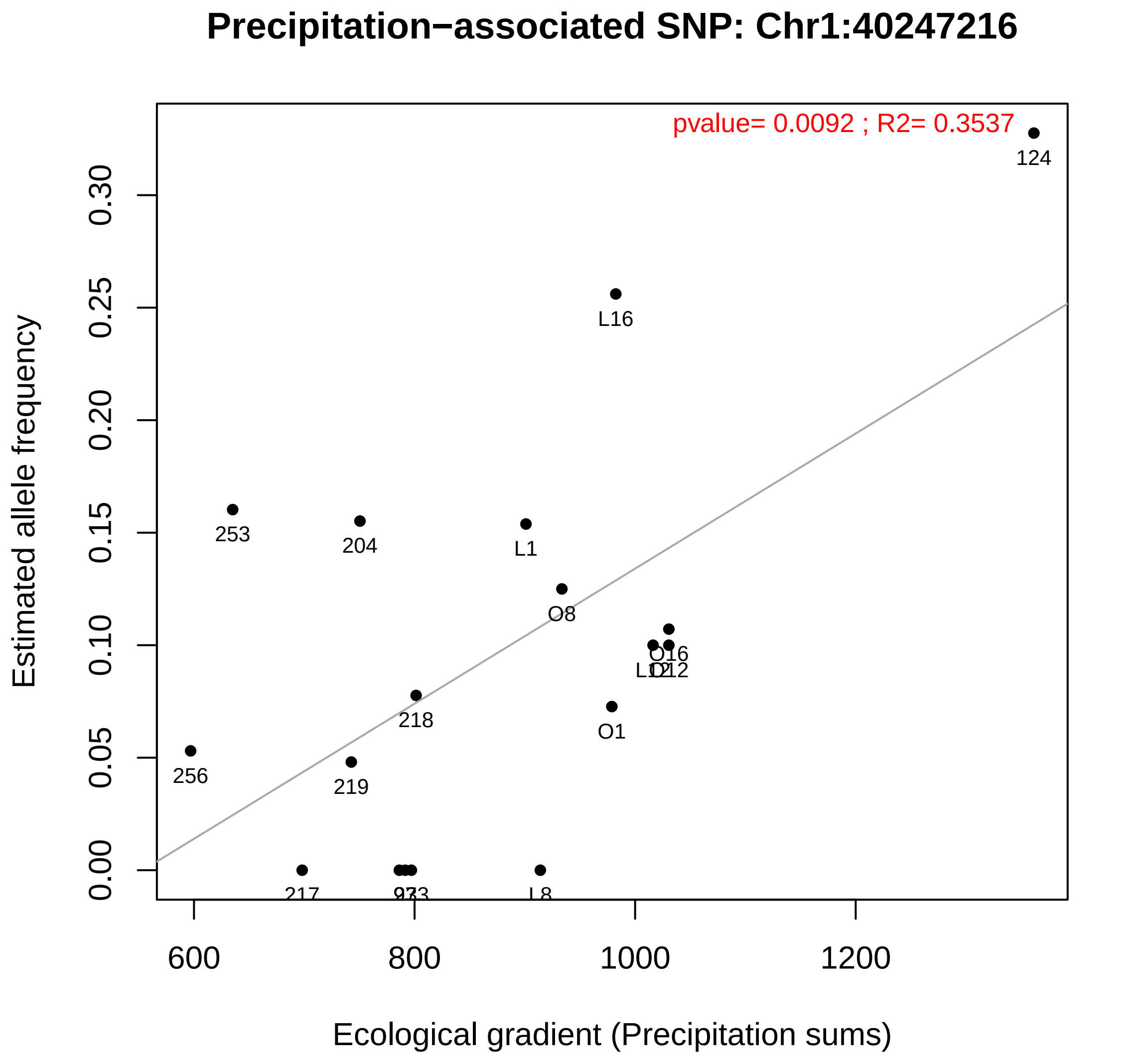

Precipitation−associated SNP: Chr1:40247216
pvalue= 0.0092 ; R2= 0.3537
Estimated allele frequency
0.00
0.05
0.10
0.15
0.20
0.25
0.30
600
800
1000
1200
Ecological gradient (Precipitation sums)
124
L16
253
204
L1
O8
O16
O12
218
O1
256
219
217
L8

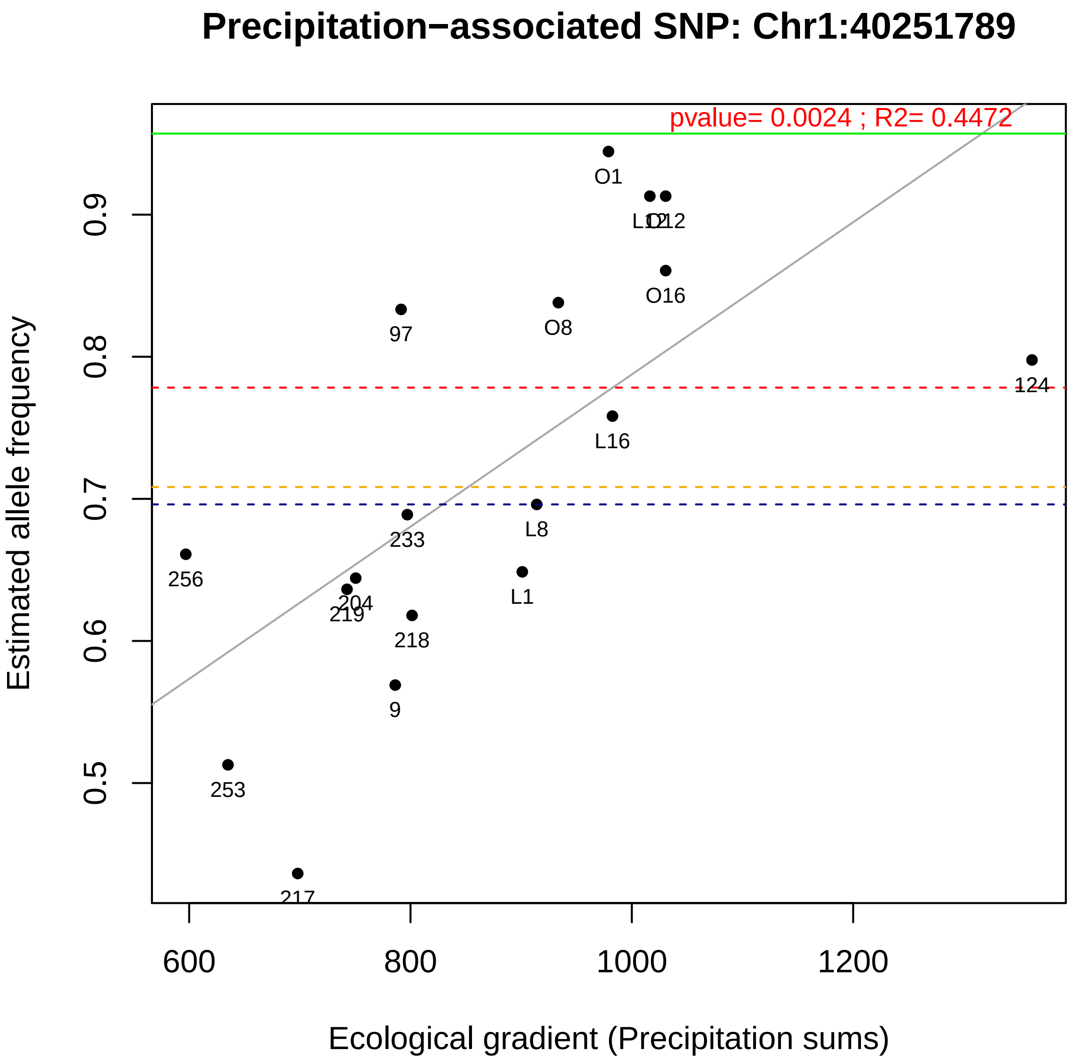

Precipitation−associated SNP: Chr1:40251789
pvalue= 0.0024 ; R2= 0.4472
Estimated allele frequency
0.5
0.6
0.7
0.8
0.9
600
800
1000
1200
Ecological gradient (Precipitation sums)
O1
O8
O16
97
124
L16
L8
233
256
204
219
218
L1
9
253
217

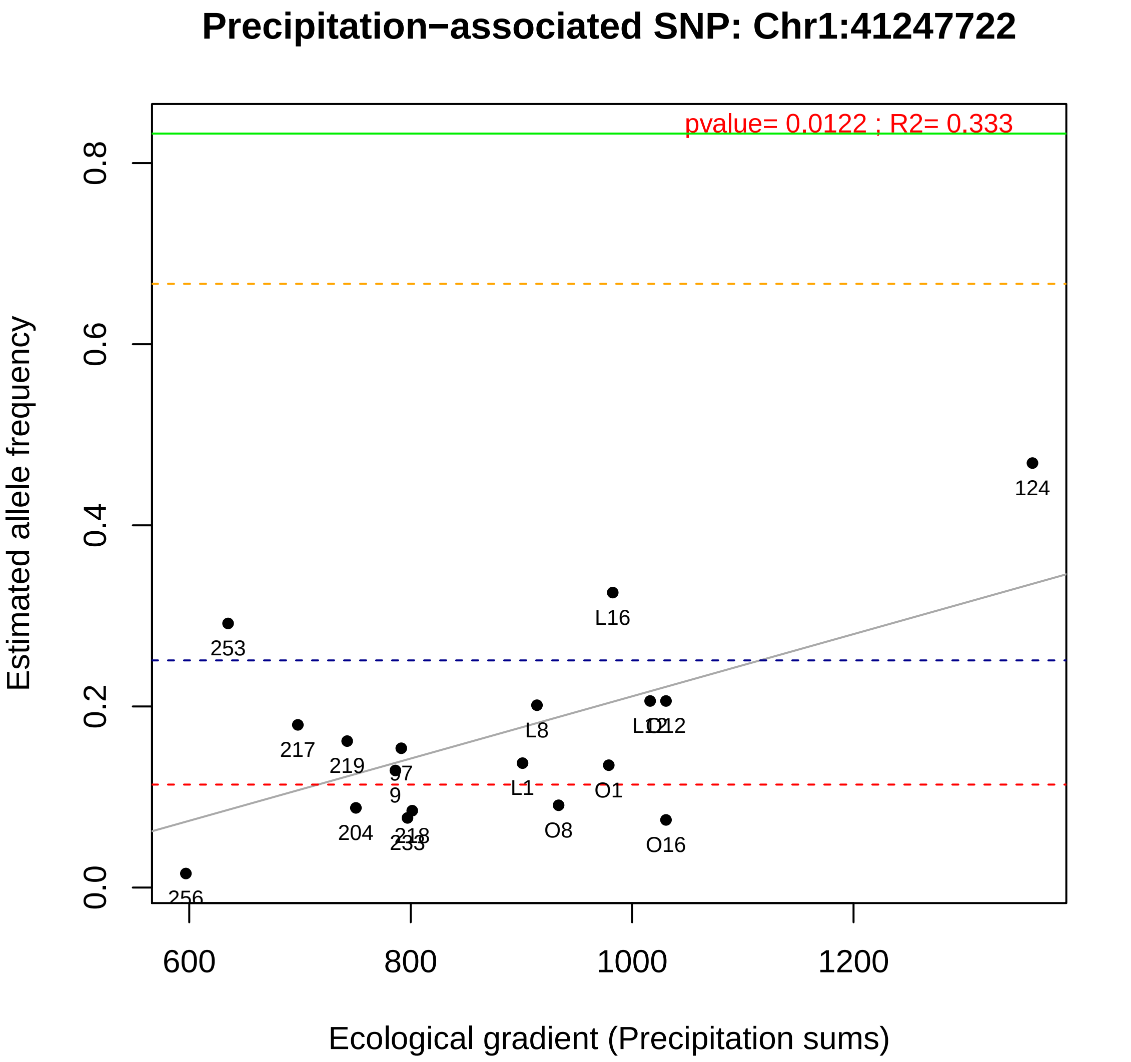

Precipitation−associated SNP: Chr1:41247722
pvalue= 0.0122 ; R2= 0.333
Estimated allele frequency
Ecological gradient (Precipitation sums)
0.0
0.2
0.4
0.6
0.8
600
800
1000
1200
124
L16
253
L8
217
219
97
9
L1
O1
204
218
233
O8
O16
256

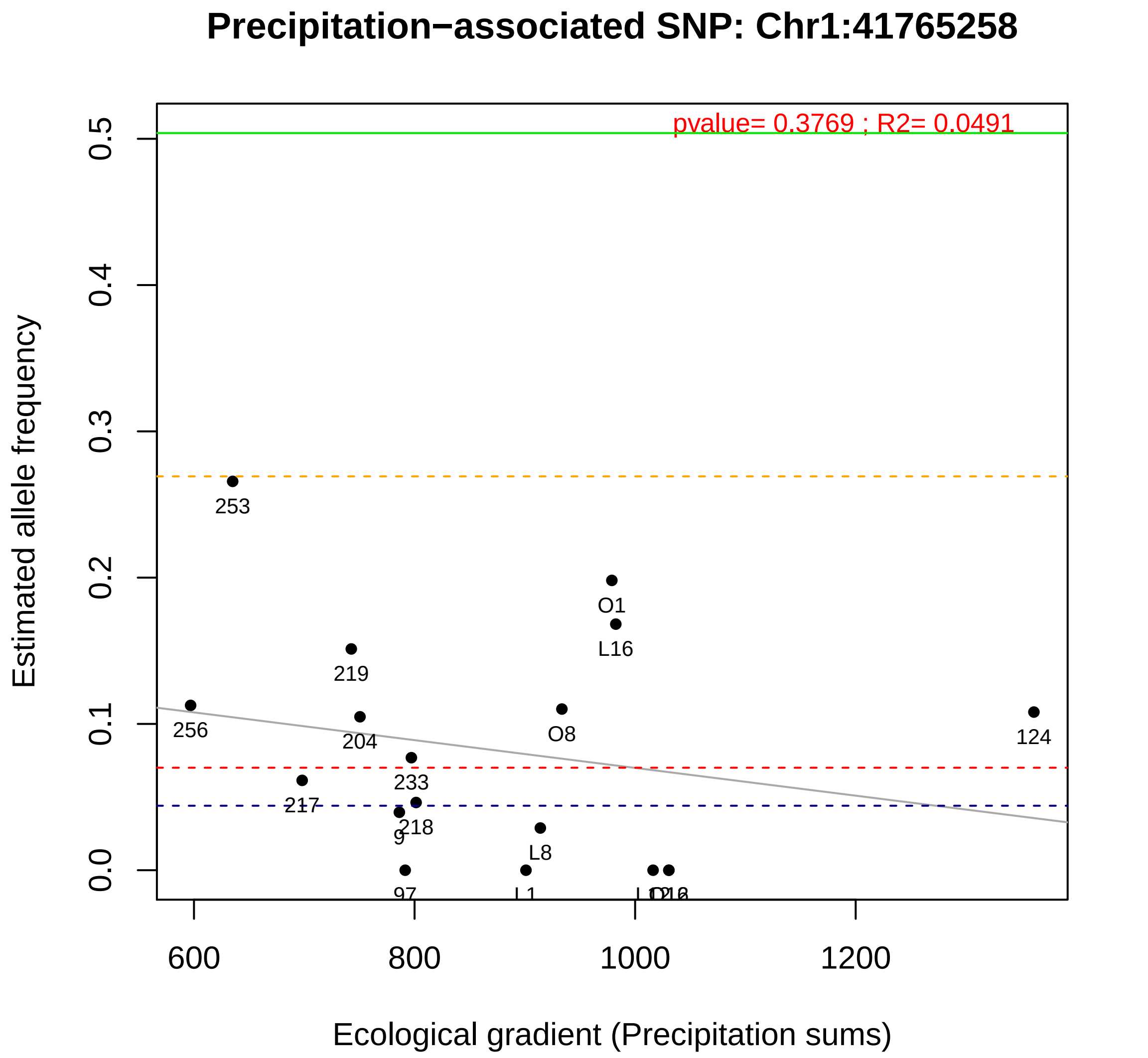
Precipitation−associated SNP: Chr1:41765258
pvalue= 0.3769 ; R2= 0.0491
Estimated allele frequency
Ecological gradient (Precipitation sums)
0.0
0.1
0.2
0.3
0.4
0.5
600
800
1000
1200
253
O1
L16
219
256
204
O8
124
233
217
218
9
L8
97

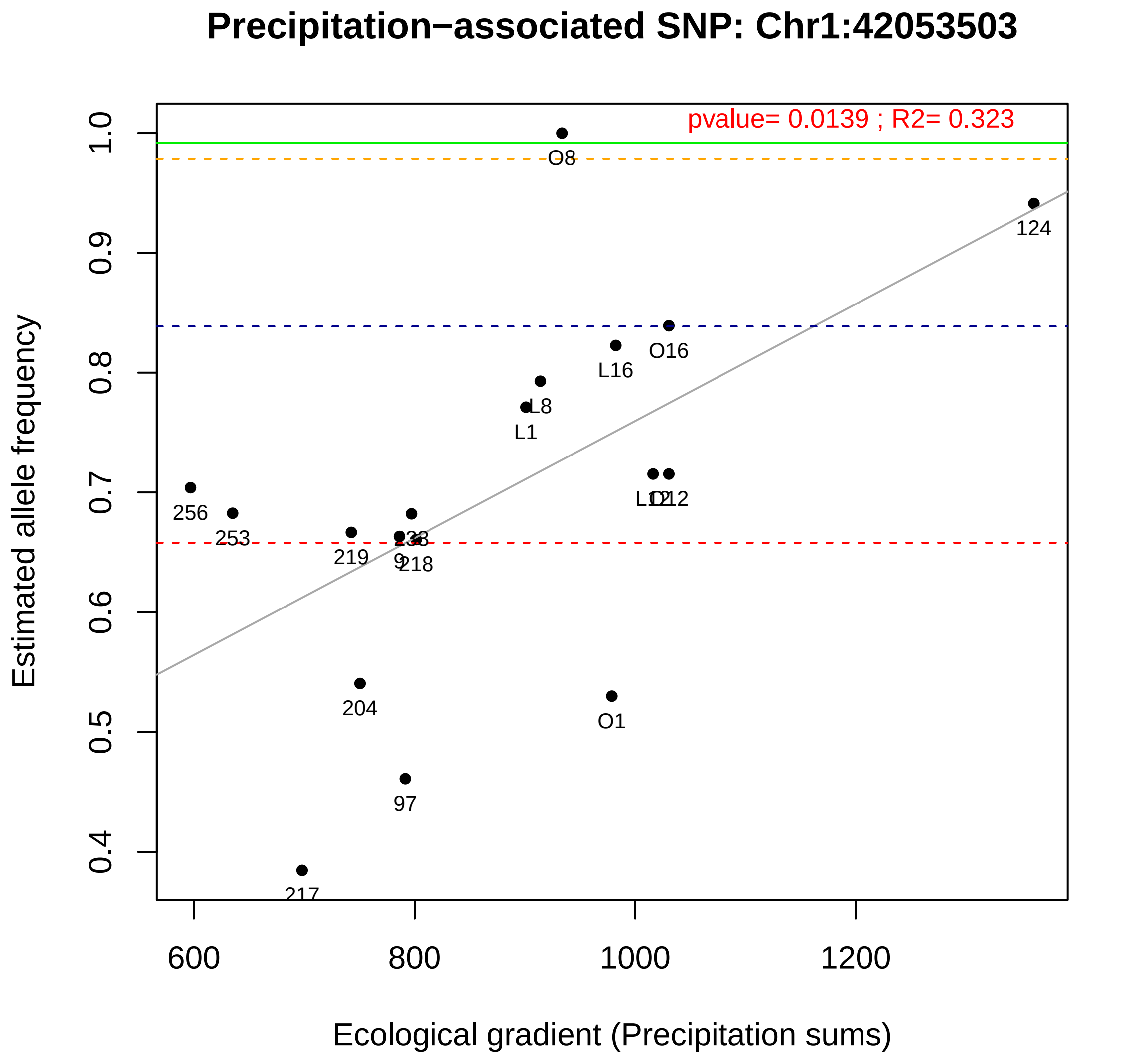

Precipitation−associated SNP: Chr1:42053503
pvalue= 0.0139 ; R2= 0.323
Estimated allele frequency
Ecological gradient (Precipitation sums)
1.0
0.9
0.8
0.7
0.6
0.5
0.4
600
800
1000
1200
O8
124
O16
L16
L8
L1
256
253
219
218
204
O1
97
217

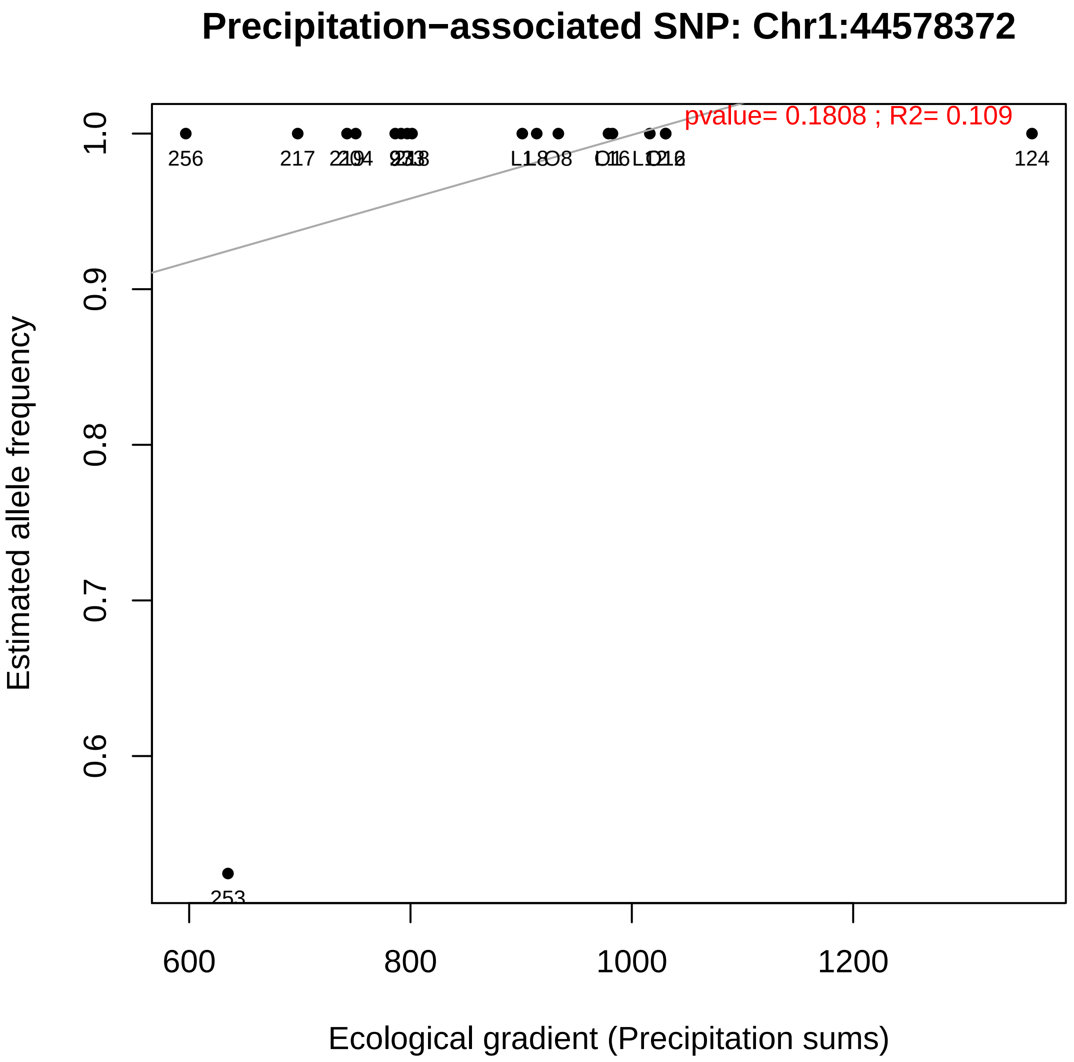

Precipitation−associated SNP: Chr1:44578372
pvalue= 0.1808 ; R2= 0.109
256
217
124
253
1.0
0.9
0.8
0.7
0.6
600
800
1000
1200
Estimated allele frequency
Ecological gradient (Precipitation sums)

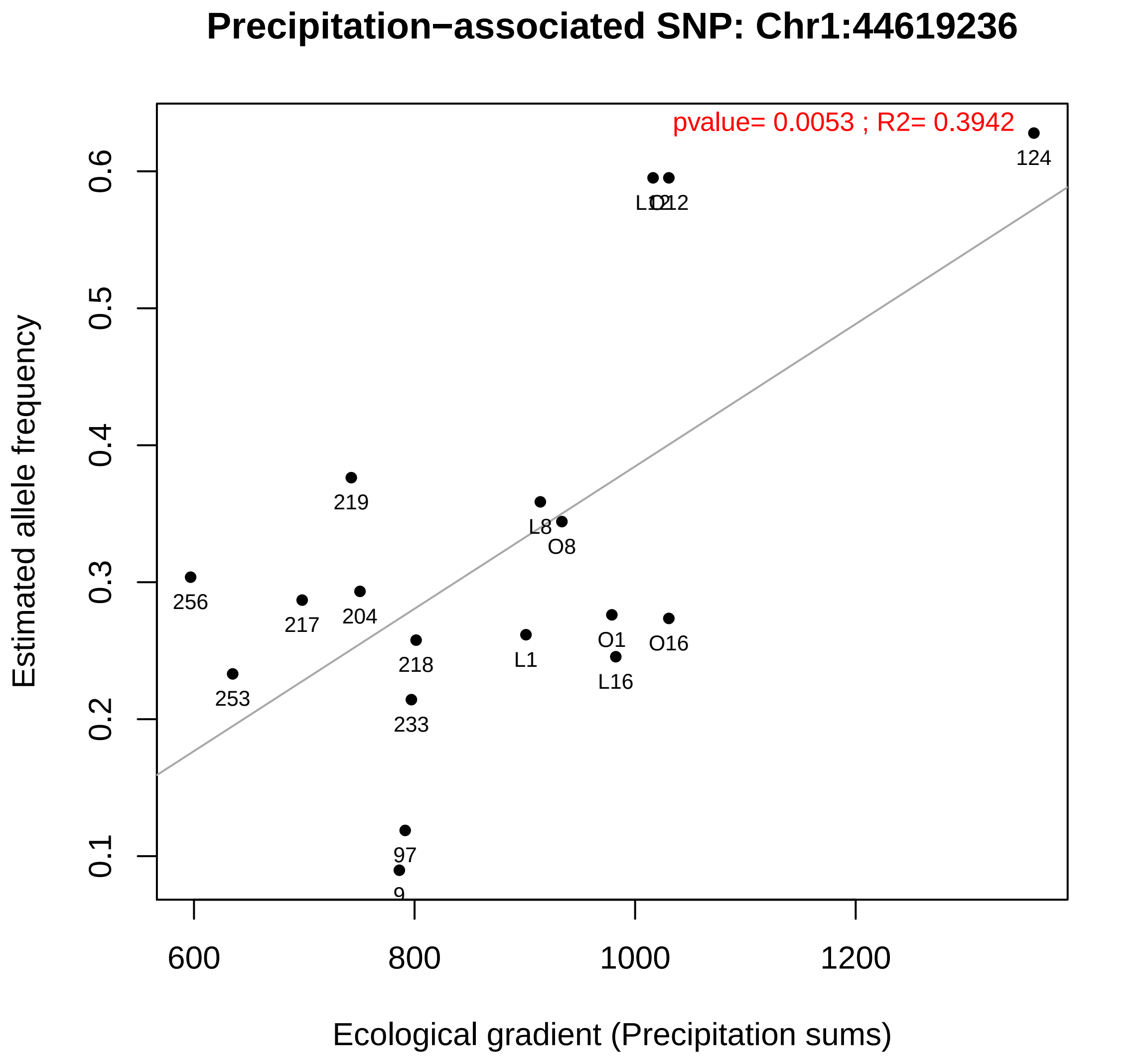
Precipitation−associated SNP: Chr1:44619236
pvalue= 0.0053 ; R2= 0.3942
Estimated allele frequency
Ecological gradient (Precipitation sums)
0.1
0.2
0.3
0.4
0.5
0.6
600
800
1000
1200
124
L12
O12
219
L8
O8
256
217
204
218
L1
O1
O16
L16
253
233
97
9

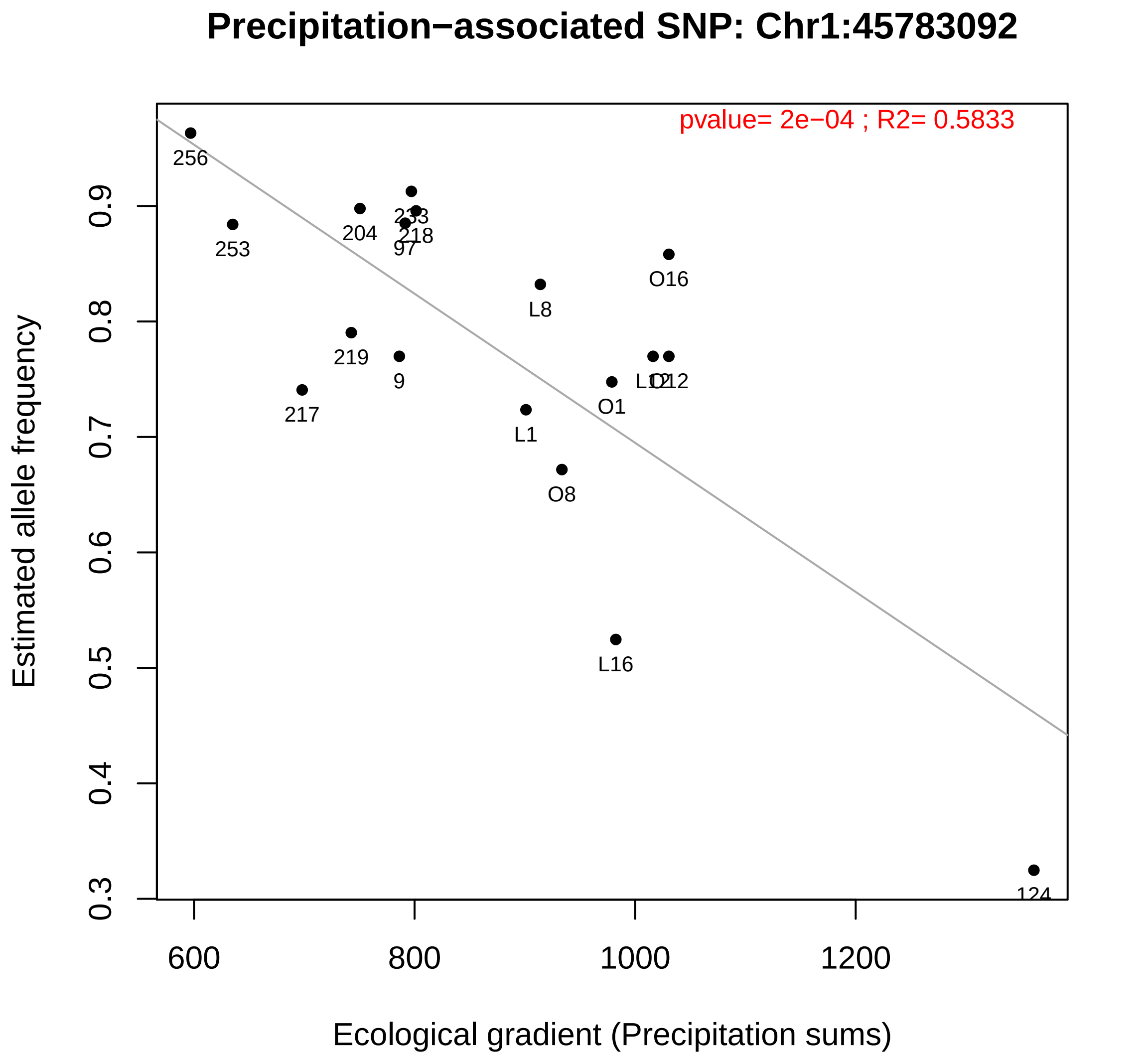
Precipitation−associated SNP: Chr1:45783092
pvalue= 2e−04 ; R2= 0.5833
256
253
204
233
218
97
O16
L8
219
9
L12
O12
O1
217
L1
O8
L16
124
0.3
0.4
0.5
0.6
0.7
0.8
0.9
600
800
1000
1200
Estimated allele frequency
Ecological gradient (Precipitation sums)

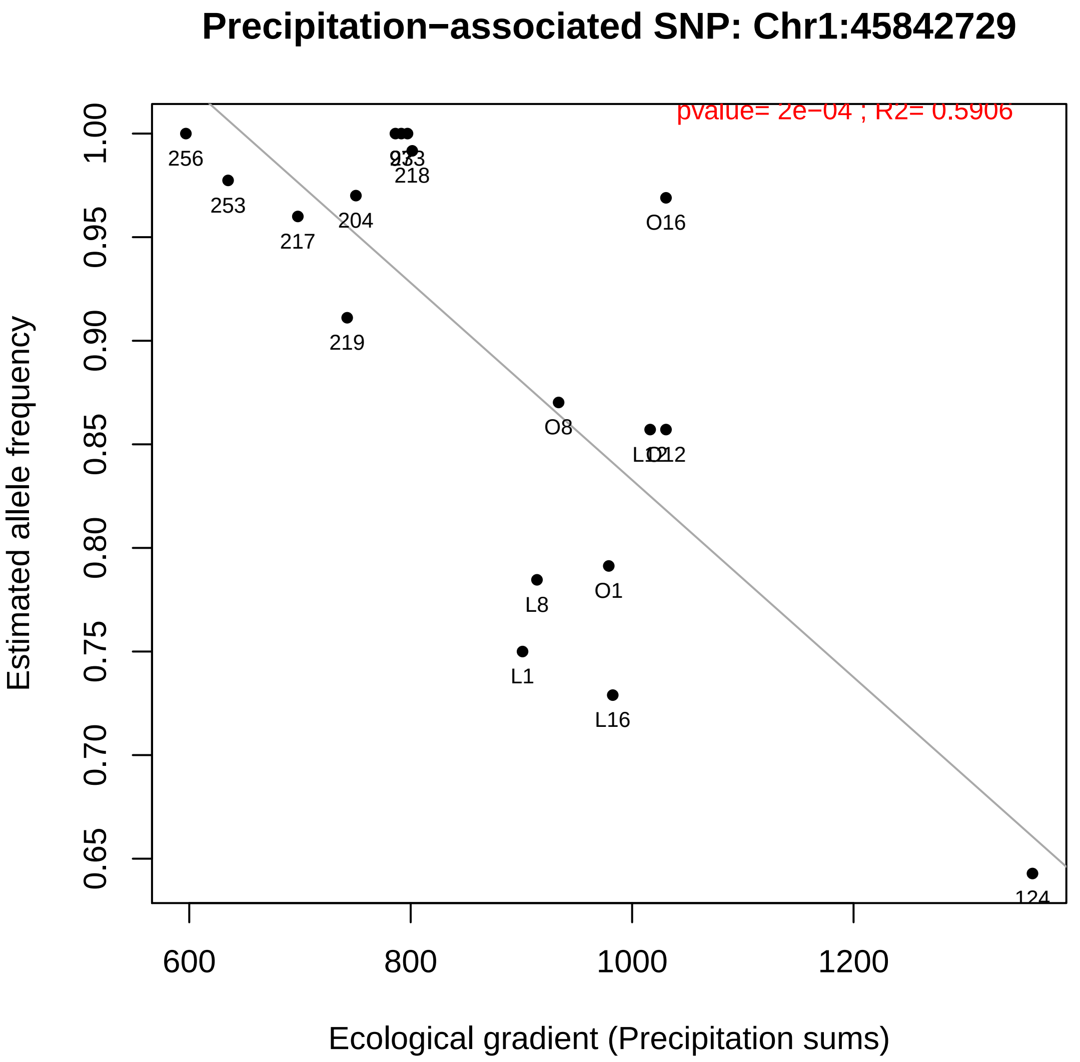

Precipitation−associated SNP: Chr1:45842729
pvalue= 2e−04 ; R2= 0.5906
Estimated allele frequency
Ecological gradient (Precipitation sums)
1.00
0.95
0.90
0.85
0.80
0.75
0.70
0.65
600
800
1000
1200
256
253
217
204
219
218
O16
O8
O12
O1
L8
L1
L16
124

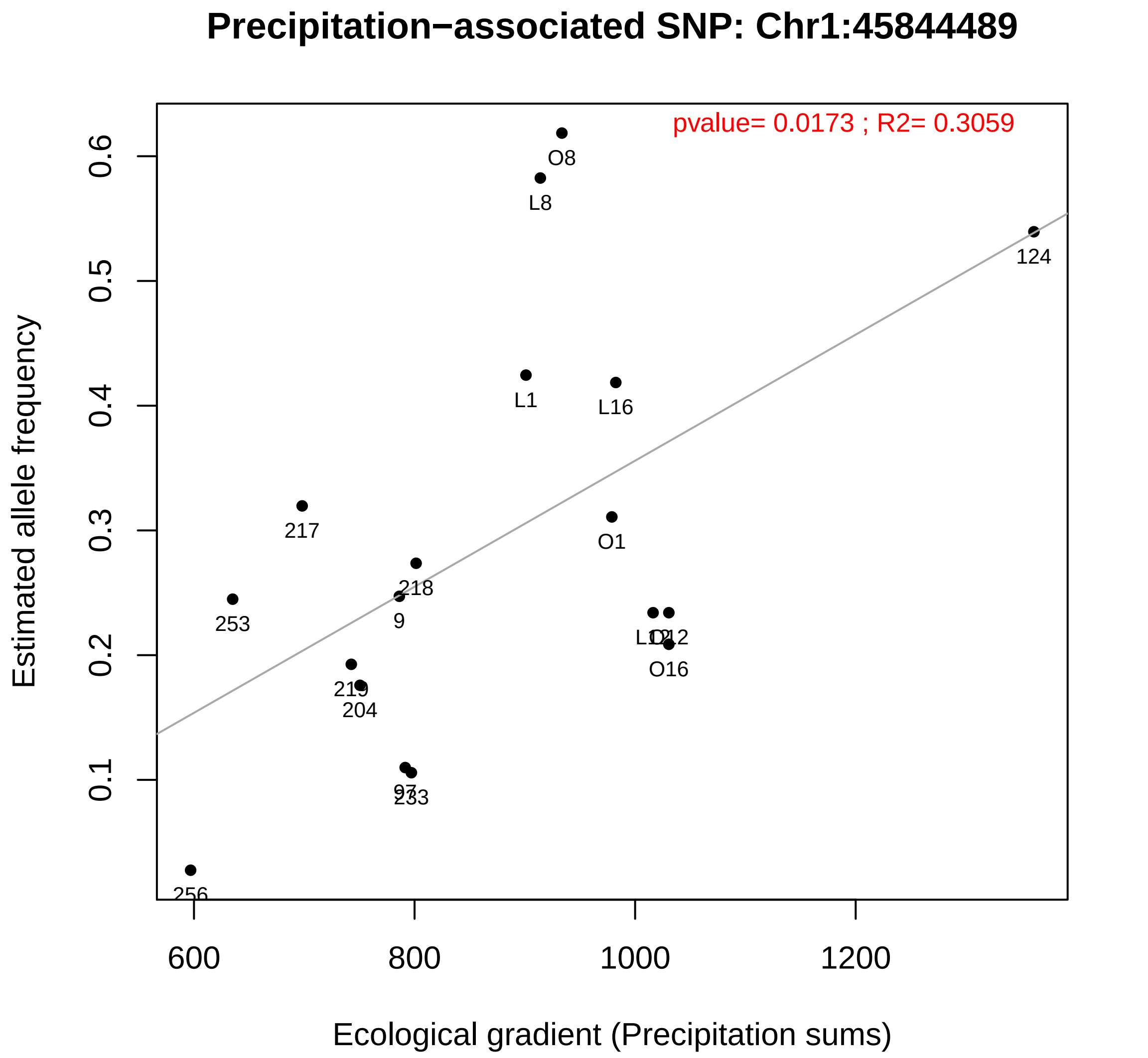

Precipitation−associated SNP: Chr1:45844489
pvalue= 0.0173 ; R2= 0.3059
Estimated allele frequency
Ecological gradient (Precipitation sums)
0.1
0.2
0.3
0.4
0.5
0.6
600
800
1000
1200
O8
L8
124
L1
L16
217
O1
218
253
9
O12
O16
219
204
97
233
256

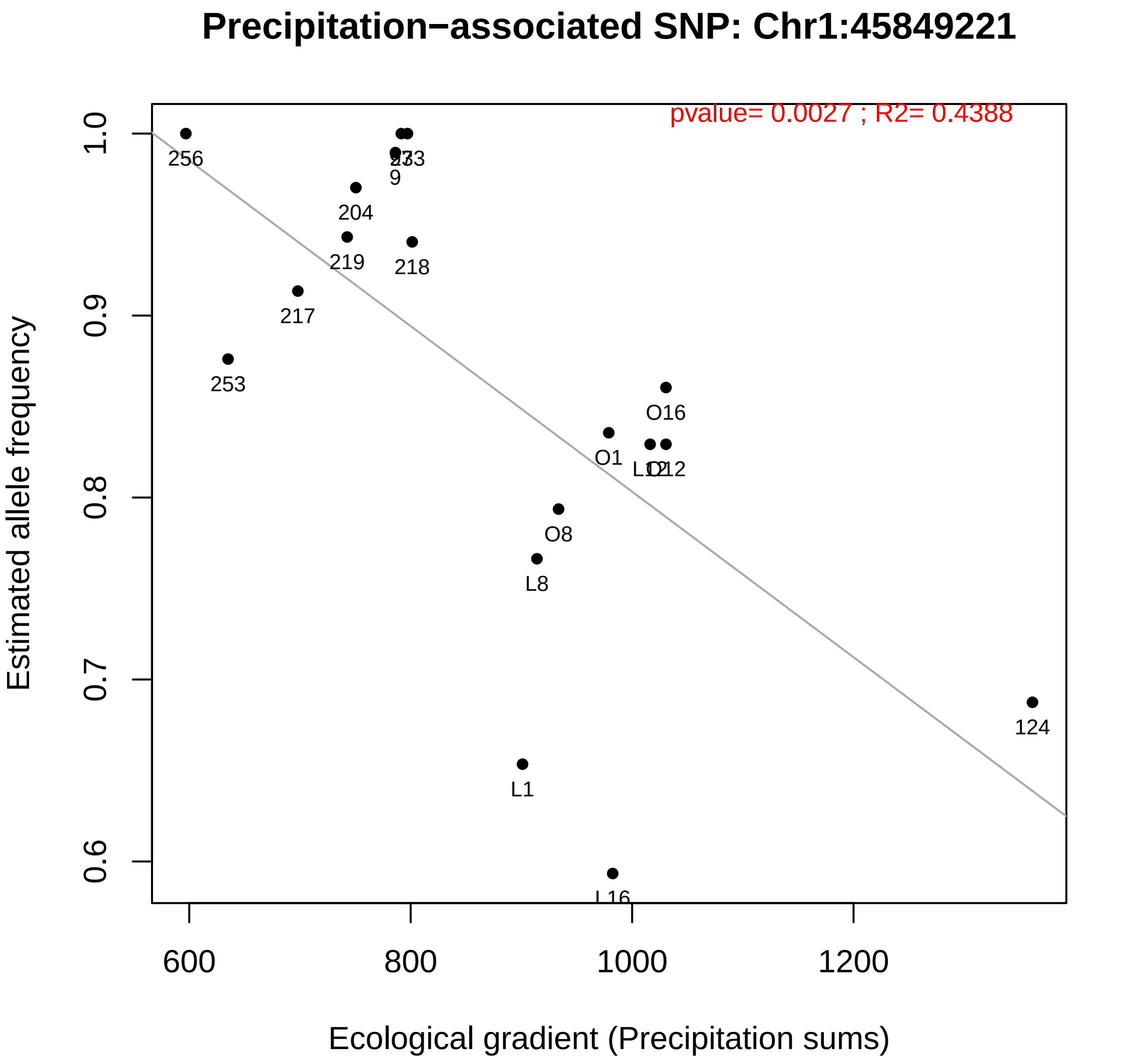

Precipitation−associated SNP: Chr1:45849221
pvalue= 0.0027 ; R2= 0.4388
Estimated allele frequency
Ecological gradient (Precipitation sums)
0.6
0.7
0.8
0.9
1.0
600
800
1000
1200
256
204
219
218
217
253
O16
O1
O8
L8
L1
124

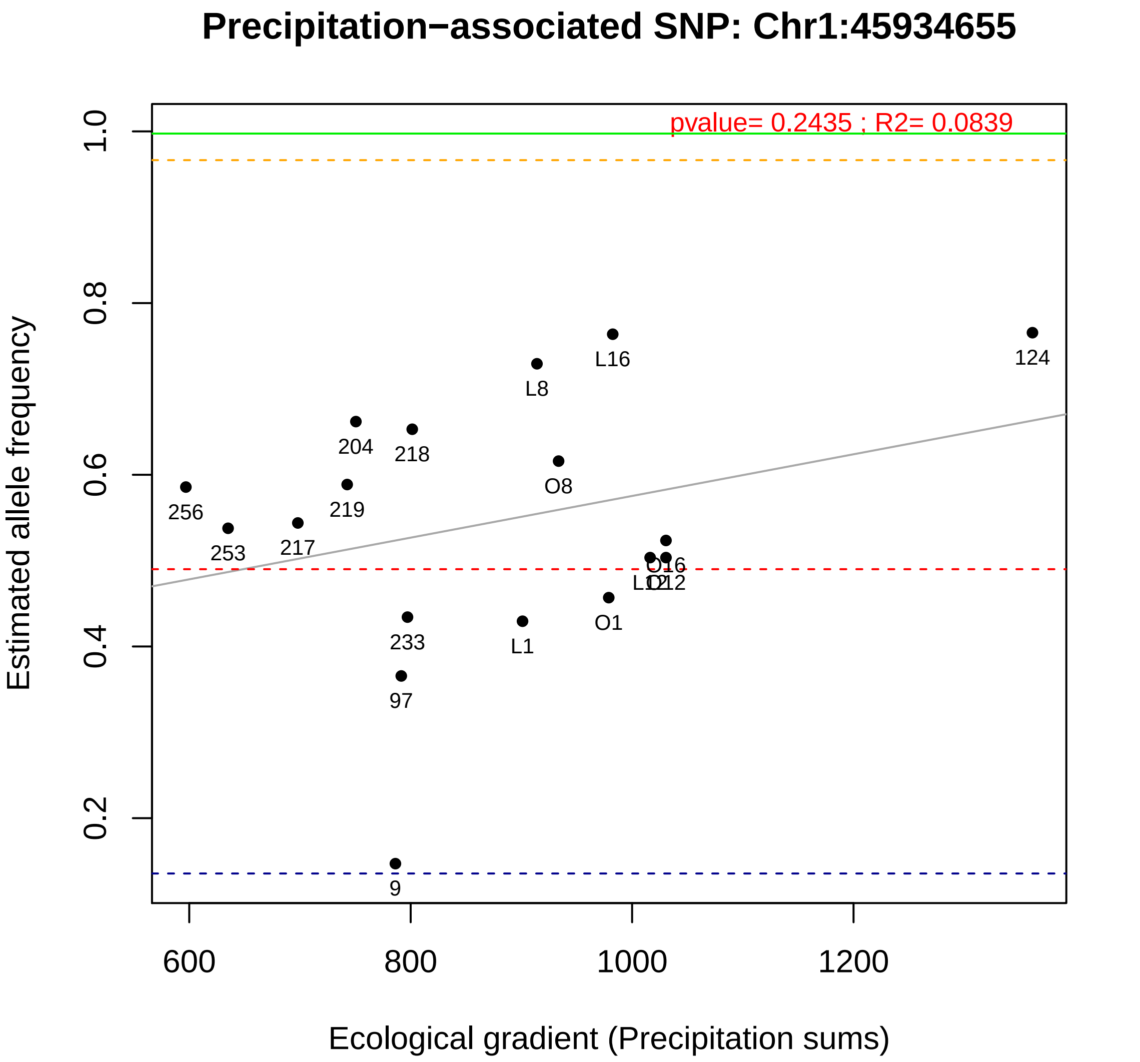
Precipitation−associated SNP: Chr1:45934655
pvalue= 0.2435 ; R2= 0.0839
Estimated allele frequency
Ecological gradient (Precipitation sums)
0.2
0.4
0.6
0.8
1.0
600
800
1000
1200
256
253
217
219
204
218
233
97
9
L1
L8
L16
O8
O1
O16
O12
124

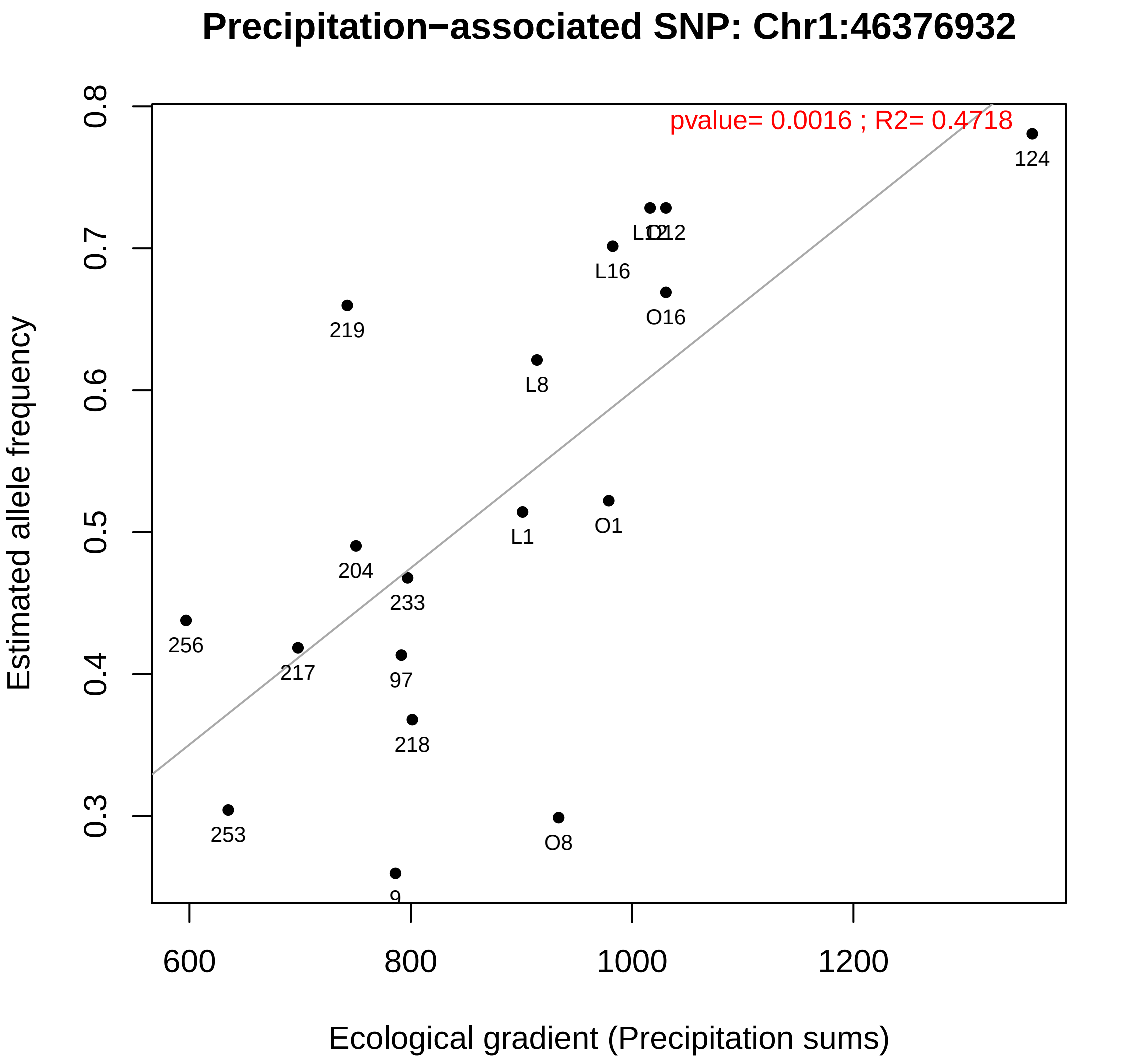
Precipitation−associated SNP: Chr1:46376932
pvalue= 0.0016 ; R2= 0.4718
Estimated allele frequency
Ecological gradient (Precipitation sums)
0.3
0.4
0.5
0.6
0.7
0.8
600
800
1000
1200
124
L12
O12
L16
O16
219
L8
L1
O1
204
233
256
217
97
218
253
O8
9

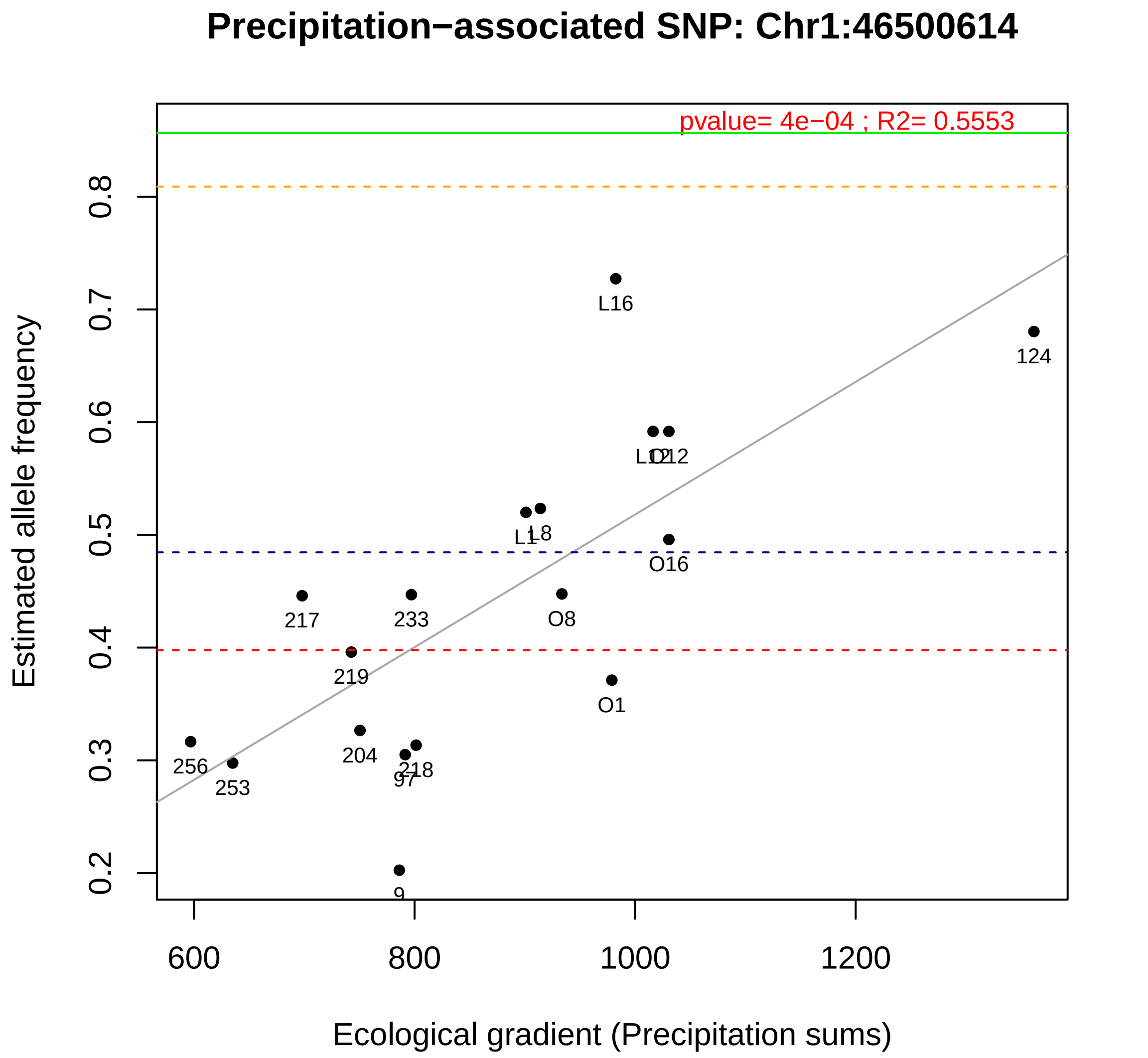

Precipitation−associated SNP: Chr1:46500614
pvalue= 4e−04 ; R2= 0.5553
Estimated allele frequency
0.2
0.3
0.4
0.5
0.6
0.7
0.8
600
800
1000
1200
Ecological gradient (Precipitation sums)
L16
124
L12
O12
L1
L8
O16
217
233
O8
219
O1
256
253
204
218
97
9

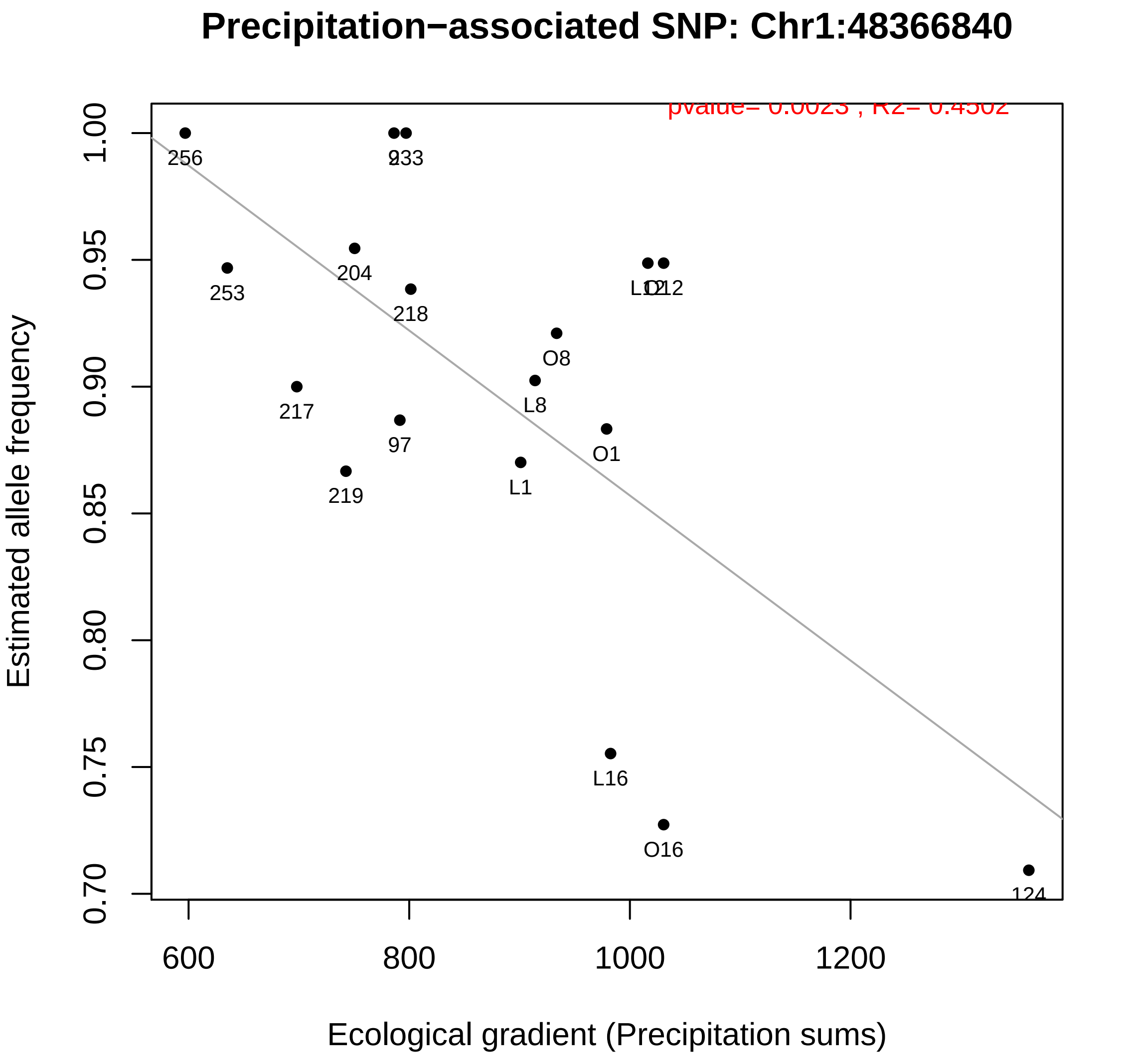

Precipitation−associated SNP: Chr1:48366840
pvalue= 0.0023 , R2= 0.4502
Estimated allele frequency
Ecological gradient (Precipitation sums)
0.70
0.75
0.80
0.85
0.90
0.95
1.00
600
800
1000
1200
256
233
253
204
218
O12
O8
217
L8
97
O1
219
L1
L16
O16
124

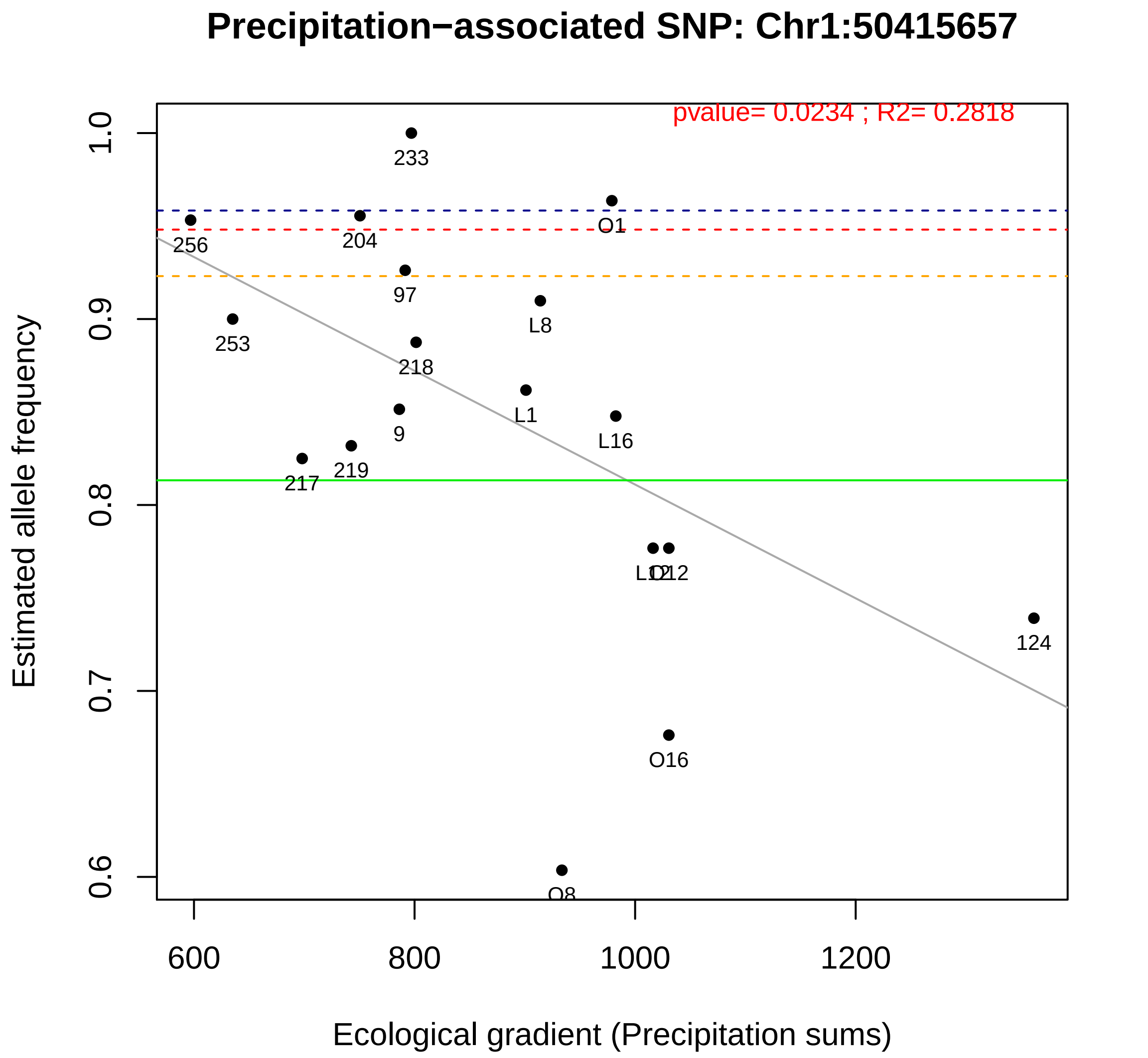

Precipitation−associated SNP: Chr1:50415657
pvalue= 0.0234 ; R2= 0.2818
Estimated allele frequency
Ecological gradient (Precipitation sums)
0.6
0.7
0.8
0.9
1.0
600
800
1000
1200
233
O1
256
204
97
L8
253
218
L1
9
L16
219
217
L12
O12
124
O16
O8

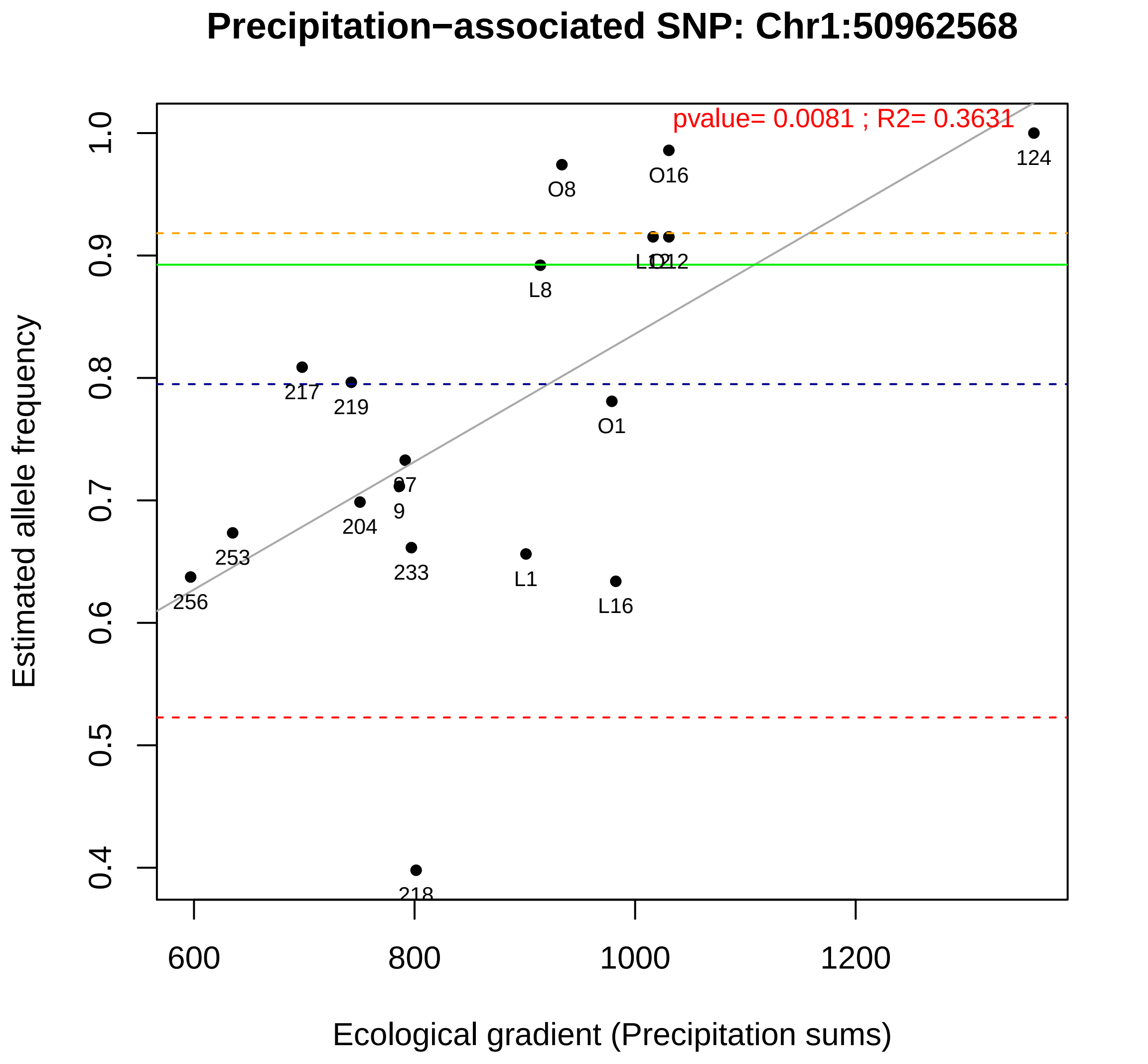

Precipitation−associated SNP: Chr1:50962568
pvalue= 0.0081 ; R2= 0.3631
Estimated allele frequency
Ecological gradient (Precipitation sums)
0.4
0.5
0.6
0.7
0.8
0.9
1.0
600
800
1000
1200
124
O16
O8
L12
O12
L8
217
219
O1
97
9
204
253
233
L1
L16
256
218

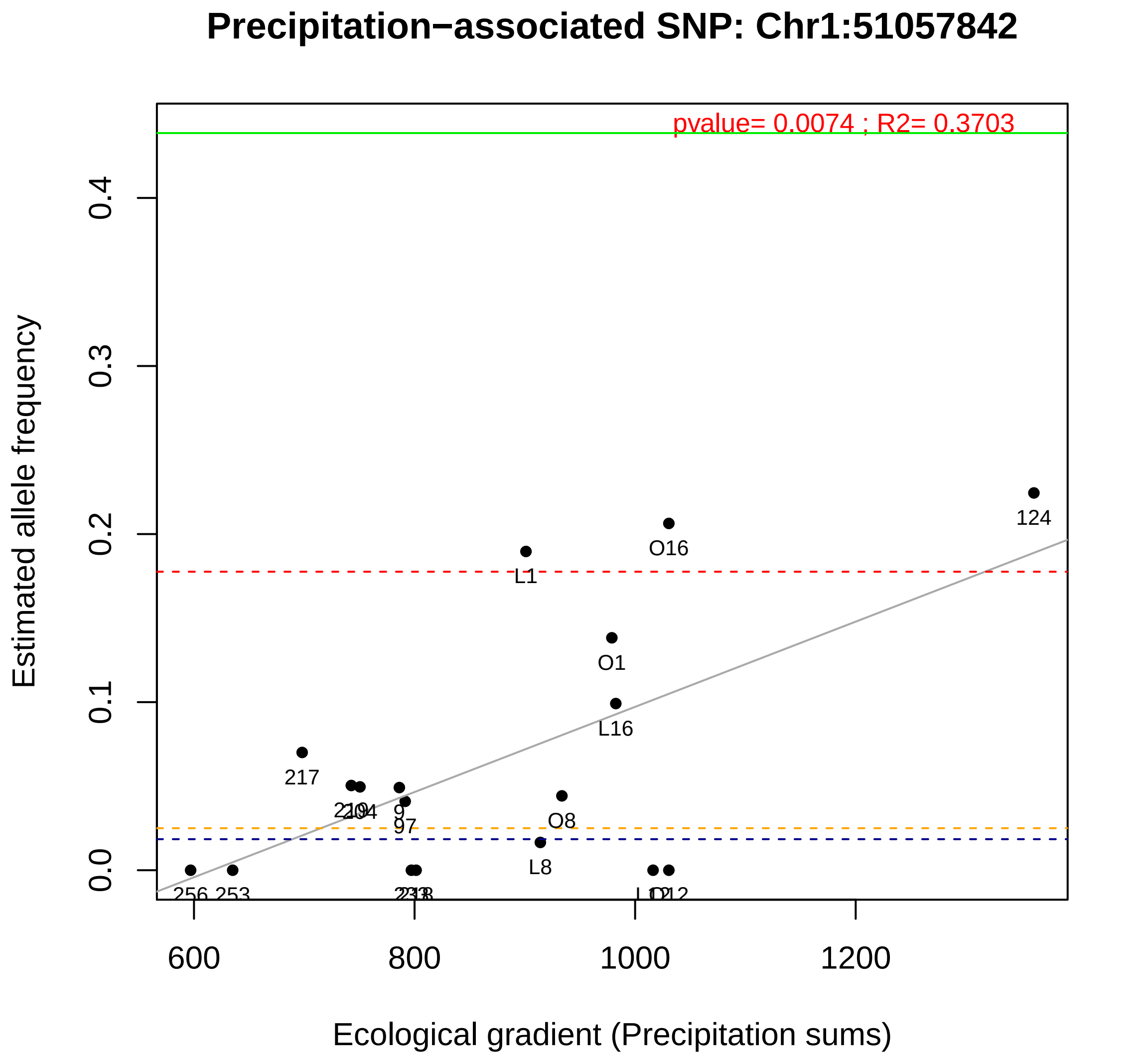

Precipitation−associated SNP: Chr1:51057842
pvalue= 0.0074 ; R2= 0.3703
Estimated allele frequency
0.0
0.1
0.2
0.3
0.4
600
800
1000
1200
Ecological gradient (Precipitation sums)
124
O16
L1
O1
L16
217
9
97
O8
L8
256
253

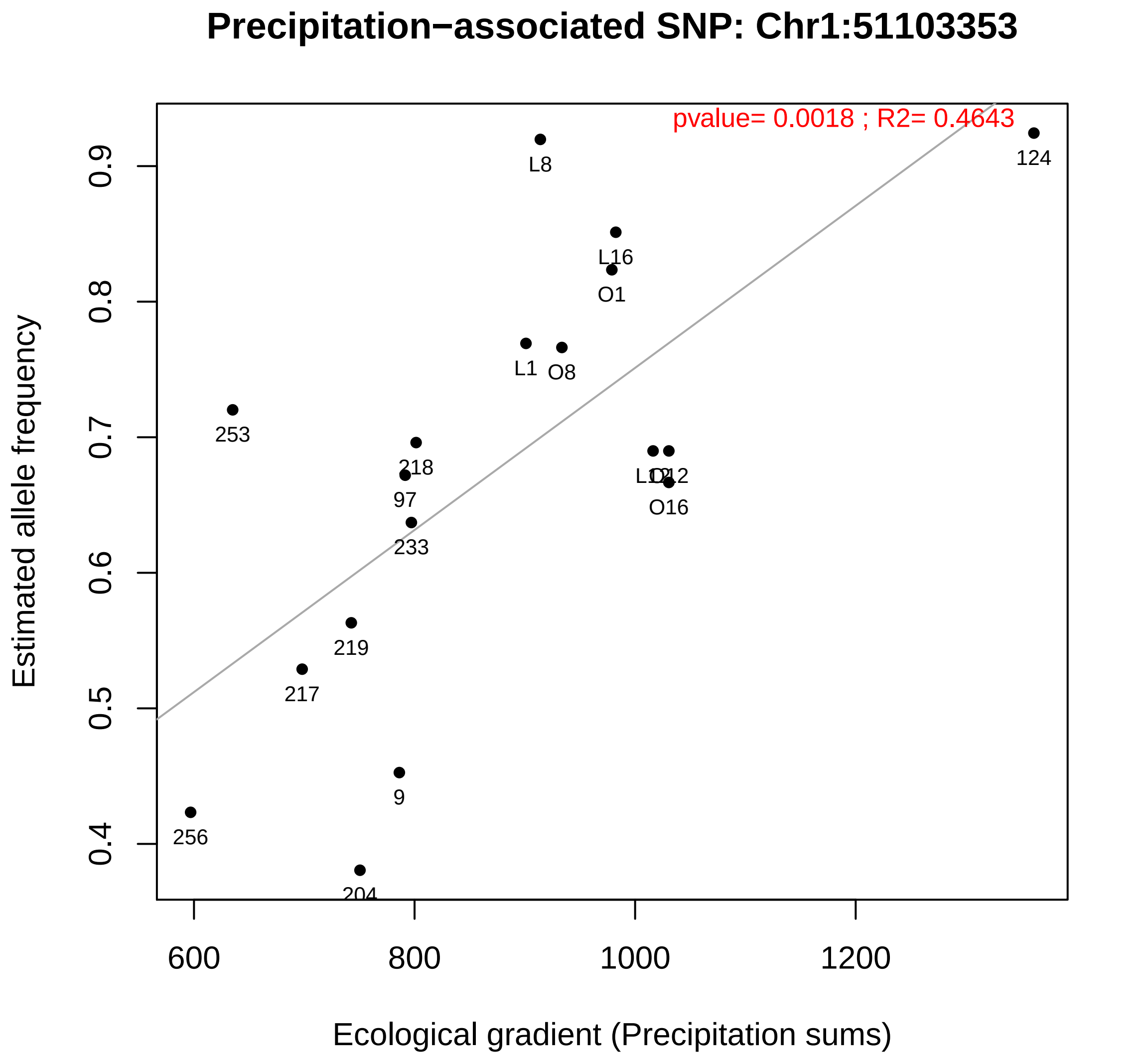

Precipitation−associated SNP: Chr1:51103353
pvalue= 0.0018 ; R2= 0.4643
Estimated allele frequency
Ecological gradient (Precipitation sums)
0.4
0.5
0.6
0.7
0.8
0.9
600
800
1000
1200
L8
124
L16
O1
L1
O8
253
218
97
233
O16
219
217
9
256
204

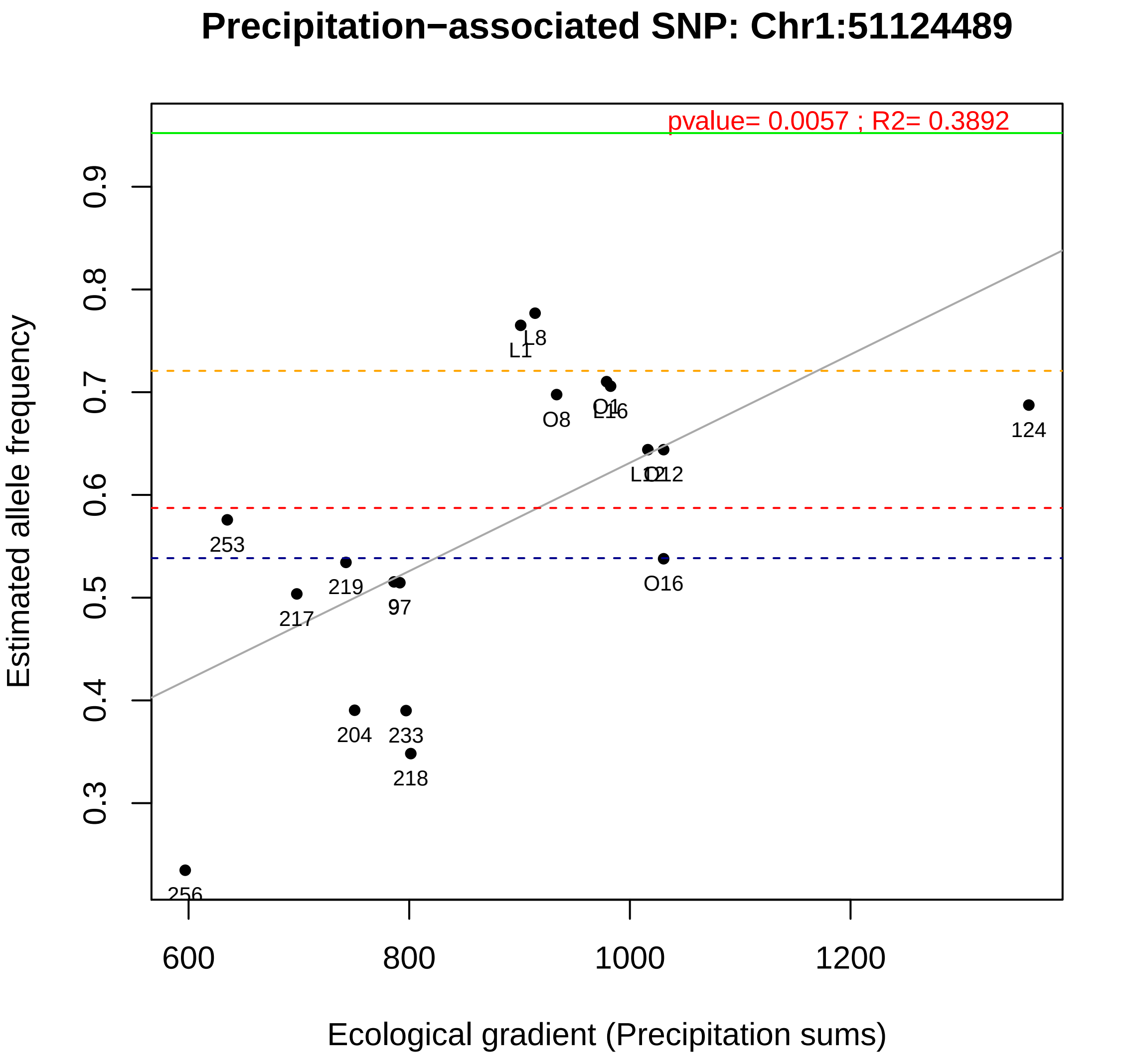
Precipitation−associated SNP: Chr1:51124489
pvalue= 0.0057 ; R2= 0.3892
Estimated allele frequency
0.3
0.4
0.5
0.6
0.7
0.8
0.9
600
800
1000
1200
Ecological gradient (Precipitation sums)
L8
L1
O8
124
253
219
217
97
O16
204
233
218
256

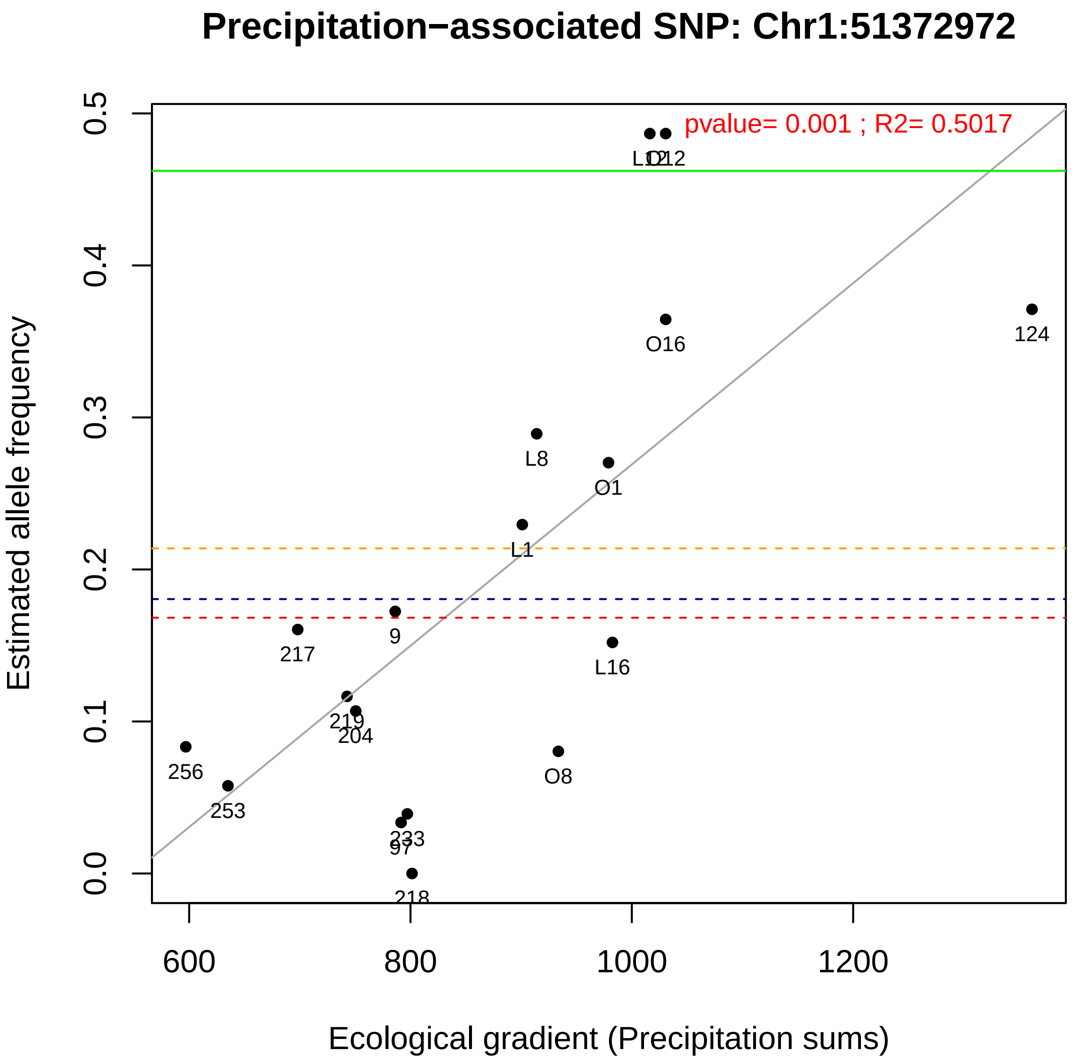

Precipitation−associated SNP: Chr1:51372972
pvalue= 0.001 ; R2= 0.5017
Estimated allele frequency
Ecological gradient (Precipitation sums)
0.0
0.1
0.2
0.3
0.4
0.5
600
800
1000
1200
L12
O12
O16
124
L8
O1
L1
9
217
L16
219
204
256
253
O8
233
97
218

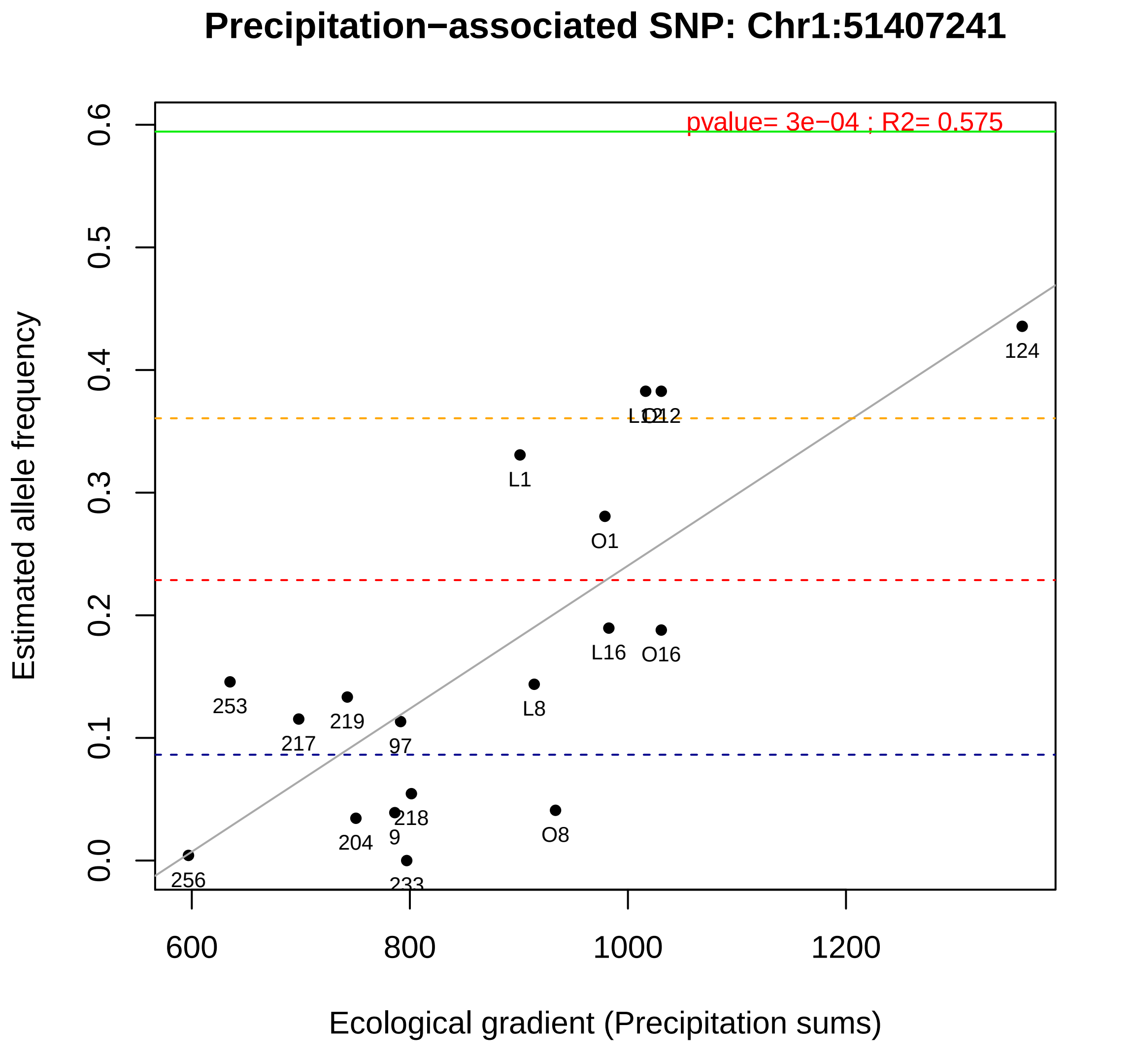

Precipitation−associated SNP: Chr1:51407241
pvalue= 3e−04 ; R2= 0.575
Estimated allele frequency
0.0
0.1
0.2
0.3
0.4
0.5
0.6
124
L1
O1
L16
O16
L8
253
217
219
97
218
204
9
O8
256
233
600
800
1000
1200
Ecological gradient (Precipitation sums)

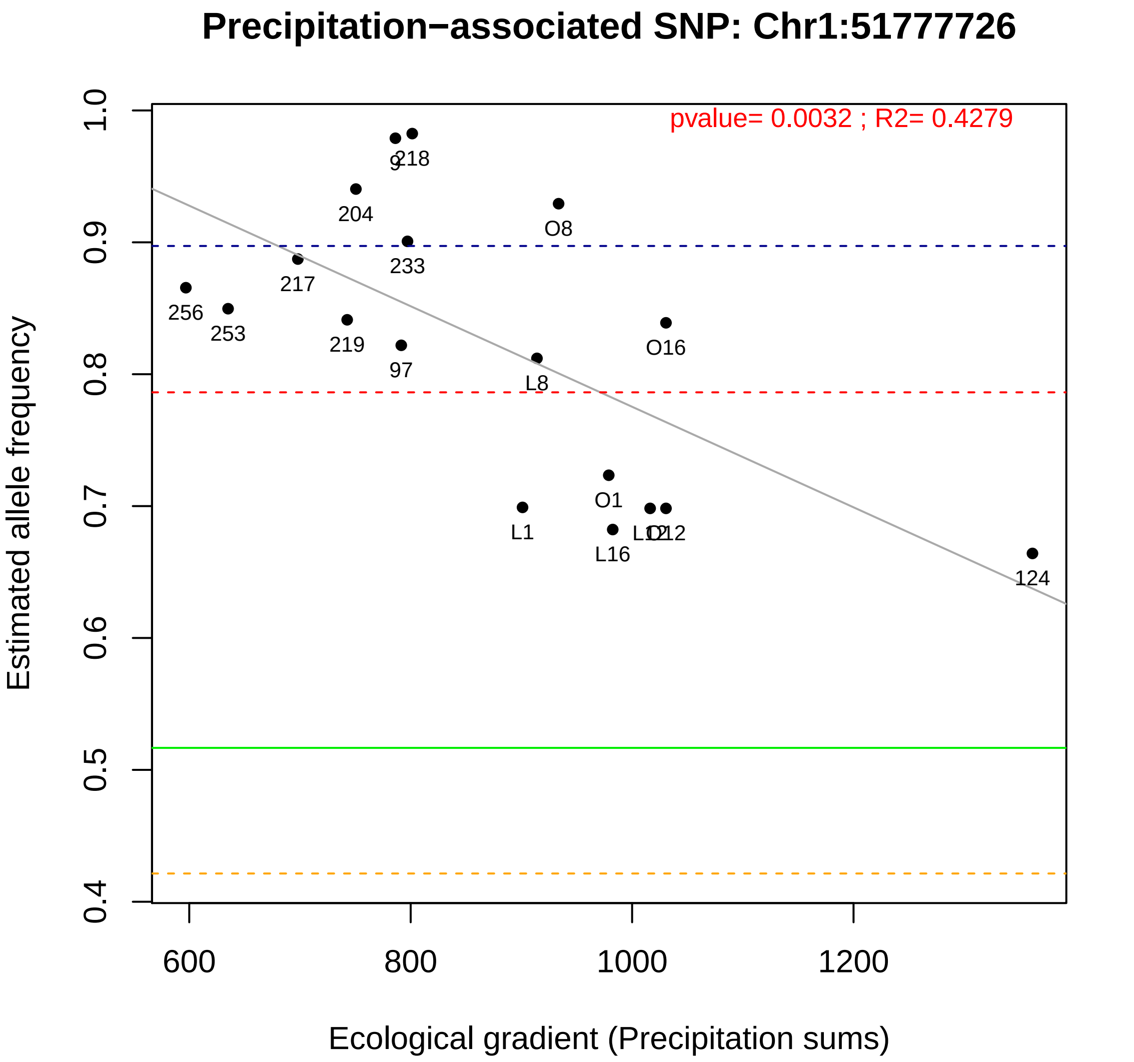

Precipitation−associated SNP: Chr1:51777726
pvalue= 0.0032 ; R2= 0.4279
Estimated allele frequency
Ecological gradient (Precipitation sums)
1.0
0.9
0.8
0.7
0.6
0.5
0.4
600
800
1000
1200
9
218
204
O8
233
217
256
253
219
97
O16
L8
O1
L1
L16
L12
O12
124

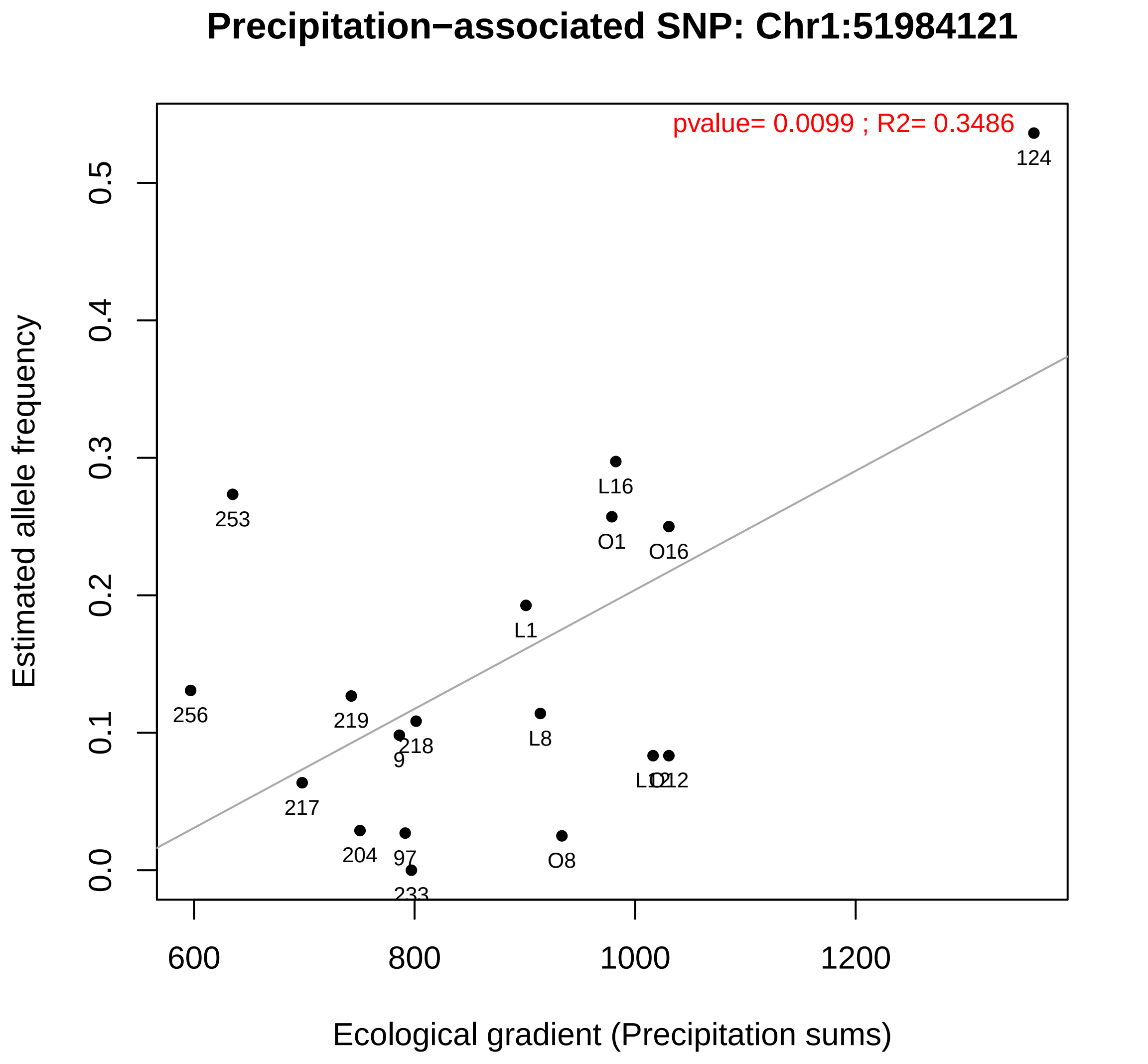
Precipitation−associated SNP: Chr1:51984121
pvalue= 0.0099 ; R2= 0.3486
Estimated allele frequency
0.0
0.1
0.2
0.3
0.4
0.5
600
800
1000
1200
Ecological gradient (Precipitation sums)
124
L16
253
O1
O16
L1
256
219
218
9
L8
L12
O12
217
204
97
233
O8

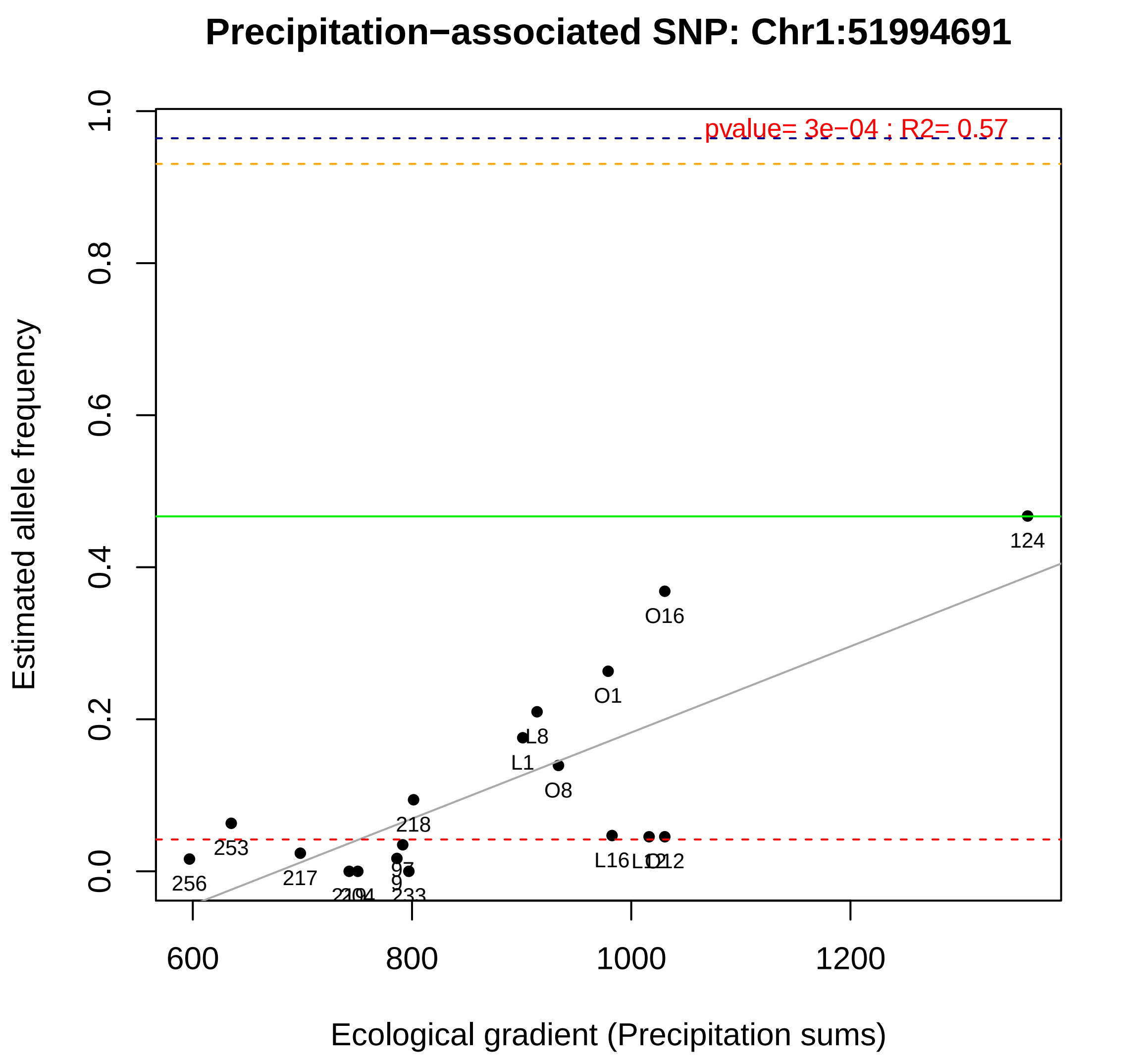

Precipitation−associated SNP: Chr1:51994691
pvalue= 3e−04 ; R2= 0.57
Estimated allele frequency
0.0
0.2
0.4
0.6
0.8
1.0
600
800
1000
1200
Ecological gradient (Precipitation sums)
124
O16
O1
L8
L1
O8
218
253
256
217
L16
97
9
233

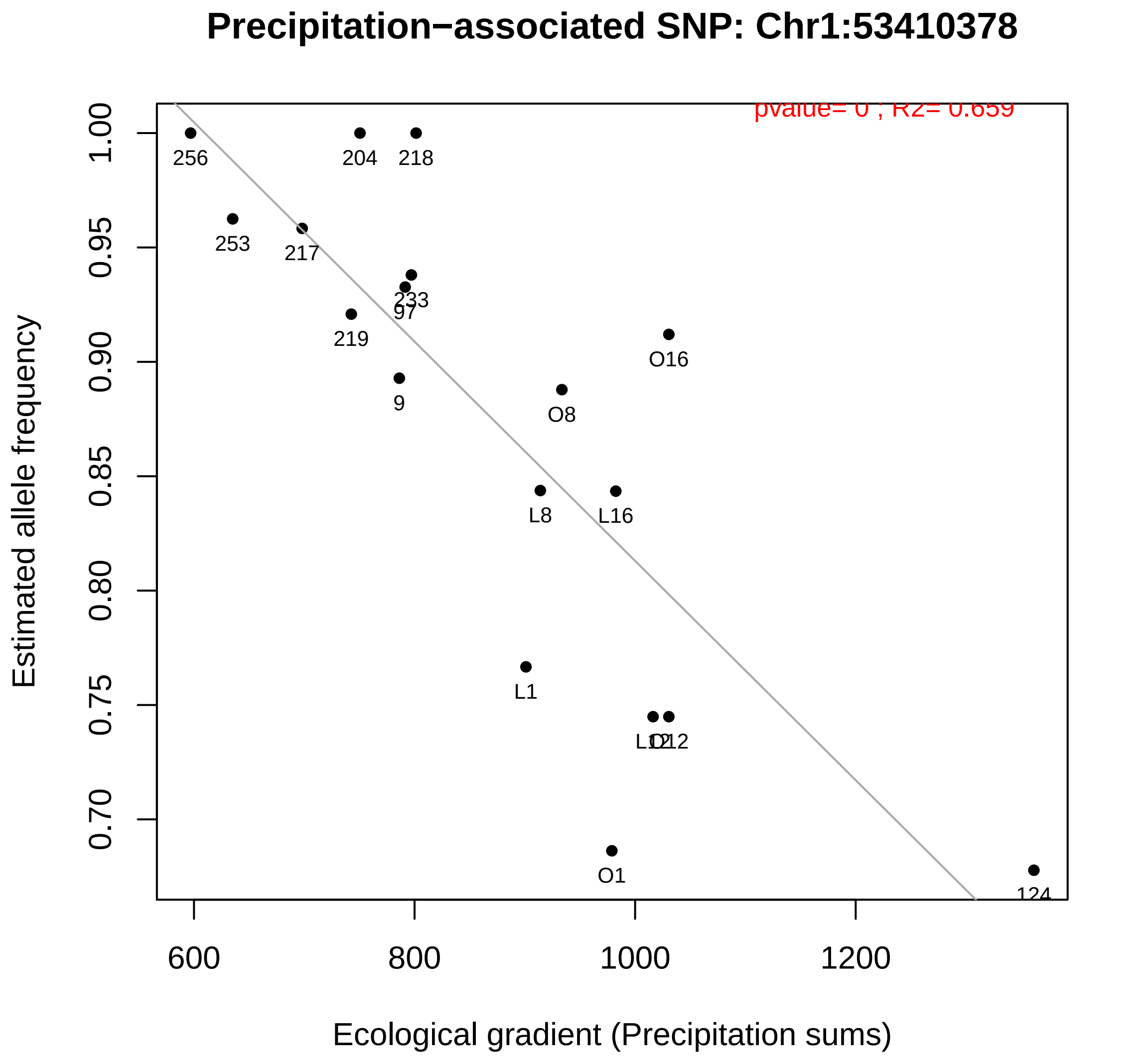

Precipitation−associated SNP: Chr1:53410378
pvalue= 0 ; R2= 0.659
Estimated allele frequency
Ecological gradient (Precipitation sums)
1.00
0.95
0.90
0.85
0.80
0.75
0.70
600
800
1000
1200
256
204
218
253
217
233
97
219
9
O16
O8
L8
L16
L1
L12
O12
O1
124

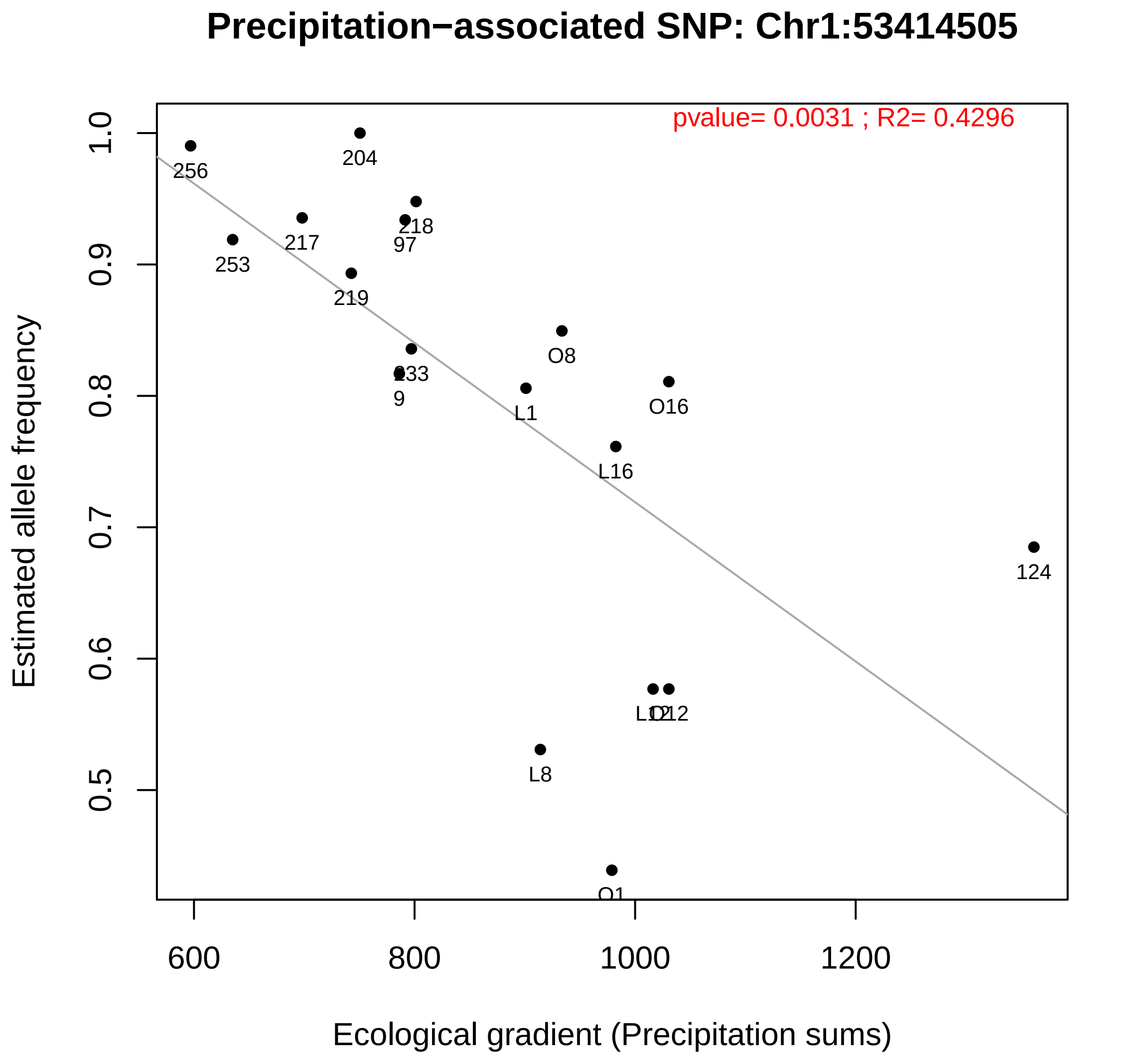

Precipitation−associated SNP: Chr1:53414505
pvalue= 0.0031 ; R2= 0.4296
Estimated allele frequency
Ecological gradient (Precipitation sums)
1.0
0.9
0.8
0.7
0.6
0.5
600
800
1000
1200
256
204
253
217
218
97
219
233
9
O8
L1
O16
L16
124
L12
O12
L8
O1

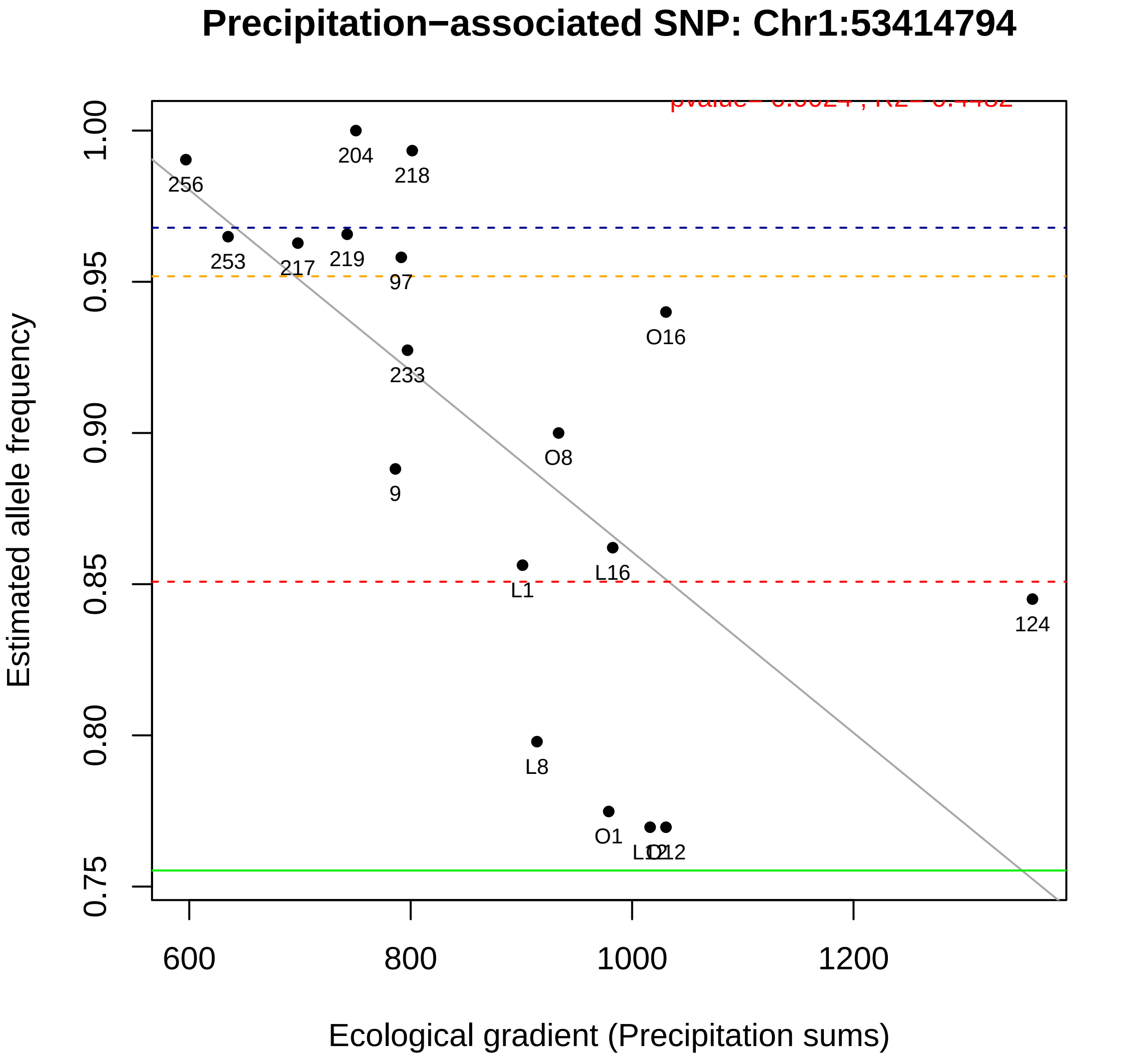
Precipitation−associated SNP: Chr1:53414794
Estimated allele frequency
Ecological gradient (Precipitation sums)
1.00
0.95
0.90
0.85
0.80
0.75
600
800
1000
1200
256
204
218
253
217
219
97
O16
233
O8
9
L1
L16
124
L8
O1
L12
O12

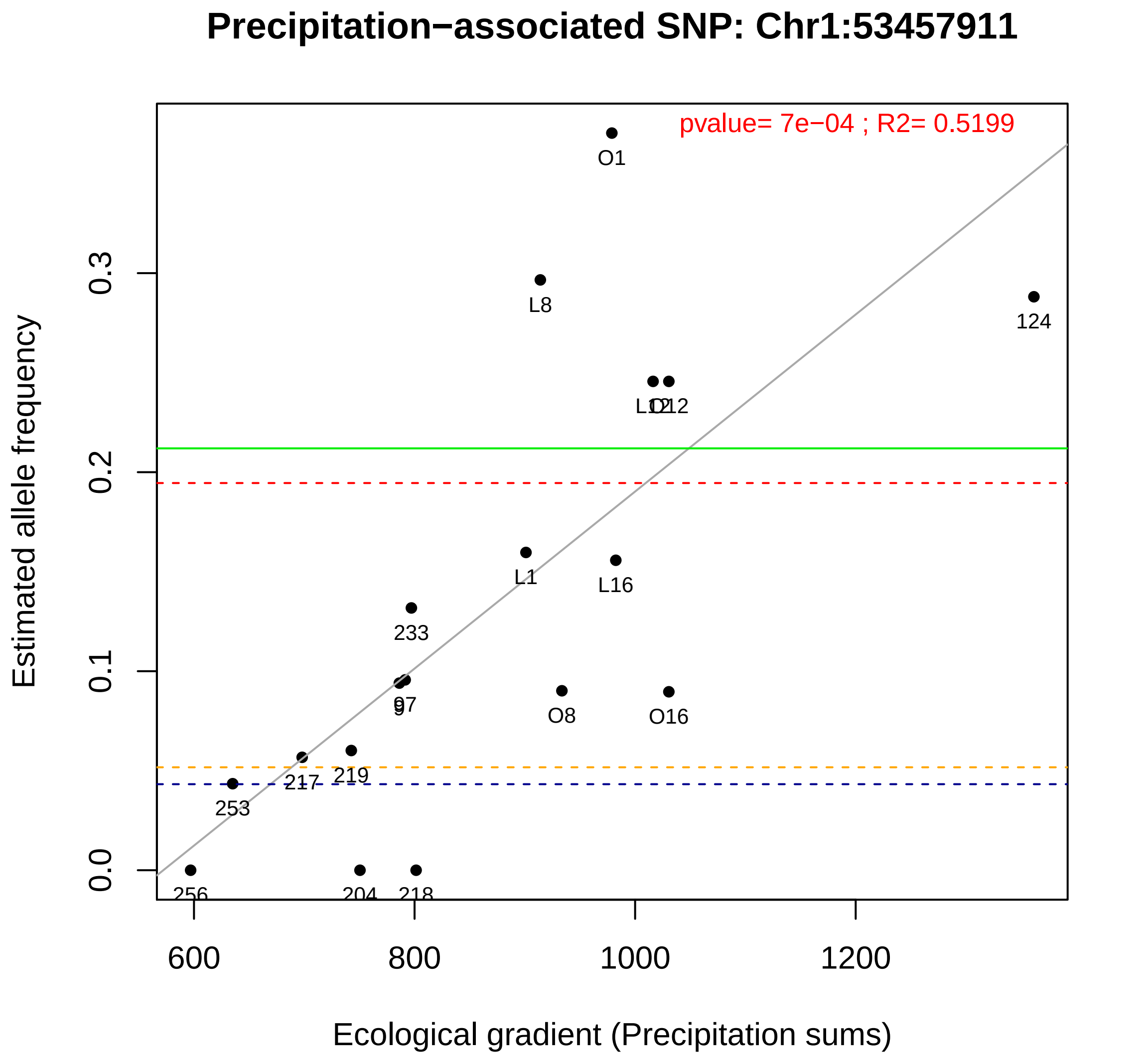

Precipitation−associated SNP: Chr1:53457911
pvalue= 7e−04 ; R2= 0.5199
O1
L8
124
L10
O12
L1
L16
233
97
9
O8
O16
219
217
253
256
204
218
0.0
0.1
0.2
0.3
600
800
1000
1200
Estimated allele frequency
Ecological gradient (Precipitation sums)

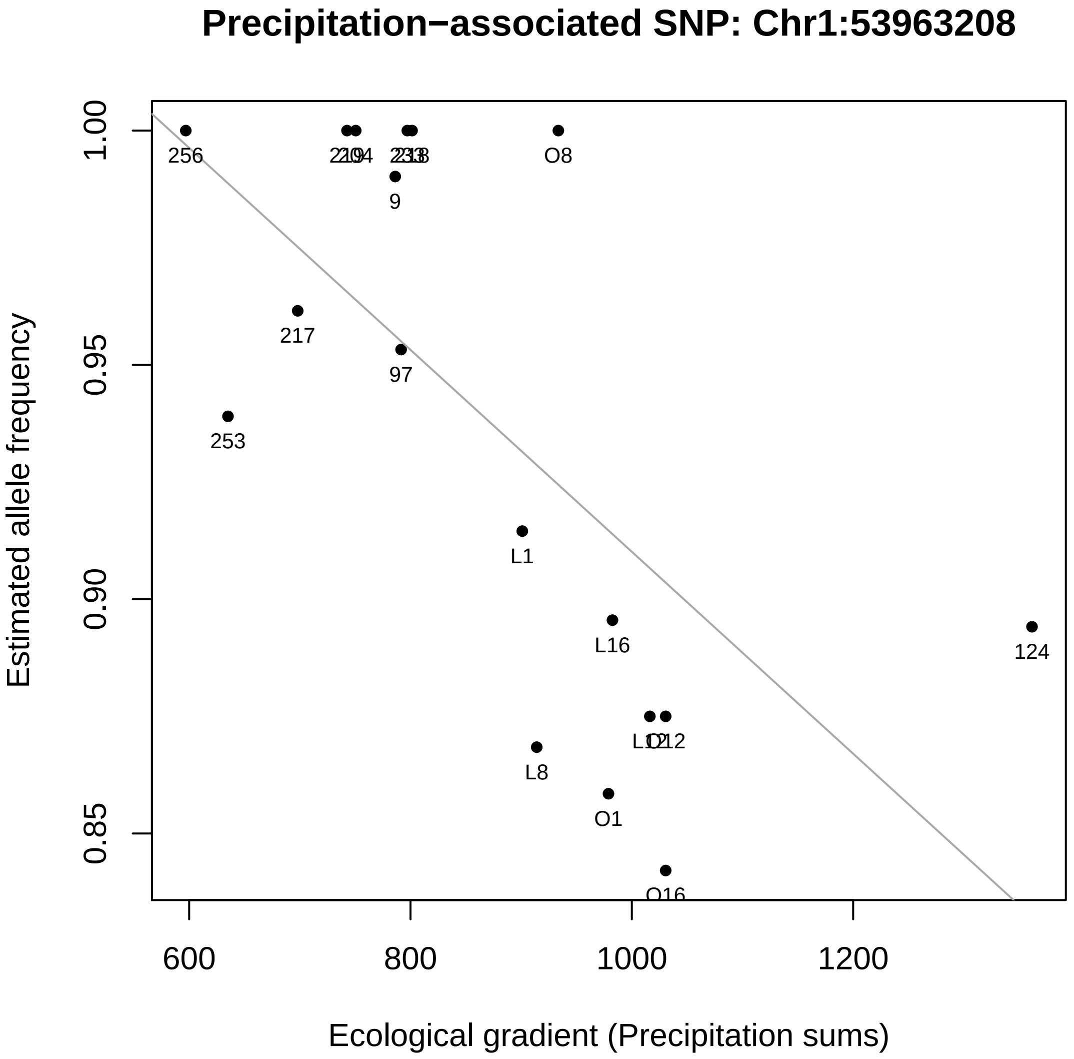
Precipitation−associated SNP: Chr1:53963208
Estimated allele frequency
Ecological gradient (Precipitation sums)
1.00
0.95
0.90
0.85
600
800
1000
1200
256
O8
9
217
97
253
L1
L16
124
L8
O1
O16

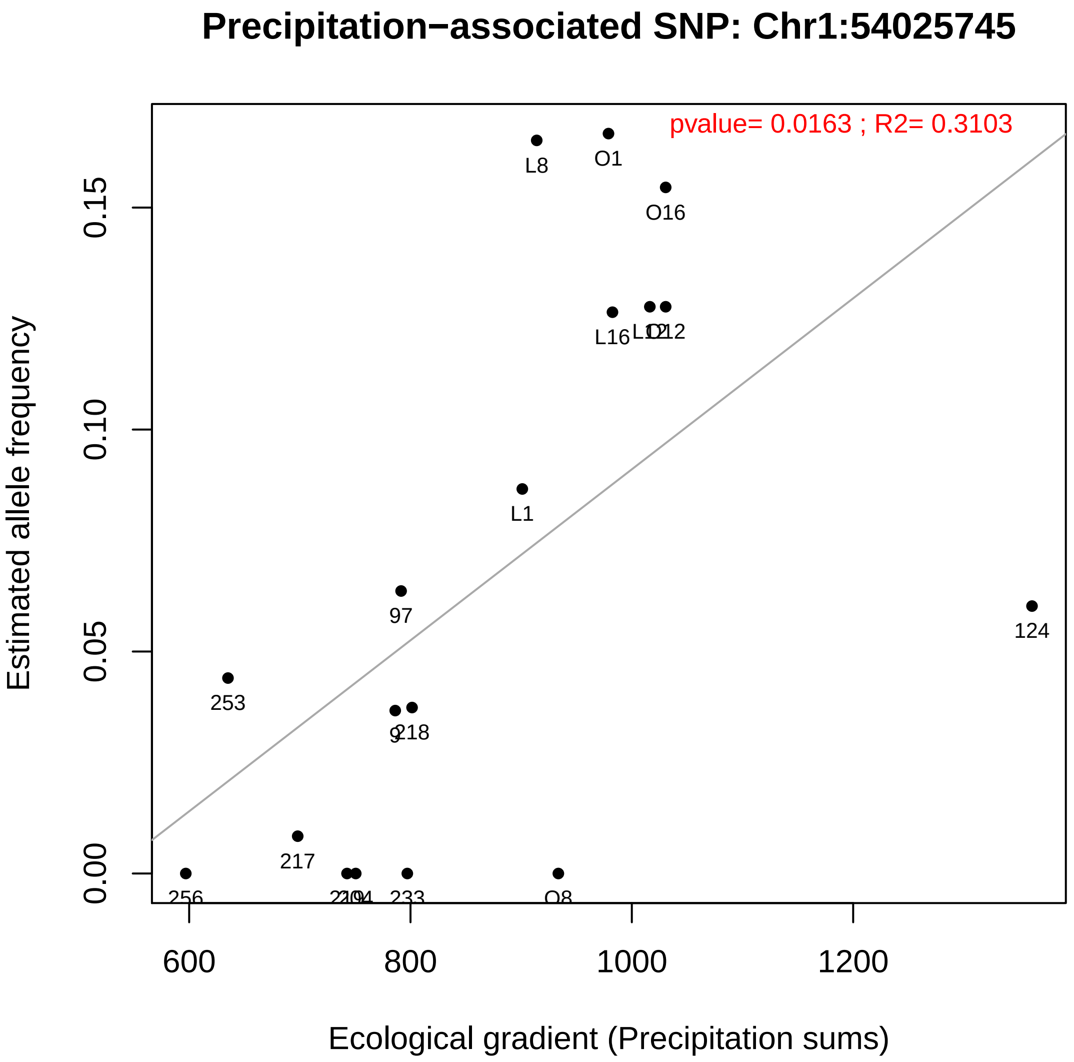

Precipitation−associated SNP: Chr1:54025745
pvalue= 0.0163 ; R2= 0.3103
Estimated allele frequency
Ecological gradient (Precipitation sums)
0.00
0.05
0.10
0.15
600
800
1000
1200
L8
O1
O16
L16
O12
L1
97
124
253
218
217
233
O8

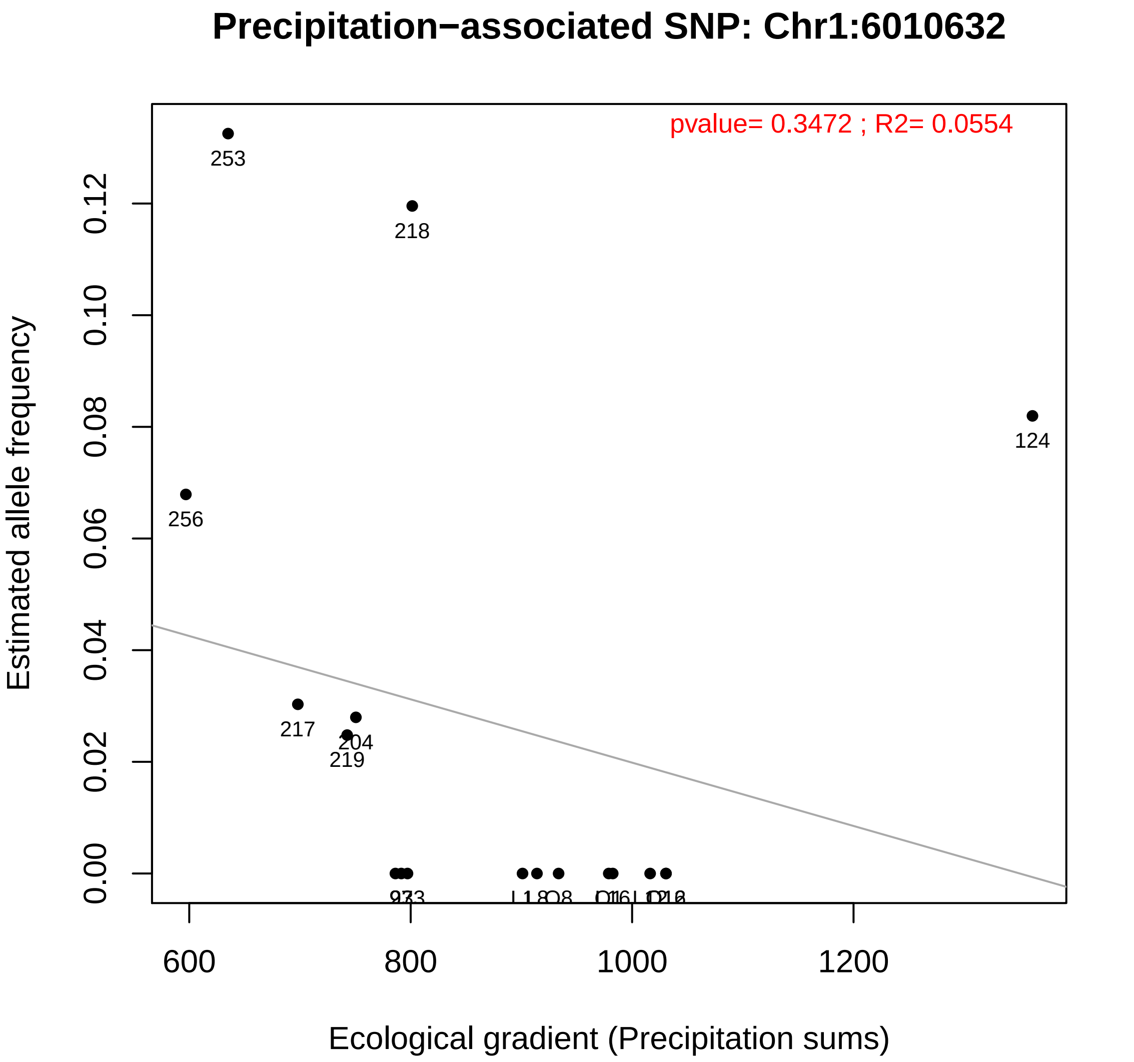
Precipitation−associated SNP: Chr1:6010632
pvalue= 0.3472 ; R2= 0.0554
253
218
124
256
217
204
219
Estimated allele frequency
0.00
0.02
0.04
0.06
0.08
0.10
0.12
600
800
1000
1200
Ecological gradient (Precipitation sums)

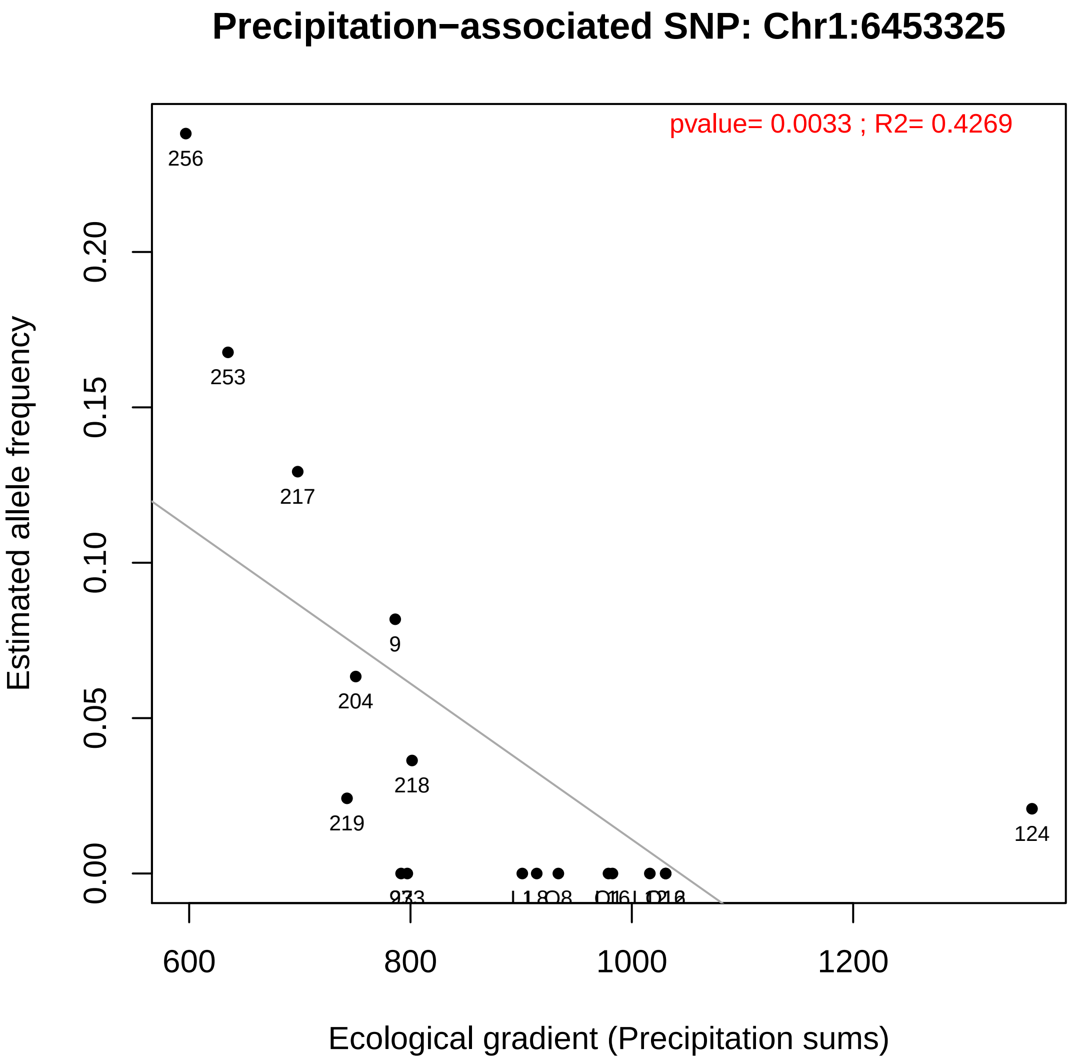
Precipitation−associated SNP: Chr1:6453325
pvalue= 0.0033 ; R2= 0.4269
Estimated allele frequency
Ecological gradient (Precipitation sums)
0.00
0.05
0.10
0.15
0.20
600
800
1000
1200
256
253
217
9
204
218
219
124

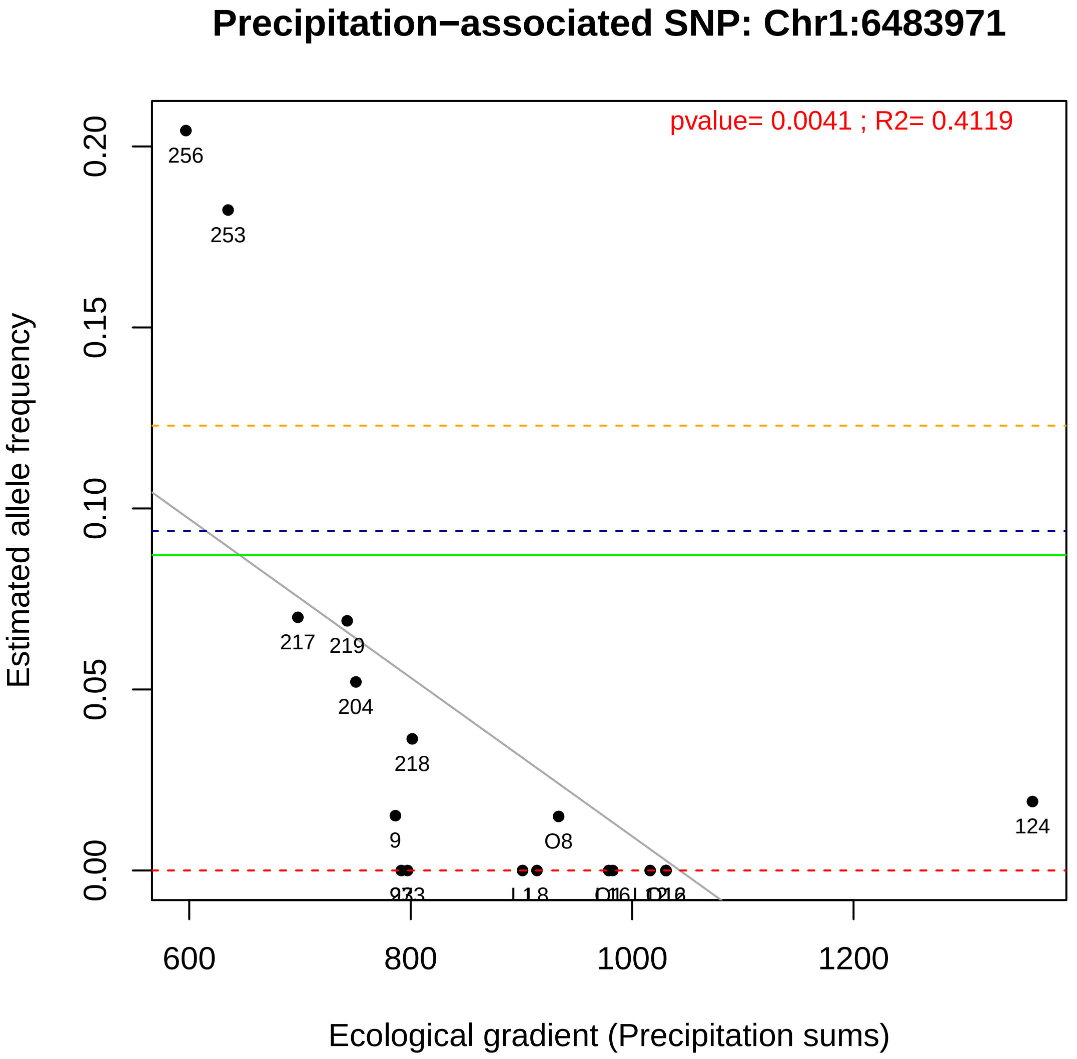
Precipitation−associated SNP: Chr1:6483971
pvalue= 0.0041 ; R2= 0.4119
Estimated allele frequency
0.00
0.05
0.10
0.15
0.20
Ecological gradient (Precipitation sums)
600
800
1000
1200
256
253
217
219
204
218
9
O8
124

**Precipitation−associated SNP: Chr1:6484629**

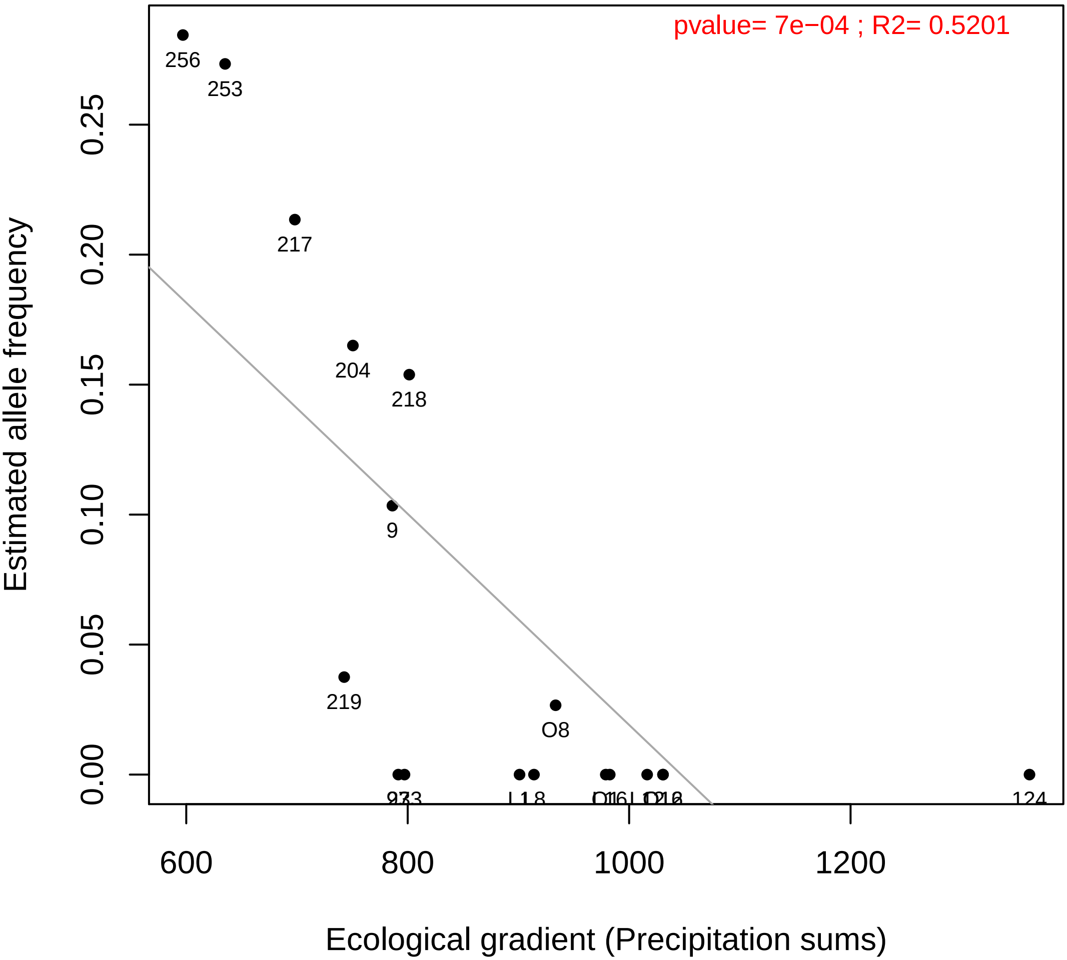

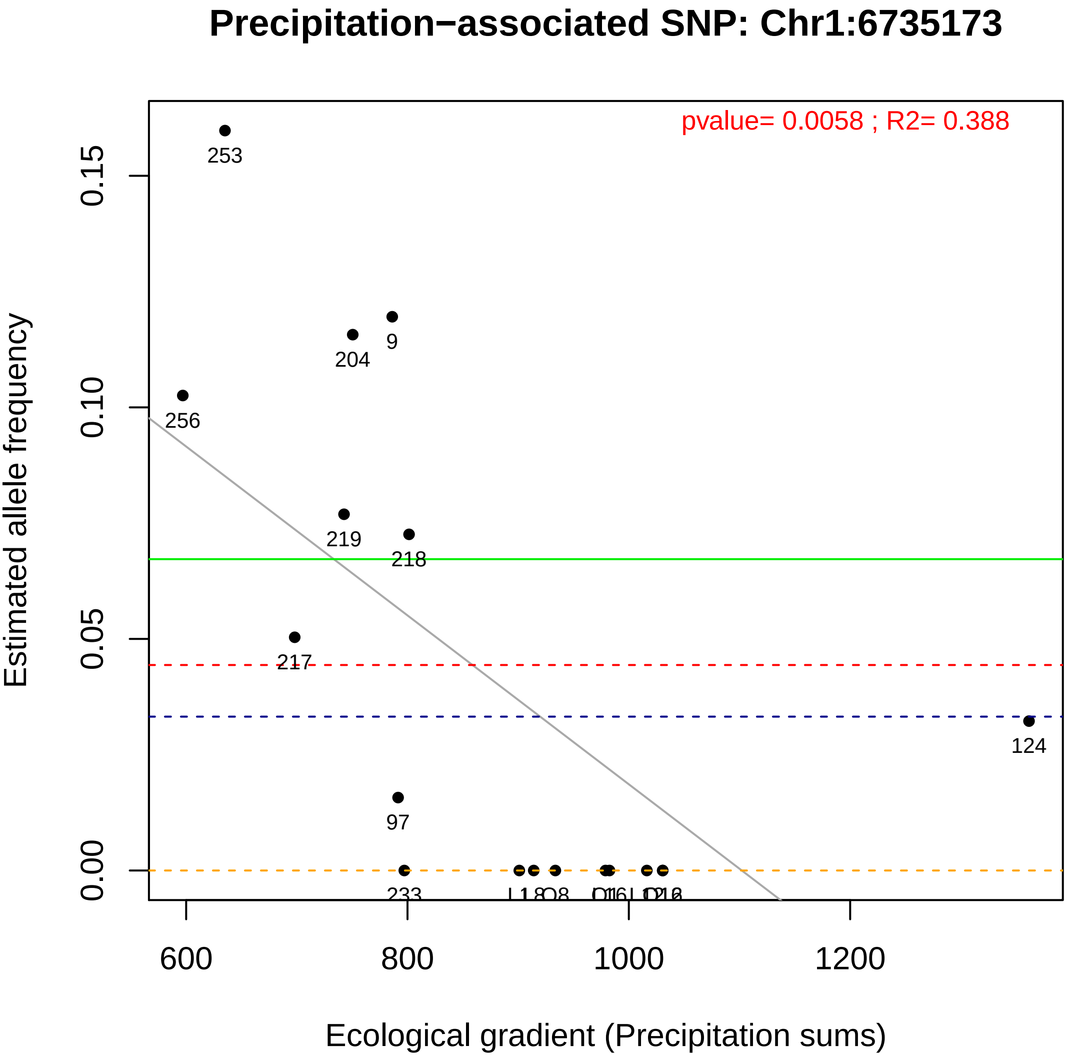

Precipitation−associated SNP: Chr1:6735173
pvalue= 0.0058 ; R2= 0.388
Estimated allele frequency
0.00
0.05
0.10
0.15
Ecological gradient (Precipitation sums)
600
800
1000
1200
253
256
204
9
219
218
217
124
97
233

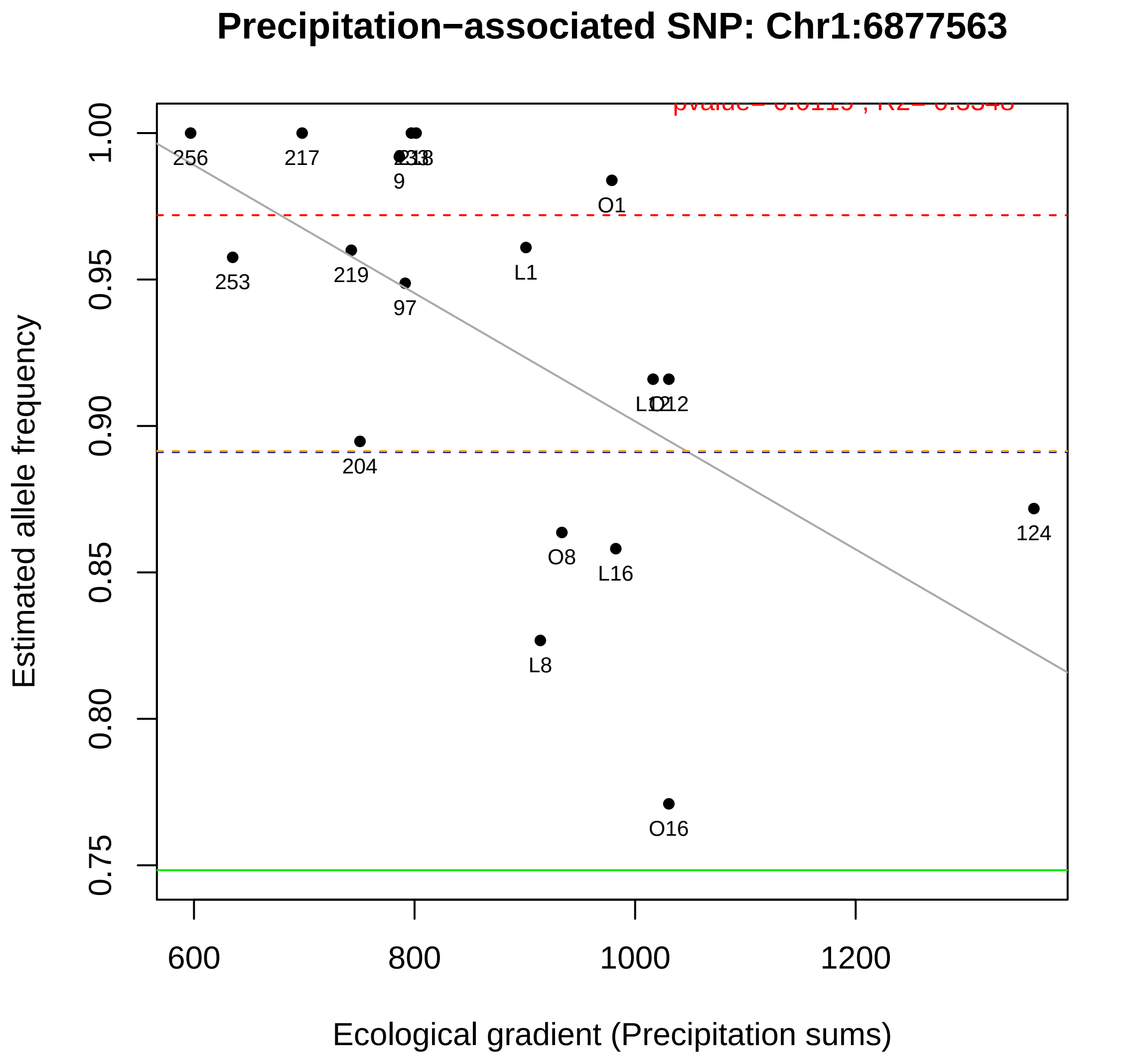

Precipitation−associated SNP: Chr1:6877563
pvalue=
Estimated allele frequency
Ecological gradient (Precipitation sums)
1.00
0.95
0.90
0.85
0.80
0.75
600
800
1000
1200
256
217
9
O1
253
219
L1
97
O12
204
124
O8
L16
L8
O16

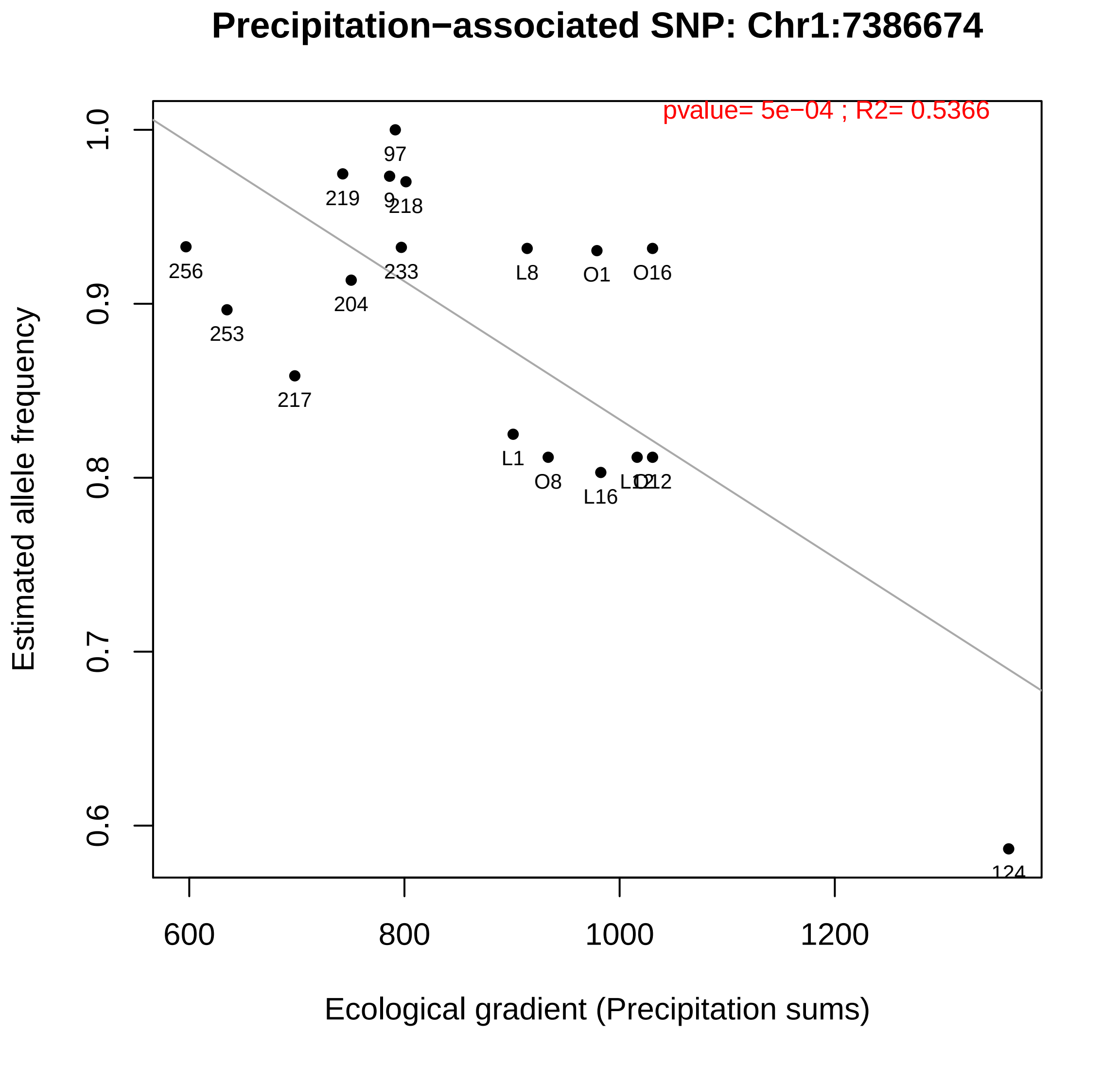

Precipitation−associated SNP: Chr1:7386674
pvalue= 5e−04 ; R2= 0.5366
Estimated allele frequency
Ecological gradient (Precipitation sums)
0.6
0.7
0.8
0.9
1.0
600
800
1000
1200
97
219
9
218
256
233
L8
O1
O16
204
253
217
L1
O8
L16
O12
124

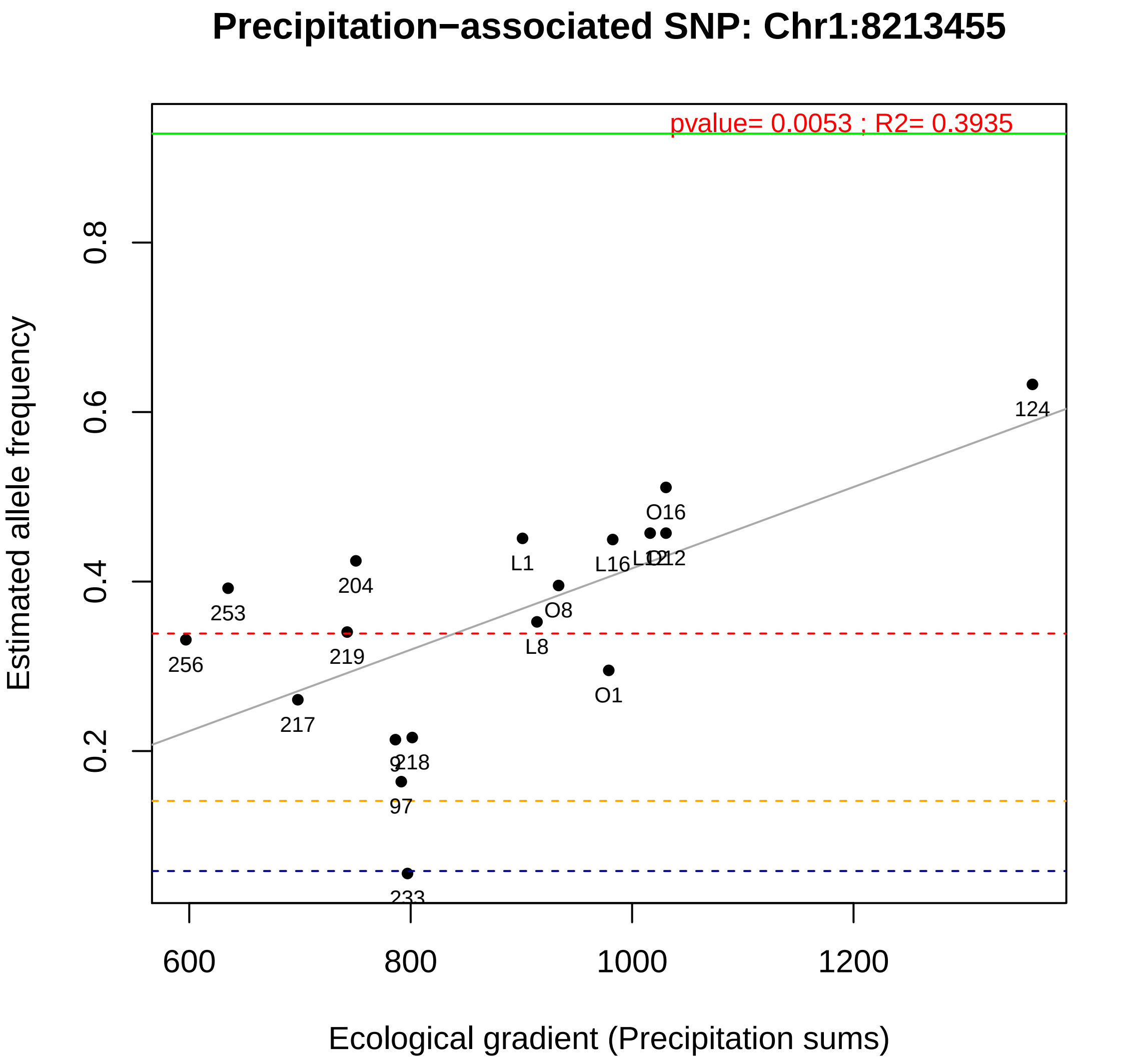
Precipitation−associated SNP: Chr1:8213455
pvalue= 0.0053 ; R2= 0.3935
Estimated allele frequency
0.2
0.4
0.6
0.8
600
800
1000
1200
Ecological gradient (Precipitation sums)
124
O16
L1
L16
O12
204
253
O8
L8
256
219
O1
217
218
97
233

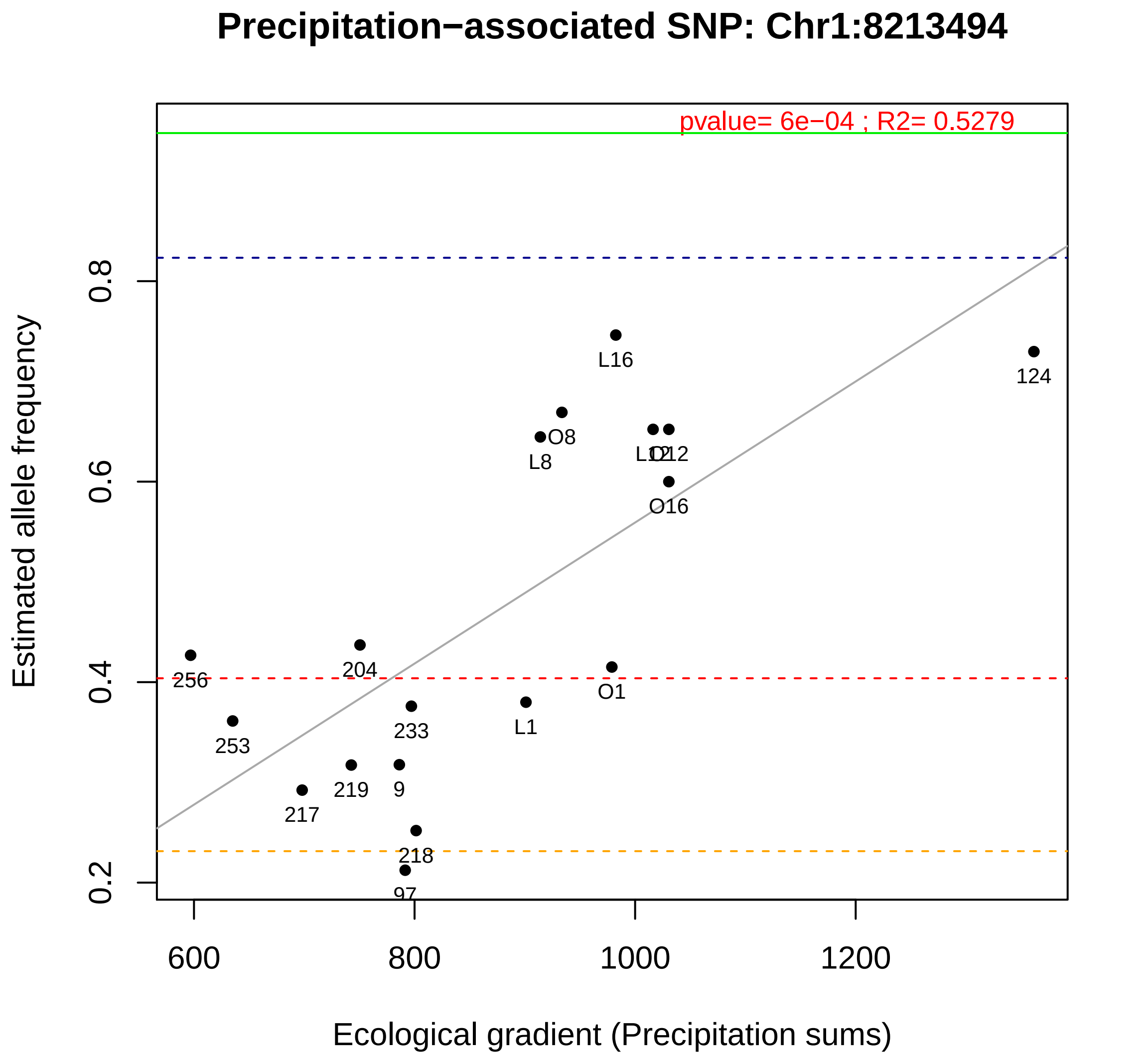
Precipitation−associated SNP: Chr1:8213494
pvalue= 6e−04 ; R2= 0.5279
Estimated allele frequency
0.2
0.4
0.6
0.8
600
800
1000
1200
Ecological gradient (Precipitation sums)
L16
124
O8
L8
O12
O16
256
204
O1
253
233
L1
219
9
217
218
97

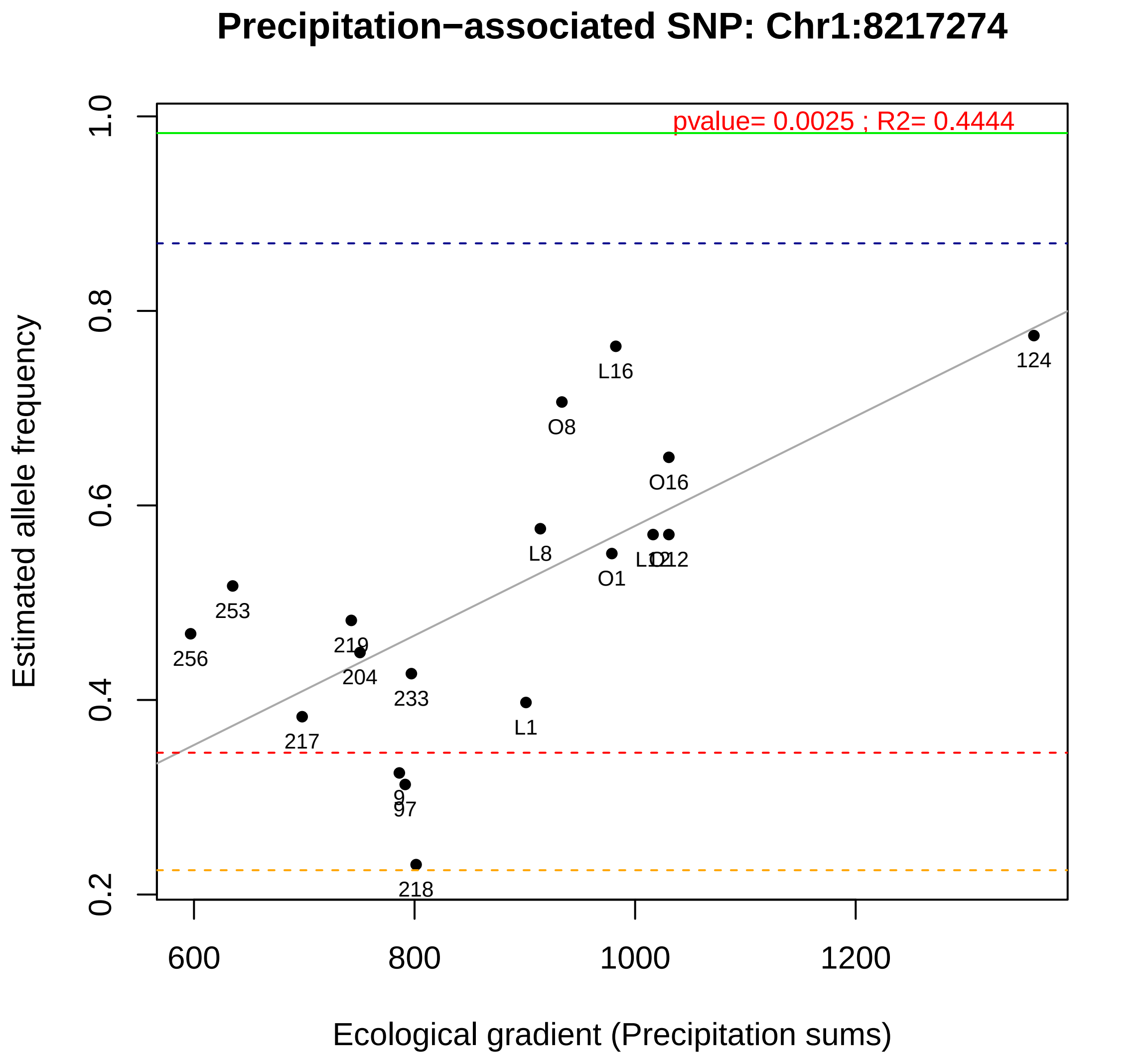
Precipitation−associated SNP: Chr1:8217274
pvalue= 0.0025 ; R2= 0.4444
Estimated allele frequency
Ecological gradient (Precipitation sums)
0.2
0.4
0.6
0.8
1.0
600
800
1000
1200
L16
124
O8
O16
L8
L12
O12
O1
253
256
219
204
233
217
L1
9
97
218

#### Precipitation−associated SNP: Chr1:8238346

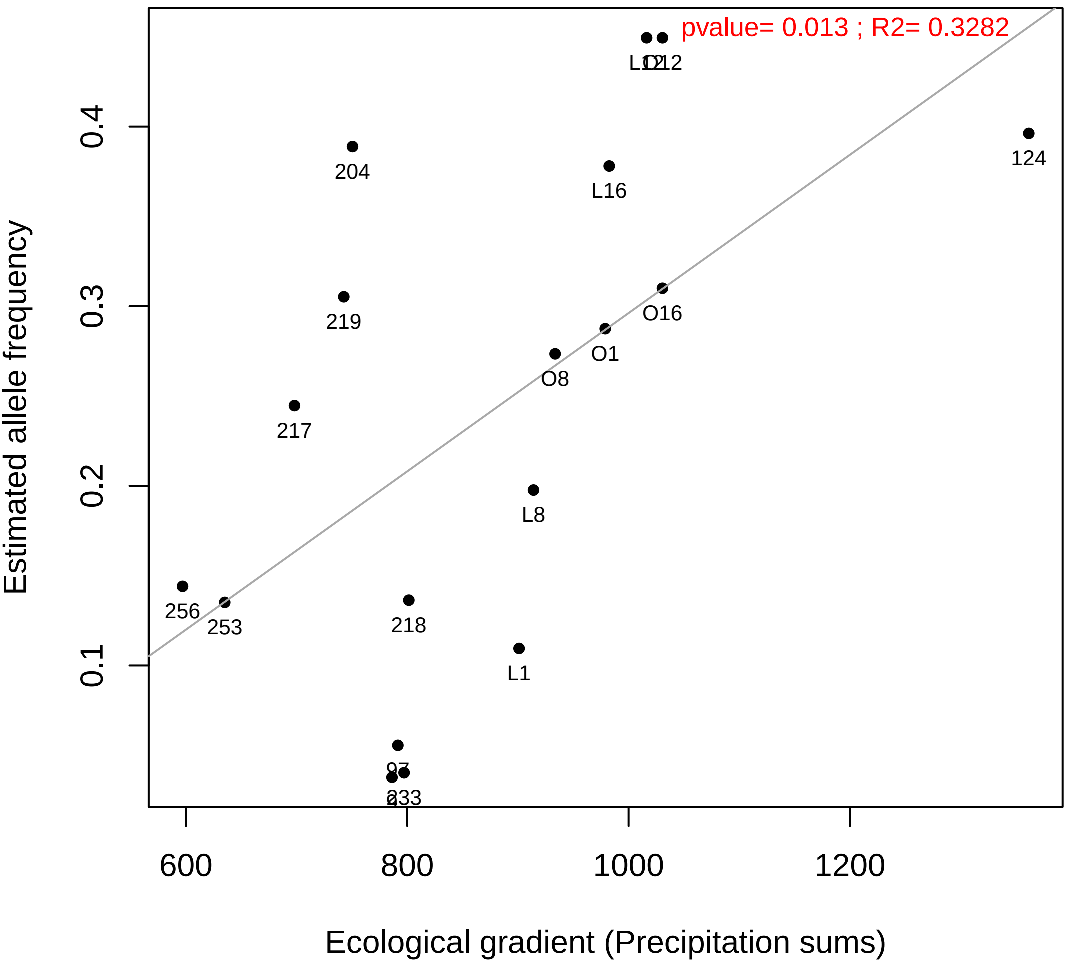

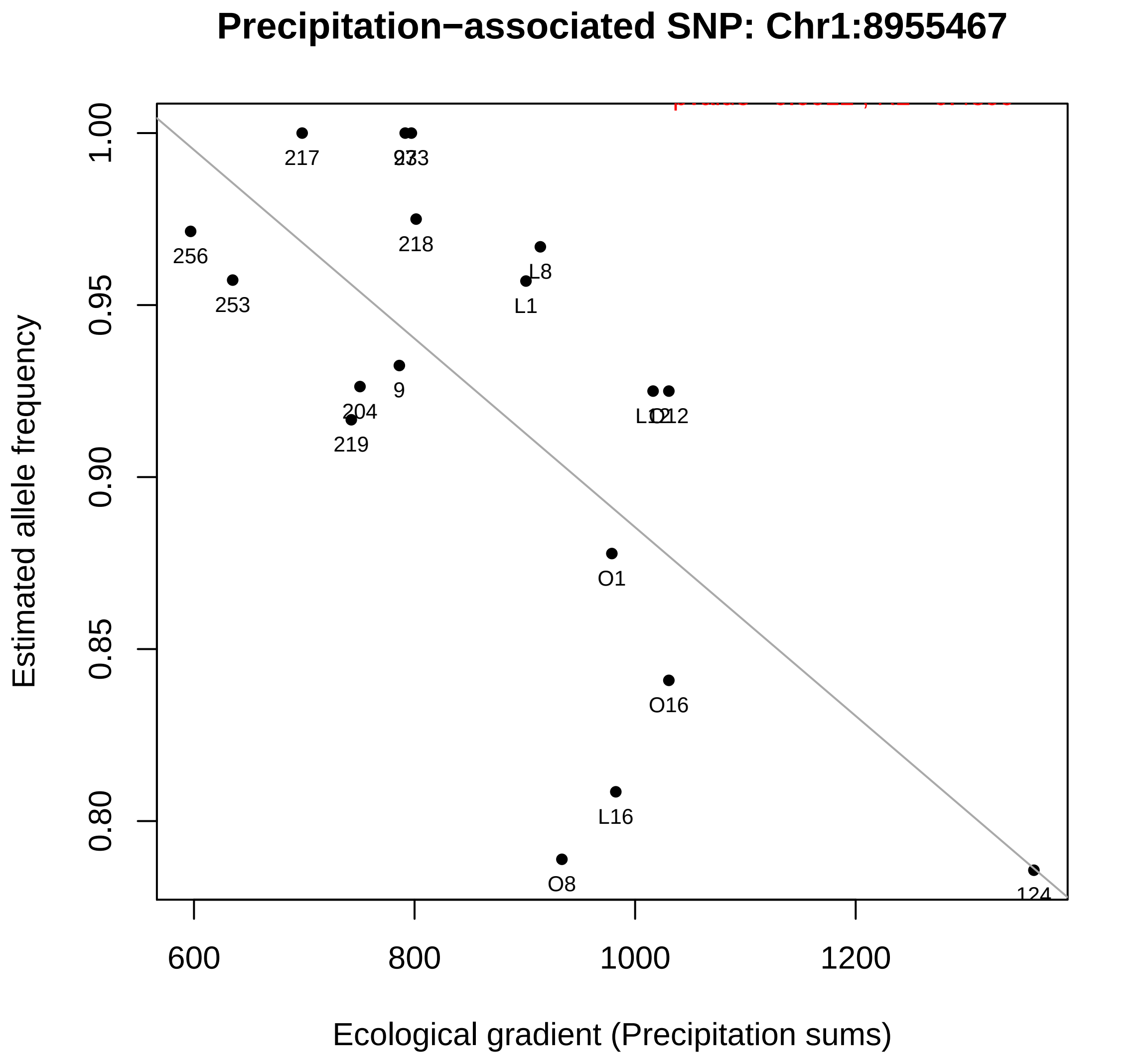
Precipitation−associated SNP: Chr1:8955467
Estimated allele frequency
Ecological gradient (Precipitation sums)
1.00
0.95
0.90
0.85
0.80
600
800
1000
1200
217
218
256
253
L8
L1
9
204
219
O1
O16
L16
O8
124

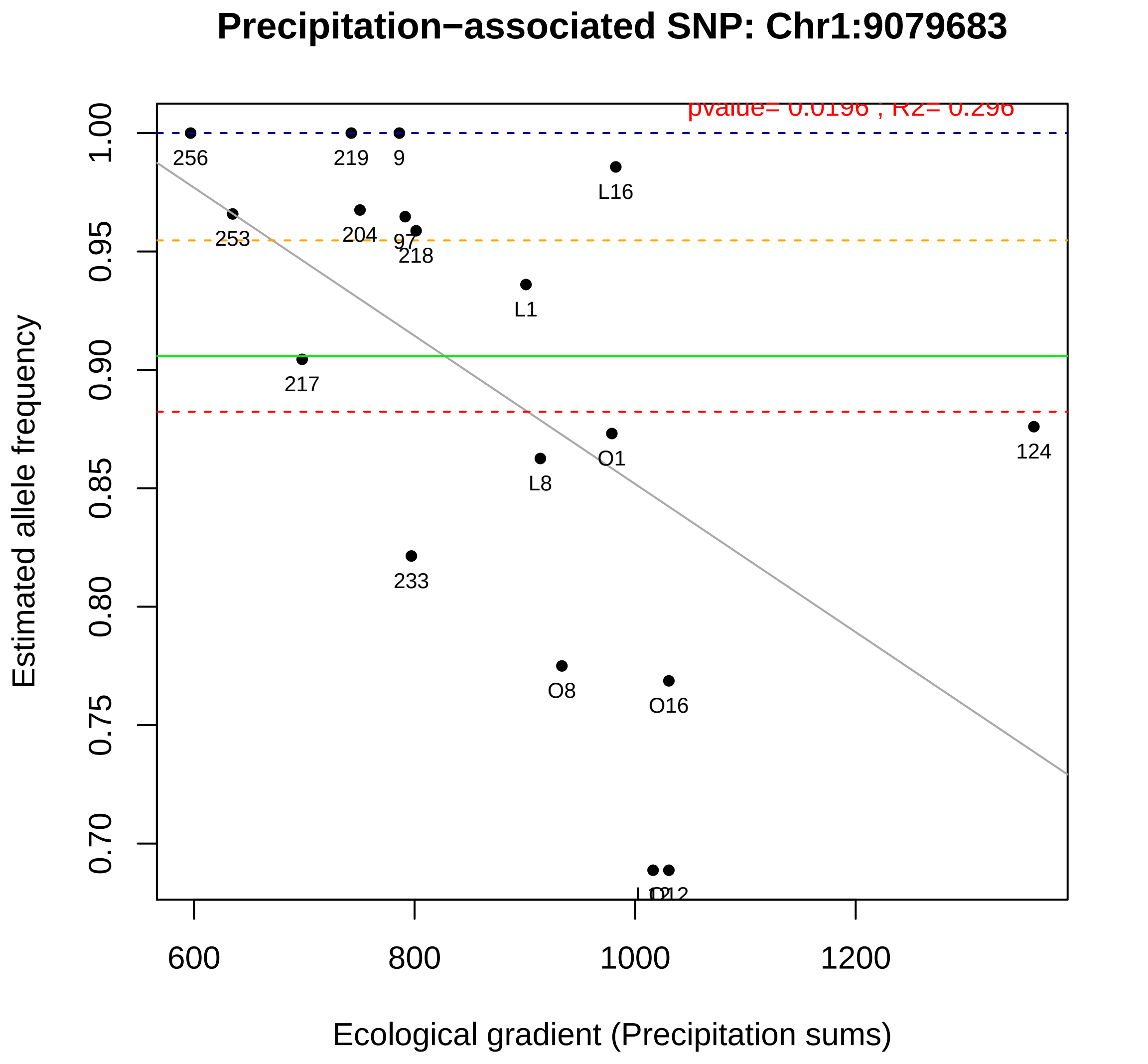

Precipitation−associated SNP: Chr1:9079683
pvalue= 0.0196 ; R2= 0.296
Estimated allele frequency
Ecological gradient (Precipitation sums)
1.00
0.95
0.90
0.85
0.80
0.75
0.70
600
800
1000
1200
256
219
9
L16
253
204
97
218
L1
217
124
O1
L8
233
O8
O16
O12

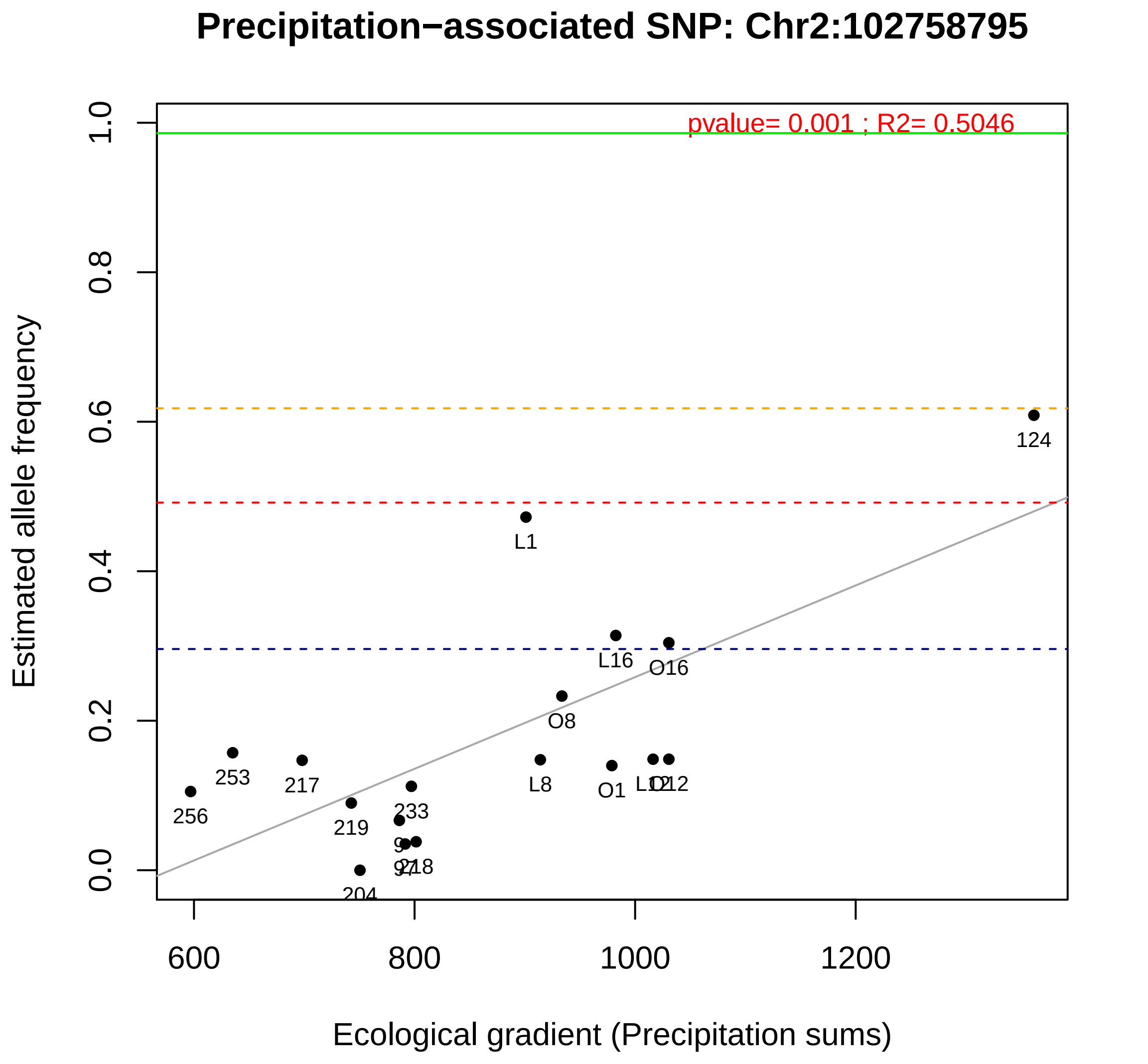

Precipitation−associated SNP: Chr2:102758795
pvalue= 0.001 ; R2= 0.5046
Estimated allele frequency
0.0
0.2
0.4
0.6
0.8
1.0
600
800
1000
1200
Ecological gradient (Precipitation sums)
124
L1
L16
O16
O8
253
217
256
219
233
L8
O1
204

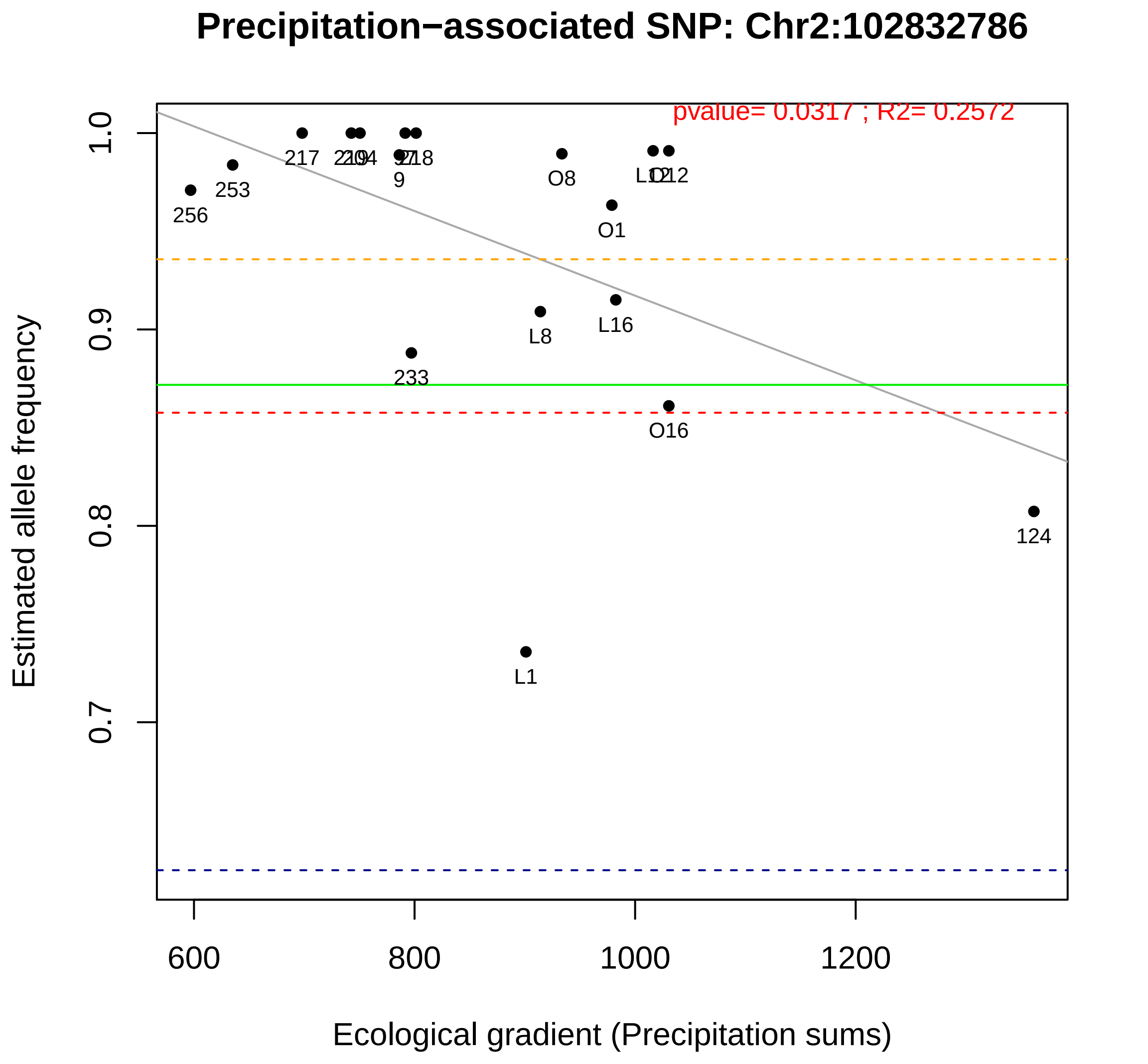
Precipitation−associated SNP: Chr2:102832786
pvalue= 0.0317 ; R2= 0.2572
Estimated allele frequency
Ecological gradient (Precipitation sums)
1.0
0.9
0.8
0.7
600
800
1000
1200
256
253
217
233
9
O8
O1
L8
L16
O16
L1
124

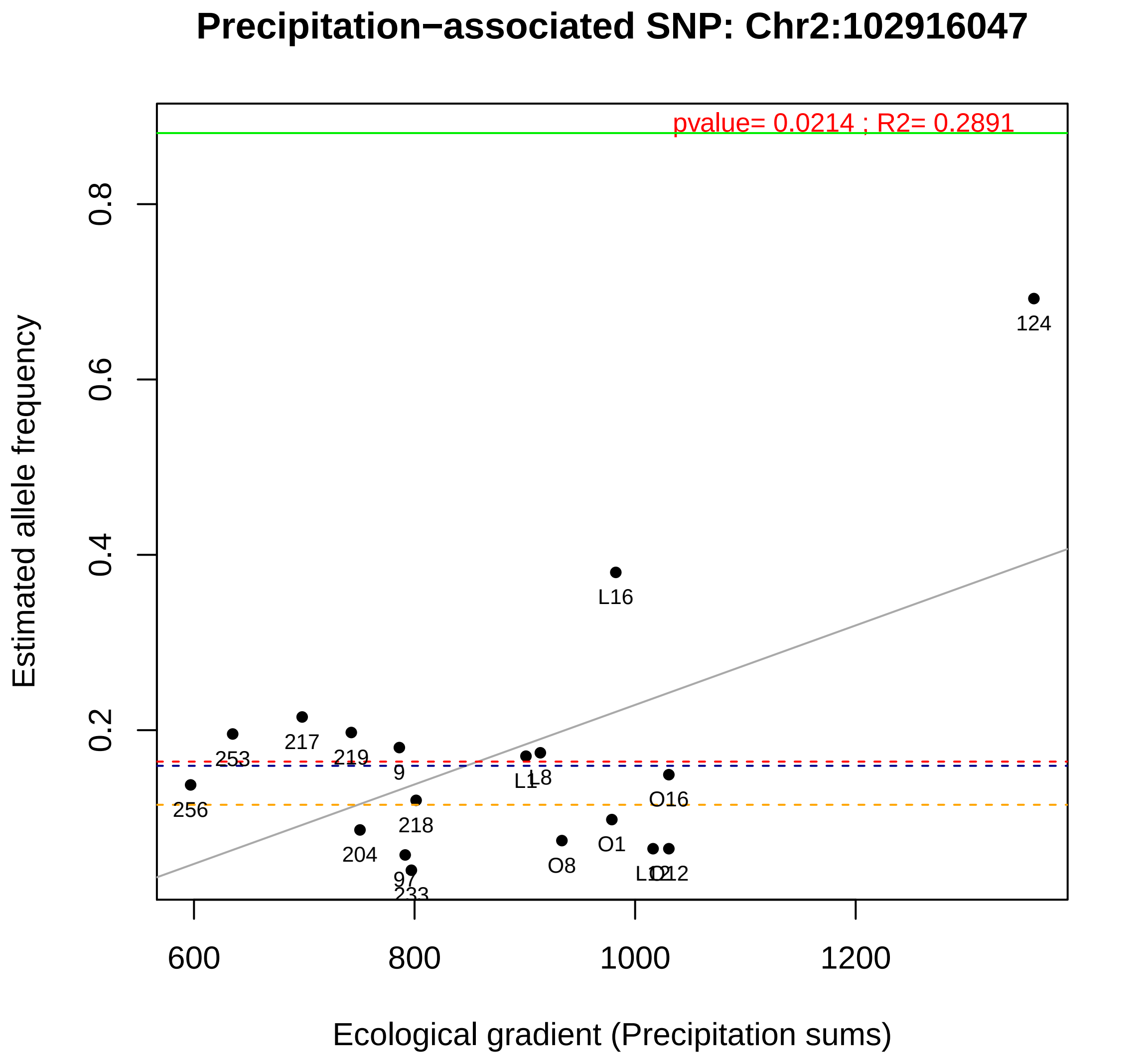

Precipitation−associated SNP: Chr2:102916047
pvalue= 0.0214 ; R2= 0.2891
Estimated allele frequency
0.2
0.4
0.6
0.8
600
800
1000
1200
Ecological gradient (Precipitation sums)
124
L16
217
253
219
9
256
218
204
97
233
L1
L8
O16
O1
O8
L12
O12

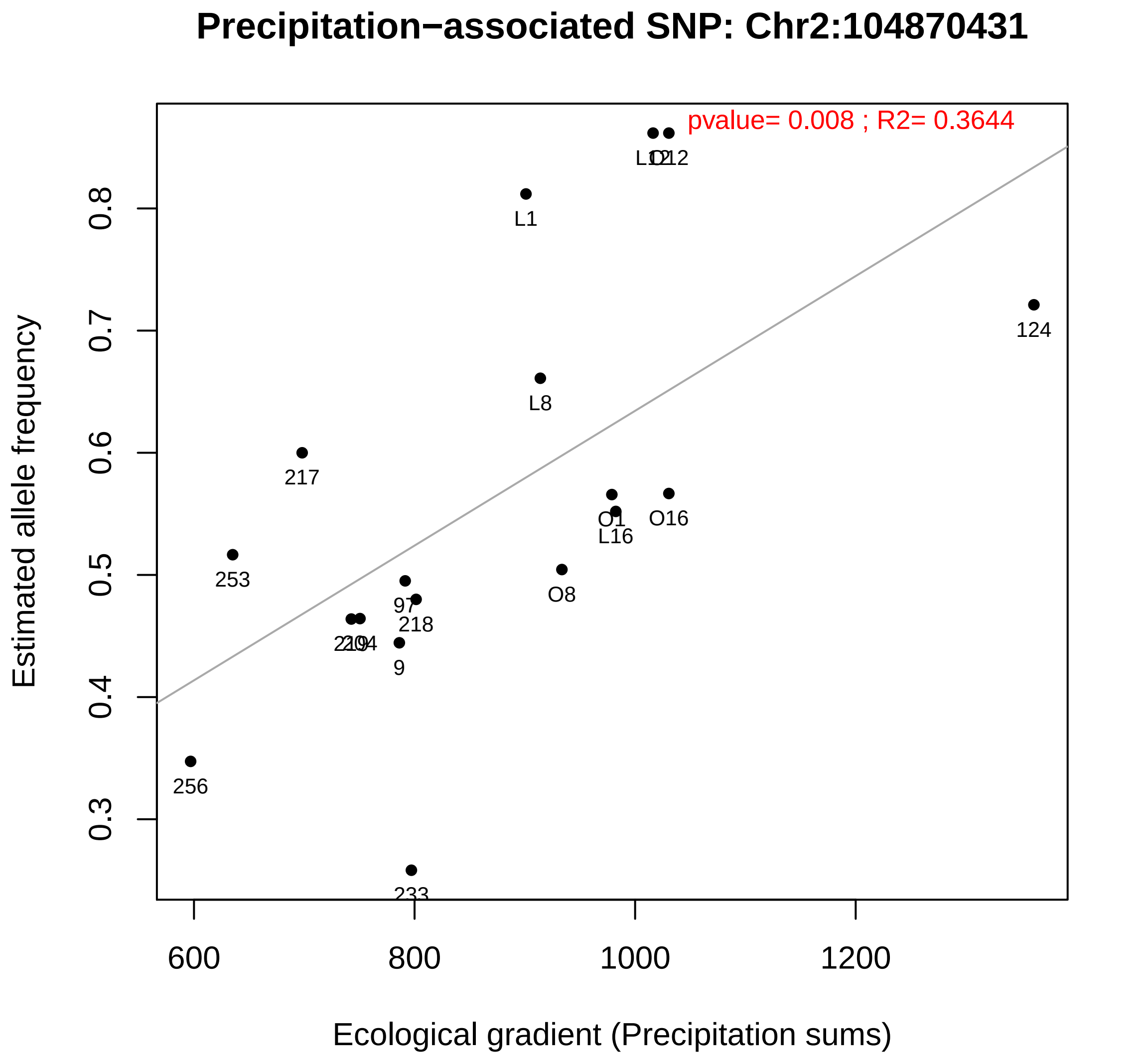
Precipitation−associated SNP: Chr2:104870431
pvalue= 0.008 ; R2= 0.3644
Estimated allele frequency
Ecological gradient (Precipitation sums)
0.3
0.4
0.5
0.6
0.7
0.8
600
800
1000
1200
L1
L8
124
217
253
256
97
218
9
233
O8
O1
L16
O16

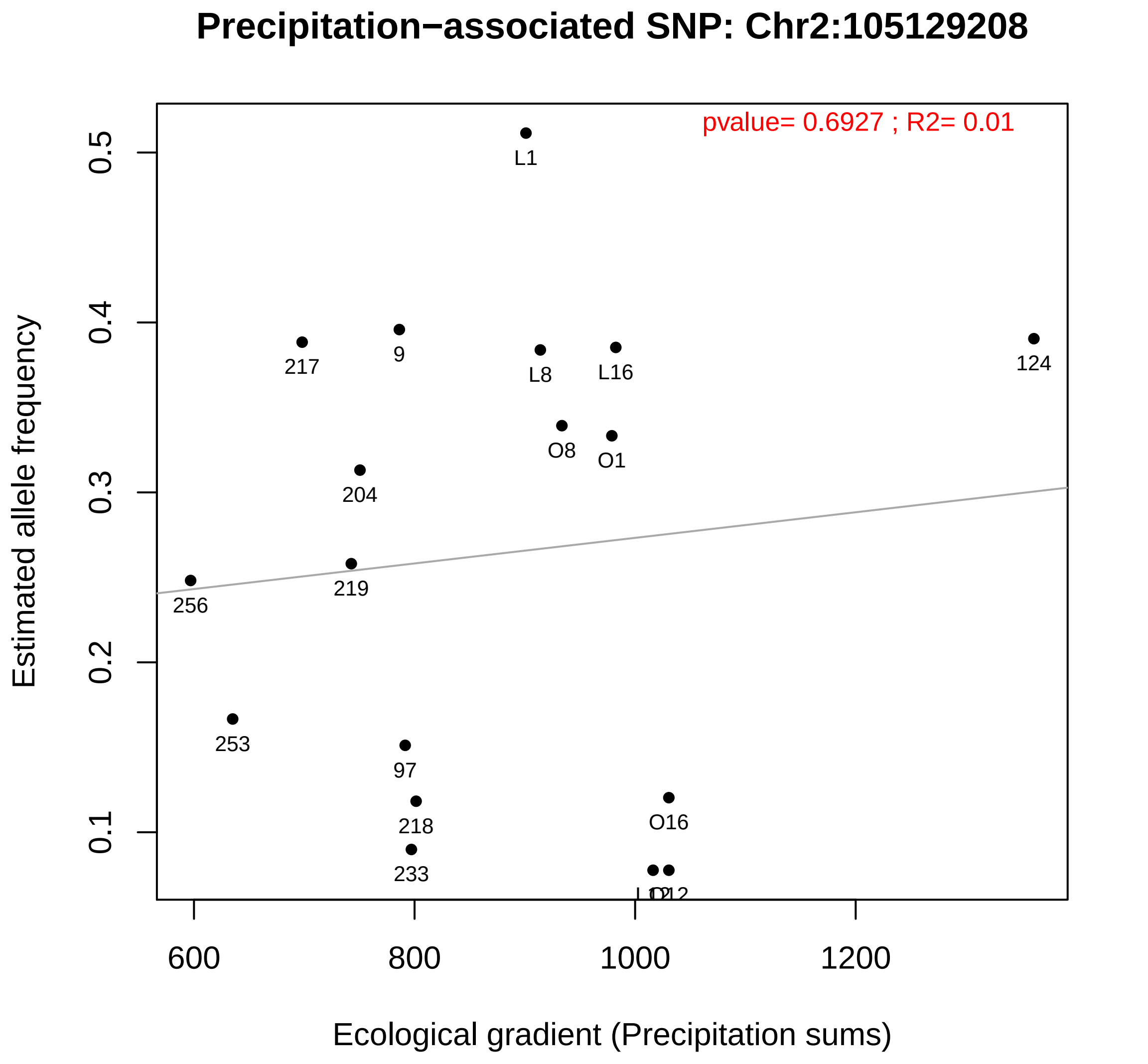
Precipitation−associated SNP: Chr2:105129208
pvalue= 0.6927 ; R2= 0.01
Estimated allele frequency
0.1
0.2
0.3
0.4
0.5
Ecological gradient (Precipitation sums)
600
800
1000
1200
L1
217
9
L8
L16
124
O8
O1
204
219
256
253
97
218
O16
233

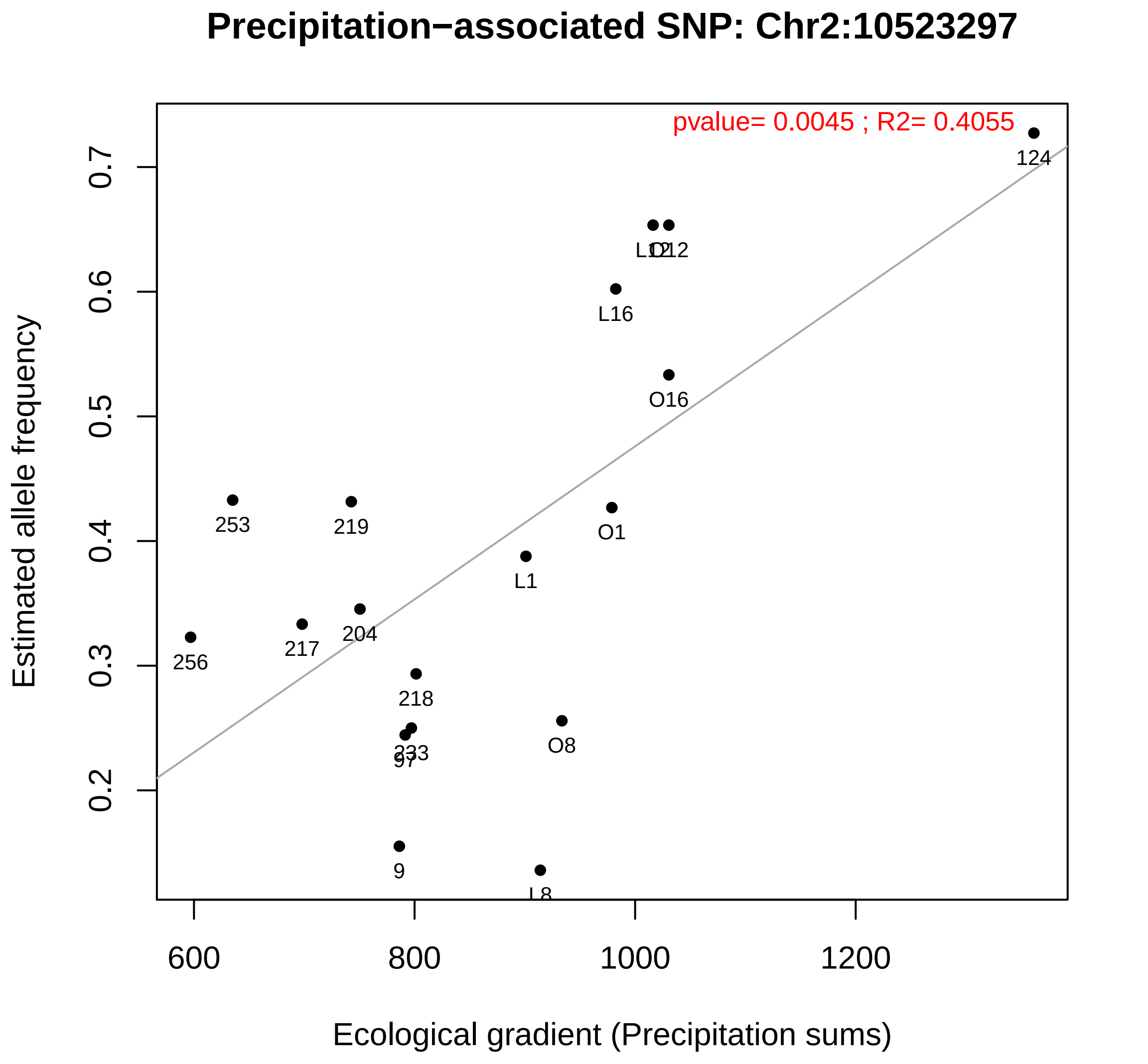

Precipitation−associated SNP: Chr2:10523297
pvalue= 0.0045 ; R2= 0.4055
124
L12
O12
L16
O16
253
219
O1
L1
204
217
256
218
O8
233
97
9
L8
0.2
0.3
0.4
0.5
0.6
0.7
600
800
1000
1200
Estimated allele frequency
Ecological gradient (Precipitation sums)

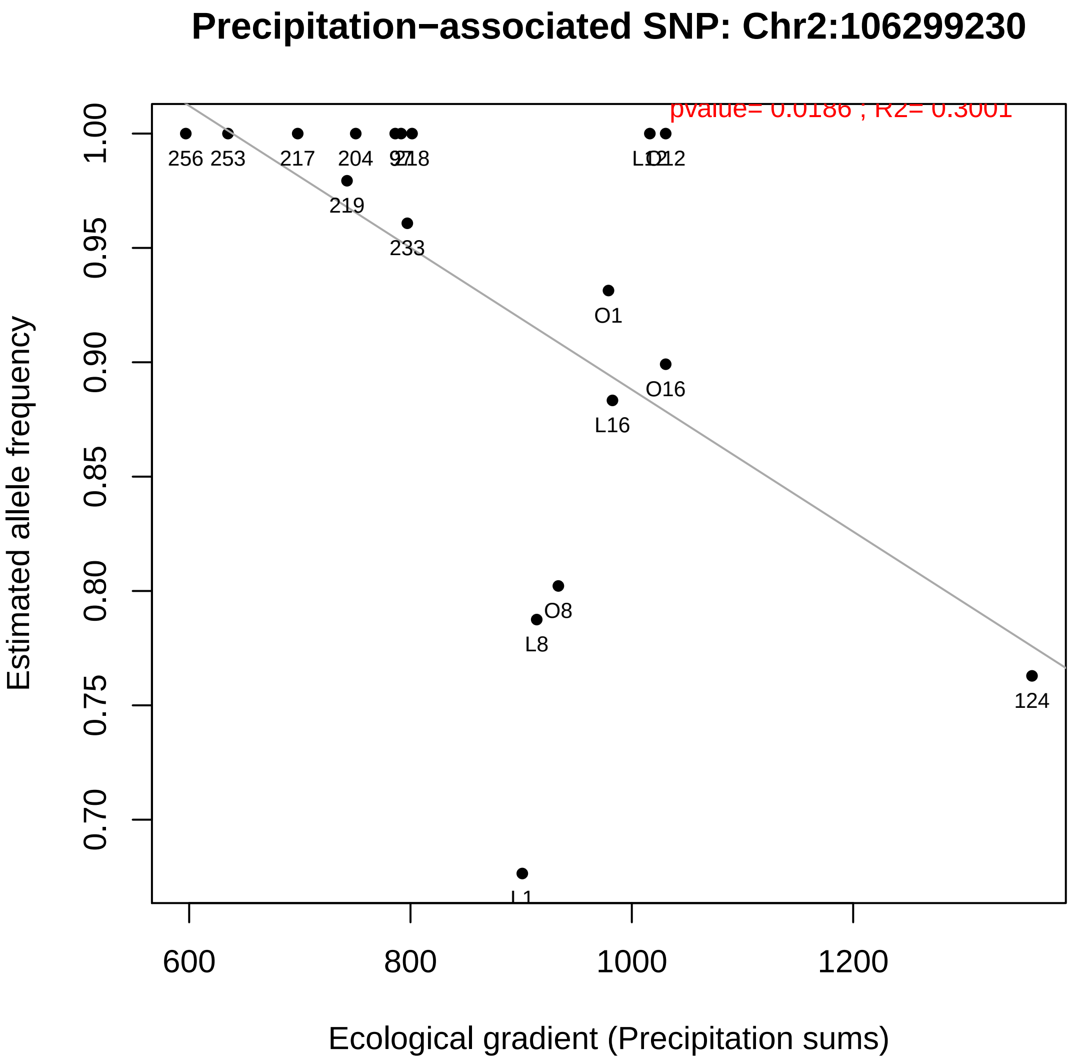

Precipitation−associated SNP: Chr2:106299230
pvalue= 0.0186 ; R2= 0.3001
Estimated allele frequency
Ecological gradient (Precipitation sums)
1.00
0.95
0.90
0.85
0.80
0.75
0.70
600
800
1000
1200
256
253
217
204
219
233
O1
O16
L16
O8
L8
124

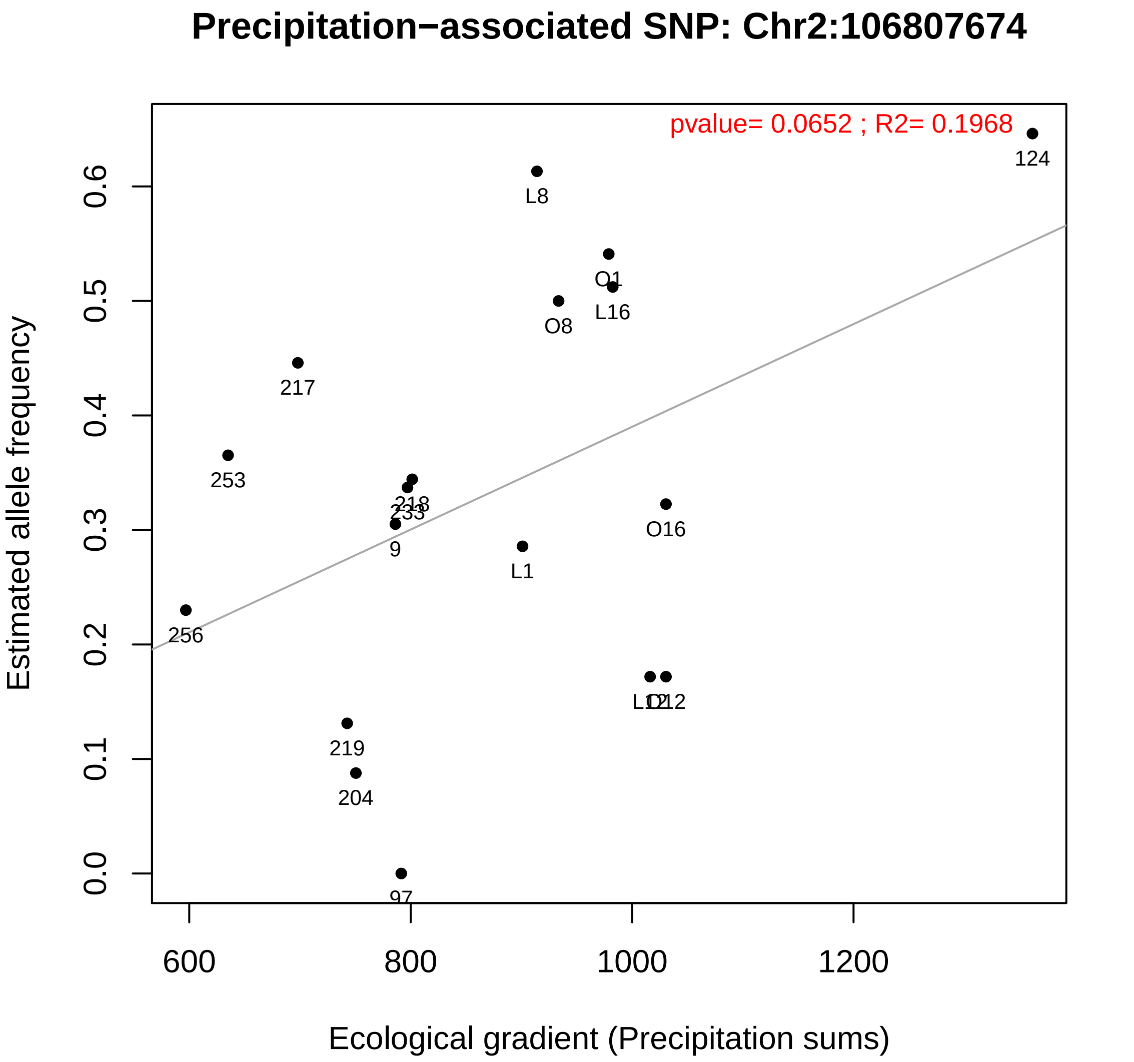
Precipitation−associated SNP: Chr2:106807674
pvalue= 0.0652 ; R2= 0.1968
Estimated allele frequency
Ecological gradient (Precipitation sums)
0.0
0.1
0.2
0.3
0.4
0.5
0.6
600
800
1000
1200
124
L8
O1
L16
O8
217
253
218
233
9
O16
L1
256
L12
O12
219
204
97

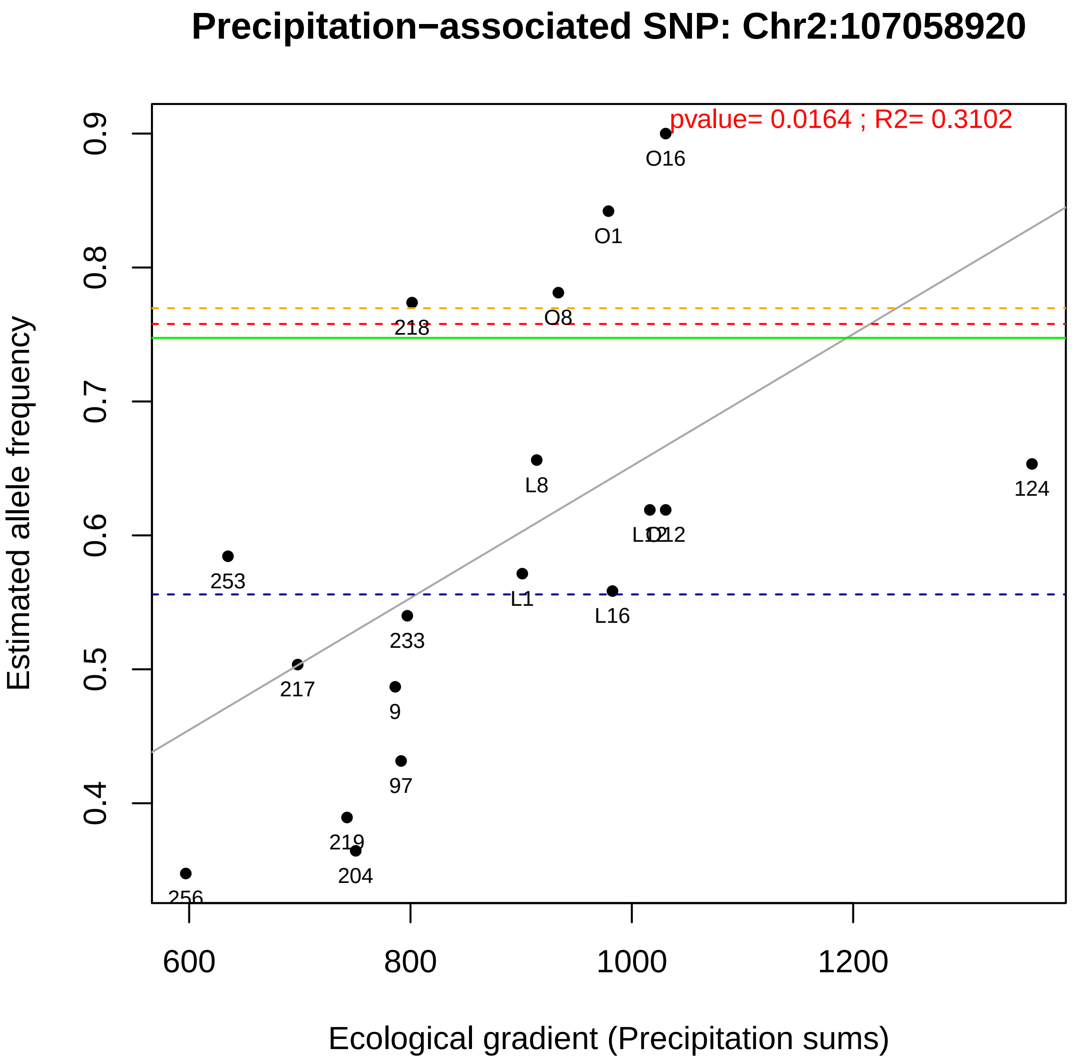

Precipitation−associated SNP: Chr2:107058920
pvalue= 0.0164 ; R2= 0.3102
Estimated allele frequency
Ecological gradient (Precipitation sums)
0.4
0.5
0.6
0.7
0.8
0.9
600
800
1000
1200
O16
O1
O8
218
L8
124
L12
O12
253
L1
L16
233
217
9
97
219
204
256

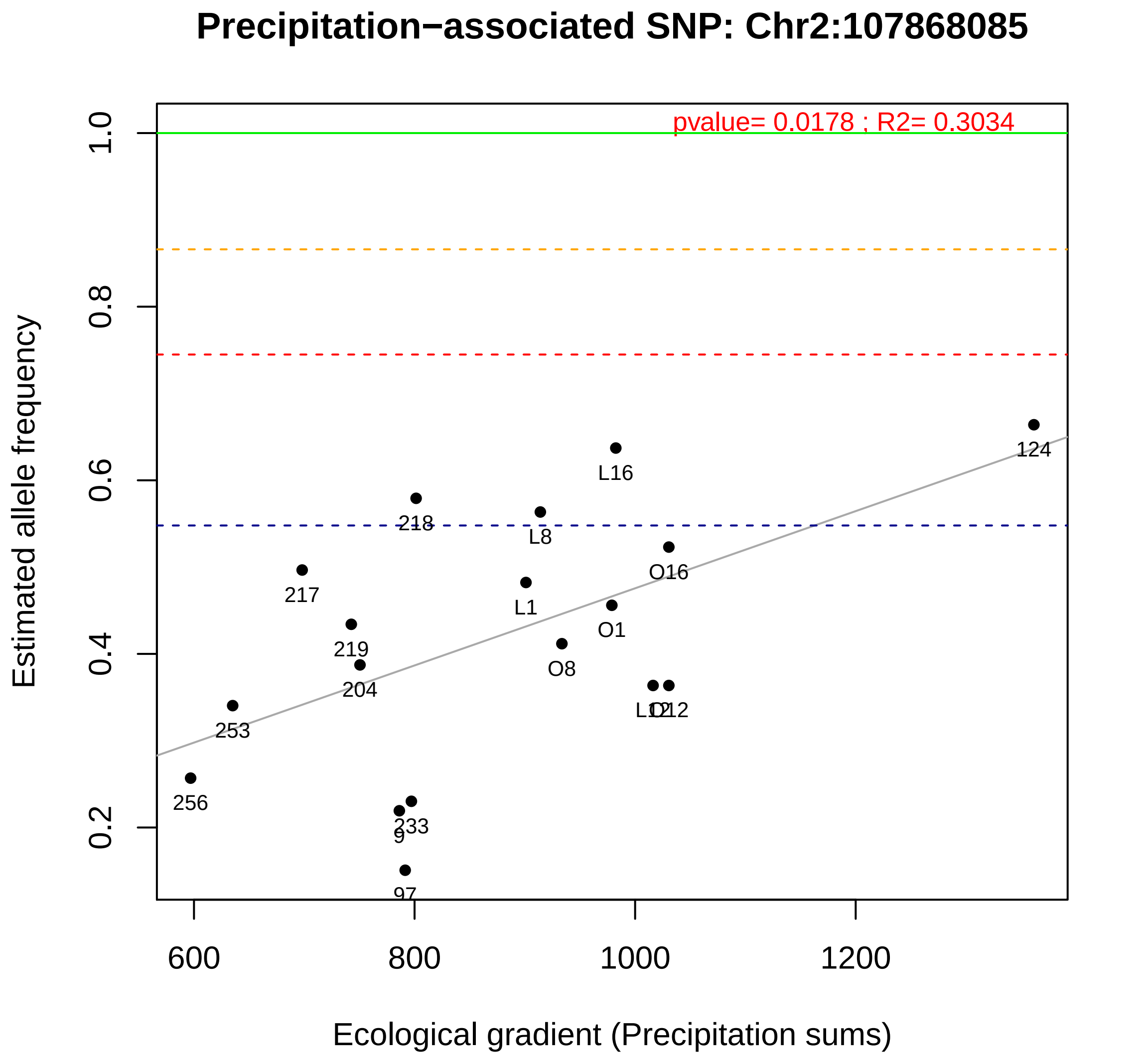

Precipitation−associated SNP: Chr2:107868085
pvalue= 0.0178 ; R2= 0.3034
Estimated allele frequency
0.2
0.4
0.6
0.8
1.0
600
800
1000
1200
Ecological gradient (Precipitation sums)
124
L16
218
L8
O16
217
L1
O1
219
O8
204
L12
O12
253
256
233
9
97

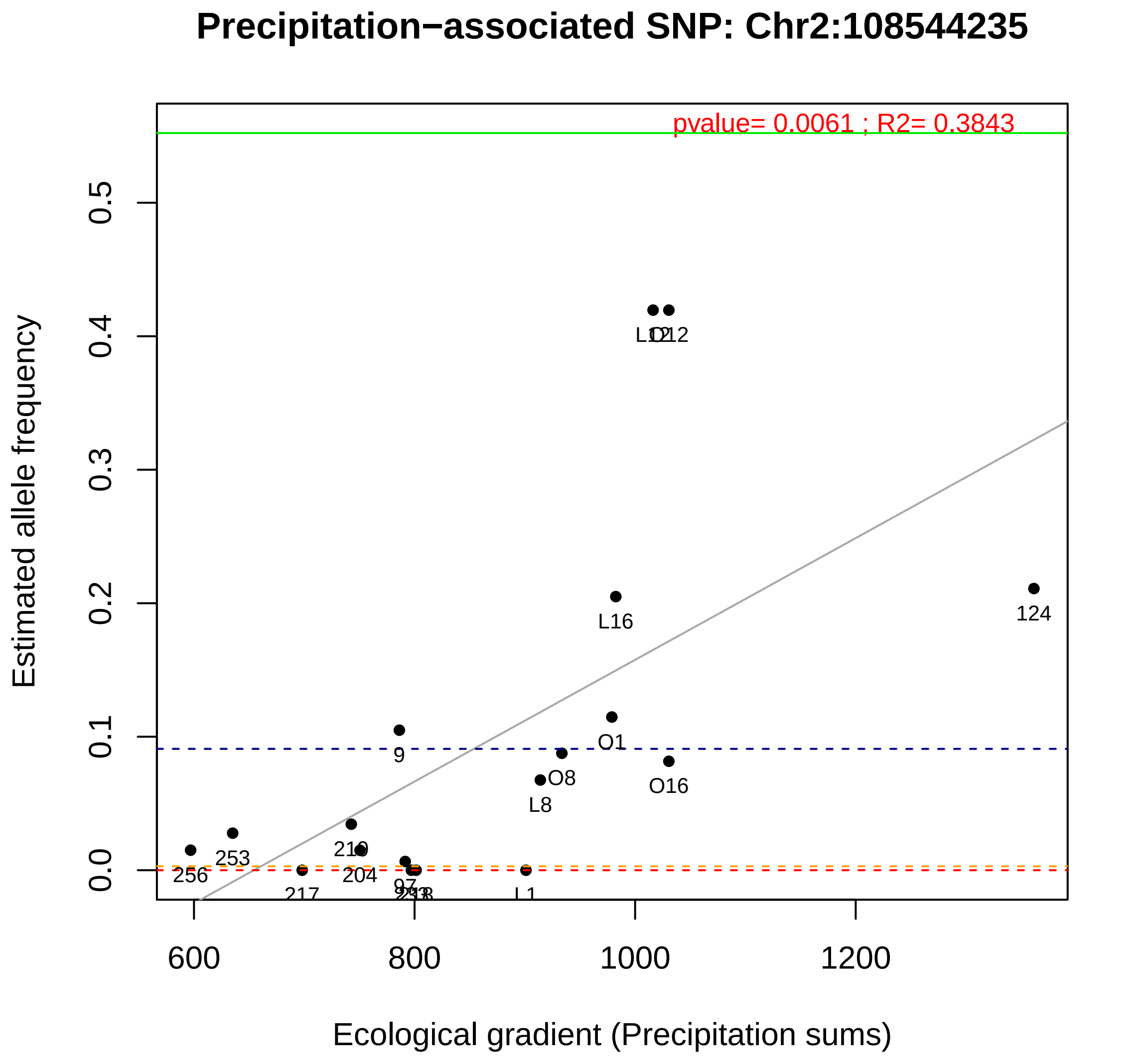

Precipitation−associated SNP: Chr2:108544235
pvalue= 0.0061 ; R2= 0.3843
Estimated allele frequency
0.0
0.1
0.2
0.3
0.4
0.5
600
800
1000
1200
Ecological gradient (Precipitation sums)
256
253
217
210
204
9
97
L1
L8
O8
O1
O16
L16
124

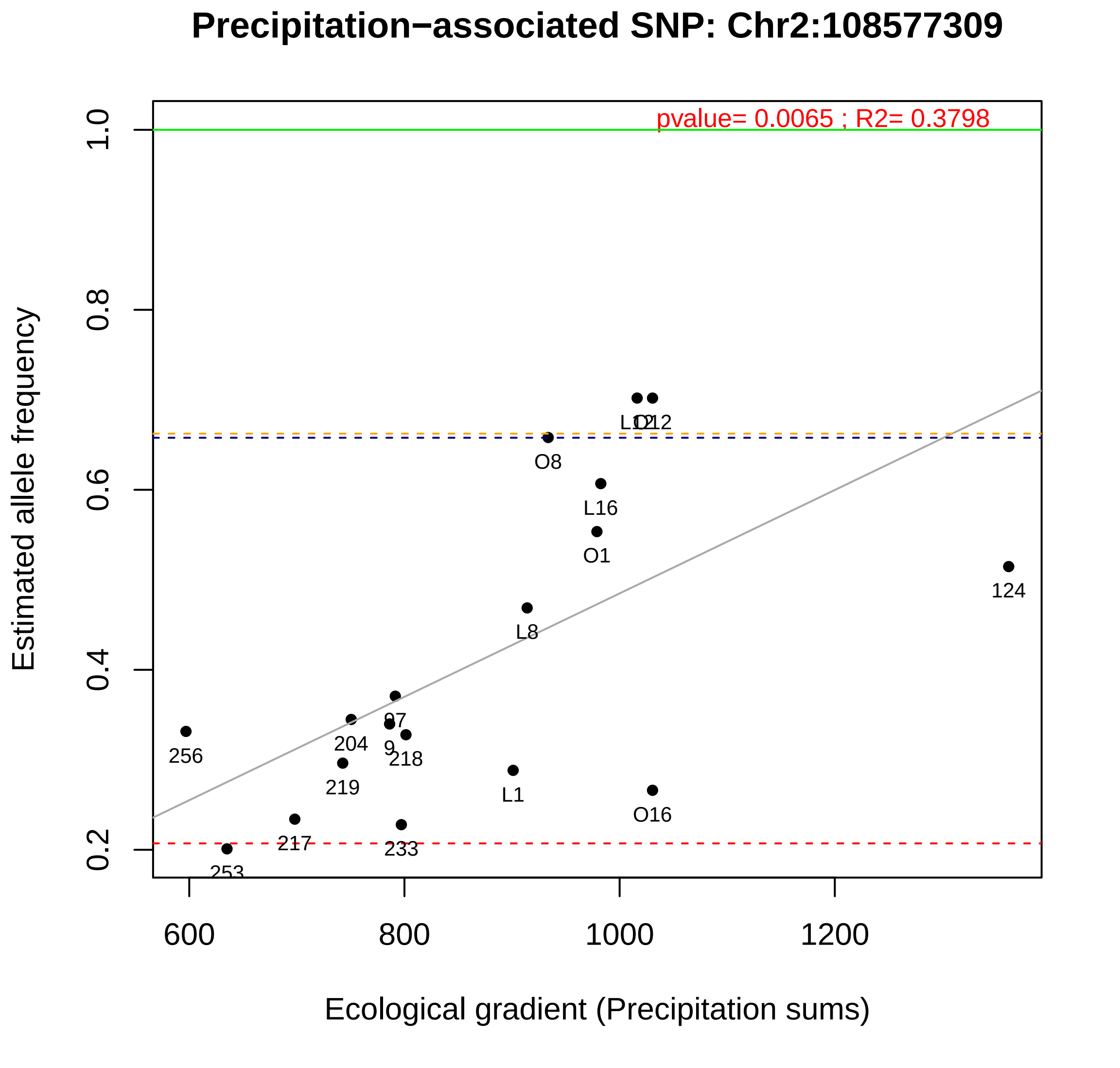

Precipitation−associated SNP: Chr2:108577309
pvalue= 0.0065 ; R2= 0.3798
Estimated allele frequency
0.2
0.4
0.6
0.8
1.0
600
800
1000
1200
Ecological gradient (Precipitation sums)
L12
O12
O8
L16
O1
124
L8
97
9
204
218
256
219
L1
O16
217
233
253

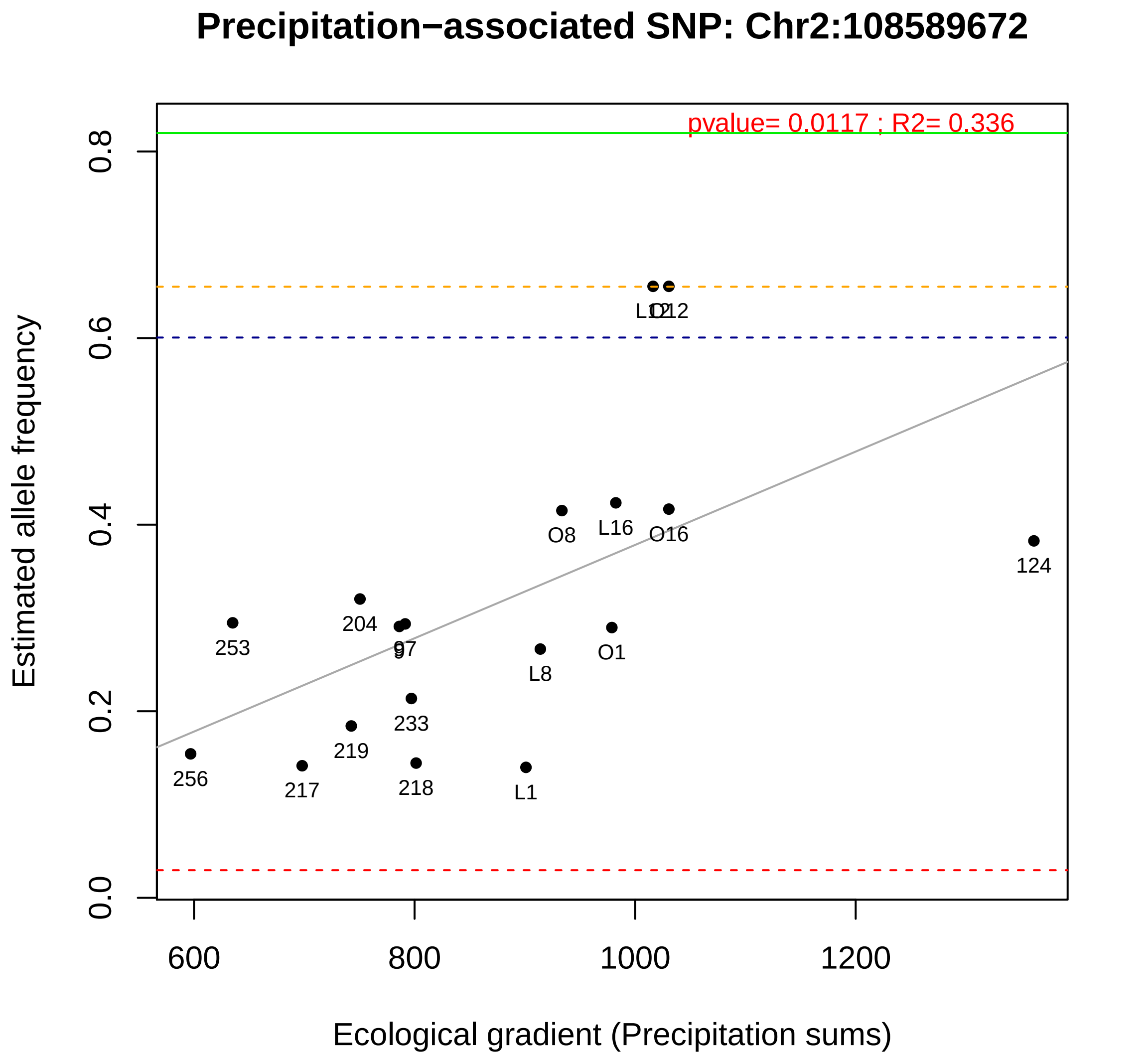

Precipitation−associated SNP: Chr2:108589672
pvalue= 0.0117 ; R2= 0.336
Estimated allele frequency
0.0
0.2
0.4
0.6
0.8
600
800
1000
1200
Ecological gradient (Precipitation sums)
256
253
217
219
204
233
218
L1
L8
O8
L16
O16
O1
124

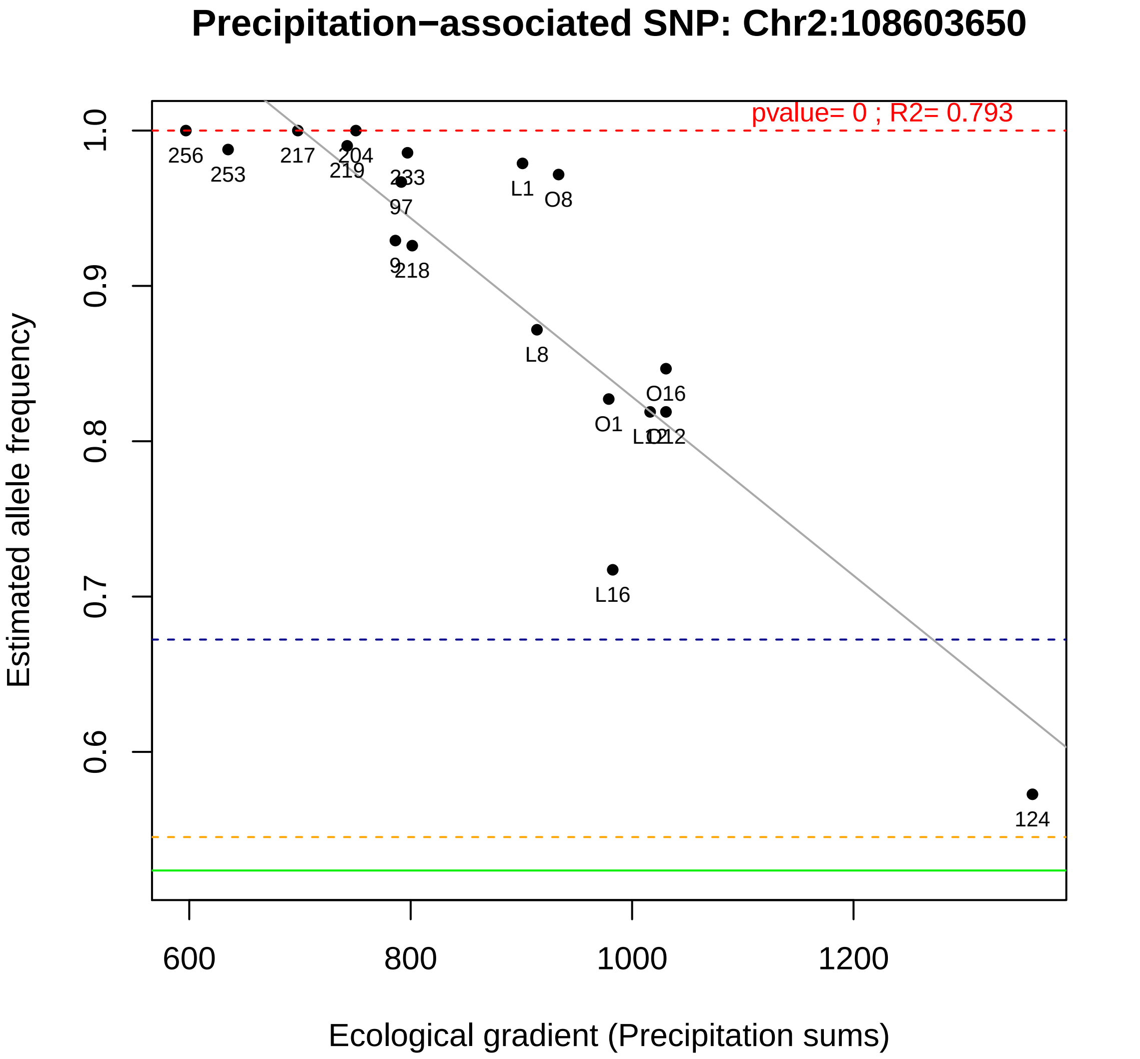
Precipitation−associated SNP: Chr2:108603650
pvalue= 0 ; R2= 0.793
Estimated allele frequency
1.0
0.9
0.8
0.7
0.6
600
800
1000
1200
Ecological gradient (Precipitation sums)
256
253
217
219
204
233
97
9
218
L1
O8
L8
O16
O1
L12
O12
L16
124

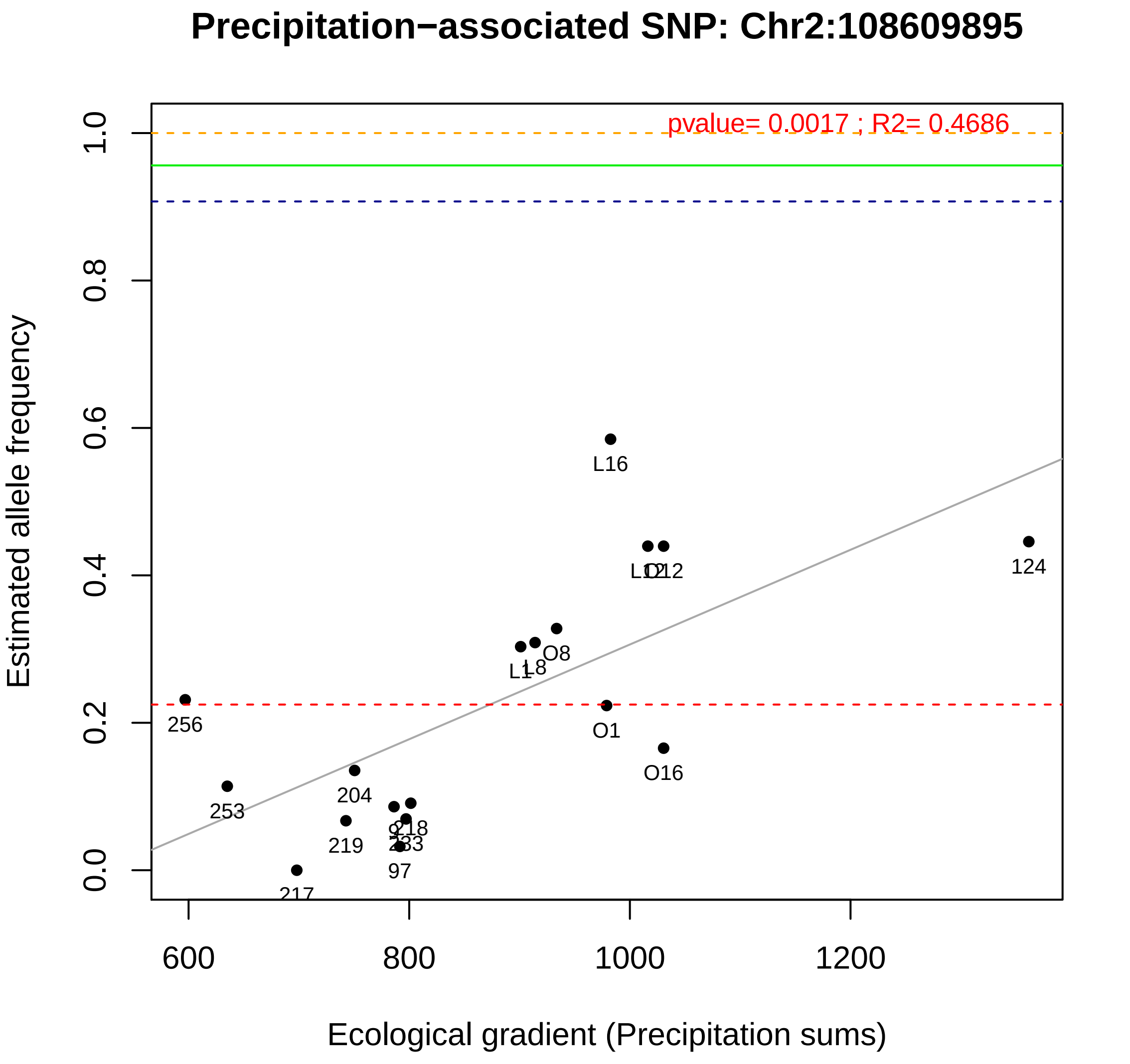
Precipitation−associated SNP: Chr2:108609895
pvalue= 0.0017 ; R2= 0.4686
Estimated allele frequency
Ecological gradient (Precipitation sums)
0.0
0.2
0.4
0.6
0.8
1.0
600
800
1000
1200
L16
L12
O12
124
O8
L1
L8
256
O1
O16
204
253
219
218
233
97
217

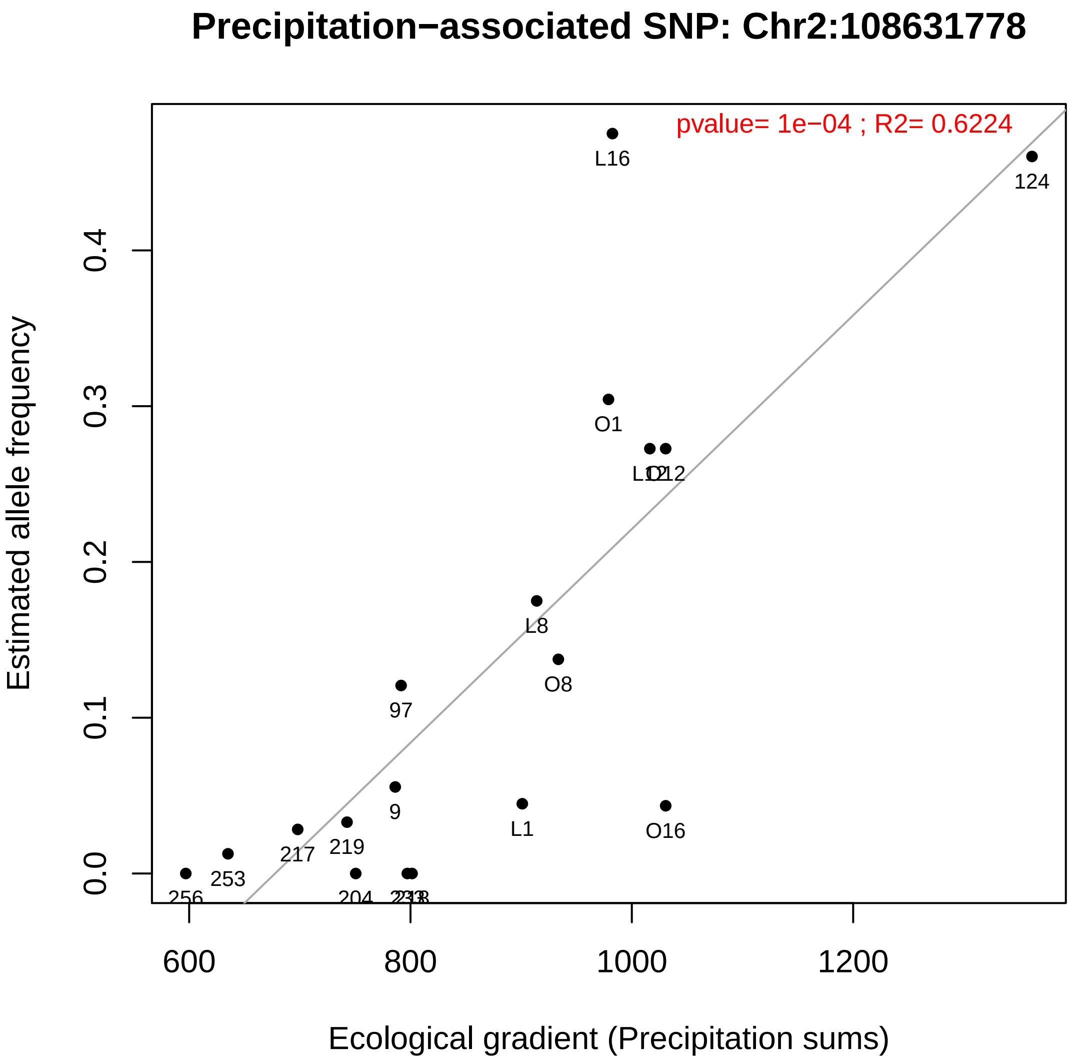

Precipitation−associated SNP: Chr2:108631778
pvalue= 1e−04 ; R2= 0.6224
L16
124
O1
L8
O8
97
9
L1
O16
217
219
253
256
204
Estimated allele frequency
Ecological gradient (Precipitation sums)
0.0
0.1
0.2
0.3
0.4
600
800
1000
1200

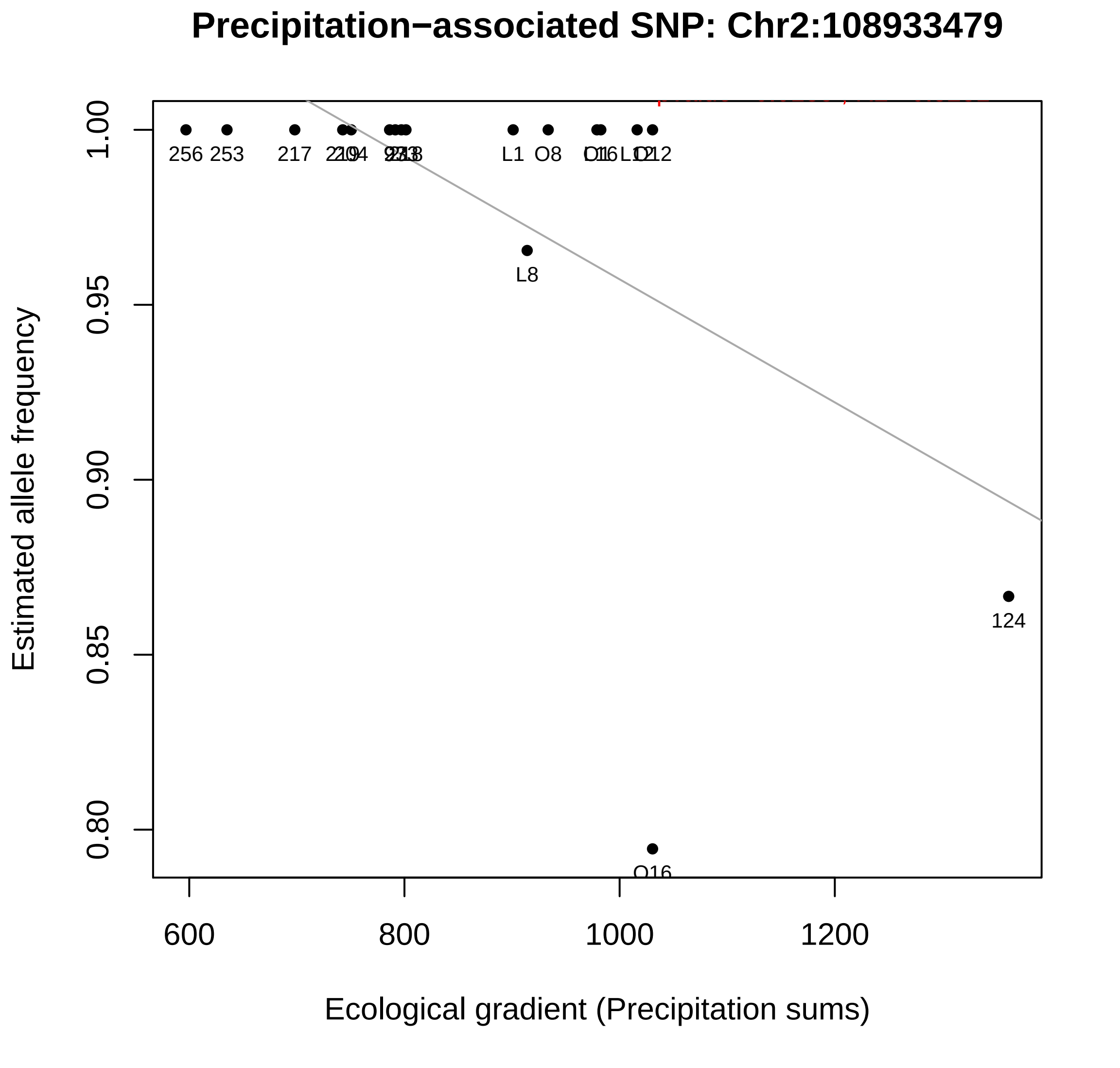
Precipitation−associated SNP: Chr2:108933479
Estimated allele frequency
Ecological gradient (Precipitation sums)
1.00
0.95
0.90
0.85
0.80
600
800
1000
1200
256
253
217
L8
124
O16
L1
O8

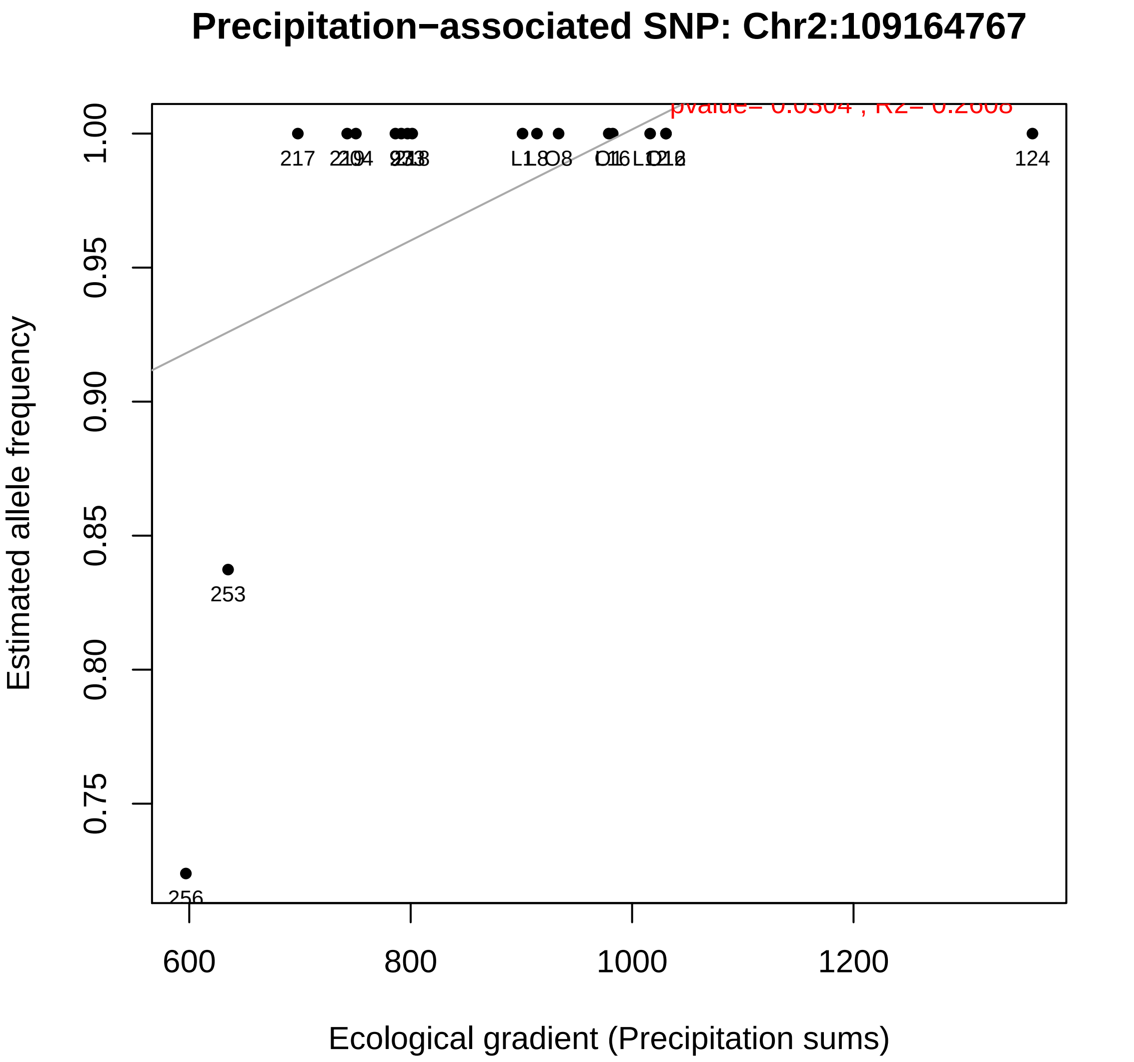

Precipitation−associated SNP: Chr2:109164767
Estimated allele frequency
Ecological gradient (Precipitation sums)
0.75
0.80
0.85
0.90
0.95
1.00
600
800
1000
1200
217
124
253
256

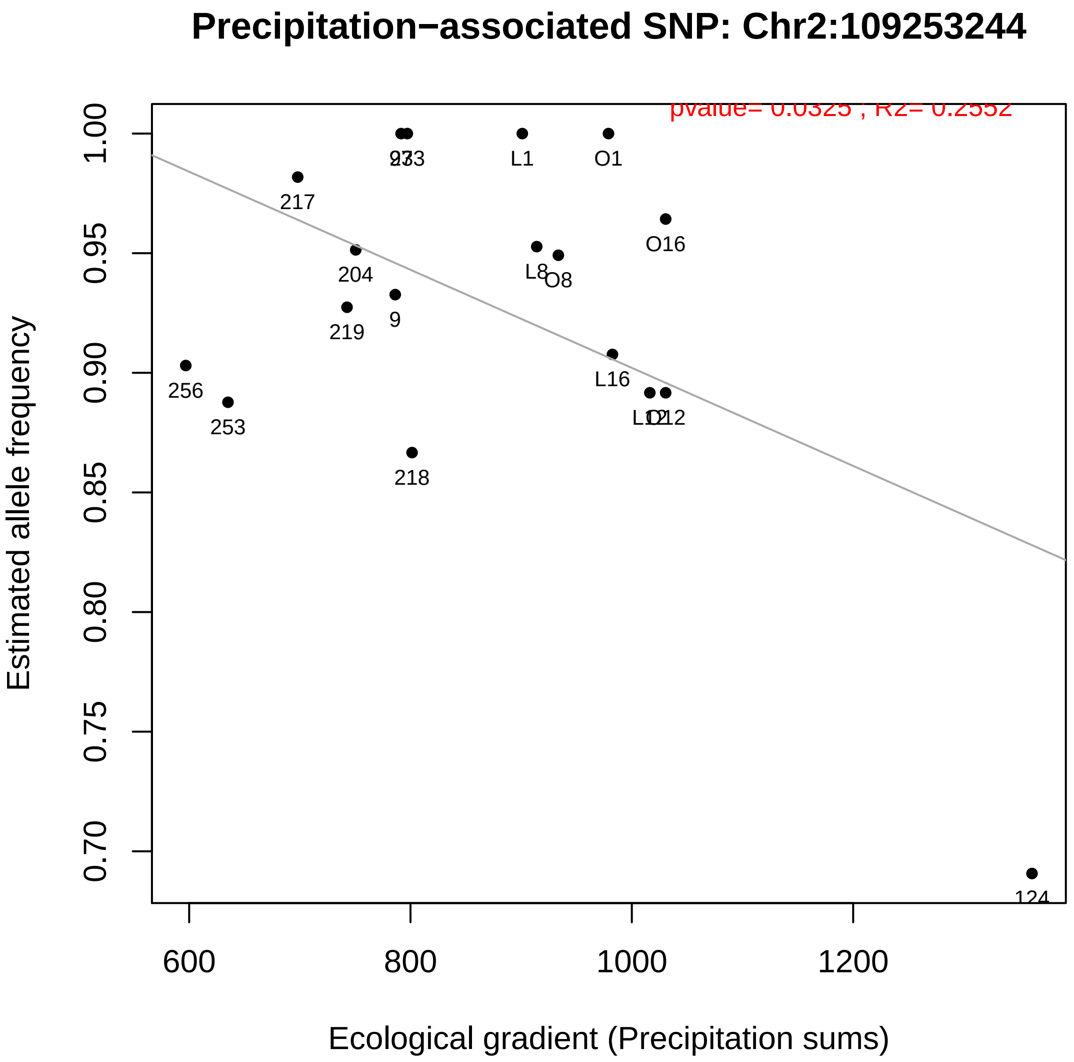
Precipitation−associated SNP: Chr2:109253244
pvalue= 0.0325 ; R2= 0.2552
Estimated allele frequency
Ecological gradient (Precipitation sums)
1.00
0.95
0.90
0.85
0.80
0.75
0.70
600
800
1000
1200
217
204
219
9
256
253
218
L1
O1
O16
L8
O8
L16
124

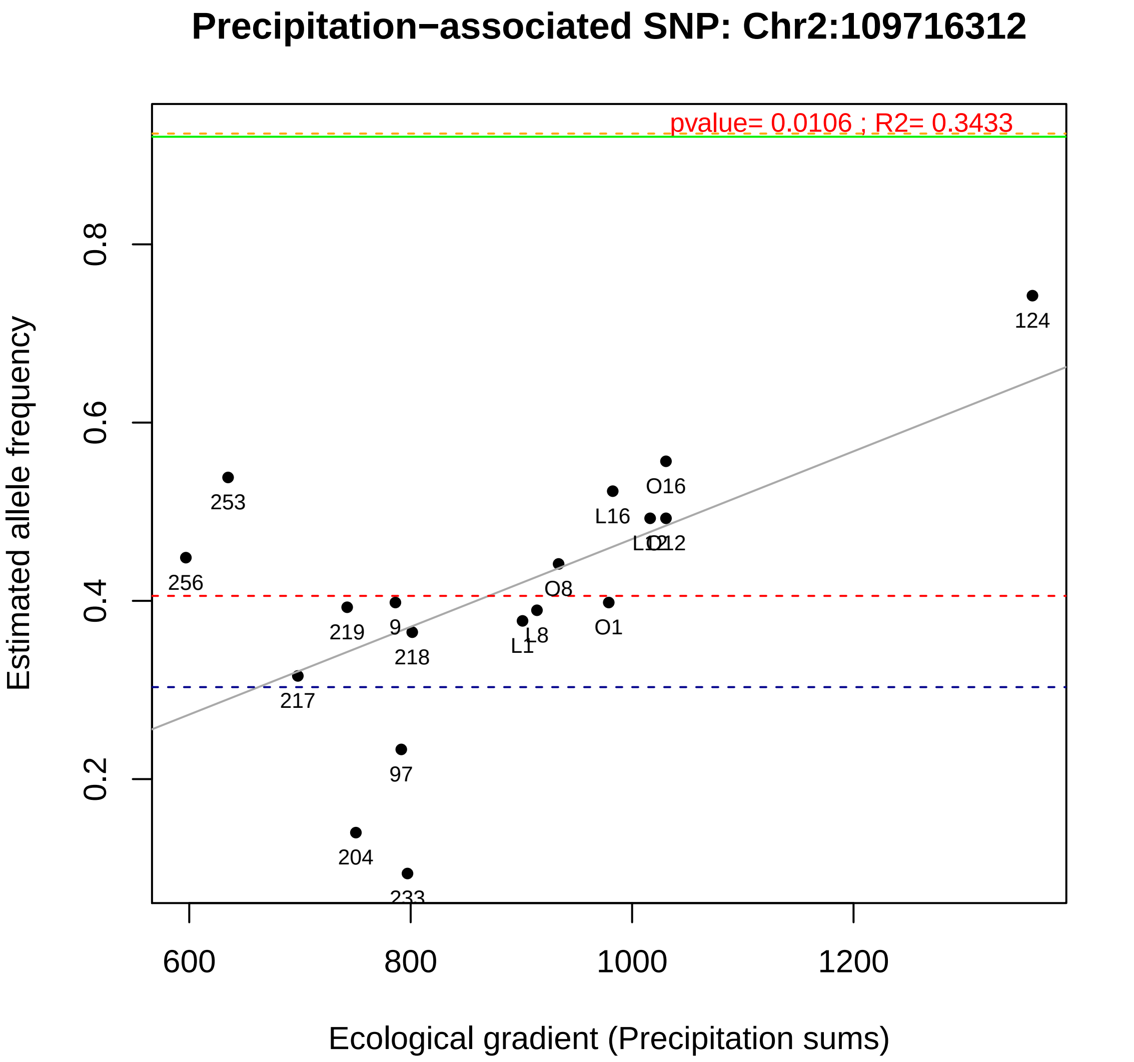

Precipitation−associated SNP: Chr2:109716312
pvalue= 0.0106 ; R2= 0.3433
Estimated allele frequency
0.2
0.4
0.6
0.8
600
800
1000
1200
Ecological gradient (Precipitation sums)
124
253
256
219
9
218
217
97
204
233
L1
L8
O8
O1
L16
O16
L12
O12

#### Precipitation−associated SNP: Chr2:111242142

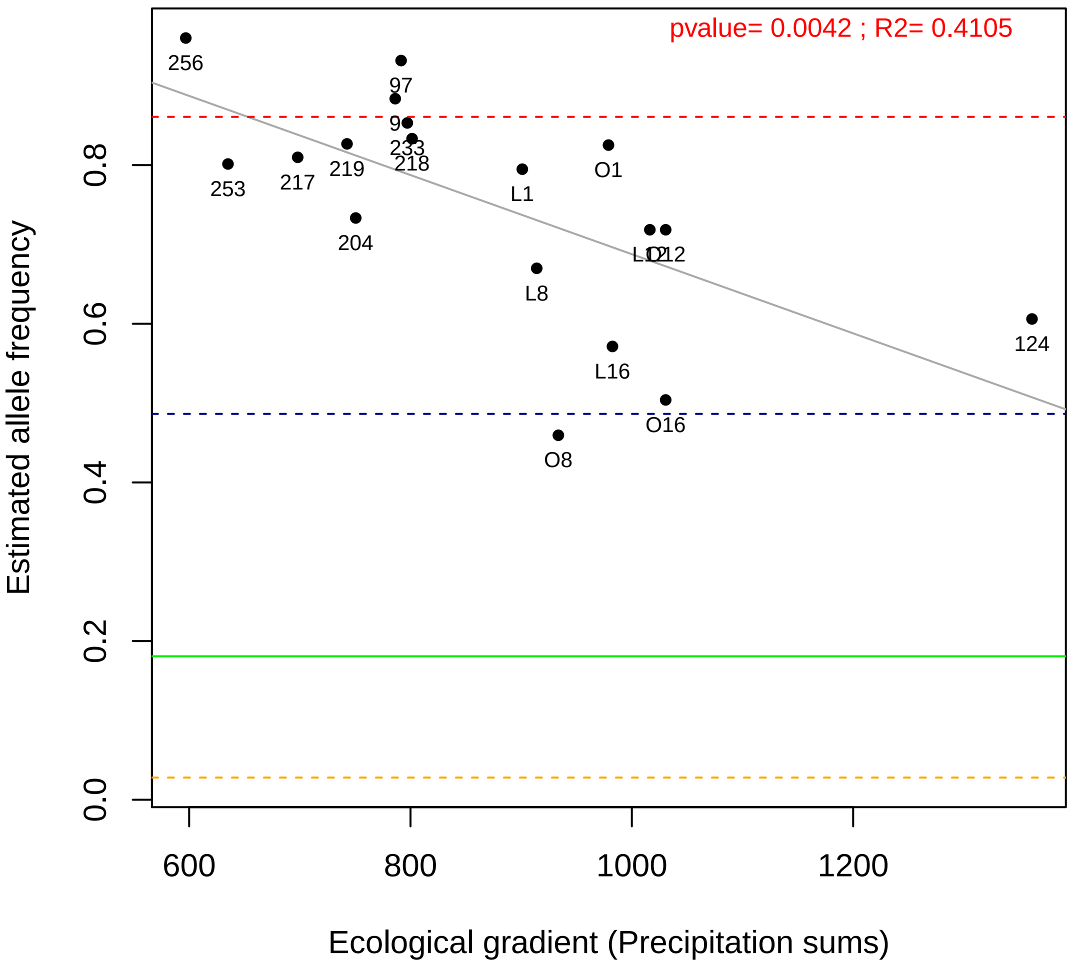

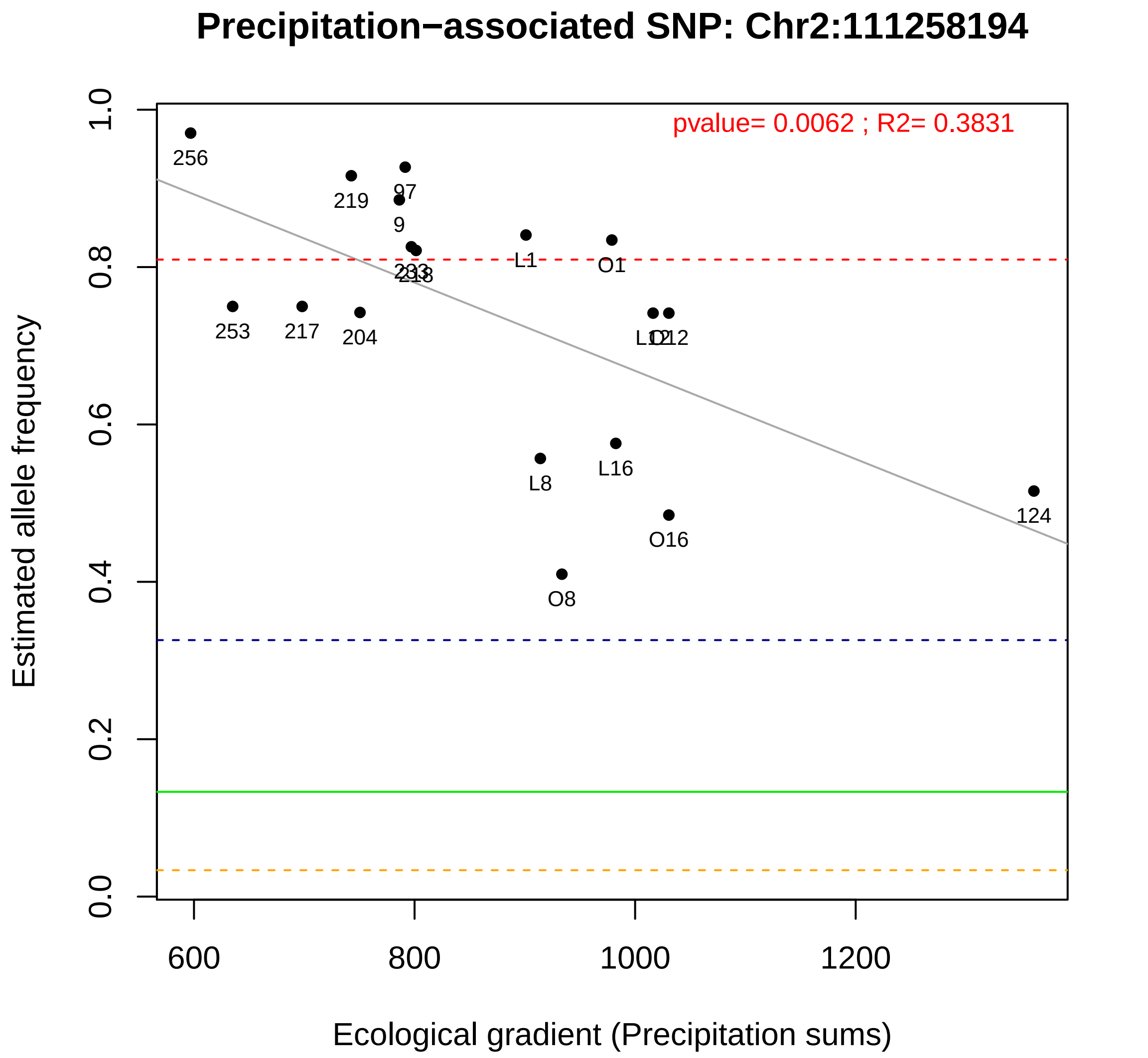

Precipitation−associated SNP: Chr2:111258194
pvalue= 0.0062 ; R2= 0.3831
256
219
97
9
L1
O1
233
218
253
217
204
L12
O12
L8
L16
124
O16
O8
Estimated allele frequency
0.0
0.2
0.4
0.6
0.8
1.0
600
800
1000
1200
Ecological gradient (Precipitation sums)

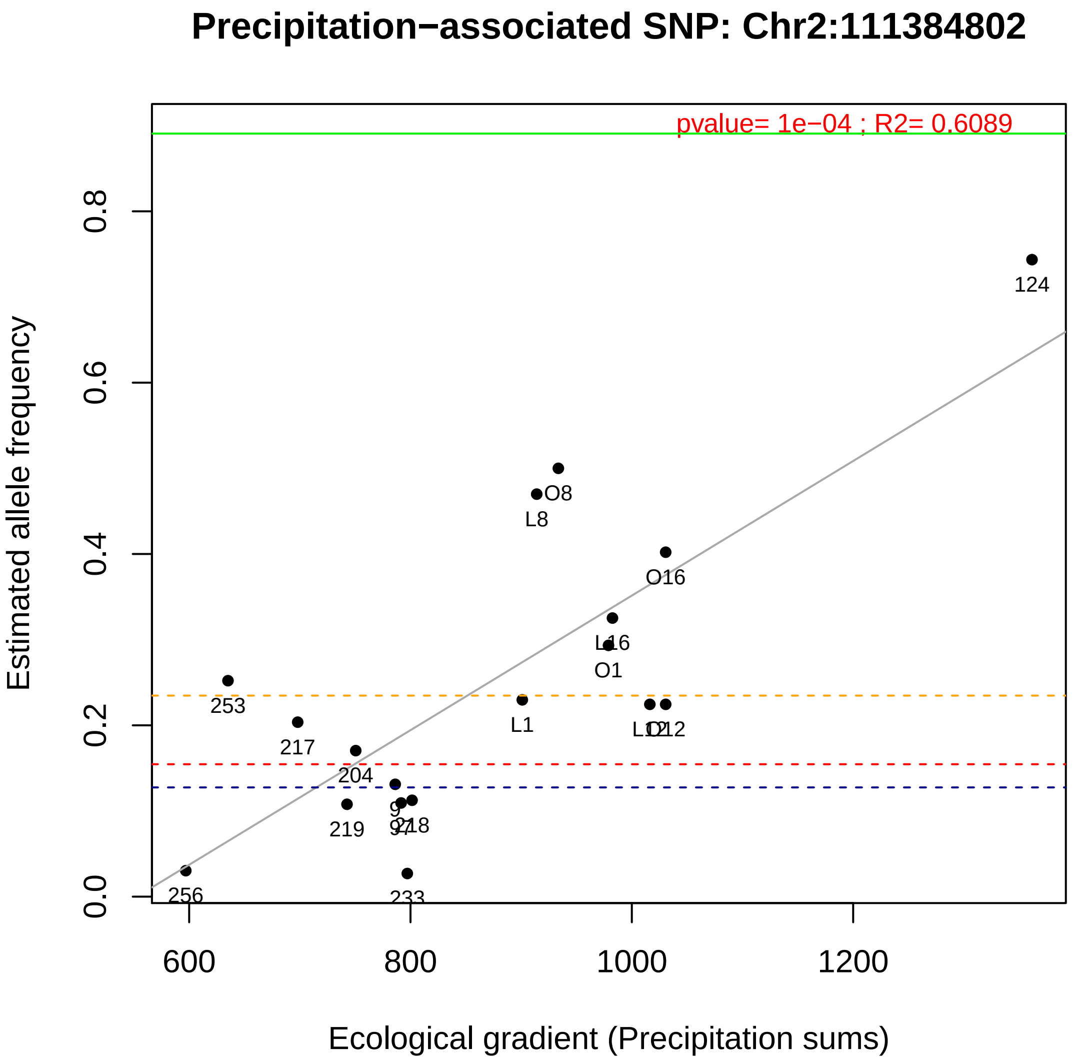

Precipitation−associated SNP: Chr2:111384802
pvalue= 1e−04 ; R2= 0.6089
Estimated allele frequency
Ecological gradient (Precipitation sums)
0.0
0.2
0.4
0.6
0.8
600
800
1000
1200
124
O8
L8
O16
L16
O1
253
217
204
L1
L12
O12
219
9
97
218
256
233

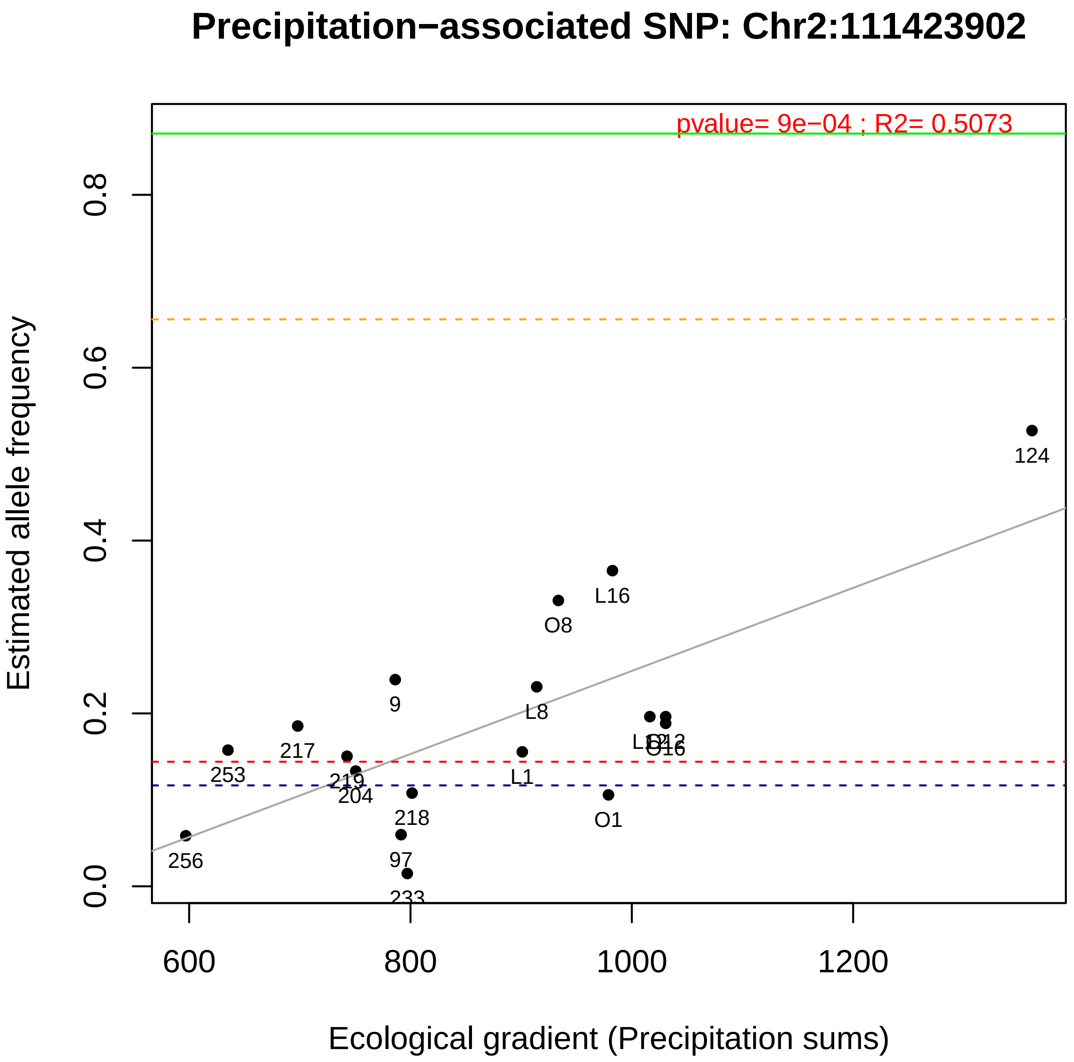
Precipitation−associated SNP: Chr2:111423902
pvalue= 9e−04 ; R2= 0.5073
Estimated allele frequency
0.0
0.2
0.4
0.6
0.8
600
800
1000
1200
Ecological gradient (Precipitation sums)
124
L16
O8
9
L8
217
253
219
204
L1
218
O1
256
97
233
O16

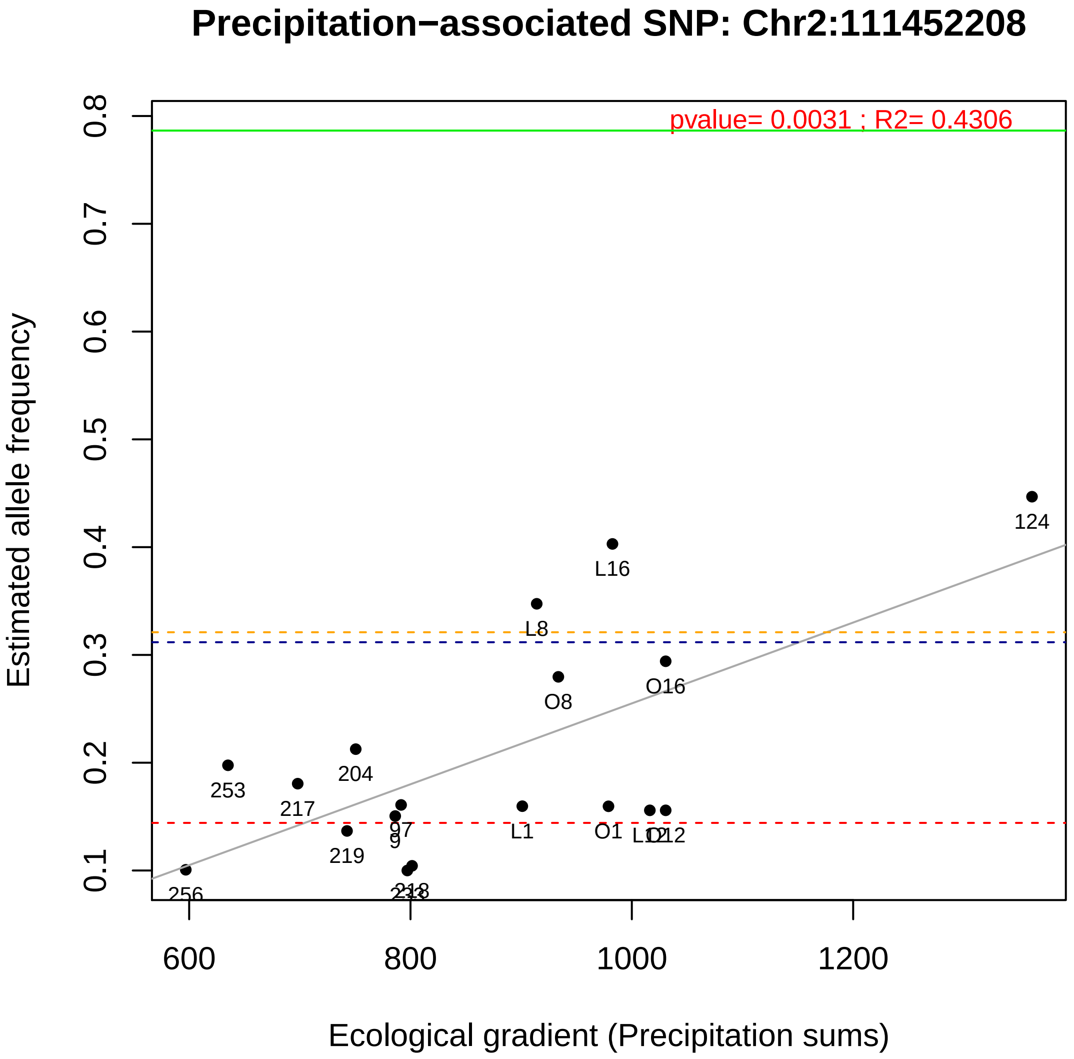
Precipitation−associated SNP: Chr2:111452208
pvalue= 0.0031 ; R2= 0.4306
Estimated allele frequency
0.1
0.2
0.3
0.4
0.5
0.6
0.7
0.8
600
800
1000
1200
Ecological gradient (Precipitation sums)
124
L16
L8
O16
O8
204
253
217
219
L1
O1
256

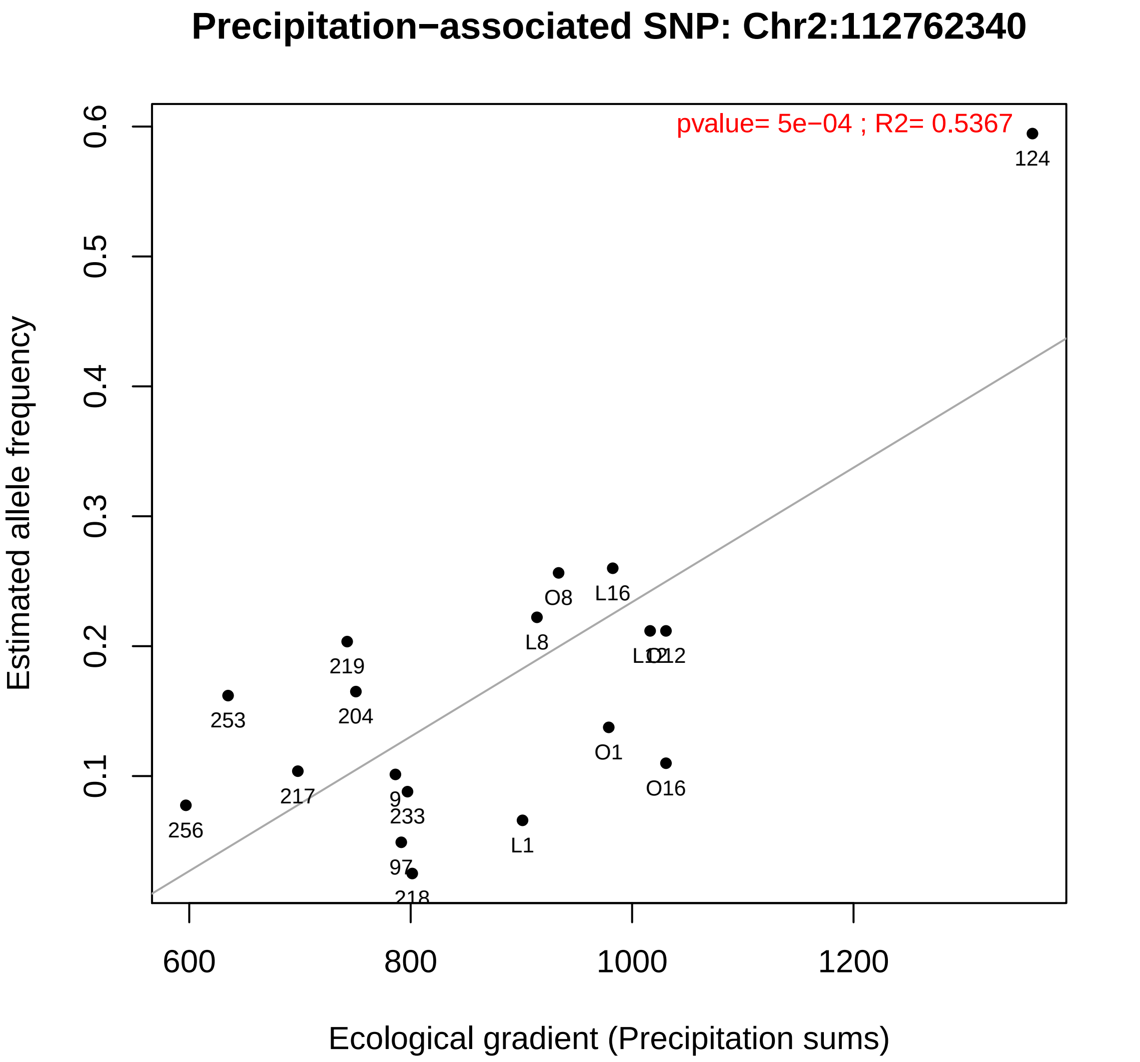
Precipitation−associated SNP: Chr2:112762340
pvalue= 5e−04 ; R2= 0.5367
Estimated allele frequency
0.1
0.2
0.3
0.4
0.5
0.6
600
800
1000
1200
Ecological gradient (Precipitation sums)
124
O8
L16
L8
219
L12
O12
253
204
O1
O16
217
9
233
256
L1
97
218

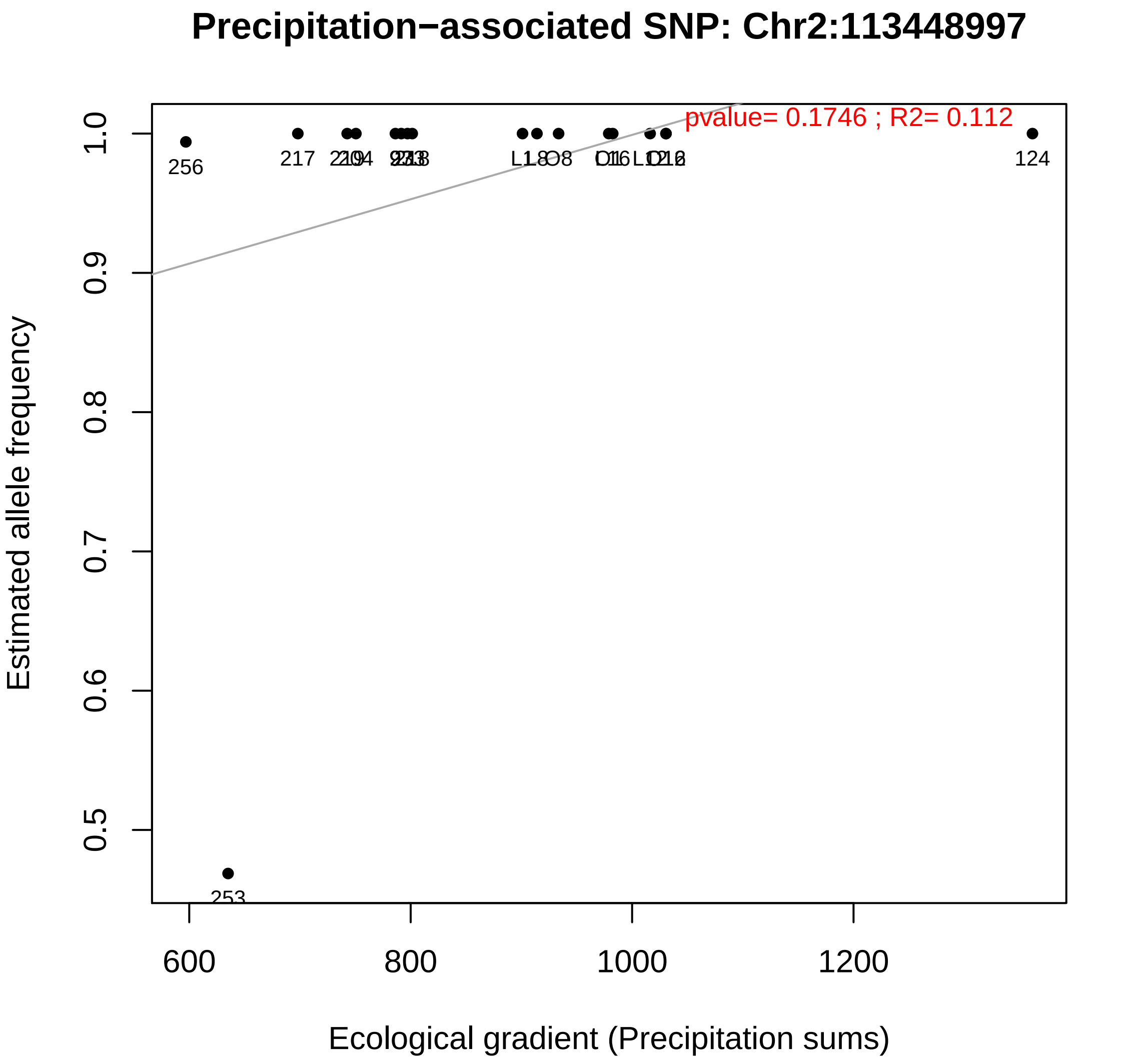
Precipitation−associated SNP: Chr2:113448997
pvalue= 0.1746 ; R2= 0.112
Estimated allele frequency
Ecological gradient (Precipitation sums)
0.5
0.6
0.7
0.8
0.9
1.0
600
800
1000
1200
256
217
124
253

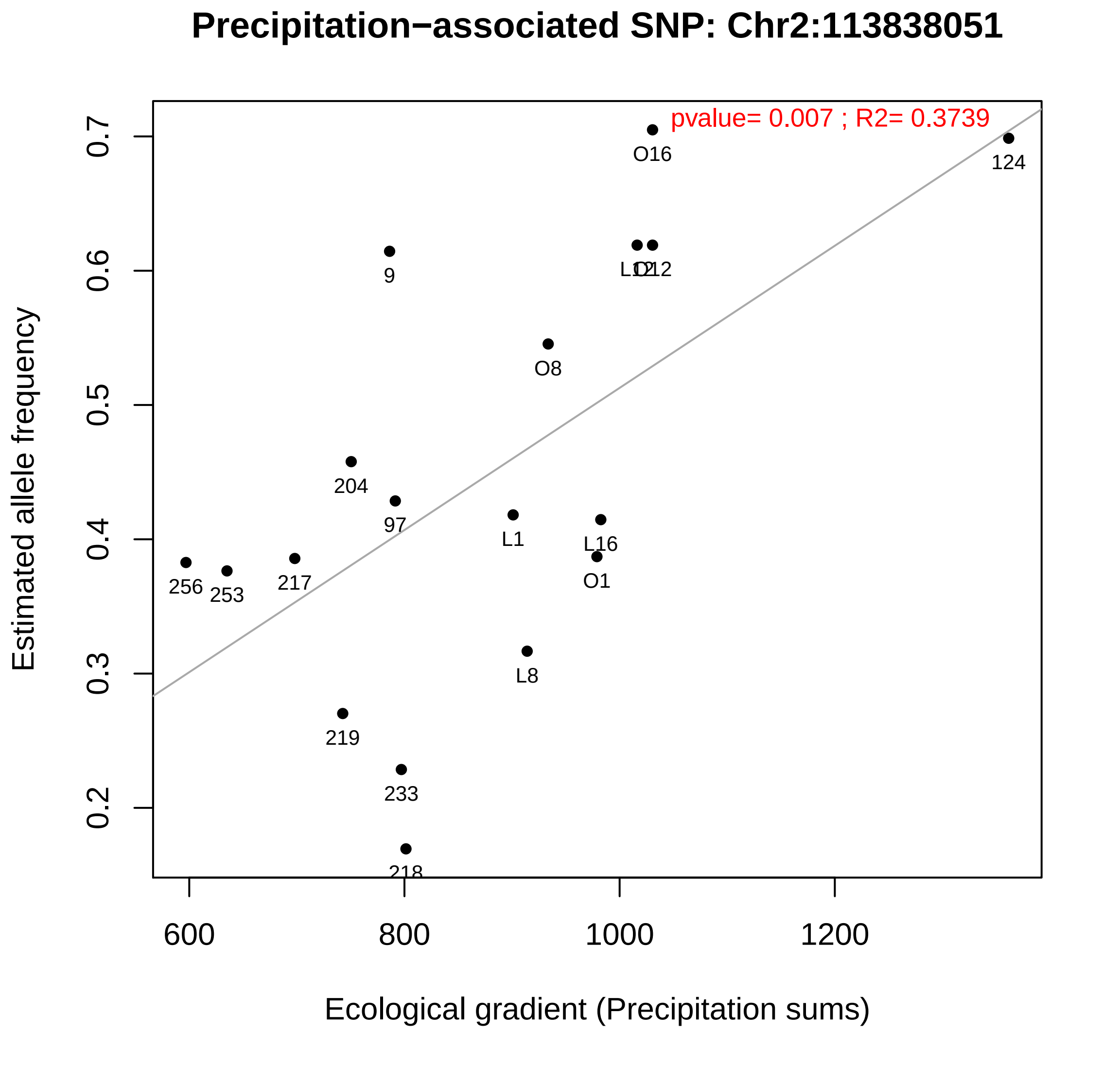

Precipitation−associated SNP: Chr2:113838051
pvalue= 0.007 ; R2= 0.3739
Estimated allele frequency
Ecological gradient (Precipitation sums)
0.2
0.3
0.4
0.5
0.6
0.7
600
800
1000
1200
O16
124
9
L12
O12
O8
204
97
L1
L16
O1
256
253
217
L8
219
233
218

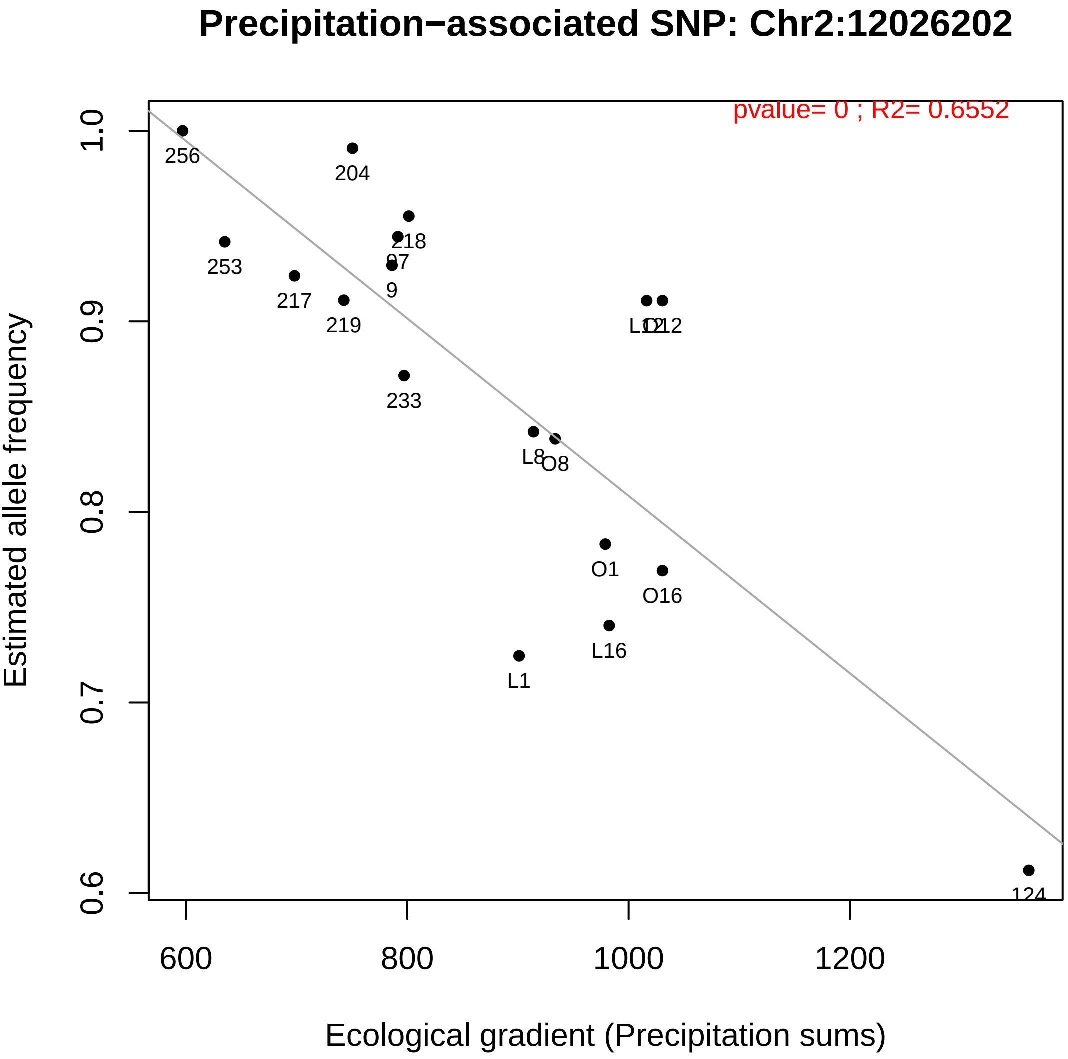

Precipitation−associated SNP: Chr2:12026202
pvalue= 0 ; R2= 0.6552
Estimated allele frequency
1.0
0.9
0.8
0.7
0.6
600
800
1000
1200
Ecological gradient (Precipitation sums)
256
204
253
218
97
9
217
219
233
L12
O12
L8
O8
O1
O16
L16
L1
124

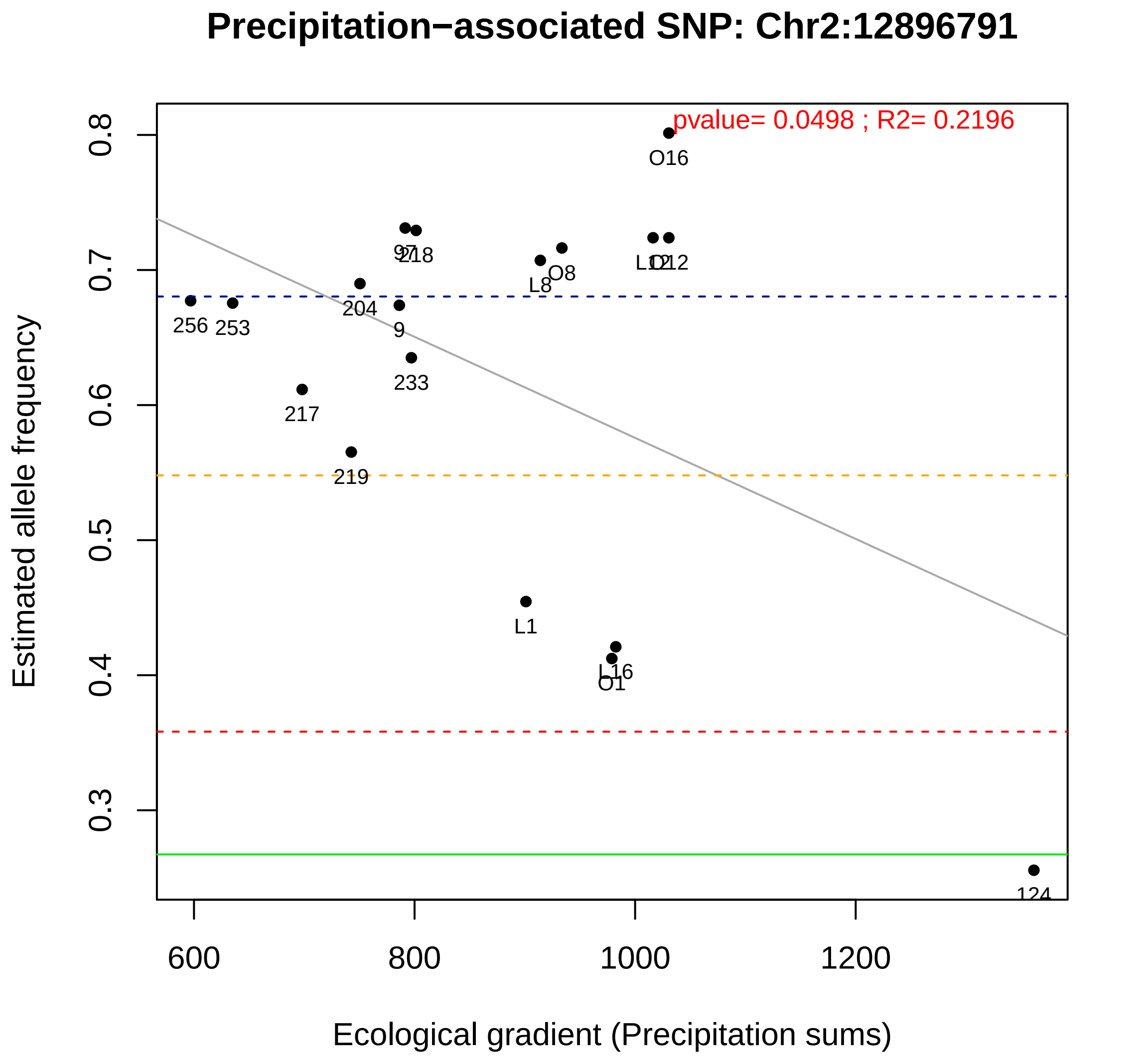

Precipitation−associated SNP: Chr2:12896791
pvalue= 0.0498 ; R2= 0.2196
Estimated allele frequency
0.3
0.4
0.5
0.6
0.7
0.8
600
800
1000
1200
Ecological gradient (Precipitation sums)
O16
97
218
L12
O12
O8
L8
204
256
253
9
233
217
219
L1
L16
O1
124

**Precipitation−associated SNP: Chr2:12955563**

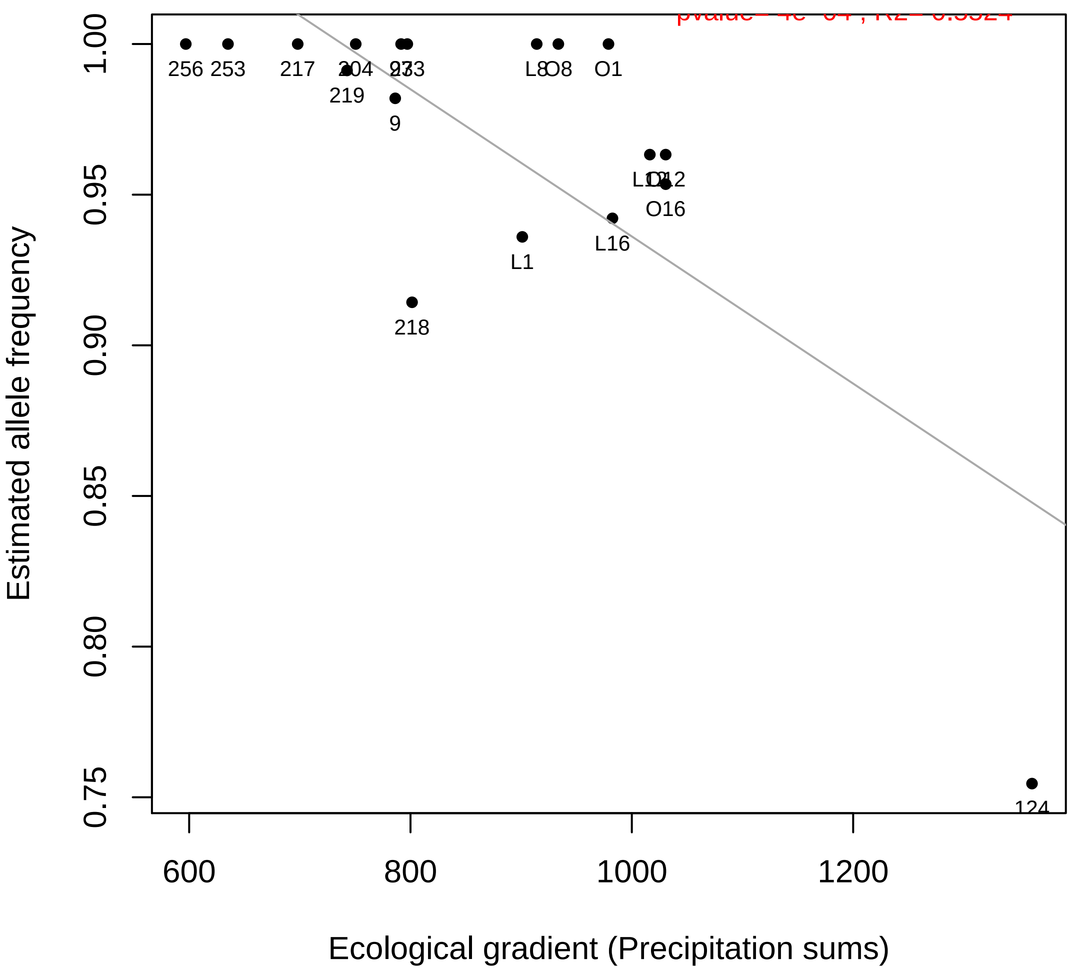

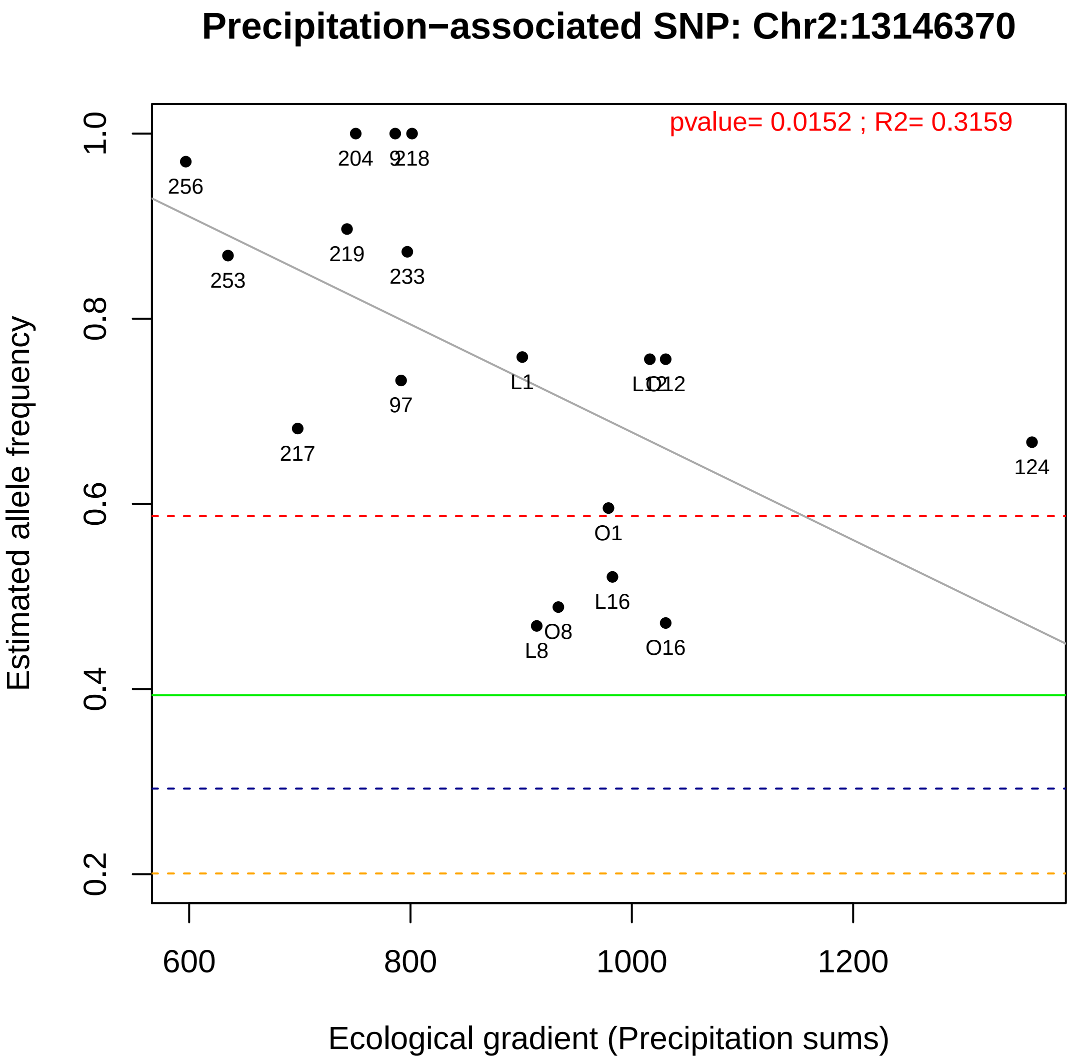

Precipitation−associated SNP: Chr2:13146370
pvalue= 0.0152 ; R2= 0.3159
Estimated allele frequency
Ecological gradient (Precipitation sums)
1.0
0.8
0.6
0.4
0.2
600
800
1000
1200
256
204
218
219
253
233
L1
97
217
124
O1
L16
O8
L8
O16

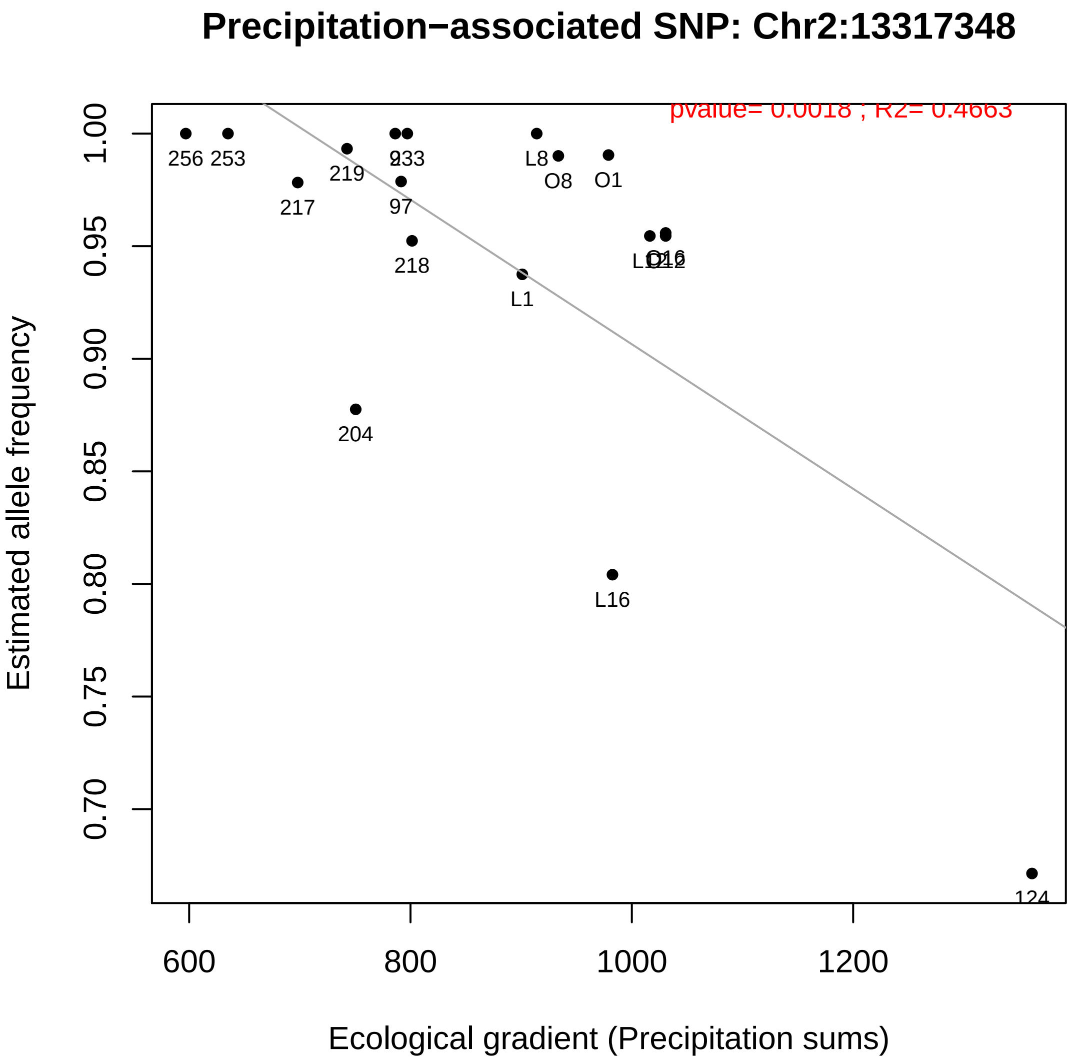

Precipitation−associated SNP: Chr2:13317348
pvalue= 0.0018 ; R2= 0.4663
Estimated allele frequency
Ecological gradient (Precipitation sums)
1.00
0.95
0.90
0.85
0.80
0.75
0.70
600
800
1000
1200
256
253
219
233
L8
O8
O1
217
97
218
L1
O16
204
L16
124

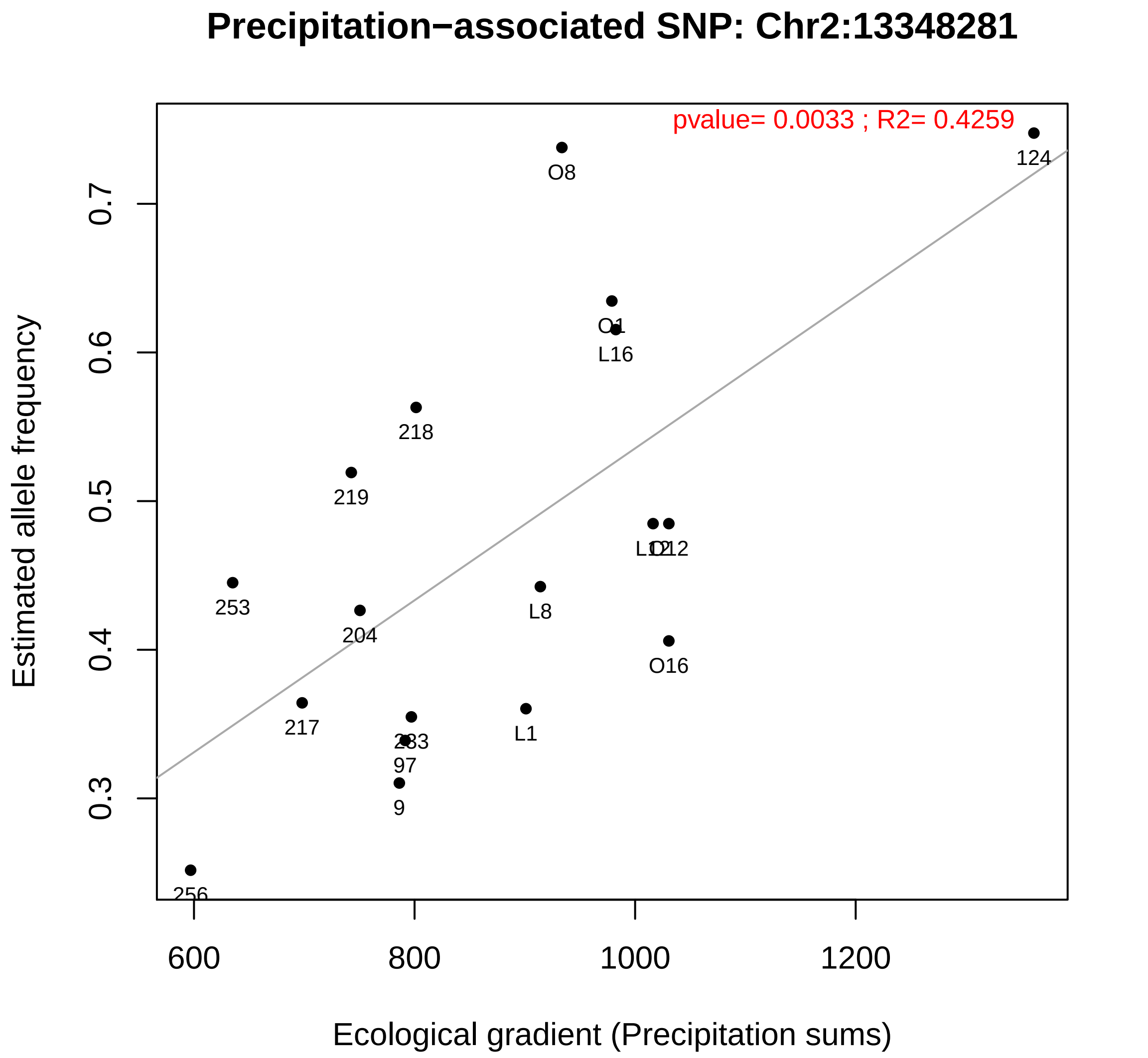

Precipitation−associated SNP: Chr2:13348281
pvalue= 0.0033 ; R2= 0.4259
Estimated allele frequency
0.3
0.4
0.5
0.6
0.7
600
800
1000
1200
Ecological gradient (Precipitation sums)
O8
124
O1
L16
218
219
L12
O12
253
204
L8
O16
217
283
97
L1
9
256

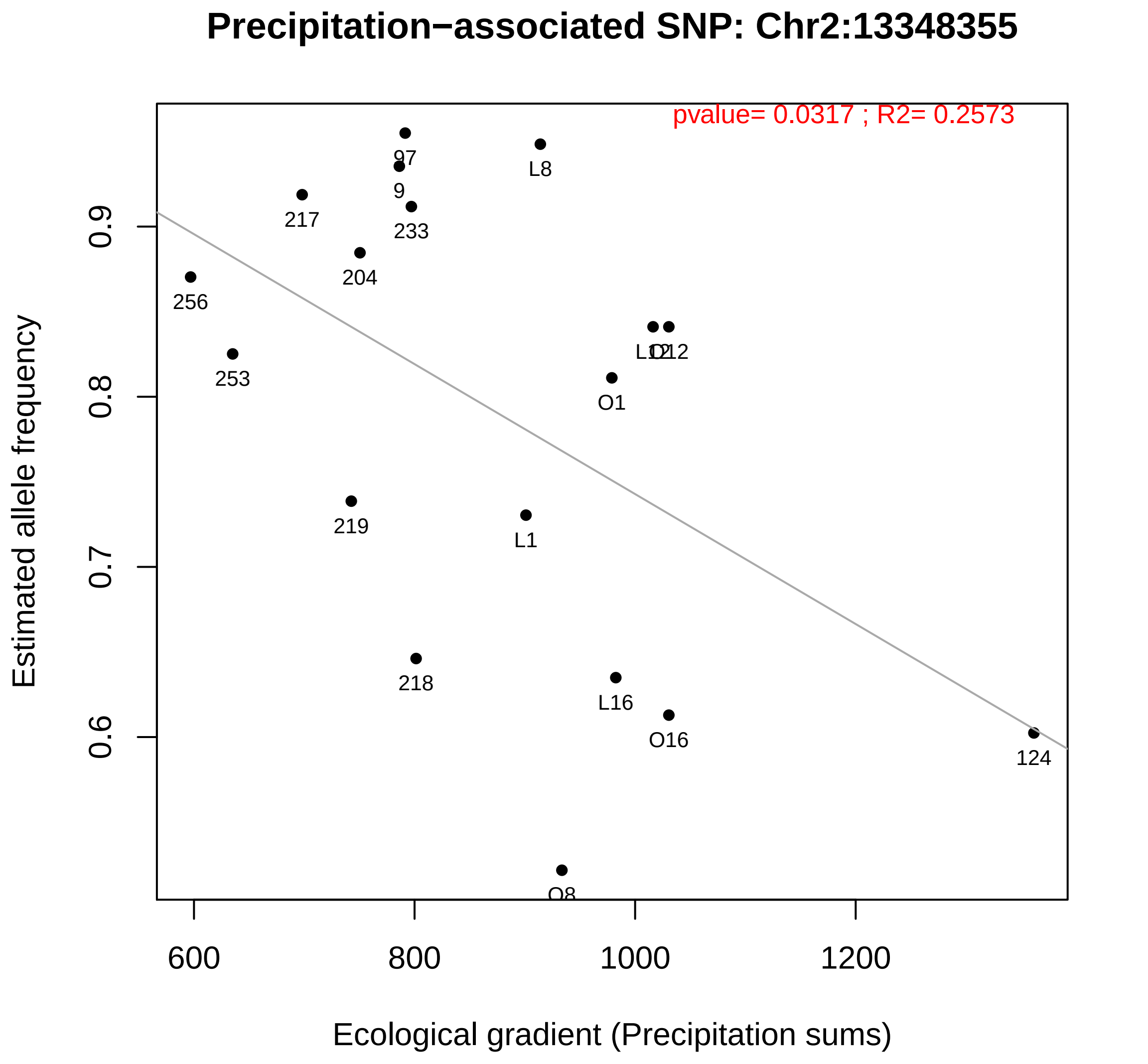
Precipitation−associated SNP: Chr2:13348355
pvalue= 0.0317 ; R2= 0.2573
97
9
L8
217
233
204
256
L10
O12
253
O1
219
L1
218
L16
O16
124
O8
0.6
0.7
0.8
0.9
Estimated allele frequency
600
800
1000
1200
Ecological gradient (Precipitation sums)

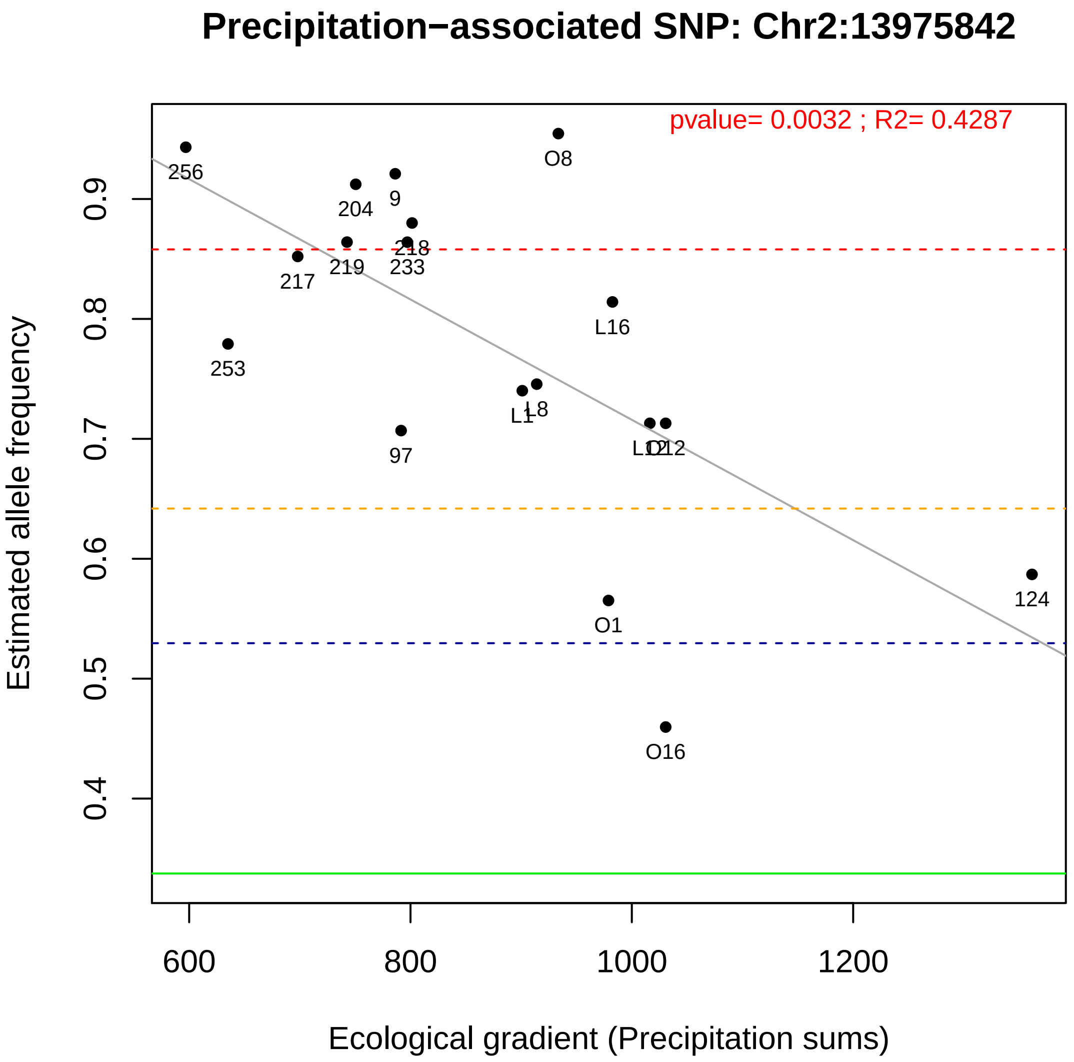

Precipitation−associated SNP: Chr2:13975842
pvalue= 0.0032 ; R2= 0.4287
Estimated allele frequency
Ecological gradient (Precipitation sums)
0.4
0.5
0.6
0.7
0.8
0.9
600
800
1000
1200
256
O8
204
9
218
219
233
217
L16
253
L1
L8
97
L12
O12
124
O1
O16

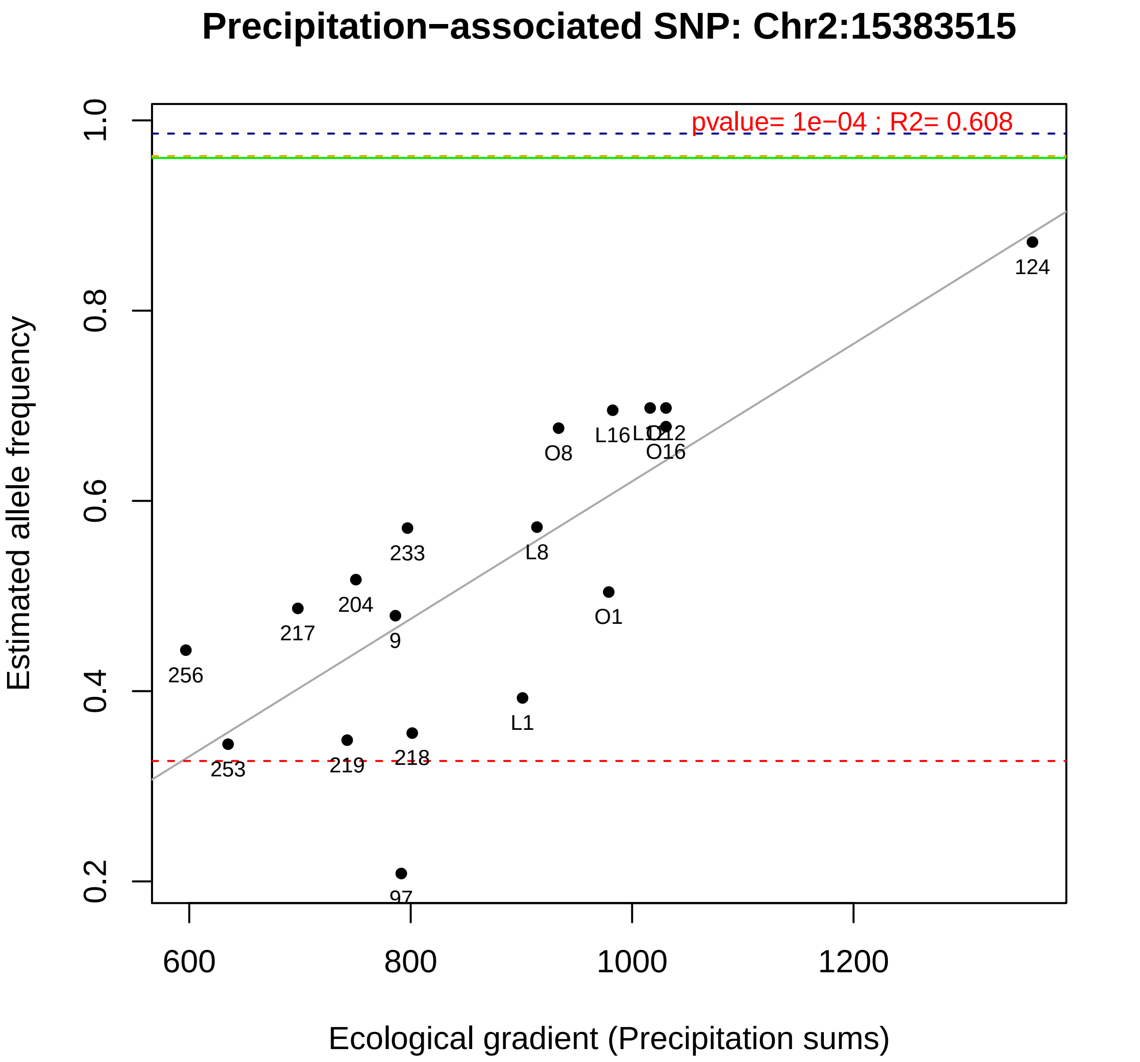

Precipitation−associated SNP: Chr2:15383515
pvalue= 1e−04 ; R2= 0.608
124
L16
O12
O16
O8
233
L8
204
O1
217
9
256
L1
253
219
218
97
0.2
0.4
0.6
0.8
1.0
600
800
1000
1200
Estimated allele frequency
Ecological gradient (Precipitation sums)

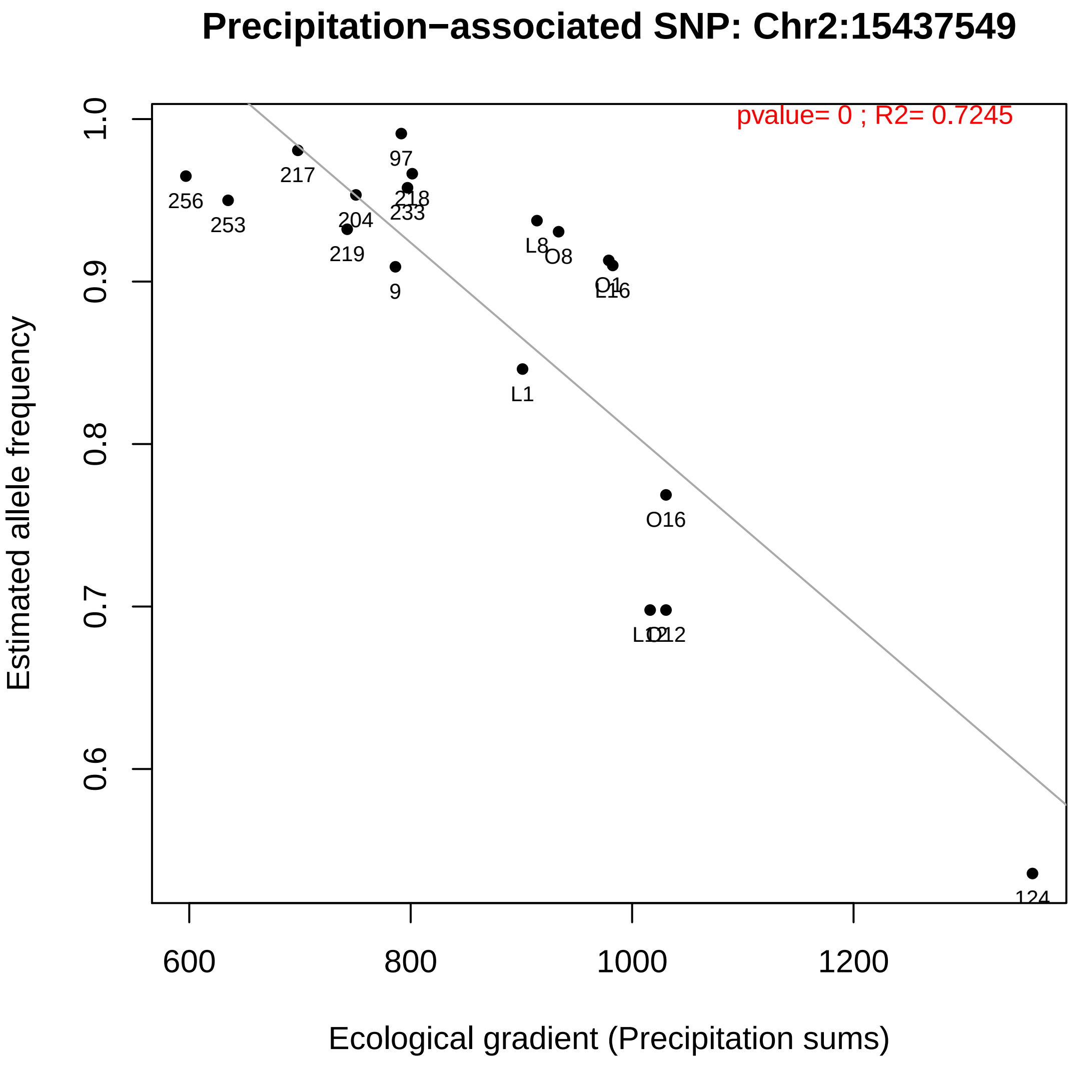
Precipitation−associated SNP: Chr2:15437549
pvalue= 0 ; R2= 0.7245
Estimated allele frequency
Ecological gradient (Precipitation sums)
1.0
0.9
0.8
0.7
0.6
600
800
1000
1200
97
217
256
253
204
219
218
233
9
L8
O8
L16
L1
O16
O12
124

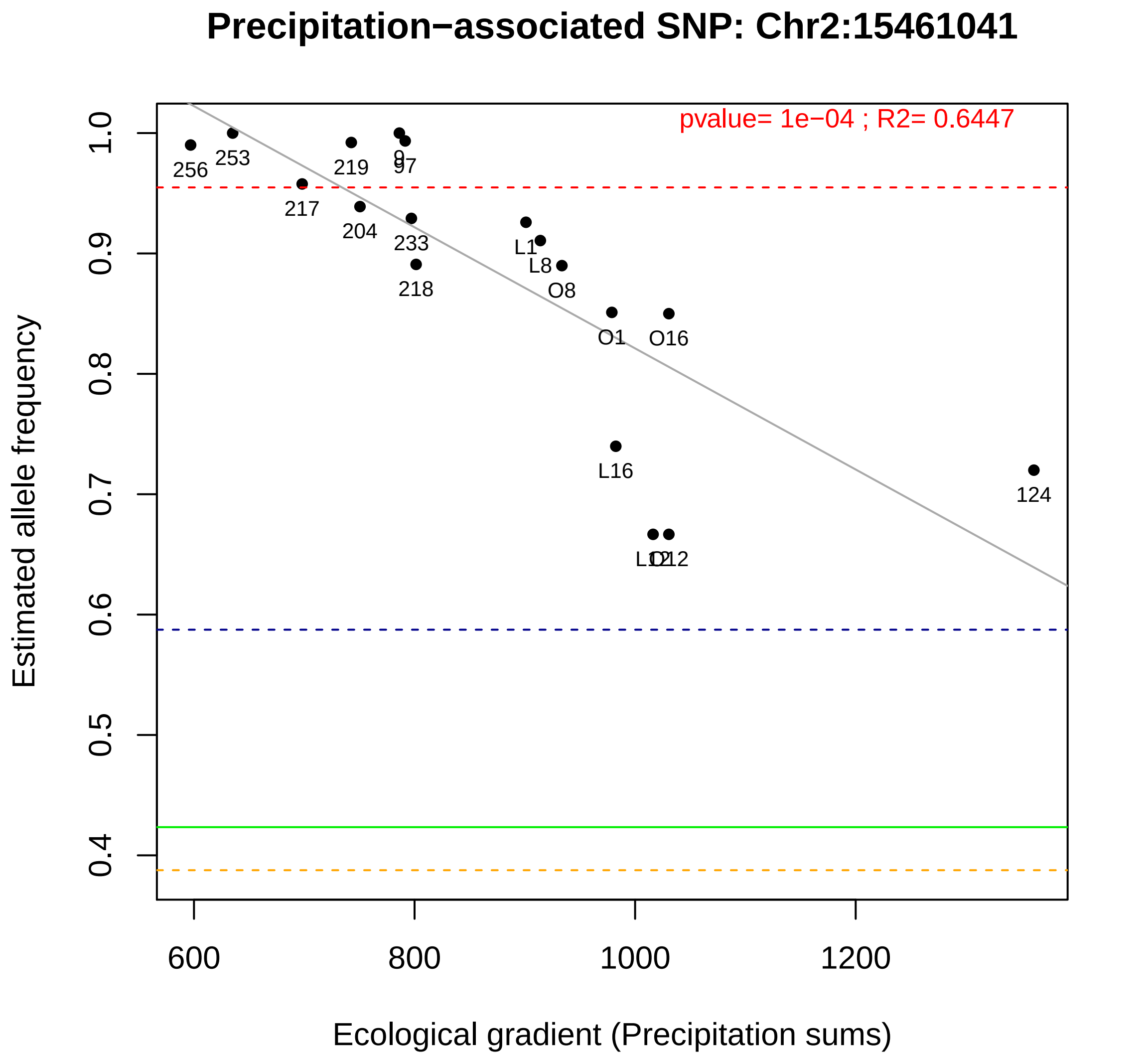
Precipitation−associated SNP: Chr2:15461041
pvalue= 1e−04 ; R2= 0.6447
Estimated allele frequency
Ecological gradient (Precipitation sums)
1.0
0.9
0.8
0.7
0.6
0.5
0.4
600
800
1000
1200
256
253
219
9
97
217
204
233
218
L1
L8
O8
O1
O16
L16
124
L12
O12

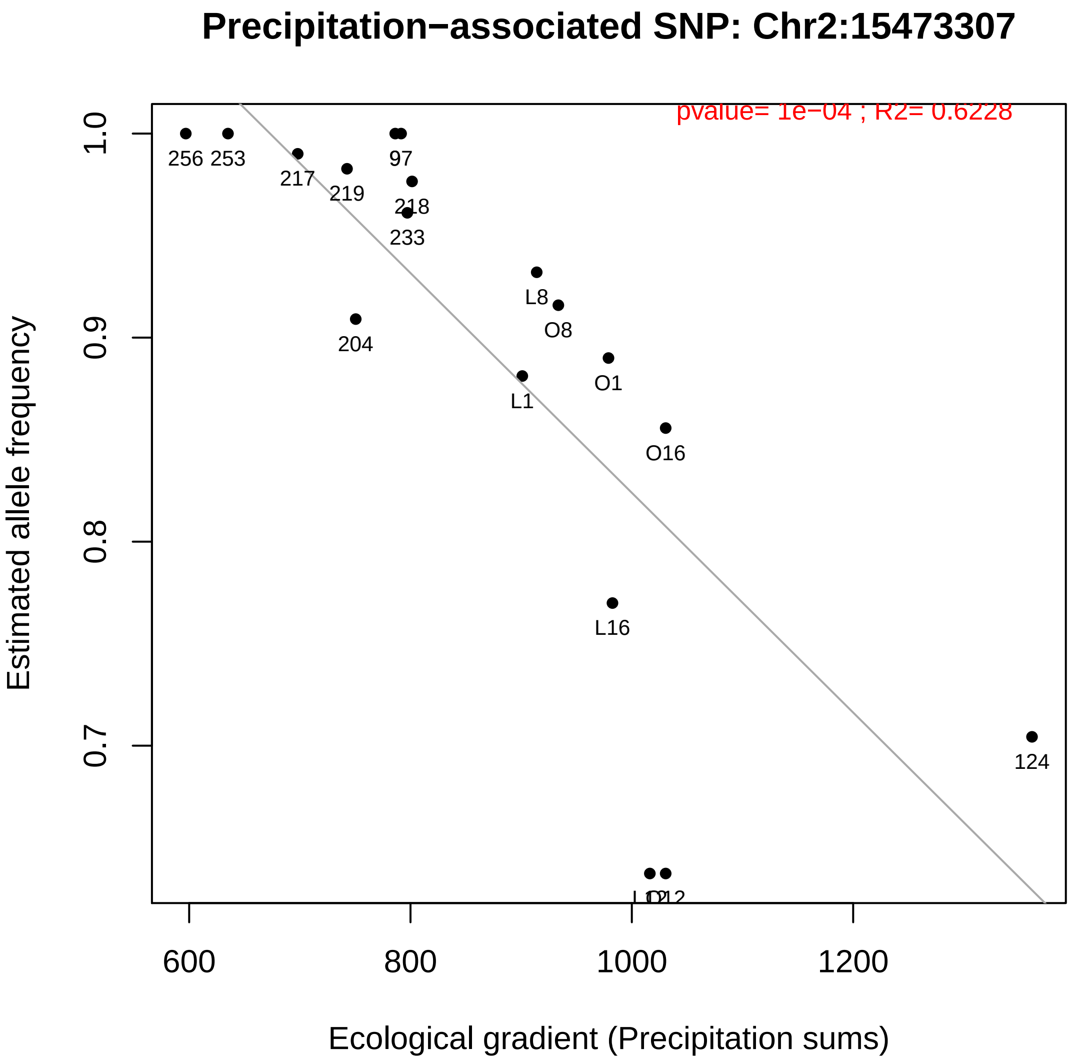

Precipitation−associated SNP: Chr2:15473307
pvalue= 1e−04 ; R2= 0.6228
Estimated allele frequency
1.0
0.9
0.8
0.7
600
800
1000
1200
Ecological gradient (Precipitation sums)
256
253
217
219
97
218
233
204
L8
O8
L1
O1
O16
L16
124

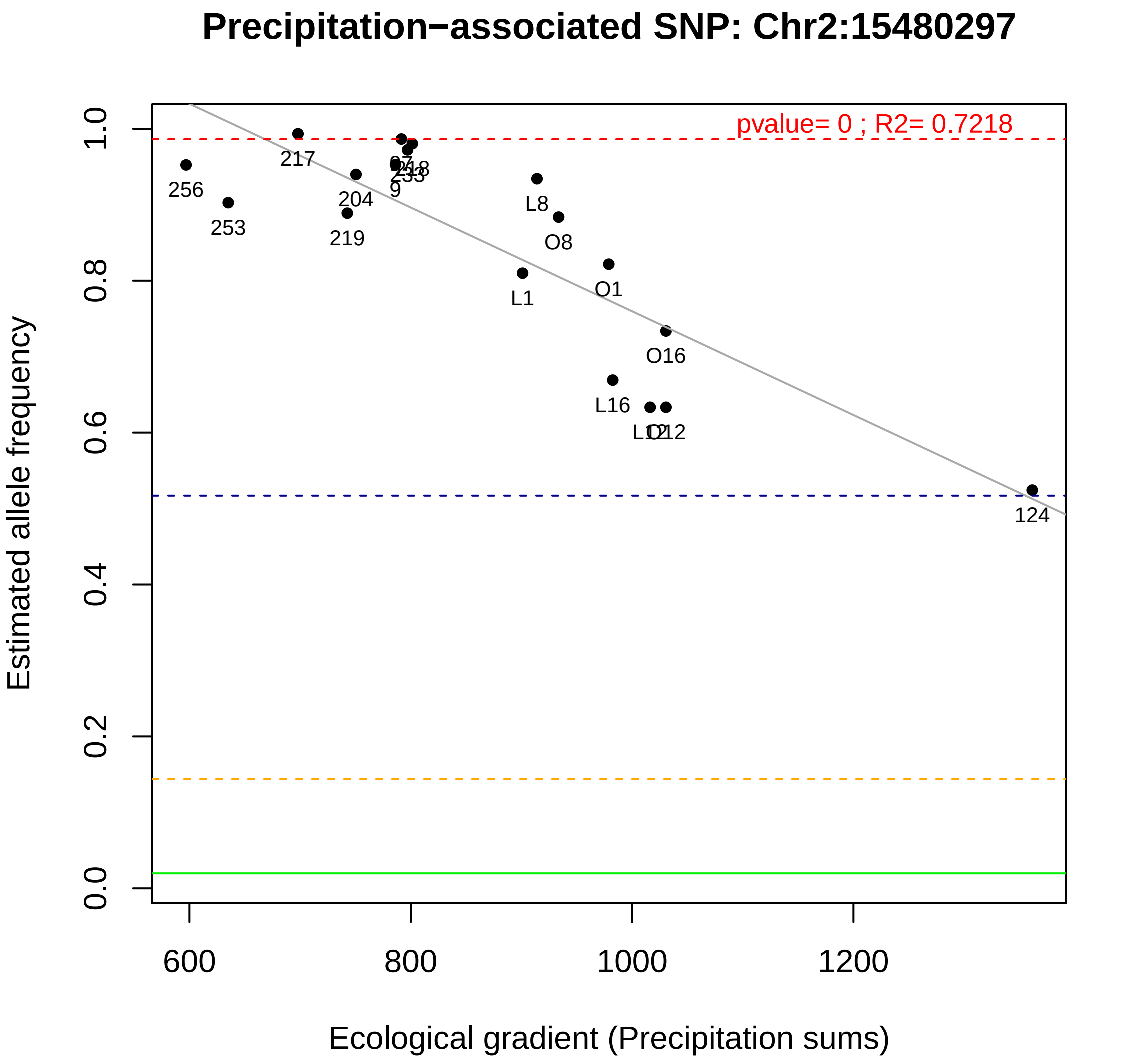
Precipitation−associated SNP: Chr2:15480297
pvalue= 0 ; R2= 0.7218
Estimated allele frequency
Ecological gradient (Precipitation sums)
0.0
0.2
0.4
0.6
0.8
1.0
600
800
1000
1200
256
253
217
204
219
L8
O8
L1
O1
O16
L16
124

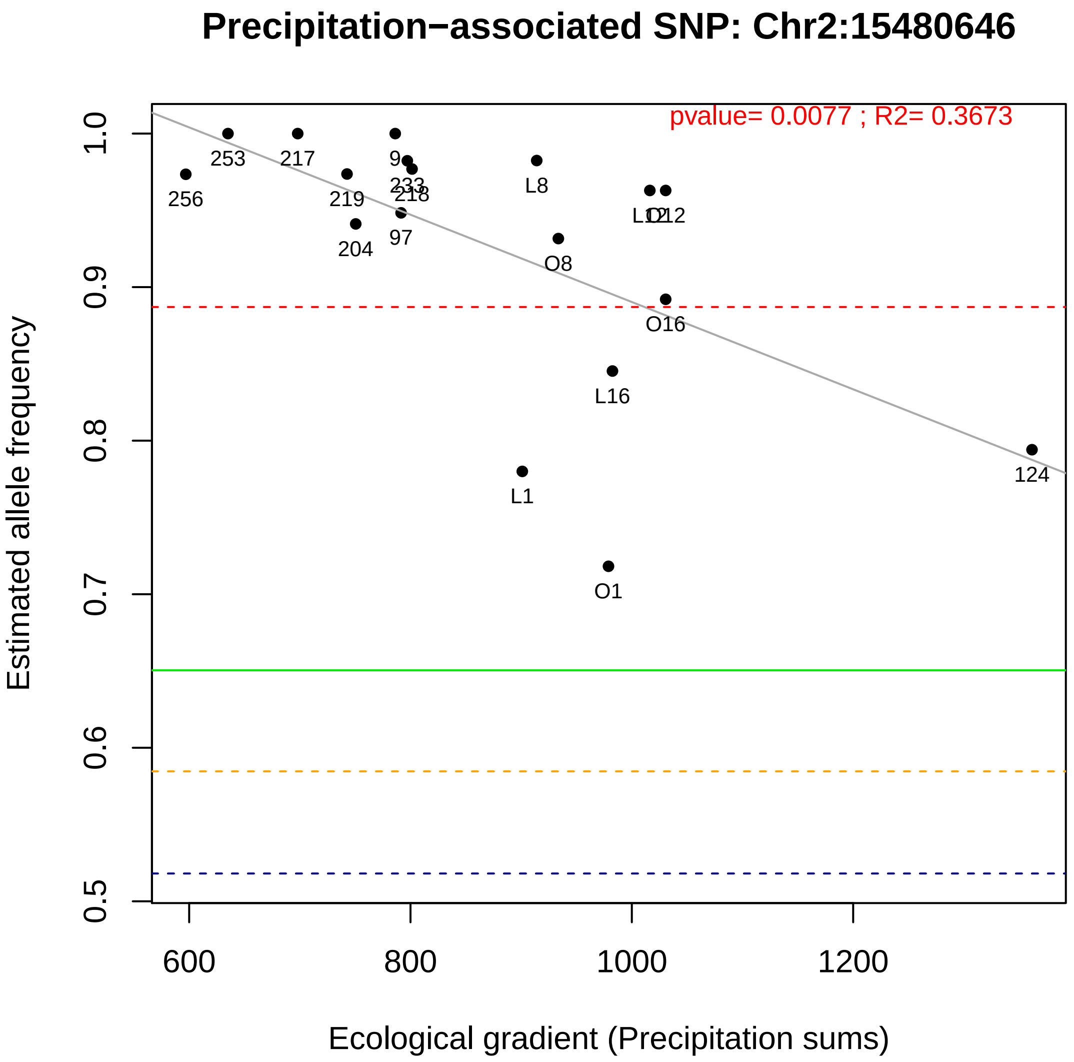

Precipitation−associated SNP: Chr2:15480646
pvalue= 0.0077 ; R2= 0.3673
Estimated allele frequency
0.5
0.6
0.7
0.8
0.9
1.0
600
800
1000
1200
Ecological gradient (Precipitation sums)
253
217
256
219
204
97
233
218
L8
O8
O16
L16
L1
O1
124

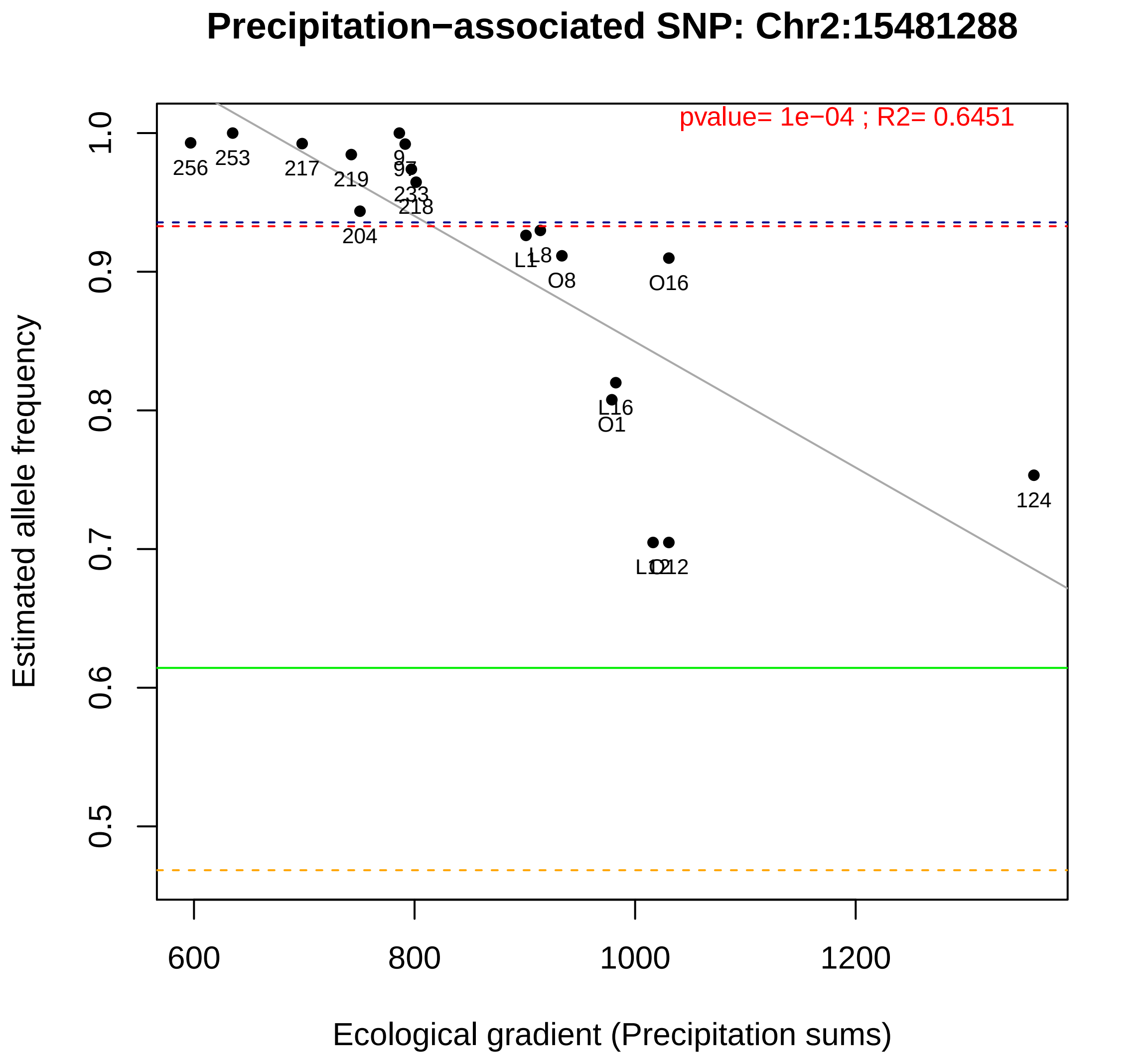

Precipitation−associated SNP: Chr2:15481288
pvalue= 1e−04 ; R2= 0.6451
Estimated allele frequency
Ecological gradient (Precipitation sums)
0.5
0.6
0.7
0.8
0.9
1.0
600
800
1000
1200
256
253
217
219
9
97
233
218
204
L1
L8
O8
O16
L16
O1
124
L12
O12

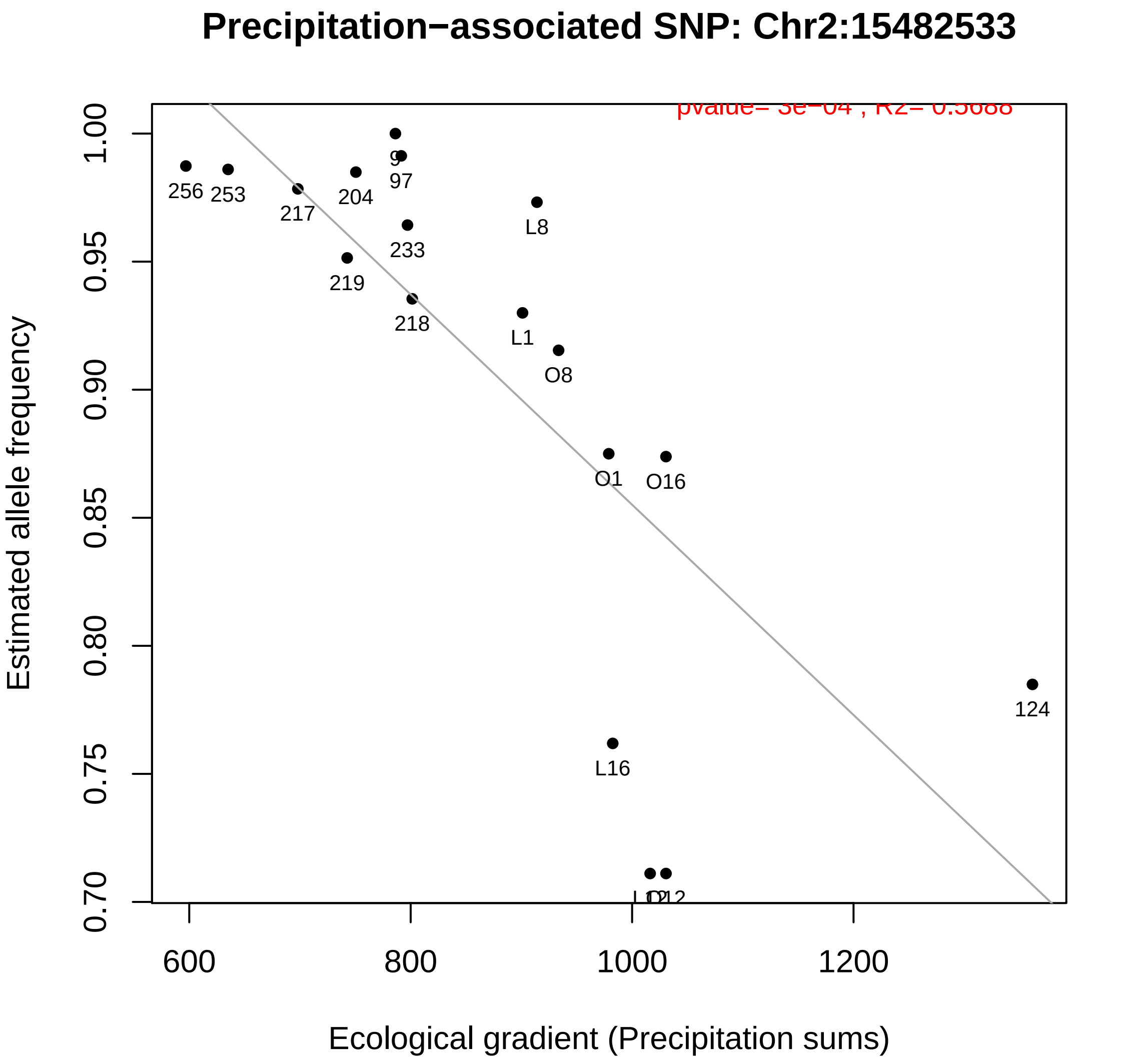

Precipitation−associated SNP: Chr2:15482533
pvalue= 3e−04 , R2= 0.5688
Estimated allele frequency
1.00
0.95
0.90
0.85
0.80
0.75
0.70
600
800
1000
1200
Ecological gradient (Precipitation sums)
256
253
217
204
97
233
219
218
L8
L1
O8
O1
O16
124
L16
O12

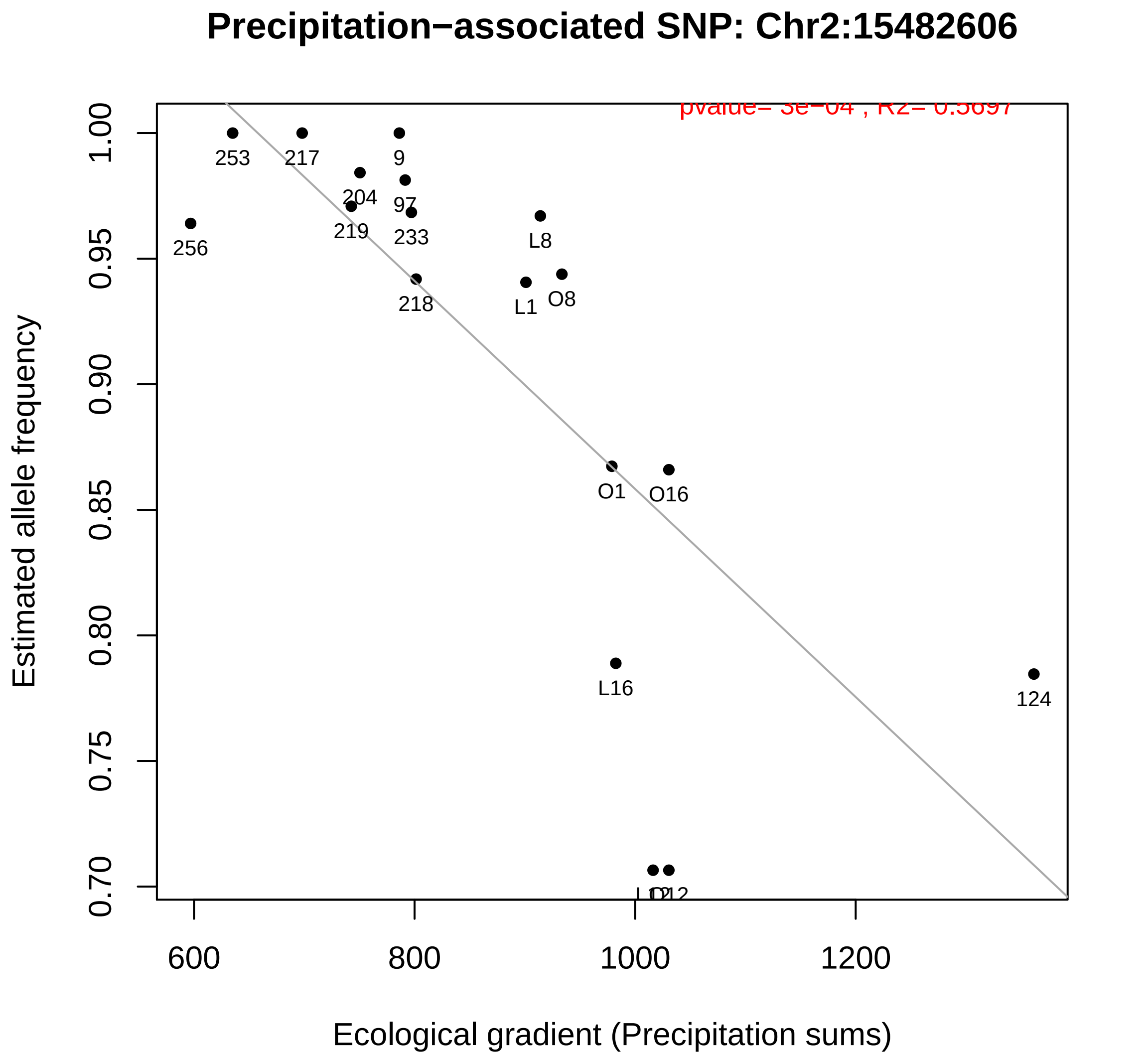

Precipitation−associated SNP: Chr2:15482606
pvalue= 3e−04 , R2= 0.5697
Estimated allele frequency
Ecological gradient (Precipitation sums)
1.00
0.95
0.90
0.85
0.80
0.75
0.70
600
800
1000
1200
253
217
9
204
97
219
233
256
L8
218
L1
O8
O1
O16
L16
124

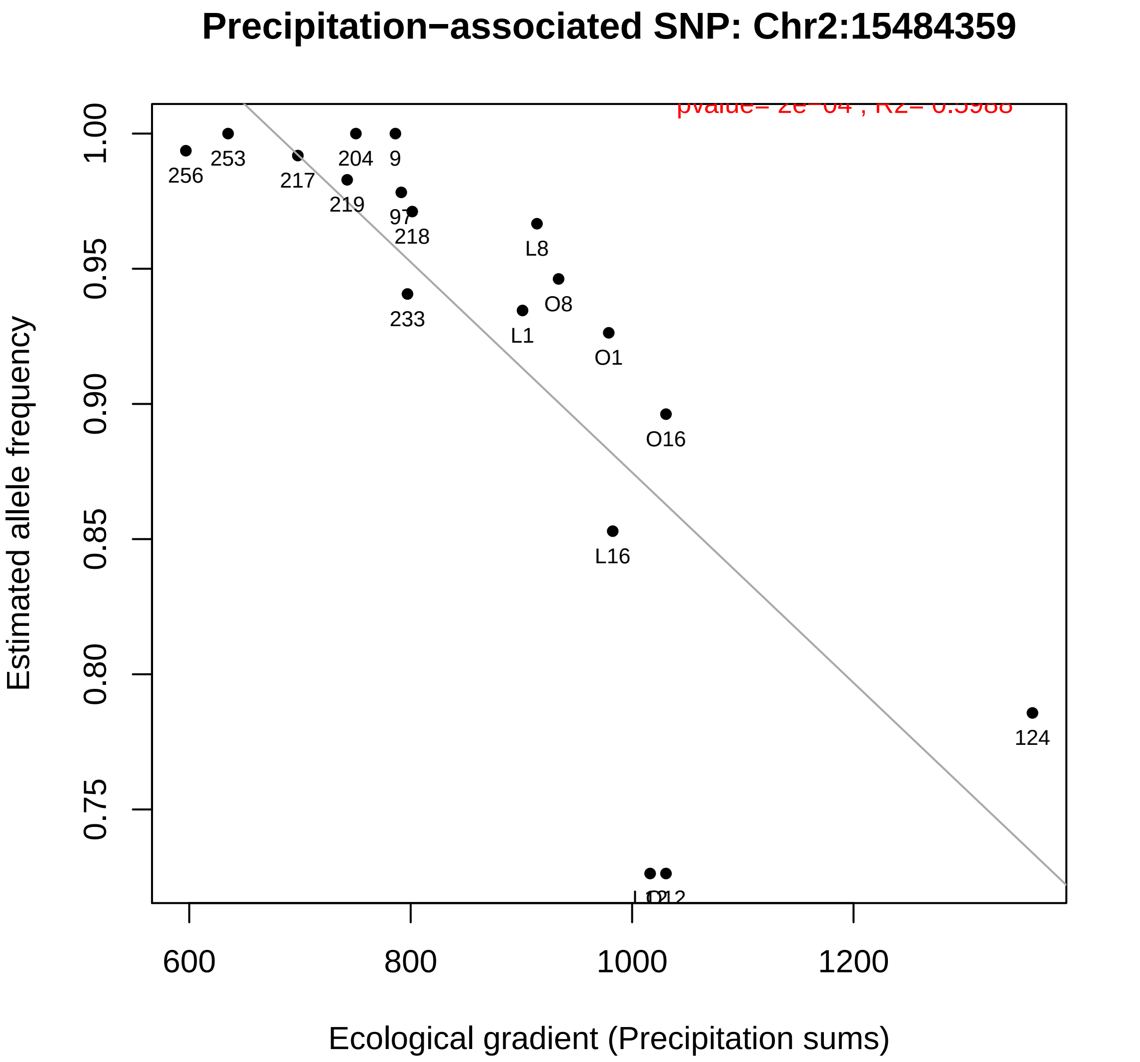

Precipitation−associated SNP: Chr2:15484359
pvalue= 2e−04 , R2= 0.5988
Estimated allele frequency
1.00
0.95
0.90
0.85
0.80
0.75
600
800
1000
1200
Ecological gradient (Precipitation sums)
256
253
217
204
9
219
97
218
L8
O8
233
L1
O1
O16
L16
124

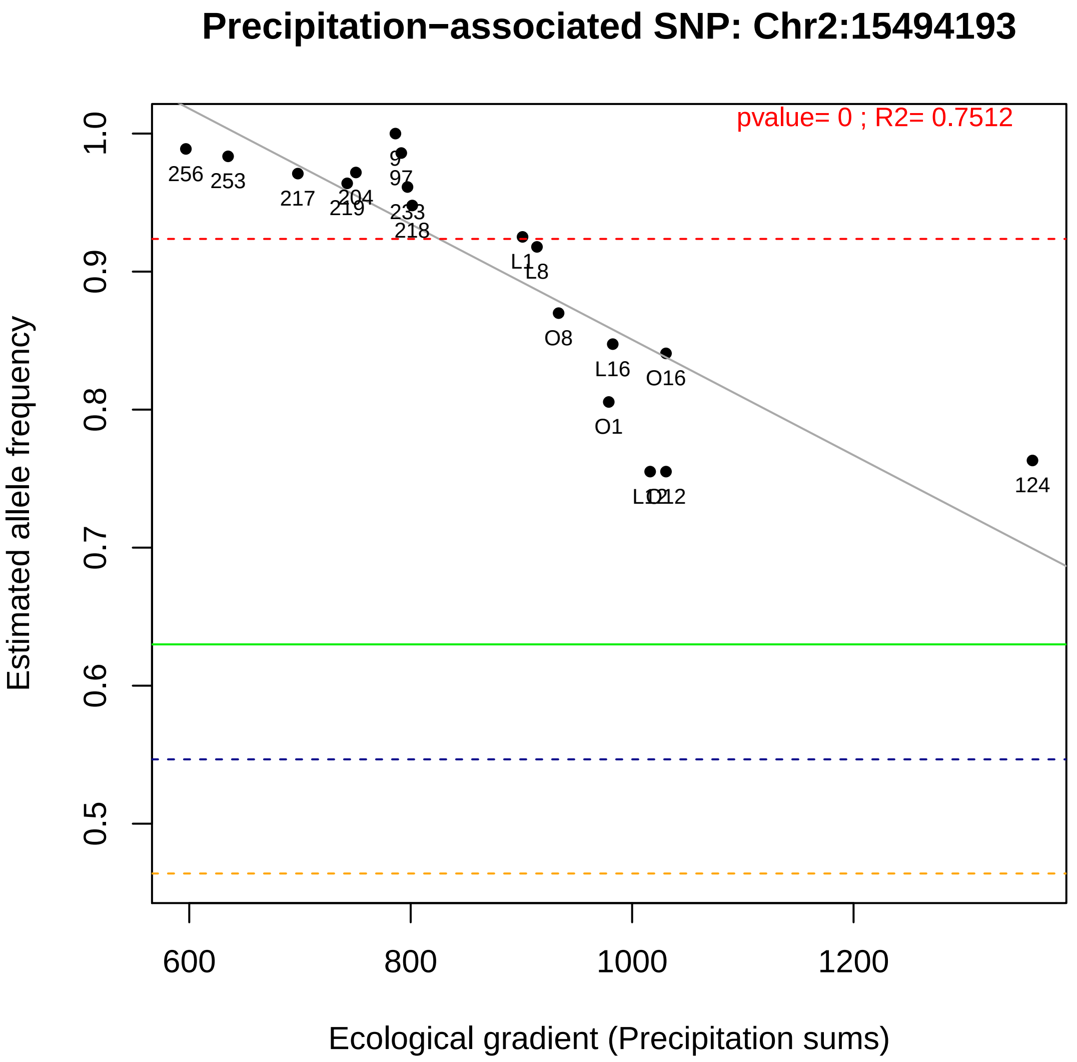
Precipitation−associated SNP: Chr2:15494193
pvalue= 0 ; R2= 0.7512
Estimated allele frequency
Ecological gradient (Precipitation sums)
1.0
0.9
0.8
0.7
0.6
0.5
600
800
1000
1200
256
253
217
204
219
9
97
233
218
L1
L8
O8
L16
O16
O1
L12
O12
124

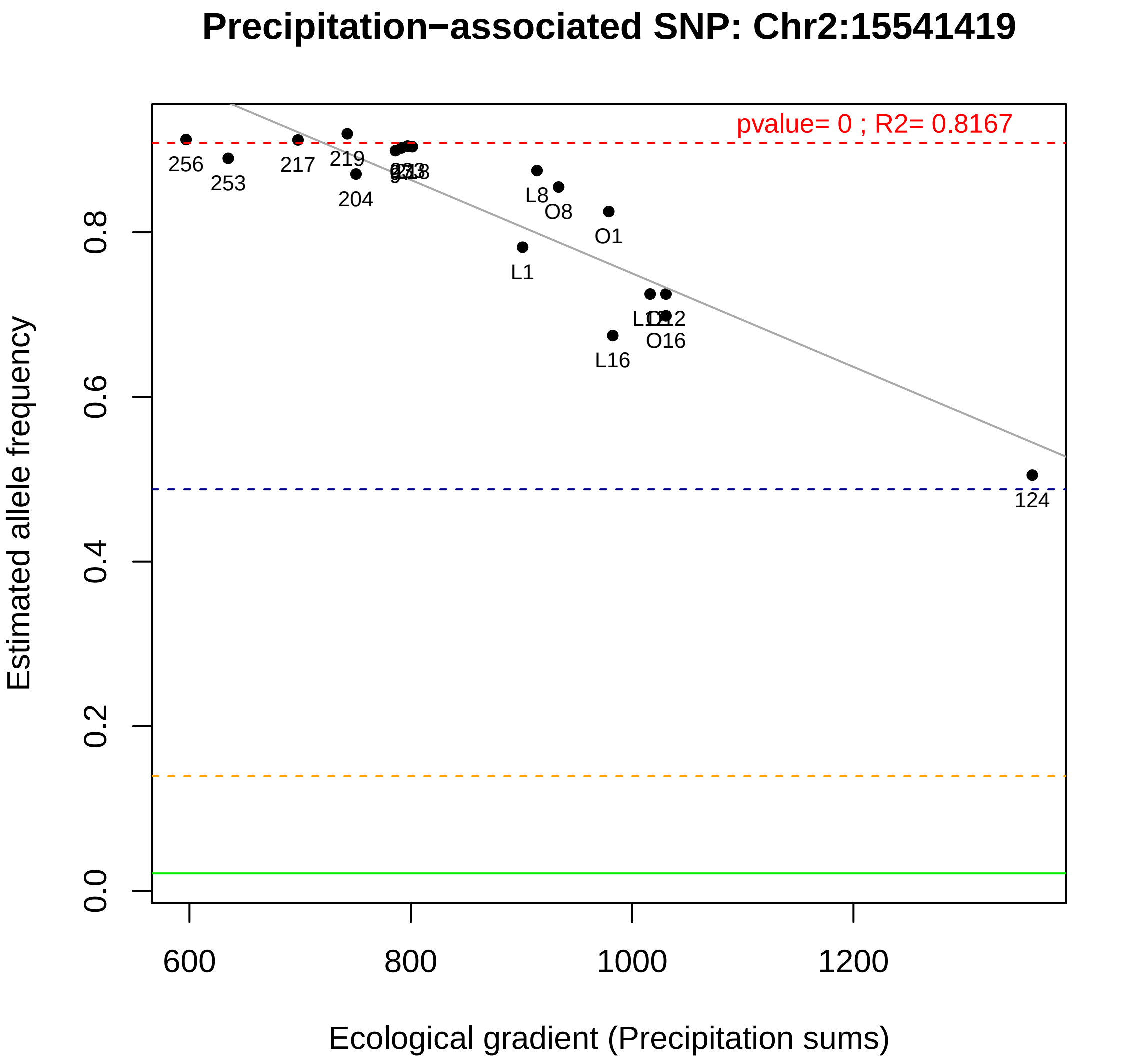
Precipitation−associated SNP: Chr2:15541419
pvalue= 0 ; R2= 0.8167
Estimated allele frequency
Ecological gradient (Precipitation sums)
0.0
0.2
0.4
0.6
0.8
600
800
1000
1200
256
253
217
219
204
L8
O8
O1
L1
O16
L16
124

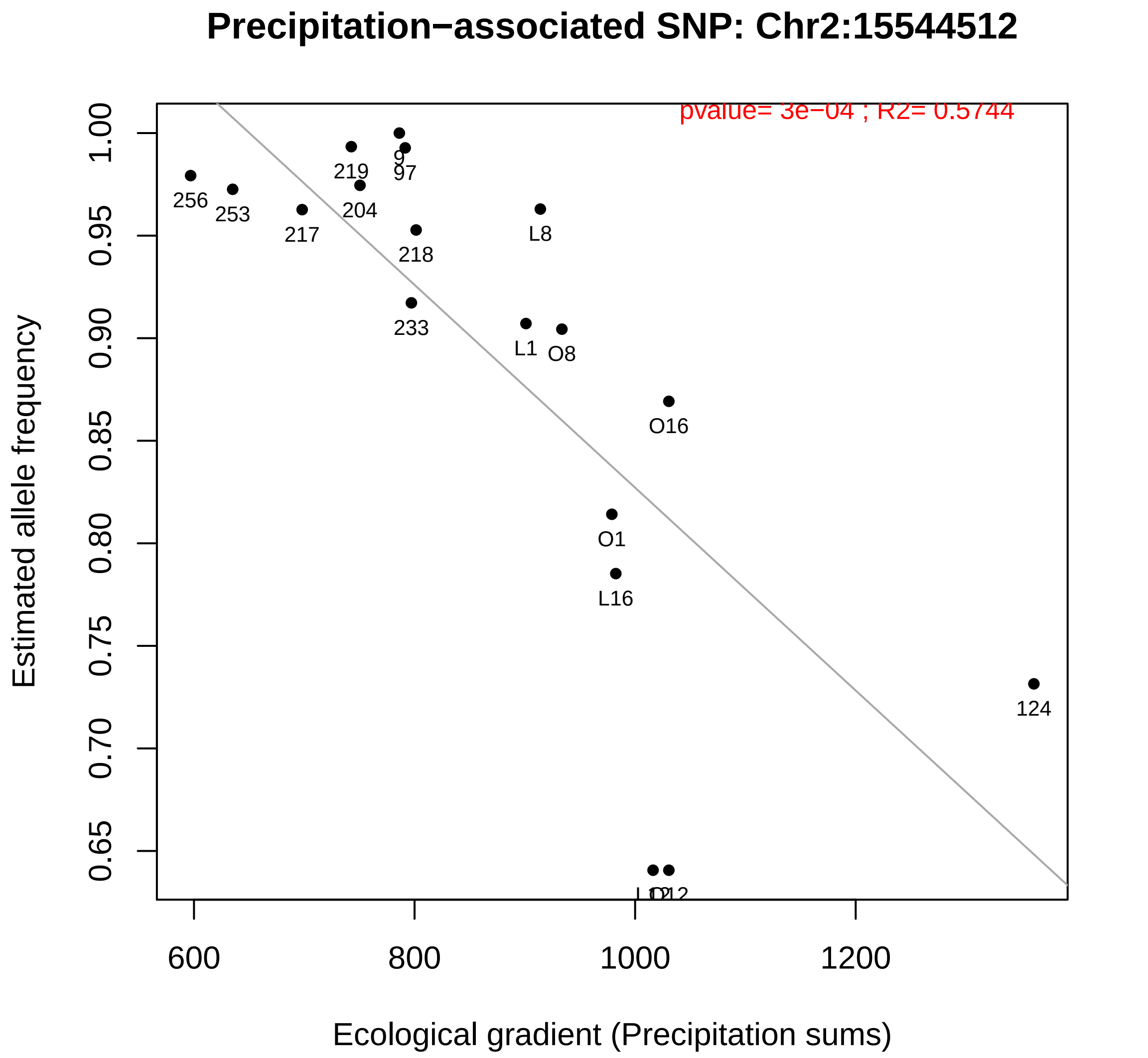

Precipitation−associated SNP: Chr2:15544512
pvalue= 3e−04 ; R2= 0.5744
Estimated allele frequency
Ecological gradient (Precipitation sums)
1.00
0.95
0.90
0.85
0.80
0.75
0.70
0.65
600
800
1000
1200
256
253
217
219
204
9
97
218
233
L8
L1
O8
O16
O1
L16
124

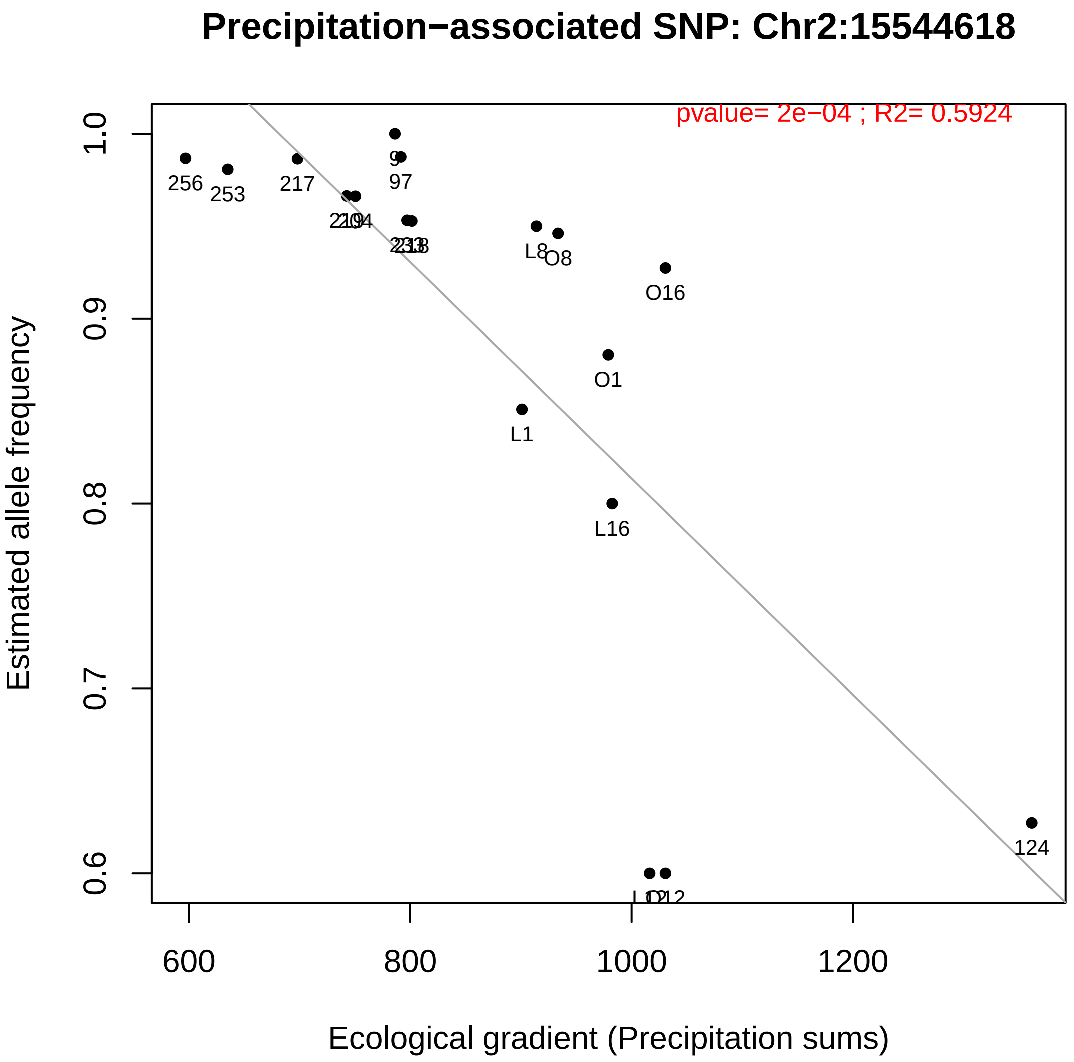
Precipitation−associated SNP: Chr2:15544618
pvalue= 2e−04 ; R2= 0.5924
Estimated allele frequency
Ecological gradient (Precipitation sums)
1.0
0.9
0.8
0.7
0.6
600
800
1000
1200
256
253
217
97
L8
O8
O16
O1
L1
L16
124

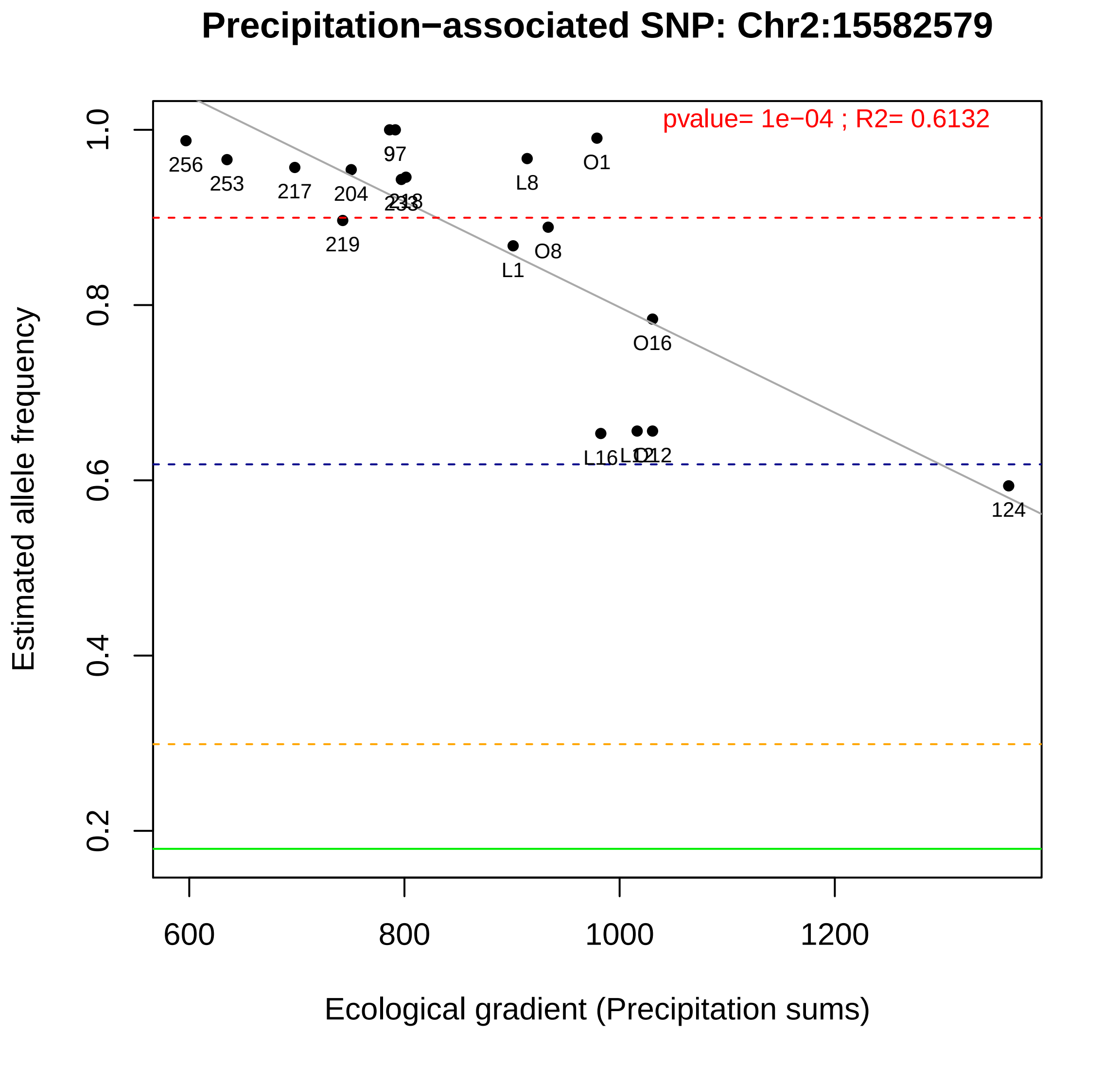
Precipitation−associated SNP: Chr2:15582579
pvalue= 1e−04 ; R2= 0.6132
Estimated allele frequency
Ecological gradient (Precipitation sums)
1.0
0.8
0.6
0.4
0.2
600
800
1000
1200
256
253
217
204
97
219
L8
O1
O8
L1
O16
L16
124

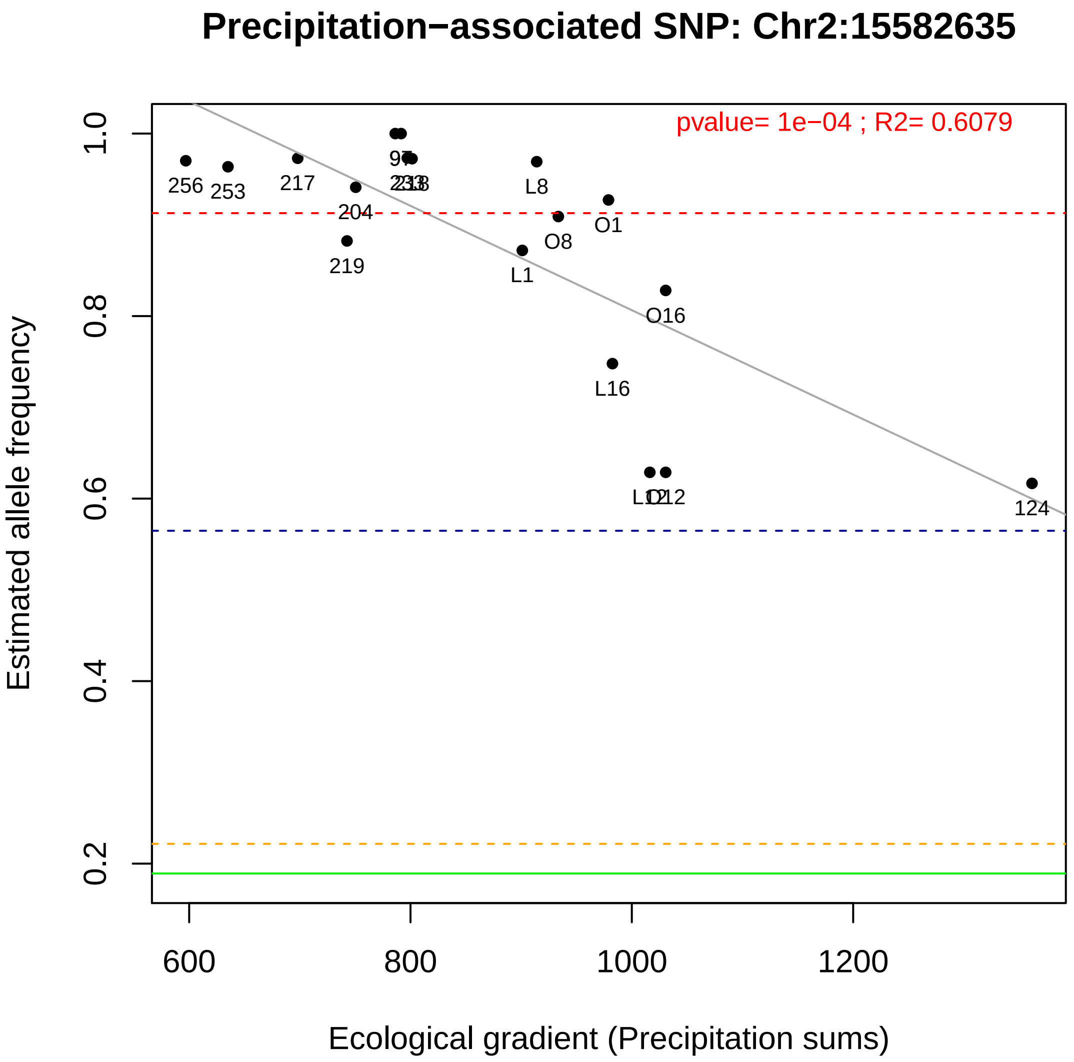

Precipitation−associated SNP: Chr2:15582635
pvalue= 1e−04 ; R2= 0.6079
Estimated allele frequency
Ecological gradient (Precipitation sums)
0.2
0.4
0.6
0.8
1.0
600
800
1000
1200
256
253
217
204
219
L8
O1
O8
L1
O16
L16
124

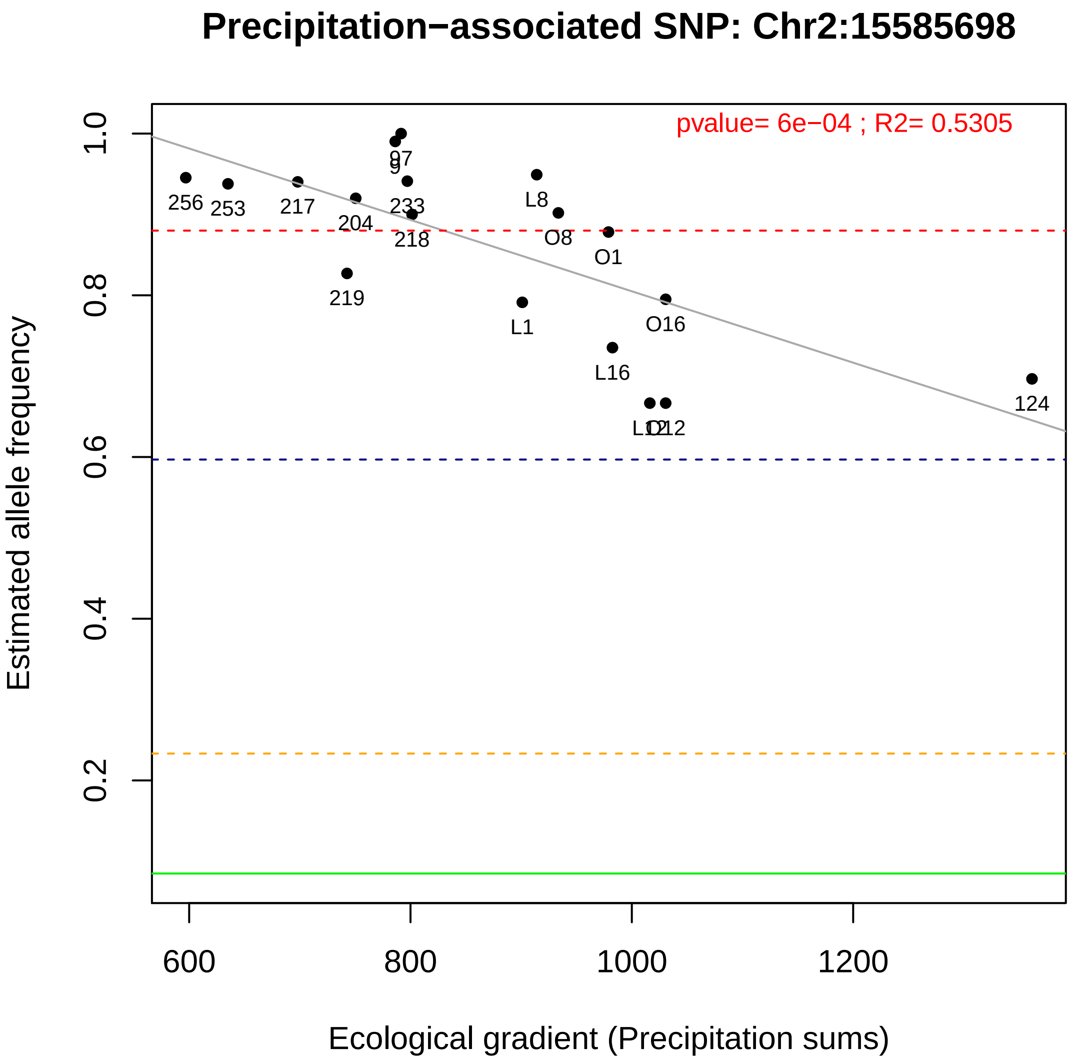
Precipitation−associated SNP: Chr2:15585698
pvalue= 6e−04 ; R2= 0.5305
Estimated allele frequency
Ecological gradient (Precipitation sums)
1.0
0.8
0.6
0.4
0.2
600
800
1000
1200
256
253
217
204
219
97
9
233
218
L8
O8
O1
L1
O16
L16
124

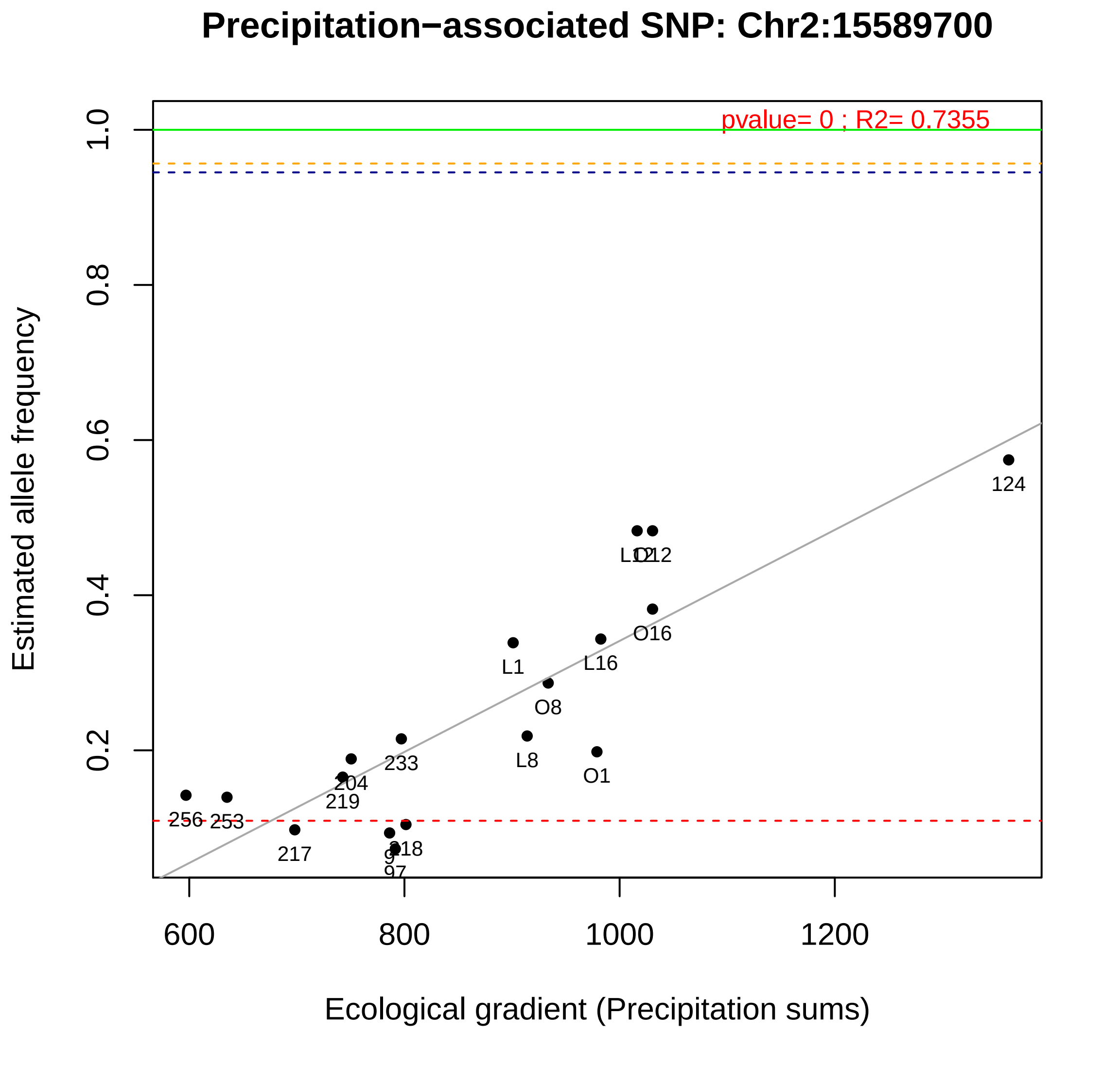

Precipitation−associated SNP: Chr2:15589700
pvalue= 0 ; R2= 0.7355
Estimated allele frequency
0.2
0.4
0.6
0.8
1.0
600
800
1000
1200
Ecological gradient (Precipitation sums)
124
L12
O12
O16
L16
L1
O8
L8
O1
233
204
219
256
253
217
218
9
97

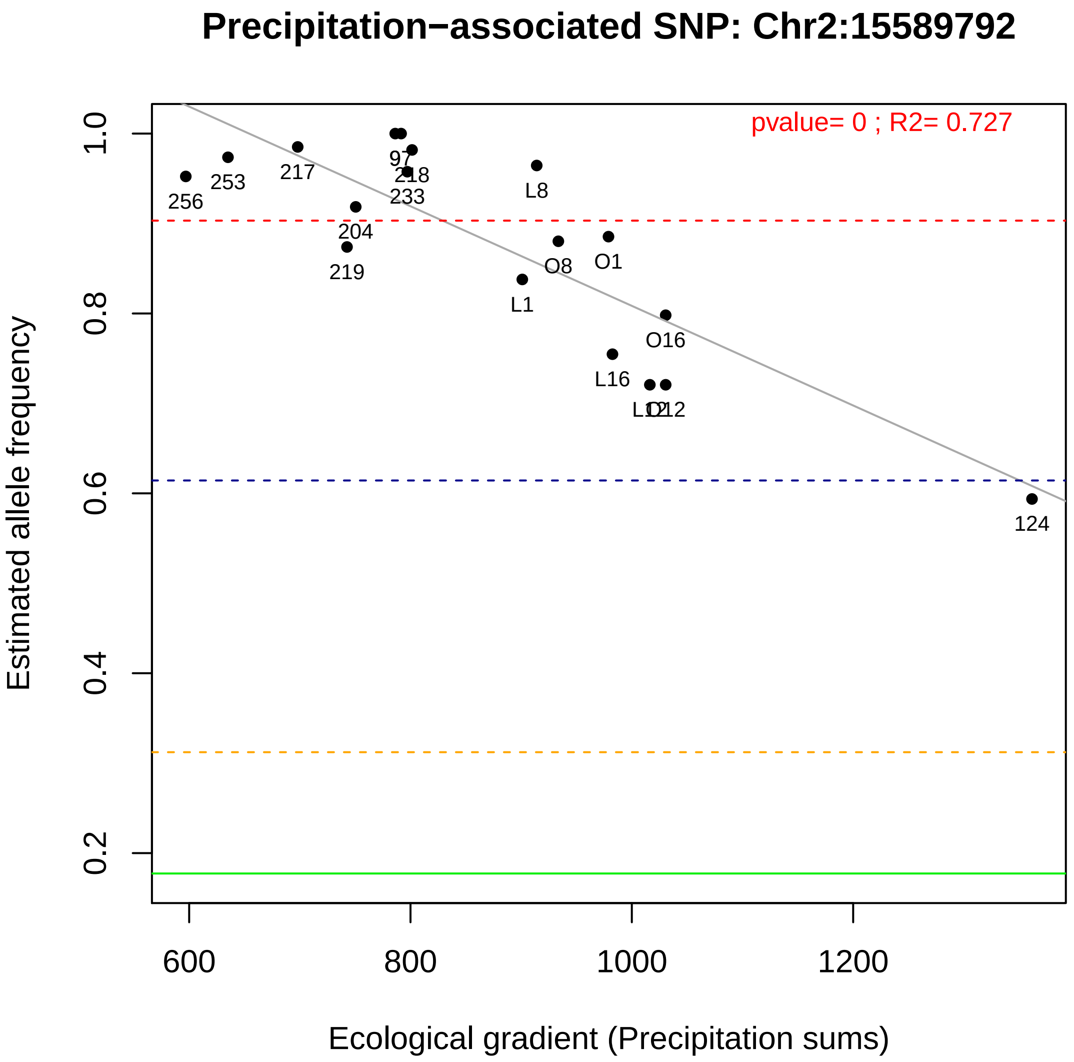

Precipitation−associated SNP: Chr2:15589792
pvalue= 0 ; R2= 0.727
256
253
217
97
218
233
204
219
L8
O8
O1
L1
O16
L16
L12
O12
124
1.0
0.8
0.6
0.4
0.2
600
800
1000
1200
Estimated allele frequency
Ecological gradient (Precipitation sums)

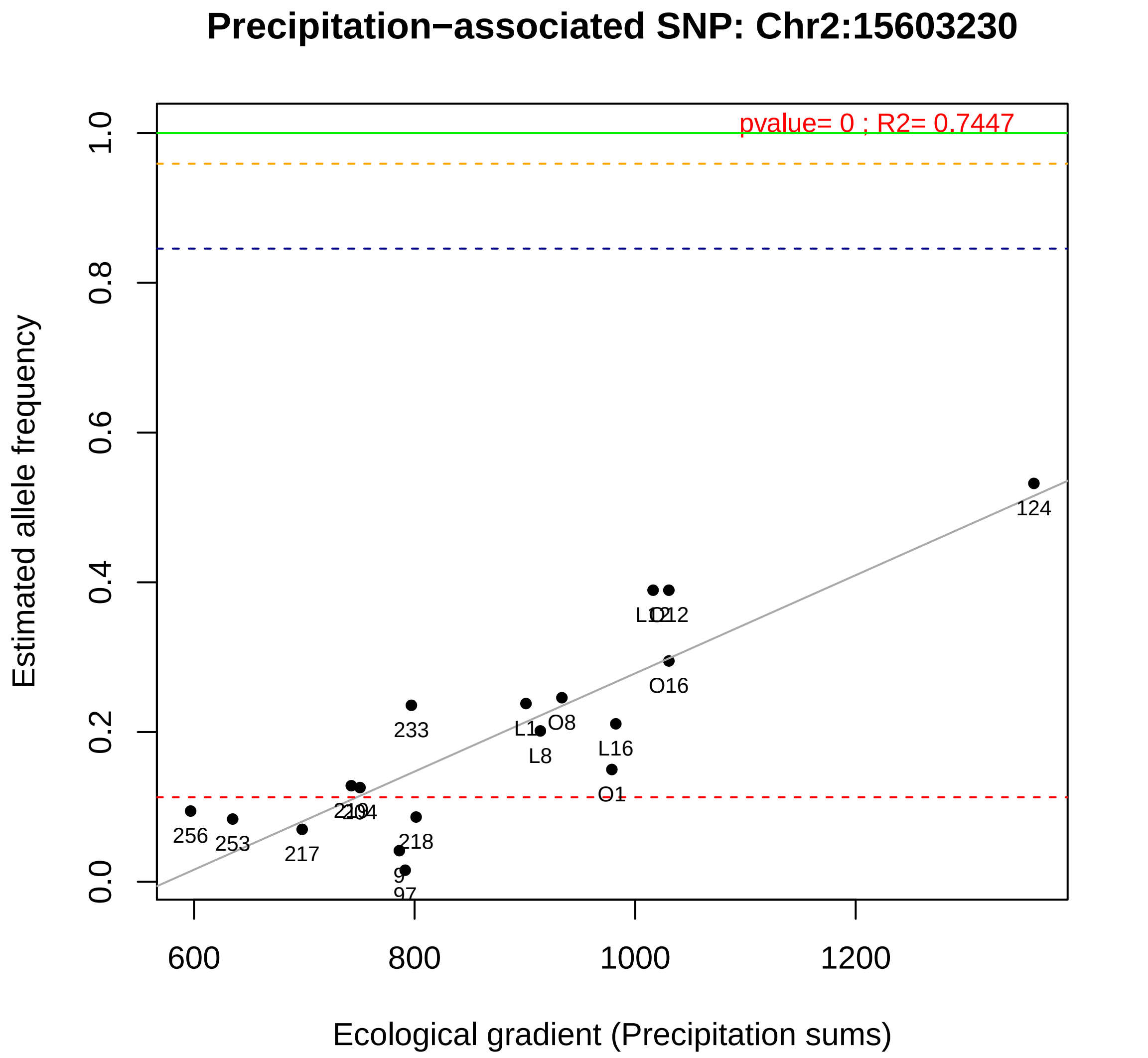

Precipitation−associated SNP: Chr2:15603230
pvalue= 0 ; R2= 0.7447
Estimated allele frequency
0.0
0.2
0.4
0.6
0.8
1.0
Ecological gradient (Precipitation sums)
600
800
1000
1200
124
L10
O12
O16
233
L1
O8
L8
L16
O1
256
253
217
218
9
97

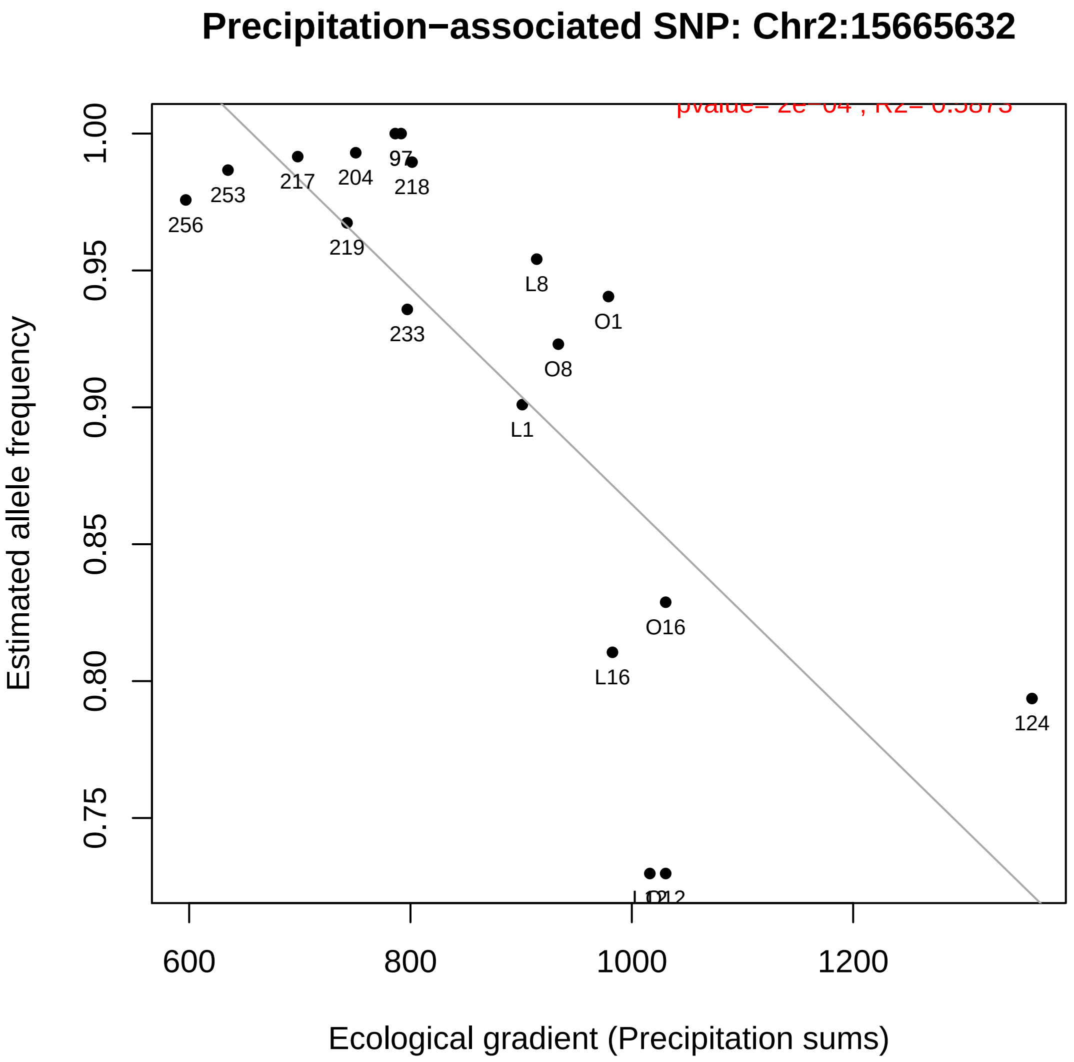
Precipitation−associated SNP: Chr2:15665632
Estimated allele frequency
Ecological gradient (Precipitation sums)
0.75
0.80
0.85
0.90
0.95
1.00
600
800
1000
1200
256
253
217
204
97
218
219
233
L8
O1
O8
L1
O16
L16
124

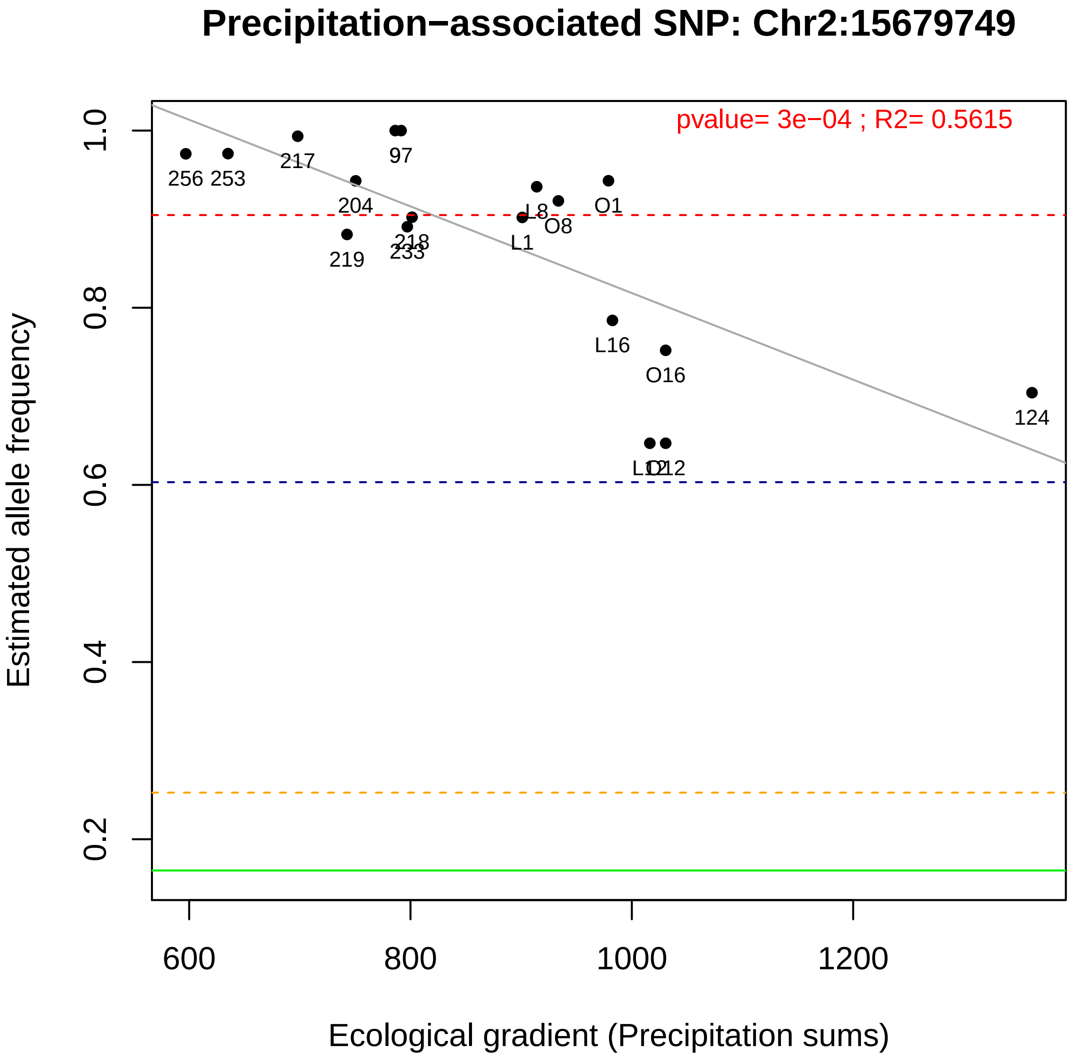

Precipitation−associated SNP: Chr2:15679749
pvalue= 3e−04 ; R2= 0.5615
Estimated allele frequency
Ecological gradient (Precipitation sums)
0.2
0.4
0.6
0.8
1.0
600
800
1000
1200
256
253
217
97
204
219
218
233
L1
L8
O8
O1
L16
O16
124
L12
O12

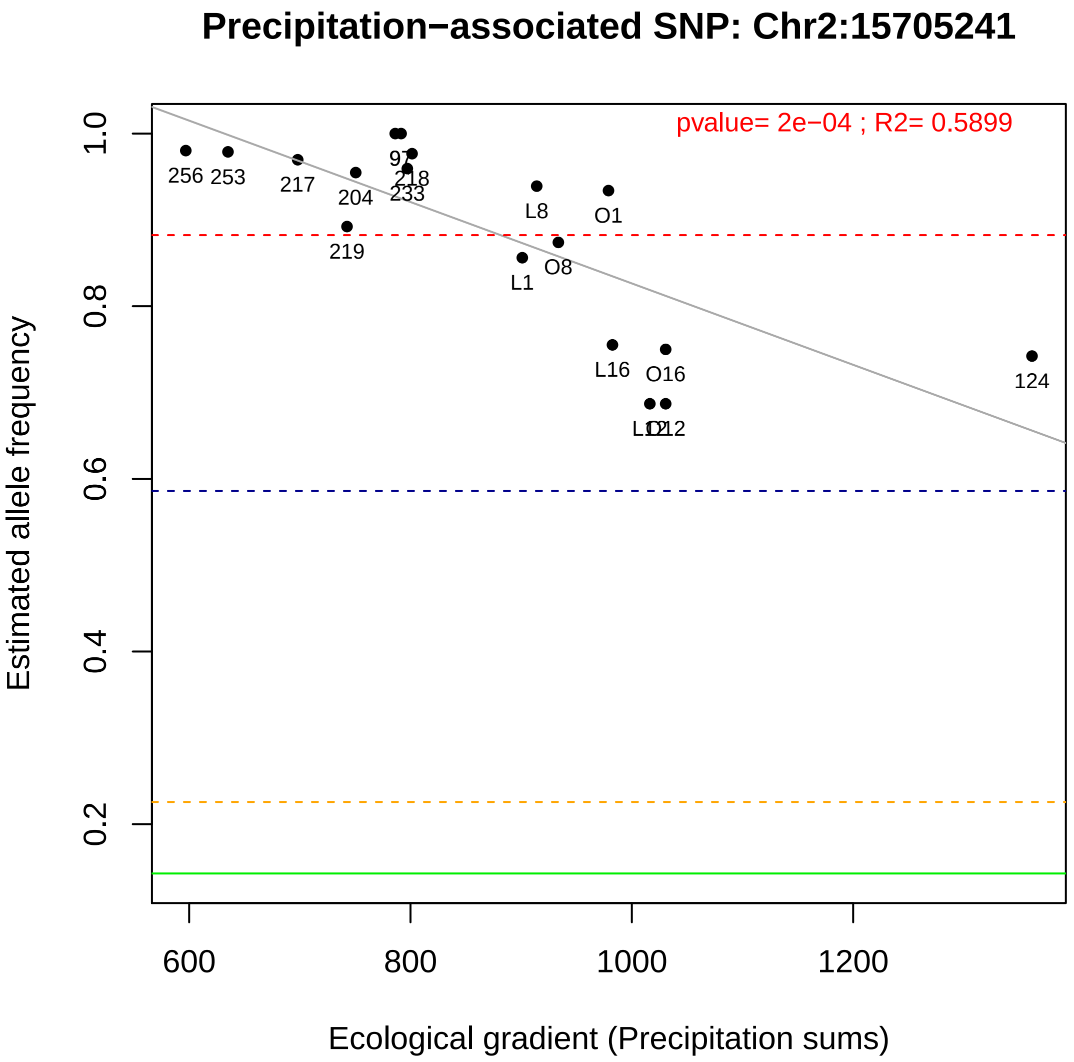
Precipitation−associated SNP: Chr2:15705241
pvalue= 2e−04 ; R2= 0.5899
Estimated allele frequency
Ecological gradient (Precipitation sums)
0.2
0.4
0.6
0.8
1.0
600
800
1000
1200
256
253
217
204
219
97
218
233
L8
O1
L1
O8
L16
O16
124
L12
O12

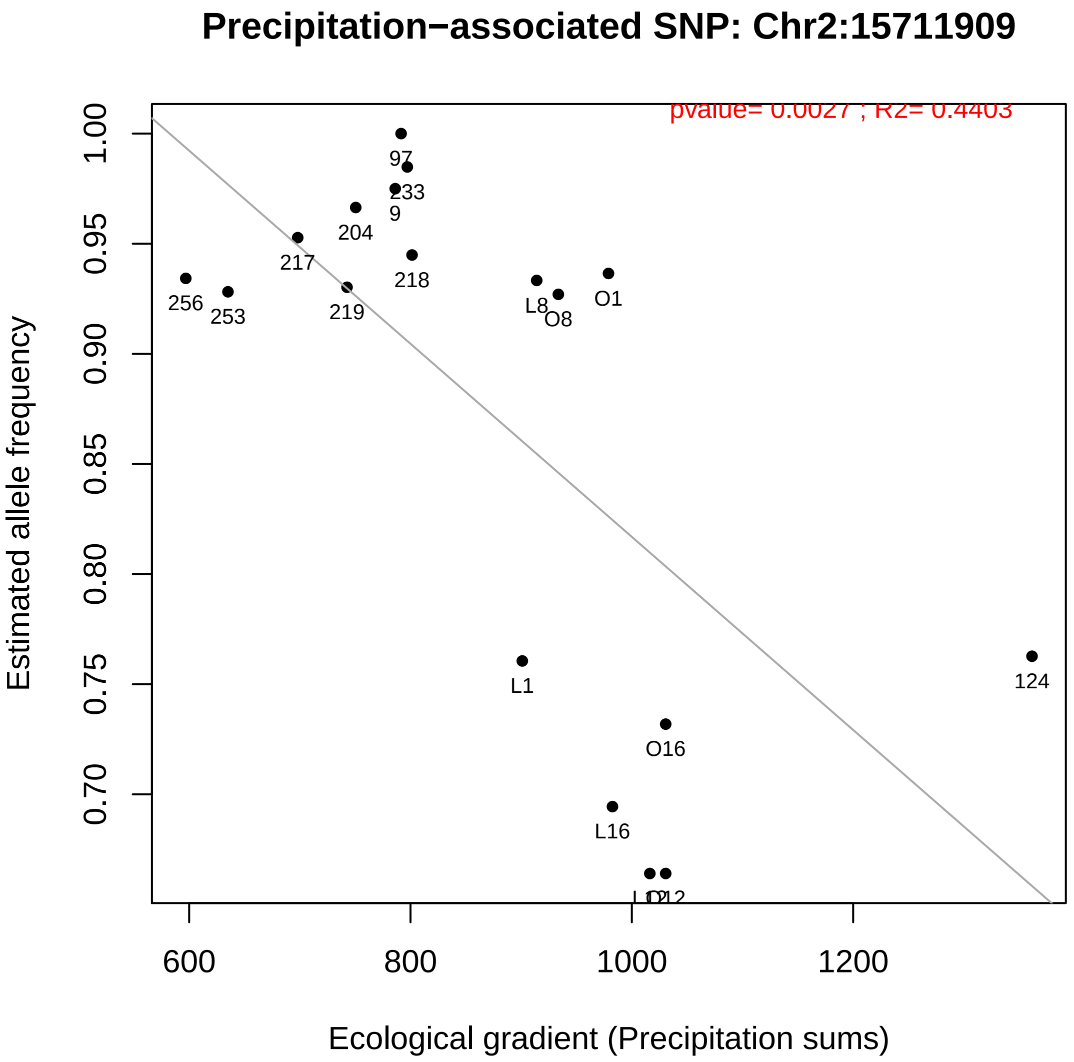

Precipitation−associated SNP: Chr2:15711909
pvalue= 0.0027 ; R2= 0.4403
Estimated allele frequency
1.00
0.95
0.90
0.85
0.80
0.75
0.70
97
233
9
204
217
218
256
253
219
L8
O8
O1
L1
124
O16
L16
600
800
1000
1200
Ecological gradient (Precipitation sums)

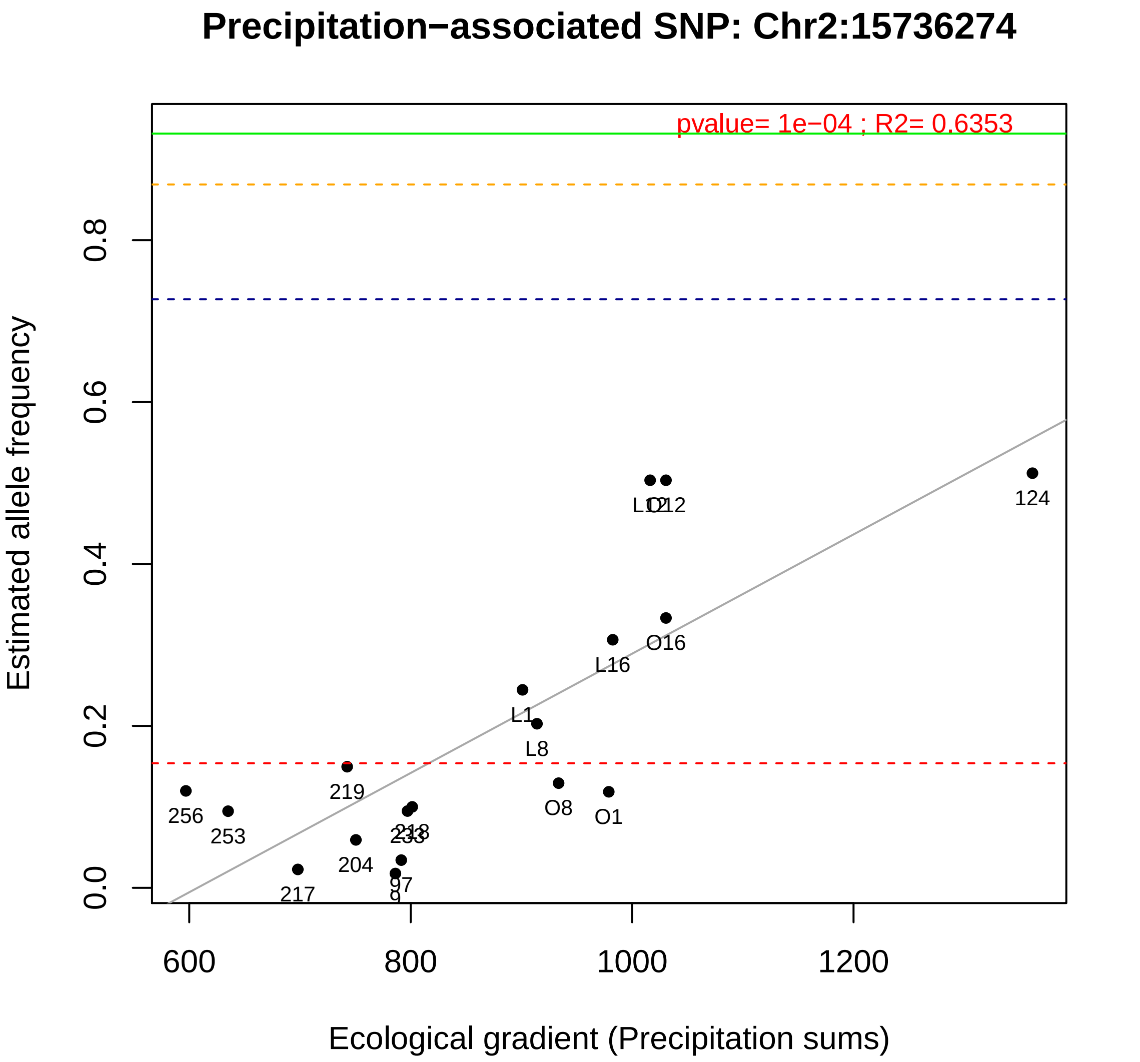

Precipitation−associated SNP: Chr2:15736274
pvalue= 1e−04 ; R2= 0.6353
Estimated allele frequency
0.0
0.2
0.4
0.6
0.8
600
800
1000
1200
Ecological gradient (Precipitation sums)
124
L12
O12
O16
L16
L1
L8
219
256
253
O8
O1
204
217
97

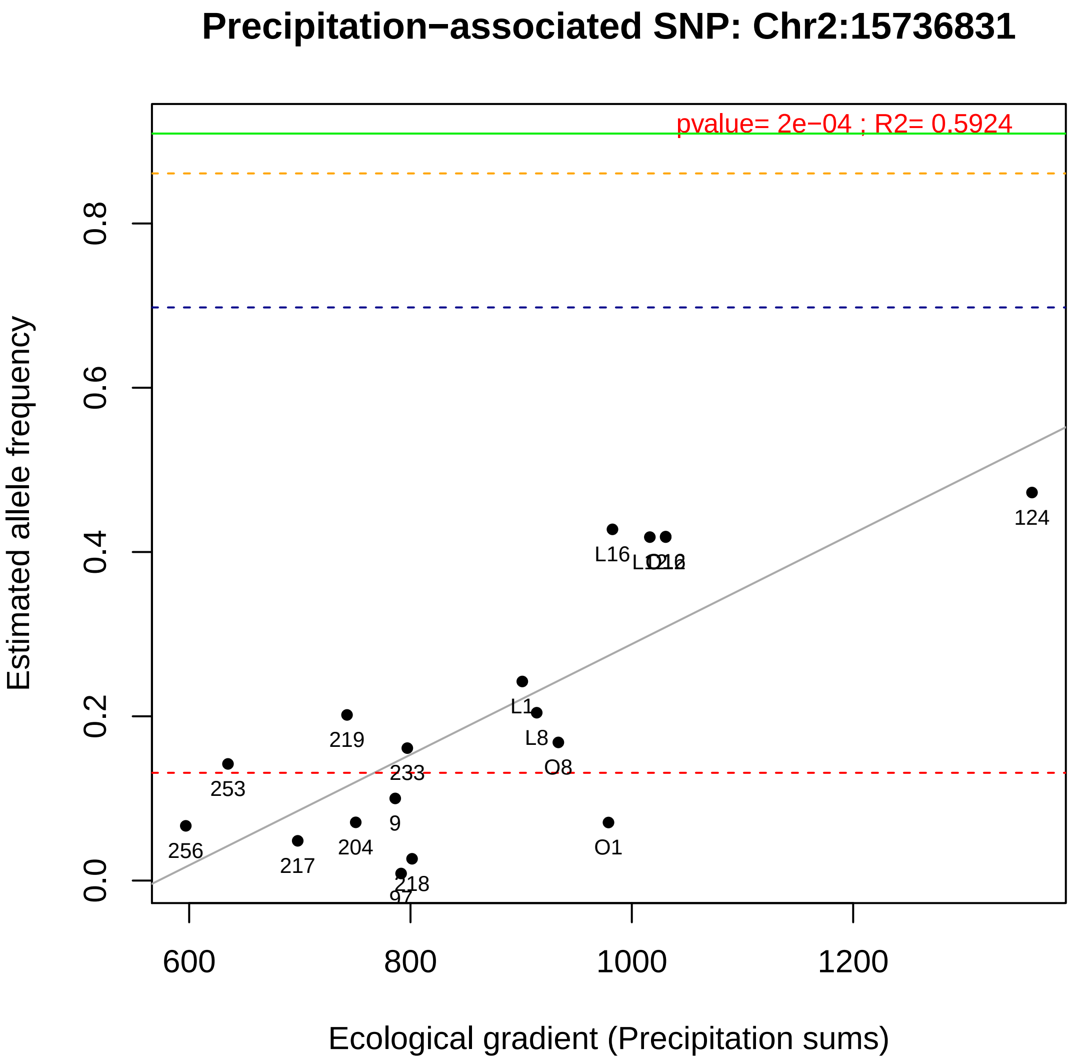
Precipitation−associated SNP: Chr2:15736831
pvalue= 2e−04 ; R2= 0.5924
Estimated allele frequency
0.0
0.2
0.4
0.6
0.8
600
800
1000
1200
Ecological gradient (Precipitation sums)
124
L16
219
253
233
L1
L8
O8
9
256
204
217
O1
218

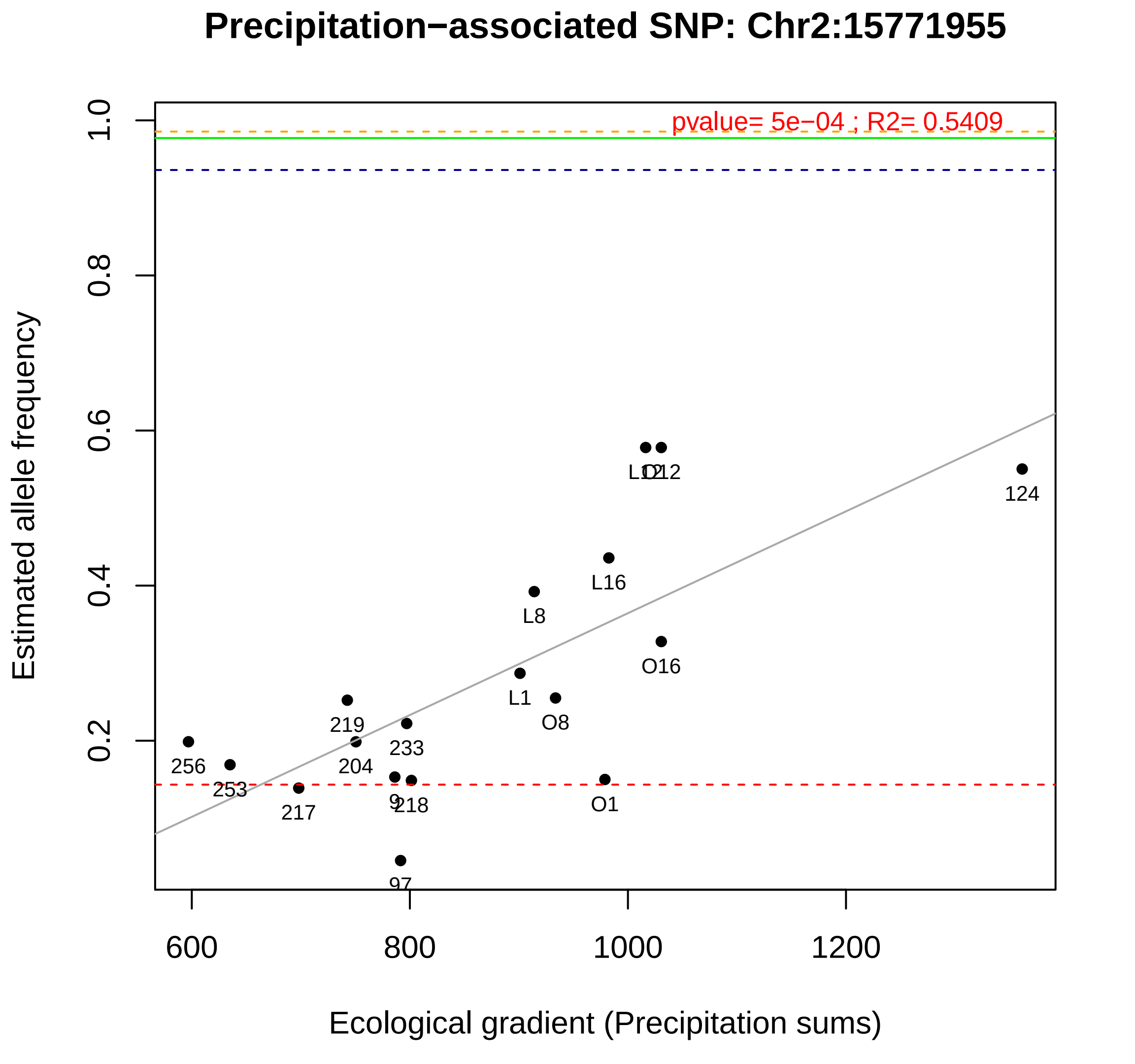

Precipitation−associated SNP: Chr2:15771955
pvalue= 5e−04 ; R2= 0.5409
Estimated allele frequency
0.2
0.4
0.6
0.8
1.0
600
800
1000
1200
Ecological gradient (Precipitation sums)
L12
O12
124
L16
L8
O16
L1
O8
219
233
204
256
253
217
9
218
O1
97

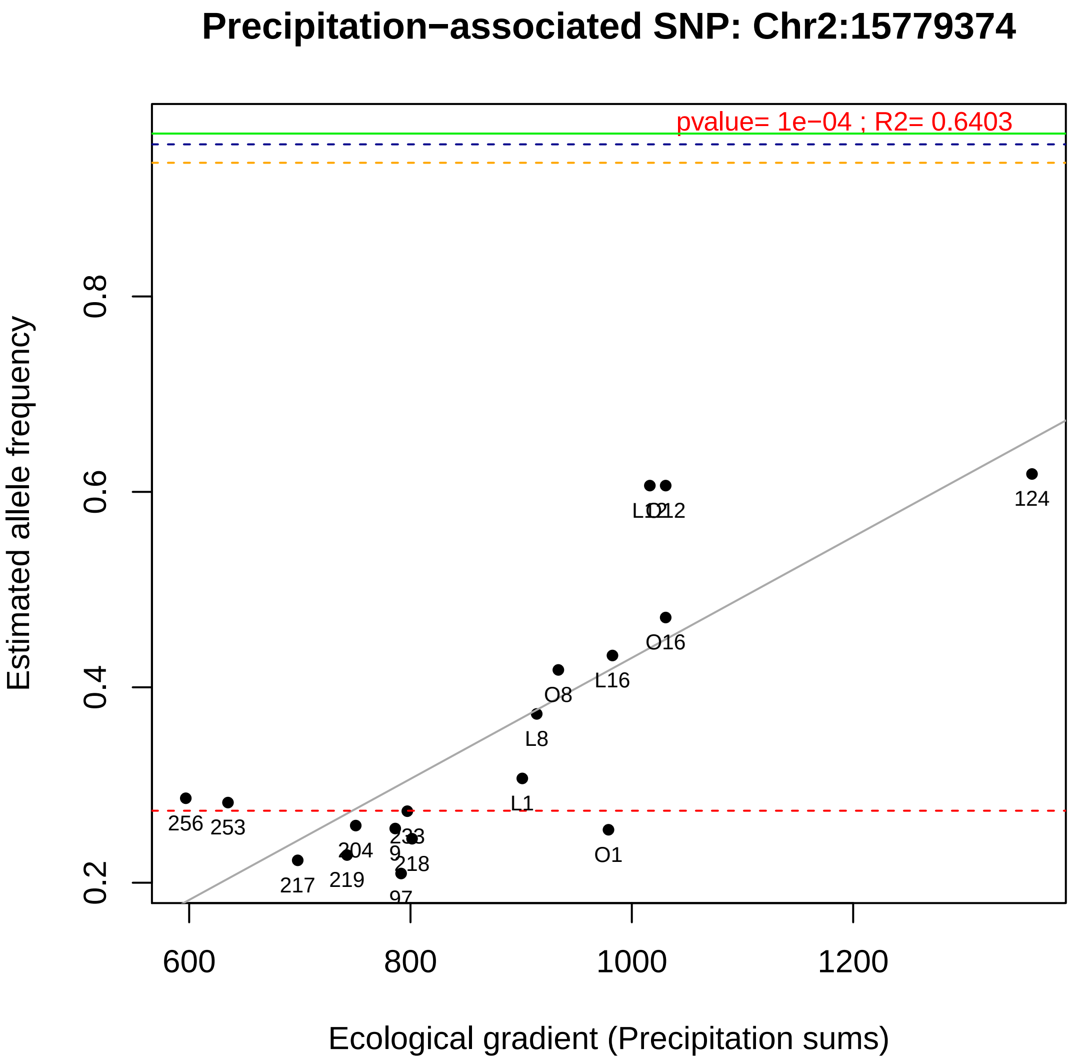
Precipitation−associated SNP: Chr2:15779374
pvalue= 1e−04 ; R2= 0.6403
Estimated allele frequency
0.2
0.4
0.6
0.8
600
800
1000
1200
Ecological gradient (Precipitation sums)
256
253
217
219
204
218
97
L1
O1
L8
O8
L16
O16
124

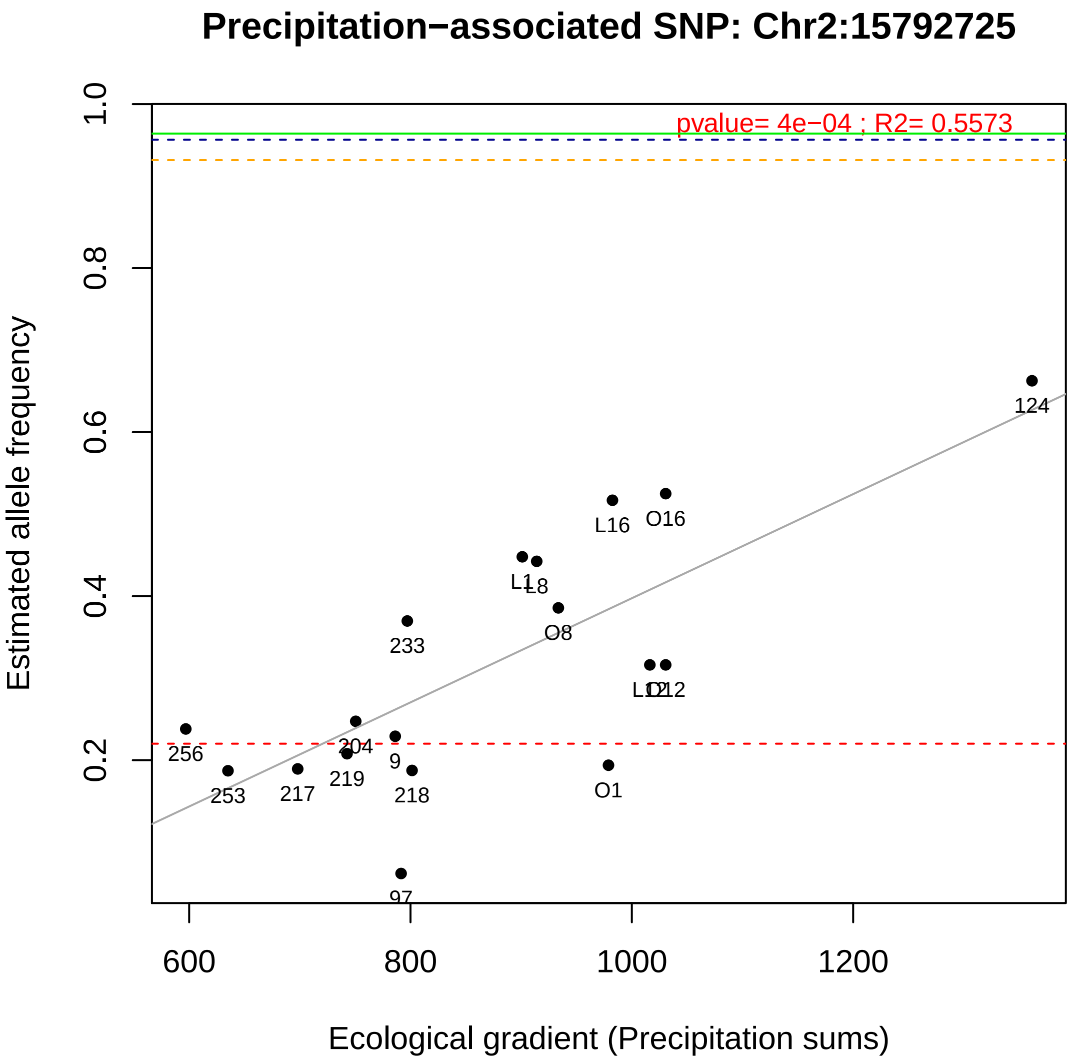
Precipitation−associated SNP: Chr2:15792725
pvalue= 4e−04 ; R2= 0.5573
Estimated allele frequency
Ecological gradient (Precipitation sums)
1.0
0.8
0.6
0.4
0.2
600
800
1000
1200
124
L16
O16
L1
L8
O8
233
L12
O12
256
204
9
219
253
217
218
O1
97

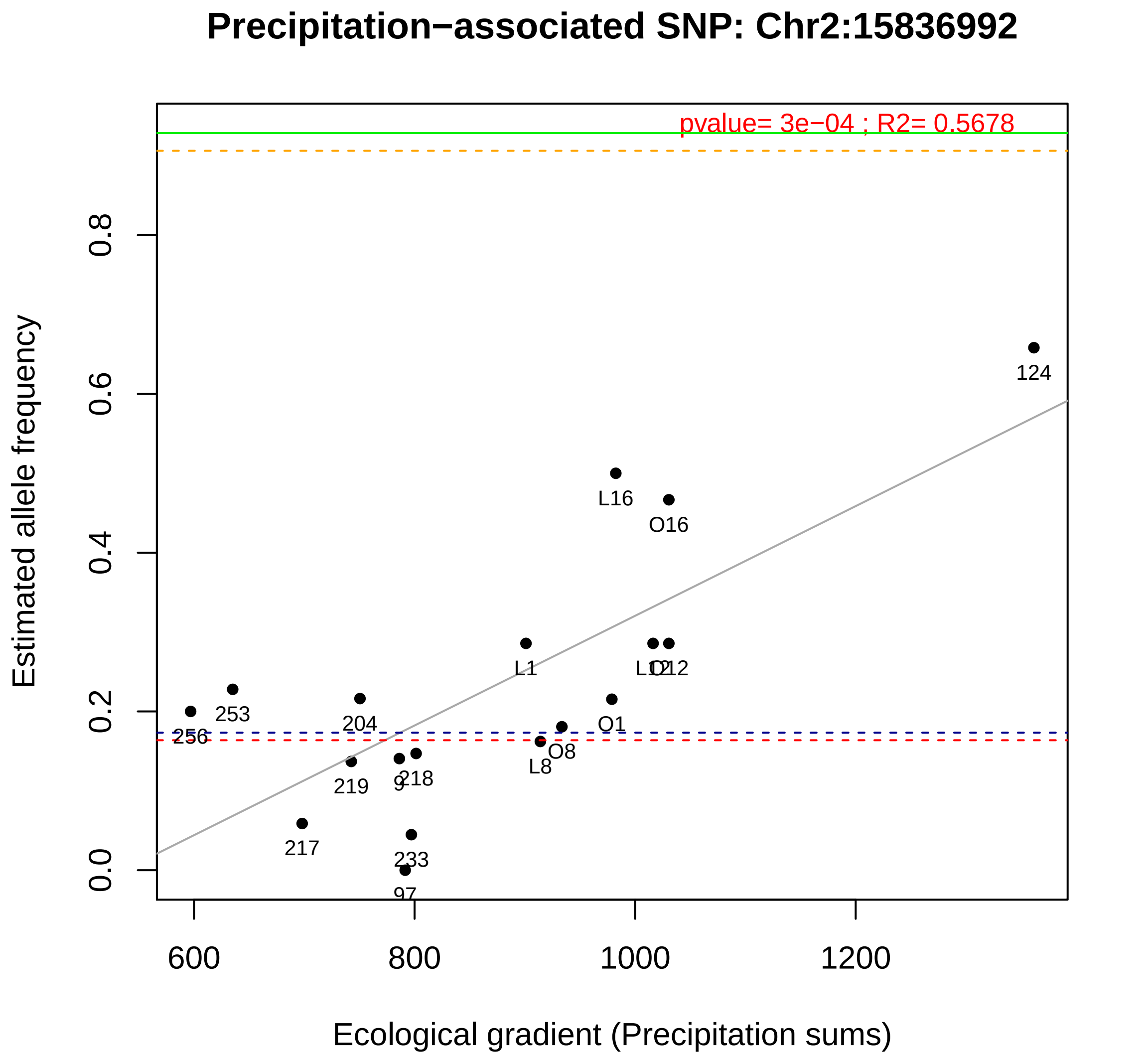

Precipitation−associated SNP: Chr2:15836992
pvalue= 3e−04 ; R2= 0.5678
Estimated allele frequency
0.0
0.2
0.4
0.6
0.8
600
800
1000
1200
Ecological gradient (Precipitation sums)
124
L16
O16
L1
L12
O12
256
253
204
O1
O8
L8
219
9
218
217
233
97

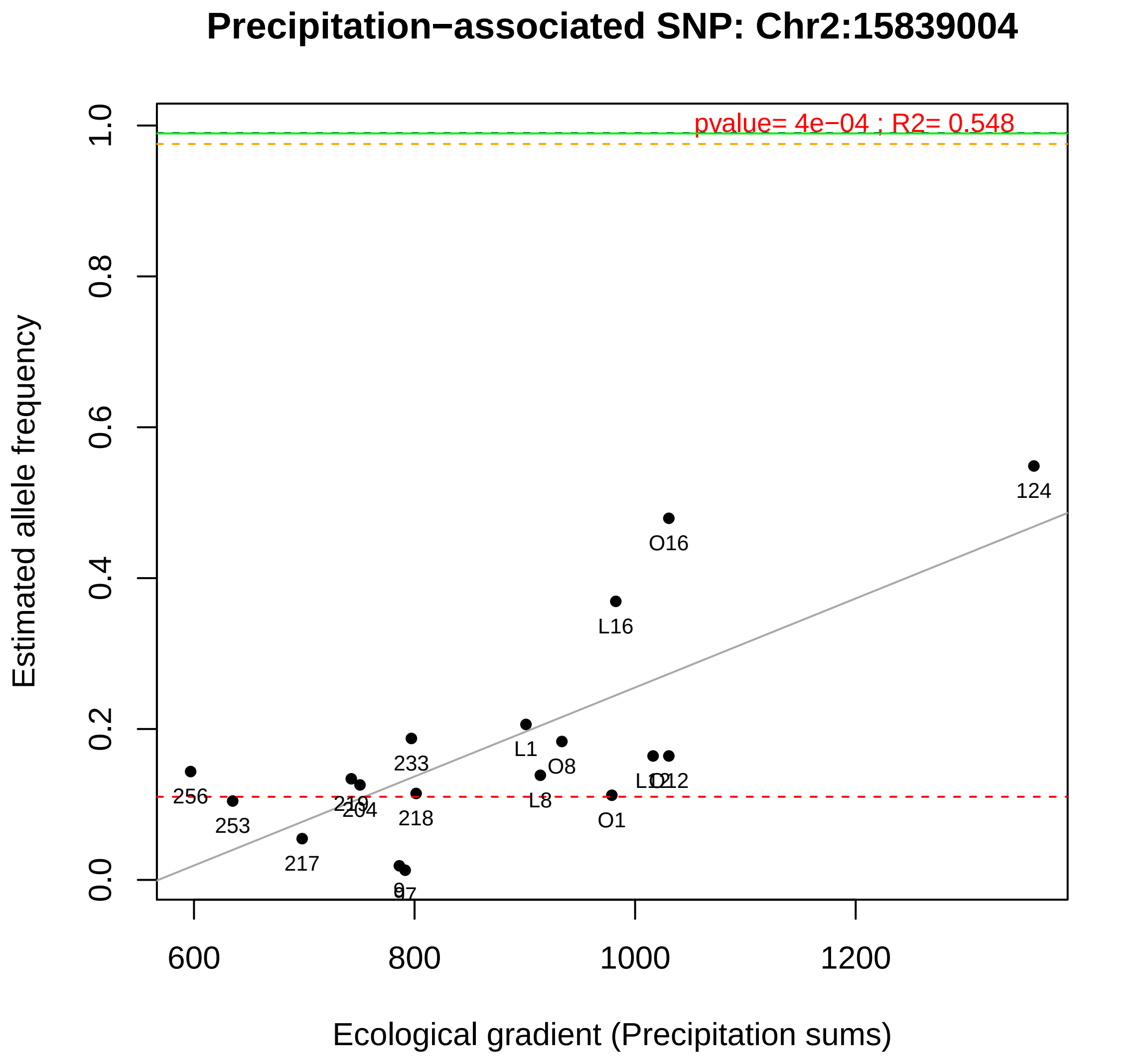

Precipitation−associated SNP: Chr2:15839004
pvalue= 4e−04 ; R2= 0.548
Estimated allele frequency
Ecological gradient (Precipitation sums)
0.0
0.2
0.4
0.6
0.8
1.0
600
800
1000
1200
124
O16
L16
L1
O8
233
L8
O1
256
253
217
218
204

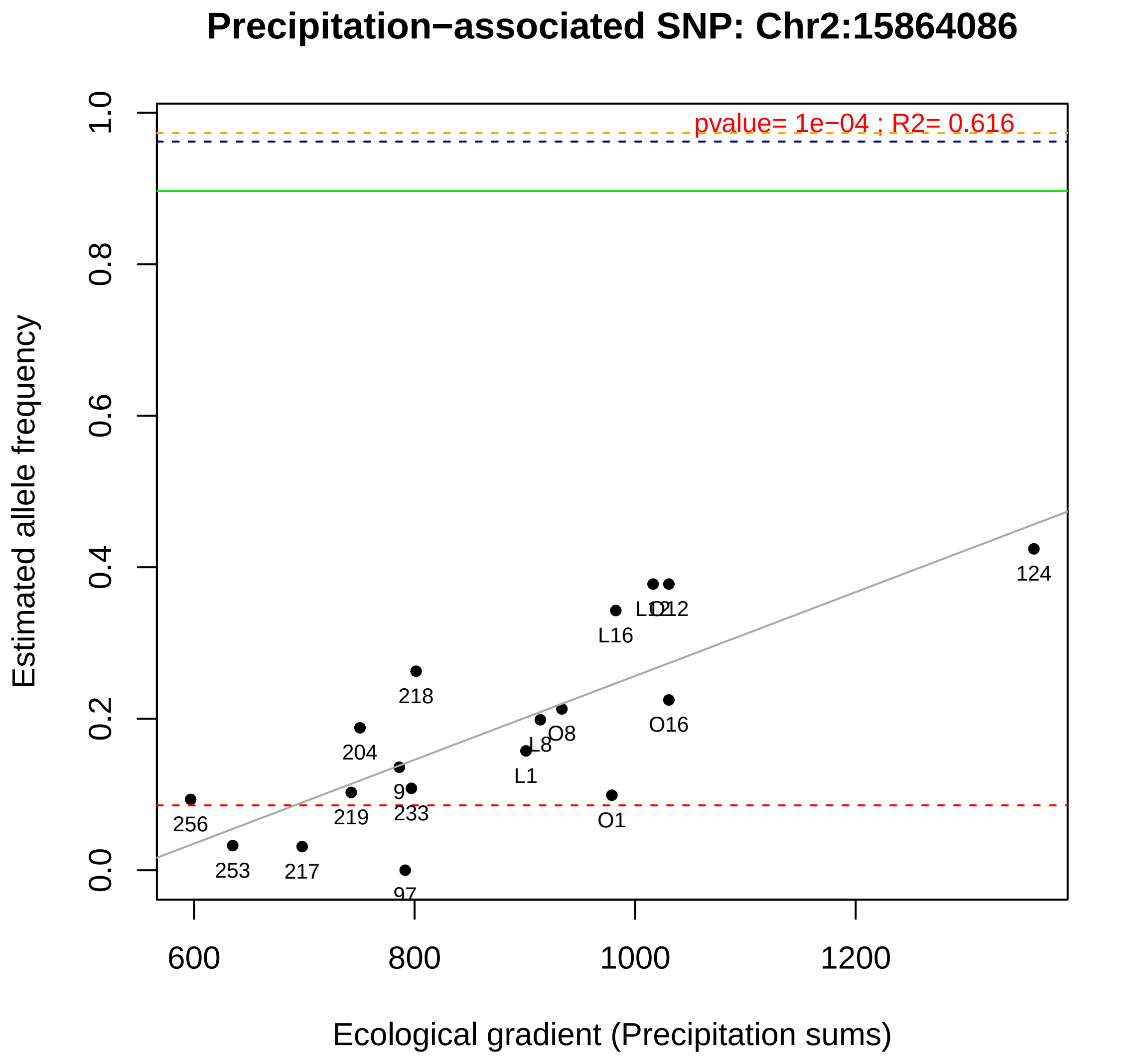

Precipitation−associated SNP: Chr2:15864086
pvalue= 1e−04 ; R2= 0.616
Estimated allele frequency
Ecological gradient (Precipitation sums)
0.0
0.2
0.4
0.6
0.8
1.0
600
800
1000
1200
124
L12
O12
L16
218
O16
O8
L8
204
L1
9
233
219
256
O1
253
217
97

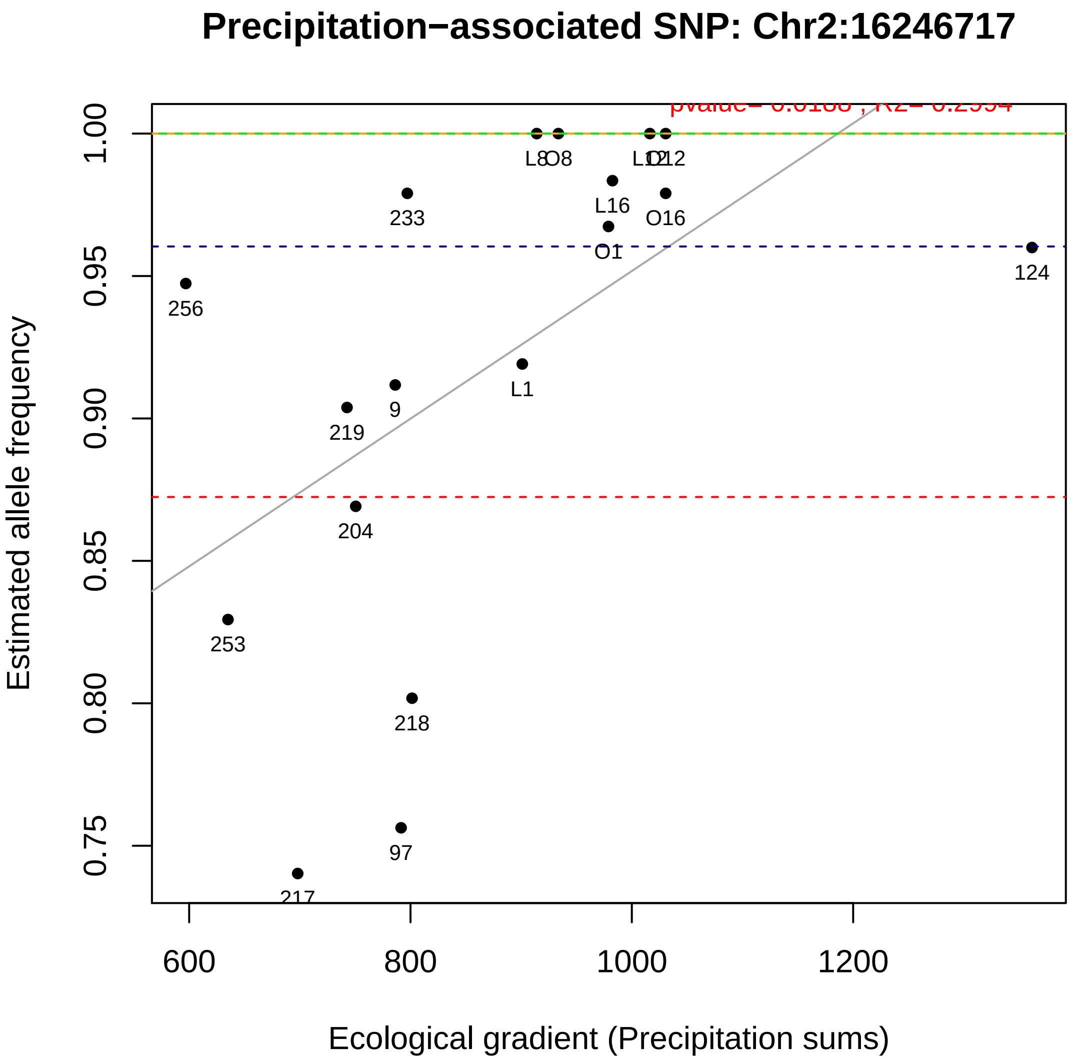
Precipitation−associated SNP: Chr2:16246717
Estimated allele frequency
Ecological gradient (Precipitation sums)
0.75
0.80
0.85
0.90
0.95
1.00
600
800
1000
1200
L8
O8
L16
O16
O1
233
124
256
L1
9
219
204
253
218
97
217

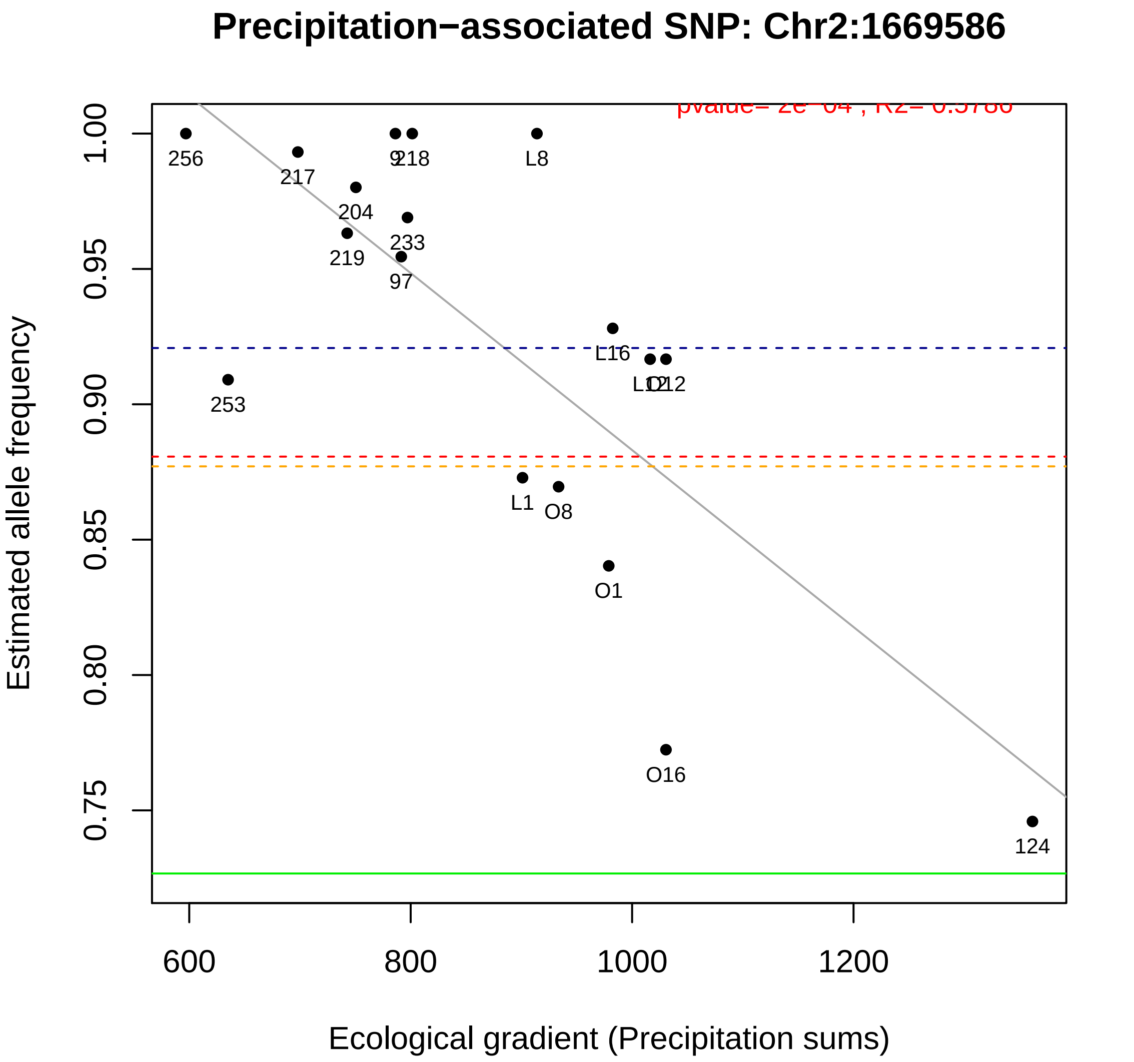
Precipitation−associated SNP: Chr2:1669586
pvalue= 2e−04 ; R2= 0.5786
Estimated allele frequency
Ecological gradient (Precipitation sums)
1.00
0.95
0.90
0.85
0.80
0.75
600
800
1000
1200
256
217
204
219
233
97
9
218
L8
L16
L10
O12
253
L1
O8
O1
O16
124

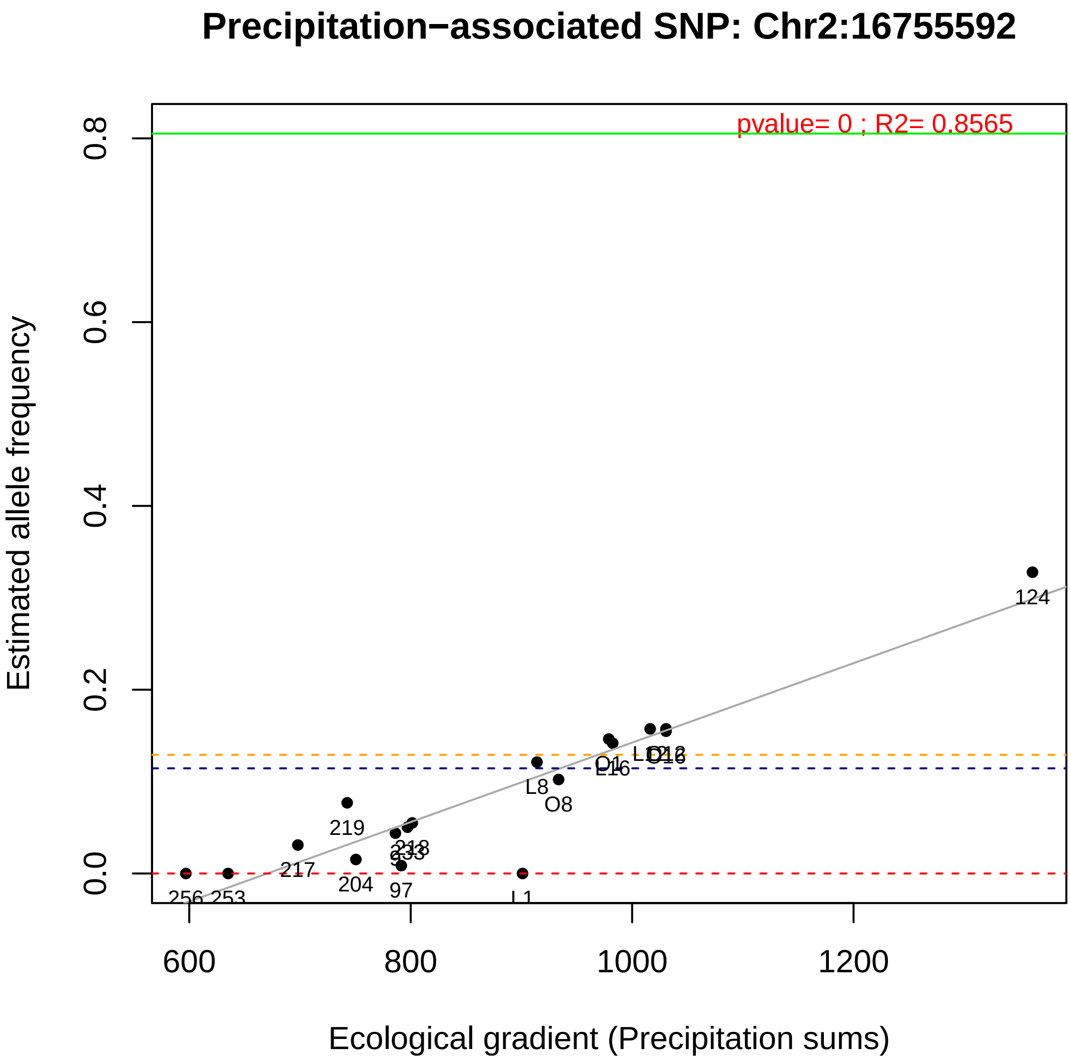

Precipitation−associated SNP: Chr2:16755592
pvalue= 0 ; R2= 0.8565
Estimated allele frequency
Ecological gradient (Precipitation sums)
0.0
0.2
0.4
0.6
0.8
600
800
1000
1200
124
219
217
204
97
L8
O8

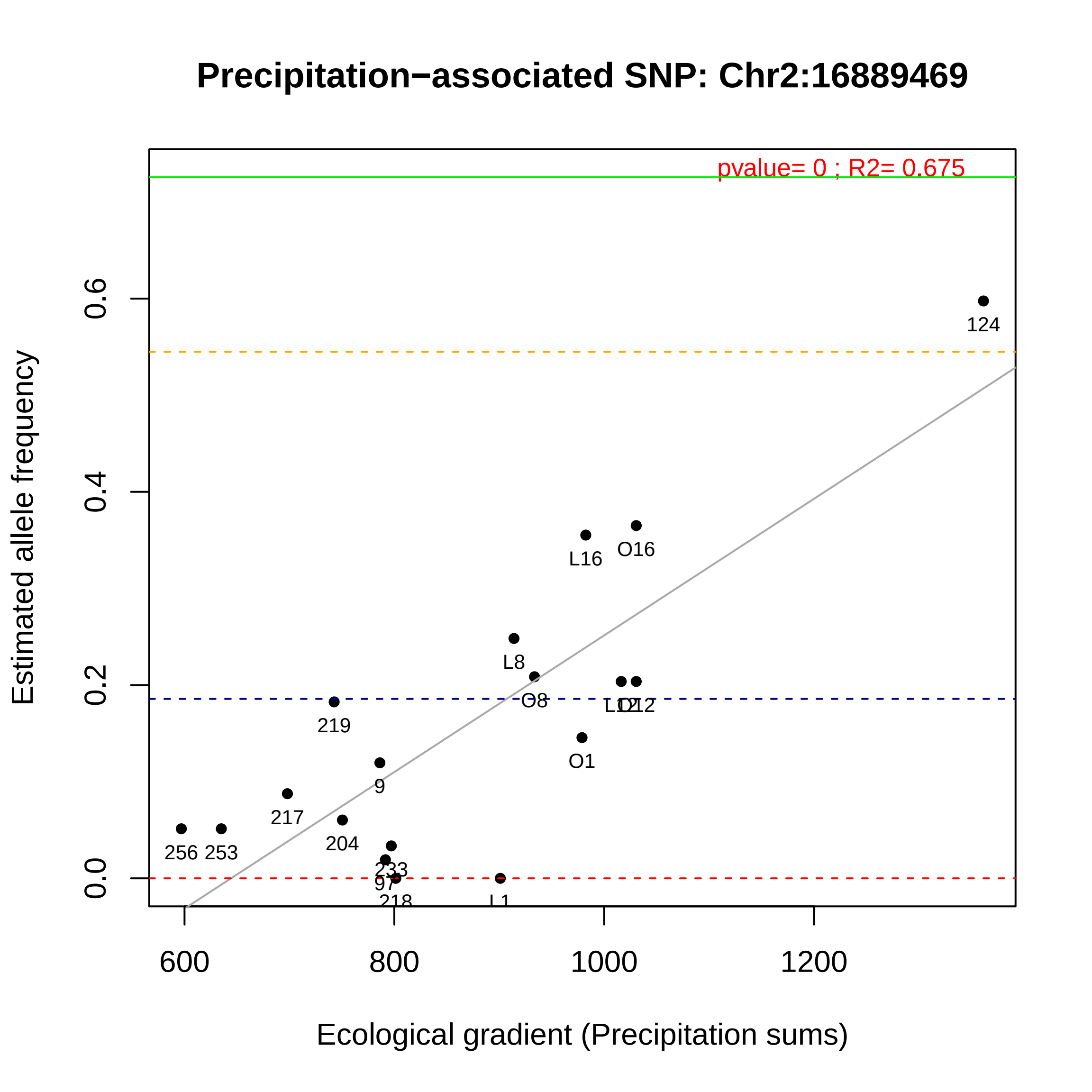

Precipitation−associated SNP: Chr2:16889469
pvalue= 0 ; R2= 0.675
Estimated allele frequency
0.0
0.2
0.4
0.6
600
800
1000
1200
Ecological gradient (Precipitation sums)
124
L16
O16
L8
O8
O1
219
9
217
204
256
253
233
97
218
L1

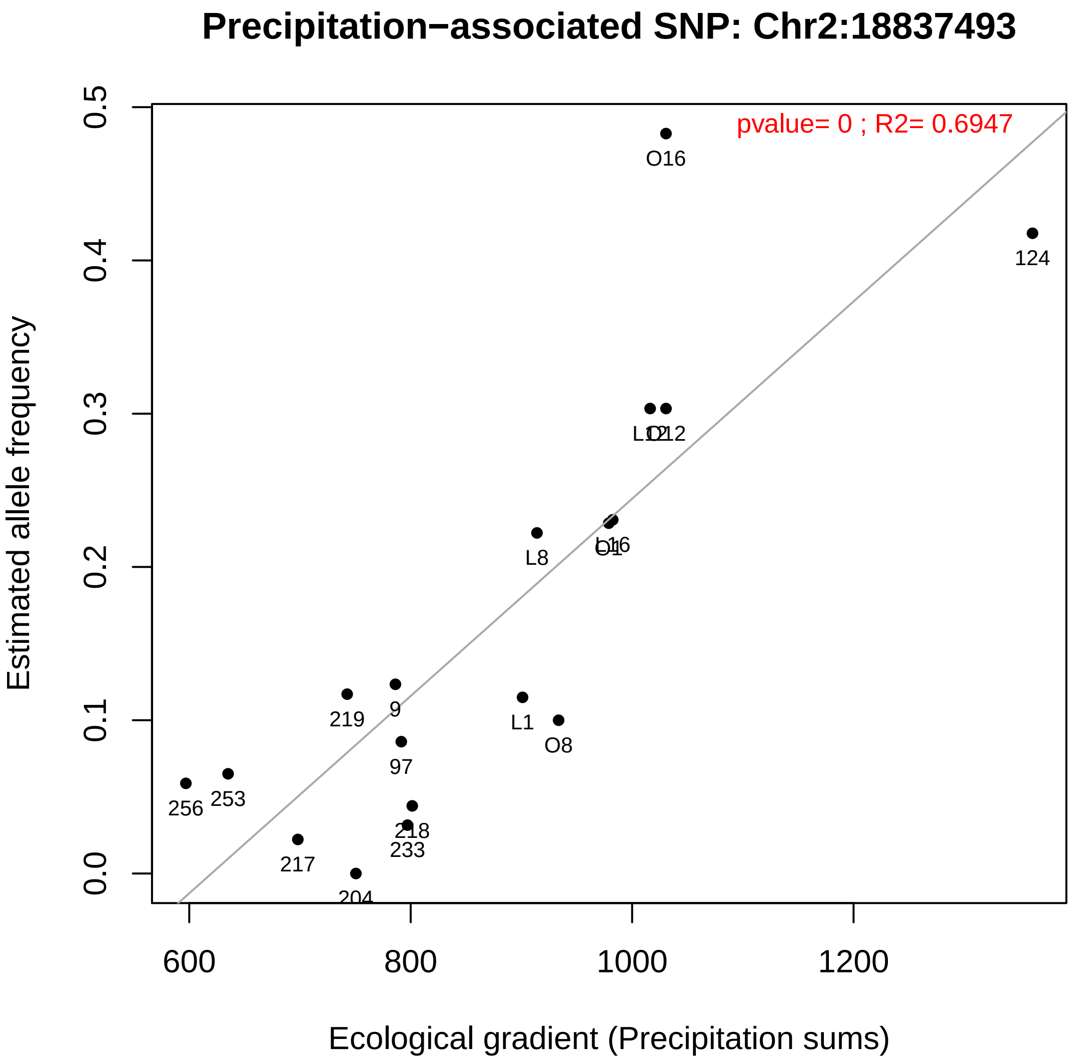
Precipitation−associated SNP: Chr2:18837493
pvalue= 0 ; R2= 0.6947
Estimated allele frequency
Ecological gradient (Precipitation sums)
0.0
0.1
0.2
0.3
0.4
0.5
600
800
1000
1200
O16
124
L10
O12
L16
O1
L8
219
9
97
L1
O8
256
253
217
218
233
204

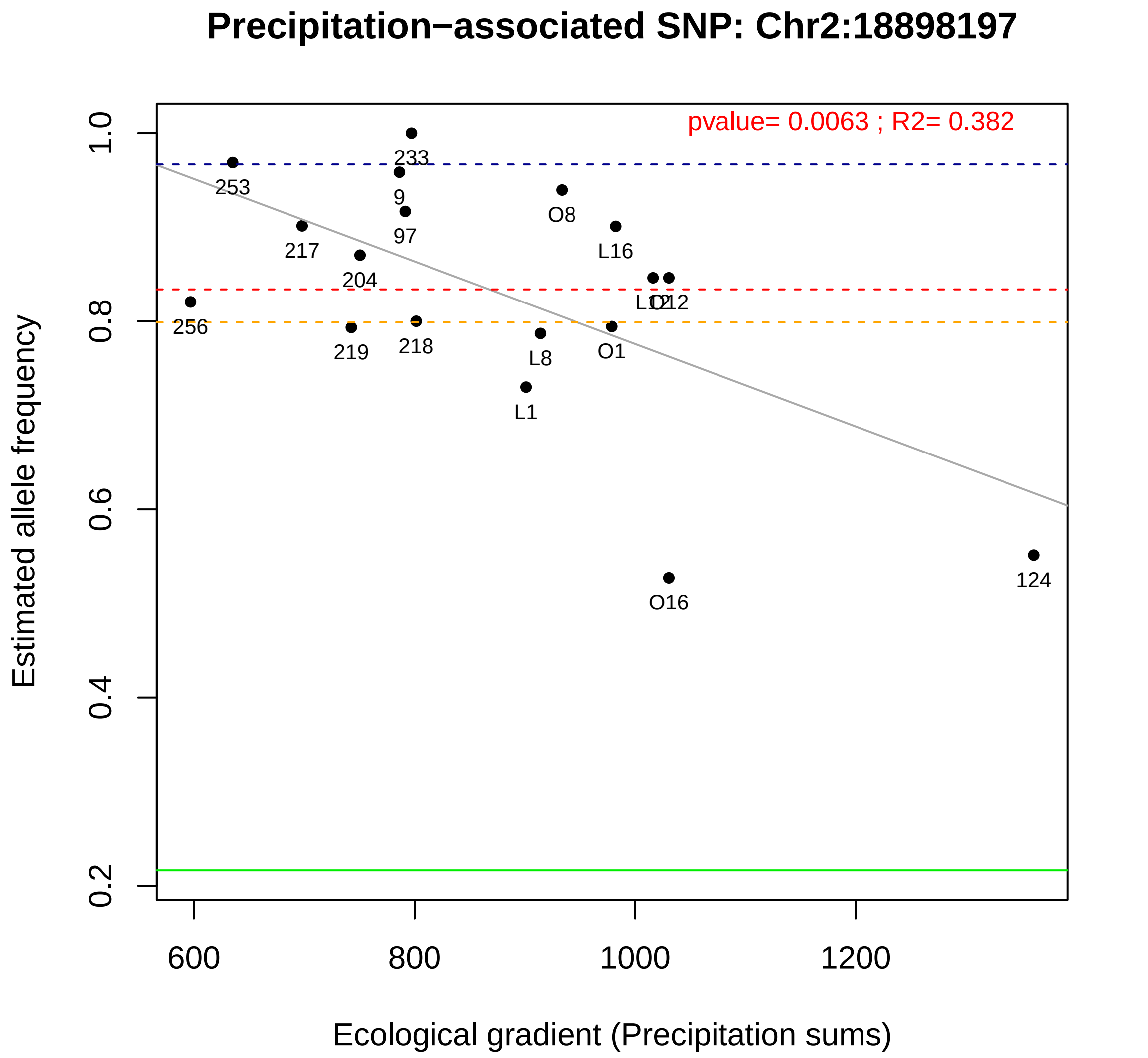
Precipitation−associated SNP: Chr2:18898197
pvalue= 0.0063 ; R2= 0.382
Estimated allele frequency
Ecological gradient (Precipitation sums)
1.0
0.8
0.6
0.4
0.2
600
800
1000
1200
233
253
9
97
O8
217
L16
204
L12
O12
256
219
218
L8
O1
L1
124
O16

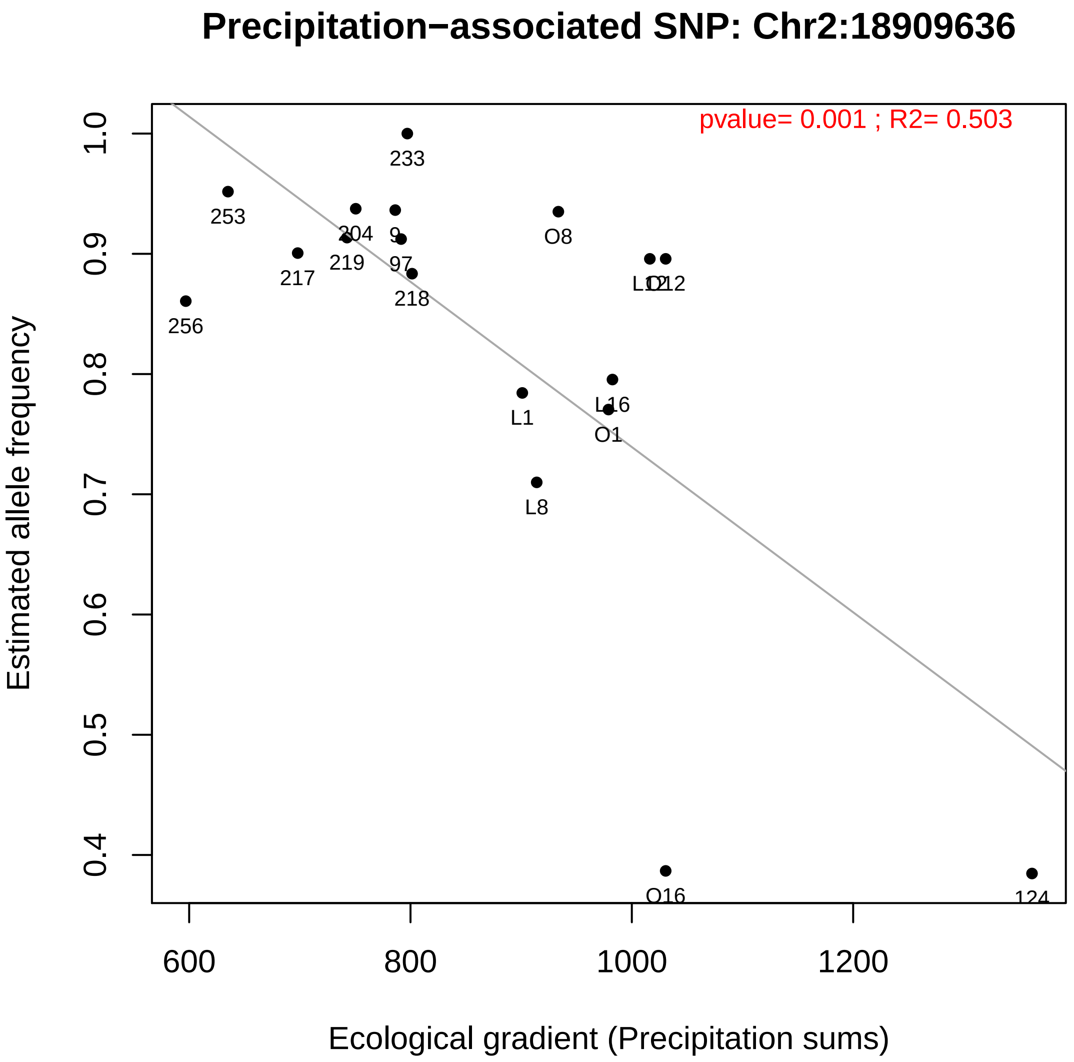

Precipitation−associated SNP: Chr2:18909636
pvalue= 0.001 ; R2= 0.503
233
253
204
9
O8
219
97
217
218
L12
O12
256
L16
L1
O1
L8
O16
124
1.0
0.9
0.8
0.7
0.6
0.5
0.4
600
800
1000
1200
Estimated allele frequency
Ecological gradient (Precipitation sums)

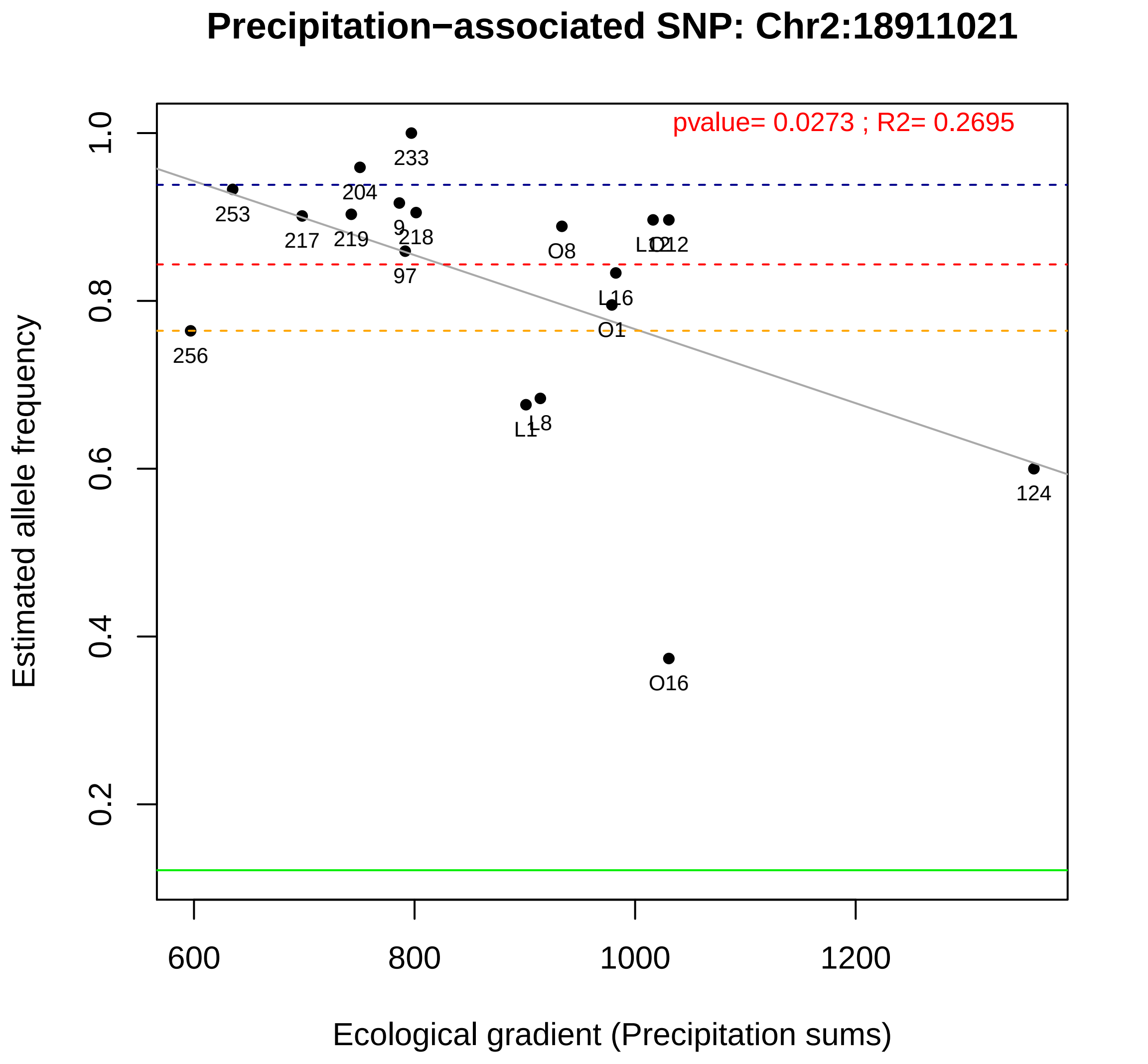

Precipitation−associated SNP: Chr2:18911021
pvalue= 0.0273 ; R2= 0.2695
Estimated allele frequency
Ecological gradient (Precipitation sums)
0.2
0.4
0.6
0.8
1.0
600
800
1000
1200
233
204
253
217
219
9
218
97
O8
L12
O12
L16
O1
256
L1
L8
124
O16

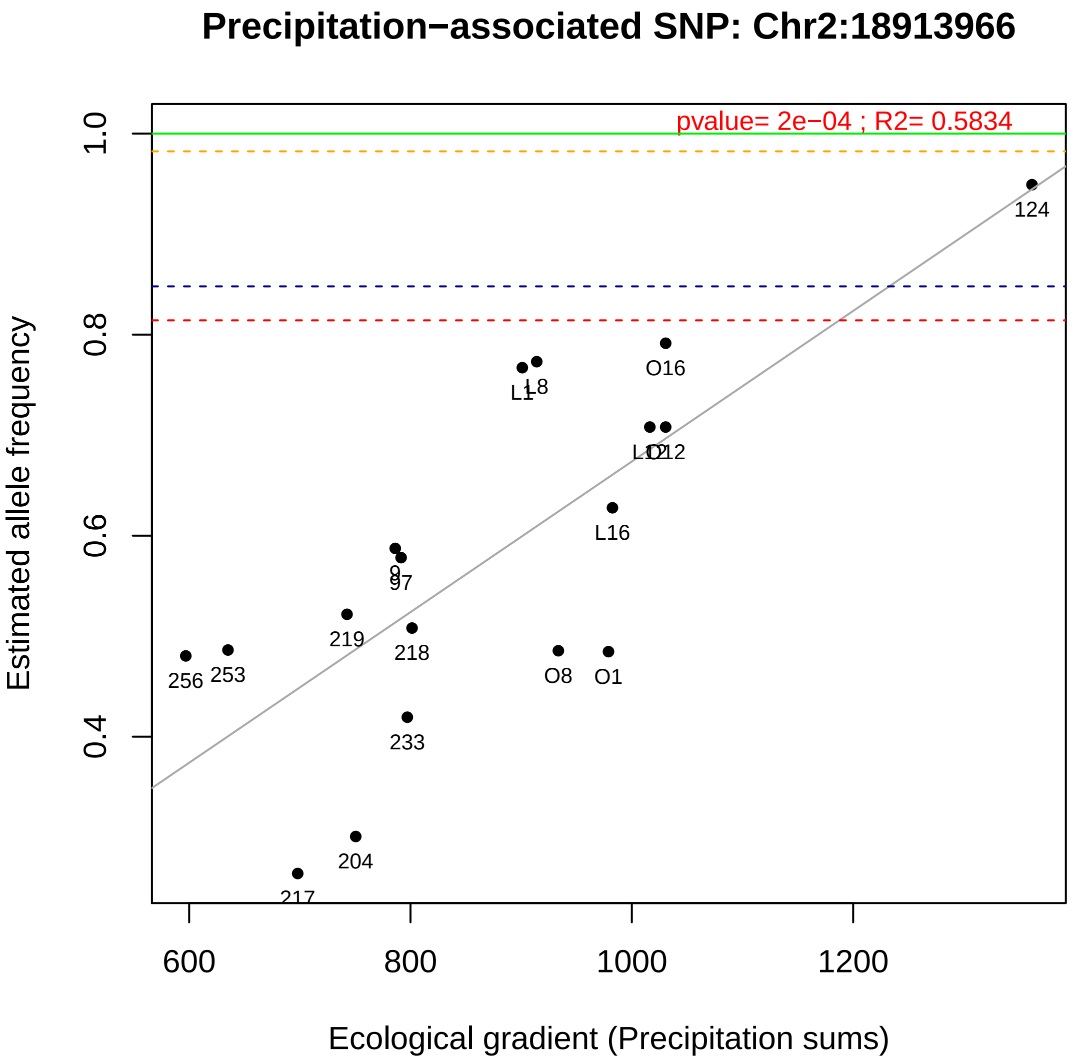

Precipitation−associated SNP: Chr2:18913966
pvalue= 2e−04 ; R2= 0.5834
Estimated allele frequency
0.4
0.6
0.8
1.0
600
800
1000
1200
Ecological gradient (Precipitation sums)
124
O16
L1
L8
L12
O12
L16
9
97
219
218
256
253
O8
O1
233
204
217

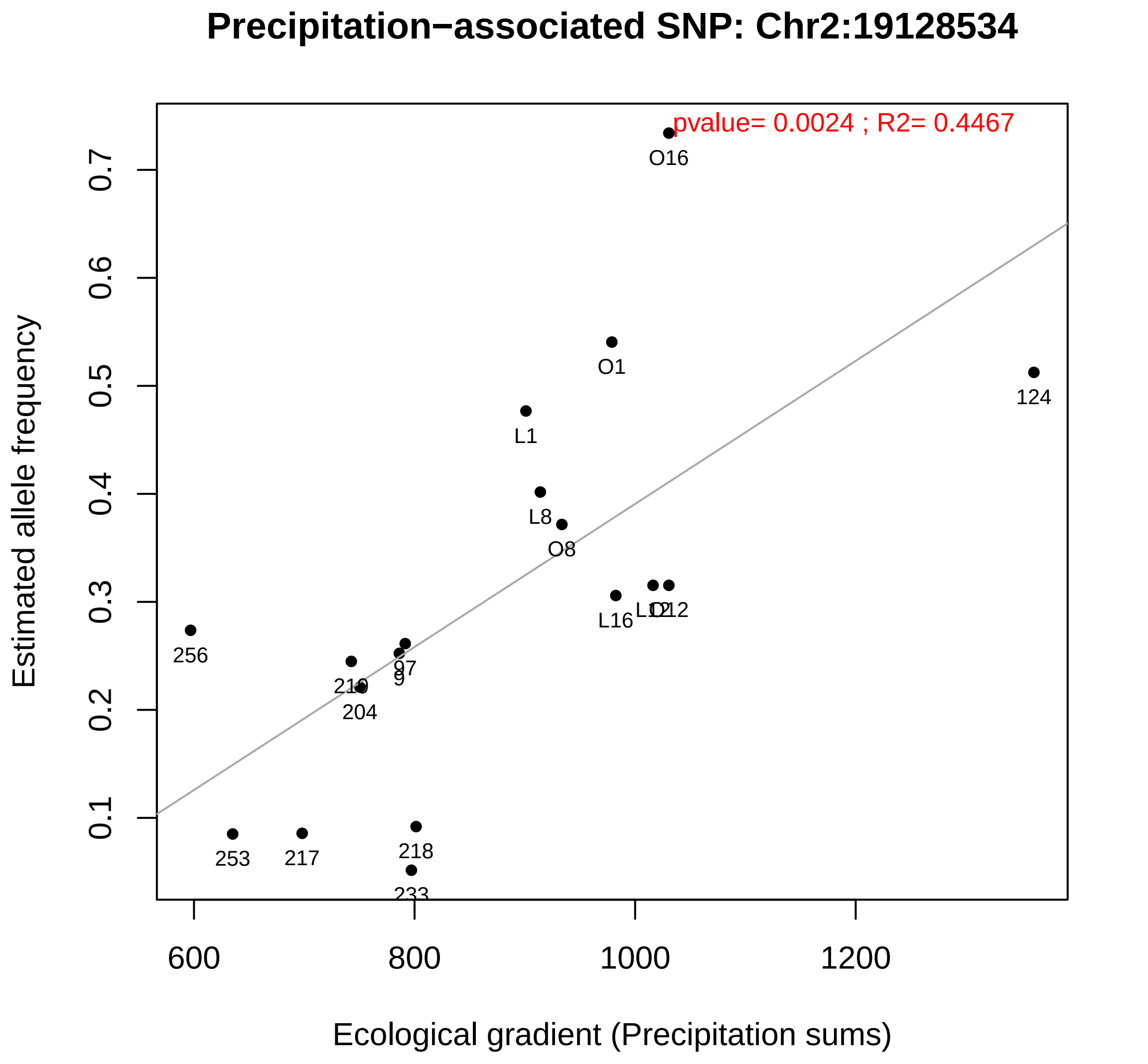
Precipitation−associated SNP: Chr2:19128534
pvalue= 0.0024 ; R2= 0.4467
O16
O1
124
L1
L8
O8
L16
L12
O12
256
210
204
97
9
253
217
218
233
0.1
0.2
0.3
0.4
0.5
0.6
0.7
600
800
1000
1200
Estimated allele frequency
Ecological gradient (Precipitation sums)

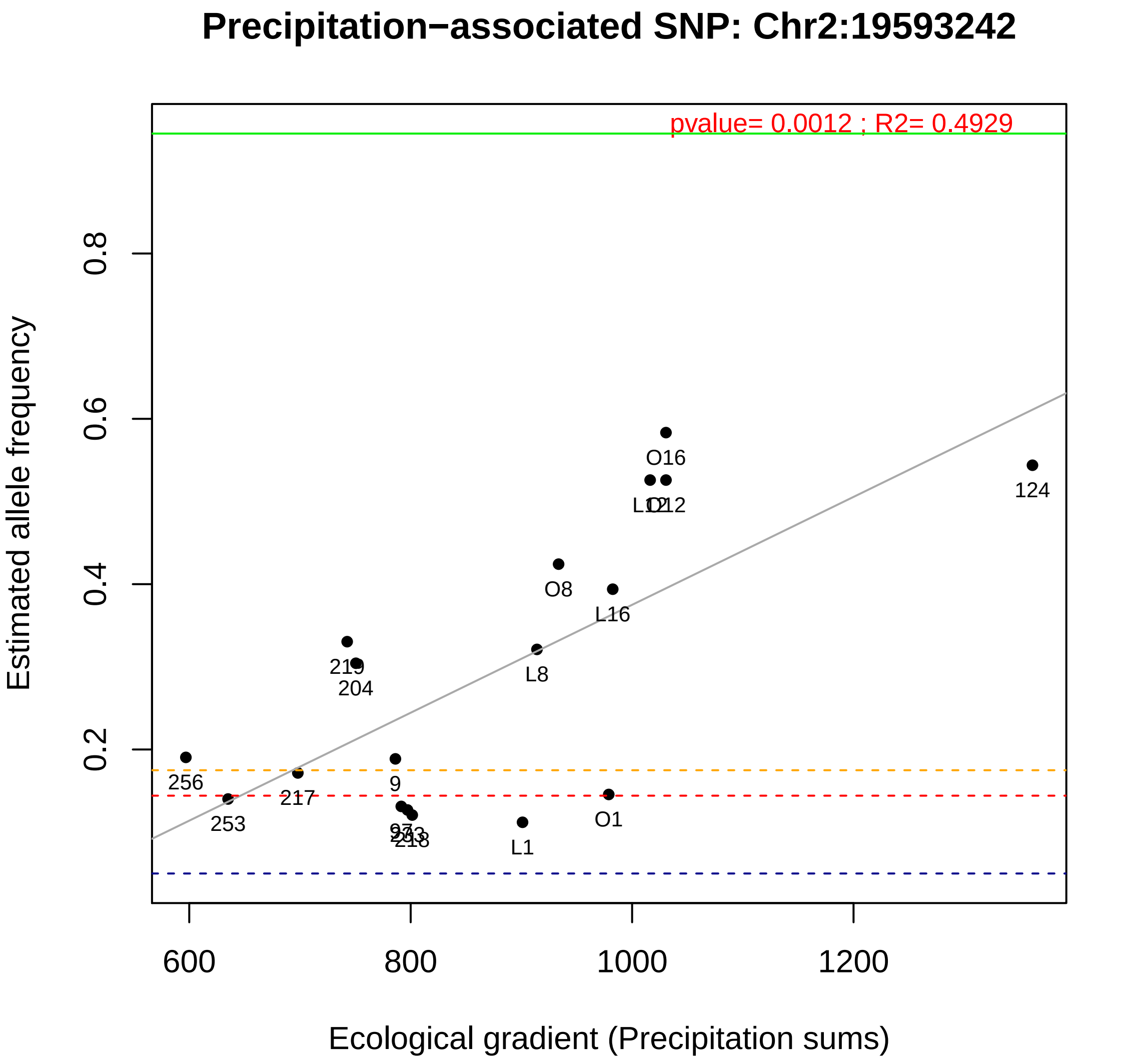

Precipitation−associated SNP: Chr2:19593242
pvalue= 0.0012 ; R2= 0.4929
Estimated allele frequency
0.2
0.4
0.6
0.8
600
800
1000
1200
Ecological gradient (Precipitation sums)
O16
L12
O2
124
O8
L16
219
204
L8
256
253
217
9
97
233
218
L1
O1

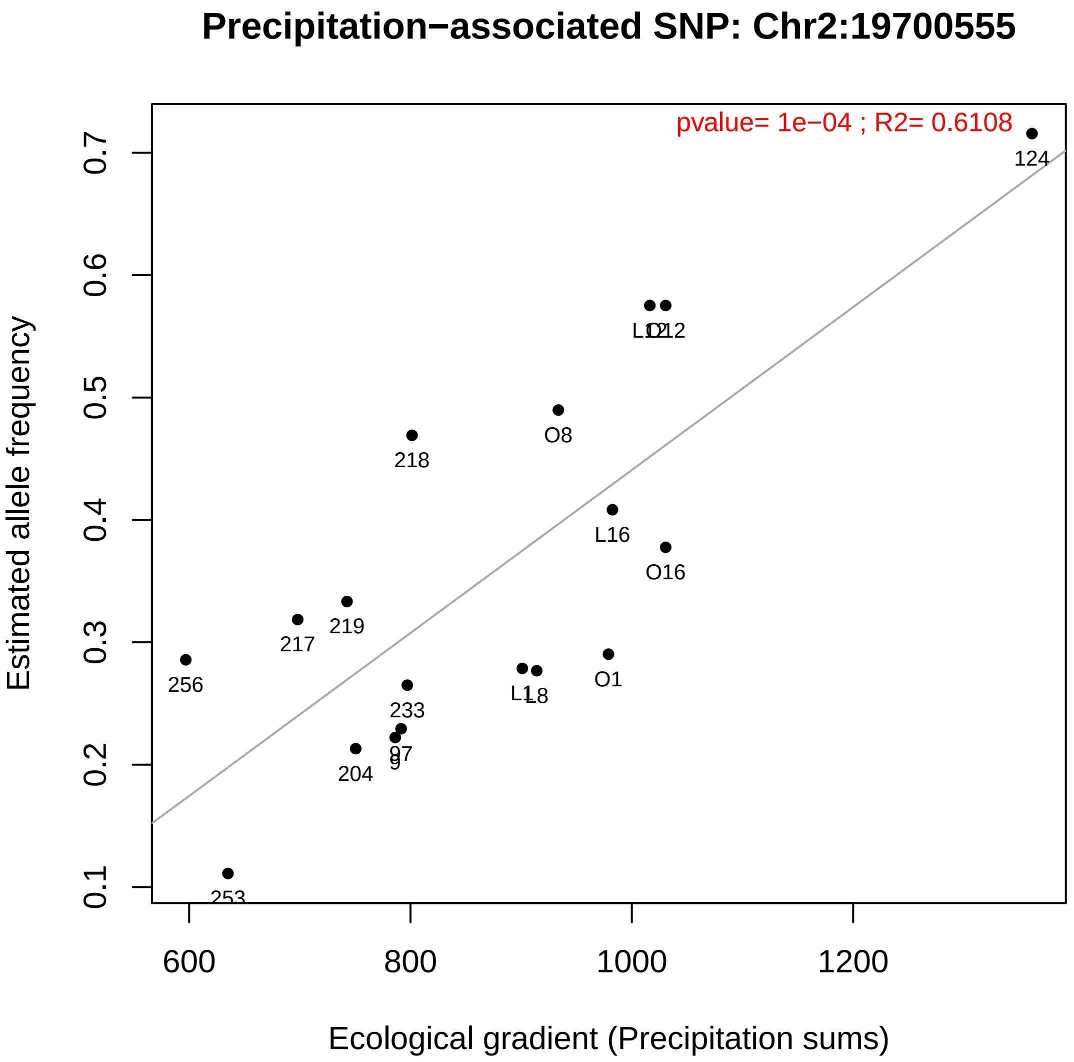
Precipitation−associated SNP: Chr2:19700555
pvalue= 1e−04 ; R2= 0.6108
Estimated allele frequency
0.1
0.2
0.3
0.4
0.5
0.6
0.7
Ecological gradient (Precipitation sums)
600
800
1000
1200
124
O8
218
L16
O16
219
217
256
233
O1
204
253

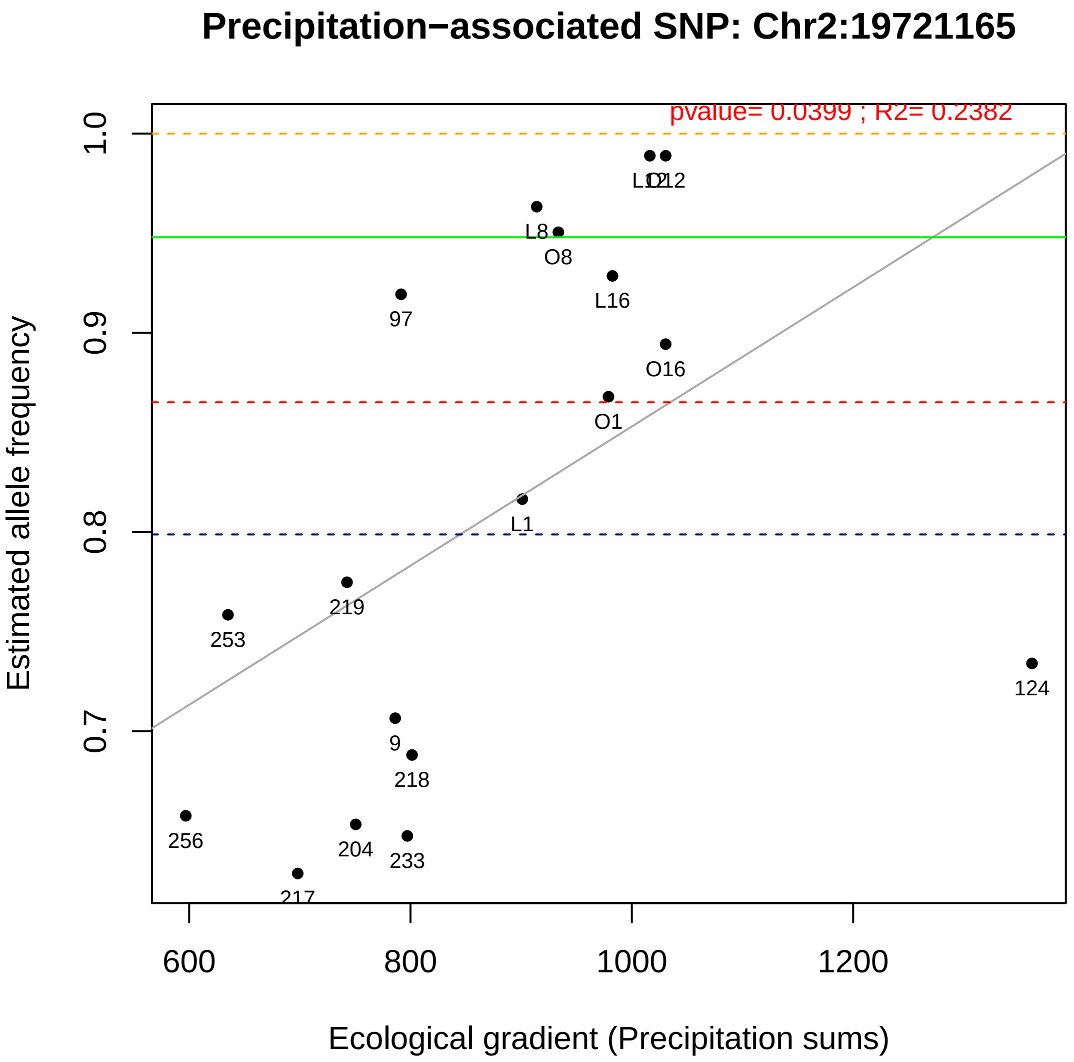

Precipitation−associated SNP: Chr2:19721165
pvalue= 0.0399 ; R2= 0.2382
Estimated allele frequency
1.0
0.9
0.8
0.7
600
800
1000
1200
Ecological gradient (Precipitation sums)
L12
O12
L8
O8
L16
97
O16
O1
L1
219
253
124
9
218
256
204
233
217

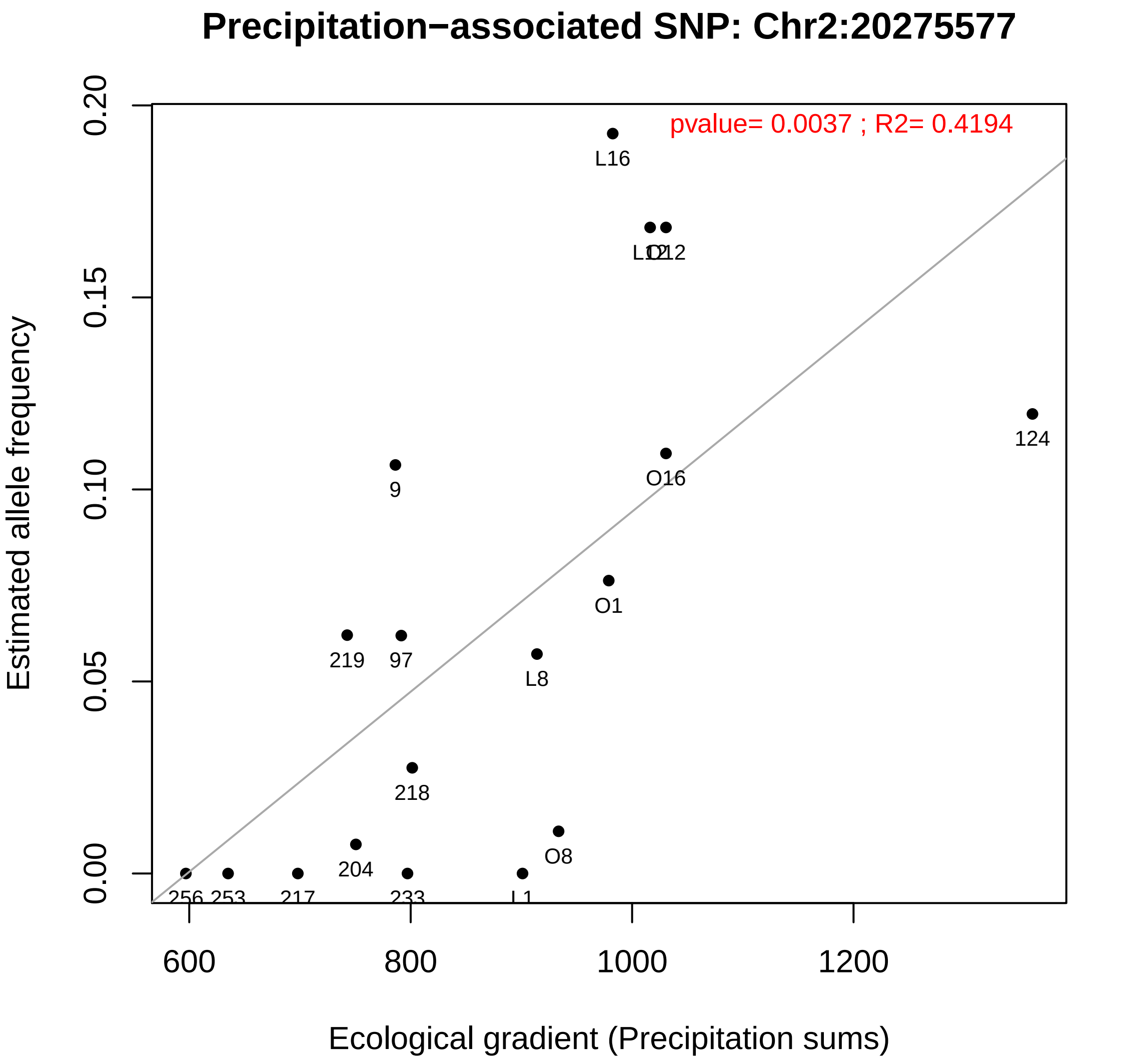

Precipitation−associated SNP: Chr2:20275577
pvalue= 0.0037 ; R2= 0.4194
Estimated allele frequency
0.00
0.05
0.10
0.15
0.20
Ecological gradient (Precipitation sums)
600
800
1000
1200
L16
124
O16
9
O1
219
97
L8
218
204
O8
256
253
217
233
L1

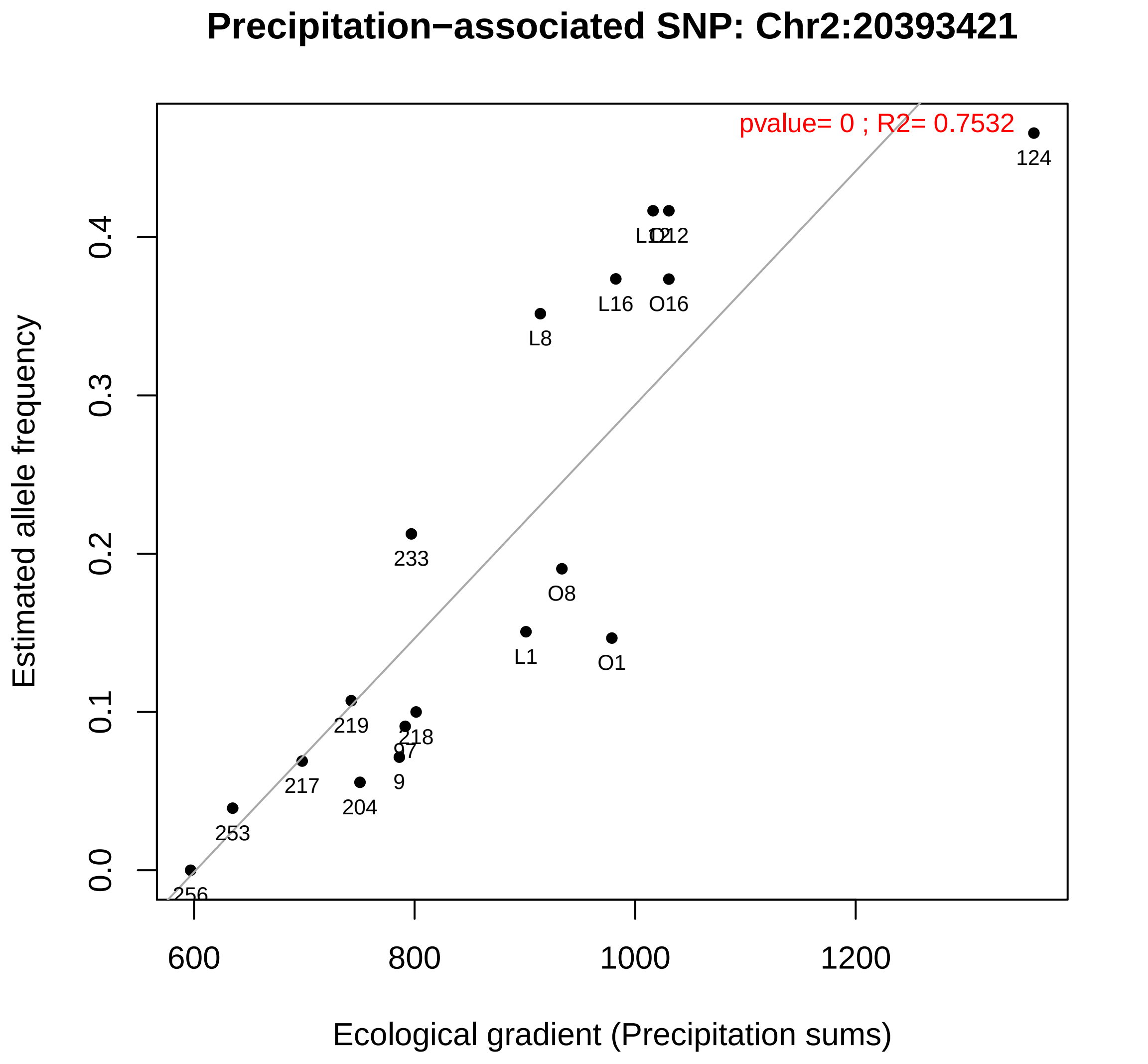

Precipitation−associated SNP: Chr2:20393421
pvalue= 0 ; R2= 0.7532
124
L12
O12
L16
O16
L8
233
O8
L1
O1
219
218
97
217
9
204
253
256
0.0
0.1
0.2
0.3
0.4
600
800
1000
1200
Estimated allele frequency
Ecological gradient (Precipitation sums)

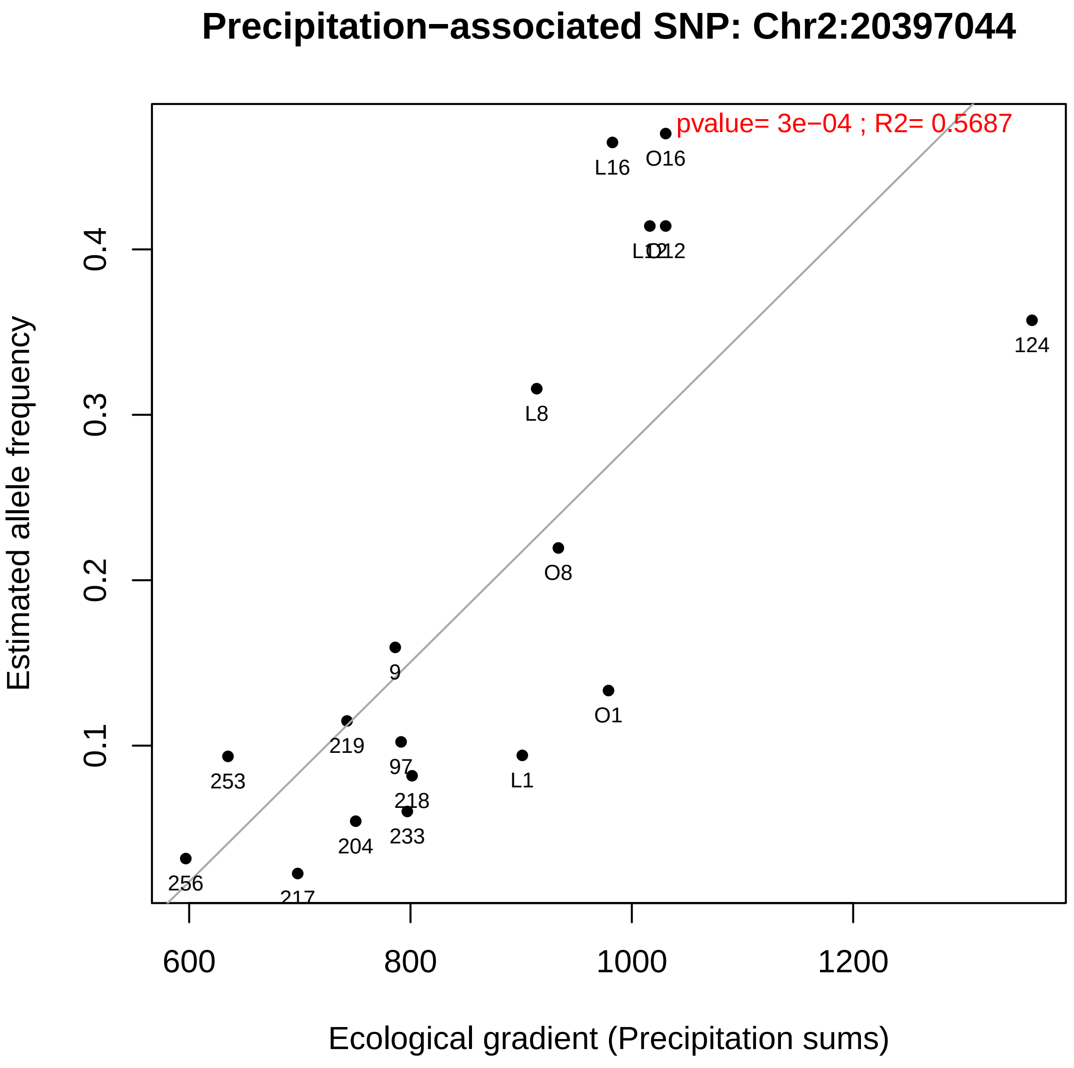

Precipitation−associated SNP: Chr2:20397044
pvalue= 3e−04 ; R2= 0.5687
L16
O16
L12
O12
124
L8
O8
9
O1
219
97
L1
253
218
233
204
256
217
0.1
0.2
0.3
0.4
600
800
1000
1200
Estimated allele frequency
Ecological gradient (Precipitation sums)

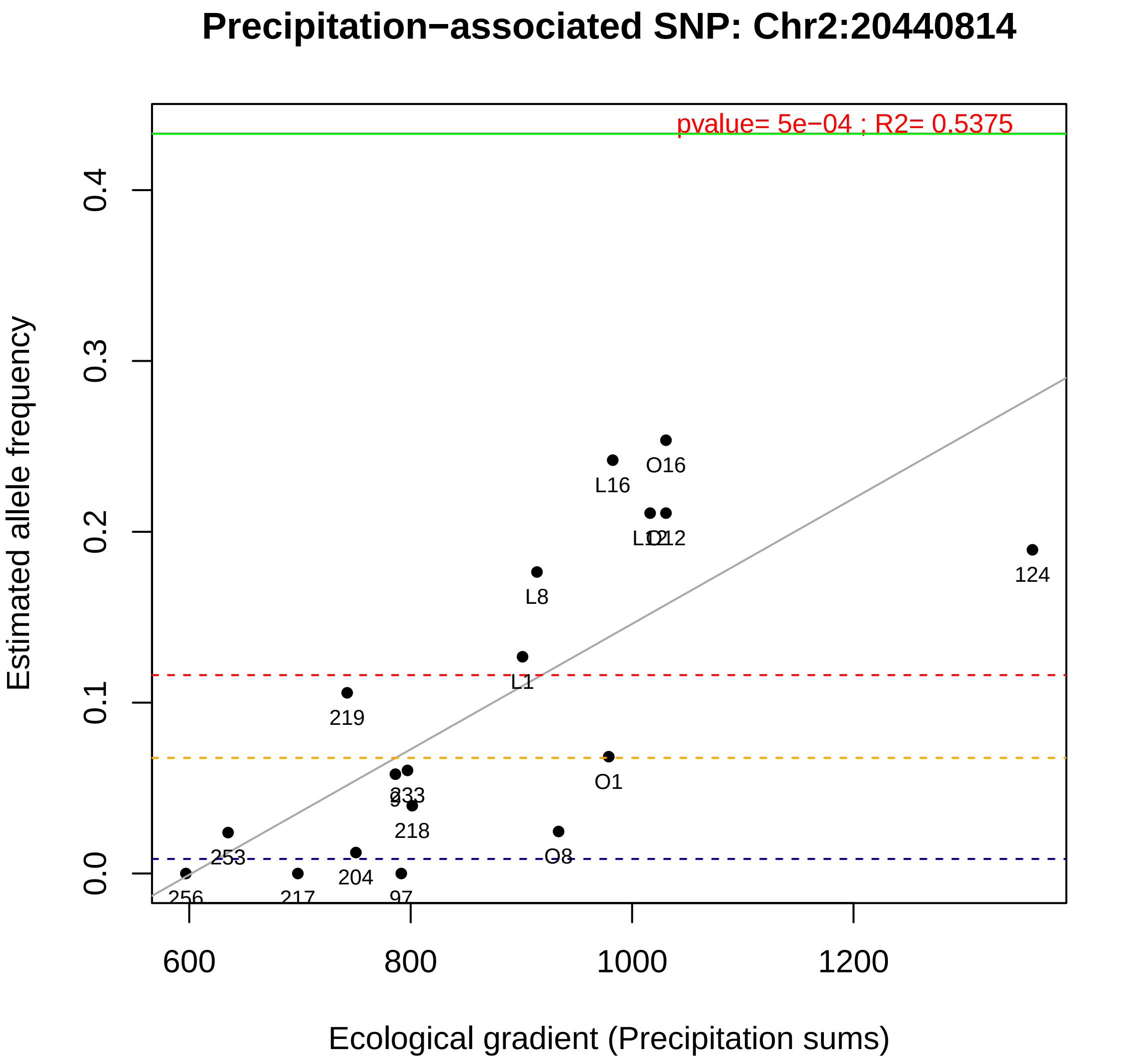
Precipitation−associated SNP: Chr2:20440814
pvalue= 5e−04 ; R2= 0.5375
Estimated allele frequency
0.0
0.1
0.2
0.3
0.4
600
800
1000
1200
Ecological gradient (Precipitation sums)
O16
L16
L12
O12
124
L8
L1
219
O1
233
218
253
O8
204
256
217
97

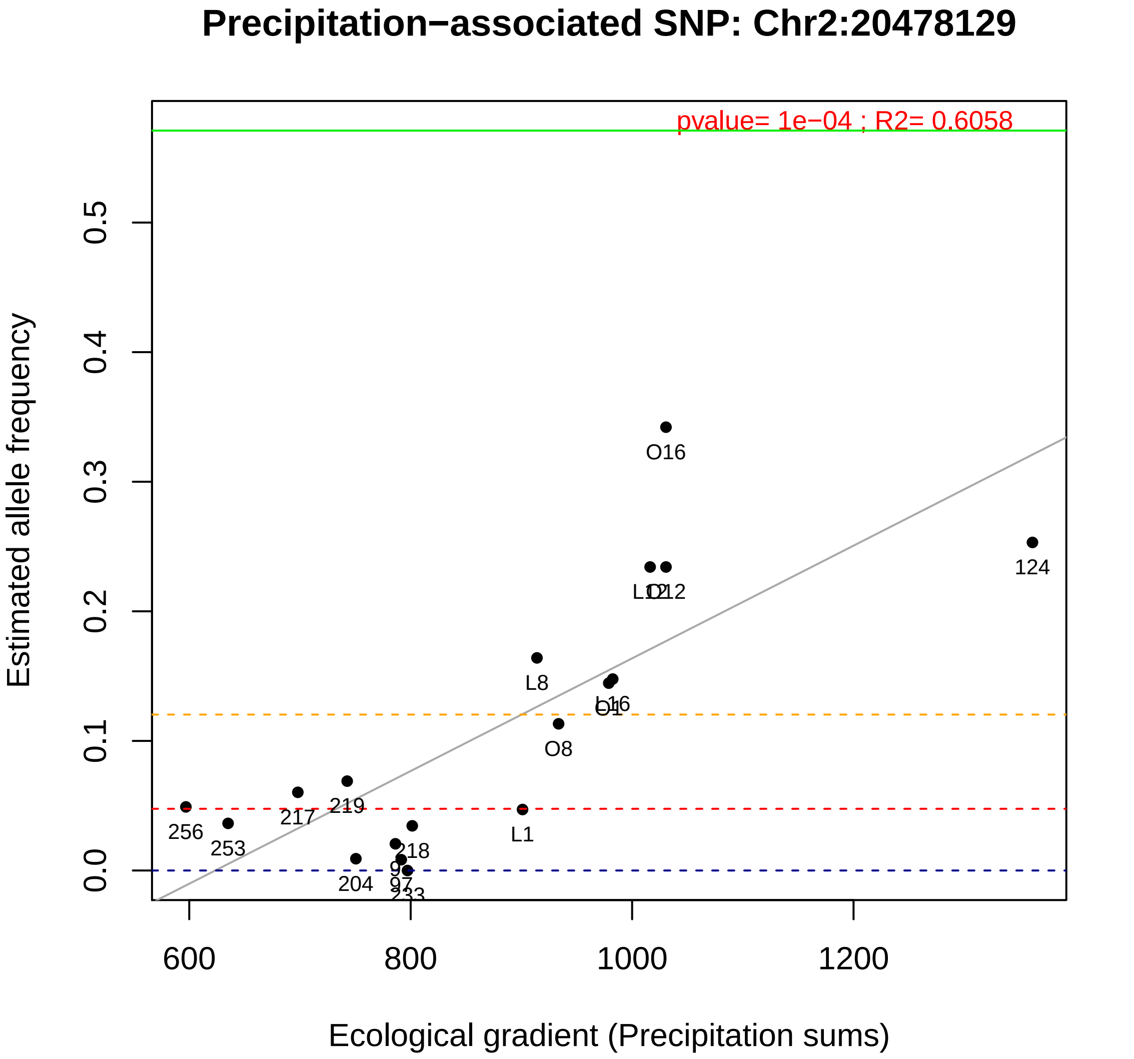

Precipitation−associated SNP: Chr2:20478129
pvalue= 1e−04 ; R2= 0.6058
Estimated allele frequency
0.0
0.1
0.2
0.3
0.4
0.5
Ecological gradient (Precipitation sums)
600
800
1000
1200
O16
124
O12
L8
O8
219
217
256
253
L1
218
204

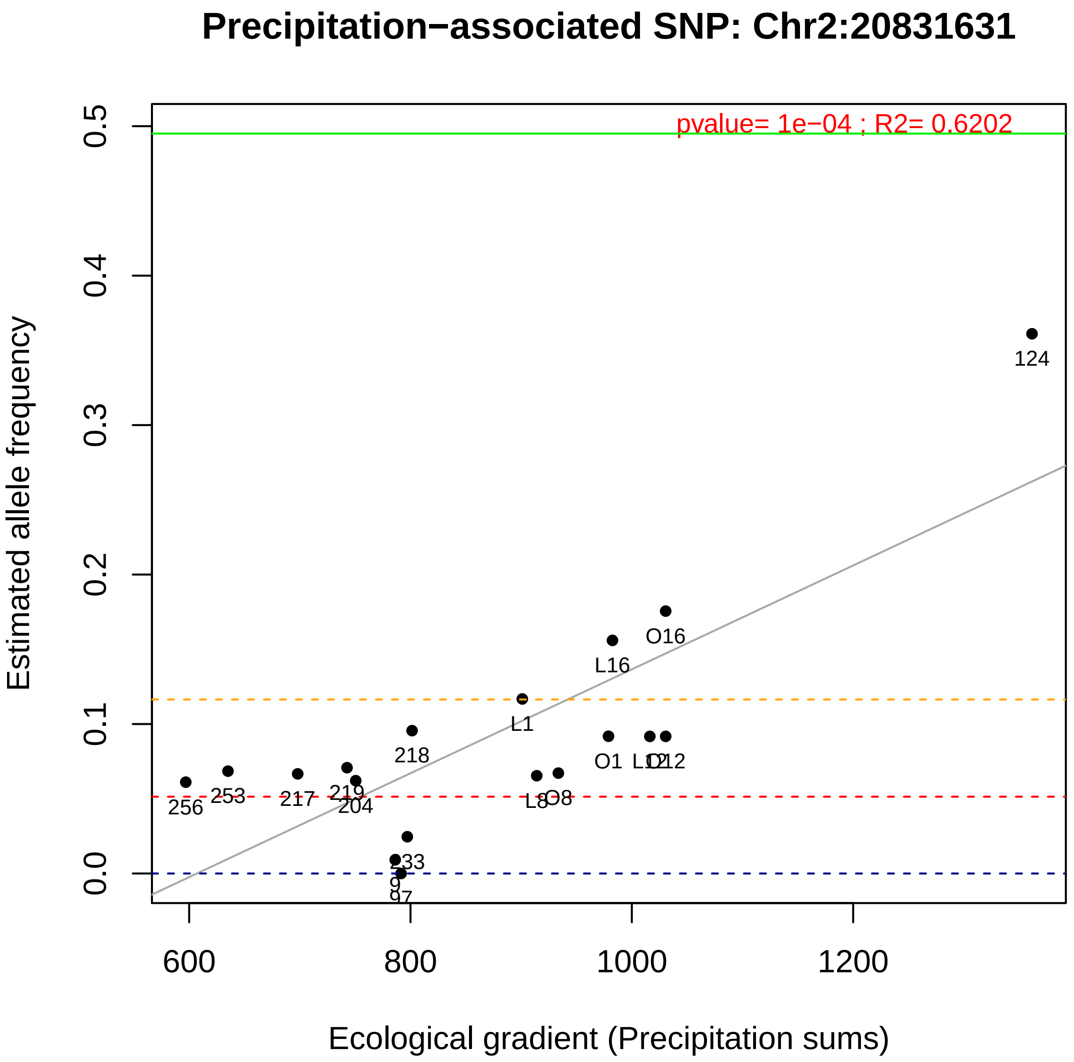

Precipitation−associated SNP: Chr2:20831631
pvalue= 1e−04 ; R2= 0.6202
Estimated allele frequency
0.0
0.1
0.2
0.3
0.4
0.5
600
800
1000
1200
Ecological gradient (Precipitation sums)
124
O16
L16
L1
218
O1
O12
256
253
217
219
204
L8
O8
233
97

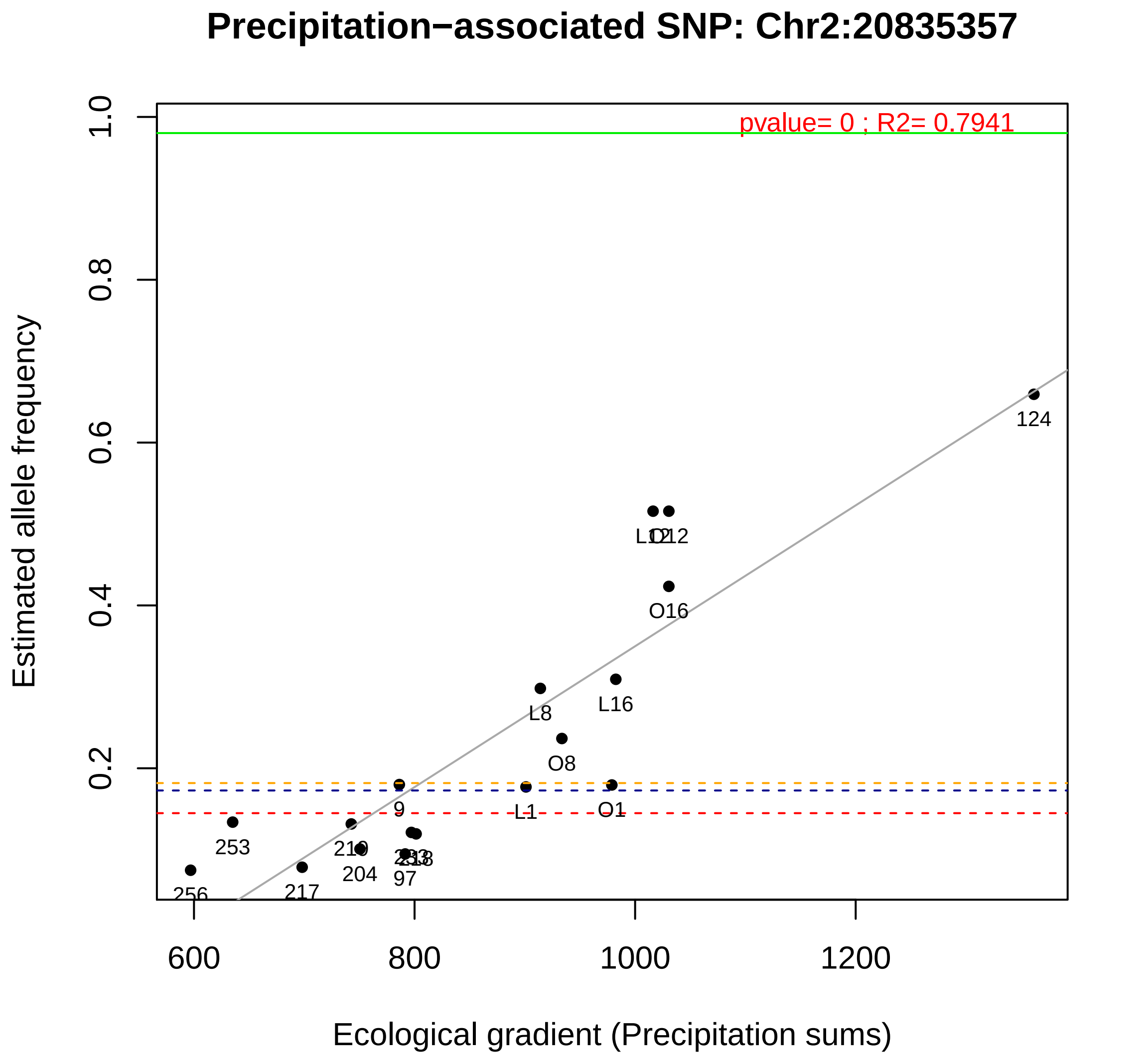

Precipitation−associated SNP: Chr2:20835357
pvalue= 0 ; R2= 0.7941
Estimated allele frequency
Ecological gradient (Precipitation sums)
0.2
0.4
0.6
0.8
1.0
600
800
1000
1200
124
O16
L16
L8
O8
9
L1
O1
253
204
97
217

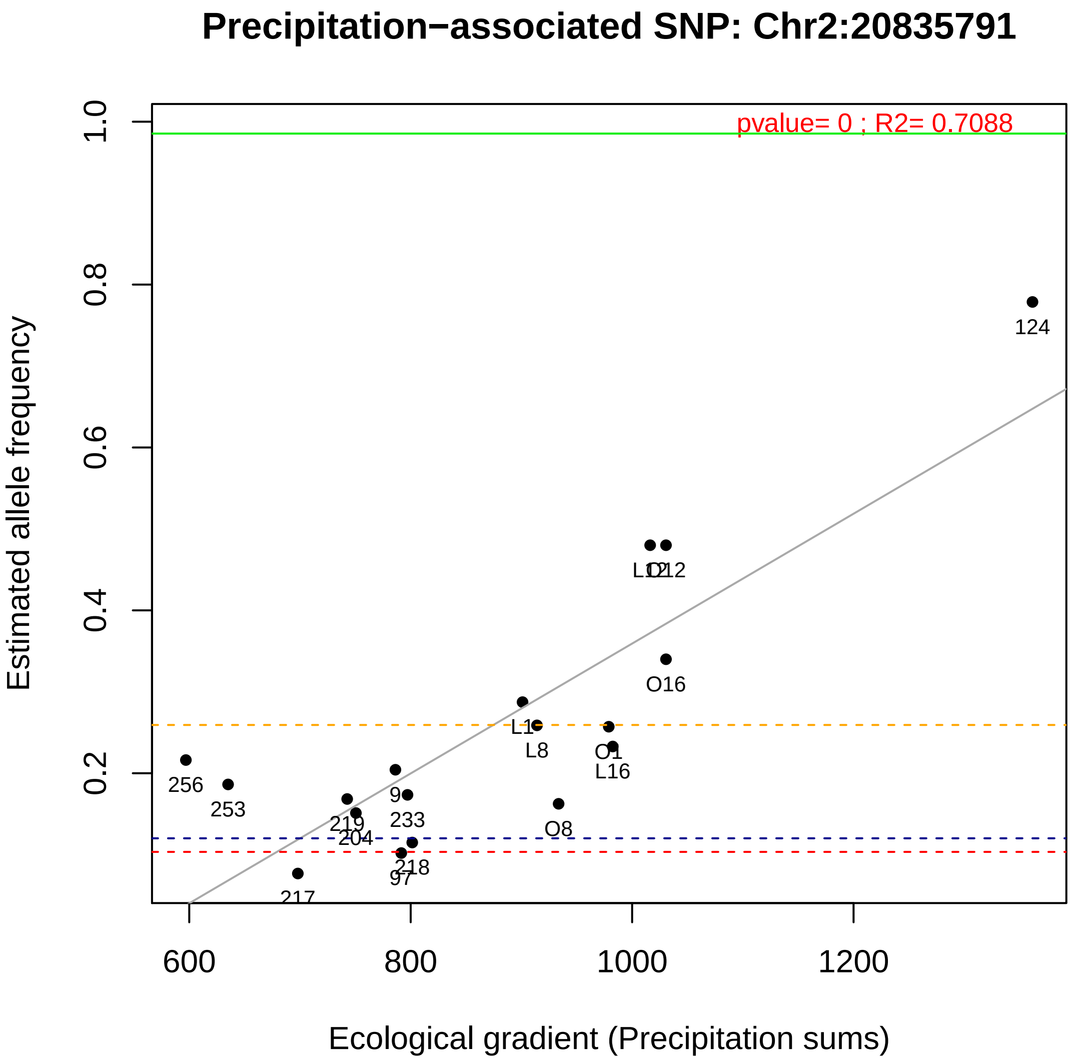

Precipitation−associated SNP: Chr2:20835791
pvalue= 0 ; R2= 0.7088
Estimated allele frequency
1.0
0.8
0.6
0.4
0.2
124
L12
O12
O16
L1
L8
O1
L16
256
253
9
233
219
204
O8
218
97
217
600
800
1000
1200
Ecological gradient (Precipitation sums)

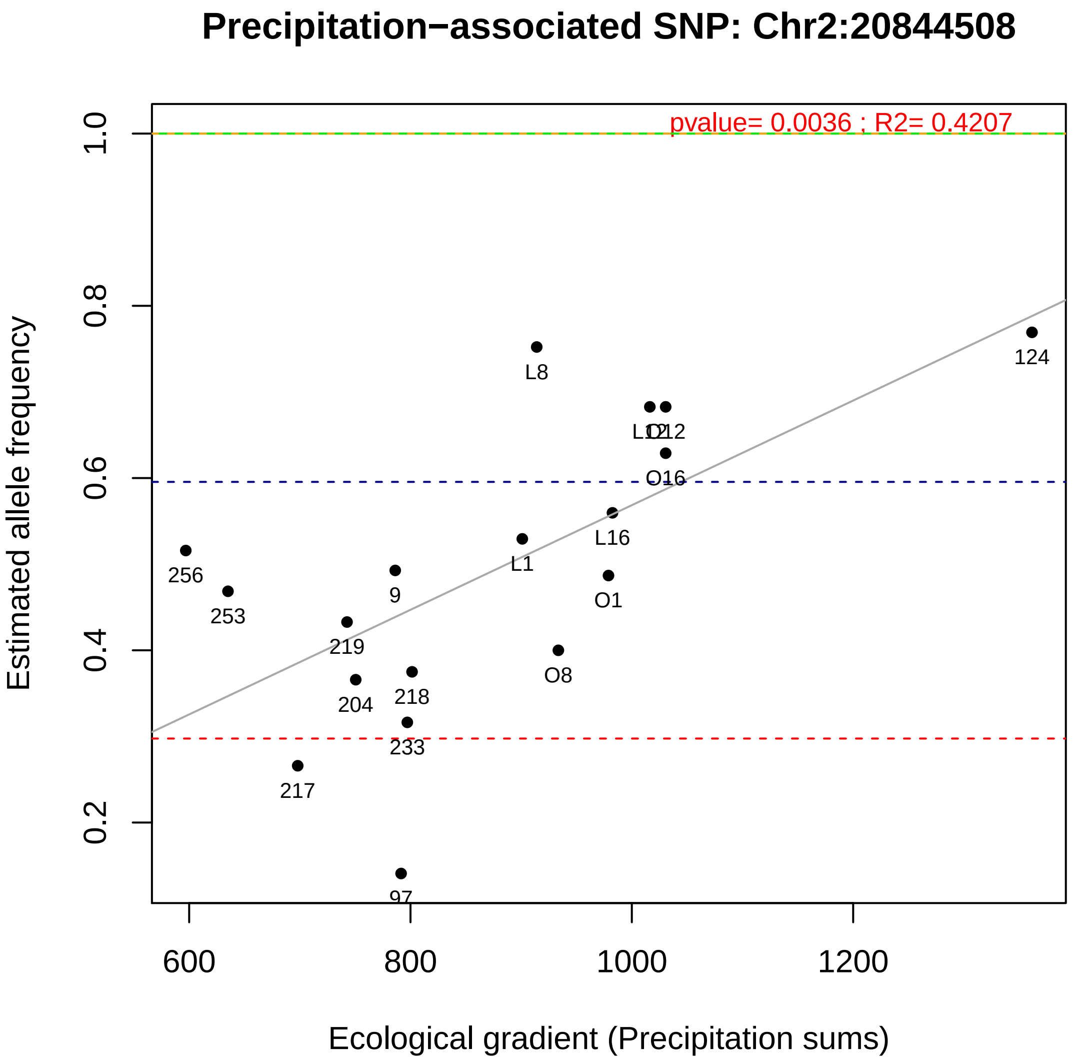
Precipitation−associated SNP: Chr2:20844508
pvalue= 0.0036 ; R2= 0.4207
Estimated allele frequency
Ecological gradient (Precipitation sums)
1.0
0.8
0.6
0.4
0.2
600
800
1000
1200
L8
124
L12
O12
O16
L16
L1
256
9
O1
253
219
O8
218
204
233
217
97

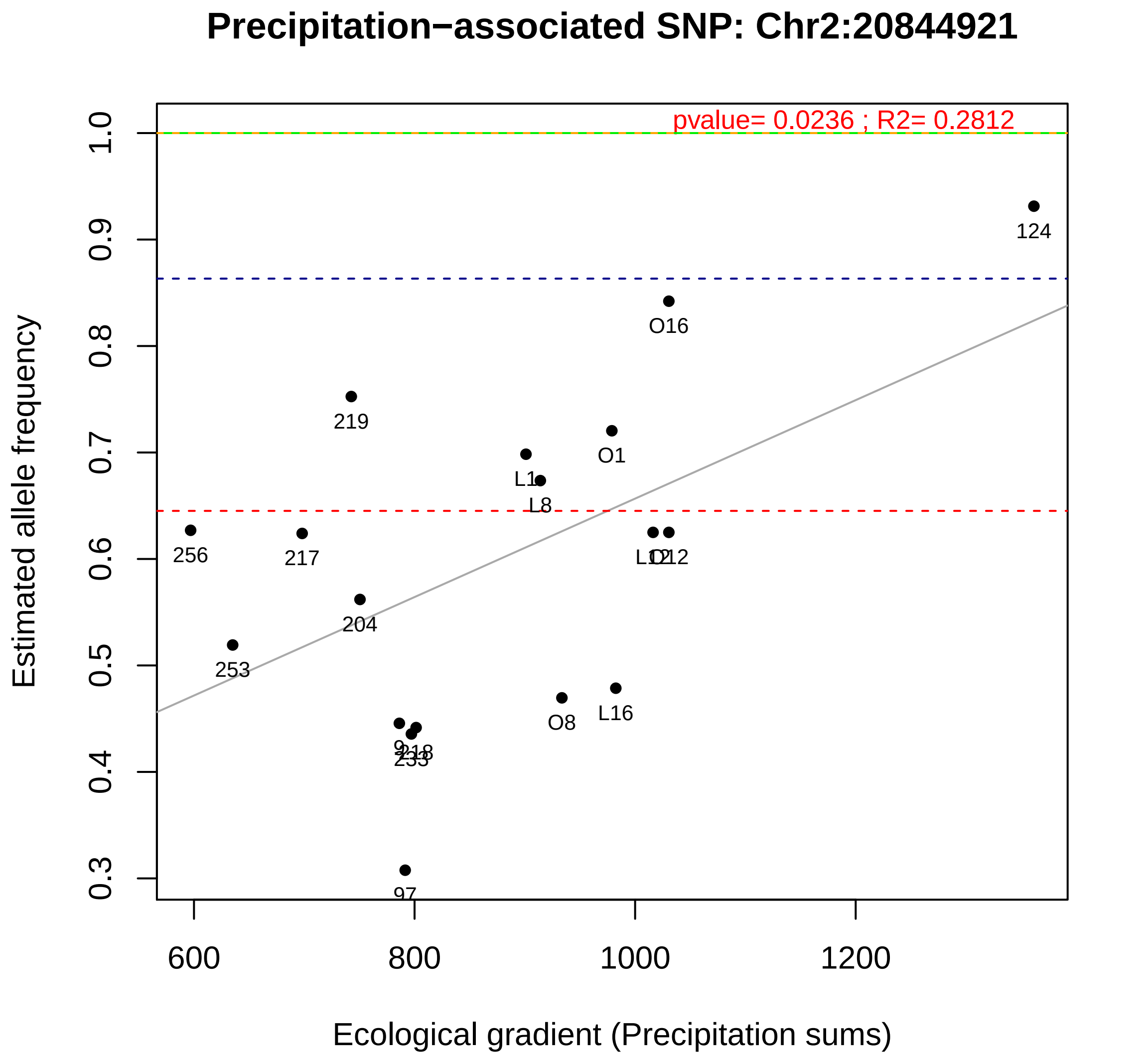
Precipitation−associated SNP: Chr2:20844921
pvalue= 0.0236 ; R2= 0.2812
Estimated allele frequency
Ecological gradient (Precipitation sums)
0.3
0.4
0.5
0.6
0.7
0.8
0.9
1.0
600
800
1000
1200
124
O16
219
O1
L1
L8
256
217
L10
O12
204
253
O8
L16
9
218
233
97

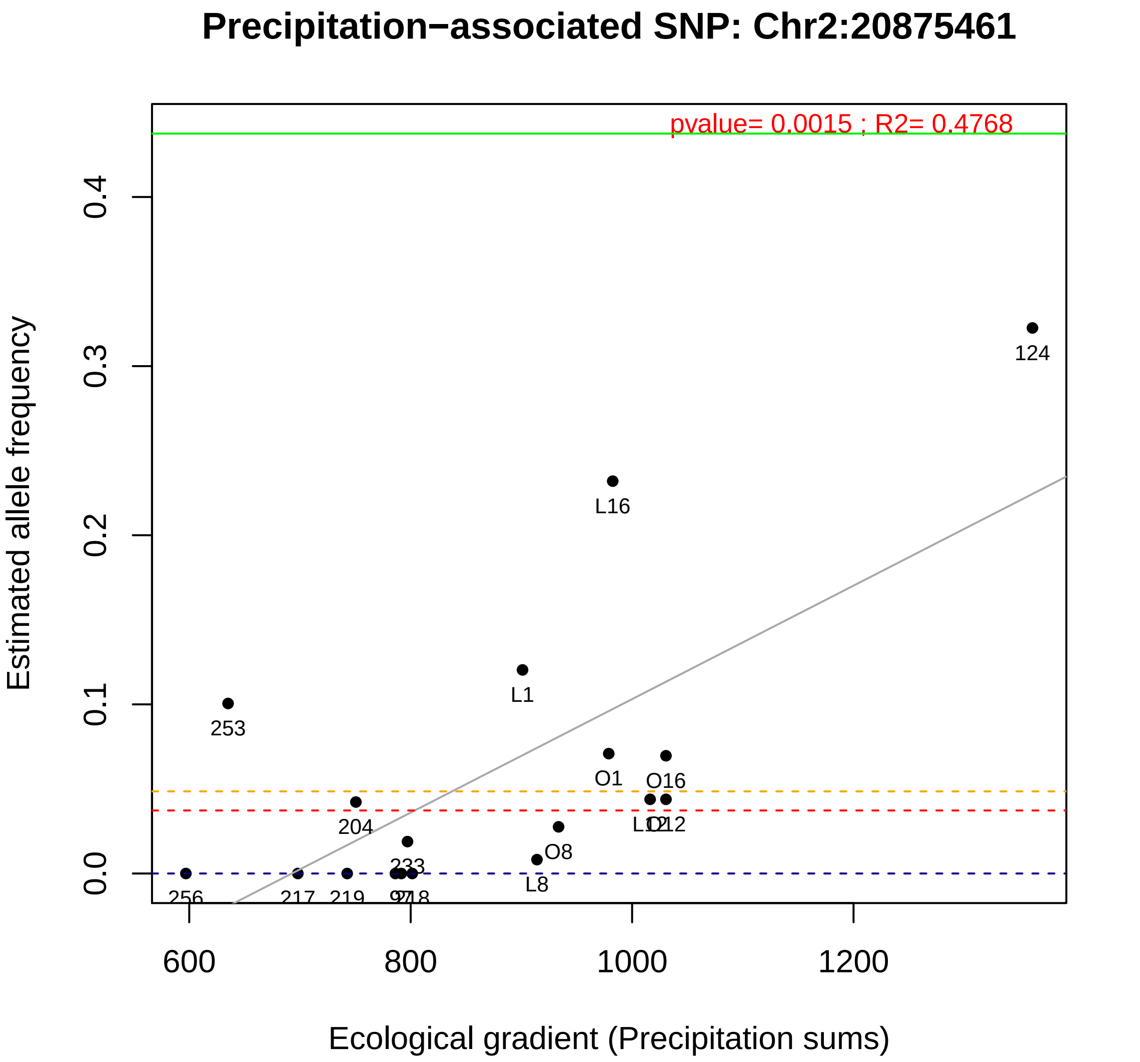

Precipitation−associated SNP: Chr2:20875461
pvalue= 0.0015 ; R2= 0.4768
Estimated allele frequency
0.0
0.1
0.2
0.3
0.4
124
L16
L1
253
O1
O16
204
L12
O12
O8
233
L8
256
217
219
218
600
800
1000
1200
Ecological gradient (Precipitation sums)

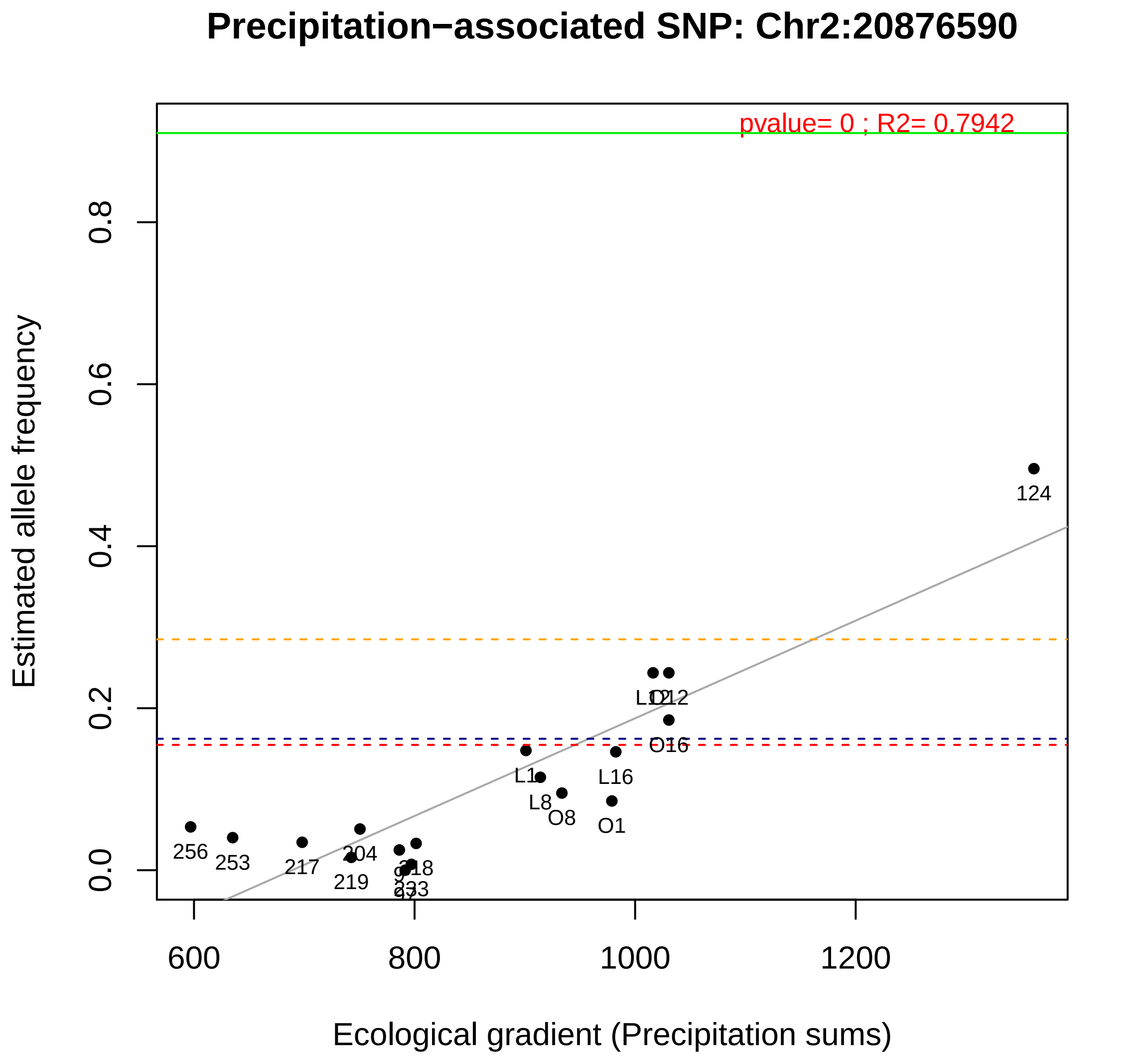

Precipitation−associated SNP: Chr2:20876590
pvalue= 0 ; R2= 0.7942
Estimated allele frequency
0.0
0.2
0.4
0.6
0.8
600
800
1000
1200
Ecological gradient (Precipitation sums)
124
256
253
217
219
204
218
233
L1
L8
O8
O1
L16
O16
L12
O12

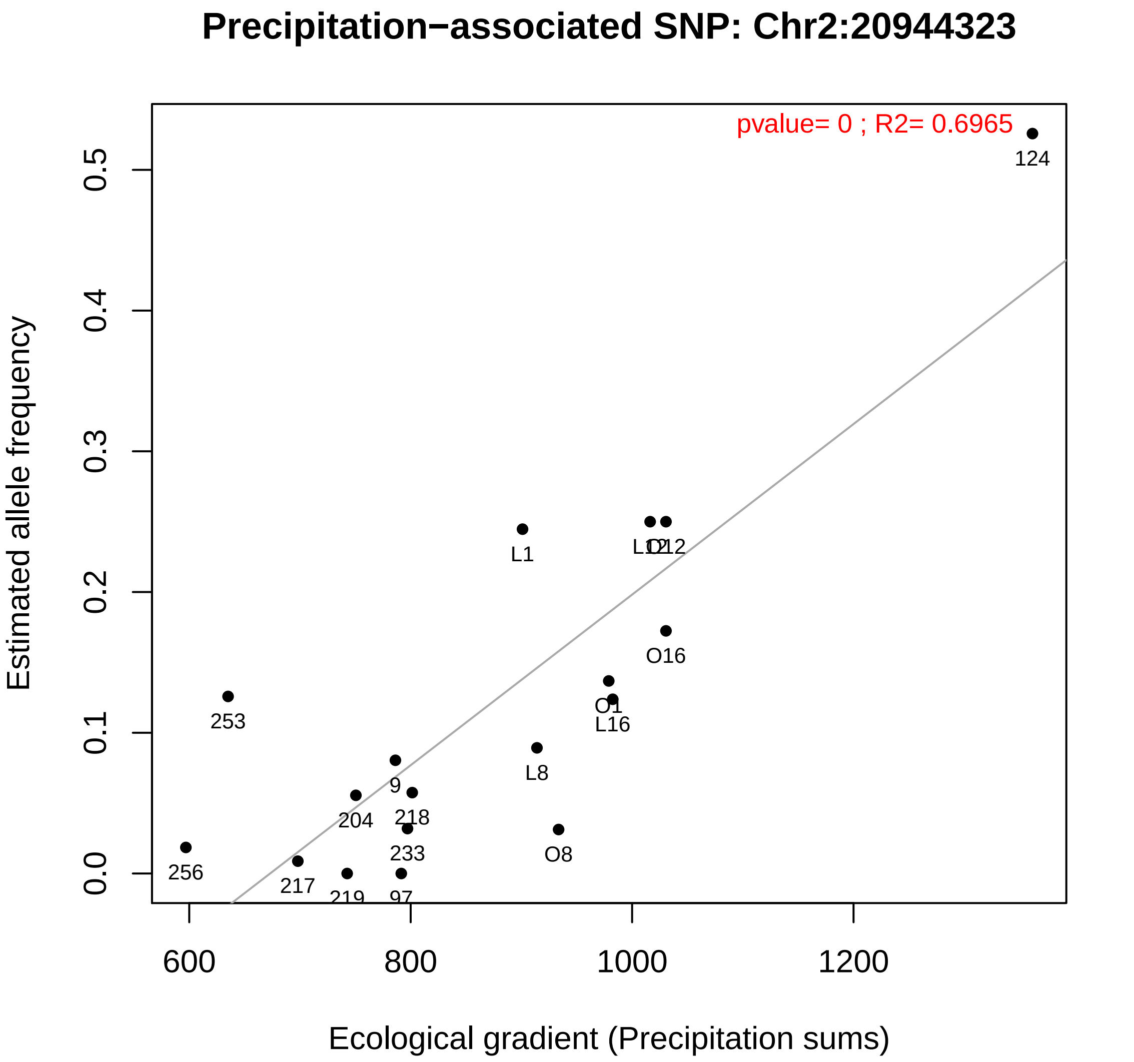

Precipitation−associated SNP: Chr2:20944323
pvalue= 0 ; R2= 0.6965
Estimated allele frequency
0.0
0.1
0.2
0.3
0.4
0.5
600
800
1000
1200
Ecological gradient (Precipitation sums)
124
L1
O16
253
L8
9
204
218
233
O8
256
217
219
97
L16

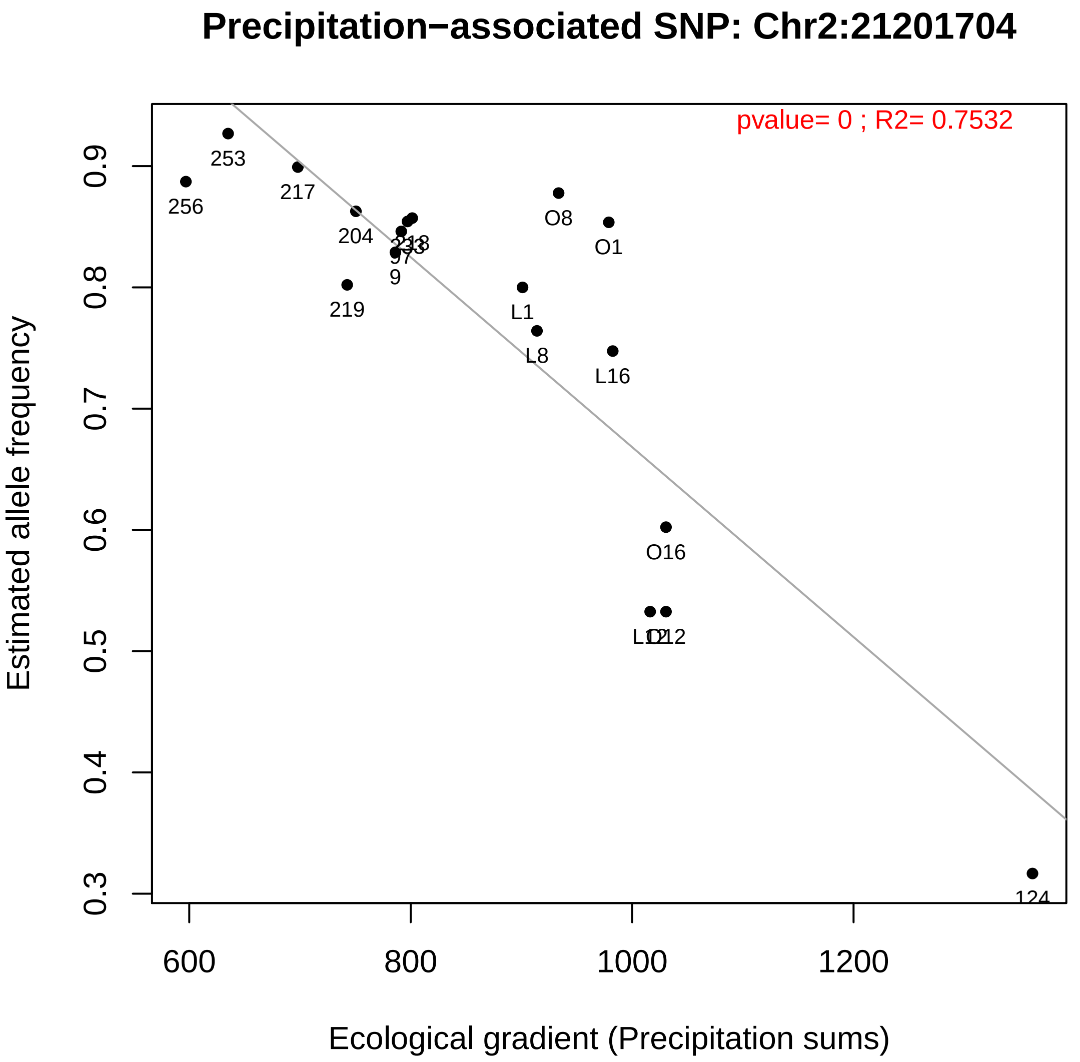

Precipitation−associated SNP: Chr2:21201704
pvalue= 0 ; R2= 0.7532
Estimated allele frequency
0.3
0.4
0.5
0.6
0.7
0.8
0.9
600
800
1000
1200
Ecological gradient (Precipitation sums)
253
256
217
204
219
9
O8
O1
L1
L8
L16
O16
O12
124

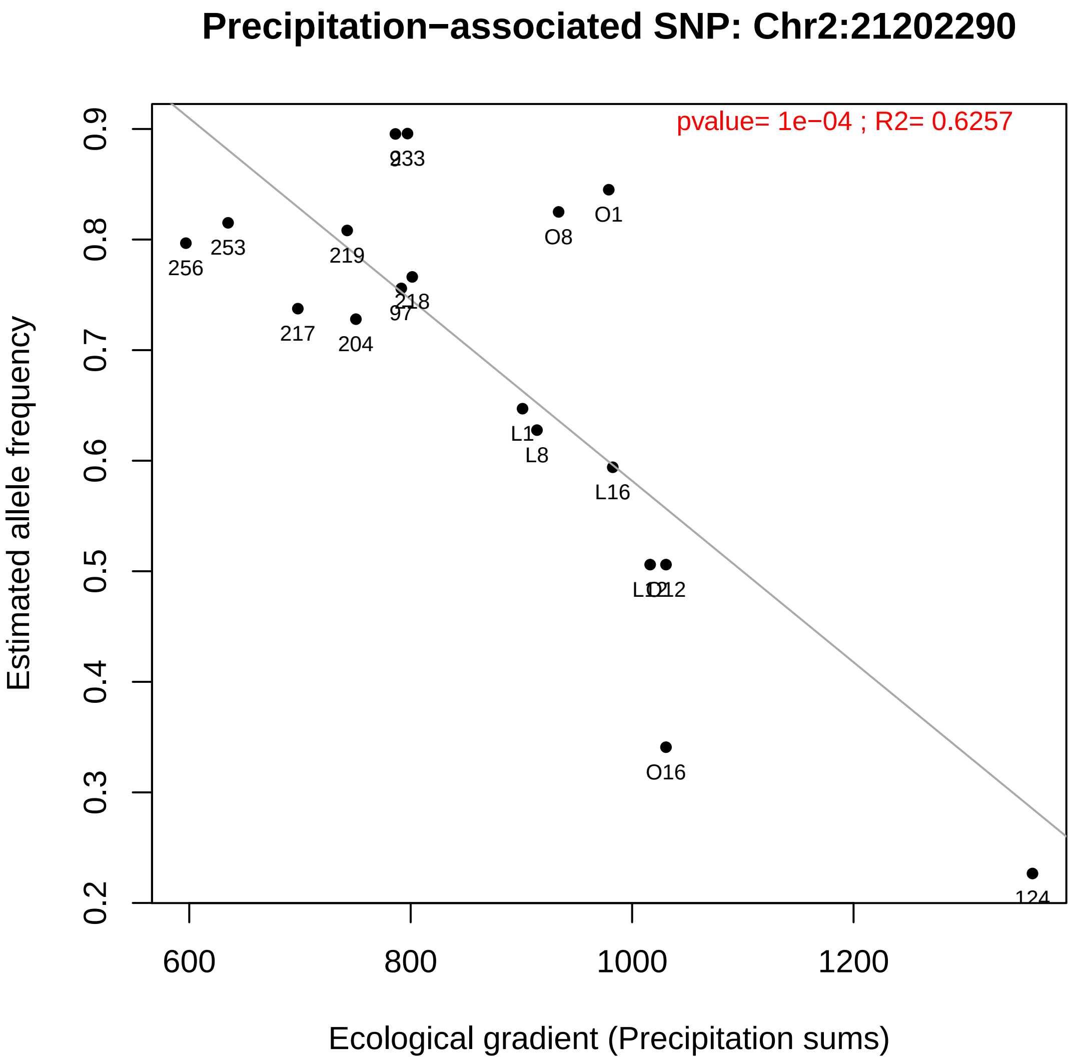

Precipitation−associated SNP: Chr2:21202290
pvalue= 1e−04 ; R2= 0.6257
Estimated allele frequency
Ecological gradient (Precipitation sums)
0.2
0.3
0.4
0.5
0.6
0.7
0.8
0.9
600
800
1000
1200
2
233
O1
O8
253
256
219
218
97
217
204
L1
L8
L16
L12
O12
O16
124

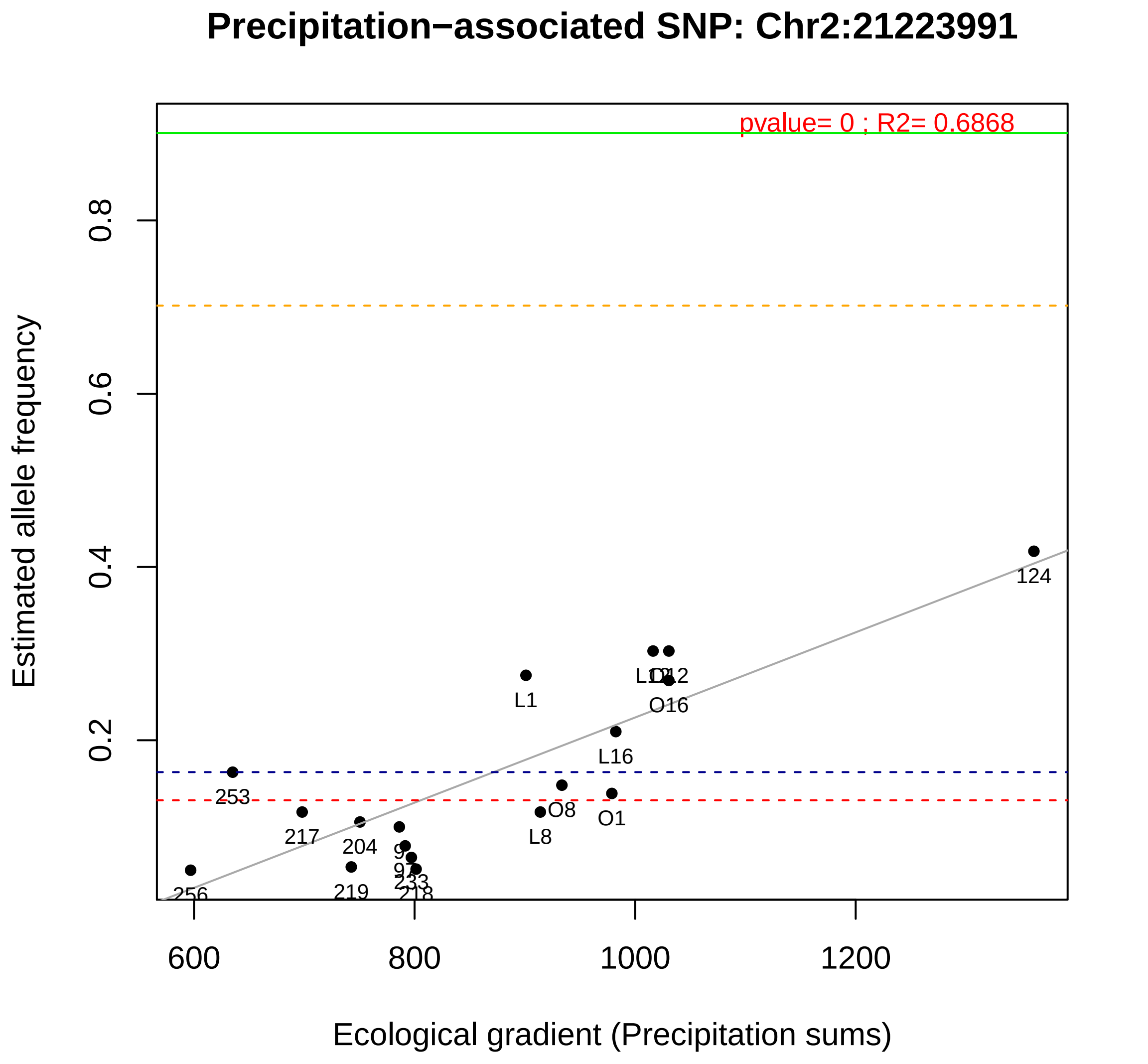

Precipitation−associated SNP: Chr2:21223991
pvalue= 0 ; R2= 0.6868
Estimated allele frequency
0.2
0.4
0.6
0.8
600
800
1000
1200
Ecological gradient (Precipitation sums)
124
L1
O16
L16
253
O8
O1
L8
217
204
219
256

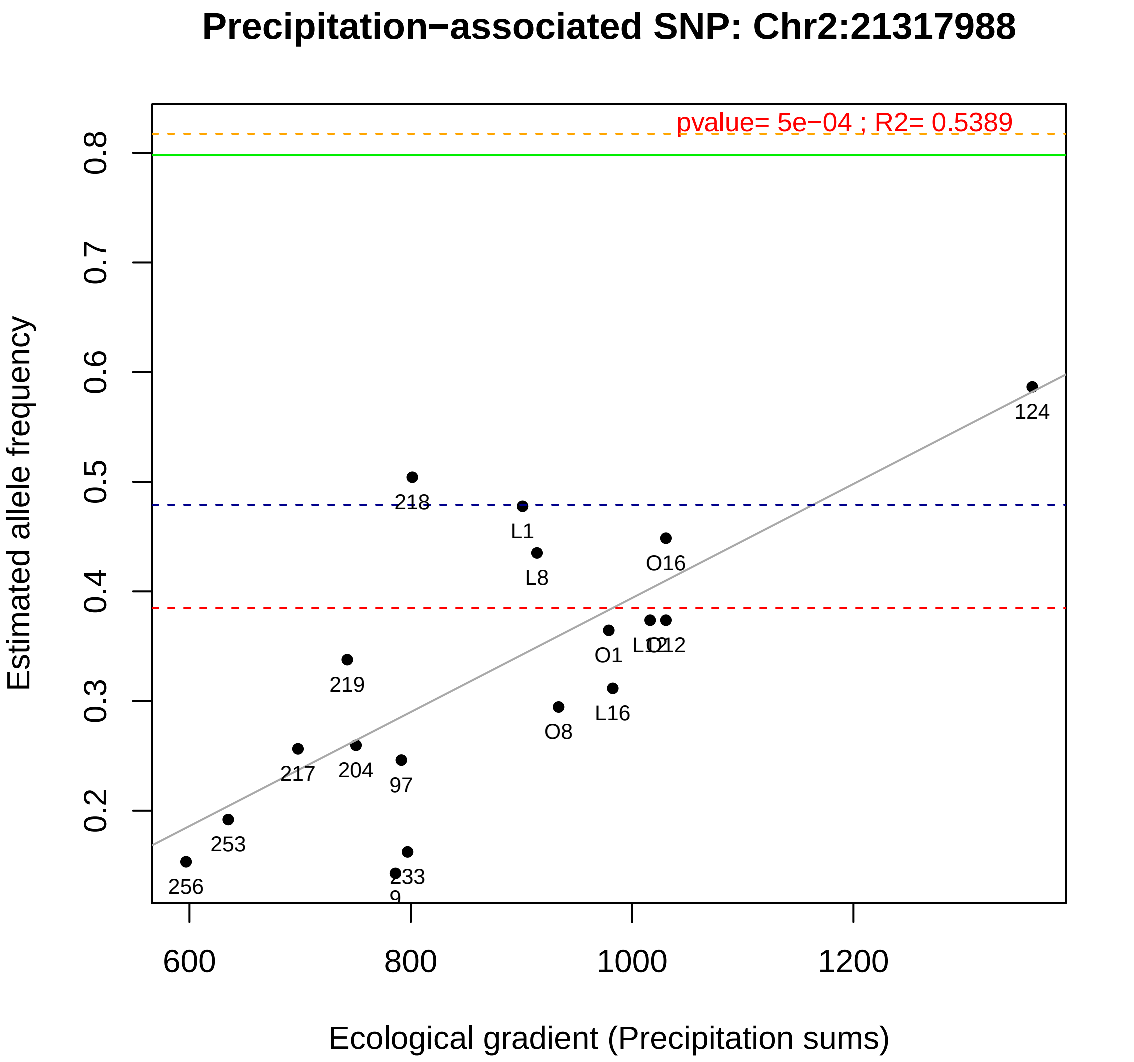
Precipitation−associated SNP: Chr2:21317988
pvalue= 5e−04 ; R2= 0.5389
Estimated allele frequency
0.2
0.3
0.4
0.5
0.6
0.7
0.8
Ecological gradient (Precipitation sums)
600
800
1000
1200
218
L1
L8
O16
124
O1
L10
O12
219
L16
O8
217
204
97
253
256
233
9

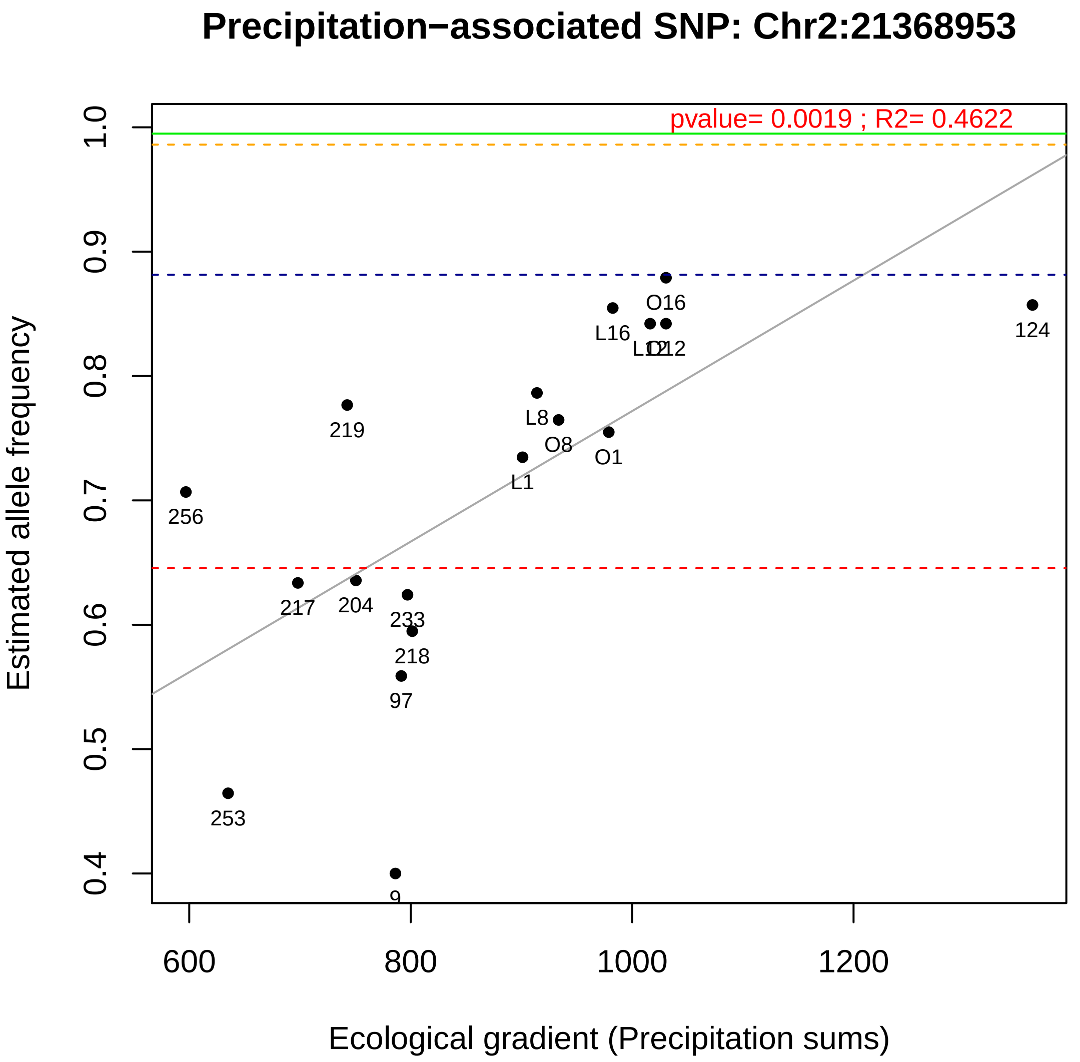
Precipitation−associated SNP: Chr2:21368953
pvalue= 0.0019 ; R2= 0.4622
Estimated allele frequency
Ecological gradient (Precipitation sums)
0.4
0.5
0.6
0.7
0.8
0.9
1.0
600
800
1000
1200
O16
124
L16
L12
O12
219
L8
O8
O1
L1
256
217
204
233
218
97
253
9

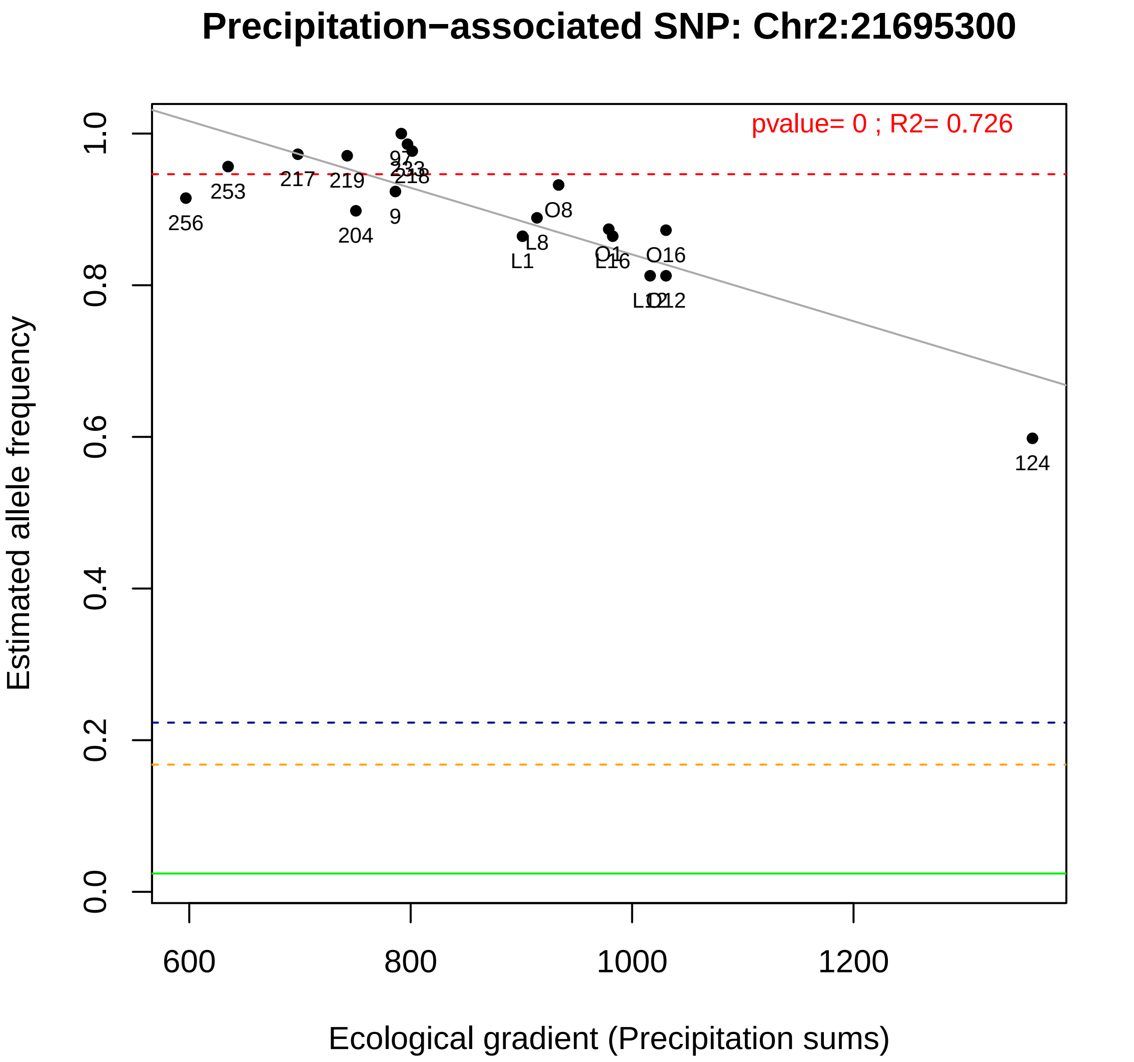

Precipitation−associated SNP: Chr2:21695300
pvalue= 0 ; R2= 0.726
Estimated allele frequency
Ecological gradient (Precipitation sums)
0.0
0.2
0.4
0.6
0.8
1.0
600
800
1000
1200
256
253
217
219
204
97
233
218
9
O8
L8
L1
O1
L16
O16
O12
124

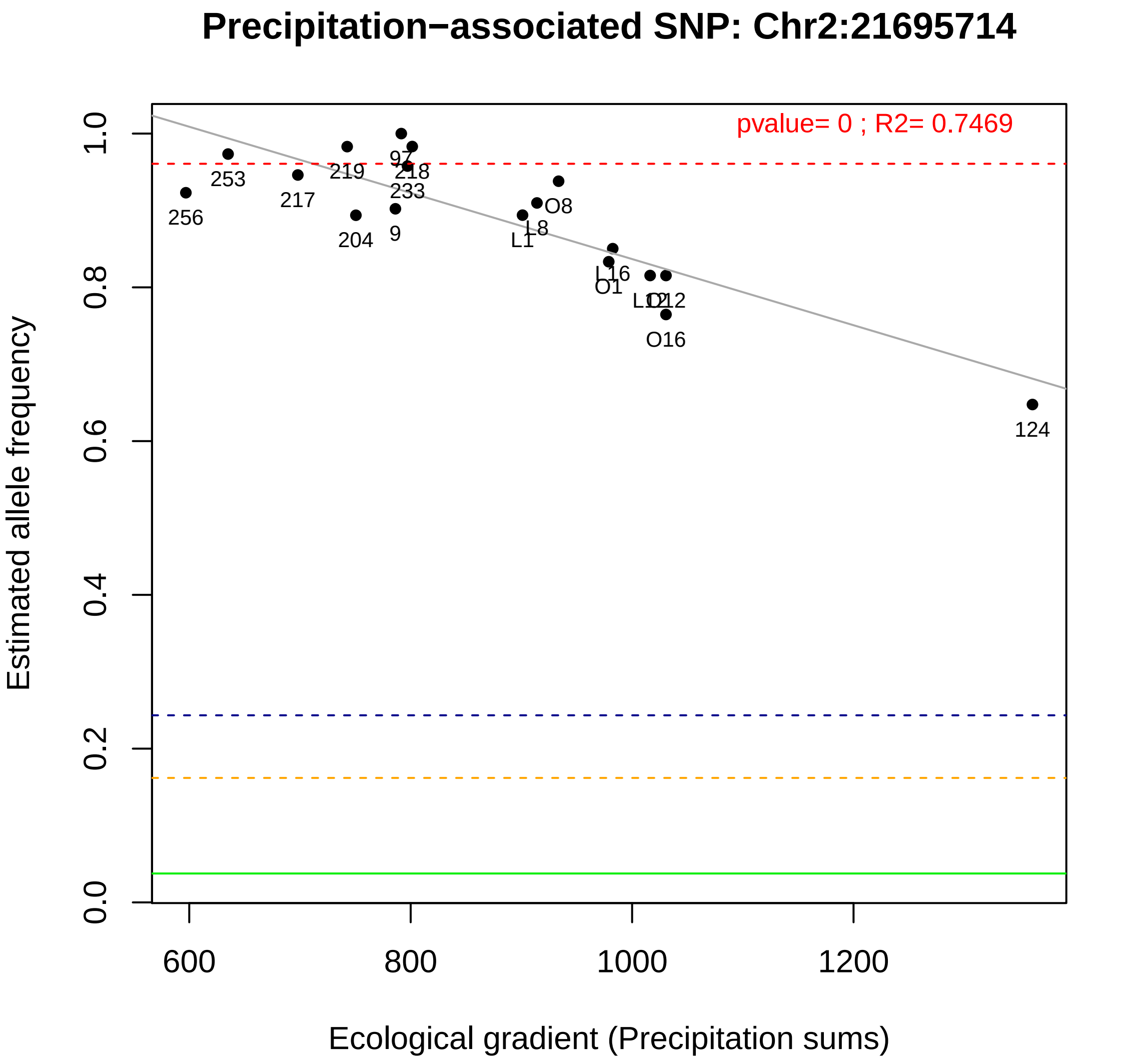

Precipitation−associated SNP: Chr2:21695714
pvalue= 0 ; R2= 0.7469
Estimated allele frequency
Ecological gradient (Precipitation sums)
0.0
0.2
0.4
0.6
0.8
1.0
600
800
1000
1200
256
253
217
219
204
97
218
233
9
O8
L8
L1
L16
O1
O12
O16
124

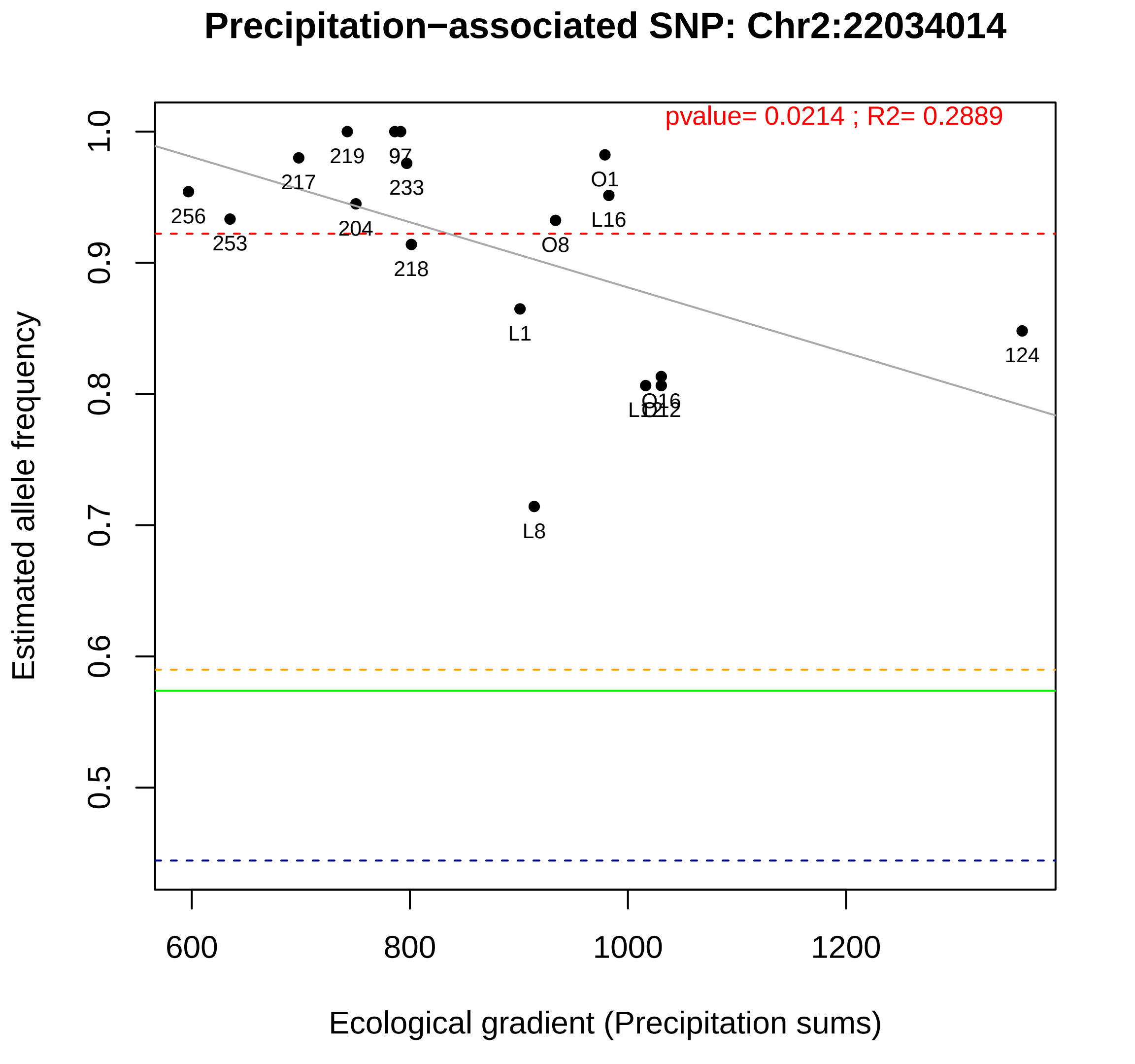
Precipitation−associated SNP: Chr2:22034014
pvalue= 0.0214 ; R2= 0.2889
Estimated allele frequency
Ecological gradient (Precipitation sums)
1.0
0.9
0.8
0.7
0.6
0.5
600
800
1000
1200
219
97
217
233
256
204
253
218
O1
L16
O8
L1
124
O16
O12
L8

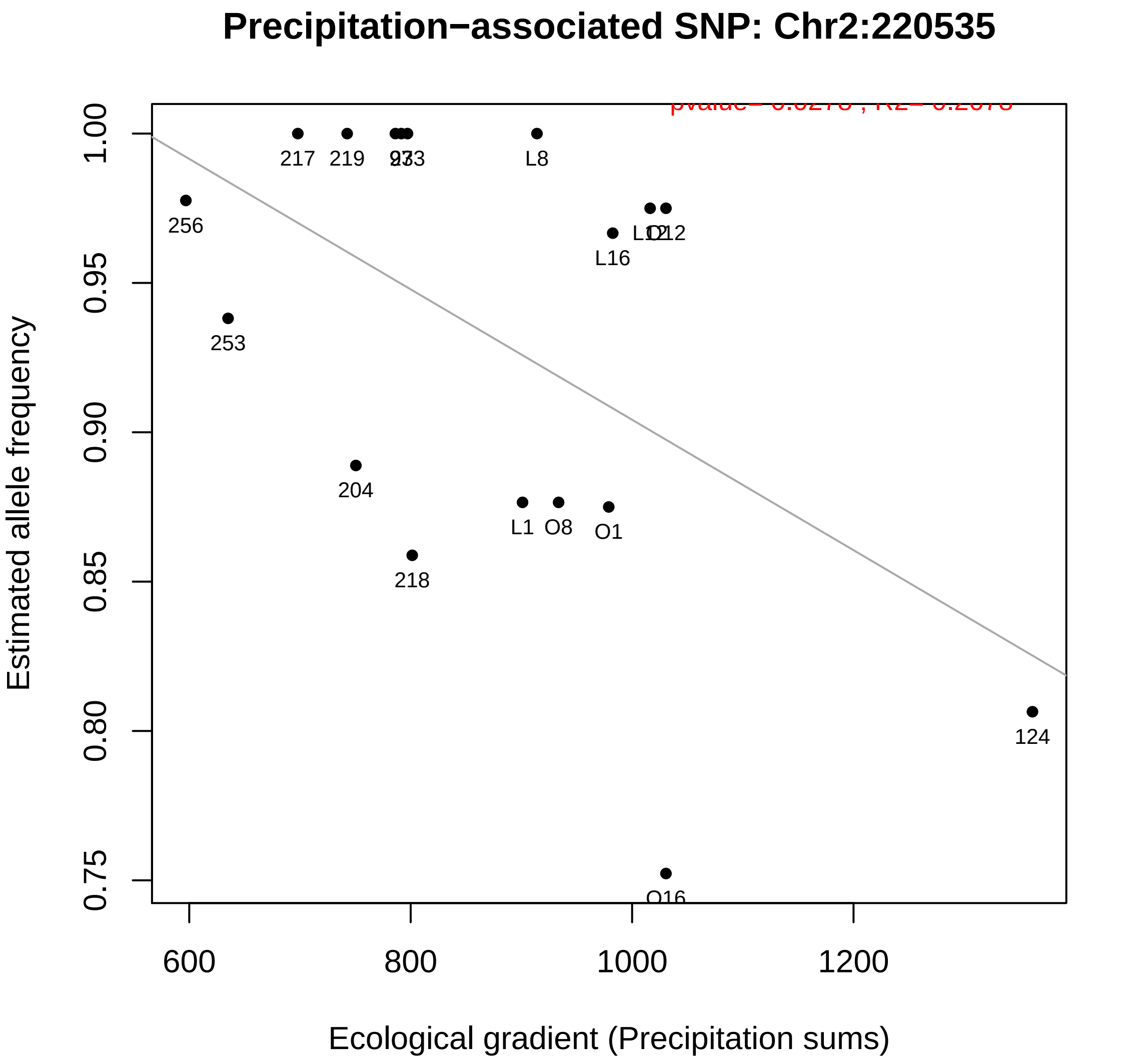
Precipitation−associated SNP: Chr2:220535
Estimated allele frequency
Ecological gradient (Precipitation sums)
1.00
0.95
0.90
0.85
0.80
0.75
600
800
1000
1200
217
219
L8
256
253
L16
204
L1
O8
O1
218
124
O16

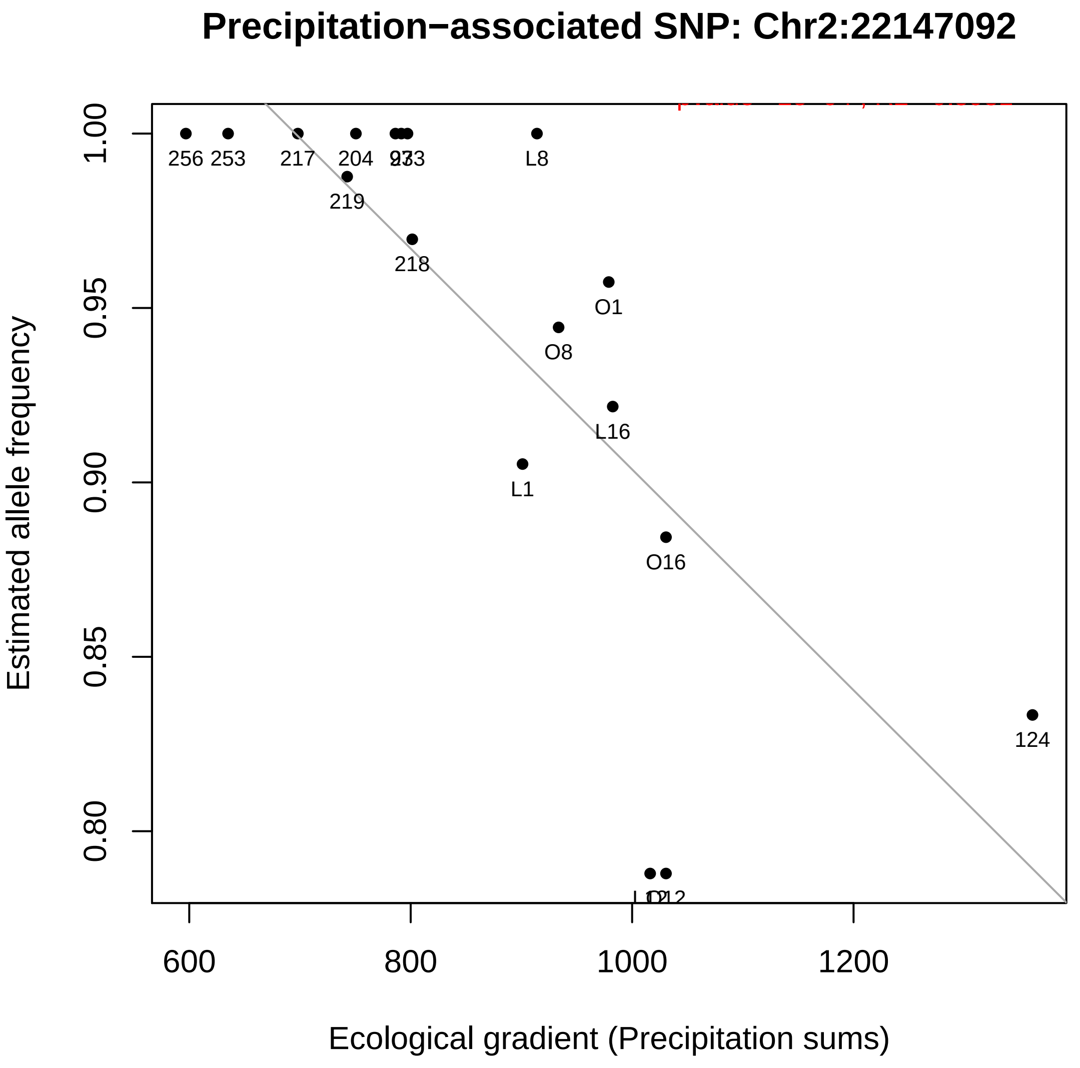
Precipitation−associated SNP: Chr2:22147092
Estimated allele frequency
1.00
0.95
0.90
0.85
0.80
600
800
1000
1200
Ecological gradient (Precipitation sums)
256
253
217
204
L8
219
218
O1
O8
L16
L1
O16
124

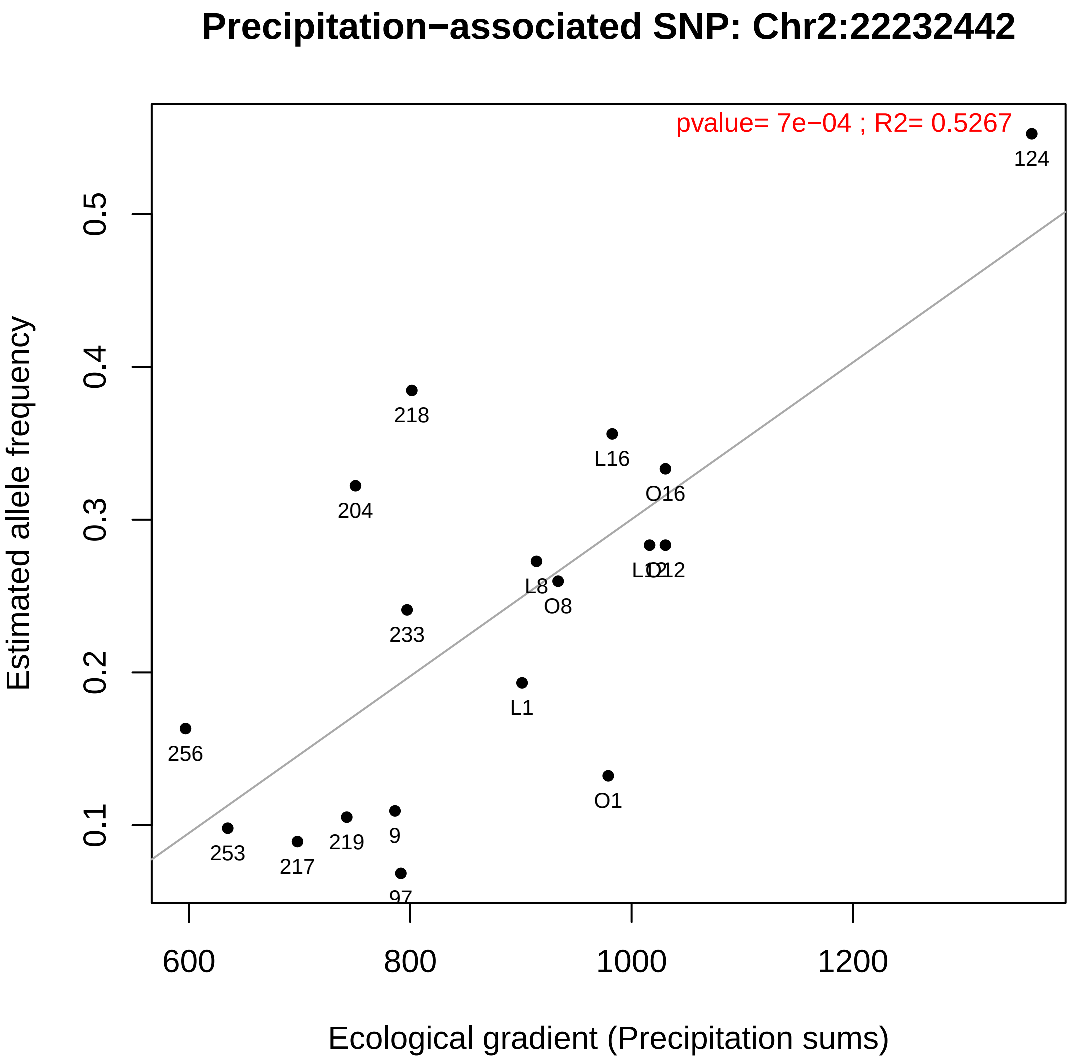

Precipitation−associated SNP: Chr2:22232442
pvalue= 7e−04 ; R2= 0.5267
124
218
L16
O16
204
L12
O12
L8
O8
233
L1
256
O1
253
217
219
9
97
0.1
0.2
0.3
0.4
0.5
600
800
1000
1200
Estimated allele frequency
Ecological gradient (Precipitation sums)

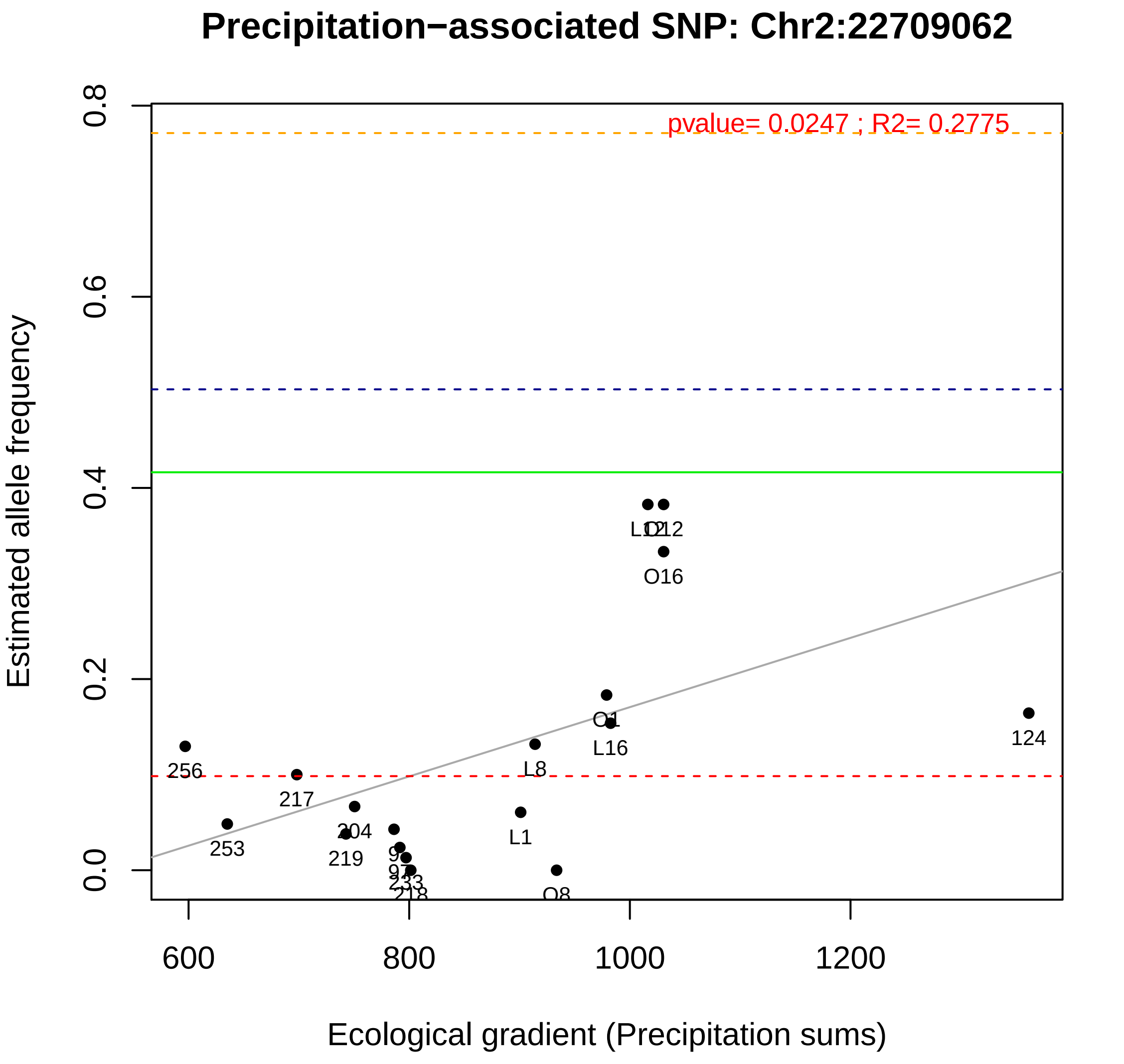
Precipitation−associated SNP: Chr2:22709062
pvalue= 0.0247 ; R2= 0.2775
Estimated allele frequency
0.0
0.2
0.4
0.6
0.8
600
800
1000
1200
Ecological gradient (Precipitation sums)
O16
L16
L8
L1
124
256
217
253
219
204
O1
O8

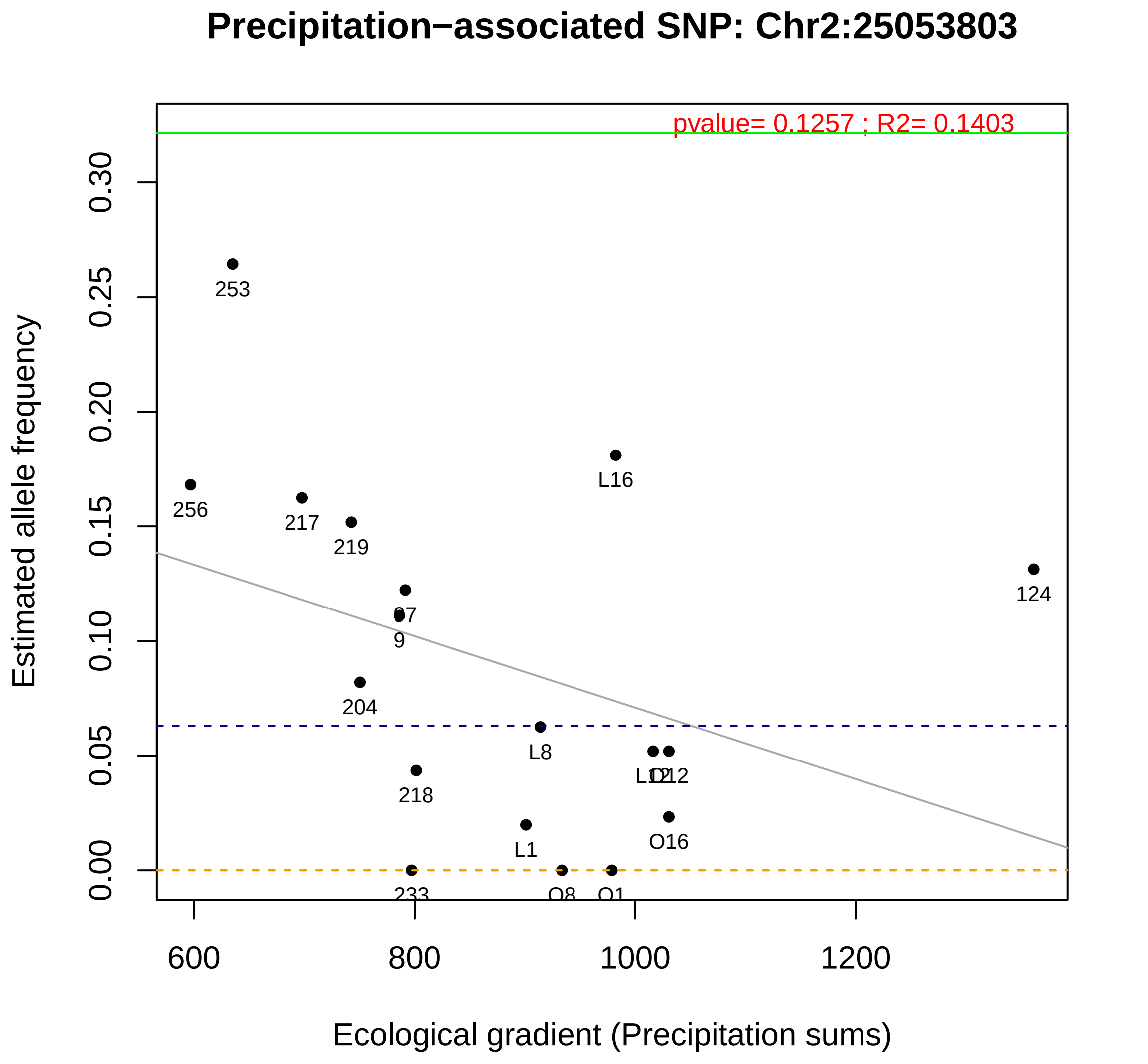

Precipitation−associated SNP: Chr2:25053803
pvalue= 0.1257 ; R2= 0.1403
Estimated allele frequency
0.00
0.05
0.10
0.15
0.20
0.25
0.30
253
256
217
219
L16
124
97
9
204
L8
L12
O12
218
O16
L1
233
O8
O1
600
800
1000
1200
Ecological gradient (Precipitation sums)

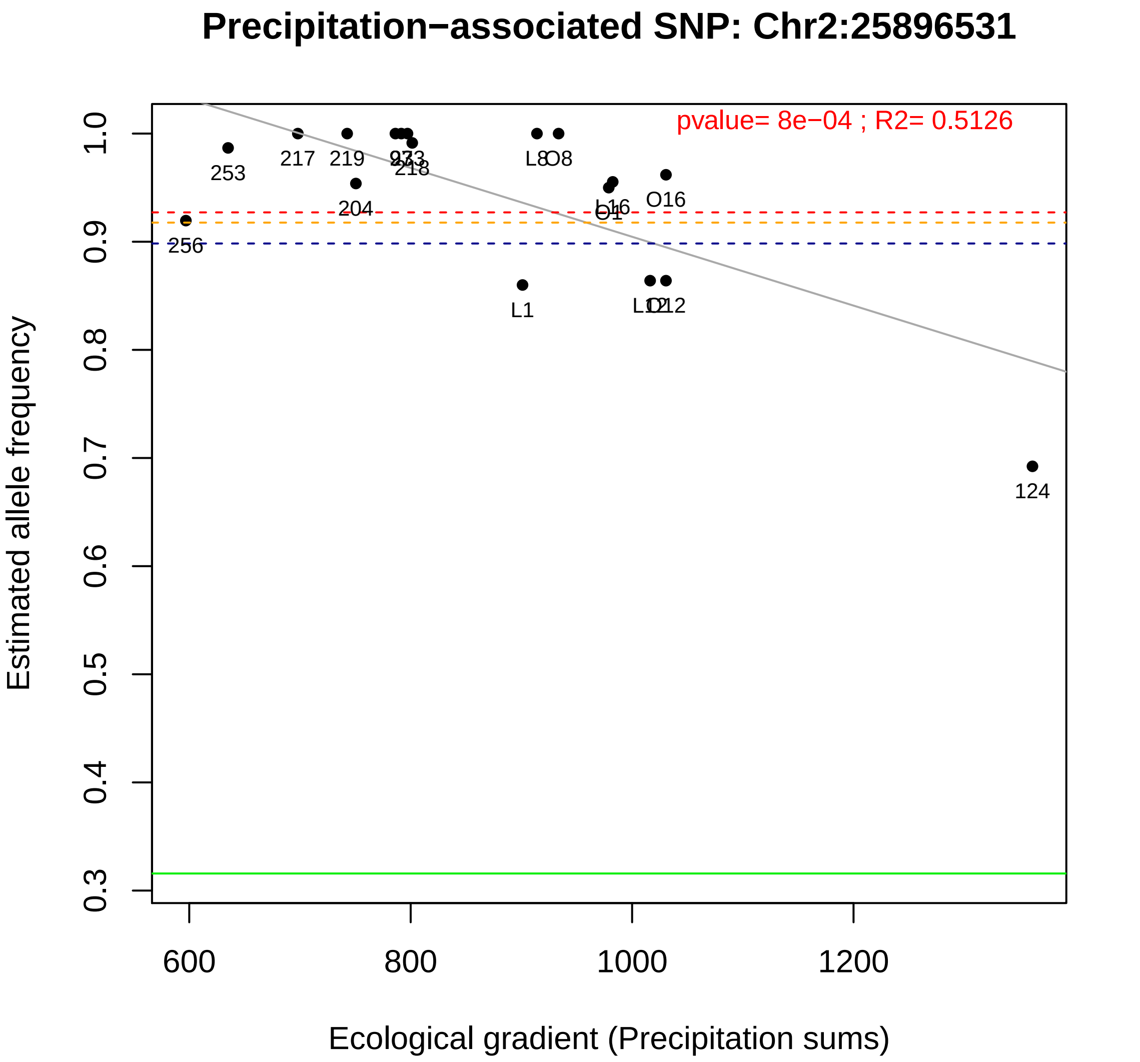
Precipitation−associated SNP: Chr2:25896531
pvalue= 8e−04 ; R2= 0.5126
Estimated allele frequency
Ecological gradient (Precipitation sums)
1.0
0.9
0.8
0.7
0.6
0.5
0.4
0.3
600
800
1000
1200
253
217
219
218
L8
O8
204
256
O16
L1
124

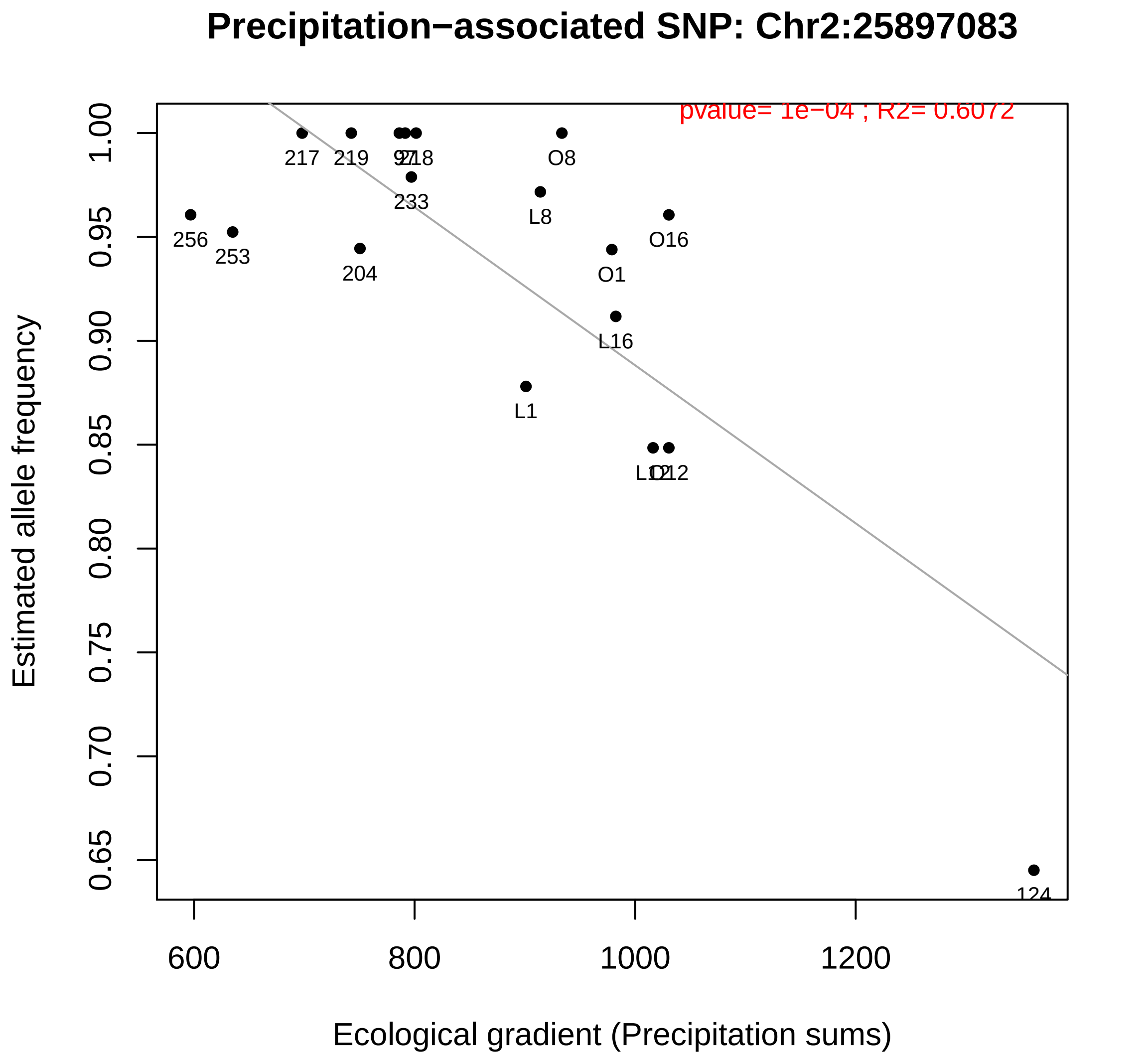
Precipitation−associated SNP: Chr2:25897083
pvalue= 1e−04 ; R2= 0.6072
Estimated allele frequency
1.00
0.95
0.90
0.85
0.80
0.75
0.70
0.65
217
219
97
218
O8
233
L8
256
253
204
O16
O1
L16
L1
L10
O12
124
600
800
1000
1200
Ecological gradient (Precipitation sums)

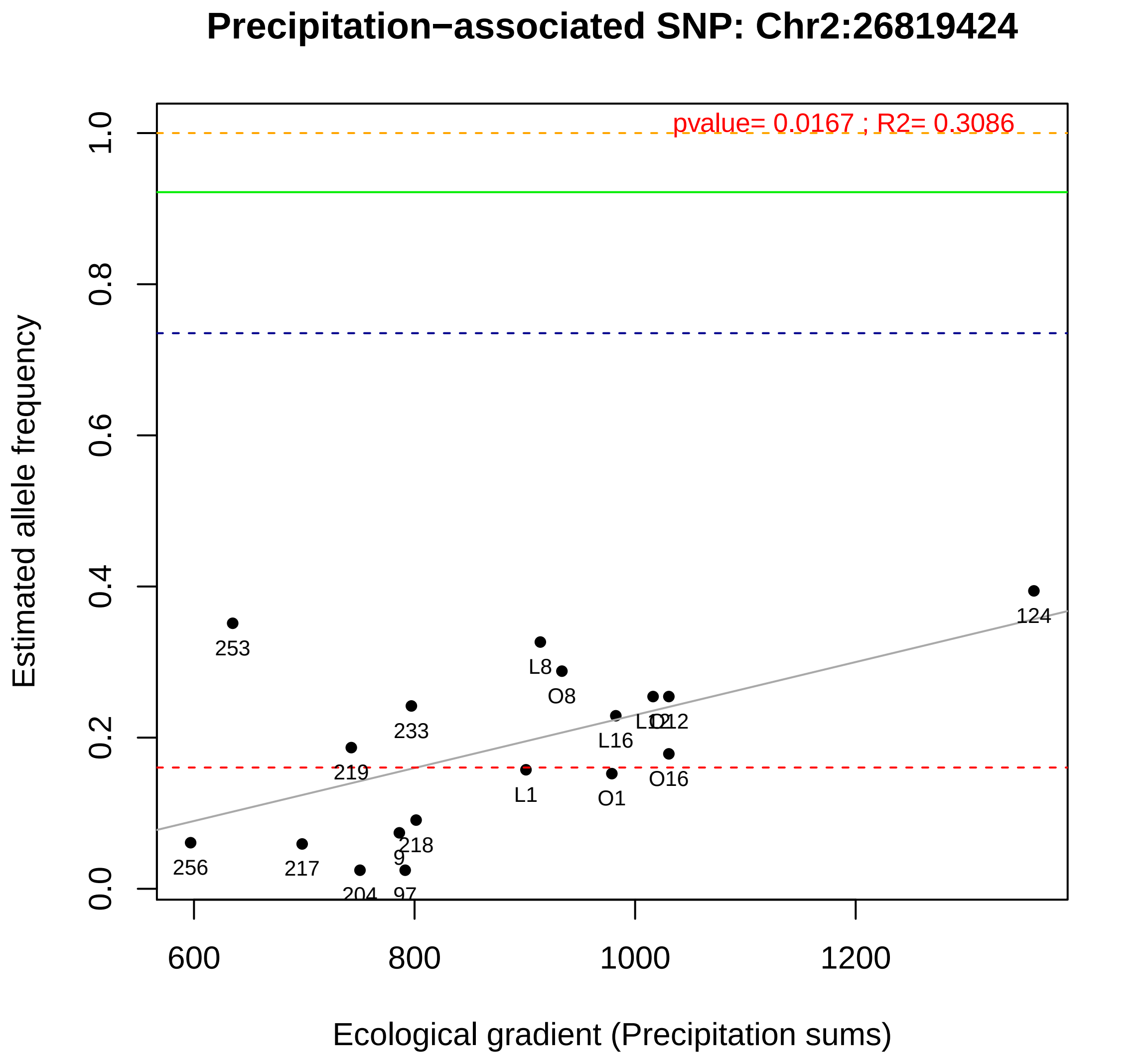

Precipitation−associated SNP: Chr2:26819424
pvalue= 0.0167 ; R2= 0.3086
Estimated allele frequency
Ecological gradient (Precipitation sums)
0.0
0.2
0.4
0.6
0.8
1.0
600
800
1000
1200
253
124
L8
O8
233
L12
O12
L16
219
O16
L1
O1
218
9
256
217
204
97

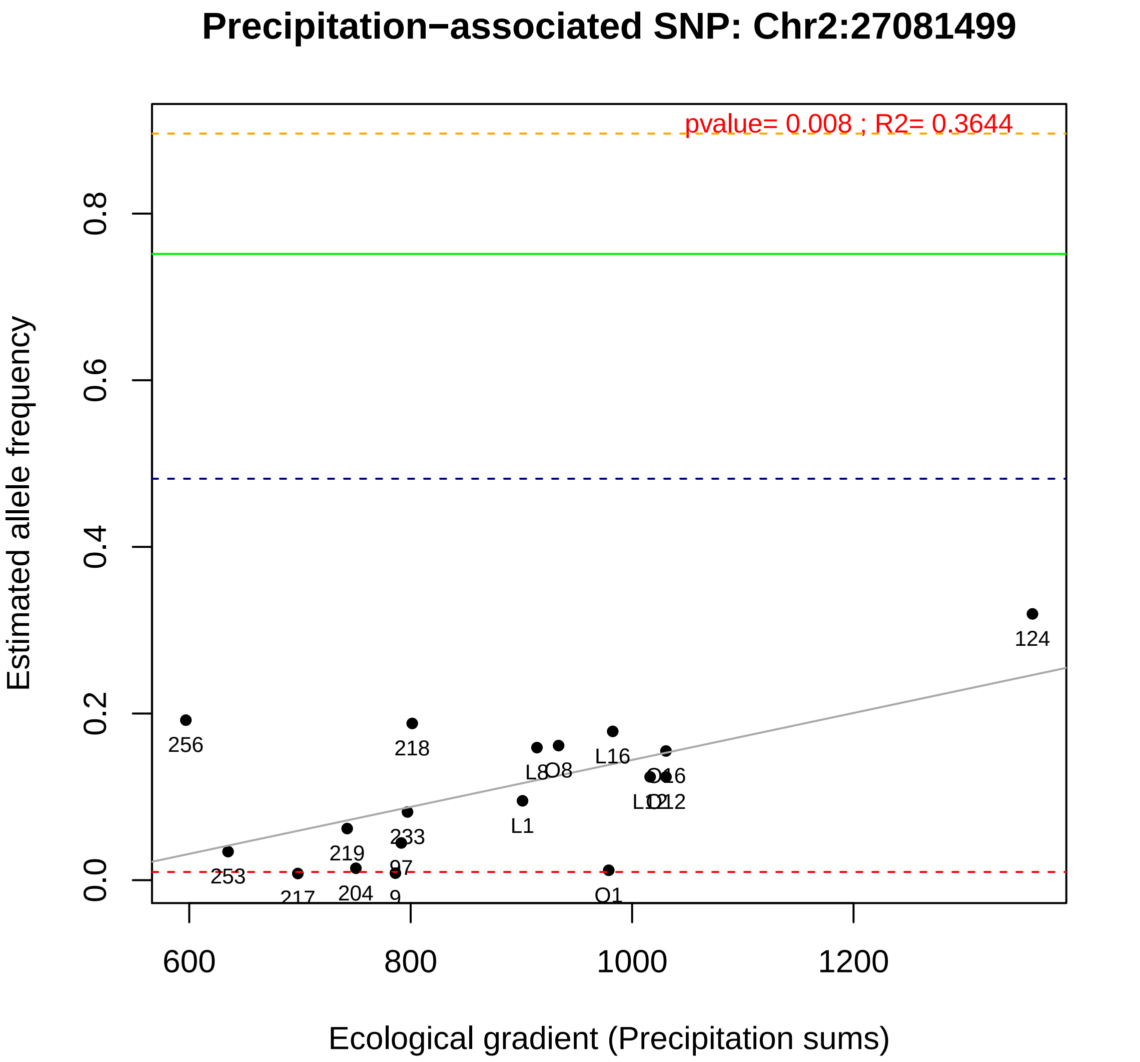

Precipitation−associated SNP: Chr2:27081499
pvalue= 0.008 ; R2= 0.3644
Estimated allele frequency
0.0
0.2
0.4
0.6
0.8
600
800
1000
1200
Ecological gradient (Precipitation sums)
256
253
217
219
204
218
233
97
9
L1
L8
O8
L16
O1
O16
L12
O12
124

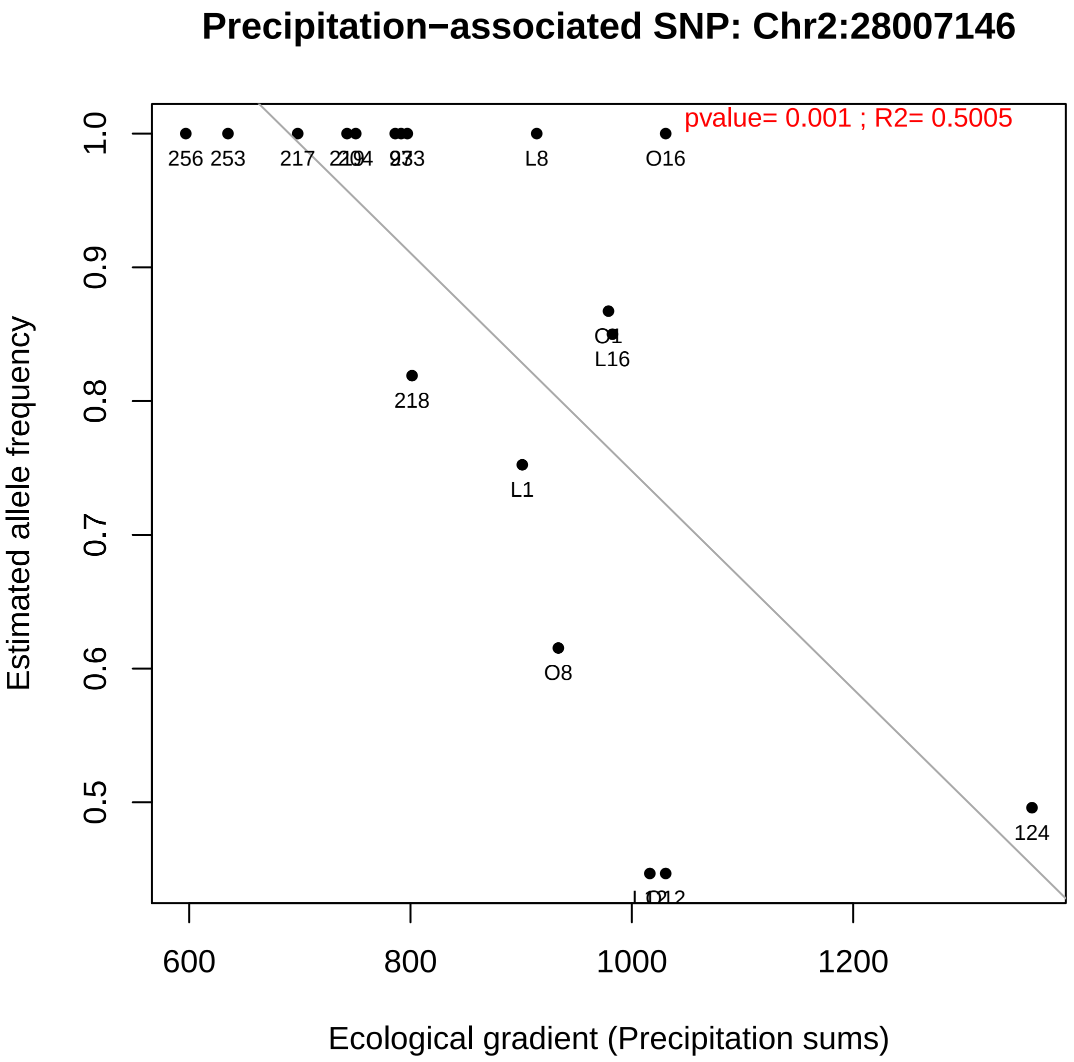
Precipitation−associated SNP: Chr2:28007146
pvalue= 0.001 ; R2= 0.5005
Estimated allele frequency
Ecological gradient (Precipitation sums)
1.0
0.9
0.8
0.7
0.6
0.5
600
800
1000
1200
256
253
217
L8
O16
218
L1
L16
O8
124

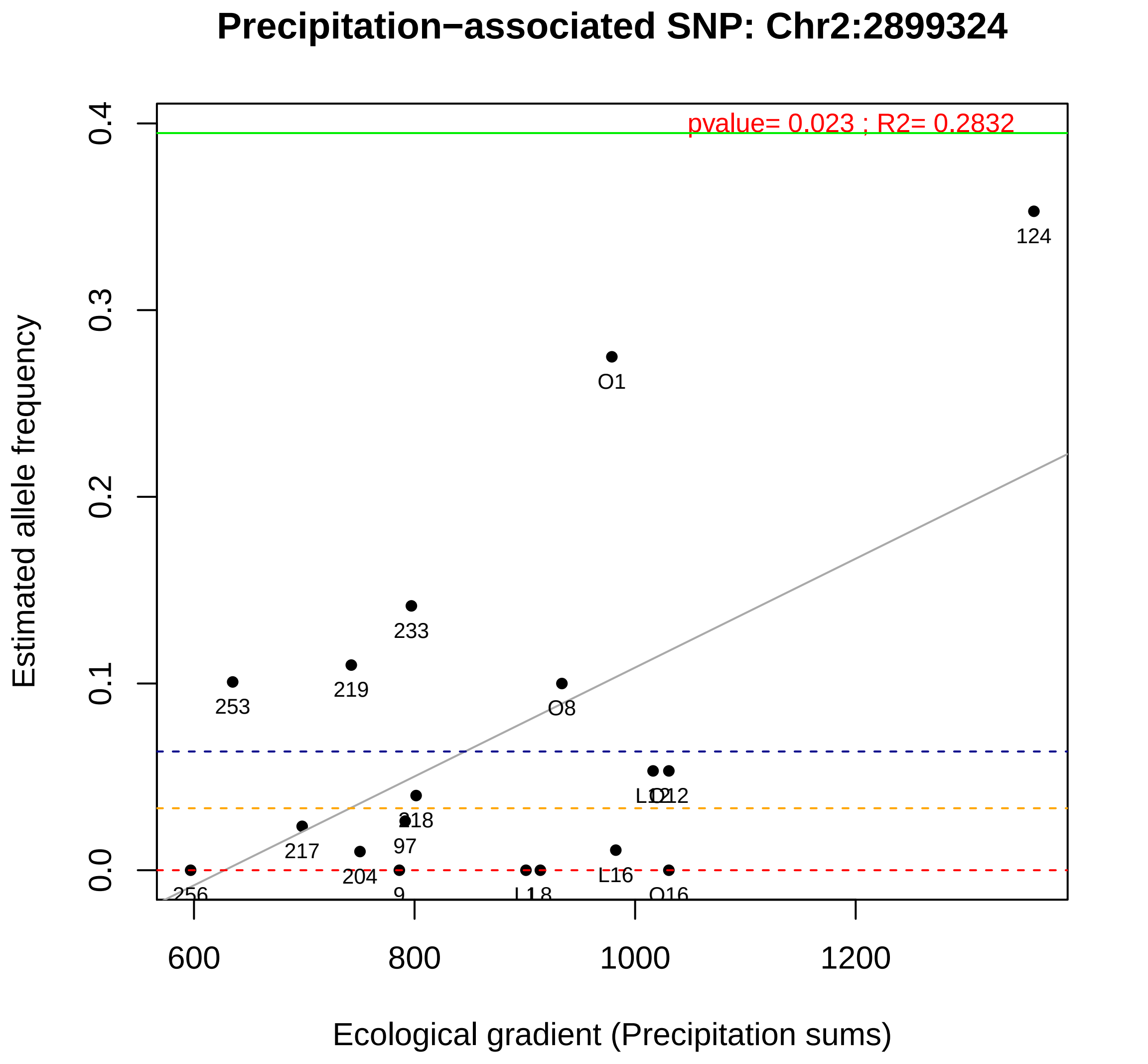

Precipitation−associated SNP: Chr2:2899324
pvalue= 0.023 ; R2= 0.2832
Estimated allele frequency
0.0
0.1
0.2
0.3
0.4
600
800
1000
1200
Ecological gradient (Precipitation sums)
124
O1
233
219
253
O8
217
204
97
9
256
L16
O16

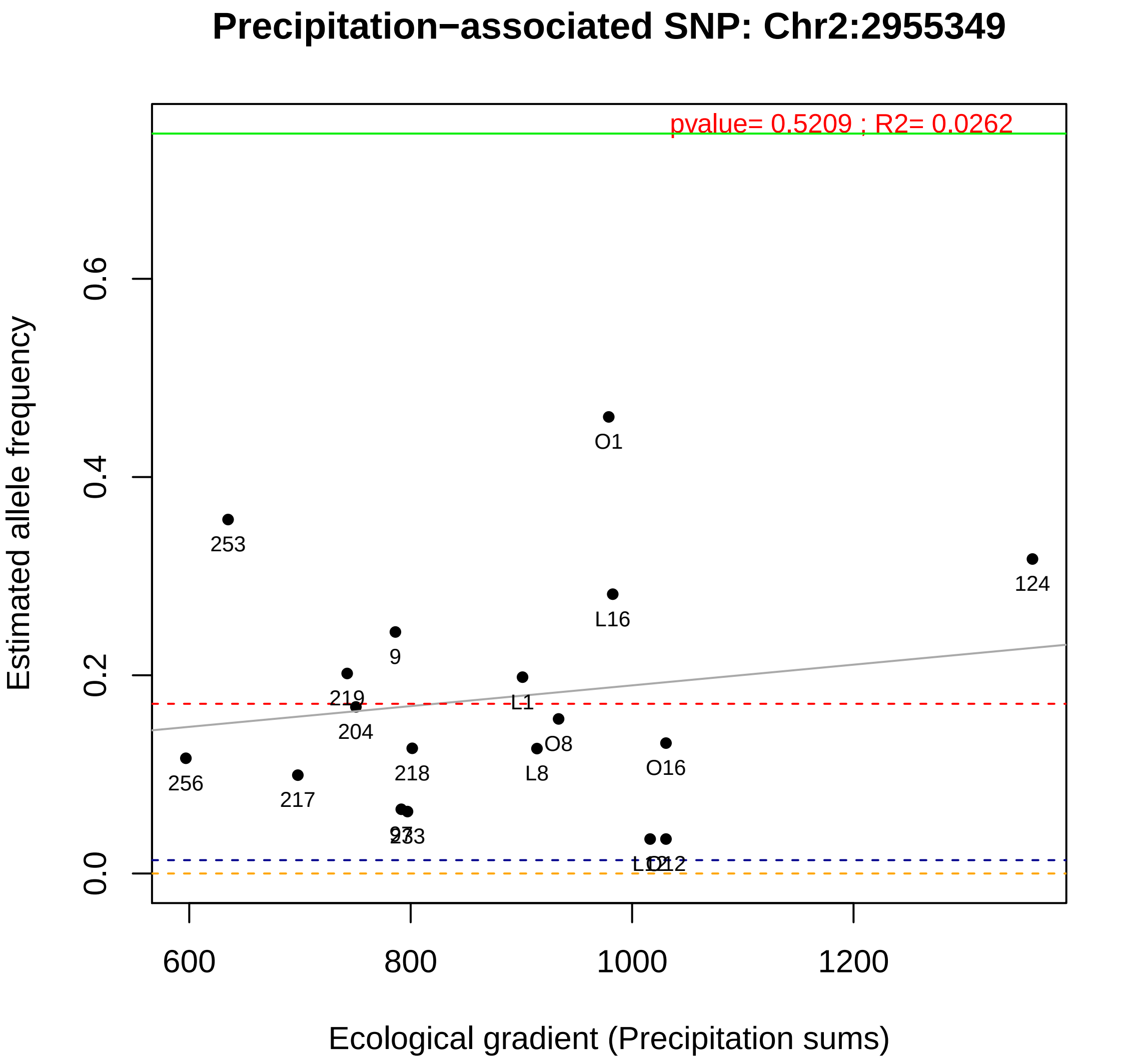
Precipitation−associated SNP: Chr2:2955349
pvalue= 0.5209 ; R2= 0.0262
Estimated allele frequency
0.0
0.2
0.4
0.6
600
800
1000
1200
Ecological gradient (Precipitation sums)
O1
253
124
L16
9
219
L1
204
O8
218
L8
O16
256
217

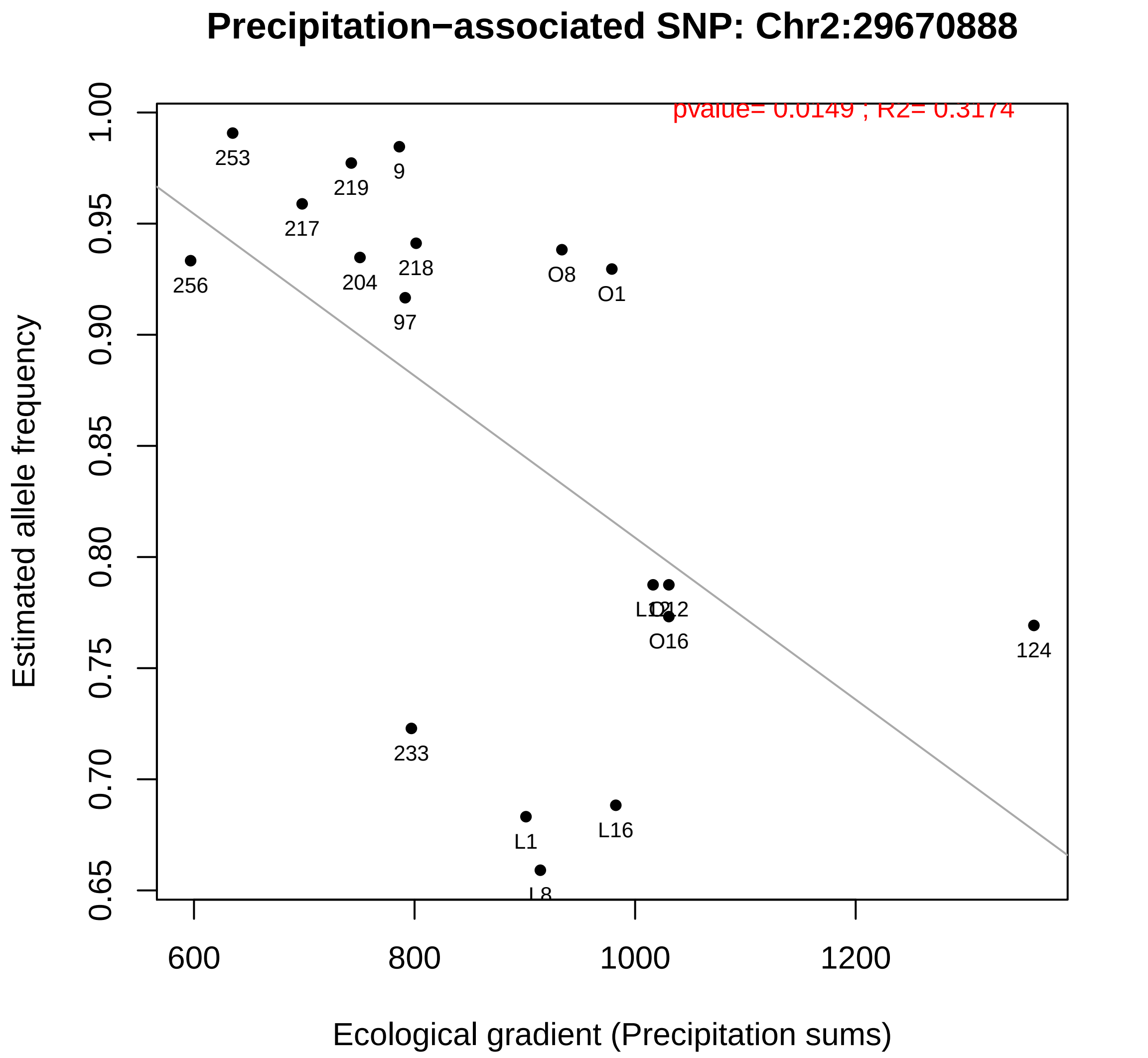

Precipitation−associated SNP: Chr2:29670888
pvalue= 0.0149 ; R2= 0.3174
Estimated allele frequency
Ecological gradient (Precipitation sums)
0.65
0.70
0.75
0.80
0.85
0.90
0.95
1.00
600
800
1000
1200
253
219
9
217
256
204
218
97
O8
O1
L12
O12
O16
124
233
L1
L16
L8

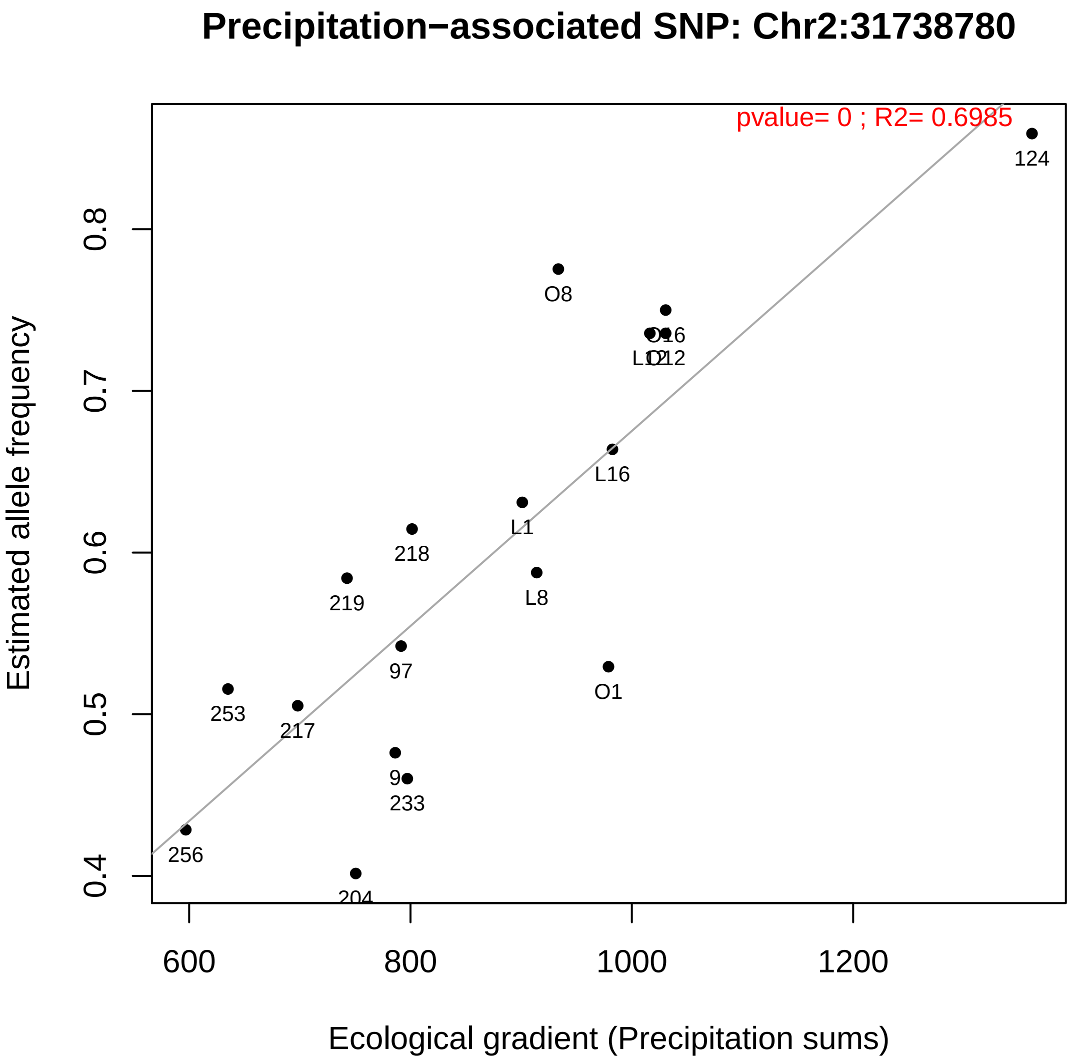

Precipitation−associated SNP: Chr2:31738780
pvalue= 0 ; R2= 0.6985
124
O8
O16
L12
O12
L16
L1
218
L8
219
97
O1
253
217
9
233
256
204
0.4
0.5
0.6
0.7
0.8
600
800
1000
1200
Estimated allele frequency
Ecological gradient (Precipitation sums)

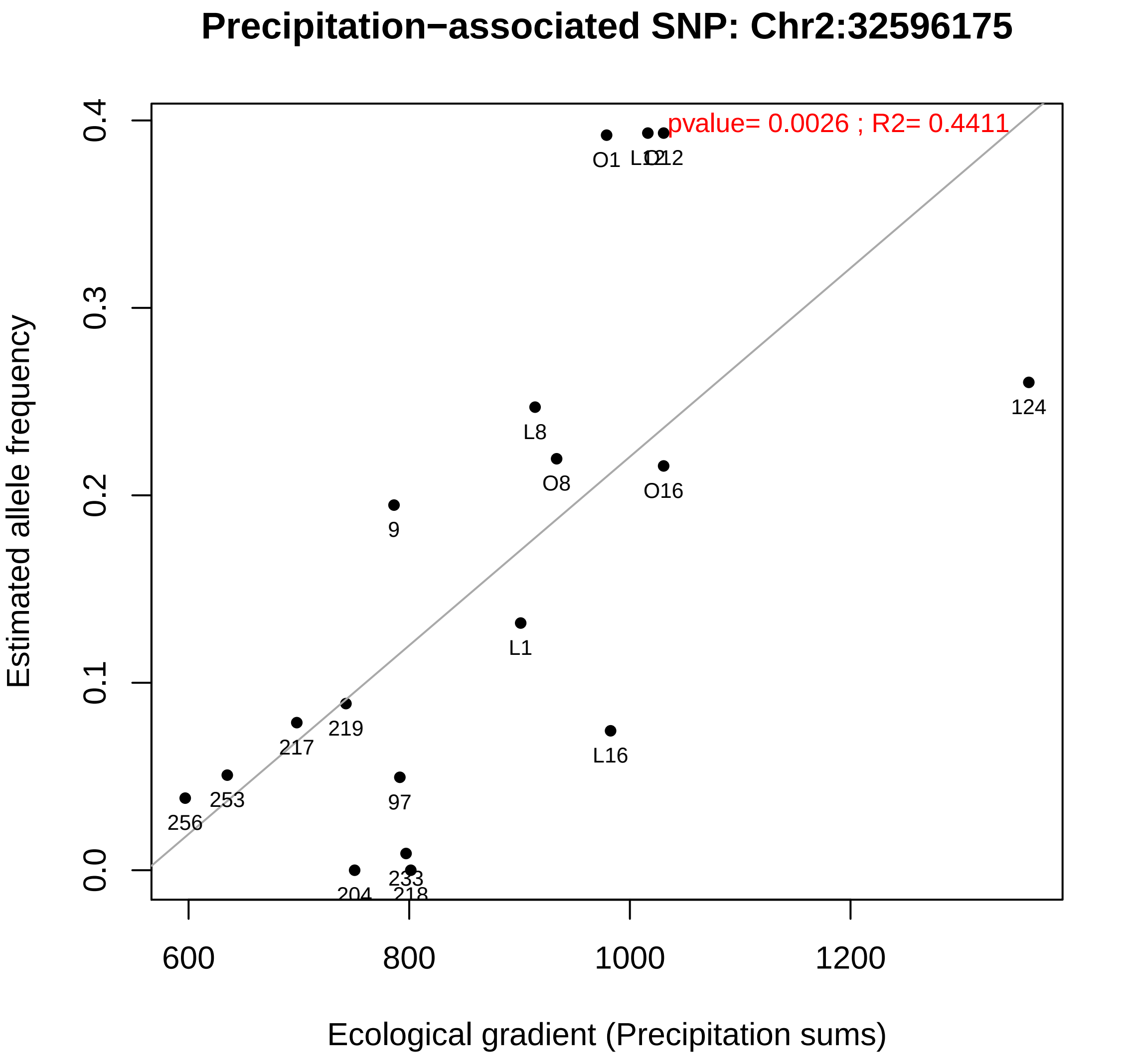
Precipitation−associated SNP: Chr2:32596175
pvalue= 0.0026 ; R2= 0.4411
Estimated allele frequency
0.0
0.1
0.2
0.3
0.4
600
800
1000
1200
Ecological gradient (Precipitation sums)
O1
L12
O12
L8
O8
O16
9
124
L1
L16
219
217
253
256
97
233
204
218

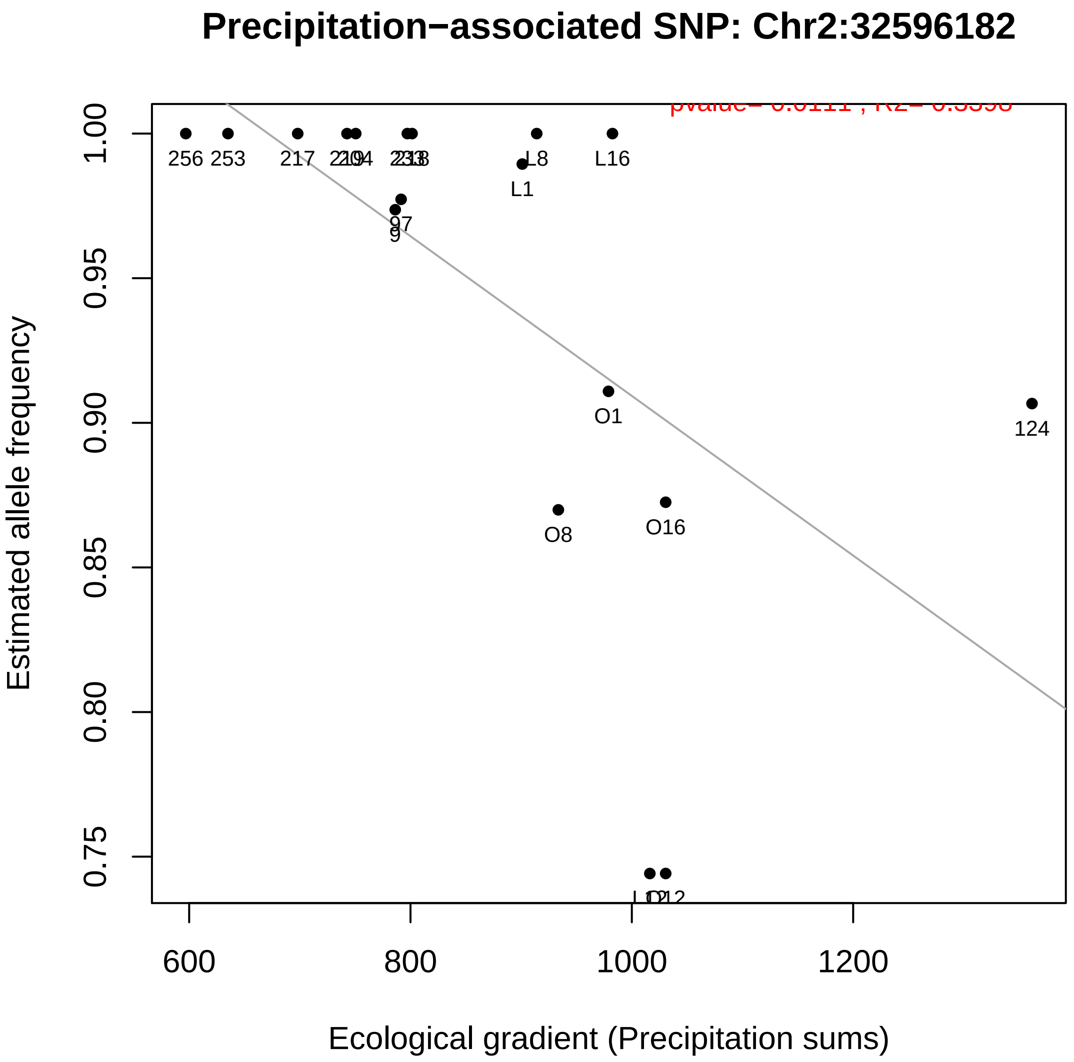

Precipitation−associated SNP: Chr2:32596182
Estimated allele frequency
1.00
0.95
0.90
0.85
0.80
0.75
600
800
1000
1200
Ecological gradient (Precipitation sums)
256
253
217
L8
L16
L1
97
9
O1
124
O8
O16

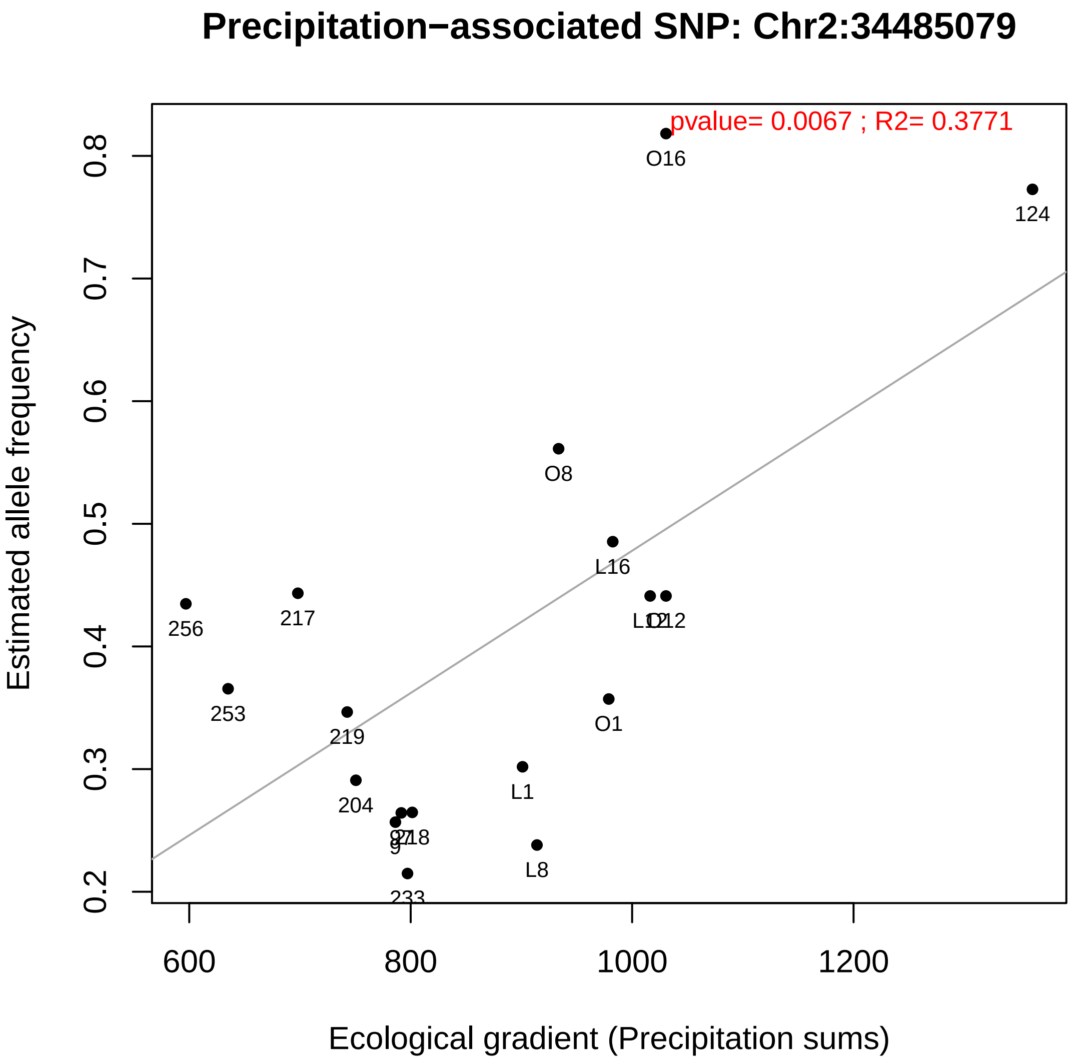
Precipitation−associated SNP: Chr2:34485079
pvalue= 0.0067 ; R2= 0.3771
O16
124
O8
L16
L12
O12
256
217
253
219
O1
L1
204
97
218
9
L8
233
0.2
0.3
0.4
0.5
0.6
0.7
0.8
600
800
1000
1200
Estimated allele frequency
Ecological gradient (Precipitation sums)

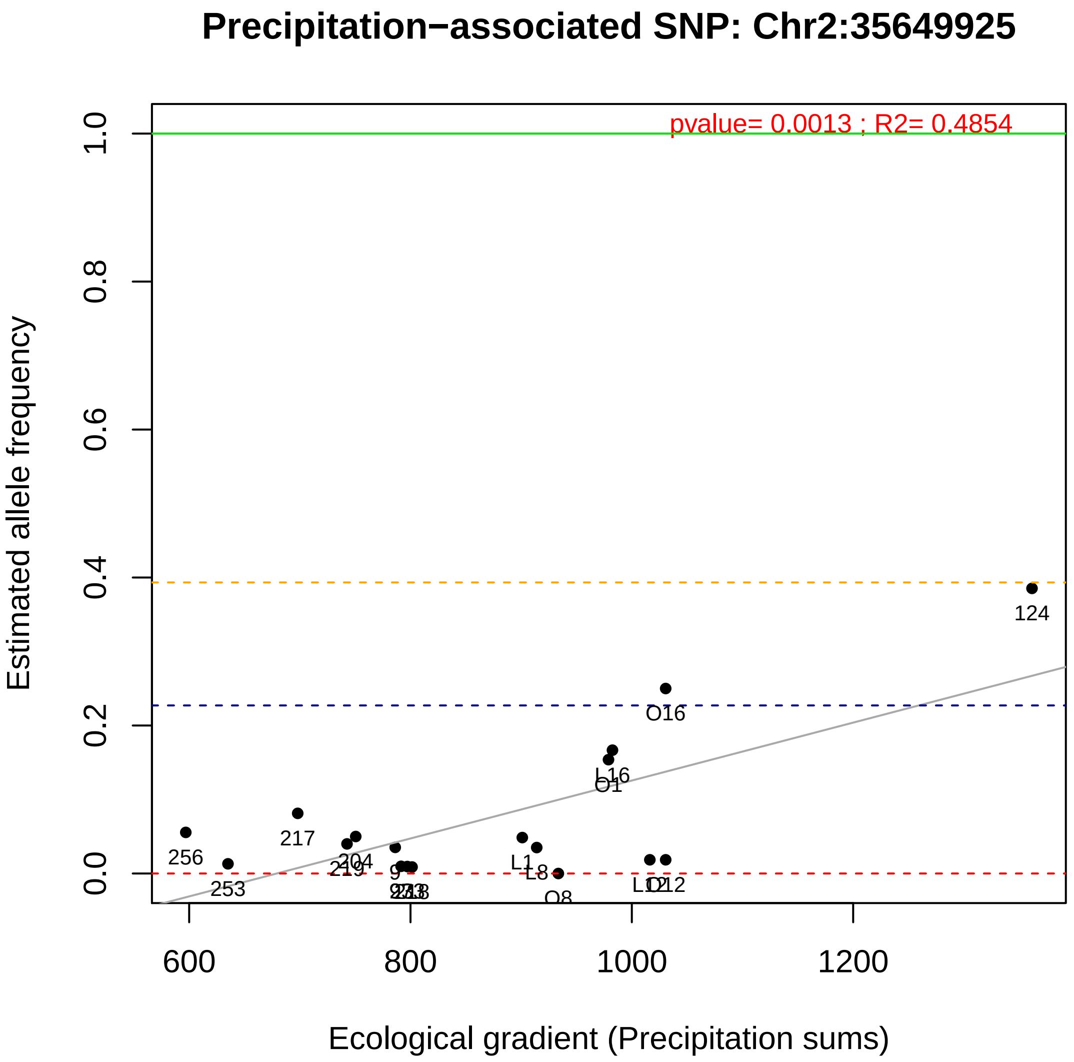

Precipitation−associated SNP: Chr2:35649925
pvalue= 0.0013 ; R2= 0.4854
Estimated allele frequency
0.0
0.2
0.4
0.6
0.8
1.0
600
800
1000
1200
Ecological gradient (Precipitation sums)
124
O16
L16
O1
217
256
253
204
219
9
L1
L8
O8
O12

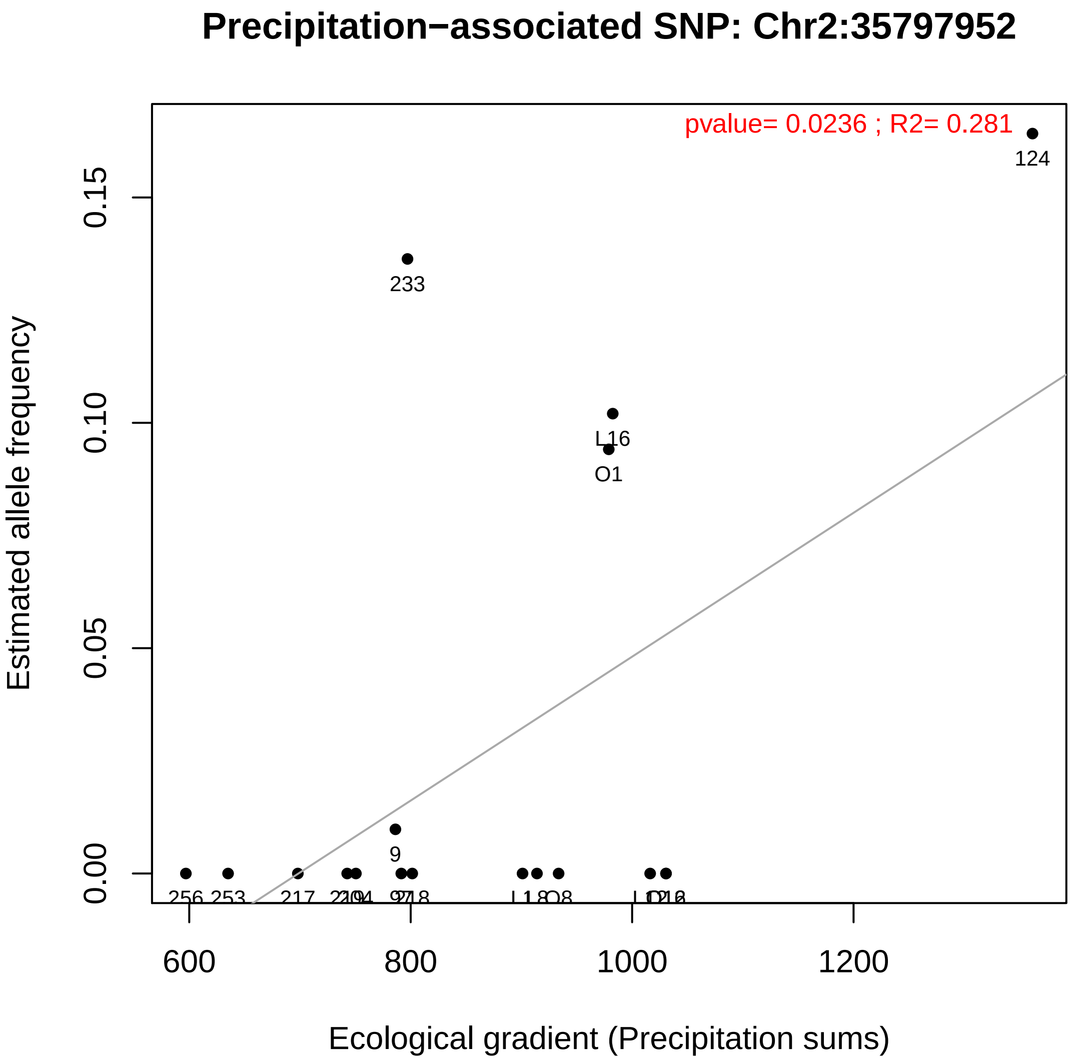

Precipitation−associated SNP: Chr2:35797952
pvalue= 0.0236 ; R2= 0.281
Estimated allele frequency
0.00
0.05
0.10
0.15
Ecological gradient (Precipitation sums)
600
800
1000
1200
124
233
L16
O1
9
256
253
217
O8

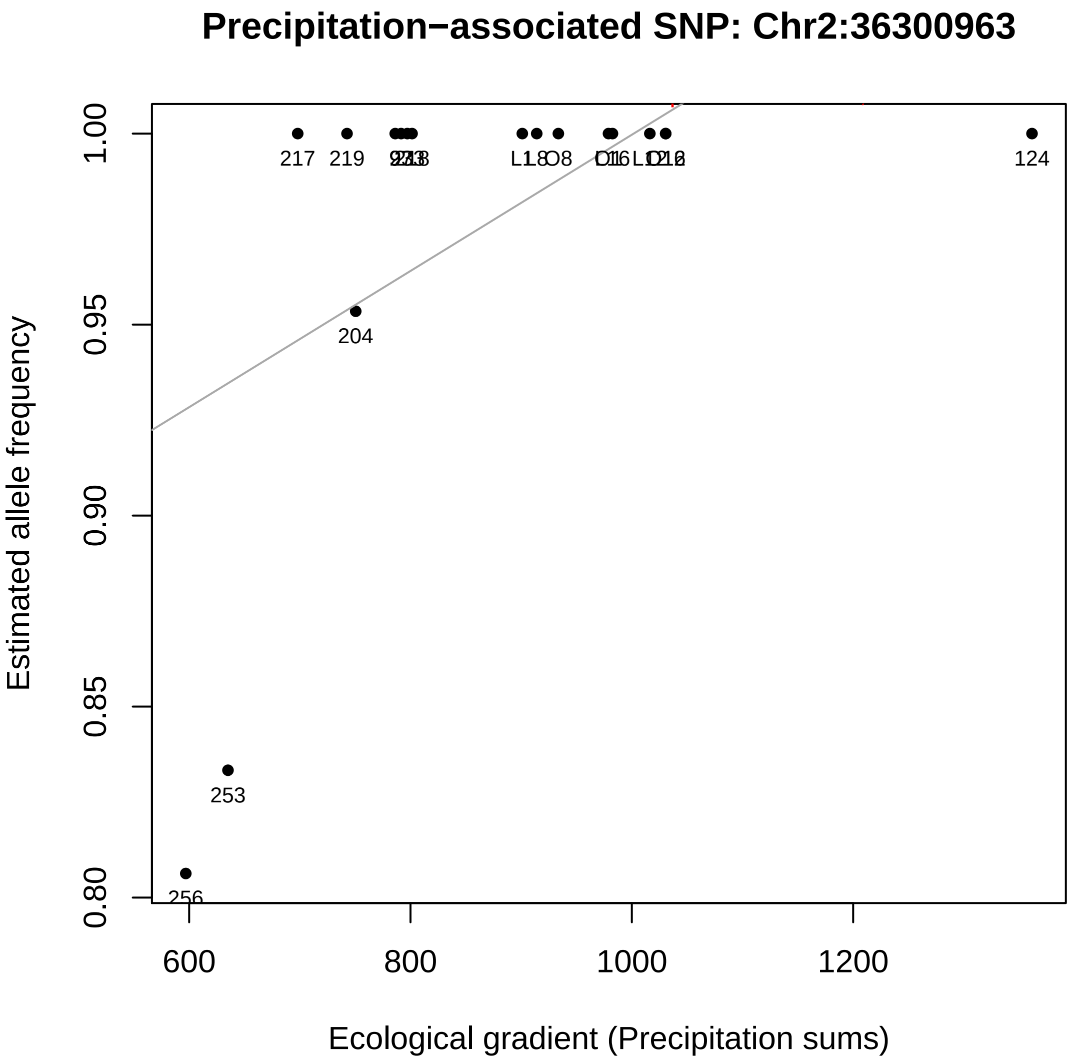

Precipitation−associated SNP: Chr2:36300963
Estimated allele frequency
Ecological gradient (Precipitation sums)
1.00
0.95
0.90
0.85
0.80
600
800
1000
1200
217
219
204
253
256
124
O8

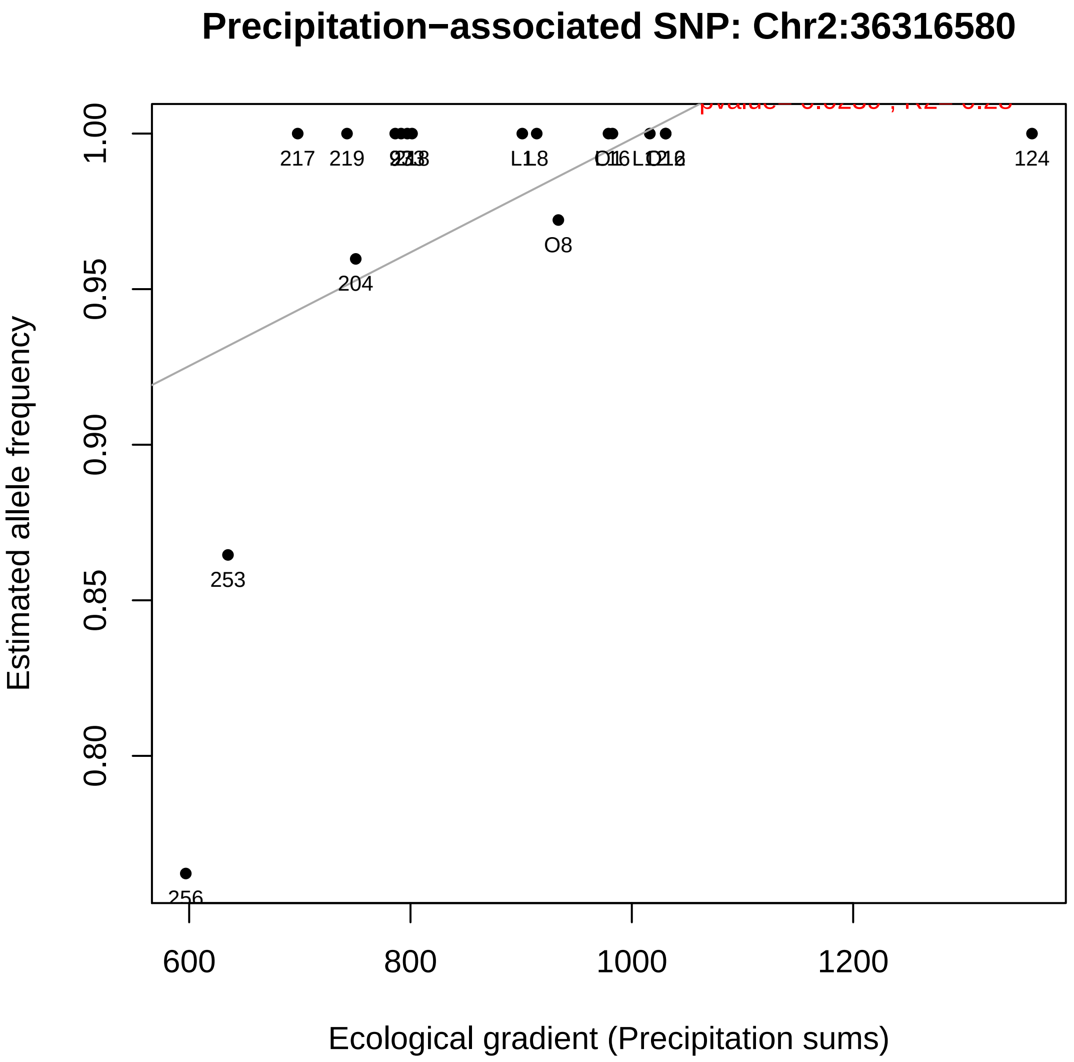
Precipitation−associated SNP: Chr2:36316580
Estimated allele frequency
Ecological gradient (Precipitation sums)
1.00
0.95
0.90
0.85
0.80
600
800
1000
1200
217
219
204
O8
253
256
124

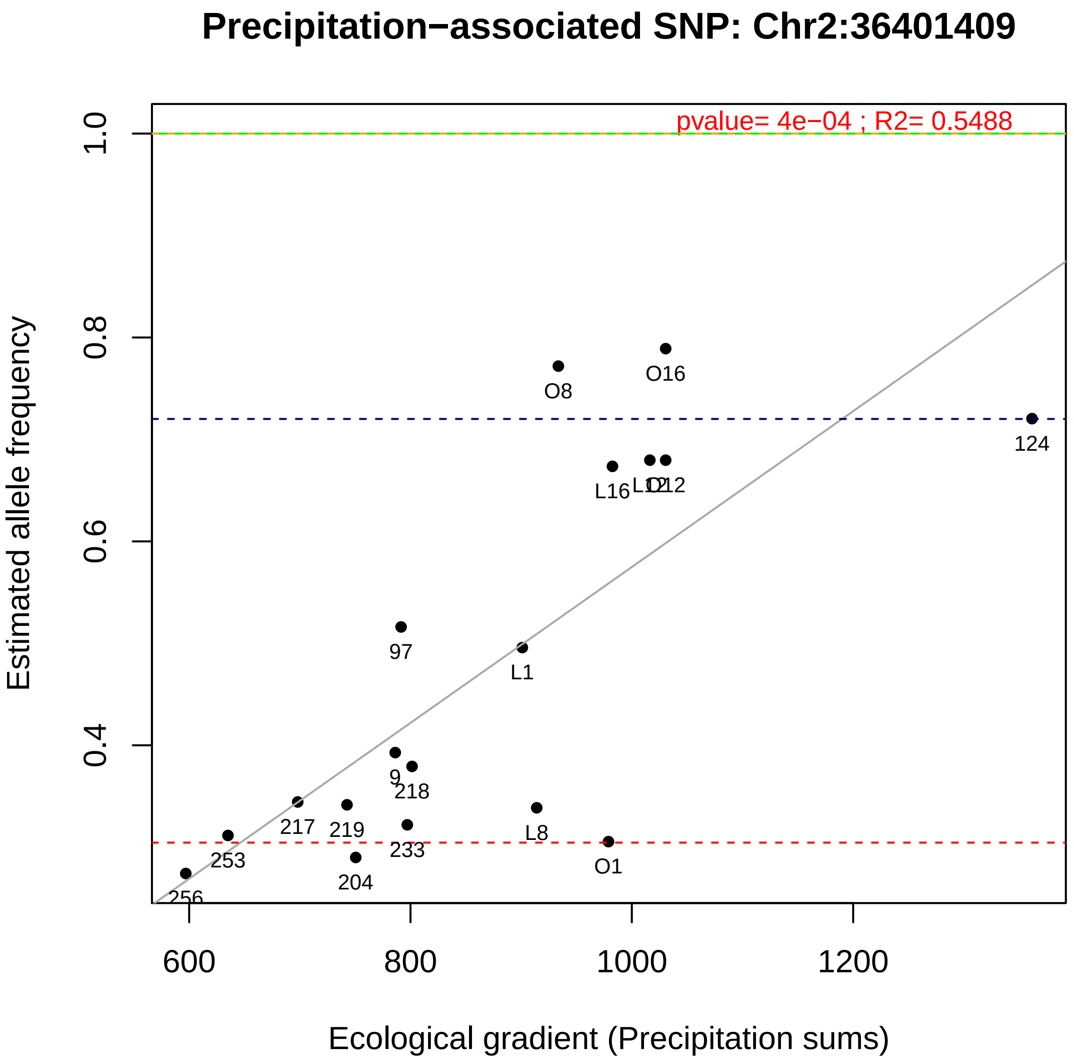

Precipitation−associated SNP: Chr2:36401409
pvalue= 4e−04 ; R2= 0.5488
Estimated allele frequency
1.0
0.8
0.6
0.4
O16
O8
124
L16
L12
O12
97
L1
9
218
217
219
233
L8
O1
253
204
256
600
800
1000
1200
Ecological gradient (Precipitation sums)

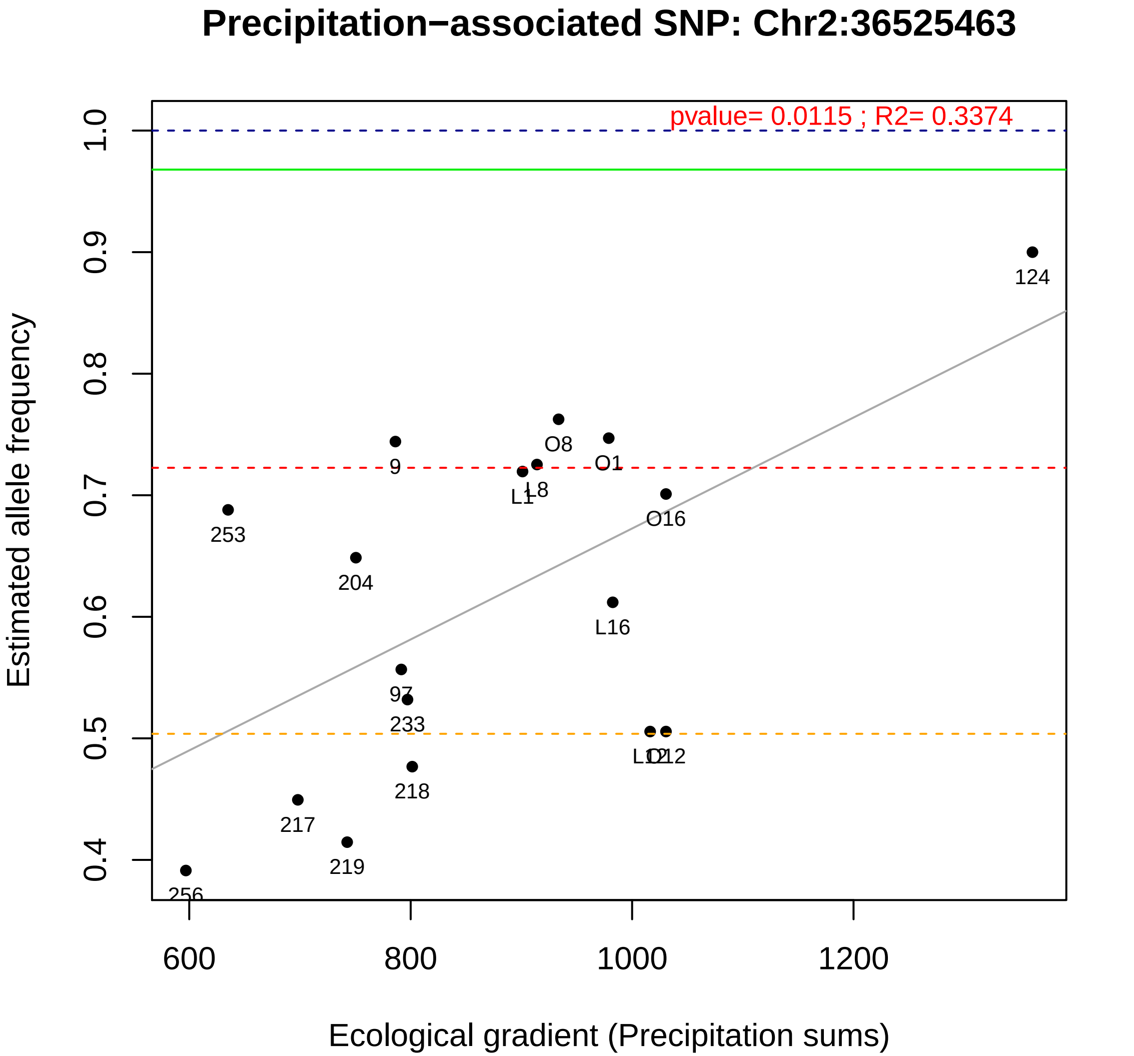
Precipitation−associated SNP: Chr2:36525463
pvalue= 0.0115 ; R2= 0.3374
Estimated allele frequency
Ecological gradient (Precipitation sums)
1.0
0.9
0.8
0.7
0.6
0.5
0.4
600
800
1000
1200
124
O8
O1
9
L1
L8
O16
253
204
L16
97
233
L12
O12
218
217
219
256

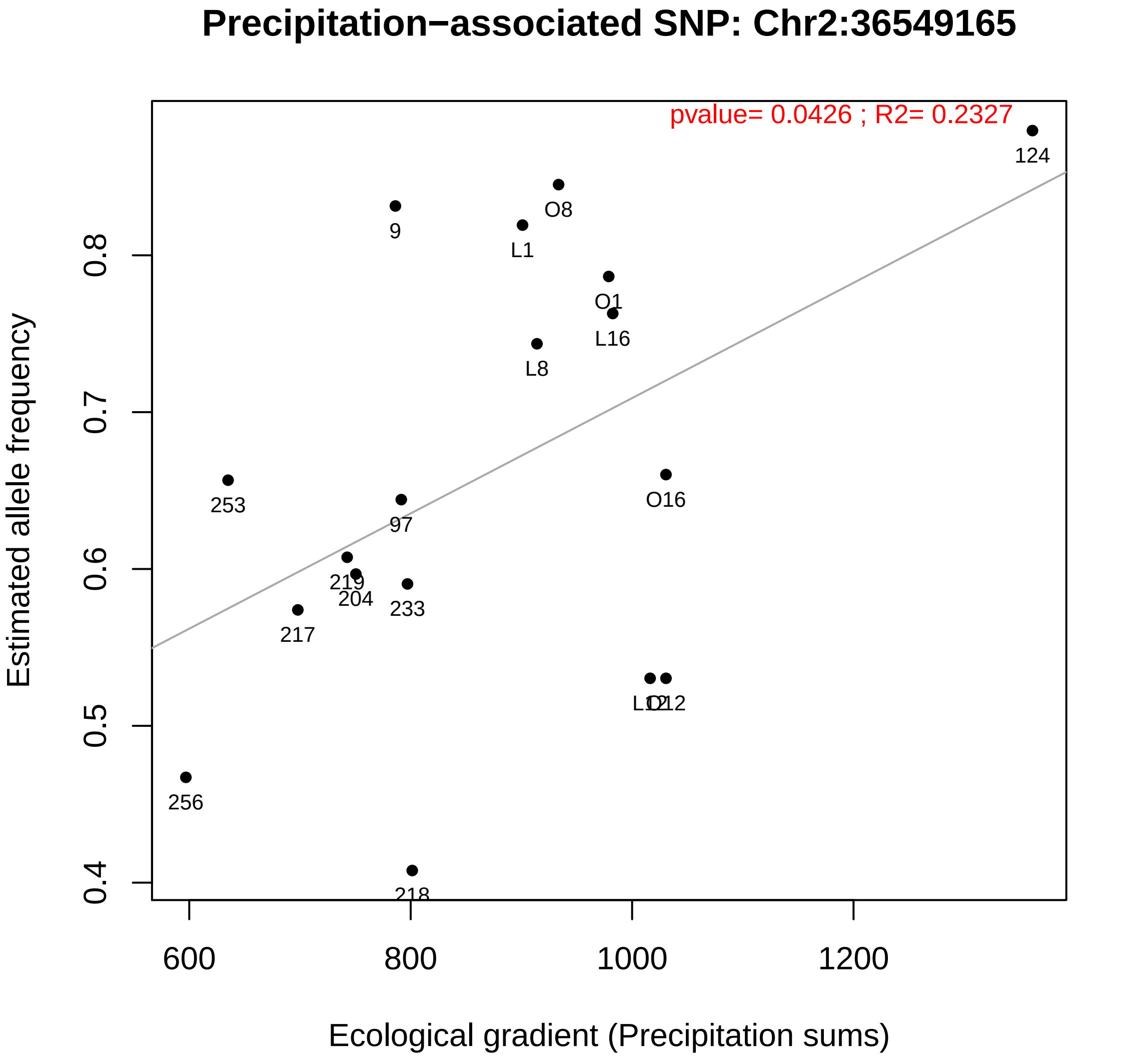
Precipitation−associated SNP: Chr2:36549165
pvalue= 0.0426 ; R2= 0.2327
Estimated allele frequency
Ecological gradient (Precipitation sums)
0.4
0.5
0.6
0.7
0.8
600
800
1000
1200
124
O8
9
L1
O1
L16
L8
253
O16
97
219
204
233
217
256
218

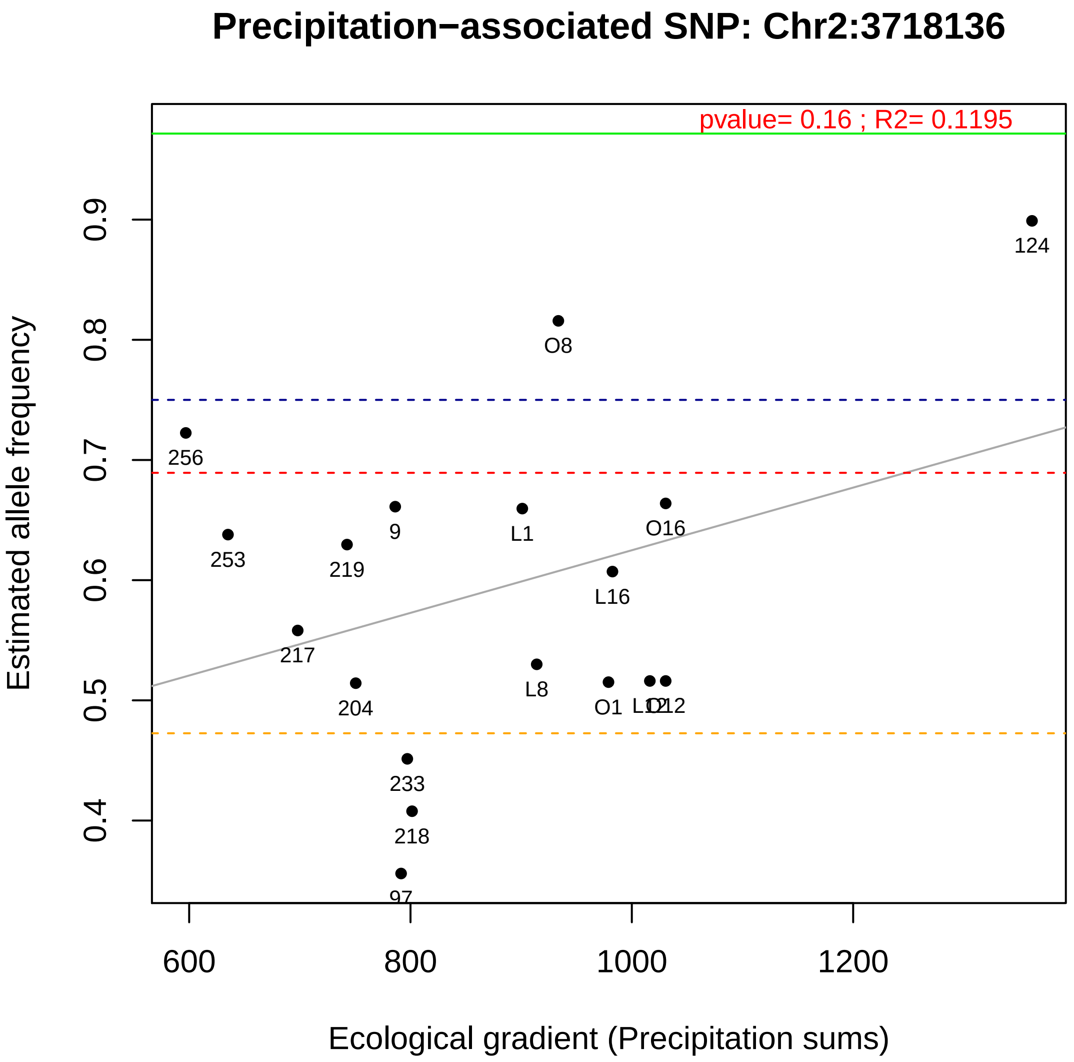

Precipitation−associated SNP: Chr2:3718136
pvalue= 0.16 ; R2= 0.1195
Estimated allele frequency
0.4
0.5
0.6
0.7
0.8
0.9
600
800
1000
1200
Ecological gradient (Precipitation sums)
124
O8
256
253
219
9
L1
O16
L16
217
204
L8
O1
O12
233
218
97

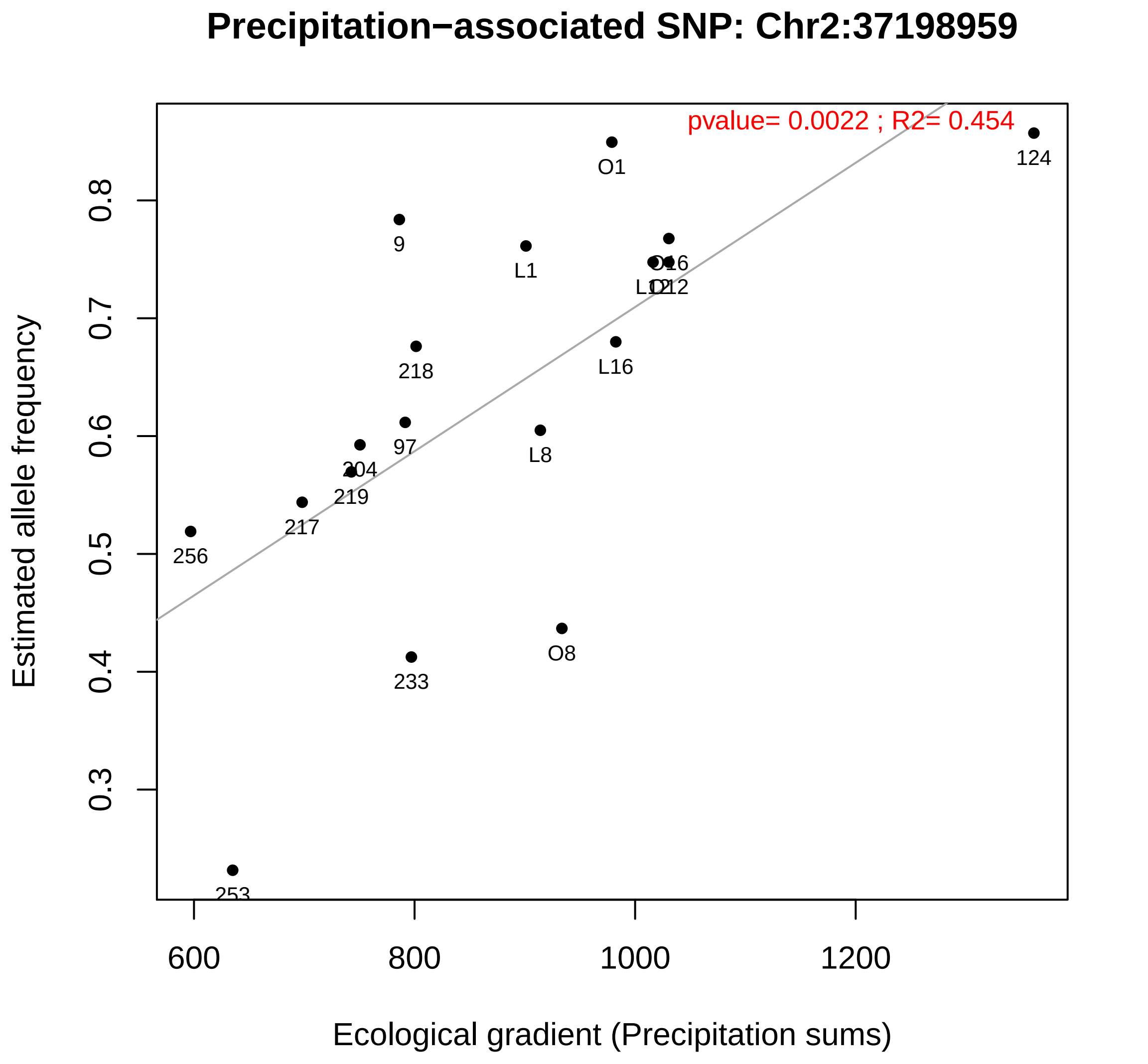
Precipitation−associated SNP: Chr2:37198959
pvalue= 0.0022 ; R2= 0.454
Estimated allele frequency
Ecological gradient (Precipitation sums)
0.3
0.4
0.5
0.6
0.7
0.8
600
800
1000
1200
O1
124
9
L1
O16
L12
O12
218
L16
97
L8
204
219
217
256
O8
233
253

**Precipitation−associated SNP: Chr2:3720275**

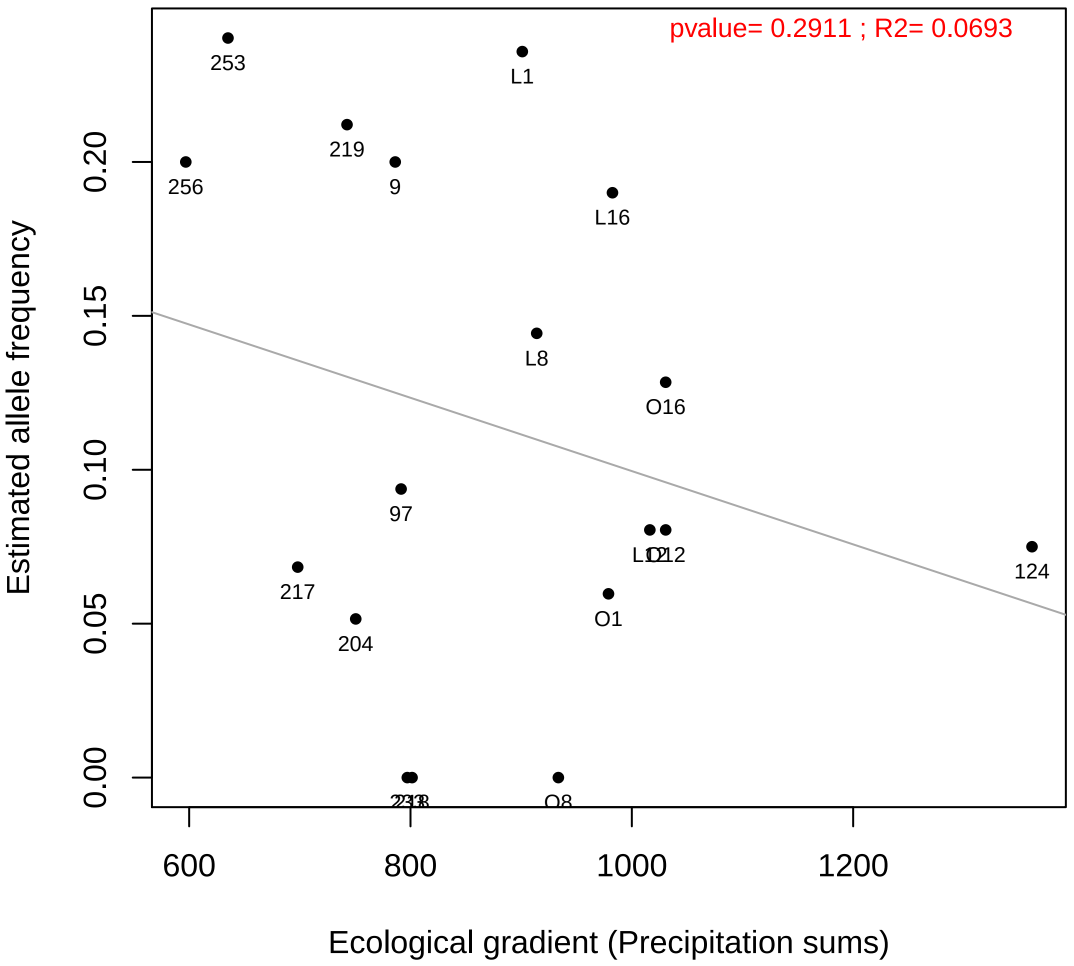

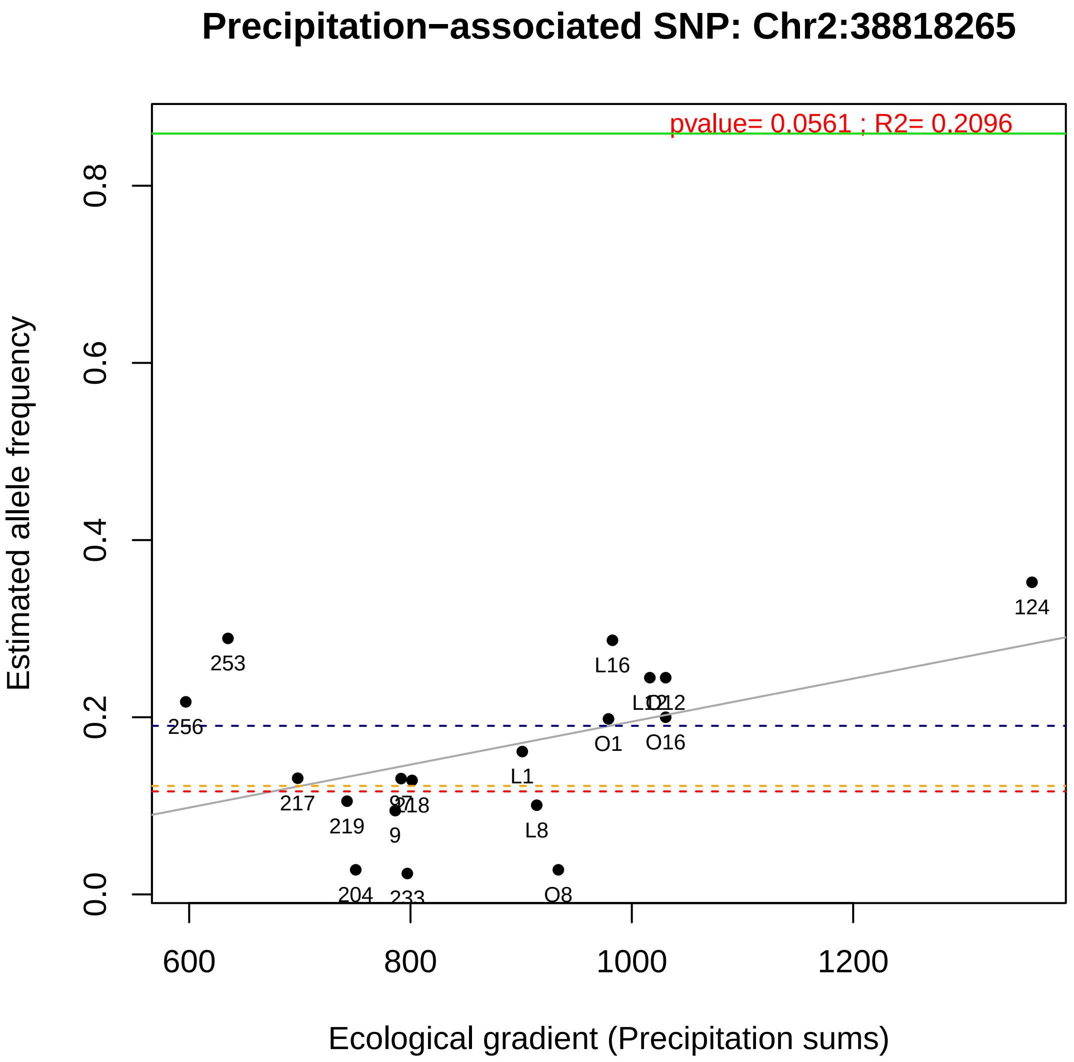

Precipitation−associated SNP: Chr2:38818265
pvalue= 0.0561 ; R2= 0.2096
Estimated allele frequency
0.0
0.2
0.4
0.6
0.8
600
800
1000
1200
Ecological gradient (Precipitation sums)
124
253
L16
256
O12
O1
O16
L1
217
219
218
9
L8
204
233
O8

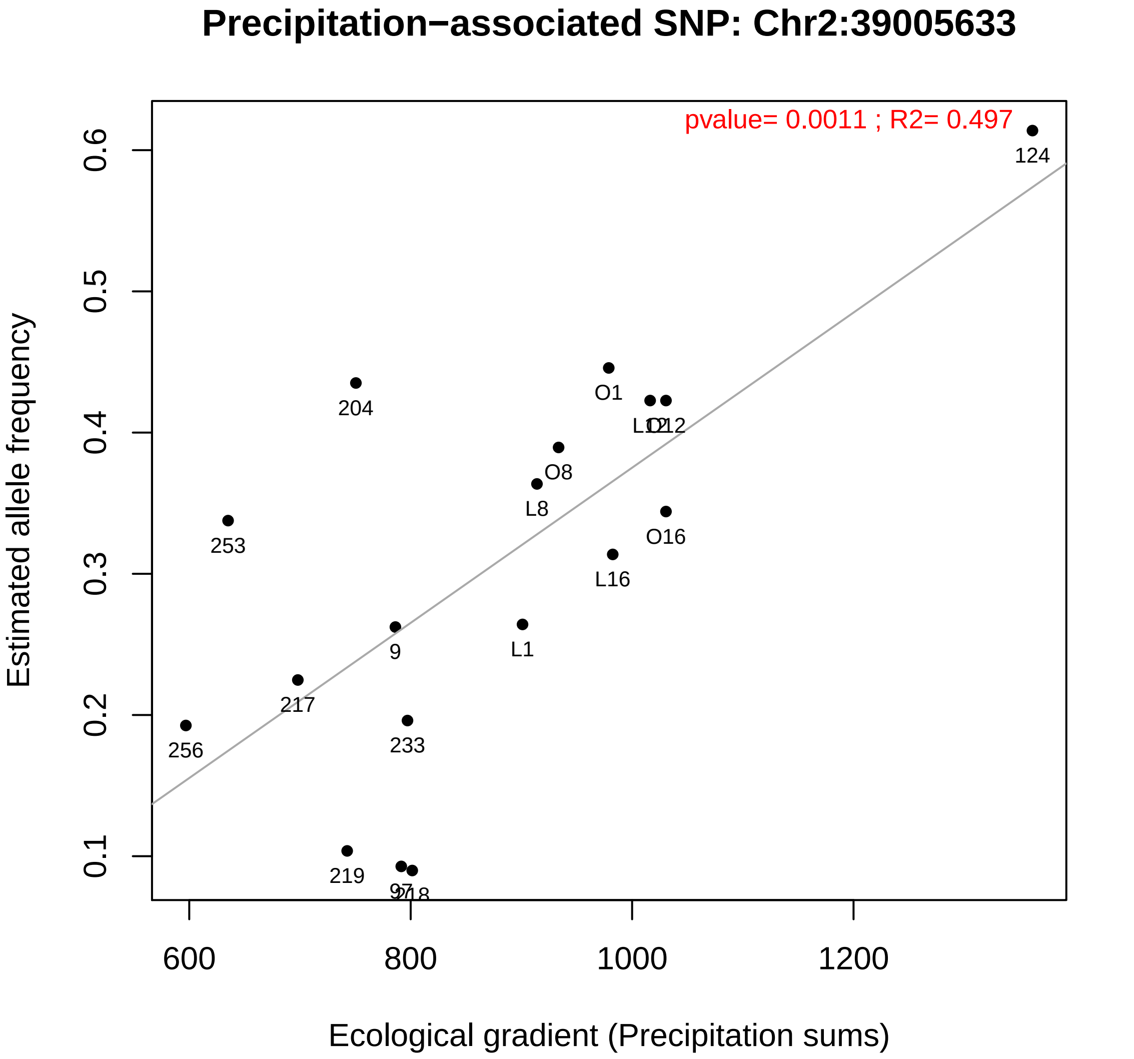
Precipitation−associated SNP: Chr2:39005633
pvalue= 0.0011 ; R2= 0.497
Estimated allele frequency
Ecological gradient (Precipitation sums)
0.1
0.2
0.3
0.4
0.5
0.6
600
800
1000
1200
124
204
O1
O8
L8
253
O16
L16
9
L1
217
256
233
219

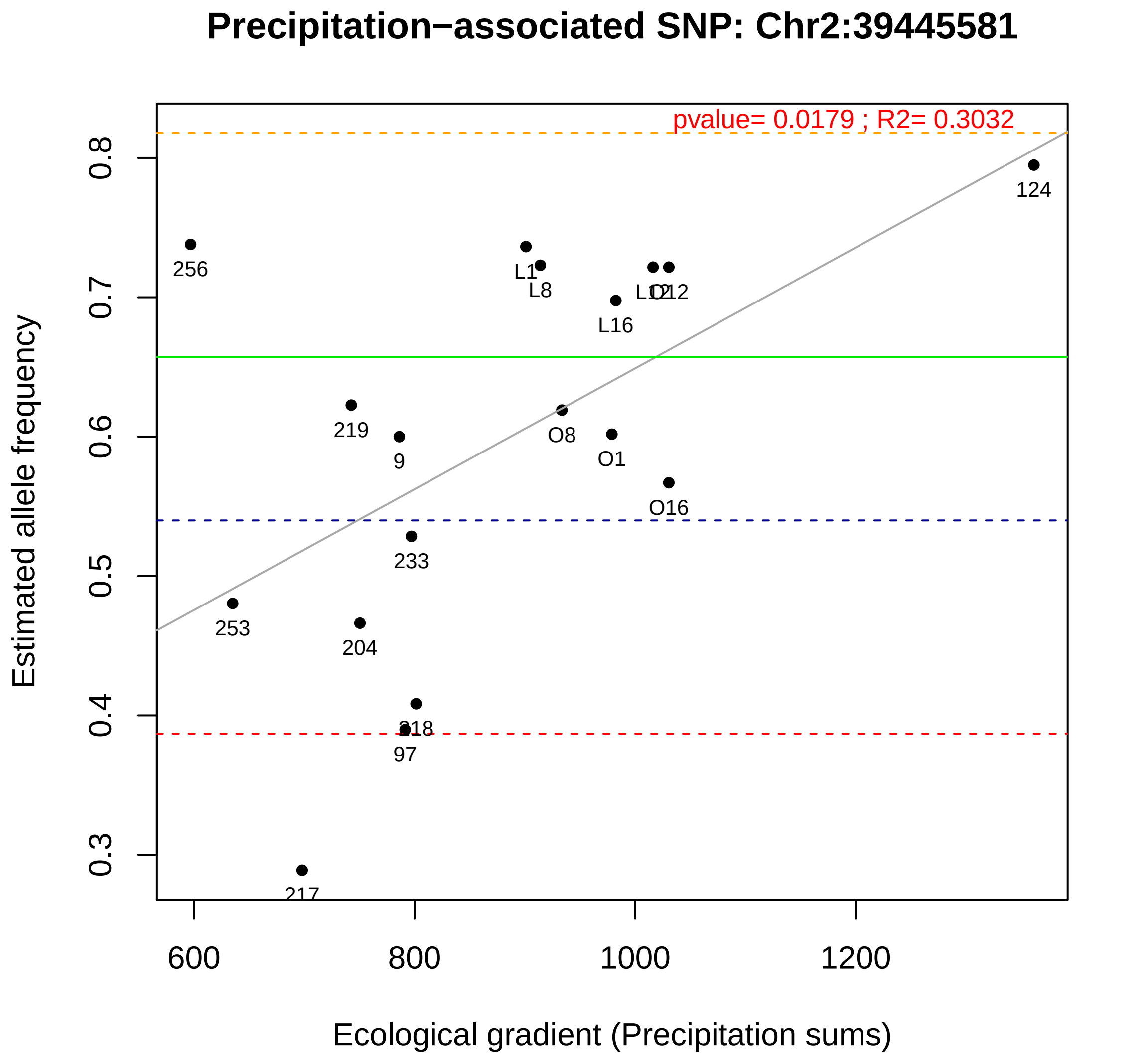

Precipitation−associated SNP: Chr2:39445581
pvalue= 0.0179 ; R2= 0.3032
Estimated allele frequency
Ecological gradient (Precipitation sums)
0.3
0.4
0.5
0.6
0.7
0.8
600
800
1000
1200
124
256
L1
L8
L12
O12
L16
219
9
O8
O1
O16
233
253
204
218
97
217

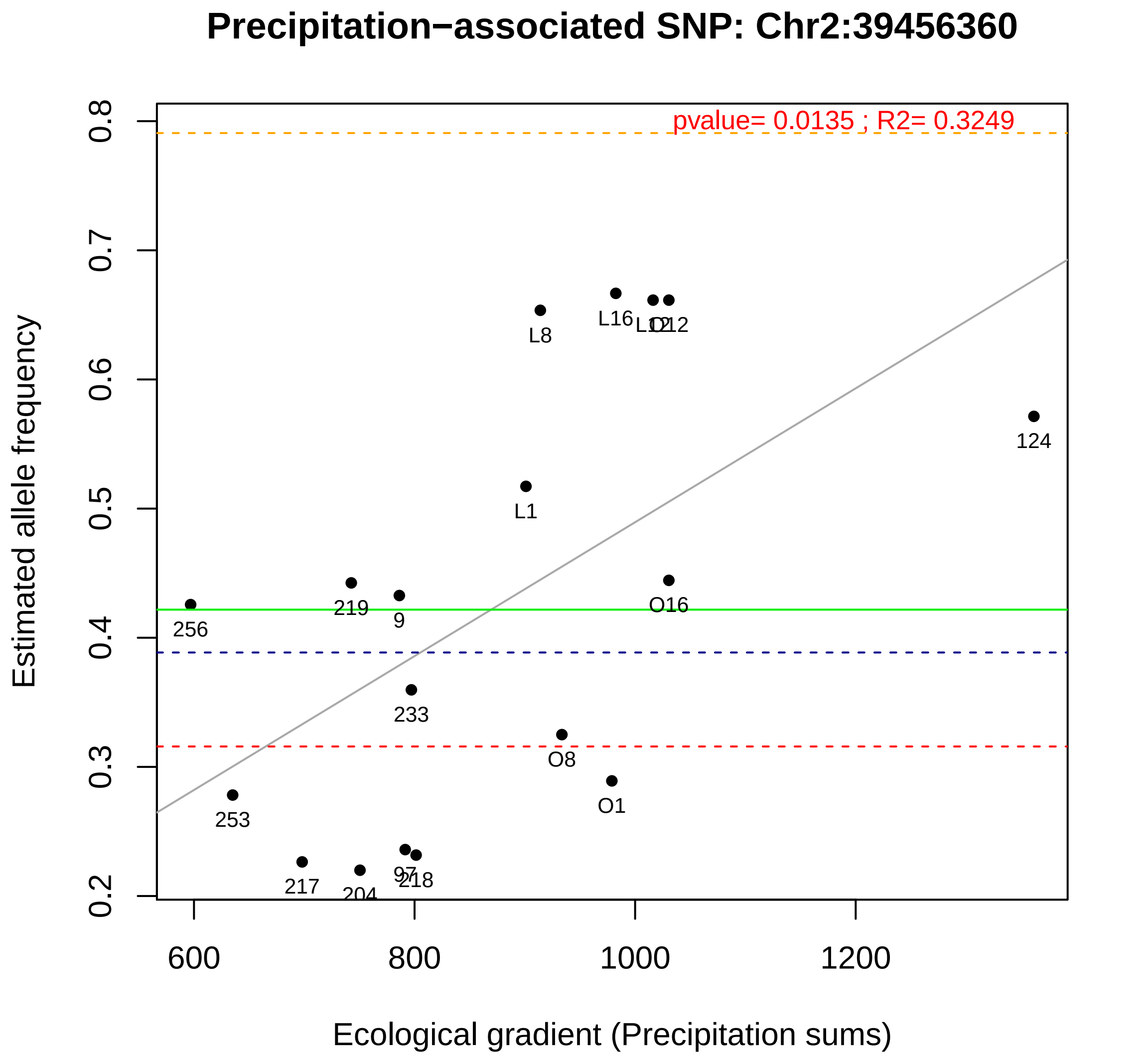
Precipitation−associated SNP: Chr2:39456360
pvalue= 0.0135 ; R2= 0.3249
Estimated allele frequency
0.2
0.3
0.4
0.5
0.6
0.7
0.8
600
800
1000
1200
Ecological gradient (Precipitation sums)
L8
L16
L12
O12
124
L1
256
219
9
O16
233
O8
O1
253
217
204
97
218

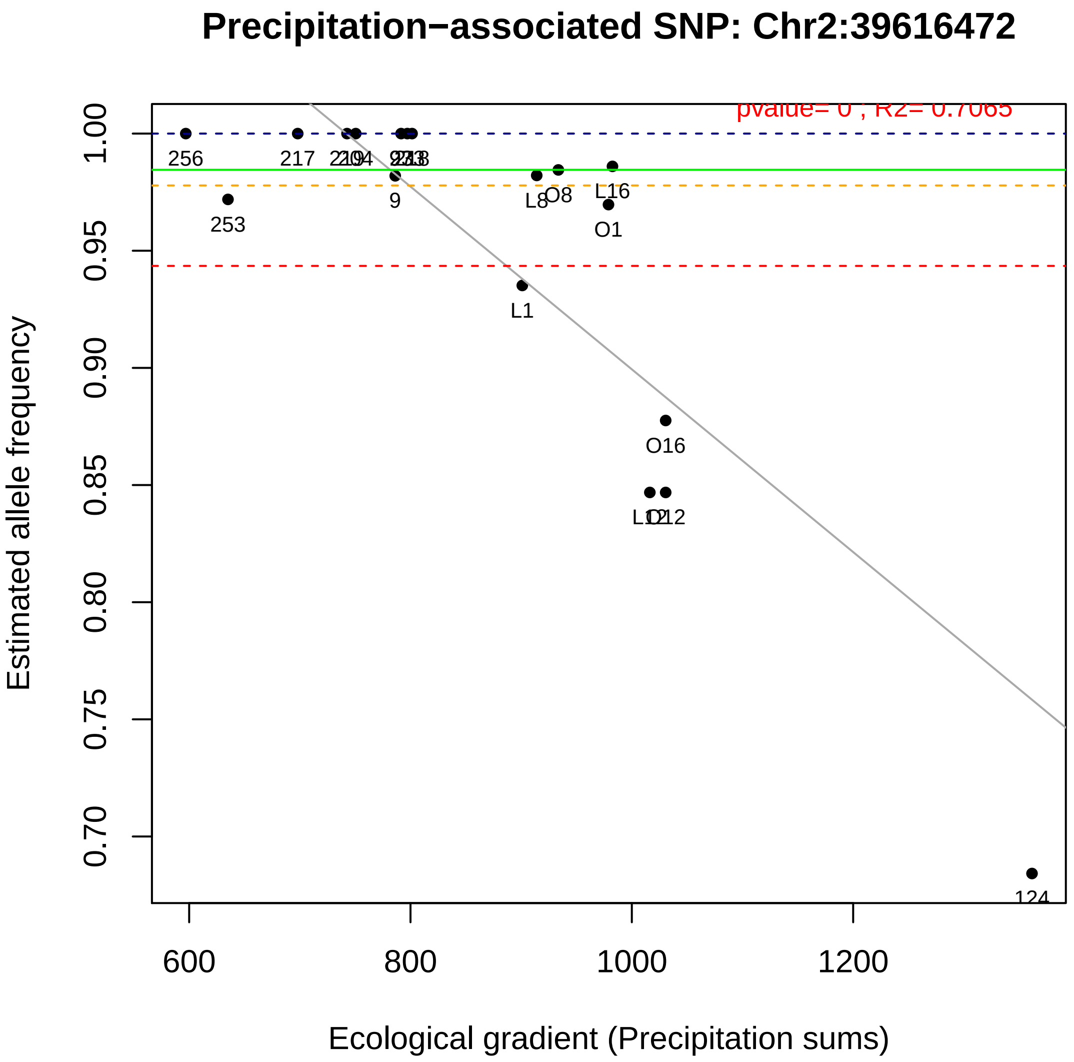

Precipitation−associated SNP: Chr2:39616472
pvalue= 0 ; R2= 0.7065
Estimated allele frequency
Ecological gradient (Precipitation sums)
1.00
0.95
0.90
0.85
0.80
0.75
0.70
600
800
1000
1200
256
217
253
9
L8
O8
L16
O1
L1
O16
O12
124

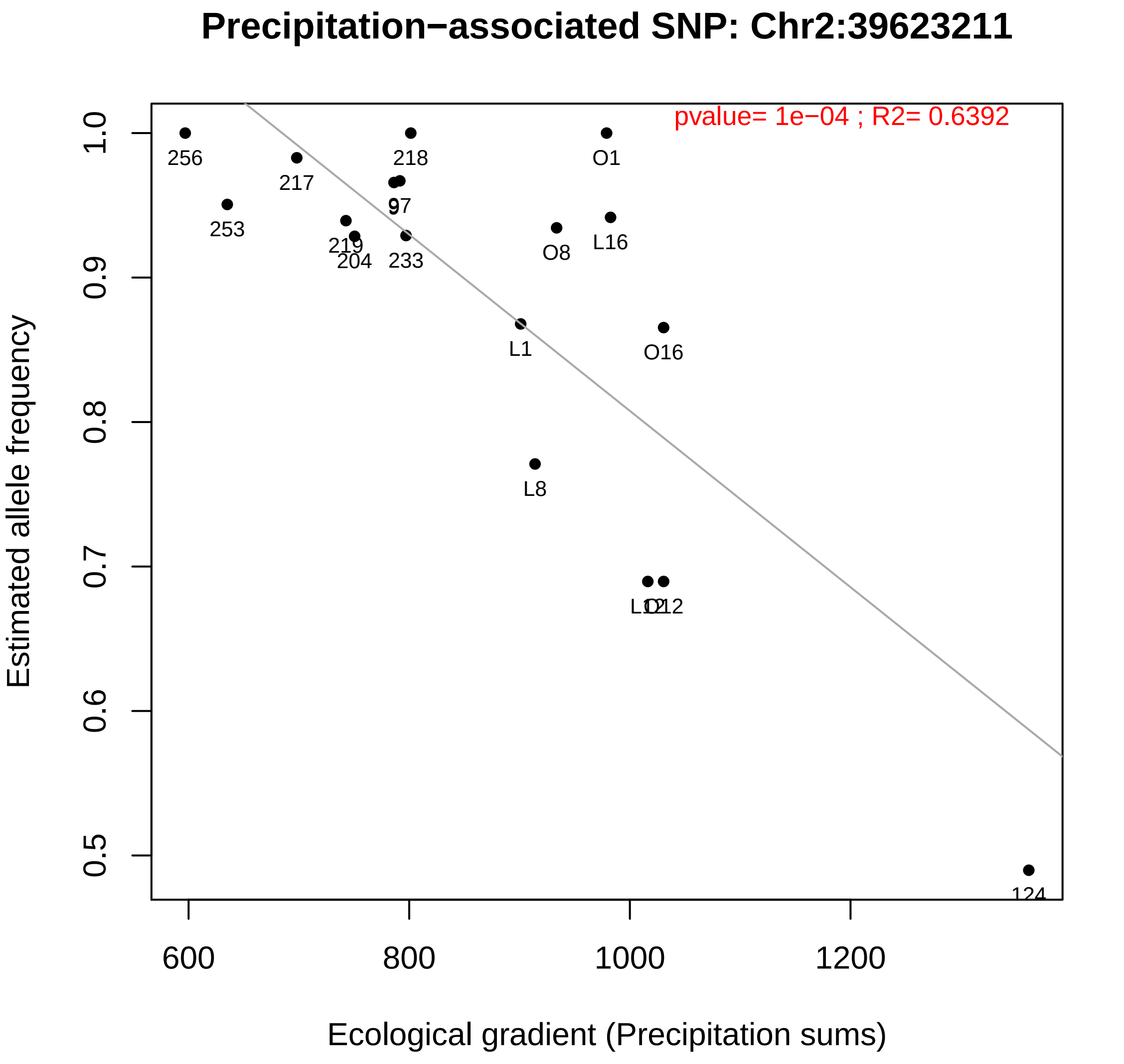

Precipitation−associated SNP: Chr2:39623211
pvalue= 1e−04 ; R2= 0.6392
256
217
218
O1
253
97
219
204
233
O8
L16
L1
O16
L8
L12
O12
124
1.0
0.9
0.8
0.7
0.6
0.5
600
800
1000
1200
Estimated allele frequency
Ecological gradient (Precipitation sums)

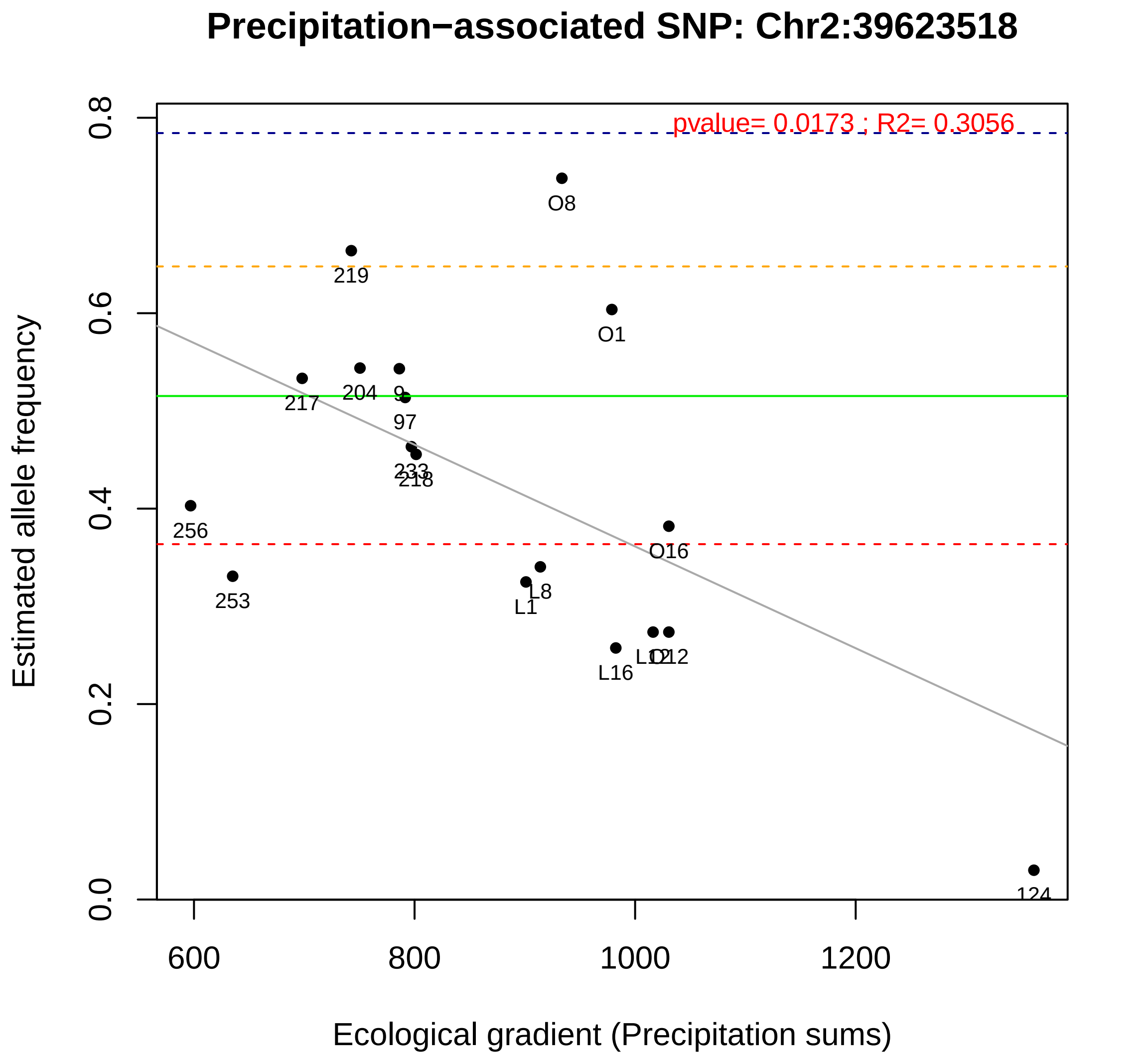
Precipitation−associated SNP: Chr2:39623518
pvalue= 0.0173 ; R2= 0.3056
O8
219
O1
217
204
9
97
233
218
256
O16
253
L8
L1
L16
L12
O12
124
0.0
0.2
0.4
0.6
0.8
600
800
1000
1200
Estimated allele frequency
Ecological gradient (Precipitation sums)

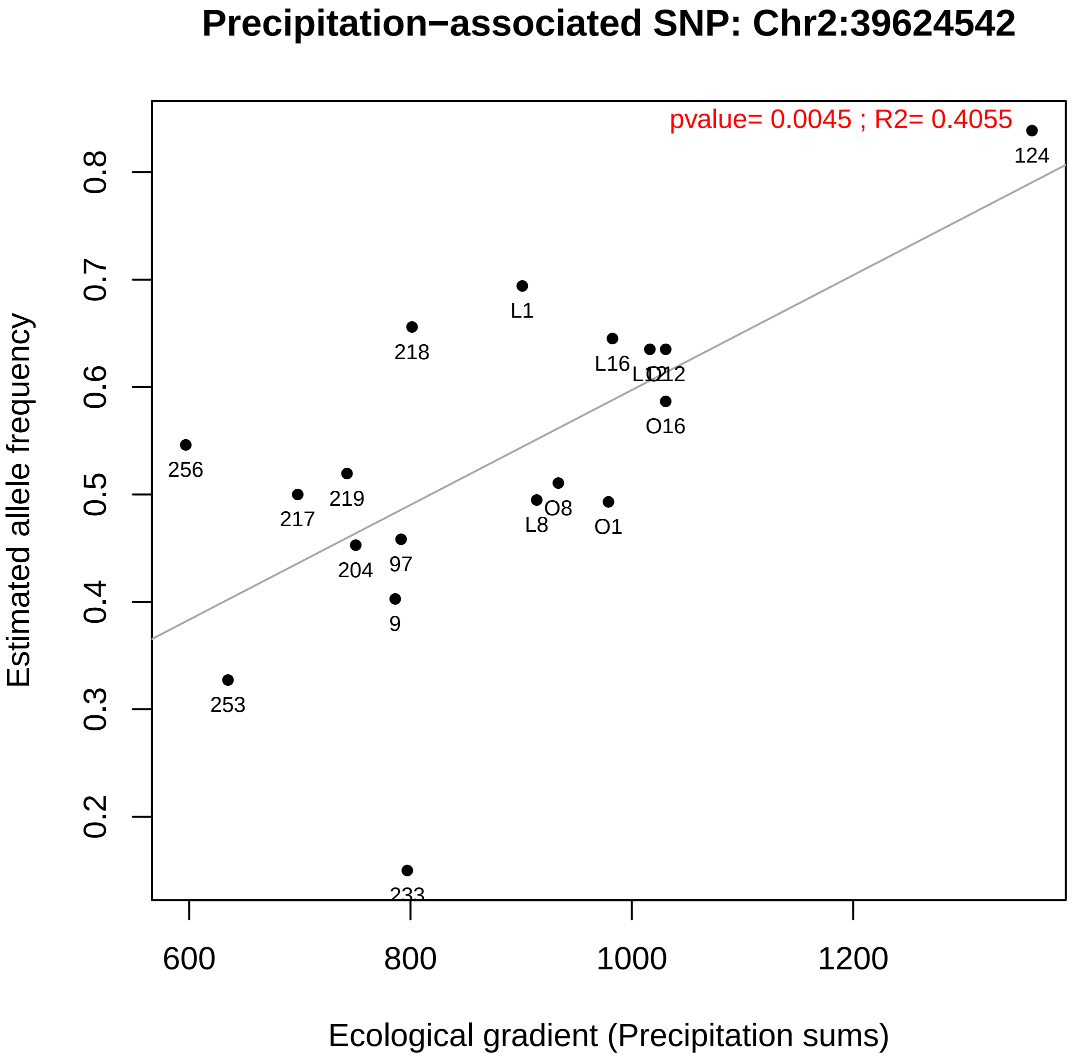
Precipitation−associated SNP: Chr2:39624542
pvalue= 0.0045 ; R2= 0.4055
Estimated allele frequency
0.2
0.3
0.4
0.5
0.6
0.7
0.8
600
800
1000
1200
Ecological gradient (Precipitation sums)
124
L1
218
L16
O12
O16
256
217
219
204
97
9
L8
O8
O1
253
233

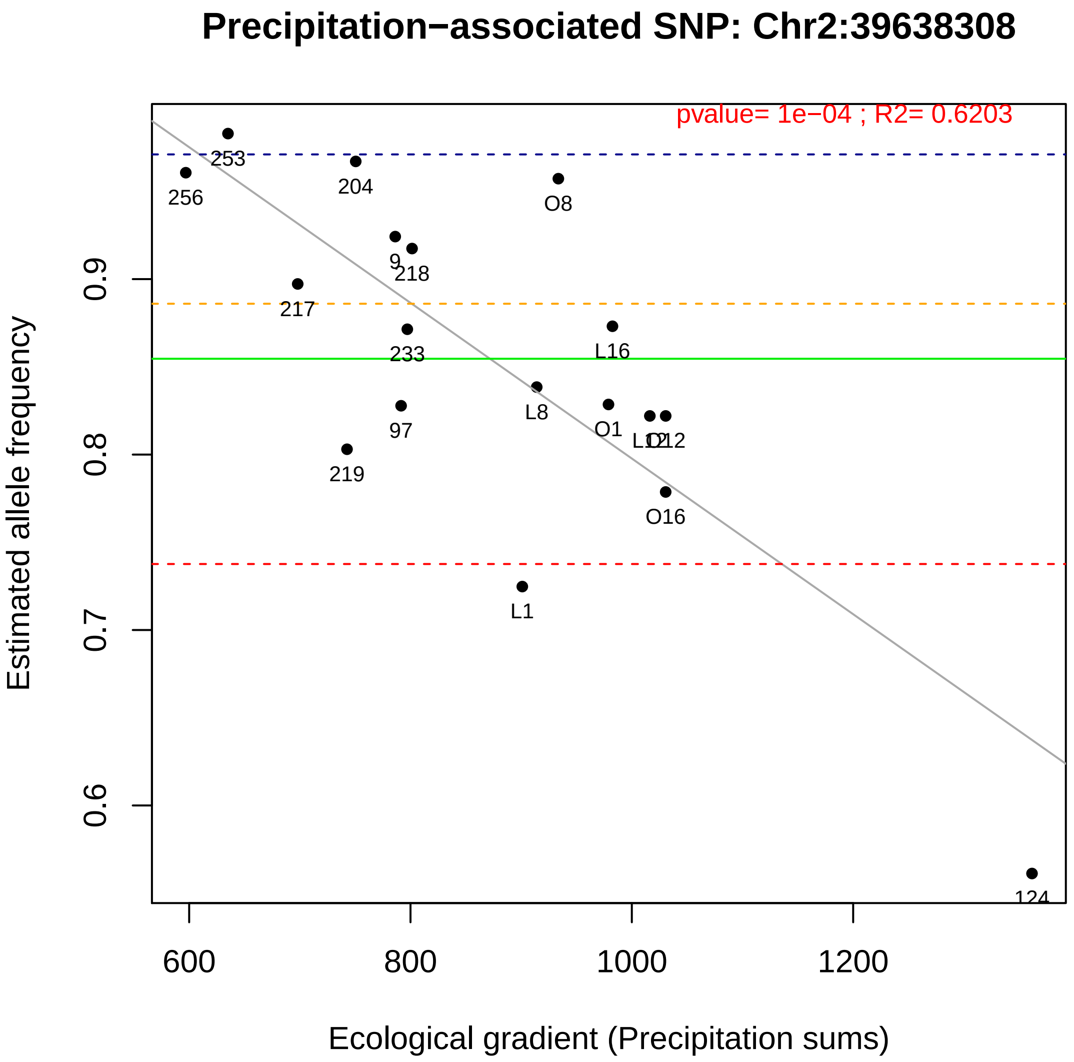
Precipitation−associated SNP: Chr2:39638308
pvalue= 1e−04 ; R2= 0.6203
Estimated allele frequency
Ecological gradient (Precipitation sums)
0.6
0.7
0.8
0.9
600
800
1000
1200
253
256
204
O8
9
218
217
233
L16
L8
97
O1
219
O16
L1
124

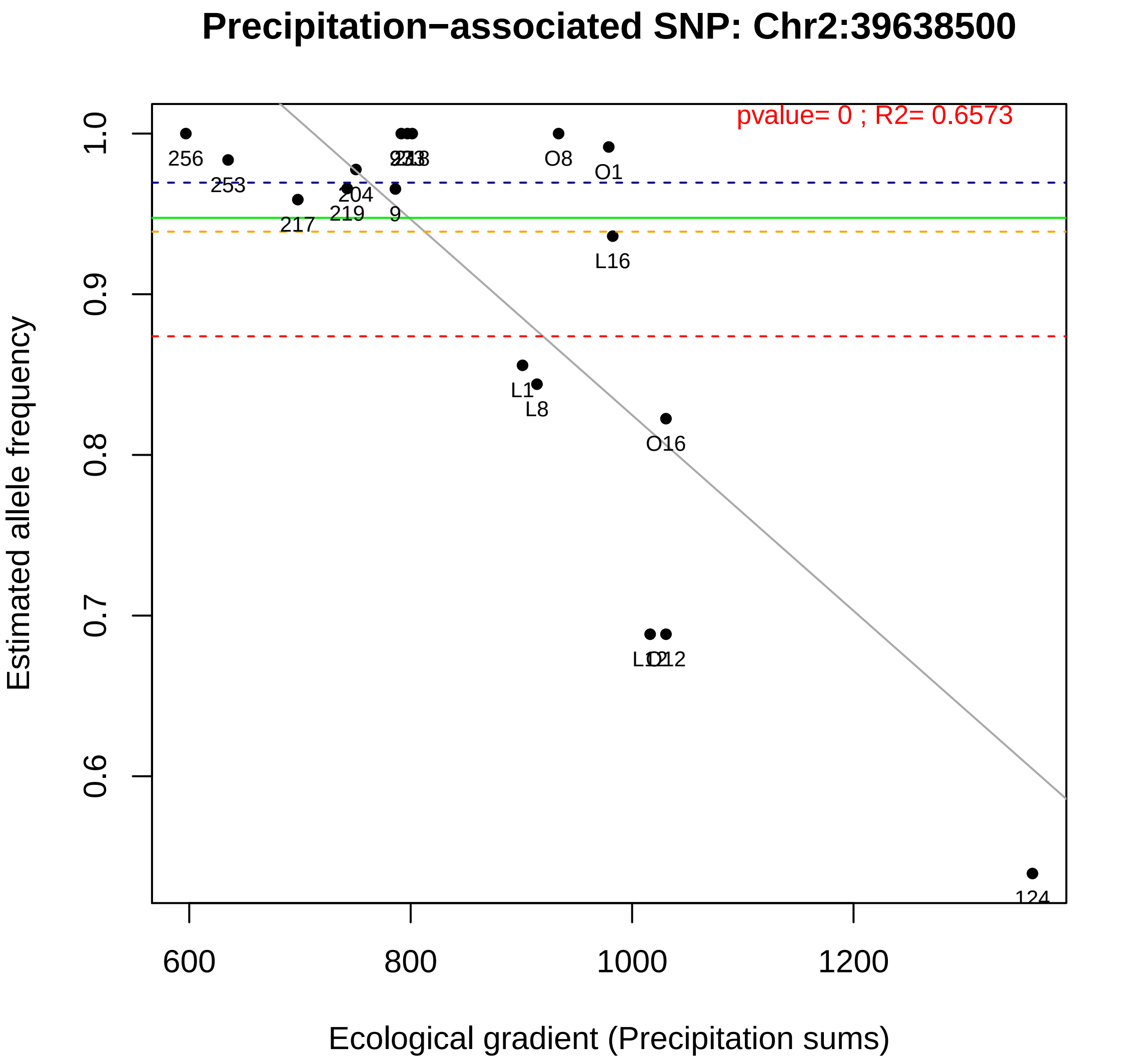

Precipitation−associated SNP: Chr2:39638500
pvalue= 0 ; R2= 0.6573
256
253
217
219
204
9
O8
O1
L16
L1
L8
O16
O12
124
1.0
0.9
0.8
0.7
0.6
600
800
1000
1200
Estimated allele frequency
Ecological gradient (Precipitation sums)

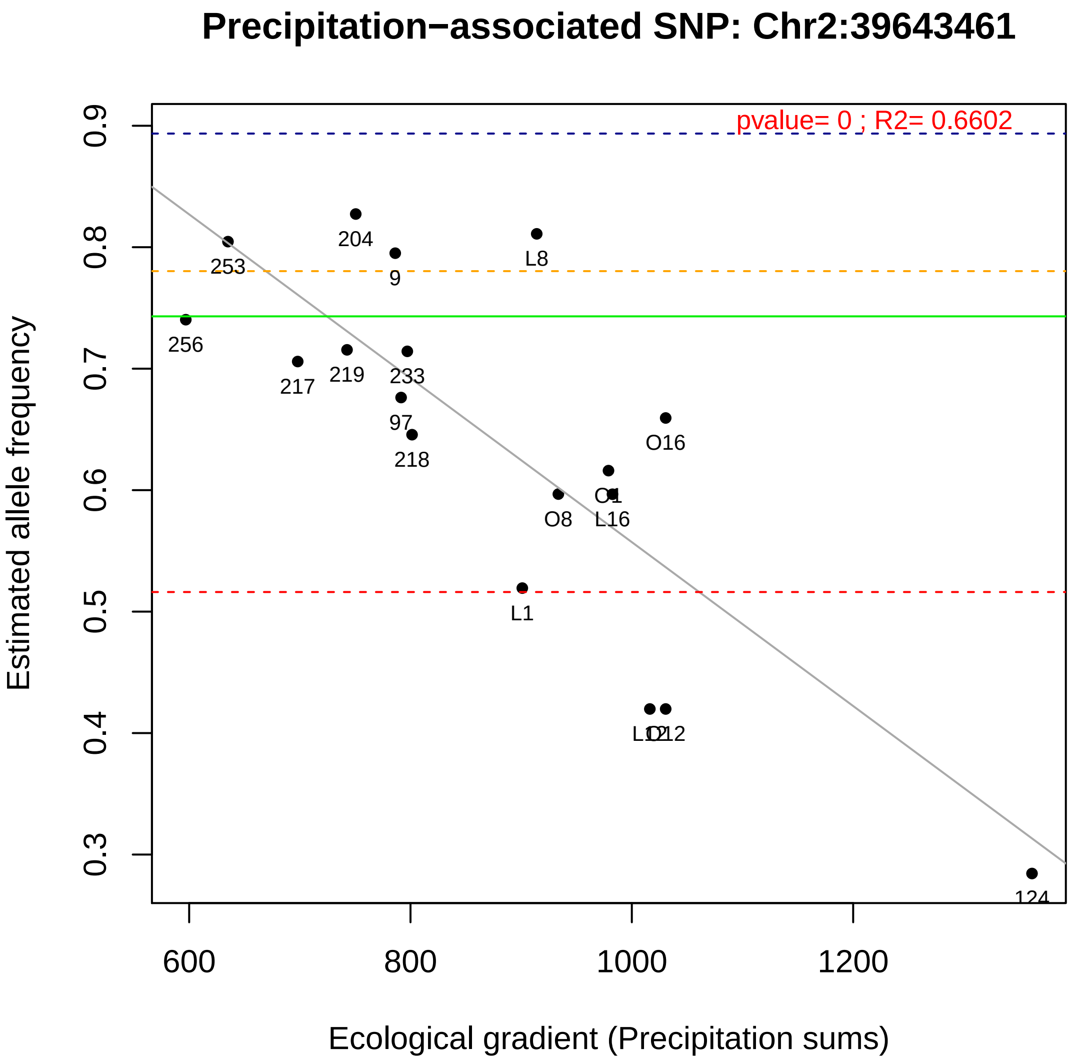
Precipitation−associated SNP: Chr2:39643461
pvalue= 0 ; R2= 0.6602
Estimated allele frequency
Ecological gradient (Precipitation sums)
0.3
0.4
0.5
0.6
0.7
0.8
0.9
600
800
1000
1200
253
204
9
L8
256
217
219
233
97
218
O16
O1
L16
O8
L1
L12
O12
124

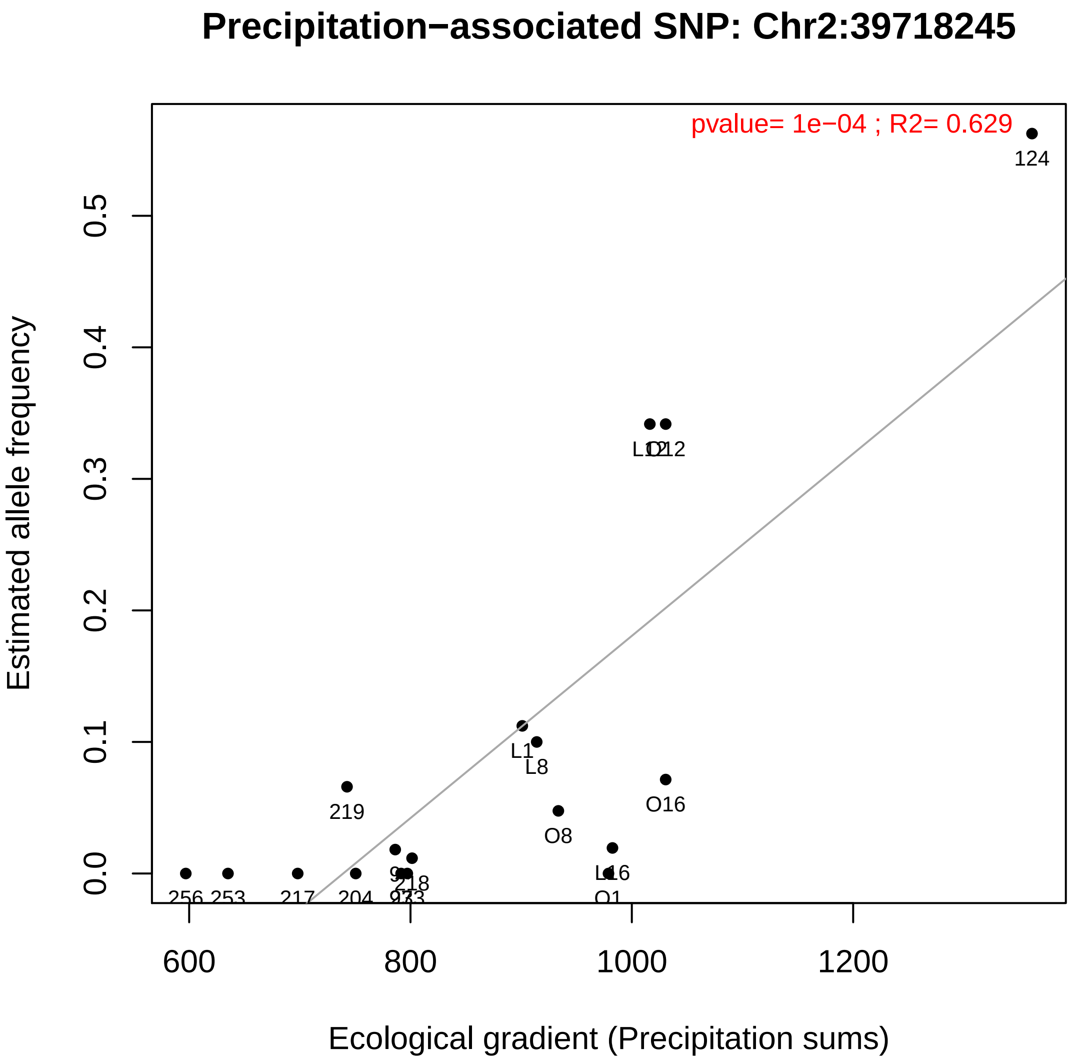

Precipitation−associated SNP: Chr2:39718245
pvalue= 1e−04 ; R2= 0.629
124
L12
O12
L1
L8
O16
219
O8
L16
O1
218
256
253
217
204
0.0
0.1
0.2
0.3
0.4
0.5
600
800
1000
1200
Estimated allele frequency
Ecological gradient (Precipitation sums)

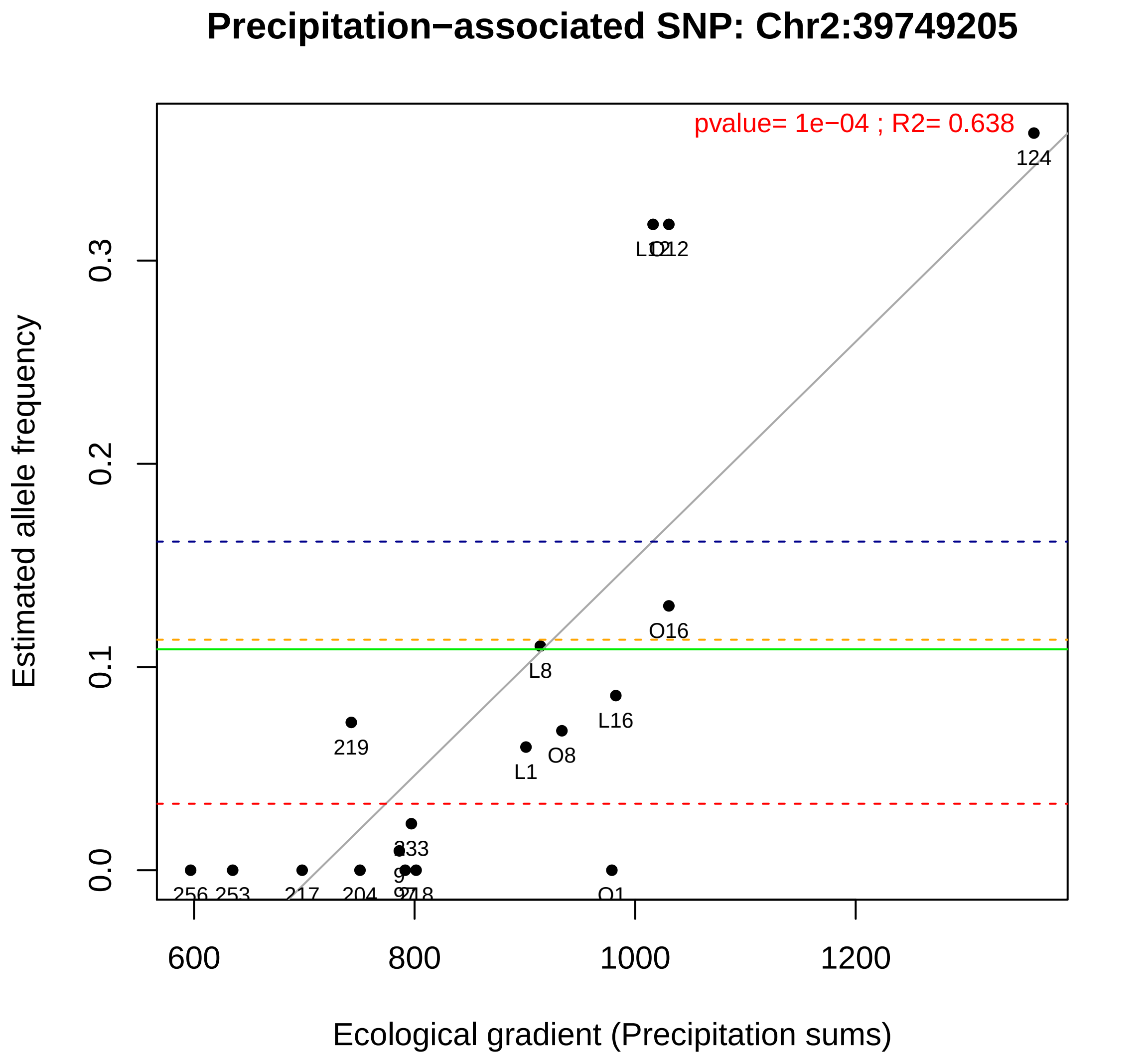

Precipitation−associated SNP: Chr2:39749205
pvalue= 1e−04 ; R2= 0.638
124
L12
O12
O16
L8
L16
219
O8
L1
233
9
97
218
256
253
217
204
O1
0.0
0.1
0.2
0.3
600
800
1000
1200
Estimated allele frequency
Ecological gradient (Precipitation sums)

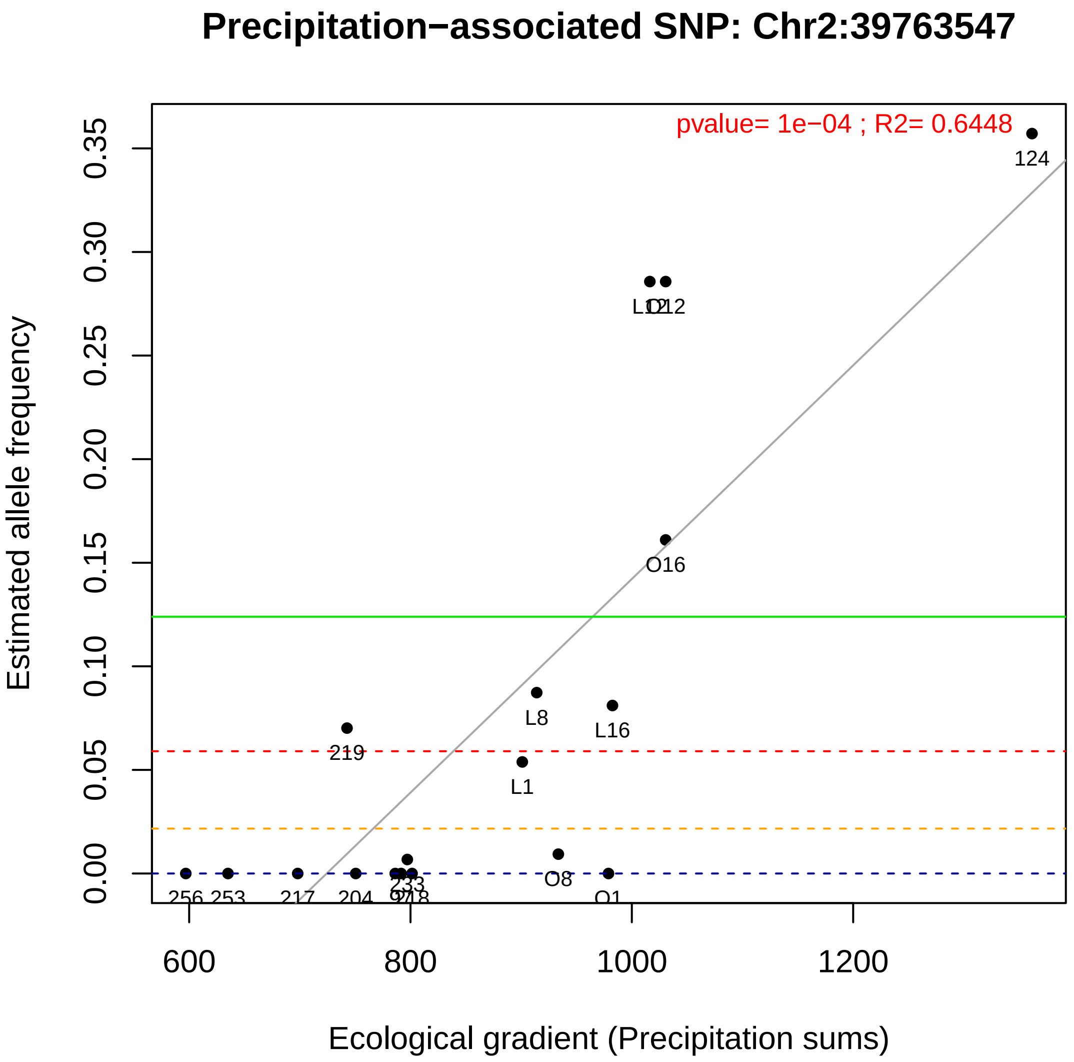
Precipitation−associated SNP: Chr2:39763547
pvalue= 1e−04 ; R2= 0.6448
Estimated allele frequency
Ecological gradient (Precipitation sums)
0.00
0.05
0.10
0.15
0.20
0.25
0.30
0.35
600
800
1000
1200
124
O16
L8
L16
219
L1
O8
233

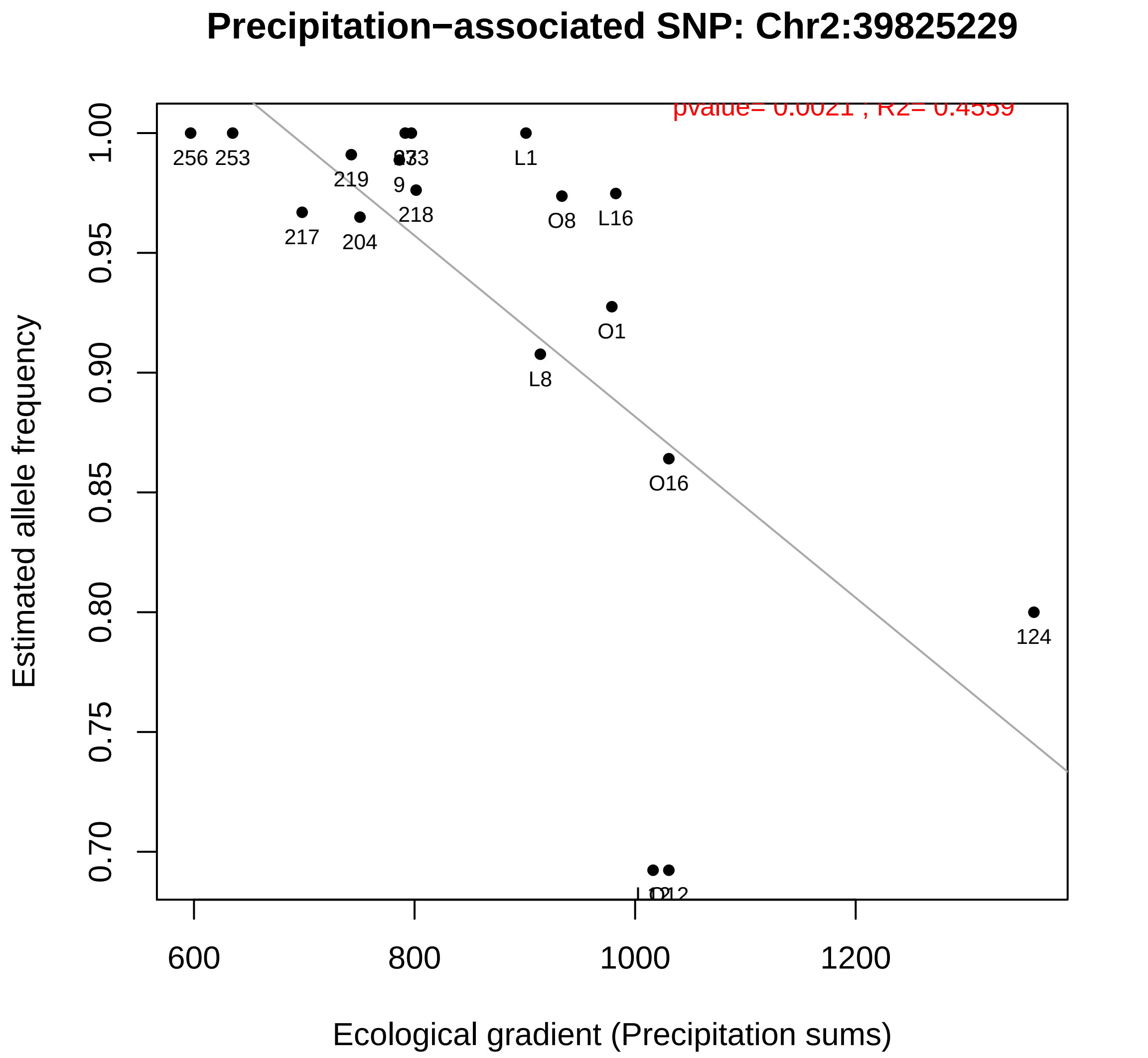

Precipitation−associated SNP: Chr2:39825229
pvalue= 0.0021 ; R2= 0.4559
256
253
219
217
204
218
9
L1
O8
L16
O1
L8
O16
124
1.00
0.95
0.90
0.85
0.80
0.75
0.70
600
800
1000
1200
Estimated allele frequency
Ecological gradient (Precipitation sums)

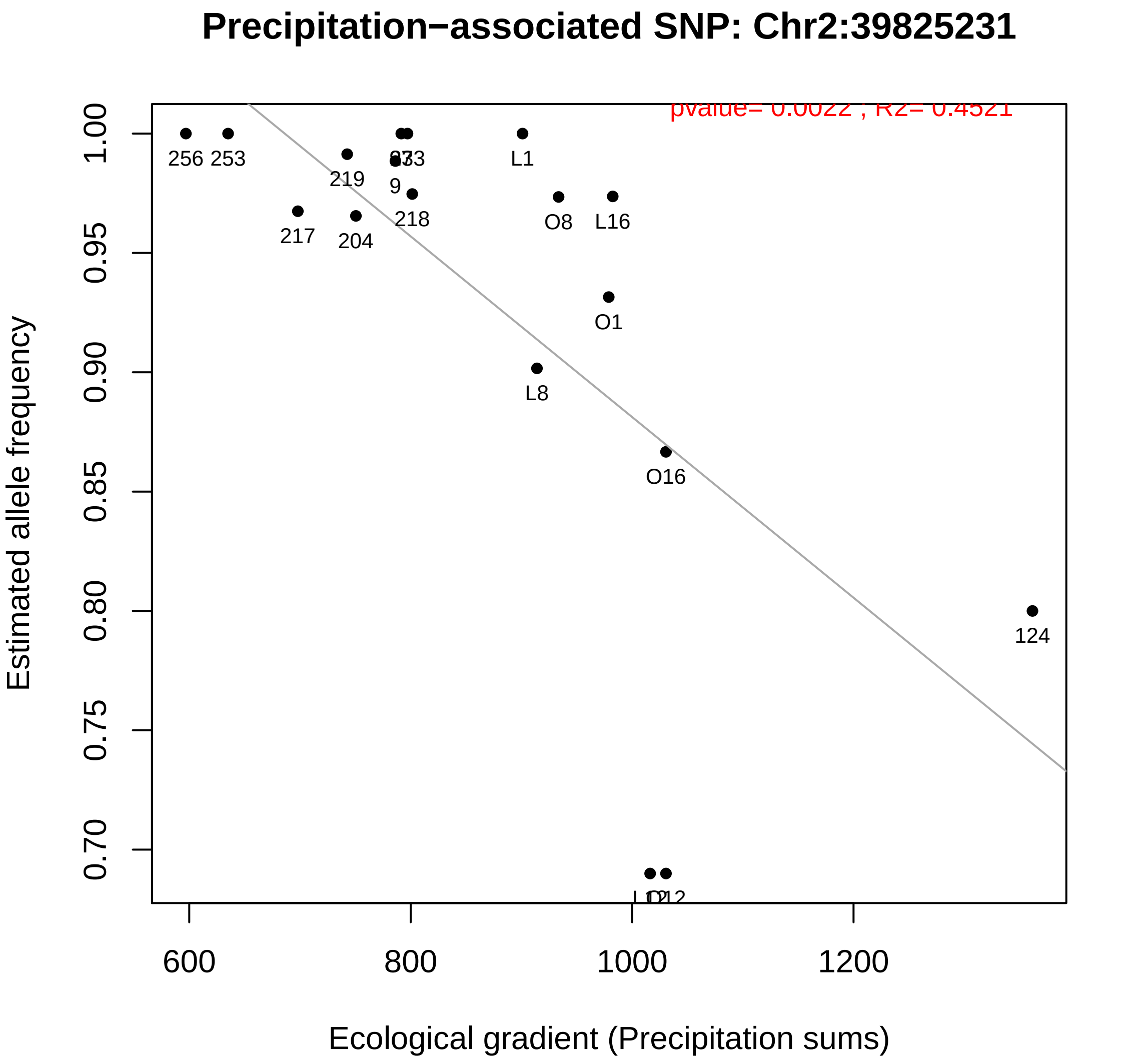

Precipitation−associated SNP: Chr2:39825231
pvalue= 0.0022 ; R2= 0.4521
Estimated allele frequency
1.00
0.95
0.90
0.85
0.80
0.75
0.70
256
253
219
9
218
217
204
L1
O8
L16
O1
L8
O16
124
600
800
1000
1200
Ecological gradient (Precipitation sums)

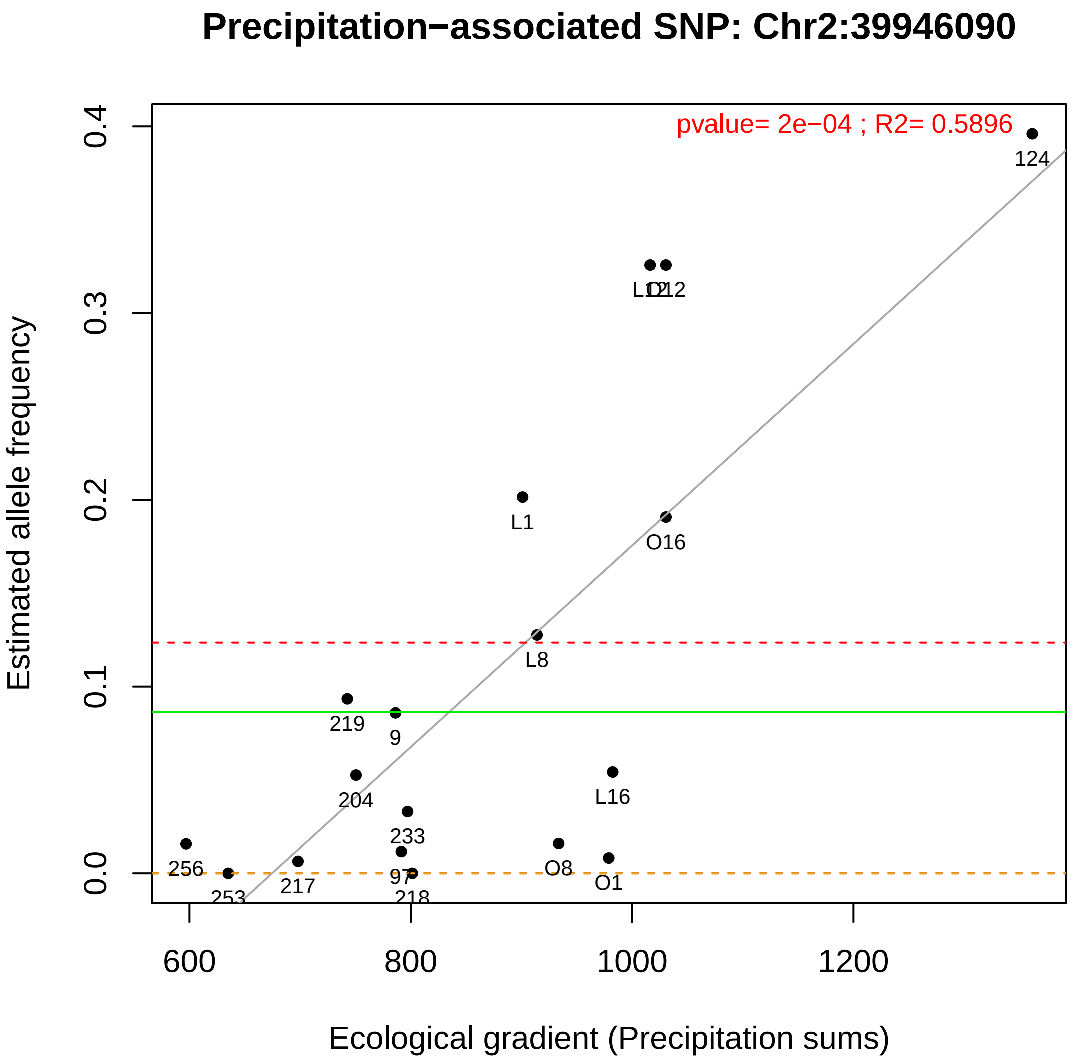

Precipitation−associated SNP: Chr2:39946090
pvalue= 2e−04 ; R2= 0.5896
Estimated allele frequency
0.0
0.1
0.2
0.3
0.4
Ecological gradient (Precipitation sums)
600
800
1000
1200
124
L12
O12
L1
O16
L8
219
9
204
L16
233
256
253
217
97
218
O8
O1

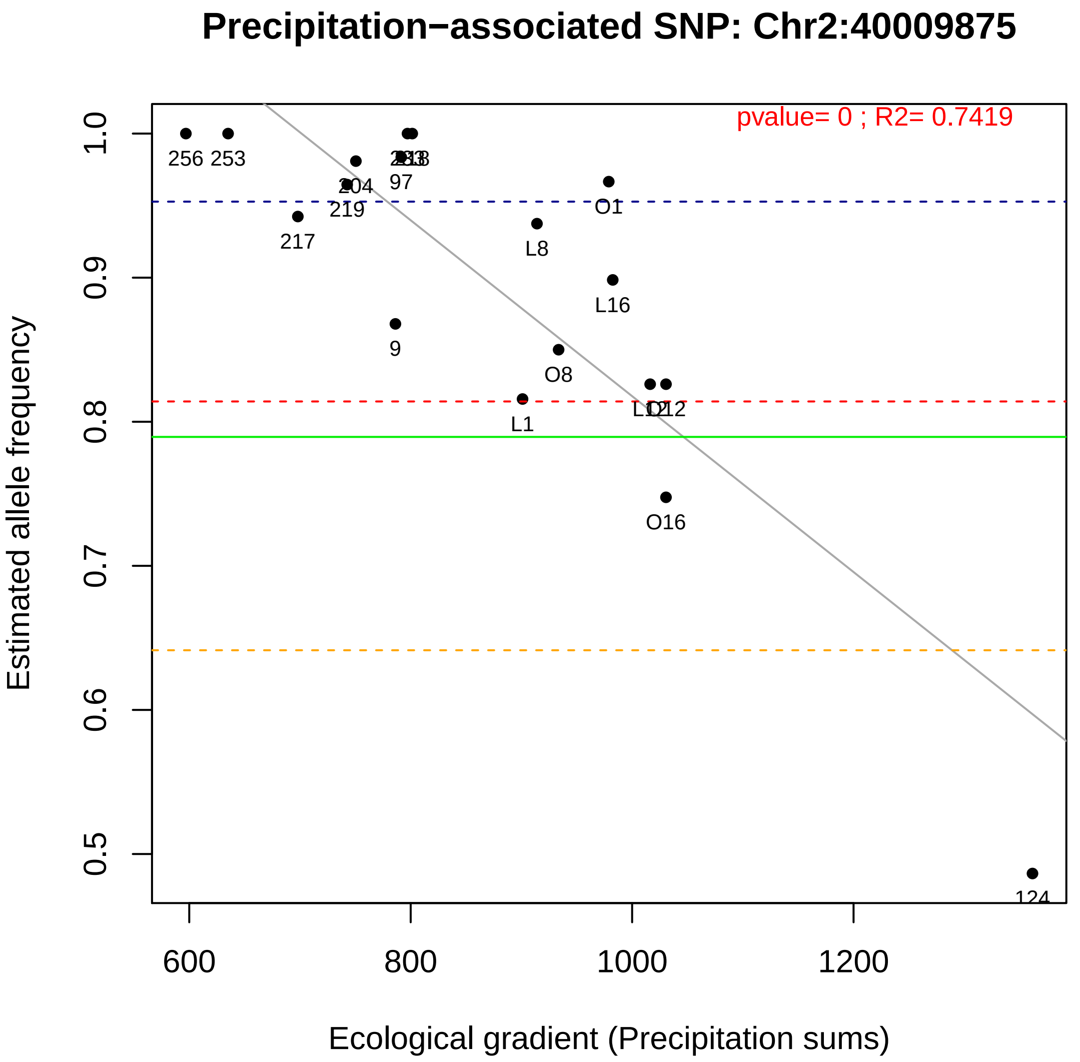
Precipitation−associated SNP: Chr2:40009875
pvalue= 0 ; R2= 0.7419
Estimated allele frequency
Ecological gradient (Precipitation sums)
1.0
0.9
0.8
0.7
0.6
0.5
600
800
1000
1200
256
253
204
219
97
217
9
L8
O1
L16
O8
L1
O16
124

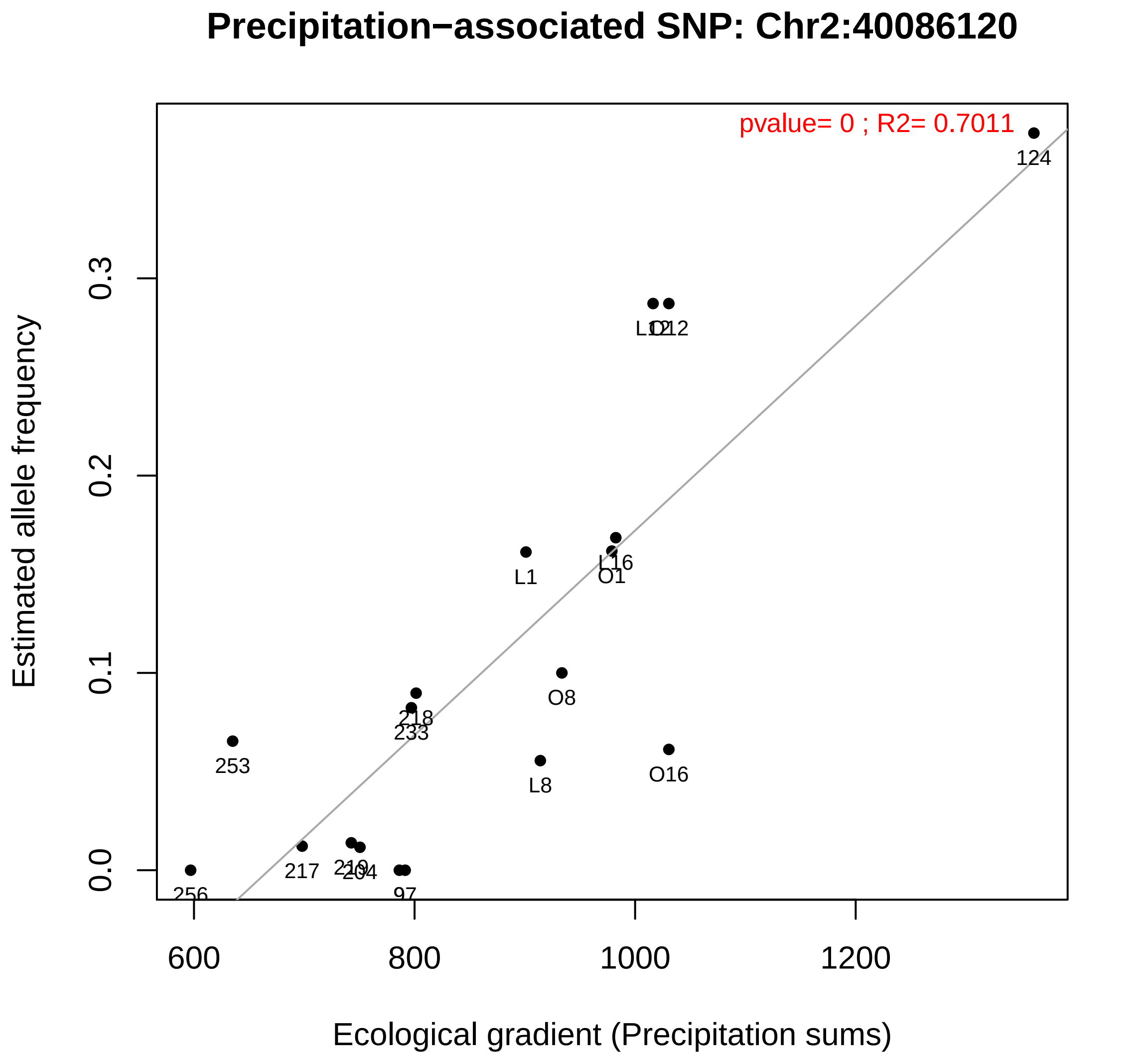

Precipitation−associated SNP: Chr2:40086120
pvalue= 0 ; R2= 0.7011
Estimated allele frequency
0.0
0.1
0.2
0.3
600
800
1000
1200
Ecological gradient (Precipitation sums)
124
L1
O1
L16
O8
L8
O16
218
233
253
217
204
97
256

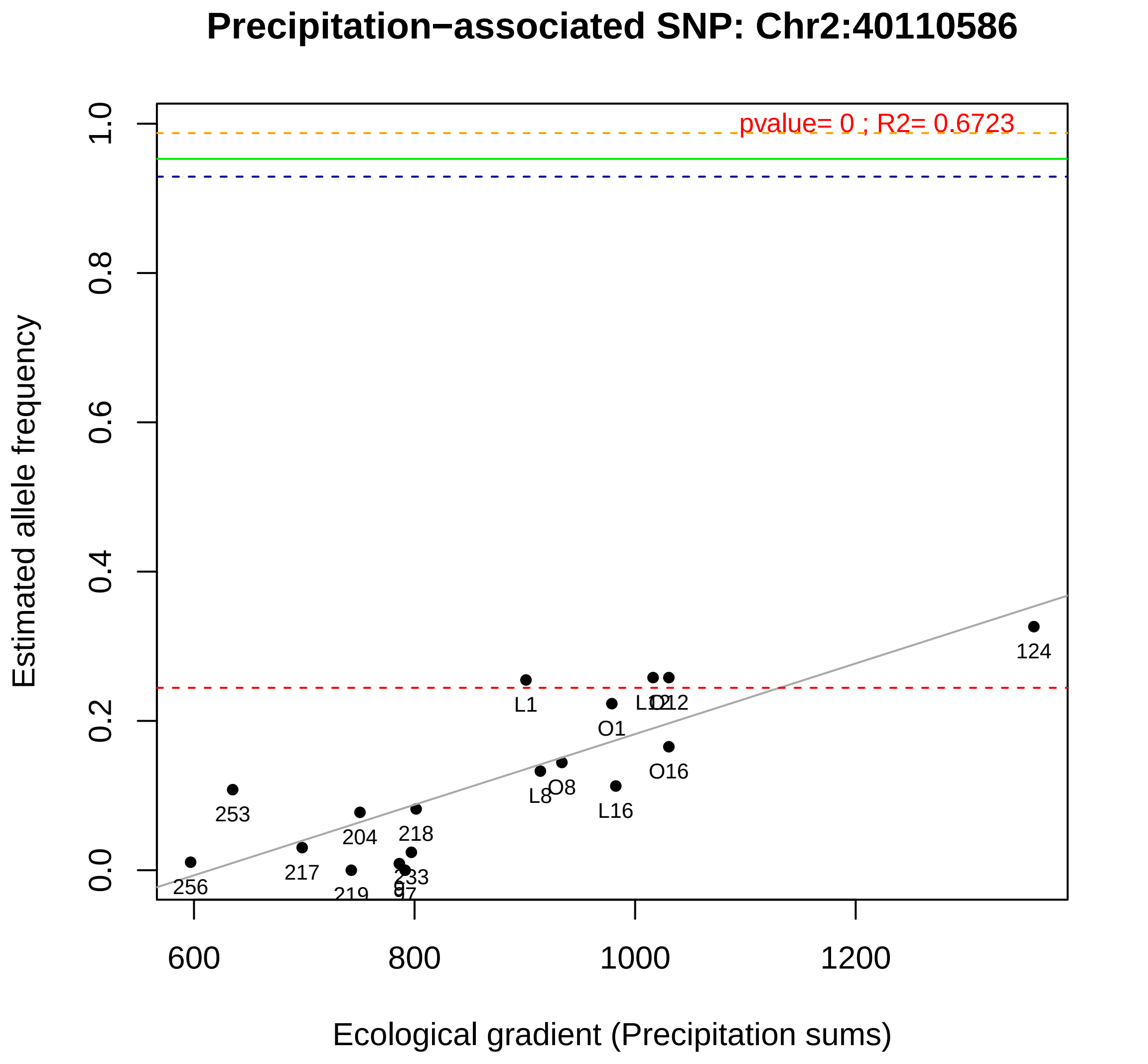

Precipitation−associated SNP: Chr2:40110586
pvalue= 0 ; R2= 0.6723
Estimated allele frequency
0.0
0.2
0.4
0.6
0.8
1.0
600
800
1000
1200
Ecological gradient (Precipitation sums)
124
L1
L12
O12
O1
O16
L8
O8
L16
253
204
218
217
233
256
219

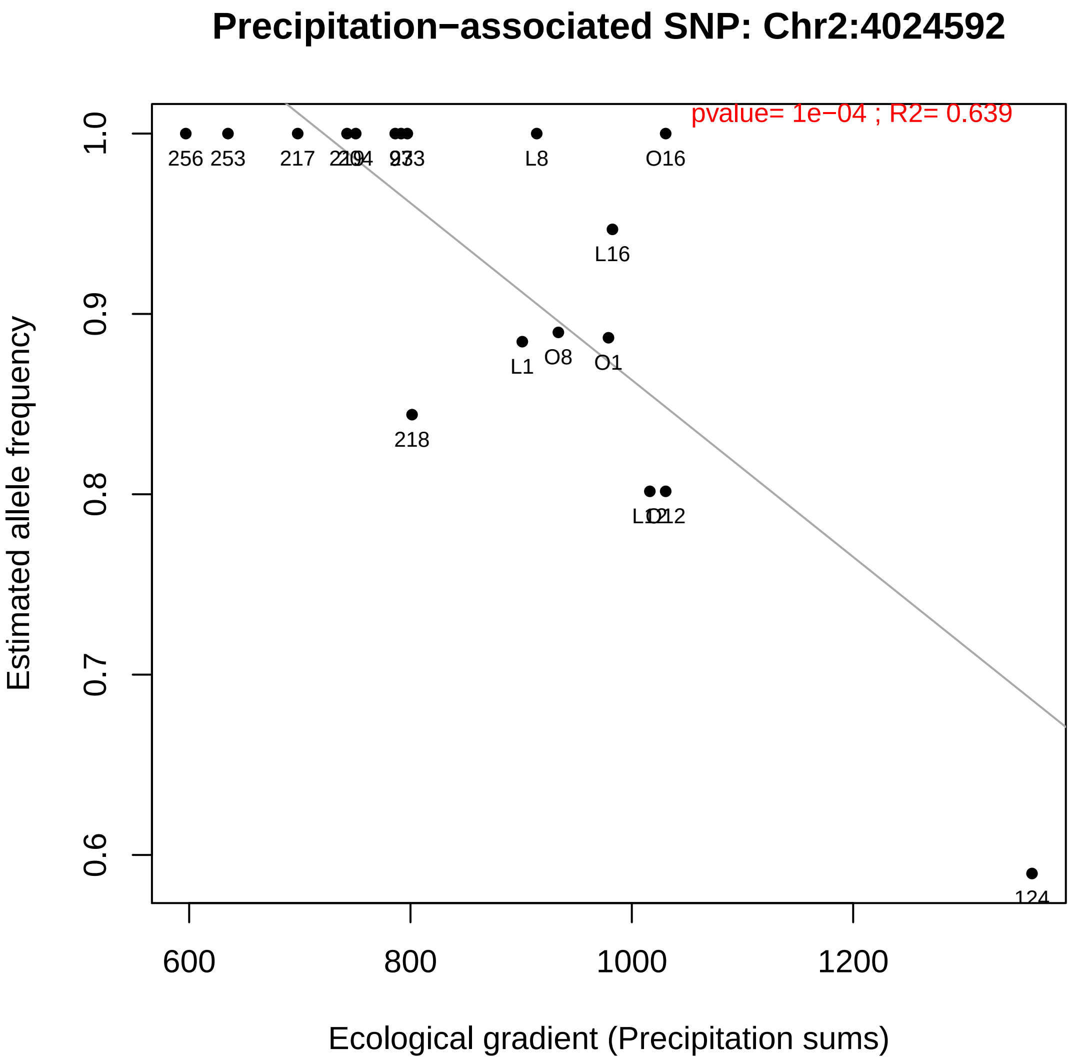

Precipitation−associated SNP: Chr2:4024592
pvalue= 1e−04 ; R2= 0.639
Estimated allele frequency
Ecological gradient (Precipitation sums)
1.0
0.9
0.8
0.7
0.6
600
800
1000
1200
256
253
217
L8
O16
L16
L1
O8
O1
218
124

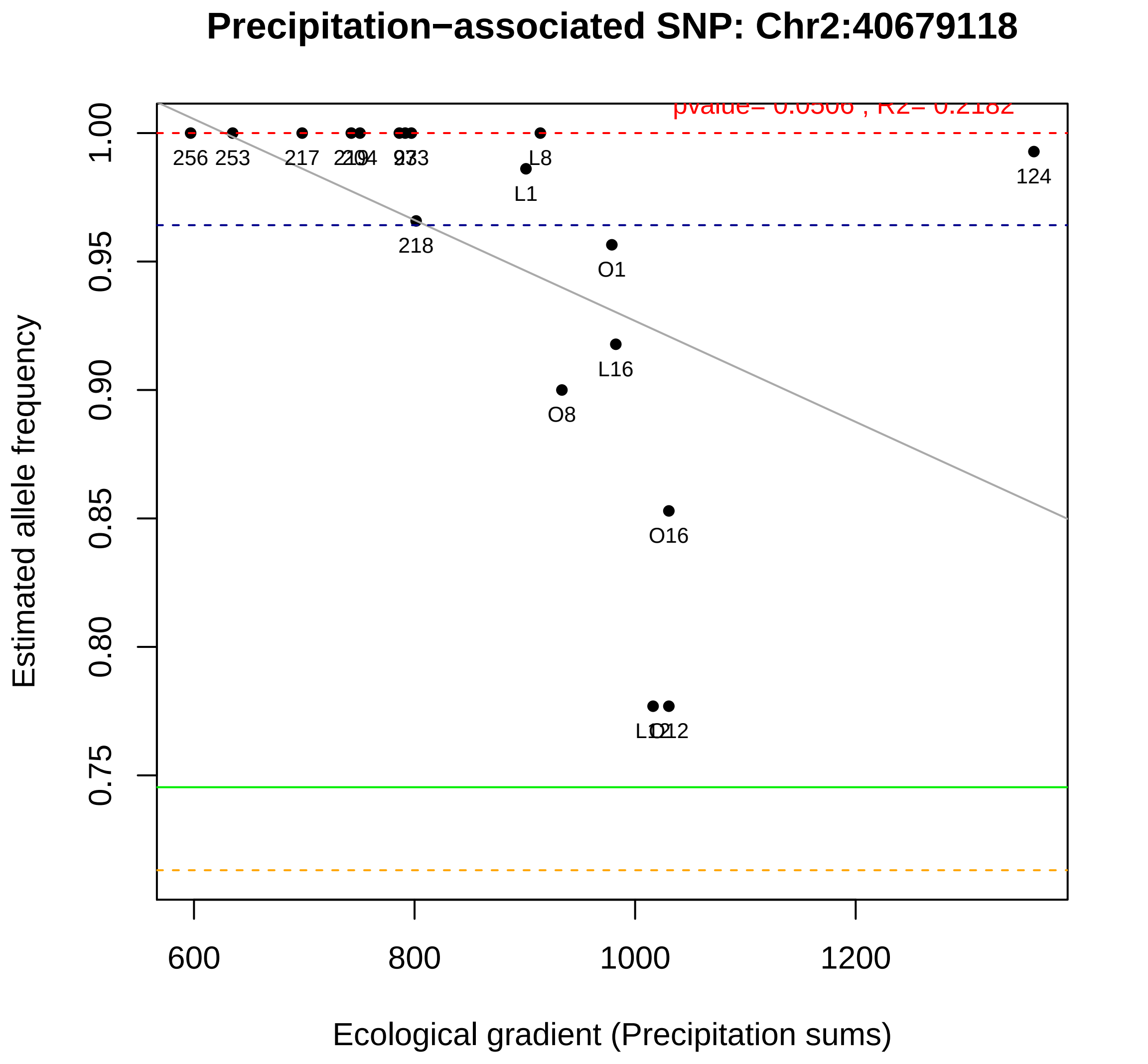

Precipitation−associated SNP: Chr2:40679118
pvalue= 0.0506 , R2= 0.2182
Estimated allele frequency
1.00
0.95
0.90
0.85
0.80
0.75
256
253
217
219
204
97
233
L8
L1
218
O1
L16
O8
O16
L12
O12
124
600
800
1000
1200
Ecological gradient (Precipitation sums)

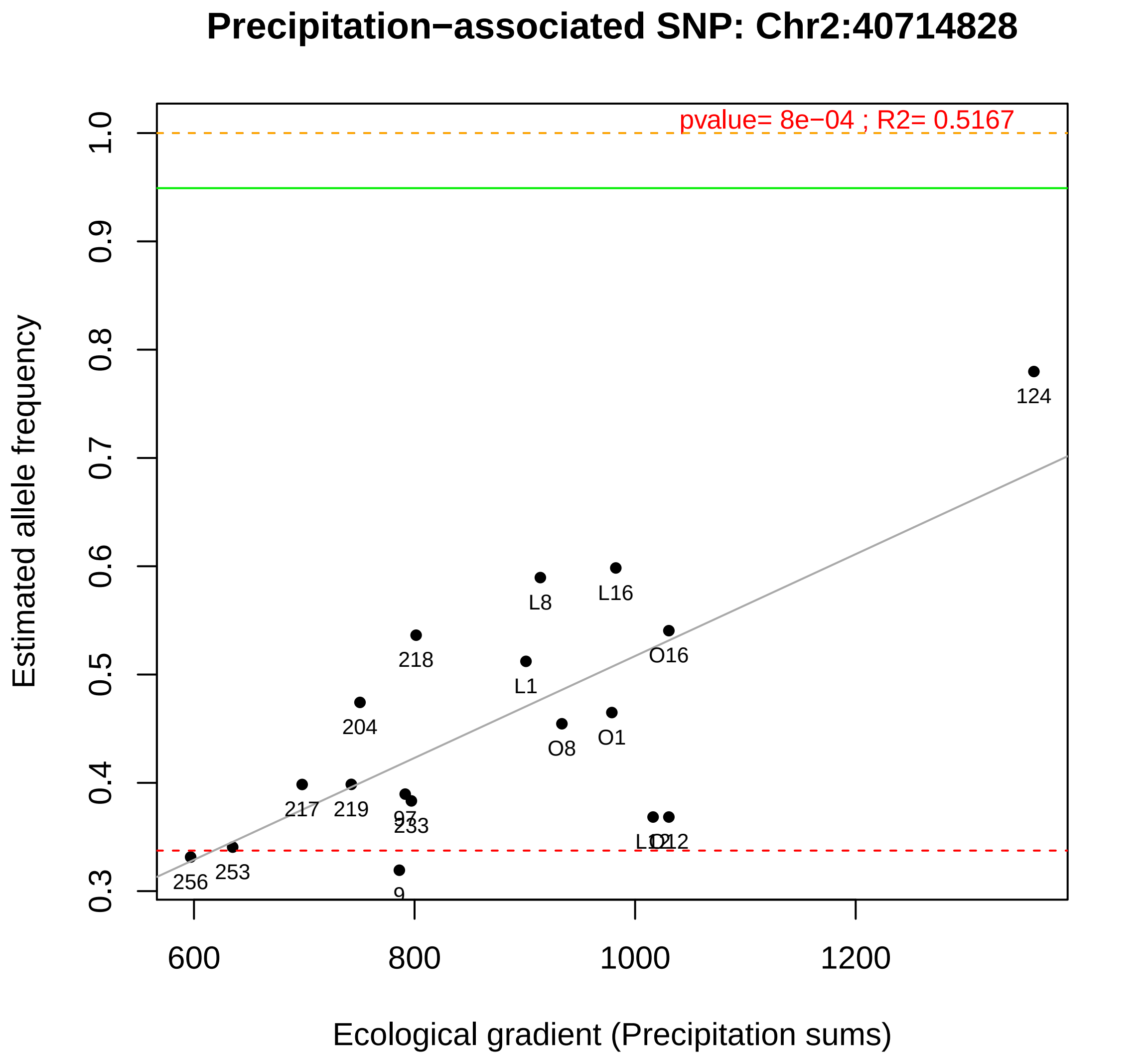

Precipitation−associated SNP: Chr2:40714828
pvalue= 8e−04 ; R2= 0.5167
Estimated allele frequency
0.3
0.4
0.5
0.6
0.7
0.8
0.9
1.0
600
800
1000
1200
Ecological gradient (Precipitation sums)
124
L8
L16
O16
218
L1
204
O8
O1
217
219
97
233
L10
O12
256
253
9

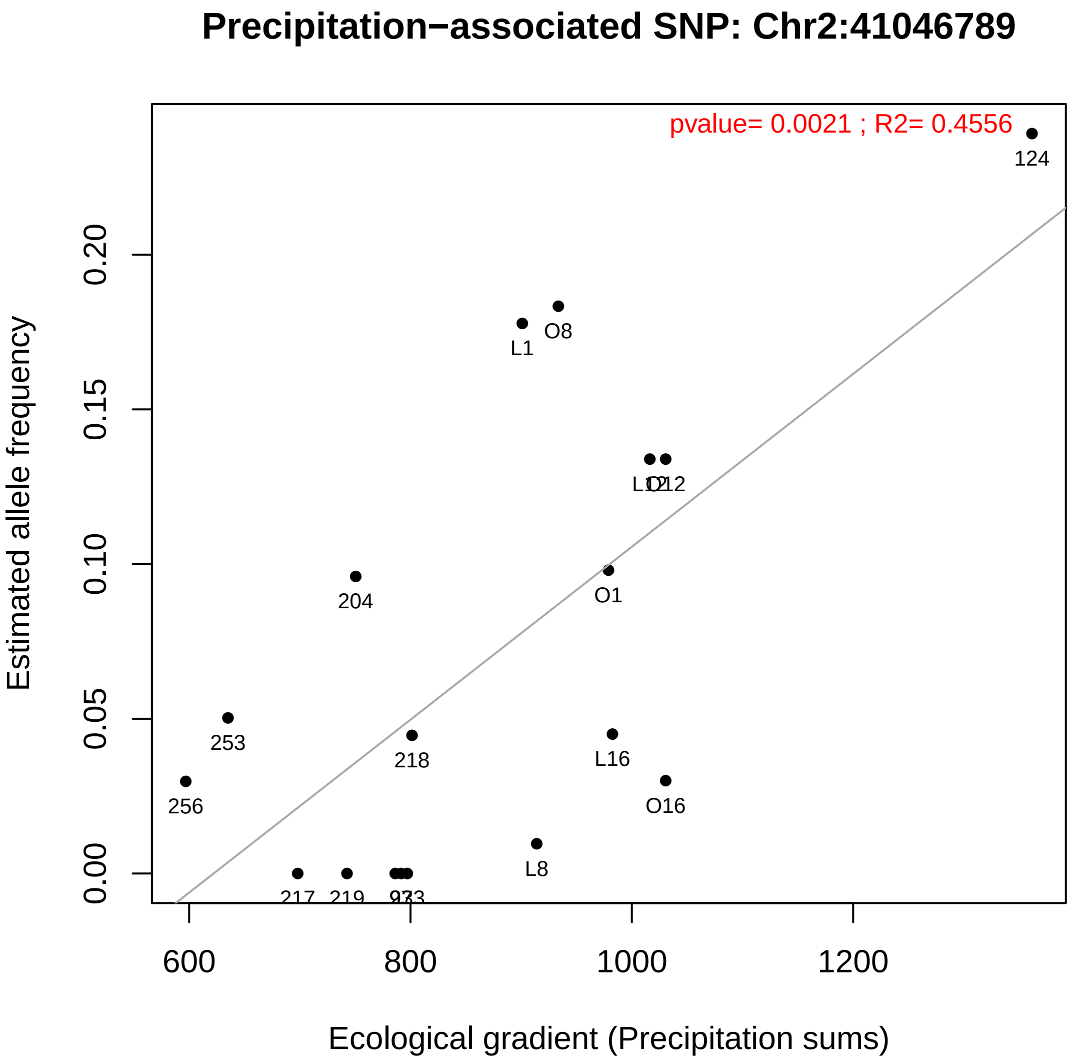
Precipitation−associated SNP: Chr2:41046789
pvalue= 0.0021 ; R2= 0.4556
Estimated allele frequency
0.00
0.05
0.10
0.15
0.20
600
800
1000
1200
Ecological gradient (Precipitation sums)
124
O8
L1
O12
O1
204
253
218
L16
256
O16
L8
217
219

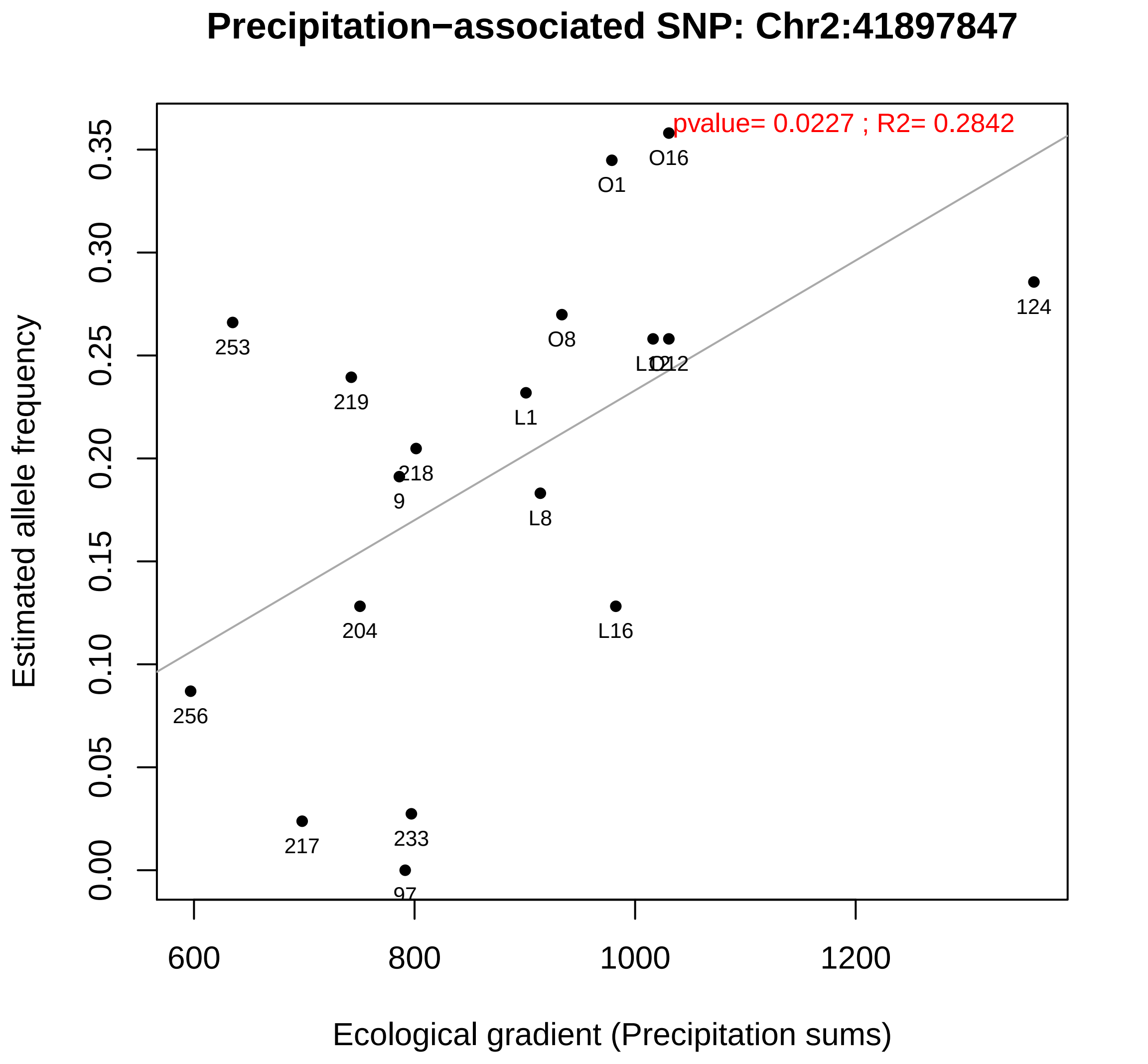

Precipitation−associated SNP: Chr2:41897847
pvalue= 0.0227 ; R2= 0.2842
Estimated allele frequency
Ecological gradient (Precipitation sums)
0.00
0.05
0.10
0.15
0.20
0.25
0.30
0.35
600
800
1000
1200
O16
O1
124
O8
253
219
L1
218
9
L8
204
L16
256
217
233
97

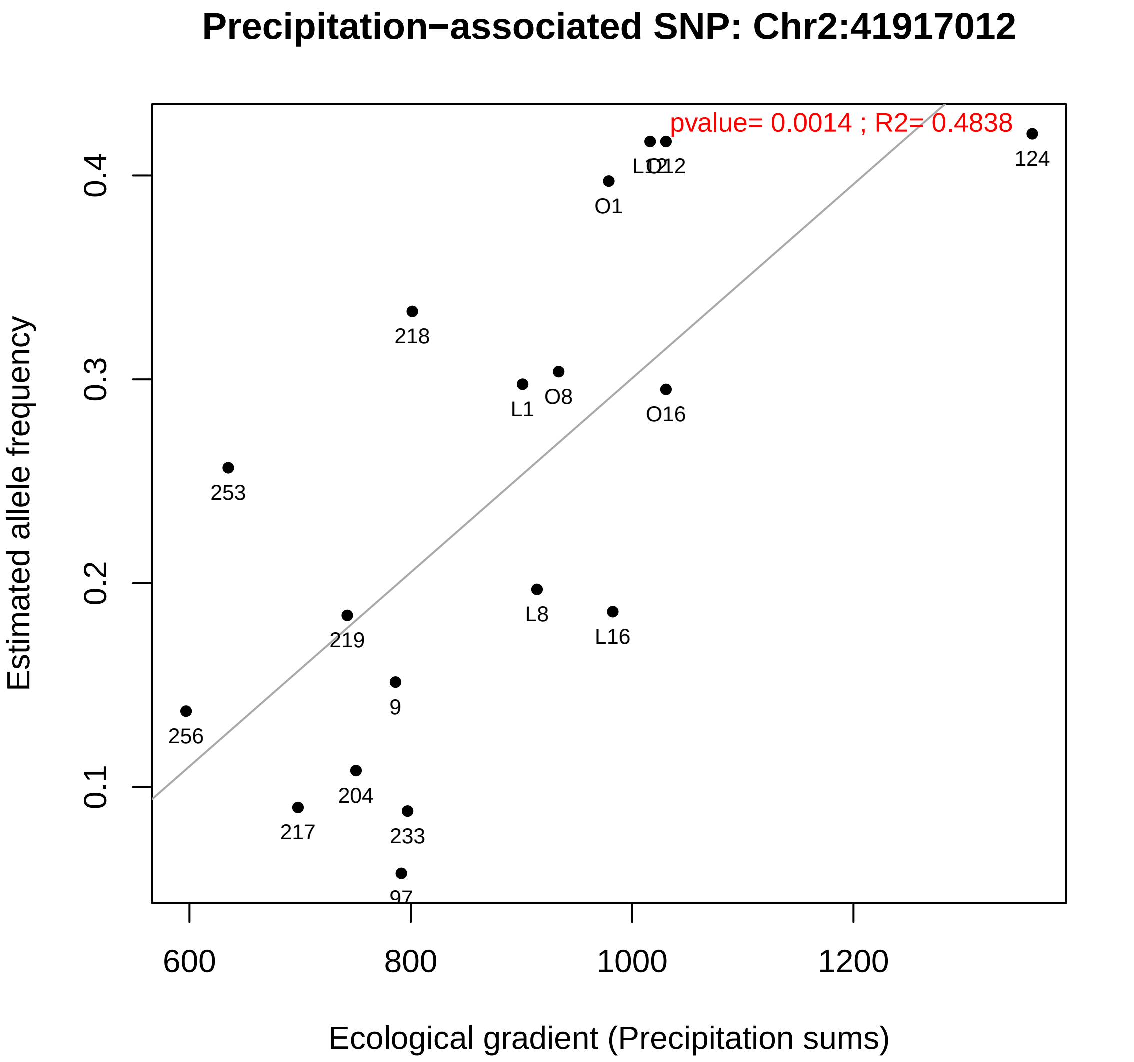
Precipitation−associated SNP: Chr2:41917012
pvalue= 0.0014 ; R2= 0.4838
124
L12
O12
O1
218
O8
L1
O16
253
219
L8
L16
9
256
204
217
233
97
0.1
0.2
0.3
0.4
600
800
1000
1200
Estimated allele frequency
Ecological gradient (Precipitation sums)

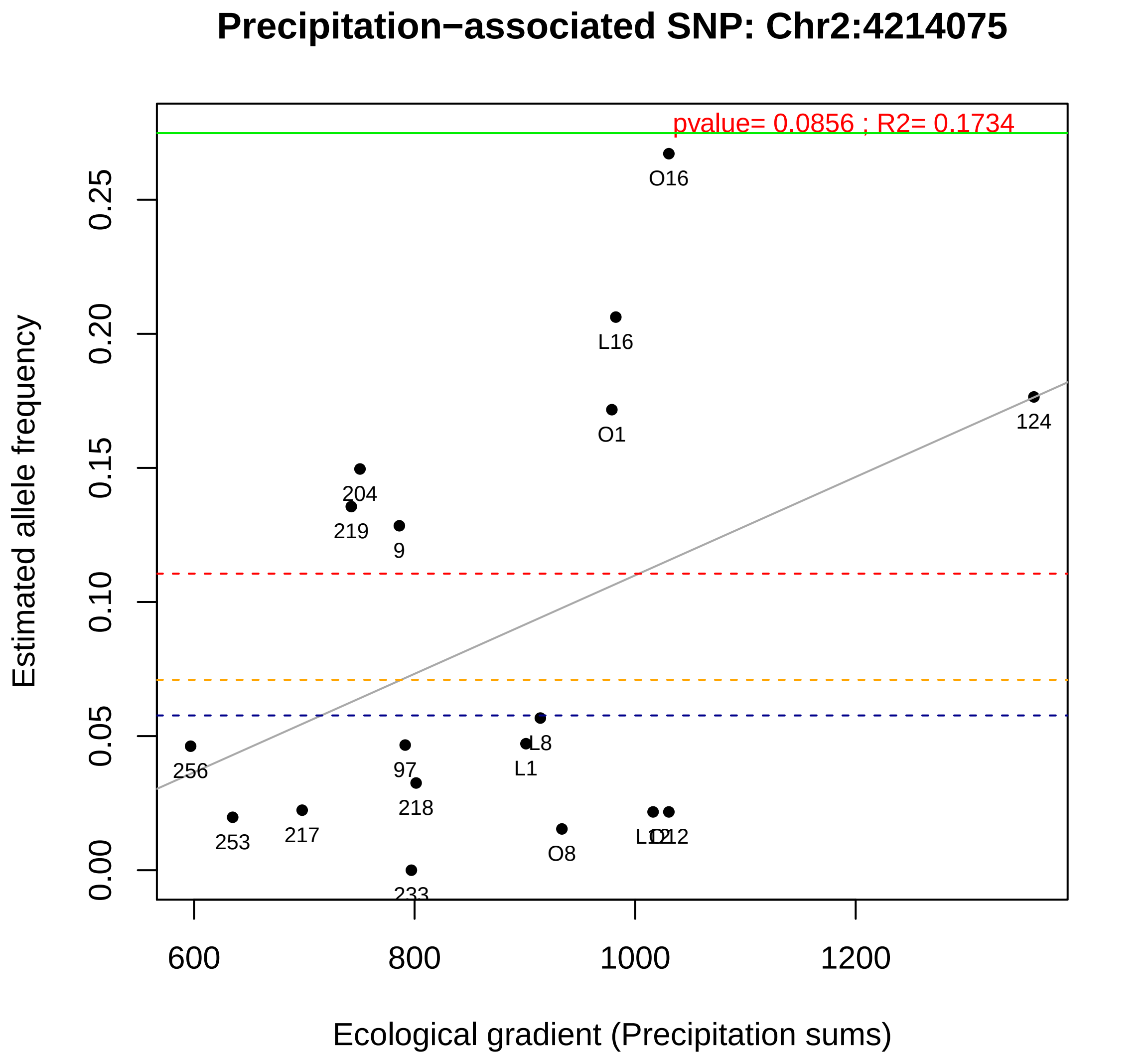
Precipitation−associated SNP: Chr2:4214075
pvalue= 0.0856 ; R2= 0.1734
Estimated allele frequency
Ecological gradient (Precipitation sums)
0.00
0.05
0.10
0.15
0.20
0.25
600
800
1000
1200
O16
L16
O1
124
204
219
9
256
97
L8
L1
218
253
217
O8
L10
O12
233

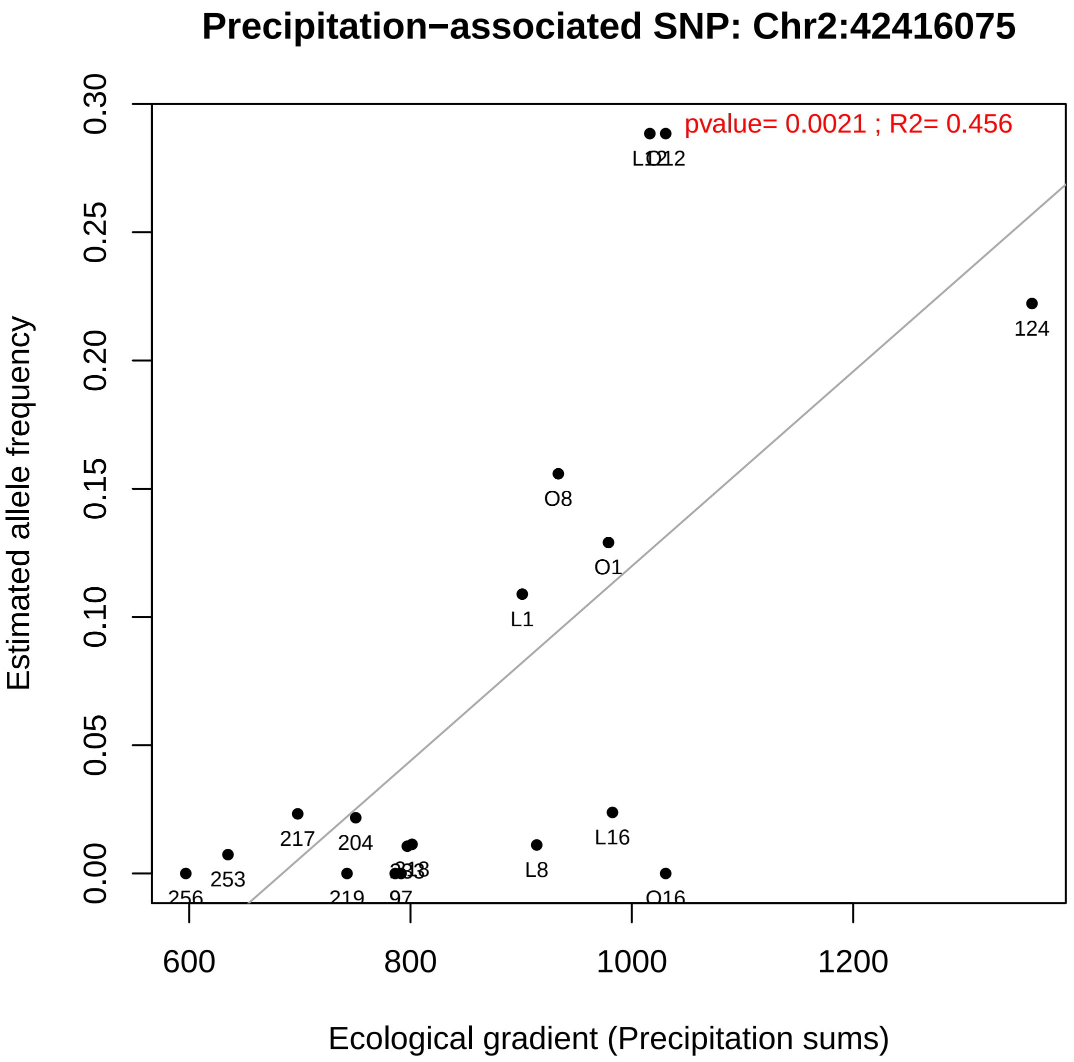
Precipitation−associated SNP: Chr2:42416075
pvalue= 0.0021 ; R2= 0.456
Estimated allele frequency
0.00
0.05
0.10
0.15
0.20
0.25
0.30
Ecological gradient (Precipitation sums)
600
800
1000
1200
124
O8
O1
L1
217
204
253
219
97
L8
L16
O16

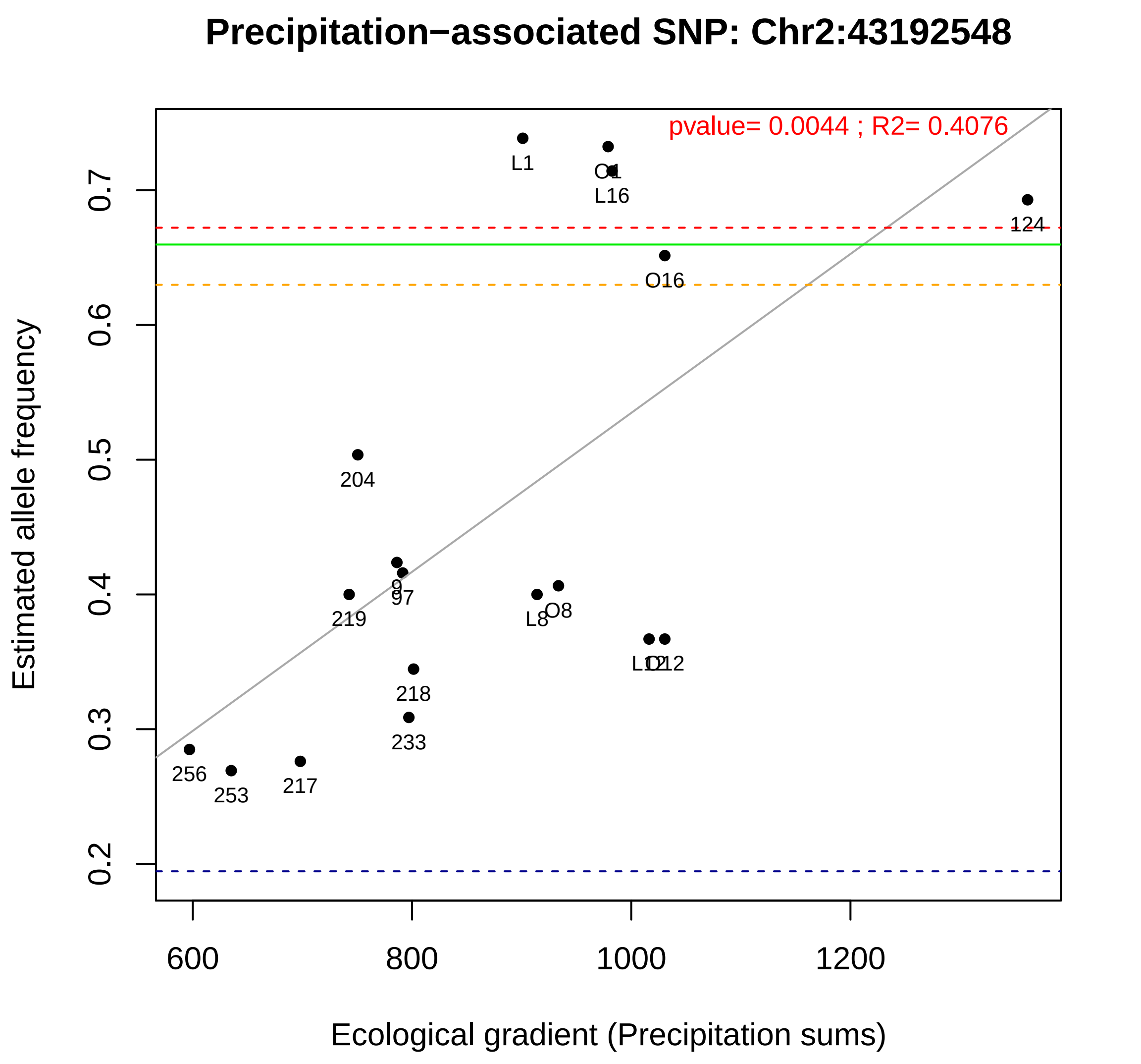

Precipitation−associated SNP: Chr2:43192548
pvalue= 0.0044 ; R2= 0.4076
Estimated allele frequency
Ecological gradient (Precipitation sums)
0.2
0.3
0.4
0.5
0.6
0.7
600
800
1000
1200
L1
O1
L16
124
O16
204
9
97
219
L8
O8
L12
O12
218
233
256
253
217

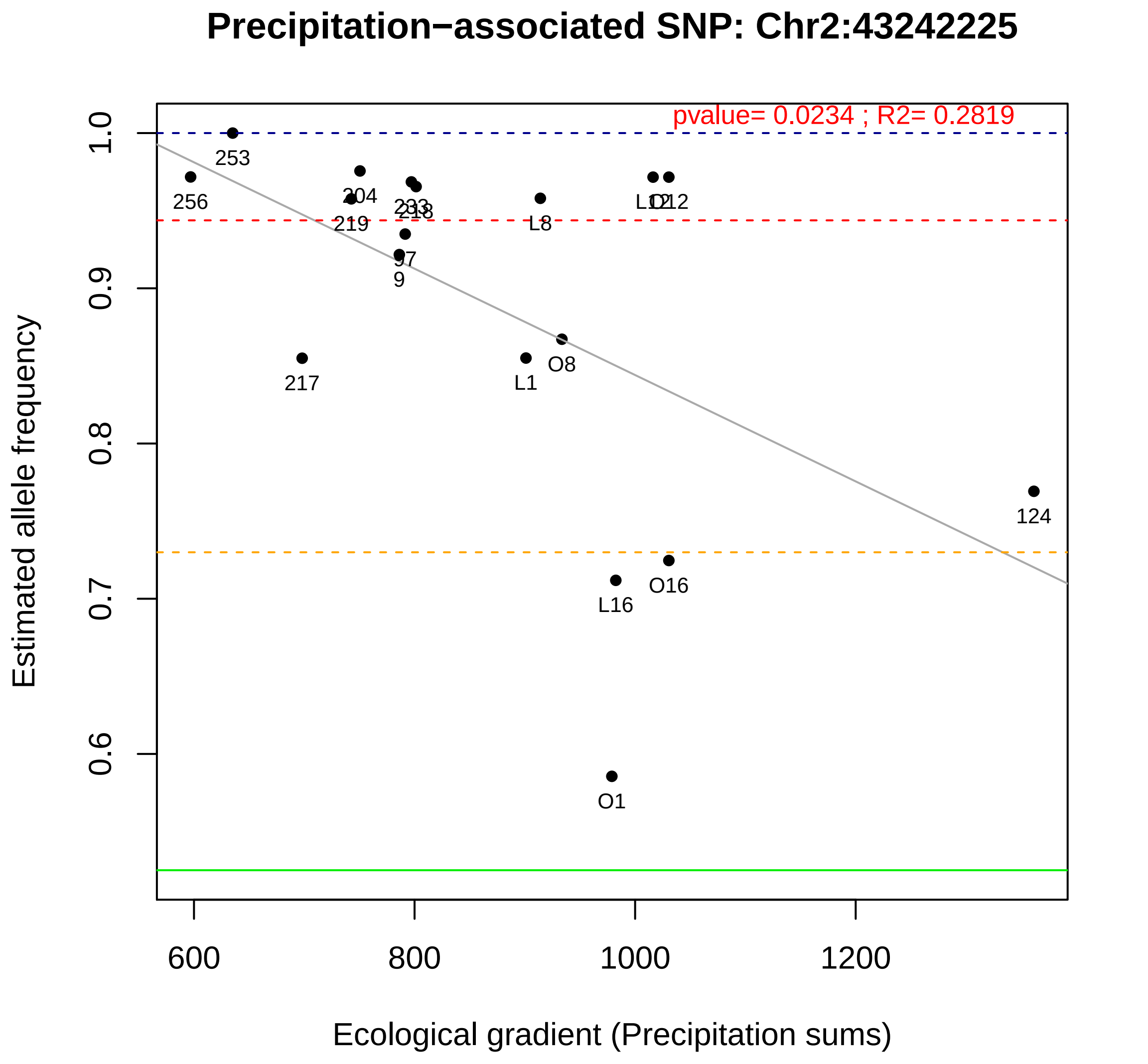
Precipitation−associated SNP: Chr2:43242225
pvalue= 0.0234 ; R2= 0.2819
Estimated allele frequency
Ecological gradient (Precipitation sums)
1.0
0.9
0.8
0.7
0.6
600
800
1000
1200
253
256
204
219
233
218
97
9
L8
L12
O12
217
L1
O8
124
O16
L16
O1

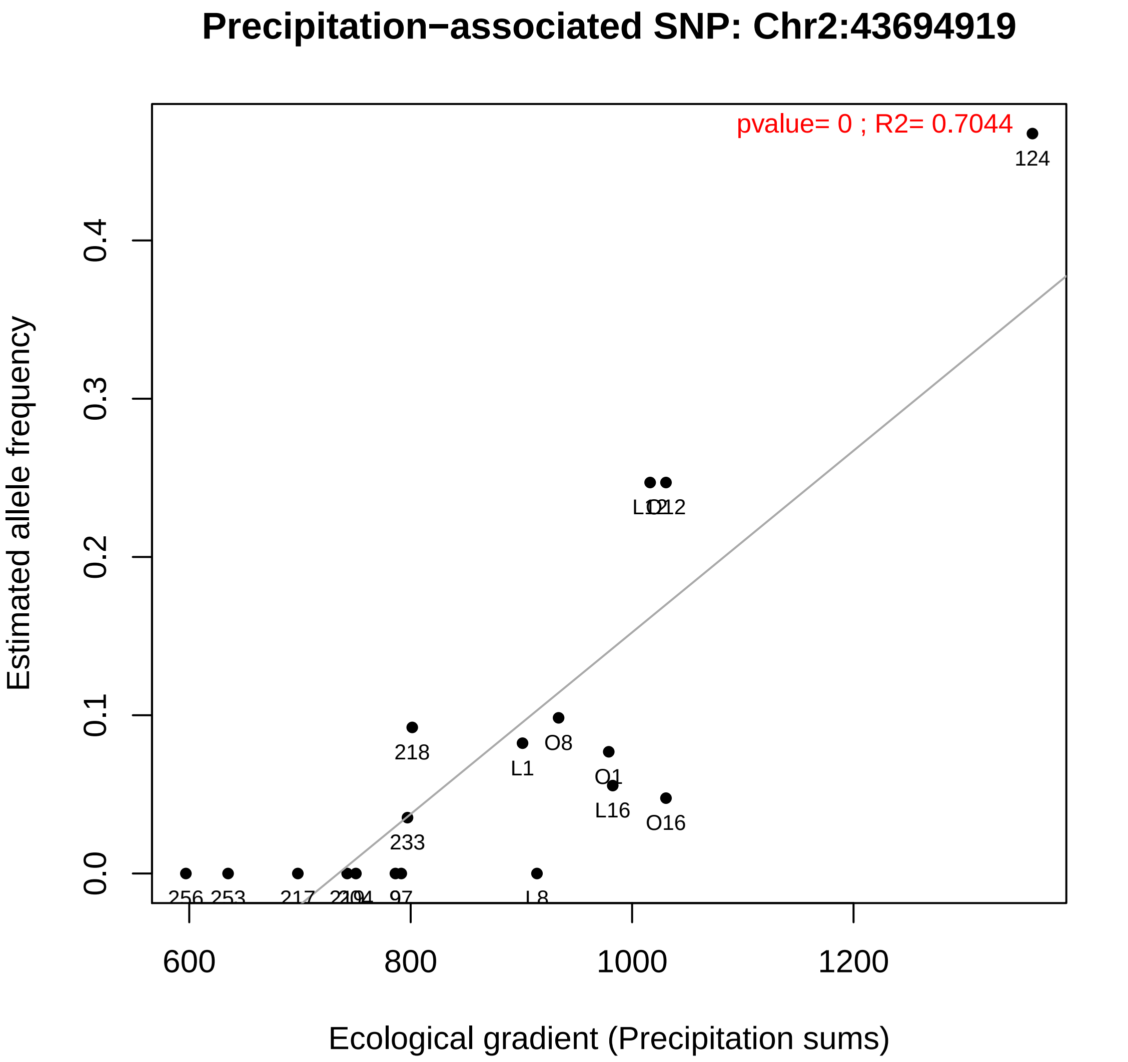
Precipitation−associated SNP: Chr2:43694919
pvalue= 0 ; R2= 0.7044
124
L12
O12
218
O8
L1
O1
L16
O16
233
256
253
217
L8
97
0.0
0.1
0.2
0.3
0.4
600
800
1000
1200
Estimated allele frequency
Ecological gradient (Precipitation sums)

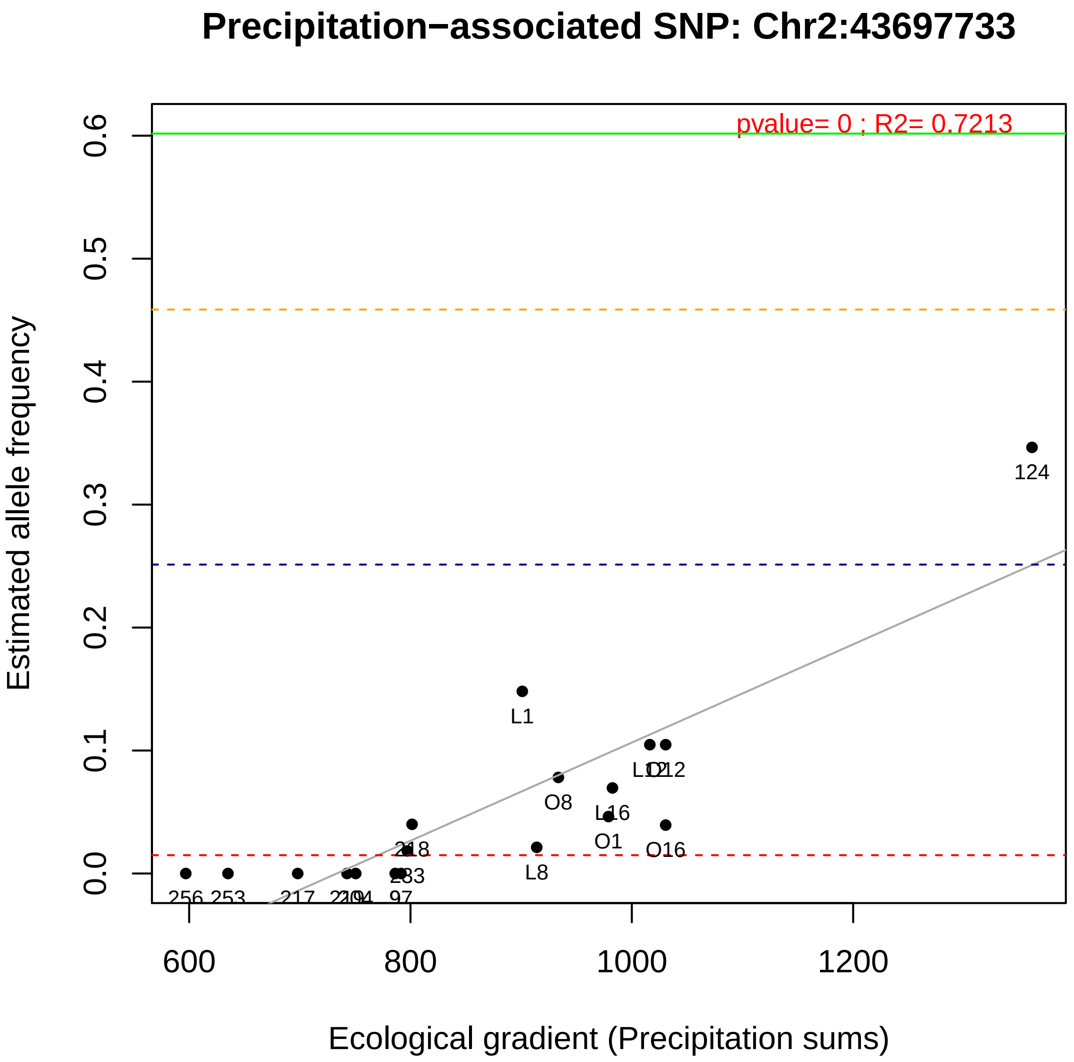
Precipitation−associated SNP: Chr2:43697733
pvalue= 0 ; R2= 0.7213
Estimated allele frequency
Ecological gradient (Precipitation sums)
0.0
0.1
0.2
0.3
0.4
0.5
0.6
600
800
1000
1200
124
L1
O8
L16
O1
O16
L8
218
283
97
256
253
217

**Precipitation−associated SNP: Chr2:43701032**

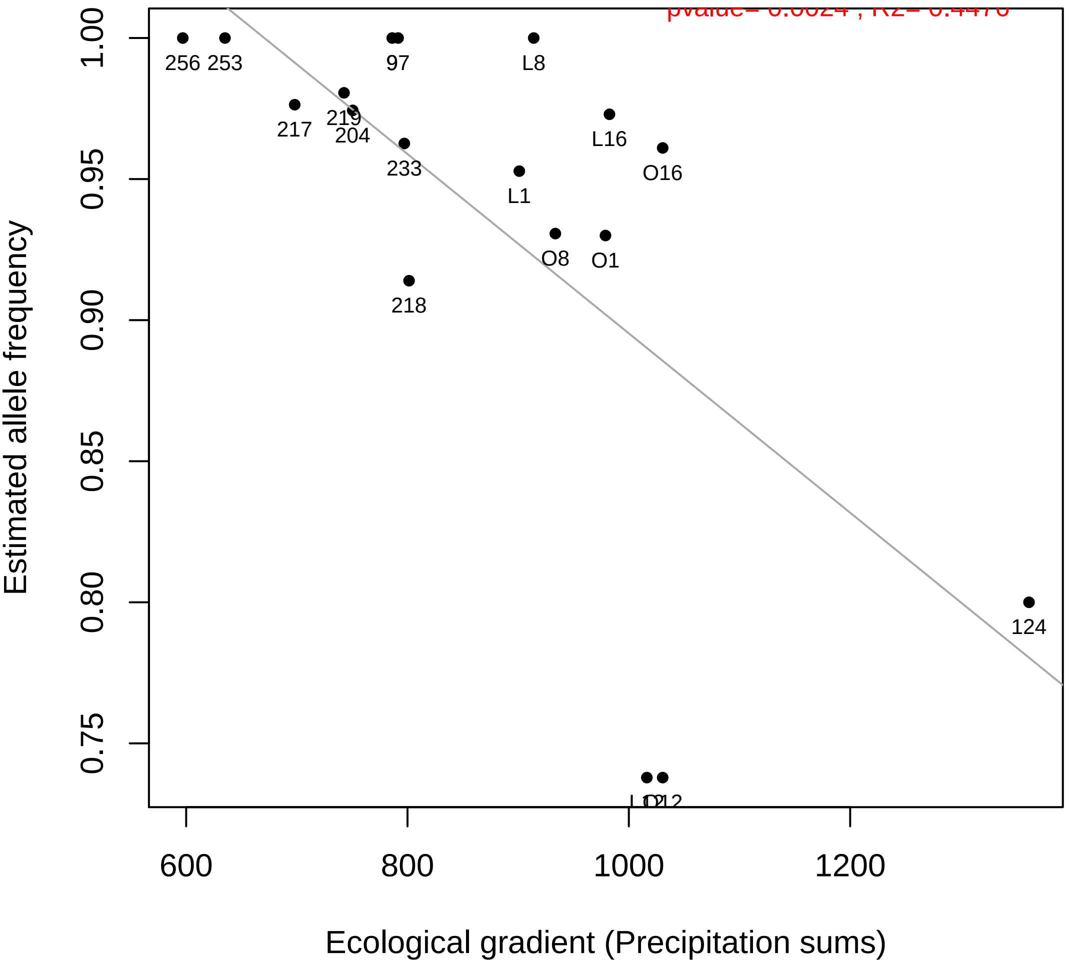

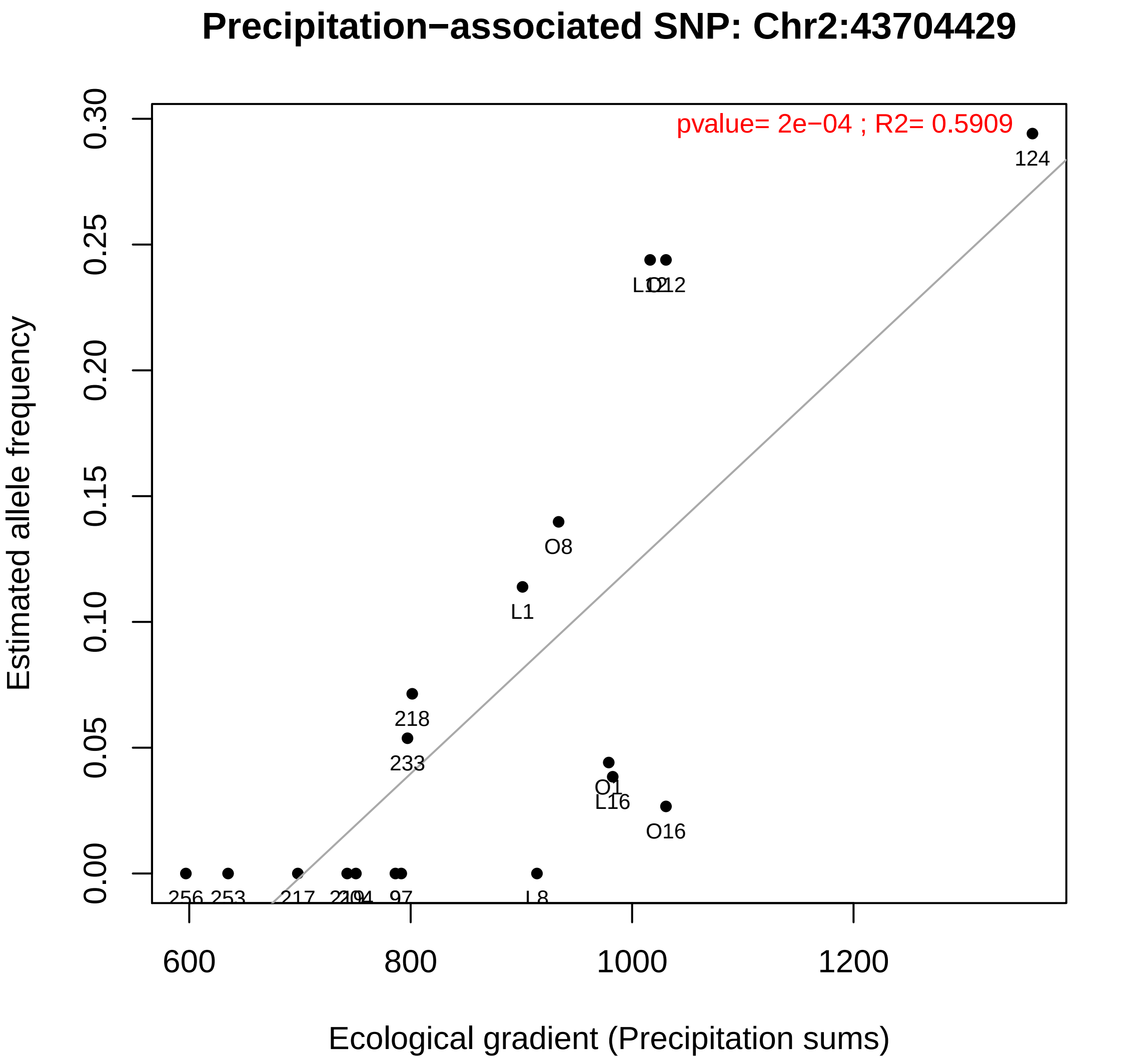

Precipitation−associated SNP: Chr2:43704429
pvalue= 2e−04 ; R2= 0.5909
Estimated allele frequency
0.00
0.05
0.10
0.15
0.20
0.25
0.30
Ecological gradient (Precipitation sums)
600
800
1000
1200
124
O8
L1
218
233
O1
L16
O16

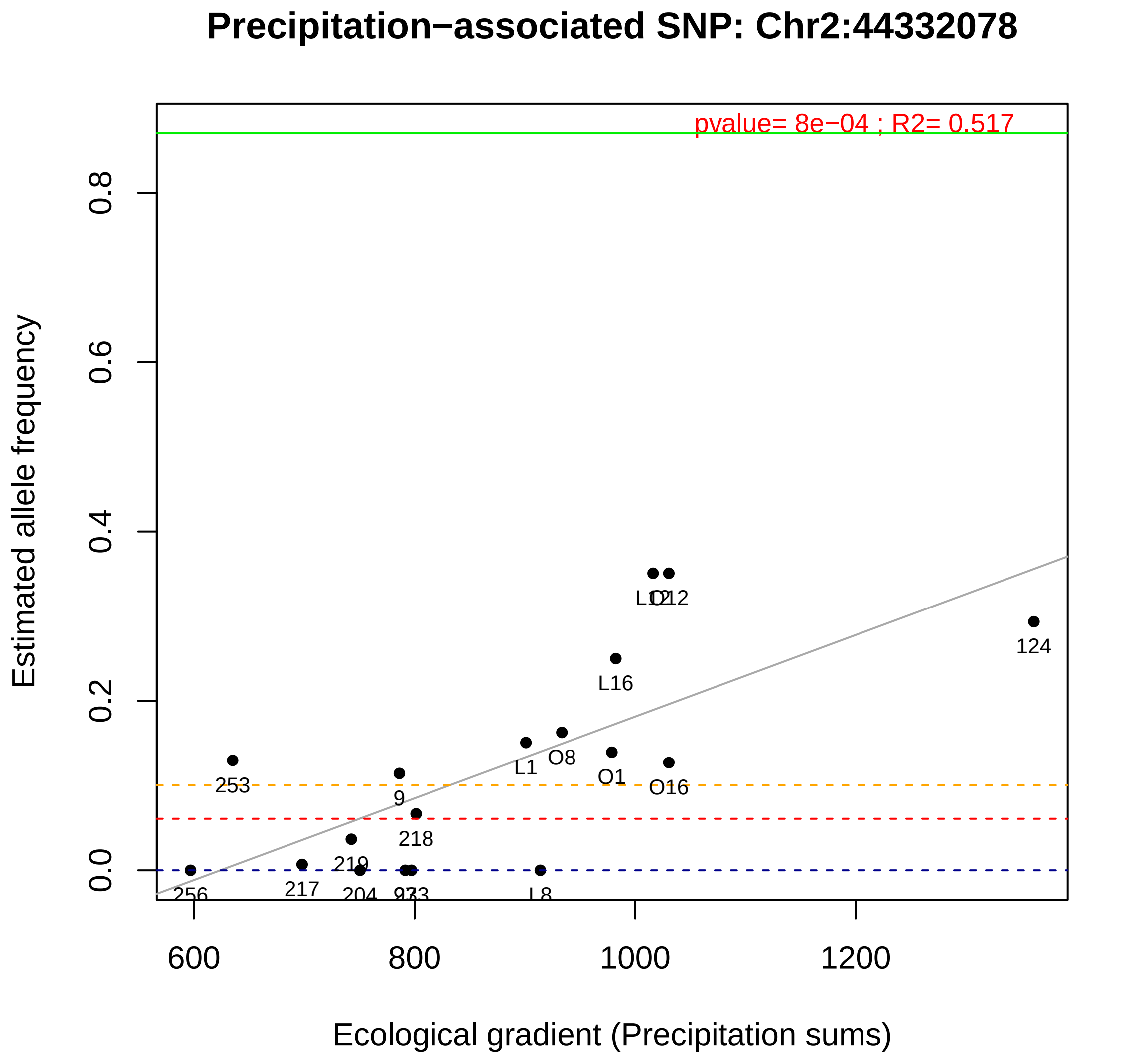
Precipitation−associated SNP: Chr2:44332078
pvalue= 8e−04 ; R2= 0.517
Estimated allele frequency
0.0
0.2
0.4
0.6
0.8
600
800
1000
1200
Ecological gradient (Precipitation sums)
253
9
218
219
217
L1
O8
O1
O16
L16
124

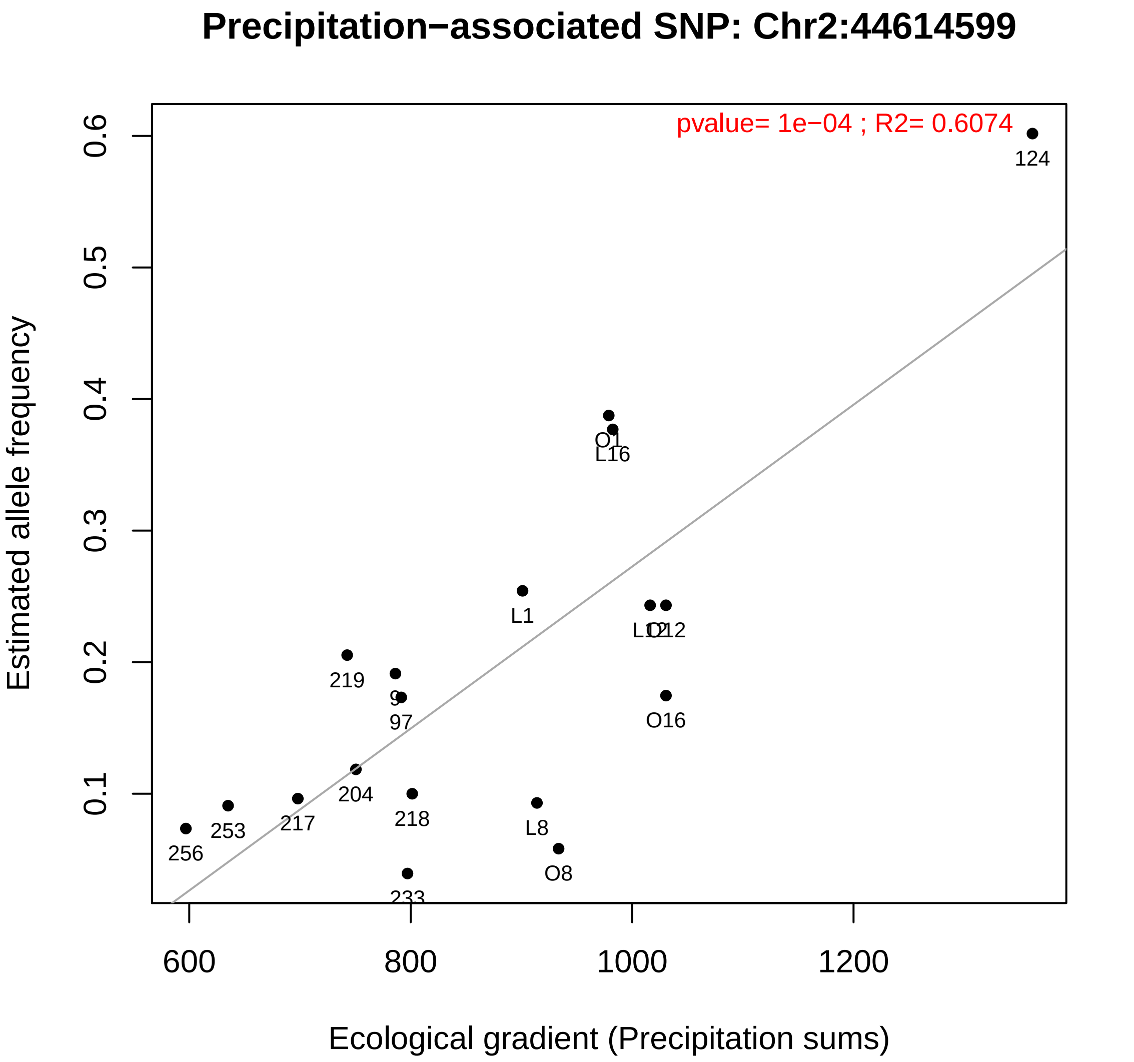
Precipitation−associated SNP: Chr2:44614599
pvalue= 1e−04 ; R2= 0.6074
Estimated allele frequency
Ecological gradient (Precipitation sums)
0.1
0.2
0.3
0.4
0.5
0.6
600
800
1000
1200
124
O1
L16
L1
L12
O12
219
9
97
O16
204
218
217
253
256
L8
O8
233

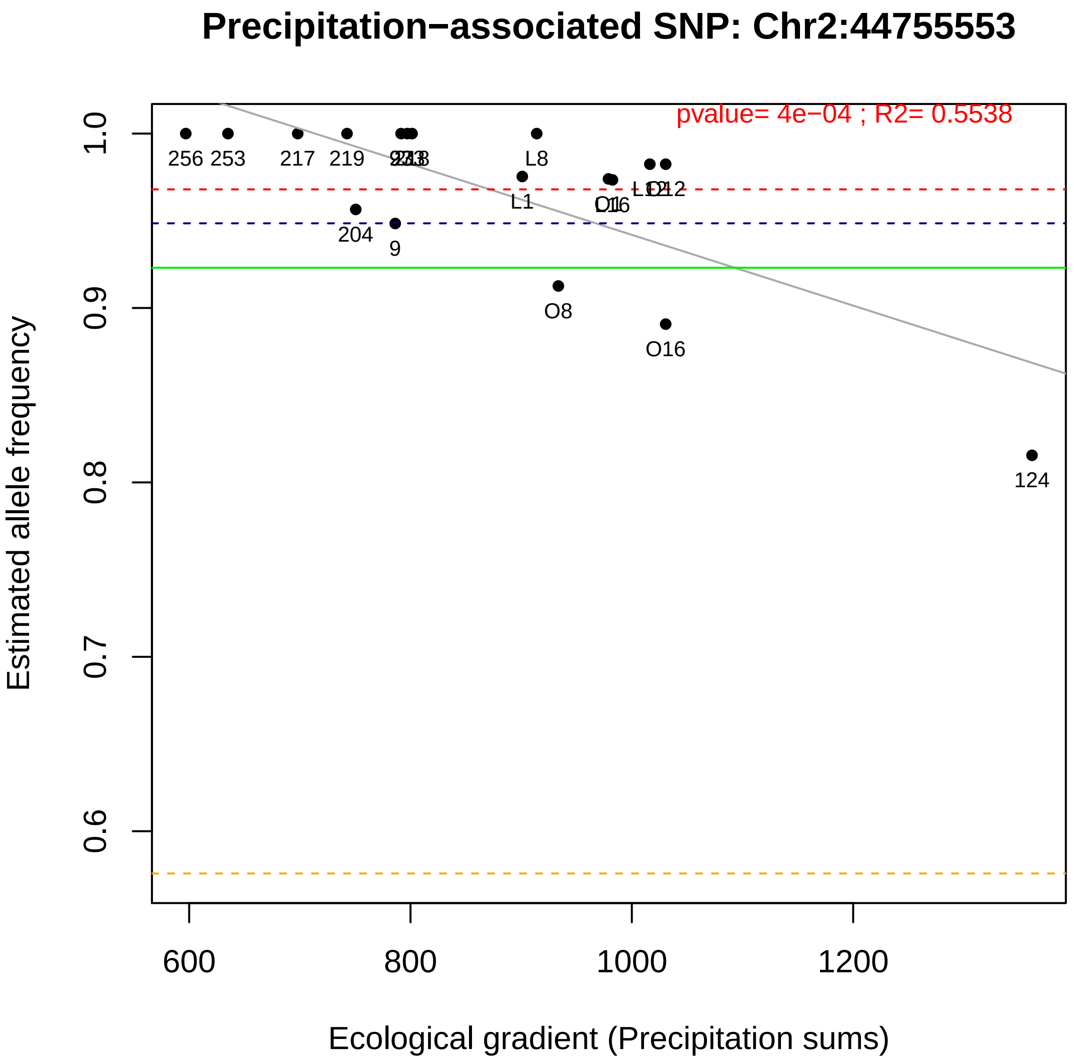

Precipitation−associated SNP: Chr2:44755553
pvalue= 4e−04 ; R2= 0.5538
Estimated allele frequency
Ecological gradient (Precipitation sums)
0.6
0.7
0.8
0.9
1.0
600
800
1000
1200
256
253
217
219
L8
L1
204
9
O8
O16
O12
124

**Precipitation−associated SNP: Chr2:47185998**

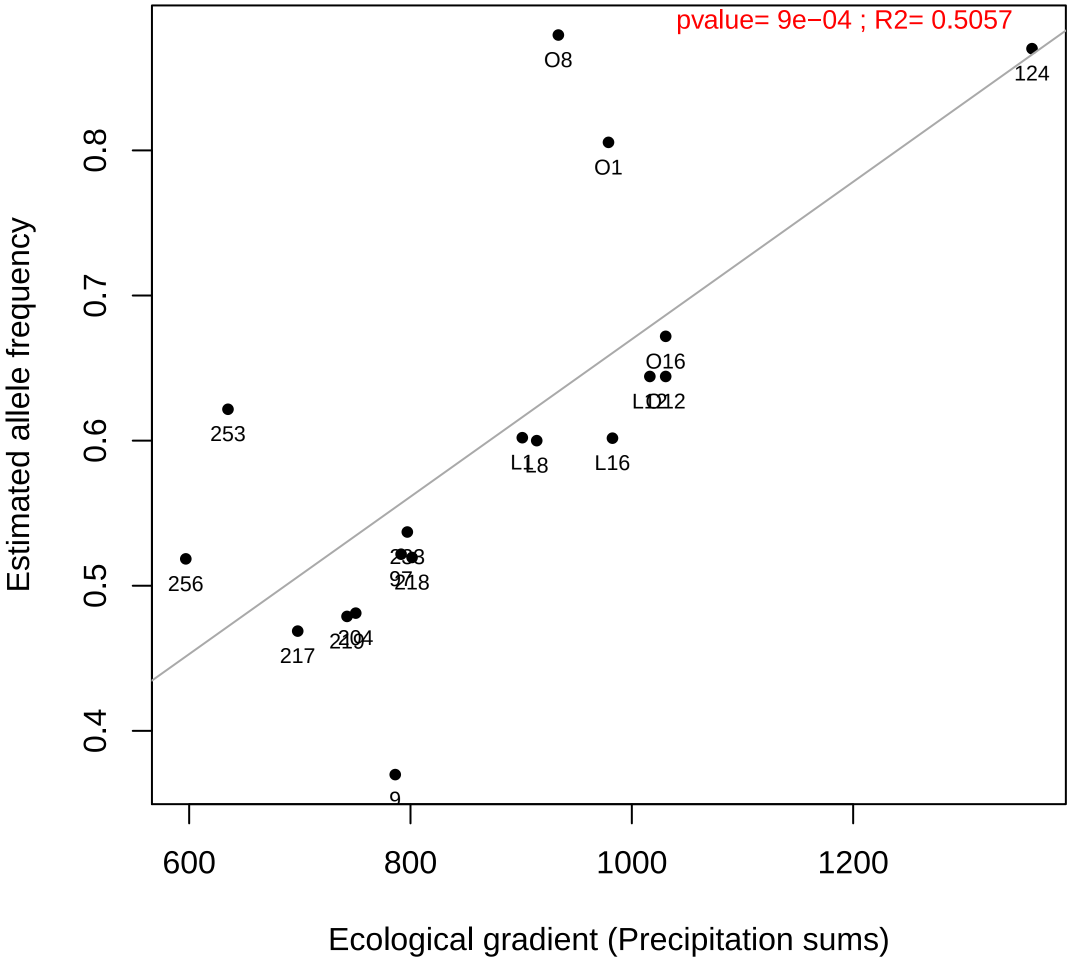

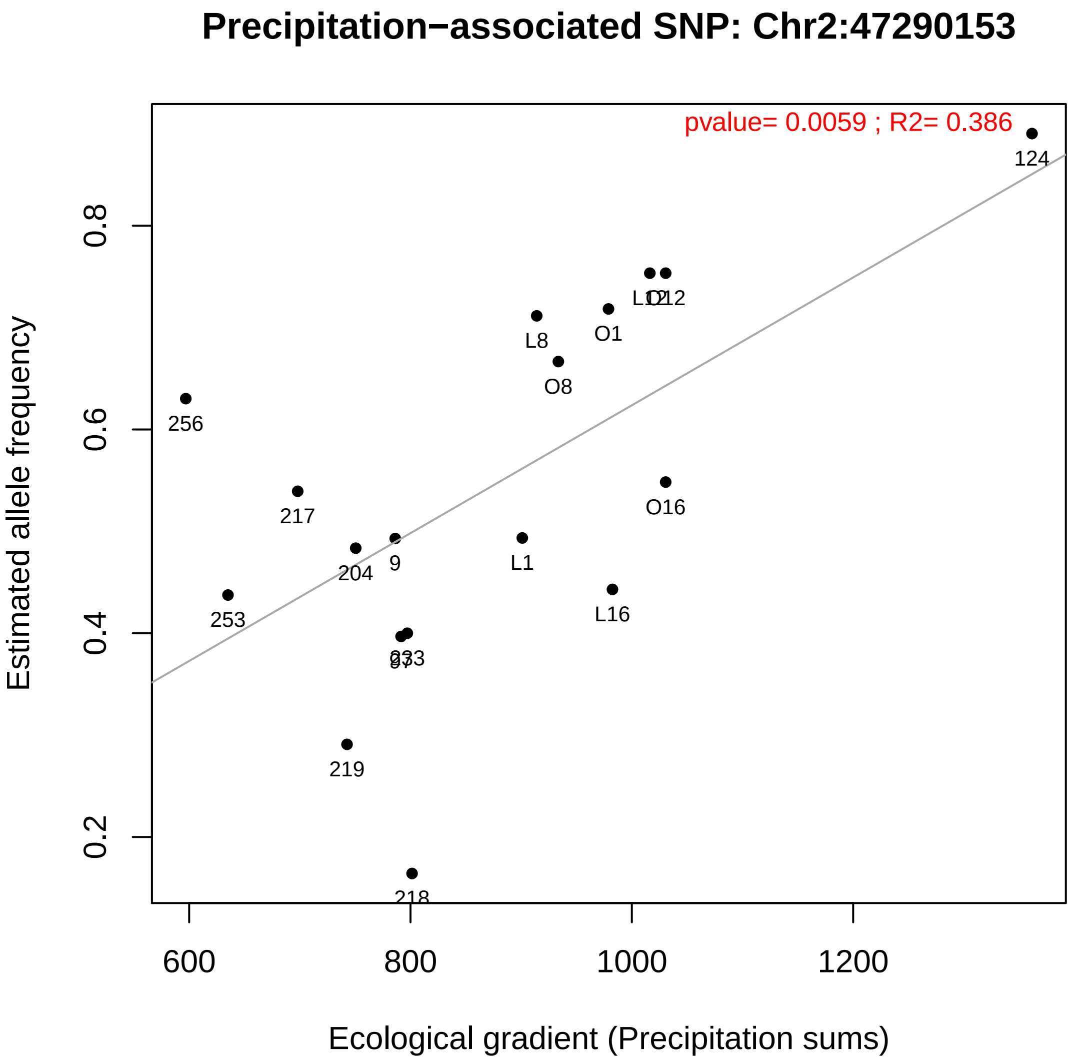
Precipitation−associated SNP: Chr2:47290153
pvalue= 0.0059 ; R2= 0.386
Estimated allele frequency
Ecological gradient (Precipitation sums)
0.2
0.4
0.6
0.8
600
800
1000
1200
124
O12
L8
O1
O8
256
O16
217
9
L1
204
253
L16
233
219
218

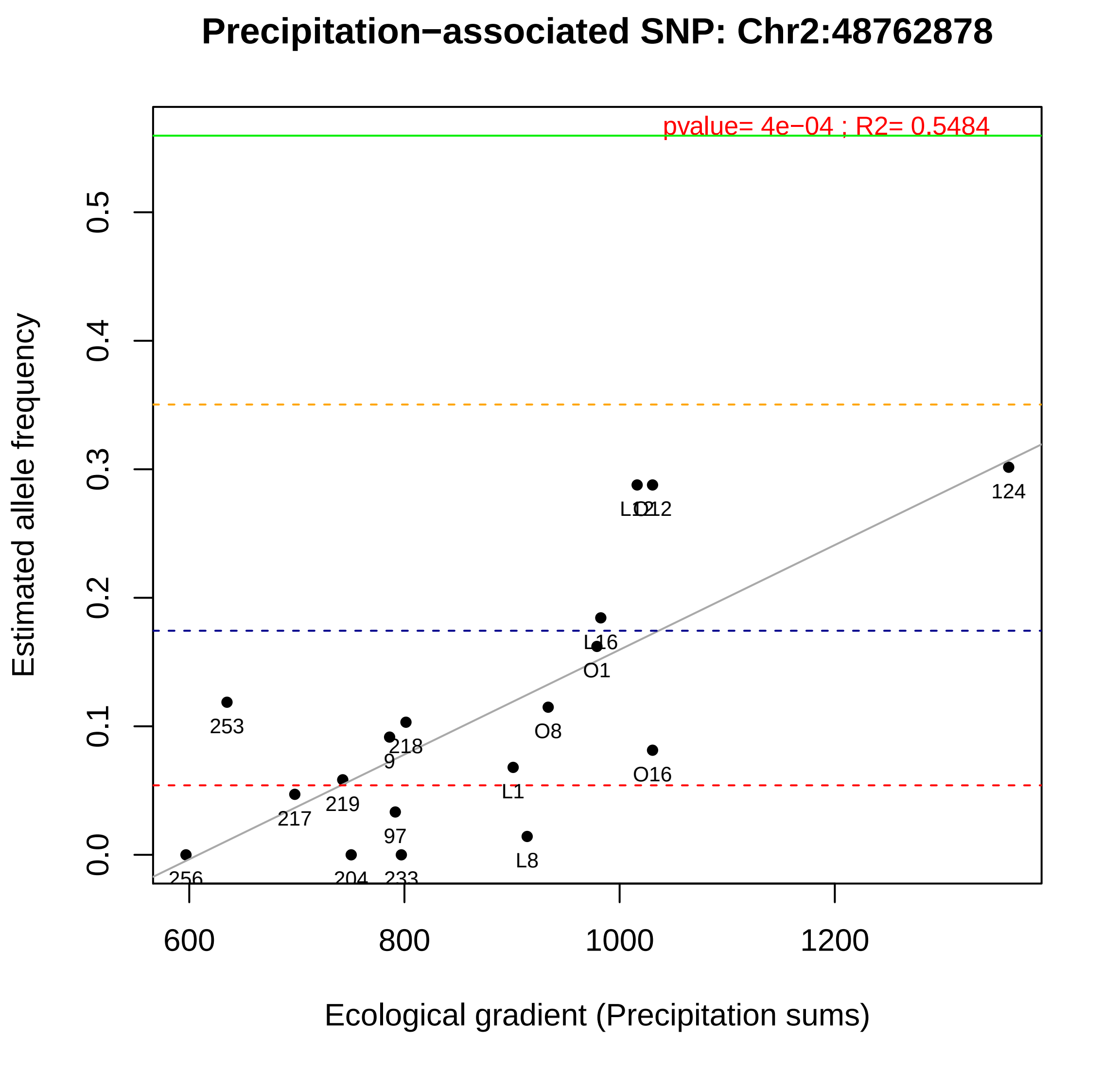
Precipitation−associated SNP: Chr2:48762878
pvalue= 4e−04 ; R2= 0.5484
Estimated allele frequency
0.0
0.1
0.2
0.3
0.4
0.5
600
800
1000
1200
Ecological gradient (Precipitation sums)
124
L12
O12
L16
O1
253
O8
218
9
O16
L1
219
217
97
L8
256
204
233

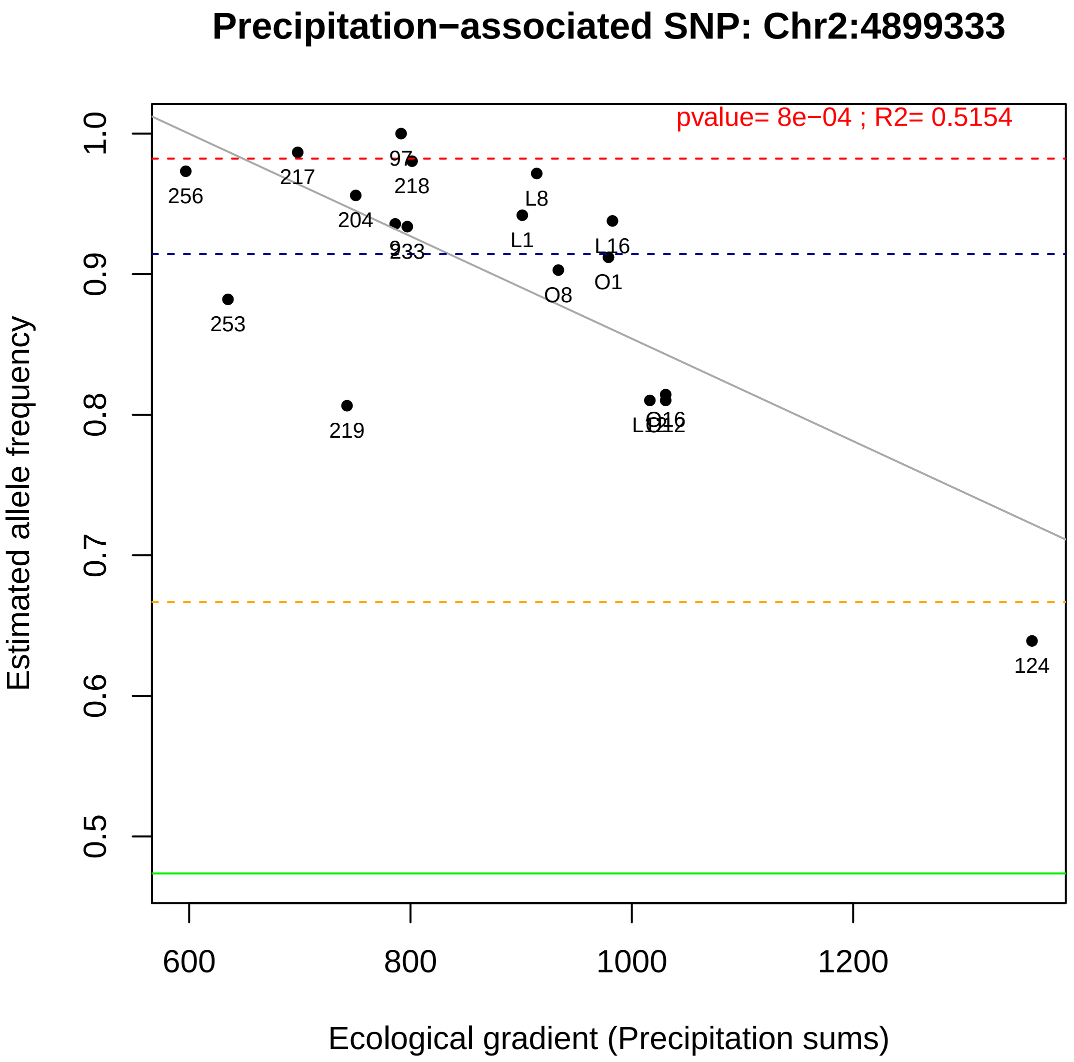
Precipitation−associated SNP: Chr2:4899333
pvalue= 8e−04 ; R2= 0.5154
Estimated allele frequency
1.0
0.9
0.8
0.7
0.6
0.5
256
217
97
218
204
233
253
219
L8
L1
L16
O1
O8
O16
124
600
800
1000
1200
Ecological gradient (Precipitation sums)

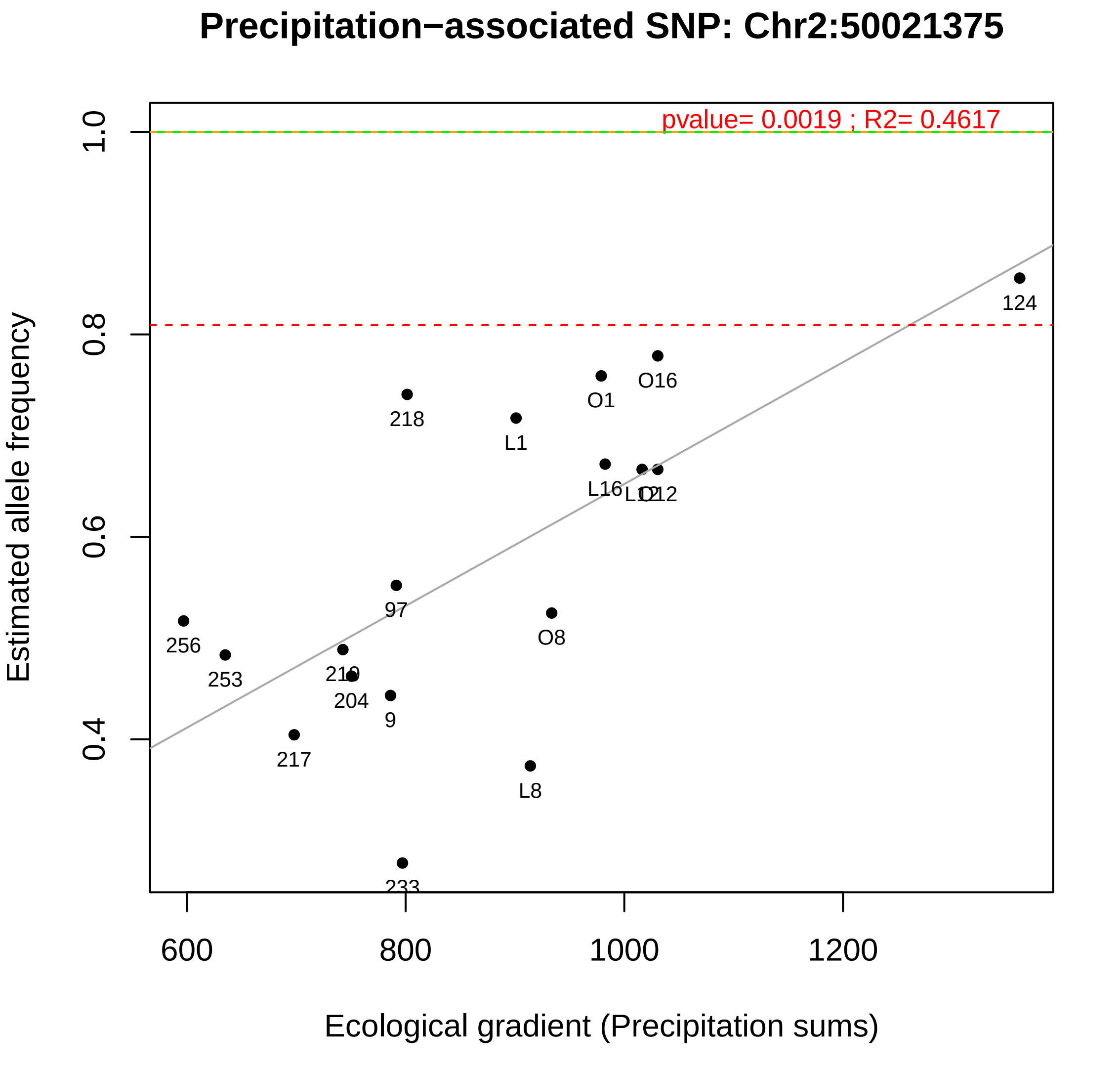
Precipitation−associated SNP: Chr2:50021375
pvalue= 0.0019 ; R2= 0.4617
Estimated allele frequency
Ecological gradient (Precipitation sums)
0.4
0.6
0.8
1.0
600
800
1000
1200
124
O16
O1
218
L1
L16
L12
O12
97
O8
256
253
210
204
9
217
L8
233

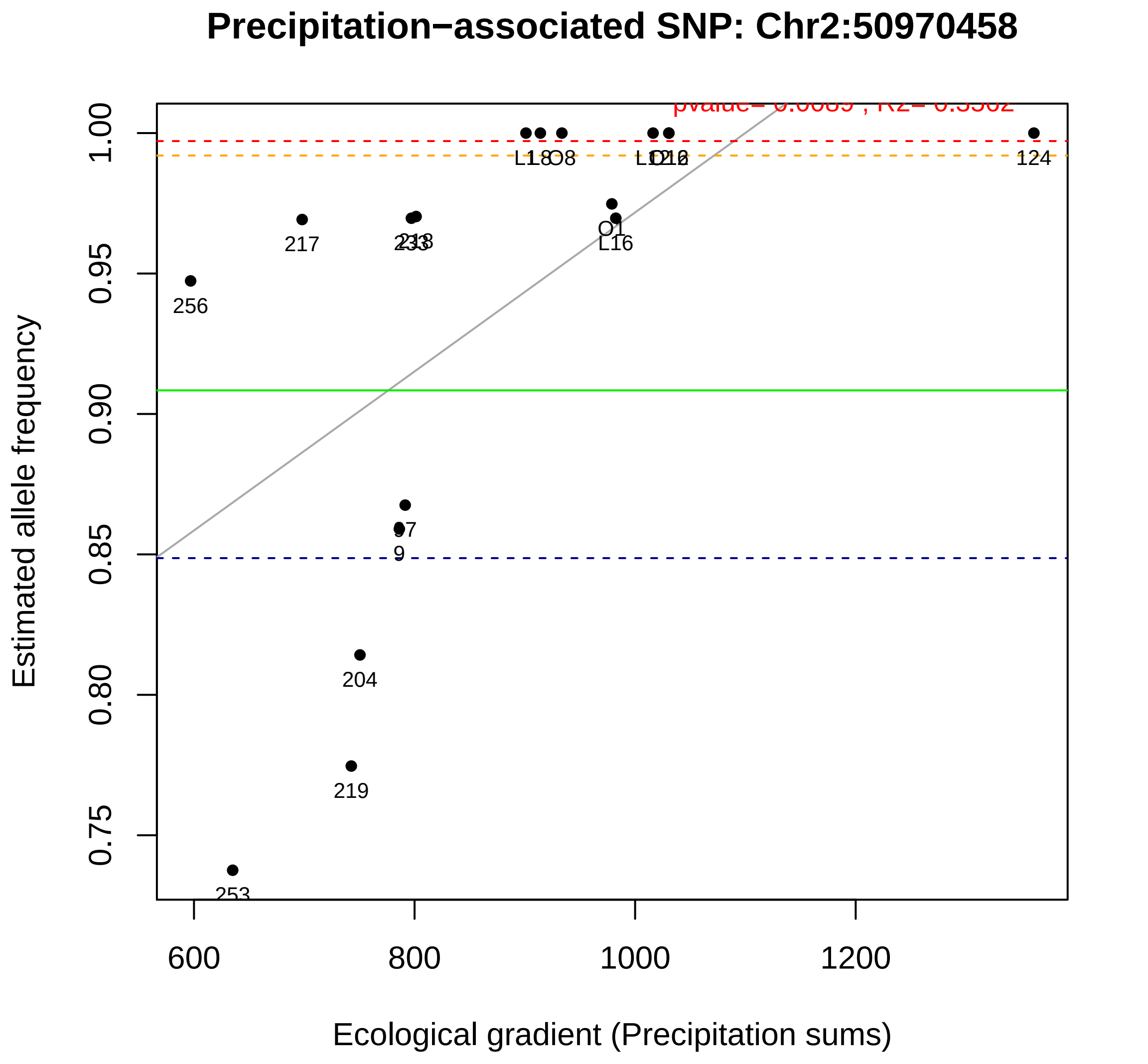
Precipitation−associated SNP: Chr2:50970458
Estimated allele frequency
Ecological gradient (Precipitation sums)
0.75
0.80
0.85
0.90
0.95
1.00
600
800
1000
1200
256
217
253
204
219
9
124
L16
O8

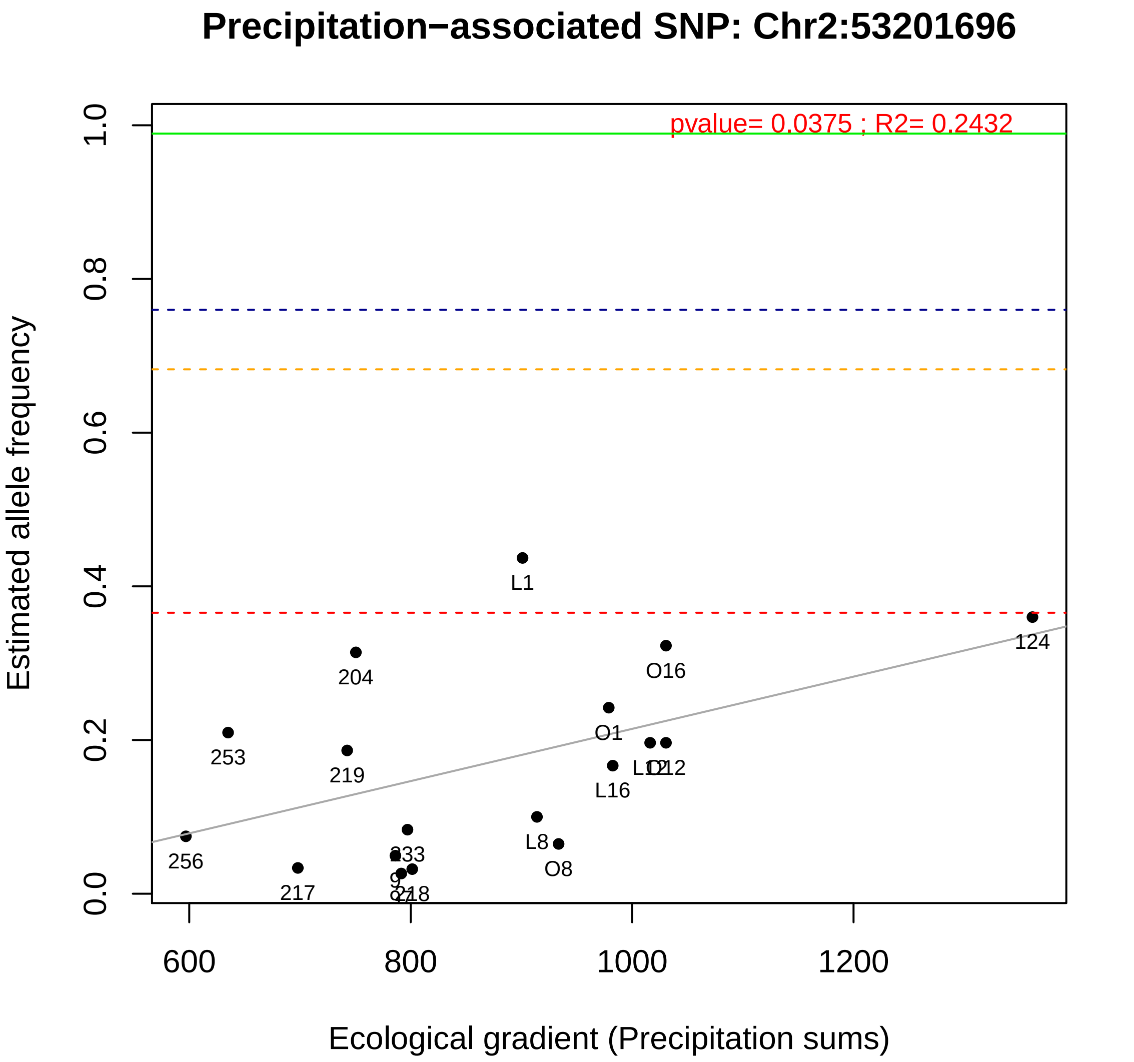
Precipitation−associated SNP: Chr2:53201696
pvalue= 0.0375 ; R2= 0.2432
Estimated allele frequency
0.0
0.2
0.4
0.6
0.8
1.0
600
800
1000
1200
Ecological gradient (Precipitation sums)
L1
124
204
O16
O1
253
219
L16
L8
O8
256
217
218

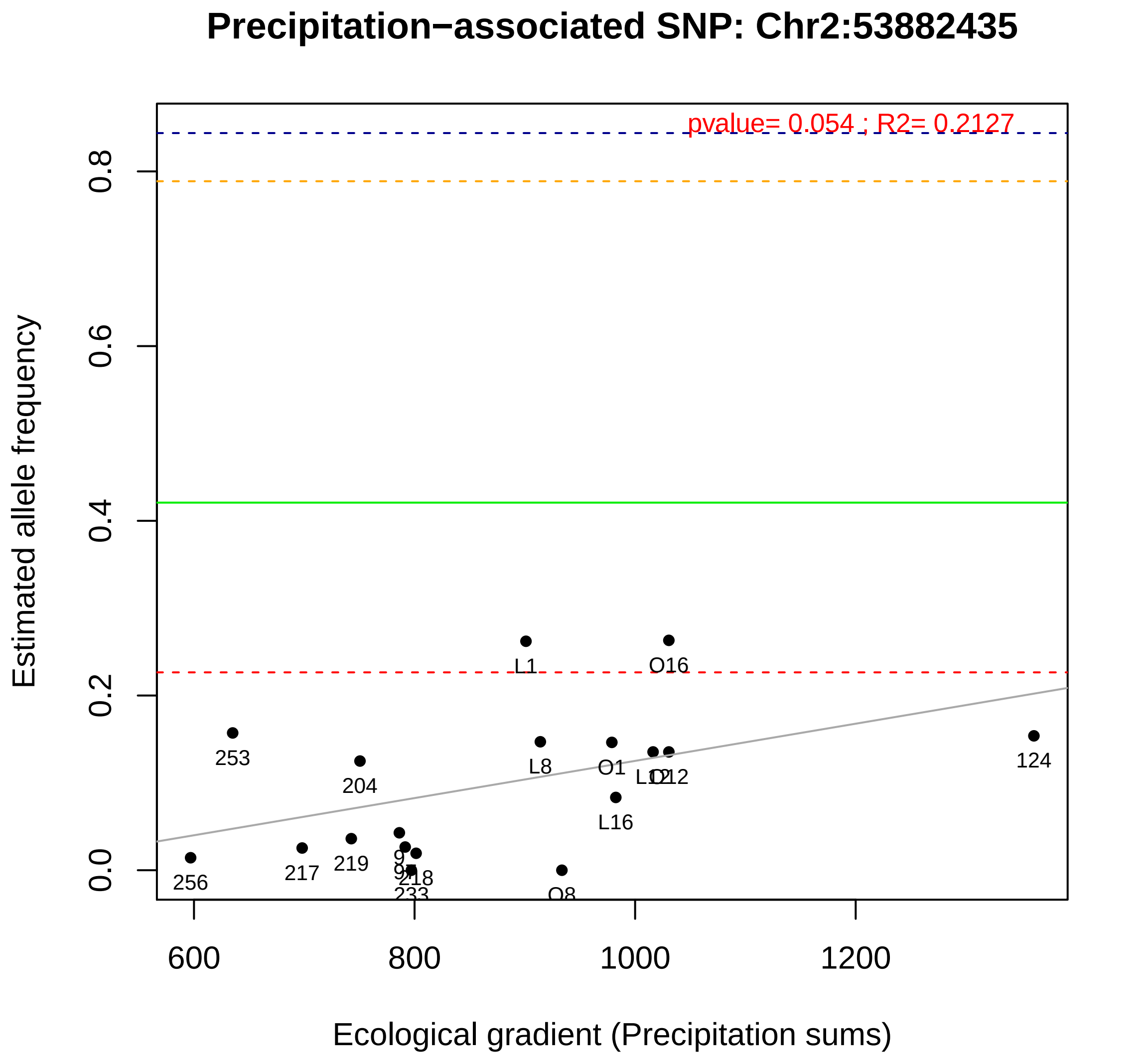

Precipitation−associated SNP: Chr2:53882435
pvalue= 0.054 ; R2= 0.2127
Estimated allele frequency
0.0
0.2
0.4
0.6
0.8
600
800
1000
1200
Ecological gradient (Precipitation sums)
L1
O16
253
204
L8
O1
124
L16
256
217
219
218
233
O8

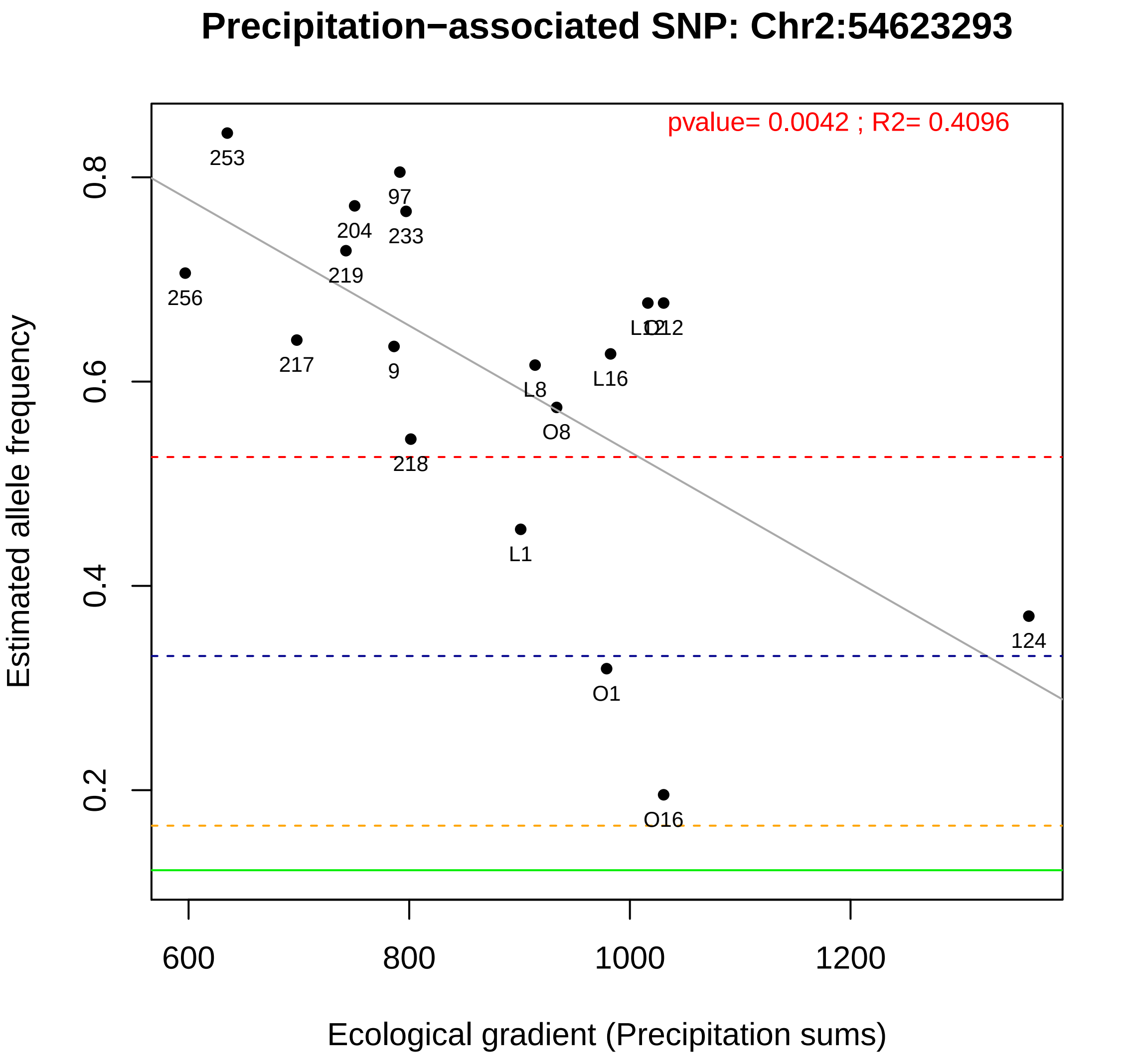

Precipitation−associated SNP: Chr2:54623293
pvalue= 0.0042 ; R2= 0.4096
Estimated allele frequency
0.2
0.4
0.6
0.8
600
800
1000
1200
Ecological gradient (Precipitation sums)
253
97
204
233
219
256
217
9
L12
O12
L16
L8
O8
218
L1
124
O1
O16

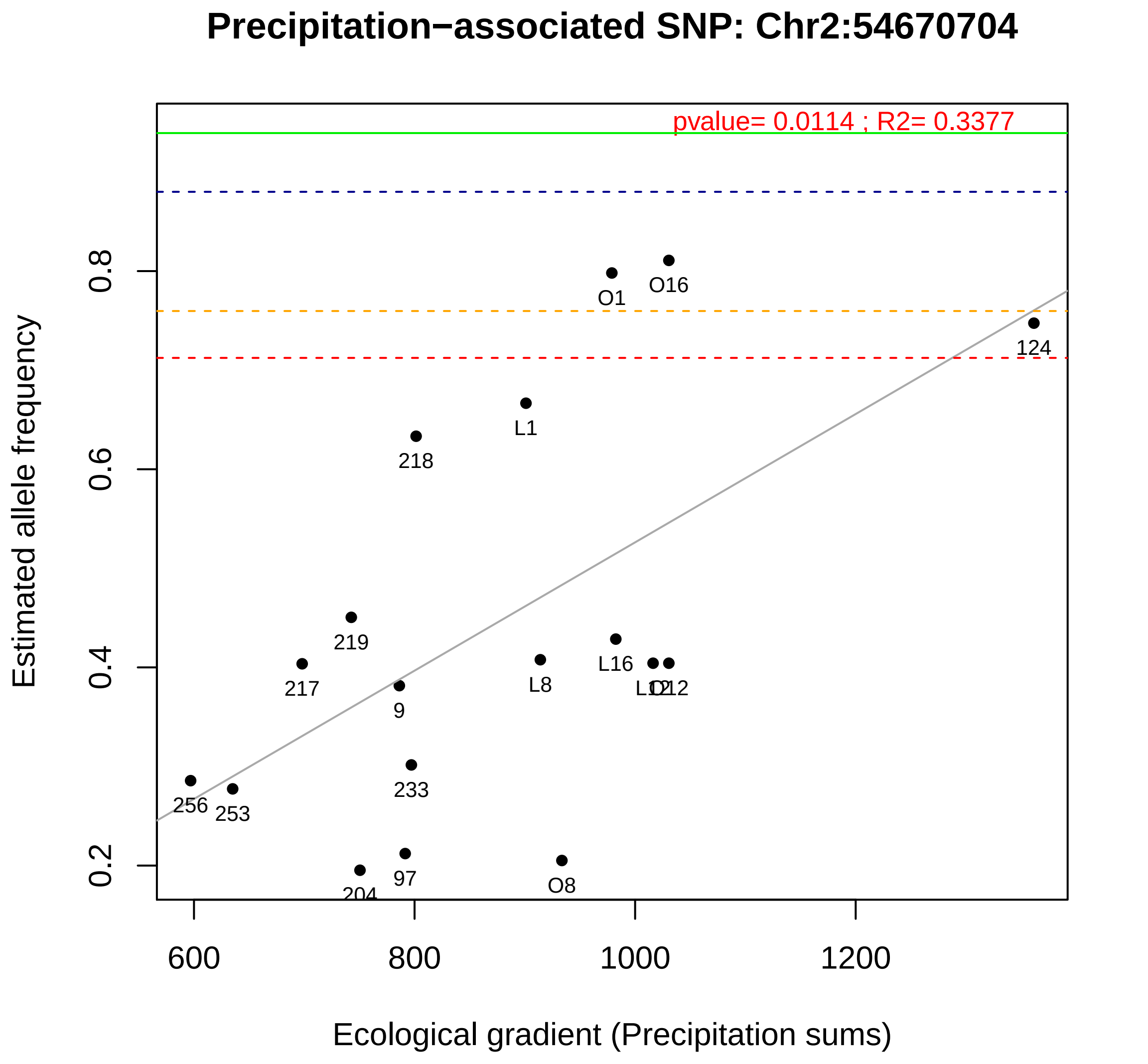

Precipitation−associated SNP: Chr2:54670704
pvalue= 0.0114 ; R2= 0.3377
Estimated allele frequency
0.2
0.4
0.6
0.8
600
800
1000
1200
Ecological gradient (Precipitation sums)
O1
O16
124
L1
218
219
217
9
L8
L16
L12
O12
233
256
253
204
97
O8

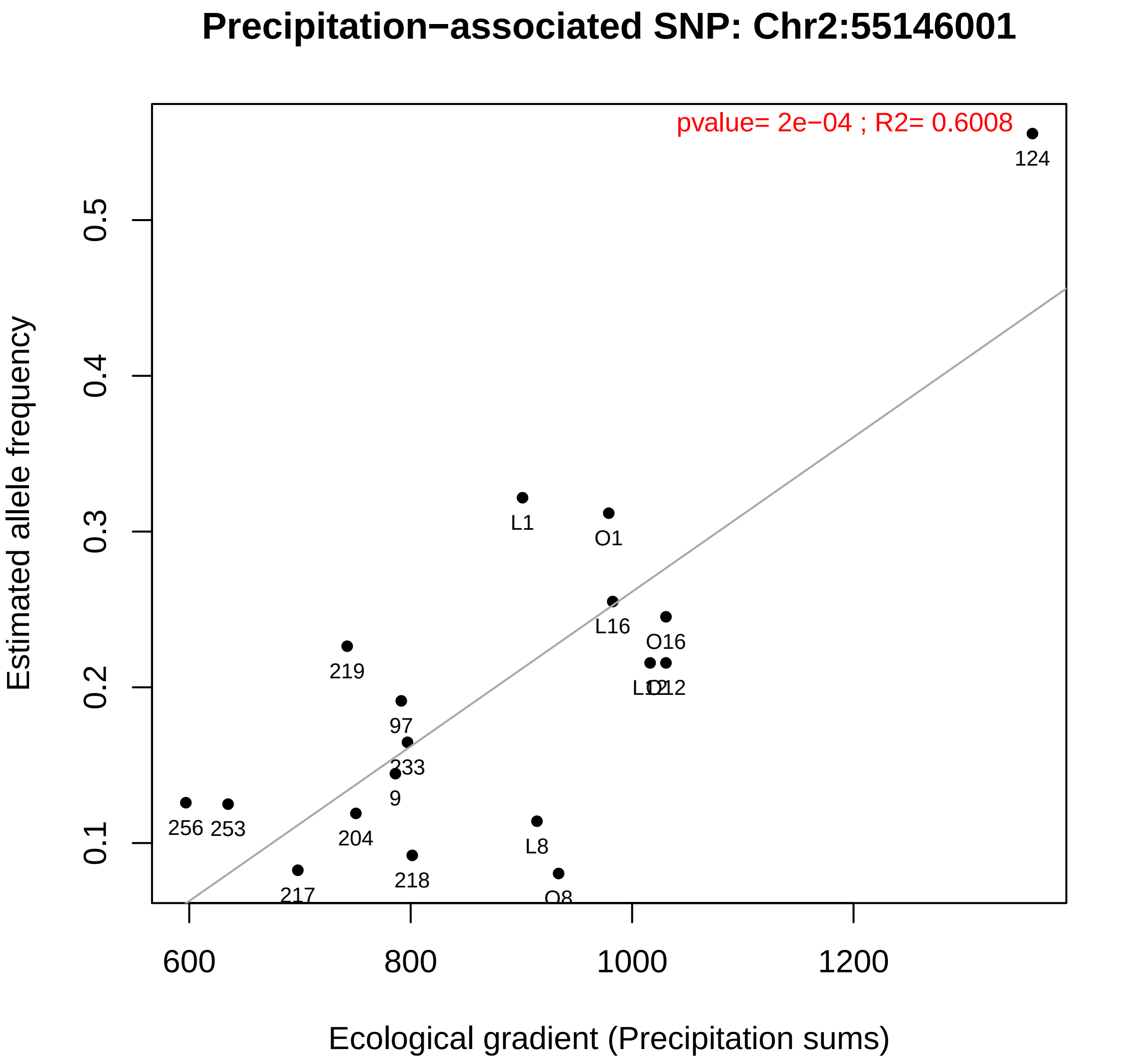

Precipitation−associated SNP: Chr2:55146001
pvalue= 2e−04 ; R2= 0.6008
124
L1
O1
L16
O16
L12
O12
219
97
233
9
256
253
204
217
218
L8
O8
0.1
0.2
0.3
0.4
0.5
600
800
1000
1200
Estimated allele frequency
Ecological gradient (Precipitation sums)

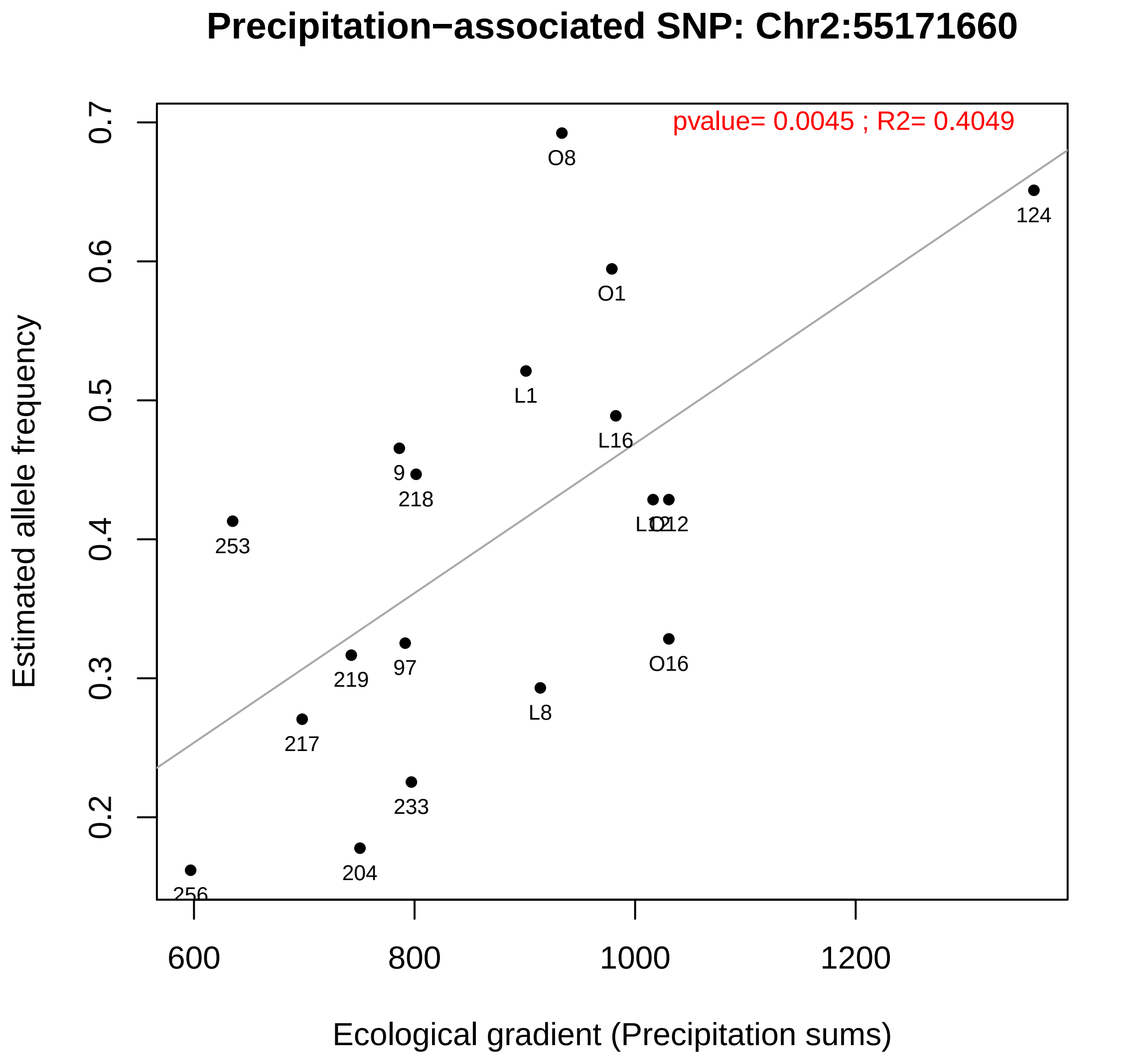

Precipitation−associated SNP: Chr2:55171660
pvalue= 0.0045 ; R2= 0.4049
Estimated allele frequency
Ecological gradient (Precipitation sums)
0.2
0.3
0.4
0.5
0.6
0.7
600
800
1000
1200
O8
124
O1
L1
L16
9
218
253
L12
O12
O16
219
97
L8
217
233
204
256

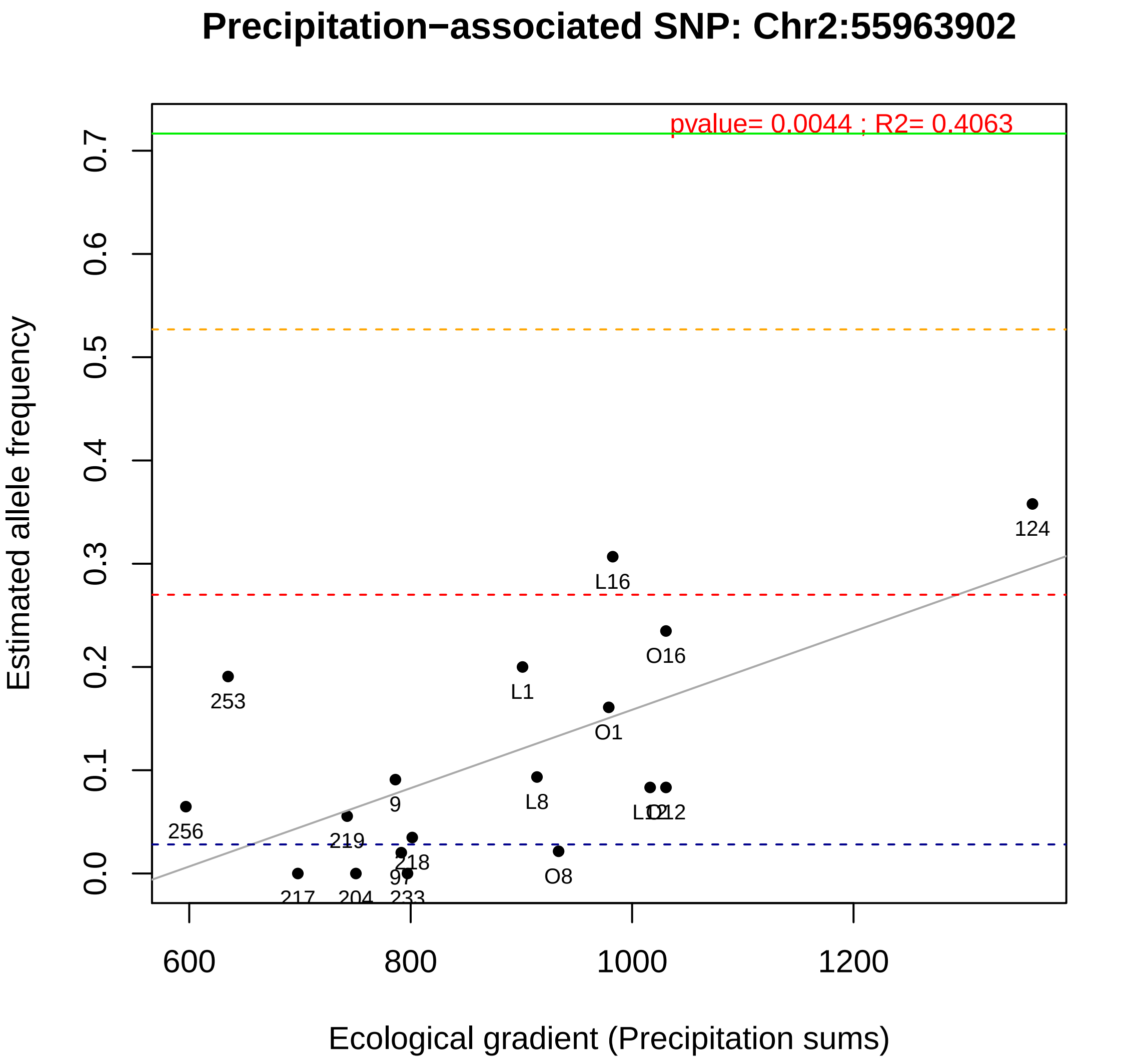

Precipitation−associated SNP: Chr2:55963902
pvalue= 0.0044 ; R2= 0.4063
Estimated allele frequency
0.0
0.1
0.2
0.3
0.4
0.5
0.6
0.7
600
800
1000
1200
Ecological gradient (Precipitation sums)
124
L16
O16
253
L1
O1
9
L8
256
219
218
O8
217
204
97
233

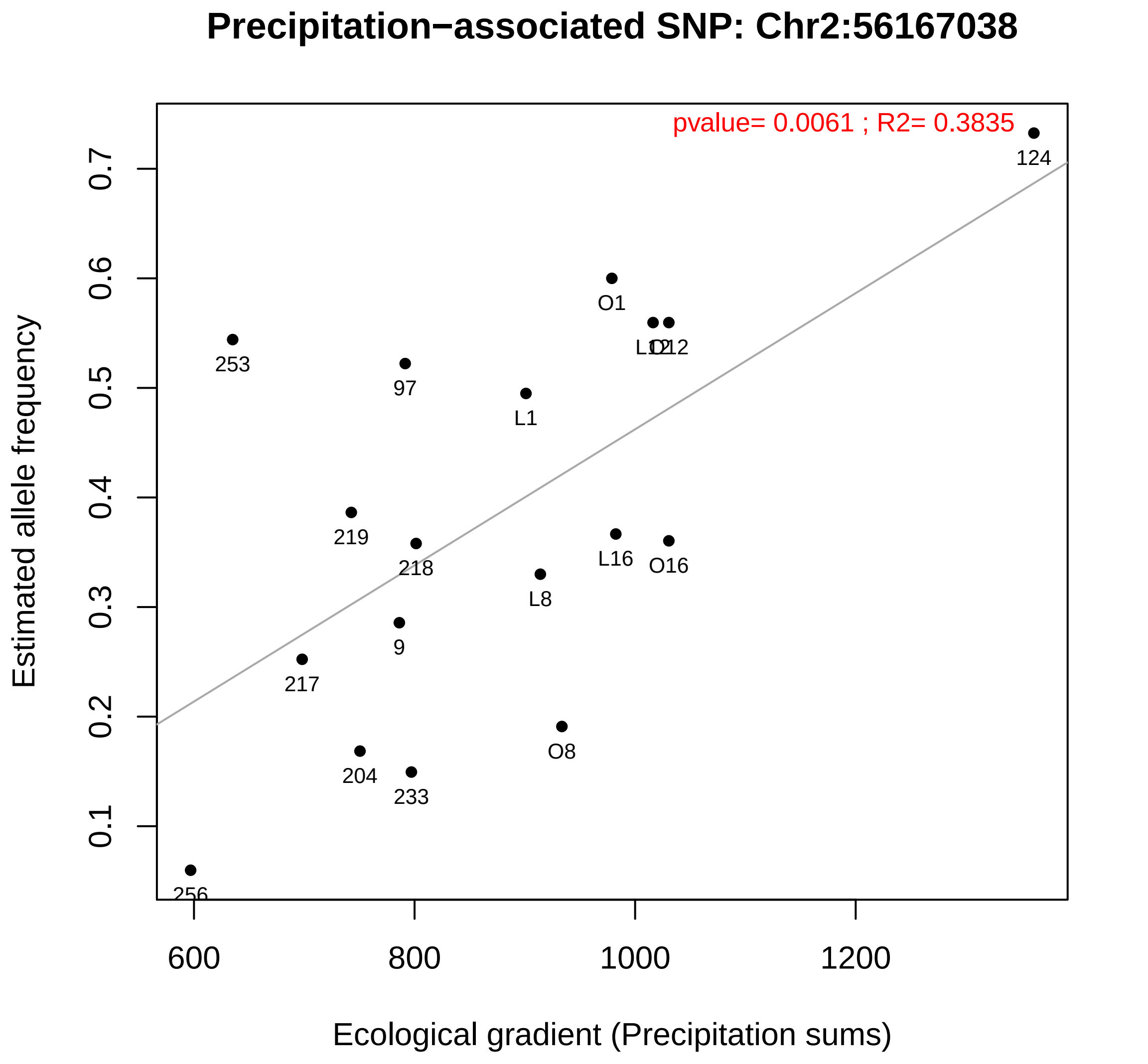
Precipitation−associated SNP: Chr2:56167038
pvalue= 0.0061 ; R2= 0.3835
Estimated allele frequency
Ecological gradient (Precipitation sums)
0.1
0.2
0.3
0.4
0.5
0.6
0.7
600
800
1000
1200
124
O1
L12
O12
253
97
L1
219
218
L16
O16
L8
9
217
O8
204
233
256

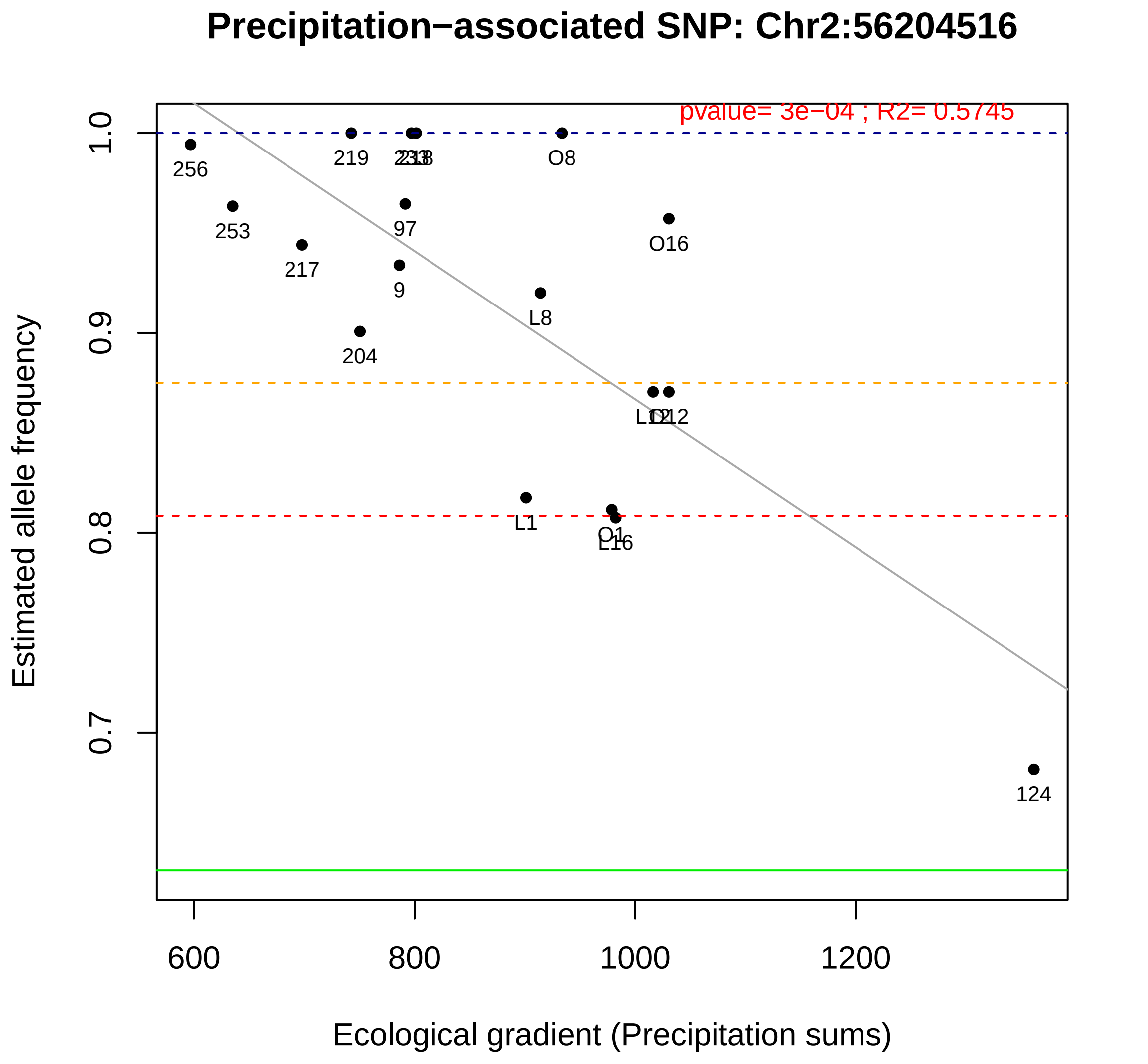

Precipitation−associated SNP: Chr2:56204516
pvalue= 3e−04 ; R2= 0.5745
256
219
O8
253
97
O16
217
9
L8
204
L1
124
Estimated allele frequency
1.0
0.9
0.8
0.7
600
800
1000
1200
Ecological gradient (Precipitation sums)

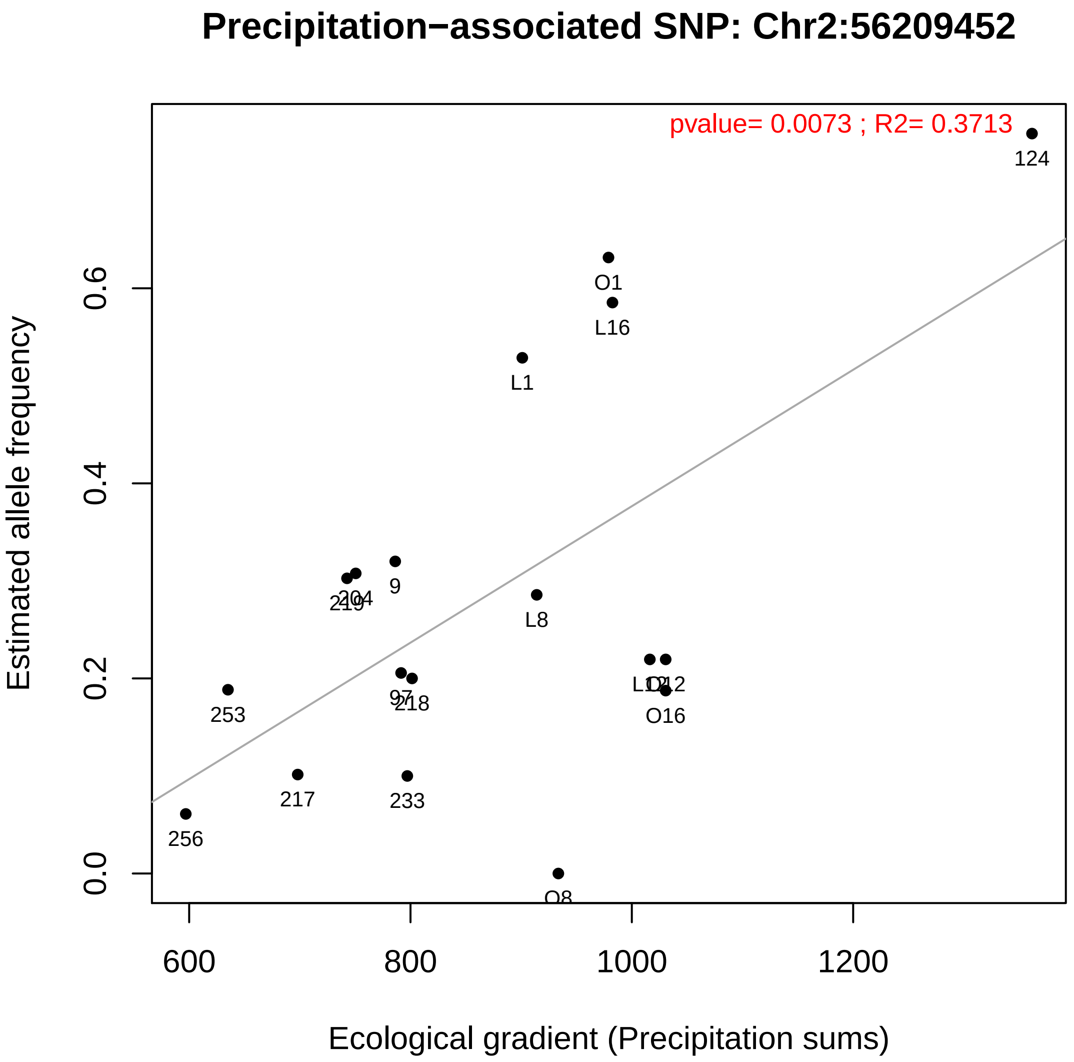

Precipitation−associated SNP: Chr2:56209452
pvalue= 0.0073 ; R2= 0.3713
Estimated allele frequency
0.0
0.2
0.4
0.6
600
800
1000
1200
Ecological gradient (Precipitation sums)
124
O1
L16
L1
9
L8
253
97
218
O16
217
233
256

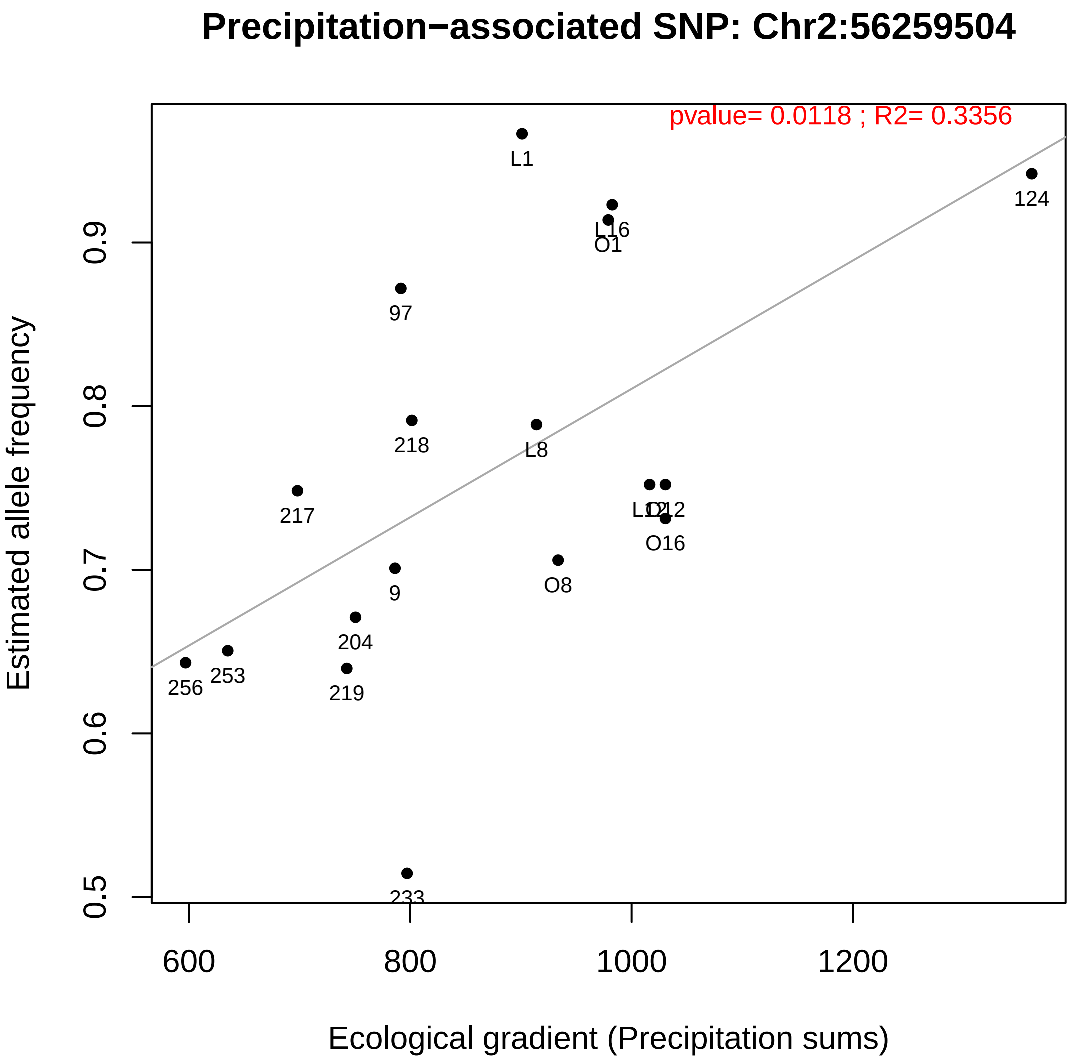

Precipitation−associated SNP: Chr2:56259504
pvalue= 0.0118 ; R2= 0.3356
L1
124
L16
O1
97
218
L8
217
L12
O12
O16
9
O8
204
253
256
219
233
0.5
0.6
0.7
0.8
0.9
600
800
1000
1200
Estimated allele frequency
Ecological gradient (Precipitation sums)

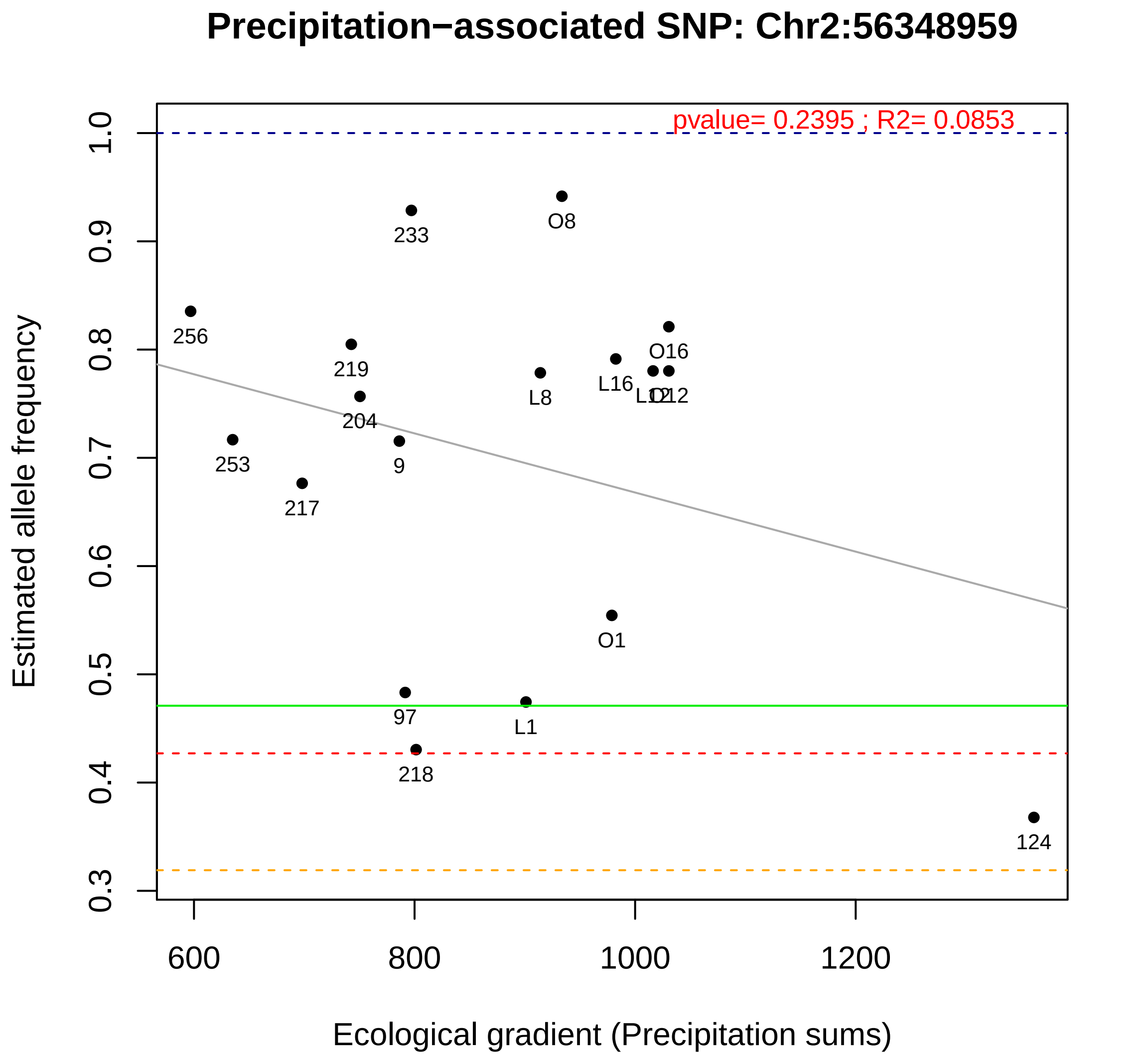

Precipitation−associated SNP: Chr2:56348959
pvalue= 0.2395 ; R2= 0.0853
Estimated allele frequency
Ecological gradient (Precipitation sums)
0.3
0.4
0.5
0.6
0.7
0.8
0.9
1.0
600
800
1000
1200
256
233
O8
219
204
253
217
9
L8
L16
O16
O12
O1
97
L1
218
124

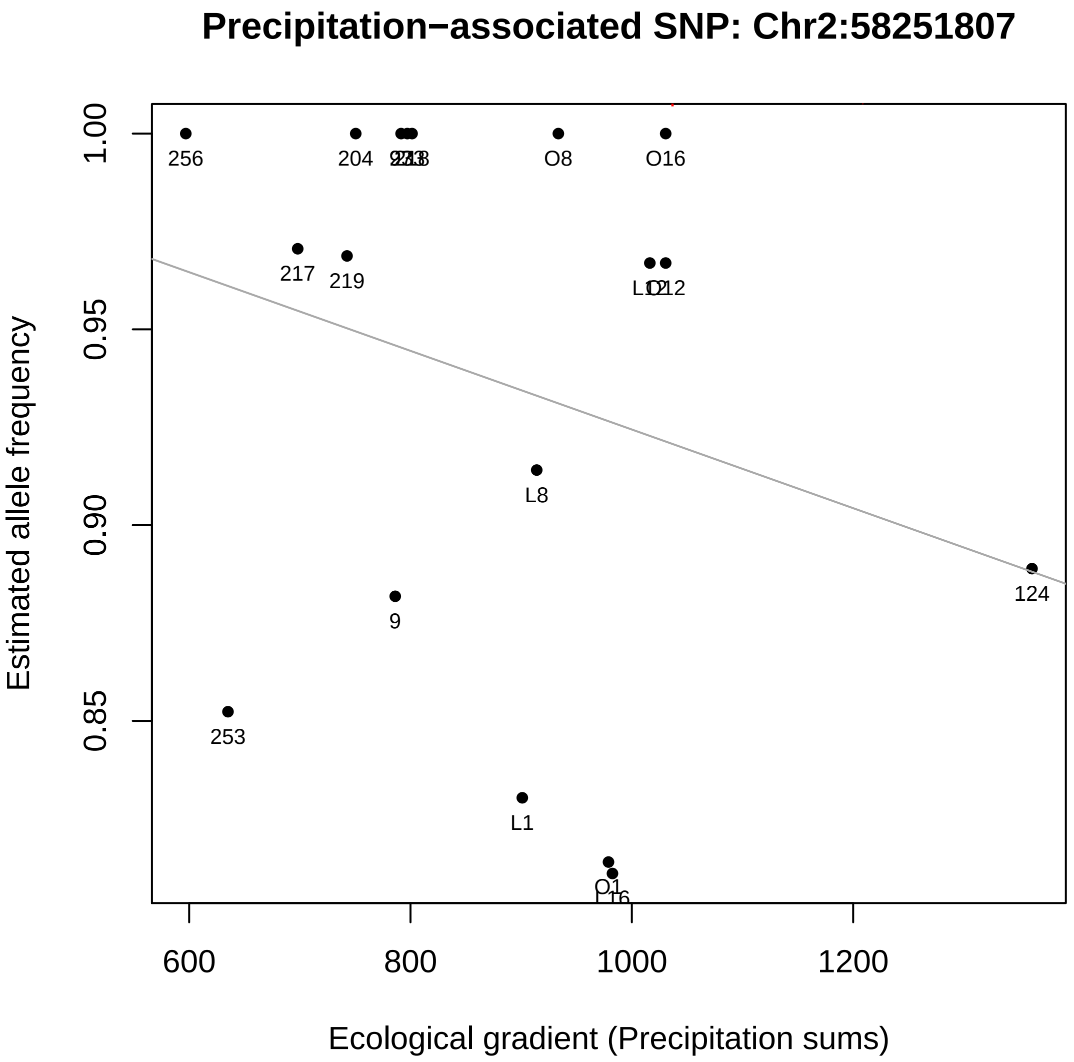
Precipitation−associated SNP: Chr2:58251807
Estimated allele frequency
Ecological gradient (Precipitation sums)
1.00
0.95
0.90
0.85
600
800
1000
1200
256
204
O8
O16
217
219
O12
L8
124
9
253
L1
O1
L16

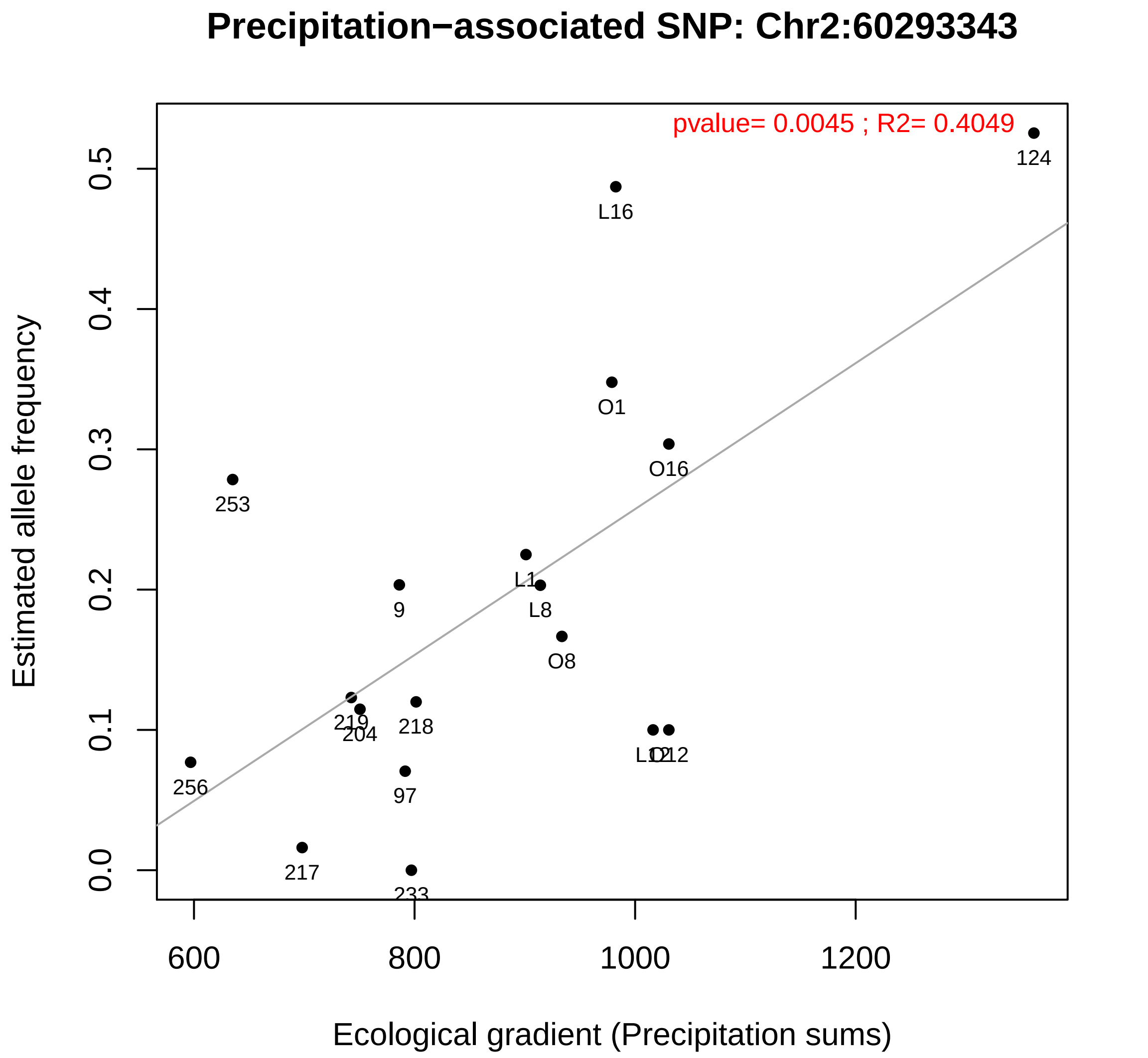

Precipitation−associated SNP: Chr2:60293343
pvalue= 0.0045 ; R2= 0.4049
Estimated allele frequency
Ecological gradient (Precipitation sums)
0.0
0.1
0.2
0.3
0.4
0.5
600
800
1000
1200
124
L16
O1
O16
253
L1
L8
9
O8
219
204
218
L12
O12
256
97
217
233

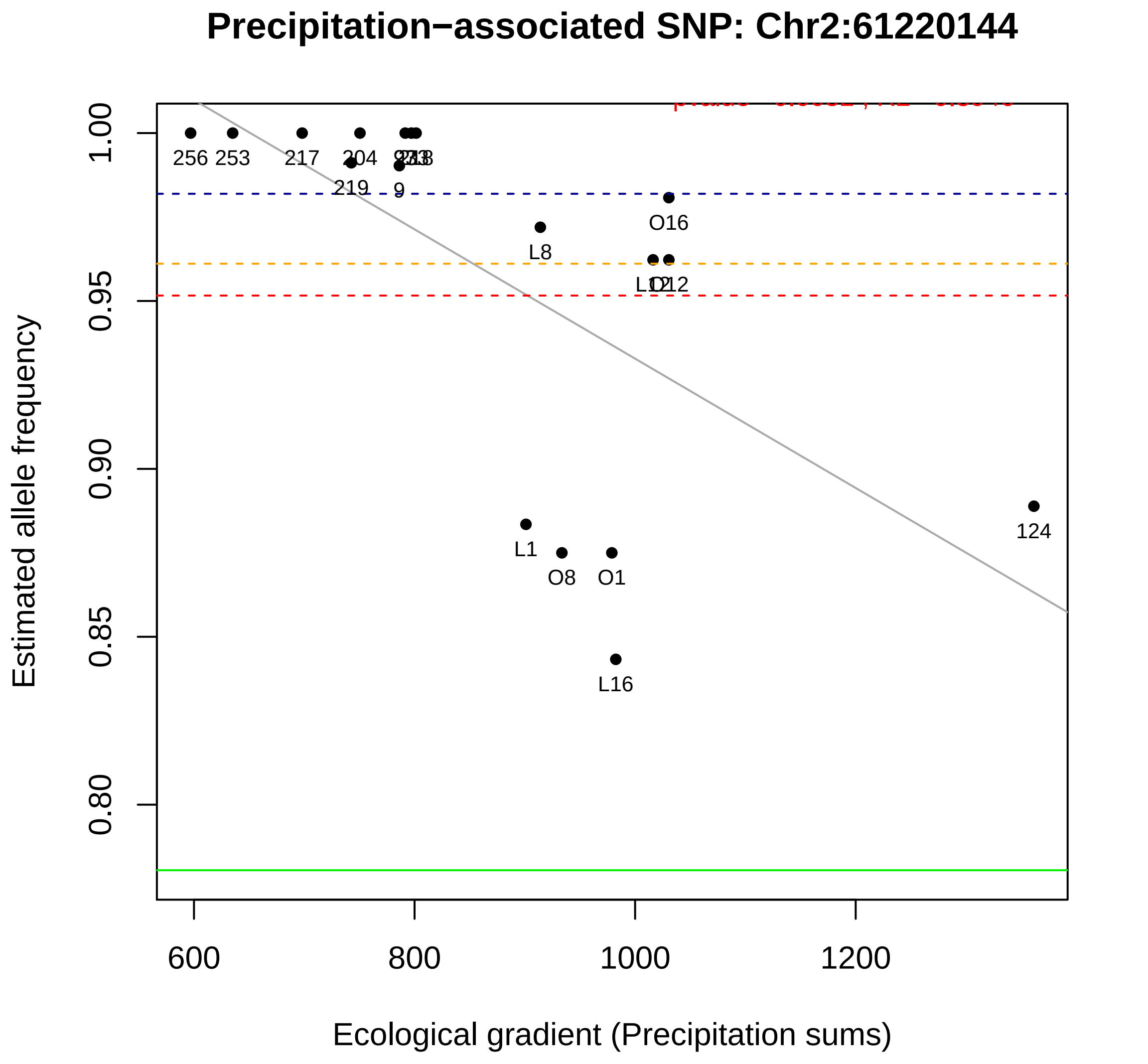

Precipitation−associated SNP: Chr2:61220144
Estimated allele frequency
Ecological gradient (Precipitation sums)
1.00
0.95
0.90
0.85
0.80
600
800
1000
1200
256
253
217
204
219
9
O16
L8
124
L1
O8
O1
L16

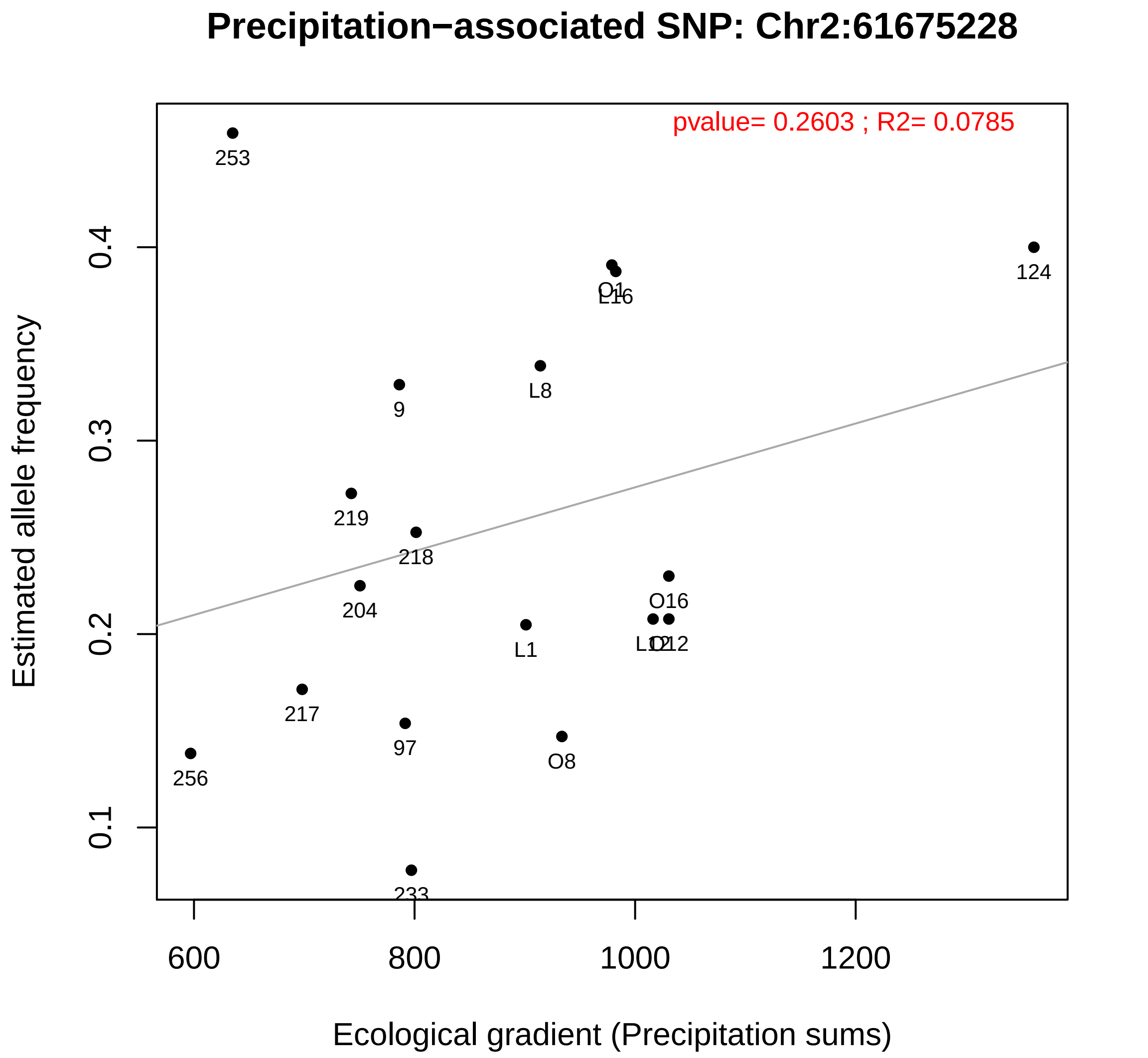

Precipitation−associated SNP: Chr2:61675228
pvalue= 0.2603 ; R2= 0.0785
Estimated allele frequency
0.1
0.2
0.3
0.4
600
800
1000
1200
Ecological gradient (Precipitation sums)
253
124
O1
L16
L8
9
219
218
204
O16
L1
L12
O12
217
97
O8
256
233

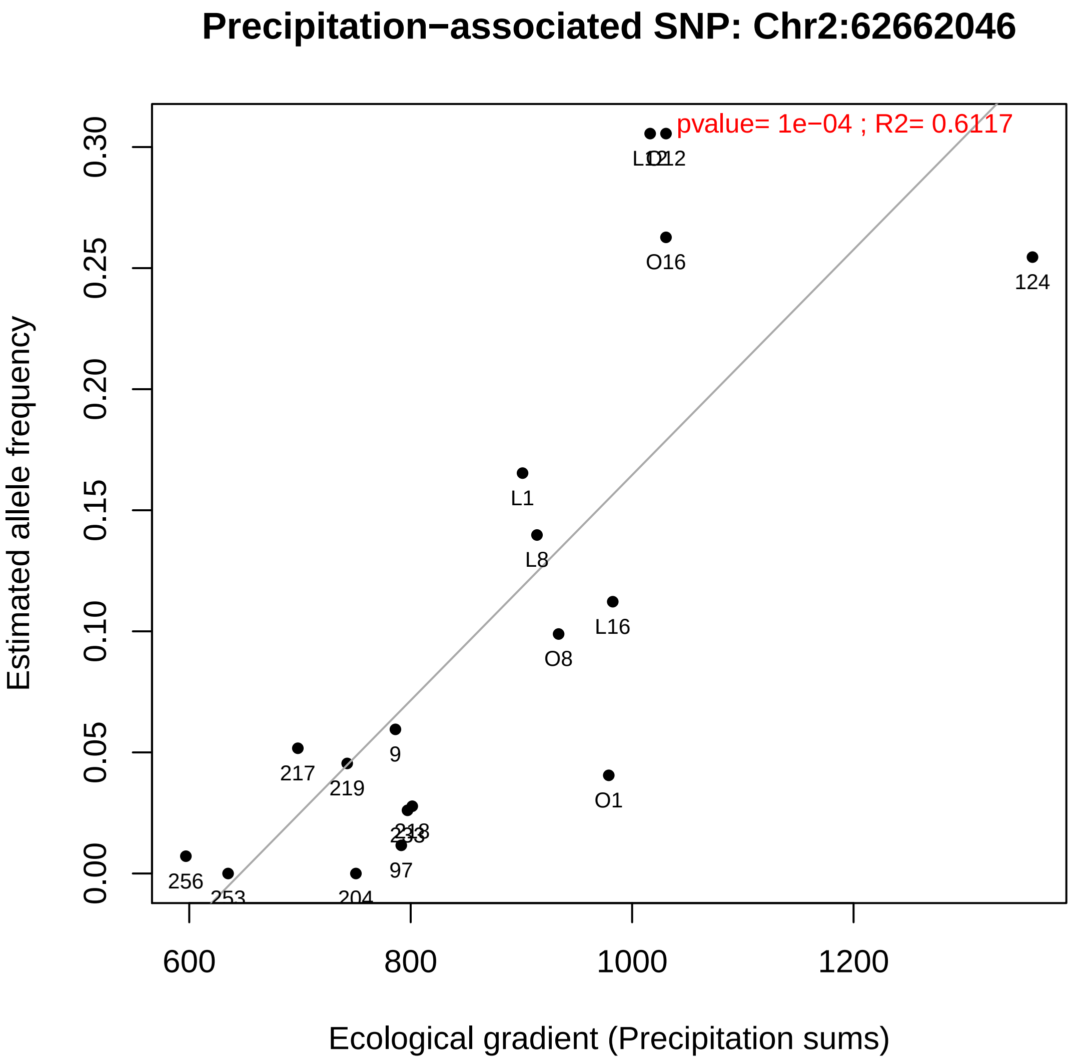
Precipitation−associated SNP: Chr2:62662046
pvalue= 1e−04 ; R2= 0.6117
Estimated allele frequency
Ecological gradient (Precipitation sums)
0.00
0.05
0.10
0.15
0.20
0.25
0.30
600
800
1000
1200
L12
O12
O16
124
L1
L8
L16
O8
9
217
219
O1
218
233
97
256
253
204

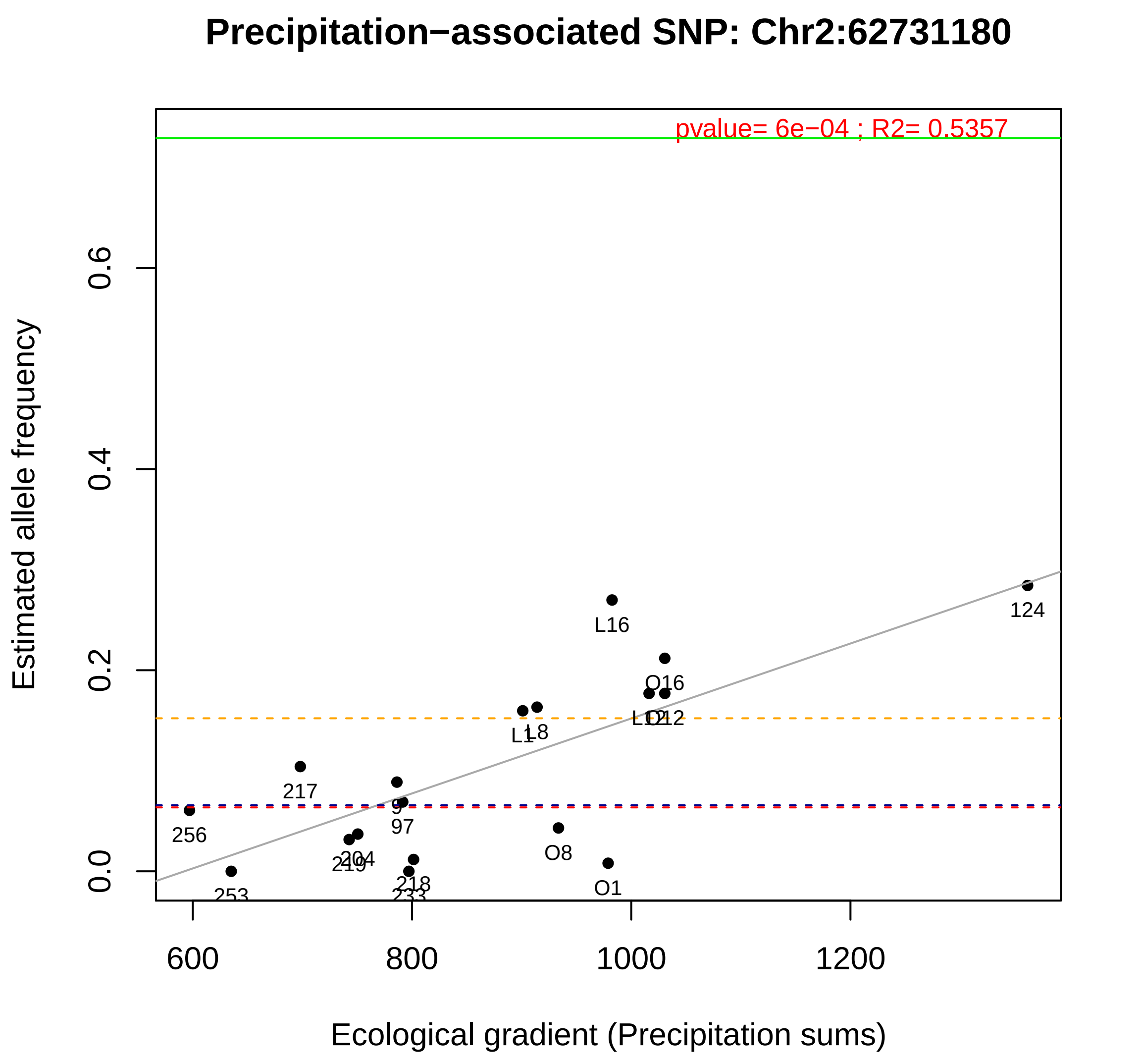

Precipitation−associated SNP: Chr2:62731180
pvalue= 6e−04 ; R2= 0.5357
Estimated allele frequency
0.0
0.2
0.4
0.6
600
800
1000
1200
Ecological gradient (Precipitation sums)
124
L16
O16
O12
L18
217
256
97
204
O8
O1
218
253

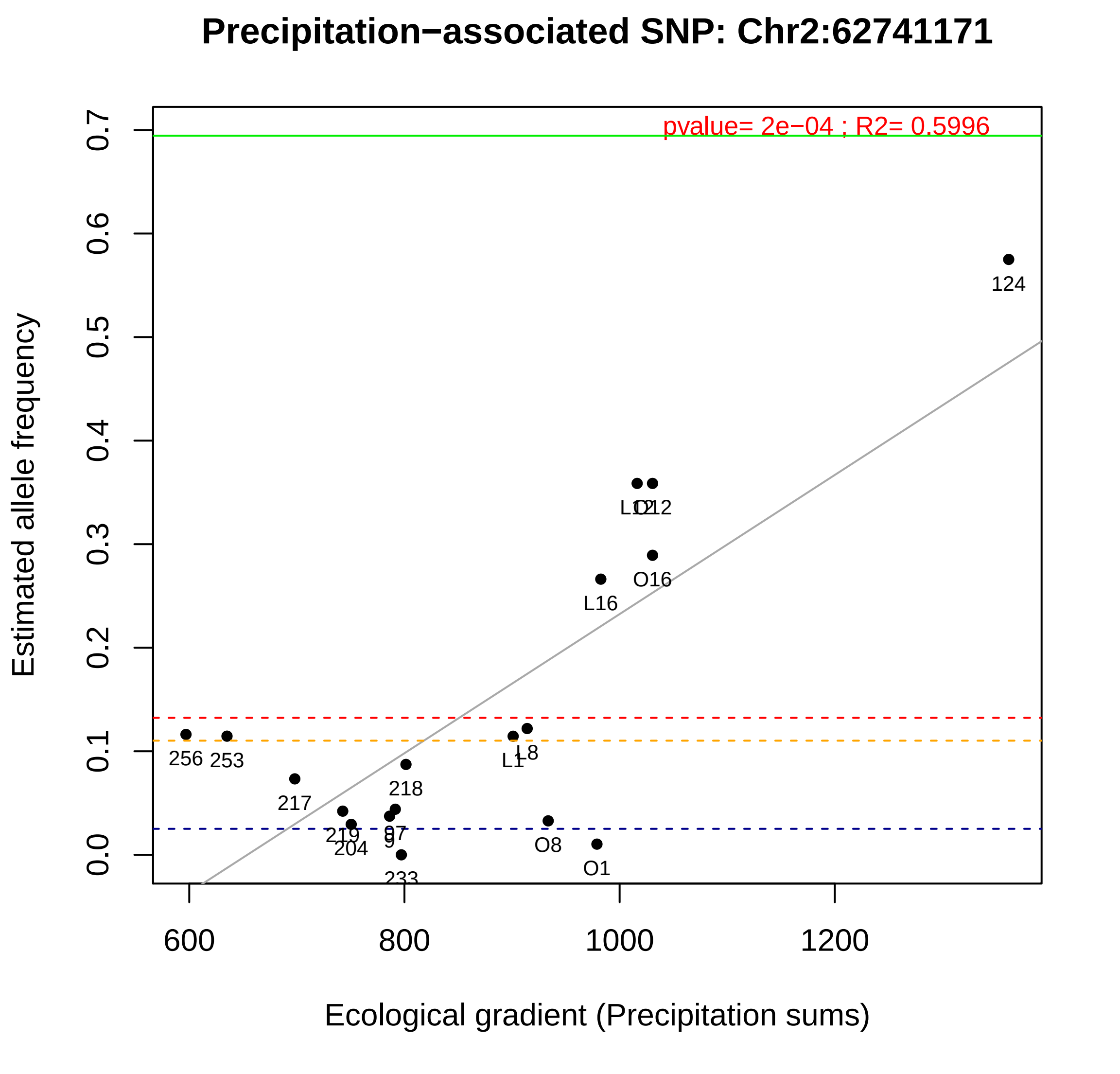
Precipitation−associated SNP: Chr2:62741171
pvalue= 2e−04 ; R2= 0.5996
Estimated allele frequency
0.0
0.1
0.2
0.3
0.4
0.5
0.6
0.7
600
800
1000
1200
Ecological gradient (Precipitation sums)
124
L12
O12
O16
L16
256
253
217
218
L1
L8
219
204
97
233
O8
O1

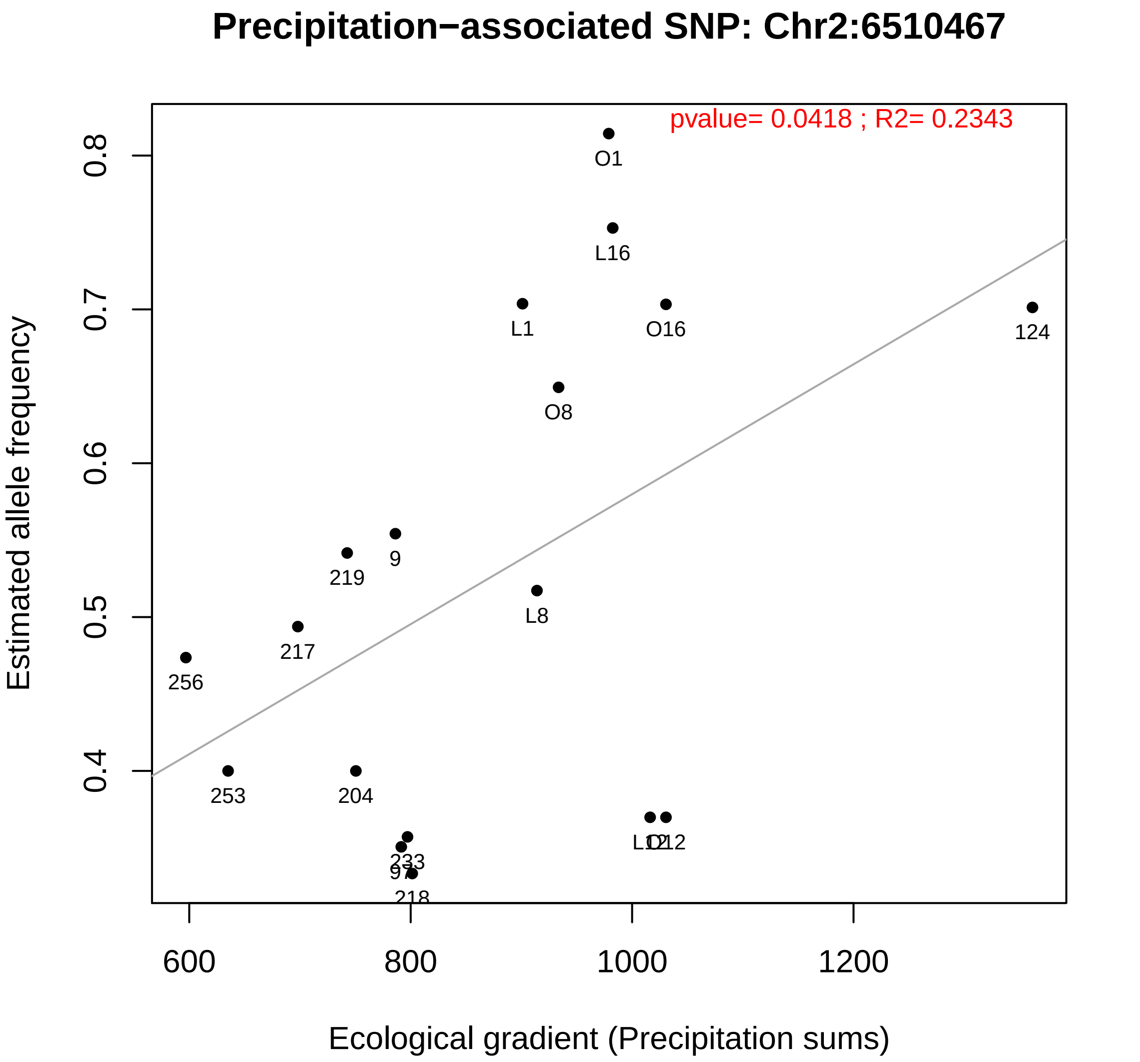
Precipitation−associated SNP: Chr2:6510467
pvalue= 0.0418 ; R2= 0.2343
Estimated allele frequency
Ecological gradient (Precipitation sums)
0.4
0.5
0.6
0.7
0.8
600
800
1000
1200
O1
L16
L1
O16
124
O8
219
9
L8
217
256
253
204
233
97
218
L10
O12

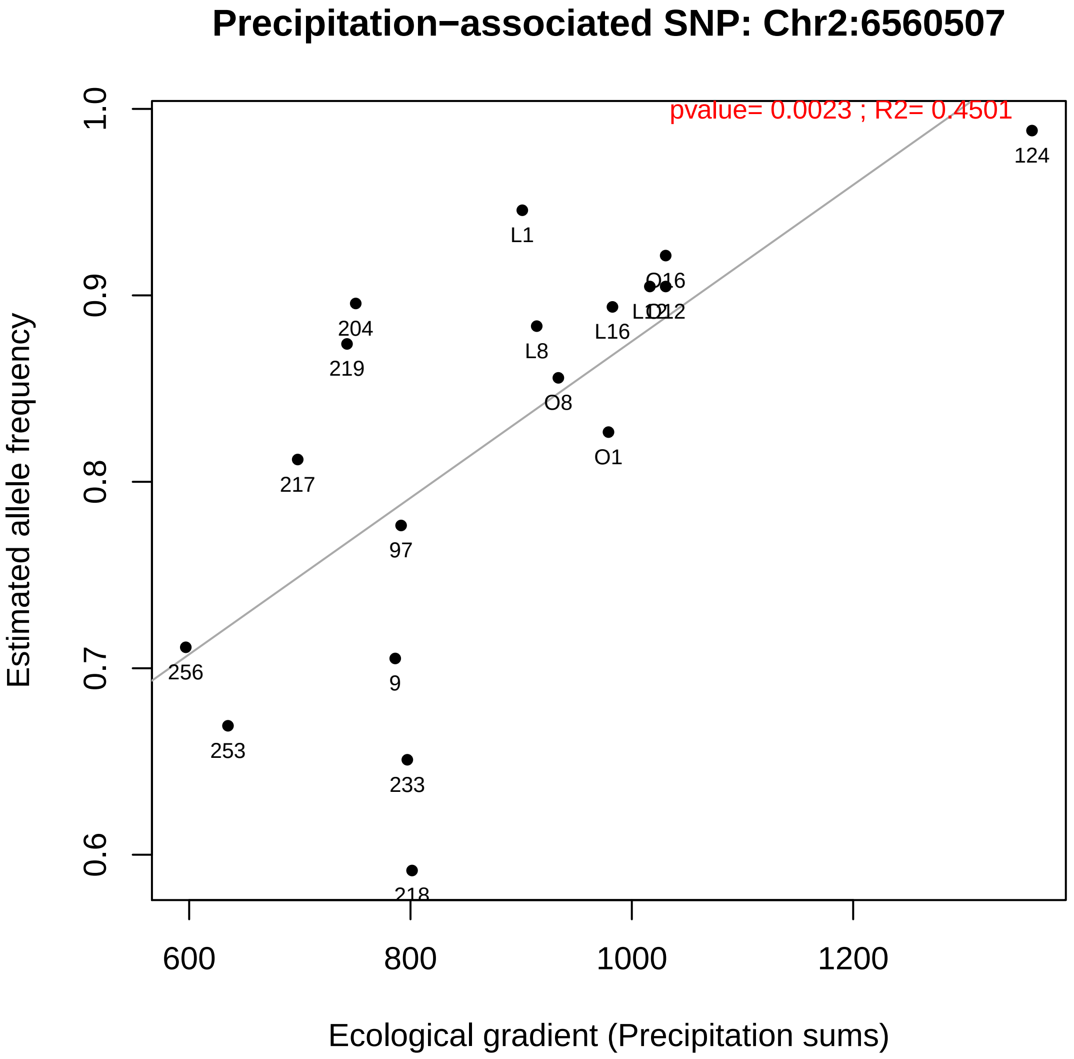
Precipitation−associated SNP: Chr2:6560507
pvalue= 0.0023 ; R2= 0.4501
Estimated allele frequency
Ecological gradient (Precipitation sums)
0.6
0.7
0.8
0.9
1.0
600
800
1000
1200
124
L1
O16
L12
O12
L16
204
219
L8
O8
O1
217
97
256
9
253
233
218

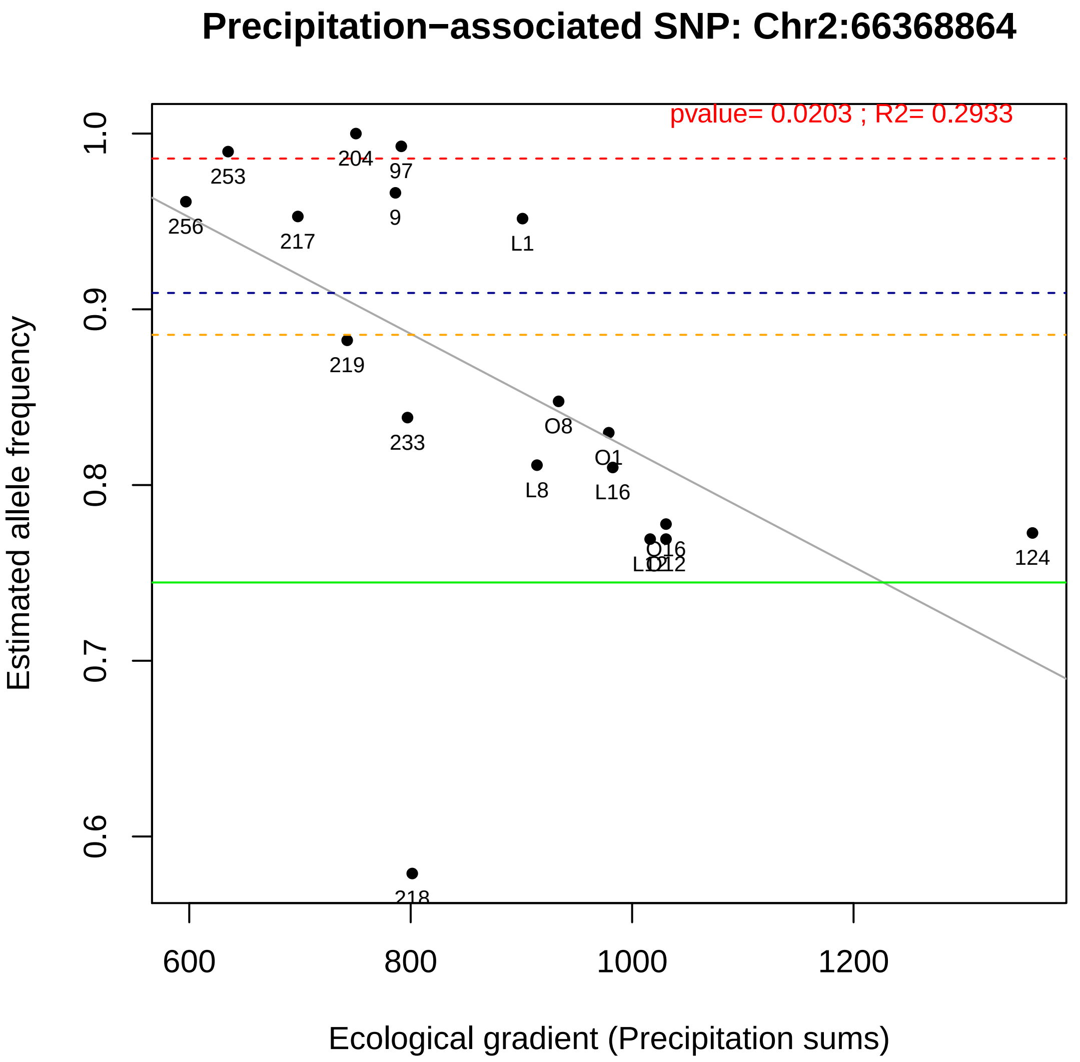

Precipitation−associated SNP: Chr2:66368864
pvalue= 0.0203 ; R2= 0.2933
Estimated allele frequency
Ecological gradient (Precipitation sums)
0.6
0.7
0.8
0.9
1.0
600
800
1000
1200
256
253
204
97
9
217
L1
219
233
O8
O1
L8
L16
O16
L12
O12
124
218

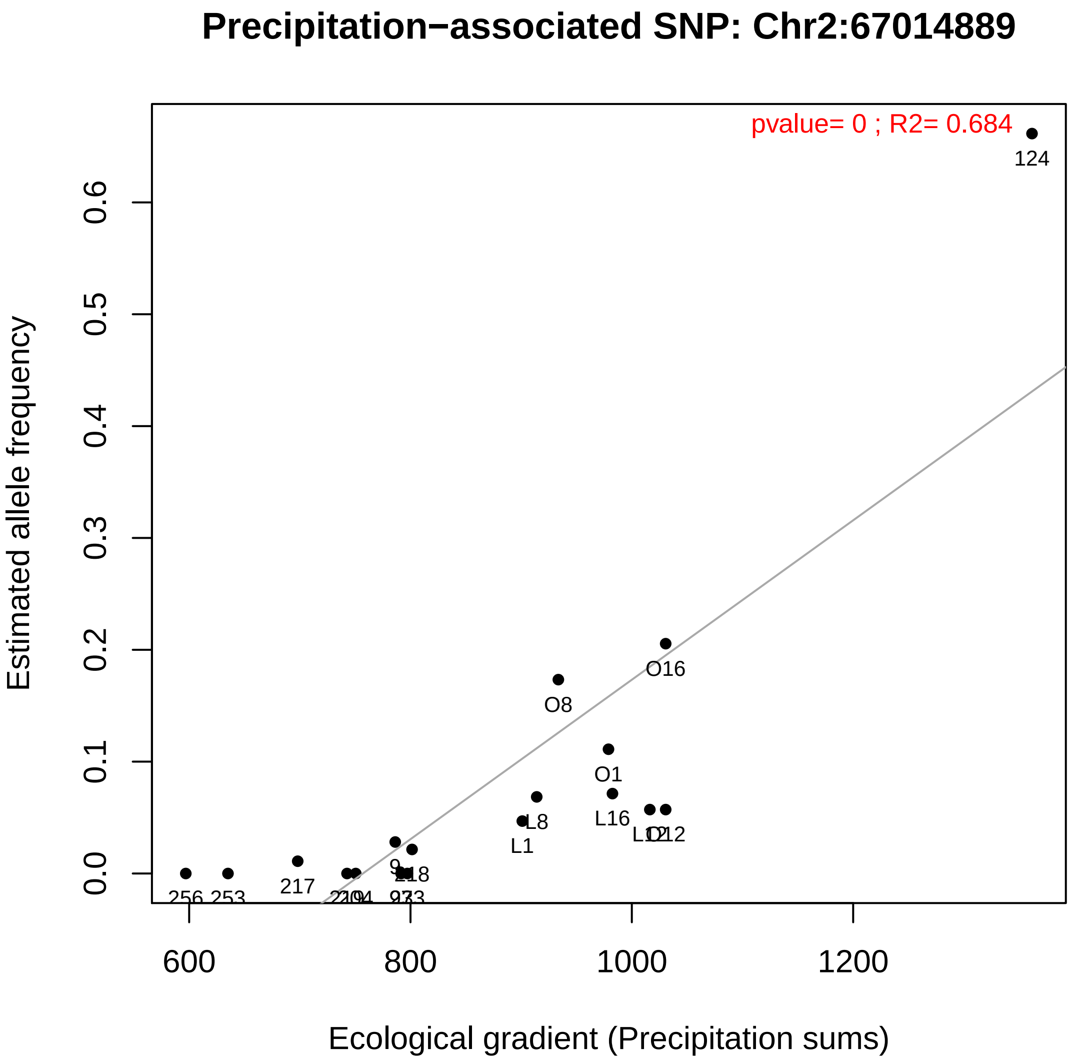
Precipitation−associated SNP: Chr2:67014889
pvalue= 0 ; R2= 0.684
Estimated allele frequency
Ecological gradient (Precipitation sums)
0.0
0.1
0.2
0.3
0.4
0.5
0.6
600
800
1000
1200
124
O16
O8
O1
L16
L8
L1
217
218
9
256
253

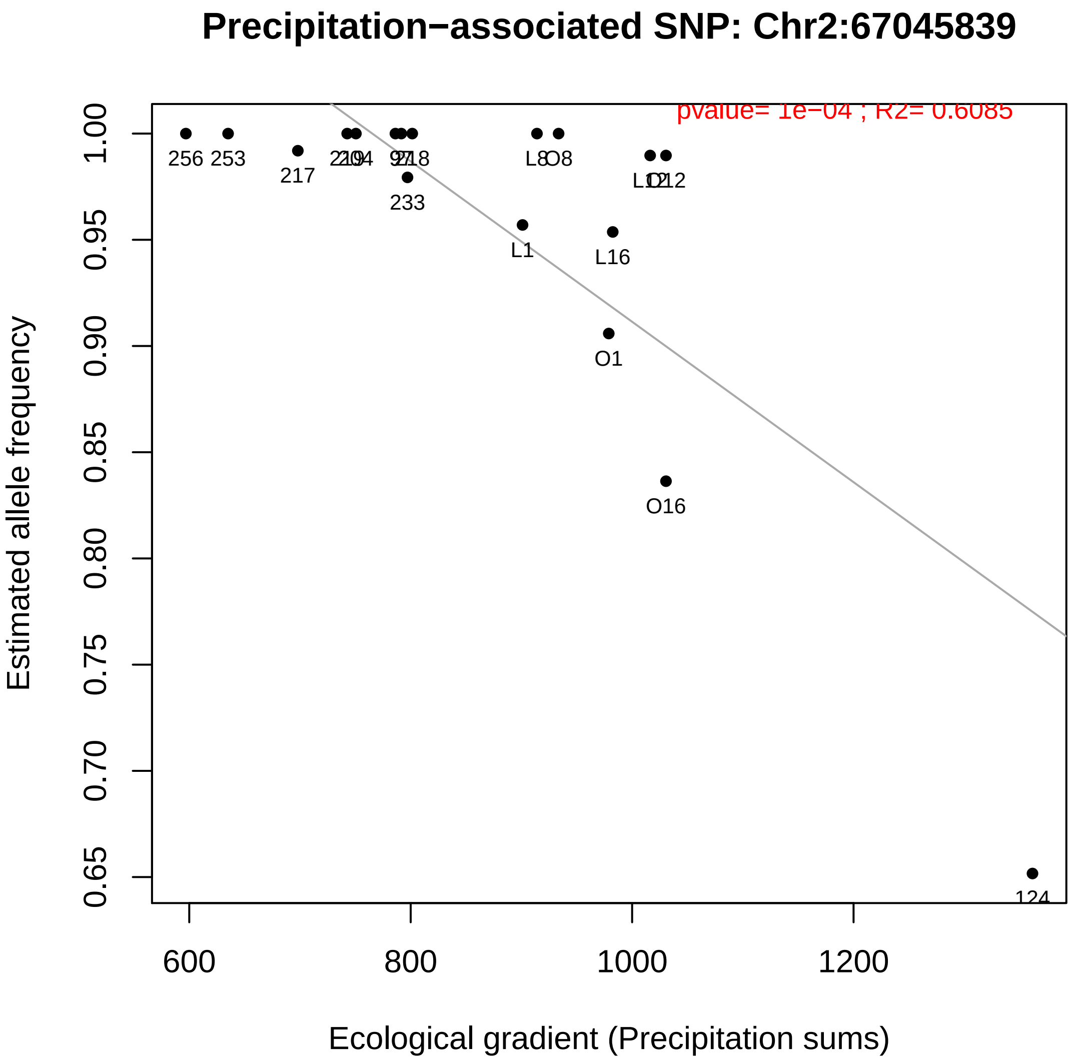

Precipitation−associated SNP: Chr2:67045839
pvalue= 1e−04 ; R2= 0.6085
Estimated allele frequency
Ecological gradient (Precipitation sums)
1.00
0.95
0.90
0.85
0.80
0.75
0.70
0.65
600
800
1000
1200
256
253
217
233
L1
L8
O8
L16
O1
O16
124

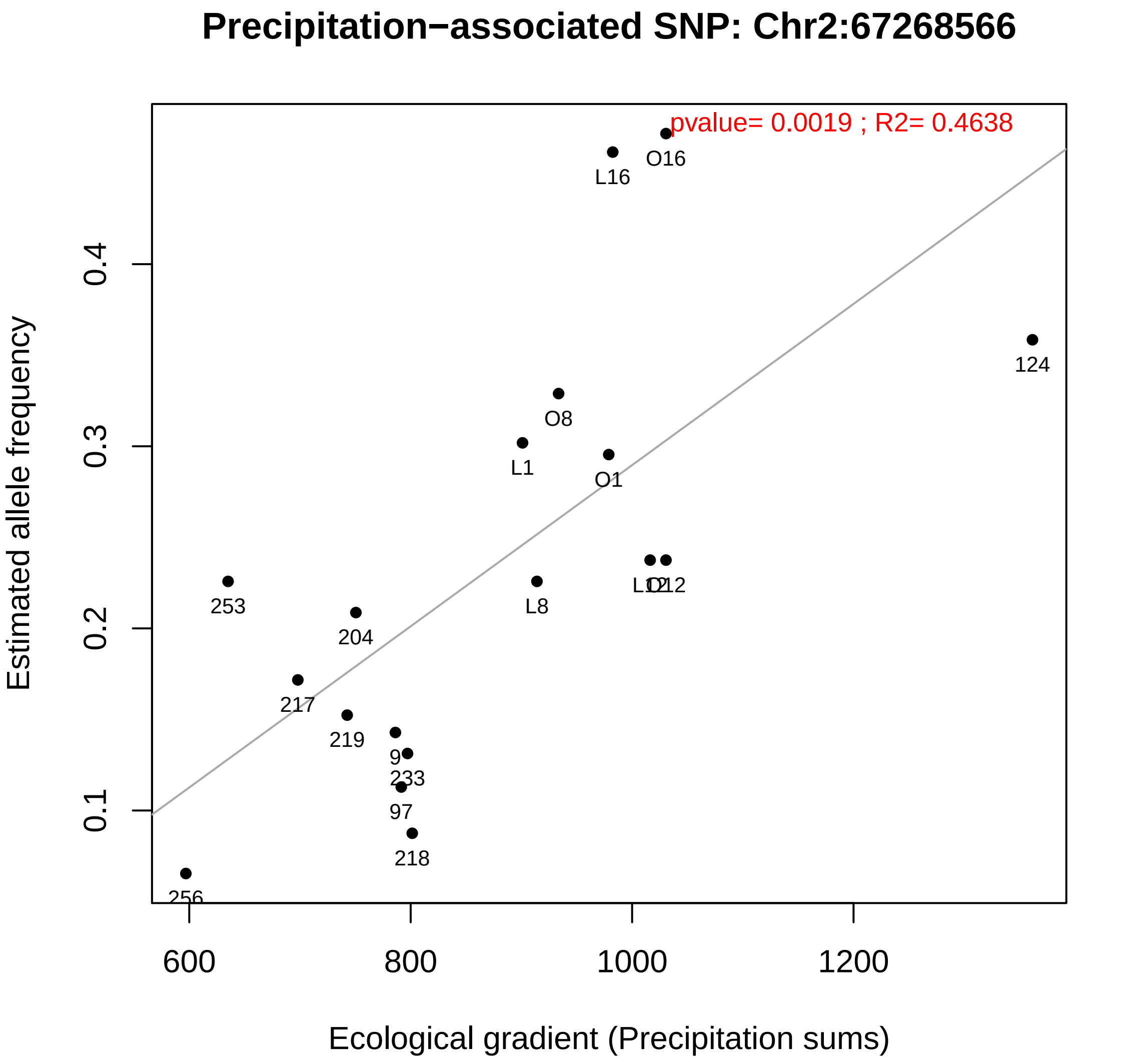

Precipitation−associated SNP: Chr2:67268566
pvalue= 0.0019 ; R2= 0.4638
O16
L16
124
O8
L1
O1
L12
O12
253
L8
204
217
219
9
233
97
218
256
0.1
0.2
0.3
0.4
600
800
1000
1200
Estimated allele frequency
Ecological gradient (Precipitation sums)

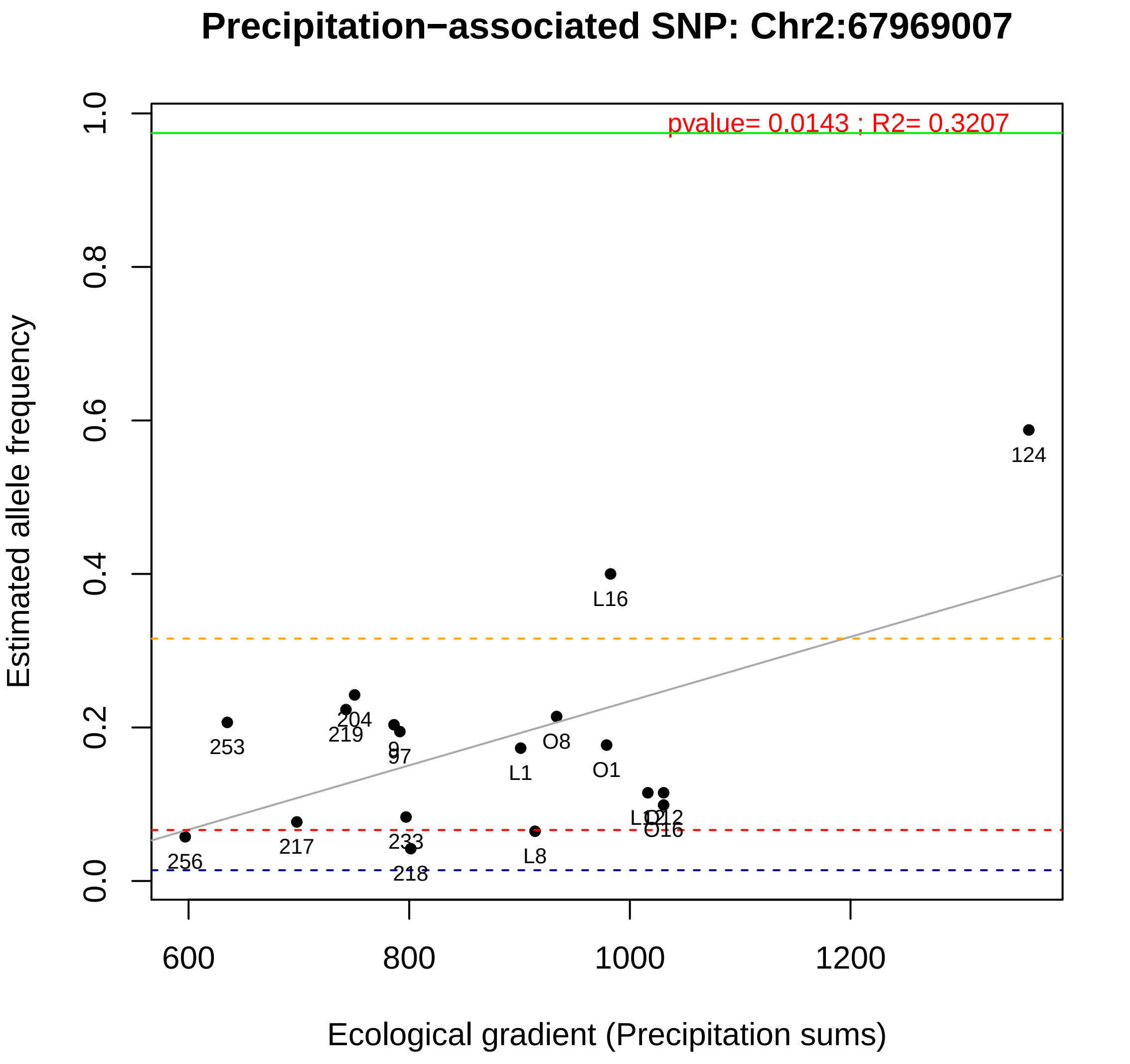
Precipitation−associated SNP: Chr2:67969007
pvalue= 0.0143 ; R2= 0.3207
Estimated allele frequency
Ecological gradient (Precipitation sums)
0.0
0.2
0.4
0.6
0.8
1.0
600
800
1000
1200
124
L16
253
204
219
97
L1
O8
O1
256
217
233
218
L8
O16

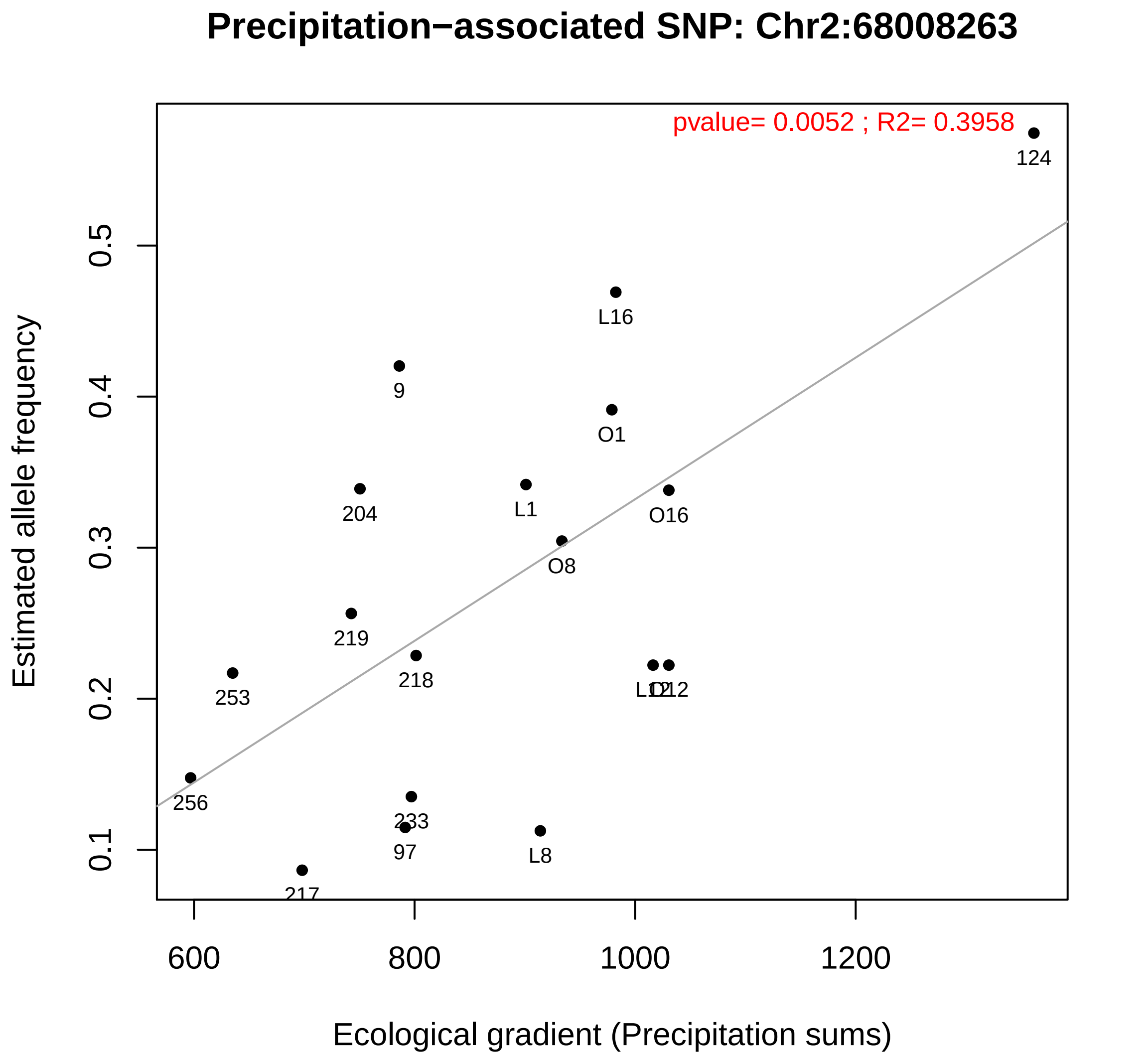

Precipitation−associated SNP: Chr2:68008263
pvalue= 0.0052 ; R2= 0.3958
124
L16
9
O1
204
L1
O16
O8
219
218
253
L12
O12
256
233
97
L8
217
0.1
0.2
0.3
0.4
0.5
600
800
1000
1200
Estimated allele frequency
Ecological gradient (Precipitation sums)

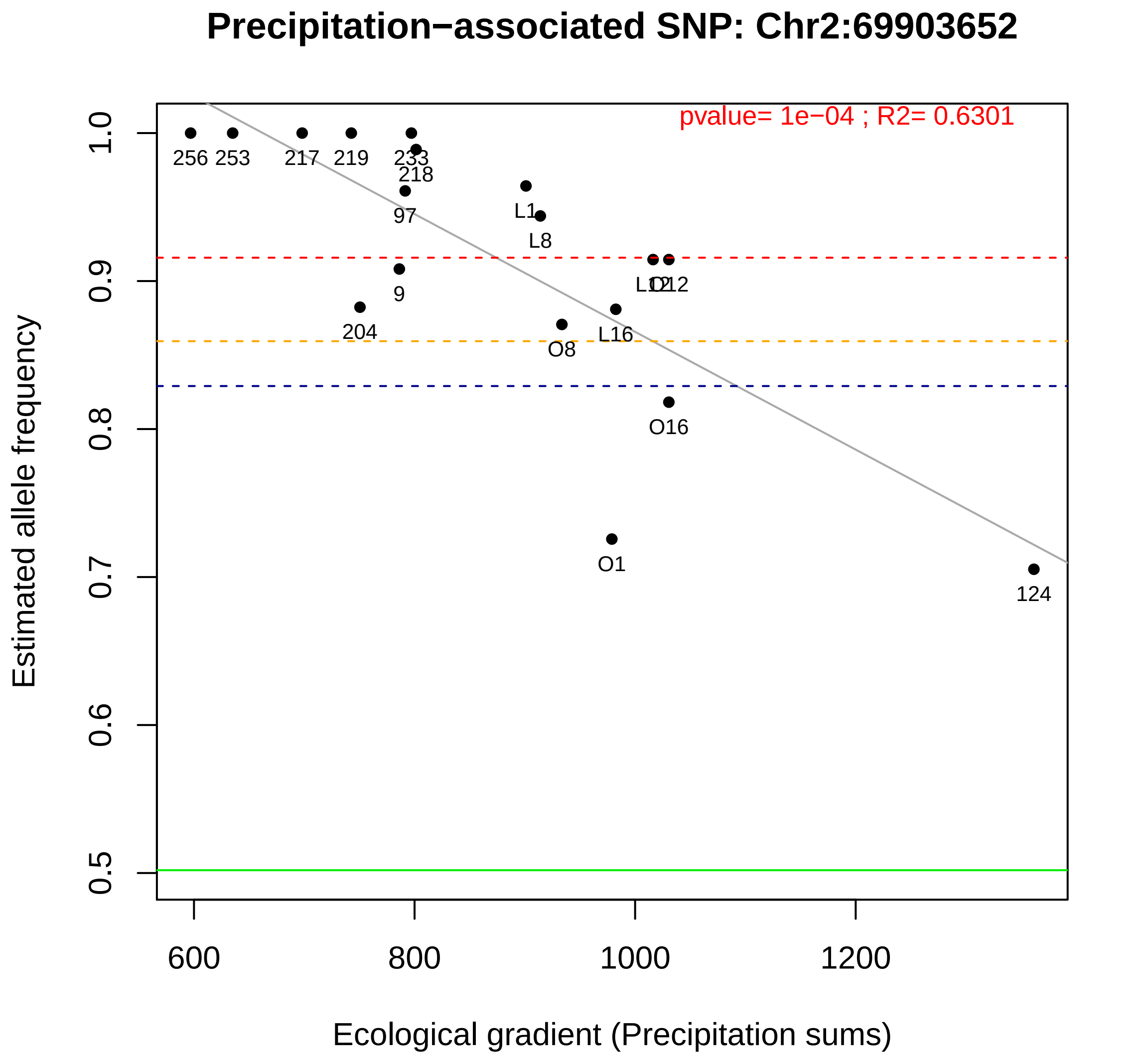

Precipitation−associated SNP: Chr2:69903652
pvalue= 1e−04 ; R2= 0.6301
256
253
217
219
233
218
97
L1
L8
9
204
L10
O12
L16
O8
O16
O1
124
Estimated allele frequency
0.5
0.6
0.7
0.8
0.9
1.0
600
800
1000
1200
Ecological gradient (Precipitation sums)

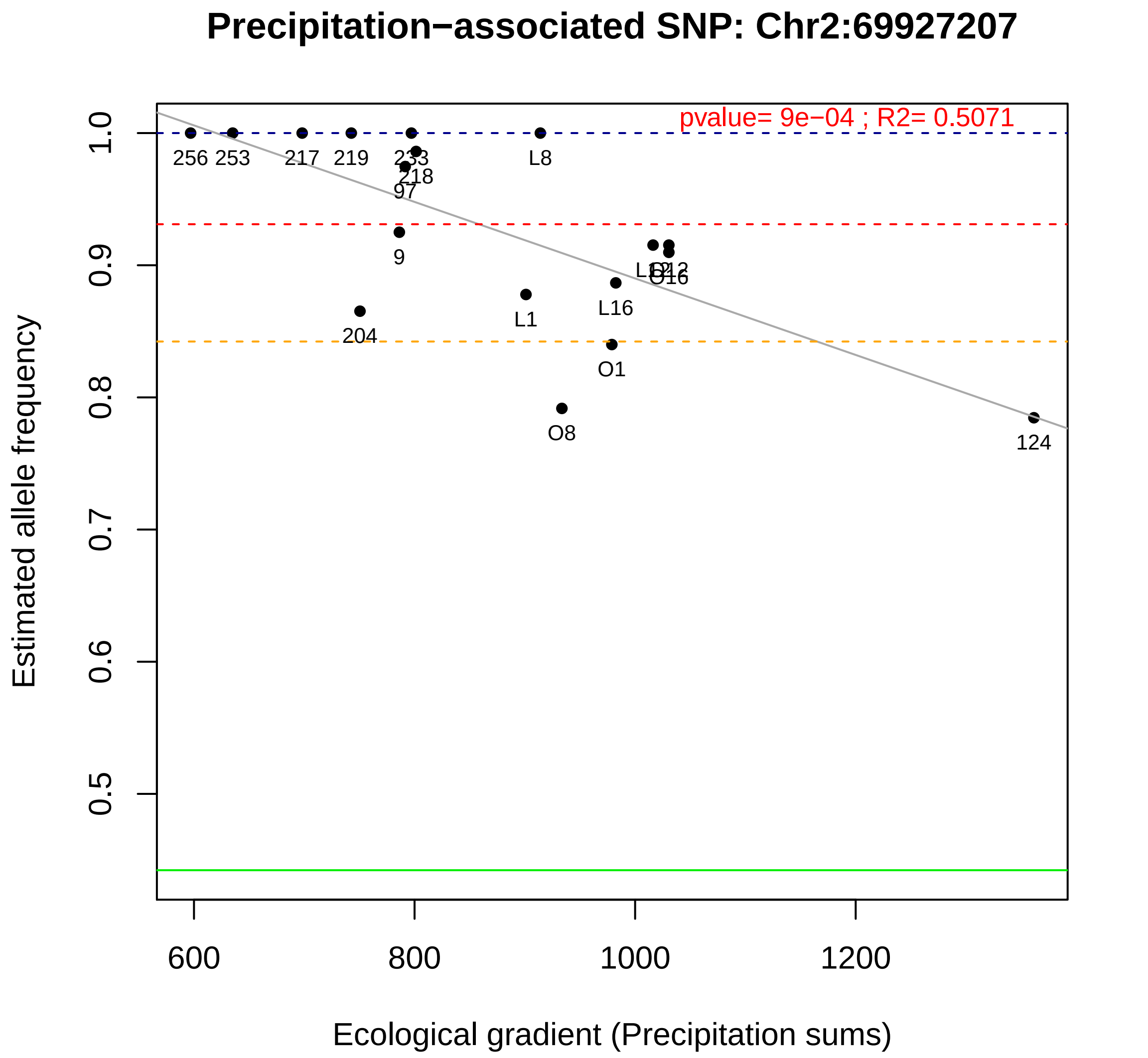
Precipitation−associated SNP: Chr2:69927207
pvalue= 9e−04 ; R2= 0.5071
Estimated allele frequency
Ecological gradient (Precipitation sums)
0.5
0.6
0.7
0.8
0.9
1.0
600
800
1000
1200
256
253
217
219
233
218
97
L8
9
204
L1
L16
O1
O8
O16
124

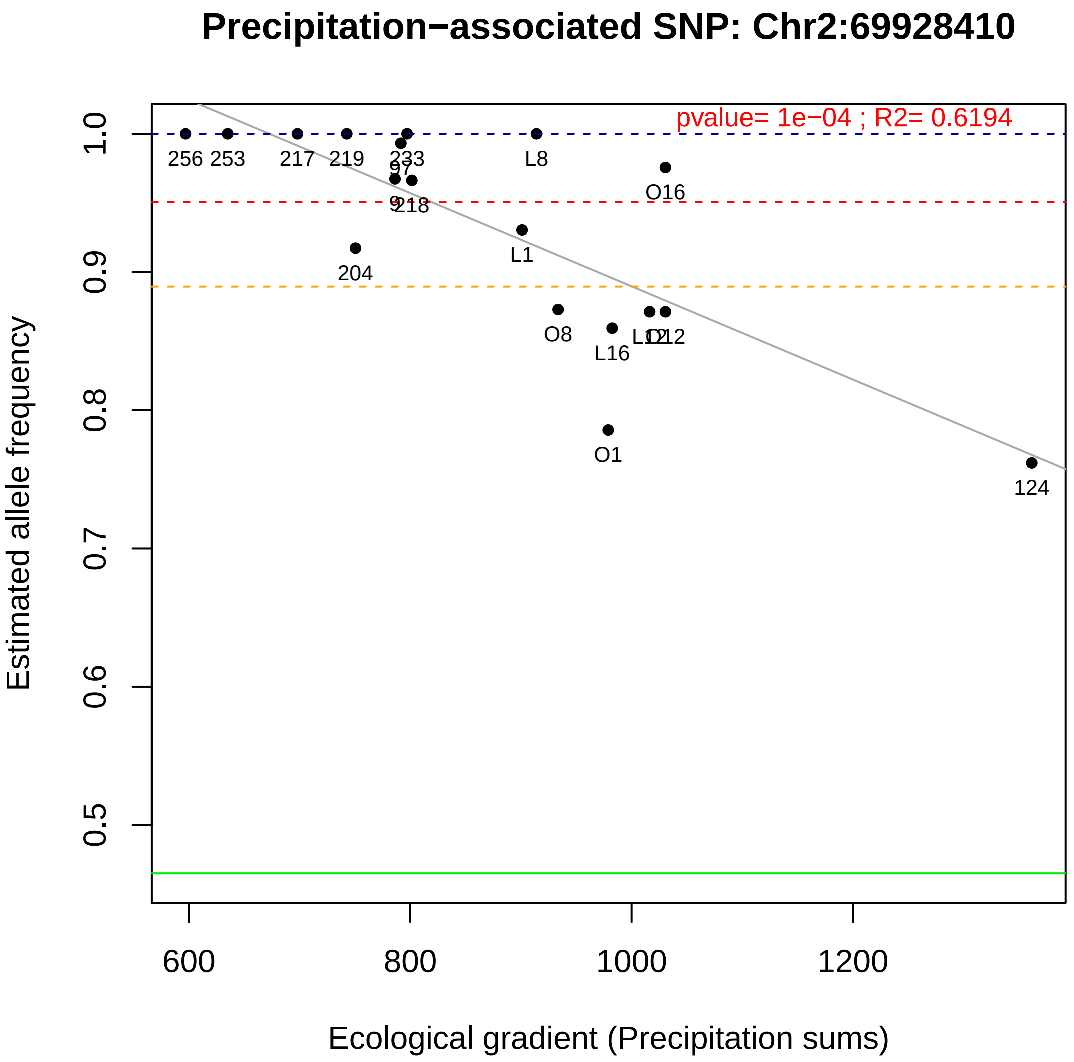

Precipitation−associated SNP: Chr2:69928410
pvalue= 1e−04 ; R2= 0.6194
Estimated allele frequency
Ecological gradient (Precipitation sums)
0.5
0.6
0.7
0.8
0.9
1.0
600
800
1000
1200
256
253
217
219
233
97
92
218
L8
O16
L1
204
O8
L16
L12
O12
O1
124

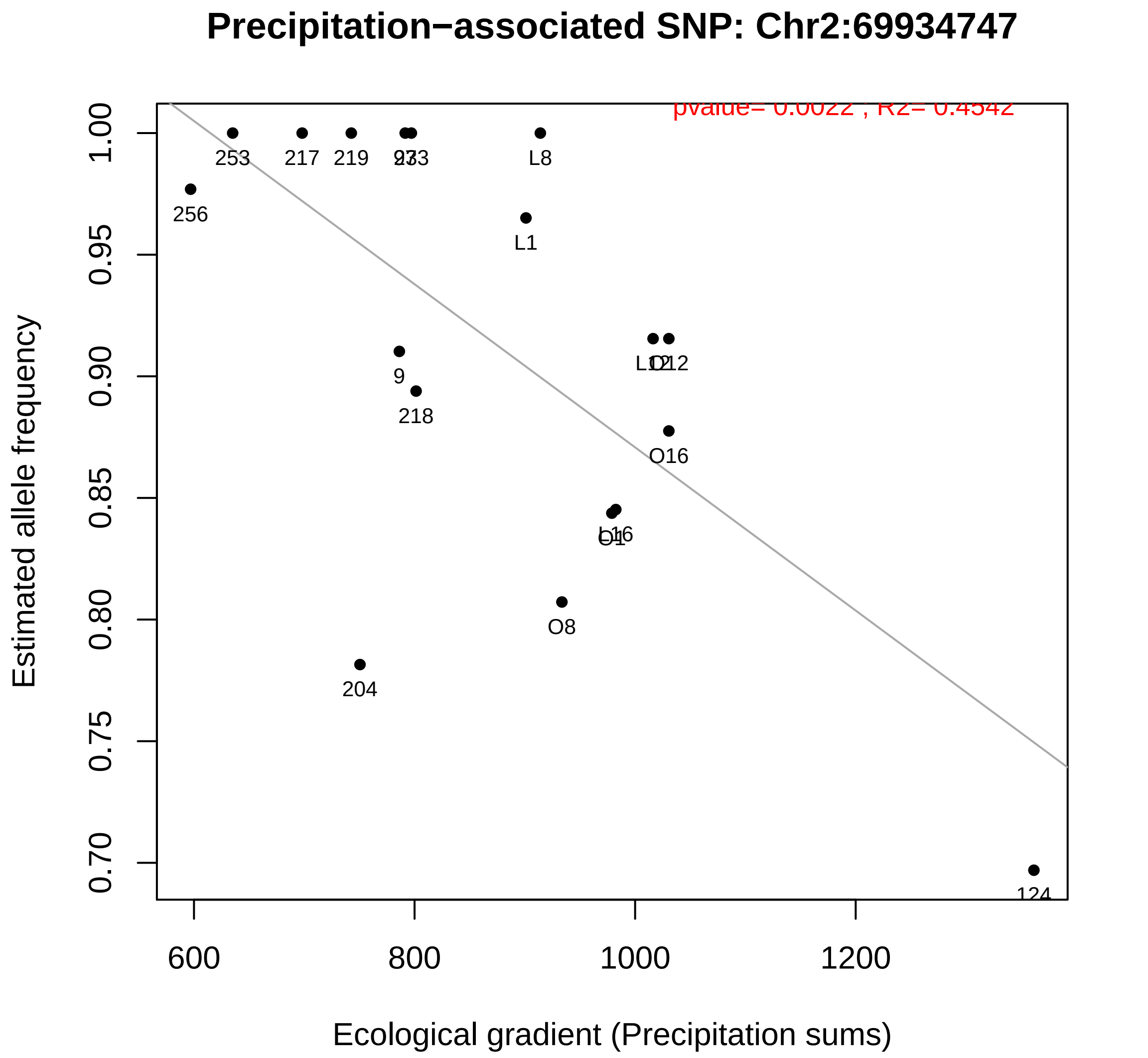

Precipitation−associated SNP: Chr2:69934747
pvalue= 0.0022 ; R2= 0.4542
Estimated allele frequency
Ecological gradient (Precipitation sums)
1.00
0.95
0.90
0.85
0.80
0.75
0.70
600
800
1000
1200
253
217
219
233
L8
256
L1
9
218
L12
O12
O16
L16
O1
O8
204
124

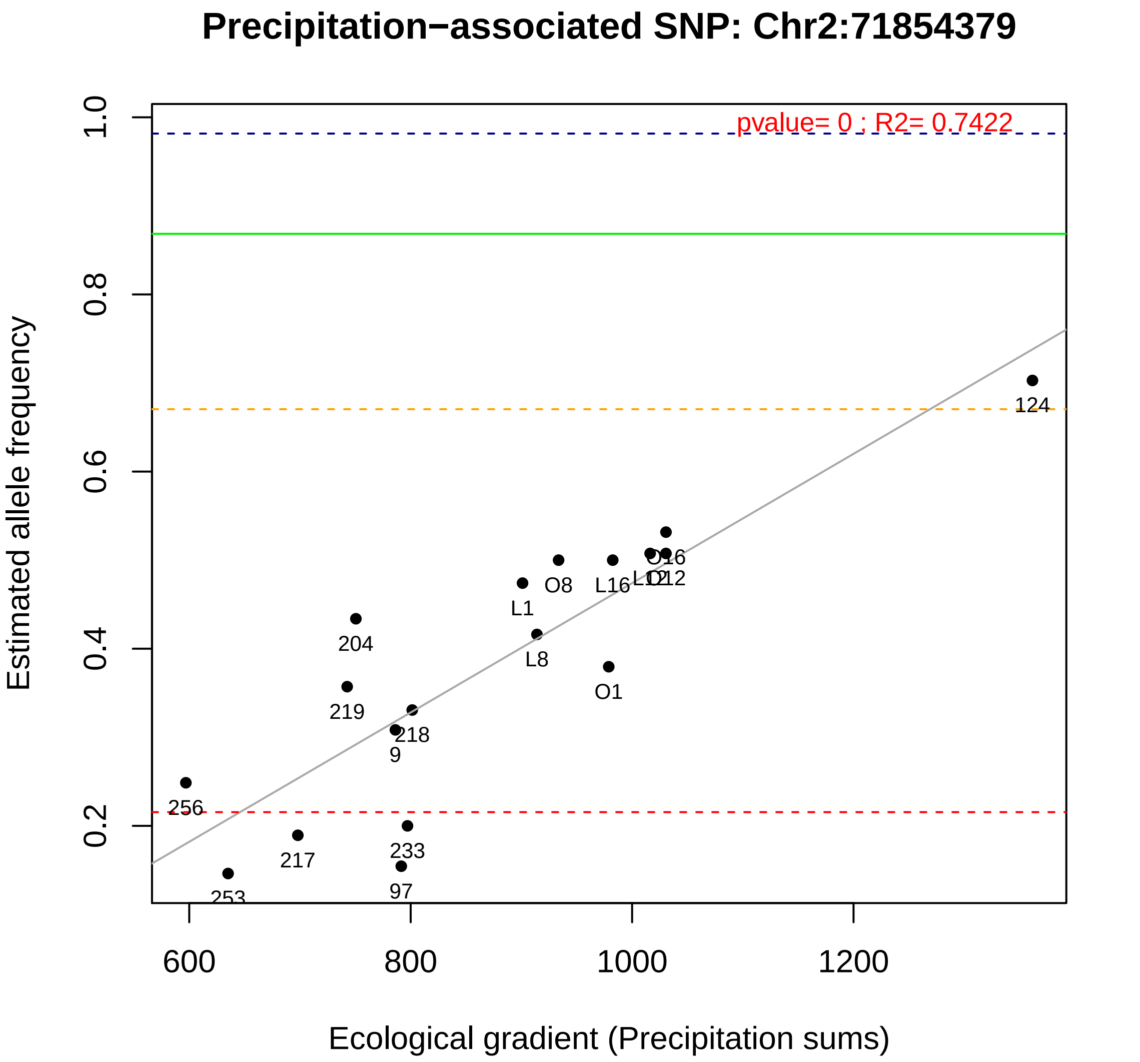

Precipitation−associated SNP: Chr2:71854379
pvalue= 0 ; R2= 0.7422
Estimated allele frequency
Ecological gradient (Precipitation sums)
0.2
0.4
0.6
0.8
1.0
600
800
1000
1200
124
O16
O12
L10
L16
O8
L1
L8
O1
204
219
218
9
256
217
233
97
253

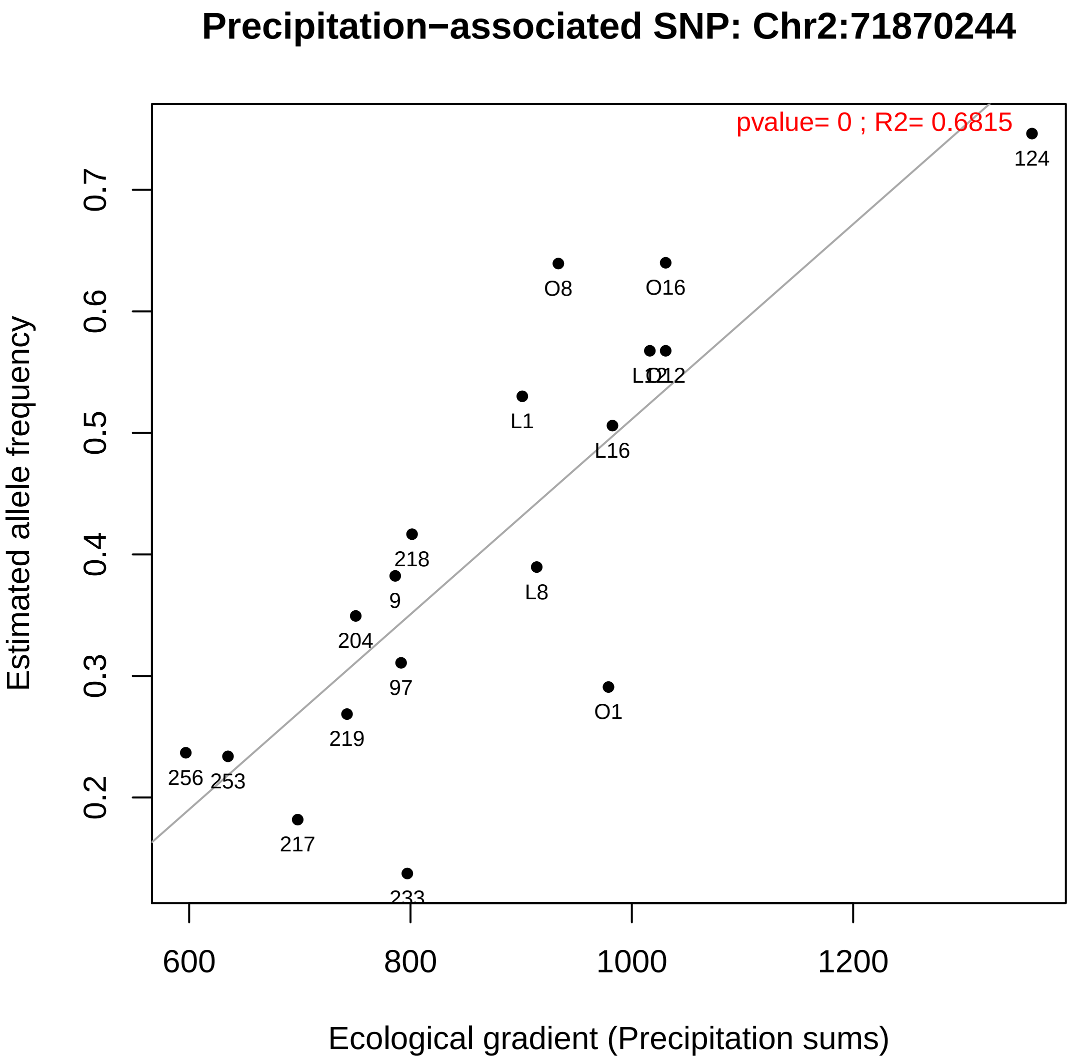
Precipitation−associated SNP: Chr2:71870244
pvalue= 0 ; R2= 0.6815
Estimated allele frequency
0.2
0.3
0.4
0.5
0.6
0.7
600
800
1000
1200
Ecological gradient (Precipitation sums)
124
O8
O16
L12
O12
L1
L16
218
9
L8
204
97
O1
219
256
253
217
233

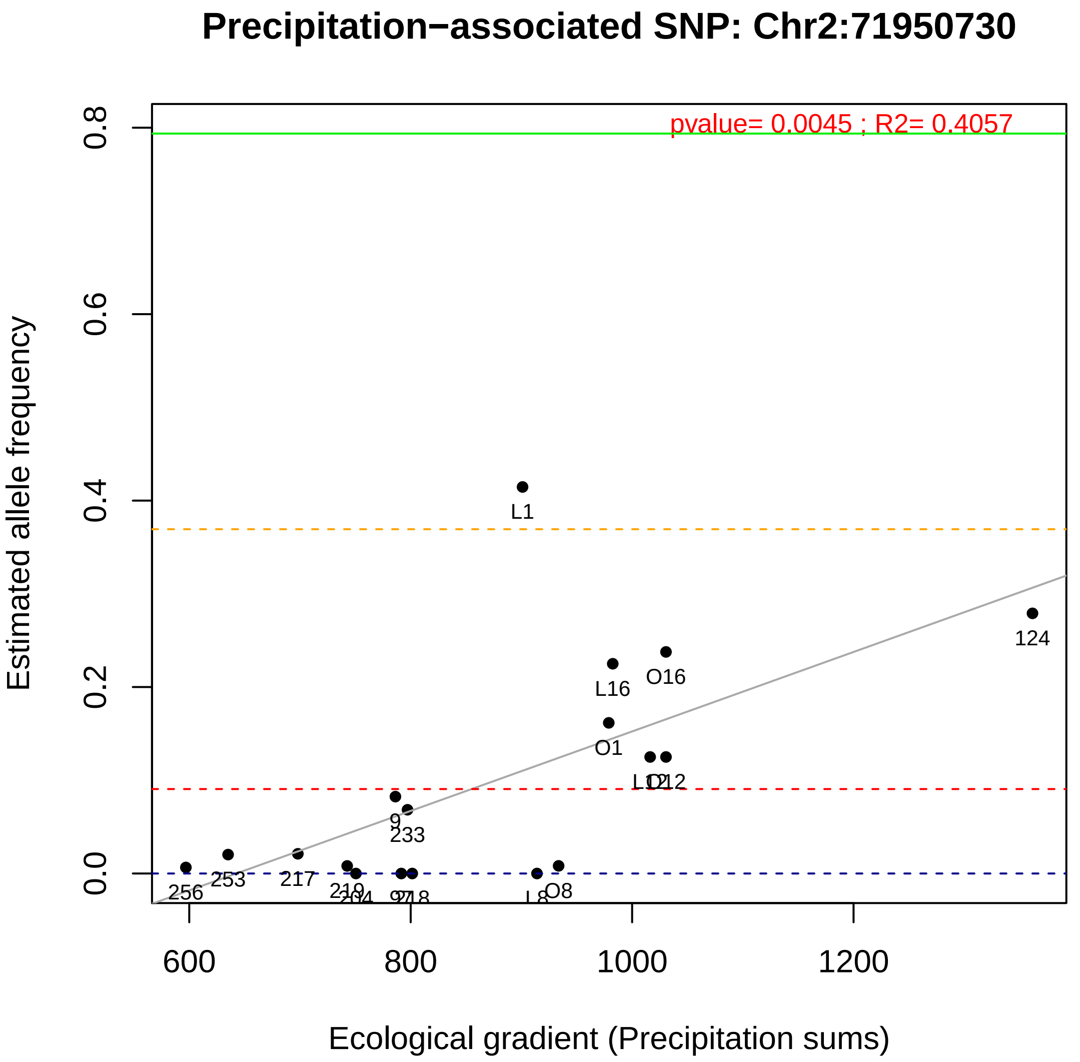
Precipitation−associated SNP: Chr2:71950730
pvalue= 0.0045 ; R2= 0.4057
Estimated allele frequency
0.0
0.2
0.4
0.6
0.8
Ecological gradient (Precipitation sums)
600
800
1000
1200
L1
124
L16
O16
O1
9
233
256
253
217
219
O8

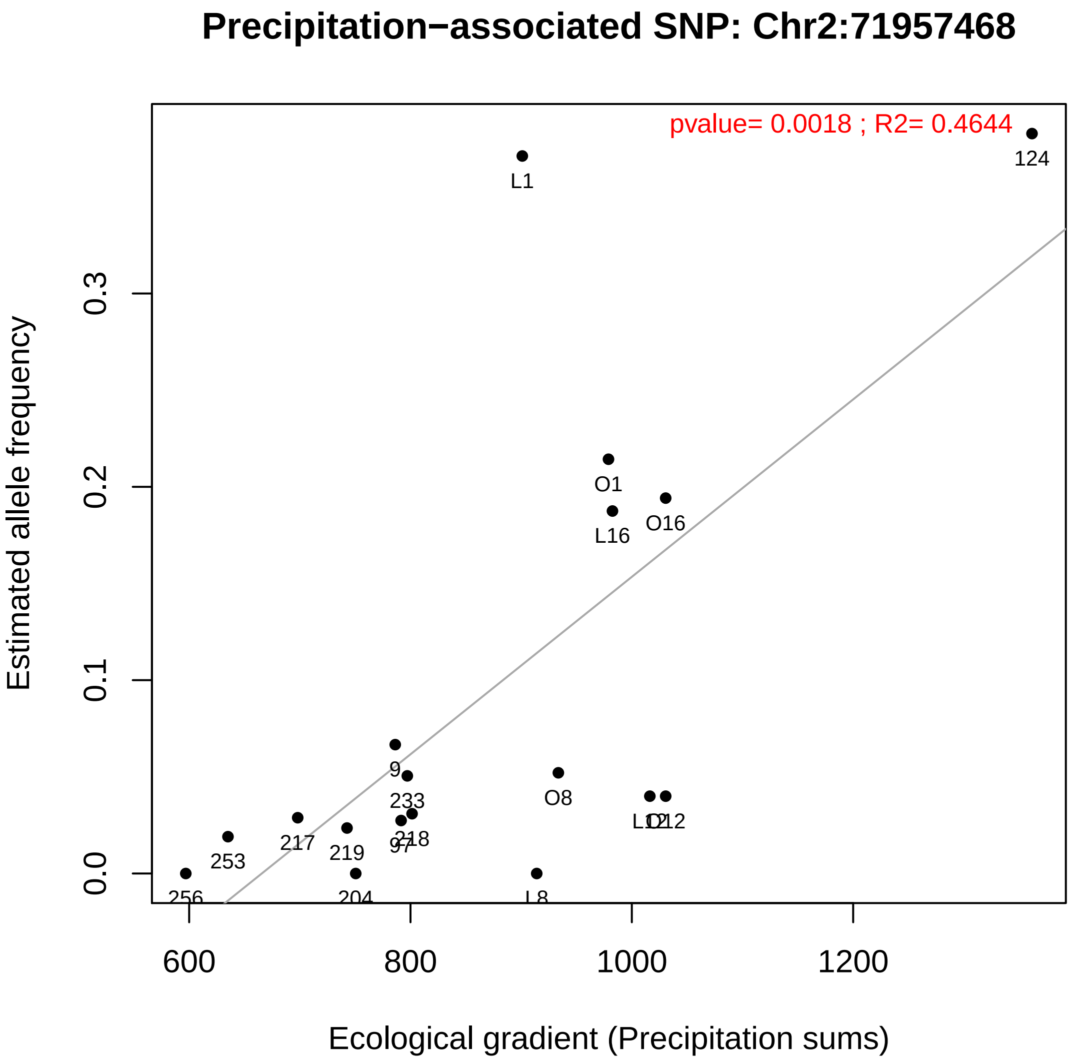
Precipitation−associated SNP: Chr2:71957468
pvalue= 0.0018 ; R2= 0.4644
124
L1
O1
O16
L16
9
233
O8
L12
O12
217
219
218
97
253
256
204
L8
Estimated allele frequency
0.0
0.1
0.2
0.3
600
800
1000
1200
Ecological gradient (Precipitation sums)

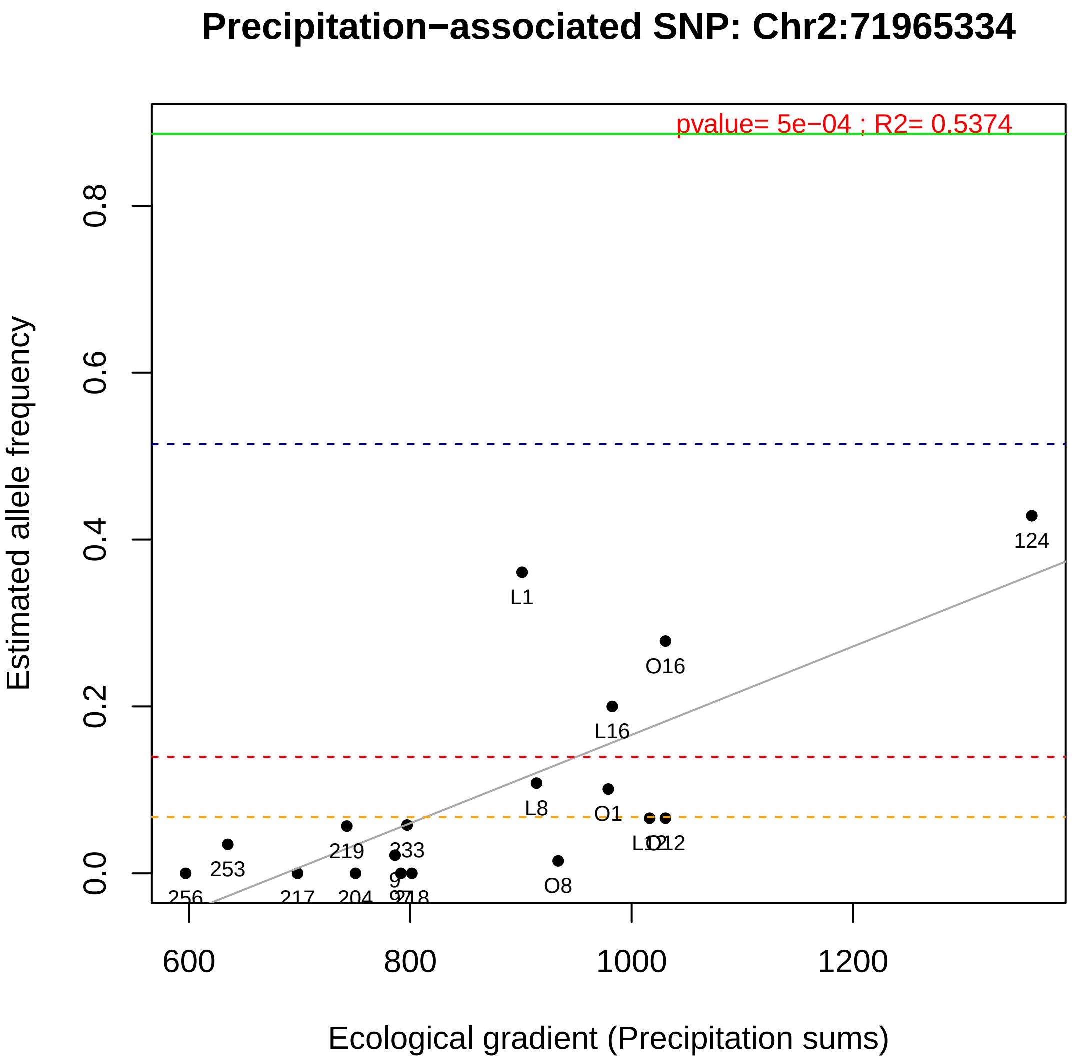

Precipitation−associated SNP: Chr2:71965334
pvalue= 5e−04 ; R2= 0.5374
Estimated allele frequency
0.0
0.2
0.4
0.6
0.8
600
800
1000
1200
Ecological gradient (Precipitation sums)
124
L1
O16
L16
L8
O1
O8
219
233
253
256
217
204

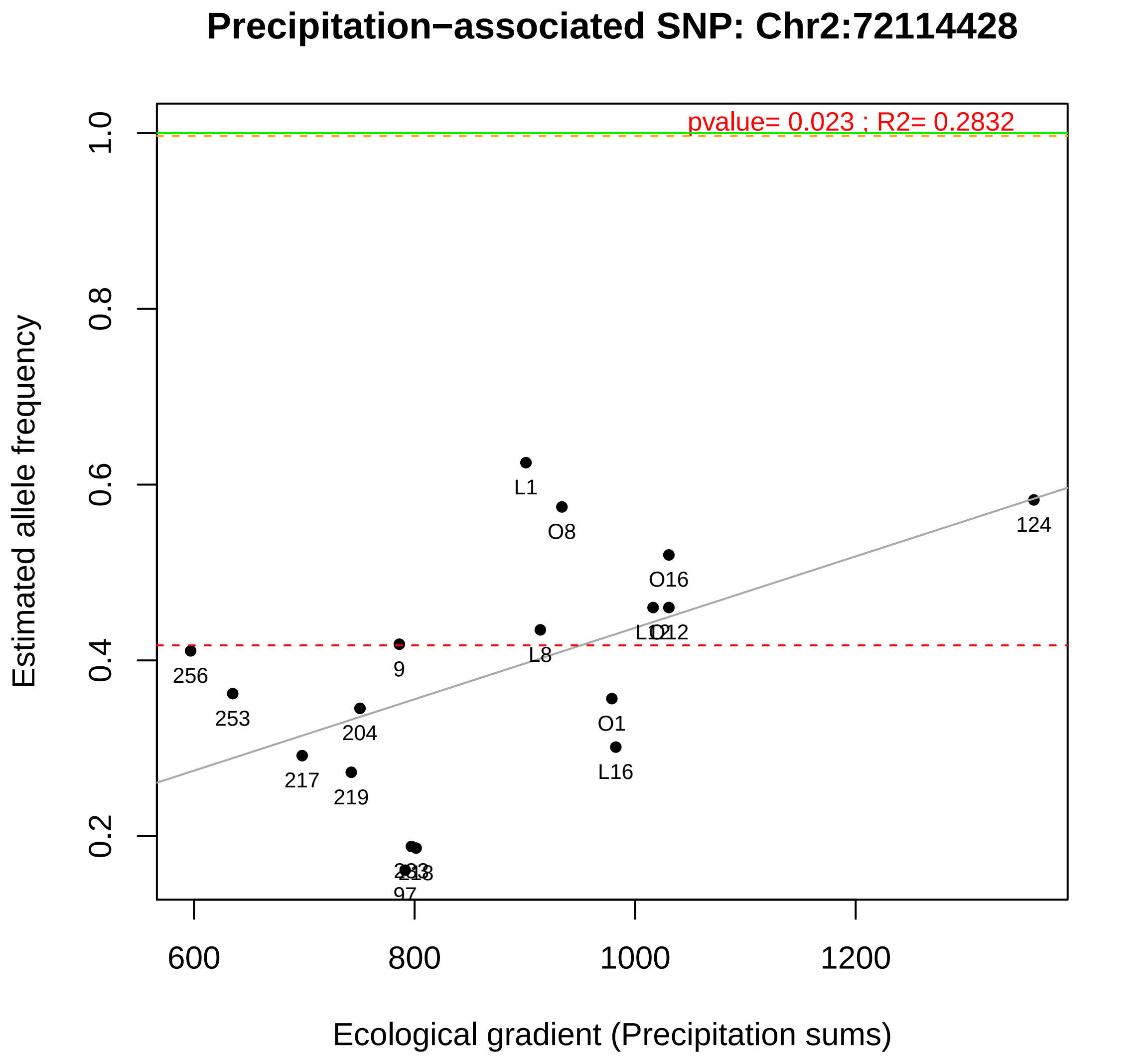
Precipitation−associated SNP: Chr2:72114428
pvalue= 0.023 ; R2= 0.2832
Estimated allele frequency
Ecological gradient (Precipitation sums)
0.2
0.4
0.6
0.8
1.0
600
800
1000
1200
L1
O8
124
O16
L12
O12
L8
9
256
253
204
217
219
O1
L16
97

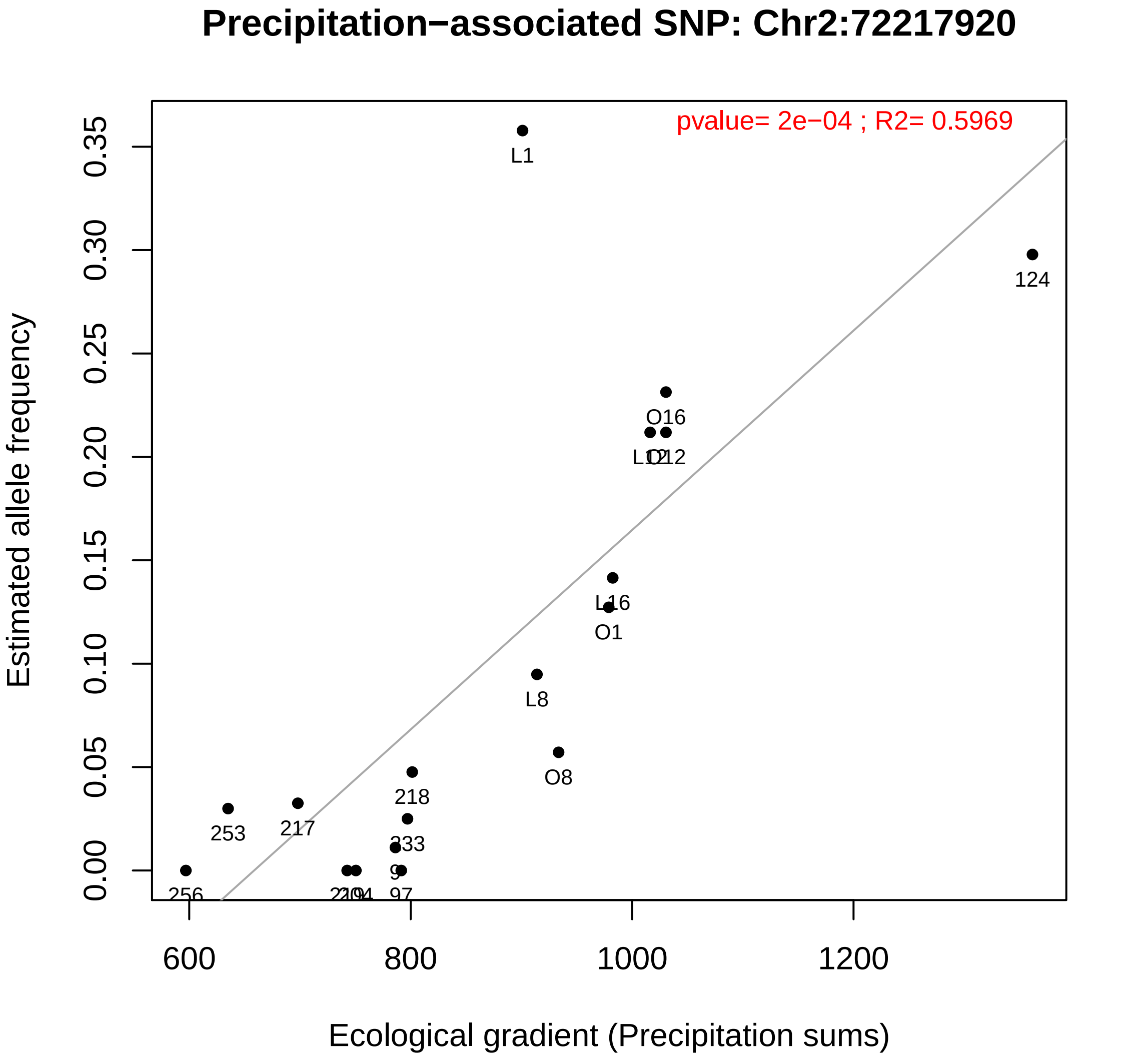
Precipitation−associated SNP: Chr2:72217920
pvalue= 2e−04 ; R2= 0.5969
Estimated allele frequency
Ecological gradient (Precipitation sums)
0.00
0.05
0.10
0.15
0.20
0.25
0.30
0.35
600
800
1000
1200
L1
124
O16
L16
O1
L8
O8
218
333
253
217
256
97

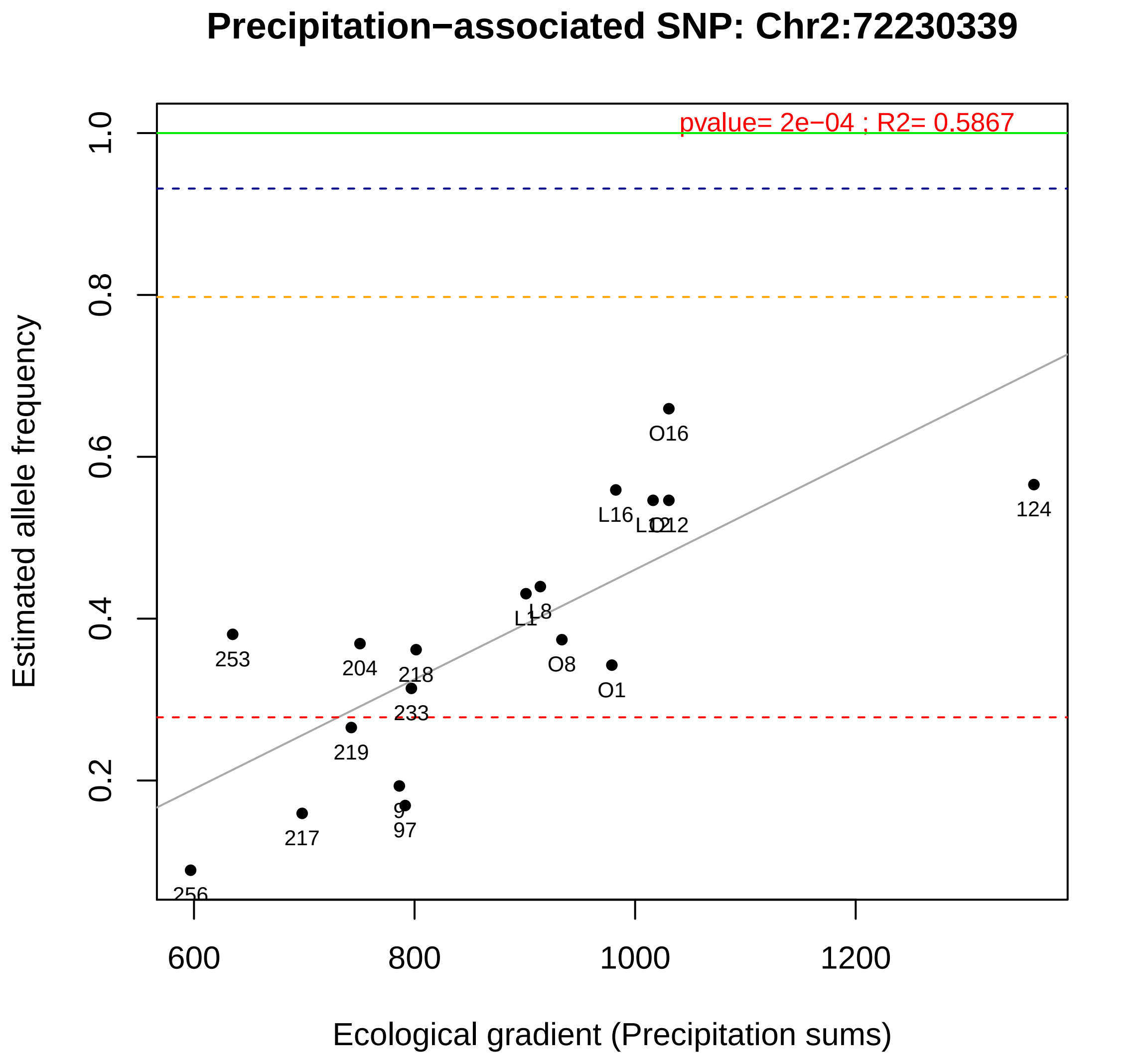
Precipitation−associated SNP: Chr2:72230339
pvalue= 2e−04 ; R2= 0.5867
Estimated allele frequency
0.2
0.4
0.6
0.8
1.0
600
800
1000
1200
Ecological gradient (Precipitation sums)
O16
L16
124
L8
L1
O8
O1
253
204
218
233
219
9
97
217
256

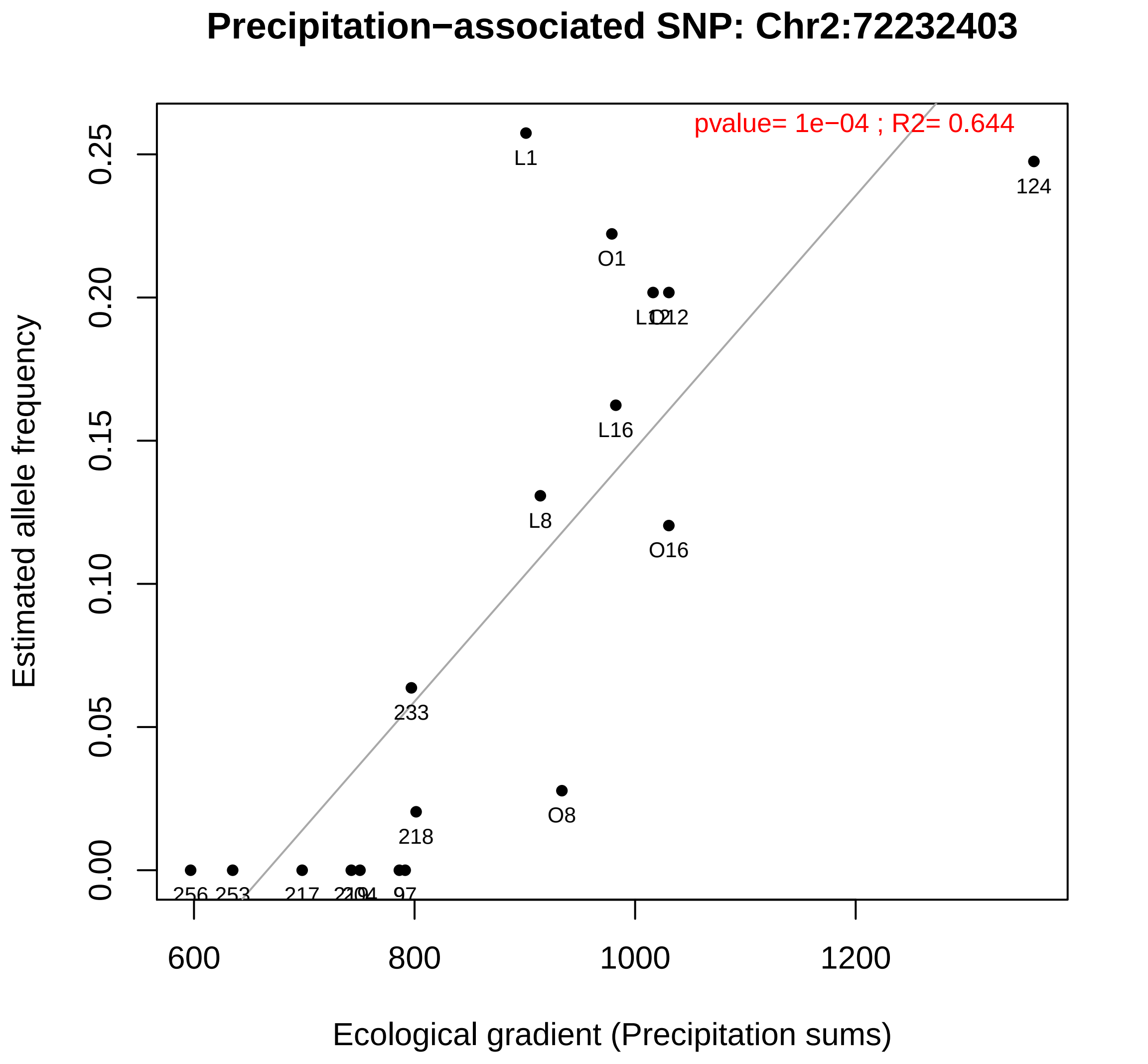
Precipitation−associated SNP: Chr2:72232403
pvalue= 1e−04 ; R2= 0.644
Estimated allele frequency
Ecological gradient (Precipitation sums)
0.00
0.05
0.10
0.15
0.20
0.25
600
800
1000
1200
L1
124
O1
L12
O12
L16
L8
O16
233
O8
218
256
253
217
97

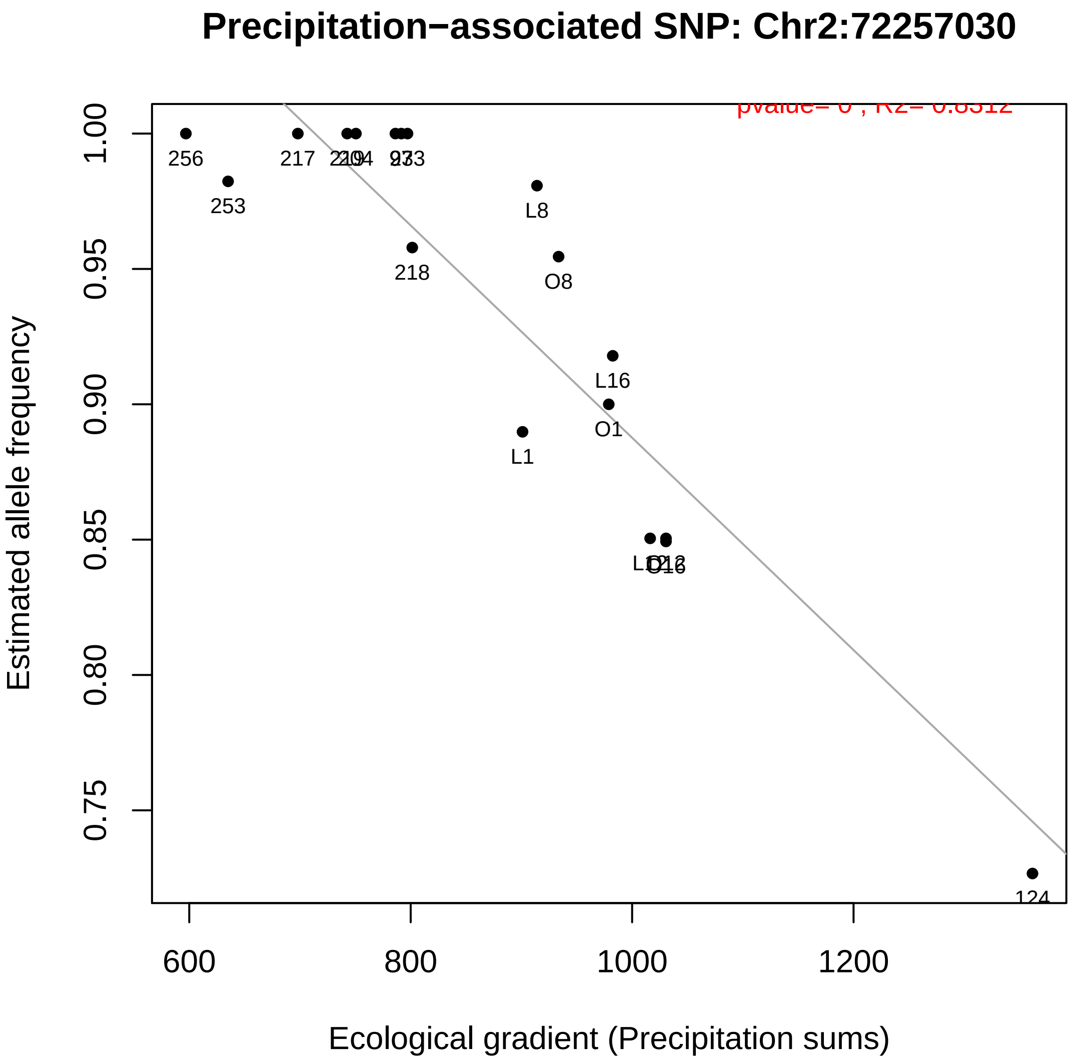

Precipitation−associated SNP: Chr2:72257030
pvalue= 0 , R2= 0.8312
Estimated allele frequency
1.00
0.95
0.90
0.85
0.80
0.75
256
217
253
218
L8
O8
L16
O1
L1
124
600
800
1000
1200
Ecological gradient (Precipitation sums)

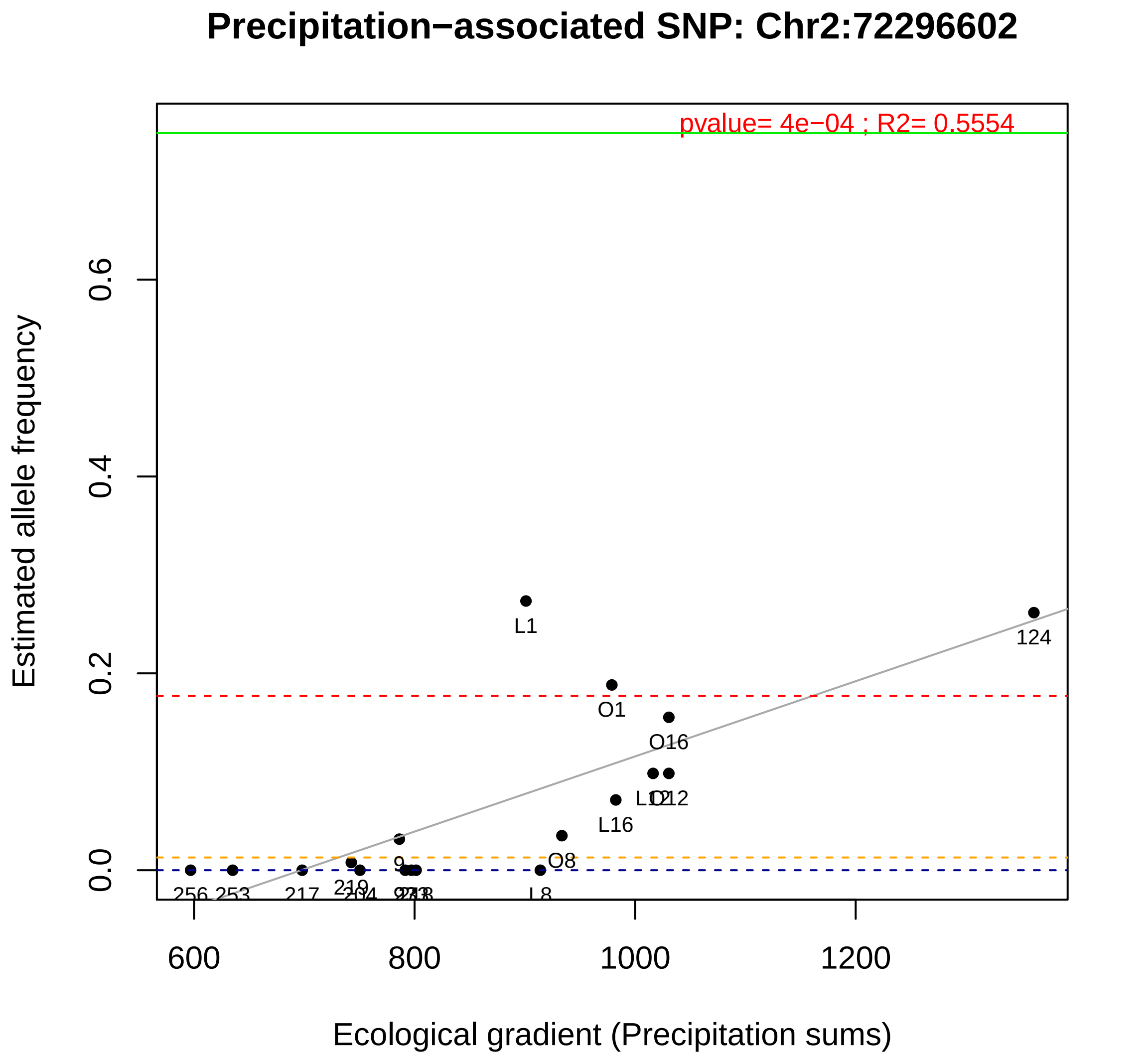
Precipitation−associated SNP: Chr2:72296602
pvalue= 4e−04 ; R2= 0.5554
Estimated allele frequency
0.0
0.2
0.4
0.6
600
800
1000
1200
Ecological gradient (Precipitation sums)
L1
124
O1
O16
L16
O8
L8
9
219
256
253
217

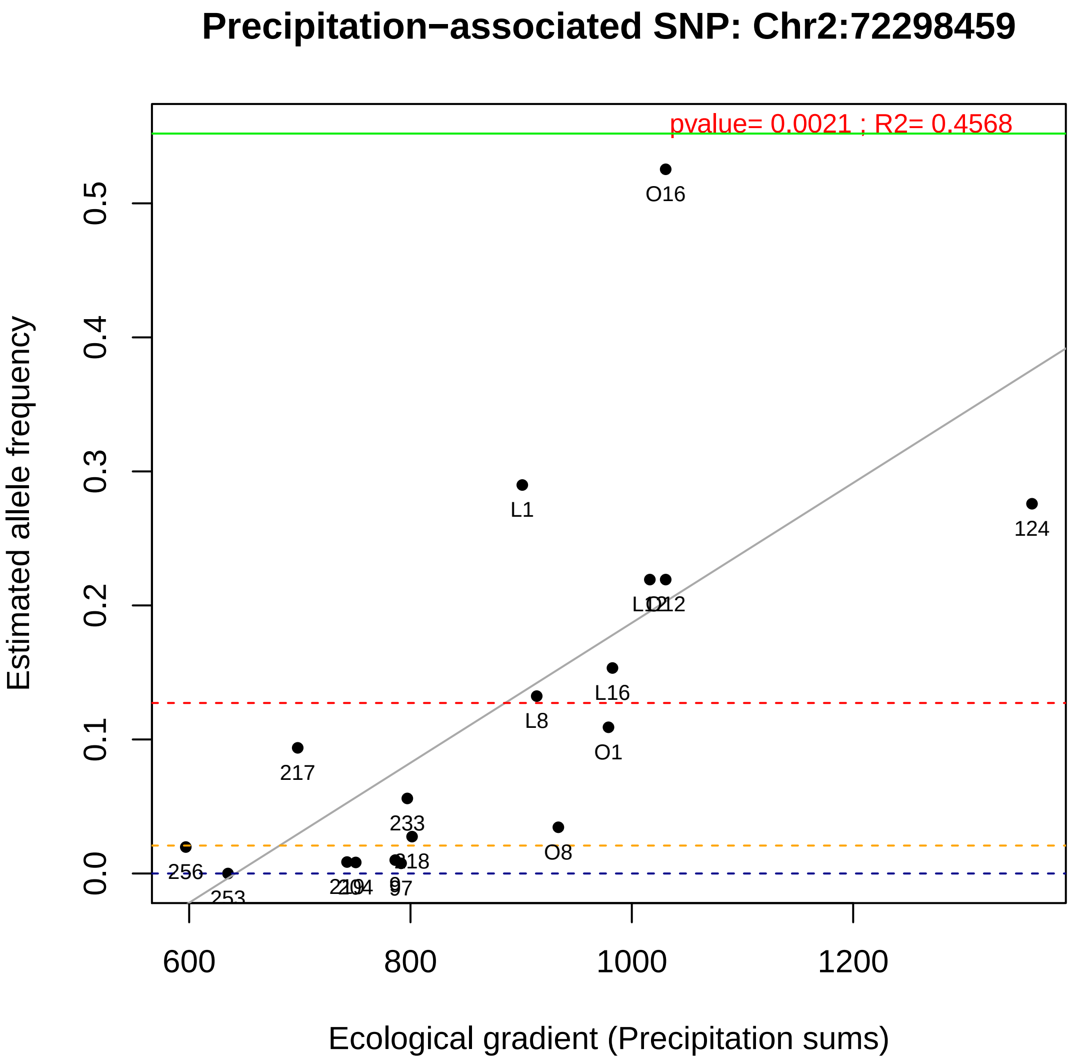
Precipitation−associated SNP: Chr2:72298459
pvalue= 0.0021 ; R2= 0.4568
Estimated allele frequency
Ecological gradient (Precipitation sums)
0.0
0.1
0.2
0.3
0.4
0.5
600
800
1000
1200
O16
L1
124
L16
L8
O1
217
233
O8
256
253
97

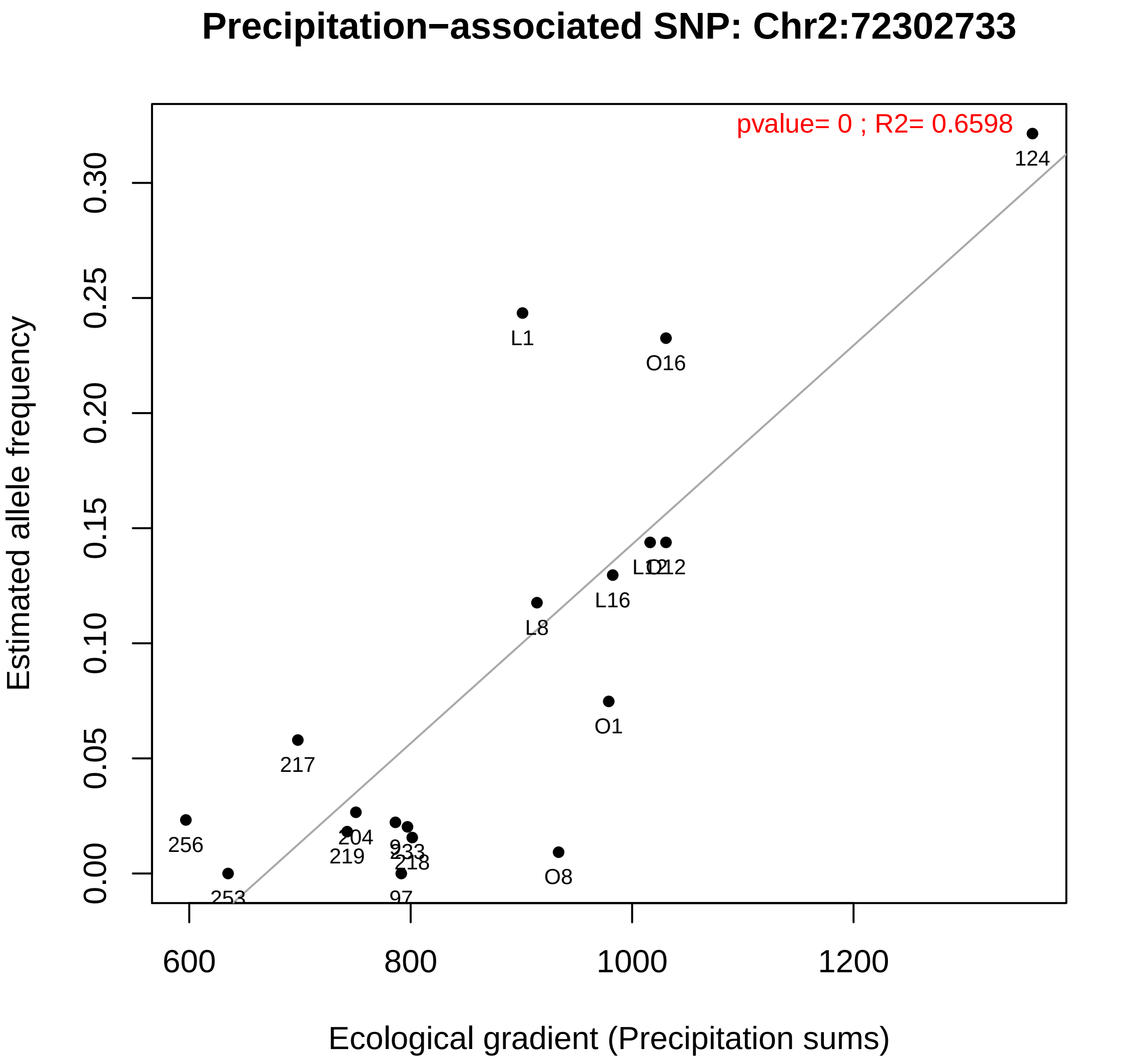

Precipitation−associated SNP: Chr2:72302733
pvalue= 0 ; R2= 0.6598
124
L1
O16
L12
O12
L16
L8
O1
217
256
204
219
9
233
218
O8
253
97
0.00
0.05
0.10
0.15
0.20
0.25
0.30
600
800
1000
1200
Estimated allele frequency
Ecological gradient (Precipitation sums)

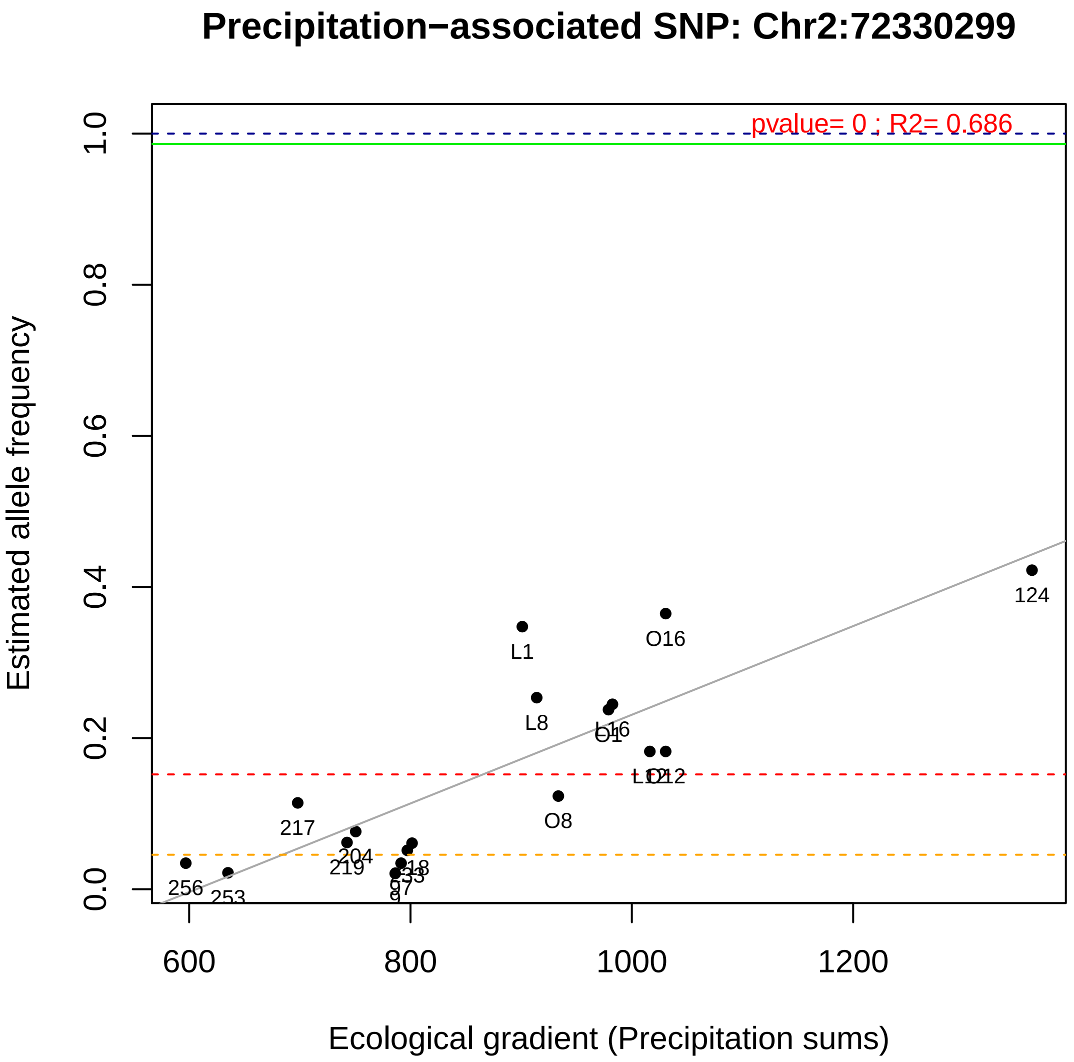

Precipitation−associated SNP: Chr2:72330299
pvalue= 0 ; R2= 0.686
Estimated allele frequency
0.0
0.2
0.4
0.6
0.8
1.0
600
800
1000
1200
Ecological gradient (Precipitation sums)
124
O16
L1
L8
O8
217
204
219
256
253

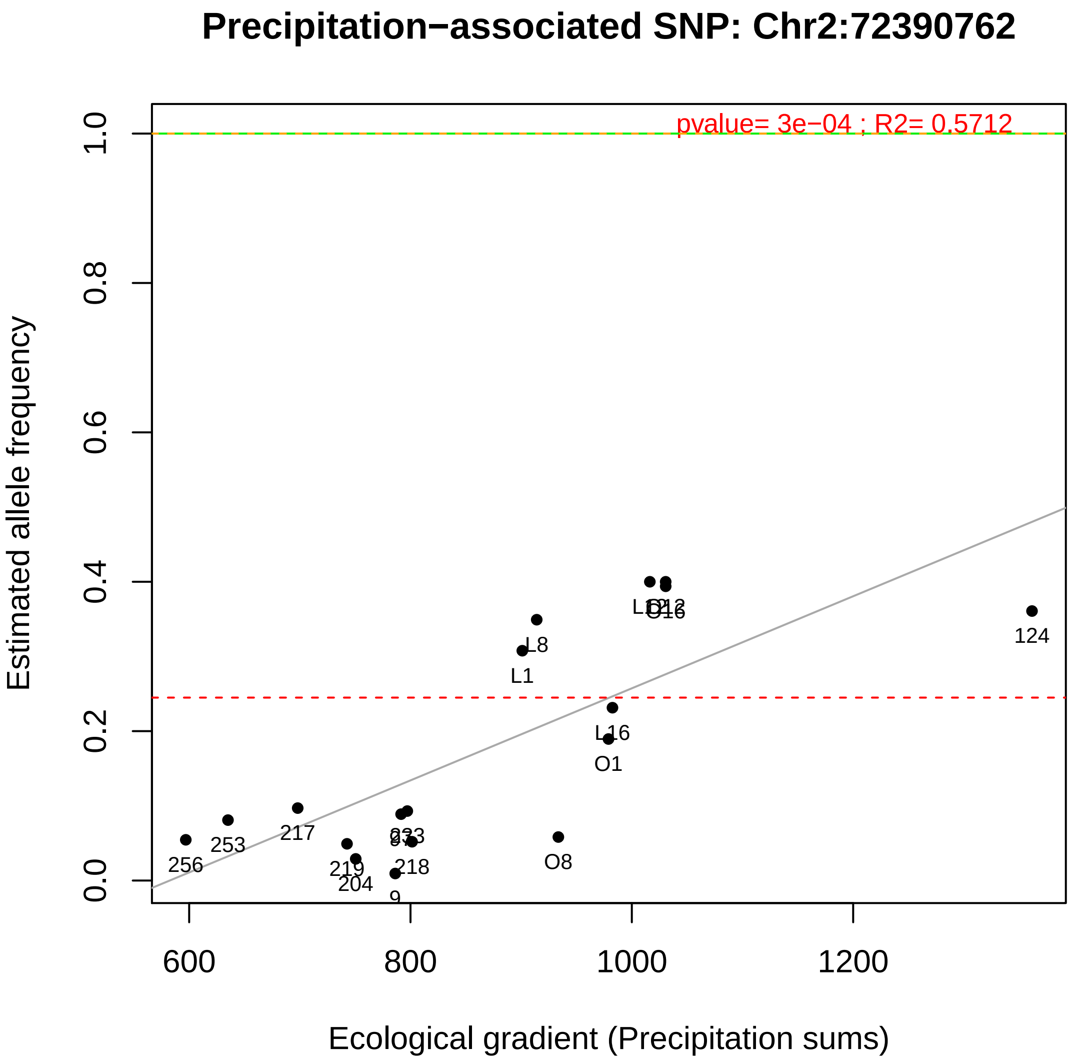
Precipitation−associated SNP: Chr2:72390762
pvalue= 3e−04 ; R2= 0.5712
Estimated allele frequency
Ecological gradient (Precipitation sums)
0.0
0.2
0.4
0.6
0.8
1.0
600
800
1000
1200
256
253
217
219
204
218
O8
L16
O1
L8
L1
124
9

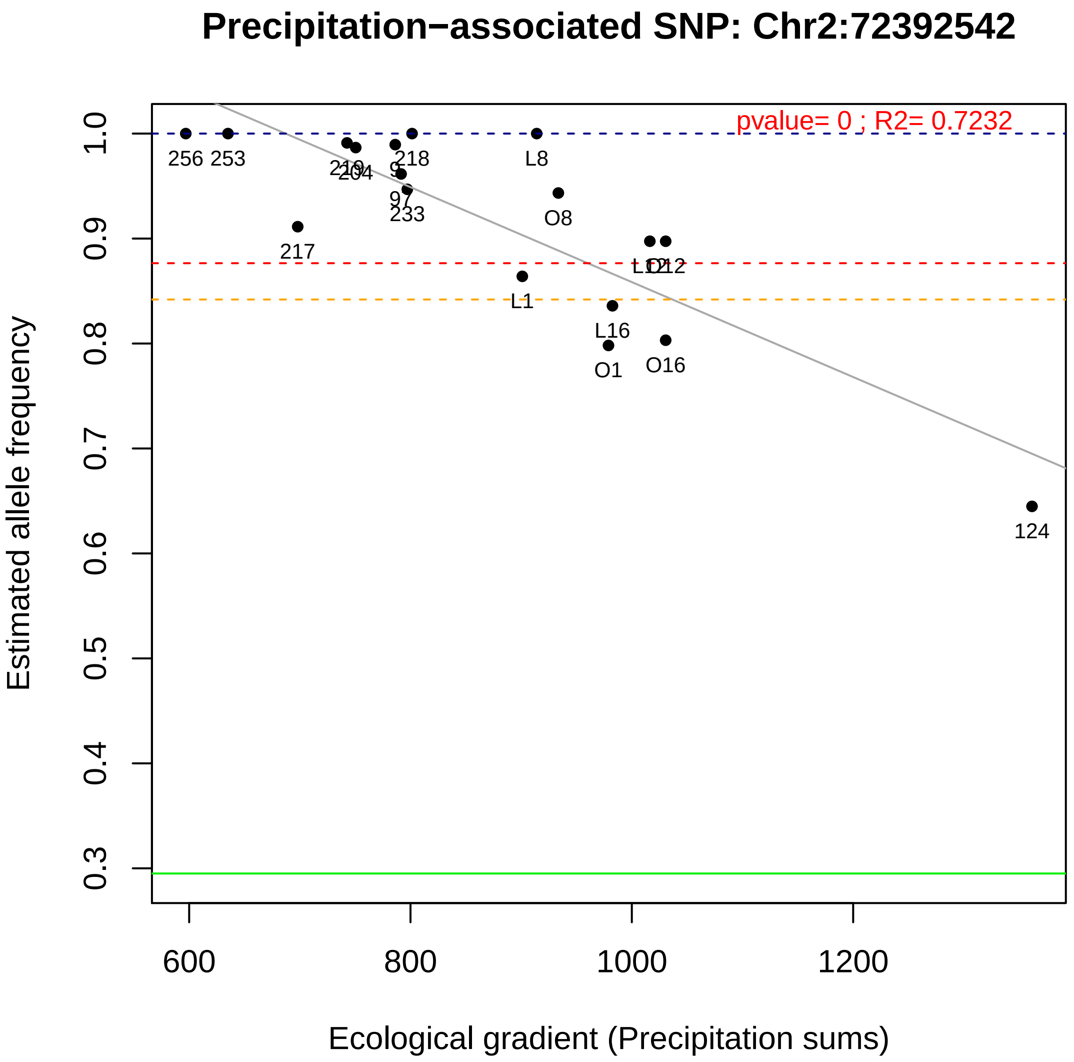

Precipitation−associated SNP: Chr2:72392542
pvalue= 0 ; R2= 0.7232
Estimated allele frequency
Ecological gradient (Precipitation sums)
0.3
0.4
0.5
0.6
0.7
0.8
0.9
1.0
600
800
1000
1200
256
253
210
204
9
218
97
233
L8
O8
217
L12
O12
L1
L16
O1
O16
124

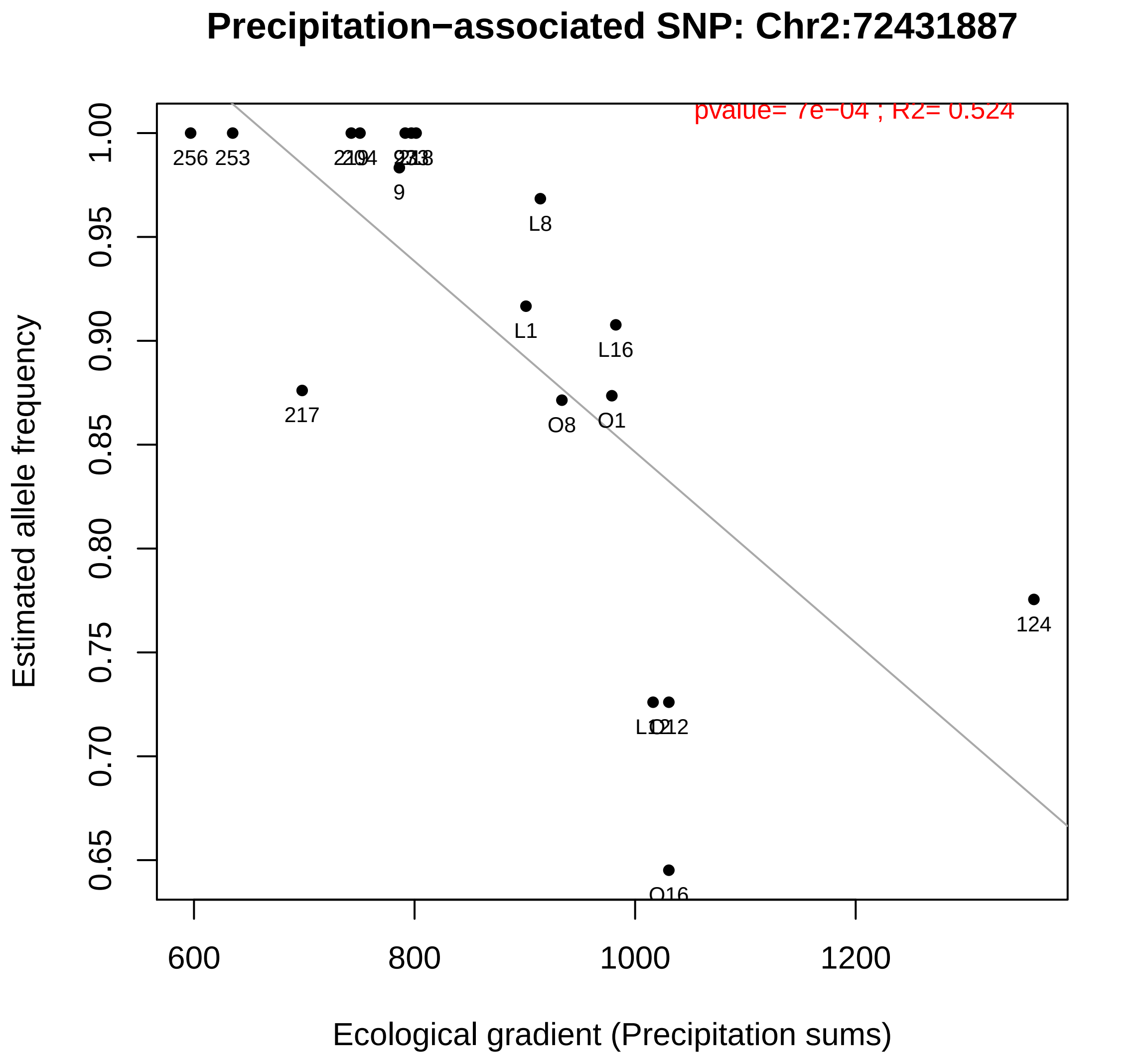

Precipitation−associated SNP: Chr2:72431887
pvalue= 7e−04 ; R2= 0.524
Estimated allele frequency
1.00
0.95
0.90
0.85
0.80
0.75
0.70
0.65
256
253
217
9
L8
L1
L16
O8
O1
124
O16
600
800
1000
1200
Ecological gradient (Precipitation sums)

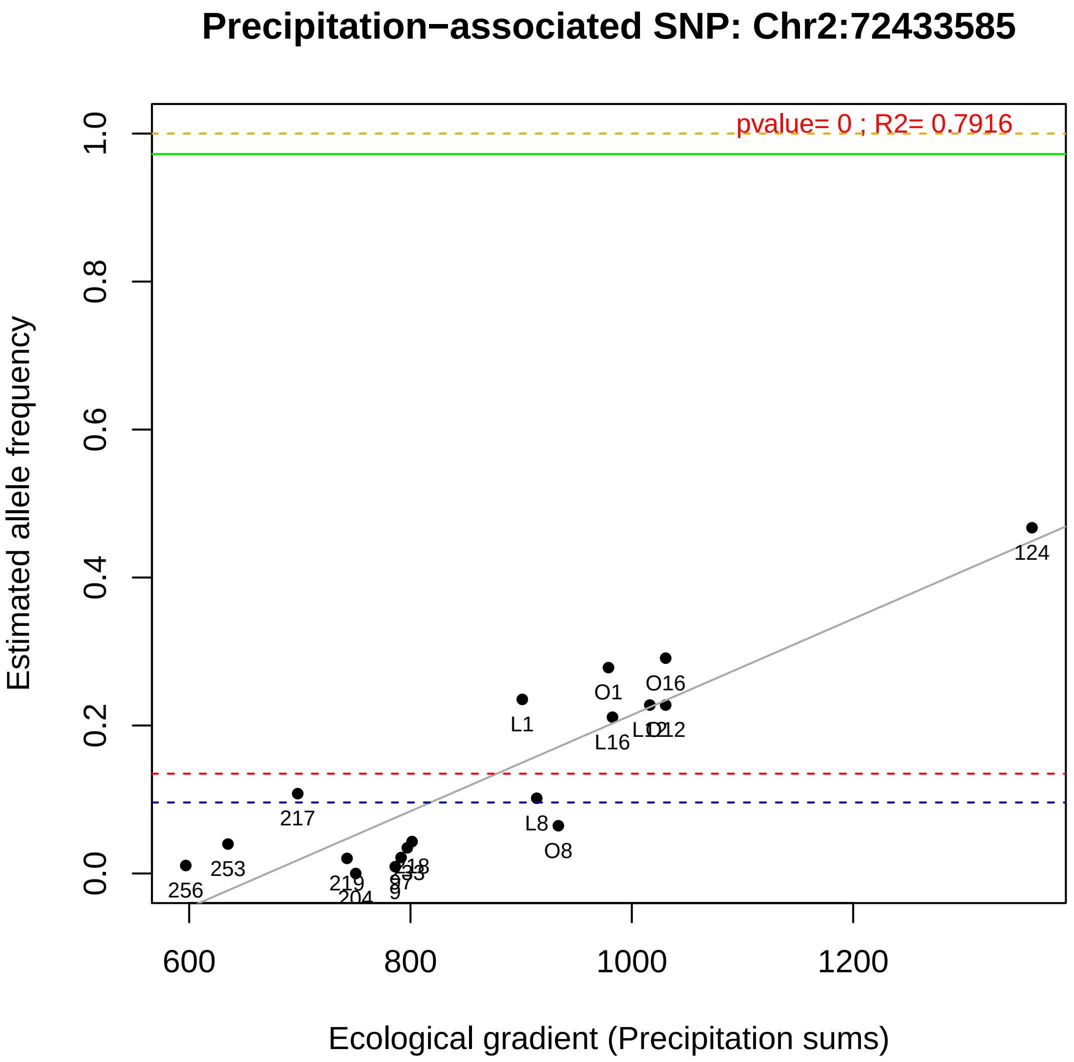

Precipitation−associated SNP: Chr2:72433585
pvalue= 0 ; R2= 0.7916
Estimated allele frequency
0.0
0.2
0.4
0.6
0.8
1.0
600
800
1000
1200
Ecological gradient (Precipitation sums)
124
O16
O1
L1
L16
O12
L8
O8
217
253
256
219
204

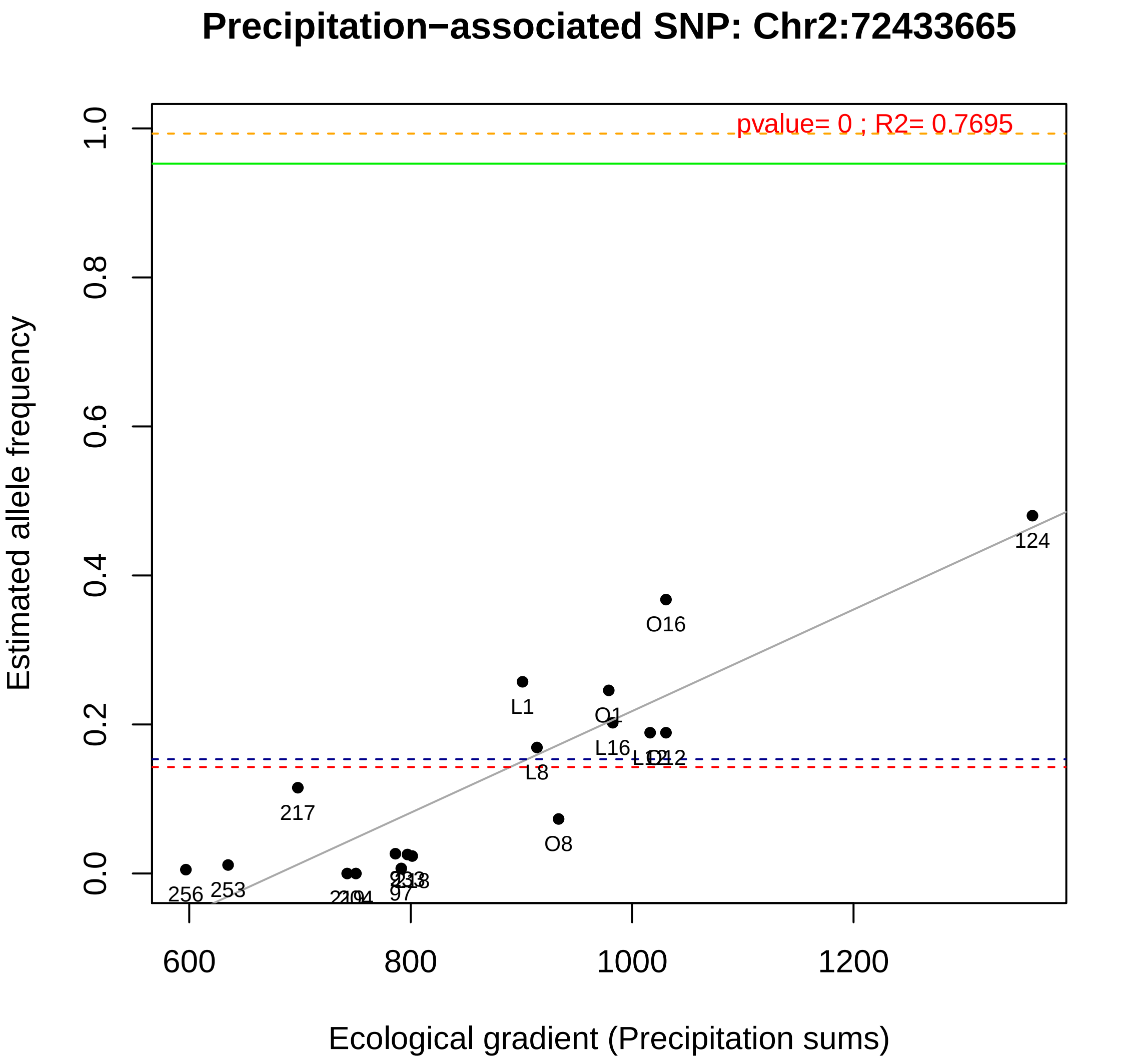
Precipitation−associated SNP: Chr2:72433665
pvalue= 0 ; R2= 0.7695
Estimated allele frequency
0.0
0.2
0.4
0.6
0.8
1.0
600
800
1000
1200
Ecological gradient (Precipitation sums)
256
253
217
97
L1
O1
L16
L8
O8
O16
124

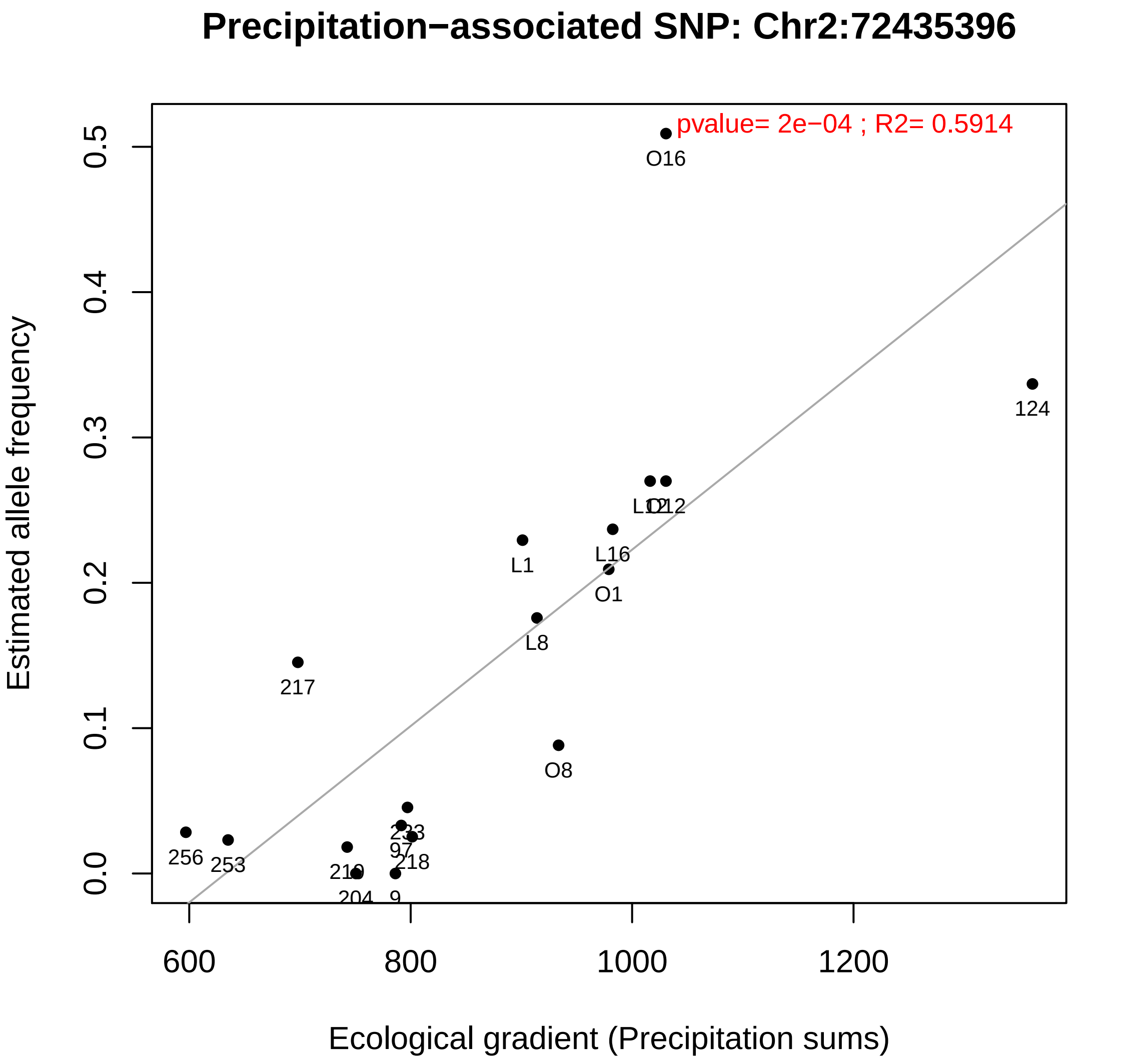

Precipitation−associated SNP: Chr2:72435396
pvalue= 2e−04 ; R2= 0.5914
Estimated allele frequency
Ecological gradient (Precipitation sums)
0.0
0.1
0.2
0.3
0.4
0.5
600
800
1000
1200
O16
124
O12
L16
L1
O1
L8
217
O8
256
253
218
97
204
9

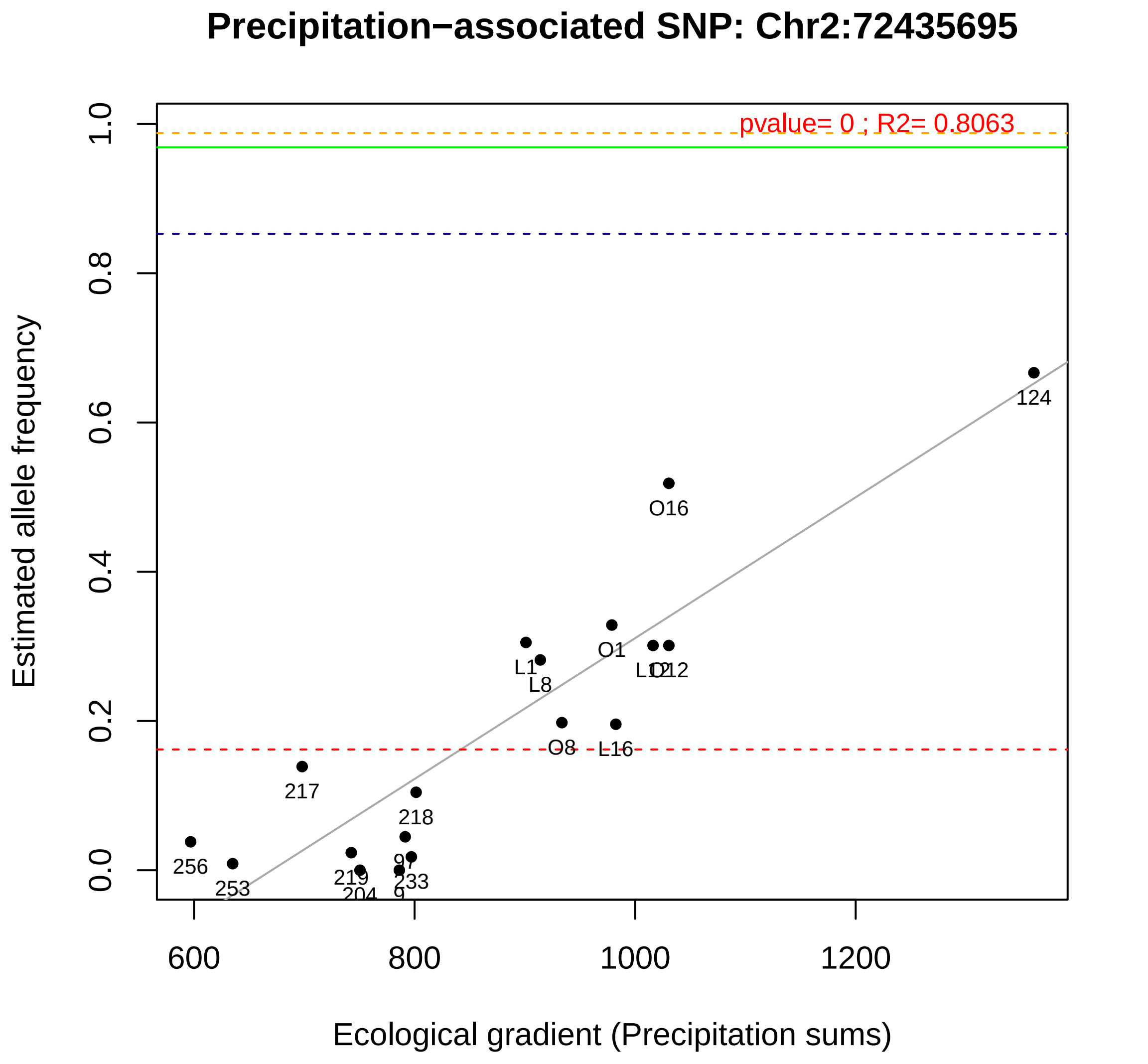

Precipitation−associated SNP: Chr2:72435695
pvalue= 0 ; R2= 0.8063
Estimated allele frequency
0.0
0.2
0.4
0.6
0.8
1.0
600
800
1000
1200
Ecological gradient (Precipitation sums)
124
O16
O1
L1
L8
O8
L16
O12
217
218
256
253
219
204
233

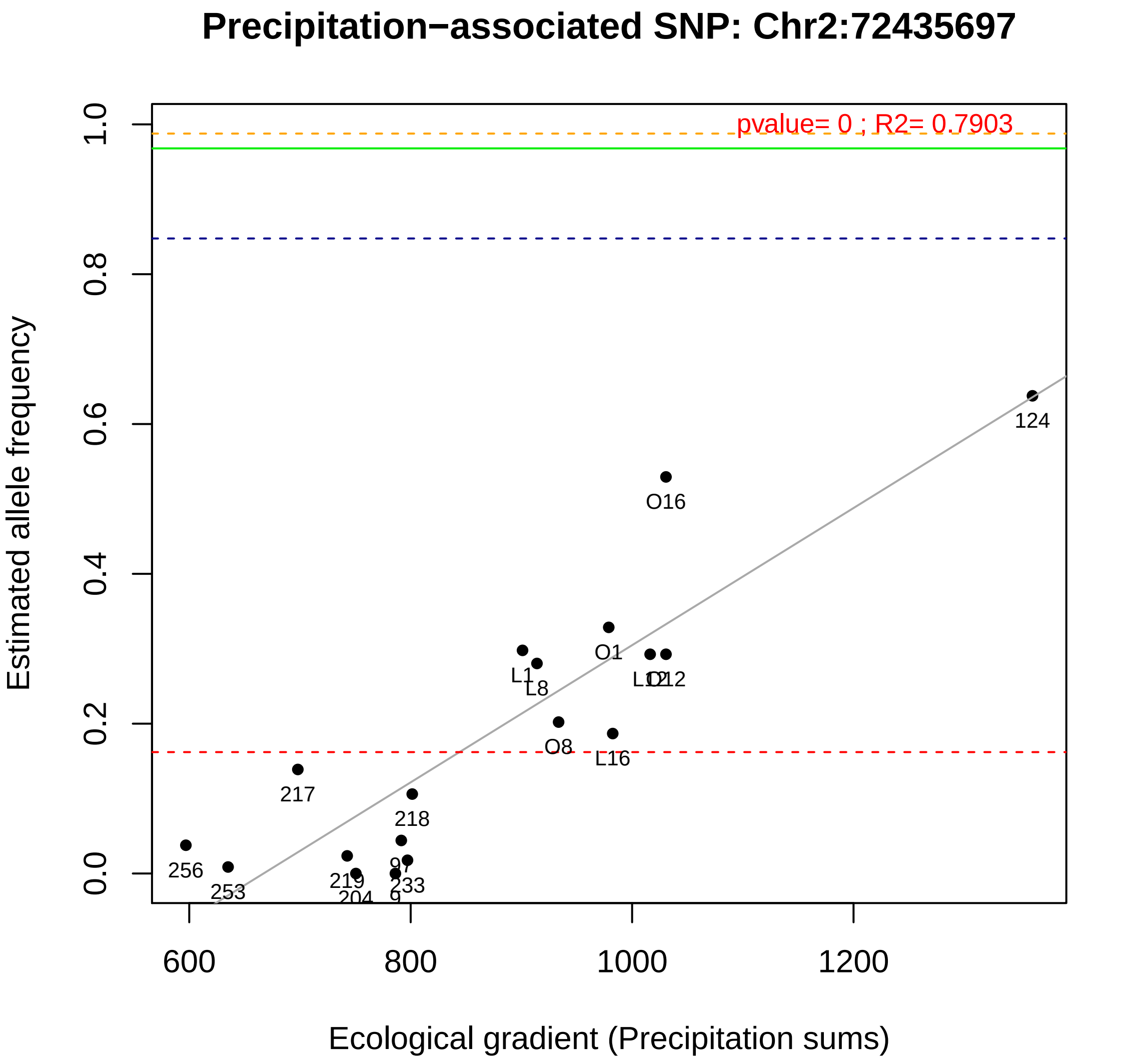

Precipitation−associated SNP: Chr2:72435697
pvalue= 0 ; R2= 0.7903
Estimated allele frequency
0.0
0.2
0.4
0.6
0.8
1.0
600
800
1000
1200
Ecological gradient (Precipitation sums)
124
O16
O1
L1
L8
O12
O8
L16
217
218
256
253
219
204
233

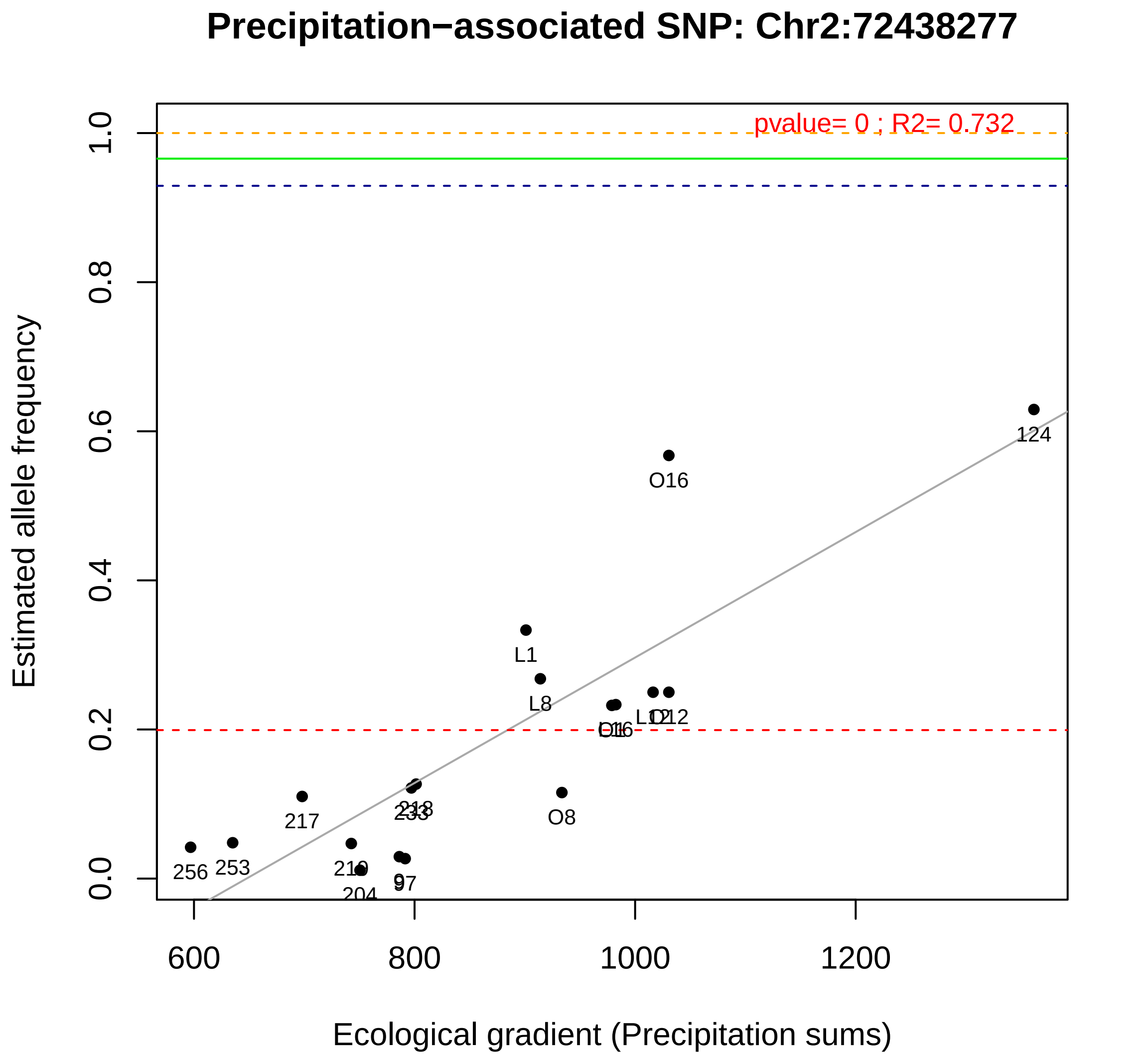
Precipitation−associated SNP: Chr2:72438277
pvalue= 0 ; R2= 0.732
Estimated allele frequency
Ecological gradient (Precipitation sums)
0.0
0.2
0.4
0.6
0.8
1.0
600
800
1000
1200
124
O16
L1
L8
L12
O12
O8
217
256
253
204
97

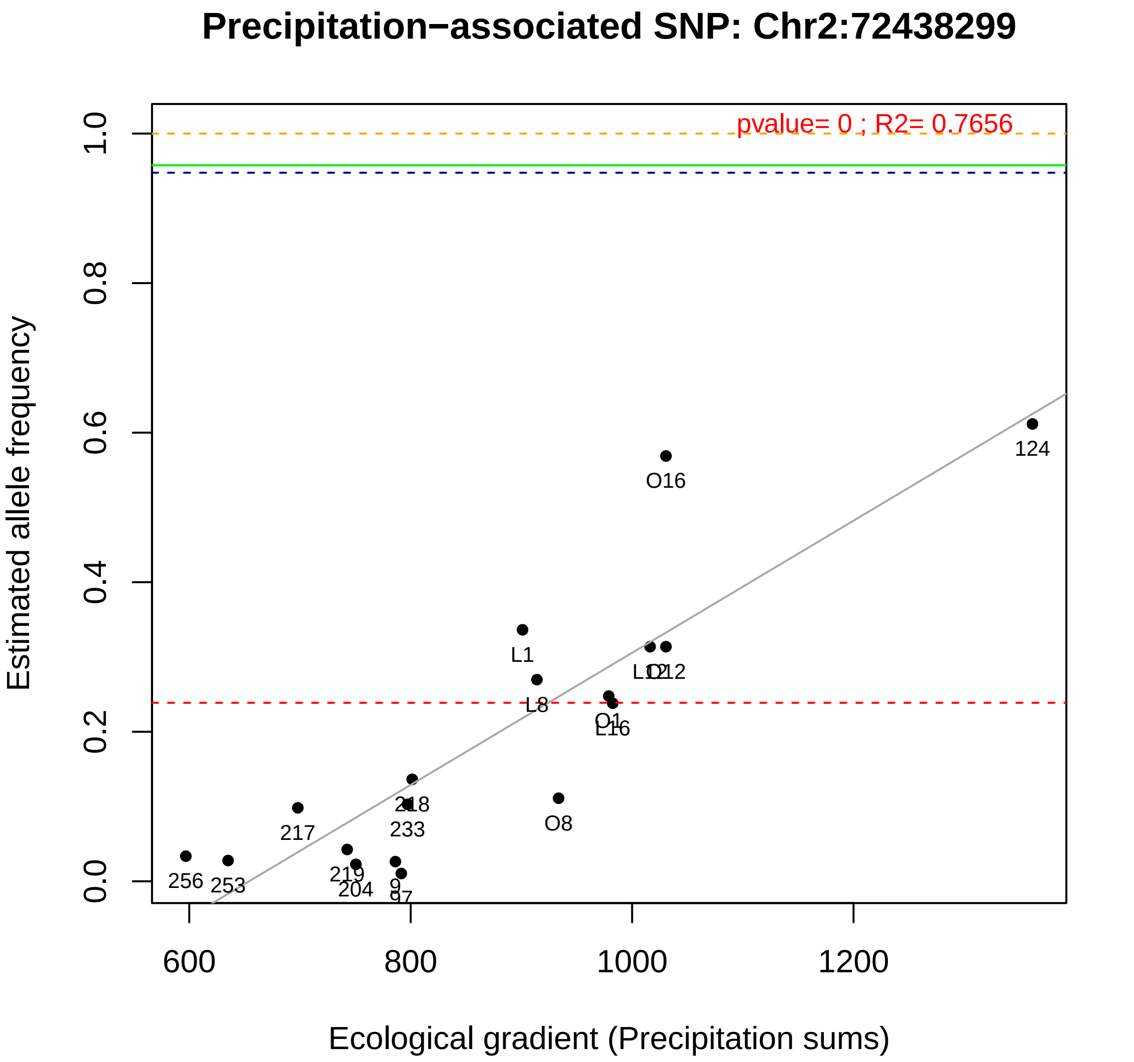
Precipitation−associated SNP: Chr2:72438299
pvalue= 0 ; R2= 0.7656
Estimated allele frequency
Ecological gradient (Precipitation sums)
0.0
0.2
0.4
0.6
0.8
1.0
600
800
1000
1200
124
O16
L1
L8
O12
O1
L16
218
233
217
O8
256
253
219
204
9

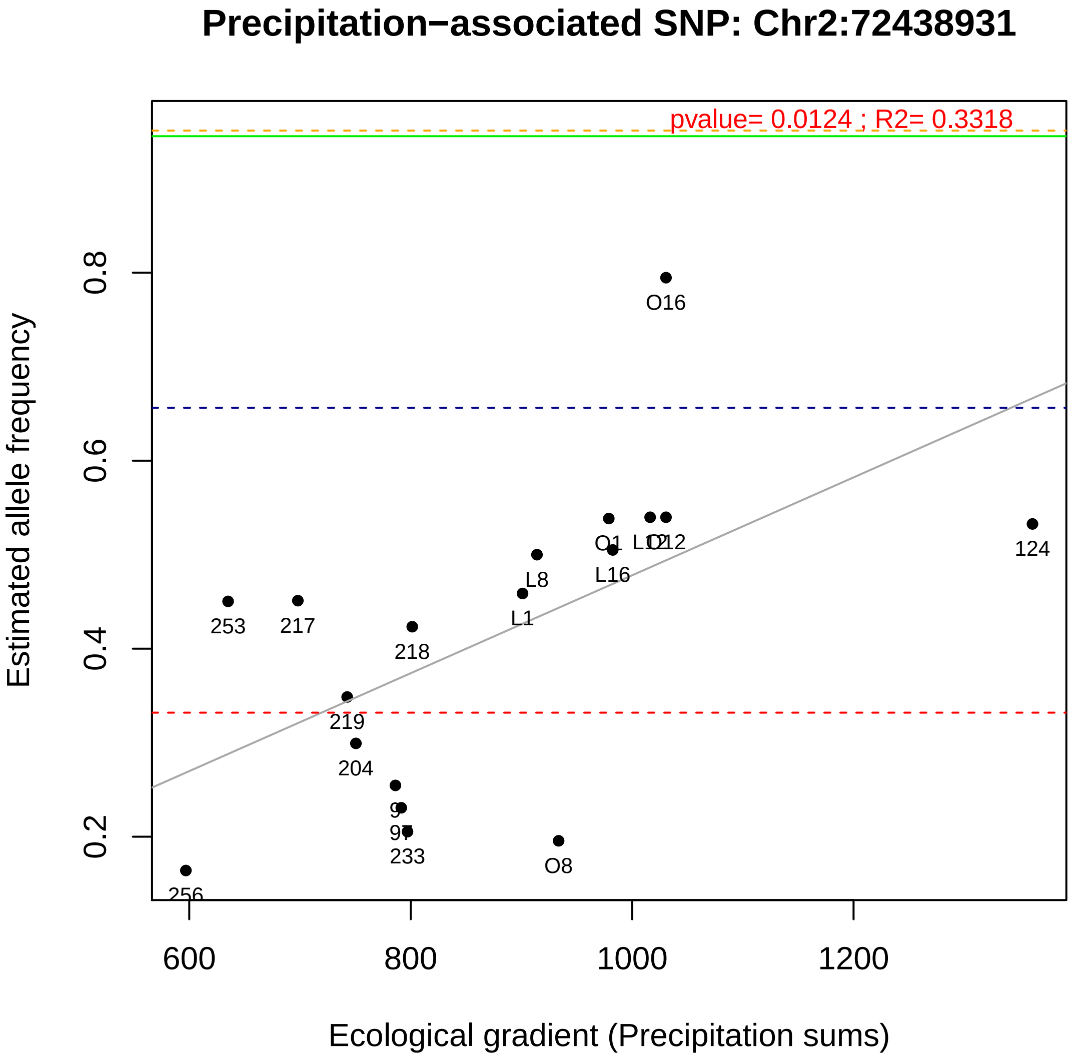
Precipitation−associated SNP: Chr2:72438931
pvalue= 0.0124 ; R2= 0.3318
Estimated allele frequency
0.2
0.4
0.6
0.8
600
800
1000
1200
Ecological gradient (Precipitation sums)
O16
O1
L16
L8
L1
124
253
217
218
219
204
233
O8
256

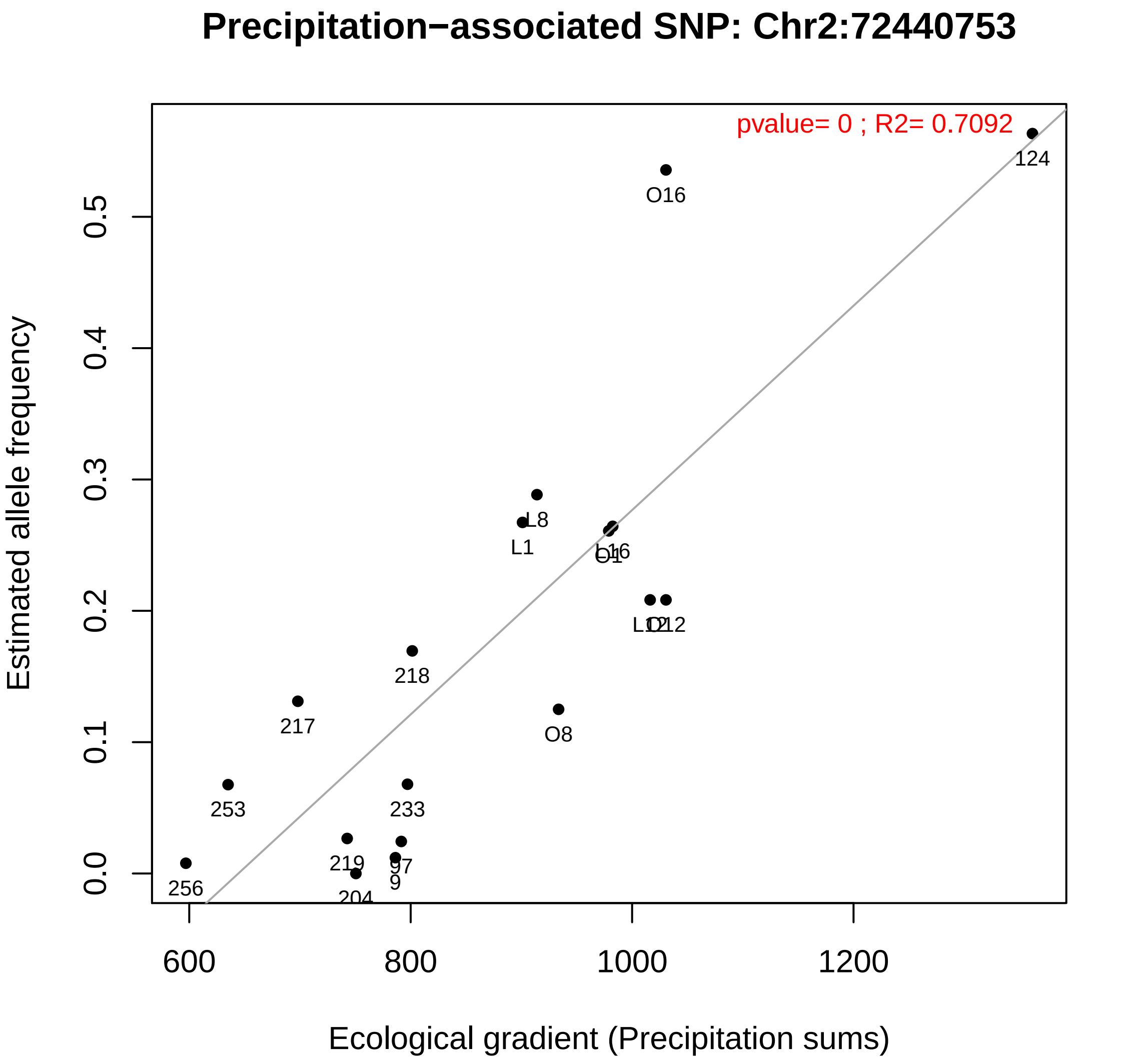

Precipitation−associated SNP: Chr2:72440753
pvalue= 0 ; R2= 0.7092
124
O16
L8
L1
L16
O1
L12
O12
218
217
O8
253
233
219
97
9
256
204
0.0
0.1
0.2
0.3
0.4
0.5
600
800
1000
1200
Estimated allele frequency
Ecological gradient (Precipitation sums)

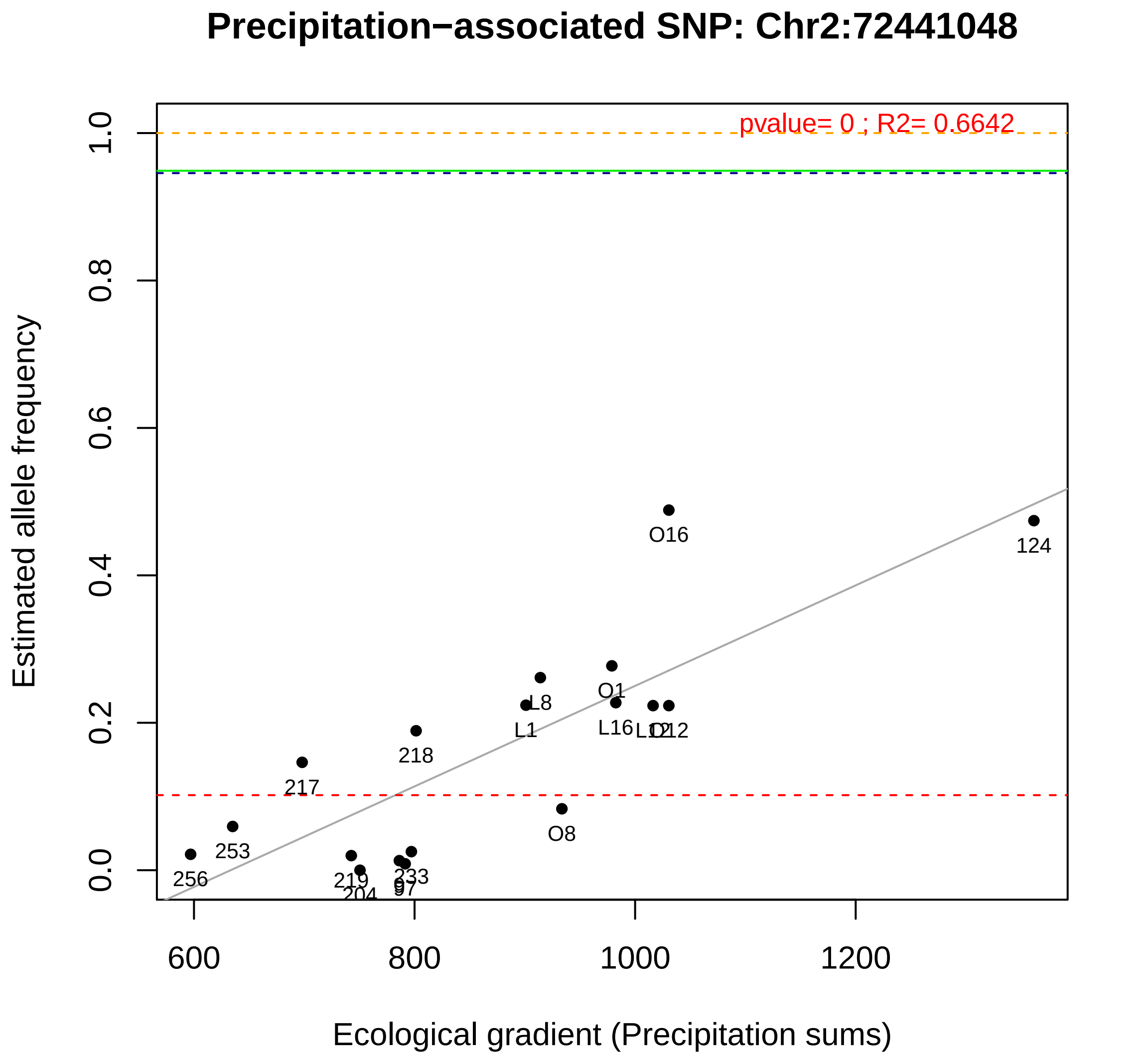
Precipitation−associated SNP: Chr2:72441048
pvalue= 0 ; R2= 0.6642
Estimated allele frequency
Ecological gradient (Precipitation sums)
0.0
0.2
0.4
0.6
0.8
1.0
600
800
1000
1200
O16
124
O1
L8
L1
L16
218
217
O8
253
256
219
204
233

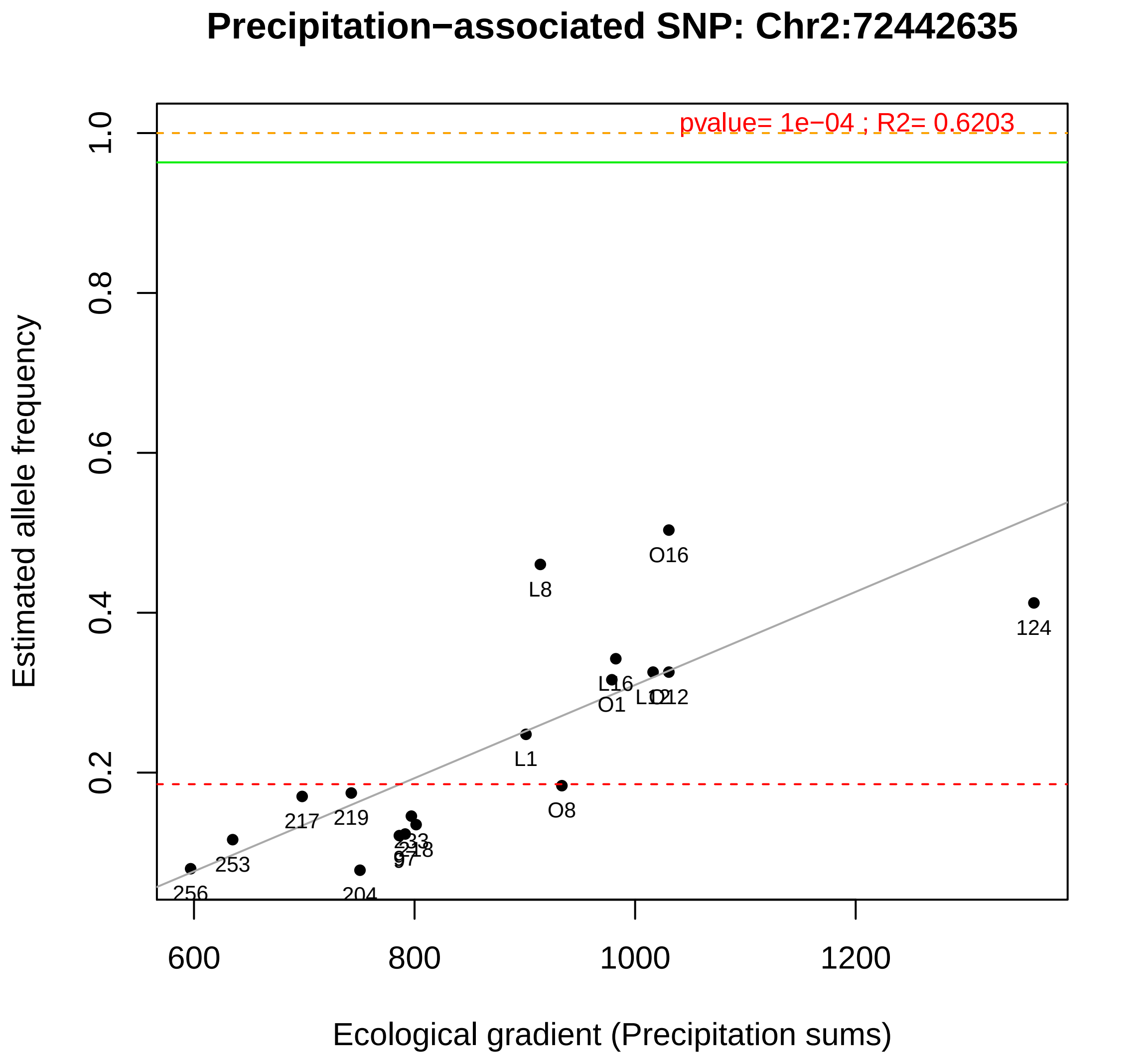

Precipitation−associated SNP: Chr2:72442635
pvalue= 1e−04 ; R2= 0.6203
Estimated allele frequency
0.2
0.4
0.6
0.8
1.0
600
800
1000
1200
Ecological gradient (Precipitation sums)
O16
L8
124
L16
O1
L12
O12
L1
O8
217
219
253
256
204
233
218
97

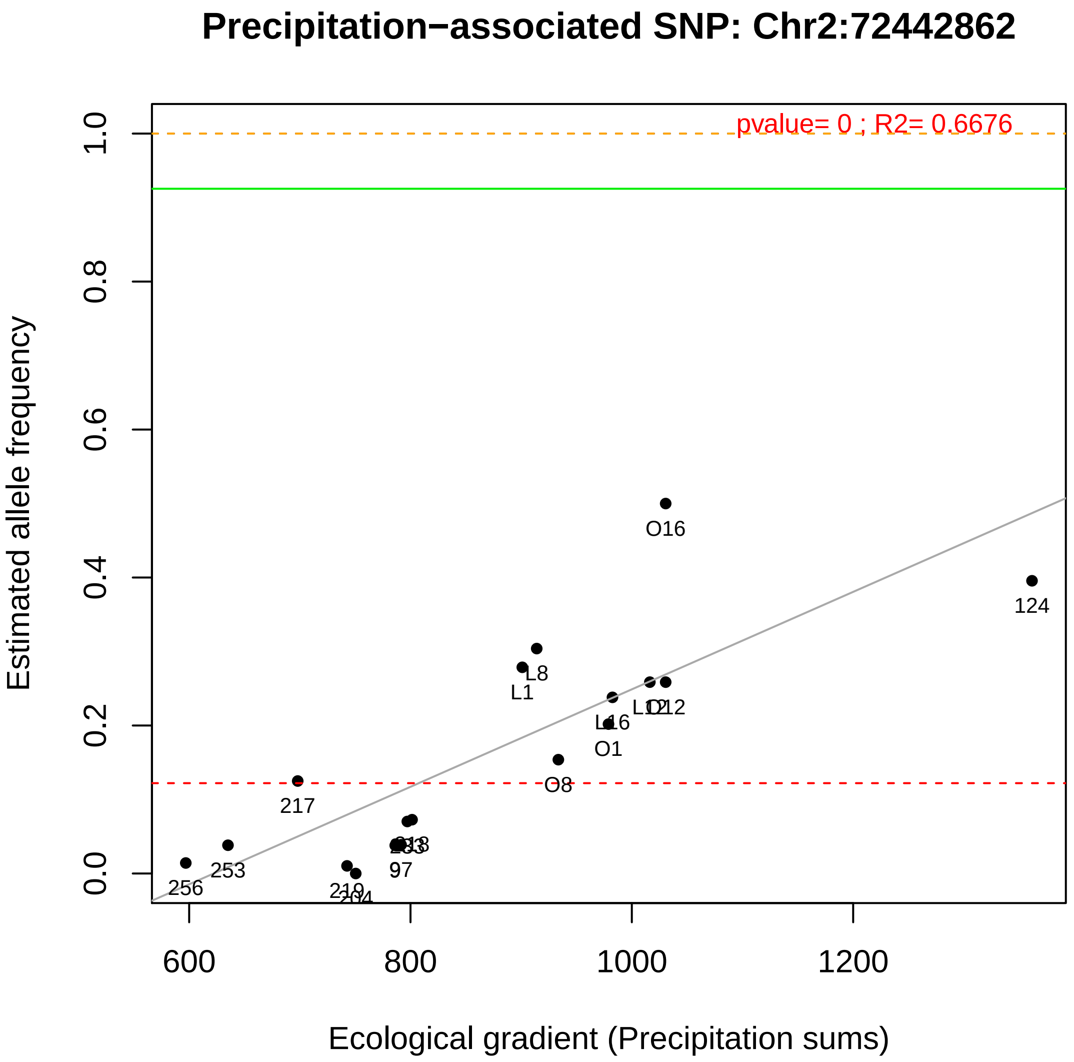

Precipitation−associated SNP: Chr2:72442862
pvalue= 0 ; R2= 0.6676
Estimated allele frequency
Ecological gradient (Precipitation sums)
0.0
0.2
0.4
0.6
0.8
1.0
600
800
1000
1200
O16
124
L8
L1
L12
O12
L16
O1
O8
217
253
256
97

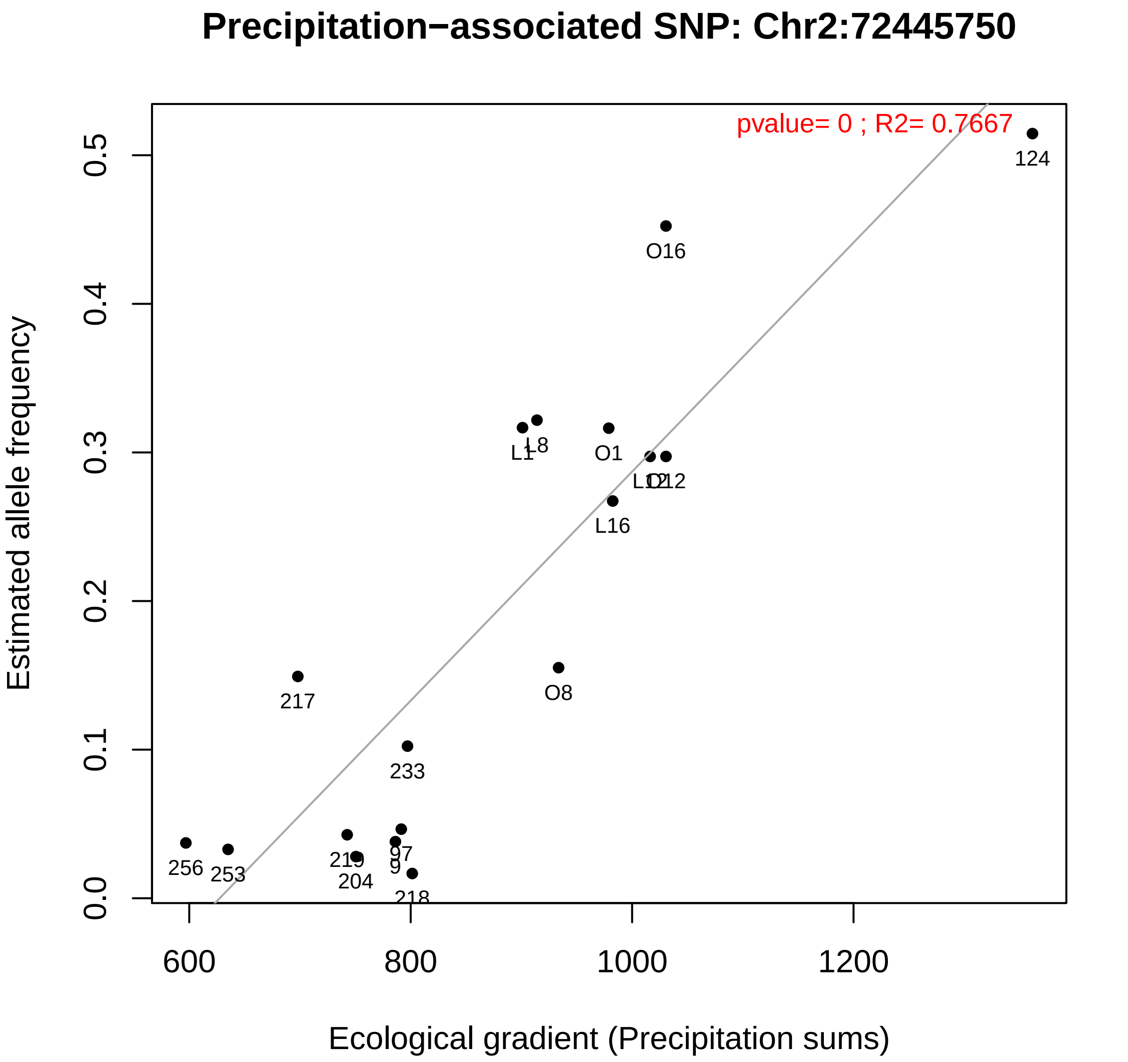

Precipitation−associated SNP: Chr2:72445750
pvalue= 0 ; R2= 0.7667
124
O16
L1
L8
O1
L12
O12
L16
O8
217
233
256
253
219
204
97
9
218
0.0
0.1
0.2
0.3
0.4
0.5
600
800
1000
1200
Estimated allele frequency
Ecological gradient (Precipitation sums)

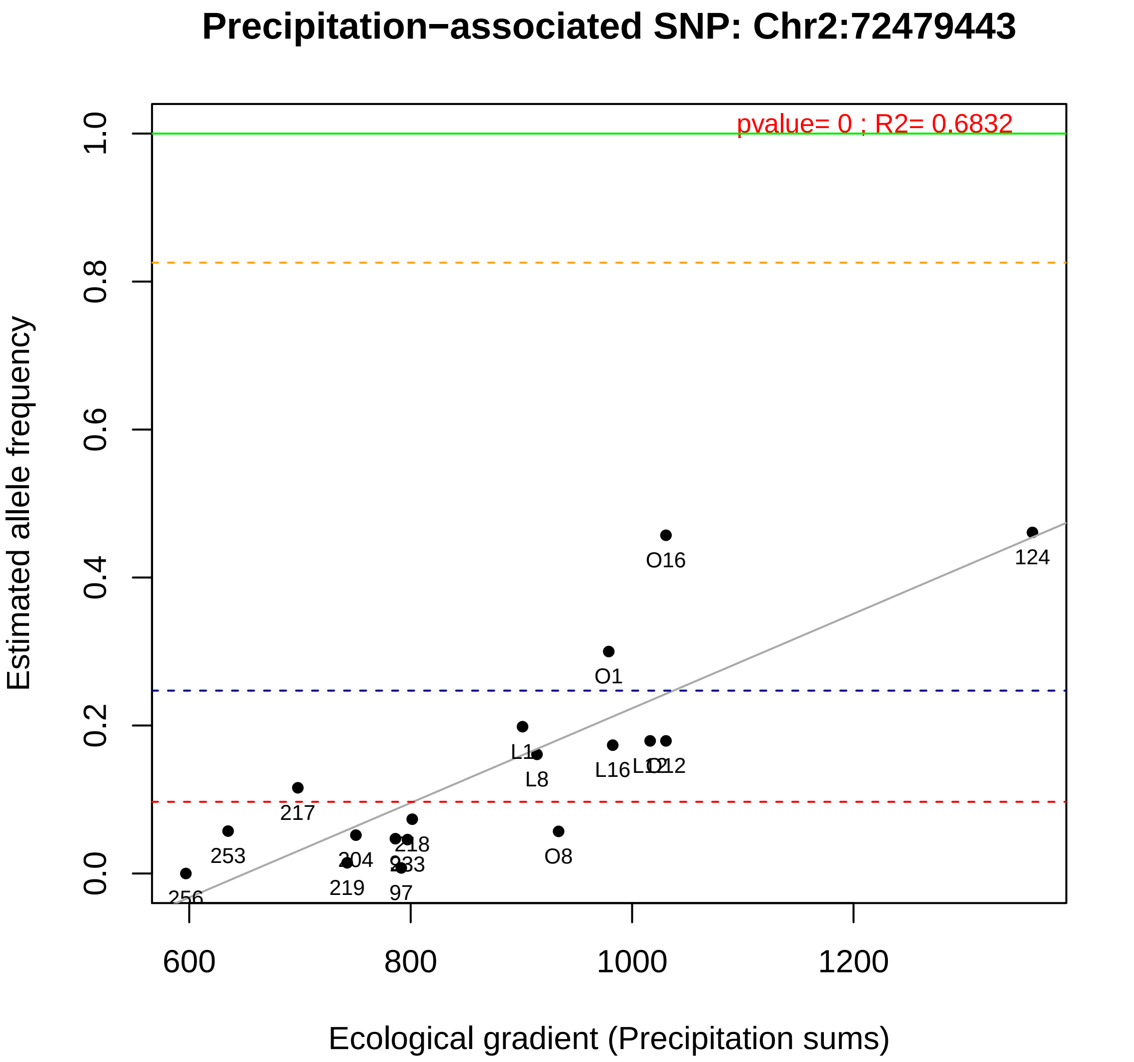

Precipitation−associated SNP: Chr2:72479443
pvalue= 0 ; R2= 0.6832
Estimated allele frequency
0.0
0.2
0.4
0.6
0.8
1.0
600
800
1000
1200
Ecological gradient (Precipitation sums)
O16
124
O1
L1
L8
L16
217
253
204
218
233
219
97
O8

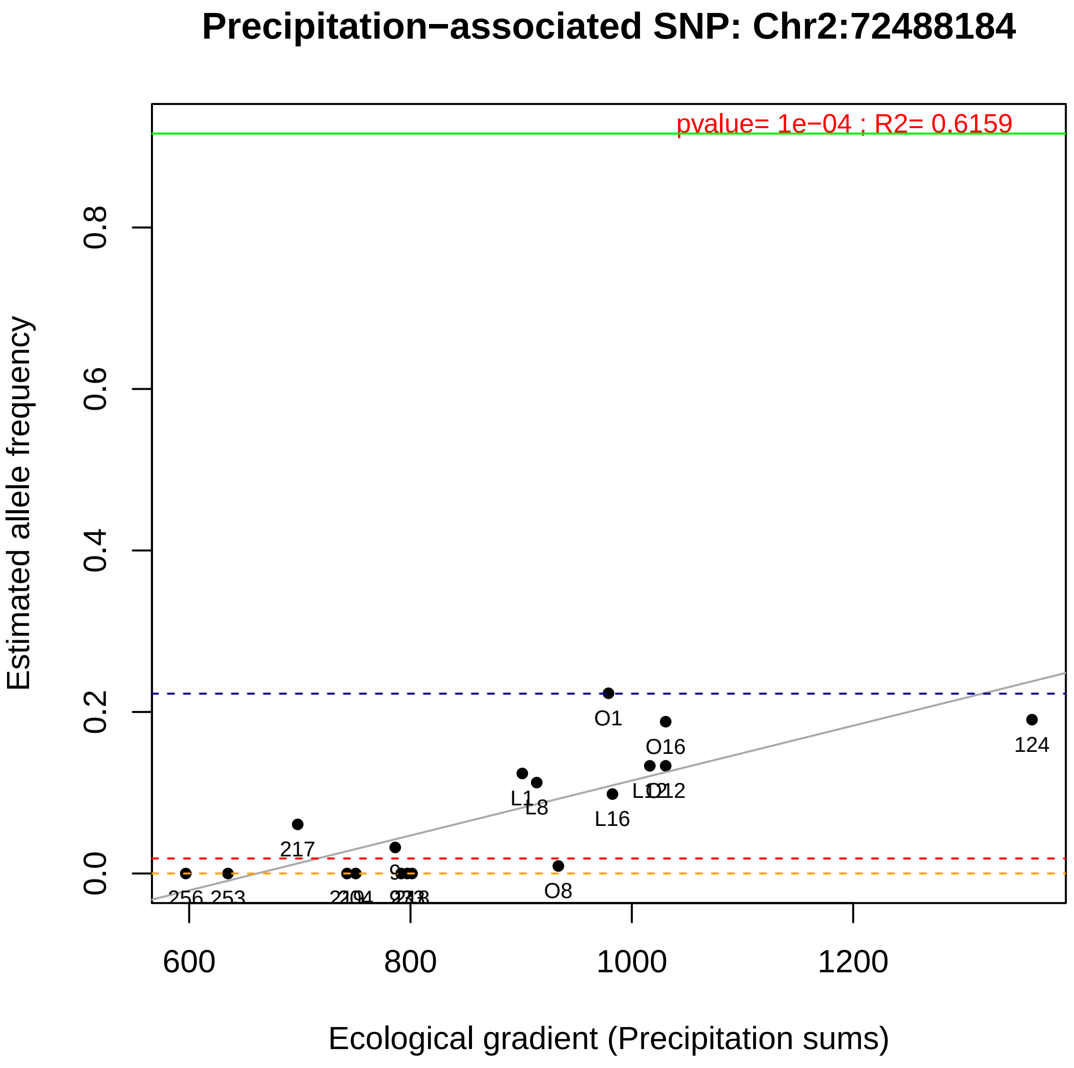

Precipitation−associated SNP: Chr2:72488184
pvalue= 1e−04 ; R2= 0.6159
Estimated allele frequency
0.0
0.2
0.4
0.6
0.8
600
800
1000
1200
Ecological gradient (Precipitation sums)
O1
O16
124
L1
L8
L16
217
O8
256
253

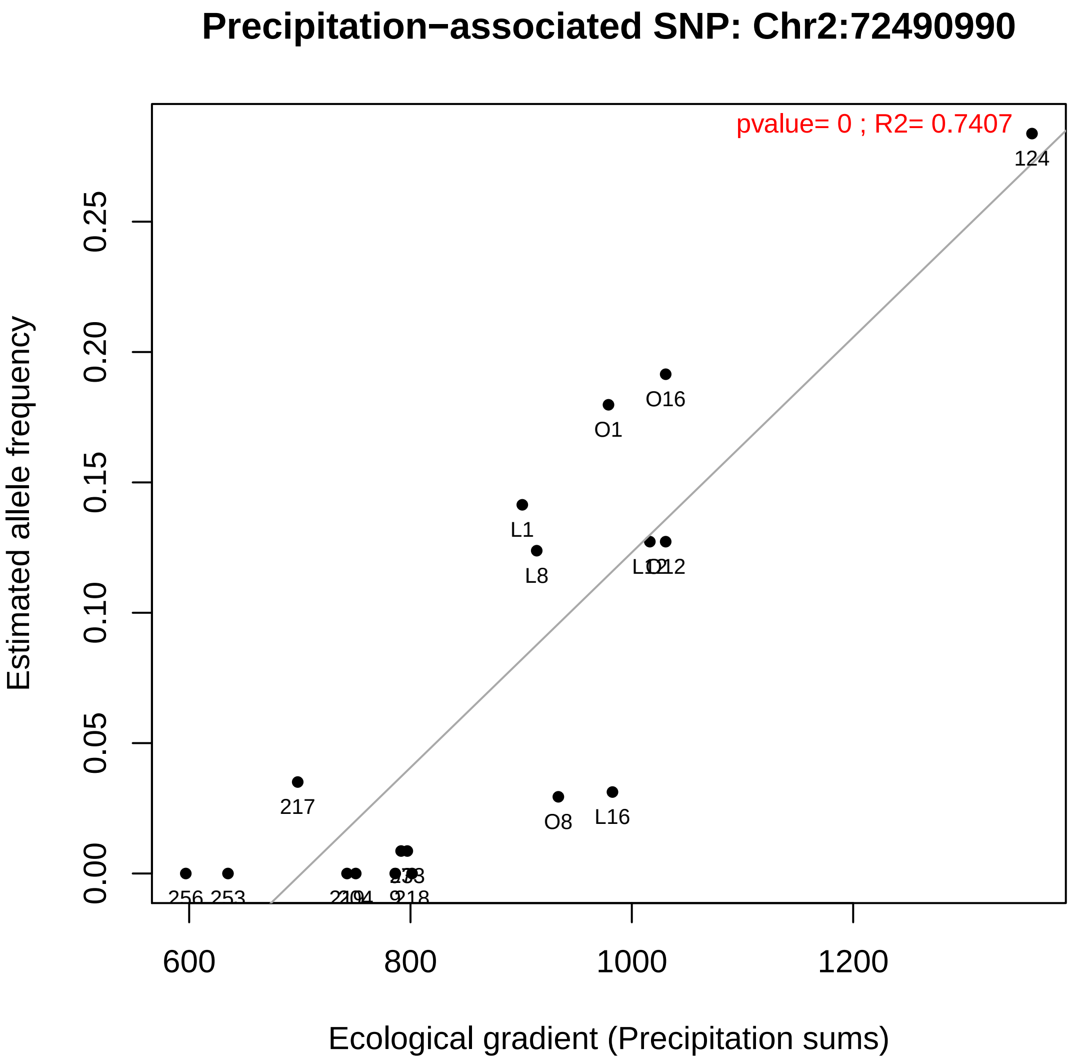
Precipitation−associated SNP: Chr2:72490990
pvalue= 0 ; R2= 0.7407
124
O16
O1
L1
L8
O12
217
O8
L16
256
253
218
0.00
0.05
0.10
0.15
0.20
0.25
600
800
1000
1200
Estimated allele frequency
Ecological gradient (Precipitation sums)

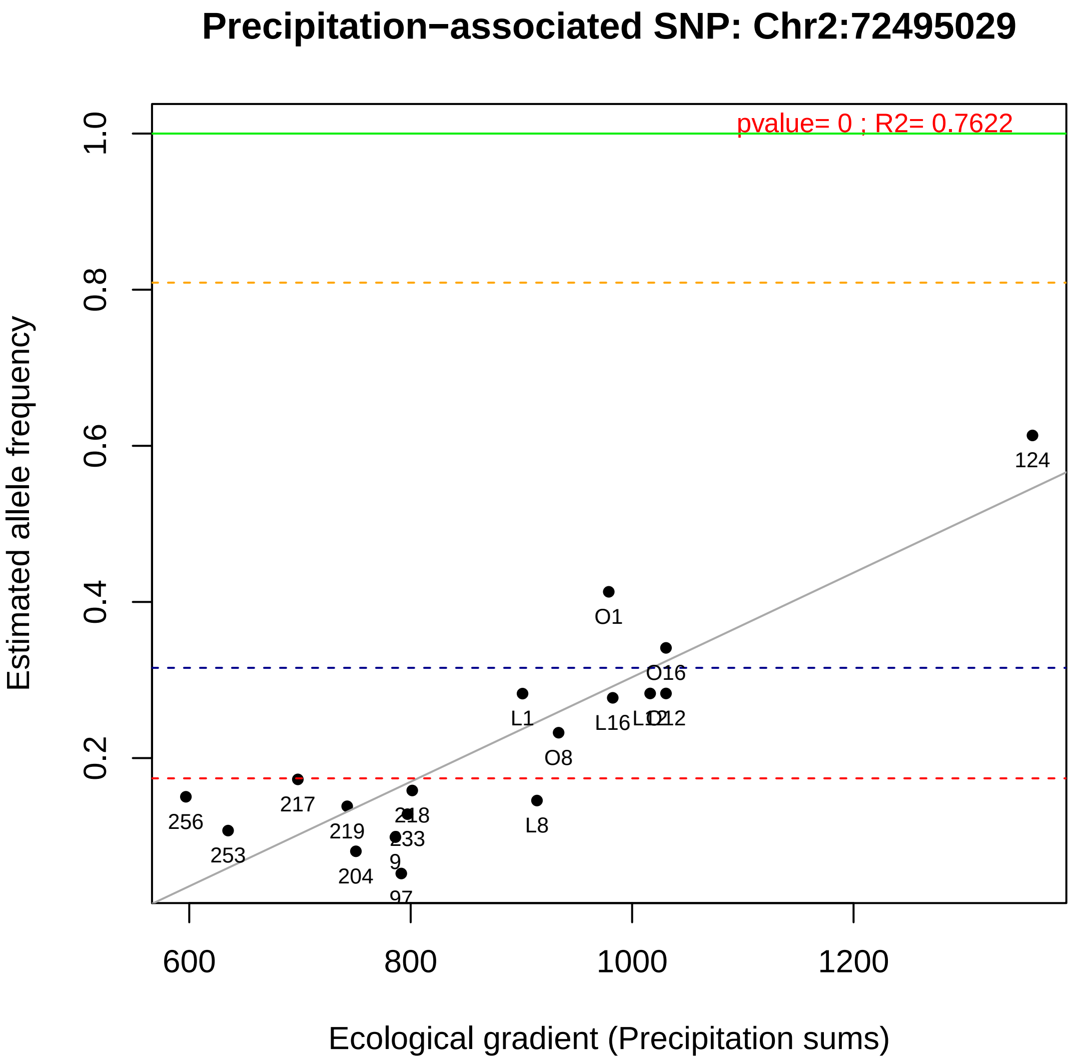
Precipitation−associated SNP: Chr2:72495029
pvalue= 0 ; R2= 0.7622
Estimated allele frequency
Ecological gradient (Precipitation sums)
0.2
0.4
0.6
0.8
1.0
600
800
1000
1200
124
O1
O16
L1
L16
O12
O8
217
256
253
219
218
233
9
204
97
L8

### Precipitation−associated SNP: Chr2:72500438

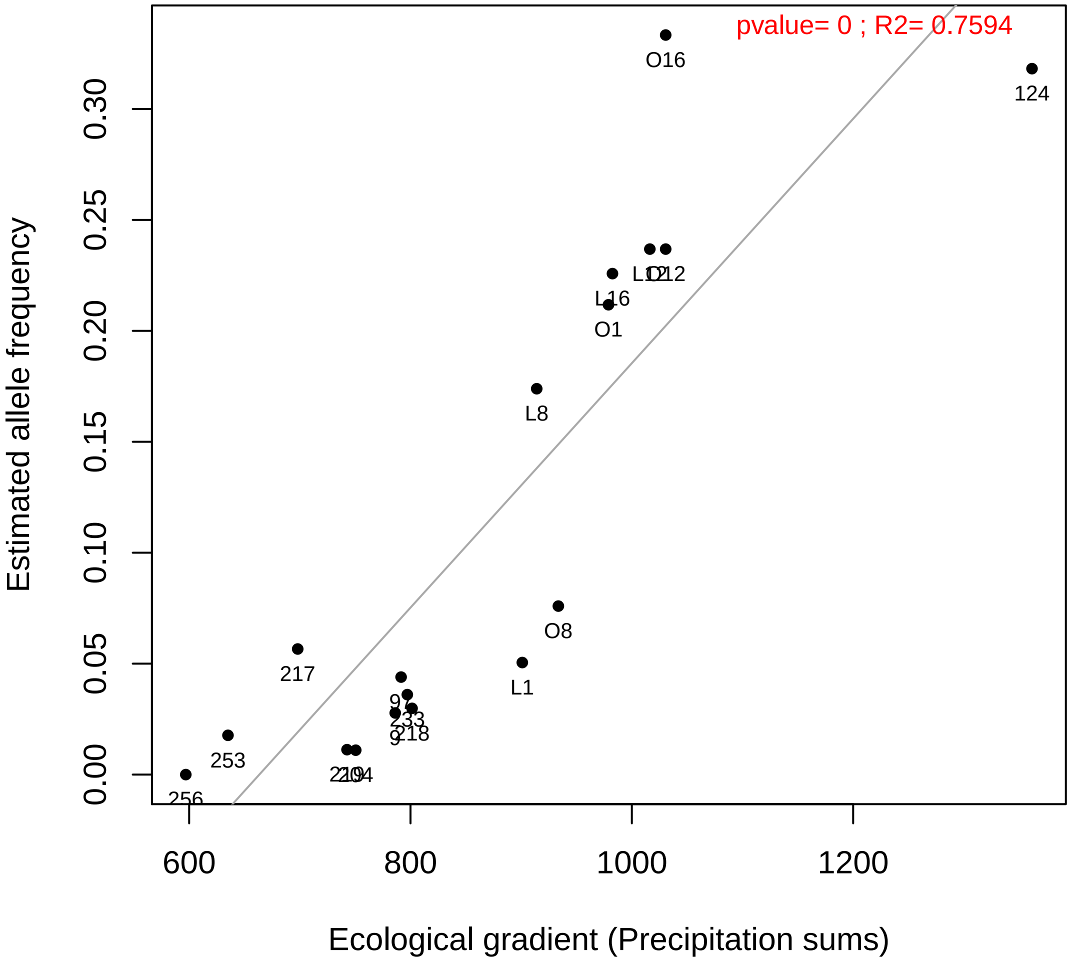

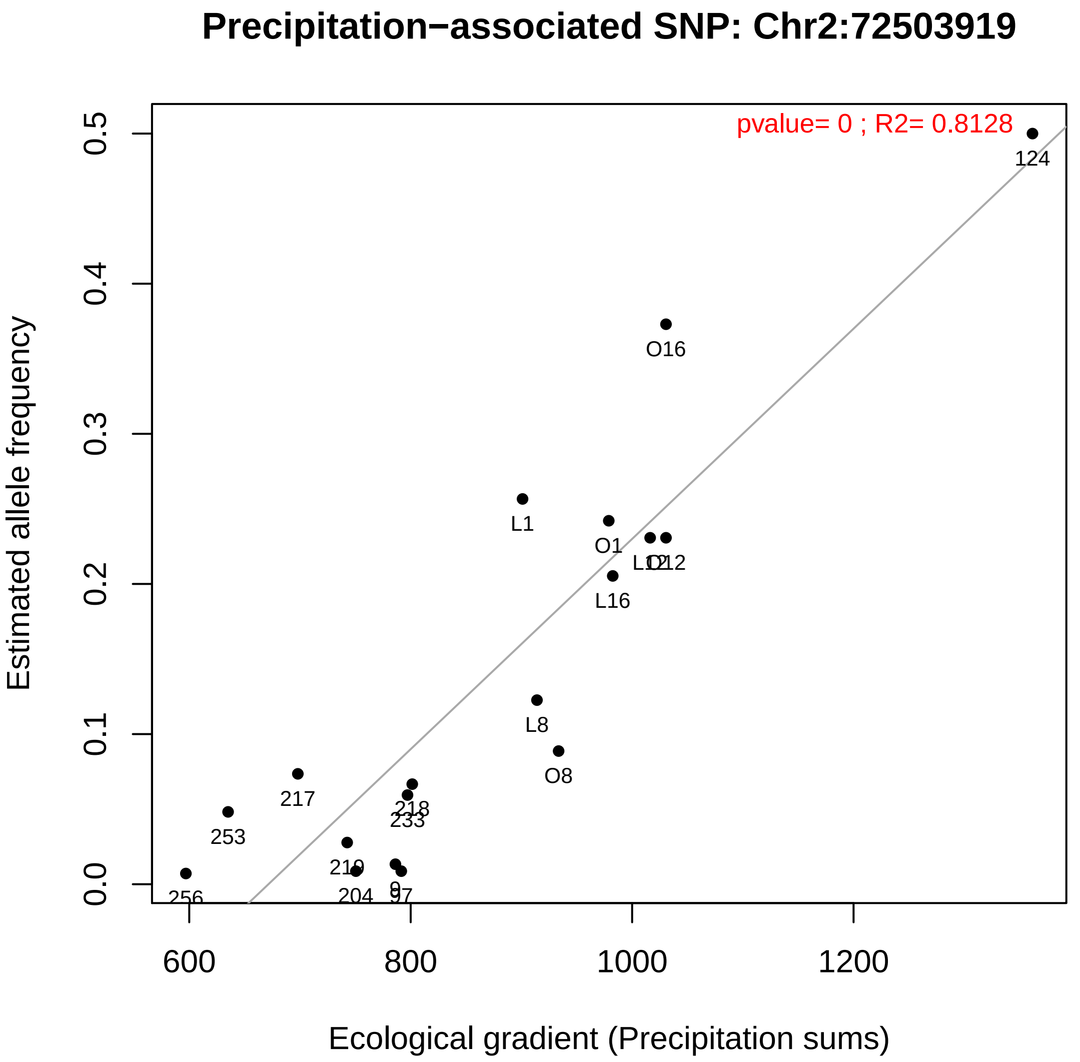
Precipitation−associated SNP: Chr2:72503919
pvalue= 0 ; R2= 0.8128
Estimated allele frequency
Ecological gradient (Precipitation sums)
0.0
0.1
0.2
0.3
0.4
0.5
600
800
1000
1200
124
O16
L1
O1
L12
O12
L16
L8
O8
217
218
233
253
210
204
9
97
256

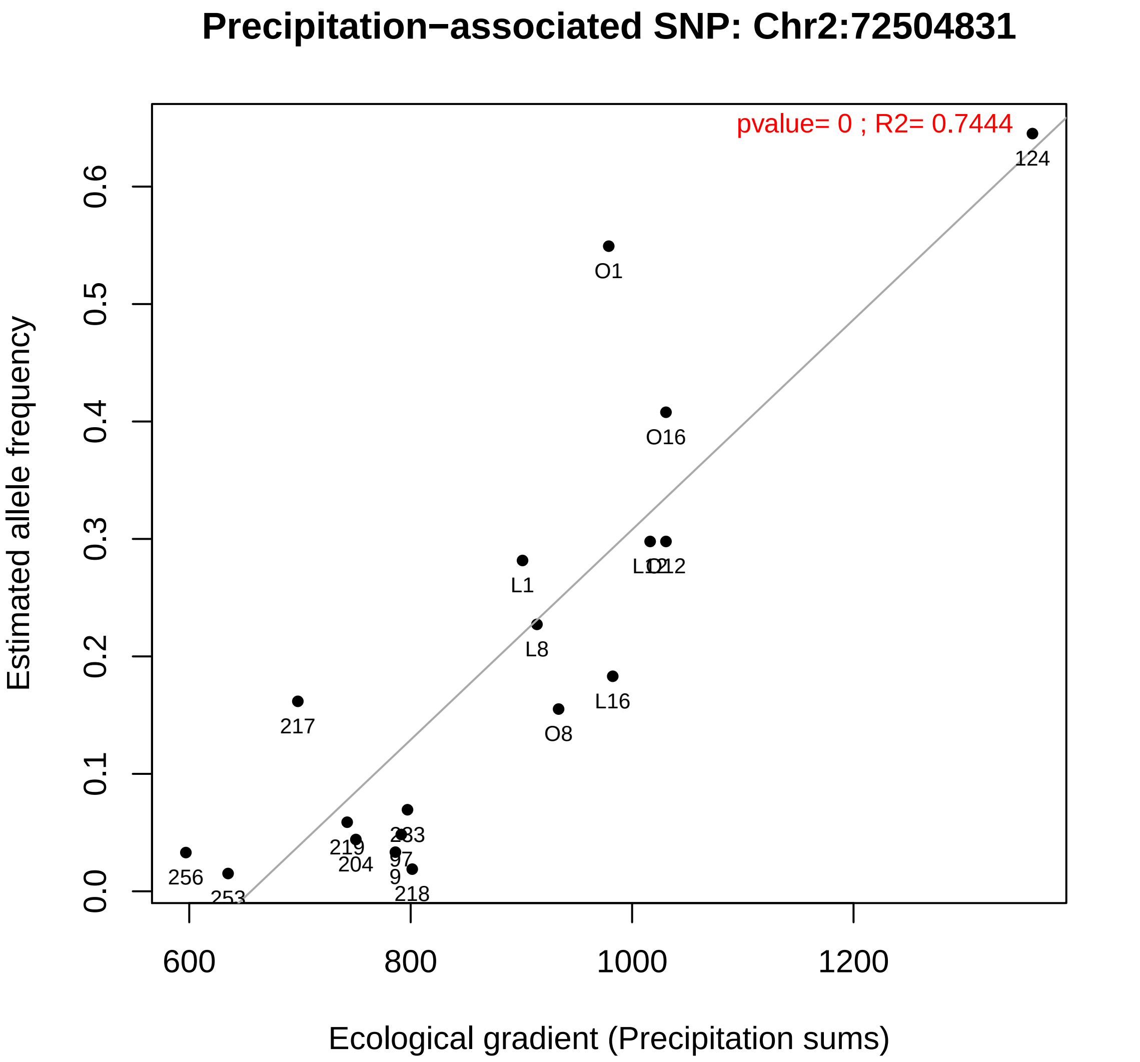

Precipitation−associated SNP: Chr2:72504831
pvalue= 0 ; R2= 0.7444
Estimated allele frequency
0.0
0.1
0.2
0.3
0.4
0.5
0.6
600
800
1000
1200
Ecological gradient (Precipitation sums)
124
O1
O16
L1
L8
L16
O8
217
256
219
204
218

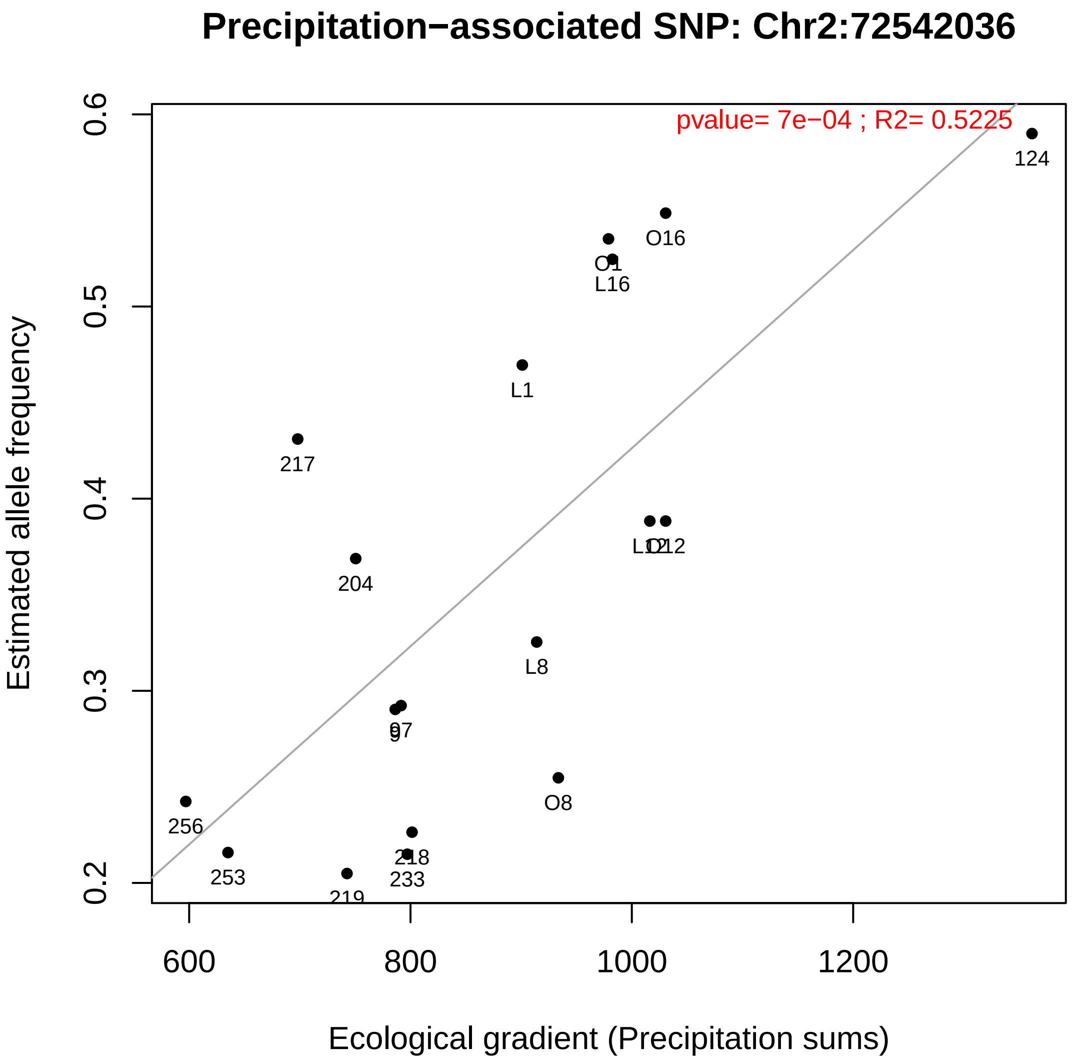

Precipitation−associated SNP: Chr2:72542036
pvalue= 7e−04 ; R2= 0.5225
Estimated allele frequency
0.2
0.3
0.4
0.5
0.6
600
800
1000
1200
Ecological gradient (Precipitation sums)
124
O16
O1
L16
L1
217
L12
O12
204
L8
97
9
O8
256
253
218
233
219

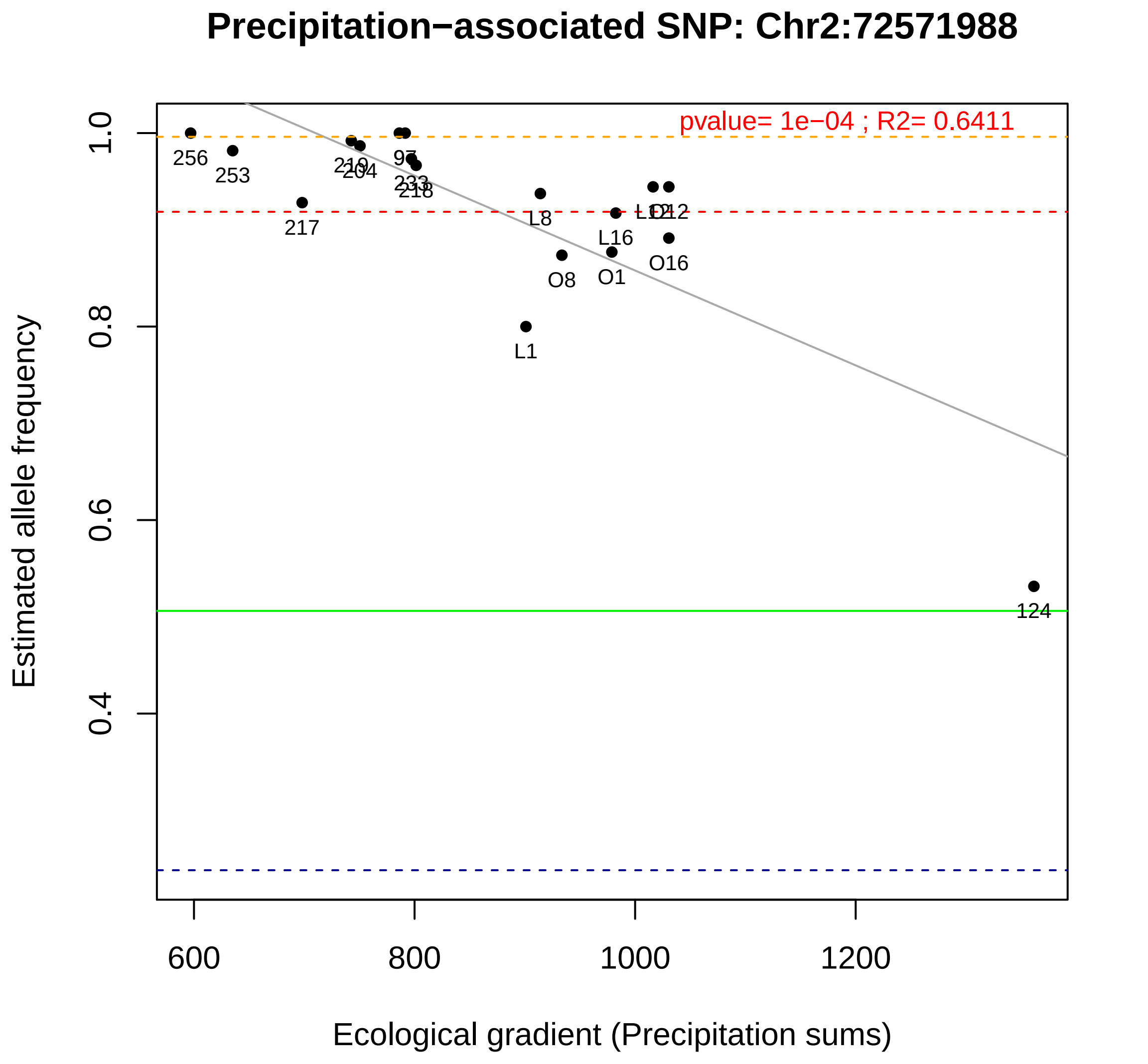

Precipitation−associated SNP: Chr2:72571988
pvalue= 1e−04 ; R2= 0.6411
256
253
210
204
97
233
218
217
L8
L12
O12
L16
O16
O8
O1
L1
124
1.0
0.8
0.6
0.4
600
800
1000
1200
Estimated allele frequency
Ecological gradient (Precipitation sums)

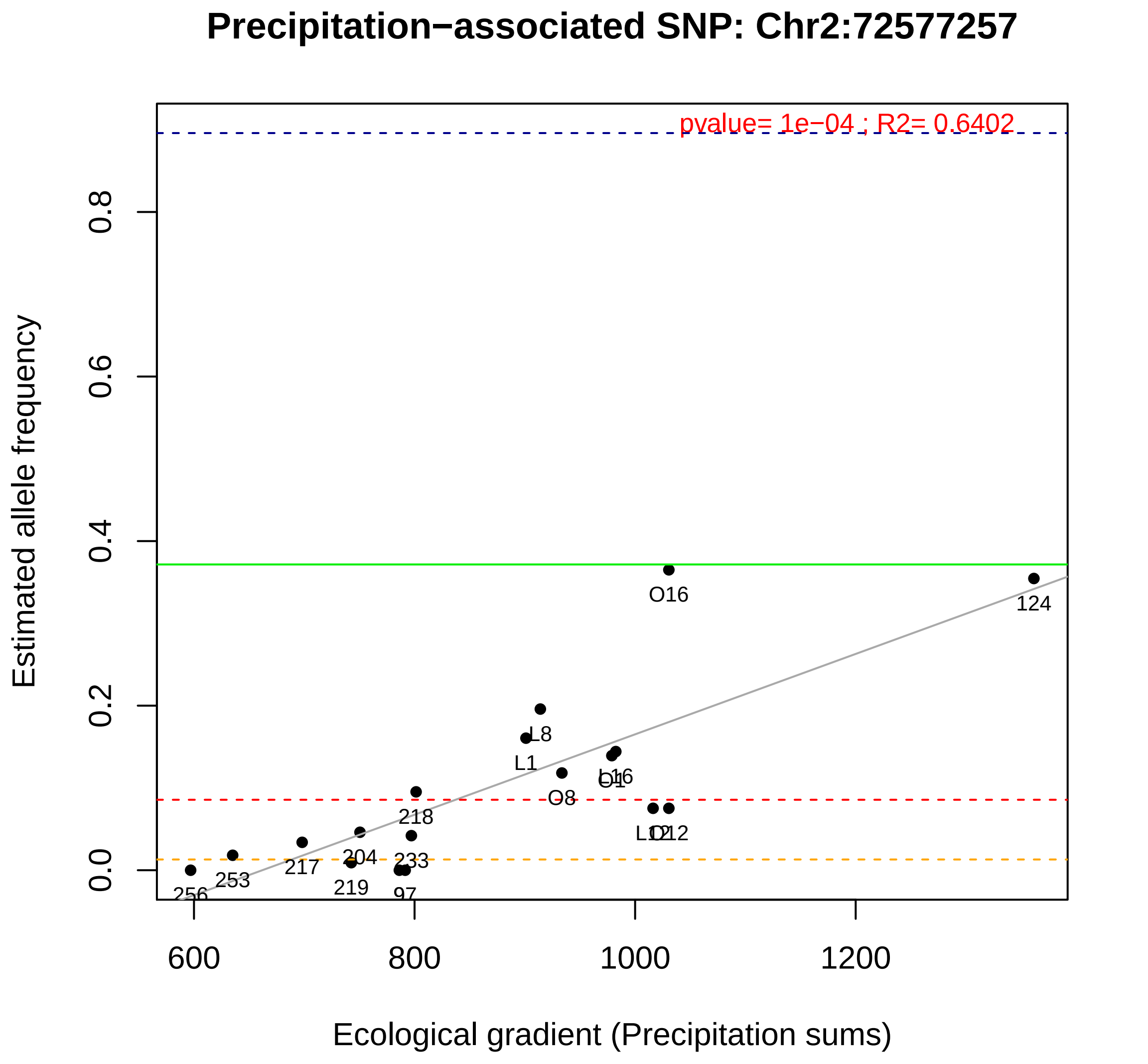

Precipitation−associated SNP: Chr2:72577257
pvalue= 1e−04 ; R2= 0.6402
Estimated allele frequency
0.0
0.2
0.4
0.6
0.8
Ecological gradient (Precipitation sums)
600
800
1000
1200
O16
124
L8
L1
L16
O1
O8
218
L12
O12
204
233
217
253
219
256
97

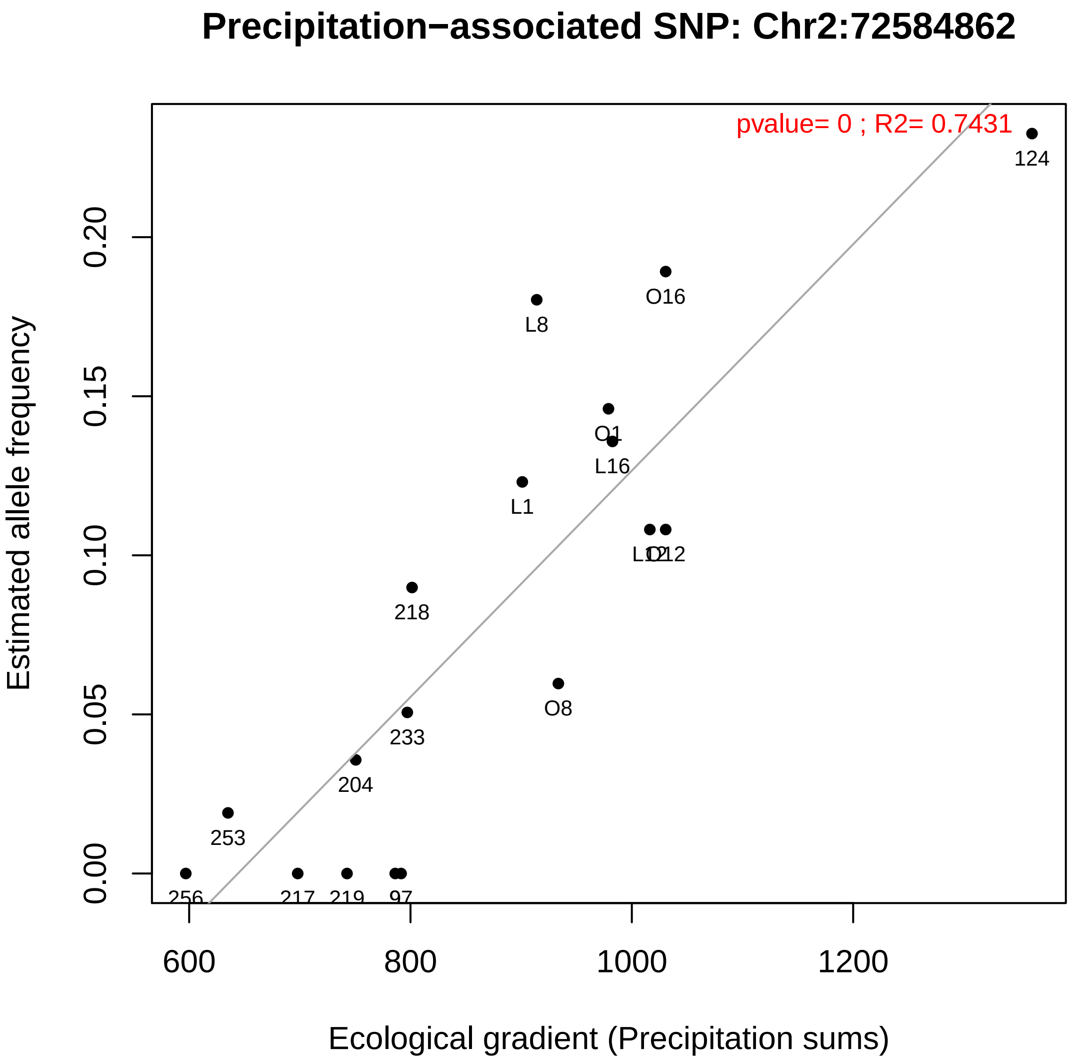
Precipitation−associated SNP: Chr2:72584862
pvalue= 0 ; R2= 0.7431
Estimated allele frequency
Ecological gradient (Precipitation sums)
0.00
0.05
0.10
0.15
0.20
600
800
1000
1200
124
O16
L8
O1
L16
L1
L12
O12
218
O8
233
204
253
256
217
219
97

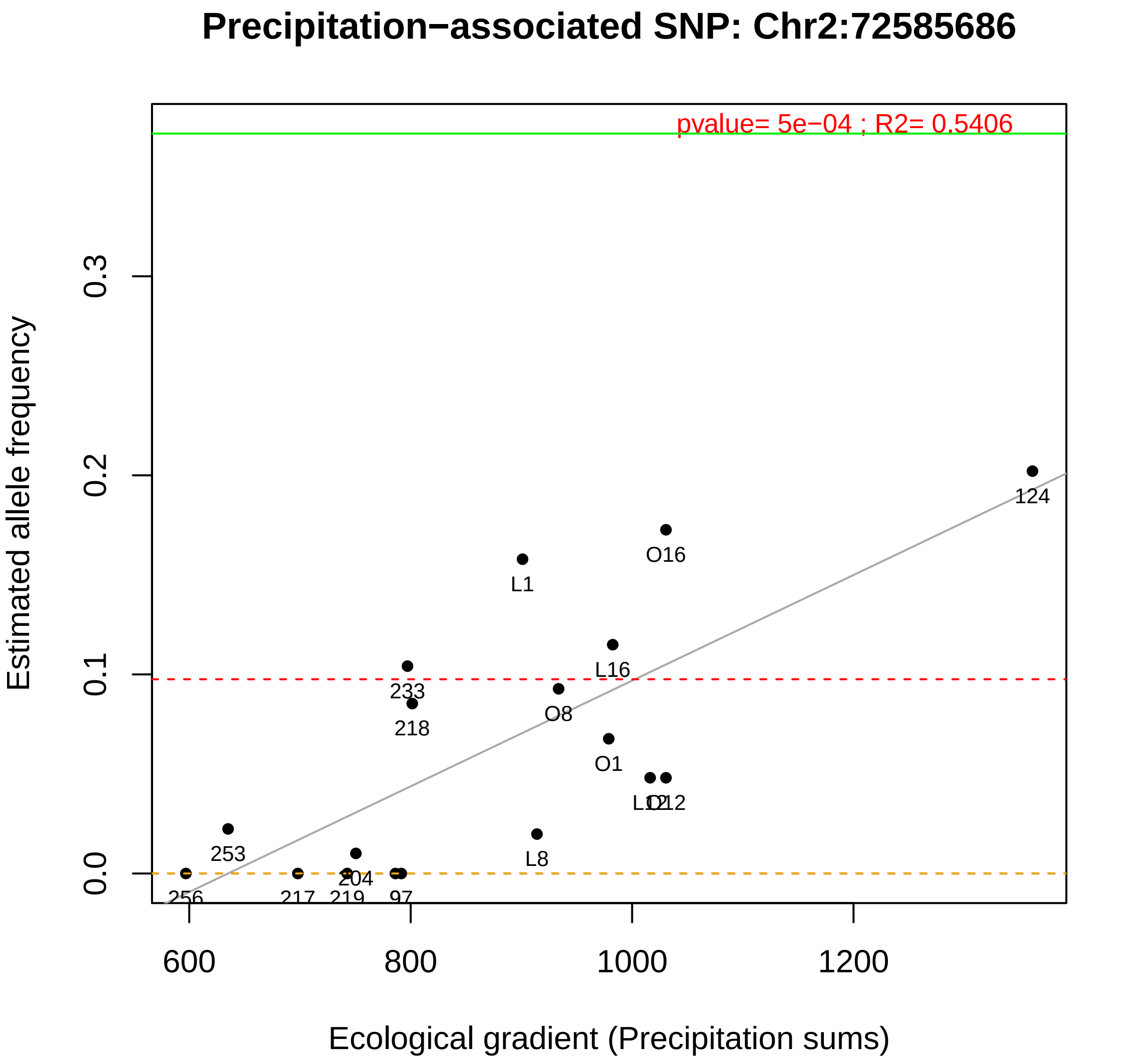
Precipitation−associated SNP: Chr2:72585686
pvalue= 5e−04 ; R2= 0.5406
Estimated allele frequency
0.0
0.1
0.2
0.3
600
800
1000
1200
Ecological gradient (Precipitation sums)
124
O16
L1
L16
233
218
O8
O1
L10
O12
253
L8
204
256
217
219
97

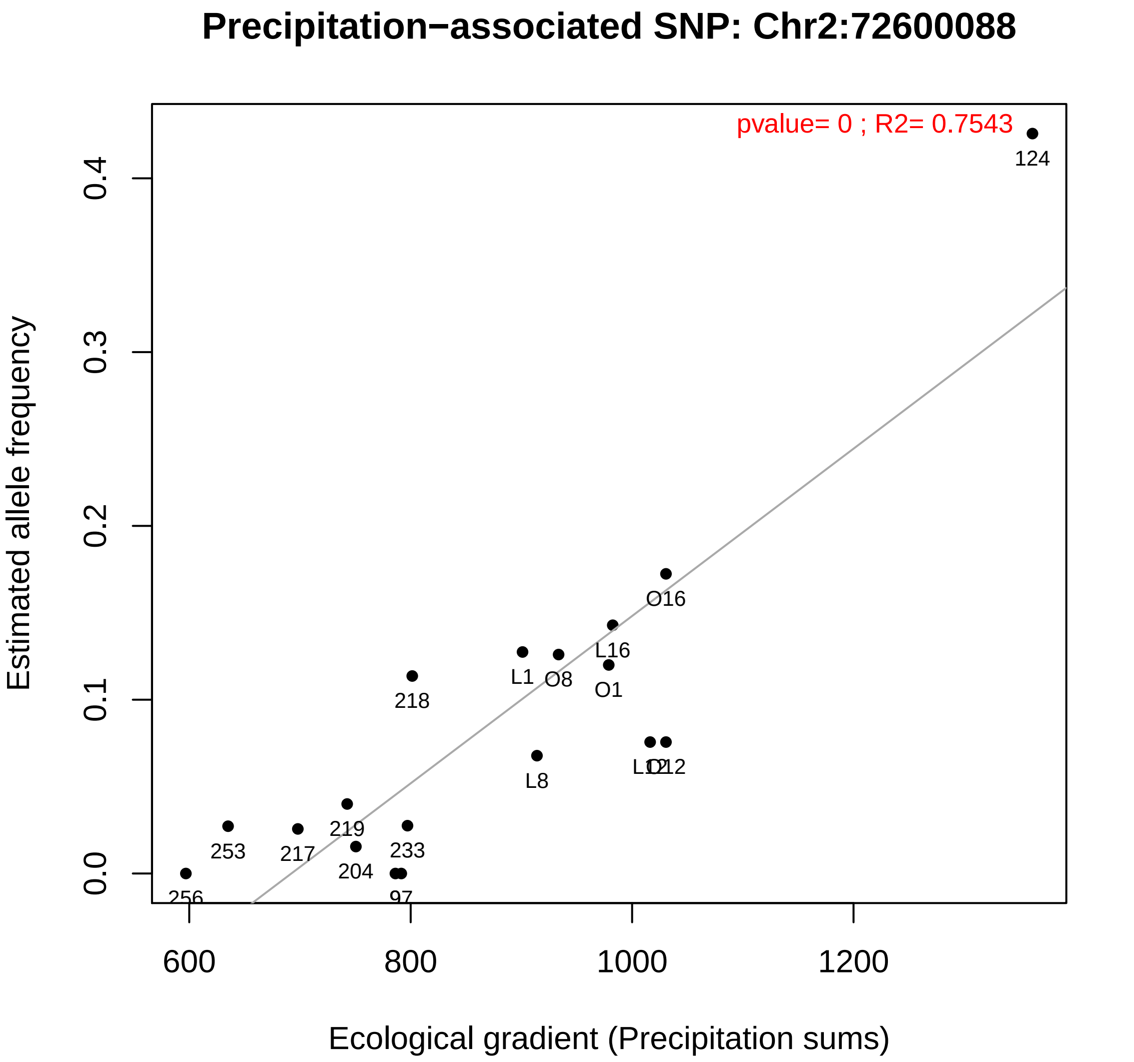

Precipitation−associated SNP: Chr2:72600088
pvalue= 0 ; R2= 0.7543
Estimated allele frequency
0.0
0.1
0.2
0.3
0.4
600
800
1000
1200
Ecological gradient (Precipitation sums)
124
O16
L16
L1
O8
O1
218
L8
219
253
217
233
204

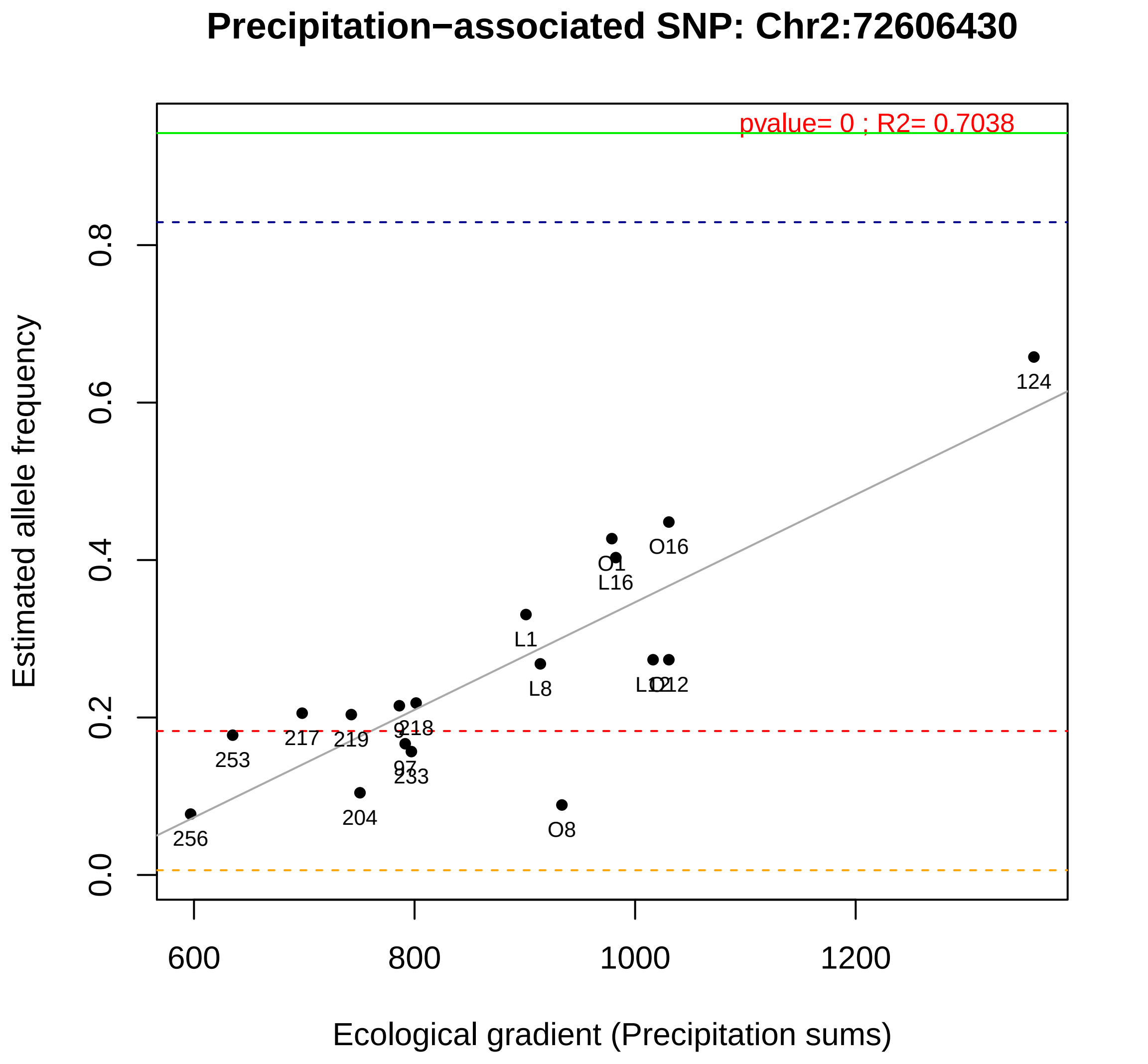

Precipitation−associated SNP: Chr2:72606430
pvalue= 0 ; R2= 0.7038
Estimated allele frequency
0.0
0.2
0.4
0.6
0.8
600
800
1000
1200
Ecological gradient (Precipitation sums)
124
O16
O1
L16
L1
L8
L10
O12
218
9
217
219
253
97
233
204
256
O8

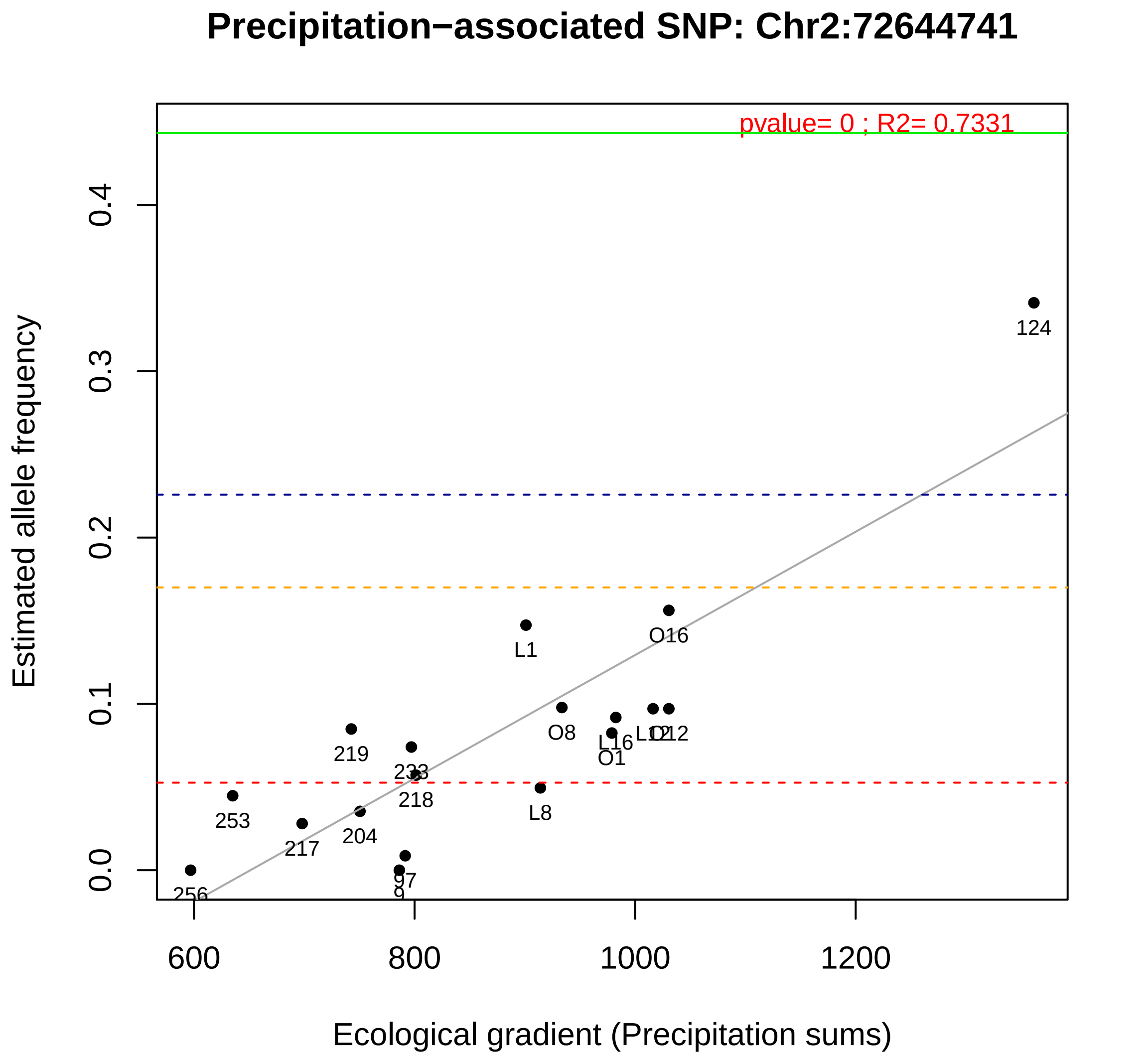

Precipitation−associated SNP: Chr2:72644741
pvalue= 0 ; R2= 0.7331
Estimated allele frequency
0.0
0.1
0.2
0.3
0.4
600
800
1000
1200
Ecological gradient (Precipitation sums)
124
O16
L1
O8
L16
O1
L12
O12
219
233
218
L8
253
217
204
97
9
256

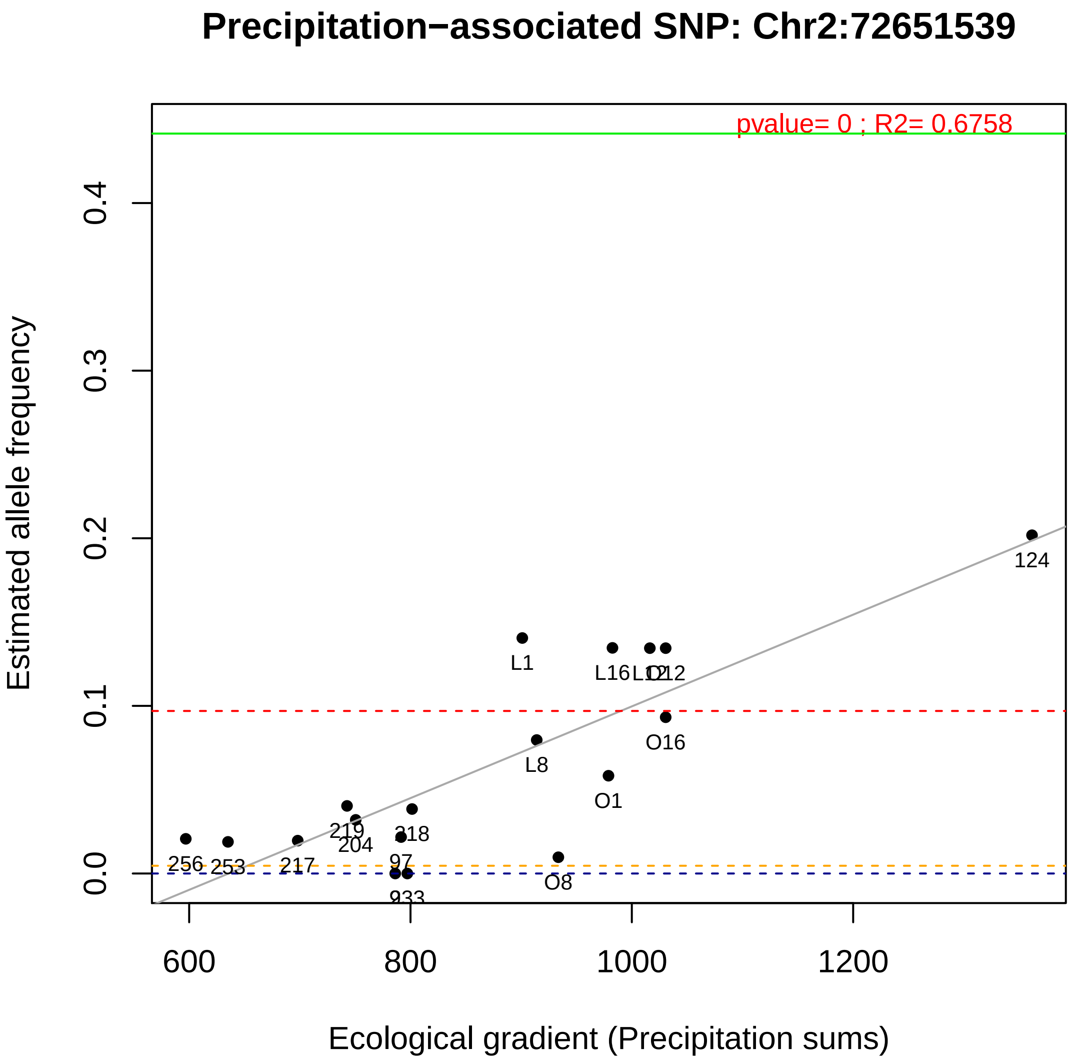

Precipitation−associated SNP: Chr2:72651539
pvalue= 0 ; R2= 0.6758
Estimated allele frequency
0.0
0.1
0.2
0.3
0.4
600
800
1000
1200
Ecological gradient (Precipitation sums)
124
L1
L16
O12
L8
O16
O1
O8
256
253
217
219
204
218
97
233

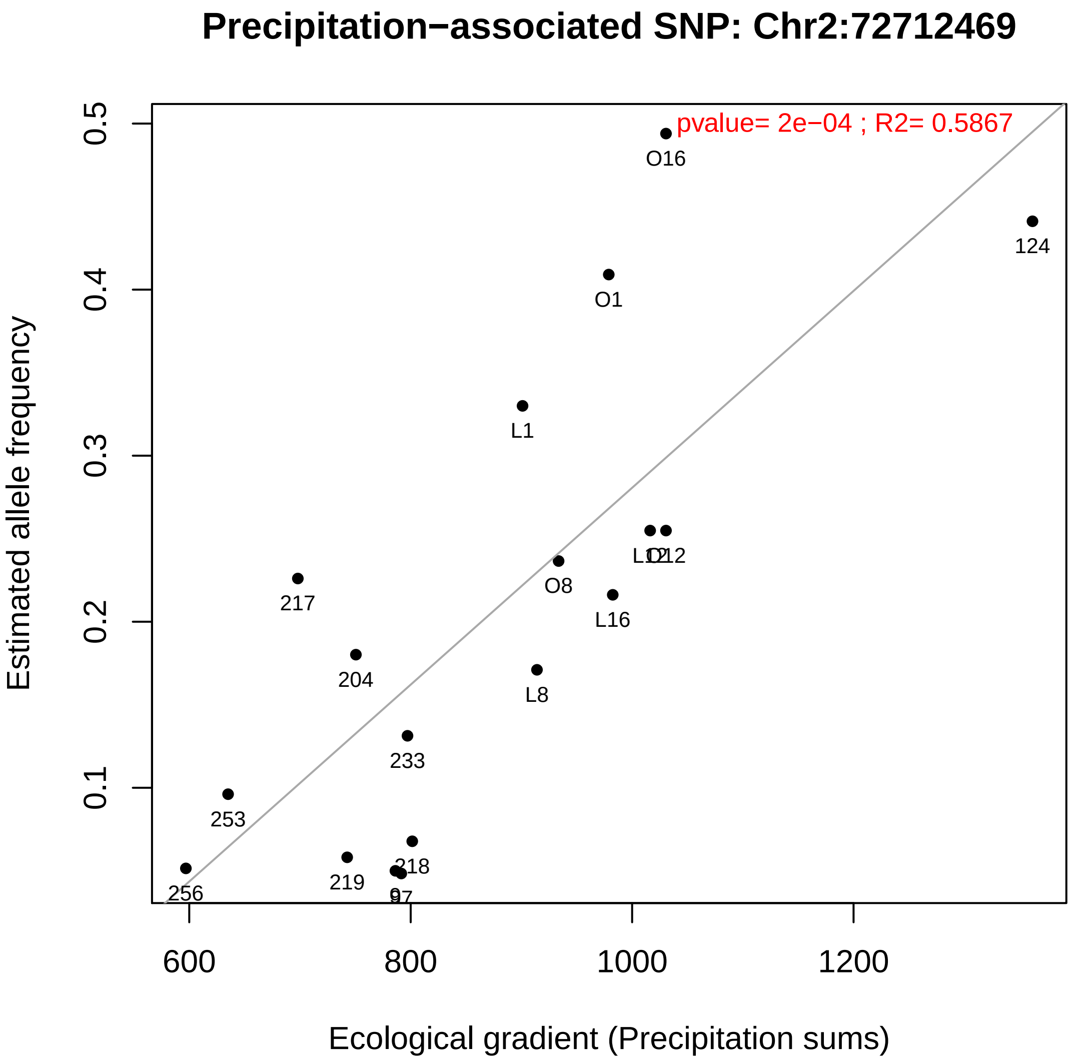

Precipitation−associated SNP: Chr2:72712469
pvalue= 2e−04 ; R2= 0.5867
Estimated allele frequency
0.1
0.2
0.3
0.4
0.5
Ecological gradient (Precipitation sums)
600
800
1000
1200
O16
124
O1
L1
O8
L16
217
204
L8
233
253
256
219
218

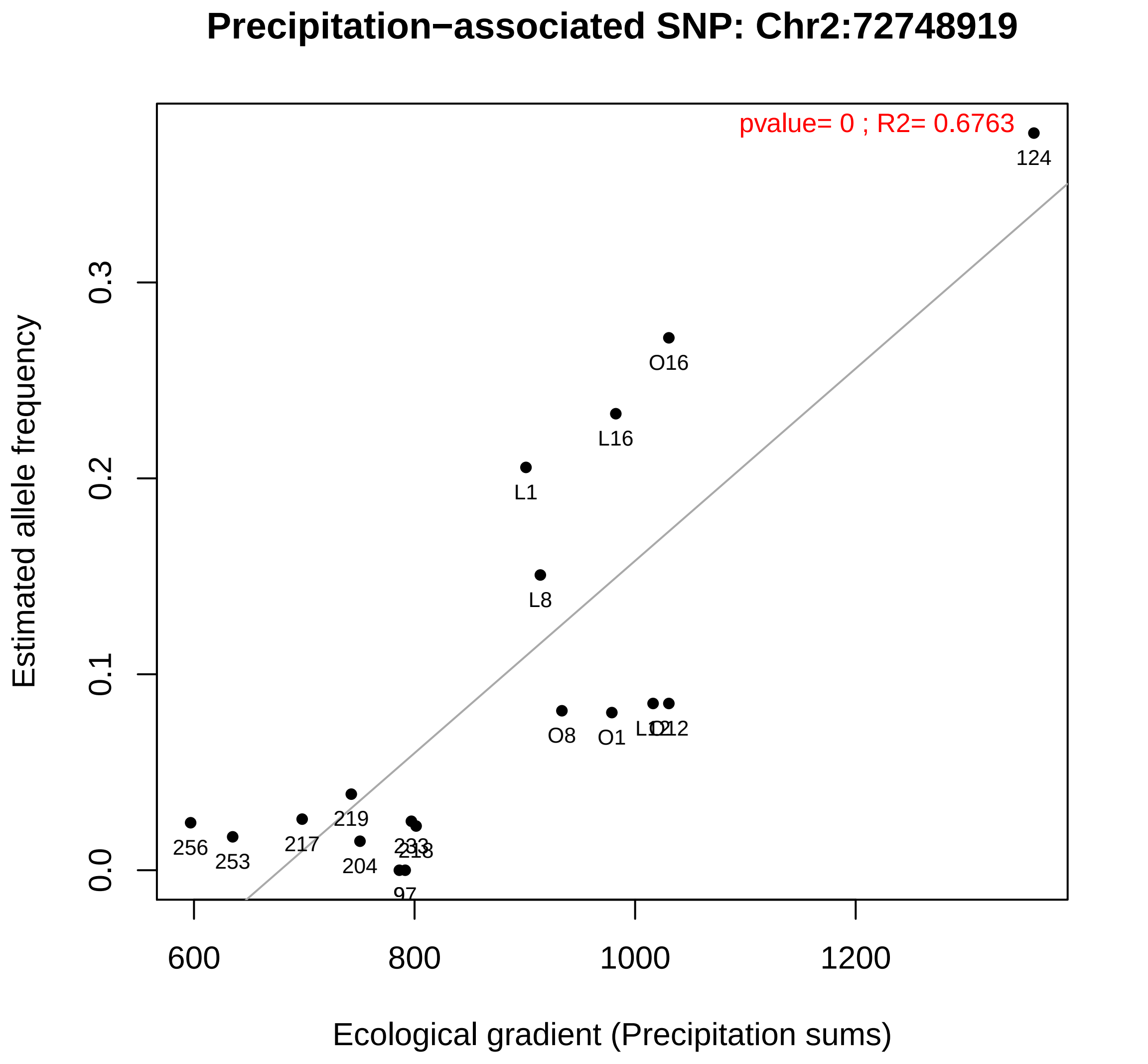
Precipitation−associated SNP: Chr2:72748919
pvalue= 0 ; R2= 0.6763
124
O16
L16
L1
L8
O8
O1
L12
O12
219
256
253
217
204
233
218
97
0.0
0.1
0.2
0.3
600
800
1000
1200
Estimated allele frequency
Ecological gradient (Precipitation sums)

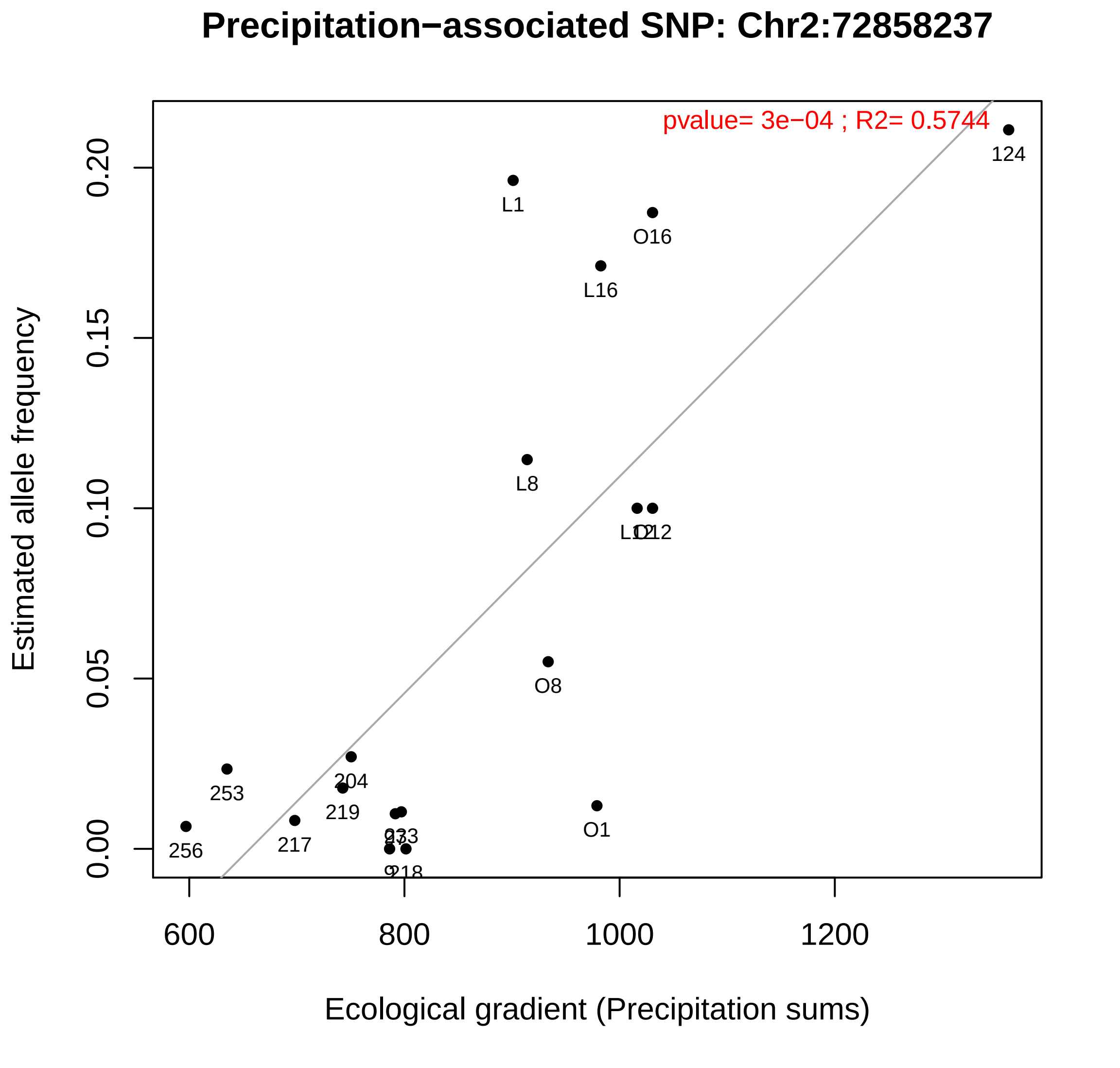

Precipitation−associated SNP: Chr2:72858237
pvalue= 3e−04 ; R2= 0.5744
Estimated allele frequency
Ecological gradient (Precipitation sums)
0.00
0.05
0.10
0.15
0.20
600
800
1000
1200
124
L1
O16
L16
L8
O8
O1
253
204
219
217
256
218

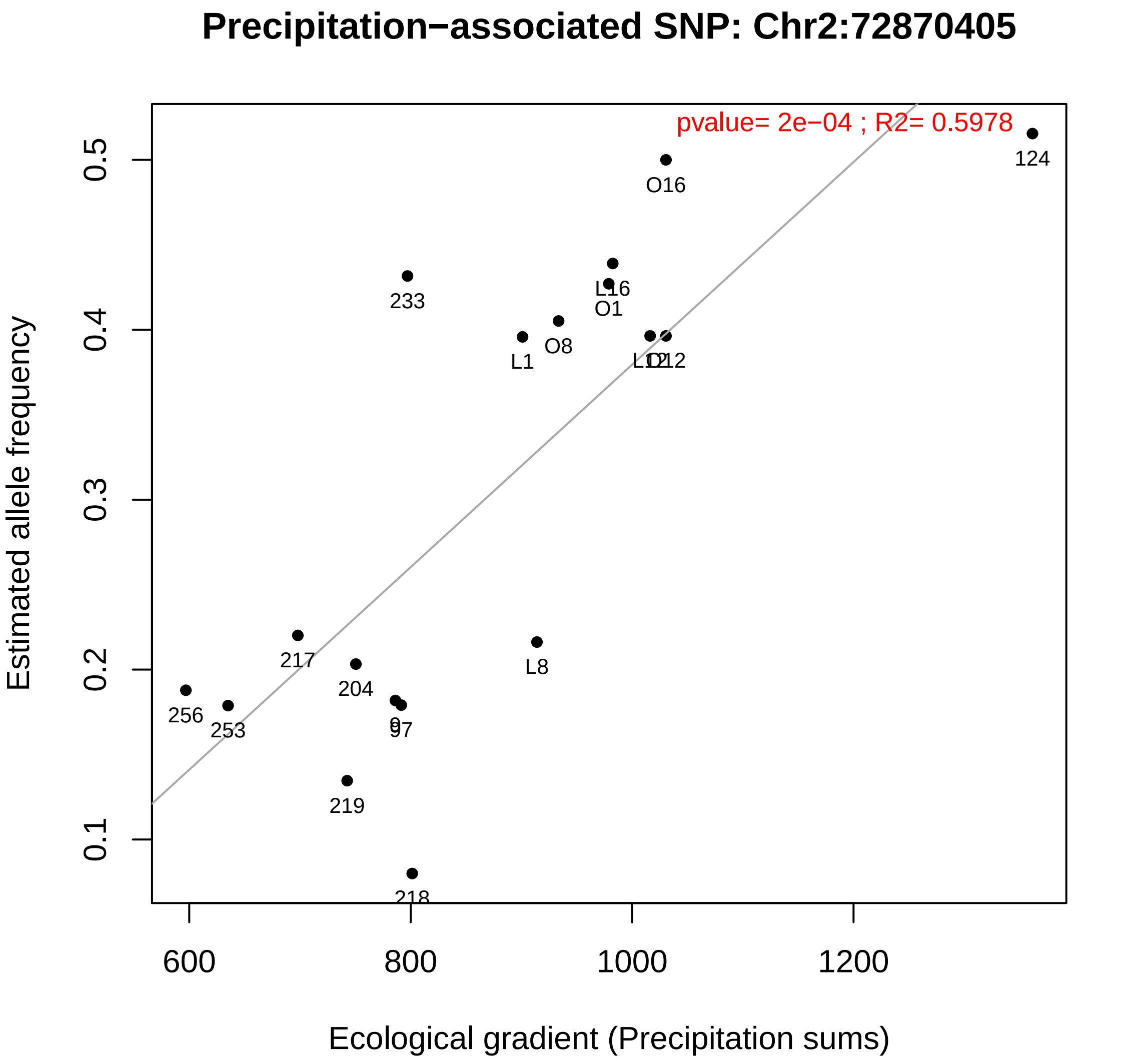
Precipitation−associated SNP: Chr2:72870405
pvalue= 2e−04 ; R2= 0.5978
124
O16
233
L16
O1
O8
L1
O12
L8
217
204
256
253
97
219
218
0.1
0.2
0.3
0.4
0.5
Estimated allele frequency
600
800
1000
1200
Ecological gradient (Precipitation sums)

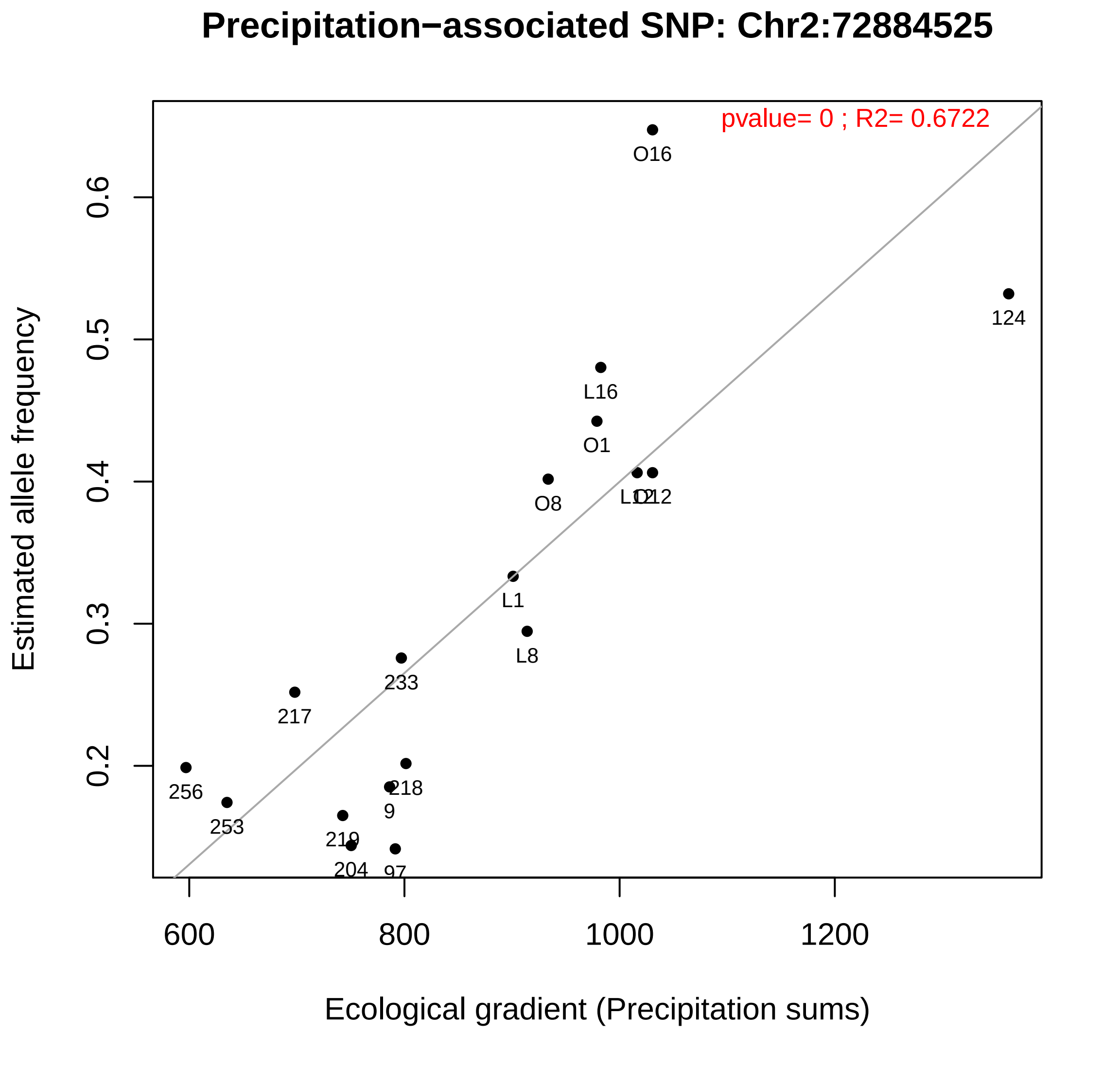
Precipitation−associated SNP: Chr2:72884525
pvalue= 0 ; R2= 0.6722
Estimated allele frequency
0.2
0.3
0.4
0.5
0.6
600
800
1000
1200
Ecological gradient (Precipitation sums)
O16
124
L16
O1
O8
L1
L8
233
217
256
253
218
9
219
204
97

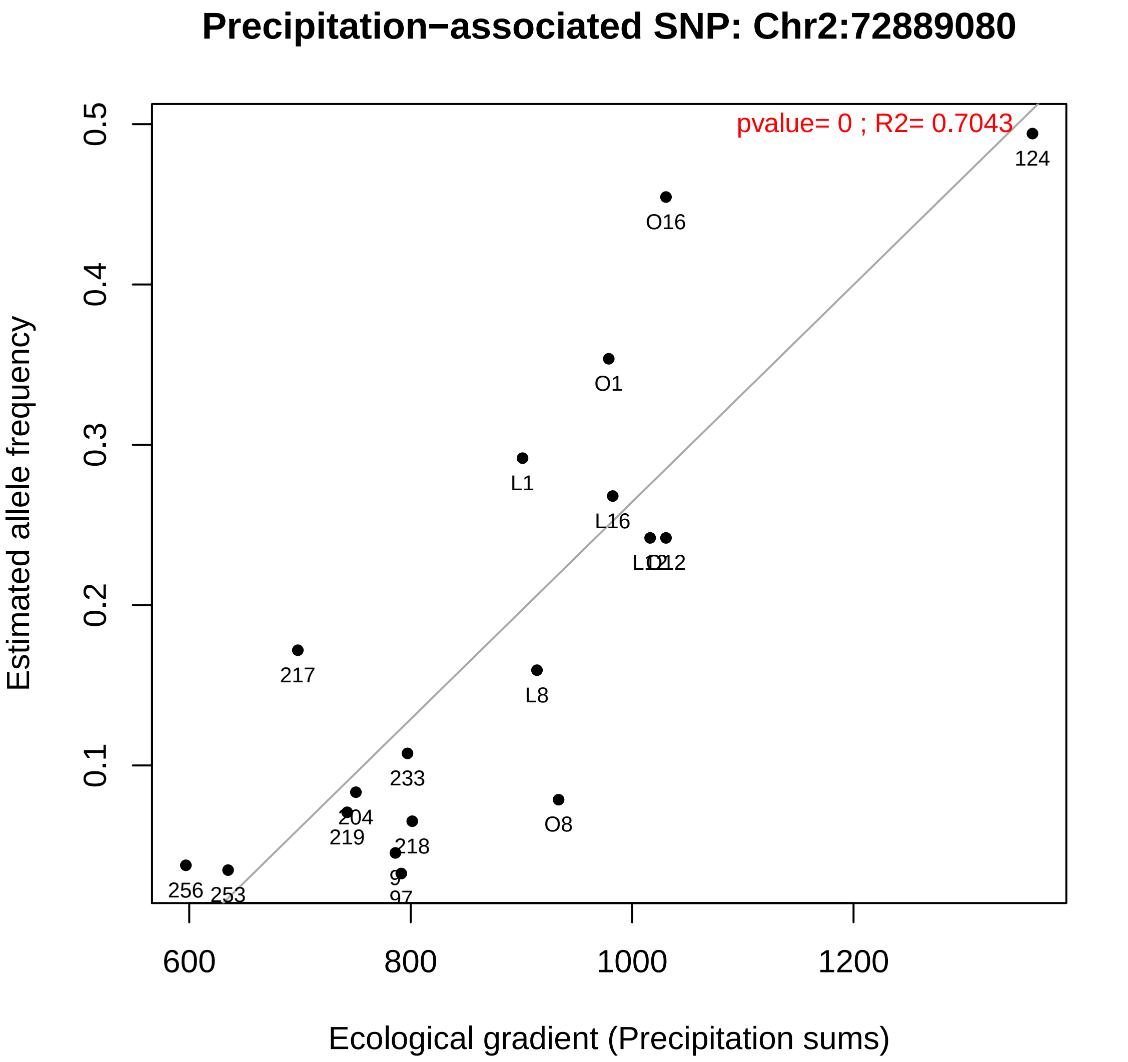

Precipitation−associated SNP: Chr2:72889080
pvalue= 0 ; R2= 0.7043
124
O16
O1
L1
L16
L12
O12
217
L8
233
204
219
218
O8
9
97
256
253
0.1
0.2
0.3
0.4
0.5
600
800
1000
1200
Estimated allele frequency
Ecological gradient (Precipitation sums)

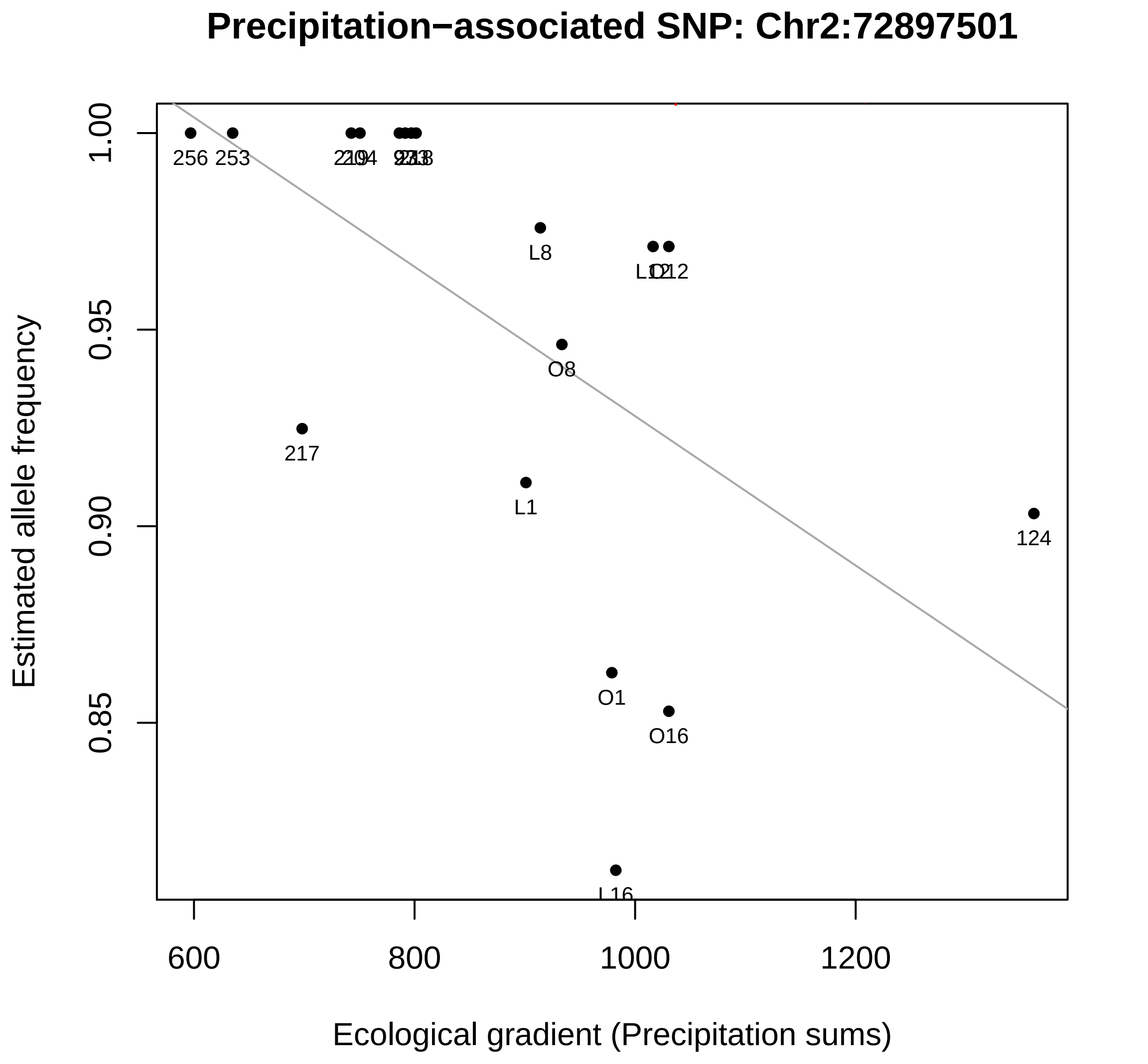
Precipitation−associated SNP: Chr2:72897501
Estimated allele frequency
Ecological gradient (Precipitation sums)
1.00
0.95
0.90
0.85
600
800
1000
1200
256
253
217
L8
O8
L1
124
O1
O16

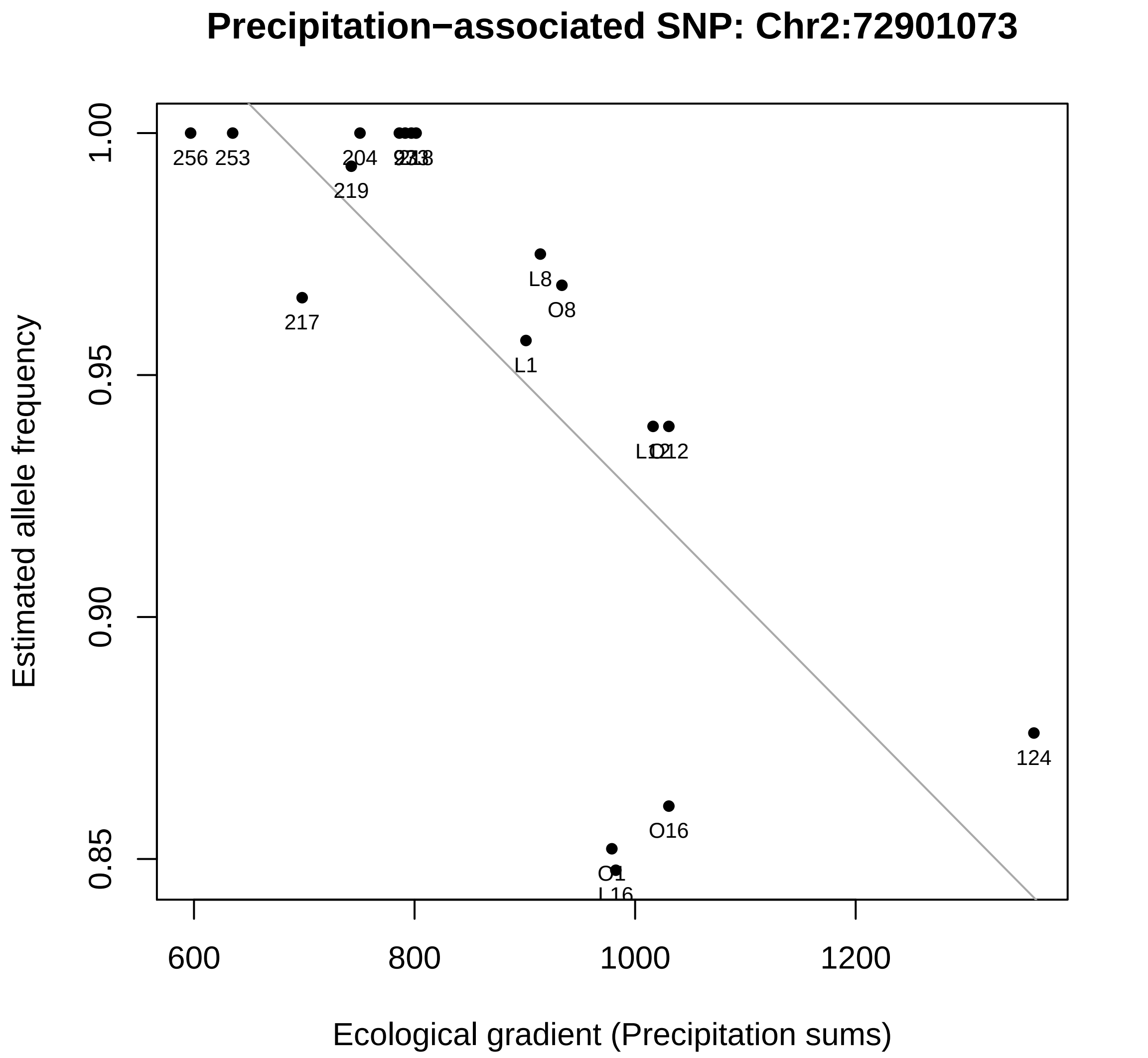
Precipitation−associated SNP: Chr2:72901073
Estimated allele frequency
Ecological gradient (Precipitation sums)
1.00
0.95
0.90
0.85
600
800
1000
1200
256
253
204
219
217
L8
O8
L1
O16
124

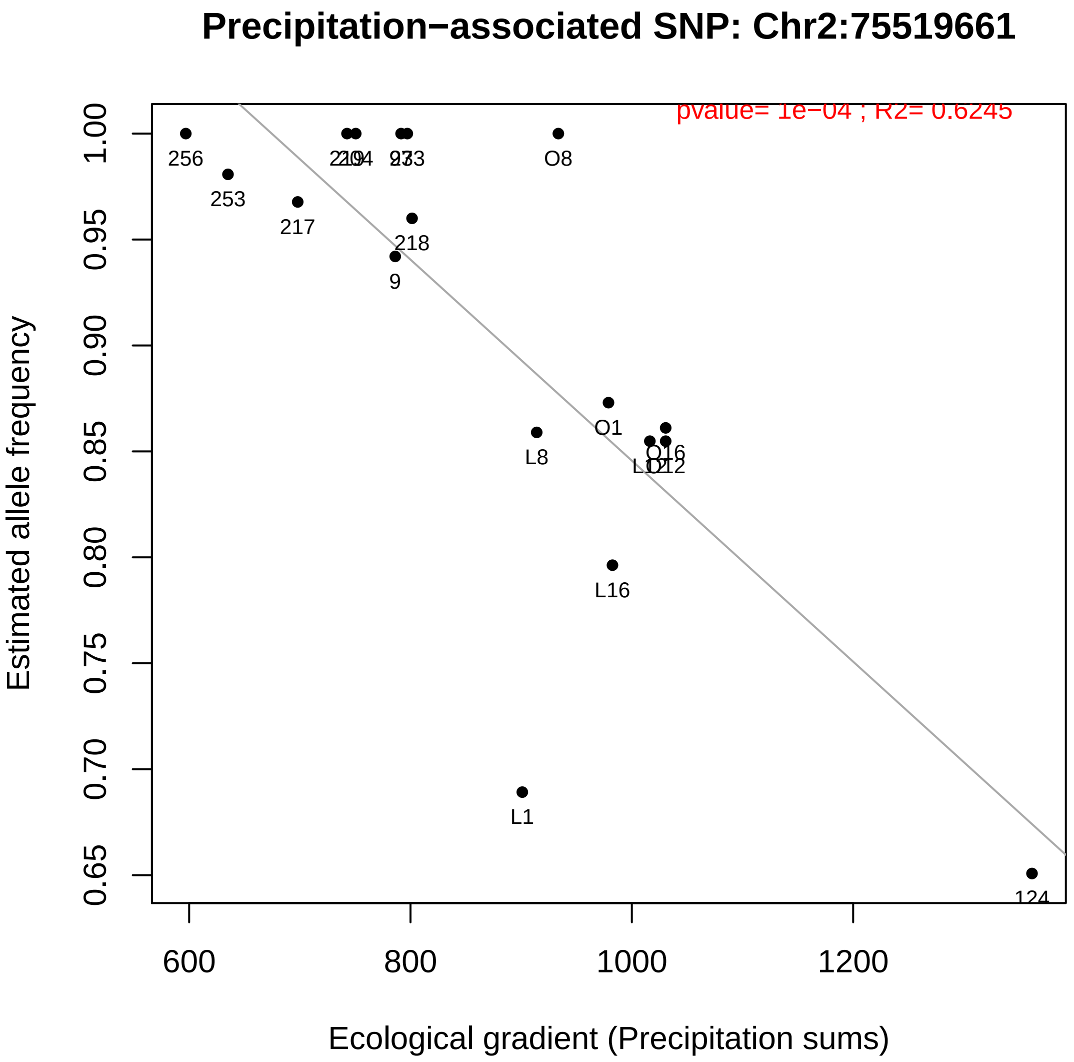

Precipitation−associated SNP: Chr2:75519661
pvalue= 1e−04 ; R2= 0.6245
Estimated allele frequency
Ecological gradient (Precipitation sums)
1.00
0.95
0.90
0.85
0.80
0.75
0.70
0.65
600
800
1000
1200
256
253
217
218
9
O8
O1
L8
L16
L1
124

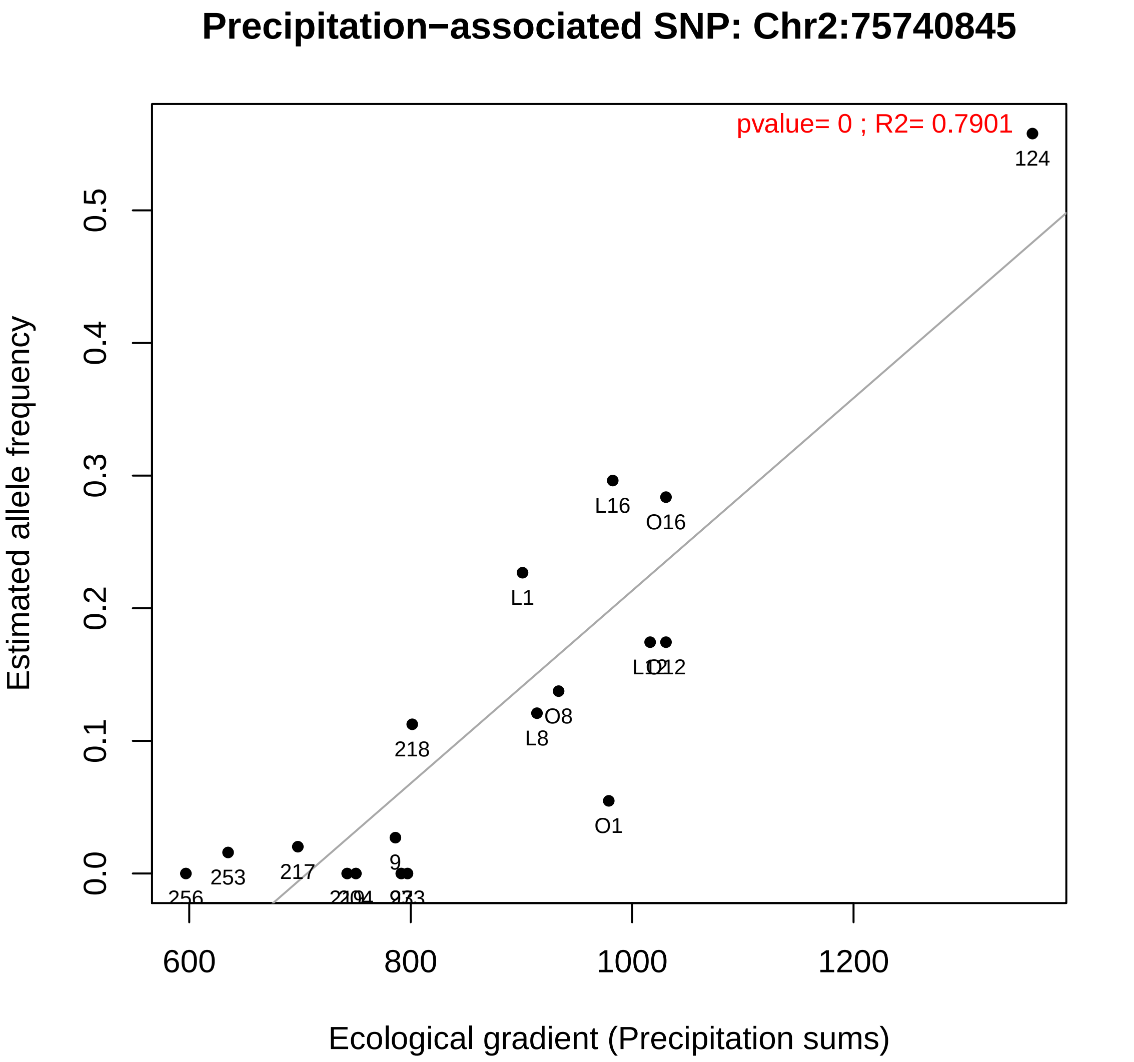
Precipitation−associated SNP: Chr2:75740845
pvalue= 0 ; R2= 0.7901
Estimated allele frequency
Ecological gradient (Precipitation sums)
0.0
0.1
0.2
0.3
0.4
0.5
600
800
1000
1200
124
L16
O16
L1
L12
O12
O8
L8
218
O1
9
217
253
256

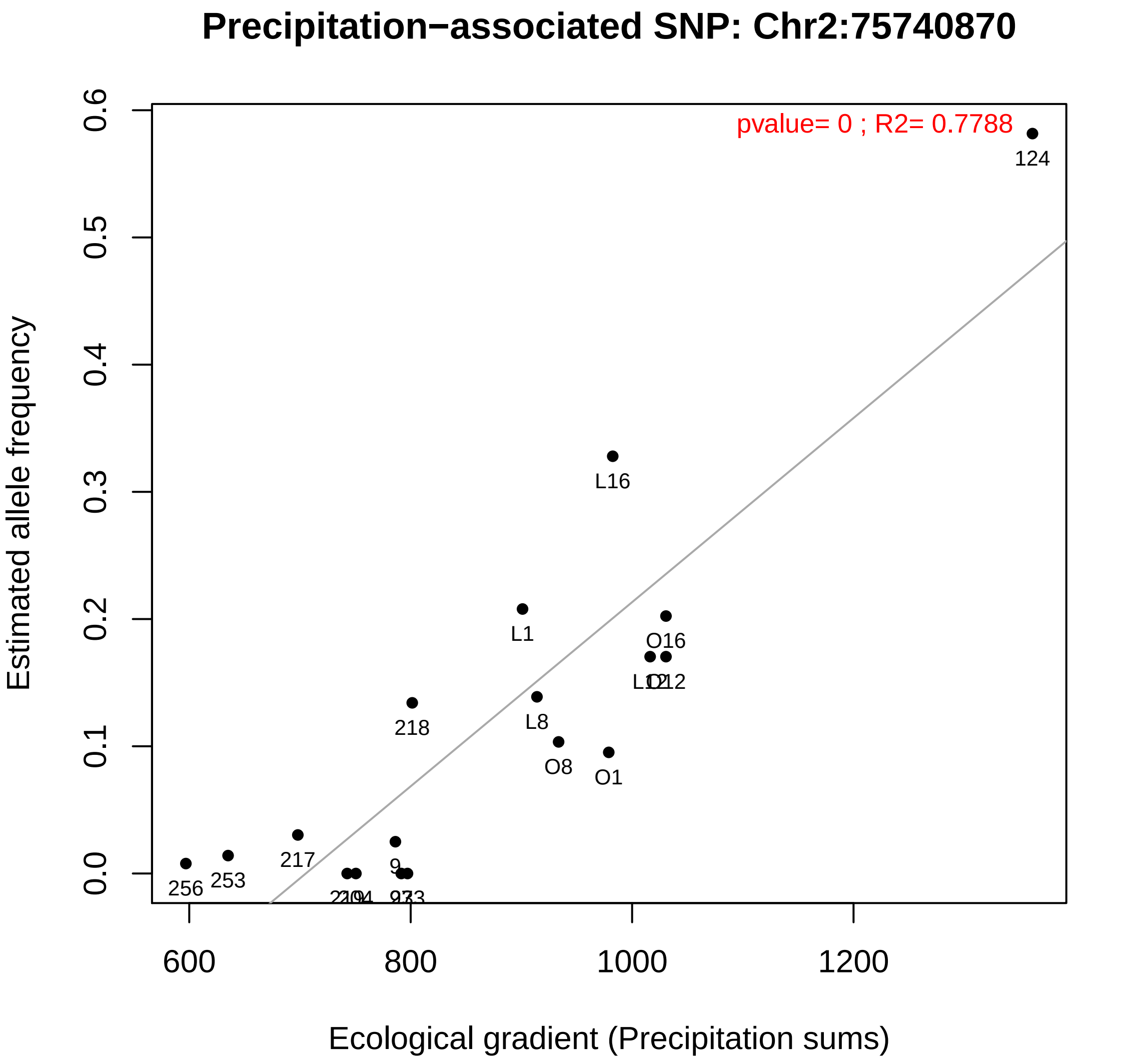

Precipitation−associated SNP: Chr2:75740870
pvalue= 0 ; R2= 0.7788
Estimated allele frequency
0.0
0.1
0.2
0.3
0.4
0.5
0.6
Ecological gradient (Precipitation sums)
600
800
1000
1200
124
L16
L1
O16
O12
L8
218
O8
O1
217
9
256
253

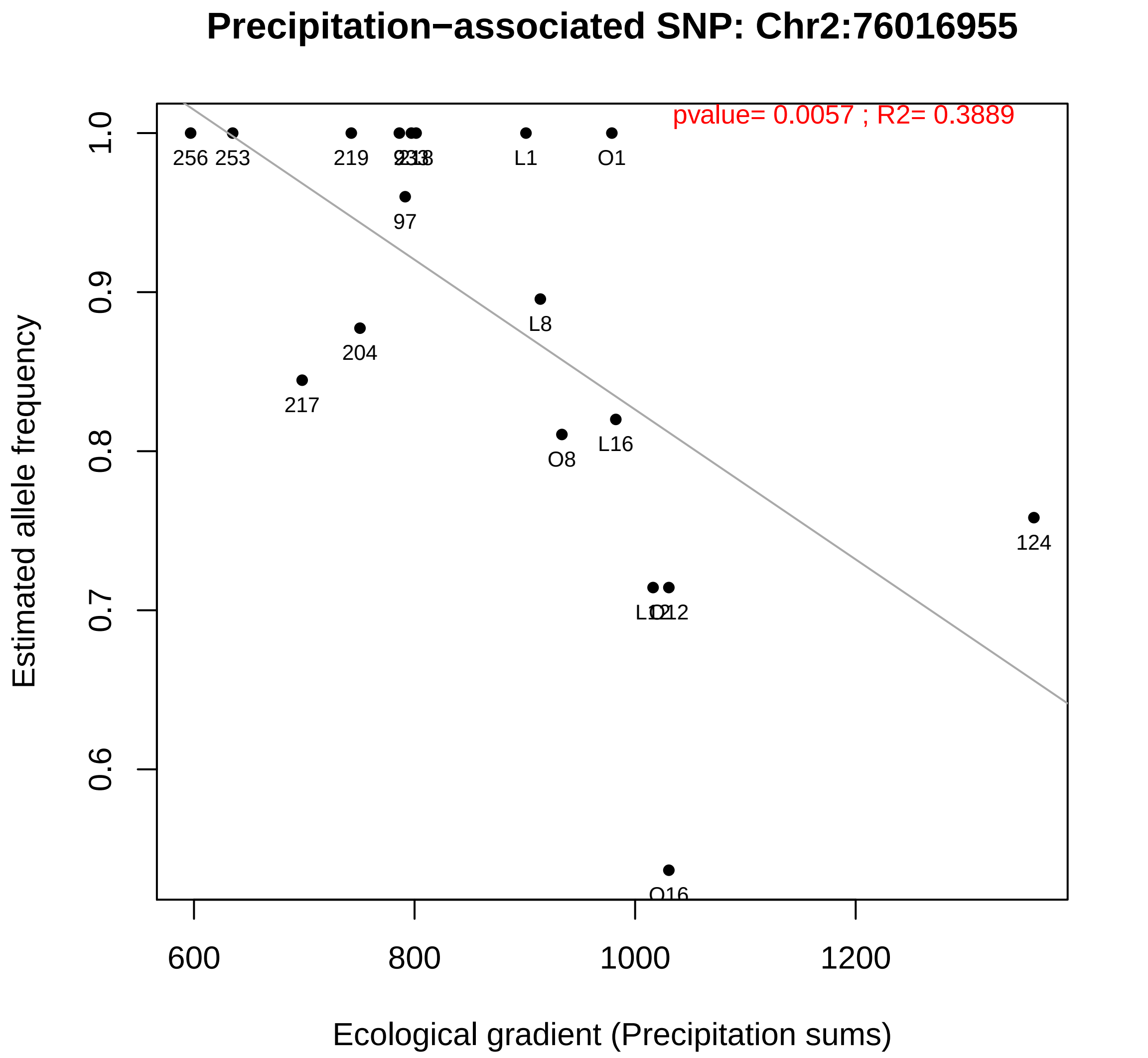

Precipitation−associated SNP: Chr2:76016955
pvalue= 0.0057 ; R2= 0.3889
Estimated allele frequency
Ecological gradient (Precipitation sums)
1.0
0.9
0.8
0.7
0.6
600
800
1000
1200
256
253
219
L1
O1
97
L8
204
217
L16
O8
124
O12
O16

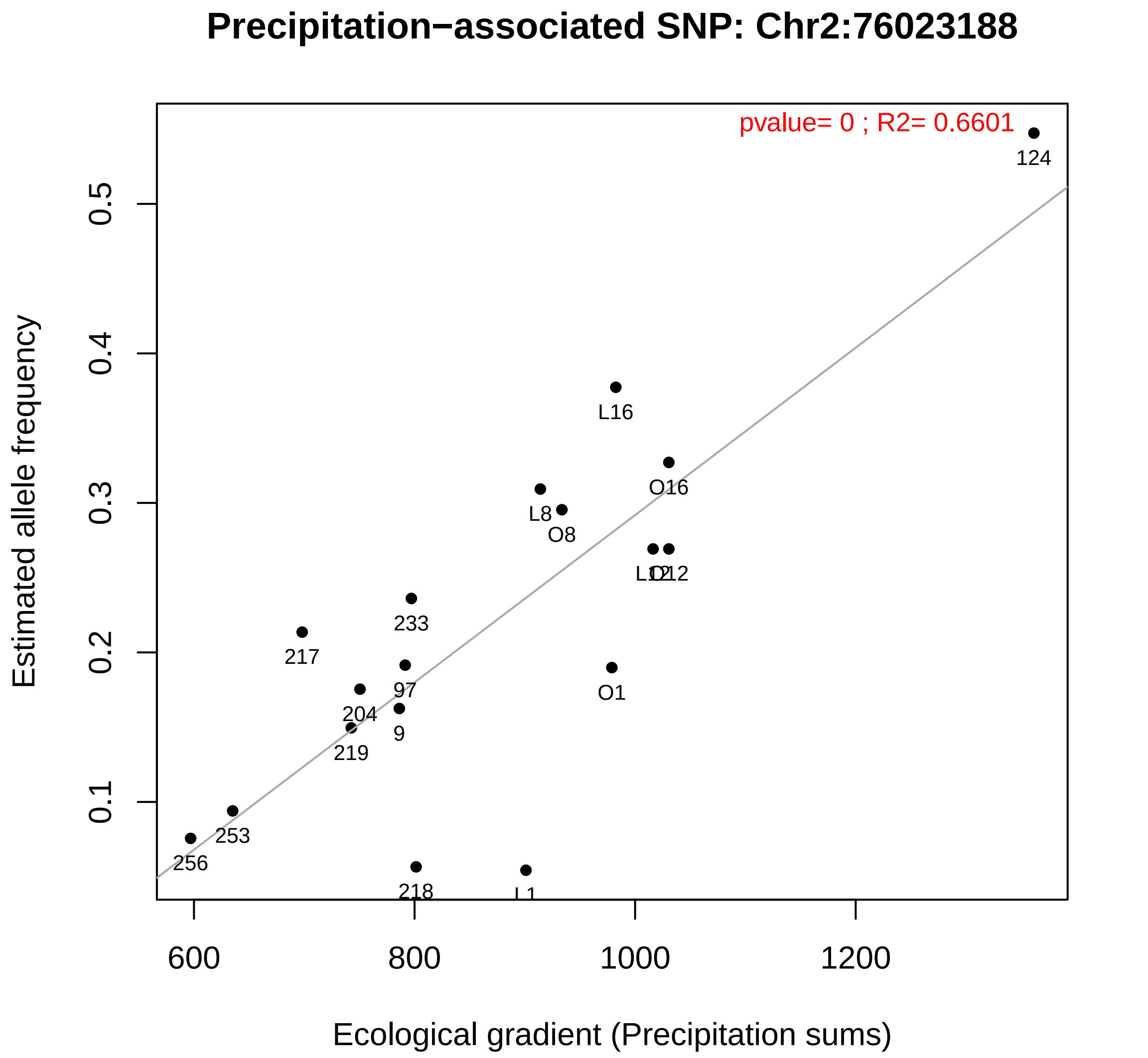
Precipitation−associated SNP: Chr2:76023188
pvalue= 0 ; R2= 0.6601
124
L16
O16
L8
O8
L12
O12
233
217
97
O1
204
9
219
253
256
218
L1
0.1
0.2
0.3
0.4
0.5
600
800
1000
1200
Estimated allele frequency
Ecological gradient (Precipitation sums)

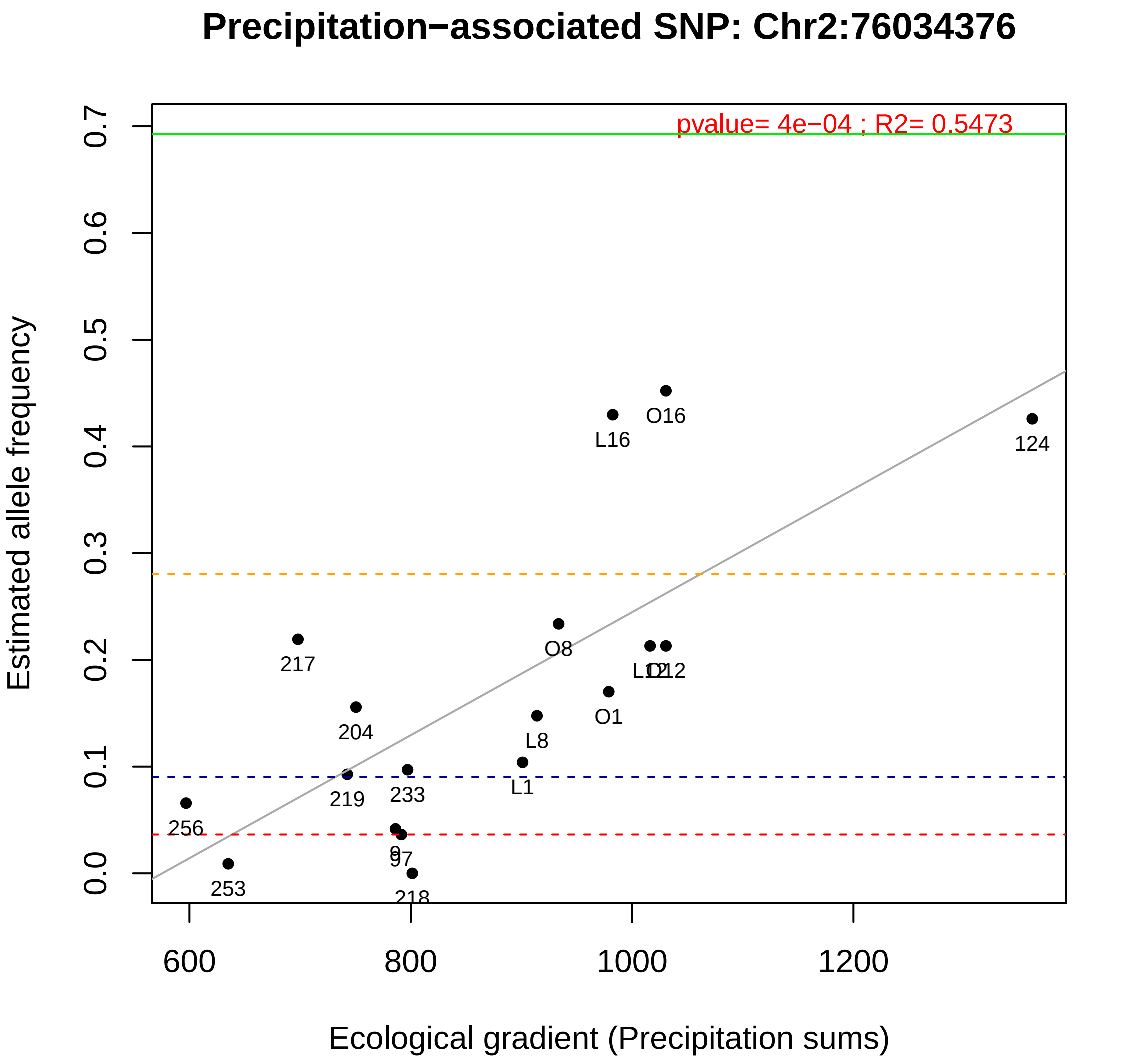
Precipitation−associated SNP: Chr2:76034376
pvalue= 4e−04 ; R2= 0.5473
Estimated allele frequency
0.0
0.1
0.2
0.3
0.4
0.5
0.6
0.7
O16
L16
124
O8
217
L12
O12
O1
204
L8
219
233
L1
256
97
253
218
600
800
1000
1200
Ecological gradient (Precipitation sums)

#### Precipitation−associated SNP: Chr2:76039684

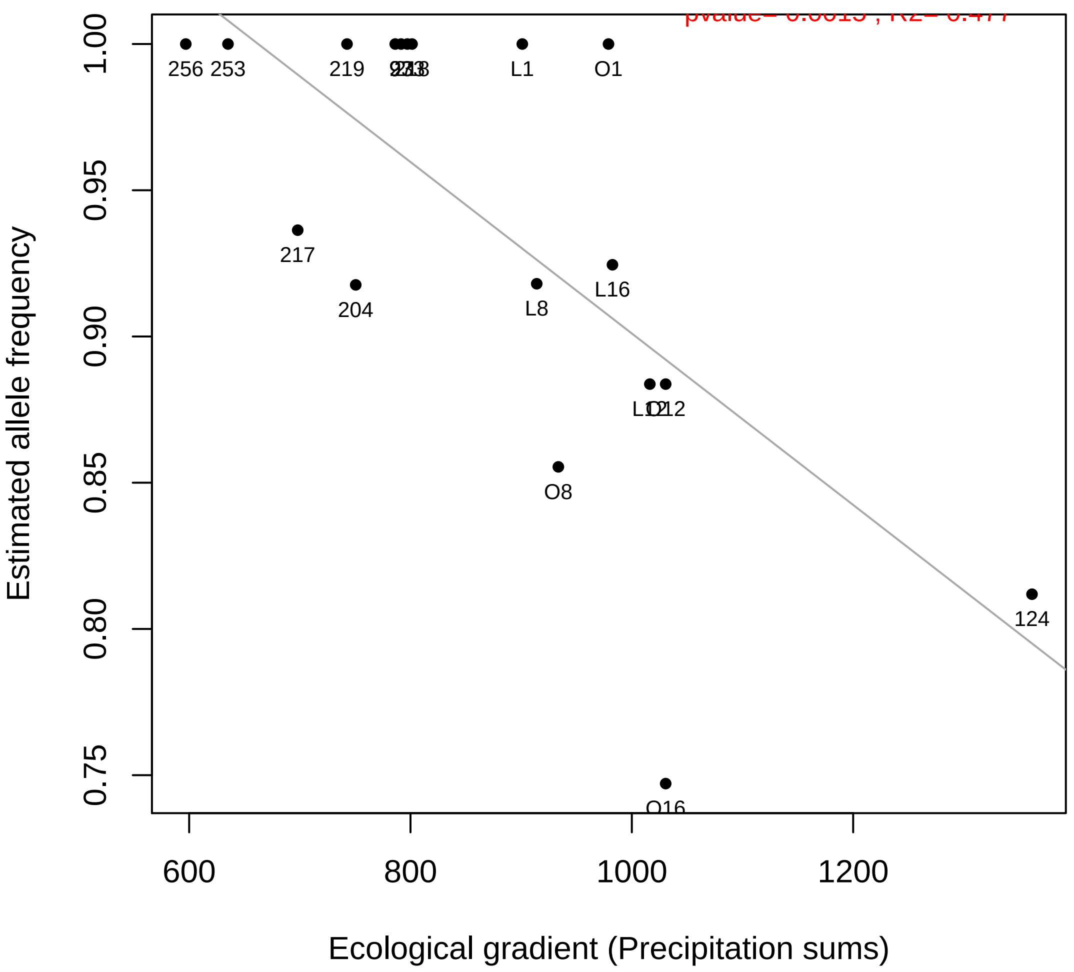

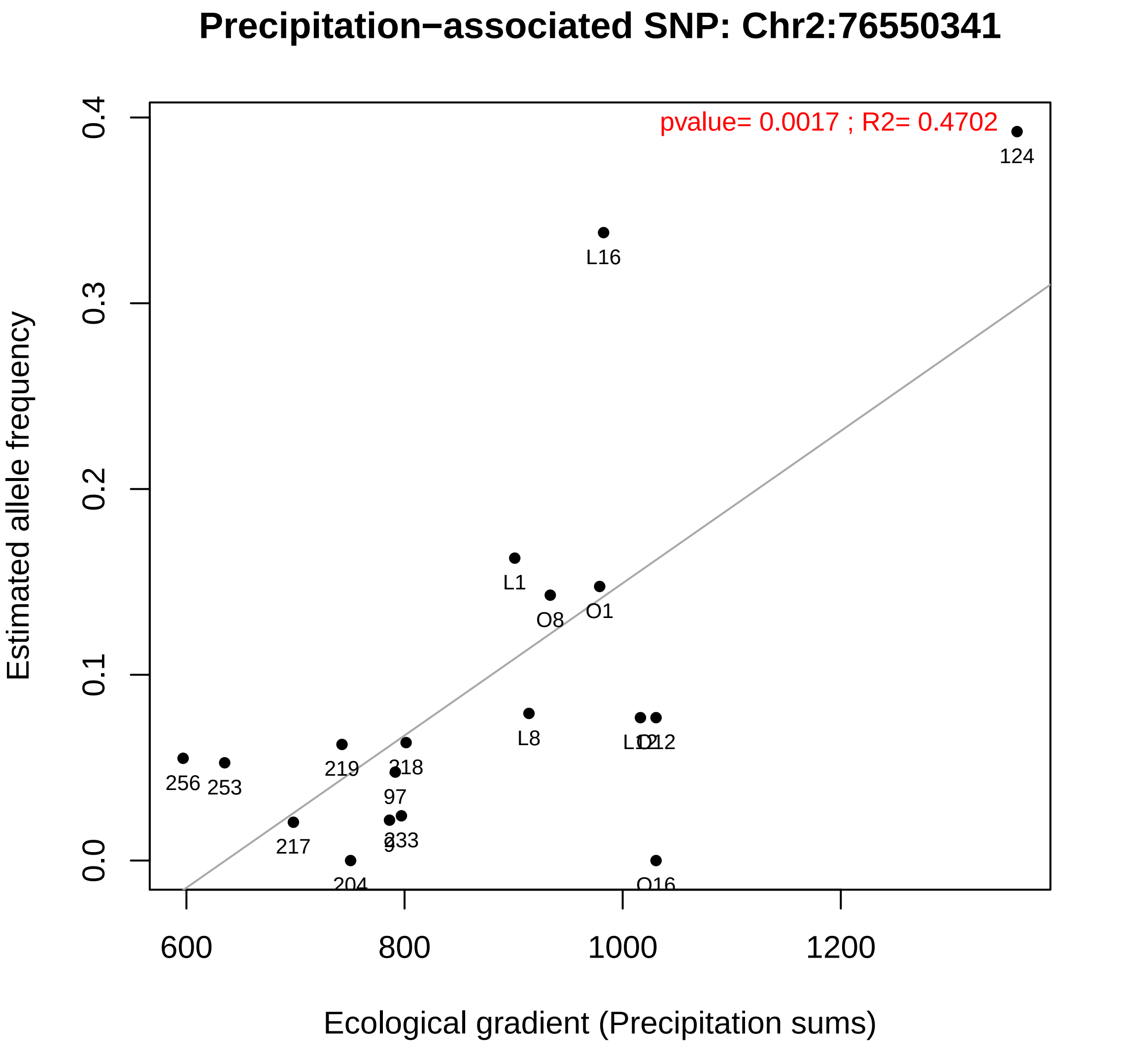
Precipitation−associated SNP: Chr2:76550341
pvalue= 0.0017 ; R2= 0.4702
124
L16
L1
O8
O1
L8
L12
O12
219
218
256
253
97
217
233
204
O16
0.0
0.1
0.2
0.3
0.4
600
800
1000
1200
Estimated allele frequency
Ecological gradient (Precipitation sums)

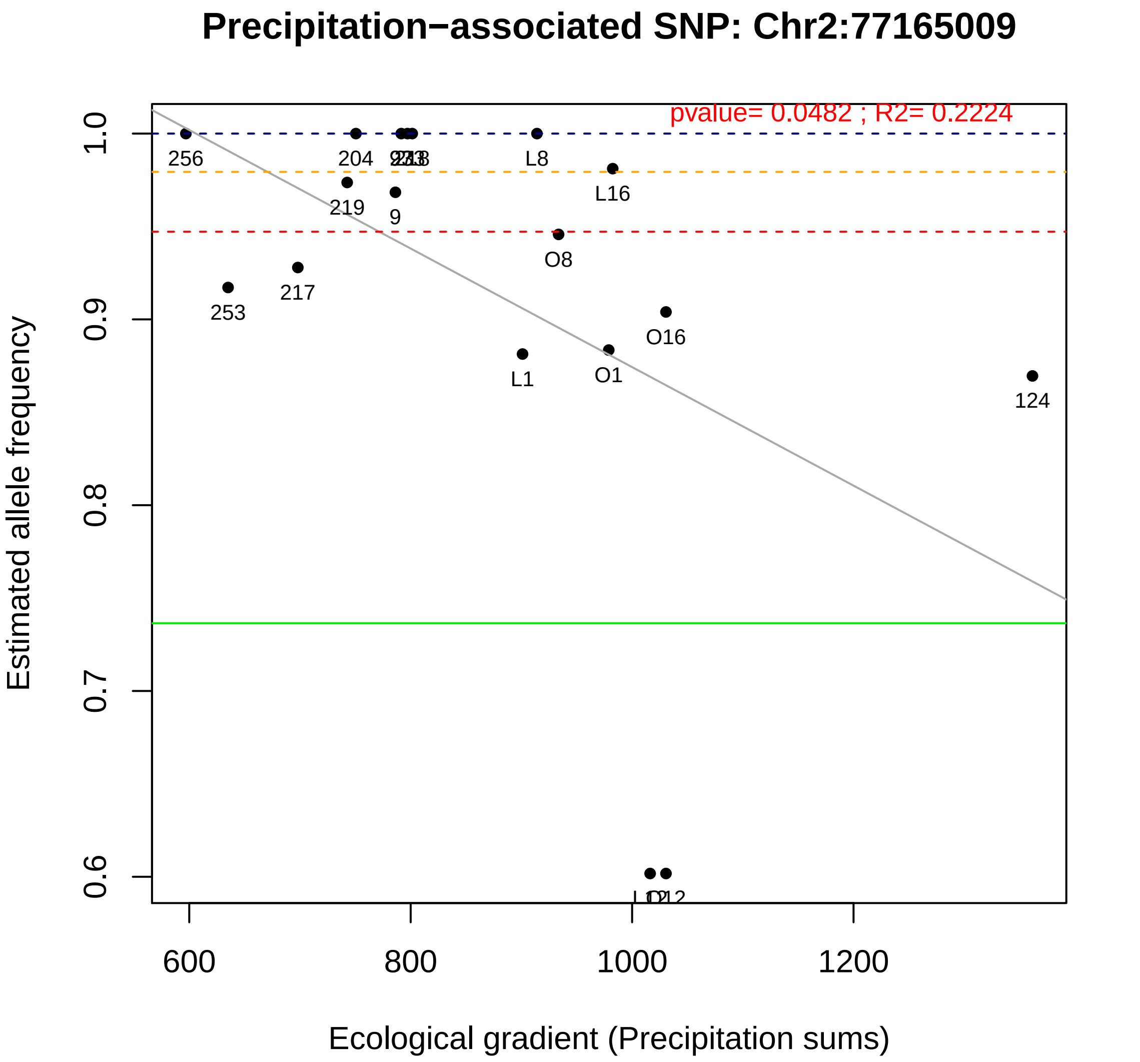
Precipitation−associated SNP: Chr2:77165009
pvalue= 0.0482 ; R2= 0.2224
Estimated allele frequency
Ecological gradient (Precipitation sums)
1.0
0.9
0.8
0.7
0.6
600
800
1000
1200
256
204
L8
L16
219
9
O8
217
253
O16
L1
O1
124

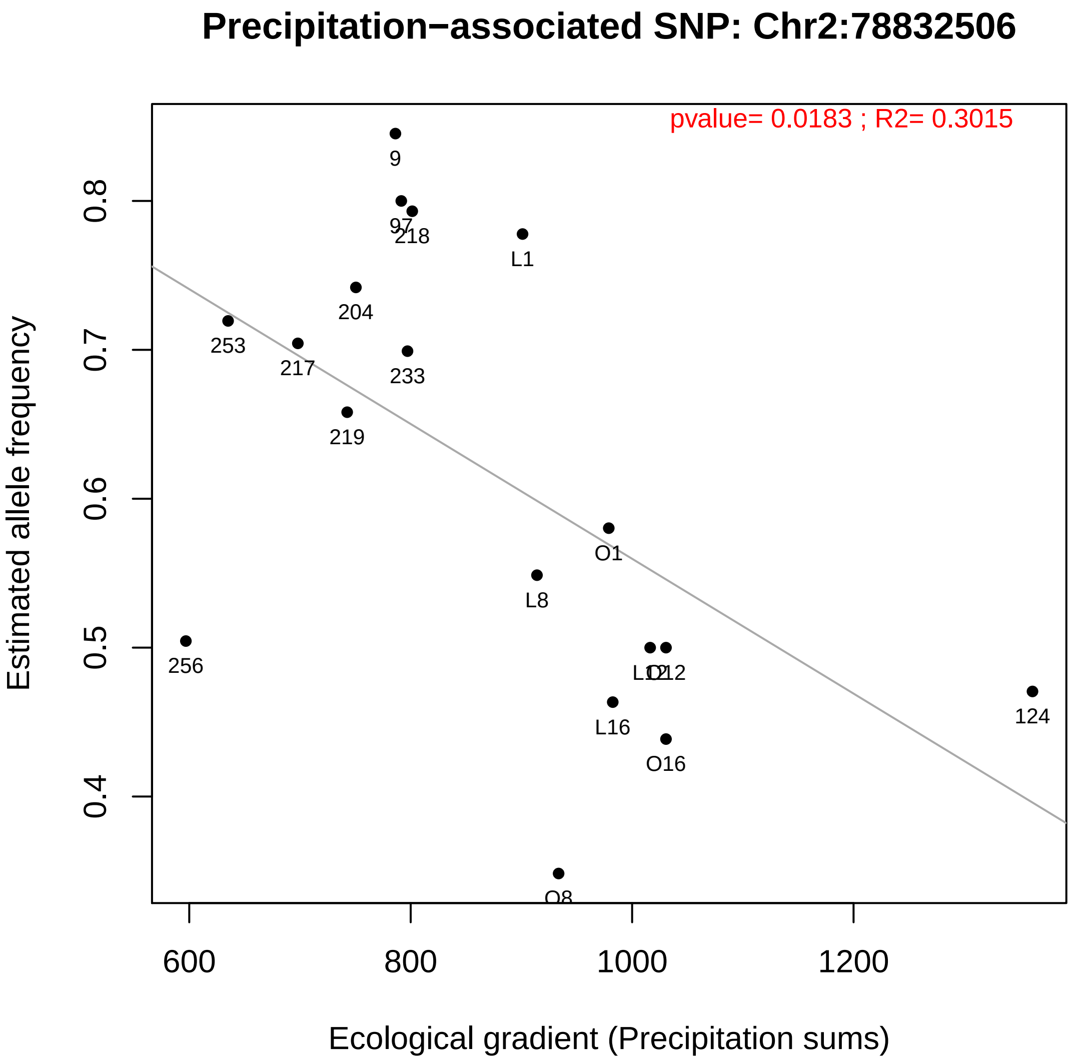

Precipitation−associated SNP: Chr2:78832506
pvalue= 0.0183 ; R2= 0.3015
Estimated allele frequency
Ecological gradient (Precipitation sums)
0.4
0.5
0.6
0.7
0.8
600
800
1000
1200
9
97
218
L1
204
253
217
233
219
O1
L8
256
L12
O12
L16
124
O16
O8

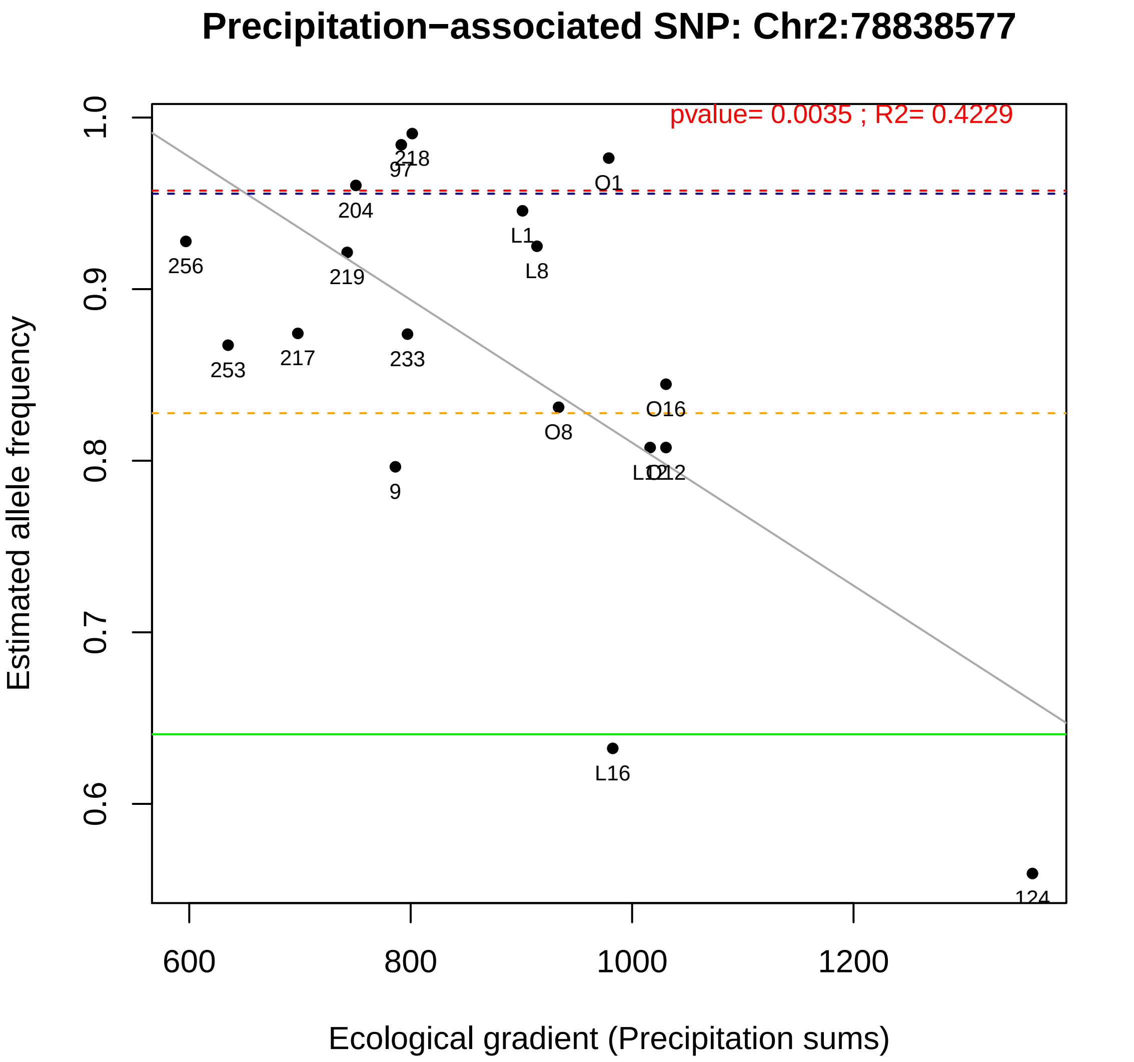

Precipitation−associated SNP: Chr2:78838577
pvalue= 0.0035 ; R2= 0.4229
218
97
O1
204
L1
L8
256
219
253
217
233
O16
O8
9
L12
O12
L16
124
1.0
0.9
0.8
0.7
0.6
600
800
1000
1200
Estimated allele frequency
Ecological gradient (Precipitation sums)

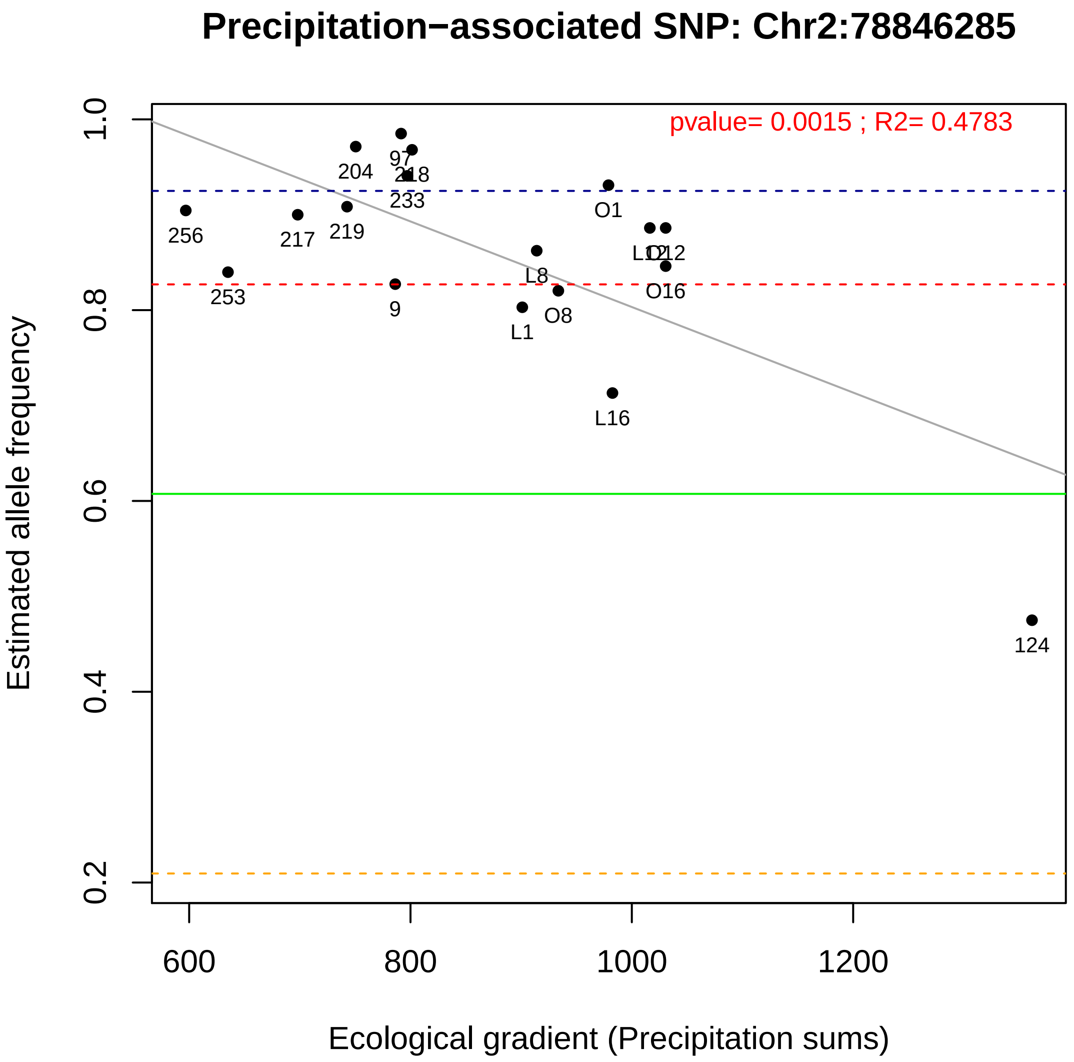
Precipitation−associated SNP: Chr2:78846285
pvalue= 0.0015 ; R2= 0.4783
Estimated allele frequency
Ecological gradient (Precipitation sums)
0.2
0.4
0.6
0.8
1.0
600
800
1000
1200
204
97
218
233
256
217
219
253
9
L8
O8
L1
O1
L12
O12
O16
L16
124

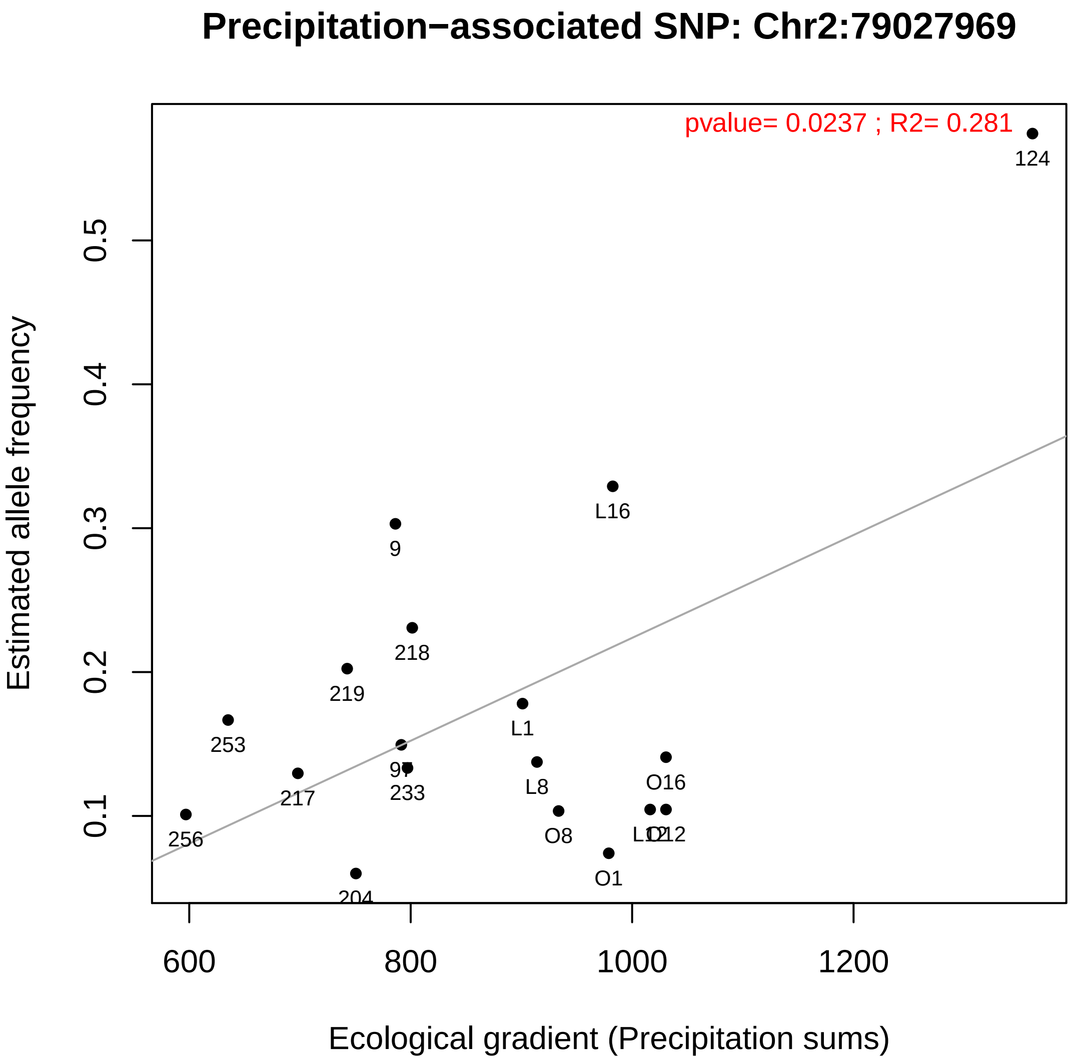

Precipitation−associated SNP: Chr2:79027969
pvalue= 0.0237 ; R2= 0.281
Estimated allele frequency
Ecological gradient (Precipitation sums)
0.1
0.2
0.3
0.4
0.5
600
800
1000
1200
124
L16
9
218
219
253
L1
97
233
217
L8
O16
256
O8
L12
O12
O1
204

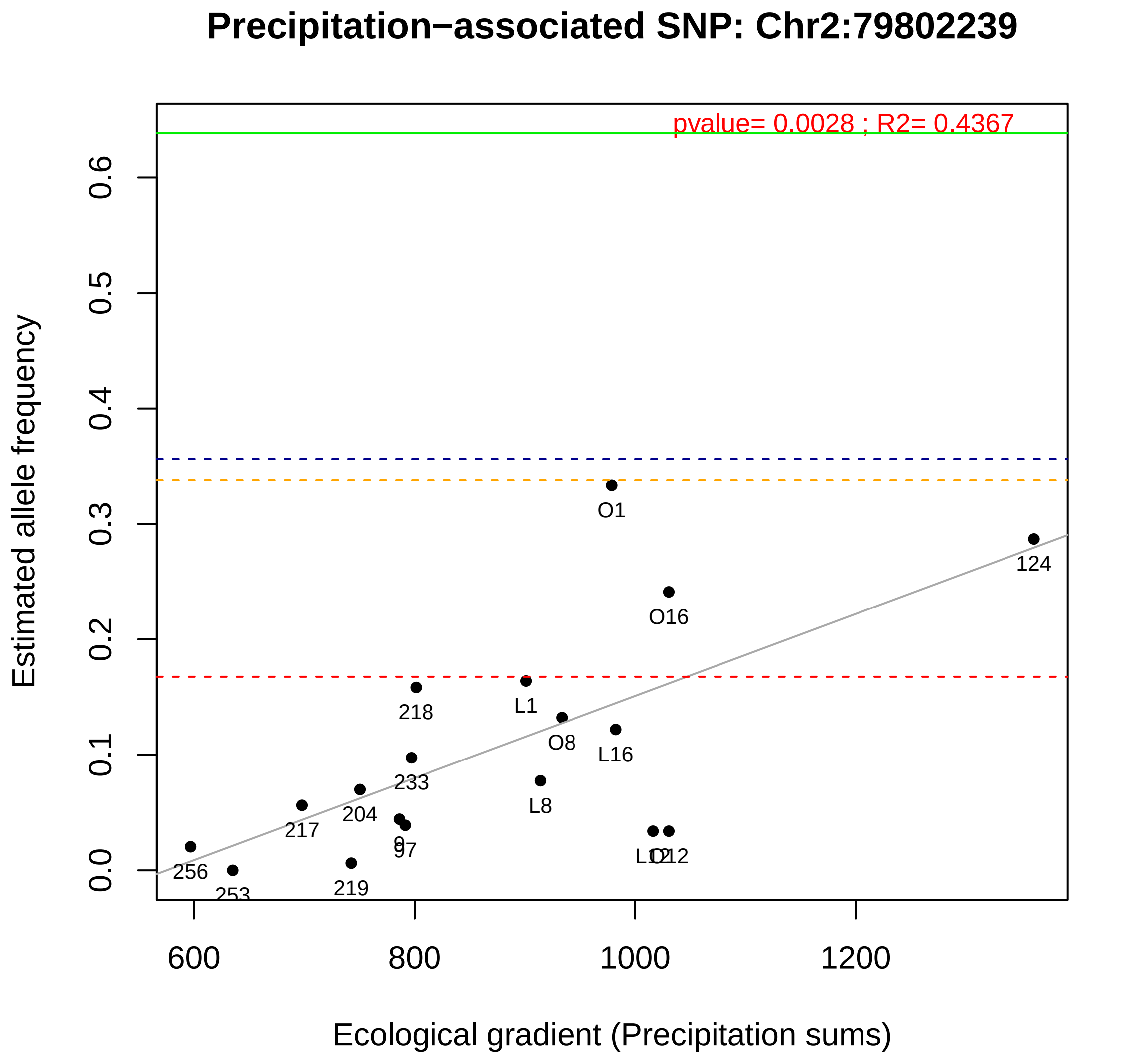

Precipitation−associated SNP: Chr2:79802239
pvalue= 0.0028 ; R2= 0.4367
Estimated allele frequency
0.0
0.1
0.2
0.3
0.4
0.5
0.6
600
800
1000
1200
Ecological gradient (Precipitation sums)
O1
124
O16
218
L1
O8
L16
233
L8
204
217
97
256
253
219

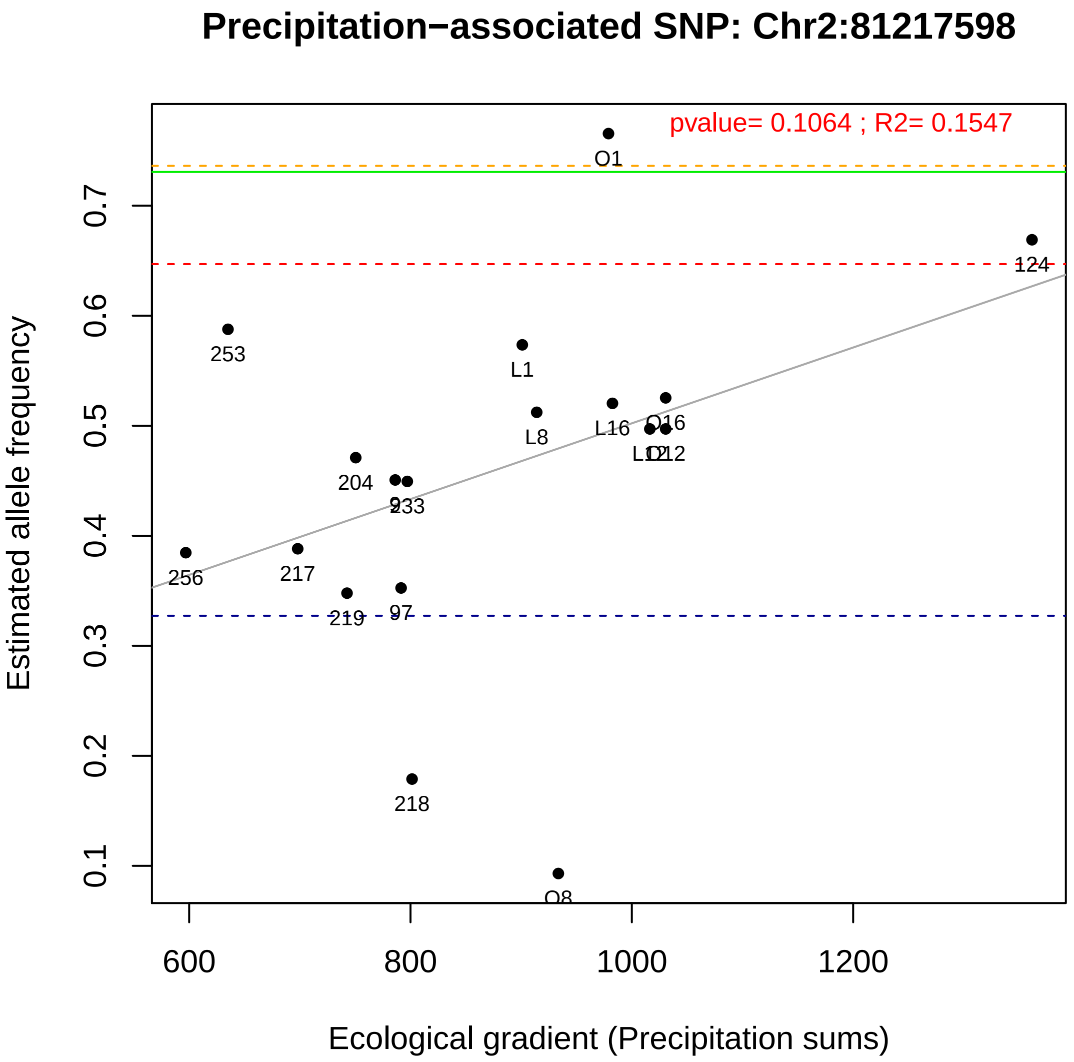
Precipitation−associated SNP: Chr2:81217598
pvalue= 0.1064 ; R2= 0.1547
O1
124
253
L1
L8
L16
O16
L12
O12
204
9
233
256
217
219
97
218
O8
0.1
0.2
0.3
0.4
0.5
0.6
0.7
600
800
1000
1200
Estimated allele frequency
Ecological gradient (Precipitation sums)

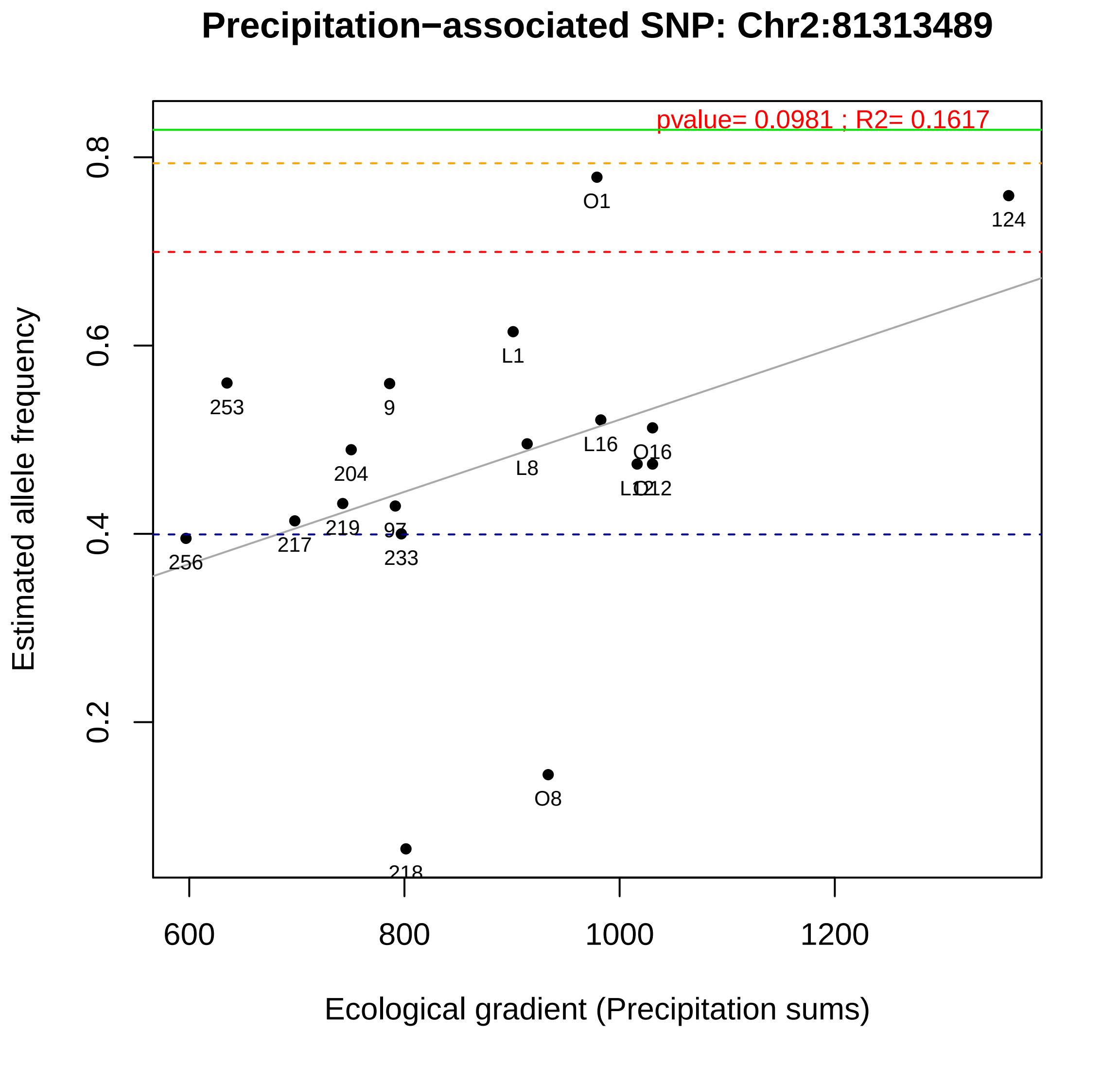

Precipitation−associated SNP: Chr2:81313489
pvalue= 0.0981 ; R2= 0.1617
Estimated allele frequency
Ecological gradient (Precipitation sums)
0.2
0.4
0.6
0.8
600
800
1000
1200
O1
124
L1
253
9
L16
O16
204
L8
L12
O12
219
97
217
233
256
O8
218

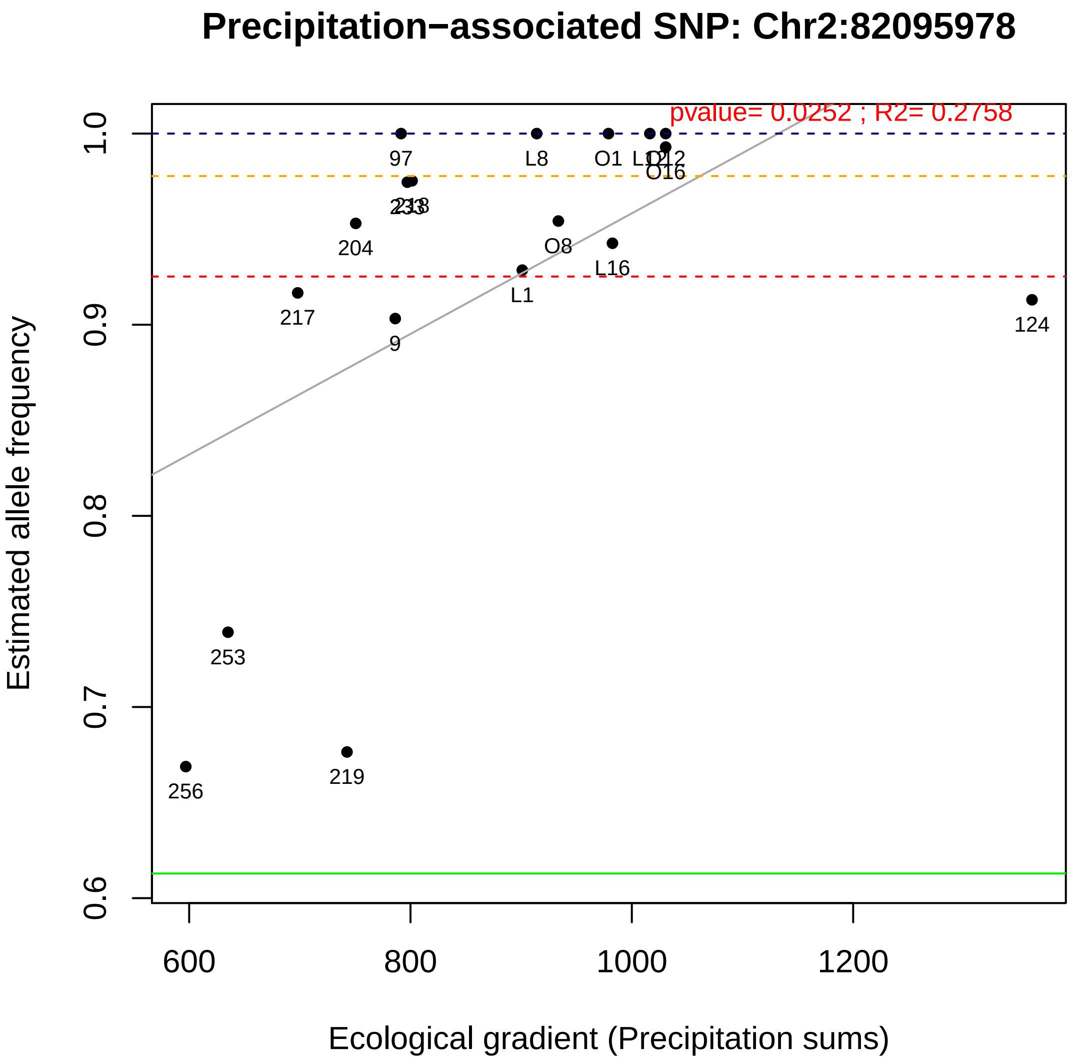

Precipitation−associated SNP: Chr2:82095978
pvalue= 0.0252 ; R2= 0.2758
Estimated allele frequency
Ecological gradient (Precipitation sums)
0.6
0.7
0.8
0.9
1.0
600
800
1000
1200
97
L8
O1
O16
233
204
O8
L16
L1
217
9
124
253
256
219

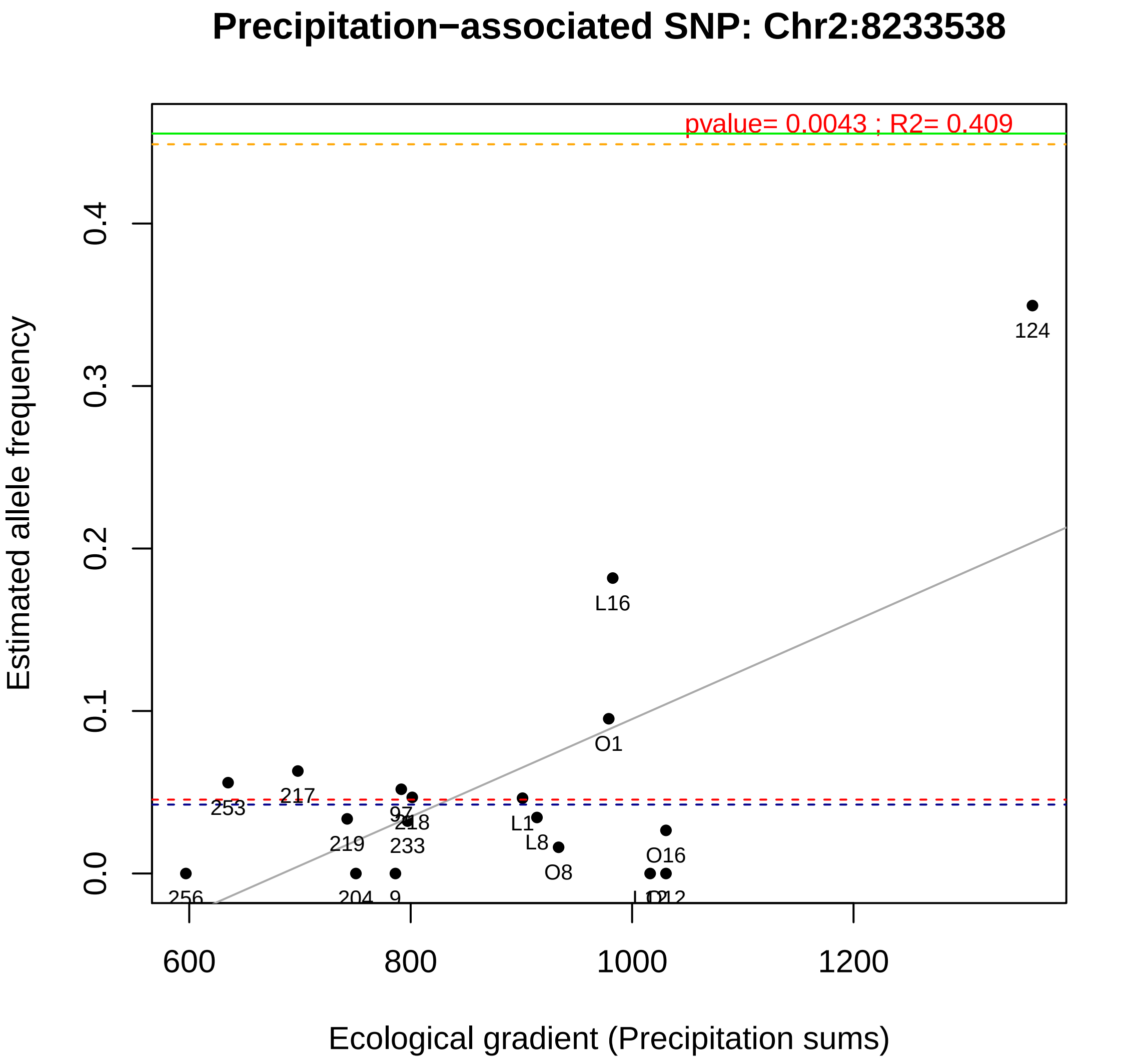

Precipitation−associated SNP: Chr2:8233538
pvalue= 0.0043 ; R2= 0.409
Estimated allele frequency
Ecological gradient (Precipitation sums)
0.0
0.1
0.2
0.3
0.4
600
800
1000
1200
124
L16
O1
217
253
97
218
219
233
L1
L8
O8
O16
256
204
9

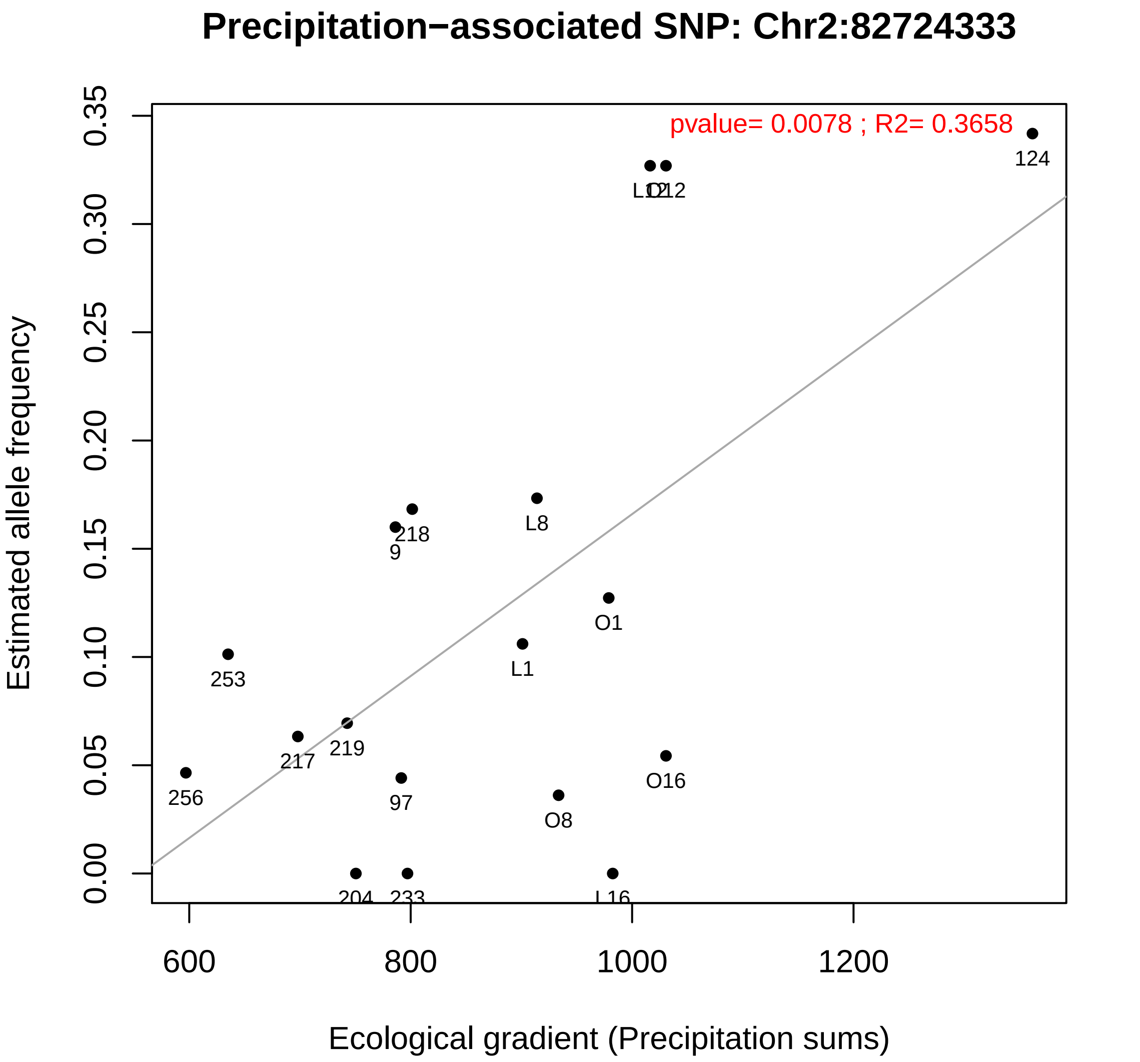

Precipitation−associated SNP: Chr2:82724333
pvalue= 0.0078 ; R2= 0.3658
Estimated allele frequency
Ecological gradient (Precipitation sums)
0.00
0.05
0.10
0.15
0.20
0.25
0.30
0.35
600
800
1000
1200
124
L10
O12
L8
218
9
O1
L1
253
219
217
256
97
O8
O16
204
233
L16

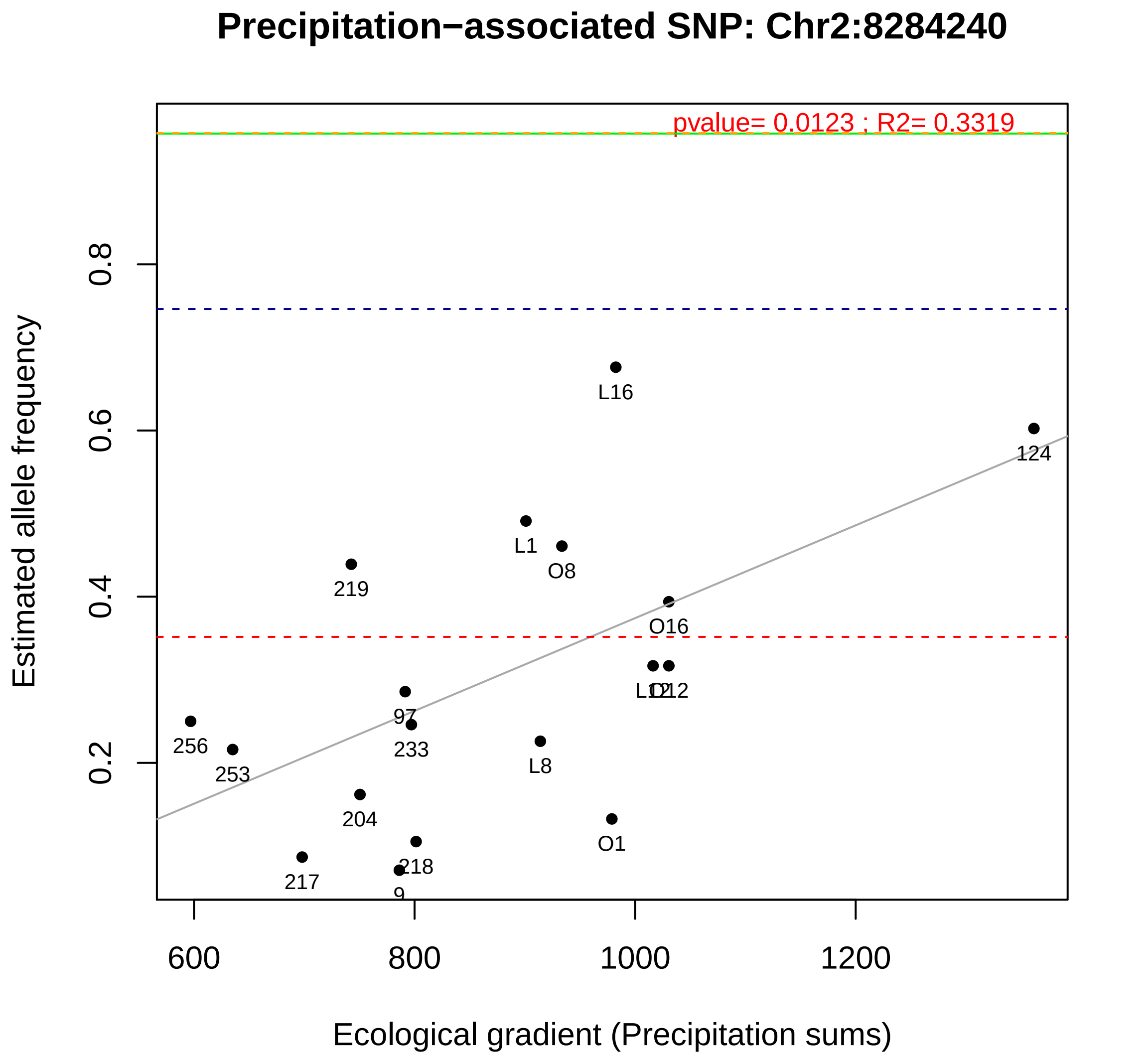

Precipitation−associated SNP: Chr2:8284240
pvalue= 0.0123 ; R2= 0.3319
Estimated allele frequency
0.2
0.4
0.6
0.8
600
800
1000
1200
Ecological gradient (Precipitation sums)
L16
124
L1
O8
219
O16
L12
O12
97
256
233
253
L8
204
O1
218
217
9

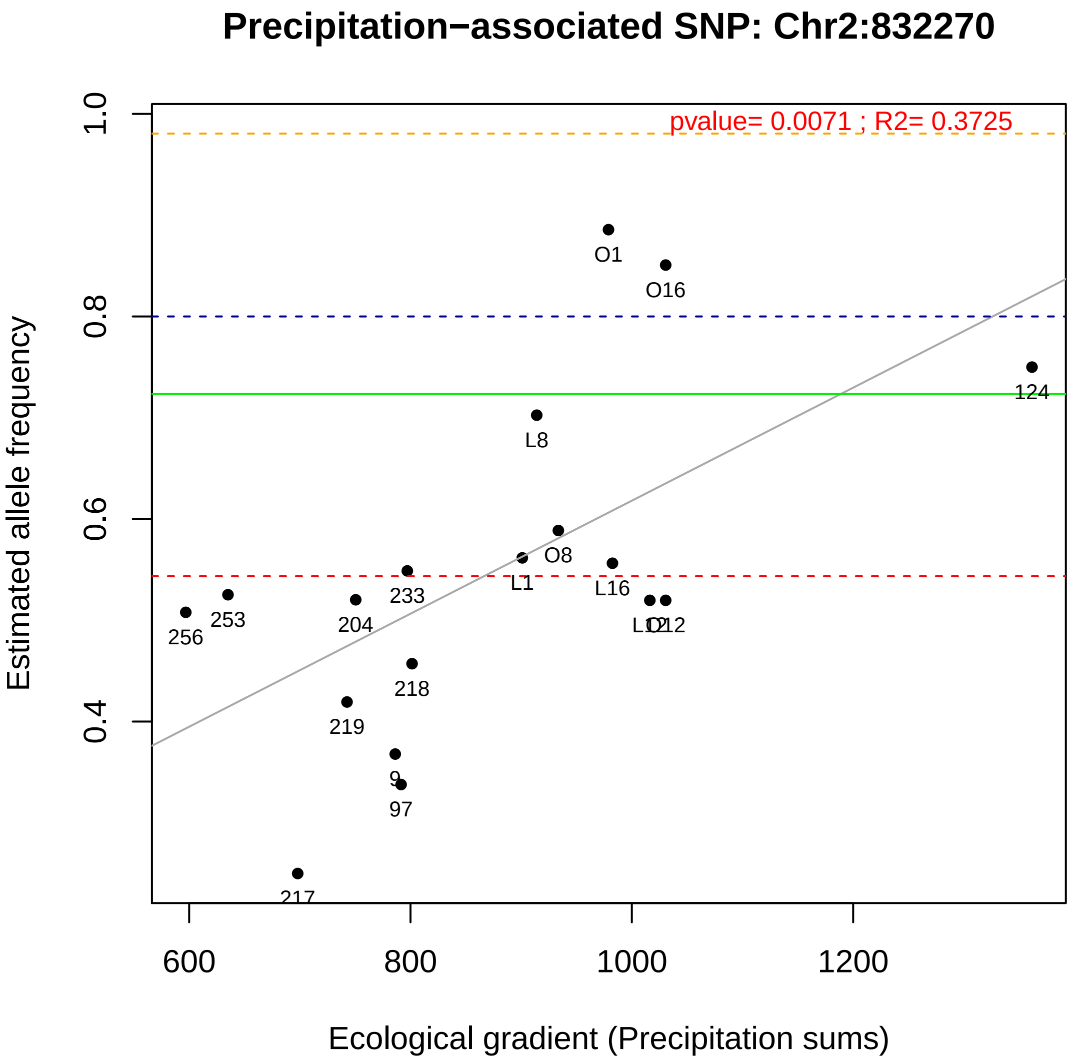

Precipitation−associated SNP: Chr2:832270
pvalue= 0.0071 ; R2= 0.3725
Estimated allele frequency
Ecological gradient (Precipitation sums)
0.4
0.6
0.8
1.0
600
800
1000
1200
O1
O16
124
L8
O8
L1
L16
233
253
256
204
218
219
9
97
217

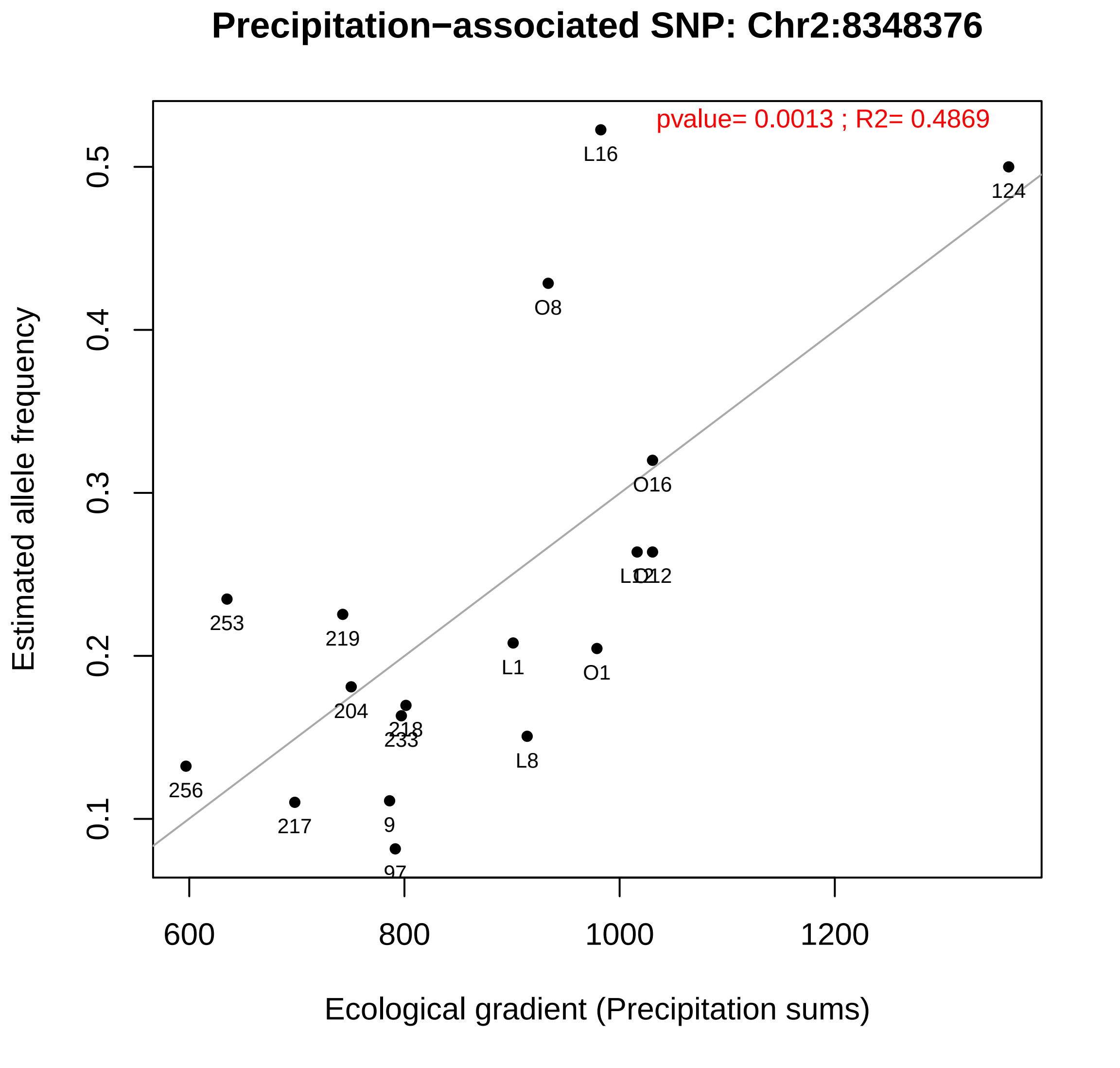

Precipitation−associated SNP: Chr2:8348376
pvalue= 0.0013 ; R2= 0.4869
Estimated allele frequency
0.1
0.2
0.3
0.4
0.5
600
800
1000
1200
Ecological gradient (Precipitation sums)
L16
124
O8
O16
L12
O12
253
219
L1
O1
204
218
233
L8
256
217
9
97

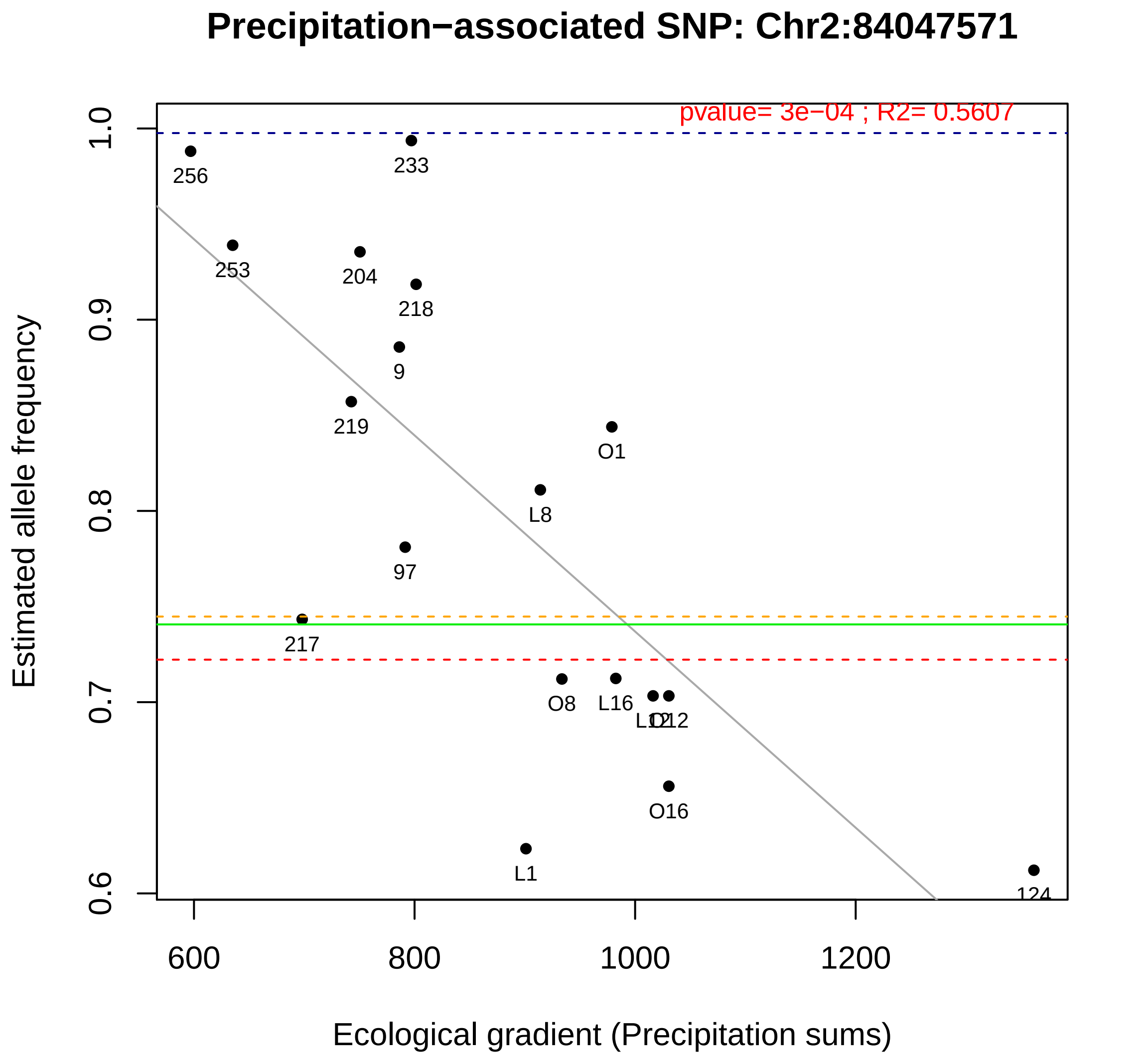
Precipitation−associated SNP: Chr2:84047571
pvalue= 3e−04 ; R2= 0.5607
Estimated allele frequency
Ecological gradient (Precipitation sums)
0.6
0.7
0.8
0.9
1.0
600
800
1000
1200
256
233
253
204
218
9
219
O1
L8
97
217
O8
L16
L12
O12
O16
L1
124

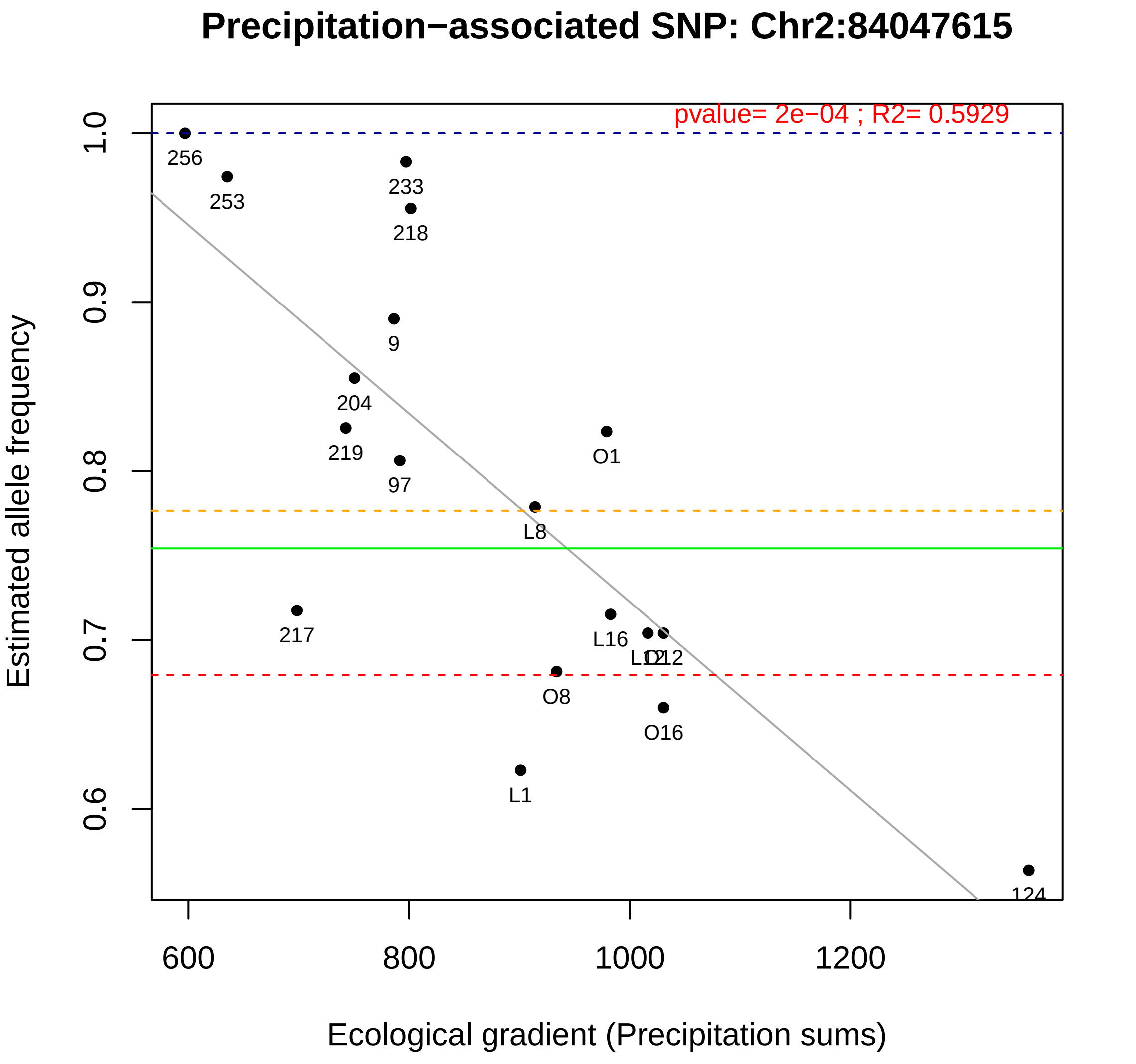
Precipitation−associated SNP: Chr2:84047615
pvalue= 2e−04 ; R2= 0.5929
Estimated allele frequency
Ecological gradient (Precipitation sums)
0.6
0.7
0.8
0.9
1.0
600
800
1000
1200
256
253
233
218
9
204
219
97
O1
L8
217
L16
L12
O12
O8
O16
L1
124

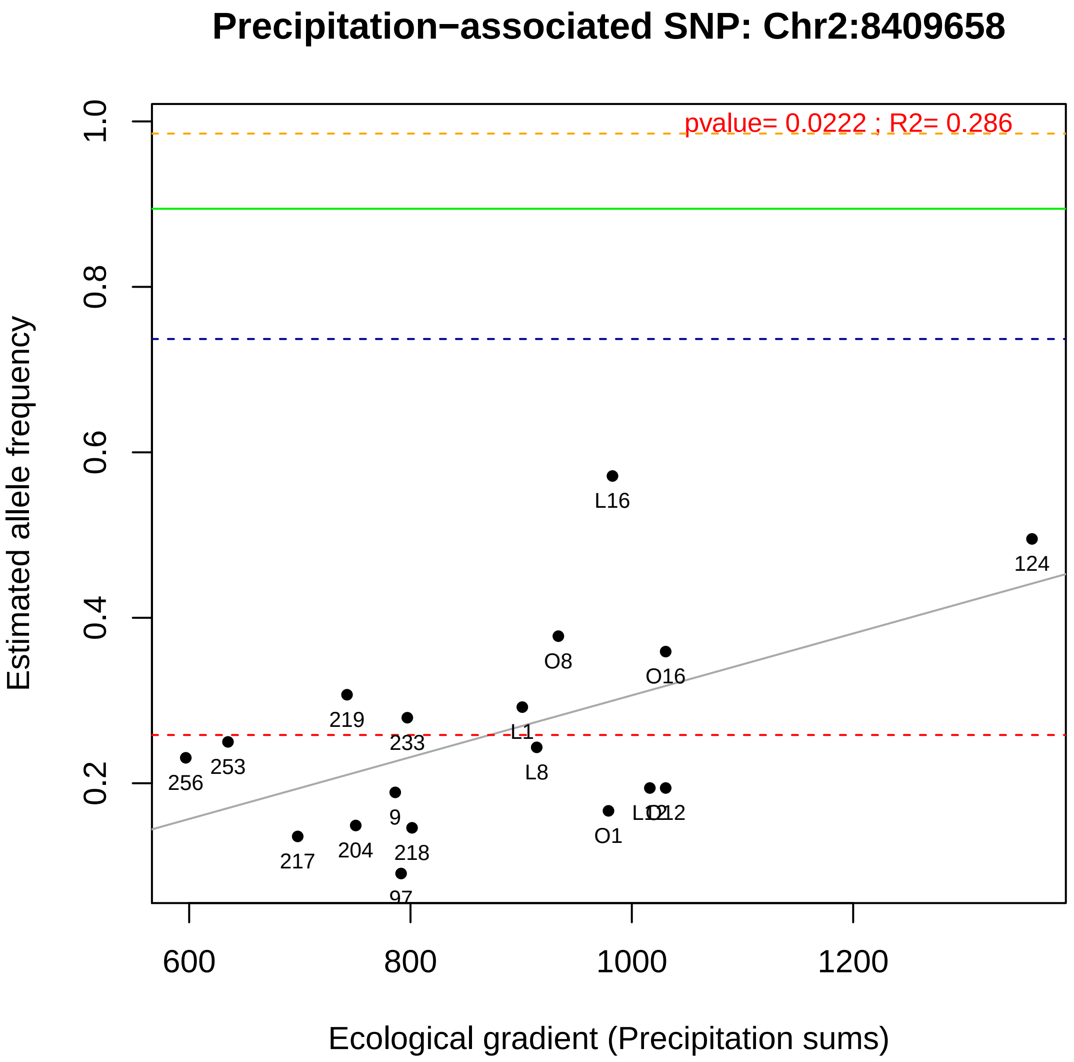

Precipitation−associated SNP: Chr2:8409658
pvalue= 0.0222 ; R2= 0.286
Estimated allele frequency
Ecological gradient (Precipitation sums)
0.2
0.4
0.6
0.8
1.0
600
800
1000
1200
L16
124
O8
O16
219
233
L1
L8
253
256
9
O1
L12
O12
217
204
218
97

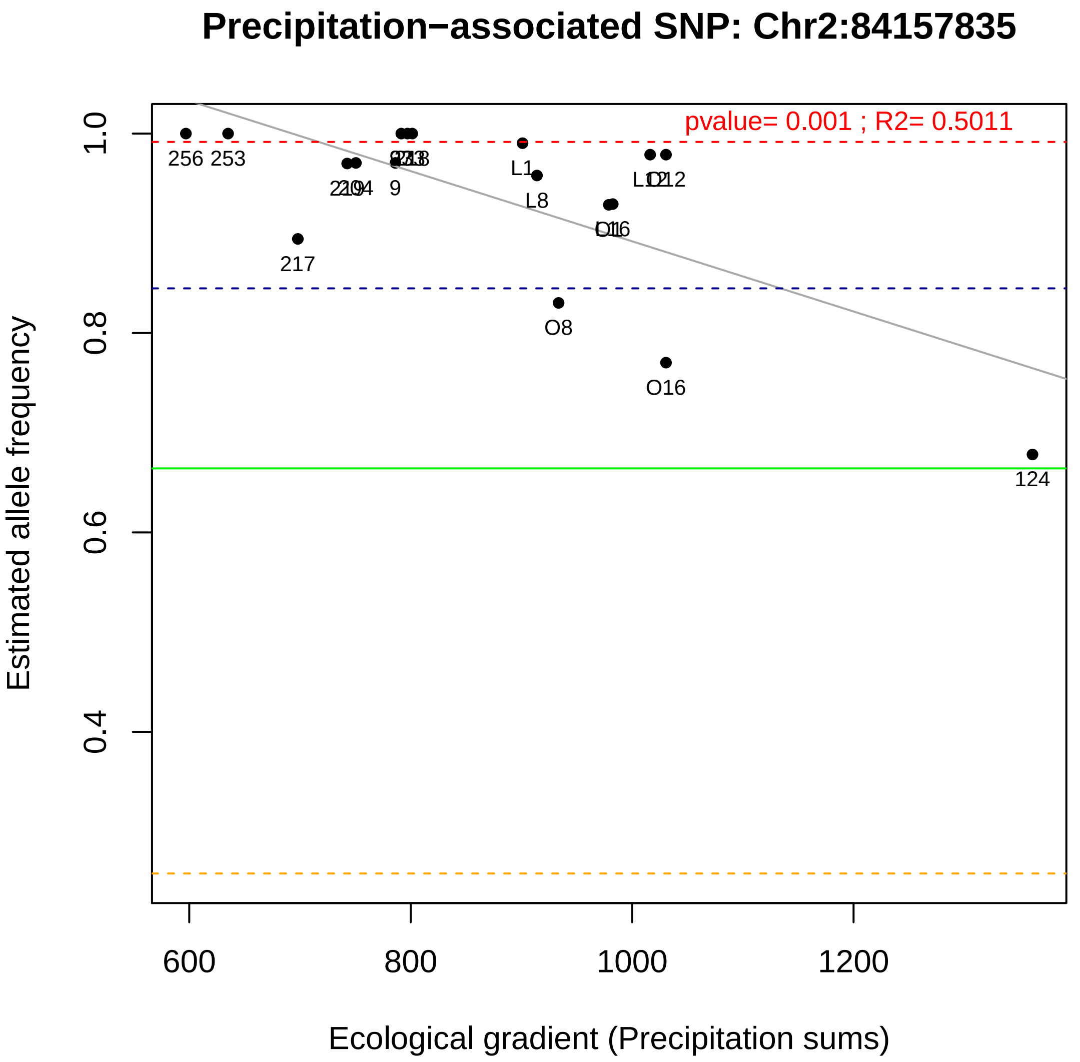
Precipitation−associated SNP: Chr2:84157835
pvalue= 0.001 ; R2= 0.5011
Estimated allele frequency
Ecological gradient (Precipitation sums)
1.0
0.8
0.6
0.4
600
800
1000
1200
256
253
217
9
L1
L8
O8
O16
124

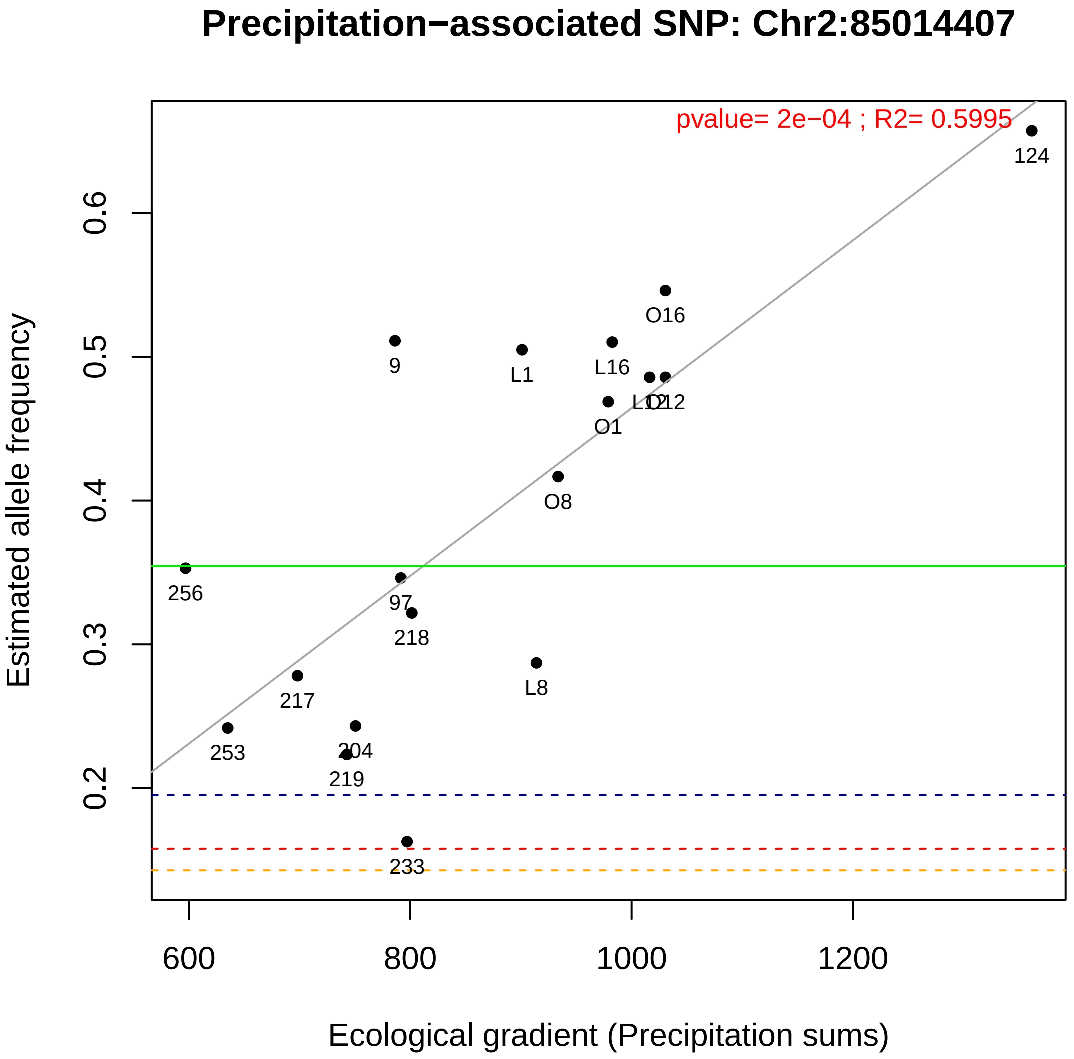
Precipitation−associated SNP: Chr2:85014407
pvalue= 2e−04 ; R2= 0.5995
Estimated allele frequency
Ecological gradient (Precipitation sums)
0.2
0.3
0.4
0.5
0.6
600
800
1000
1200
124
O16
9
L1
L16
O12
O1
O8
256
97
218
L8
217
253
204
219
233

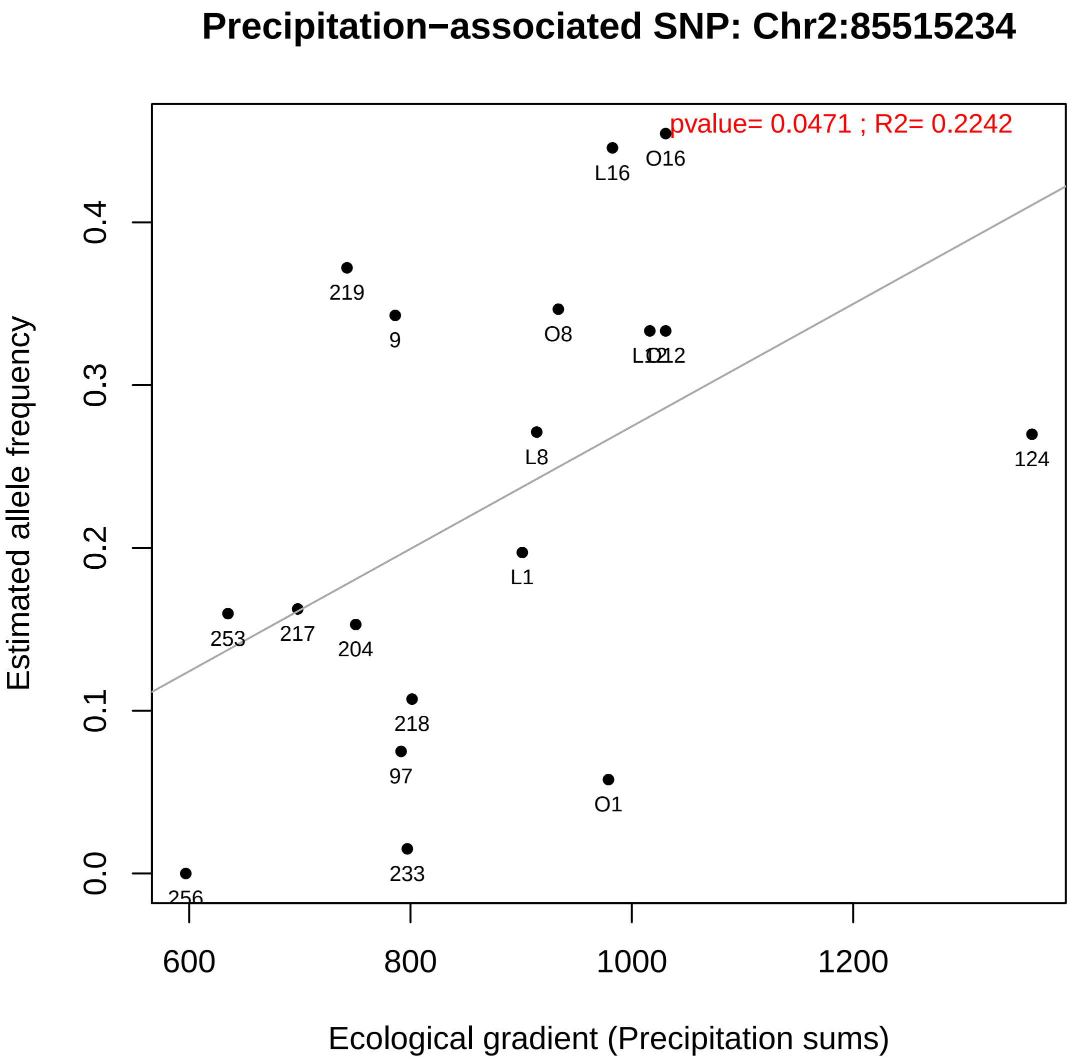
Precipitation−associated SNP: Chr2:85515234
pvalue= 0.0471 ; R2= 0.2242
Estimated allele frequency
0.0
0.1
0.2
0.3
0.4
600
800
1000
1200
Ecological gradient (Precipitation sums)
O16
L16
219
9
O8
L12
O12
L8
124
L1
253
217
204
218
97
O1
233
256

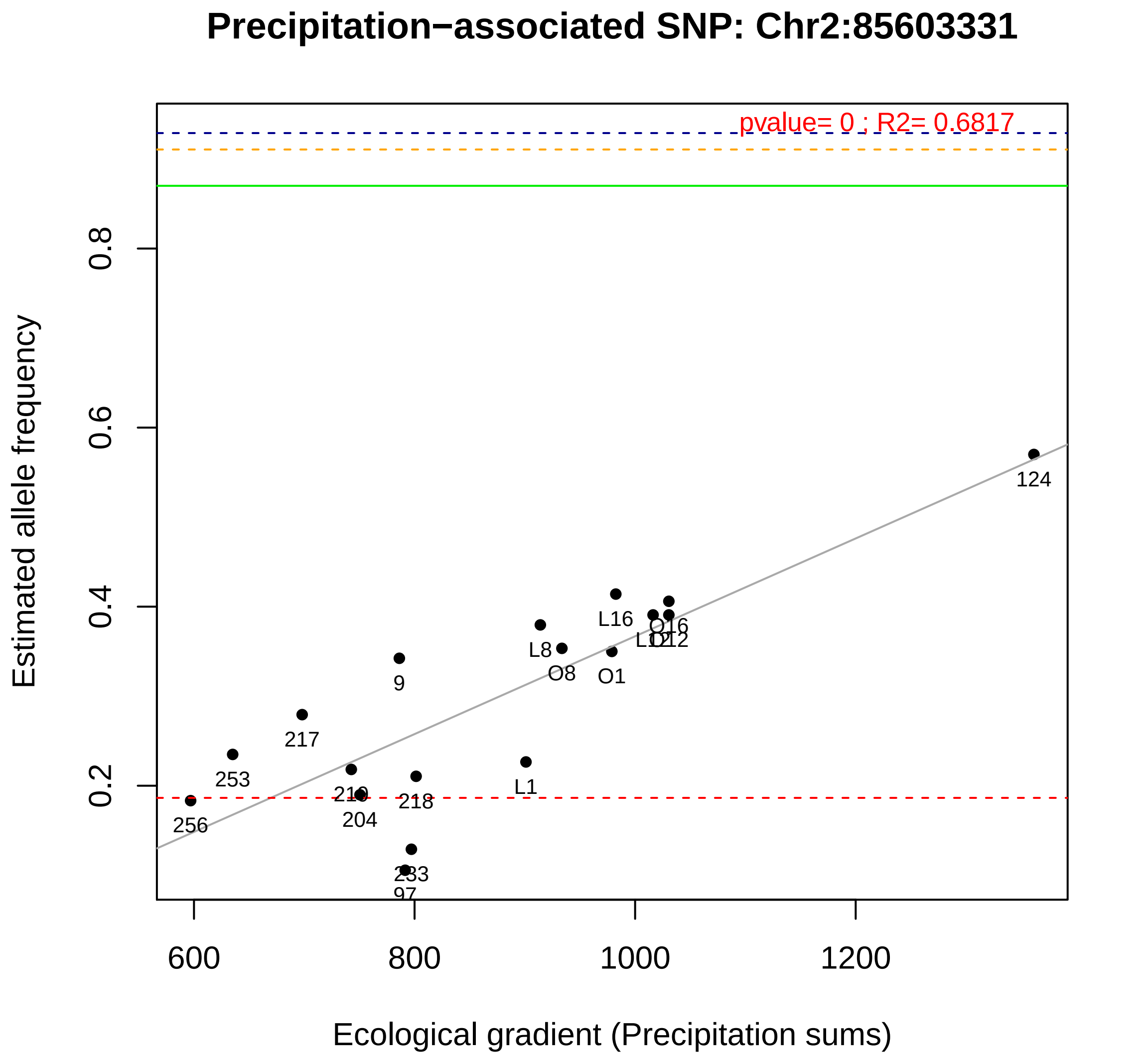

Precipitation−associated SNP: Chr2:85603331
pvalue= 0 ; R2= 0.6817
Estimated allele frequency
0.2
0.4
0.6
0.8
600
800
1000
1200
Ecological gradient (Precipitation sums)
124
L16
O16
L12
O12
L8
O8
O1
9
217
253
256
210
204
218
L1
233
97

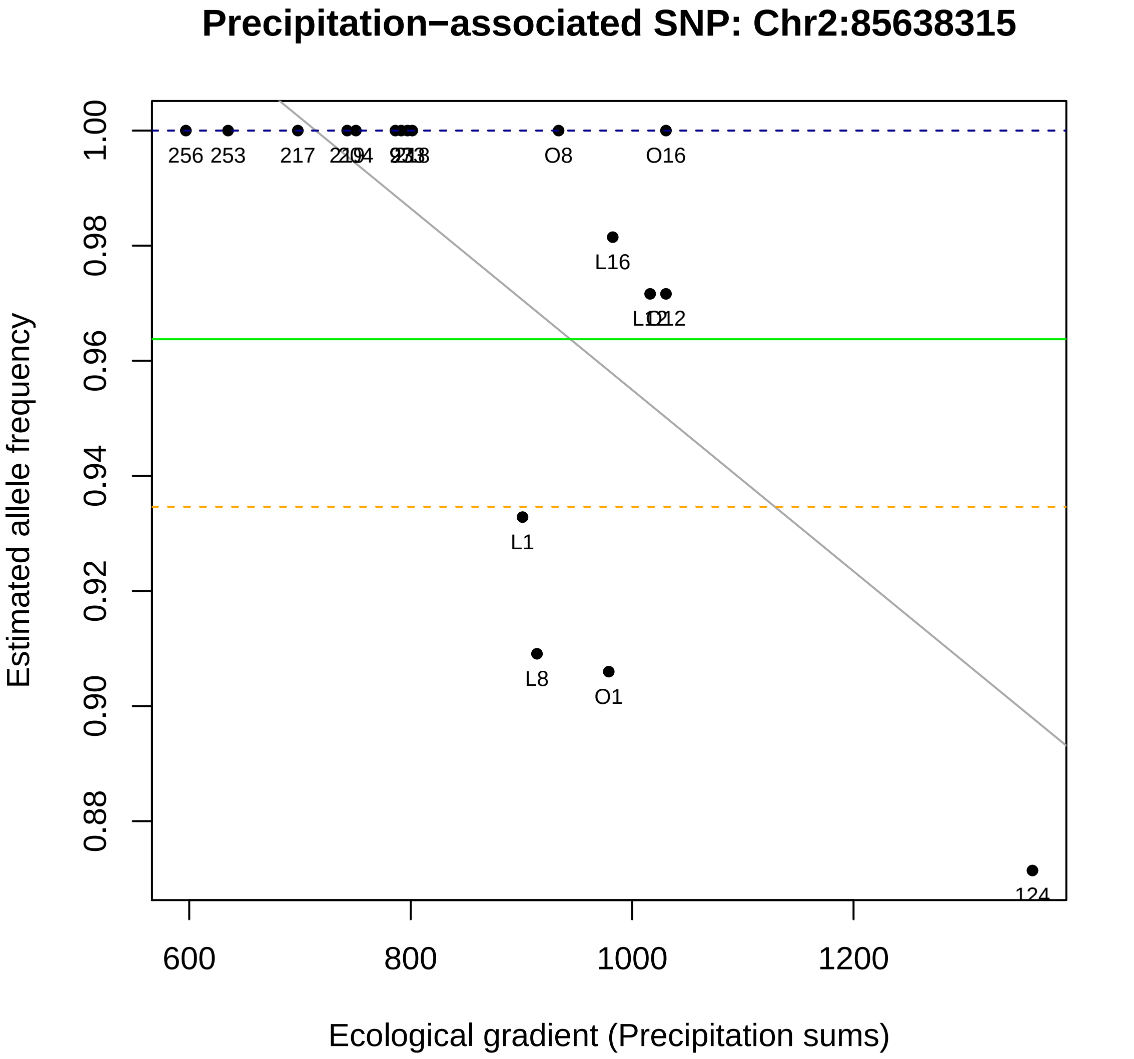
Precipitation−associated SNP: Chr2:85638315
Estimated allele frequency
Ecological gradient (Precipitation sums)
1.00
0.98
0.96
0.94
0.92
0.90
0.88
600
800
1000
1200
256
253
217
O8
O16
L16
L1
L8
O1
124

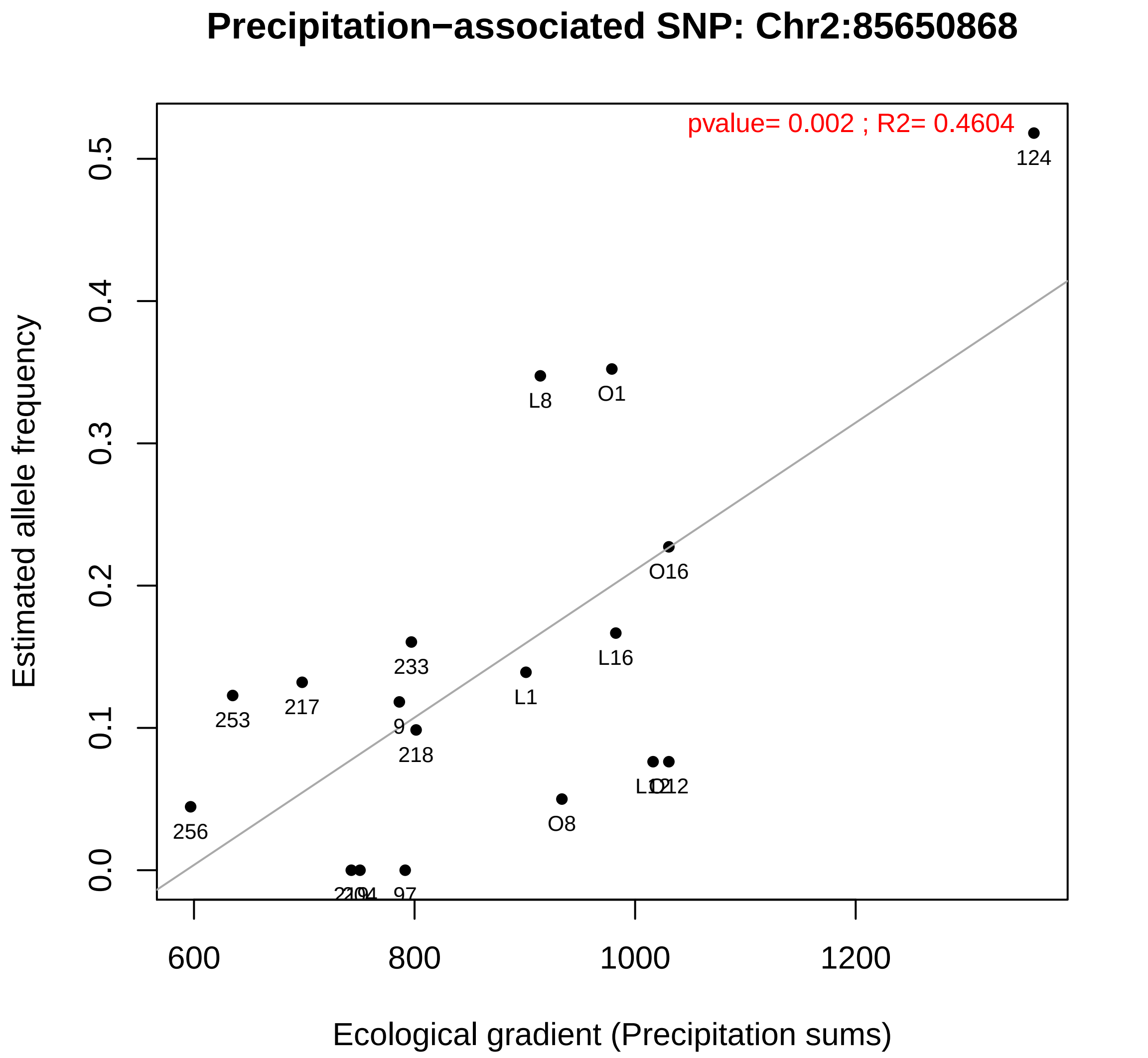

Precipitation−associated SNP: Chr2:85650868
pvalue= 0.002 ; R2= 0.4604
Estimated allele frequency
Ecological gradient (Precipitation sums)
0.0
0.1
0.2
0.3
0.4
0.5
600
800
1000
1200
124
L8
O1
O16
L16
233
L1
217
253
9
218
O8
256
97

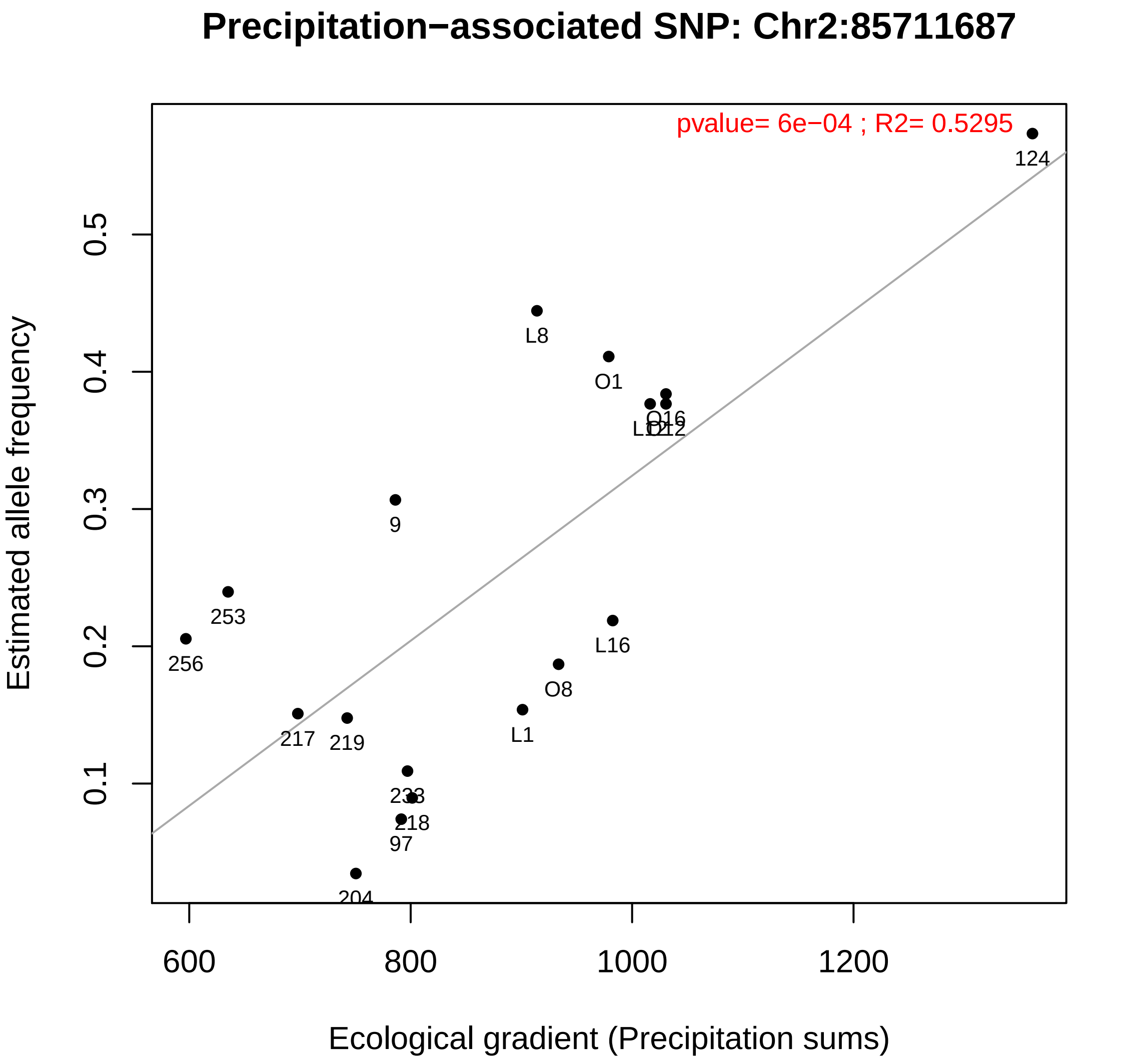

Precipitation−associated SNP: Chr2:85711687
pvalue= 6e−04 ; R2= 0.5295
Estimated allele frequency
Ecological gradient (Precipitation sums)
0.1
0.2
0.3
0.4
0.5
600
800
1000
1200
124
L8
O1
O16
L12
9
253
256
L16
O8
217
219
L1
218
97
204

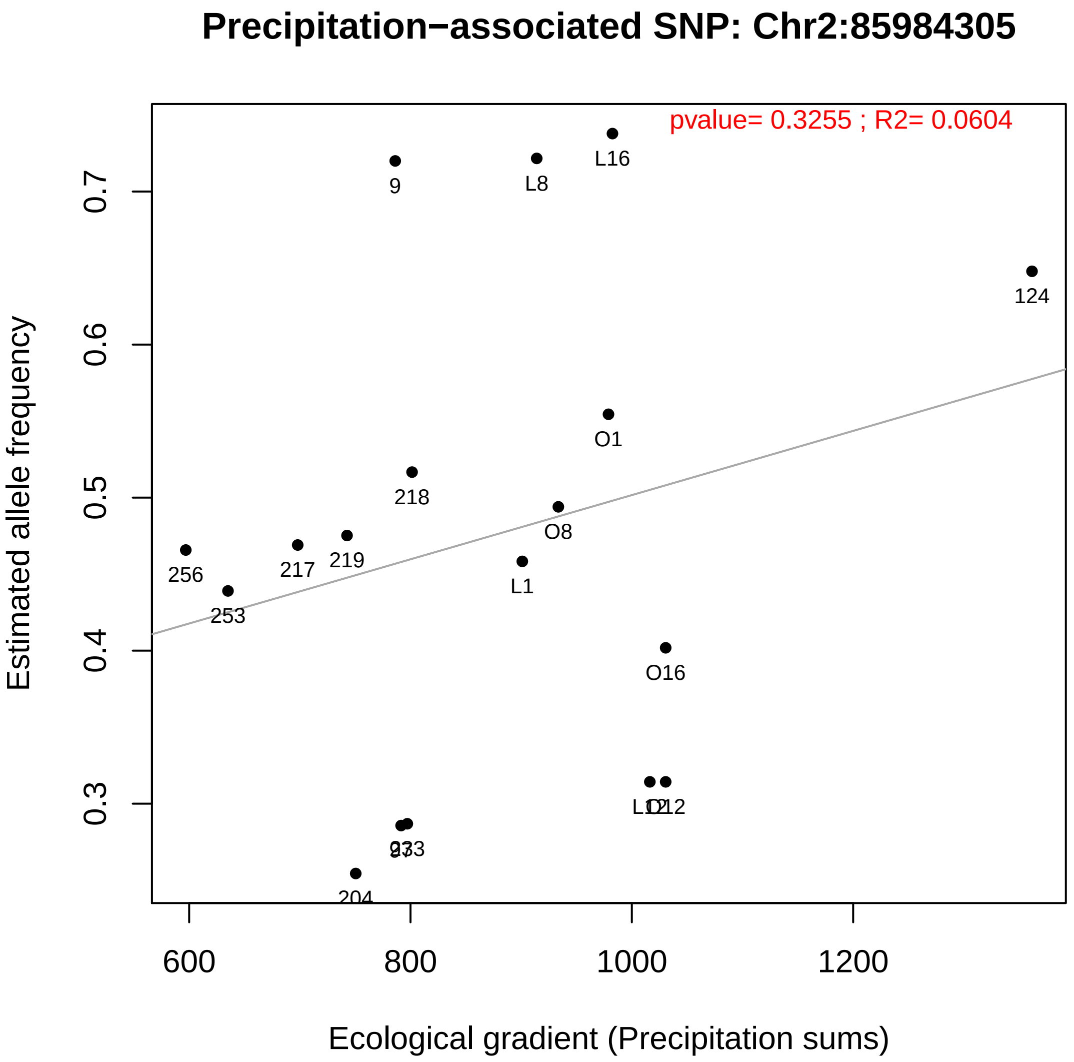
Precipitation−associated SNP: Chr2:85984305
pvalue= 0.3255 ; R2= 0.0604
Estimated allele frequency
0.3
0.4
0.5
0.6
0.7
600
800
1000
1200
Ecological gradient (Precipitation sums)
9
L8
L16
124
O1
218
O8
256
217
219
L1
253
O16
204

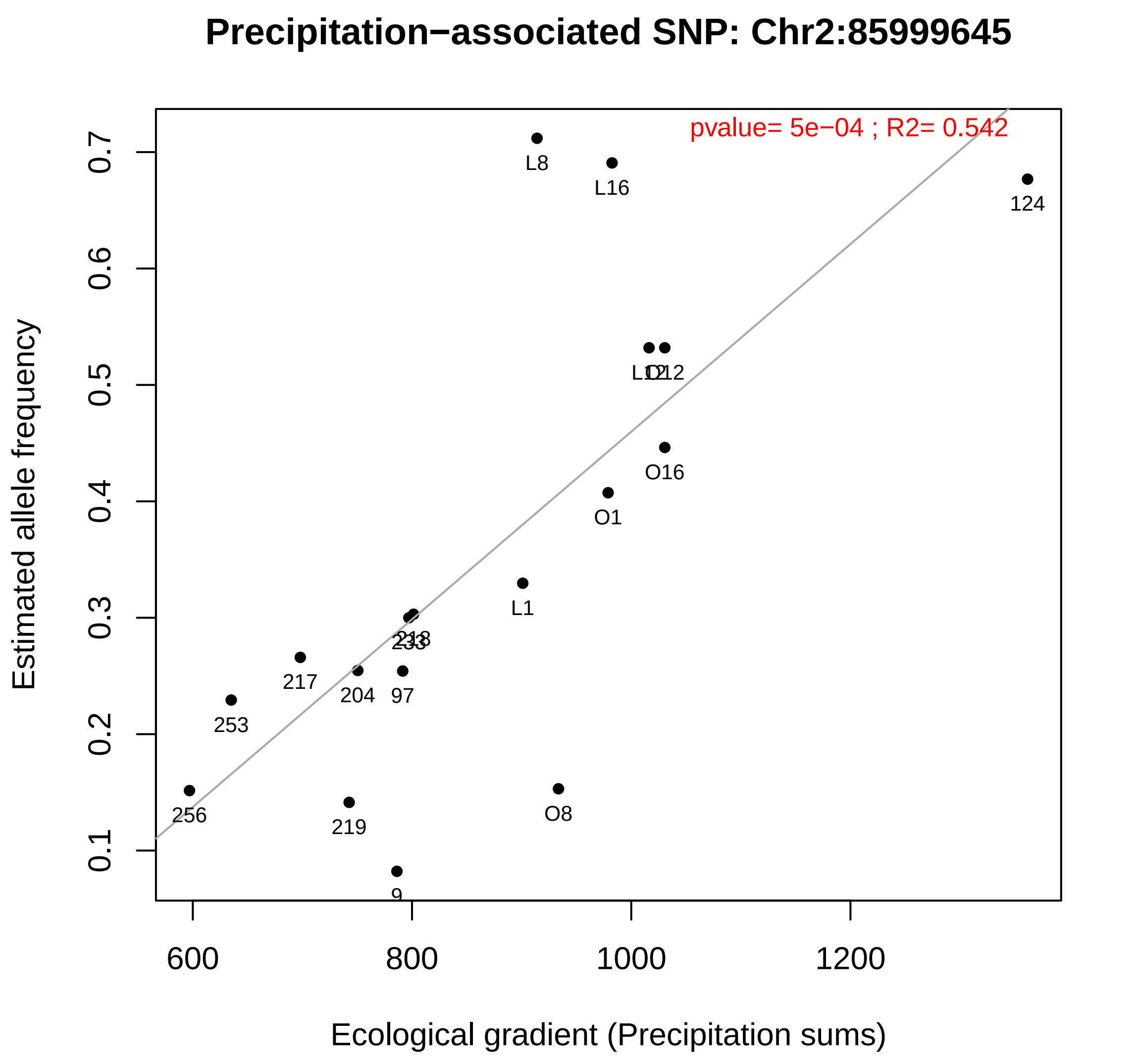
Precipitation−associated SNP: Chr2:85999645
pvalue= 5e−04 ; R2= 0.542
Estimated allele frequency
Ecological gradient (Precipitation sums)
0.1
0.2
0.3
0.4
0.5
0.6
0.7
600
800
1000
1200
L8
L16
124
L12
O12
O16
O1
L1
218
233
217
204
97
253
256
219
O8
9

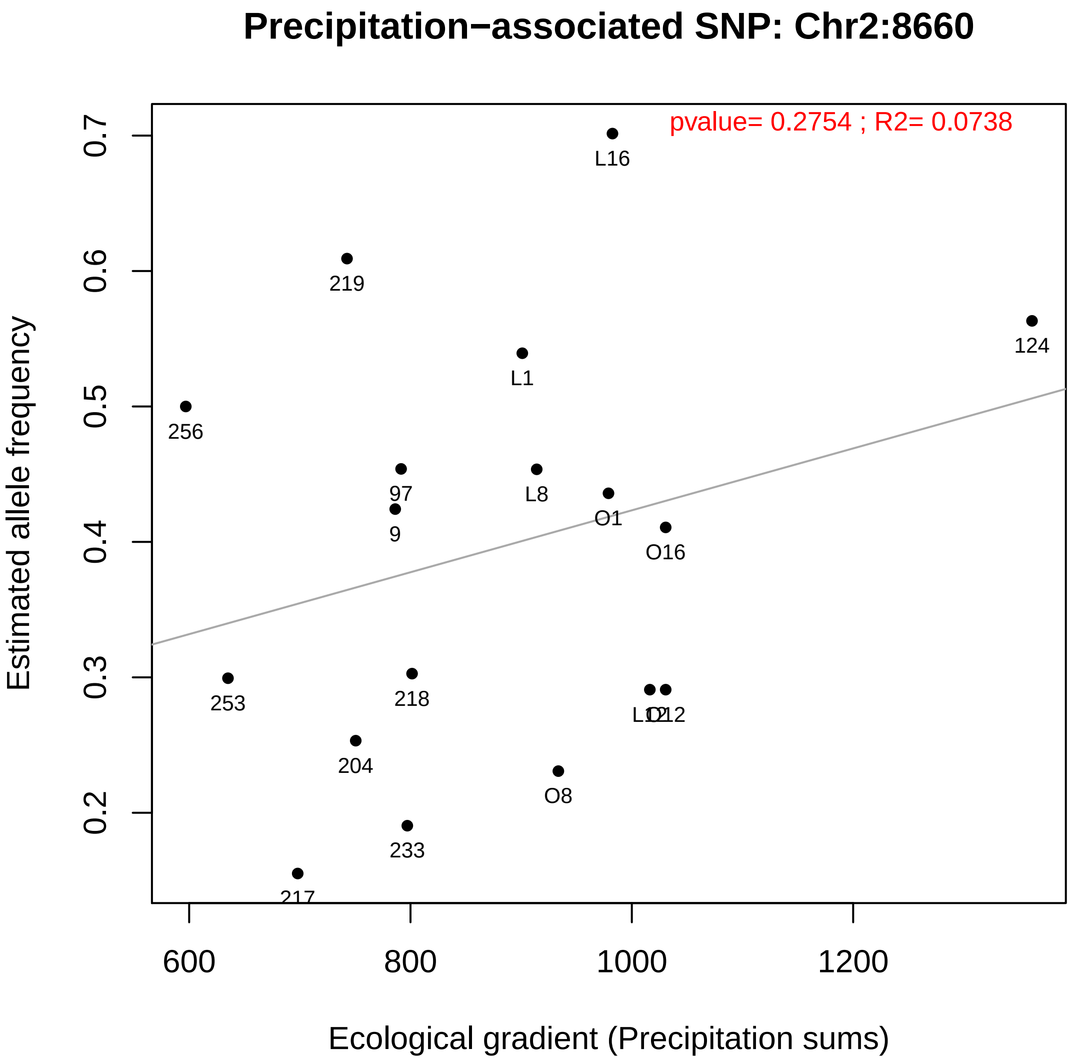

Precipitation−associated SNP: Chr2:8660
pvalue= 0.2754 ; R2= 0.0738
Estimated allele frequency
Ecological gradient (Precipitation sums)
0.2
0.3
0.4
0.5
0.6
0.7
600
800
1000
1200
L16
219
124
L1
256
97
L8
9
O1
O16
253
218
O12
204
O8
233
217

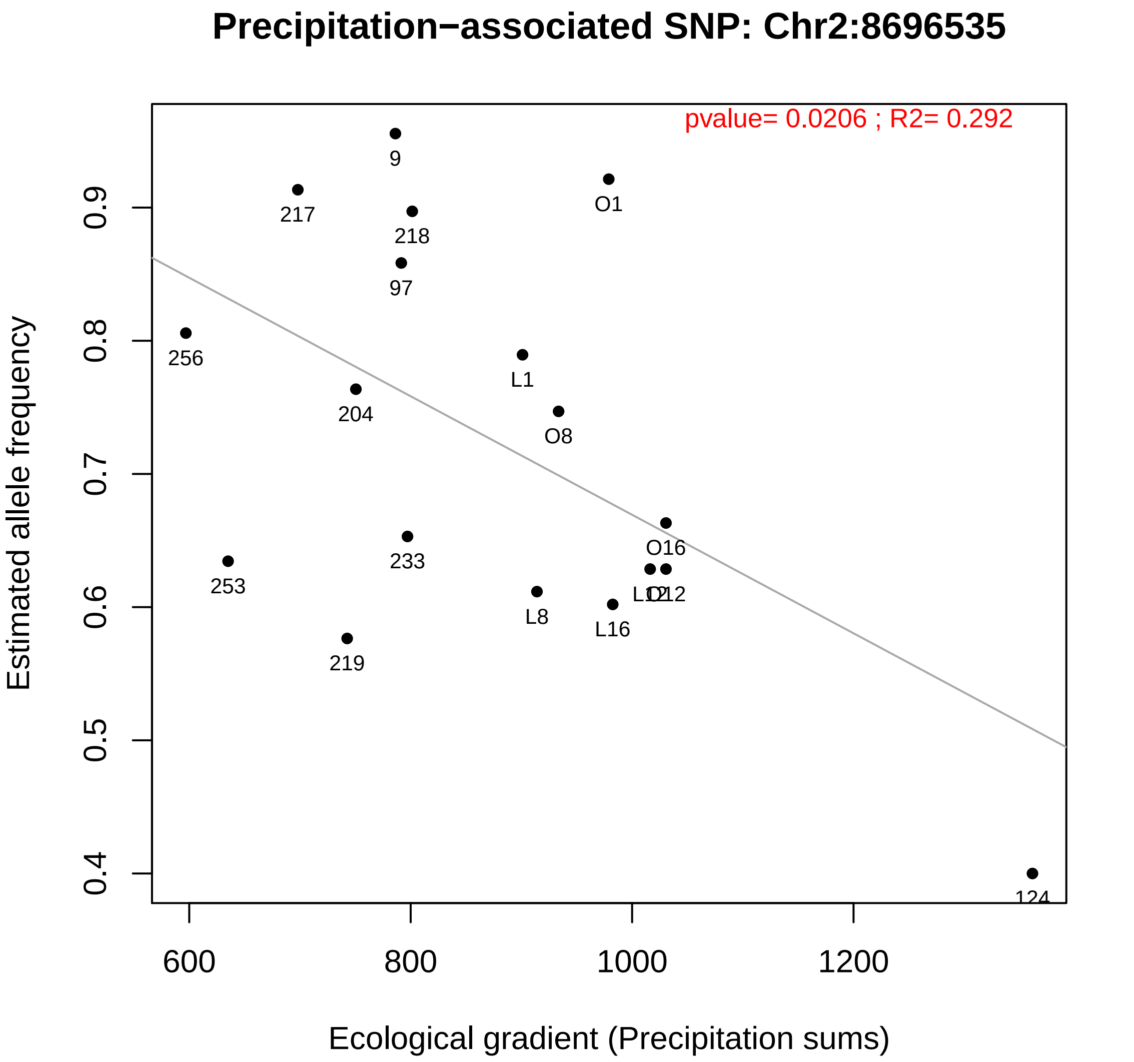
Precipitation−associated SNP: Chr2:8696535
pvalue= 0.0206 ; R2= 0.292
9
217
218
O1
97
256
L1
204
O8
O16
233
253
L12
O12
L8
L16
219
124
0.4
0.5
0.6
0.7
0.8
0.9
600
800
1000
1200
Estimated allele frequency
Ecological gradient (Precipitation sums)

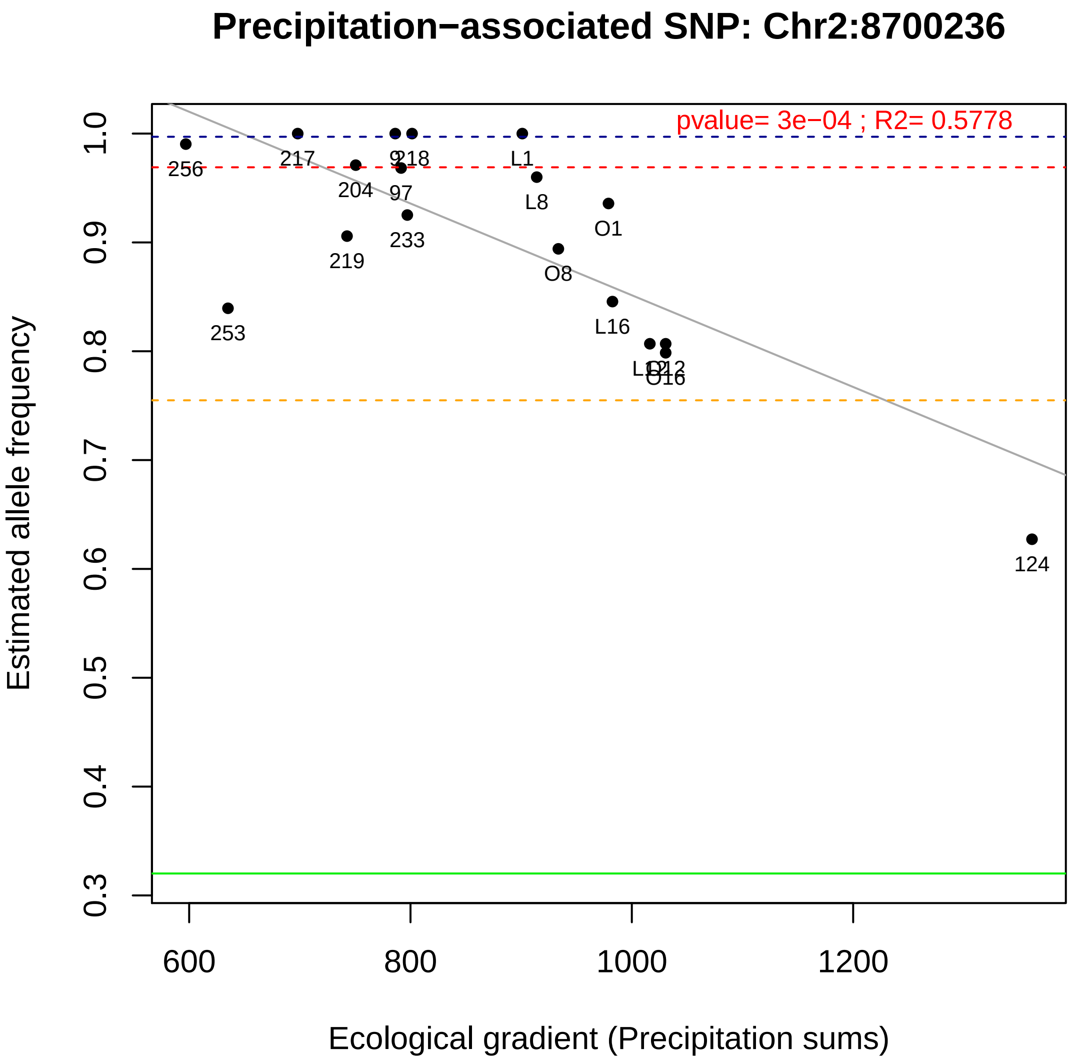

Precipitation−associated SNP: Chr2:8700236
pvalue= 3e−04 ; R2= 0.5778
Estimated allele frequency
Ecological gradient (Precipitation sums)
0.3
0.4
0.5
0.6
0.7
0.8
0.9
1.0
600
800
1000
1200
256
217
218
204
97
233
219
253
L1
L8
O1
O8
L16
O16
124

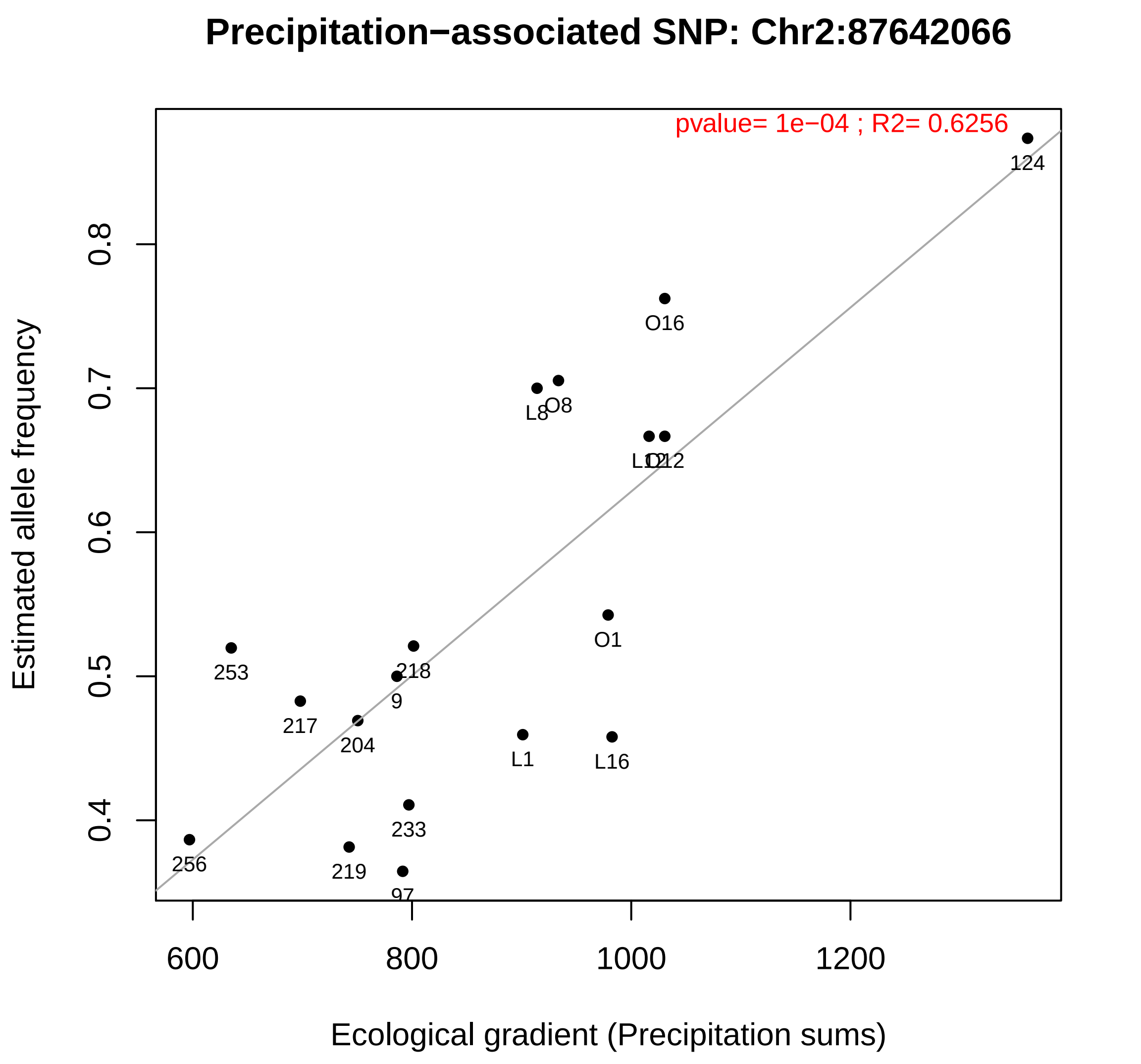
Precipitation−associated SNP: Chr2:87642066
pvalue= 1e−04 ; R2= 0.6256
Estimated allele frequency
0.4
0.5
0.6
0.7
0.8
600
800
1000
1200
Ecological gradient (Precipitation sums)
124
O16
L8
O8
O12
O1
253
218
9
217
204
L1
L16
233
256
219
97

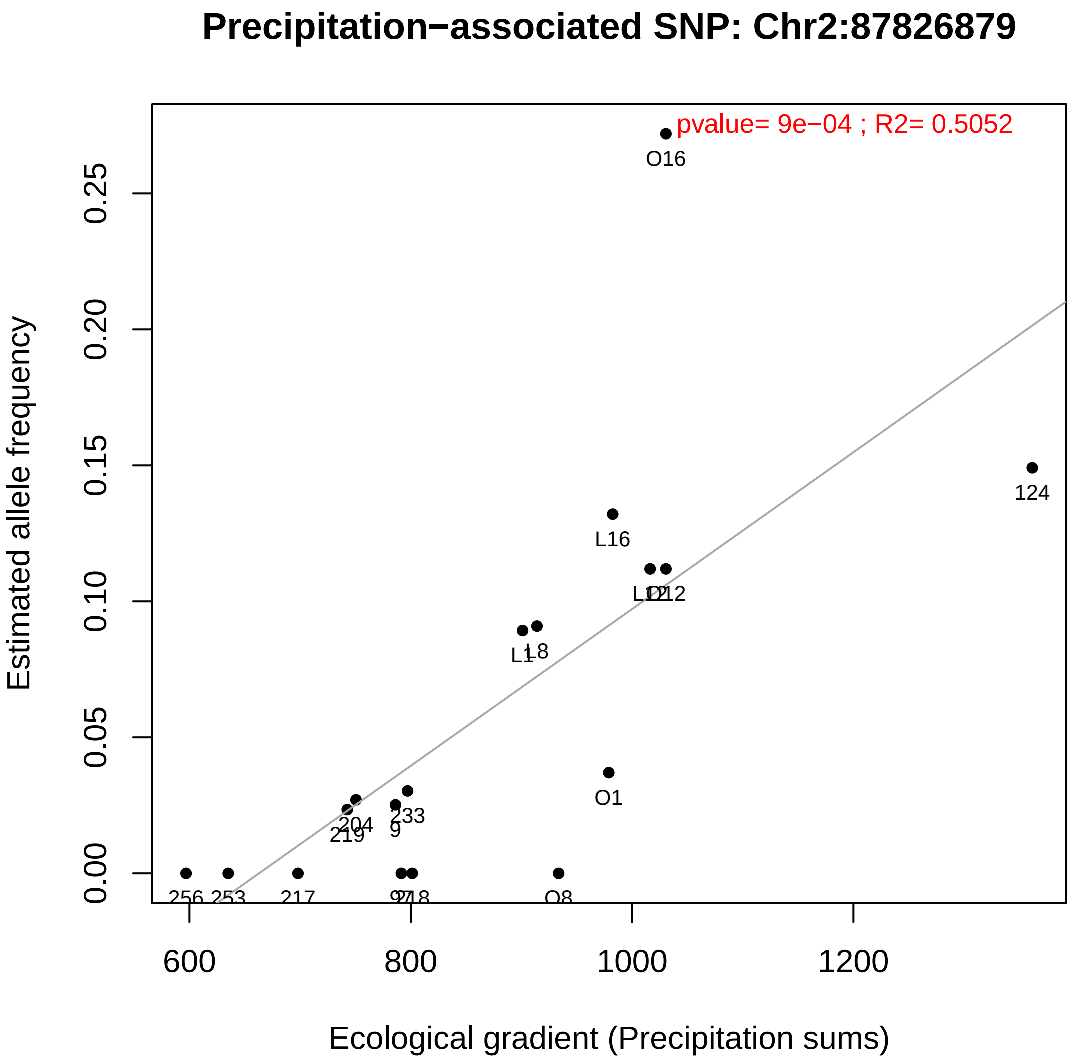

Precipitation−associated SNP: Chr2:87826879
pvalue= 9e−04 ; R2= 0.5052
Estimated allele frequency
Ecological gradient (Precipitation sums)
0.00
0.05
0.10
0.15
0.20
0.25
600
800
1000
1200
O16
124
L16
O12
O1
O8

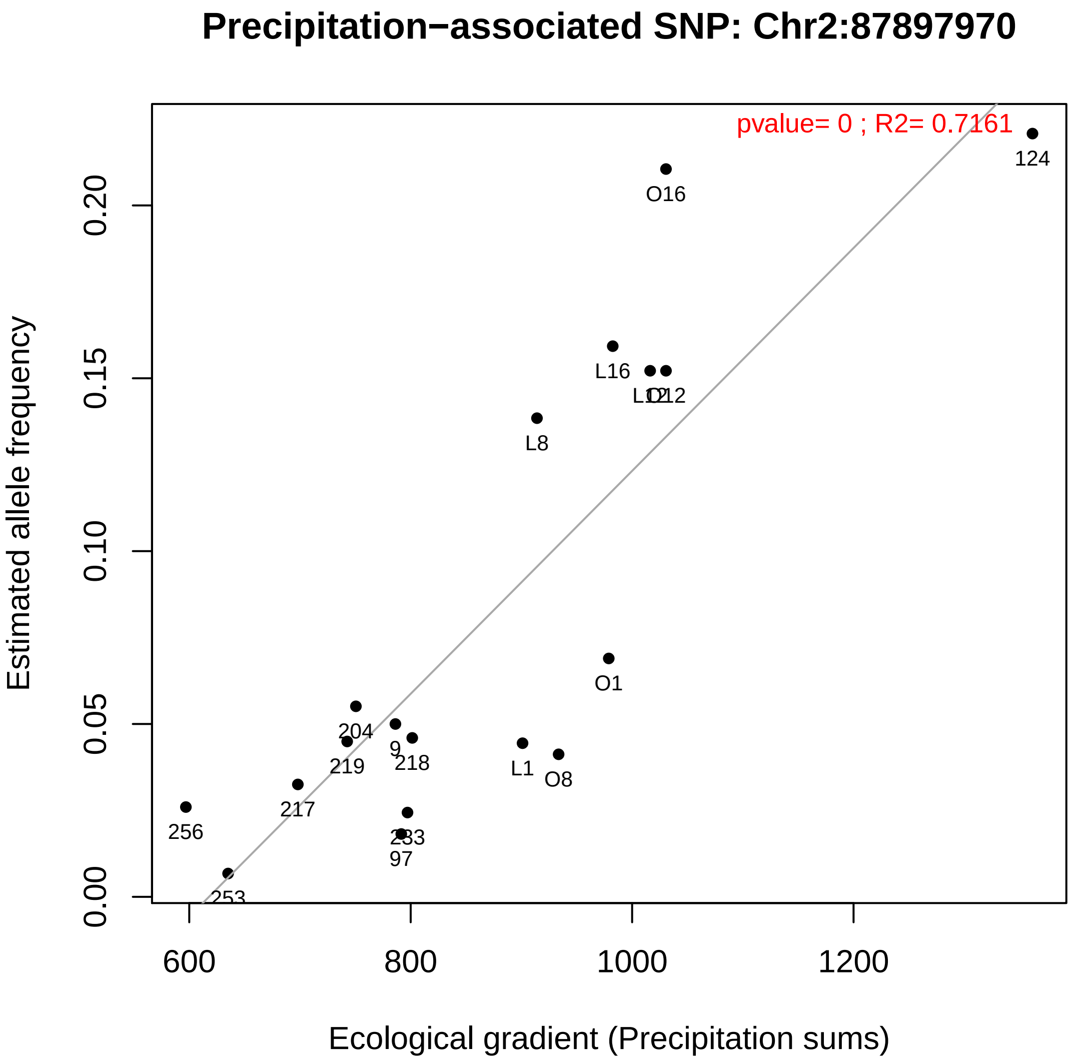
Precipitation−associated SNP: Chr2:87897970
pvalue= 0 ; R2= 0.7161
124
O16
L16
L12
O12
L8
O1
204
9
218
219
L1
O8
217
256
233
97
253
0.00
0.05
0.10
0.15
0.20
600
800
1000
1200
Estimated allele frequency
Ecological gradient (Precipitation sums)

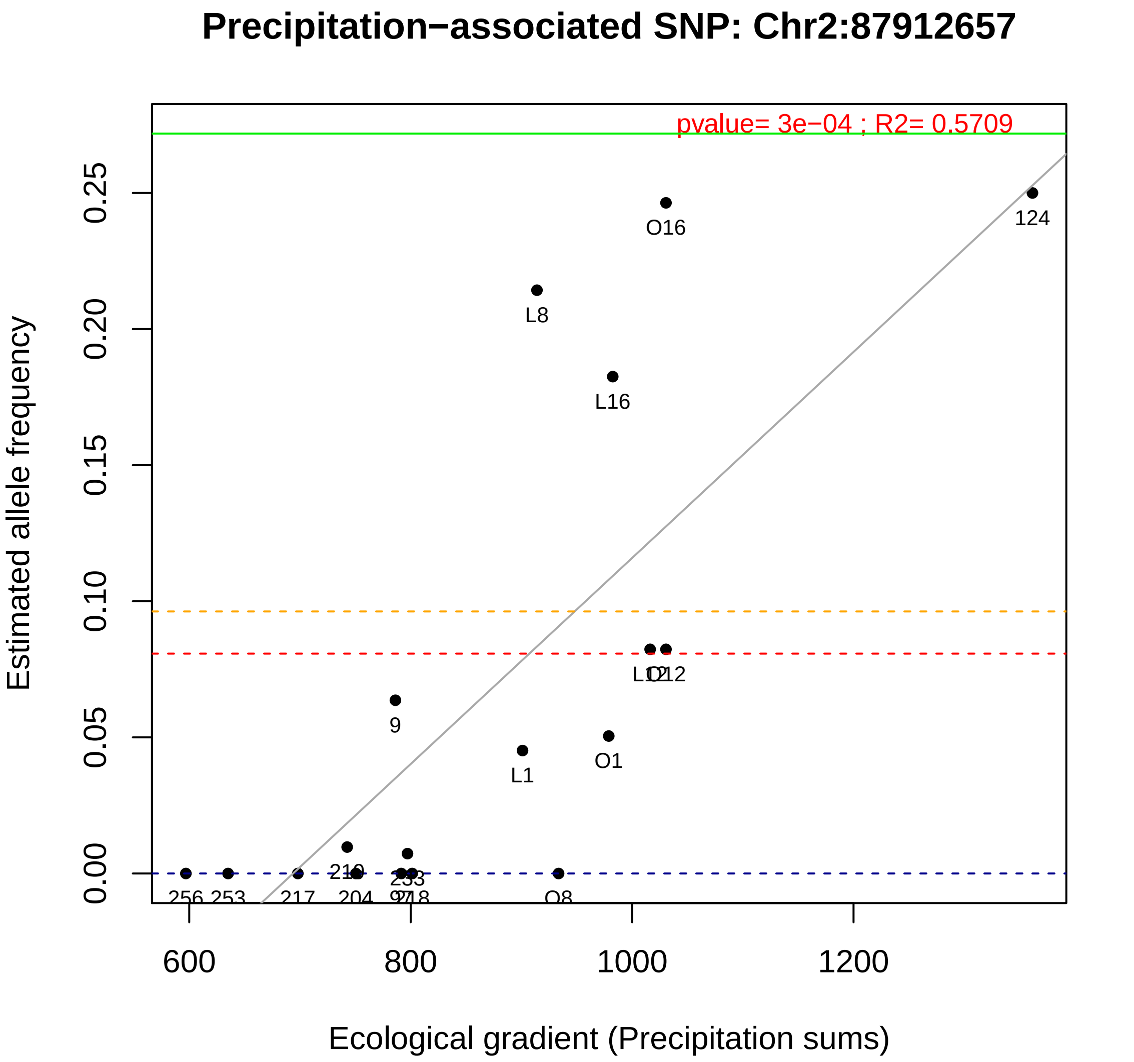

Precipitation−associated SNP: Chr2:87912657
pvalue= 3e−04 ; R2= 0.5709
Estimated allele frequency
0.00
0.05
0.10
0.15
0.20
0.25
600
800
1000
1200
Ecological gradient (Precipitation sums)
O16
124
L8
L16
9
L1
O1
256
253
217
204
233
218
O8

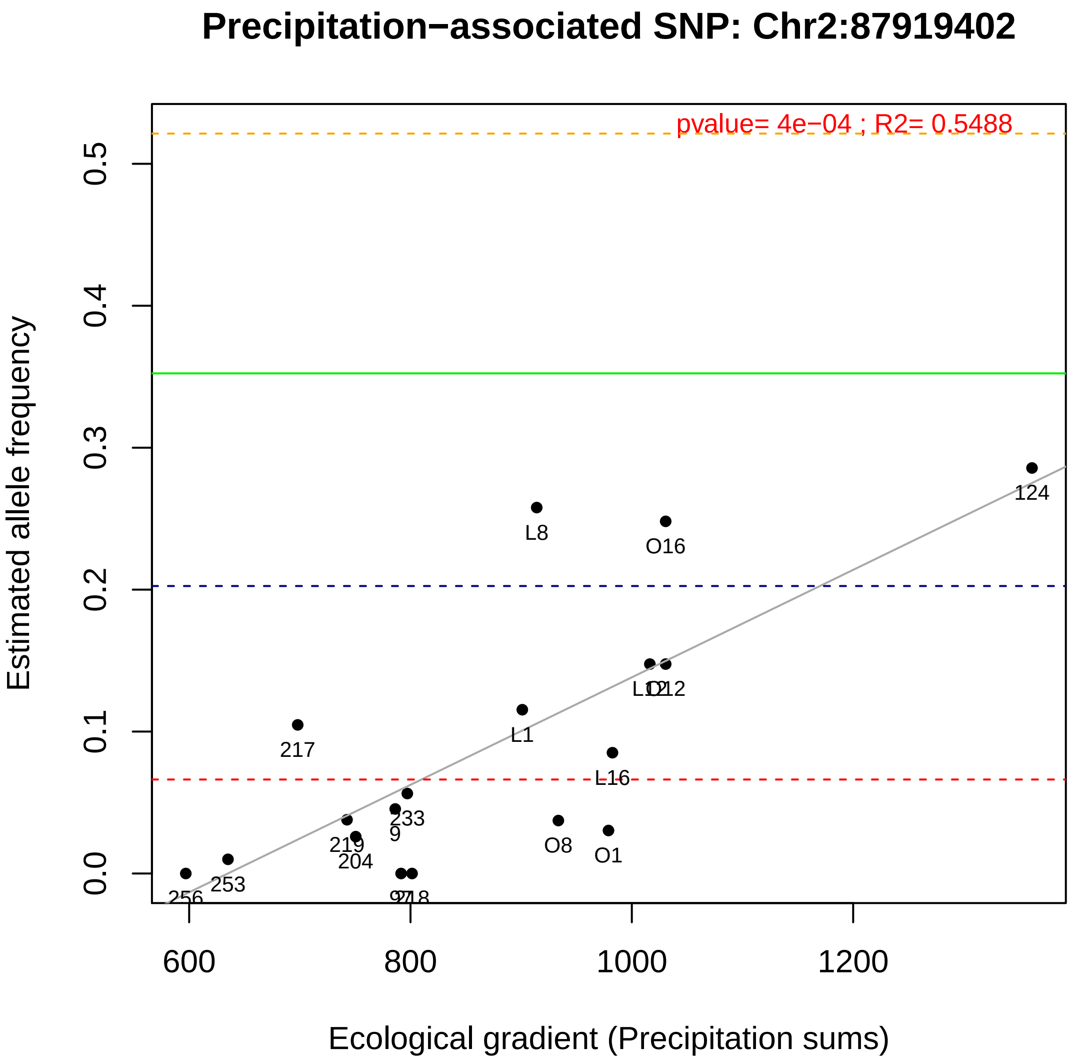

Precipitation−associated SNP: Chr2:87919402
pvalue= 4e−04 ; R2= 0.5488
Estimated allele frequency
0.0
0.1
0.2
0.3
0.4
0.5
600
800
1000
1200
Ecological gradient (Precipitation sums)
124
L8
O16
L12
O12
L1
217
L16
233
9
219
204
O8
O1
253
256
218

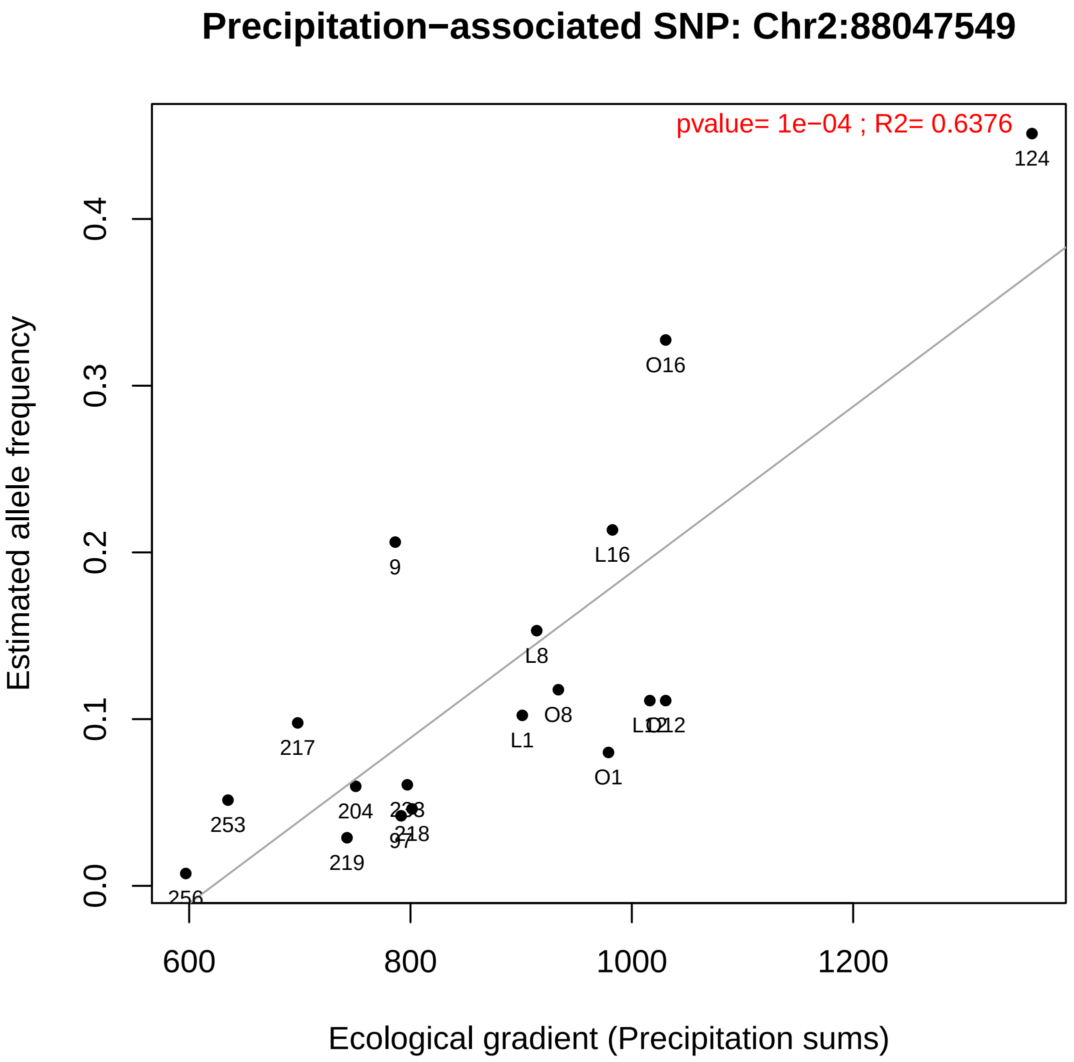
Precipitation−associated SNP: Chr2:88047549
pvalue= 1e−04 ; R2= 0.6376
Estimated allele frequency
Ecological gradient (Precipitation sums)
0.0
0.1
0.2
0.3
0.4
600
800
1000
1200
124
O16
L16
9
L8
O8
L1
O1
217
253
204
218
219
97
256

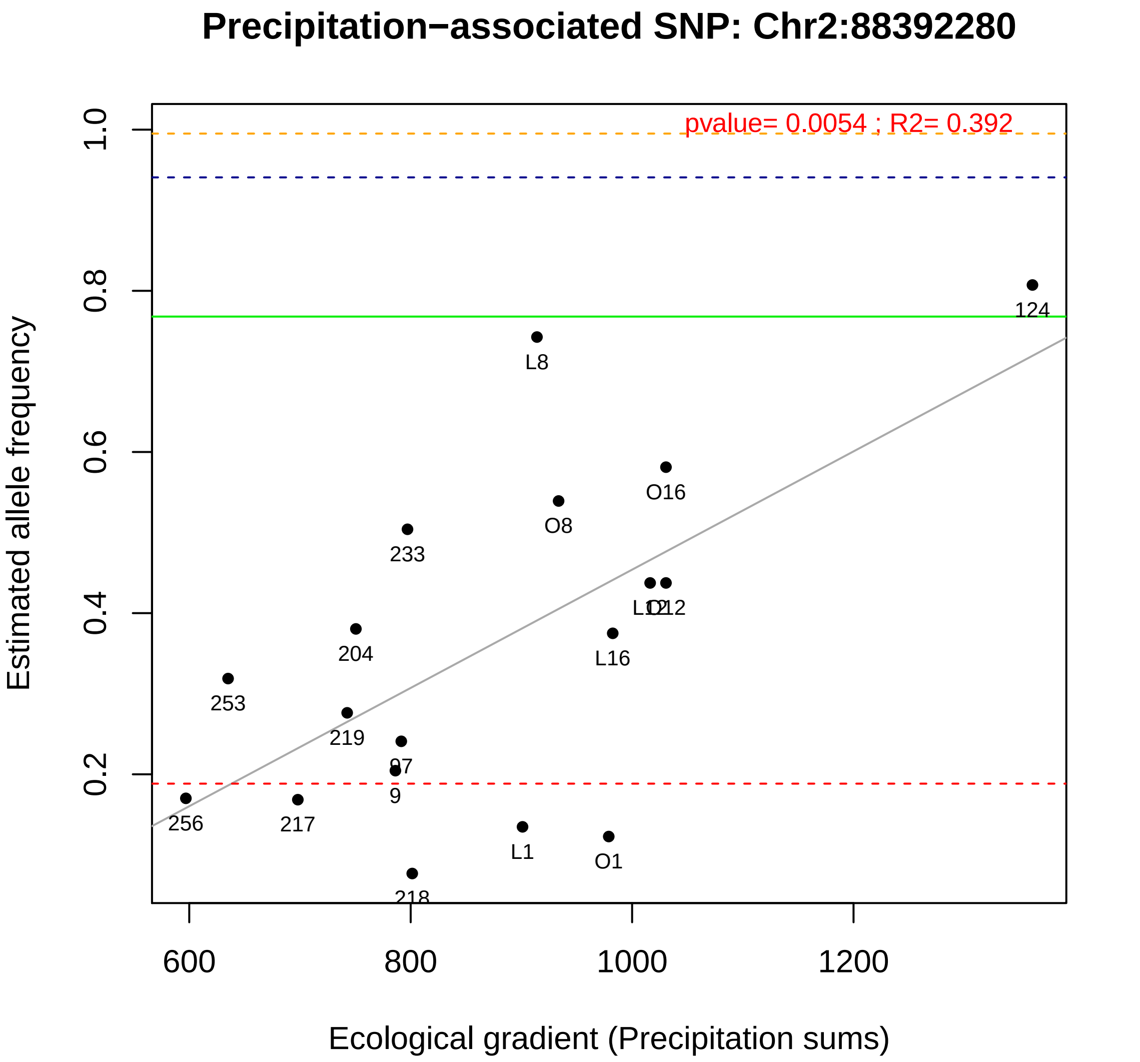
Precipitation−associated SNP: Chr2:88392280
pvalue= 0.0054 ; R2= 0.392
Estimated allele frequency
Ecological gradient (Precipitation sums)
0.2
0.4
0.6
0.8
1.0
600
800
1000
1200
124
L8
O16
O8
233
L12
O12
204
L16
253
219
97
9
256
217
L1
O1
218

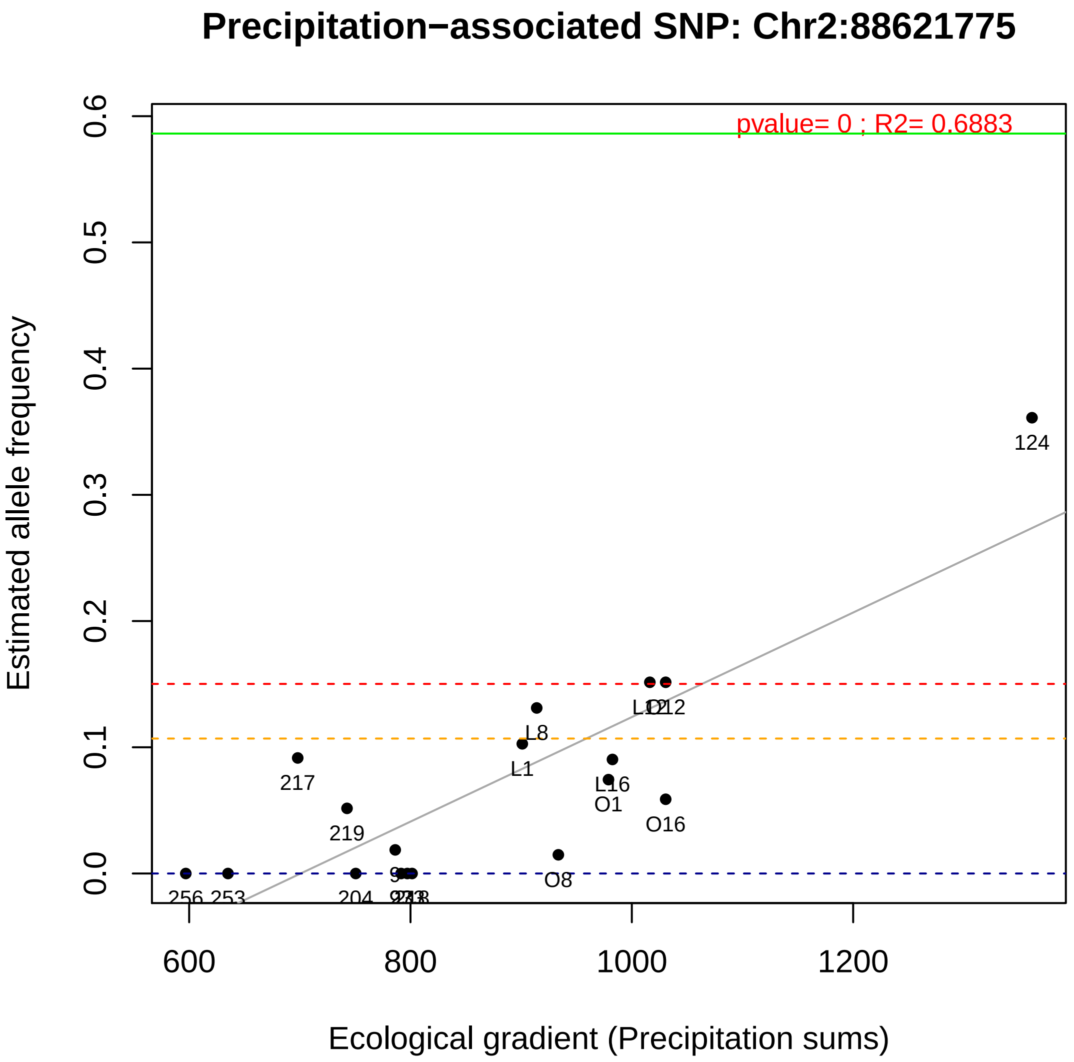

Precipitation−associated SNP: Chr2:88621775
pvalue= 0 ; R2= 0.6883
Estimated allele frequency
Ecological gradient (Precipitation sums)
0.0
0.1
0.2
0.3
0.4
0.5
0.6
600
800
1000
1200
124
L8
L1
217
219
L16
O1
O16
O8
204
256
253

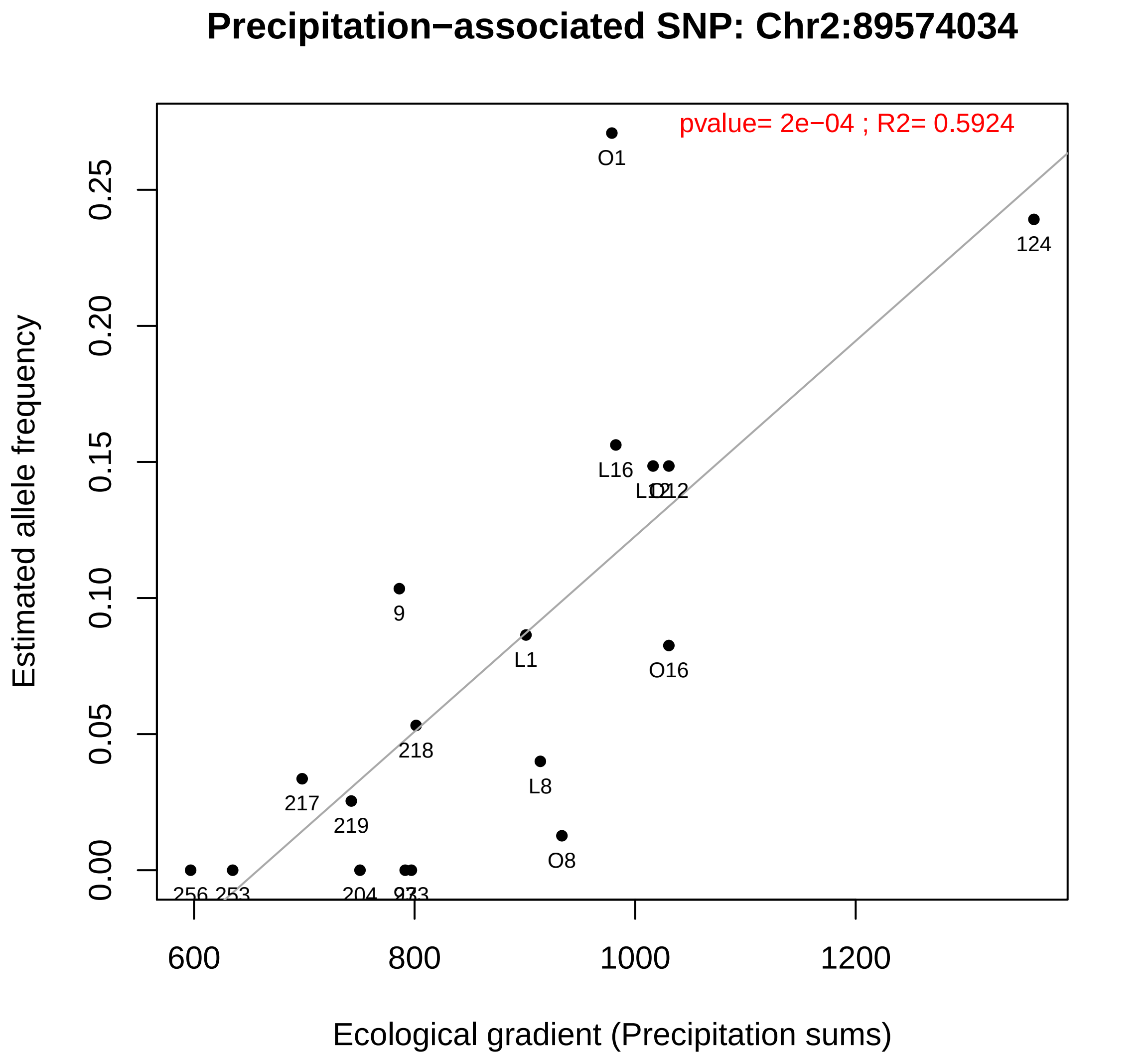

Precipitation−associated SNP: Chr2:89574034
pvalue= 2e−04 ; R2= 0.5924
Estimated allele frequency
Ecological gradient (Precipitation sums)
0.00
0.05
0.10
0.15
0.20
0.25
600
800
1000
1200
O1
124
L16
O12
9
L1
O16
218
L8
217
219
O8
256
253
204

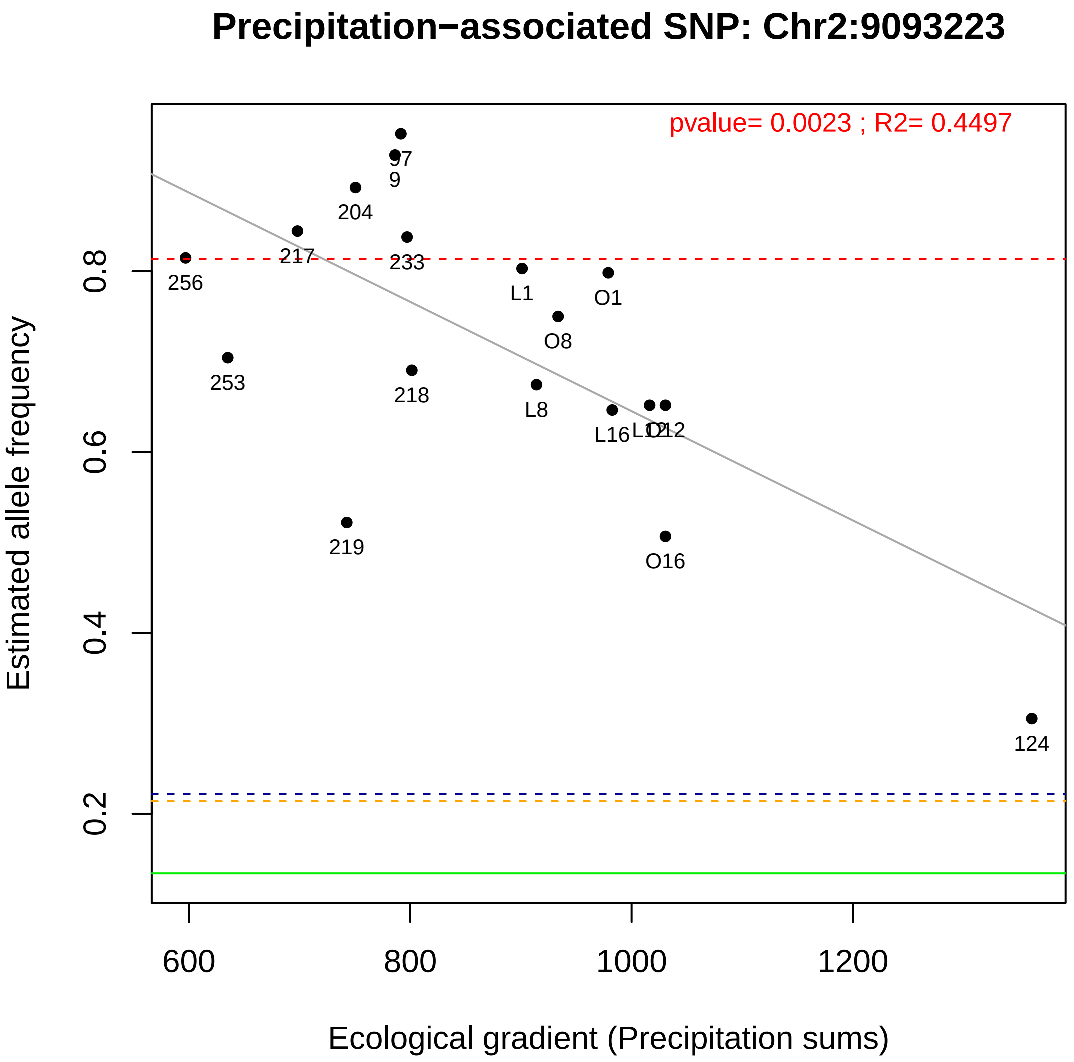

Precipitation−associated SNP: Chr2:9093223
pvalue= 0.0023 ; R2= 0.4497
Estimated allele frequency
Ecological gradient (Precipitation sums)
0.2
0.4
0.6
0.8
600
800
1000
1200
97
9
204
217
233
256
L1
O1
O8
253
218
L8
L16
L12
O12
219
O16
124

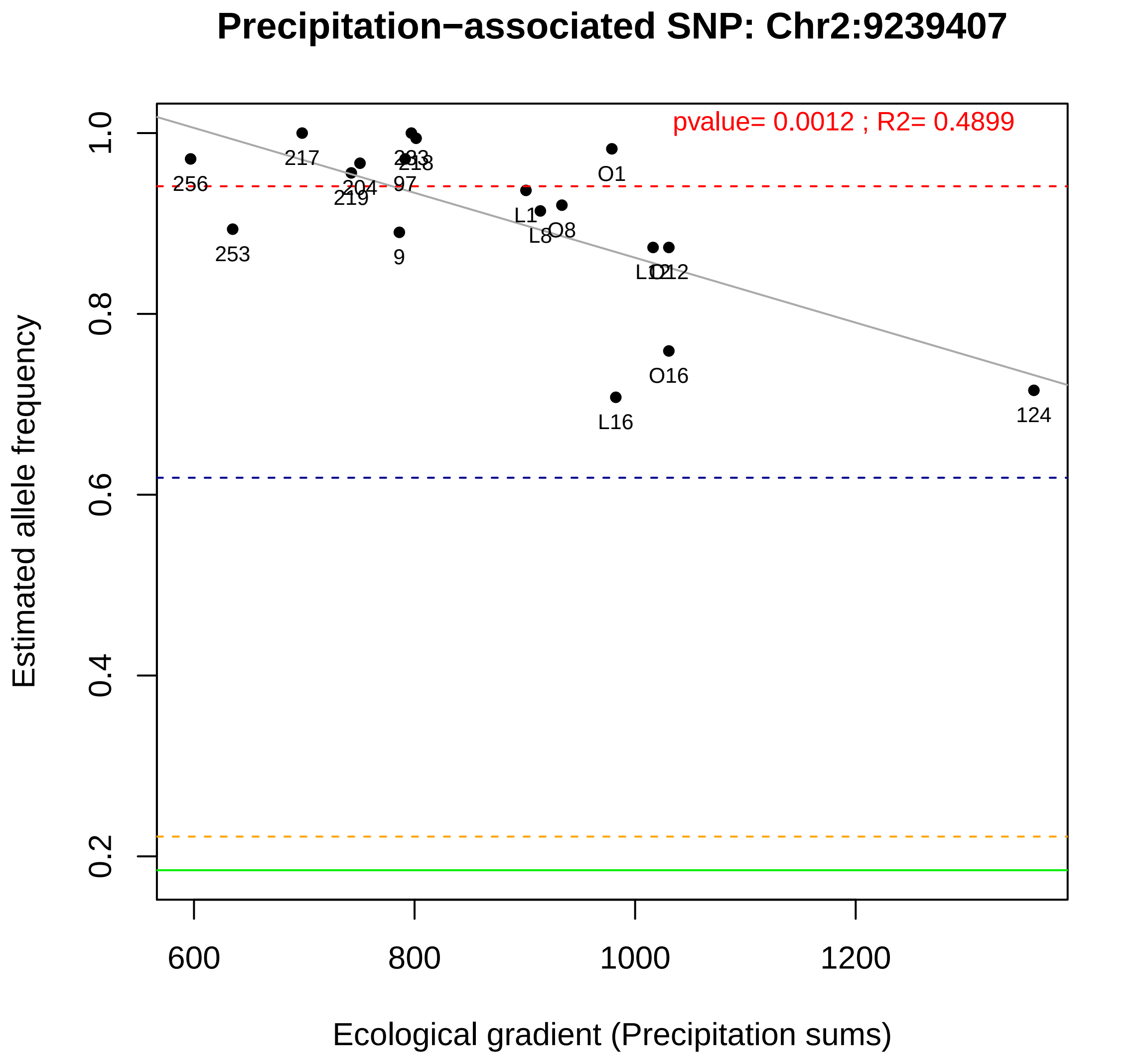

Precipitation−associated SNP: Chr2:9239407
pvalue= 0.0012 ; R2= 0.4899
Estimated allele frequency
Ecological gradient (Precipitation sums)
0.2
0.4
0.6
0.8
1.0
600
800
1000
1200
256
253
217
204
219
283
218
97
9
L1
L8
O8
O1
L12
O12
O16
L16
124

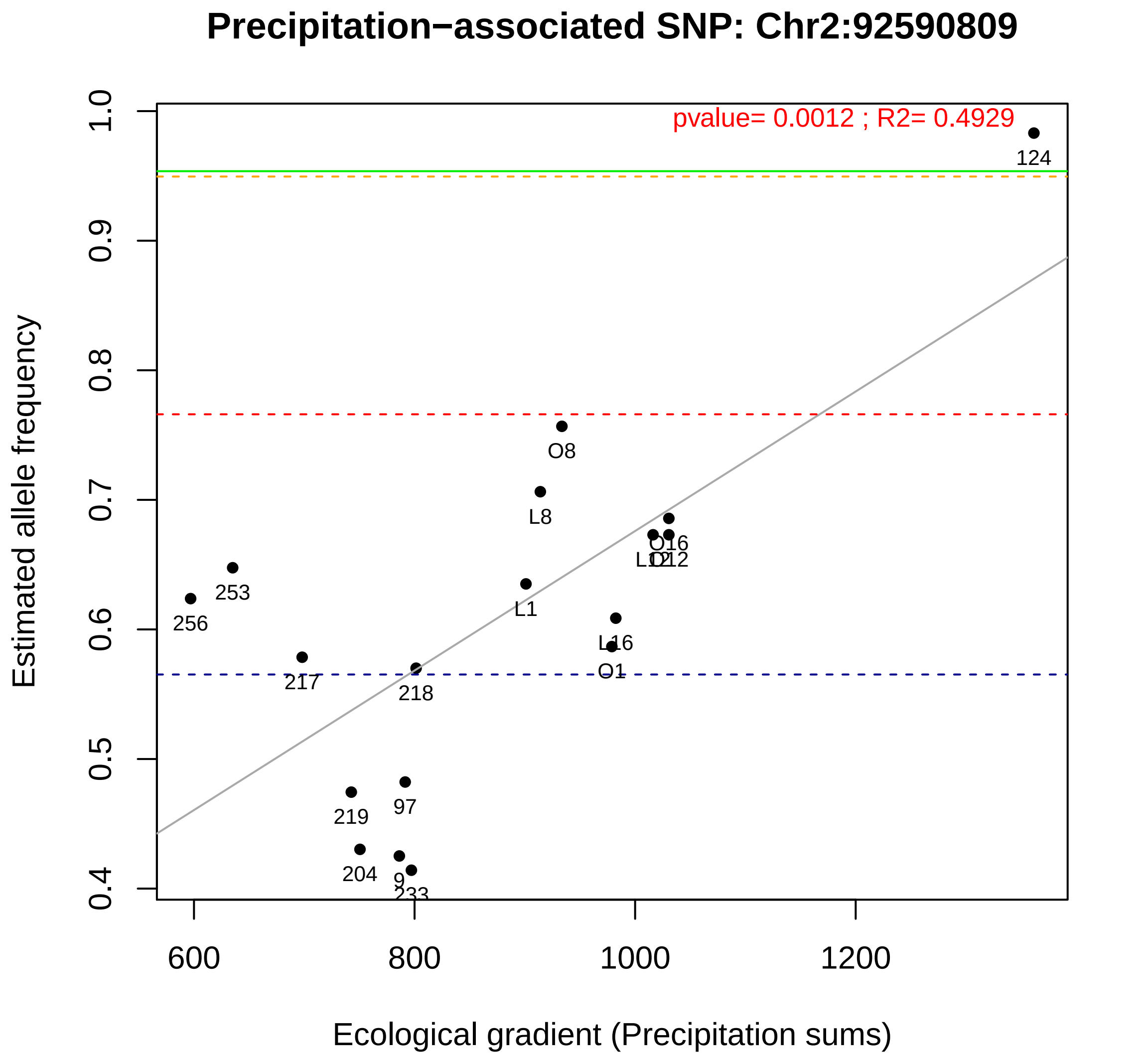
Precipitation−associated SNP: Chr2:92590809
pvalue= 0.0012 ; R2= 0.4929
Estimated allele frequency
Ecological gradient (Precipitation sums)
0.4
0.5
0.6
0.7
0.8
0.9
1.0
600
800
1000
1200
124
O8
L8
O16
L10
O12
253
256
L1
L16
O1
217
218
219
97
204
9
233

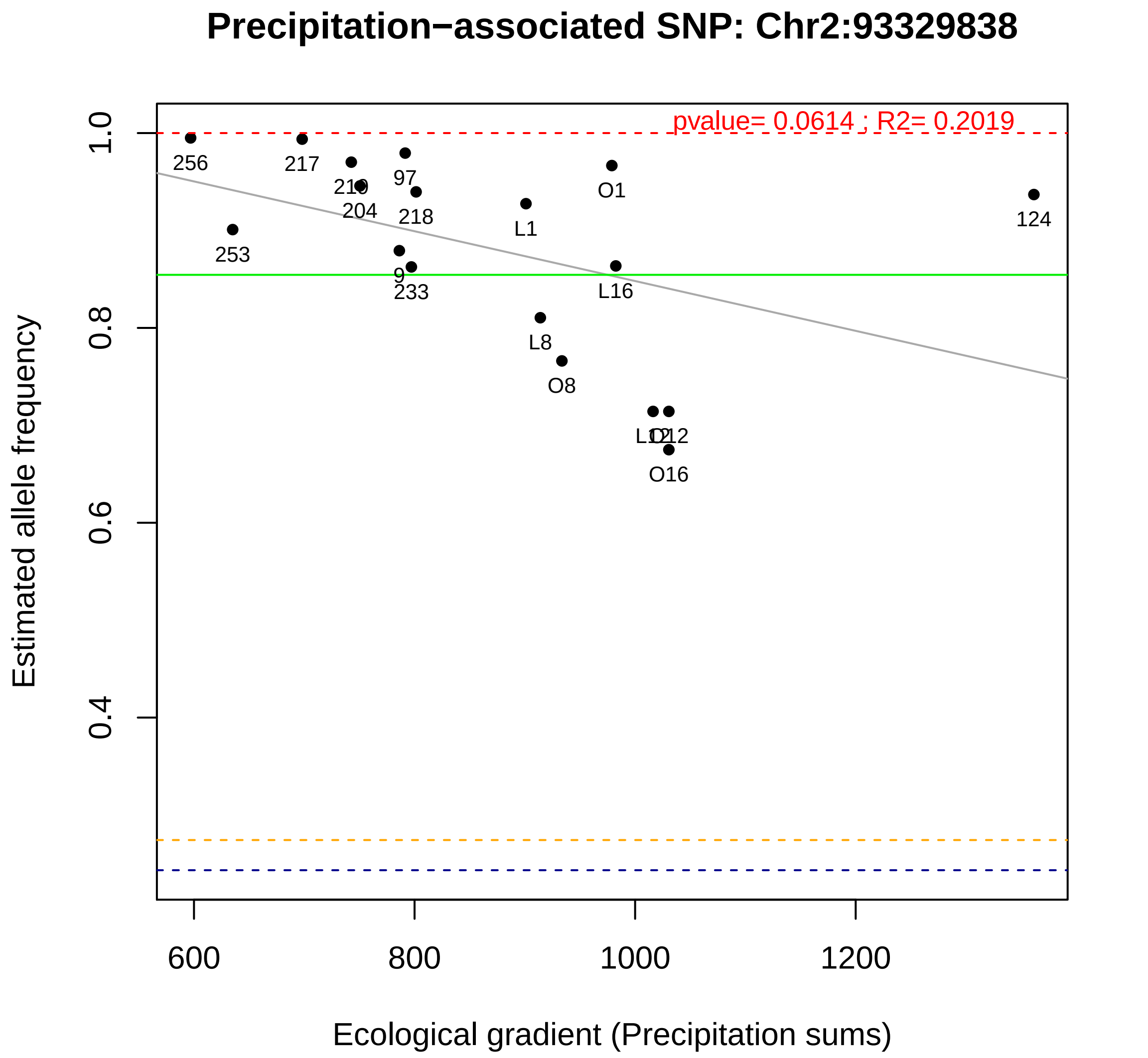
Precipitation−associated SNP: Chr2:93329838
pvalue= 0.0614 ; R2= 0.2019
Estimated allele frequency
0.4
0.6
0.8
1.0
600
800
1000
1200
Ecological gradient (Precipitation sums)
256
217
210
204
97
218
253
9
233
L1
O1
L16
L8
O8
L12
O12
O16
124

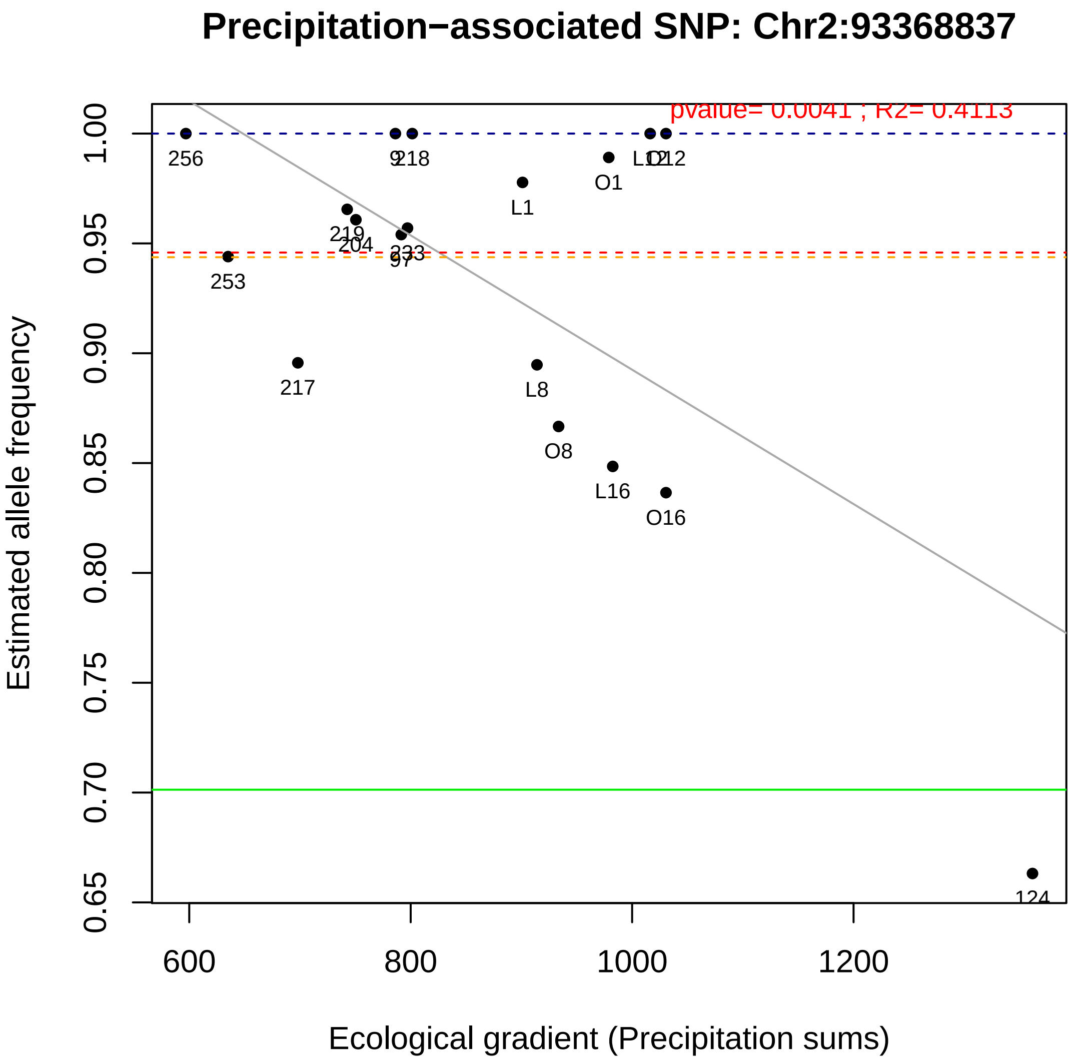

Precipitation−associated SNP: Chr2:93368837
pvalue= 0.0041 ; R2= 0.4113
Estimated allele frequency
Ecological gradient (Precipitation sums)
0.65
0.70
0.75
0.80
0.85
0.90
0.95
1.00
600
800
1000
1200
256
9
218
L12
O12
O1
L1
219
204
233
97
253
217
L8
O8
L16
O16
124

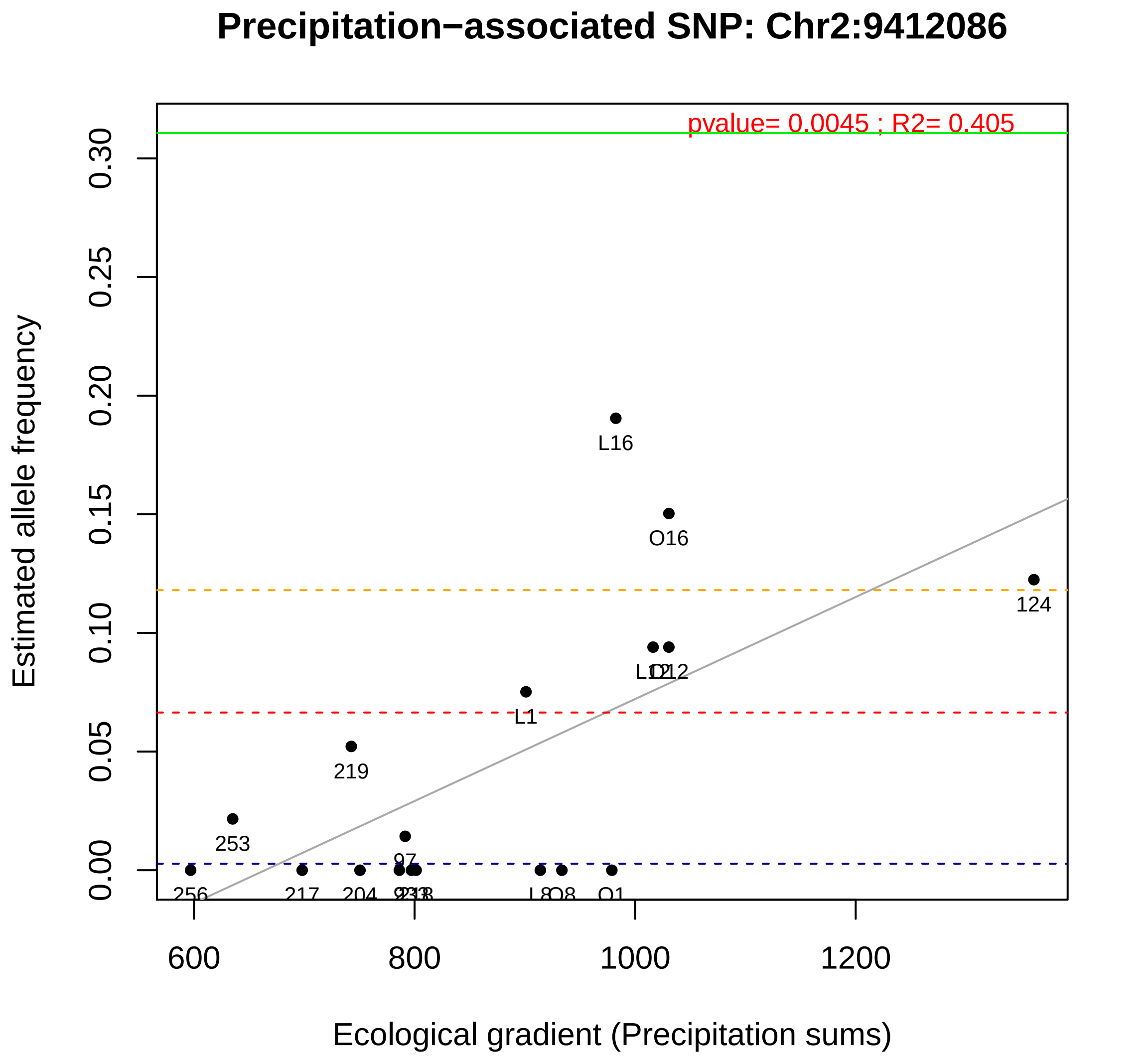
Precipitation−associated SNP: Chr2:9412086
pvalue= 0.0045 ; R2= 0.405
Estimated allele frequency
0.00
0.05
0.10
0.15
0.20
0.25
0.30
600
800
1000
1200
Ecological gradient (Precipitation sums)
L16
O16
124
L1
219
253
97
256
217
204
O1

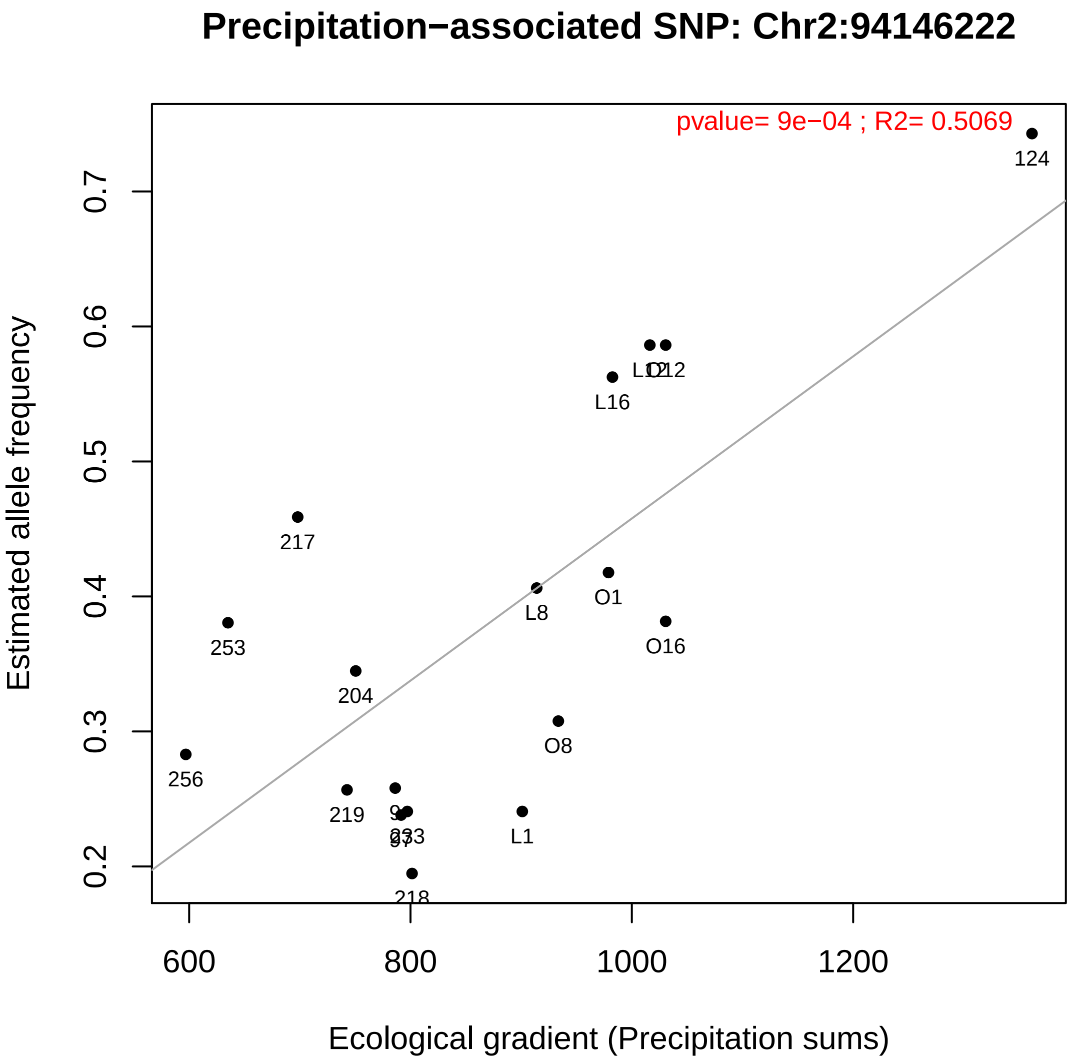
Precipitation−associated SNP: Chr2:94146222
pvalue= 9e−04 ; R2= 0.5069
Estimated allele frequency
Ecological gradient (Precipitation sums)
0.2
0.3
0.4
0.5
0.6
0.7
600
800
1000
1200
124
L16
217
L8
O1
O16
253
204
O8
256
219
L1
218

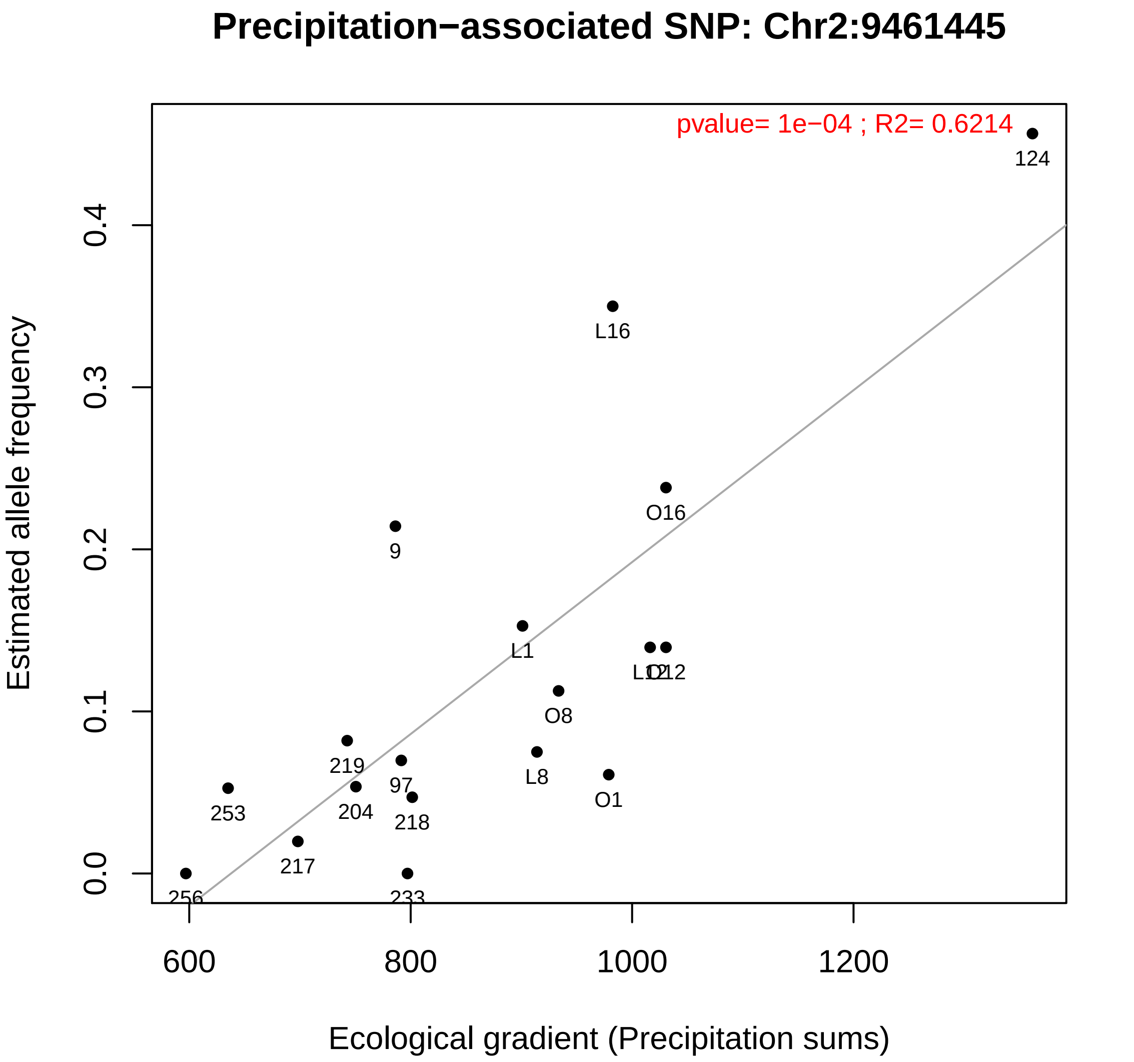
Precipitation−associated SNP: Chr2:9461445
pvalue= 1e−04 ; R2= 0.6214
Estimated allele frequency
0.0
0.1
0.2
0.3
0.4
600
800
1000
1200
Ecological gradient (Precipitation sums)
124
L16
O16
9
L1
O8
219
97
L8
O1
253
204
218
217
256
233

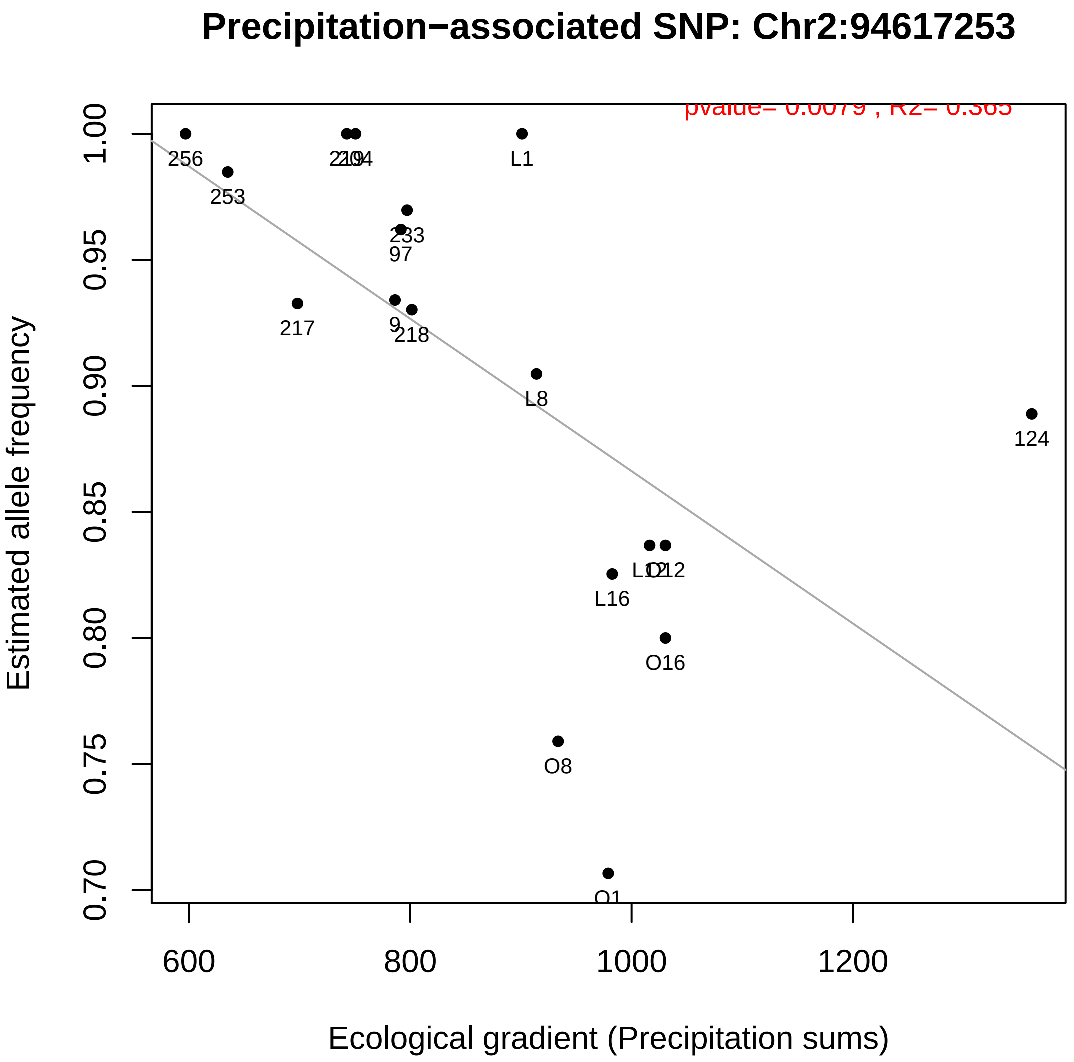
Precipitation−associated SNP: Chr2:94617253
pvalue= 0.0079 , R2= 0.365
Estimated allele frequency
Ecological gradient (Precipitation sums)
1.00
0.95
0.90
0.85
0.80
0.75
0.70
600
800
1000
1200
256
253
210
194
L1
233
97
217
9
218
L8
124
L10
O12
L16
O16
O8
O1

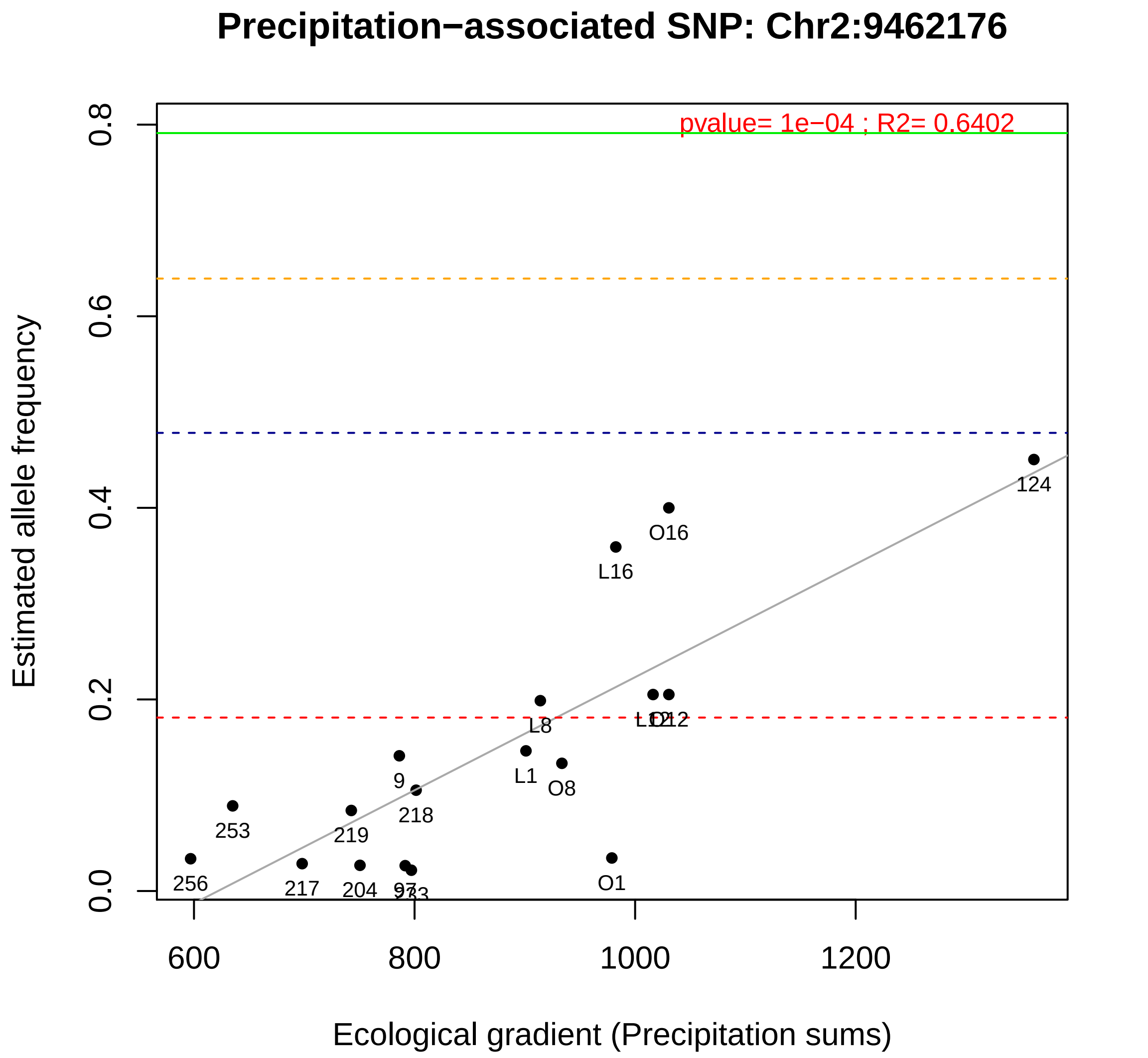

Precipitation−associated SNP: Chr2:9462176
pvalue= 1e−04 ; R2= 0.6402
Estimated allele frequency
Ecological gradient (Precipitation sums)
0.0
0.2
0.4
0.6
0.8
600
800
1000
1200
124
O16
L16
L8
L1
O8
O1
9
218
219
253
256
217
204

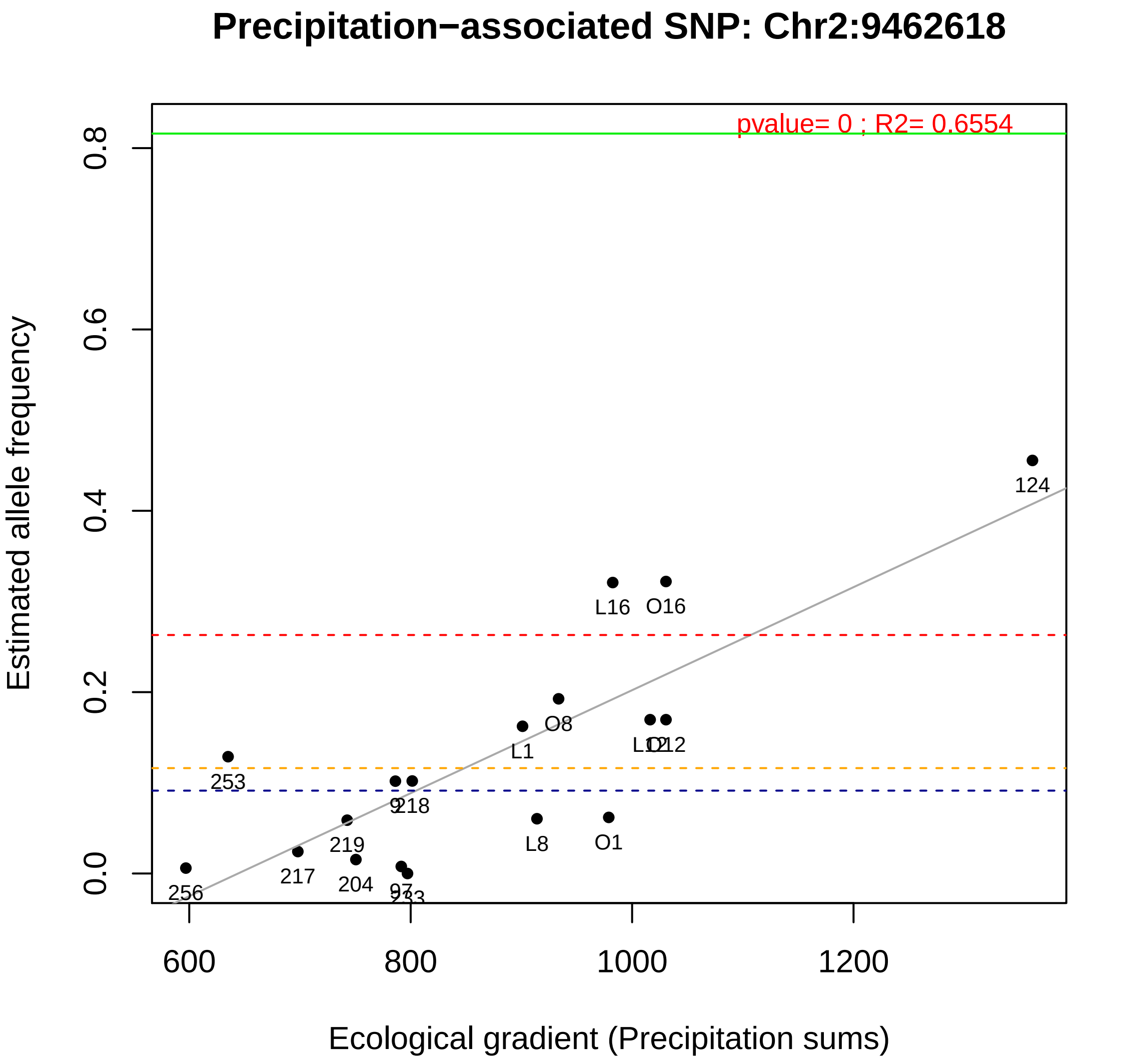
Precipitation−associated SNP: Chr2:9462618
pvalue= 0 ; R2= 0.6554
Estimated allele frequency
Ecological gradient (Precipitation sums)
0.0
0.2
0.4
0.6
0.8
600
800
1000
1200
124
L16
O16
O8
L1
L12
O12
253
218
219
L8
O1
217
204
256
97
233

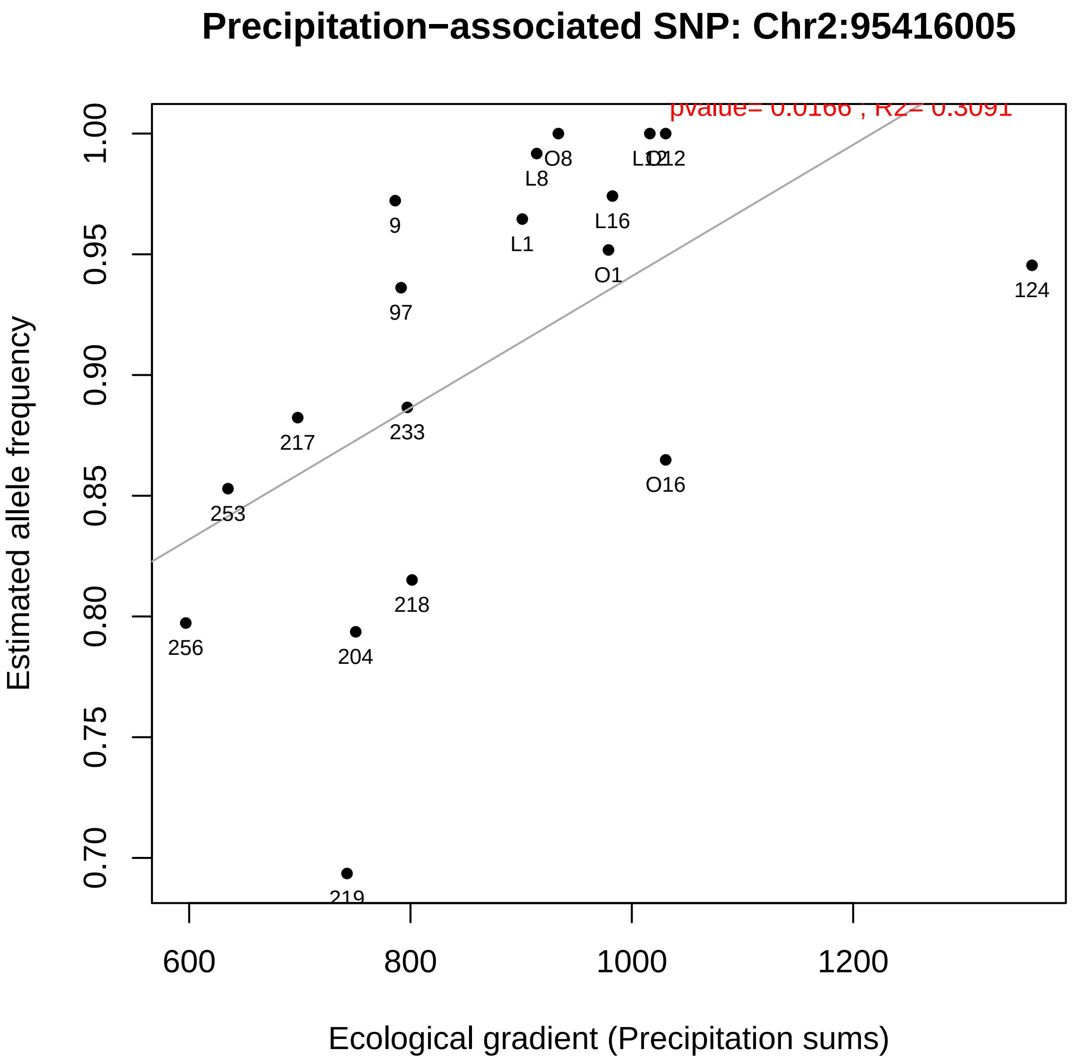

Precipitation−associated SNP: Chr2:95416005
pvalue= 0.0166 ; R2= 0.3091
Estimated allele frequency
Ecological gradient (Precipitation sums)
0.70
0.75
0.80
0.85
0.90
0.95
1.00
600
800
1000
1200
O8
L8
L12
O12
9
L1
L16
O1
97
124
217
233
253
O16
218
256
204
219

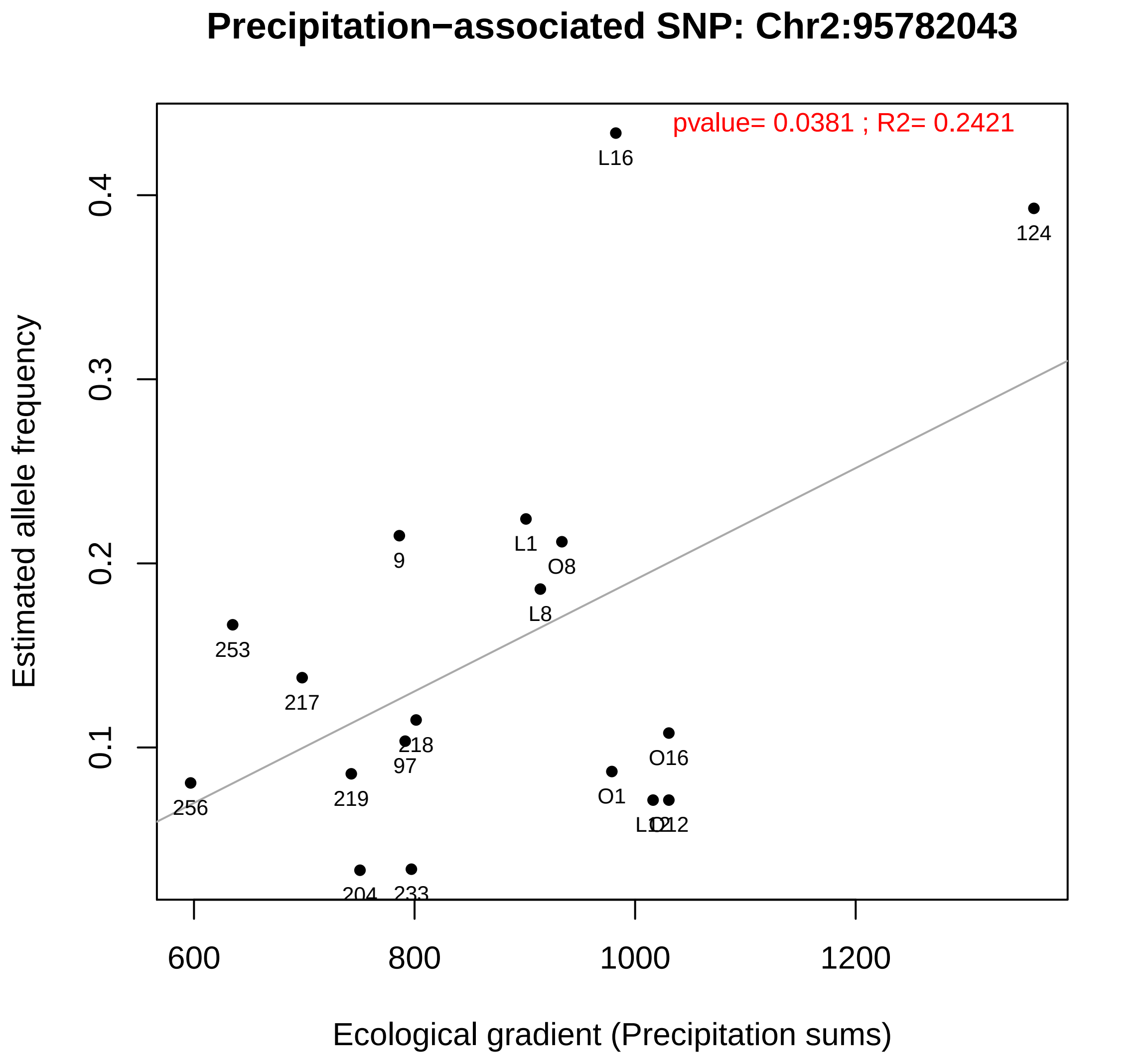

Precipitation−associated SNP: Chr2:95782043
pvalue= 0.0381 ; R2= 0.2421
Estimated allele frequency
0.1
0.2
0.3
0.4
600
800
1000
1200
Ecological gradient (Precipitation sums)
L16
124
9
L1
O8
L8
253
217
218
97
O16
256
219
O1
L12
O12
204
233

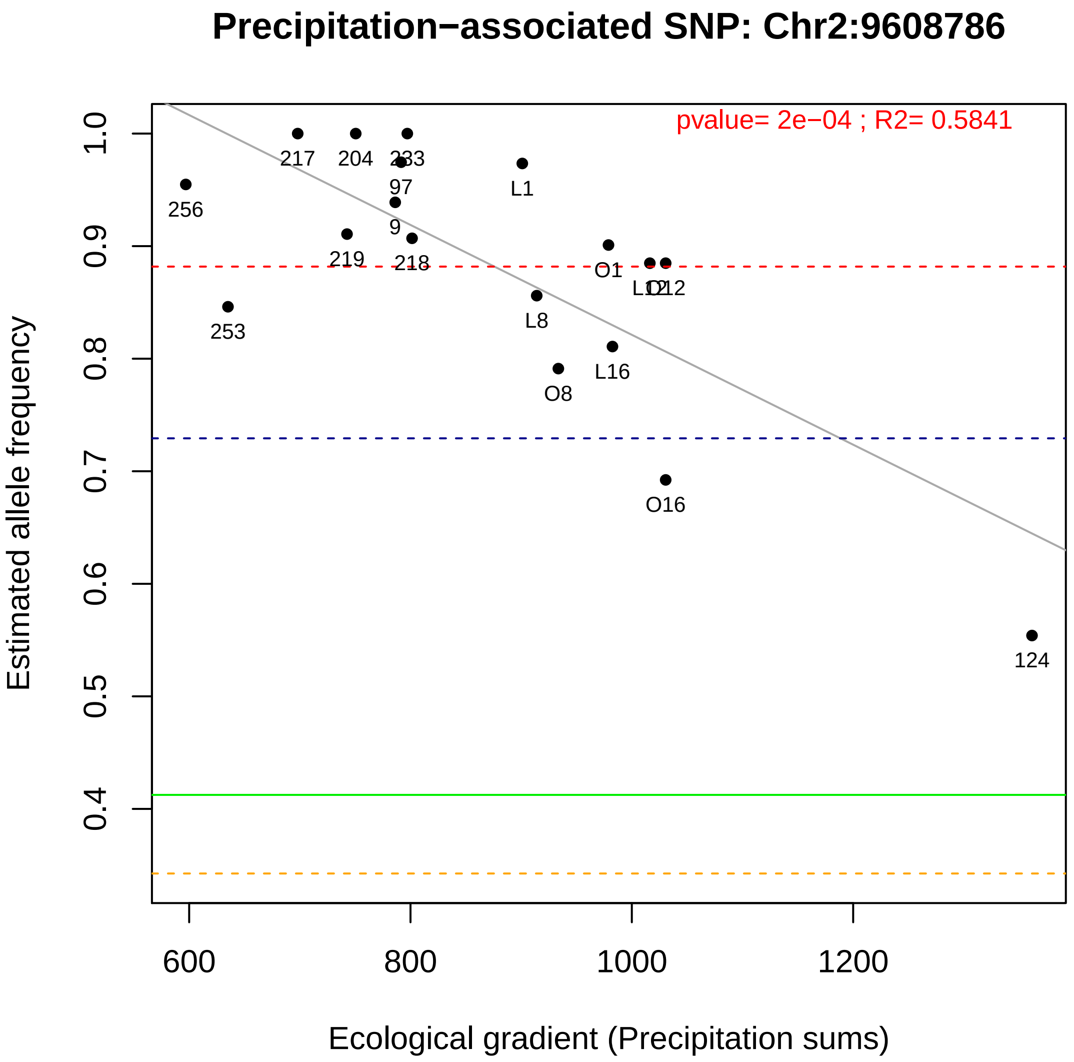

Precipitation−associated SNP: Chr2:9608786
pvalue= 2e−04 ; R2= 0.5841
Estimated allele frequency
Ecological gradient (Precipitation sums)
1.0
0.9
0.8
0.7
0.6
0.5
0.4
600
800
1000
1200
217
204
233
97
256
9
219
218
L1
O1
L12
O12
253
L8
L16
O8
O16
124

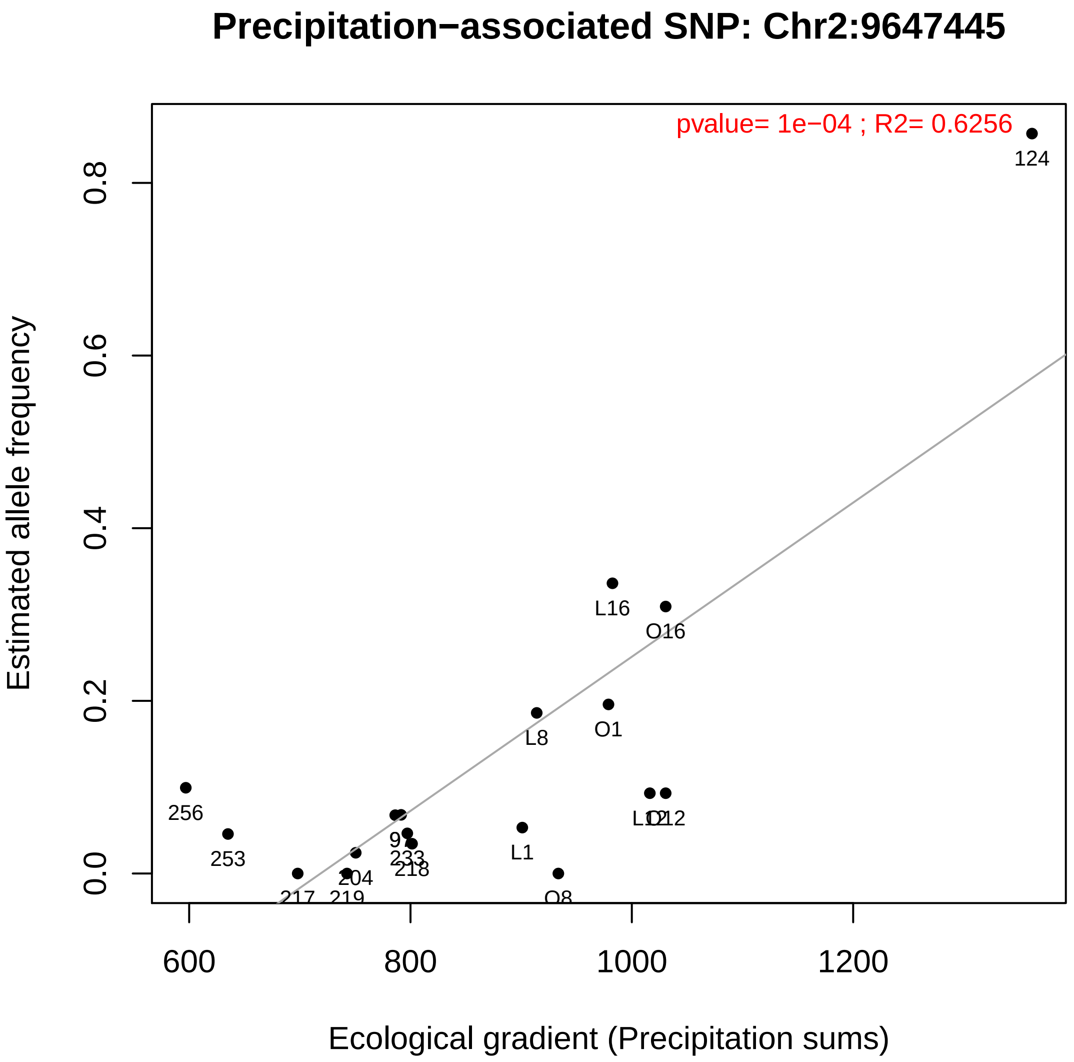

Precipitation−associated SNP: Chr2:9647445
pvalue= 1e−04 ; R2= 0.6256
124
L16
O16
L8
O1
256
253
217
219
204
97
233
218
L1
O8
L12
O12
0.0
0.2
0.4
0.6
0.8
600
800
1000
1200
Estimated allele frequency
Ecological gradient (Precipitation sums)

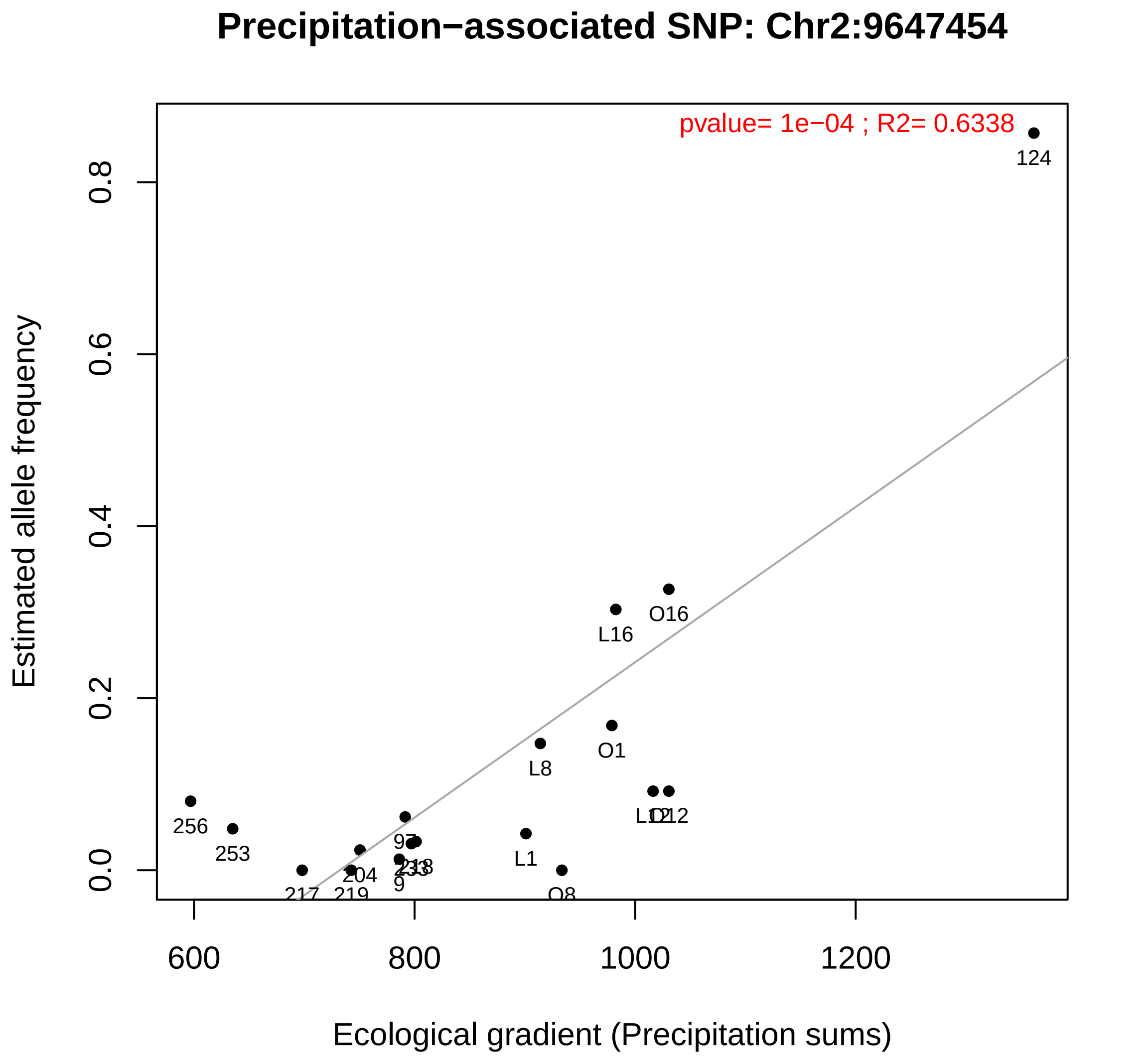
Precipitation−associated SNP: Chr2:9647454
pvalue= 1e−04 ; R2= 0.6338
124
O16
L16
O1
L8
L12
O12
256
253
97
L1
204
213
218
9
217
219
O8
0.0
0.2
0.4
0.6
0.8
600
800
1000
1200
Estimated allele frequency
Ecological gradient (Precipitation sums)

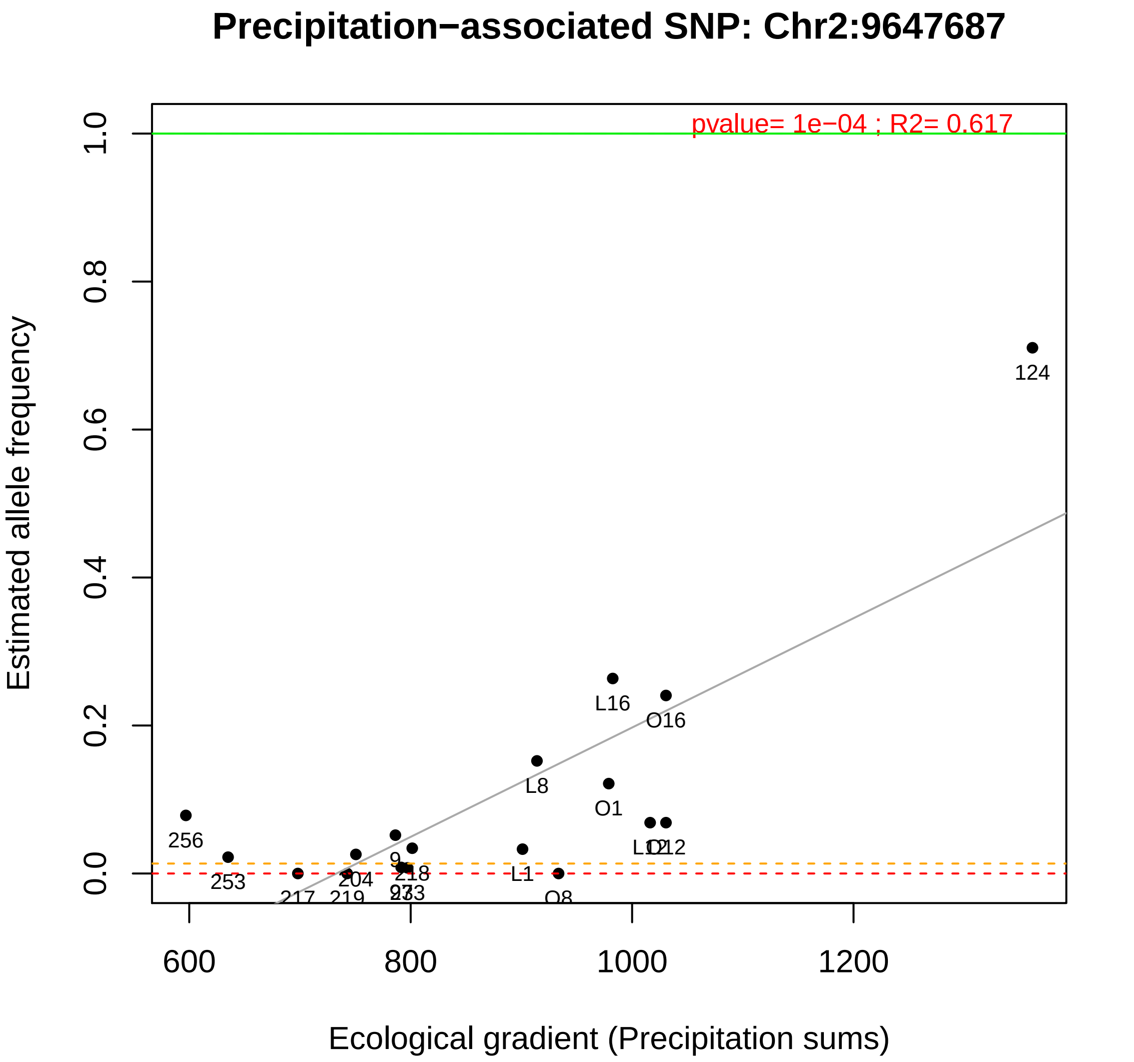

Precipitation−associated SNP: Chr2:9647687
pvalue= 1e−04 ; R2= 0.617
Estimated allele frequency
0.0
0.2
0.4
0.6
0.8
1.0
600
800
1000
1200
Ecological gradient (Precipitation sums)
124
L16
O16
L8
O1
256
253
204
218
L1

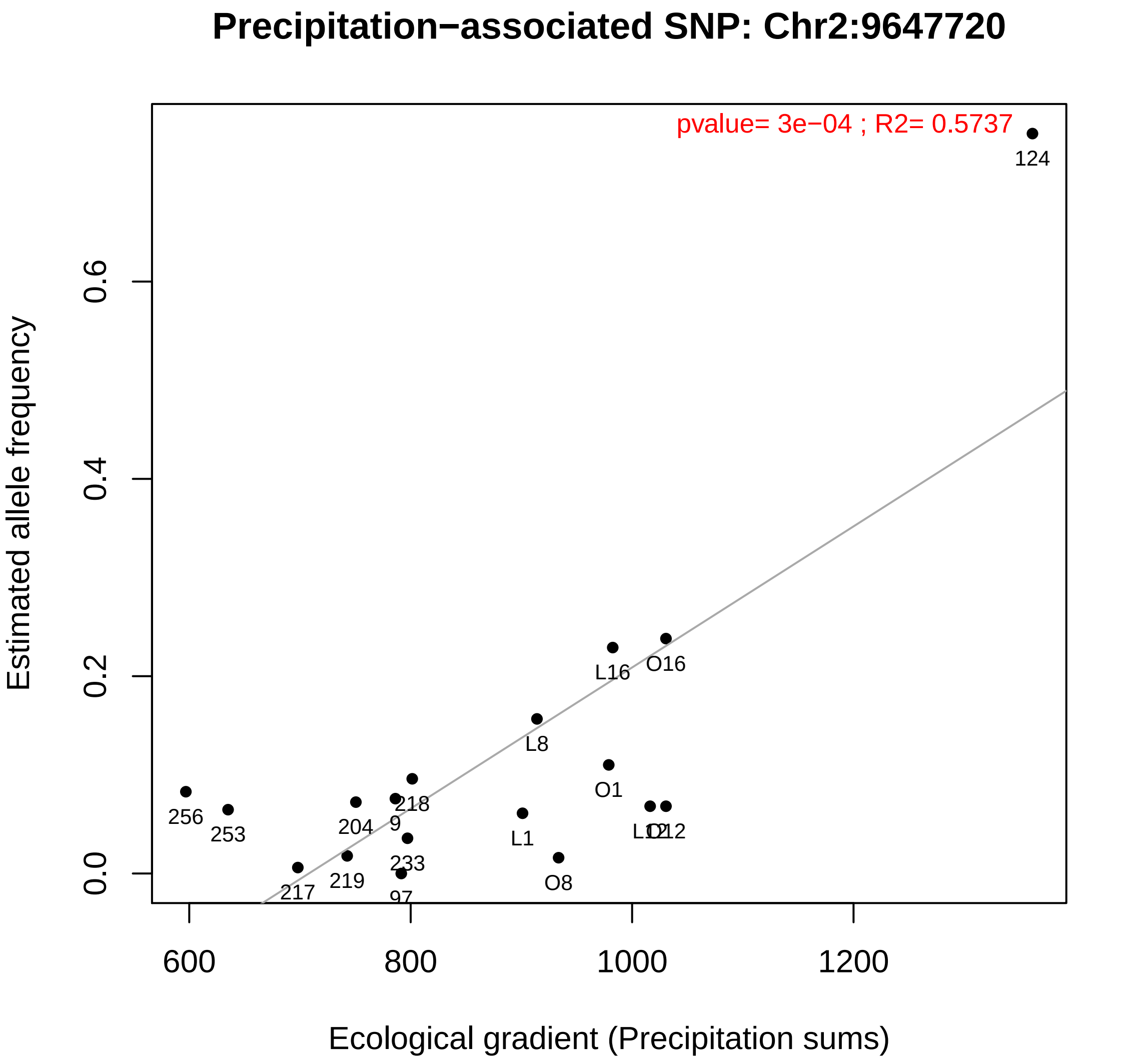
Precipitation−associated SNP: Chr2:9647720
pvalue= 3e−04 ; R2= 0.5737
124
L16
O16
L8
O1
256
253
204
9
218
233
217
219
97
L1
O8
L12
O12
0.0
0.2
0.4
0.6
600
800
1000
1200
Estimated allele frequency
Ecological gradient (Precipitation sums)

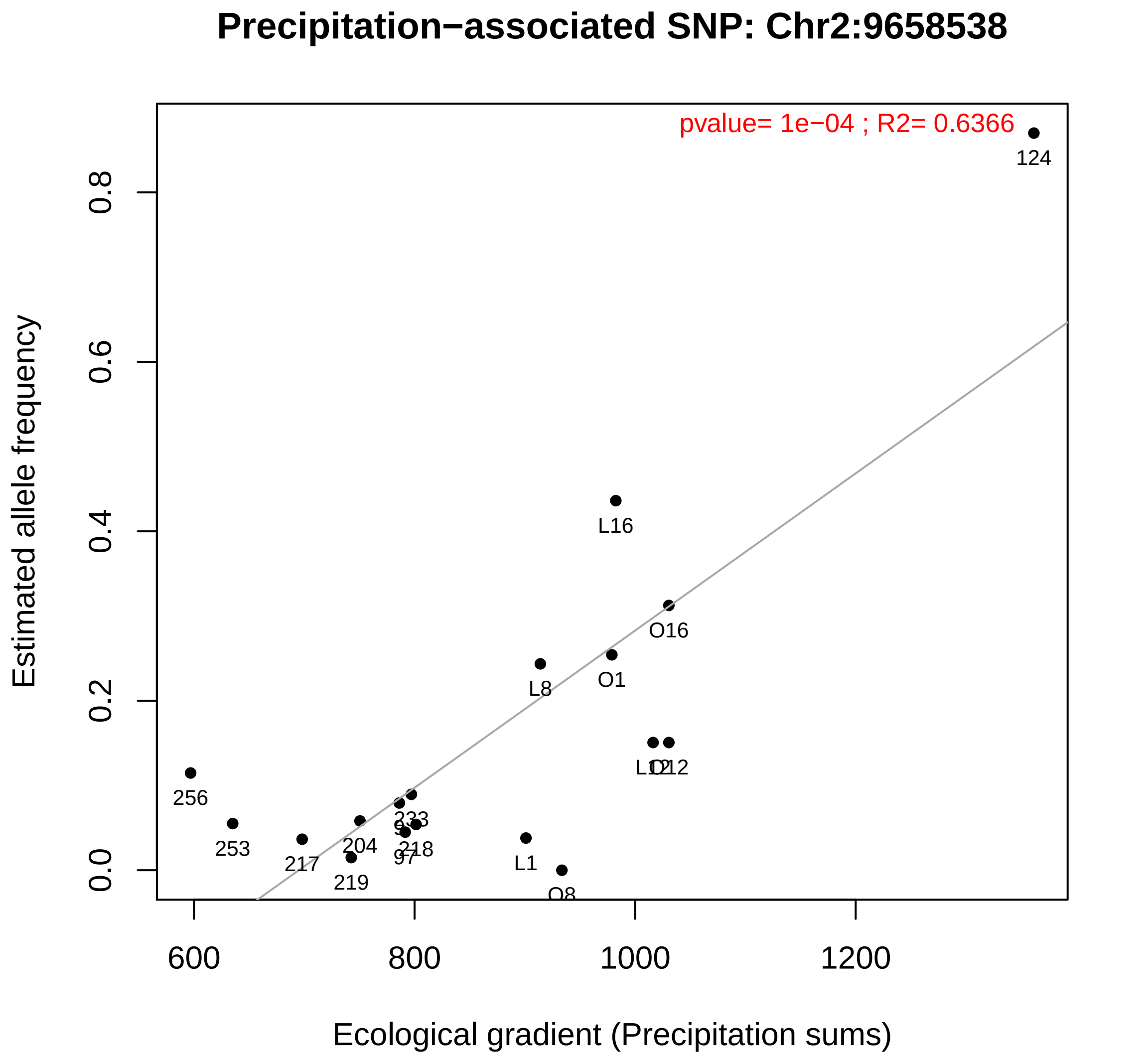

Precipitation−associated SNP: Chr2:9658538
pvalue= 1e−04 ; R2= 0.6366
Estimated allele frequency
0.0
0.2
0.4
0.6
0.8
600
800
1000
1200
Ecological gradient (Precipitation sums)
124
L16
O16
O1
L8
256
253
217
219
204
233
218
97
L1
O8

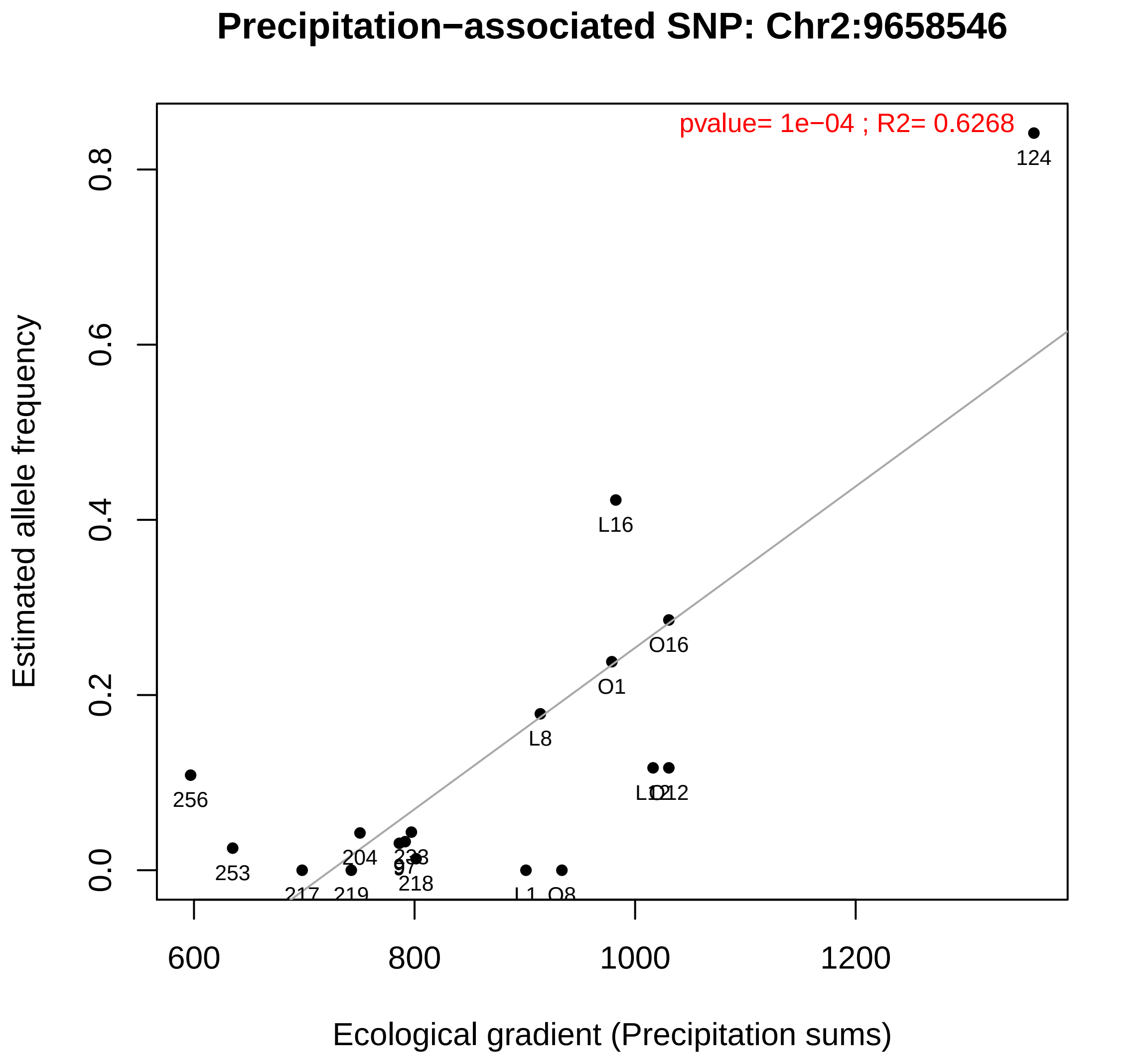

Precipitation−associated SNP: Chr2:9658546
pvalue= 1e−04 ; R2= 0.6268
Estimated allele frequency
Ecological gradient (Precipitation sums)
0.0
0.2
0.4
0.6
0.8
600
800
1000
1200
124
L16
O16
O1
L8
256
253
204
217
219
218
L1
O8
O12

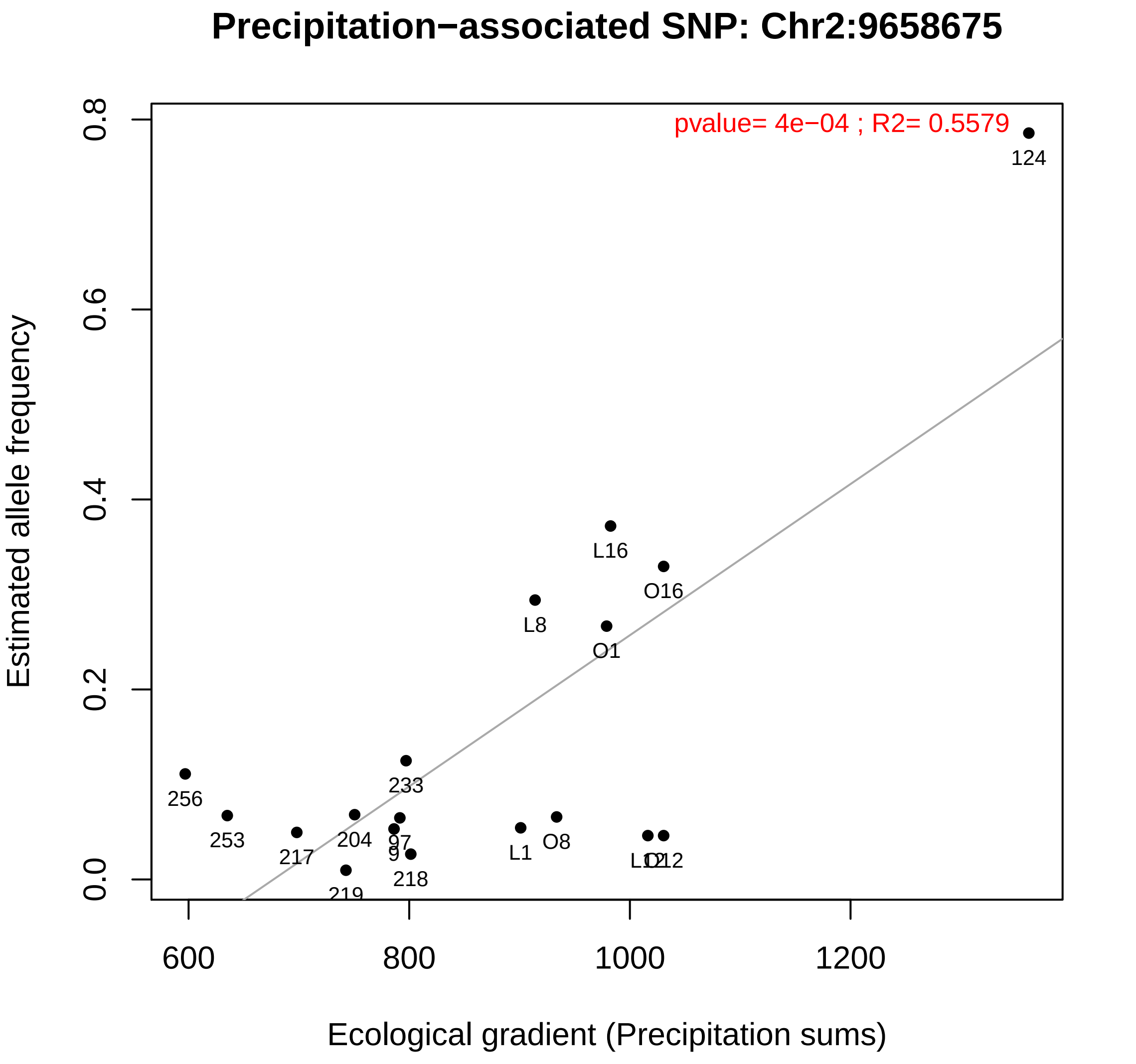

Precipitation−associated SNP: Chr2:9658675
pvalue= 4e−04 ; R2= 0.5579
Estimated allele frequency
0.0
0.2
0.4
0.6
0.8
600
800
1000
1200
Ecological gradient (Precipitation sums)
124
L16
O16
L8
O1
233
256
253
217
204
97
9
218
219
L1
O8
L12
O12

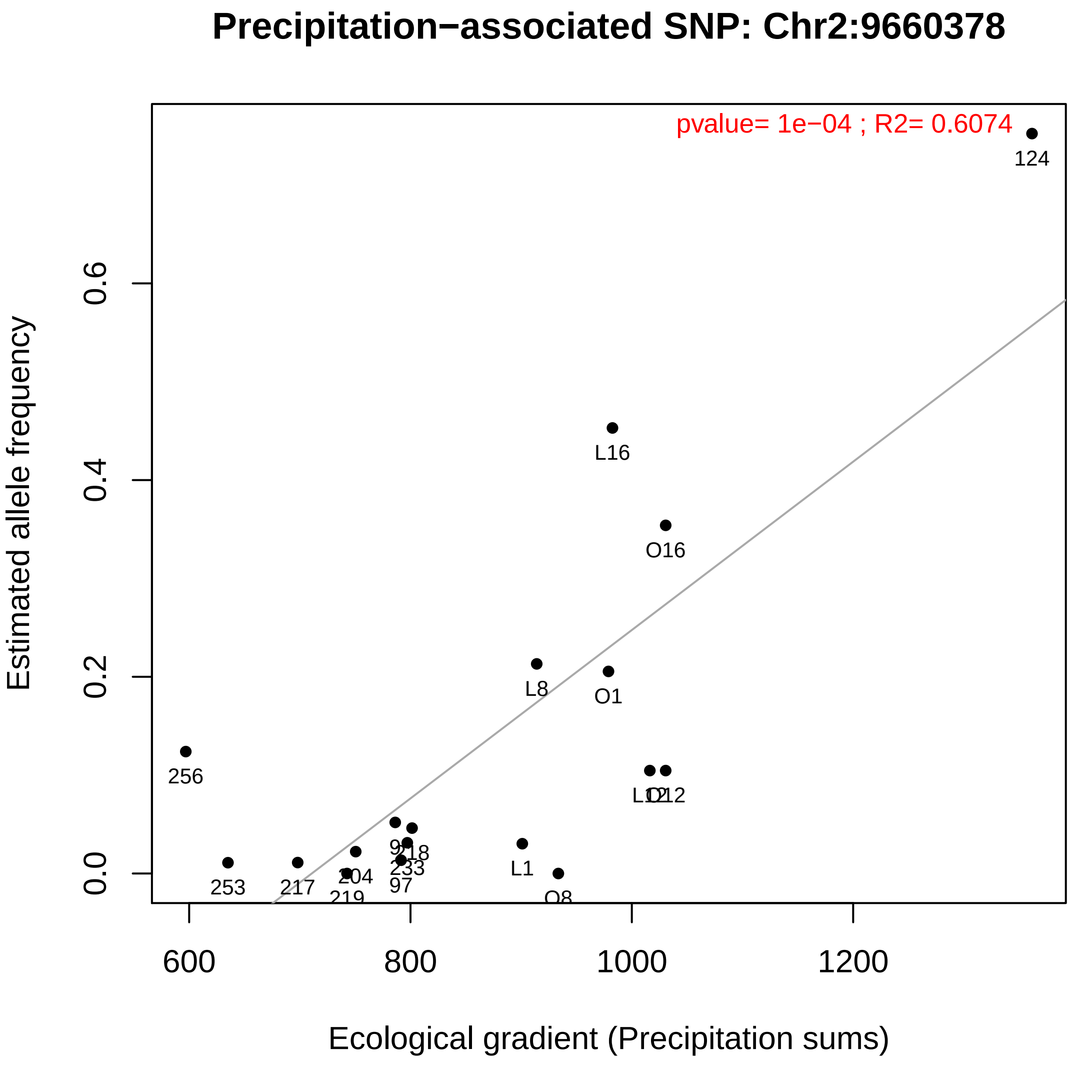

Precipitation−associated SNP: Chr2:9660378
pvalue= 1e−04 ; R2= 0.6074
124
L16
O16
L8
O1
256
L12
O12
253
217
204
219
218
233
97
L1
O8
0.0
0.2
0.4
0.6
600
800
1000
1200
Estimated allele frequency
Ecological gradient (Precipitation sums)

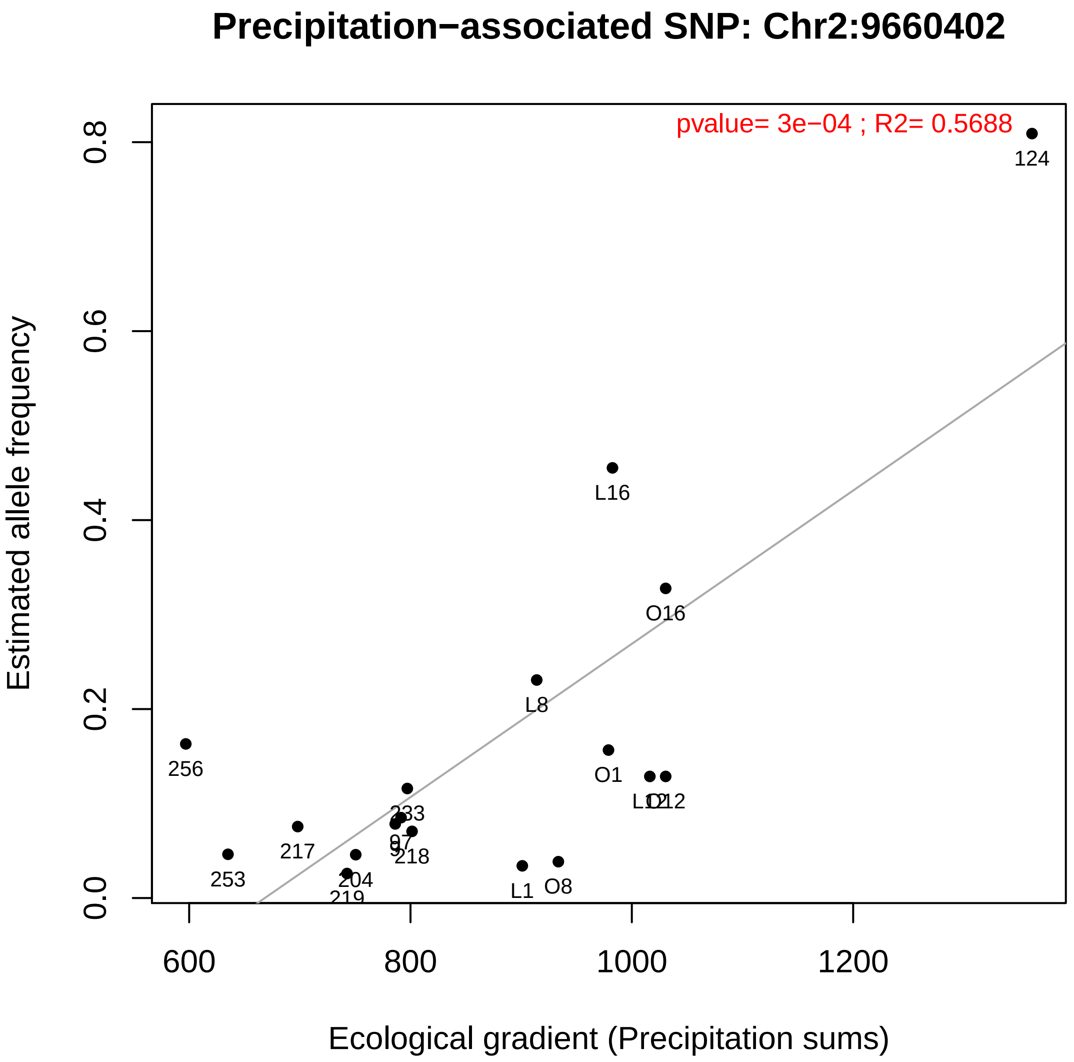
Precipitation−associated SNP: Chr2:9660402
pvalue= 3e−04 ; R2= 0.5688
Estimated allele frequency
Ecological gradient (Precipitation sums)
0.0
0.2
0.4
0.6
0.8
600
800
1000
1200
124
L16
O16
L8
256
O1
L12
O12
233
217
97
218
253
204
219
L1
O8

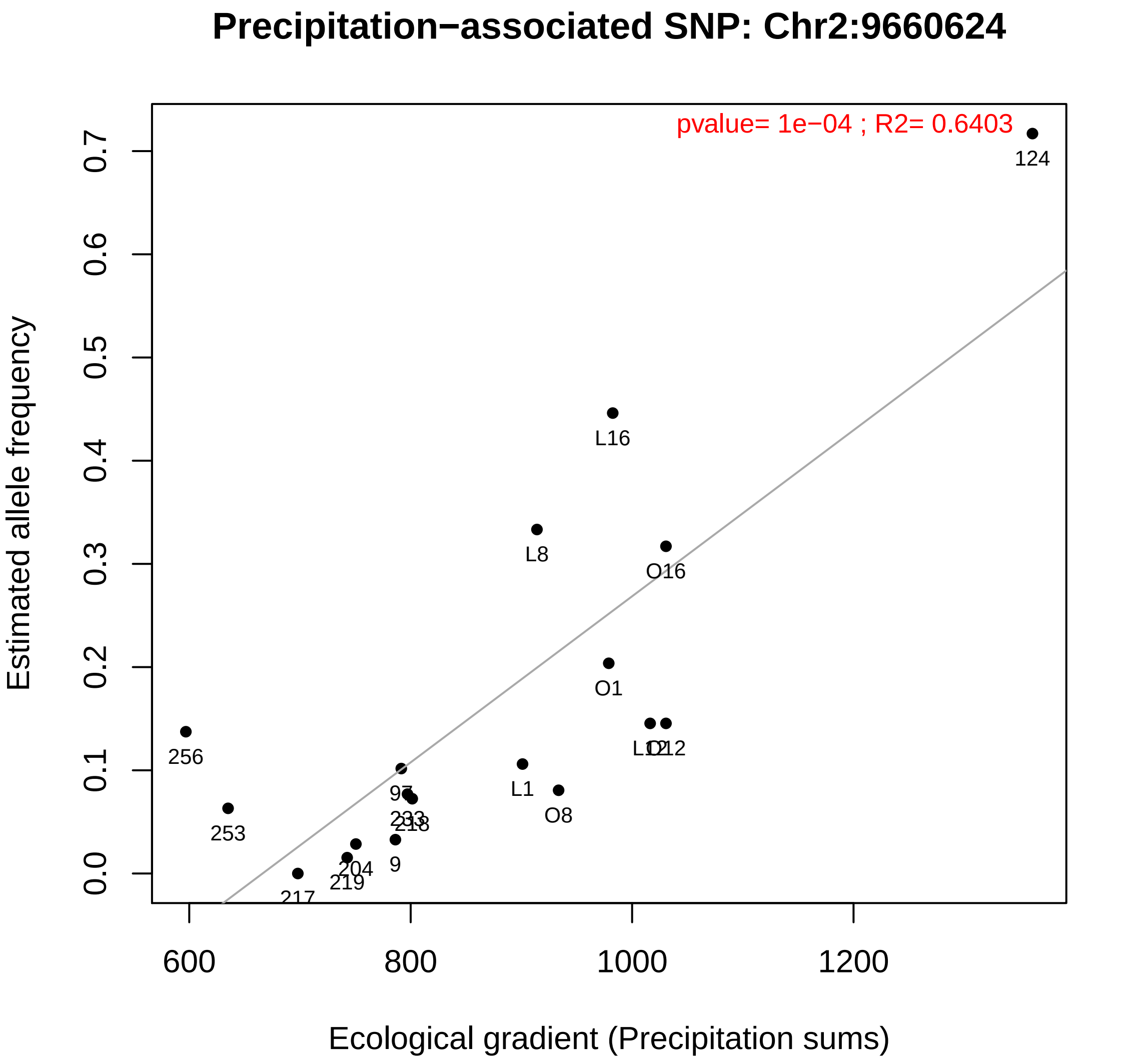
Precipitation−associated SNP: Chr2:9660624
pvalue= 1e−04 ; R2= 0.6403
Estimated allele frequency
Ecological gradient (Precipitation sums)
0.0
0.1
0.2
0.3
0.4
0.5
0.6
0.7
600
800
1000
1200
124
L16
L8
O16
O1
O12
256
253
L1
O8
97
233
218
9
204
219
217

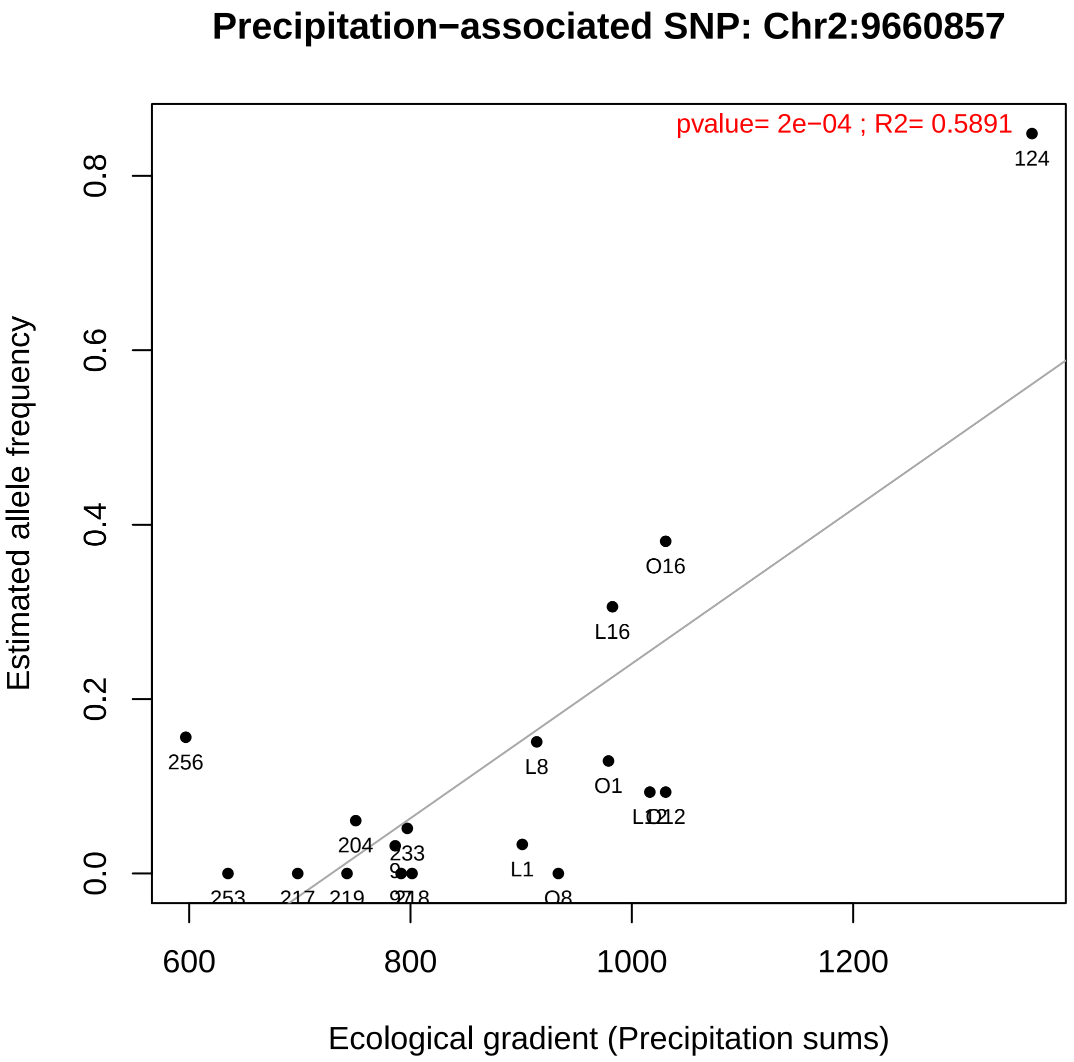
Precipitation−associated SNP: Chr2:9660857
pvalue= 2e−04 ; R2= 0.5891
Estimated allele frequency
Ecological gradient (Precipitation sums)
0.0
0.2
0.4
0.6
0.8
600
800
1000
1200
124
O16
L16
256
L8
O1
204
233
L1
253
217
219
O8

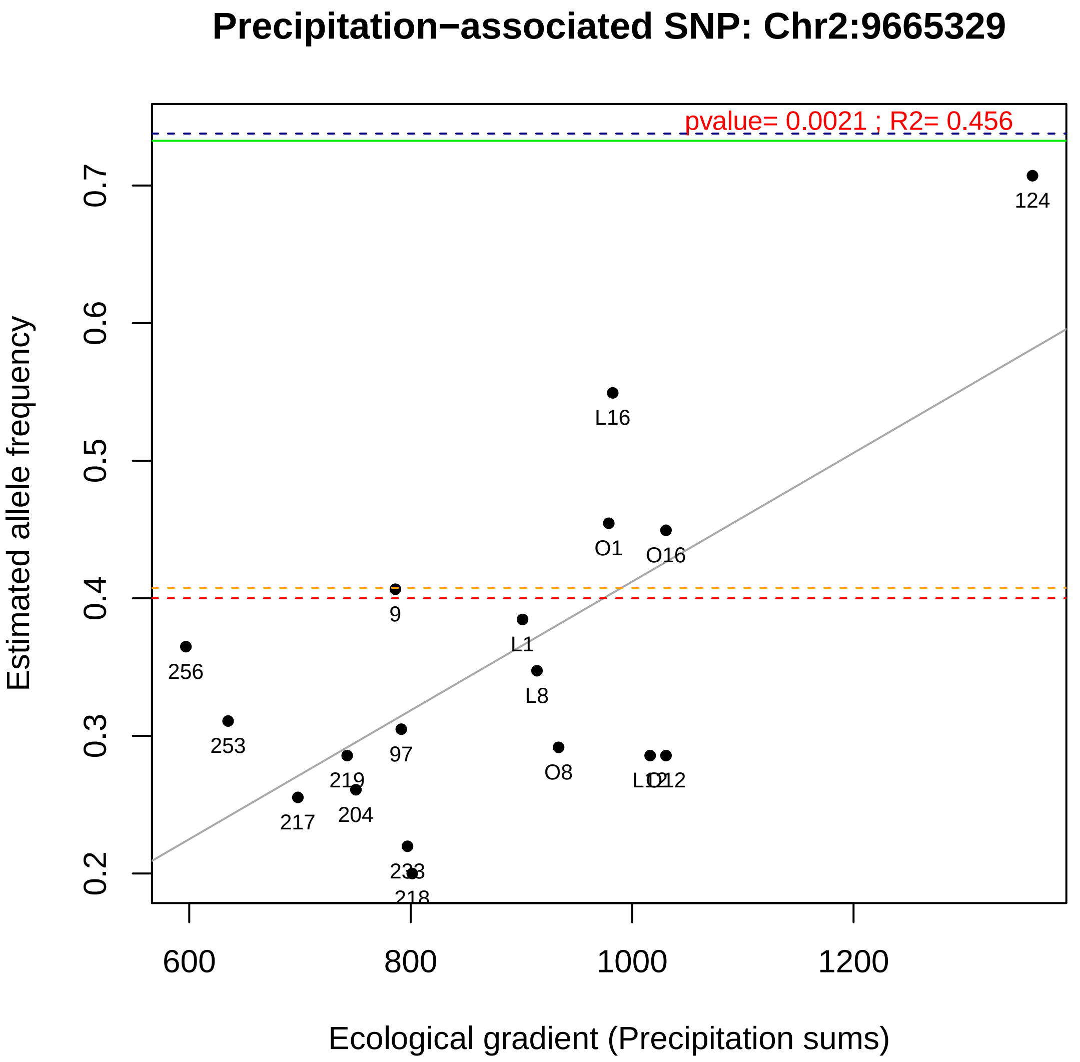
Precipitation−associated SNP: Chr2:9665329
pvalue= 0.0021 ; R2= 0.456
Estimated allele frequency
Ecological gradient (Precipitation sums)
0.2
0.3
0.4
0.5
0.6
0.7
600
800
1000
1200
124
L16
O1
O16
9
L1
256
L8
253
97
O8
219
204
217
218

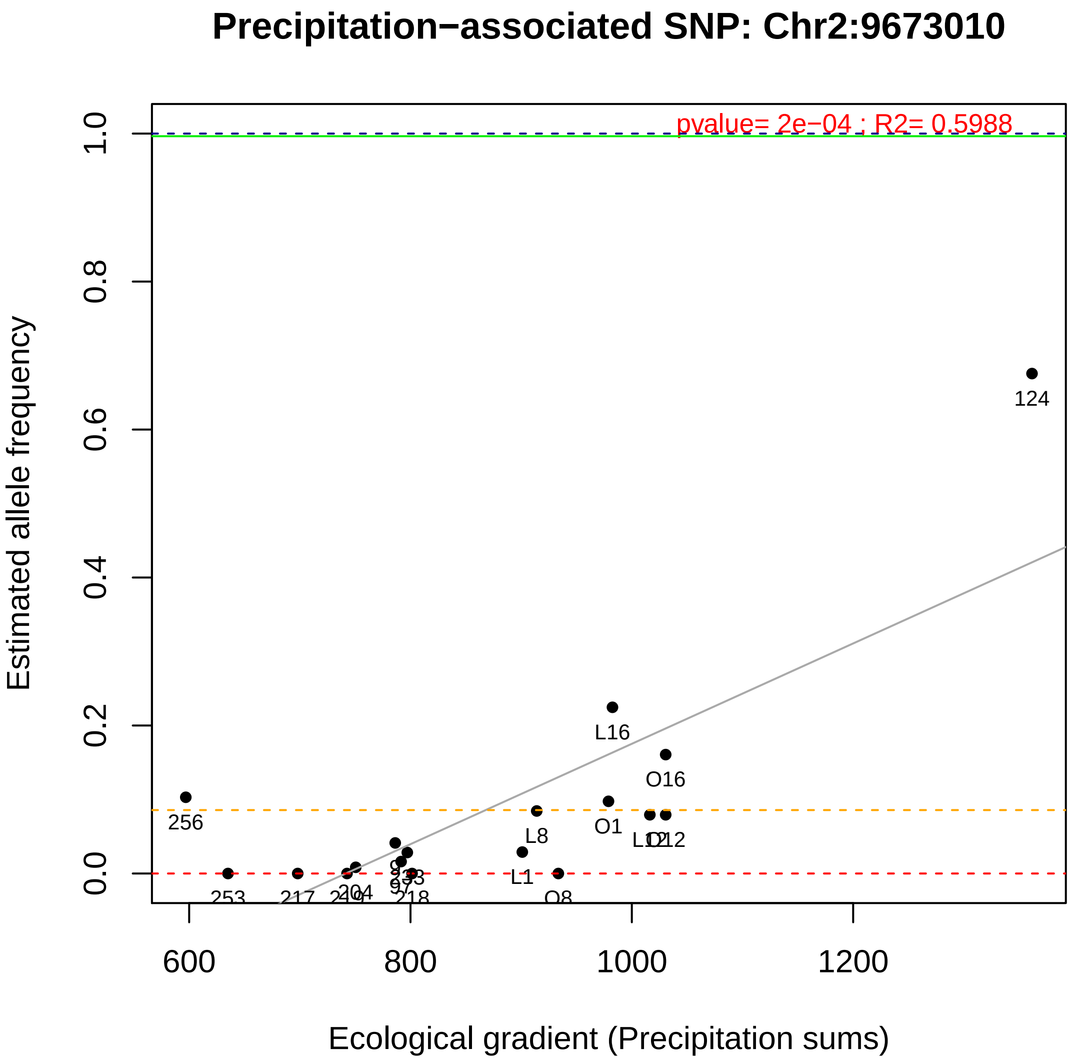

Precipitation−associated SNP: Chr2:9673010
pvalue= 2e−04 ; R2= 0.5988
Estimated allele frequency
0.0
0.2
0.4
0.6
0.8
1.0
600
800
1000
1200
Ecological gradient (Precipitation sums)
124
L16
O16
256
O1
L8
L1
O8
204
253
217
218

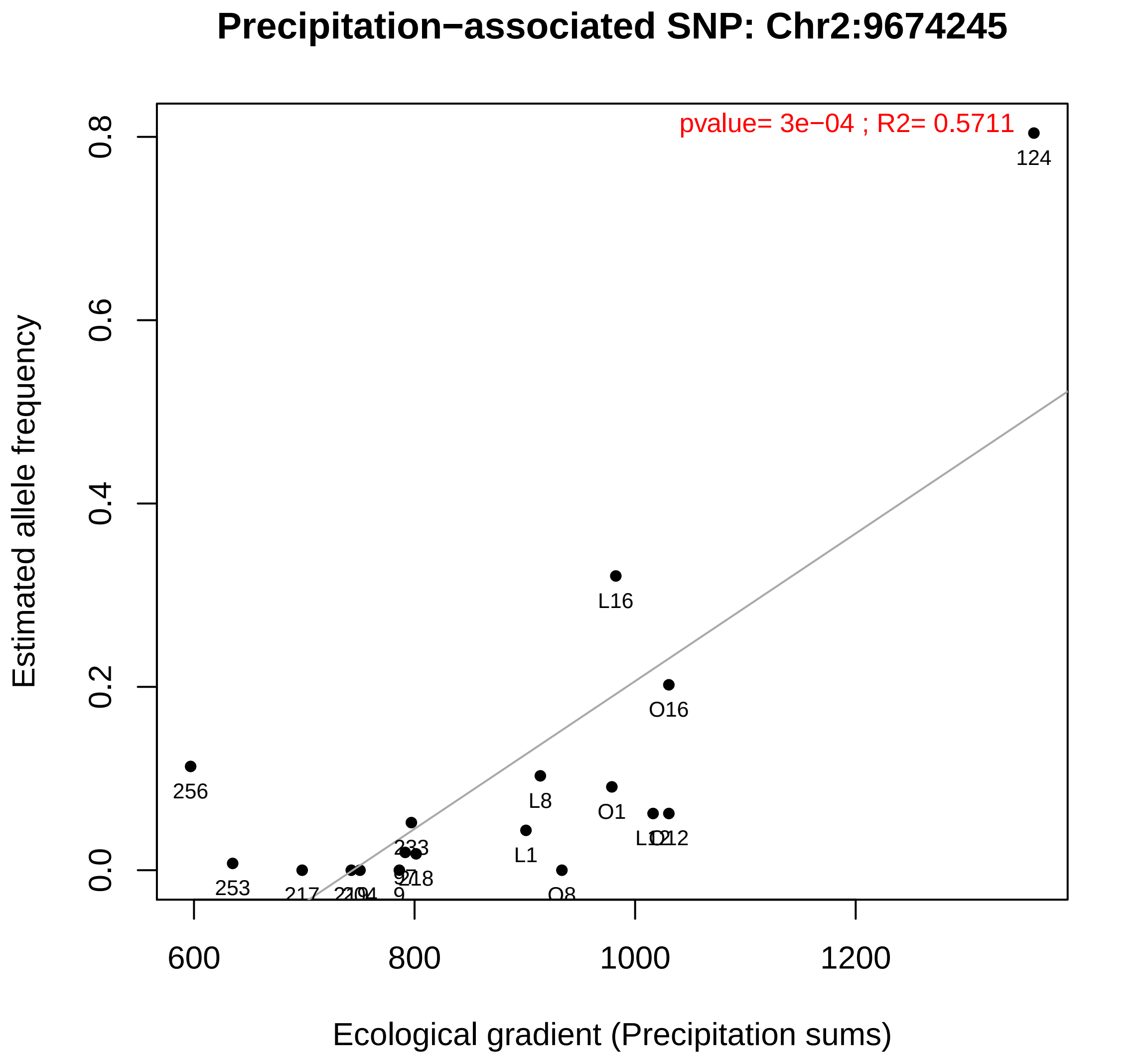
Precipitation−associated SNP: Chr2:9674245
pvalue= 3e−04 ; R2= 0.5711
Estimated allele frequency
Ecological gradient (Precipitation sums)
0.0
0.2
0.4
0.6
0.8
600
800
1000
1200
124
L16
O16
256
L8
O1
L1
233
218
253
217
O8

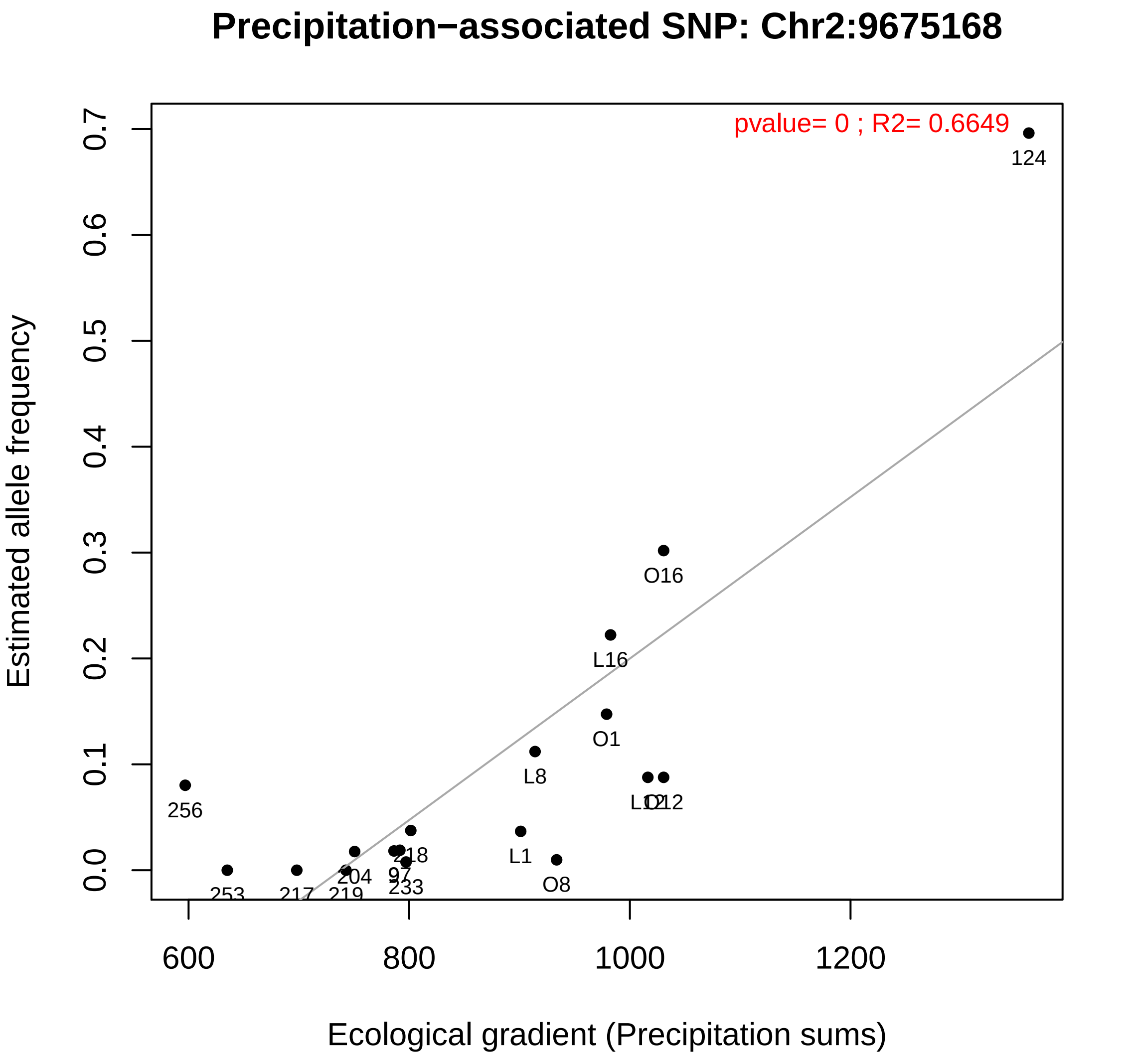

Precipitation−associated SNP: Chr2:9675168
pvalue= 0 ; R2= 0.6649
Estimated allele frequency
Ecological gradient (Precipitation sums)
0.0
0.1
0.2
0.3
0.4
0.5
0.6
0.7
600
800
1000
1200
124
O16
L16
O1
L8
256
L1
O8
233
253
217
219
204
97
218

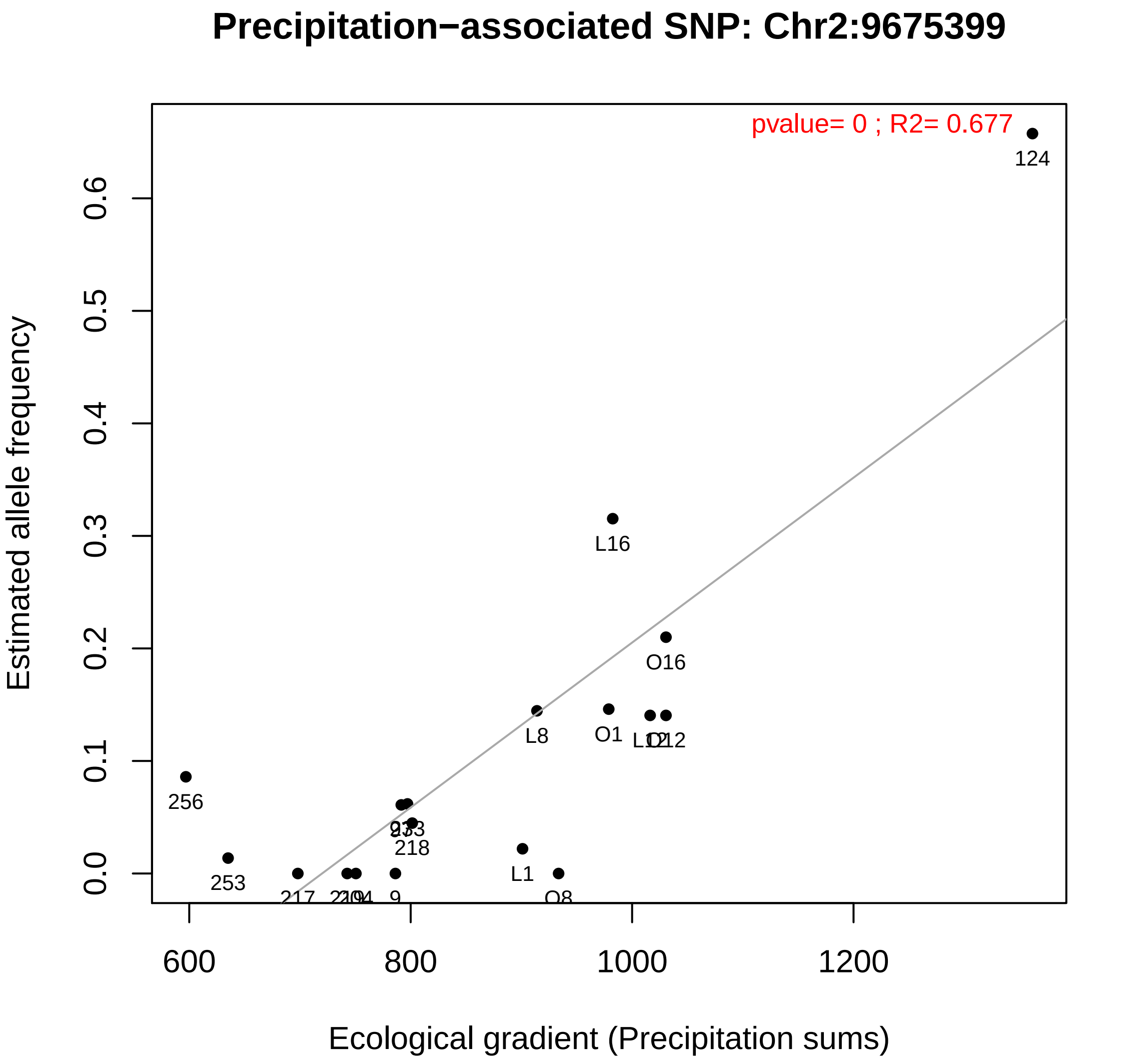
Precipitation−associated SNP: Chr2:9675399
pvalue= 0 ; R2= 0.677
124
L16
O16
L8
O1
L12
O12
256
253
217
218
L1
O8
9
0.0
0.1
0.2
0.3
0.4
0.5
0.6
600
800
1000
1200
Estimated allele frequency
Ecological gradient (Precipitation sums)

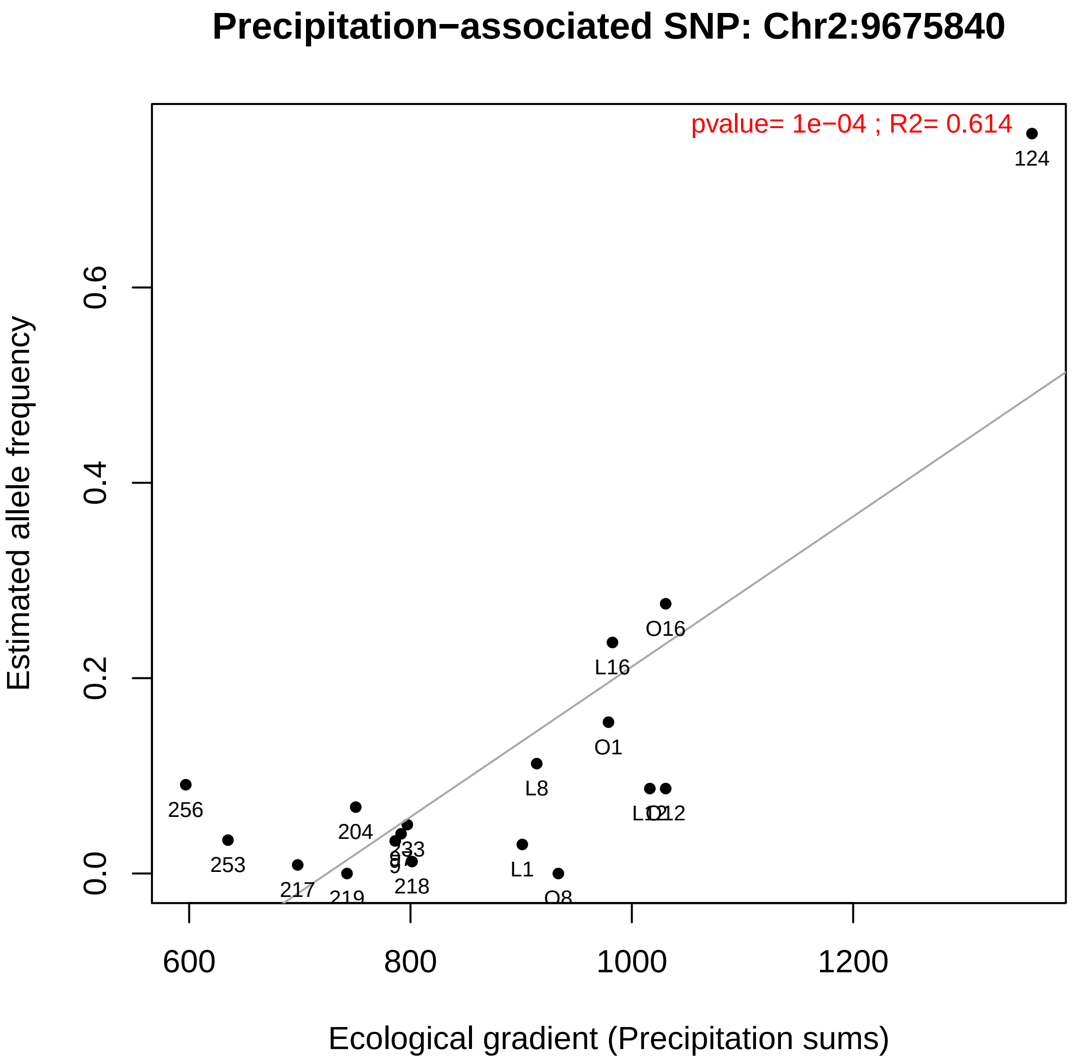

Precipitation−associated SNP: Chr2:9675840
pvalue= 1e−04 ; R2= 0.614
124
O16
L16
O1
L8
L12
O12
256
204
253
217
219
233
9
218
L1
O8
0.0
0.2
0.4
0.6
600
800
1000
1200
Estimated allele frequency
Ecological gradient (Precipitation sums)

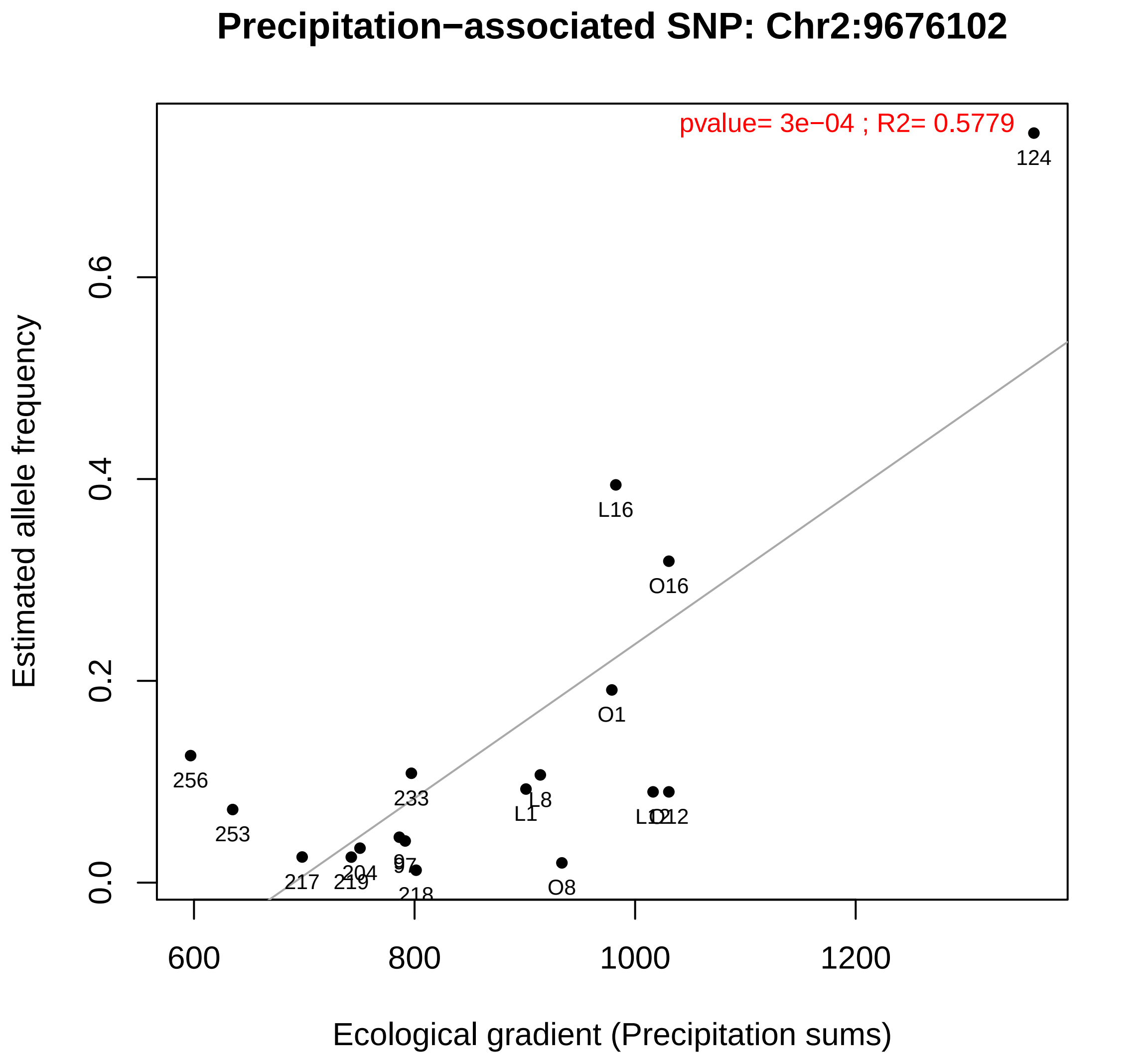

Precipitation−associated SNP: Chr2:9676102
pvalue= 3e−04 ; R2= 0.5779
124
L16
O16
O1
256
233
L8
L1
L12
O12
253
217
219
204
9
97
218
O8
0.0
0.2
0.4
0.6
600
800
1000
1200
Estimated allele frequency
Ecological gradient (Precipitation sums)

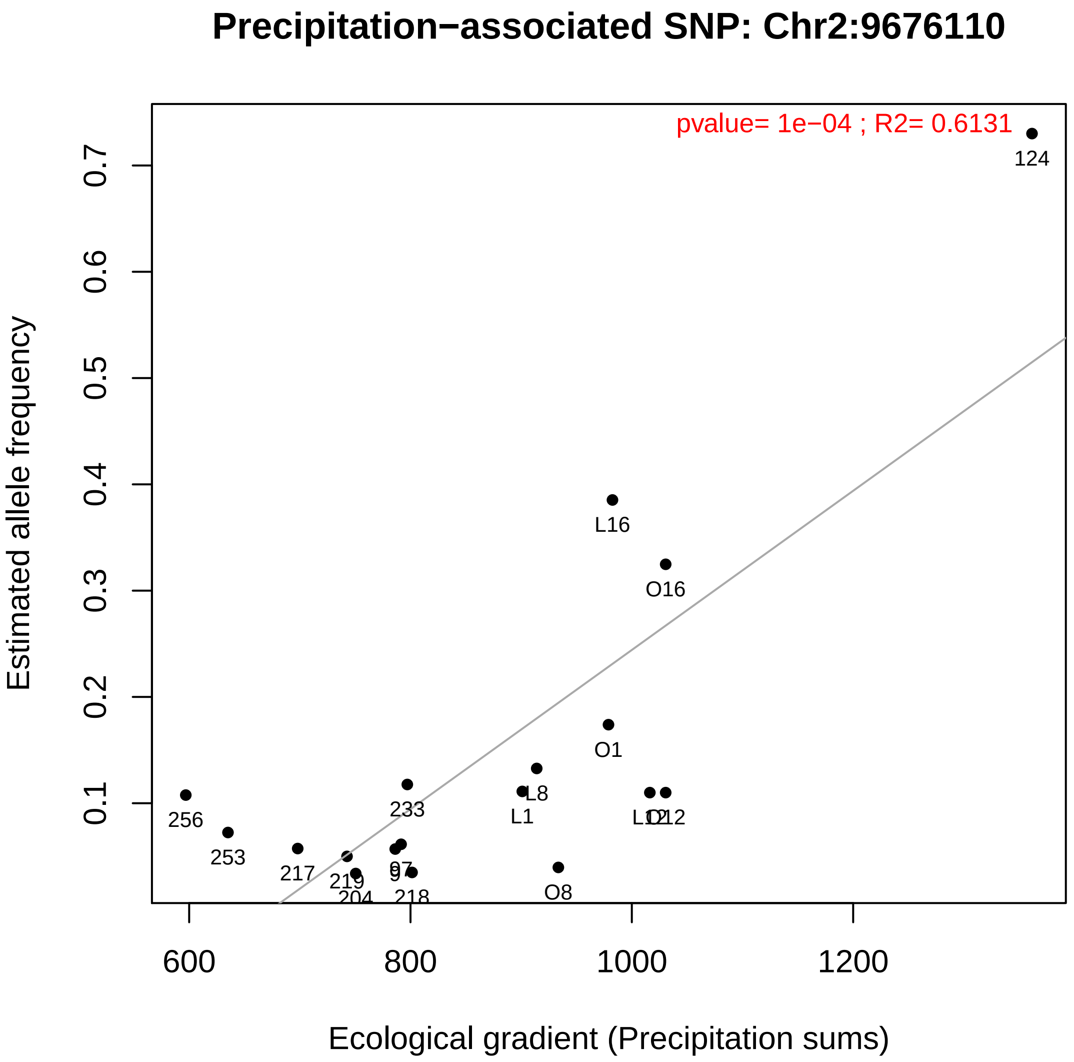

Precipitation−associated SNP: Chr2:9676110
pvalue= 1e−04 ; R2= 0.6131
Estimated allele frequency
Ecological gradient (Precipitation sums)
0.1
0.2
0.3
0.4
0.5
0.6
0.7
600
800
1000
1200
124
L16
O16
O1
L8
L1
233
256
253
217
219
204
97
218
O8
L12
O12

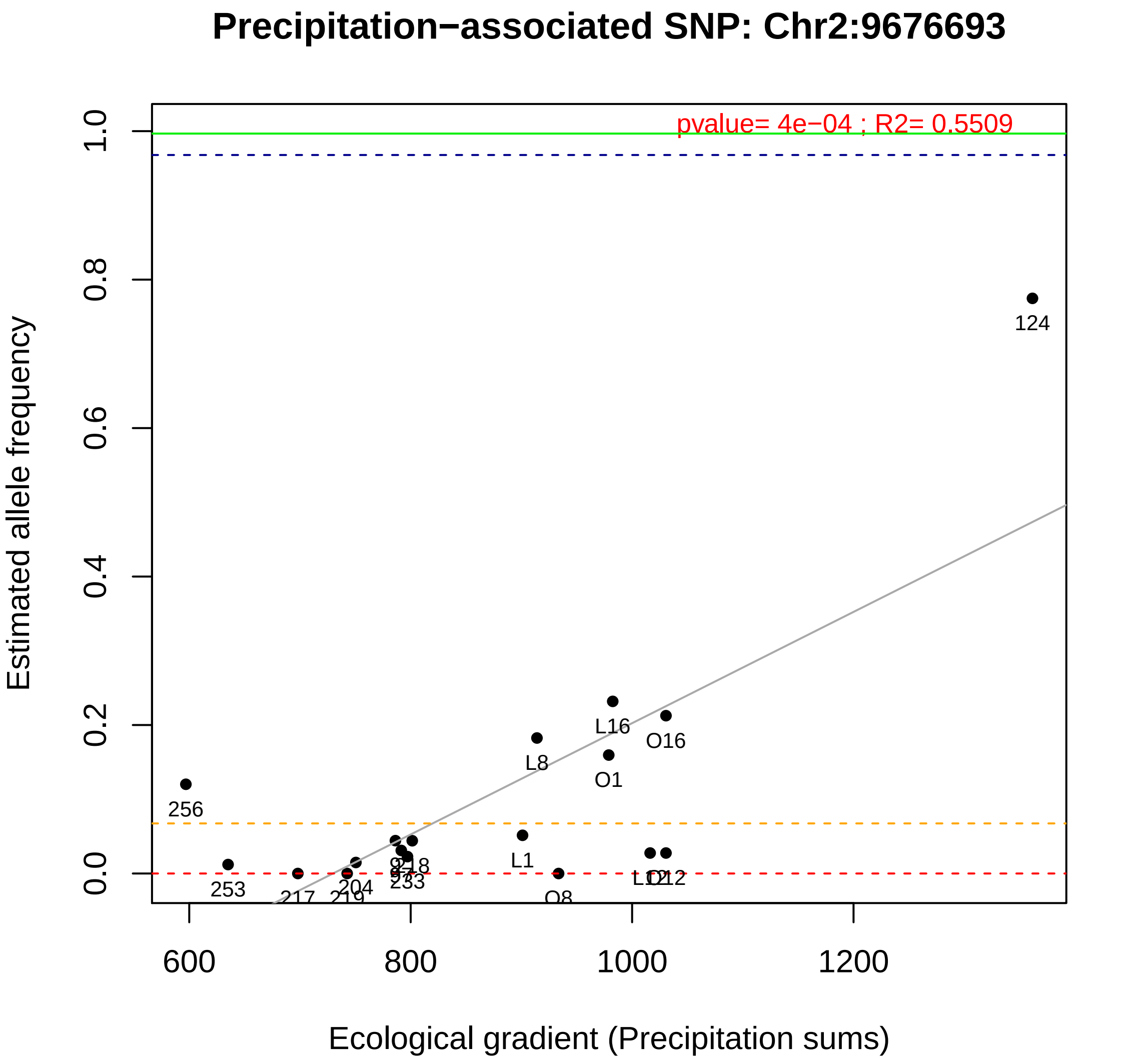
Precipitation−associated SNP: Chr2:9676693
pvalue= 4e−04 ; R2= 0.5509
Estimated allele frequency
Ecological gradient (Precipitation sums)
0.0
0.2
0.4
0.6
0.8
1.0
600
800
1000
1200
124
L16
O16
L8
O1
256
253
204
233
L1
O12

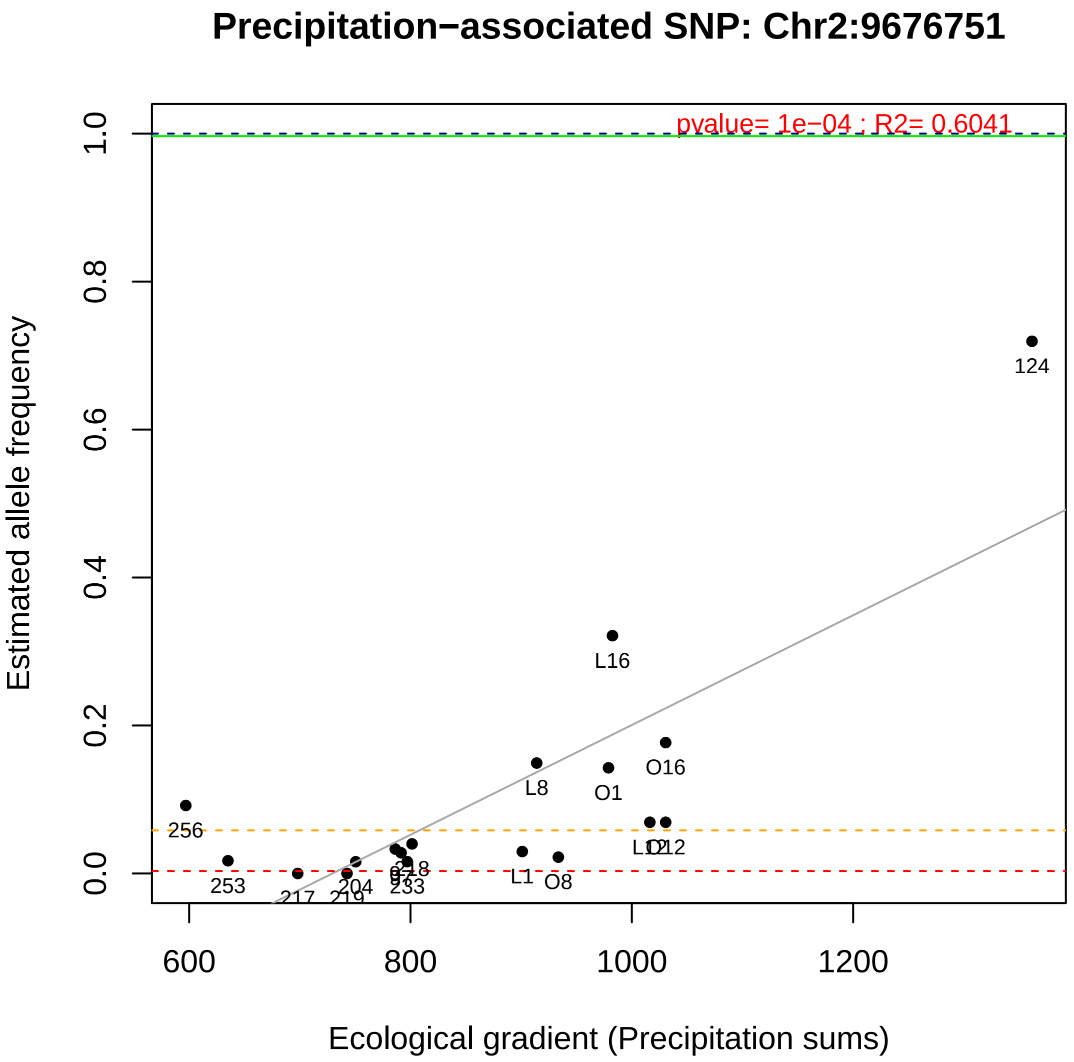

Precipitation−associated SNP: Chr2:9676751
pvalue= 1e−04 ; R2= 0.6041
Estimated allele frequency
0.0
0.2
0.4
0.6
0.8
1.0
Ecological gradient (Precipitation sums)
600
800
1000
1200
124
L16
O16
L8
O1
256
253
L1
O8

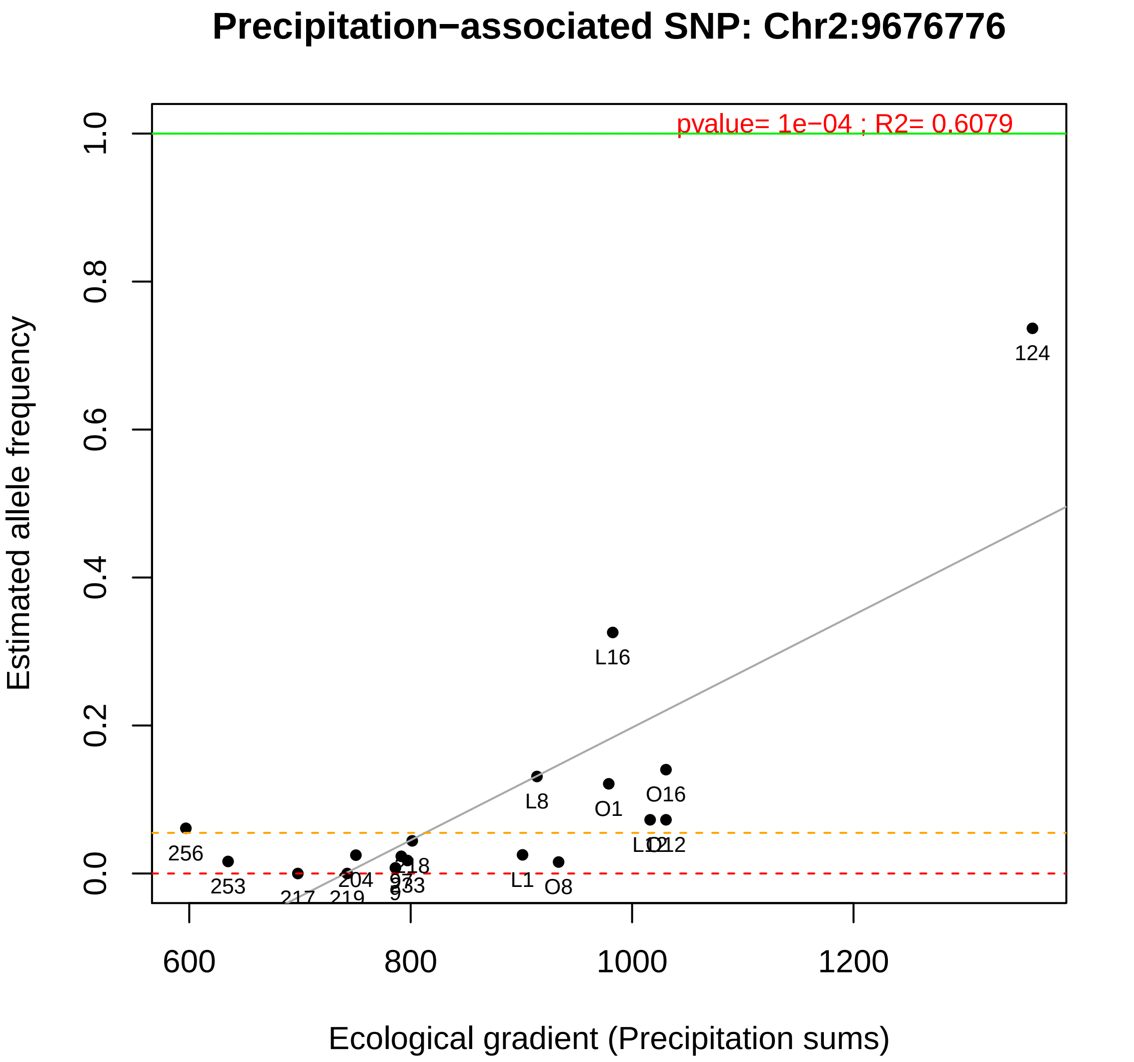

Precipitation−associated SNP: Chr2:9676776
pvalue= 1e−04 ; R2= 0.6079
Estimated allele frequency
0.0
0.2
0.4
0.6
0.8
1.0
Ecological gradient (Precipitation sums)
600
800
1000
1200
124
L16
L8
O1
O16
L12
O12
256
253
217
219
204
L18
97
233
9
L1
O8

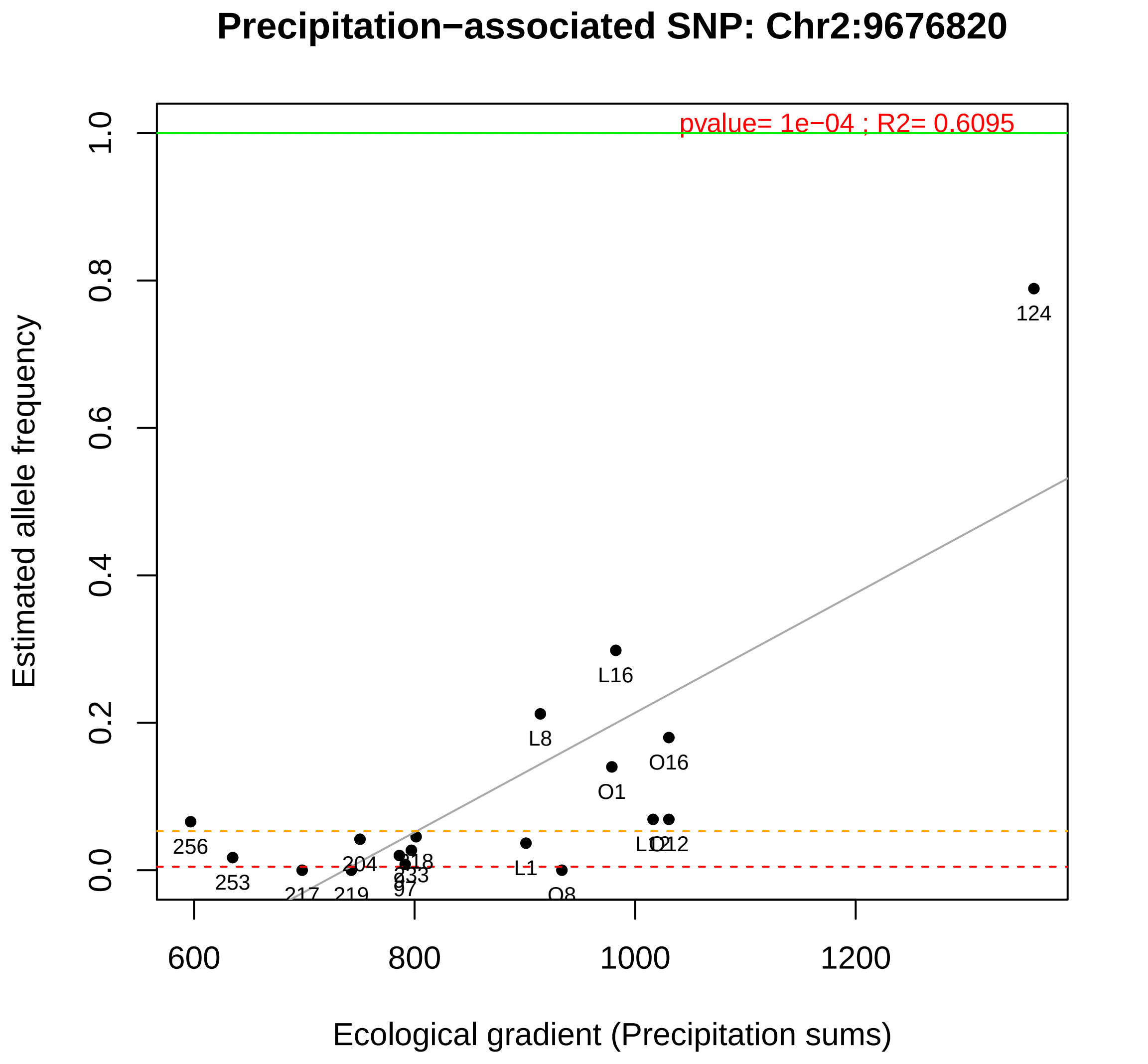
Precipitation−associated SNP: Chr2:9676820
pvalue= 1e−04 ; R2= 0.6095
Estimated allele frequency
0.0
0.2
0.4
0.6
0.8
1.0
Ecological gradient (Precipitation sums)
600
800
1000
1200
124
L16
L8
O16
O1
256
253
217
219
204
218
233
97
L1
O8
L12
O12

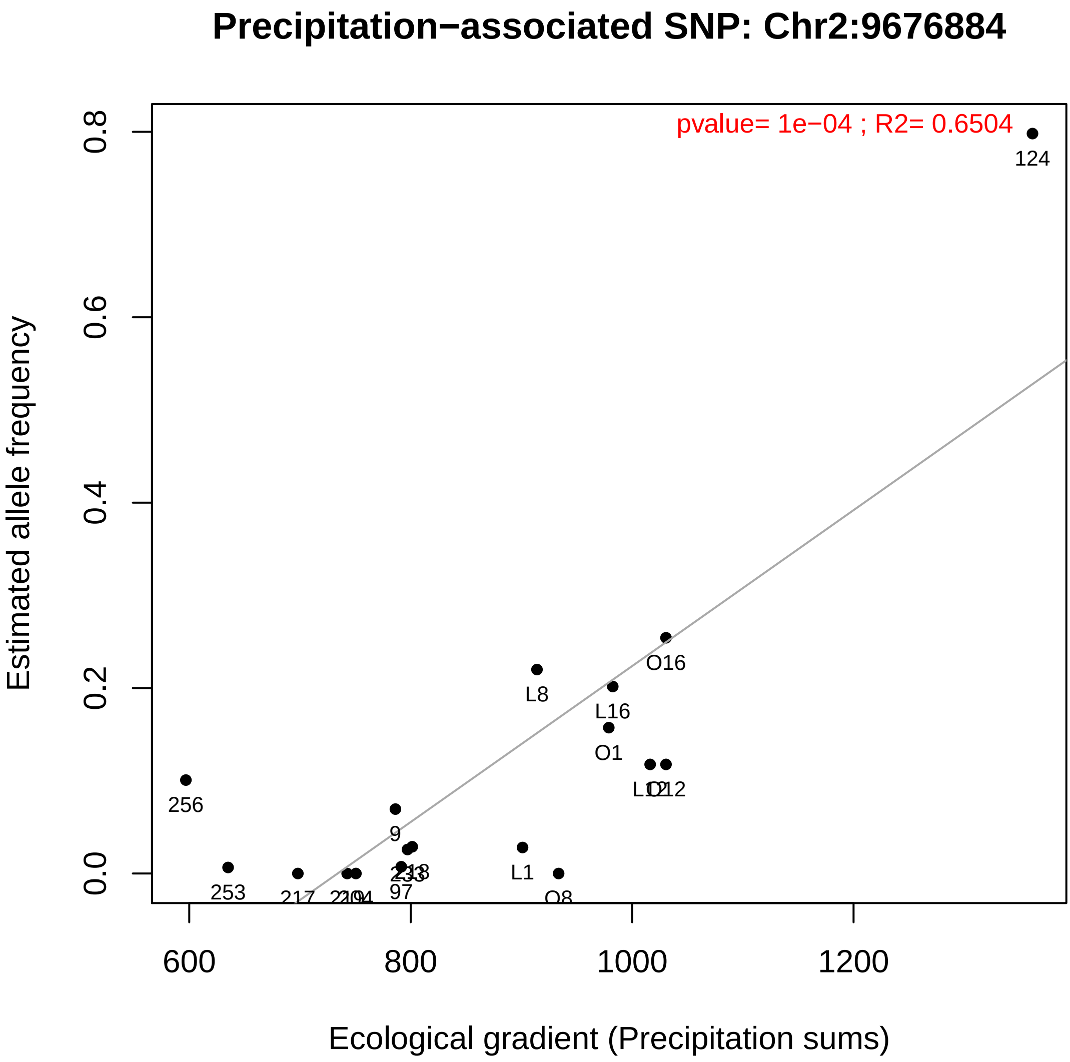

Precipitation−associated SNP: Chr2:9676884
pvalue= 1e−04 ; R2= 0.6504
Estimated allele frequency
0.0
0.2
0.4
0.6
0.8
600
800
1000
1200
Ecological gradient (Precipitation sums)
124
O16
L8
L16
O1
256
9
L1
253
217
97

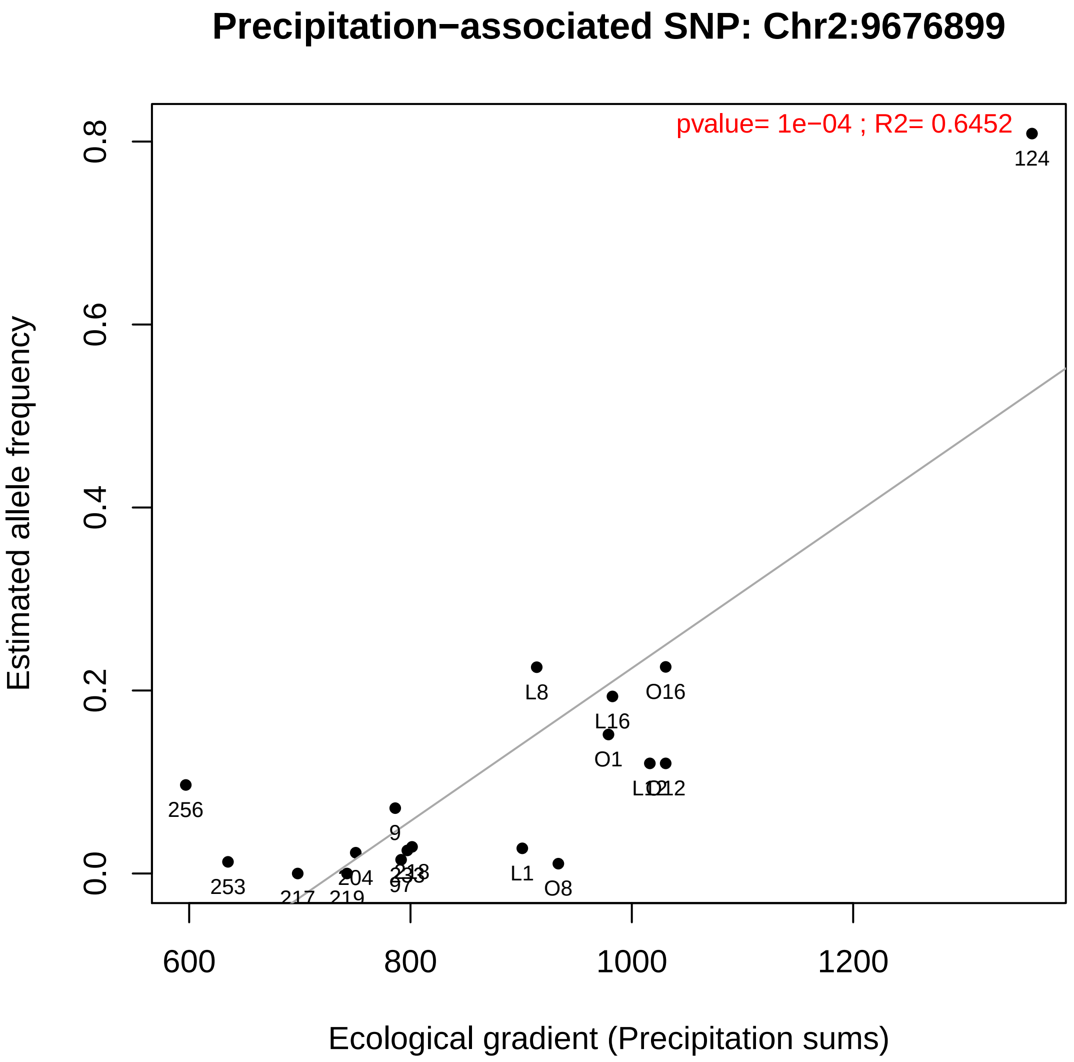

Precipitation−associated SNP: Chr2:9676899
pvalue= 1e−04 ; R2= 0.6452
Estimated allele frequency
Ecological gradient (Precipitation sums)
0.0
0.2
0.4
0.6
0.8
600
800
1000
1200
124
L8
O16
L16
O1
256
9
253
204
217
219
97
L1
O8

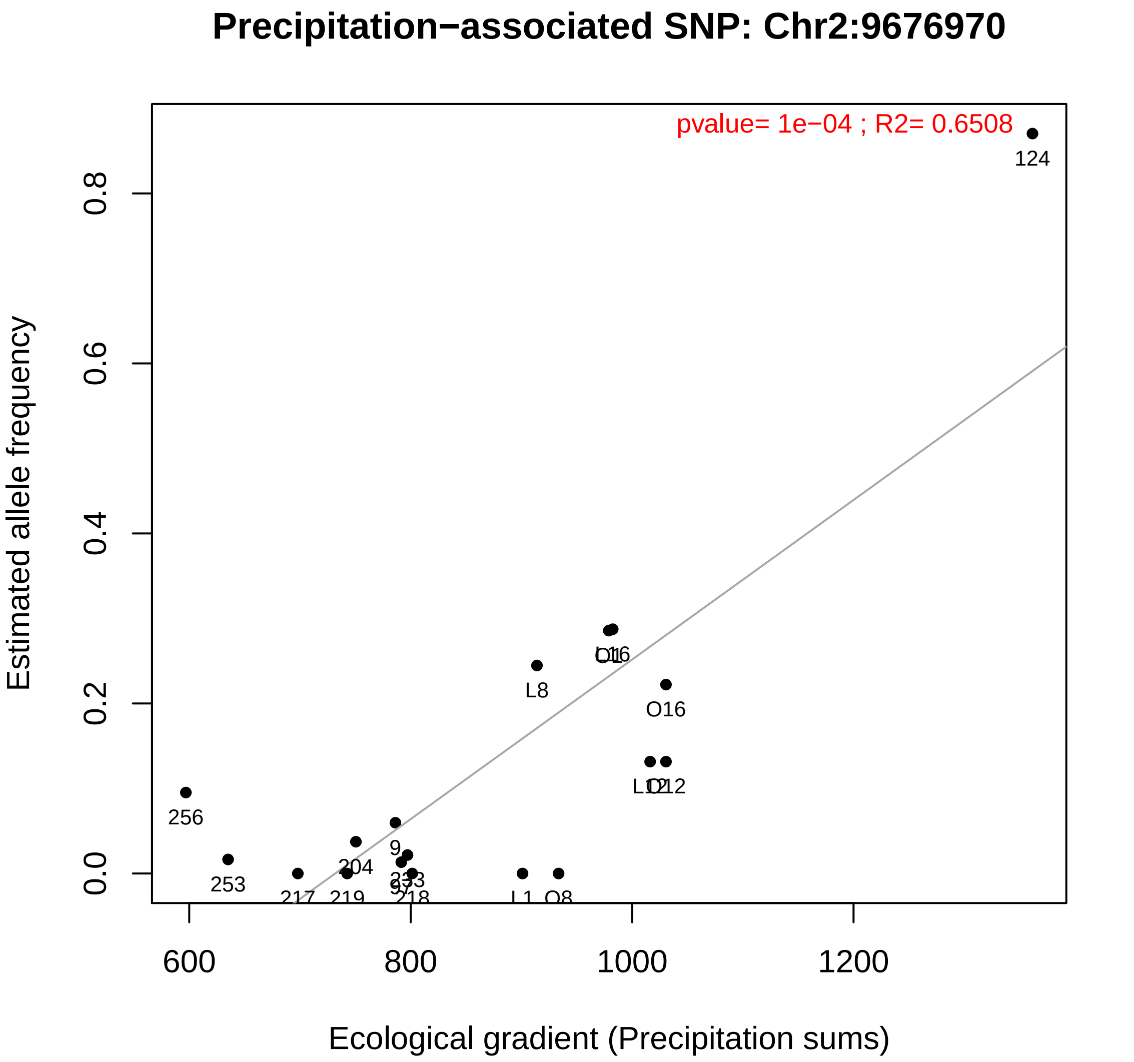

Precipitation−associated SNP: Chr2:9676970
pvalue= 1e−04 ; R2= 0.6508
Estimated allele frequency
Ecological gradient (Precipitation sums)
0.0
0.2
0.4
0.6
0.8
600
800
1000
1200
124
L8
O16
256
253
204
9
217
219
218
L1
O8

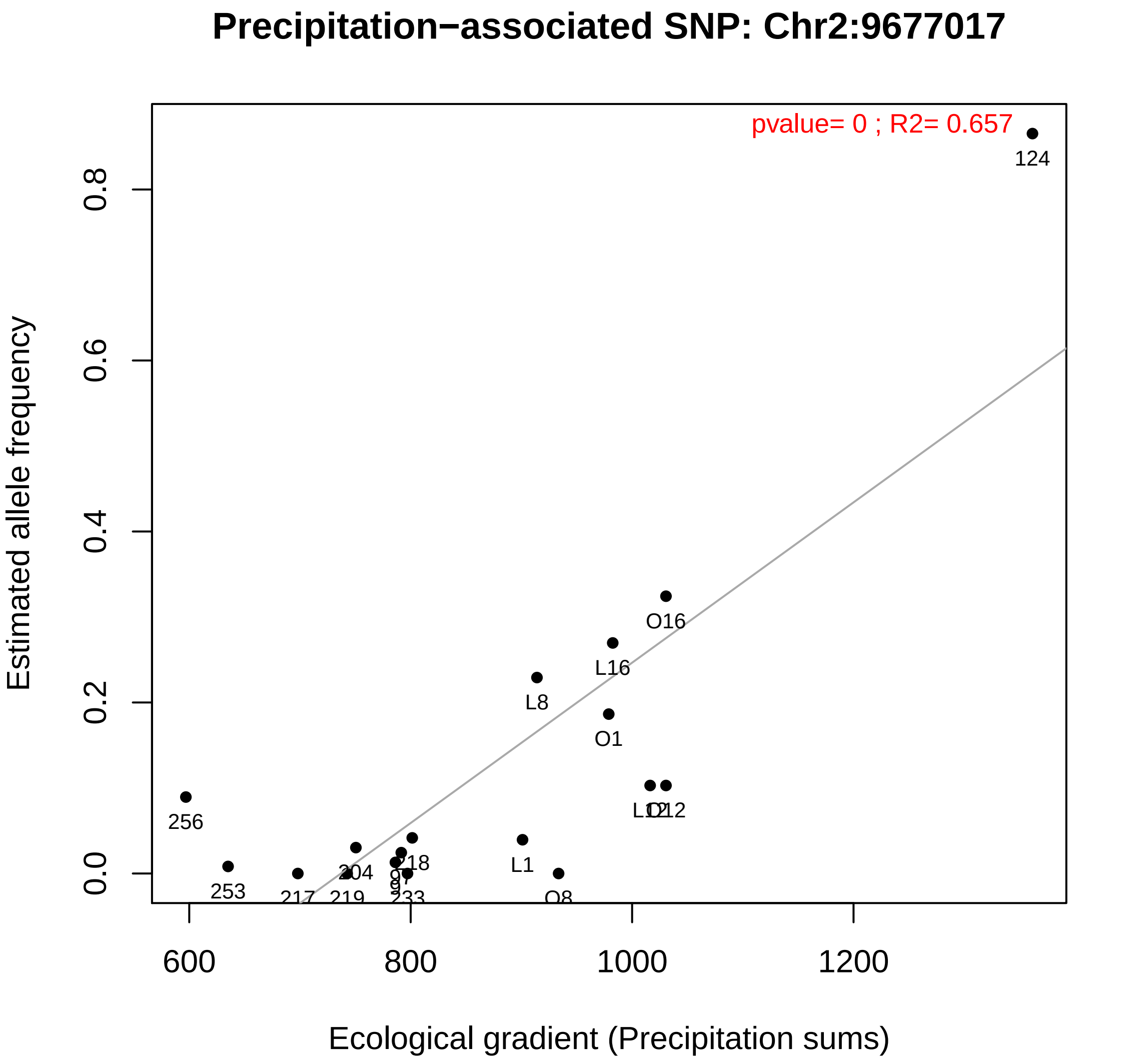
Precipitation−associated SNP: Chr2:9677017
pvalue= 0 ; R2= 0.657
124
O16
L16
L8
O1
L12
O12
256
253
217
219
204
218
97
233
L1
O8
0.0
0.2
0.4
0.6
0.8
600
800
1000
1200
Estimated allele frequency
Ecological gradient (Precipitation sums)

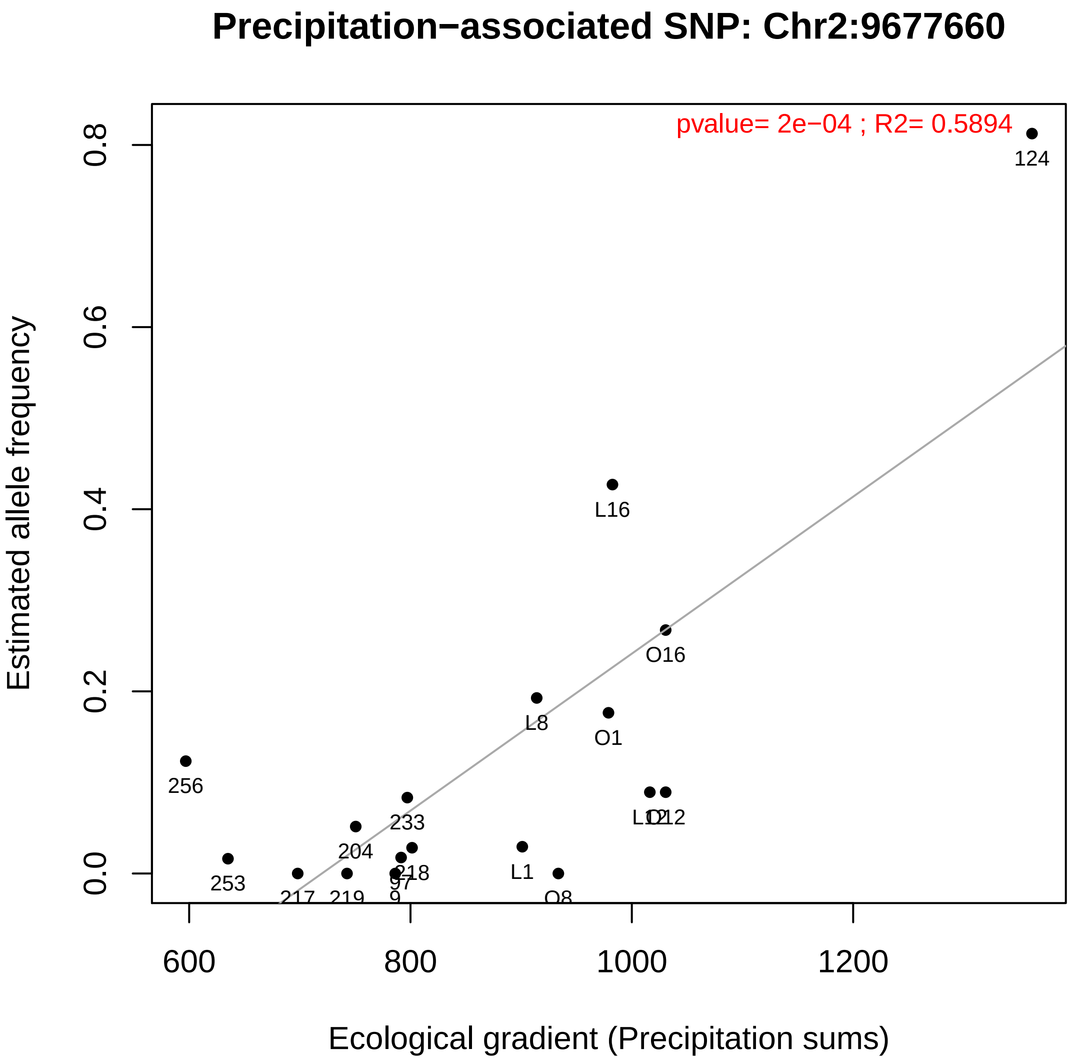

Precipitation−associated SNP: Chr2:9677660
pvalue= 2e−04 ; R2= 0.5894
124
L16
O16
L8
O1
256
233
204
253
218
97
L1
217
219
9
O8
0.0
0.2
0.4
0.6
0.8
600
800
1000
1200
Estimated allele frequency
Ecological gradient (Precipitation sums)

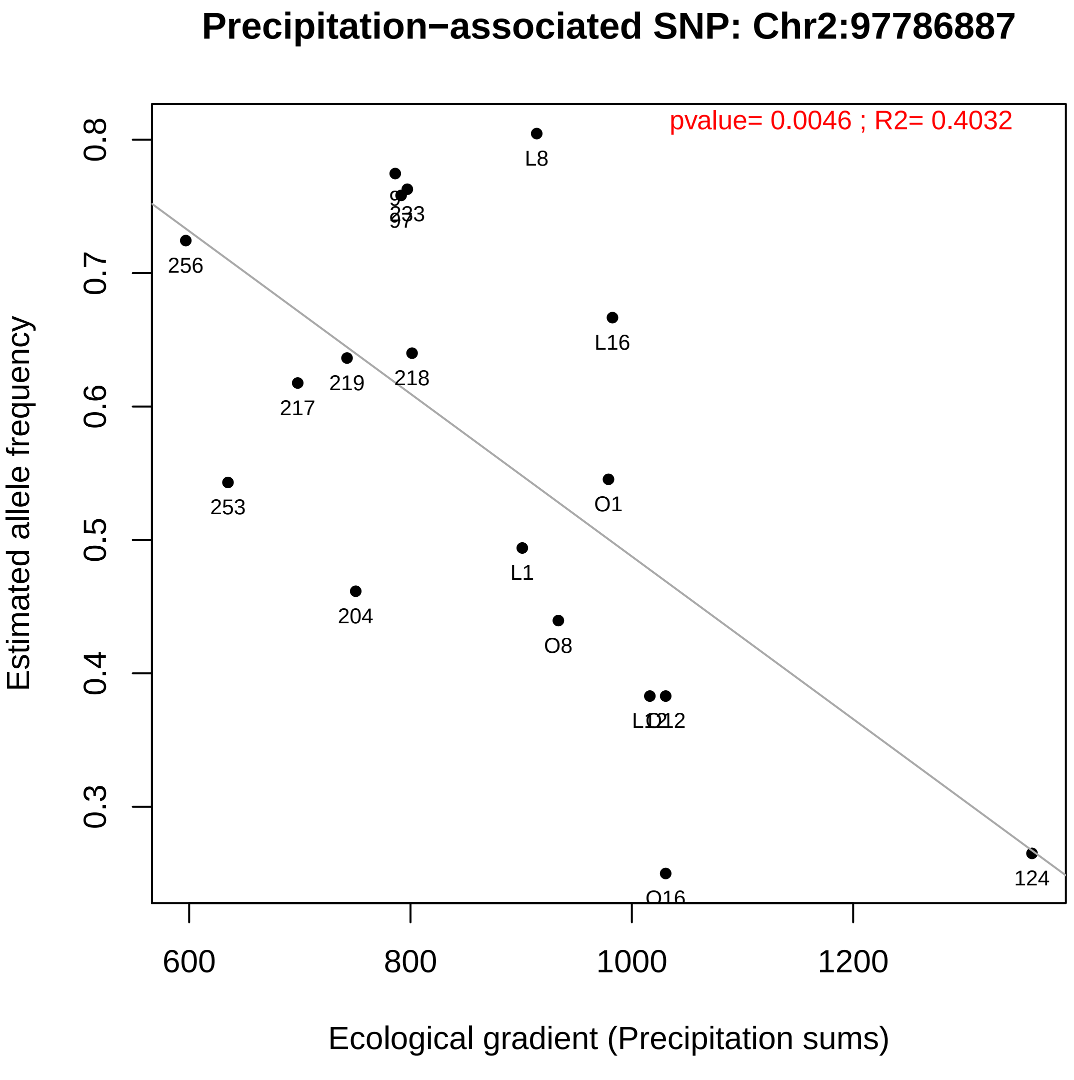
Precipitation−associated SNP: Chr2:97786887
pvalue= 0.0046 ; R2= 0.4032
Estimated allele frequency
Ecological gradient (Precipitation sums)
0.3
0.4
0.5
0.6
0.7
0.8
600
800
1000
1200
L8
233
97
256
L16
219
218
217
253
O1
L1
204
O8
L12
O12
124
O16

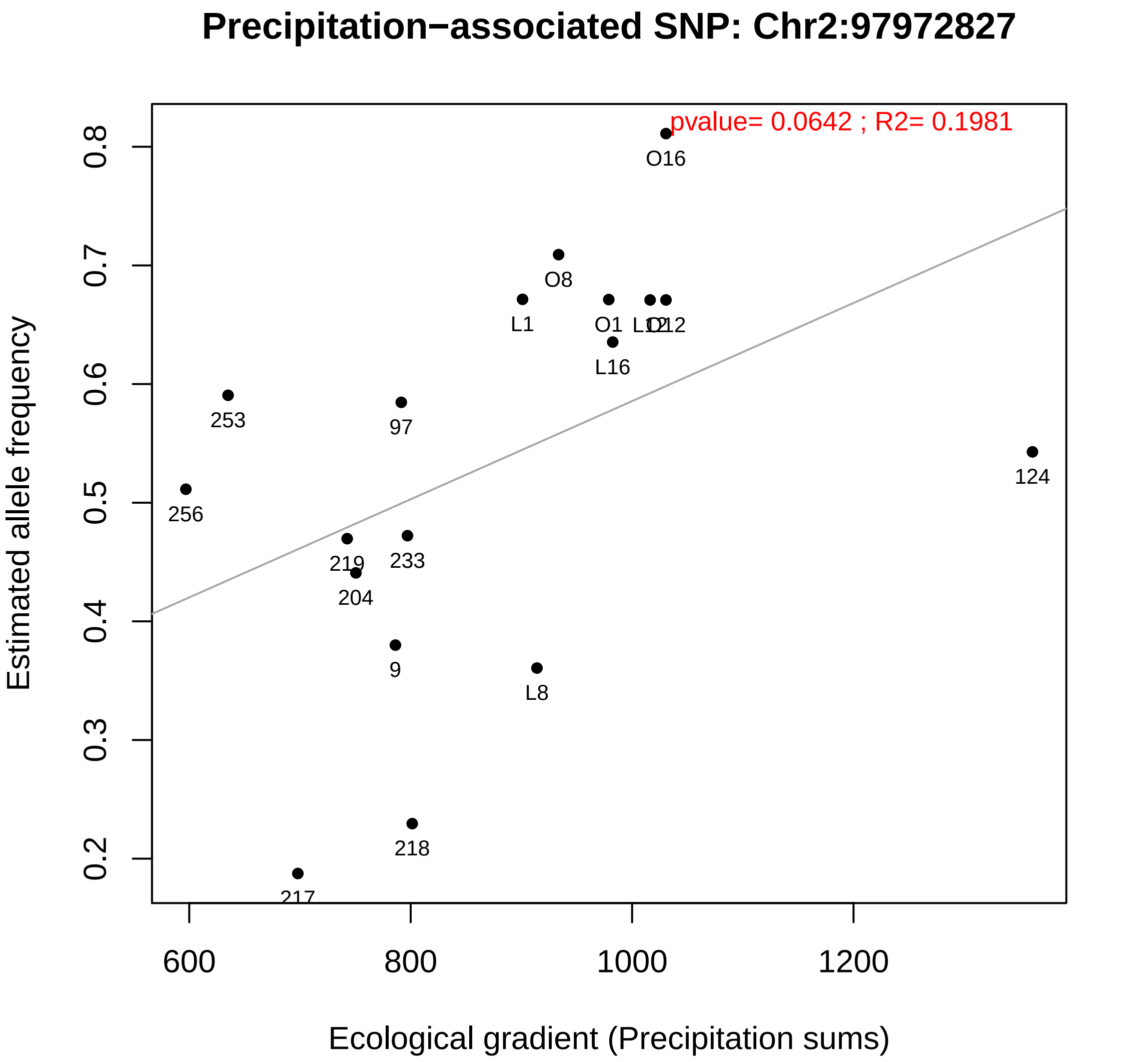
Precipitation−associated SNP: Chr2:97972827
pvalue= 0.0642 ; R2= 0.1981
O16
O8
L1
O1
L12
O12
L16
253
97
124
256
219
233
204
9
L8
218
217
0.2
0.3
0.4
0.5
0.6
0.7
0.8
600
800
1000
1200
Estimated allele frequency
Ecological gradient (Precipitation sums)

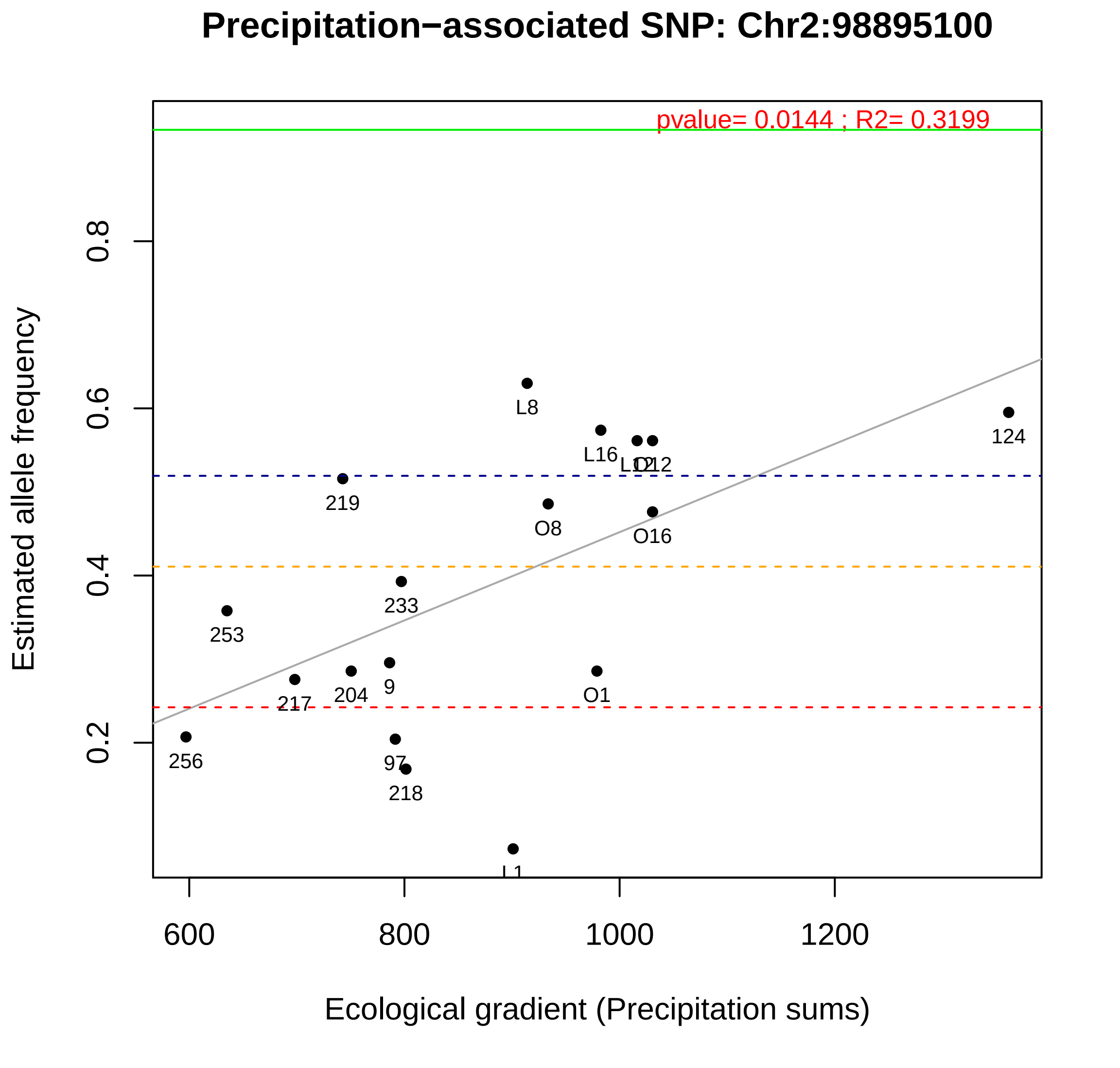

Precipitation−associated SNP: Chr2:98895100
pvalue= 0.0144 ; R2= 0.3199
Estimated allele frequency
0.2
0.4
0.6
0.8
600
800
1000
1200
Ecological gradient (Precipitation sums)
L8
124
L16
O12
219
O8
O16
233
253
9
204
217
O1
256
97
218

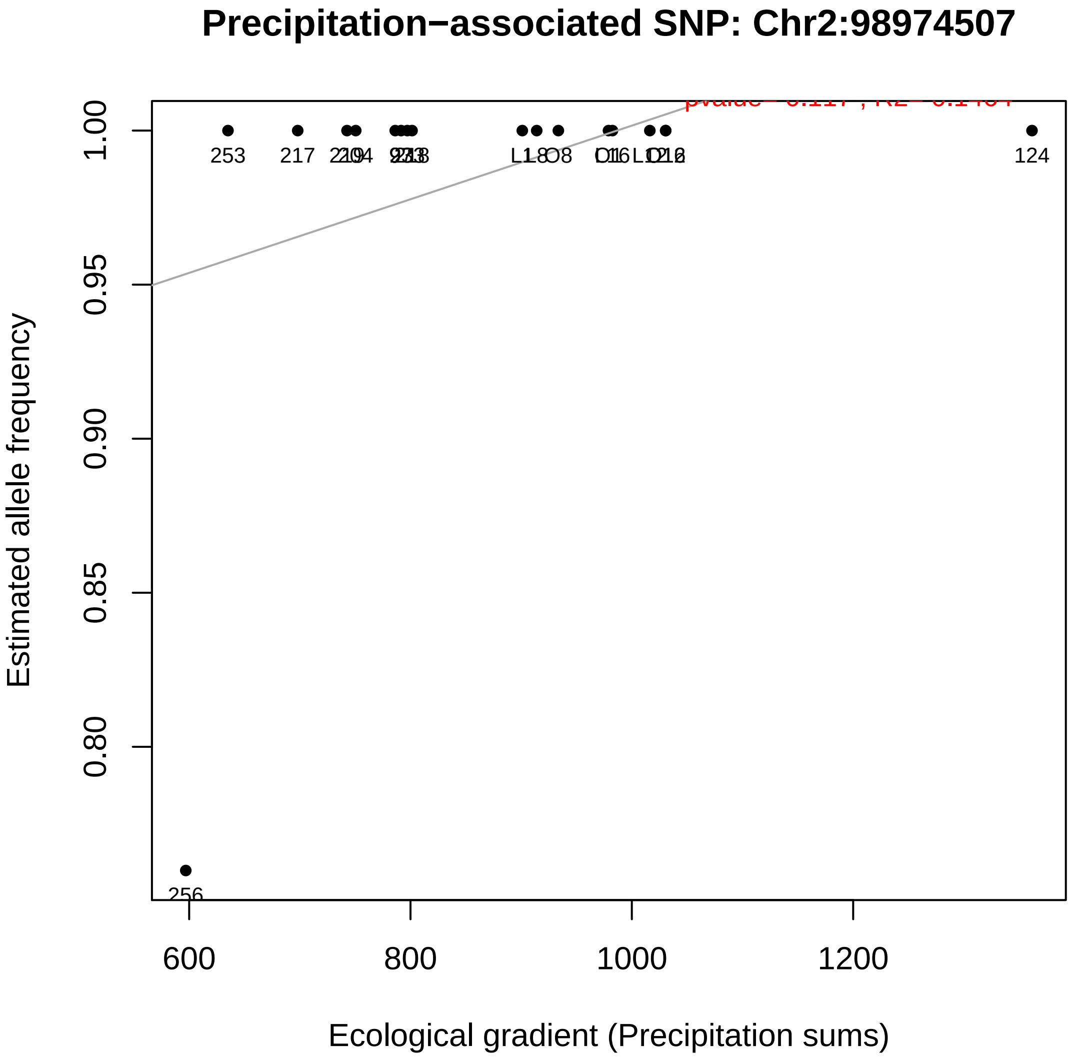
Precipitation−associated SNP: Chr2:98974507
Estimated allele frequency
1.00
0.95
0.90
0.85
0.80
253
217
124
256
600
800
1000
1200
Ecological gradient (Precipitation sums)

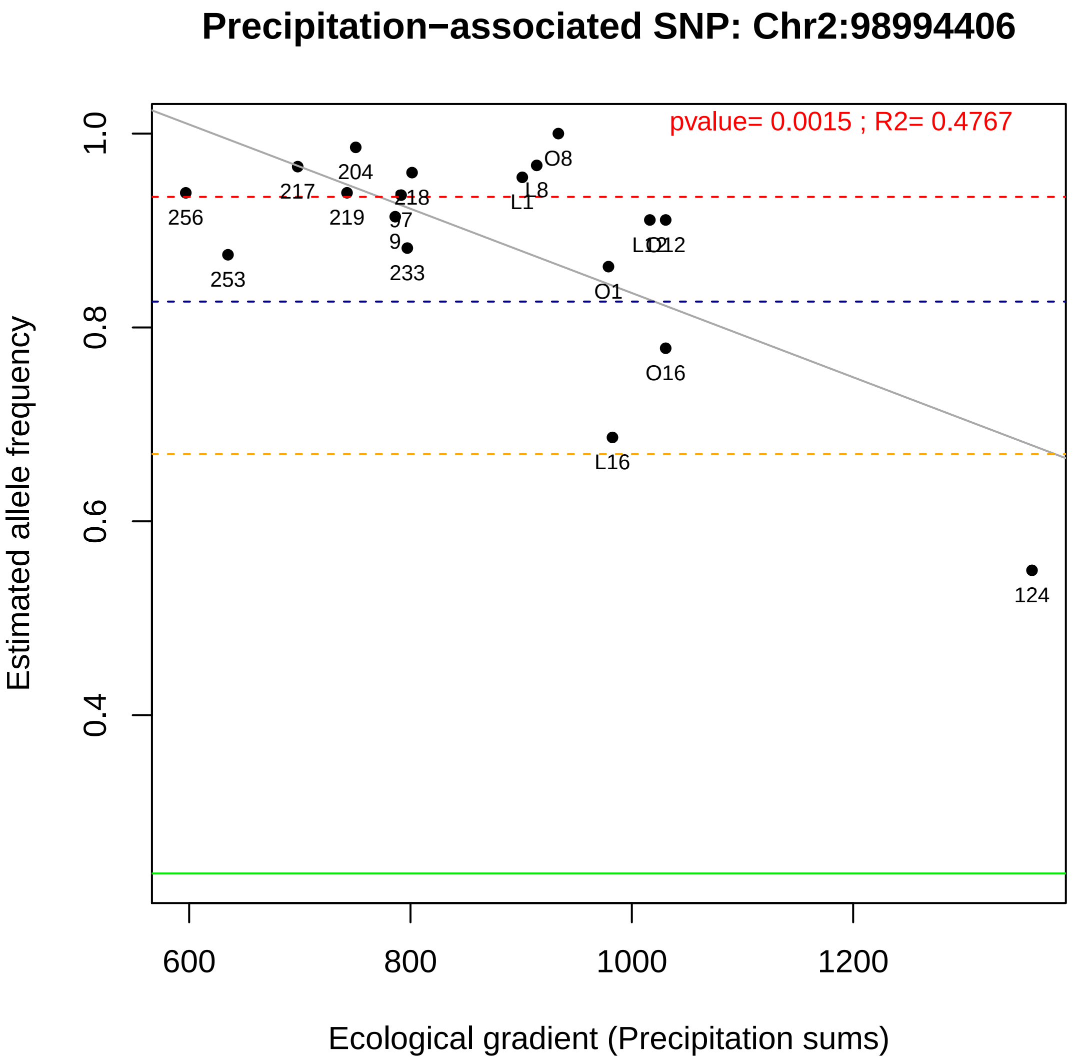

Precipitation−associated SNP: Chr2:98994406
pvalue= 0.0015 ; R2= 0.4767
Estimated allele frequency
Ecological gradient (Precipitation sums)
1.0
0.8
0.6
0.4
600
800
1000
1200
256
253
217
204
219
218
97
9
233
L1
L8
O8
L12
O12
O1
O16
L16
124

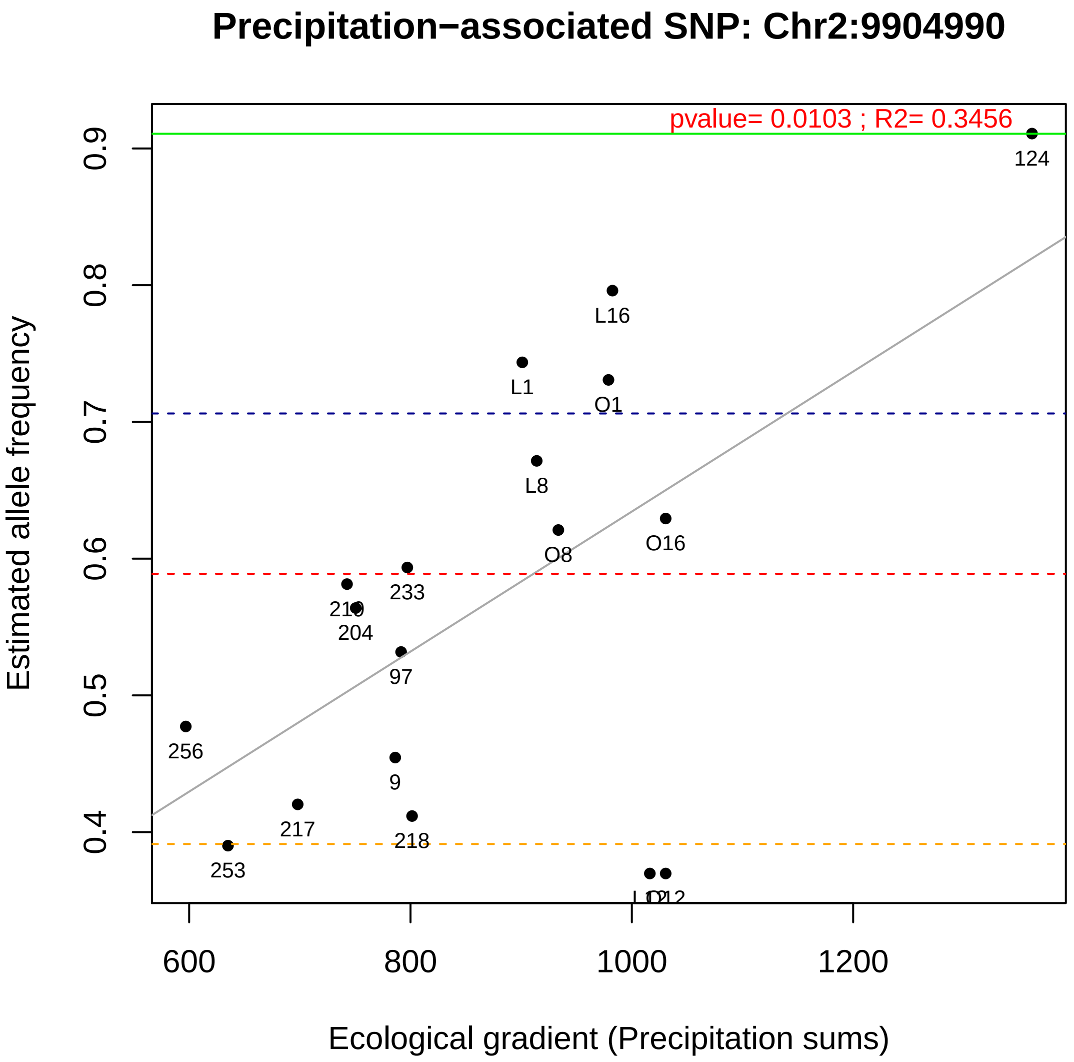
Precipitation−associated SNP: Chr2:9904990
pvalue= 0.0103 ; R2= 0.3456
Estimated allele frequency
Ecological gradient (Precipitation sums)
0.4
0.5
0.6
0.7
0.8
0.9
600
800
1000
1200
124
L16
L1
O1
L8
O16
O8
233
204
97
256
9
217
218
253

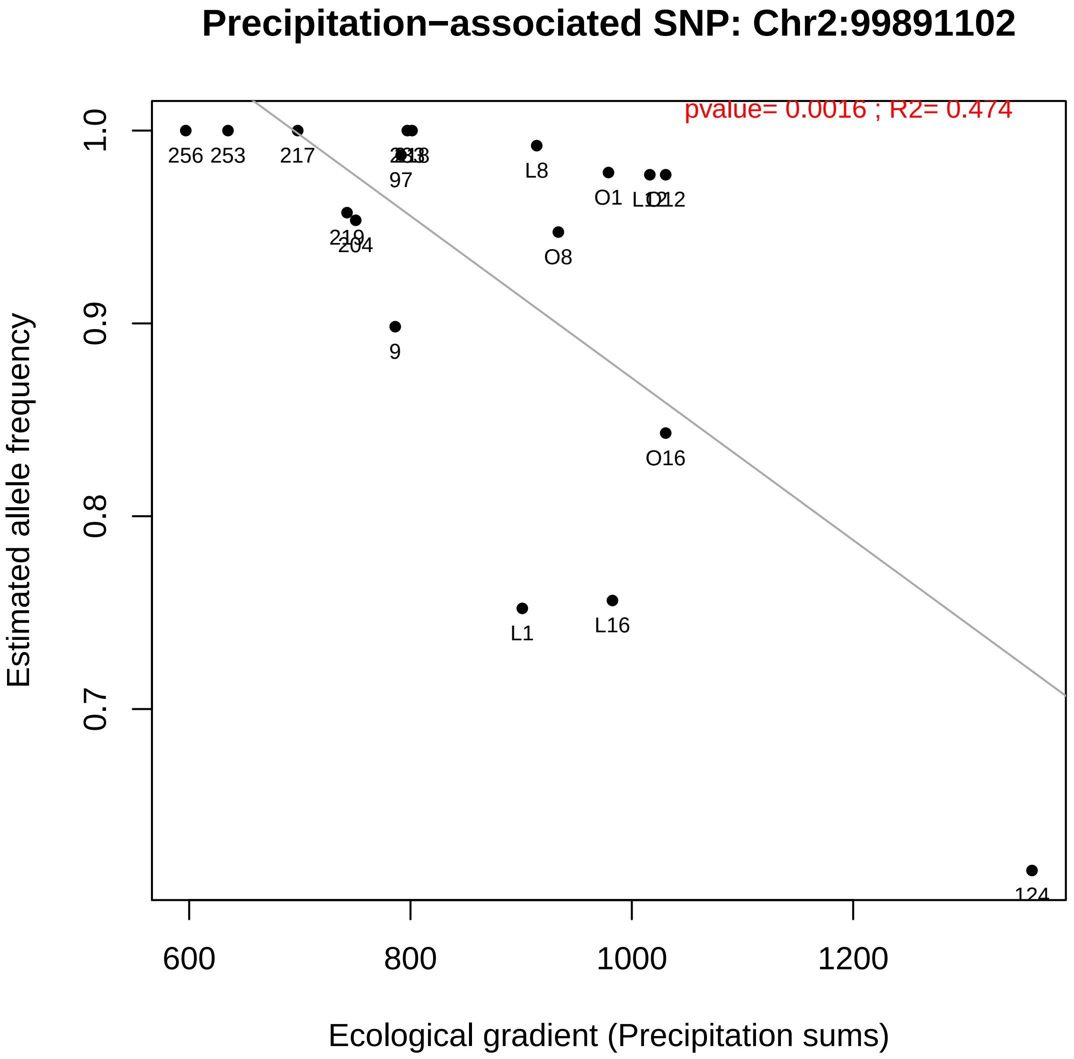
Precipitation−associated SNP: Chr2:99891102
pvalue= 0.0016 ; R2= 0.474
Estimated allele frequency
1.0
0.9
0.8
0.7
Ecological gradient (Precipitation sums)
600
800
1000
1200
256
253
217
97
L8
O1
219
204
O8
9
O16
L1
L16
124

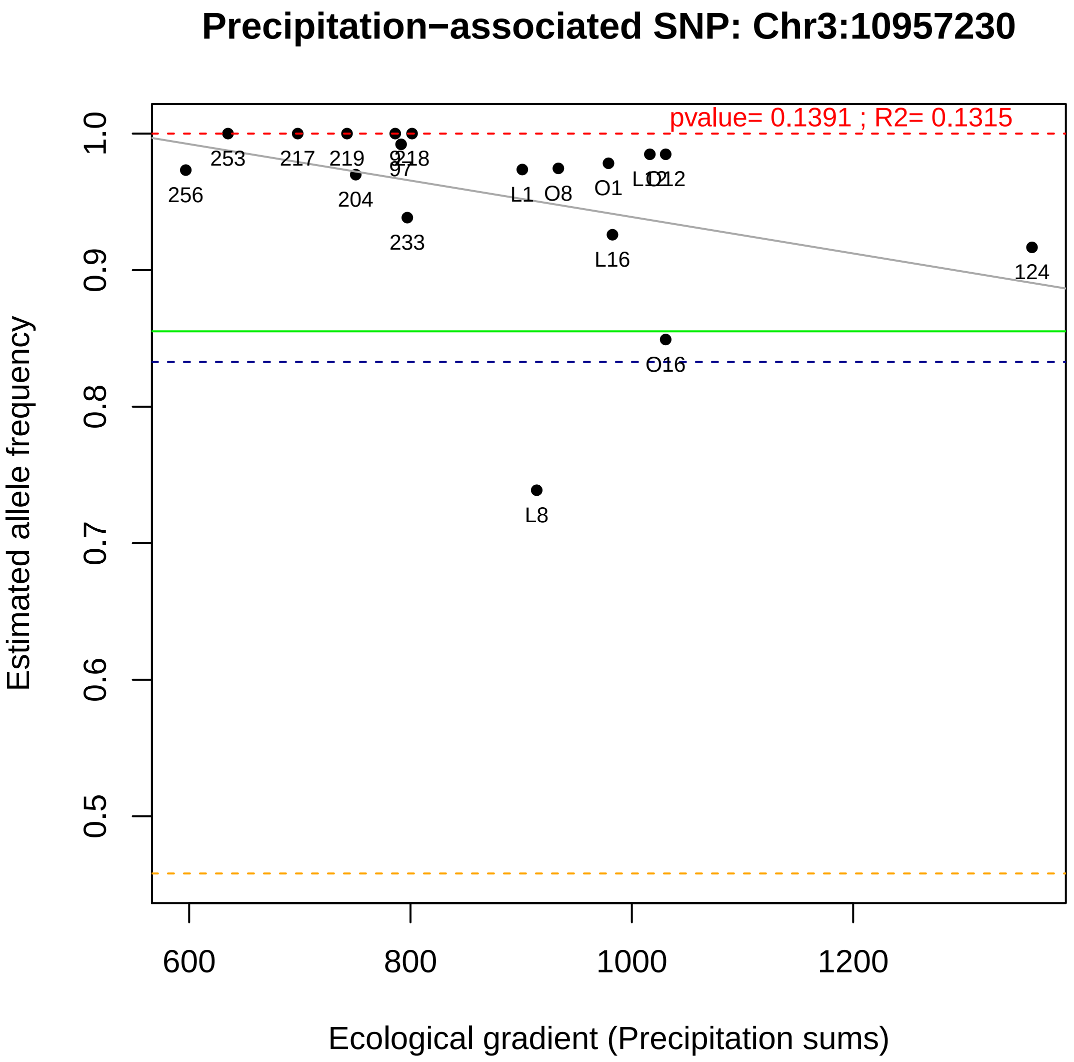

Precipitation−associated SNP: Chr3:10957230
pvalue= 0.1391 ; R2= 0.1315
Estimated allele frequency
Ecological gradient (Precipitation sums)
1.0
0.9
0.8
0.7
0.6
0.5
600
800
1000
1200
256
253
217
219
204
97
218
233
L1
O8
O1
L12
O12
L16
124
O16
L8

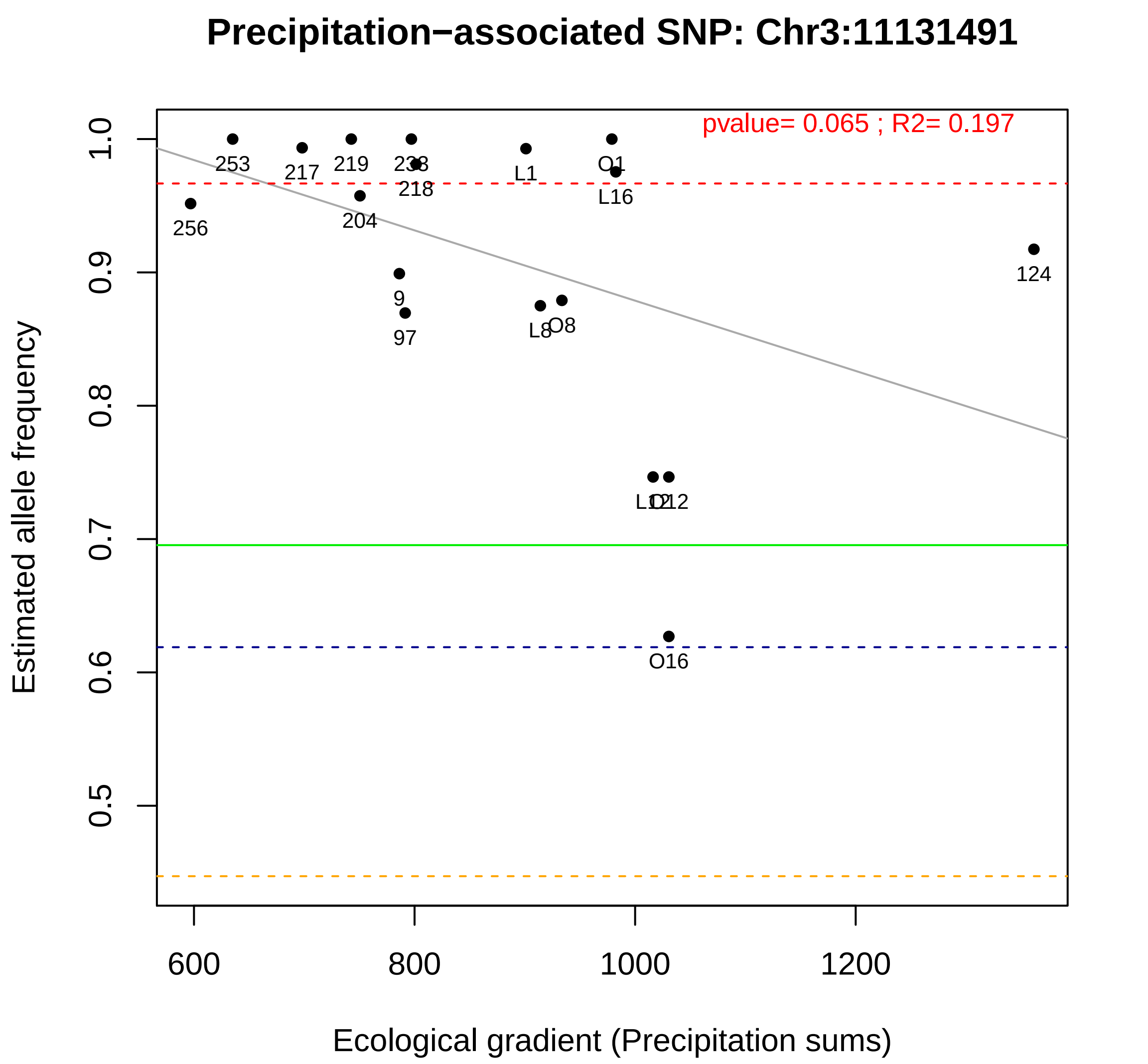
Precipitation−associated SNP: Chr3:11131491
pvalue= 0.065 ; R2= 0.197
Estimated allele frequency
Ecological gradient (Precipitation sums)
0.5
0.6
0.7
0.8
0.9
1.0
600
800
1000
1200
253
217
219
218
204
256
9
97
L1
L8
O8
O1
L16
124
O16

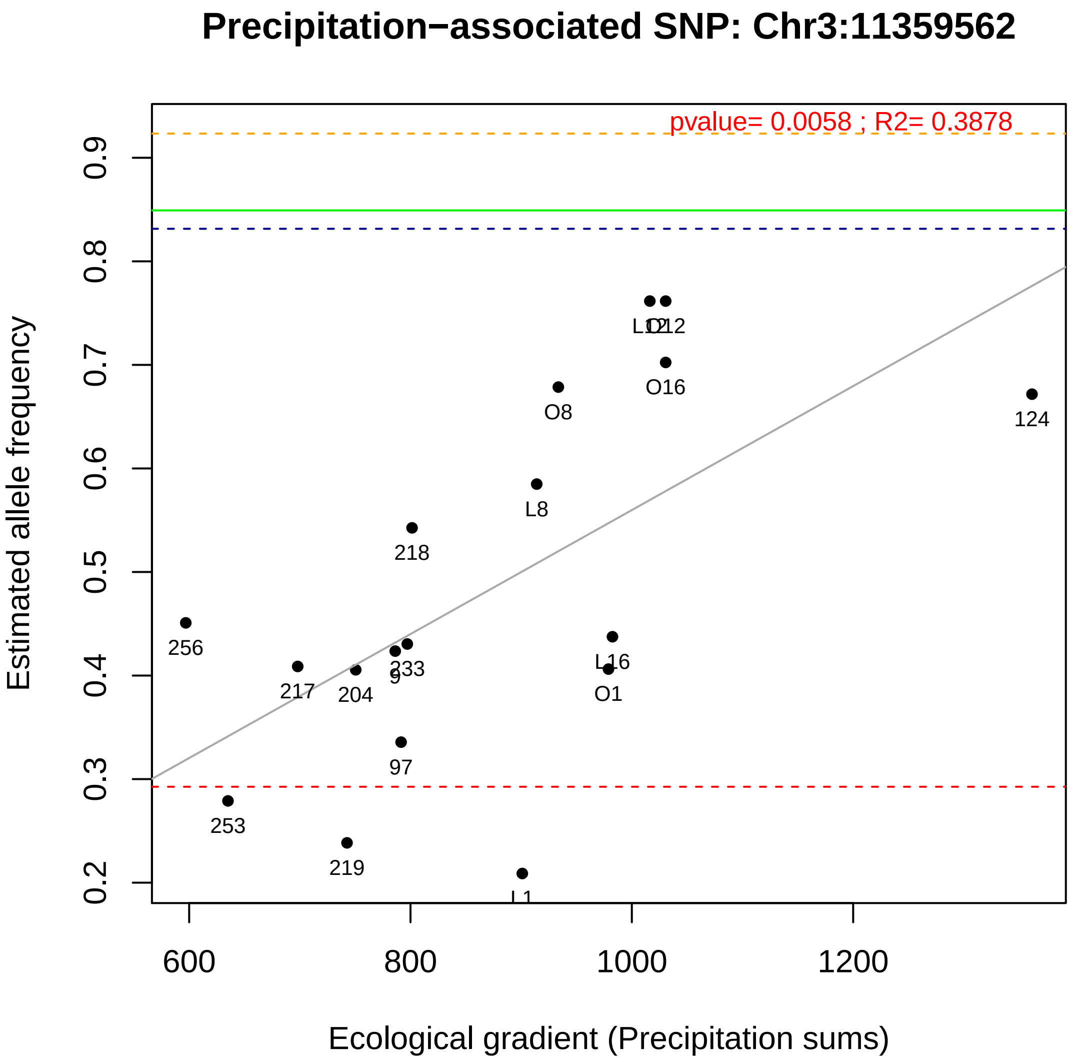

Precipitation−associated SNP: Chr3:11359562
pvalue= 0.0058 ; R2= 0.3878
Estimated allele frequency
0.2
0.3
0.4
0.5
0.6
0.7
0.8
0.9
600
800
1000
1200
Ecological gradient (Precipitation sums)
256
217
204
233
9
97
253
219
218
L8
O8
L12
O12
O16
L16
O1
124

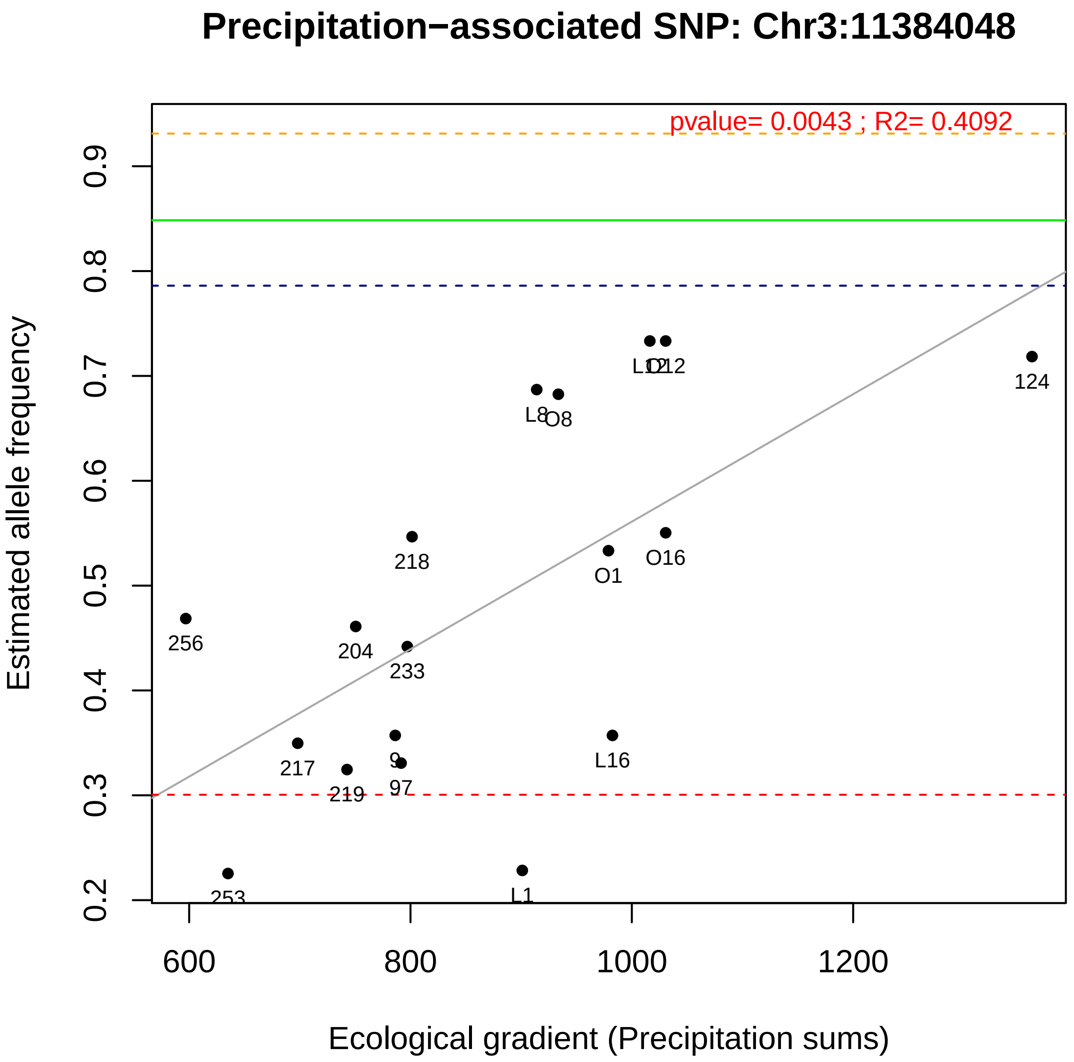

Precipitation−associated SNP: Chr3:11384048
pvalue= 0.0043 ; R2= 0.4092
Estimated allele frequency
Ecological gradient (Precipitation sums)
0.2
0.3
0.4
0.5
0.6
0.7
0.8
0.9
600
800
1000
1200
124
L12
O12
L8
O8
218
O16
O1
256
204
233
217
219
9
97
L16
253
L1

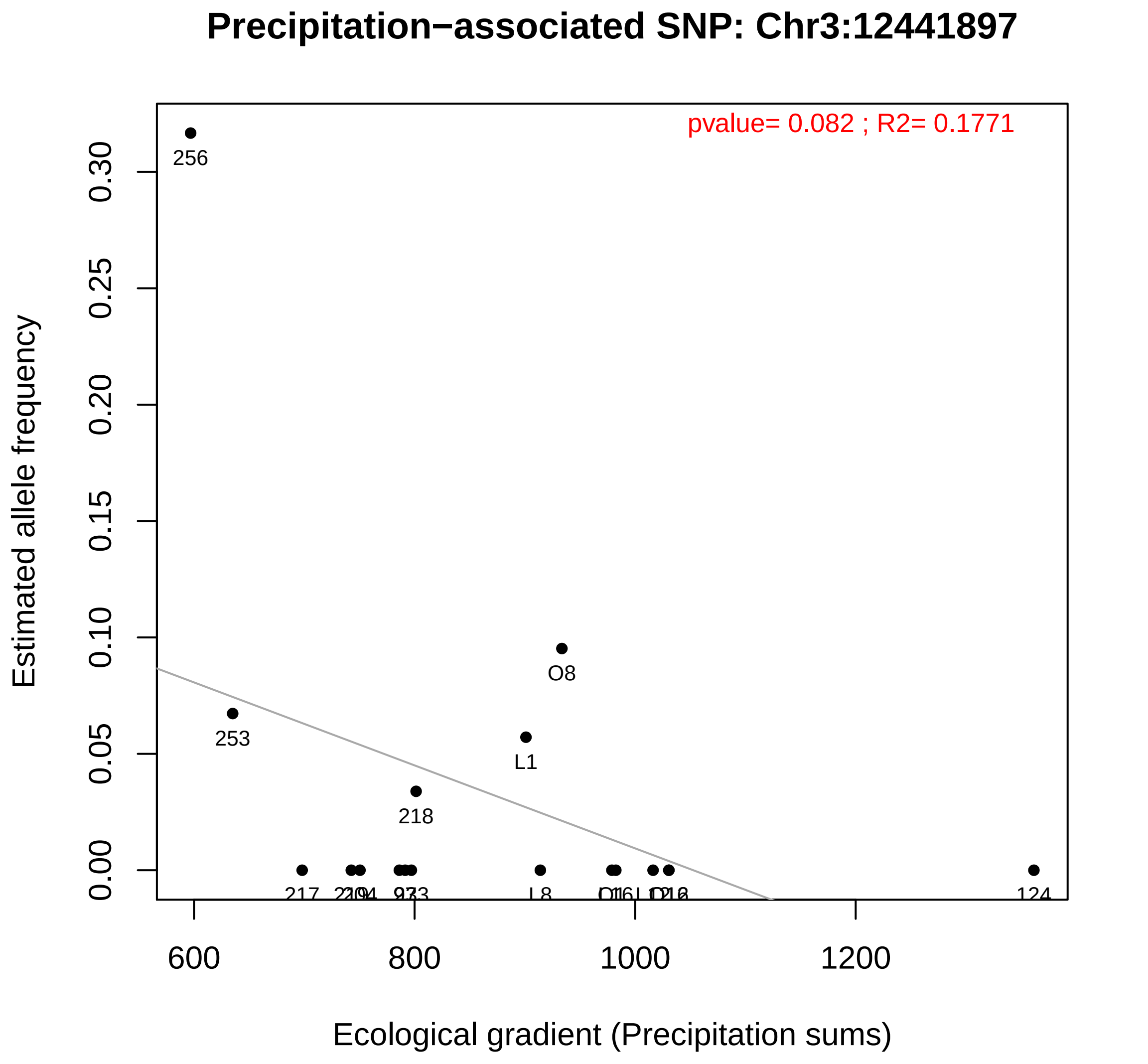

Precipitation−associated SNP: Chr3:12441897
pvalue= 0.082 ; R2= 0.1771
Estimated allele frequency
Ecological gradient (Precipitation sums)
0.00
0.05
0.10
0.15
0.20
0.25
0.30
600
800
1000
1200
256
253
O8
L1
218
217
L8
124

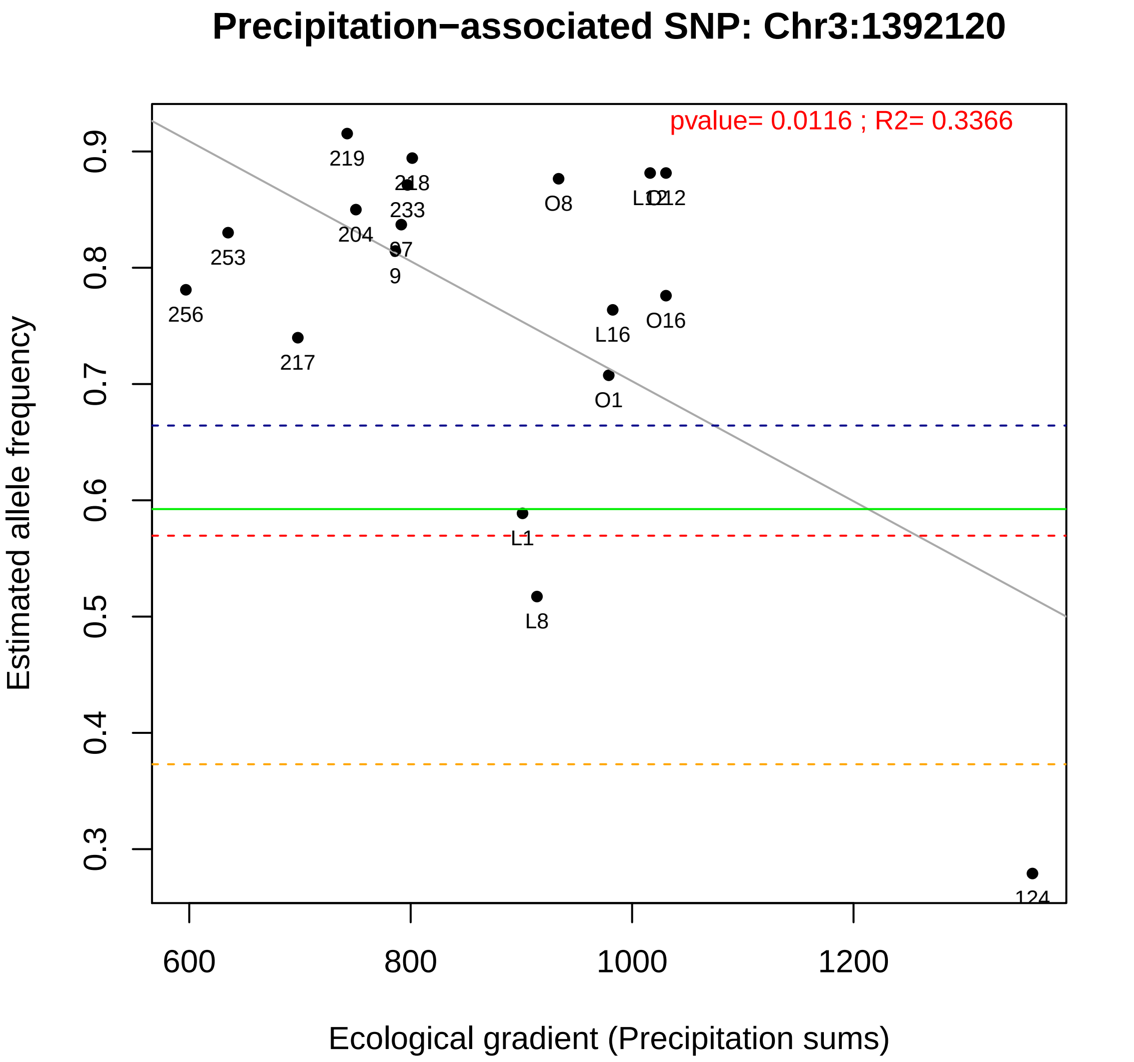
Precipitation−associated SNP: Chr3:1392120
pvalue= 0.0116 ; R2= 0.3366
Estimated allele frequency
0.3
0.4
0.5
0.6
0.7
0.8
0.9
600
800
1000
1200
Ecological gradient (Precipitation sums)
219
218
233
204
253
97
9
256
217
O8
L12
O12
L16
O16
O1
L1
L8
124

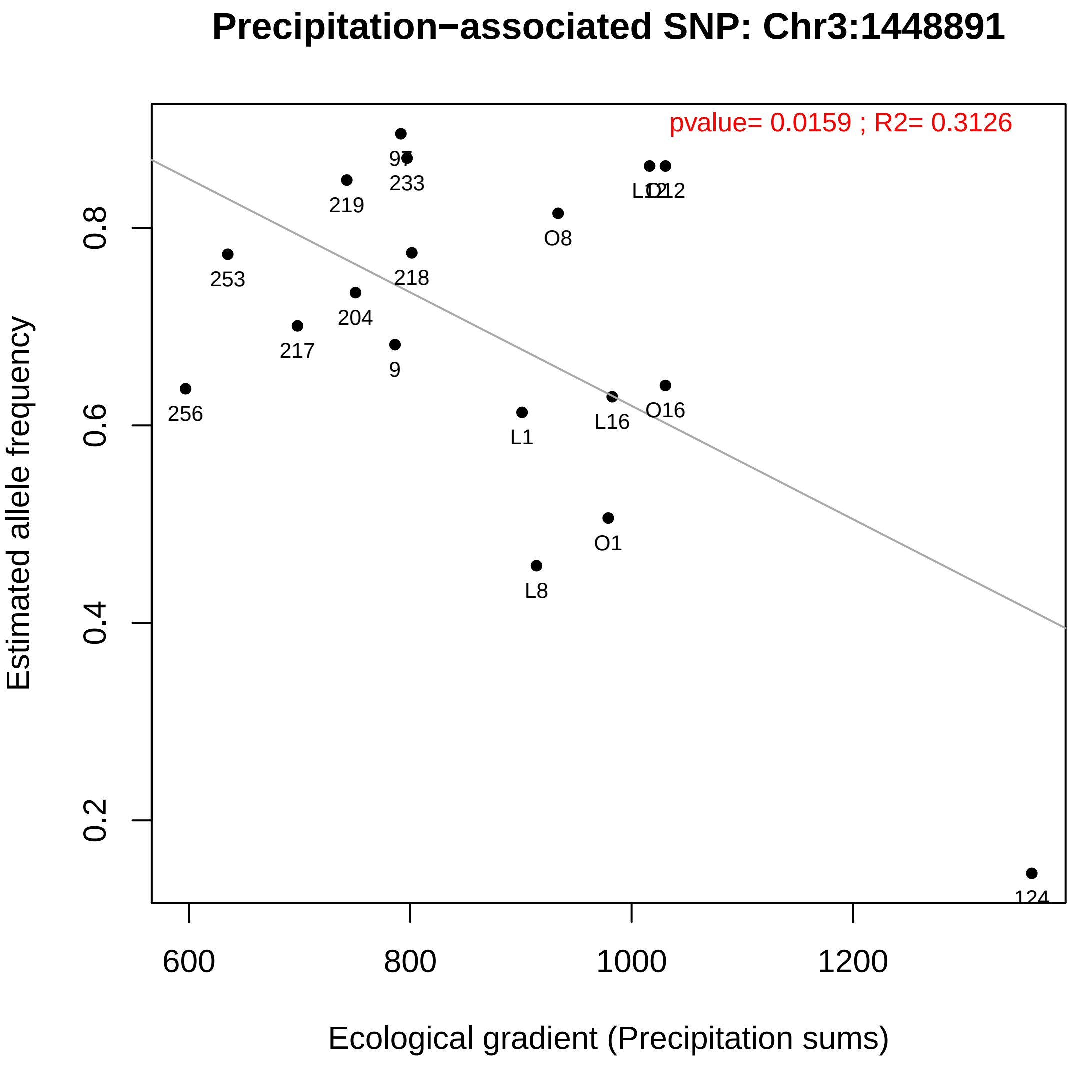

Precipitation−associated SNP: Chr3:1448891
pvalue= 0.0159 ; R2= 0.3126
Estimated allele frequency
0.2
0.4
0.6
0.8
600
800
1000
1200
Ecological gradient (Precipitation sums)
233
219
O8
253
218
204
217
9
256
L1
L16
O16
O1
L8
124

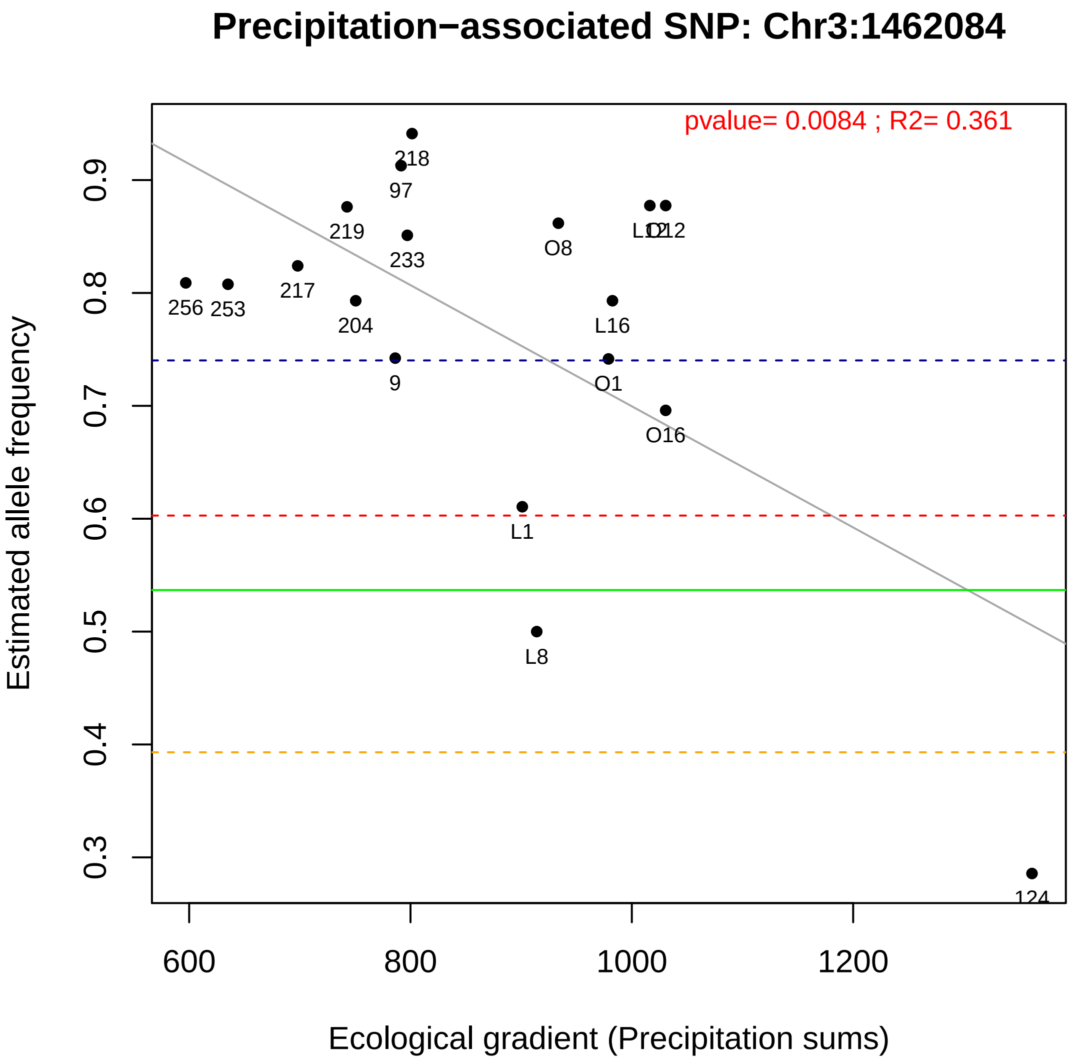
Precipitation−associated SNP: Chr3:1462084
pvalue= 0.0084 ; R2= 0.361
Estimated allele frequency
0.3
0.4
0.5
0.6
0.7
0.8
0.9
600
800
1000
1200
Ecological gradient (Precipitation sums)
218
97
219
233
217
256
253
204
9
O8
L12
O12
L16
O1
O16
L1
L8
124

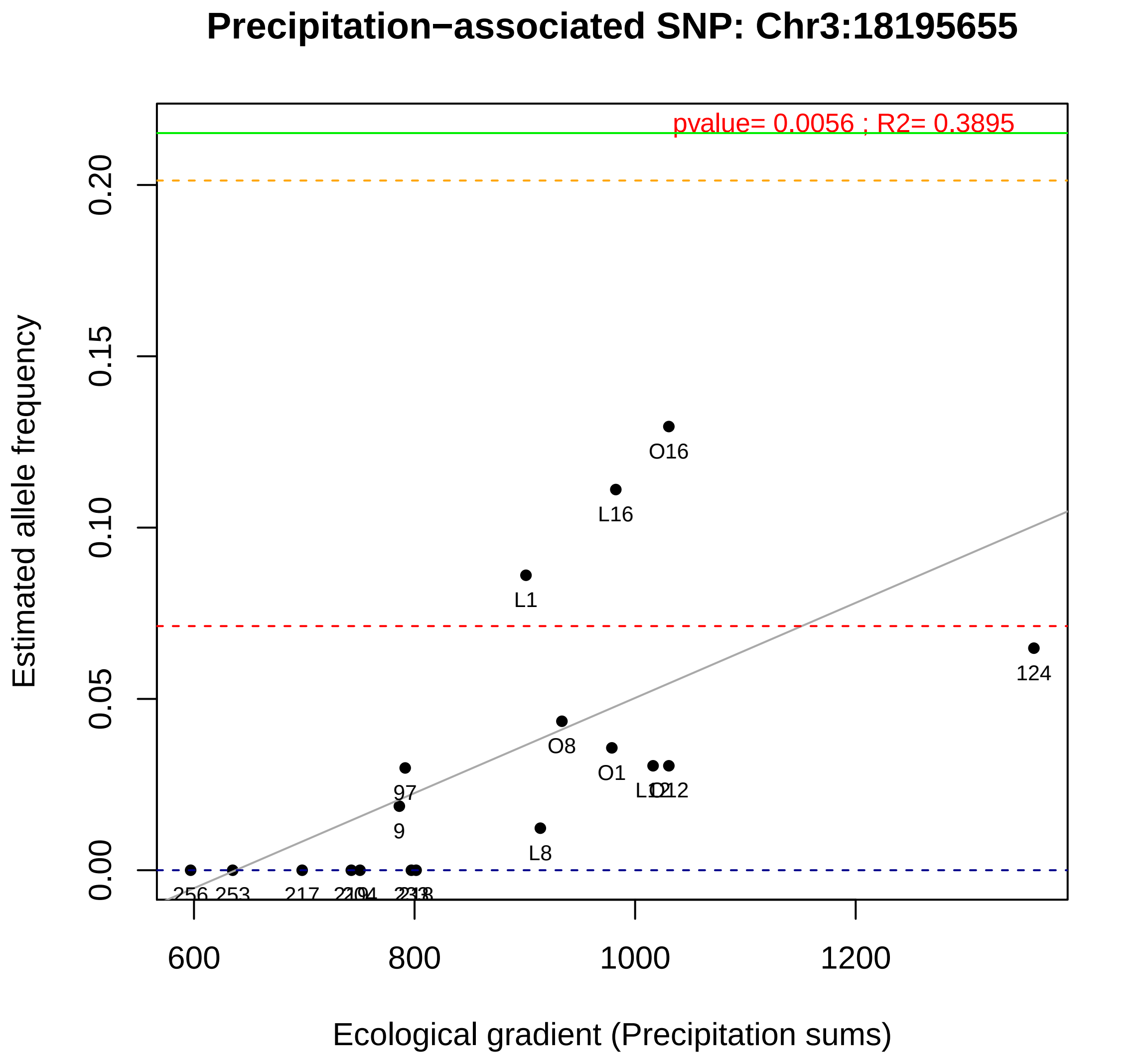

Precipitation−associated SNP: Chr3:18195655
pvalue= 0.0056 ; R2= 0.3895
Estimated allele frequency
0.00
0.05
0.10
0.15
0.20
600
800
1000
1200
Ecological gradient (Precipitation sums)
O16
L16
L1
124
O8
O1
97
9
L8
256
253
217

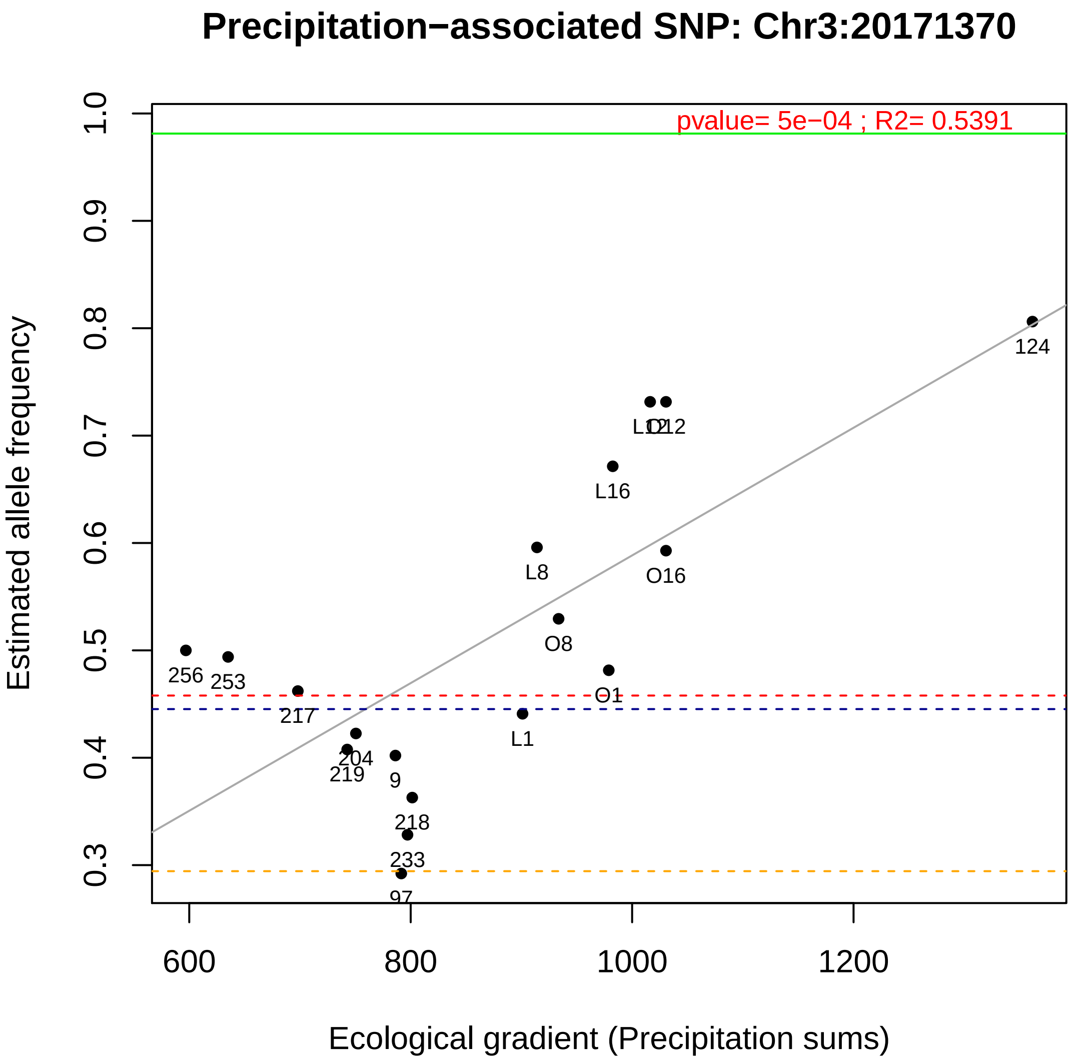

Precipitation−associated SNP: Chr3:20171370
pvalue= 5e−04 ; R2= 0.5391
Estimated allele frequency
Ecological gradient (Precipitation sums)
1.0
0.9
0.8
0.7
0.6
0.5
0.4
0.3
600
800
1000
1200
124
L12
O12
L16
L8
O16
O8
O1
L1
256
253
217
204
219
9
218
233
97

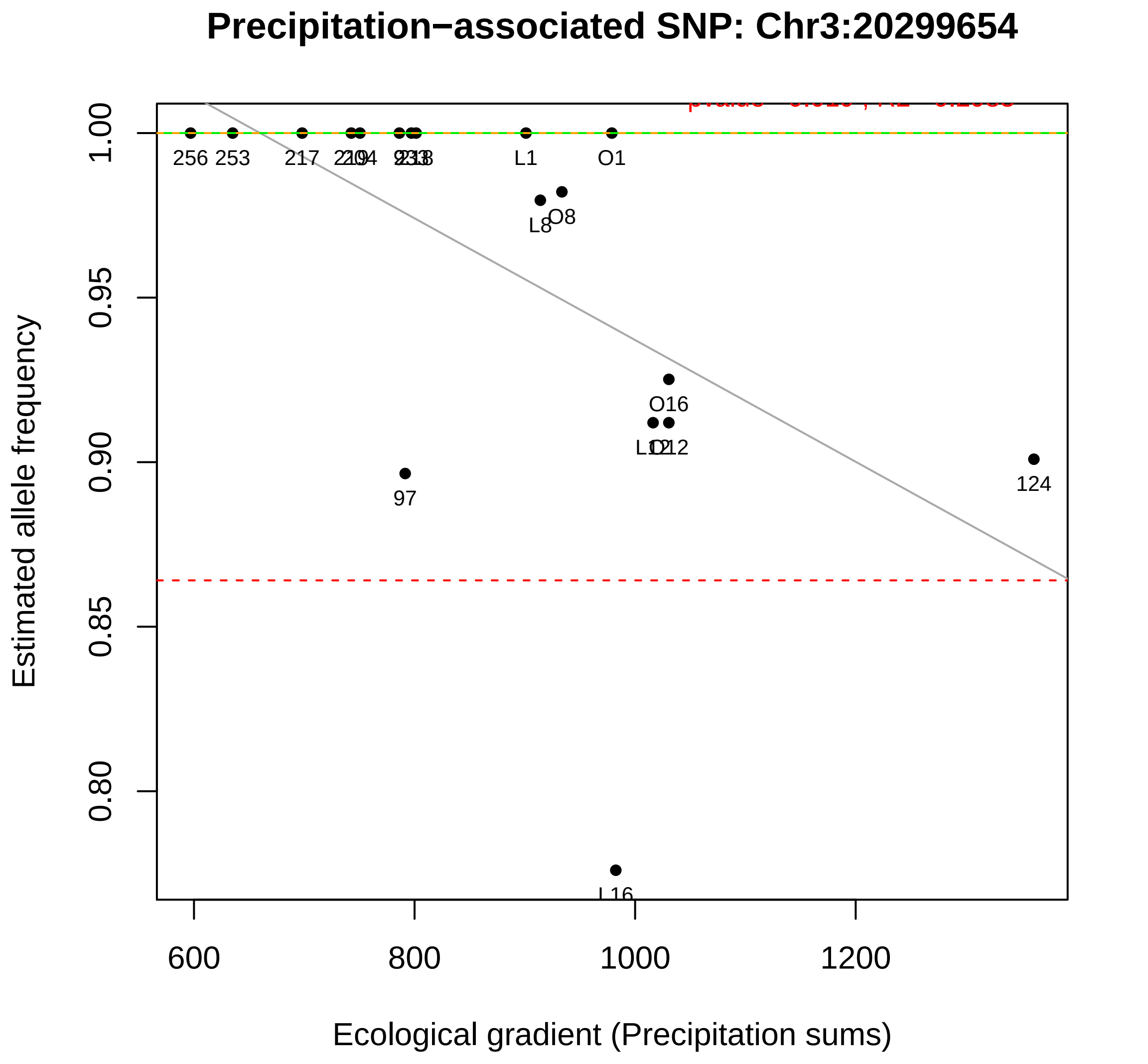
Precipitation−associated SNP: Chr3:20299654
Estimated allele frequency
1.00
0.95
0.90
0.85
0.80
256
253
217
L1
O1
L8
O8
O16
L12
O12
97
124
L16
600
800
1000
1200
Ecological gradient (Precipitation sums)

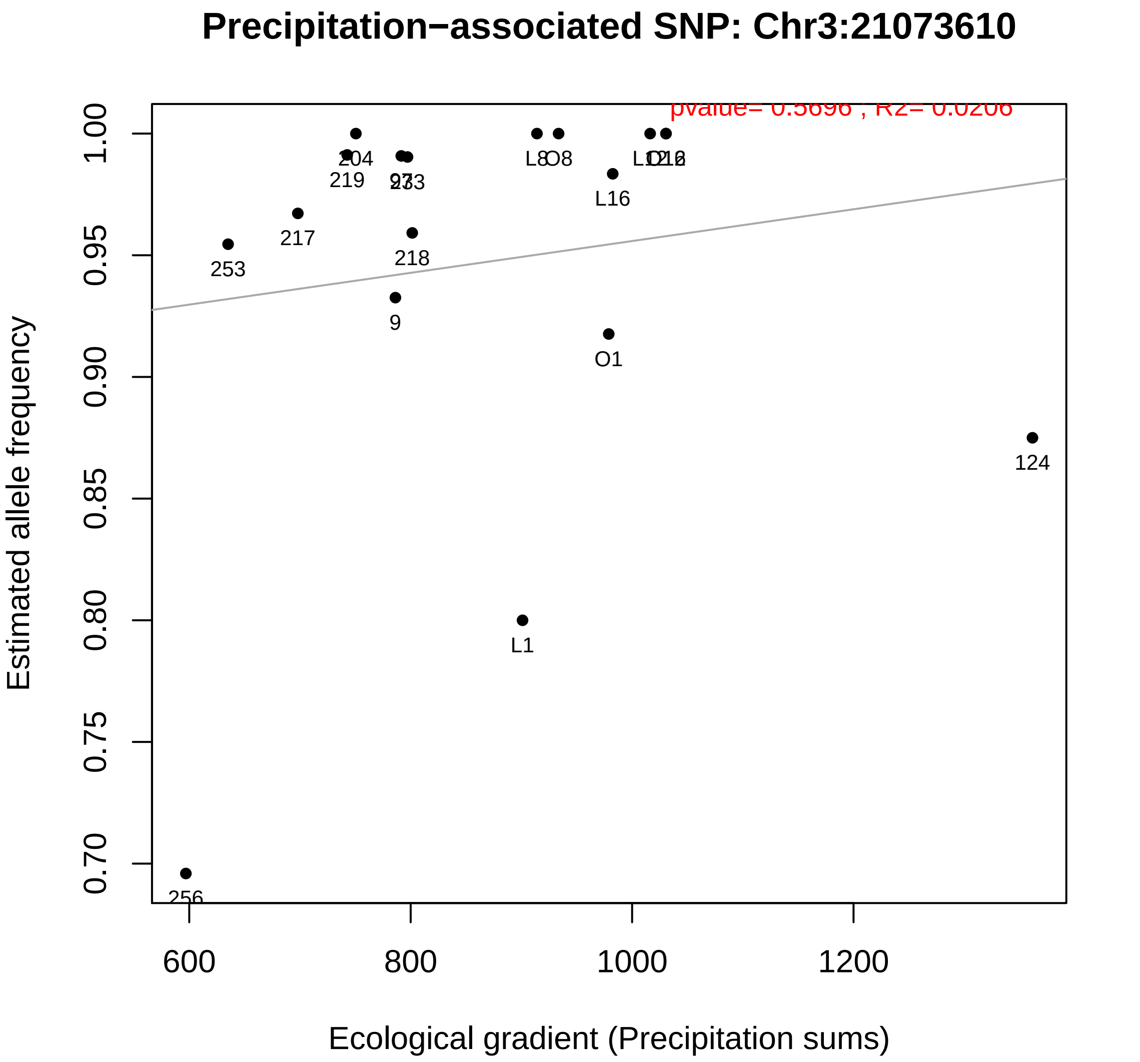

Precipitation−associated SNP: Chr3:21073610
pvalue= 0.5696 ; R2= 0.0206
Estimated allele frequency
Ecological gradient (Precipitation sums)
1.00
0.95
0.90
0.85
0.80
0.75
0.70
600
800
1000
1200
204
219
97
233
L8
O8
L16
217
253
218
9
O1
124
L1
256

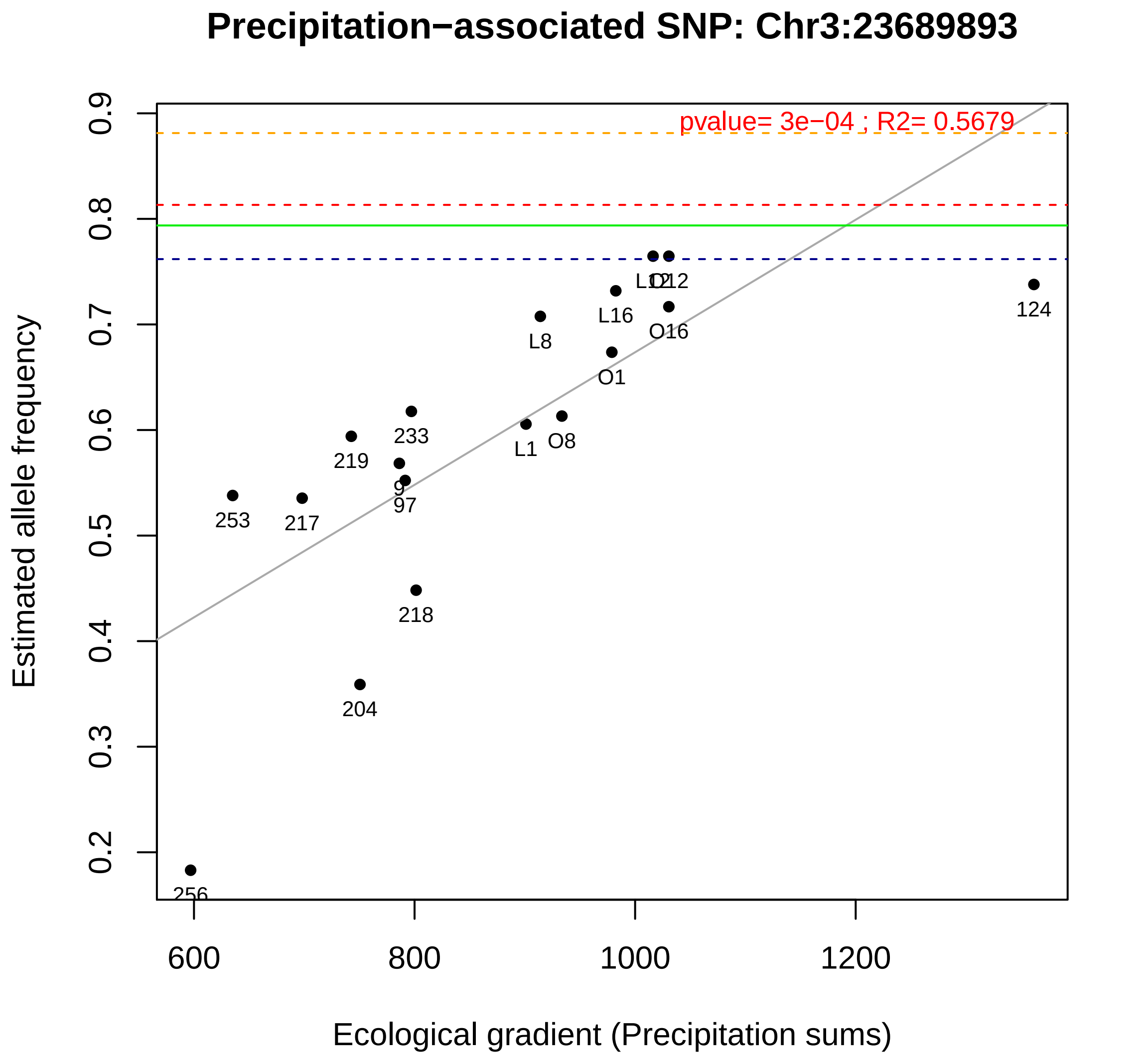
Precipitation−associated SNP: Chr3:23689893
pvalue= 3e−04 ; R2= 0.5679
Estimated allele frequency
0.2
0.3
0.4
0.5
0.6
0.7
0.8
0.9
600
800
1000
1200
Ecological gradient (Precipitation sums)
L12
O12
L16
124
O16
L8
O1
233
L1
O8
219
9
97
253
217
218
204
256

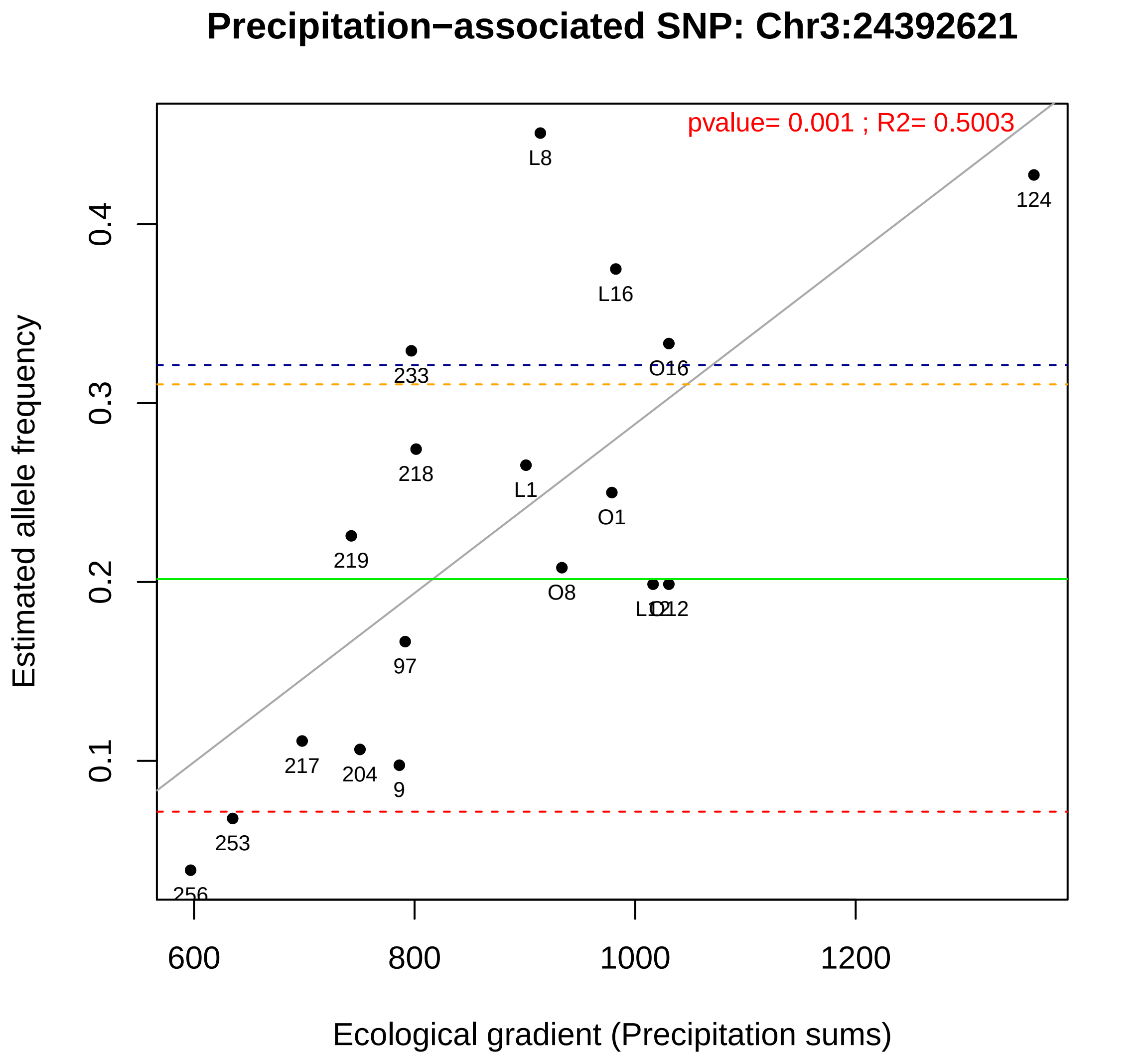

Precipitation−associated SNP: Chr3:24392621
pvalue= 0.001 ; R2= 0.5003
Estimated allele frequency
0.1
0.2
0.3
0.4
600
800
1000
1200
Ecological gradient (Precipitation sums)
L8
124
L16
O16
233
218
L1
O1
219
O8
L12
O12
97
217
204
9
253
256

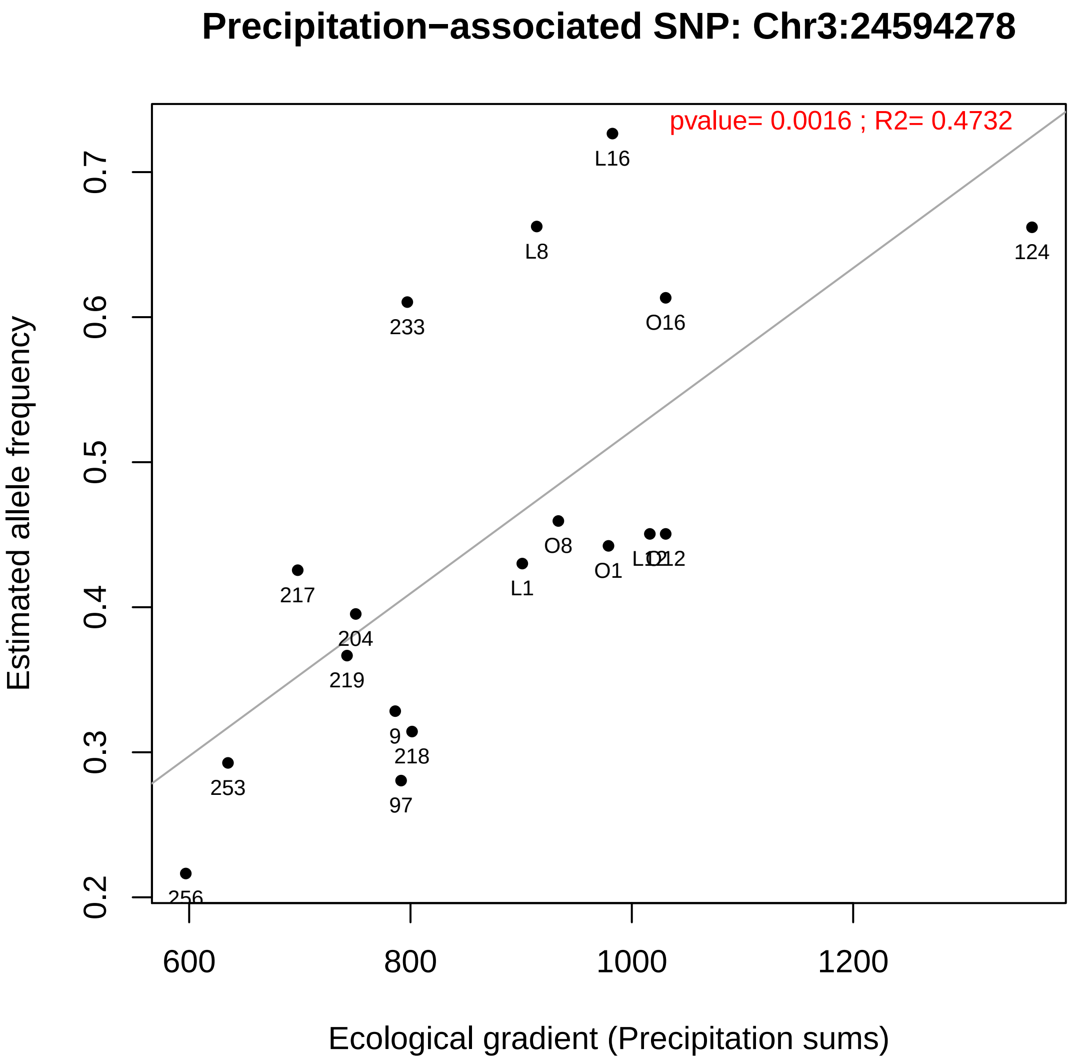
Precipitation−associated SNP: Chr3:24594278
pvalue= 0.0016 ; R2= 0.4732
L16
L8
124
233
O16
O8
O1
L1
217
204
219
9
218
253
97
256
0.2
0.3
0.4
0.5
0.6
0.7
600
800
1000
1200
Estimated allele frequency
Ecological gradient (Precipitation sums)

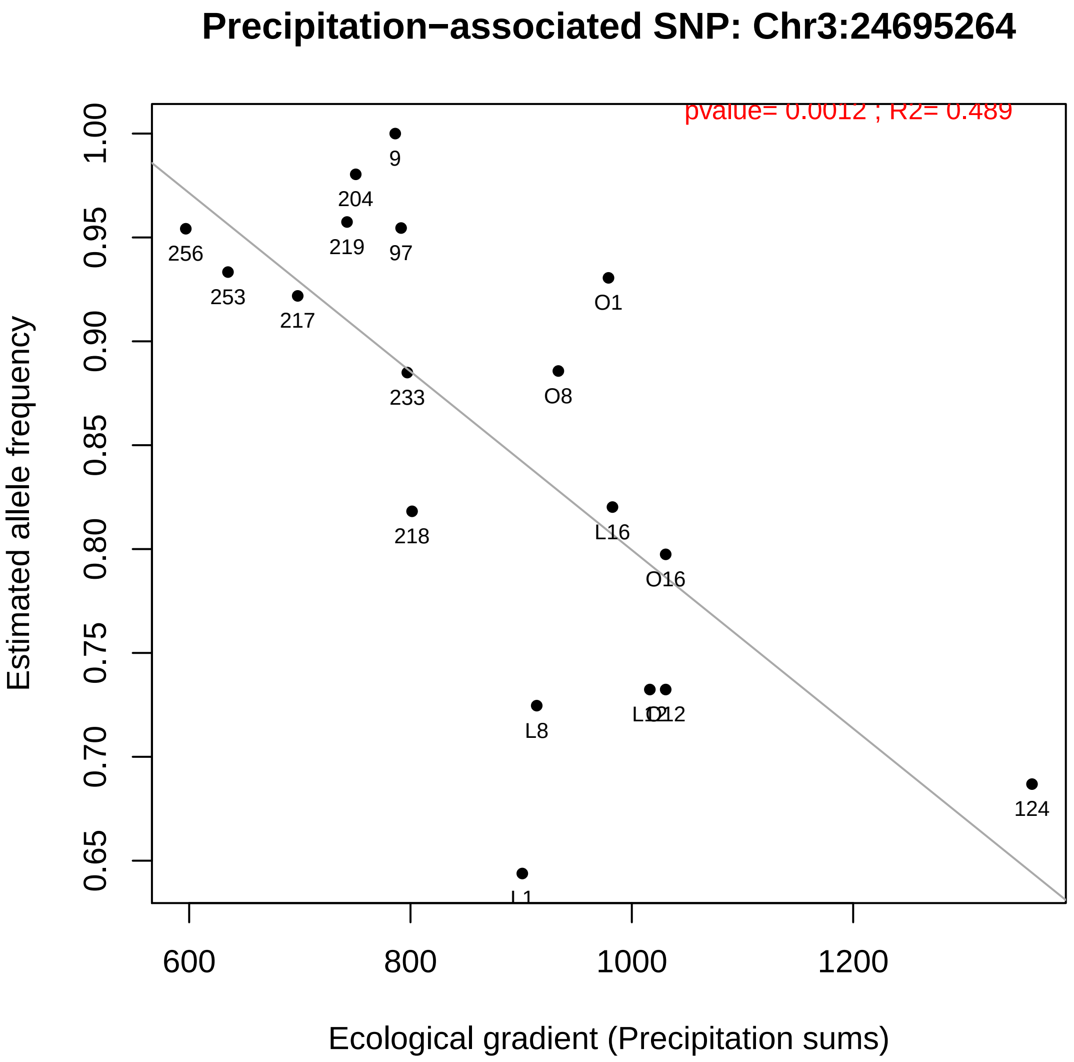
Precipitation−associated SNP: Chr3:24695264
pvalue= 0.0012 ; R2= 0.489
Estimated allele frequency
Ecological gradient (Precipitation sums)
0.65
0.70
0.75
0.80
0.85
0.90
0.95
1.00
600
800
1000
1200
9
204
256
219
97
253
217
O1
233
O8
218
L16
O16
L8
O12
124
L1

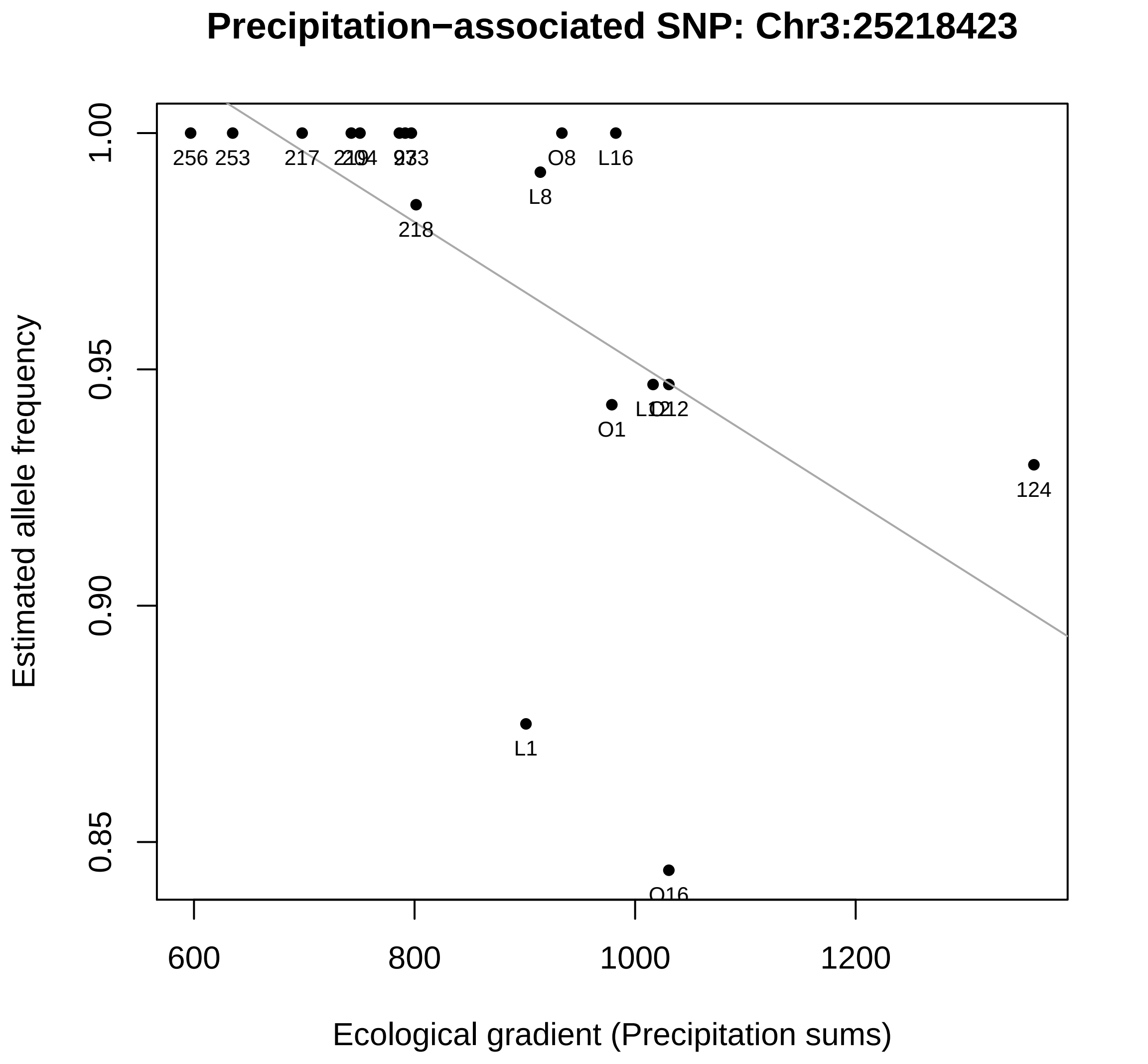

Precipitation−associated SNP: Chr3:25218423
Estimated allele frequency
Ecological gradient (Precipitation sums)
1.00
0.95
0.90
0.85
600
800
1000
1200
256
253
217
O8
L16
L8
218
O1
124
L1
O16

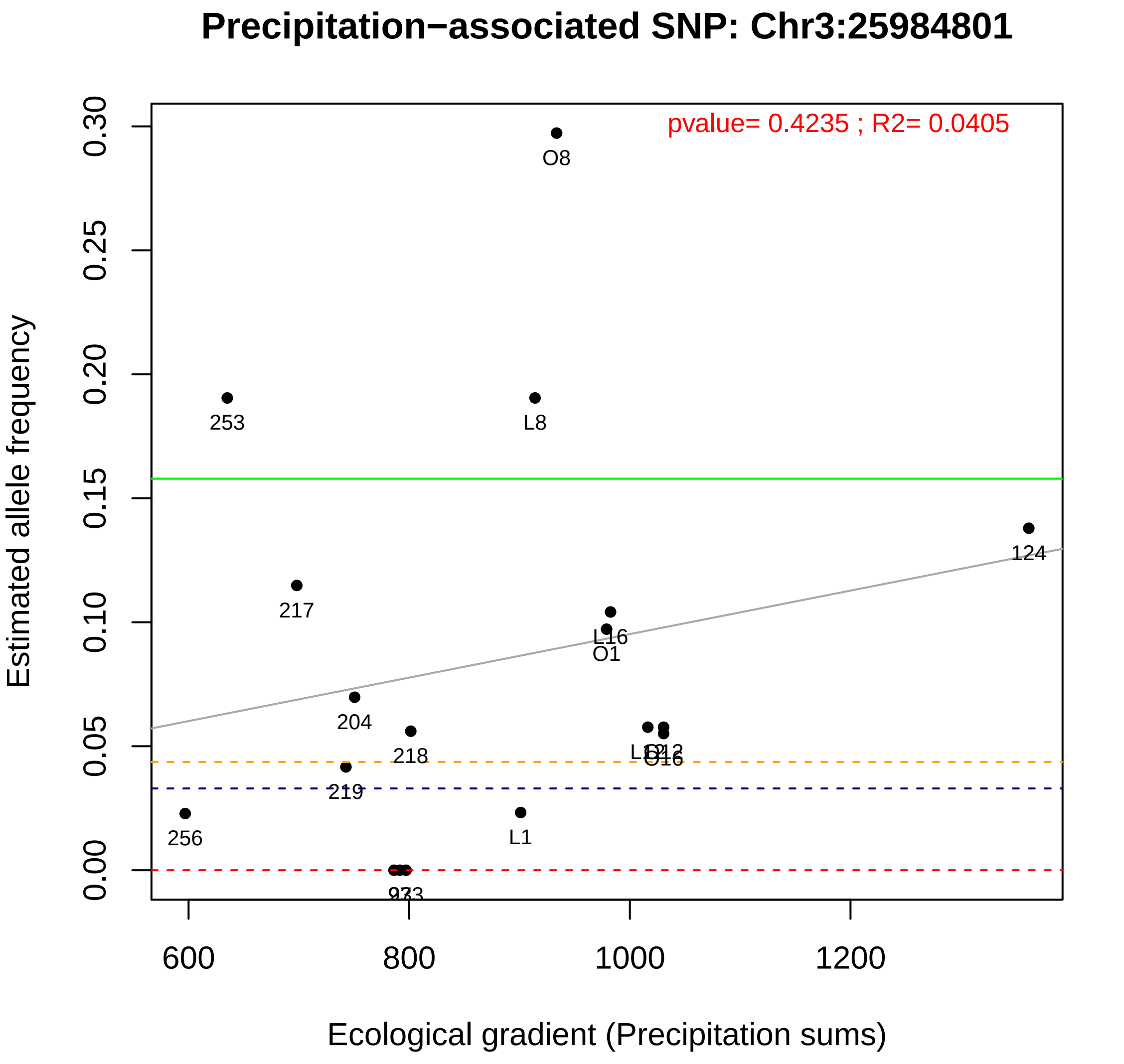

Precipitation−associated SNP: Chr3:25984801
pvalue= 0.4235 ; R2= 0.0405
Estimated allele frequency
Ecological gradient (Precipitation sums)
0.00
0.05
0.10
0.15
0.20
0.25
0.30
600
800
1000
1200
O8
253
L8
124
217
L16
O1
204
218
219
256
L1

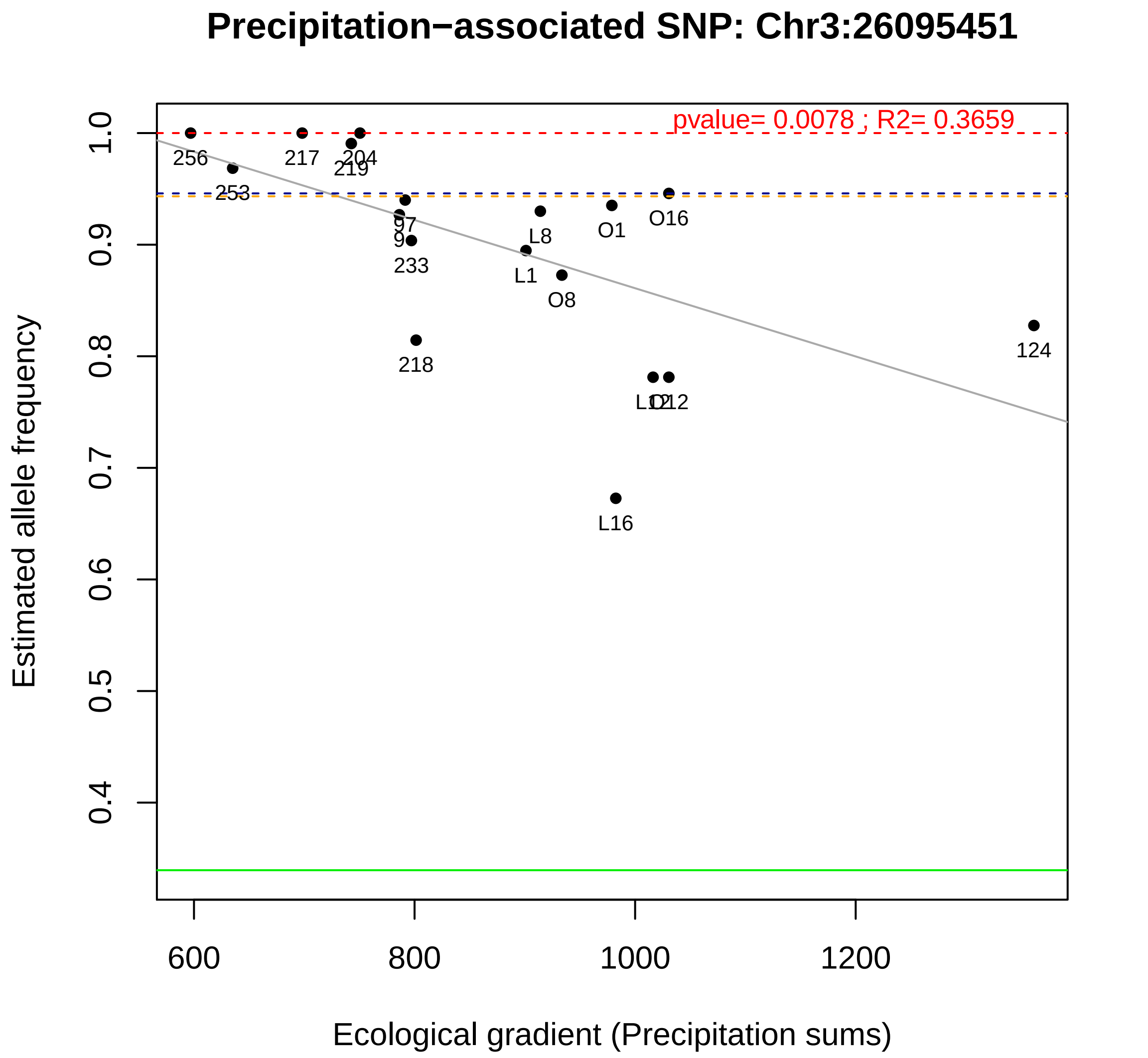
Precipitation−associated SNP: Chr3:26095451
pvalue= 0.0078 ; R2= 0.3659
Estimated allele frequency
Ecological gradient (Precipitation sums)
0.4
0.5
0.6
0.7
0.8
0.9
1.0
600
800
1000
1200
256
253
217
204
219
97
233
218
L8
L1
O8
O1
O16
L16
O12
124

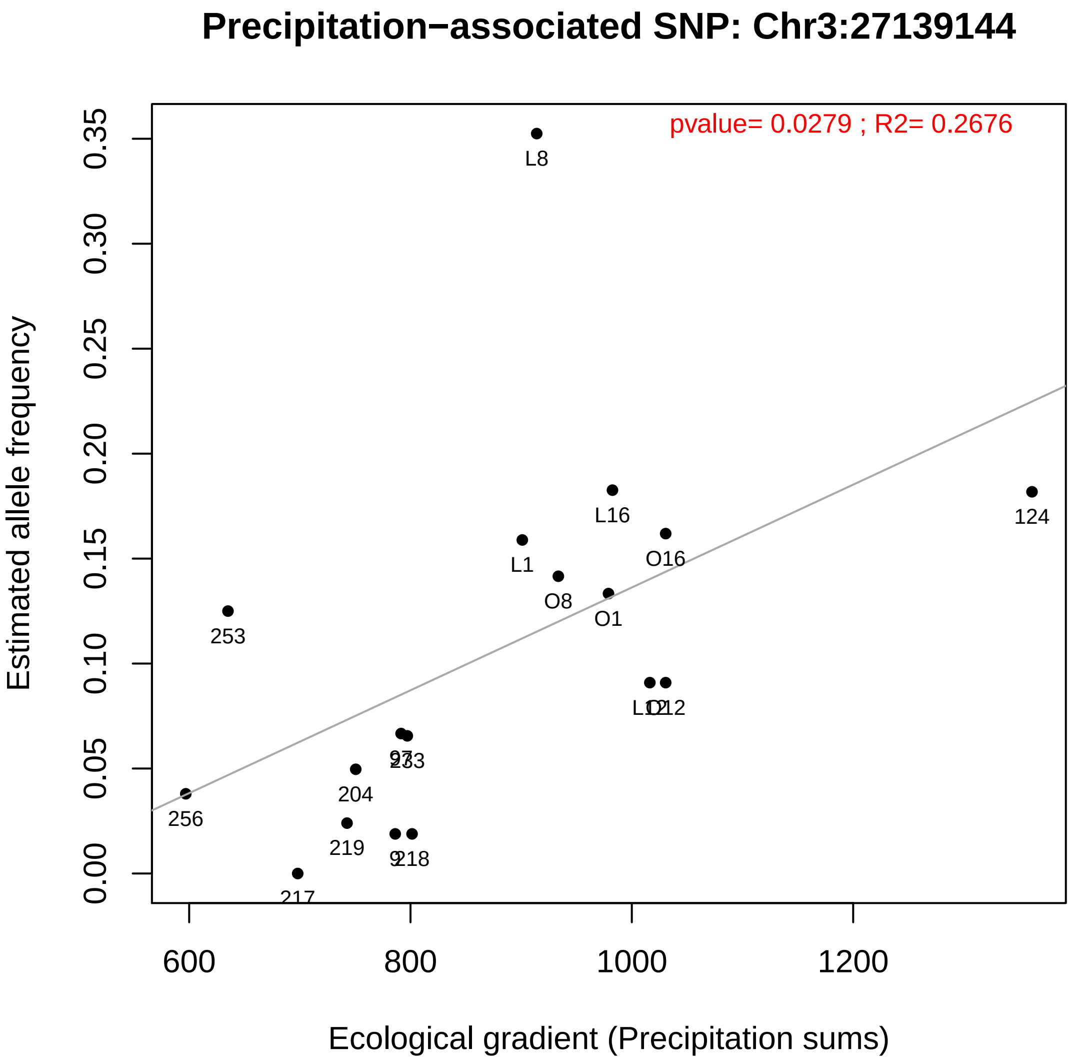
Precipitation−associated SNP: Chr3:27139144
pvalue= 0.0279 ; R2= 0.2676
Estimated allele frequency
0.00
0.05
0.10
0.15
0.20
0.25
0.30
0.35
Ecological gradient (Precipitation sums)
600
800
1000
1200
L8
L16
124
O16
L1
O8
O1
253
204
256
219
218
217

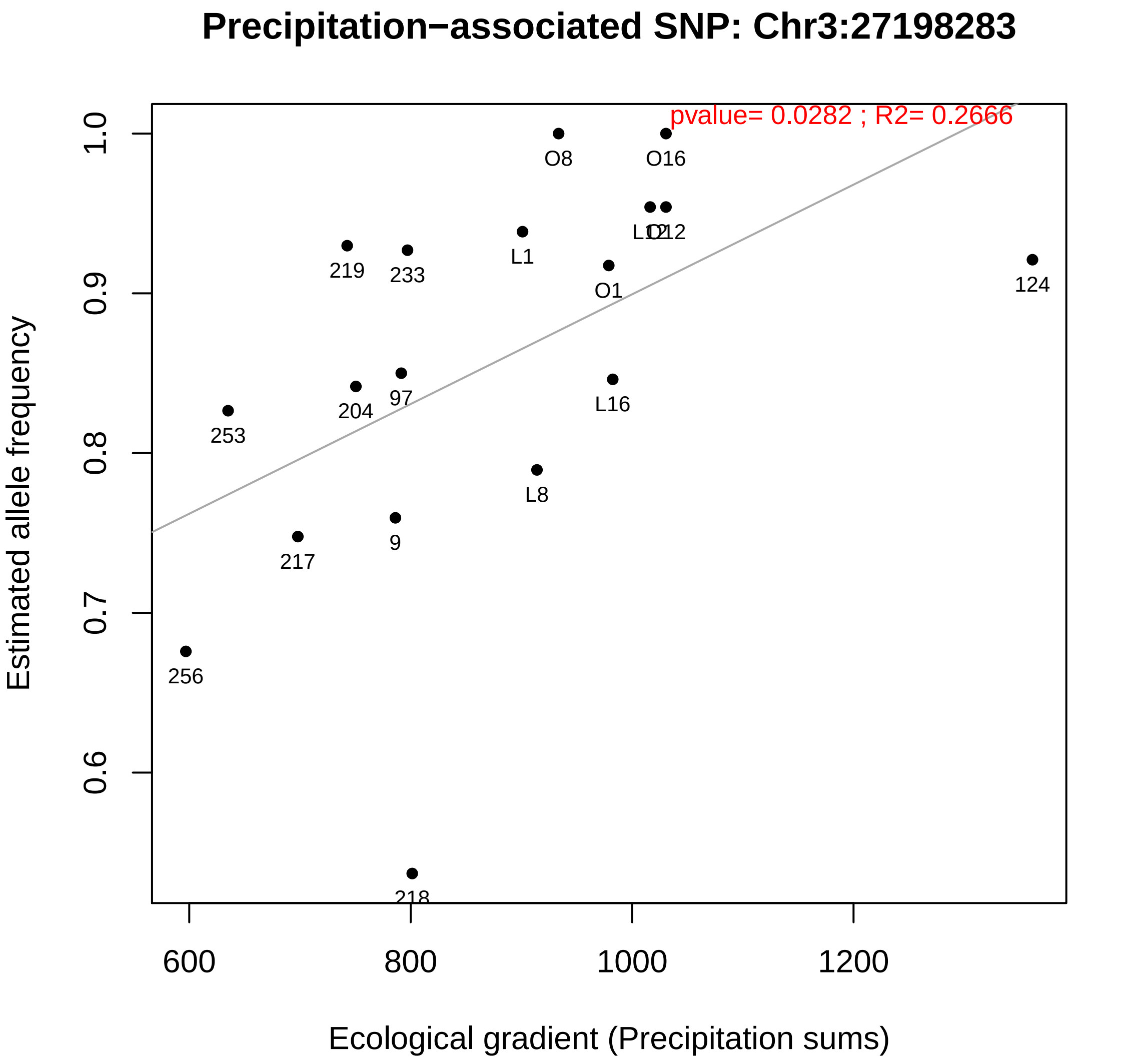

Precipitation−associated SNP: Chr3:27198283
pvalue= 0.0282 ; R2= 0.2666
Estimated allele frequency
Ecological gradient (Precipitation sums)
0.6
0.7
0.8
0.9
1.0
600
800
1000
1200
O8
O16
L12
O12
L1
219
233
O1
124
97
204
L16
253
L8
9
217
256
218

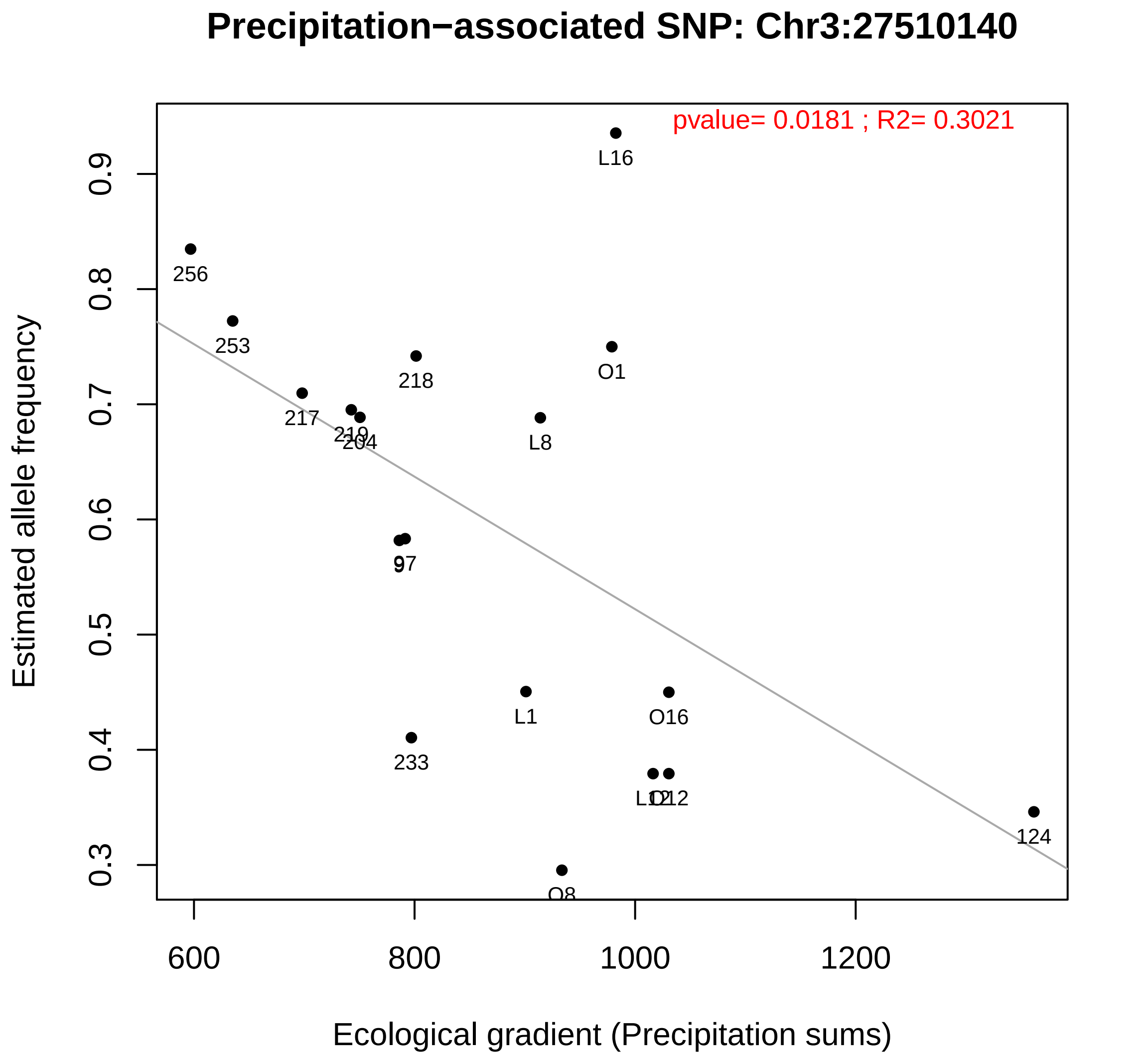
Precipitation−associated SNP: Chr3:27510140
pvalue= 0.0181 ; R2= 0.3021
Estimated allele frequency
Ecological gradient (Precipitation sums)
0.3
0.4
0.5
0.6
0.7
0.8
0.9
600
800
1000
1200
L16
256
253
218
O1
217
219
204
L8
97
L1
O16
233
L12
O12
124
O8

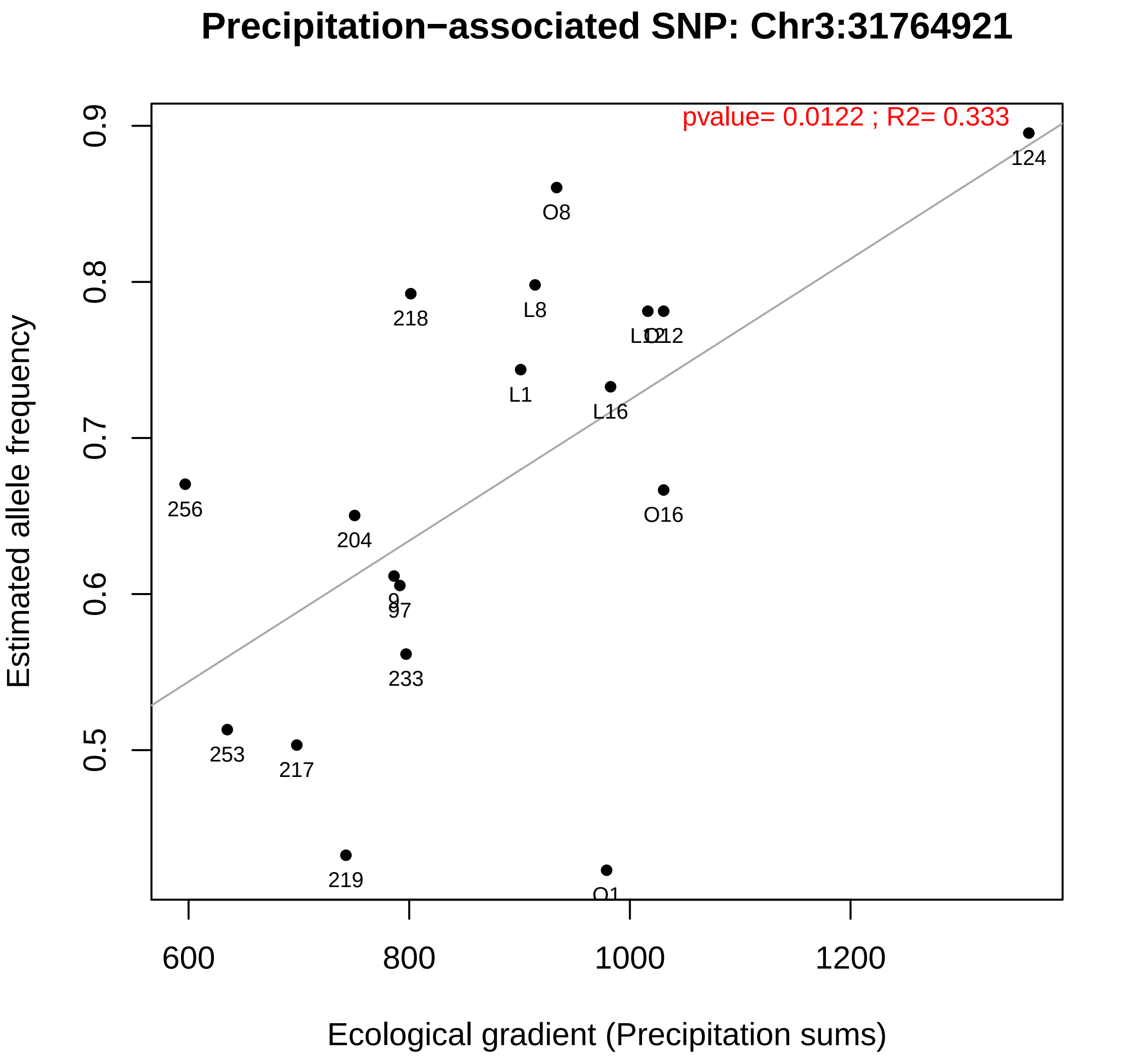
Precipitation−associated SNP: Chr3:31764921
pvalue= 0.0122 ; R2= 0.333
Estimated allele frequency
0.5
0.6
0.7
0.8
0.9
600
800
1000
1200
Ecological gradient (Precipitation sums)
124
O8
218
L8
L1
L16
O16
256
204
9
97
233
253
217
219
O1

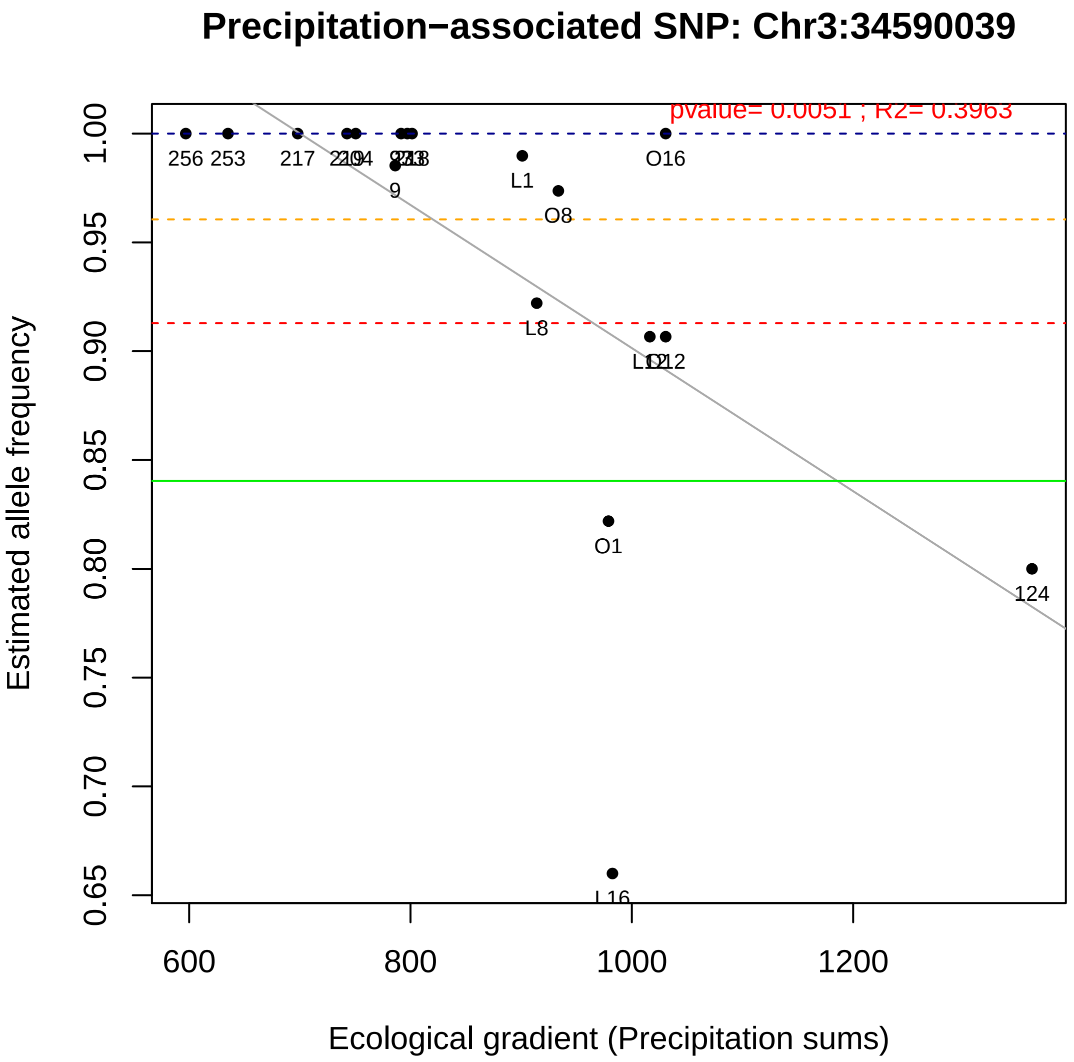

Precipitation−associated SNP: Chr3:34590039
pvalue= 0.0051 ; R2= 0.3963
Estimated allele frequency
Ecological gradient (Precipitation sums)
0.65
0.70
0.75
0.80
0.85
0.90
0.95
1.00
600
800
1000
1200
256
253
217
9
L1
O8
O16
L8
O12
O1
124
L16

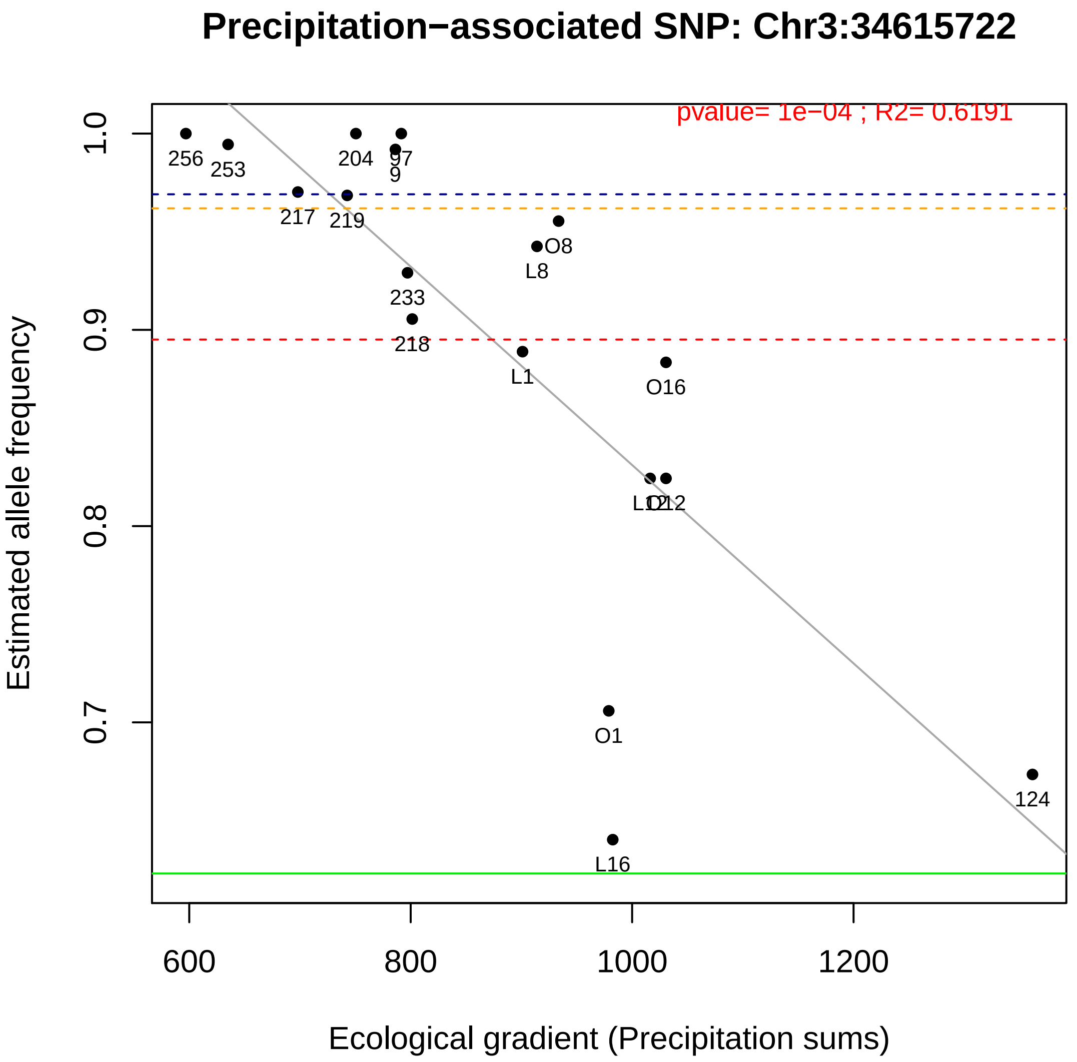

Precipitation−associated SNP: Chr3:34615722
pvalue= 1e−04 ; R2= 0.6191
Estimated allele frequency
1.0
0.9
0.8
0.7
600
800
1000
1200
Ecological gradient (Precipitation sums)
256
253
204
97
9
217
219
O8
L8
233
218
L1
O16
L12
O12
O1
124
L16

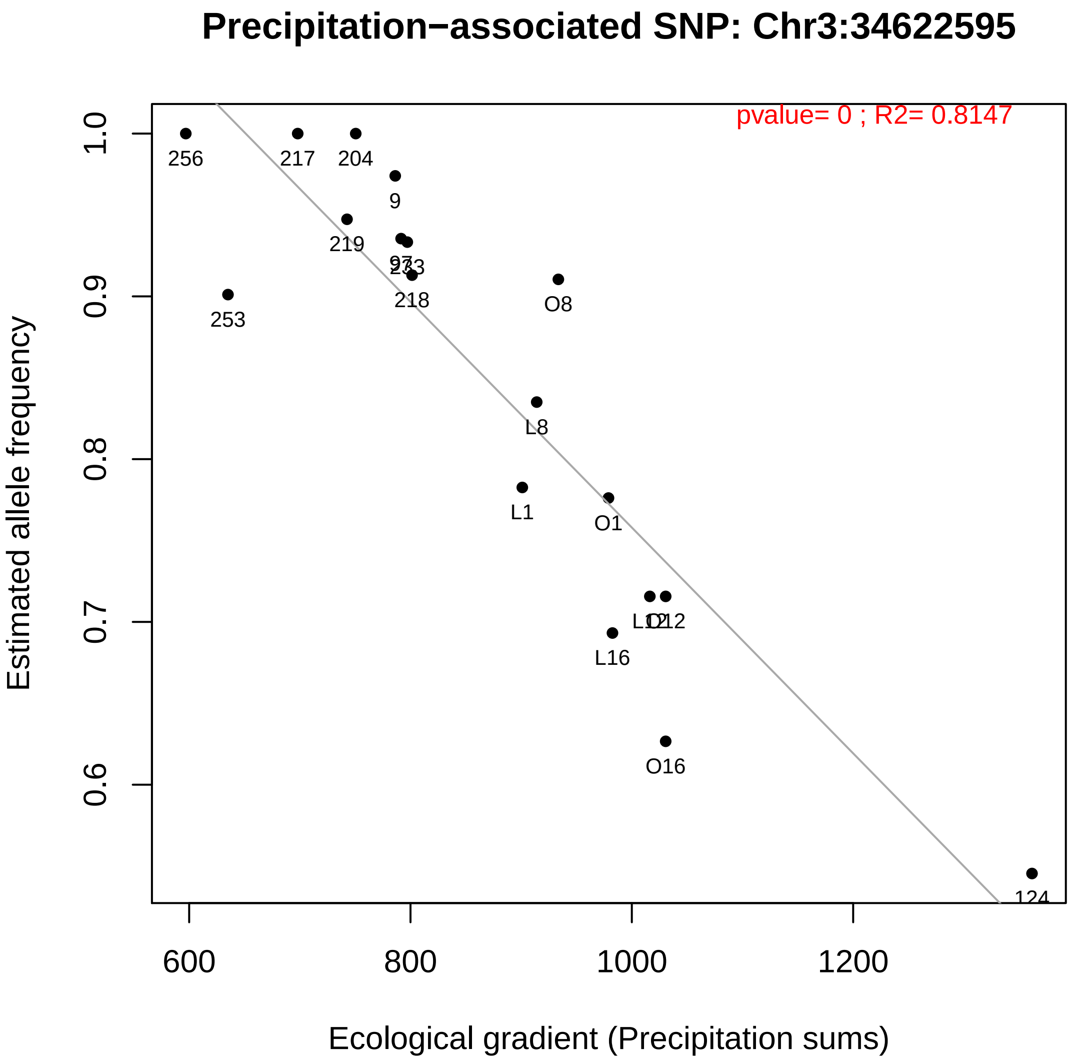
Precipitation−associated SNP: Chr3:34622595
pvalue= 0 ; R2= 0.8147
Estimated allele frequency
Ecological gradient (Precipitation sums)
1.0
0.9
0.8
0.7
0.6
600
800
1000
1200
256
217
204
9
219
97
233
218
253
O8
L8
L1
O1
L12
O12
L16
O16
124

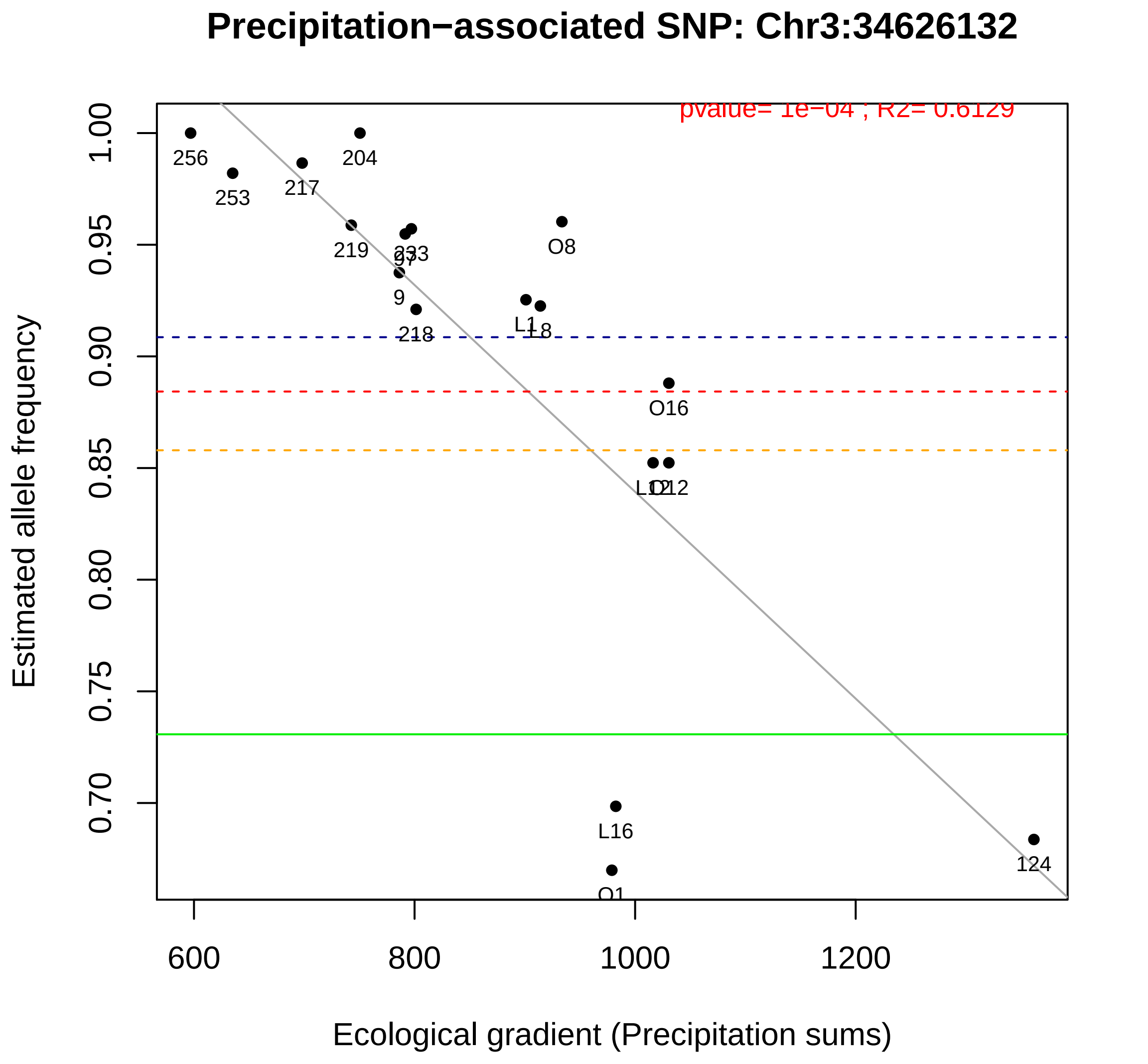
Precipitation−associated SNP: Chr3:34626132
pvalue= 1e−04 ; R2= 0.6129
Estimated allele frequency
Ecological gradient (Precipitation sums)
1.00
0.95
0.90
0.85
0.80
0.75
0.70
600
800
1000
1200
256
204
253
217
219
233
9
218
O8
L1
L18
O16
L12
O12
L16
O1
124

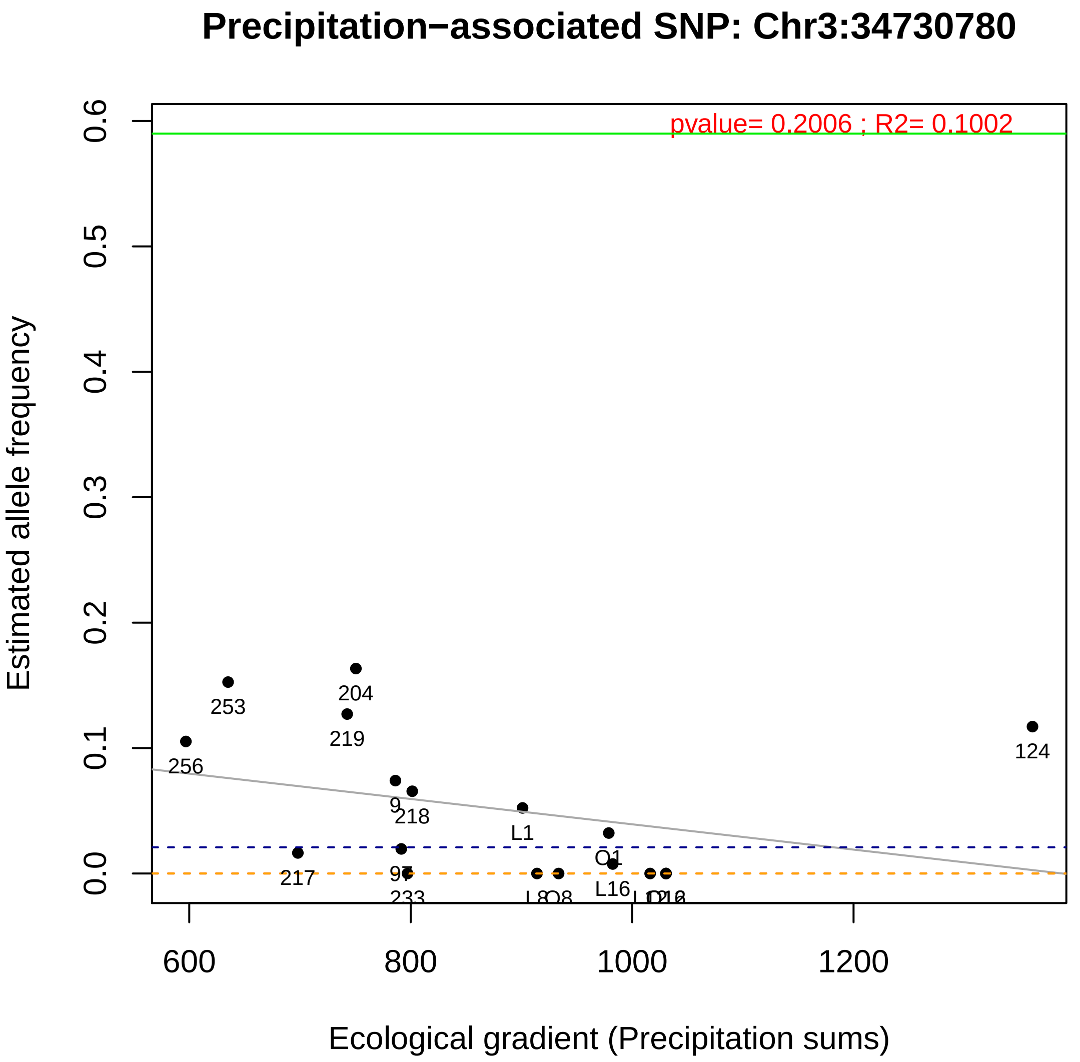

Precipitation−associated SNP: Chr3:34730780
pvalue= 0.2006 ; R2= 0.1002
Estimated allele frequency
0.0
0.1
0.2
0.3
0.4
0.5
0.6
600
800
1000
1200
Ecological gradient (Precipitation sums)
253
204
219
256
124
9
218
L1
217
97
233
O1
L16
L8
O8

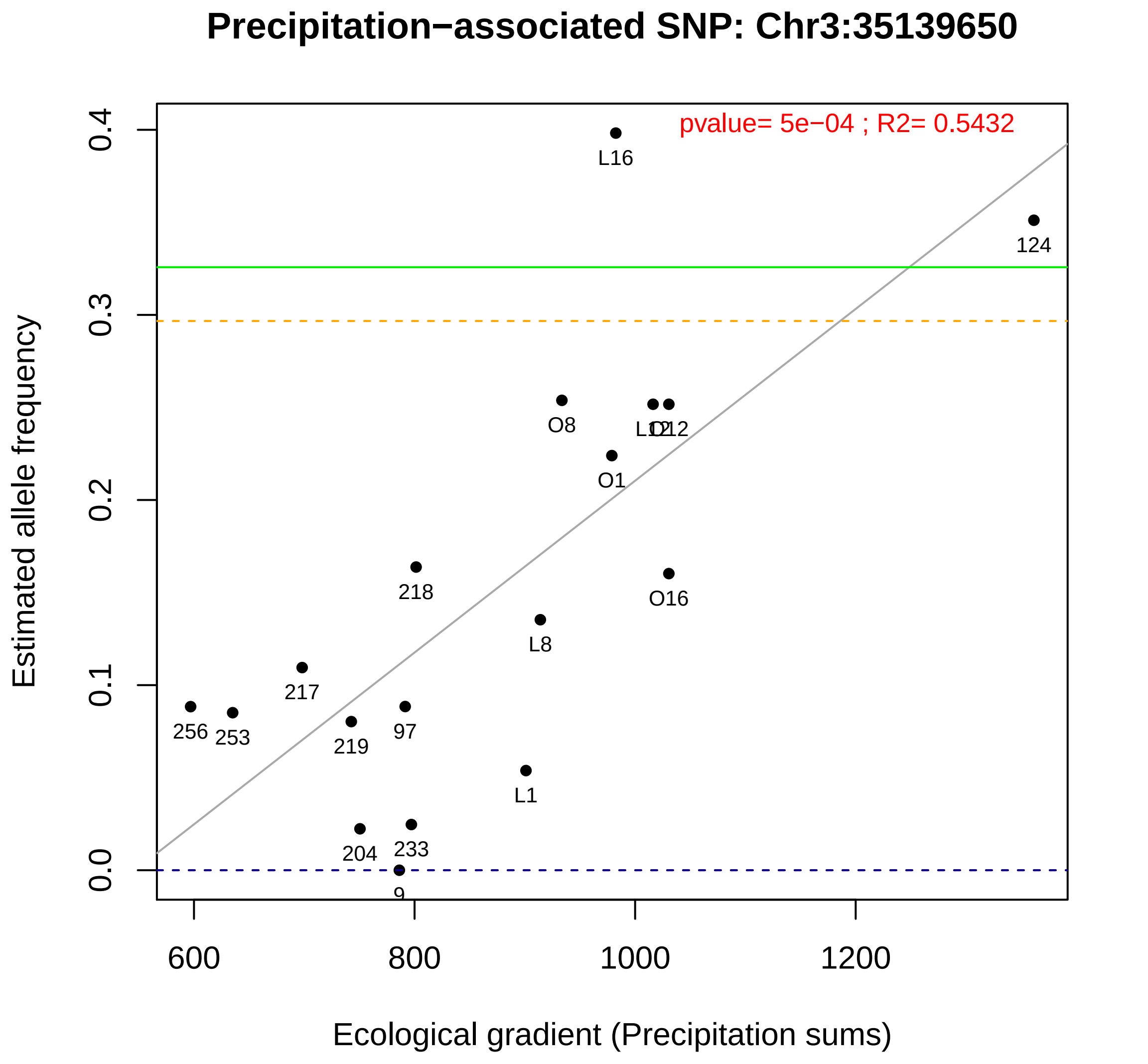
Precipitation−associated SNP: Chr3:35139650
pvalue= 5e−04 ; R2= 0.5432
Estimated allele frequency
Ecological gradient (Precipitation sums)
0.0
0.1
0.2
0.3
0.4
600
800
1000
1200
L16
124
O8
L12
O12
O1
218
O16
L8
217
256
253
219
97
L1
204
233
9

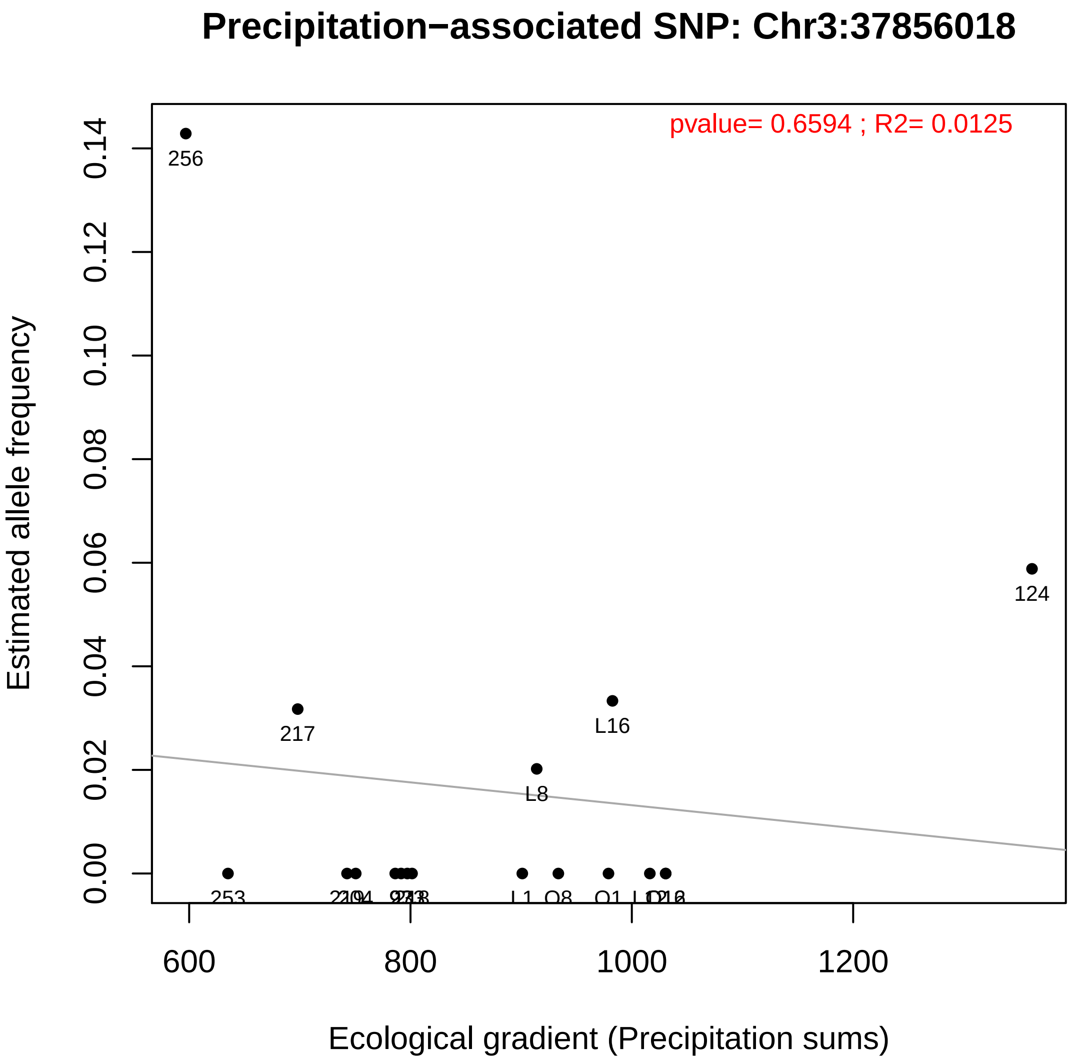
Precipitation−associated SNP: Chr3:37856018
pvalue= 0.6594 ; R2= 0.0125
Estimated allele frequency
0.00
0.02
0.04
0.06
0.08
0.10
0.12
0.14
600
800
1000
1200
Ecological gradient (Precipitation sums)
256
217
L16
L8
124
253
L1
O8
O1

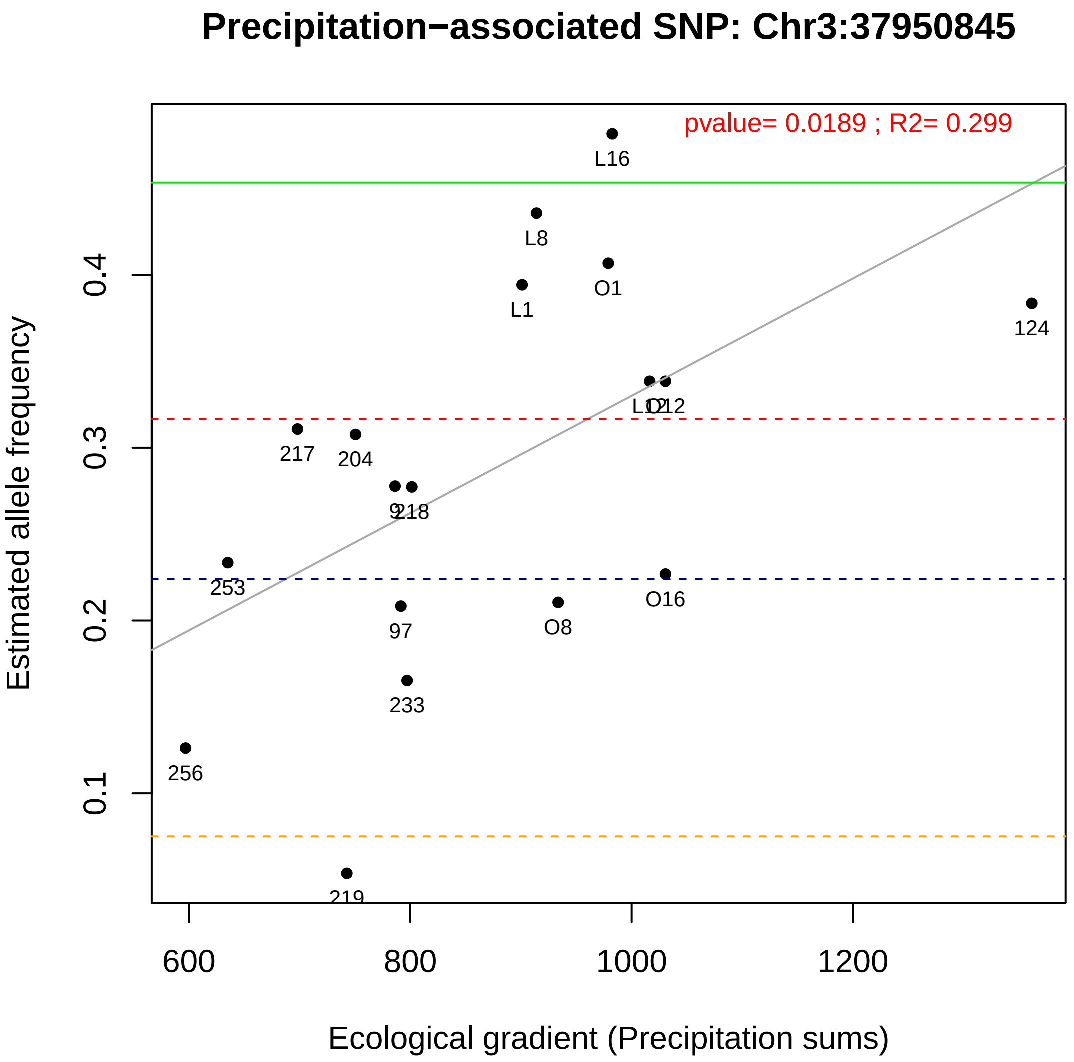

Precipitation−associated SNP: Chr3:37950845
pvalue= 0.0189 ; R2= 0.299
L16
L8
O1
L1
124
L10
O12
217
204
9
218
253
O16
97
O8
233
256
219
0.1
0.2
0.3
0.4
600
800
1000
1200
Estimated allele frequency
Ecological gradient (Precipitation sums)

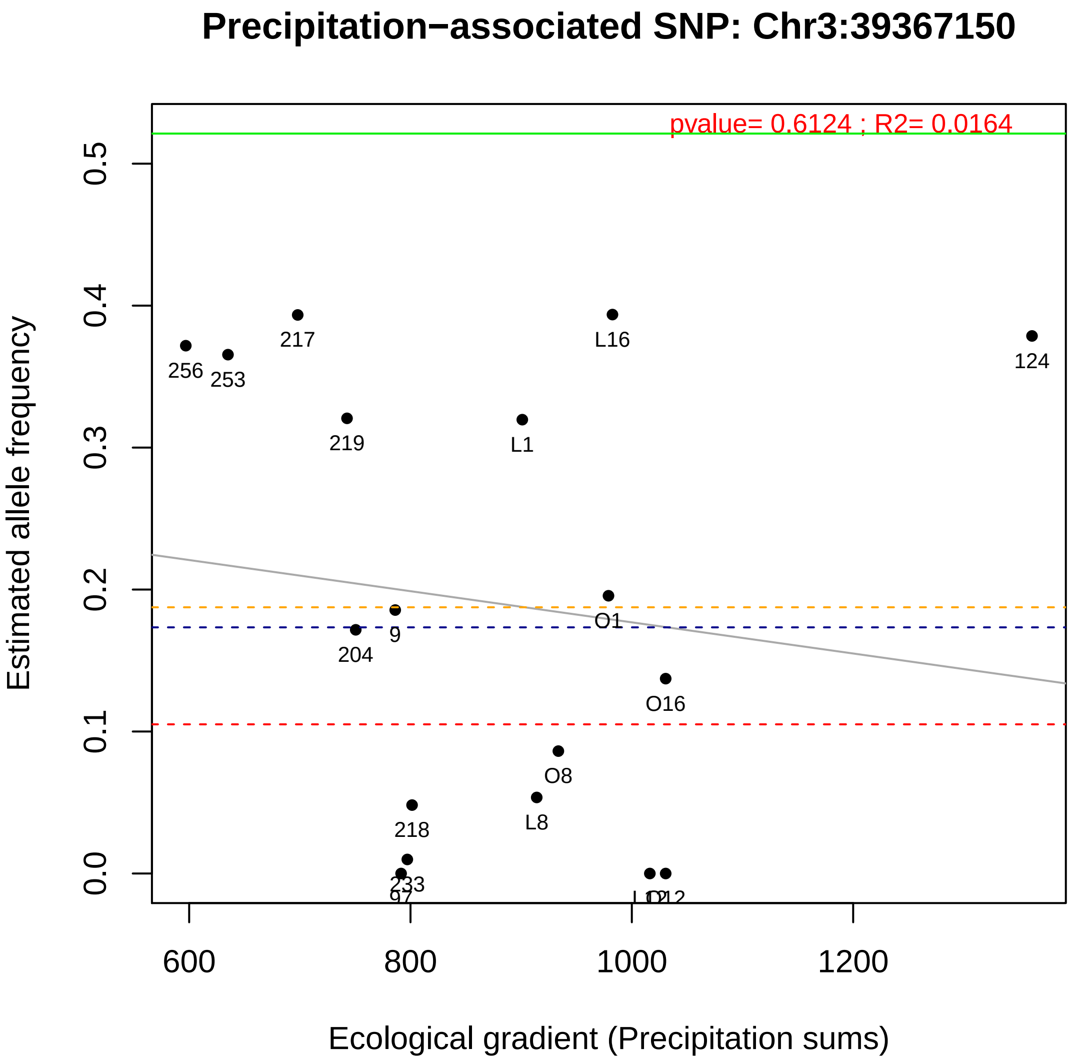

Precipitation−associated SNP: Chr3:39367150
pvalue= 0.6124 ; R2= 0.0164
Estimated allele frequency
0.0
0.1
0.2
0.3
0.4
0.5
600
800
1000
1200
Ecological gradient (Precipitation sums)
256
253
217
219
L1
L16
124
O1
204
9
O16
O8
L8
218
233
97

**Precipitation−associated SNP: Chr3:39545090**

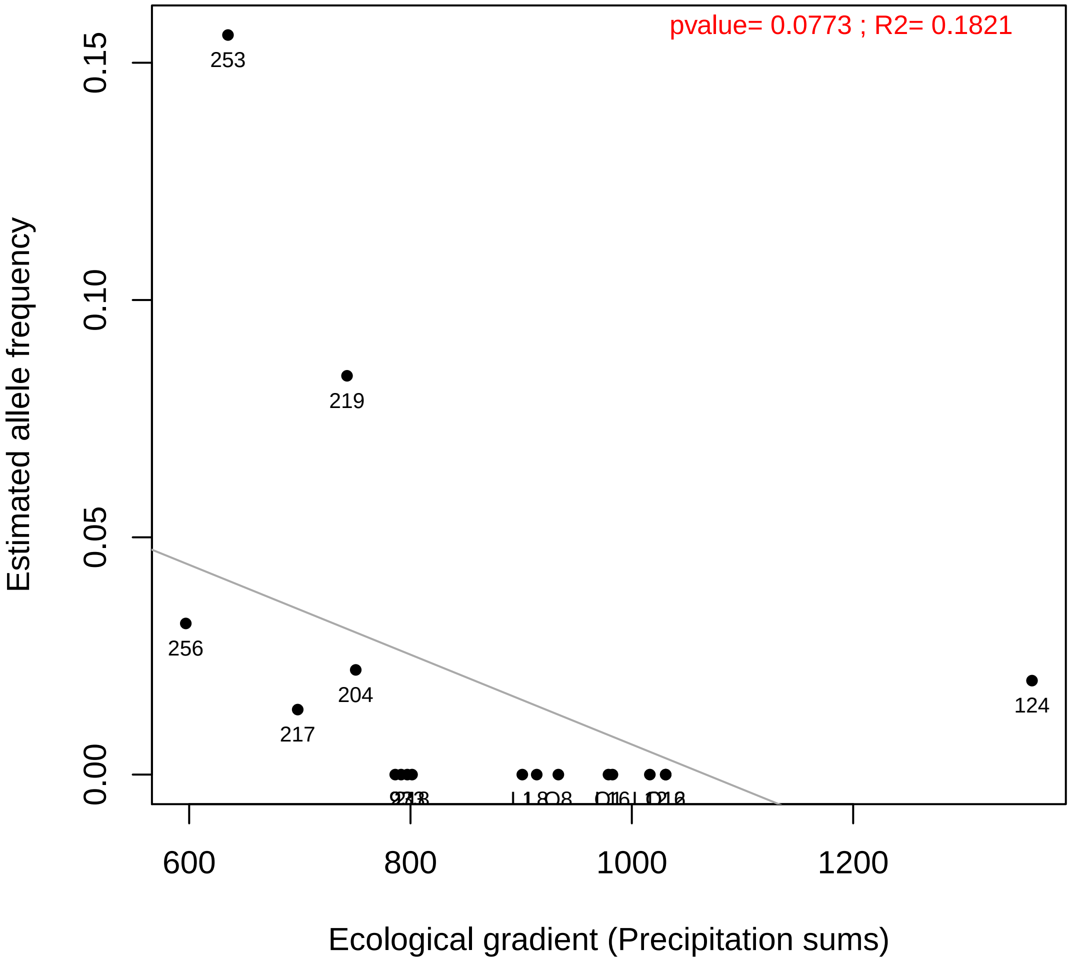

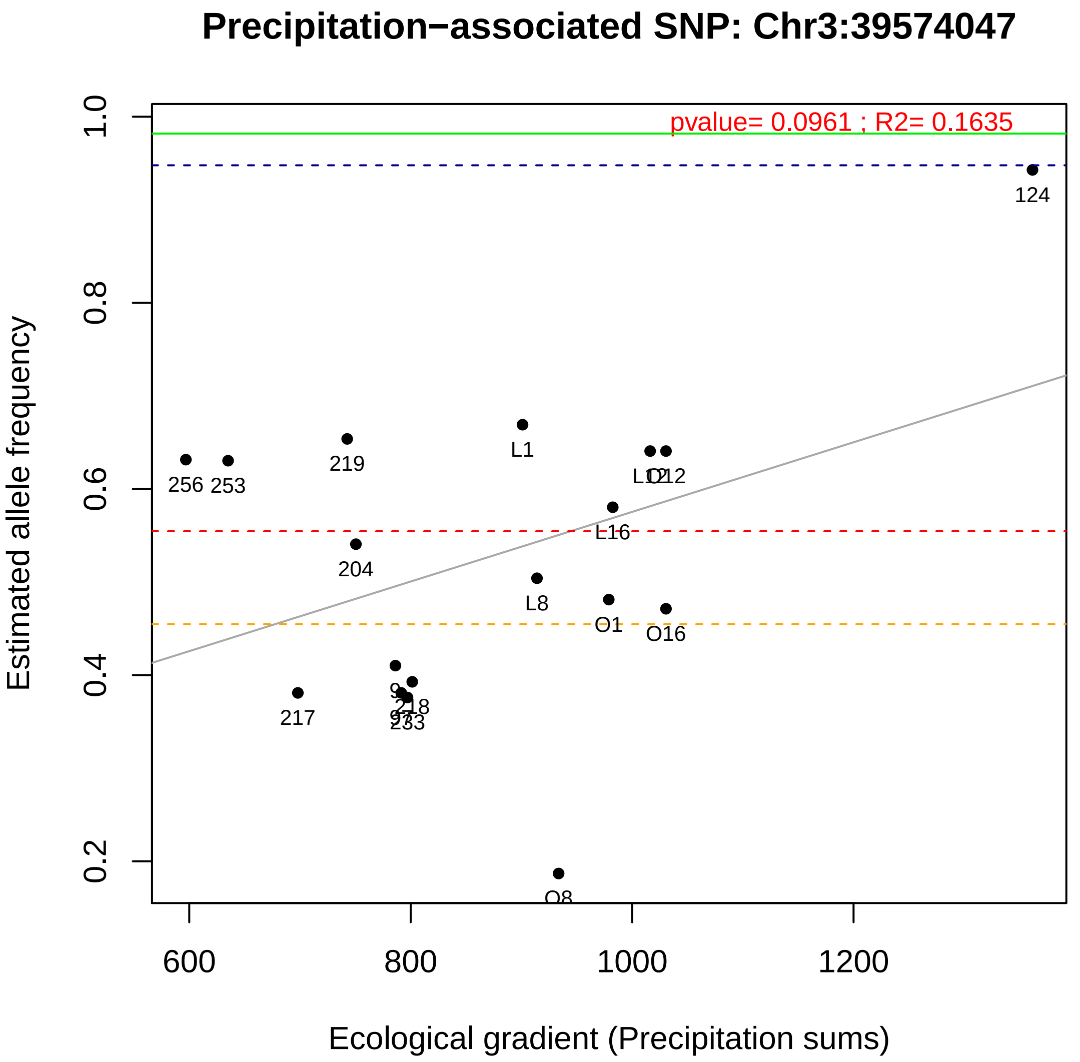

Precipitation−associated SNP: Chr3:39574047
pvalue= 0.0961 ; R2= 0.1635
Estimated allele frequency
Ecological gradient (Precipitation sums)
0.2
0.4
0.6
0.8
1.0
600
800
1000
1200
124
L1
219
256
253
L16
204
L8
O1
O16
217
218
O8

**Precipitation−associated SNP: Chr3:39609169**

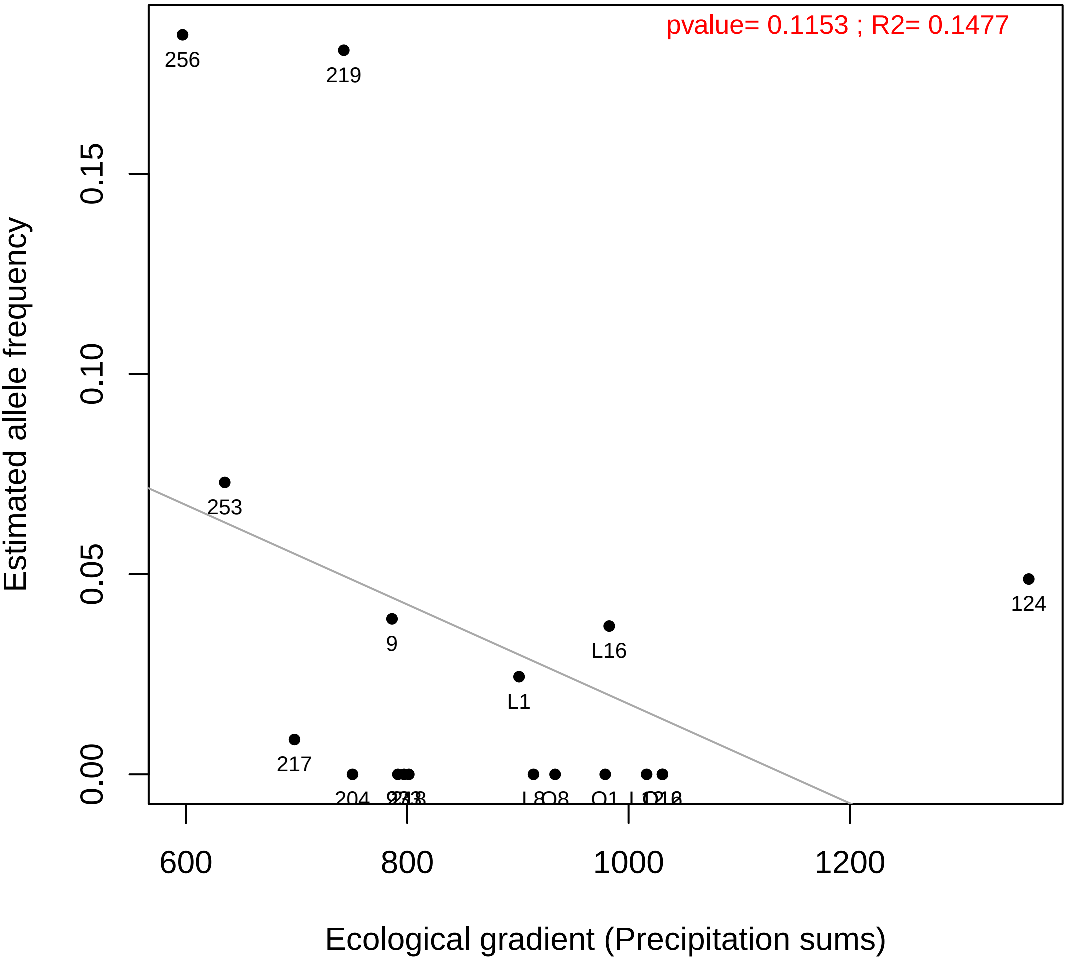

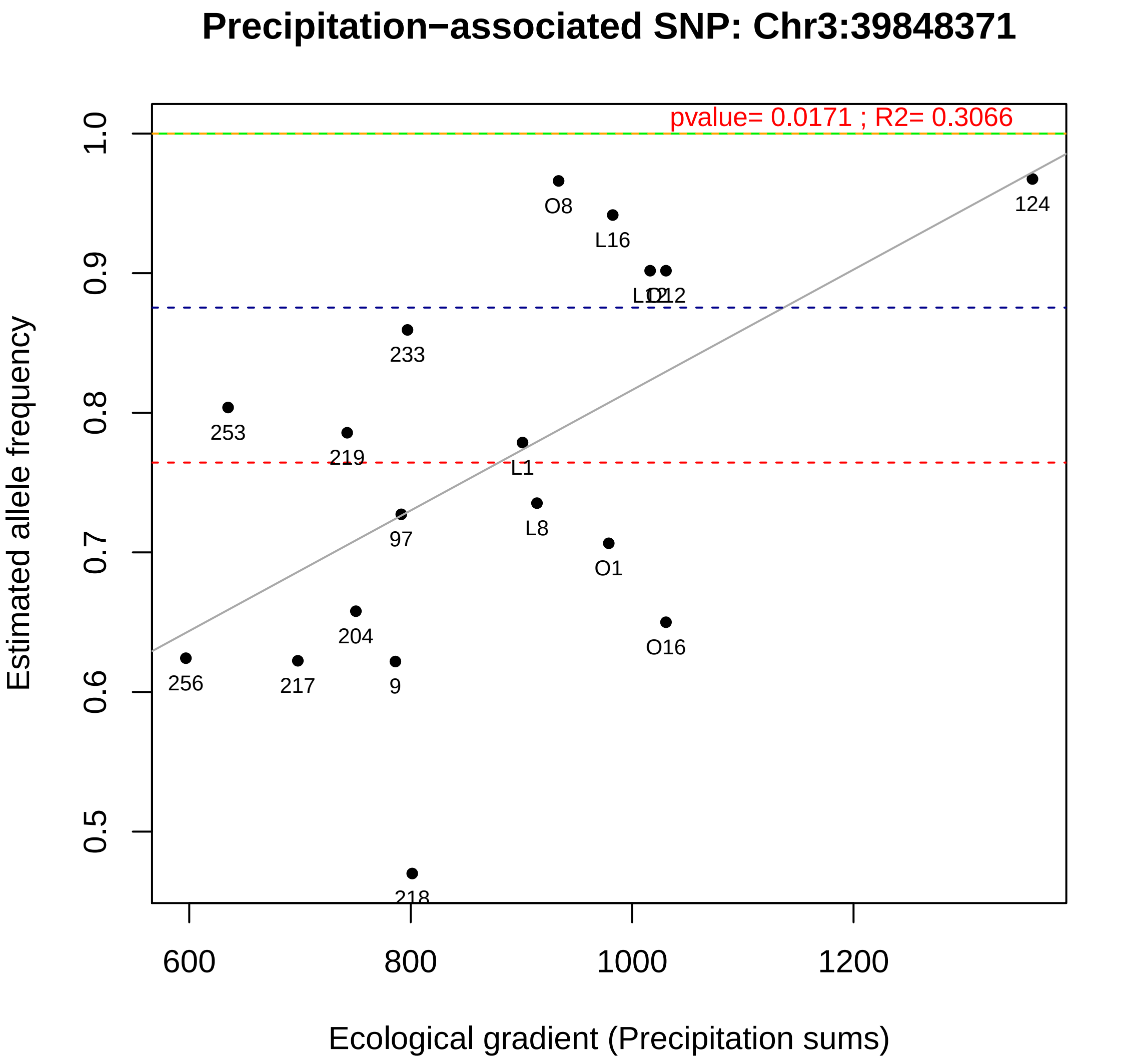
Precipitation−associated SNP: Chr3:39848371
pvalue= 0.0171 ; R2= 0.3066
O8
L16
124
L12
O12
233
253
219
L1
L8
97
O1
204
O16
256
217
9
218
0.5
0.6
0.7
0.8
0.9
1.0
600
800
1000
1200
Estimated allele frequency
Ecological gradient (Precipitation sums)

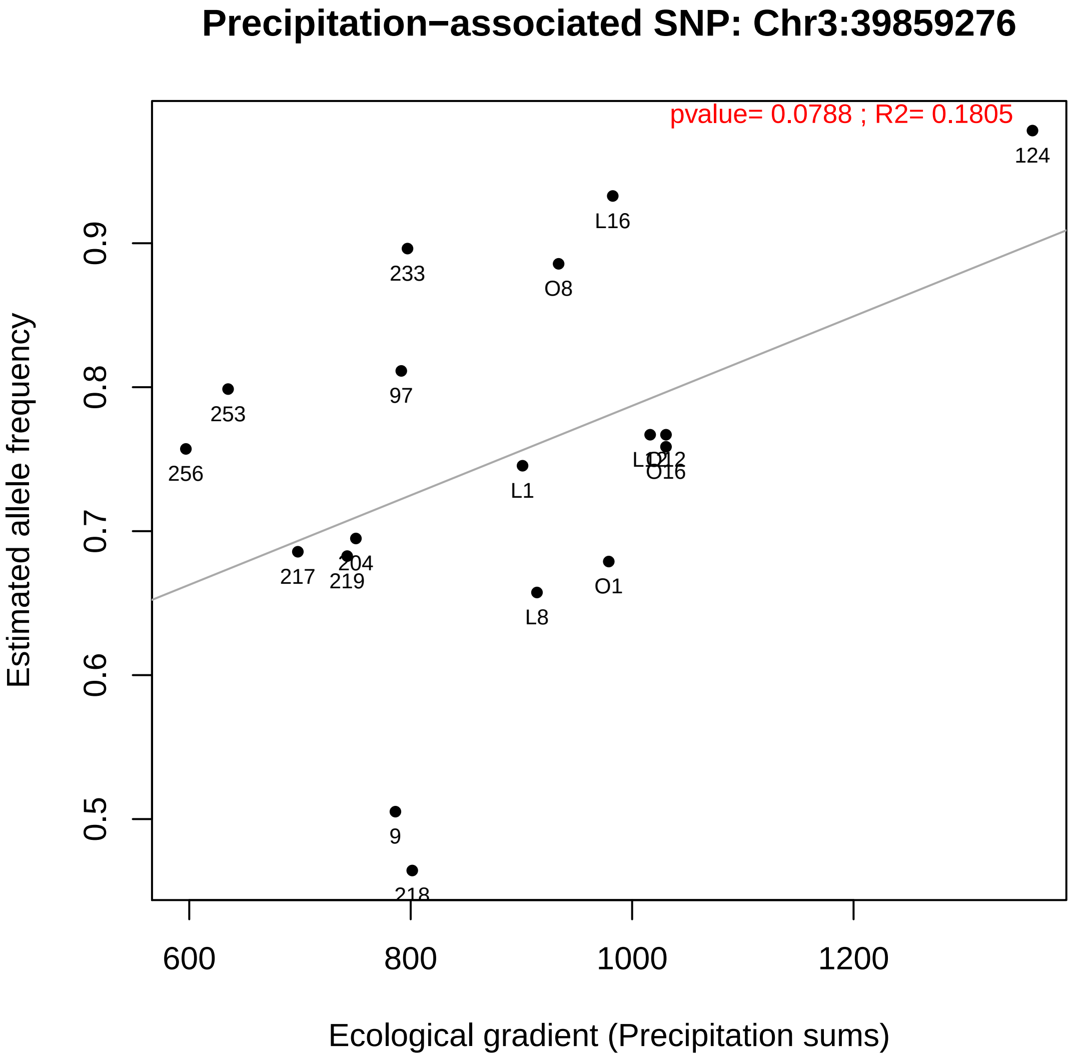
Precipitation−associated SNP: Chr3:39859276
pvalue= 0.0788 ; R2= 0.1805
124
L16
233
O8
97
253
256
L1
L12
O12
O16
217
204
219
O1
L8
9
218
0.5
0.6
0.7
0.8
0.9
Estimated allele frequency
600
800
1000
1200
Ecological gradient (Precipitation sums)

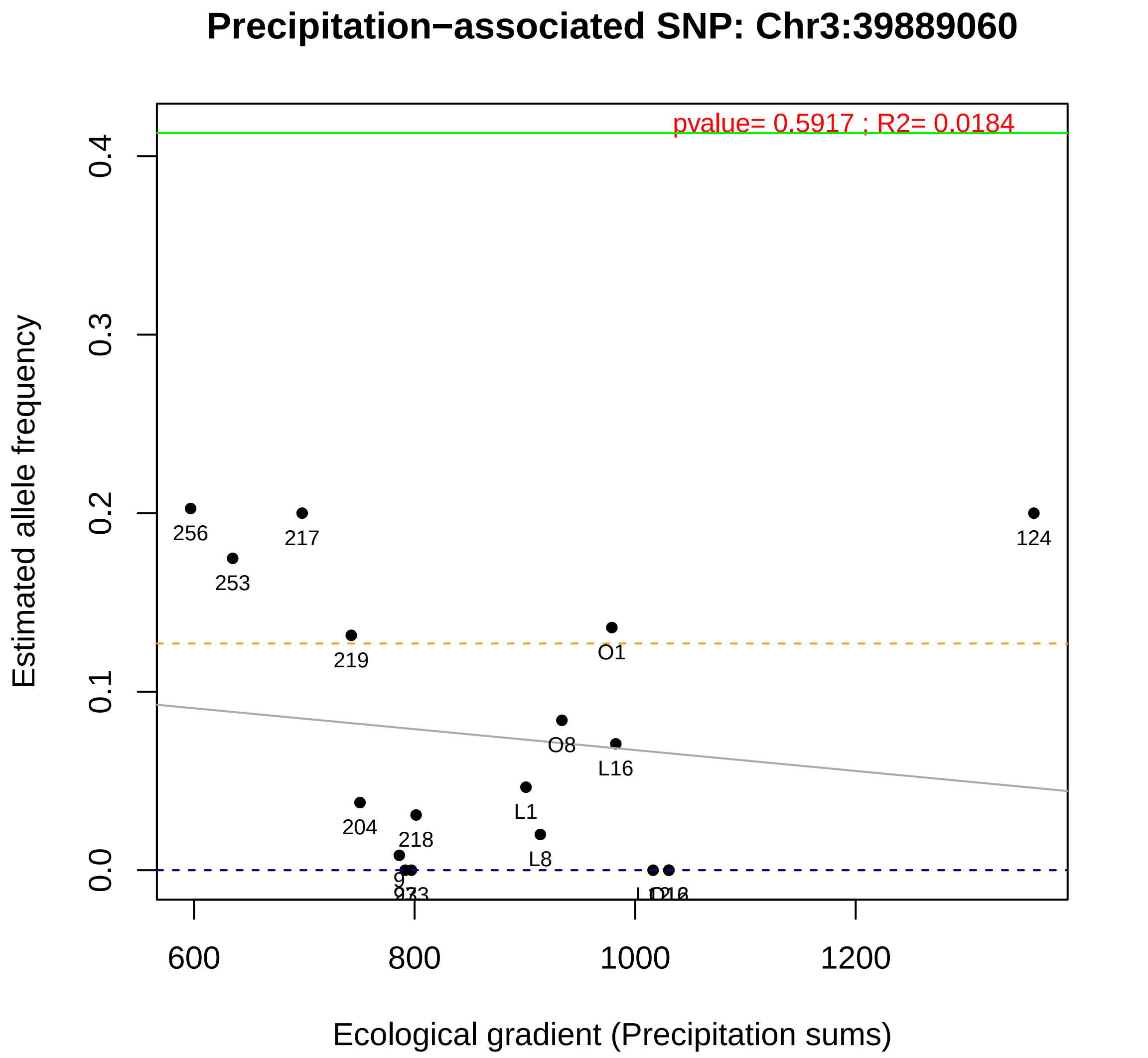
Precipitation−associated SNP: Chr3:39889060
pvalue= 0.5917 ; R2= 0.0184
Estimated allele frequency
0.0
0.1
0.2
0.3
0.4
256
253
217
124
219
O1
O8
L16
L1
204
218
L8
600
800
1000
1200
Ecological gradient (Precipitation sums)

#### Precipitation−associated SNP: Chr3:39955031

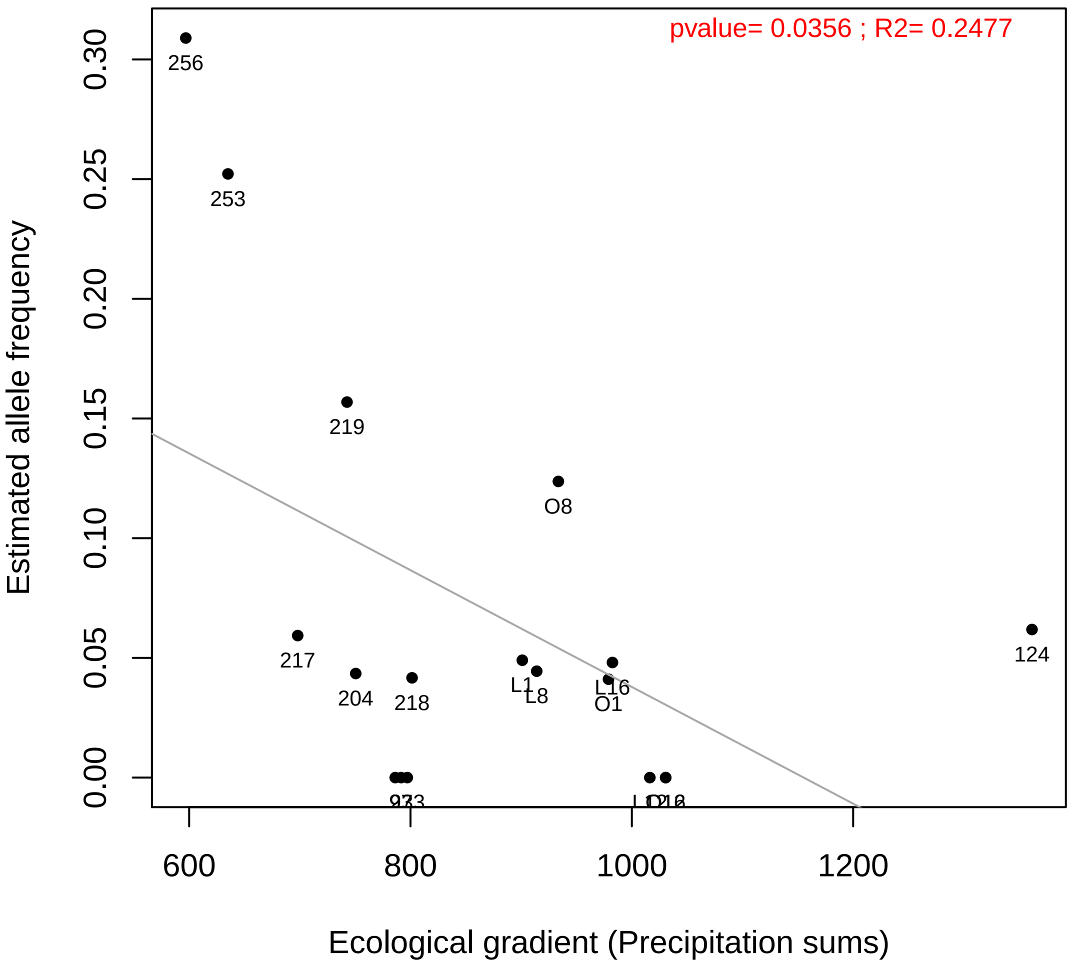

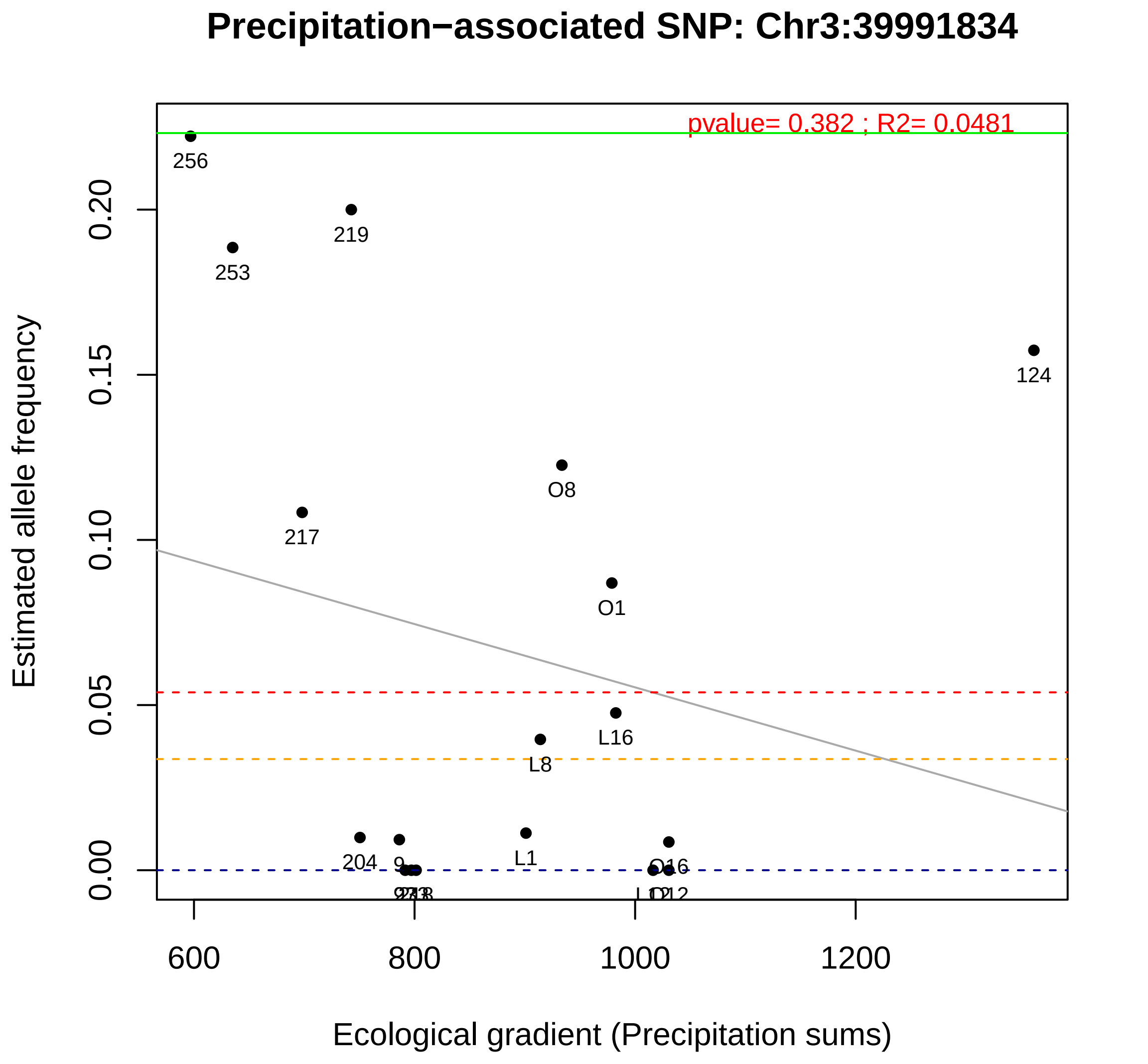

Precipitation−associated SNP: Chr3:39991834
pvalue= 0.382 ; R2= 0.0481
Estimated allele frequency
Ecological gradient (Precipitation sums)
0.00
0.05
0.10
0.15
0.20
600
800
1000
1200
256
253
219
217
124
O8
O1
L16
L8
L1
204
O16

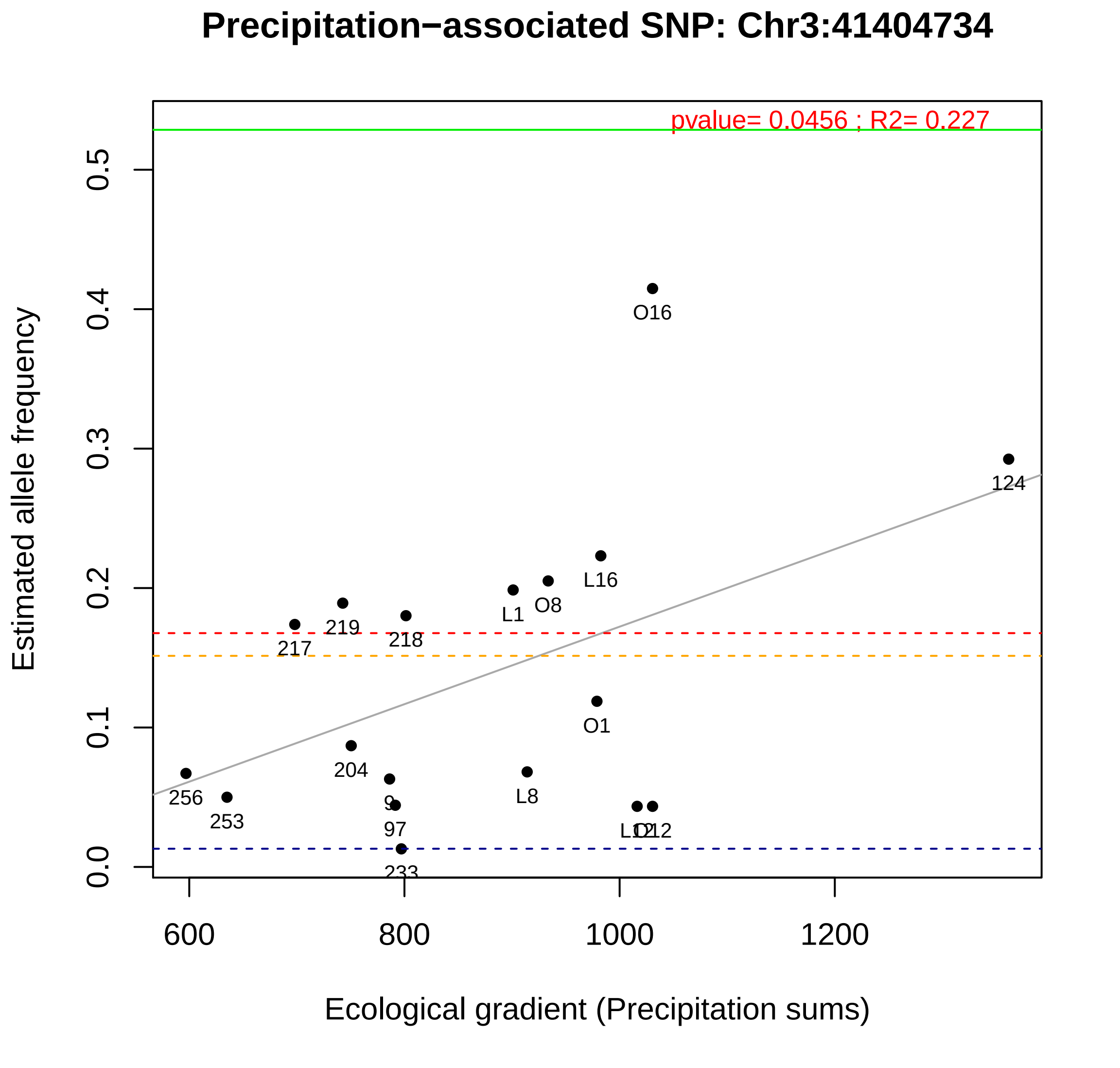
Precipitation−associated SNP: Chr3:41404734
pvalue= 0.0456 ; R2= 0.227
Estimated allele frequency
0.0
0.1
0.2
0.3
0.4
0.5
600
800
1000
1200
Ecological gradient (Precipitation sums)
O16
124
L16
O8
L1
219
218
217
O1
204
256
253
L8
97
233

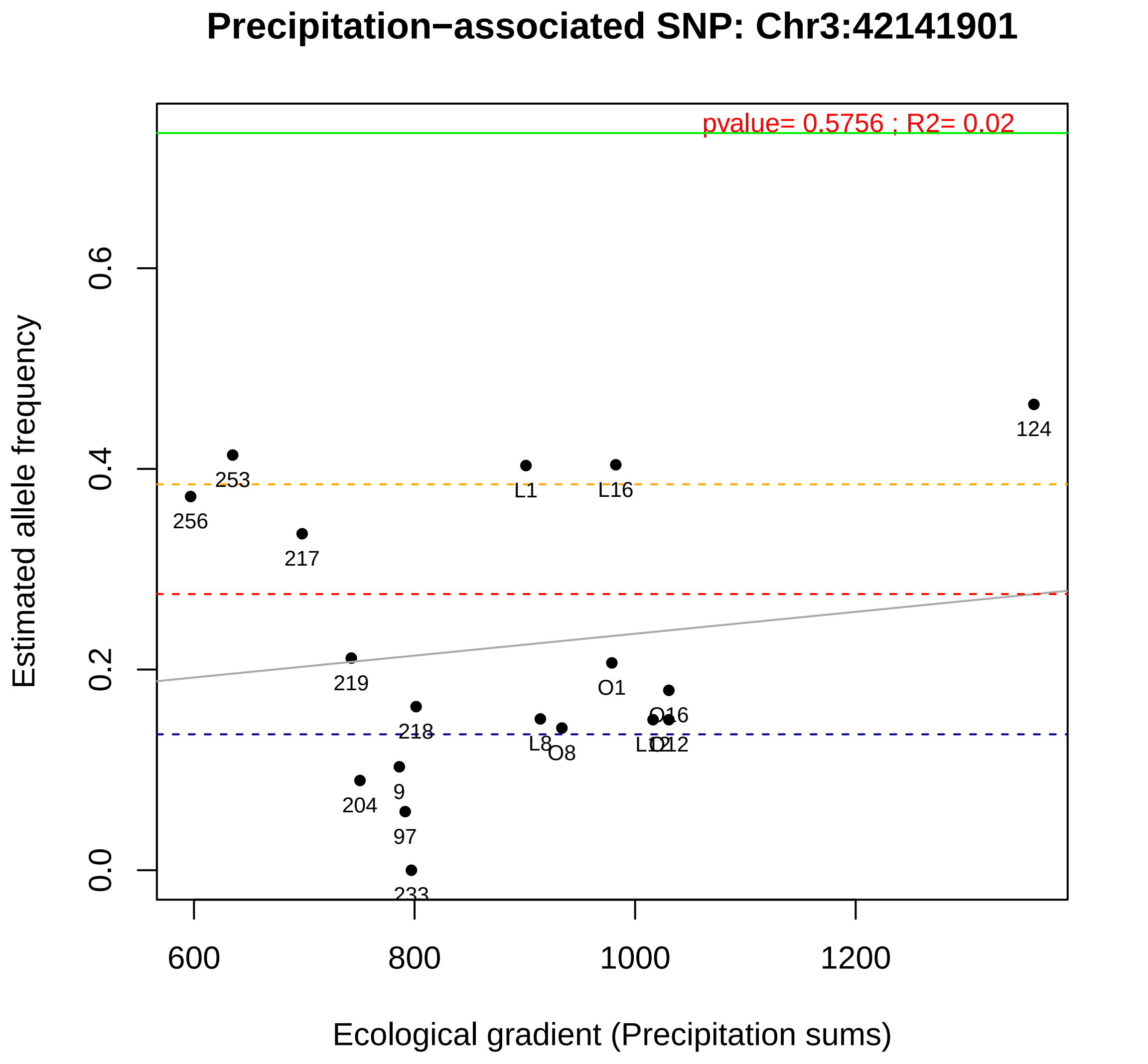

Precipitation−associated SNP: Chr3:42141901
pvalue= 0.5756 ; R2= 0.02
Estimated allele frequency
0.0
0.2
0.4
0.6
600
800
1000
1200
Ecological gradient (Precipitation sums)
256
253
217
219
218
204
9
97
233
L1
L16
L8
O8
O1
O16
L12
O12
124

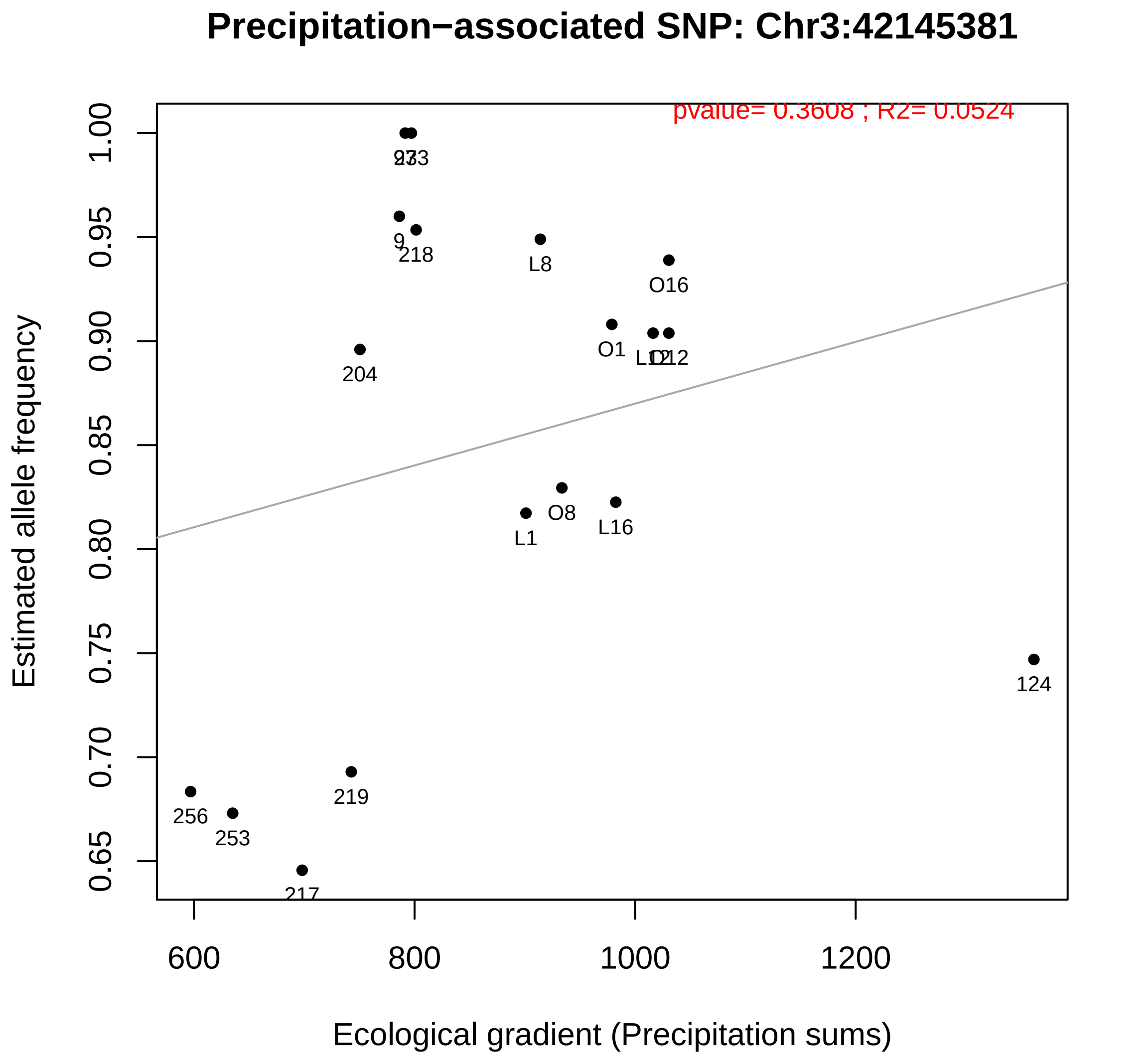

Precipitation−associated SNP: Chr3:42145381
pvalue= 0.3608 ; R2= 0.0524
Estimated allele frequency
Ecological gradient (Precipitation sums)
1.00
0.95
0.90
0.85
0.80
0.75
0.70
0.65
600
800
1000
1200
233
9
218
L8
O16
O1
204
O8
L16
L1
124
219
256
253
217

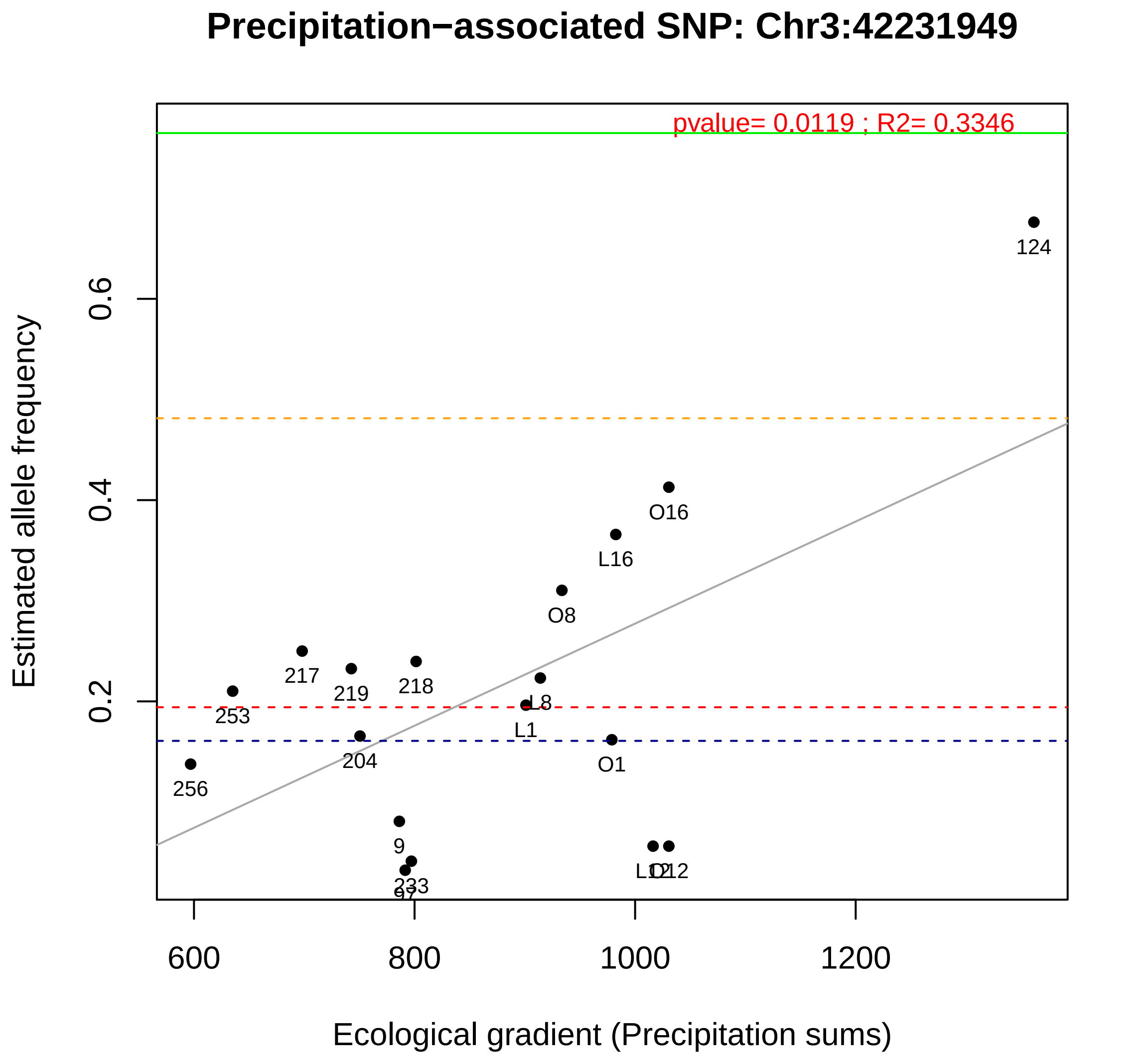

Precipitation−associated SNP: Chr3:42231949
pvalue= 0.0119 ; R2= 0.3346
Estimated allele frequency
0.2
0.4
0.6
600
800
1000
1200
Ecological gradient (Precipitation sums)
124
O16
L16
O8
217
219
218
253
L8
L1
204
O1
256
9
233
L12
O12

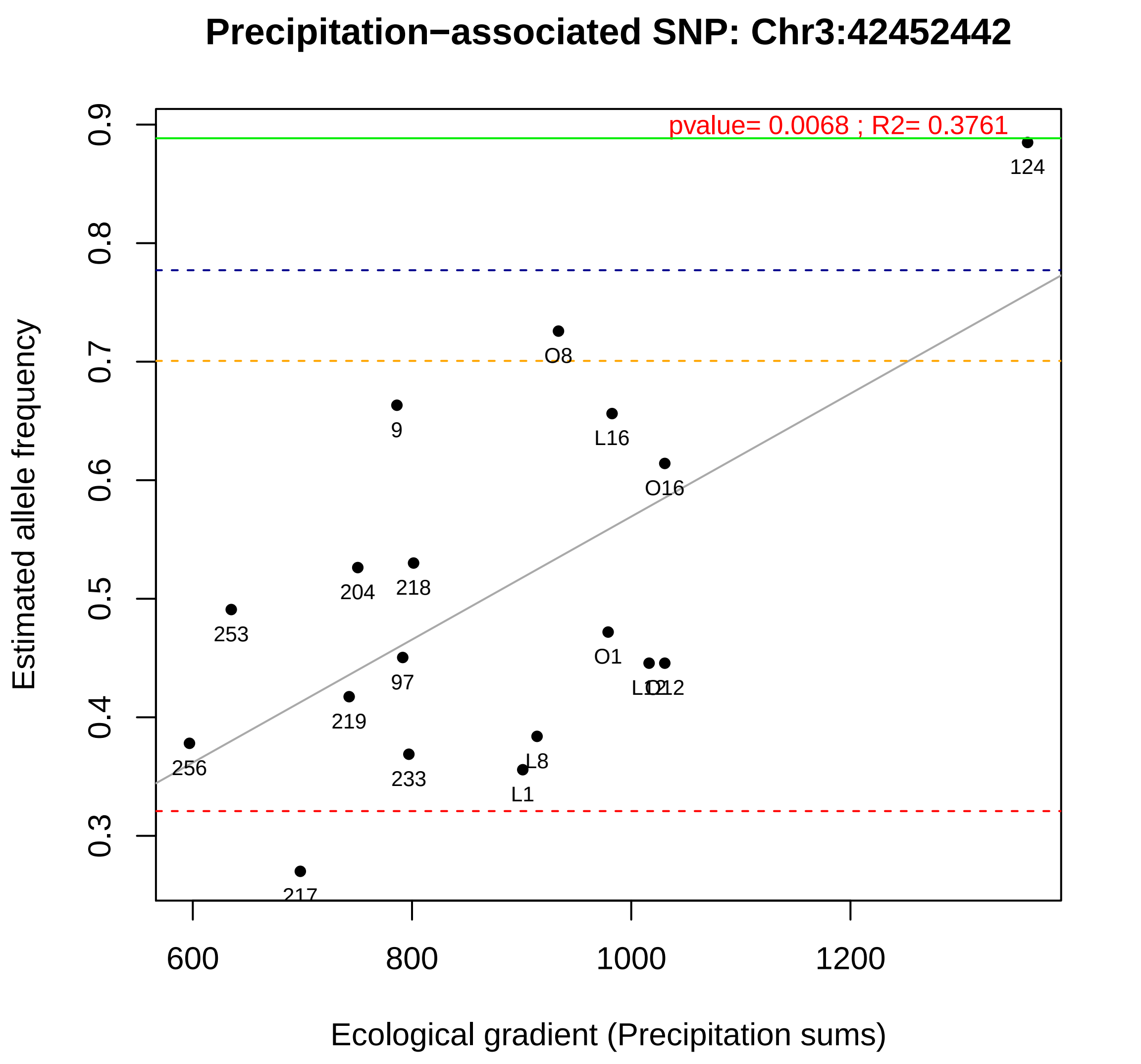
Precipitation−associated SNP: Chr3:42452442
pvalue= 0.0068 ; R2= 0.3761
Estimated allele frequency
Ecological gradient (Precipitation sums)
0.3
0.4
0.5
0.6
0.7
0.8
0.9
600
800
1000
1200
124
O8
9
L16
O16
204
218
253
O1
97
L12
O12
219
256
233
L8
L1
217

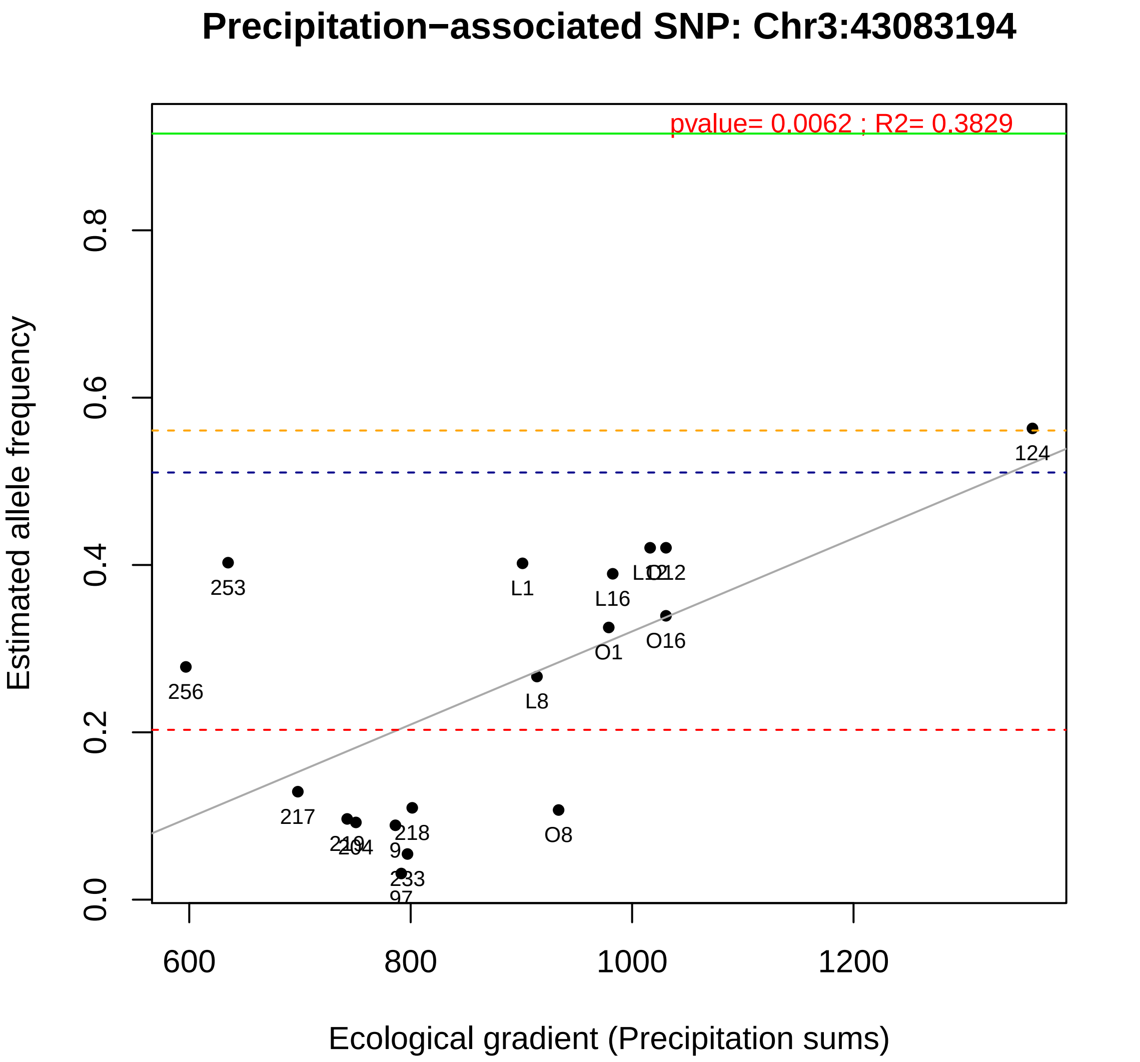

Precipitation−associated SNP: Chr3:43083194
pvalue= 0.0062 ; R2= 0.3829
Estimated allele frequency
0.0
0.2
0.4
0.6
0.8
600
800
1000
1200
Ecological gradient (Precipitation sums)
124
253
L1
L16
O16
O1
256
L8
217
218
O8
233
97

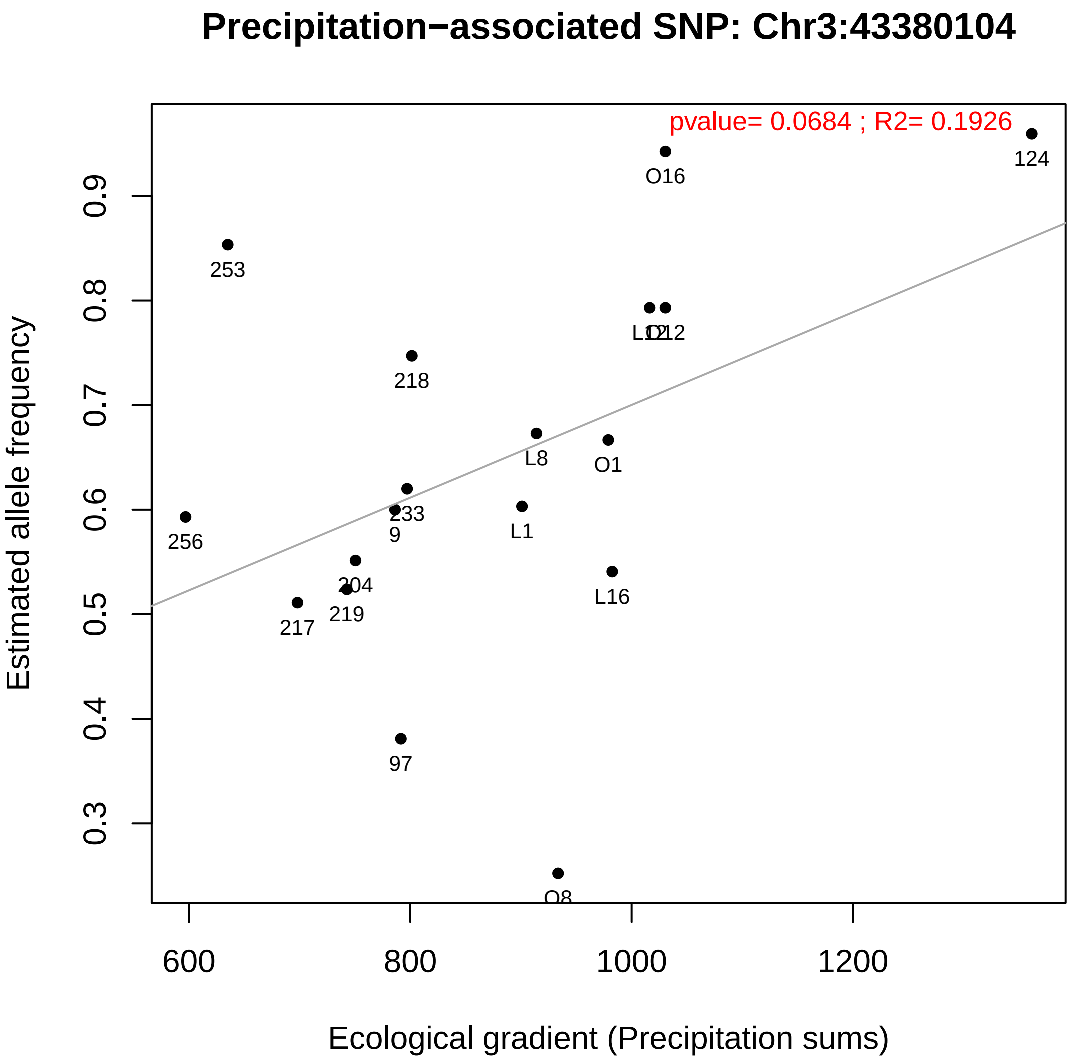
Precipitation−associated SNP: Chr3:43380104
pvalue= 0.0684 ; R2= 0.1926
124
O16
253
L10
O12
218
L8
O1
233
9
L1
256
204
219
217
L16
97
O8
0.3
0.4
0.5
0.6
0.7
0.8
0.9
600
800
1000
1200
Estimated allele frequency
Ecological gradient (Precipitation sums)

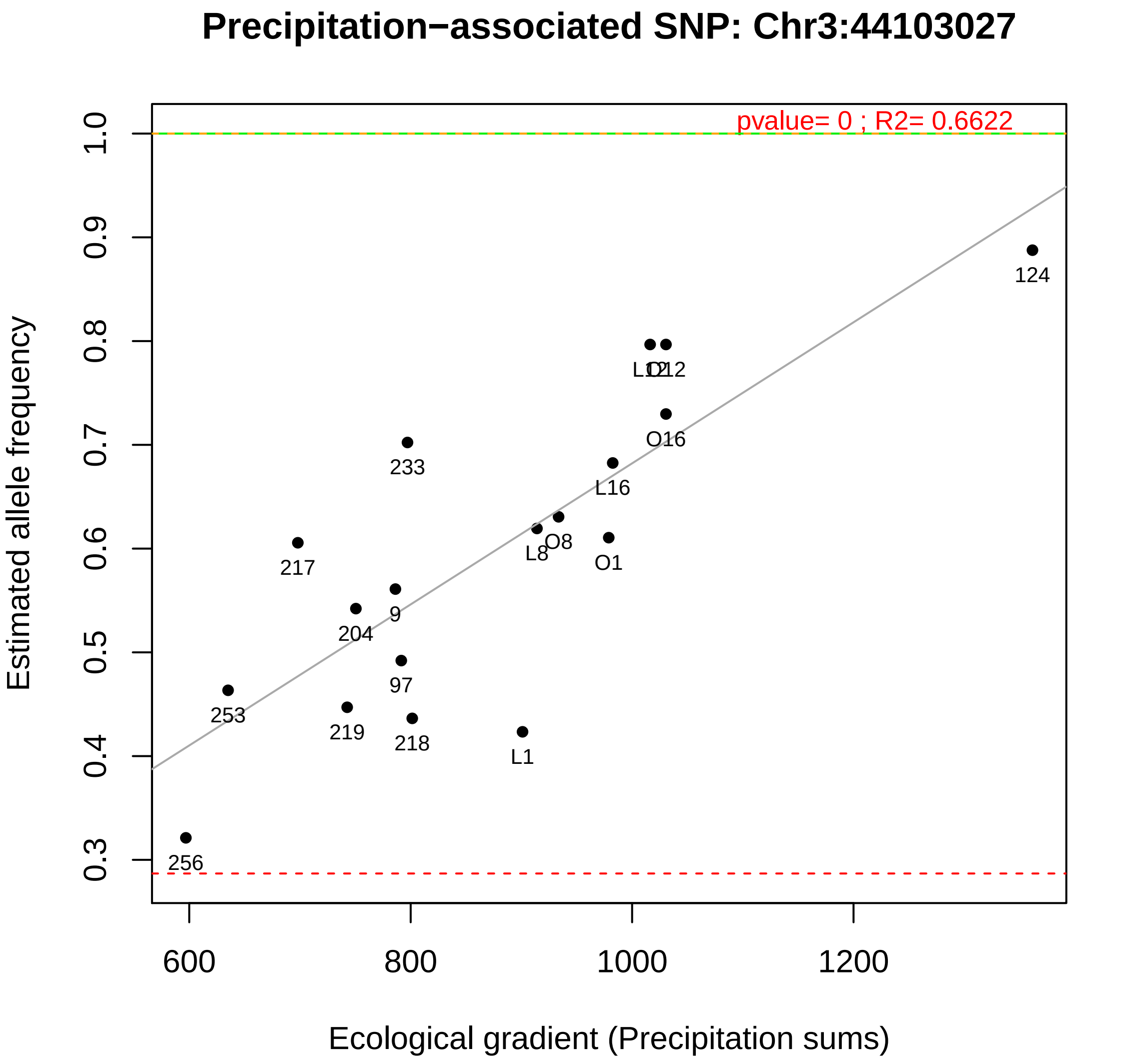
Precipitation−associated SNP: Chr3:44103027
pvalue= 0 ; R2= 0.6622
Estimated allele frequency
Ecological gradient (Precipitation sums)
0.3
0.4
0.5
0.6
0.7
0.8
0.9
1.0
600
800
1000
1200
124
O12
O16
233
L16
L8
O8
O1
217
9
204
97
253
219
218
L1
256

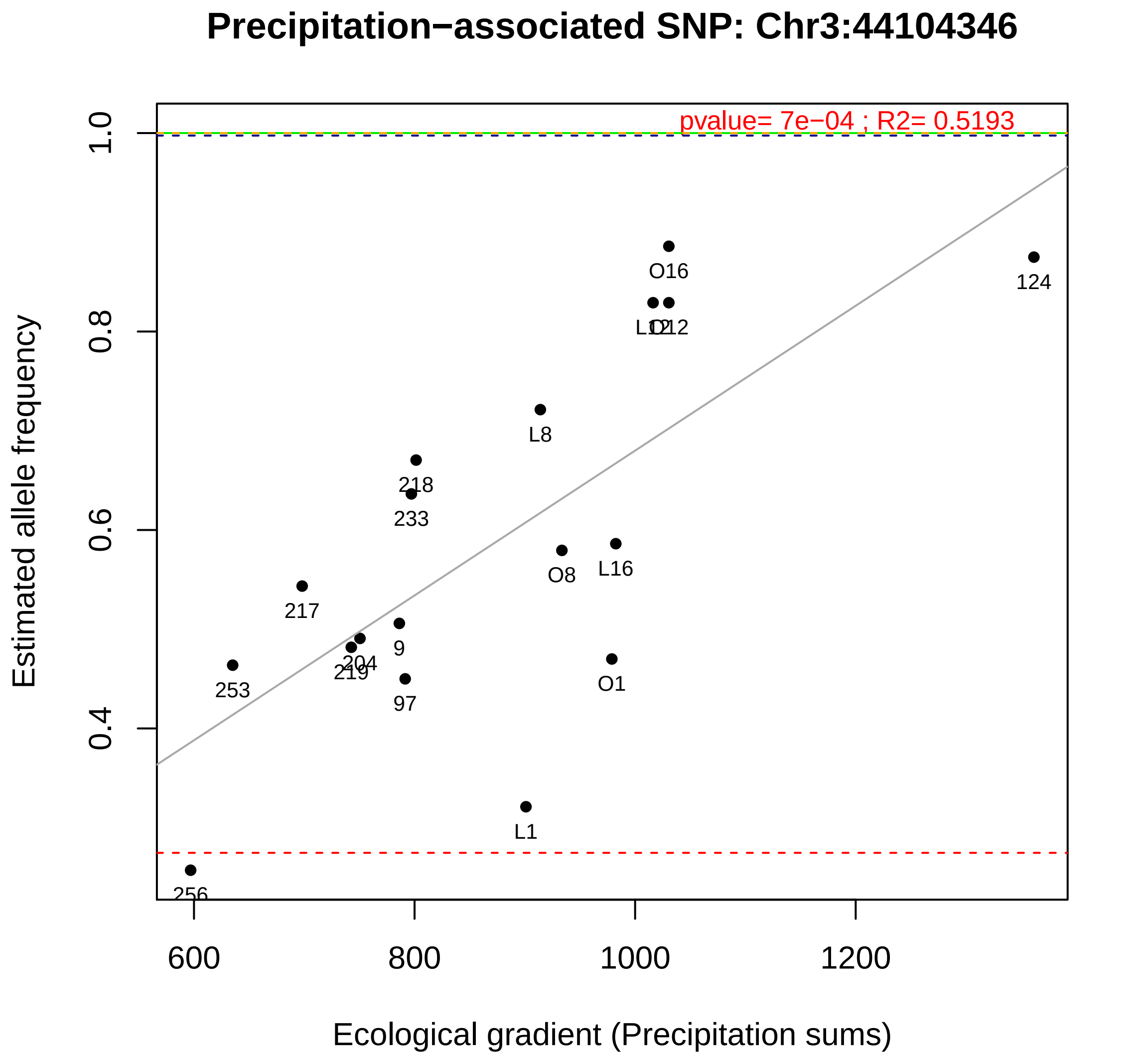

Precipitation−associated SNP: Chr3:44104346
pvalue= 7e−04 ; R2= 0.5193
Estimated allele frequency
Ecological gradient (Precipitation sums)
0.4
0.6
0.8
1.0
600
800
1000
1200
O16
124
L10
O12
L8
218
233
O8
L16
217
9
204
219
253
97
O1
L1
256

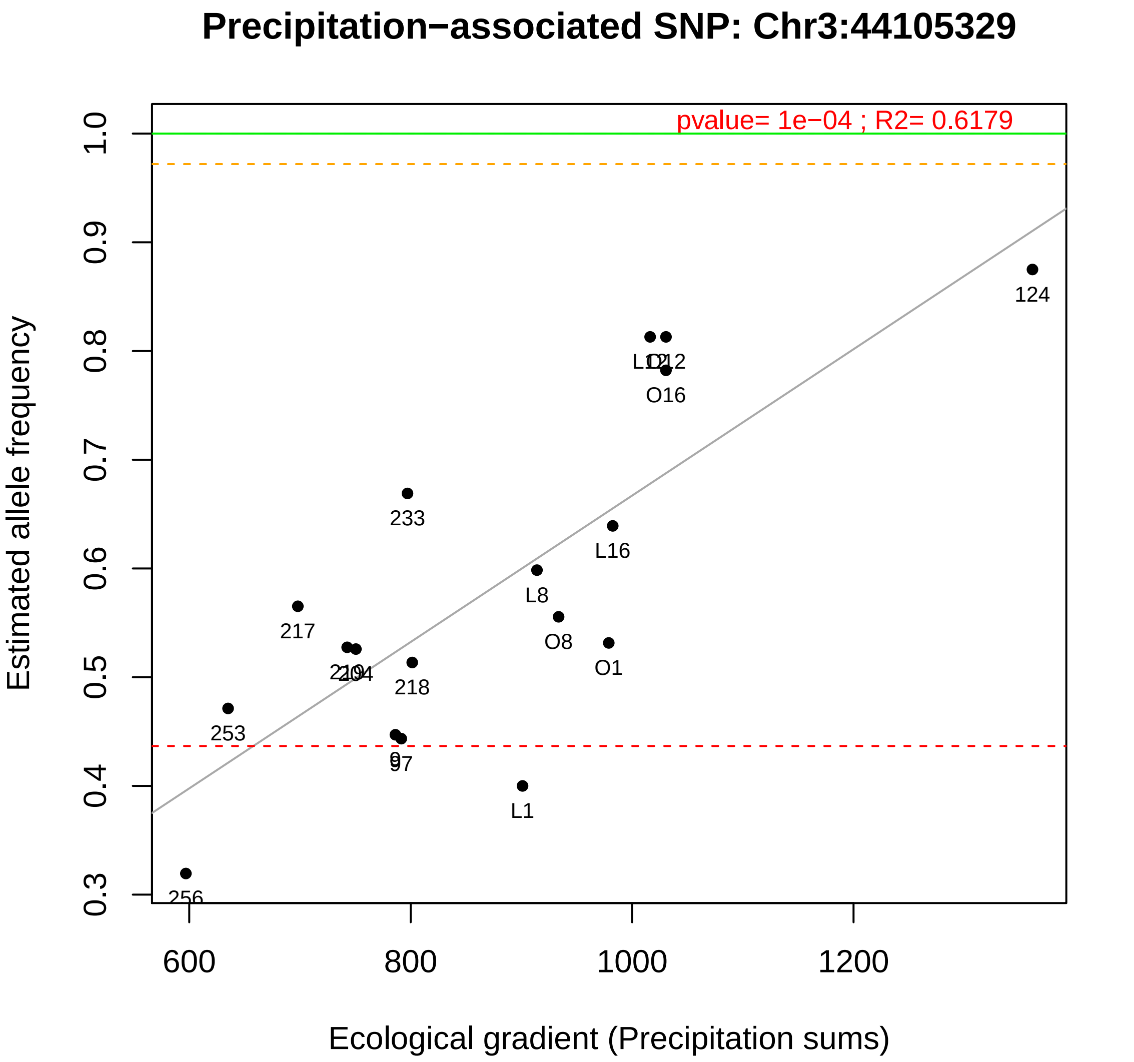
Precipitation−associated SNP: Chr3:44105329
pvalue= 1e−04 ; R2= 0.6179
Estimated allele frequency
0.3
0.4
0.5
0.6
0.7
0.8
0.9
1.0
600
800
1000
1200
Ecological gradient (Precipitation sums)
124
O16
233
L16
L8
O8
O1
217
218
253
97
L1
256

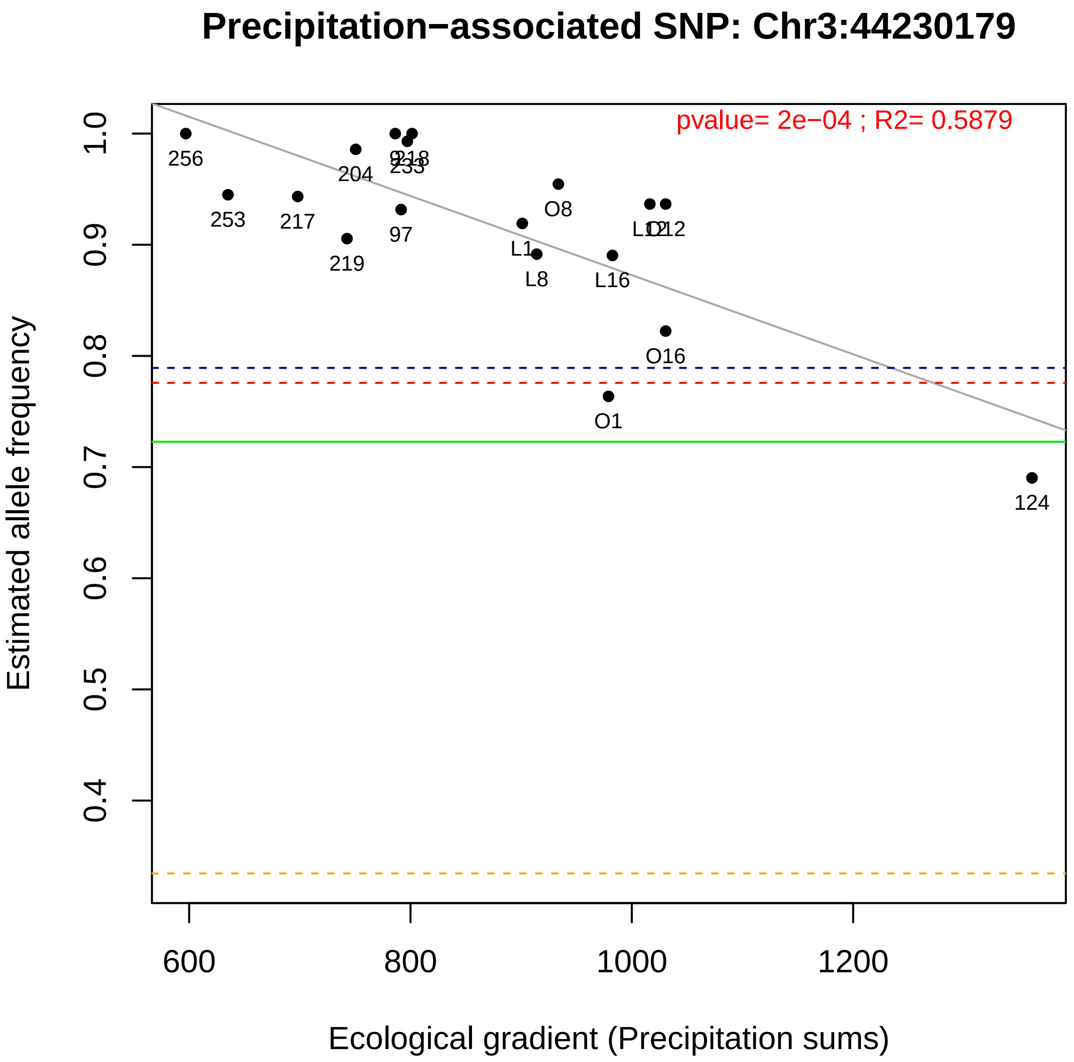

Precipitation−associated SNP: Chr3:44230179
pvalue= 2e−04 ; R2= 0.5879
Estimated allele frequency
Ecological gradient (Precipitation sums)
0.4
0.5
0.6
0.7
0.8
0.9
1.0
600
800
1000
1200
256
253
217
204
219
97
218
233
O8
L1
L8
L16
O12
O16
O1
124

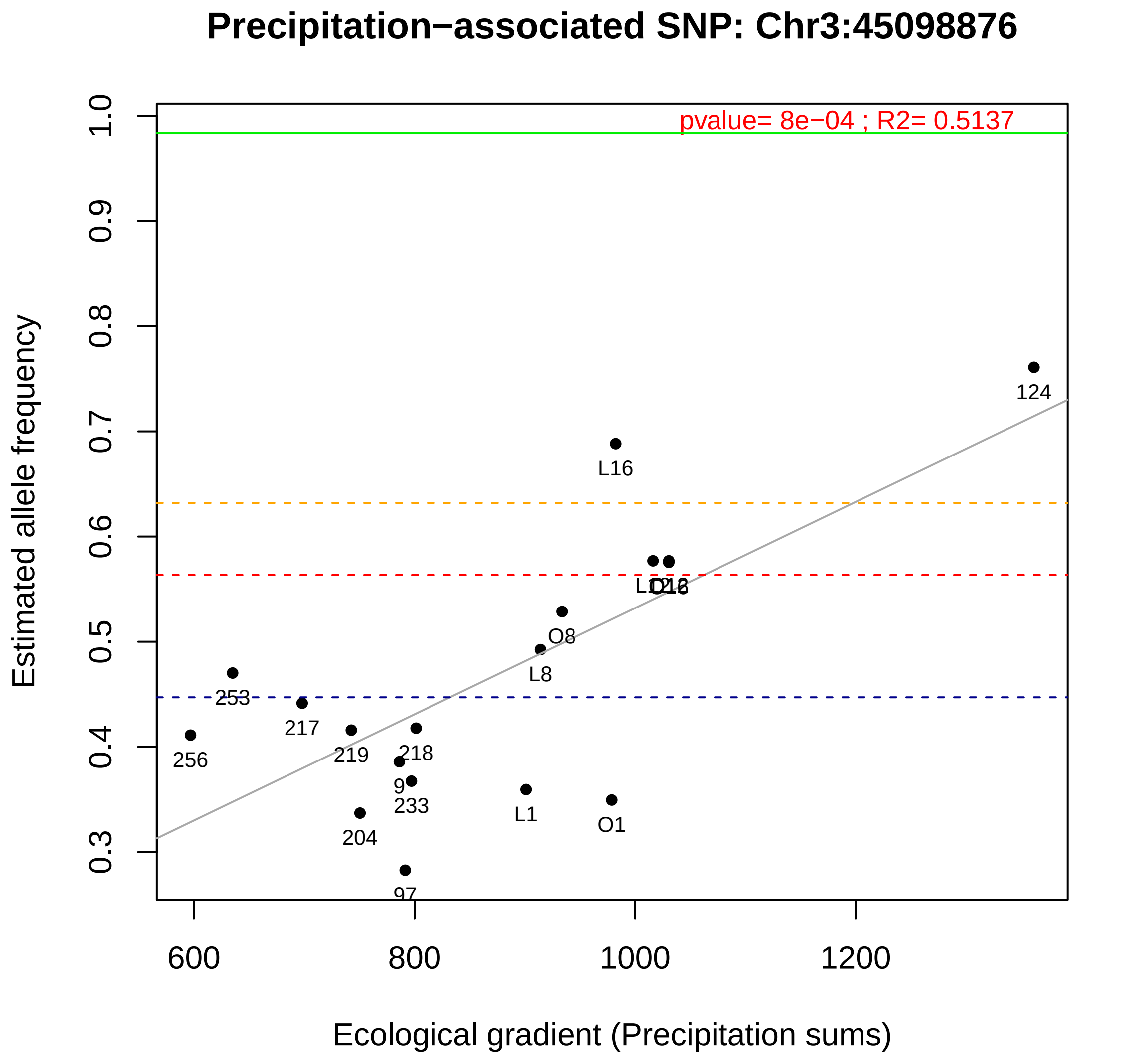

Precipitation−associated SNP: Chr3:45098876
pvalue= 8e−04 ; R2= 0.5137
Estimated allele frequency
Ecological gradient (Precipitation sums)
0.3
0.4
0.5
0.6
0.7
0.8
0.9
1.0
600
800
1000
1200
124
L16
O8
L8
253
217
256
219
218
9
233
204
L1
O1
97

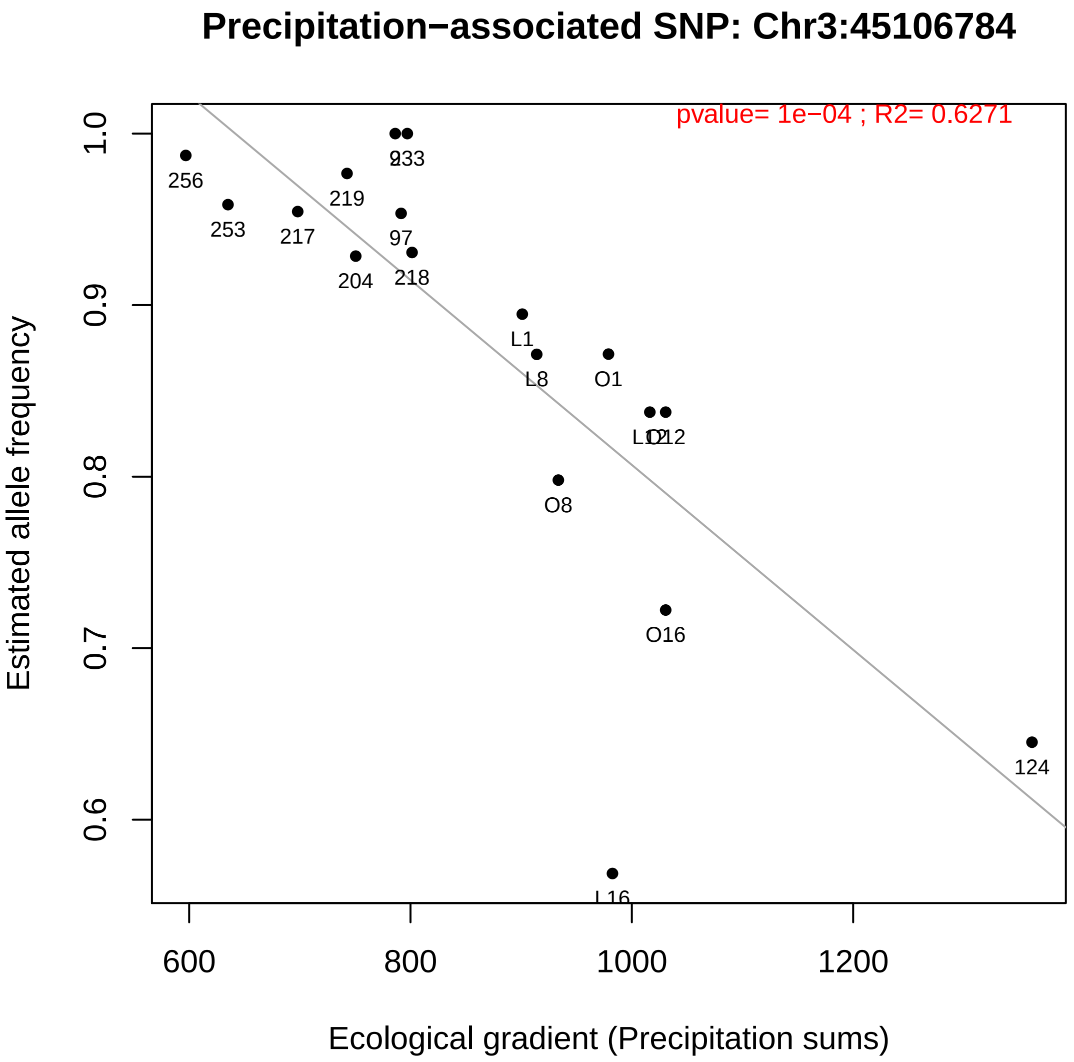
Precipitation−associated SNP: Chr3:45106784
pvalue= 1e−04 ; R2= 0.6271
Estimated allele frequency
Ecological gradient (Precipitation sums)
0.6
0.7
0.8
0.9
1.0
600
800
1000
1200
256
253
217
219
204
97
218
233
L1
L8
O1
O8
O12
O16
L16
124

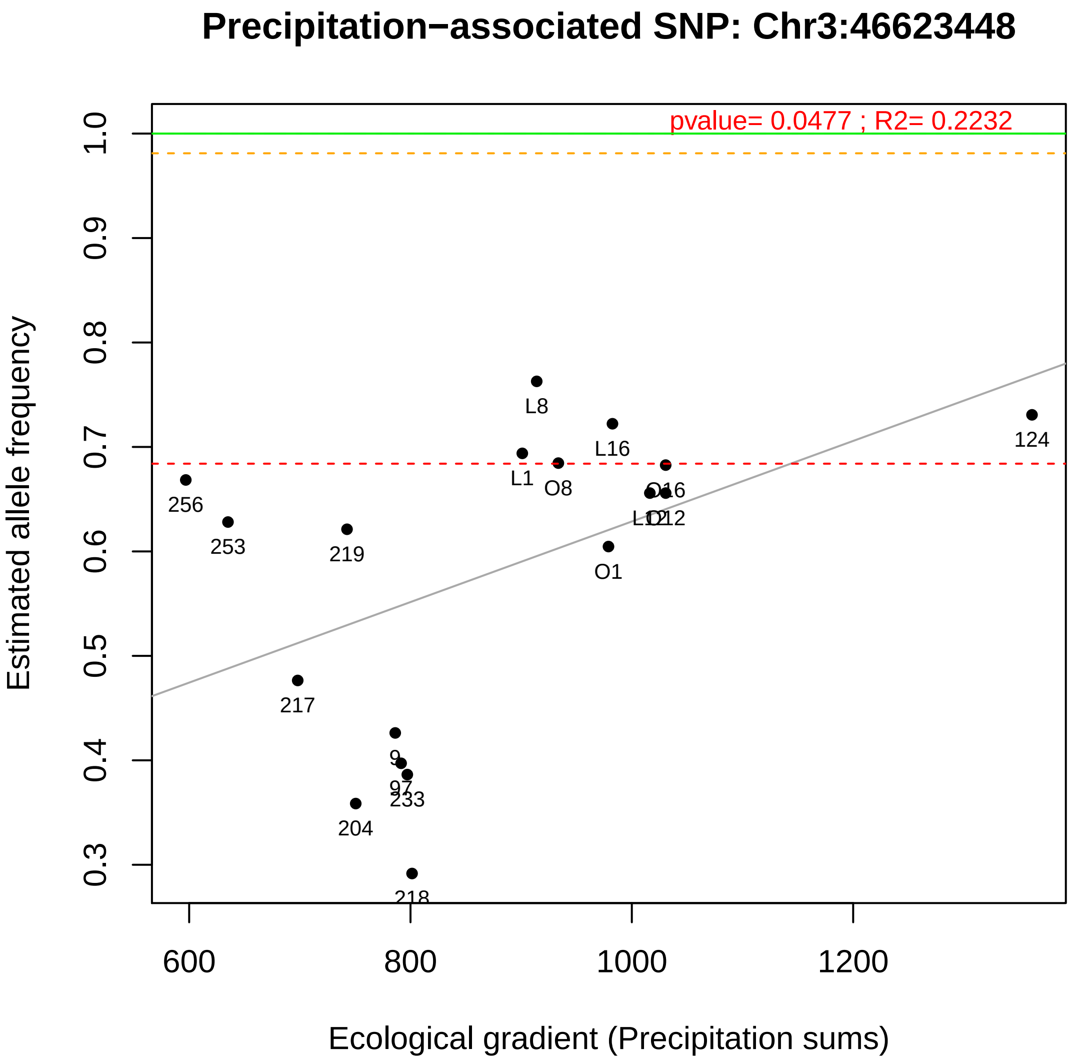

Precipitation−associated SNP: Chr3:46623448
pvalue= 0.0477 ; R2= 0.2232
Estimated allele frequency
Ecological gradient (Precipitation sums)
1.0
0.9
0.8
0.7
0.6
0.5
0.4
0.3
600
800
1000
1200
L8
L16
124
L1
O8
256
O16
L12
O12
253
219
O1
217
9
97
233
204
218

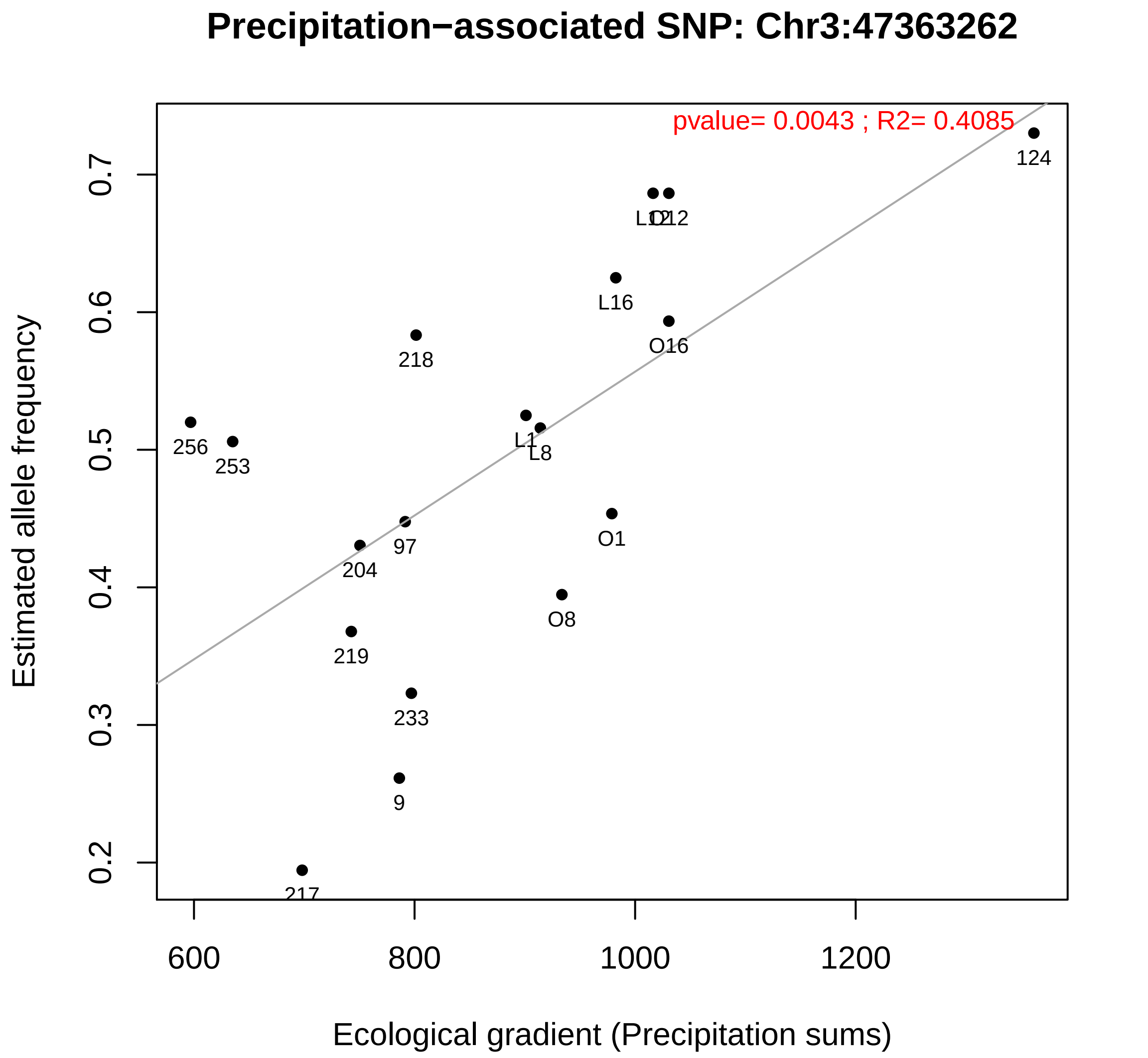

Precipitation−associated SNP: Chr3:47363262
pvalue= 0.0043 ; R2= 0.4085
Estimated allele frequency
Ecological gradient (Precipitation sums)
0.2
0.3
0.4
0.5
0.6
0.7
600
800
1000
1200
124
L10
O12
L16
O16
218
256
253
L1
L8
97
204
O1
O8
219
233
9
217

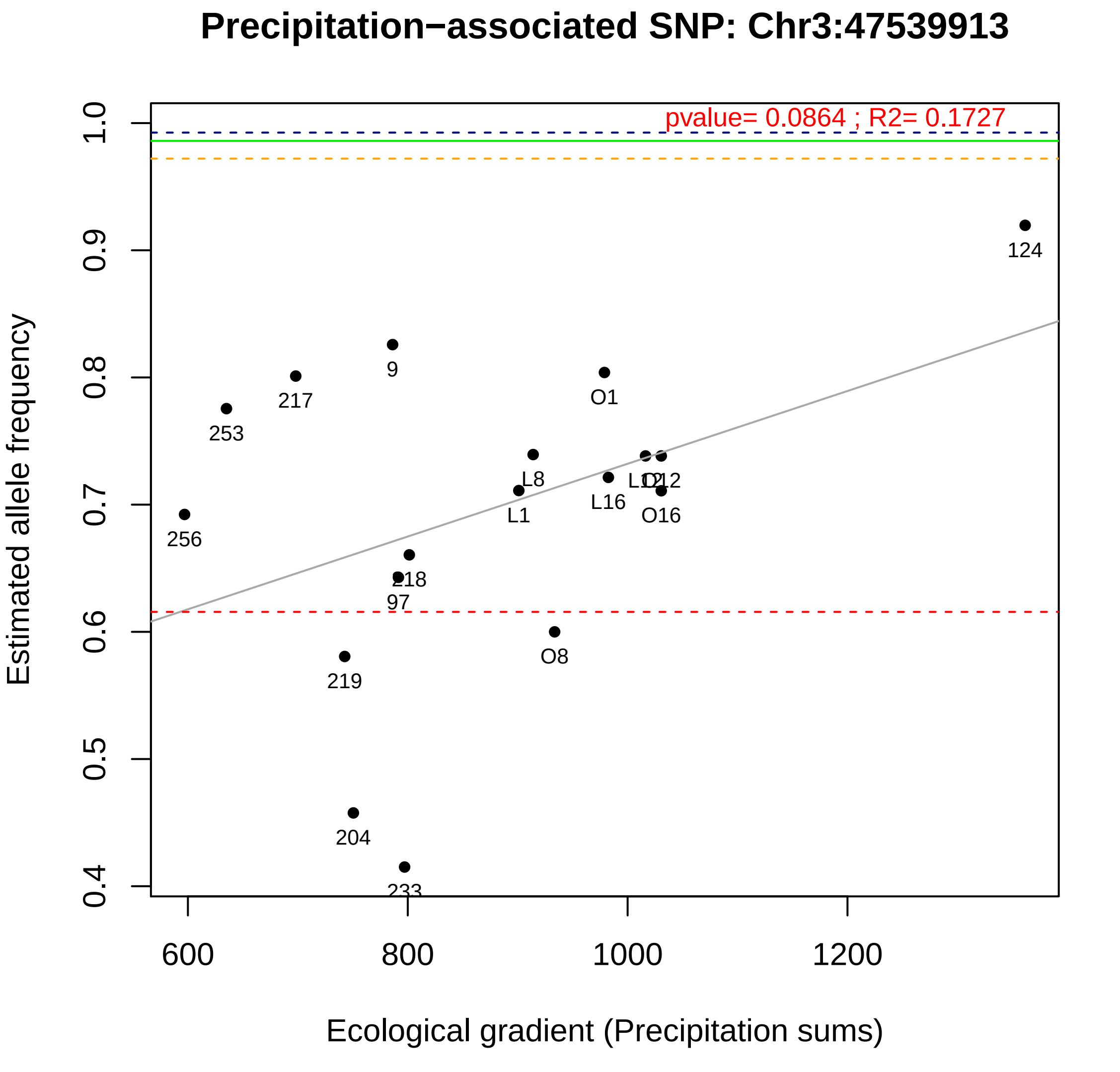

Precipitation−associated SNP: Chr3:47539913
pvalue= 0.0864 ; R2= 0.1727
Estimated allele frequency
Ecological gradient (Precipitation sums)
0.4
0.5
0.6
0.7
0.8
0.9
1.0
600
800
1000
1200
124
9
217
253
O1
L8
L12
O12
L16
L1
O16
256
218
97
O8
219
204
233

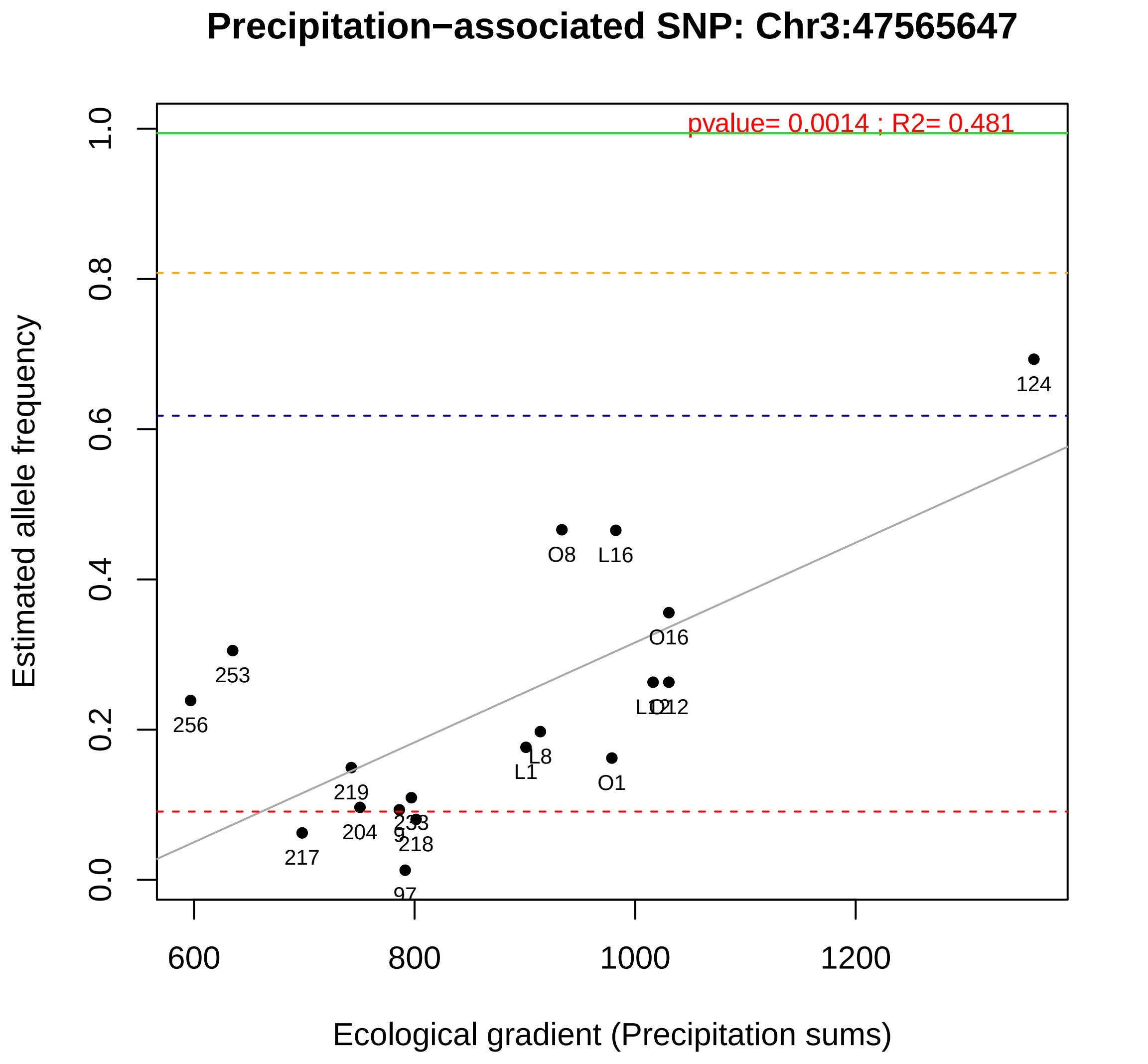

Precipitation−associated SNP: Chr3:47565647
pvalue= 0.0014 ; R2= 0.481
Estimated allele frequency
Ecological gradient (Precipitation sums)
0.0
0.2
0.4
0.6
0.8
1.0
600
800
1000
1200
124
O8
L16
O16
253
256
219
204
217
218
97
L8
L1
O1

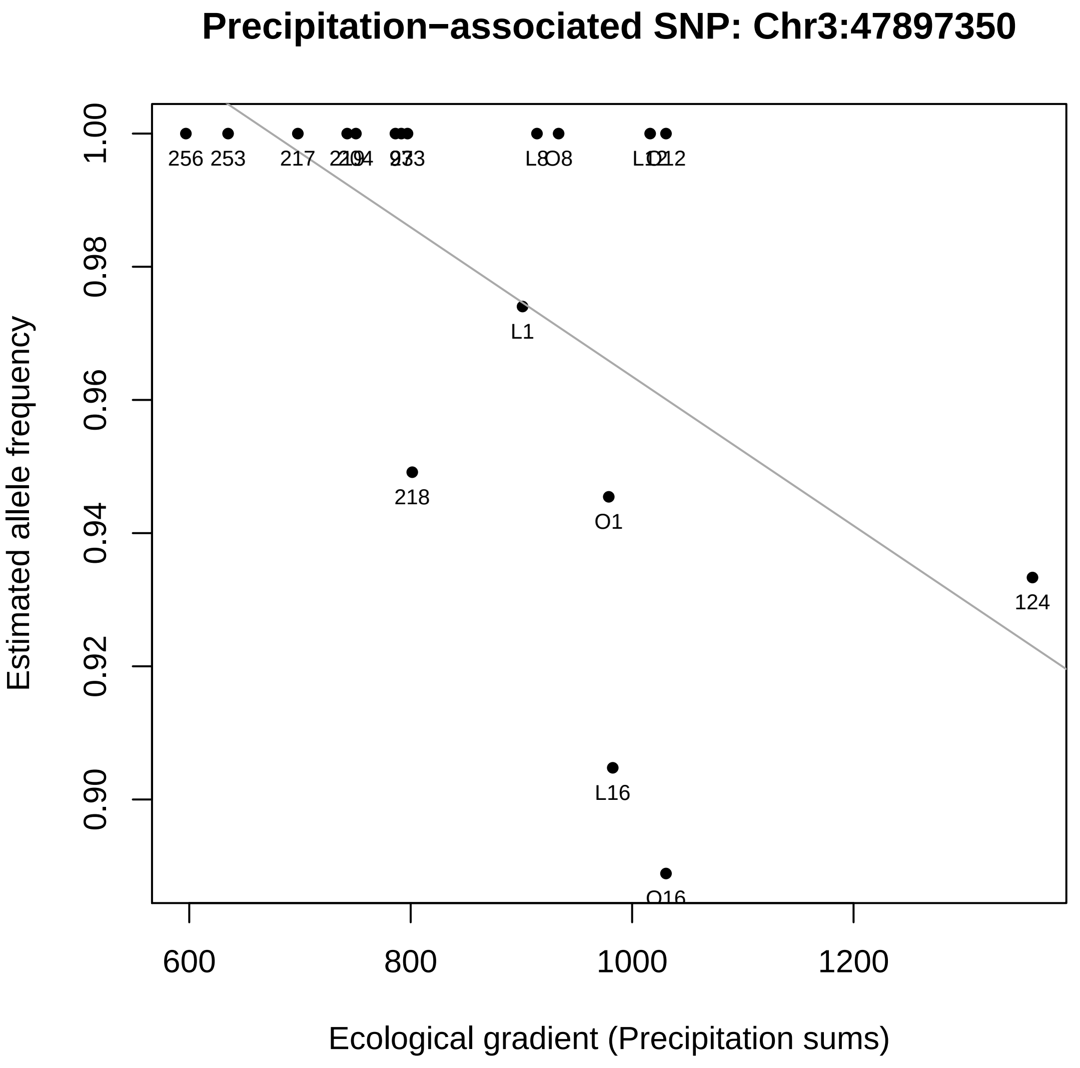
Precipitation−associated SNP: Chr3:47897350
Estimated allele frequency
Ecological gradient (Precipitation sums)
0.90
0.92
0.94
0.96
0.98
1.00
600
800
1000
1200
256
253
217
124
218
L1
O1
L16
L8
O8

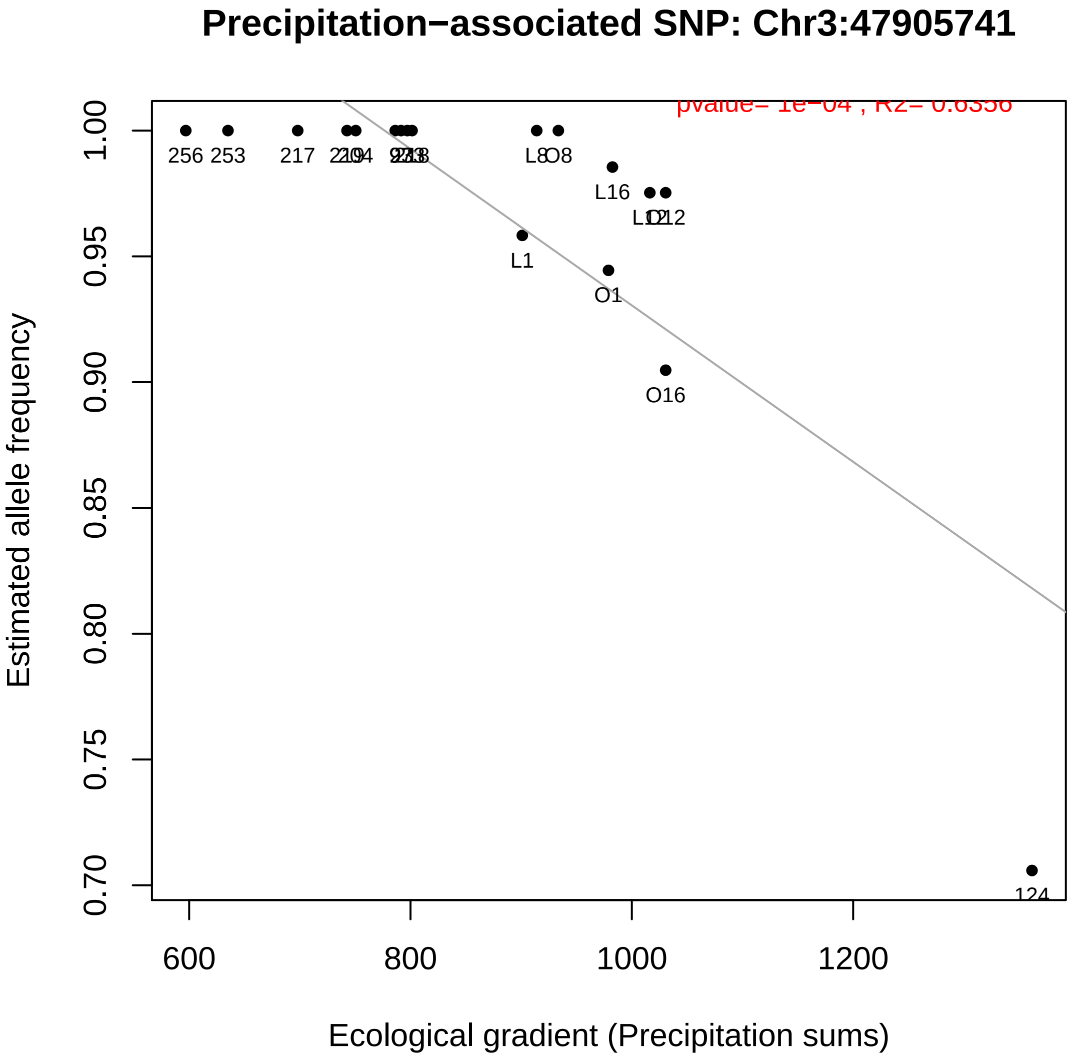
Precipitation−associated SNP: Chr3:47905741
pvalue= 1e−04 , R2= 0.6356
Estimated allele frequency
1.00
0.95
0.90
0.85
0.80
0.75
0.70
600
800
1000
1200
Ecological gradient (Precipitation sums)
256
253
217
L8
O8
L16
L1
O1
O16
124

**Precipitation−associated SNP: Chr3:48316250**

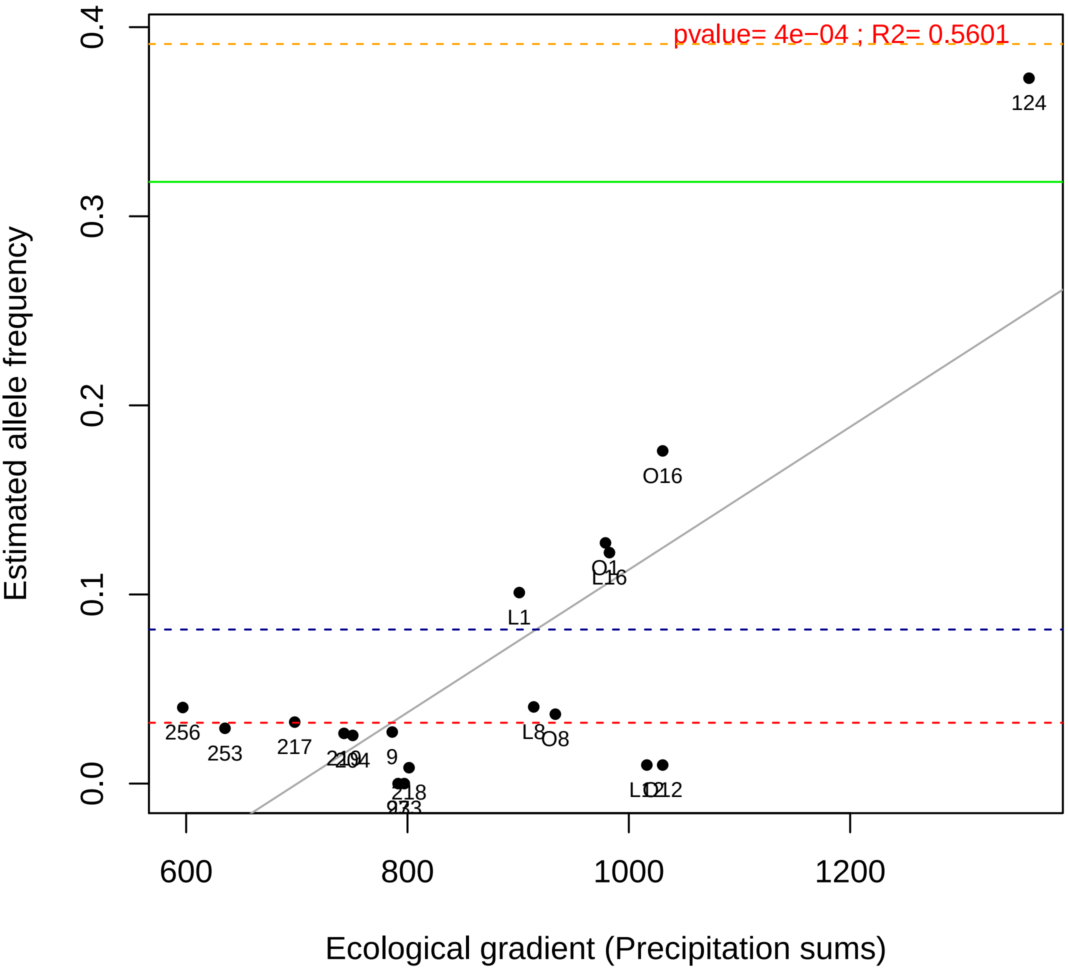

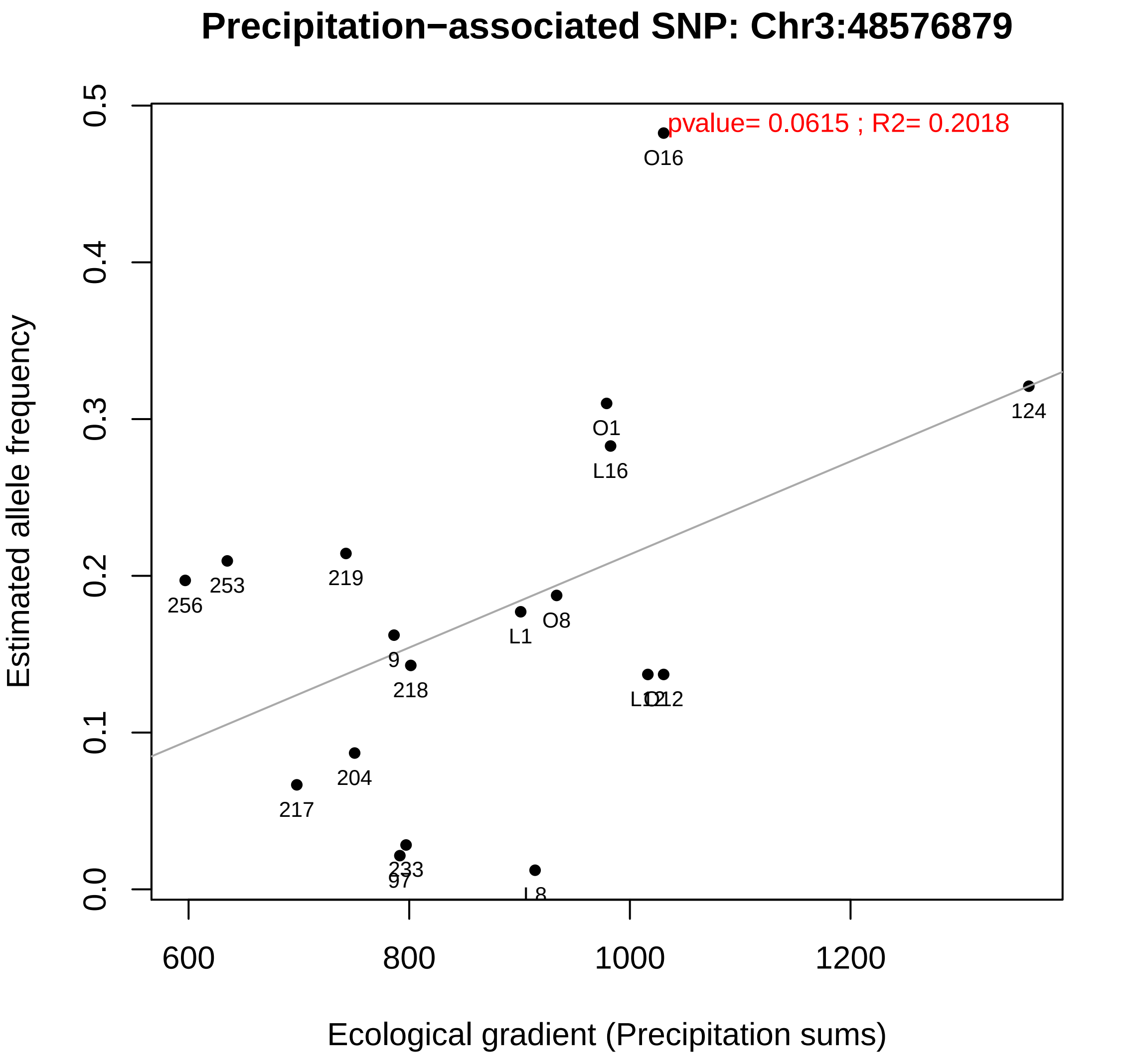

Precipitation−associated SNP: Chr3:48576879
pvalue= 0.0615 ; R2= 0.2018
Estimated allele frequency
Ecological gradient (Precipitation sums)
0.0
0.1
0.2
0.3
0.4
0.5
600
800
1000
1200
O16
124
O1
L16
256
253
219
O8
L1
9
218
L12
O12
204
217
233
97
L8

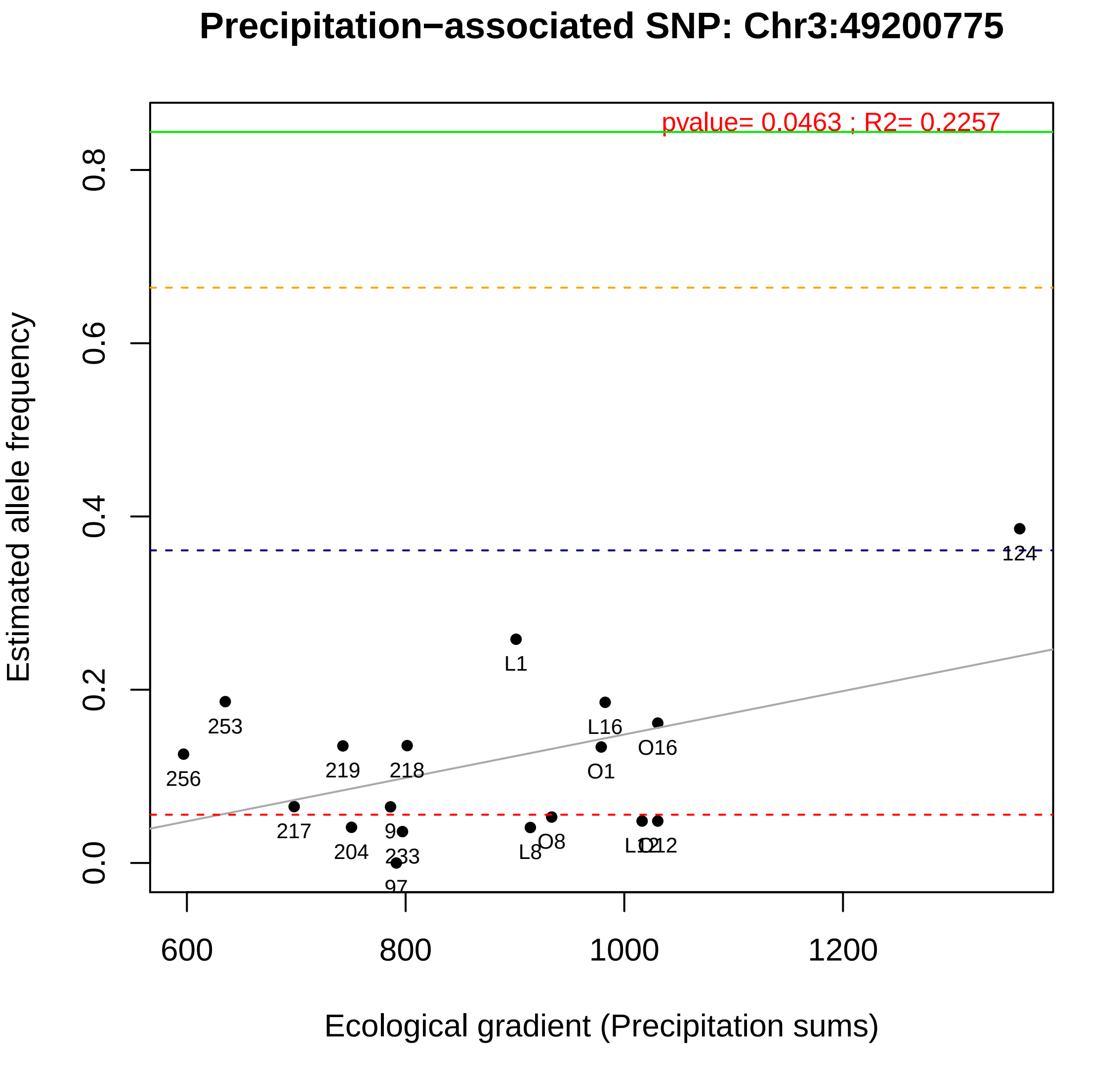
Precipitation−associated SNP: Chr3:49200775
pvalue= 0.0463 ; R2= 0.2257
Estimated allele frequency
0.0
0.2
0.4
0.6
0.8
600
800
1000
1200
Ecological gradient (Precipitation sums)
124
L1
253
L16
O16
256
219
218
O1
217
204
233
97
L8
O8
O12

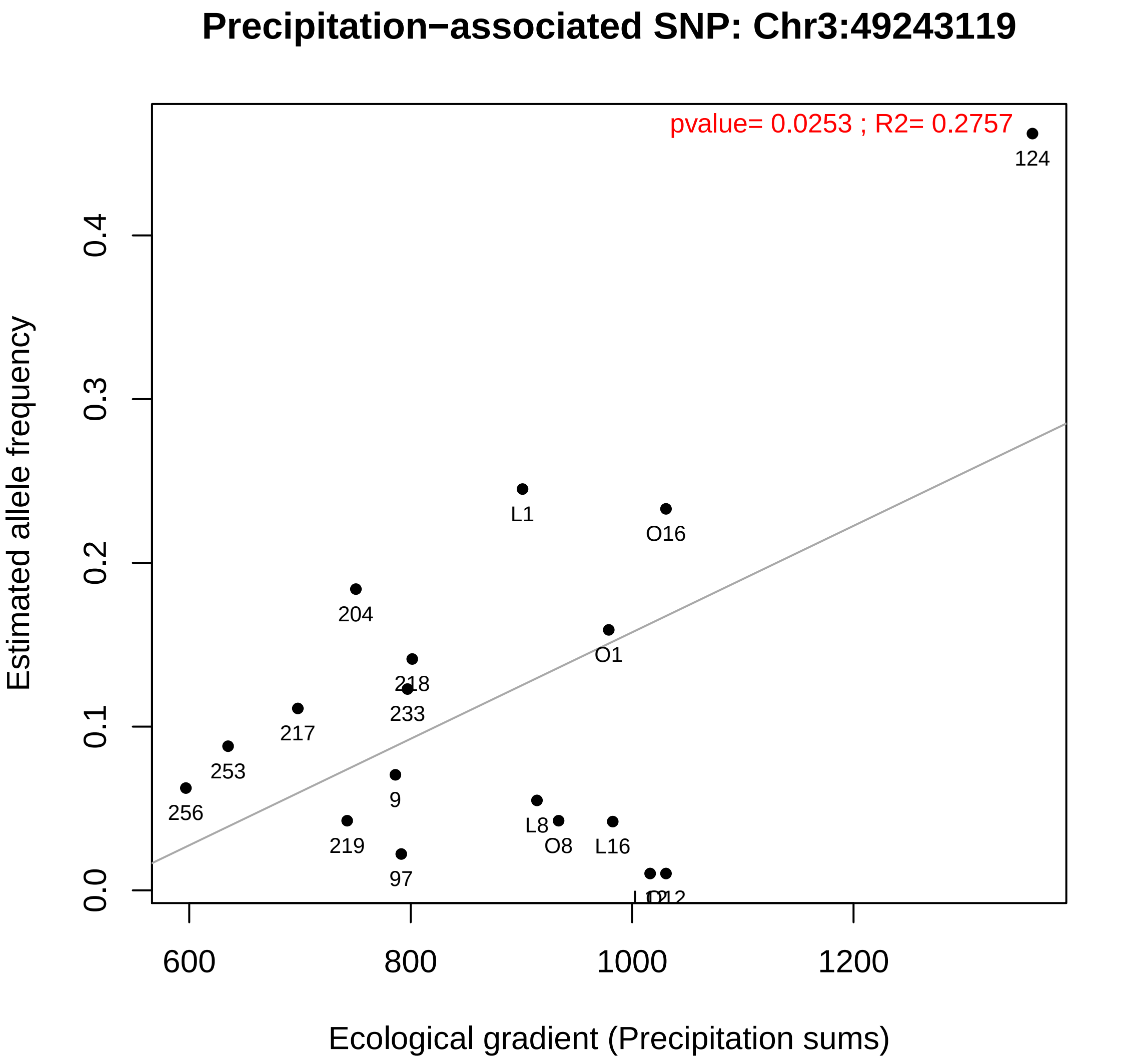

Precipitation−associated SNP: Chr3:49243119
pvalue= 0.0253 ; R2= 0.2757
Estimated allele frequency
0.0
0.1
0.2
0.3
0.4
600
800
1000
1200
Ecological gradient (Precipitation sums)
124
L1
O16
204
O1
218
233
217
253
256
9
219
97
L8
O8
L16

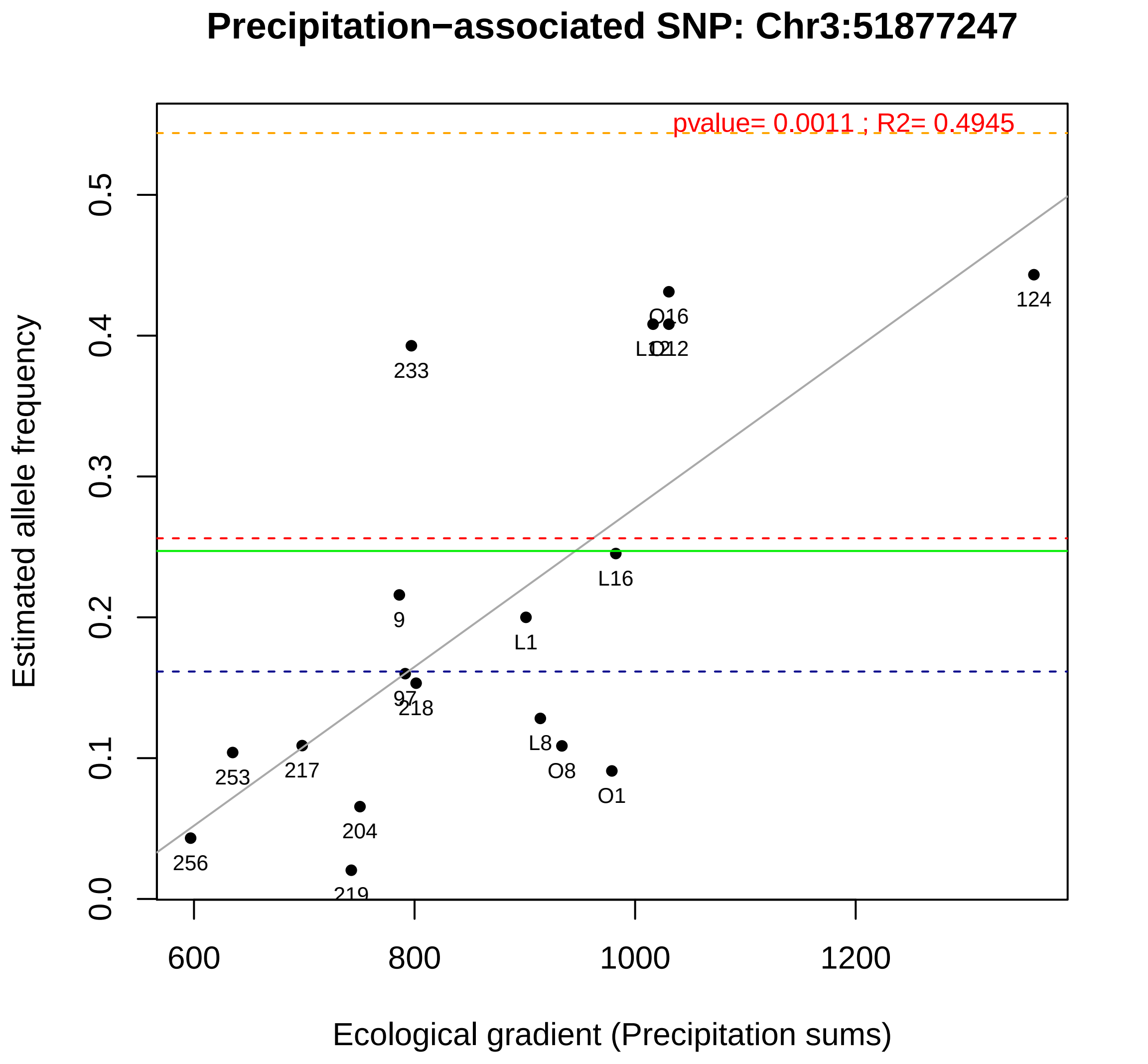
Precipitation−associated SNP: Chr3:51877247
pvalue= 0.0011 ; R2= 0.4945
Estimated allele frequency
0.0
0.1
0.2
0.3
0.4
0.5
600
800
1000
1200
Ecological gradient (Precipitation sums)
124
O16
233
L16
9
L1
97
218
L8
O8
O1
253
217
204
256
219

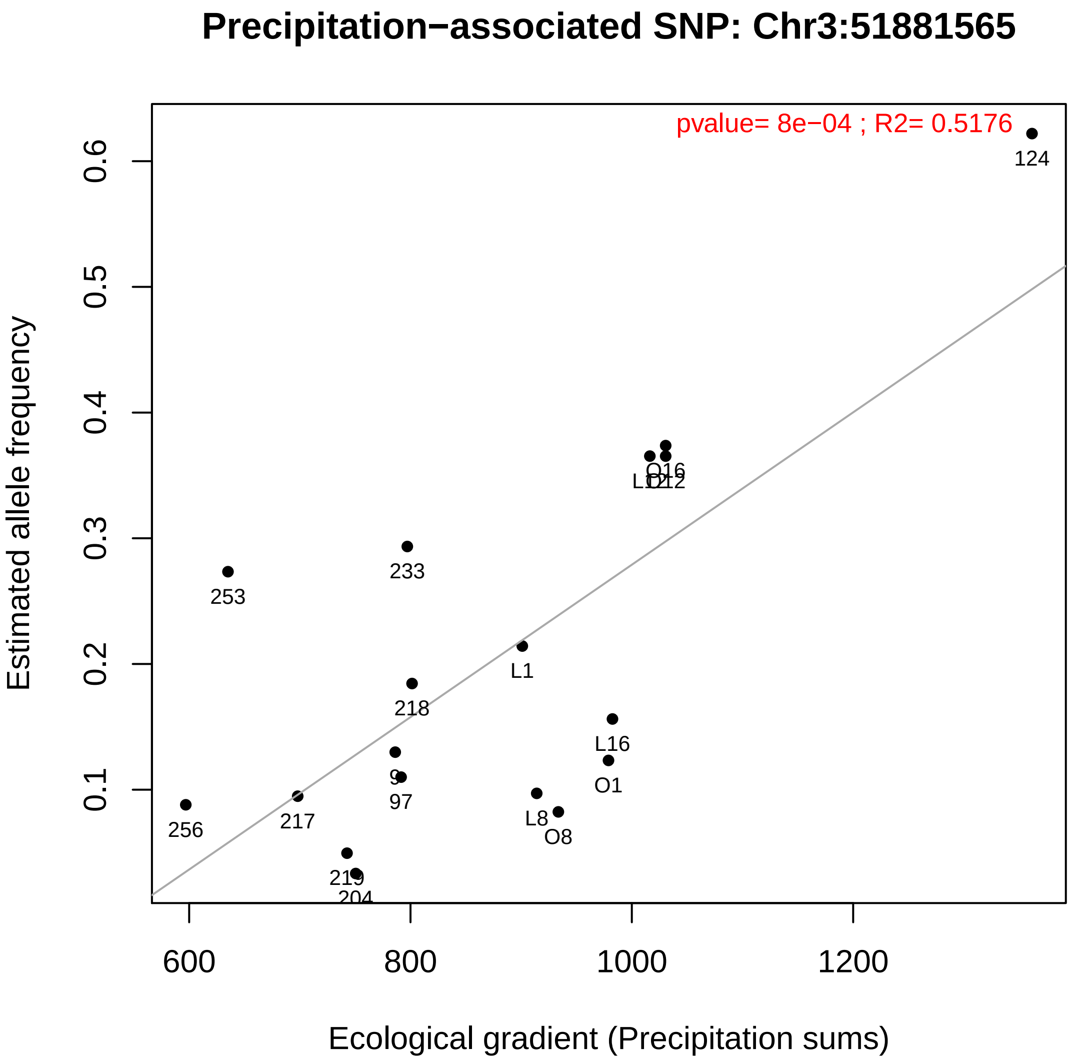
Precipitation−associated SNP: Chr3:51881565
pvalue= 8e−04 ; R2= 0.5176
Estimated allele frequency
0.1
0.2
0.3
0.4
0.5
0.6
600
800
1000
1200
Ecological gradient (Precipitation sums)
124
O16
L12
O12
233
253
L1
218
L16
O1
97
L8
O8
256
217
219
204

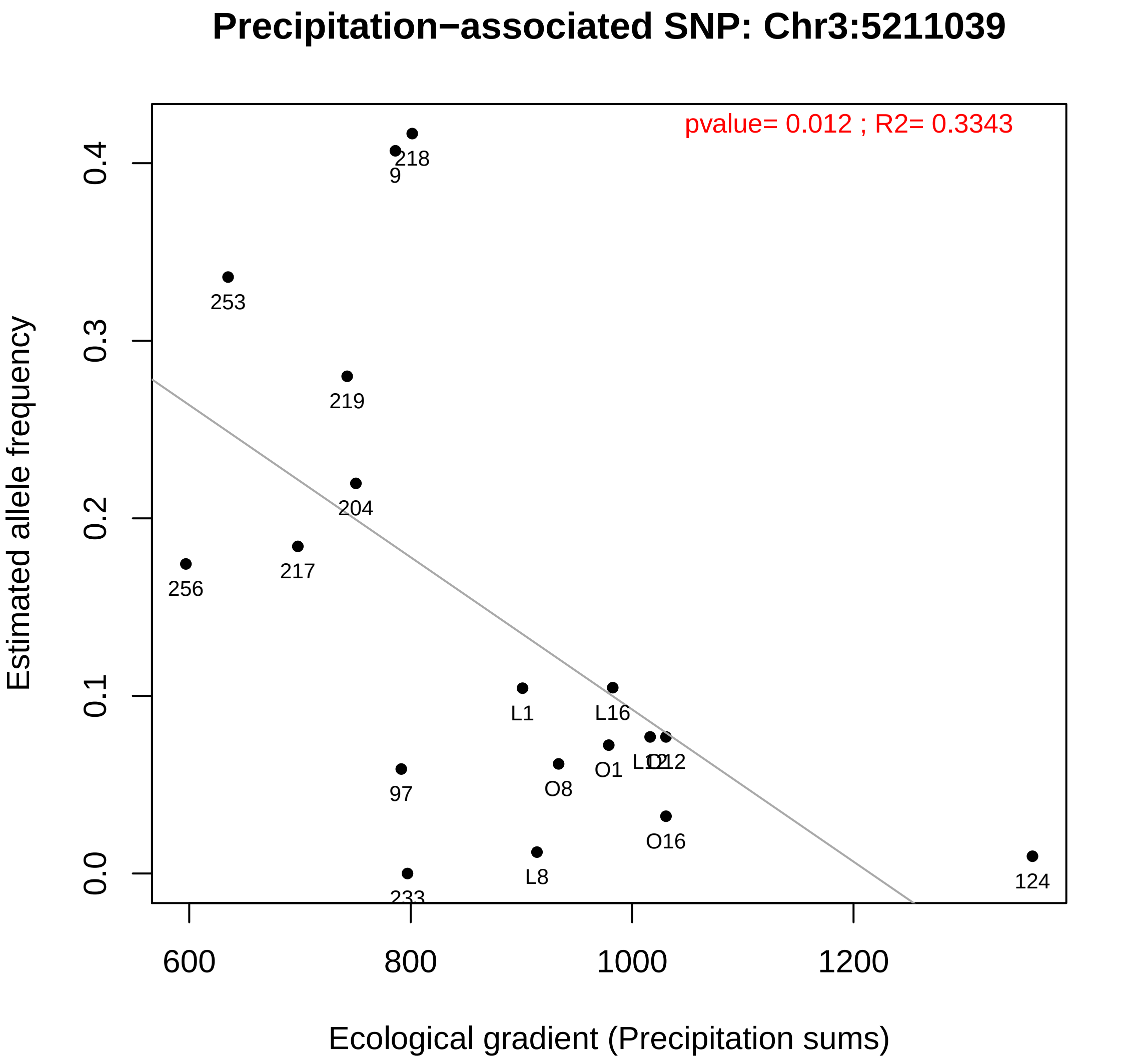
Precipitation−associated SNP: Chr3:5211039
pvalue= 0.012 ; R2= 0.3343
Estimated allele frequency
0.0
0.1
0.2
0.3
0.4
600
800
1000
1200
Ecological gradient (Precipitation sums)
218
9
253
219
204
217
256
L1
L16
L12
O12
O1
O8
97
O16
L8
233
124

#### Precipitation−associated SNP: Chr3:54839537

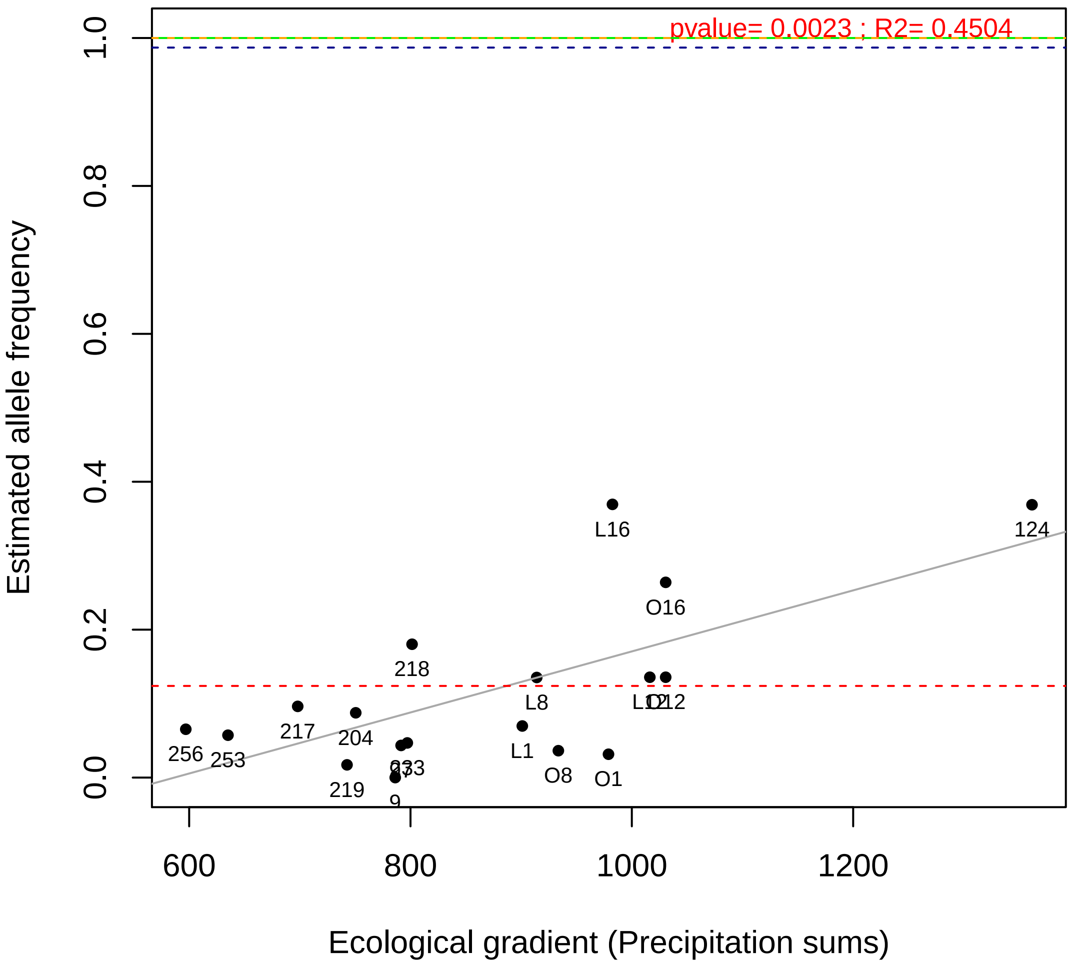

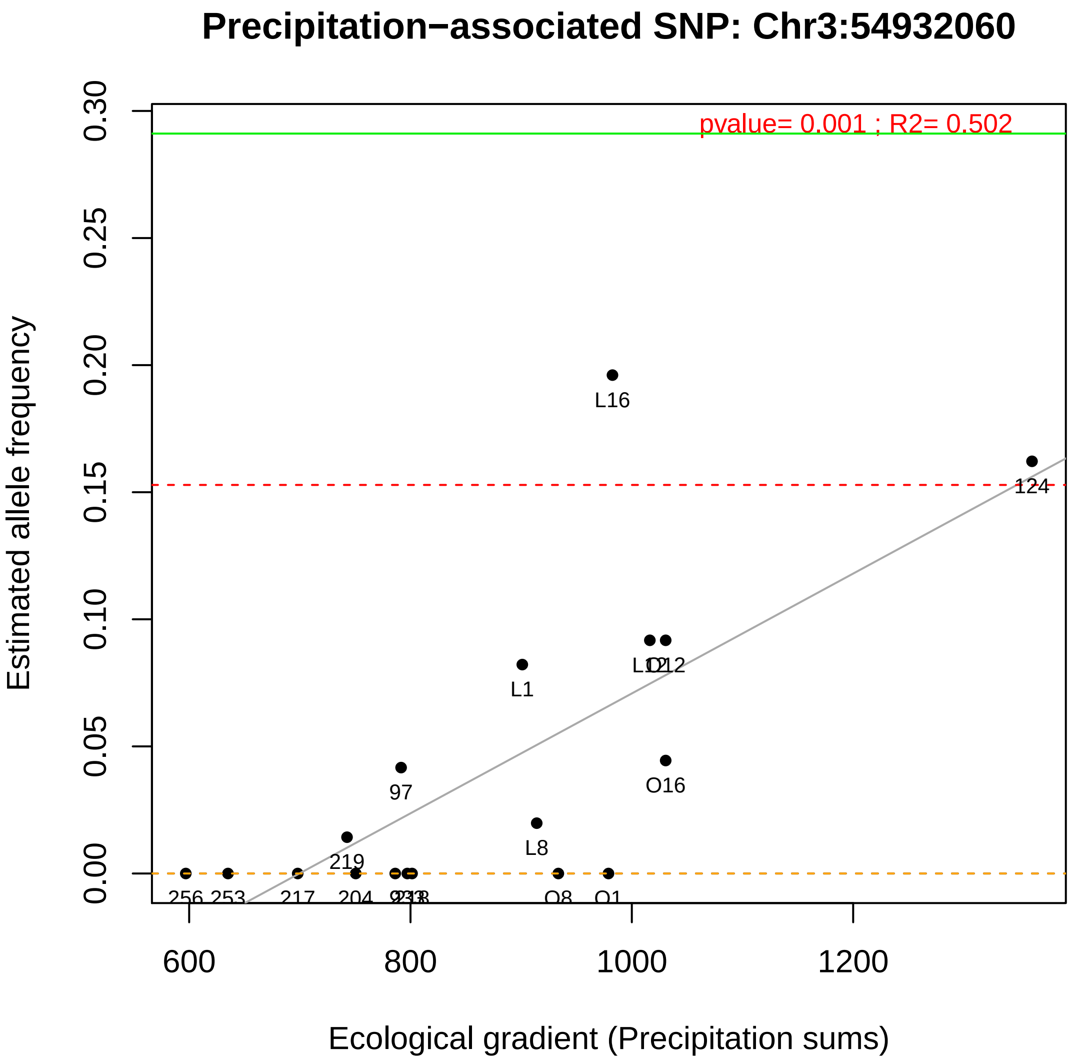
Precipitation−associated SNP: Chr3:54932060
pvalue= 0.001 ; R2= 0.502
Estimated allele frequency
0.30
0.25
0.20
0.15
0.10
0.05
0.00
L16
124
L12
O12
L1
O16
97
L8
219
256
253
217
204
O8
O1
600
800
1000
1200
Ecological gradient (Precipitation sums)

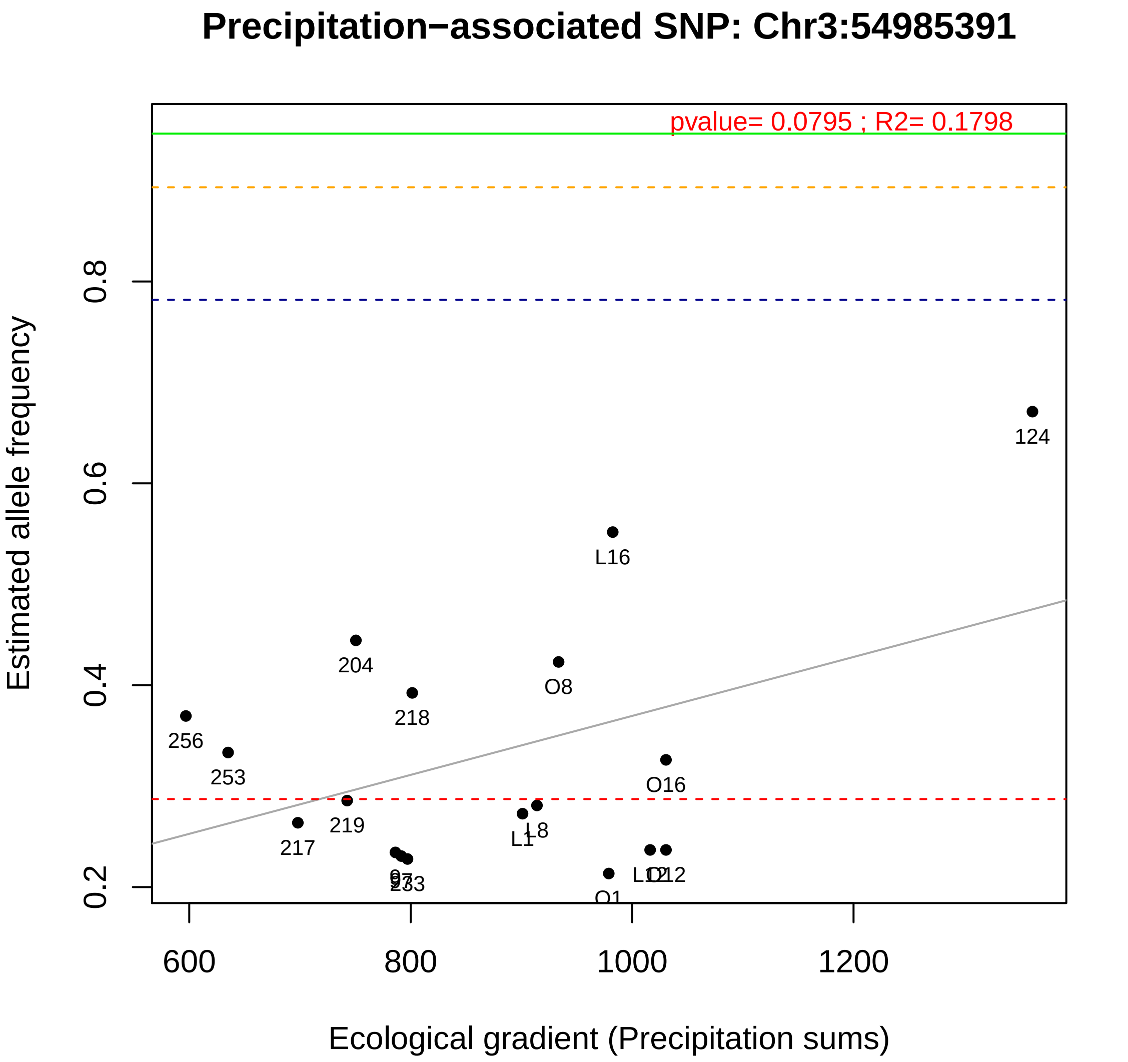

Precipitation−associated SNP: Chr3:54985391
pvalue= 0.0795 ; R2= 0.1798
Estimated allele frequency
0.2
0.4
0.6
0.8
600
800
1000
1200
Ecological gradient (Precipitation sums)
124
L16
204
O8
218
256
253
O16
219
217
L8
L1
O1

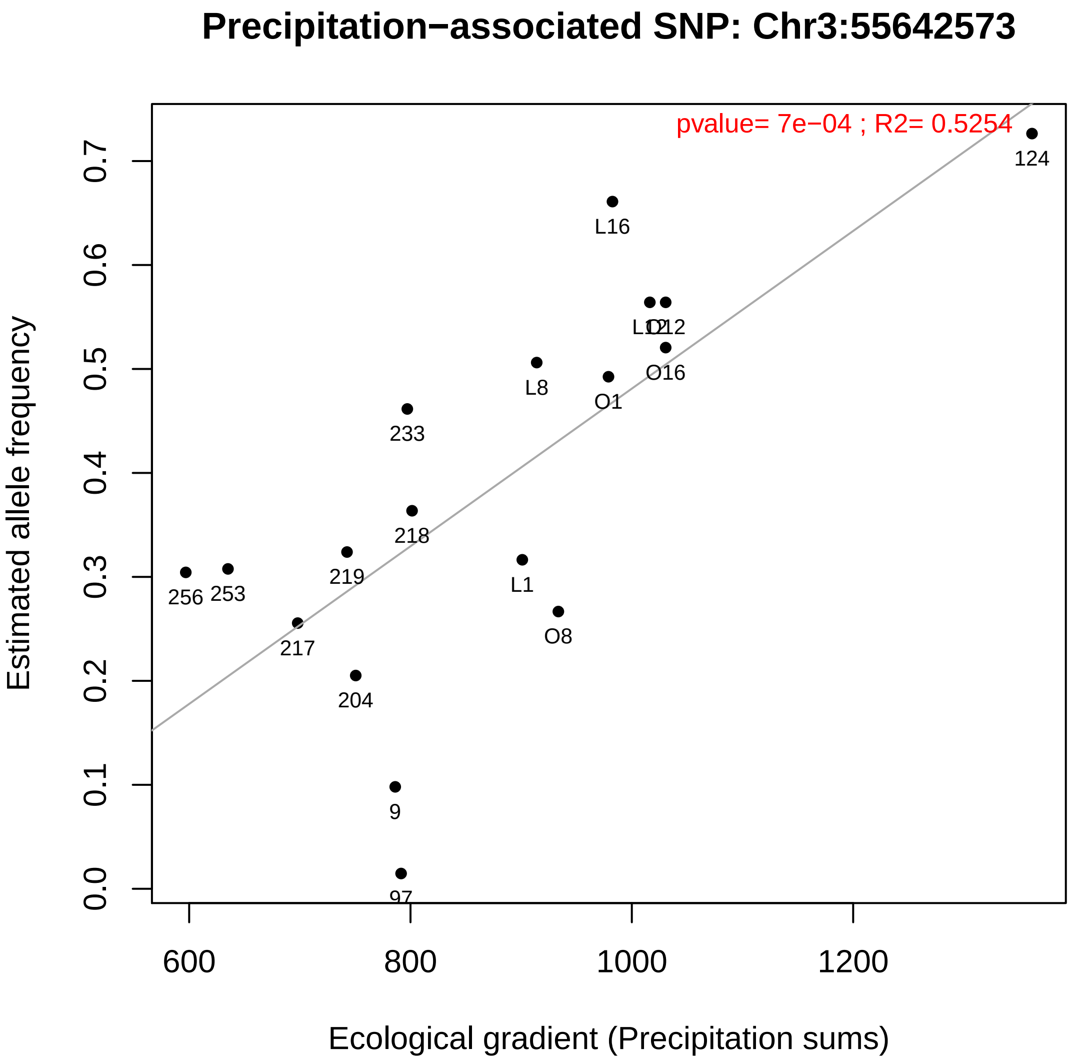
Precipitation−associated SNP: Chr3:55642573
pvalue= 7e−04 ; R2= 0.5254
Estimated allele frequency
Ecological gradient (Precipitation sums)
0.0
0.1
0.2
0.3
0.4
0.5
0.6
0.7
600
800
1000
1200
124
L16
L12
O12
O16
L8
O1
233
218
219
L1
256
253
217
O8
204
9
97

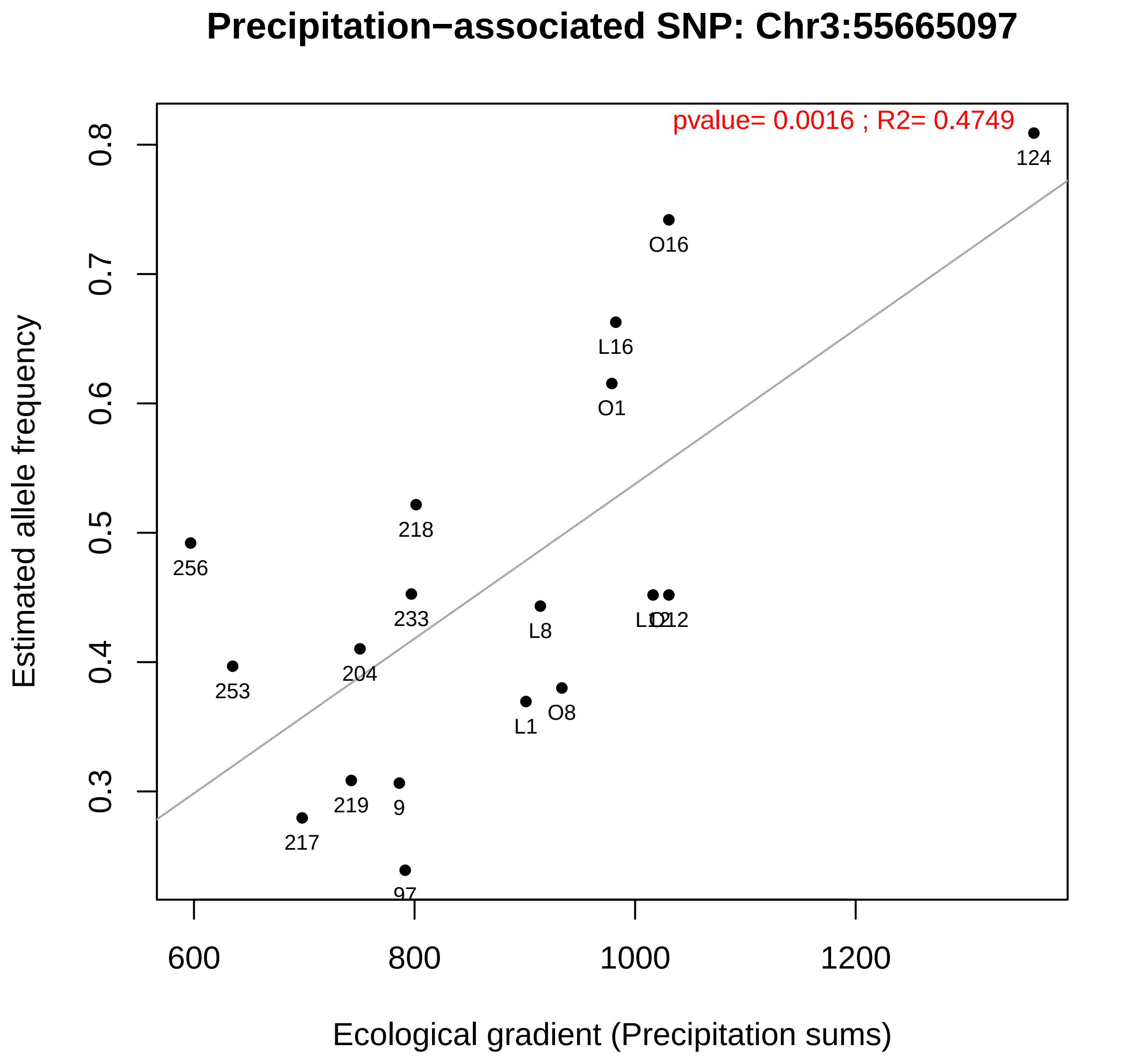
Precipitation−associated SNP: Chr3:55665097
pvalue= 0.0016 ; R2= 0.4749
124
O16
L16
O1
218
256
233
L8
L12
O12
204
253
O8
L1
219
9
217
97
0.3
0.4
0.5
0.6
0.7
0.8
600
800
1000
1200
Estimated allele frequency
Ecological gradient (Precipitation sums)

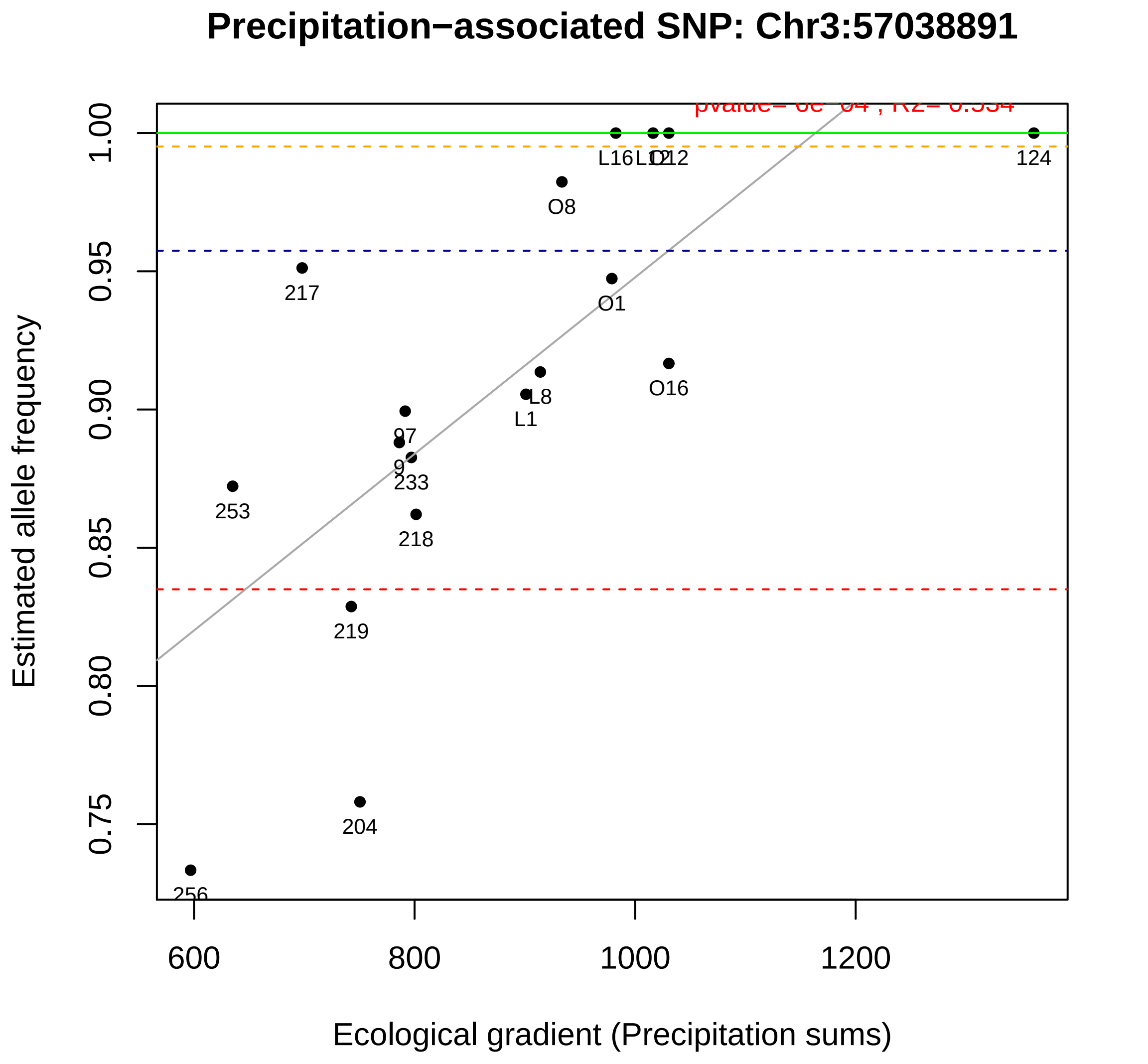

Precipitation−associated SNP: Chr3:57038891
Estimated allele frequency
Ecological gradient (Precipitation sums)
0.75
0.80
0.85
0.90
0.95
1.00
600
800
1000
1200
L16
124
O8
217
O1
O16
L8
L1
253
233
218
219
204

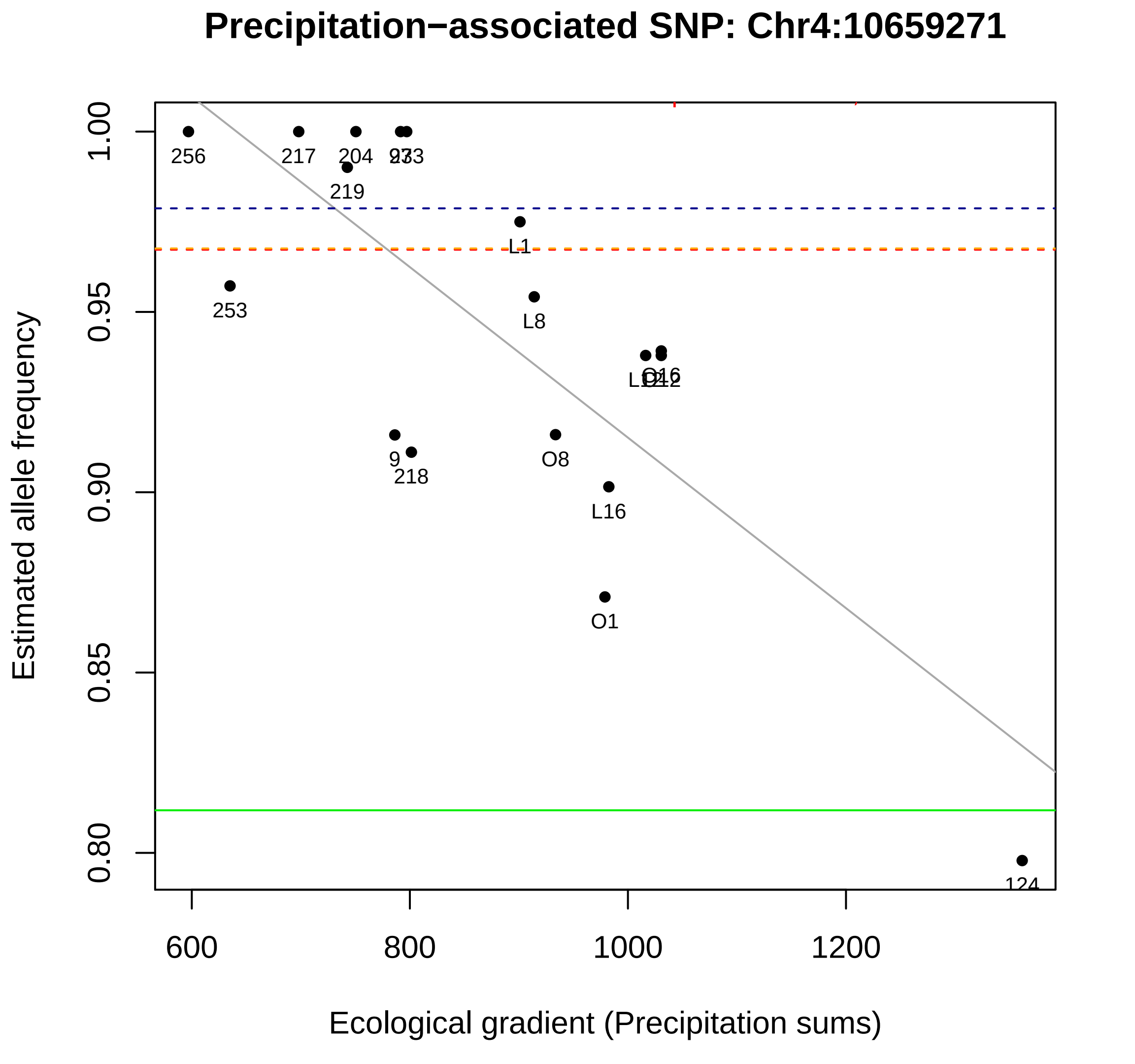

Precipitation−associated SNP: Chr4:10659271
Estimated allele frequency
Ecological gradient (Precipitation sums)
1.00
0.95
0.90
0.85
0.80
600
800
1000
1200
256
217
204
219
L1
253
L8
O16
9
218
O8
L16
O1
124

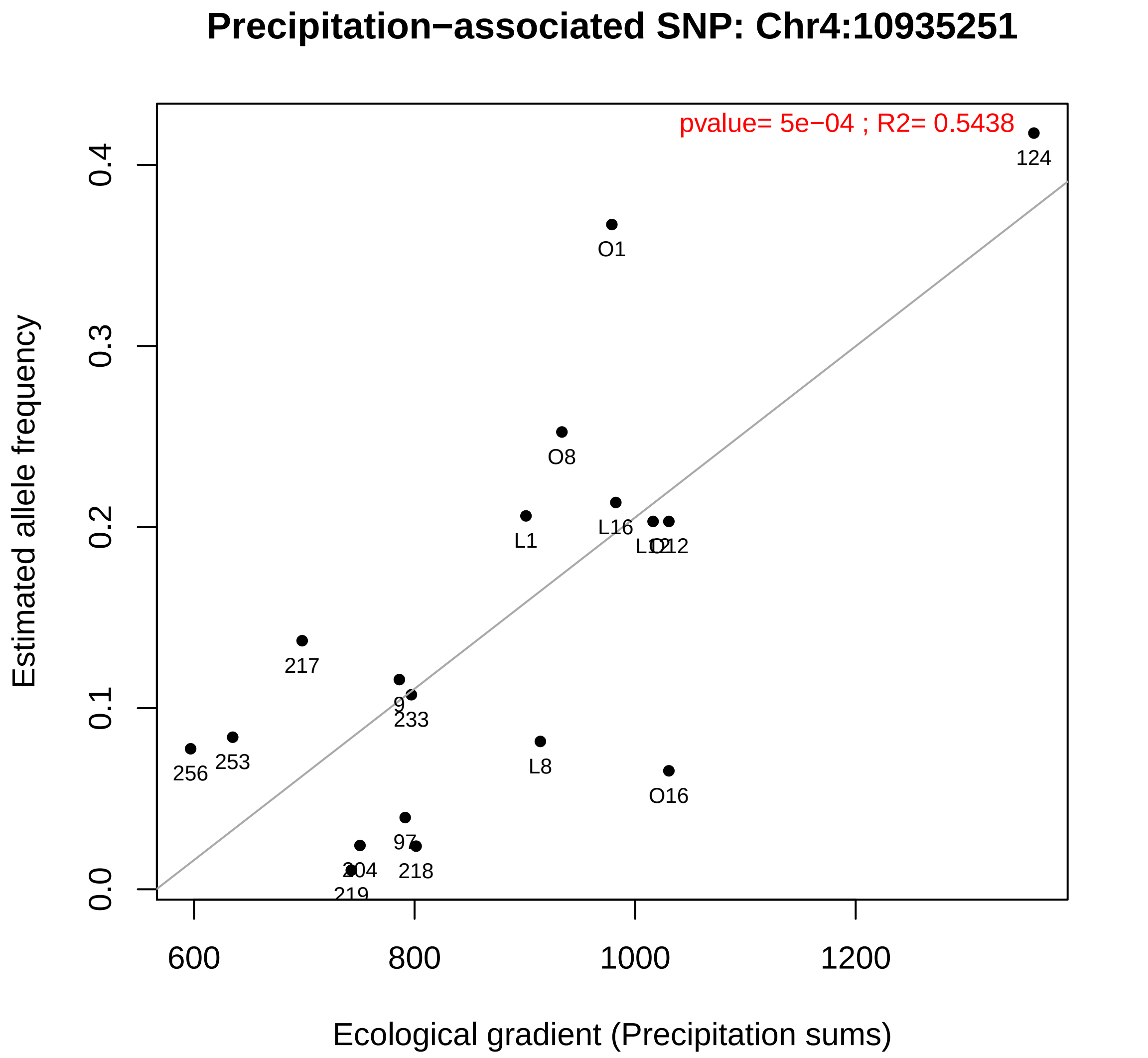

Precipitation−associated SNP: Chr4:10935251
pvalue= 5e−04 ; R2= 0.5438
124
O1
O8
L16
L1
L12
O12
217
9
233
253
256
L8
O16
97
204
218
219
0.0
0.1
0.2
0.3
0.4
600
800
1000
1200
Estimated allele frequency
Ecological gradient (Precipitation sums)

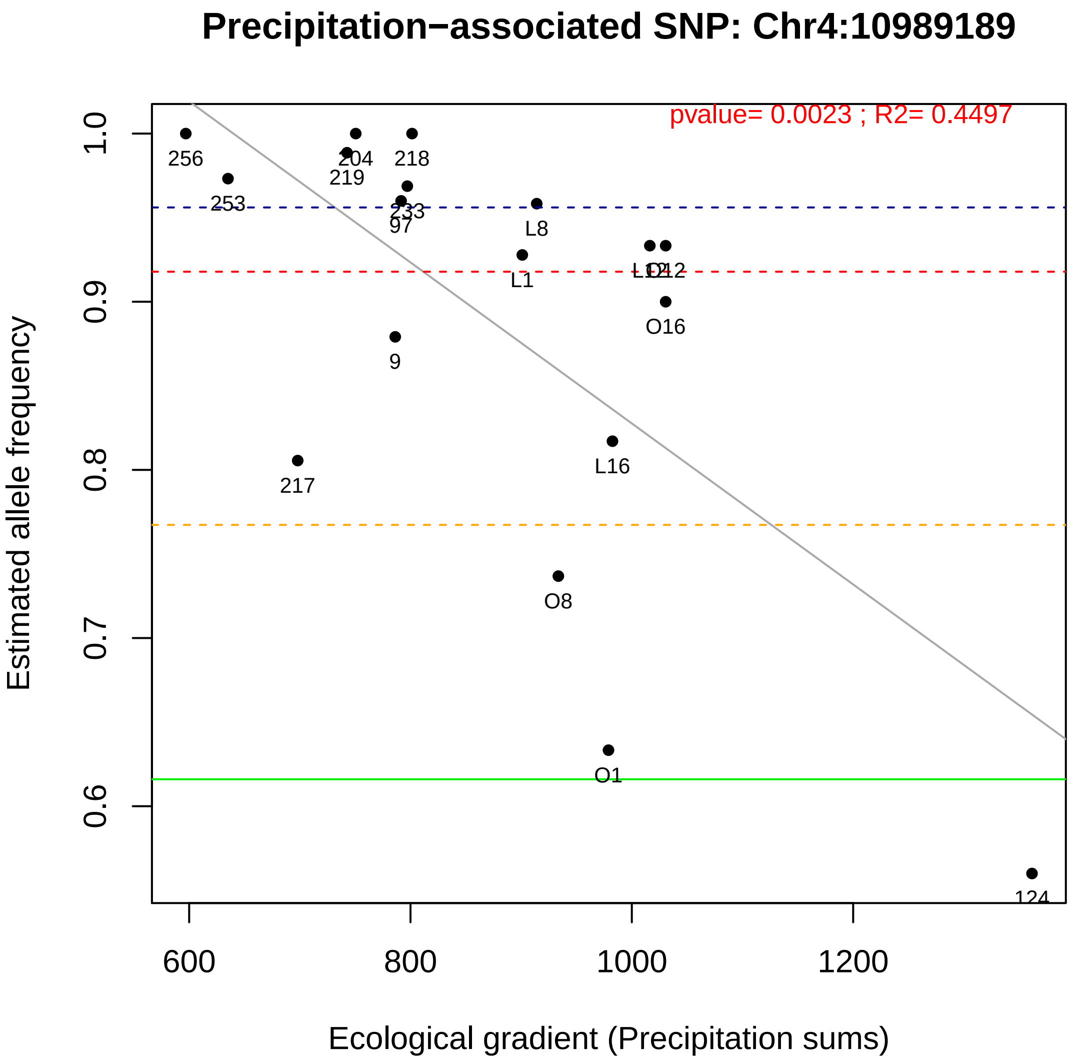

Precipitation−associated SNP: Chr4:10989189
pvalue= 0.0023 ; R2= 0.4497
Estimated allele frequency
Ecological gradient (Precipitation sums)
1.0
0.9
0.8
0.7
0.6
600
800
1000
1200
256
204
218
219
253
233
97
L8
L1
O16
9
217
L16
O8
O1
124

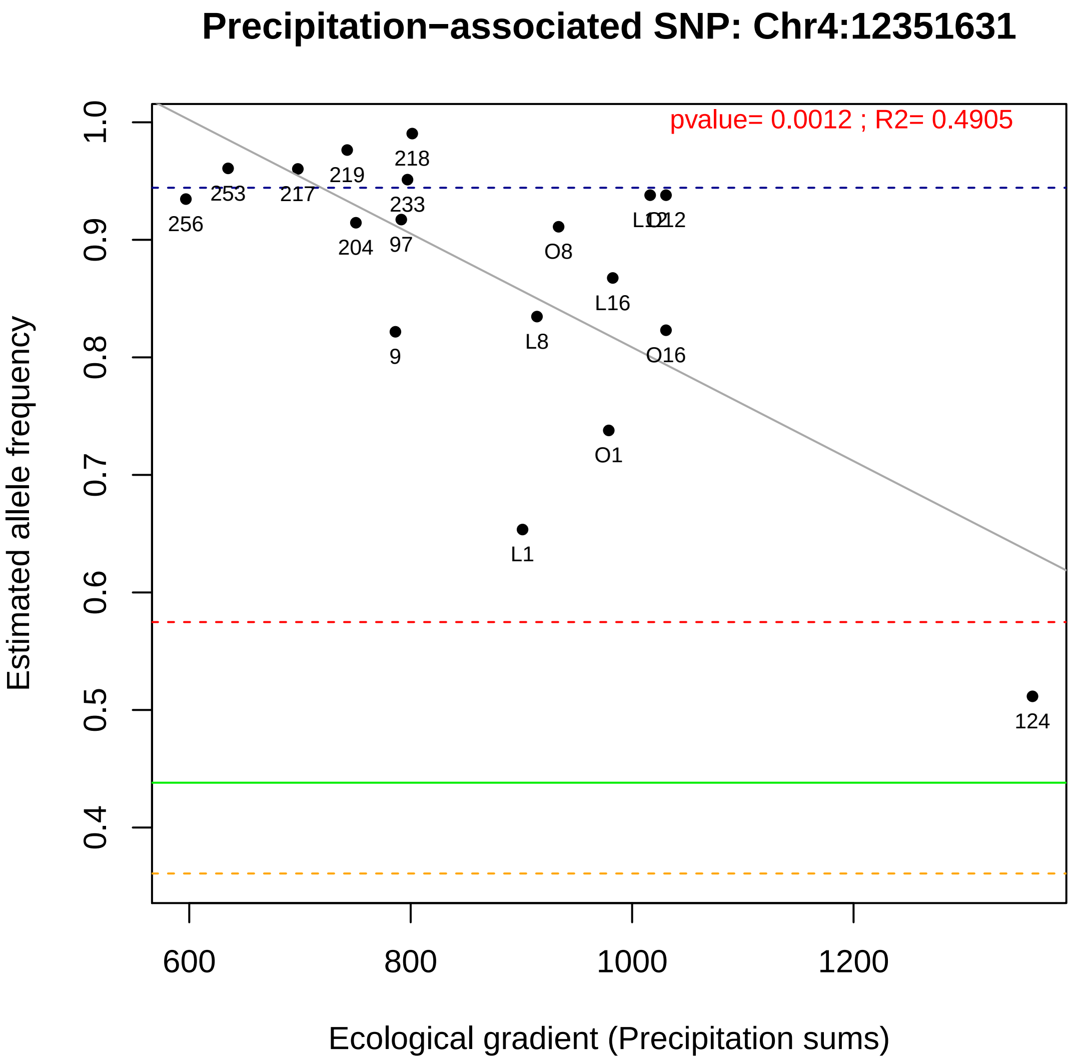
Precipitation−associated SNP: Chr4:12351631
pvalue= 0.0012 ; R2= 0.4905
Estimated allele frequency
Ecological gradient (Precipitation sums)
0.4
0.5
0.6
0.7
0.8
0.9
1.0
600
800
1000
1200
256
253
217
219
218
233
204
97
9
O8
L16
L8
O16
O1
L1
124

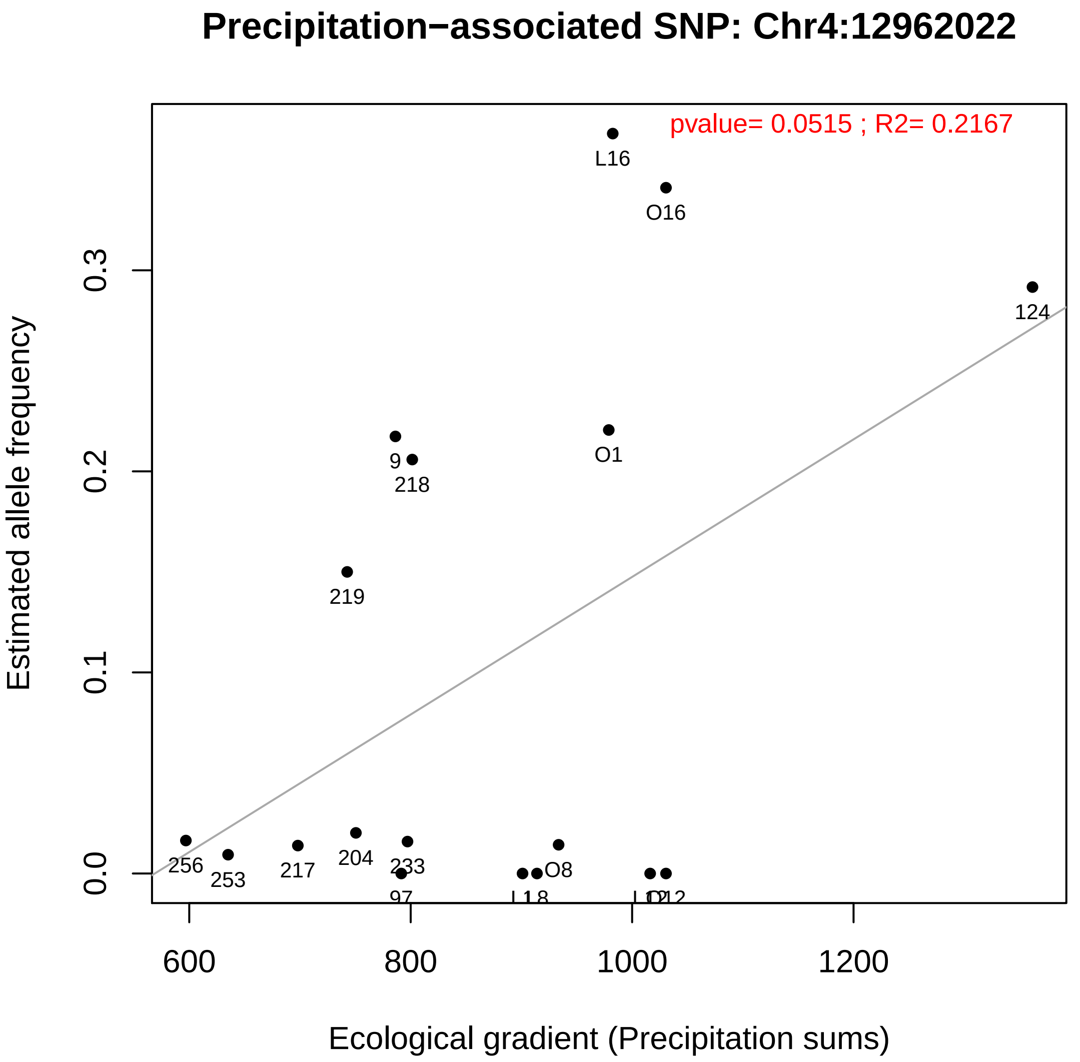
Precipitation−associated SNP: Chr4:12962022
pvalue= 0.0515 ; R2= 0.2167
Estimated allele frequency
0.0
0.1
0.2
0.3
Ecological gradient (Precipitation sums)
600
800
1000
1200
L16
O16
124
O1
9
218
219
256
253
217
204
233
97
O8

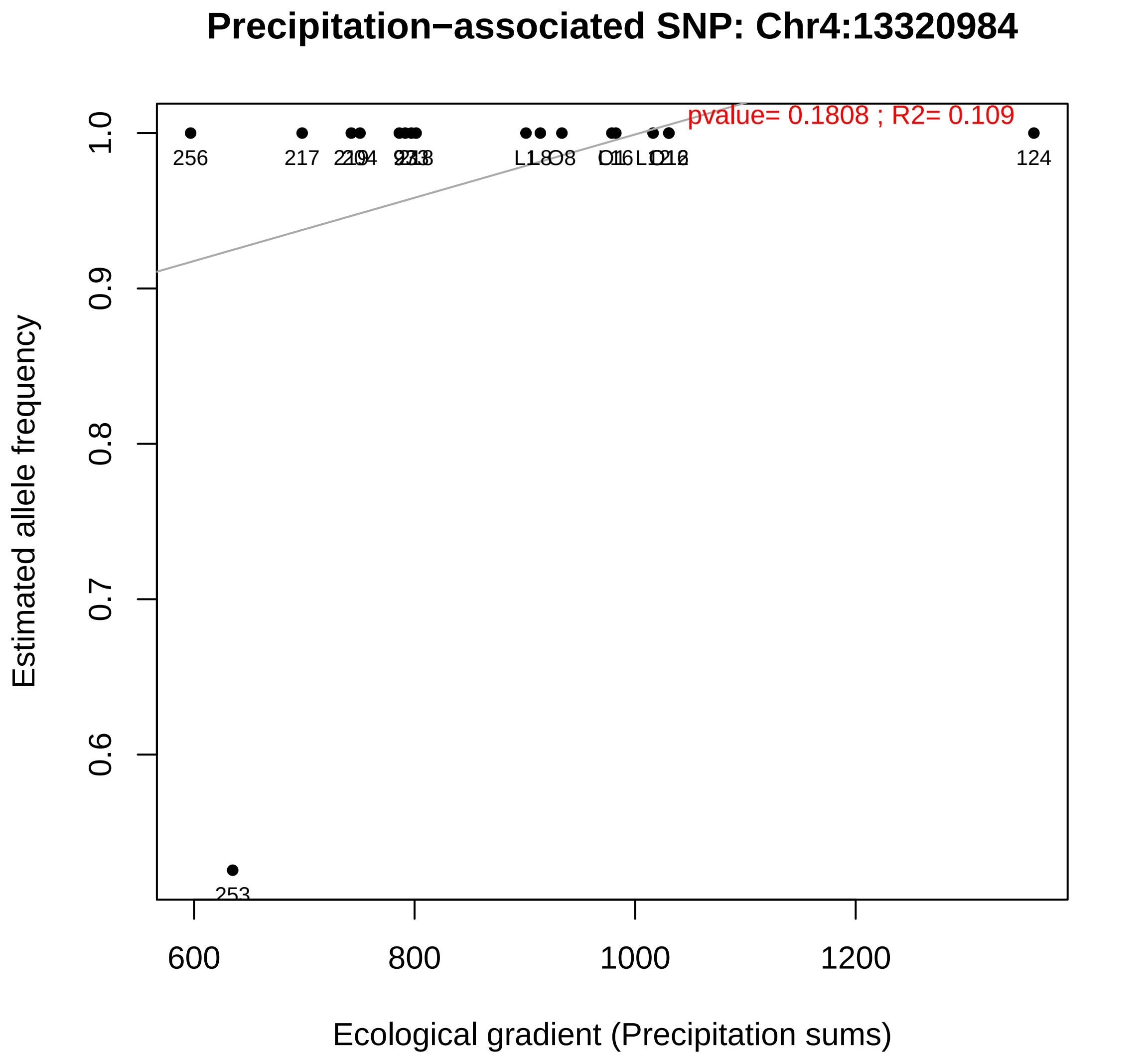
Precipitation−associated SNP: Chr4:13320984
pvalue= 0.1808 ; R2= 0.109
Estimated allele frequency
Ecological gradient (Precipitation sums)
1.0
0.9
0.8
0.7
0.6
600
800
1000
1200
256
217
124
253

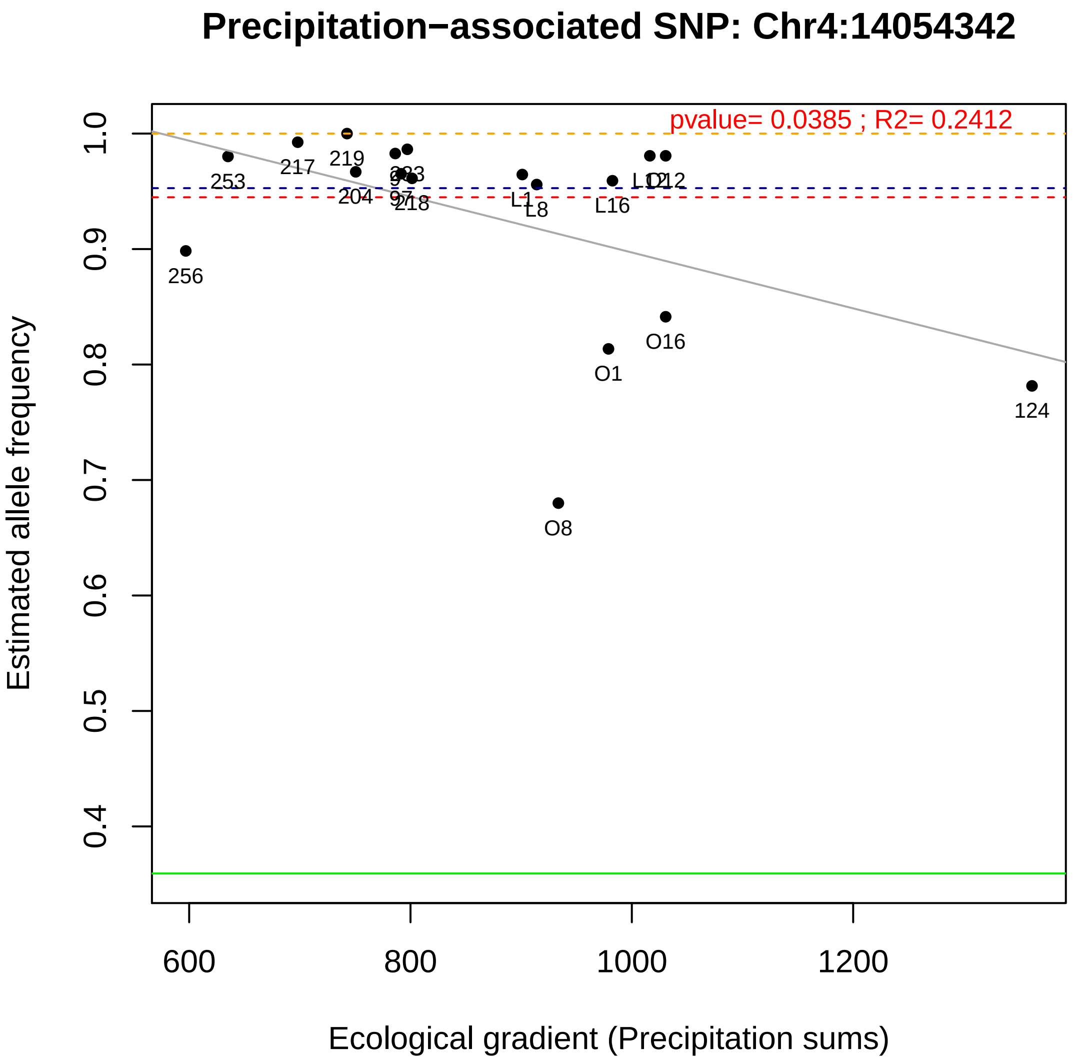
Precipitation−associated SNP: Chr4:14054342
pvalue= 0.0385 ; R2= 0.2412
Estimated allele frequency
Ecological gradient (Precipitation sums)
0.4
0.5
0.6
0.7
0.8
0.9
1.0
600
800
1000
1200
256
253
217
219
204
218
L1
L8
L16
O16
O1
O8
124

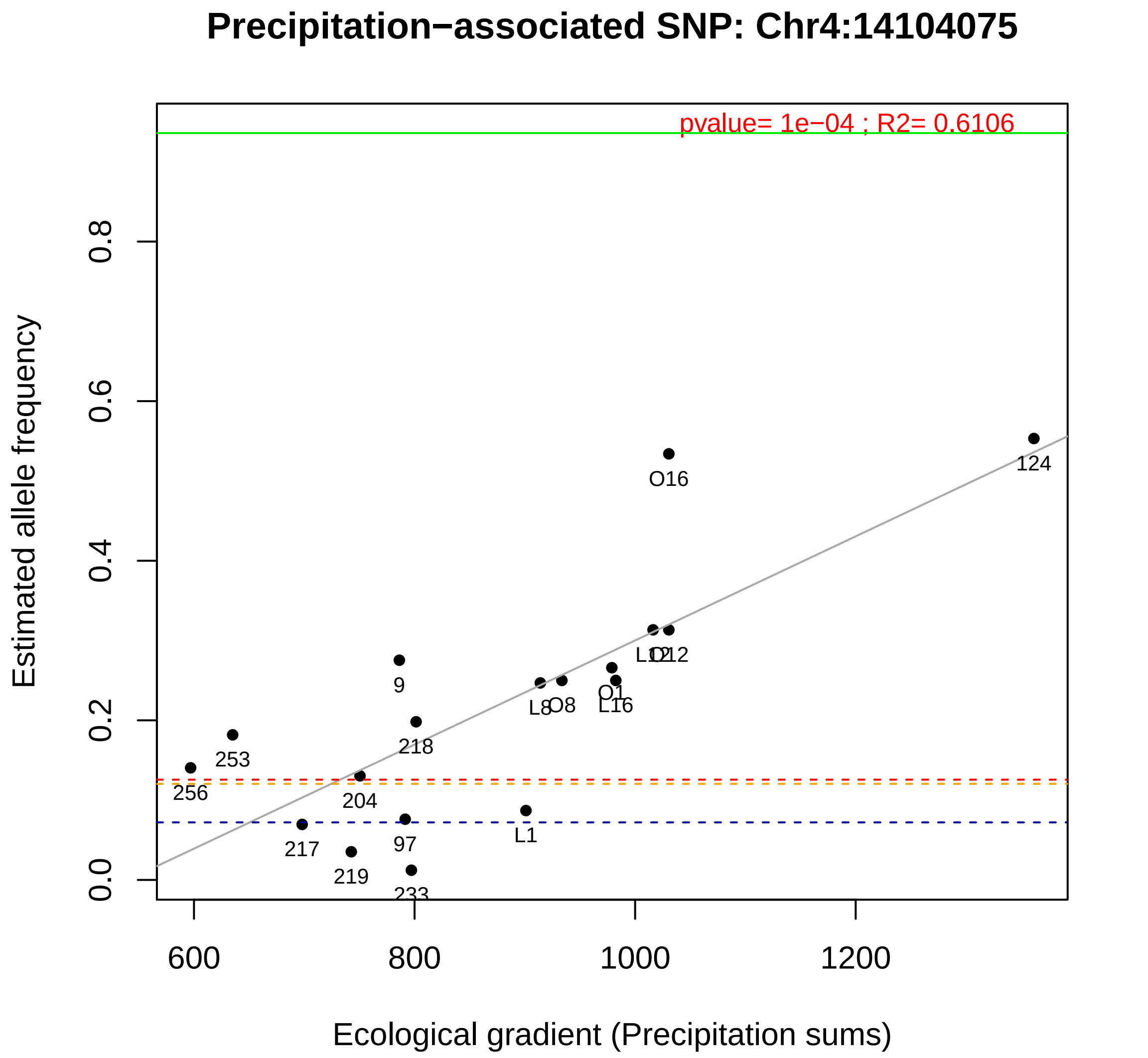

Precipitation−associated SNP: Chr4:14104075
pvalue= 1e−04 ; R2= 0.6106
Estimated allele frequency
0.0
0.2
0.4
0.6
0.8
600
800
1000
1200
Ecological gradient (Precipitation sums)
124
O16
L12
O12
9
O1
L16
L8
O8
218
253
256
204
217
97
L1
219
233

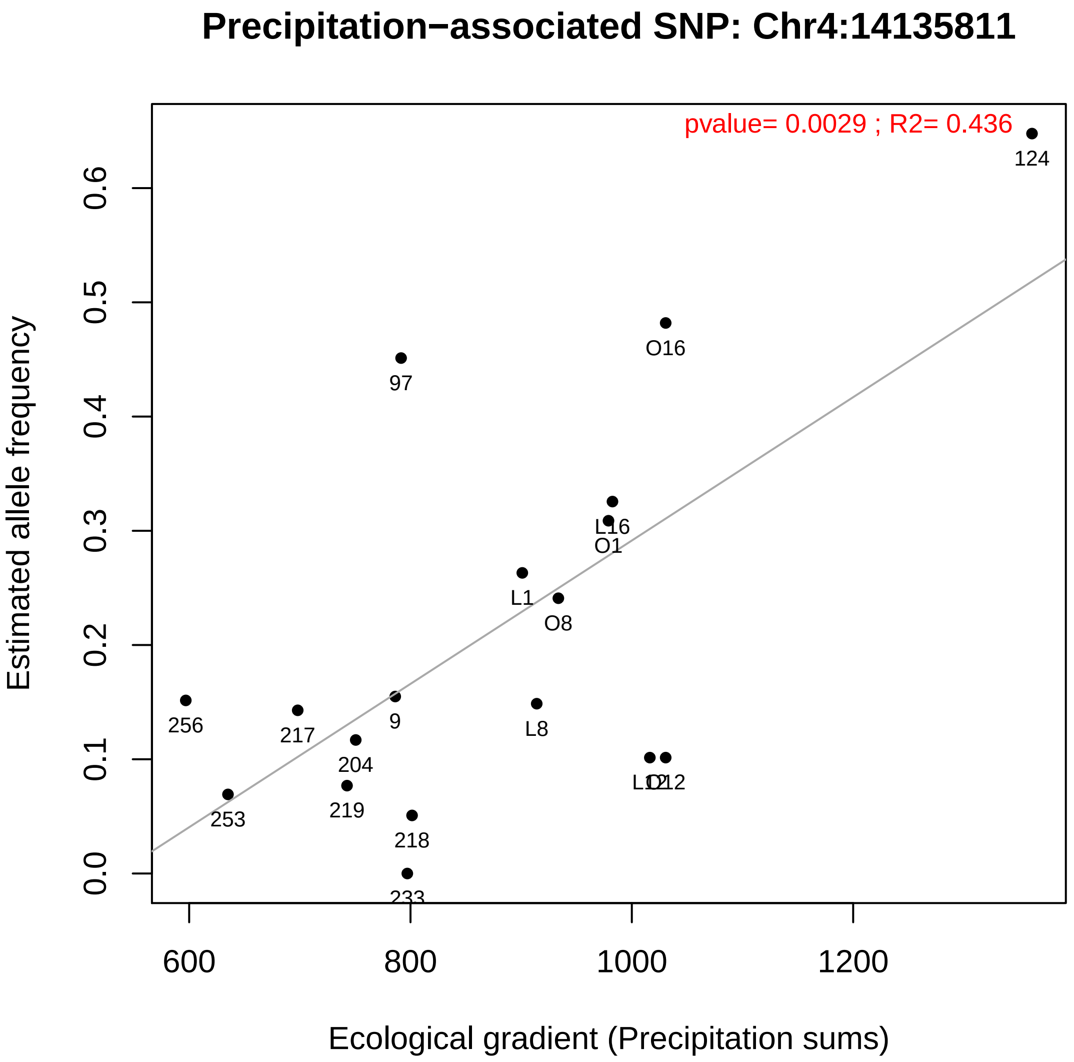

Precipitation−associated SNP: Chr4:14135811
pvalue= 0.0029 ; R2= 0.436
Estimated allele frequency
Ecological gradient (Precipitation sums)
0.0
0.1
0.2
0.3
0.4
0.5
0.6
600
800
1000
1200
124
O16
97
L16
O1
L1
O8
256
217
9
L8
204
219
218
233
253
L12
O12

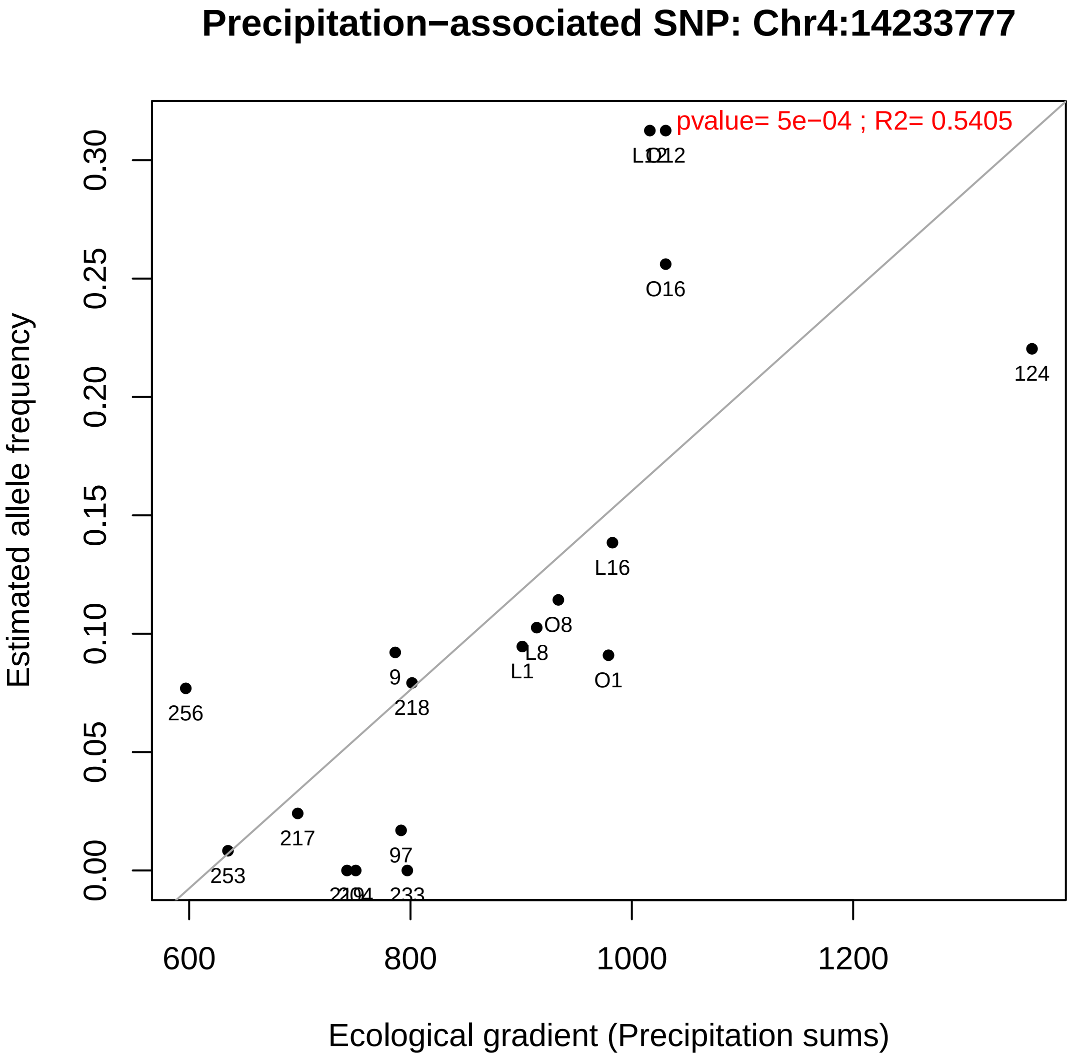
Precipitation−associated SNP: Chr4:14233777
pvalue= 5e−04 ; R2= 0.5405
Estimated allele frequency
Ecological gradient (Precipitation sums)
0.00
0.05
0.10
0.15
0.20
0.25
0.30
600
800
1000
1200
O12
O16
124
L16
O8
L8
L1
O1
9
218
256
217
253
97
233

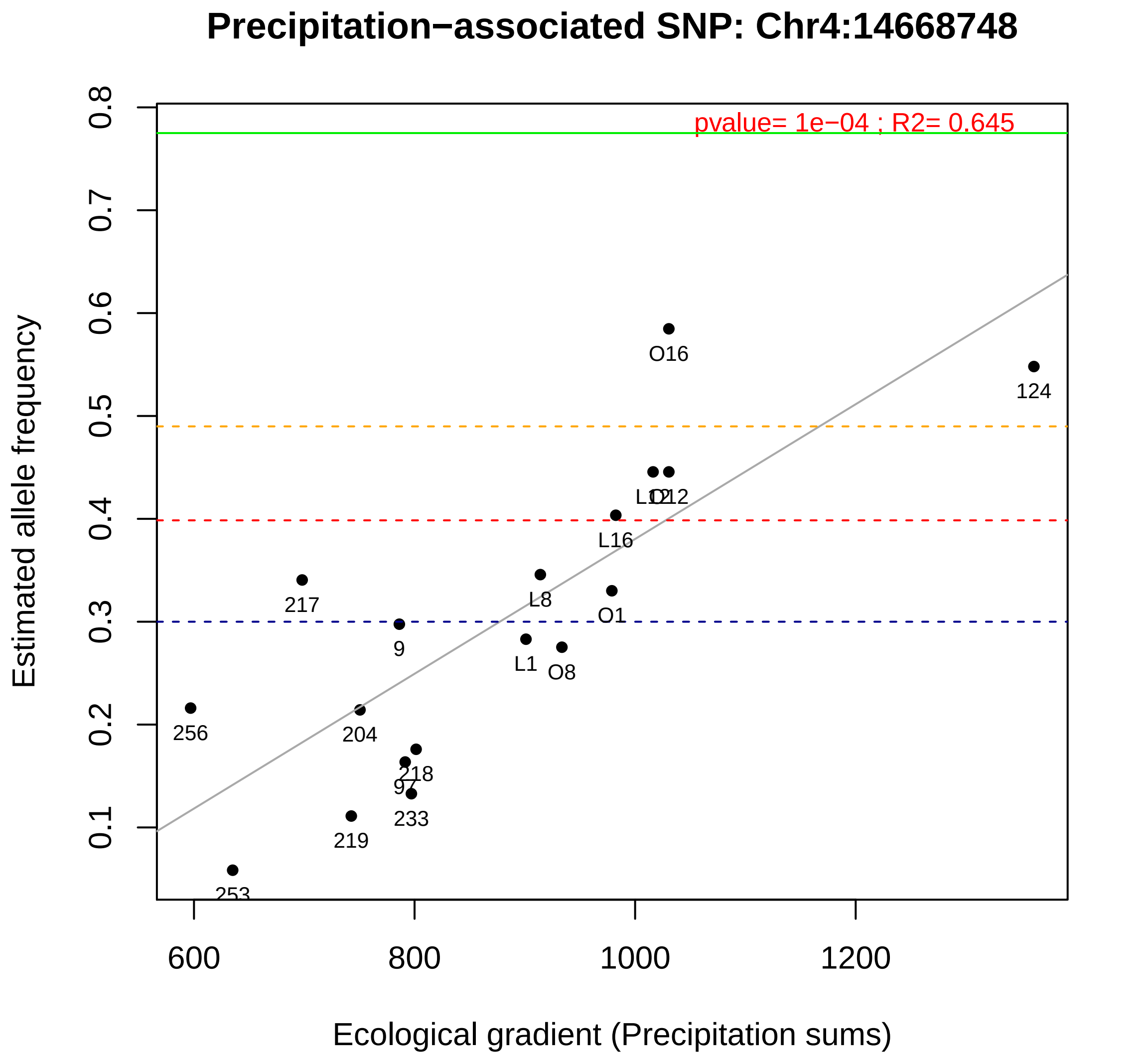

Precipitation−associated SNP: Chr4:14668748
pvalue= 1e−04 ; R2= 0.645
Estimated allele frequency
0.1
0.2
0.3
0.4
0.5
0.6
0.7
0.8
600
800
1000
1200
Ecological gradient (Precipitation sums)
O16
124
L12
O12
L16
217
L8
O1
9
L1
O8
256
204
218
97
233
219
253

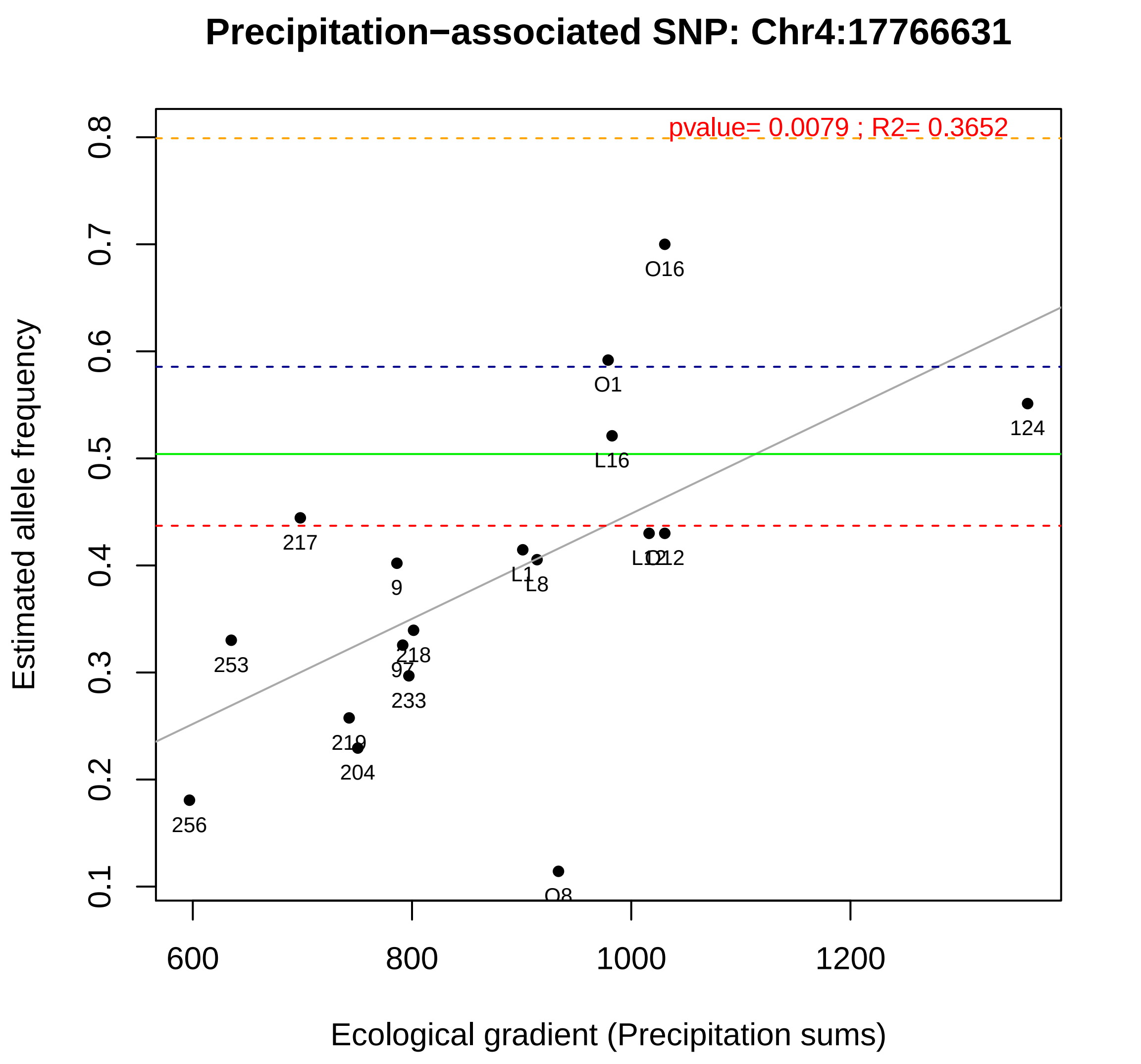

Precipitation−associated SNP: Chr4:17766631
pvalue= 0.0079 ; R2= 0.3652
Estimated allele frequency
Ecological gradient (Precipitation sums)
0.1
0.2
0.3
0.4
0.5
0.6
0.7
0.8
600
800
1000
1200
O16
O1
124
L16
217
L12
O12
9
L1
L8
253
218
97
233
219
204
256
O8

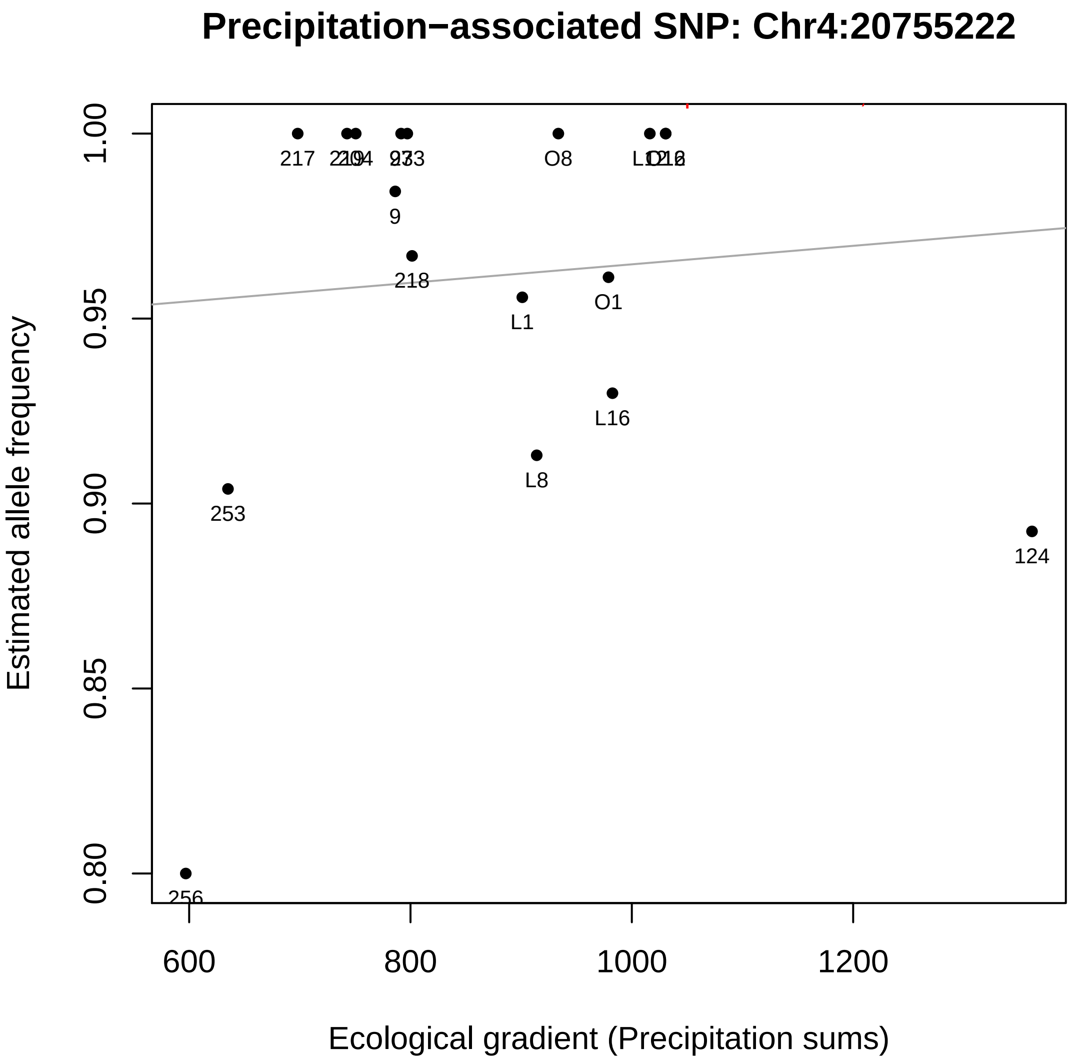

Precipitation−associated SNP: Chr4:20755222
Estimated allele frequency
Ecological gradient (Precipitation sums)
0.80
0.85
0.90
0.95
1.00
600
800
1000
1200
217
9
218
O8
O1
L1
L16
L8
253
124
256

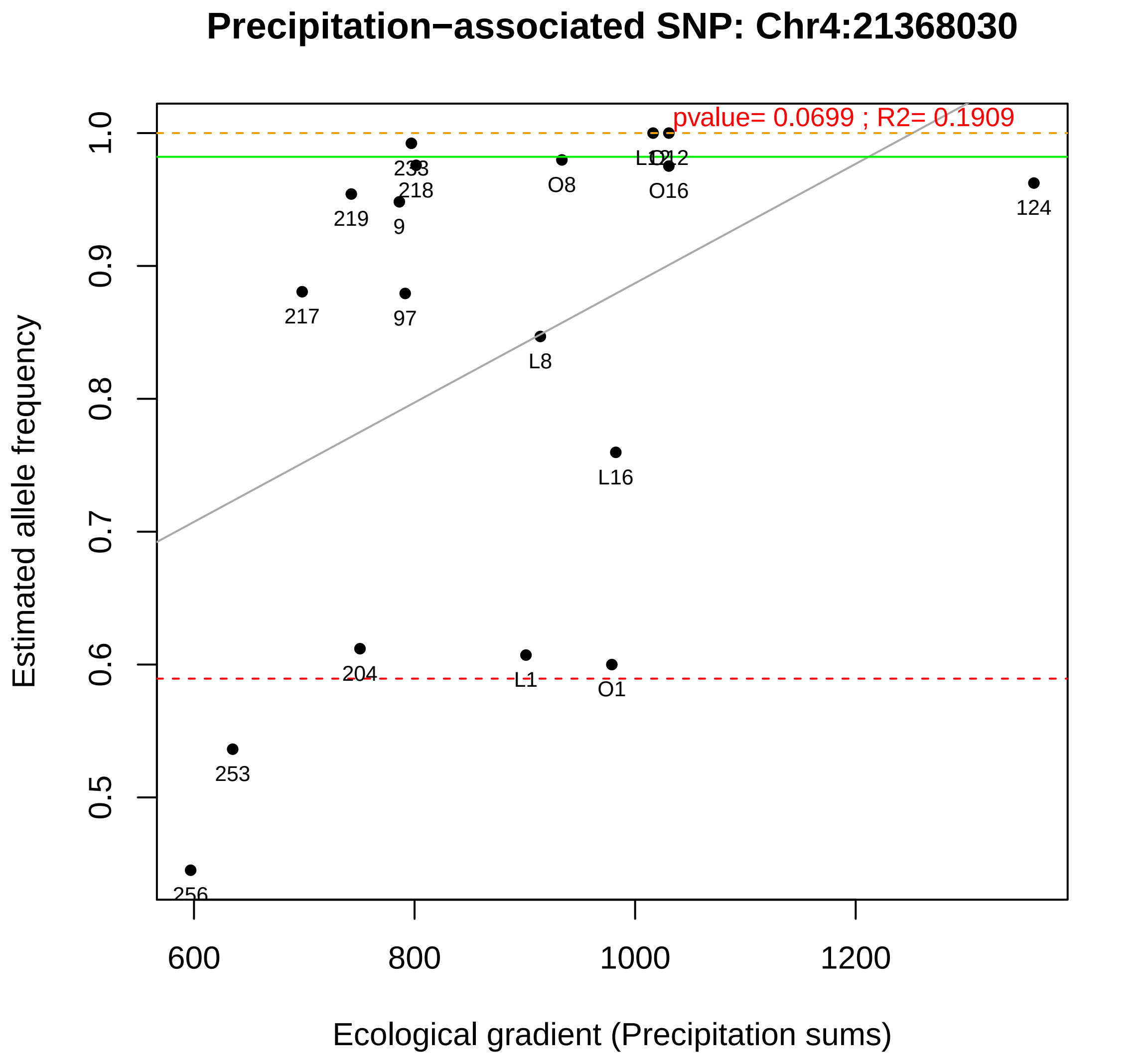

Precipitation−associated SNP: Chr4:21368030
pvalue= 0.0699 ; R2= 0.1909
Estimated allele frequency
Ecological gradient (Precipitation sums)
0.5
0.6
0.7
0.8
0.9
1.0
600
800
1000
1200
233
218
219
9
217
97
O8
L12
O12
O16
124
L8
L16
204
L1
O1
253
256

### Precipitation−associated SNP: Chr4:21497721

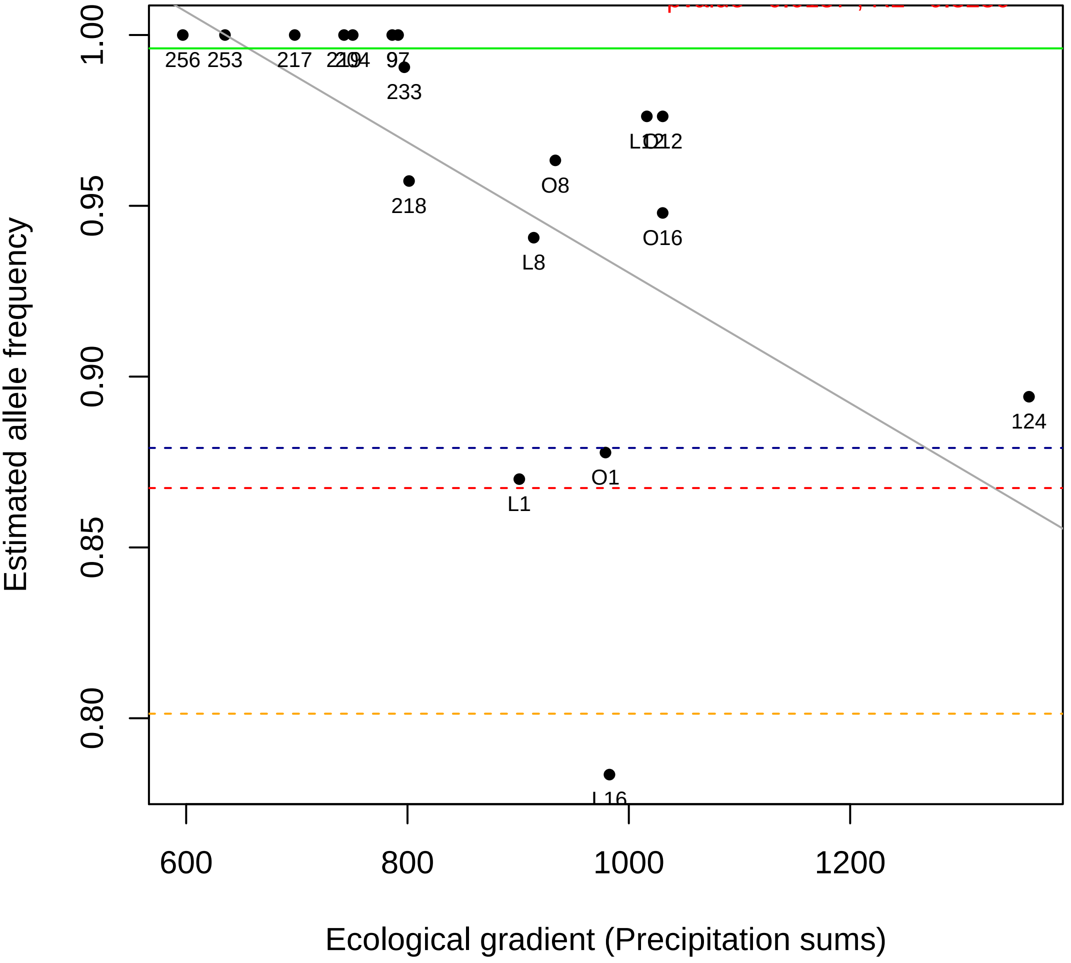

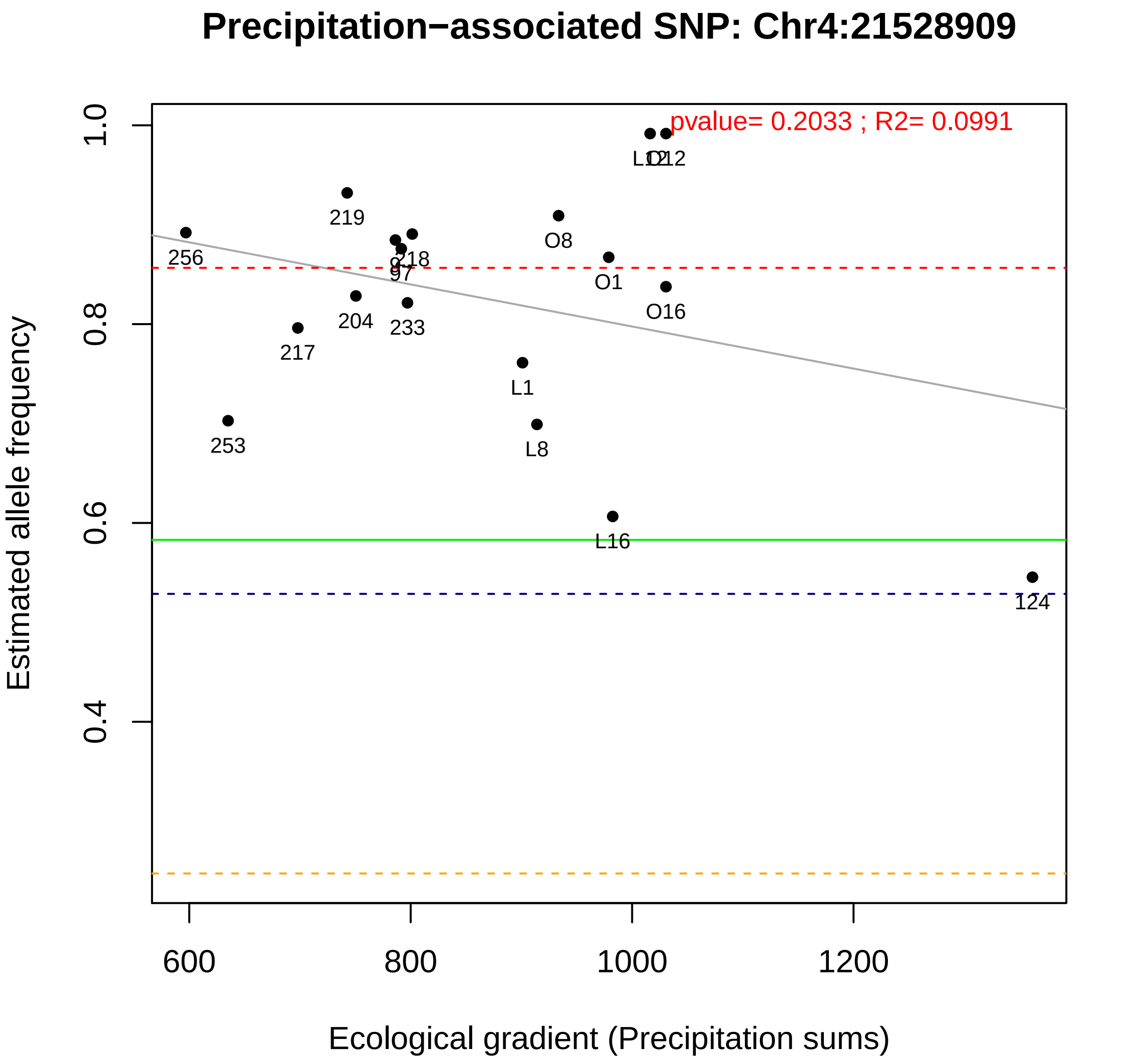

Precipitation−associated SNP: Chr4:21528909
pvalue= 0.2033 ; R2= 0.0991
Estimated allele frequency
Ecological gradient (Precipitation sums)
1.0
0.8
0.6
0.4
600
800
1000
1200
256
253
217
219
204
218
97
233
L1
L8
O8
O1
O16
L16
124

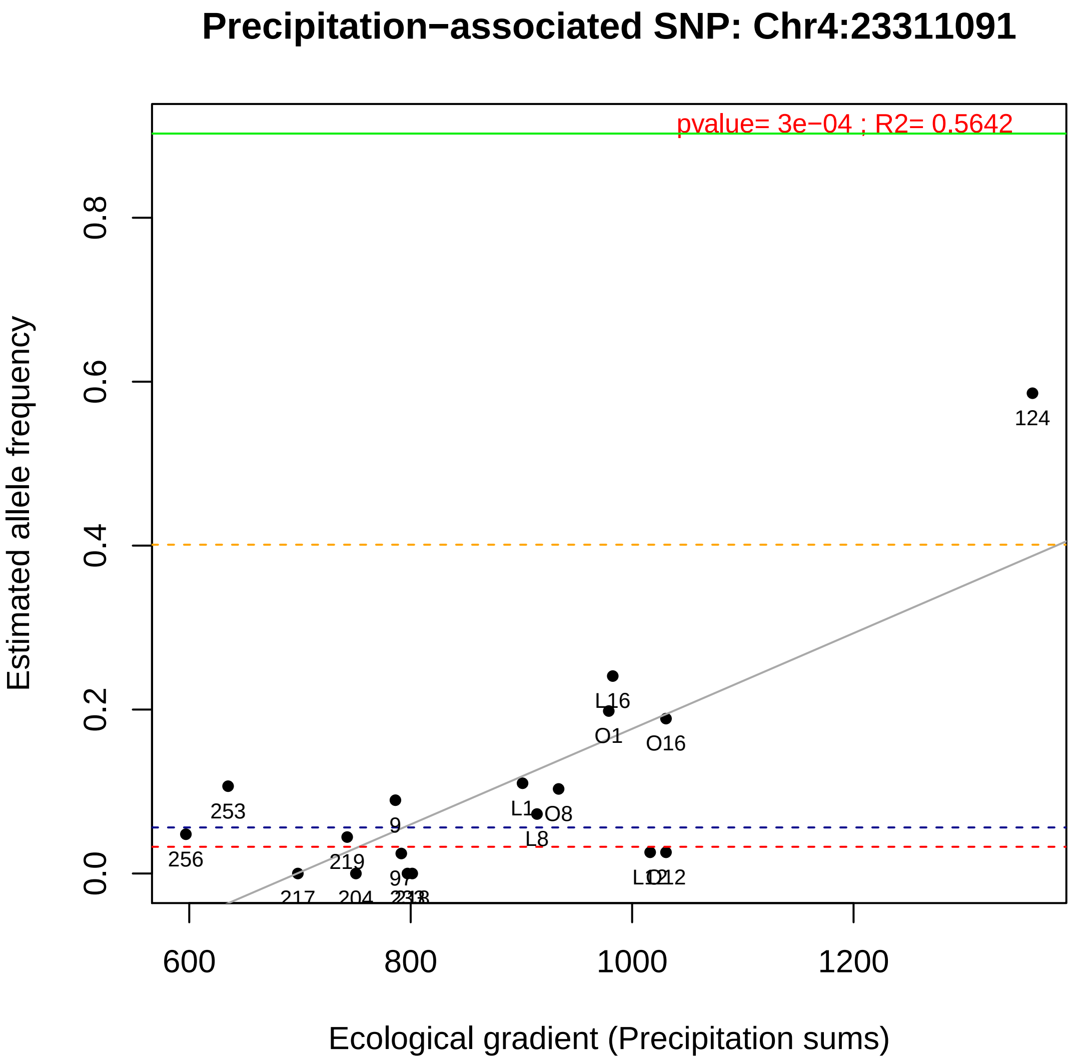
Precipitation−associated SNP: Chr4:23311091
pvalue= 3e−04 ; R2= 0.5642
Estimated allele frequency
0.0
0.2
0.4
0.6
0.8
600
800
1000
1200
Ecological gradient (Precipitation sums)
124
L16
O1
O16
L1
O8
L8
253
256
9
219
217
204
97
L12
O12

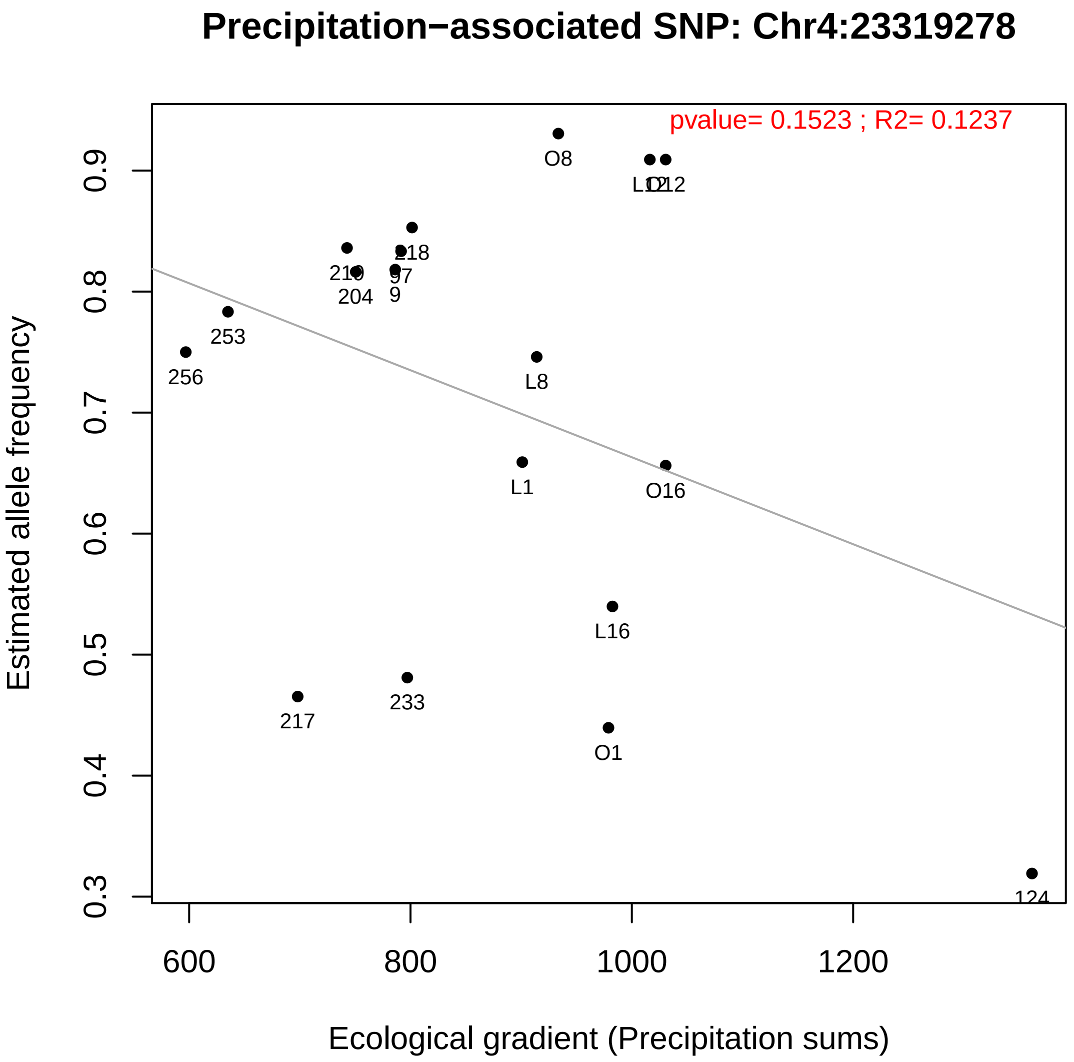
Precipitation−associated SNP: Chr4:23319278
pvalue= 0.1523 ; R2= 0.1237
Estimated allele frequency
Ecological gradient (Precipitation sums)
0.3
0.4
0.5
0.6
0.7
0.8
0.9
600
800
1000
1200
O8
L12
O12
218
210
204
97
9
253
256
L8
L1
O16
L16
217
233
O1
124

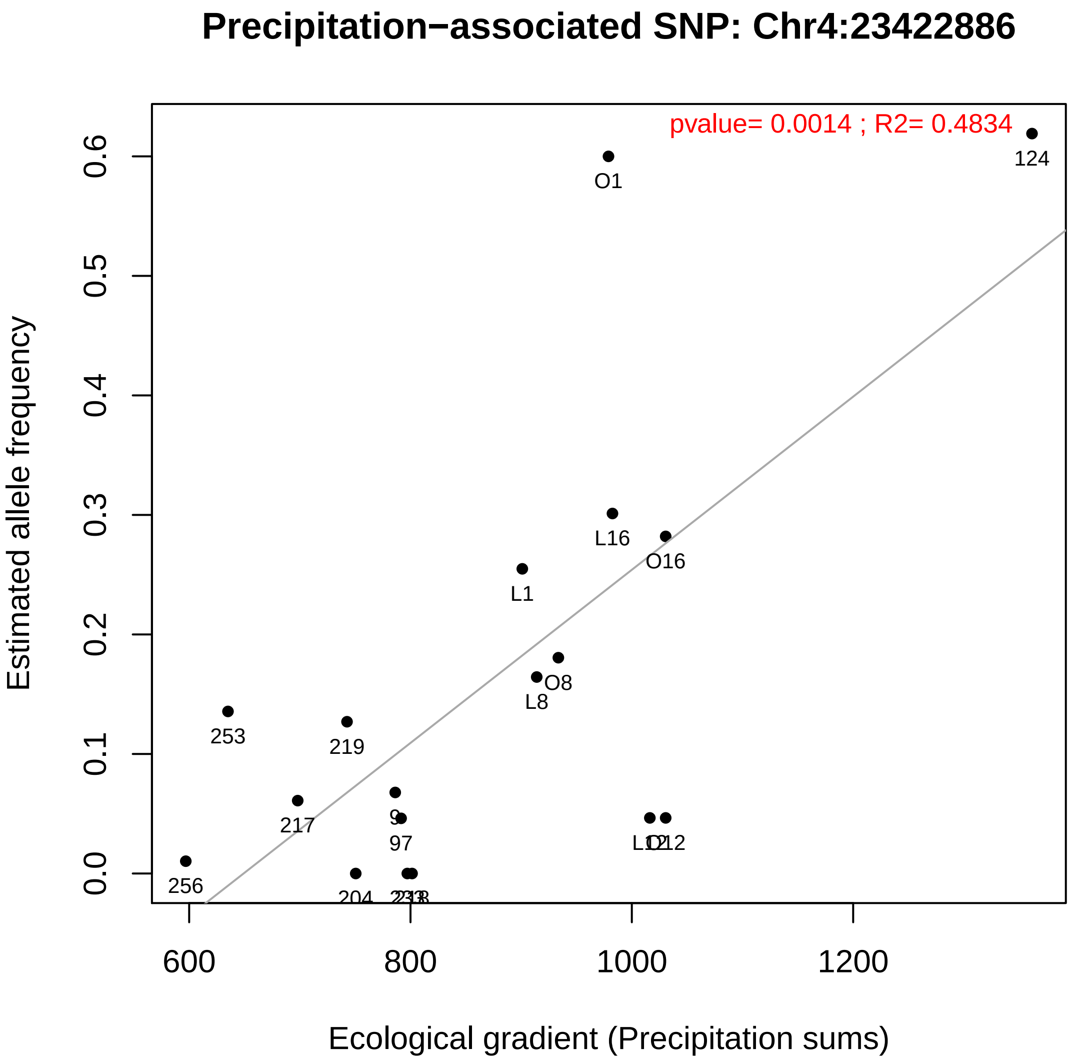
Precipitation−associated SNP: Chr4:23422886
pvalue= 0.0014 ; R2= 0.4834
Estimated allele frequency
Ecological gradient (Precipitation sums)
0.0
0.1
0.2
0.3
0.4
0.5
0.6
600
800
1000
1200
124
O1
L16
O16
L1
O8
L8
253
219
217
9
97
256
204

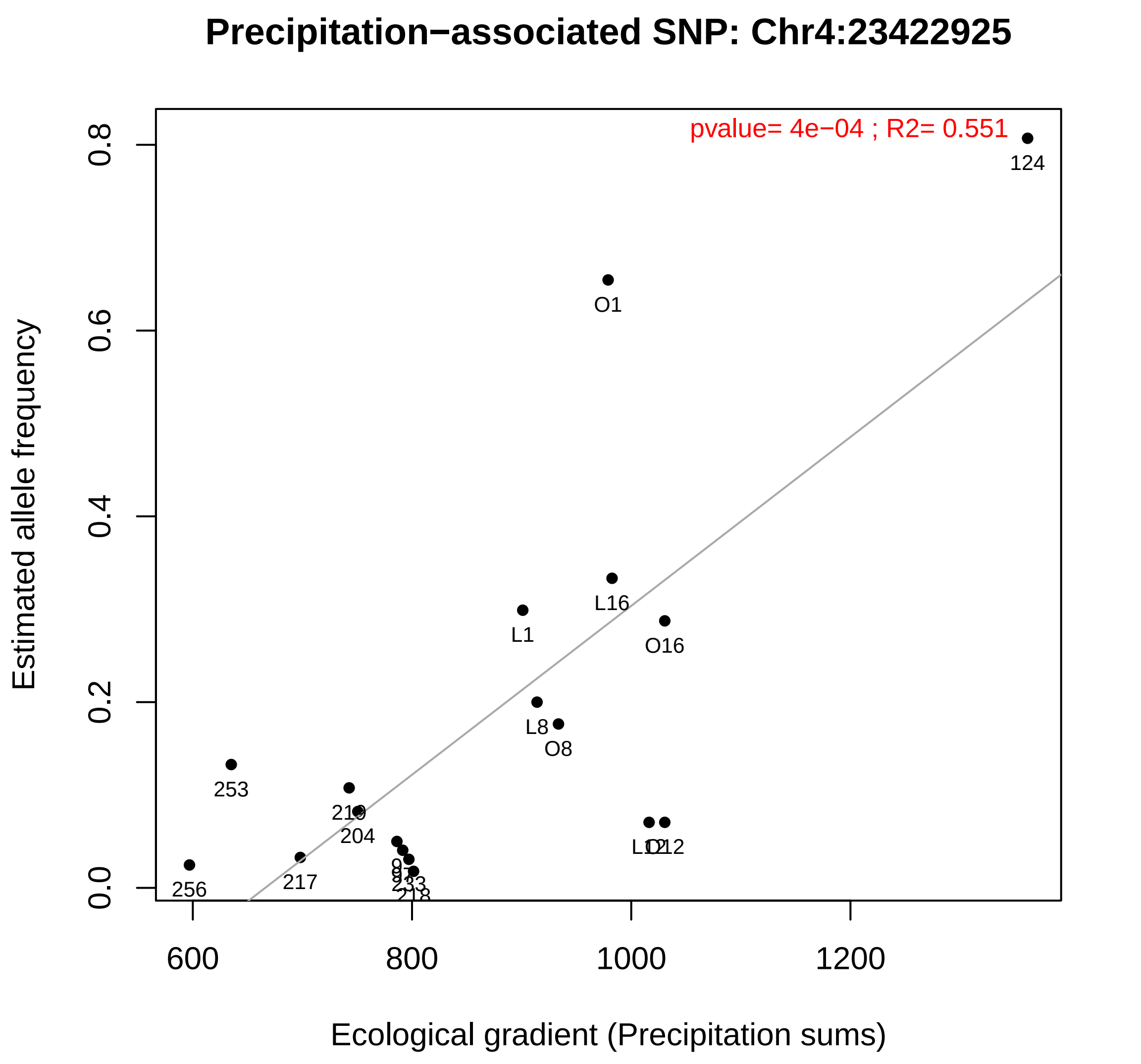
Precipitation−associated SNP: Chr4:23422925
pvalue= 4e−04 ; R2= 0.551
124
O1
L16
L1
O16
L8
O8
253
204
256
217
0.0
0.2
0.4
0.6
0.8
600
800
1000
1200
Estimated allele frequency
Ecological gradient (Precipitation sums)

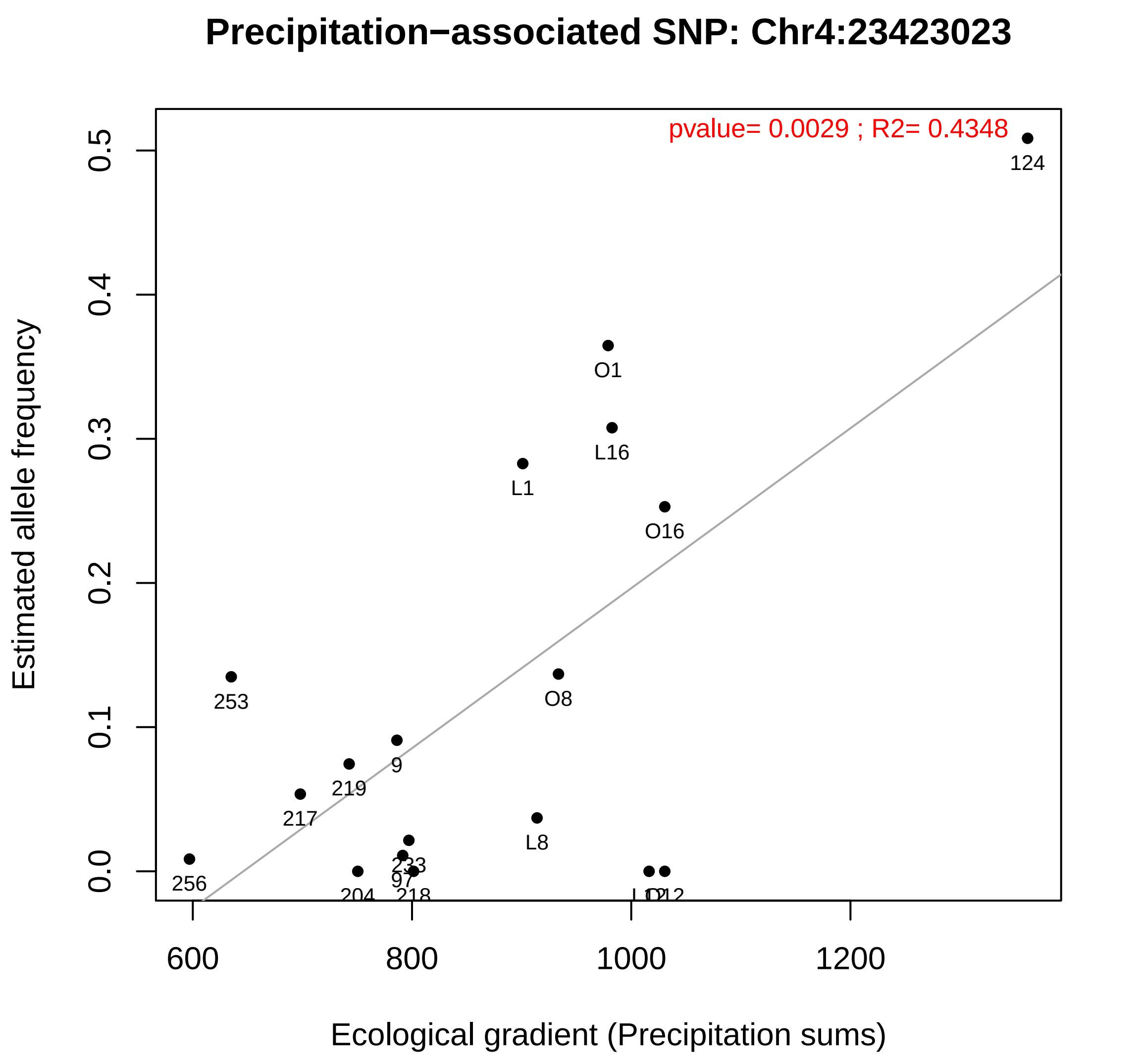

Precipitation−associated SNP: Chr4:23423023
pvalue= 0.0029 ; R2= 0.4348
Estimated allele frequency
0.0
0.1
0.2
0.3
0.4
0.5
Ecological gradient (Precipitation sums)
600
800
1000
1200
124
O1
L16
L1
O16
253
O8
9
219
217
L8
256
233
97
204
218

**Precipitation−associated SNP: Chr4:23427867**

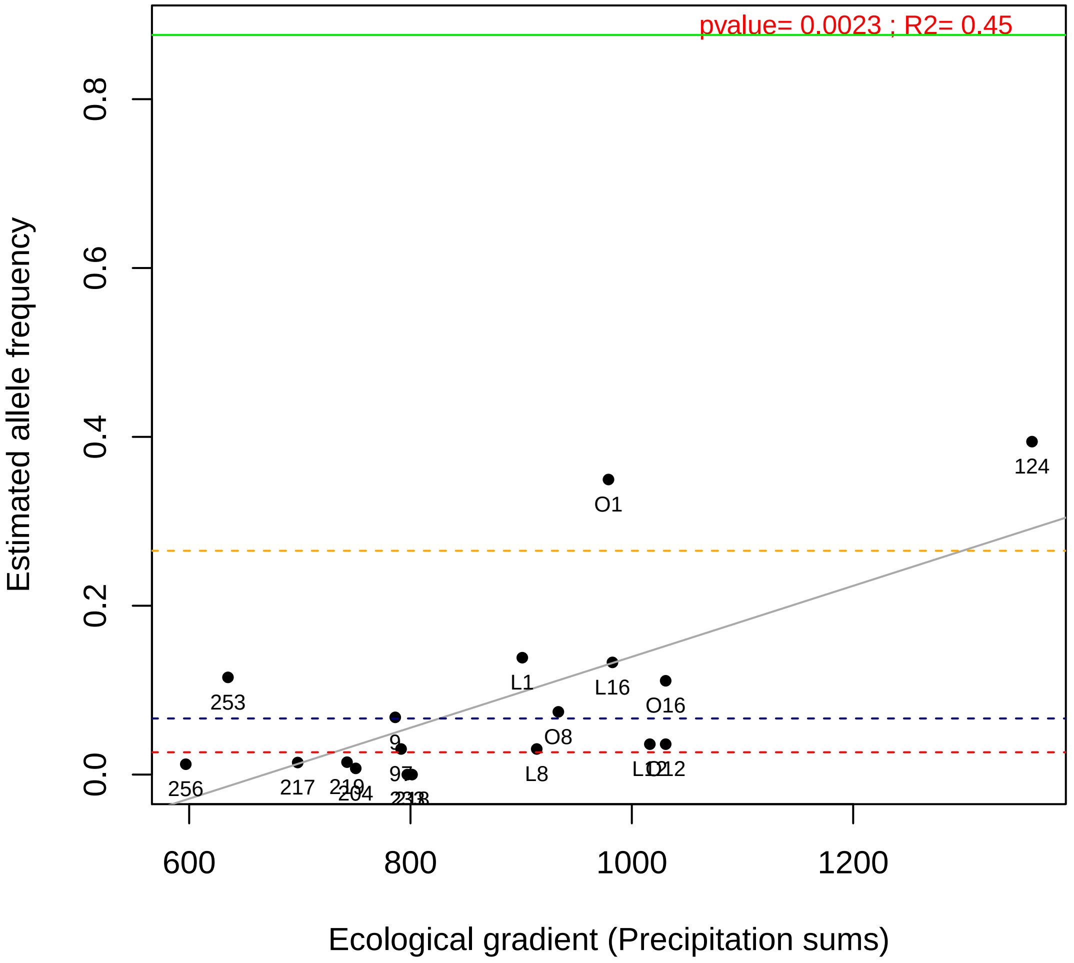

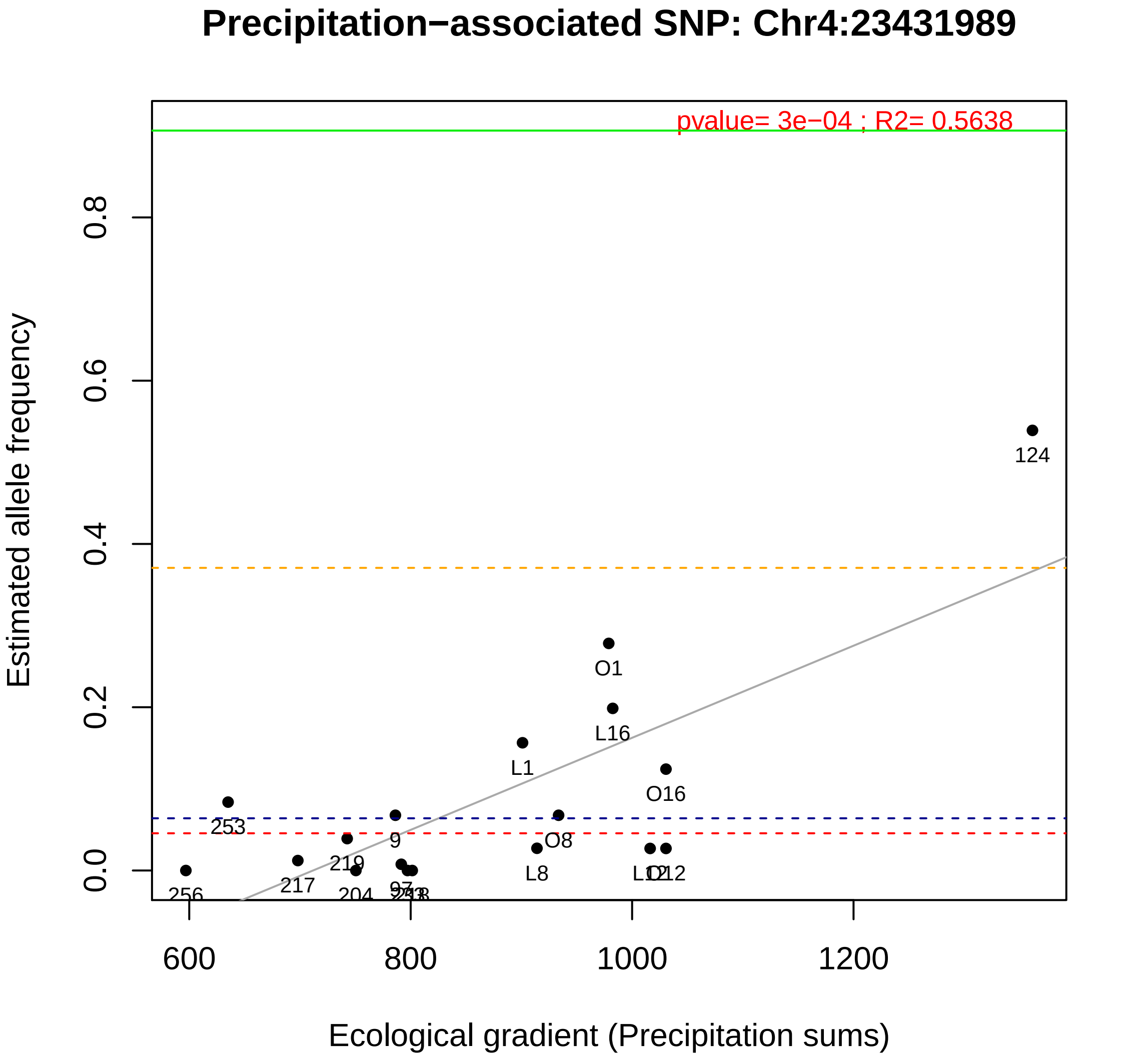
Precipitation−associated SNP: Chr4:23431989
pvalue= 3e−04 ; R2= 0.5638
Estimated allele frequency
0.0
0.2
0.4
0.6
0.8
Ecological gradient (Precipitation sums)
600
800
1000
1200
124
O1
L16
L1
O16
253
9
O8
L8
219
217
204
97

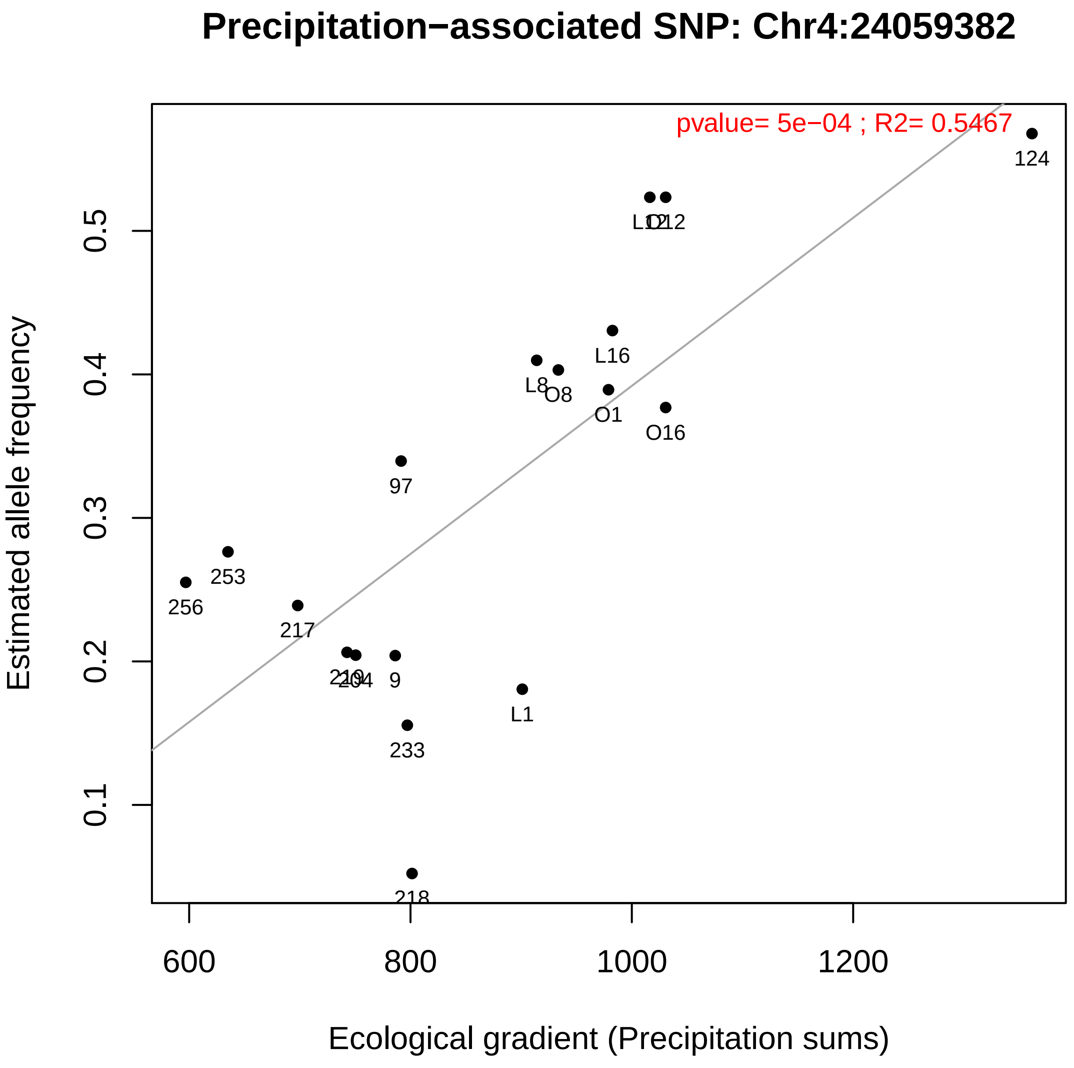
Precipitation−associated SNP: Chr4:24059382
pvalue= 5e−04 ; R2= 0.5467
Estimated allele frequency
Ecological gradient (Precipitation sums)
0.1
0.2
0.3
0.4
0.5
600
800
1000
1200
124
L12
O12
L16
L8
O8
O1
O16
97
253
256
217
210
204
9
L1
233
218

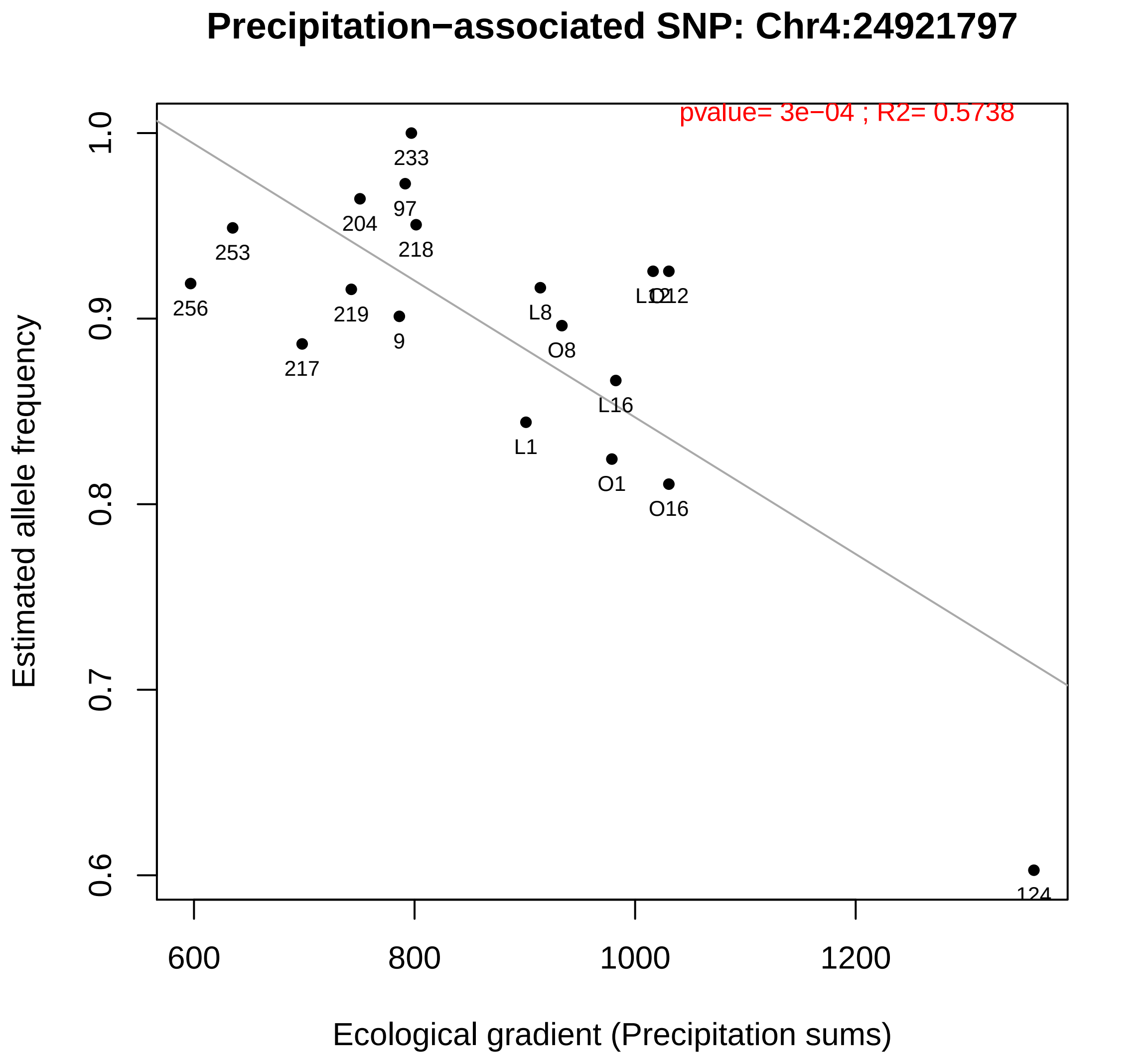
Precipitation−associated SNP: Chr4:24921797
pvalue= 3e−04 ; R2= 0.5738
Estimated allele frequency
Ecological gradient (Precipitation sums)
0.6
0.7
0.8
0.9
1.0
600
800
1000
1200
233
97
204
218
253
256
219
9
217
L8
O8
L12
O12
L16
L1
O1
O16
124

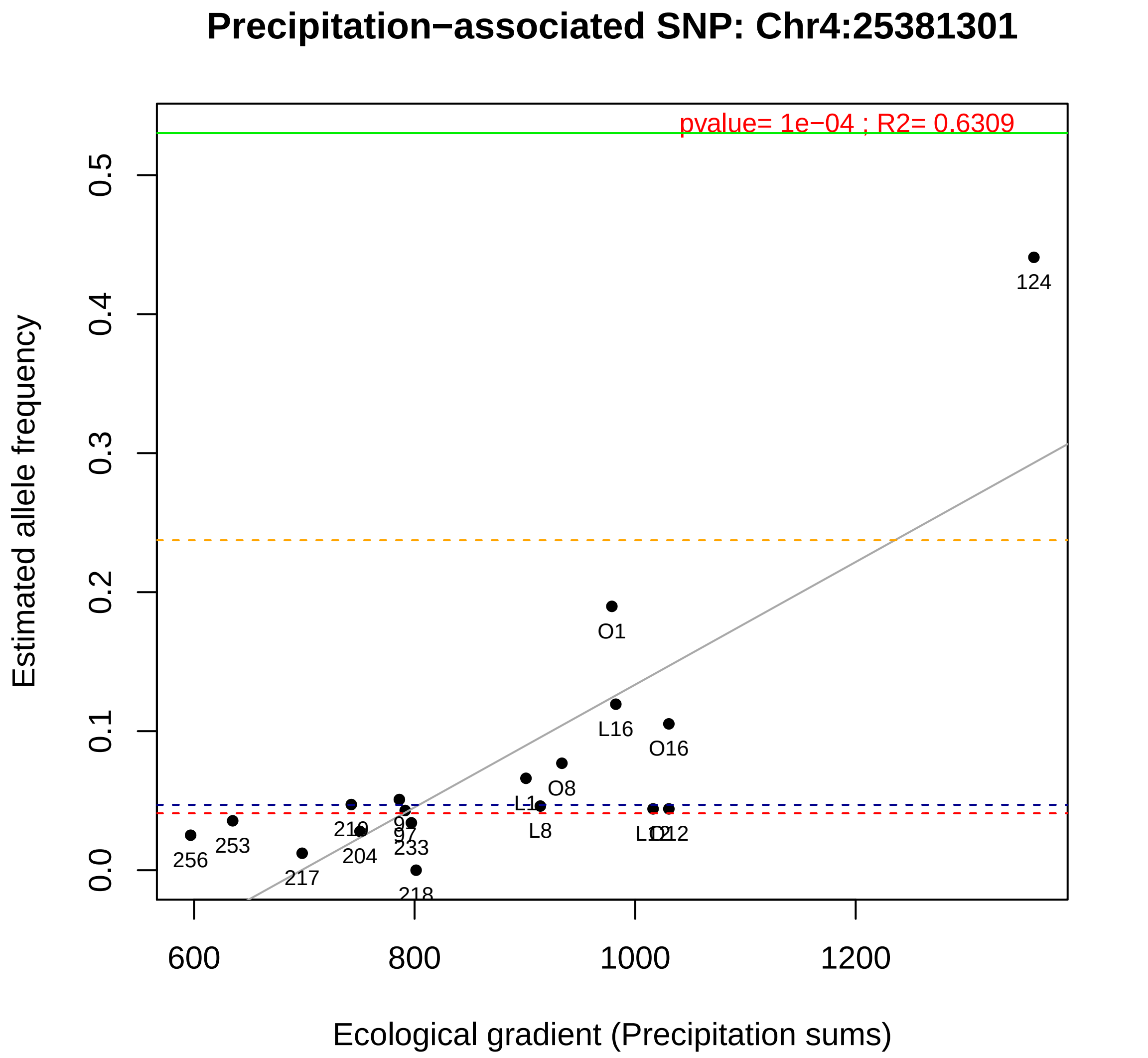

Precipitation−associated SNP: Chr4:25381301
pvalue= 1e−04 ; R2= 0.6309
Estimated allele frequency
0.0
0.1
0.2
0.3
0.4
0.5
600
800
1000
1200
Ecological gradient (Precipitation sums)
124
O1
L16
O16
O8
L1
L8
L12
O12
210
204
233
218
217
253
256

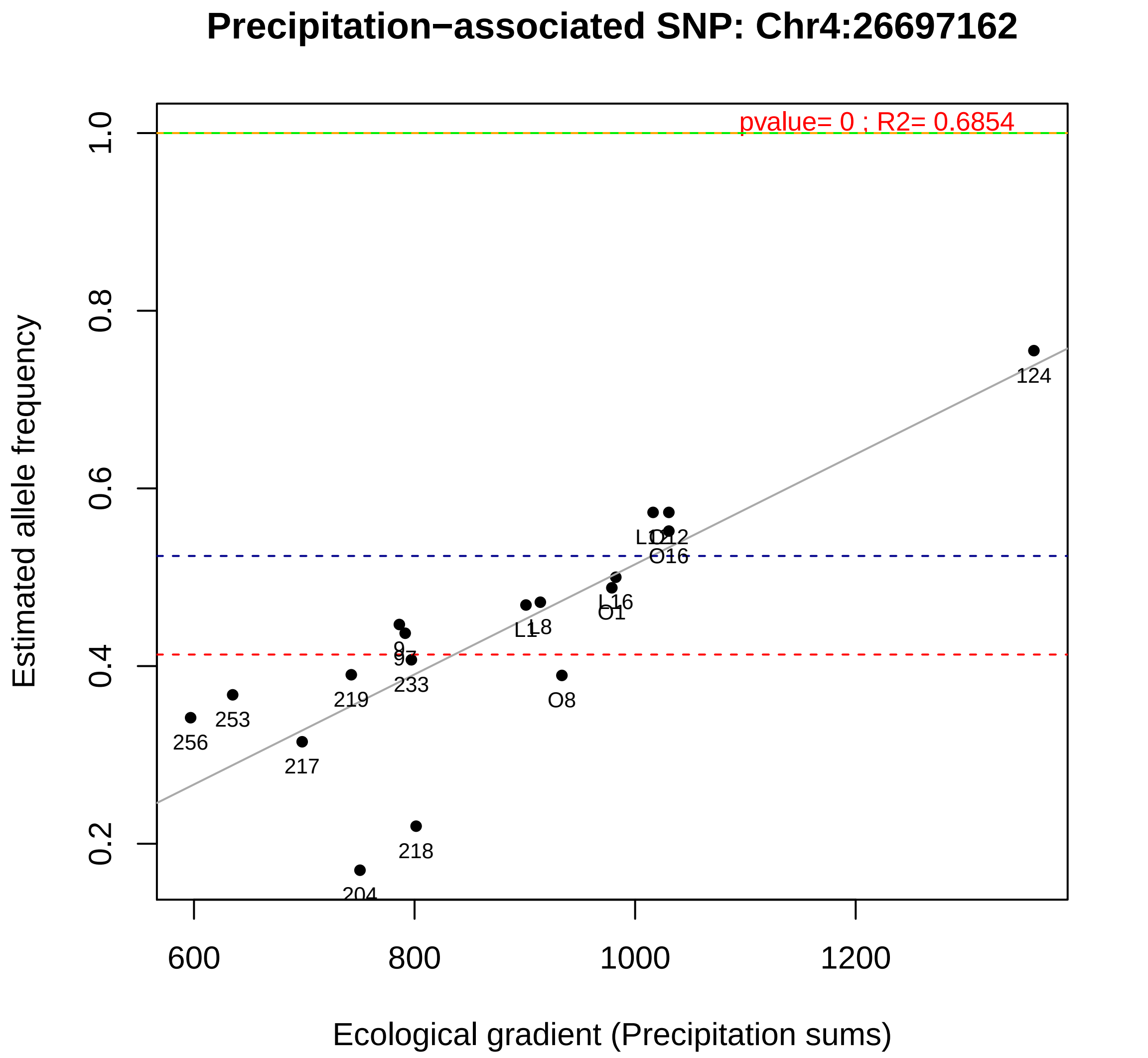

Precipitation−associated SNP: Chr4:26697162
pvalue= 0 ; R2= 0.6854
Estimated allele frequency
Ecological gradient (Precipitation sums)
0.2
0.4
0.6
0.8
1.0
600
800
1000
1200
124
L12
O12
O16
L16
O1
L1
L8
9
97
233
219
O8
253
256
217
218
204

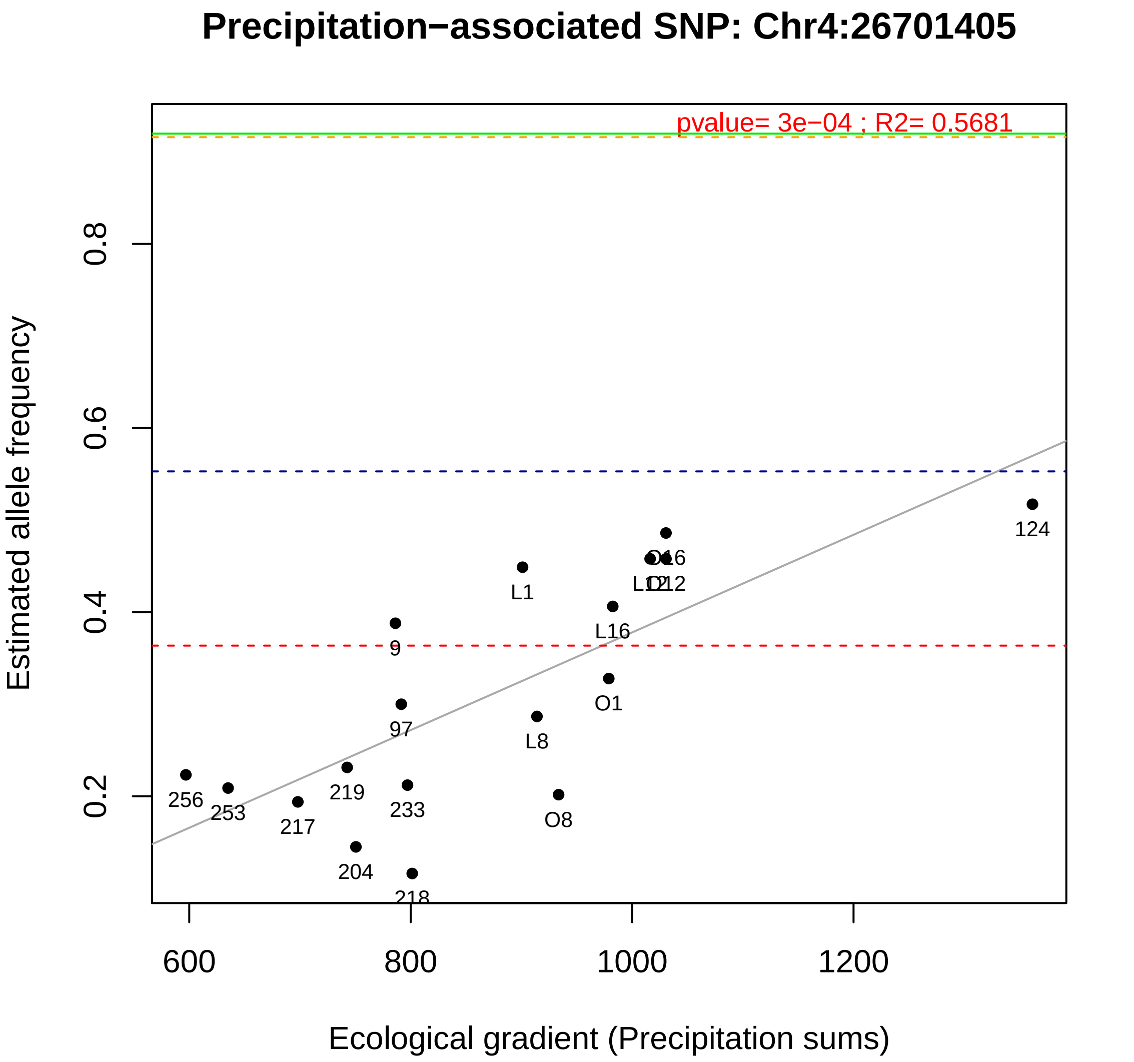

Precipitation−associated SNP: Chr4:26701405
pvalue= 3e−04 ; R2= 0.5681
Estimated allele frequency
0.2
0.4
0.6
0.8
600
800
1000
1200
Ecological gradient (Precipitation sums)
124
O16
L12
O12
L1
L16
9
O1
97
L8
219
233
256
253
217
O8
204
218

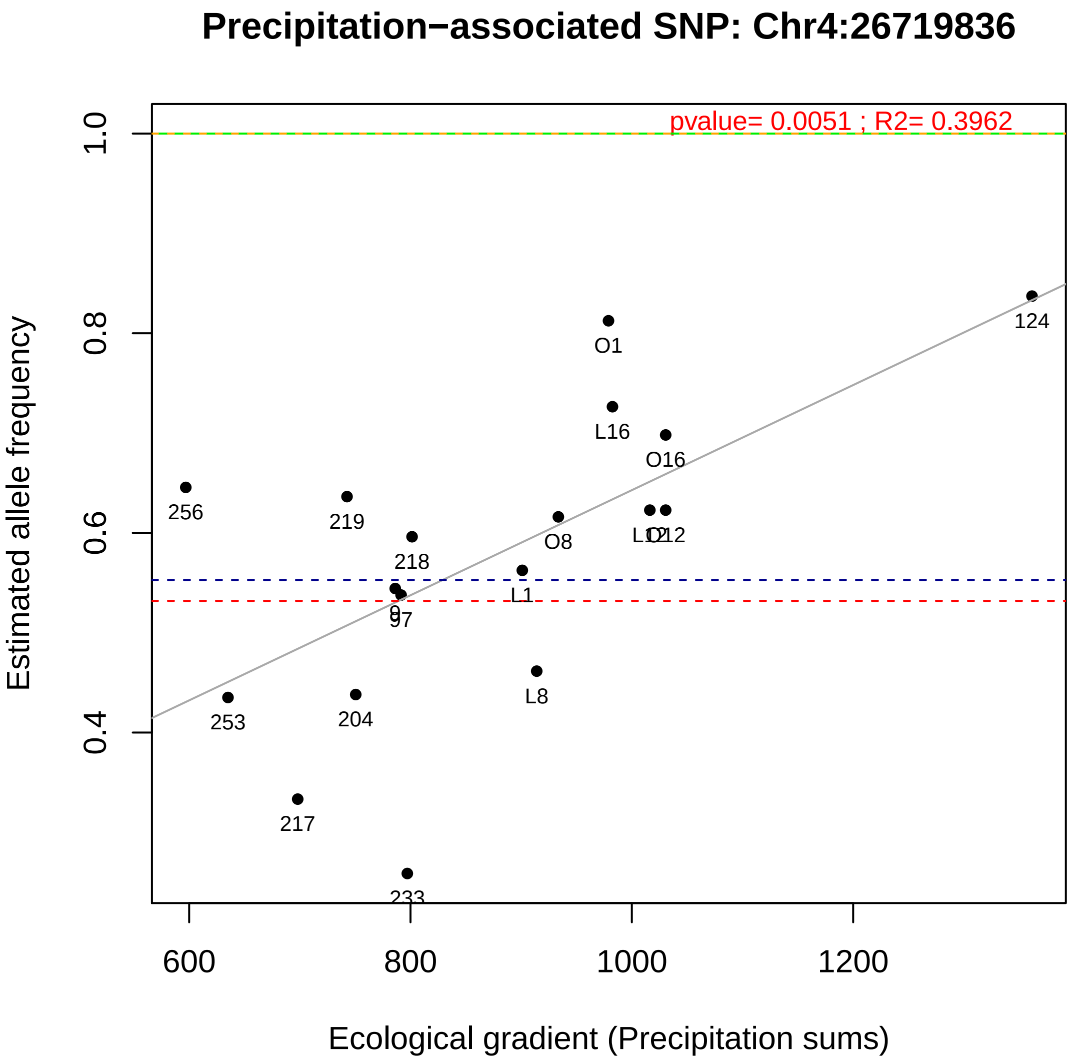
Precipitation−associated SNP: Chr4:26719836
pvalue= 0.0051 ; R2= 0.3962
Estimated allele frequency
Ecological gradient (Precipitation sums)
0.4
0.6
0.8
1.0
600
800
1000
1200
124
O1
L16
O16
256
219
218
O8
L1
L8
253
204
217
233

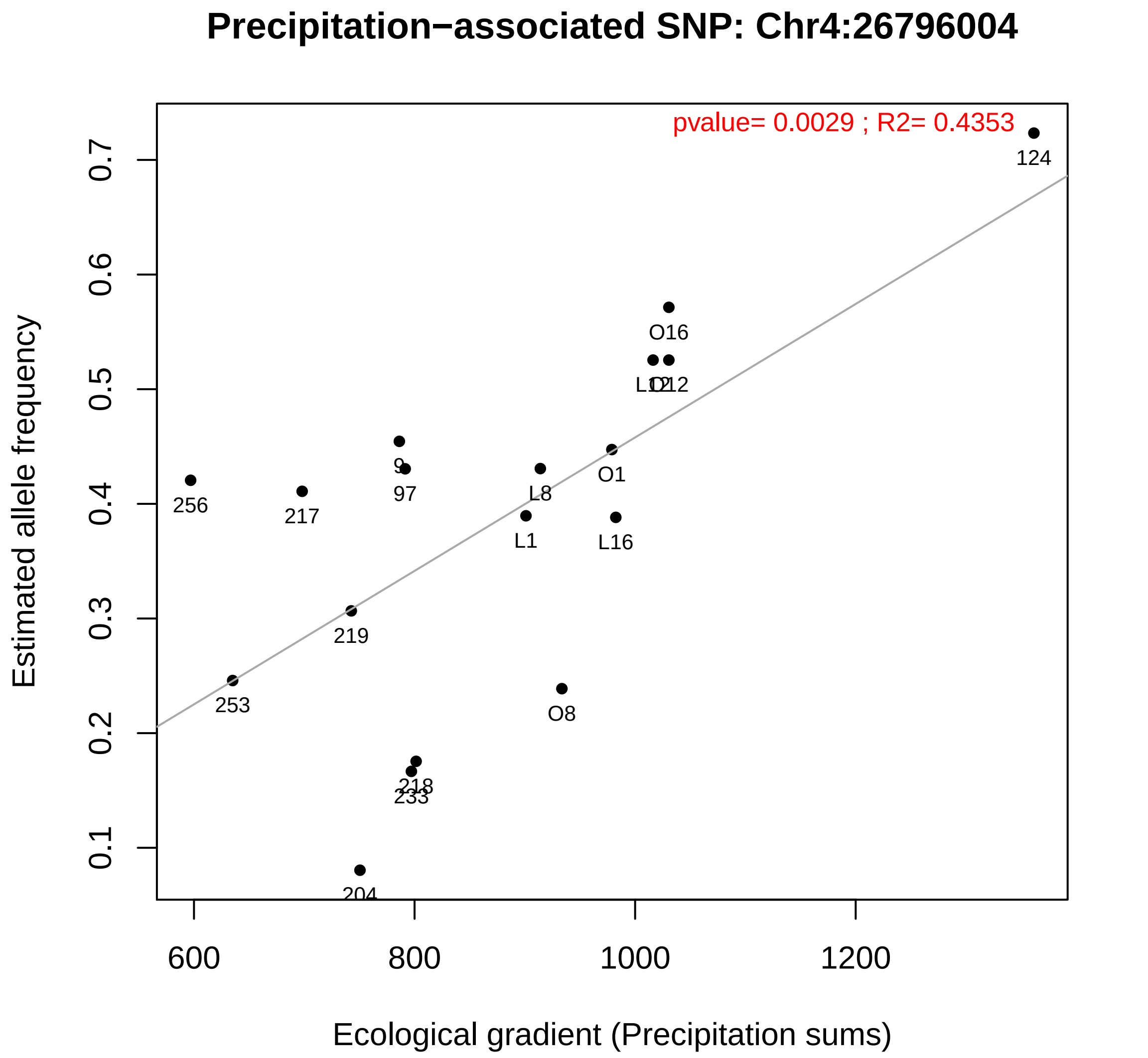
Precipitation−associated SNP: Chr4:26796004
pvalue= 0.0029 ; R2= 0.4353
Estimated allele frequency
Ecological gradient (Precipitation sums)
0.1
0.2
0.3
0.4
0.5
0.6
0.7
600
800
1000
1200
124
O16
L12
O12
9
97
256
217
L8
O1
L1
L16
219
253
O8
218
233
204

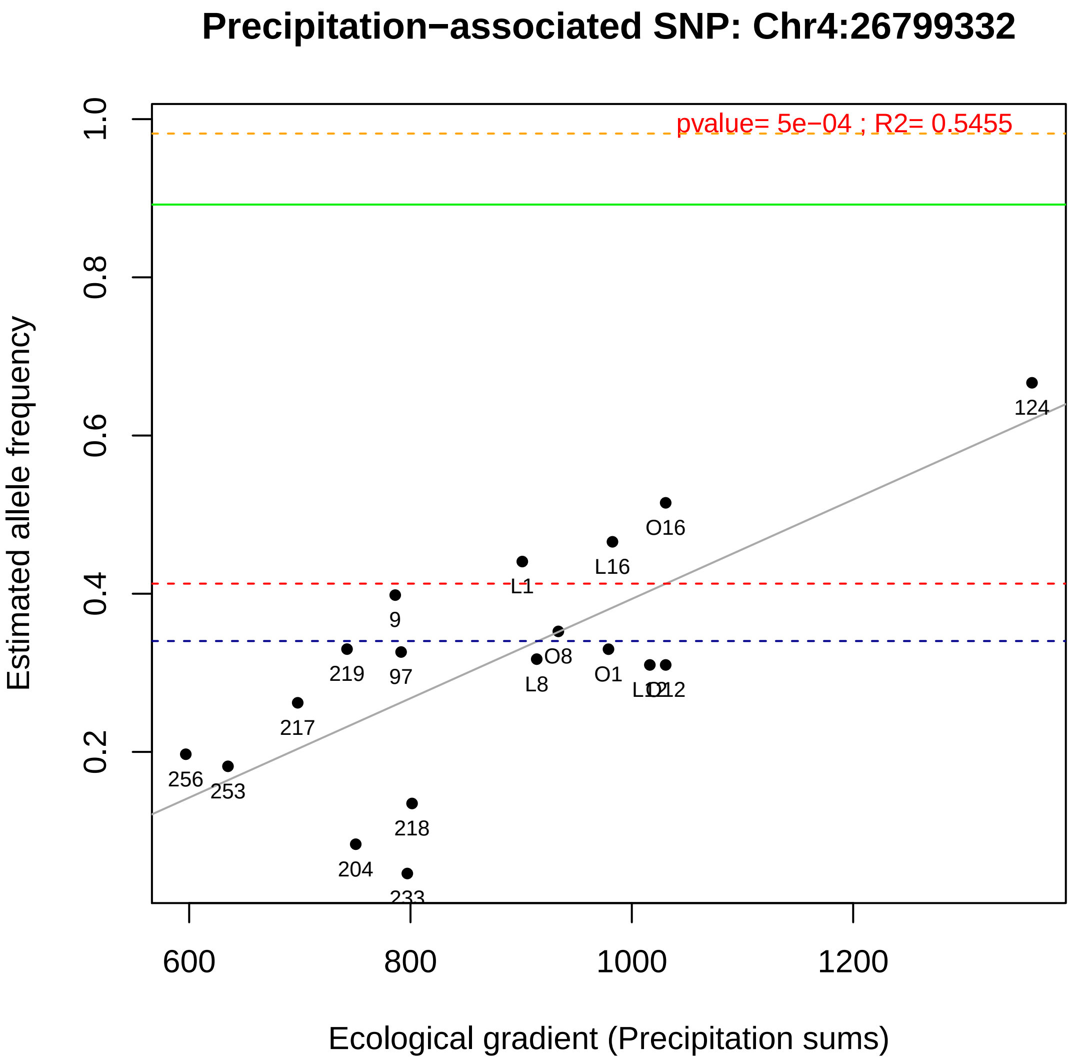

Precipitation−associated SNP: Chr4:26799332
pvalue= 5e−04 ; R2= 0.5455
Estimated allele frequency
Ecological gradient (Precipitation sums)
0.2
0.4
0.6
0.8
1.0
600
800
1000
1200
124
O16
L16
L1
9
O8
219
97
O1
L8
L12
O12
217
256
253
218
204
233

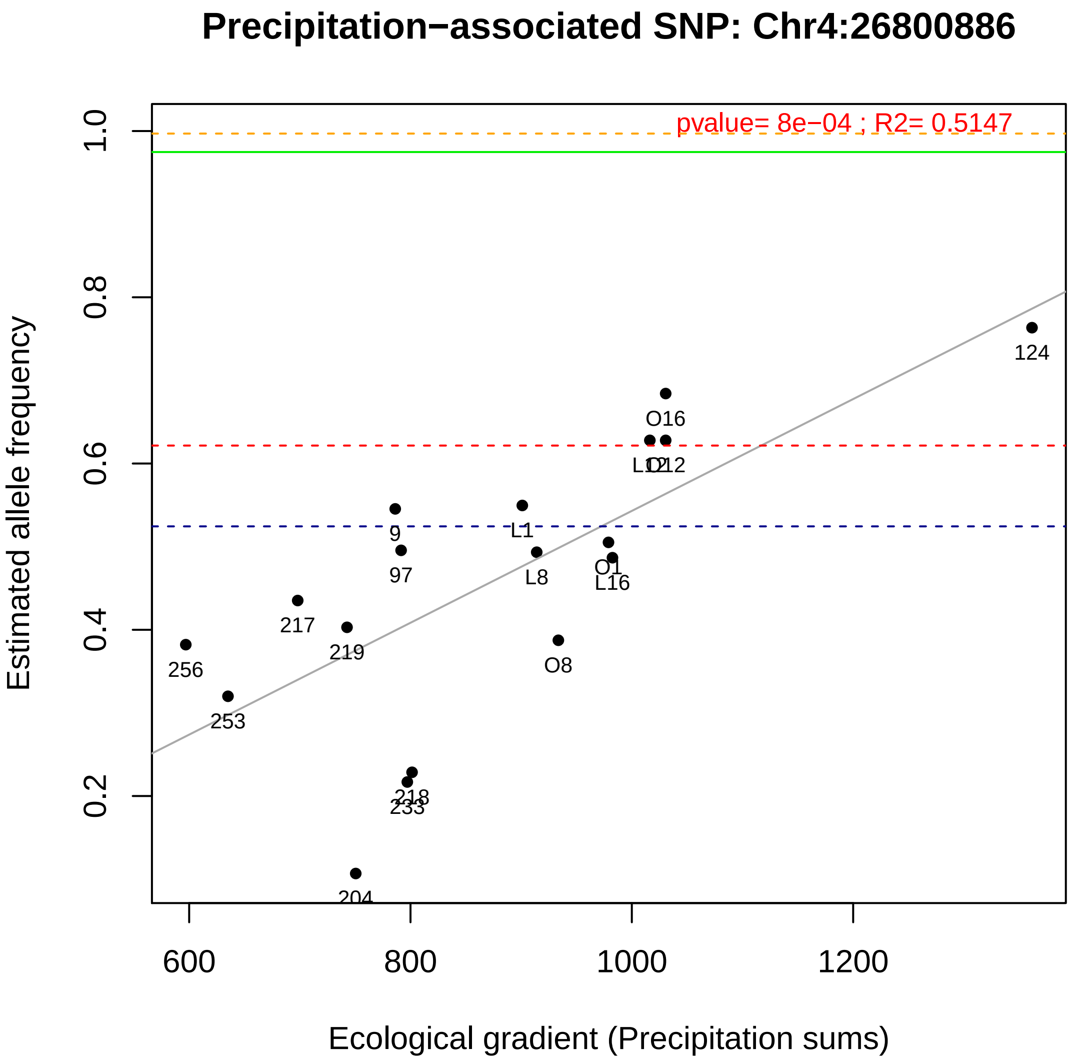
Precipitation−associated SNP: Chr4:26800886
pvalue= 8e−04 ; R2= 0.5147
Estimated allele frequency
Ecological gradient (Precipitation sums)
0.2
0.4
0.6
0.8
1.0
600
800
1000
1200
124
O16
L12
O12
9
L1
97
L8
O1
L16
217
219
256
O8
253
218
233
204

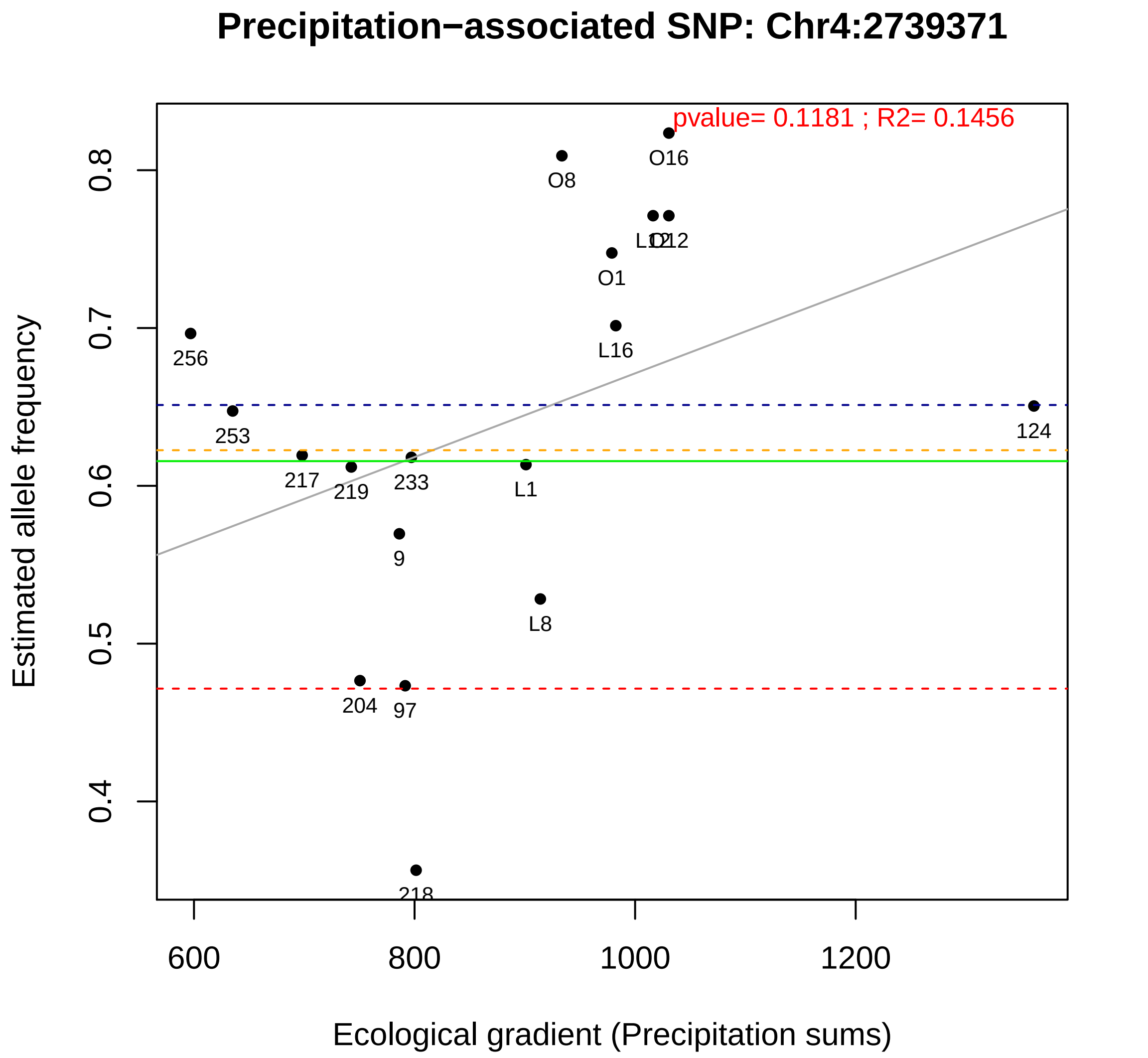

Precipitation−associated SNP: Chr4:2739371
pvalue= 0.1181 ; R2= 0.1456
Estimated allele frequency
Ecological gradient (Precipitation sums)
0.4
0.5
0.6
0.7
0.8
600
800
1000
1200
O16
O8
L12
O12
O1
L16
256
253
124
217
219
233
L1
9
L8
204
97
218

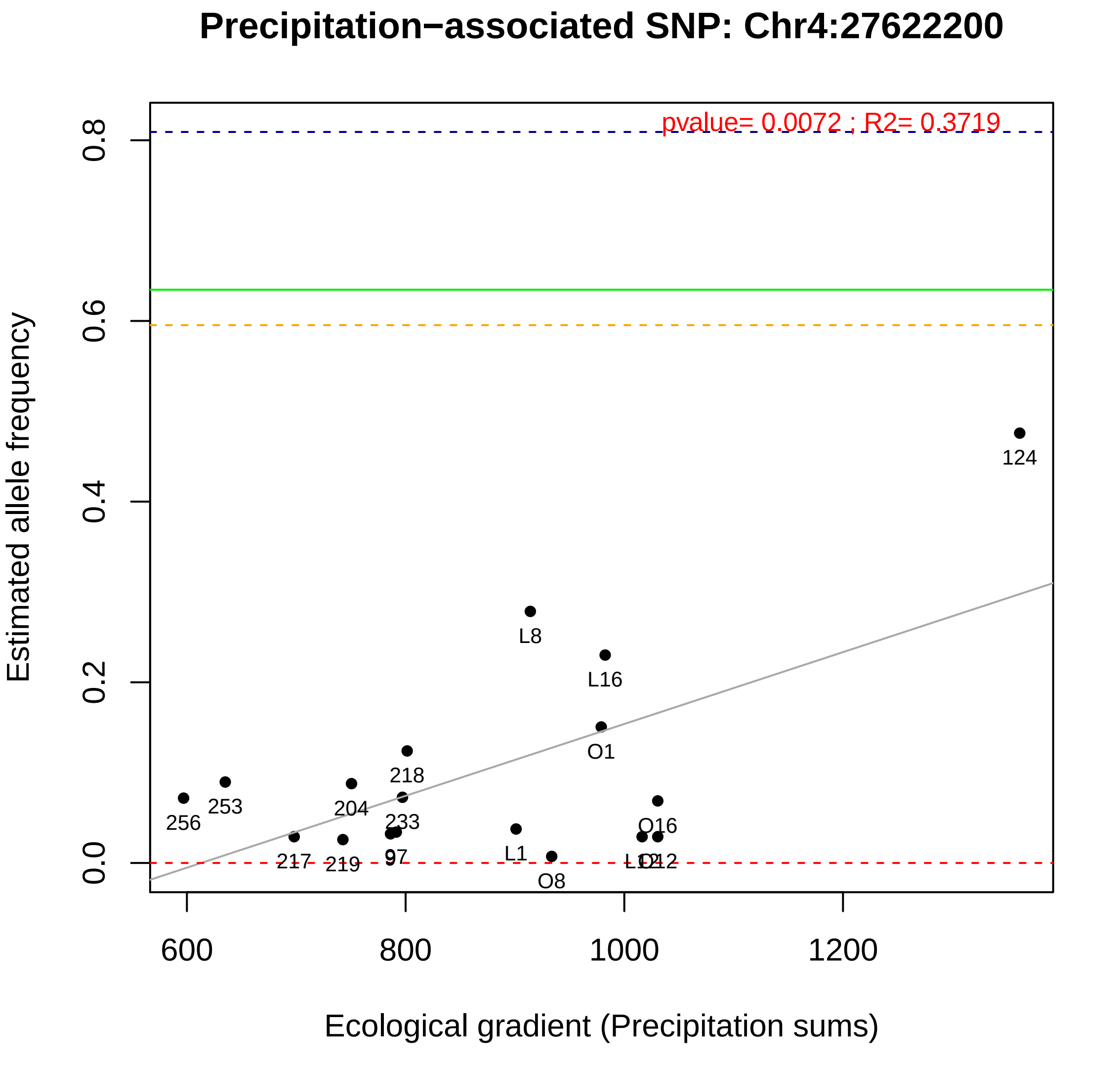

Precipitation−associated SNP: Chr4:27622200
pvalue= 0.0072 ; R2= 0.3719
Estimated allele frequency
0.0
0.2
0.4
0.6
0.8
600
800
1000
1200
Ecological gradient (Precipitation sums)
124
L8
L16
O1
218
253
256
204
233
217
219
L1
O8
O16

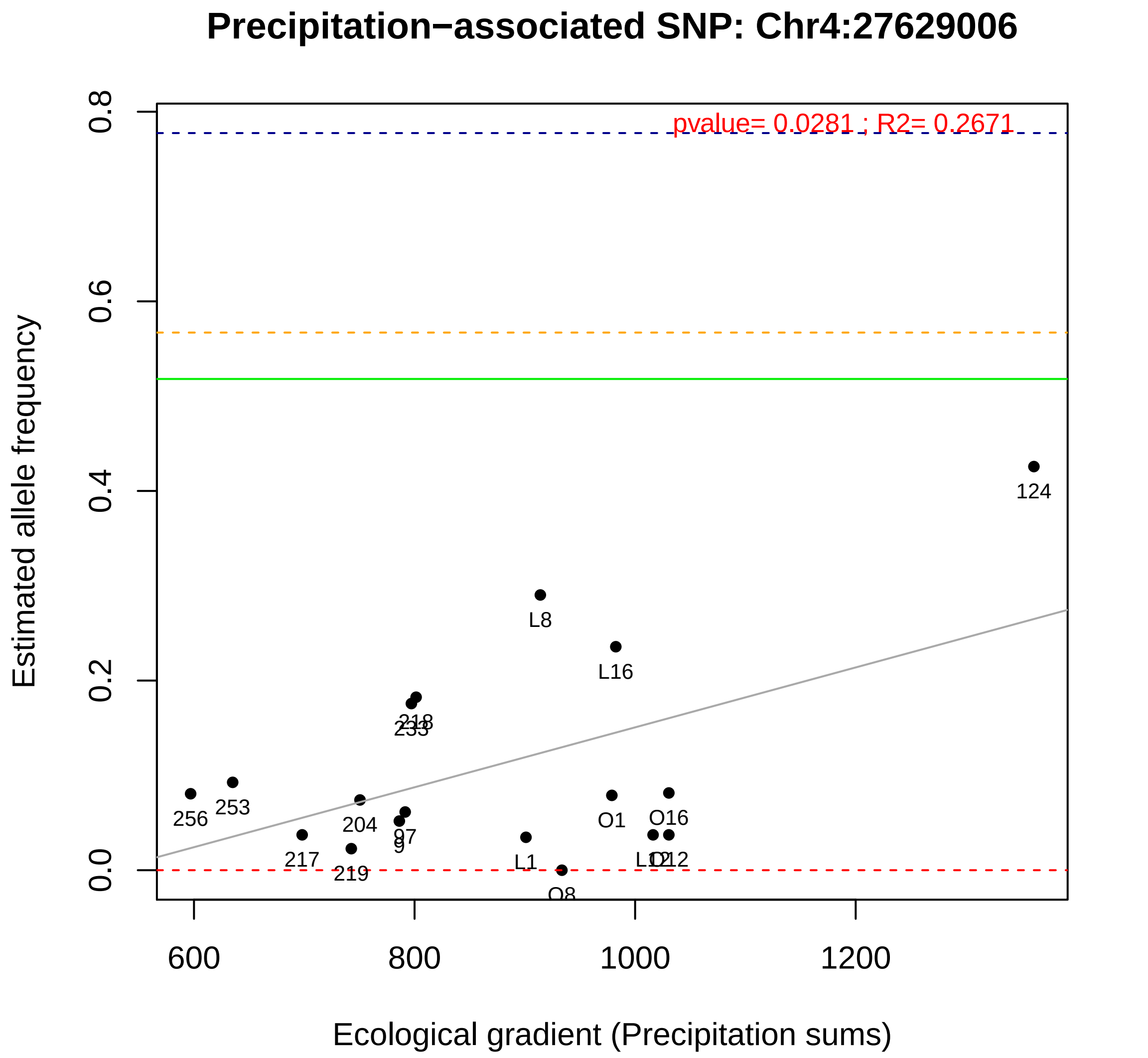

Precipitation−associated SNP: Chr4:27629006
pvalue= 0.0281 ; R2= 0.2671
Estimated allele frequency
0.0
0.2
0.4
0.6
0.8
600
800
1000
1200
Ecological gradient (Precipitation sums)
124
L8
L16
218
233
256
253
204
97
9
217
219
L1
O8
O1
O16
L10
O12

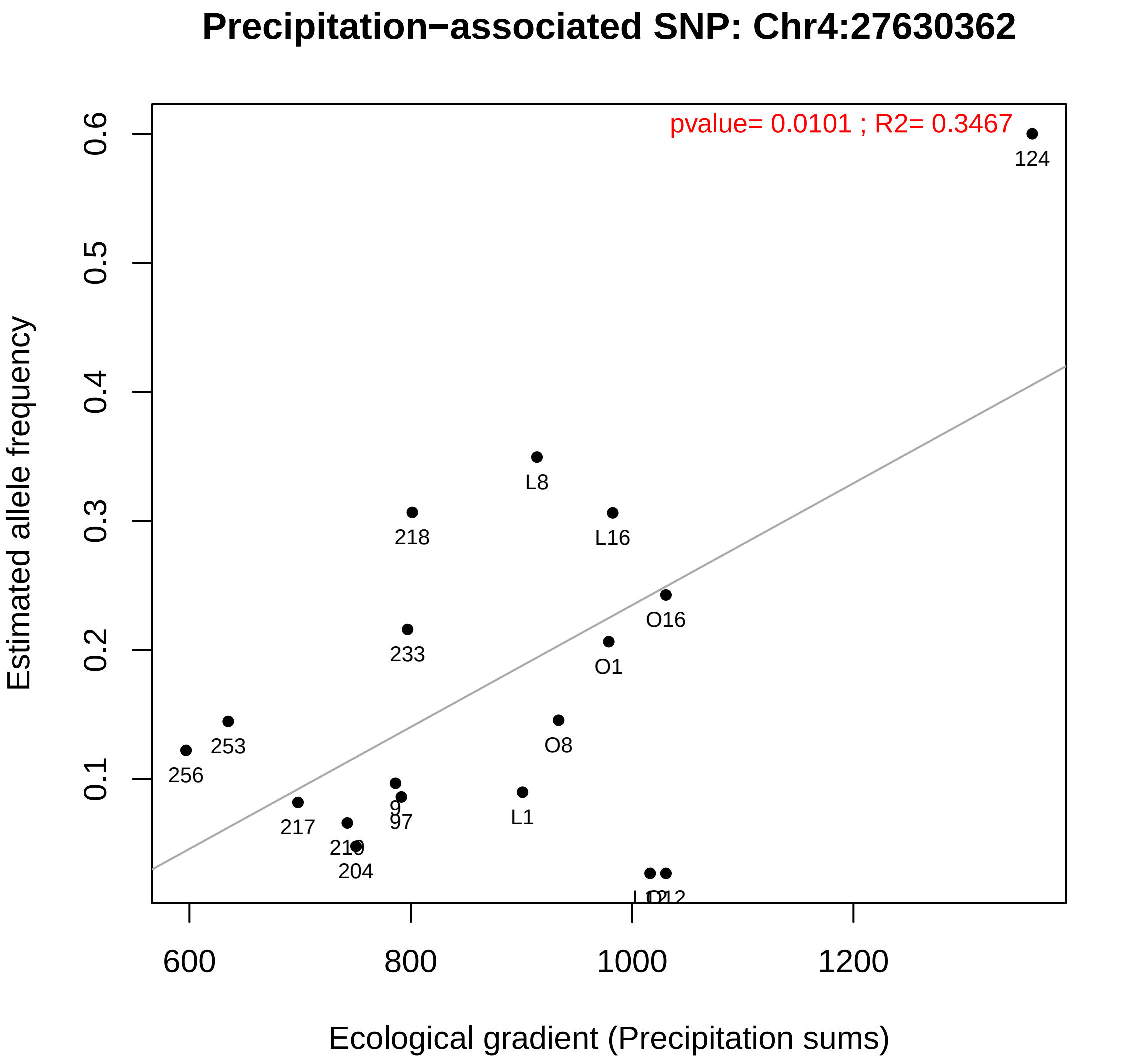

Precipitation−associated SNP: Chr4:27630362
pvalue= 0.0101 ; R2= 0.3467
Estimated allele frequency
Ecological gradient (Precipitation sums)
0.1
0.2
0.3
0.4
0.5
0.6
600
800
1000
1200
124
L8
218
L16
O16
233
O1
253
O8
256
217
97
L1
204

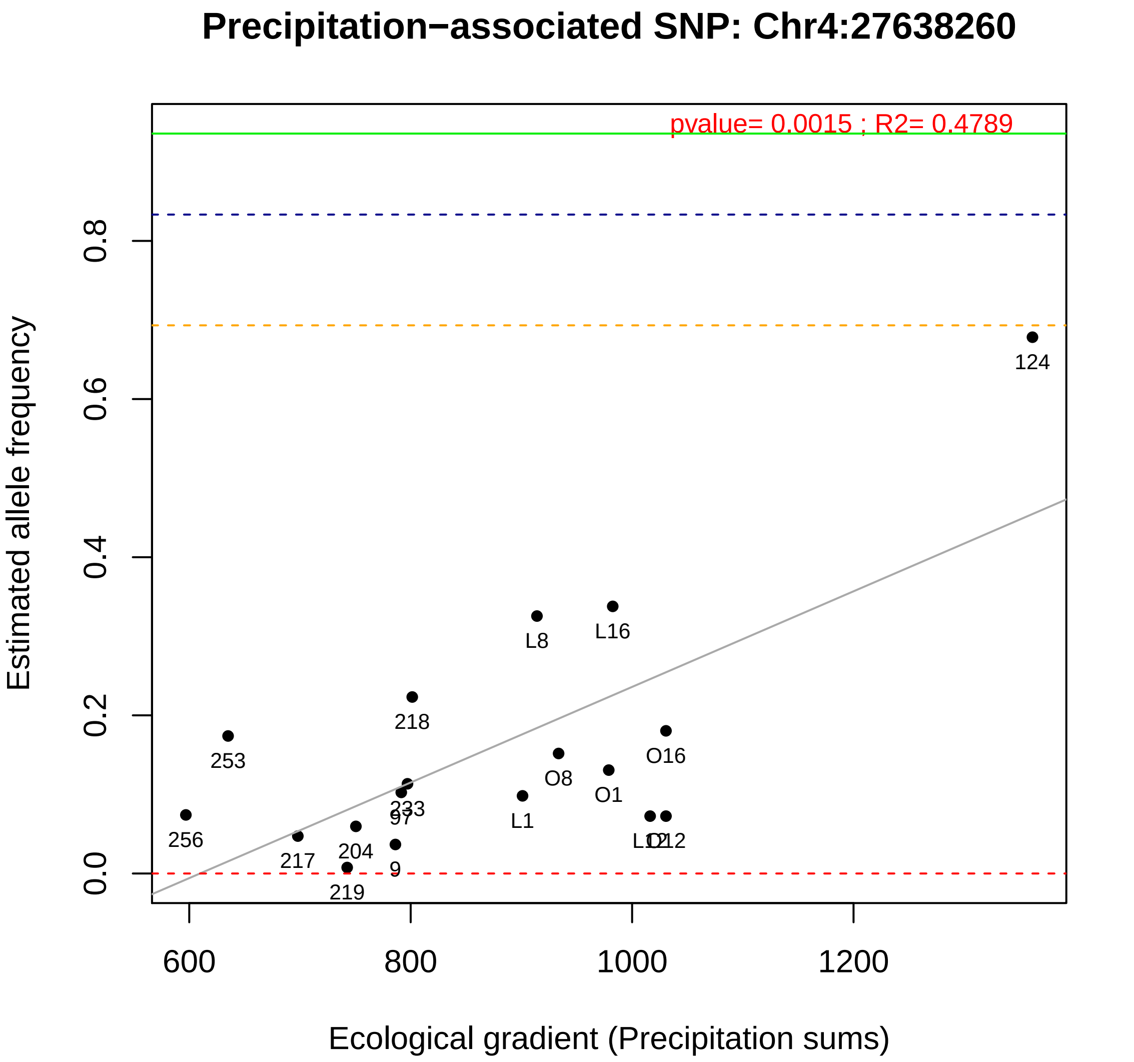
Precipitation−associated SNP: Chr4:27638260
pvalue= 0.0015 ; R2= 0.4789
Estimated allele frequency
0.0
0.2
0.4
0.6
0.8
600
800
1000
1200
Ecological gradient (Precipitation sums)
124
L8
L16
218
253
O16
O8
O1
233
97
L1
256
217
204
9
219

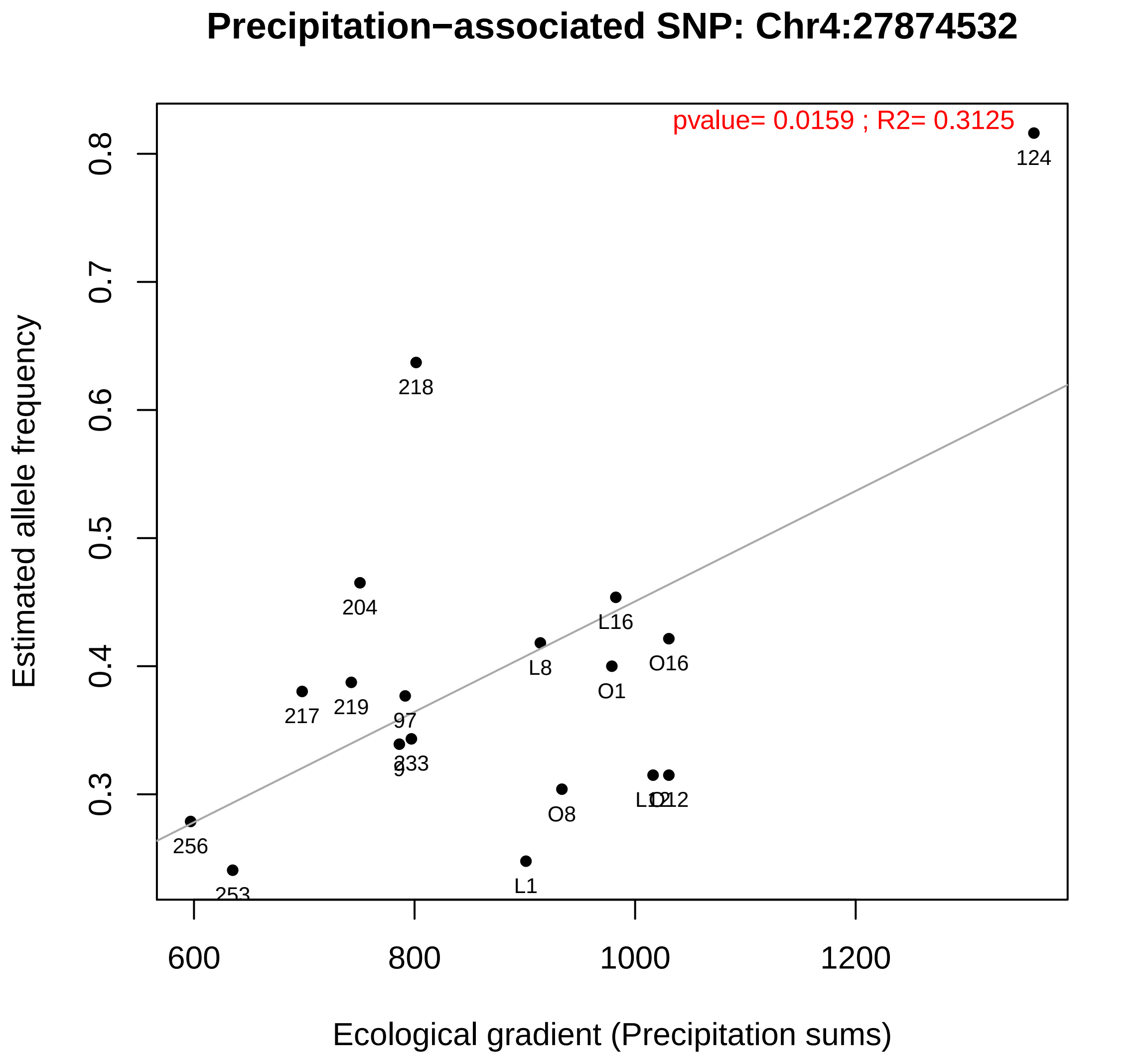
Precipitation−associated SNP: Chr4:27874532
pvalue= 0.0159 ; R2= 0.3125
Estimated allele frequency
0.3
0.4
0.5
0.6
0.7
0.8
600
800
1000
1200
Ecological gradient (Precipitation sums)
124
218
204
L16
O16
L8
O1
217
219
97
233
O8
256
253
L1

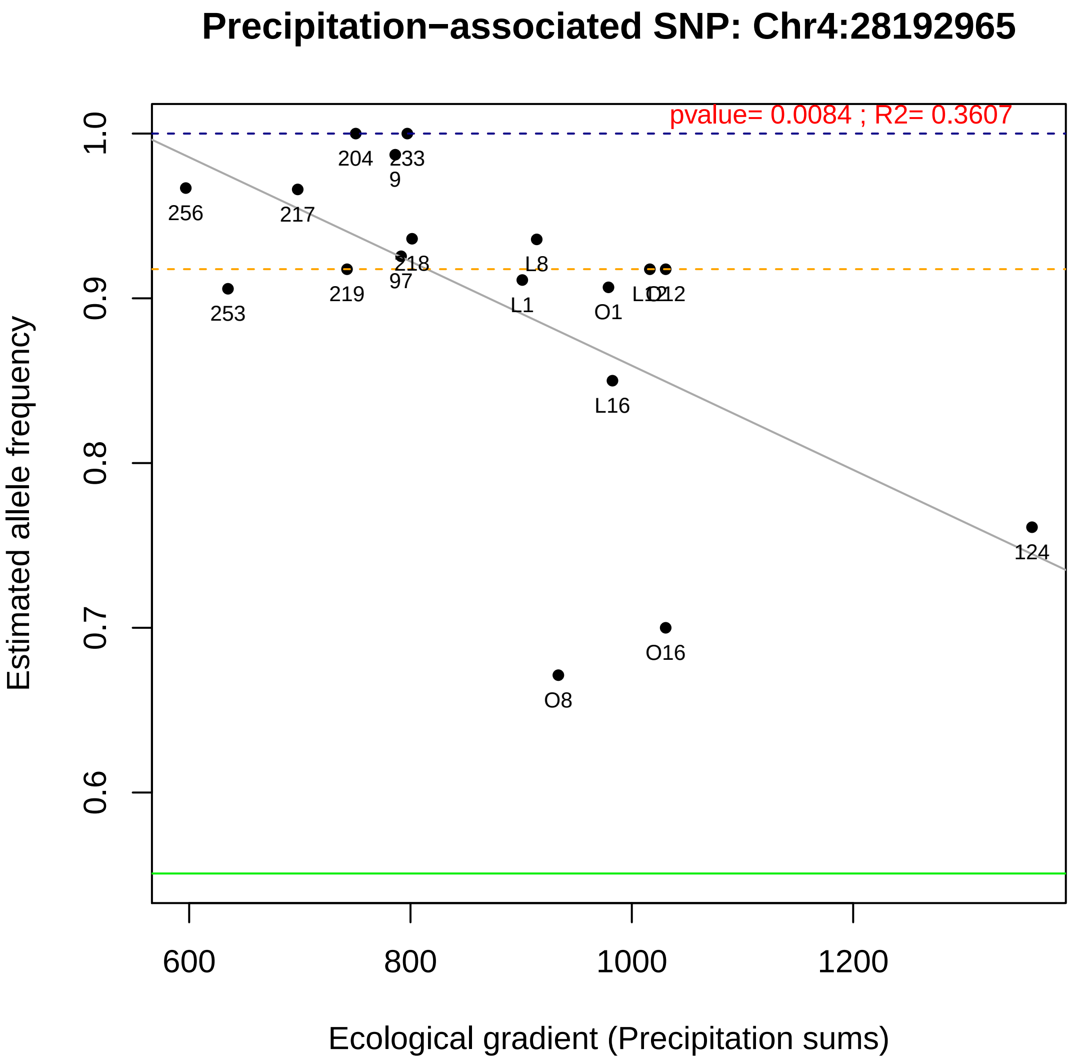
Precipitation−associated SNP: Chr4:28192965
pvalue= 0.0084 ; R2= 0.3607
Estimated allele frequency
Ecological gradient (Precipitation sums)
1.0
0.9
0.8
0.7
0.6
600
800
1000
1200
204
233
9
256
217
218
97
219
253
L8
L1
O1
L12
O12
L16
124
O16
O8

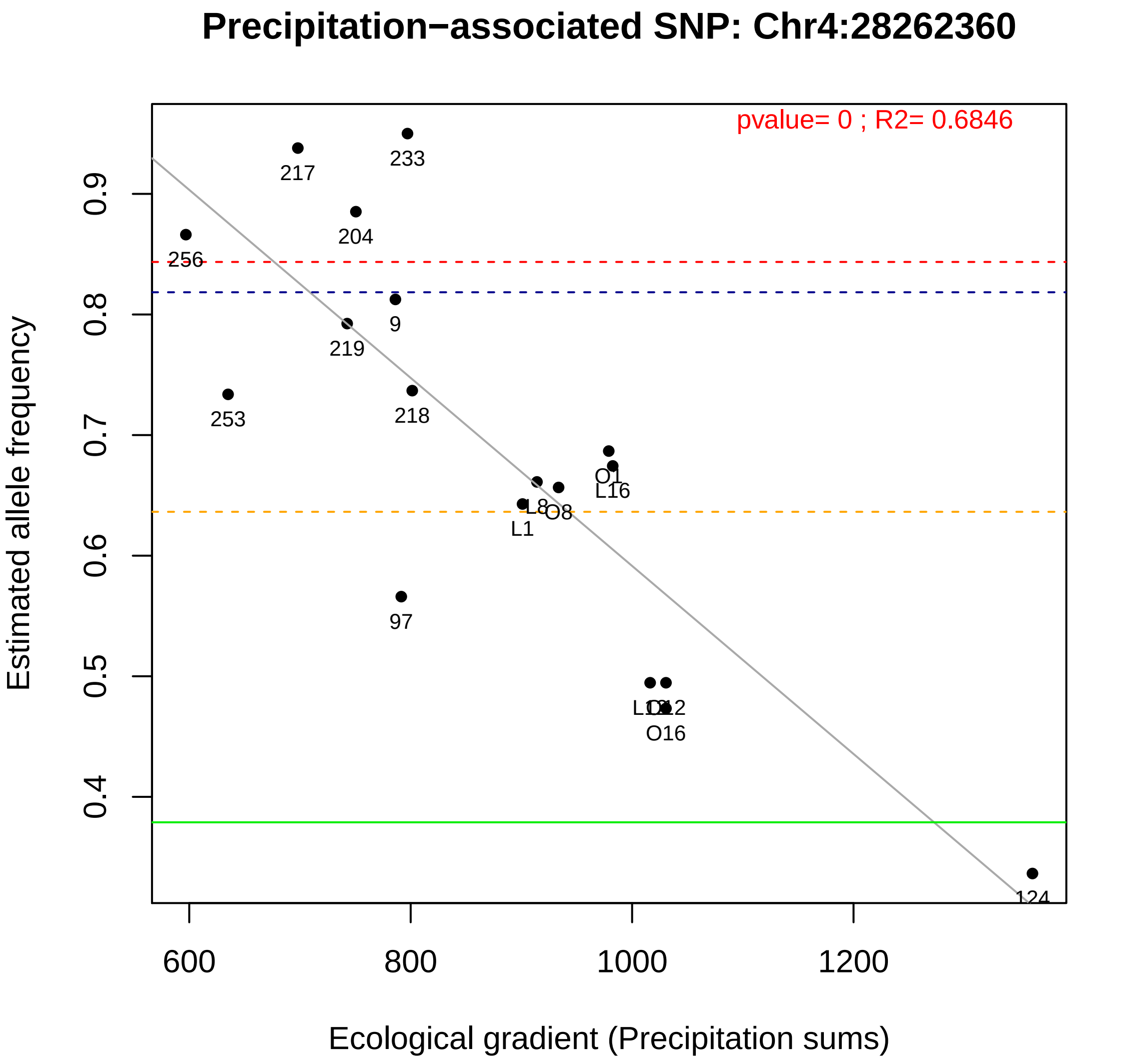

Precipitation−associated SNP: Chr4:28262360
pvalue= 0 ; R2= 0.6846
Estimated allele frequency
Ecological gradient (Precipitation sums)
0.4
0.5
0.6
0.7
0.8
0.9
600
800
1000
1200
217
233
204
256
9
219
253
218
O1
L16
L8
O8
L1
97
O16
124

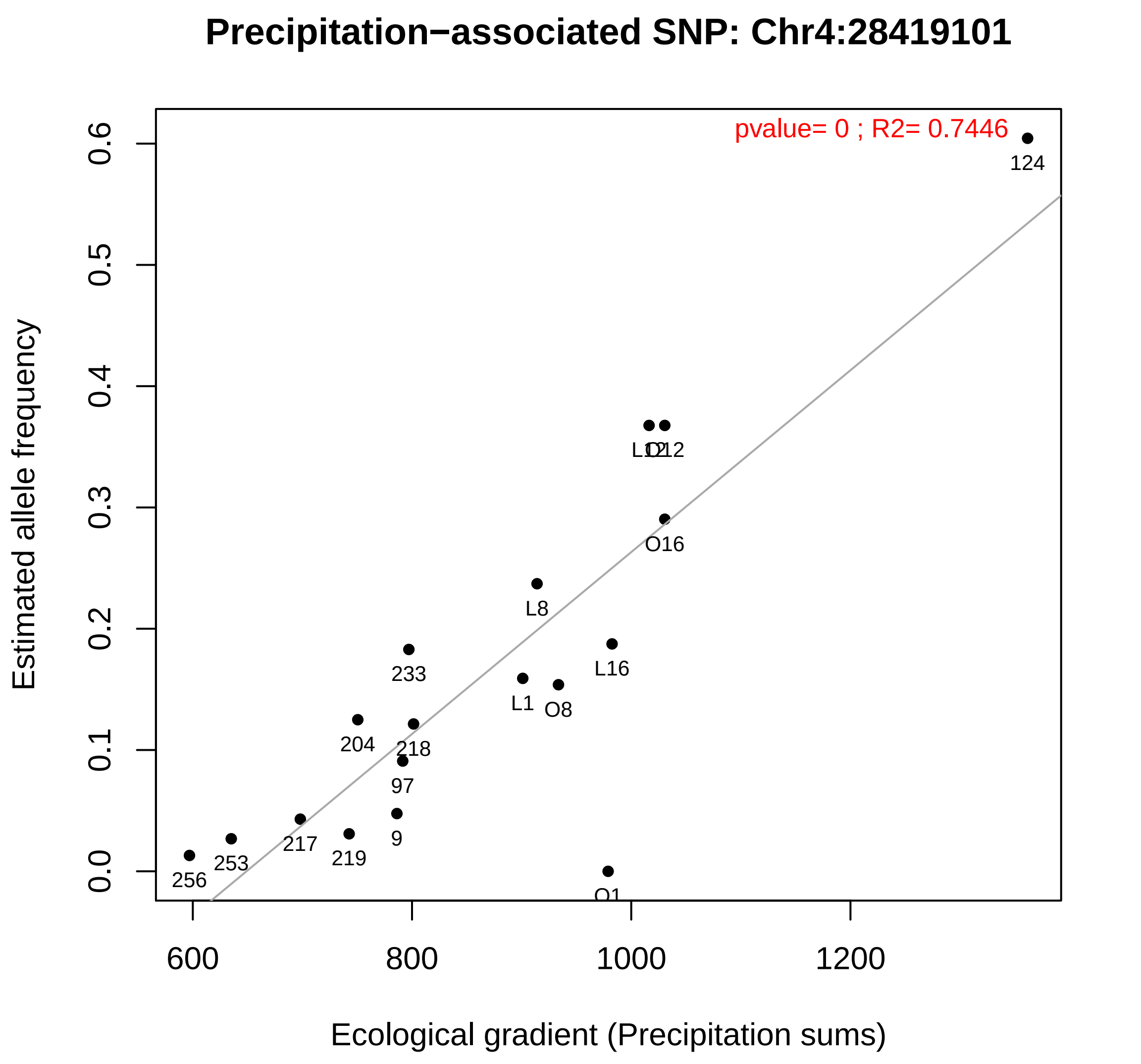
Precipitation−associated SNP: Chr4:28419101
pvalue= 0 ; R2= 0.7446
Estimated allele frequency
0.0
0.1
0.2
0.3
0.4
0.5
0.6
600
800
1000
1200
Ecological gradient (Precipitation sums)
124
L12
O12
O16
L8
233
L16
L1
O8
204
218
97
217
9
219
253
256
O1

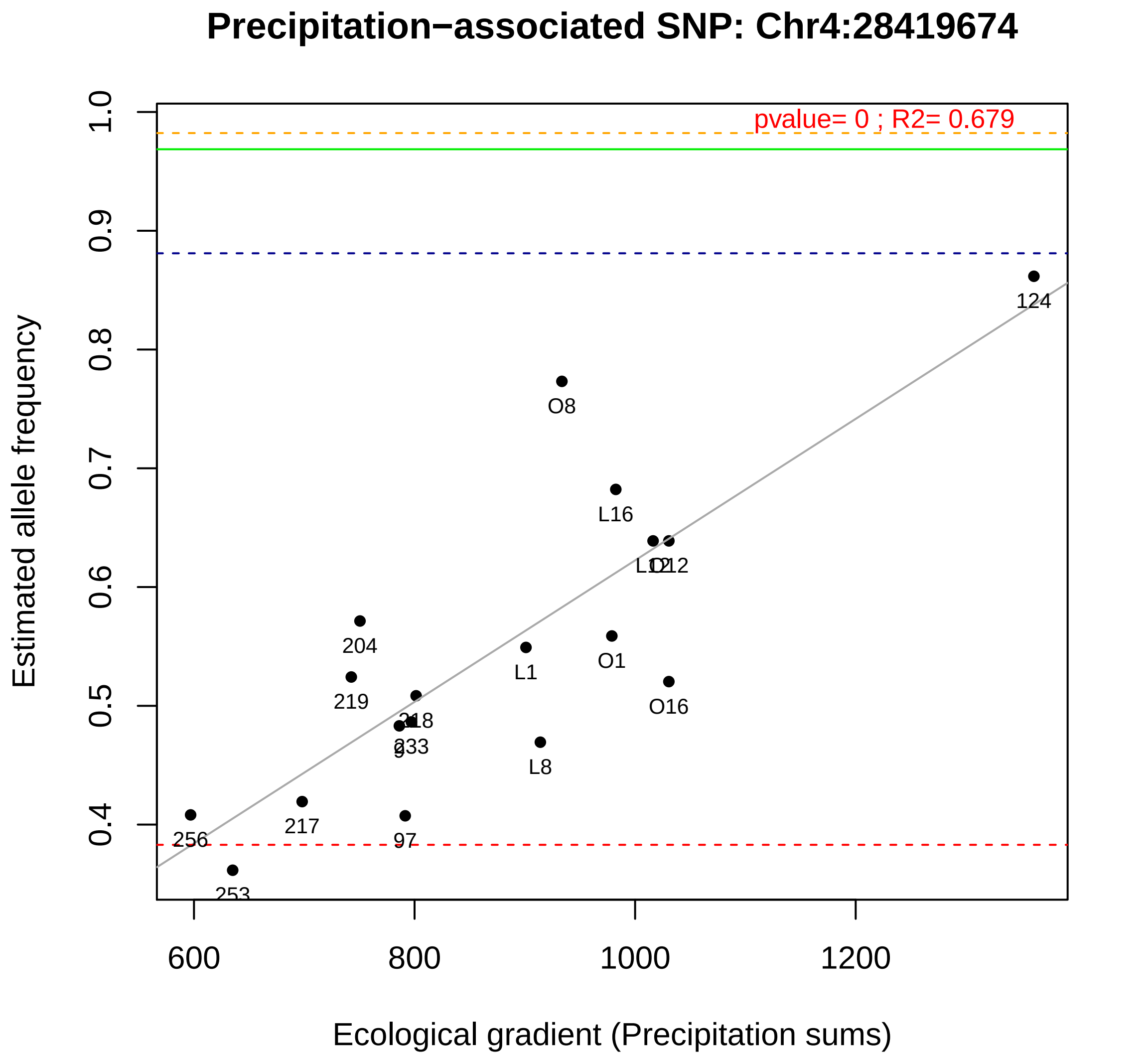

Precipitation−associated SNP: Chr4:28419674
pvalue= 0 ; R2= 0.679
Estimated allele frequency
Ecological gradient (Precipitation sums)
0.4
0.5
0.6
0.7
0.8
0.9
1.0
600
800
1000
1200
124
O8
L16
L12
O12
204
O1
L1
219
218
233
5
O16
L8
256
217
97
253

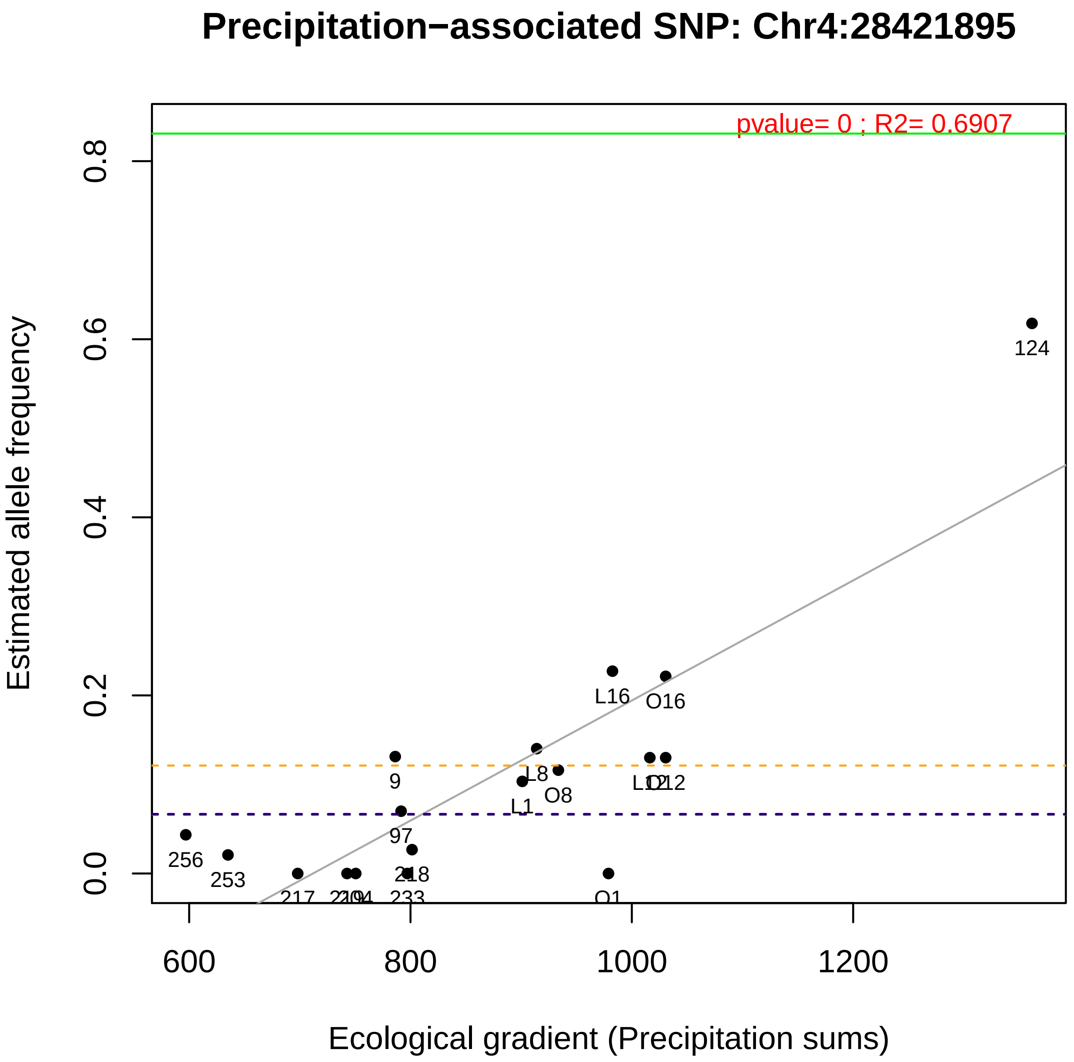
Precipitation−associated SNP: Chr4:28421895
pvalue= 0 ; R2= 0.6907
Estimated allele frequency
0.0
0.2
0.4
0.6
0.8
600
800
1000
1200
Ecological gradient (Precipitation sums)
124
L16
O16
9
L8
O8
L1
97
256
253
217
218
233
O1

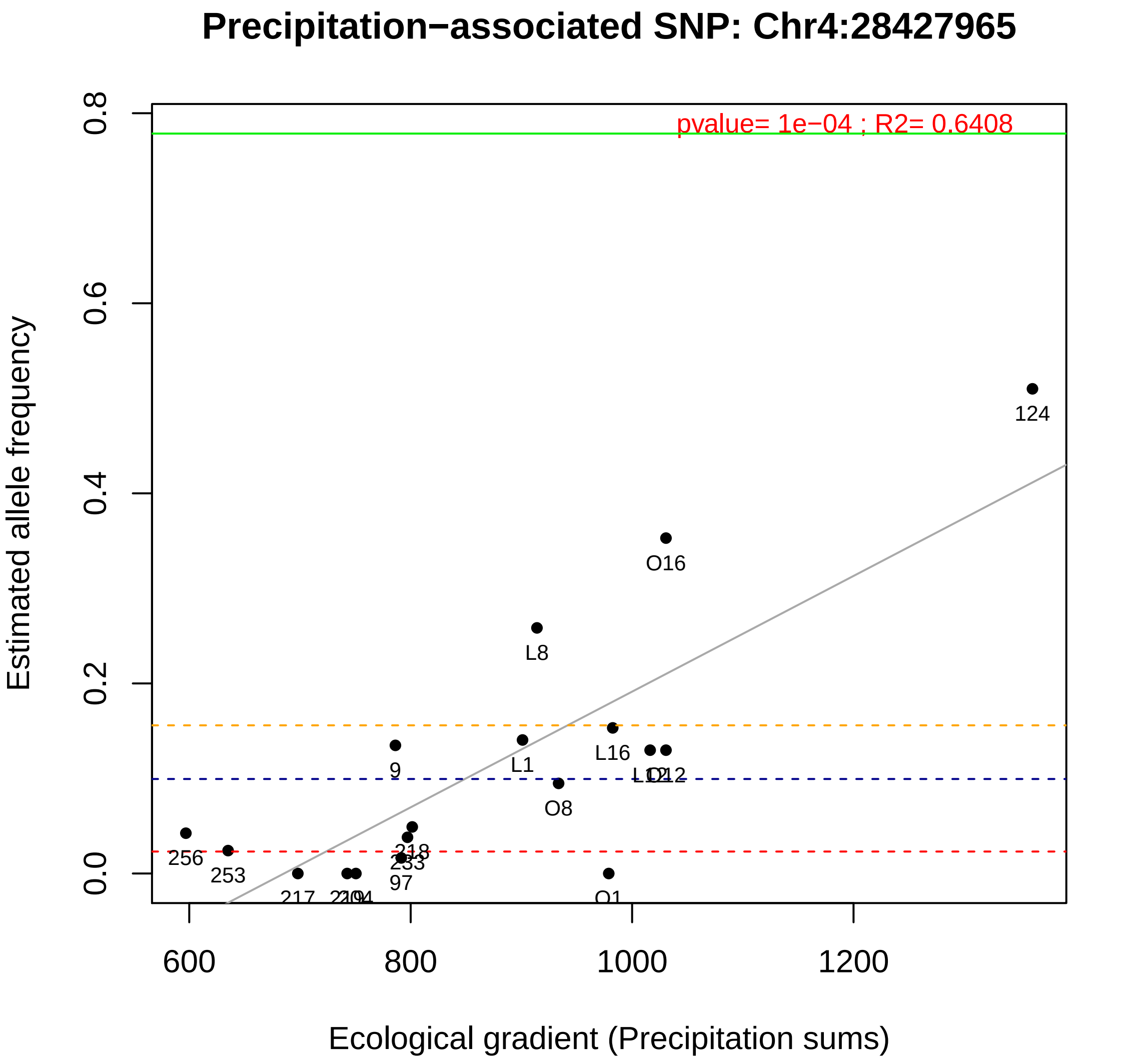
Precipitation−associated SNP: Chr4:28427965
pvalue= 1e−04 ; R2= 0.6408
Estimated allele frequency
0.0
0.2
0.4
0.6
0.8
600
800
1000
1200
Ecological gradient (Precipitation sums)
124
O16
L8
L16
L1
9
O8
256
253
217
218
233
97
O1

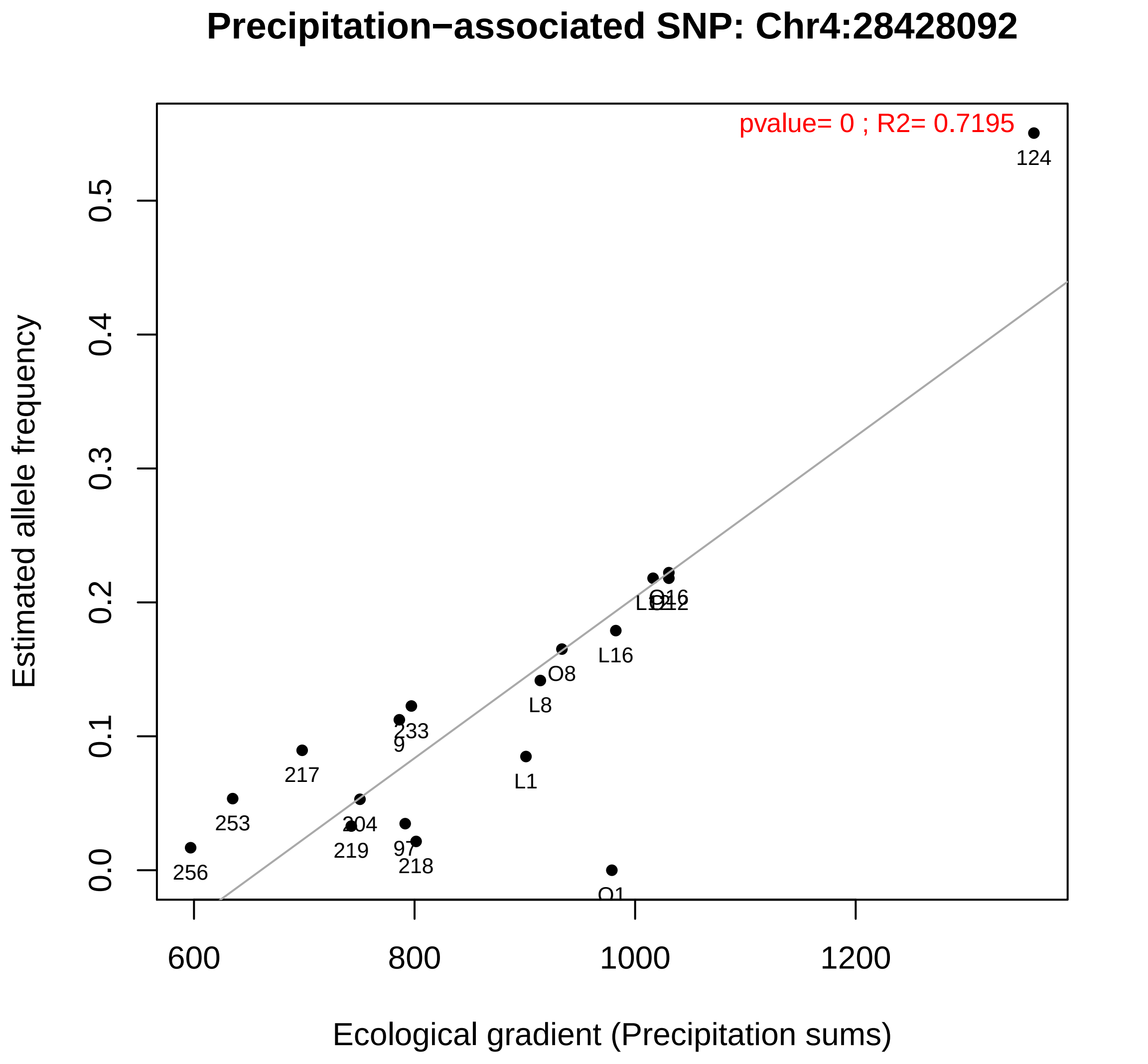
Precipitation−associated SNP: Chr4:28428092
pvalue= 0 ; R2= 0.7195
Estimated allele frequency
Ecological gradient (Precipitation sums)
0.0
0.1
0.2
0.3
0.4
0.5
600
800
1000
1200
124
O16
L12
L16
O8
L8
233
9
217
L1
253
204
219
97
218
256
O1

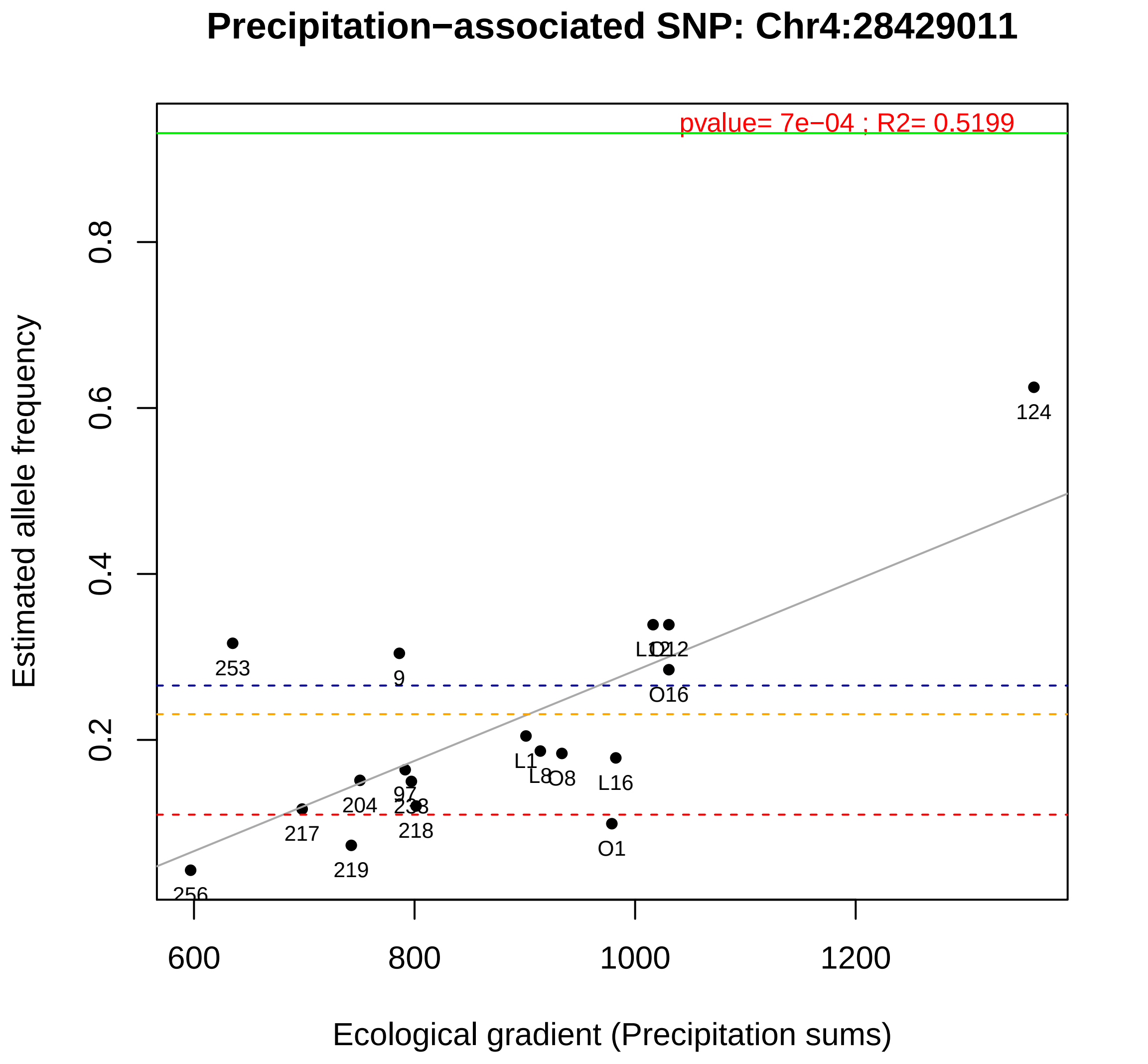
Precipitation−associated SNP: Chr4:28429011
pvalue= 7e−04 ; R2= 0.5199
Estimated allele frequency
0.2
0.4
0.6
0.8
600
800
1000
1200
Ecological gradient (Precipitation sums)
124
253
9
L1
L8
O8
L16
O16
O1
204
217
219
218
97
256

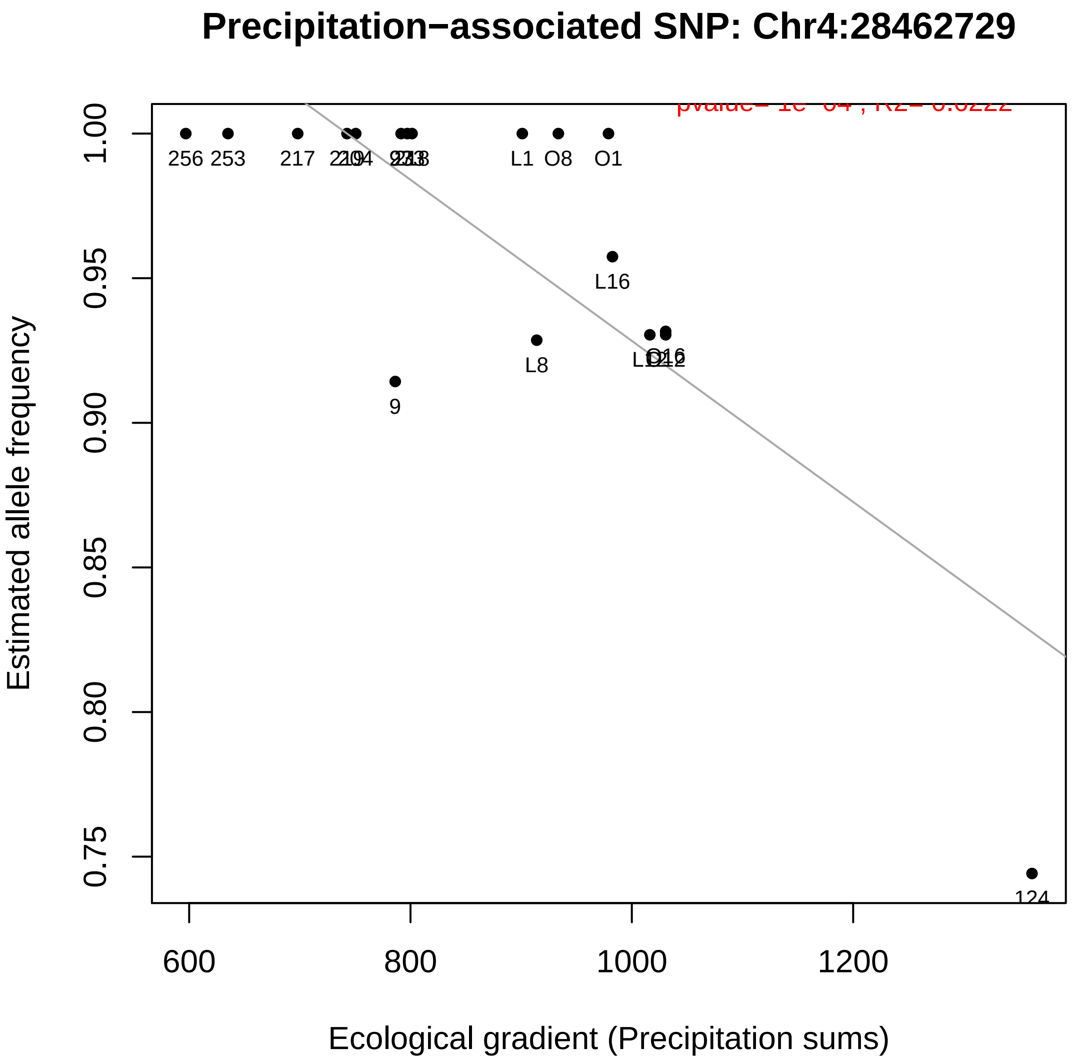
Precipitation−associated SNP: Chr4:28462729
pvalue= 1e−04 ; R2= 0.6222
Estimated allele frequency
1.00
0.95
0.90
0.85
0.80
0.75
256
253
217
L1
O8
O1
L16
L8
9
124
600
800
1000
1200
Ecological gradient (Precipitation sums)

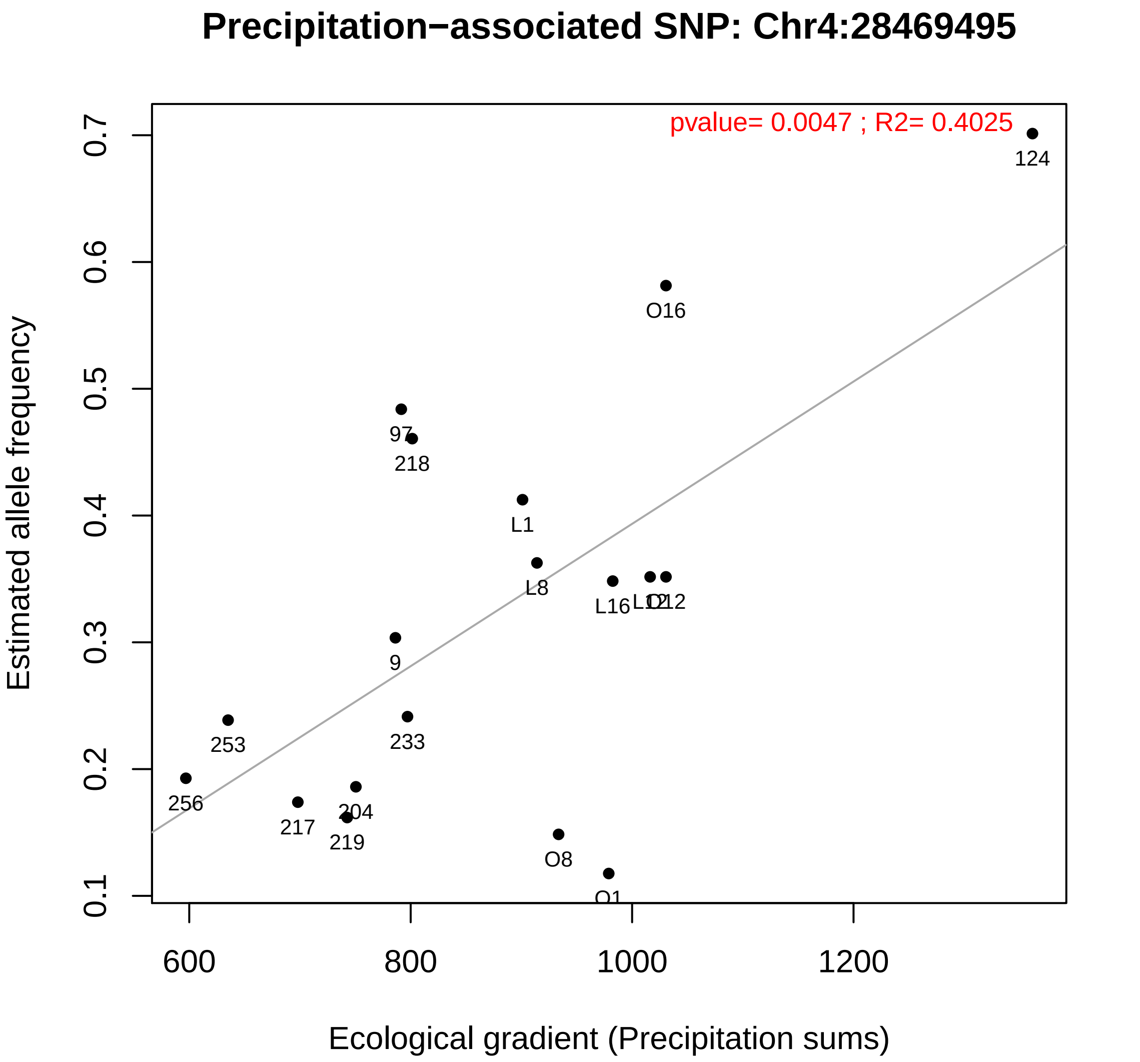
Precipitation−associated SNP: Chr4:28469495
pvalue= 0.0047 ; R2= 0.4025
Estimated allele frequency
Ecological gradient (Precipitation sums)
0.1
0.2
0.3
0.4
0.5
0.6
0.7
600
800
1000
1200
124
O16
97
218
L1
L8
L16
O12
9
253
233
256
217
204
219
O8

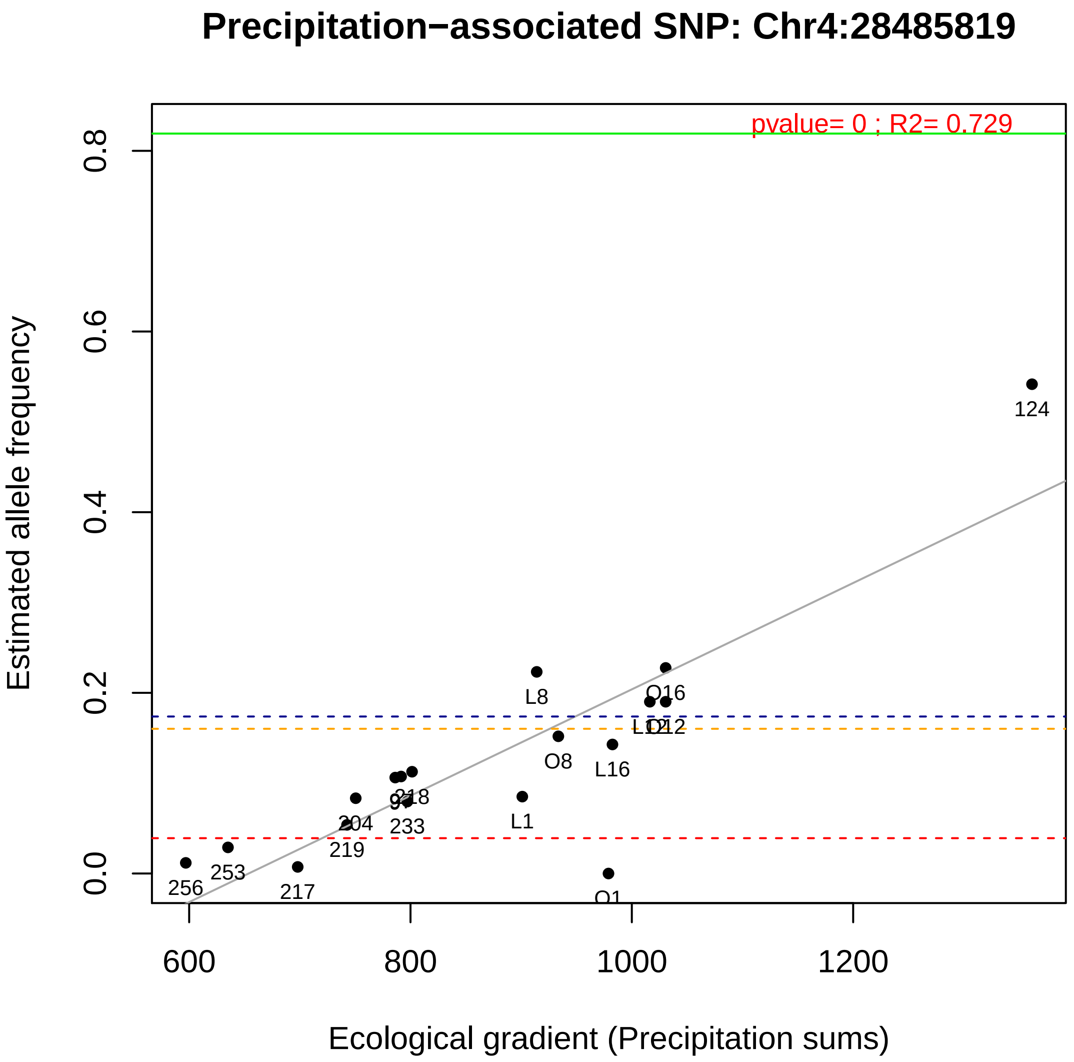

Precipitation−associated SNP: Chr4:28485819
pvalue= 0 ; R2= 0.729
Estimated allele frequency
0.0
0.2
0.4
0.6
0.8
Ecological gradient (Precipitation sums)
600
800
1000
1200
124
L8
O16
O8
L16
L1
204
219
233
253
256
217

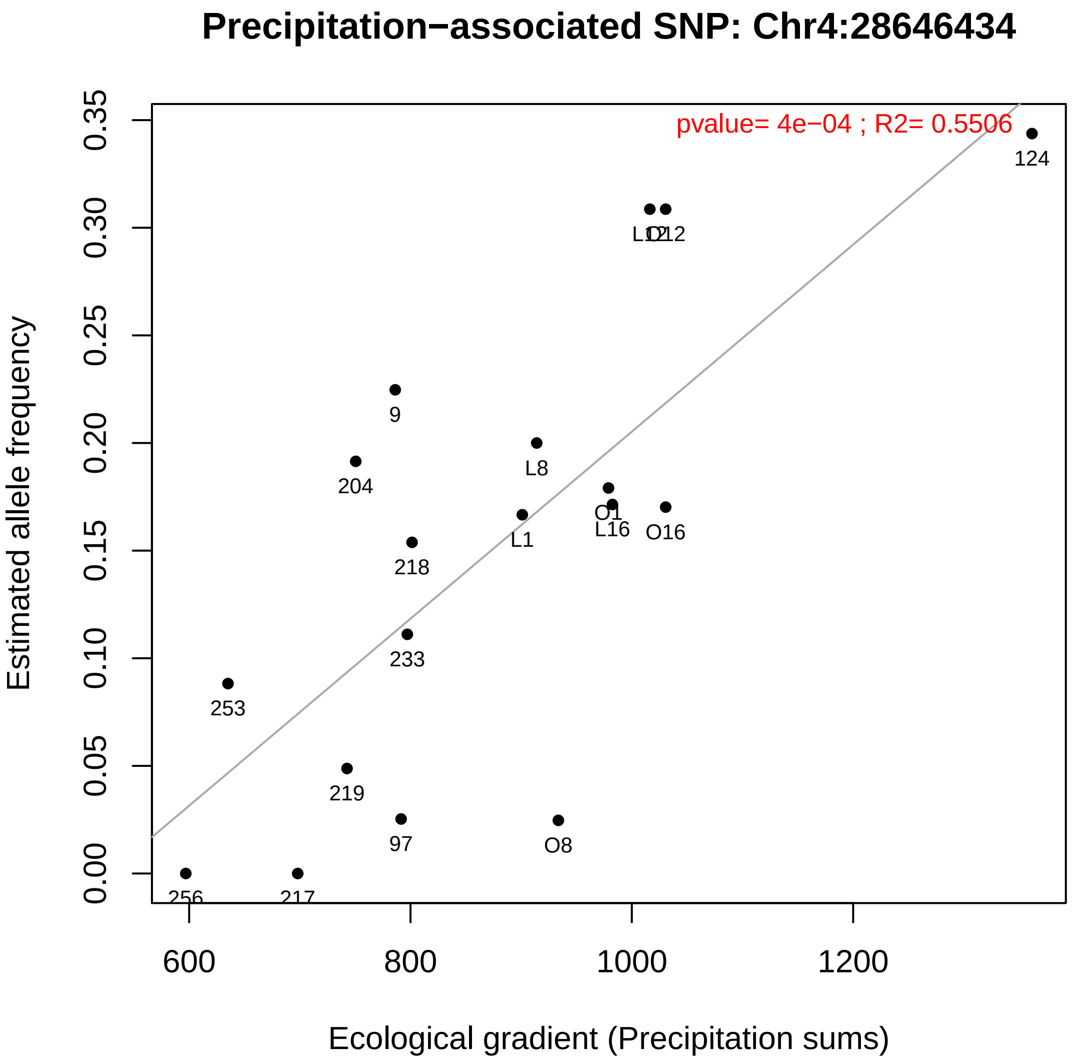
Precipitation−associated SNP: Chr4:28646434
pvalue= 4e−04 ; R2= 0.5506
124
L12
O12
9
L8
204
O1
L16
O16
L1
218
233
253
219
97
O8
256
217
0.00
0.05
0.10
0.15
0.20
0.25
0.30
0.35
Estimated allele frequency
600
800
1000
1200
Ecological gradient (Precipitation sums)

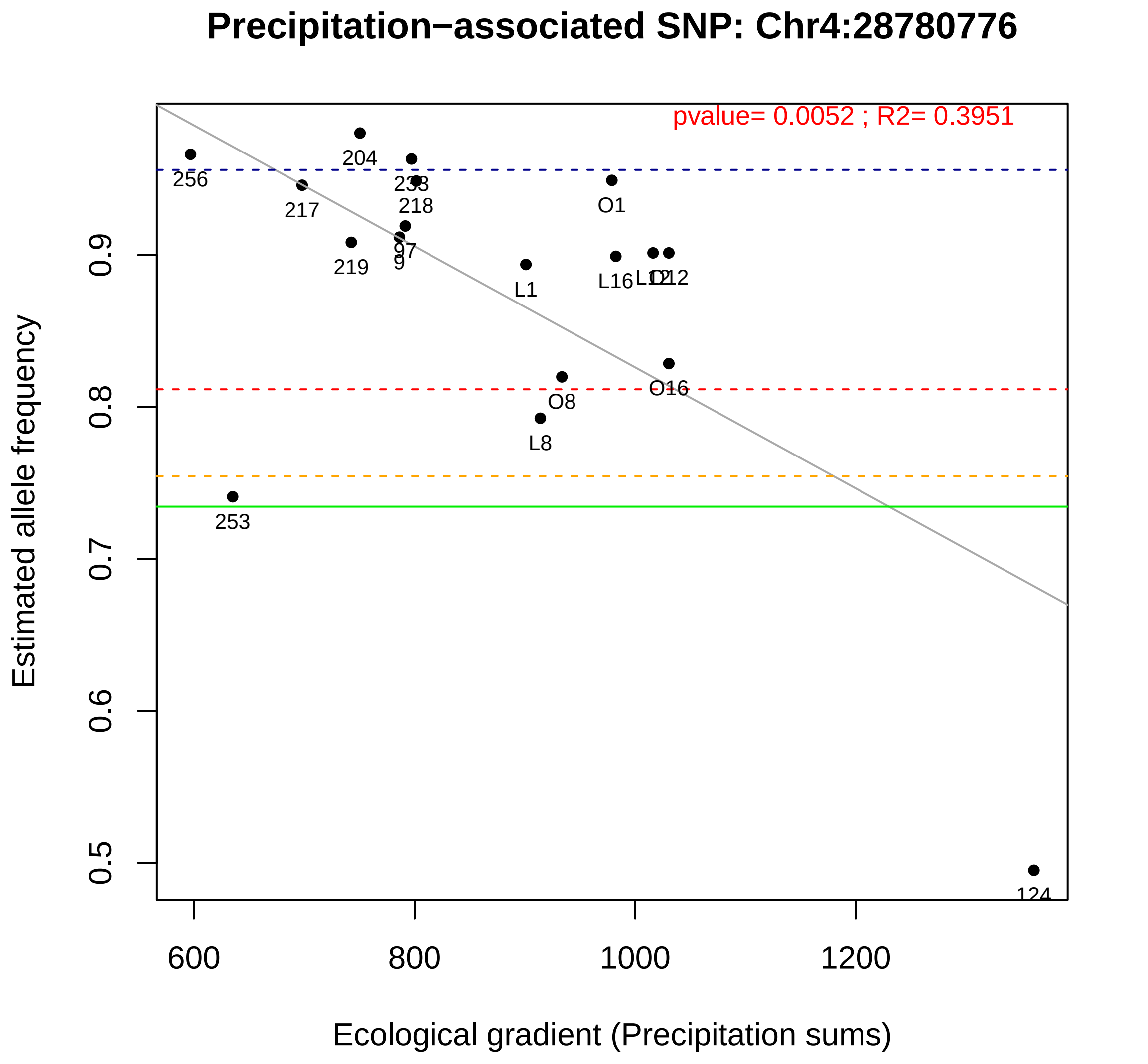

Precipitation−associated SNP: Chr4:28780776
pvalue= 0.0052 ; R2= 0.3951
Estimated allele frequency
Ecological gradient (Precipitation sums)
0.5
0.6
0.7
0.8
0.9
600
800
1000
1200
256
204
233
217
218
219
97
9
O1
L1
L16
L12
O12
O16
O8
L8
253
124

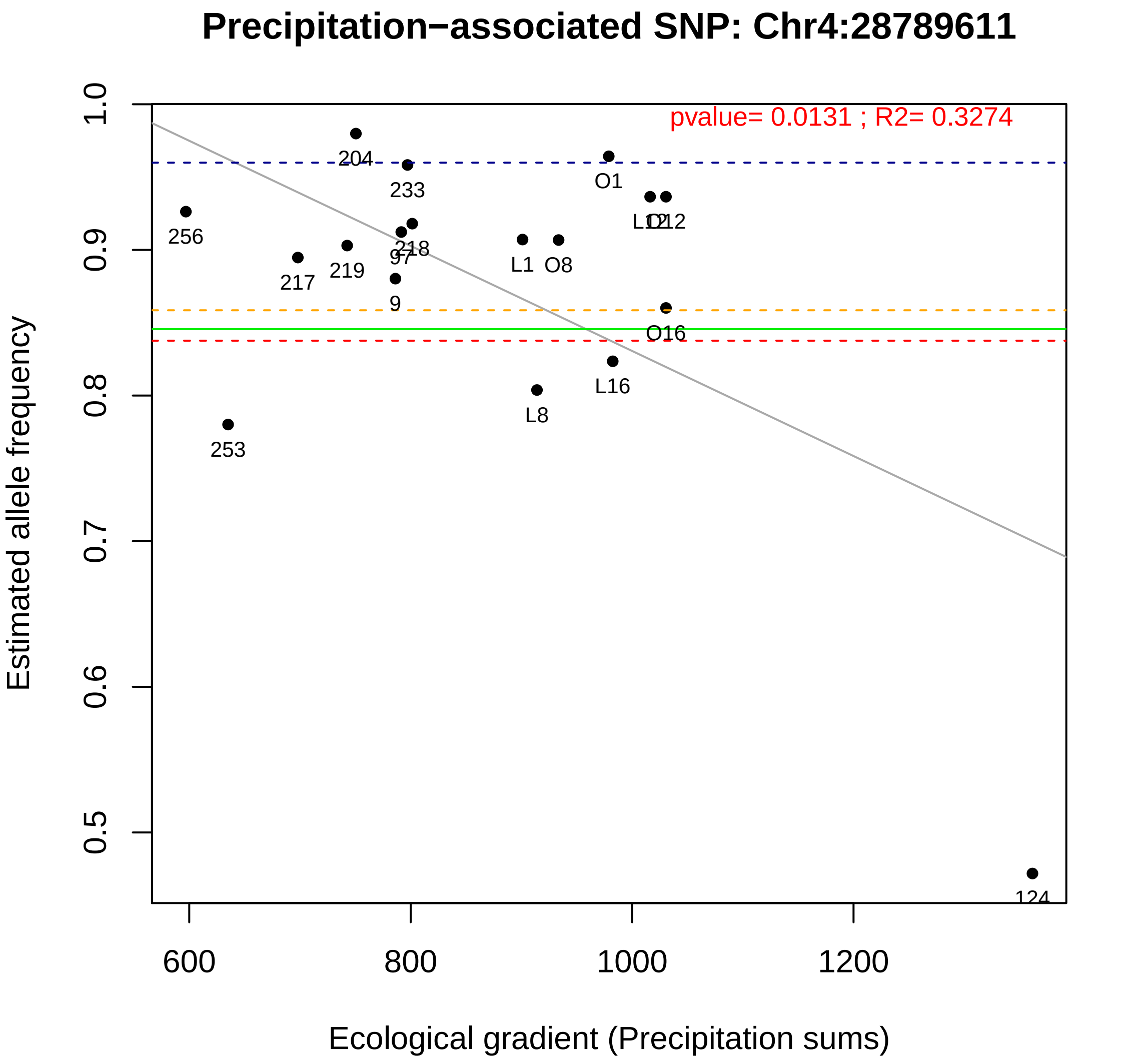

Precipitation−associated SNP: Chr4:28789611
pvalue= 0.0131 ; R2= 0.3274
Estimated allele frequency
1.0
0.9
0.8
0.7
0.6
0.5
600
800
1000
1200
Ecological gradient (Precipitation sums)
204
233
256
217
219
218
97
9
L1
O8
O1
L12
O12
O16
L16
L8
253
124

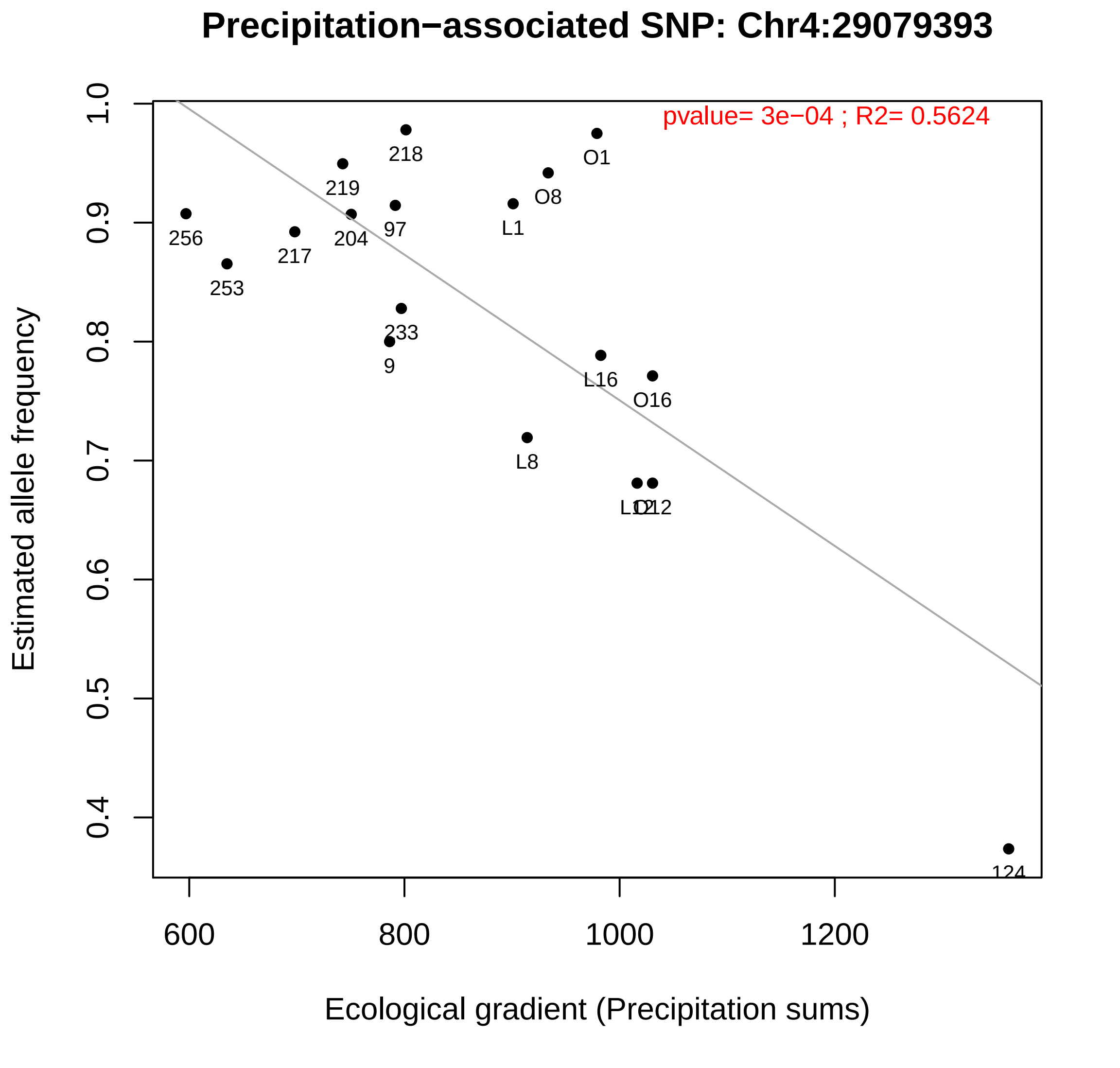
Precipitation−associated SNP: Chr4:29079393
pvalue= 3e−04 ; R2= 0.5624
Estimated allele frequency
1.0
0.9
0.8
0.7
0.6
0.5
0.4
600
800
1000
1200
Ecological gradient (Precipitation sums)
218
O1
219
O8
256
204
97
L1
217
253
233
9
L16
O16
L8
L12
O12
124

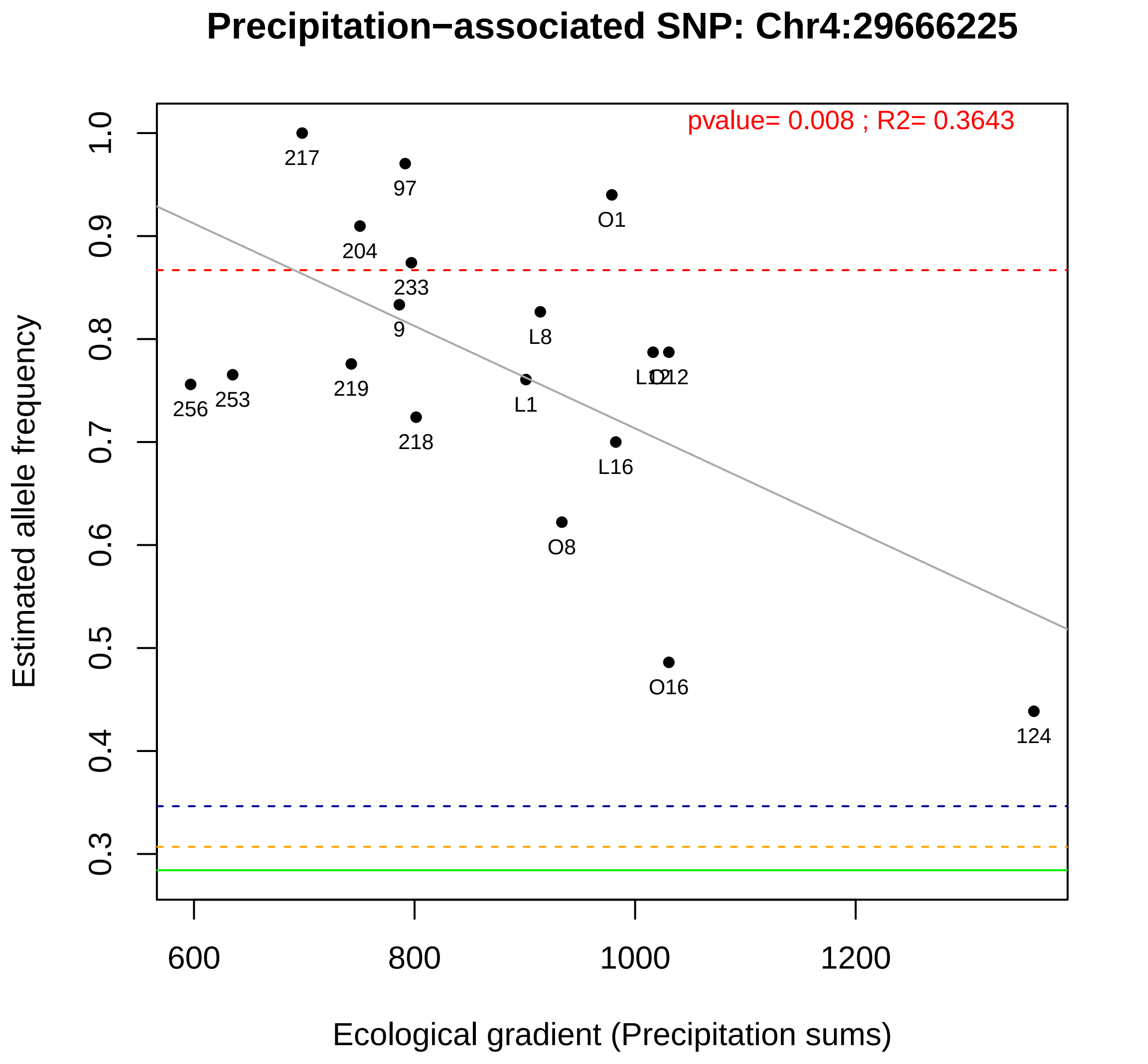

Precipitation−associated SNP: Chr4:29666225
pvalue= 0.008 ; R2= 0.3643
Estimated allele frequency
Ecological gradient (Precipitation sums)
0.3
0.4
0.5
0.6
0.7
0.8
0.9
1.0
600
800
1000
1200
217
97
O1
204
233
9
L8
L10
O12
256
253
219
L1
218
L16
O8
O16
124

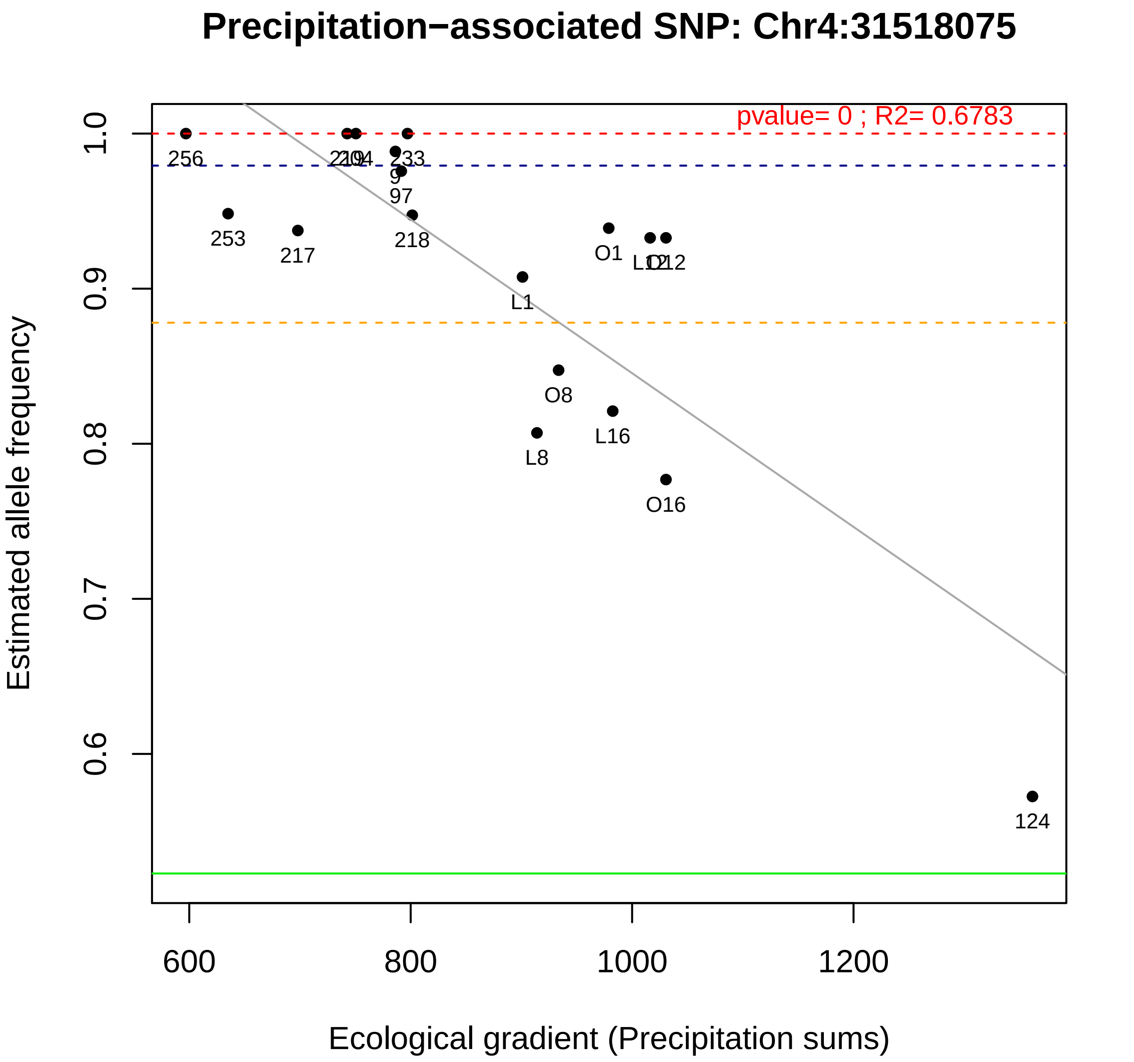
Precipitation−associated SNP: Chr4:31518075
pvalue= 0 ; R2= 0.6783
Estimated allele frequency
Ecological gradient (Precipitation sums)
1.0
0.9
0.8
0.7
0.6
600
800
1000
1200
256
233
9
97
253
217
218
O1
L1
O8
L16
L8
O16
124

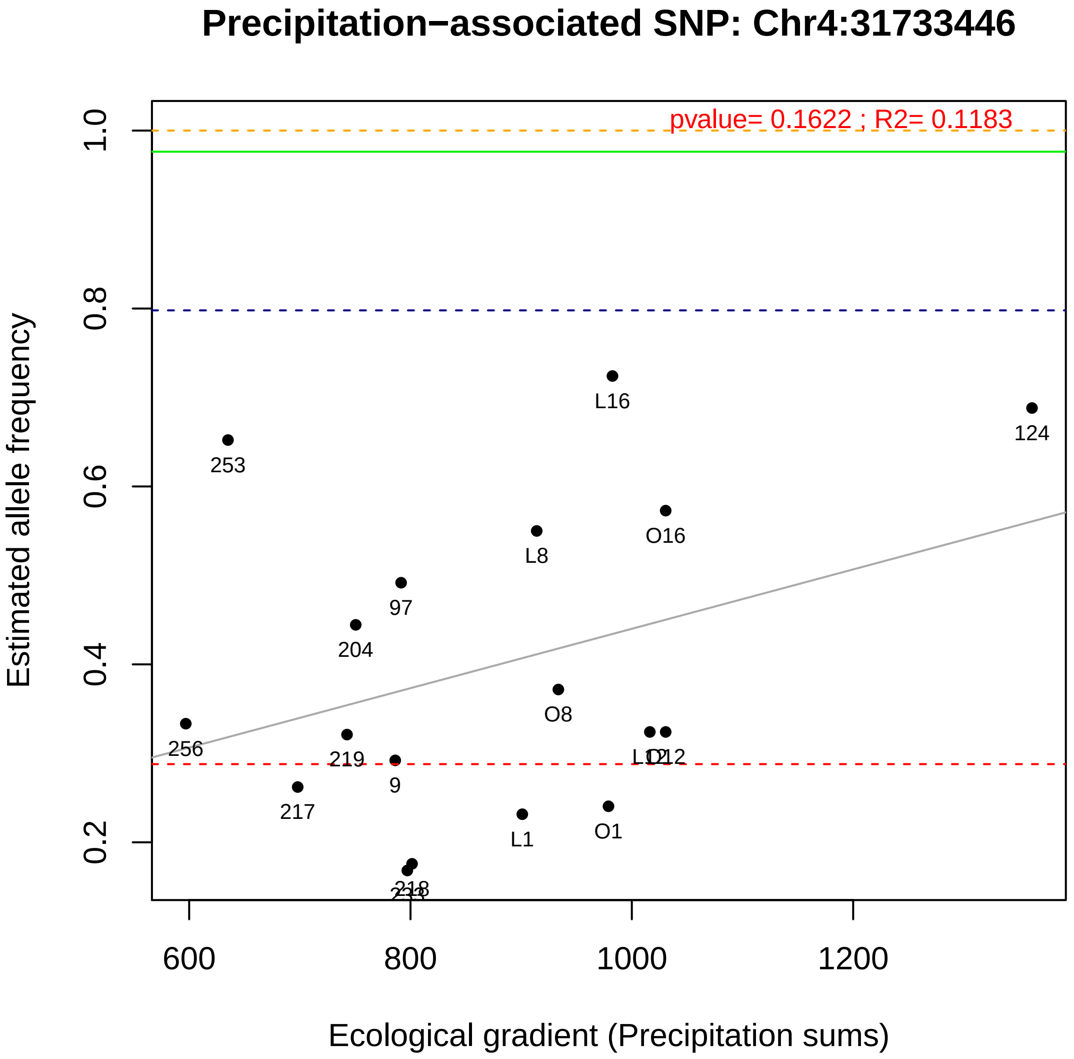
Precipitation−associated SNP: Chr4:31733446
pvalue= 0.1622 ; R2= 0.1183
Estimated allele frequency
0.2
0.4
0.6
0.8
1.0
Ecological gradient (Precipitation sums)
600
800
1000
1200
L16
124
253
O16
L8
97
204
O8
256
219
9
217
L1
O1
218

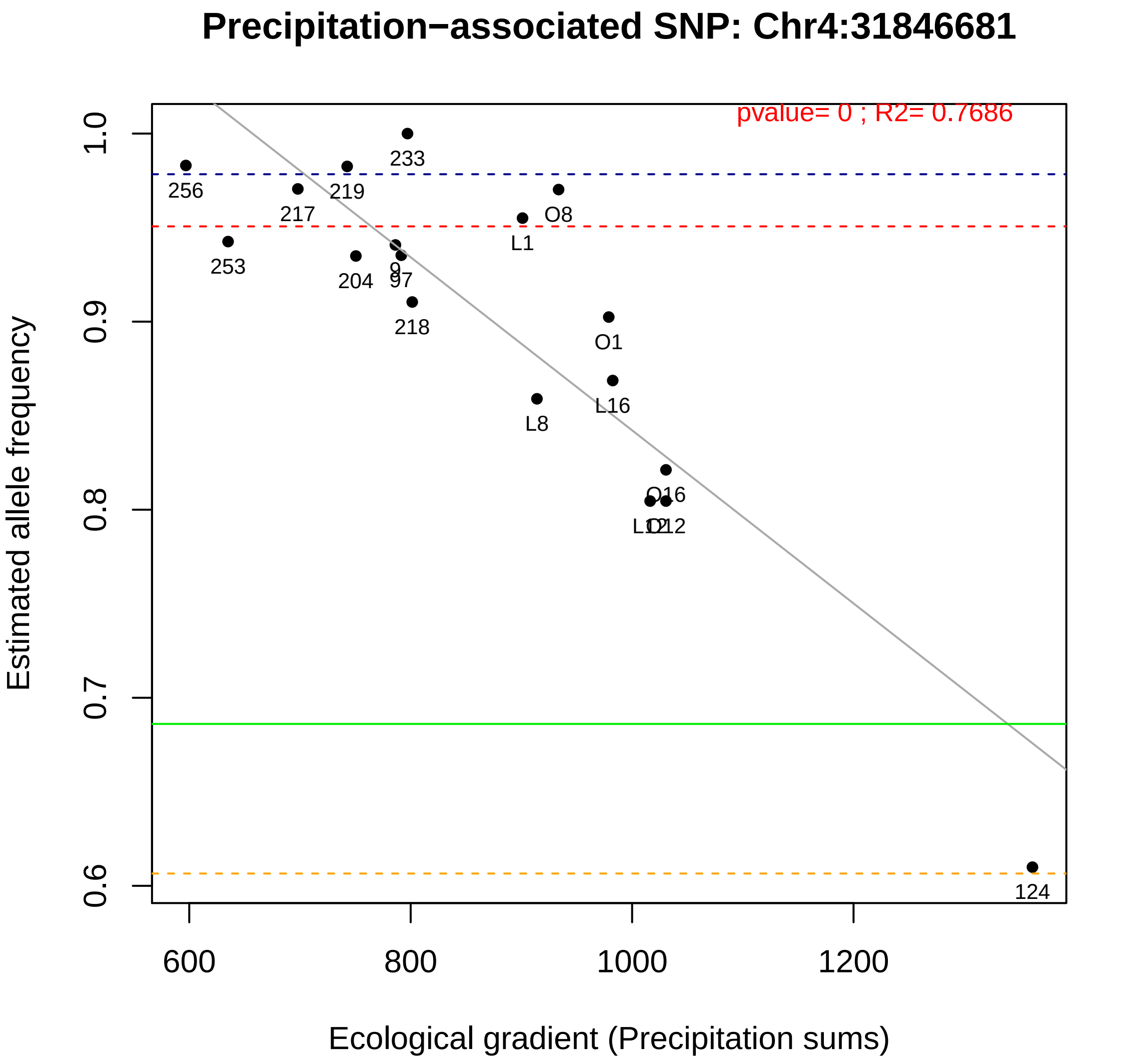
Precipitation−associated SNP: Chr4:31846681
pvalue= 0 ; R2= 0.7686
Estimated allele frequency
1.0
0.9
0.8
0.7
0.6
600
800
1000
1200
Ecological gradient (Precipitation sums)
256
253
217
219
233
204
9
97
218
L1
O8
O1
L16
L8
O16
L10
O12
124

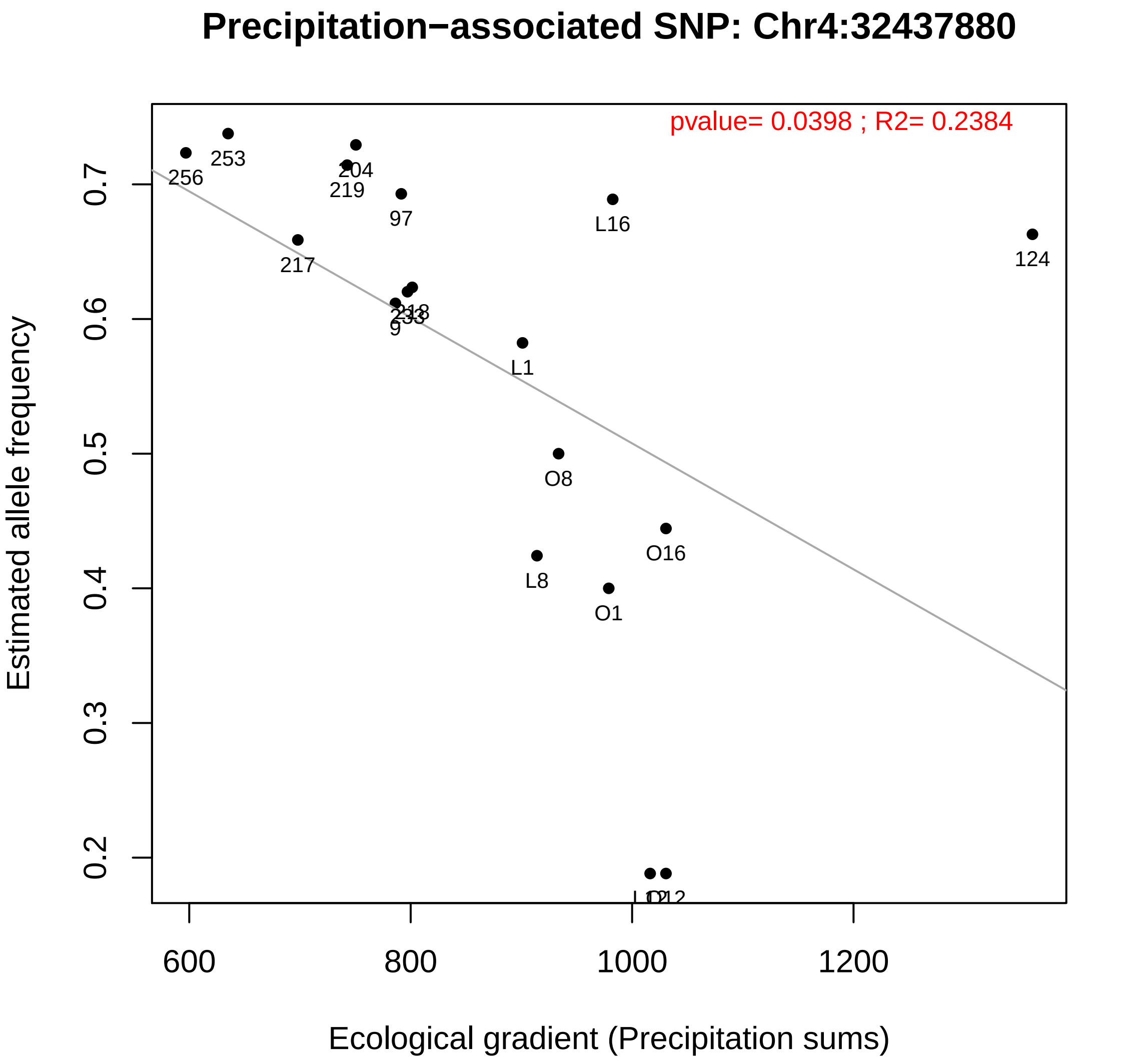

Precipitation−associated SNP: Chr4:32437880
pvalue= 0.0398 ; R2= 0.2384
Estimated allele frequency
Ecological gradient (Precipitation sums)
0.2
0.3
0.4
0.5
0.6
0.7
600
800
1000
1200
256
253
204
219
97
L16
124
217
L1
O8
O16
L8
O1

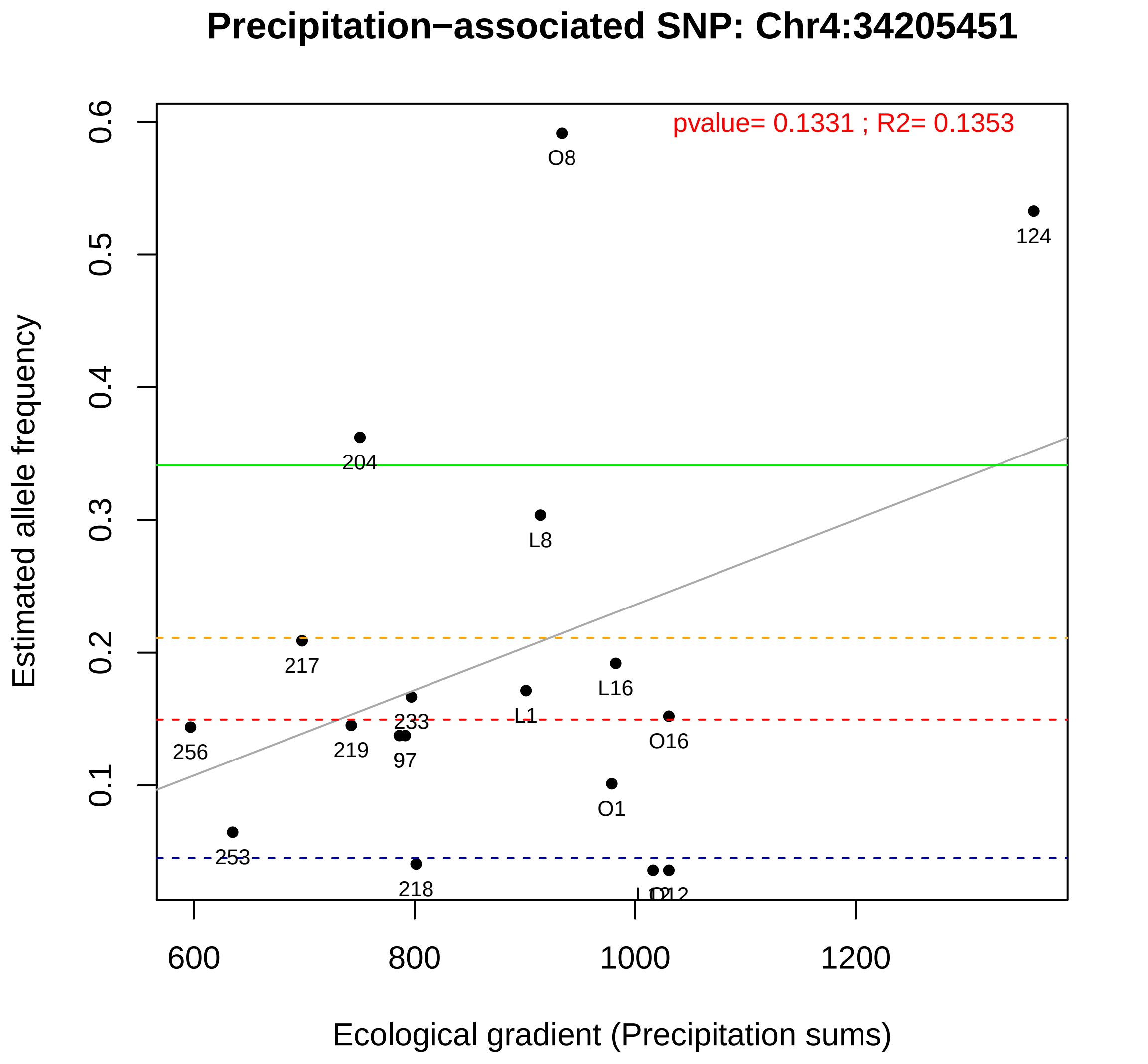

Precipitation−associated SNP: Chr4:34205451
pvalue= 0.1331 ; R2= 0.1353
O8
124
204
L8
217
L16
233
L1
O16
256
219
97
O1
253
218
L12
O12
0.1
0.2
0.3
0.4
0.5
0.6
Estimated allele frequency
600
800
1000
1200
Ecological gradient (Precipitation sums)

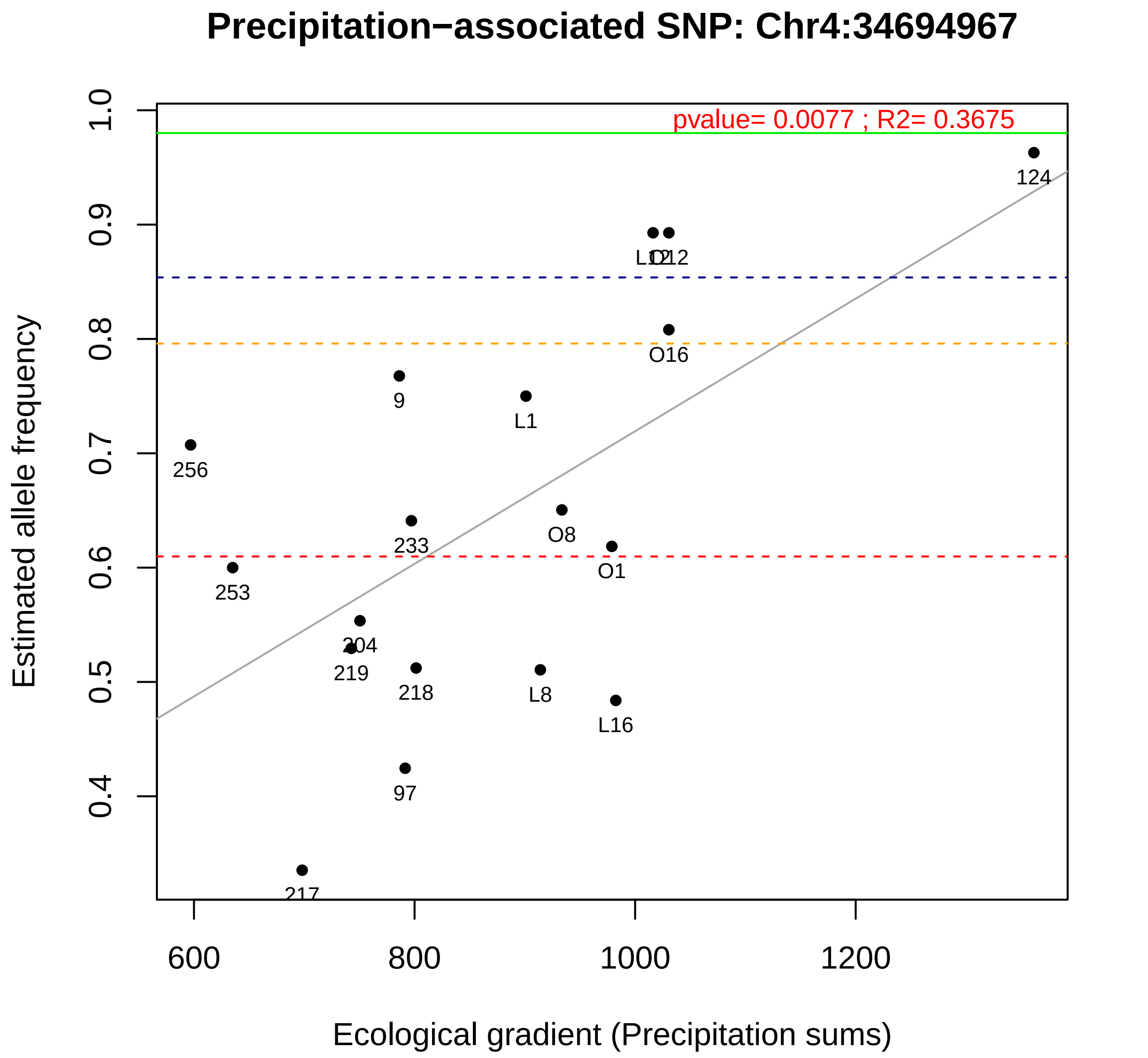

Precipitation−associated SNP: Chr4:34694967
pvalue= 0.0077 ; R2= 0.3675
Estimated allele frequency
Ecological gradient (Precipitation sums)
0.4
0.5
0.6
0.7
0.8
0.9
1.0
600
800
1000
1200
124
L12
O12
O16
9
L1
256
233
O8
O1
253
204
219
218
L8
L16
97
217

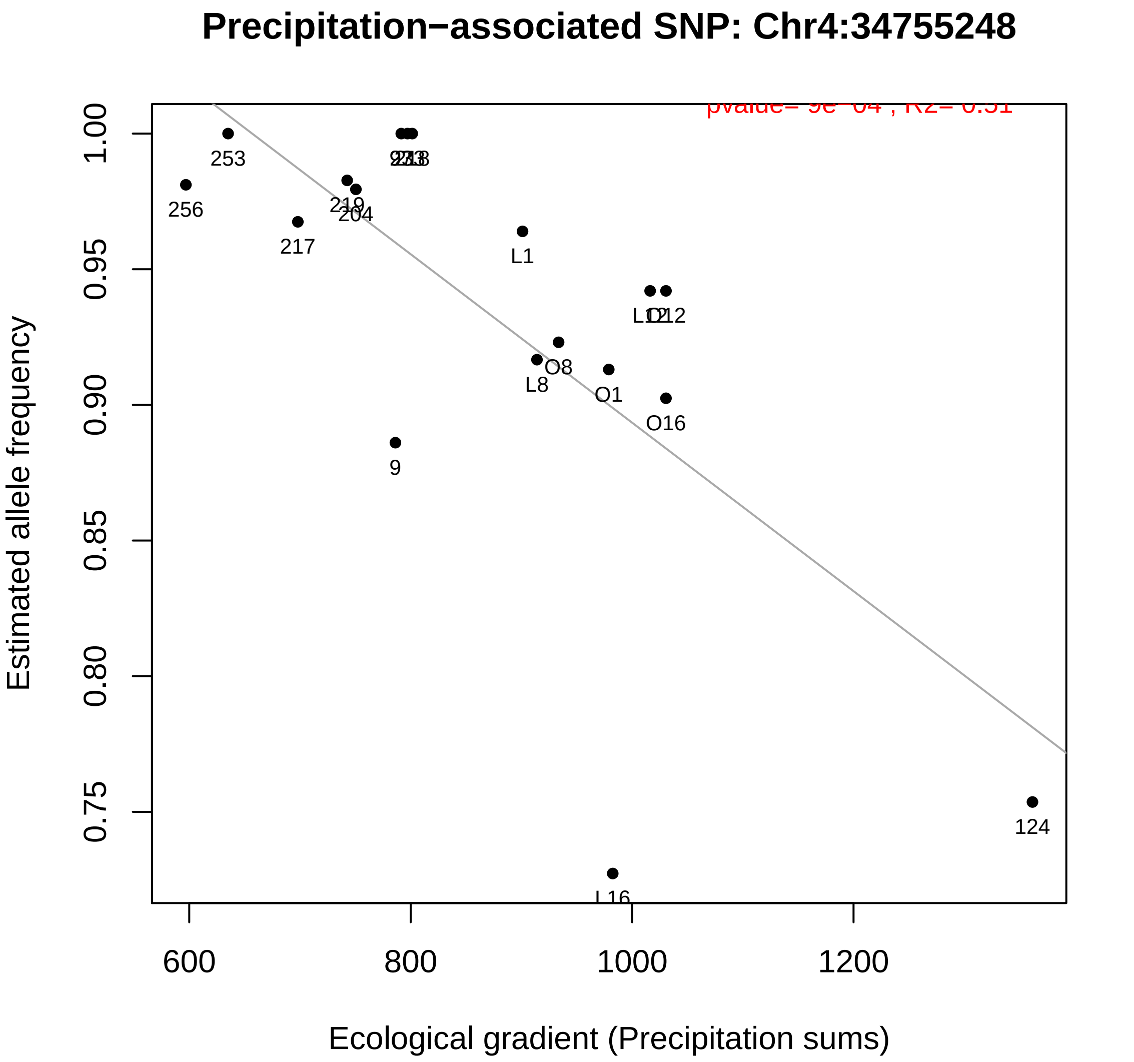

Precipitation−associated SNP: Chr4:34755248
pvalue= 9e−04 ; R2= 0.51
Estimated allele frequency
Ecological gradient (Precipitation sums)
1.00
0.95
0.90
0.85
0.80
0.75
600
800
1000
1200
253
256
217
219
204
L1
L12
O12
O8
L8
O1
O16
9
124
L16

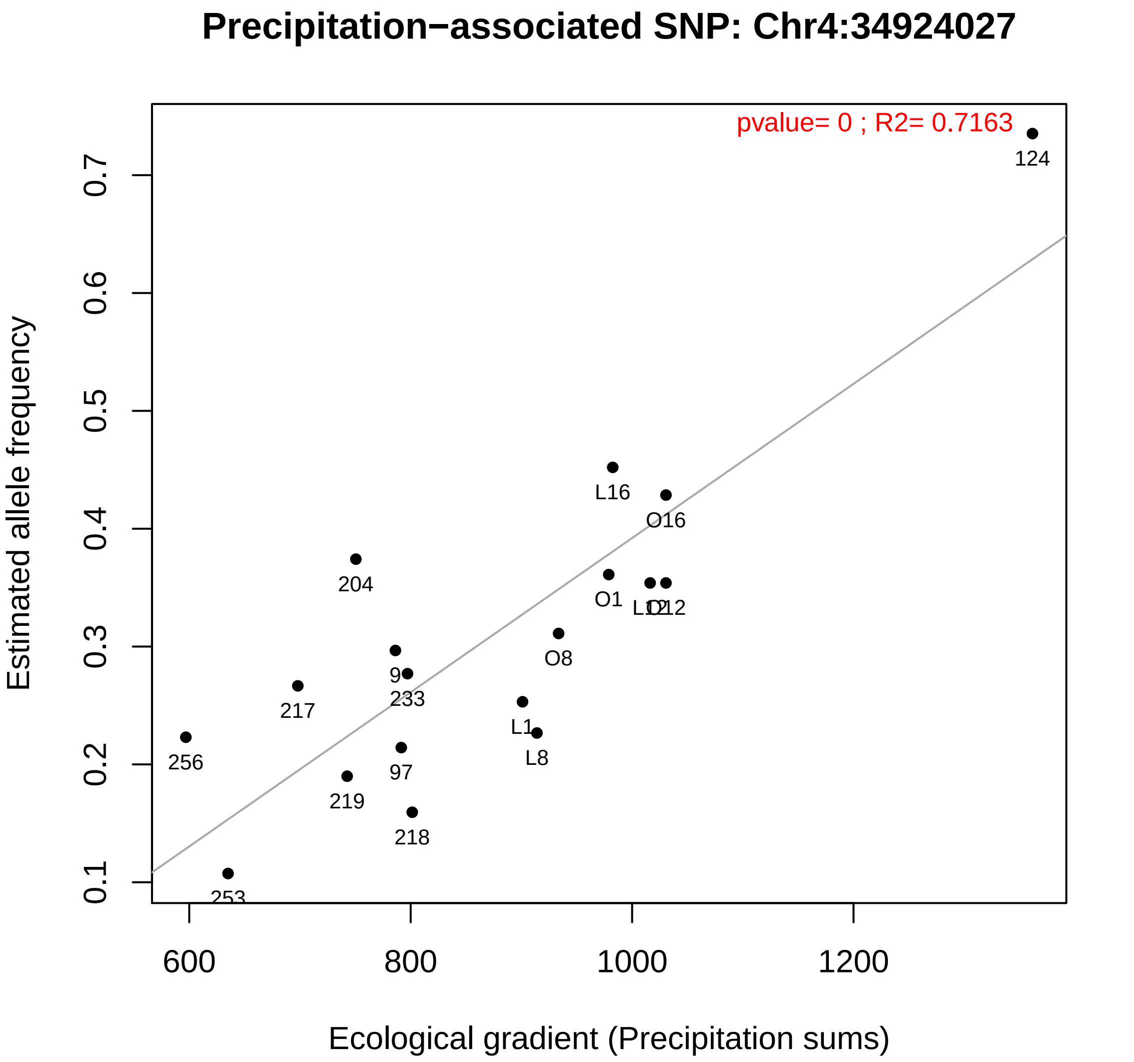
Precipitation−associated SNP: Chr4:34924027
pvalue= 0 ; R2= 0.7163
Estimated allele frequency
Ecological gradient (Precipitation sums)
0.1
0.2
0.3
0.4
0.5
0.6
0.7
600
800
1000
1200
124
L16
O16
204
O1
L12
O12
O8
9
233
217
L1
L8
256
97
219
218
253

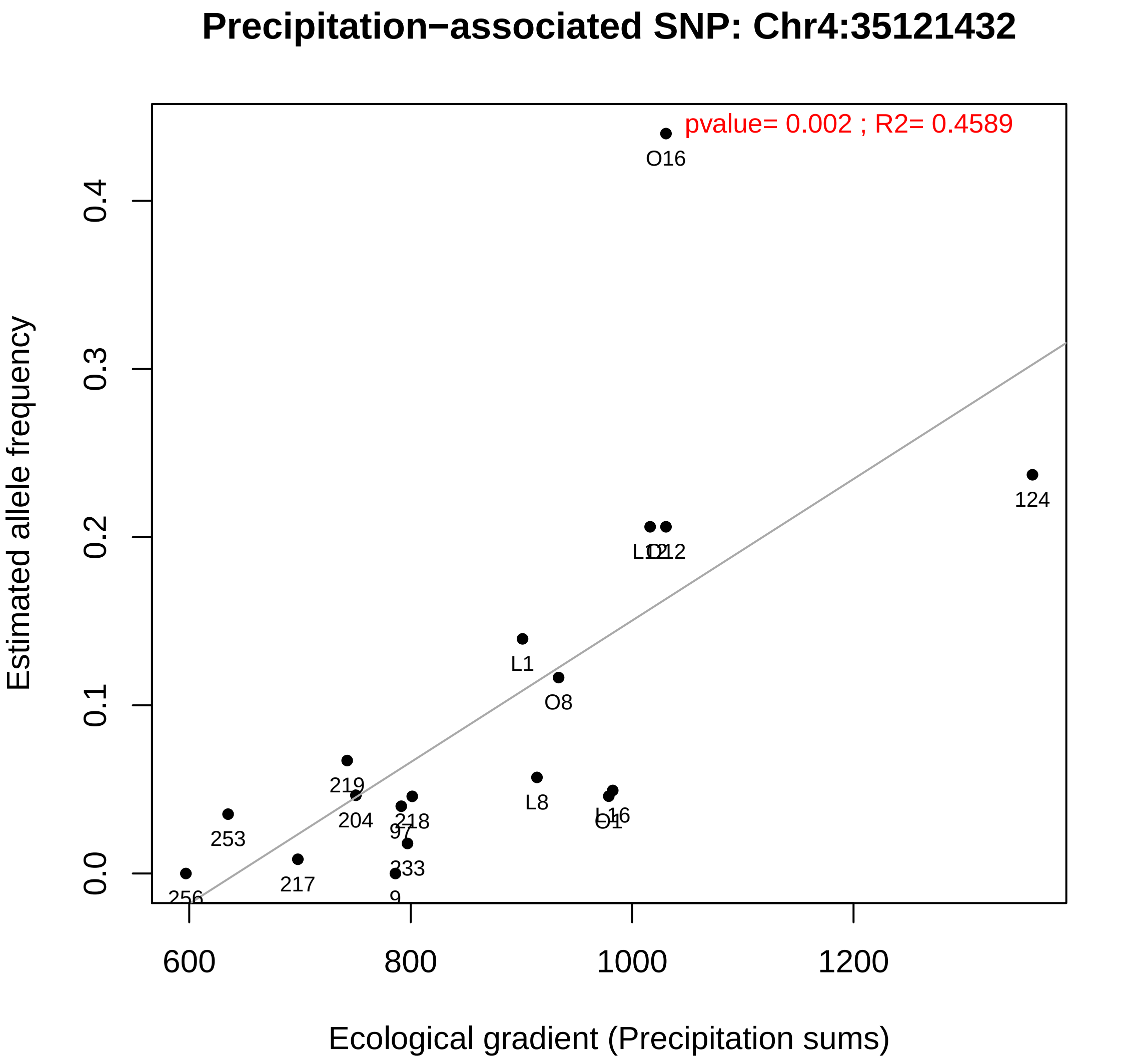

Precipitation−associated SNP: Chr4:35121432
pvalue= 0.002 ; R2= 0.4589
O16
124
L10
O12
L1
O8
219
204
218
97
233
9
253
217
256
L8
L16
O1
0.0
0.1
0.2
0.3
0.4
600
800
1000
1200
Estimated allele frequency
Ecological gradient (Precipitation sums)

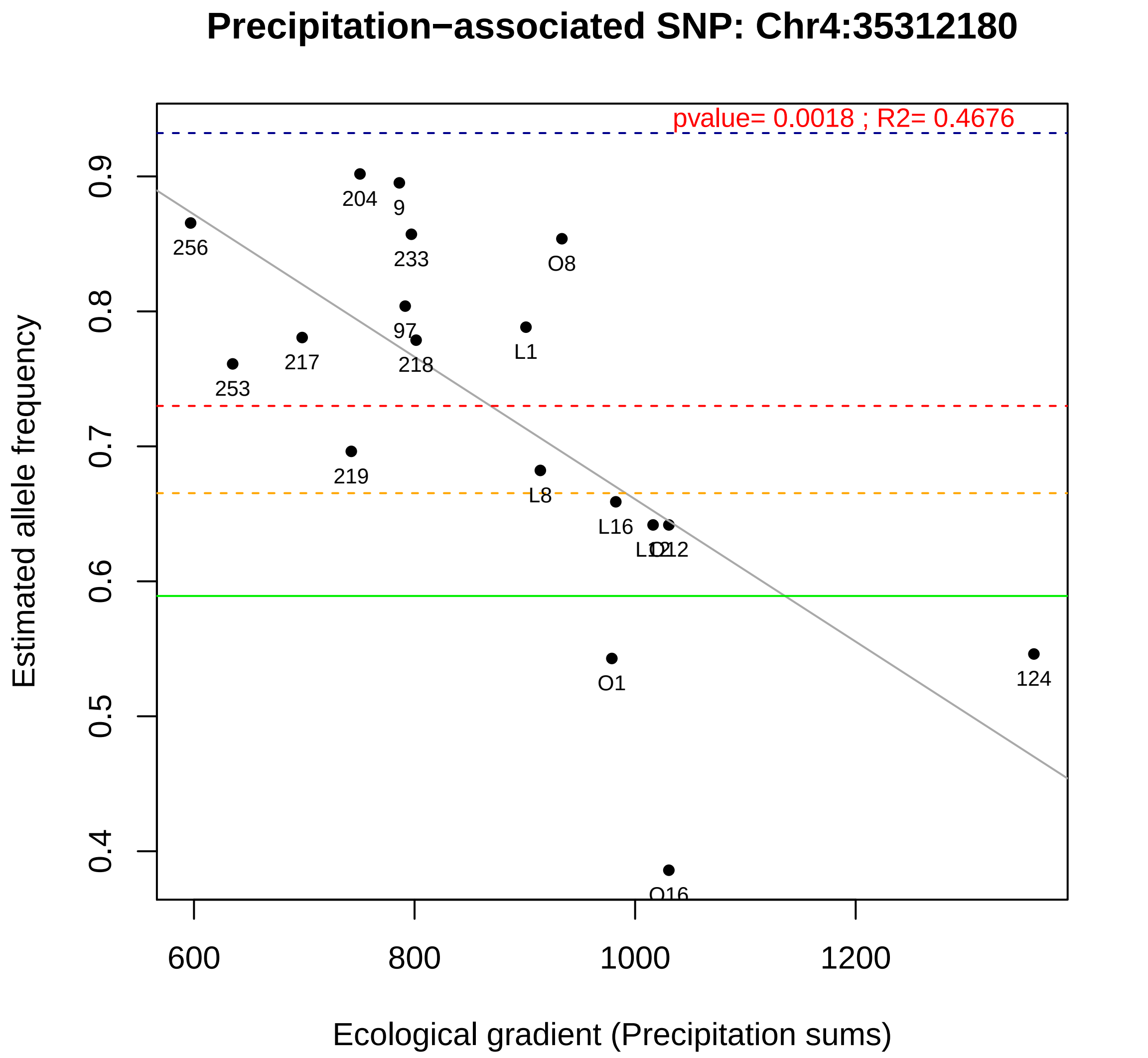
Precipitation−associated SNP: Chr4:35312180
pvalue= 0.0018 ; R2= 0.4676
Estimated allele frequency
Ecological gradient (Precipitation sums)
0.4
0.5
0.6
0.7
0.8
0.9
600
800
1000
1200
204
9
256
233
O8
97
L1
217
218
253
219
L8
L16
O1
124

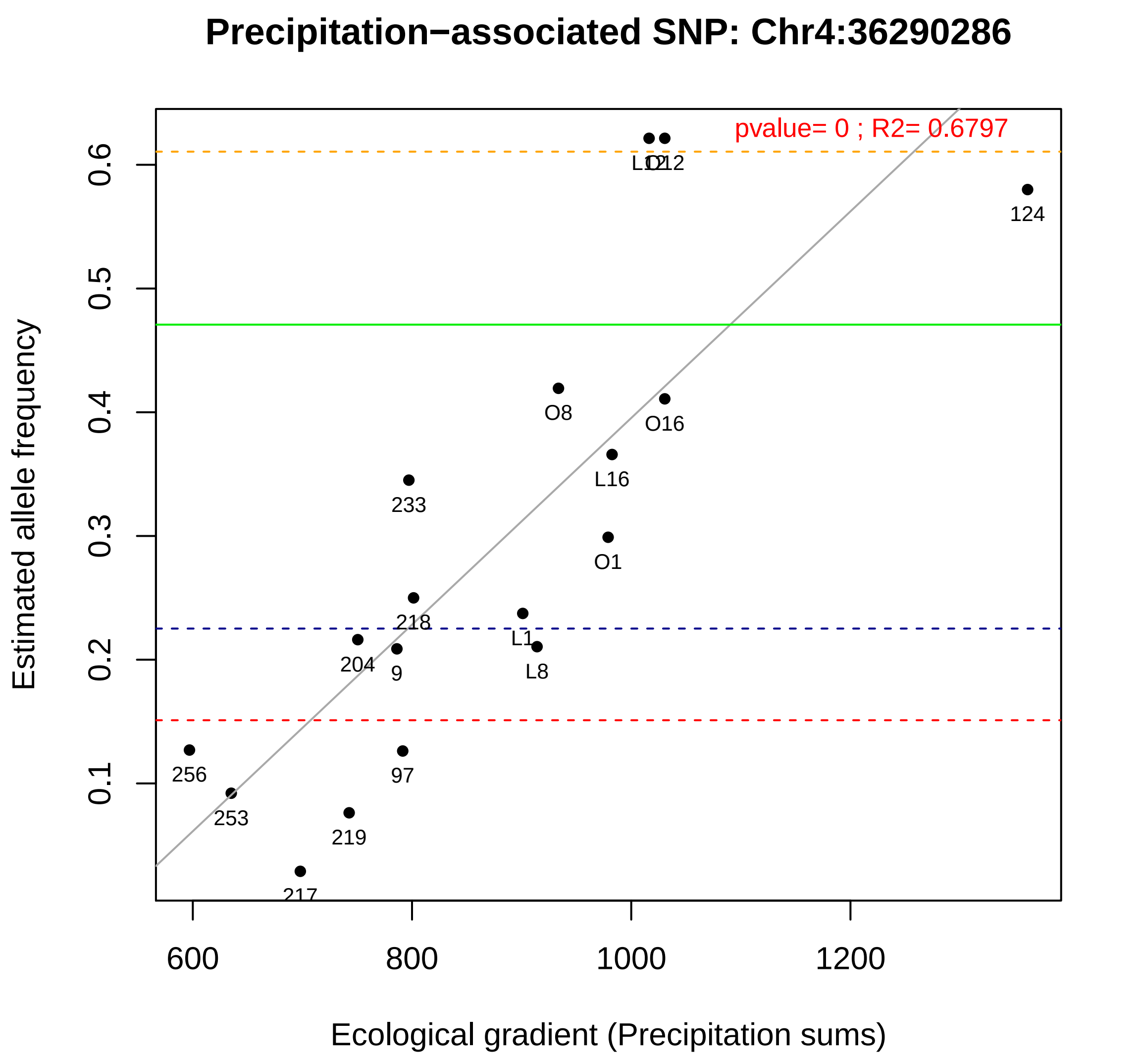

Precipitation−associated SNP: Chr4:36290286
pvalue= 0 ; R2= 0.6797
Estimated allele frequency
0.1
0.2
0.3
0.4
0.5
0.6
600
800
1000
1200
Ecological gradient (Precipitation sums)
124
O8
O16
L16
233
O1
218
L1
204
9
L8
256
97
253
219
217

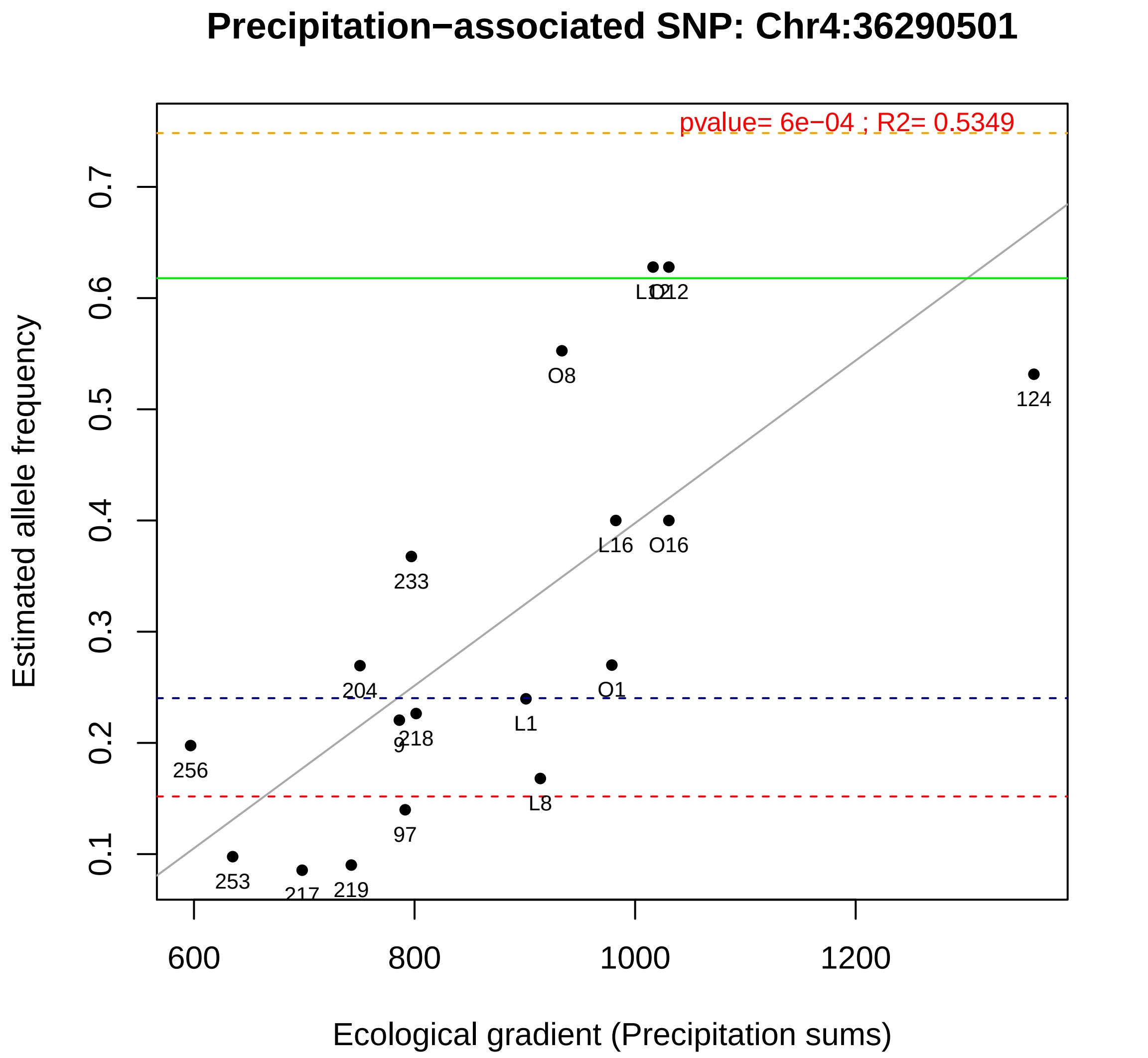

Precipitation−associated SNP: Chr4:36290501
pvalue= 6e−04 ; R2= 0.5349
Estimated allele frequency
Ecological gradient (Precipitation sums)
0.1
0.2
0.3
0.4
0.5
0.6
0.7
600
800
1000
1200
L12
O12
O8
124
L16
O16
233
204
O1
L1
218
9
256
L8
97
253
217
219

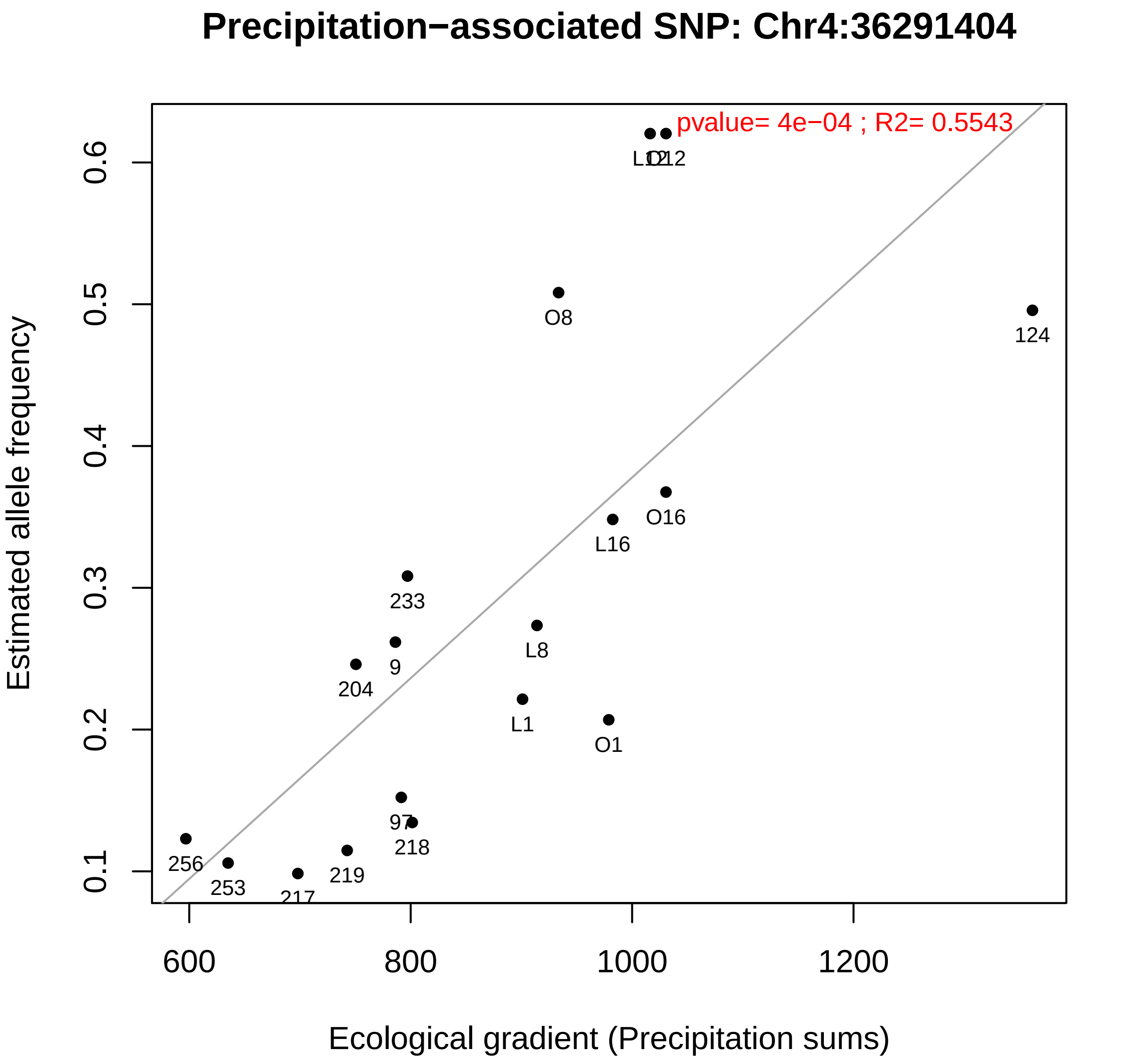

Precipitation−associated SNP: Chr4:36291404
pvalue= 4e−04 ; R2= 0.5543
Estimated allele frequency
Ecological gradient (Precipitation sums)
0.1
0.2
0.3
0.4
0.5
0.6
600
800
1000
1200
L12
O12
O8
124
O16
L16
233
L8
9
204
L1
O1
97
218
256
253
217
219

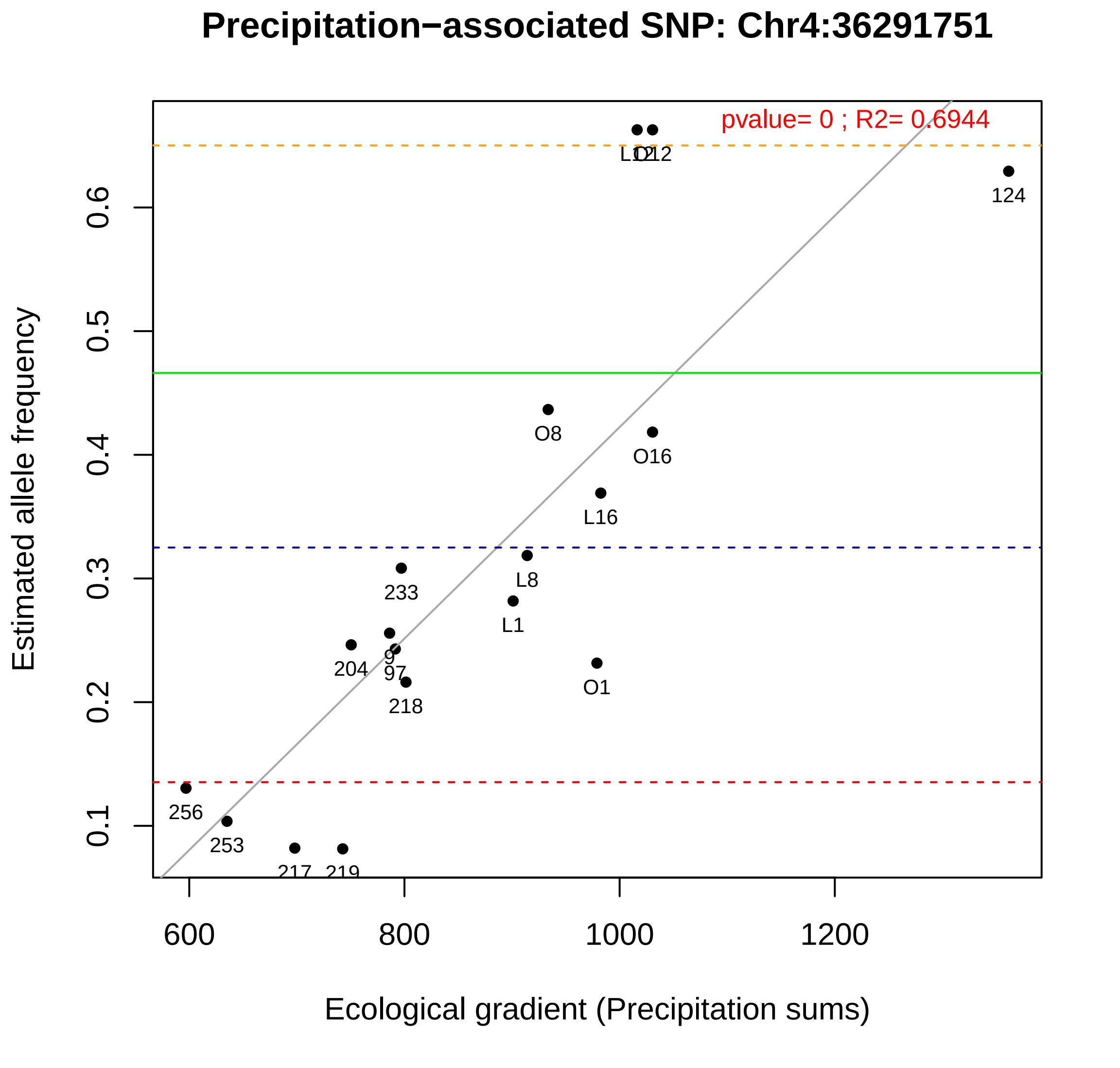
Precipitation−associated SNP: Chr4:36291751
pvalue= 0 ; R2= 0.6944
Estimated allele frequency
Ecological gradient (Precipitation sums)
0.1
0.2
0.3
0.4
0.5
0.6
600
800
1000
1200
L10
O12
124
O8
O16
L16
L8
233
L1
204
9
97
218
O1
256
253
217
219

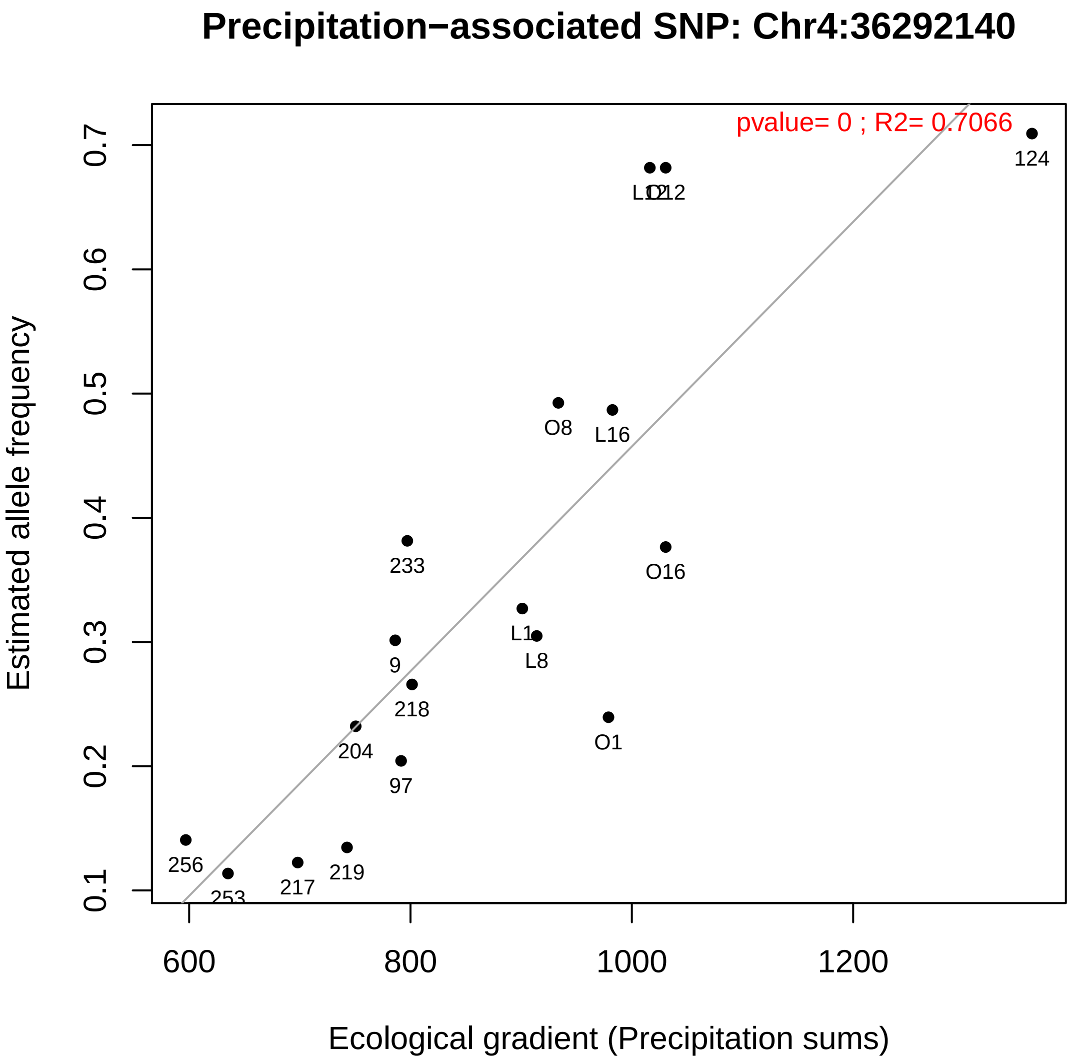
Precipitation−associated SNP: Chr4:36292140
pvalue= 0 ; R2= 0.7066
Estimated allele frequency
Ecological gradient (Precipitation sums)
0.1
0.2
0.3
0.4
0.5
0.6
0.7
600
800
1000
1200
124
L12
O12
O8
L16
233
O16
L1
L8
9
218
204
97
O1
256
253
217
219

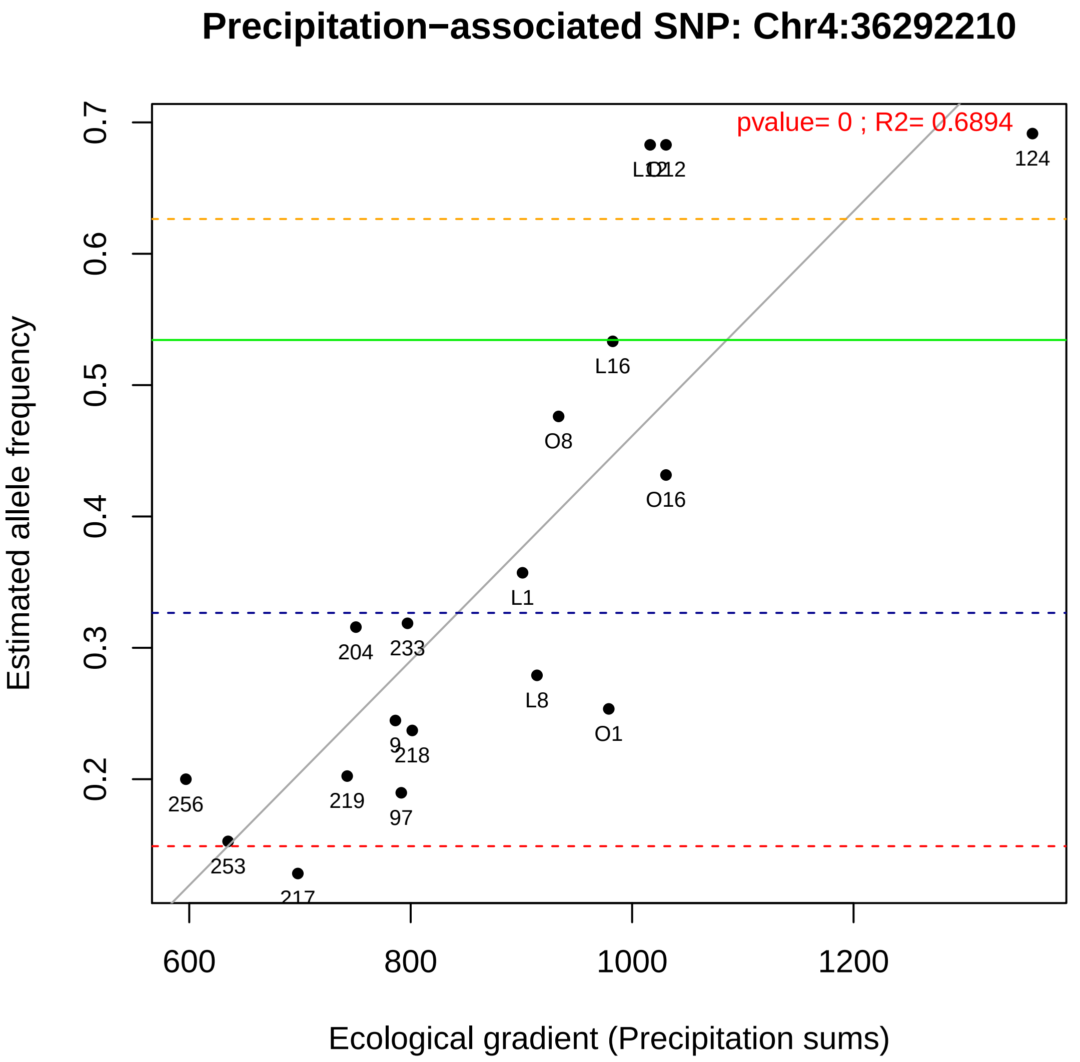

Precipitation−associated SNP: Chr4:36292210
pvalue= 0 ; R2= 0.6894
Estimated allele frequency
Ecological gradient (Precipitation sums)
0.2
0.3
0.4
0.5
0.6
0.7
600
800
1000
1200
124
L12
O12
L16
O8
O16
L1
204
233
L8
O1
9
218
256
219
97
253
217

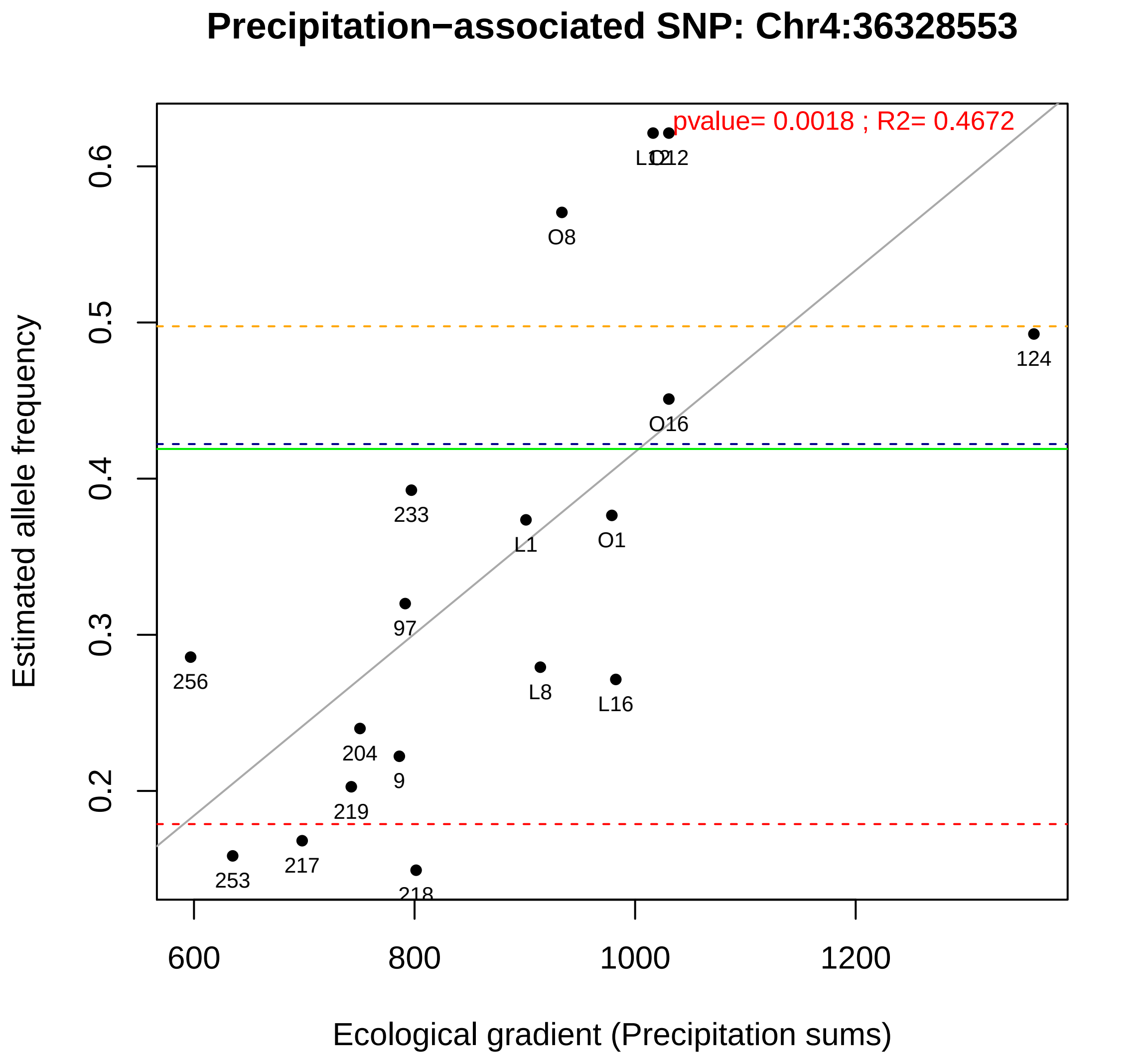
Precipitation−associated SNP: Chr4:36328553
pvalue= 0.0018 ; R2= 0.4672
Estimated allele frequency
Ecological gradient (Precipitation sums)
0.2
0.3
0.4
0.5
0.6
600
800
1000
1200
L12
O12
O8
124
O16
233
L1
O1
97
256
L8
L16
204
9
219
217
253
218

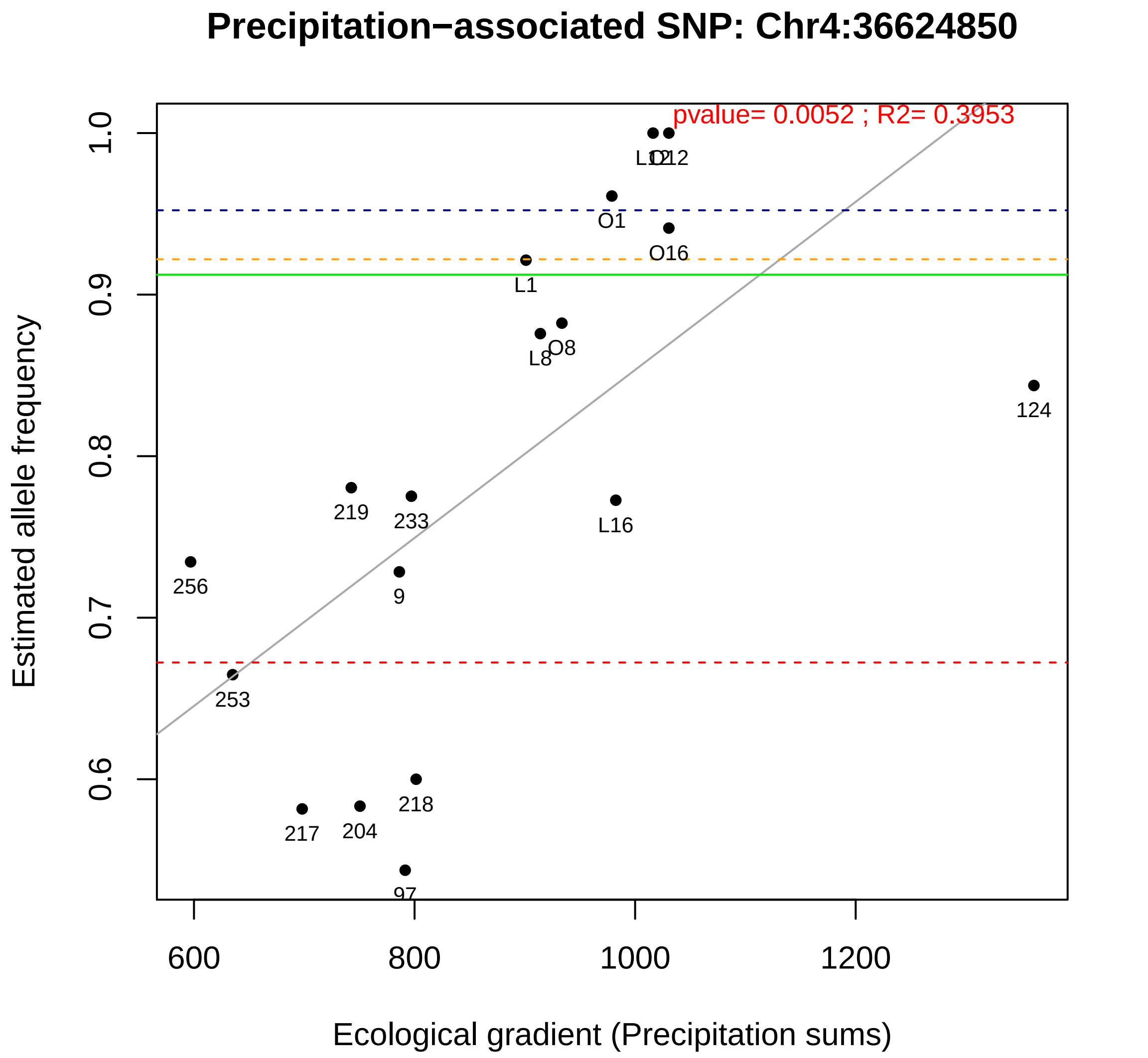
Precipitation−associated SNP: Chr4:36624850
pvalue= 0.0052 ; R2= 0.3953
Estimated allele frequency
Ecological gradient (Precipitation sums)
1.0
0.9
0.8
0.7
0.6
600
800
1000
1200
O12
O1
O16
L1
L8
O8
124
219
233
L16
256
9
253
218
217
204
97

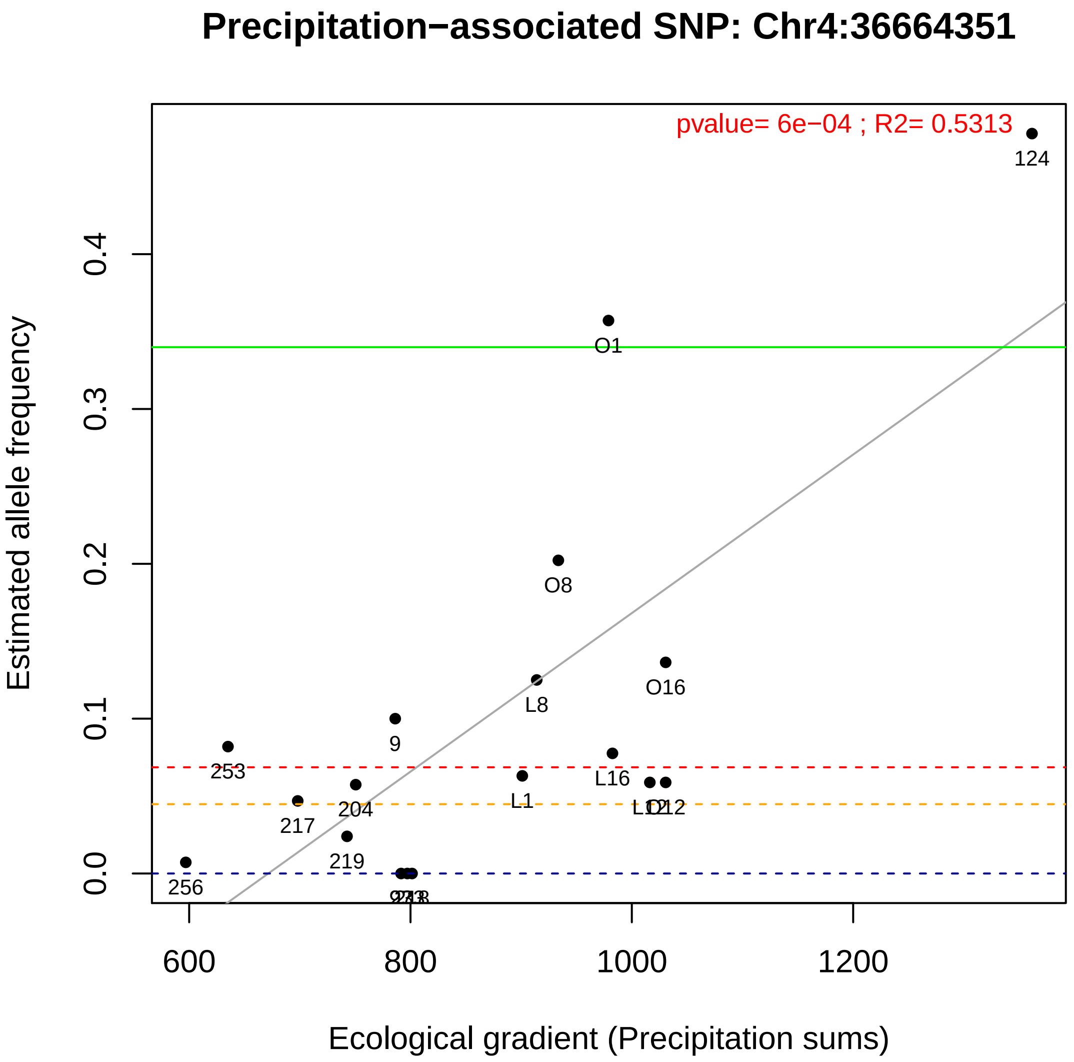

Precipitation−associated SNP: Chr4:36664351
pvalue= 6e−04 ; R2= 0.5313
Estimated allele frequency
Ecological gradient (Precipitation sums)
0.0
0.1
0.2
0.3
0.4
600
800
1000
1200
124
O1
O8
O16
L8
9
253
L16
L1
204
217
219
256

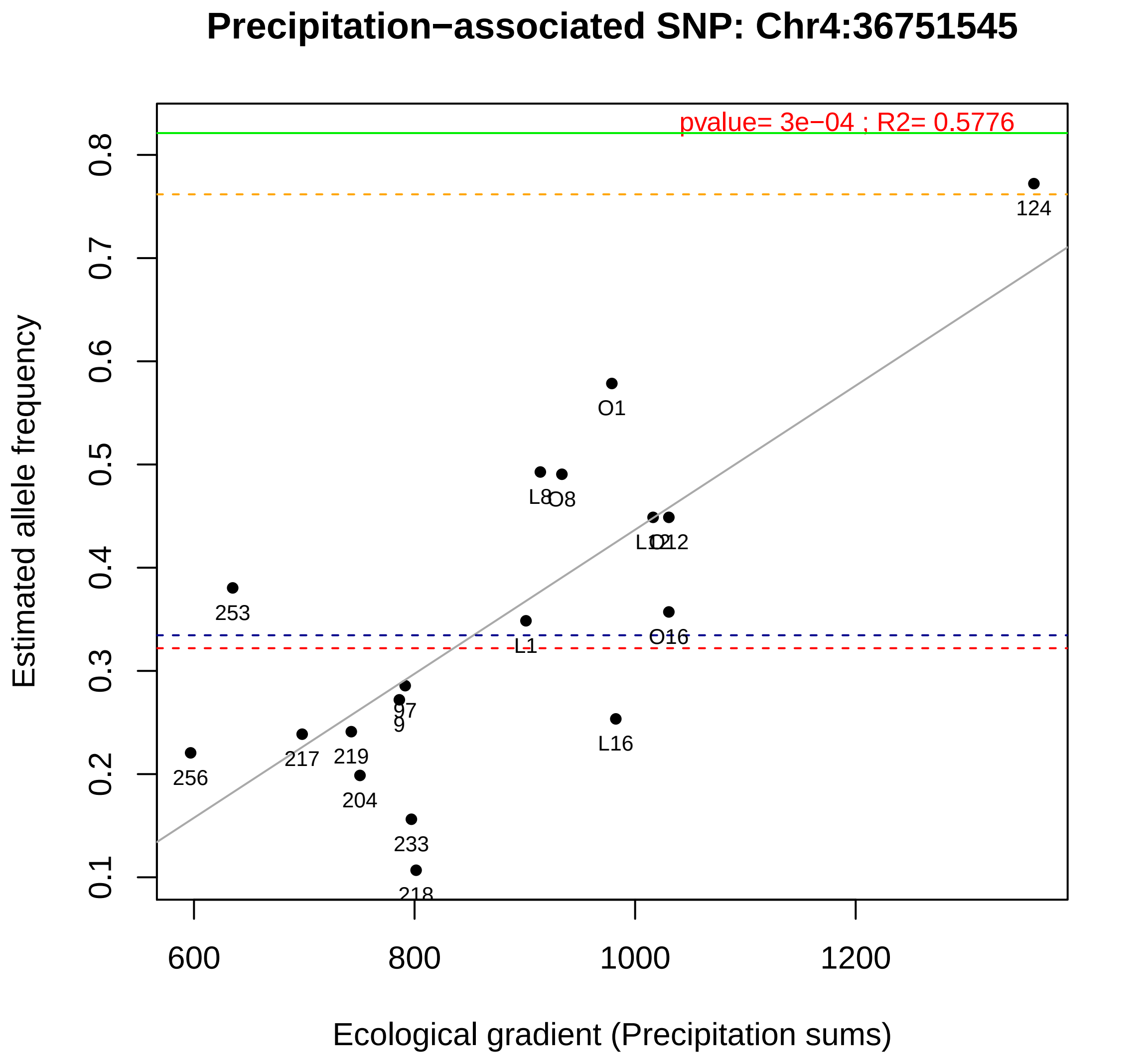

Precipitation−associated SNP: Chr4:36751545
pvalue= 3e−04 ; R2= 0.5776
124
O1
L8
O8
L12
O12
253
L1
O16
97
9
L16
217
219
256
204
233
218
0.1
0.2
0.3
0.4
0.5
0.6
0.7
0.8
600
800
1000
1200
Estimated allele frequency
Ecological gradient (Precipitation sums)

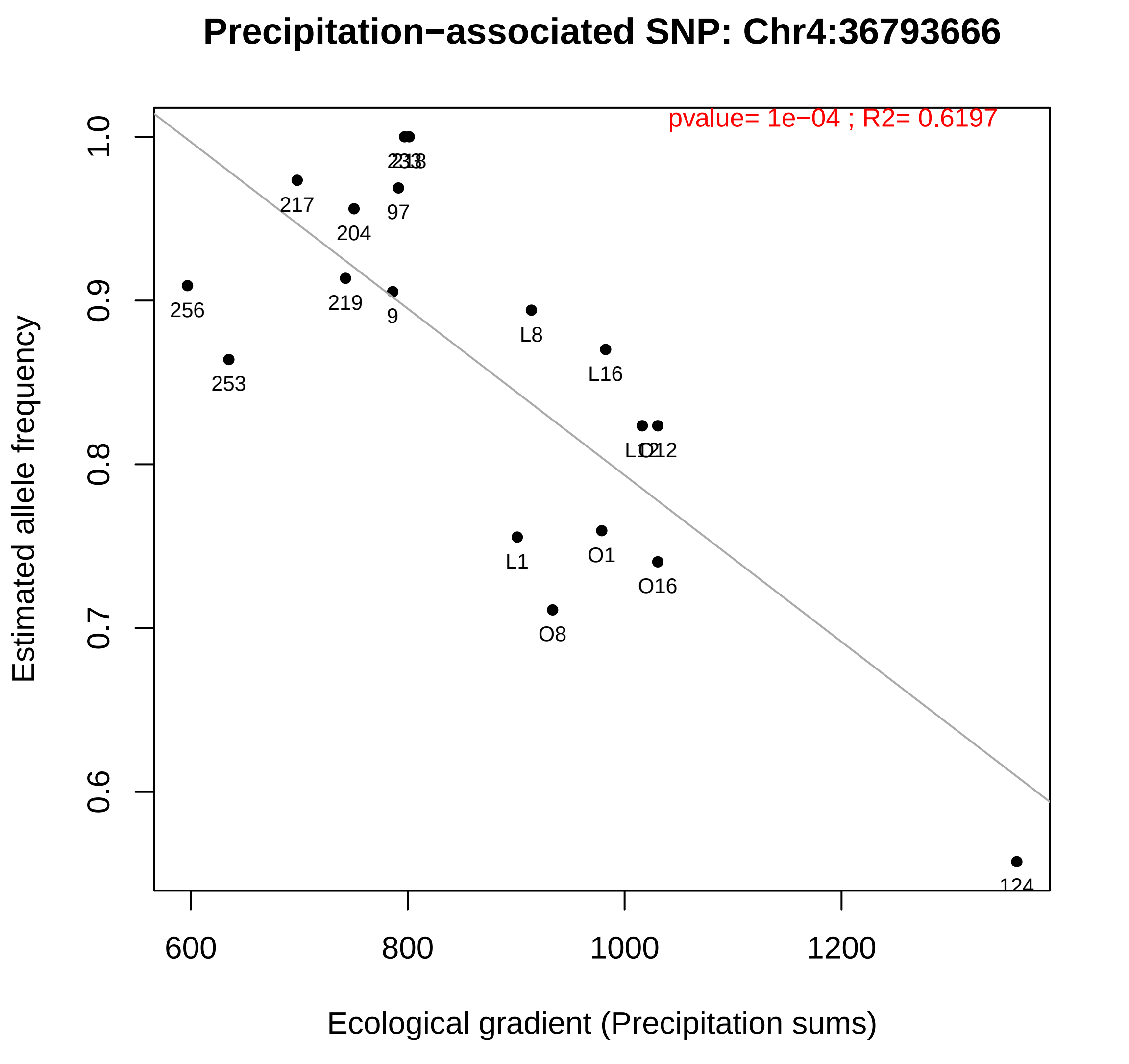

Precipitation−associated SNP: Chr4:36793666
pvalue= 1e−04 ; R2= 0.6197
Estimated allele frequency
Ecological gradient (Precipitation sums)
0.6
0.7
0.8
0.9
1.0
600
800
1000
1200
217
204
97
256
219
9
L8
253
L16
L1
O1
O16
O8
124

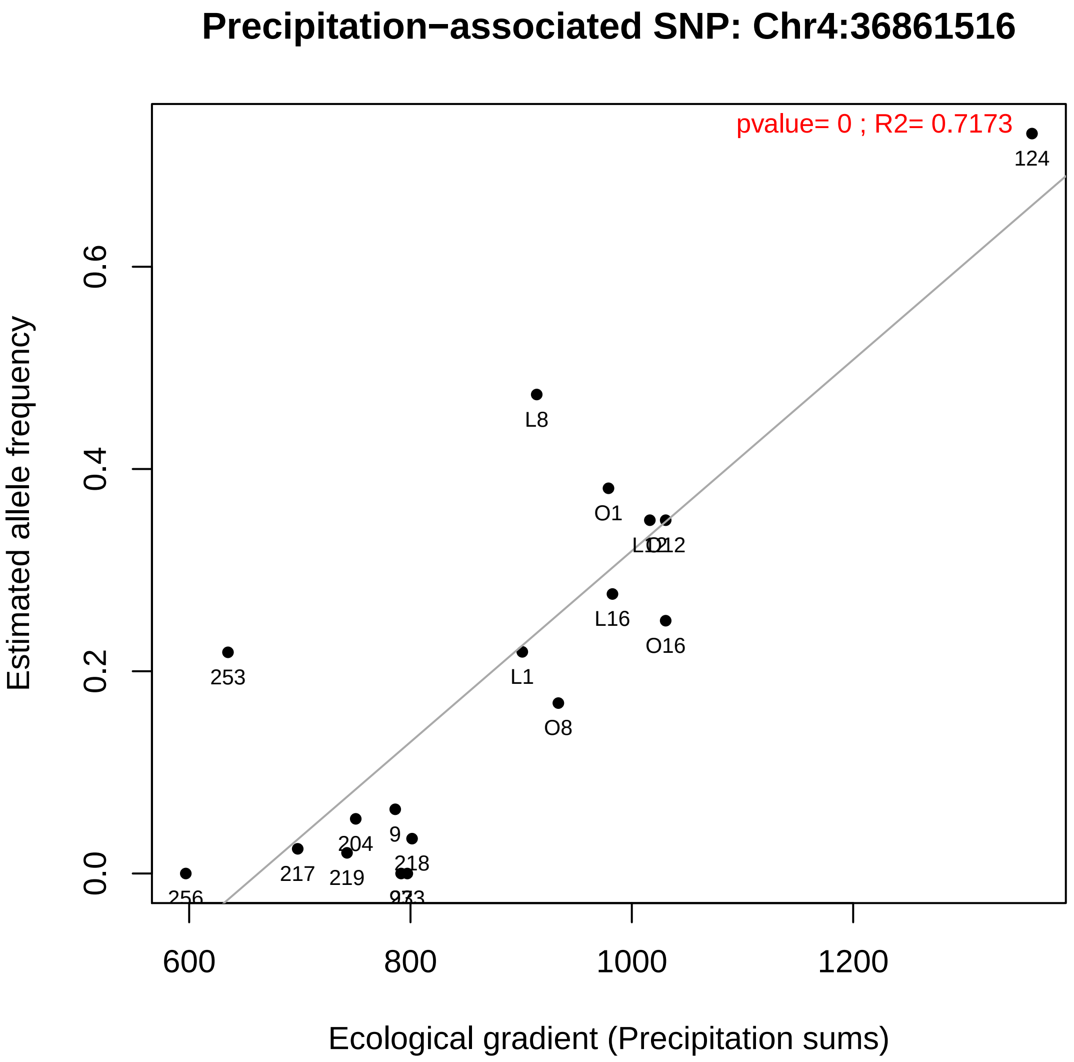
Precipitation−associated SNP: Chr4:36861516
pvalue= 0 ; R2= 0.7173
124
L8
O1
L10
O12
L16
O16
253
L1
O8
204
9
217
219
218
256
0.0
0.2
0.4
0.6
600
800
1000
1200
Estimated allele frequency
Ecological gradient (Precipitation sums)

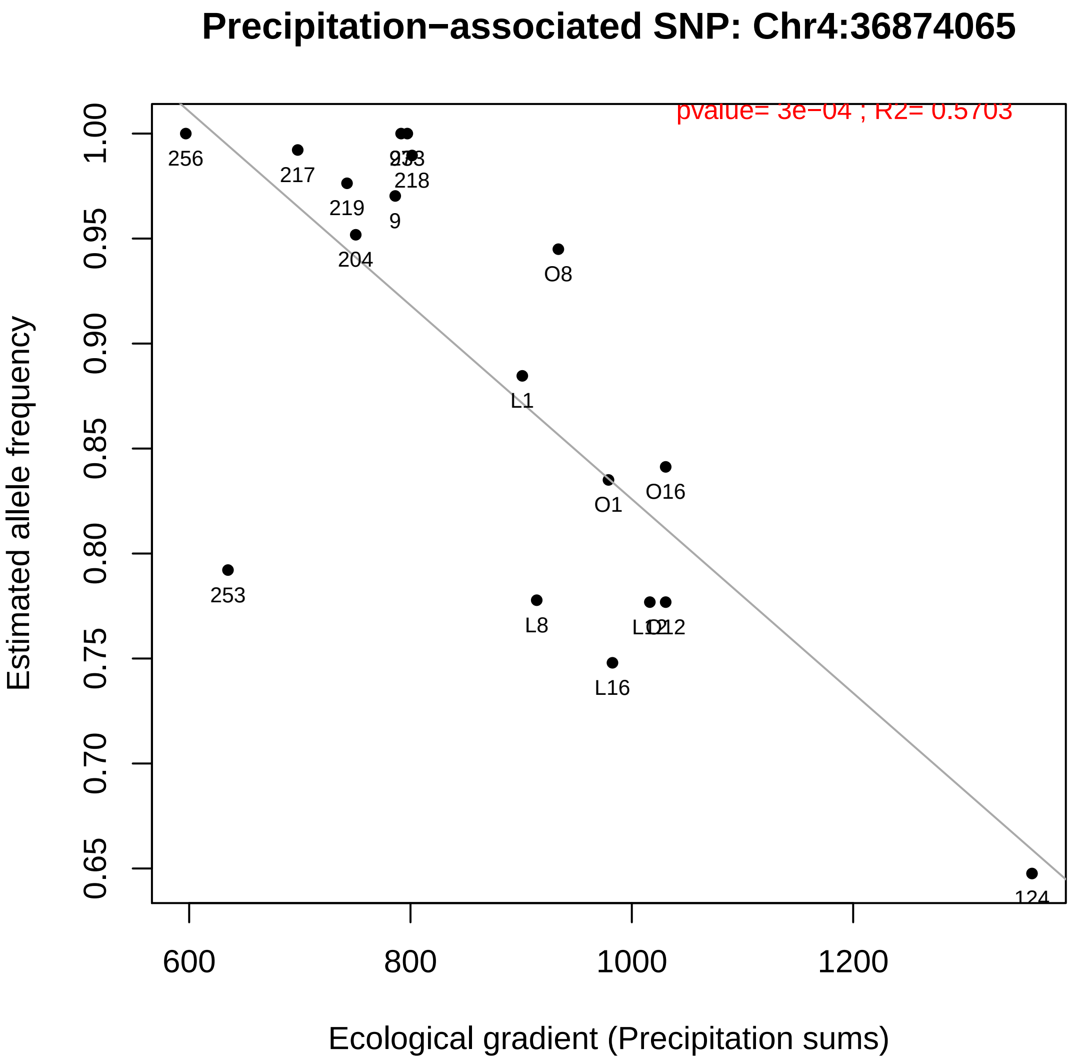
Precipitation−associated SNP: Chr4:36874065
pvalue= 3e−04 ; R2= 0.5703
Estimated allele frequency
Ecological gradient (Precipitation sums)
1.00
0.95
0.90
0.85
0.80
0.75
0.70
0.65
600
800
1000
1200
256
217
219
204
9
218
O8
L1
O1
O16
253
L8
L16
O12
124

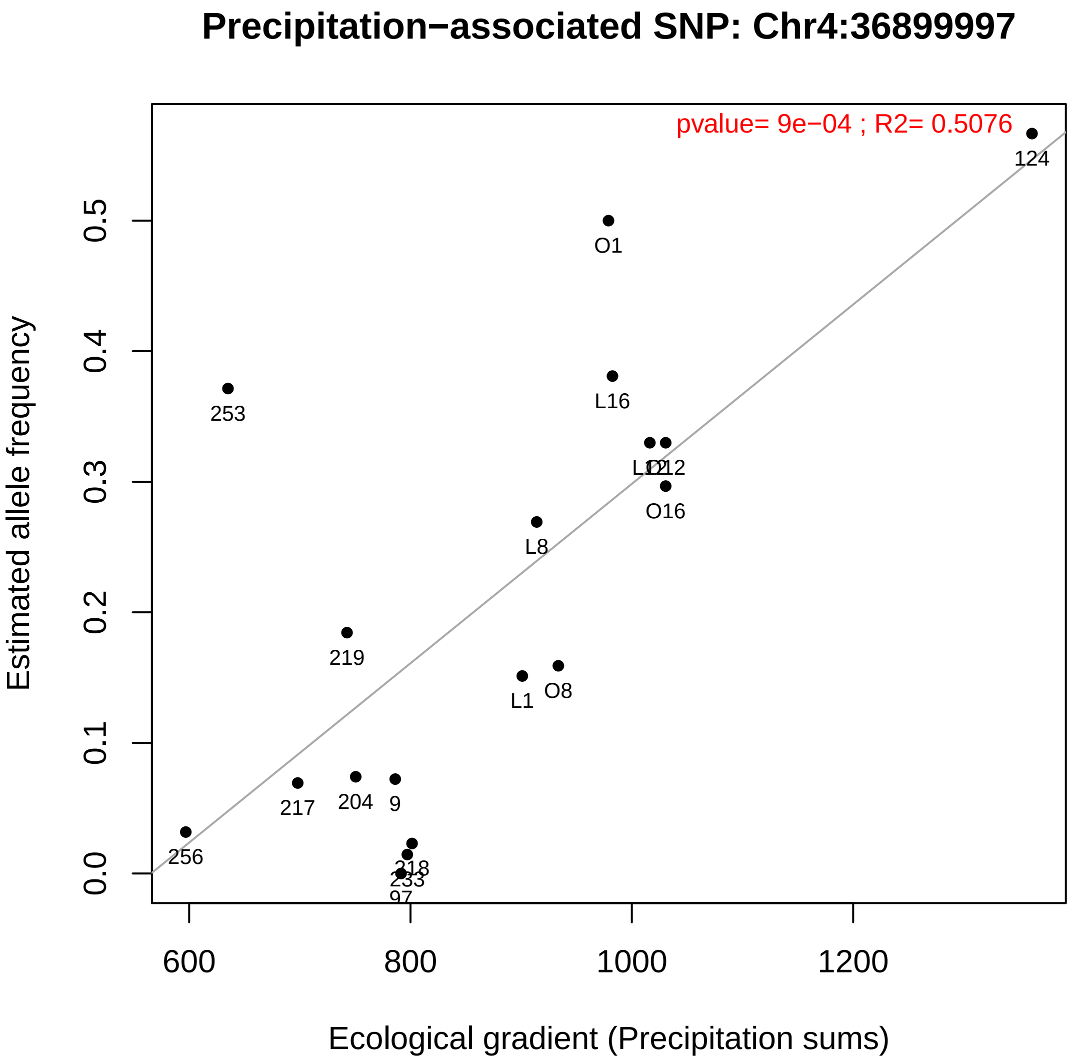

Precipitation−associated SNP: Chr4:36899997
pvalue= 9e−04 ; R2= 0.5076
Estimated allele frequency
Ecological gradient (Precipitation sums)
0.0
0.1
0.2
0.3
0.4
0.5
600
800
1000
1200
124
O1
L16
253
L12
O12
O16
L8
219
L1
O8
217
204
9
256
218
233
97

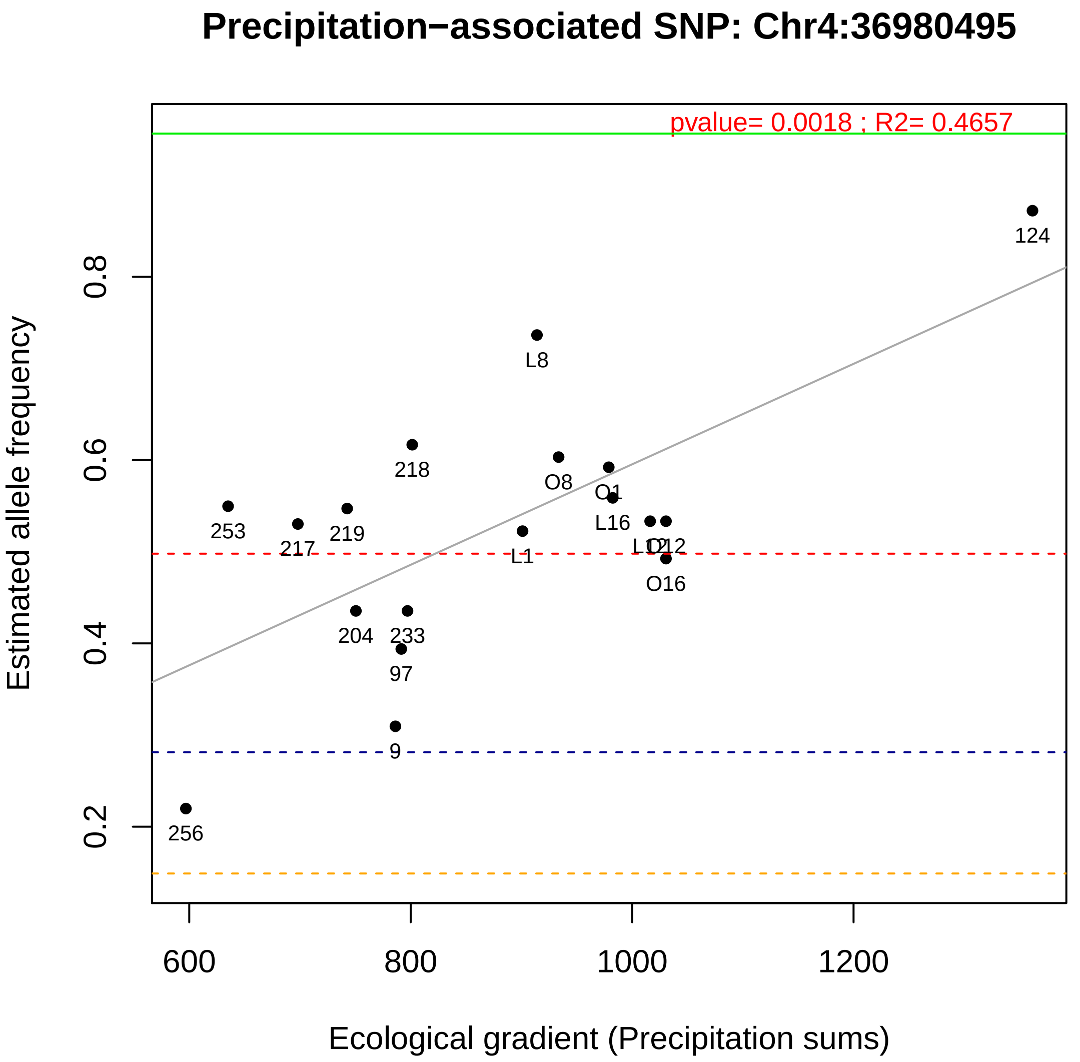

Precipitation−associated SNP: Chr4:36980495
pvalue= 0.0018 ; R2= 0.4657
Estimated allele frequency
0.2
0.4
0.6
0.8
600
800
1000
1200
Ecological gradient (Precipitation sums)
124
L8
218
O8
O1
L16
253
217
219
L1
L12
O12
O16
204
233
97
9
256

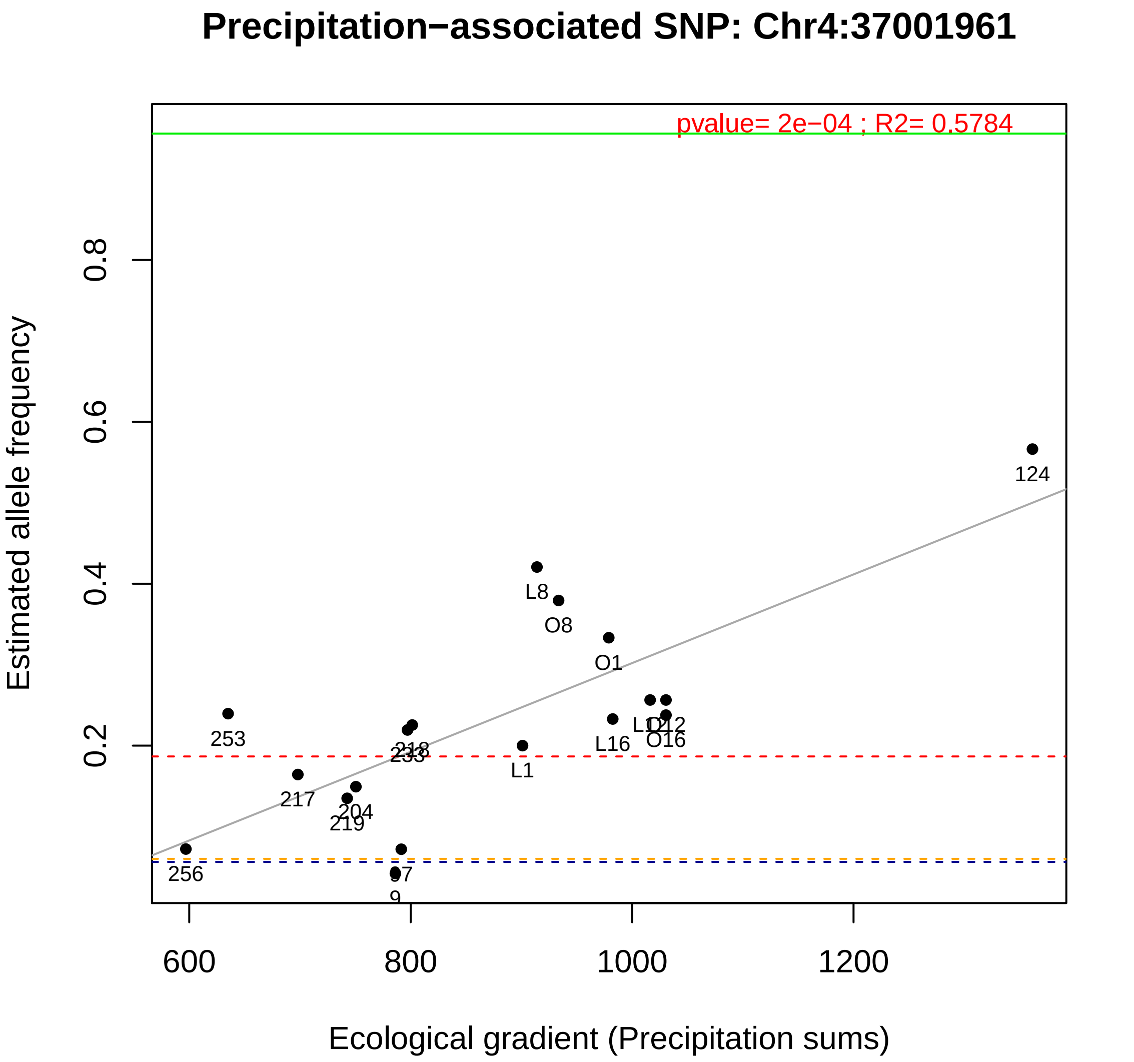
Precipitation−associated SNP: Chr4:37001961
pvalue= 2e−04 ; R2= 0.5784
Estimated allele frequency
0.2
0.4
0.6
0.8
600
800
1000
1200
Ecological gradient (Precipitation sums)
124
L8
O8
O1
L16
O16
L1
253
217
204
219
256
9

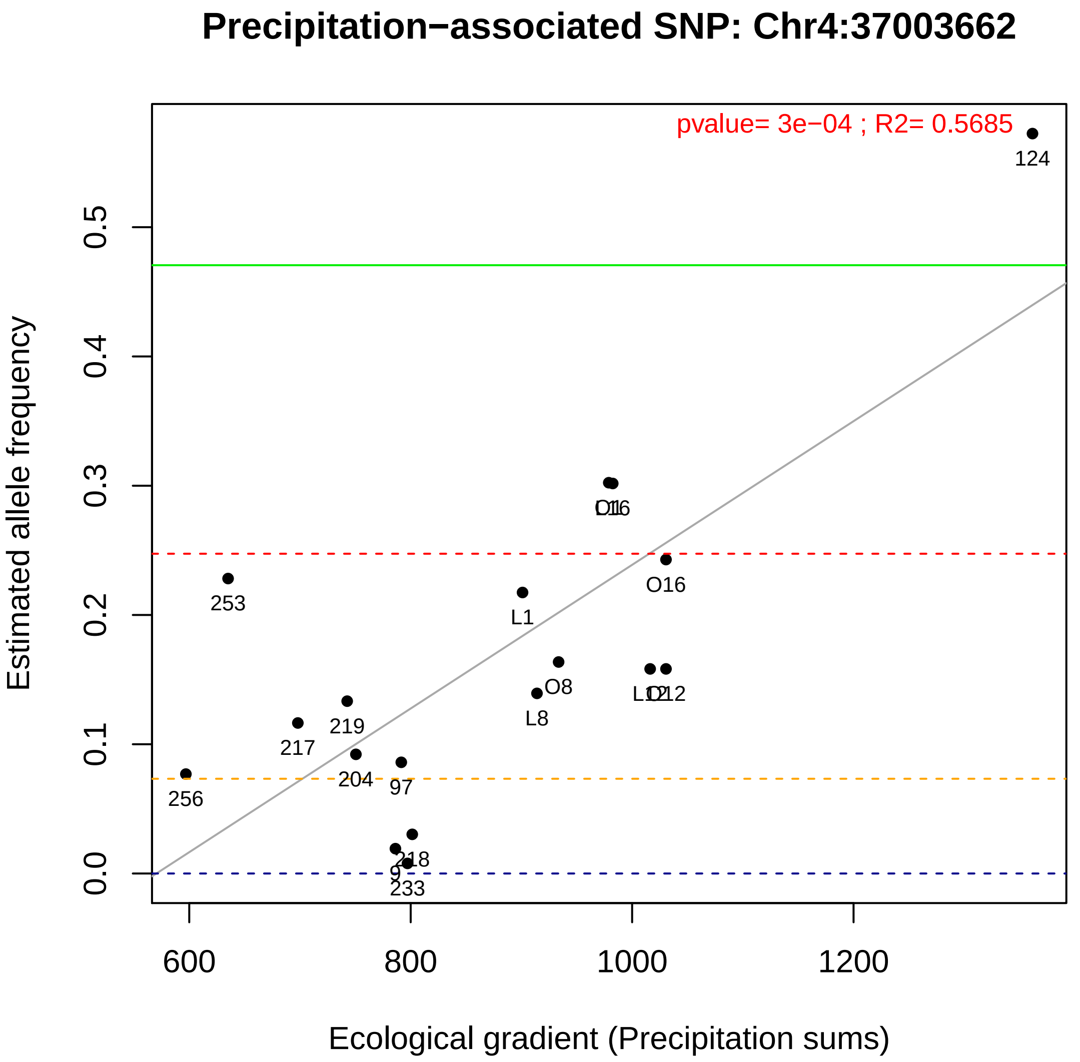
Precipitation−associated SNP: Chr4:37003662
pvalue= 3e−04 ; R2= 0.5685
Estimated allele frequency
Ecological gradient (Precipitation sums)
0.0
0.1
0.2
0.3
0.4
0.5
600
800
1000
1200
124
253
L1
O16
O8
L8
219
217
204
97
256
218
9
233

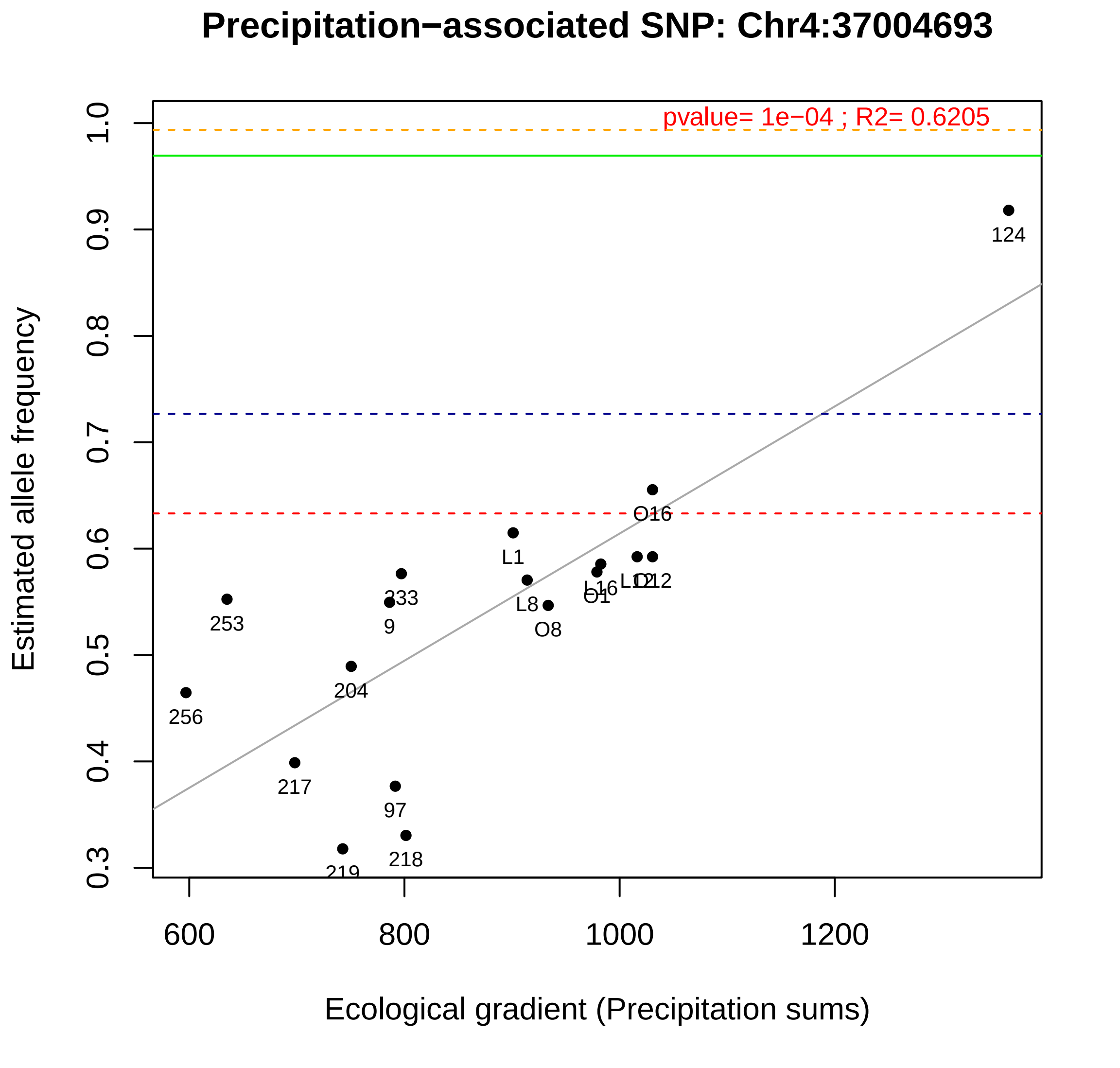

Precipitation−associated SNP: Chr4:37004693
pvalue= 1e−04 ; R2= 0.6205
Estimated allele frequency
Ecological gradient (Precipitation sums)
124
O16
L1
L12
O12
L16
O1
L8
O8
233
9
253
204
256
217
97
218
219

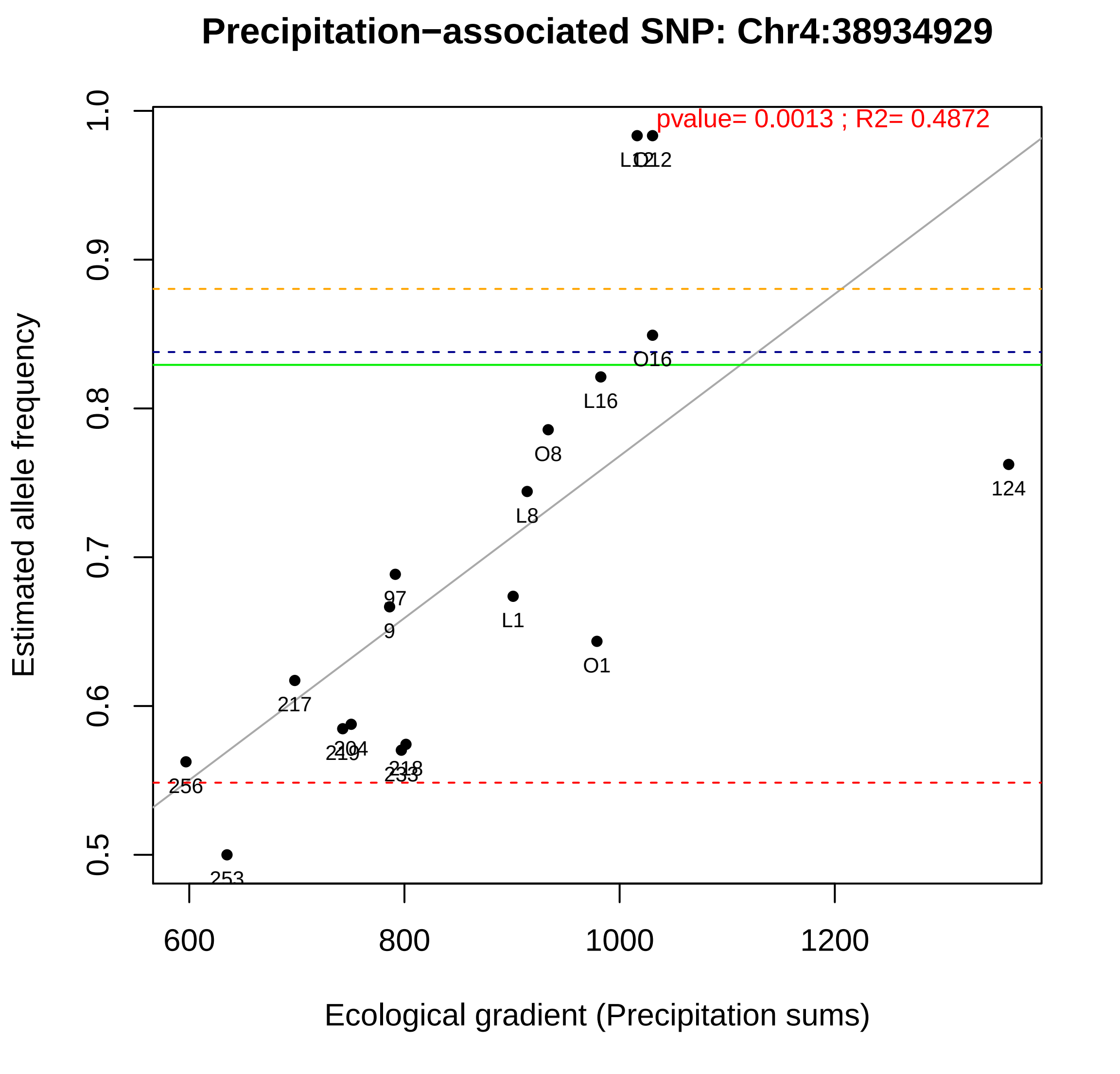
Precipitation−associated SNP: Chr4:38934929
pvalue= 0.0013 ; R2= 0.4872
Estimated allele frequency
Ecological gradient (Precipitation sums)
1.0
0.9
0.8
0.7
0.6
0.5
600
800
1000
1200
L10
O12
O16
L16
O8
L8
124
97
9
L1
O1
217
204
219
218
233
256
253

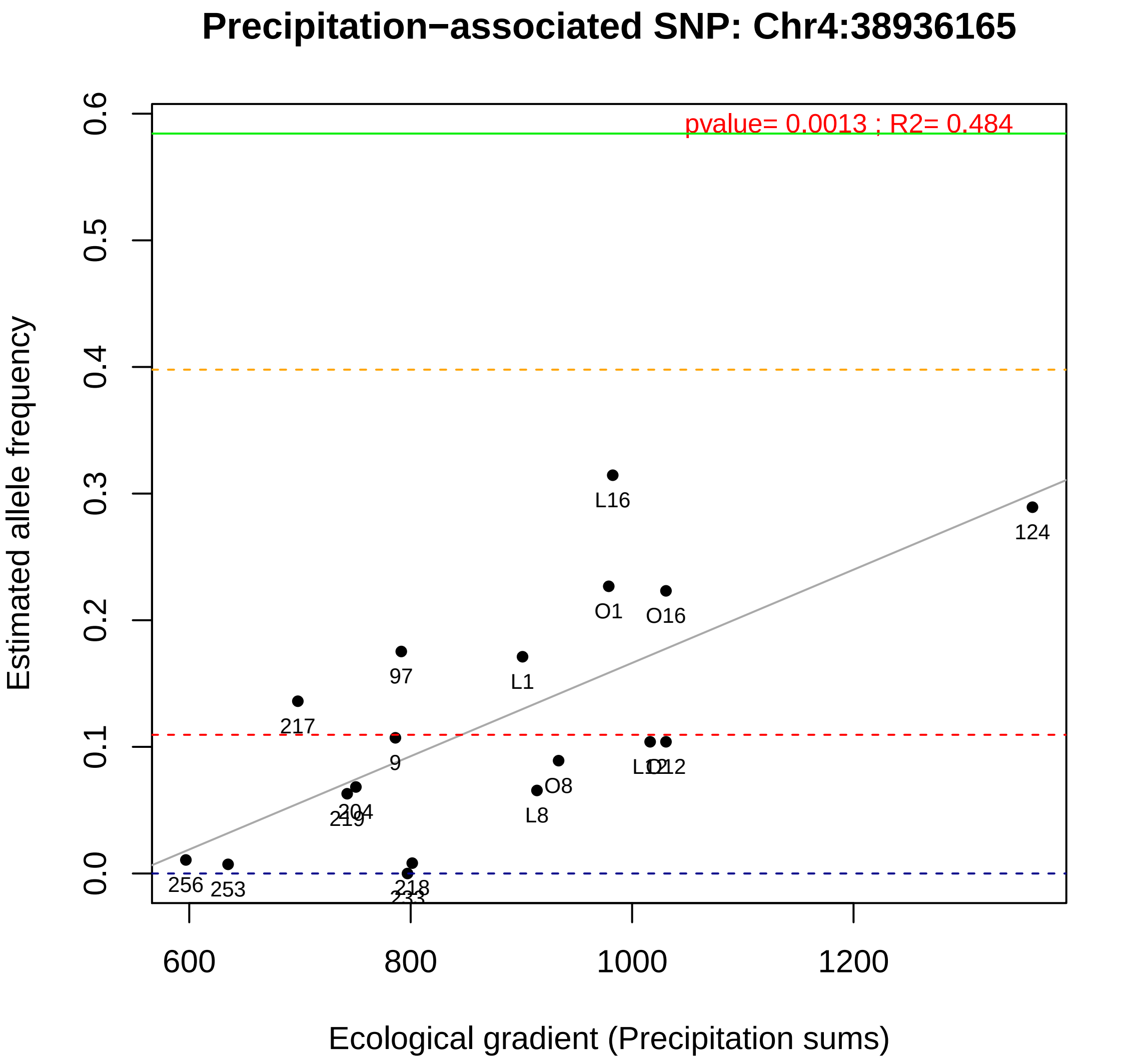
Precipitation−associated SNP: Chr4:38936165
pvalue= 0.0013 ; R2= 0.484
Estimated allele frequency
0.0
0.1
0.2
0.3
0.4
0.5
0.6
600
800
1000
1200
Ecological gradient (Precipitation sums)
L16
124
O1
O16
97
L1
217
9
O8
L8
256
253
218

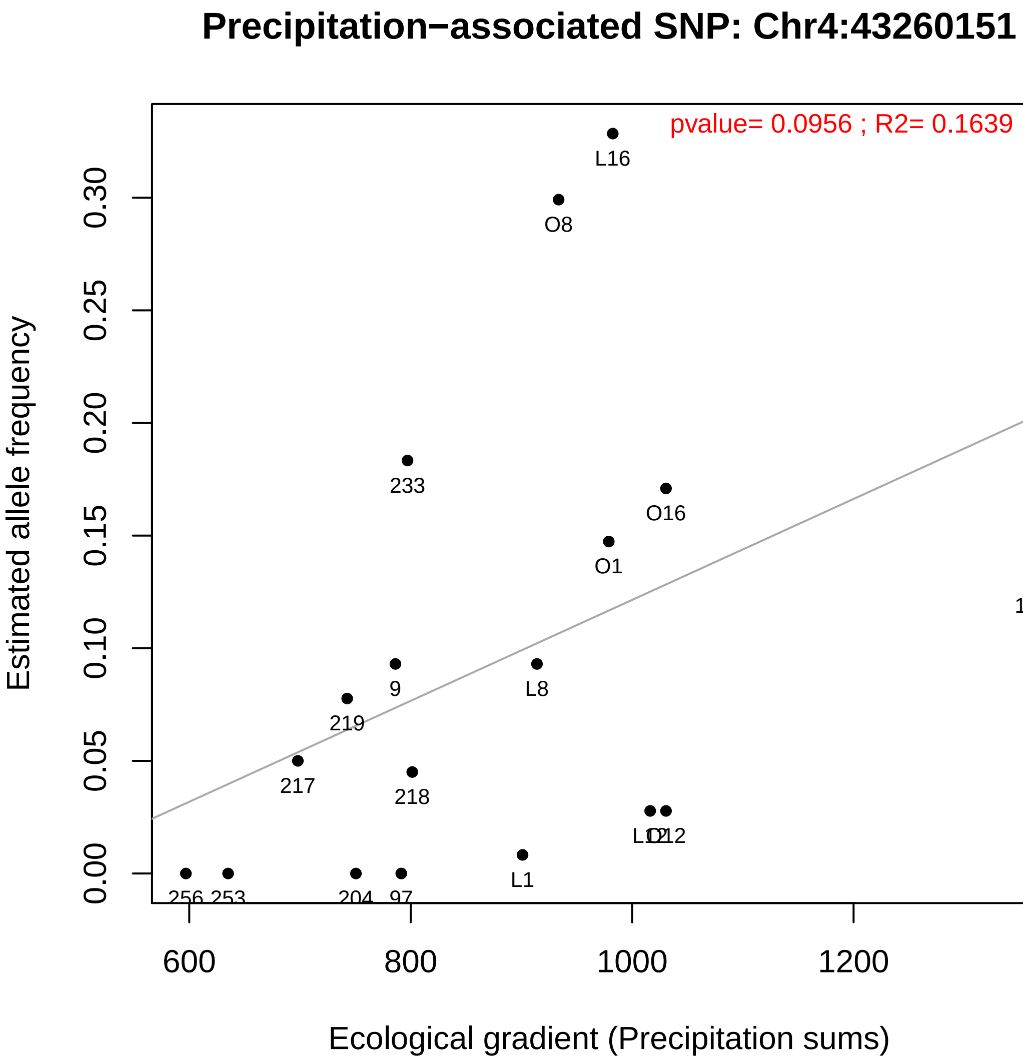
Precipitation−associated SNP: Chr4:43260151
pvalue= 0.0956 ; R2= 0.1639
L16
O8
233
O16
O1
124
9
L8
219
217
218
L1
256
253
204
97
0.00
0.05
0.10
0.15
0.20
0.25
0.30
Estimated allele frequency
600
800
1000
1200
Ecological gradient (Precipitation sums)

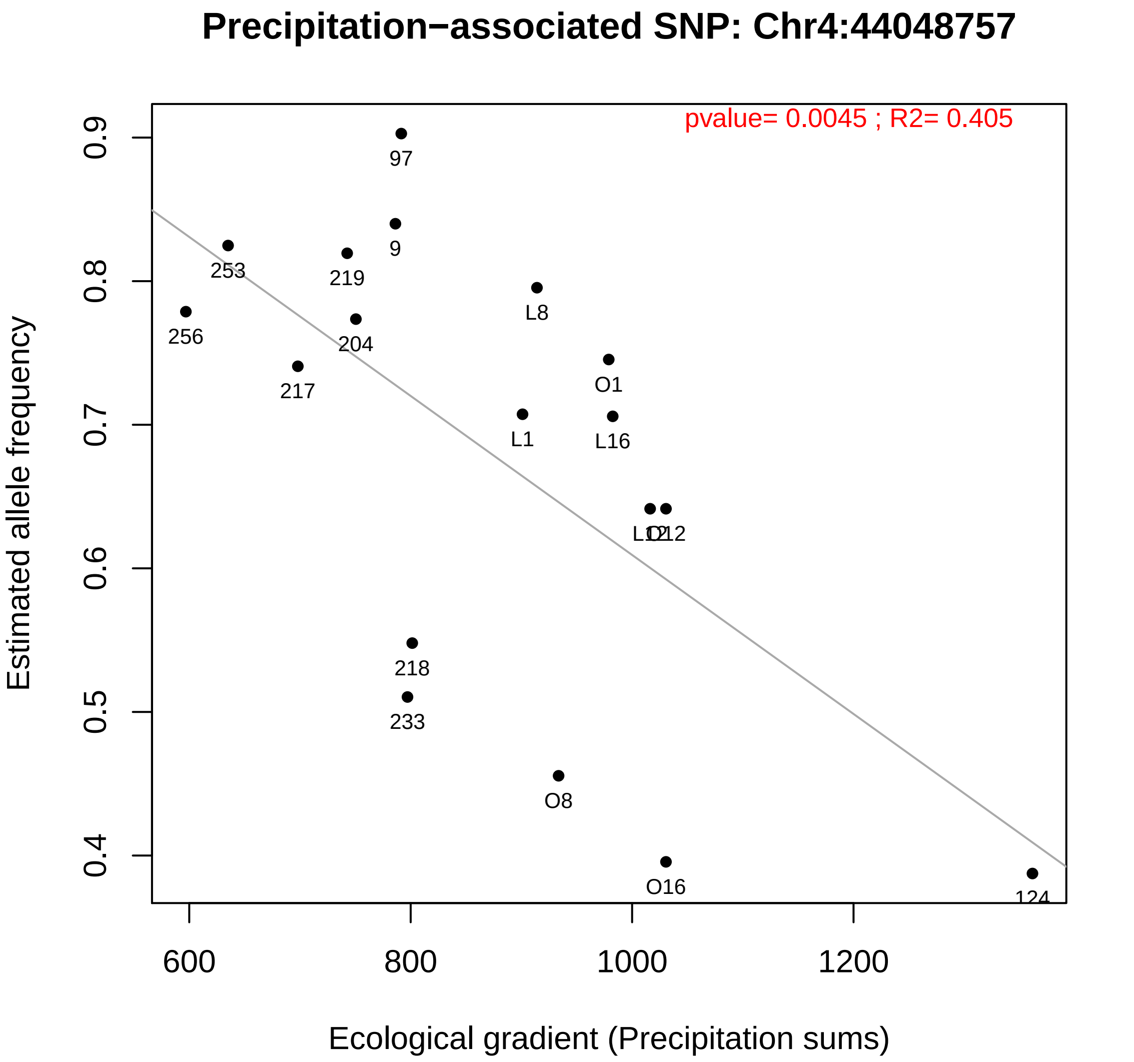
Precipitation−associated SNP: Chr4:44048757
pvalue= 0.0045 ; R2= 0.405
Estimated allele frequency
0.9
0.8
0.7
0.6
0.5
0.4
600
800
1000
1200
Ecological gradient (Precipitation sums)
97
9
253
219
256
204
217
L8
O1
L1
L16
218
233
O8
O16
124

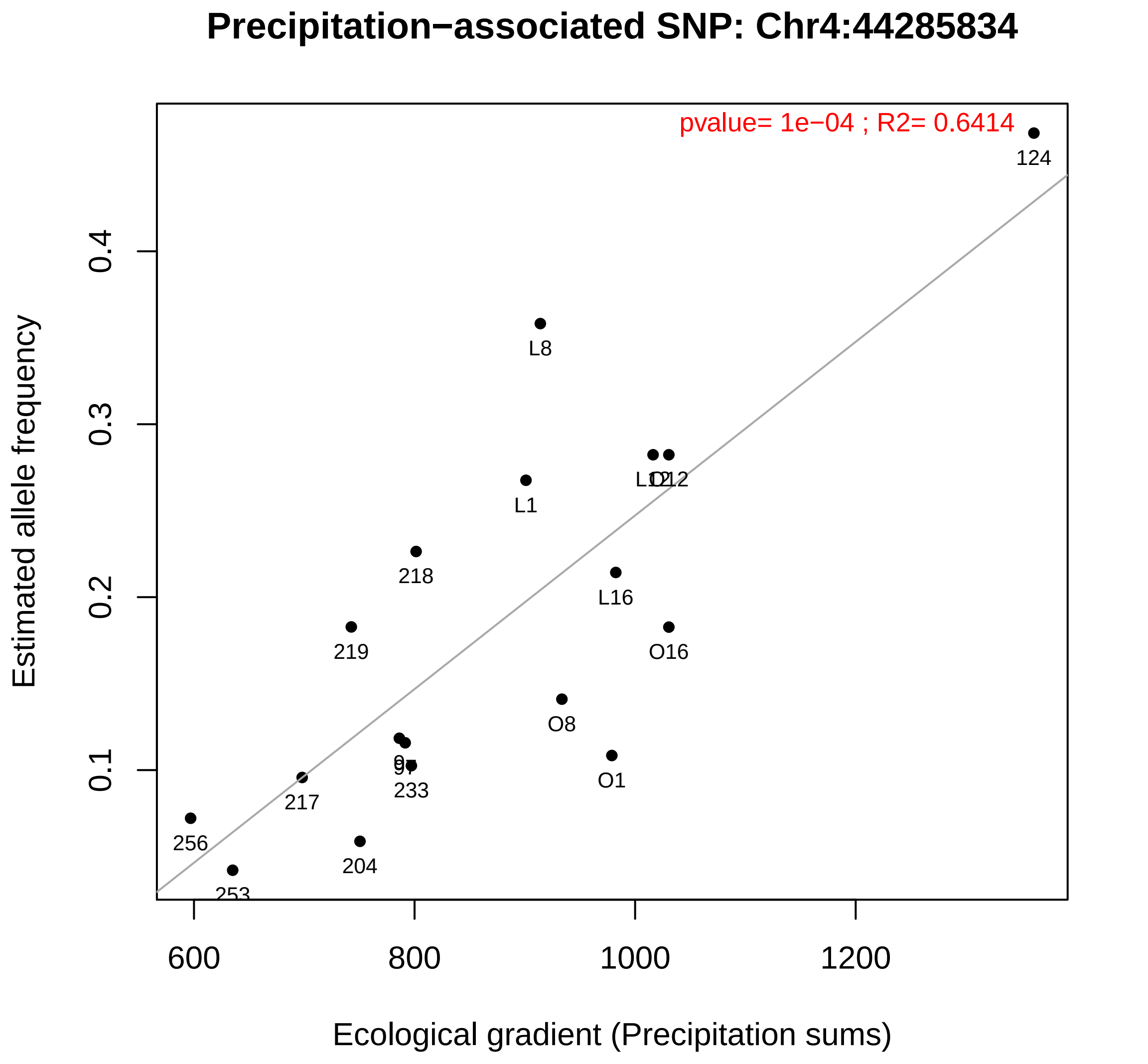

Precipitation−associated SNP: Chr4:44285834
pvalue= 1e−04 ; R2= 0.6414
124
L8
L12
O12
L1
218
L16
219
O16
O8
9
97
233
O1
217
256
204
253
0.1
0.2
0.3
0.4
600
800
1000
1200
Estimated allele frequency
Ecological gradient (Precipitation sums)

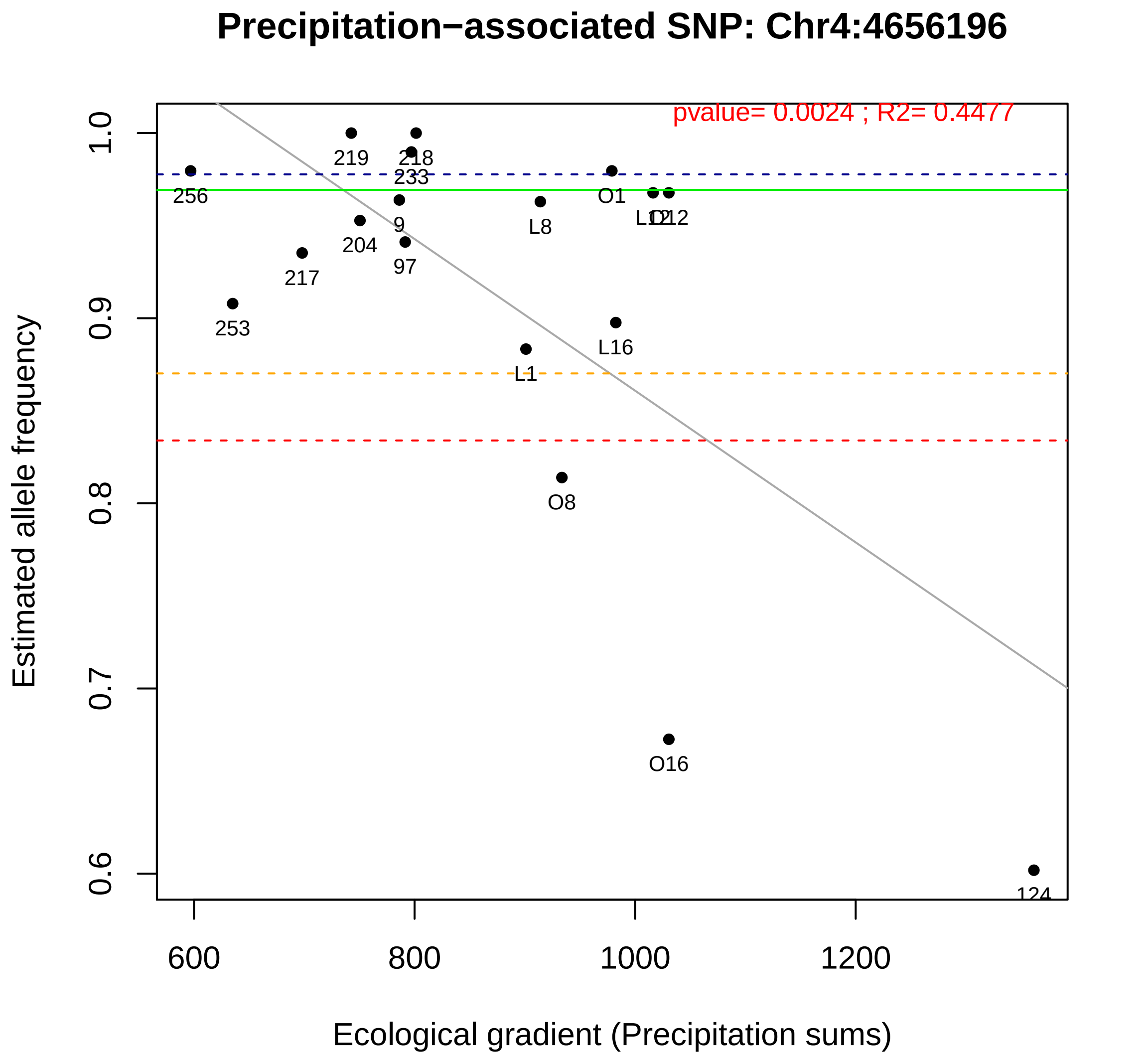

Precipitation−associated SNP: Chr4:4656196
pvalue= 0.0024 ; R2= 0.4477
Estimated allele frequency
Ecological gradient (Precipitation sums)
0.6
0.7
0.8
0.9
1.0
600
800
1000
1200
256
253
217
219
204
233
218
9
97
L1
L8
O8
O1
L16
L12
O12
O16
124

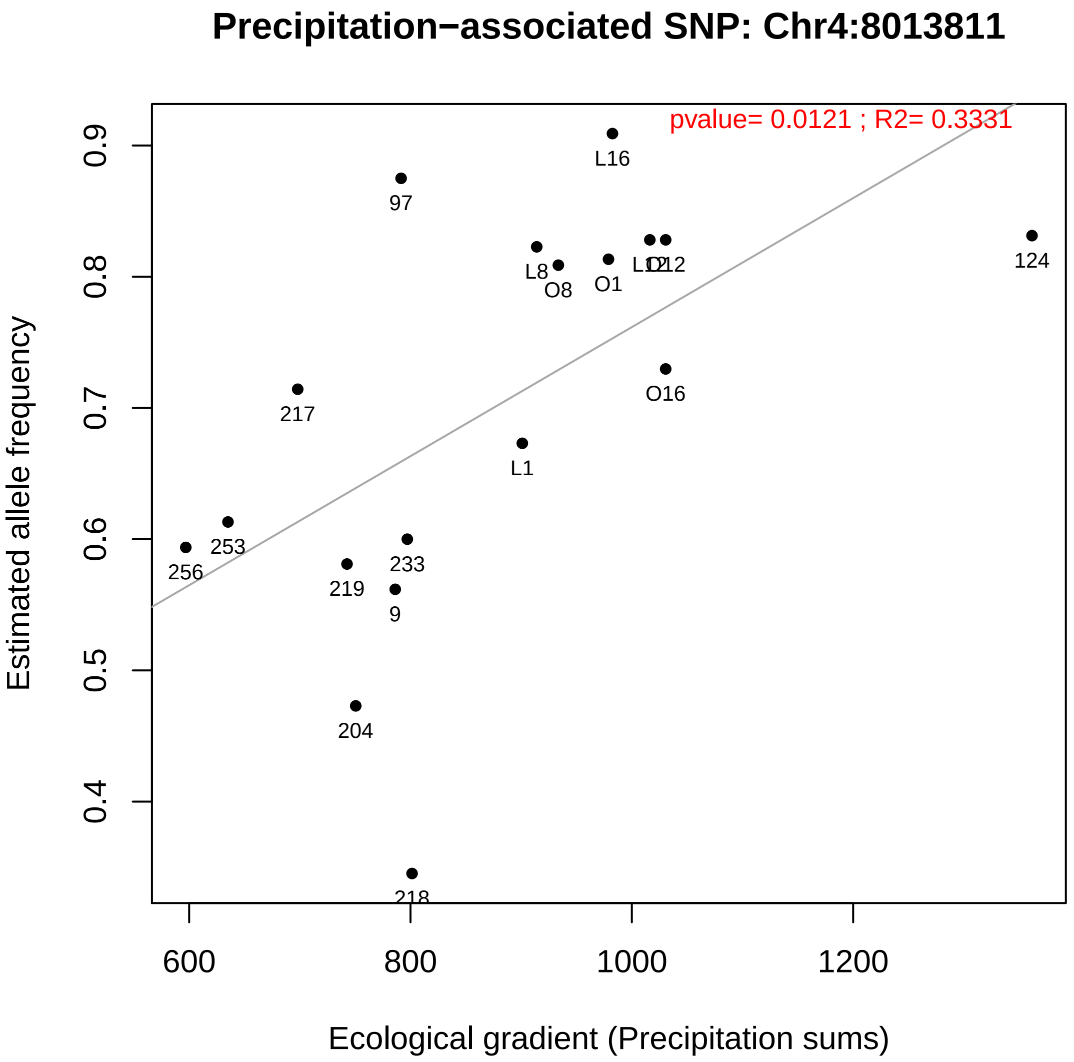

Precipitation−associated SNP: Chr4:8013811
pvalue= 0.0121 ; R2= 0.3331
Estimated allele frequency
0.4
0.5
0.6
0.7
0.8
0.9
600
800
1000
1200
Ecological gradient (Precipitation sums)
L16
97
L8
O8
O1
L12
O12
124
O16
217
L1
253
256
219
233
9
204
218

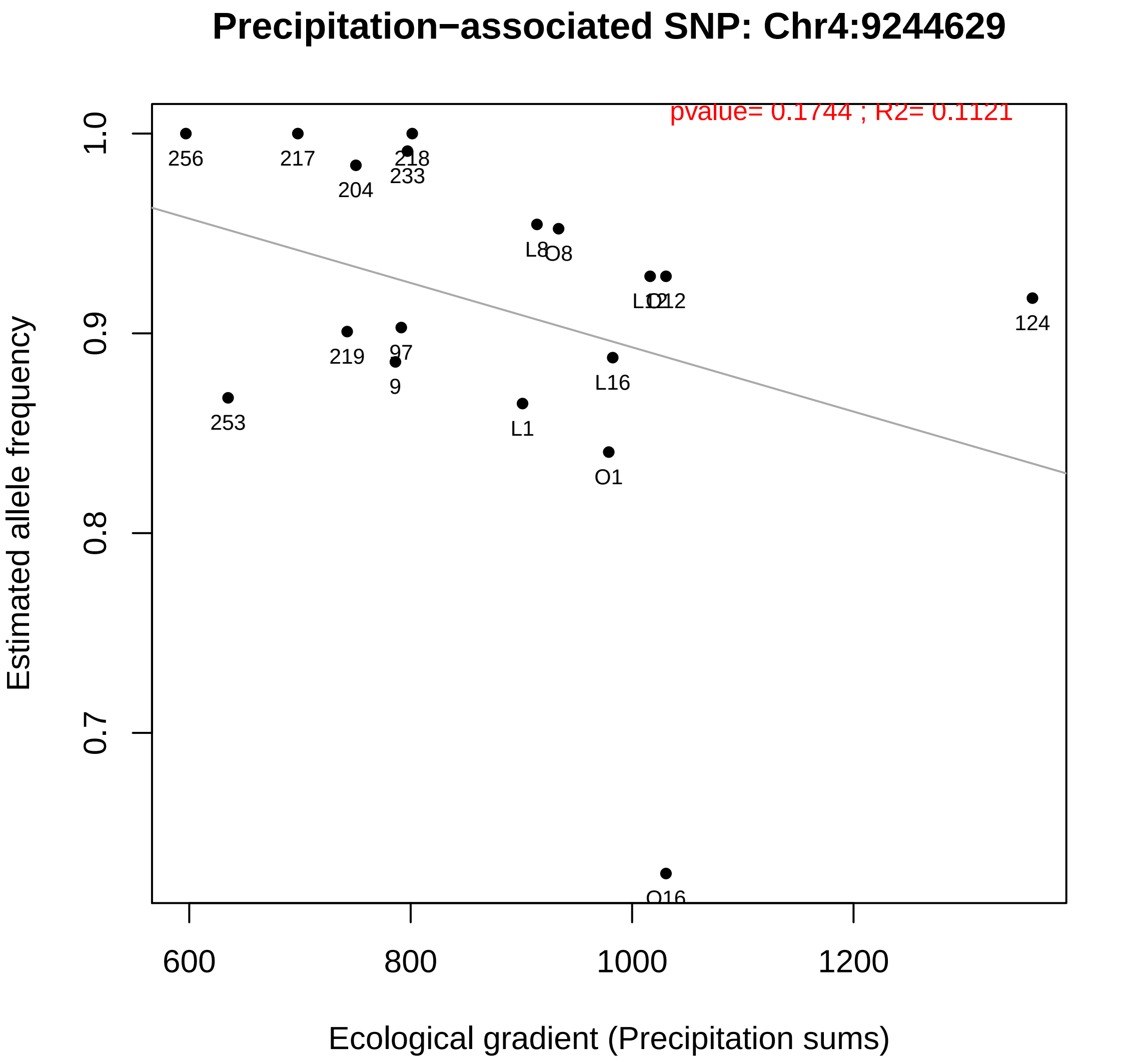

Precipitation−associated SNP: Chr4:9244629
pvalue= 0.1744 ; R2= 0.1121
Estimated allele frequency
1.0
0.9
0.8
0.7
600
800
1000
1200
Ecological gradient (Precipitation sums)
256
217
218
233
204
L8
O8
L12
O12
124
219
97
9
253
L1
L16
O1
O16

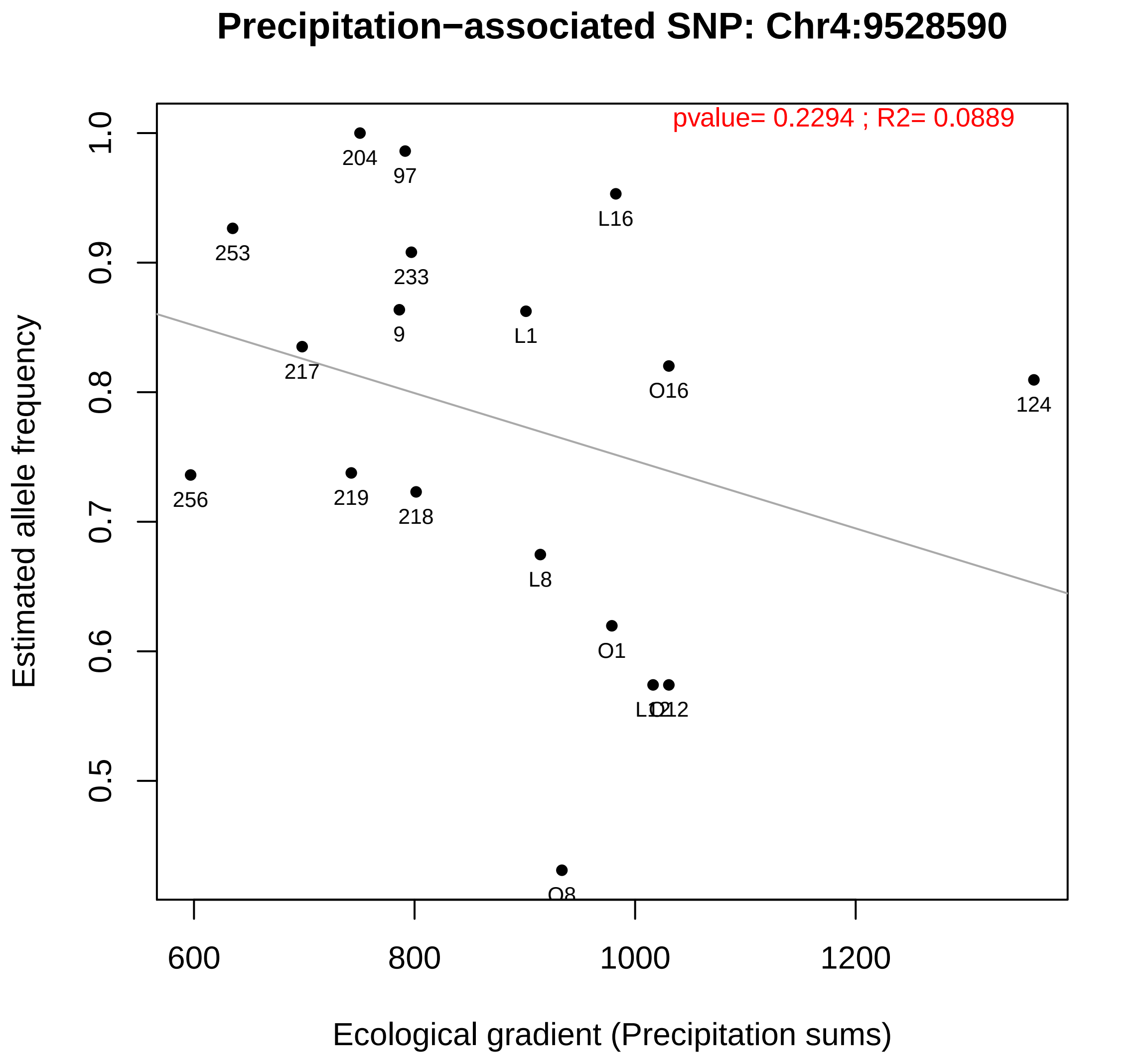
Precipitation−associated SNP: Chr4:9528590
pvalue= 0.2294 ; R2= 0.0889
Estimated allele frequency
Ecological gradient (Precipitation sums)
0.5
0.6
0.7
0.8
0.9
1.0
600
800
1000
1200
204
97
L16
253
233
9
L1
217
O16
124
256
219
218
L8
O1
L12
O12
O8

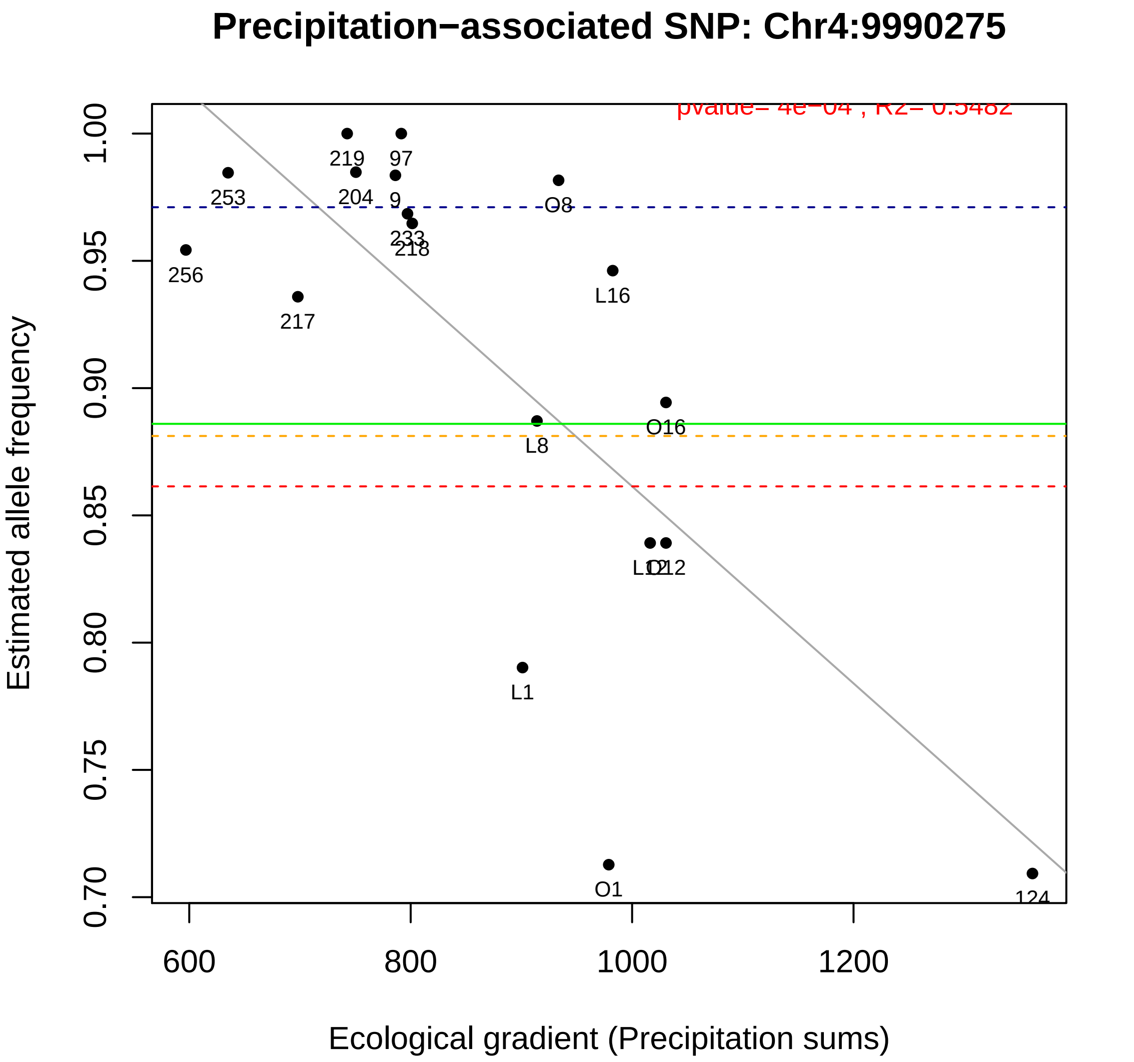

Precipitation−associated SNP: Chr4:9990275
pvalue= 4e−04 , R2= 0.5482
Estimated allele frequency
Ecological gradient (Precipitation sums)
0.70
0.75
0.80
0.85
0.90
0.95
1.00
600
800
1000
1200
219
97
253
204
9
O8
233
218
256
217
L16
O16
L8
L12
O12
L1
O1
124

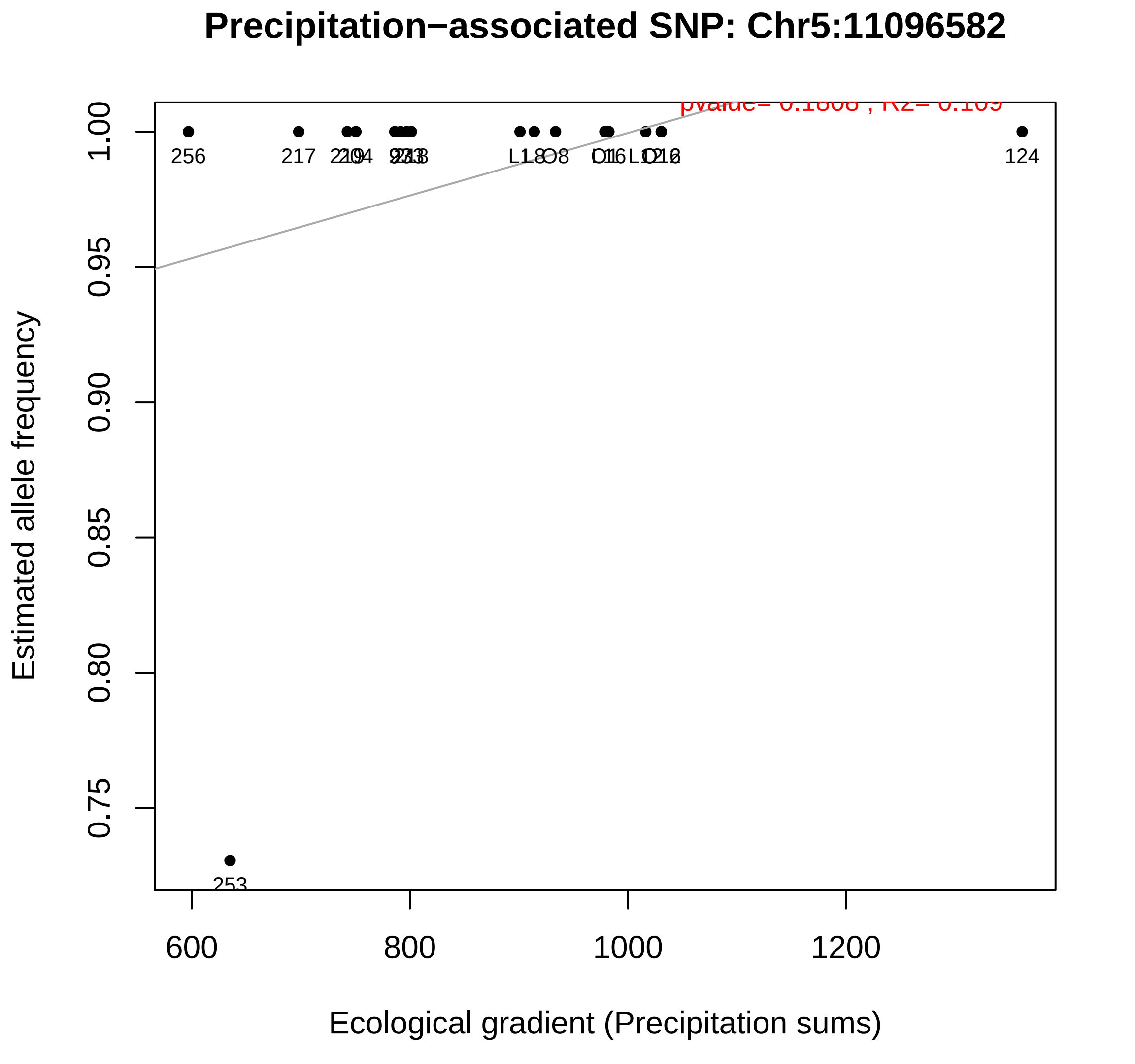
Precipitation−associated SNP: Chr5:11096582
pvalue= 0.1808 ; R2= 0.109
Estimated allele frequency
Ecological gradient (Precipitation sums)
0.75
0.80
0.85
0.90
0.95
1.00
600
800
1000
1200
256
217
124
253

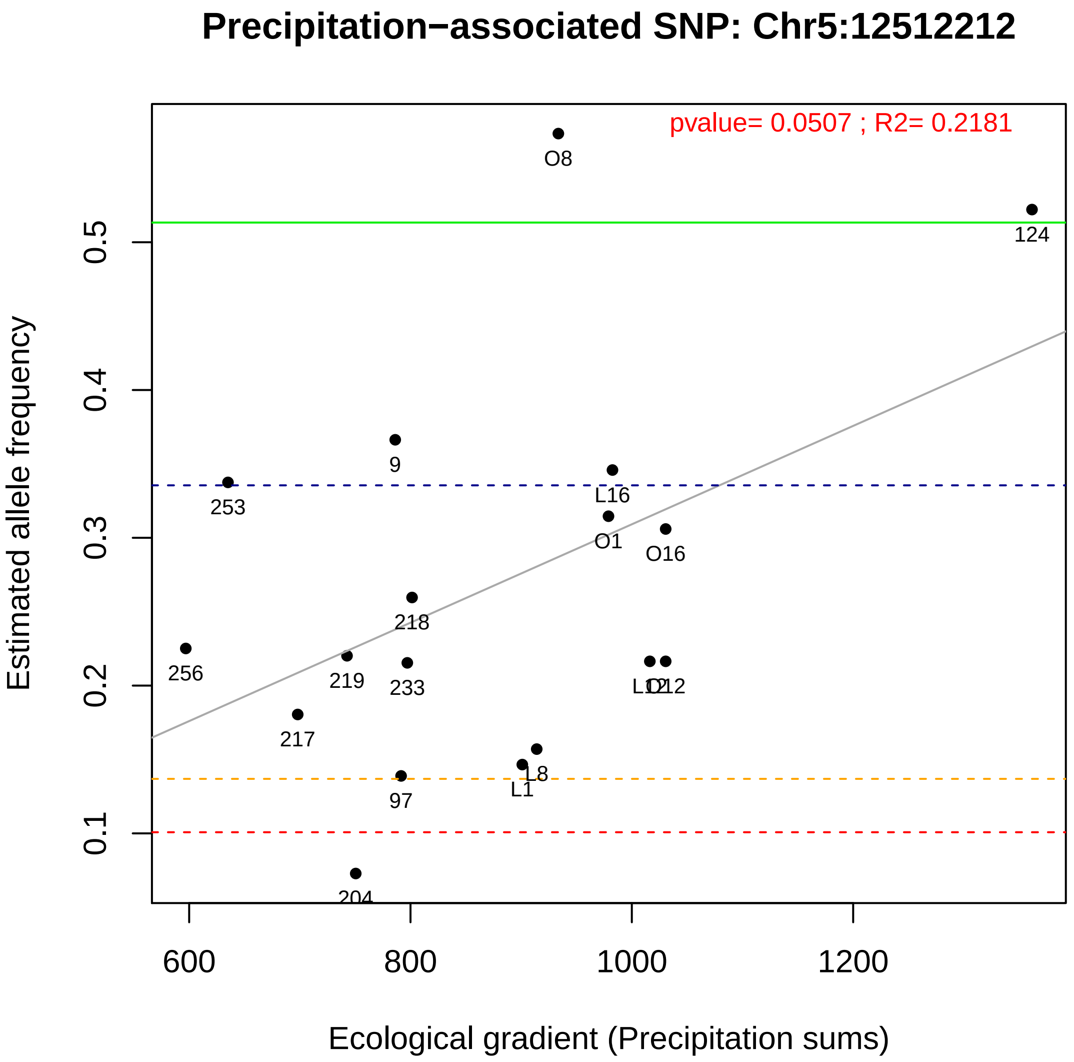

Precipitation−associated SNP: Chr5:12512212
pvalue= 0.0507 ; R2= 0.2181
Estimated allele frequency
0.1
0.2
0.3
0.4
0.5
600
800
1000
1200
Ecological gradient (Precipitation sums)
O8
124
9
253
L16
O1
O16
218
256
219
233
L12
O12
217
L8
L1
97
204

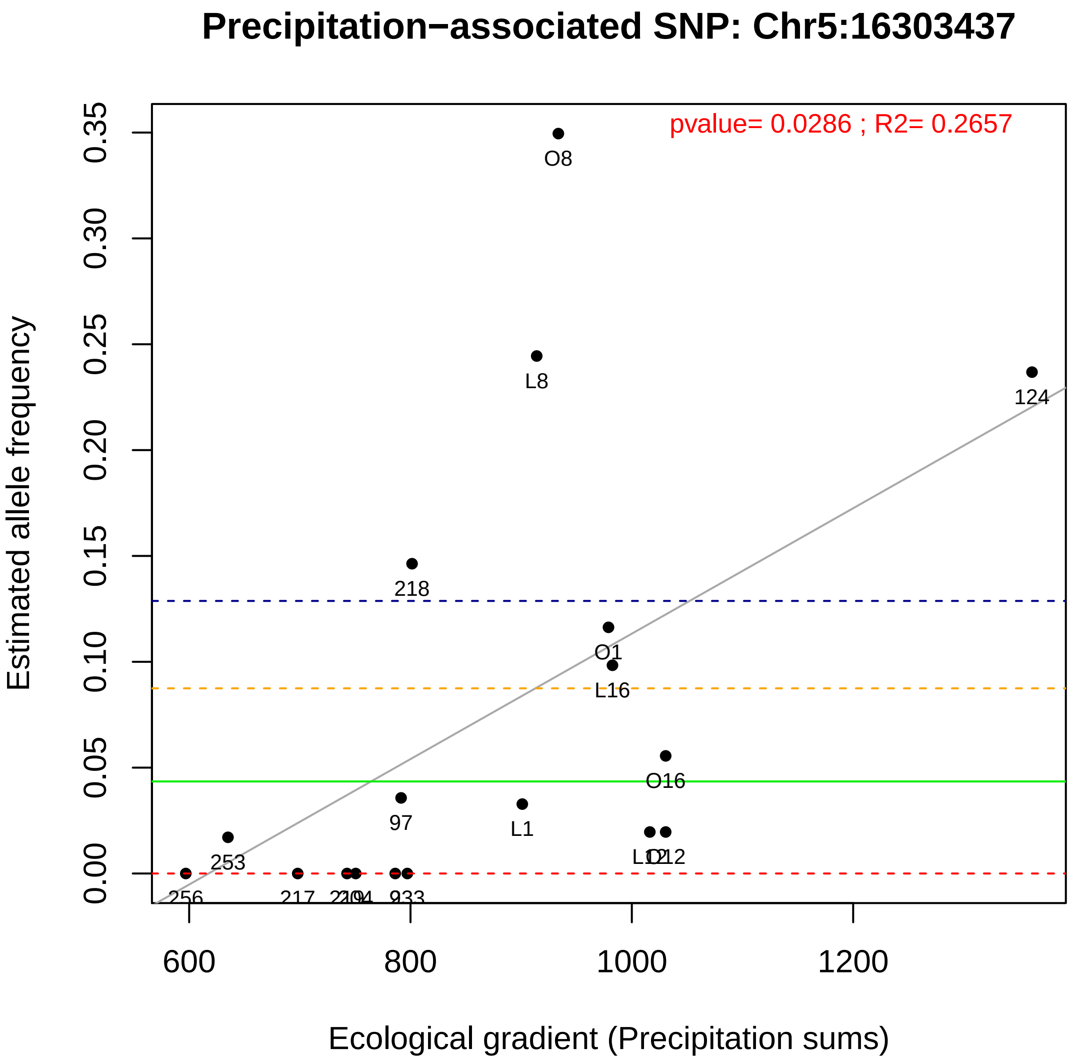
Precipitation−associated SNP: Chr5:16303437
pvalue= 0.0286 ; R2= 0.2657
Estimated allele frequency
Ecological gradient (Precipitation sums)
0.00
0.05
0.10
0.15
0.20
0.25
0.30
0.35
600
800
1000
1200
O8
L8
124
218
O1
L16
O16
97
L1
253
O12
256
217

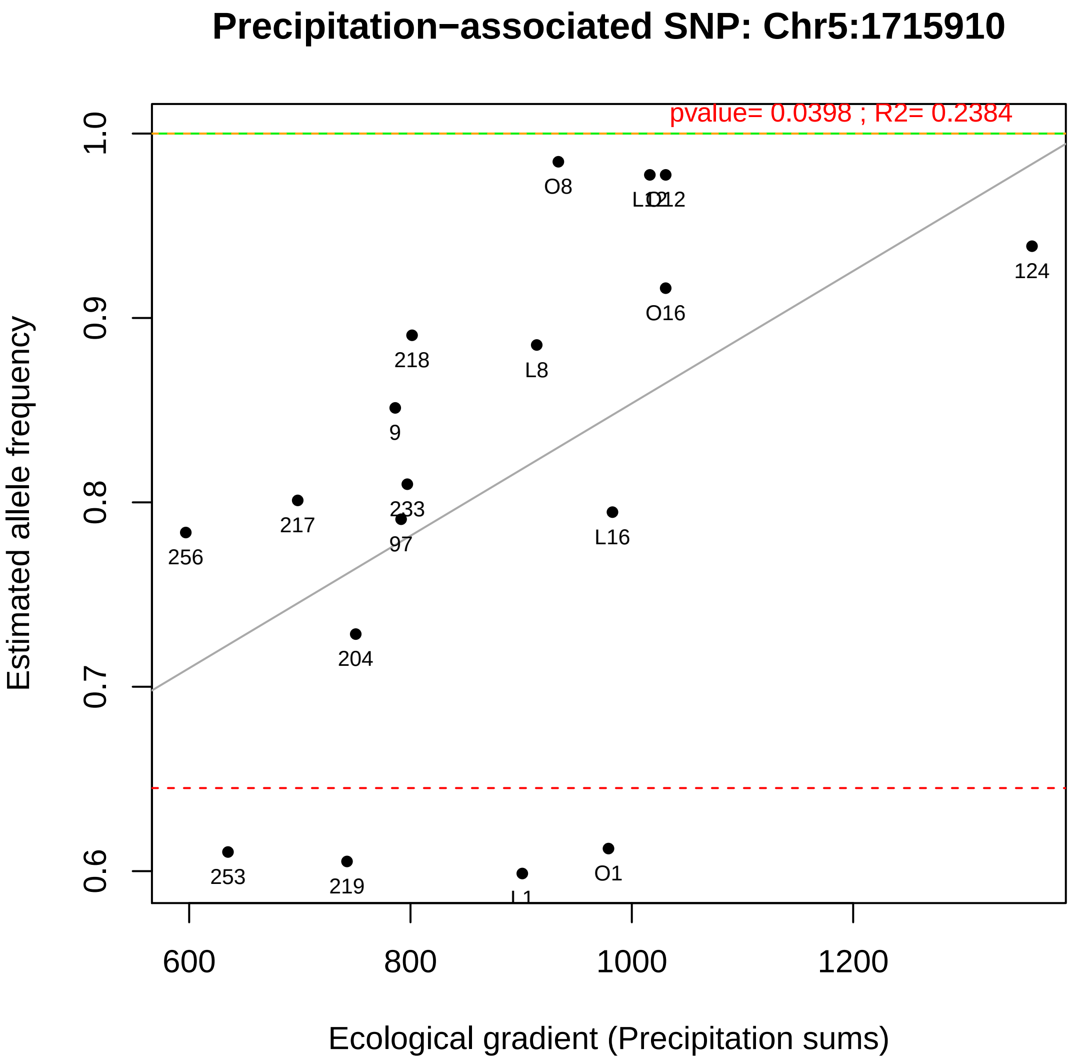

Precipitation−associated SNP: Chr5:1715910
pvalue= 0.0398 ; R2= 0.2384
Estimated allele frequency
0.6
0.7
0.8
0.9
1.0
600
800
1000
1200
Ecological gradient (Precipitation sums)
O8
124
O16
218
L8
9
233
217
97
L16
256
204
253
219
O1

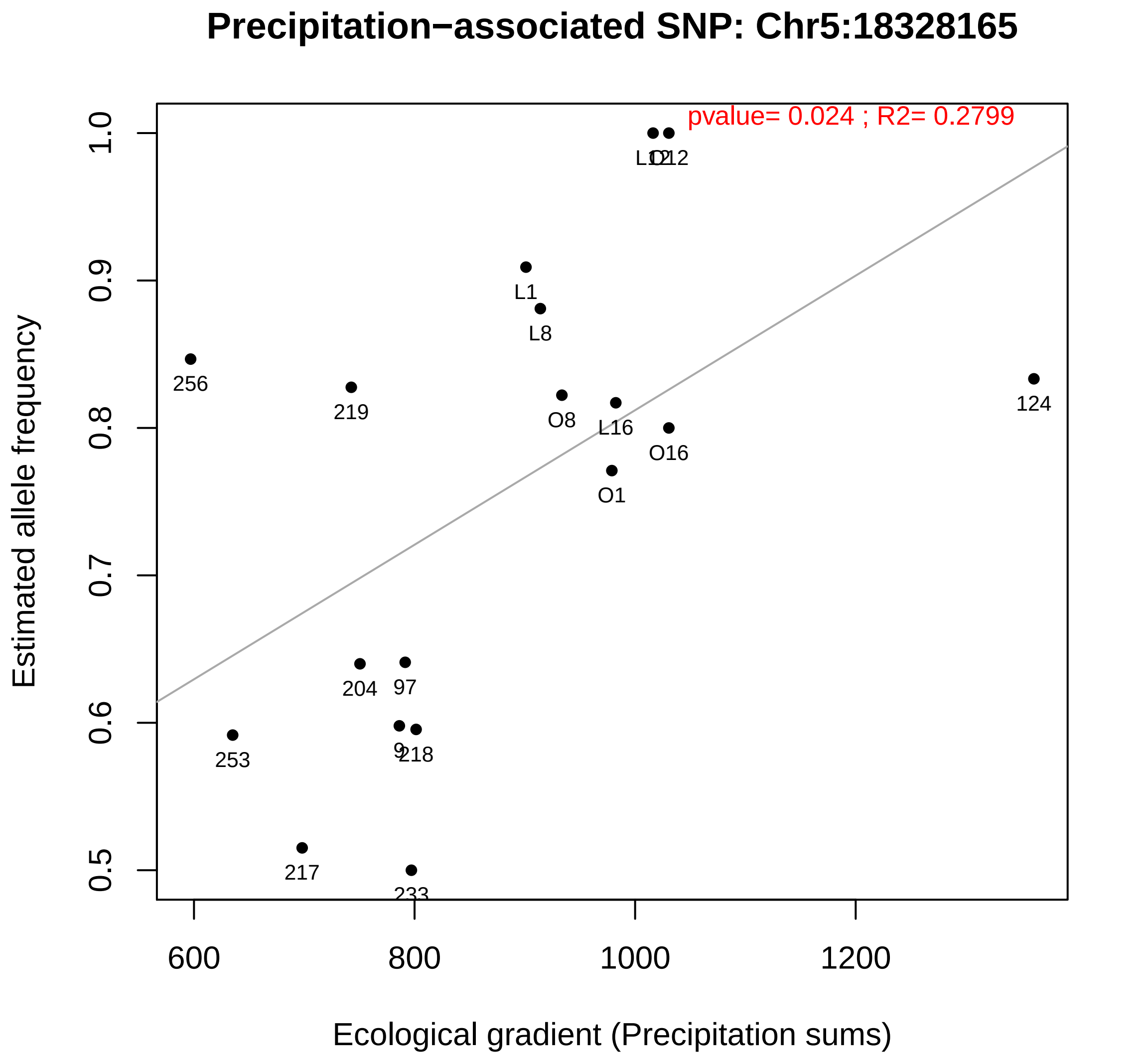

Precipitation−associated SNP: Chr5:18328165
pvalue= 0.024 ; R2= 0.2799
Estimated allele frequency
Ecological gradient (Precipitation sums)
1.0
0.9
0.8
0.7
0.6
0.5
600
800
1000
1200
L12
O12
L1
L8
256
219
O8
L16
O16
124
O1
204
97
253
9
218
217
233

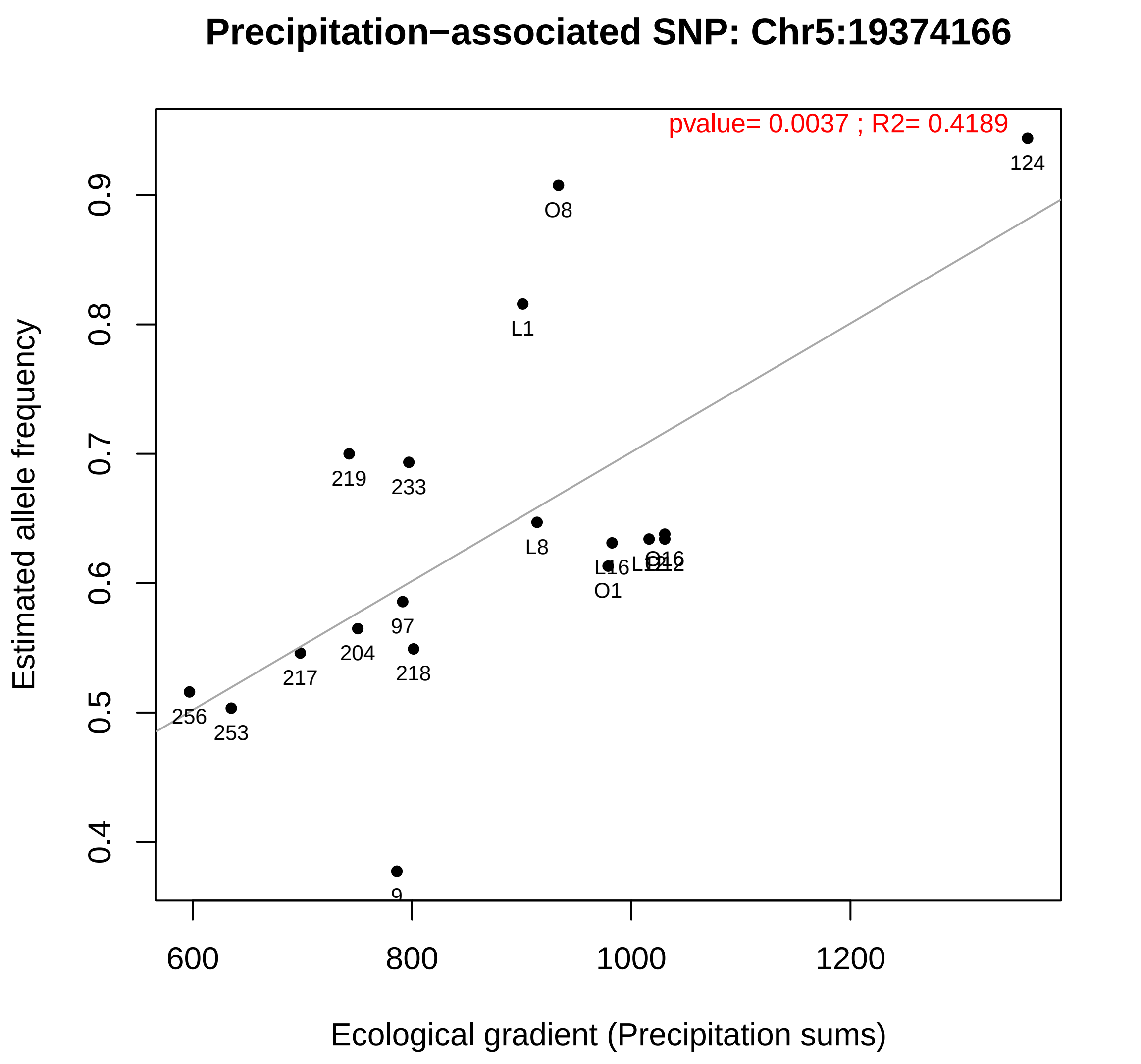
Precipitation−associated SNP: Chr5:19374166
pvalue= 0.0037 ; R2= 0.4189
Estimated allele frequency
Ecological gradient (Precipitation sums)
0.4
0.5
0.6
0.7
0.8
0.9
600
800
1000
1200
124
O8
L1
219
233
L8
L16
O1
O16
L12
97
204
218
217
256
253
9

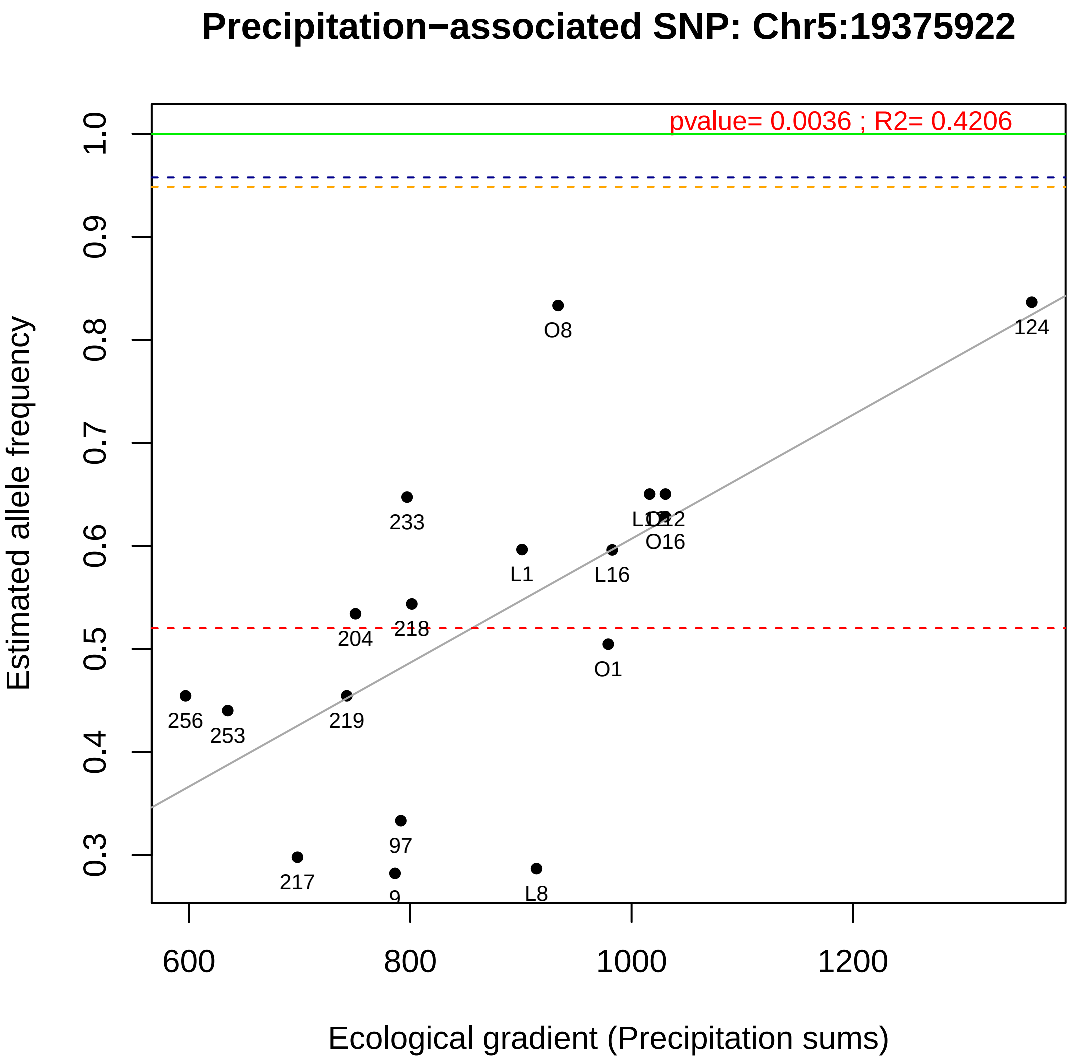
Precipitation−associated SNP: Chr5:19375922
pvalue= 0.0036 ; R2= 0.4206
Estimated allele frequency
0.3
0.4
0.5
0.6
0.7
0.8
0.9
1.0
600
800
1000
1200
Ecological gradient (Precipitation sums)
O8
124
233
L1
L16
O16
218
204
O1
256
253
219
97
217
L8

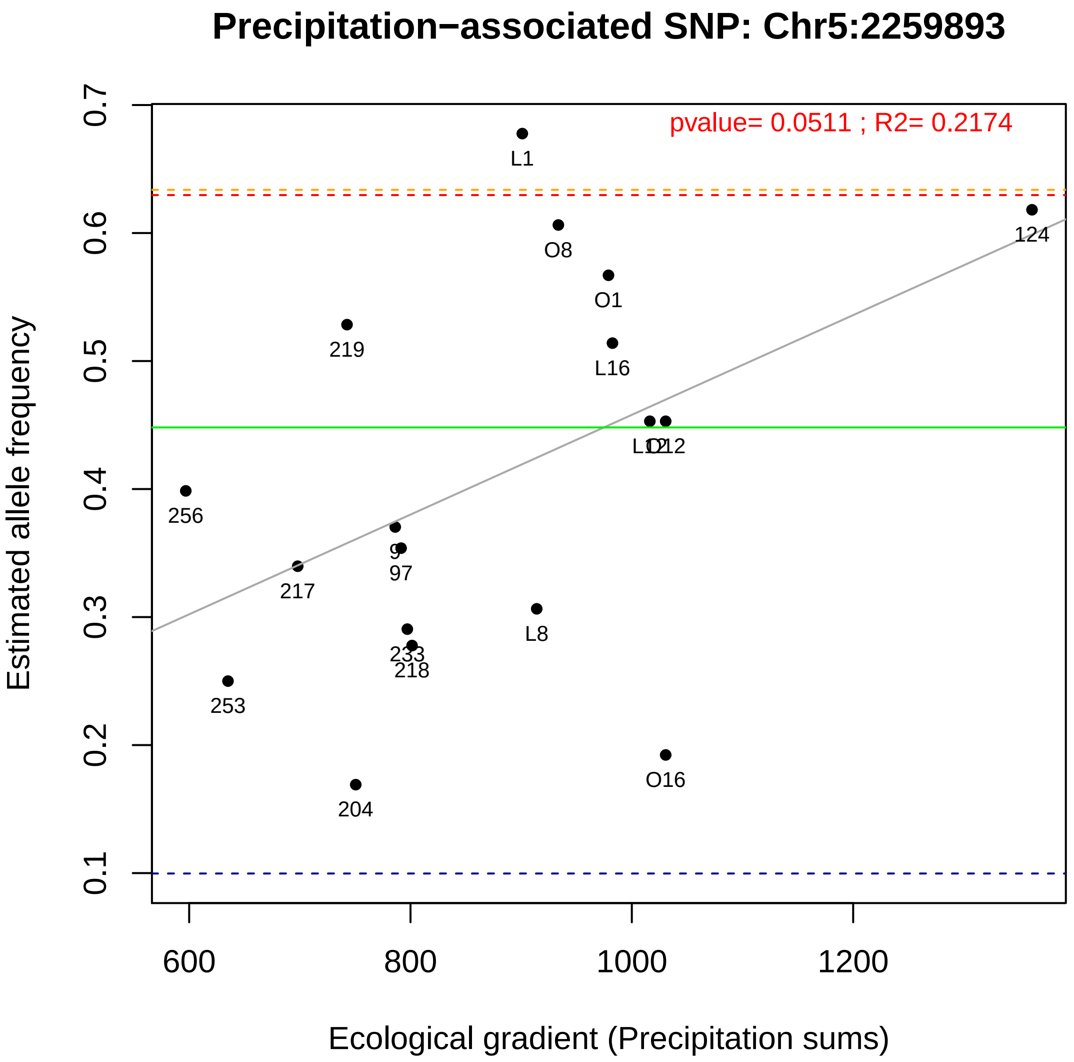

Precipitation−associated SNP: Chr5:2259893
pvalue= 0.0511 ; R2= 0.2174
Estimated allele frequency
0.1
0.2
0.3
0.4
0.5
0.6
0.7
600
800
1000
1200
Ecological gradient (Precipitation sums)
L1
O8
O1
219
L16
124
256
217
97
233
218
L8
253
204
O16

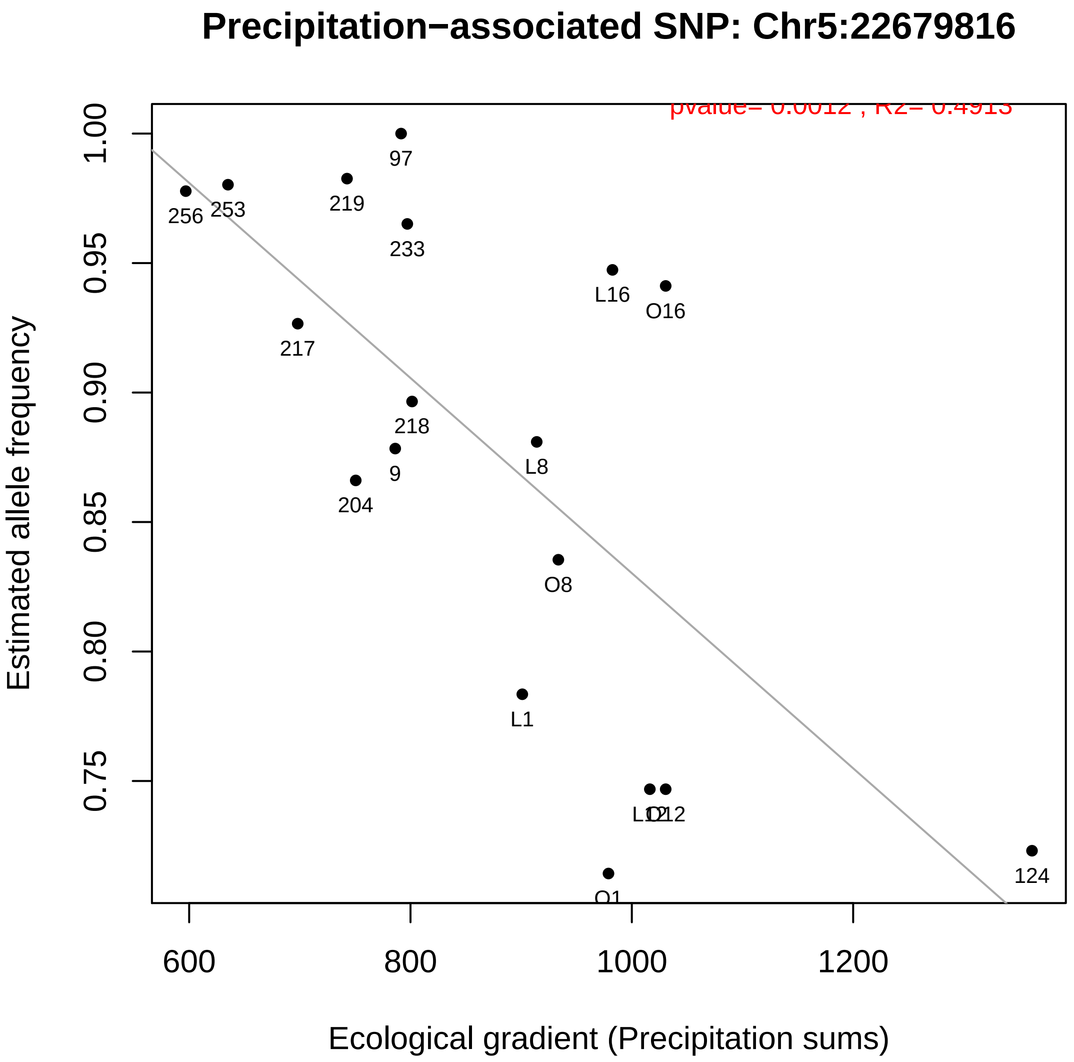

Precipitation−associated SNP: Chr5:22679816
pvalue= 0.0012 , R2= 0.4913
Estimated allele frequency
Ecological gradient (Precipitation sums)
0.75
0.80
0.85
0.90
0.95
1.00
600
800
1000
1200
97
256
253
219
233
L16
O16
217
218
L8
9
204
O8
L1
124

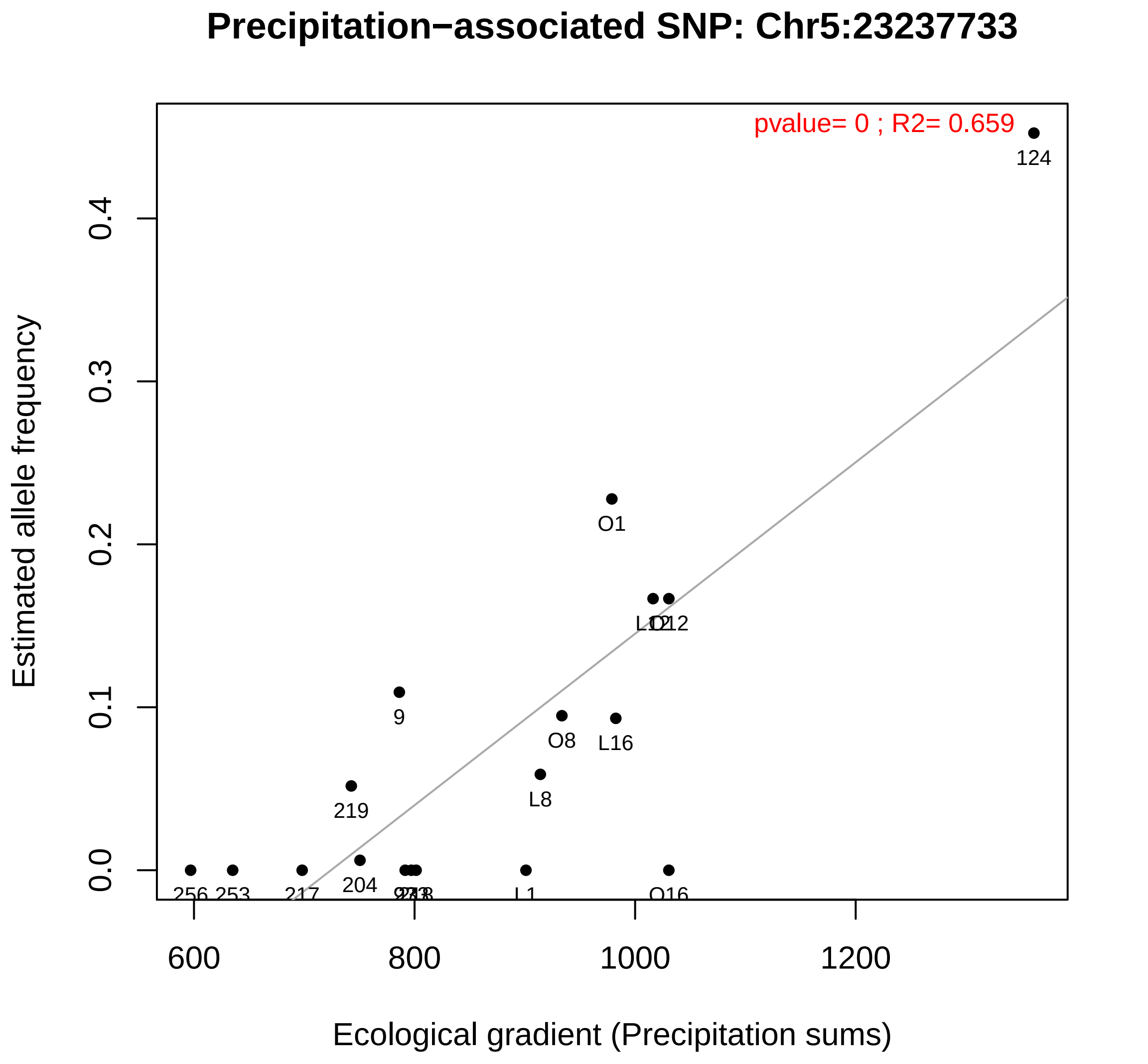

Precipitation−associated SNP: Chr5:23237733
pvalue= 0 ; R2= 0.659
124
O1
9
O8
L16
219
L8
204
256
253
217
L1
O16
0.0
0.1
0.2
0.3
0.4
Estimated allele frequency
600
800
1000
1200
Ecological gradient (Precipitation sums)

**Precipitation−associated SNP: Chr5:24517770**

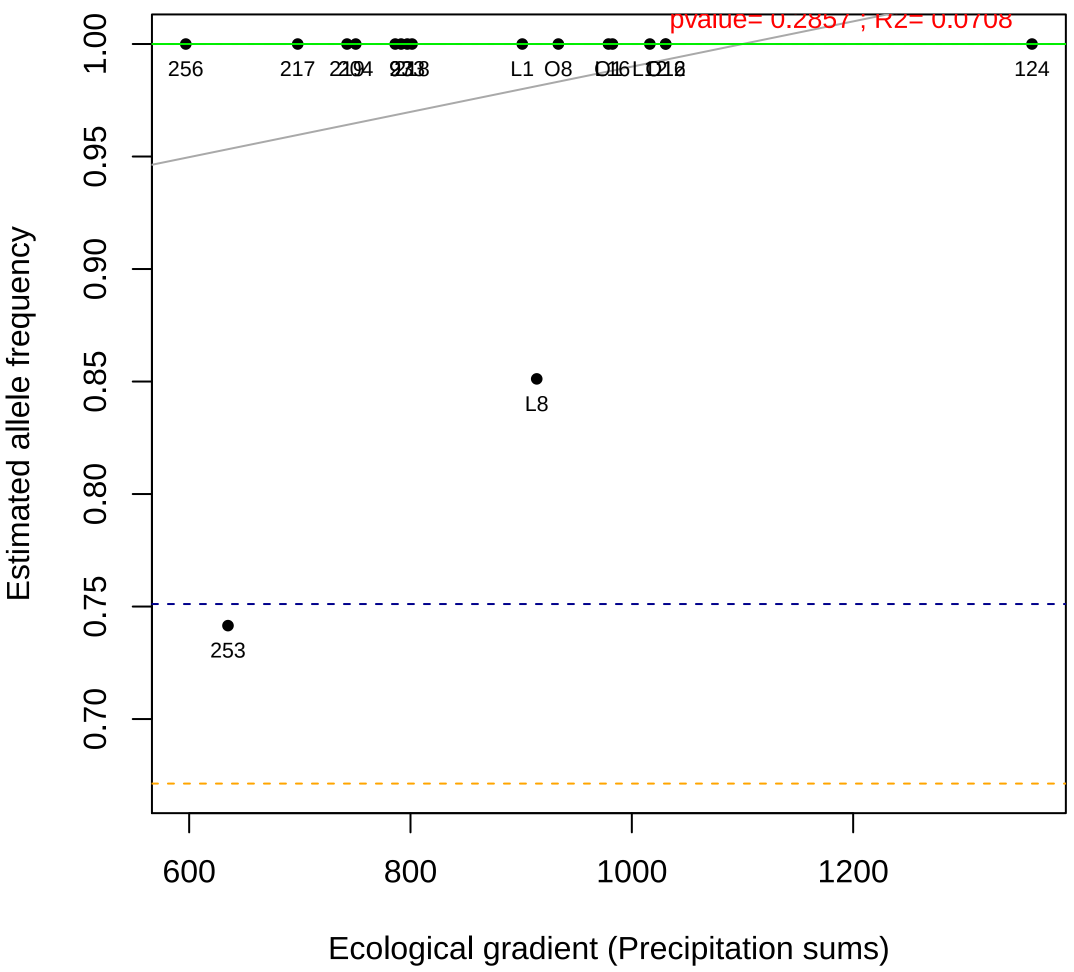

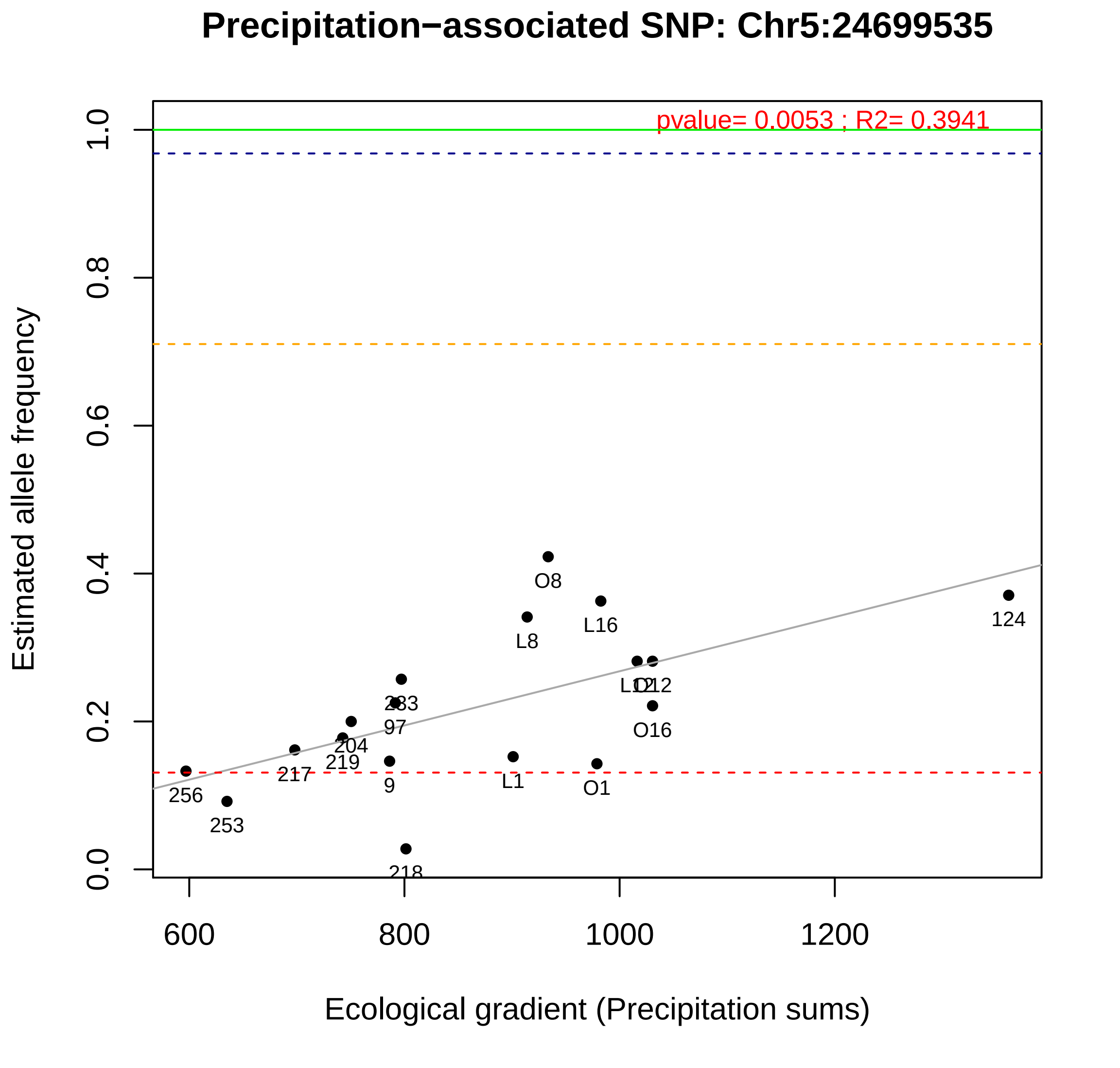

Precipitation−associated SNP: Chr5:24699535
pvalue= 0.0053 ; R2= 0.3941
Estimated allele frequency
0.0
0.2
0.4
0.6
0.8
1.0
600
800
1000
1200
Ecological gradient (Precipitation sums)
O8
L16
124
L8
L12
O12
233
97
O16
204
219
217
9
L1
O1
256
253
218

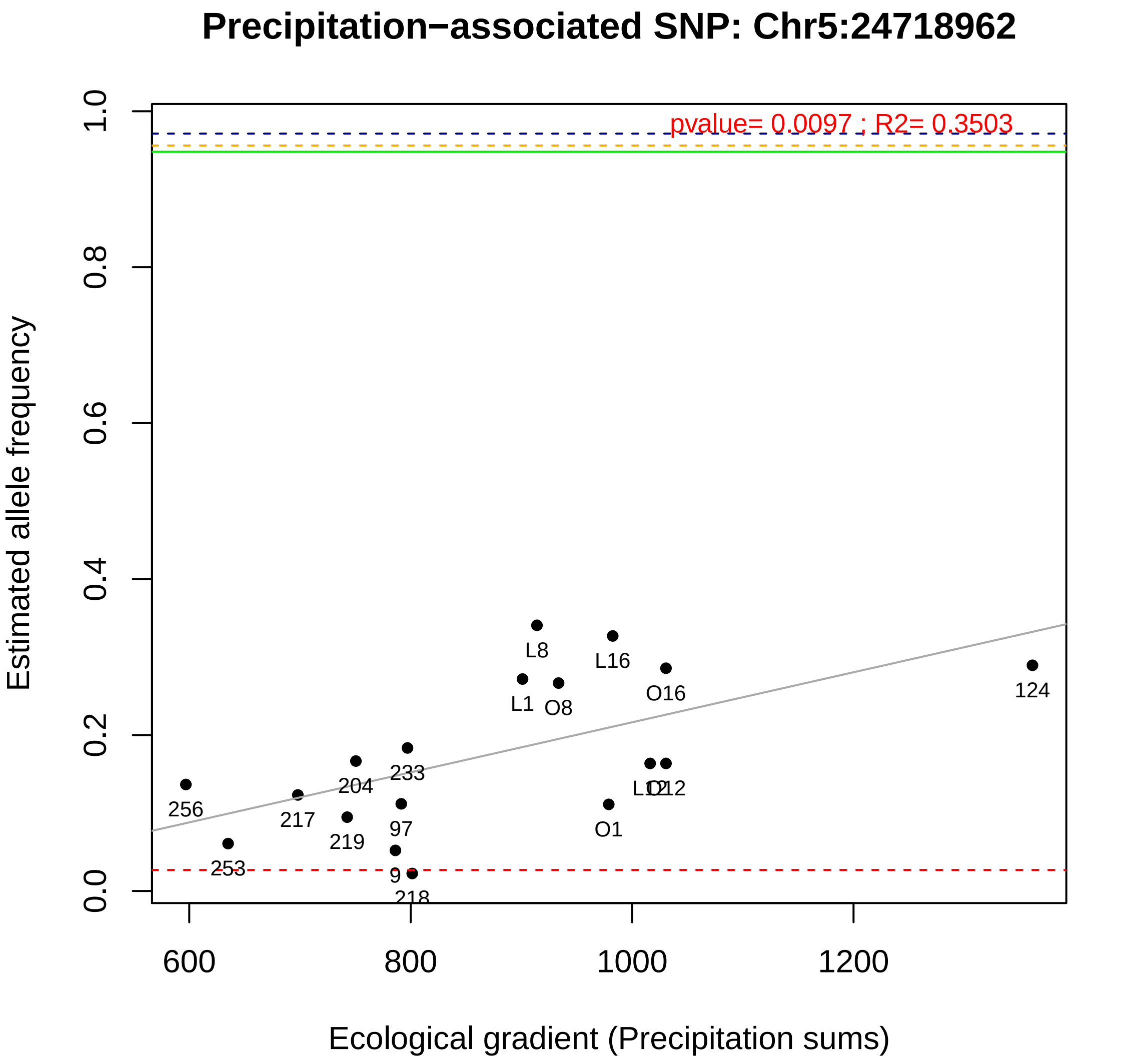

Precipitation−associated SNP: Chr5:24718962
pvalue= 0.0097 ; R2= 0.3503
Estimated allele frequency
0.0
0.2
0.4
0.6
0.8
1.0
600
800
1000
1200
Ecological gradient (Precipitation sums)
L8
L16
L1
O8
O16
124
233
204
256
217
97
219
253
9
218
O1
L12
O12

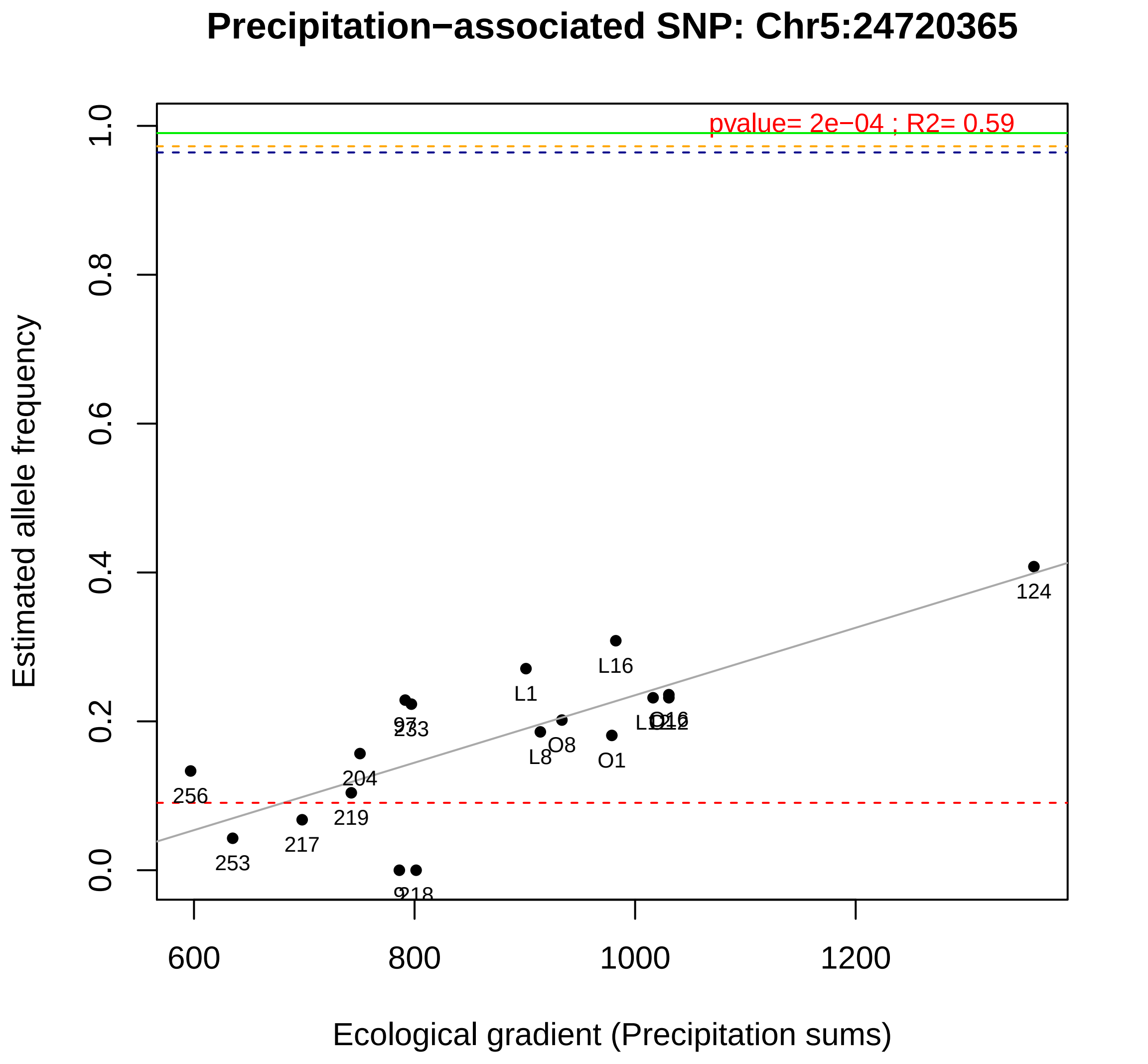

Precipitation−associated SNP: Chr5:24720365
pvalue= 2e−04 ; R2= 0.59
Estimated allele frequency
0.0
0.2
0.4
0.6
0.8
1.0
600
800
1000
1200
Ecological gradient (Precipitation sums)
124
L16
L1
O8
L8
O1
204
256
219
217
253

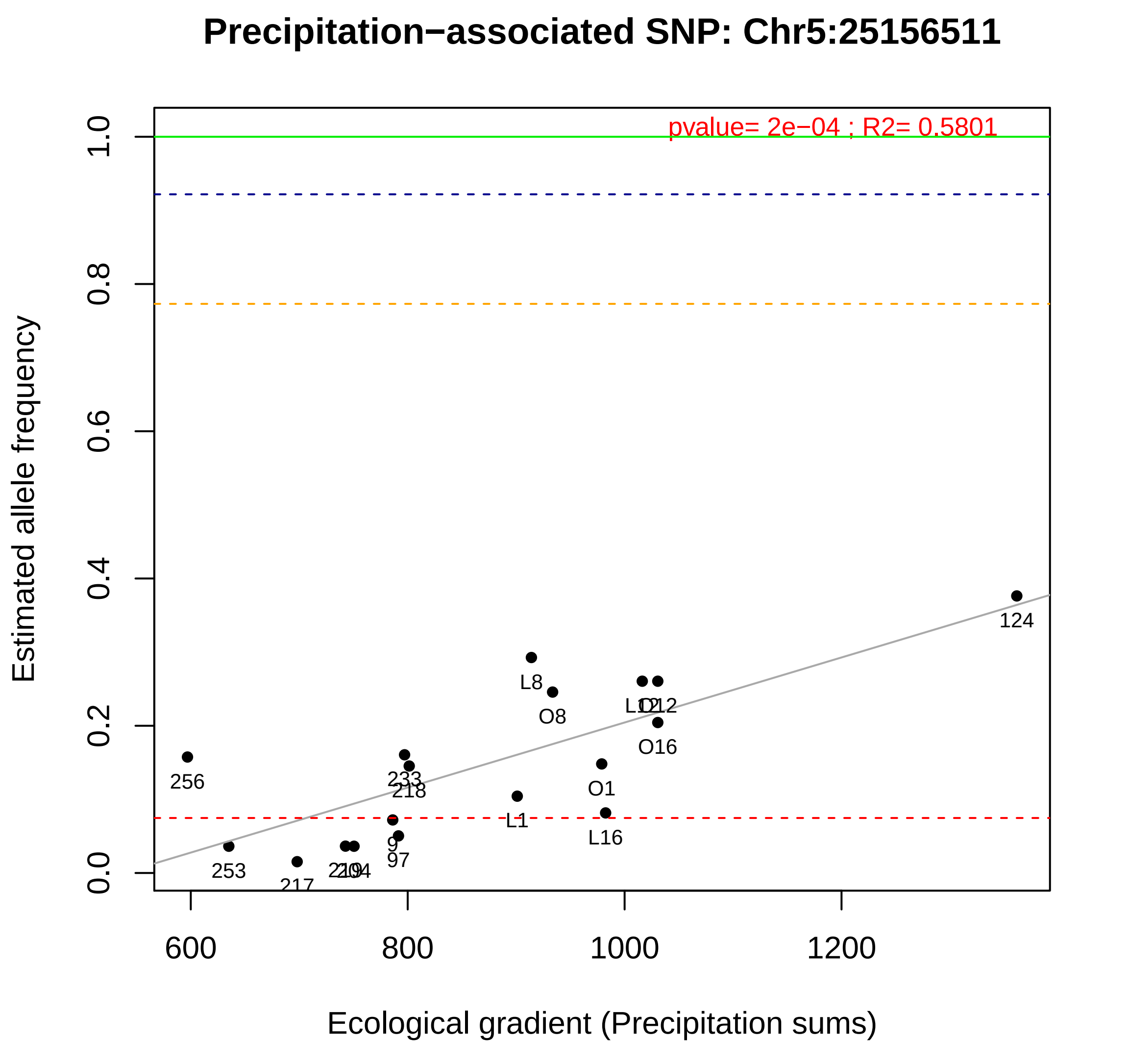
Precipitation−associated SNP: Chr5:25156511
pvalue= 2e−04 ; R2= 0.5801
Estimated allele frequency
0.0
0.2
0.4
0.6
0.8
1.0
600
800
1000
1200
Ecological gradient (Precipitation sums)
256
253
217
233
218
9
97
L1
L8
O8
O1
L16
O16
124

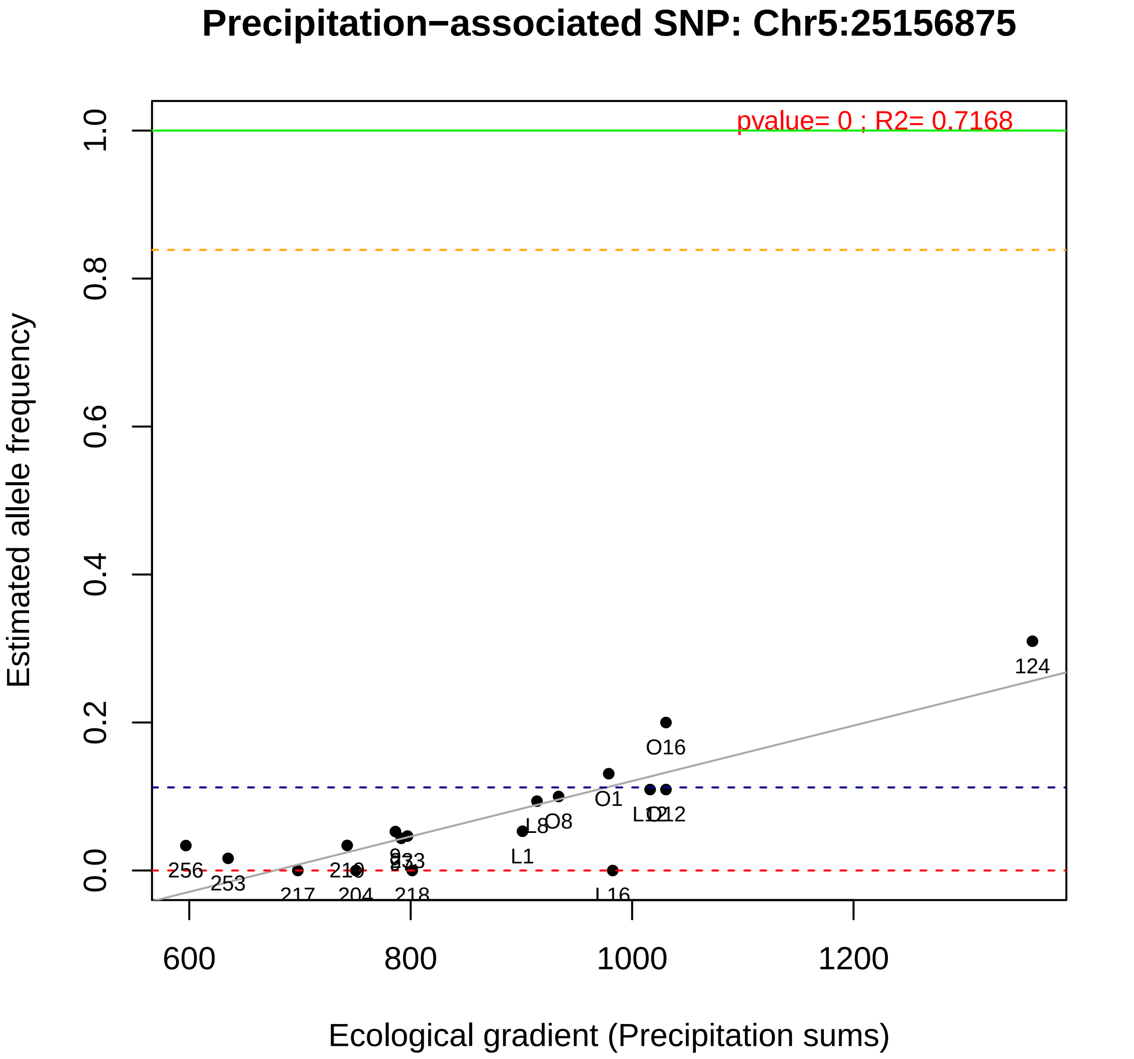
Precipitation−associated SNP: Chr5:25156875
pvalue= 0 ; R2= 0.7168
Estimated allele frequency
0.0
0.2
0.4
0.6
0.8
1.0
Ecological gradient (Precipitation sums)
600
800
1000
1200
124
O16
O1
O8
L8
L1
O12
256
253
217
204
218
L16

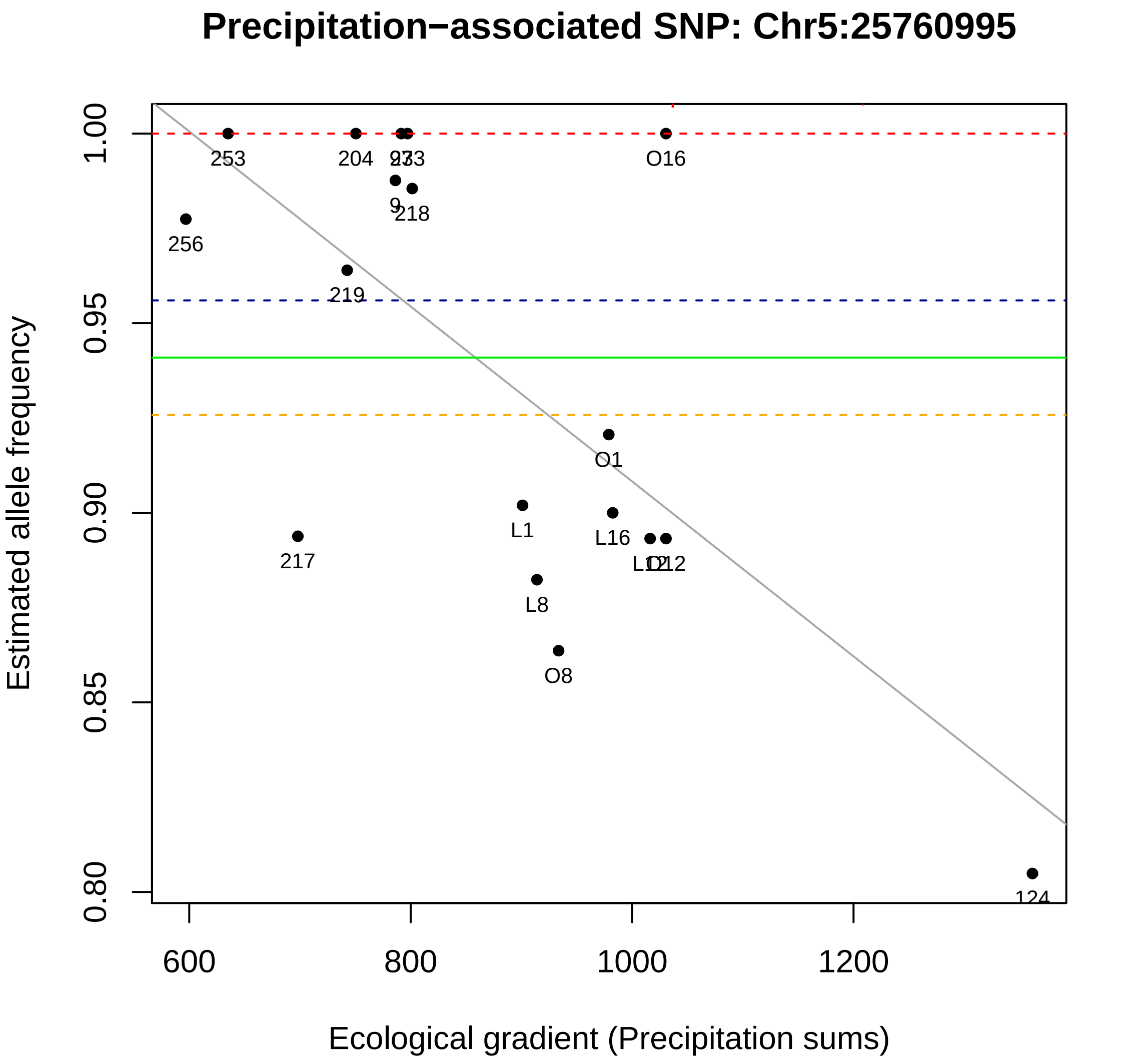

Precipitation−associated SNP: Chr5:25760995
Estimated allele frequency
Ecological gradient (Precipitation sums)
1.00
0.95
0.90
0.85
0.80
600
800
1000
1200
253
204
O16
9
218
256
219
O1
L1
L16
217
L8
O8
124

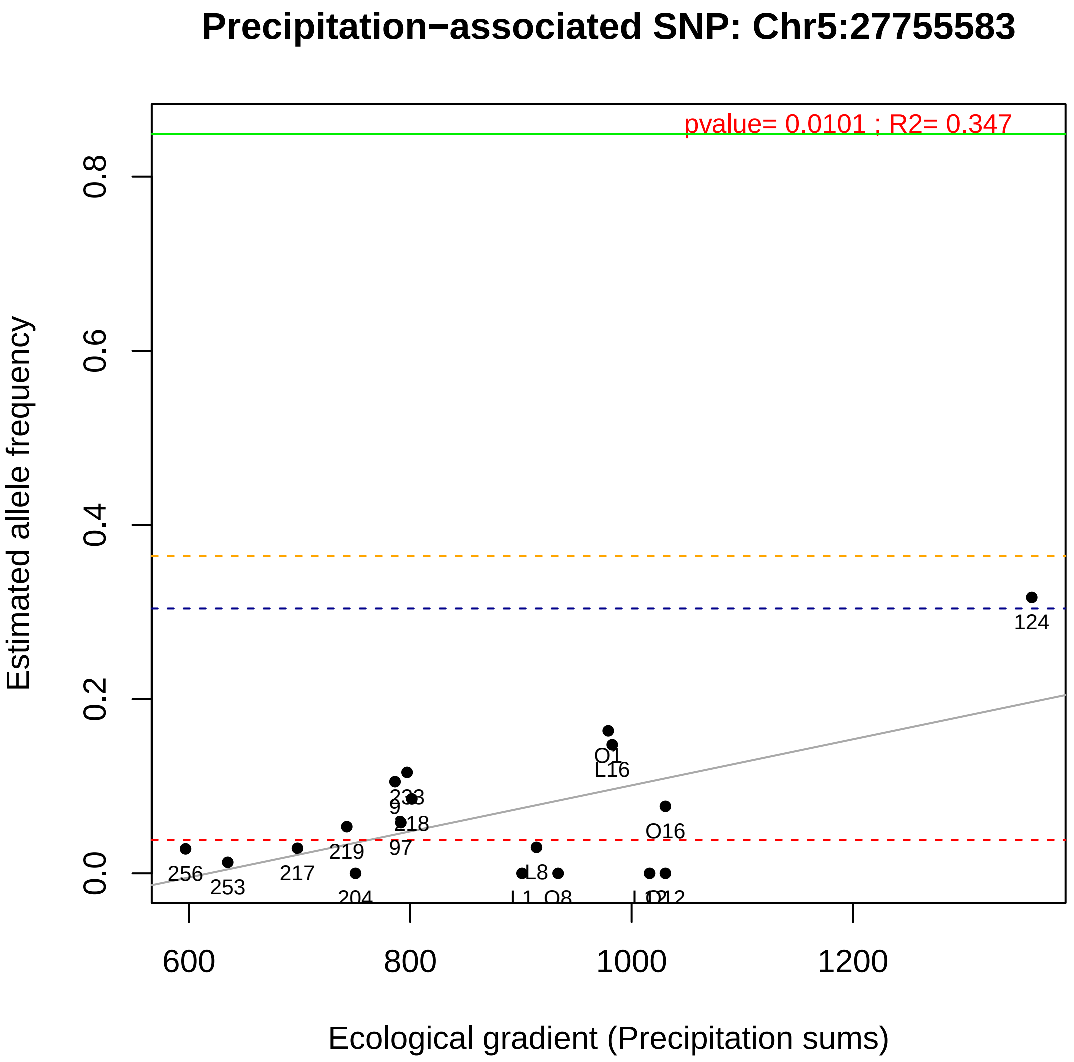

Precipitation−associated SNP: Chr5:27755583
pvalue= 0.0101 ; R2= 0.347
Estimated allele frequency
0.0
0.2
0.4
0.6
0.8
600
800
1000
1200
Ecological gradient (Precipitation sums)
124
O1
L16
O16
218
97
219
256
253
217
204
L8
O8

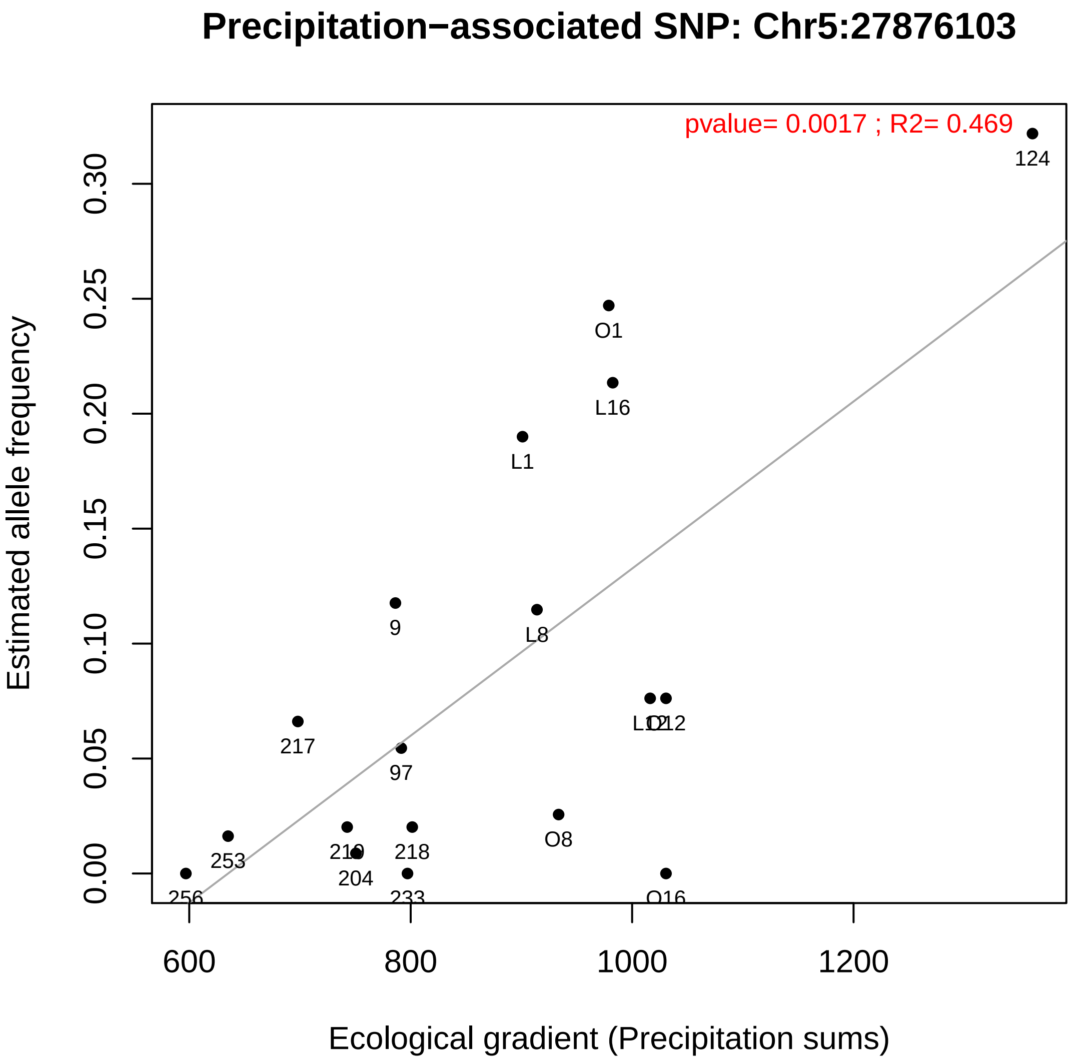

Precipitation−associated SNP: Chr5:27876103
pvalue= 0.0017 ; R2= 0.469
Estimated allele frequency
0.00
0.05
0.10
0.15
0.20
0.25
0.30
600
800
1000
1200
Ecological gradient (Precipitation sums)
124
O1
L16
L1
9
L8
L12
O12
217
97
O8
253
210
218
204
256
233
O16

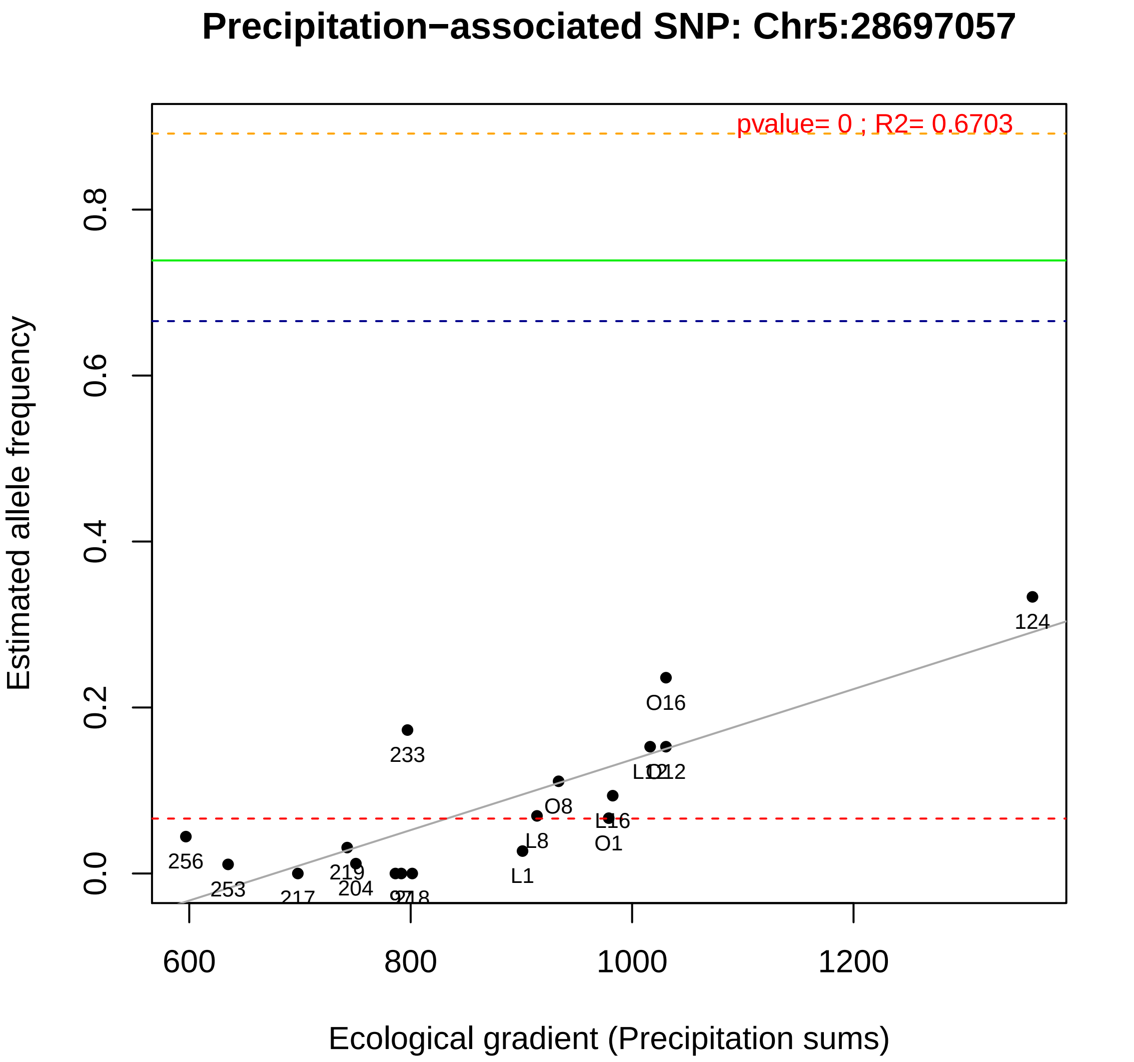
Precipitation−associated SNP: Chr5:28697057
pvalue= 0 ; R2= 0.6703
Estimated allele frequency
0.0
0.2
0.4
0.6
0.8
Ecological gradient (Precipitation sums)
600
800
1000
1200
124
O16
233
O8
L8
L16
O1
L1
256
253
217
219
204

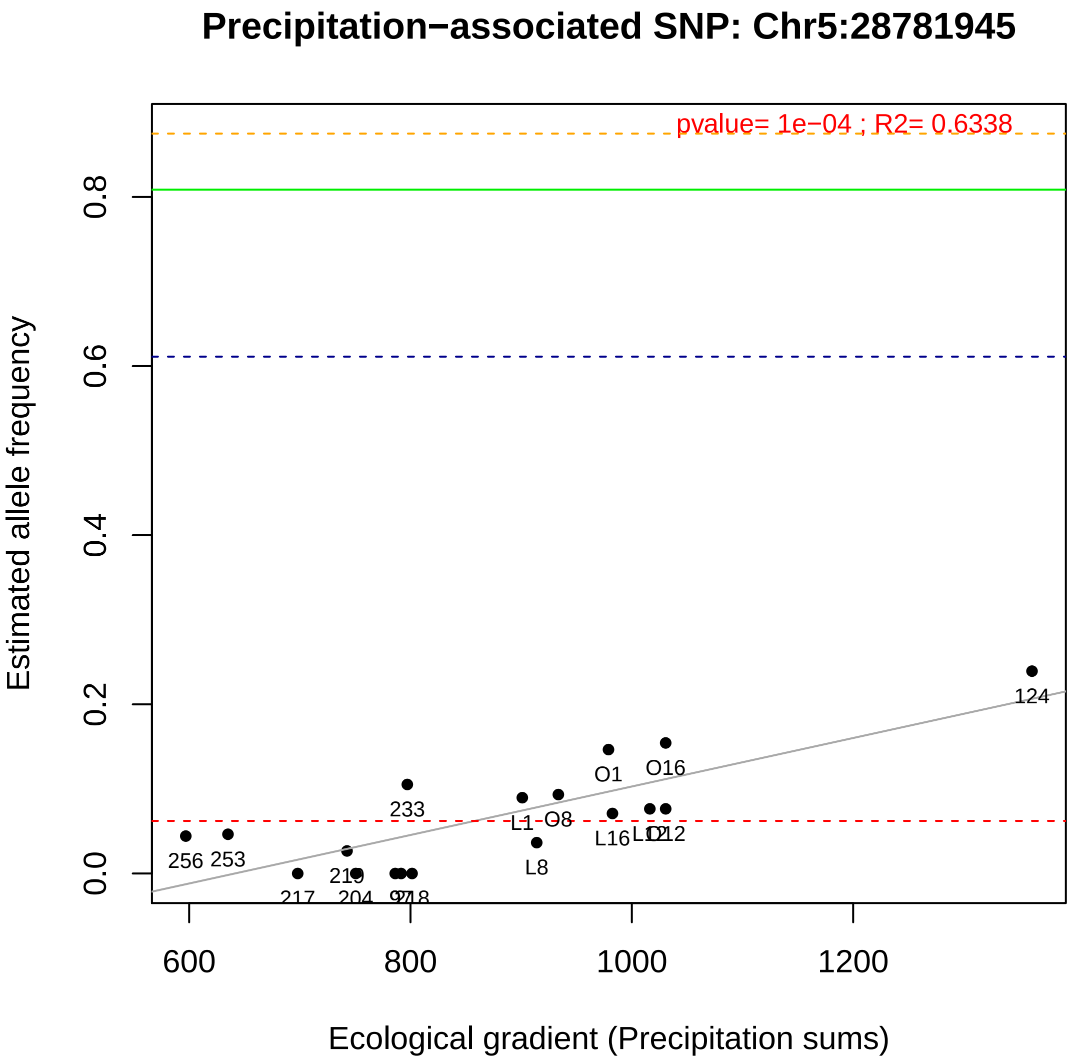
Precipitation−associated SNP: Chr5:28781945
pvalue= 1e−04 ; R2= 0.6338
Estimated allele frequency
0.0
0.2
0.4
0.6
0.8
600
800
1000
1200
Ecological gradient (Precipitation sums)
124
O1
O16
233
L1
O8
L16
L8
256
253
217
204

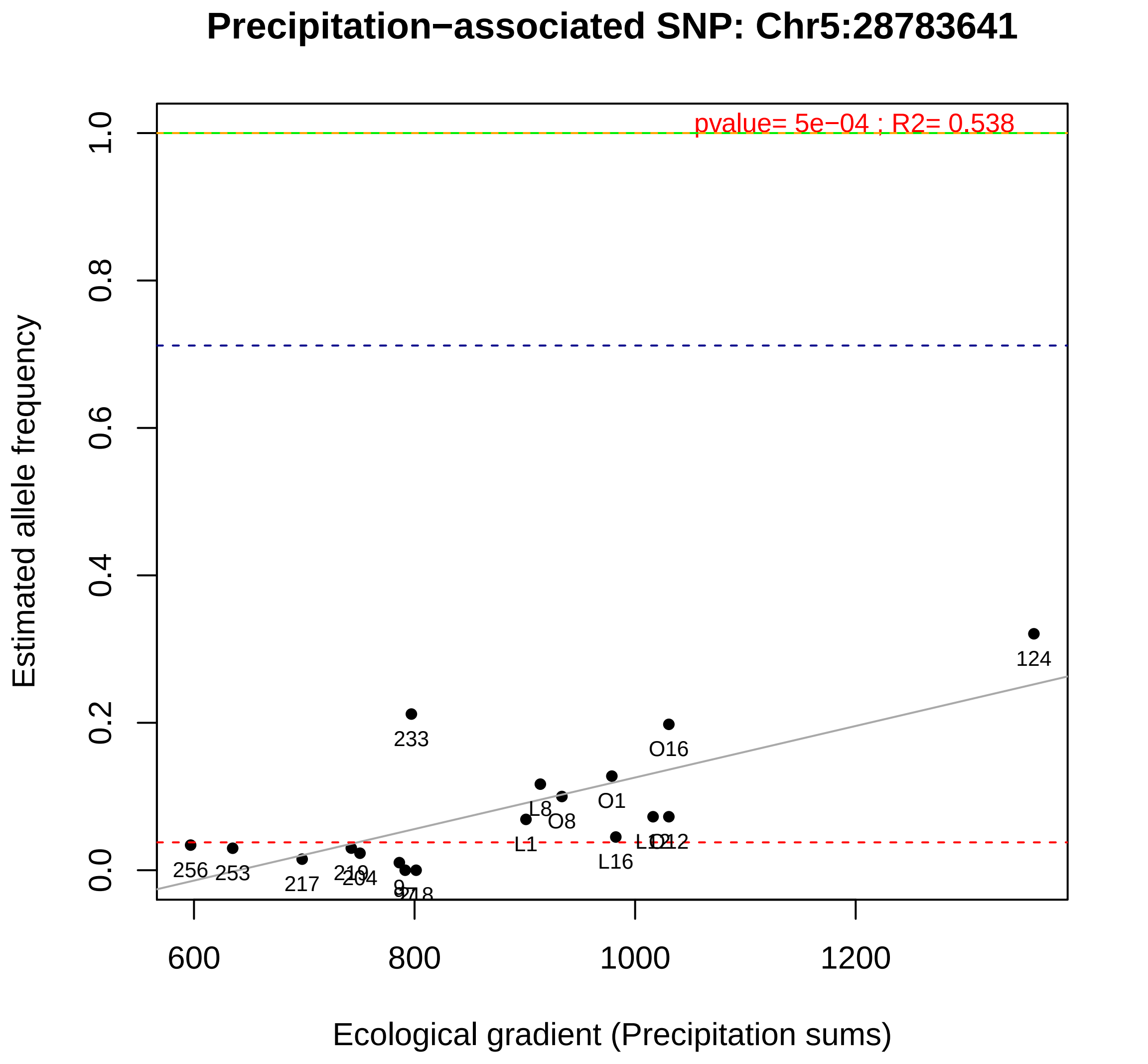

Precipitation−associated SNP: Chr5:28783641
pvalue= 5e−04 ; R2= 0.538
Estimated allele frequency
Ecological gradient (Precipitation sums)
0.0
0.2
0.4
0.6
0.8
1.0
600
800
1000
1200
124
233
O16
O1
L8
O8
L1
L16
256
253
217
204

**Precipitation−associated SNP: Chr5:2894896**

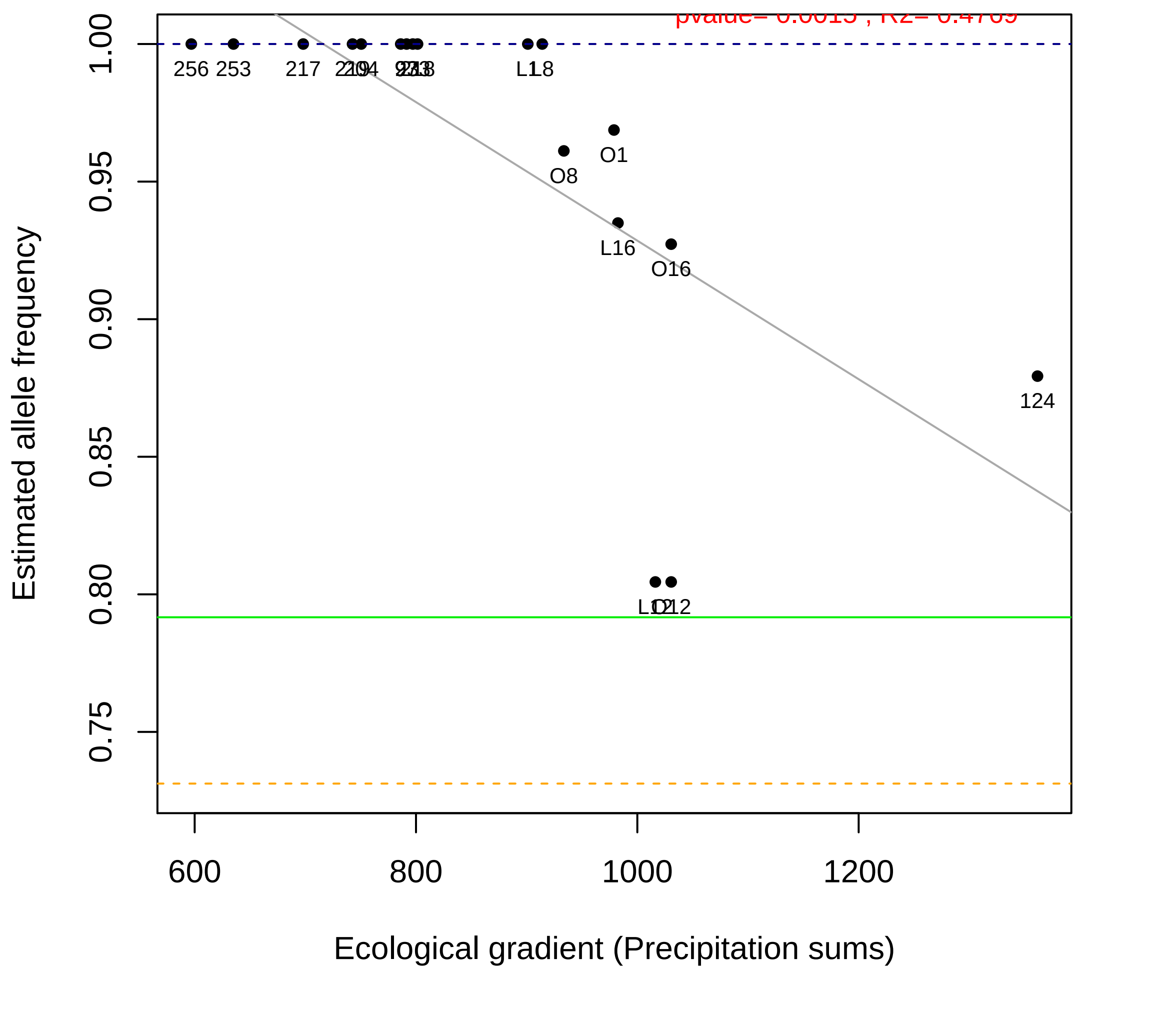

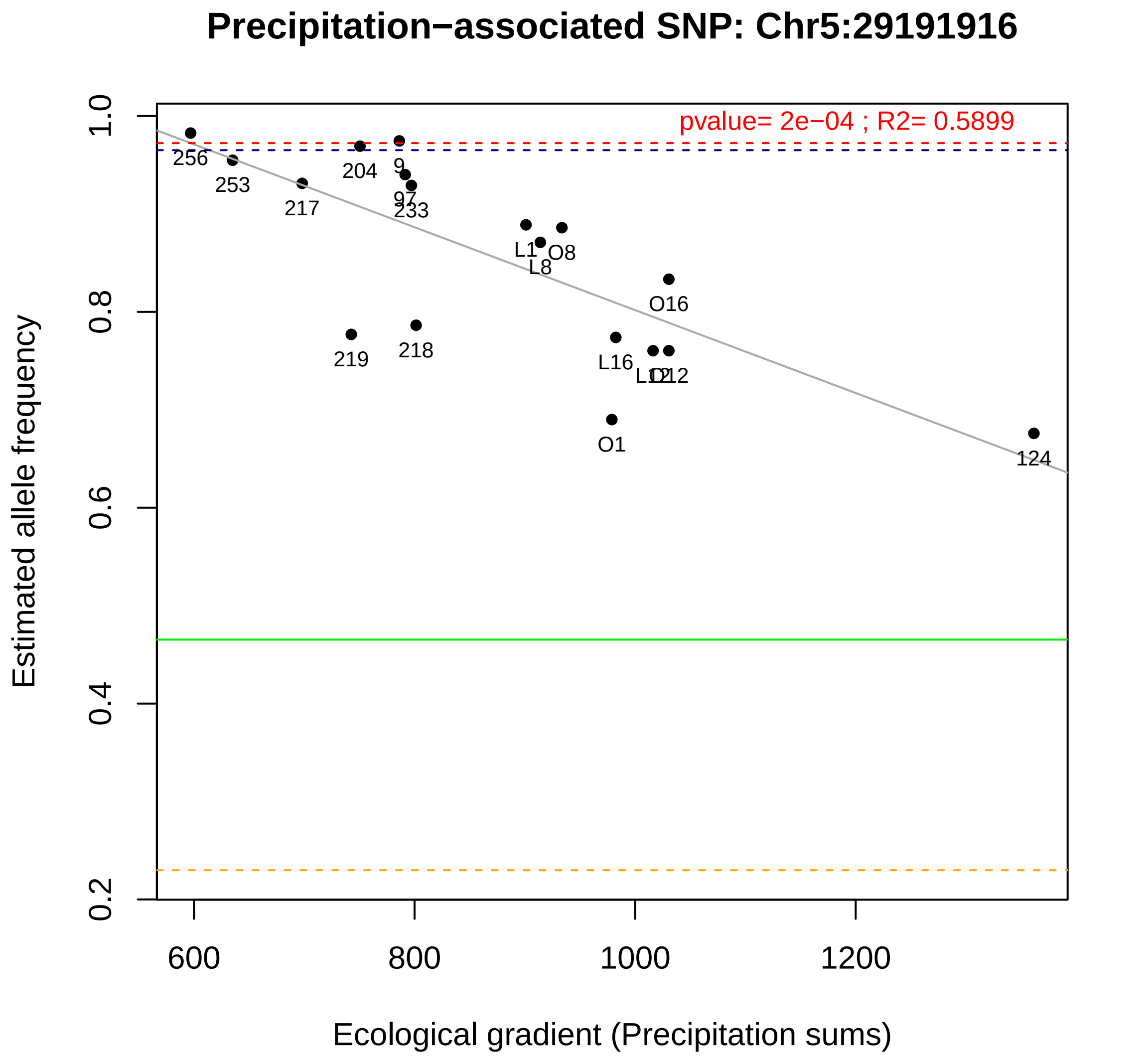

Precipitation−associated SNP: Chr5:29191916
pvalue= 2e−04 ; R2= 0.5899
Estimated allele frequency
Ecological gradient (Precipitation sums)
1.0
0.8
0.6
0.4
0.2
600
800
1000
1200
256
253
217
204
9
97
233
219
218
L1
L8
O8
O16
L16
L12
O12
O1
124

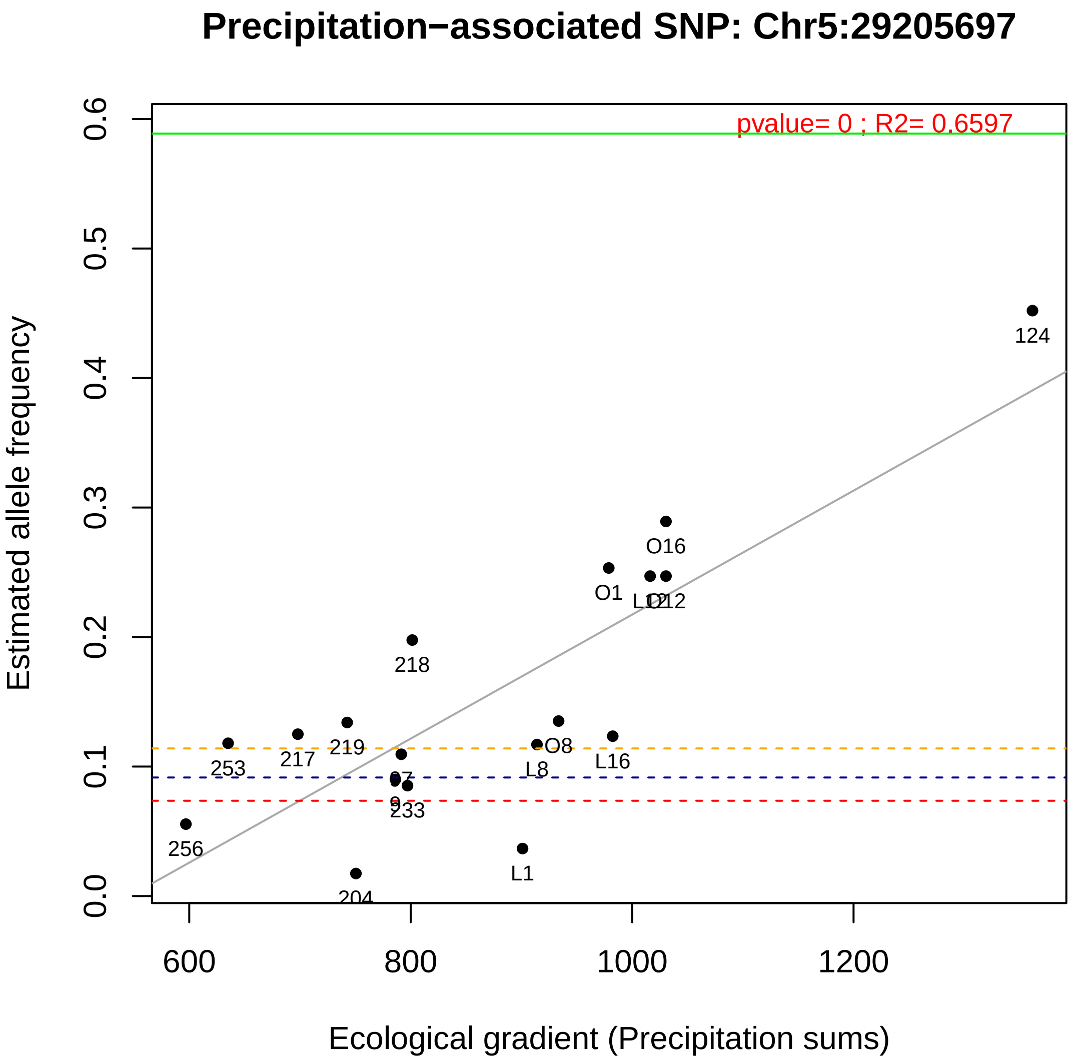

Precipitation−associated SNP: Chr5:29205697
pvalue= 0 ; R2= 0.6597
Estimated allele frequency
0.0
0.1
0.2
0.3
0.4
0.5
0.6
600
800
1000
1200
Ecological gradient (Precipitation sums)
124
O16
O1
O12
218
219
253
217
O8
L8
L16
233
256
204
L1

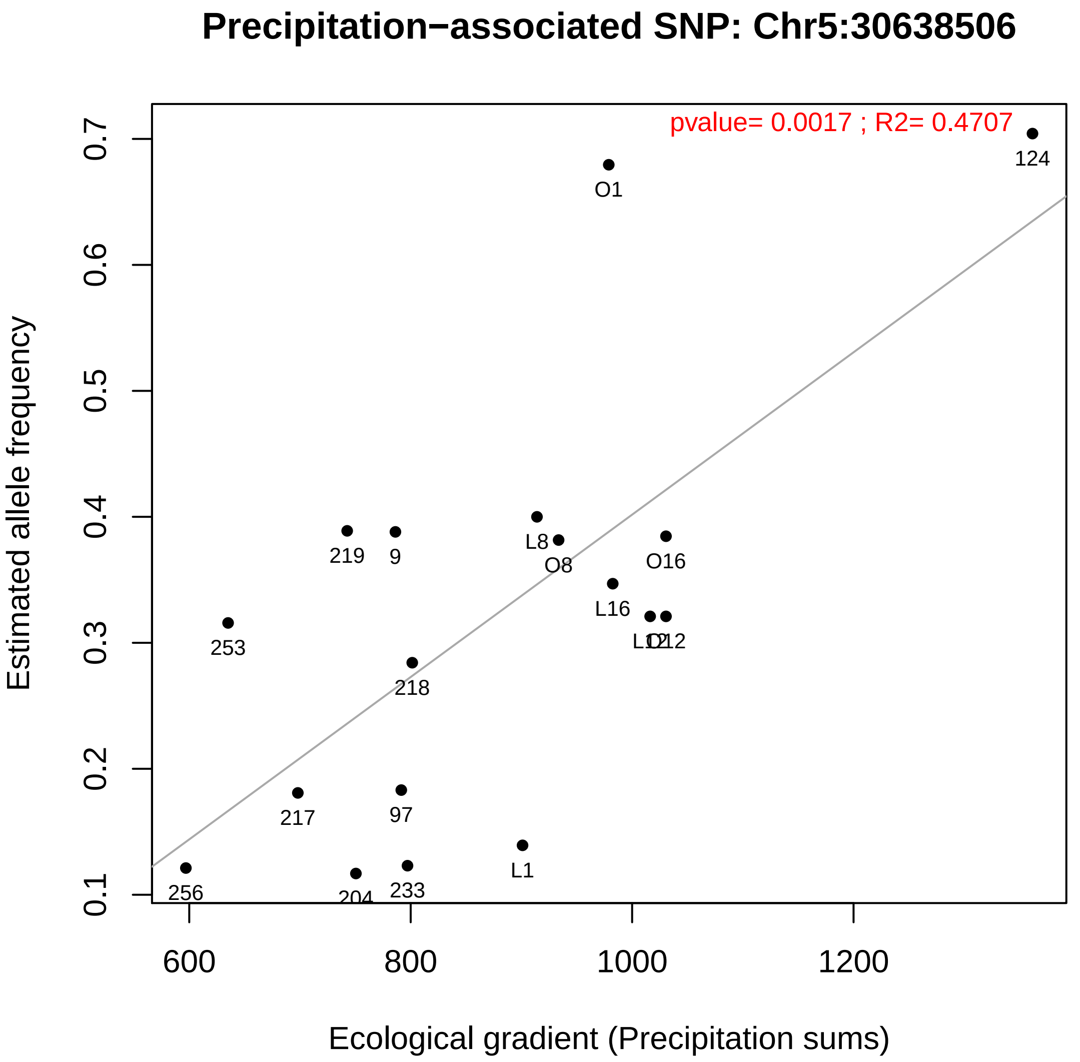
Precipitation−associated SNP: Chr5:30638506
pvalue= 0.0017 ; R2= 0.4707
Estimated allele frequency
Ecological gradient (Precipitation sums)
0.1
0.2
0.3
0.4
0.5
0.6
0.7
600
800
1000
1200
124
O1
219
9
L8
O8
O16
L16
253
218
217
97
L1
256
204
233

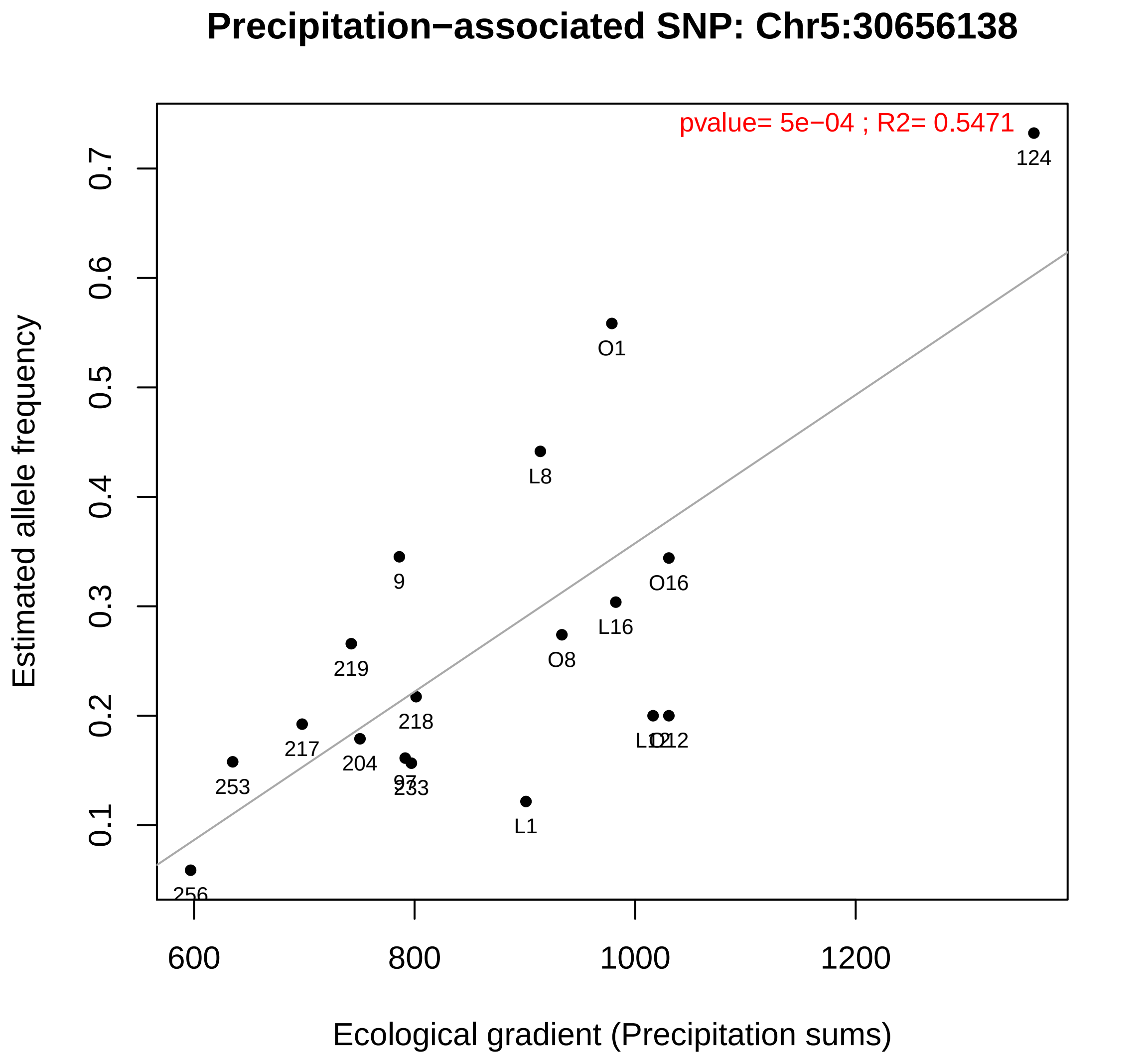
Precipitation−associated SNP: Chr5:30656138
pvalue= 5e−04 ; R2= 0.5471
Estimated allele frequency
0.1
0.2
0.3
0.4
0.5
0.6
0.7
600
800
1000
1200
Ecological gradient (Precipitation sums)
124
O1
L8
9
O16
L16
O8
219
218
217
204
253
97
233
L1
L12
O12
256

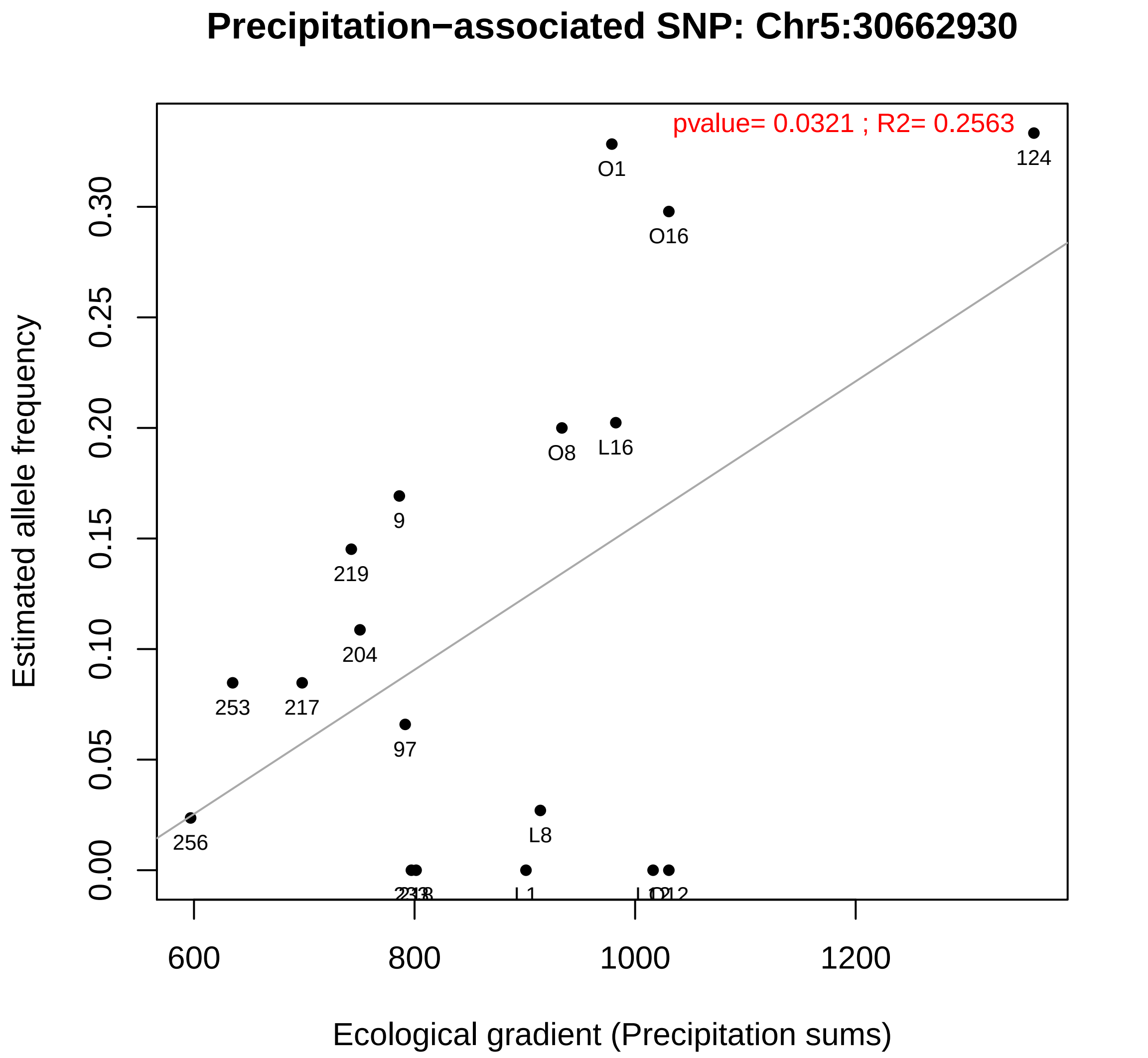
Precipitation−associated SNP: Chr5:30662930
pvalue= 0.0321 ; R2= 0.2563
Estimated allele frequency
0.00
0.05
0.10
0.15
0.20
0.25
0.30
600
800
1000
1200
Ecological gradient (Precipitation sums)
O1
124
O16
O8
L16
9
219
204
253
217
97
L8
256
L1

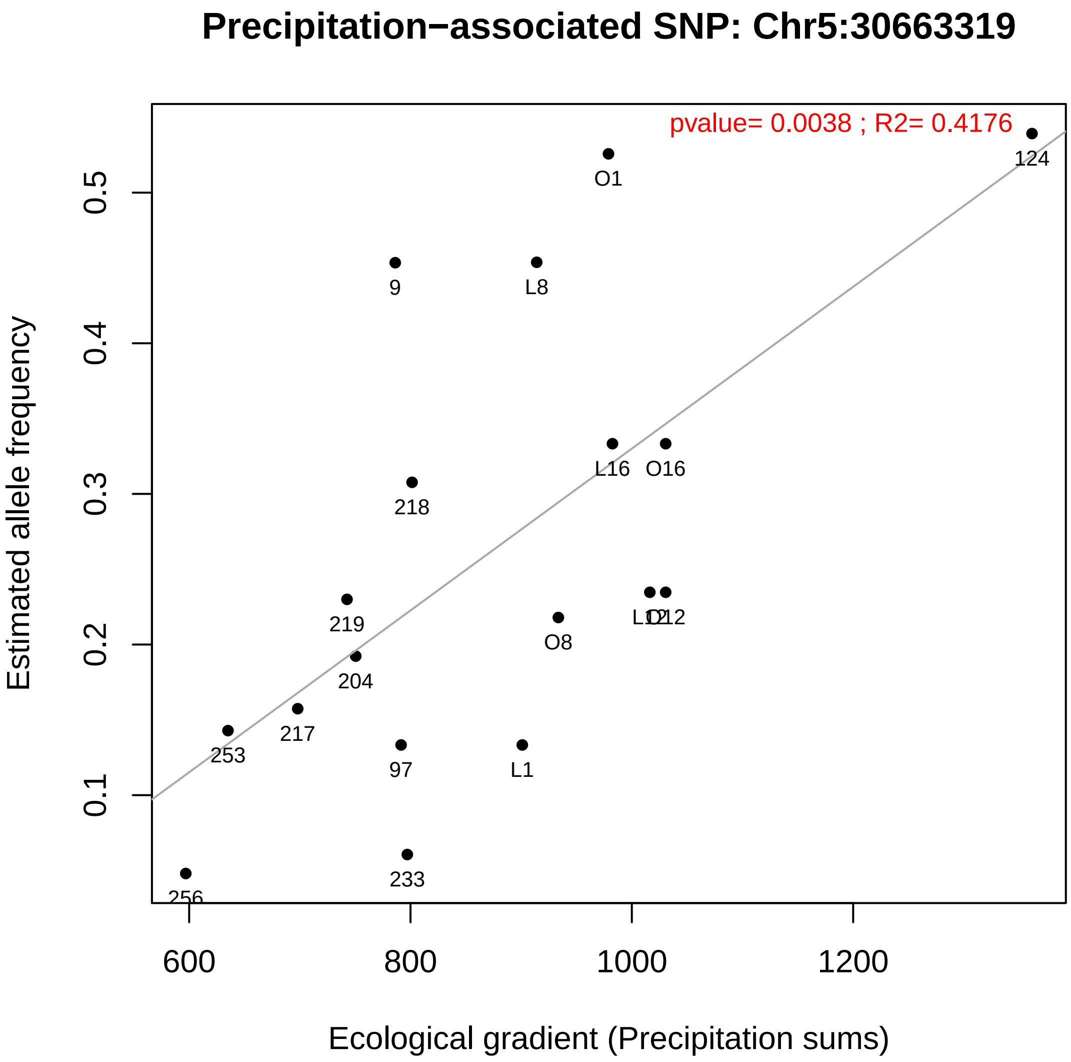
Precipitation−associated SNP: Chr5:30663319
pvalue= 0.0038 ; R2= 0.4176
Estimated allele frequency
0.1
0.2
0.3
0.4
0.5
600
800
1000
1200
Ecological gradient (Precipitation sums)
124
O1
9
L8
L16
O16
218
219
O8
204
217
253
97
L1
233
256

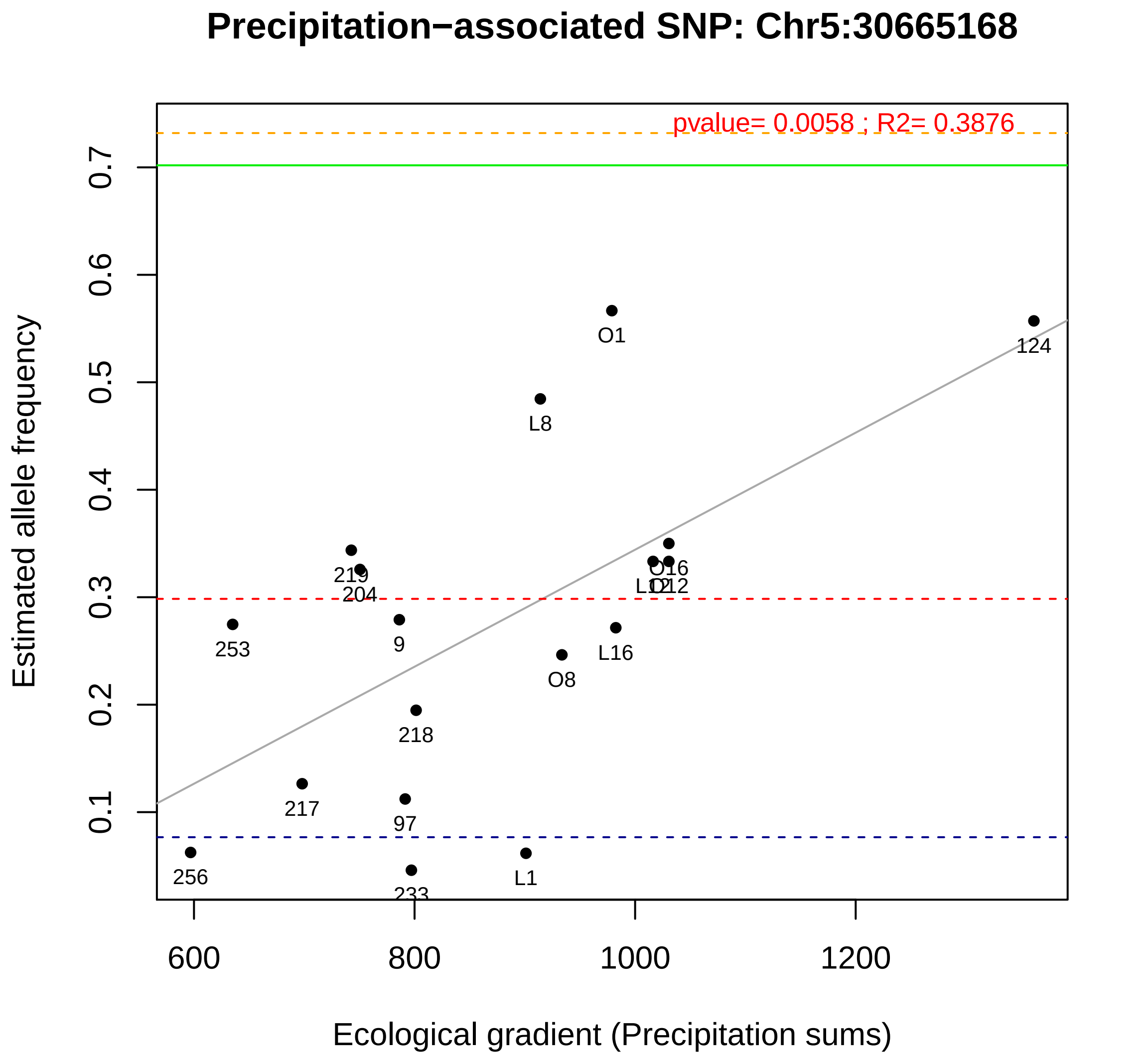

Precipitation−associated SNP: Chr5:30665168
pvalue= 0.0058 ; R2= 0.3876
Estimated allele frequency
0.1
0.2
0.3
0.4
0.5
0.6
0.7
600
800
1000
1200
Ecological gradient (Precipitation sums)
O1
124
L8
219
204
O16
L12
O12
253
9
L16
O8
218
217
97
256
233
L1

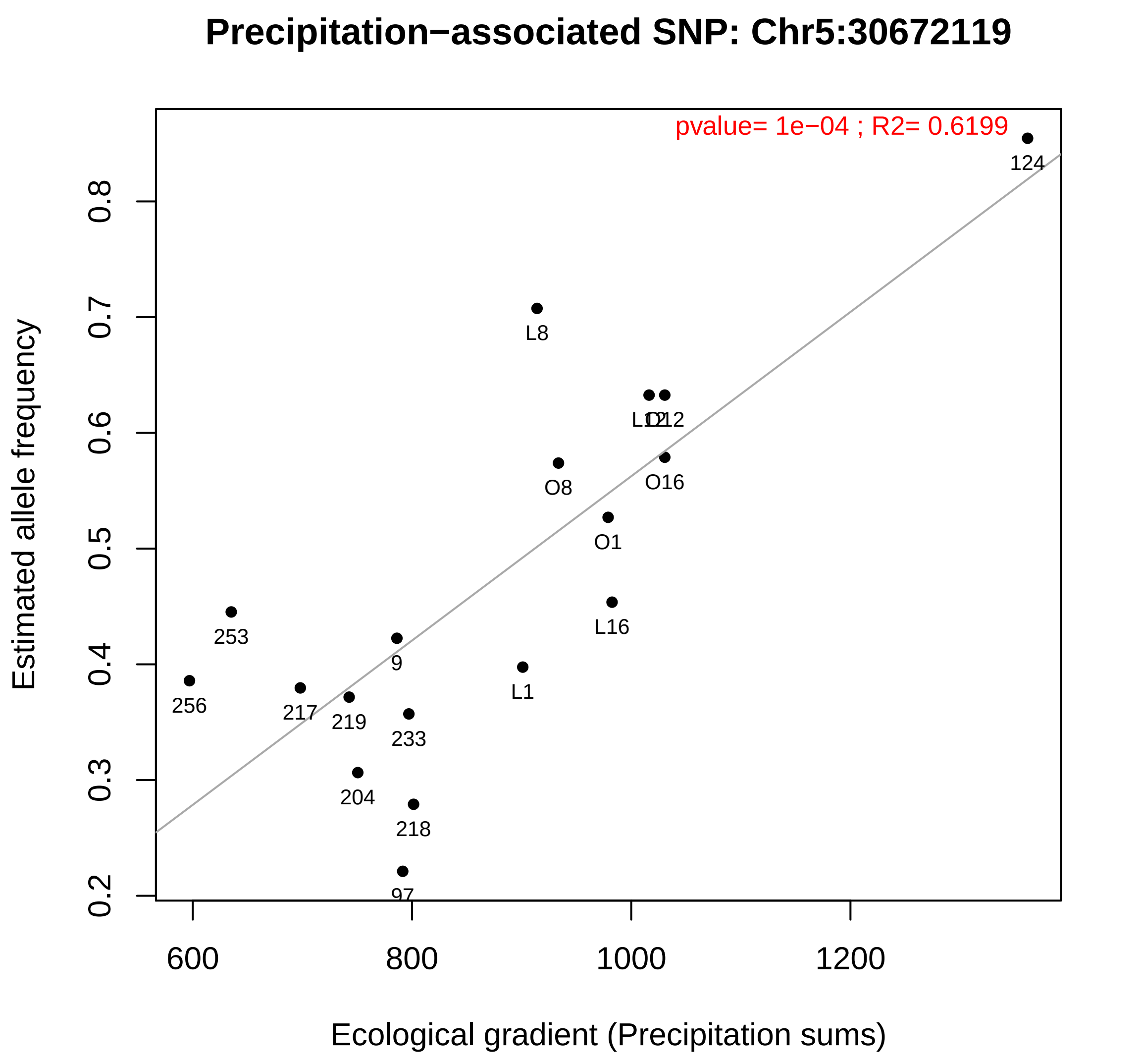

Precipitation−associated SNP: Chr5:30672119
pvalue= 1e−04 ; R2= 0.6199
Estimated allele frequency
Ecological gradient (Precipitation sums)
0.2
0.3
0.4
0.5
0.6
0.7
0.8
600
800
1000
1200
124
L8
L12
O12
O16
O8
O1
L16
253
9
L1
256
217
219
233
204
218
97

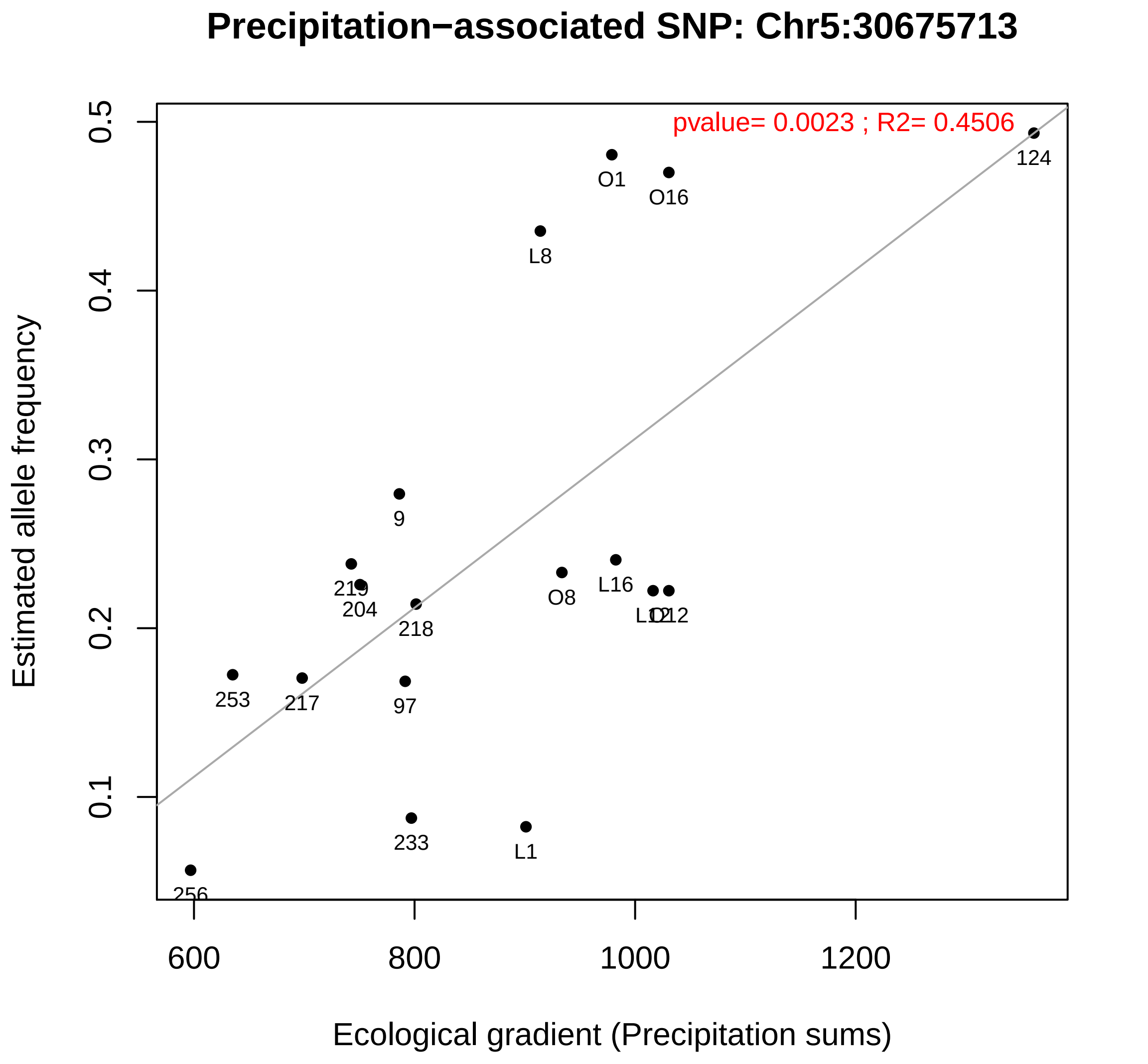

Precipitation−associated SNP: Chr5:30675713
pvalue= 0.0023 ; R2= 0.4506
124
O1
O16
L8
9
219
204
218
O8
L16
L12
O12
253
217
97
233
L1
256
0.1
0.2
0.3
0.4
0.5
600
800
1000
1200
Estimated allele frequency
Ecological gradient (Precipitation sums)

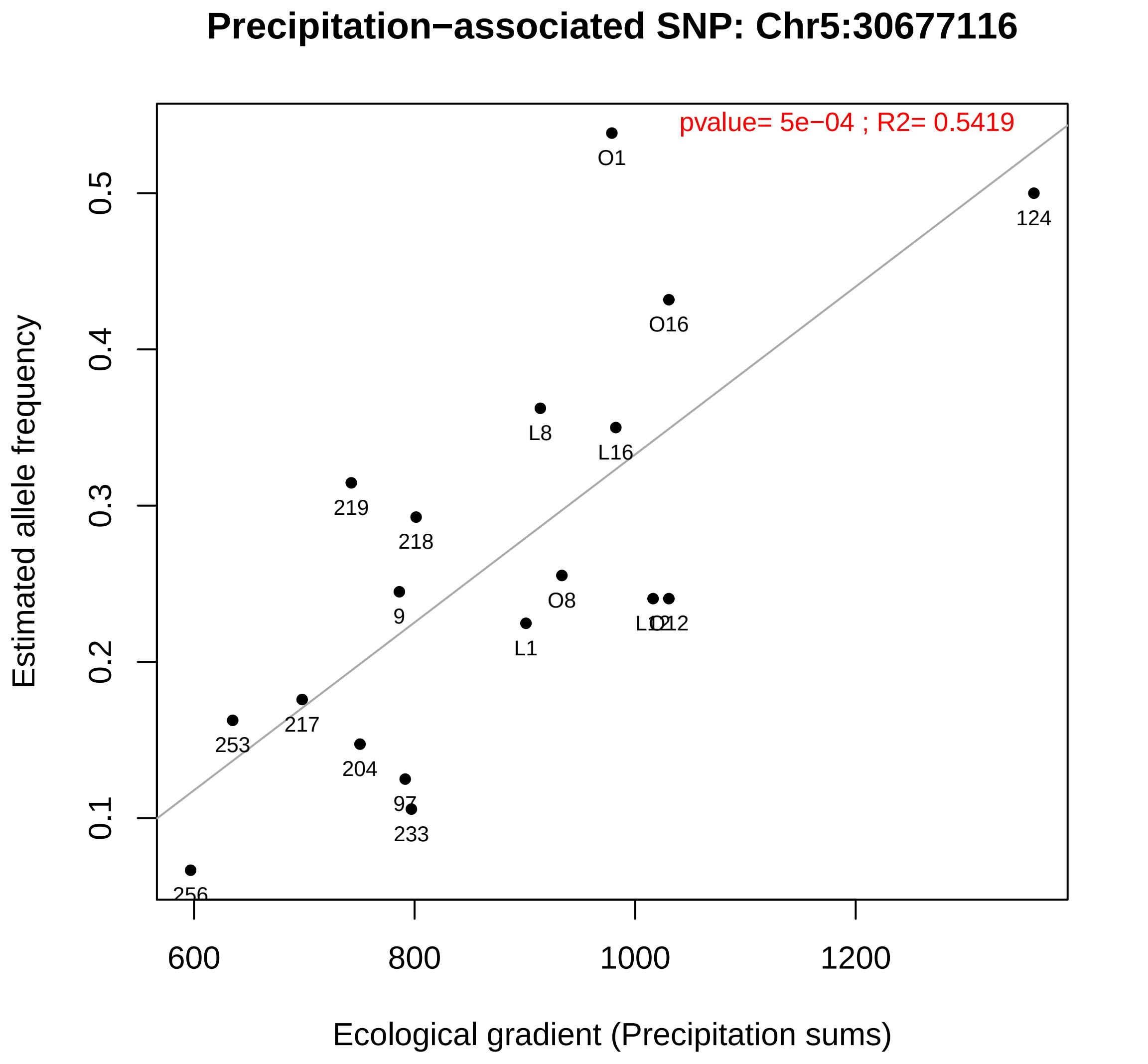

Precipitation−associated SNP: Chr5:30677116
pvalue= 5e−04 ; R2= 0.5419
O1
124
O16
L8
L16
219
218
O8
9
L1
217
253
204
97
233
256
0.1
0.2
0.3
0.4
0.5
600
800
1000
1200
Estimated allele frequency
Ecological gradient (Precipitation sums)

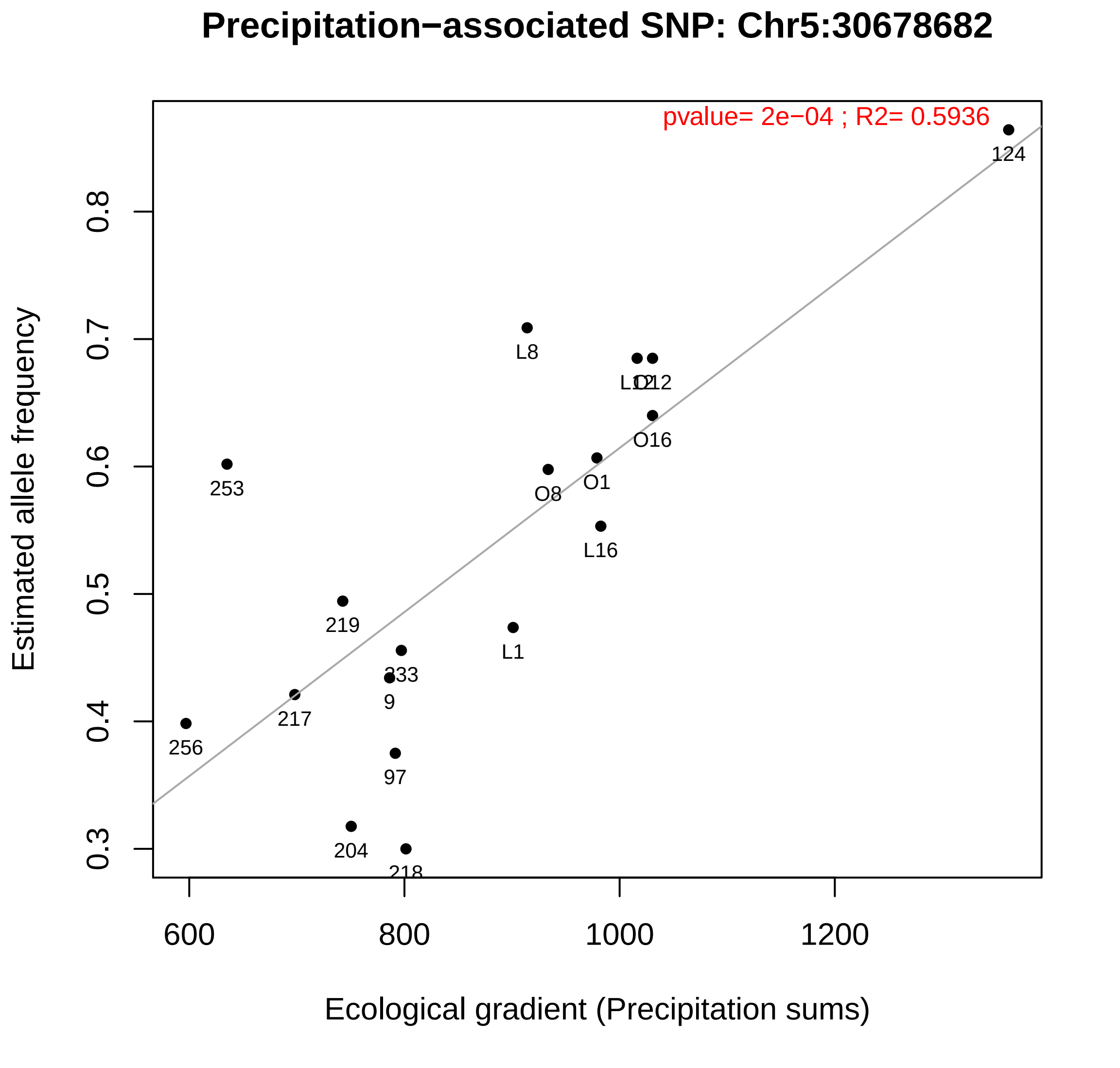

Precipitation−associated SNP: Chr5:30678682
pvalue= 2e−04 ; R2= 0.5936
Estimated allele frequency
0.3
0.4
0.5
0.6
0.7
0.8
600
800
1000
1200
Ecological gradient (Precipitation sums)
124
L8
L12
O12
O16
253
O1
O8
L16
219
L1
233
9
217
256
97
204
218

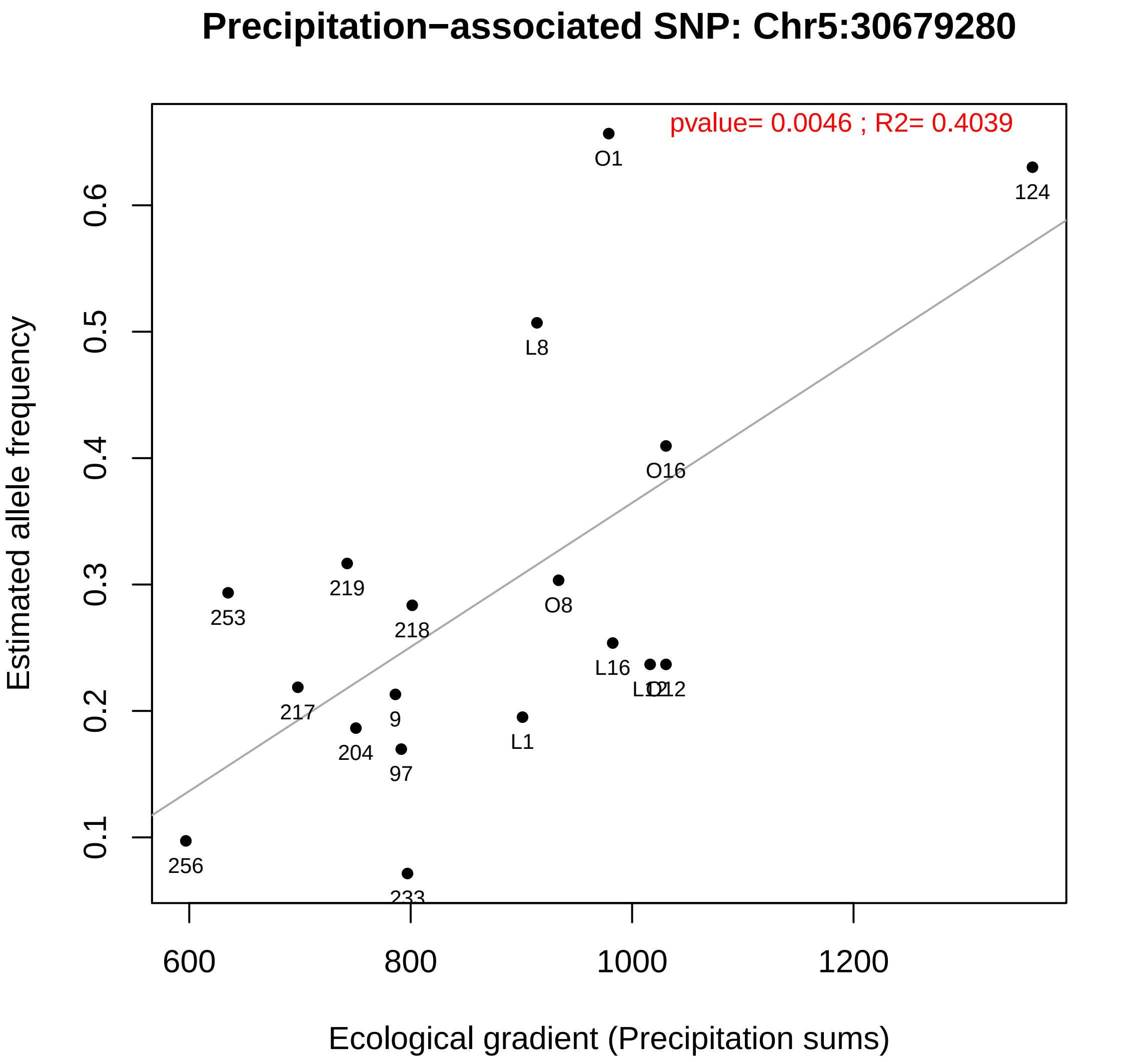
Precipitation−associated SNP: Chr5:30679280
pvalue= 0.0046 ; R2= 0.4039
O1
124
L8
O16
219
253
218
O8
L16
L12
O12
217
9
204
L1
97
256
233
0.1
0.2
0.3
0.4
0.5
0.6
600
800
1000
1200
Estimated allele frequency
Ecological gradient (Precipitation sums)

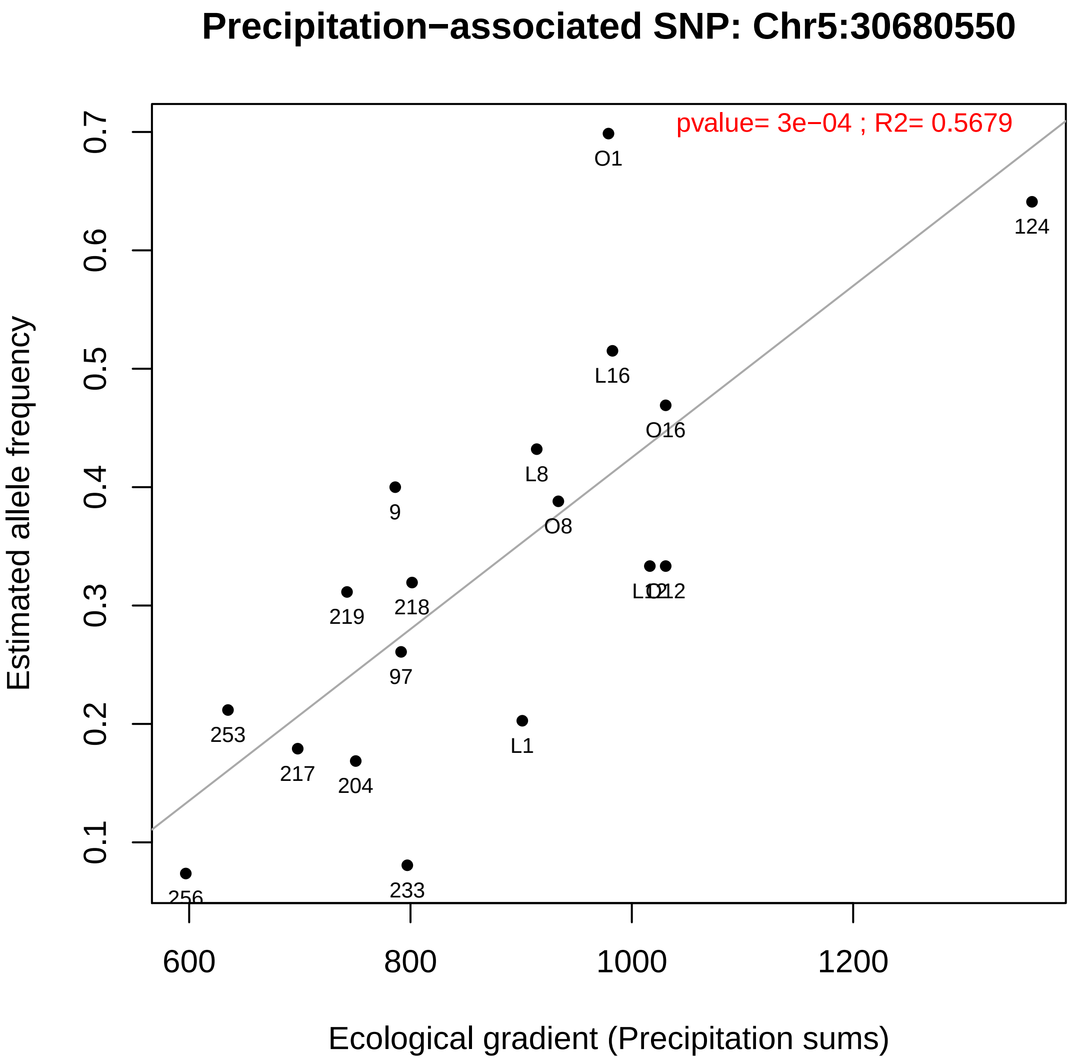
Precipitation−associated SNP: Chr5:30680550
pvalue= 3e−04 ; R2= 0.5679
O1
124
L16
O16
L8
9
O8
L12
O12
219
218
97
253
L1
217
204
256
233
0.1
0.2
0.3
0.4
0.5
0.6
0.7
600
800
1000
1200
Estimated allele frequency
Ecological gradient (Precipitation sums)

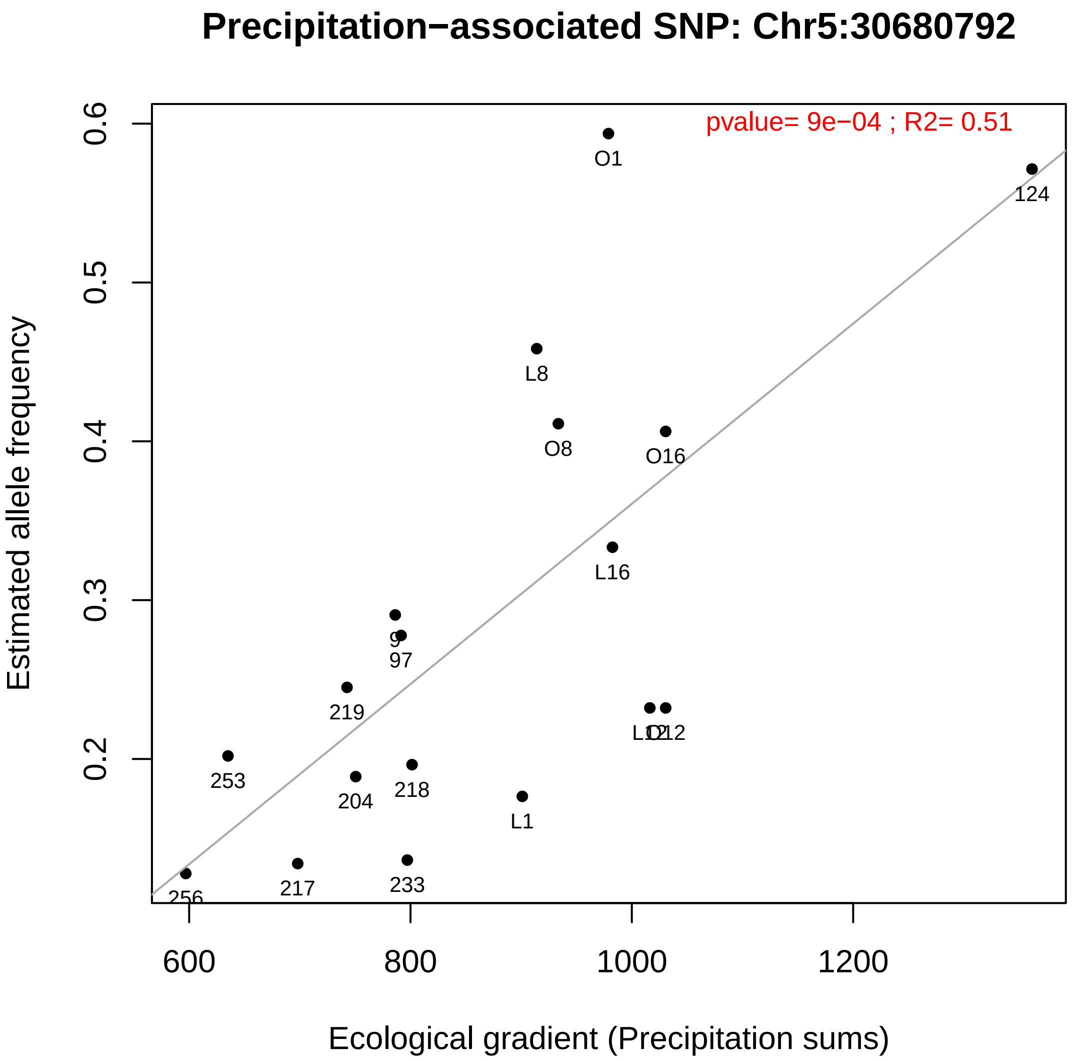

Precipitation−associated SNP: Chr5:30680792
pvalue= 9e−04 ; R2= 0.51
O1
124
L8
O8
O16
L16
9
97
219
L12
O12
253
204
218
L1
256
217
233
0.2
0.3
0.4
0.5
0.6
600
800
1000
1200
Estimated allele frequency
Ecological gradient (Precipitation sums)

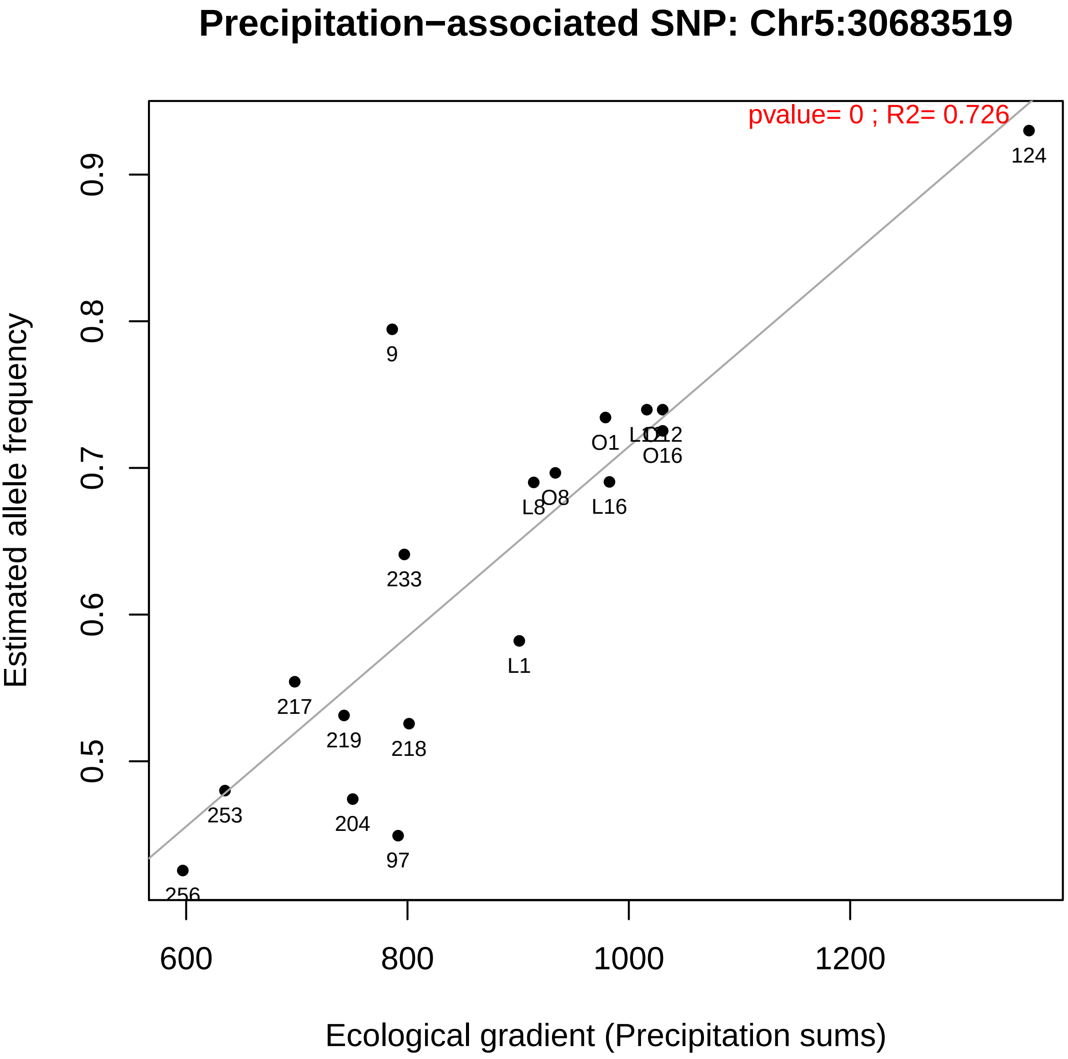
Precipitation−associated SNP: Chr5:30683519
pvalue= 0 ; R2= 0.726
Estimated allele frequency
0.5
0.6
0.7
0.8
0.9
600
800
1000
1200
Ecological gradient (Precipitation sums)
124
9
O1
O16
L8
O8
L16
233
L1
217
219
218
253
204
97
256

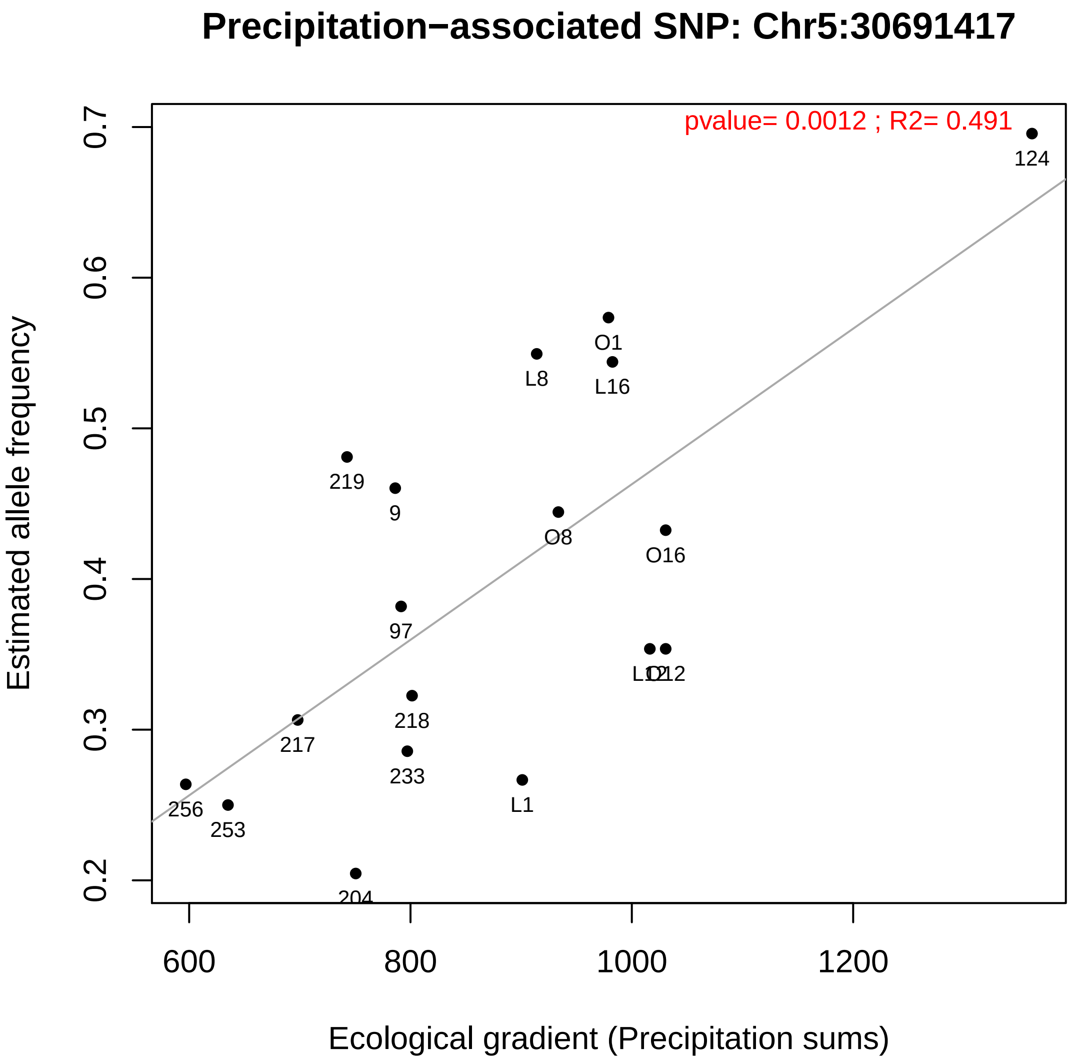

Precipitation−associated SNP: Chr5:30691417
pvalue= 0.0012 ; R2= 0.491
124
O1
L8
L16
219
9
O8
O16
97
L12
O12
218
217
233
L1
256
253
204
0.2
0.3
0.4
0.5
0.6
0.7
600
800
1000
1200
Estimated allele frequency
Ecological gradient (Precipitation sums)

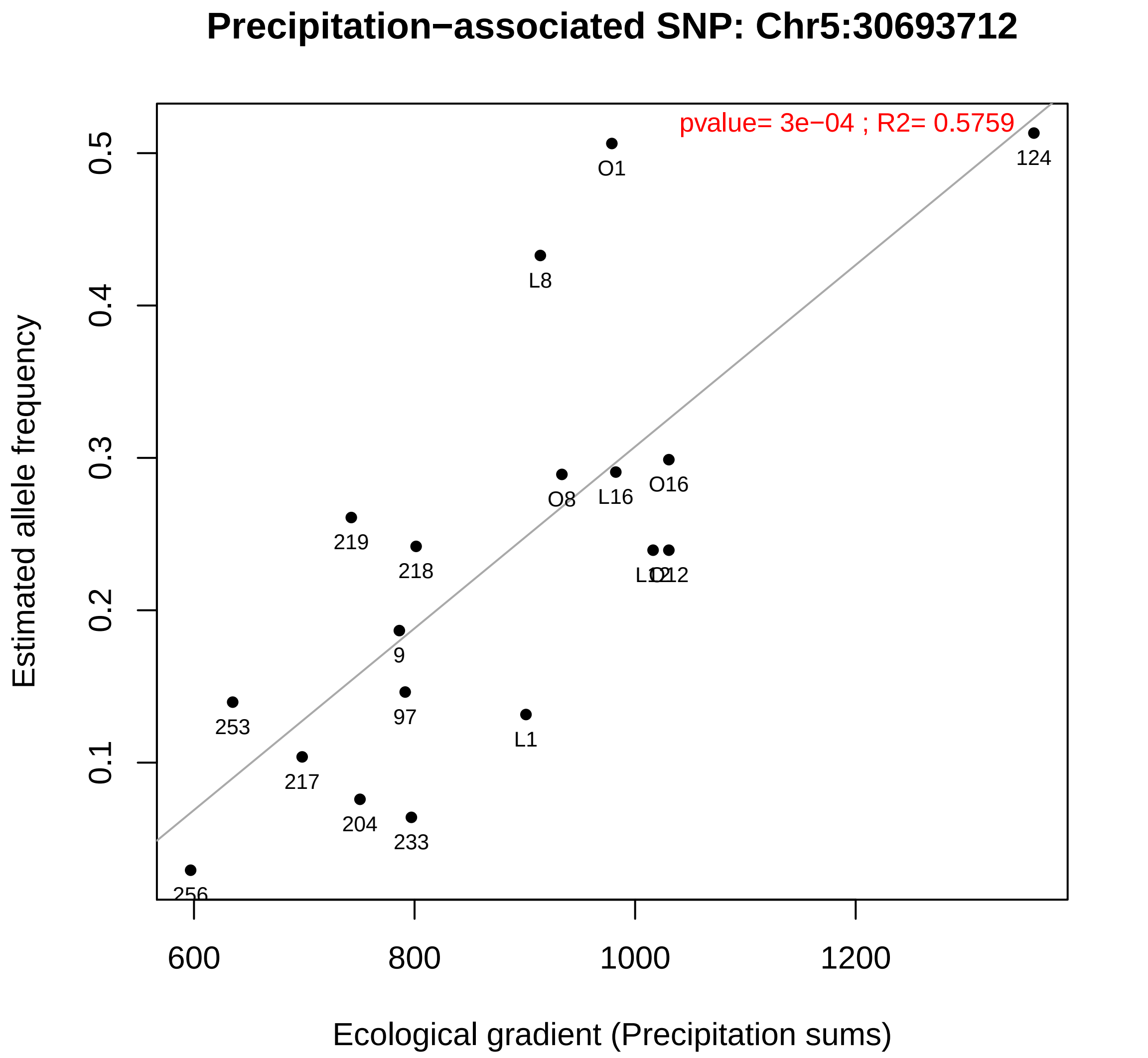

Precipitation−associated SNP: Chr5:30693712
pvalue= 3e−04 ; R2= 0.5759
Estimated allele frequency
0.1
0.2
0.3
0.4
0.5
600
800
1000
1200
Ecological gradient (Precipitation sums)
O1
124
L8
O8
L16
O16
219
218
L12
O12
9
97
253
L1
217
204
233
256

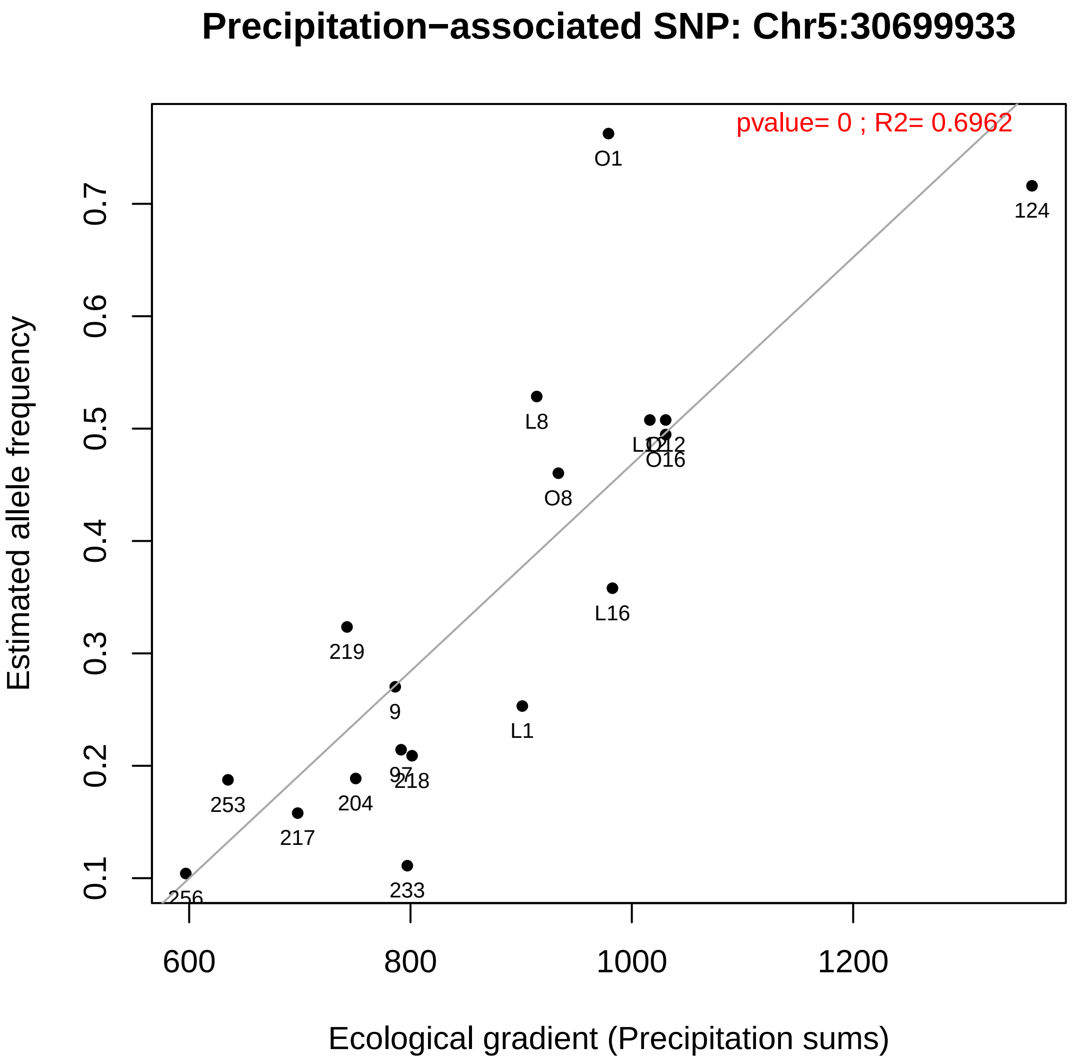

Precipitation−associated SNP: Chr5:30699933
pvalue= 0 ; R2= 0.6962
O1
124
L8
O12
O16
O8
L16
219
9
L1
97
218
253
204
217
233
256
0.1
0.2
0.3
0.4
0.5
0.6
0.7
600
800
1000
1200
Estimated allele frequency
Ecological gradient (Precipitation sums)

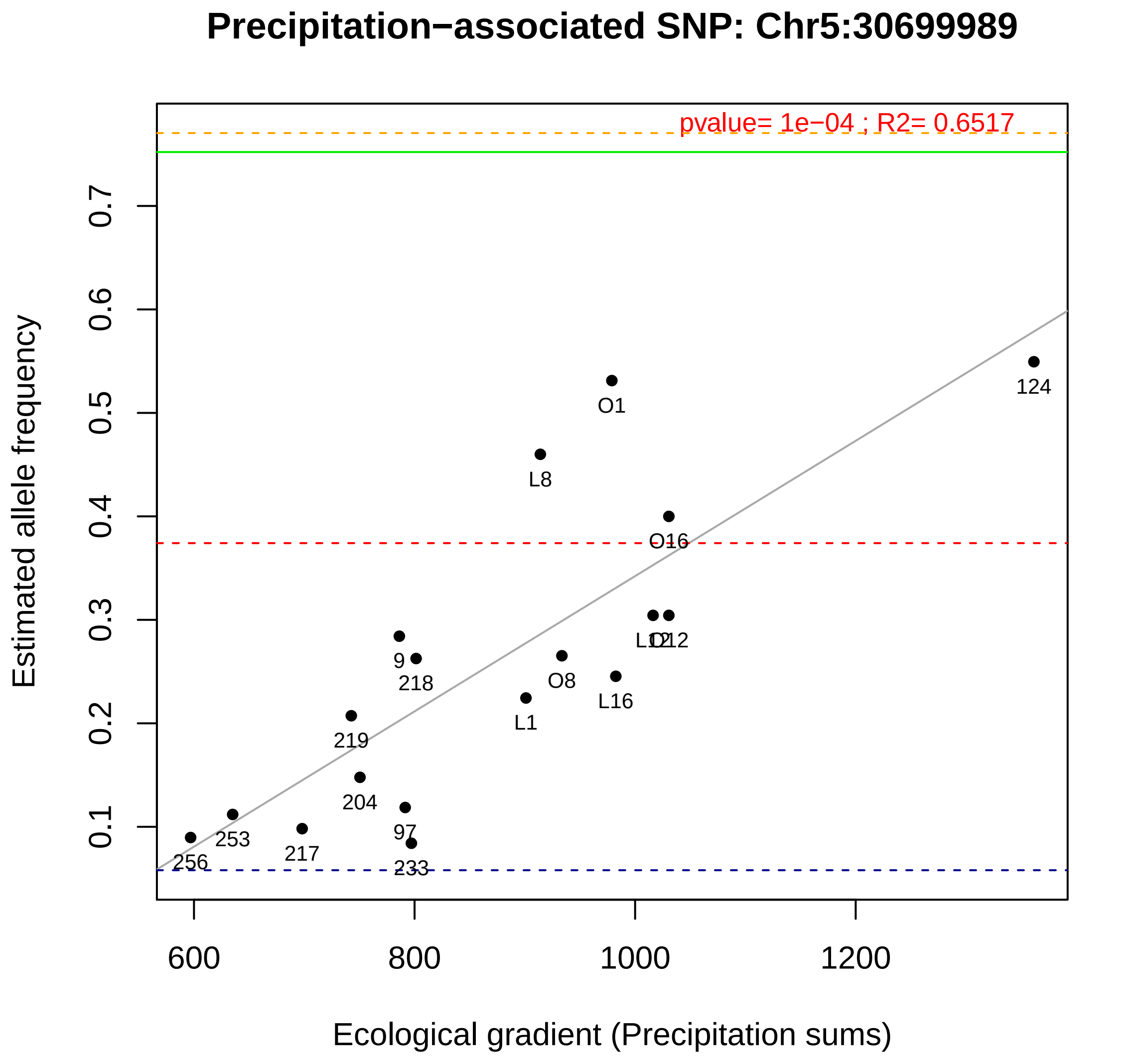
Precipitation−associated SNP: Chr5:30699989
pvalue= 1e−04 ; R2= 0.6517
Estimated allele frequency
0.1
0.2
0.3
0.4
0.5
0.6
0.7
600
800
1000
1200
Ecological gradient (Precipitation sums)
124
O1
L8
O16
O12
O8
L16
L1
9
218
219
204
97
233
253
256
217

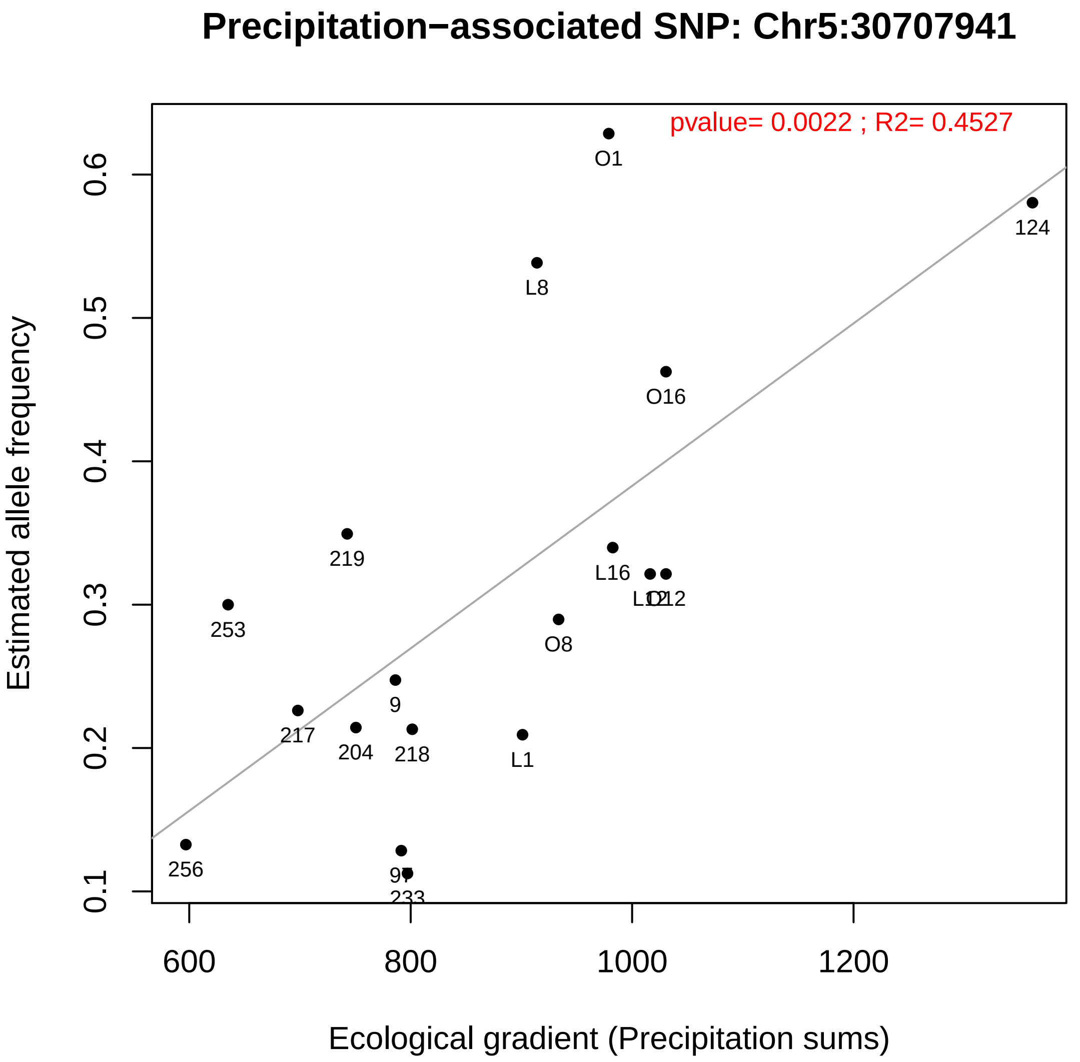

Precipitation−associated SNP: Chr5:30707941
pvalue= 0.0022 ; R2= 0.4527
Estimated allele frequency
0.1
0.2
0.3
0.4
0.5
0.6
600
800
1000
1200
Ecological gradient (Precipitation sums)
O1
124
L8
O16
219
L16
L12
O12
253
O8
9
217
204
218
L1
256
97
233

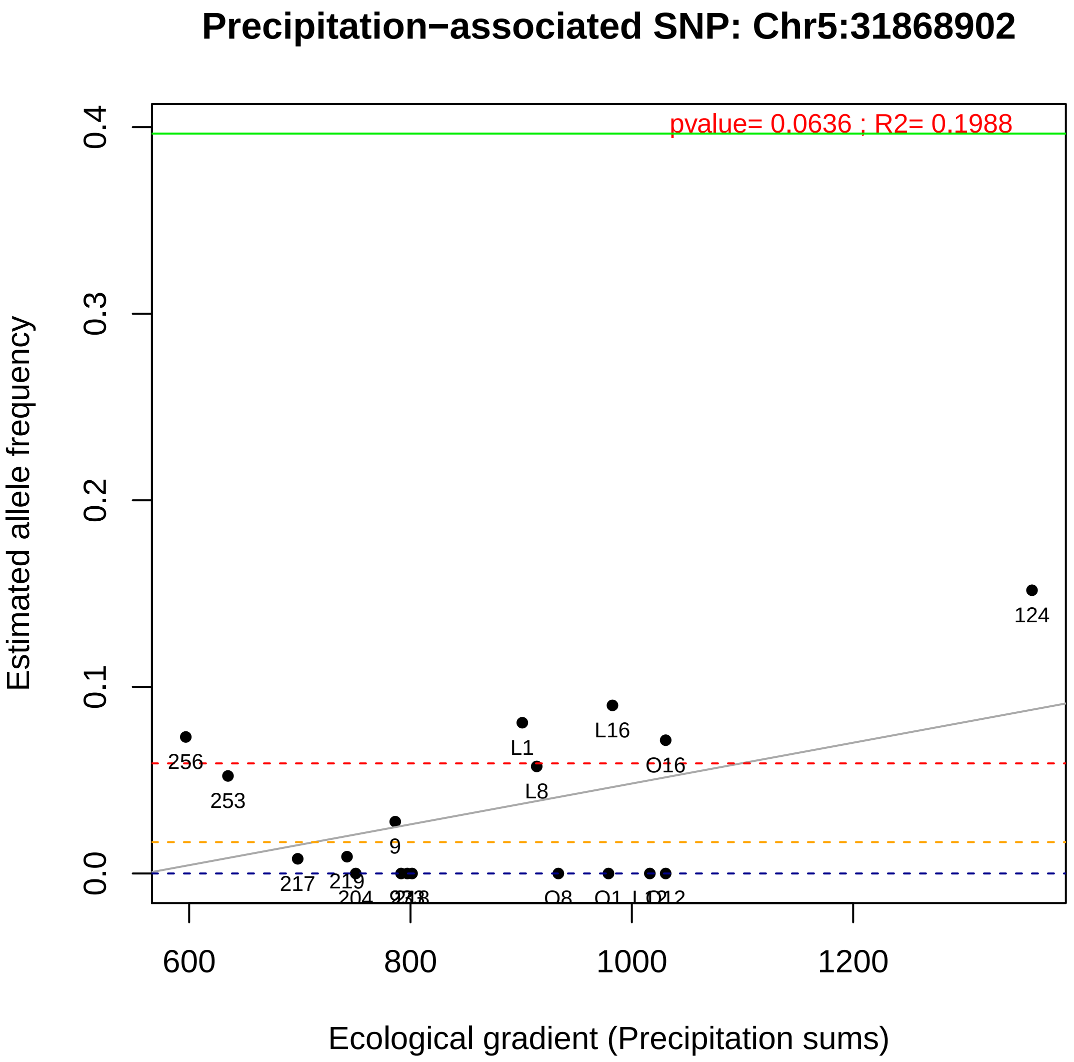
Precipitation−associated SNP: Chr5:31868902
pvalue= 0.0636 ; R2= 0.1988
Estimated allele frequency
0.0
0.1
0.2
0.3
0.4
600
800
1000
1200
Ecological gradient (Precipitation sums)
256
253
217
219
204
9
L1
L8
L16
O16
124
O8
O1

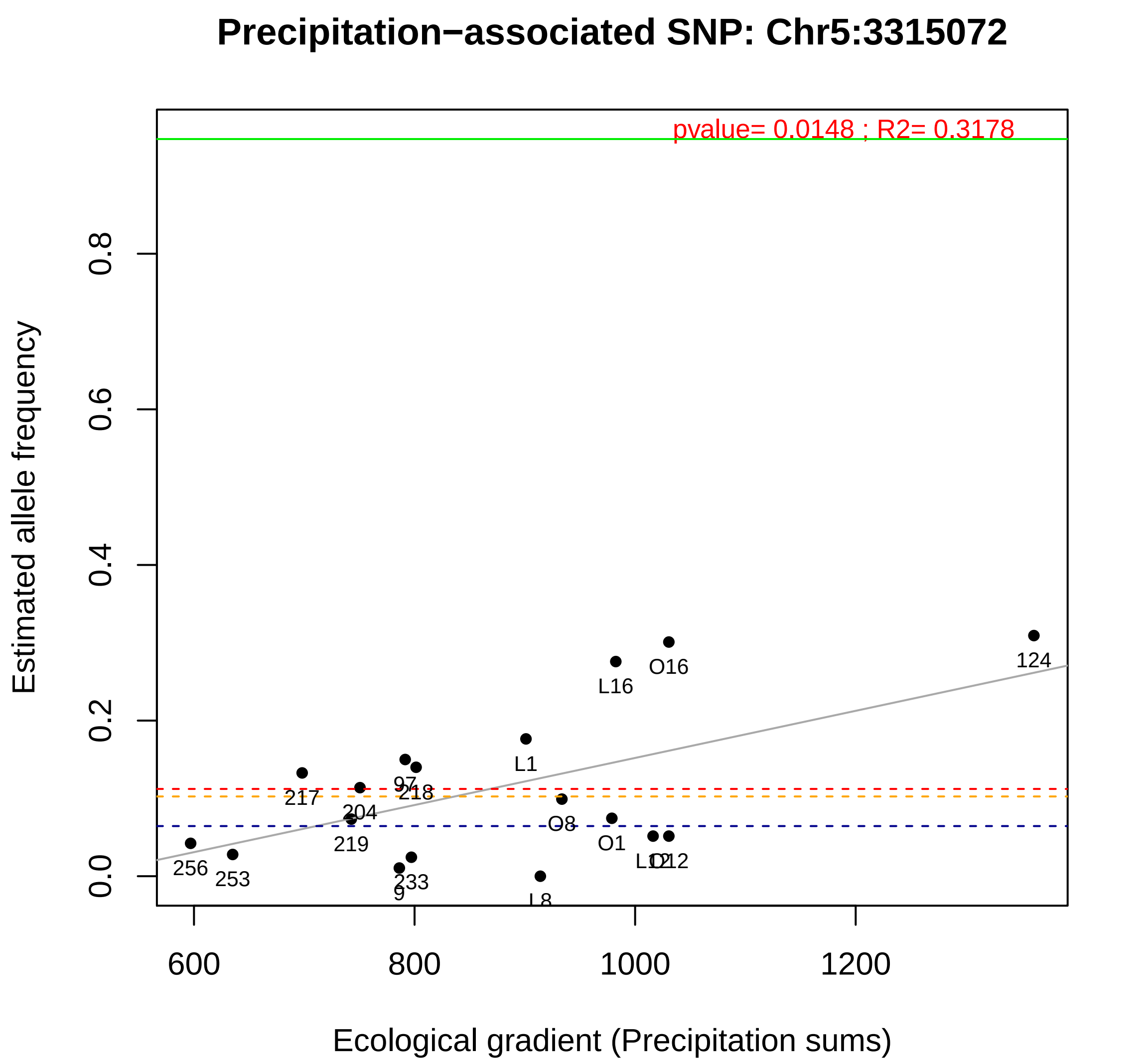

Precipitation−associated SNP: Chr5:3315072
pvalue= 0.0148 ; R2= 0.3178
Estimated allele frequency
0.0
0.2
0.4
0.6
0.8
600
800
1000
1200
Ecological gradient (Precipitation sums)
124
O16
L16
L1
97
218
217
204
219
O8
O1
256
253
233
9
L8

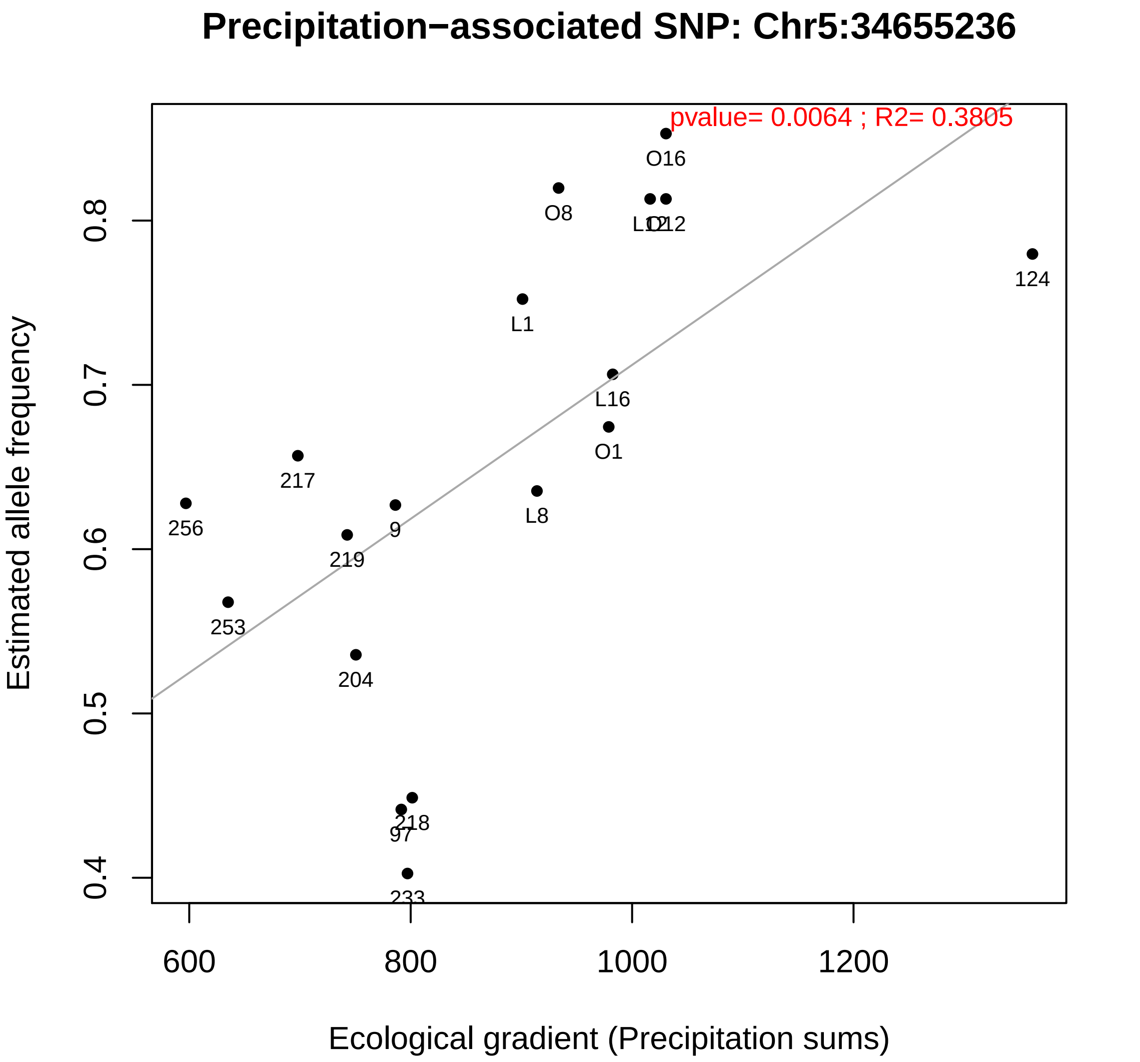

Precipitation−associated SNP: Chr5:34655236
pvalue= 0.0064 ; R2= 0.3805
Estimated allele frequency
Ecological gradient (Precipitation sums)
0.4
0.5
0.6
0.7
0.8
600
800
1000
1200
O16
O8
L10
O12
124
L1
L16
O1
217
256
9
L8
219
253
204
218
97
233

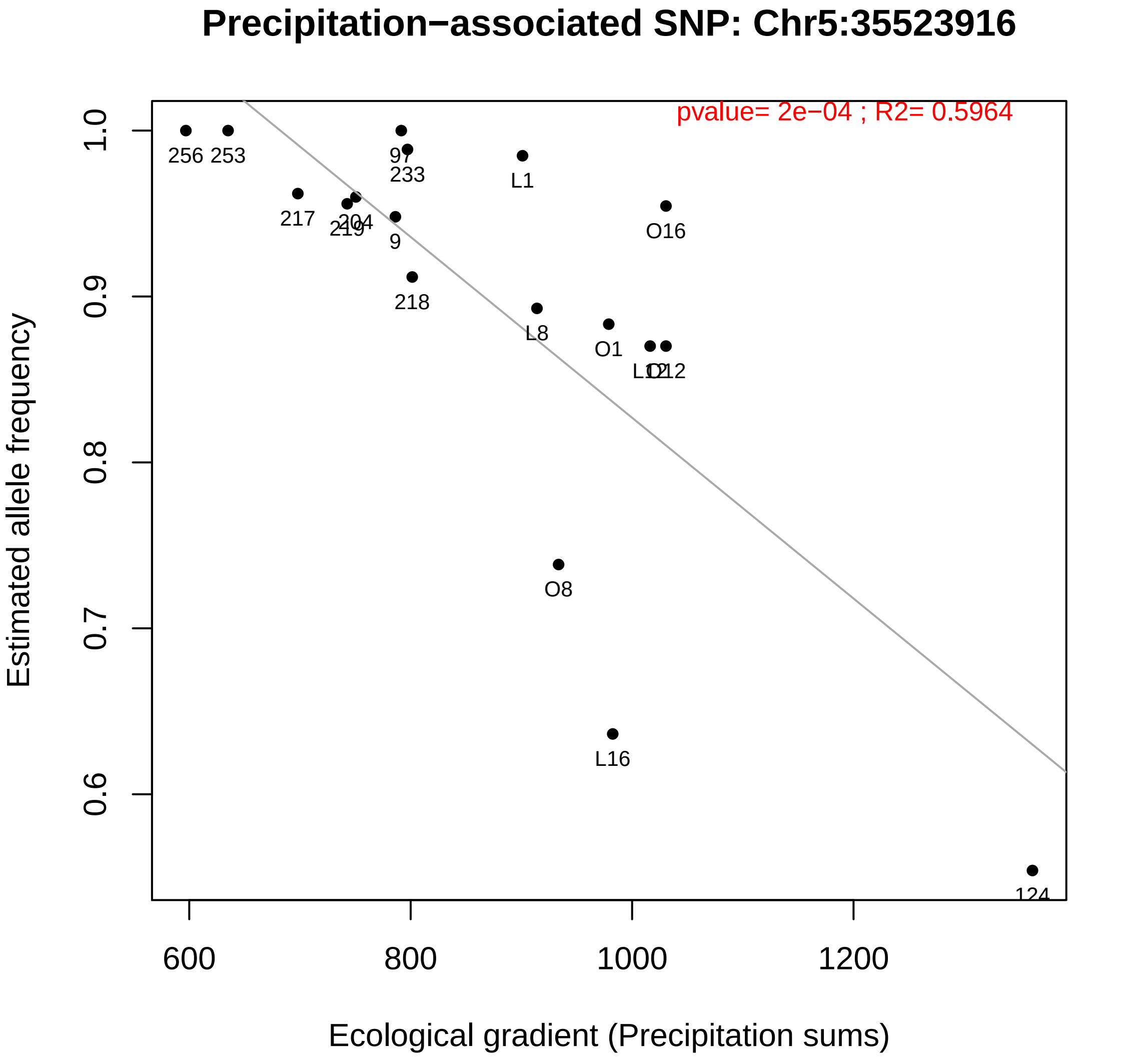
Precipitation−associated SNP: Chr5:35523916
pvalue= 2e−04 ; R2= 0.5964
Estimated allele frequency
0.6
0.7
0.8
0.9
1.0
Ecological gradient (Precipitation sums)
600
800
1000
1200
256
253
97
233
L1
217
219
204
9
O16
218
L8
O1
L12
O12
O8
L16
124

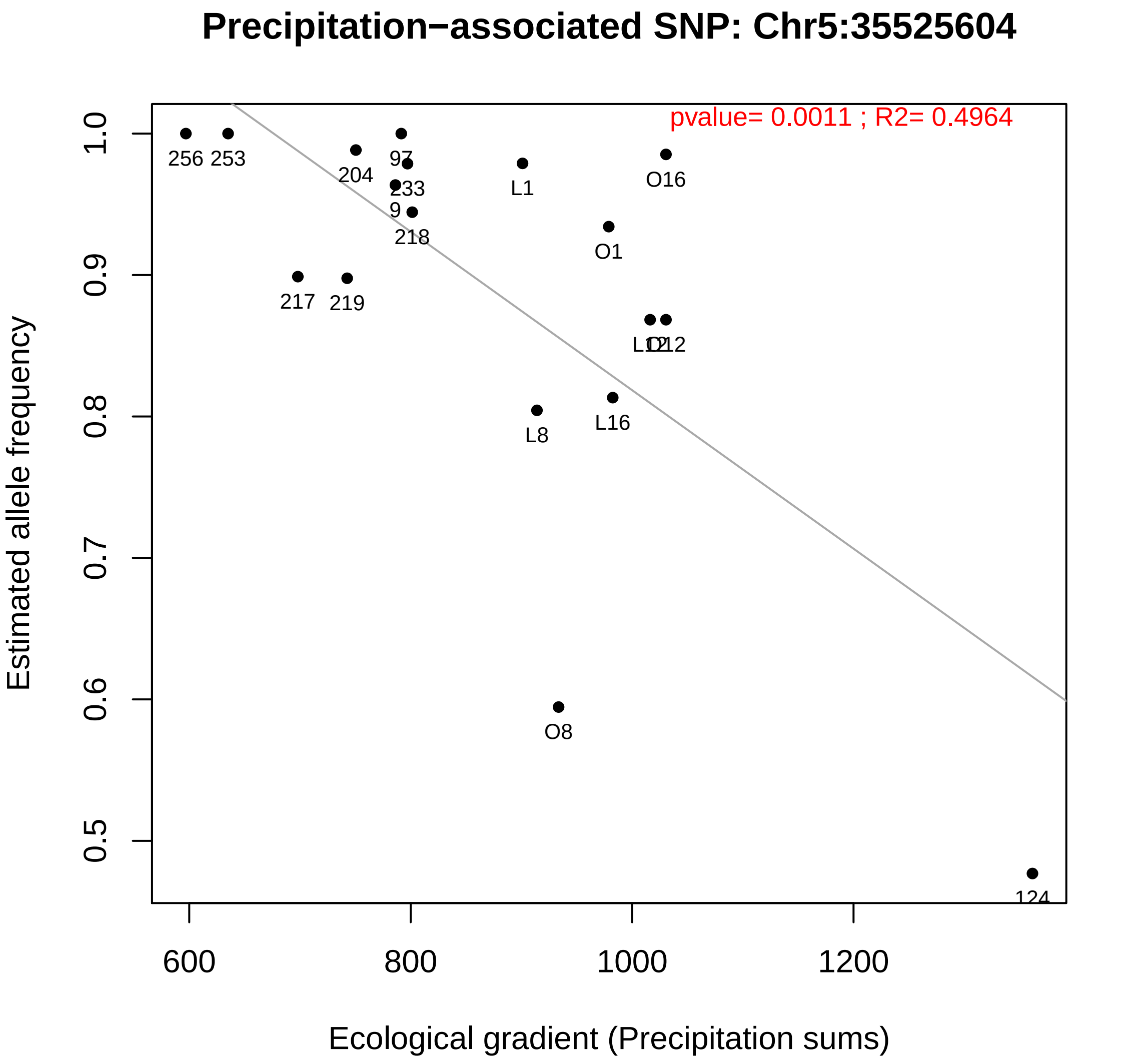

Precipitation−associated SNP: Chr5:35525604
pvalue= 0.0011 ; R2= 0.4964
Estimated allele frequency
Ecological gradient (Precipitation sums)
0.5
0.6
0.7
0.8
0.9
1.0
600
800
1000
1200
256
253
204
97
233
9
218
217
219
L1
O16
O1
L12
O12
L16
L8
O8
124

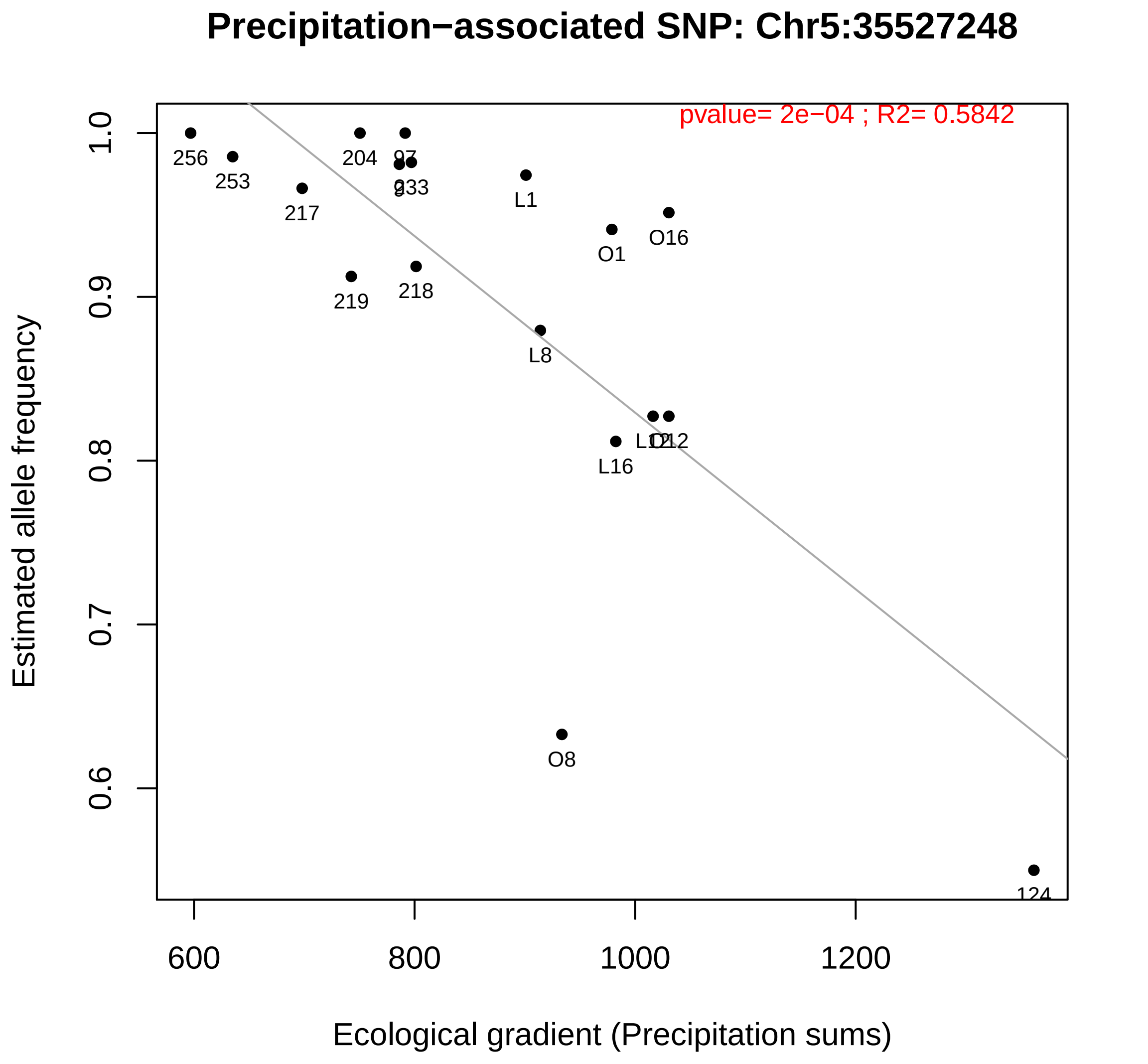

Precipitation−associated SNP: Chr5:35527248
pvalue= 2e−04 ; R2= 0.5842
256
253
217
204
97
233
219
218
L1
O1
O16
L8
L16
O12
O8
124
1.0
0.9
0.8
0.7
0.6
600
800
1000
1200
Estimated allele frequency
Ecological gradient (Precipitation sums)

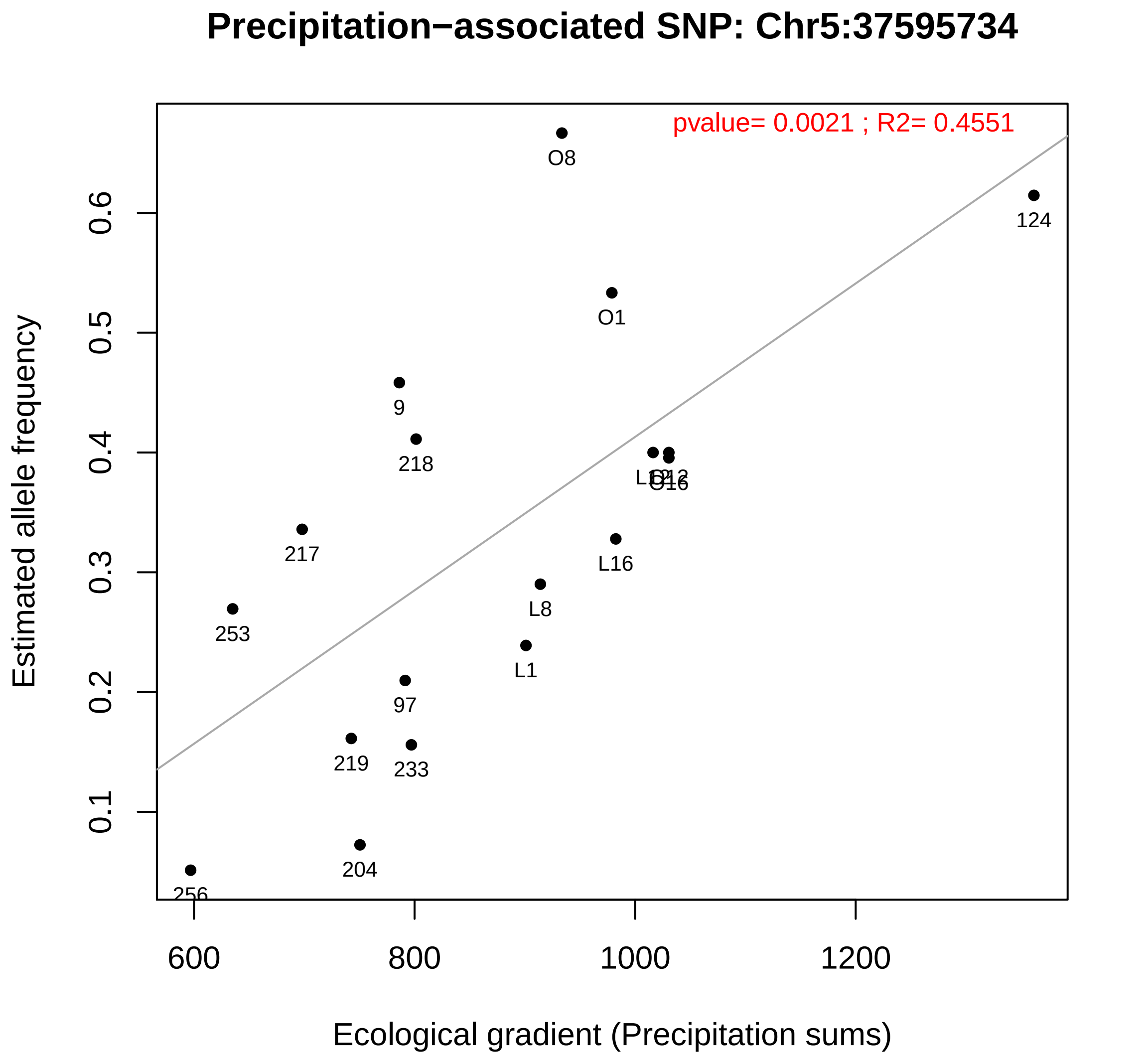

Precipitation−associated SNP: Chr5:37595734
pvalue= 0.0021 ; R2= 0.4551
Estimated allele frequency
Ecological gradient (Precipitation sums)
0.1
0.2
0.3
0.4
0.5
0.6
600
800
1000
1200
O8
124
O1
9
218
217
253
L16
L8
L1
97
219
233
204
256

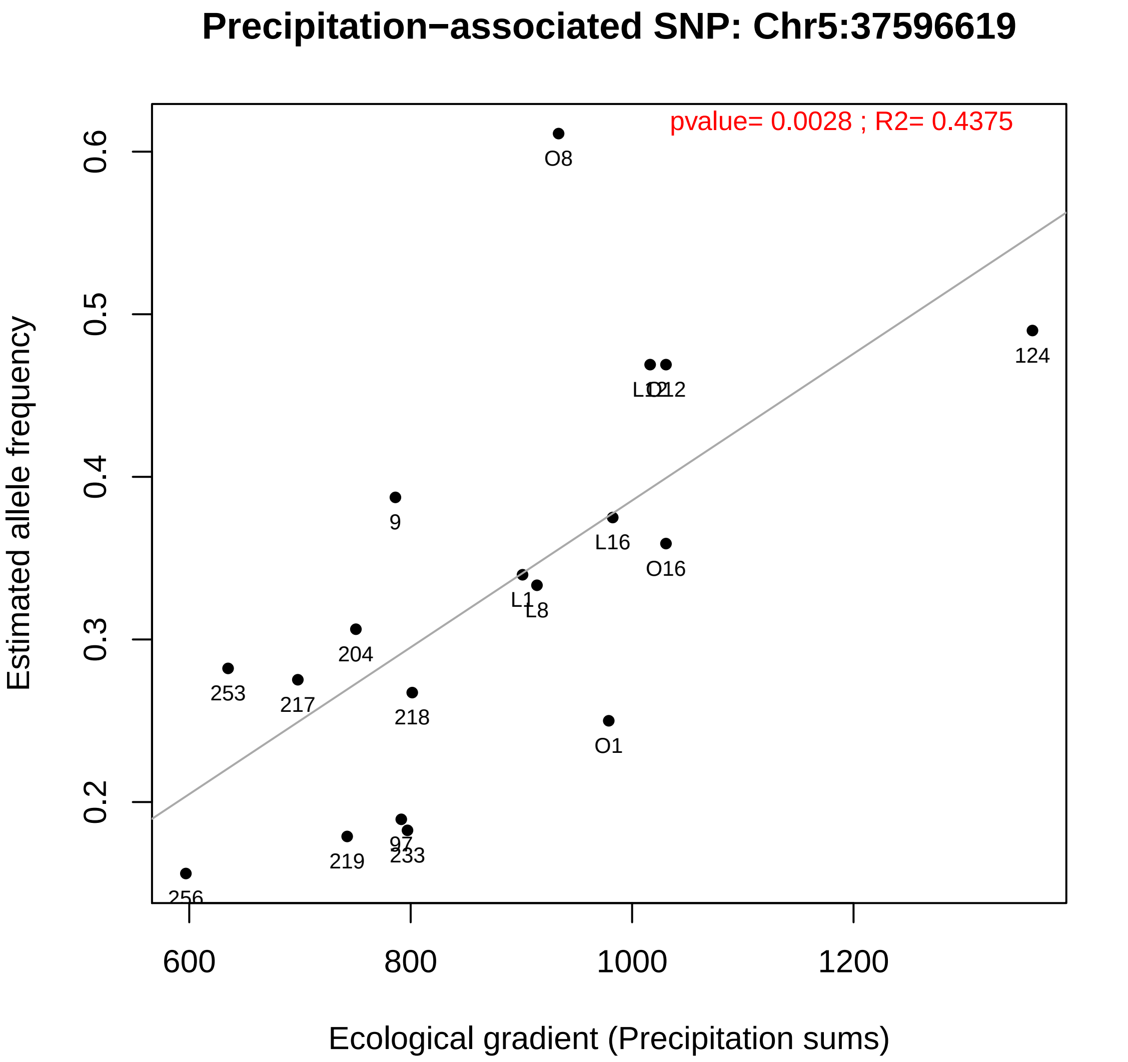

Precipitation−associated SNP: Chr5:37596619
pvalue= 0.0028 ; R2= 0.4375
Estimated allele frequency
0.2
0.3
0.4
0.5
0.6
600
800
1000
1200
Ecological gradient (Precipitation sums)
O8
124
L12
O12
9
L16
O16
L1
L8
204
253
217
218
O1
97
233
219
256

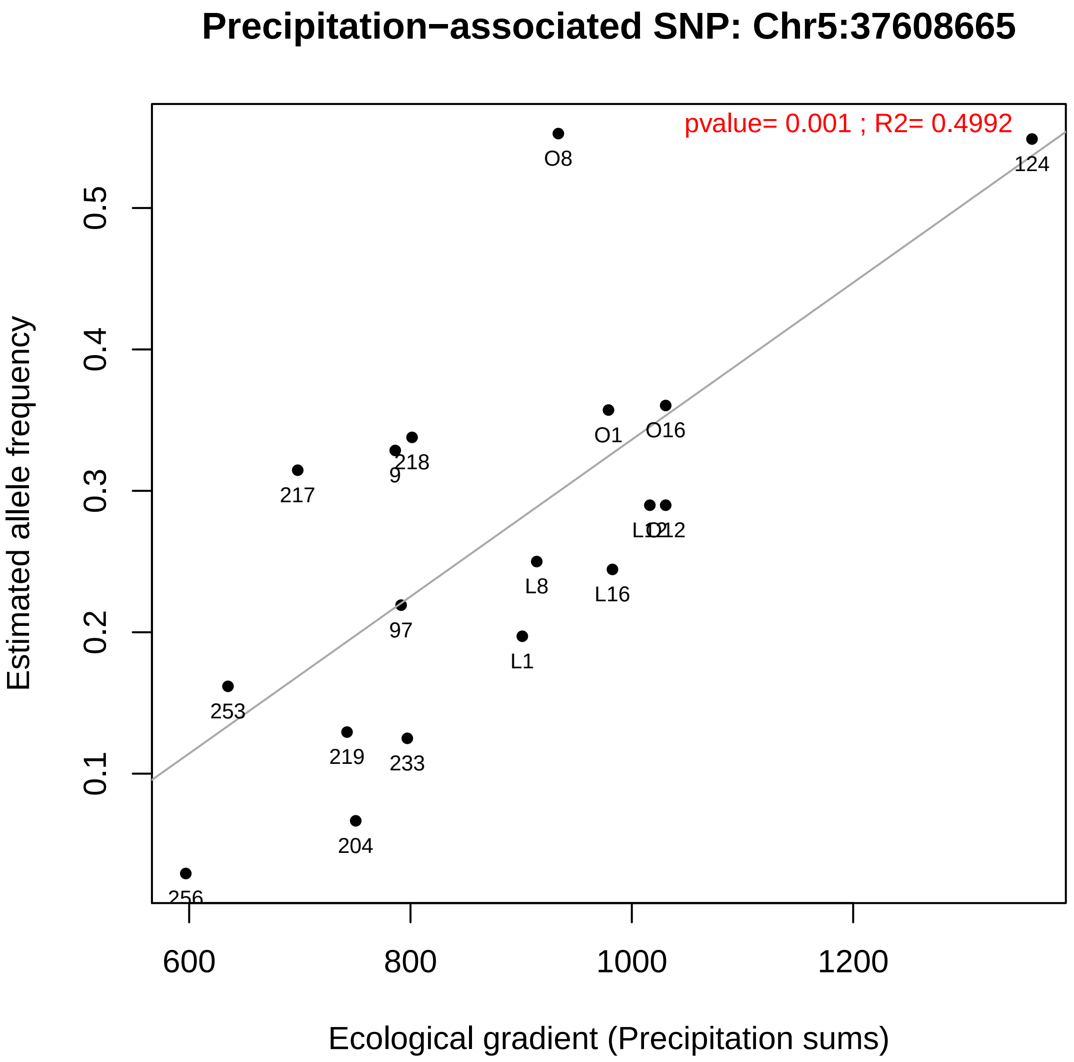

Precipitation−associated SNP: Chr5:37608665
pvalue= 0.001 ; R2= 0.4992
O8
124
O1
O16
218
9
217
L12
O12
L8
L16
97
L1
253
219
233
204
256
0.1
0.2
0.3
0.4
0.5
600
800
1000
1200
Estimated allele frequency
Ecological gradient (Precipitation sums)

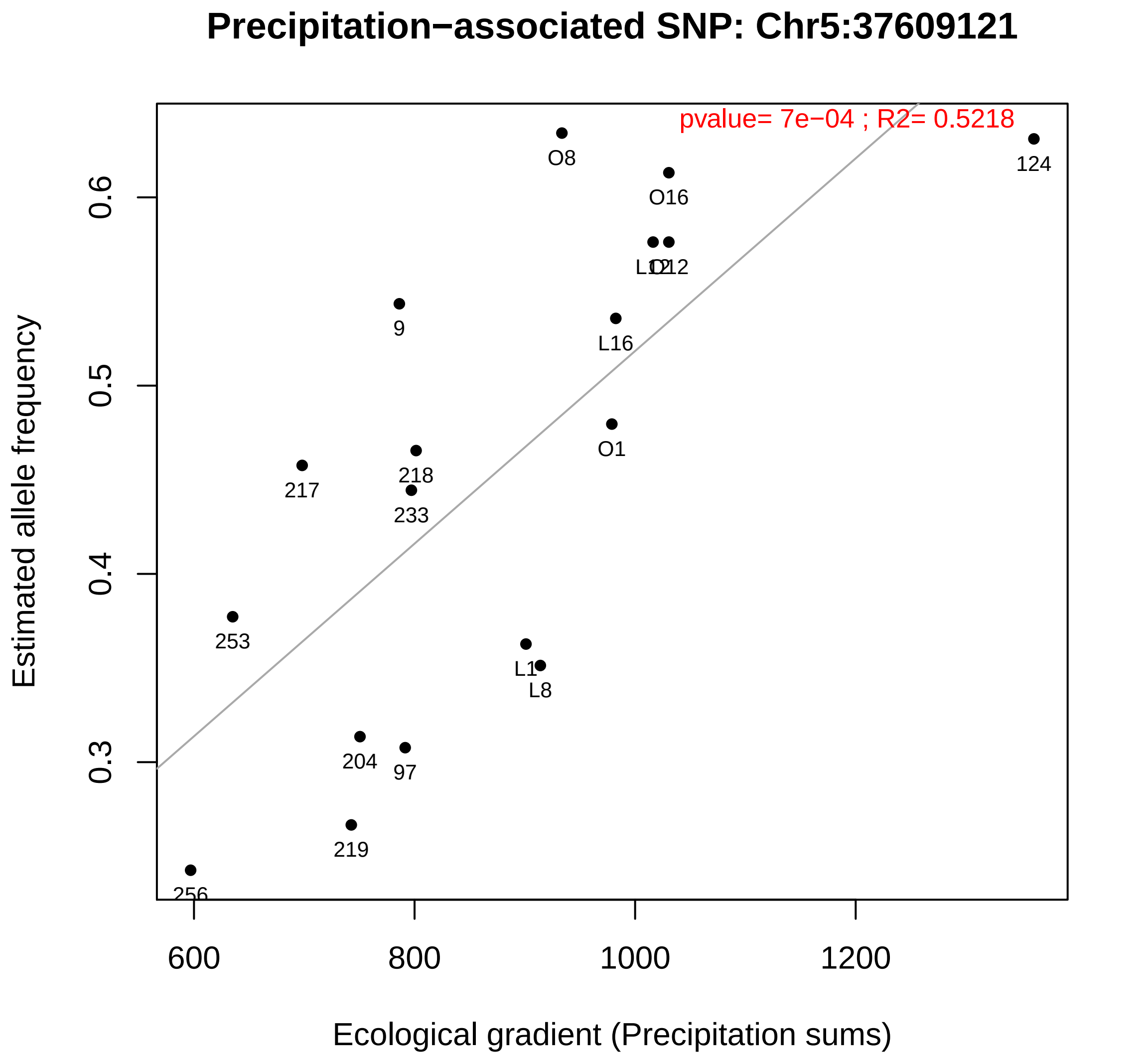
Precipitation−associated SNP: Chr5:37609121
pvalue= 7e−04 ; R2= 0.5218
Estimated allele frequency
0.3
0.4
0.5
0.6
600
800
1000
1200
Ecological gradient (Precipitation sums)
O8
124
O16
L12
O12
9
L16
O1
217
218
233
253
L1
L8
204
97
219
256

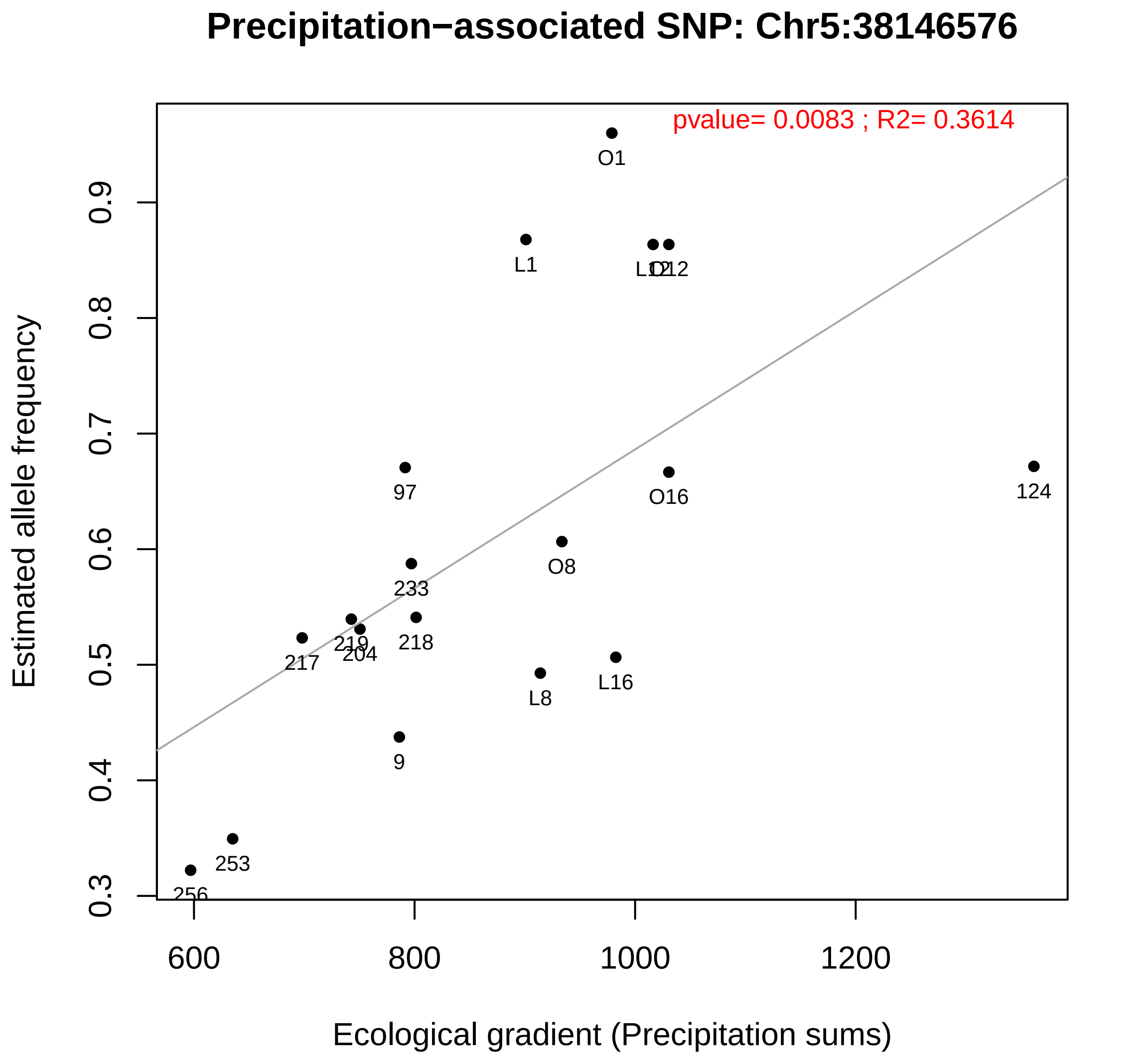

Precipitation−associated SNP: Chr5:38146576
pvalue= 0.0083 ; R2= 0.3614
O1
L1
L12
O12
97
O16
124
O8
233
218
219
204
217
L16
L8
9
253
256
Estimated allele frequency
0.3
0.4
0.5
0.6
0.7
0.8
0.9
600
800
1000
1200
Ecological gradient (Precipitation sums)

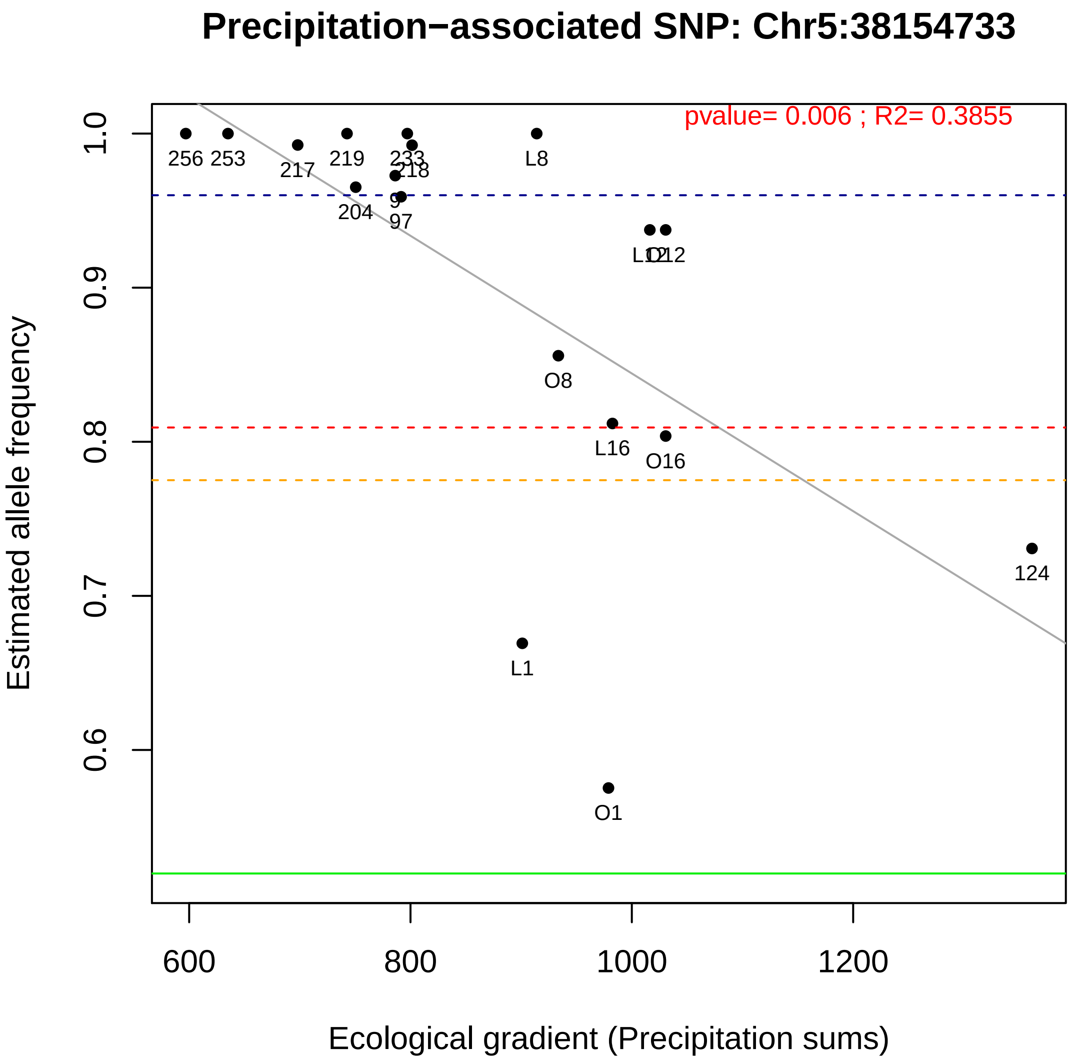
Precipitation−associated SNP: Chr5:38154733
pvalue= 0.006 ; R2= 0.3855
Estimated allele frequency
Ecological gradient (Precipitation sums)
1.0
0.9
0.8
0.7
0.6
600
800
1000
1200
256
253
217
219
233
218
204
9
97
L8
L12
O12
O8
L16
O16
124
L1
O1

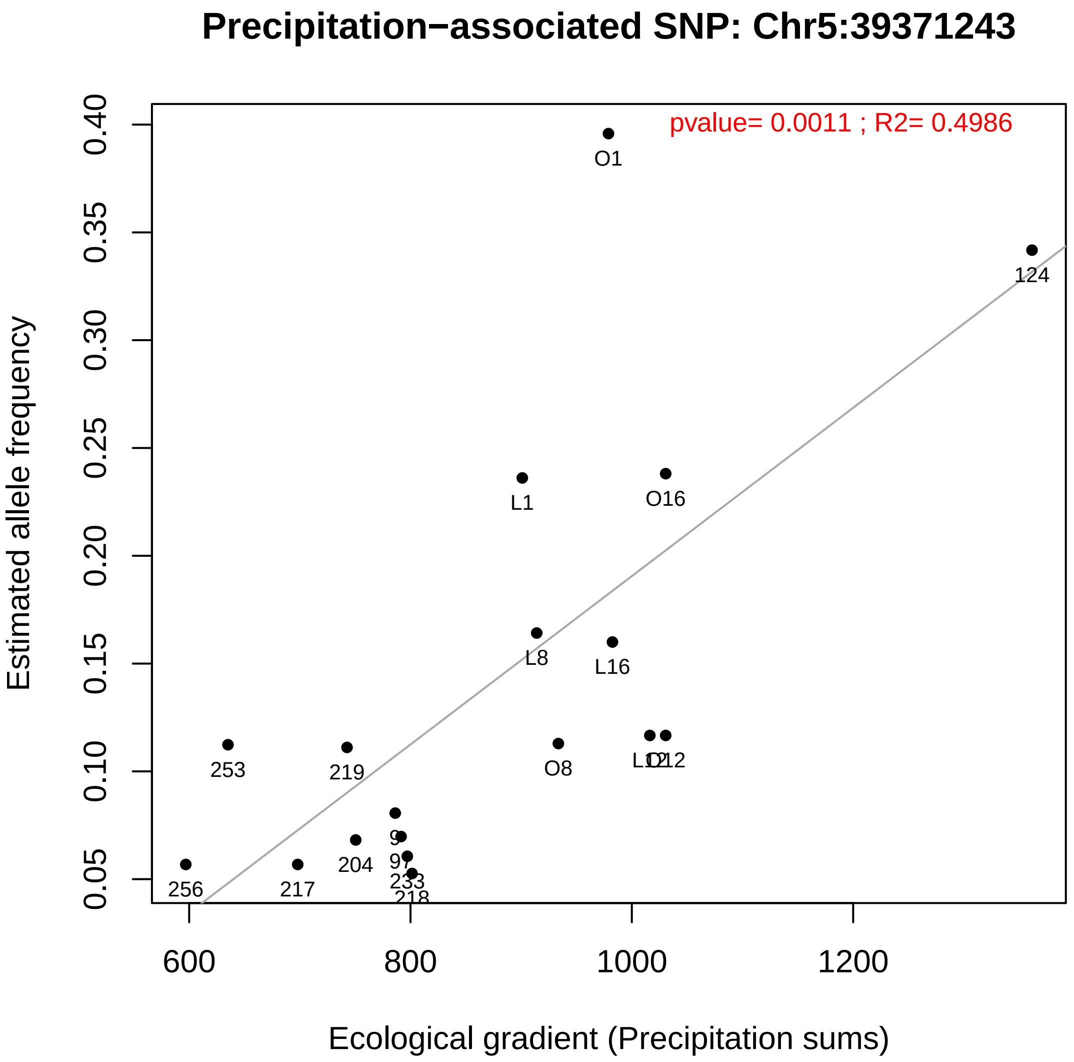

Precipitation−associated SNP: Chr5:39371243
pvalue= 0.0011 ; R2= 0.4986
Estimated allele frequency
Ecological gradient (Precipitation sums)
0.05
0.10
0.15
0.20
0.25
0.30
0.35
0.40
600
800
1000
1200
O1
124
L1
O16
L8
L16
253
219
O8
O12
204
256
217
233
218

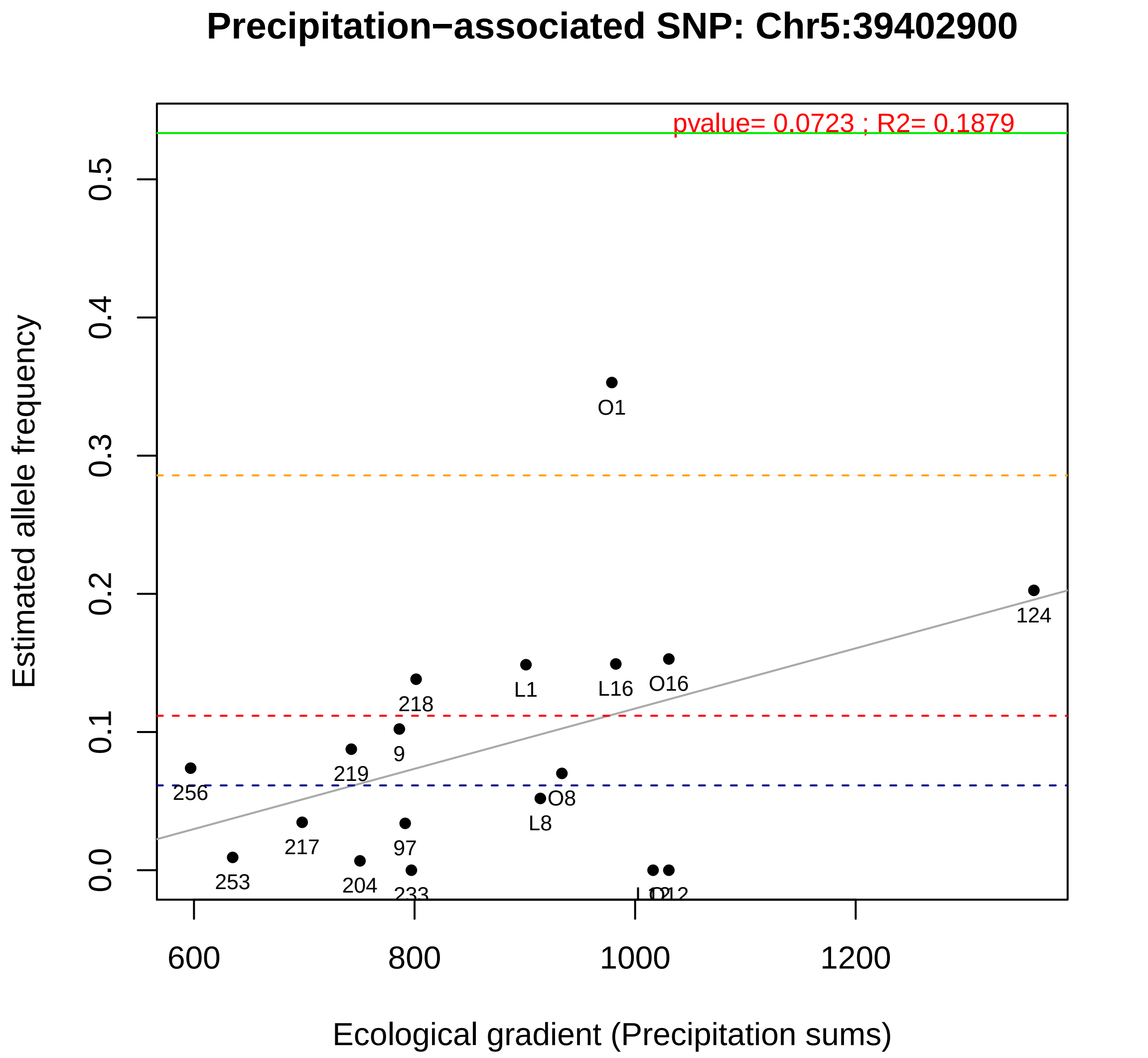
Precipitation−associated SNP: Chr5:39402900
pvalue= 0.0723 ; R2= 0.1879
Estimated allele frequency
0.0
0.1
0.2
0.3
0.4
0.5
600
800
1000
1200
Ecological gradient (Precipitation sums)
O1
124
L1
L16
O16
218
9
219
256
O8
L8
217
97
253
204
233

**Precipitation−associated SNP: Chr5:40192876**

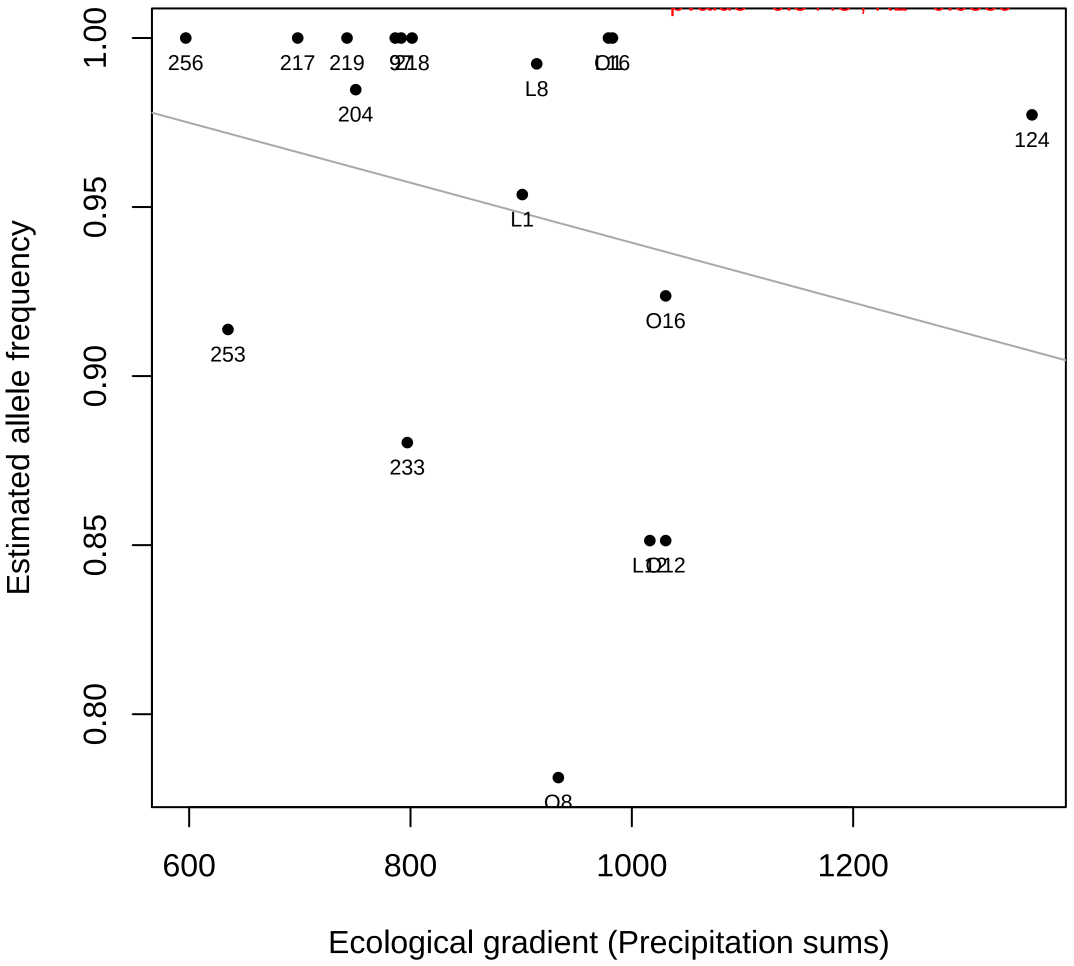

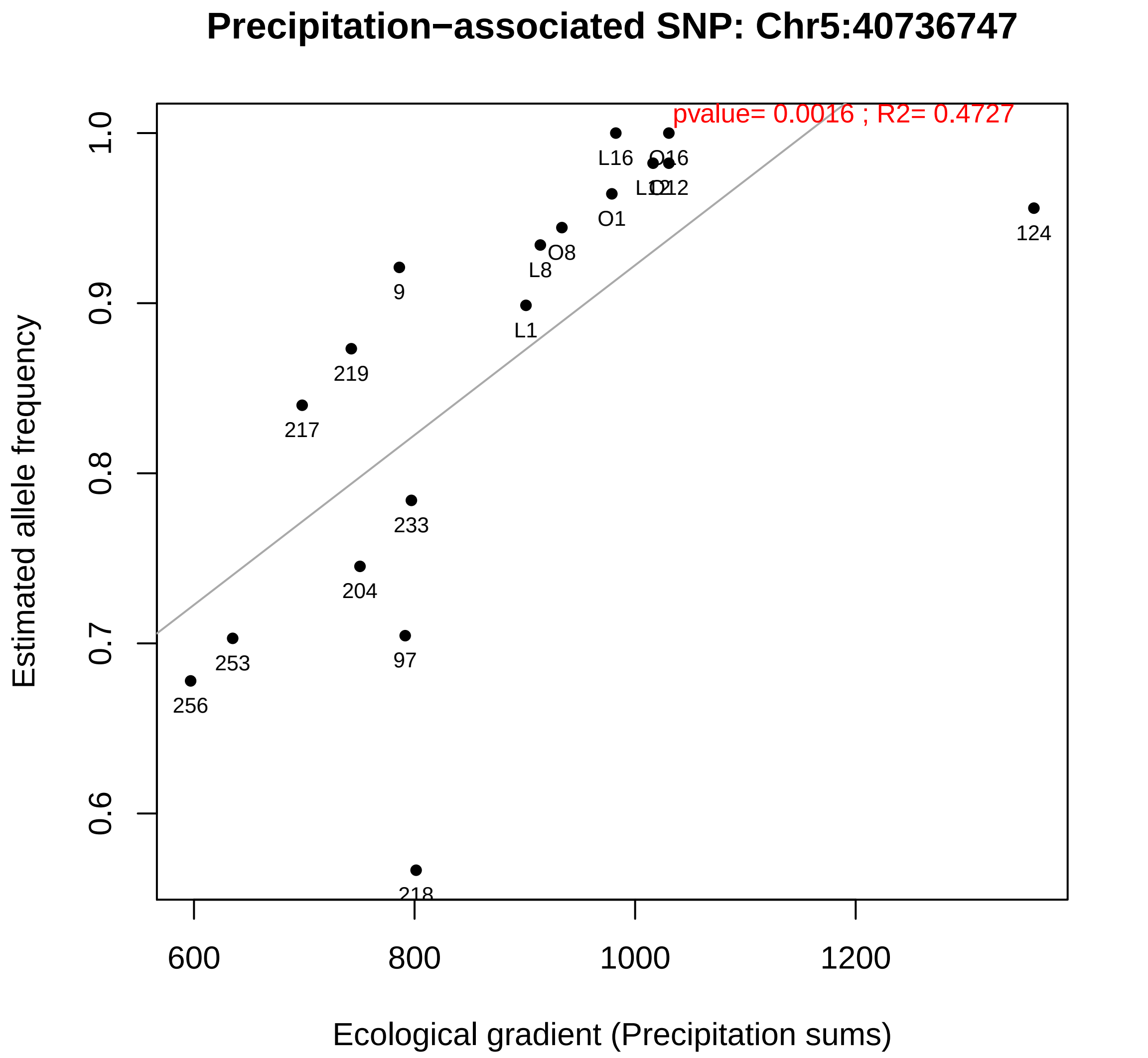
Precipitation−associated SNP: Chr5:40736747
pvalue= 0.0016 ; R2= 0.4727
Estimated allele frequency
Ecological gradient (Precipitation sums)
0.6
0.7
0.8
0.9
1.0
600
800
1000
1200
L16
O16
L12
O12
O1
124
O8
L8
9
L1
219
217
233
204
253
97
256
218

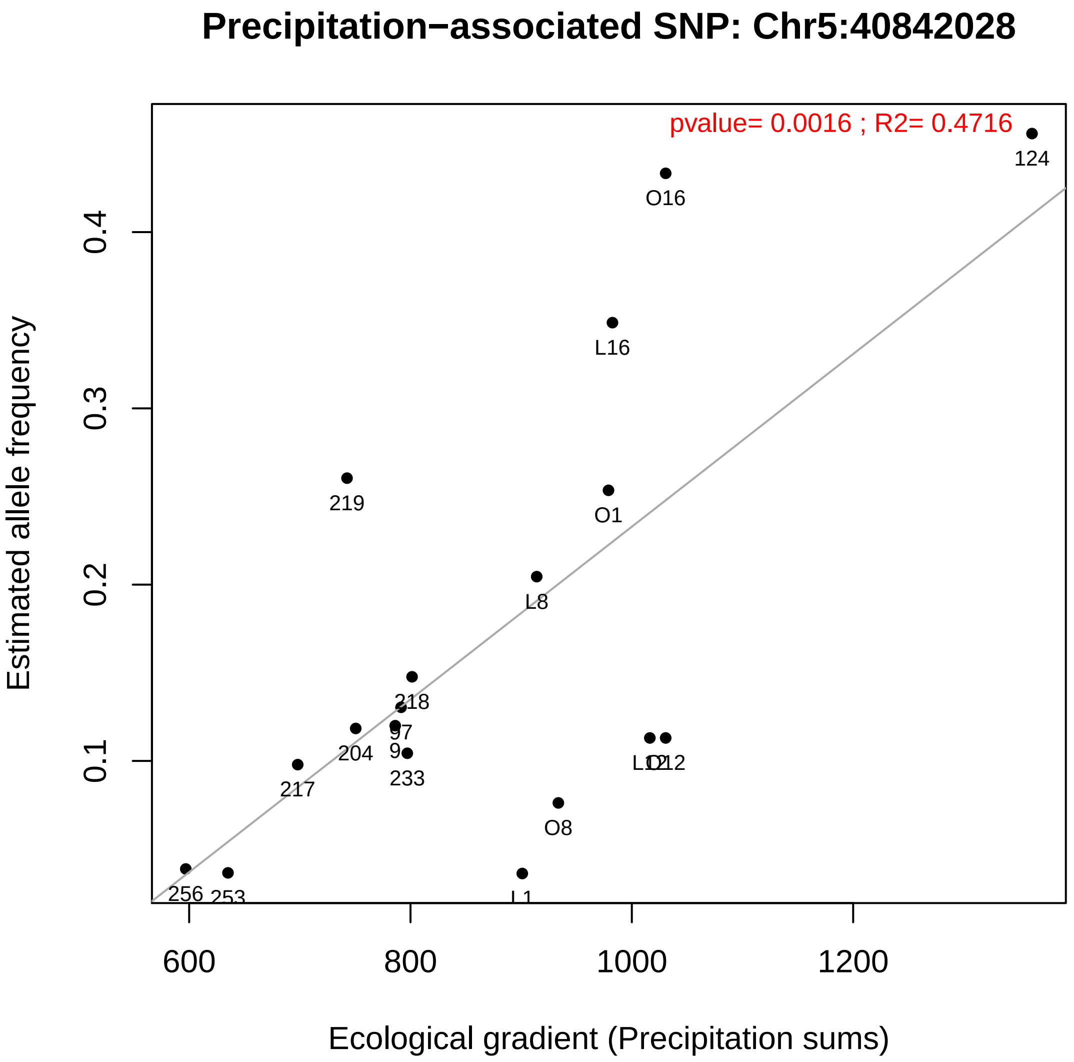

Precipitation−associated SNP: Chr5:40842028
pvalue= 0.0016 ; R2= 0.4716
Estimated allele frequency
Ecological gradient (Precipitation sums)
0.1
0.2
0.3
0.4
600
800
1000
1200
124
O16
L16
219
O1
L8
218
97
204
233
217
O8
256

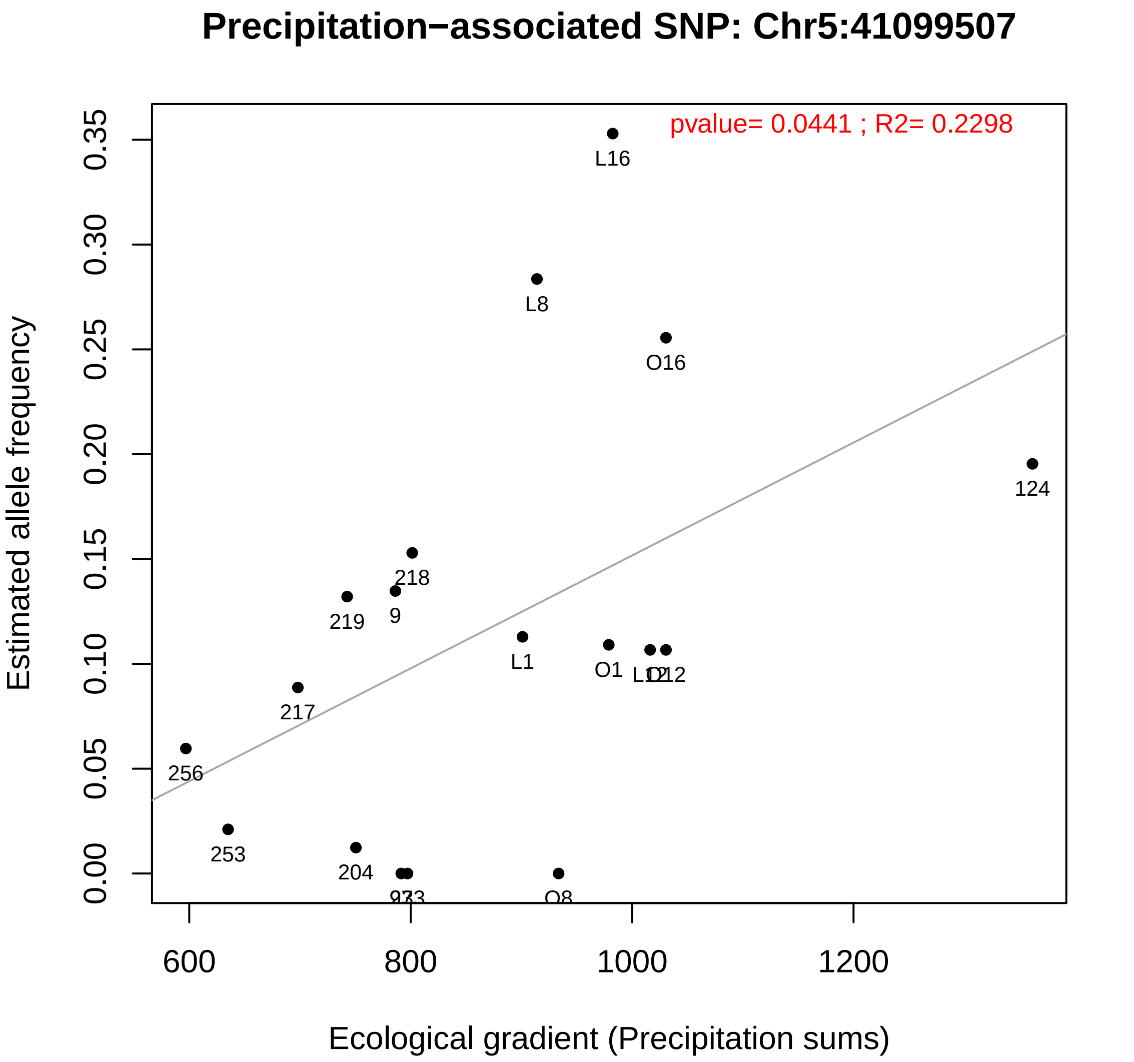

Precipitation−associated SNP: Chr5:41099507
pvalue= 0.0441 ; R2= 0.2298
Estimated allele frequency
Ecological gradient (Precipitation sums)
0.00
0.05
0.10
0.15
0.20
0.25
0.30
0.35
600
800
1000
1200
L16
L8
O16
124
218
219
9
L1
O1
217
256
253
204
O8

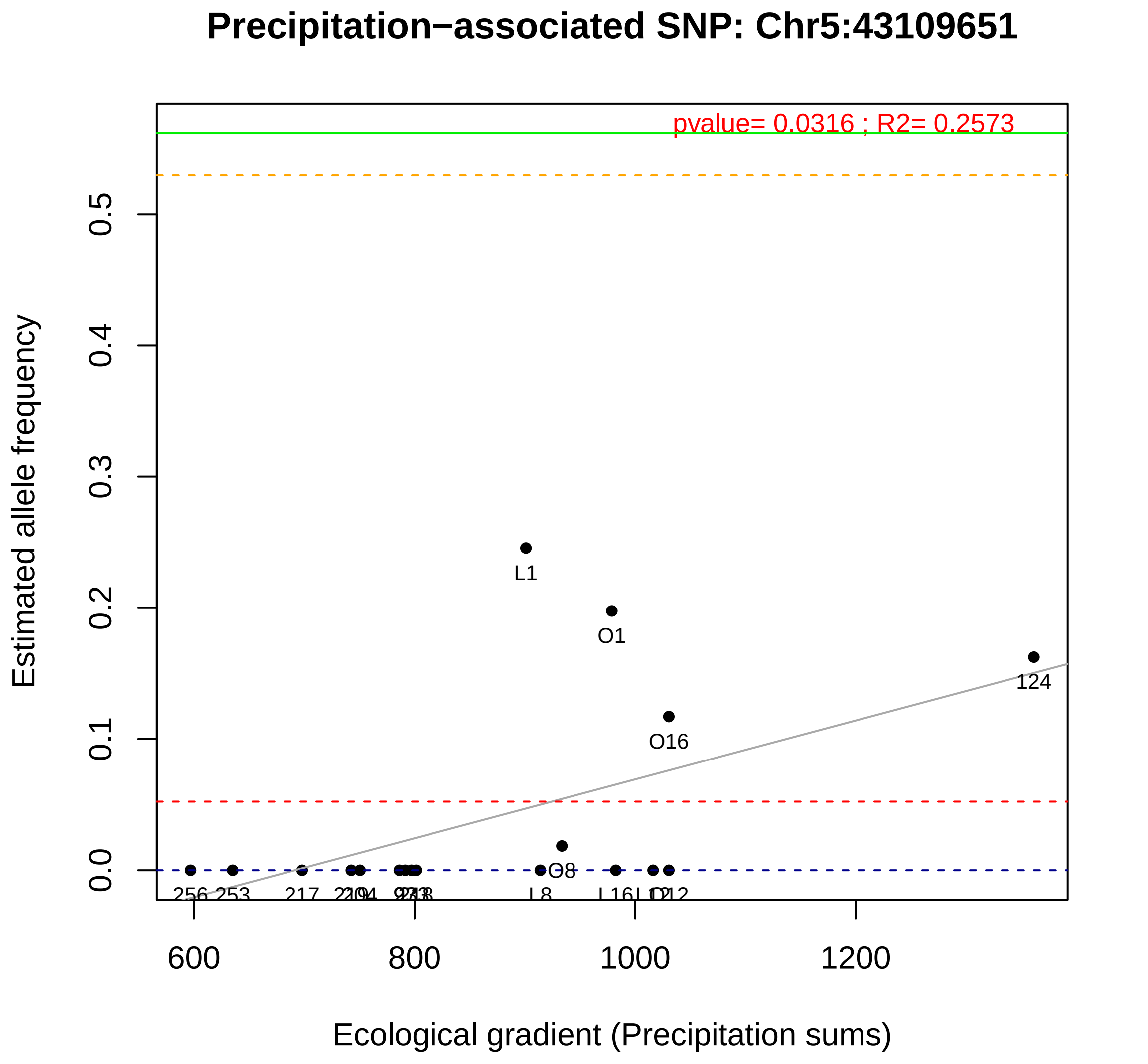

Precipitation−associated SNP: Chr5:43109651
pvalue= 0.0316 ; R2= 0.2573
Estimated allele frequency
0.0
0.1
0.2
0.3
0.4
0.5
600
800
1000
1200
Ecological gradient (Precipitation sums)
L1
O1
124
O16
O8
256
253
217
L8

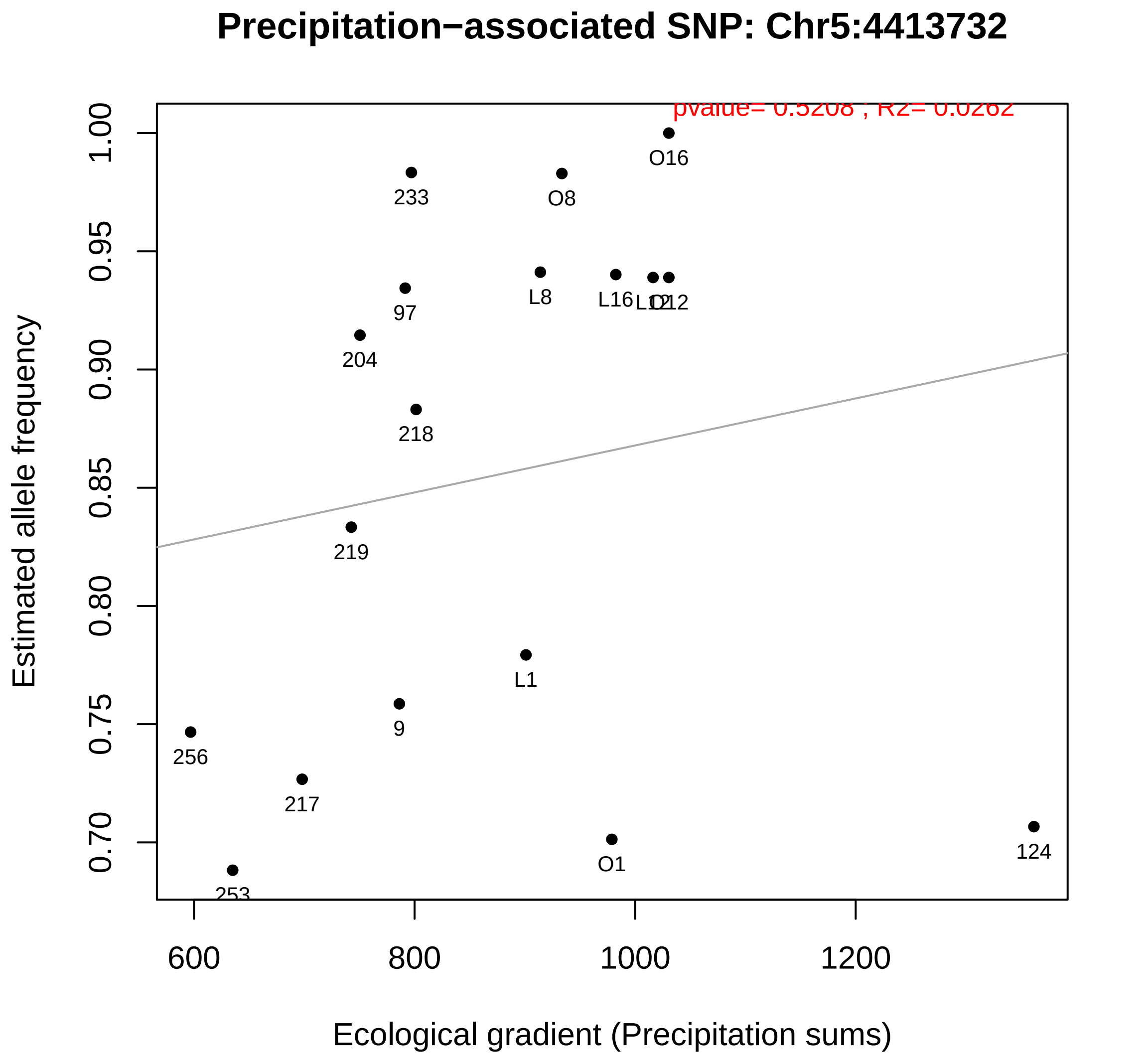

Precipitation−associated SNP: Chr5:4413732
pvalue= 0.5208 ; R2= 0.0262
Estimated allele frequency
Ecological gradient (Precipitation sums)
1.00
0.95
0.90
0.85
0.80
0.75
0.70
600
800
1000
1200
O16
233
O8
L8
L16
L12
O12
97
204
218
219
L1
9
256
217
O1
124
253

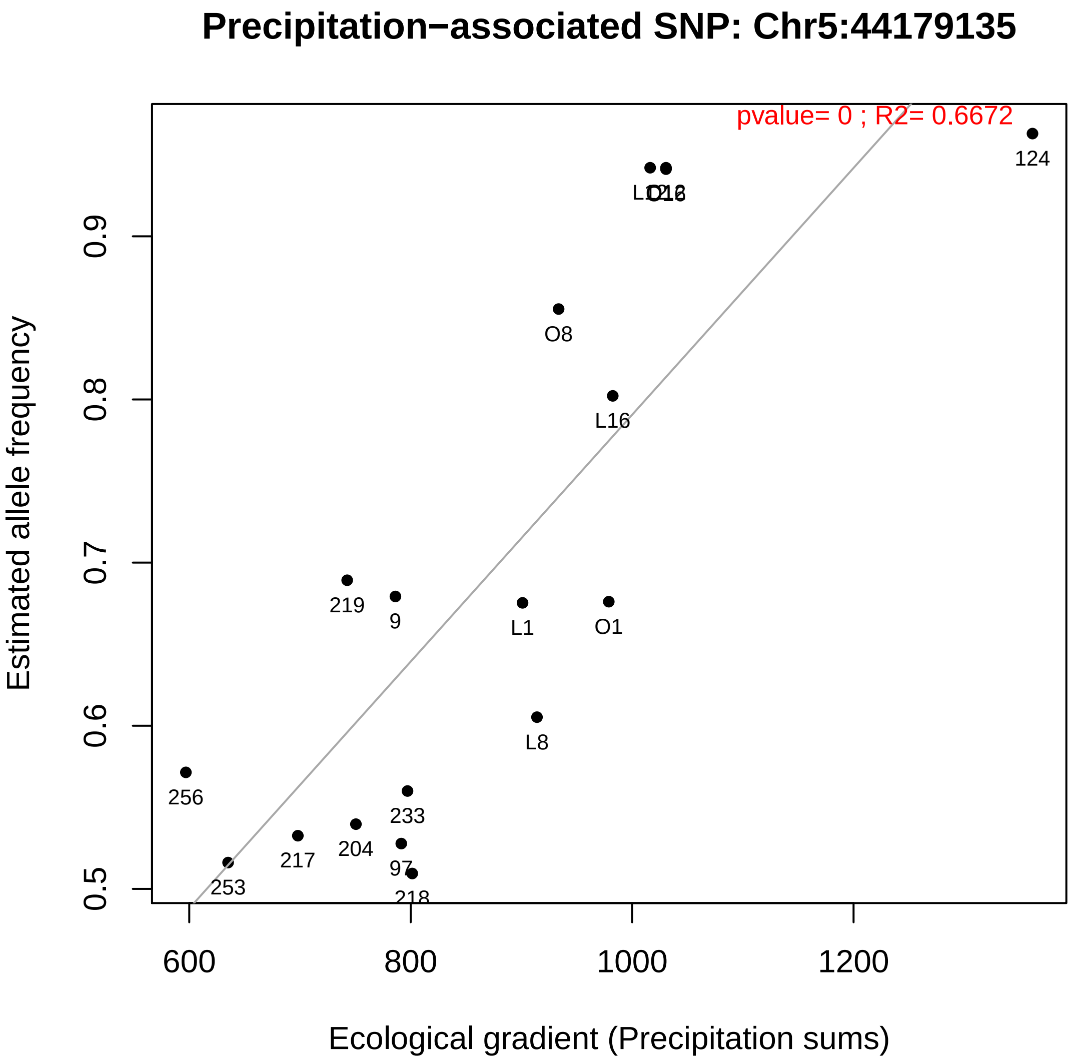

Precipitation−associated SNP: Chr5:44179135
pvalue= 0 ; R2= 0.6672
124
O8
L16
219
9
L1
O1
L8
256
233
204
217
97
253
218
0.5
0.6
0.7
0.8
0.9
Estimated allele frequency
600
800
1000
1200
Ecological gradient (Precipitation sums)

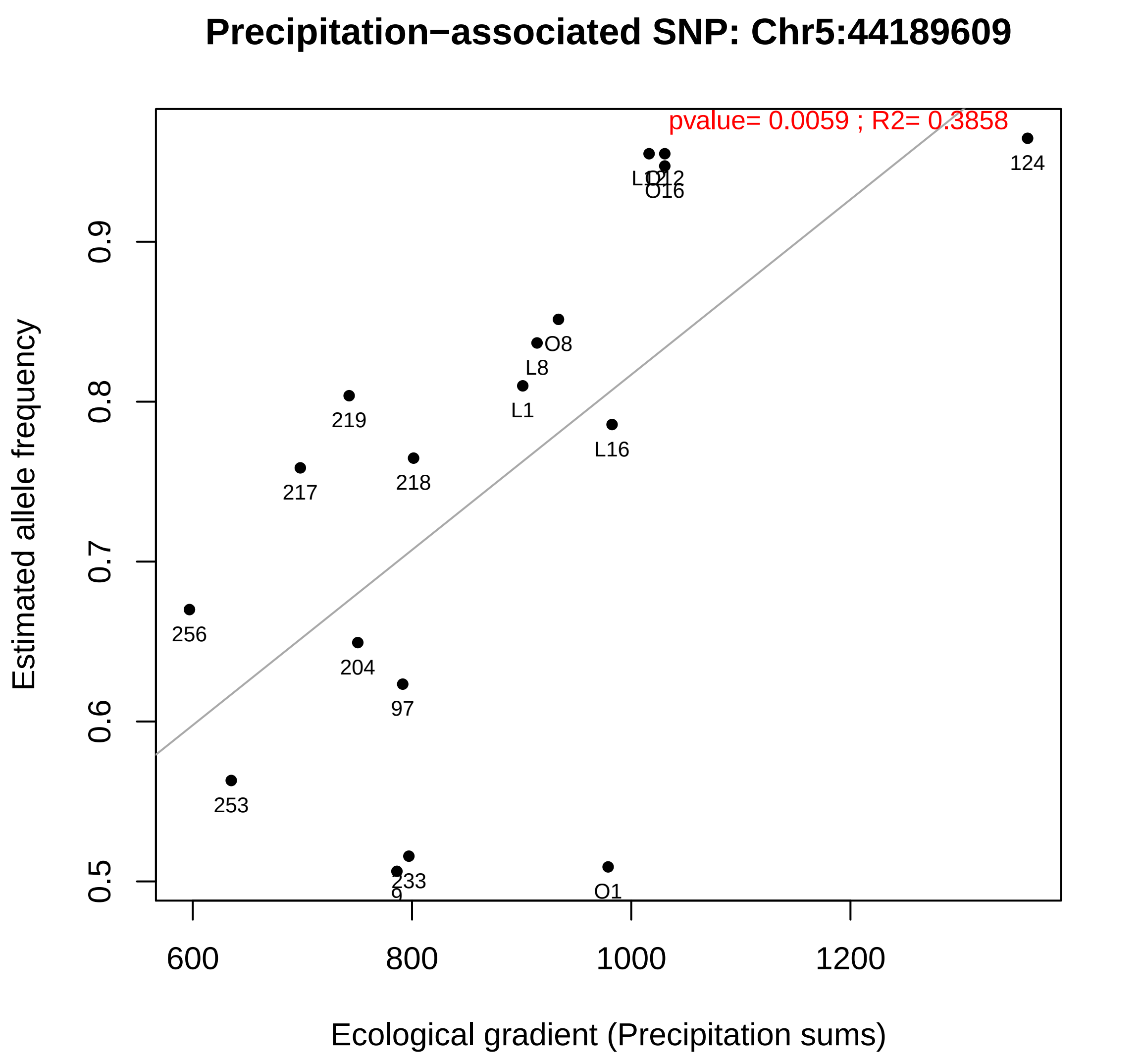
Precipitation−associated SNP: Chr5:44189609
pvalue= 0.0059 ; R2= 0.3858
Estimated allele frequency
0.5
0.6
0.7
0.8
0.9
600
800
1000
1200
Ecological gradient (Precipitation sums)
124
O12
O16
O8
L8
L1
L16
219
217
218
256
204
97
253
233
O1

Precipitation−associated SNP: Chr5:44193125

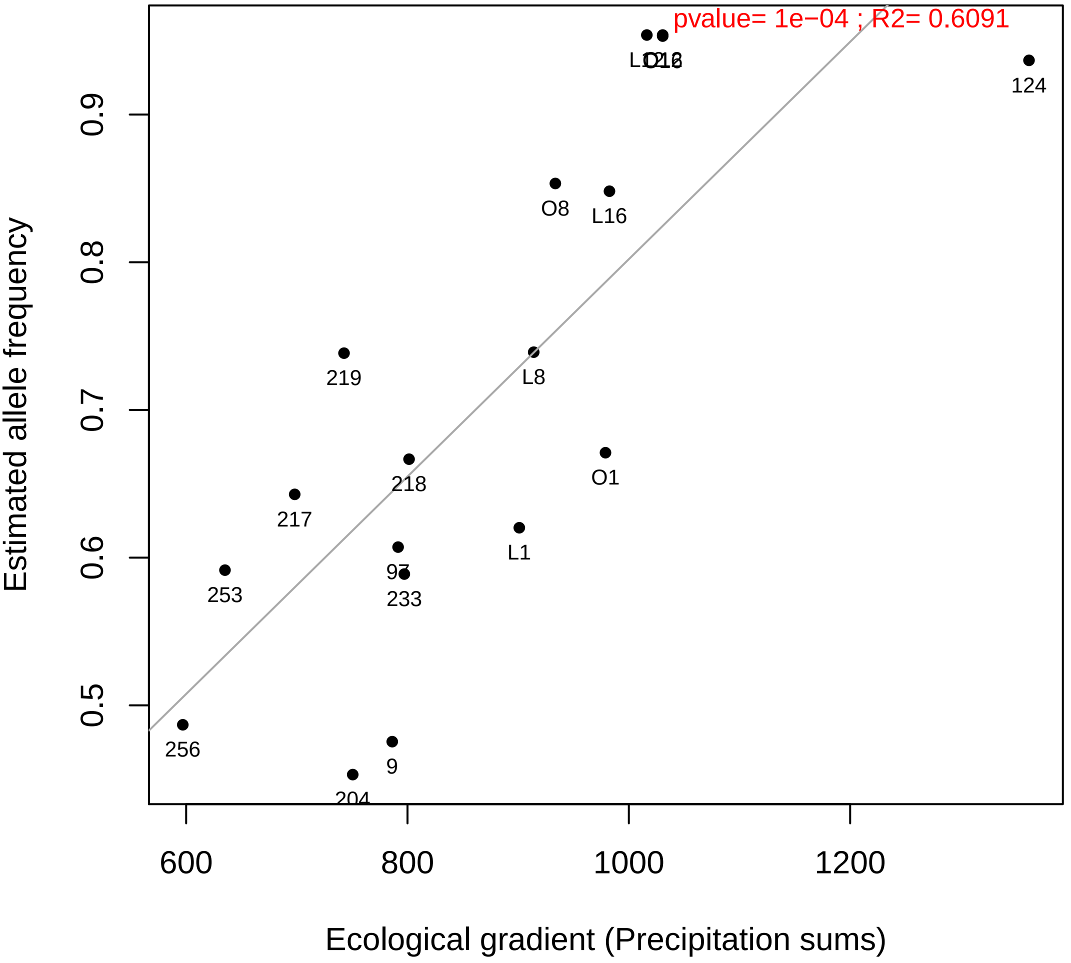

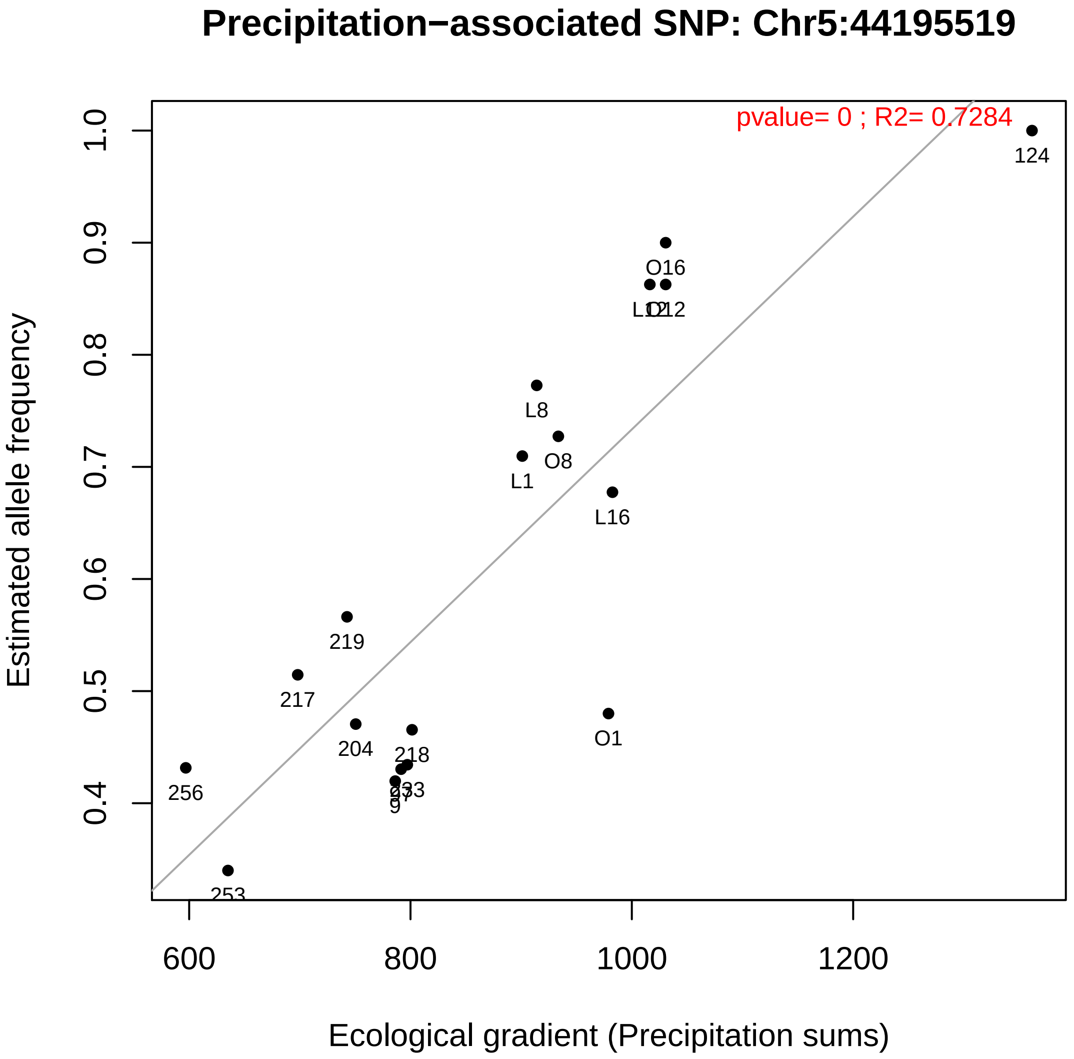
Precipitation−associated SNP: Chr5:44195519
pvalue= 0 ; R2= 0.7284
Estimated allele frequency
Ecological gradient (Precipitation sums)
0.4
0.5
0.6
0.7
0.8
0.9
1.0
600
800
1000
1200
124
O16
L8
O8
L1
L16
219
217
204
218
256
O1

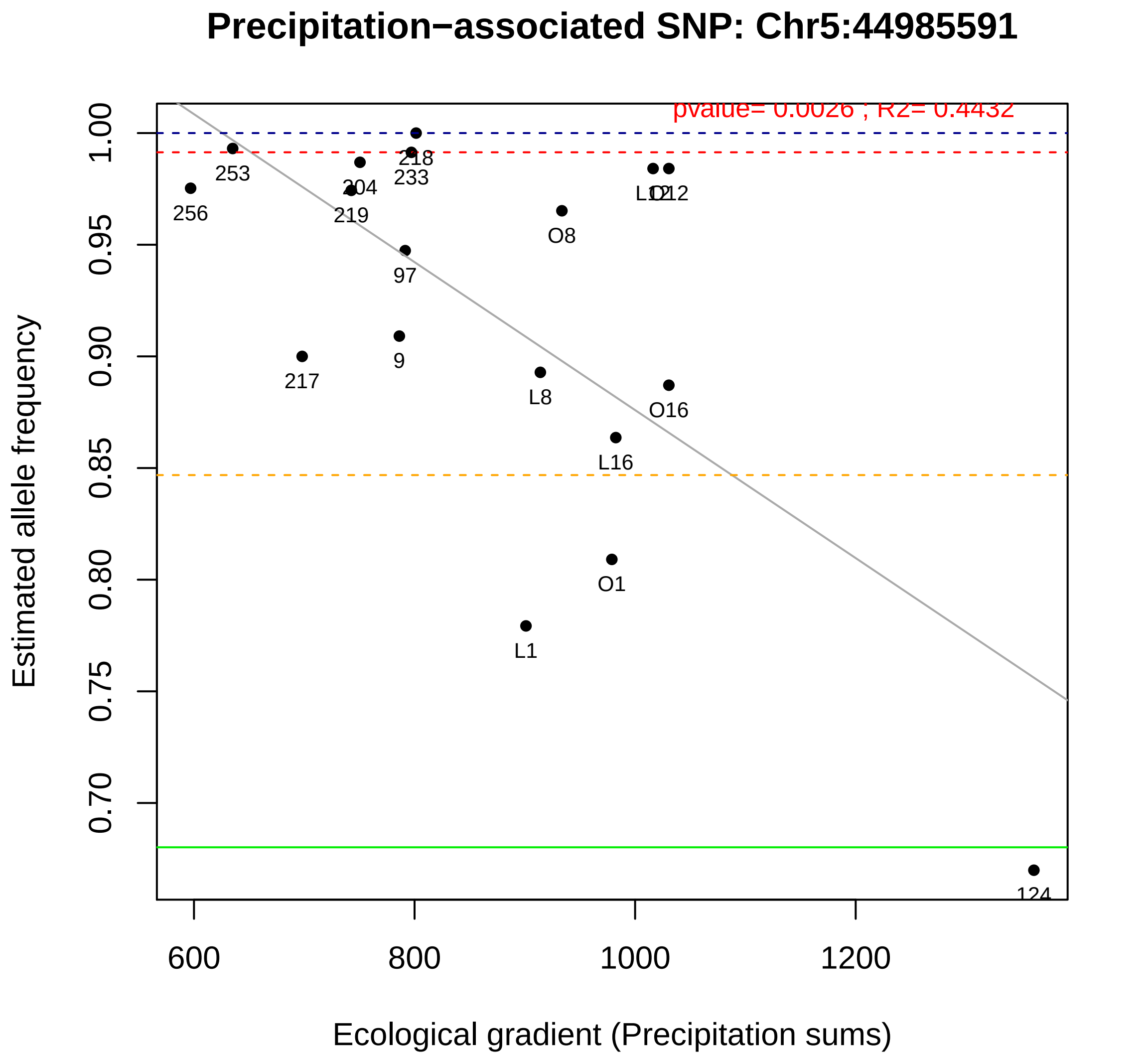
Precipitation−associated SNP: Chr5:44985591
pvalue= 0.0026 ; R2= 0.4432
Estimated allele frequency
Ecological gradient (Precipitation sums)
0.70
0.75
0.80
0.85
0.90
0.95
1.00
600
800
1000
1200
256
253
204
219
218
233
97
9
217
L8
L1
O8
L16
O1
O16
L12
O12
124

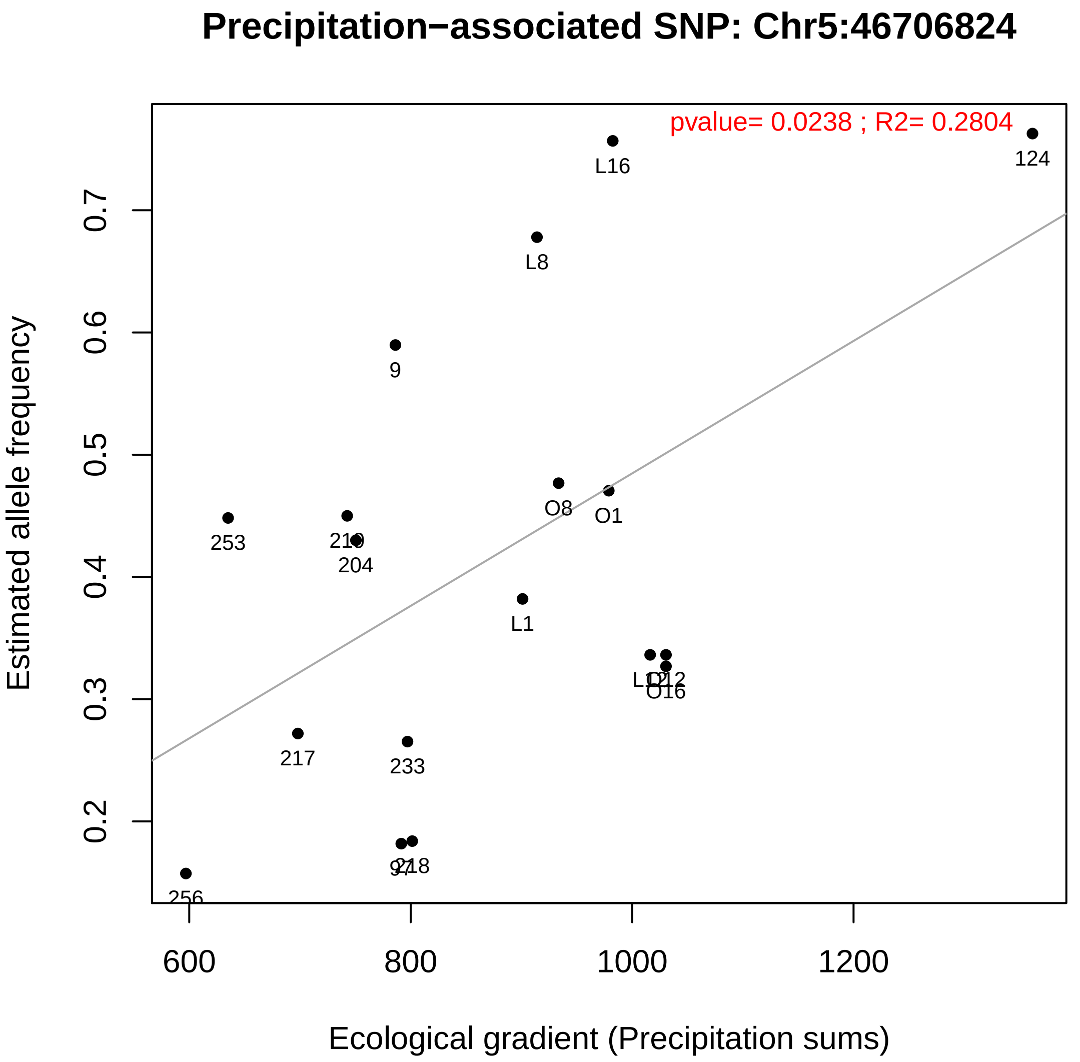
Precipitation−associated SNP: Chr5:46706824
pvalue= 0.0238 ; R2= 0.2804
Estimated allele frequency
0.2
0.3
0.4
0.5
0.6
0.7
Ecological gradient (Precipitation sums)
600
800
1000
1200
L16
124
L8
9
O8
O1
253
204
L1
O16
217
233
218
256

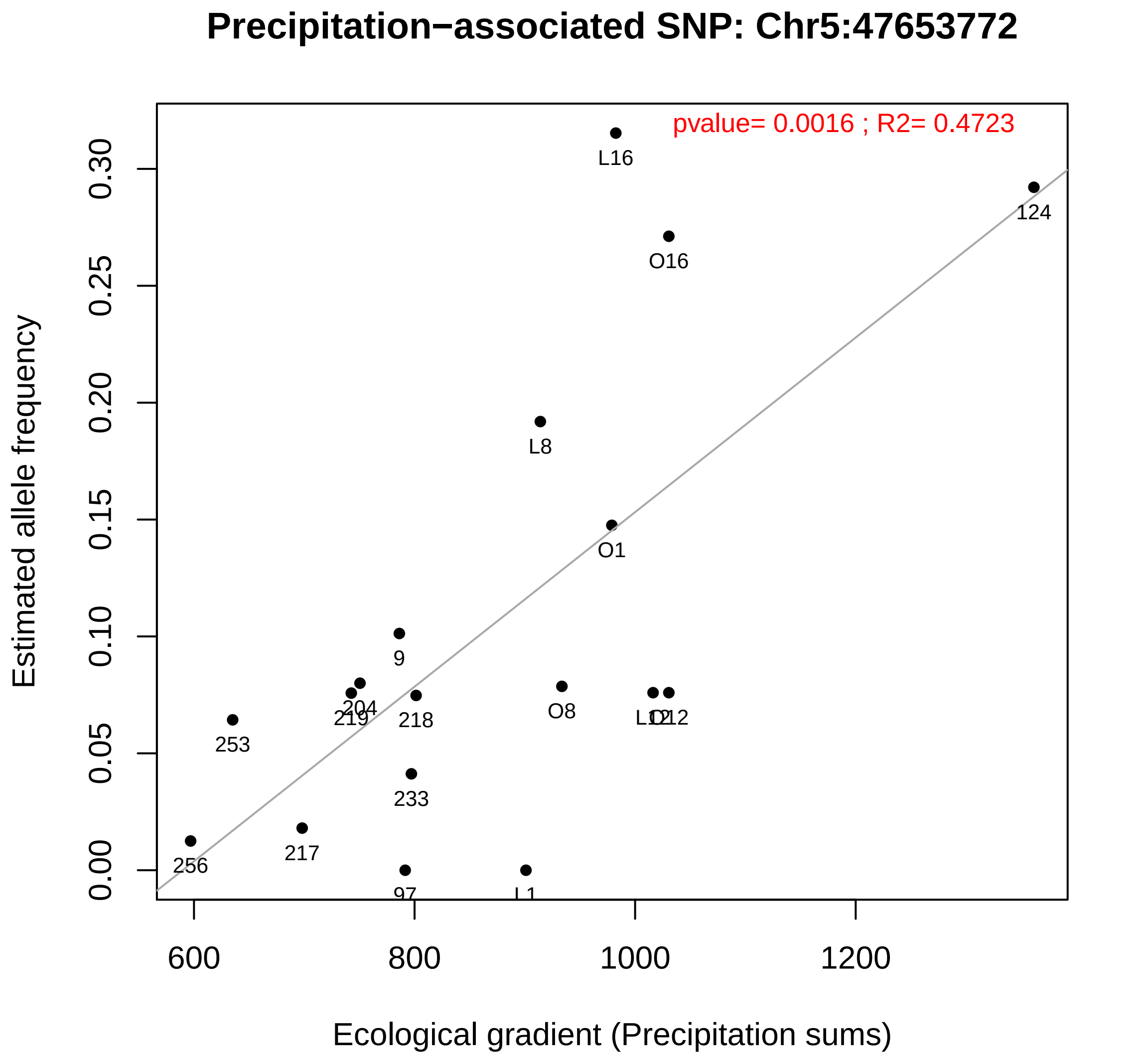
Precipitation−associated SNP: Chr5:47653772
pvalue= 0.0016 ; R2= 0.4723
Estimated allele frequency
0.00
0.05
0.10
0.15
0.20
0.25
0.30
600
800
1000
1200
Ecological gradient (Precipitation sums)
L16
124
O16
L8
O1
9
204
219
218
O8
L12
O12
253
233
217
256
97
L1

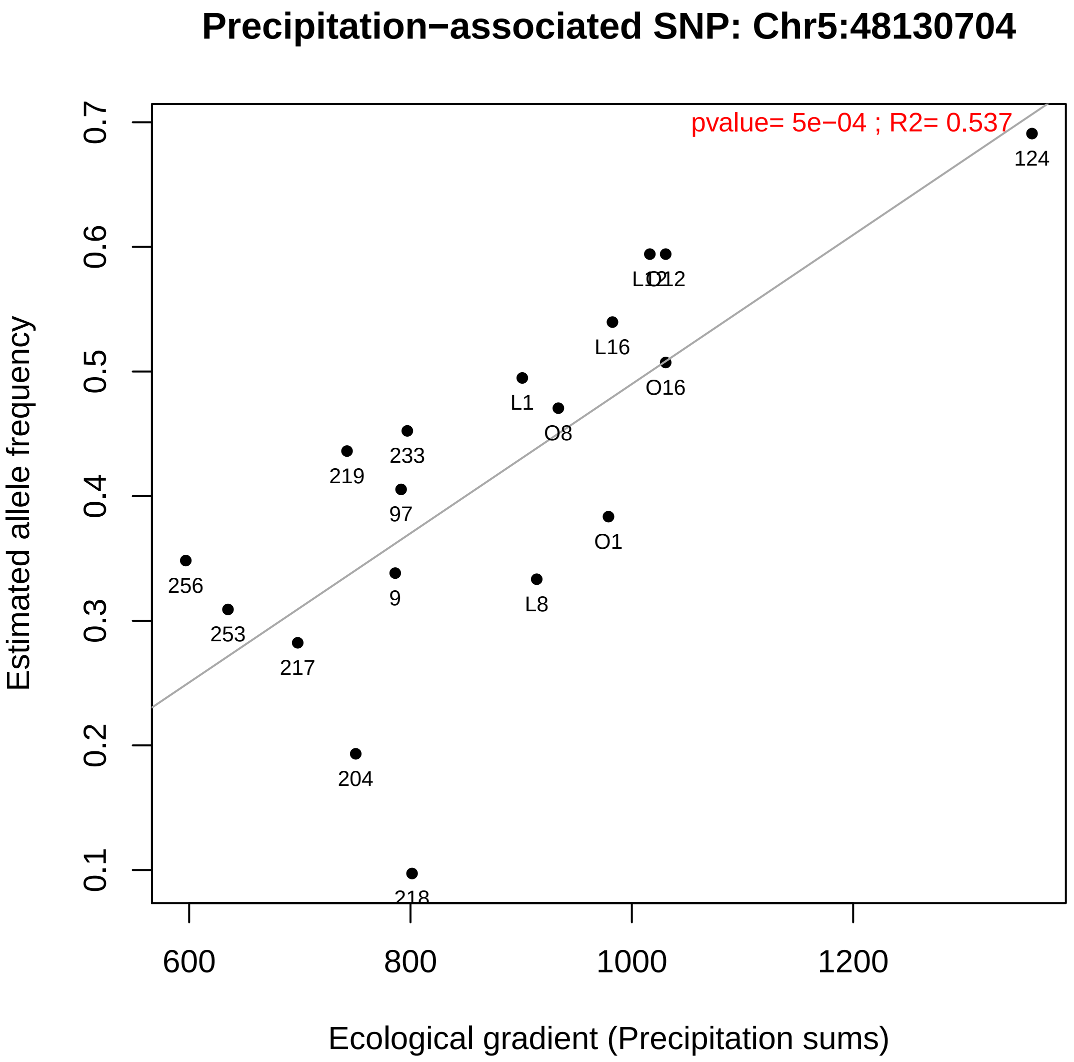
Precipitation−associated SNP: Chr5:48130704
pvalue= 5e−04 ; R2= 0.537
Estimated allele frequency
Ecological gradient (Precipitation sums)
0.1
0.2
0.3
0.4
0.5
0.6
0.7
600
800
1000
1200
124
L12
O12
L16
O16
L1
O8
233
219
97
O1
256
9
L8
253
217
204
218

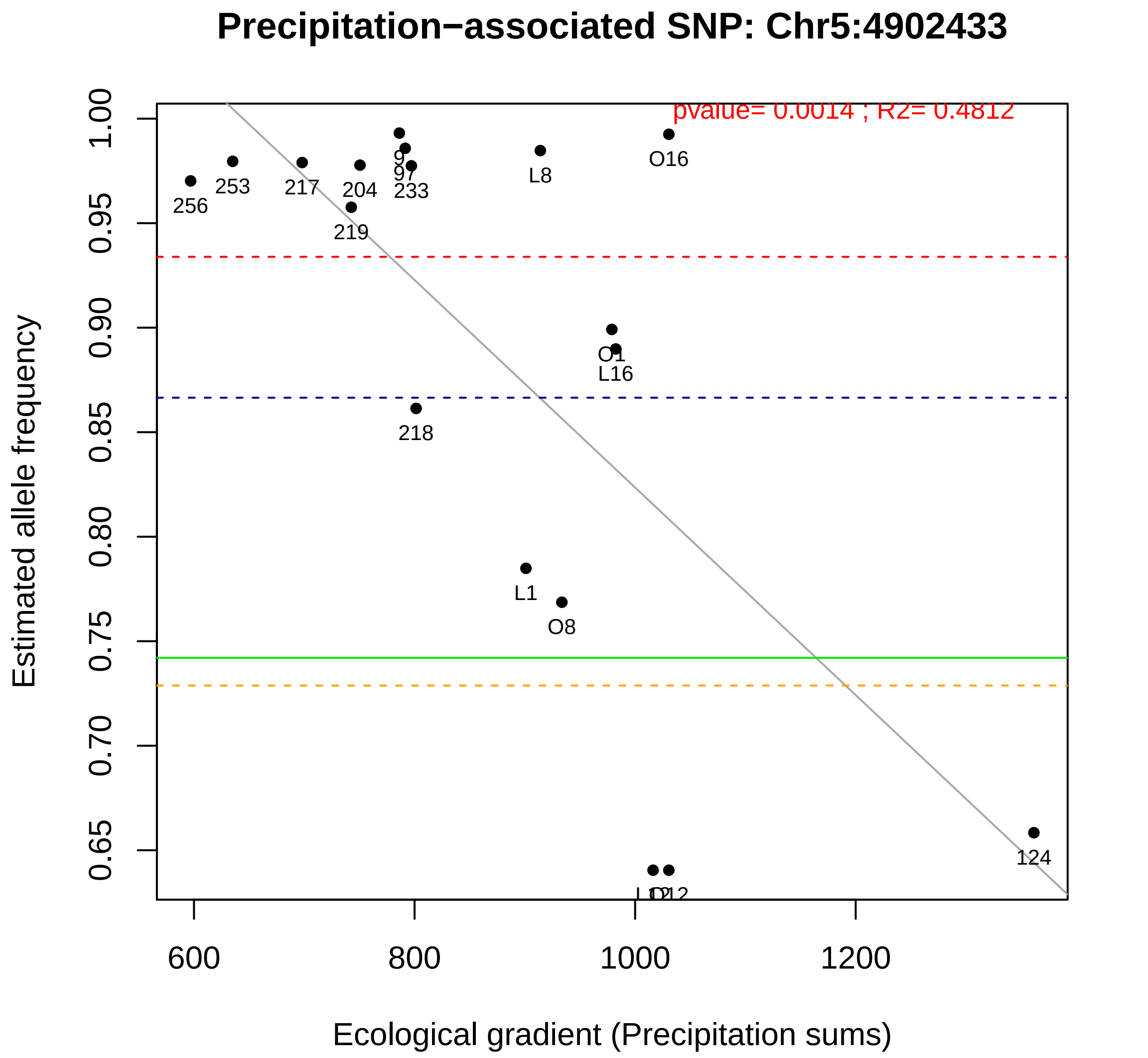

Precipitation−associated SNP: Chr5:4902433
pvalue= 0.0014 ; R2= 0.4812
Estimated allele frequency
Ecological gradient (Precipitation sums)
0.65
0.70
0.75
0.80
0.85
0.90
0.95
1.00
600
800
1000
1200
256
253
217
204
219
9
97
233
L8
O16
O1
L16
218
L1
O8
124

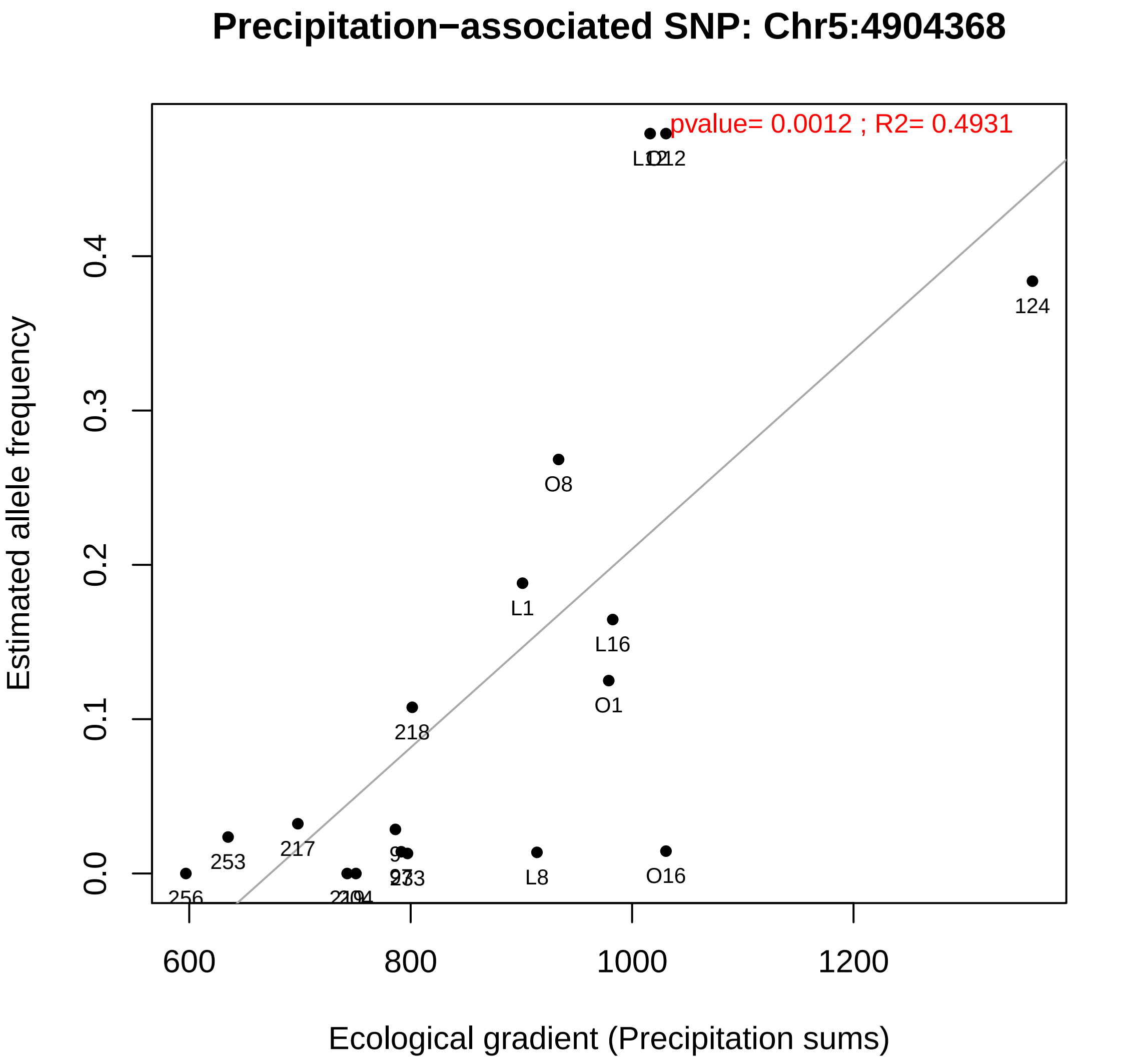

Precipitation−associated SNP: Chr5:4904368
pvalue= 0.0012 ; R2= 0.4931
124
O8
L1
L16
O1
218
217
253
256
L8
O16
0.0
0.1
0.2
0.3
0.4
600
800
1000
1200
Estimated allele frequency
Ecological gradient (Precipitation sums)

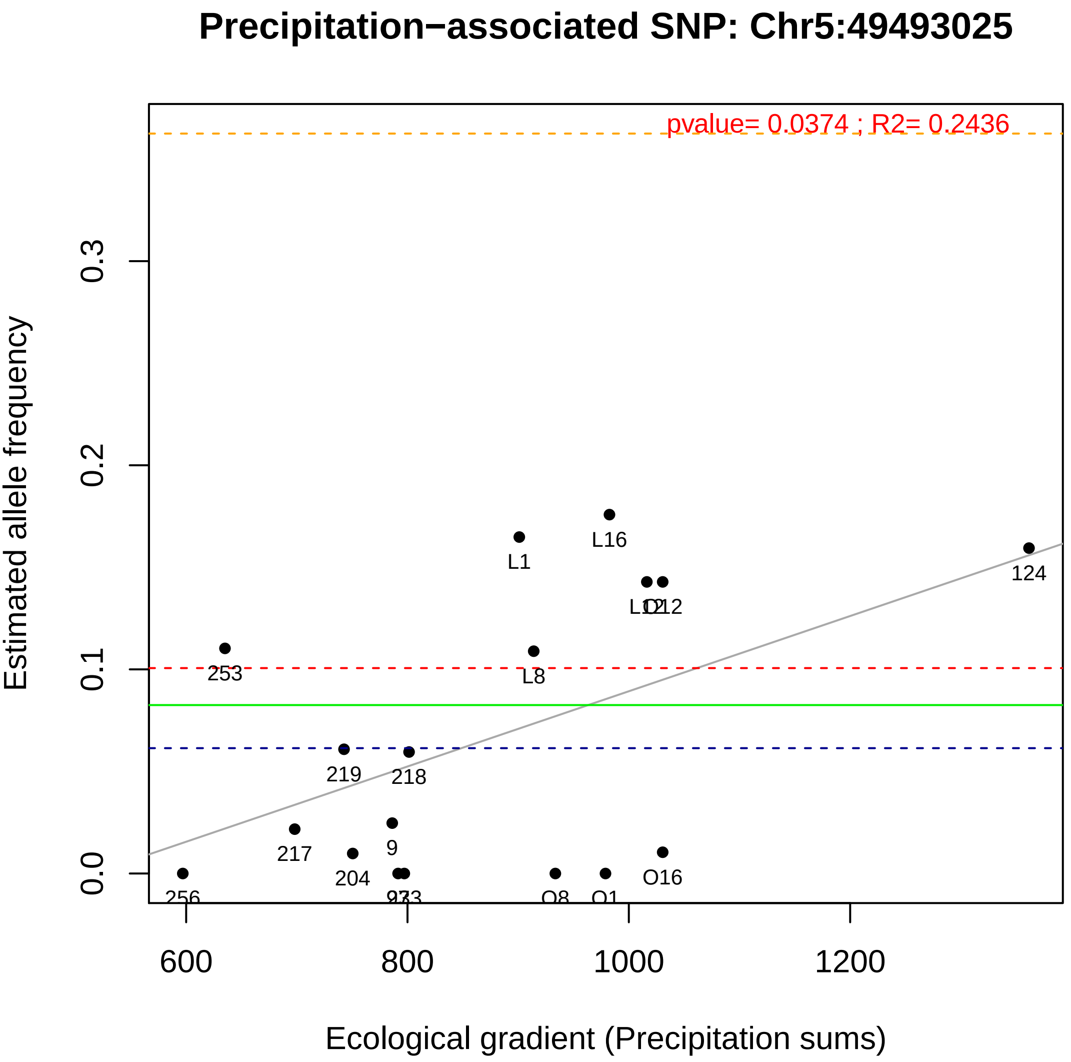

Precipitation−associated SNP: Chr5:49493025
pvalue= 0.0374 ; R2= 0.2436
Estimated allele frequency
0.0
0.1
0.2
0.3
Ecological gradient (Precipitation sums)
600
800
1000
1200
L16
L1
124
253
L8
219
218
217
9
204
O16
O8
O1
256

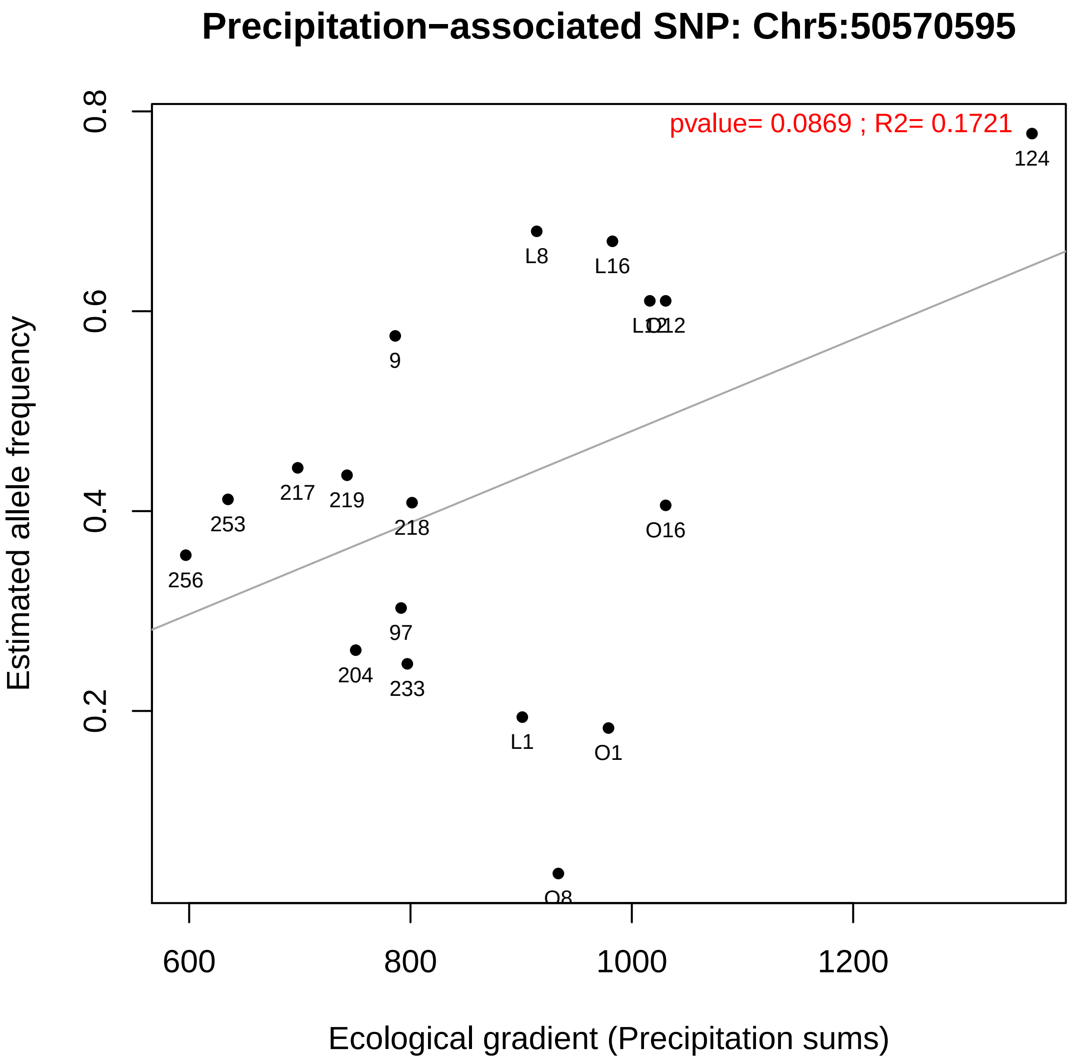

Precipitation−associated SNP: Chr5:50570595
pvalue= 0.0869 ; R2= 0.1721
Estimated allele frequency
0.2
0.4
0.6
0.8
600
800
1000
1200
Ecological gradient (Precipitation sums)
124
L8
L16
O12
9
217
219
253
218
O16
256
97
204
233
L1
O1
O8

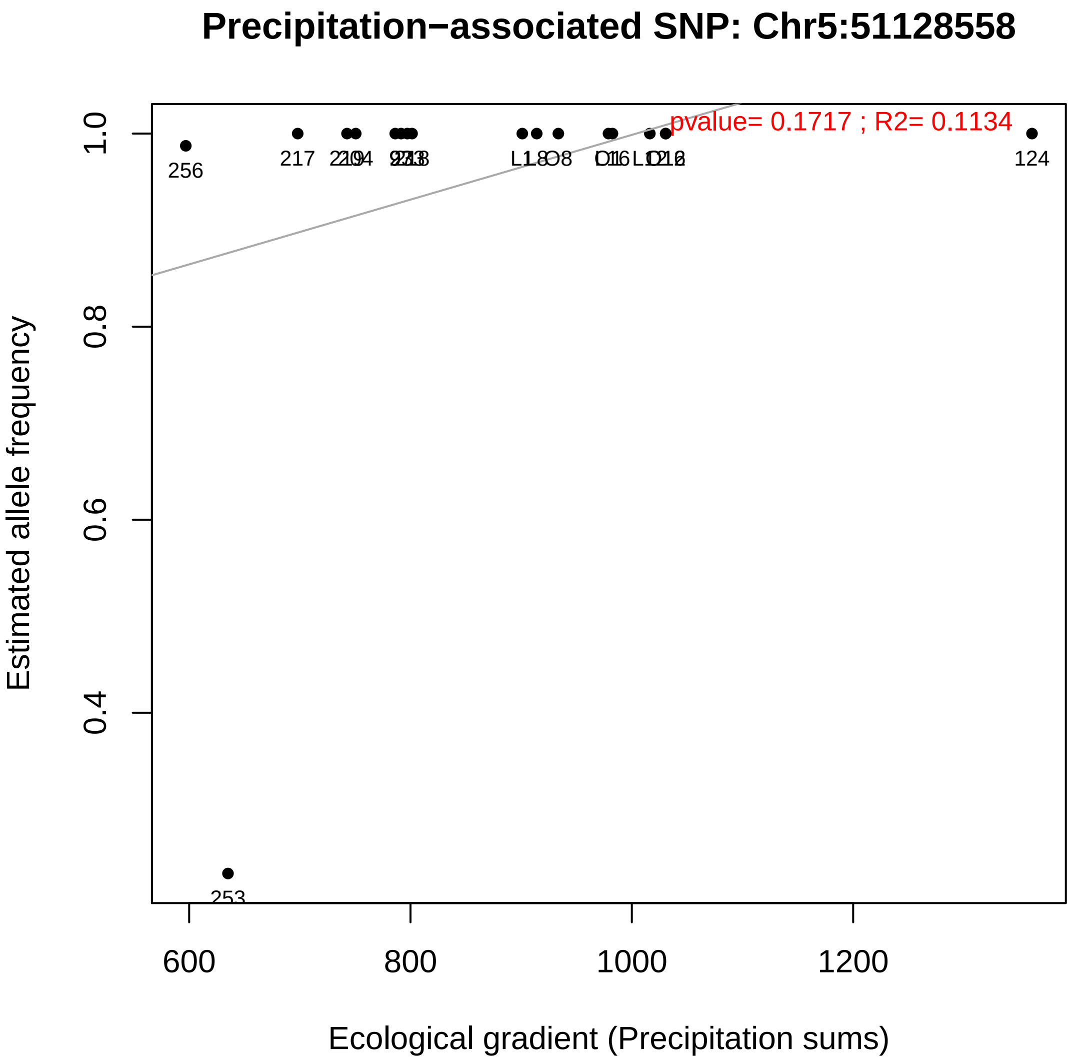

Precipitation−associated SNP: Chr5:51128558
pvalue= 0.1717 ; R2= 0.1134
256
217
124
253
1.0
0.8
0.6
0.4
600
800
1000
1200
Estimated allele frequency
Ecological gradient (Precipitation sums)

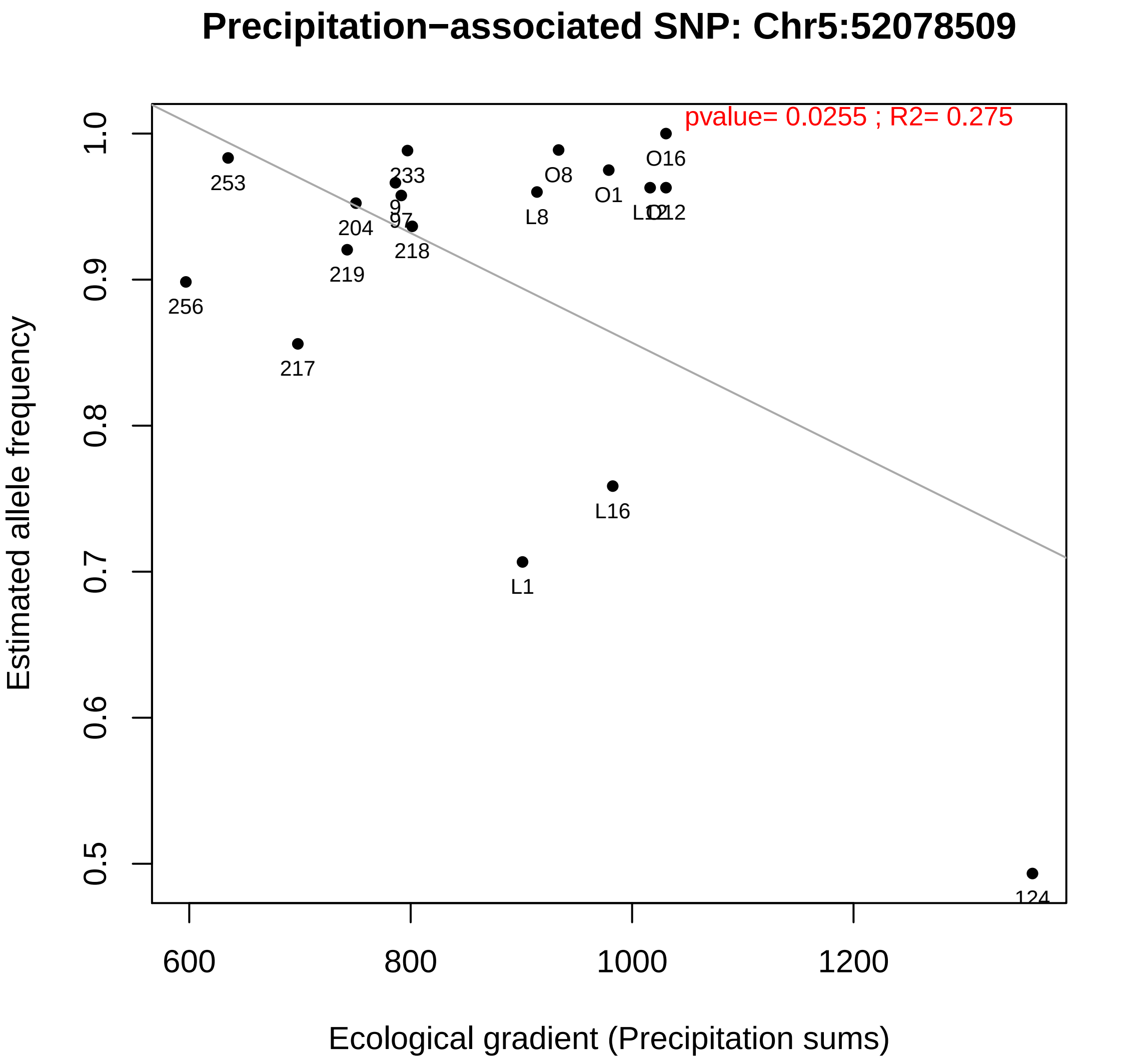

Precipitation−associated SNP: Chr5:52078509
pvalue= 0.0255 ; R2= 0.275
Estimated allele frequency
Ecological gradient (Precipitation sums)
1.0
0.9
0.8
0.7
0.6
0.5
600
800
1000
1200
253
233
O8
O16
O1
L8
L12
O12
204
9
97
218
219
256
217
L16
L1
124

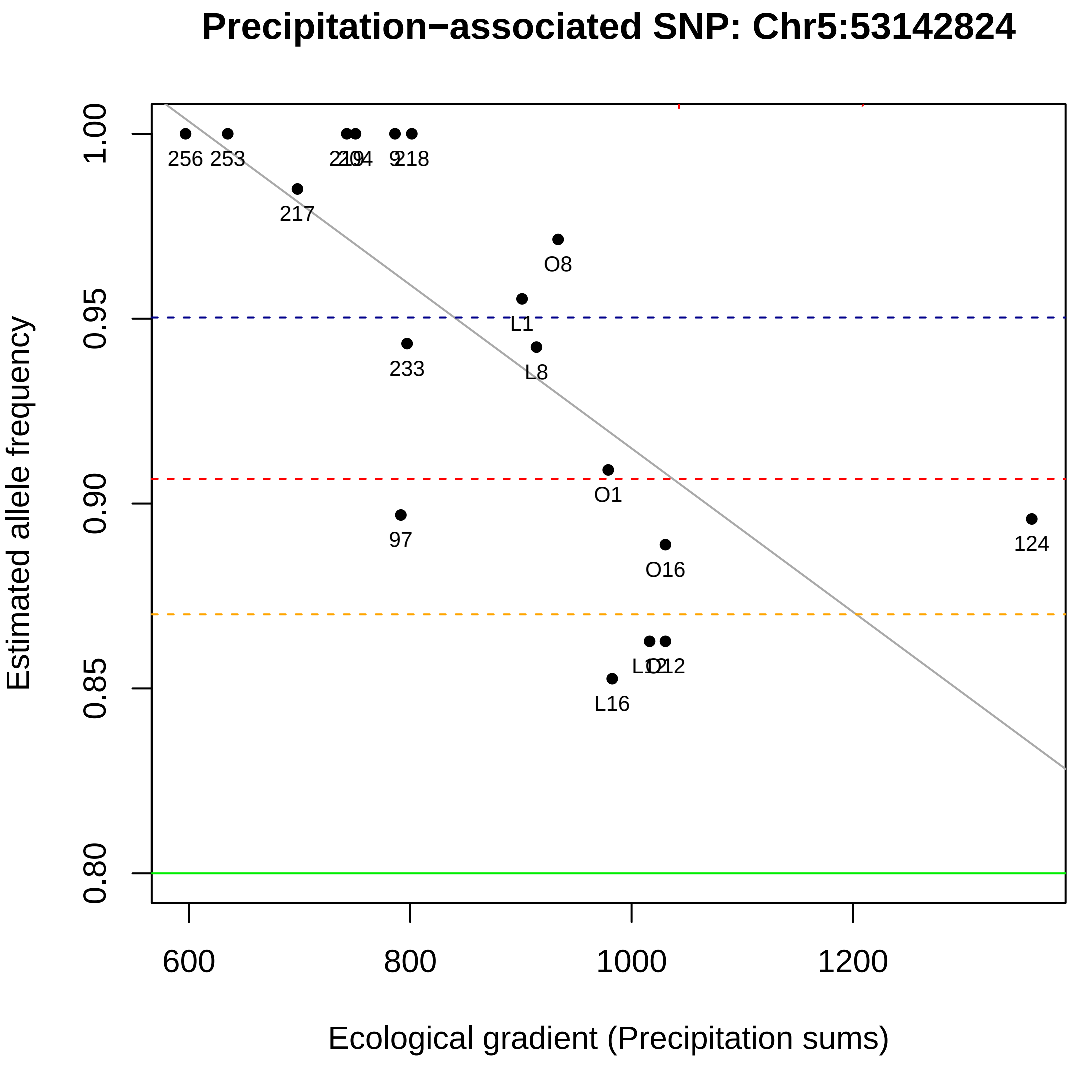
Precipitation−associated SNP: Chr5:53142824
Estimated allele frequency
Ecological gradient (Precipitation sums)
1.00
0.95
0.90
0.85
0.80
600
800
1000
1200
256
253
217
O8
L1
233
L8
O1
97
O16
124
L16
9
218

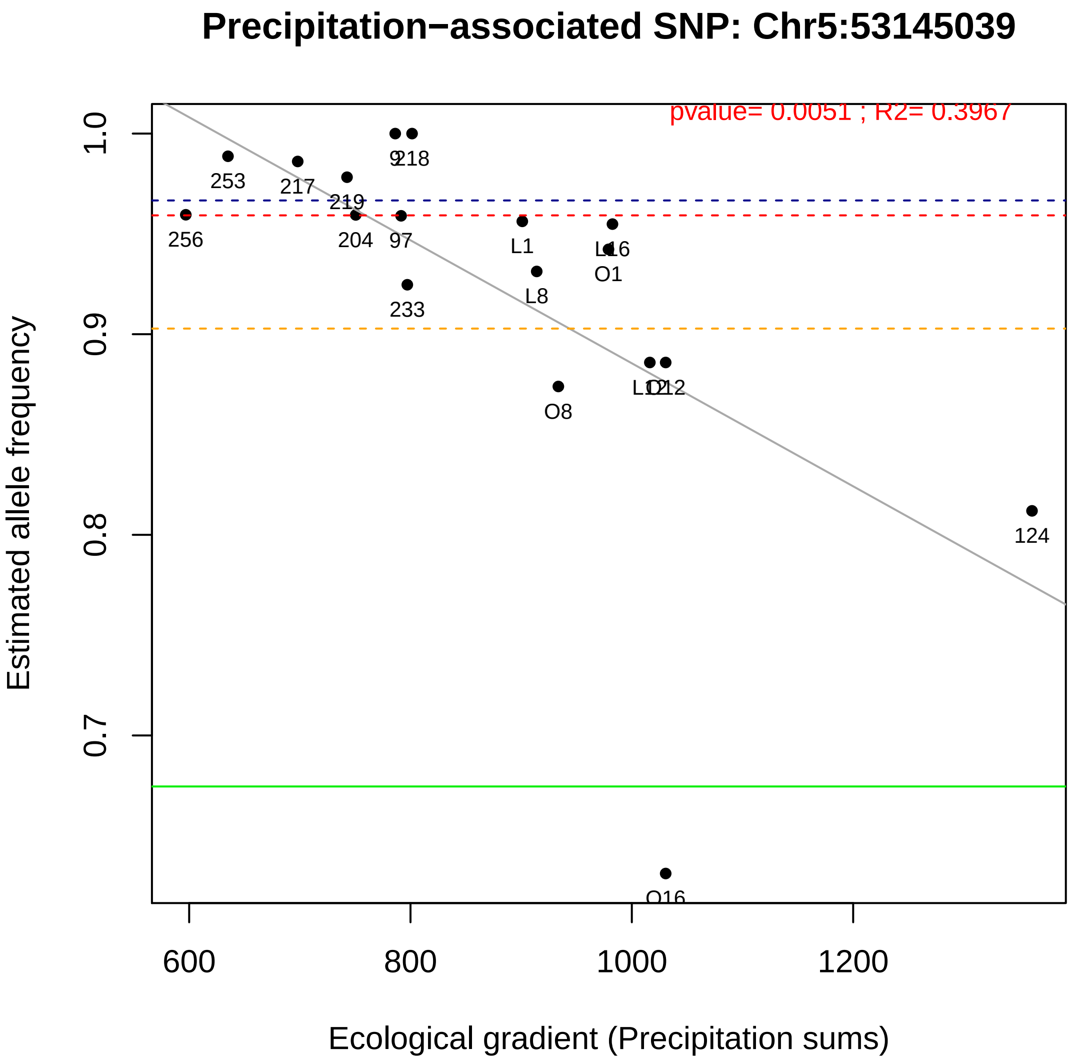

Precipitation−associated SNP: Chr5:53145039
pvalue= 0.0051 ; R2= 0.3967
Estimated allele frequency
Ecological gradient (Precipitation sums)
0.7
0.8
0.9
1.0
600
800
1000
1200
253
217
219
256
204
97
233
218
L1
L16
O1
L8
O8
O12
124
O16

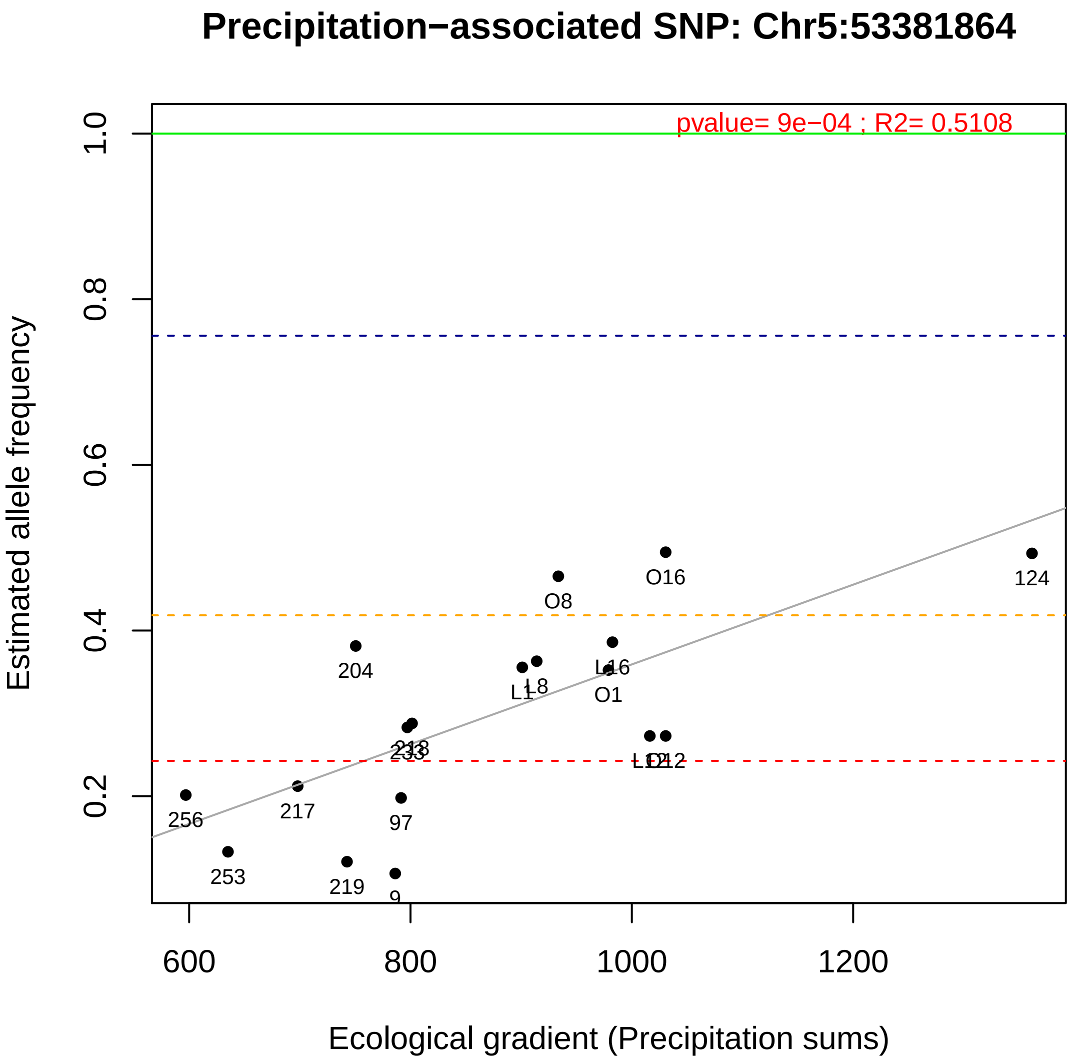

Precipitation−associated SNP: Chr5:53381864
pvalue= 9e−04 ; R2= 0.5108
Estimated allele frequency
0.2
0.4
0.6
0.8
1.0
600
800
1000
1200
Ecological gradient (Precipitation sums)
256
253
217
219
204
97
9
O8
O16
124
L1
L8
L16
O1

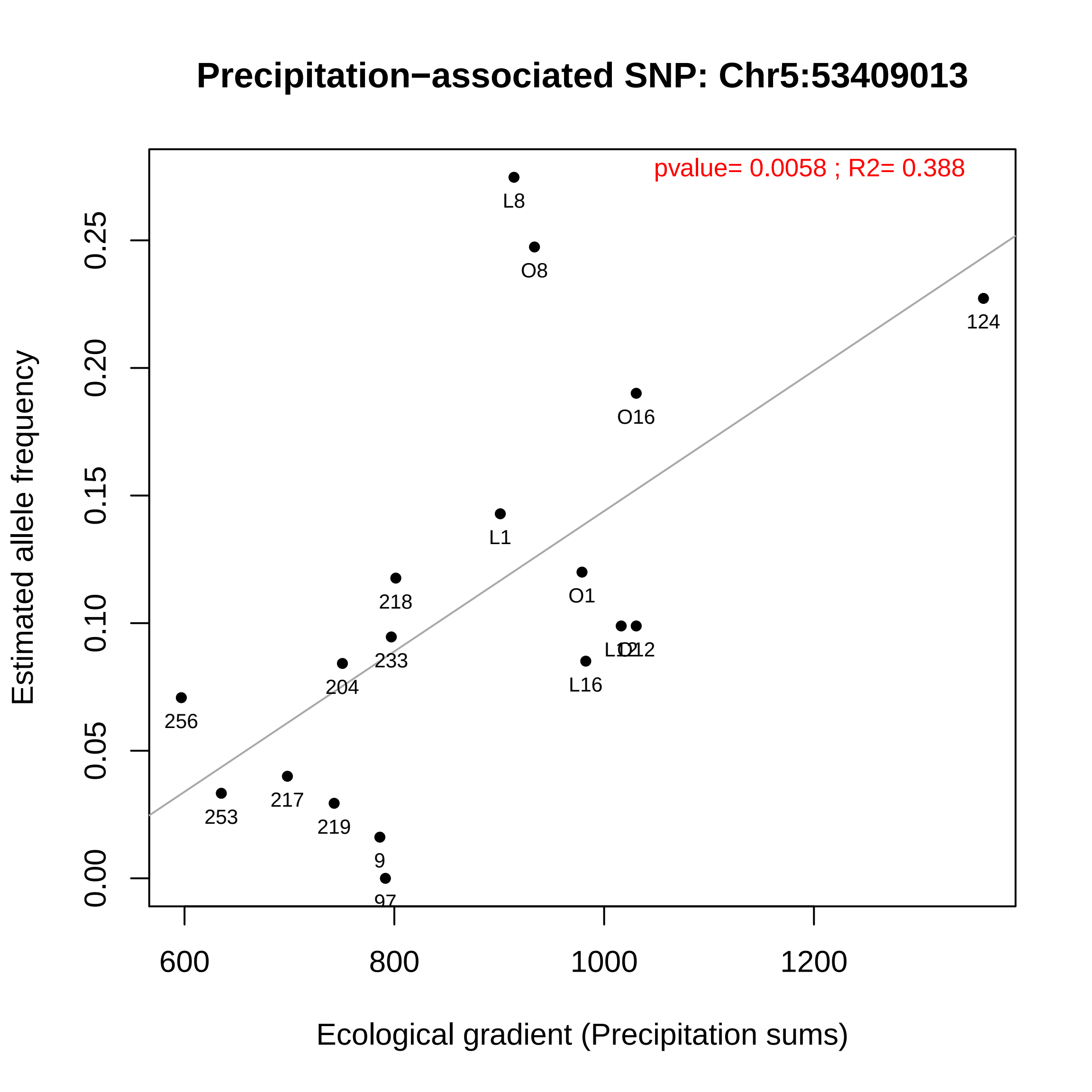

Precipitation−associated SNP: Chr5:53409013
pvalue= 0.0058 ; R2= 0.388
Estimated allele frequency
0.00
0.05
0.10
0.15
0.20
0.25
600
800
1000
1200
Ecological gradient (Precipitation sums)
L8
O8
124
O16
L1
218
O1
233
L12
O12
204
L16
256
217
253
219
9
97

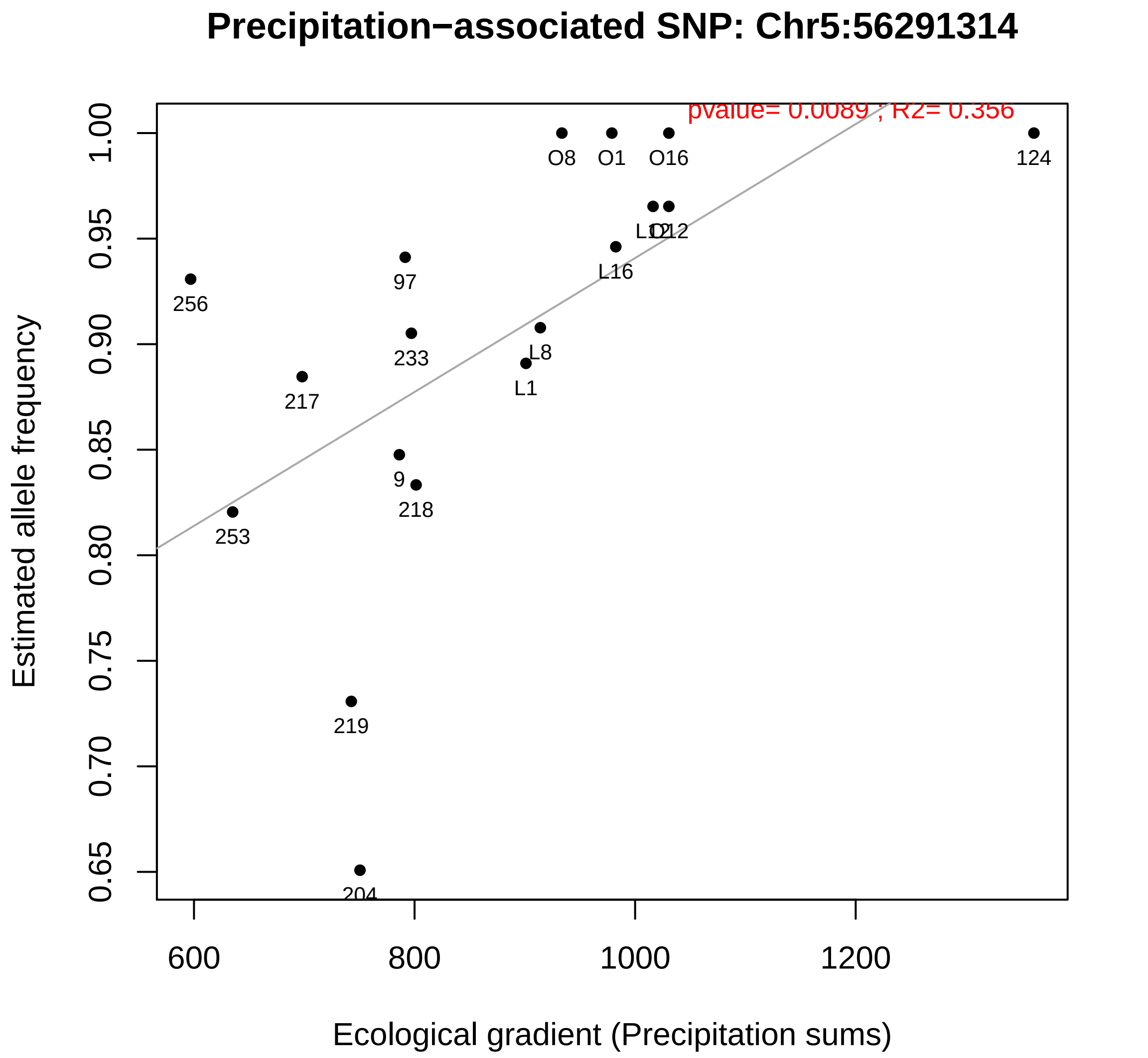

Precipitation−associated SNP: Chr5:56291314
pvalue= 0.0089 ; R2= 0.356
Estimated allele frequency
1.00
0.95
0.90
0.85
0.80
0.75
0.70
0.65
600
800
1000
1200
Ecological gradient (Precipitation sums)
O8
O1
O16
124
L12
O12
L16
97
256
233
L8
L1
217
9
218
253
219
204

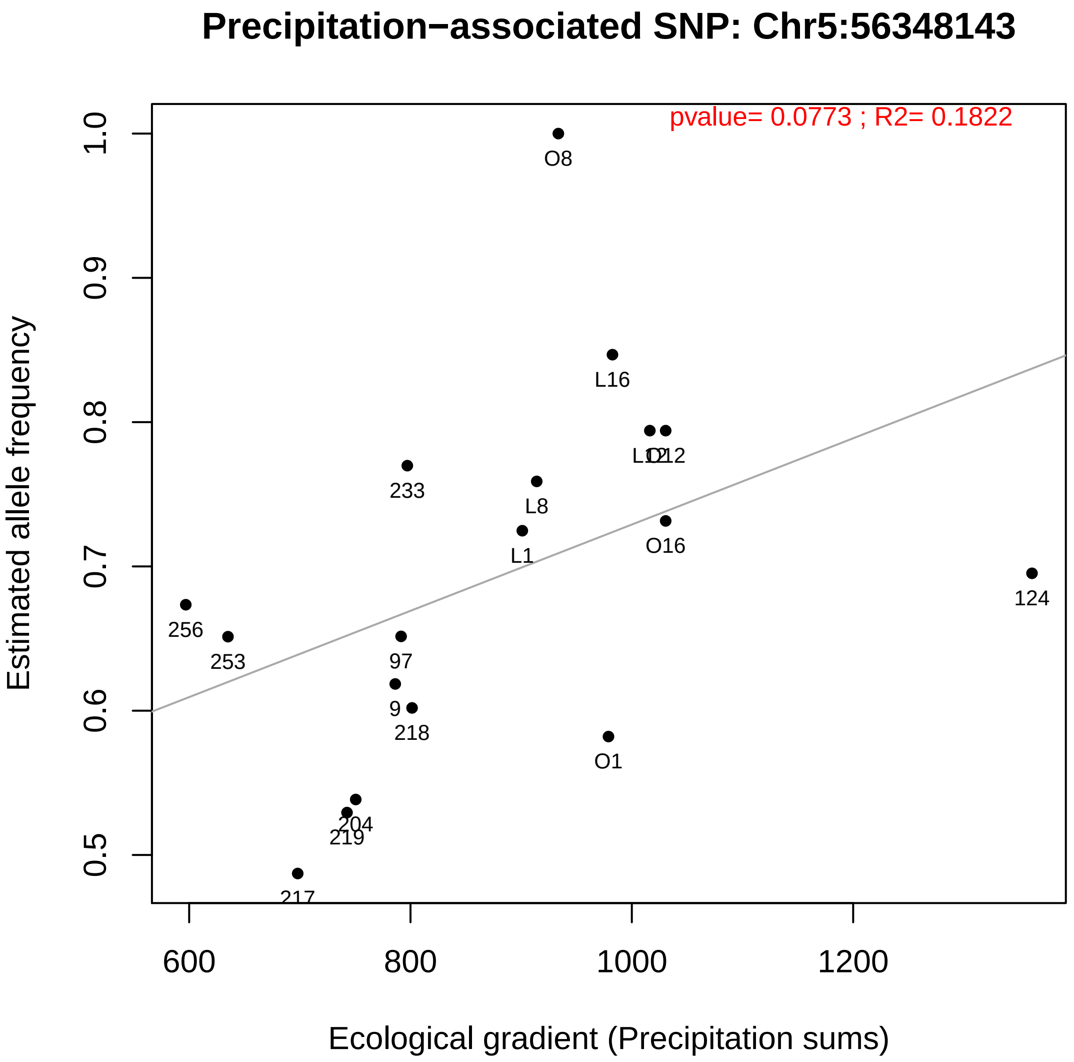
Precipitation−associated SNP: Chr5:56348143
pvalue= 0.0773 ; R2= 0.1822
Estimated allele frequency
Ecological gradient (Precipitation sums)
0.5
0.6
0.7
0.8
0.9
1.0
600
800
1000
1200
O8
L16
L12
O12
233
L8
L1
O16
124
256
253
97
9
218
O1
204
219
217

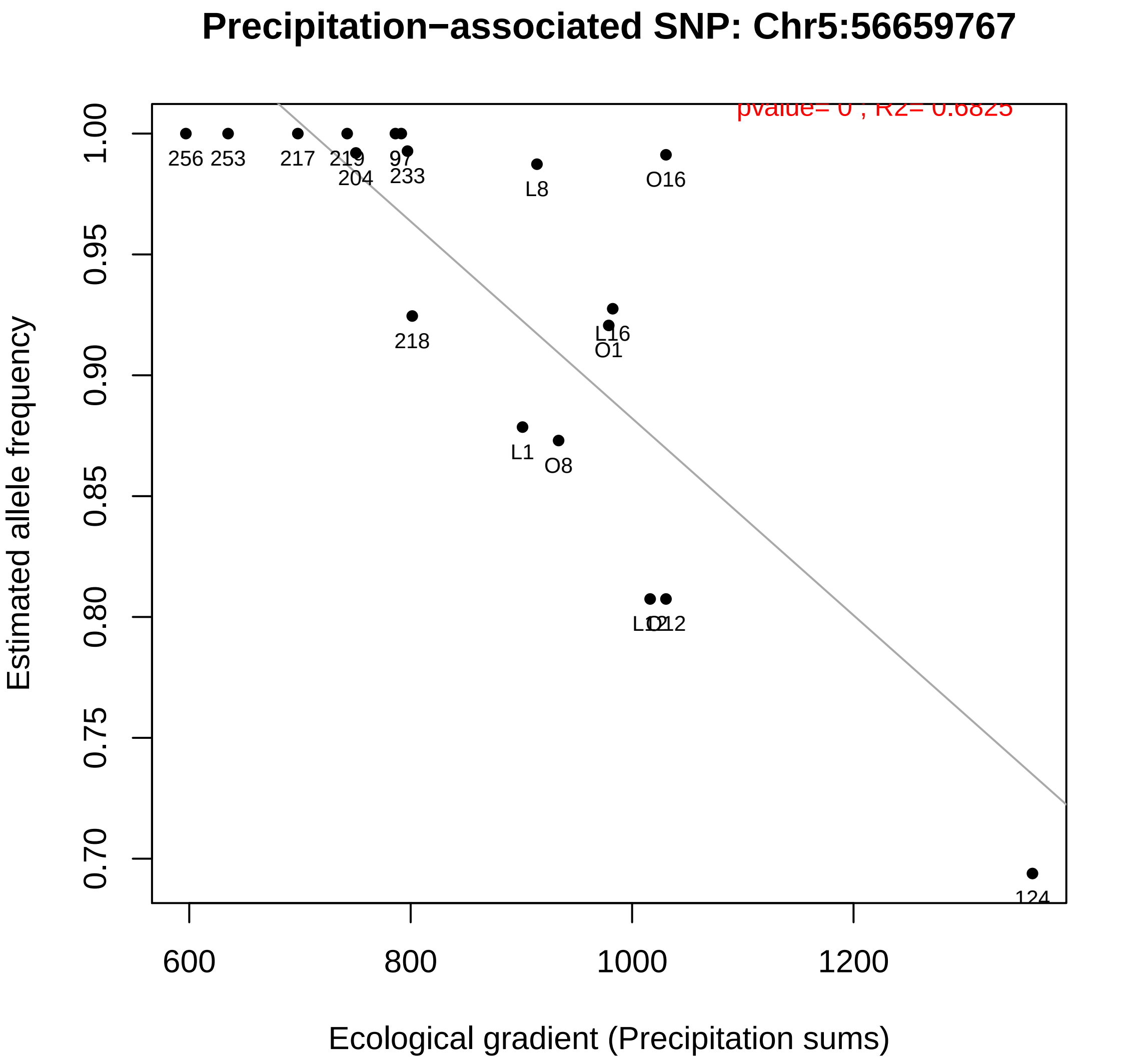
Precipitation−associated SNP: Chr5:56659767
pvalue= 0 ; R2= 0.6825
Estimated allele frequency
Ecological gradient (Precipitation sums)
1.00
0.95
0.90
0.85
0.80
0.75
0.70
600
800
1000
1200
256
253
217
219
204
97
233
L8
O16
218
L16
O1
L1
O8
L12
O12
124

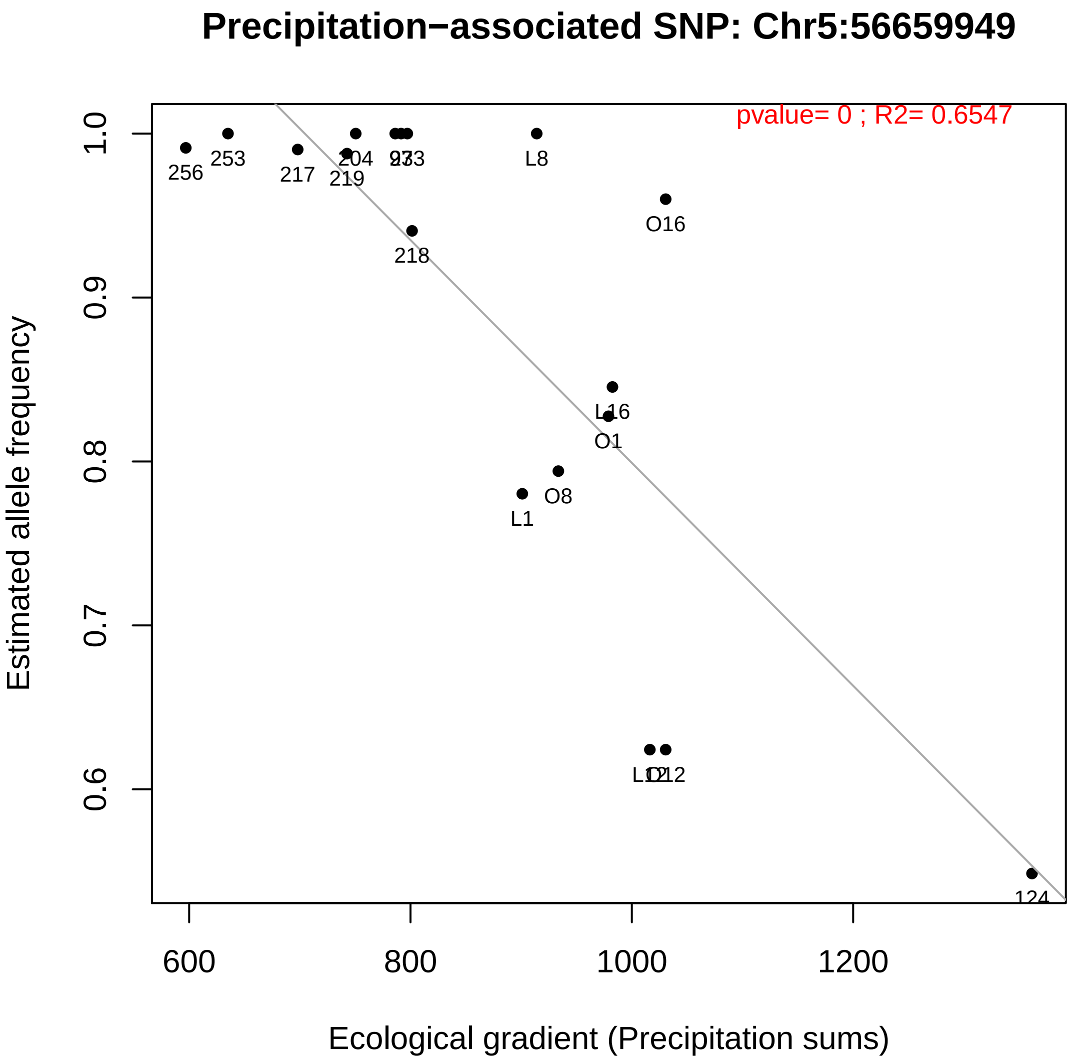
Precipitation−associated SNP: Chr5:56659949
pvalue= 0 ; R2= 0.6547
Estimated allele frequency
1.0
0.9
0.8
0.7
0.6
600
800
1000
1200
Ecological gradient (Precipitation sums)
256
253
217
219
204
218
L8
O16
L16
O1
O8
L1
124

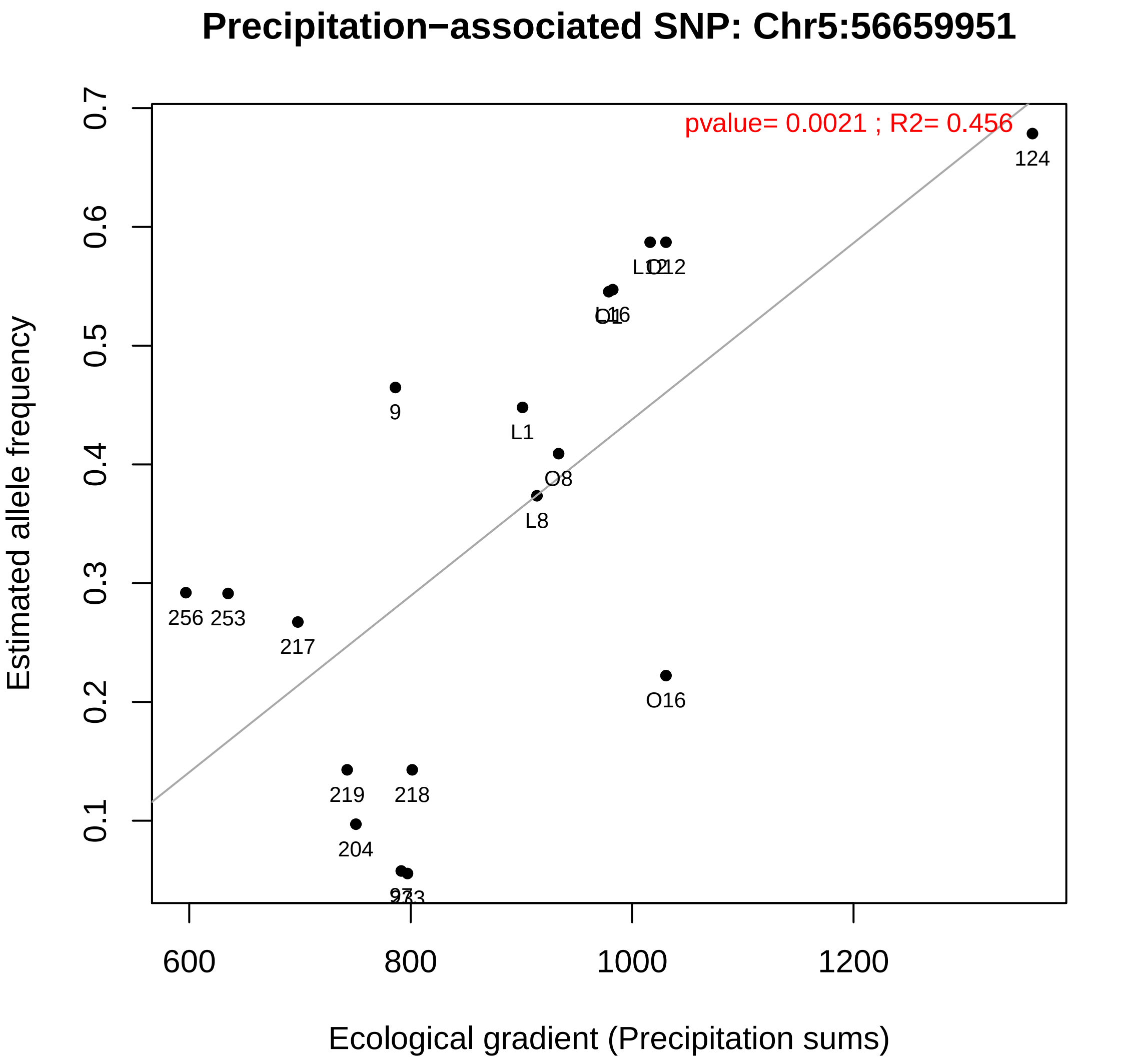

Precipitation−associated SNP: Chr5:56659951
pvalue= 0.0021 ; R2= 0.456
Estimated allele frequency
Ecological gradient (Precipitation sums)
0.1
0.2
0.3
0.4
0.5
0.6
0.7
600
800
1000
1200
124
L12
O12
L16
O1
9
L1
O8
L8
256
253
217
O16
219
218
204
97
273

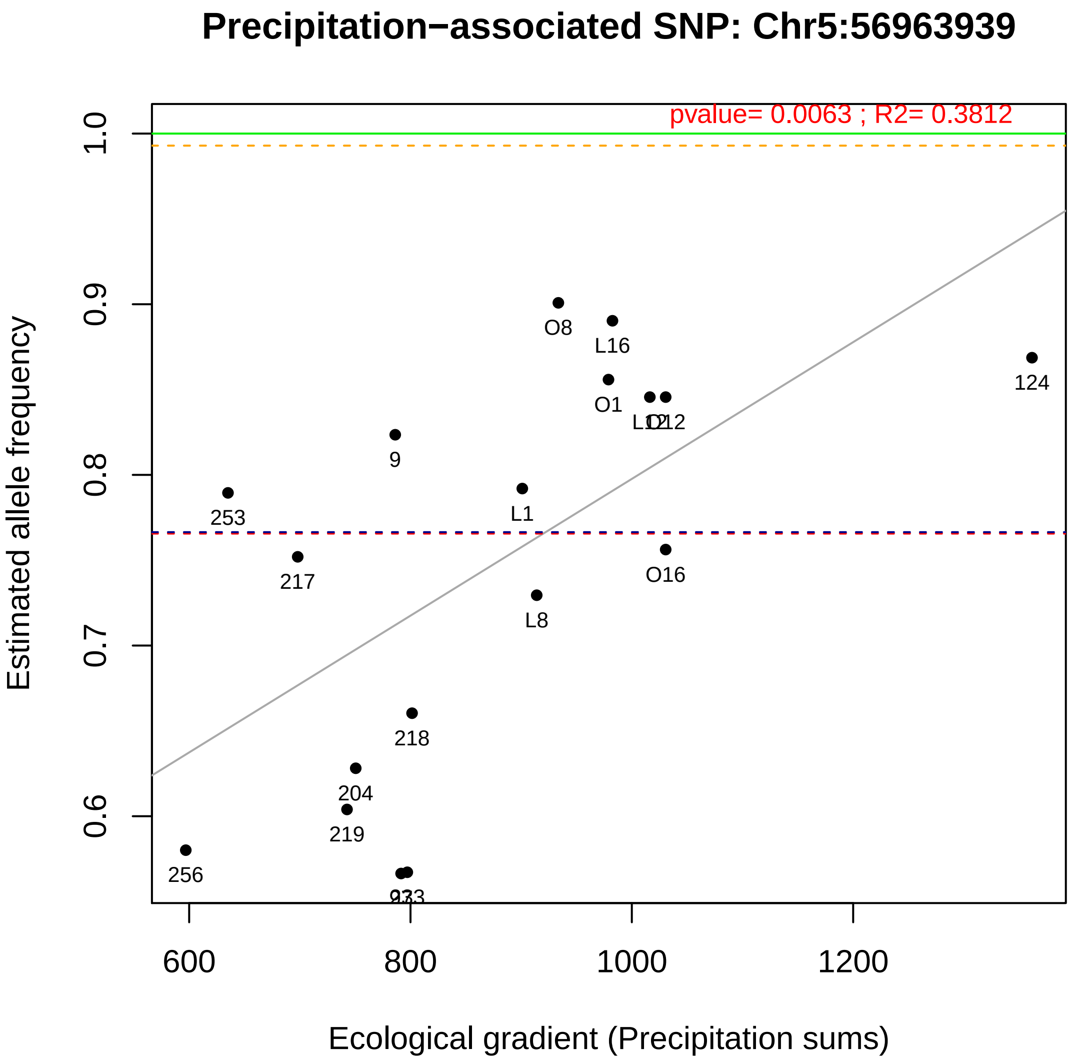

Precipitation−associated SNP: Chr5:56963939
pvalue= 0.0063 ; R2= 0.3812
Estimated allele frequency
0.6
0.7
0.8
0.9
1.0
600
800
1000
1200
Ecological gradient (Precipitation sums)
O8
L16
124
O1
9
253
L1
217
O16
L8
218
204
219
256

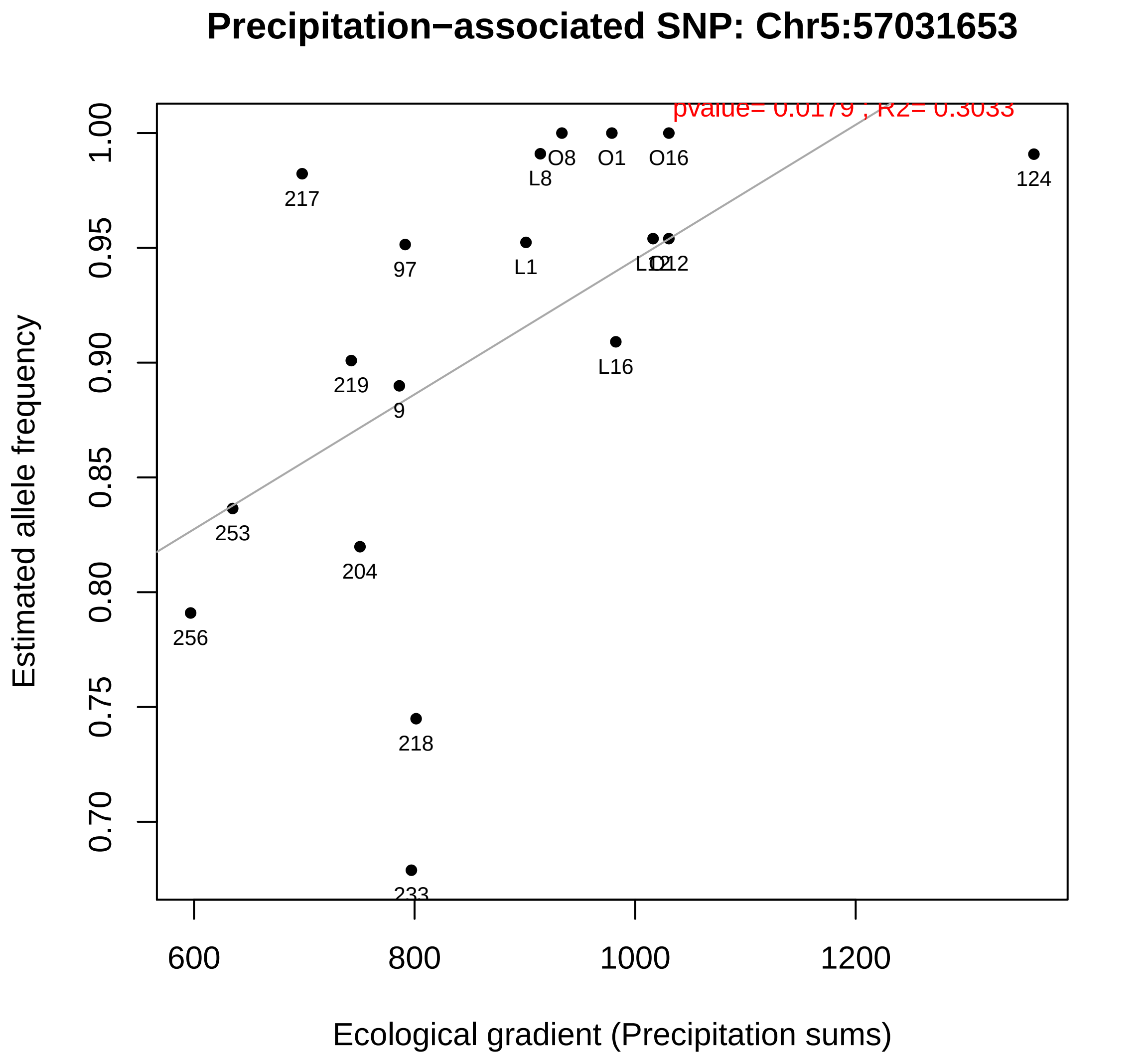
Precipitation−associated SNP: Chr5:57031653
pvalue= 0.0179 ; R2= 0.3033
Estimated allele frequency
Ecological gradient (Precipitation sums)
0.70
0.75
0.80
0.85
0.90
0.95
1.00
600
800
1000
1200
O8
O1
O16
L8
124
217
97
L1
L10
O12
L16
219
9
253
204
256
218
233

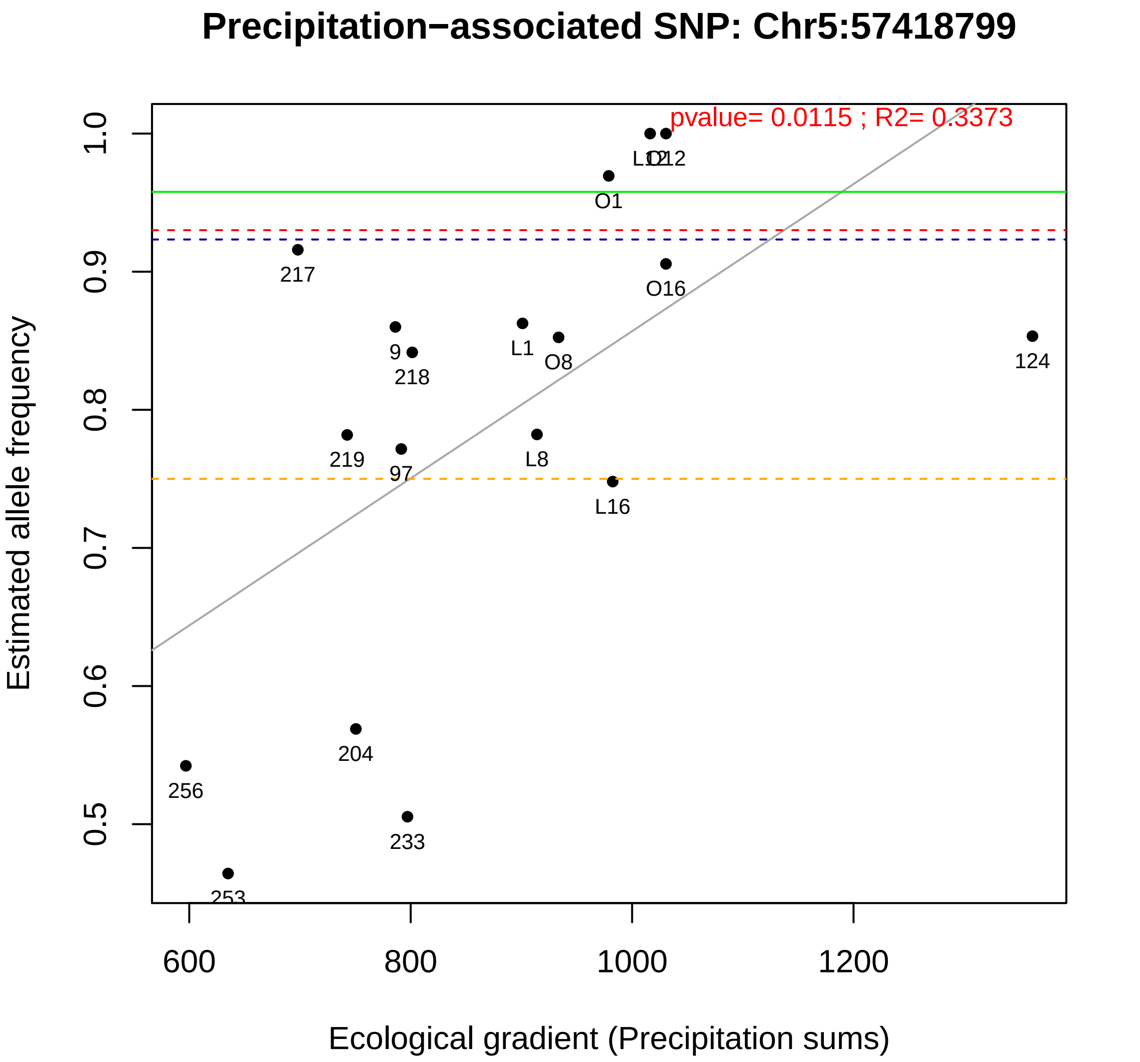

Precipitation−associated SNP: Chr5:57418799
pvalue= 0.0115 ; R2= 0.3373
Estimated allele frequency
Ecological gradient (Precipitation sums)
1.0
0.9
0.8
0.7
0.6
0.5
600
800
1000
1200
O12
O1
217
O16
9
218
L1
O8
124
219
97
L8
L16
204
256
233
253

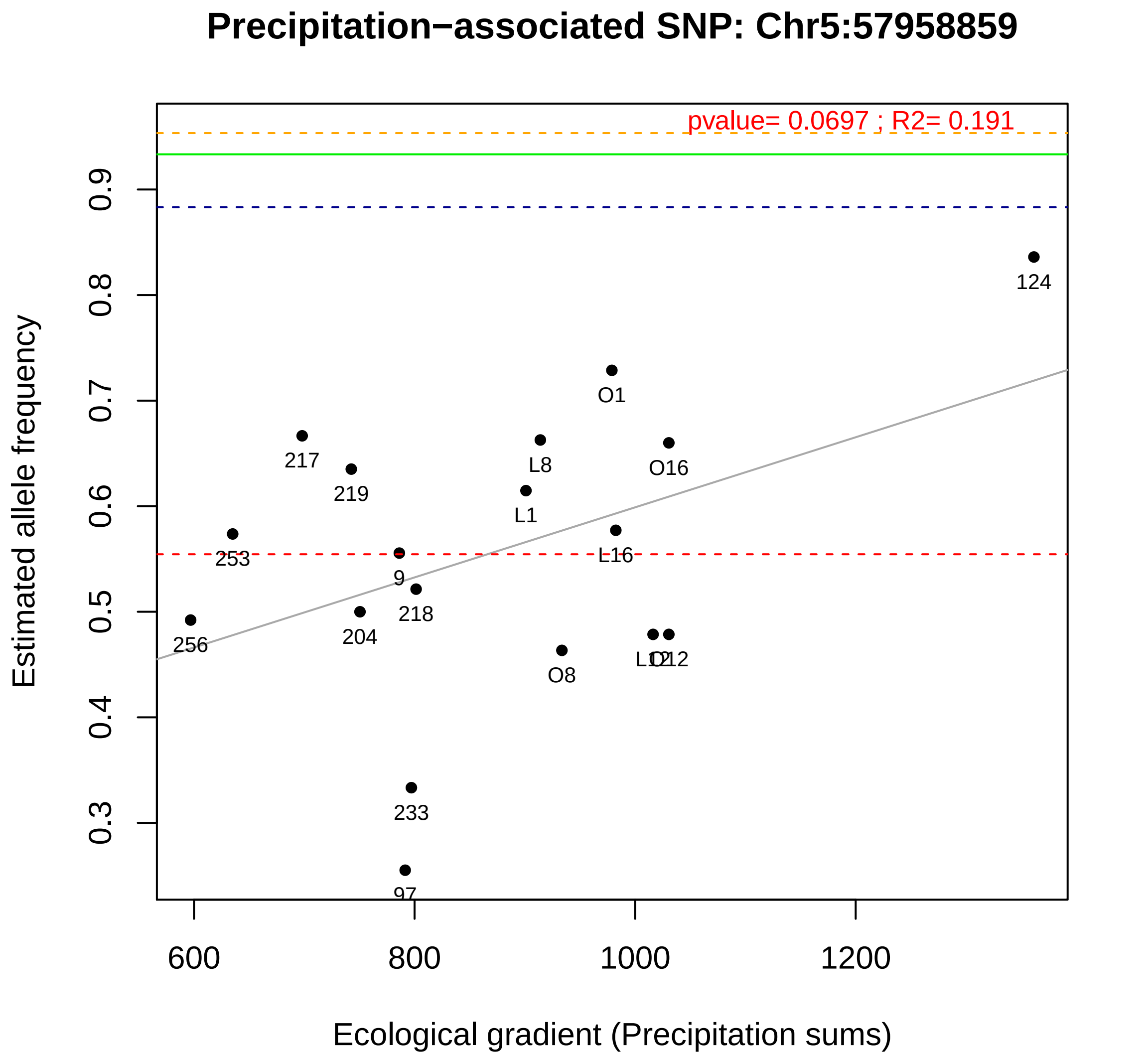
Precipitation−associated SNP: Chr5:57958859
pvalue= 0.0697 ; R2= 0.191
Estimated allele frequency
Ecological gradient (Precipitation sums)
0.3
0.4
0.5
0.6
0.7
0.8
0.9
600
800
1000
1200
124
O1
217
219
L8
O16
L1
253
L16
9
218
204
256
O8
233
97

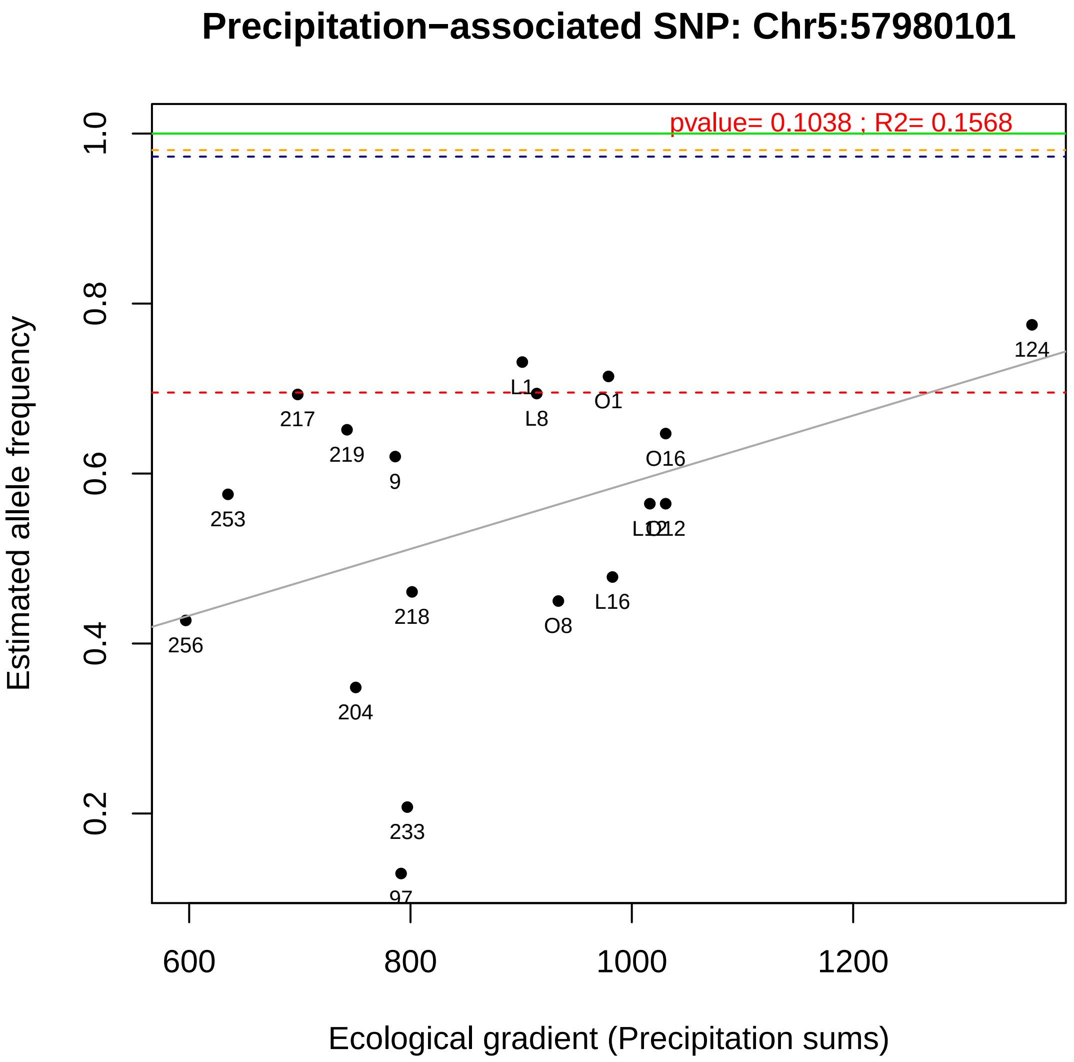

Precipitation−associated SNP: Chr5:57980101
pvalue= 0.1038 ; R2= 0.1568
Estimated allele frequency
Ecological gradient (Precipitation sums)
0.2
0.4
0.6
0.8
1.0
600
800
1000
1200
124
217
L1
L8
O1
219
O16
9
253
L12
O12
L16
218
O8
256
204
233
97

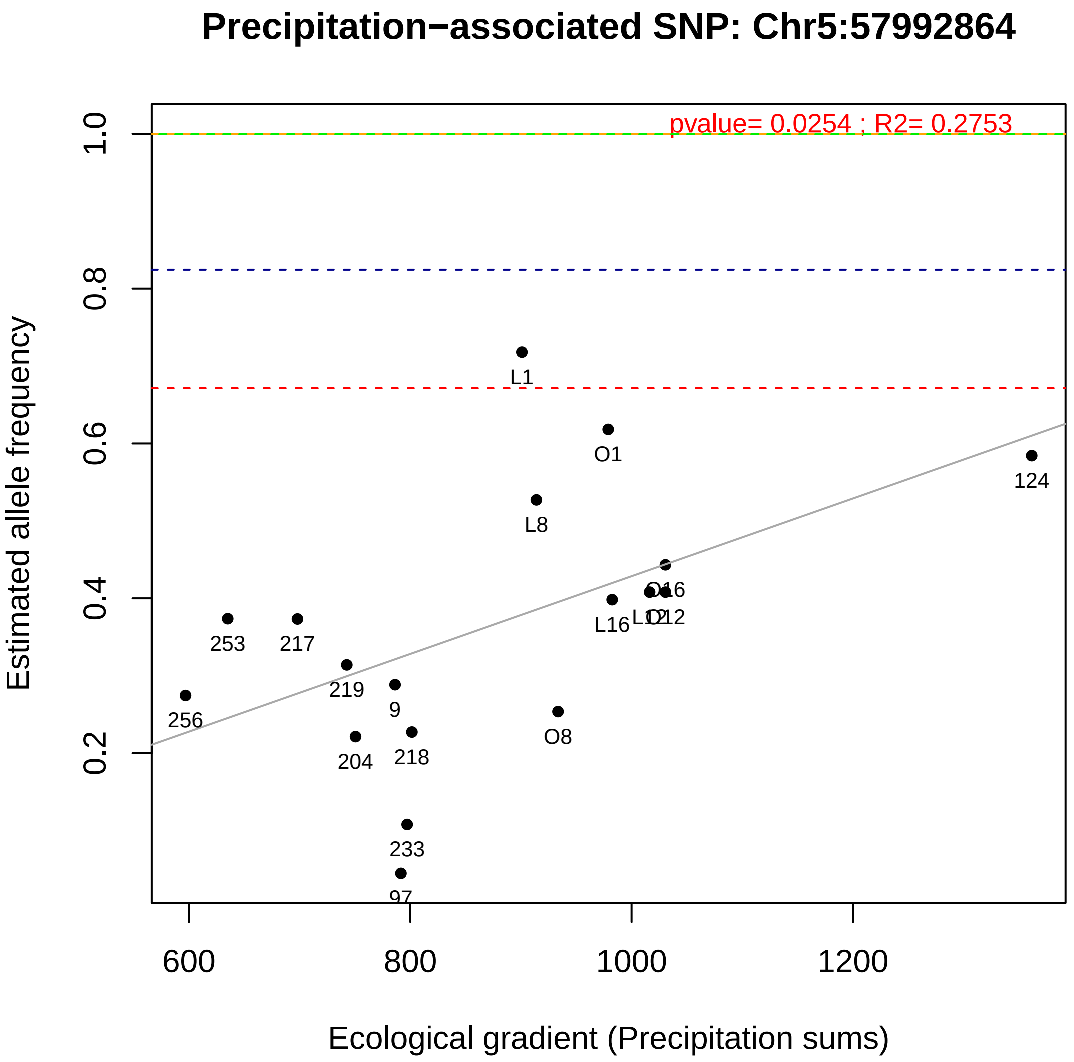

Precipitation−associated SNP: Chr5:57992864
pvalue= 0.0254 ; R2= 0.2753
Estimated allele frequency
Ecological gradient (Precipitation sums)
0.2
0.4
0.6
0.8
1.0
600
800
1000
1200
L1
O1
124
L8
O16
L16
L10
O12
253
217
219
9
256
O8
204
218
233
97

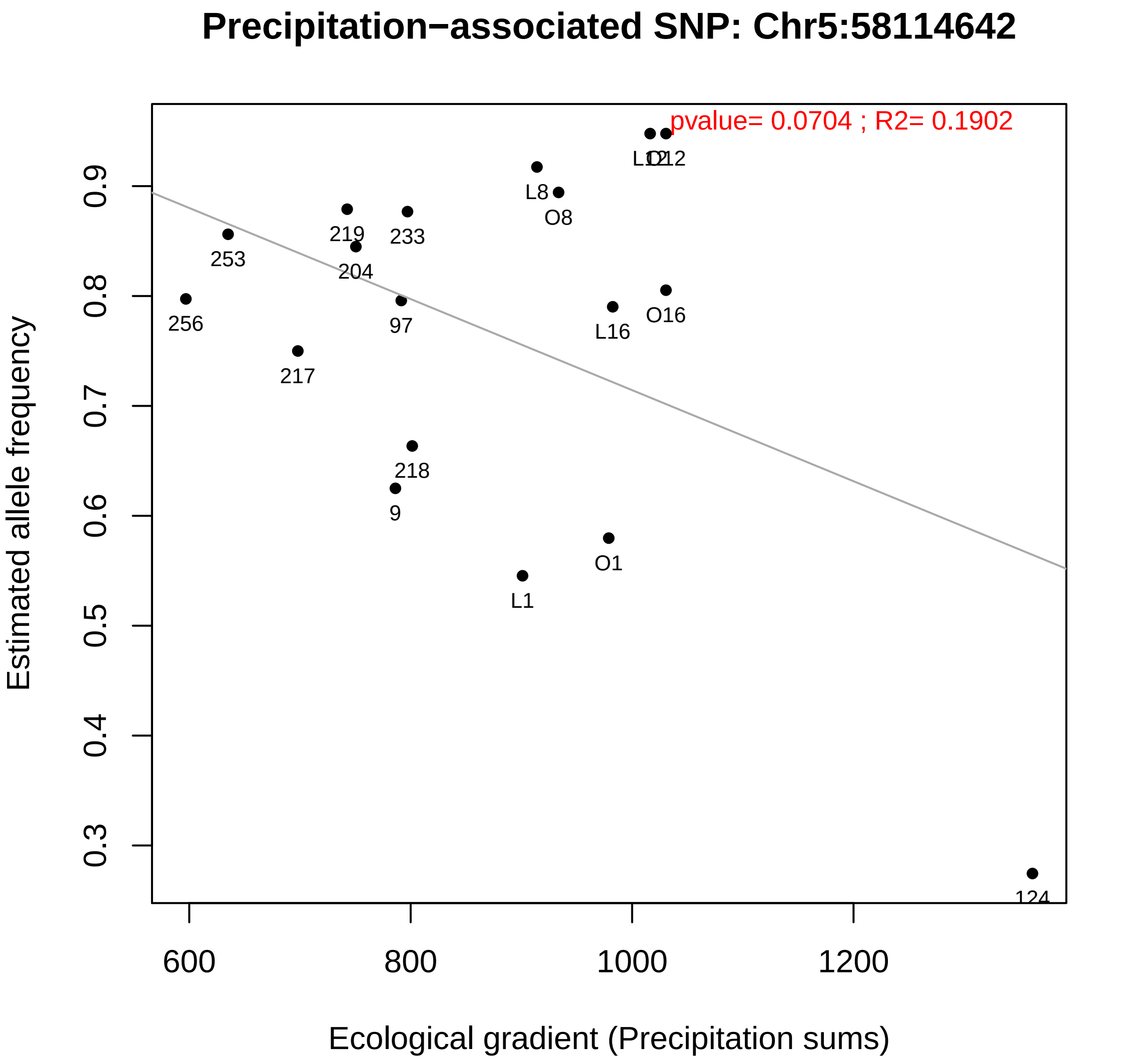
Precipitation−associated SNP: Chr5:58114642
pvalue= 0.0704 ; R2= 0.1902
Estimated allele frequency
Ecological gradient (Precipitation sums)
0.3
0.4
0.5
0.6
0.7
0.8
0.9
600
800
1000
1200
256
253
217
219
204
233
97
218
9
L8
O8
L12
O12
L16
O16
O1
L1
124

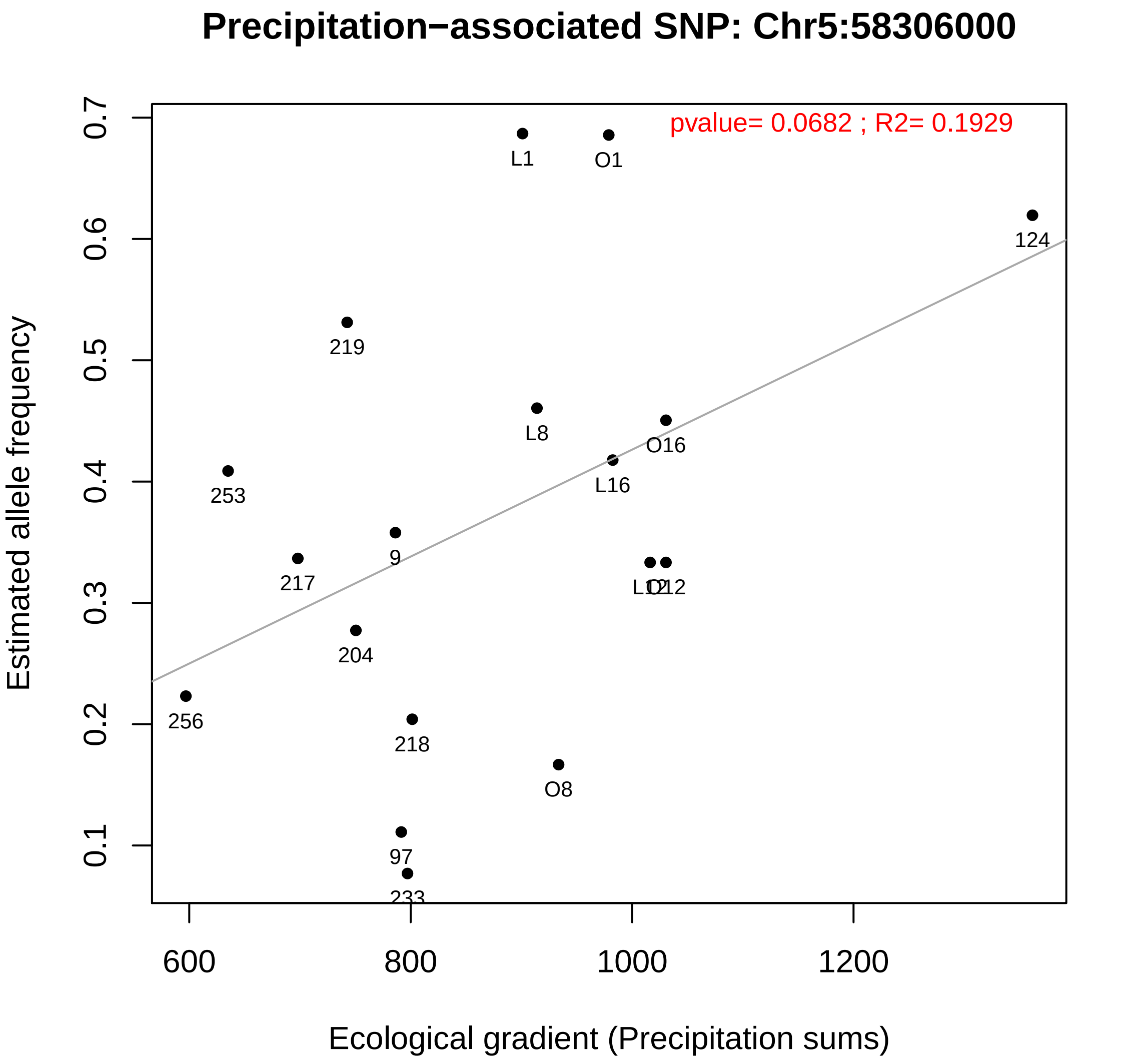
Precipitation−associated SNP: Chr5:58306000
pvalue= 0.0682 ; R2= 0.1929
L1
O1
124
219
L8
O16
L16
253
9
217
L12
O12
204
256
218
O8
97
233
0.1
0.2
0.3
0.4
0.5
0.6
0.7
600
800
1000
1200
Estimated allele frequency
Ecological gradient (Precipitation sums)

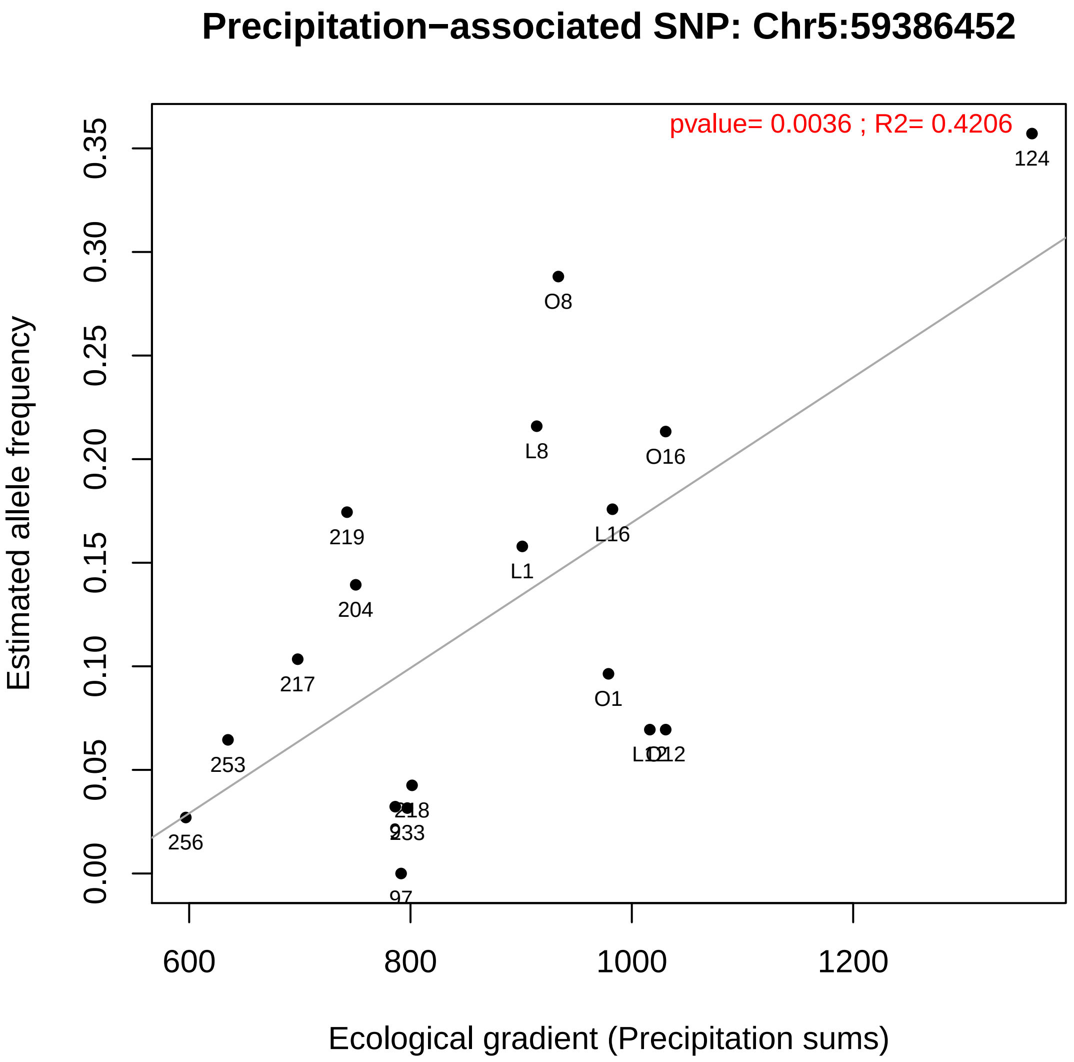
Precipitation−associated SNP: Chr5:59386452
pvalue= 0.0036 ; R2= 0.4206
124
O8
L8
O16
219
L16
L1
204
217
O1
L12
O12
253
218
233
256
97
0.00
0.05
0.10
0.15
0.20
0.25
0.30
0.35
Estimated allele frequency
600
800
1000
1200
Ecological gradient (Precipitation sums)

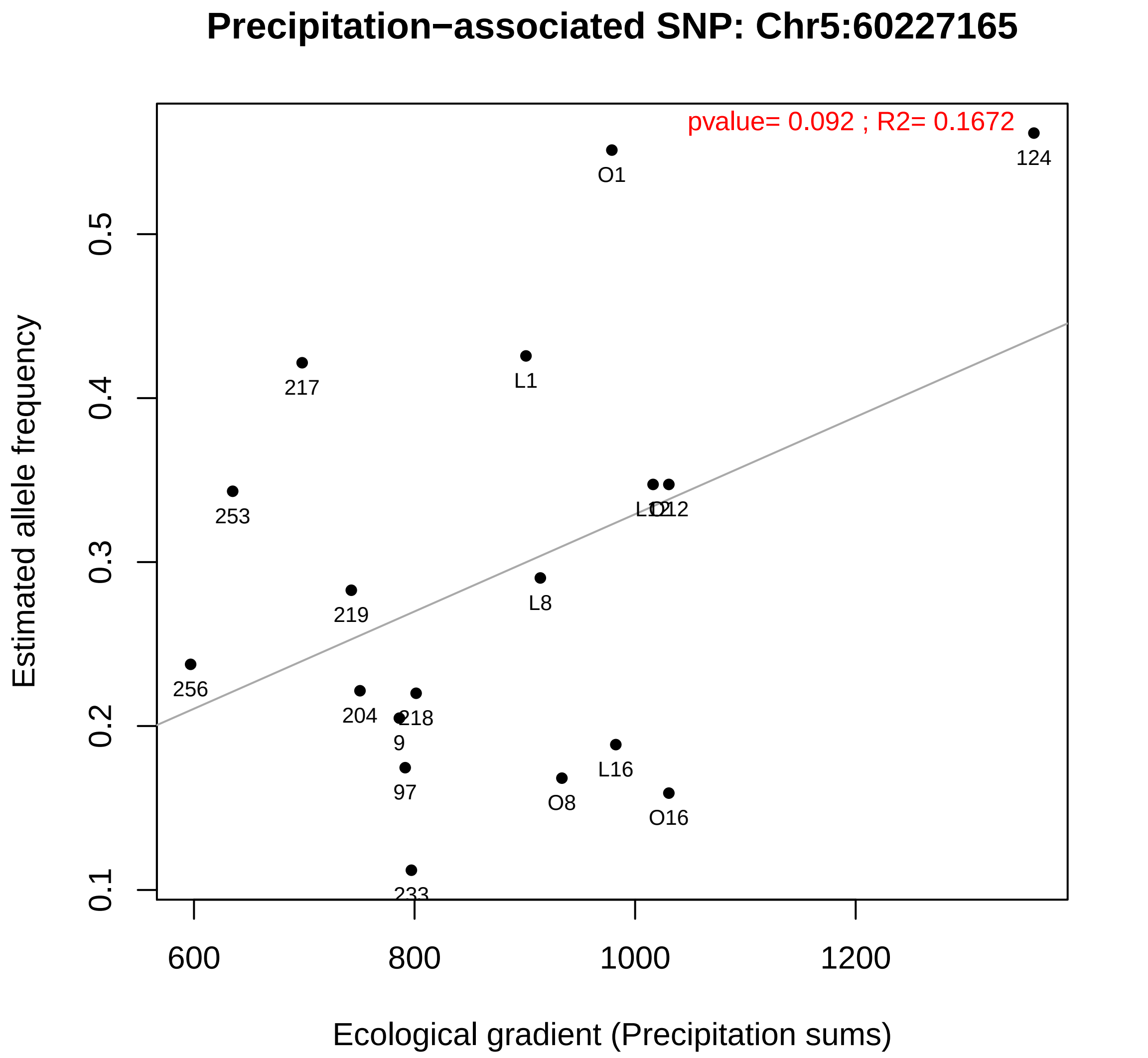
Precipitation−associated SNP: Chr5:60227165
pvalue= 0.092 ; R2= 0.1672
124
O1
217
L1
253
L10
O12
219
L8
256
204
218
9
97
L16
O8
O16
233
0.1
0.2
0.3
0.4
0.5
600
800
1000
1200
Estimated allele frequency
Ecological gradient (Precipitation sums)

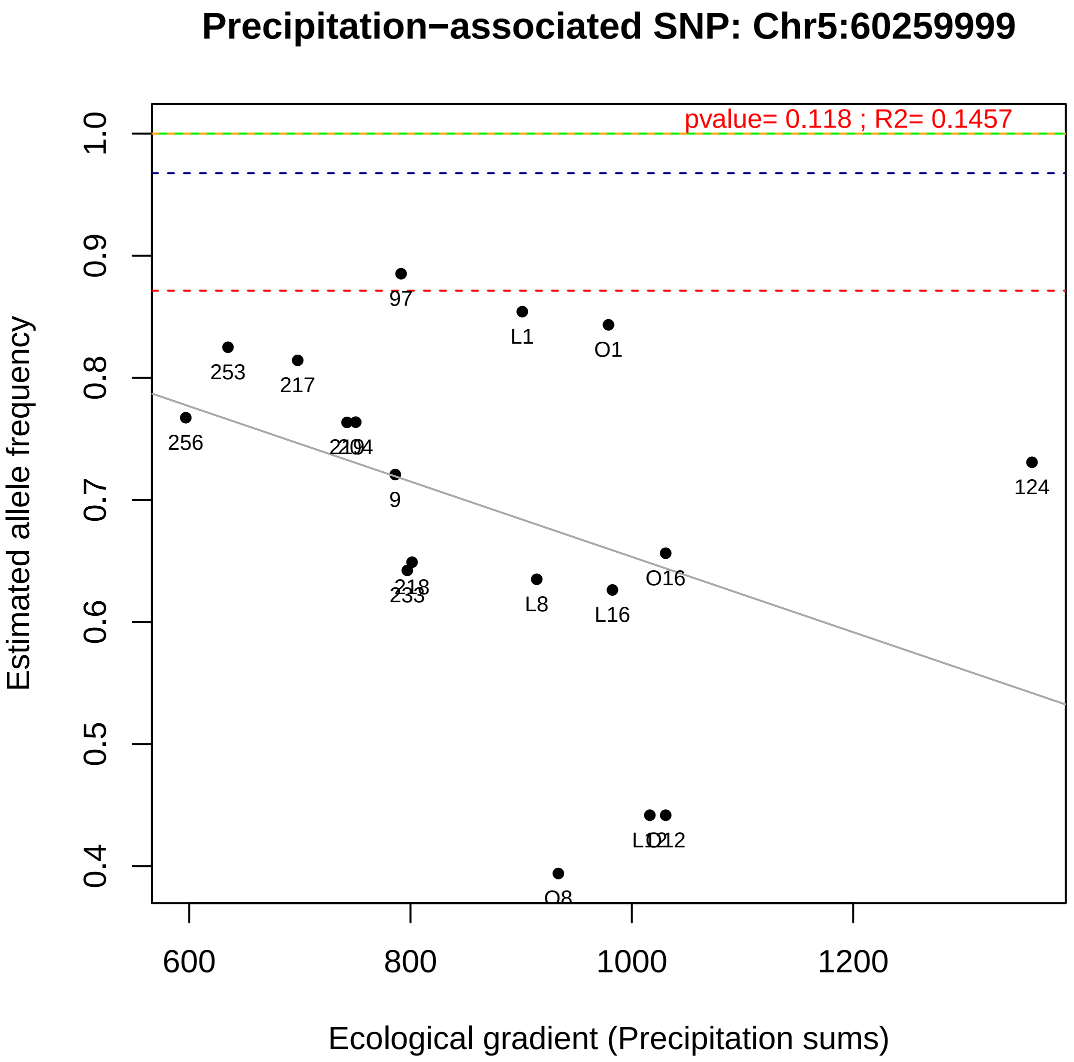

Precipitation−associated SNP: Chr5:60259999
pvalue= 0.118 ; R2= 0.1457
Estimated allele frequency
Ecological gradient (Precipitation sums)
1.0
0.9
0.8
0.7
0.6
0.5
0.4
600
800
1000
1200
97
L1
O1
253
217
256
124
9
O16
L8
L16
218
233
O8

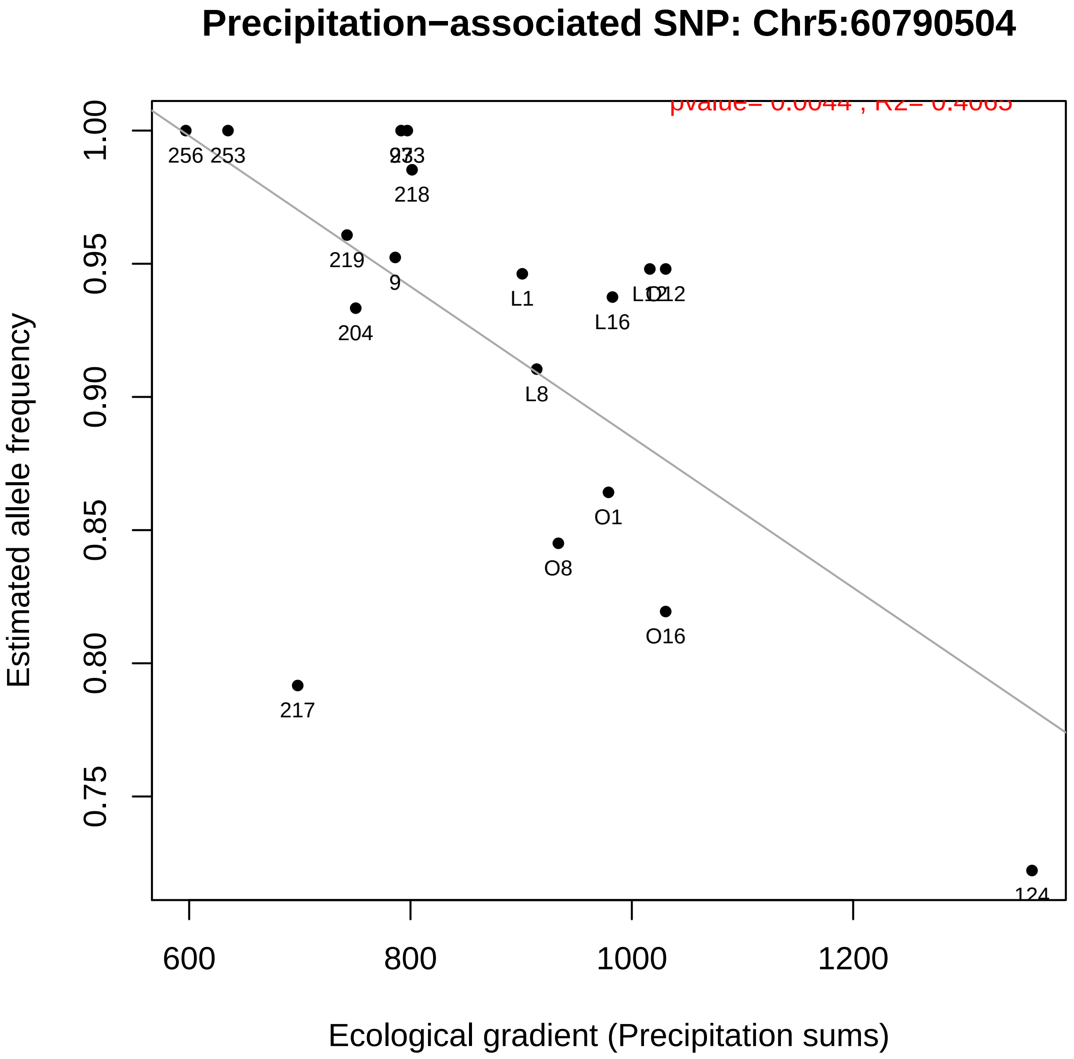
Precipitation−associated SNP: Chr5:60790504
pvalue= 0.0044 , R2= 0.4065
Estimated allele frequency
1.00
0.95
0.90
0.85
0.80
0.75
600
800
1000
1200
Ecological gradient (Precipitation sums)
256
253
218
219
9
204
L1
L16
L8
O1
O8
O16
217
124

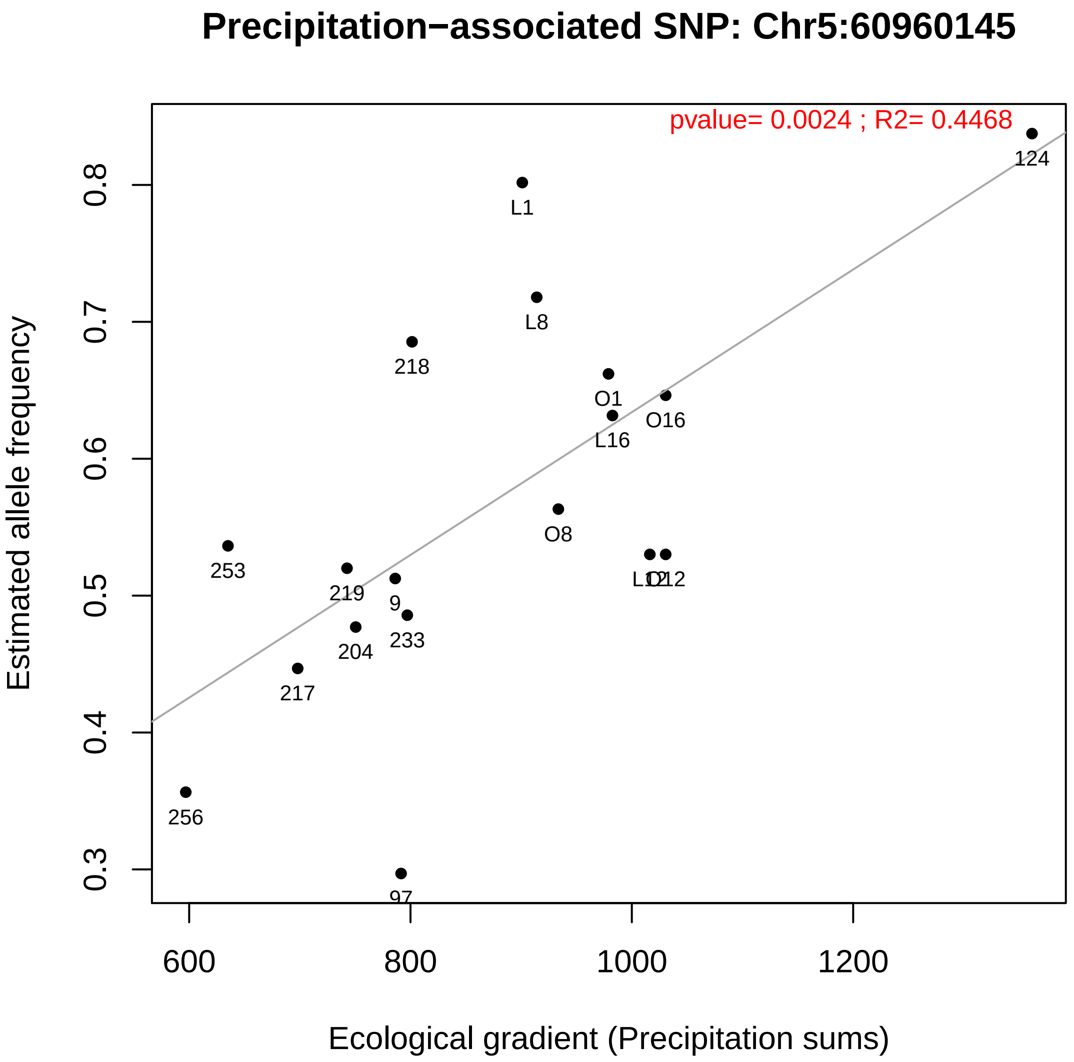

Precipitation−associated SNP: Chr5:60960145
pvalue= 0.0024 ; R2= 0.4468
Estimated allele frequency
Ecological gradient (Precipitation sums)
0.3
0.4
0.5
0.6
0.7
0.8
600
800
1000
1200
124
L1
L8
218
O1
O16
L16
O8
253
219
9
233
204
217
256
97
O12

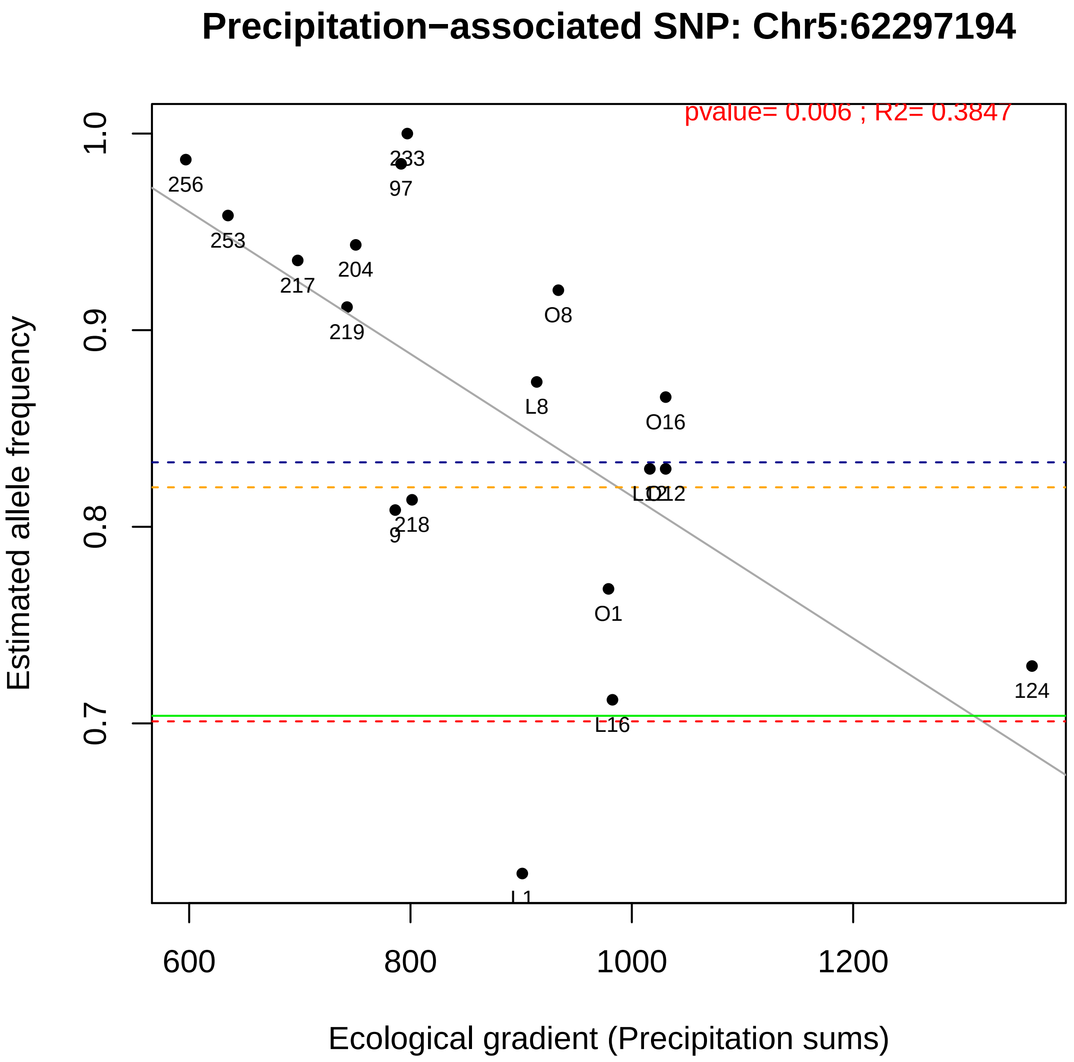

Precipitation−associated SNP: Chr5:62297194
pvalue= 0.006 ; R2= 0.3847
Estimated allele frequency
1.0
0.9
0.8
0.7
256
233
97
253
204
217
219
O8
L8
O16
L12
O12
218
9
O1
124
L16
L1
600
800
1000
1200
Ecological gradient (Precipitation sums)

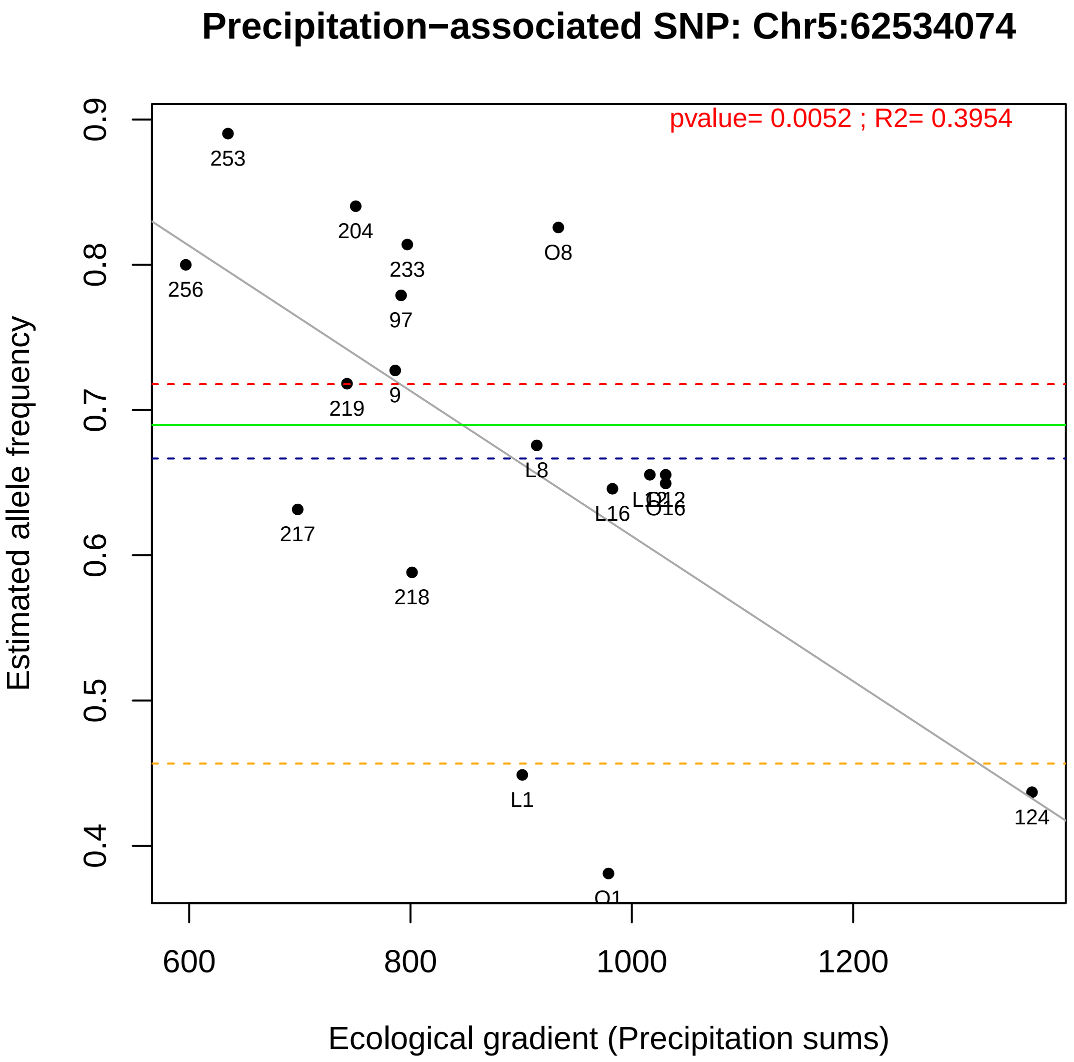
Precipitation−associated SNP: Chr5:62534074
pvalue= 0.0052 ; R2= 0.3954
Estimated allele frequency
Ecological gradient (Precipitation sums)
0.4
0.5
0.6
0.7
0.8
0.9
600
800
1000
1200
253
204
233
O8
256
97
9
219
L8
L16
217
218
L1
124

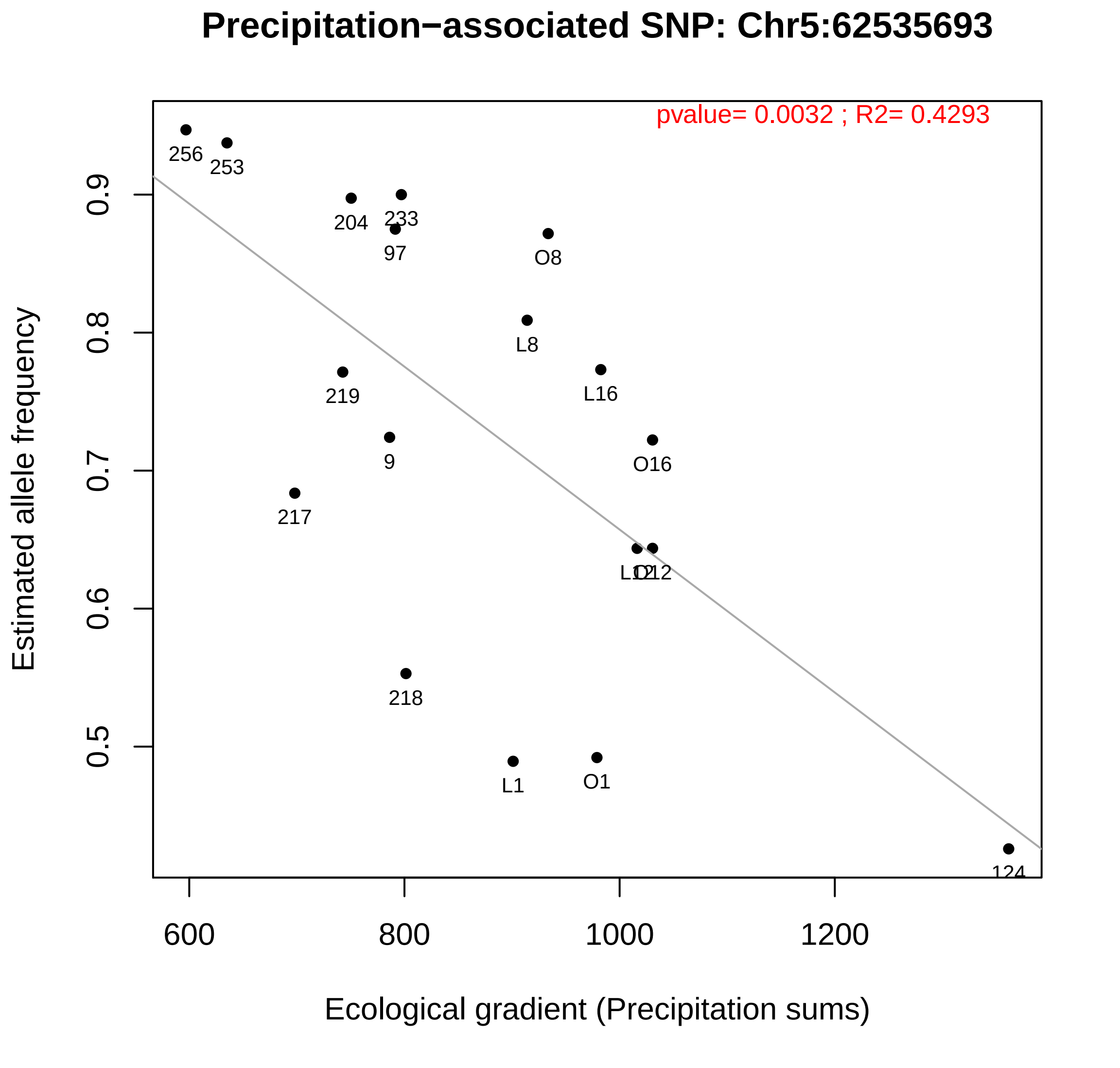
Precipitation−associated SNP: Chr5:62535693
pvalue= 0.0032 ; R2= 0.4293
Estimated allele frequency
0.5
0.6
0.7
0.8
0.9
600
800
1000
1200
Ecological gradient (Precipitation sums)
256
253
204
233
97
O8
L8
219
L16
9
O16
217
L12
O12
218
L1
O1
124

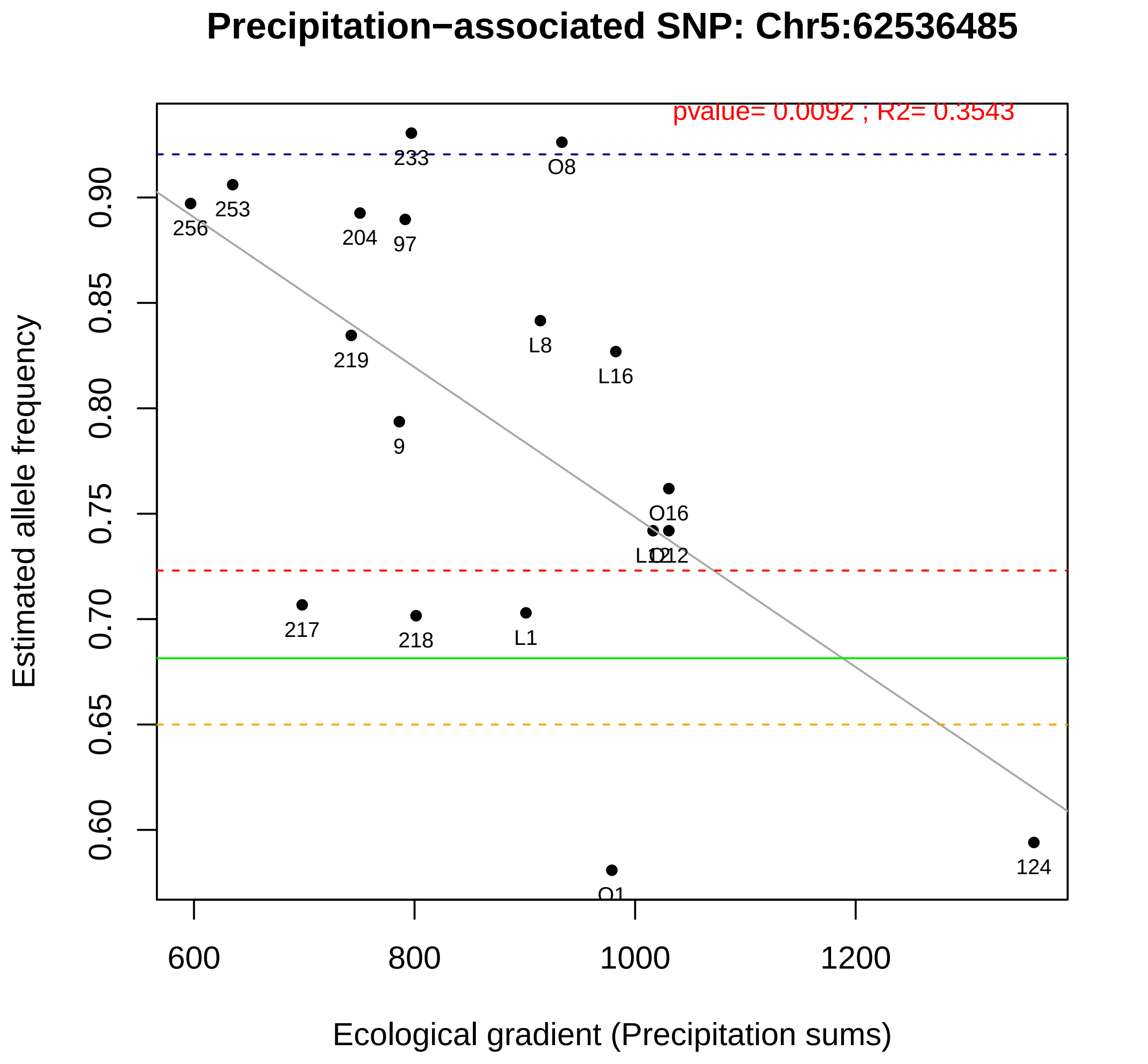
Precipitation−associated SNP: Chr5:62536485
pvalue= 0.0092 ; R2= 0.3543
Estimated allele frequency
0.60
0.65
0.70
0.75
0.80
0.85
0.90
600
800
1000
1200
Ecological gradient (Precipitation sums)
233
O8
253
256
204
97
219
L8
L16
9
O16
L10
O12
217
218
L1
124
O1

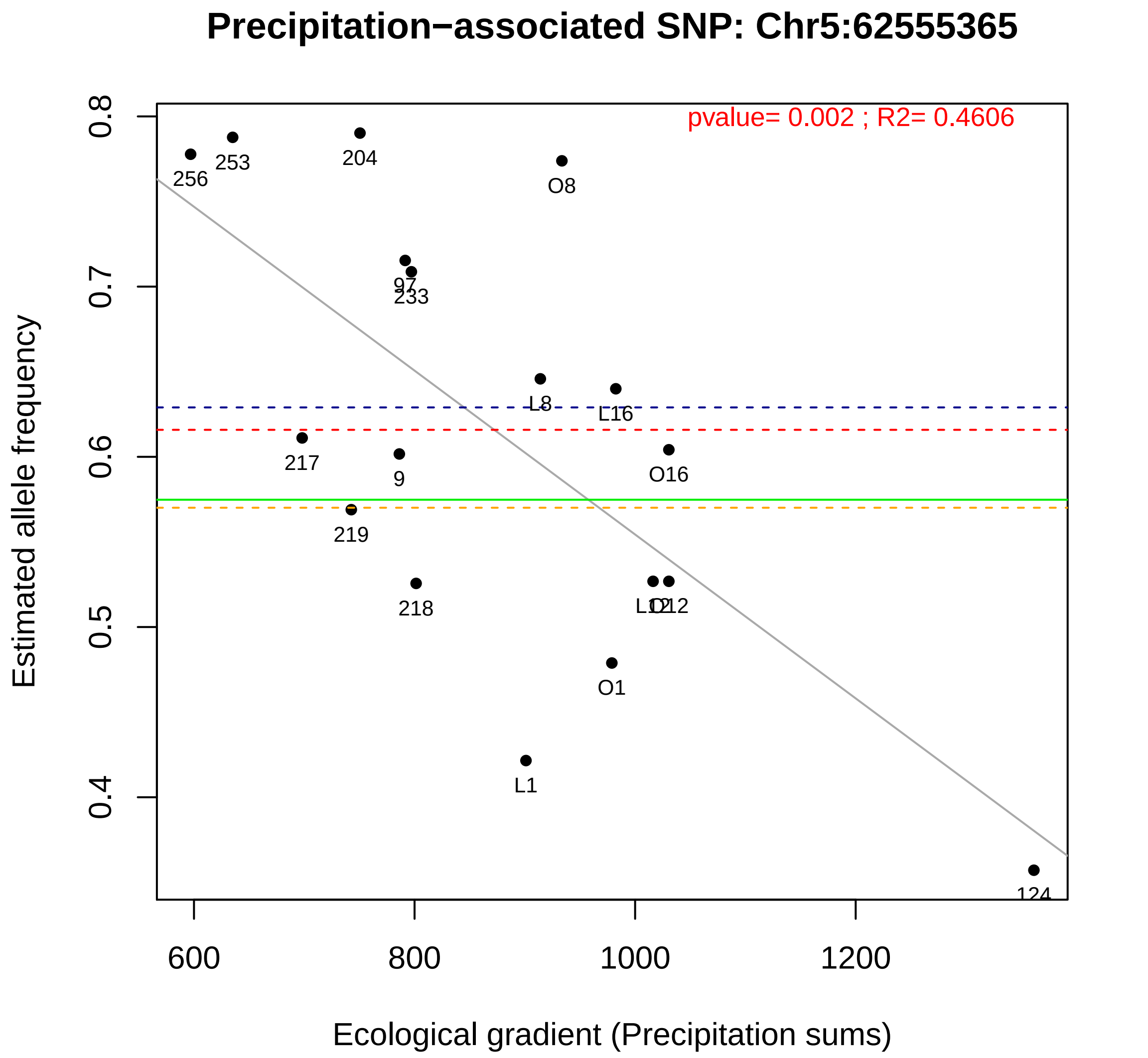
Precipitation−associated SNP: Chr5:62555365
pvalue= 0.002 ; R2= 0.4606
Estimated allele frequency
0.4
0.5
0.6
0.7
0.8
600
800
1000
1200
Ecological gradient (Precipitation sums)
256
253
204
O8
97
233
L8
L16
217
9
O16
219
218
L12
O12
O1
L1
124

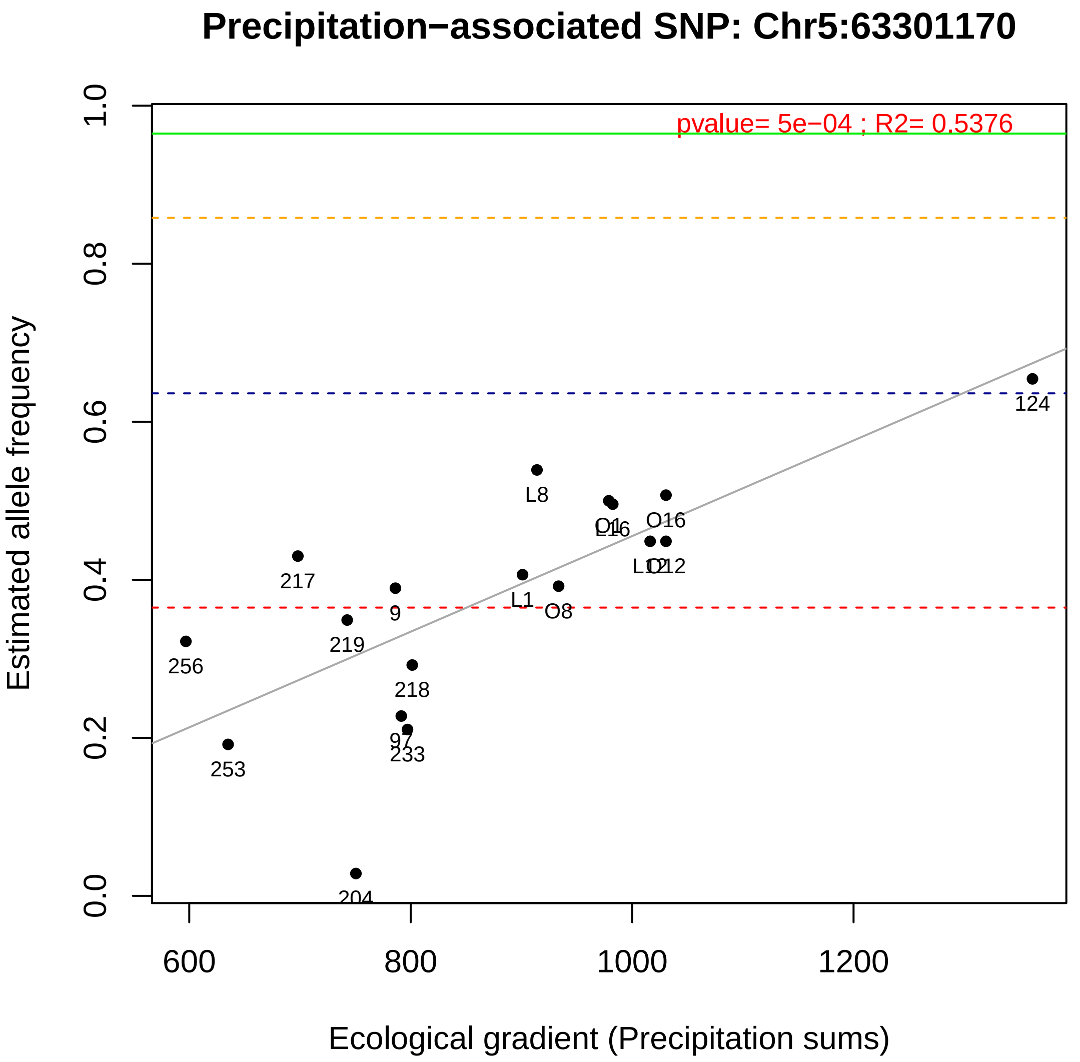

Precipitation−associated SNP: Chr5:63301170
pvalue= 5e−04 ; R2= 0.5376
Estimated allele frequency
0.0
0.2
0.4
0.6
0.8
1.0
Ecological gradient (Precipitation sums)
600
800
1000
1200
124
L8
O16
L12
O12
217
9
L1
O8
219
256
218
97
233
253
204

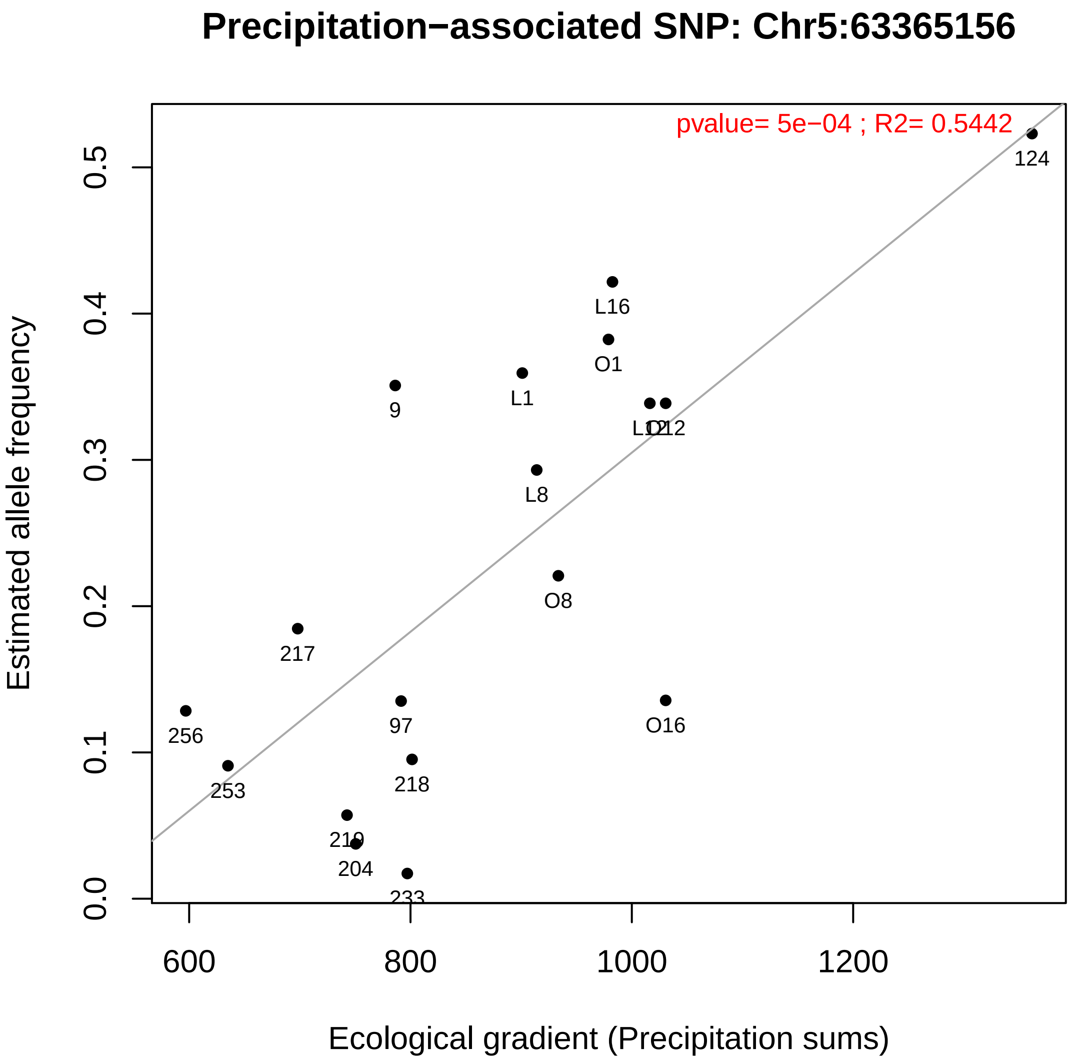
Precipitation−associated SNP: Chr5:63365156
pvalue= 5e−04 ; R2= 0.5442
Estimated allele frequency
0.0
0.1
0.2
0.3
0.4
0.5
600
800
1000
1200
Ecological gradient (Precipitation sums)
124
L16
O1
L1
9
O12
L8
O8
217
97
O16
256
253
218
204
233

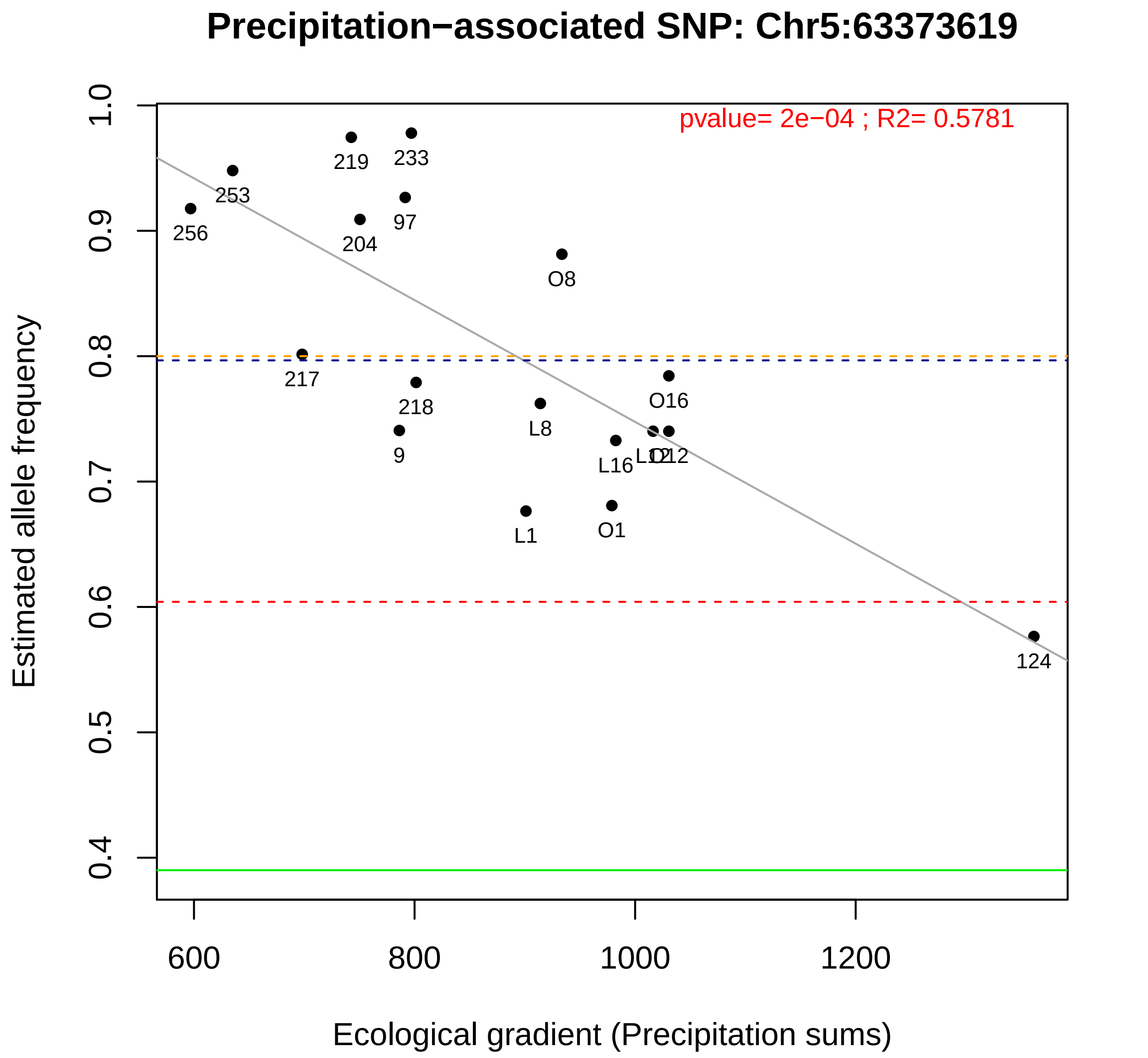
Precipitation−associated SNP: Chr5:63373619
pvalue= 2e−04 ; R2= 0.5781
Estimated allele frequency
Ecological gradient (Precipitation sums)
0.4
0.5
0.6
0.7
0.8
0.9
1.0
600
800
1000
1200
219
233
253
256
204
97
O8
217
218
O16
L8
9
L16
L12
O12
L1
O1
124

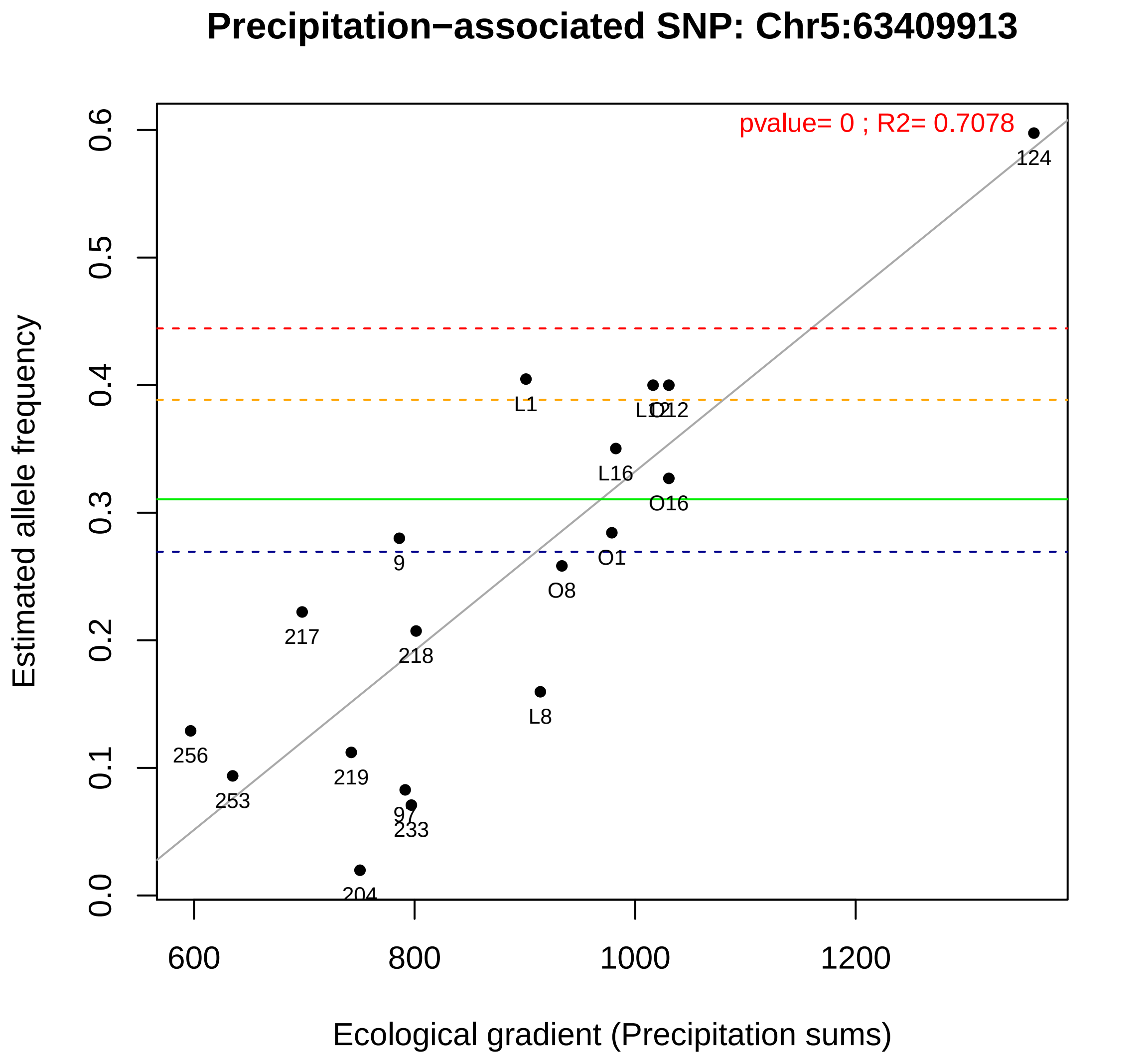

Precipitation−associated SNP: Chr5:63409913
pvalue= 0 ; R2= 0.7078
Estimated allele frequency
Ecological gradient (Precipitation sums)
0.0
0.1
0.2
0.3
0.4
0.5
0.6
600
800
1000
1200
124
L1
L10
O12
L16
O16
O1
9
O8
217
218
L8
256
253
219
97
233
204

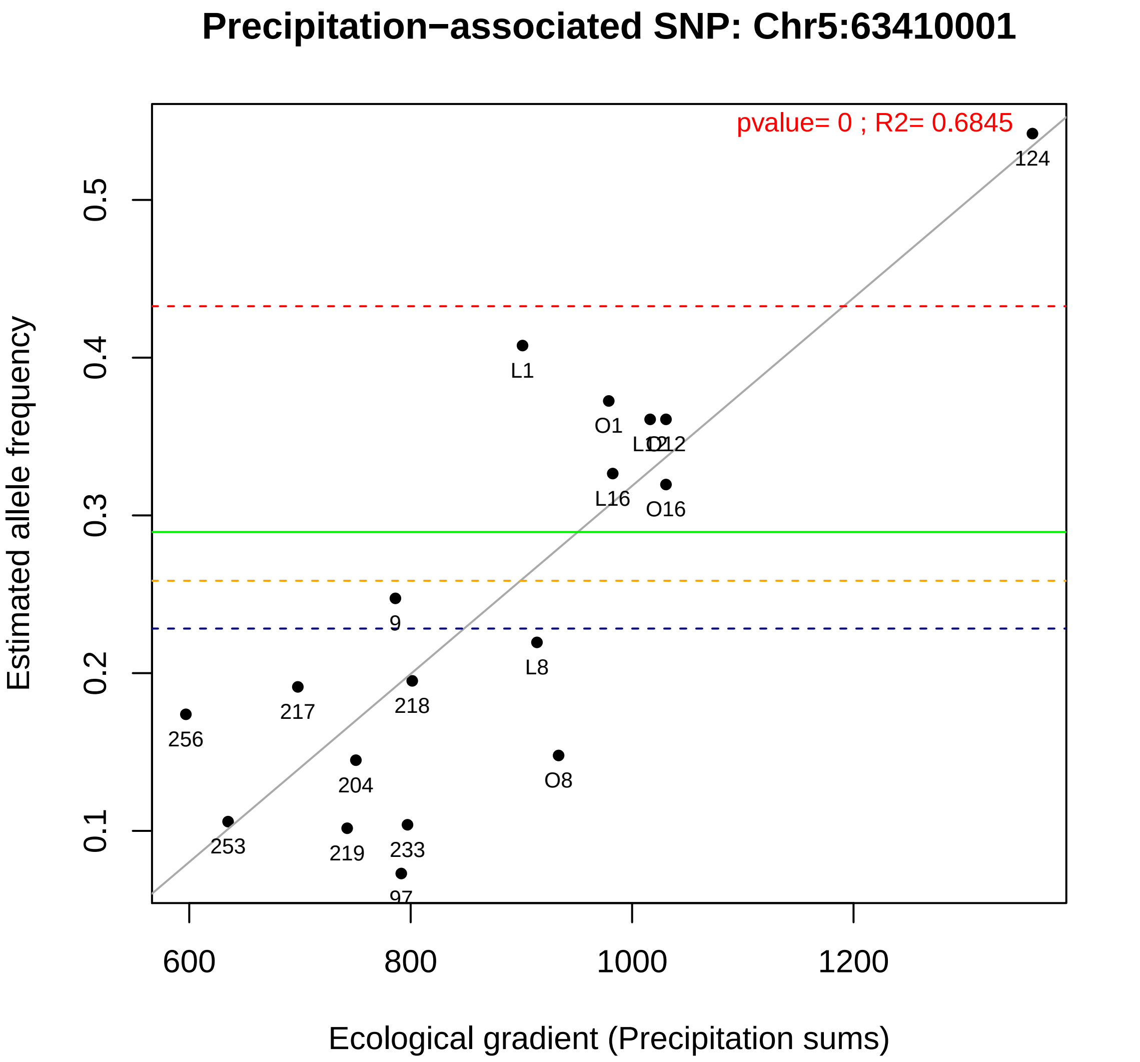

Precipitation−associated SNP: Chr5:63410001
pvalue= 0 ; R2= 0.6845
Estimated allele frequency
Ecological gradient (Precipitation sums)
0.1
0.2
0.3
0.4
0.5
600
800
1000
1200
124
L1
O1
L12
O12
L16
O16
9
L8
217
218
256
204
O8
253
219
233
97

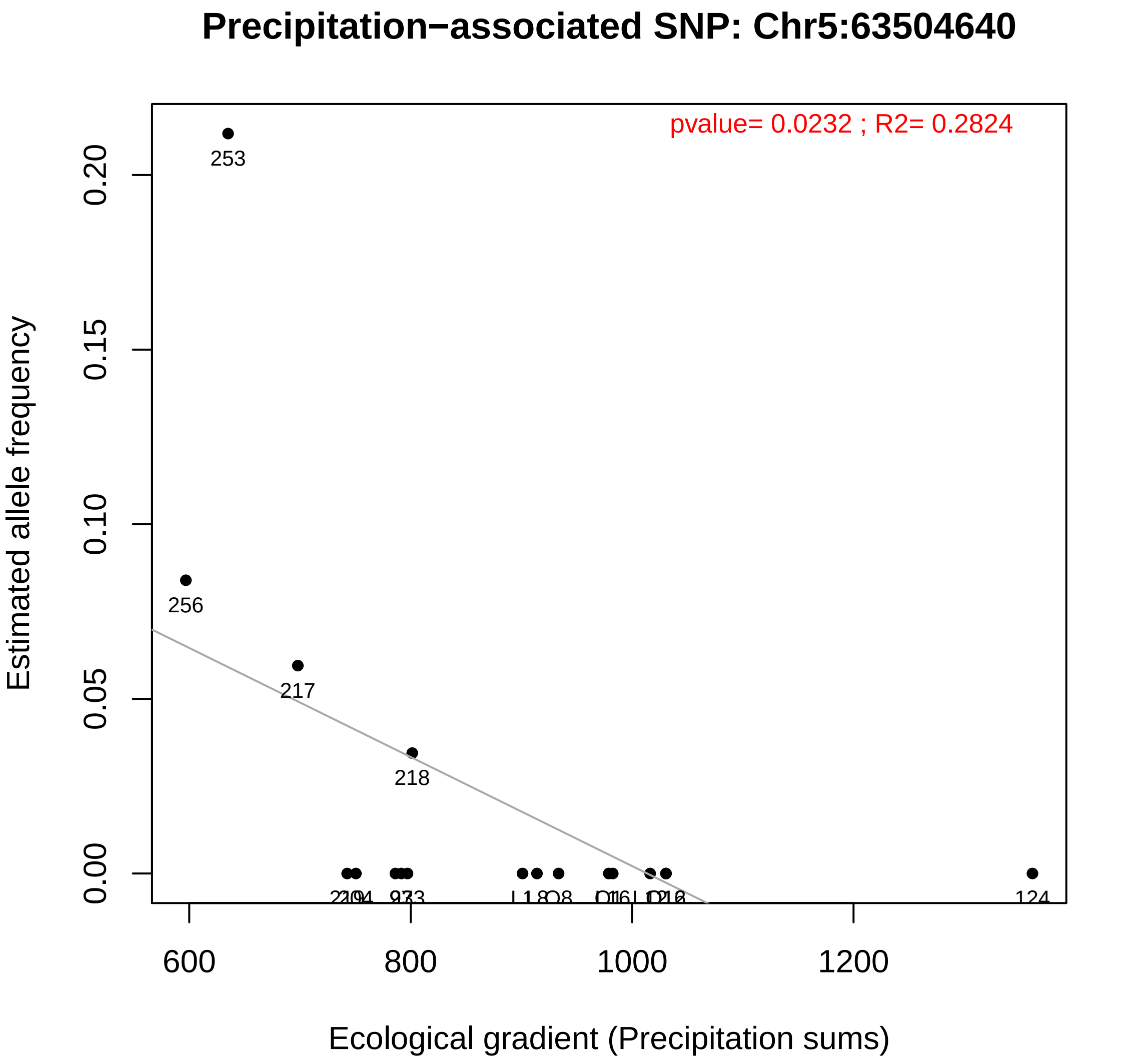

Precipitation−associated SNP: Chr5:63504640
pvalue= 0.0232 ; R2= 0.2824
Estimated allele frequency
0.00
0.05
0.10
0.15
0.20
600
800
1000
1200
Ecological gradient (Precipitation sums)
253
256
217
218
124

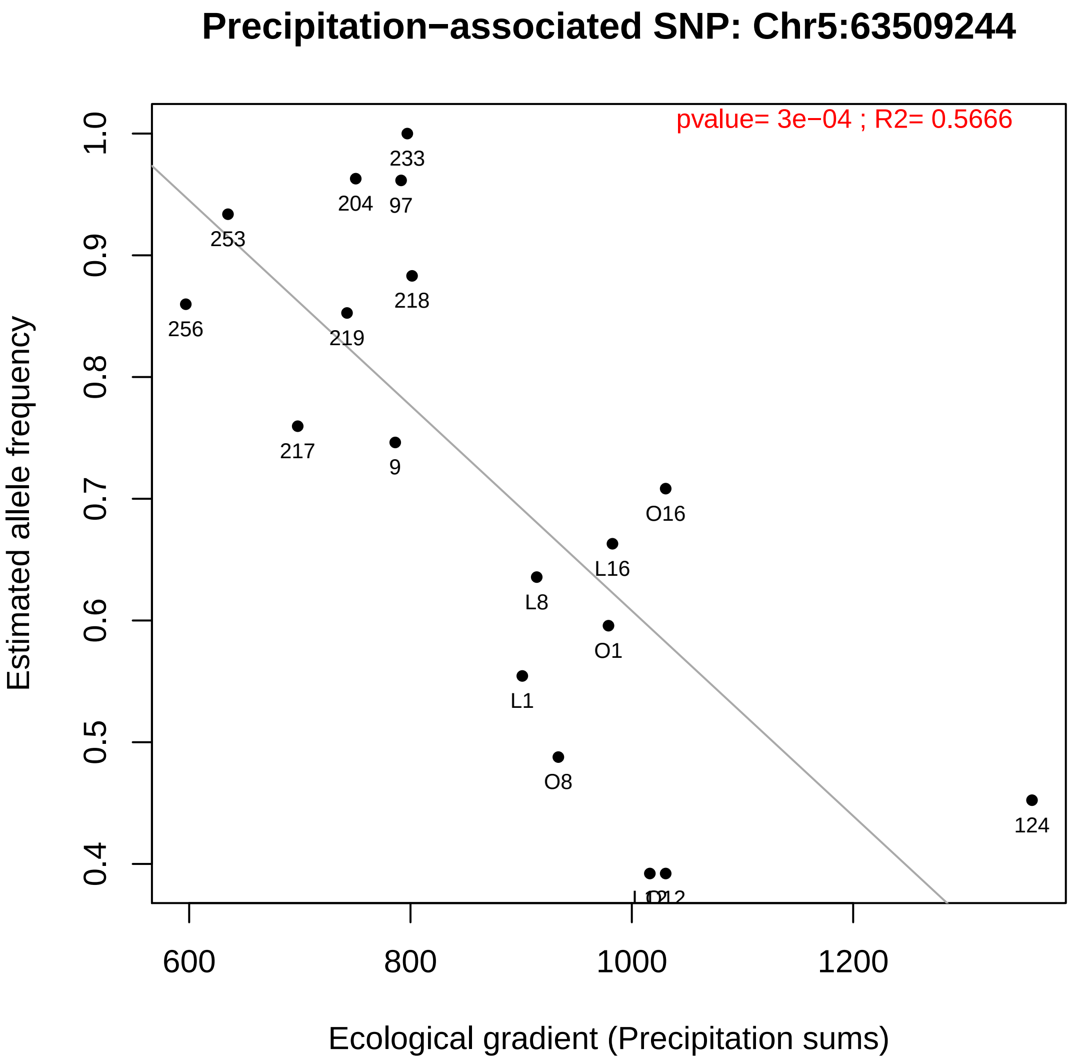

Precipitation−associated SNP: Chr5:63509244
pvalue= 3e−04 ; R2= 0.5666
Estimated allele frequency
Ecological gradient (Precipitation sums)
0.4
0.5
0.6
0.7
0.8
0.9
1.0
600
800
1000
1200
233
204
97
253
218
256
219
217
9
O16
L16
L8
O1
L1
O8
124

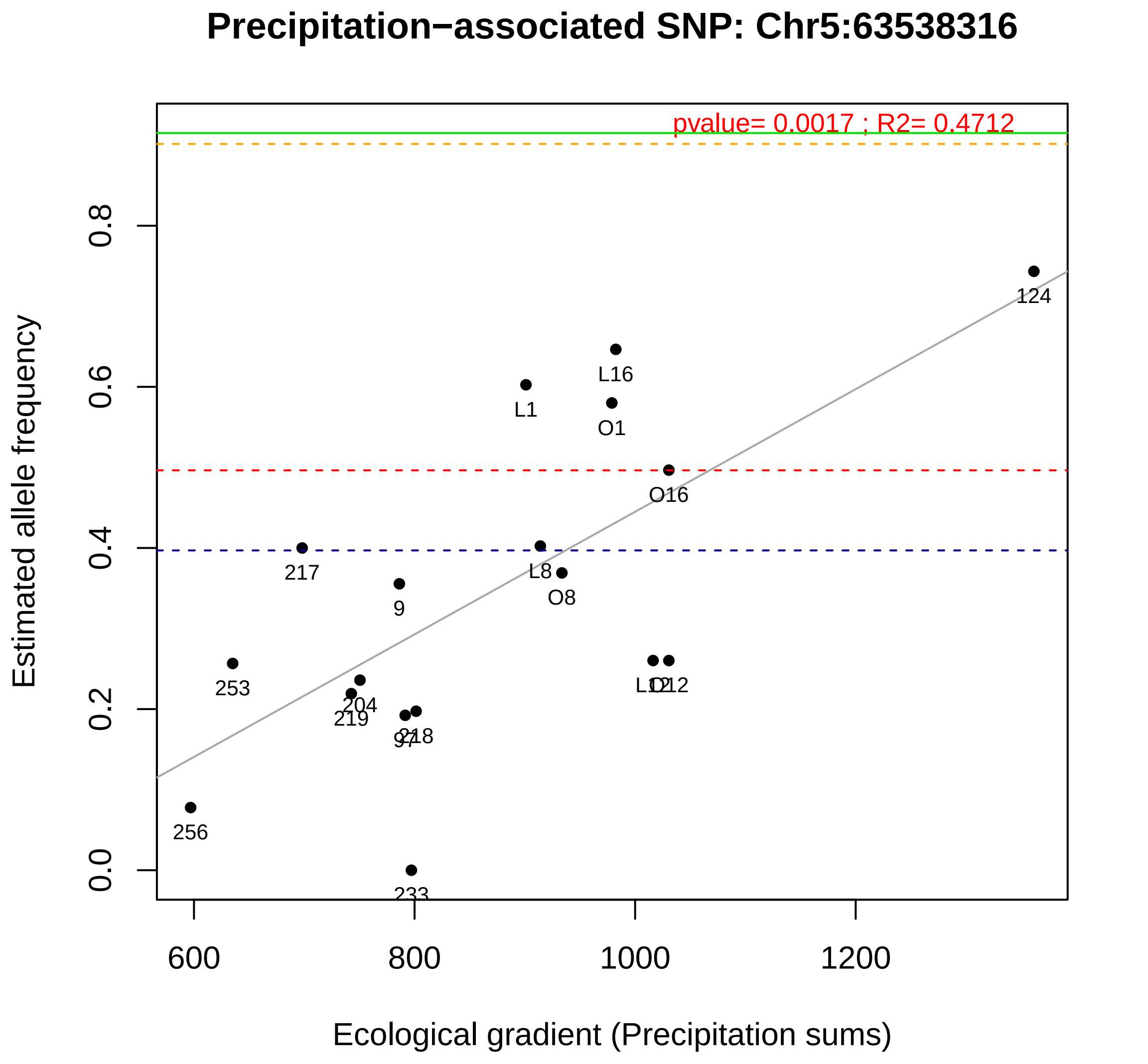

Precipitation−associated SNP: Chr5:63538316
pvalue= 0.0017 ; R2= 0.4712
Estimated allele frequency
0.0
0.2
0.4
0.6
0.8
600
800
1000
1200
Ecological gradient (Precipitation sums)
124
L16
L1
O1
O16
L8
O8
217
9
253
204
219
218
256
233

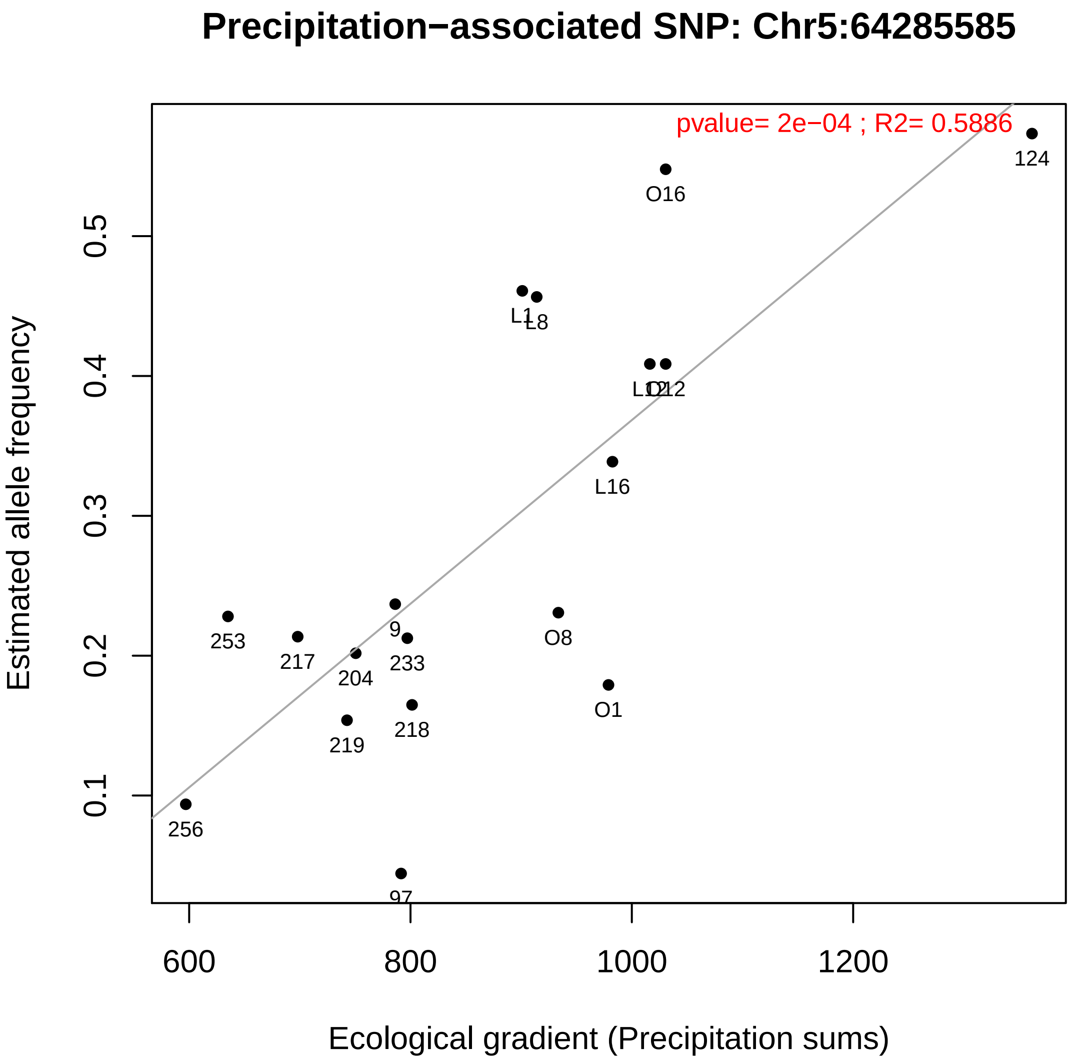

Precipitation−associated SNP: Chr5:64285585
pvalue= 2e−04 ; R2= 0.5886
Estimated allele frequency
Ecological gradient (Precipitation sums)
0.1
0.2
0.3
0.4
0.5
600
800
1000
1200
124
O16
L1
L8
L12
O12
L16
253
217
204
9
233
O8
O1
219
218
256
97

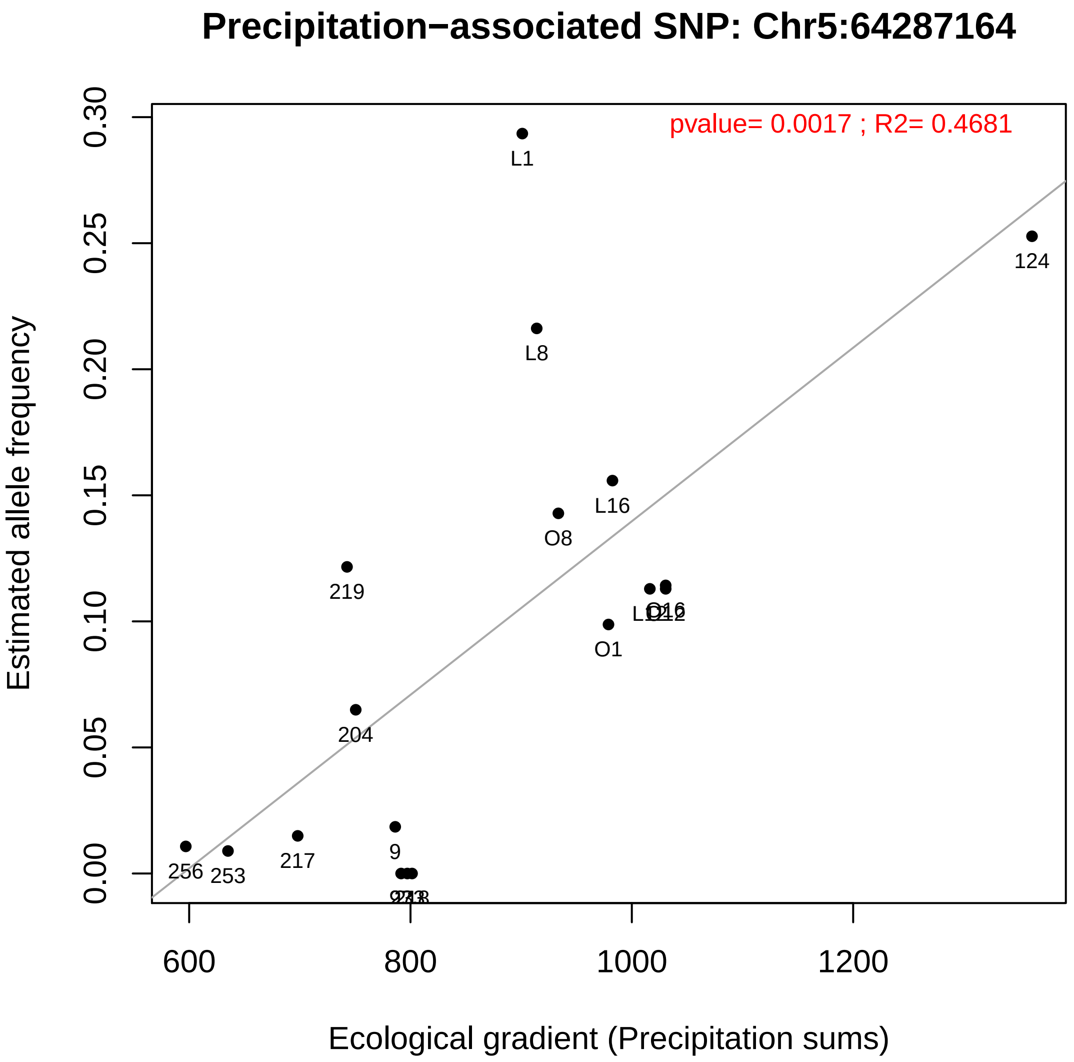

Precipitation−associated SNP: Chr5:64287164
pvalue= 0.0017 ; R2= 0.4681
Estimated allele frequency
Ecological gradient (Precipitation sums)
0.00
0.05
0.10
0.15
0.20
0.25
0.30
600
800
1000
1200
L1
124
L8
L16
O8
219
O1
204
256
253
217
9

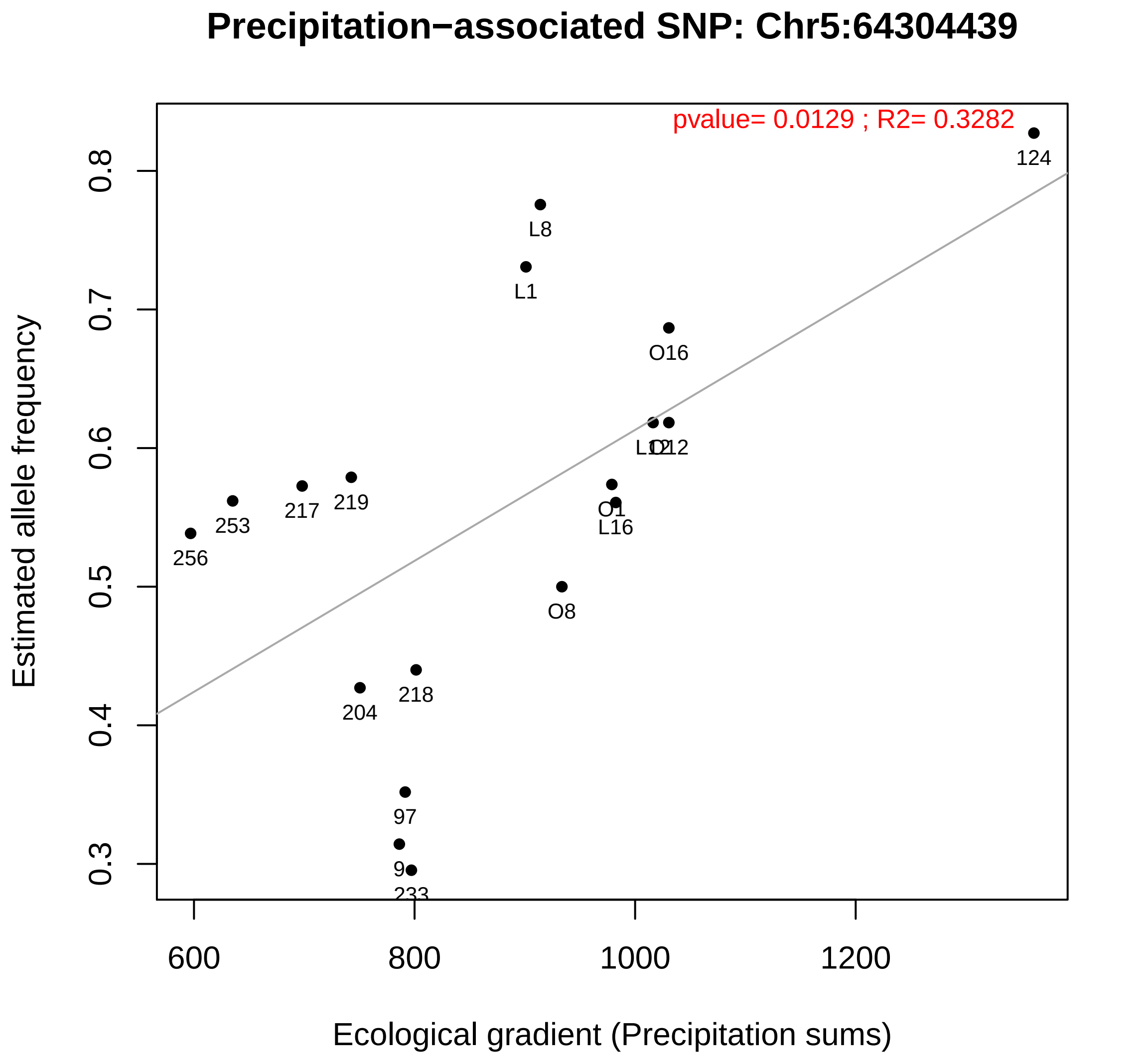
Precipitation−associated SNP: Chr5:64304439
pvalue= 0.0129 ; R2= 0.3282
Estimated allele frequency
Ecological gradient (Precipitation sums)
0.3
0.4
0.5
0.6
0.7
0.8
600
800
1000
1200
124
L8
L1
O16
L12
O12
O1
L16
O8
256
253
217
219
204
218
97
9
233

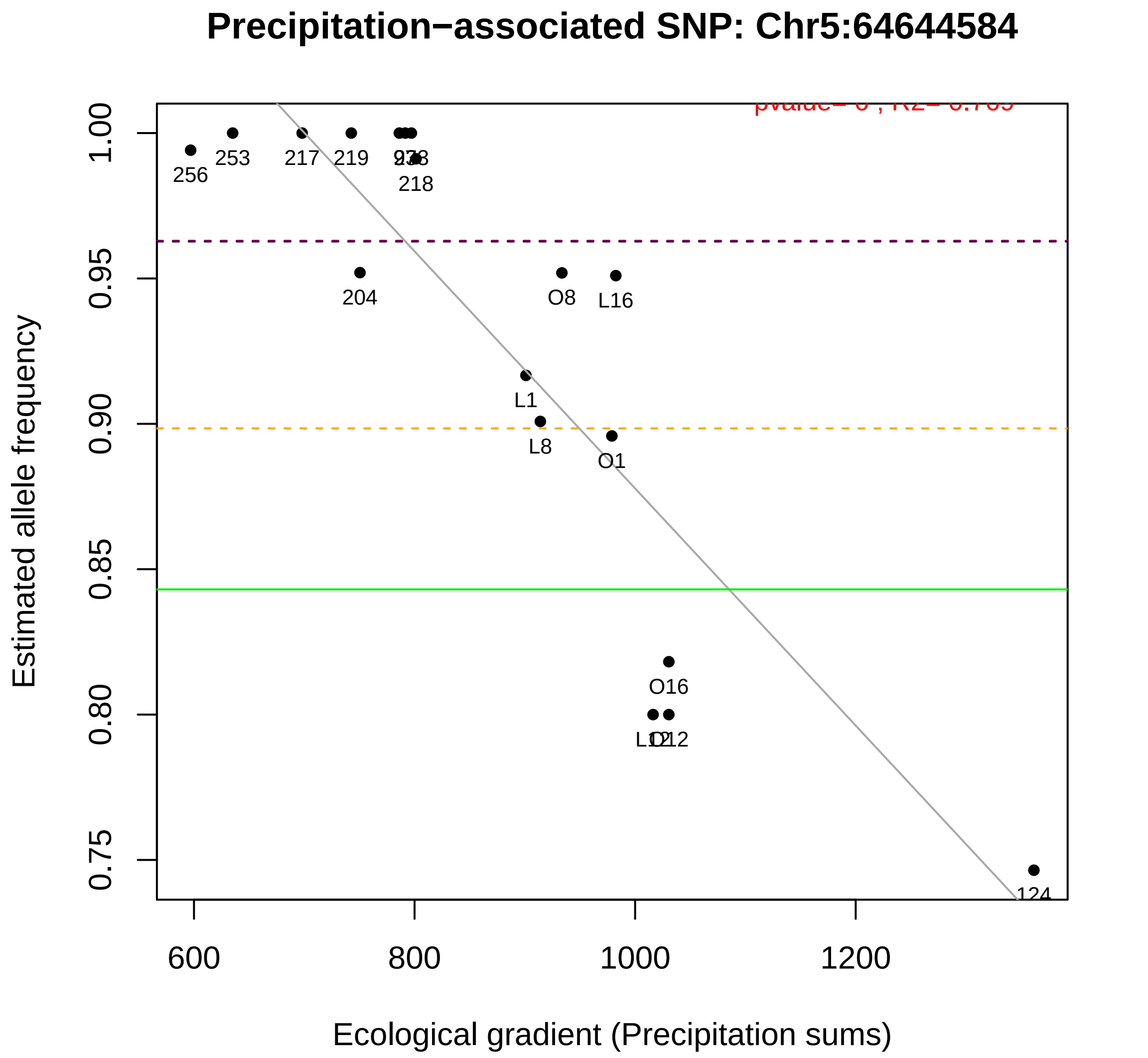
Precipitation−associated SNP: Chr5:64644584
Estimated allele frequency
Ecological gradient (Precipitation sums)
1.00
0.95
0.90
0.85
0.80
0.75
600
800
1000
1200
256
253
217
219
218
204
O8
L16
L1
L8
O1
O16
124

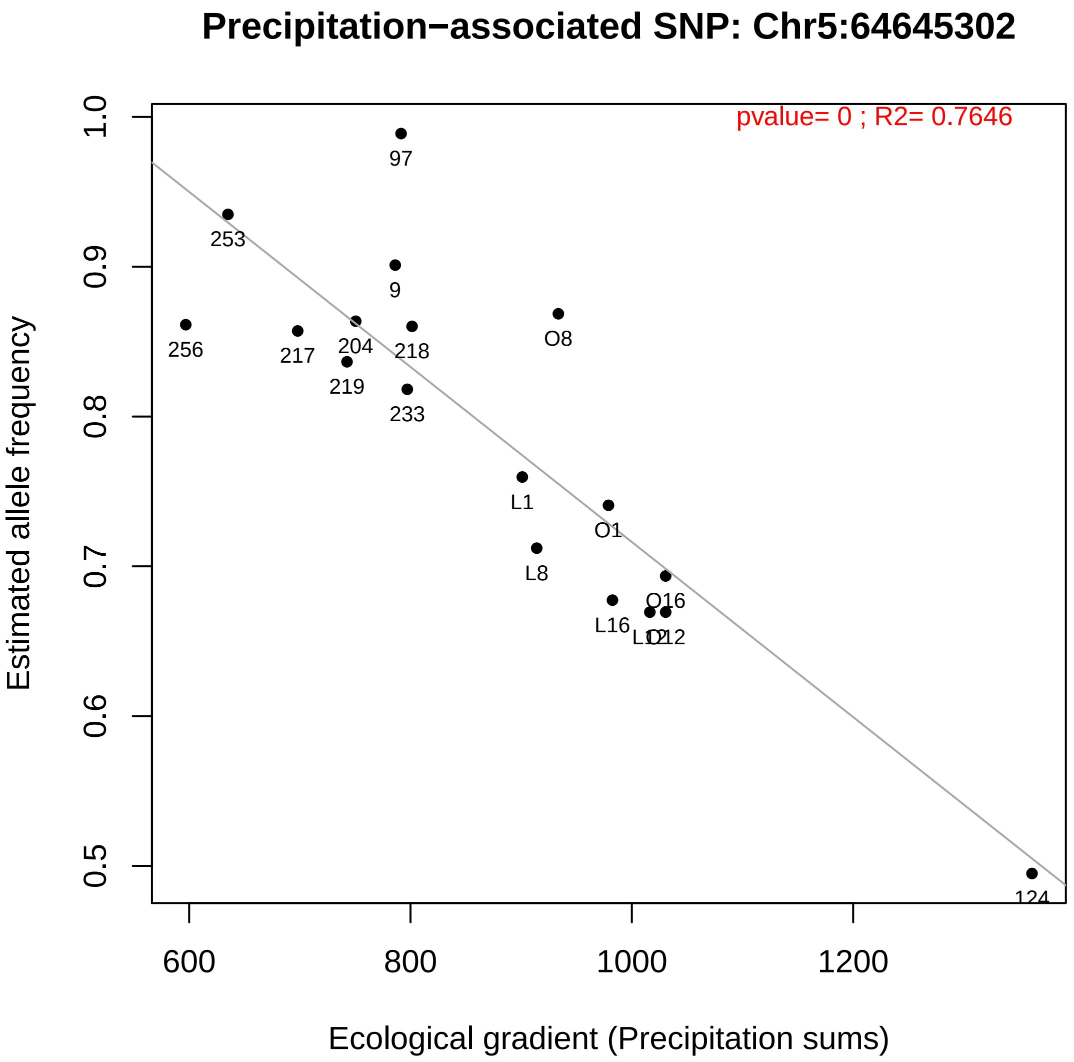
Precipitation−associated SNP: Chr5:64645302
pvalue= 0 ; R2= 0.7646
Estimated allele frequency
Ecological gradient (Precipitation sums)
1.0
0.9
0.8
0.7
0.6
0.5
600
800
1000
1200
97
253
9
256
217
204
218
219
233
O8
L1
O1
L8
O16
L16
L12
O12
124

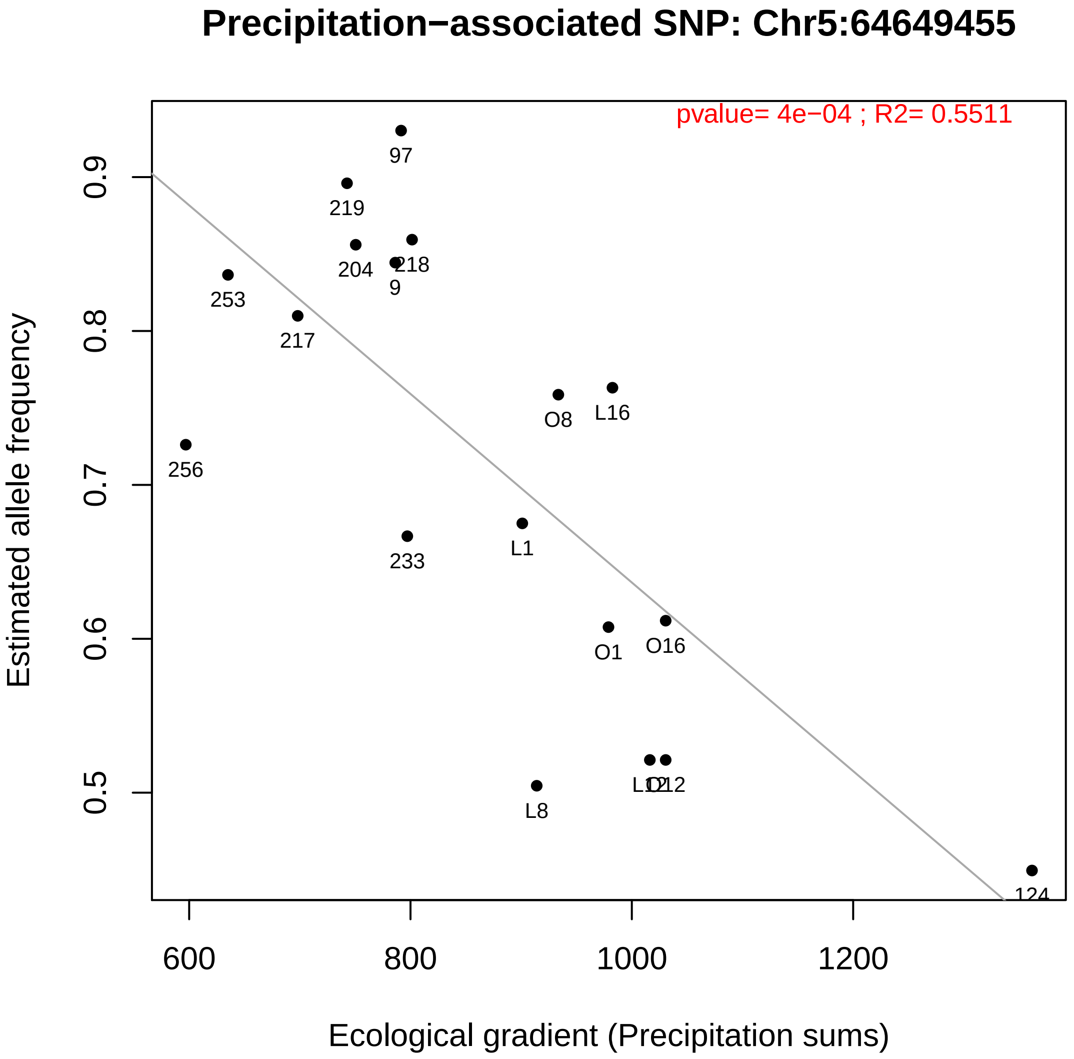

Precipitation−associated SNP: Chr5:64649455
pvalue= 4e−04 ; R2= 0.5511
97
219
204
218
9
253
217
O8
L16
256
233
L1
O1
O16
L12
O12
L8
124
0.5
0.6
0.7
0.8
0.9
600
800
1000
1200
Estimated allele frequency
Ecological gradient (Precipitation sums)

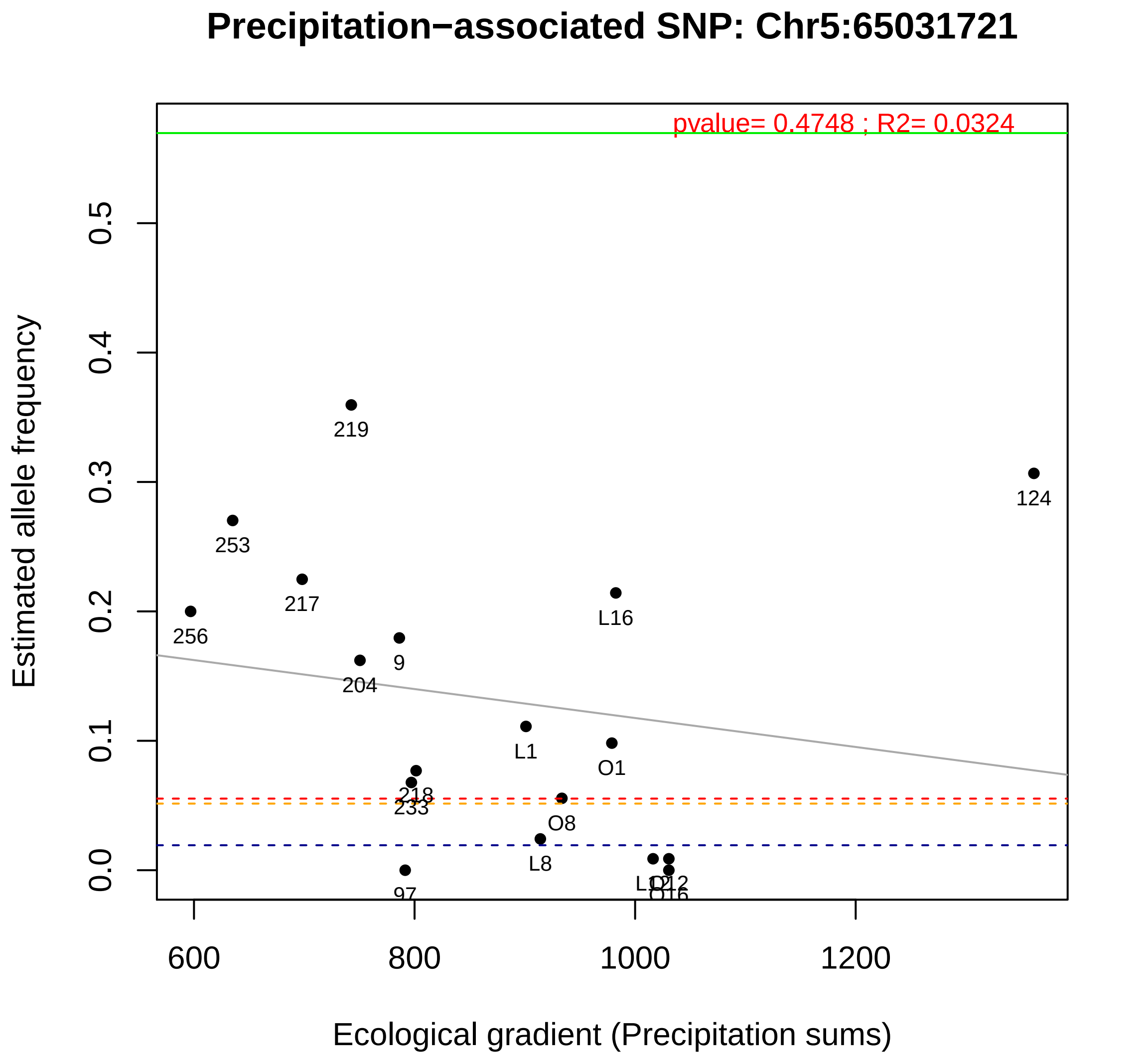

Precipitation−associated SNP: Chr5:65031721
pvalue= 0.4748 ; R2= 0.0324
Estimated allele frequency
0.0
0.1
0.2
0.3
0.4
0.5
600
800
1000
1200
Ecological gradient (Precipitation sums)
219
124
253
217
256
L16
9
204
L1
O1
218
233
O8
L8
97

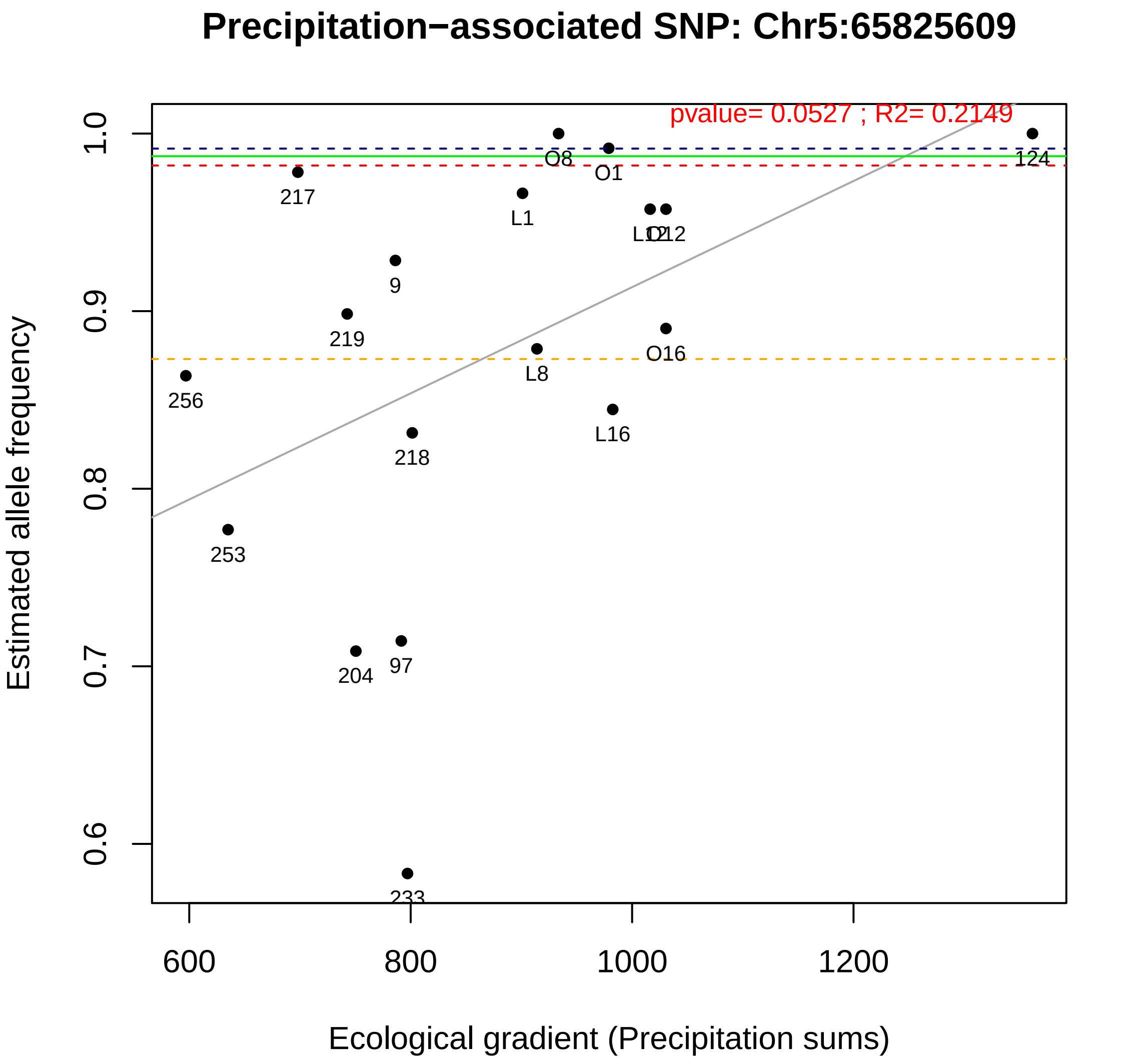
Precipitation−associated SNP: Chr5:65825609
pvalue= 0.0527 ; R2= 0.2149
Estimated allele frequency
Ecological gradient (Precipitation sums)
0.6
0.7
0.8
0.9
1.0
600
800
1000
1200
O8
O1
124
217
L1
L12
O12
9
219
O16
L8
256
L16
218
253
204
97
233

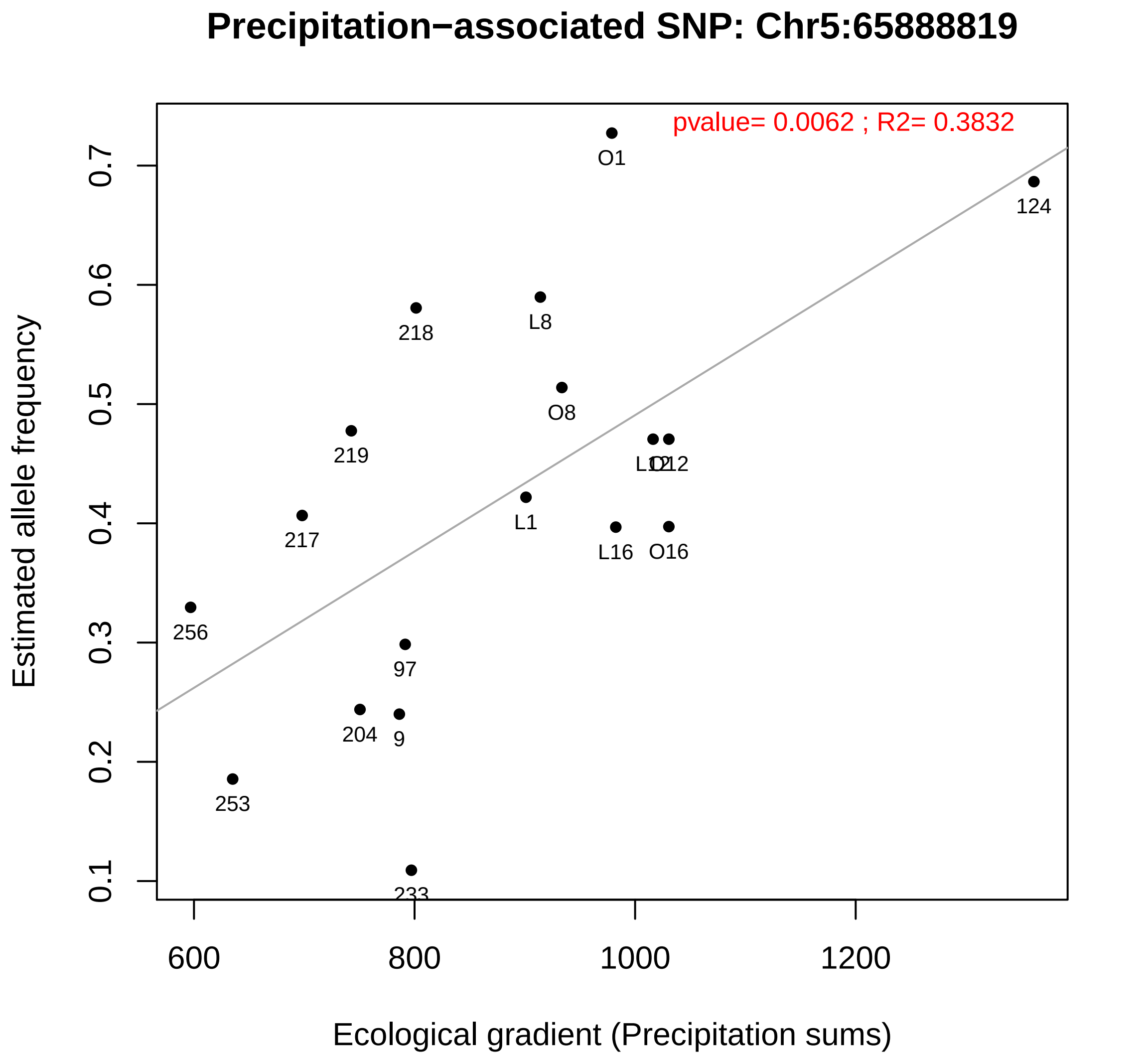
Precipitation−associated SNP: Chr5:65888819
pvalue= 0.0062 ; R2= 0.3832
Estimated allele frequency
Ecological gradient (Precipitation sums)
0.1
0.2
0.3
0.4
0.5
0.6
0.7
600
800
1000
1200
O1
124
218
L8
O8
219
L12
O12
L1
217
L16
O16
256
97
204
9
253
233

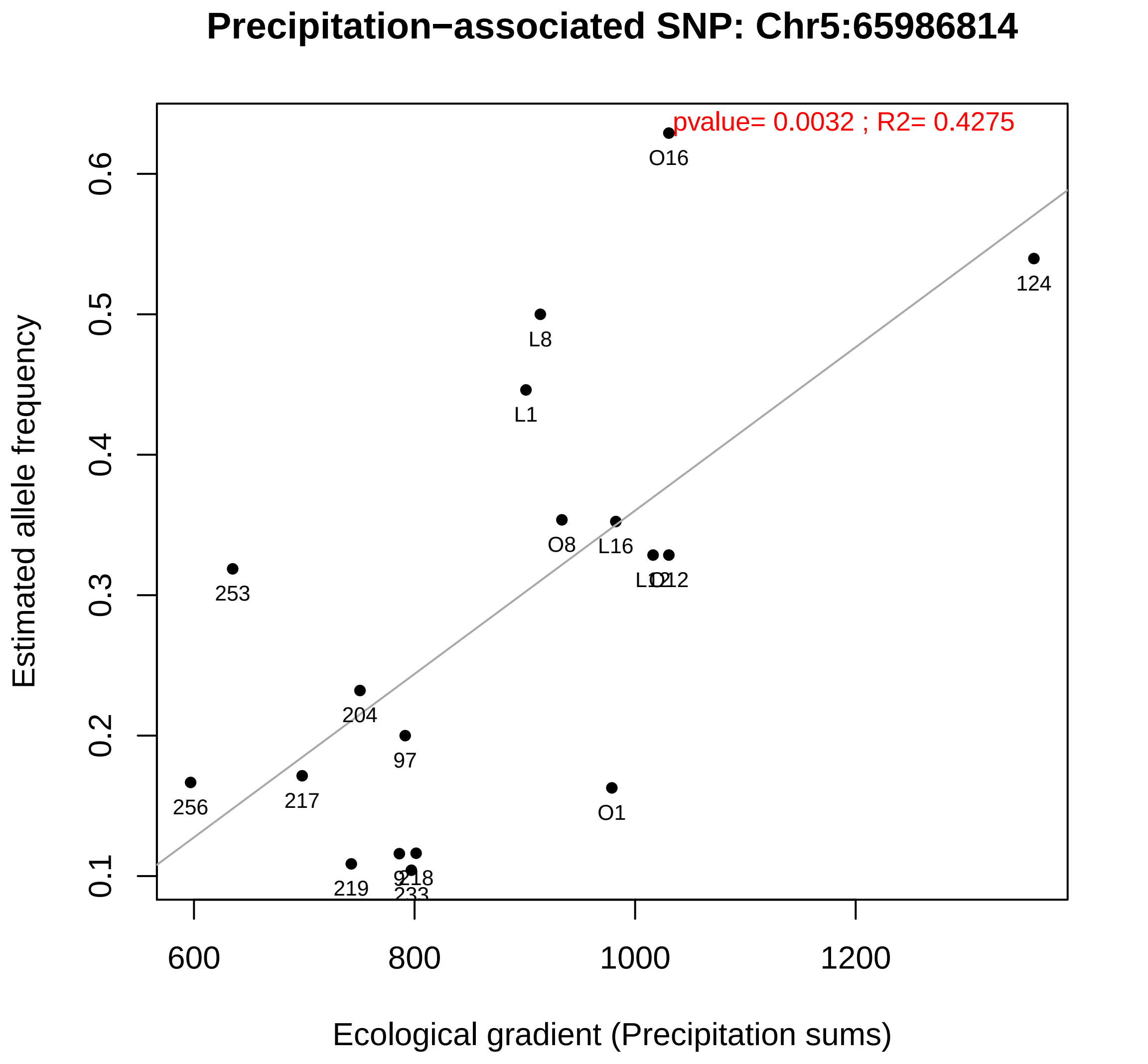

Precipitation−associated SNP: Chr5:65986814
pvalue= 0.0032 ; R2= 0.4275
Estimated allele frequency
Ecological gradient (Precipitation sums)
0.1
0.2
0.3
0.4
0.5
0.6
600
800
1000
1200
O16
124
L8
L1
O8
L16
O12
253
204
97
256
217
O1
219
218
233

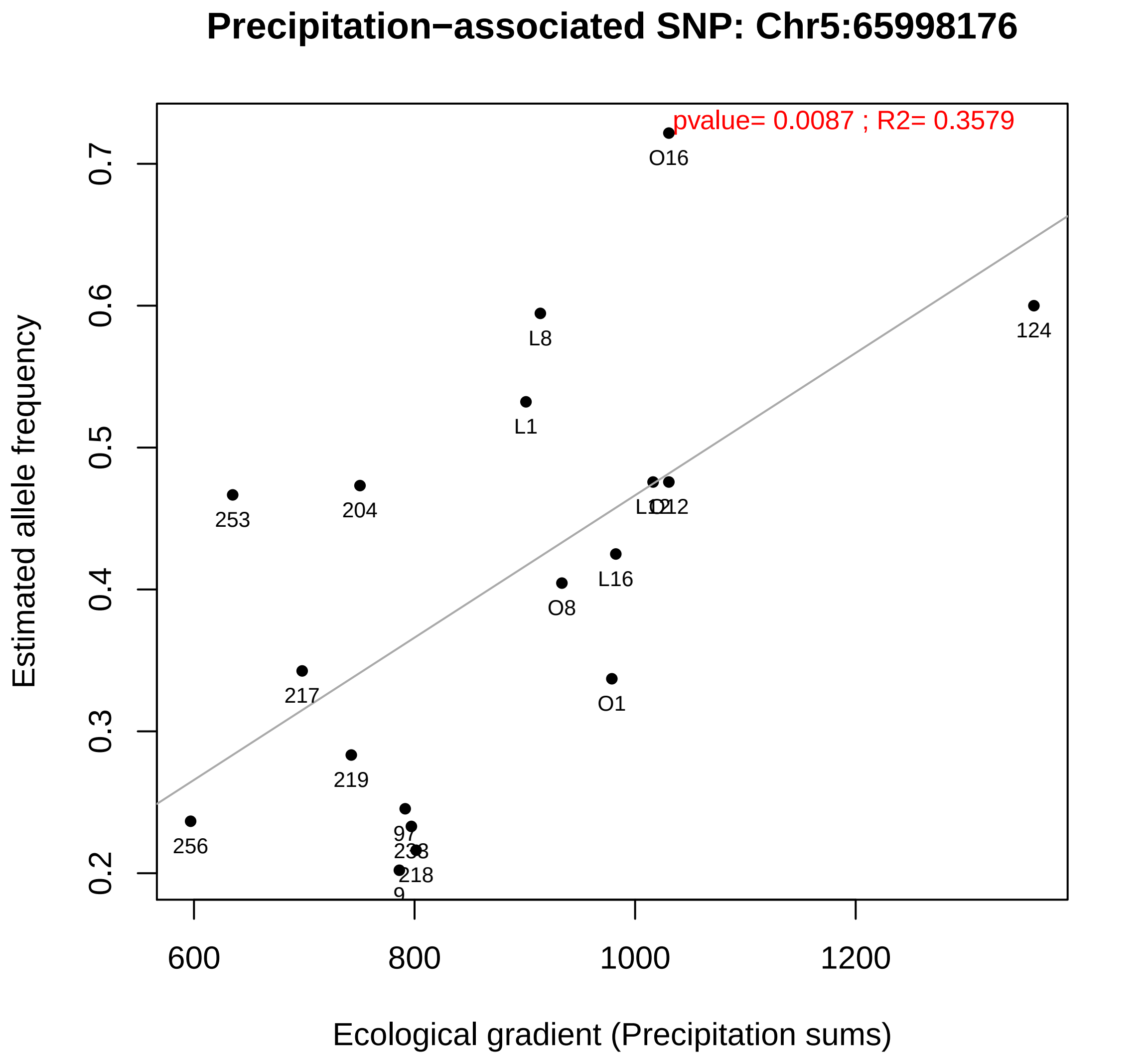
Precipitation−associated SNP: Chr5:65998176
pvalue= 0.0087 ; R2= 0.3579
Estimated allele frequency
0.2
0.3
0.4
0.5
0.6
0.7
600
800
1000
1200
Ecological gradient (Precipitation sums)
O16
L8
124
L1
253
204
L12
O12
L16
O8
217
O1
219
256
97
238
218
9

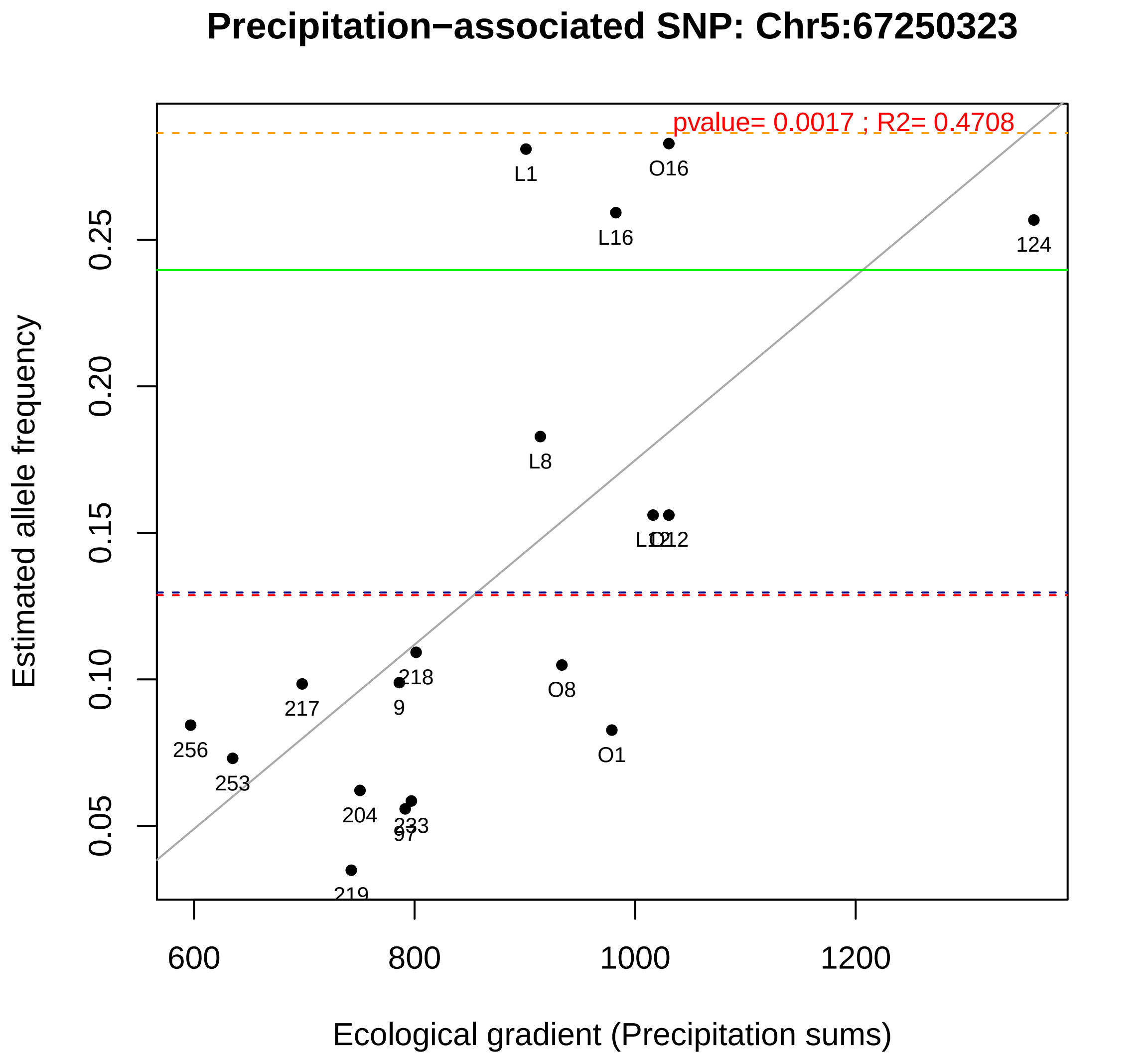

Precipitation−associated SNP: Chr5:67250323
pvalue= 0.0017 ; R2= 0.4708
Estimated allele frequency
0.05
0.10
0.15
0.20
0.25
600
800
1000
1200
Ecological gradient (Precipitation sums)
L1
O16
L16
124
L8
L12
O12
218
9
217
O8
256
O1
253
204
233
97
219

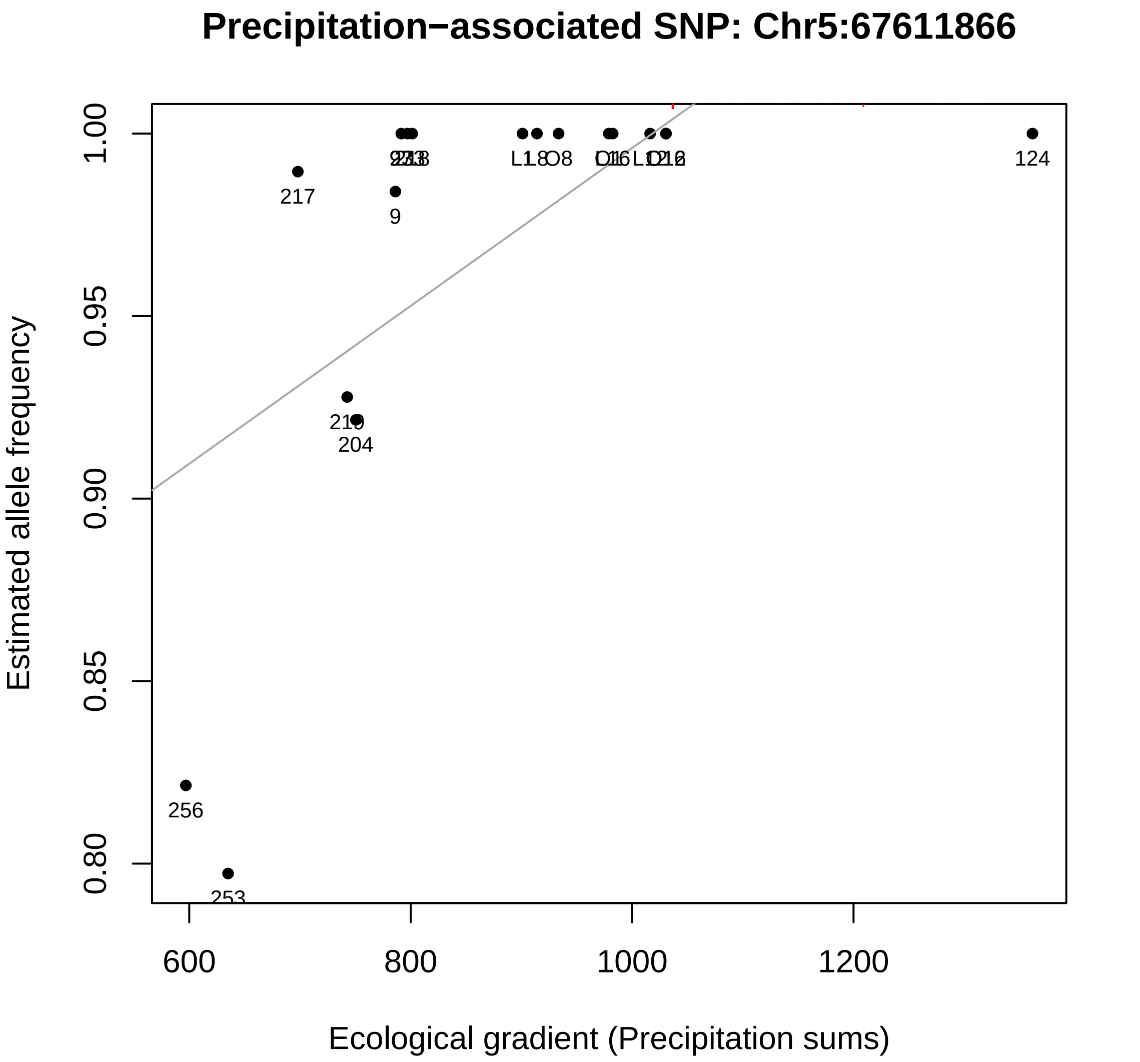

Precipitation−associated SNP: Chr5:67611866
Estimated allele frequency
Ecological gradient (Precipitation sums)
0.80
0.85
0.90
0.95
1.00
600
800
1000
1200
217
9
210
204
256
253
124

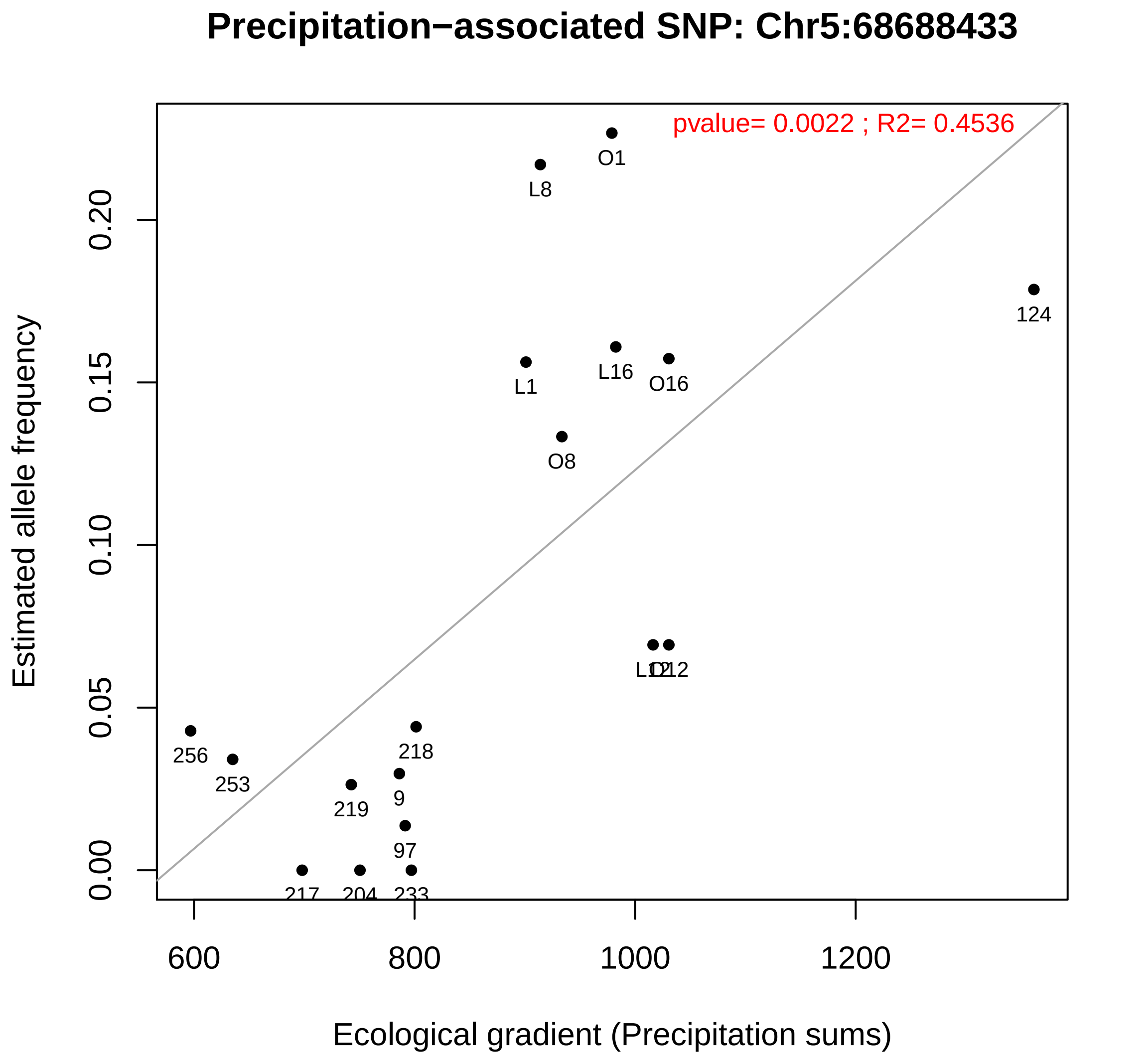

Precipitation−associated SNP: Chr5:68688433
pvalue= 0.0022 ; R2= 0.4536
Estimated allele frequency
Ecological gradient (Precipitation sums)
0.00
0.05
0.10
0.15
0.20
600
800
1000
1200
O1
L8
124
L16
L1
O16
O8
256
253
218
219
9
97
217
204
233

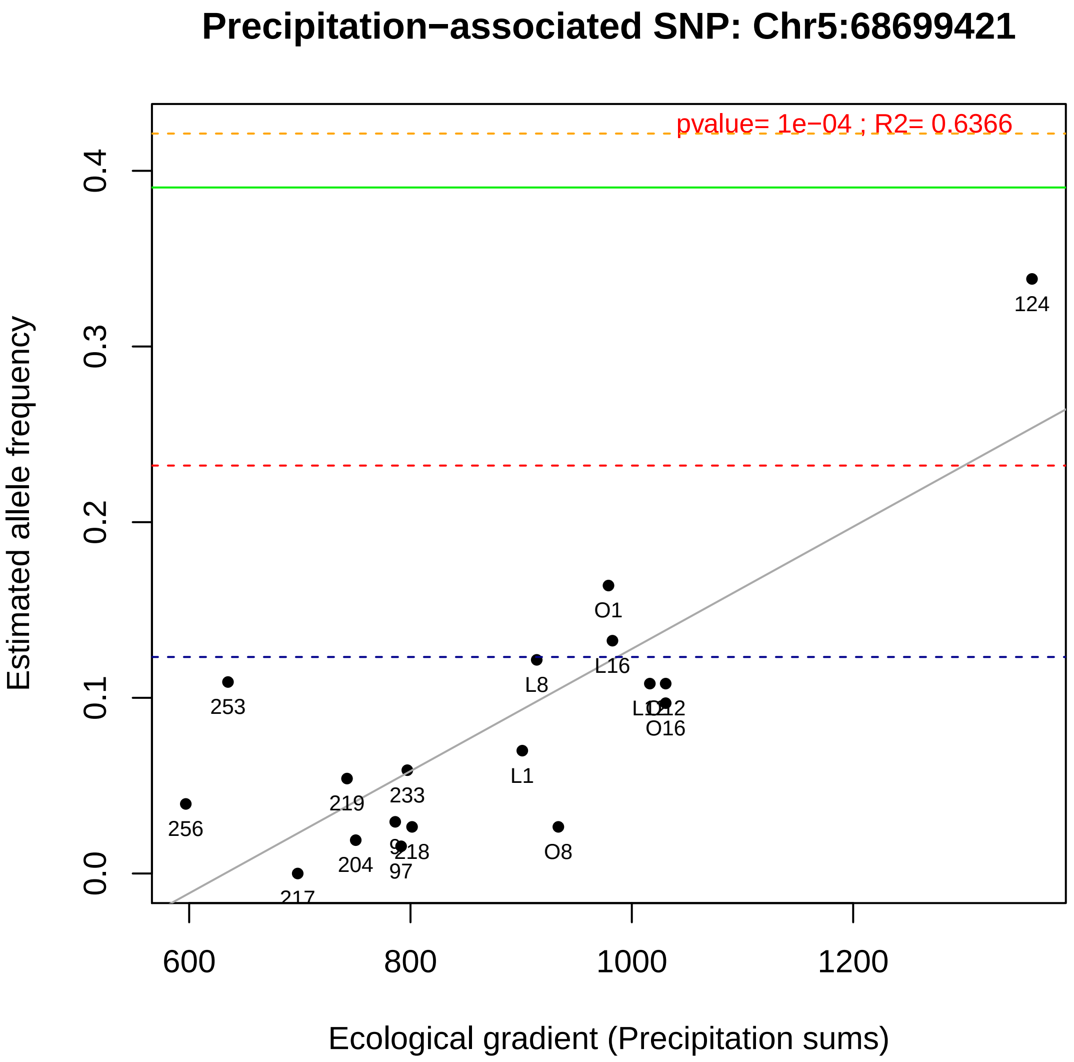
Precipitation−associated SNP: Chr5:68699421
pvalue= 1e−04 ; R2= 0.6366
Estimated allele frequency
0.0
0.1
0.2
0.3
0.4
600
800
1000
1200
Ecological gradient (Precipitation sums)
124
O1
L16
L8
253
L12
O12
O16
L1
233
219
256
9
218
204
97
O8
217

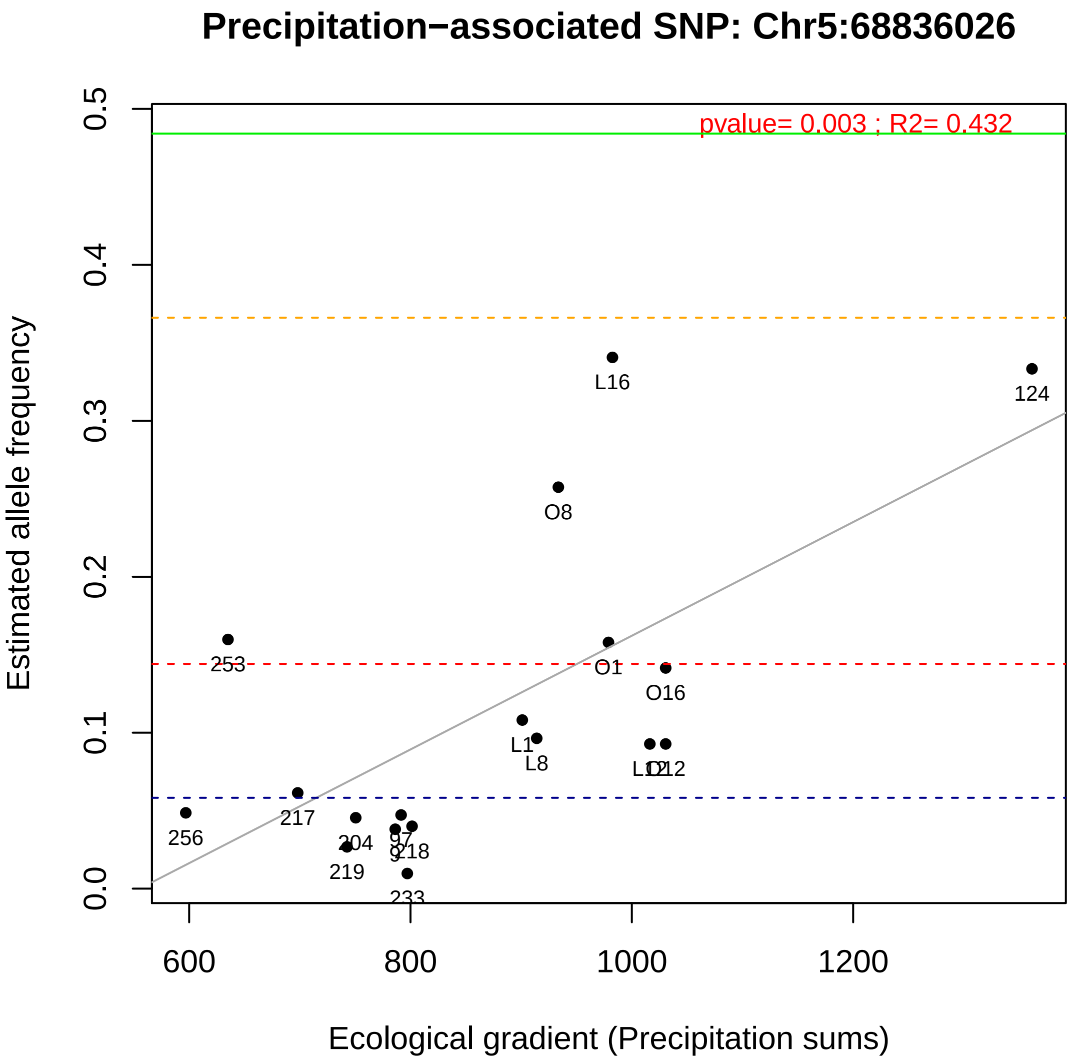

Precipitation−associated SNP: Chr5:68836026
pvalue= 0.003 ; R2= 0.432
Estimated allele frequency
Ecological gradient (Precipitation sums)
0.0
0.1
0.2
0.3
0.4
0.5
600
800
1000
1200
L16
124
O8
253
O1
O16
L1
L8
L12
O12
217
256
204
219
97
9
218
233

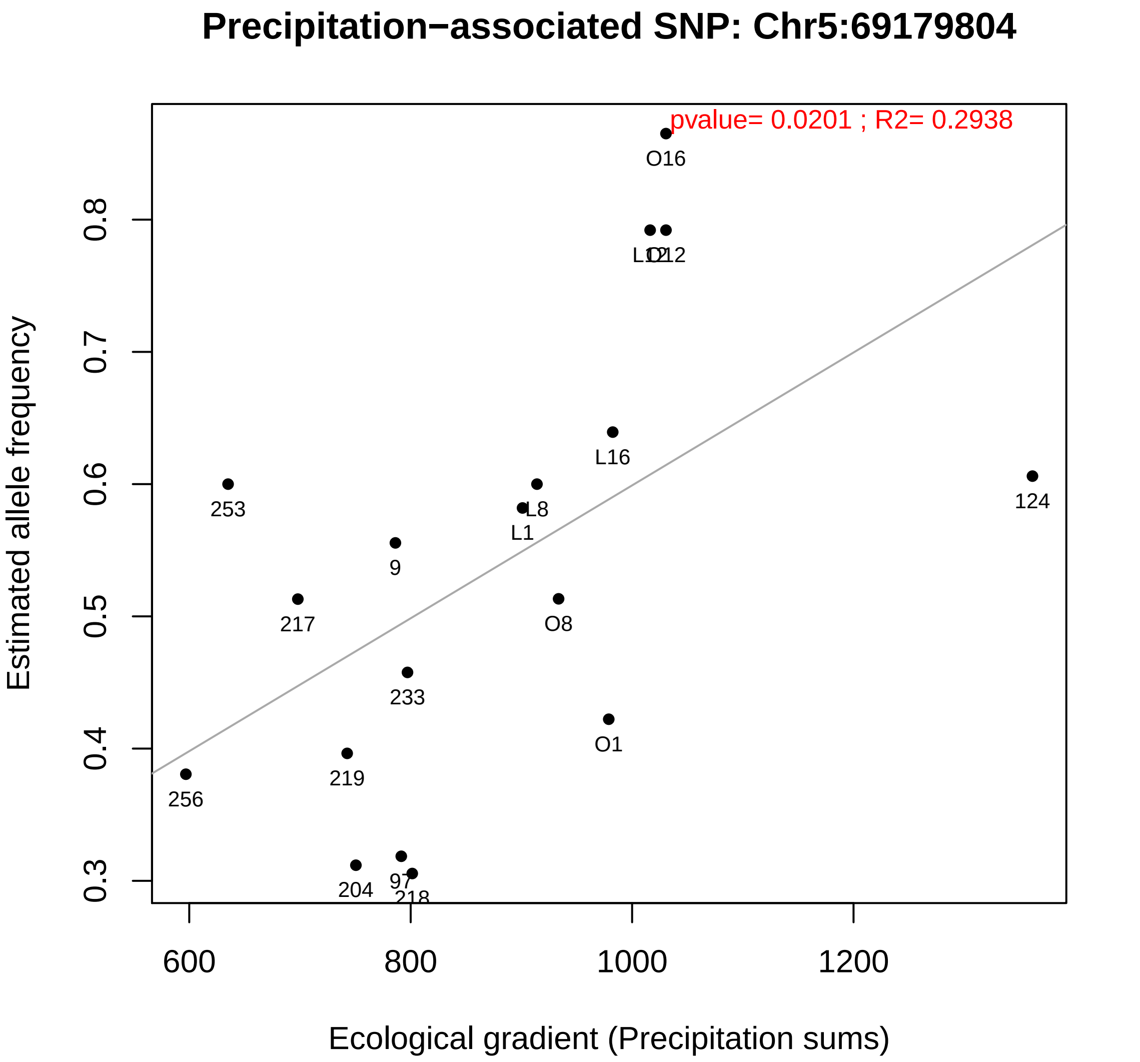
Precipitation−associated SNP: Chr5:69179804
pvalue= 0.0201 ; R2= 0.2938
O16
L12
O12
L16
253
L8
L1
124
9
217
O8
233
O1
219
256
204
97
218
0.3
0.4
0.5
0.6
0.7
0.8
Estimated allele frequency
600
800
1000
1200
Ecological gradient (Precipitation sums)

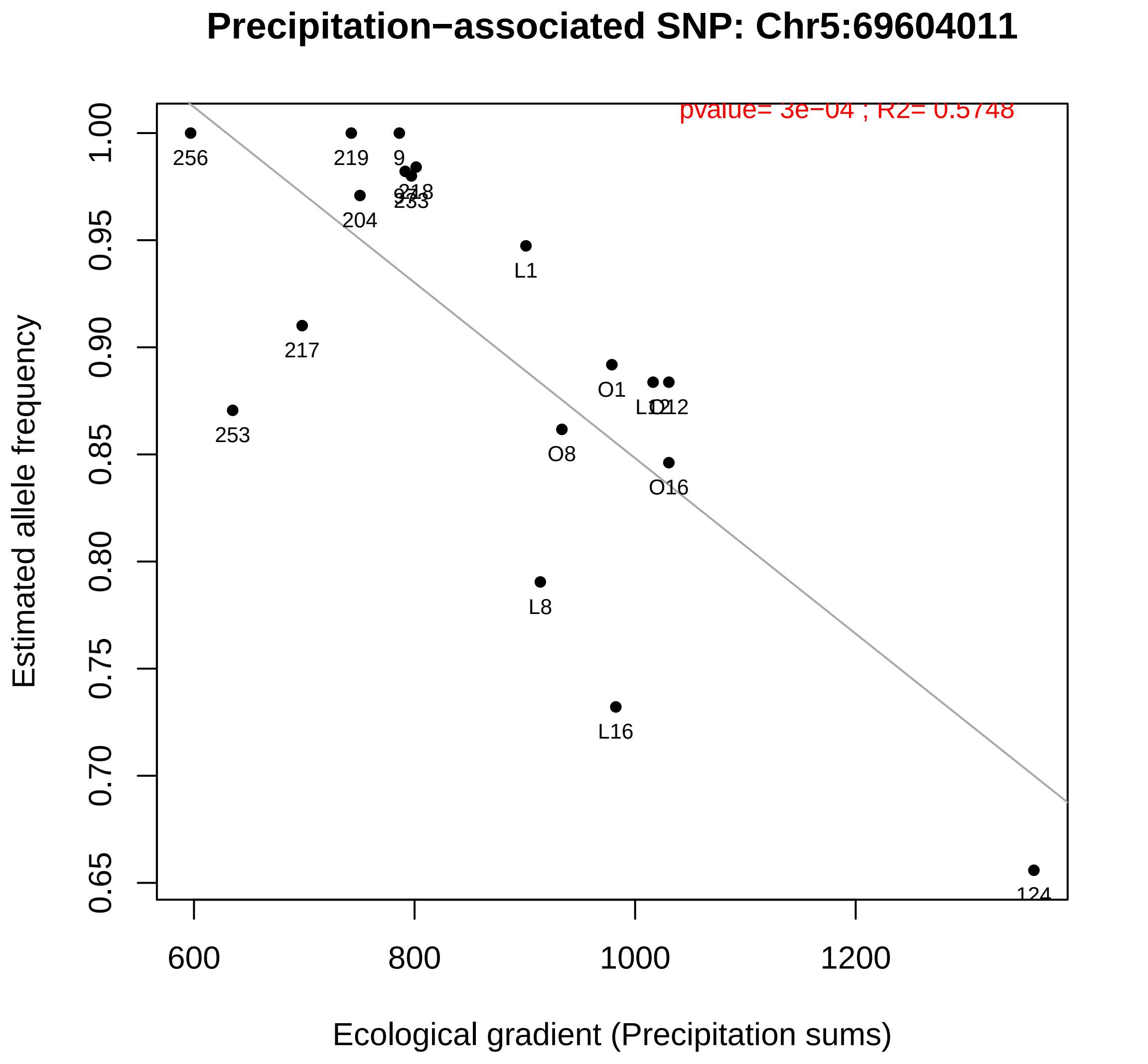
Precipitation−associated SNP: Chr5:69604011
pvalue= 3e−04 ; R2= 0.5748
Estimated allele frequency
Ecological gradient (Precipitation sums)
256
219
9
218
233
204
L1
217
253
O1
L12
O12
O8
O16
L8
L16
124
0.65
0.70
0.75
0.80
0.85
0.90
0.95
1.00
600
800
1000
1200

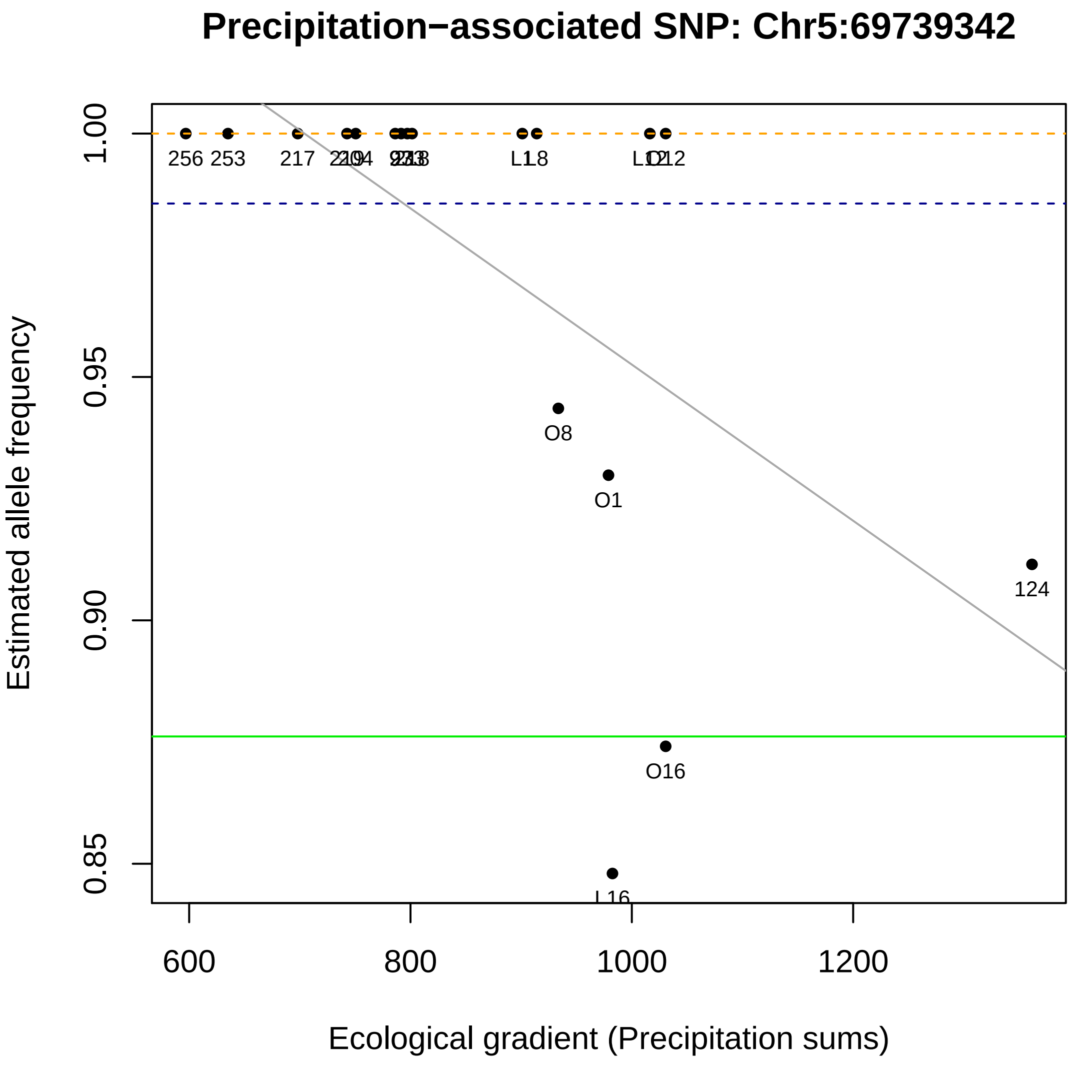
Precipitation−associated SNP: Chr5:69739342
Estimated allele frequency
Ecological gradient (Precipitation sums)
1.00
0.95
0.90
0.85
600
800
1000
1200
256
253
217
O8
O1
124
O16
L16

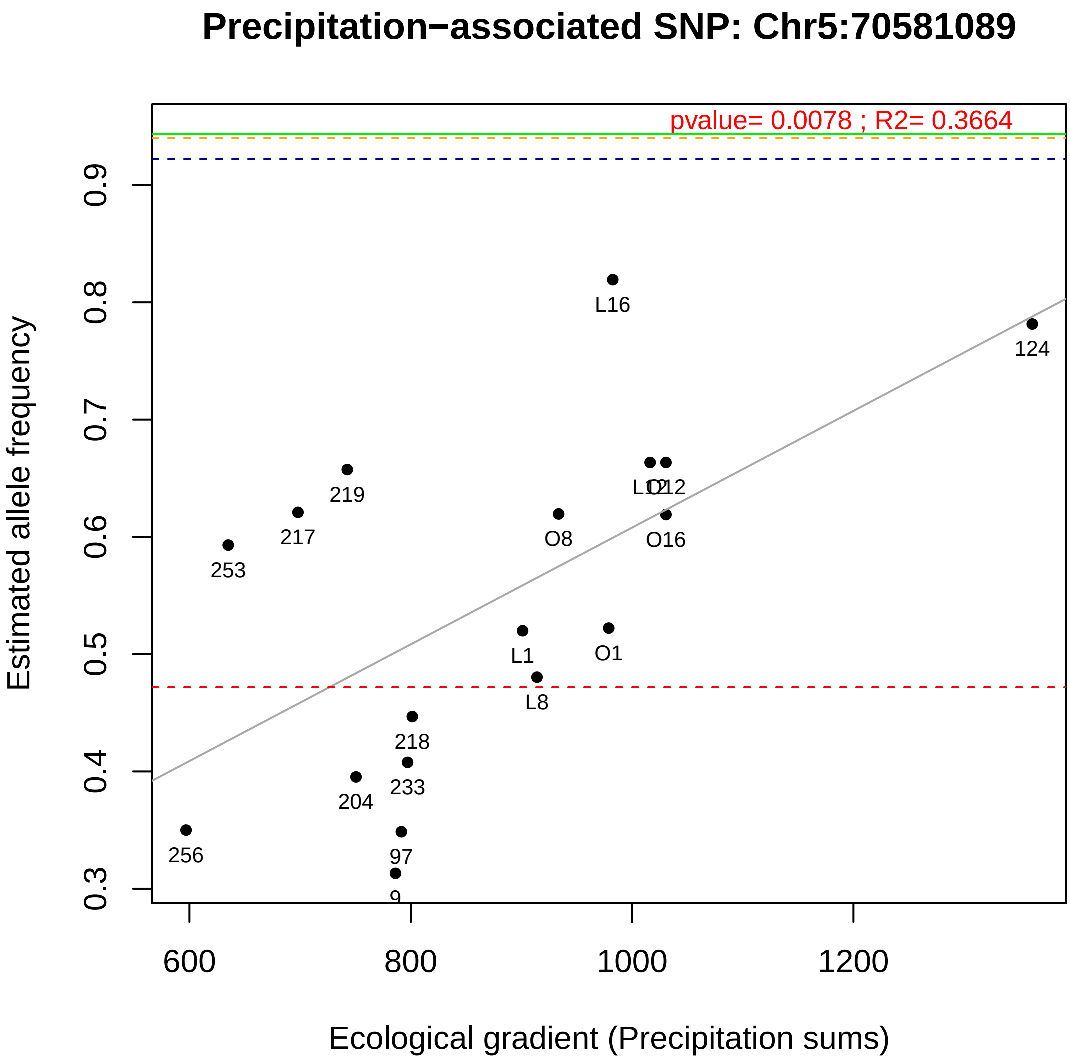

Precipitation−associated SNP: Chr5:70581089
pvalue= 0.0078 ; R2= 0.3664
Estimated allele frequency
0.3
0.4
0.5
0.6
0.7
0.8
0.9
600
800
1000
1200
Ecological gradient (Precipitation sums)
L16
124
219
217
253
O8
O16
L1
O1
L8
218
233
204
256
97
9

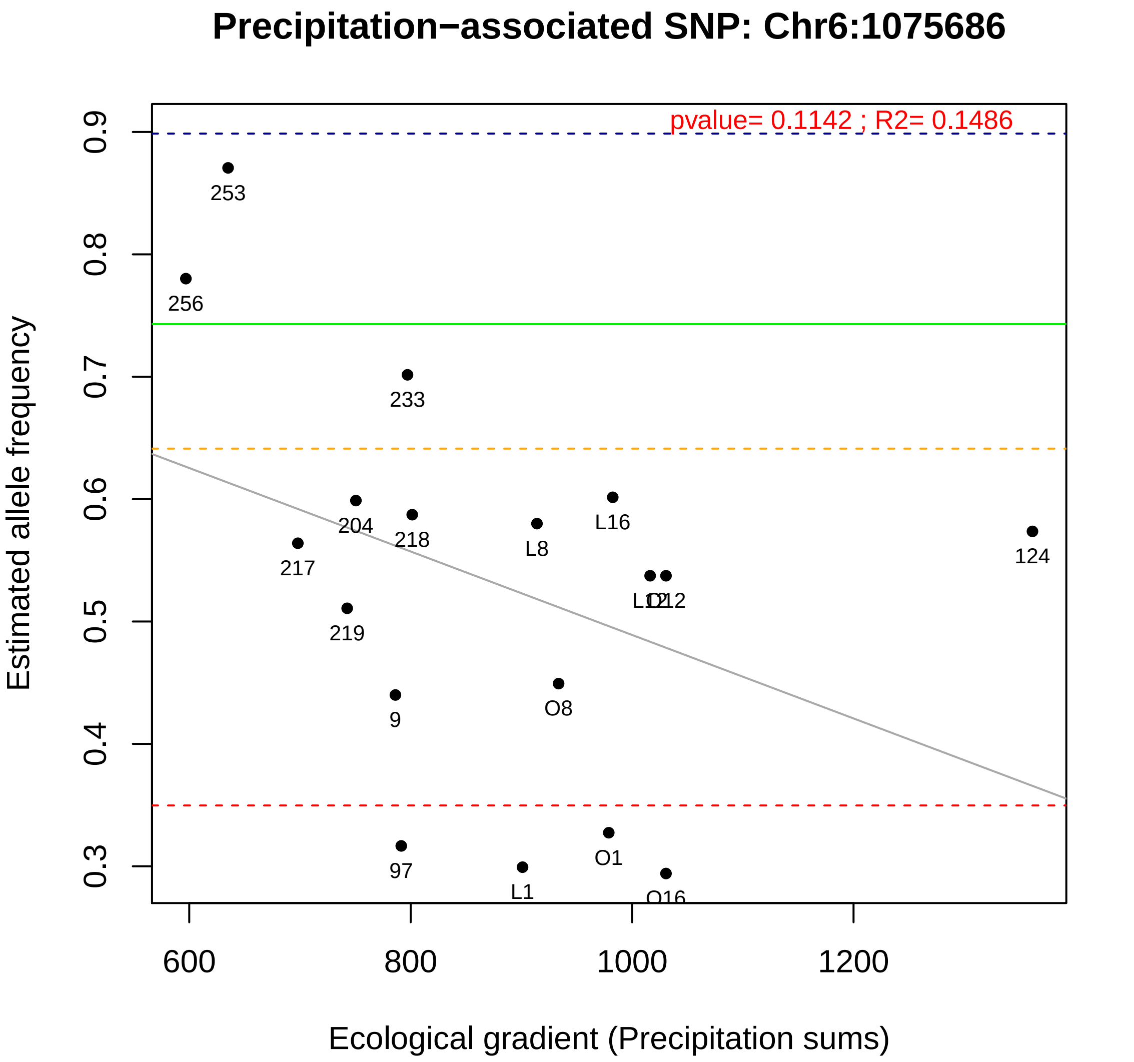

Precipitation−associated SNP: Chr6:1075686
pvalue= 0.1142 ; R2= 0.1486
Estimated allele frequency
Ecological gradient (Precipitation sums)
0.3
0.4
0.5
0.6
0.7
0.8
0.9
600
800
1000
1200
253
256
233
204
218
217
219
L8
L16
124
9
O8
97
L1
O1

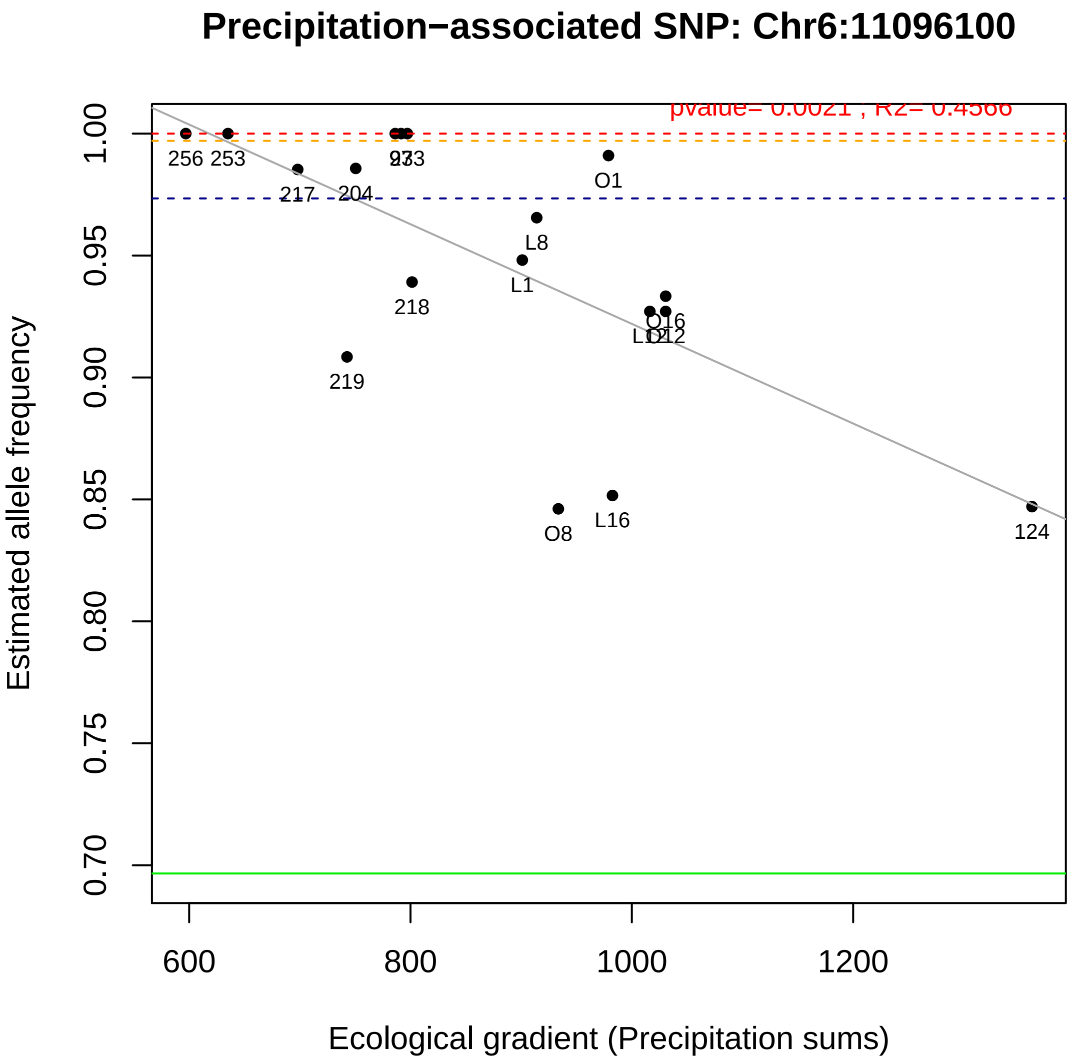

Precipitation−associated SNP: Chr6:11096100
pvalue= 0.0021 , R2= 0.4566
Estimated allele frequency
Ecological gradient (Precipitation sums)
1.00
0.95
0.90
0.85
0.80
0.75
0.70
600
800
1000
1200
256
253
217
204
219
218
L8
L1
O1
O16
O12
L16
O8
124

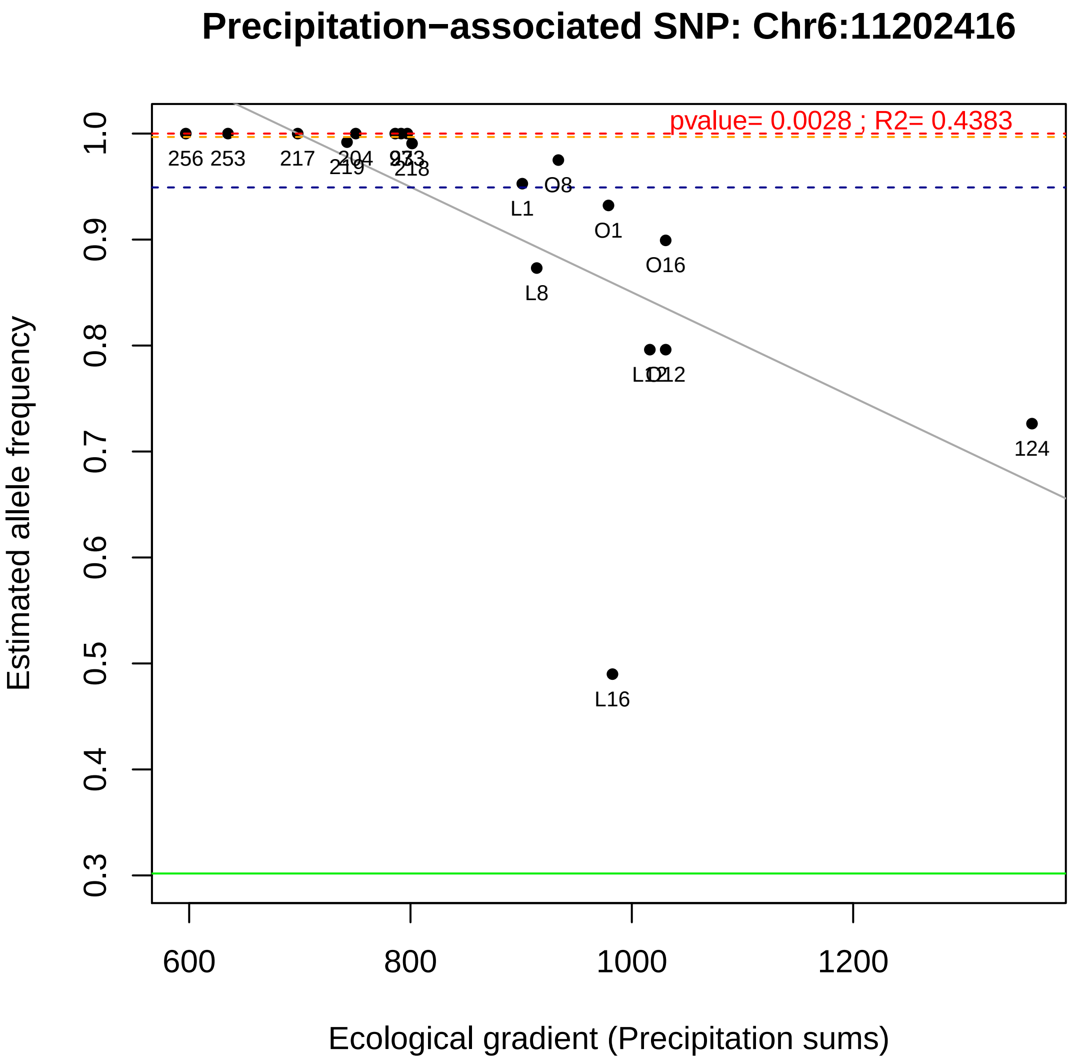
Precipitation−associated SNP: Chr6:11202416
pvalue= 0.0028 ; R2= 0.4383
Estimated allele frequency
1.0
0.9
0.8
0.7
0.6
0.5
0.4
0.3
600
800
1000
1200
Ecological gradient (Precipitation sums)
256
253
217
204
219
218
O8
L1
O1
O16
L8
O12
124
L16

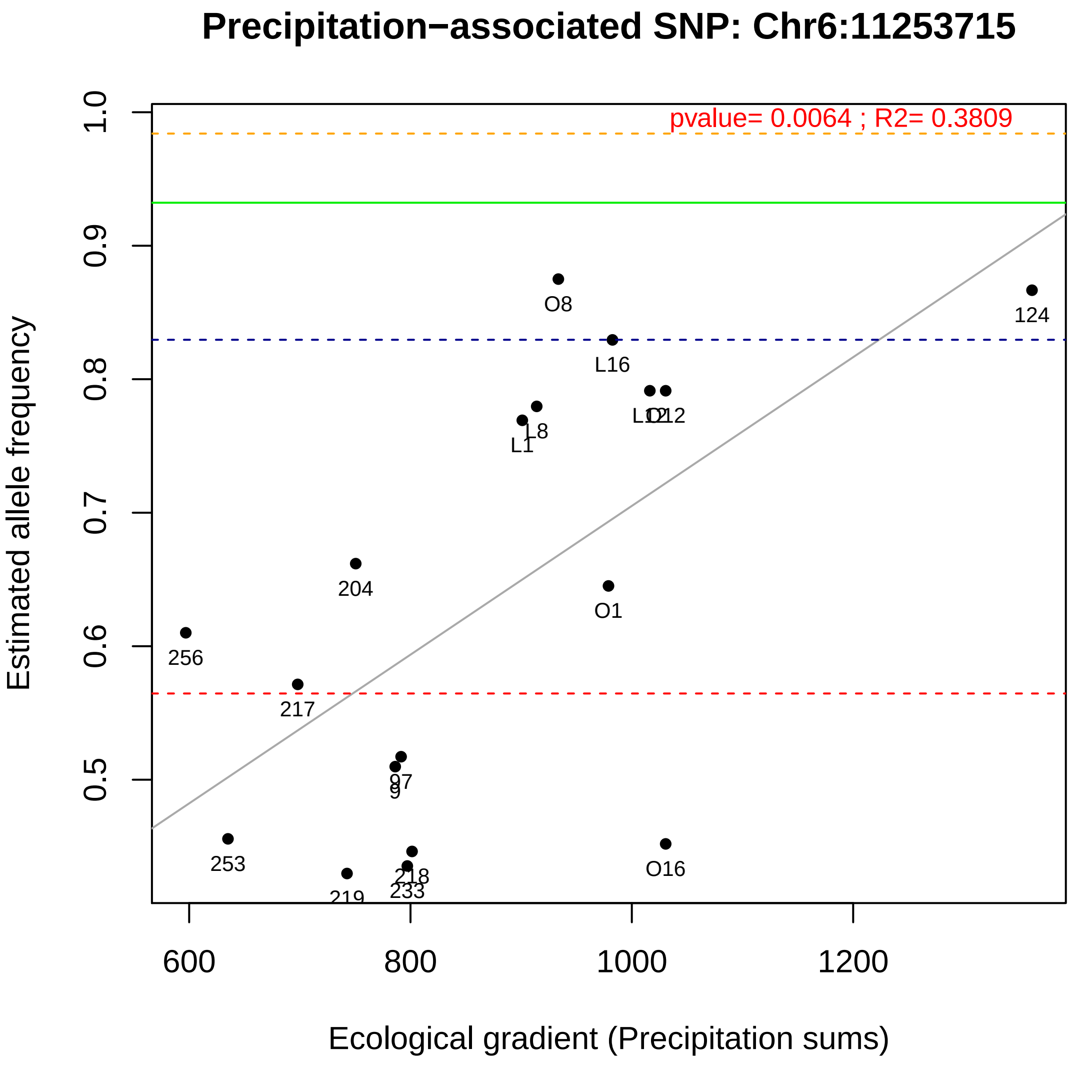
Precipitation−associated SNP: Chr6:11253715
pvalue= 0.0064 ; R2= 0.3809
Estimated allele frequency
Ecological gradient (Precipitation sums)
1.0
0.9
0.8
0.7
0.6
0.5
600
800
1000
1200
O8
124
L16
L10
O12
L8
L1
204
O1
256
217
97
9
253
219
218
233
O16

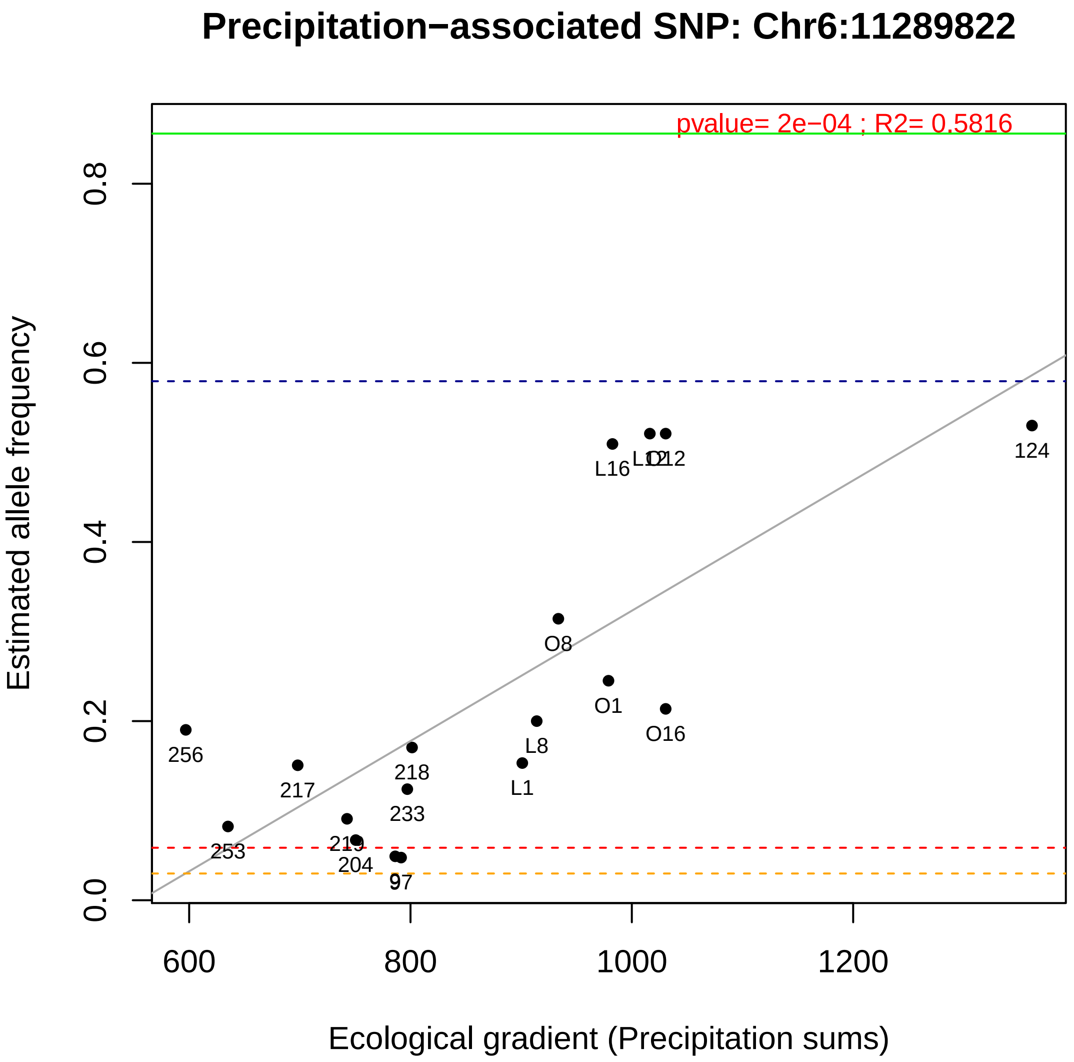

Precipitation−associated SNP: Chr6:11289822
pvalue= 2e−04 ; R2= 0.5816
Estimated allele frequency
0.0
0.2
0.4
0.6
0.8
600
800
1000
1200
Ecological gradient (Precipitation sums)
256
253
217
204
233
218
L1
L8
O8
O1
O16
L16
O12
124

#### Precipitation−associated SNP: Chr6:11309566

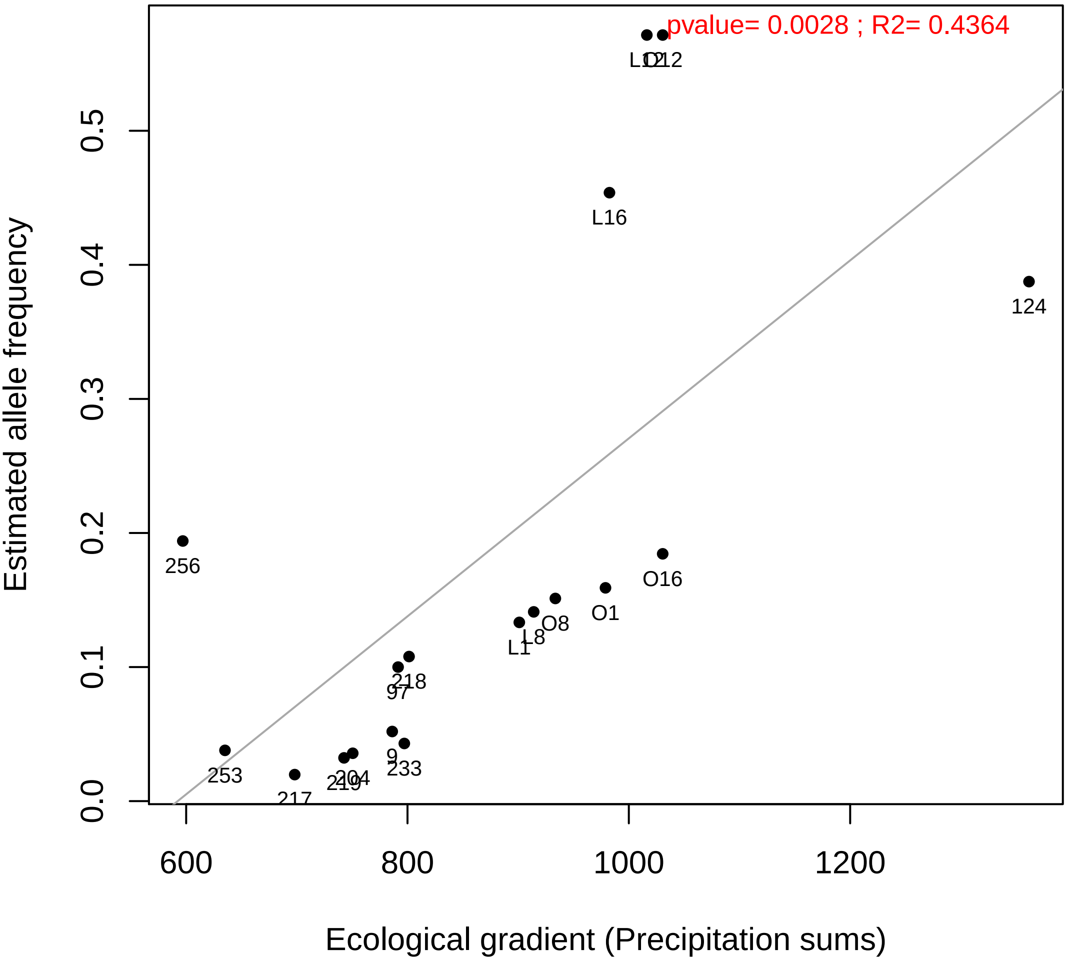

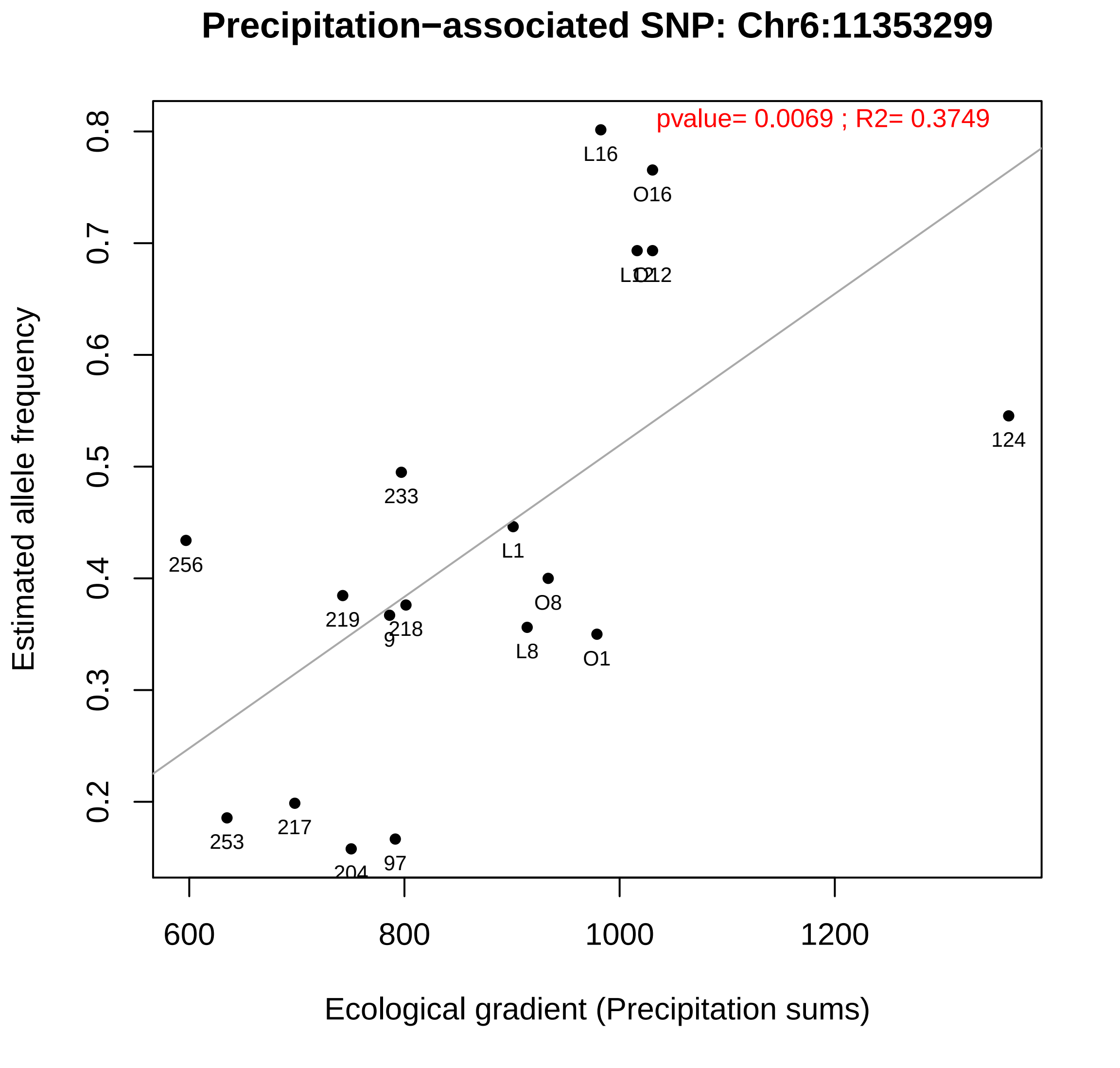

Precipitation−associated SNP: Chr6:11353299
pvalue= 0.0069 ; R2= 0.3749
L16
O16
L12
O12
124
233
256
L1
O8
219
218
9
L8
O1
253
217
204
97
0.2
0.3
0.4
0.5
0.6
0.7
0.8
600
800
1000
1200
Estimated allele frequency
Ecological gradient (Precipitation sums)

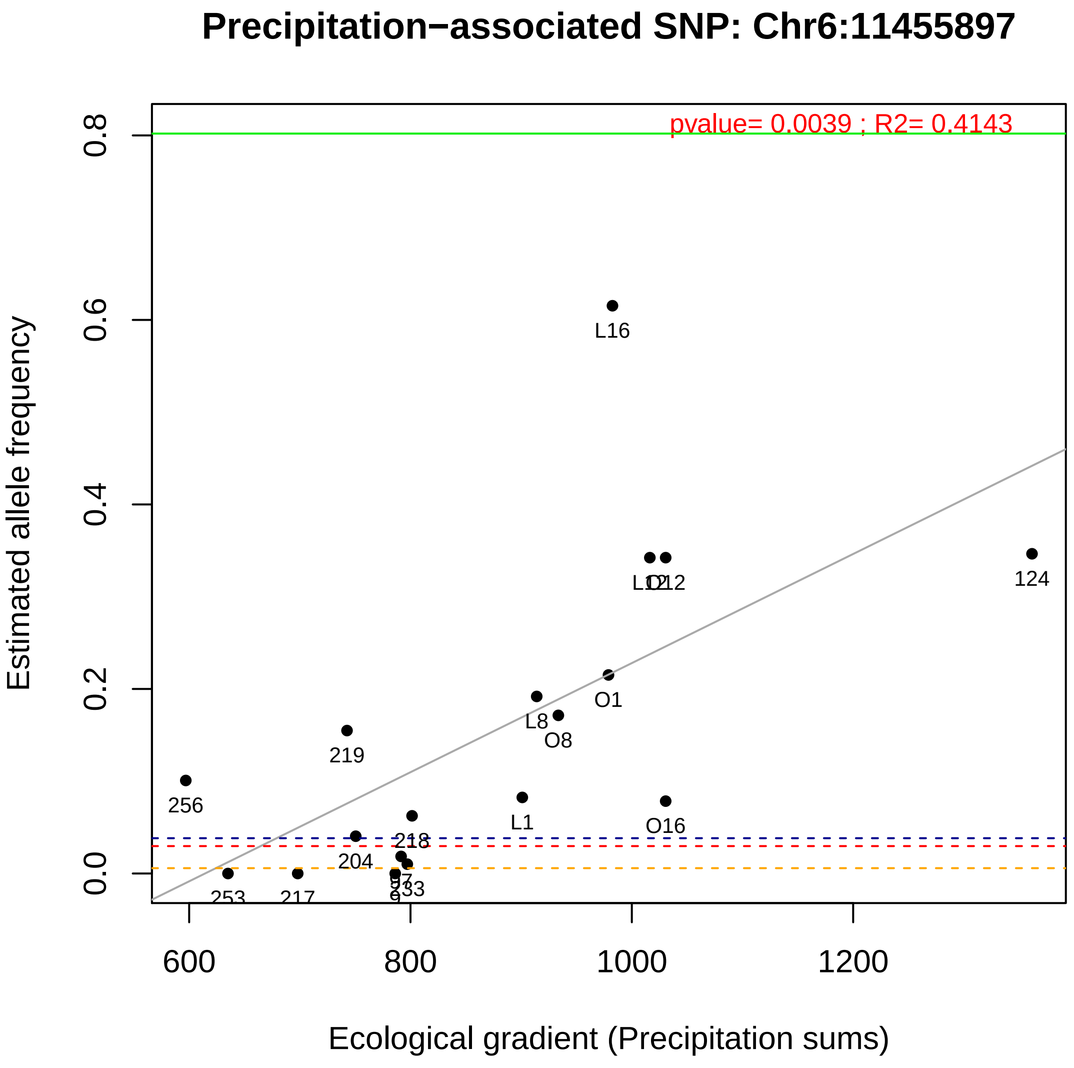

Precipitation−associated SNP: Chr6:11455897
pvalue= 0.0039 ; R2= 0.4143
Estimated allele frequency
Ecological gradient (Precipitation sums)
0.0
0.2
0.4
0.6
0.8
600
800
1000
1200
L16
L12
O12
124
O1
L8
O8
219
256
L1
O16
218
204
253
217
233

**Precipitation−associated SNP: Chr6:11459934**

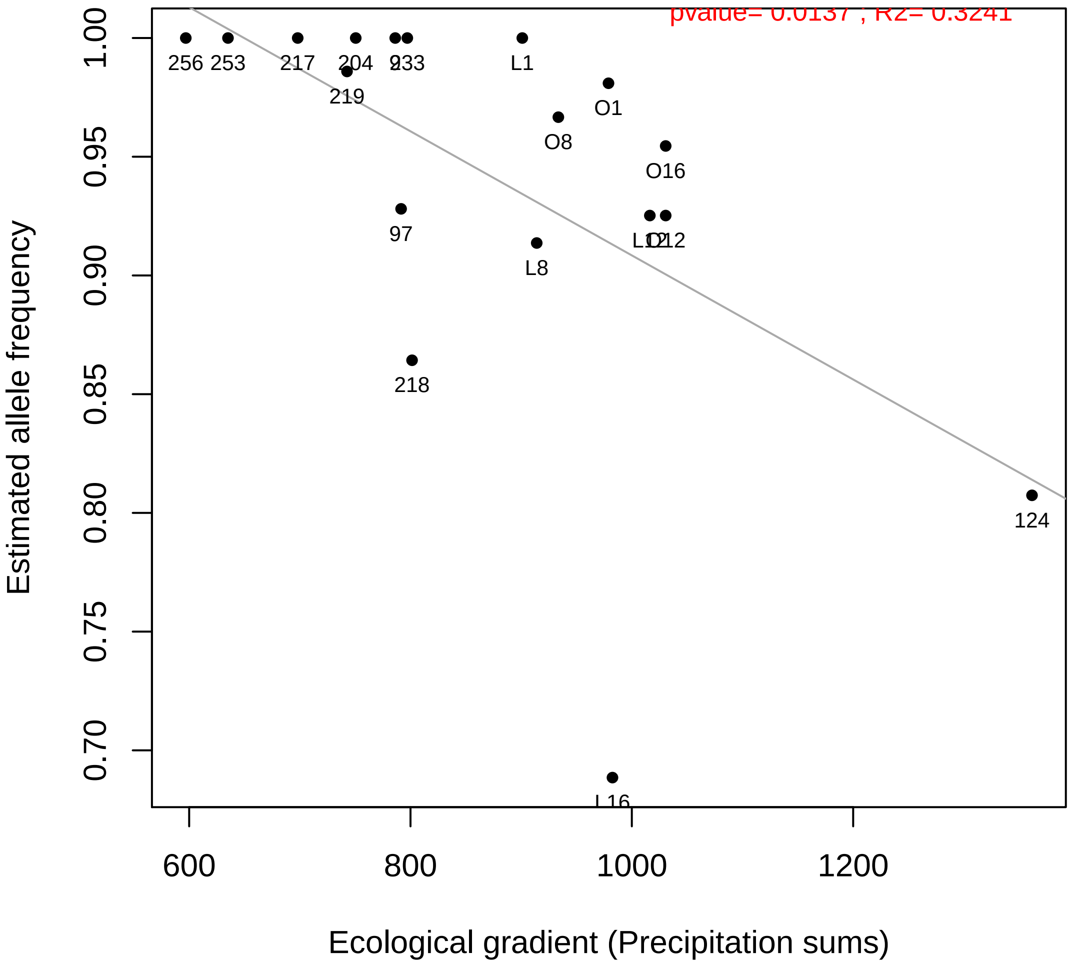

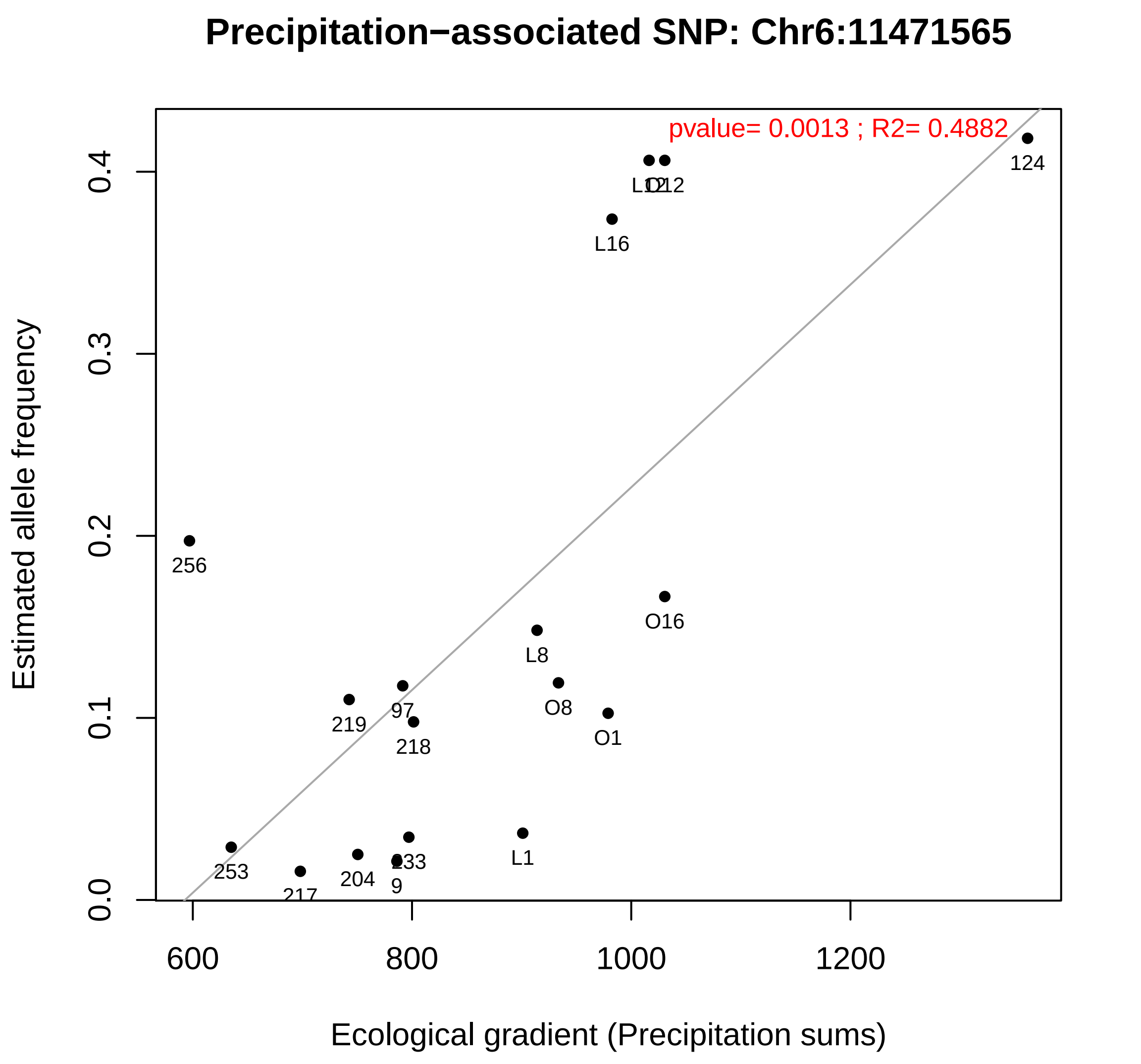
Precipitation−associated SNP: Chr6:11471565
pvalue= 0.0013 ; R2= 0.4882
Estimated allele frequency
Ecological gradient (Precipitation sums)
0.0
0.1
0.2
0.3
0.4
600
800
1000
1200
124
L12
O12
L16
256
O16
L8
O8
O1
97
219
218
253
217
204
233
9
L1

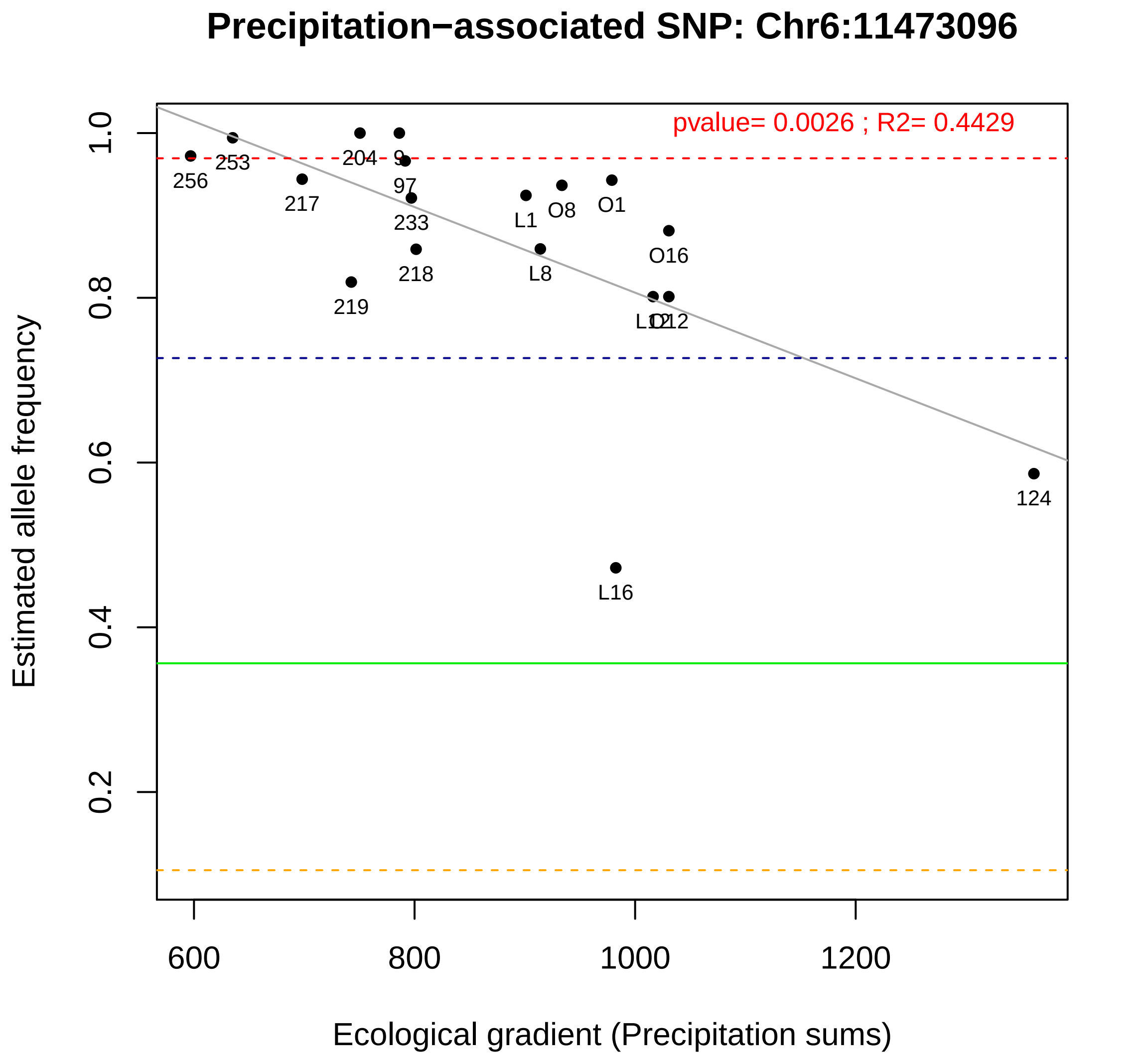
Precipitation−associated SNP: Chr6:11473096
pvalue= 0.0026 ; R2= 0.4429
Estimated allele frequency
Ecological gradient (Precipitation sums)
0.2
0.4
0.6
0.8
1.0
600
800
1000
1200
256
253
217
204
9
97
233
219
218
L1
O8
O1
L8
O16
L12
O12
124
L16

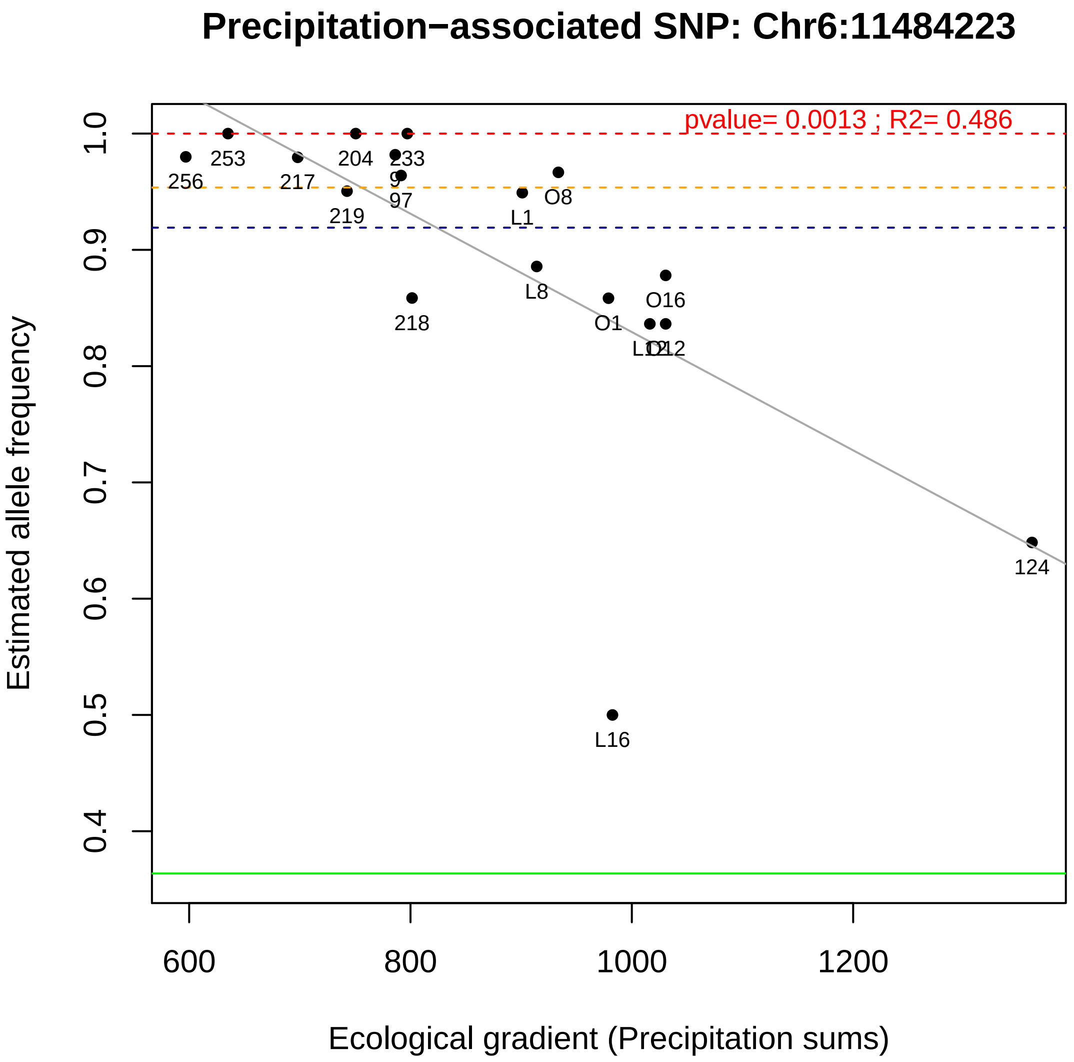
Precipitation−associated SNP: Chr6:11484223
pvalue= 0.0013 ; R2= 0.486
Estimated allele frequency
Ecological gradient (Precipitation sums)
1.0
0.9
0.8
0.7
0.6
0.5
0.4
600
800
1000
1200
256
253
217
204
233
9
97
219
L1
O8
L8
218
O1
O16
L12
O12
L16
124

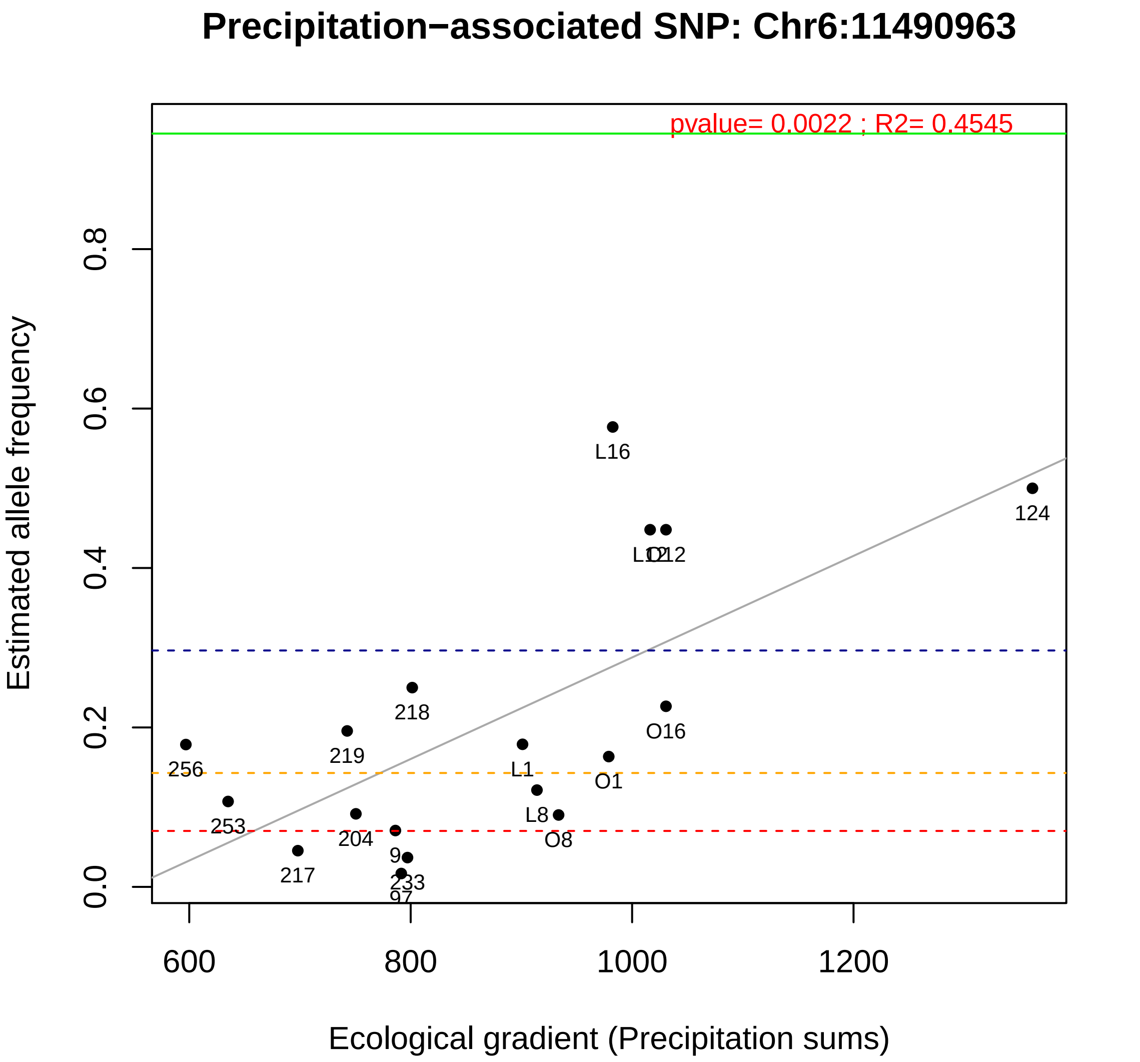

Precipitation−associated SNP: Chr6:11490963
pvalue= 0.0022 ; R2= 0.4545
Estimated allele frequency
0.0
0.2
0.4
0.6
0.8
600
800
1000
1200
Ecological gradient (Precipitation sums)
L16
124
L12
O12
218
O16
219
256
L1
O1
253
L8
204
O8
217
9
233
97

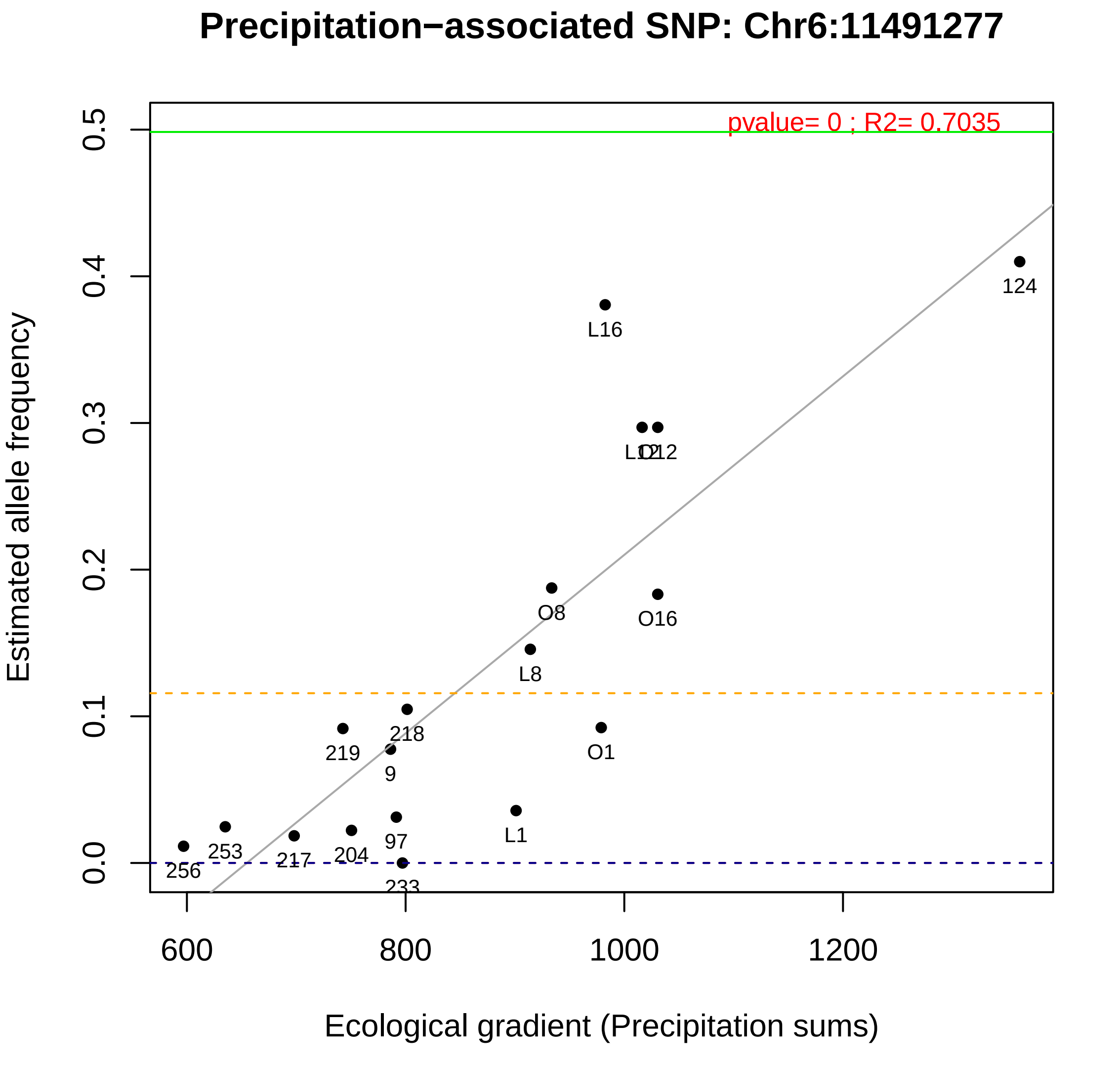

Precipitation−associated SNP: Chr6:11491277
pvalue= 0 ; R2= 0.7035
Estimated allele frequency
0.0
0.1
0.2
0.3
0.4
0.5
600
800
1000
1200
Ecological gradient (Precipitation sums)
124
L16
L12
O12
O8
O16
L8
218
219
9
O1
97
L1
253
217
204
256
233

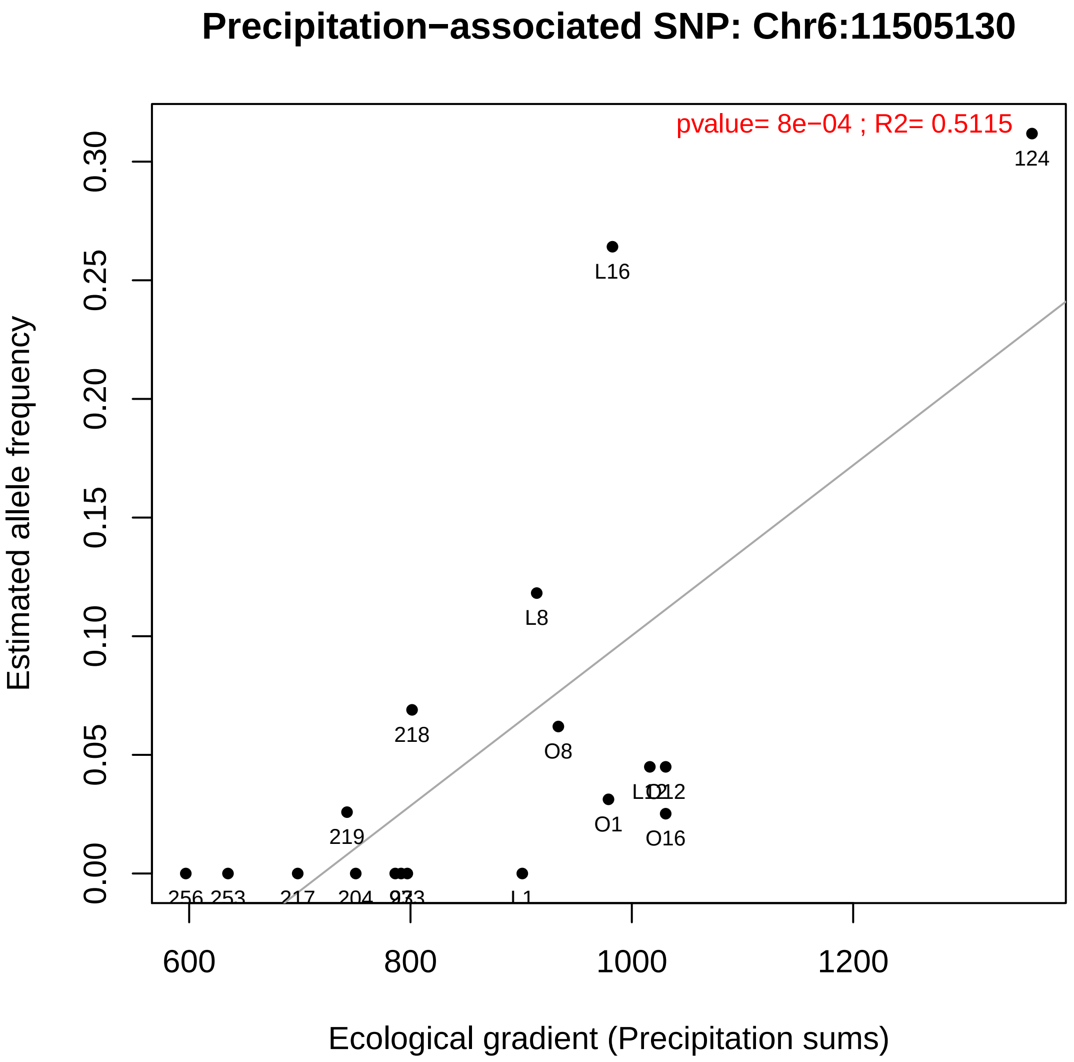
Precipitation−associated SNP: Chr6:11505130
pvalue= 8e−04 ; R2= 0.5115
Estimated allele frequency
0.00
0.05
0.10
0.15
0.20
0.25
0.30
Ecological gradient (Precipitation sums)
600
800
1000
1200
124
L16
L8
218
O8
O1
O16
219
256
253
217
204
L1

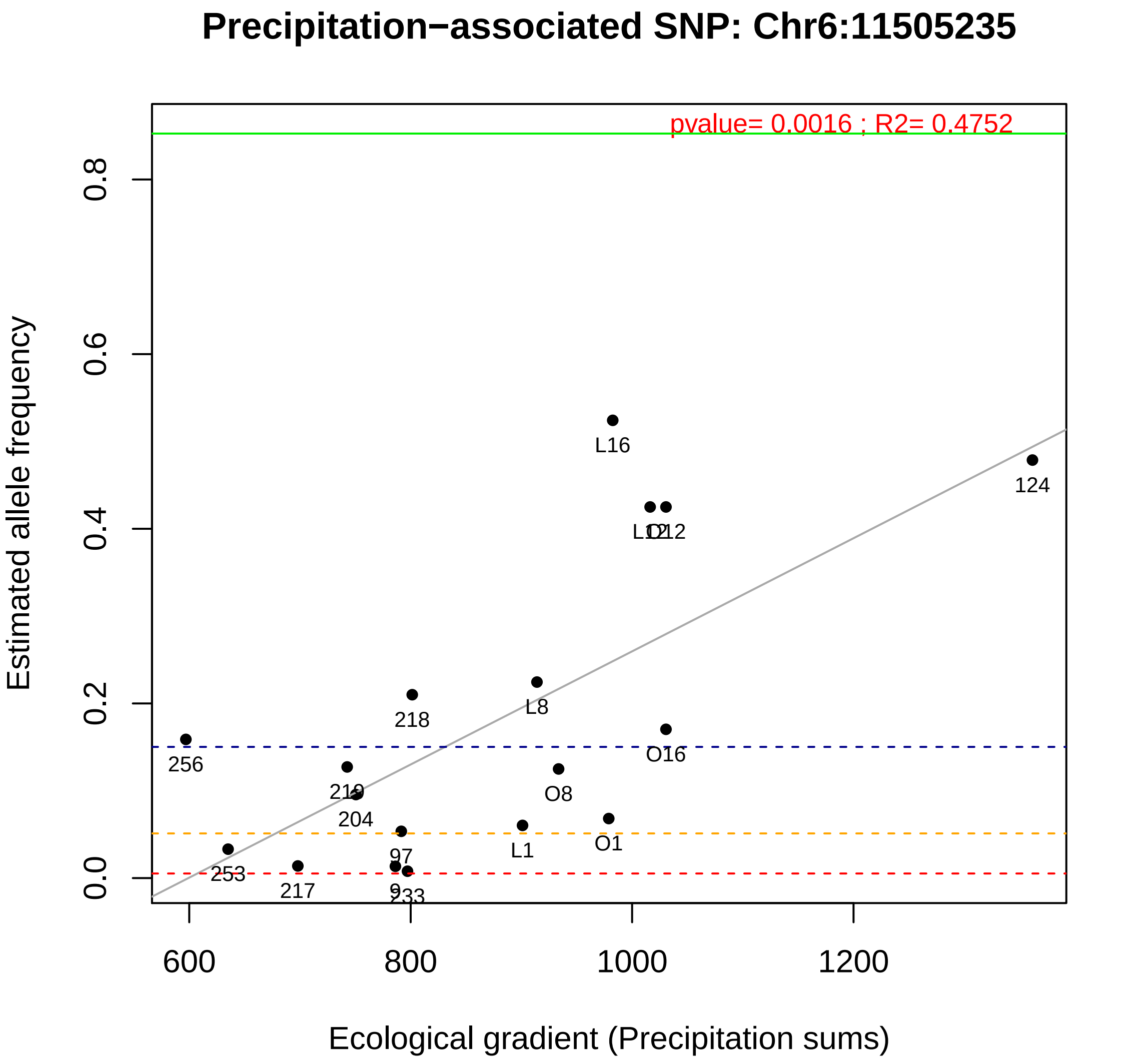
Precipitation−associated SNP: Chr6:11505235
pvalue= 0.0016 ; R2= 0.4752
Estimated allele frequency
0.0
0.2
0.4
0.6
0.8
600
800
1000
1200
Ecological gradient (Precipitation sums)
L16
124
L12
O12
218
L8
256
O16
219
204
O8
97
L1
O1
253
217
9
233

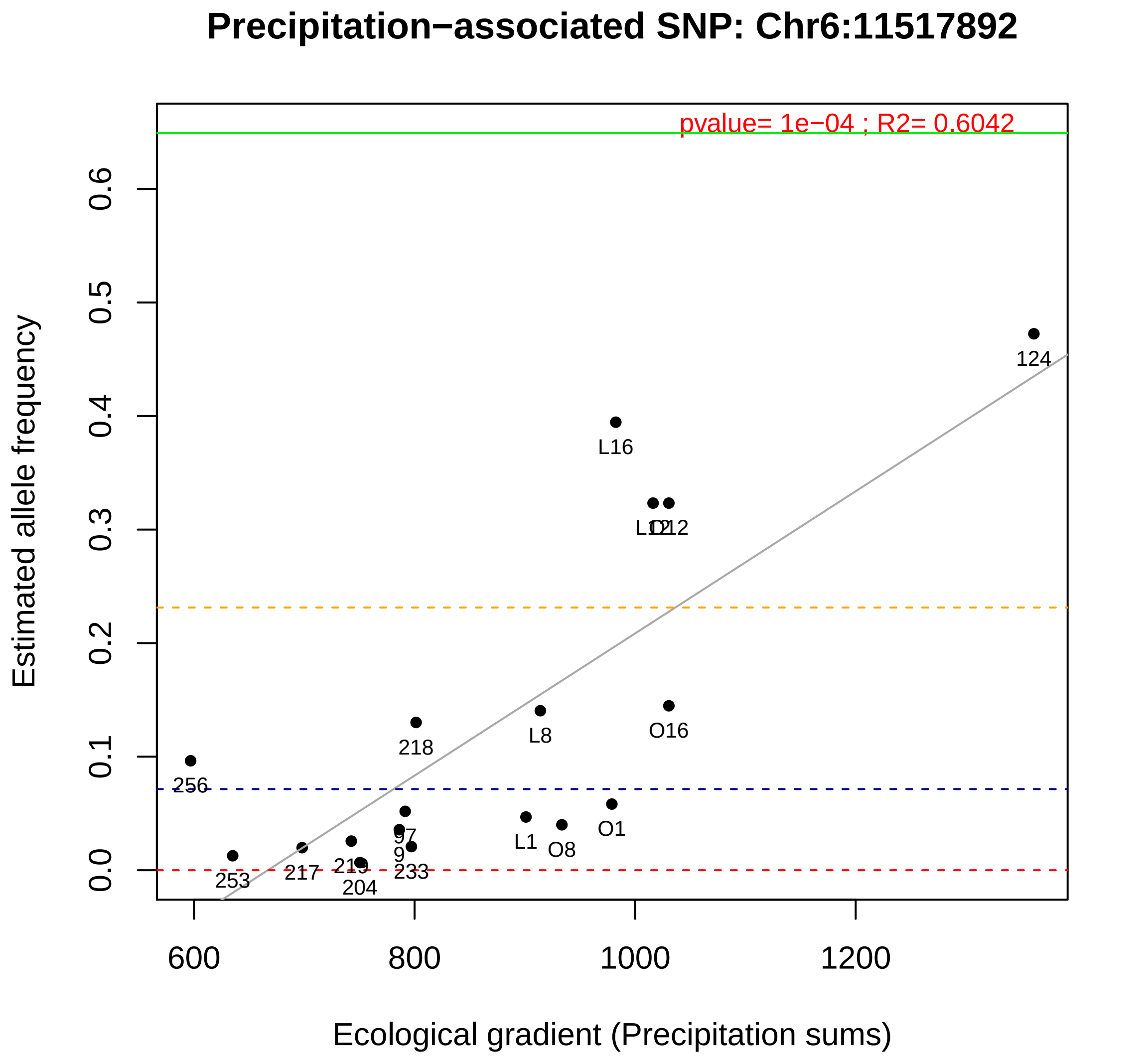
Precipitation−associated SNP: Chr6:11517892
pvalue= 1e−04 ; R2= 0.6042
Estimated allele frequency
0.0
0.1
0.2
0.3
0.4
0.5
0.6
600
800
1000
1200
Ecological gradient (Precipitation sums)
124
L16
O12
L8
O16
218
256
O1
L1
O8
253
217
204
233

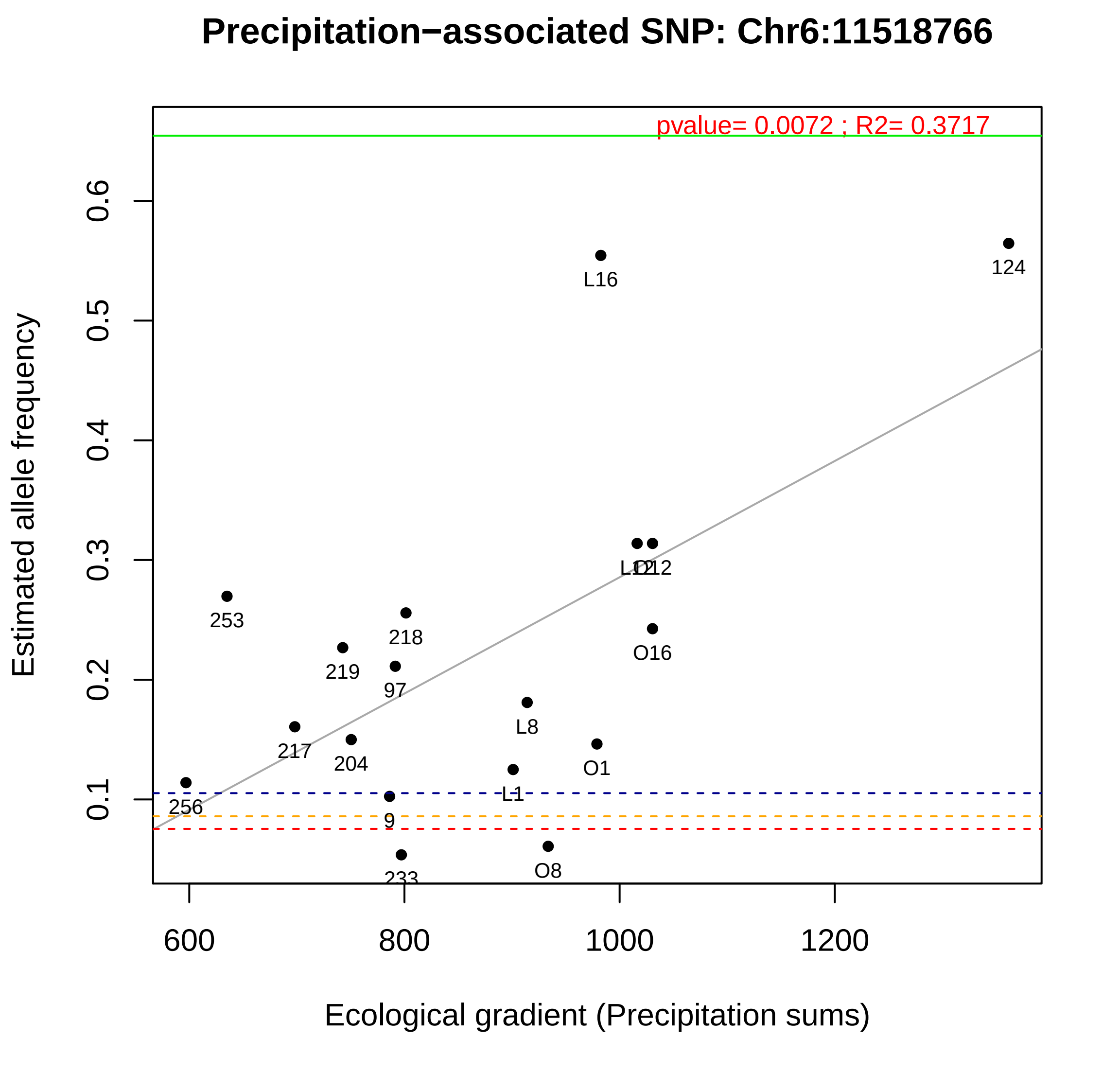
Precipitation−associated SNP: Chr6:11518766
pvalue= 0.0072 ; R2= 0.3717
Estimated allele frequency
0.1
0.2
0.3
0.4
0.5
0.6
600
800
1000
1200
Ecological gradient (Precipitation sums)
L16
124
253
218
219
97
O16
L8
217
204
O1
256
L1
9
233
O8
O12

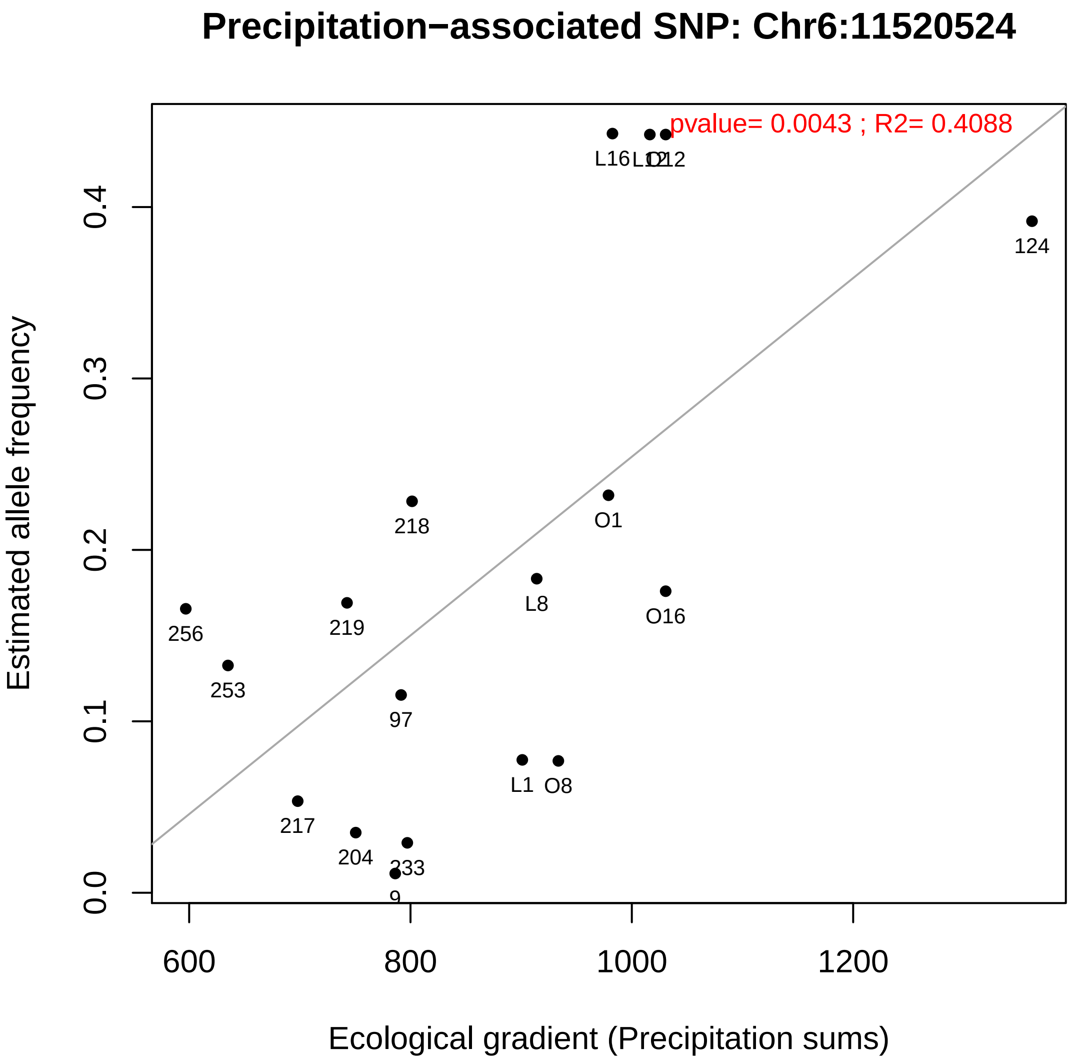

Precipitation−associated SNP: Chr6:11520524
pvalue= 0.0043 ; R2= 0.4088
Estimated allele frequency
Ecological gradient (Precipitation sums)
0.0
0.1
0.2
0.3
0.4
600
800
1000
1200
L16
124
218
O1
L8
O16
256
219
253
97
L1
O8
217
204
233
9

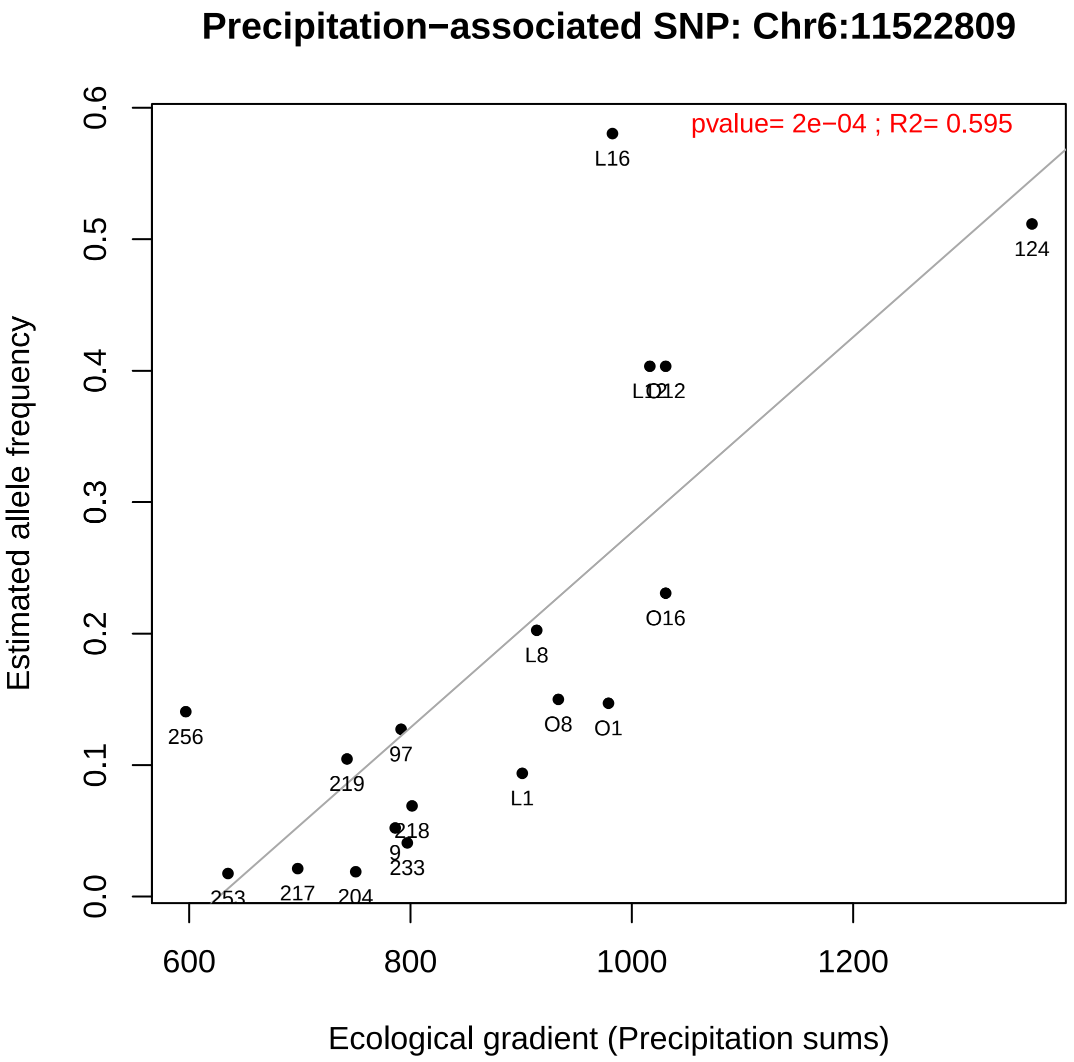

Precipitation−associated SNP: Chr6:11522809
pvalue= 2e−04 ; R2= 0.595
Estimated allele frequency
Ecological gradient (Precipitation sums)
0.0
0.1
0.2
0.3
0.4
0.5
0.6
600
800
1000
1200
L16
124
L12
O12
O16
L8
O8
O1
256
97
219
L1
218
9
233
253
217
204

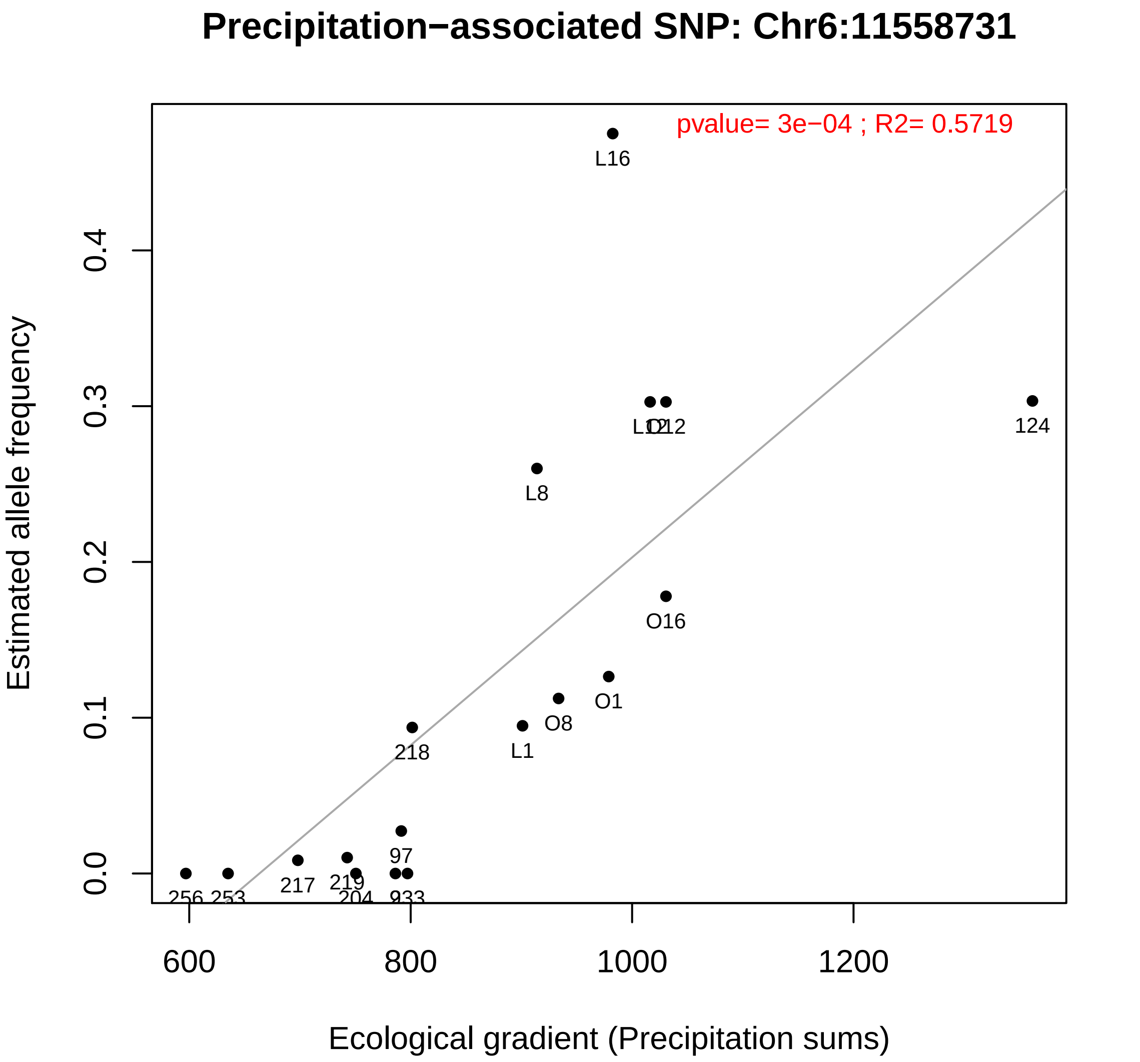

Precipitation−associated SNP: Chr6:11558731
pvalue= 3e−04 ; R2= 0.5719
L16
L12
O12
124
L8
O16
O1
O8
L1
218
97
217
219
Estimated allele frequency
0.0
0.1
0.2
0.3
0.4
600
800
1000
1200
Ecological gradient (Precipitation sums)

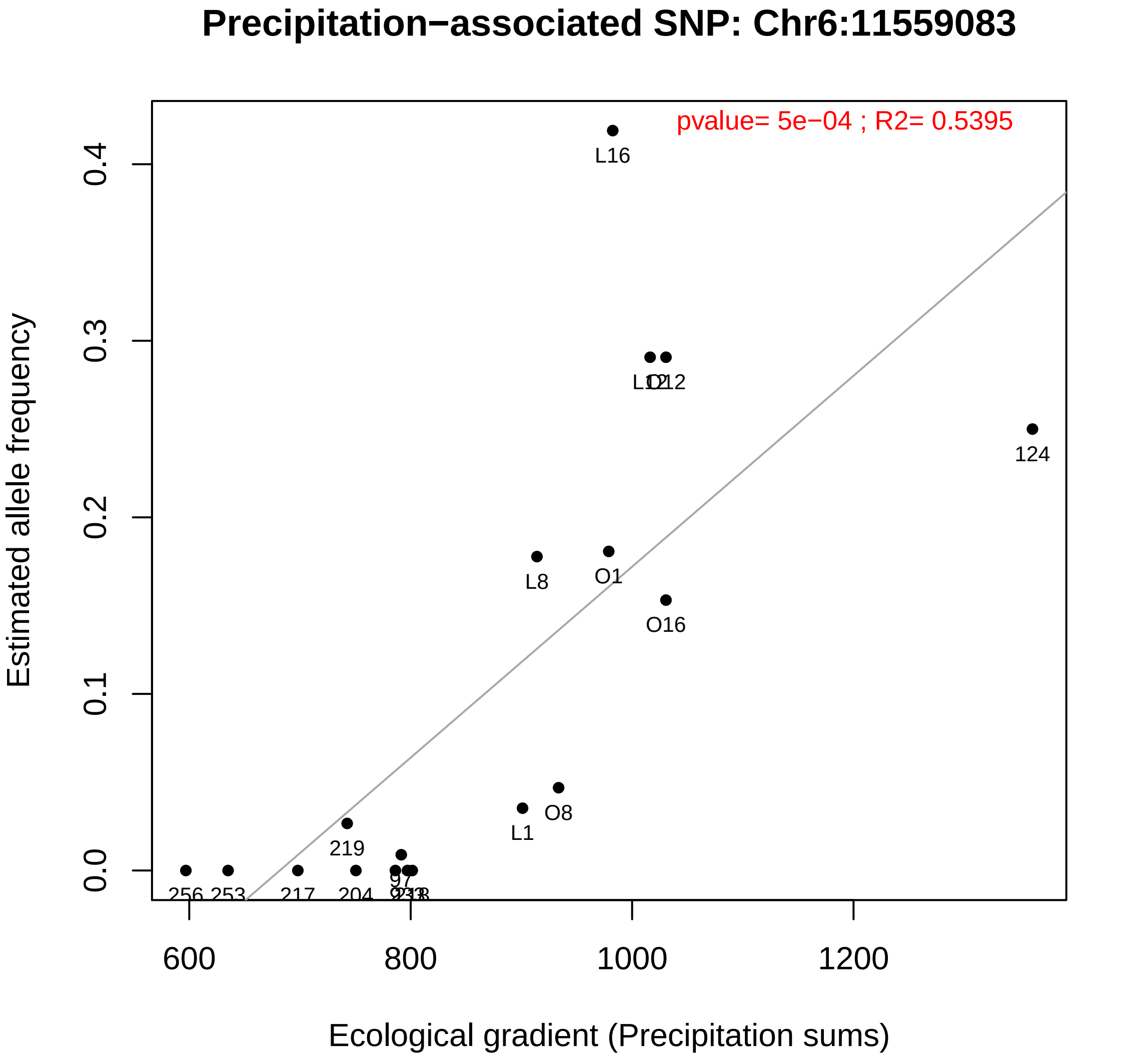
Precipitation−associated SNP: Chr6:11559083
pvalue= 5e−04 ; R2= 0.5395
L16
124
L8
O1
O16
O8
L1
219
97
0.0
0.1
0.2
0.3
0.4
600
800
1000
1200
Estimated allele frequency
Ecological gradient (Precipitation sums)

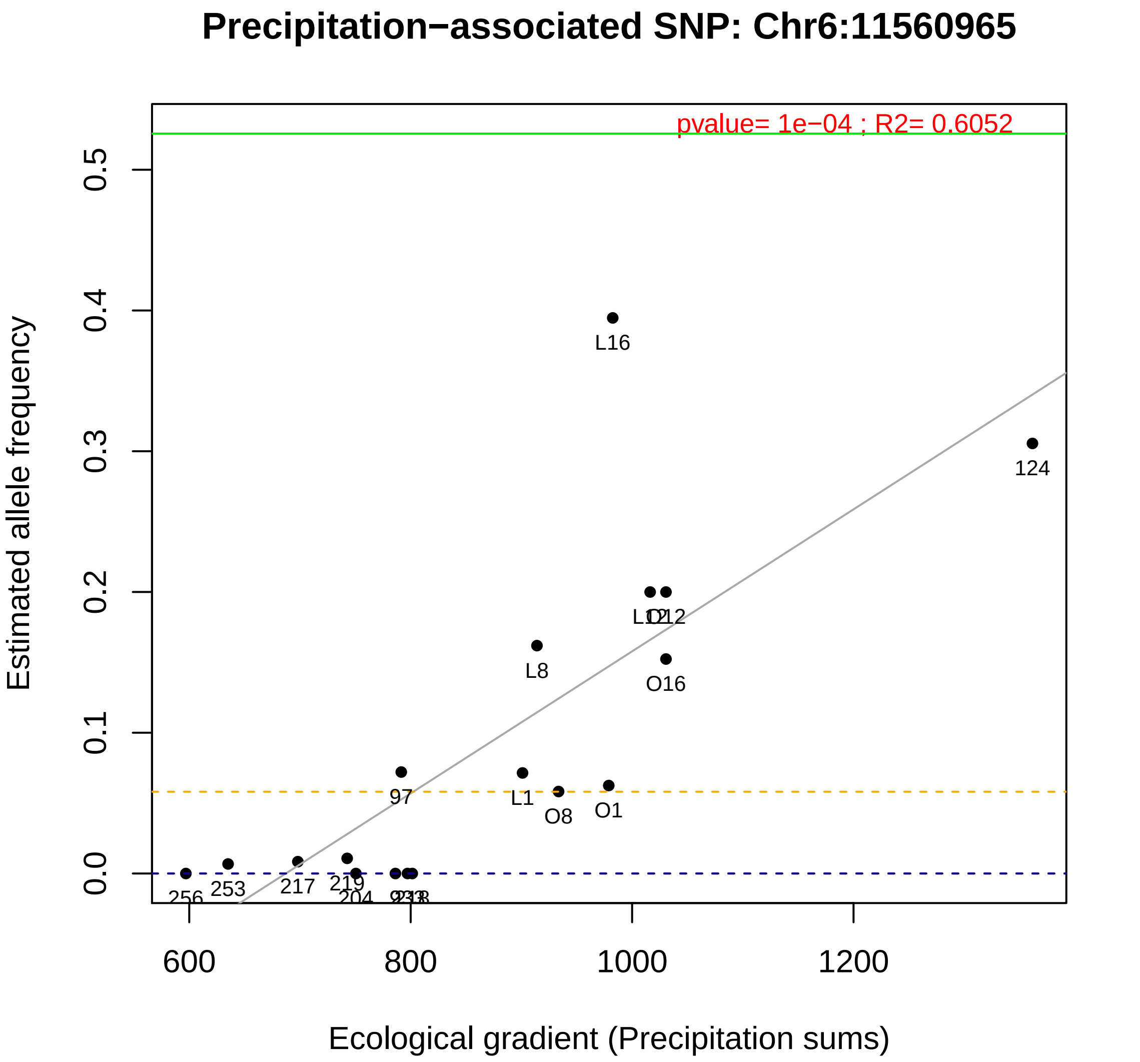

Precipitation−associated SNP: Chr6:11560965
pvalue= 1e−04 ; R2= 0.6052
Estimated allele frequency
0.0
0.1
0.2
0.3
0.4
0.5
600
800
1000
1200
Ecological gradient (Precipitation sums)
L16
124
O12
L8
O16
97
L1
O8
O1
256
253
217
219
204

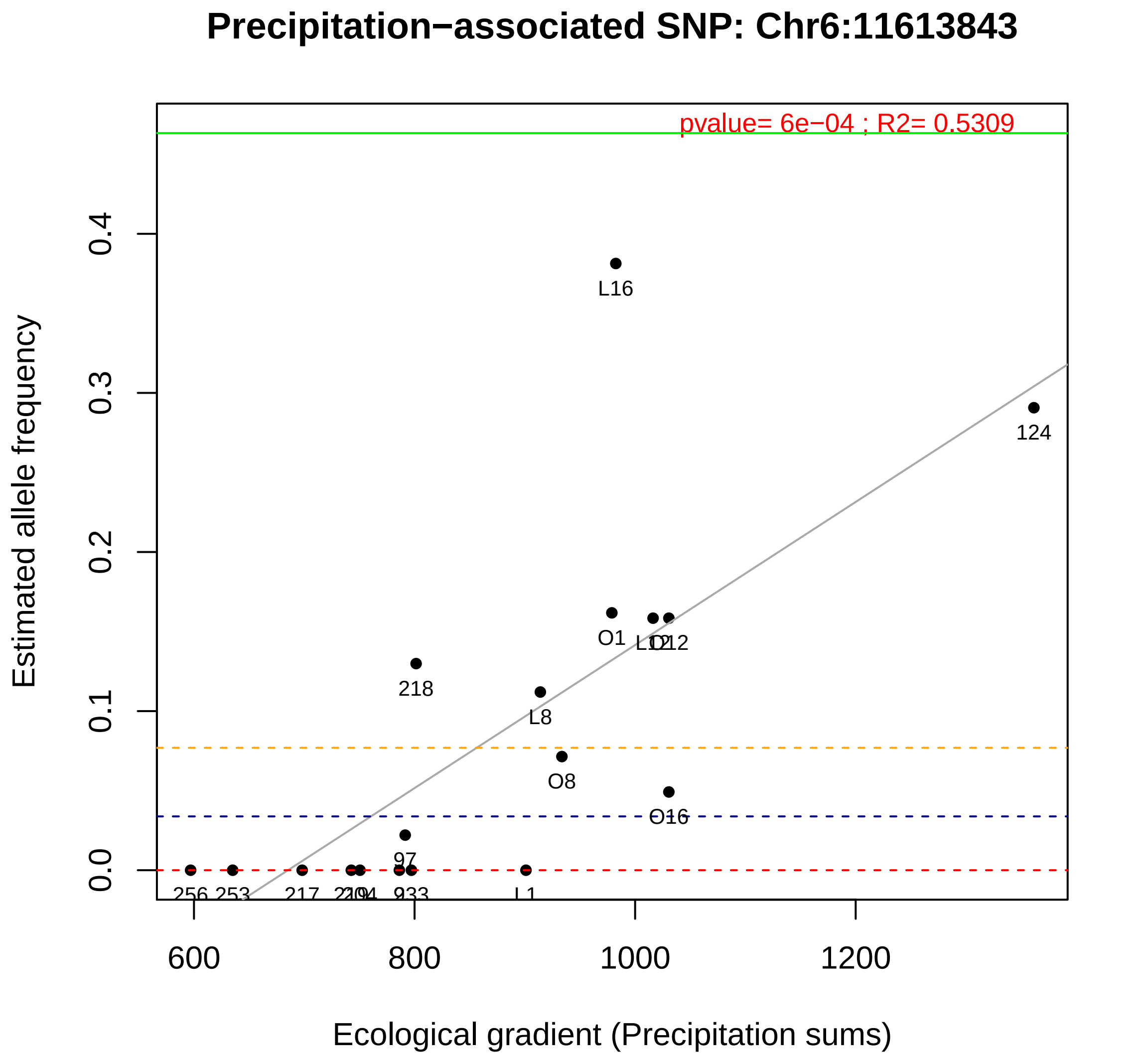

Precipitation−associated SNP: Chr6:11613843
pvalue= 6e−04 ; R2= 0.5309
Estimated allele frequency
0.0
0.1
0.2
0.3
0.4
600
800
1000
1200
Ecological gradient (Precipitation sums)
L16
124
O1
L12
O12
218
L8
O8
O16
97
256
253
217
L1

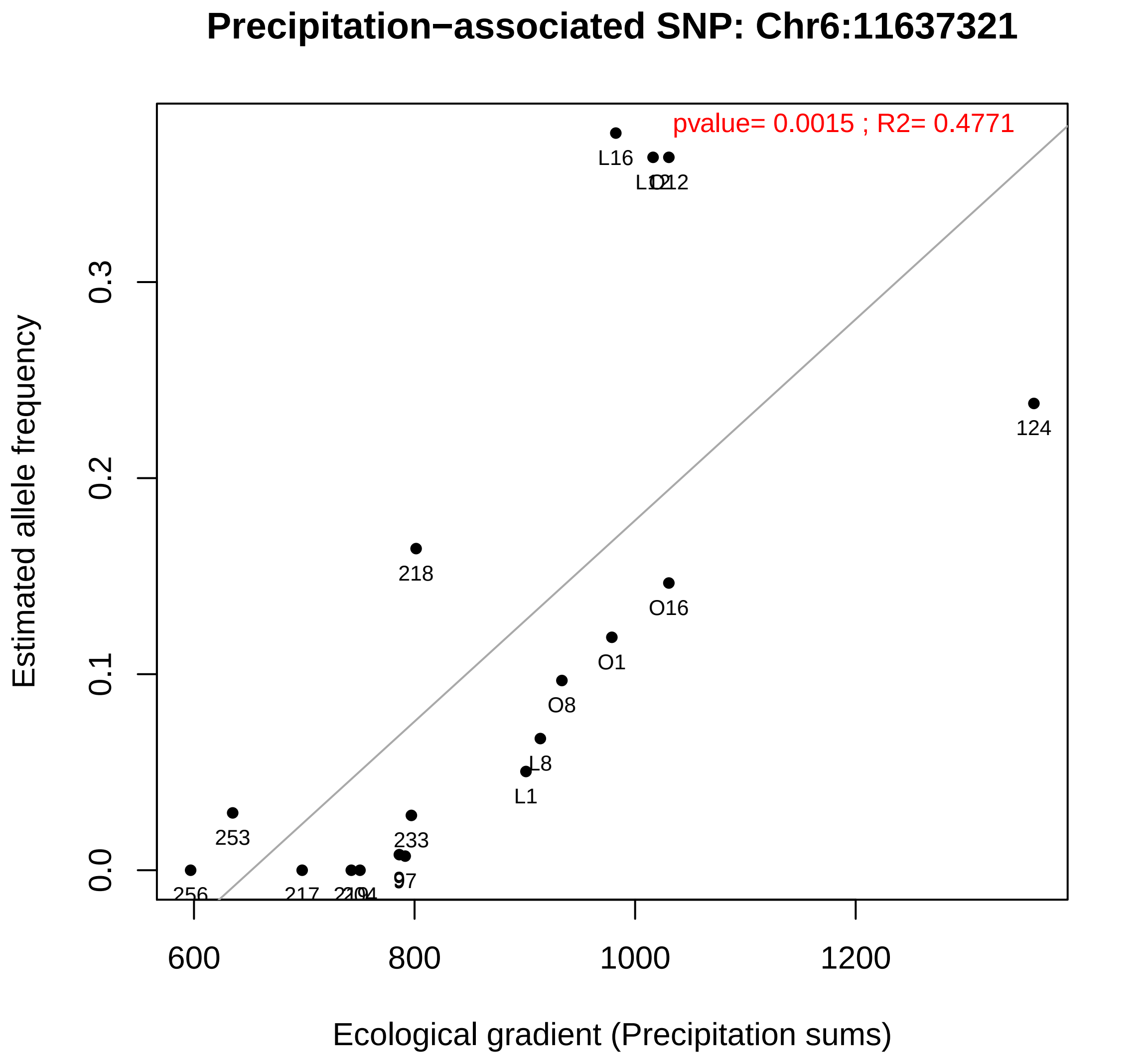

Precipitation−associated SNP: Chr6:11637321
pvalue= 0.0015 ; R2= 0.4771
Estimated allele frequency
0.0
0.1
0.2
0.3
600
800
1000
1200
Ecological gradient (Precipitation sums)
L16
O12
124
218
O16
O1
O8
L8
L1
253
233
256
217

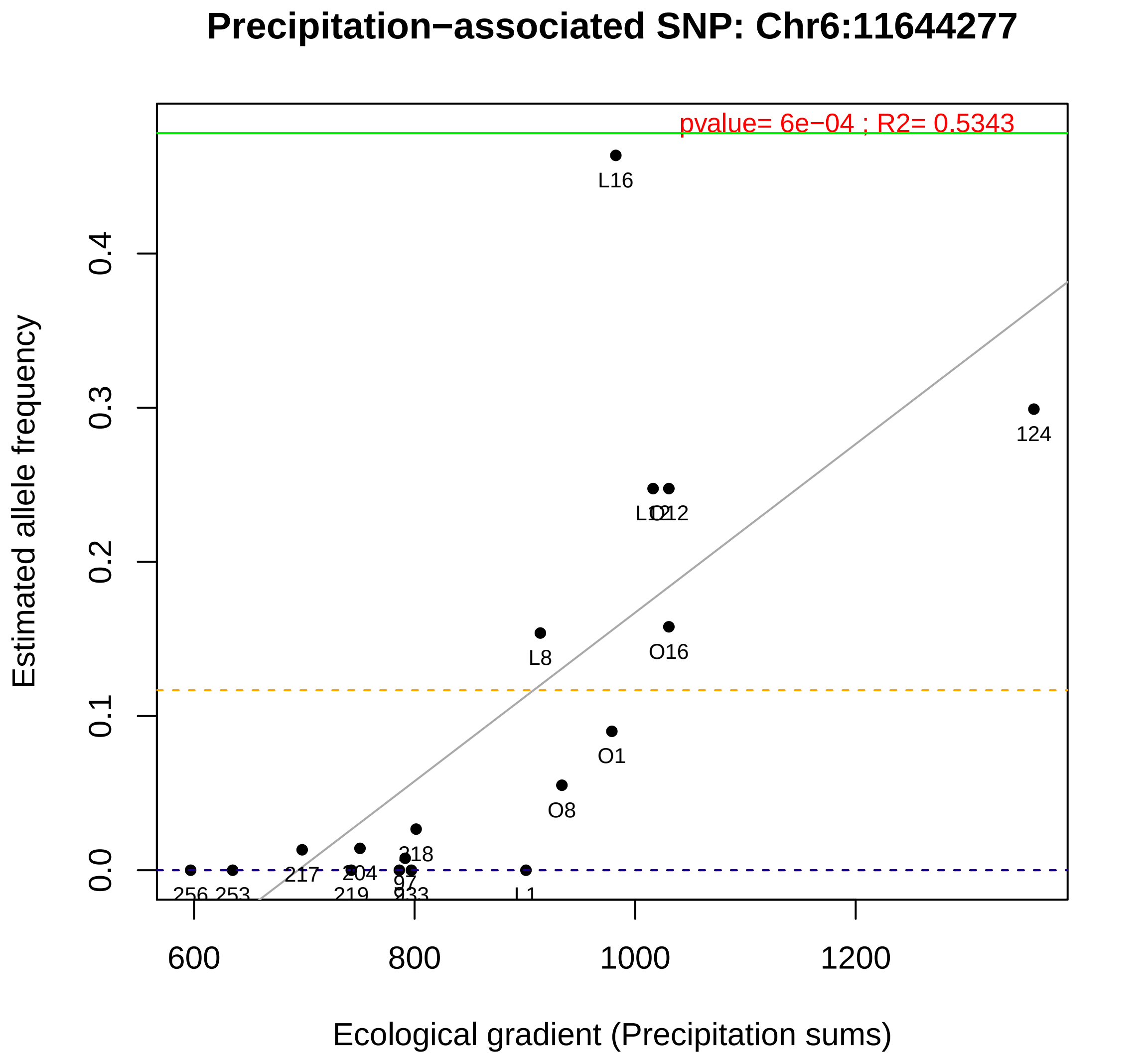

Precipitation−associated SNP: Chr6:11644277
pvalue= 6e−04 ; R2= 0.5343
L16
124
L12
O12
L8
O16
O1
O8
218
217
204
97
256
253
219
233
L1
0.0
0.1
0.2
0.3
0.4
600
800
1000
1200
Estimated allele frequency
Ecological gradient (Precipitation sums)

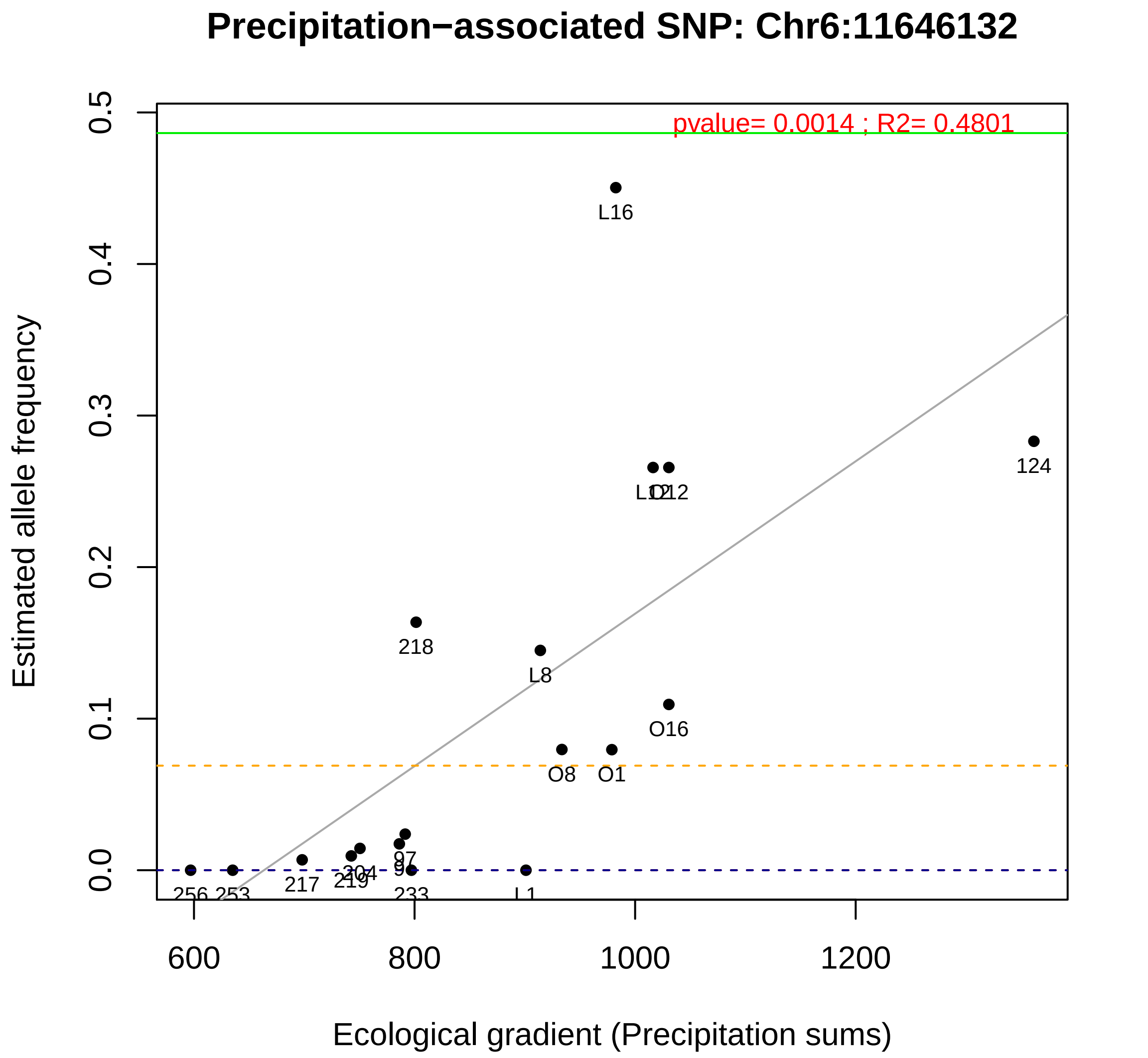

Precipitation−associated SNP: Chr6:11646132
pvalue= 0.0014 ; R2= 0.4801
Estimated allele frequency
Ecological gradient (Precipitation sums)
0.0
0.1
0.2
0.3
0.4
0.5
600
800
1000
1200
L16
124
L12
O12
218
L8
O16
O8
O1
217
219
204
97
96
233
256
253
L1

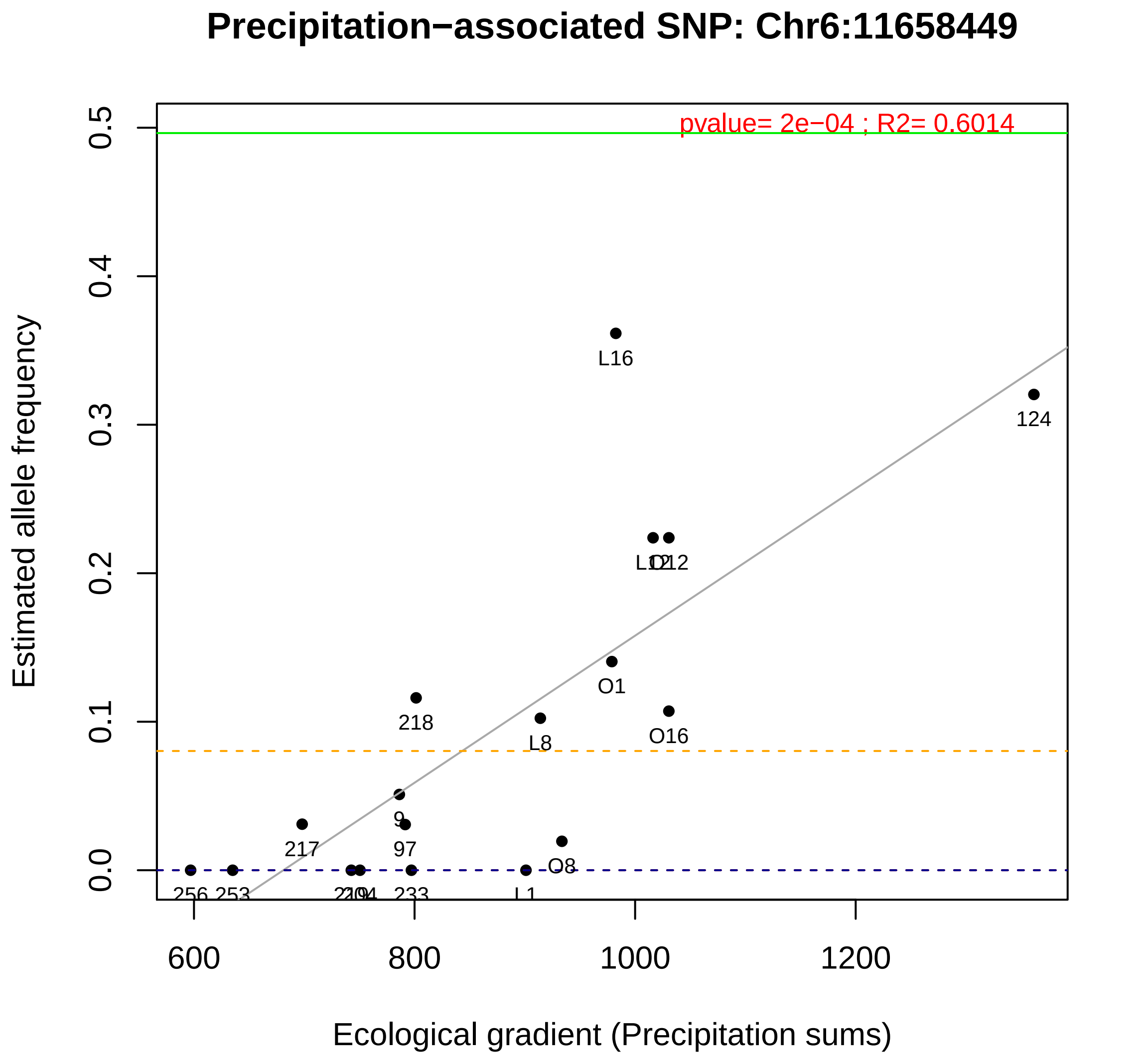

Precipitation−associated SNP: Chr6:11658449
pvalue= 2e−04 ; R2= 0.6014
Estimated allele frequency
0.0
0.1
0.2
0.3
0.4
0.5
L16
124
L12
O12
O1
218
O16
L8
9
217
97
O8
256
253
210
194
233
L1
600
800
1000
1200
Ecological gradient (Precipitation sums)

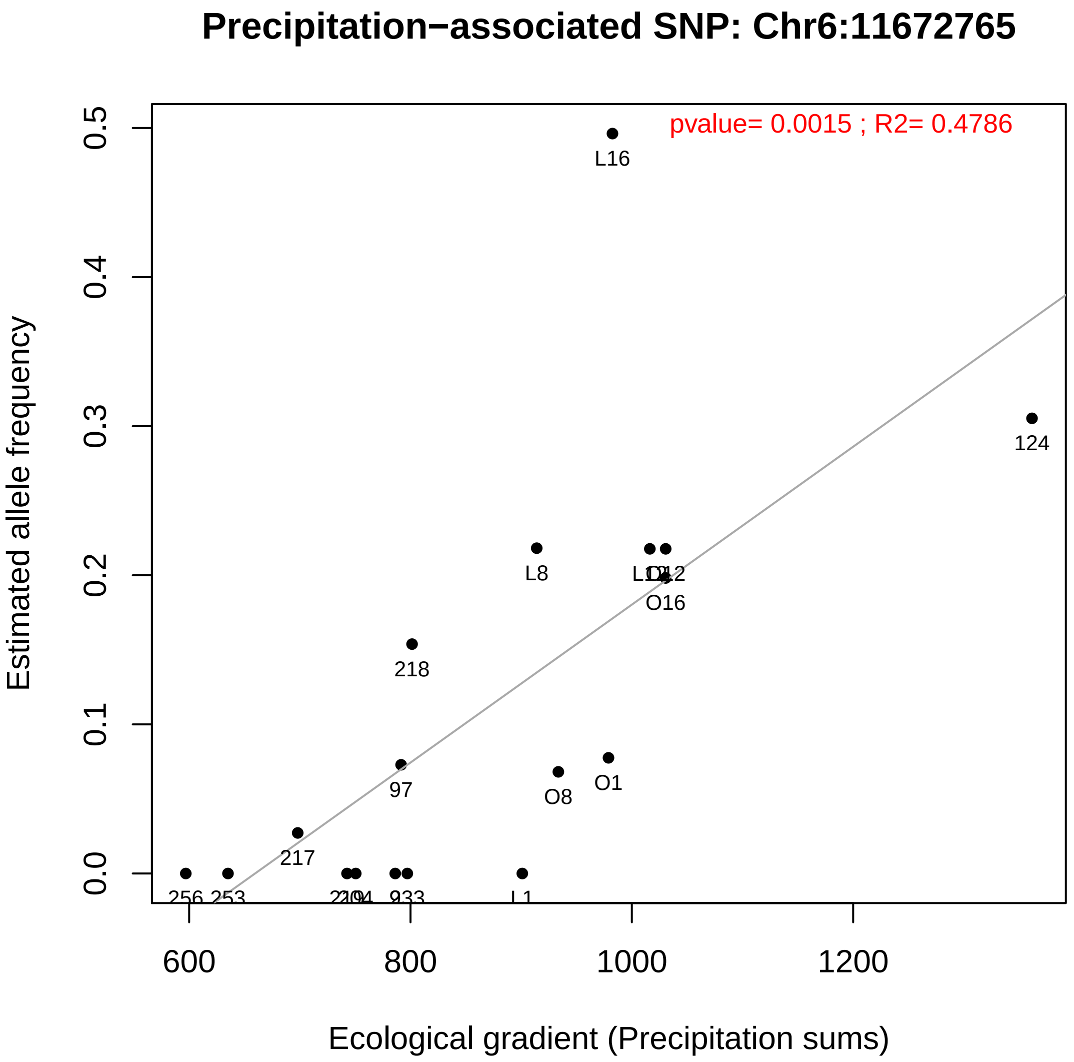
Precipitation−associated SNP: Chr6:11672765
pvalue= 0.0015 ; R2= 0.4786
Estimated allele frequency
0.0
0.1
0.2
0.3
0.4
0.5
600
800
1000
1200
Ecological gradient (Precipitation sums)
L16
124
L8
O16
218
97
O8
O1
217

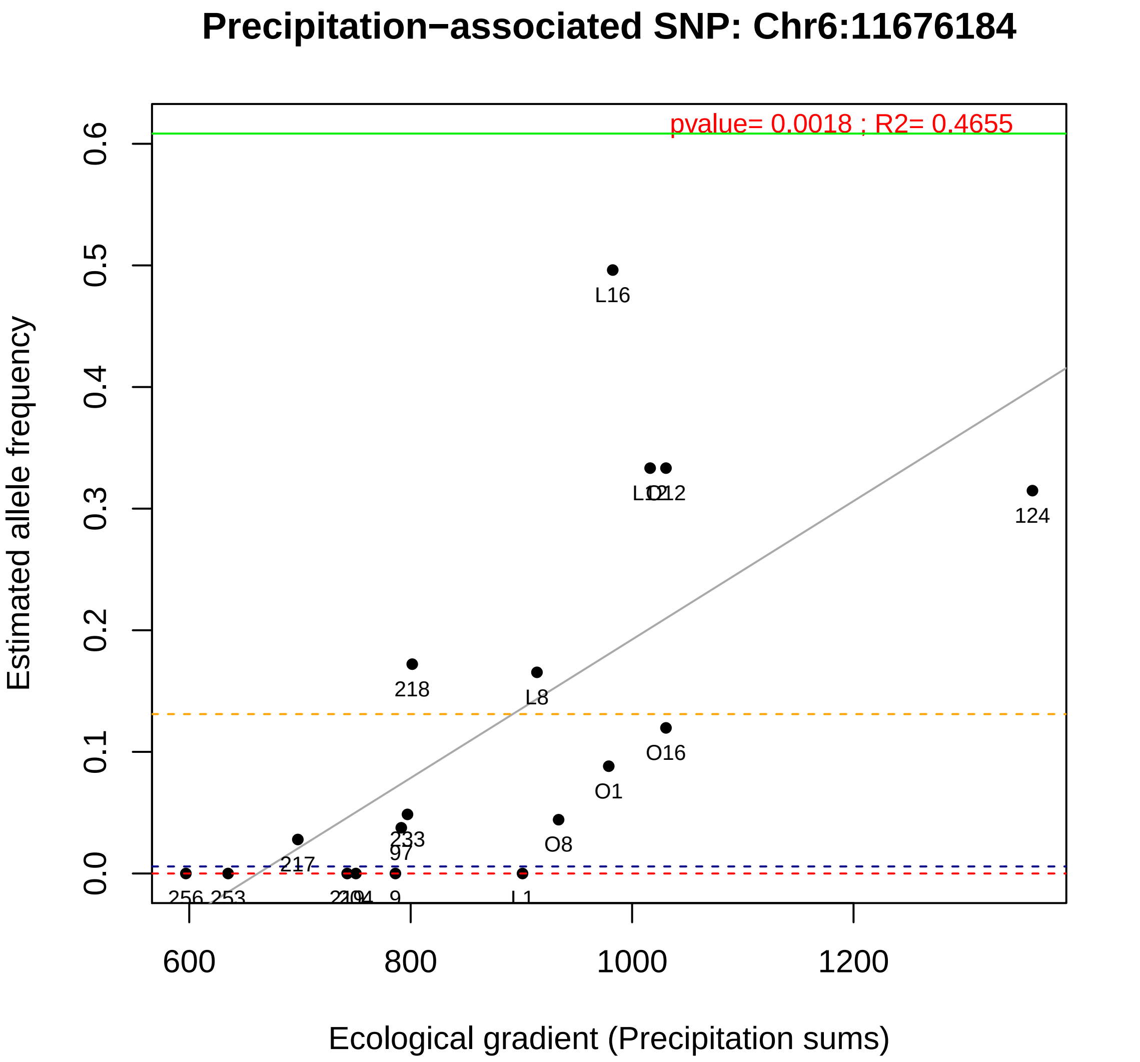

Precipitation−associated SNP: Chr6:11676184
pvalue= 0.0018 ; R2= 0.4655
Estimated allele frequency
0.0
0.1
0.2
0.3
0.4
0.5
0.6
Ecological gradient (Precipitation sums)
600
800
1000
1200
L16
124
218
L8
O16
O1
233
97
O8
217
256
253
9
L1

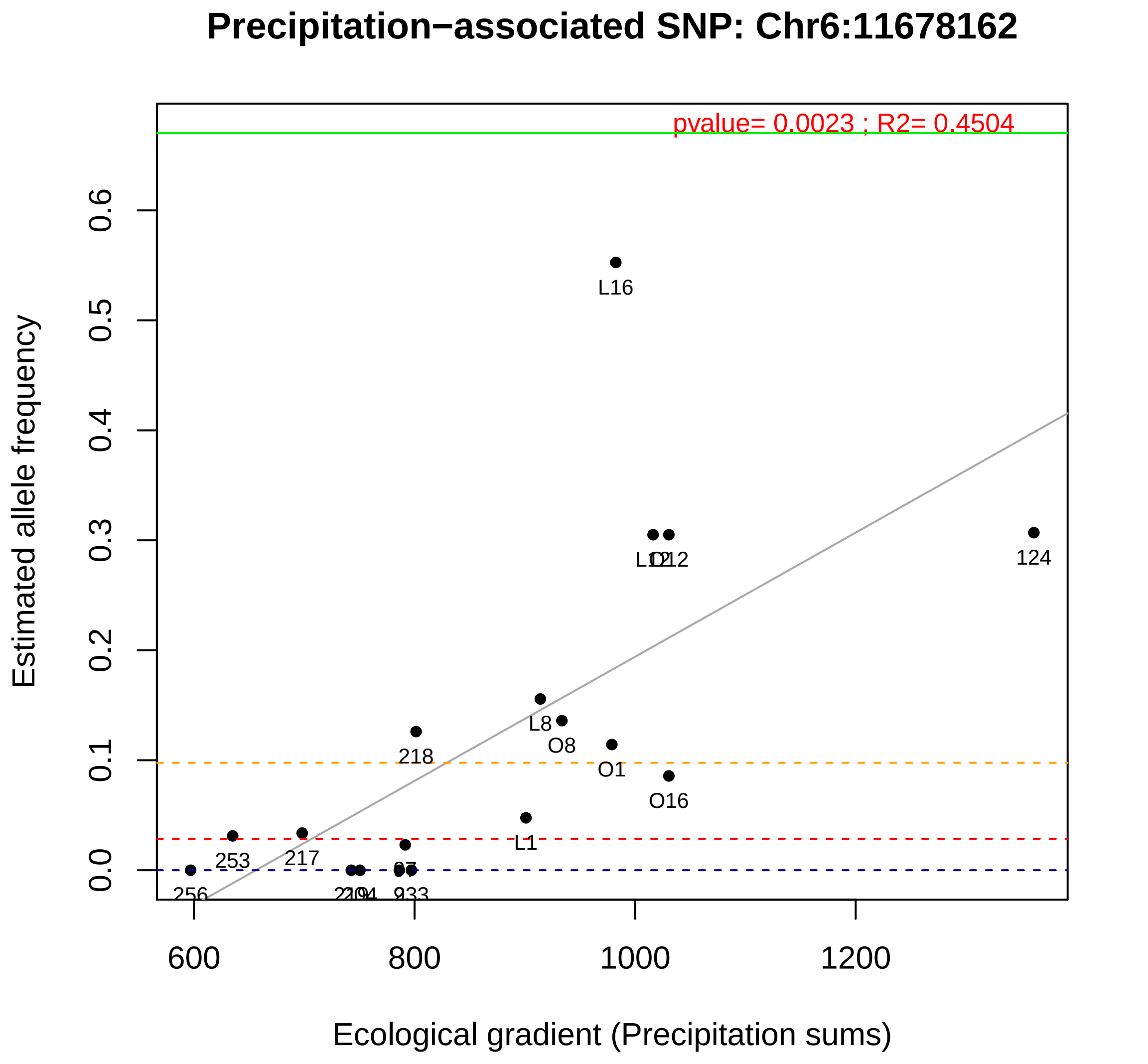

Precipitation−associated SNP: Chr6:11678162
pvalue= 0.0023 ; R2= 0.4504
Estimated allele frequency
0.0
0.1
0.2
0.3
0.4
0.5
0.6
600
800
1000
1200
Ecological gradient (Precipitation sums)
L16
L12
O12
124
L8
O8
218
O1
O16
L1
253
217
256

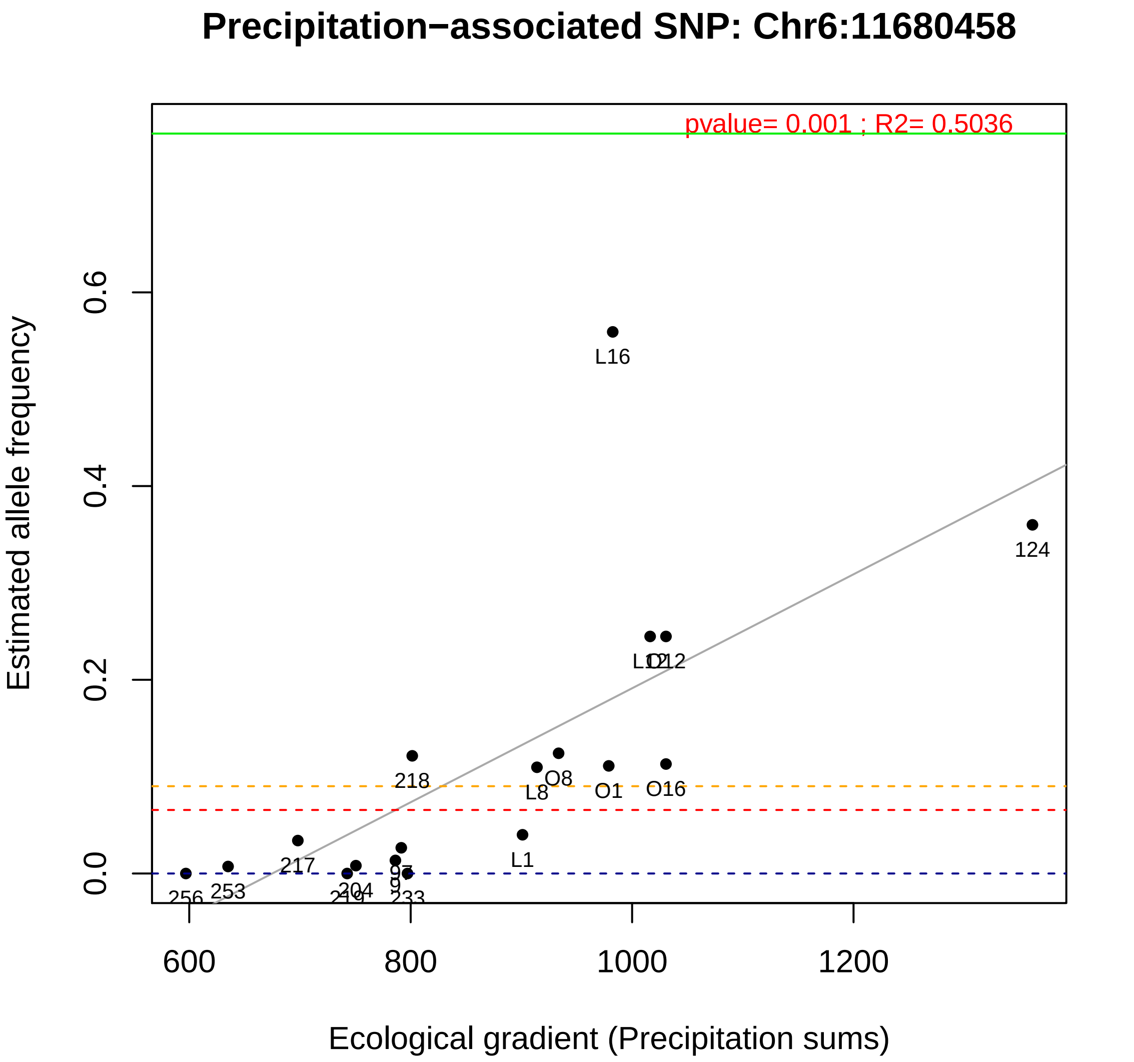

Precipitation−associated SNP: Chr6:11680458
pvalue= 0.001 ; R2= 0.5036
Estimated allele frequency
0.0
0.2
0.4
0.6
600
800
1000
1200
Ecological gradient (Precipitation sums)
L16
124
L12
O12
218
O8
L8
O1
O16
217
L1
256
253
204
233

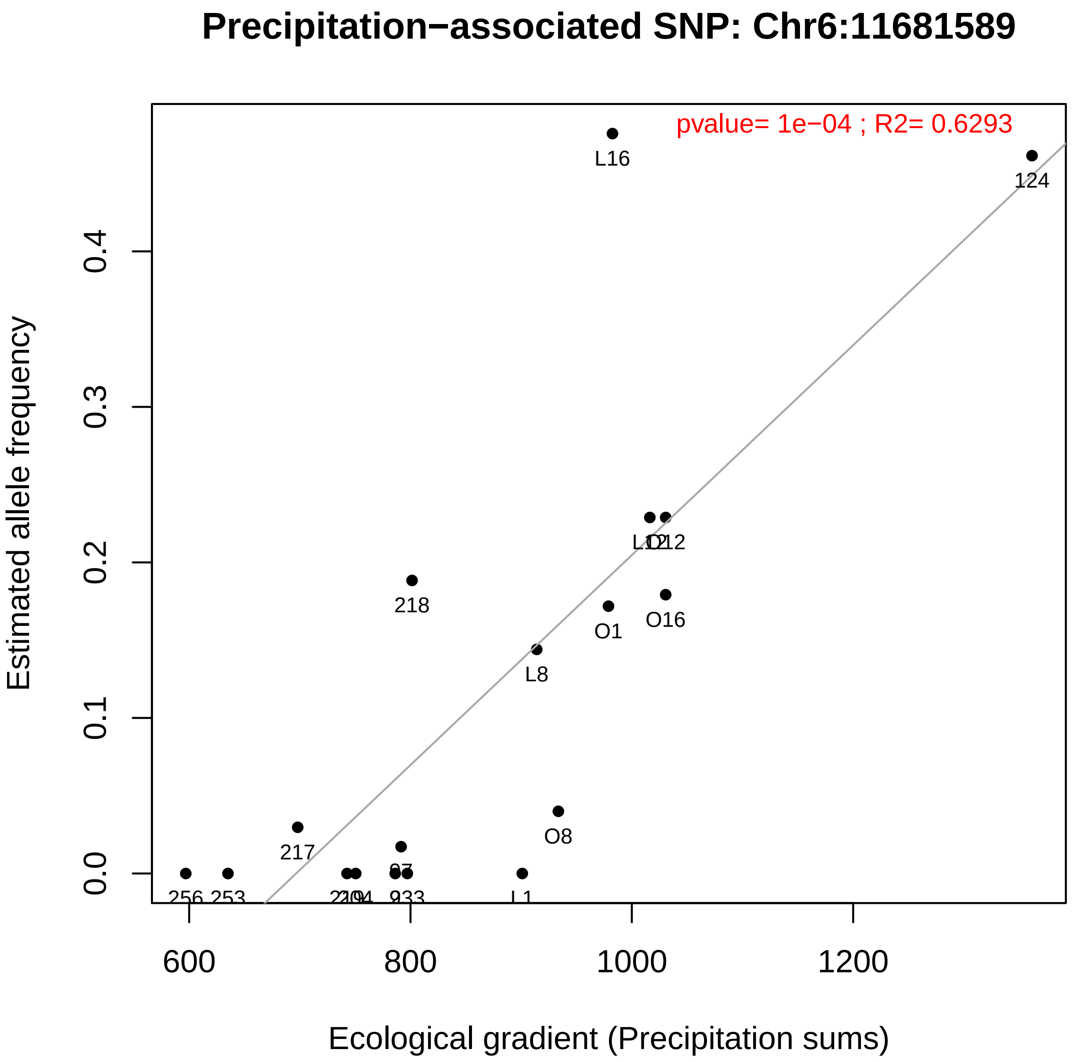
Precipitation−associated SNP: Chr6:11681589
pvalue= 1e−04 ; R2= 0.6293
Estimated allele frequency
Ecological gradient (Precipitation sums)
0.0
0.1
0.2
0.3
0.4
600
800
1000
1200
L16
124
218
L8
O1
O16
O8
217
L1
256
253

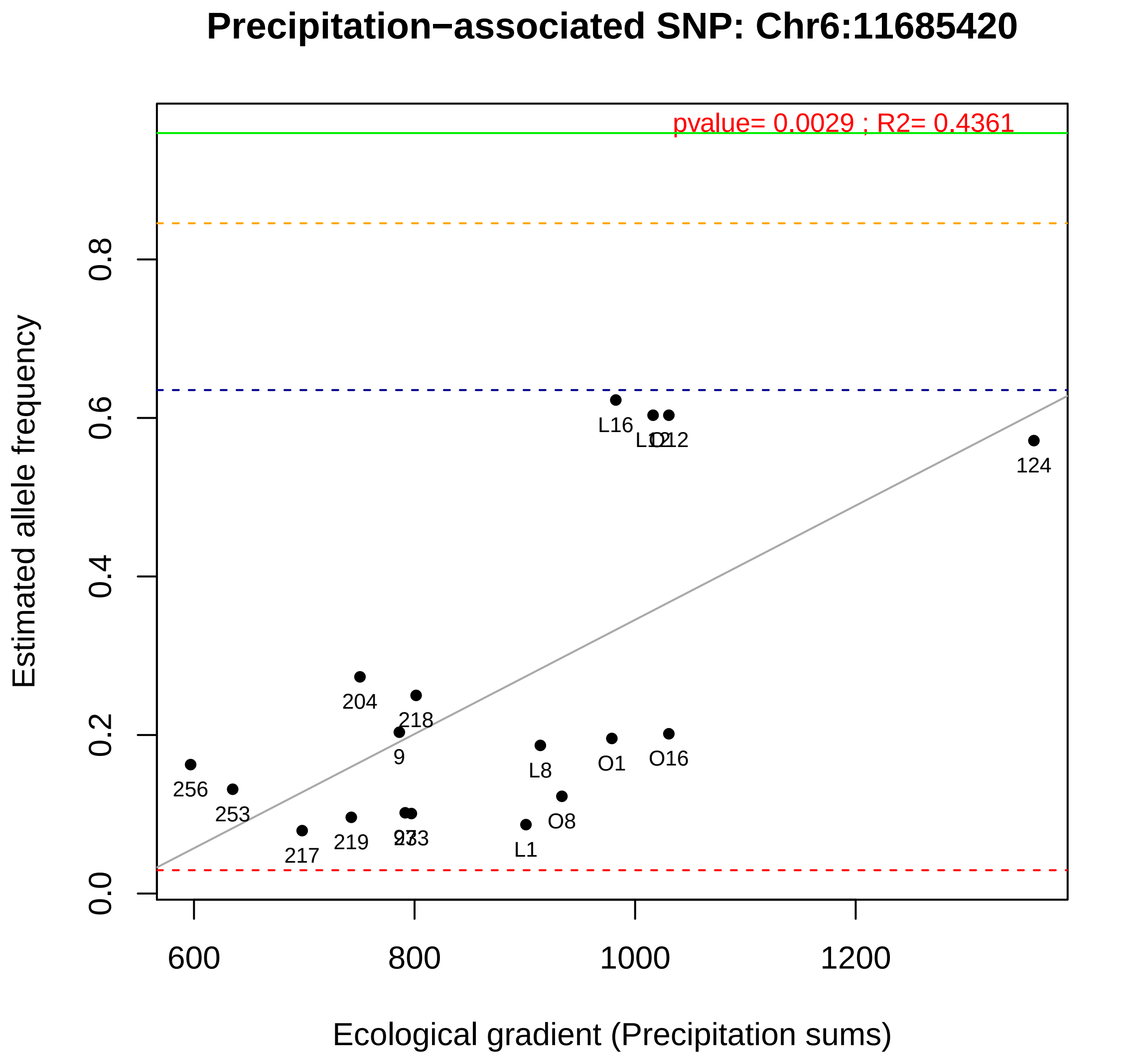
Precipitation−associated SNP: Chr6:11685420
pvalue= 0.0029 ; R2= 0.4361
Estimated allele frequency
0.0
0.2
0.4
0.6
0.8
600
800
1000
1200
Ecological gradient (Precipitation sums)
L16
L12
O12
124
204
218
9
256
253
217
219
L8
O1
O16
O8
L1

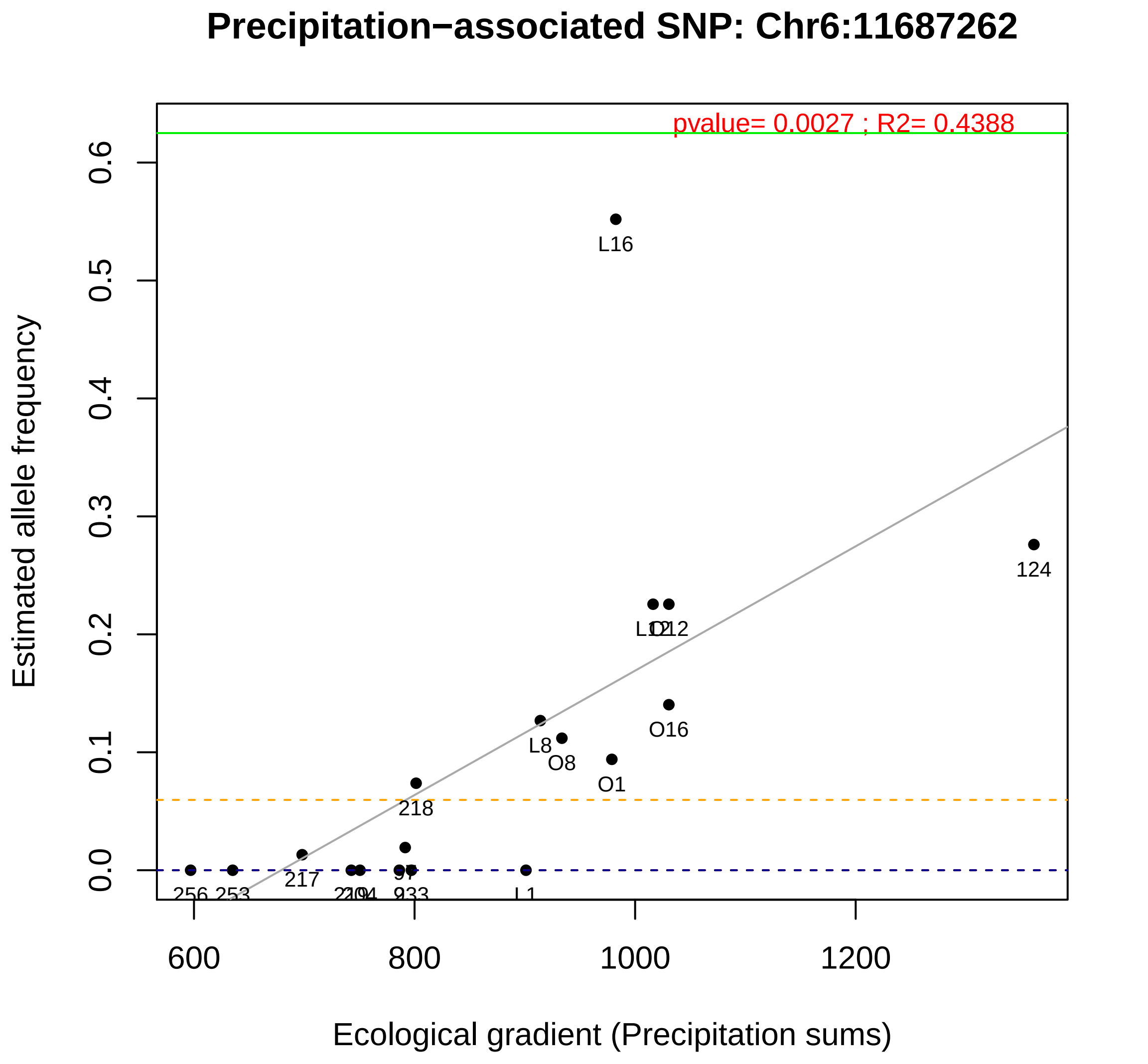

Precipitation−associated SNP: Chr6:11687262
pvalue= 0.0027 ; R2= 0.4388
Estimated allele frequency
Ecological gradient (Precipitation sums)
0.0
0.1
0.2
0.3
0.4
0.5
0.6
600
800
1000
1200
L16
124
O16
L8
O8
O1
218
217
L1

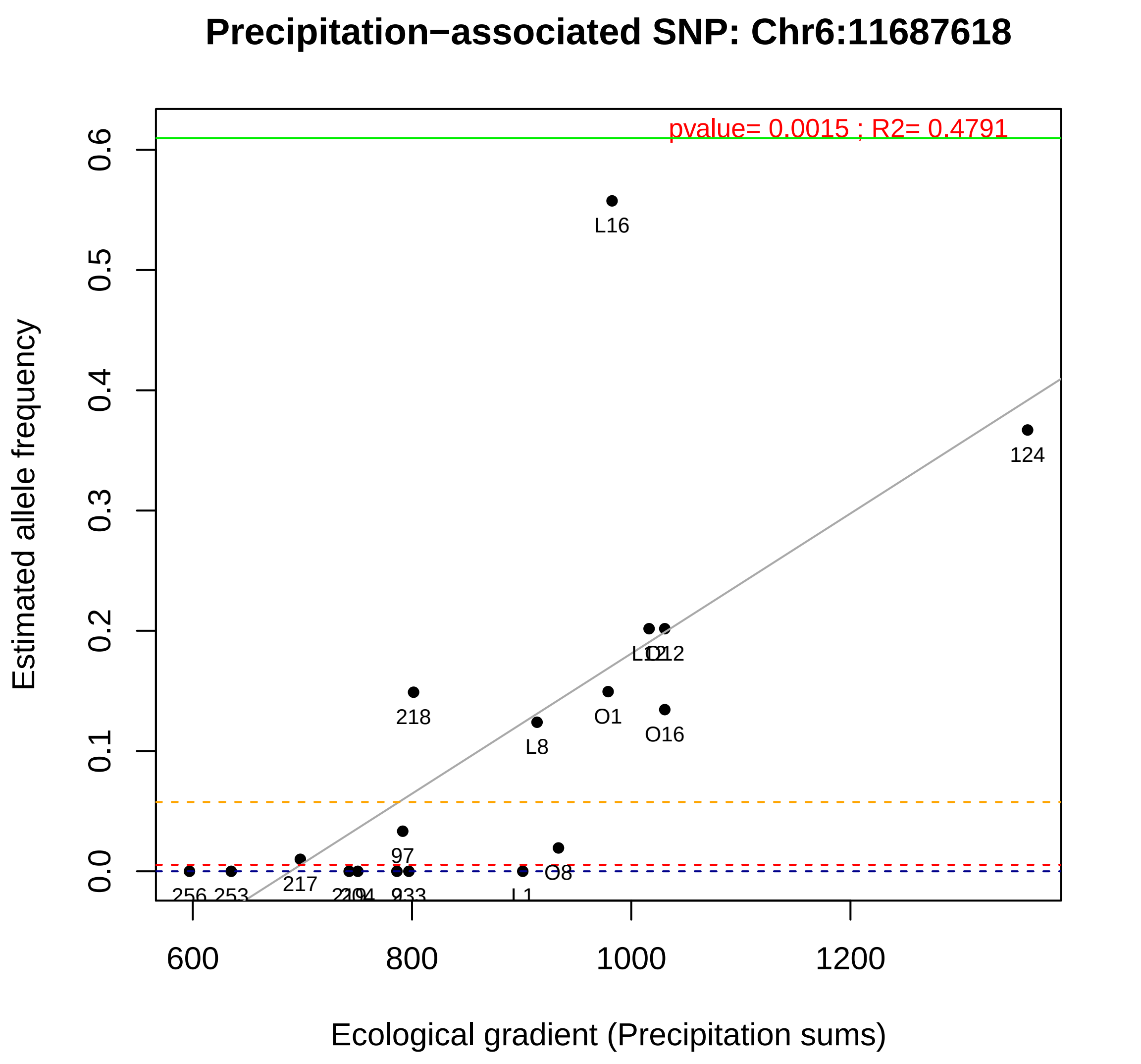

Precipitation−associated SNP: Chr6:11687618
pvalue= 0.0015 ; R2= 0.4791
Estimated allele frequency
0.0
0.1
0.2
0.3
0.4
0.5
0.6
Ecological gradient (Precipitation sums)
600
800
1000
1200
L16
124
218
L8
O1
O16
O8
97
217

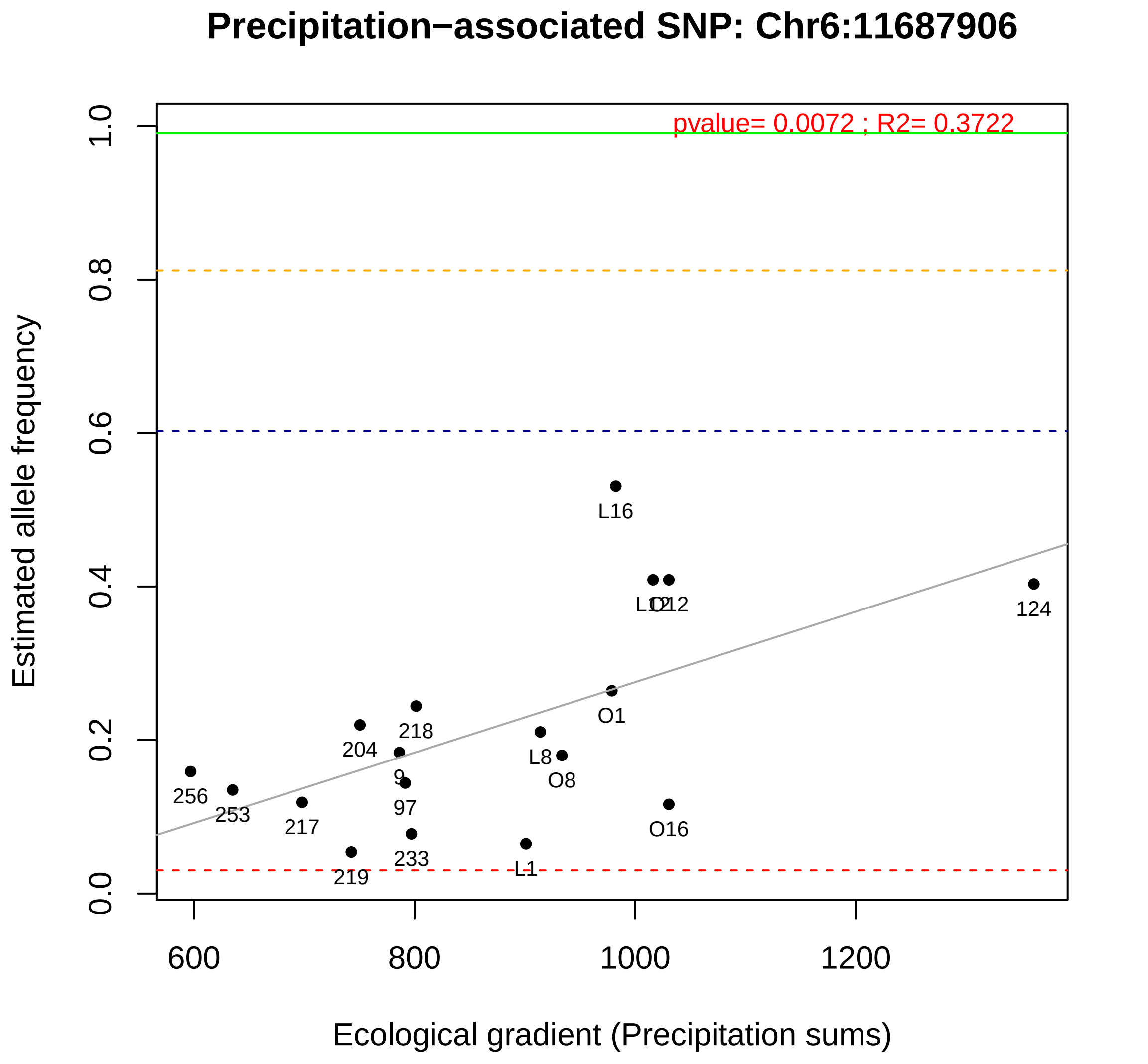
Precipitation−associated SNP: Chr6:11687906
pvalue= 0.0072 ; R2= 0.3722
Estimated allele frequency
0.0
0.2
0.4
0.6
0.8
1.0
Ecological gradient (Precipitation sums)
600
800
1000
1200
L16
L12
O12
124
O1
218
204
L8
O8
9
97
256
253
217
O16
233
219
L1

### Precipitation−associated SNP: Chr6:11692709

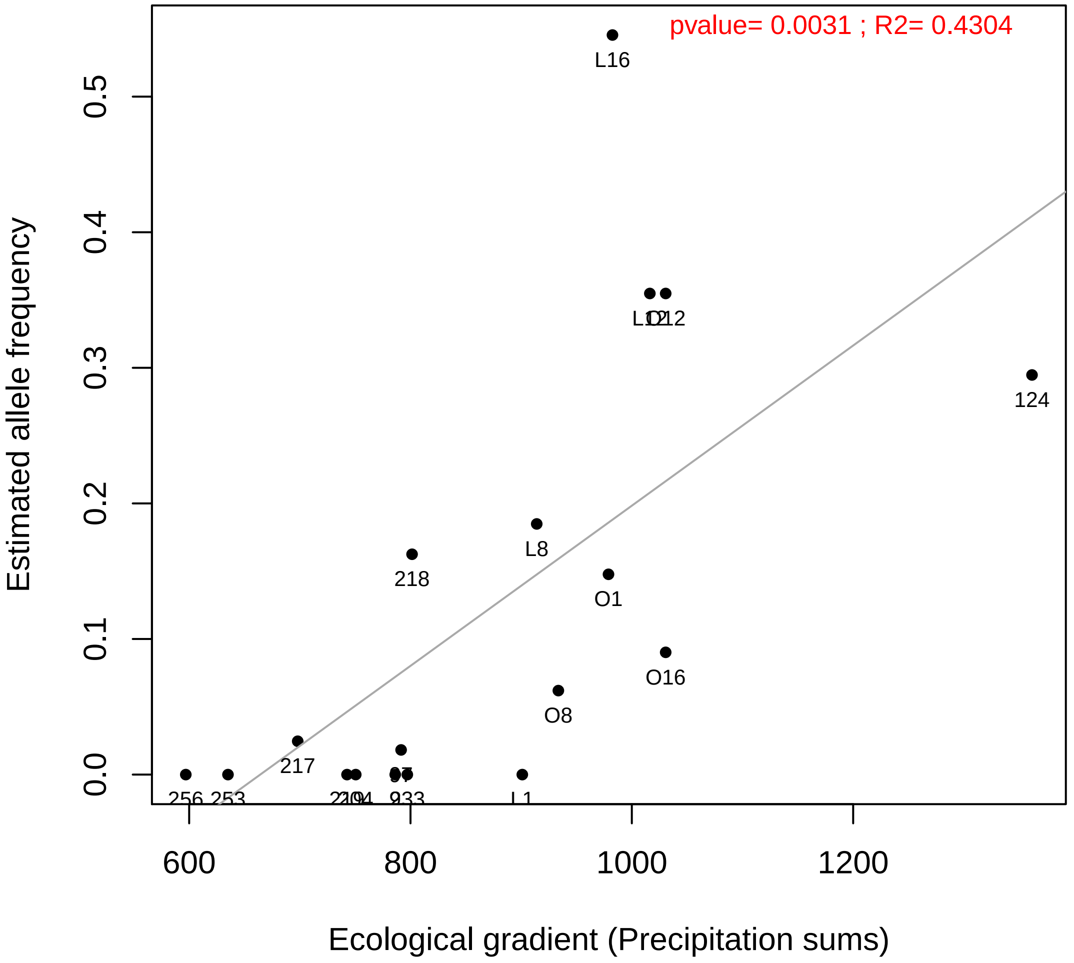

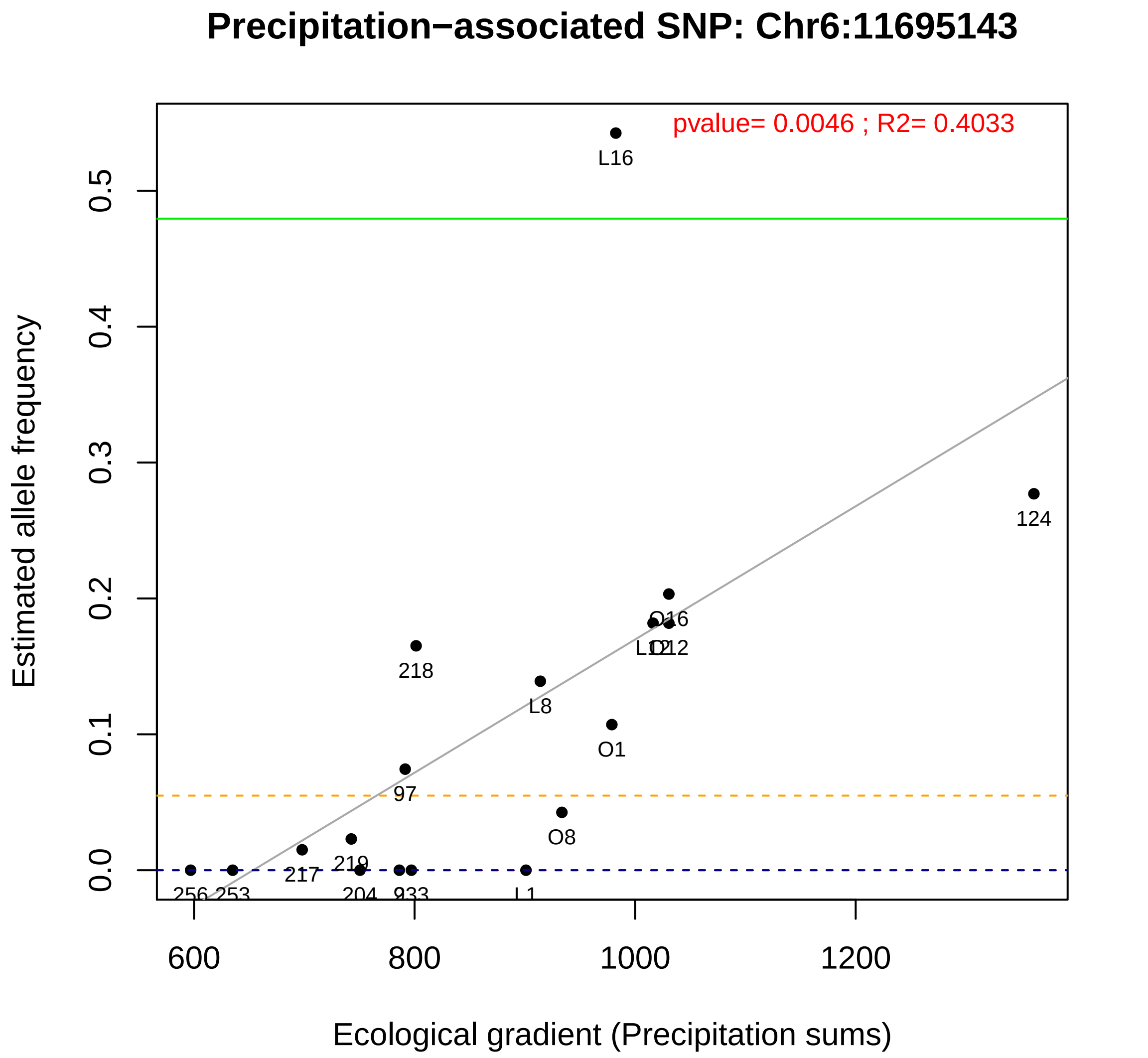
Precipitation−associated SNP: Chr6:11695143
pvalue= 0.0046 ; R2= 0.4033
L16
124
O16
L10
O12
218
L8
O1
97
O8
219
217
256
253
204
933
L1
0.0
0.1
0.2
0.3
0.4
0.5
600
800
1000
1200
Estimated allele frequency
Ecological gradient (Precipitation sums)

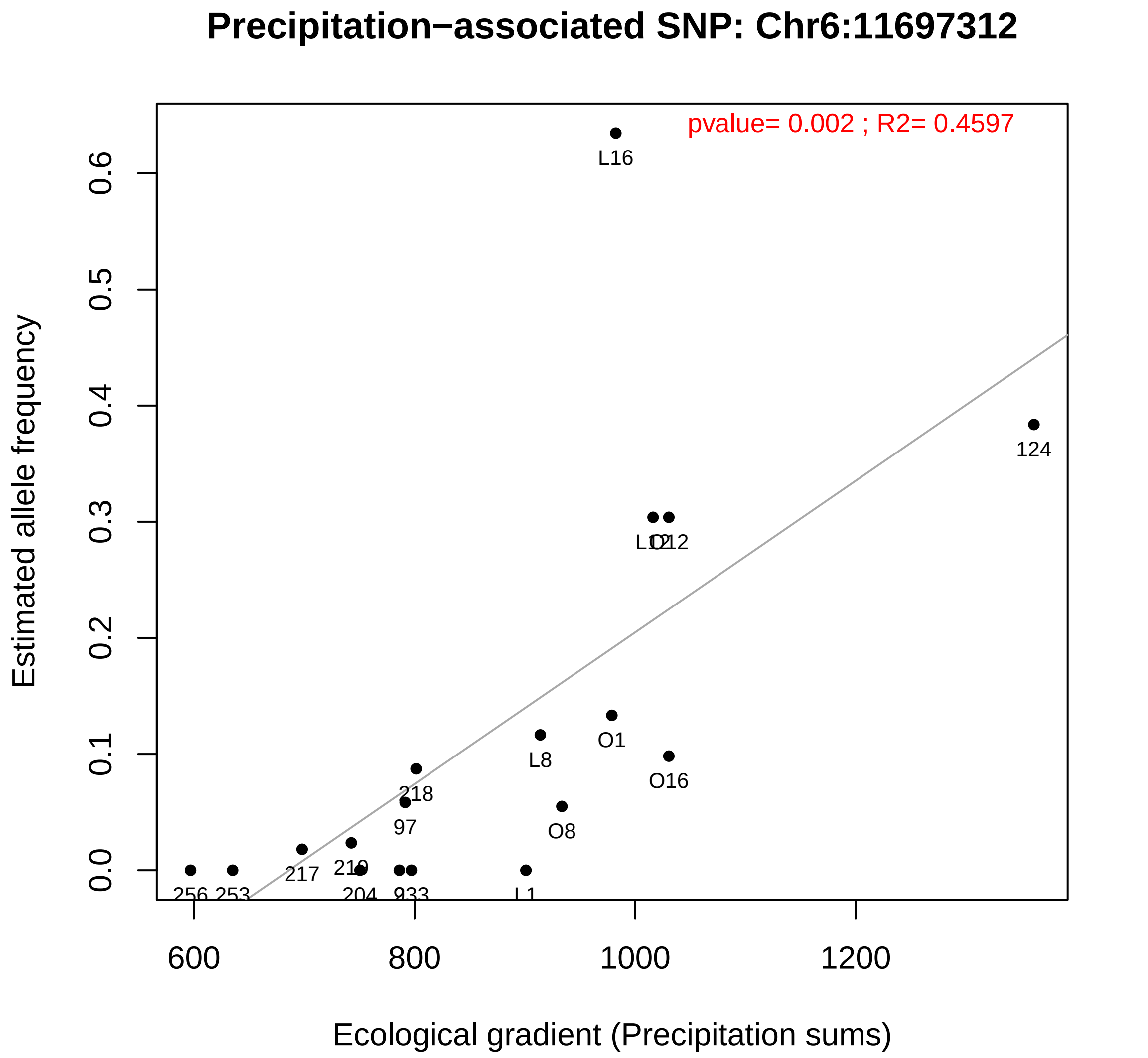

Precipitation−associated SNP: Chr6:11697312
pvalue= 0.002 ; R2= 0.4597
Estimated allele frequency
0.0
0.1
0.2
0.3
0.4
0.5
0.6
600
800
1000
1200
Ecological gradient (Precipitation sums)
L16
124
O12
O1
L8
O16
218
97
O8
217
210
204
233
L1
256
253

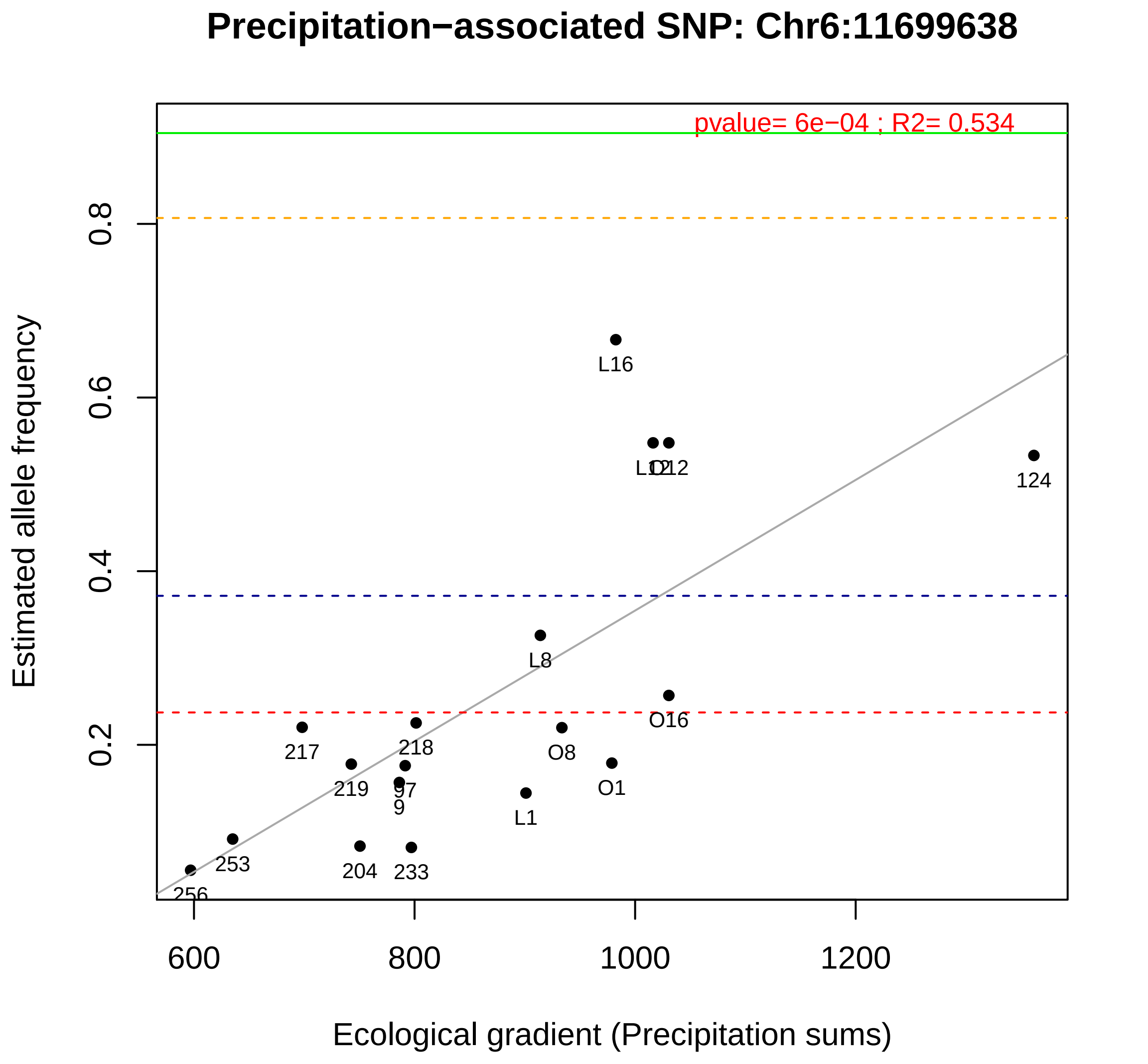

Precipitation−associated SNP: Chr6:11699638
pvalue= 6e−04 ; R2= 0.534
Estimated allele frequency
0.2
0.4
0.6
0.8
600
800
1000
1200
Ecological gradient (Precipitation sums)
L16
L12
O12
124
L8
O16
217
218
O8
219
97
9
O1
L1
253
204
233
256

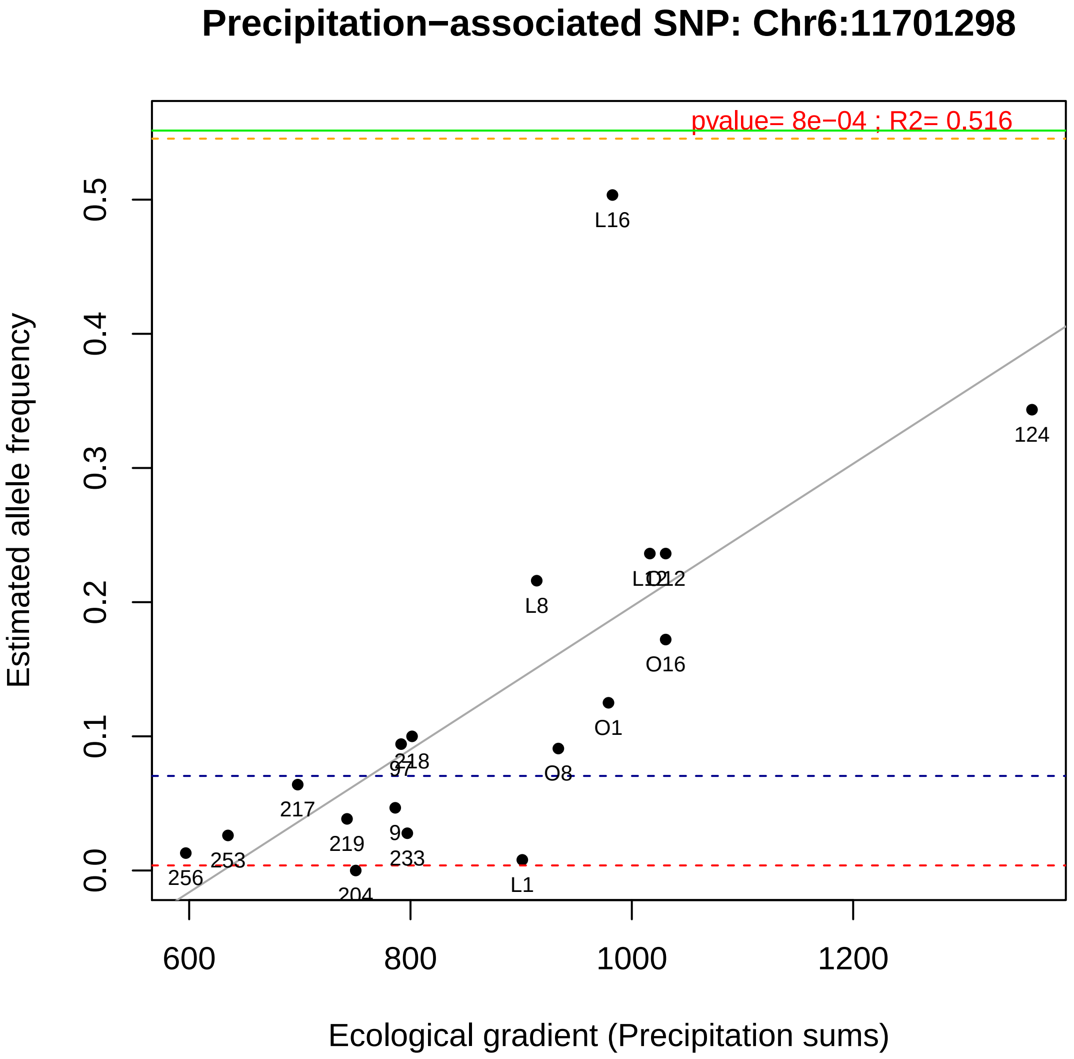

Precipitation−associated SNP: Chr6:11701298
pvalue= 8e−04 ; R2= 0.516
Estimated allele frequency
Ecological gradient (Precipitation sums)
0.0
0.1
0.2
0.3
0.4
0.5
600
800
1000
1200
L16
124
L8
O16
O1
O8
218
217
219
233
253
256
204
L1

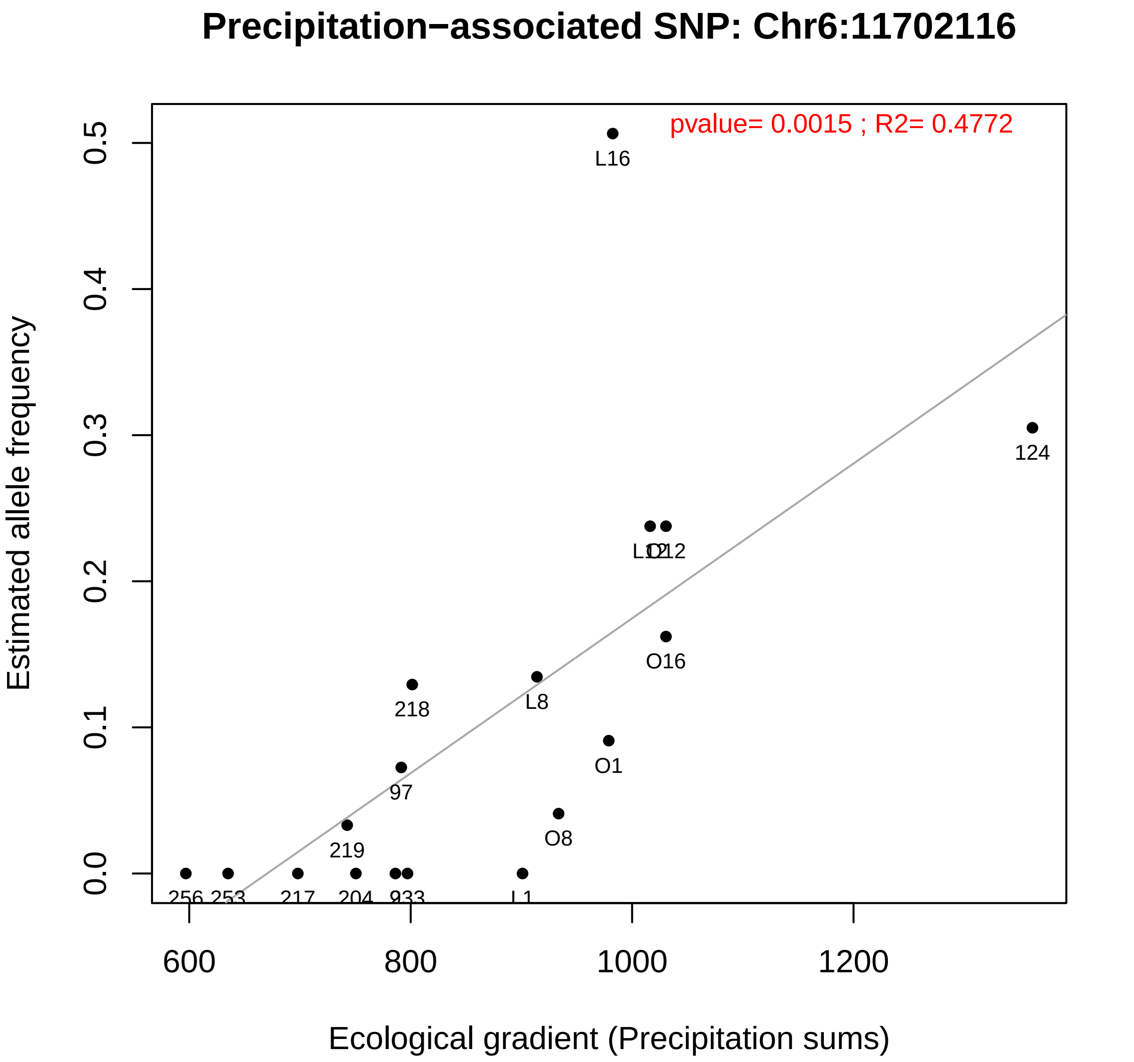

Precipitation−associated SNP: Chr6:11702116
pvalue= 0.0015 ; R2= 0.4772
L16
124
L12
O12
O16
218
L8
O1
97
219
O8
256
253
217
204
933
L1
0.0
0.1
0.2
0.3
0.4
0.5
600
800
1000
1200
Estimated allele frequency
Ecological gradient (Precipitation sums)

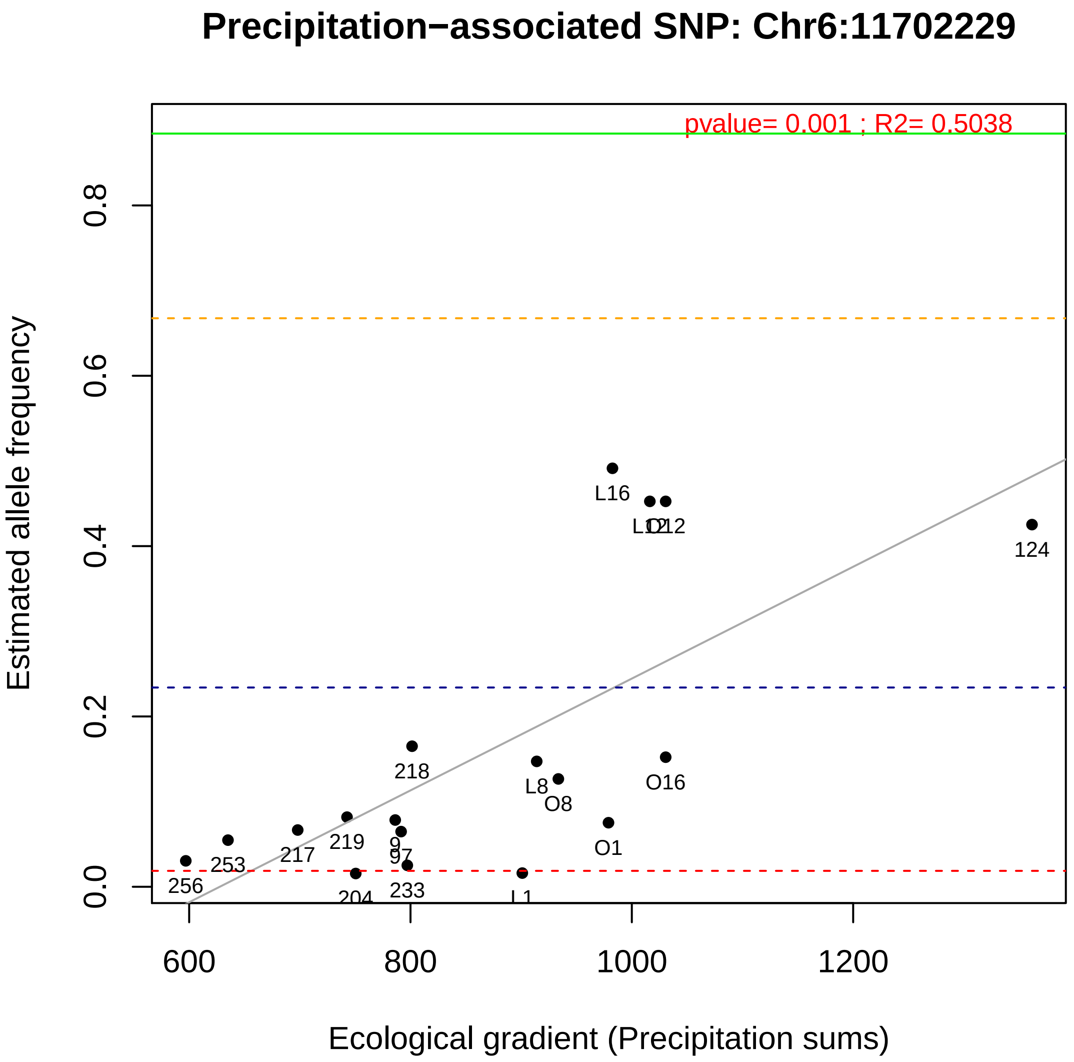
Precipitation−associated SNP: Chr6:11702229
pvalue= 0.001 ; R2= 0.5038
Estimated allele frequency
0.0
0.2
0.4
0.6
0.8
600
800
1000
1200
Ecological gradient (Precipitation sums)
L16
L12
O12
124
218
L8
O8
O16
219
217
253
256
9
97
233
204
O1
L1

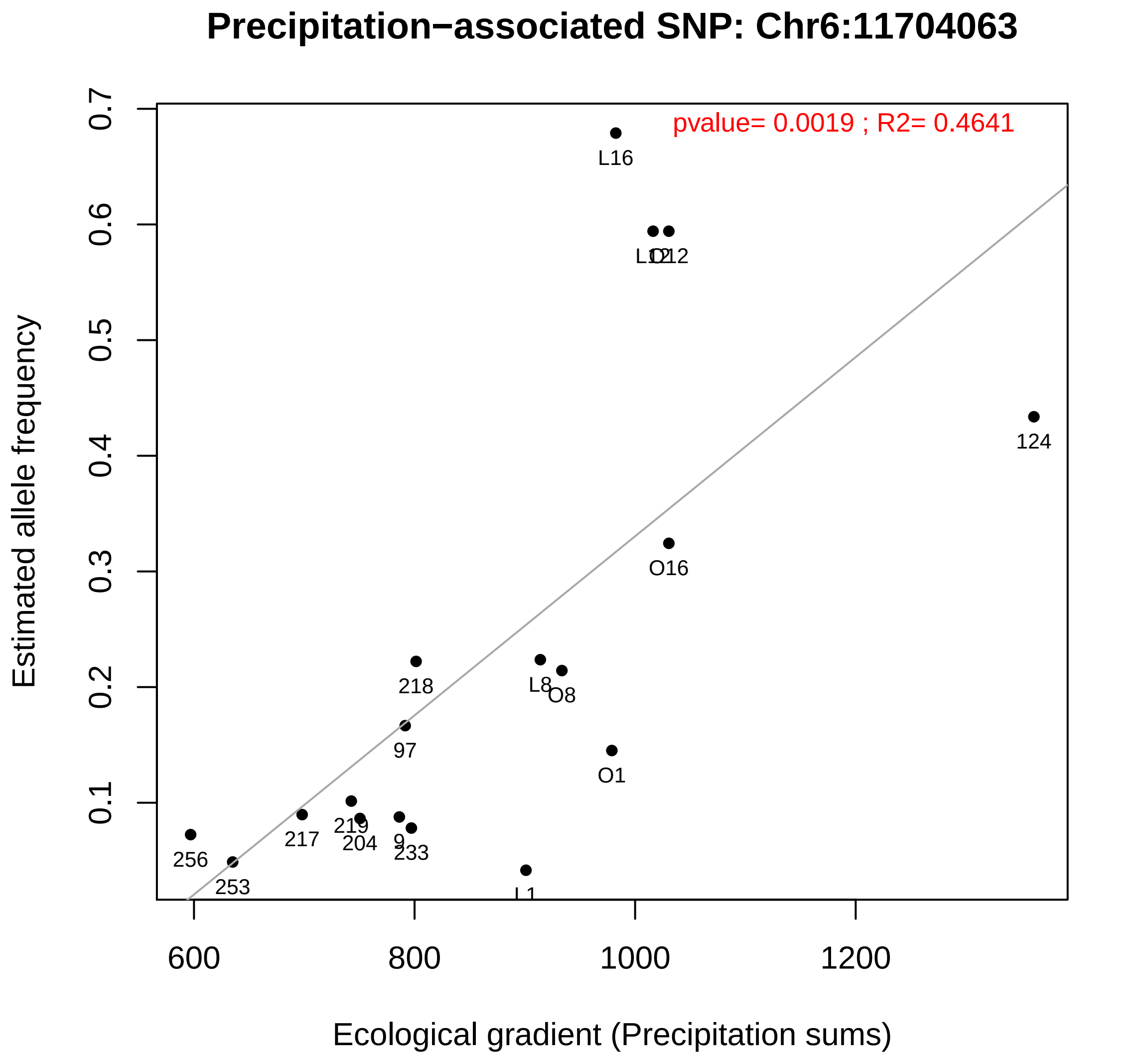

Precipitation−associated SNP: Chr6:11704063
pvalue= 0.0019 ; R2= 0.4641
Estimated allele frequency
Ecological gradient (Precipitation sums)
0.1
0.2
0.3
0.4
0.5
0.6
0.7
600
800
1000
1200
L16
L12
O12
124
O16
218
L8
O8
97
O1
219
204
217
256
253
9
233
L1

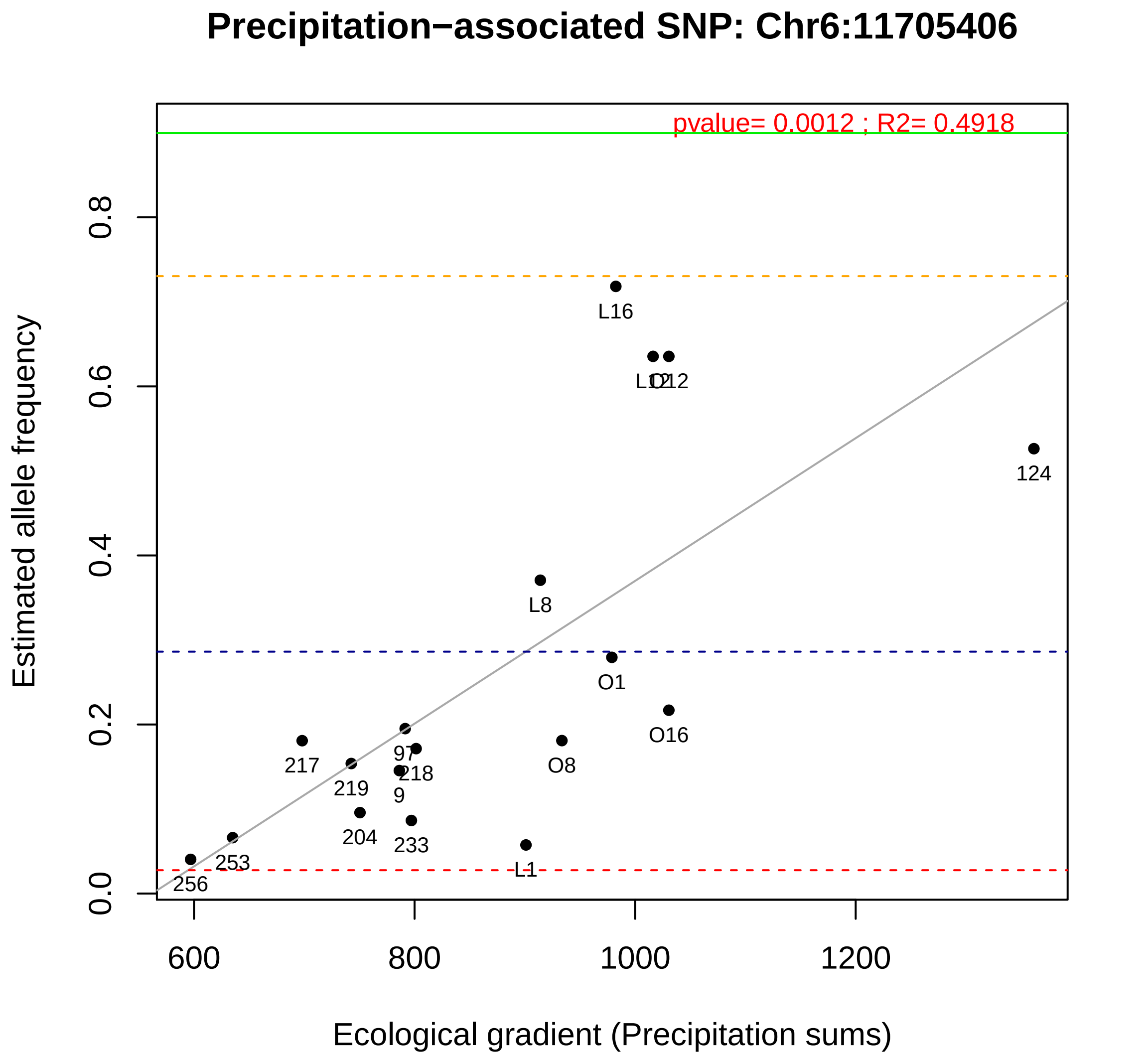

Precipitation−associated SNP: Chr6:11705406
pvalue= 0.0012 ; R2= 0.4918
Estimated allele frequency
0.0
0.2
0.4
0.6
0.8
600
800
1000
1200
Ecological gradient (Precipitation sums)
L16
L12
O12
124
L8
O1
O16
217
97
218
219
9
O8
204
233
253
256
L1

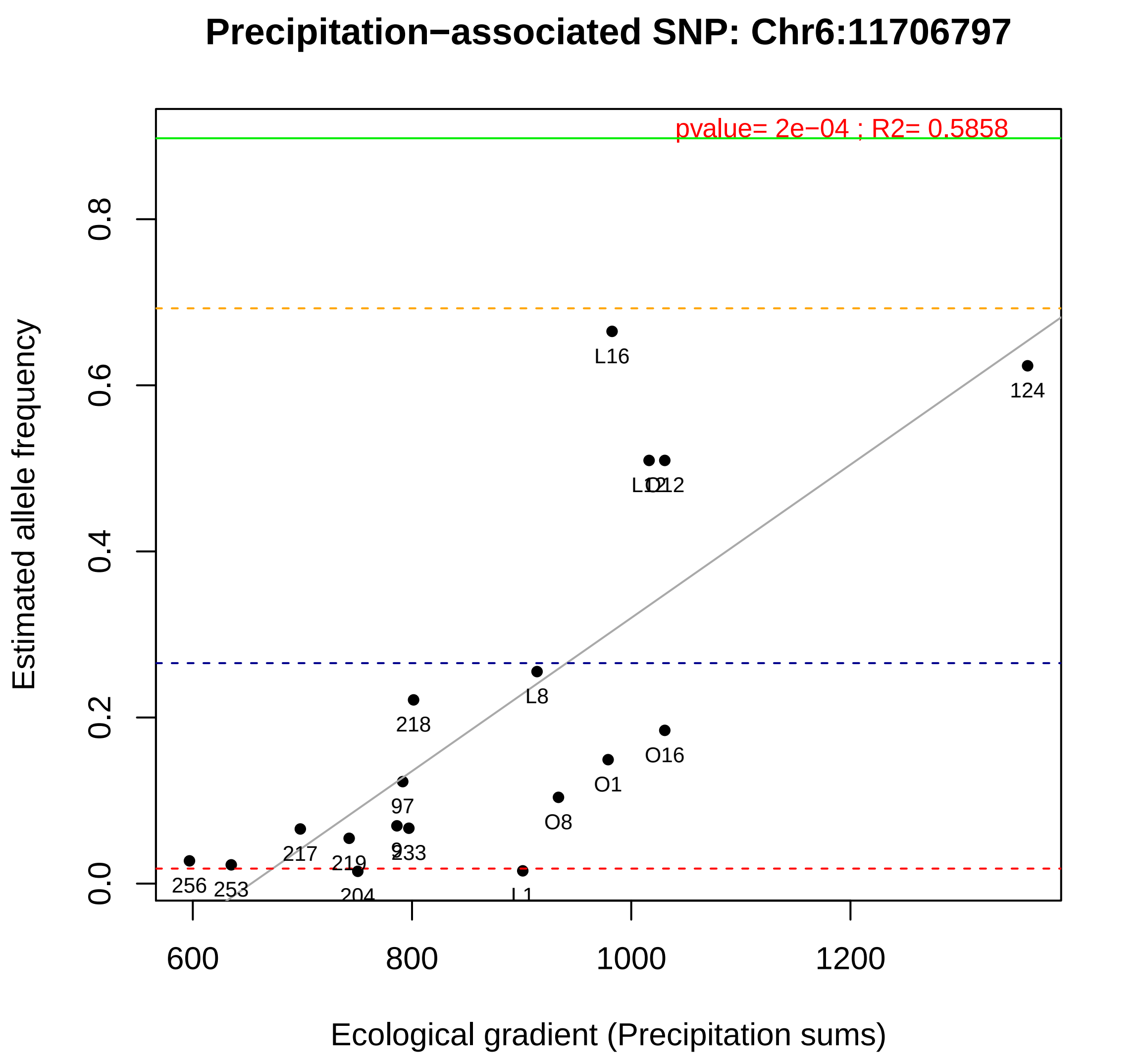

Precipitation−associated SNP: Chr6:11706797
pvalue= 2e−04 ; R2= 0.5858
Estimated allele frequency
0.0
0.2
0.4
0.6
0.8
600
800
1000
1200
Ecological gradient (Precipitation sums)
L16
124
L12
O12
L8
218
O16
O1
97
O8
217
219
9
233
256
253
204
L1

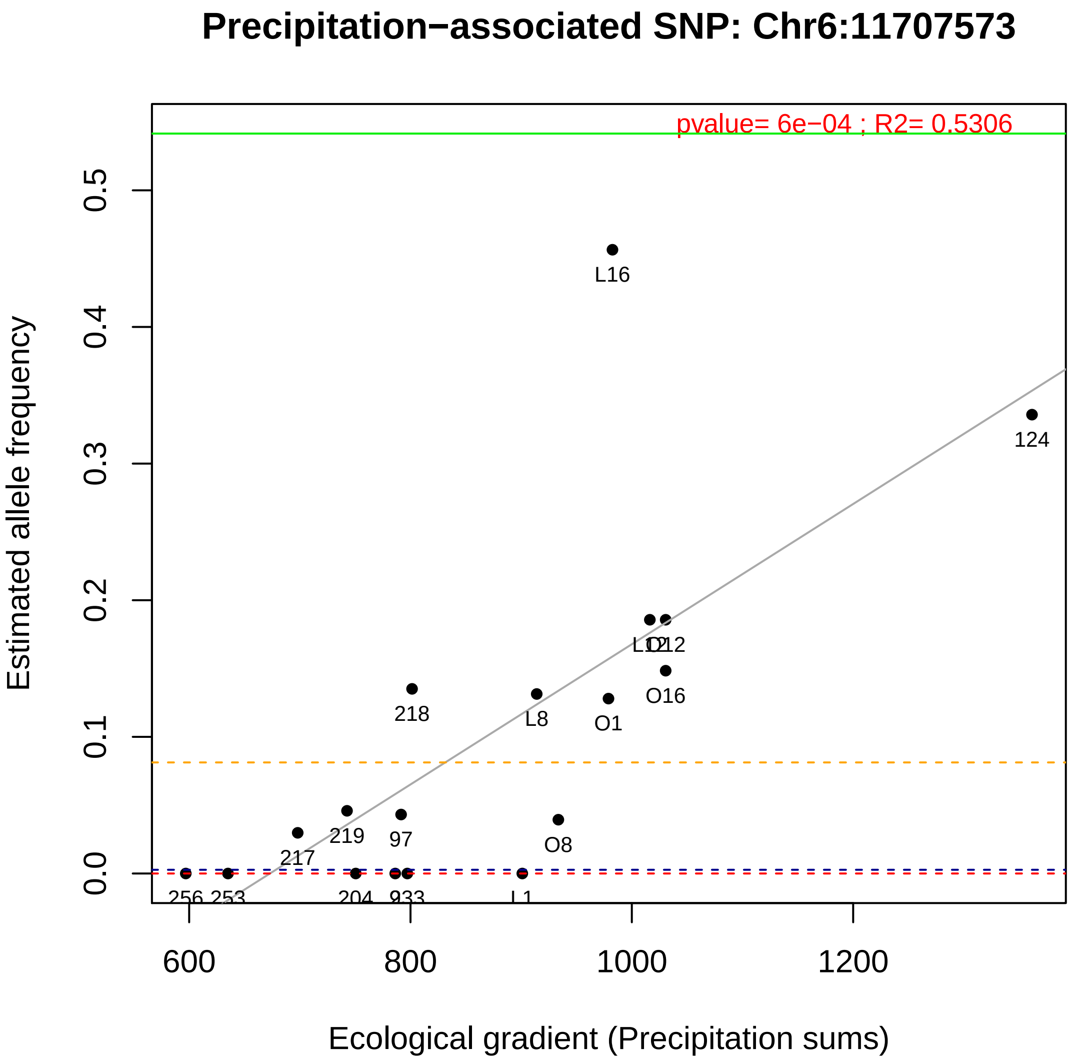

Precipitation−associated SNP: Chr6:11707573
pvalue= 6e−04 ; R2= 0.5306
Estimated allele frequency
Ecological gradient (Precipitation sums)
0.0
0.1
0.2
0.3
0.4
0.5
600
800
1000
1200
L16
124
L12
O12
O16
218
L8
O1
219
97
217
O8

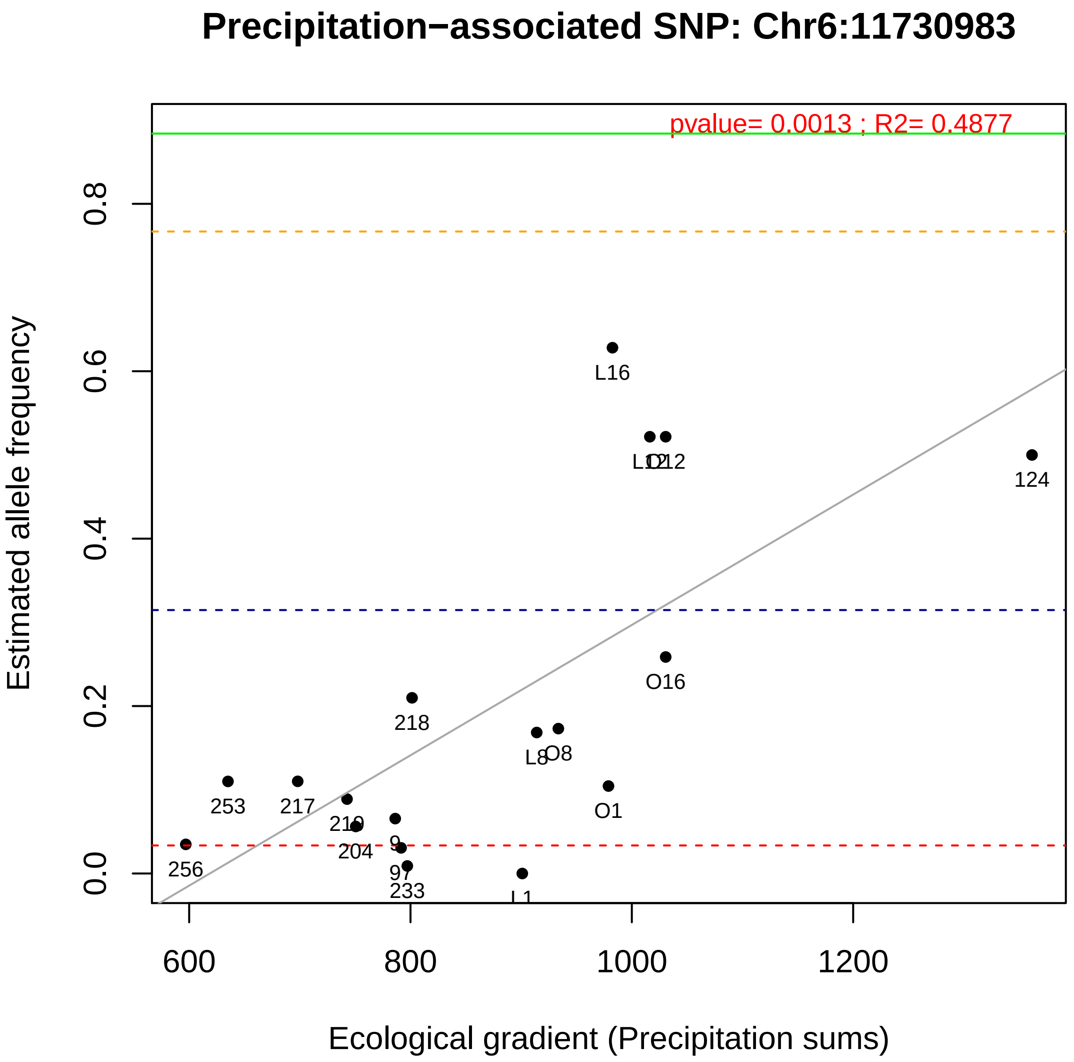

Precipitation−associated SNP: Chr6:11730983
pvalue= 0.0013 ; R2= 0.4877
Estimated allele frequency
0.0
0.2
0.4
0.6
0.8
600
800
1000
1200
Ecological gradient (Precipitation sums)
L16
L12
O12
124
O16
218
L8
O8
O1
253
217
210
204
9
97
233
256
L1

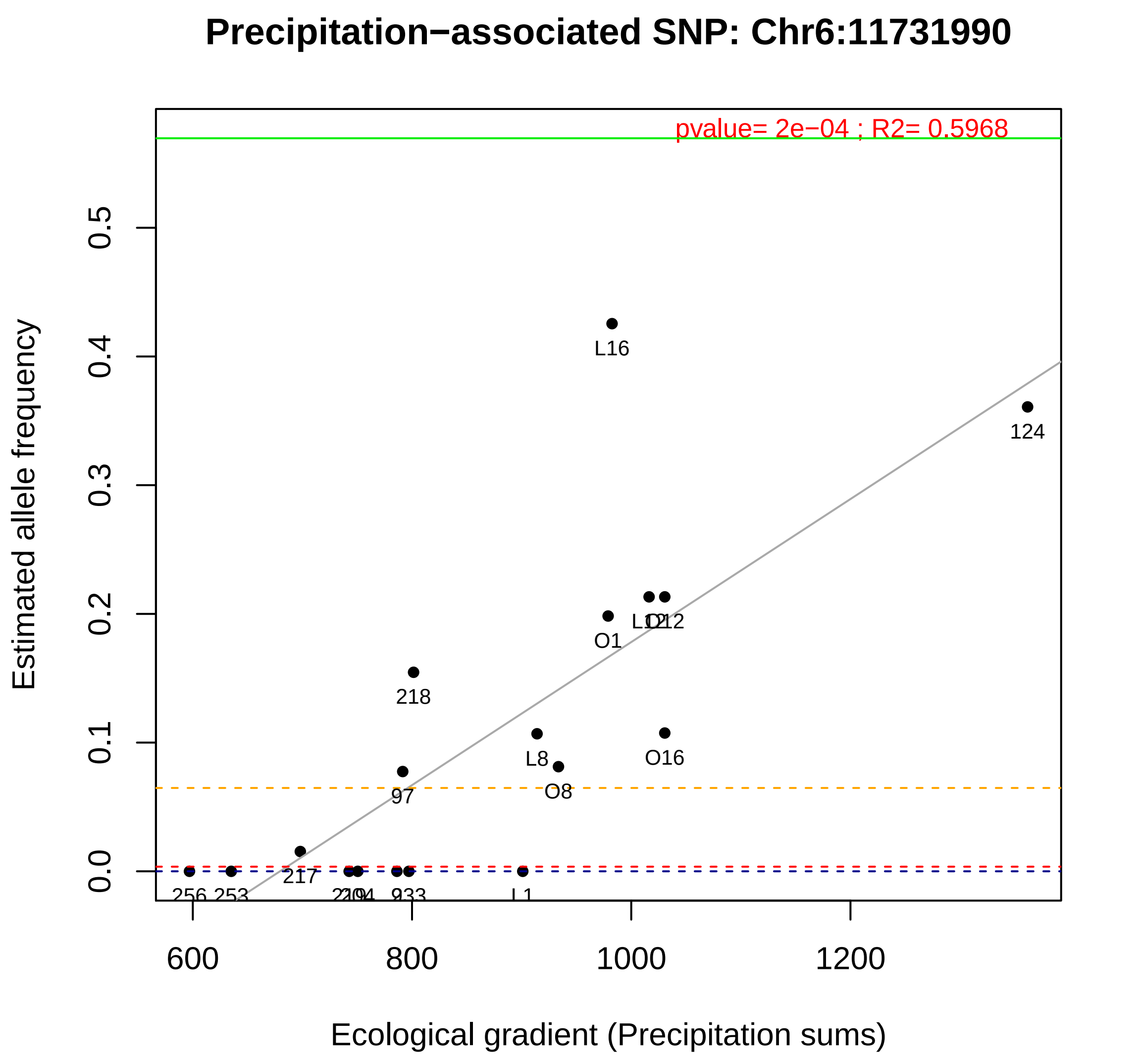

Precipitation−associated SNP: Chr6:11731990
pvalue= 2e−04 ; R2= 0.5968
Estimated allele frequency
0.0
0.1
0.2
0.3
0.4
0.5
Ecological gradient (Precipitation sums)
600
800
1000
1200
L16
124
O1
218
L8
O16
O8
97
217
256
253
L1

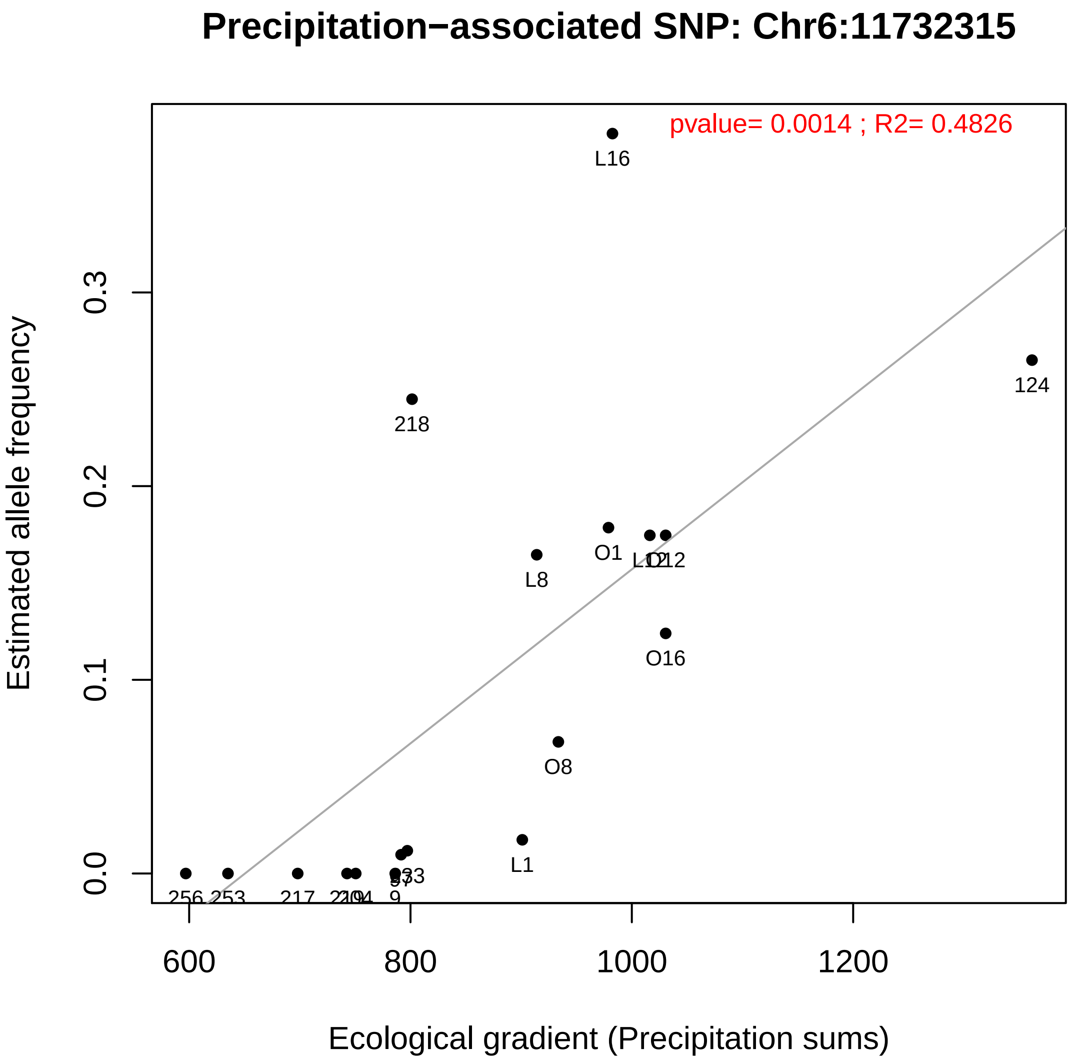

Precipitation−associated SNP: Chr6:11732315
pvalue= 0.0014 ; R2= 0.4826
Estimated allele frequency
Ecological gradient (Precipitation sums)
0.0
0.1
0.2
0.3
600
800
1000
1200
L16
124
218
O1
L8
O16
O8
L1
256
253
217
33
9

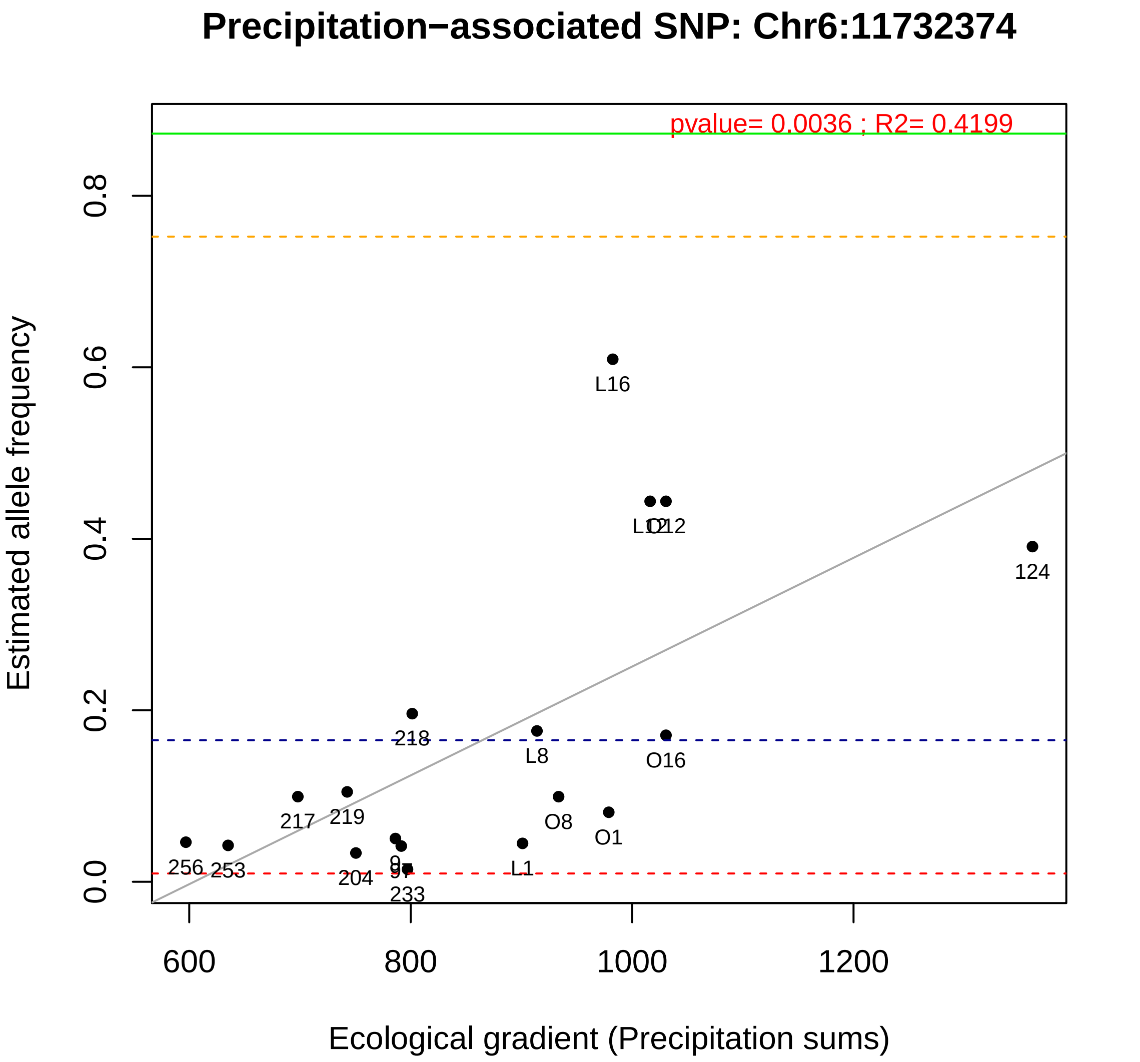
Precipitation−associated SNP: Chr6:11732374
pvalue= 0.0036 ; R2= 0.4199
Estimated allele frequency
0.0
0.2
0.4
0.6
0.8
Ecological gradient (Precipitation sums)
600
800
1000
1200
L16
124
218
L8
O16
217
219
O8
O1
256
253
204
233
L1

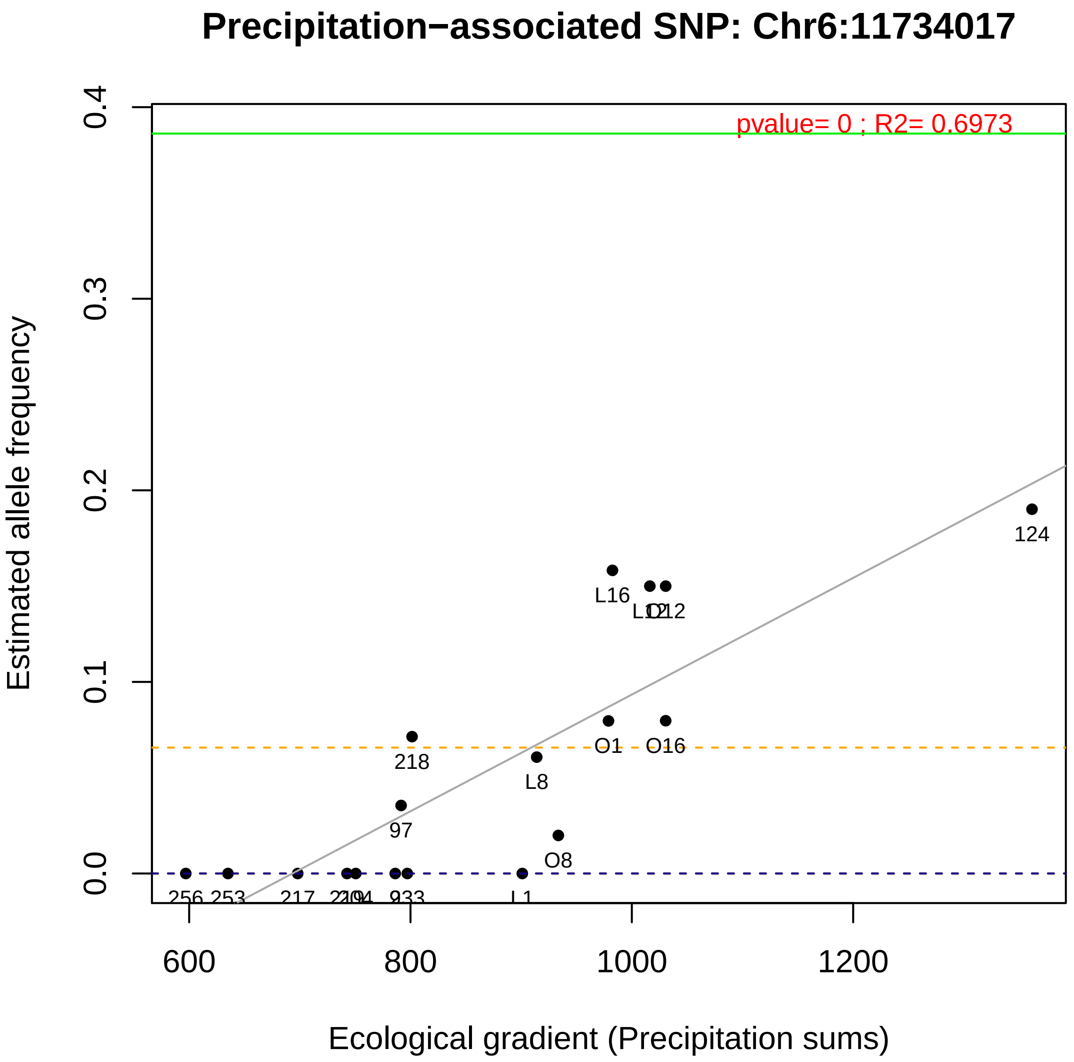

Precipitation−associated SNP: Chr6:11734017
pvalue= 0 ; R2= 0.6973
Estimated allele frequency
0.0
0.1
0.2
0.3
0.4
600
800
1000
1200
Ecological gradient (Precipitation sums)
124
L16
O16
O1
218
L8
97
O8

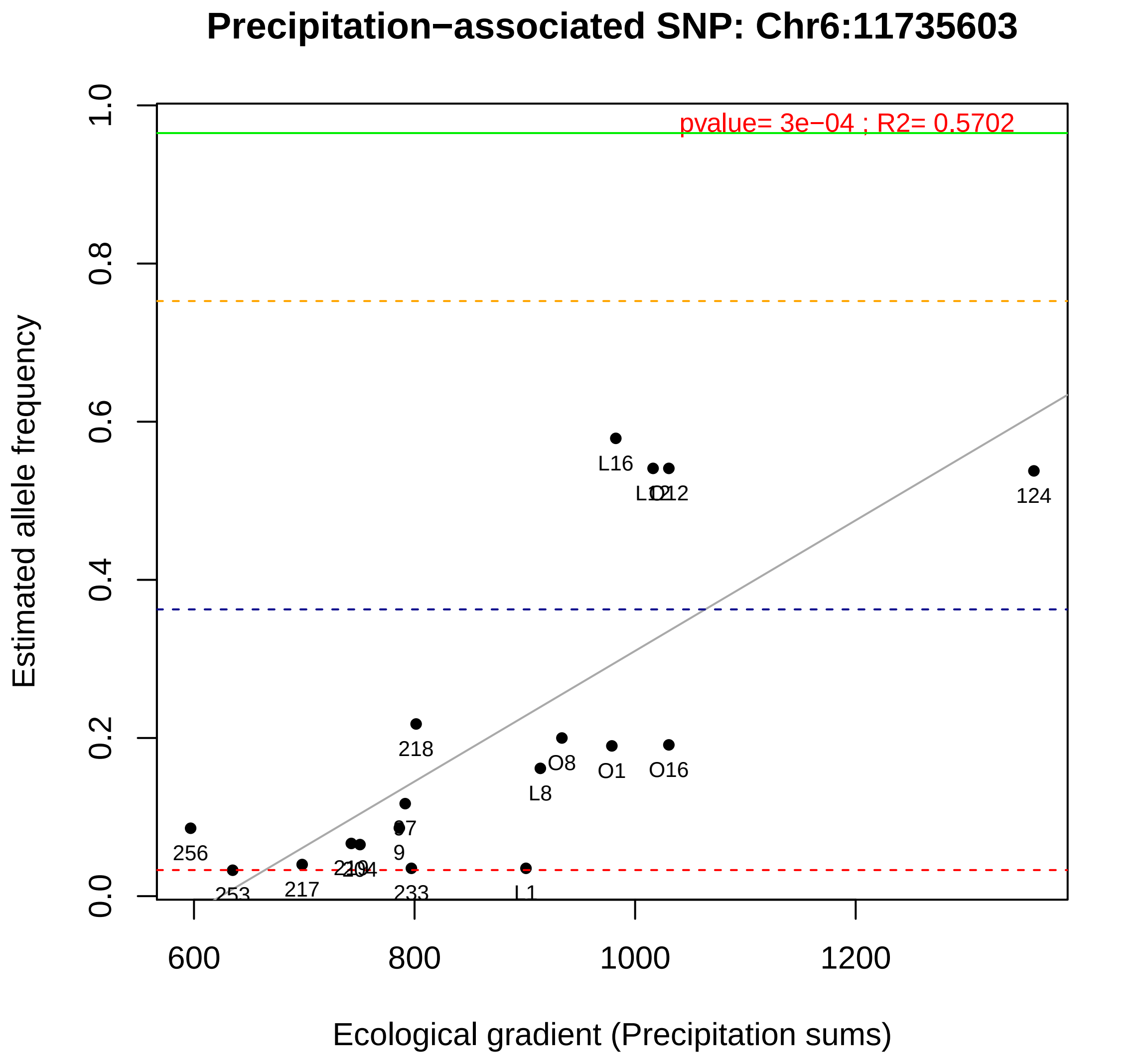

Precipitation−associated SNP: Chr6:11735603
pvalue= 3e−04 ; R2= 0.5702
Estimated allele frequency
0.0
0.2
0.4
0.6
0.8
1.0
Ecological gradient (Precipitation sums)
600
800
1000
1200
L16
L12
O12
124
218
O8
O1
O16
L8
97
9
256
253
217
204
233
L1

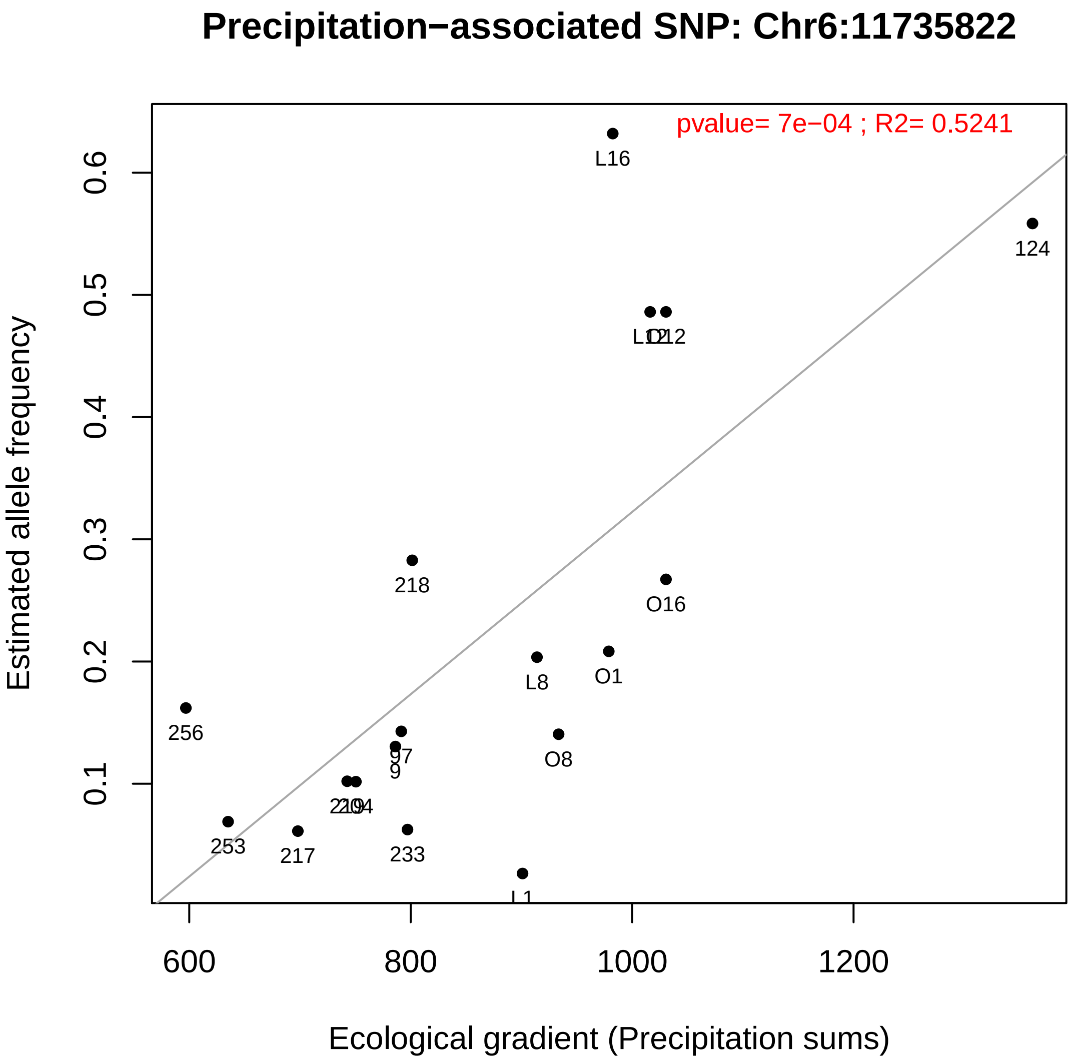
Precipitation−associated SNP: Chr6:11735822
pvalue= 7e−04 ; R2= 0.5241
Estimated allele frequency
0.1
0.2
0.3
0.4
0.5
0.6
600
800
1000
1200
Ecological gradient (Precipitation sums)
L16
124
L12
O12
218
O16
L8
O1
256
O8
97
9
210
204
253
217
233
L1

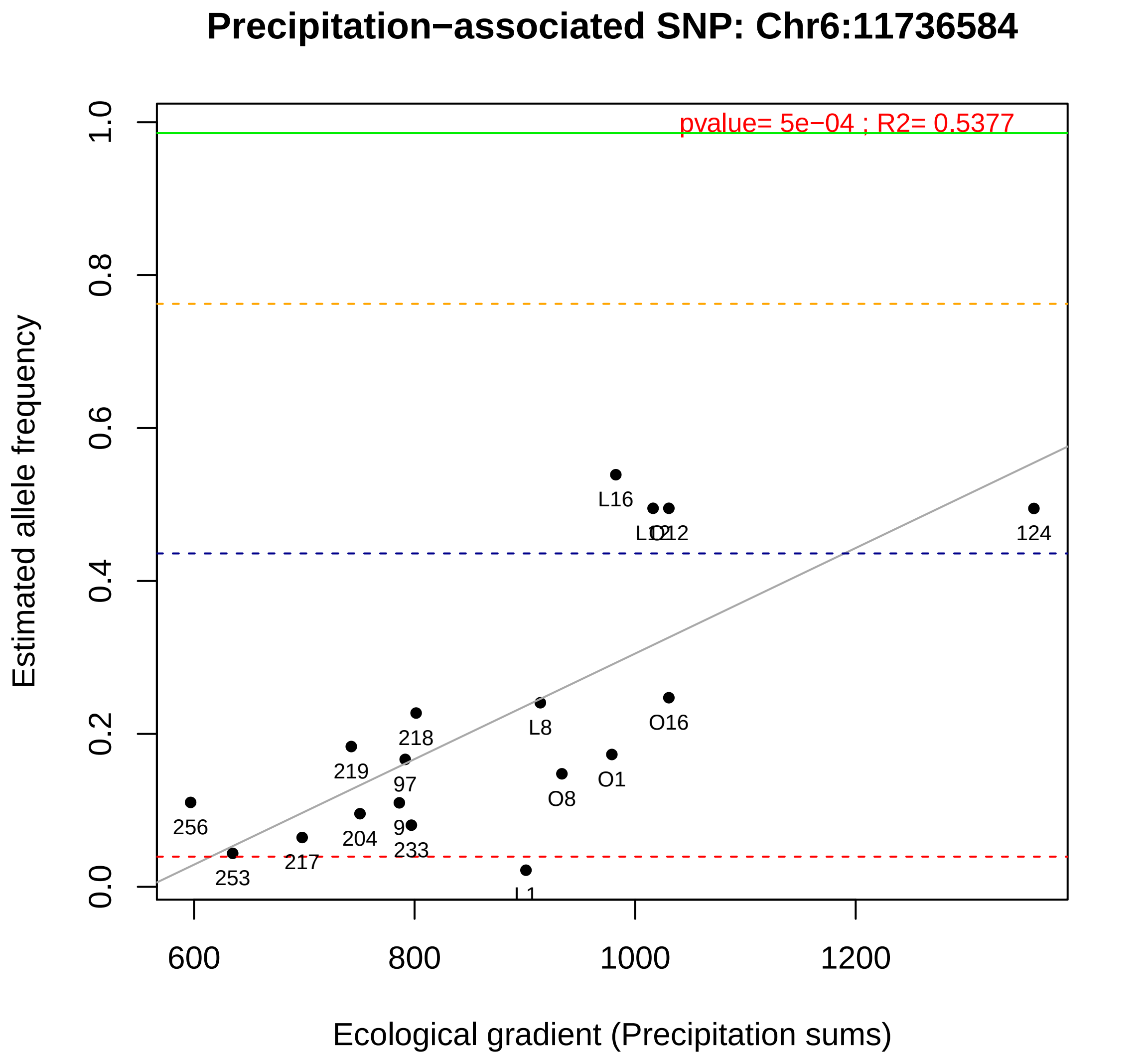

Precipitation−associated SNP: Chr6:11736584
pvalue= 5e−04 ; R2= 0.5377
Estimated allele frequency
0.0
0.2
0.4
0.6
0.8
1.0
600
800
1000
1200
Ecological gradient (Precipitation sums)
L16
124
O16
L8
218
219
97
O1
O8
256
204
217
253
233

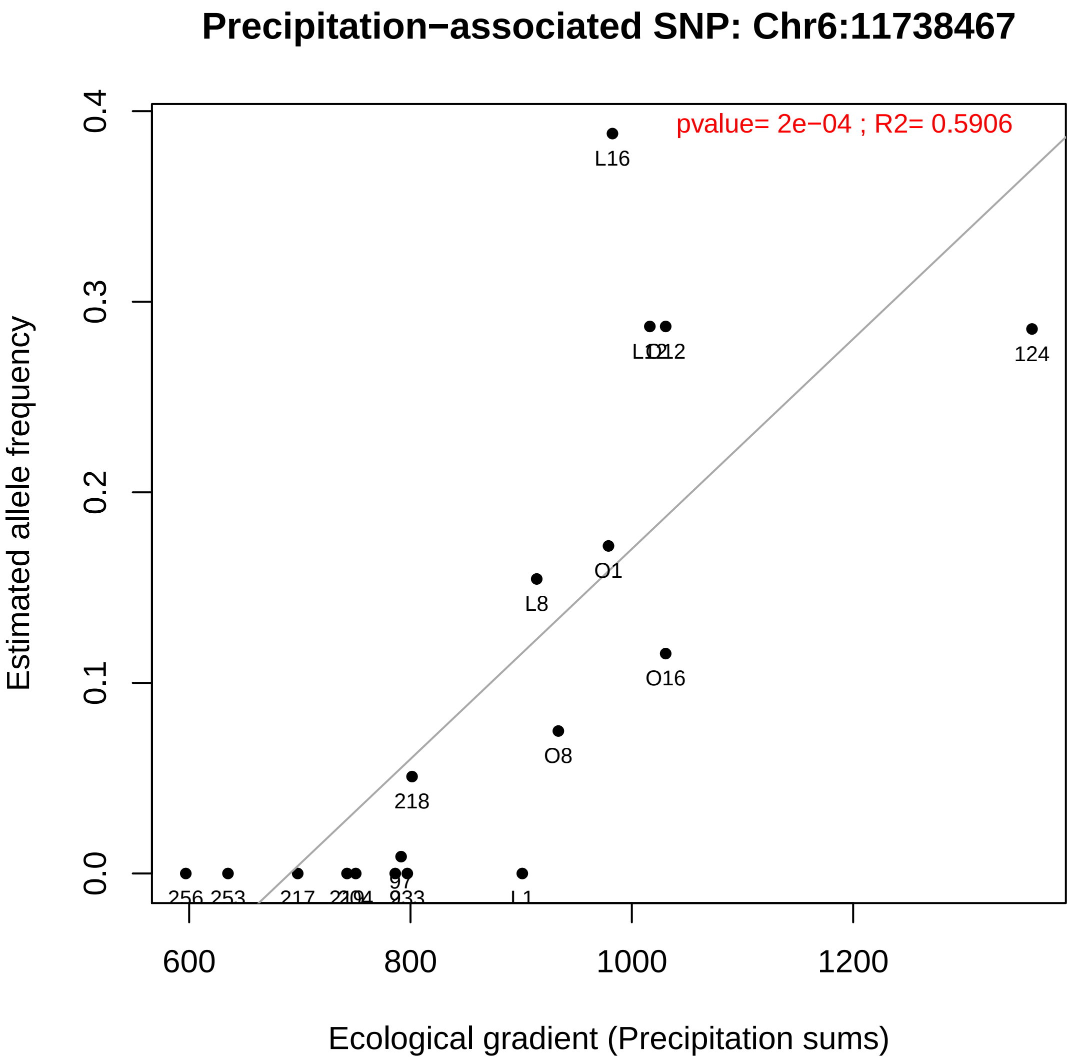

Precipitation−associated SNP: Chr6:11738467
pvalue= 2e−04 ; R2= 0.5906
L16
124
O1
L8
O16
O8
218
97
256
253
217
L1
0.0
0.1
0.2
0.3
0.4
600
800
1000
1200
Estimated allele frequency
Ecological gradient (Precipitation sums)

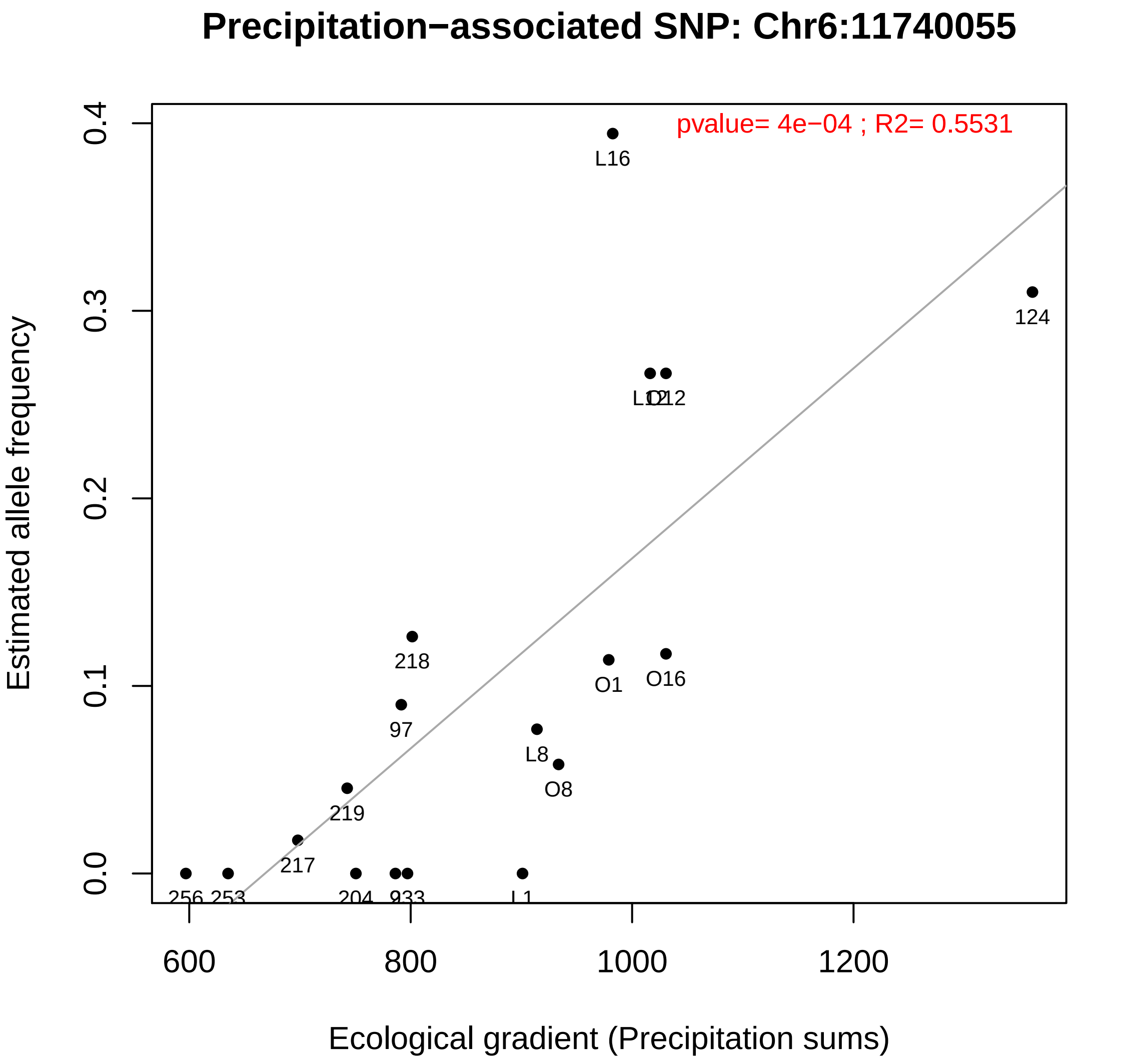
Precipitation−associated SNP: Chr6:11740055
pvalue= 4e−04 ; R2= 0.5531
Estimated allele frequency
Ecological gradient (Precipitation sums)
0.0
0.1
0.2
0.3
0.4
600
800
1000
1200
L16
124
O12
218
O1
O16
97
L8
O8
219
217
256
253
204
L1

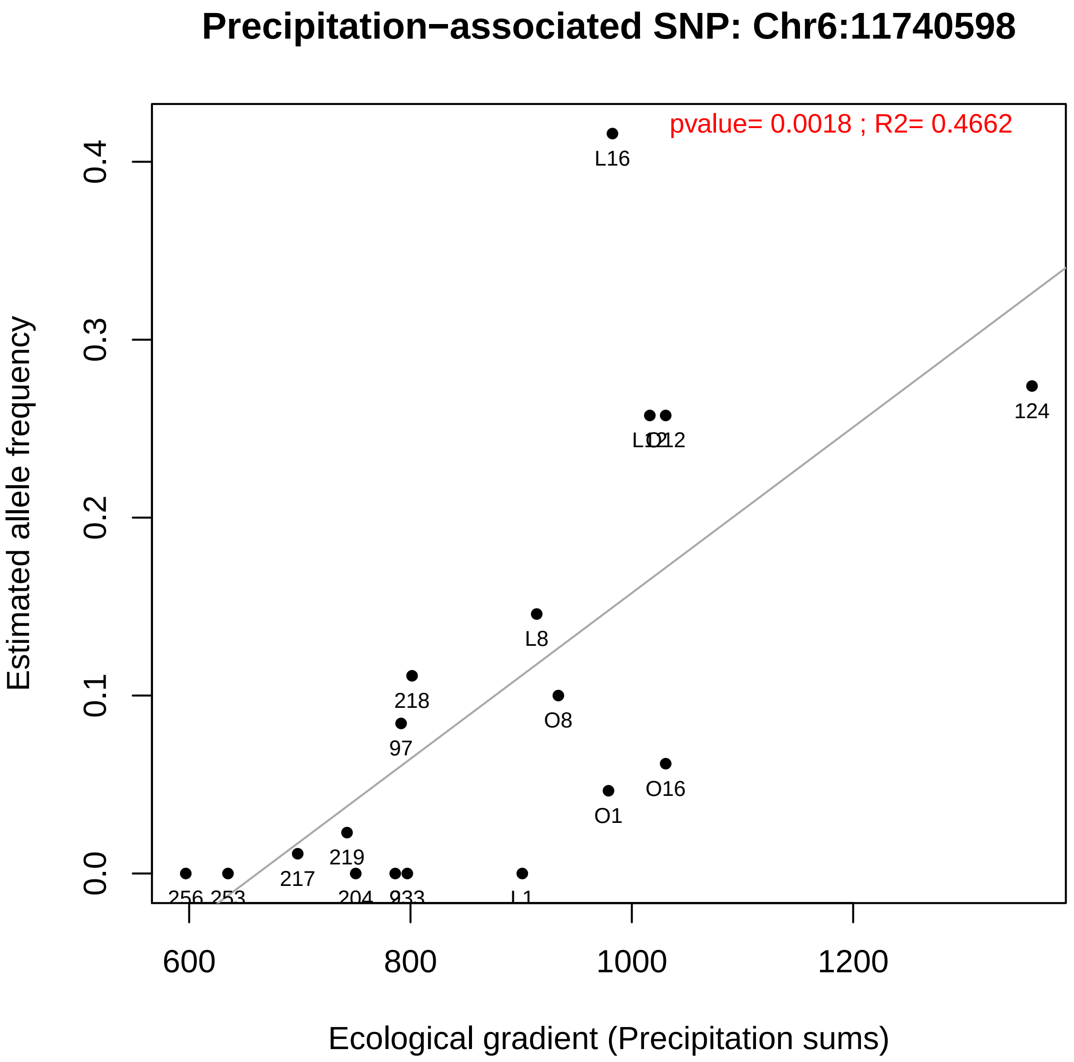
Precipitation−associated SNP: Chr6:11740598
pvalue= 0.0018 ; R2= 0.4662
L16
124
L12
O12
L8
218
O8
97
O16
O1
219
217
256
253
204
9
33
L1
0.0
0.1
0.2
0.3
0.4
600
800
1000
1200
Estimated allele frequency
Ecological gradient (Precipitation sums)

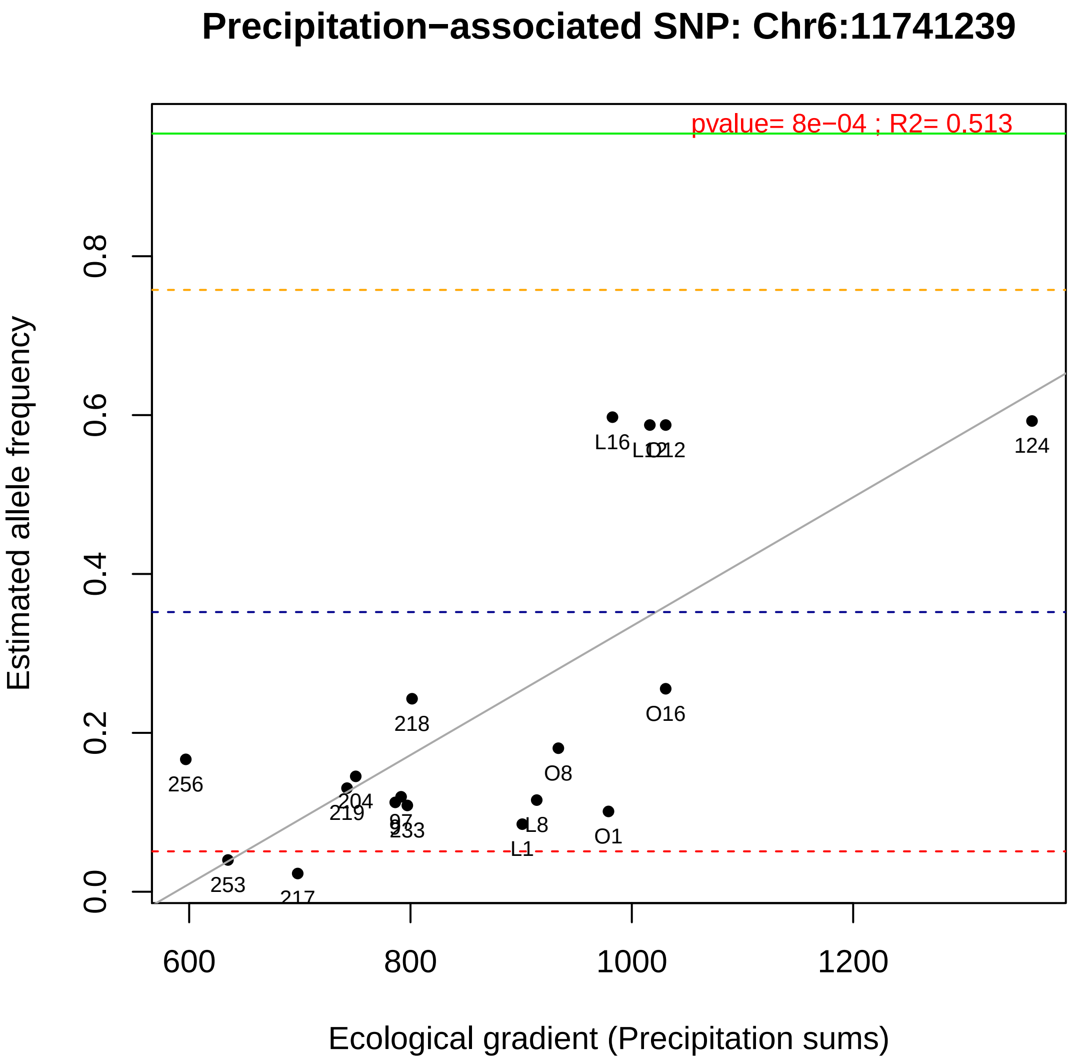

Precipitation−associated SNP: Chr6:11741239
pvalue= 8e−04 ; R2= 0.513
Estimated allele frequency
0.0
0.2
0.4
0.6
0.8
600
800
1000
1200
Ecological gradient (Precipitation sums)
L16
124
218
O16
O8
256
204
219
L8
O1
L1
253
217

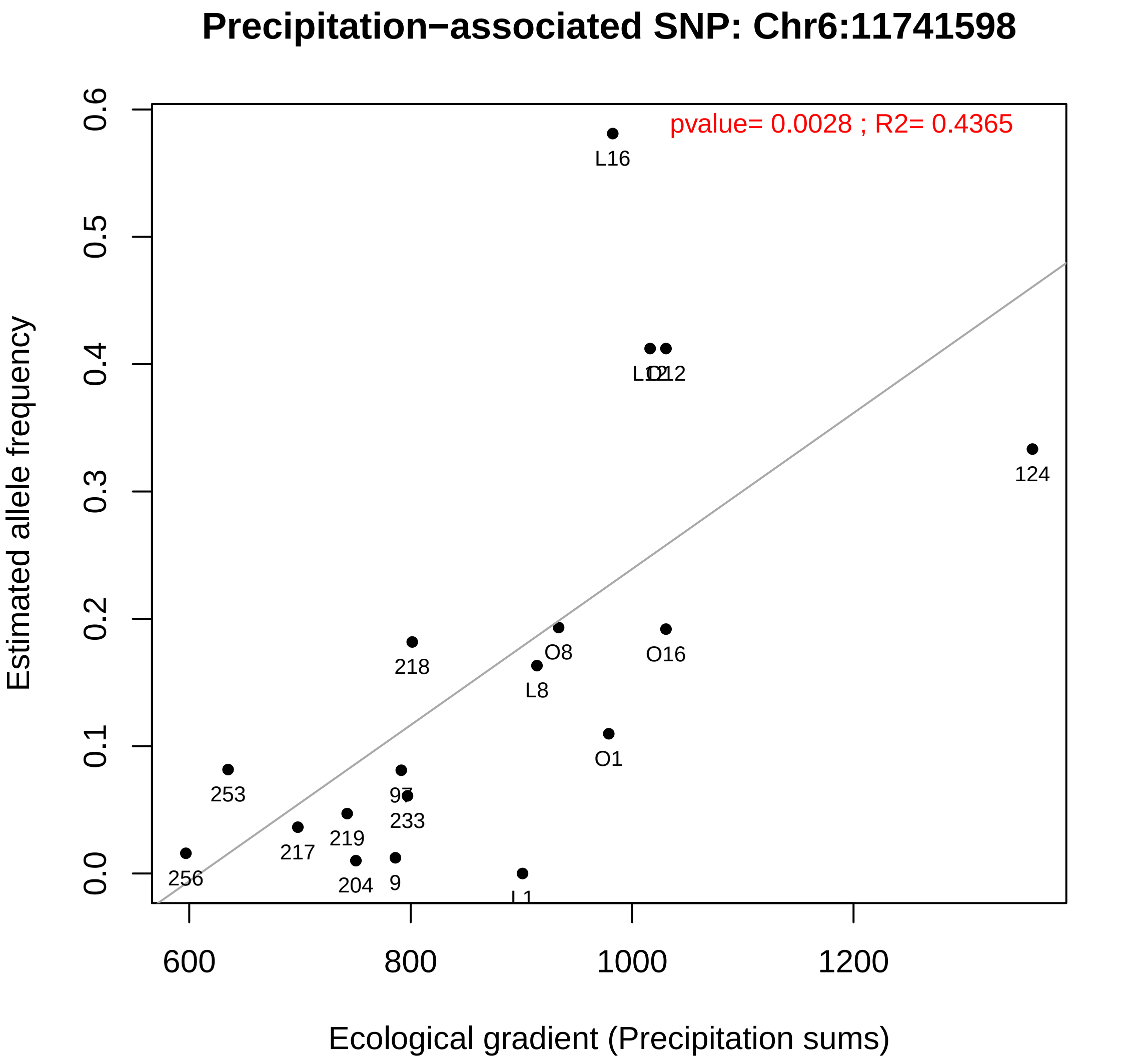
Precipitation−associated SNP: Chr6:11741598
pvalue= 0.0028 ; R2= 0.4365
Estimated allele frequency
0.0
0.1
0.2
0.3
0.4
0.5
0.6
Ecological gradient (Precipitation sums)
600
800
1000
1200
L16
L12
O12
124
218
O8
O16
L8
O1
253
97
233
219
217
256
204
9
L1

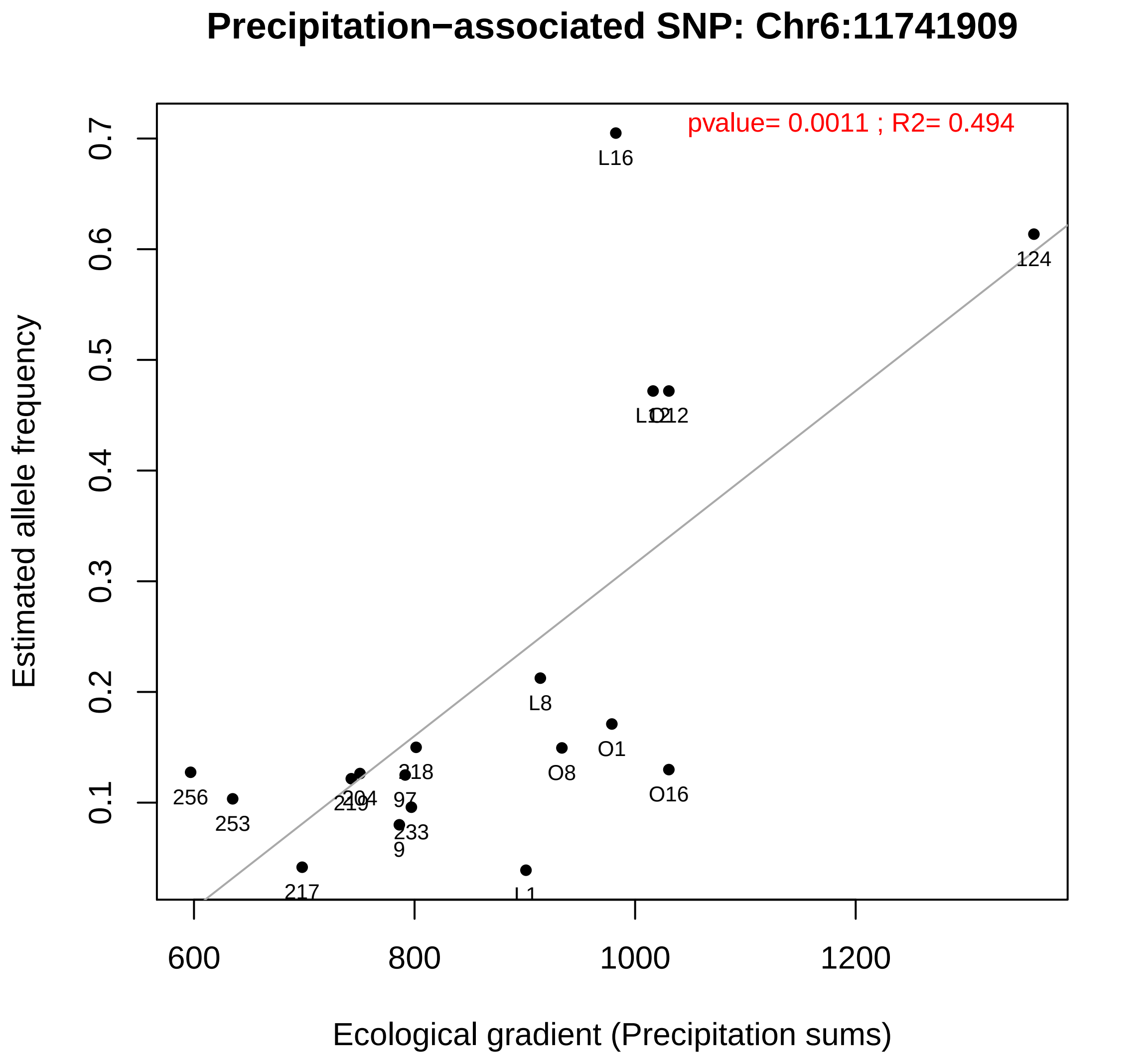
Precipitation−associated SNP: Chr6:11741909
pvalue= 0.0011 ; R2= 0.494
Estimated allele frequency
0.1
0.2
0.3
0.4
0.5
0.6
0.7
600
800
1000
1200
Ecological gradient (Precipitation sums)
L16
124
L12
O12
L8
O1
O8
O16
256
253
204
219
218
97
233
9
217
L1

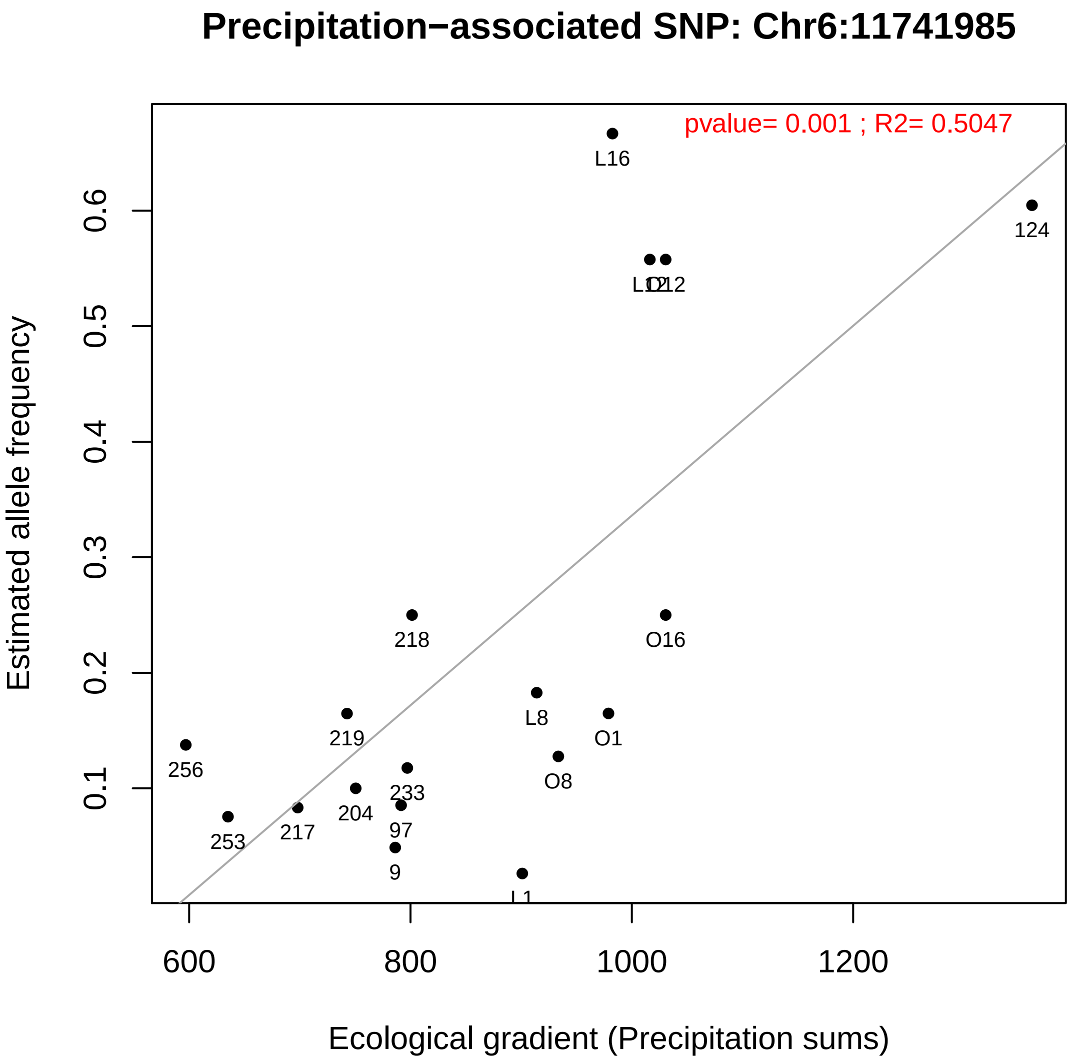
Precipitation−associated SNP: Chr6:11741985
pvalue= 0.001 ; R2= 0.5047
Estimated allele frequency
Ecological gradient (Precipitation sums)
0.1
0.2
0.3
0.4
0.5
0.6
600
800
1000
1200
L16
124
O12
O16
218
L8
O1
219
256
O8
233
204
97
217
253
9

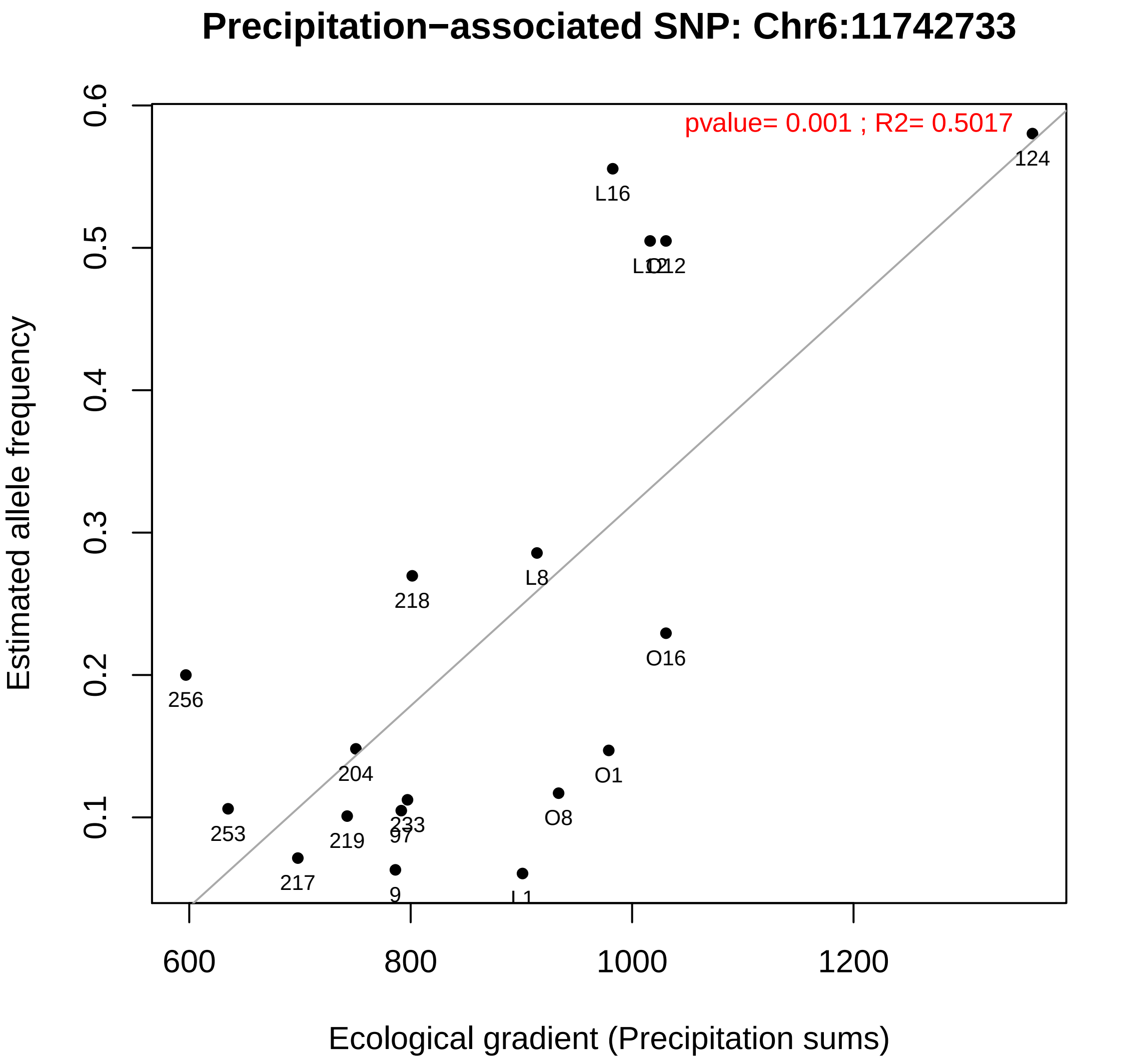
Precipitation−associated SNP: Chr6:11742733
pvalue= 0.001 ; R2= 0.5017
Estimated allele frequency
0.1
0.2
0.3
0.4
0.5
0.6
600
800
1000
1200
Ecological gradient (Precipitation sums)
124
L16
218
L8
O16
256
204
O1
O8
253
219
233
97
217
9

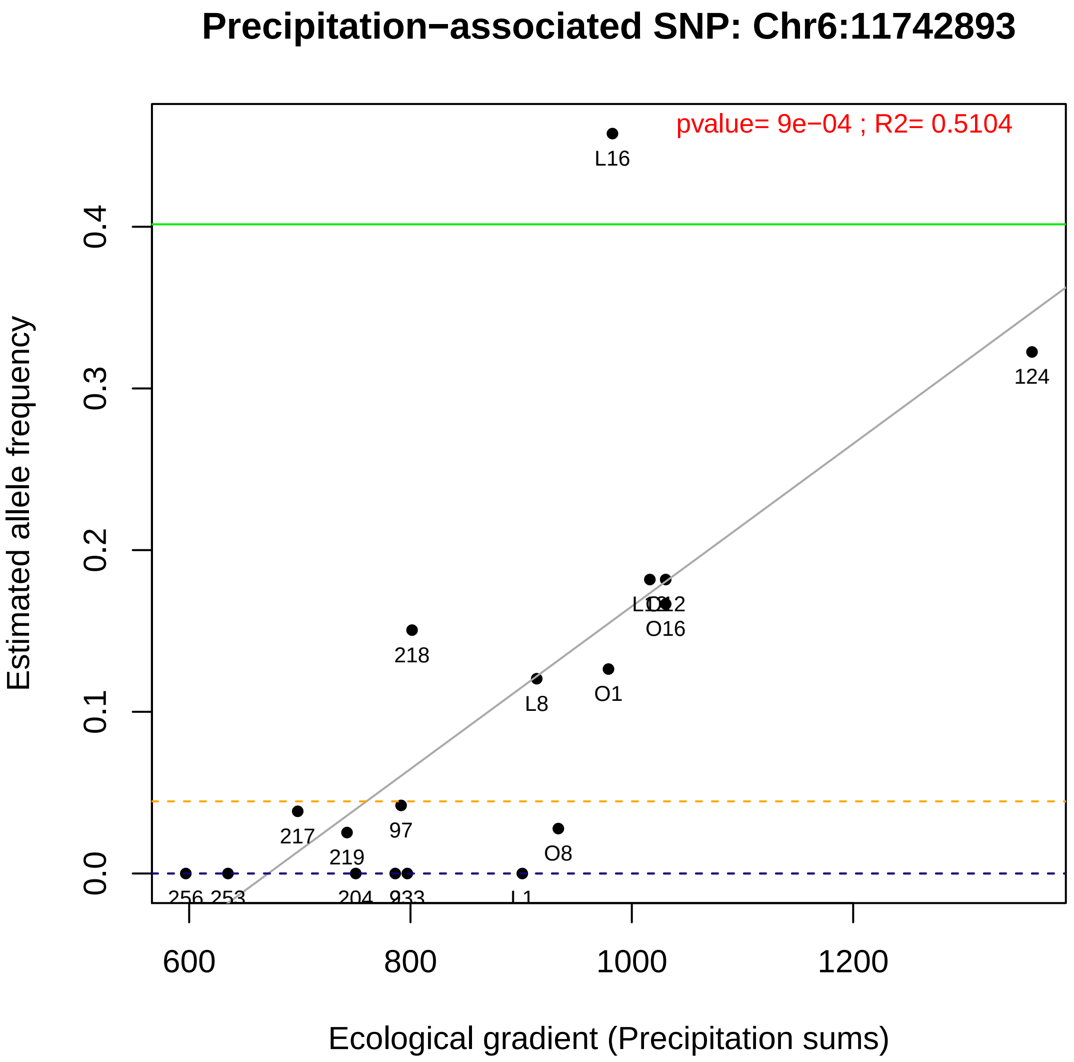

Precipitation−associated SNP: Chr6:11742893
pvalue= 9e−04 ; R2= 0.5104
L16
124
218
L8
O1
O16
217
219
97
O8
0.0
0.1
0.2
0.3
0.4
600
800
1000
1200
Estimated allele frequency
Ecological gradient (Precipitation sums)

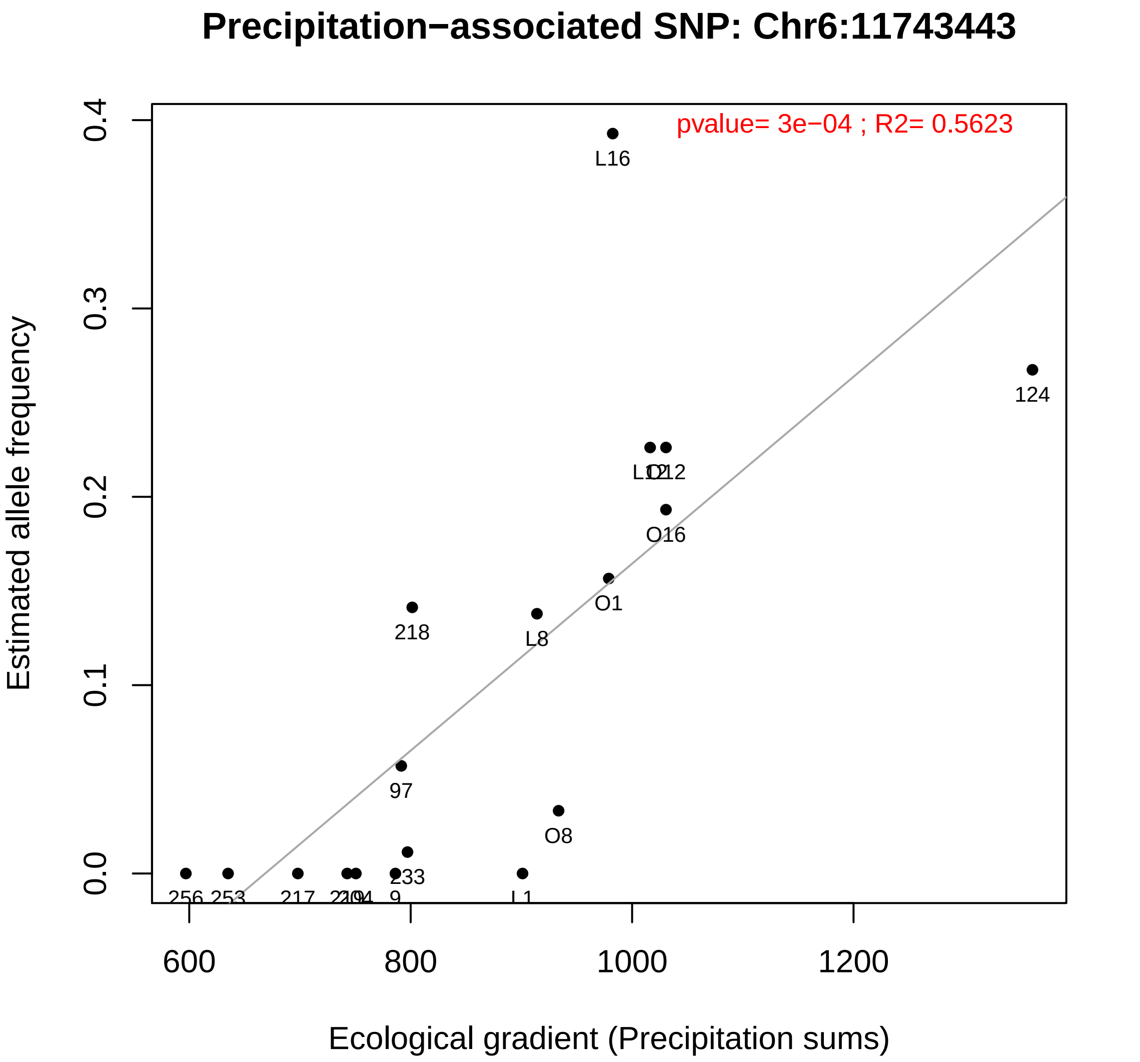
Precipitation−associated SNP: Chr6:11743443
pvalue= 3e−04 ; R2= 0.5623
Estimated allele frequency
Ecological gradient (Precipitation sums)
0.0
0.1
0.2
0.3
0.4
600
800
1000
1200
L16
124
O12
O16
O1
218
L8
97
O8
233
256
253
217
9
L1

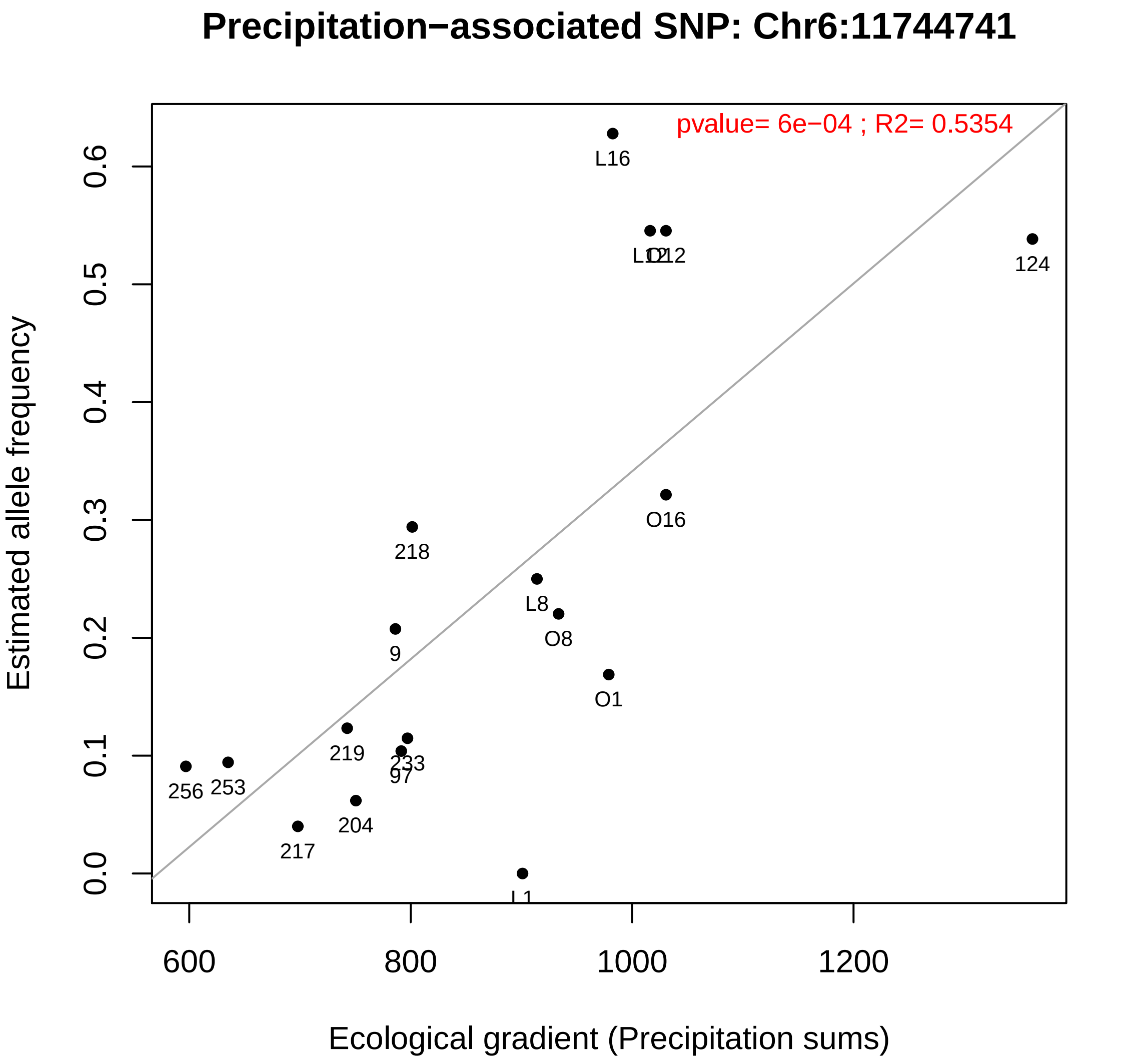
Precipitation−associated SNP: Chr6:11744741
pvalue= 6e−04 ; R2= 0.5354
L16
L12
O12
124
O16
218
L8
O8
9
O1
219
233
97
256
253
204
217
L1
0.0
0.1
0.2
0.3
0.4
0.5
0.6
600
800
1000
1200
Estimated allele frequency
Ecological gradient (Precipitation sums)

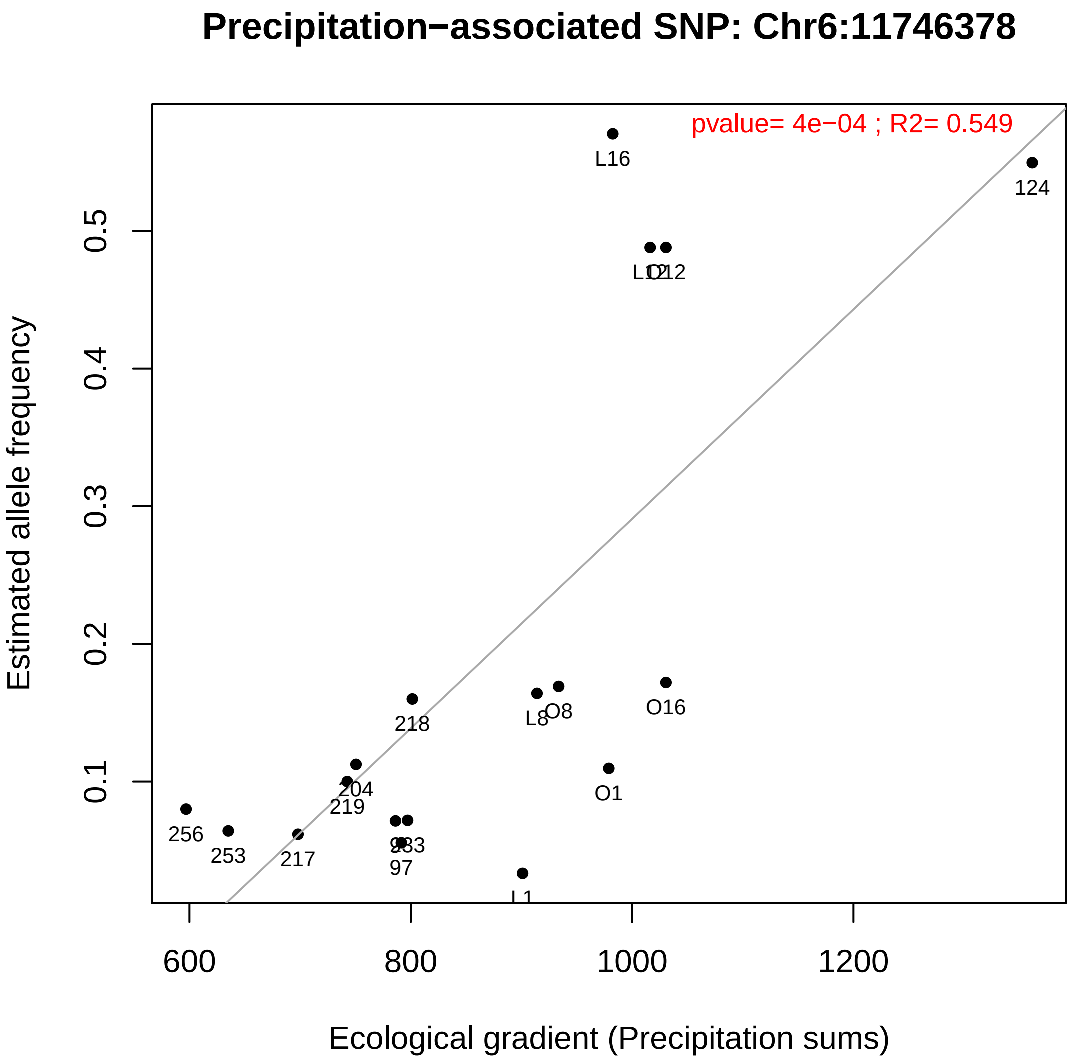

Precipitation−associated SNP: Chr6:11746378
pvalue= 4e−04 ; R2= 0.549
L16
124
L12
O12
218
L8
O8
O16
204
219
O1
256
253
217
233
97
L1
0.1
0.2
0.3
0.4
0.5
Estimated allele frequency
600
800
1000
1200
Ecological gradient (Precipitation sums)

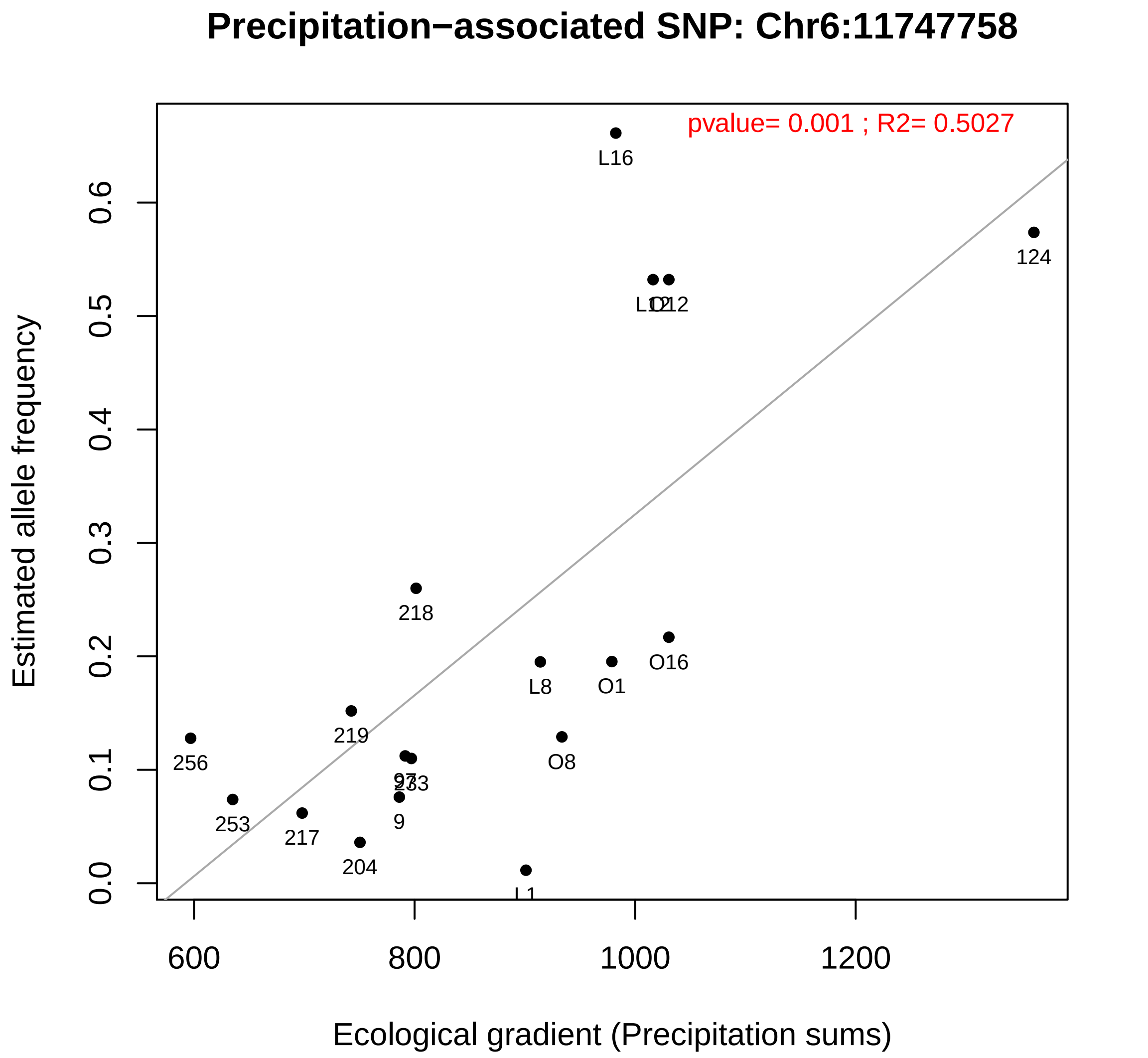

Precipitation−associated SNP: Chr6:11747758
pvalue= 0.001 ; R2= 0.5027
Estimated allele frequency
Ecological gradient (Precipitation sums)
0.0
0.1
0.2
0.3
0.4
0.5
0.6
600
800
1000
1200
L16
124
L12
O12
218
O16
L8
O1
219
256
O8
97
233
253
217
9
204
L1

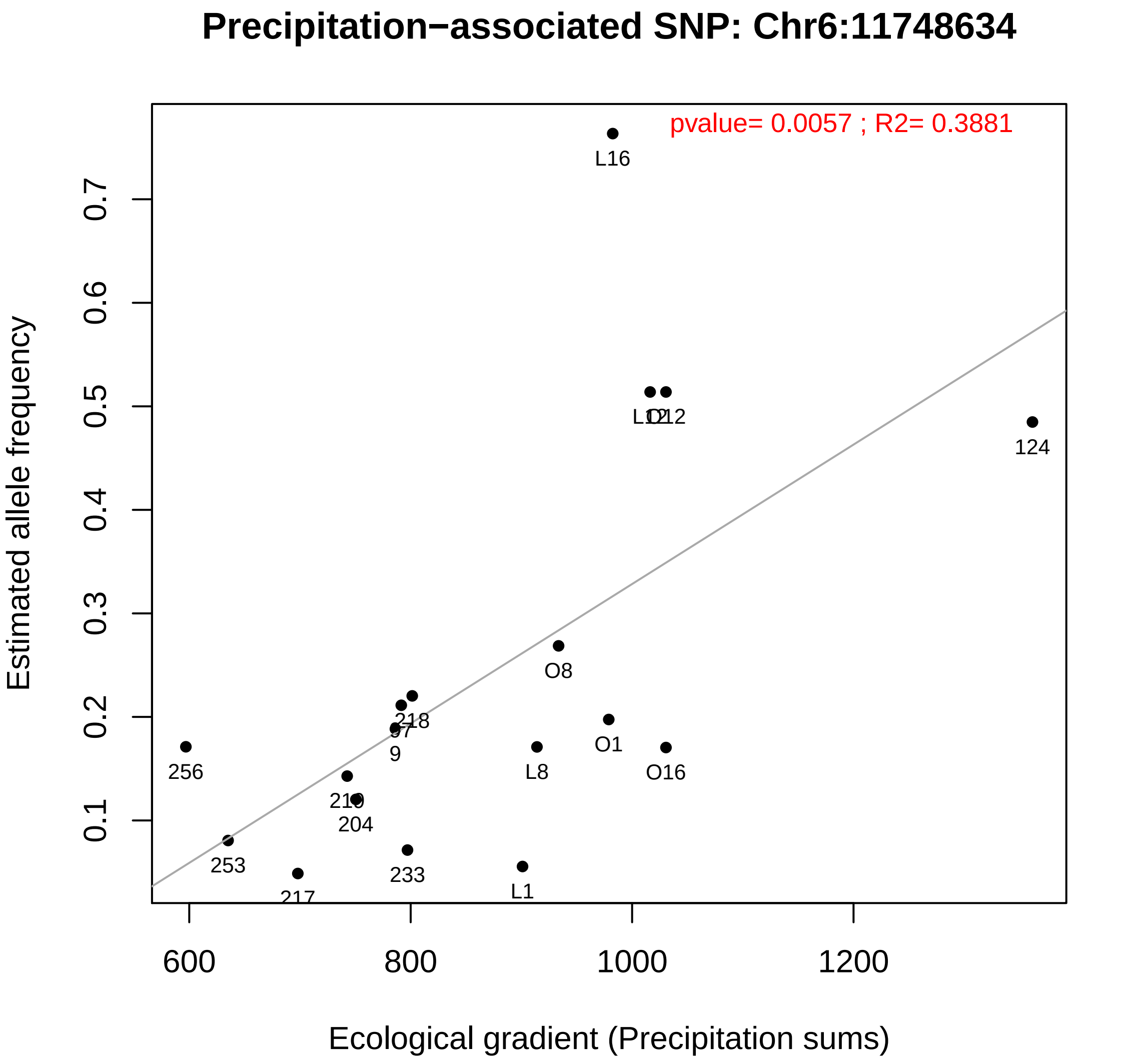
Precipitation−associated SNP: Chr6:11748634
pvalue= 0.0057 ; R2= 0.3881
Estimated allele frequency
Ecological gradient (Precipitation sums)
0.1
0.2
0.3
0.4
0.5
0.6
0.7
600
800
1000
1200
L16
124
O8
218
256
L8
O1
O16
204
253
217
233
L1

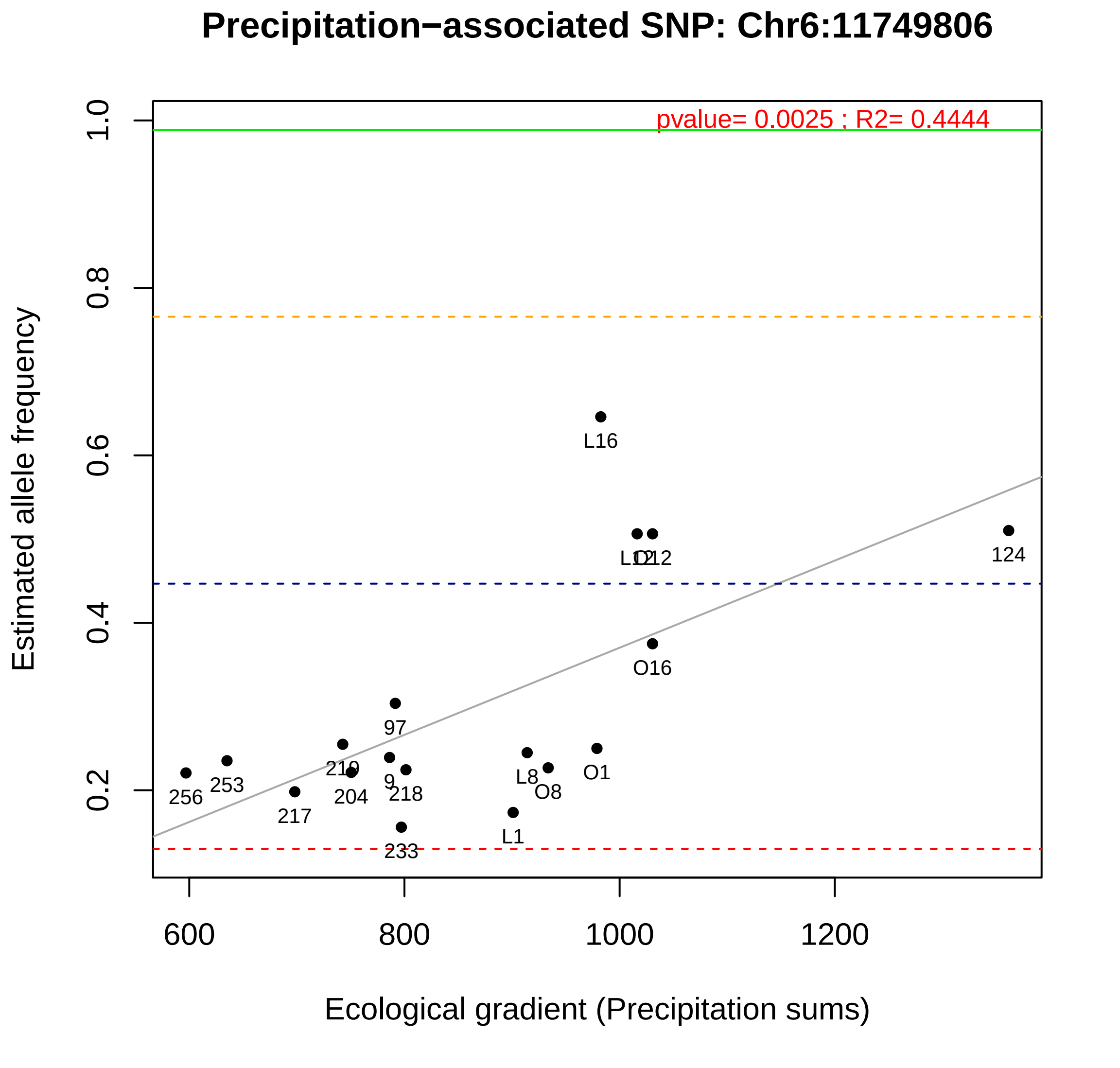
Precipitation−associated SNP: Chr6:11749806
pvalue= 0.0025 ; R2= 0.4444
Estimated allele frequency
Ecological gradient (Precipitation sums)
1.0
0.8
0.6
0.4
0.2
600
800
1000
1200
L16
L12
O12
124
O16
97
210
9
204
218
253
256
217
233
L8
O8
O1
L1

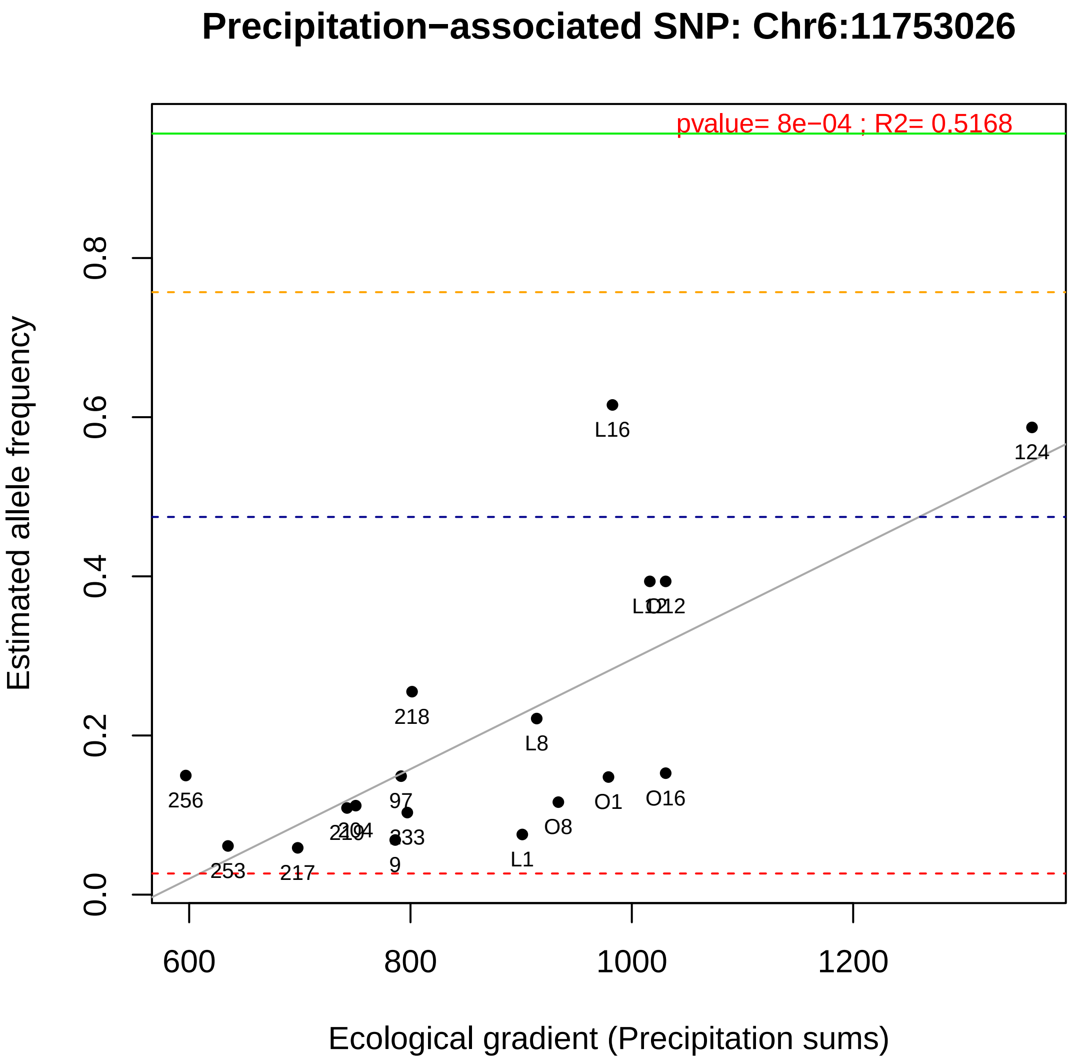

Precipitation−associated SNP: Chr6:11753026
pvalue= 8e−04 ; R2= 0.5168
Estimated allele frequency
0.0
0.2
0.4
0.6
0.8
600
800
1000
1200
Ecological gradient (Precipitation sums)
L16
124
218
L8
256
97
253
217
9
L1
O8
O1
O16

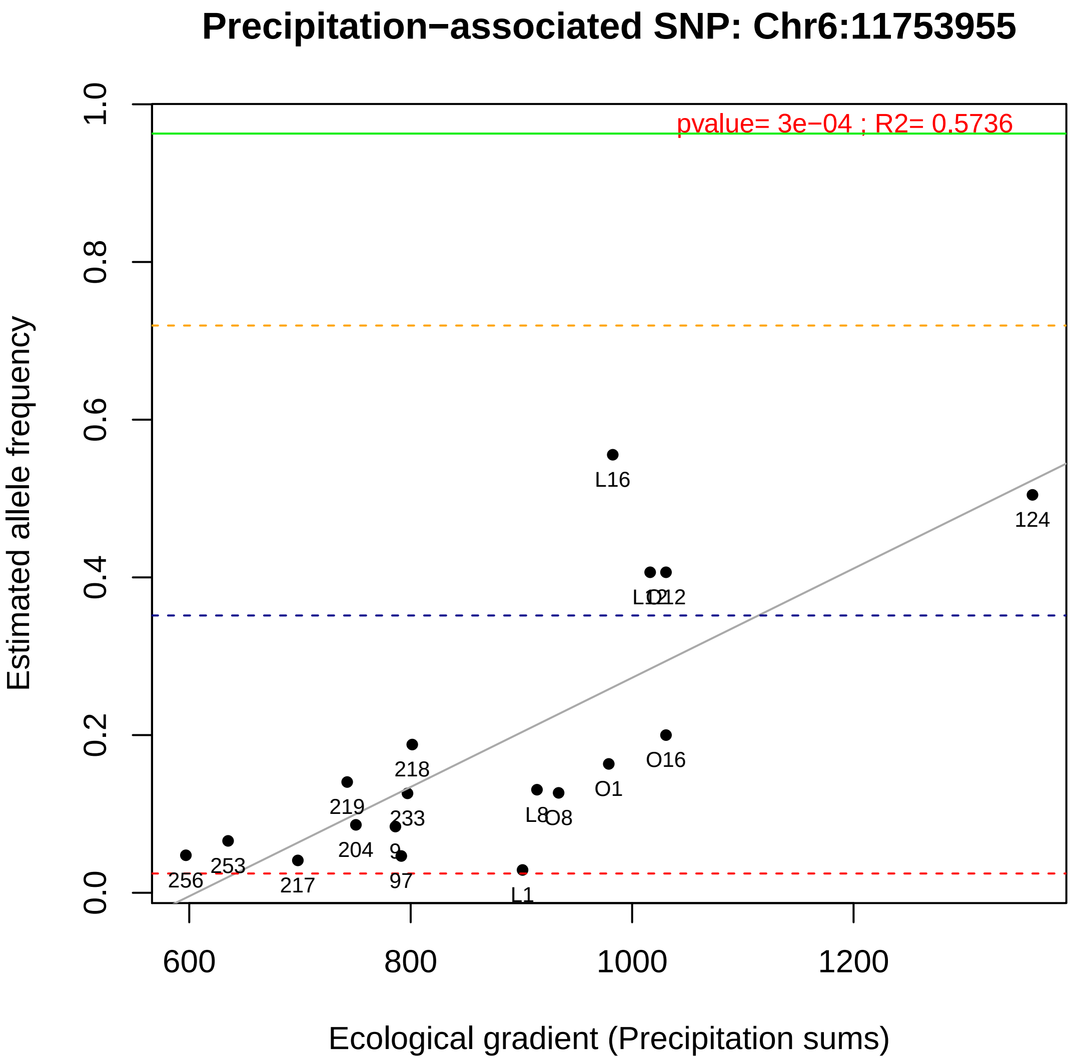
Precipitation−associated SNP: Chr6:11753955
pvalue= 3e−04 ; R2= 0.5736
Estimated allele frequency
Ecological gradient (Precipitation sums)
0.0
0.2
0.4
0.6
0.8
1.0
600
800
1000
1200
L16
124
O16
O1
218
219
233
204
9
97
253
256
217
L1
L8
O8

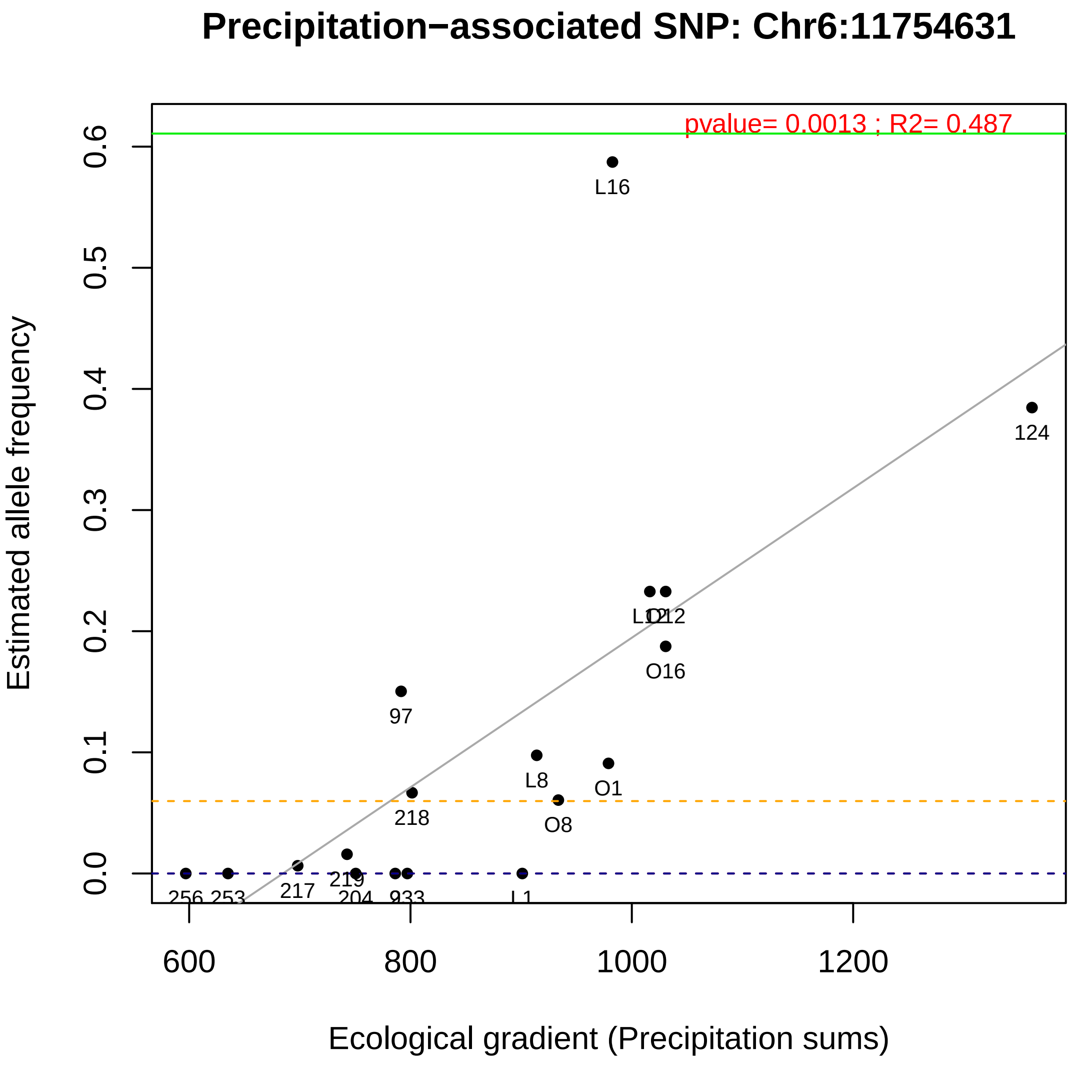

Precipitation−associated SNP: Chr6:11754631
pvalue= 0.0013 ; R2= 0.487
L16
124
L12
O12
O16
97
L8
O1
218
O8
219
217
256
253
204
L1
0.0
0.1
0.2
0.3
0.4
0.5
0.6
600
800
1000
1200
Estimated allele frequency
Ecological gradient (Precipitation sums)

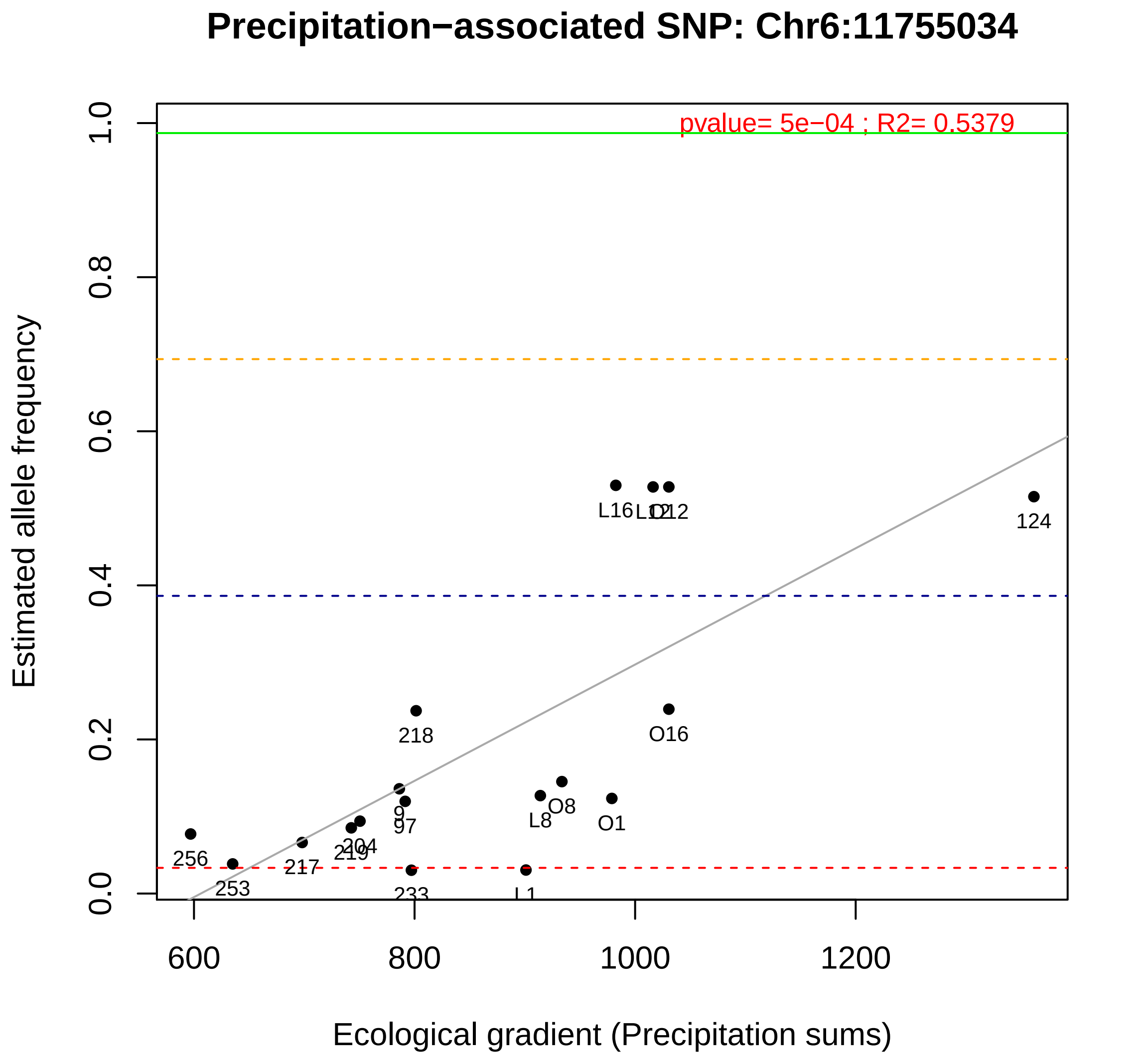

Precipitation−associated SNP: Chr6:11755034
pvalue= 5e−04 ; R2= 0.5379
Estimated allele frequency
Ecological gradient (Precipitation sums)
0.0
0.2
0.4
0.6
0.8
1.0
600
800
1000
1200
L16
L12
O12
124
218
O16
O8
L8
O1
9
97
204
219
256
217
253
233
L1

#### Precipitation−associated SNP: Chr6:11756015

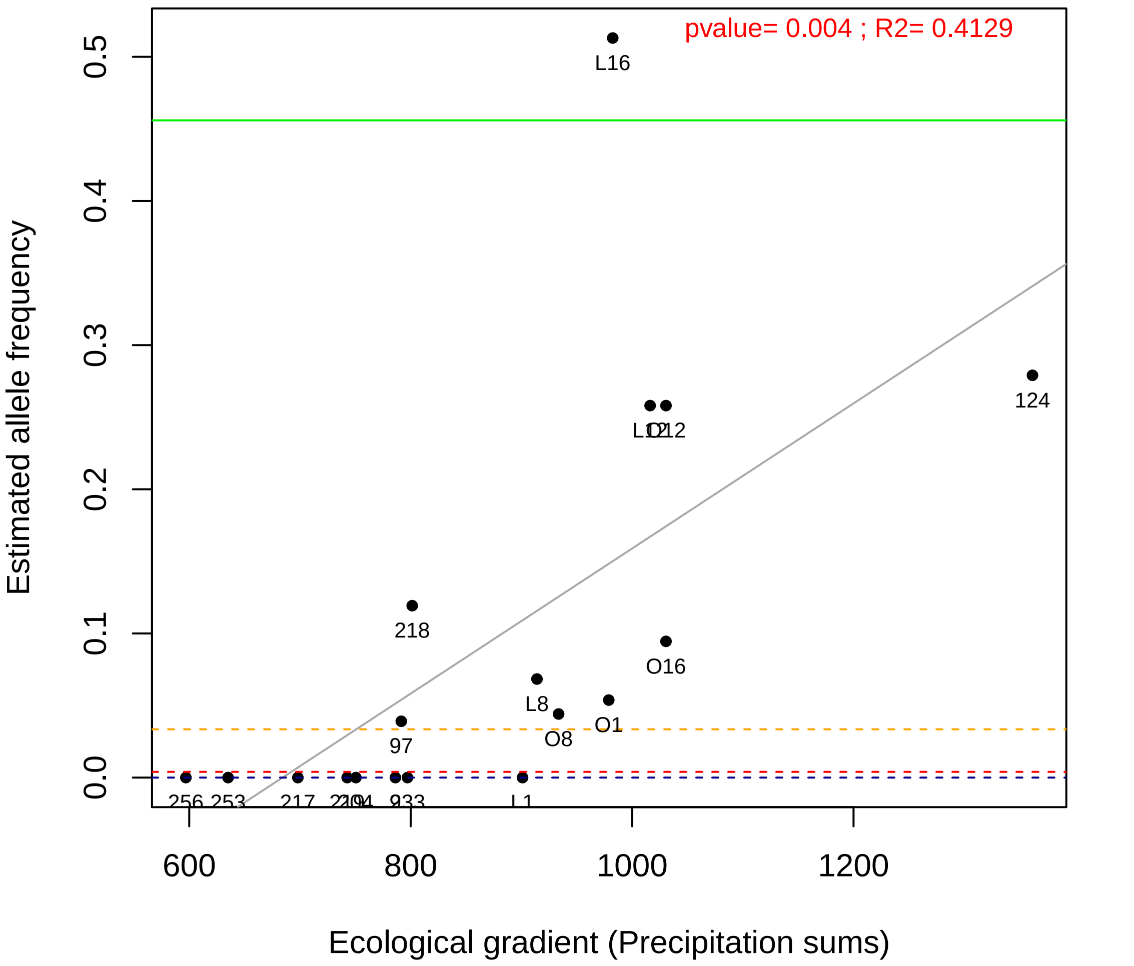

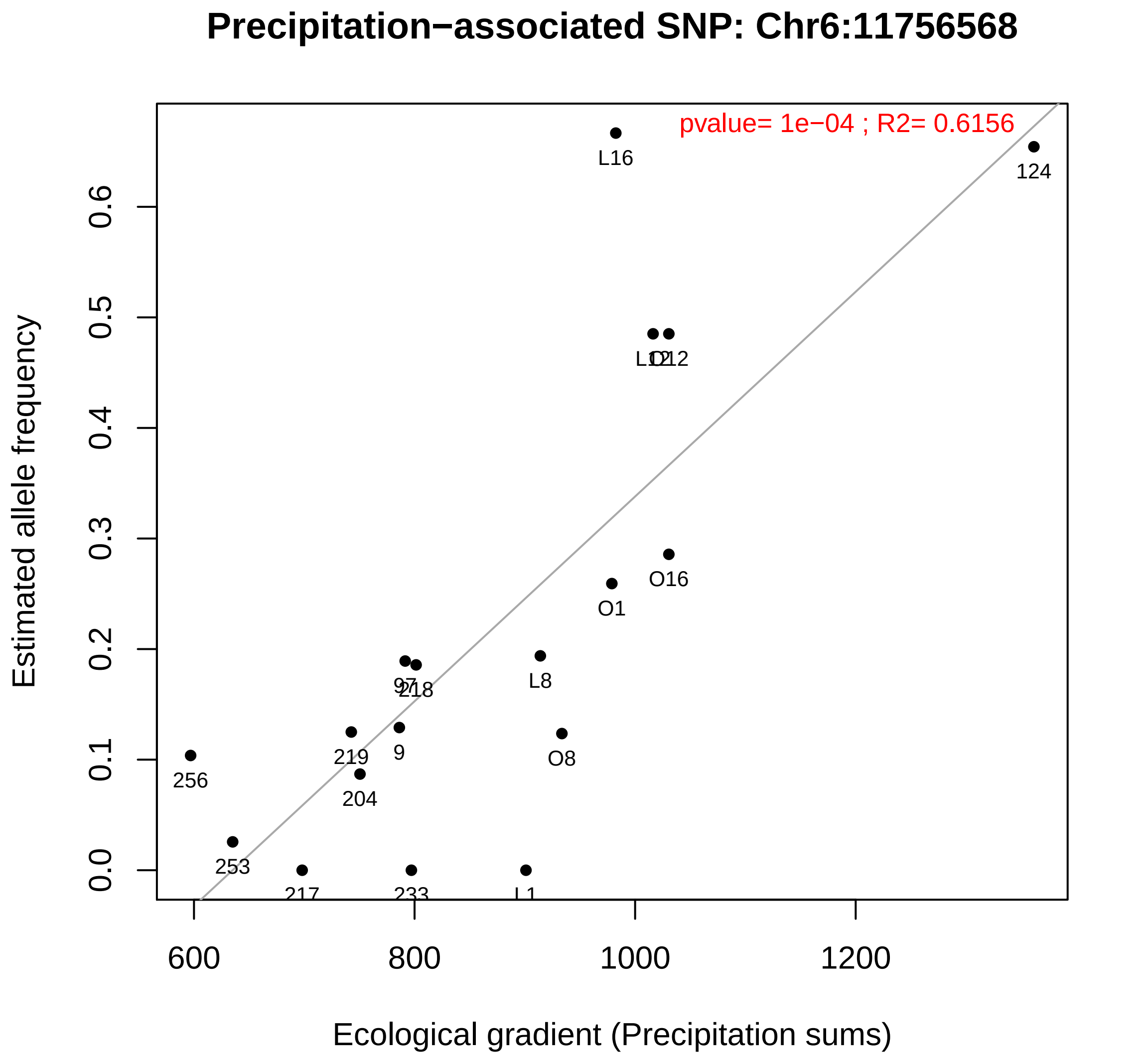

Precipitation−associated SNP: Chr6:11756568
pvalue= 1e−04 ; R2= 0.6156
L16
124
L12
O12
O16
O1
L8
97
218
219
9
O8
256
204
253
217
233
L1
0.0
0.1
0.2
0.3
0.4
0.5
0.6
600
800
1000
1200
Estimated allele frequency
Ecological gradient (Precipitation sums)

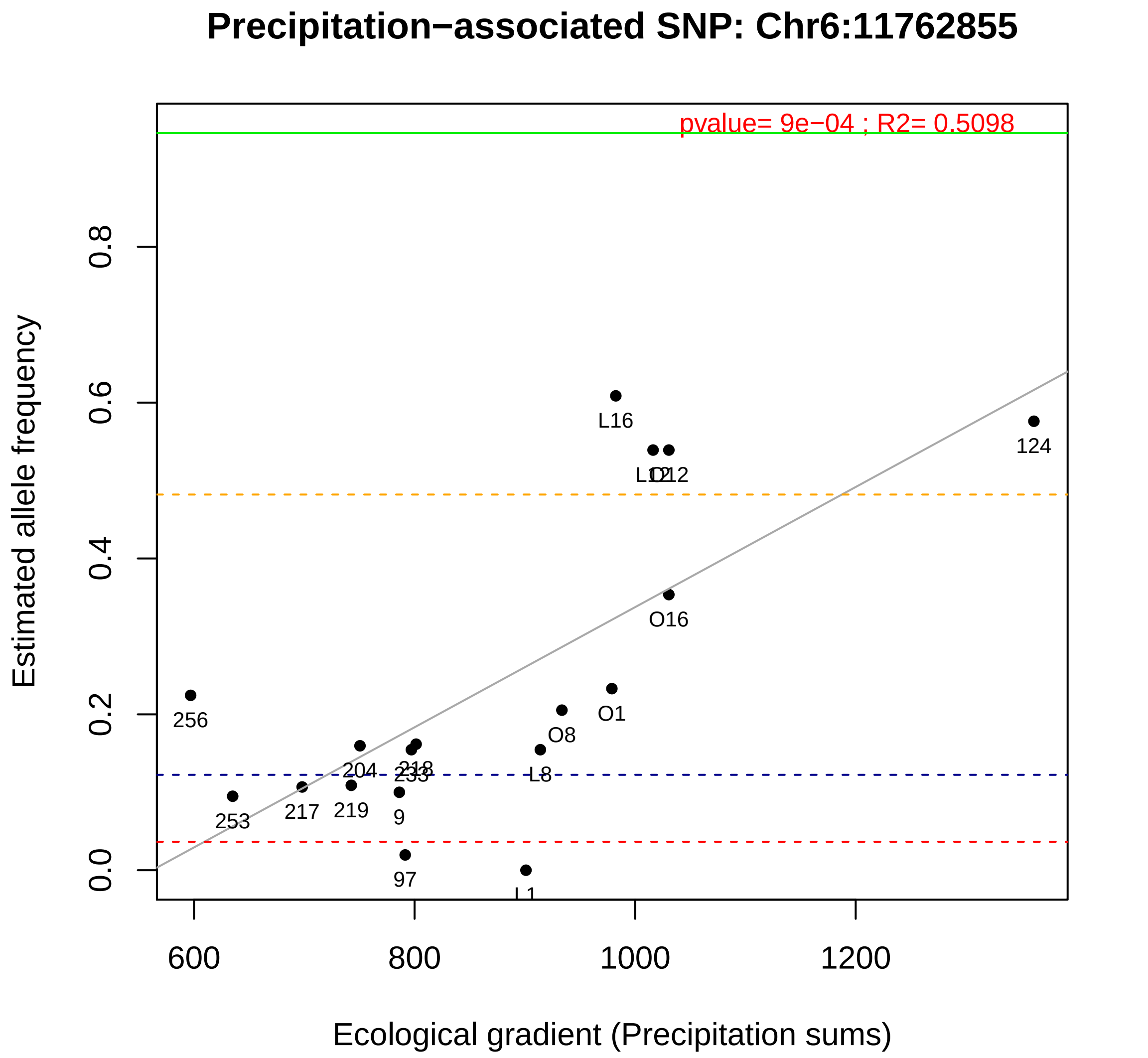

Precipitation−associated SNP: Chr6:11762855
pvalue= 9e−04 ; R2= 0.5098
Estimated allele frequency
0.0
0.2
0.4
0.6
0.8
600
800
1000
1200
Ecological gradient (Precipitation sums)
L16
124
O12
O16
256
O1
O8
L8
204
233
218
253
217
219
9
97

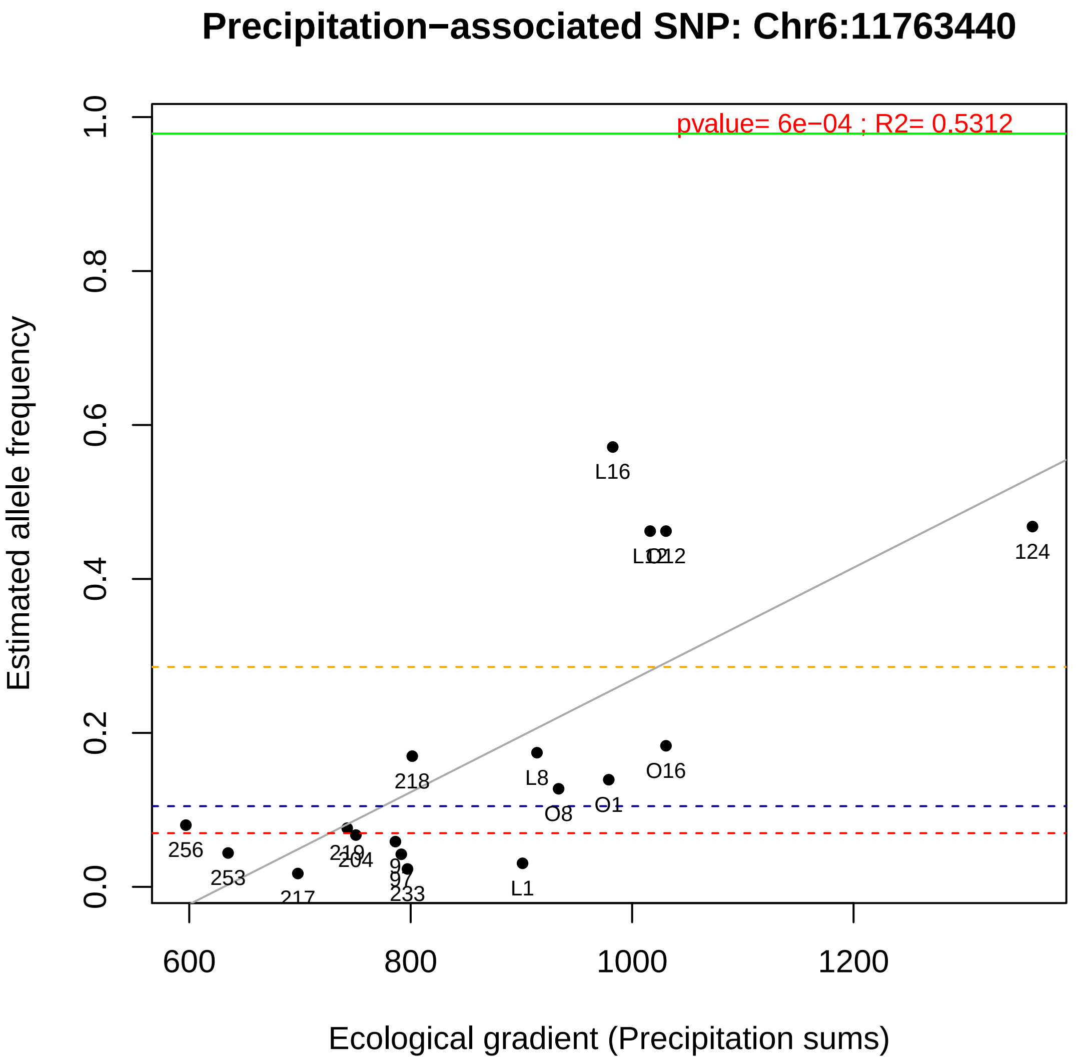
Precipitation−associated SNP: Chr6:11763440
pvalue= 6e−04 ; R2= 0.5312
Estimated allele frequency
0.0
0.2
0.4
0.6
0.8
1.0
600
800
1000
1200
Ecological gradient (Precipitation sums)
L16
124
218
L8
O8
O1
O16
256
253
217
219
204
233
L1

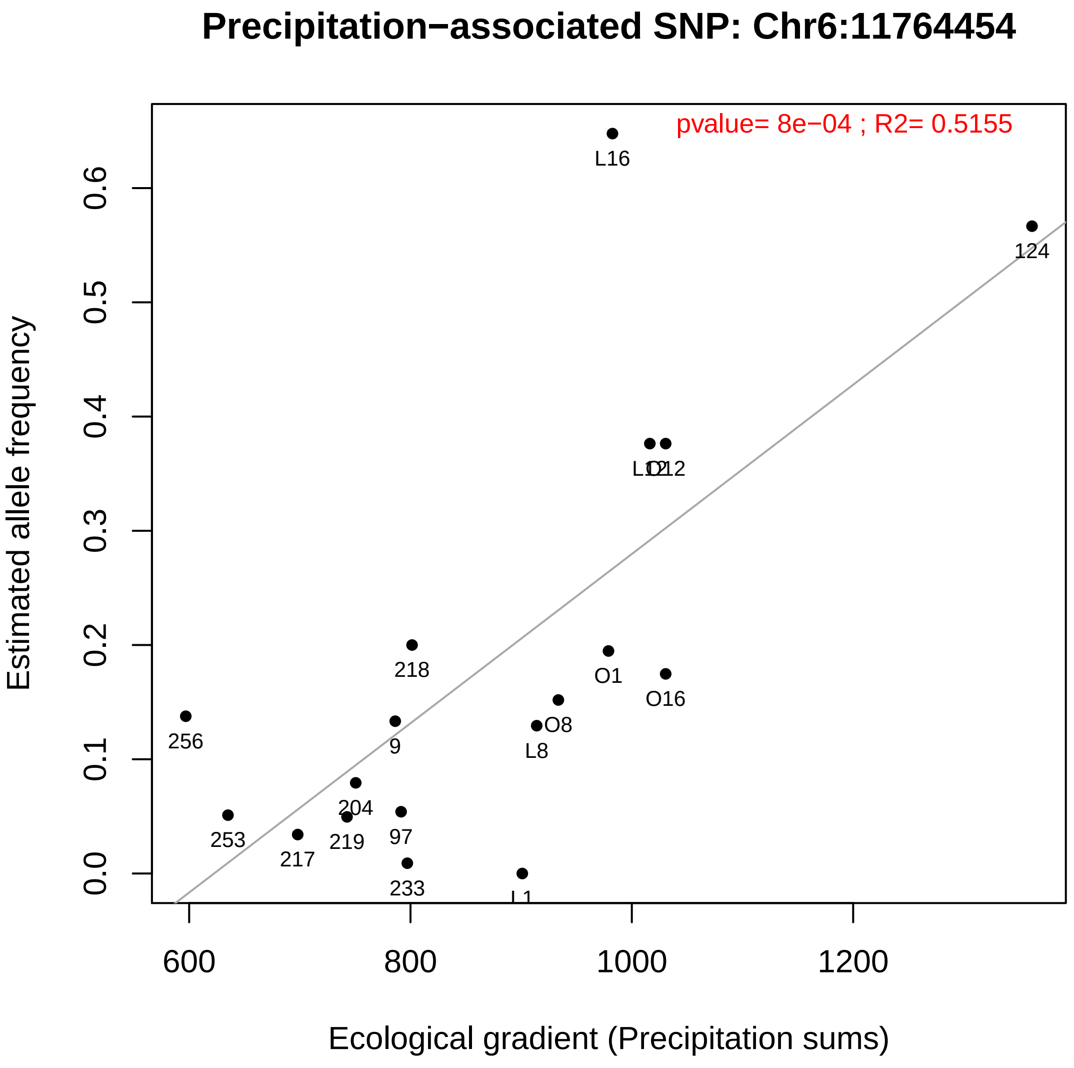
Precipitation−associated SNP: Chr6:11764454
pvalue= 8e−04 ; R2= 0.5155
L16
124
L12
O12
218
O1
O16
256
9
O8
L8
204
253
219
97
217
233
L1
0.0
0.1
0.2
0.3
0.4
0.5
0.6
600
800
1000
1200
Estimated allele frequency
Ecological gradient (Precipitation sums)

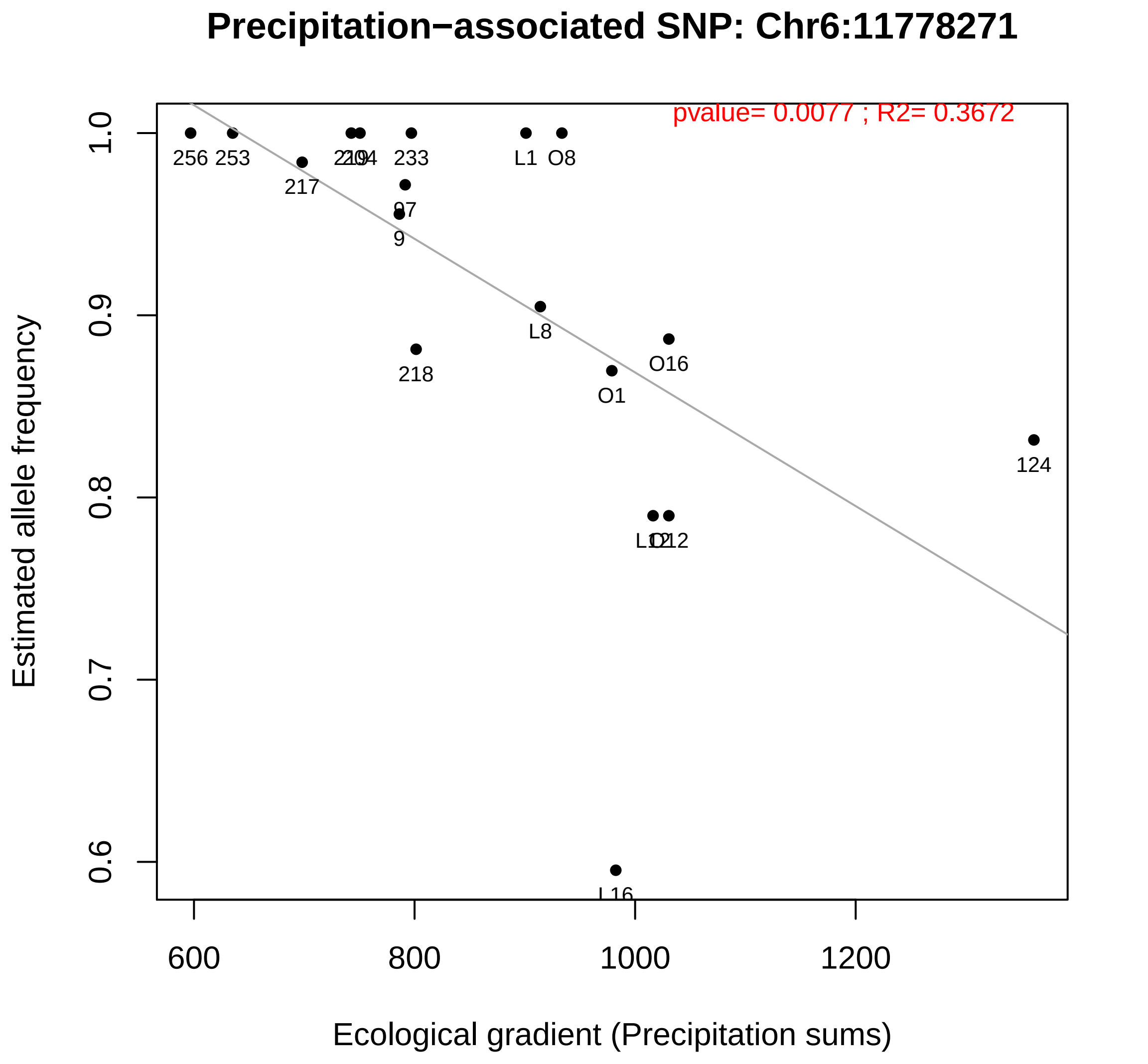

Precipitation−associated SNP: Chr6:11778271
pvalue= 0.0077 ; R2= 0.3672
Estimated allele frequency
Ecological gradient (Precipitation sums)
1.0
0.9
0.8
0.7
0.6
600
800
1000
1200
256
253
217
233
L1
O8
9
L8
218
O16
O1
124

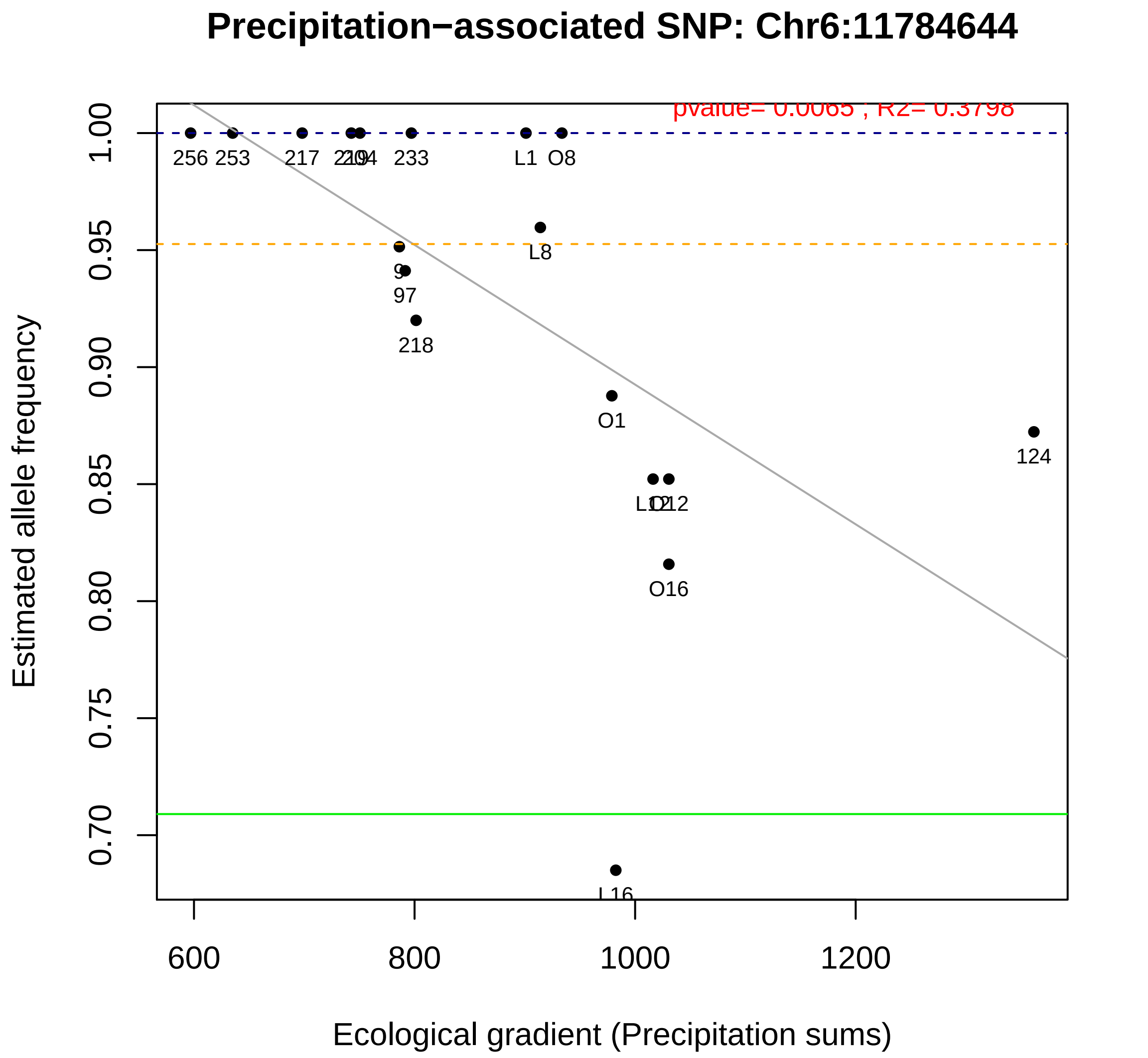

Precipitation−associated SNP: Chr6:11784644
pvalue= 0.0065 ; R2= 0.3798
Estimated allele frequency
Ecological gradient (Precipitation sums)
0.70
0.75
0.80
0.85
0.90
0.95
1.00
600
800
1000
1200
256
253
217
233
L1
O8
L8
97
218
O1
124
O16
L16

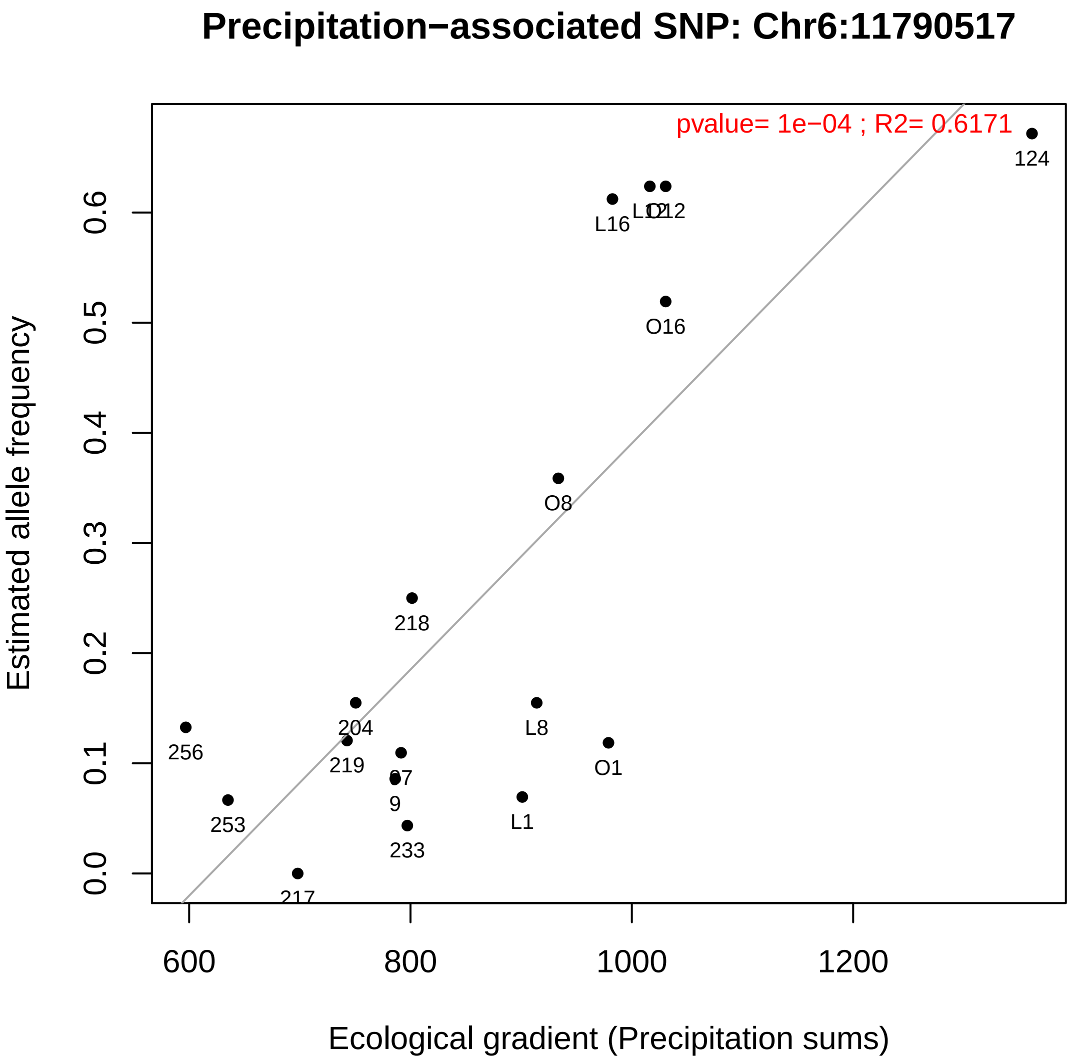
Precipitation−associated SNP: Chr6:11790517
pvalue= 1e−04 ; R2= 0.6171
Estimated allele frequency
Ecological gradient (Precipitation sums)
0.0
0.1
0.2
0.3
0.4
0.5
0.6
600
800
1000
1200
124
L12
O12
L16
O16
O8
218
204
256
219
L8
O1
9
253
233
L1
217

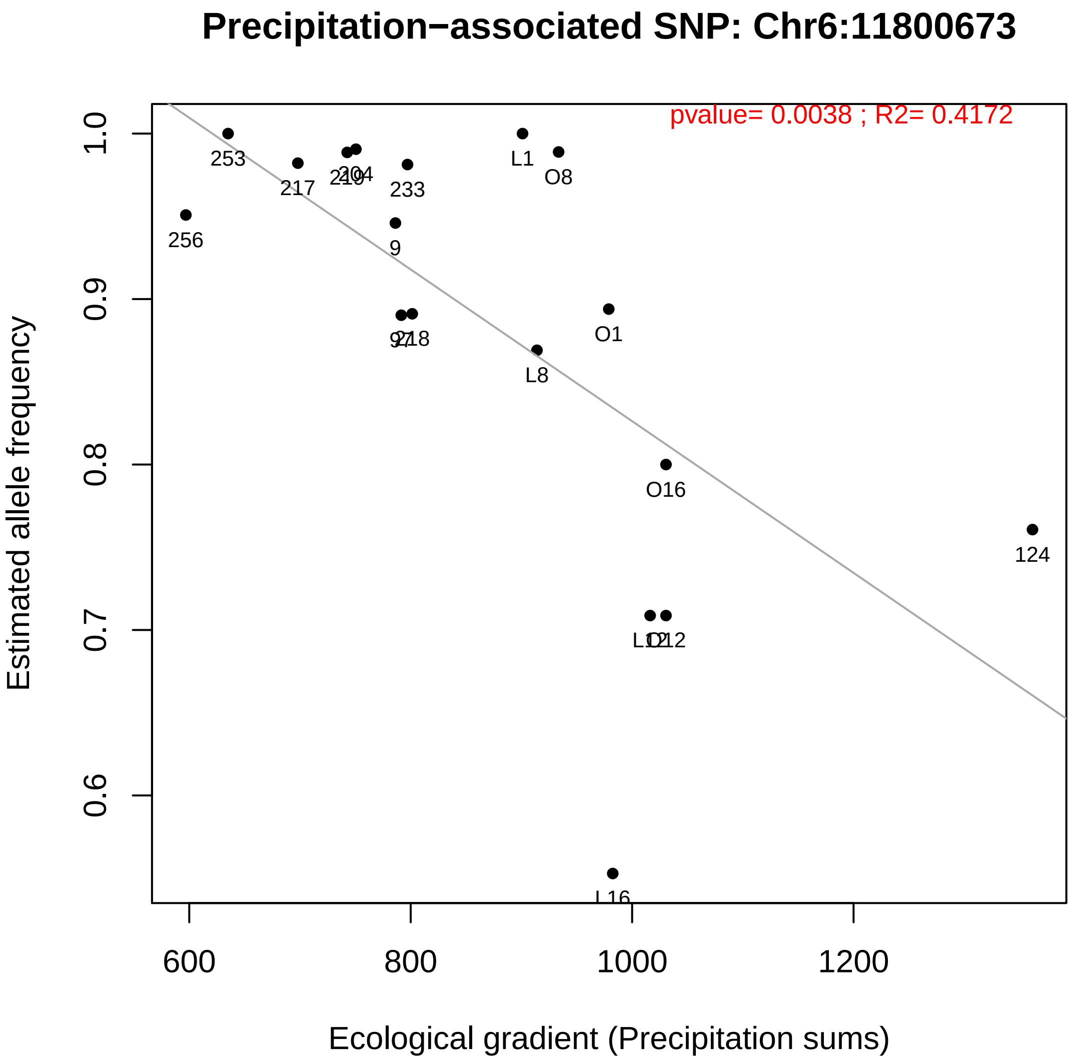

Precipitation−associated SNP: Chr6:11800673
pvalue= 0.0038 ; R2= 0.4172
Estimated allele frequency
0.6
0.7
0.8
0.9
1.0
600
800
1000
1200
Ecological gradient (Precipitation sums)
253
217
204
233
L1
O8
256
9
O1
218
L8
O16
124
L12
O12
L16

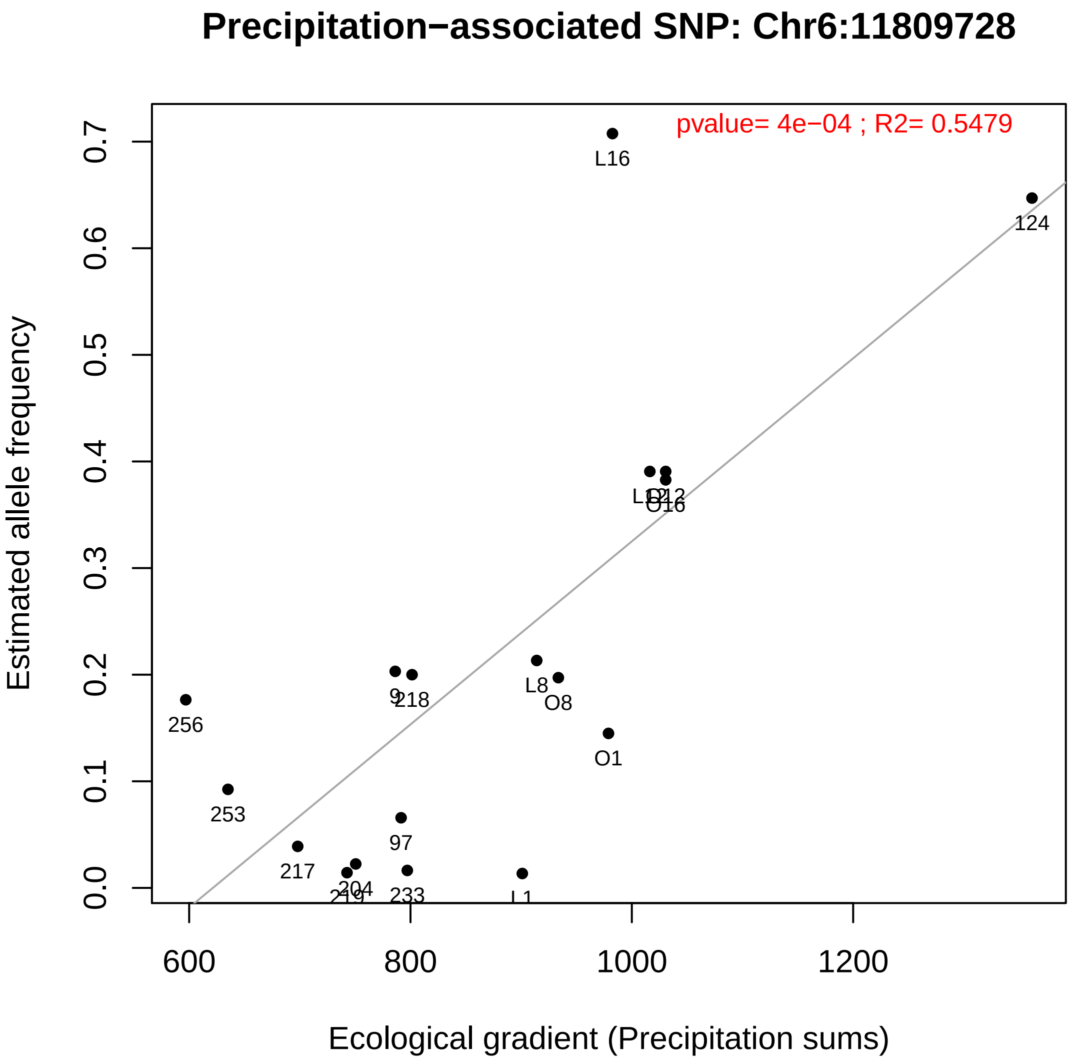

Precipitation−associated SNP: Chr6:11809728
pvalue= 4e−04 ; R2= 0.5479
Estimated allele frequency
Ecological gradient (Precipitation sums)
0.0
0.1
0.2
0.3
0.4
0.5
0.6
0.7
600
800
1000
1200
L16
124
O16
L8
O8
O1
256
253
217
204
97
233
218

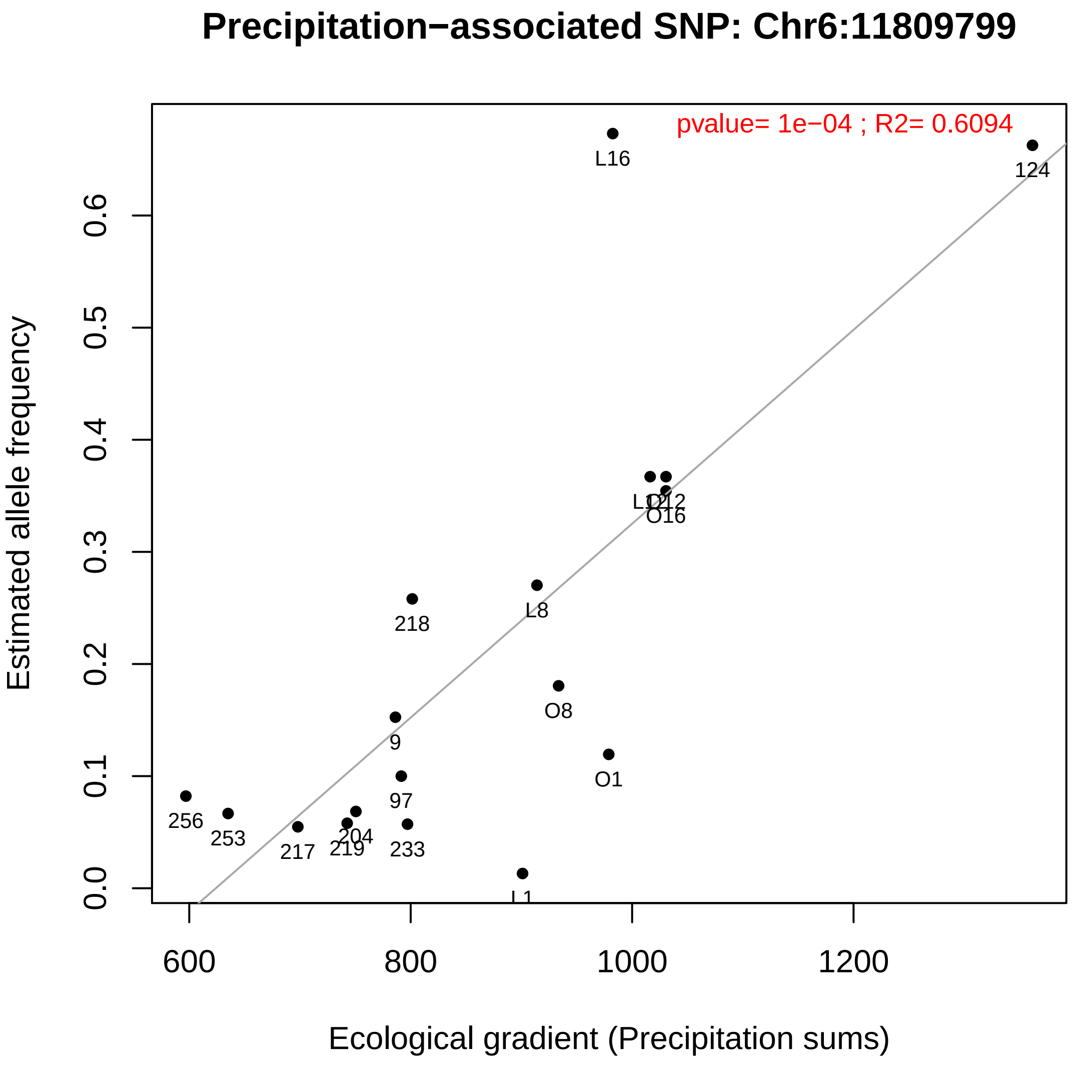
Precipitation−associated SNP: Chr6:11809799
pvalue= 1e−04 ; R2= 0.6094
Estimated allele frequency
Ecological gradient (Precipitation sums)
0.0
0.1
0.2
0.3
0.4
0.5
0.6
600
800
1000
1200
L16
124
L8
218
O8
9
O1
97
256
253
217
219
204
233
O16

#### Precipitation−associated SNP: Chr6:11842540

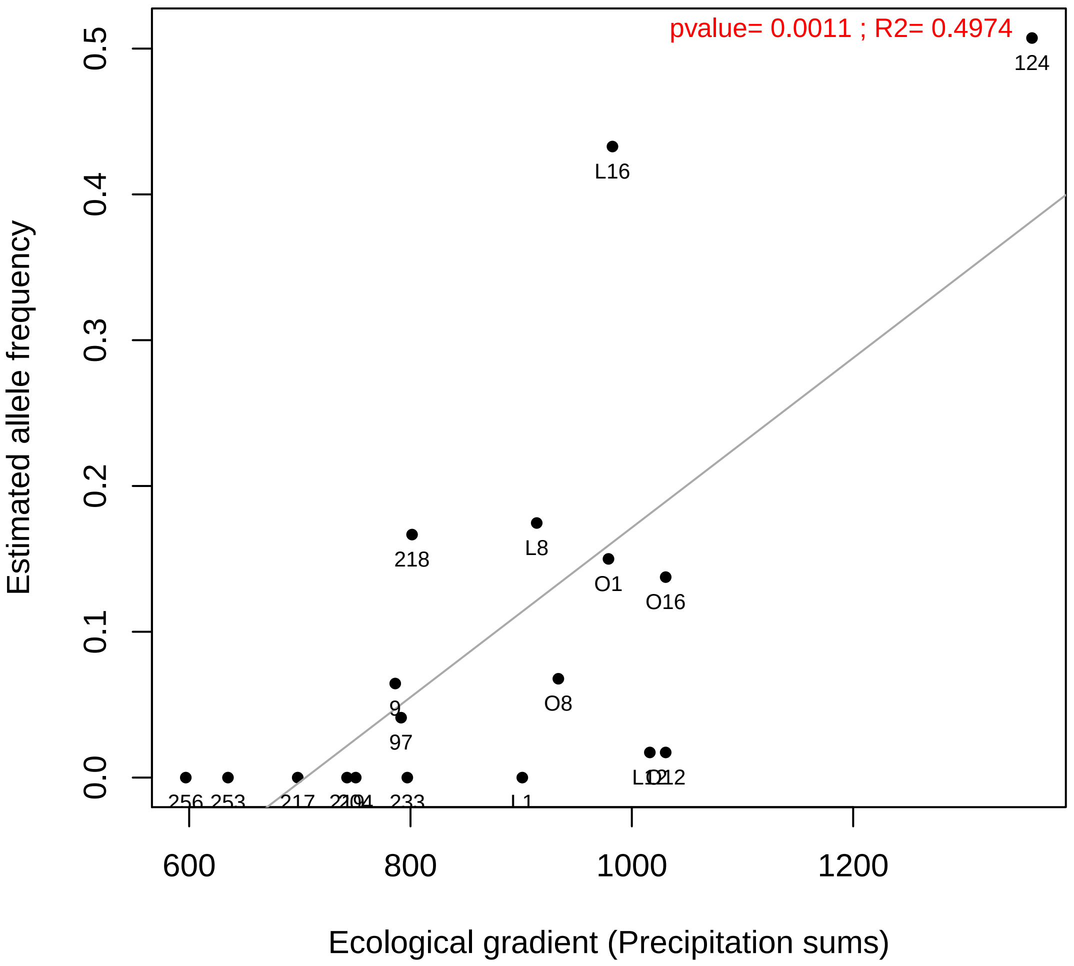

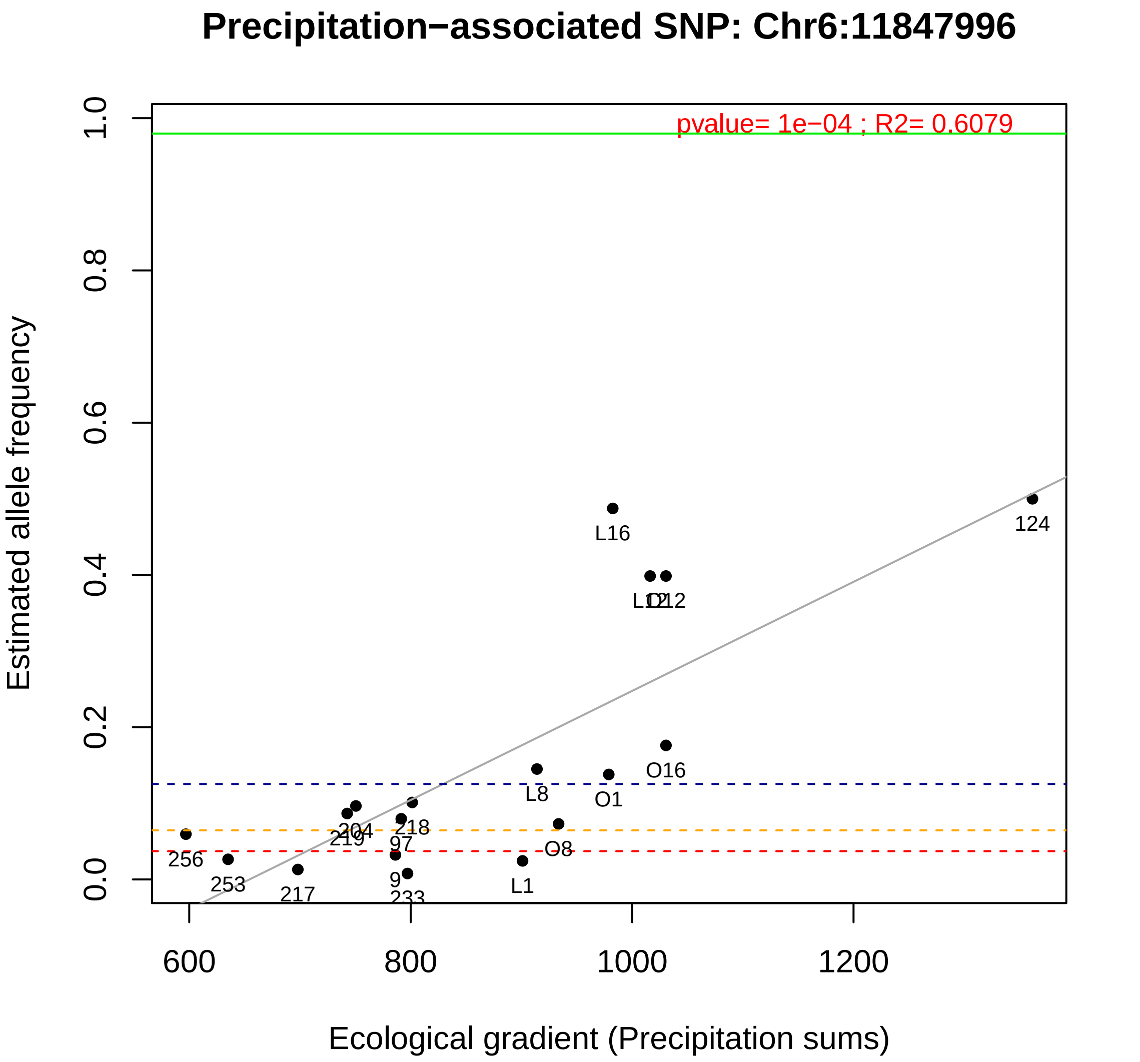

Precipitation−associated SNP: Chr6:11847996
pvalue= 1e−04 ; R2= 0.6079
Estimated allele frequency
0.0
0.2
0.4
0.6
0.8
1.0
600
800
1000
1200
Ecological gradient (Precipitation sums)
L16
124
L12
O12
O16
L8
O1
204
219
218
97
O8
256
253
217
9
233
L1

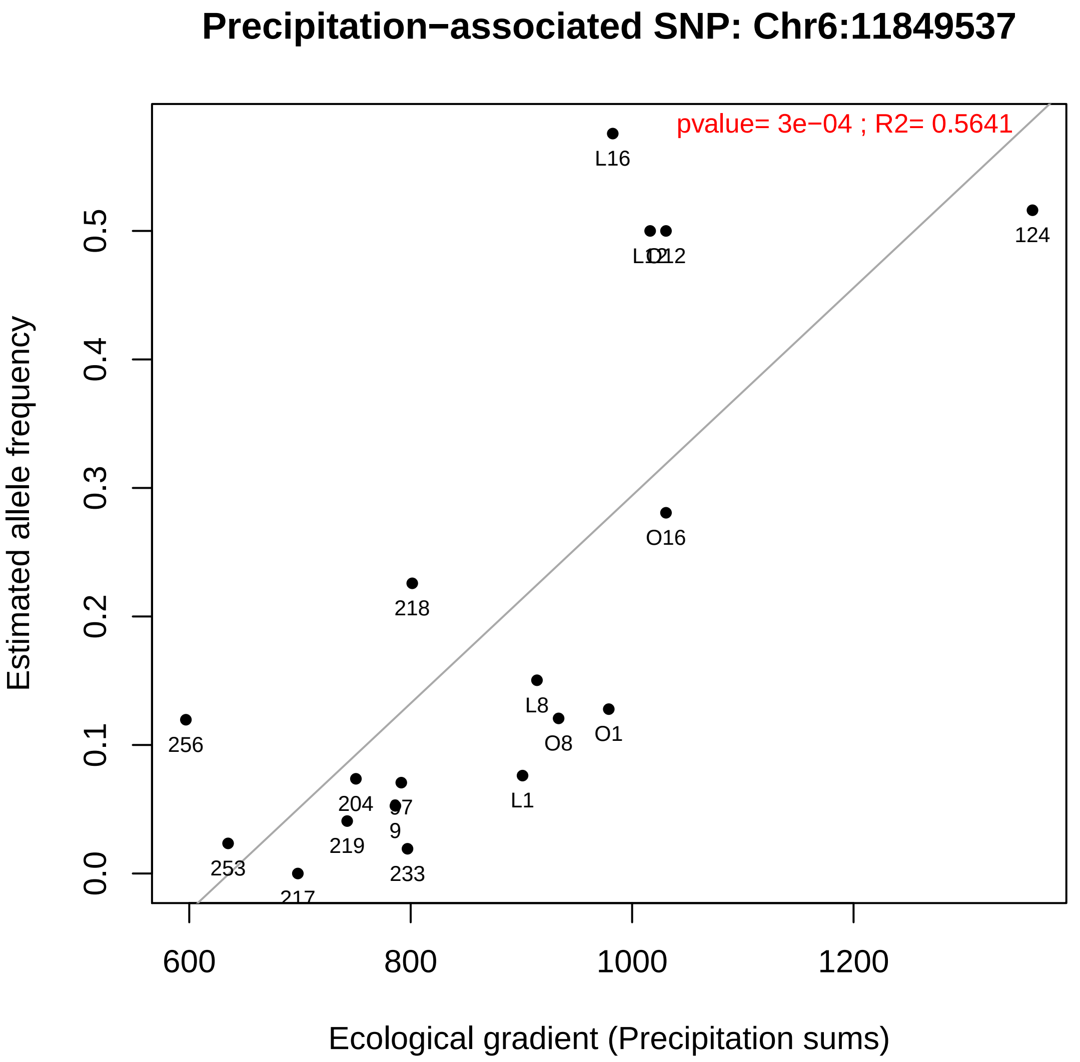
Precipitation−associated SNP: Chr6:11849537
pvalue= 3e−04 ; R2= 0.5641
L16
124
L12
O12
O16
218
L8
O1
O8
256
L1
204
97
9
219
233
253
217
0.0
0.1
0.2
0.3
0.4
0.5
600
800
1000
1200
Estimated allele frequency
Ecological gradient (Precipitation sums)

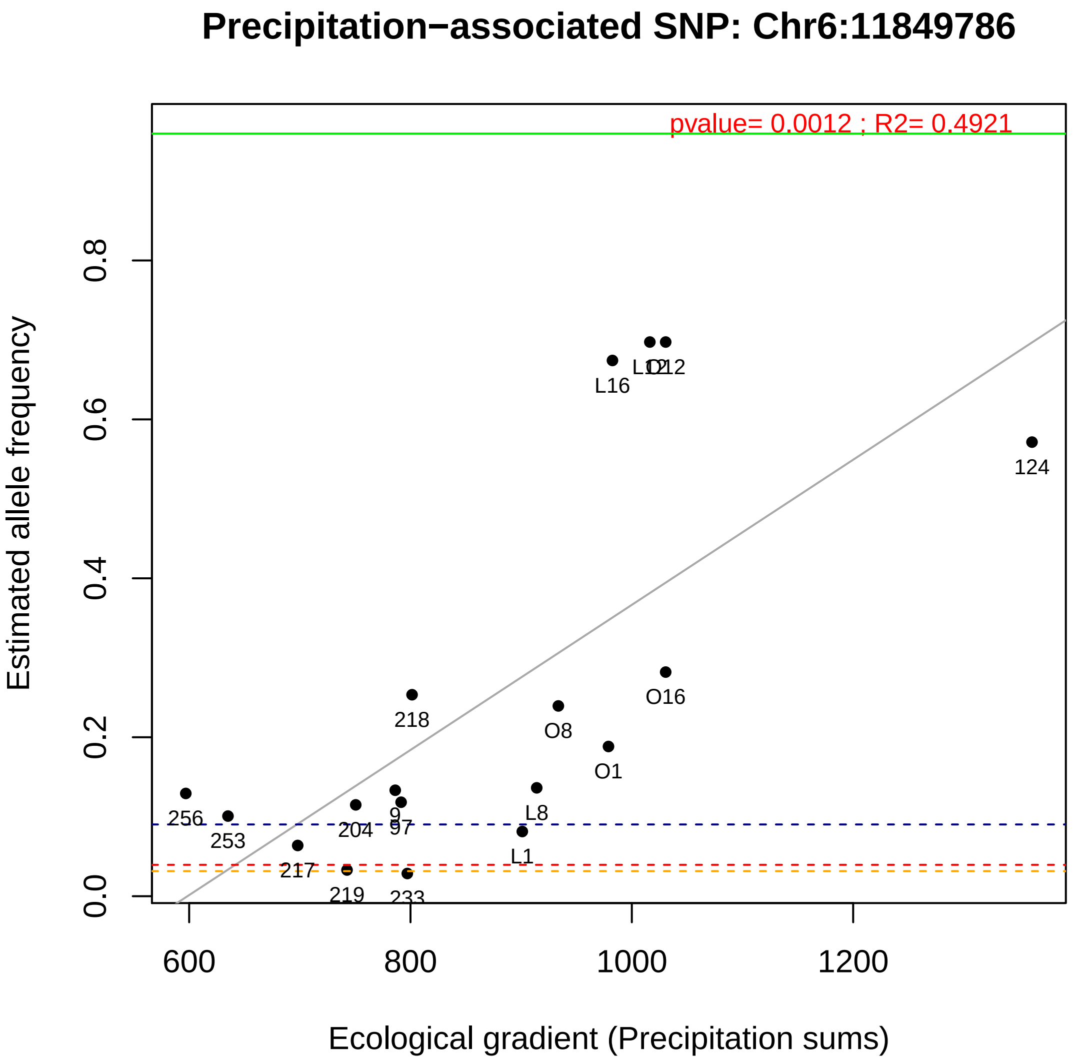

Precipitation−associated SNP: Chr6:11849786
pvalue= 0.0012 ; R2= 0.4921
Estimated allele frequency
0.0
0.2
0.4
0.6
0.8
600
800
1000
1200
Ecological gradient (Precipitation sums)
L16
L12
O12
124
O16
218
O8
O1
L8
256
253
204
9
97
217
L1
219
233

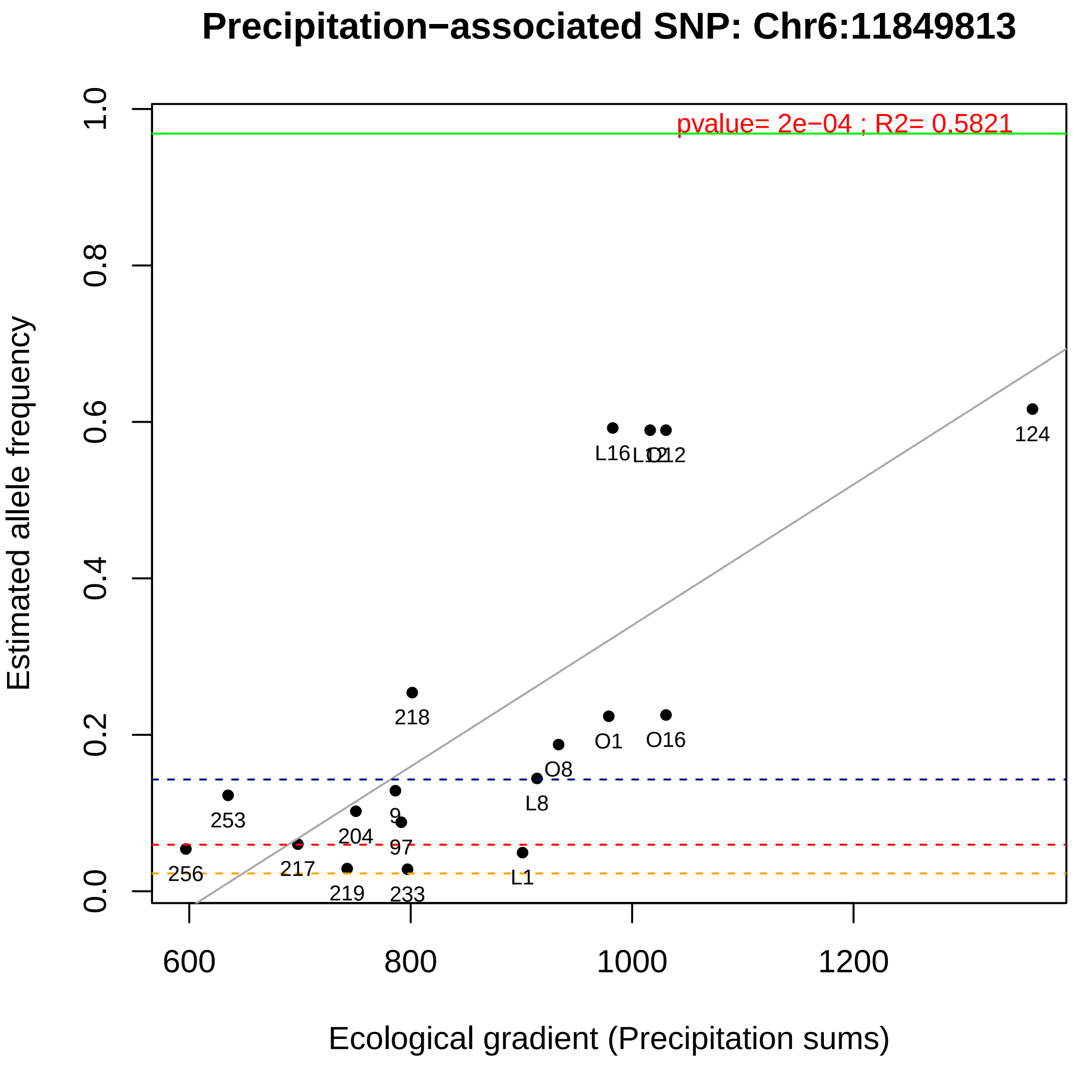
Precipitation−associated SNP: Chr6:11849813
pvalue= 2e−04 ; R2= 0.5821
Estimated allele frequency
0.0
0.2
0.4
0.6
0.8
1.0
600
800
1000
1200
Ecological gradient (Precipitation sums)
124
L16
O12
218
O1
O16
O8
L8
253
9
204
97
256
217
219
233
L1

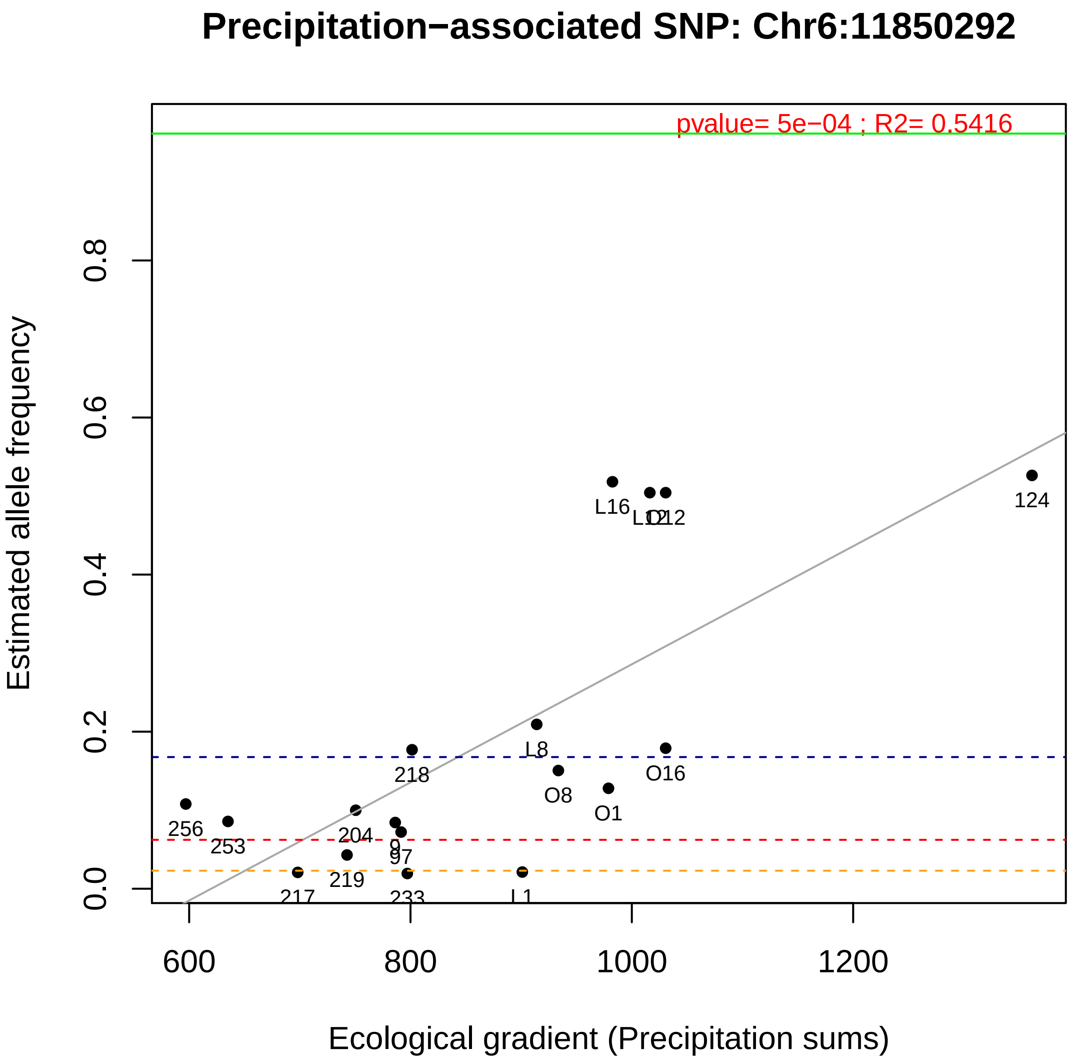

Precipitation−associated SNP: Chr6:11850292
pvalue= 5e−04 ; R2= 0.5416
Estimated allele frequency
0.0
0.2
0.4
0.6
0.8
600
800
1000
1200
Ecological gradient (Precipitation sums)
L16
124
L8
218
O16
O8
O1
256
253
204
219
217
233
L1

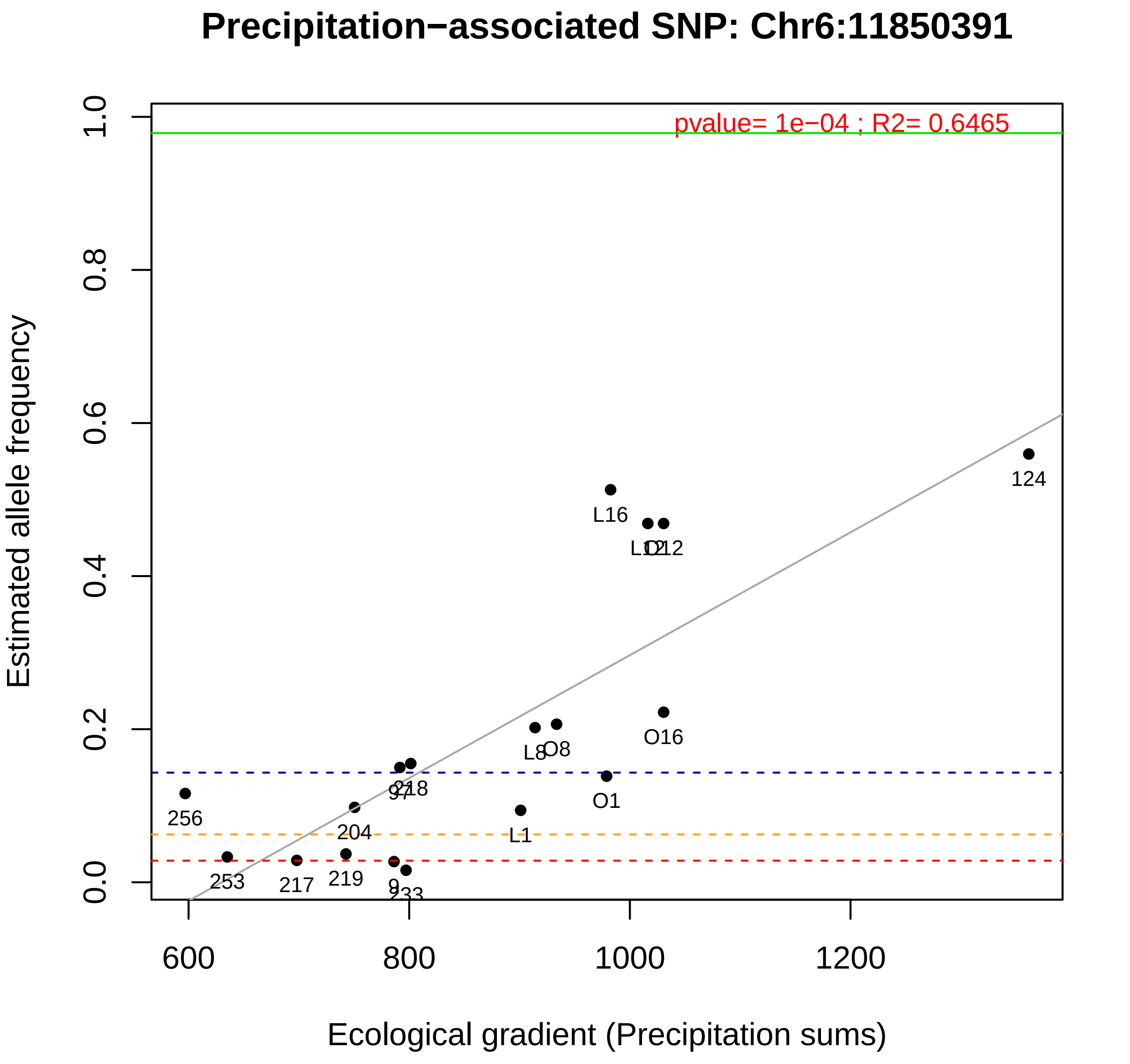

Precipitation−associated SNP: Chr6:11850391
pvalue= 1e−04 ; R2= 0.6465
Estimated allele frequency
Ecological gradient (Precipitation sums)
0.0
0.2
0.4
0.6
0.8
1.0
600
800
1000
1200
124
L16
L12
O12
O16
L8
O8
O1
L1
97
218
256
204
253
217
219
9
233

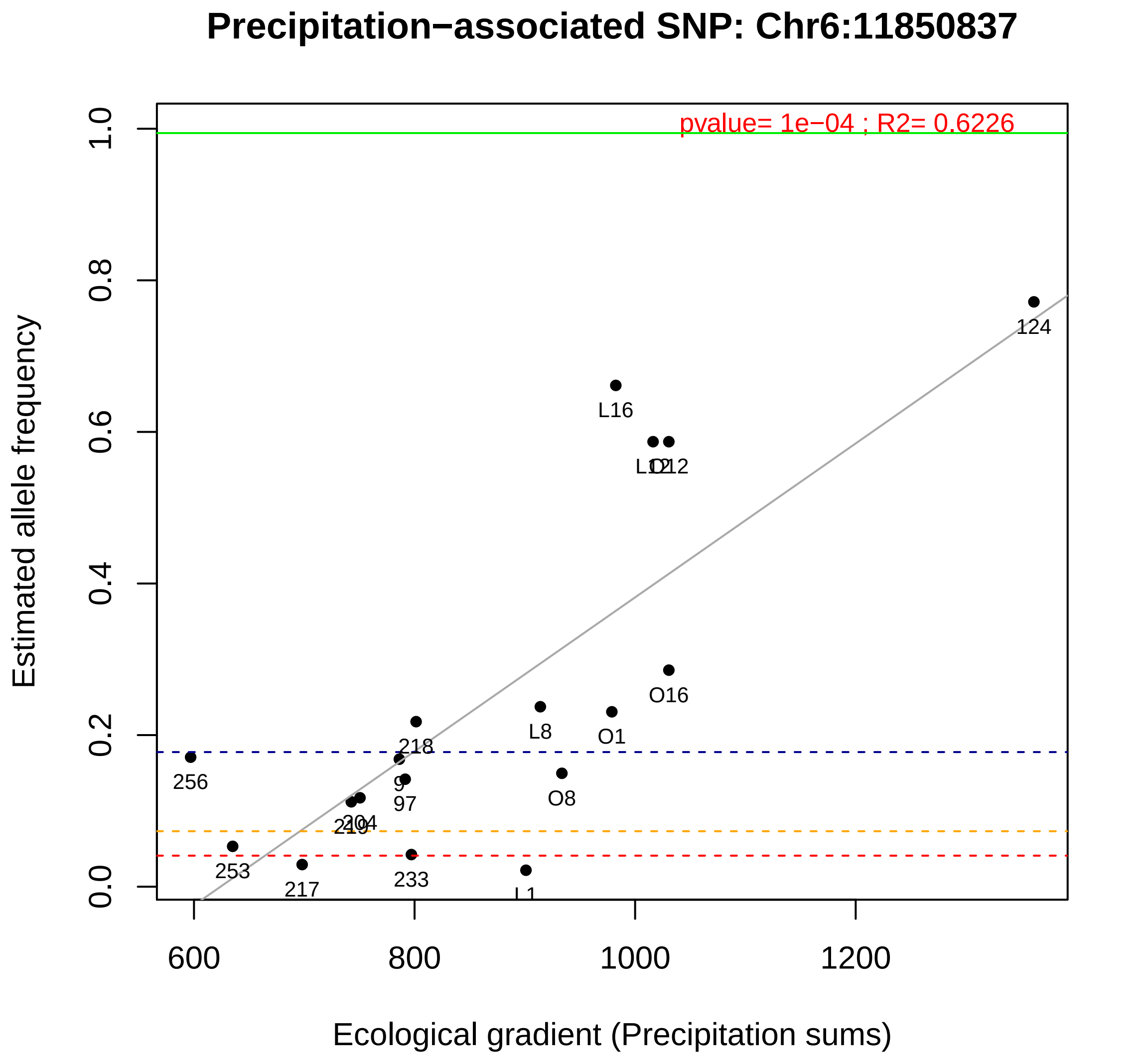

Precipitation−associated SNP: Chr6:11850837
pvalue= 1e−04 ; R2= 0.6226
Estimated allele frequency
0.0
0.2
0.4
0.6
0.8
1.0
600
800
1000
1200
Ecological gradient (Precipitation sums)
124
L16
L12
O12
O16
L8
O1
218
256
9
97
O8
219
204
253
217
233
L1

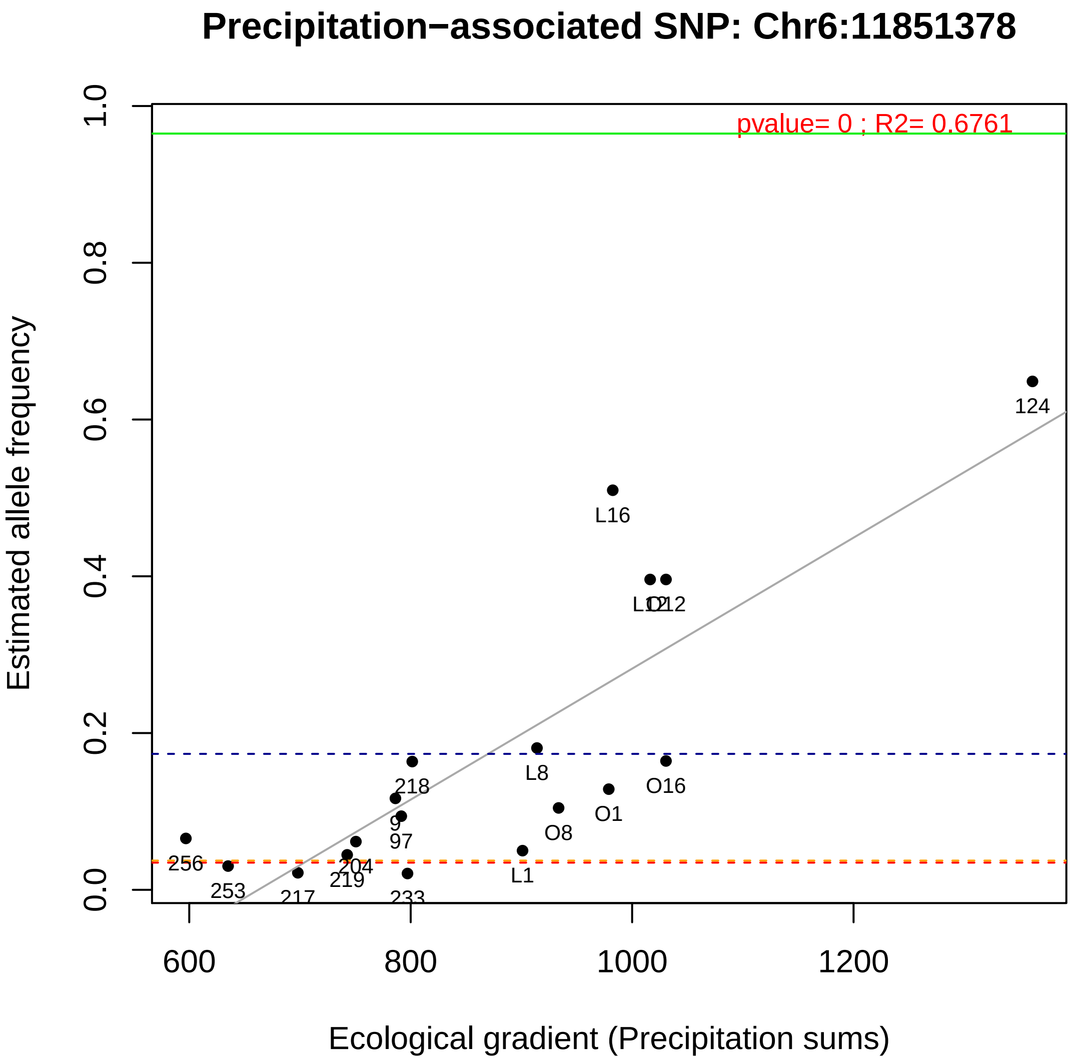
Precipitation−associated SNP: Chr6:11851378
pvalue= 0 ; R2= 0.6761
Estimated allele frequency
Ecological gradient (Precipitation sums)
0.0
0.2
0.4
0.6
0.8
1.0
600
800
1000
1200
124
L16
O12
L8
O16
218
O1
O8
9
97
256
204
219
L1
253
217
233

#### Precipitation−associated SNP: Chr6:11852079

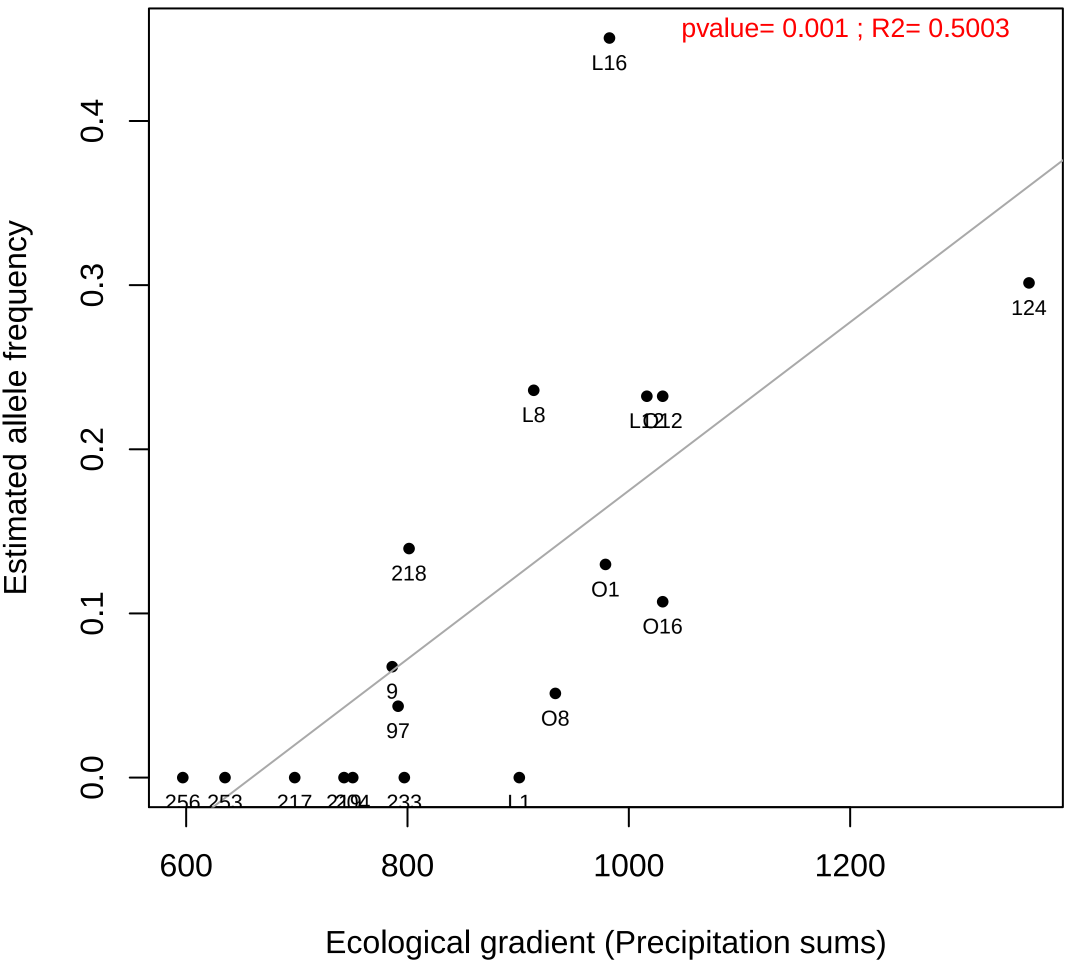

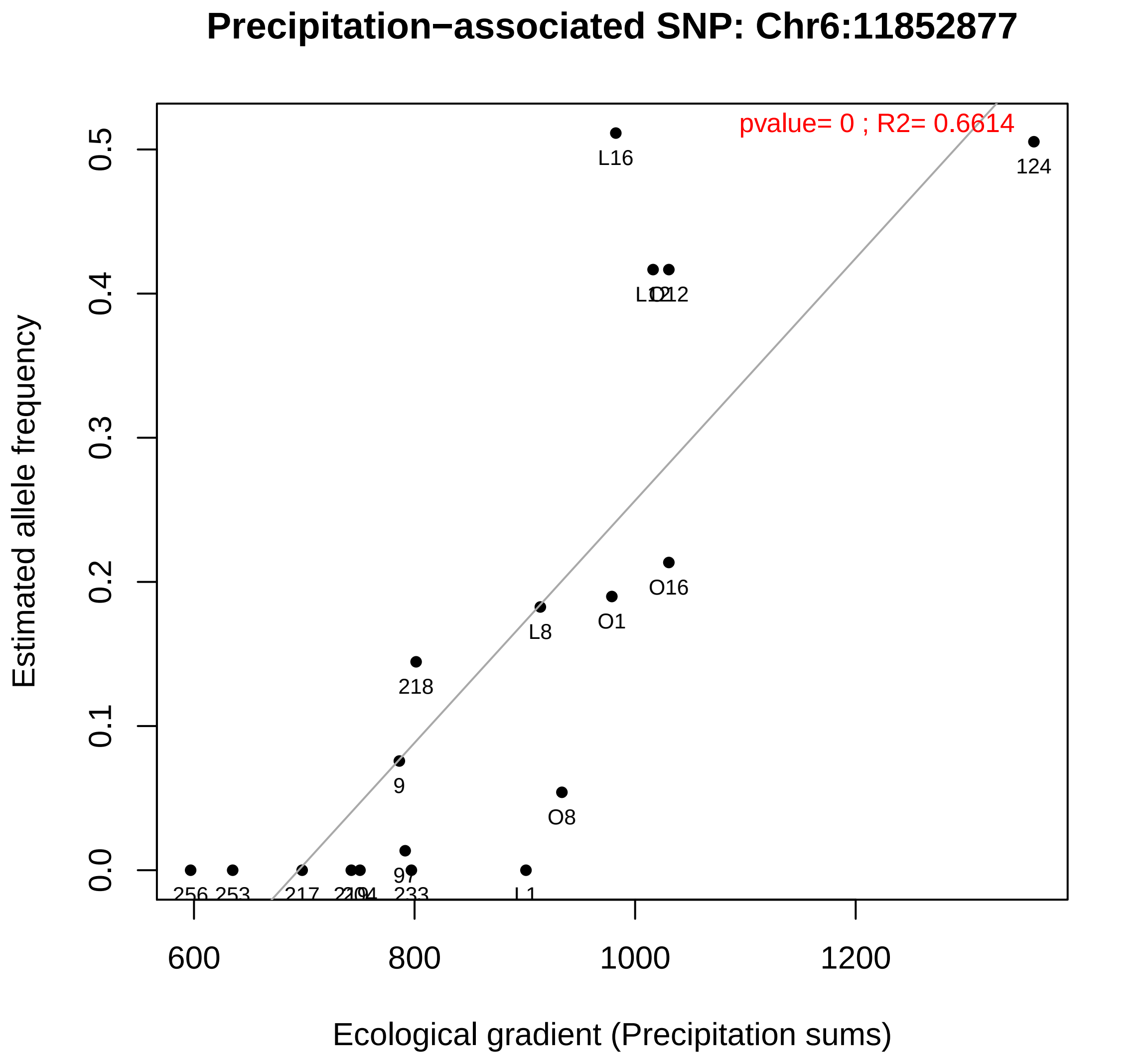

Precipitation−associated SNP: Chr6:11852877
pvalue= 0 ; R2= 0.6614
Estimated allele frequency
Ecological gradient (Precipitation sums)
0.0
0.1
0.2
0.3
0.4
0.5
600
800
1000
1200
L16
124
L12
O12
O16
O1
L8
218
9
O8
97
256
253
217
210
204
233
L1

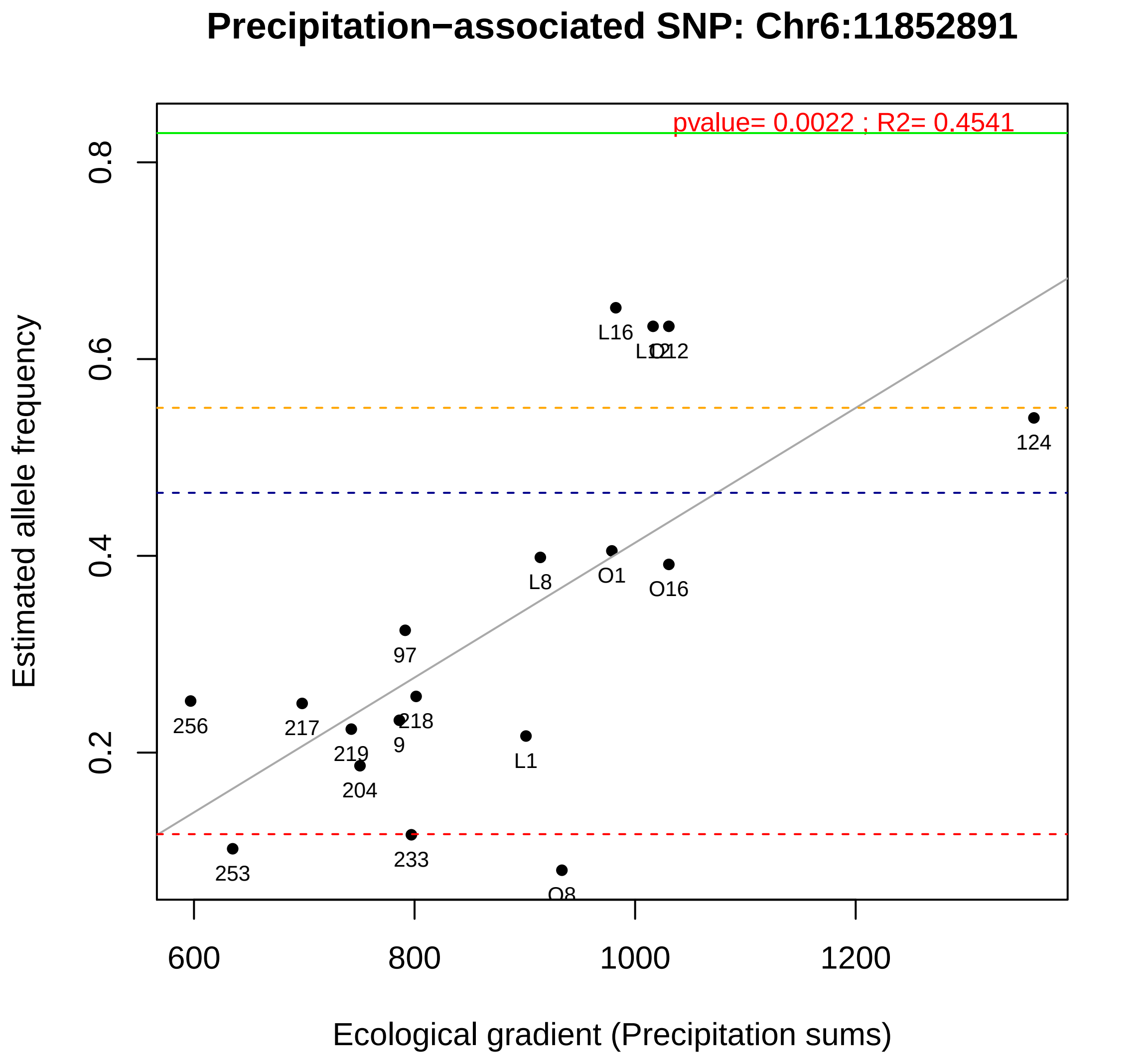
Precipitation−associated SNP: Chr6:11852891
pvalue= 0.0022 ; R2= 0.4541
Estimated allele frequency
0.2
0.4
0.6
0.8
600
800
1000
1200
Ecological gradient (Precipitation sums)
L16
L12
O12
124
L8
O1
O16
97
256
217
219
218
9
204
L1
253
233
O8

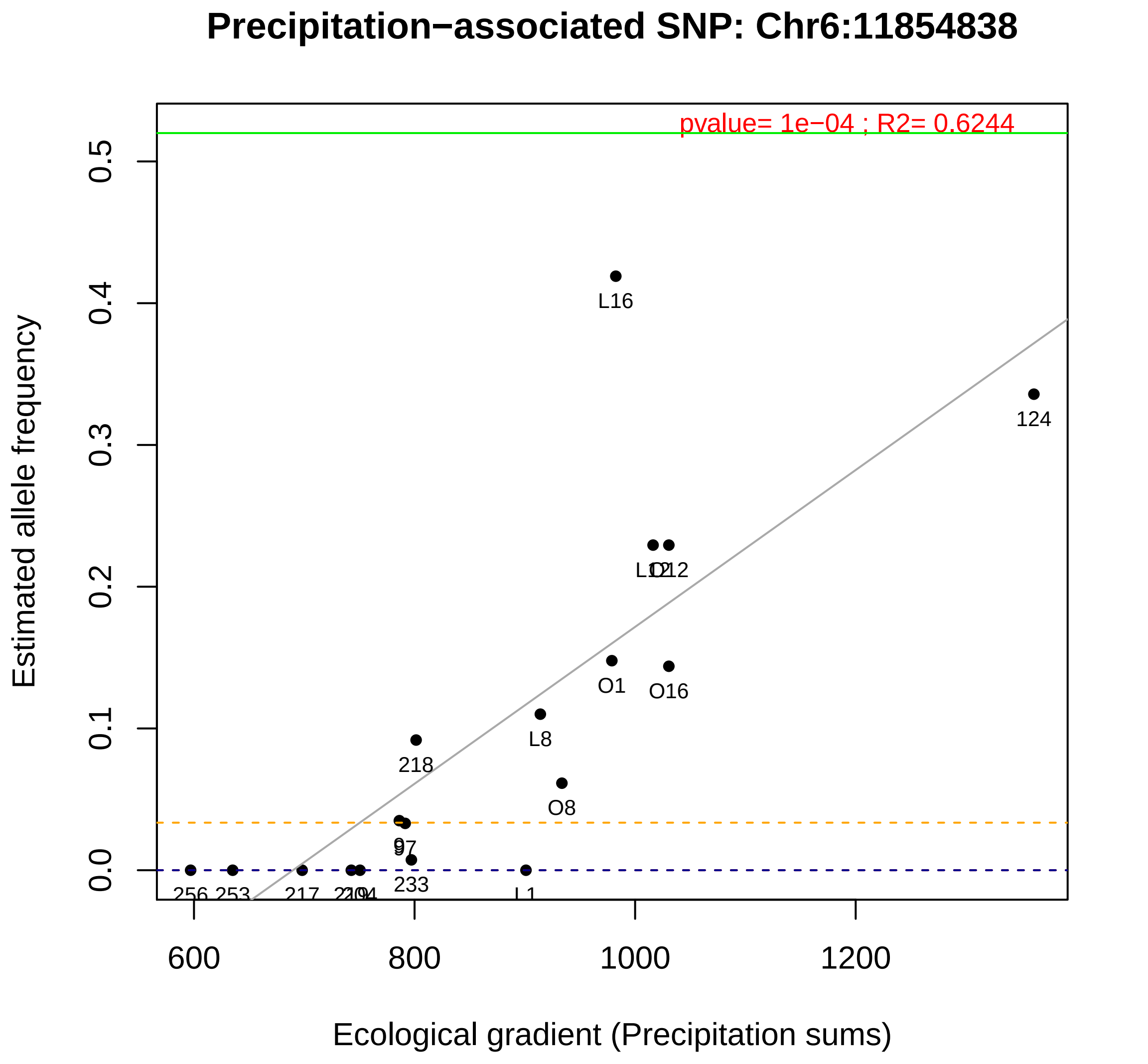

Precipitation−associated SNP: Chr6:11854838
pvalue= 1e−04 ; R2= 0.6244
Estimated allele frequency
0.0
0.1
0.2
0.3
0.4
0.5
600
800
1000
1200
Ecological gradient (Precipitation sums)
L16
124
O12
O1
O16
L8
218
O8
233
256
253
217

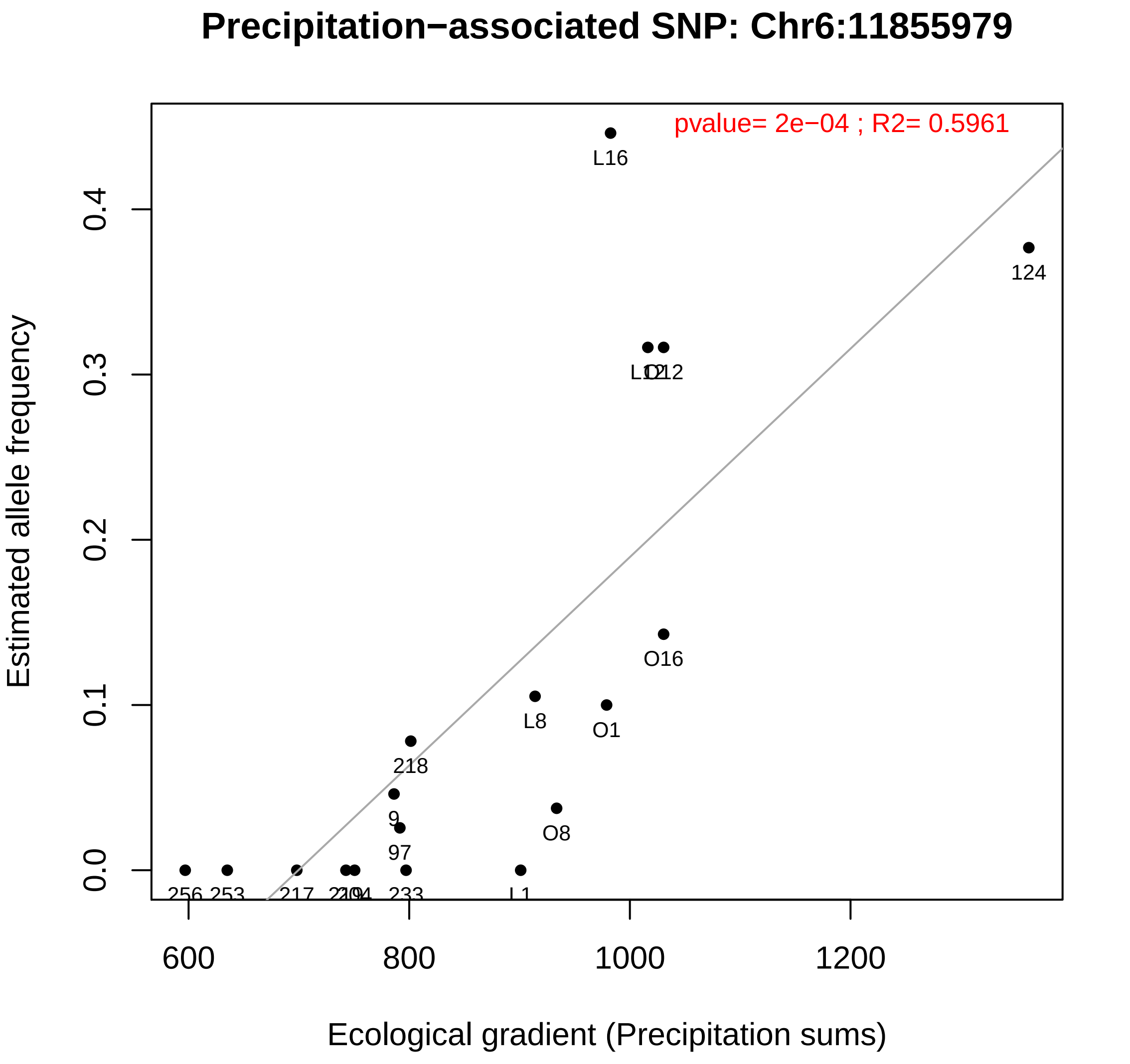

Precipitation−associated SNP: Chr6:11855979
pvalue= 2e−04 ; R2= 0.5961
Estimated allele frequency
0.0
0.1
0.2
0.3
0.4
600
800
1000
1200
Ecological gradient (Precipitation sums)
L16
124
L12
O12
O16
L8
O1
218
9
97
O8
256
253
217
233
L1

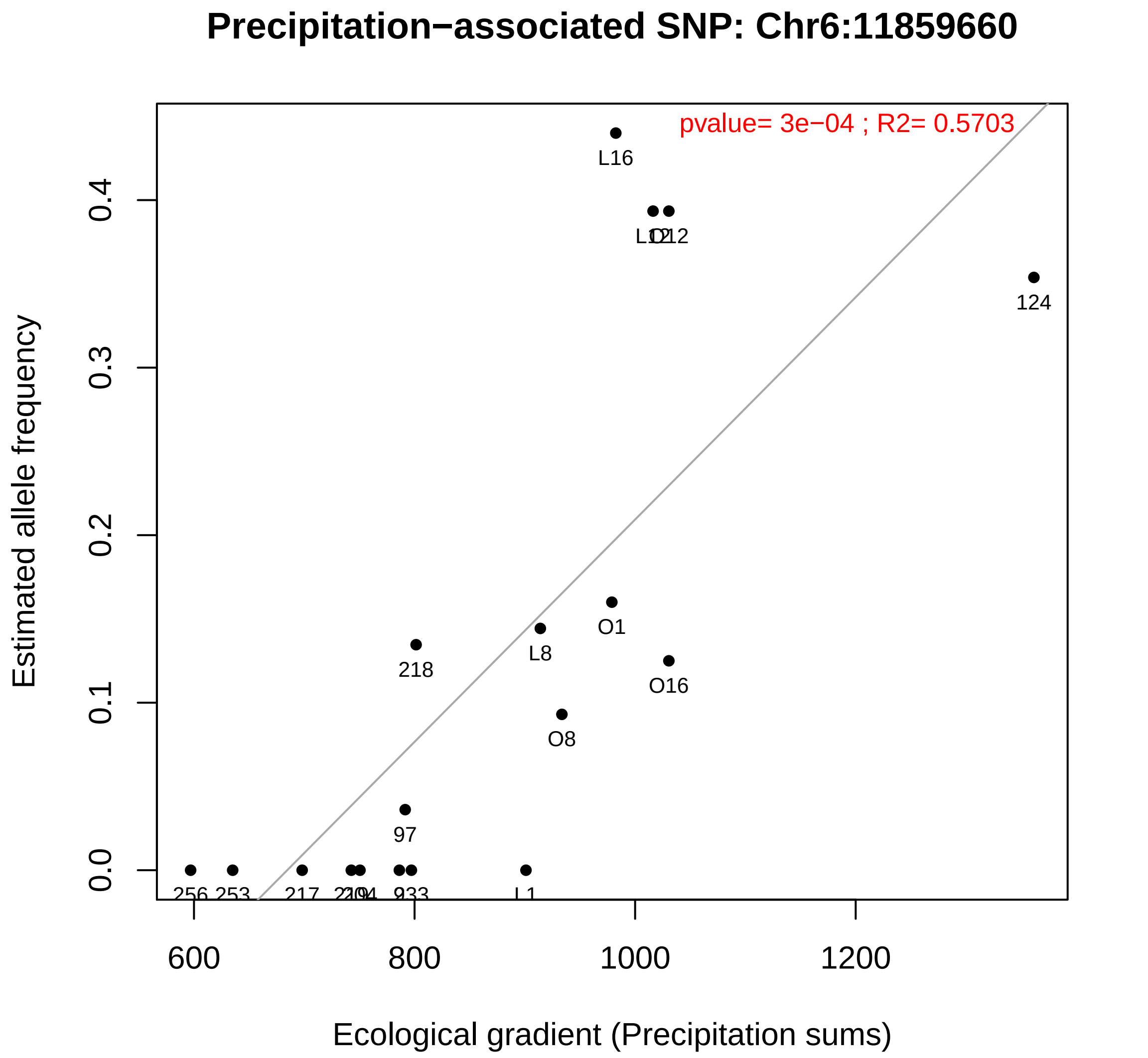

Precipitation−associated SNP: Chr6:11859660
pvalue= 3e−04 ; R2= 0.5703
L16
L12
O12
124
O1
L8
218
O16
O8
97
256
253
217
L1
0.0
0.1
0.2
0.3
0.4
600
800
1000
1200
Estimated allele frequency
Ecological gradient (Precipitation sums)

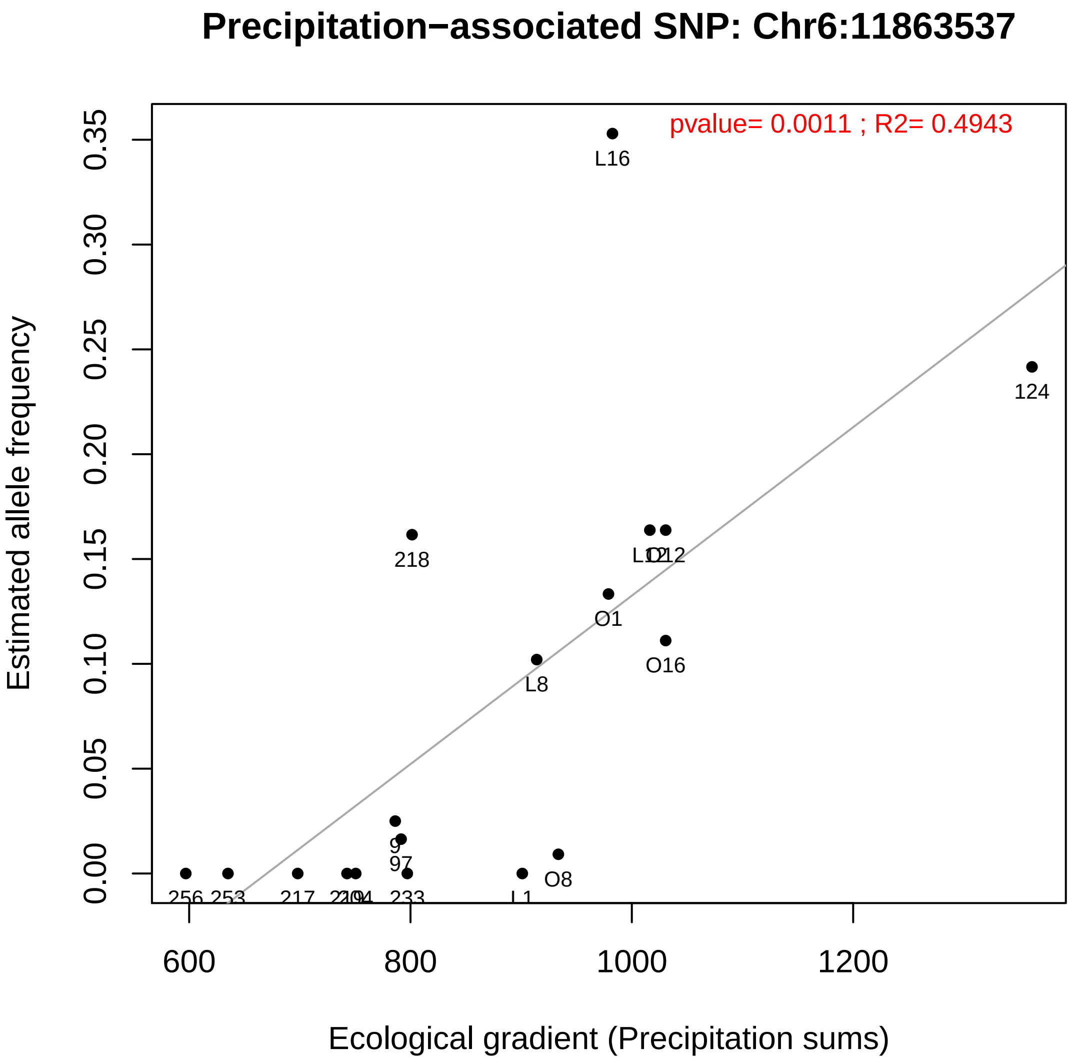

Precipitation−associated SNP: Chr6:11863537
pvalue= 0.0011 ; R2= 0.4943
Estimated allele frequency
Ecological gradient (Precipitation sums)
0.00
0.05
0.10
0.15
0.20
0.25
0.30
0.35
600
800
1000
1200
L16
124
218
O1
O16
L8
O8
9
97
233
256
253
217
L1

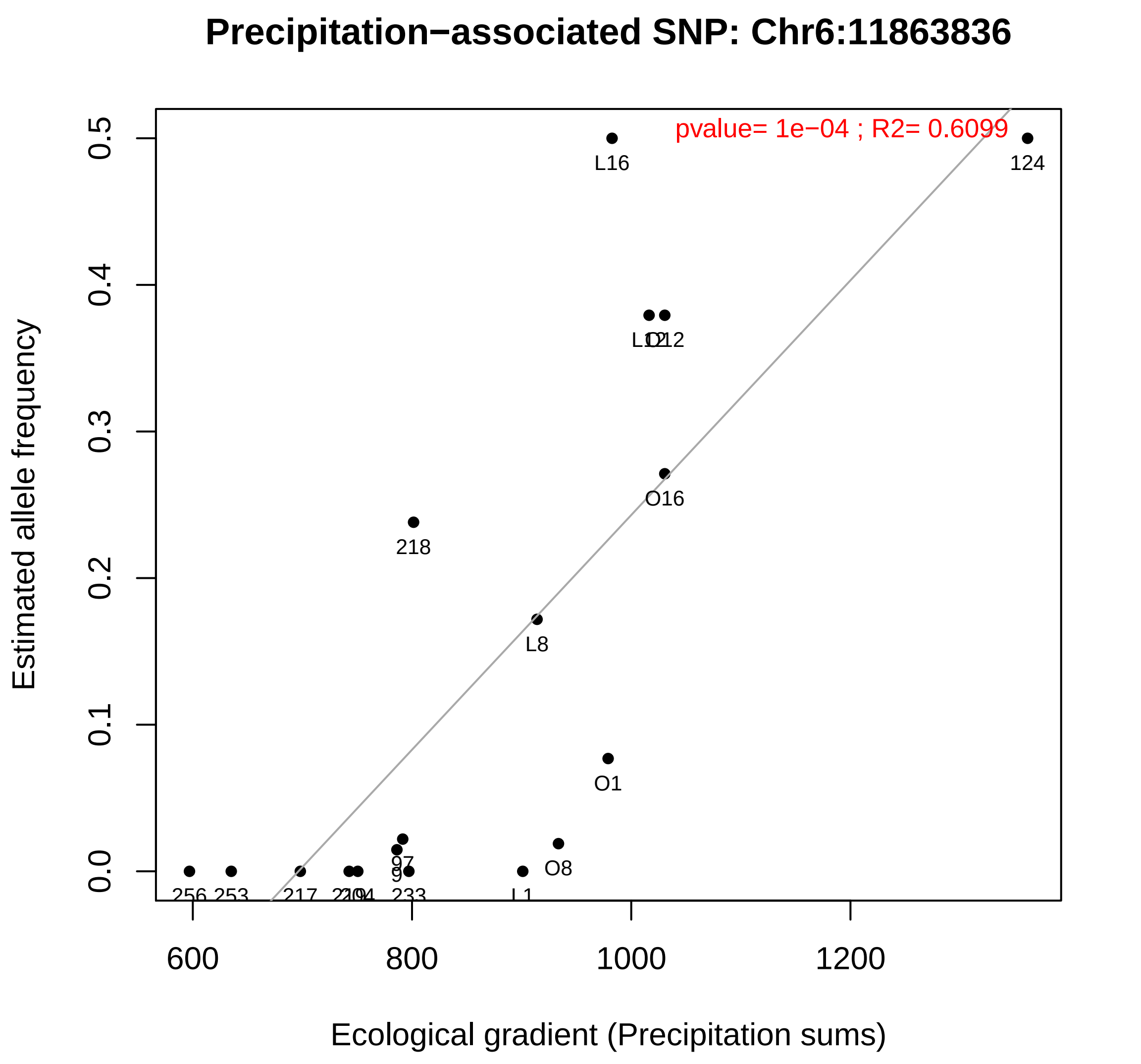

Precipitation−associated SNP: Chr6:11863836
pvalue= 1e−04 ; R2= 0.6099
Estimated allele frequency
Ecological gradient (Precipitation sums)
0.0
0.1
0.2
0.3
0.4
0.5
600
800
1000
1200
L16
124
O12
O16
218
L8
O1
O8
97
233
256
253
217

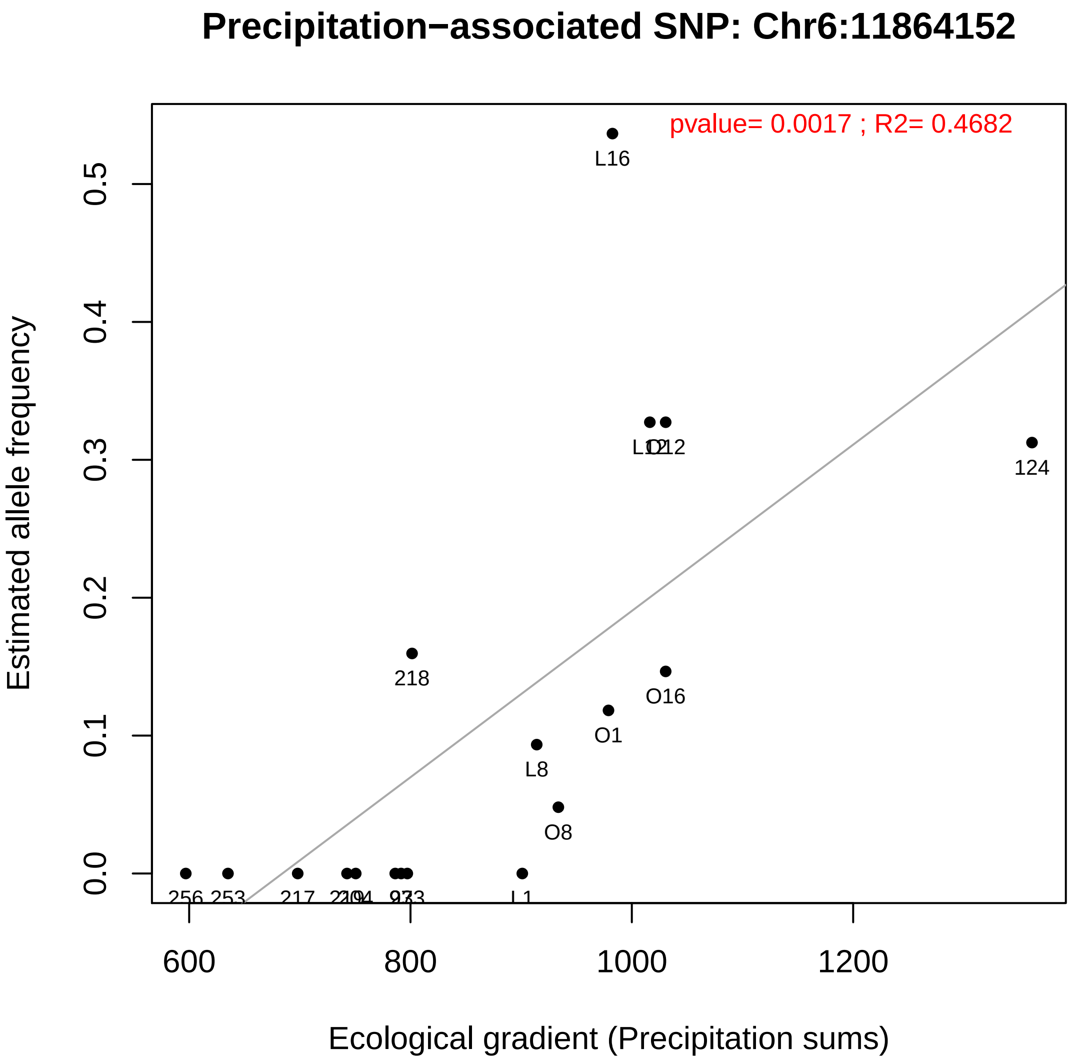

Precipitation−associated SNP: Chr6:11864152
pvalue= 0.0017 ; R2= 0.4682
Estimated allele frequency
Ecological gradient (Precipitation sums)
0.0
0.1
0.2
0.3
0.4
0.5
600
800
1000
1200
L16
124
218
O16
O1
L8
O8
256
253
217
L1

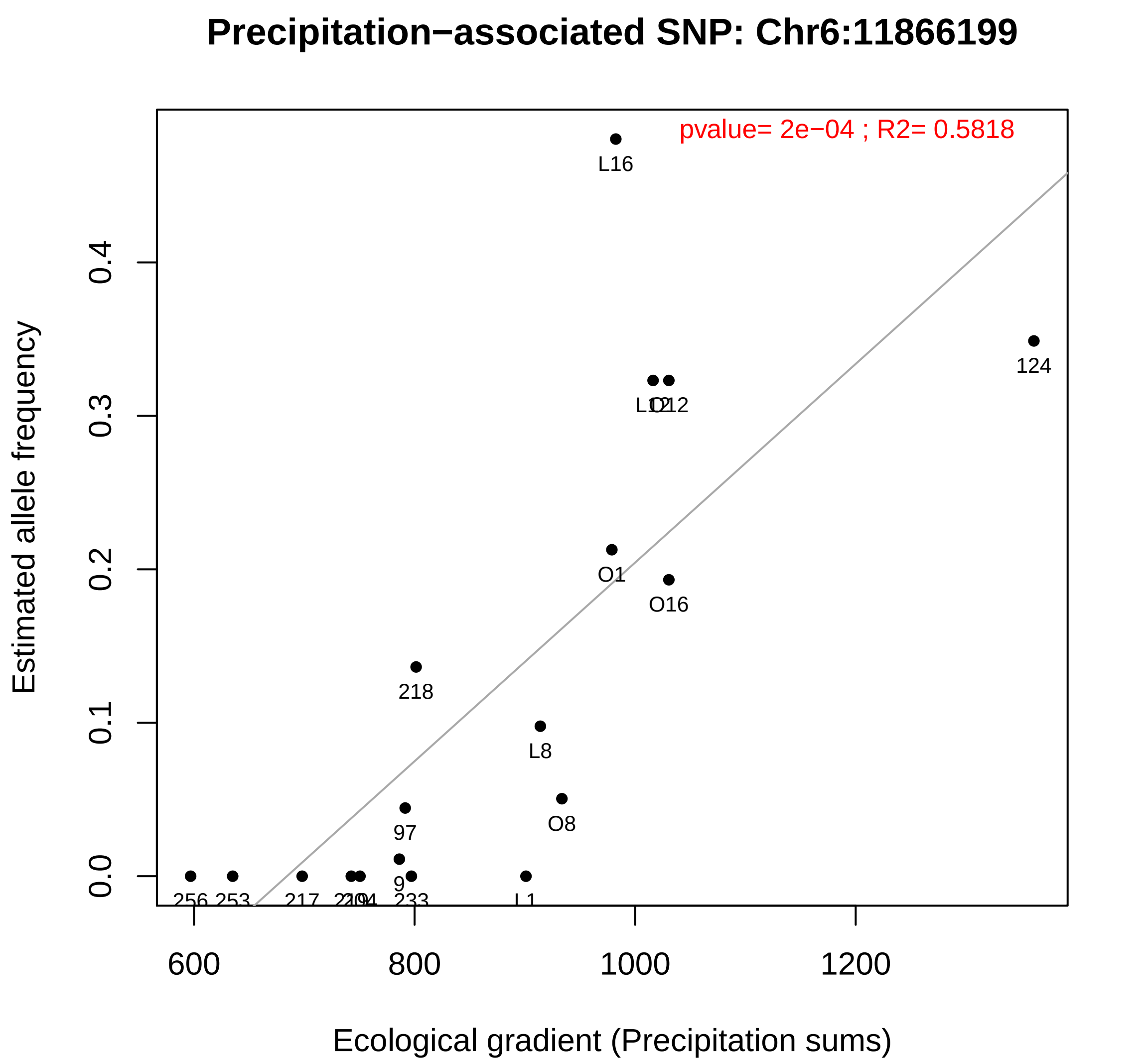

Precipitation−associated SNP: Chr6:11866199
pvalue= 2e−04 ; R2= 0.5818
Estimated allele frequency
Ecological gradient (Precipitation sums)
0.0
0.1
0.2
0.3
0.4
600
800
1000
1200
L16
124
O1
O16
218
L8
O8
97
9
256
253
217
233

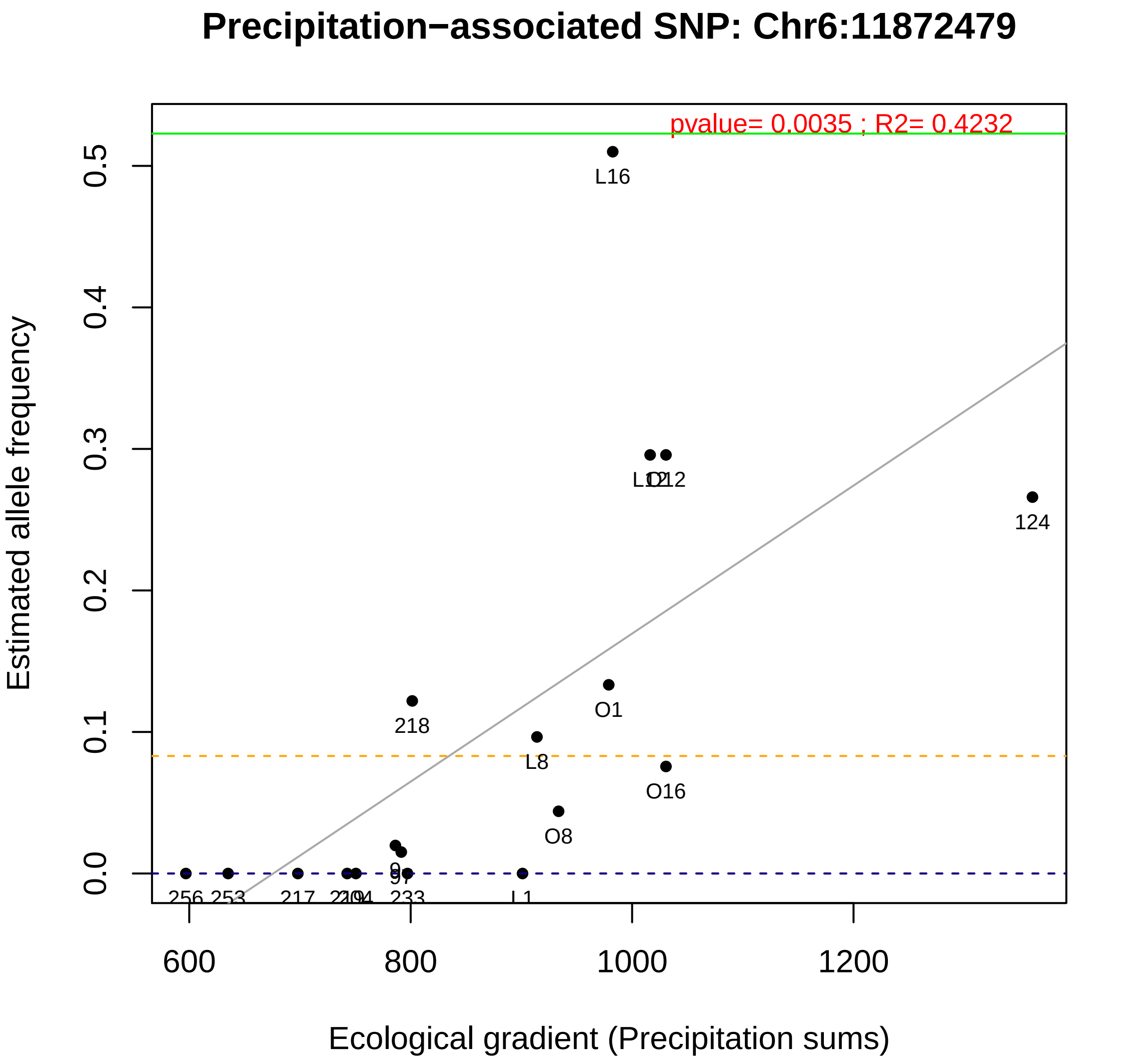

Precipitation−associated SNP: Chr6:11872479
pvalue= 0.0035 ; R2= 0.4232
Estimated allele frequency
Ecological gradient (Precipitation sums)
0.0
0.1
0.2
0.3
0.4
0.5
600
800
1000
1200
L16
124
218
O1
L8
O16
O8

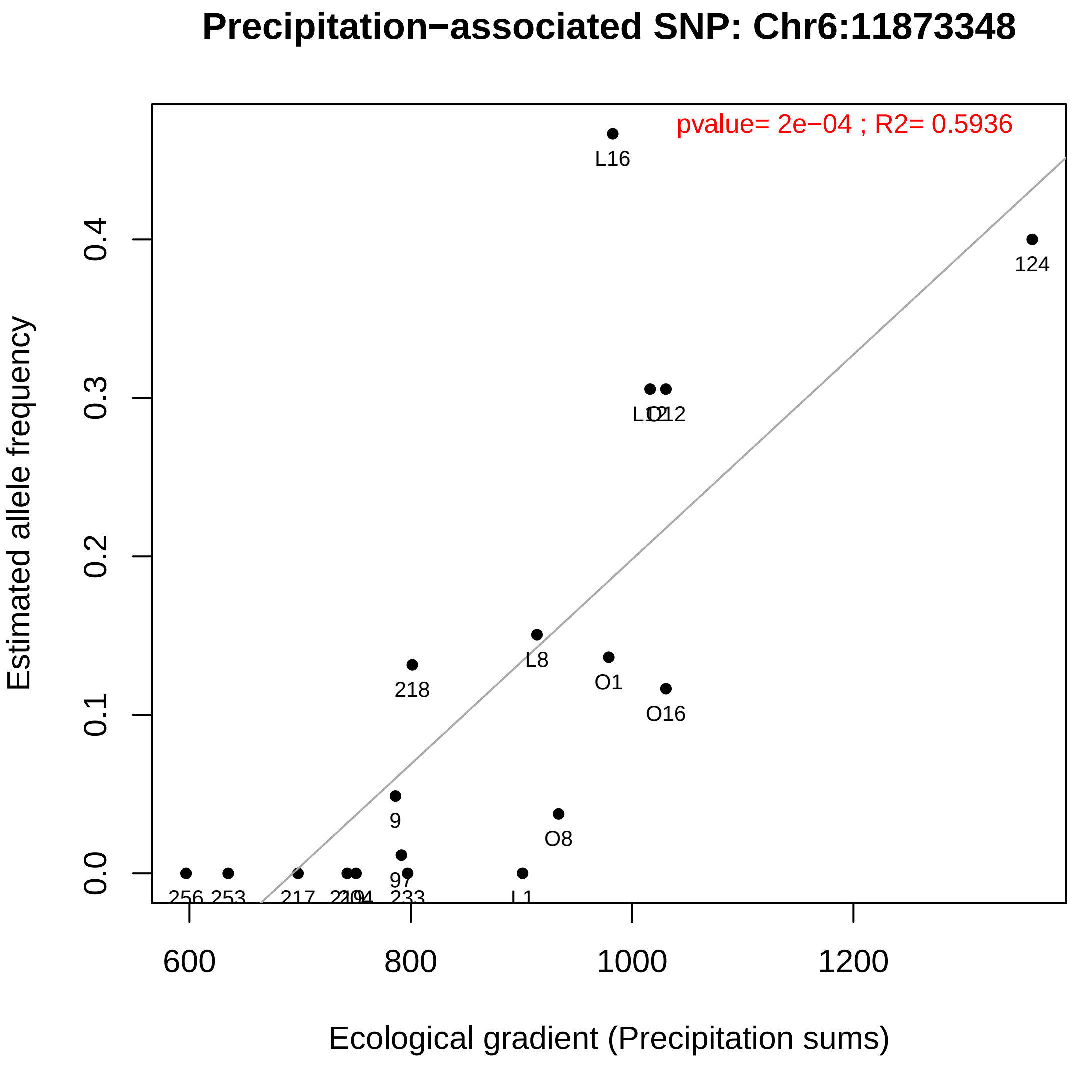
Precipitation−associated SNP: Chr6:11873348
pvalue= 2e−04 ; R2= 0.5936
Estimated allele frequency
Ecological gradient (Precipitation sums)
0.0
0.1
0.2
0.3
0.4
600
800
1000
1200
L16
124
L8
O1
O16
218
9
O8
97
256
253
217
233

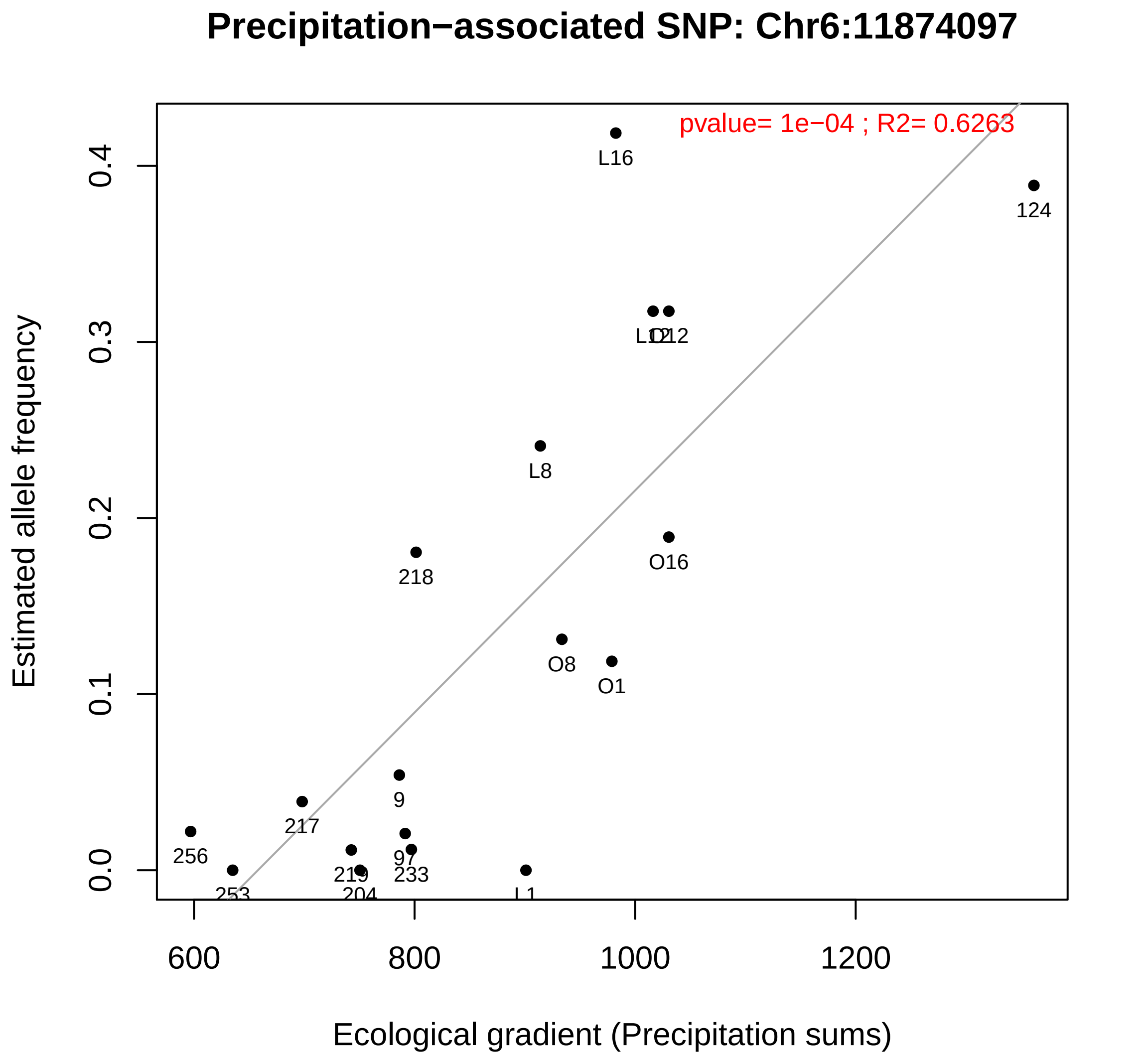
Precipitation−associated SNP: Chr6:11874097
pvalue= 1e−04 ; R2= 0.6263
Estimated allele frequency
Ecological gradient (Precipitation sums)
0.0
0.1
0.2
0.3
0.4
600
800
1000
1200
L16
124
L12
O12
L8
O16
218
O8
O1
9
217
256
97
233
219
204
253
L1

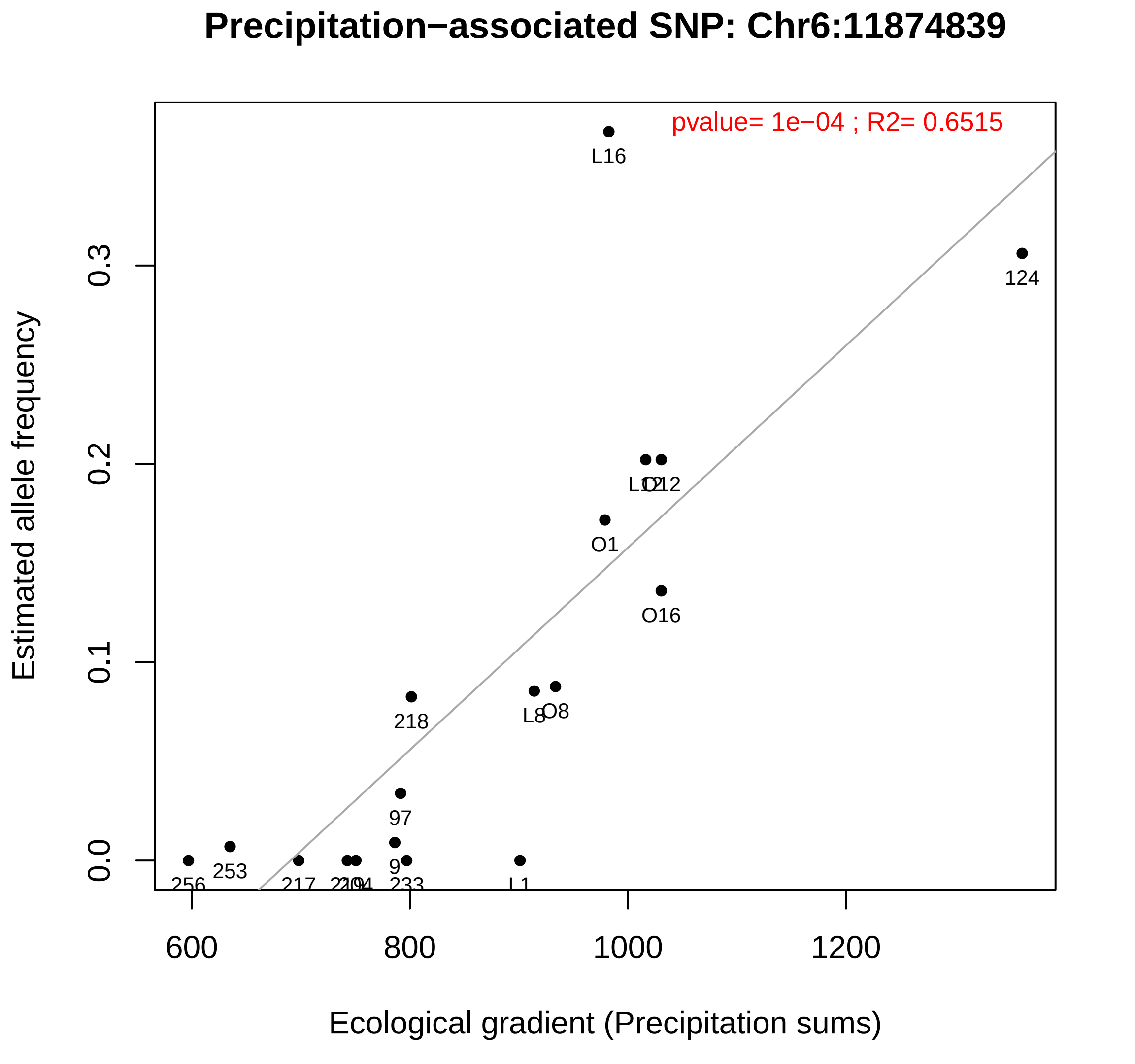

Precipitation−associated SNP: Chr6:11874839
pvalue= 1e−04 ; R2= 0.6515
L16
124
L12
O12
O1
O16
218
L8
O8
97
9
253
256
217
233
L1
0.0
0.1
0.2
0.3
Estimated allele frequency
600
800
1000
1200
Ecological gradient (Precipitation sums)

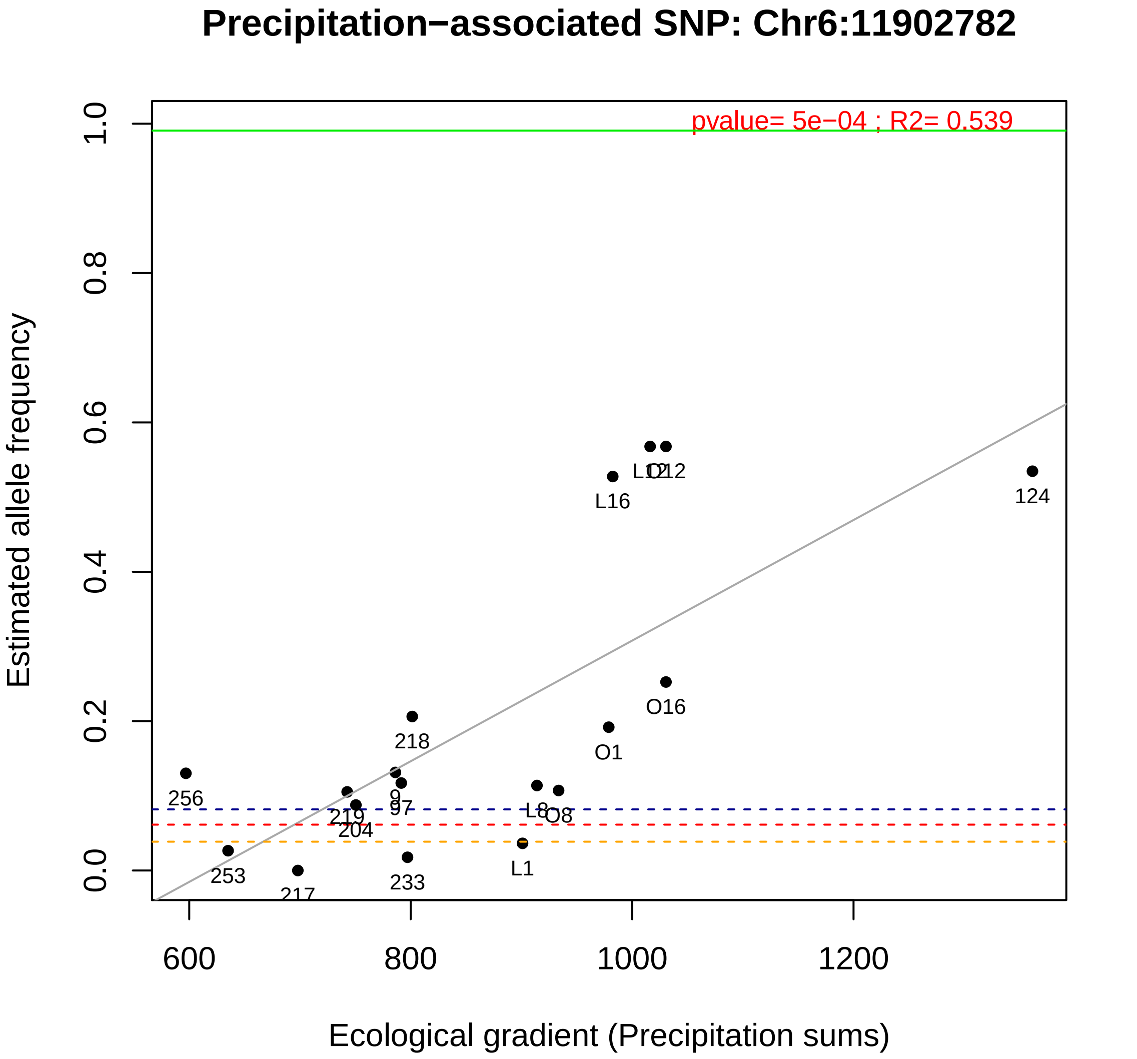
Precipitation−associated SNP: Chr6:11902782
pvalue= 5e−04 ; R2= 0.539
Estimated allele frequency
0.0
0.2
0.4
0.6
0.8
1.0
Ecological gradient (Precipitation sums)
600
800
1000
1200
L16
124
O16
218
O1
256
9
97
219
204
L8
O8
L1
253
233
217

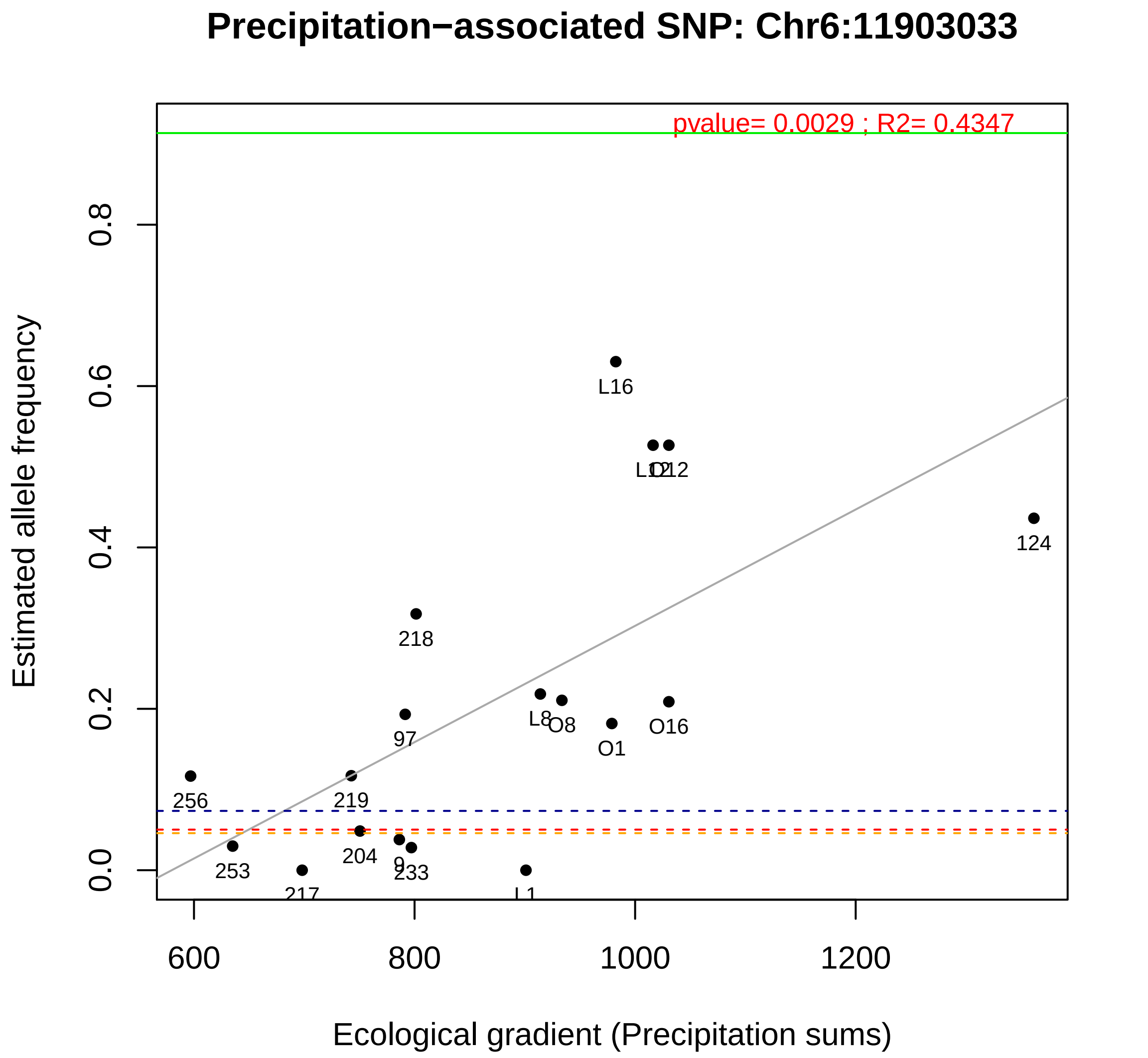
Precipitation−associated SNP: Chr6:11903033
pvalue= 0.0029 ; R2= 0.4347
Estimated allele frequency
0.0
0.2
0.4
0.6
0.8
600
800
1000
1200
Ecological gradient (Precipitation sums)
L16
L12
O12
124
218
L8
O8
O16
O1
97
256
219
253
204
9
233
217
L1

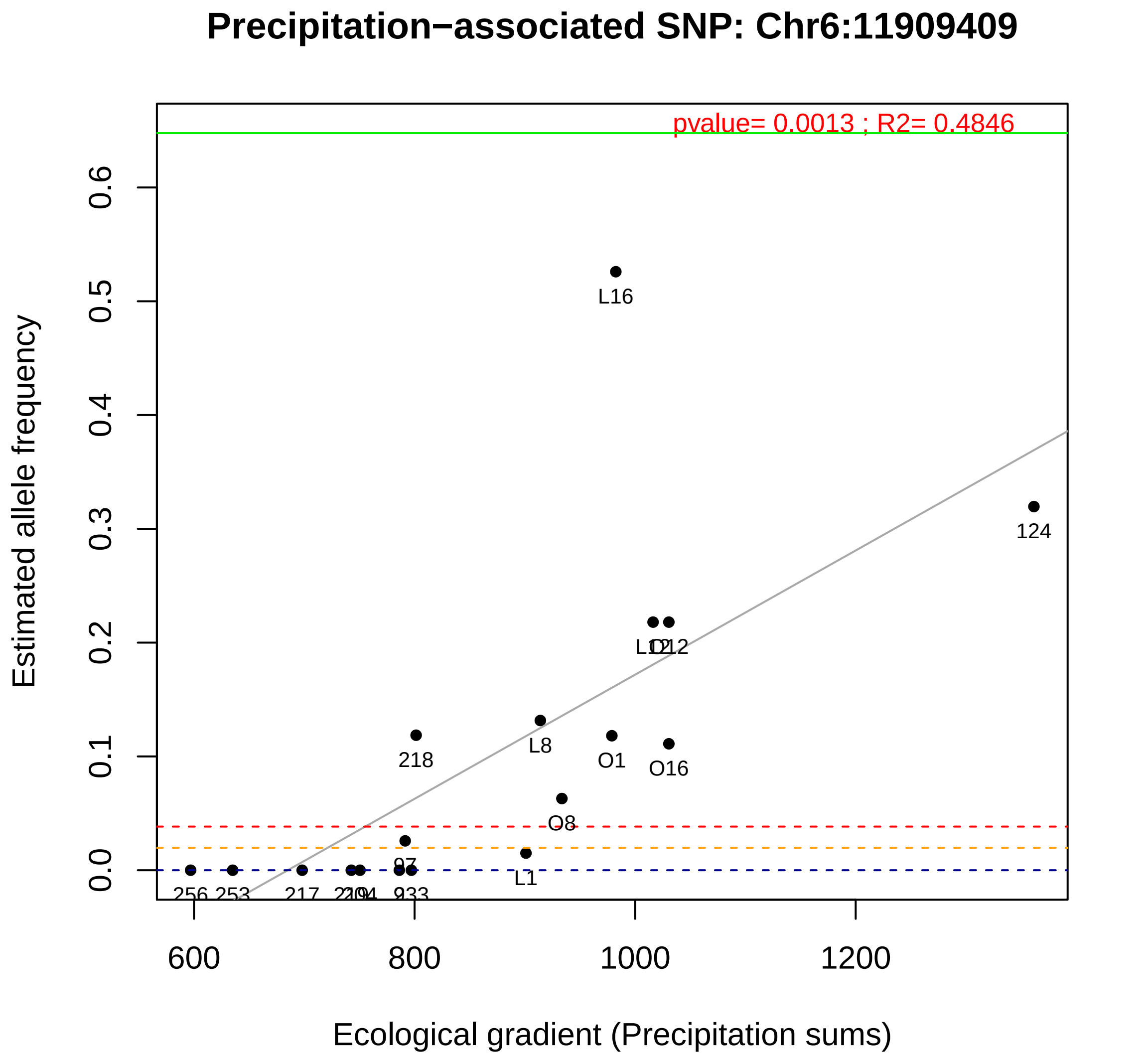
Precipitation−associated SNP: Chr6:11909409
pvalue= 0.0013 ; R2= 0.4846
Estimated allele frequency
0.0
0.1
0.2
0.3
0.4
0.5
0.6
600
800
1000
1200
Ecological gradient (Precipitation sums)
L16
124
L12
O12
L8
218
O1
O16
O8
L1
256
253
217

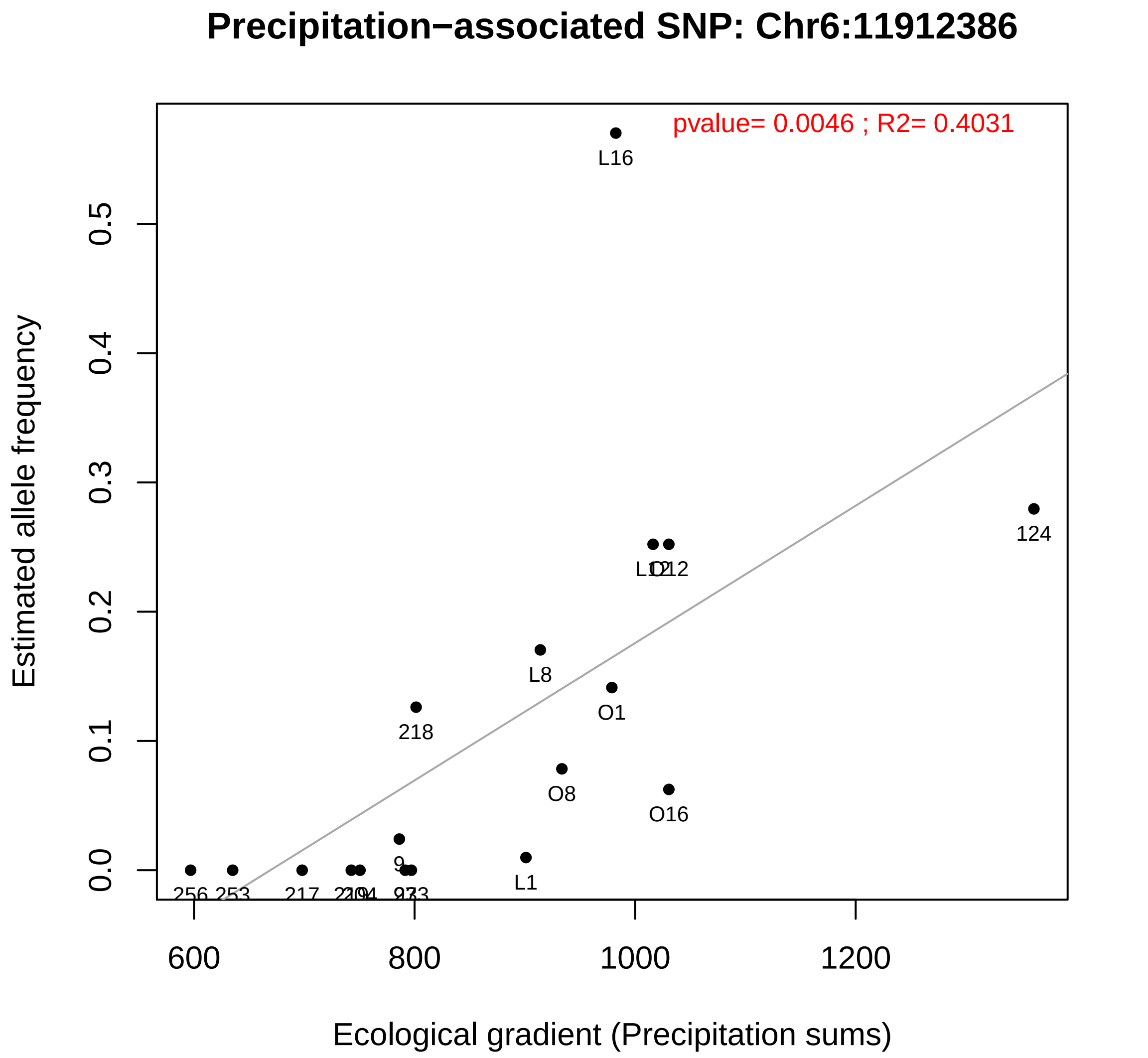
Precipitation−associated SNP: Chr6:11912386
pvalue= 0.0046 ; R2= 0.4031
Estimated allele frequency
Ecological gradient (Precipitation sums)
0.0
0.1
0.2
0.3
0.4
0.5
600
800
1000
1200
L16
124
L12
O12
L8
O1
218
O8
O16
L1
256
253
217
9

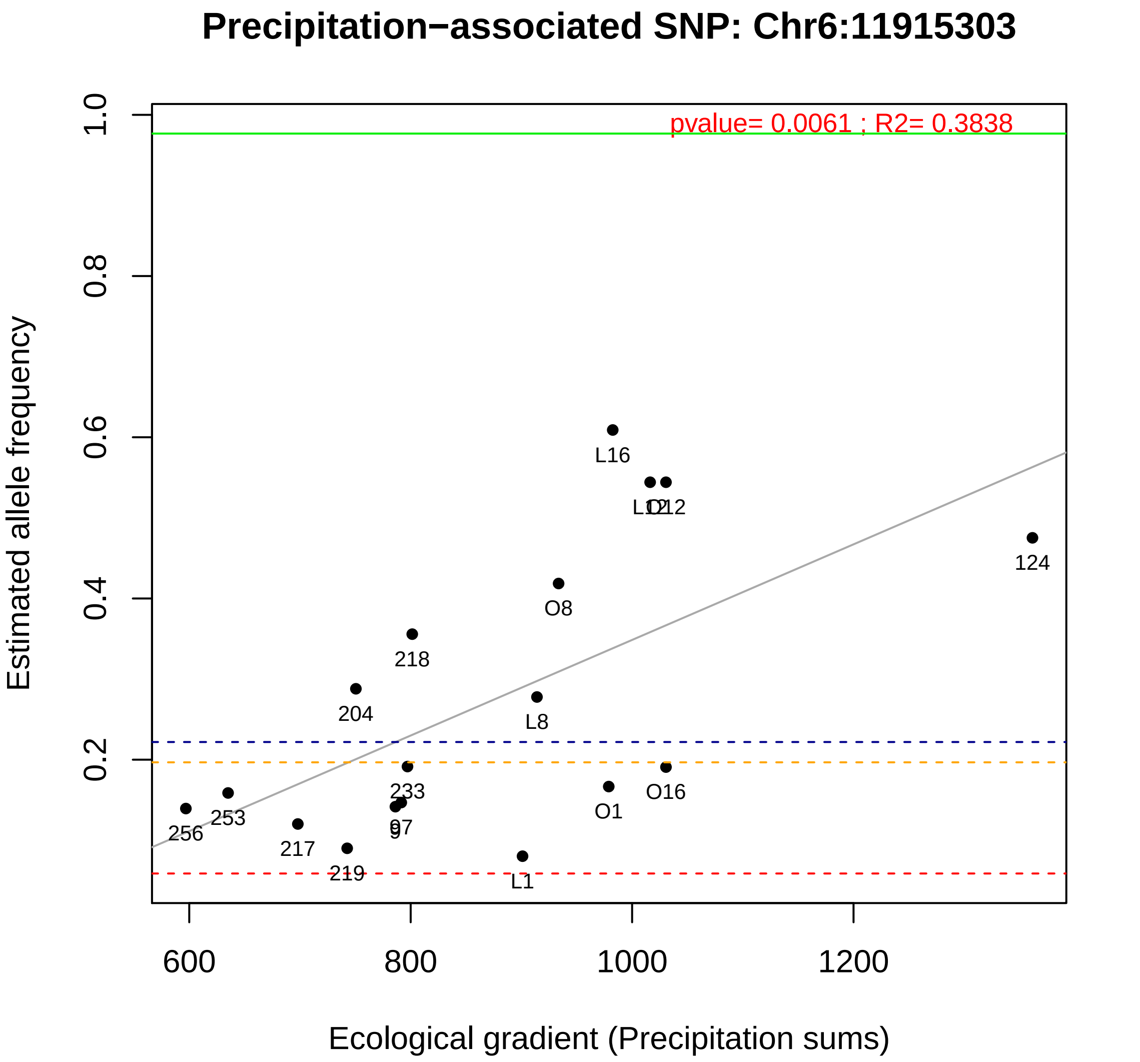

Precipitation−associated SNP: Chr6:11915303
pvalue= 0.0061 ; R2= 0.3838
Estimated allele frequency
1.0
0.8
0.6
0.4
0.2
600
800
1000
1200
Ecological gradient (Precipitation sums)
L16
L12
O12
124
O8
218
204
L8
233
O16
O1
256
253
217
219
97
9
L1

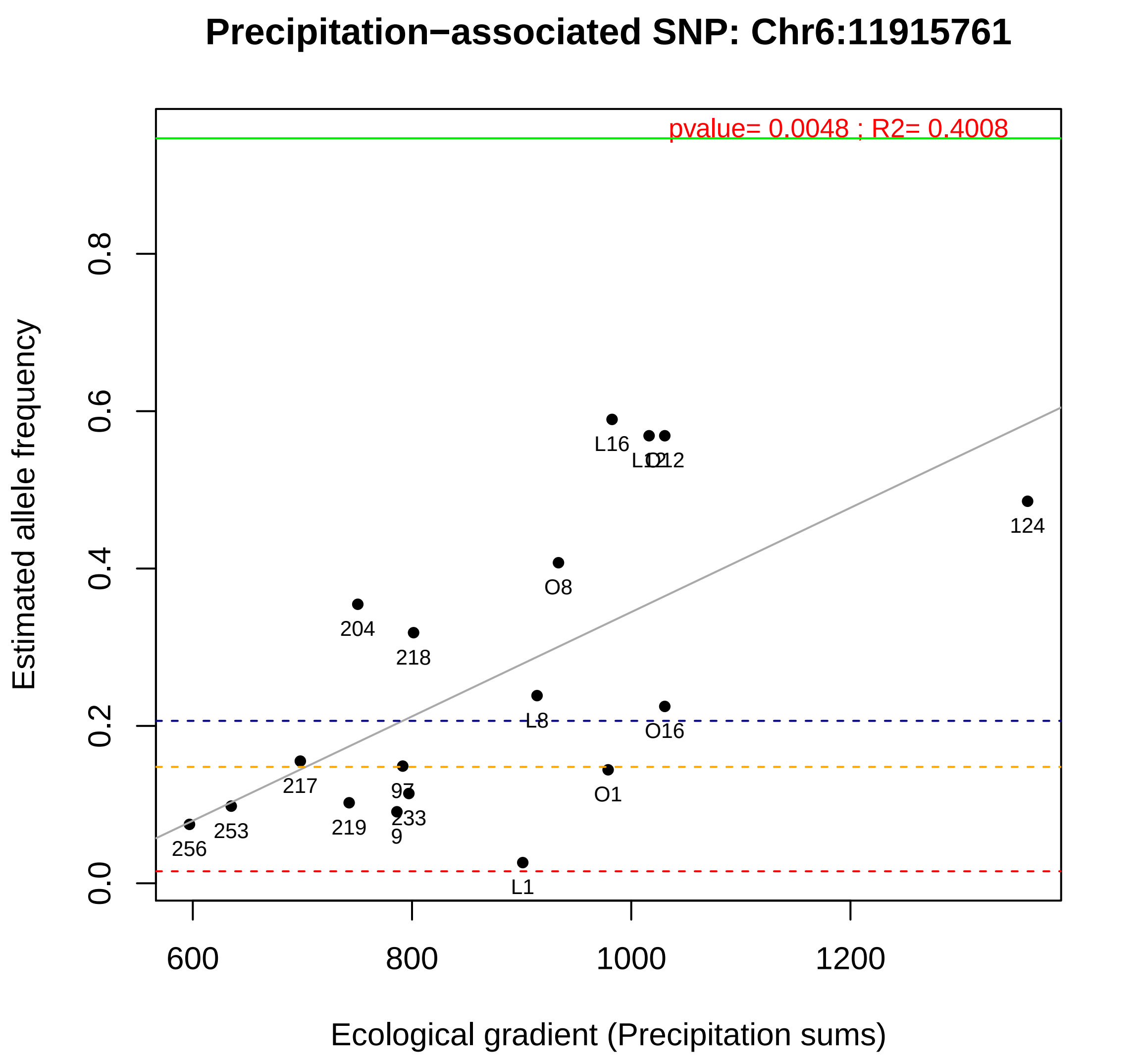

Precipitation−associated SNP: Chr6:11915761
pvalue= 0.0048 ; R2= 0.4008
Estimated allele frequency
0.0
0.2
0.4
0.6
0.8
600
800
1000
1200
Ecological gradient (Precipitation sums)
L16
L12
O12
124
O8
204
218
L8
O16
217
97
233
9
O1
219
253
256
L1

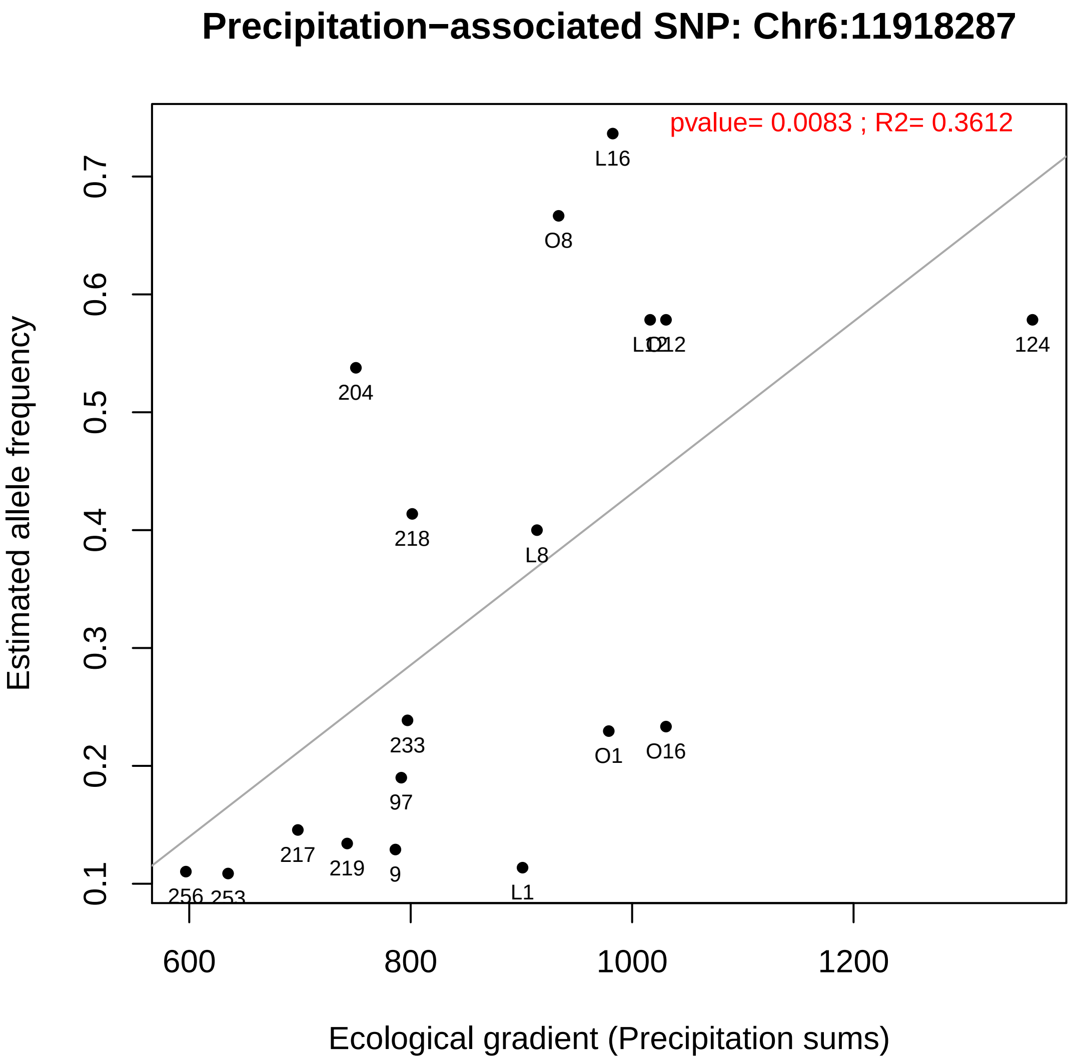

Precipitation−associated SNP: Chr6:11918287
pvalue= 0.0083 ; R2= 0.3612
Estimated allele frequency
0.1
0.2
0.3
0.4
0.5
0.6
0.7
600
800
1000
1200
Ecological gradient (Precipitation sums)
L16
O8
L12
O12
124
204
218
L8
233
O1
O16
97
217
219
9
L1
256
253

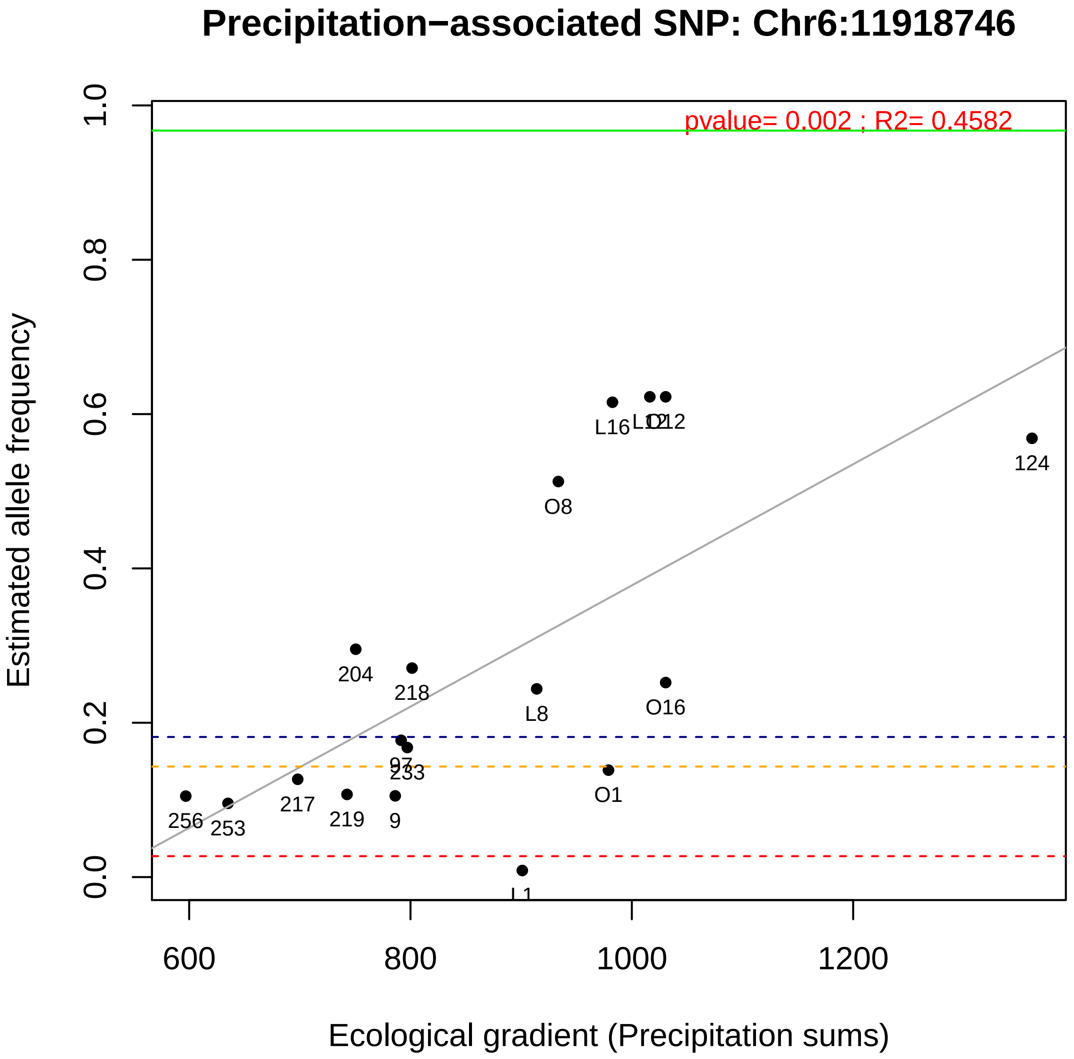
Precipitation−associated SNP: Chr6:11918746
pvalue= 0.002 ; R2= 0.4582
Estimated allele frequency
0.0
0.2
0.4
0.6
0.8
1.0
600
800
1000
1200
Ecological gradient (Precipitation sums)
L16
L12
O12
124
O8
204
218
L8
O16
97
233
O1
256
253
217
219
9
L1

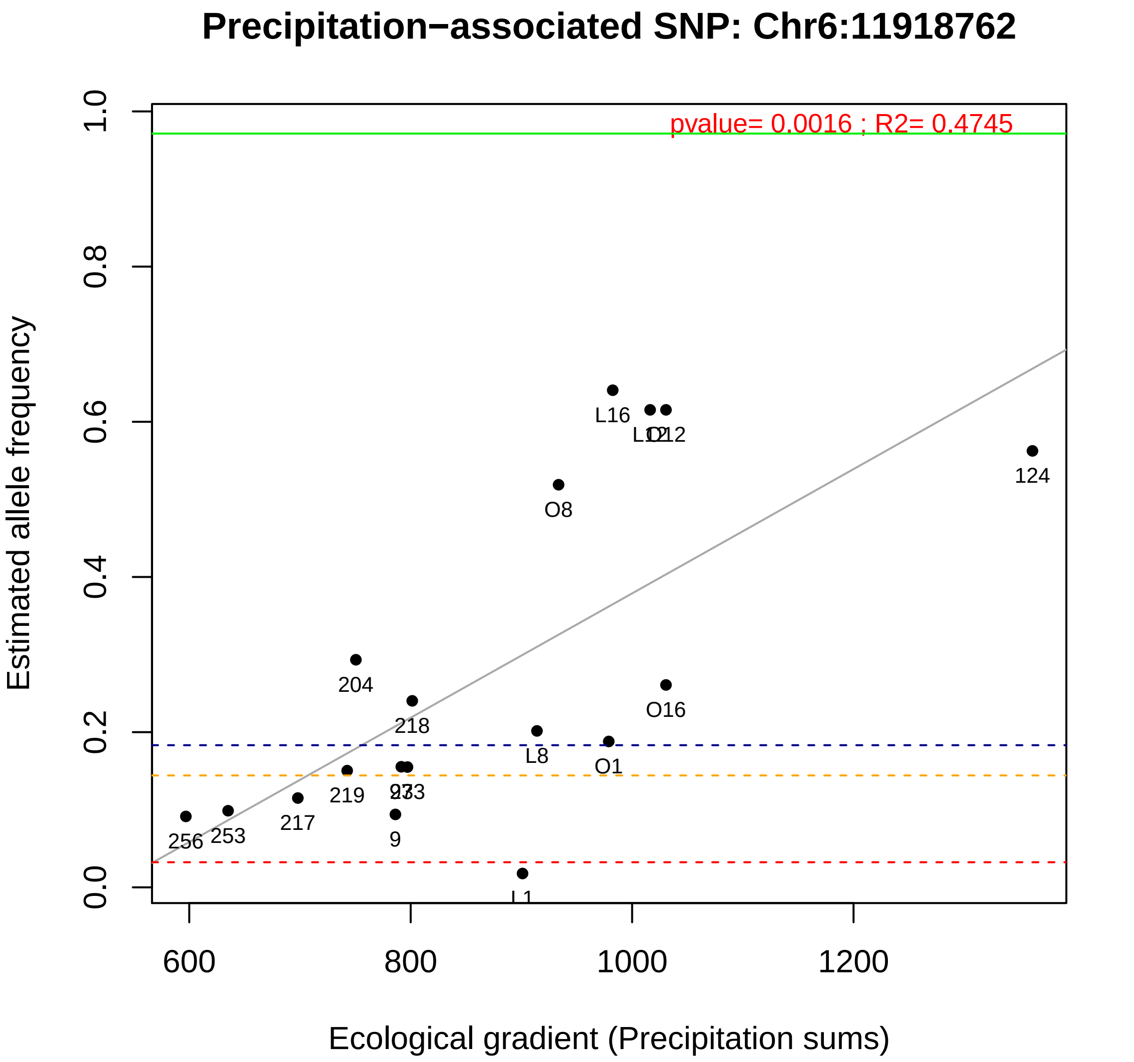

Precipitation−associated SNP: Chr6:11918762
pvalue= 0.0016 ; R2= 0.4745
Estimated allele frequency
Ecological gradient (Precipitation sums)
0.0
0.2
0.4
0.6
0.8
1.0
600
800
1000
1200
L16
O12
124
O8
204
218
O16
L8
O1
219
217
256
253
9

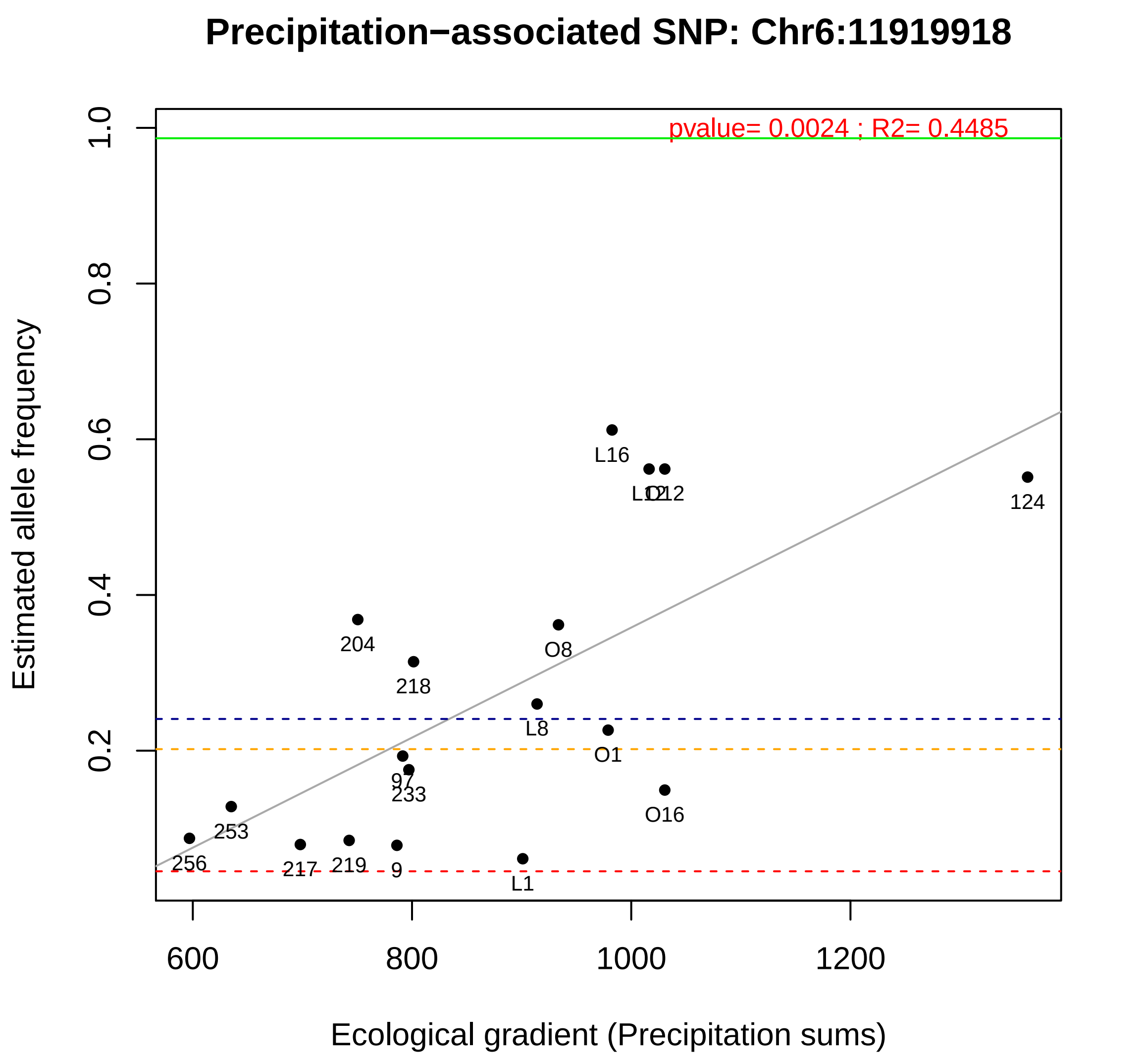

Precipitation−associated SNP: Chr6:11919918
pvalue= 0.0024 ; R2= 0.4485
Estimated allele frequency
0.2
0.4
0.6
0.8
1.0
600
800
1000
1200
Ecological gradient (Precipitation sums)
L16
L12
O12
124
204
O8
218
L8
O1
97
233
O16
253
256
217
219
9
L1

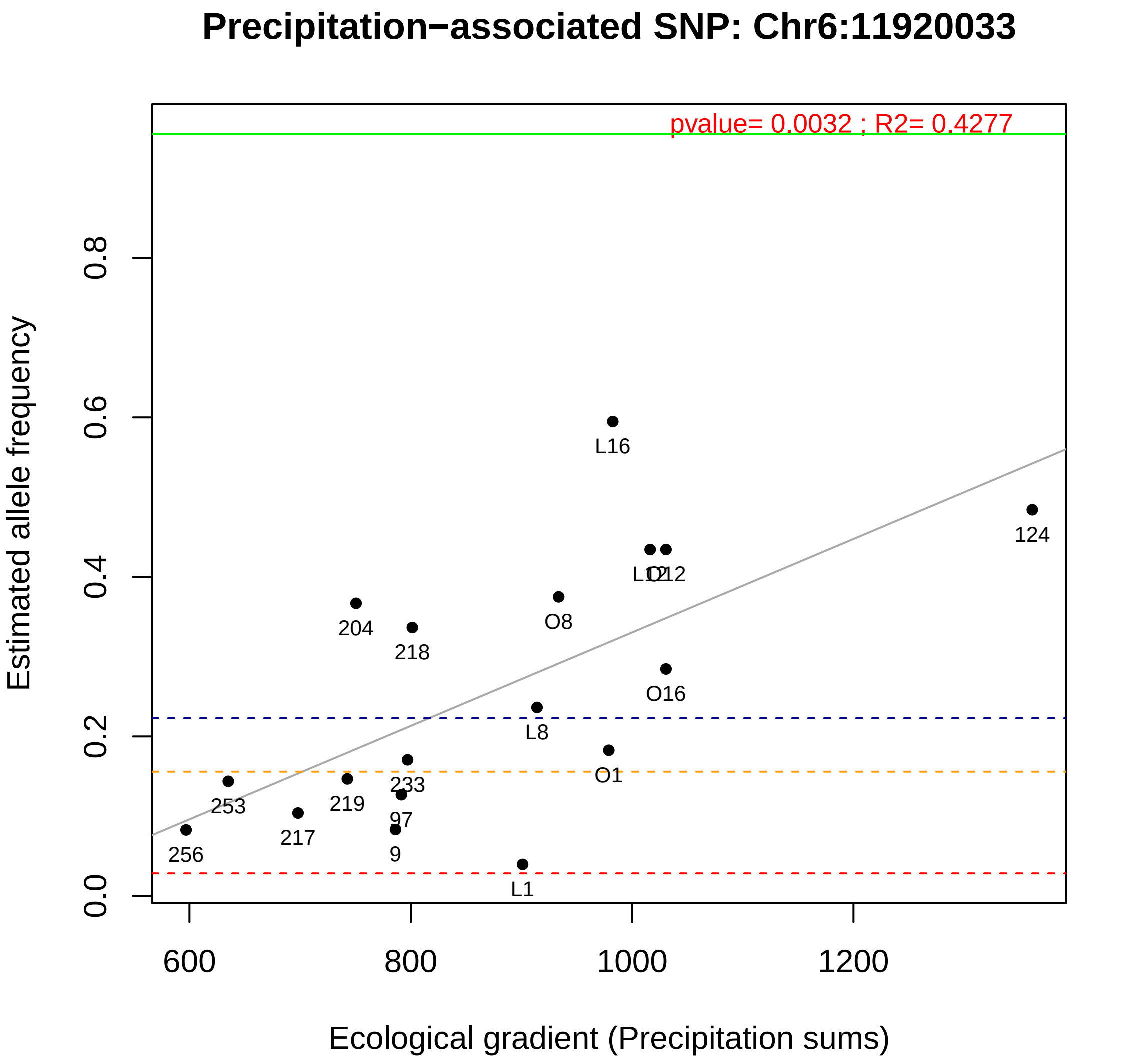

Precipitation−associated SNP: Chr6:11920033
pvalue= 0.0032 ; R2= 0.4277
Estimated allele frequency
0.0
0.2
0.4
0.6
0.8
600
800
1000
1200
Ecological gradient (Precipitation sums)
L16
124
L12
O12
O8
204
218
O16
L8
O1
233
253
219
97
217
256
9
L1

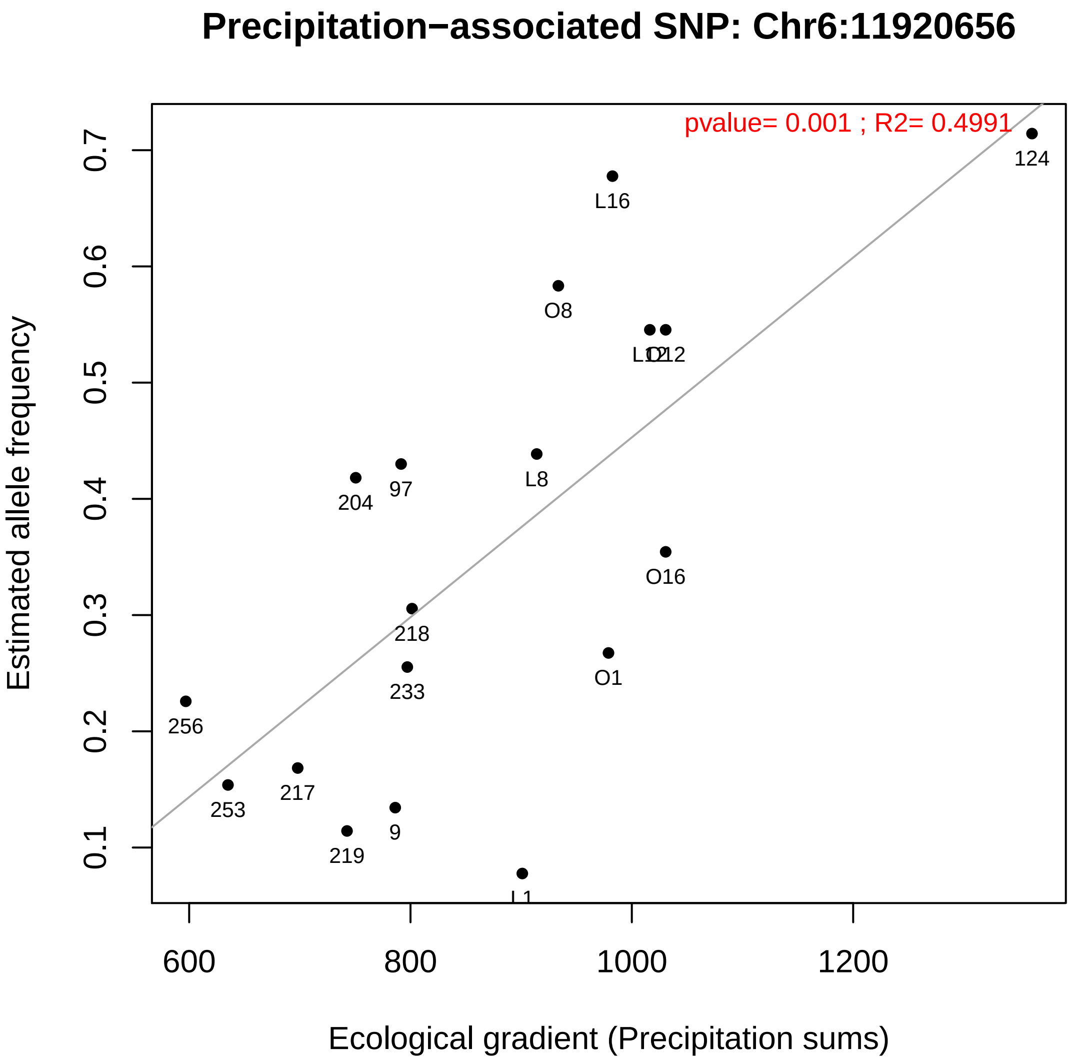
Precipitation−associated SNP: Chr6:11920656
pvalue= 0.001 ; R2= 0.4991
124
L16
O8
L12
O12
97
204
L8
O16
218
233
O1
256
217
253
9
219
L1
0.1
0.2
0.3
0.4
0.5
0.6
0.7
Estimated allele frequency
600
800
1000
1200
Ecological gradient (Precipitation sums)

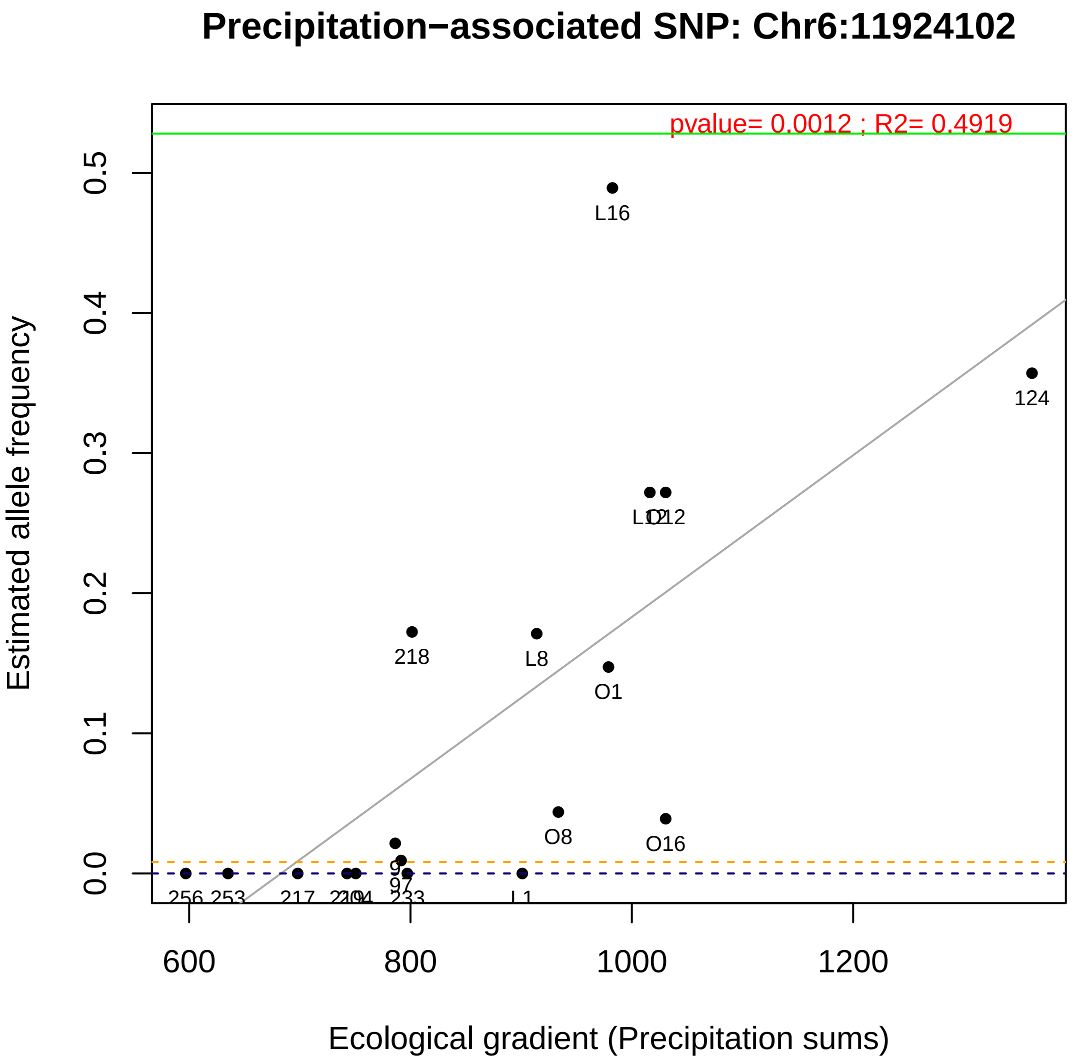
Precipitation−associated SNP: Chr6:11924102
pvalue= 0.0012 ; R2= 0.4919
Estimated allele frequency
Ecological gradient (Precipitation sums)
0.0
0.1
0.2
0.3
0.4
0.5
600
800
1000
1200
L16
124
218
L8
O1
O8
O16
97
256
253
217
233
L1

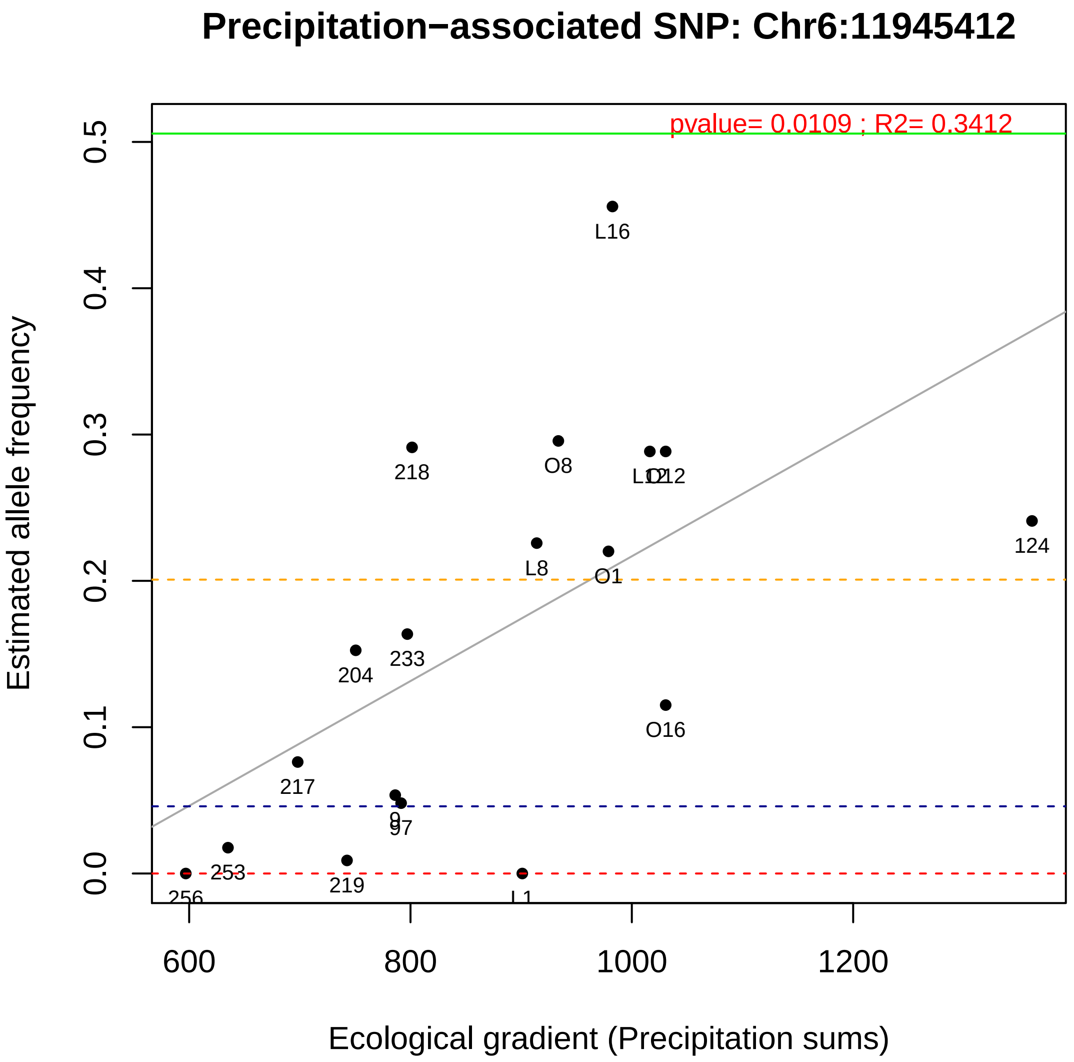

Precipitation−associated SNP: Chr6:11945412
pvalue= 0.0109 ; R2= 0.3412
Estimated allele frequency
0.0
0.1
0.2
0.3
0.4
0.5
600
800
1000
1200
Ecological gradient (Precipitation sums)
L16
218
O8
L12
O12
124
L8
O1
233
204
O16
217
9
97
253
219
256
L1

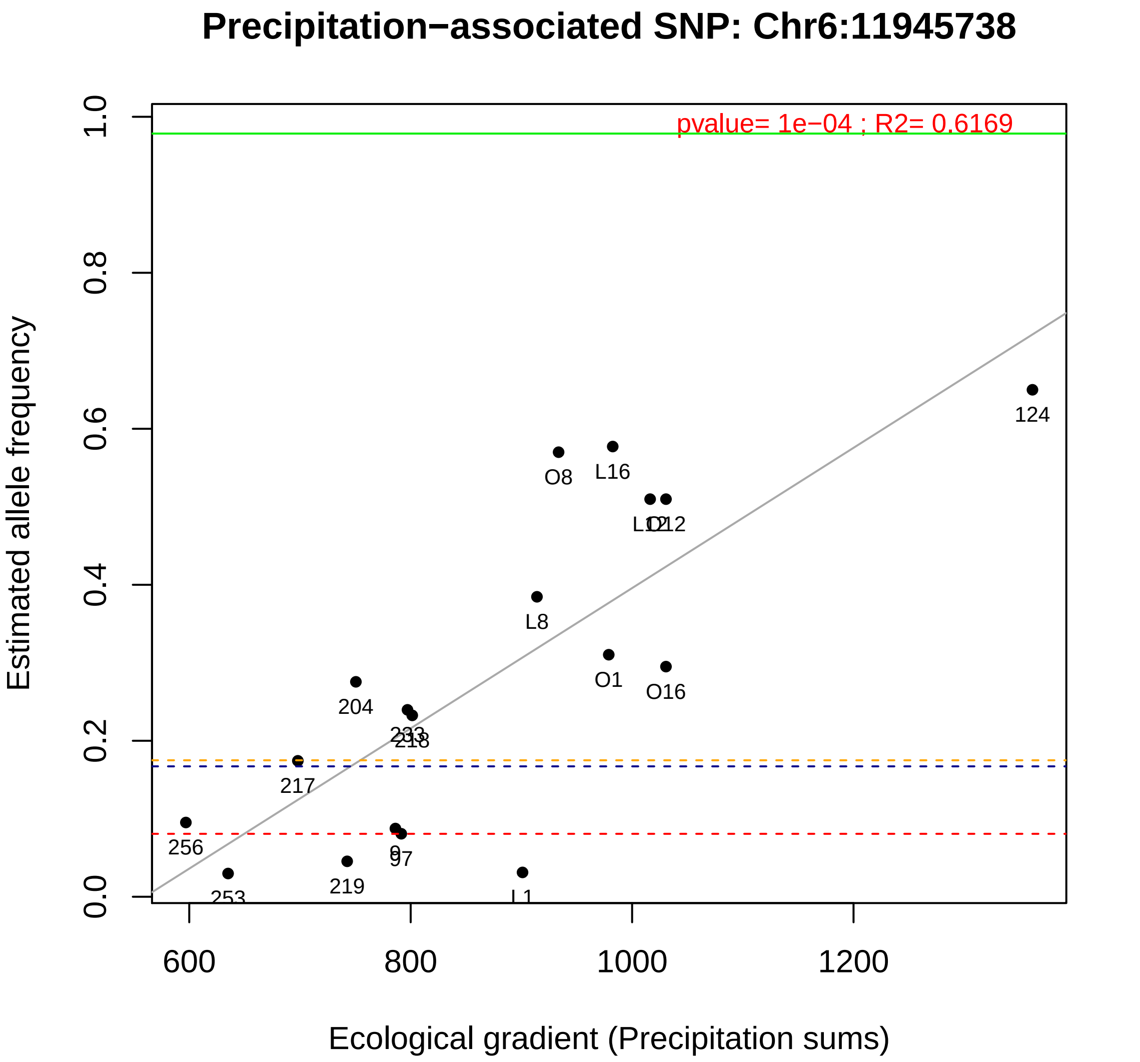
Precipitation−associated SNP: Chr6:11945738
pvalue= 1e−04 ; R2= 0.6169
Estimated allele frequency
0.0
0.2
0.4
0.6
0.8
1.0
600
800
1000
1200
Ecological gradient (Precipitation sums)
124
O8
L16
L12
O12
L8
O1
O16
204
233
218
217
256
253
219
9
97
L1

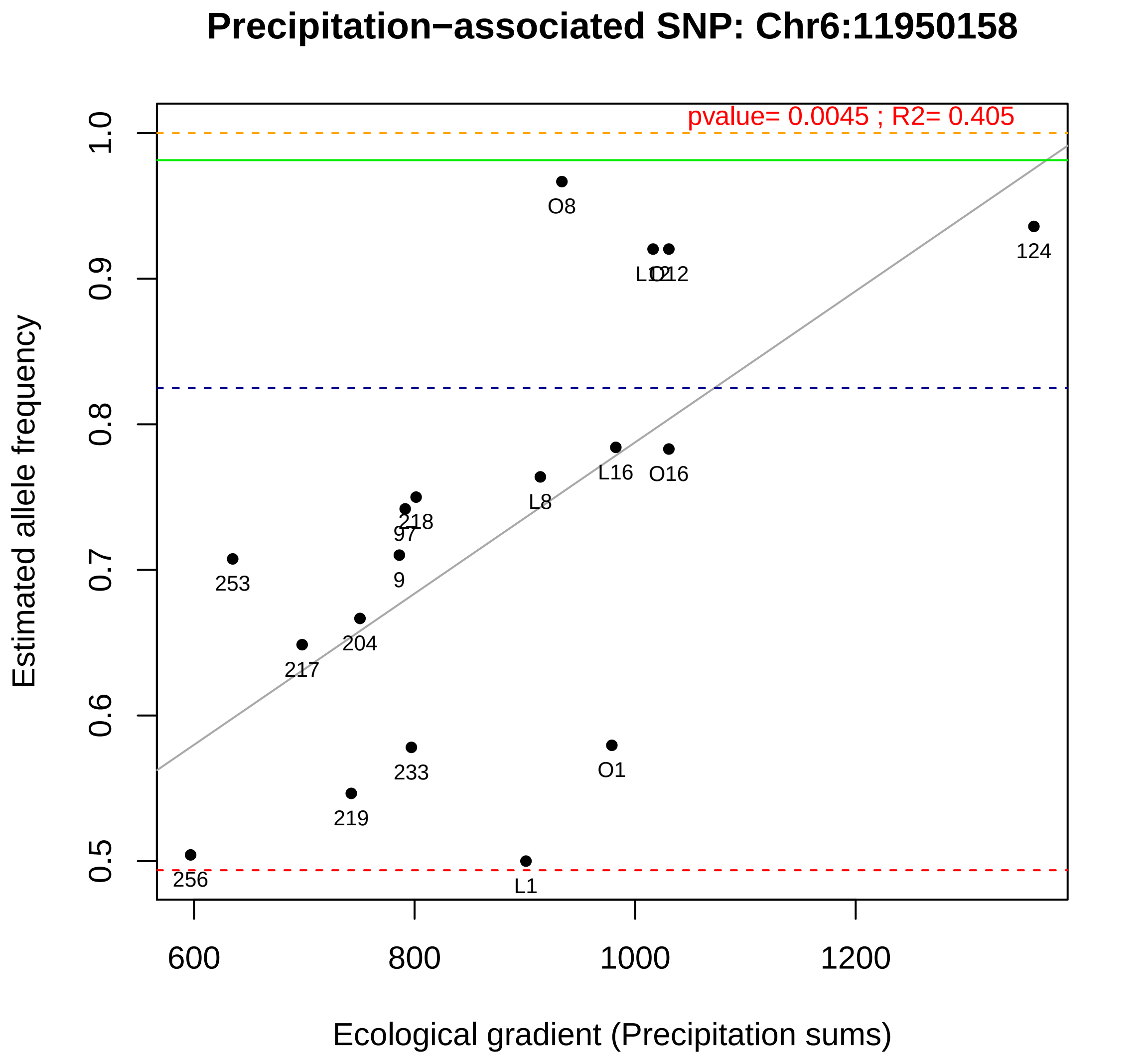
Precipitation−associated SNP: Chr6:11950158
pvalue= 0.0045 ; R2= 0.405
Estimated allele frequency
Ecological gradient (Precipitation sums)
0.5
0.6
0.7
0.8
0.9
1.0
600
800
1000
1200
O8
124
L12
O12
L16
O16
L8
218
97
9
253
204
217
233
O1
219
256
L1

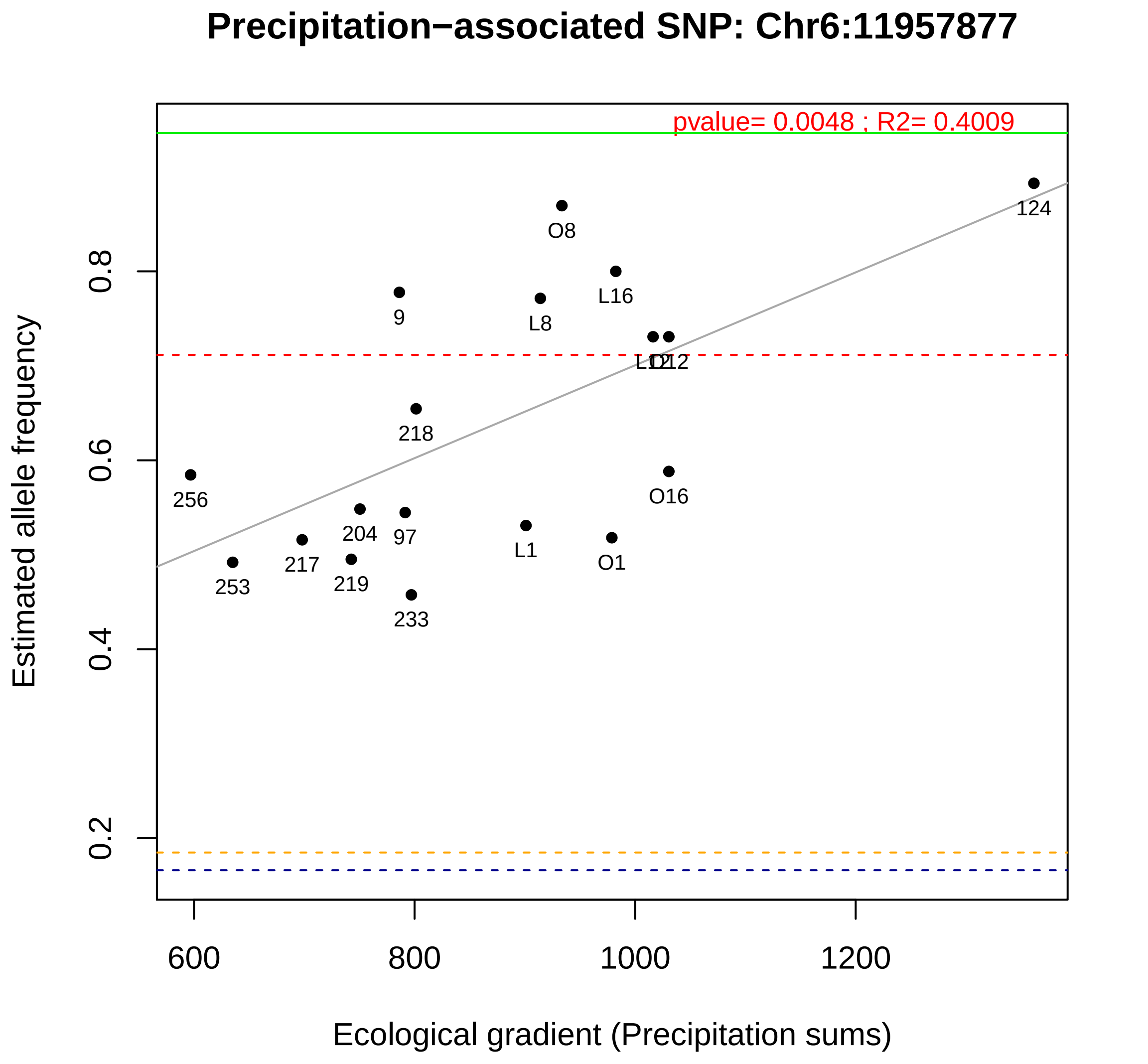

Precipitation−associated SNP: Chr6:11957877
pvalue= 0.0048 ; R2= 0.4009
Estimated allele frequency
0.2
0.4
0.6
0.8
600
800
1000
1200
Ecological gradient (Precipitation sums)
124
O8
L16
9
L8
L12
O12
218
256
O16
204
97
L1
O1
217
253
219
233

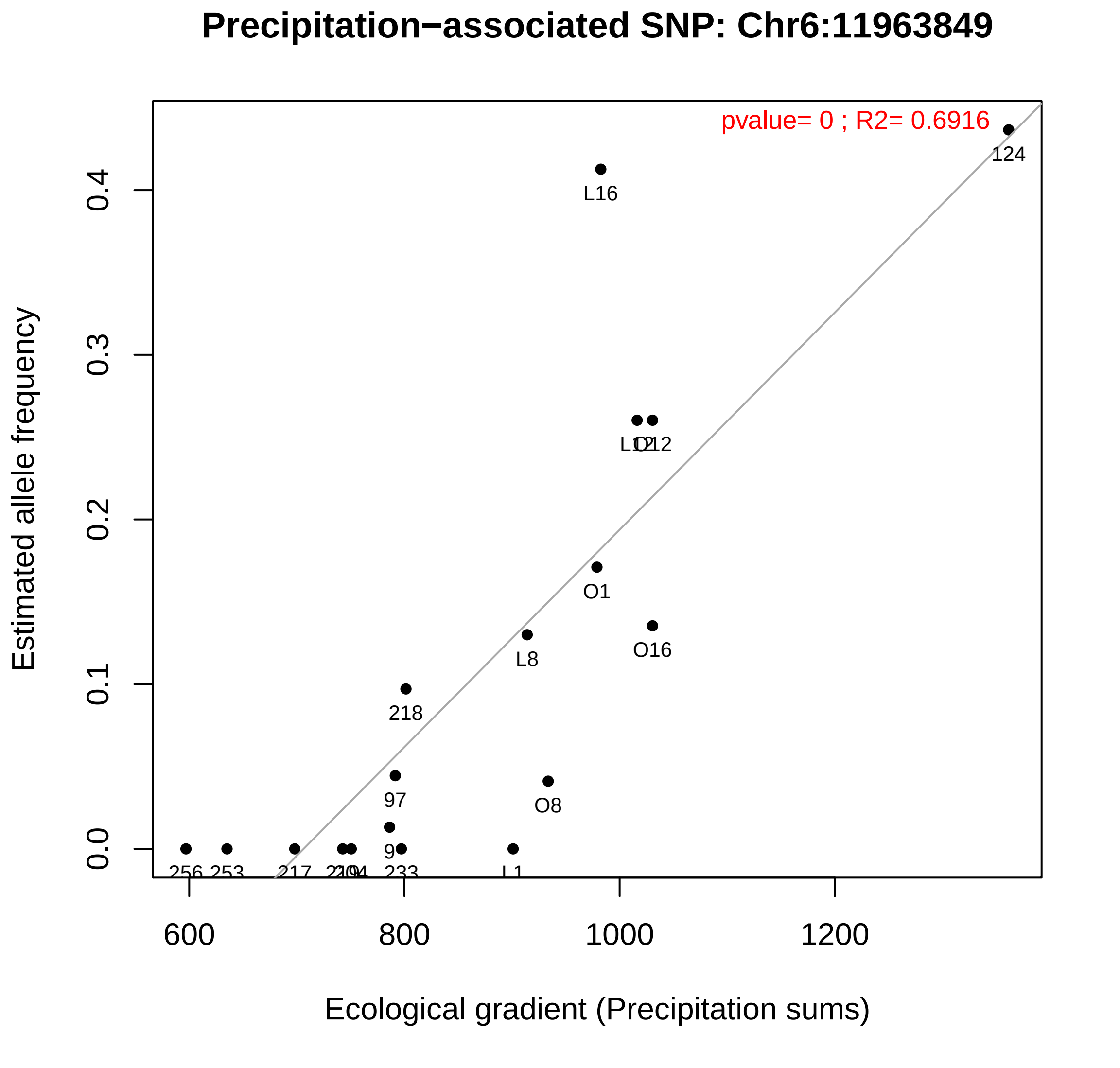
Precipitation−associated SNP: Chr6:11963849
pvalue= 0 ; R2= 0.6916
124
L16
L12
O12
O1
O16
L8
218
97
O8
9
256
253
217
233
L1
0.0
0.1
0.2
0.3
0.4
600
800
1000
1200
Estimated allele frequency
Ecological gradient (Precipitation sums)

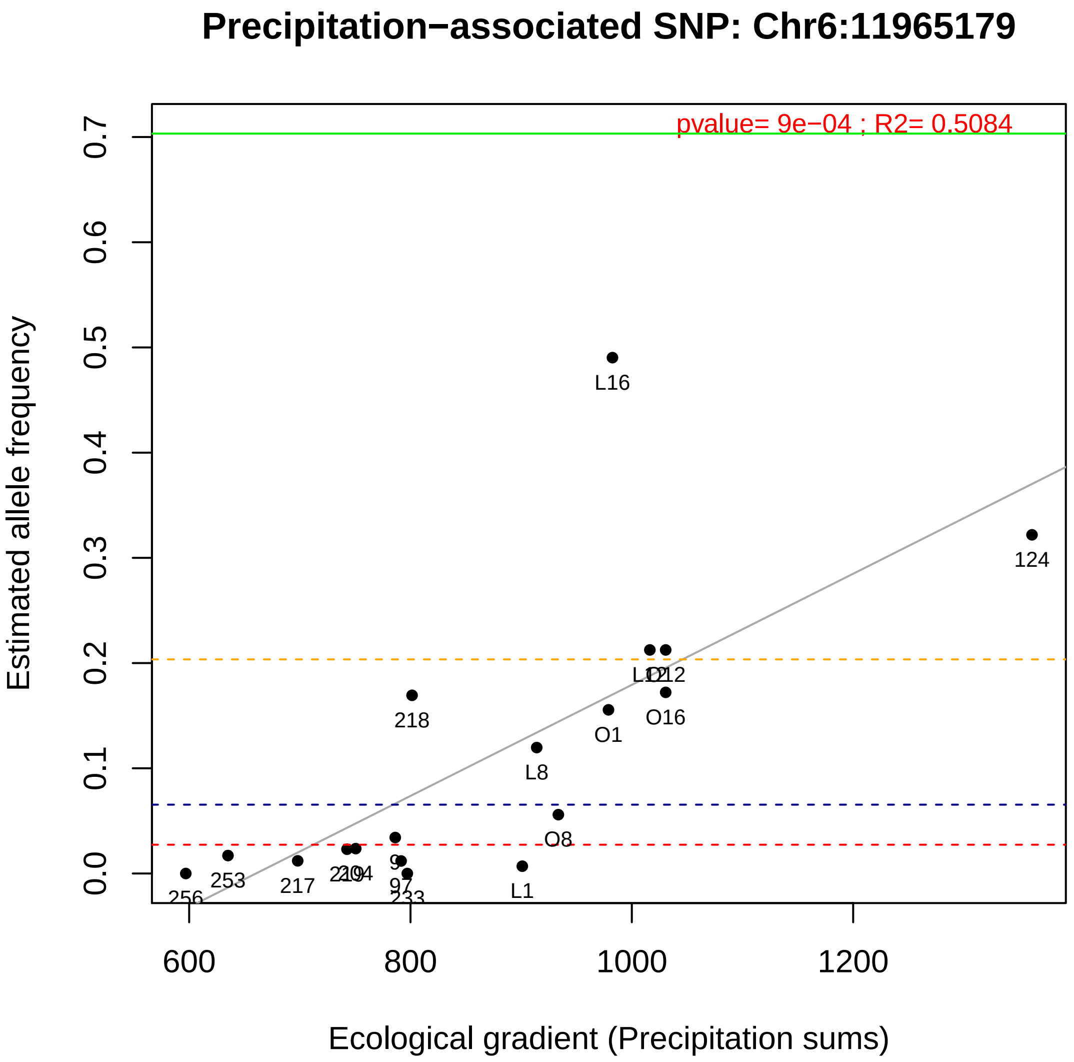
Precipitation−associated SNP: Chr6:11965179
pvalue= 9e−04 ; R2= 0.5084
Estimated allele frequency
Ecological gradient (Precipitation sums)
0.0
0.1
0.2
0.3
0.4
0.5
0.6
0.7
600
800
1000
1200
L16
124
218
O1
O16
L8
O8
L1
253
217
256

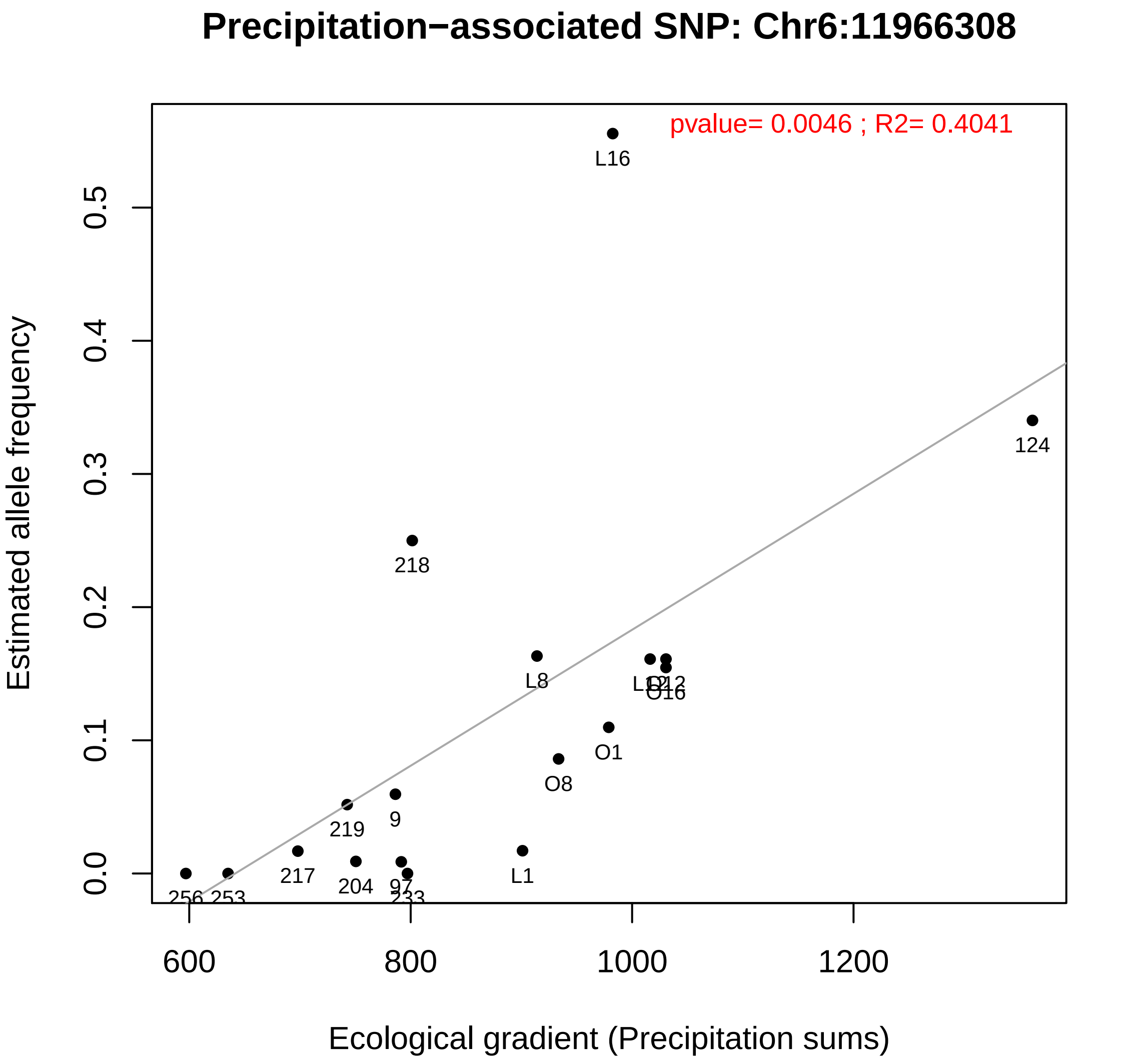
Precipitation−associated SNP: Chr6:11966308
pvalue= 0.0046 ; R2= 0.4041
L16
124
218
L8
O1
O8
219
9
217
204
97
233
L1
256
253
0.0
0.1
0.2
0.3
0.4
0.5
600
800
1000
1200
Estimated allele frequency
Ecological gradient (Precipitation sums)

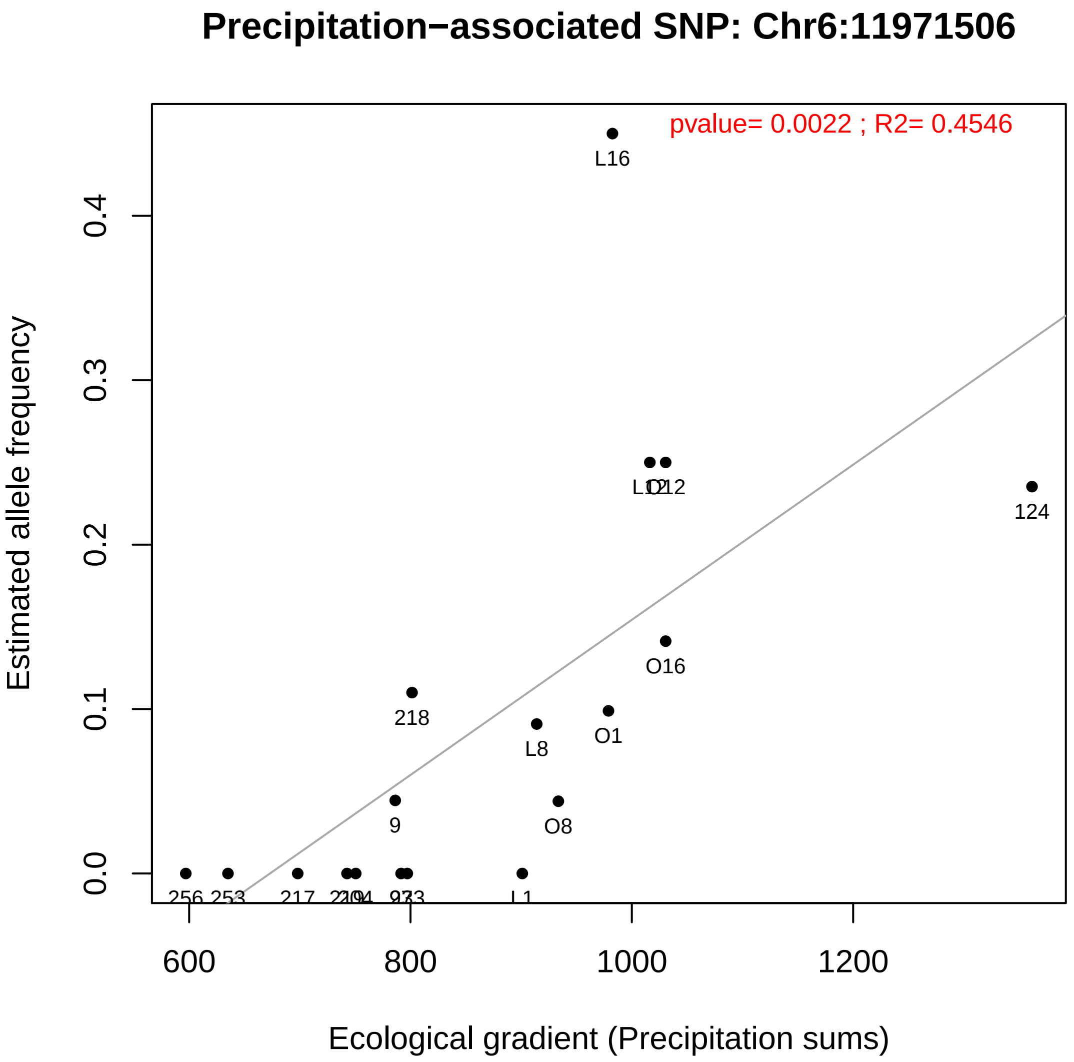

Precipitation−associated SNP: Chr6:11971506
pvalue= 0.0022 ; R2= 0.4546
L16
L12
O12
124
O16
218
O1
L8
9
O8
256
253
217
L1
0.0
0.1
0.2
0.3
0.4
600
800
1000
1200
Estimated allele frequency
Ecological gradient (Precipitation sums)

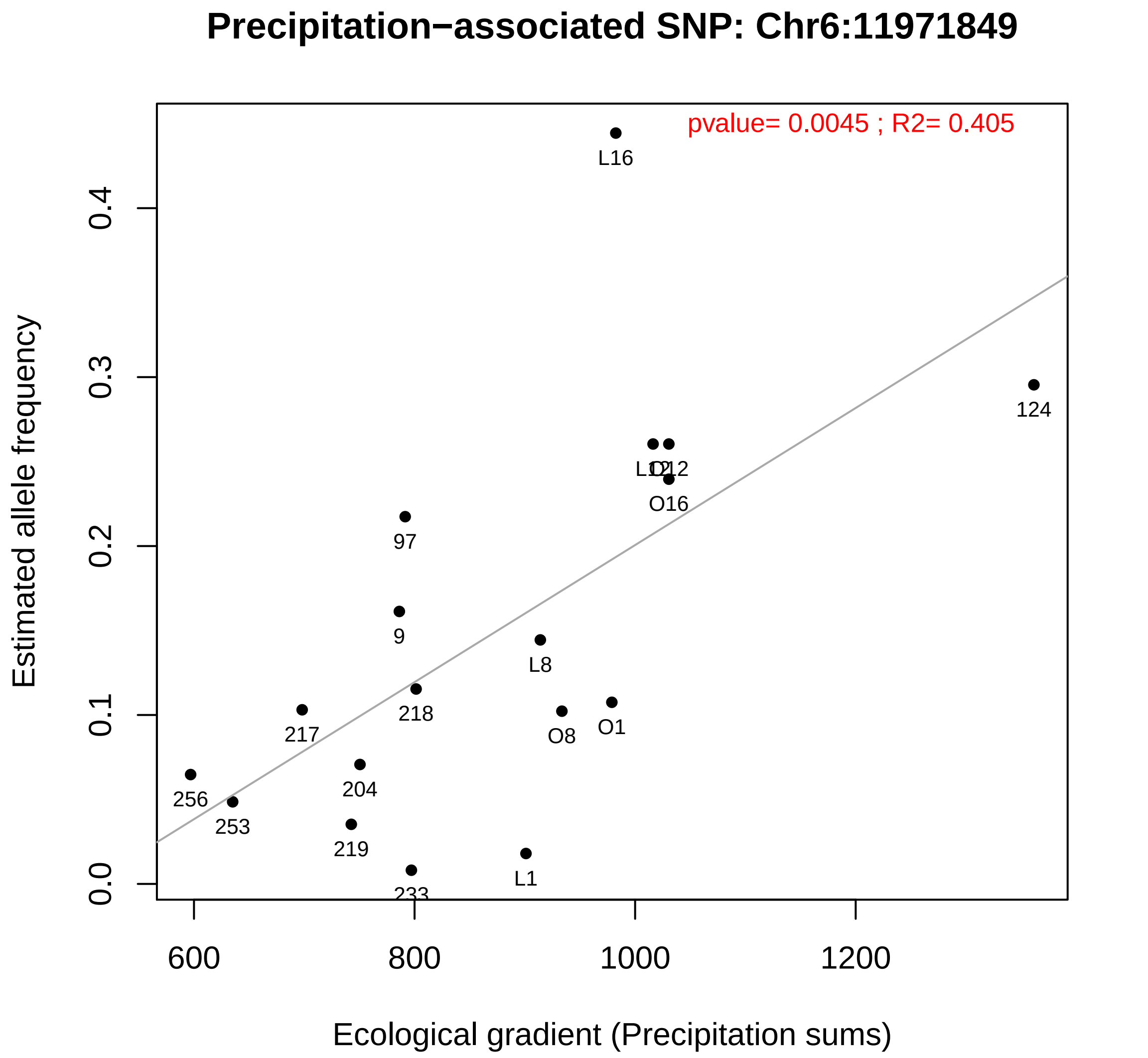
Precipitation−associated SNP: Chr6:11971849
pvalue= 0.0045 ; R2= 0.405
L16
124
L12
O12
O16
97
9
L8
218
217
O8
O1
256
204
253
219
233
L1
0.0
0.1
0.2
0.3
0.4
600
800
1000
1200
Estimated allele frequency
Ecological gradient (Precipitation sums)

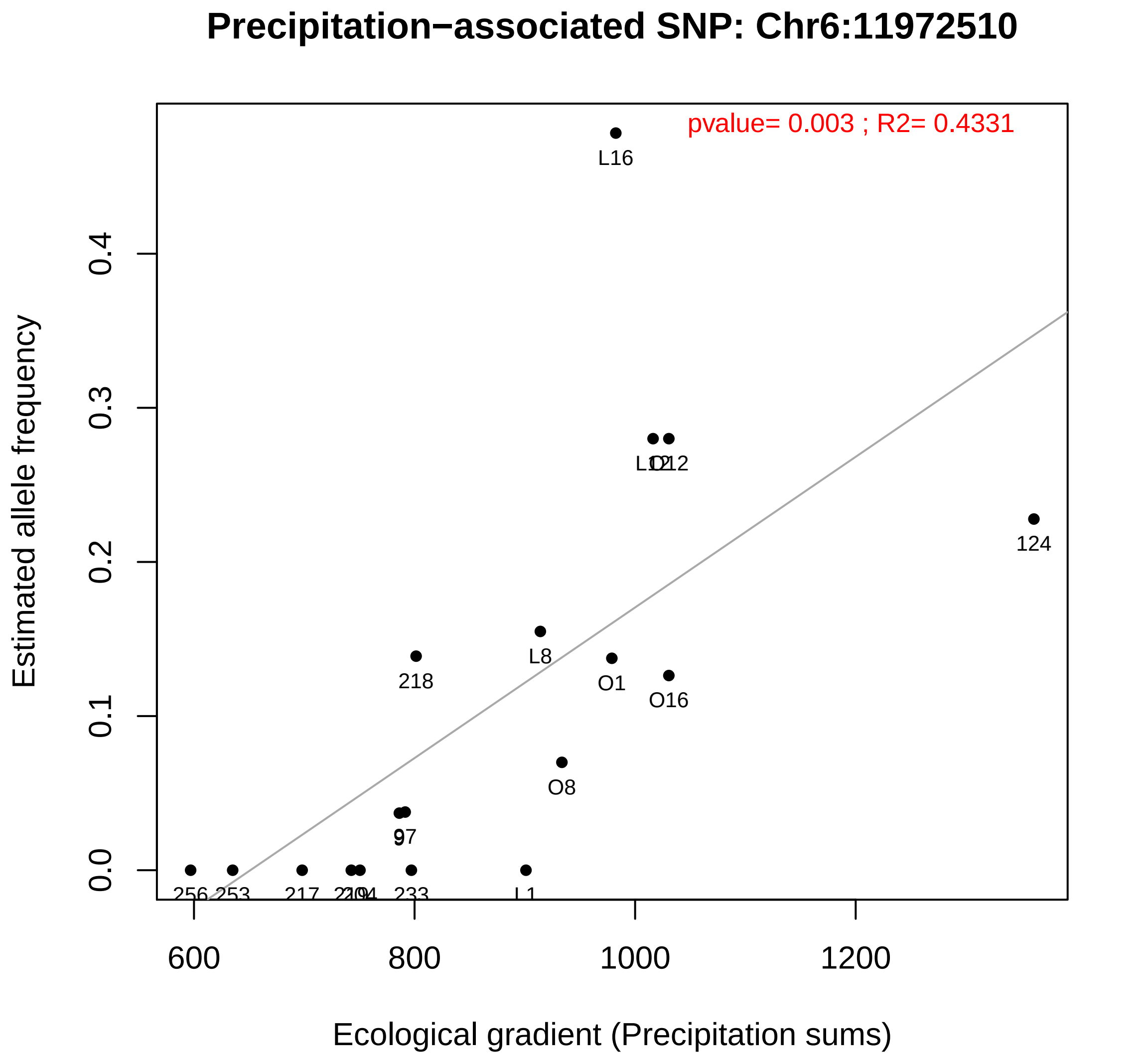

Precipitation−associated SNP: Chr6:11972510
pvalue= 0.003 ; R2= 0.4331
L16
L12
O12
124
L8
218
O1
O16
O8
9
7
256
253
217
233
L1
0.0
0.1
0.2
0.3
0.4
600
800
1000
1200
Estimated allele frequency
Ecological gradient (Precipitation sums)

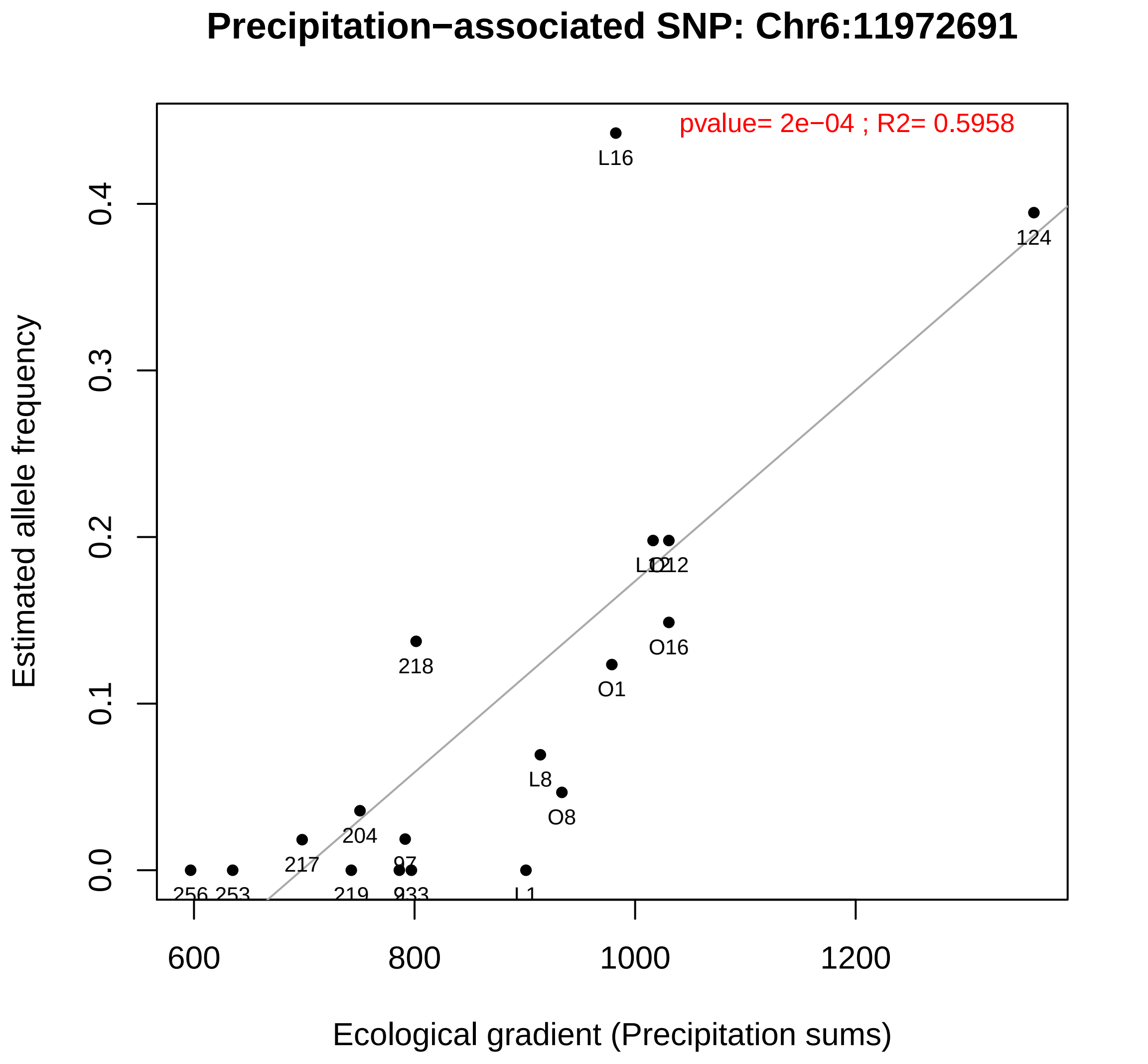

Precipitation−associated SNP: Chr6:11972691
pvalue= 2e−04 ; R2= 0.5958
Estimated allele frequency
Ecological gradient (Precipitation sums)
0.0
0.1
0.2
0.3
0.4
600
800
1000
1200
L16
124
O12
O16
218
O1
L8
O8
204
217
97
219
933
L1
256
253

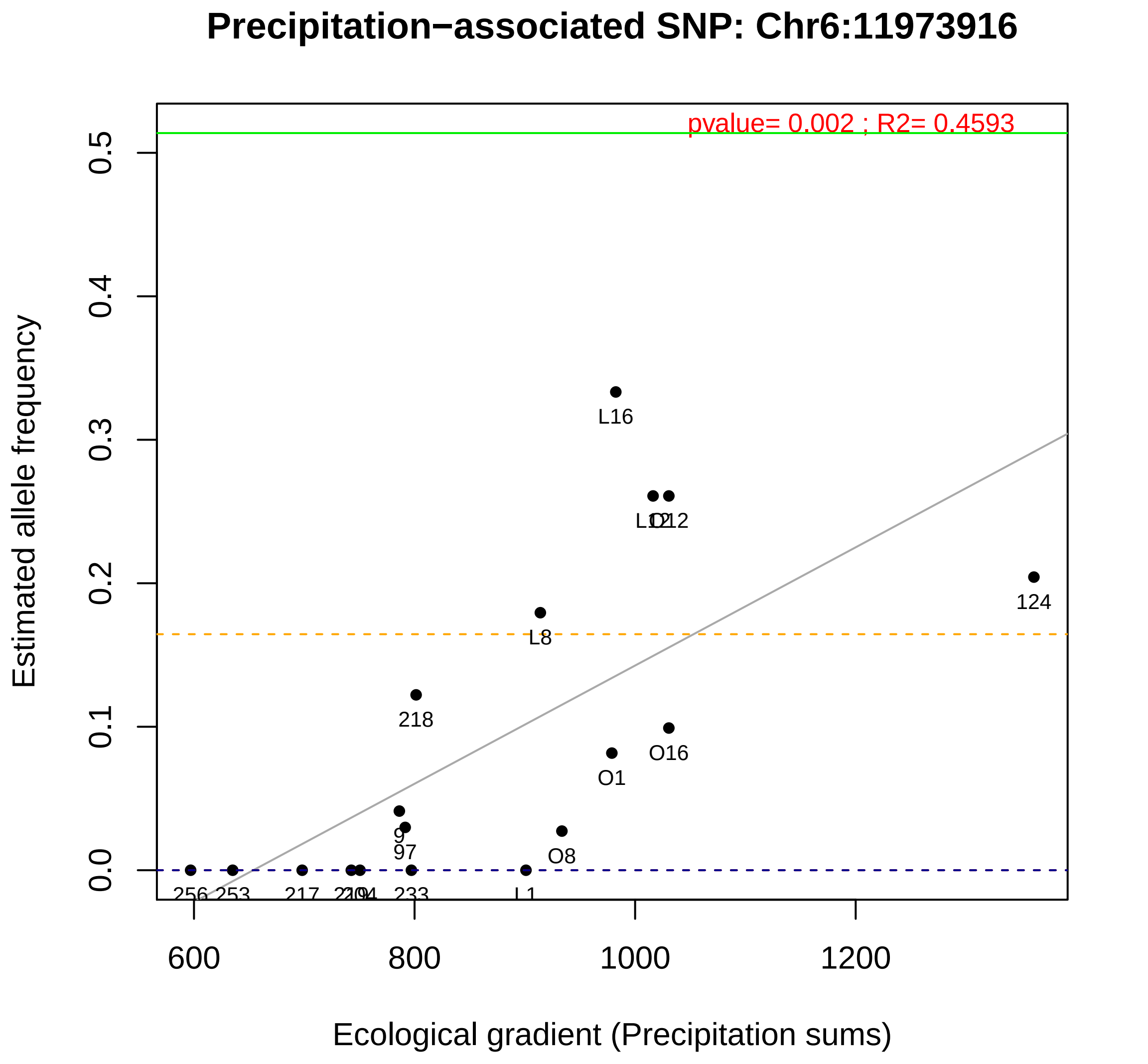

Precipitation−associated SNP: Chr6:11973916
pvalue= 0.002 ; R2= 0.4593
Estimated allele frequency
Ecological gradient (Precipitation sums)
0.0
0.1
0.2
0.3
0.4
0.5
600
800
1000
1200
L16
O12
124
L8
218
O16
O1
9
97
O8
256
253
217
233
L1

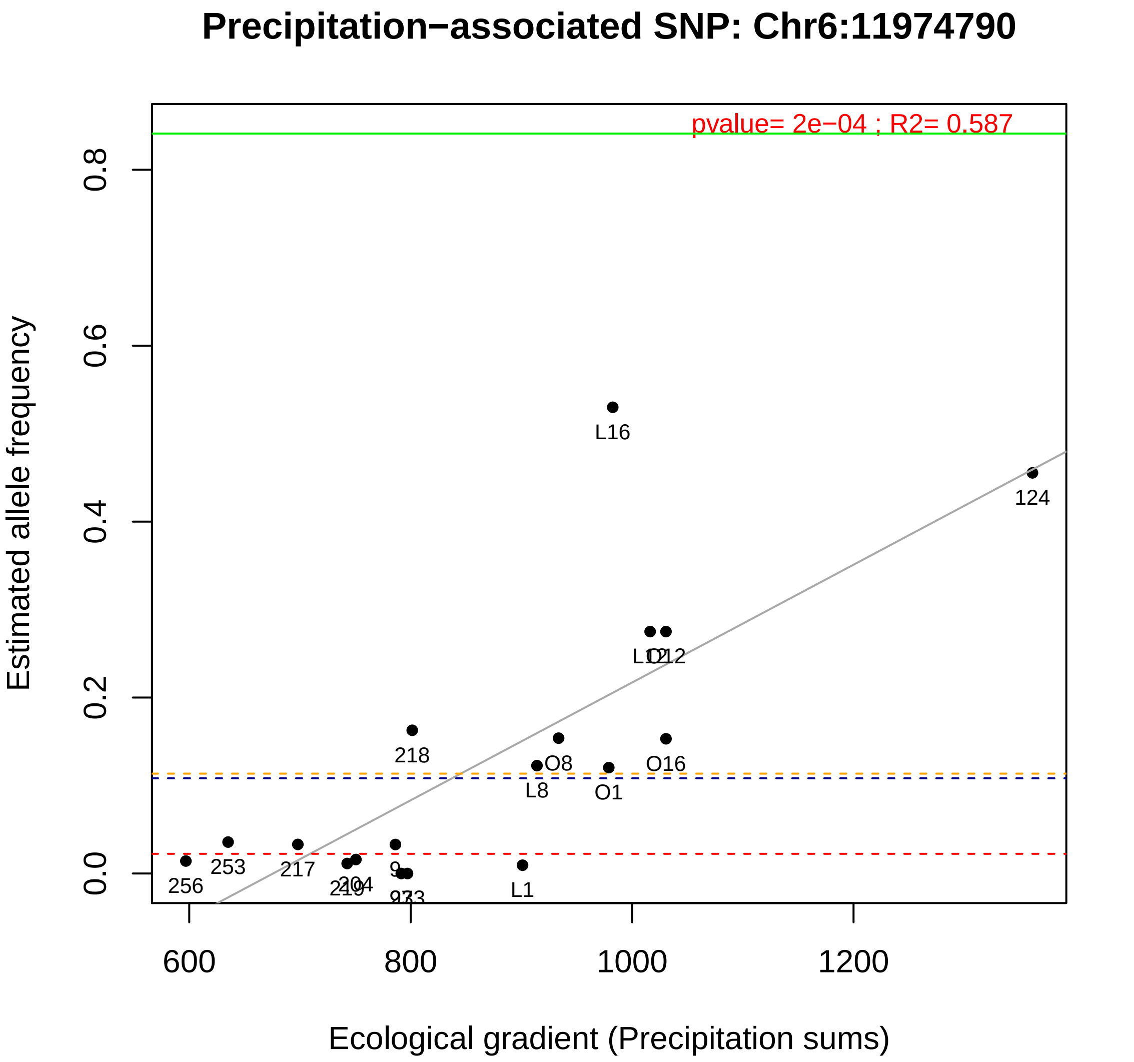

Precipitation−associated SNP: Chr6:11974790
pvalue= 2e−04 ; R2= 0.587
Estimated allele frequency
0.0
0.2
0.4
0.6
0.8
600
800
1000
1200
Ecological gradient (Precipitation sums)
L16
124
O12
218
O8
O16
L8
O1
253
217
256
9
L1

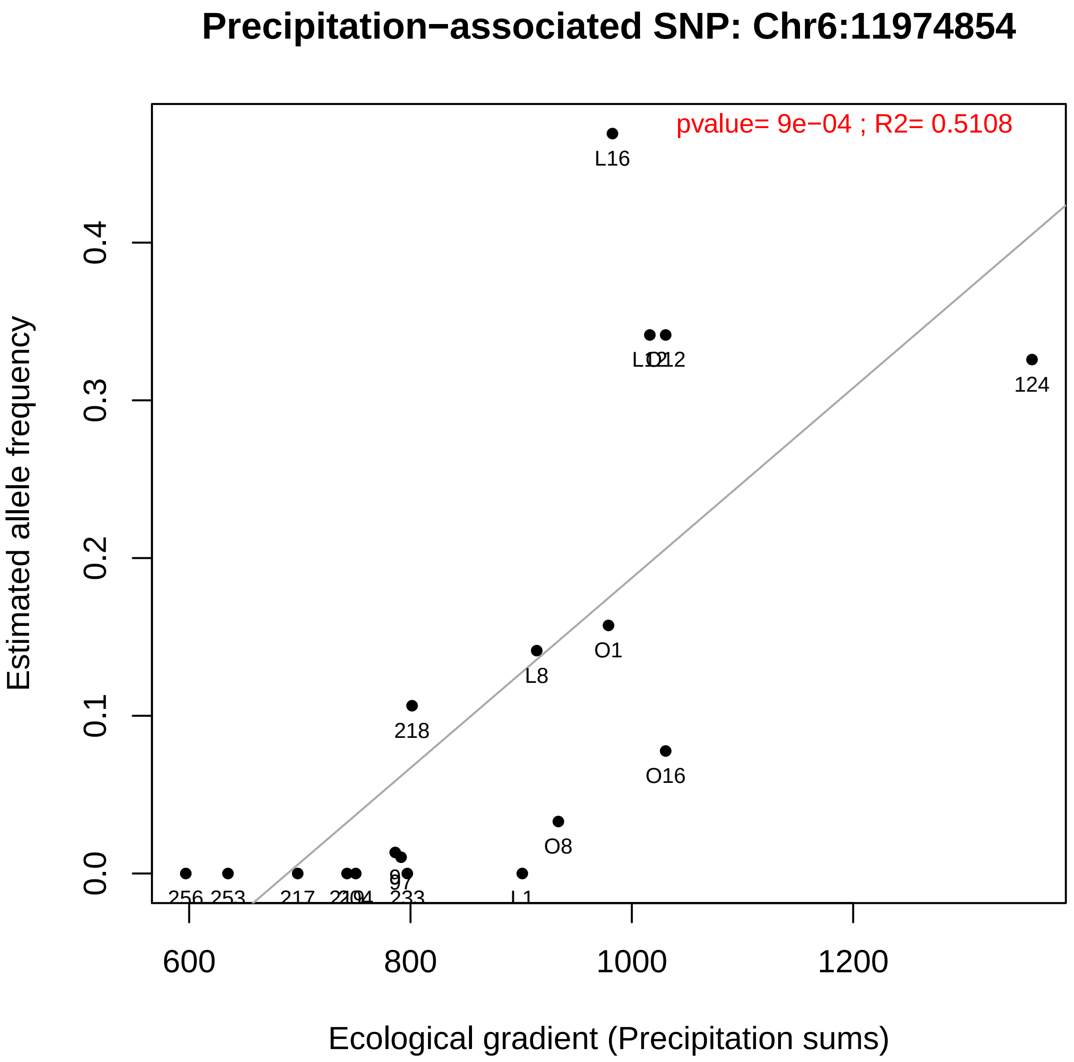

Precipitation−associated SNP: Chr6:11974854
pvalue= 9e−04 ; R2= 0.5108
Estimated allele frequency
Ecological gradient (Precipitation sums)
0.0
0.1
0.2
0.3
0.4
600
800
1000
1200
L16
124
O1
L8
218
O16
O8
256
253
217
233

### Precipitation−associated SNP: Chr6:11974941

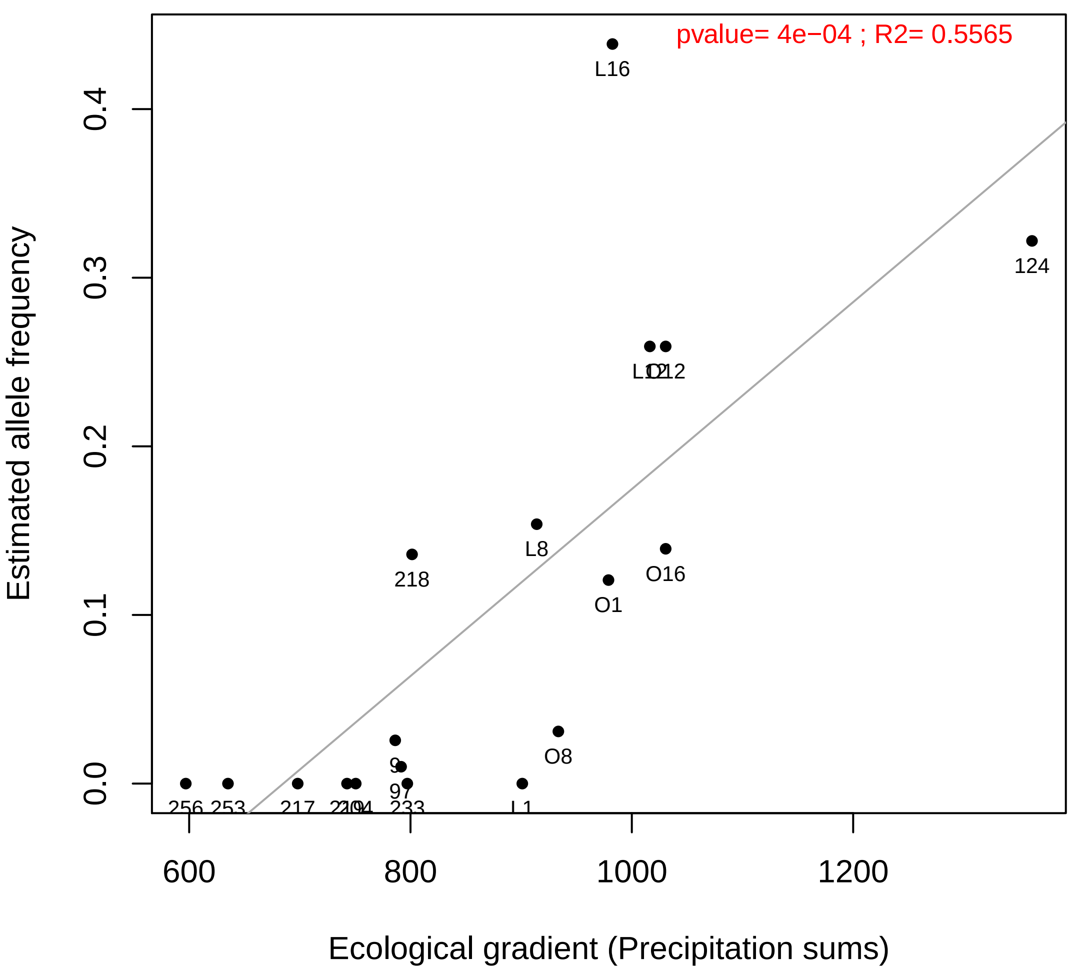

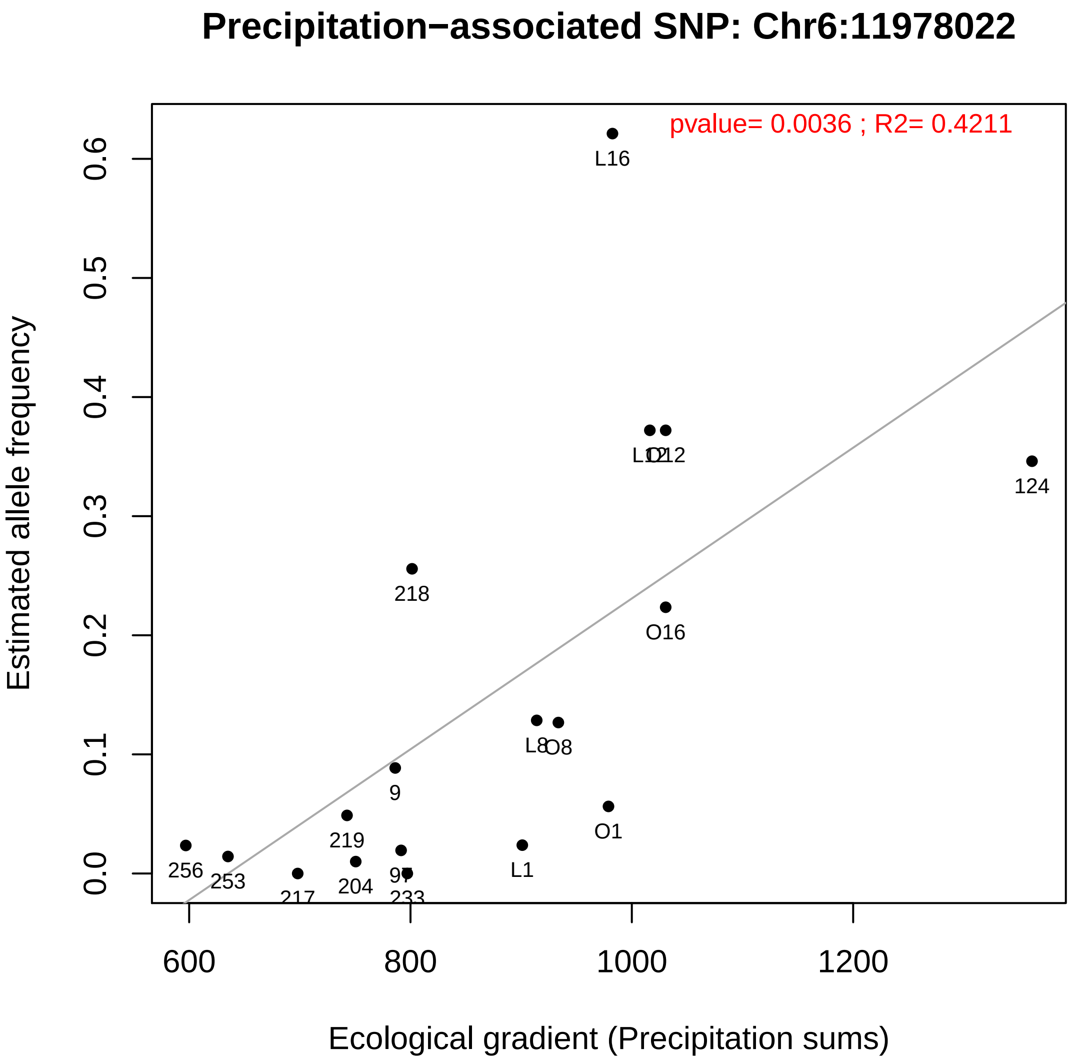

Precipitation−associated SNP: Chr6:11978022
pvalue= 0.0036 ; R2= 0.4211
Estimated allele frequency
Ecological gradient (Precipitation sums)
0.0
0.1
0.2
0.3
0.4
0.5
0.6
600
800
1000
1200
L16
124
218
O16
L8
O8
9
O1
219
L1
256
253
204
217
233

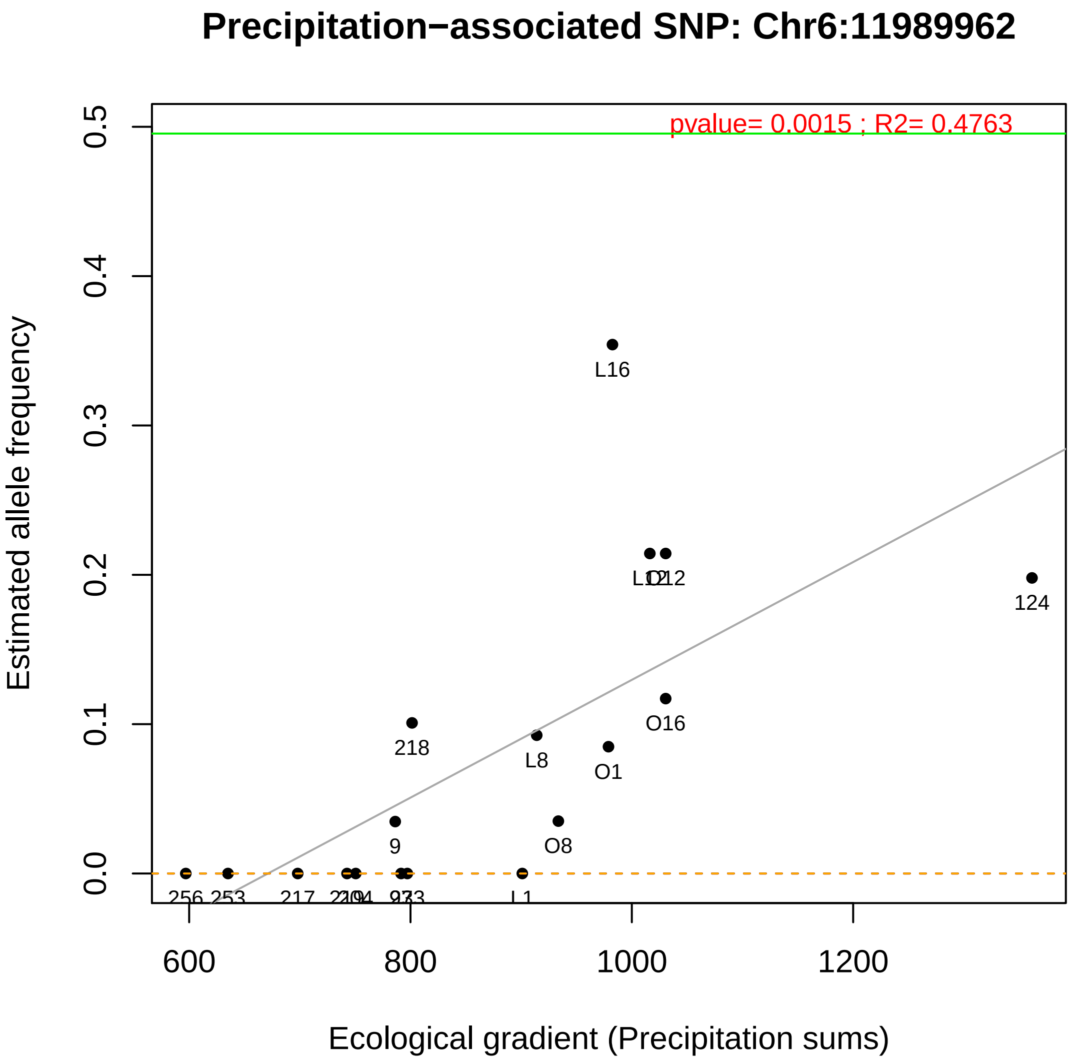

Precipitation−associated SNP: Chr6:11989962
pvalue= 0.0015 ; R2= 0.4763
Estimated allele frequency
0.0
0.1
0.2
0.3
0.4
0.5
600
800
1000
1200
Ecological gradient (Precipitation sums)
L16
124
O16
218
L8
O1
9
O8
256
253
217
L1

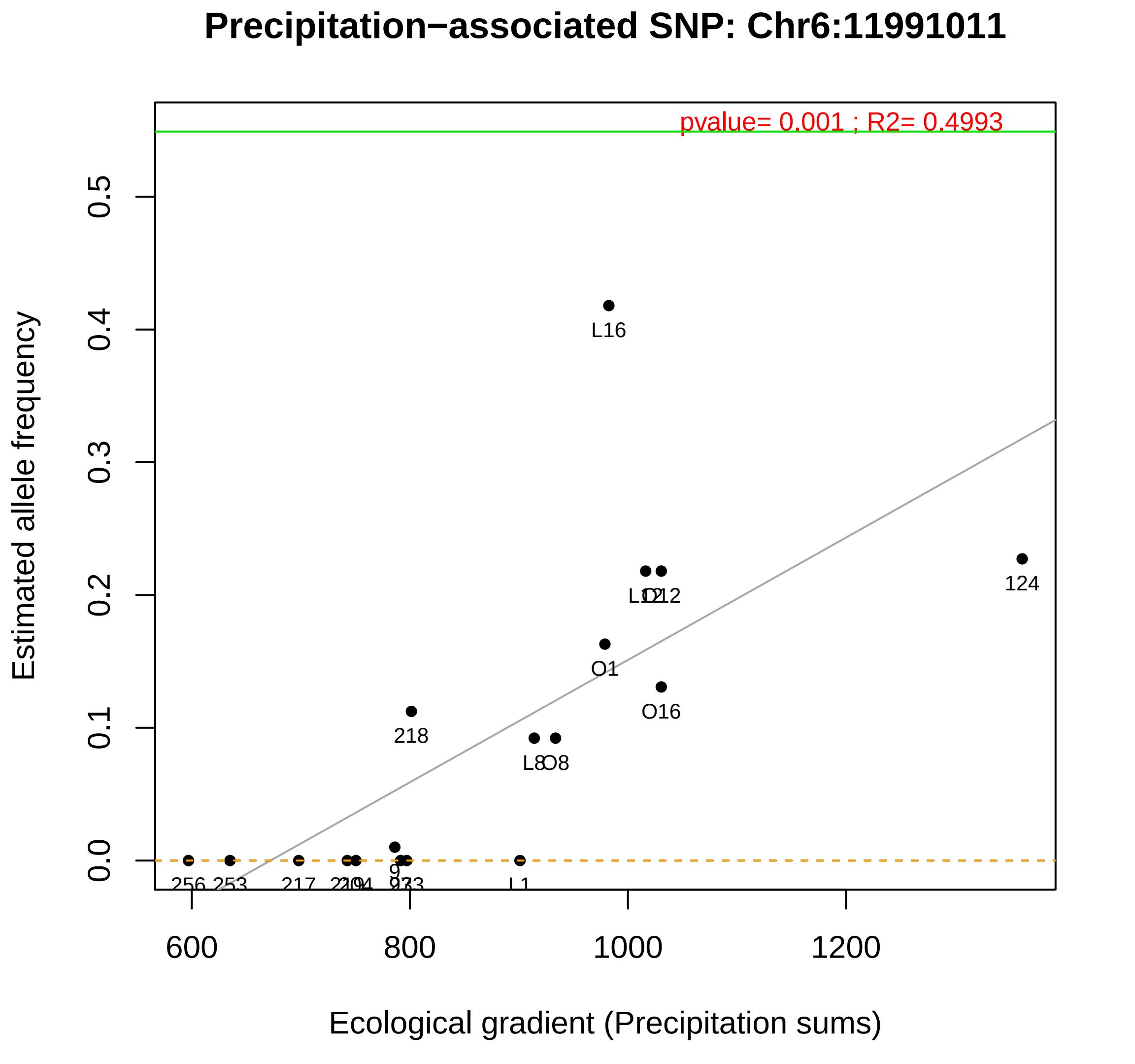
Precipitation−associated SNP: Chr6:11991011
pvalue= 0.001 ; R2= 0.4993
Estimated allele frequency
0.0
0.1
0.2
0.3
0.4
0.5
L16
L12
O2
124
O1
O16
218
L8
O8
256
253
217
210
194
9
27
73
L1
600
800
1000
1200
Ecological gradient (Precipitation sums)

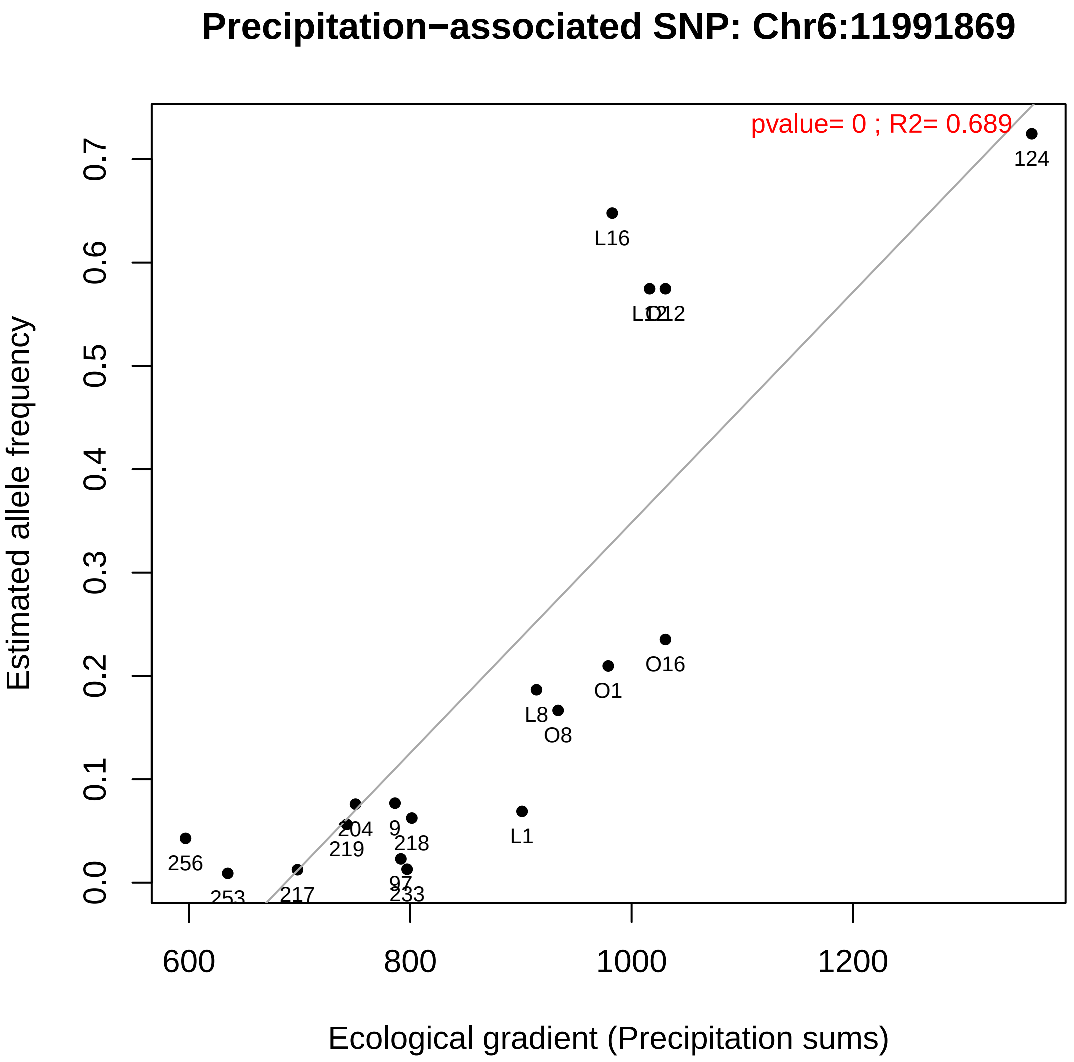
Precipitation−associated SNP: Chr6:11991869
pvalue= 0 ; R2= 0.689
Estimated allele frequency
Ecological gradient (Precipitation sums)
0.0
0.1
0.2
0.3
0.4
0.5
0.6
0.7
600
800
1000
1200
124
L16
L12
O12
O16
O1
L8
O8
L1
204
219
9
218
256
253
217
97
233

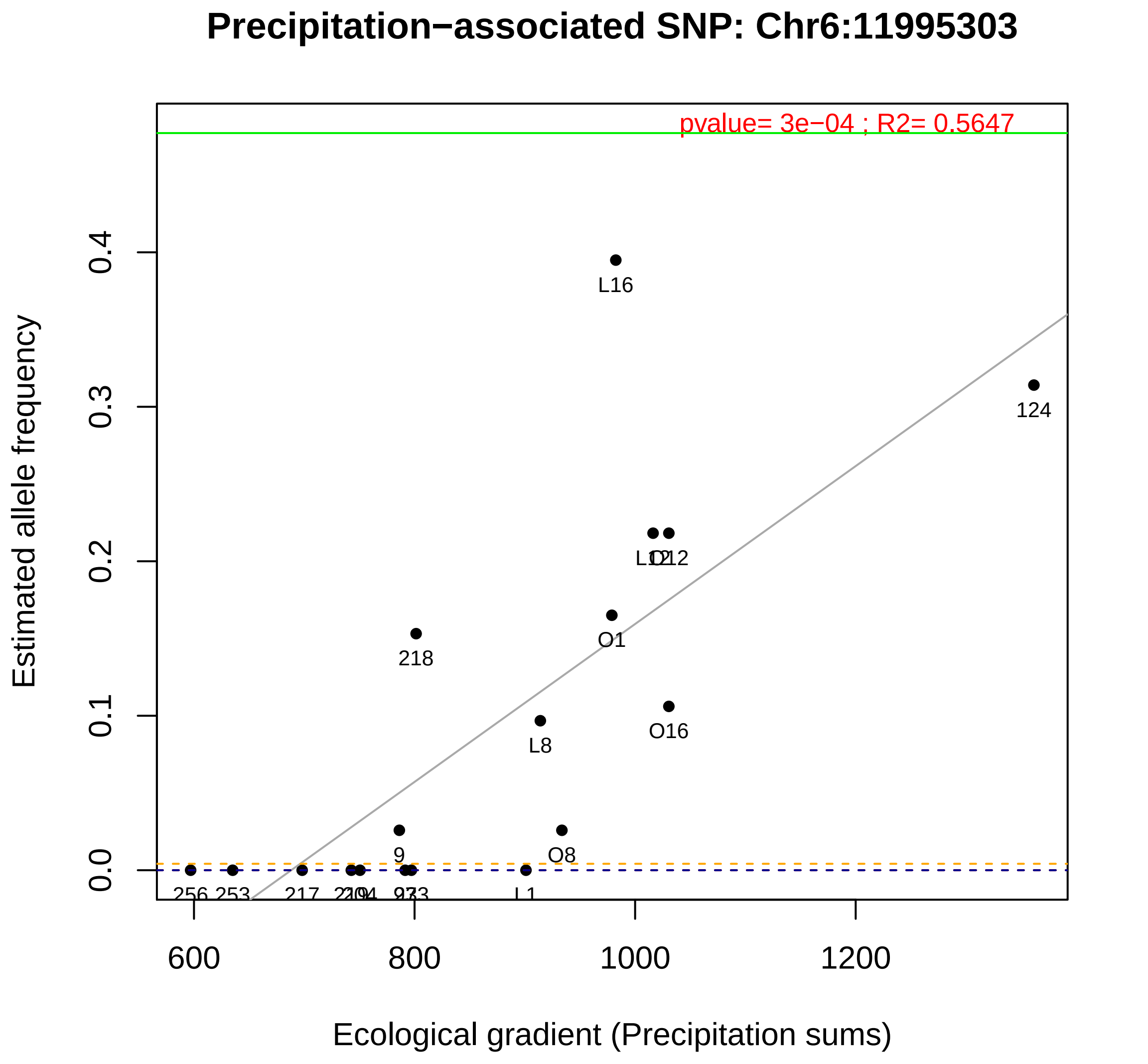
Precipitation−associated SNP: Chr6:11995303
pvalue= 3e−04 ; R2= 0.5647
Estimated allele frequency
0.0
0.1
0.2
0.3
0.4
L16
124
L12
O12
O1
218
O16
L8
9
O8
256
253
217
L1
600
800
1000
1200
Ecological gradient (Precipitation sums)

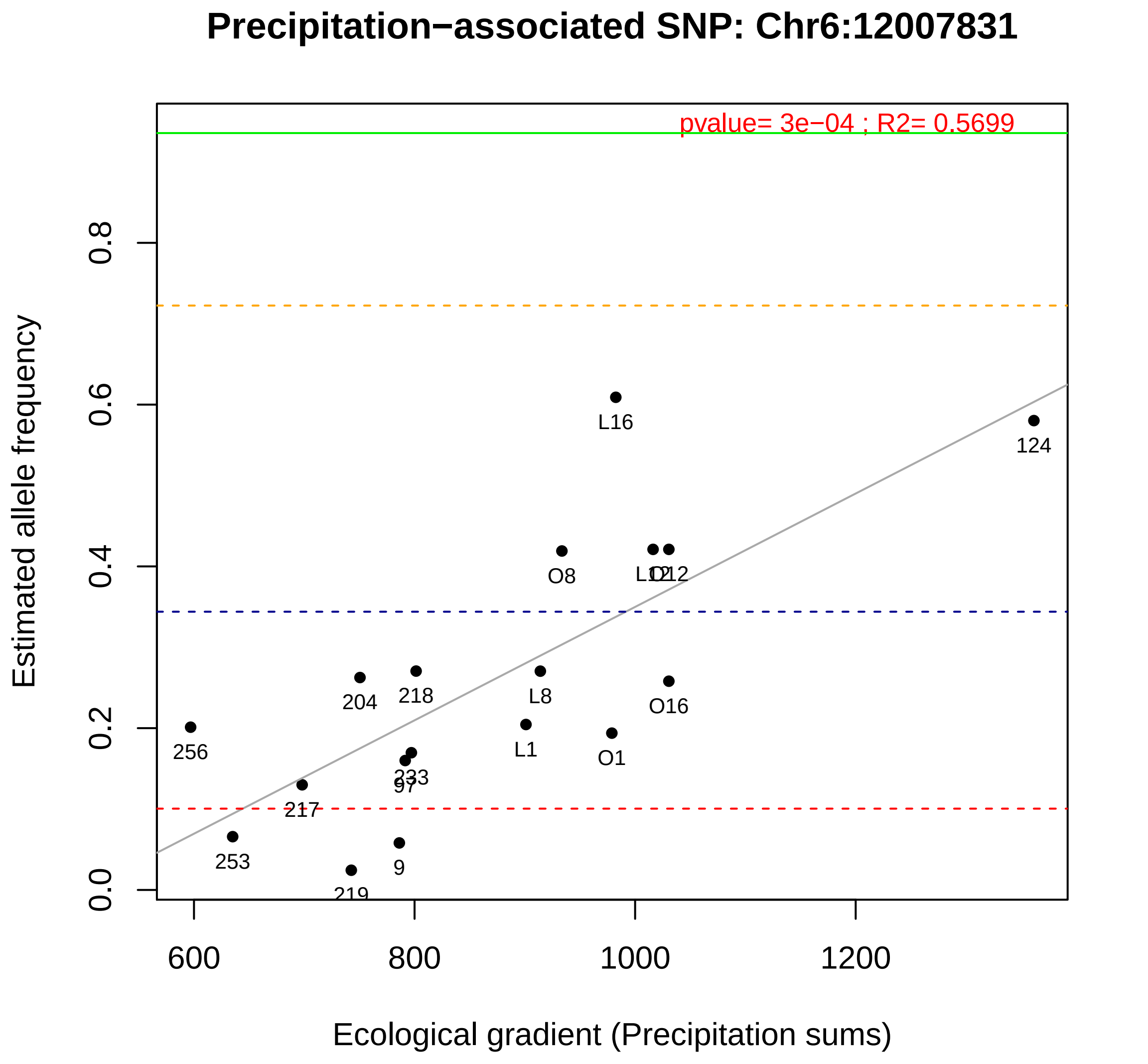

Precipitation−associated SNP: Chr6:12007831
pvalue= 3e−04 ; R2= 0.5699
Estimated allele frequency
0.0
0.2
0.4
0.6
0.8
600
800
1000
1200
Ecological gradient (Precipitation sums)
L16
124
O8
O12
O16
204
218
L8
256
L1
O1
233
97
217
253
9
219

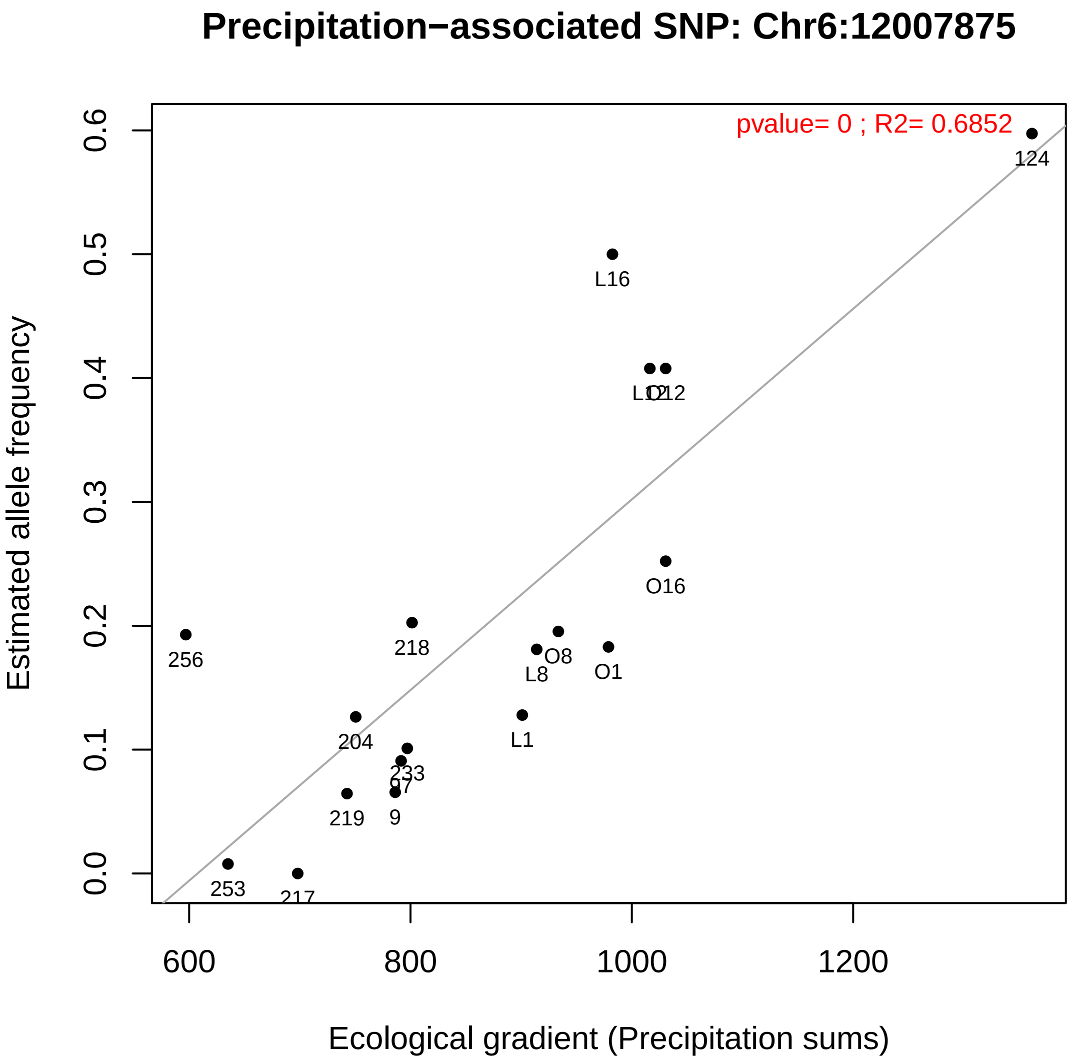

Precipitation−associated SNP: Chr6:12007875
pvalue= 0 ; R2= 0.6852
Estimated allele frequency
0.0
0.1
0.2
0.3
0.4
0.5
0.6
Ecological gradient (Precipitation sums)
600
800
1000
1200
124
L16
O12
O16
256
218
O8
L8
O1
204
L1
233
219
9
253
217

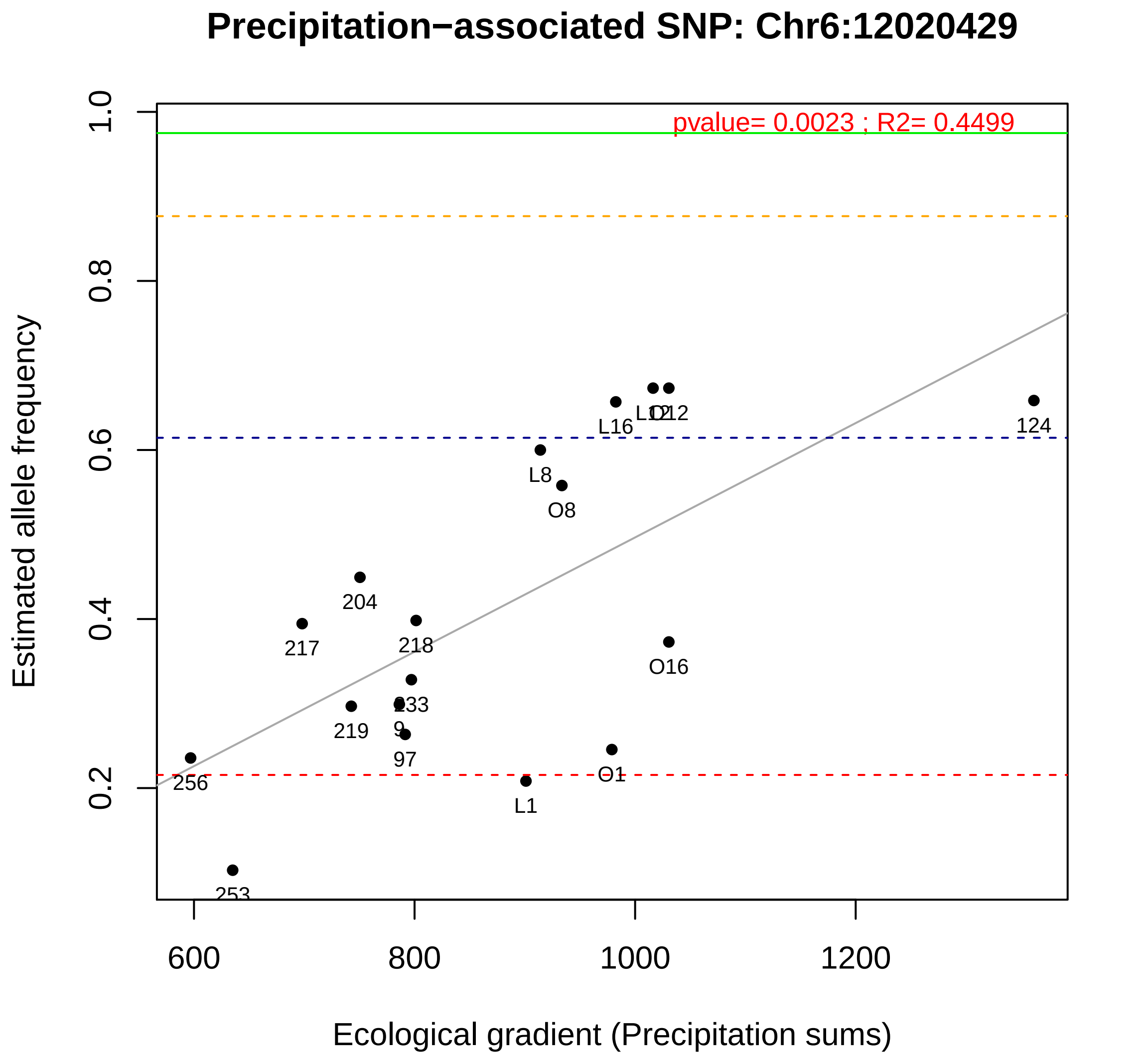

Precipitation−associated SNP: Chr6:12020429
pvalue= 0.0023 ; R2= 0.4499
Estimated allele frequency
Ecological gradient (Precipitation sums)
0.2
0.4
0.6
0.8
1.0
600
800
1000
1200
256
253
217
204
219
218
233
9
97
L1
L8
O8
L16
O1
O16
O12
124

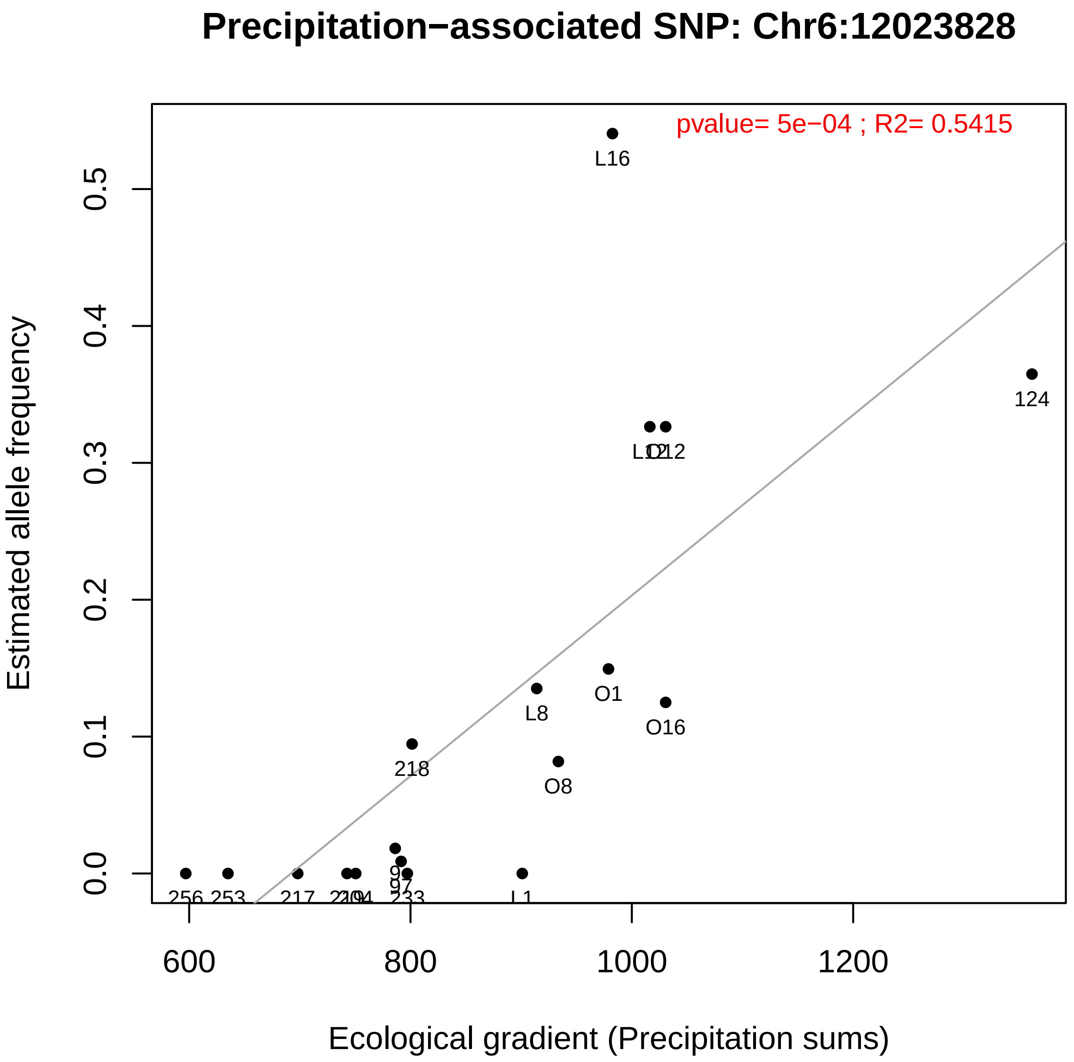
Precipitation−associated SNP: Chr6:12023828
pvalue= 5e−04 ; R2= 0.5415
L16
124
O1
L8
O16
218
O8
Estimated allele frequency
0.0
0.1
0.2
0.3
0.4
0.5
600
800
1000
1200
Ecological gradient (Precipitation sums)

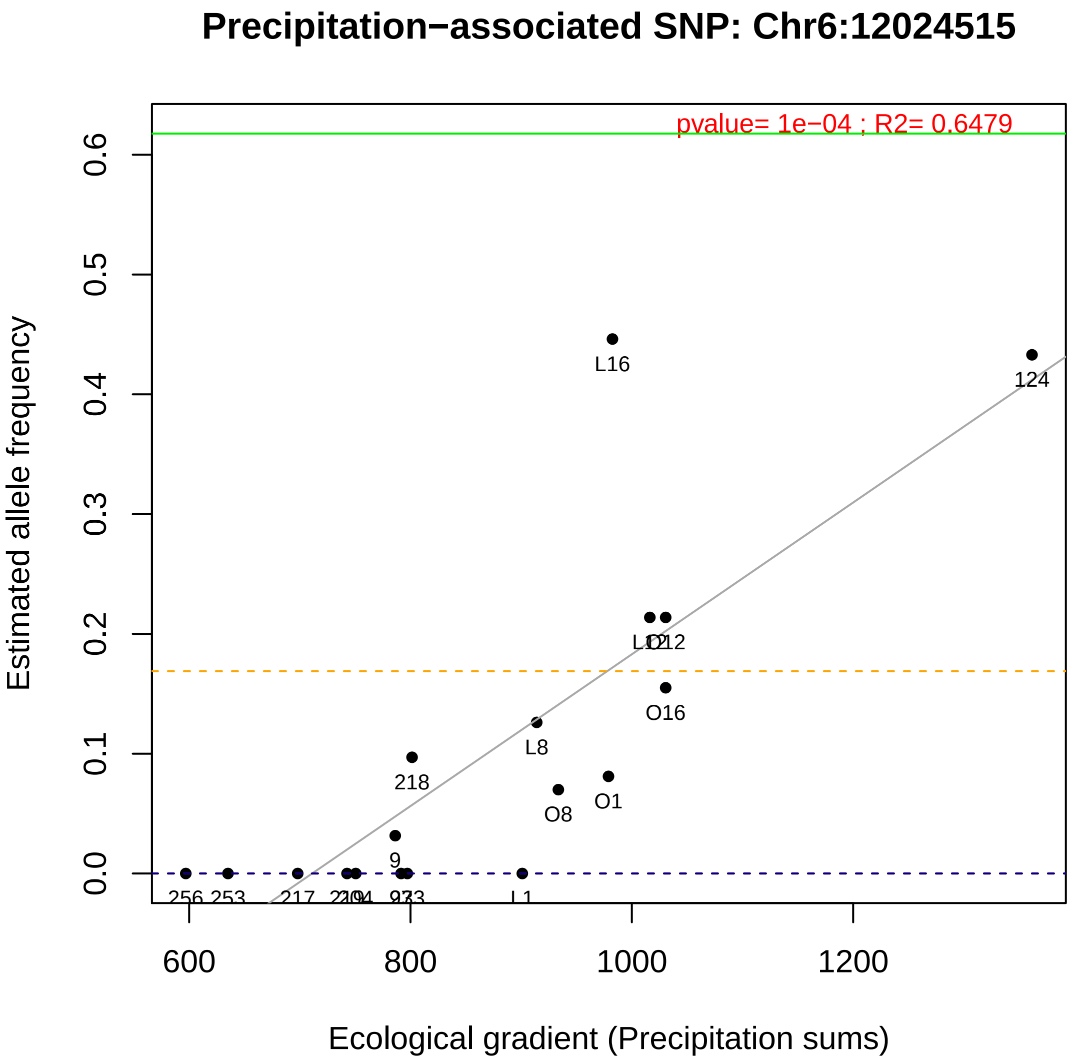
Precipitation−associated SNP: Chr6:12024515
pvalue= 1e−04 ; R2= 0.6479
Estimated allele frequency
0.0
0.1
0.2
0.3
0.4
0.5
0.6
Ecological gradient (Precipitation sums)
600
800
1000
1200
L16
124
O12
O16
L8
218
O1
O8
9
256
253
217
L1

**Precipitation−associated SNP: Chr6:12025385**

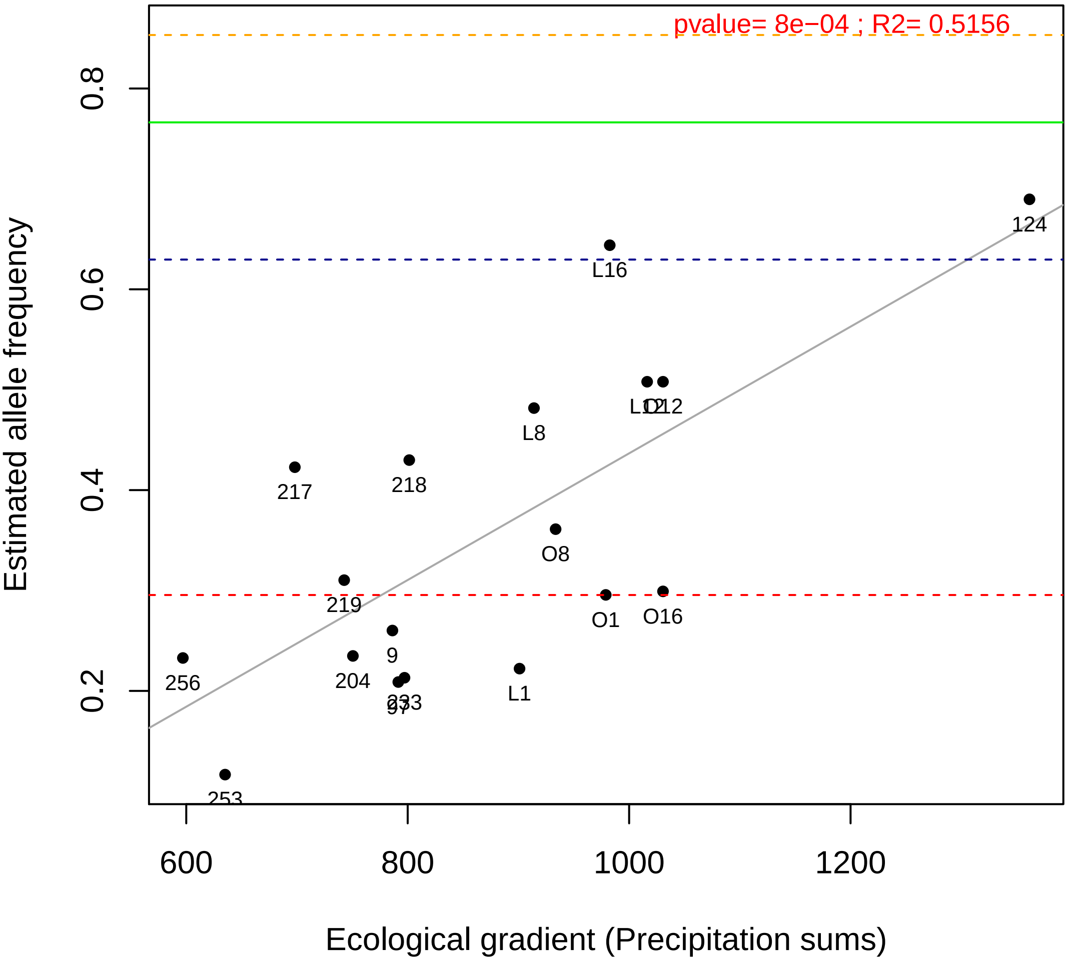

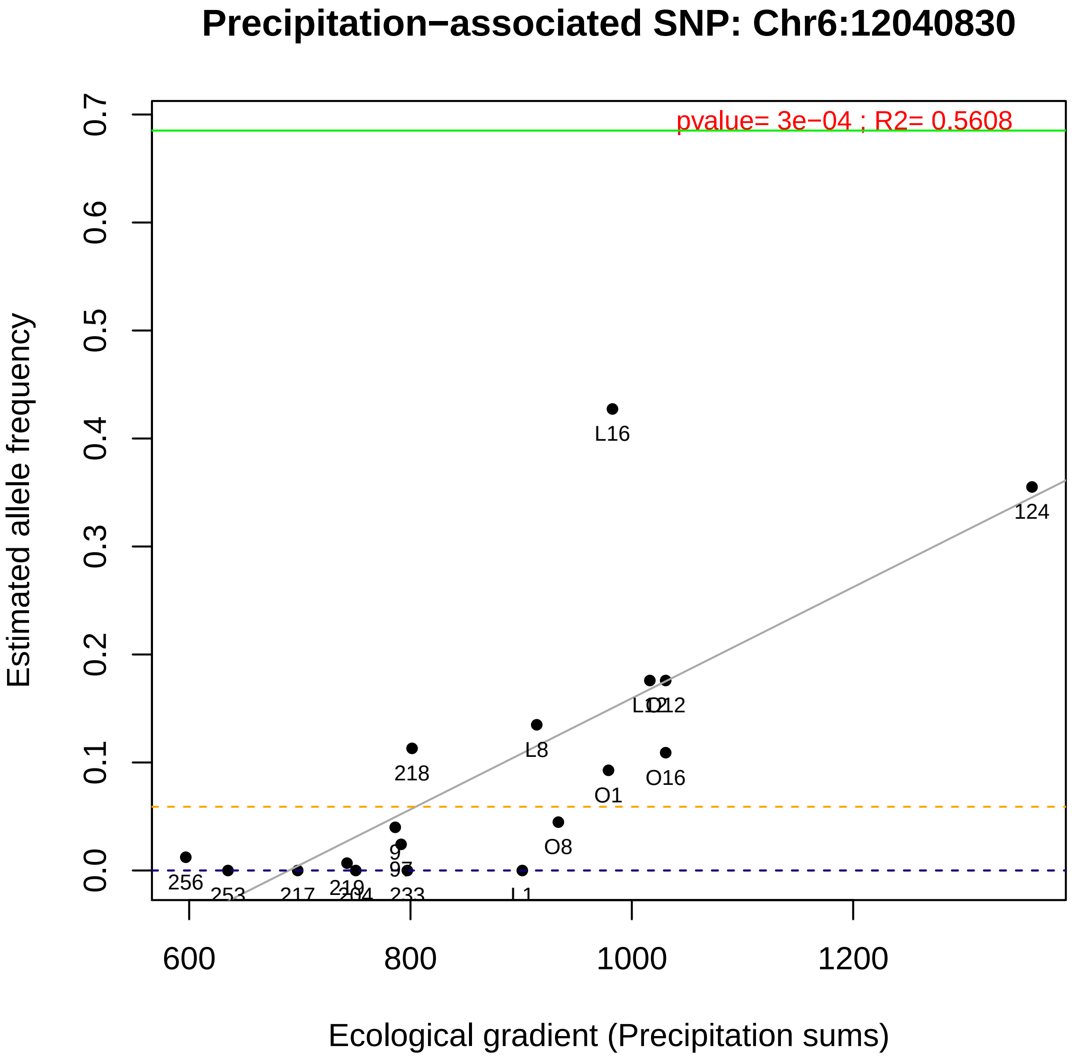
Precipitation−associated SNP: Chr6:12040830
pvalue= 3e−04 ; R2= 0.5608
Estimated allele frequency
0.0
0.1
0.2
0.3
0.4
0.5
0.6
0.7
600
800
1000
1200
Ecological gradient (Precipitation sums)
L16
124
L8
218
O1
O16
O8
256
217
219
233
L1

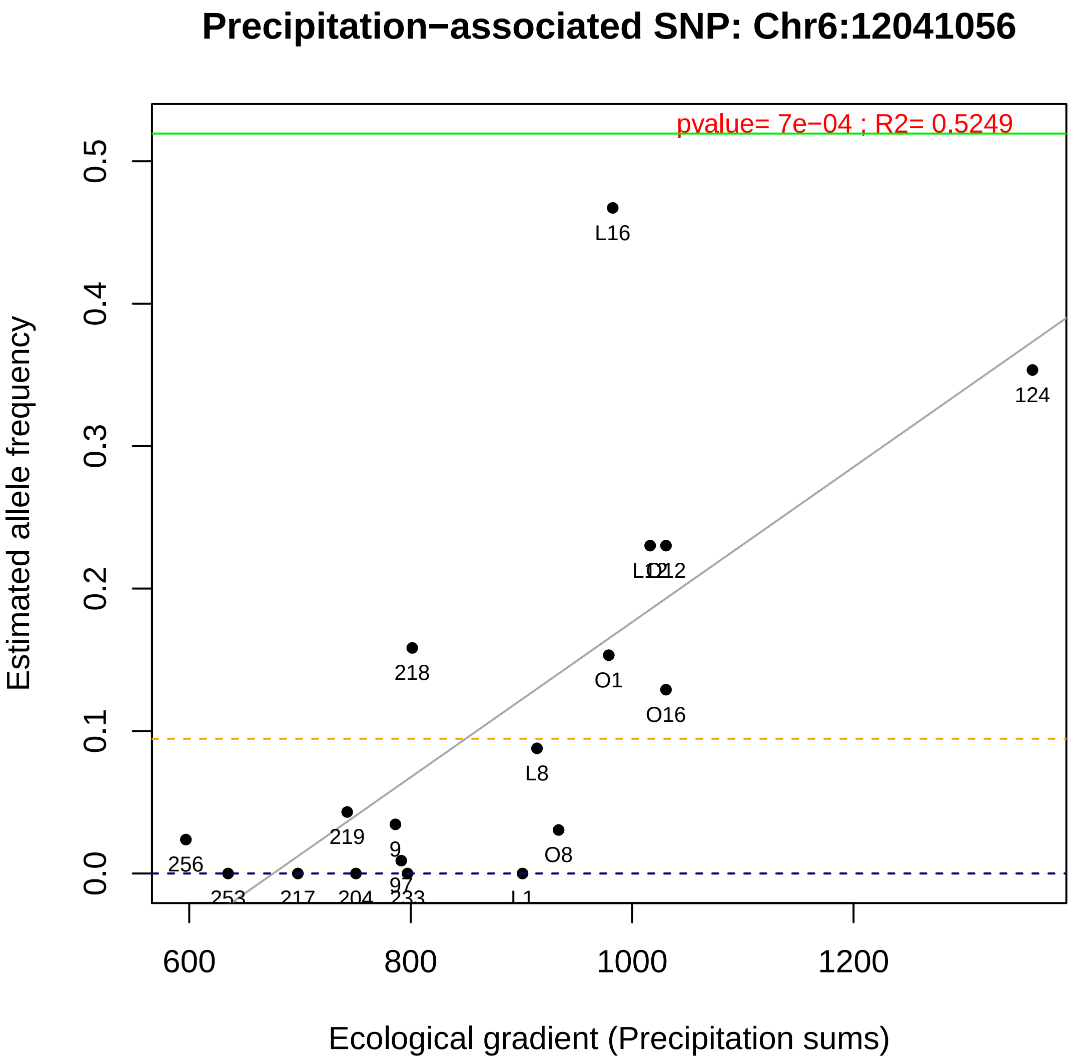

Precipitation−associated SNP: Chr6:12041056
pvalue= 7e−04 ; R2= 0.5249
Estimated allele frequency
Ecological gradient (Precipitation sums)
0.0
0.1
0.2
0.3
0.4
0.5
600
800
1000
1200
L16
124
O12
218
O1
O16
L8
219
256
9
O8
97

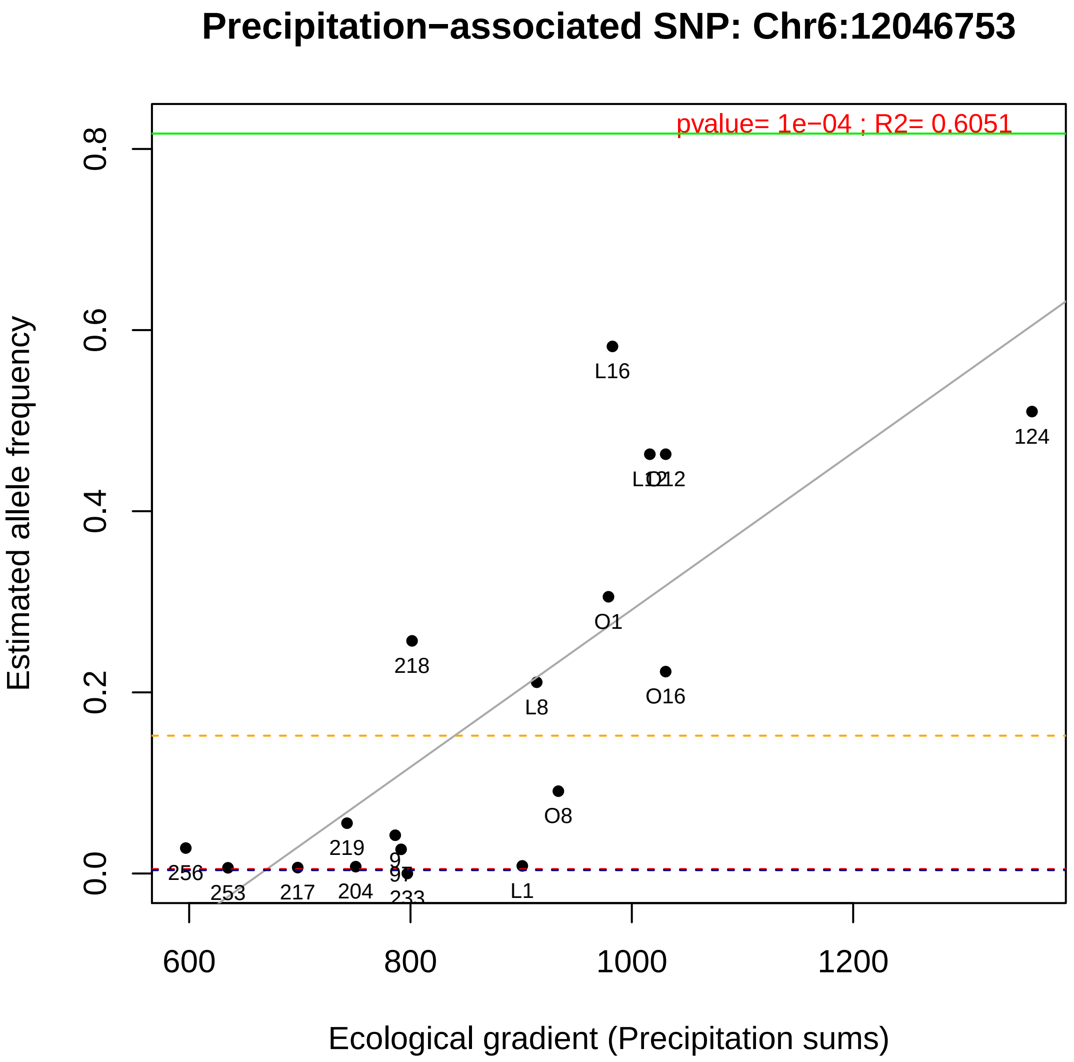

Precipitation−associated SNP: Chr6:12046753
pvalue= 1e−04 ; R2= 0.6051
Estimated allele frequency
0.0
0.2
0.4
0.6
0.8
600
800
1000
1200
Ecological gradient (Precipitation sums)
L16
124
L12
O12
O1
218
L8
O16
O8
219
256
253
217
204
9
97
233
L1

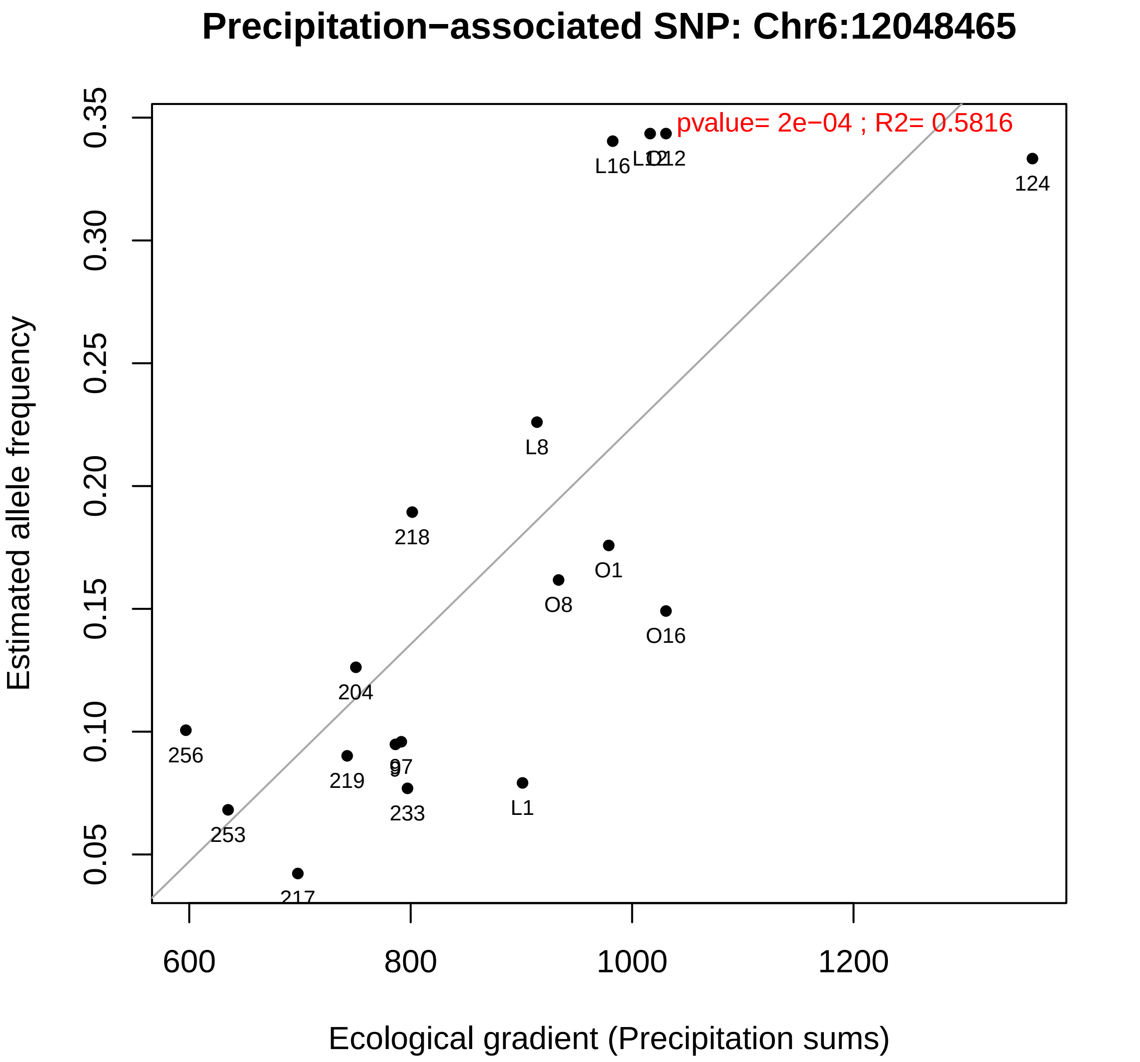

Precipitation−associated SNP: Chr6:12048465
pvalue= 2e−04 ; R2= 0.5816
Estimated allele frequency
0.05
0.10
0.15
0.20
0.25
0.30
0.35
Ecological gradient (Precipitation sums)
600
800
1000
1200
L16
124
L8
218
O1
O8
O16
204
256
219
233
L1
253
217

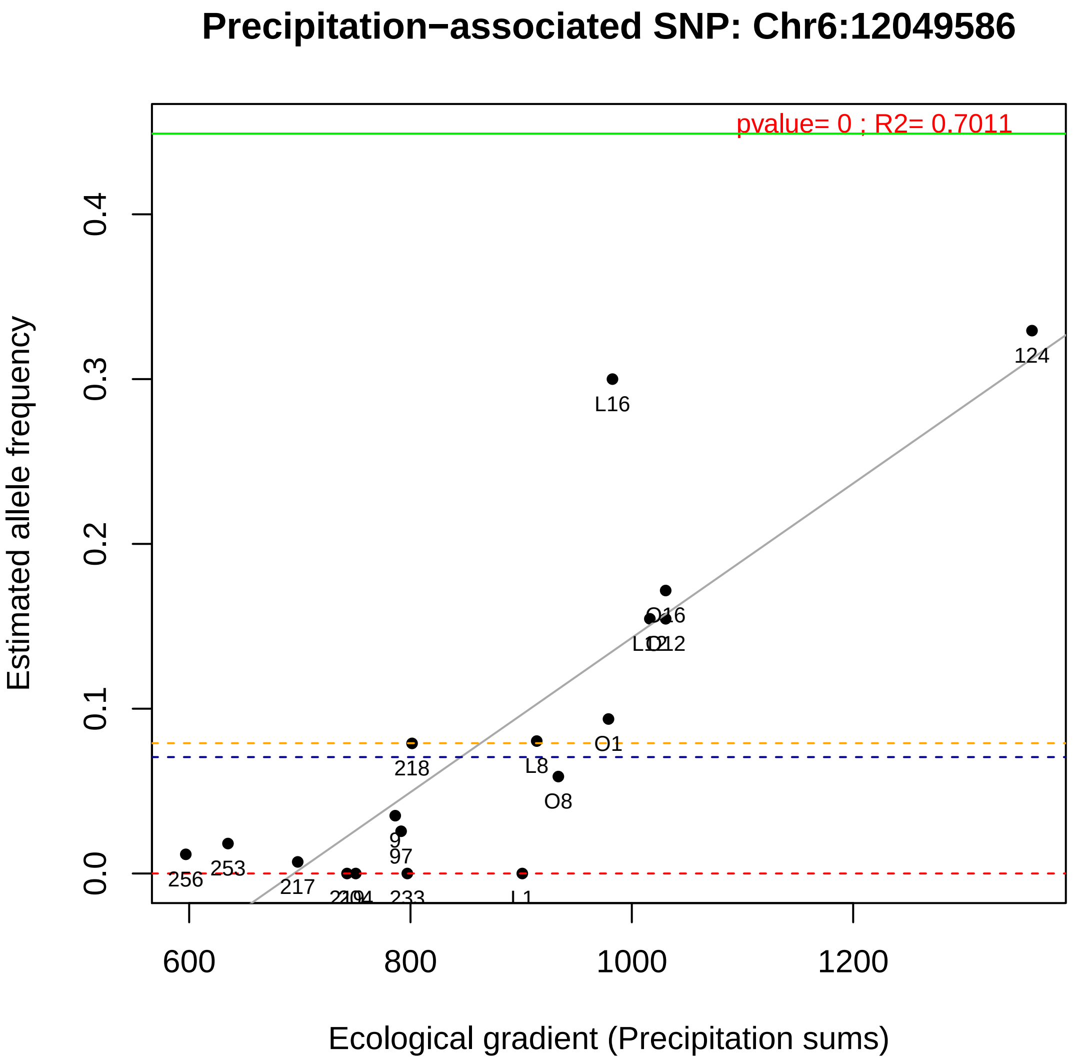
Precipitation−associated SNP: Chr6:12049586
pvalue= 0 ; R2= 0.7011
Estimated allele frequency
Ecological gradient (Precipitation sums)
0.0
0.1
0.2
0.3
0.4
600
800
1000
1200
124
L16
O16
L12
O12
O1
L8
218
O8
9
97
256
253
217
233
L1

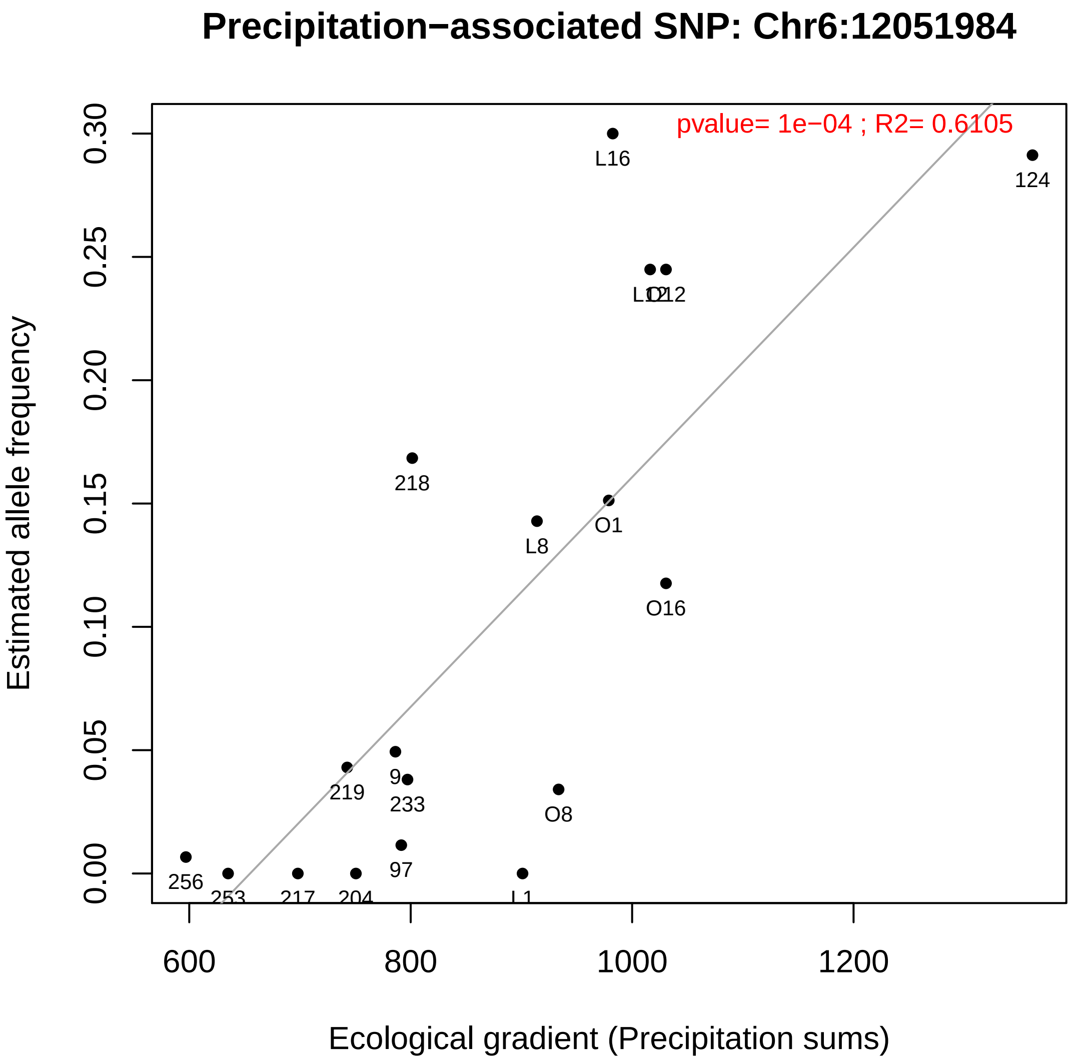

Precipitation−associated SNP: Chr6:12051984
pvalue= 1e−04 ; R2= 0.6105
Estimated allele frequency
0.00
0.05
0.10
0.15
0.20
0.25
0.30
600
800
1000
1200
Ecological gradient (Precipitation sums)
L16
124
L12
O12
218
O1
L8
O16
219
9
233
O8
97
256

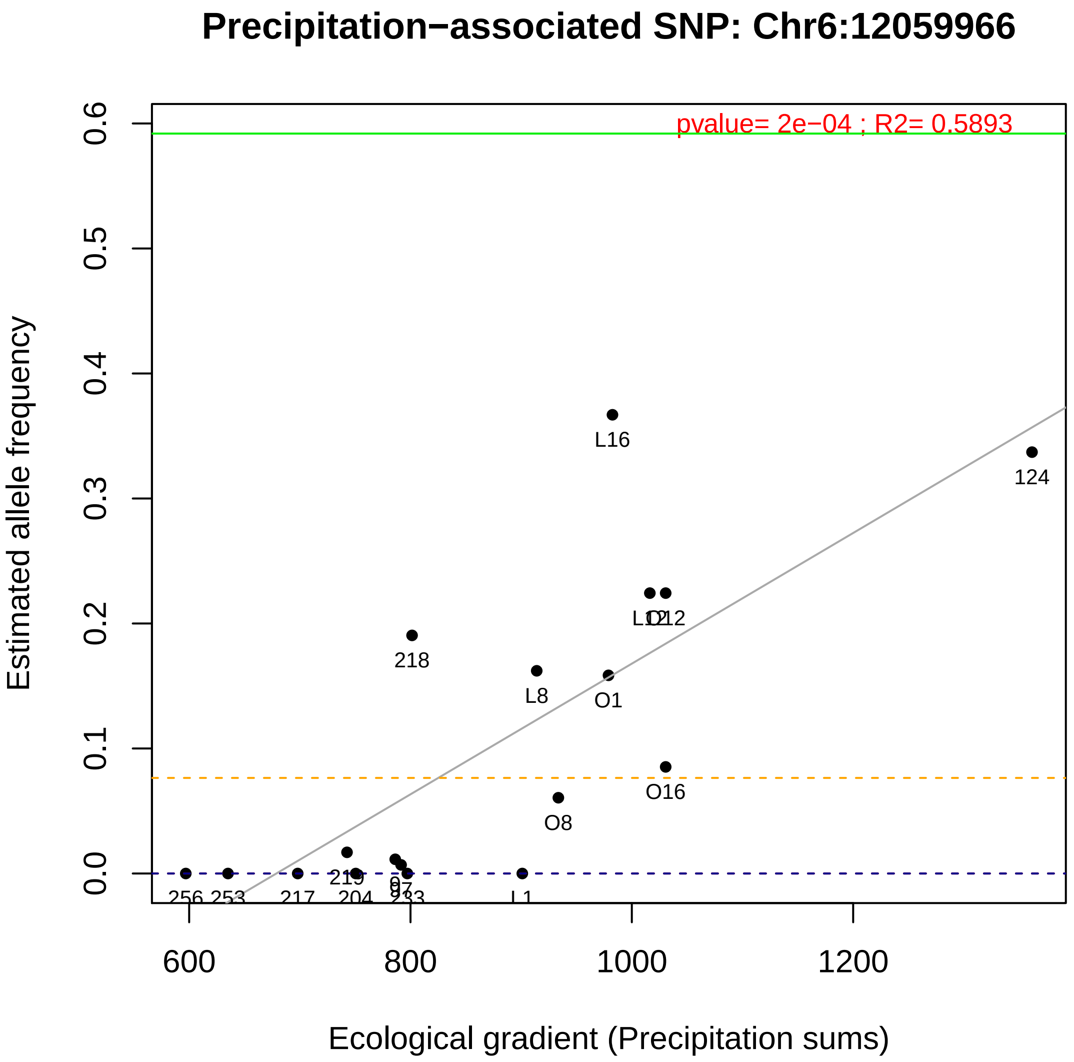

Precipitation−associated SNP: Chr6:12059966
pvalue= 2e−04 ; R2= 0.5893
Estimated allele frequency
0.0
0.1
0.2
0.3
0.4
0.5
0.6
Ecological gradient (Precipitation sums)
600
800
1000
1200
L16
124
218
L8
O1
O16
O8
219
256
253
217
204
233
L1

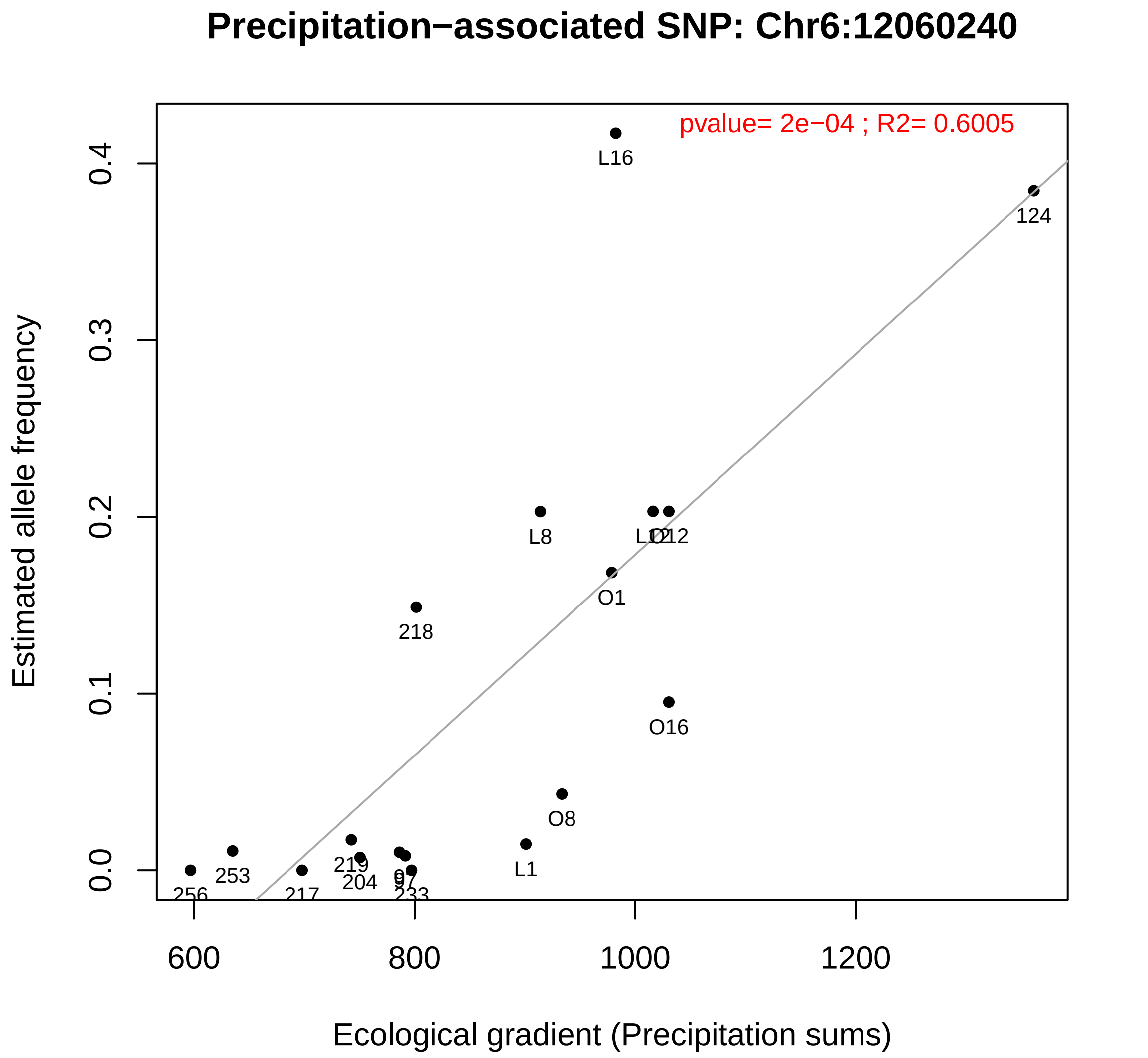

Precipitation−associated SNP: Chr6:12060240
pvalue= 2e−04 ; R2= 0.6005
Estimated allele frequency
Ecological gradient (Precipitation sums)
0.0
0.1
0.2
0.3
0.4
600
800
1000
1200
L16
124
L8
O1
218
O16
O8
L1
253
219
204
217

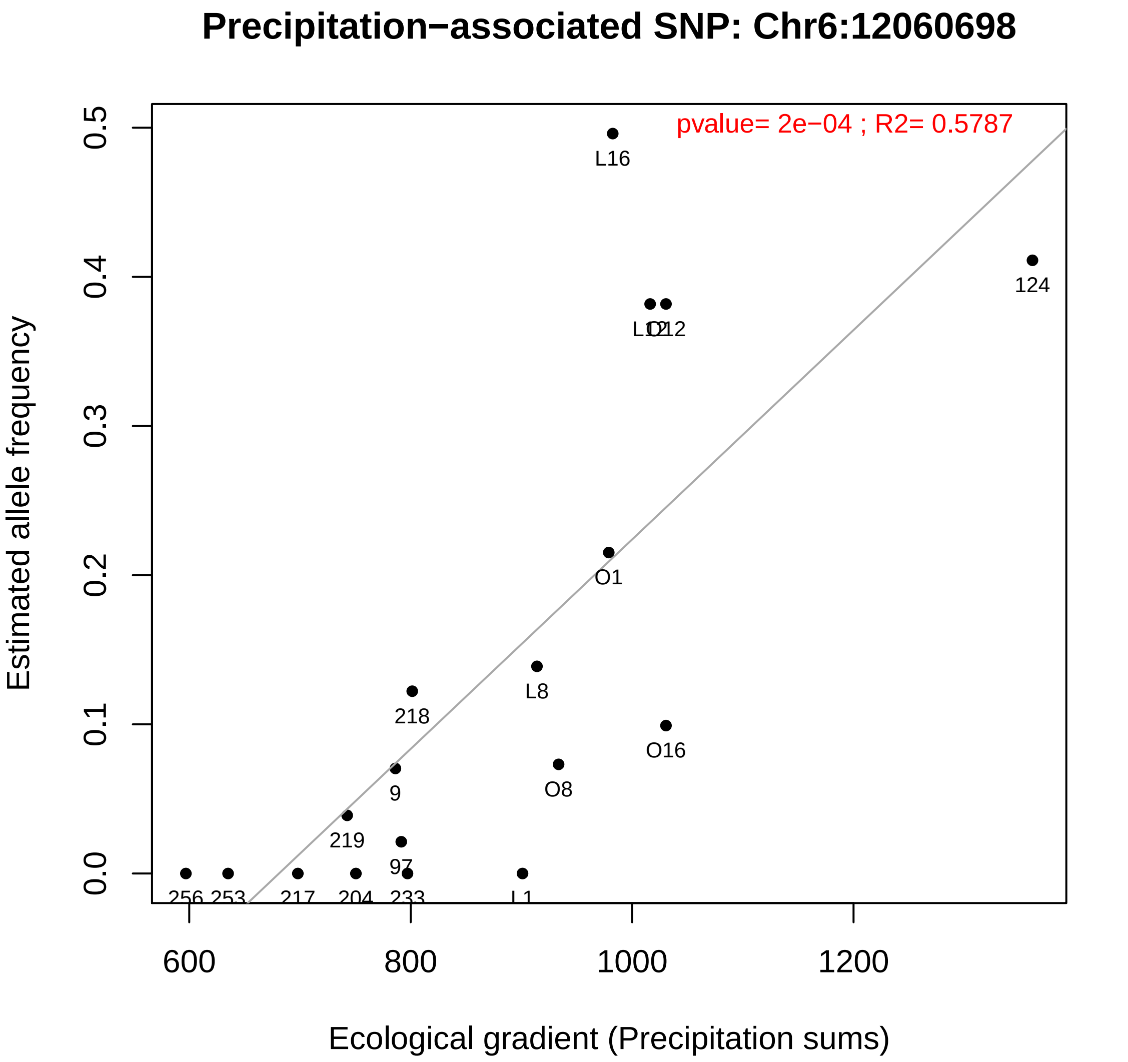
Precipitation−associated SNP: Chr6:12060698
pvalue= 2e−04 ; R2= 0.5787
Estimated allele frequency
0.0
0.1
0.2
0.3
0.4
0.5
Ecological gradient (Precipitation sums)
600
800
1000
1200
L16
124
O12
O1
L8
218
O16
9
O8
219
97
256
253
217
204
233
L1

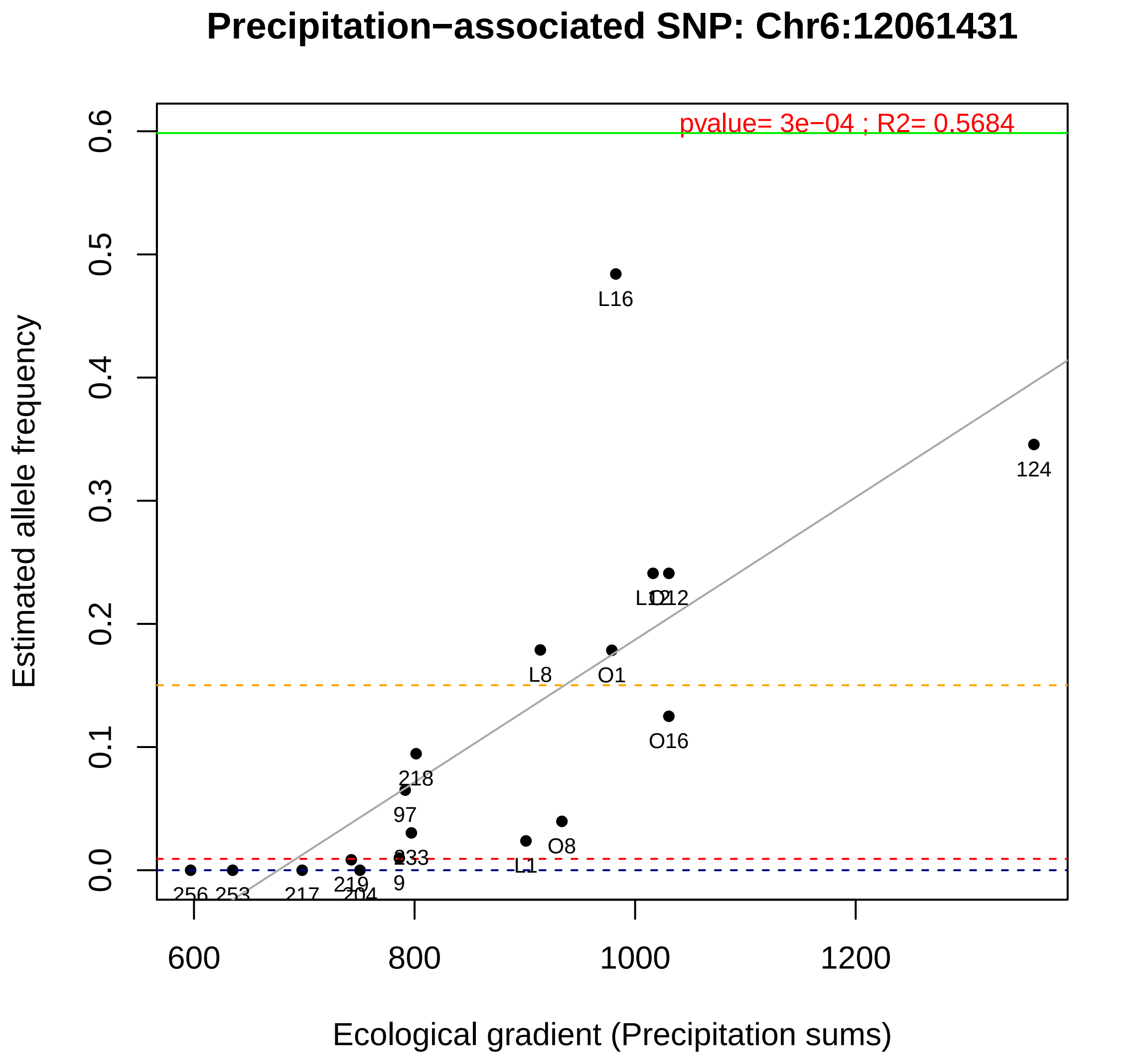

Precipitation−associated SNP: Chr6:12061431
pvalue= 3e−04 ; R2= 0.5684
Estimated allele frequency
0.0
0.1
0.2
0.3
0.4
0.5
0.6
600
800
1000
1200
Ecological gradient (Precipitation sums)
L16
124
L12
O12
L8
O1
O16
218
97
233
9
219
204
256
253
217
L1
O8

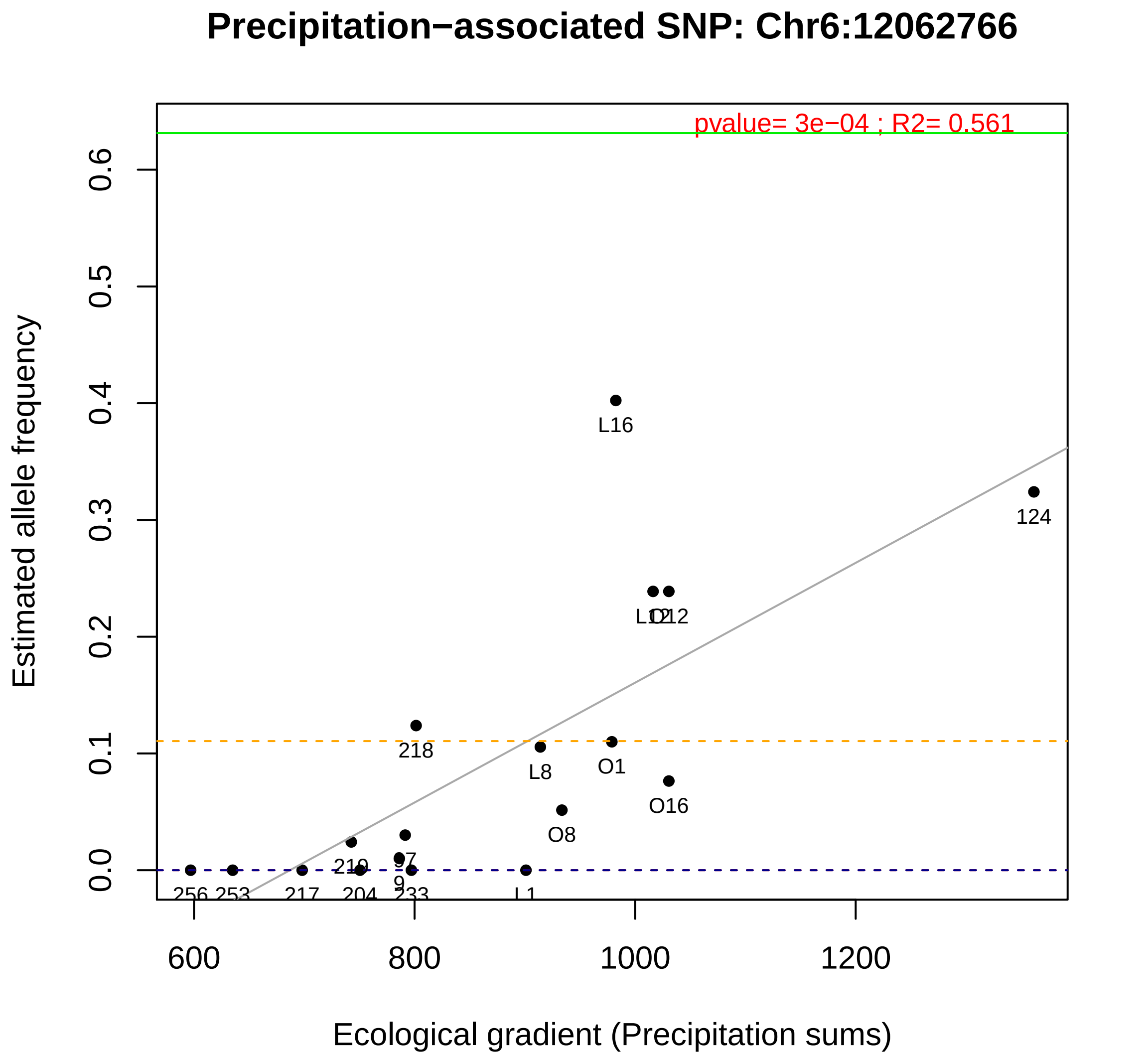
Precipitation−associated SNP: Chr6:12062766
pvalue= 3e−04 ; R2= 0.561
Estimated allele frequency
0.0
0.1
0.2
0.3
0.4
0.5
0.6
600
800
1000
1200
Ecological gradient (Precipitation sums)
L16
124
L12
O12
218
L8
O1
O16
O8
219
97
9
256
253
217
204
233
L1

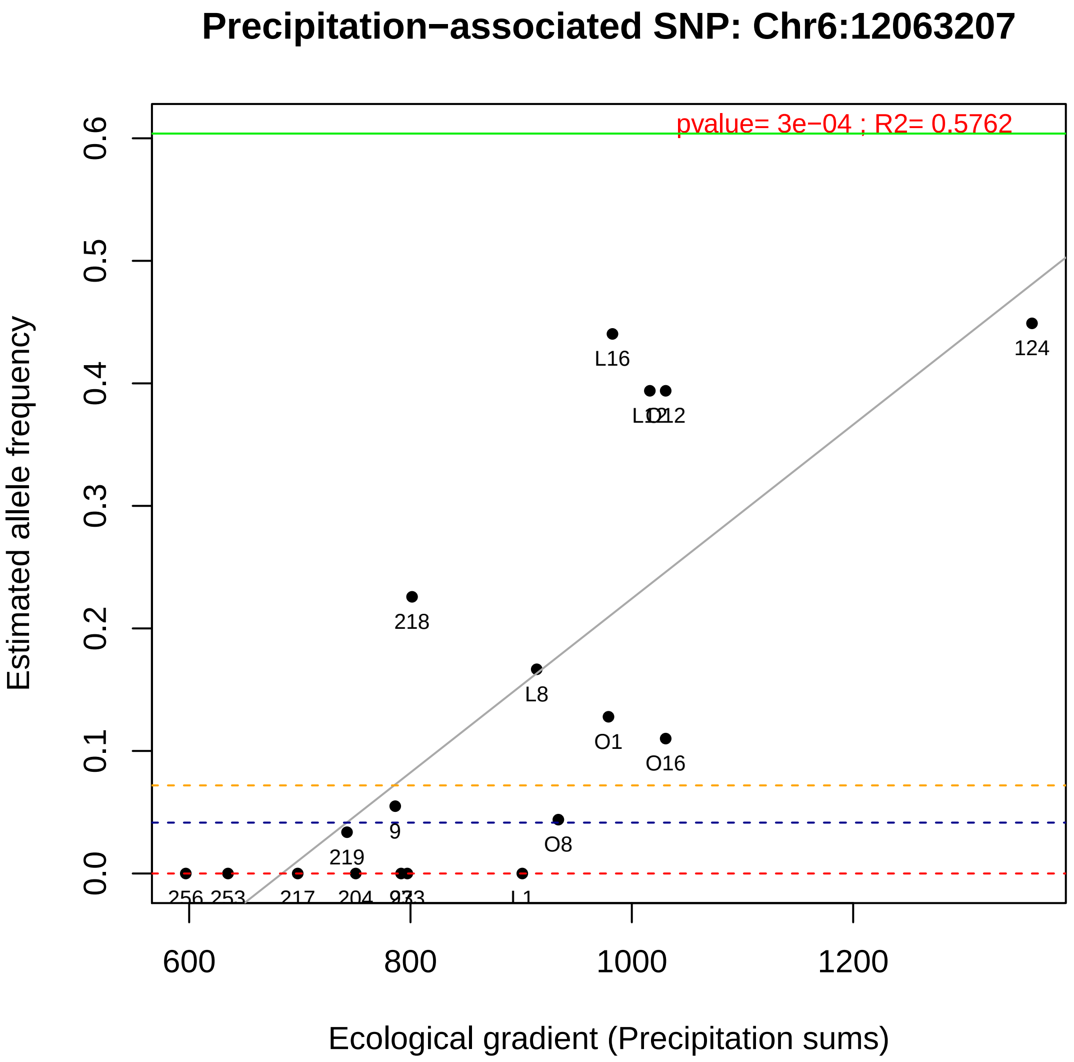

Precipitation−associated SNP: Chr6:12063207
pvalue= 3e−04 ; R2= 0.5762
Estimated allele frequency
0.0
0.1
0.2
0.3
0.4
0.5
0.6
600
800
1000
1200
Ecological gradient (Precipitation sums)
124
L16
L12
O12
218
L8
O1
O16
9
O8
219

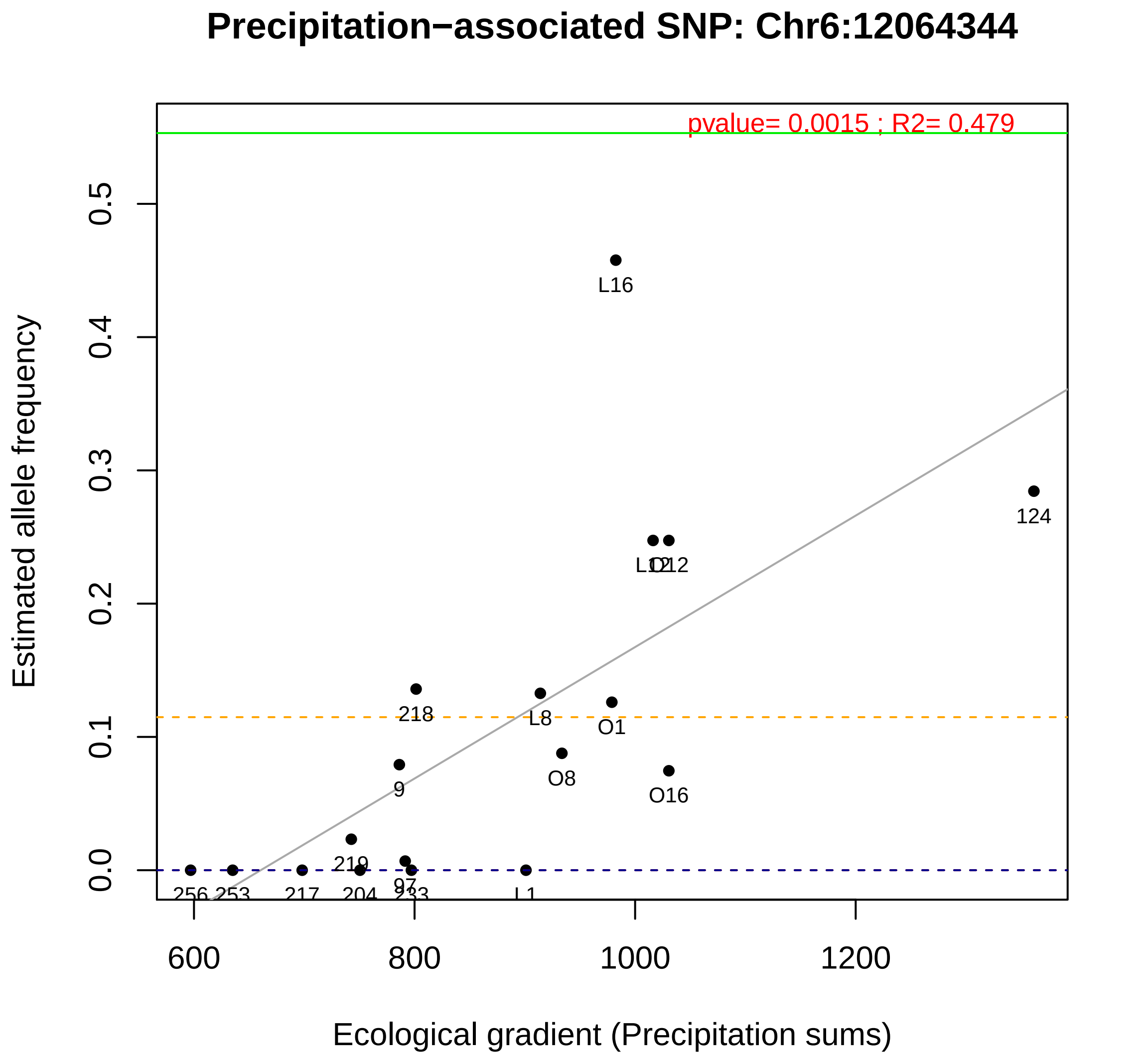
Precipitation−associated SNP: Chr6:12064344
pvalue= 0.0015 ; R2= 0.479
Estimated allele frequency
0.0
0.1
0.2
0.3
0.4
0.5
600
800
1000
1200
Ecological gradient (Precipitation sums)
L16
124
218
L8
O1
O8
9
O16
219
97
256
253
217
204
233
L1

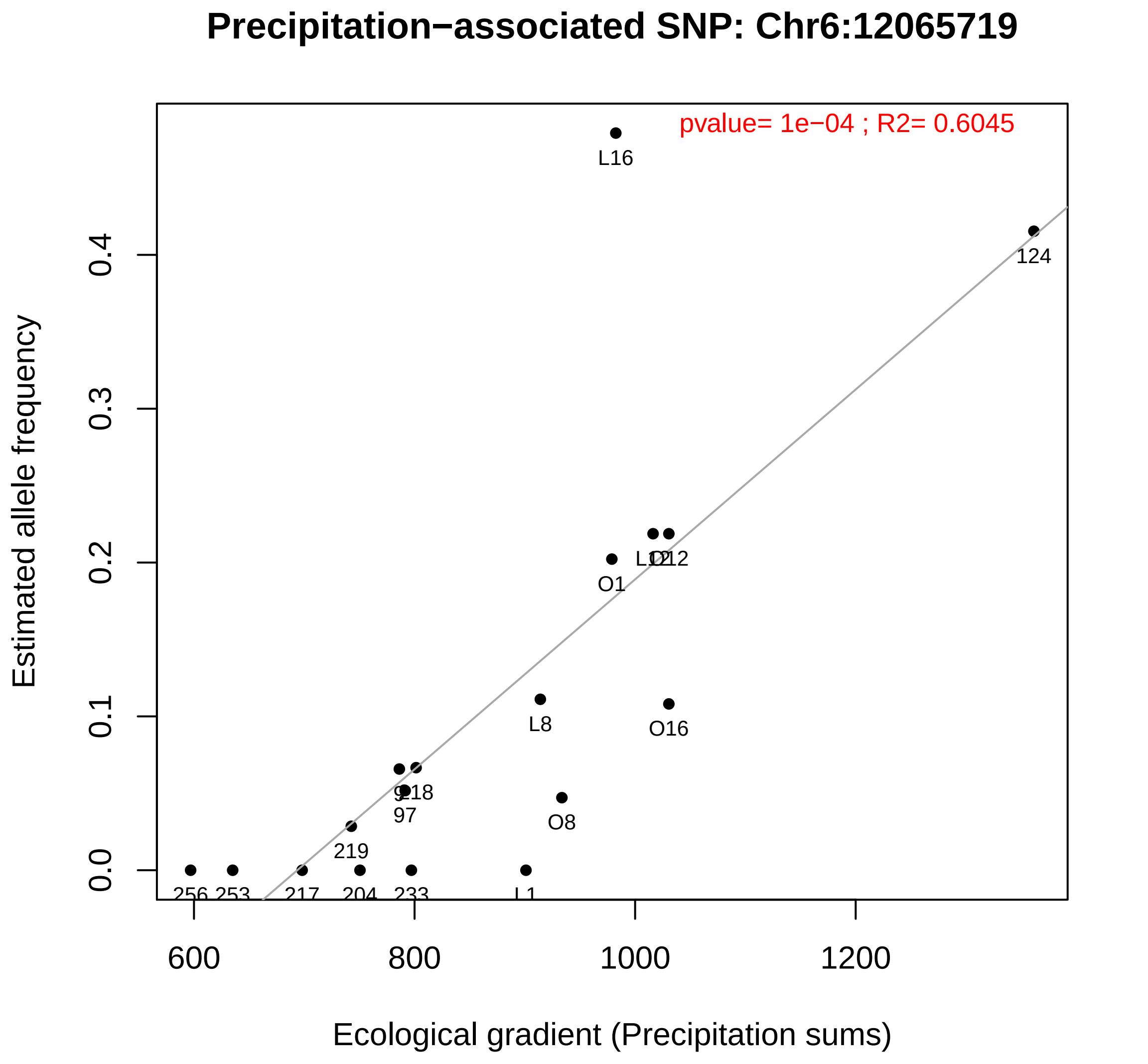

Precipitation−associated SNP: Chr6:12065719
pvalue= 1e−04 ; R2= 0.6045
L16
124
L12
O12
O1
L8
O16
918
97
O8
219
256
253
217
204
233
L1
0.0
0.1
0.2
0.3
0.4
600
800
1000
1200
Estimated allele frequency
Ecological gradient (Precipitation sums)

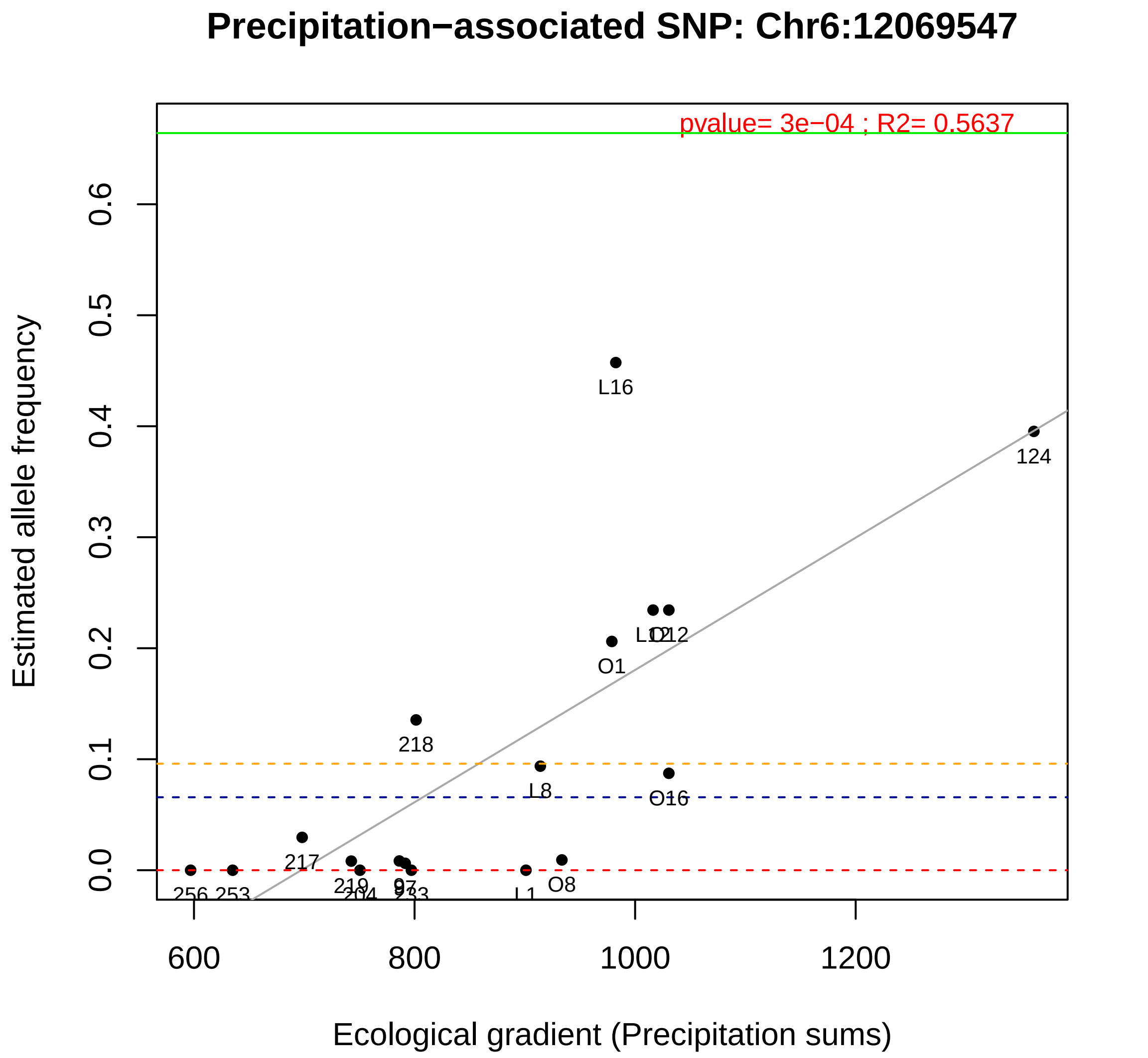

Precipitation−associated SNP: Chr6:12069547
pvalue= 3e−04 ; R2= 0.5637
Estimated allele frequency
0.0
0.1
0.2
0.3
0.4
0.5
0.6
600
800
1000
1200
Ecological gradient (Precipitation sums)
L16
124
L12
O12
O1
218
L8
O16
217
219
97
O8

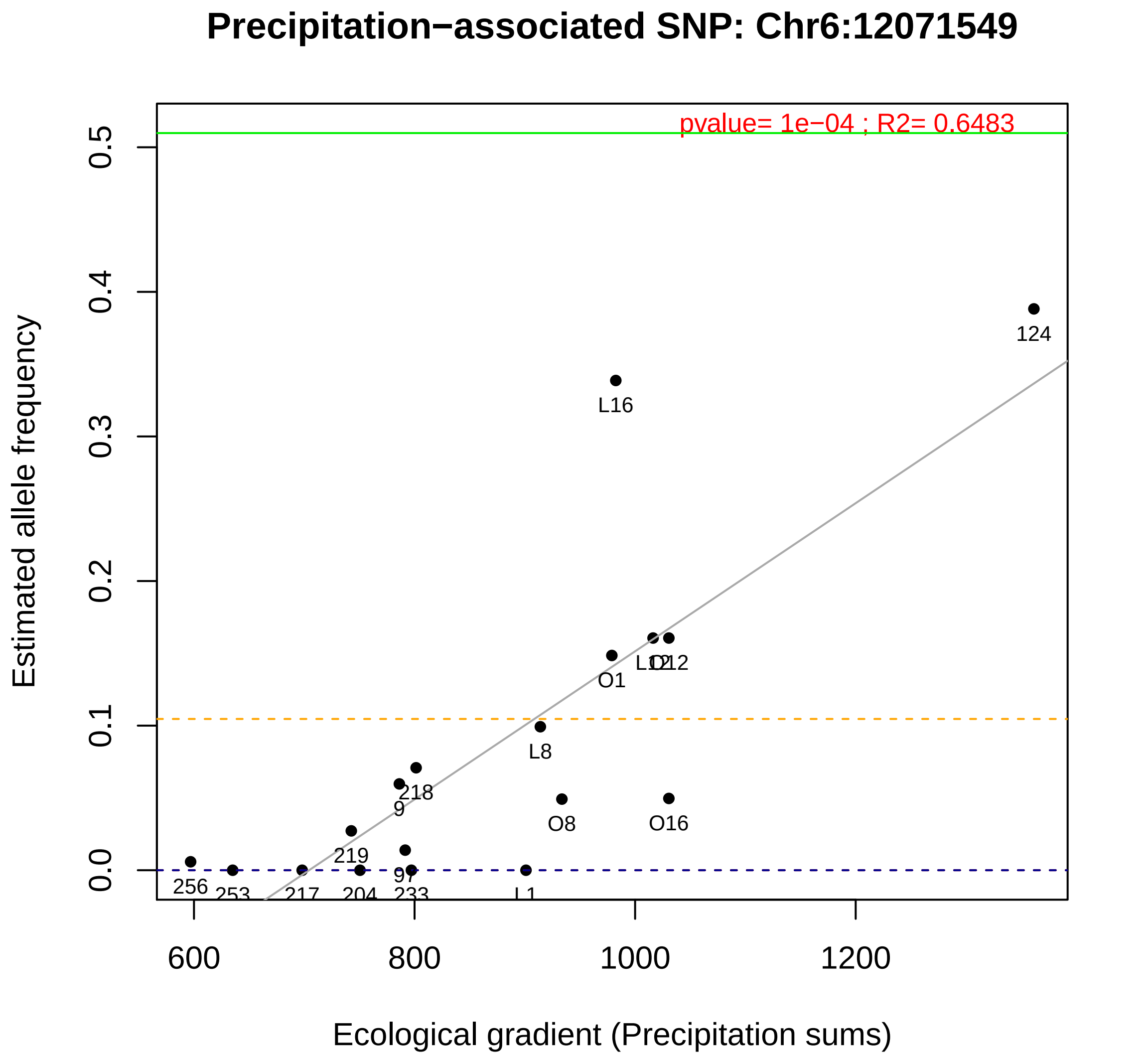

Precipitation−associated SNP: Chr6:12071549
pvalue= 1e−04 ; R2= 0.6483
Estimated allele frequency
Ecological gradient (Precipitation sums)
0.0
0.1
0.2
0.3
0.4
0.5
600
800
1000
1200
124
L16
O1
L12
O12
L8
218
9
O8
O16
219
97
256
253
217
204
233
L1

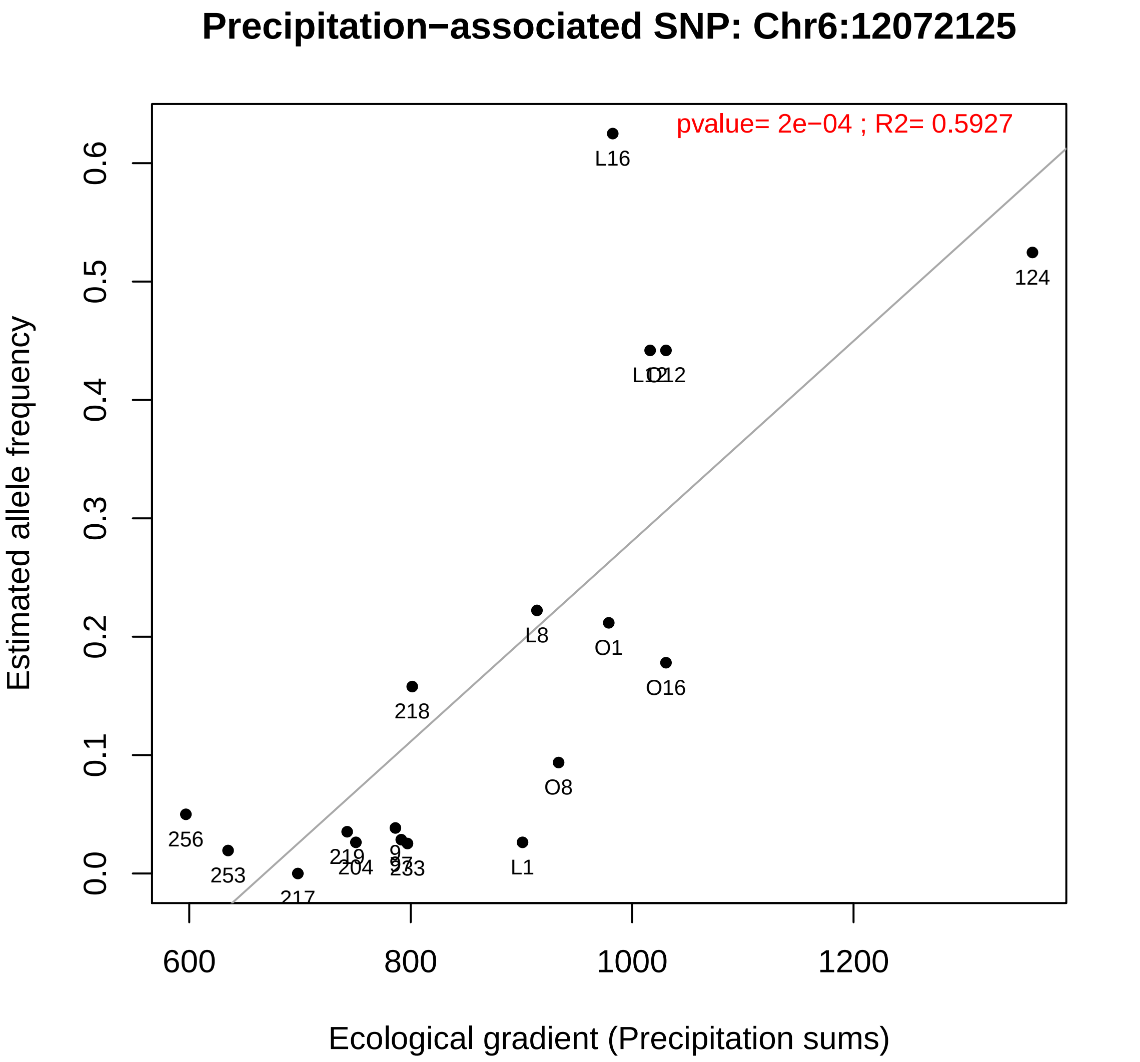

Precipitation−associated SNP: Chr6:12072125
pvalue= 2e−04 ; R2= 0.5927
Estimated allele frequency
0.0
0.1
0.2
0.3
0.4
0.5
0.6
600
800
1000
1200
Ecological gradient (Precipitation sums)
L16
124
L12
O12
L8
O1
O16
218
O8
256
253
217
219
204
9
233
L1

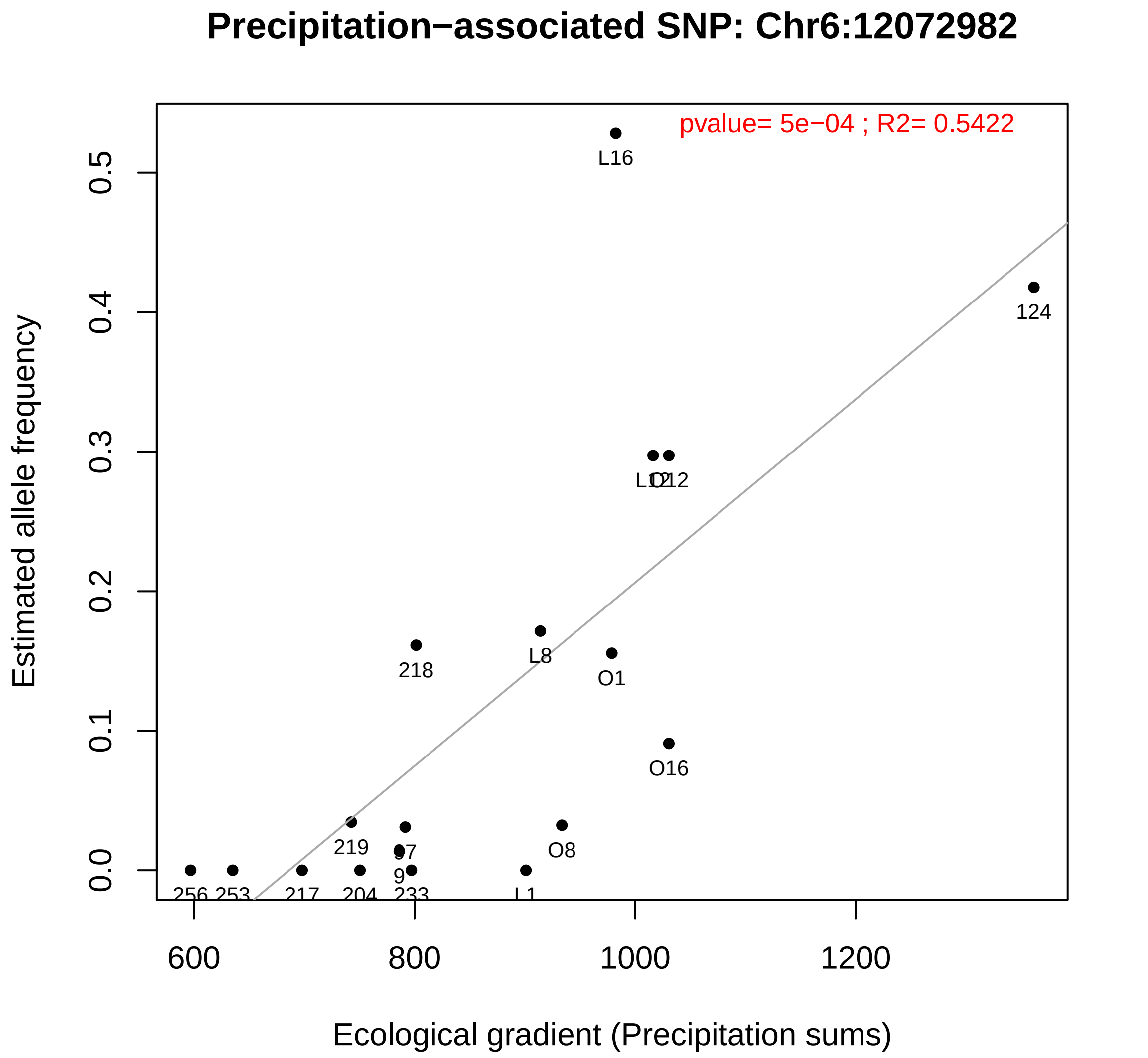
Precipitation−associated SNP: Chr6:12072982
pvalue= 5e−04 ; R2= 0.5422
Estimated allele frequency
Ecological gradient (Precipitation sums)
0.0
0.1
0.2
0.3
0.4
0.5
600
800
1000
1200
L16
124
O12
218
L8
O1
O16
219
O8
9
256
253
217
204
233

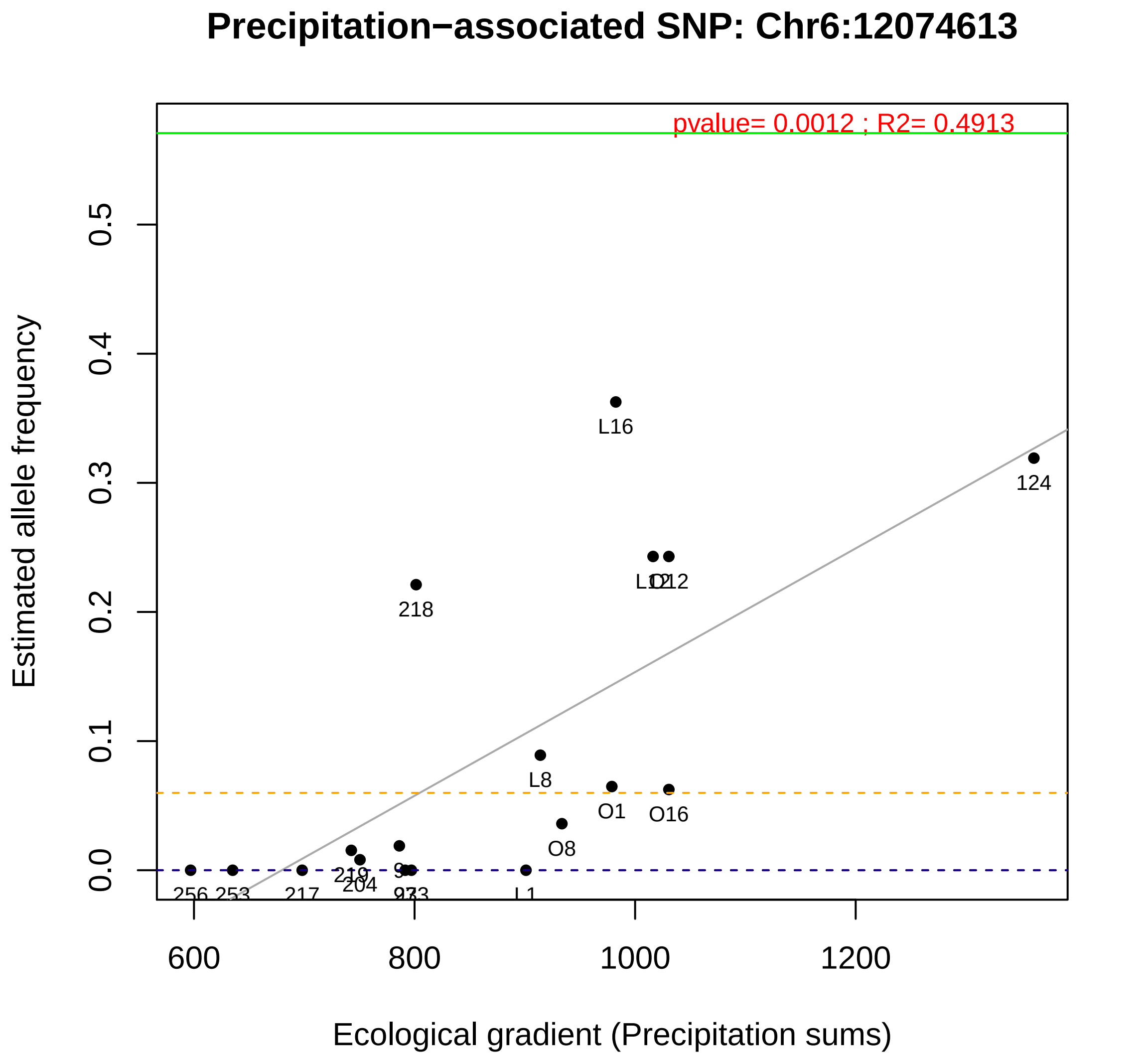
Precipitation−associated SNP: Chr6:12074613
pvalue= 0.0012 ; R2= 0.4913
Estimated allele frequency
0.0
0.1
0.2
0.3
0.4
0.5
600
800
1000
1200
Ecological gradient (Precipitation sums)
L16
124
218
L8
O1
O16
O8
219
204
217
L1
256
253

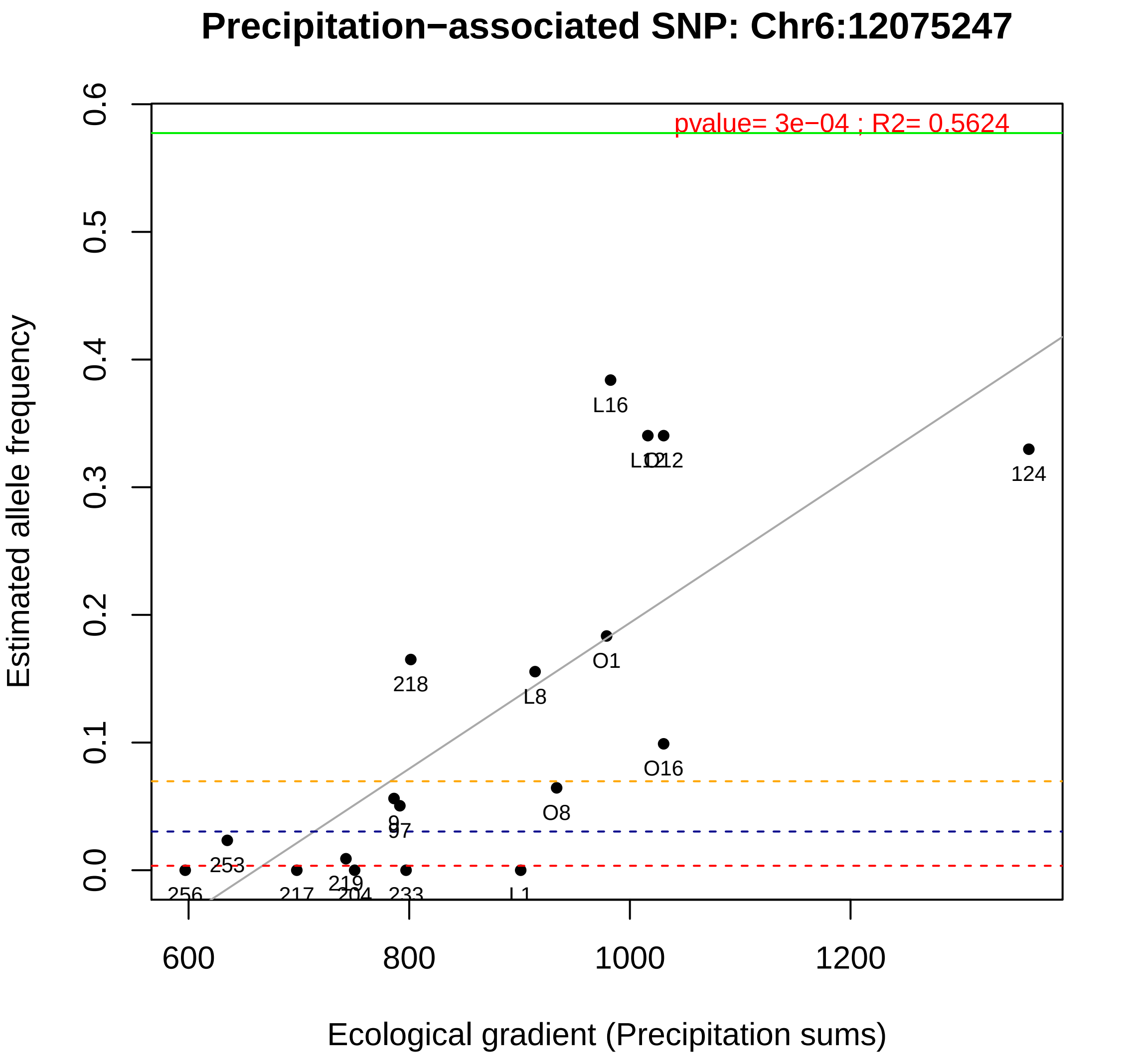

Precipitation−associated SNP: Chr6:12075247
pvalue= 3e−04 ; R2= 0.5624
Estimated allele frequency
0.0
0.1
0.2
0.3
0.4
0.5
0.6
Ecological gradient (Precipitation sums)
600
800
1000
1200
L16
L12
O12
124
218
L8
O1
O16
O8
9
97
253
219
256
217
204
233
L1

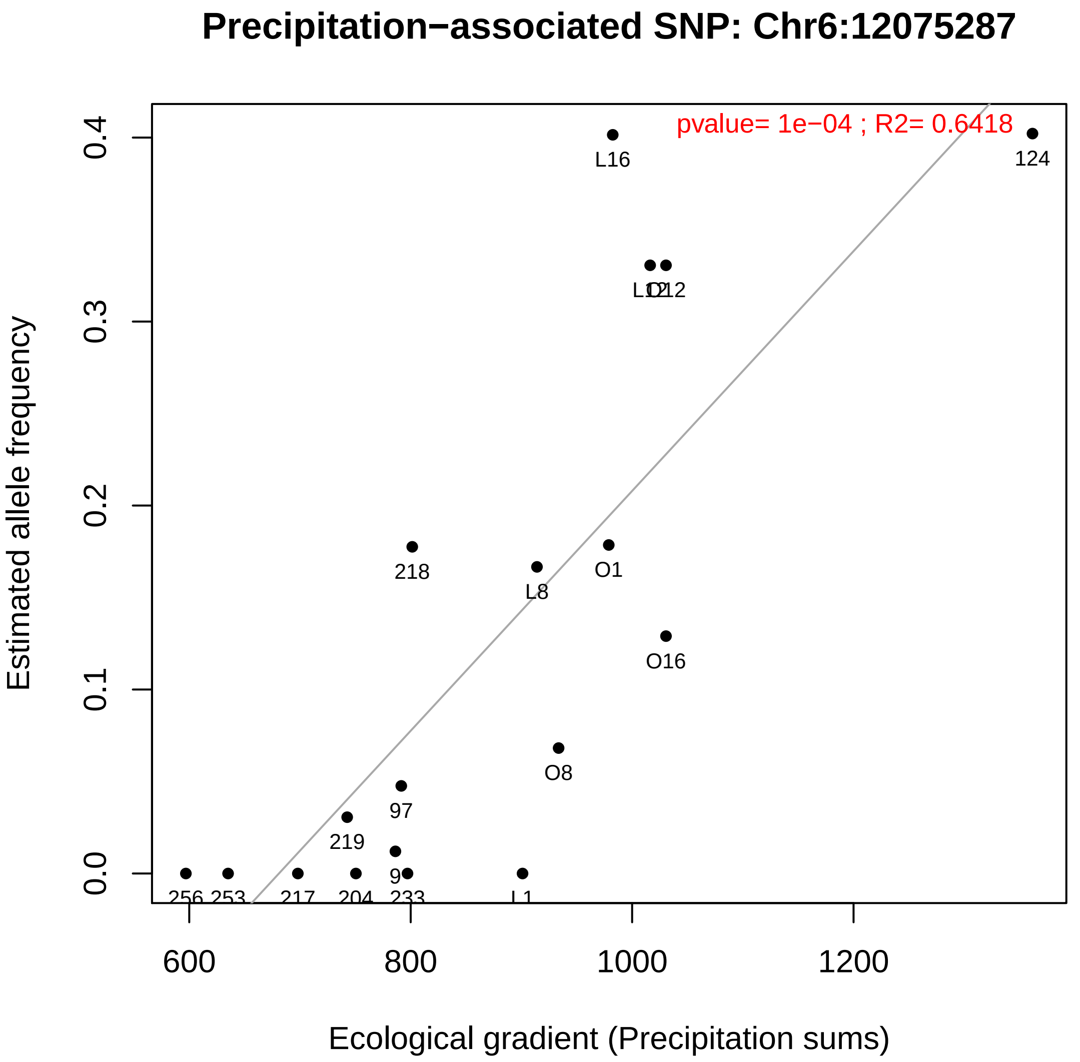
Precipitation−associated SNP: Chr6:12075287
pvalue= 1e−04 ; R2= 0.6418
Estimated allele frequency
0.0
0.1
0.2
0.3
0.4
Ecological gradient (Precipitation sums)
600
800
1000
1200
L16
124
O12
218
L8
O1
O16
O8
97
219
256
253
217
204
233
L1

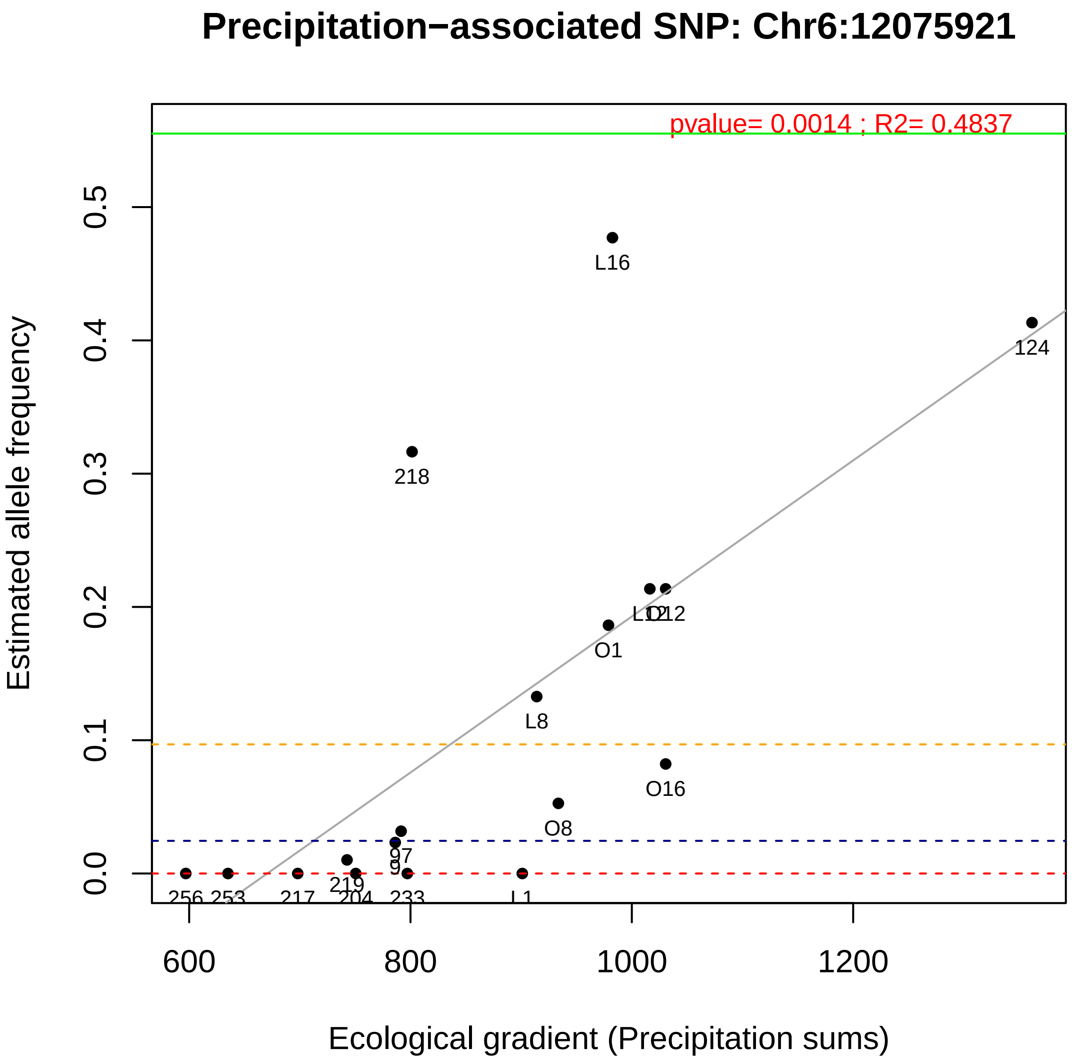

Precipitation−associated SNP: Chr6:12075921
pvalue= 0.0014 ; R2= 0.4837
Estimated allele frequency
Ecological gradient (Precipitation sums)
0.0
0.1
0.2
0.3
0.4
0.5
600
800
1000
1200
L16
124
218
O12
O1
L8
O16
O8
97
219

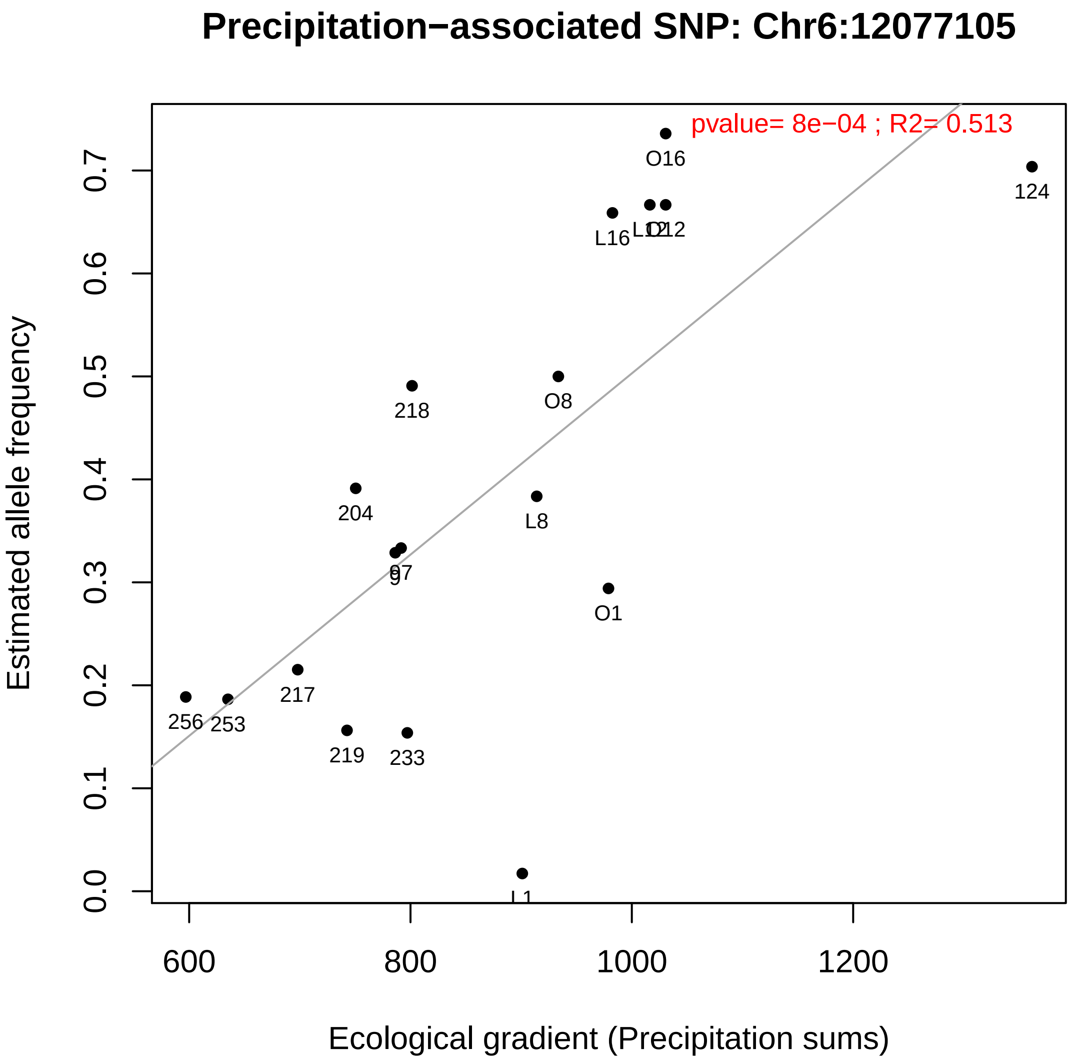
Precipitation−associated SNP: Chr6:12077105
pvalue= 8e−04 ; R2= 0.513
Estimated allele frequency
0.0
0.1
0.2
0.3
0.4
0.5
0.6
0.7
600
800
1000
1200
Ecological gradient (Precipitation sums)
O16
124
L16
218
O8
204
L8
O1
217
256
253
219
233

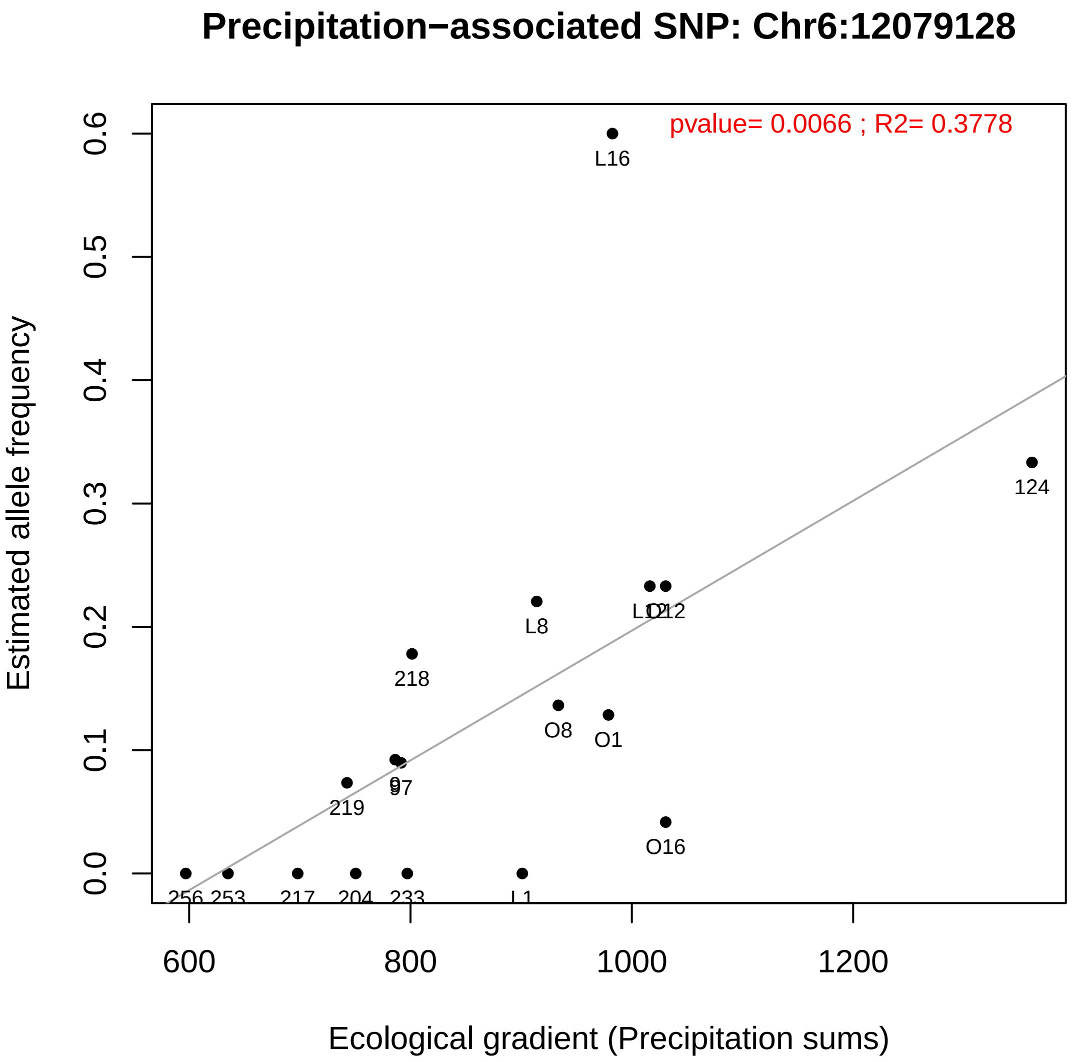

Precipitation−associated SNP: Chr6:12079128
pvalue= 0.0066 ; R2= 0.3778
Estimated allele frequency
Ecological gradient (Precipitation sums)
0.0
0.1
0.2
0.3
0.4
0.5
0.6
600
800
1000
1200
L16
124
L8
218
O8
O1
219
97
O16
256
253
217
204
233
L1

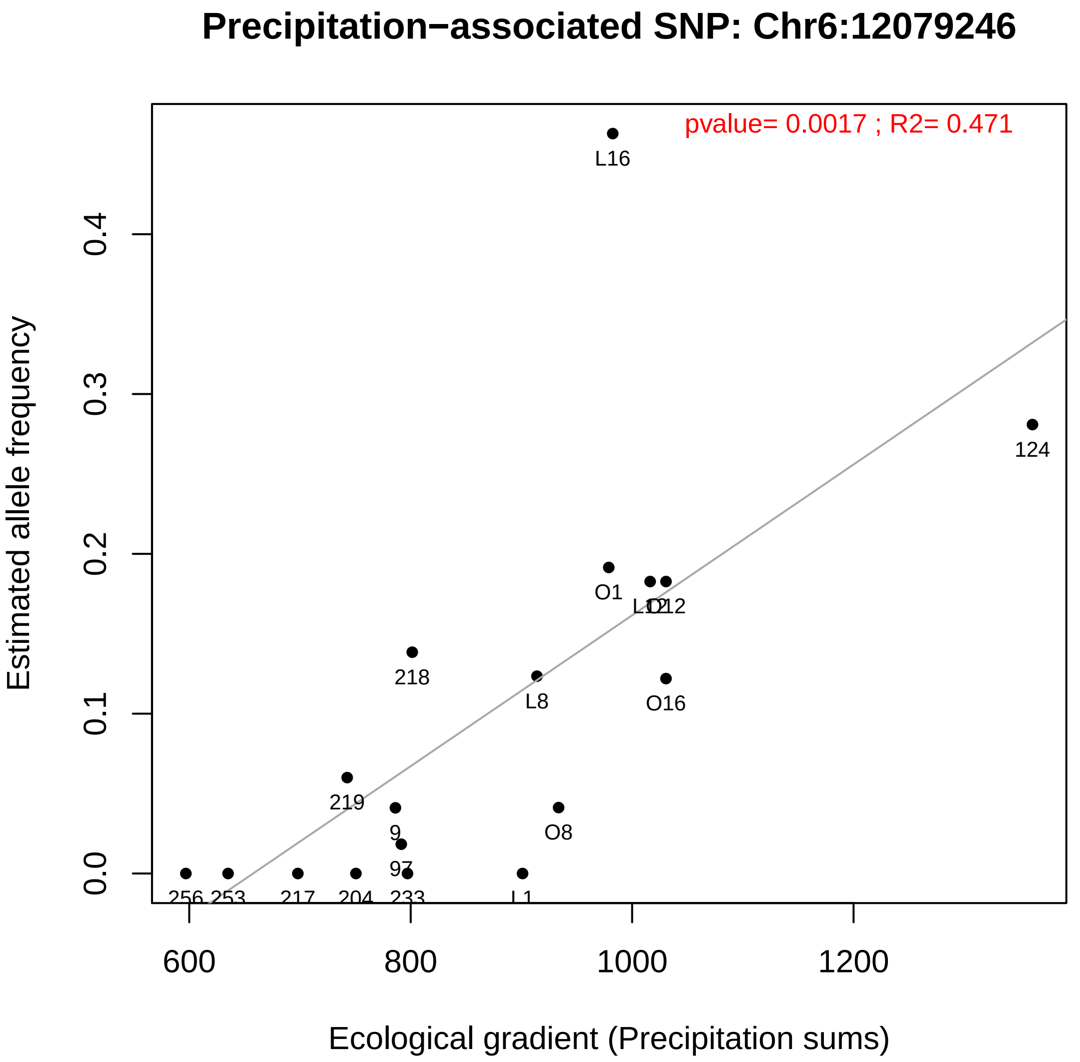

Precipitation−associated SNP: Chr6:12079246
pvalue= 0.0017 ; R2= 0.471
L16
124
O1
L12
O12
218
L8
O16
219
9
O8
97
256
253
217
204
233
L1
0.0
0.1
0.2
0.3
0.4
Estimated allele frequency
600
800
1000
1200
Ecological gradient (Precipitation sums)

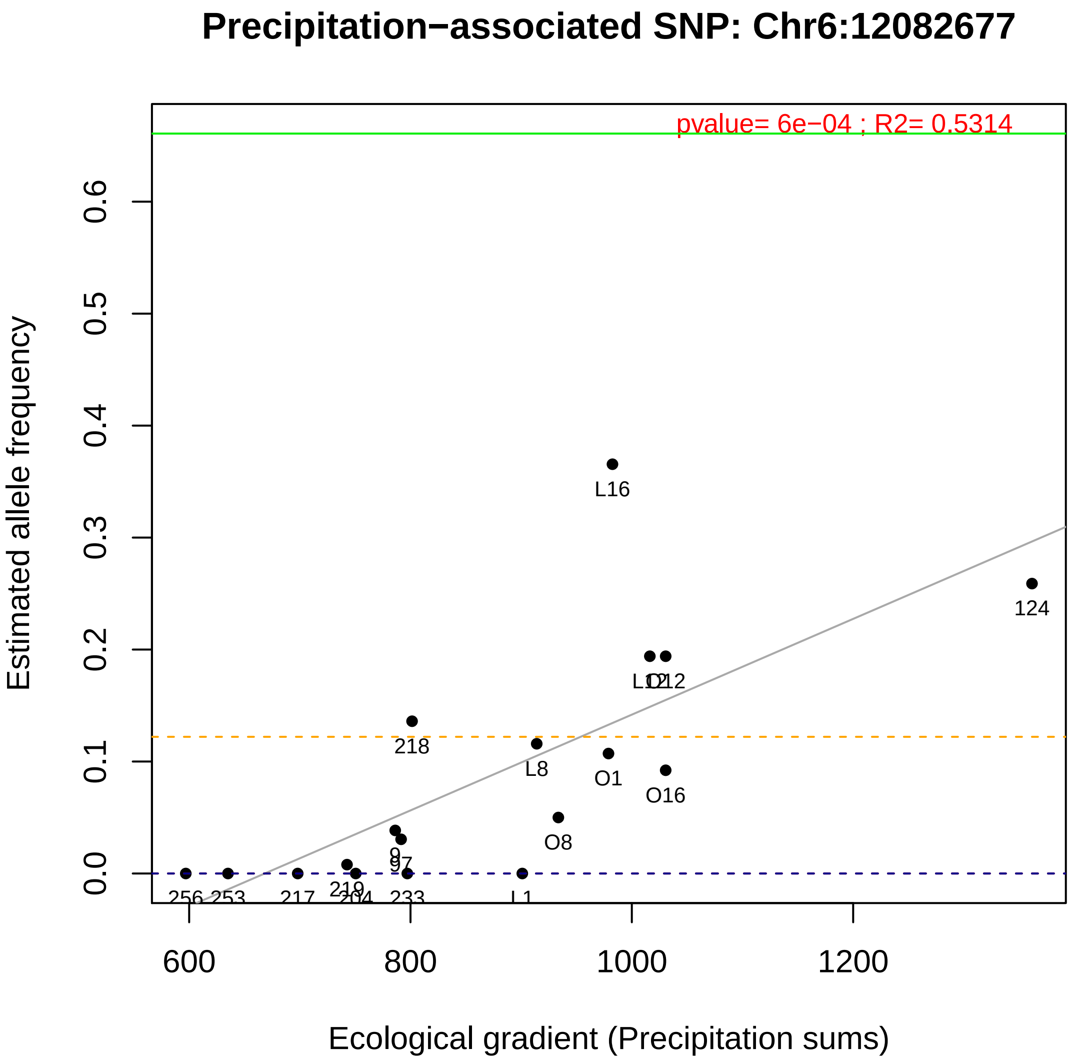
Precipitation−associated SNP: Chr6:12082677
pvalue= 6e−04 ; R2= 0.5314
Estimated allele frequency
0.0
0.1
0.2
0.3
0.4
0.5
0.6
600
800
1000
1200
Ecological gradient (Precipitation sums)
L16
124
O12
218
L8
O1
O16
O8
217
219
233
256
253

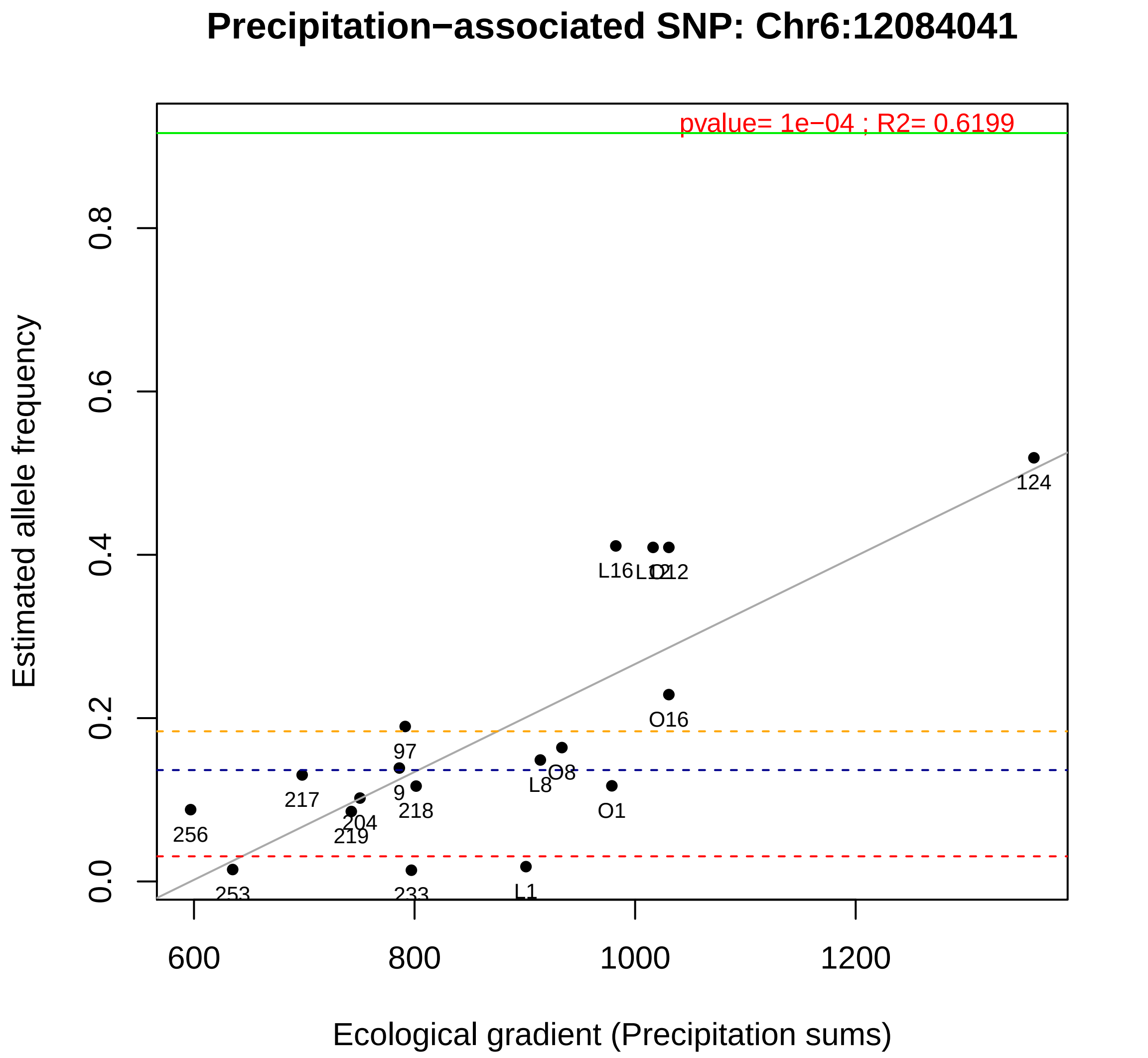

Precipitation−associated SNP: Chr6:12084041
pvalue= 1e−04 ; R2= 0.6199
Estimated allele frequency
0.0
0.2
0.4
0.6
0.8
600
800
1000
1200
Ecological gradient (Precipitation sums)
124
L16
O12
O16
97
9
218
217
204
219
256
253
233
L1
L8
O8
O1

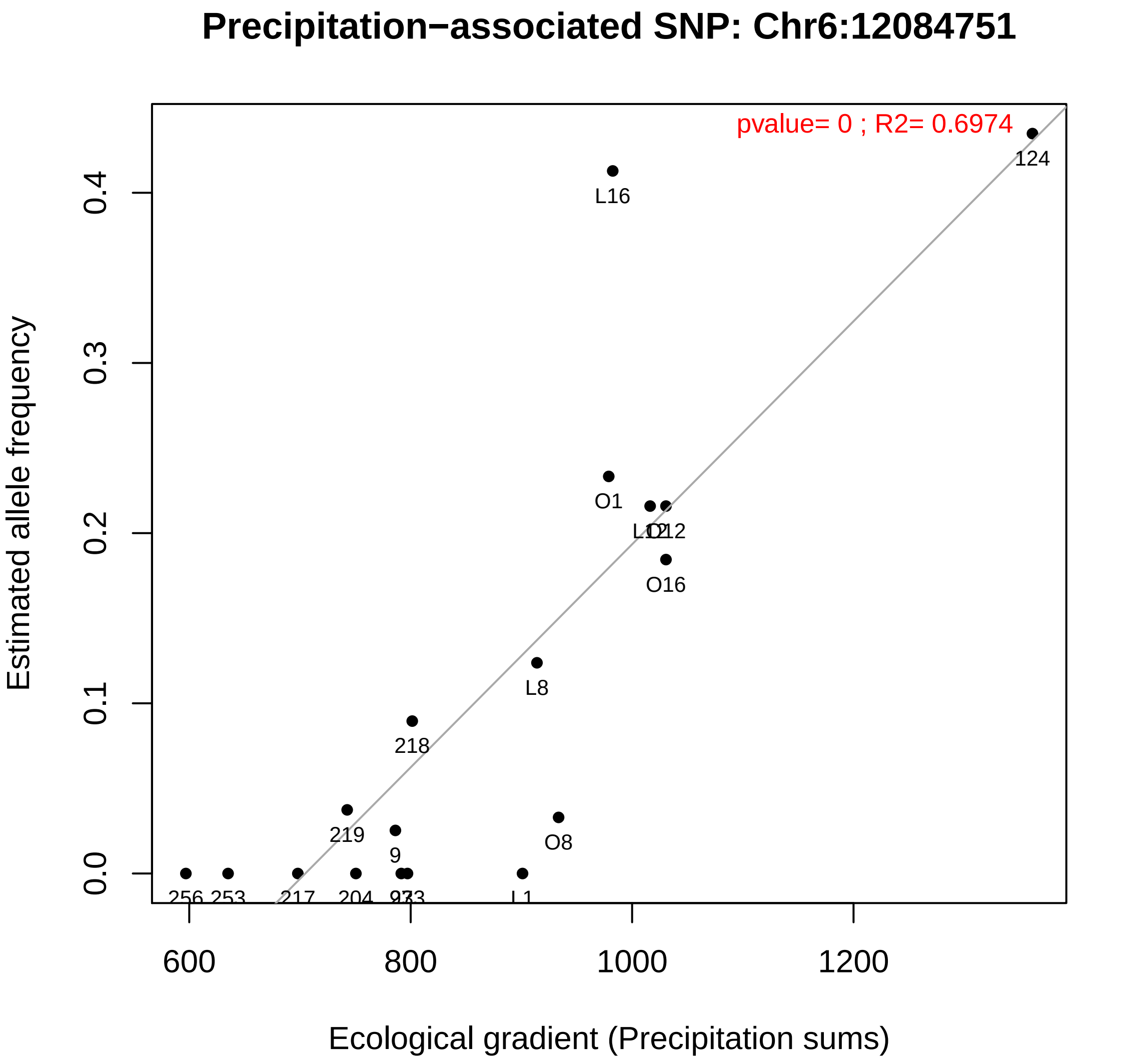

Precipitation−associated SNP: Chr6:12084751
pvalue= 0 ; R2= 0.6974
Estimated allele frequency
0.0
0.1
0.2
0.3
0.4
Ecological gradient (Precipitation sums)
600
800
1000
1200
124
L16
O1
O12
O16
L8
218
219
9
O8
L1

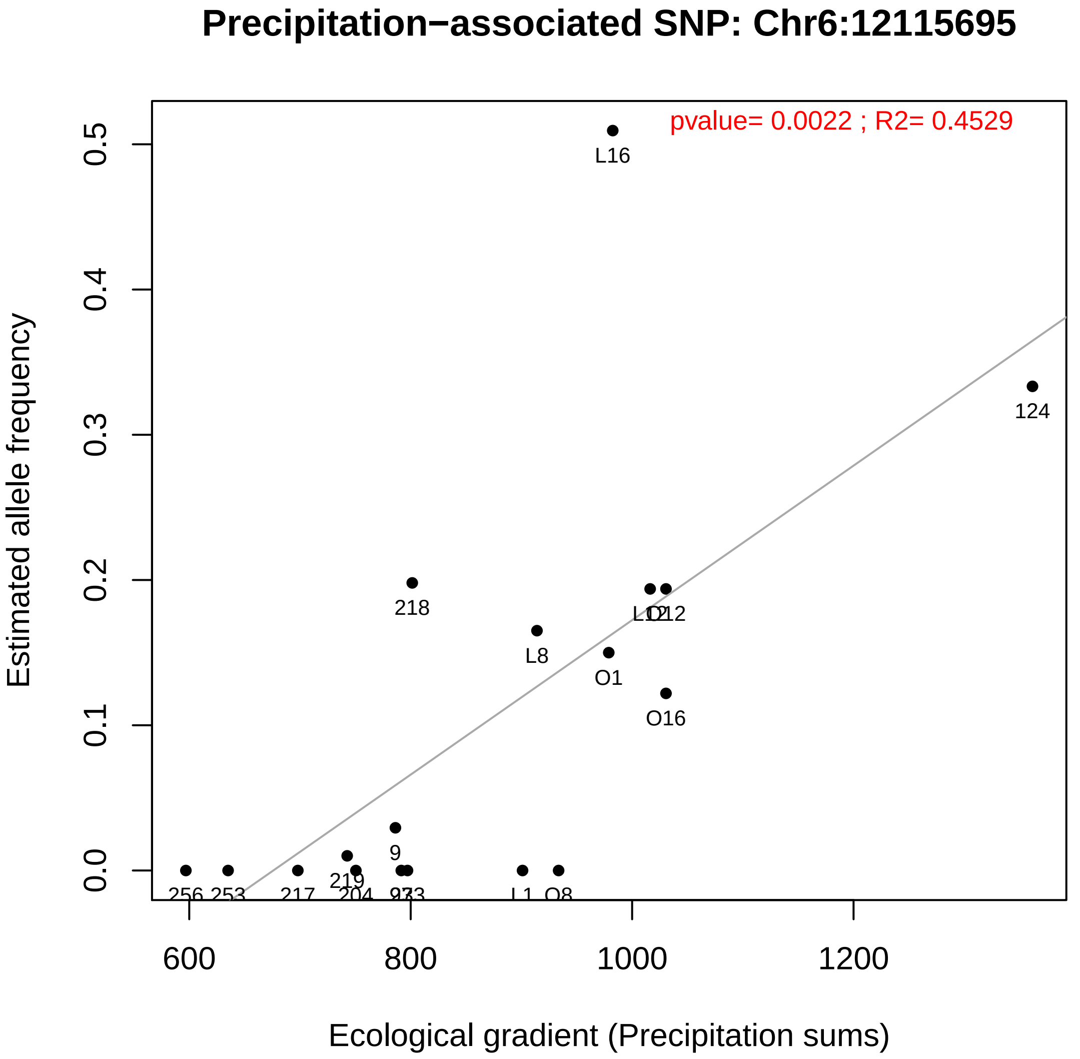
Precipitation−associated SNP: Chr6:12115695
pvalue= 0.0022 ; R2= 0.4529
Estimated allele frequency
0.0
0.1
0.2
0.3
0.4
0.5
600
800
1000
1200
Ecological gradient (Precipitation sums)
L16
124
218
L8
O1
O16
9
219

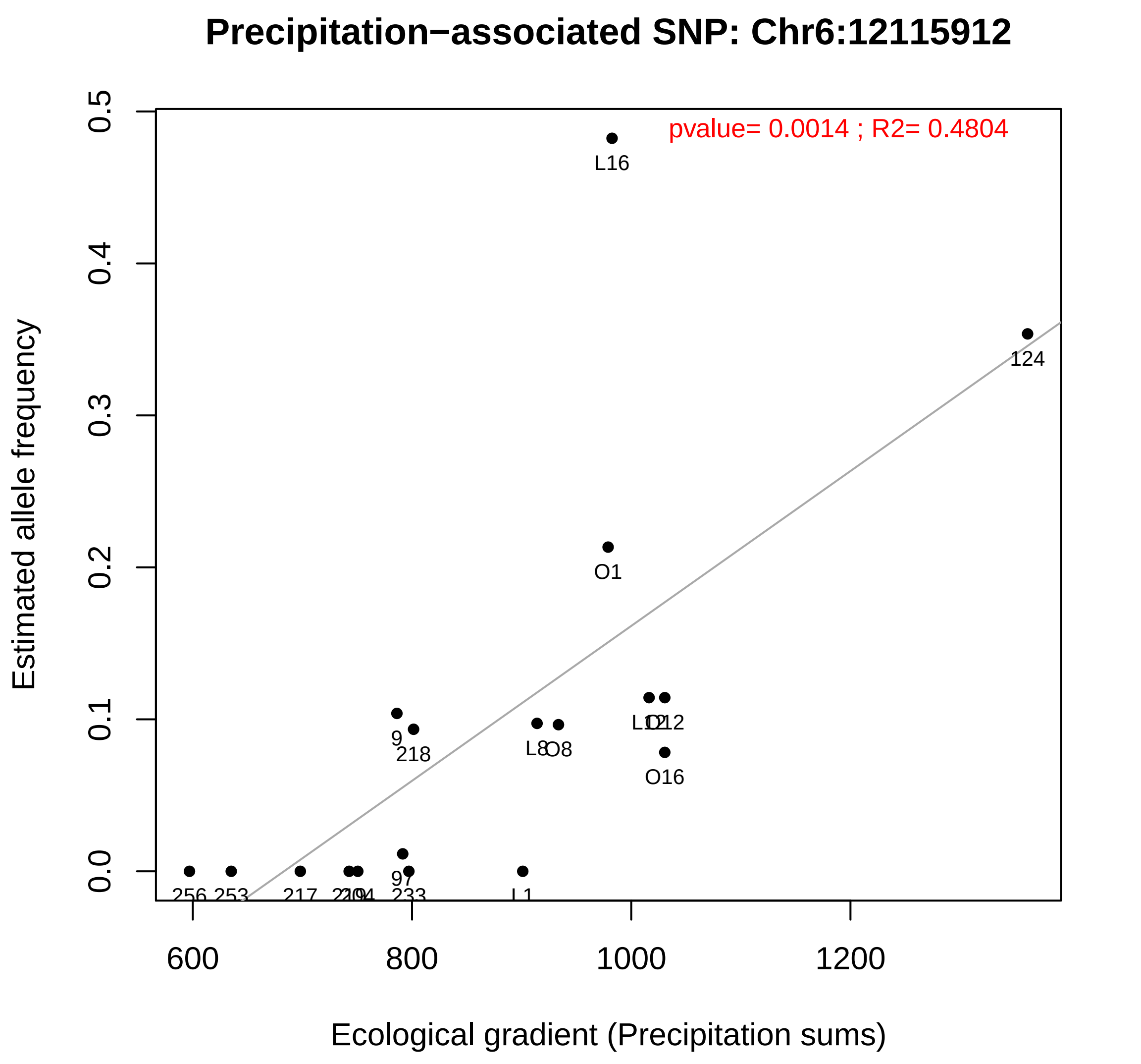

Precipitation−associated SNP: Chr6:12115912
pvalue= 0.0014 ; R2= 0.4804
Estimated allele frequency
Ecological gradient (Precipitation sums)
0.0
0.1
0.2
0.3
0.4
0.5
600
800
1000
1200
L16
124
O1
9
218
L8
O8
O16
97

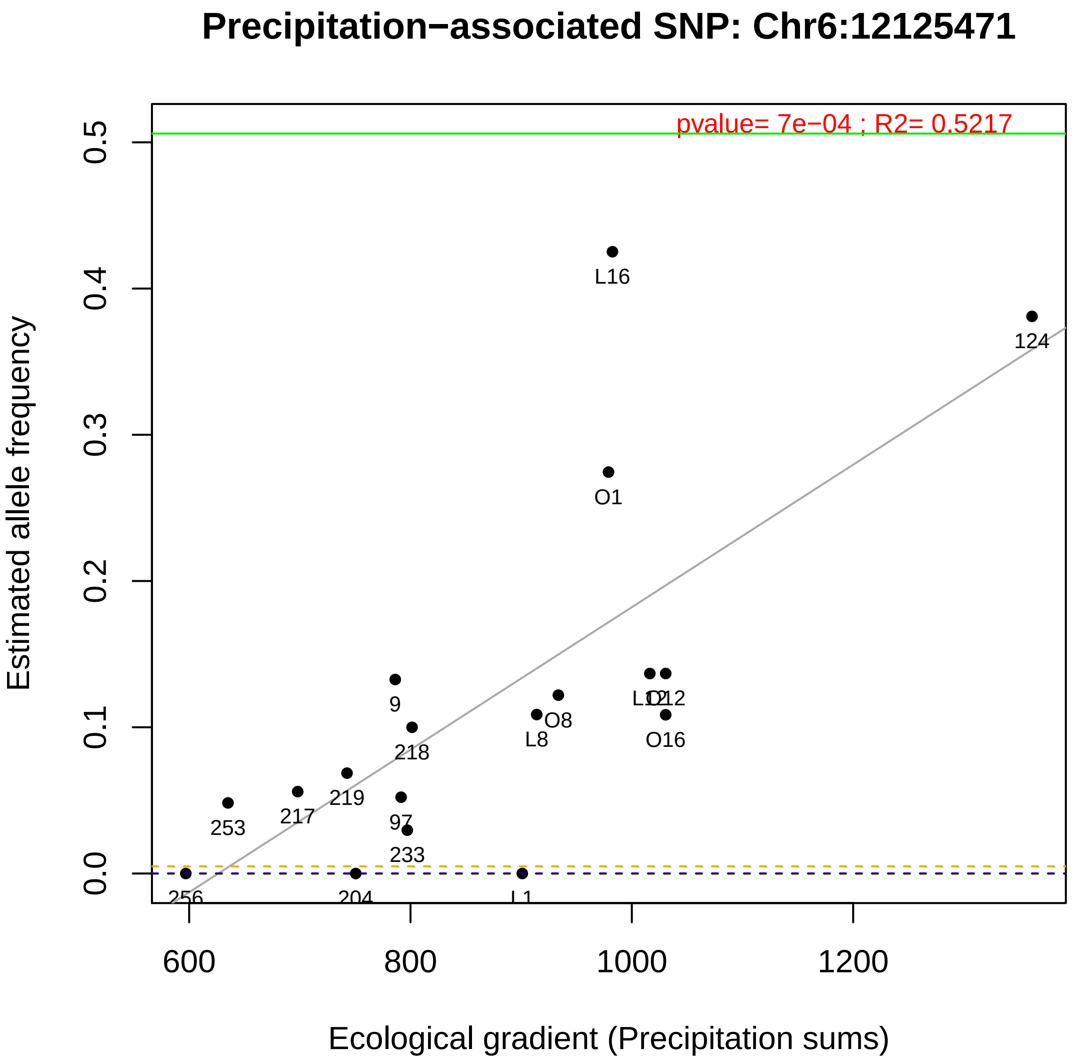
Precipitation−associated SNP: Chr6:12125471
pvalue= 7e−04 ; R2= 0.5217
L16
124
O1
9
O8
L12
O12
L8
O16
218
219
253
217
97
233
256
204
L1
Estimated allele frequency
0.0
0.1
0.2
0.3
0.4
0.5
600
800
1000
1200
Ecological gradient (Precipitation sums)

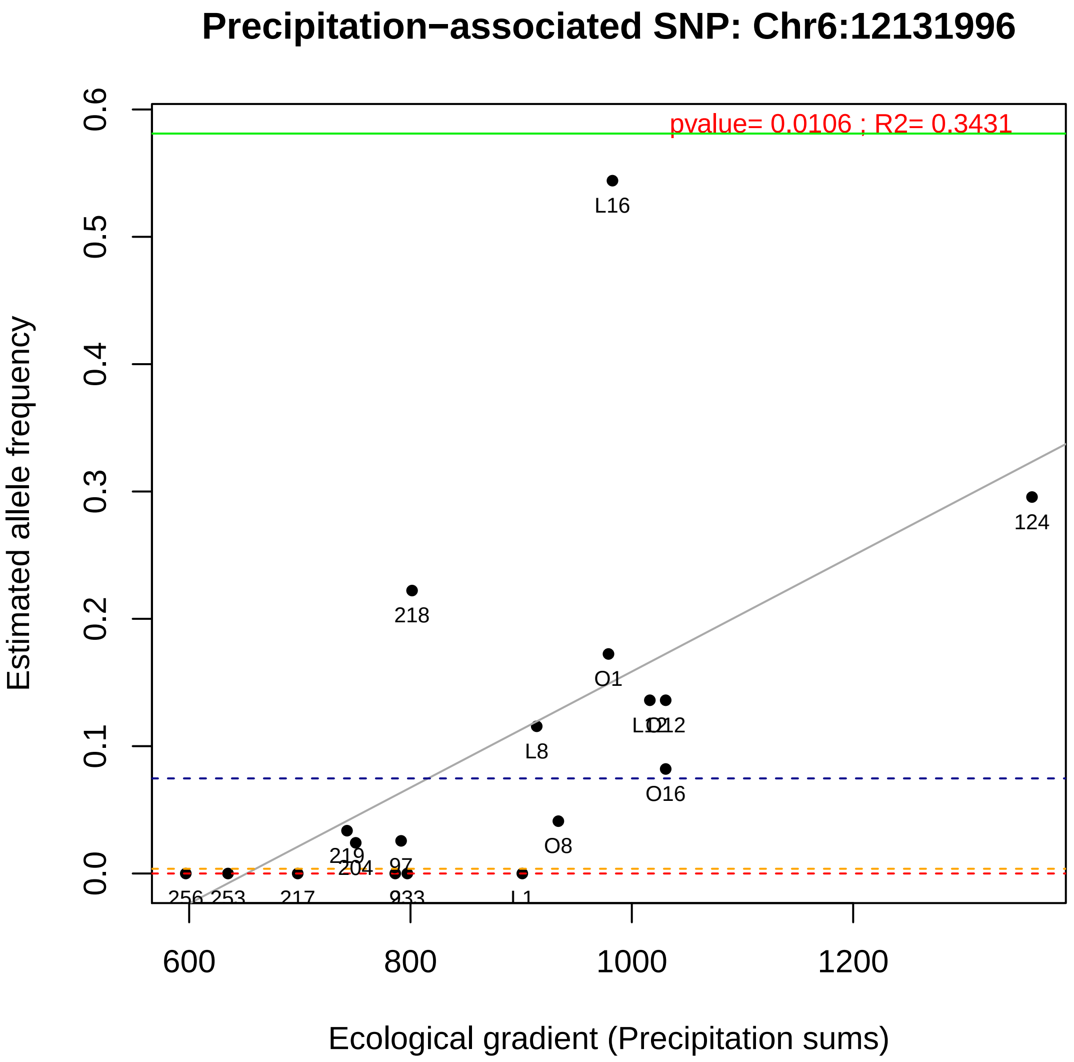

Precipitation−associated SNP: Chr6:12131996
pvalue= 0.0106 ; R2= 0.3431
Estimated allele frequency
0.0
0.1
0.2
0.3
0.4
0.5
0.6
600
800
1000
1200
Ecological gradient (Precipitation sums)
L16
124
218
O1
L12
O12
L8
O16
O8
219
204
97
256
253
217
9
33
L1

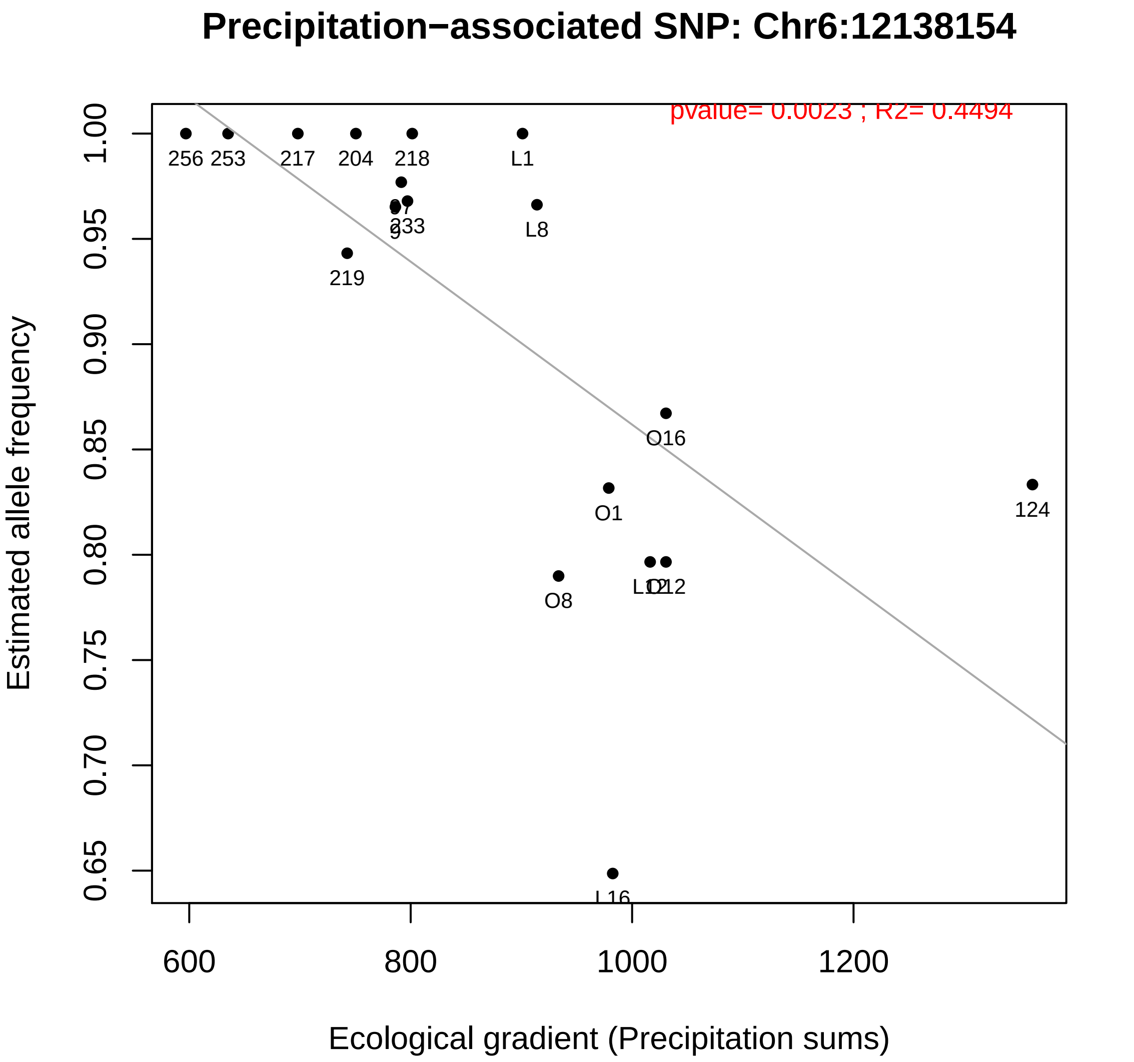

Precipitation−associated SNP: Chr6:12138154
pvalue= 0.0023 ; R2= 0.4494
Estimated allele frequency
Ecological gradient (Precipitation sums)
1.00
0.95
0.90
0.85
0.80
0.75
0.70
0.65
600
800
1000
1200
256
253
217
204
218
L1
233
L8
219
O16
O1
124
O12
O8

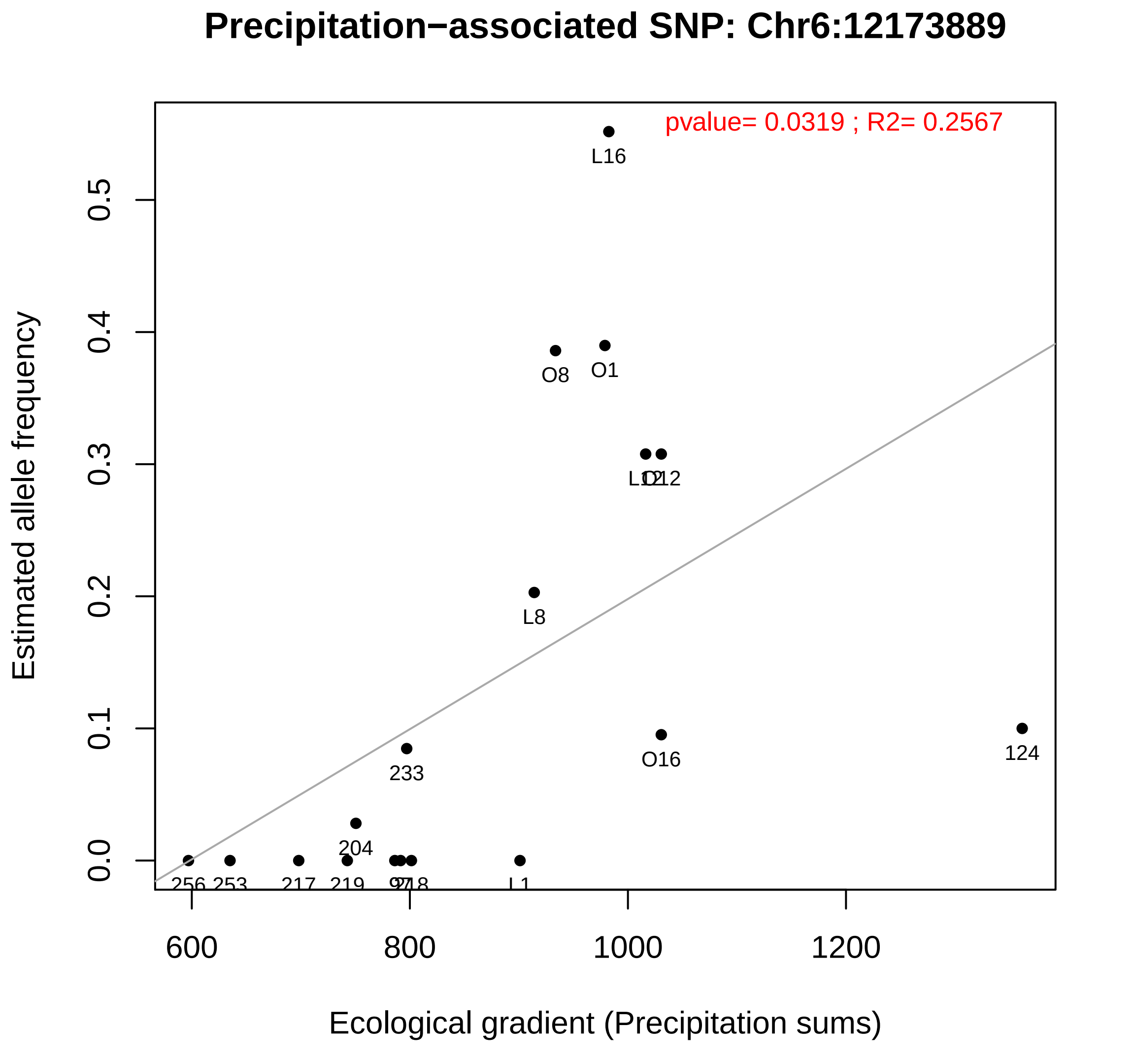

Precipitation−associated SNP: Chr6:12173889
pvalue= 0.0319 ; R2= 0.2567
Estimated allele frequency
Ecological gradient (Precipitation sums)
0.0
0.1
0.2
0.3
0.4
0.5
600
800
1000
1200
L16
O8
O1
L8
233
O16
124
204

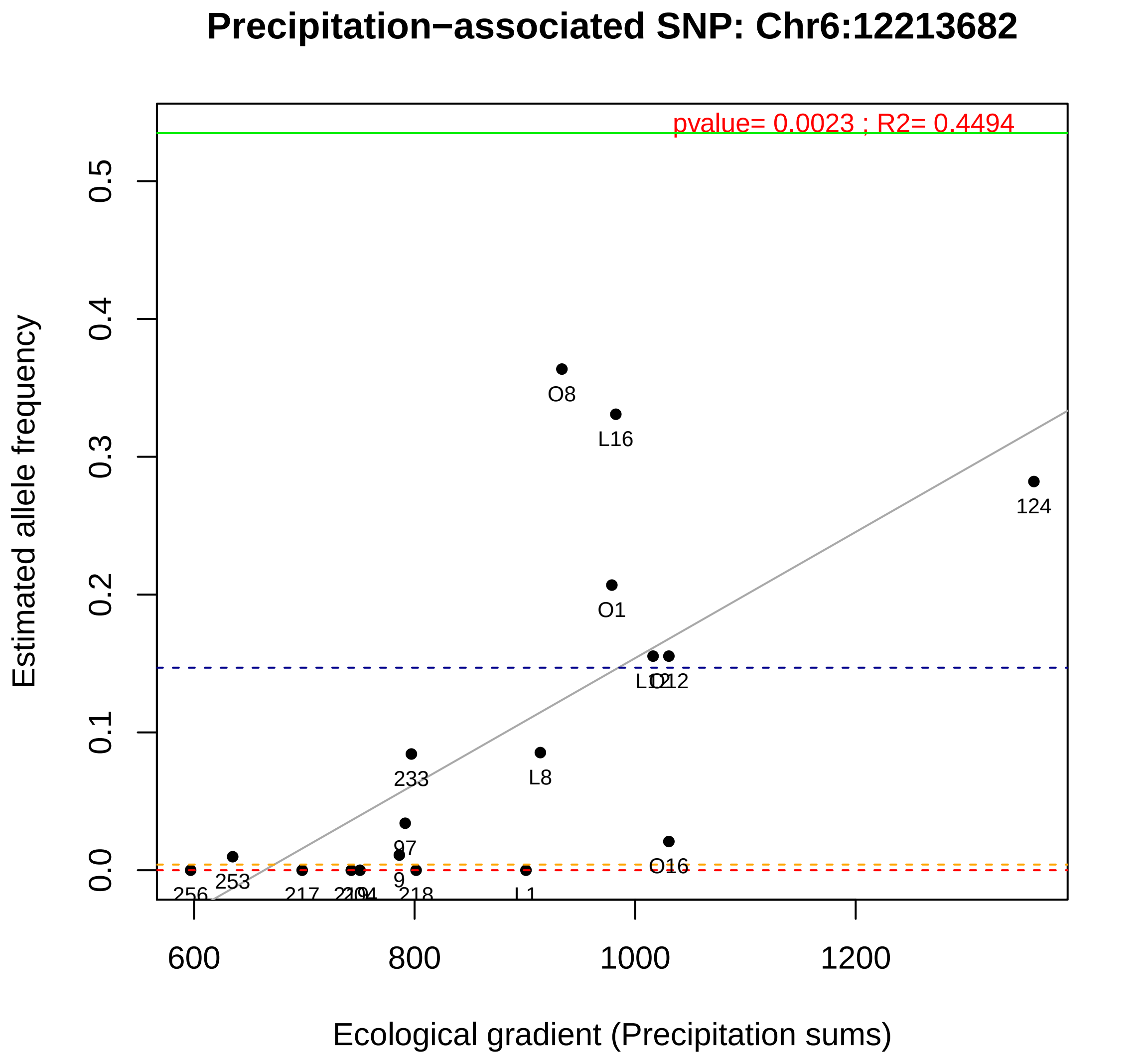
Precipitation−associated SNP: Chr6:12213682
pvalue= 0.0023 ; R2= 0.4494
Estimated allele frequency
0.0
0.1
0.2
0.3
0.4
0.5
O8
L16
124
O1
L10
O12
233
L8
97
O16
9
253
256
217
210
194
218
L1
600
800
1000
1200
Ecological gradient (Precipitation sums)

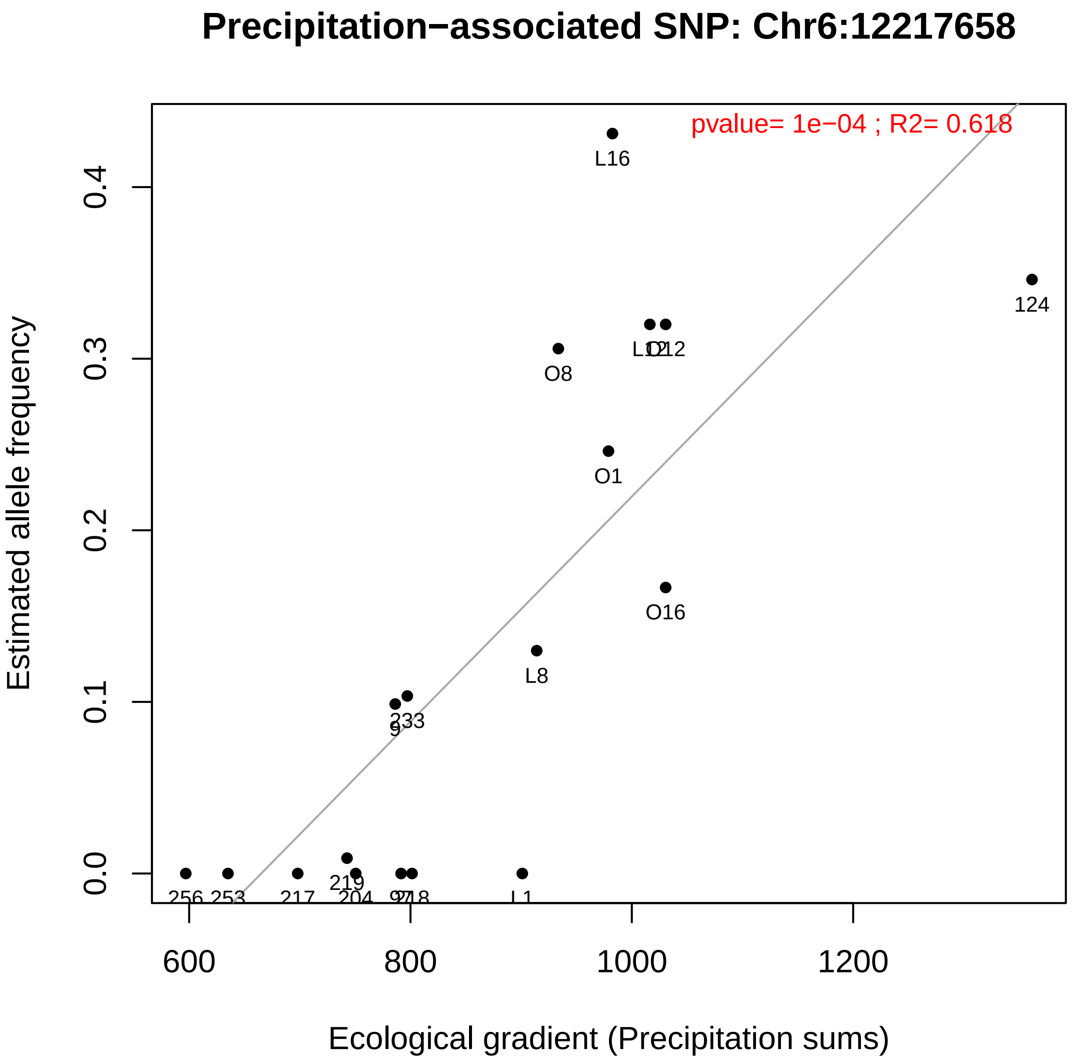
Precipitation−associated SNP: Chr6:12217658
pvalue= 1e−04 ; R2= 0.618
L16
124
L12
O12
O8
O1
O16
L8
233
9
219
256
253
217
204
218
L1
0.0
0.1
0.2
0.3
0.4
600
800
1000
1200
Estimated allele frequency
Ecological gradient (Precipitation sums)

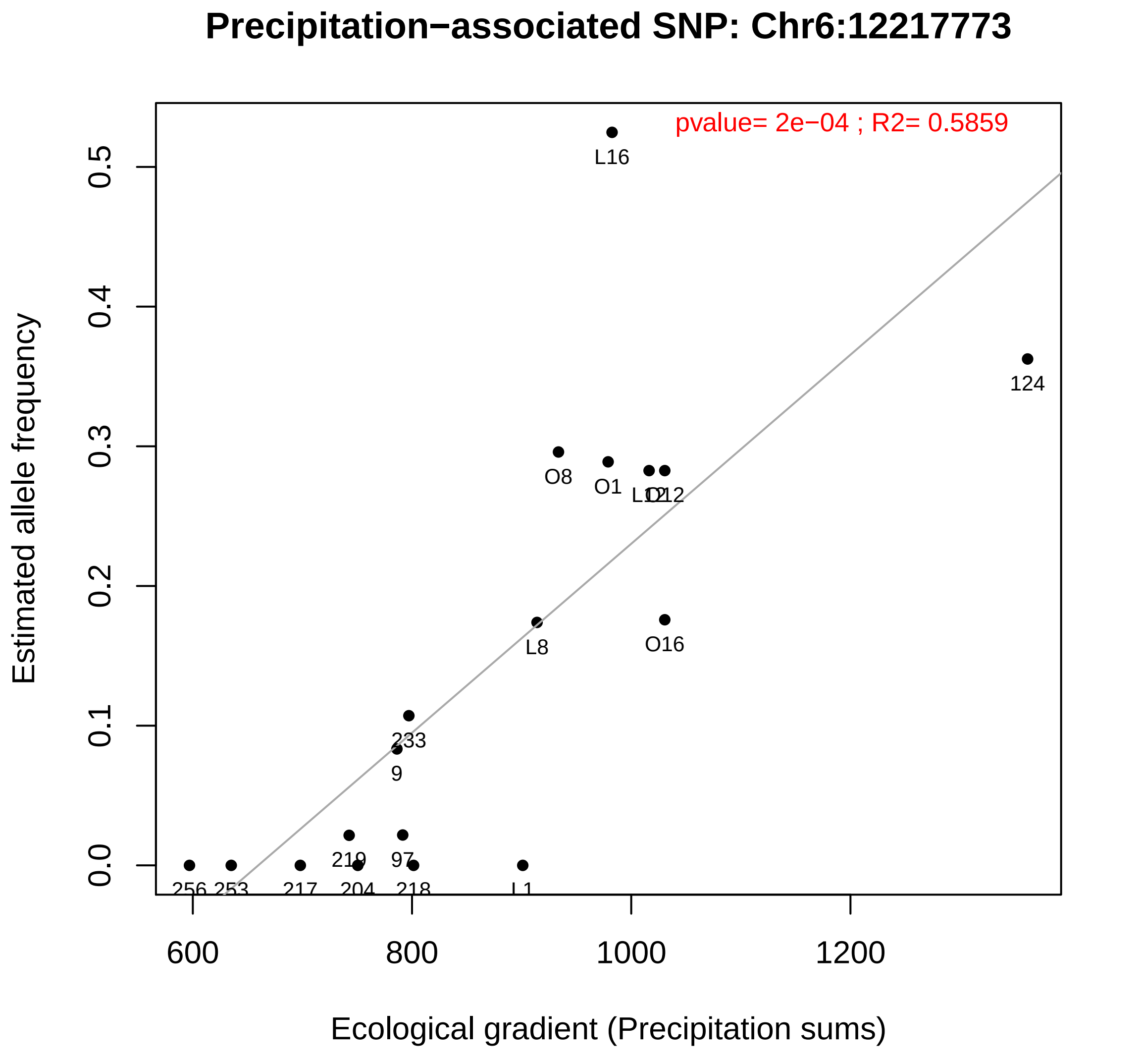
Precipitation−associated SNP: Chr6:12217773
pvalue= 2e−04 ; R2= 0.5859
Estimated allele frequency
Ecological gradient (Precipitation sums)
0.0
0.1
0.2
0.3
0.4
0.5
600
800
1000
1200
L16
124
O8
O1
L12
O12
L8
O16
233
9
219
97
256
253
217
204
218
L1

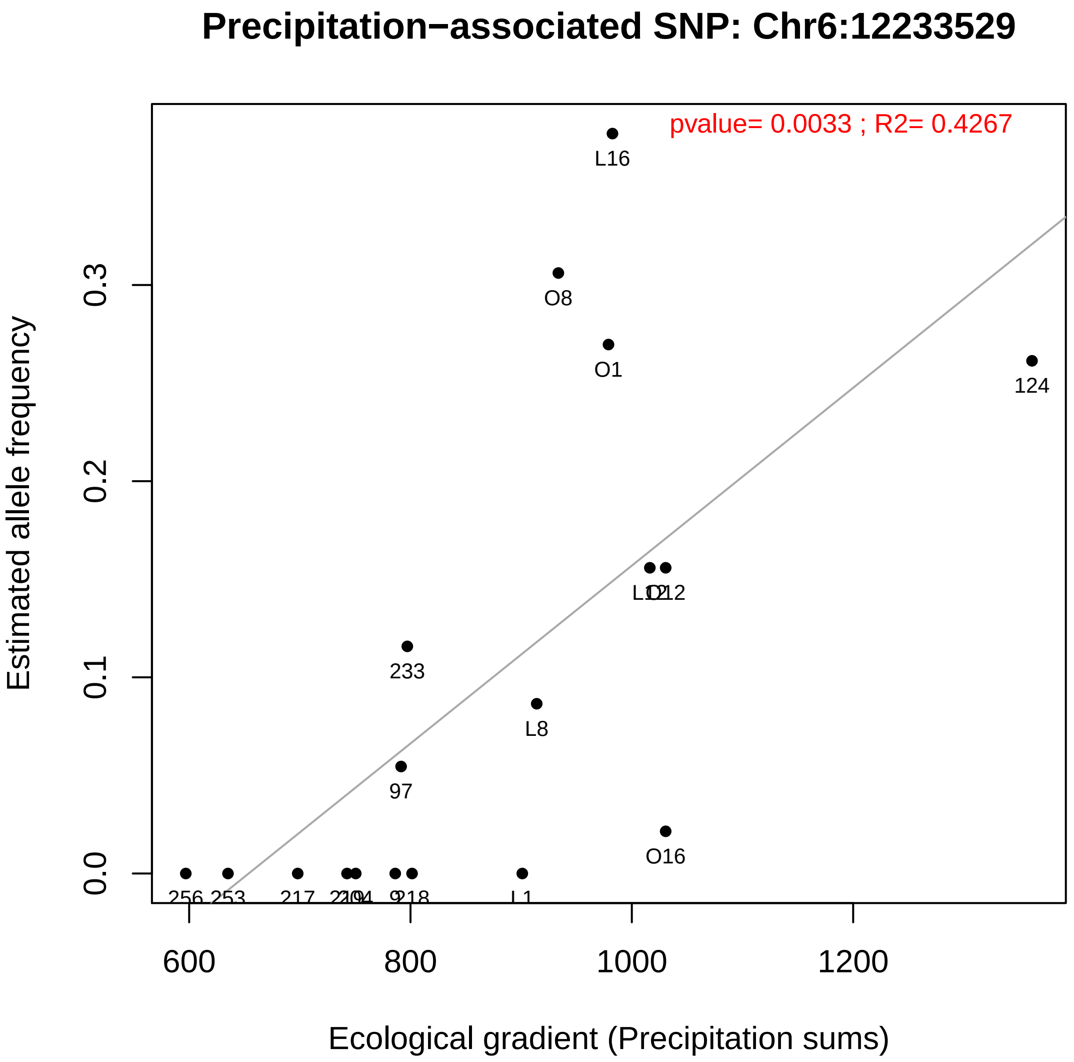
Precipitation−associated SNP: Chr6:12233529
pvalue= 0.0033 ; R2= 0.4267
Estimated allele frequency
0.0
0.1
0.2
0.3
600
800
1000
1200
Ecological gradient (Precipitation sums)
L16
O8
O1
124
233
L8
97
O16
O12
256
253
217
218
L1

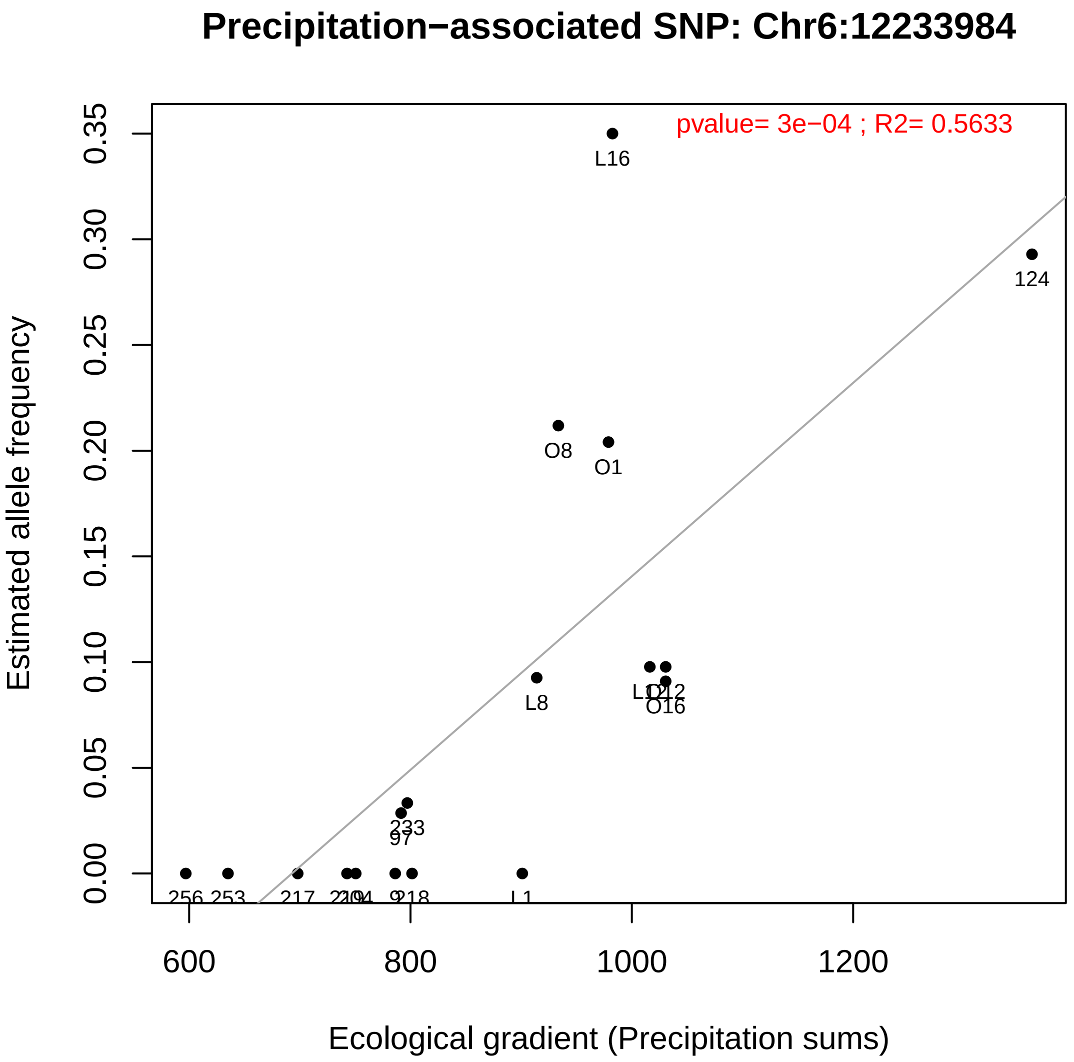

Precipitation−associated SNP: Chr6:12233984
pvalue= 3e−04 ; R2= 0.5633
Estimated allele frequency
0.00
0.05
0.10
0.15
0.20
0.25
0.30
0.35
600
800
1000
1200
Ecological gradient (Precipitation sums)
L16
124
O8
O1
L8
O16
233
97
256
253
217
218
L1

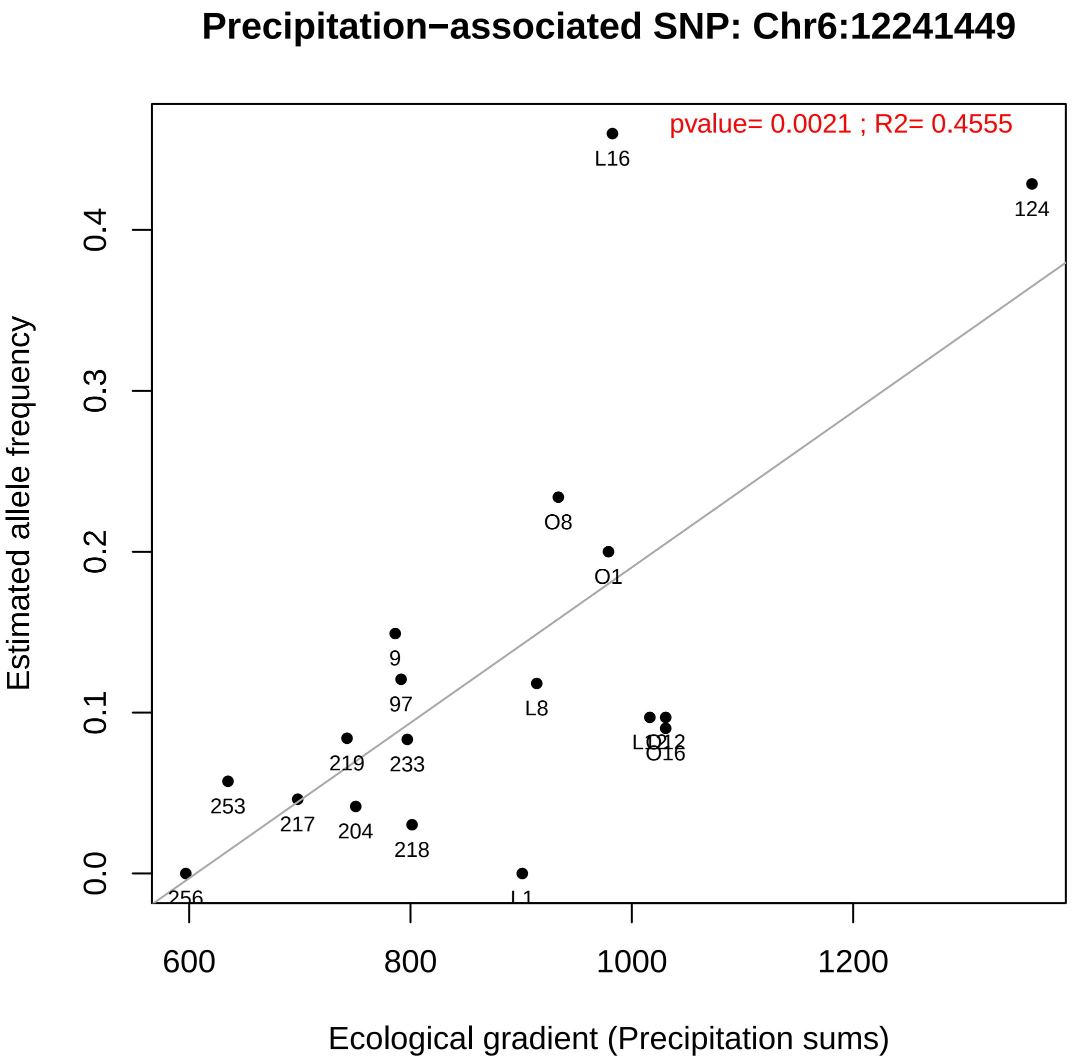
Precipitation−associated SNP: Chr6:12241449
pvalue= 0.0021 ; R2= 0.4555
Estimated allele frequency
Ecological gradient (Precipitation sums)
0.0
0.1
0.2
0.3
0.4
600
800
1000
1200
L16
124
O8
O1
9
97
L8
219
233
253
217
204
218
O16

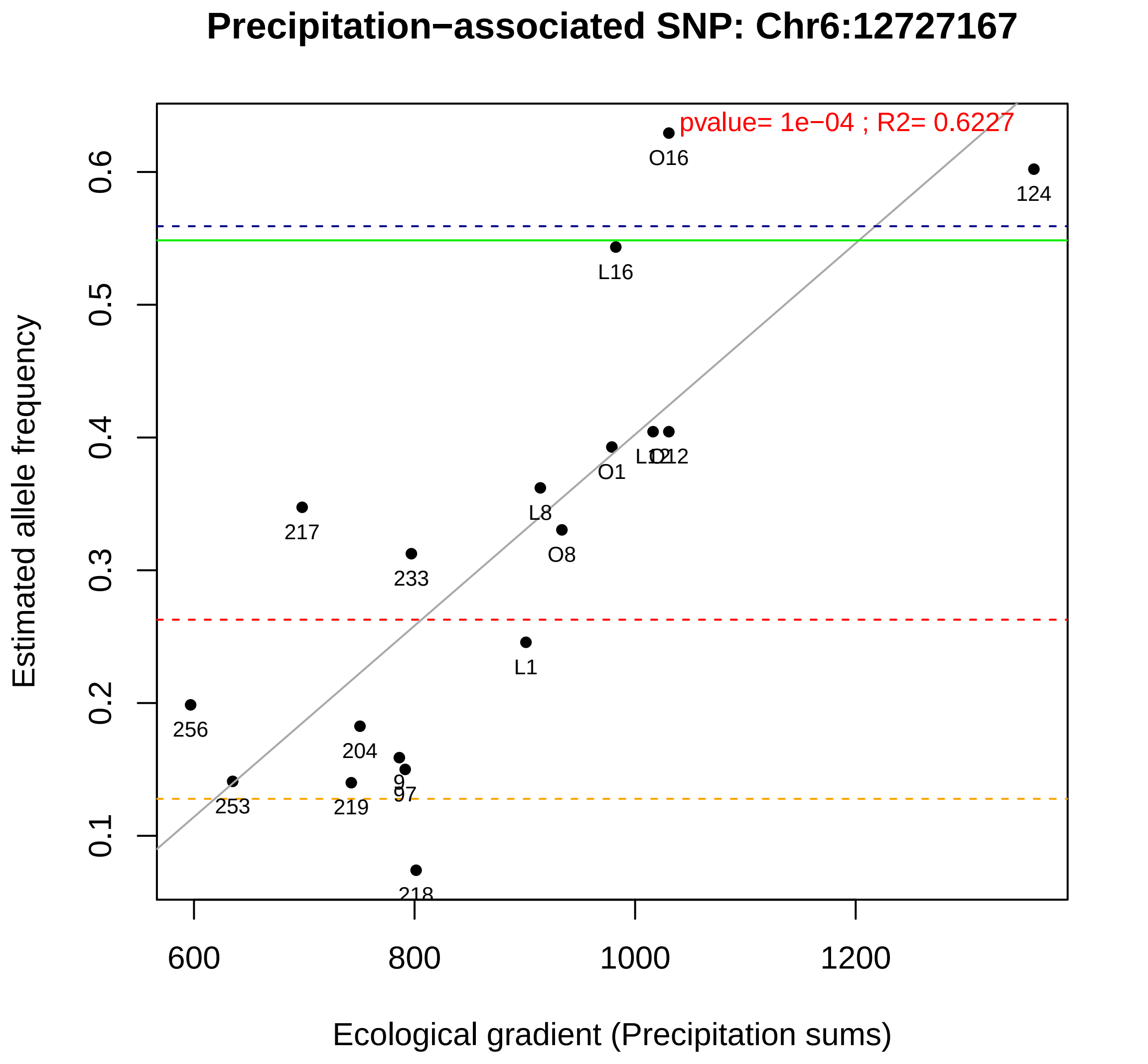

Precipitation−associated SNP: Chr6:12727167
pvalue= 1e−04 ; R2= 0.6227
Estimated allele frequency
Ecological gradient (Precipitation sums)
0.1
0.2
0.3
0.4
0.5
0.6
600
800
1000
1200
O16
124
L16
L12
O12
O1
L8
O8
217
233
L1
256
204
9
97
253
219
218

**Precipitation−associated SNP: Chr6:12776738**

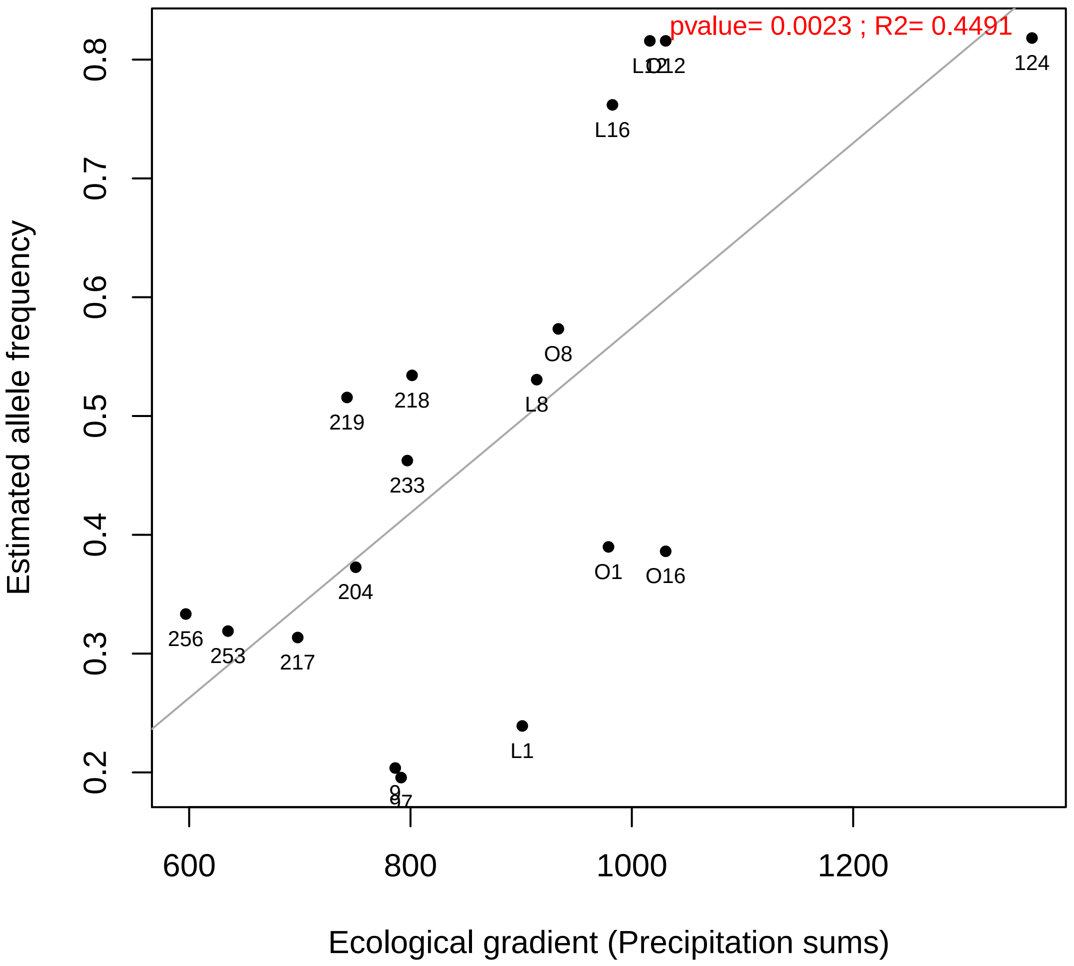

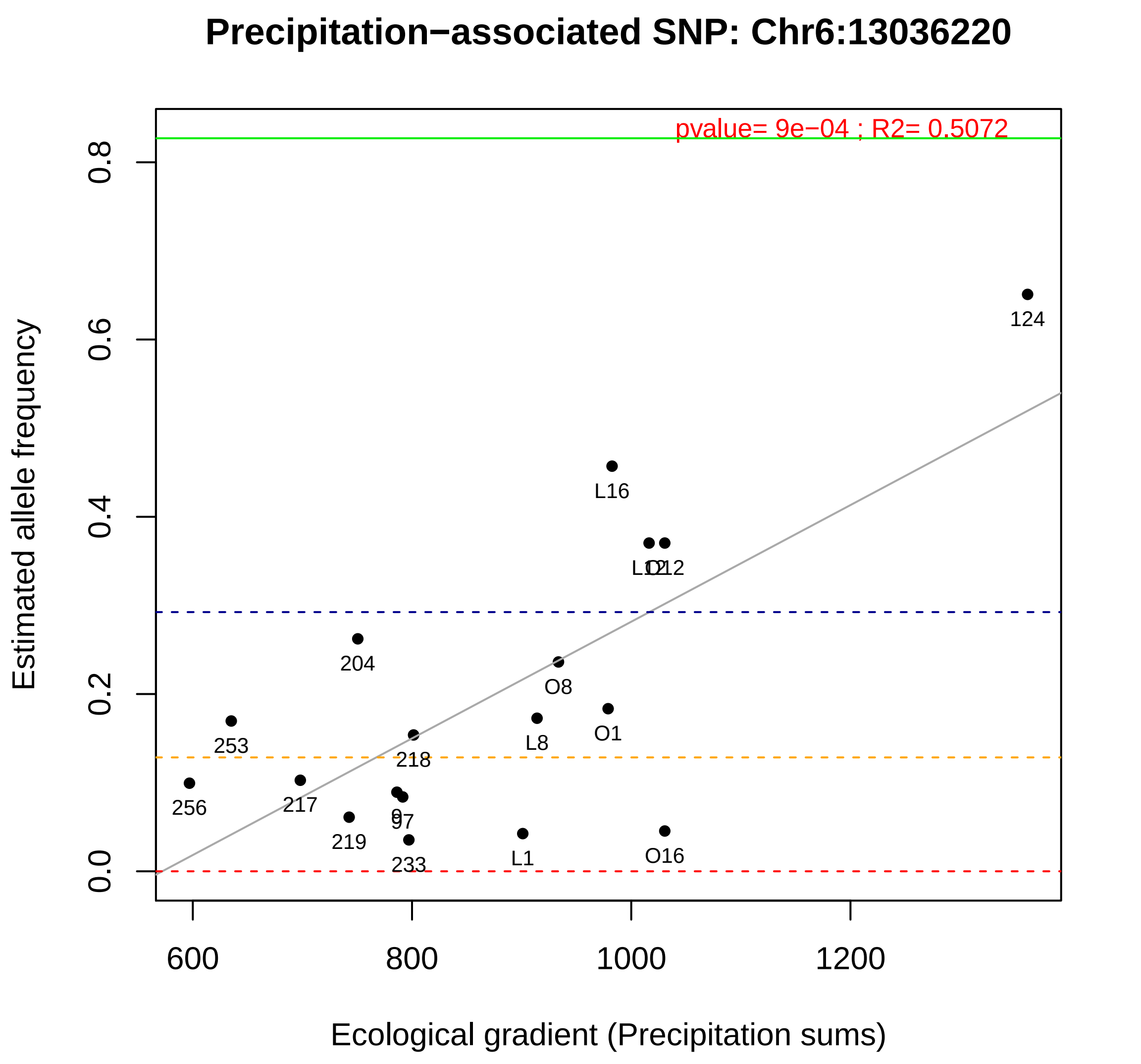
Precipitation−associated SNP: Chr6:13036220
pvalue= 9e−04 ; R2= 0.5072
Estimated allele frequency
0.0
0.2
0.4
0.6
0.8
600
800
1000
1200
Ecological gradient (Precipitation sums)
124
L16
L12
O12
204
O8
O1
L8
253
218
256
217
9
97
219
233
L1
O16

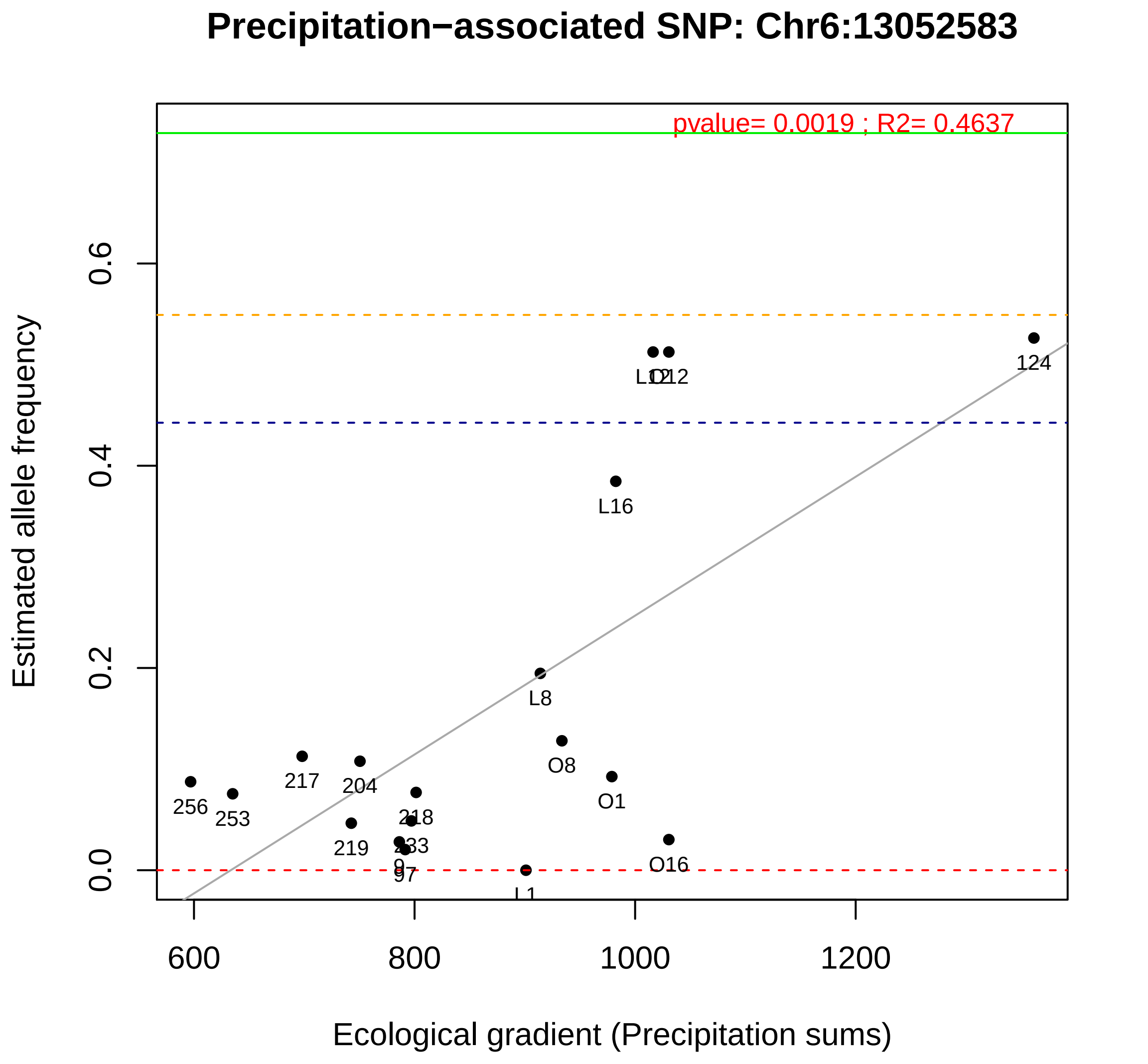

Precipitation−associated SNP: Chr6:13052583
pvalue= 0.0019 ; R2= 0.4637
Estimated allele frequency
0.0
0.2
0.4
0.6
600
800
1000
1200
Ecological gradient (Precipitation sums)
124
L16
L8
O8
O1
O16
217
204
256
253
219
218
233
97
L1

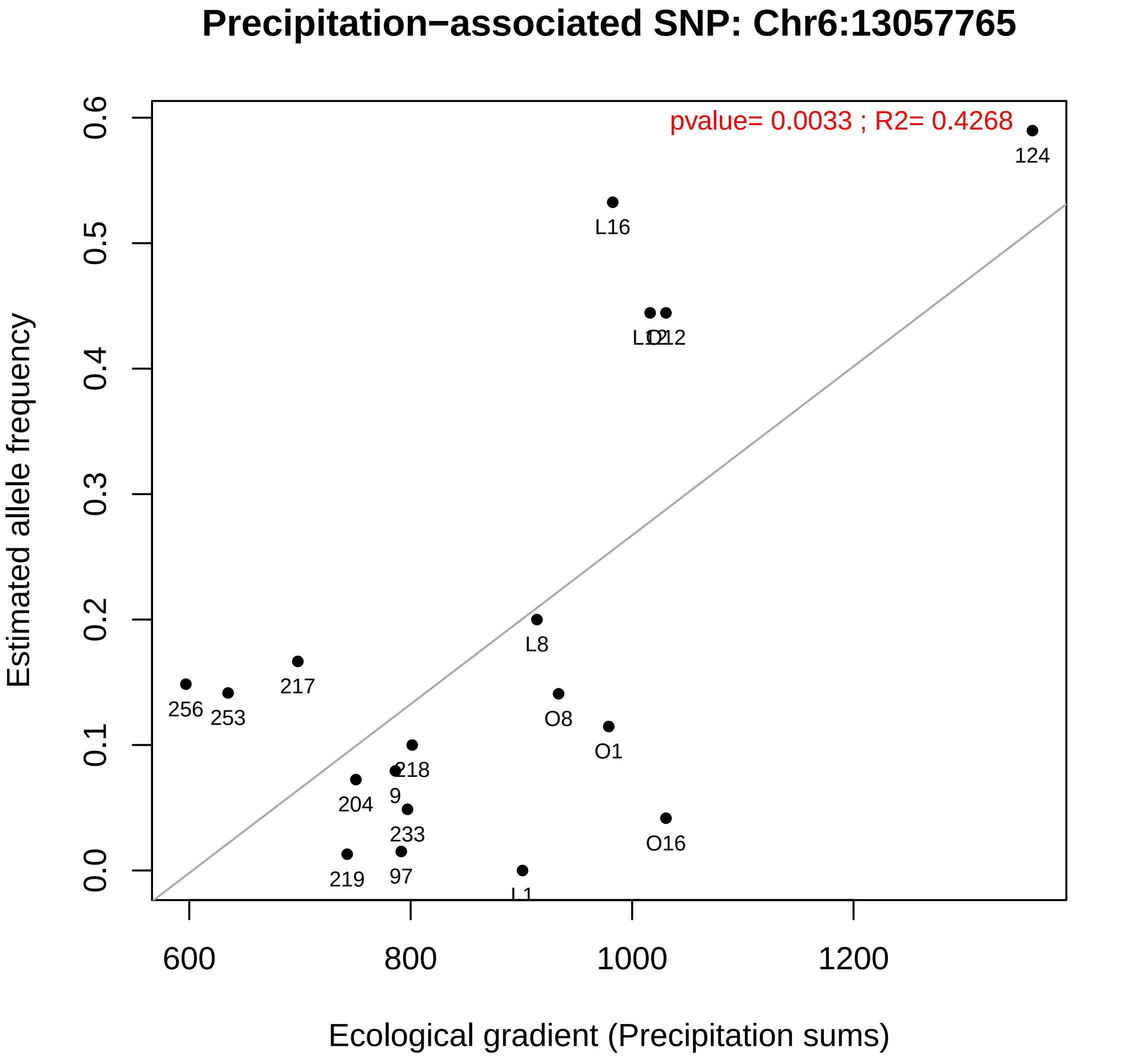
Precipitation−associated SNP: Chr6:13057765
pvalue= 0.0033 ; R2= 0.4268
Estimated allele frequency
0.0
0.1
0.2
0.3
0.4
0.5
0.6
600
800
1000
1200
Ecological gradient (Precipitation sums)
124
L16
L8
217
256
253
O8
O1
218
9
204
233
O16
219
97

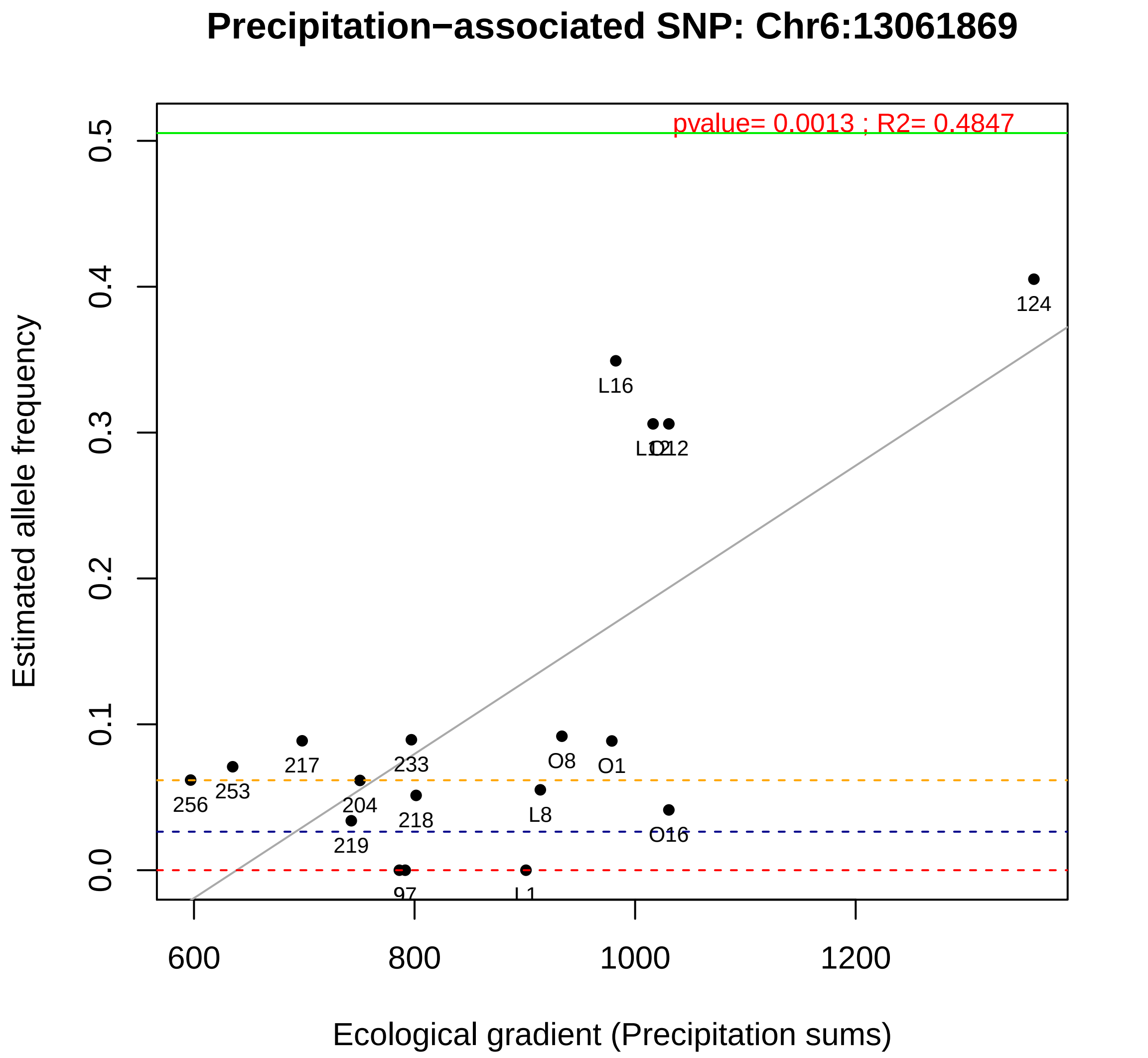

Precipitation−associated SNP: Chr6:13061869
pvalue= 0.0013 ; R2= 0.4847
Estimated allele frequency
0.0
0.1
0.2
0.3
0.4
0.5
600
800
1000
1200
Ecological gradient (Precipitation sums)
124
L16
L12
O12
217
233
O8
O1
256
253
204
218
L8
219
O16
97
L1

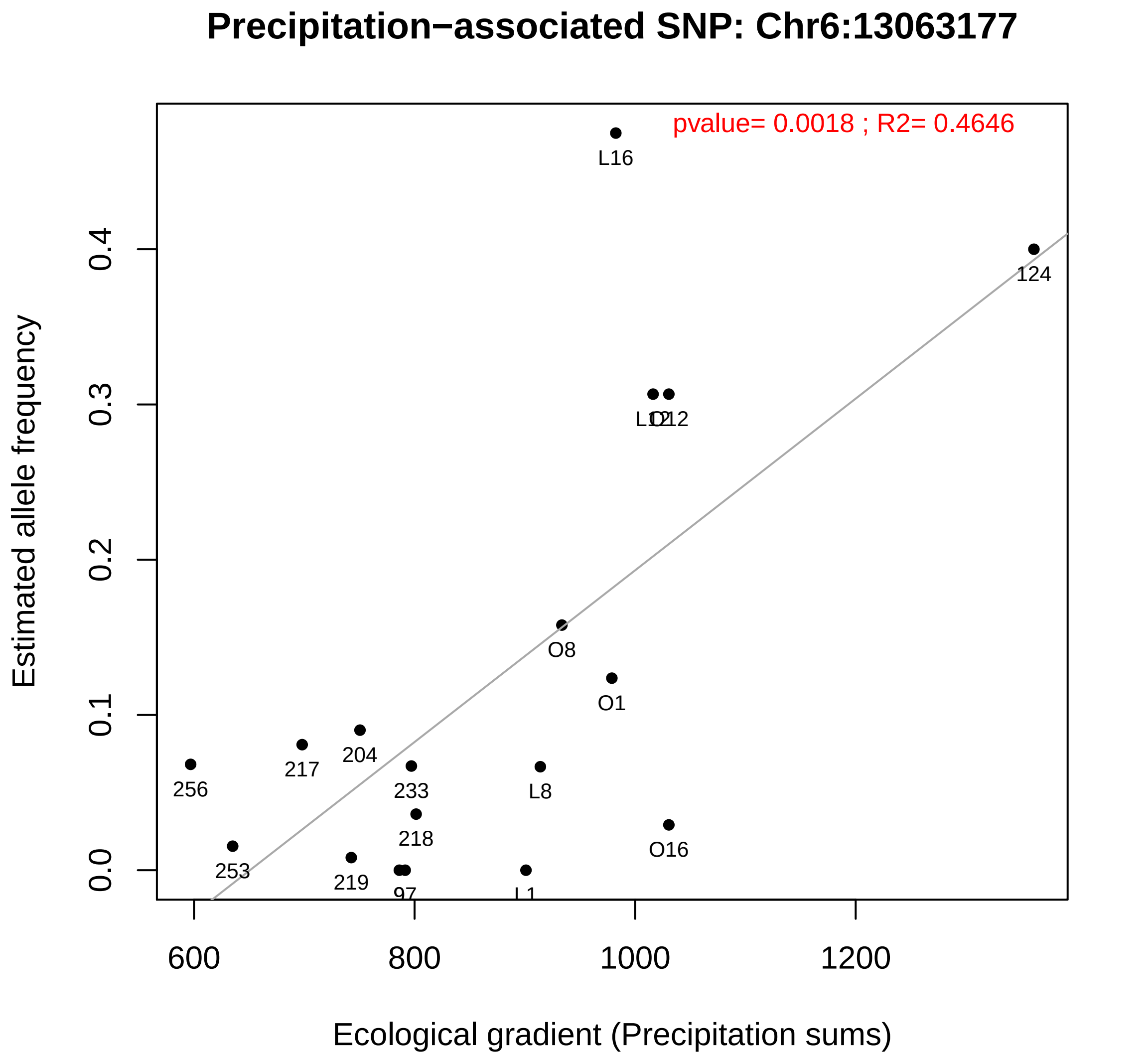

Precipitation−associated SNP: Chr6:13063177
pvalue= 0.0018 ; R2= 0.4646
Estimated allele frequency
0.0
0.1
0.2
0.3
0.4
600
800
1000
1200
Ecological gradient (Precipitation sums)
L16
124
O12
O8
O1
256
217
204
233
L8
218
O16
253
219
97
L1

Precipitation−associated SNP: Chr6:13292948

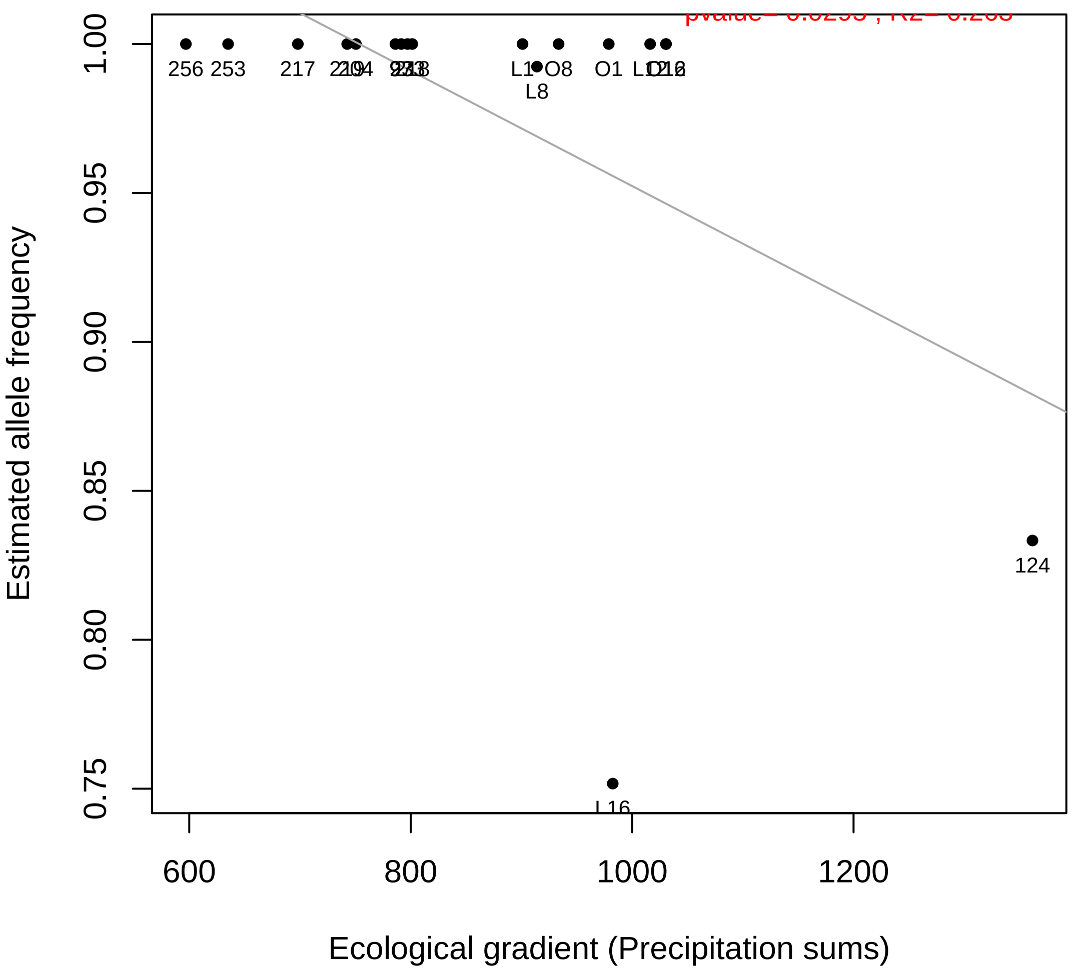

Precipitation−associated SNP: Chr6:13340795

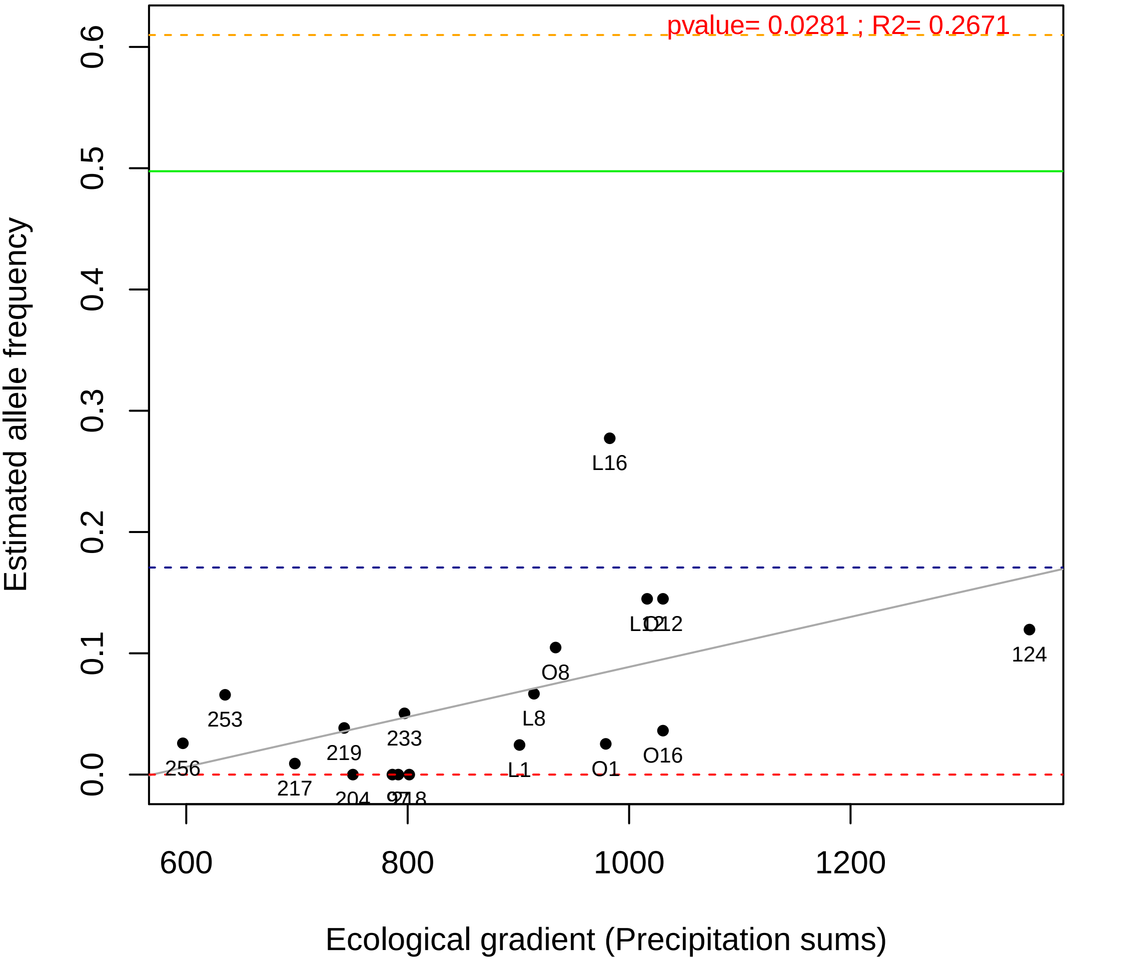

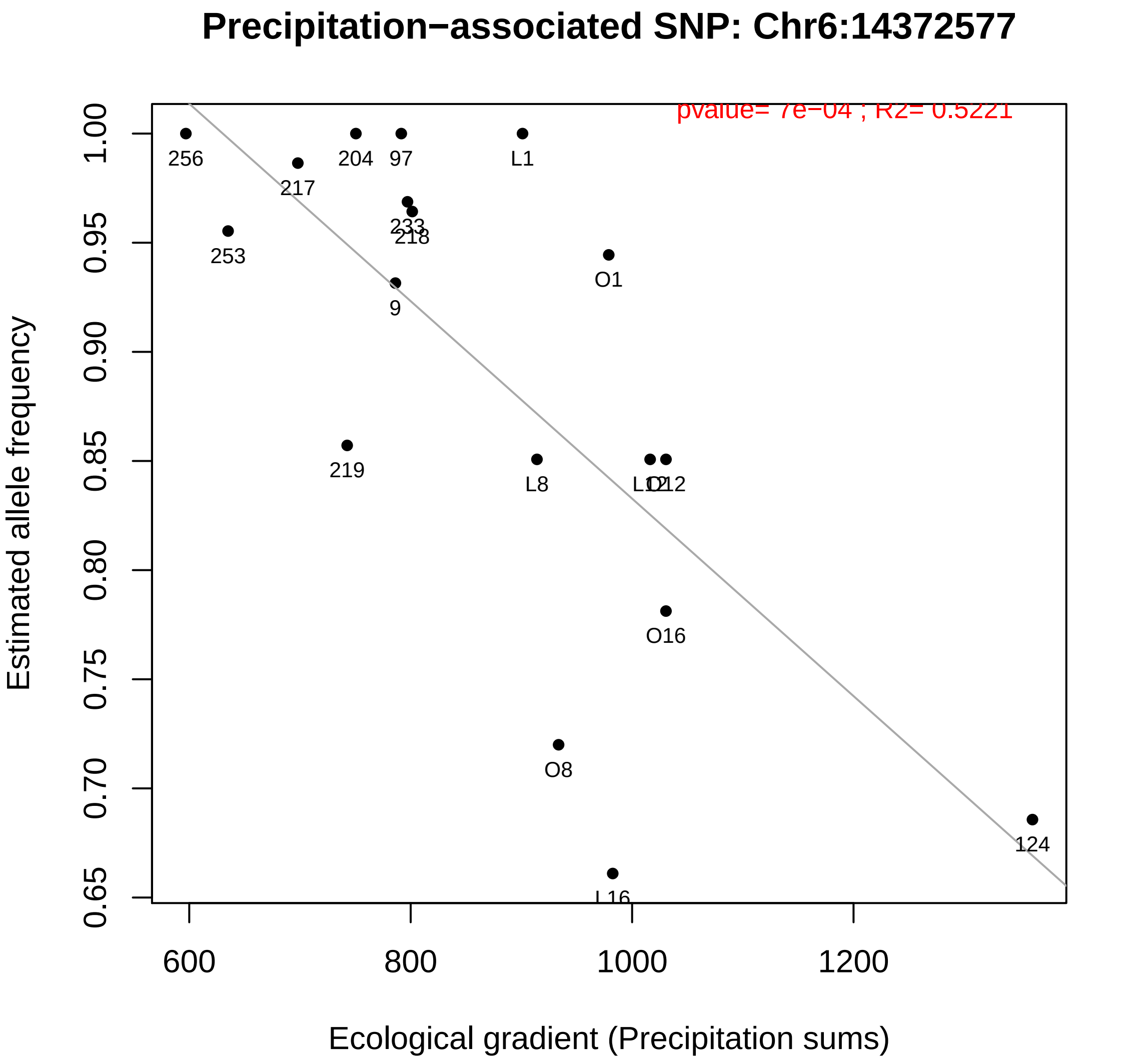
Precipitation−associated SNP: Chr6:14372577
pvalue= 7e−04 ; R2= 0.5221
Estimated allele frequency
Ecological gradient (Precipitation sums)
0.65
0.70
0.75
0.80
0.85
0.90
0.95
1.00
600
800
1000
1200
256
217
204
97
L1
233
218
253
O1
9
219
L8
L12
O12
O16
O8
124
L16

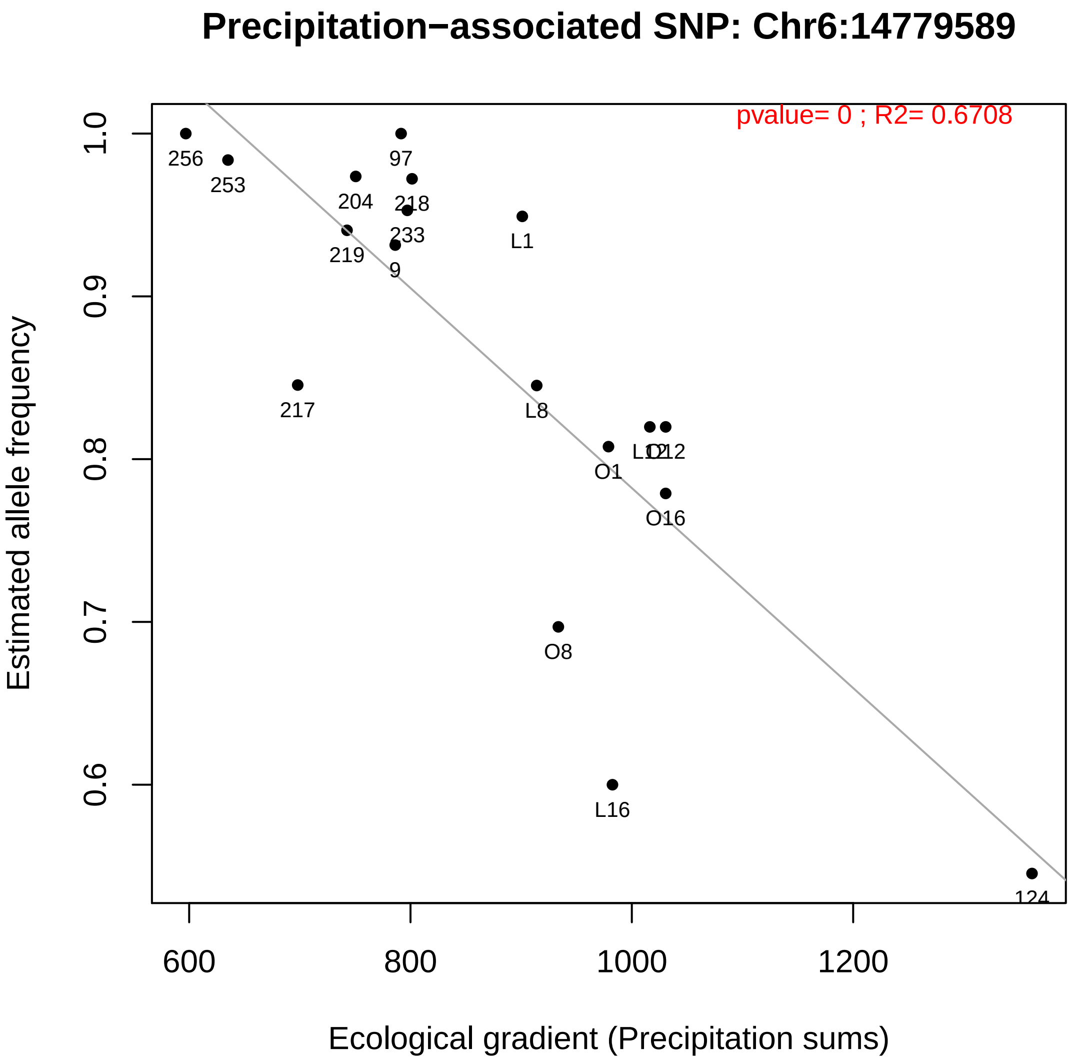
Precipitation−associated SNP: Chr6:14779589
pvalue= 0 ; R2= 0.6708
Estimated allele frequency
Ecological gradient (Precipitation sums)
1.0
0.9
0.8
0.7
0.6
600
800
1000
1200
256
253
97
204
218
233
219
9
L1
217
L8
O1
O12
O16
O8
L16
124

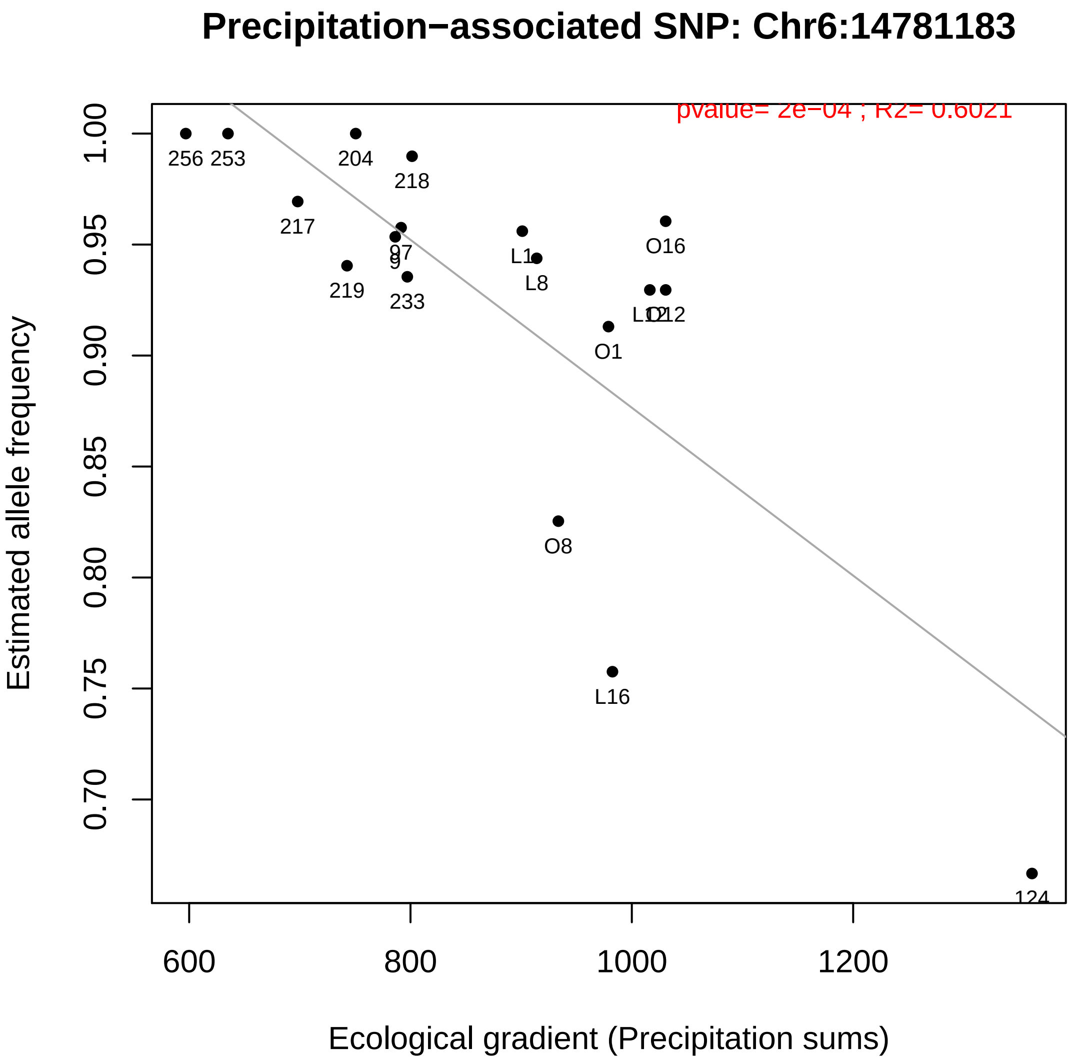

Precipitation−associated SNP: Chr6:14781183
pvalue= 2e−04 ; R2= 0.6021
Estimated allele frequency
1.00
0.95
0.90
0.85
0.80
0.75
0.70
256
253
204
218
217
97
9
219
233
L1
L8
O16
L12
O12
O1
O8
L16
124
600
800
1000
1200
Ecological gradient (Precipitation sums)

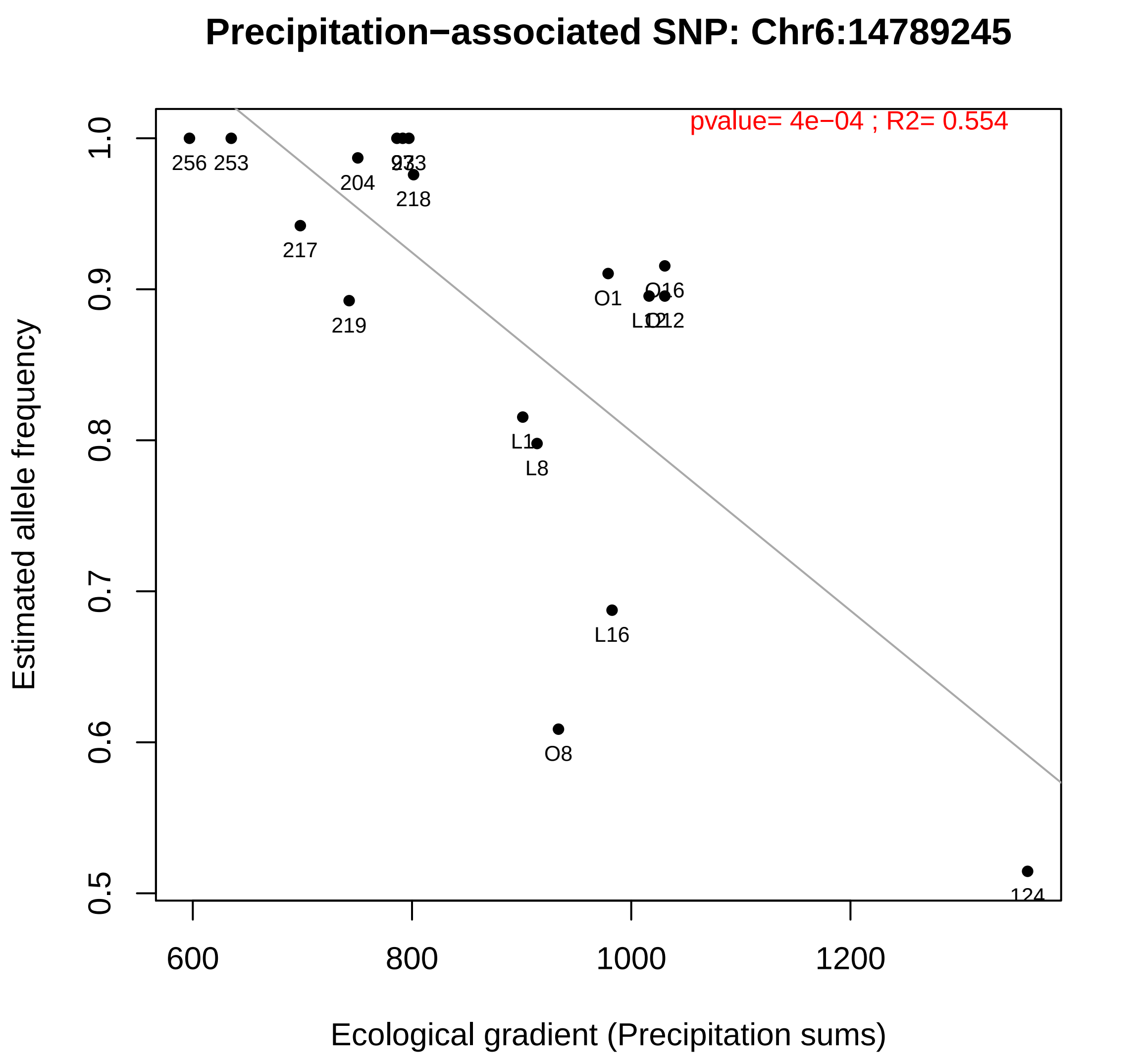

Precipitation−associated SNP: Chr6:14789245
pvalue= 4e−04 ; R2= 0.554
Estimated allele frequency
0.5
0.6
0.7
0.8
0.9
1.0
600
800
1000
1200
Ecological gradient (Precipitation sums)
256
253
204
218
217
219
O1
O16
O12
L1
L8
L16
O8
124

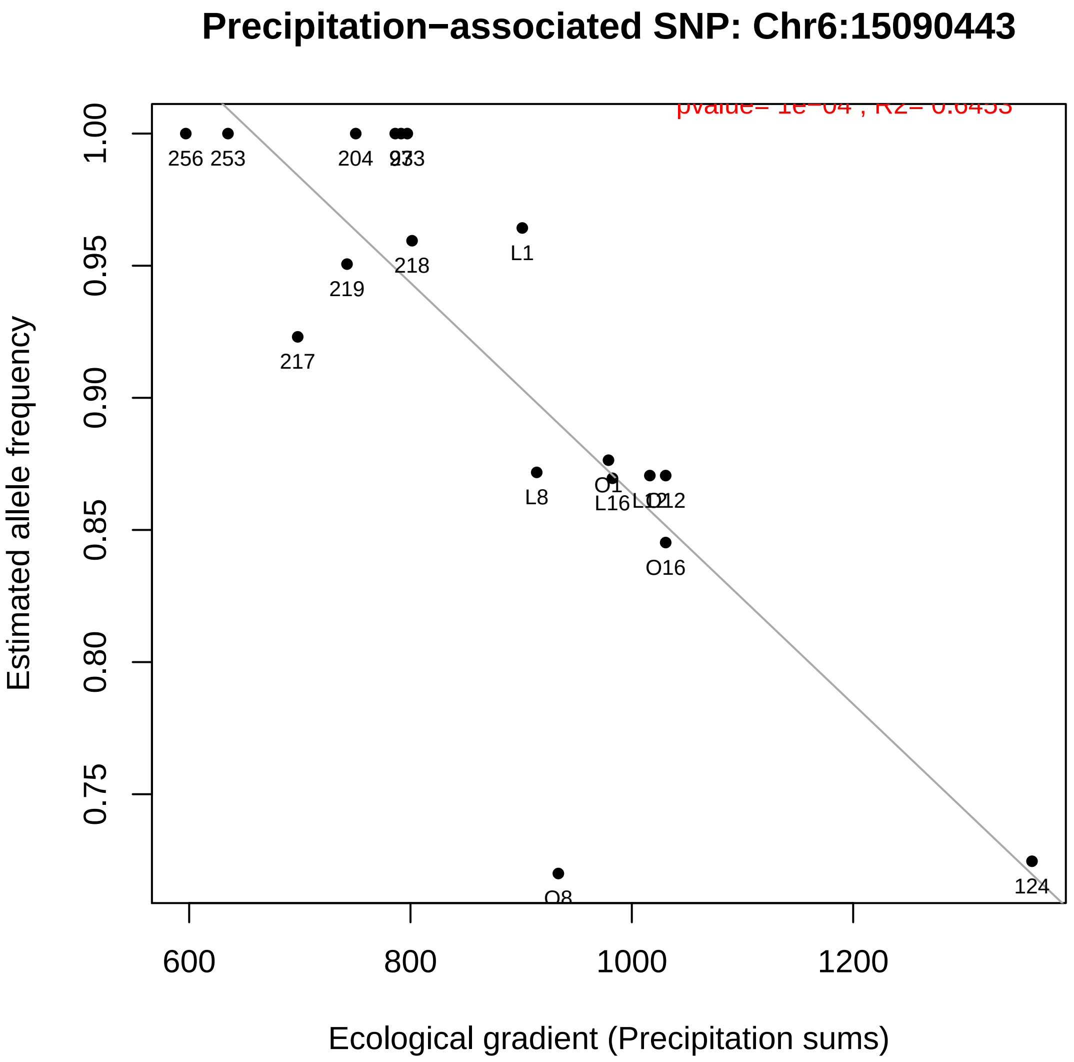

Precipitation−associated SNP: Chr6:15090443
Estimated allele frequency
Ecological gradient (Precipitation sums)
1.00
0.95
0.90
0.85
0.80
0.75
600
800
1000
1200
256
253
204
218
219
217
L1
L8
O1
L16
O16
O8
124

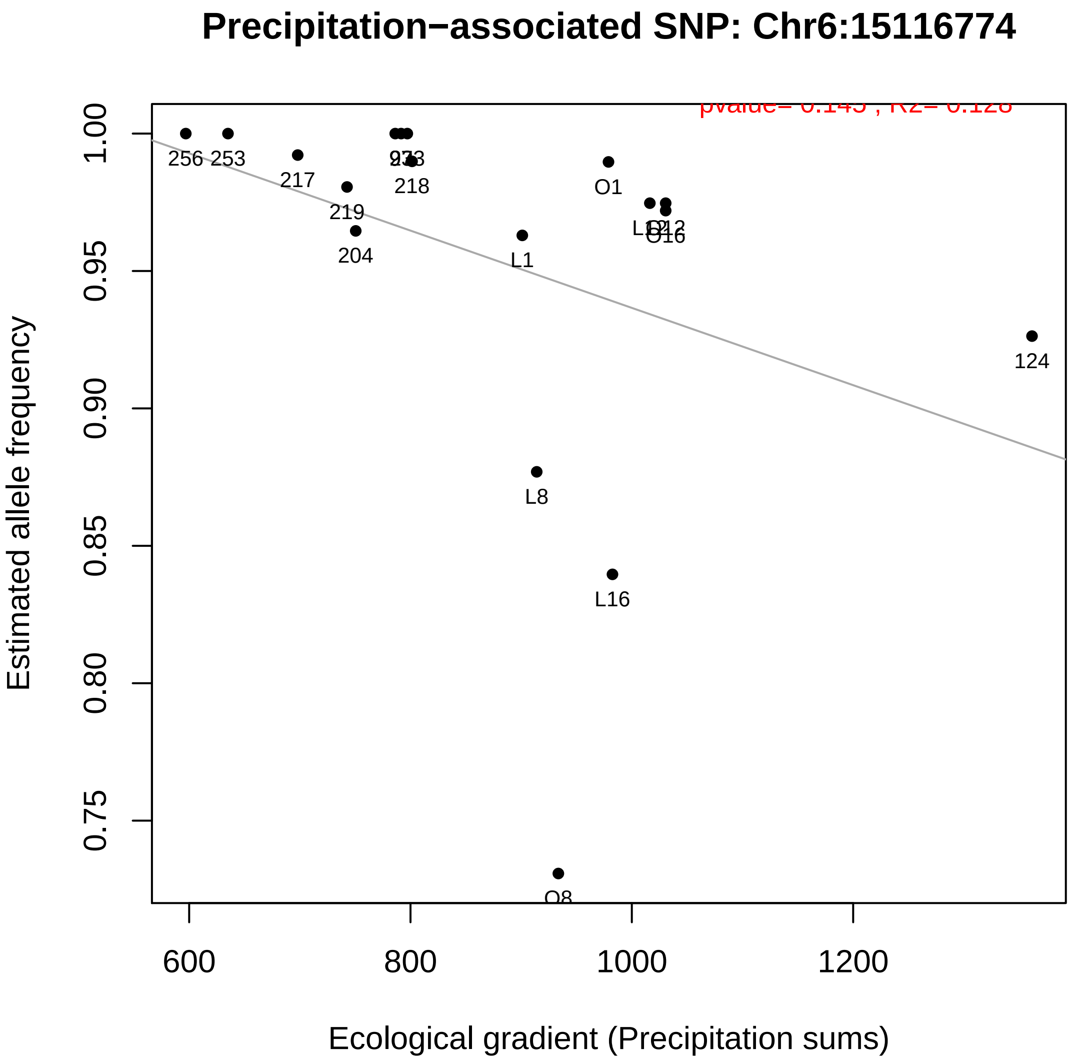
Precipitation−associated SNP: Chr6:15116774
Estimated allele frequency
Ecological gradient (Precipitation sums)
1.00
0.95
0.90
0.85
0.80
0.75
600
800
1000
1200
256
253
217
219
204
218
O1
L1
L8
L16
O8
124

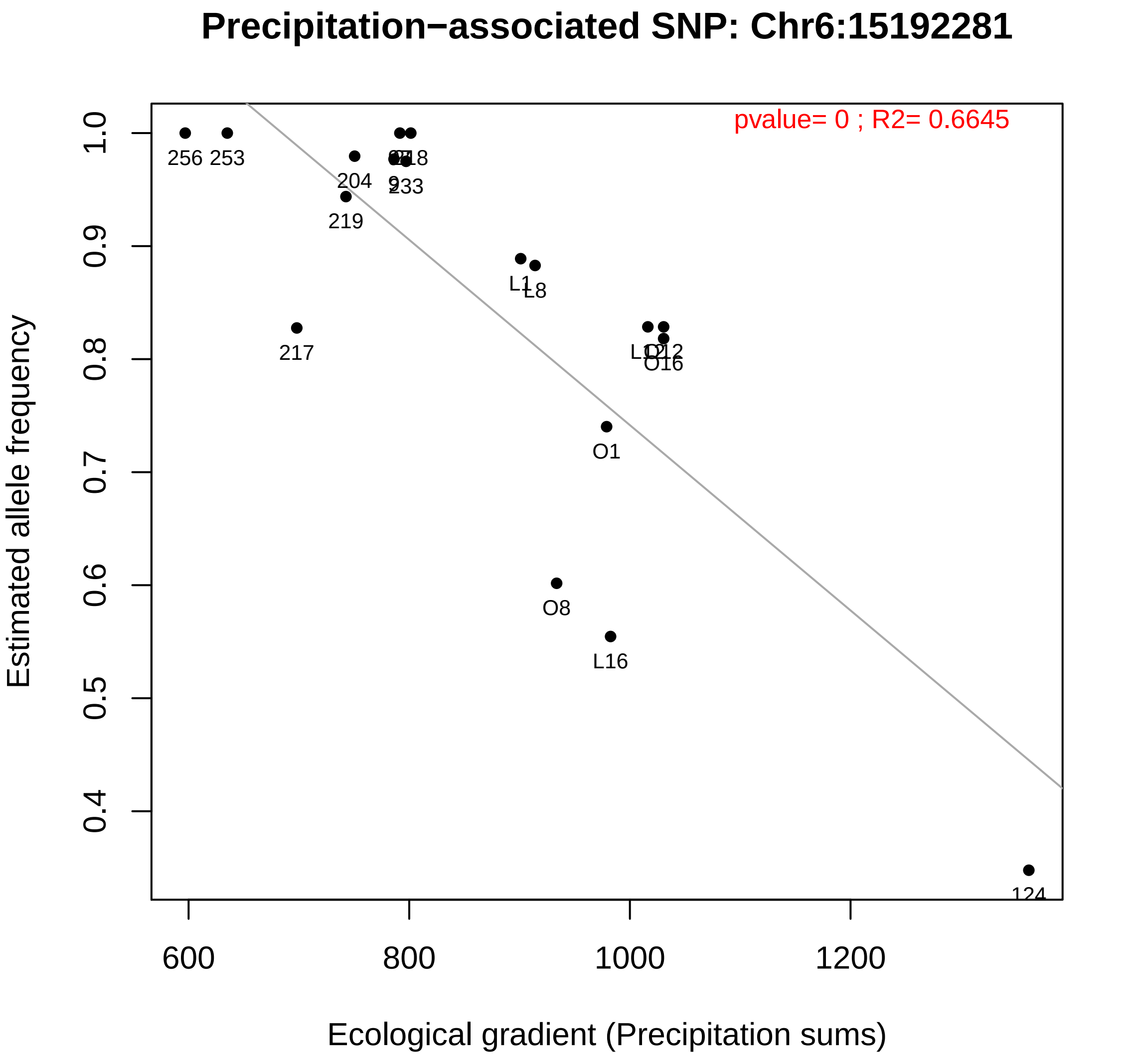
Precipitation−associated SNP: Chr6:15192281
pvalue= 0 ; R2= 0.6645
Estimated allele frequency
Ecological gradient (Precipitation sums)
0.4
0.5
0.6
0.7
0.8
0.9
1.0
600
800
1000
1200
256
253
204
219
233
217
L1
L8
O1
O8
L16
O16
124

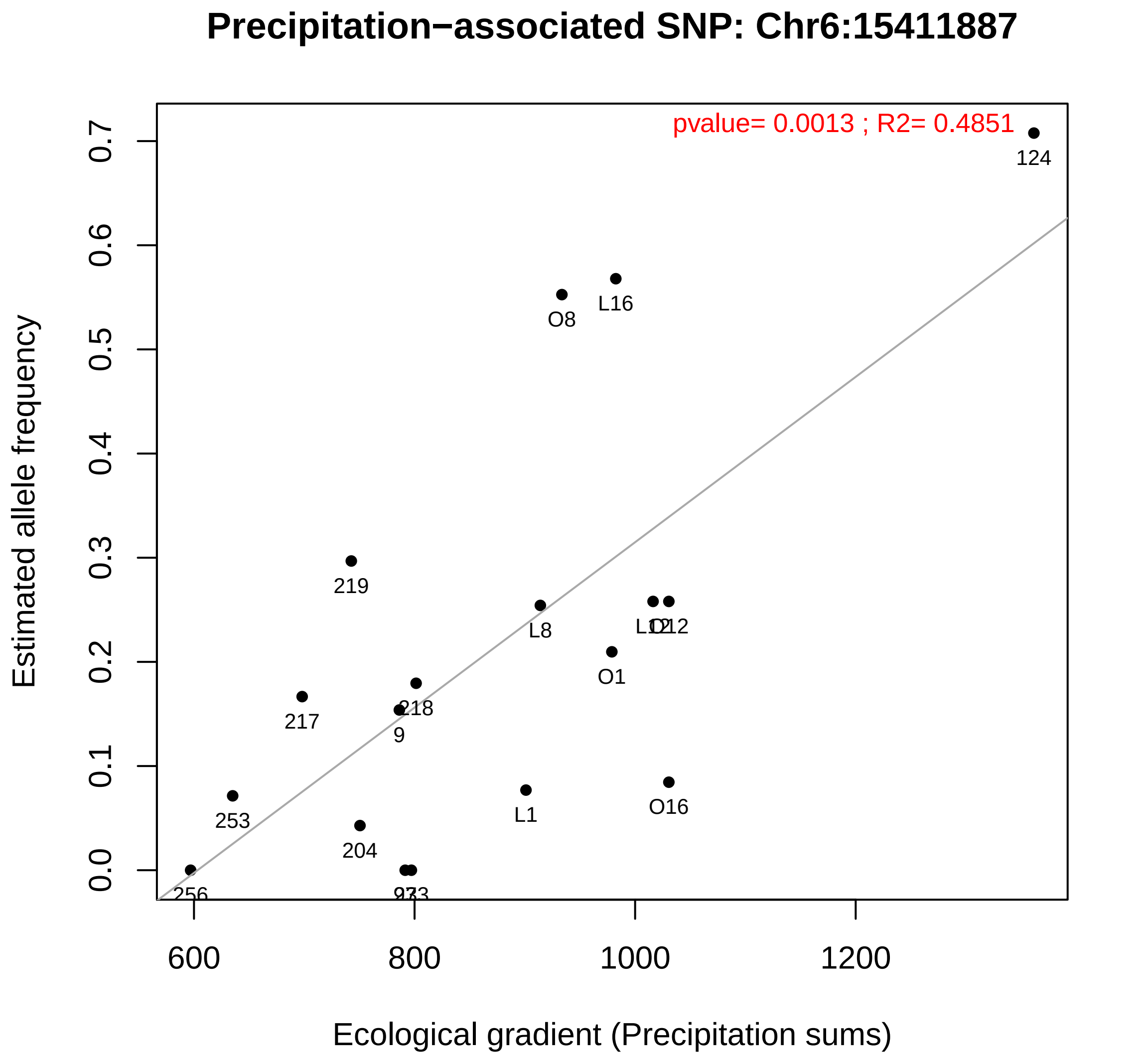
Precipitation−associated SNP: Chr6:15411887
pvalue= 0.0013 ; R2= 0.4851
Estimated allele frequency
Ecological gradient (Precipitation sums)
0.0
0.1
0.2
0.3
0.4
0.5
0.6
0.7
600
800
1000
1200
124
L16
O8
219
L8
O1
O12
217
218
9
253
204
L1
O16
256

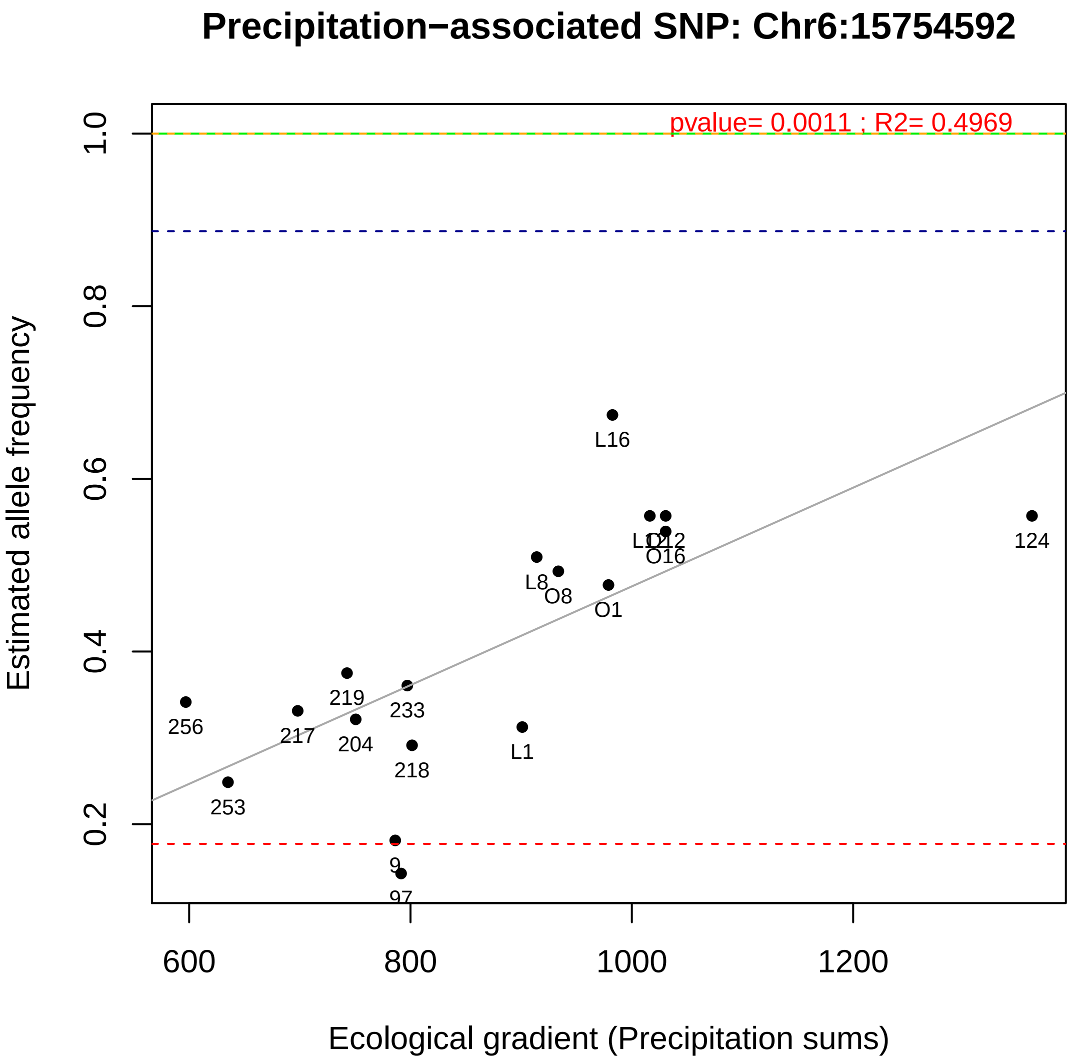

Precipitation−associated SNP: Chr6:15754592
pvalue= 0.0011 ; R2= 0.4969
Estimated allele frequency
Ecological gradient (Precipitation sums)
0.2
0.4
0.6
0.8
1.0
600
800
1000
1200
L16
L12
O12
O16
124
L8
O8
O1
219
233
256
217
204
218
L1
253
9
97

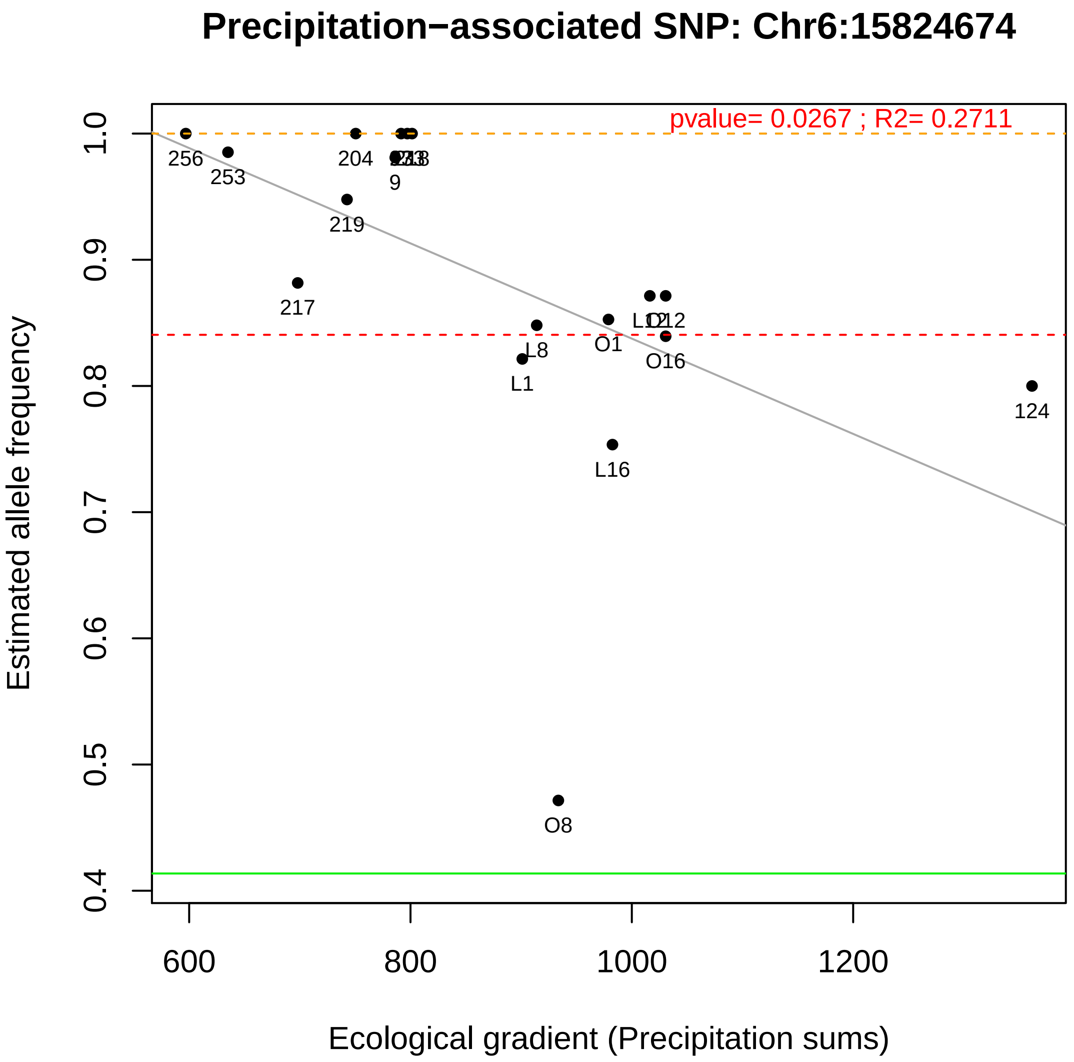
Precipitation−associated SNP: Chr6:15824674
pvalue= 0.0267 ; R2= 0.2711
Estimated allele frequency
Ecological gradient (Precipitation sums)
1.0
0.9
0.8
0.7
0.6
0.5
0.4
600
800
1000
1200
256
253
204
9
219
217
L8
L1
O1
O16
124
L16
O8

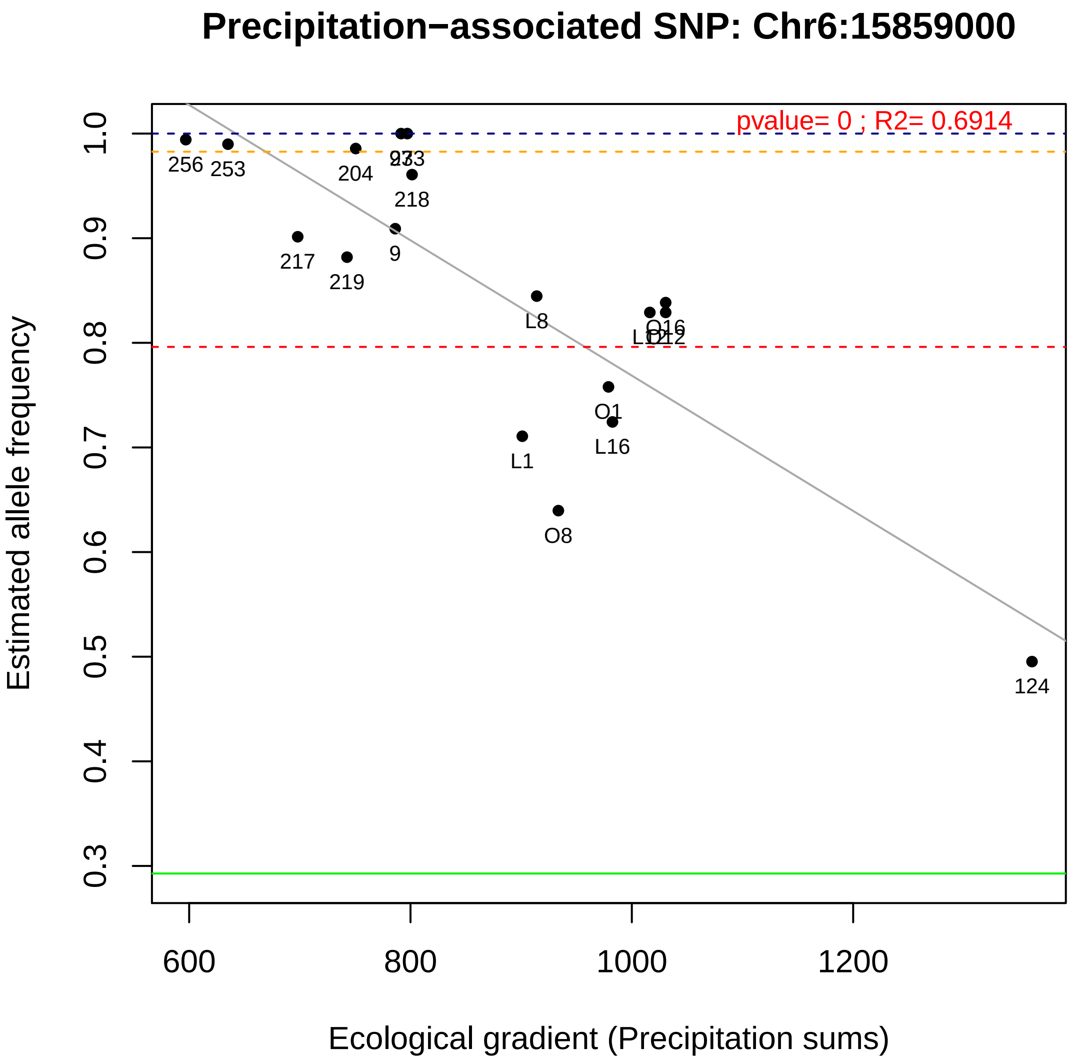

Precipitation−associated SNP: Chr6:15859000
pvalue= 0 ; R2= 0.6914
Estimated allele frequency
Ecological gradient (Precipitation sums)
0.3
0.4
0.5
0.6
0.7
0.8
0.9
1.0
600
800
1000
1200
256
253
204
218
217
219
9
L8
O16
O12
O1
L16
L1
O8
124

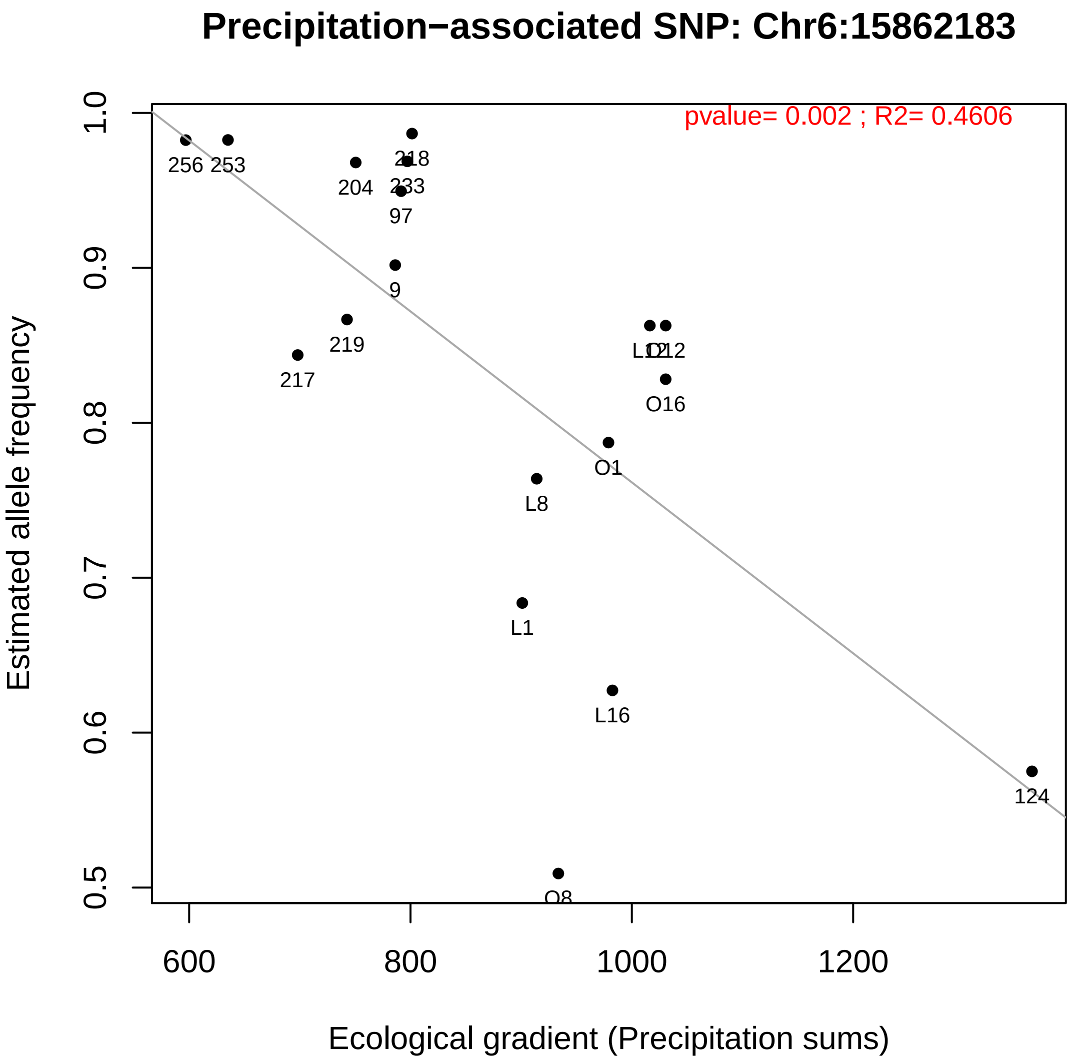
Precipitation−associated SNP: Chr6:15862183
pvalue= 0.002 ; R2= 0.4606
Estimated allele frequency
0.5
0.6
0.7
0.8
0.9
1.0
600
800
1000
1200
Ecological gradient (Precipitation sums)
256
253
204
218
233
97
9
219
217
L12
O12
O16
O1
L8
L1
L16
124
O8

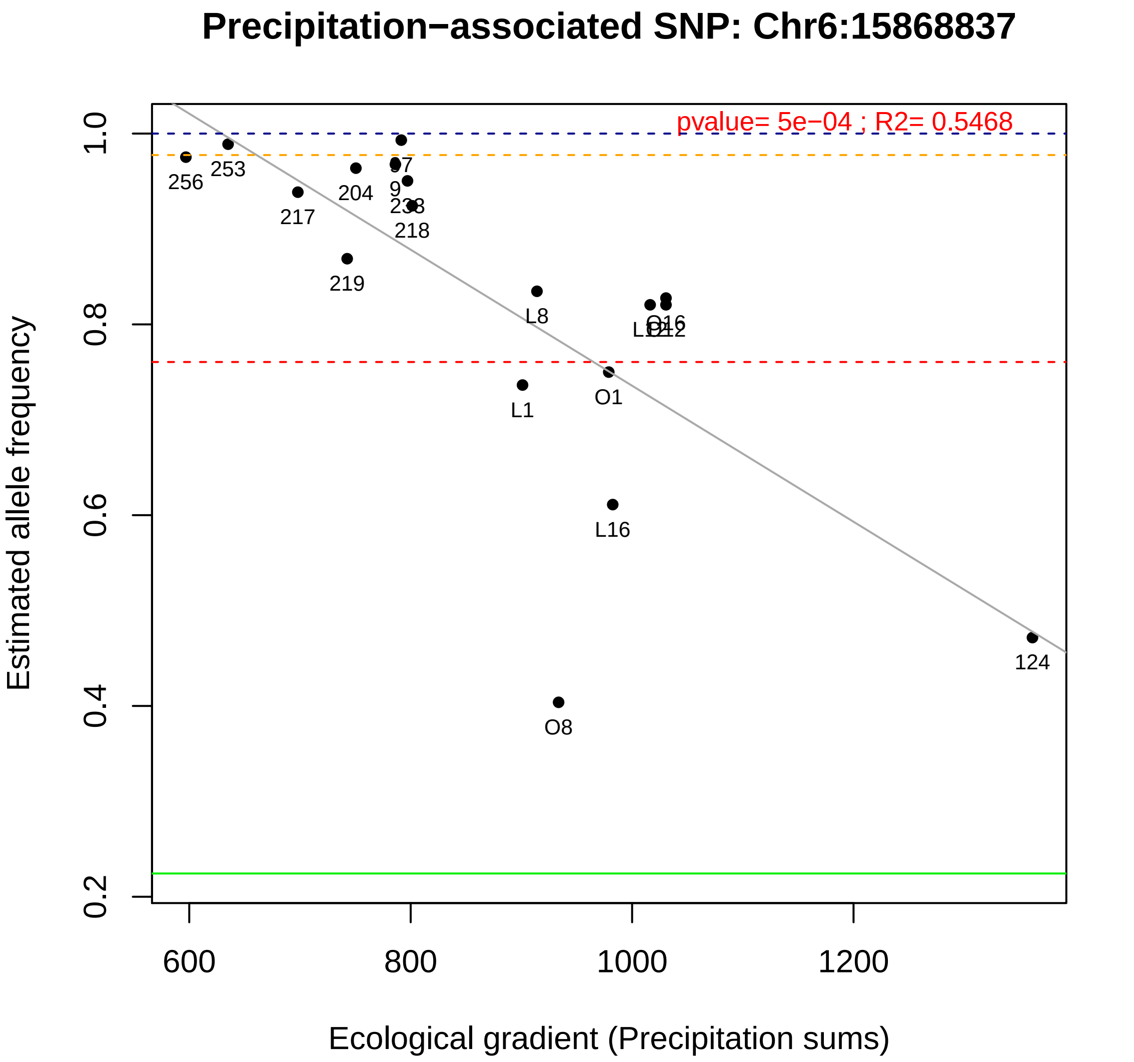
Precipitation−associated SNP: Chr6:15868837
pvalue= 5e−04 ; R2= 0.5468
Estimated allele frequency
1.0
0.8
0.6
0.4
0.2
600
800
1000
1200
Ecological gradient (Precipitation sums)
256
253
217
204
219
9
218
L8
O1
L1
L16
O8
124

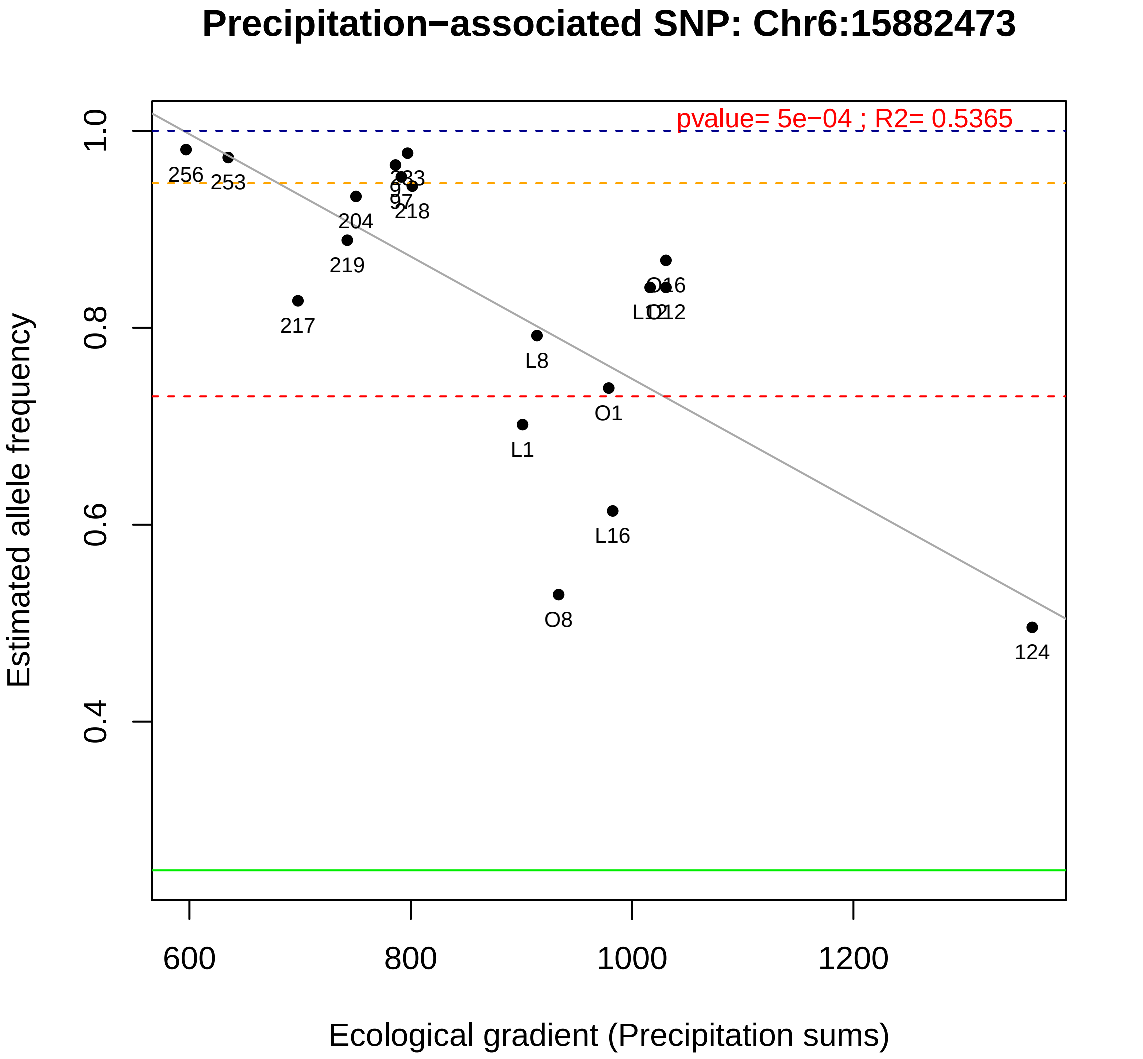

Precipitation−associated SNP: Chr6:15882473
pvalue= 5e−04 ; R2= 0.5365
Estimated allele frequency
Ecological gradient (Precipitation sums)
1.0
0.8
0.6
0.4
600
800
1000
1200
256
253
204
219
217
233
97
218
O16
O12
L12
L8
O1
L1
L16
O8
124

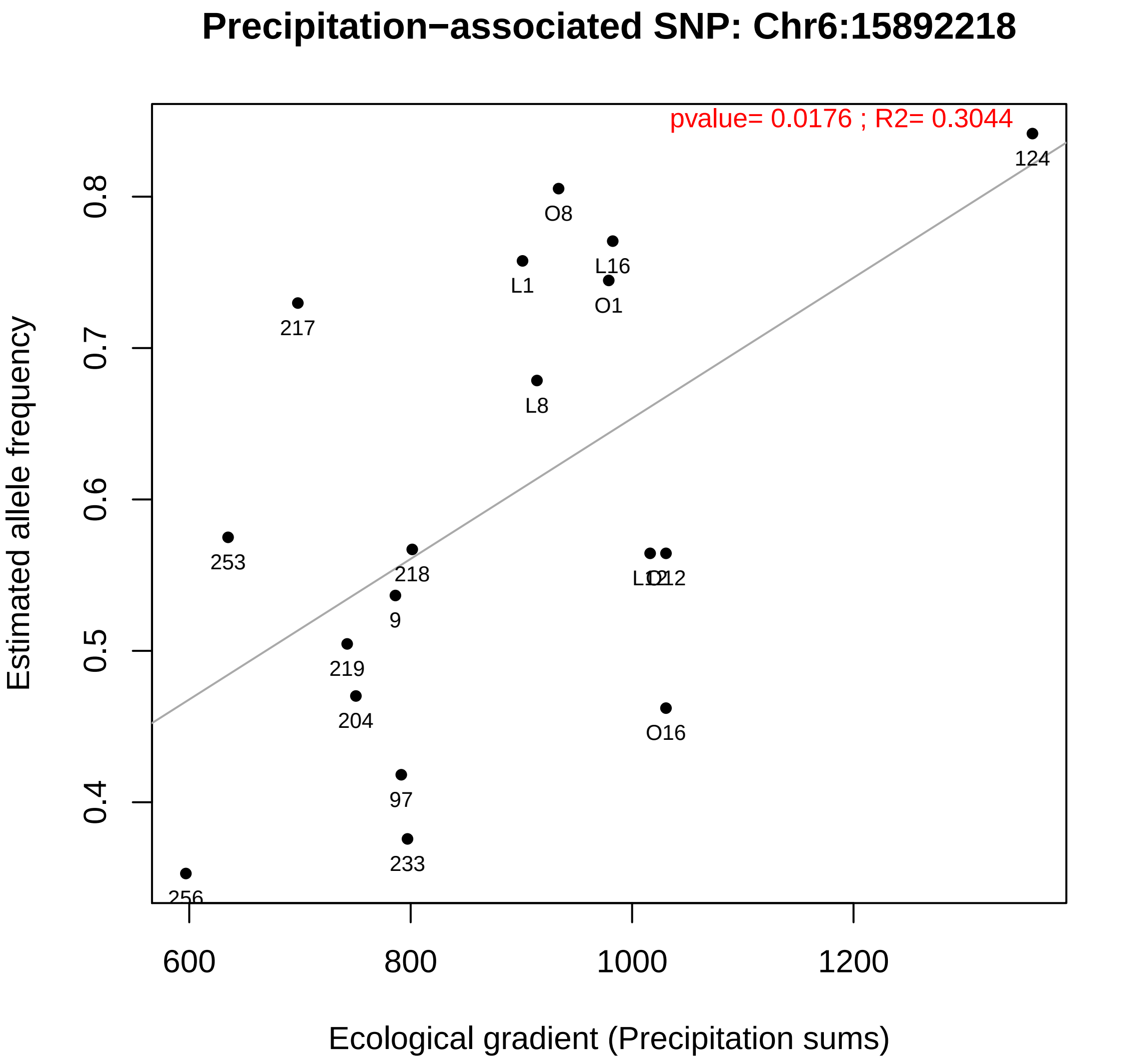
Precipitation−associated SNP: Chr6:15892218
pvalue= 0.0176 ; R2= 0.3044
Estimated allele frequency
Ecological gradient (Precipitation sums)
0.4
0.5
0.6
0.7
0.8
600
800
1000
1200
124
O8
L16
L1
O1
217
L8
253
218
L12
O12
9
219
204
O16
97
233
256

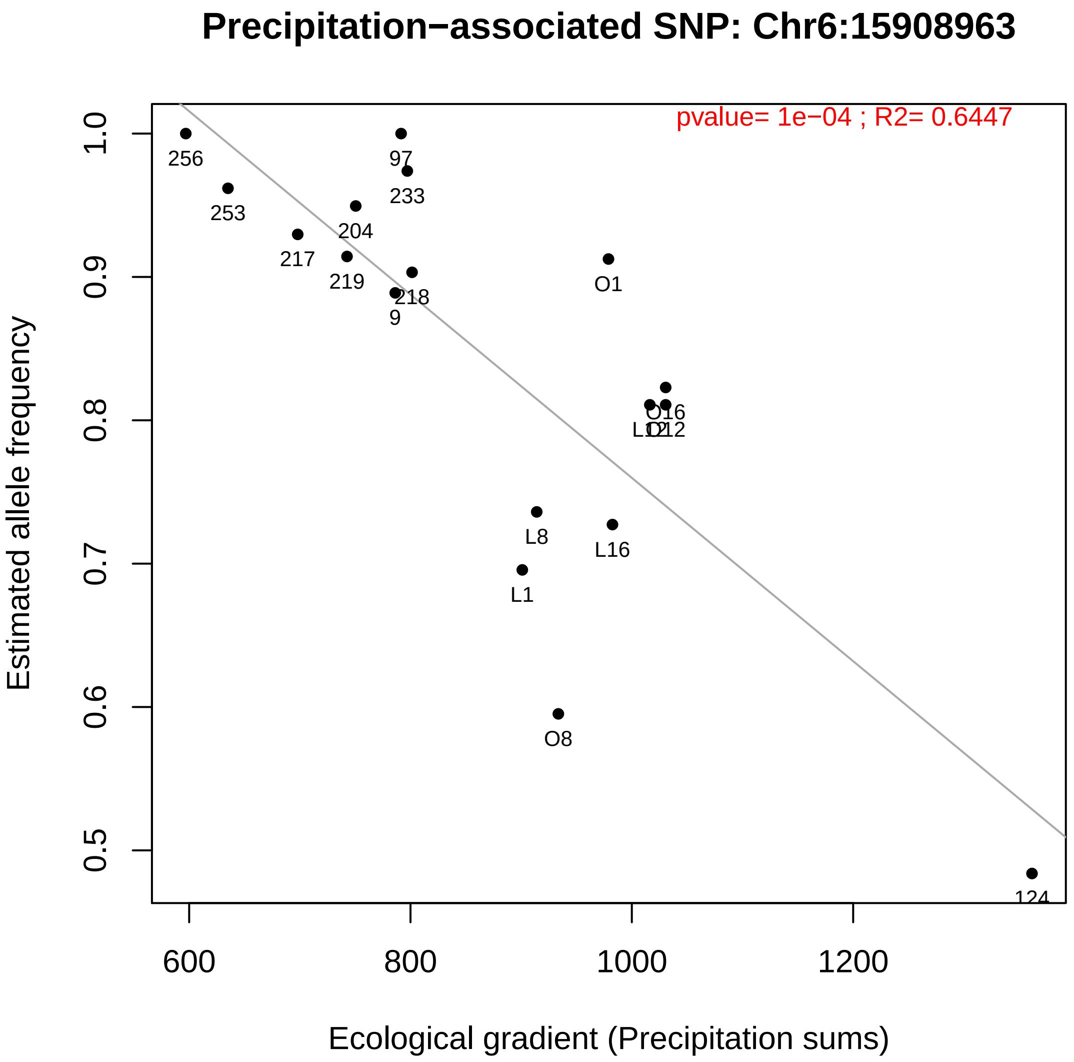
Precipitation−associated SNP: Chr6:15908963
pvalue= 1e−04 ; R2= 0.6447
Estimated allele frequency
Ecological gradient (Precipitation sums)
0.5
0.6
0.7
0.8
0.9
1.0
600
800
1000
1200
256
97
233
253
204
217
219
218
9
O1
O16
L12
O12
L8
L16
L1
O8
124

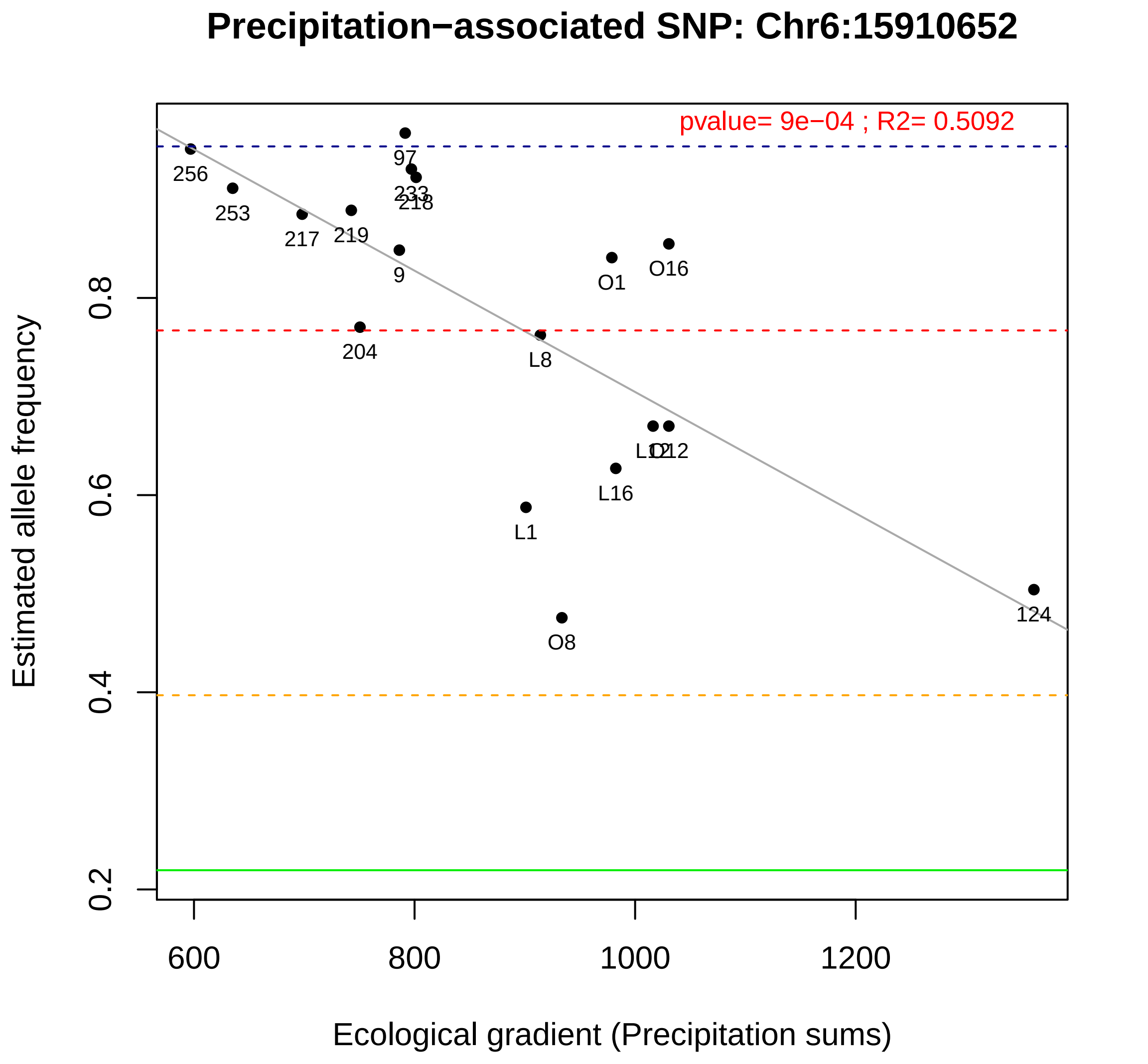
Precipitation−associated SNP: Chr6:15910652
pvalue= 9e−04 ; R2= 0.5092
Estimated allele frequency
0.2
0.4
0.6
0.8
600
800
1000
1200
Ecological gradient (Precipitation sums)
256
97
233
218
253
217
219
9
O1
O16
204
L8
L12
O12
L16
L1
O8
124

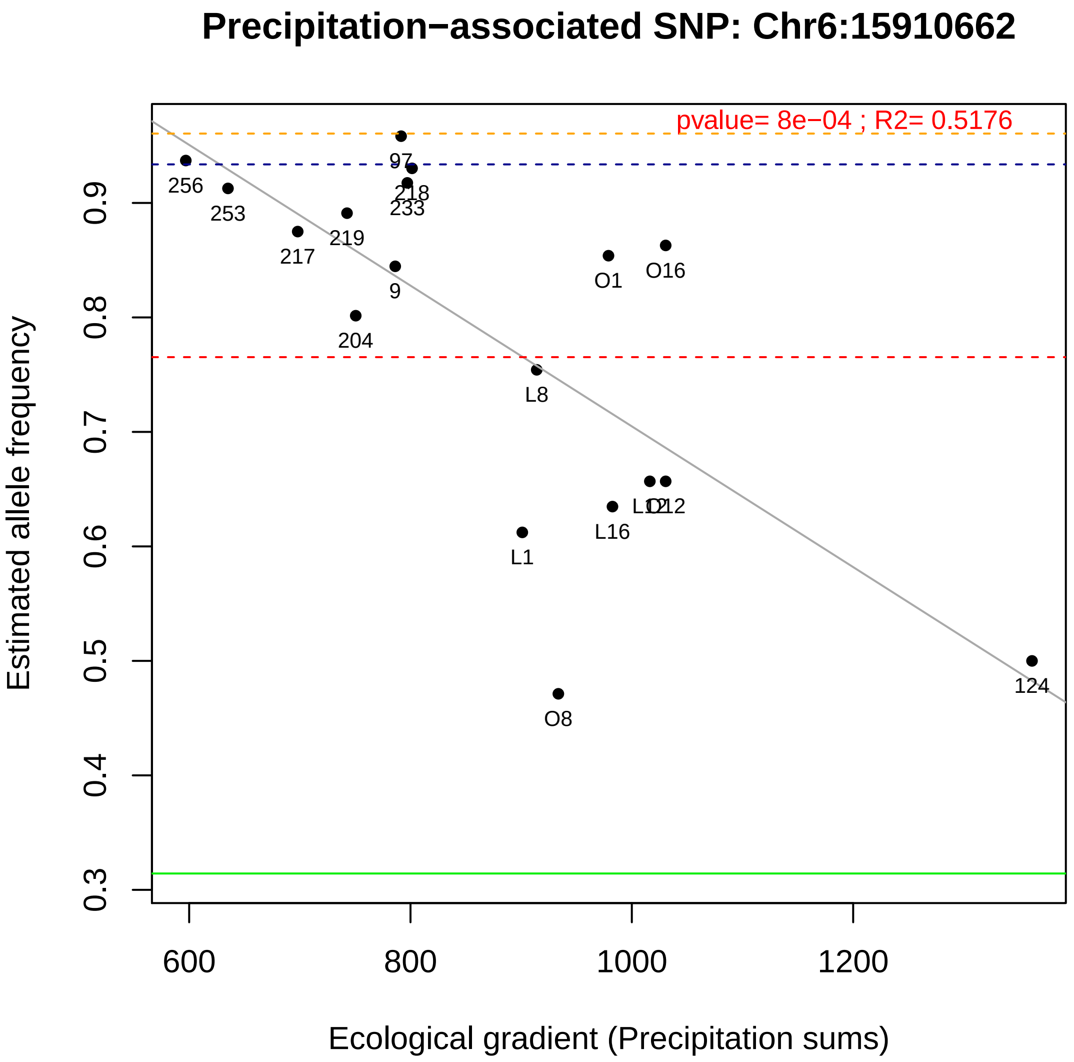
Precipitation−associated SNP: Chr6:15910662
pvalue= 8e−04 ; R2= 0.5176
Estimated allele frequency
Ecological gradient (Precipitation sums)
0.3
0.4
0.5
0.6
0.7
0.8
0.9
600
800
1000
1200
256
253
217
219
97
218
233
9
204
L8
O1
O16
L16
L12
O12
L1
O8
124

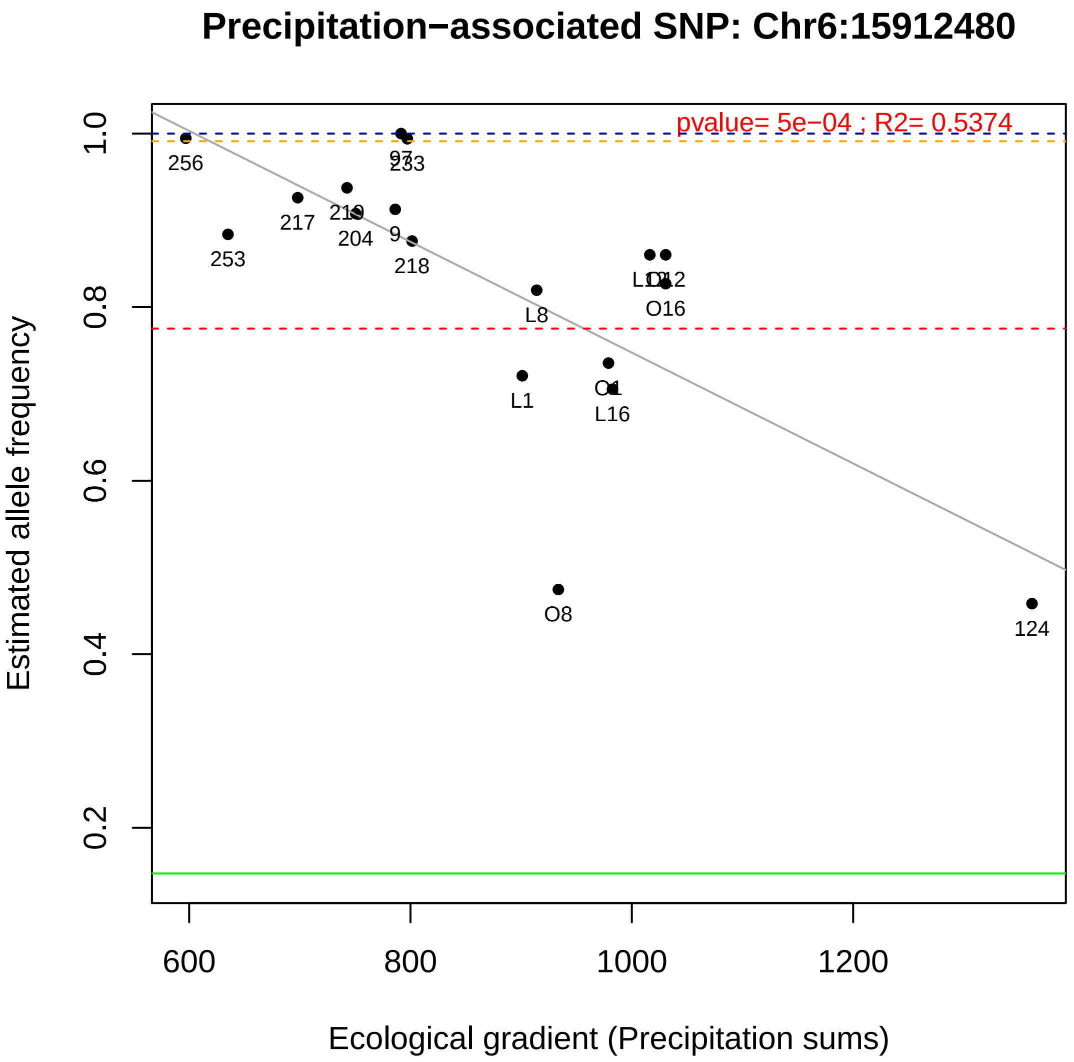

Precipitation−associated SNP: Chr6:15912480
pvalue= 5e−04 ; R2= 0.5374
Estimated allele frequency
Ecological gradient (Precipitation sums)
0.2
0.4
0.6
0.8
1.0
600
800
1000
1200
256
97
233
217
219
204
9
253
218
L12
O12
O16
L8
L1
O1
L16
O8
124

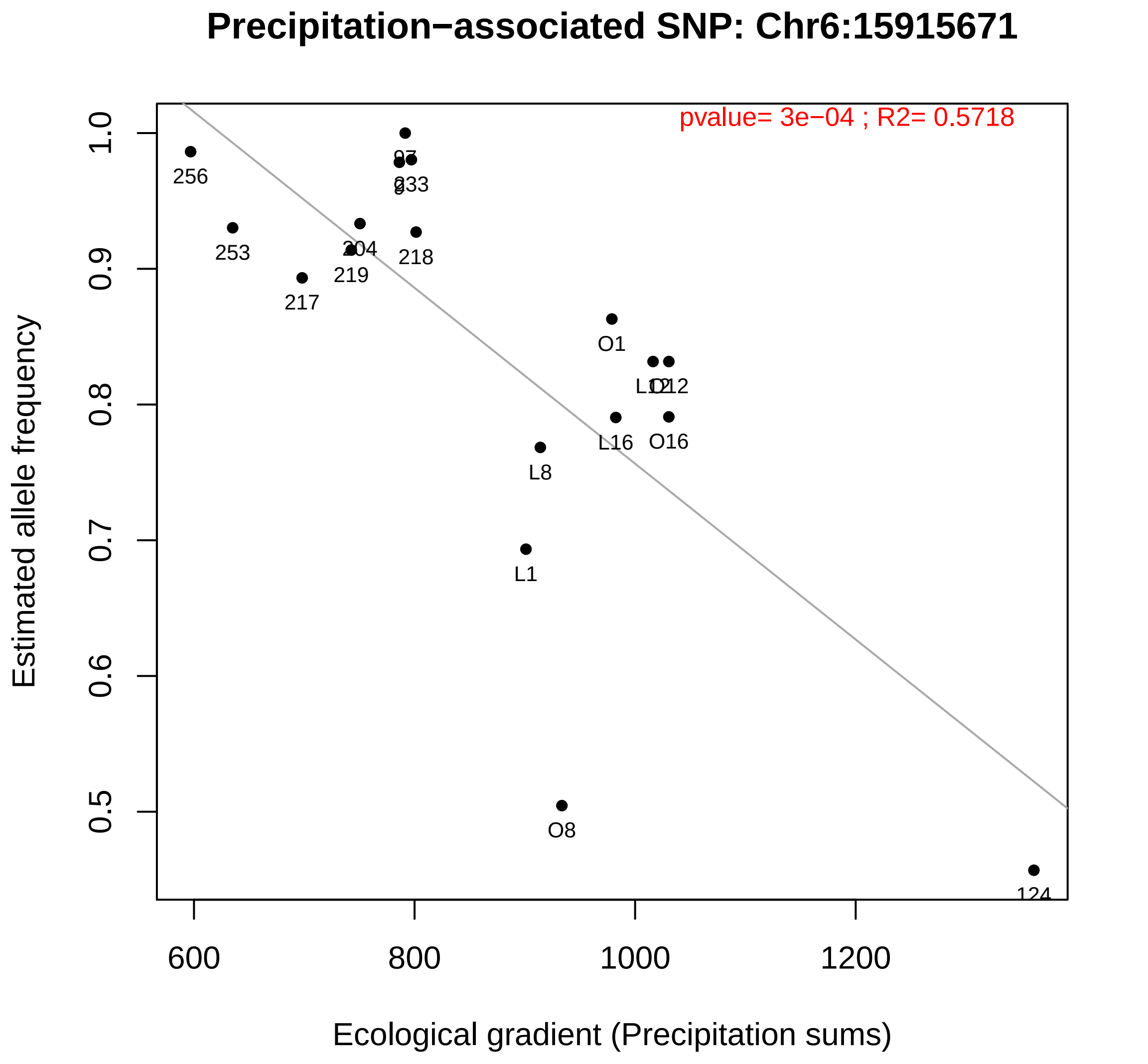
Precipitation−associated SNP: Chr6:15915671
pvalue= 3e−04 ; R2= 0.5718
Estimated allele frequency
Ecological gradient (Precipitation sums)
1.0
0.9
0.8
0.7
0.6
0.5
600
800
1000
1200
256
253
217
219
204
218
233
O1
L16
O16
L8
L1
O8
124

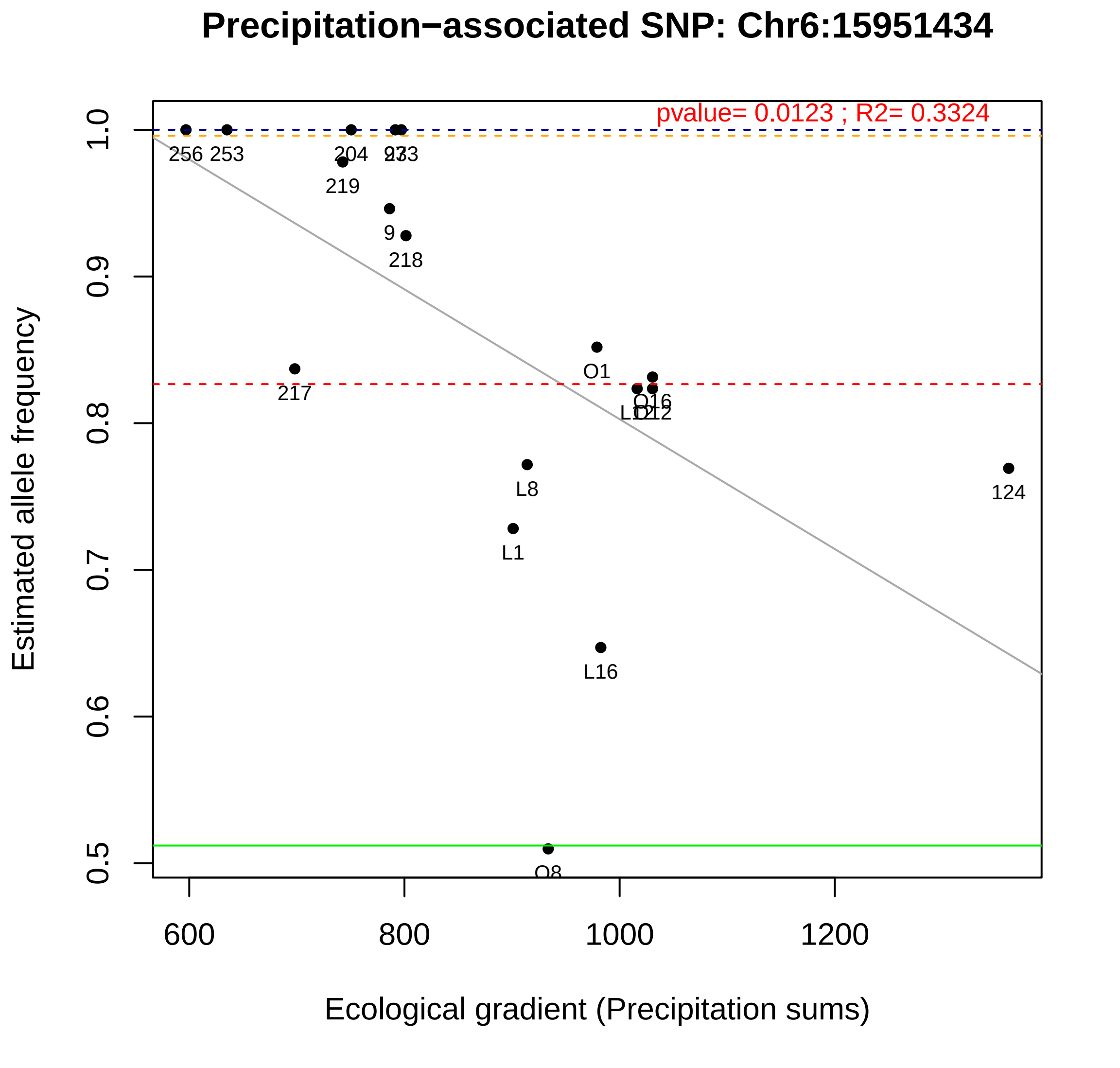

Precipitation−associated SNP: Chr6:15951434
pvalue= 0.0123 ; R2= 0.3324
Estimated allele frequency
Ecological gradient (Precipitation sums)
1.0
0.9
0.8
0.7
0.6
0.5
600
800
1000
1200
256
253
204
219
9
218
217
O1
O16
L12
L8
L1
124
L16
O8

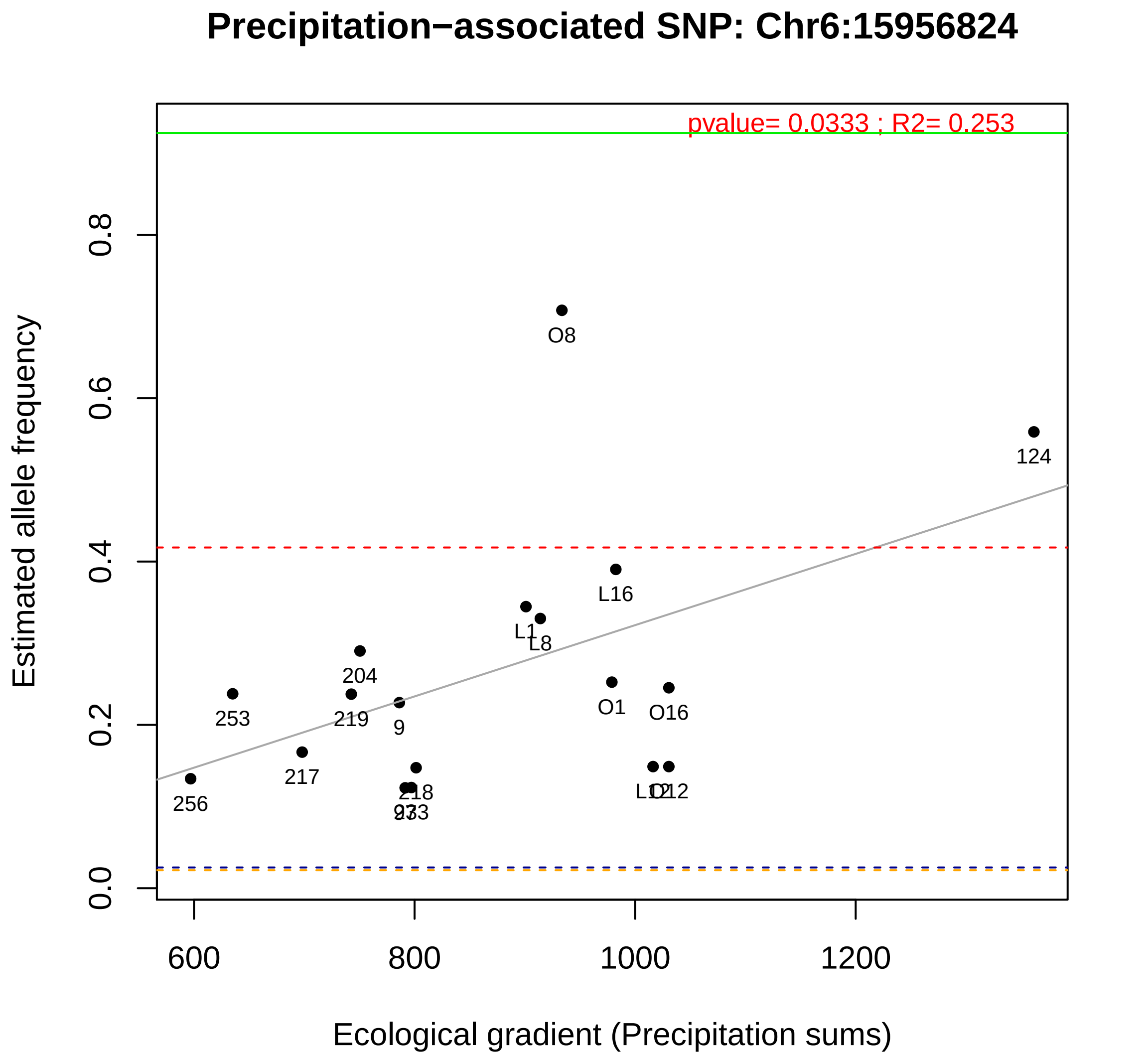

Precipitation−associated SNP: Chr6:15956824
pvalue= 0.0333 ; R2= 0.253
Estimated allele frequency
Ecological gradient (Precipitation sums)
0.0
0.2
0.4
0.6
0.8
600
800
1000
1200
O8
124
L16
L1
L8
204
253
219
9
O1
O16
217
218
256

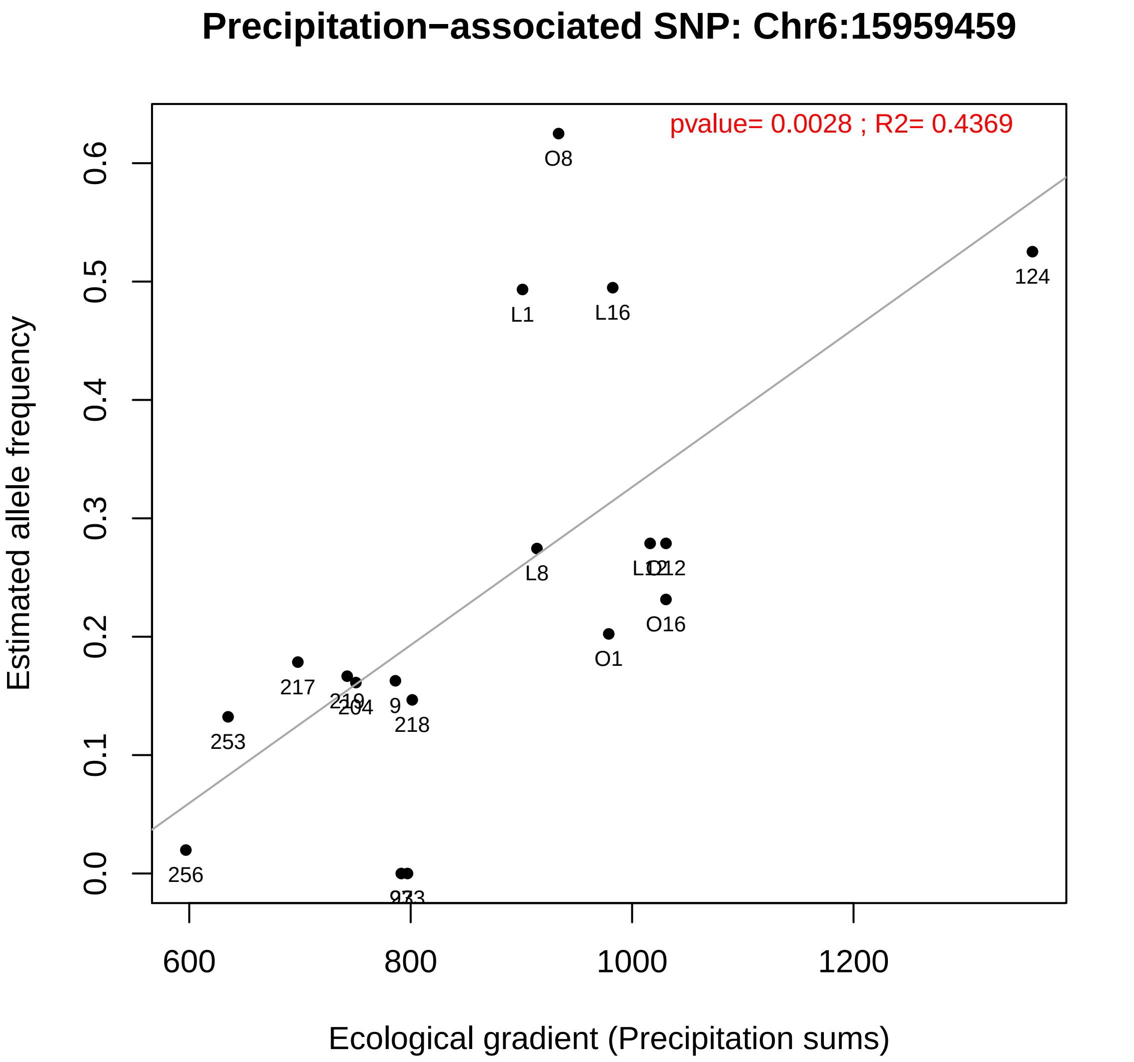

Precipitation−associated SNP: Chr6:15959459
pvalue= 0.0028 ; R2= 0.4369
Estimated allele frequency
Ecological gradient (Precipitation sums)
0.0
0.1
0.2
0.3
0.4
0.5
0.6
600
800
1000
1200
O8
L1
L16
124
L8
O16
O1
217
9
218
253
256

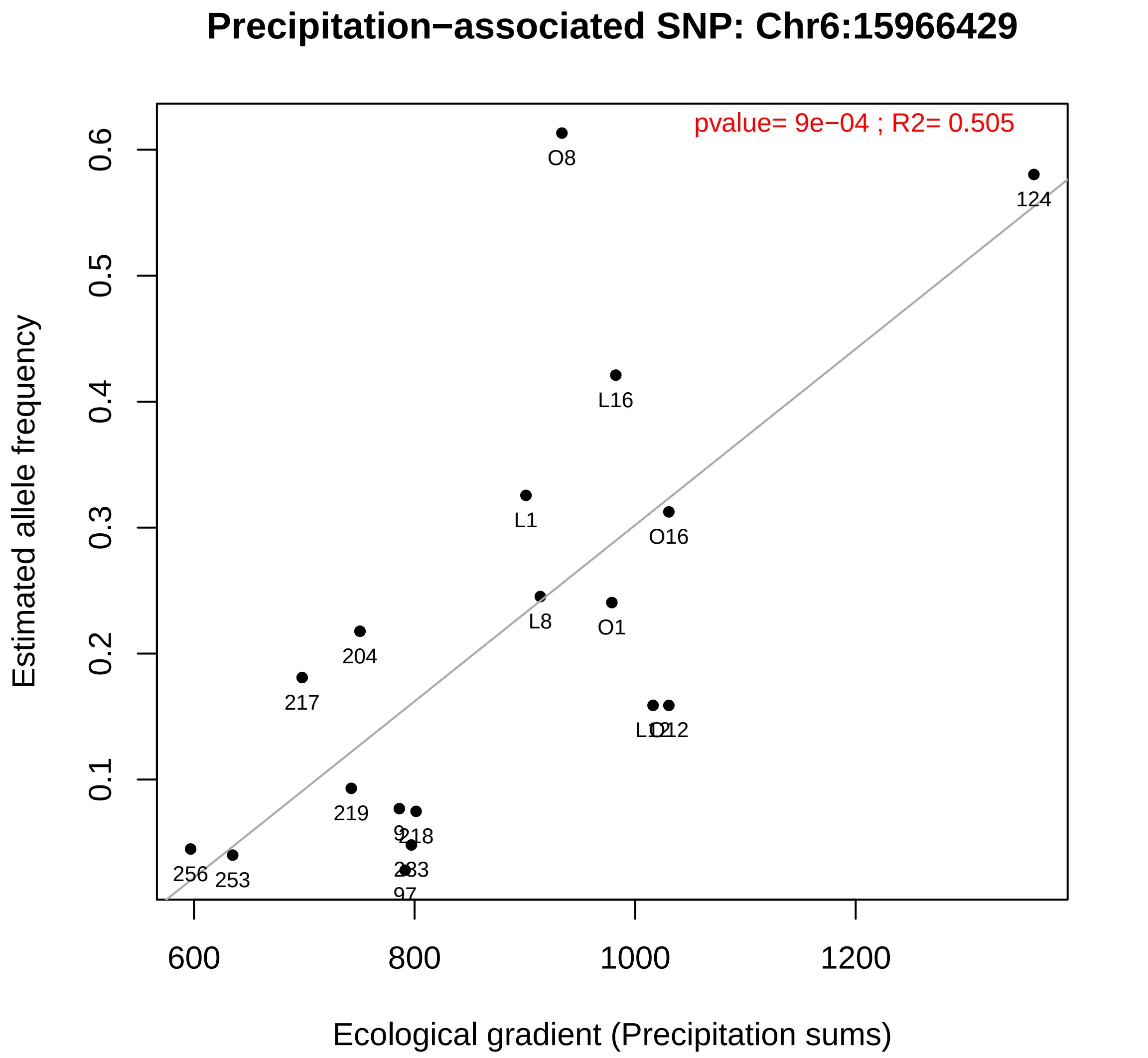
Precipitation−associated SNP: Chr6:15966429
pvalue= 9e−04 ; R2= 0.505
Estimated allele frequency
Ecological gradient (Precipitation sums)
0.1
0.2
0.3
0.4
0.5
0.6
600
800
1000
1200
O8
124
L16
L1
O16
L8
O1
204
217
219
256
253
218
233
97

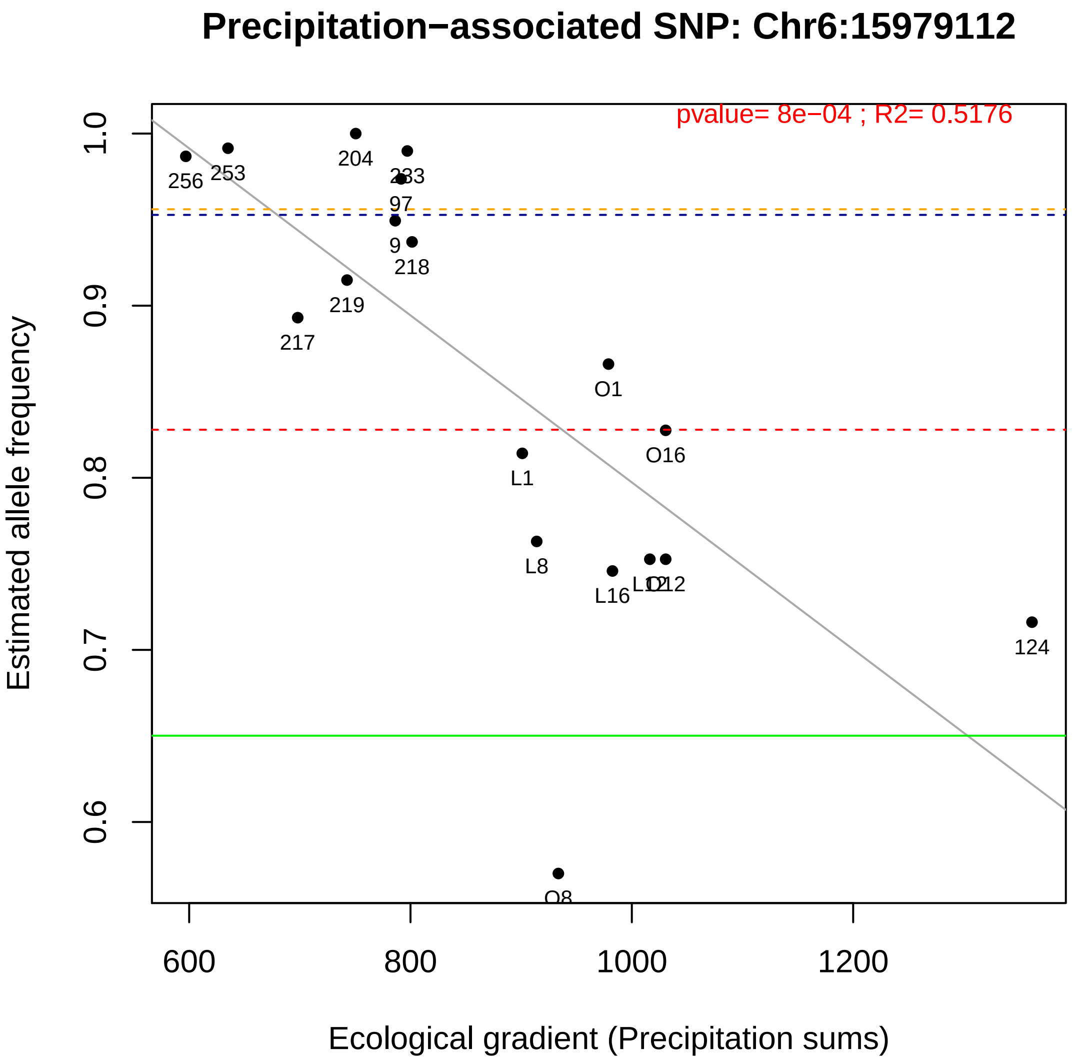

Precipitation−associated SNP: Chr6:15979112
pvalue= 8e−04 ; R2= 0.5176
Estimated allele frequency
Ecological gradient (Precipitation sums)
0.6
0.7
0.8
0.9
1.0
600
800
1000
1200
256
253
204
233
97
9
218
219
217
O1
O16
L1
L8
L16
L12
O12
124
O8

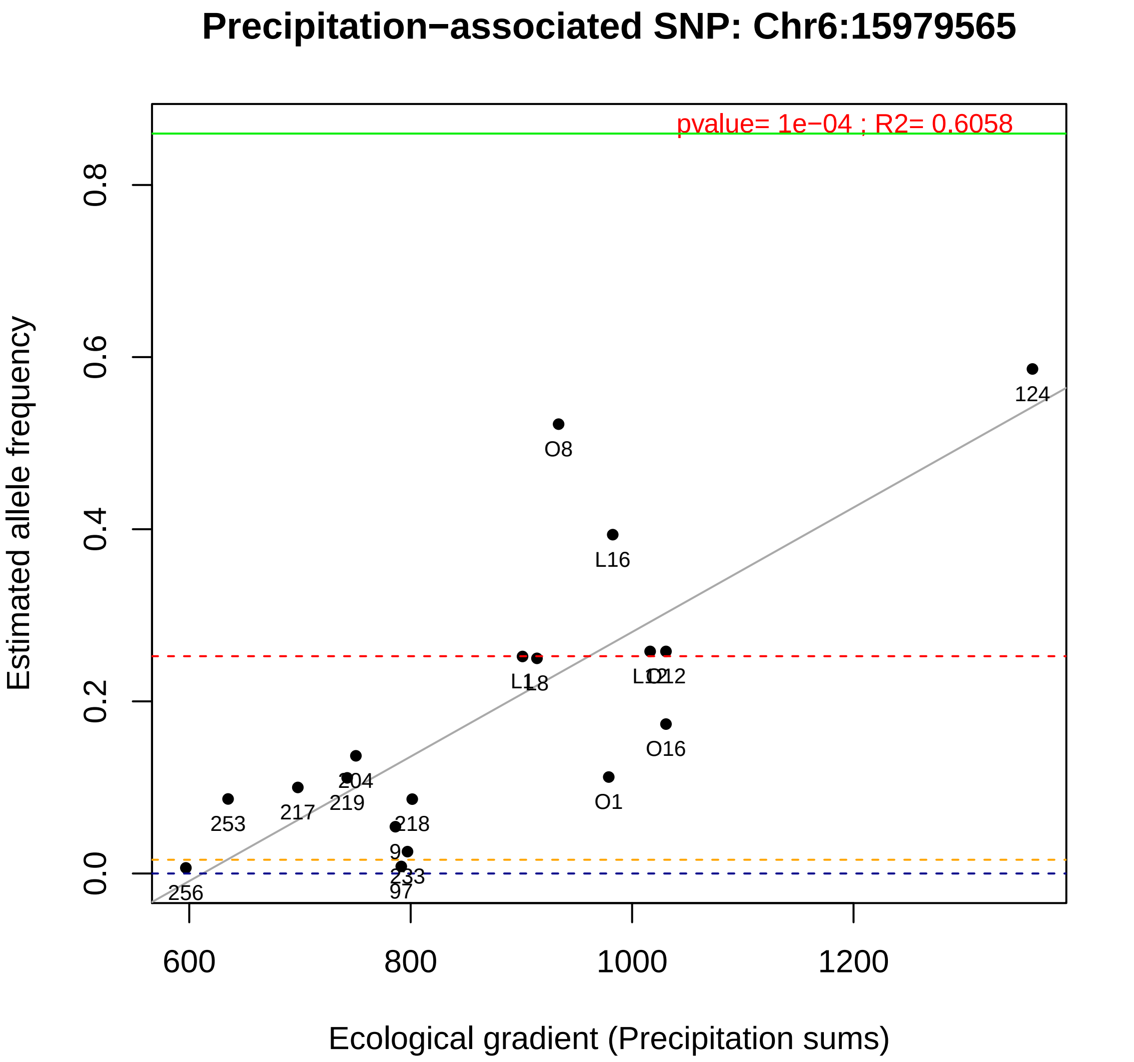
Precipitation−associated SNP: Chr6:15979565
pvalue= 1e−04 ; R2= 0.6058
Estimated allele frequency
0.0
0.2
0.4
0.6
0.8
600
800
1000
1200
Ecological gradient (Precipitation sums)
124
O8
L16
L8
O16
O1
204
219
217
253
218
233
97
256

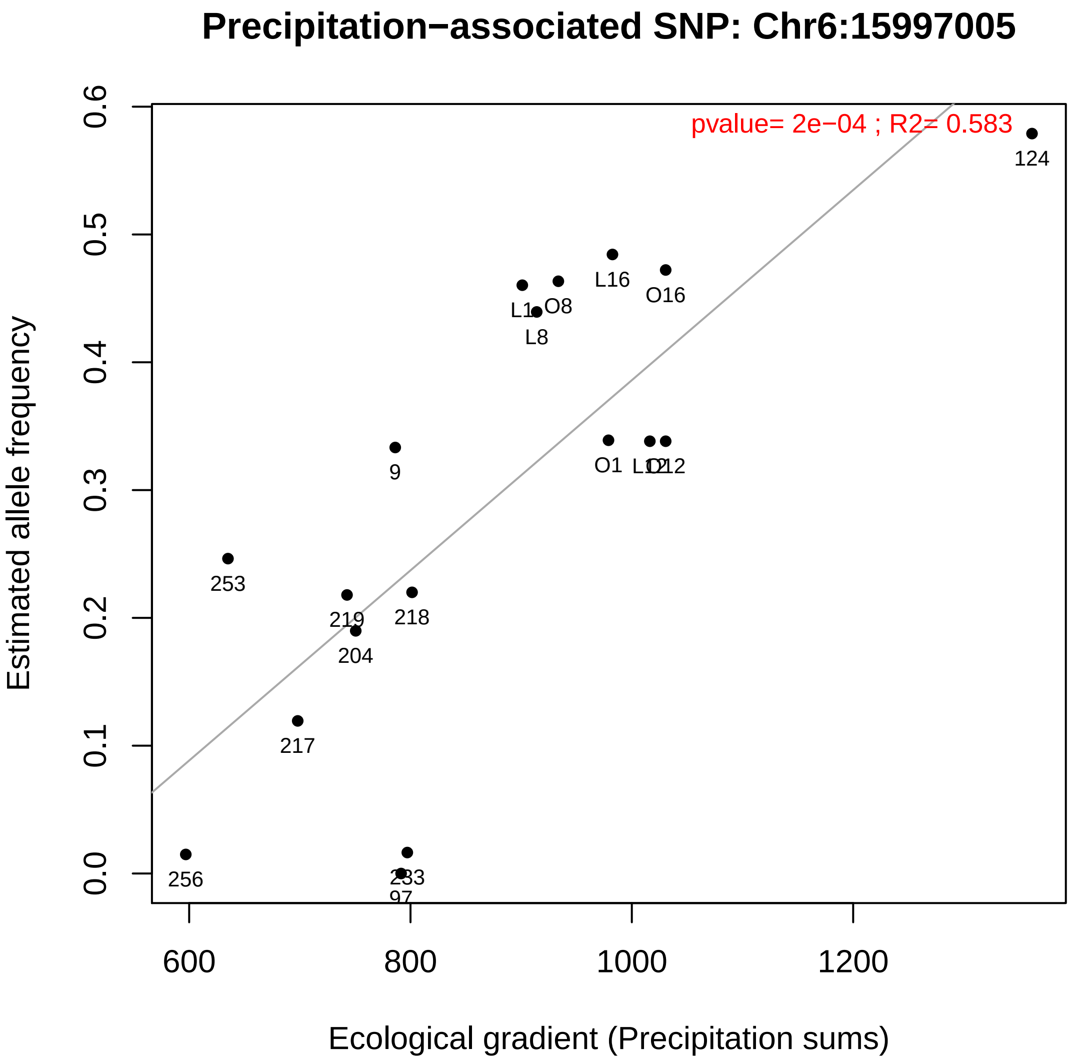
Precipitation−associated SNP: Chr6:15997005
pvalue= 2e−04 ; R2= 0.583
Estimated allele frequency
0.0
0.1
0.2
0.3
0.4
0.5
0.6
600
800
1000
1200
Ecological gradient (Precipitation sums)
124
L16
O16
L1
O8
L8
9
O1
L12
O12
253
219
218
204
217
256
233
97

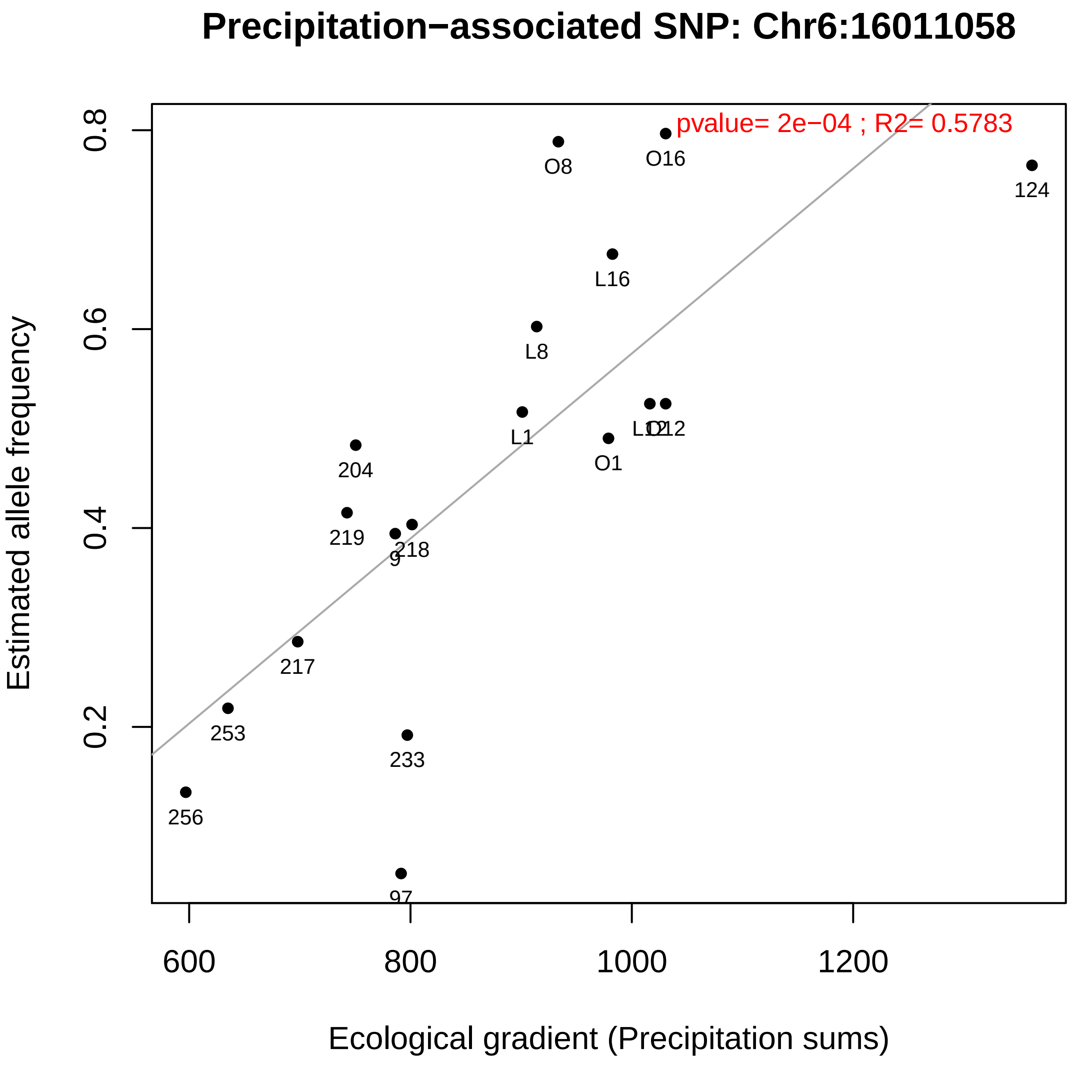
Precipitation−associated SNP: Chr6:16011058
pvalue= 2e−04 ; R2= 0.5783
O8
O16
124
L16
L8
L1
L12
O12
O1
204
219
218
9
217
253
233
256
97
0.2
0.4
0.6
0.8
600
800
1000
1200
Estimated allele frequency
Ecological gradient (Precipitation sums)

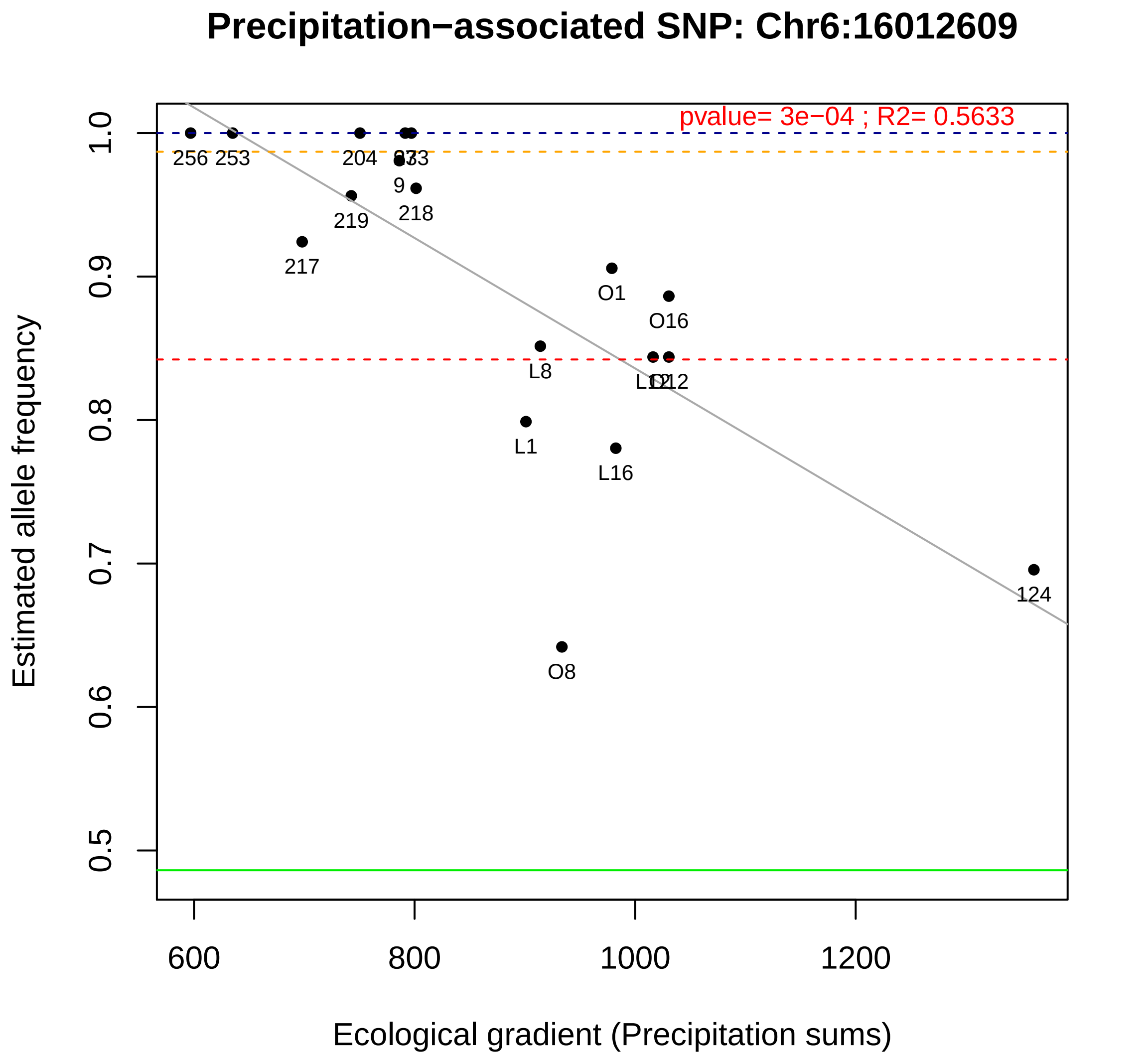

Precipitation−associated SNP: Chr6:16012609
pvalue= 3e−04 ; R2= 0.5633
Estimated allele frequency
Ecological gradient (Precipitation sums)
1.0
0.9
0.8
0.7
0.6
0.5
600
800
1000
1200
256
253
204
9
219
218
217
O1
O16
L8
L1
L16
124
O8

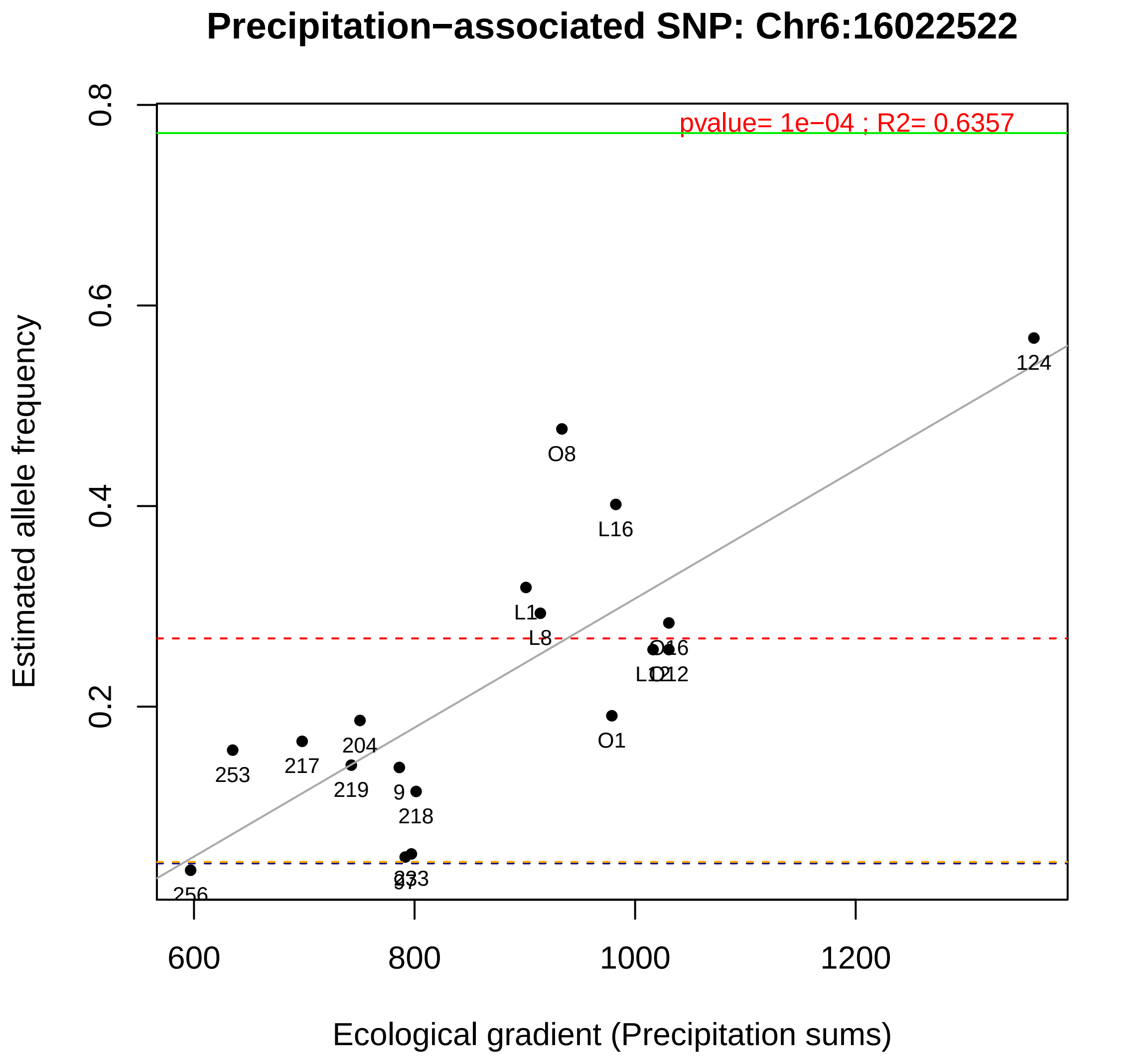

Precipitation−associated SNP: Chr6:16022522
pvalue= 1e−04 ; R2= 0.6357
Estimated allele frequency
0.2
0.4
0.6
0.8
600
800
1000
1200
Ecological gradient (Precipitation sums)
124
O8
L16
L1
L8
O16
L12
O1
204
217
253
219
9
218
256
233

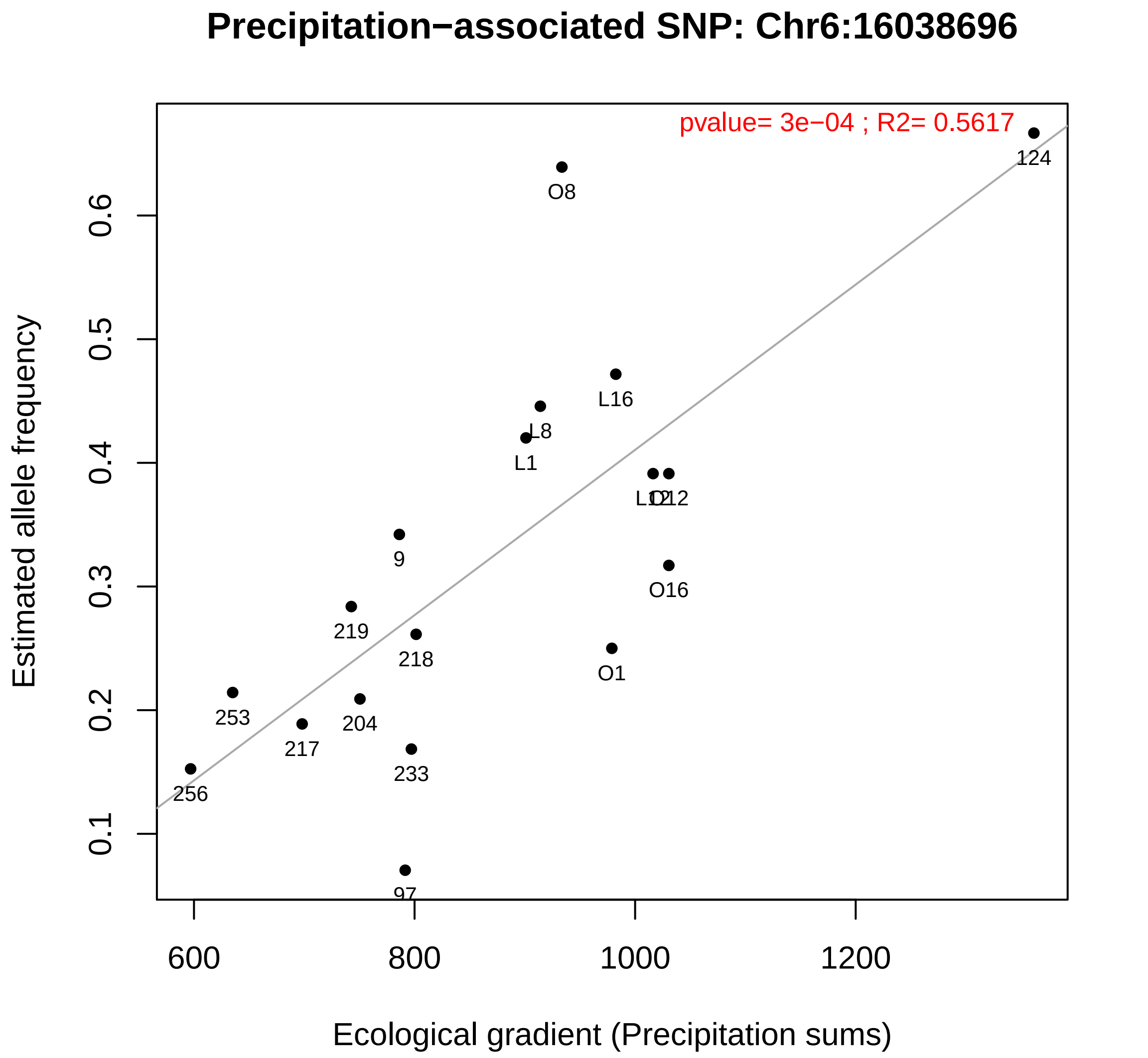
Precipitation−associated SNP: Chr6:16038696
pvalue= 3e−04 ; R2= 0.5617
124
O8
L16
L8
L1
L12
O12
9
O16
219
218
O1
253
204
217
233
256
97
0.1
0.2
0.3
0.4
0.5
0.6
600
800
1000
1200
Estimated allele frequency
Ecological gradient (Precipitation sums)

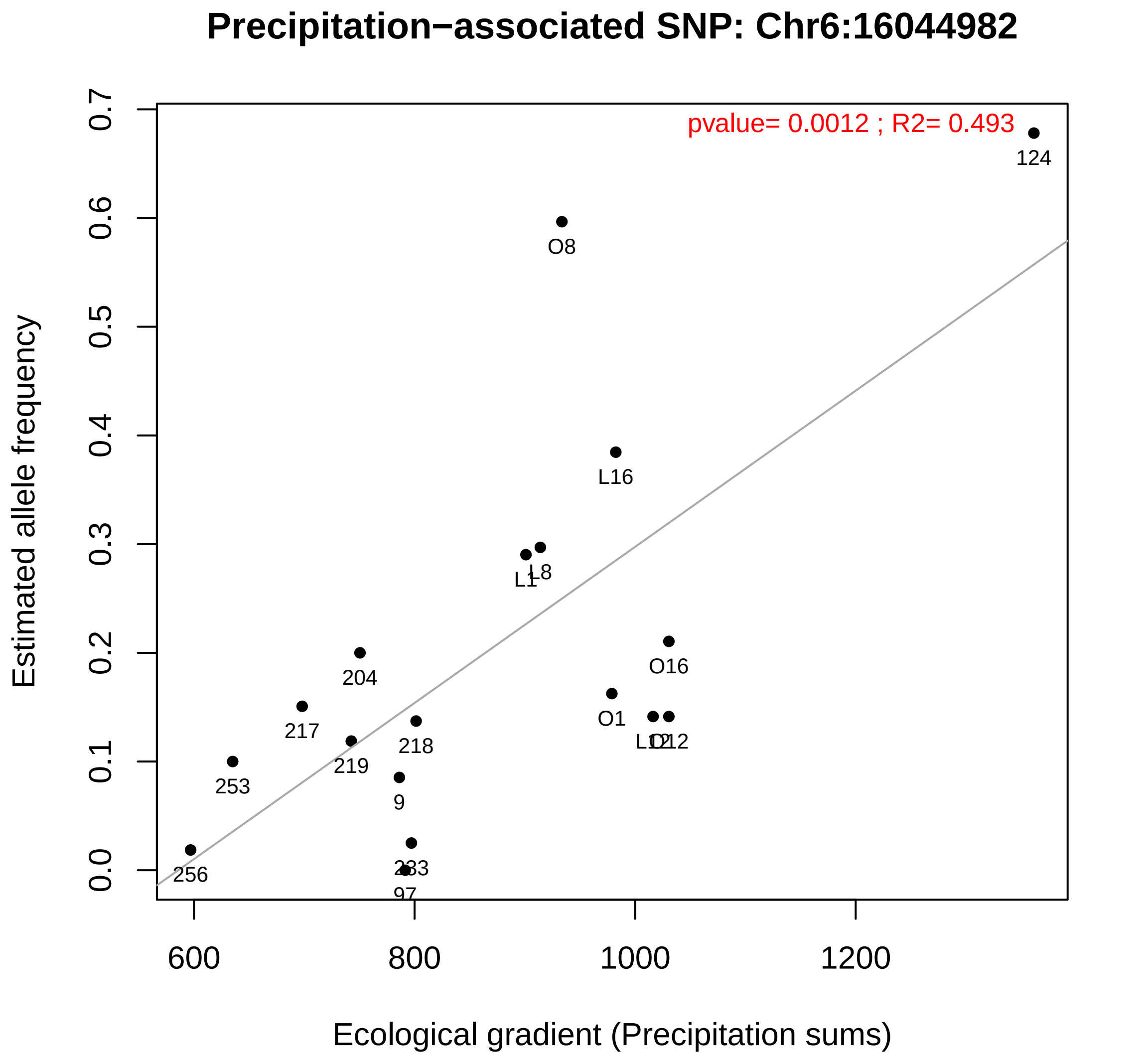
Precipitation−associated SNP: Chr6:16044982
pvalue= 0.0012 ; R2= 0.493
Estimated allele frequency
Ecological gradient (Precipitation sums)
0.0
0.1
0.2
0.3
0.4
0.5
0.6
0.7
600
800
1000
1200
124
O8
L16
L1
L8
O16
204
O1
217
218
219
253
9
256
233
97

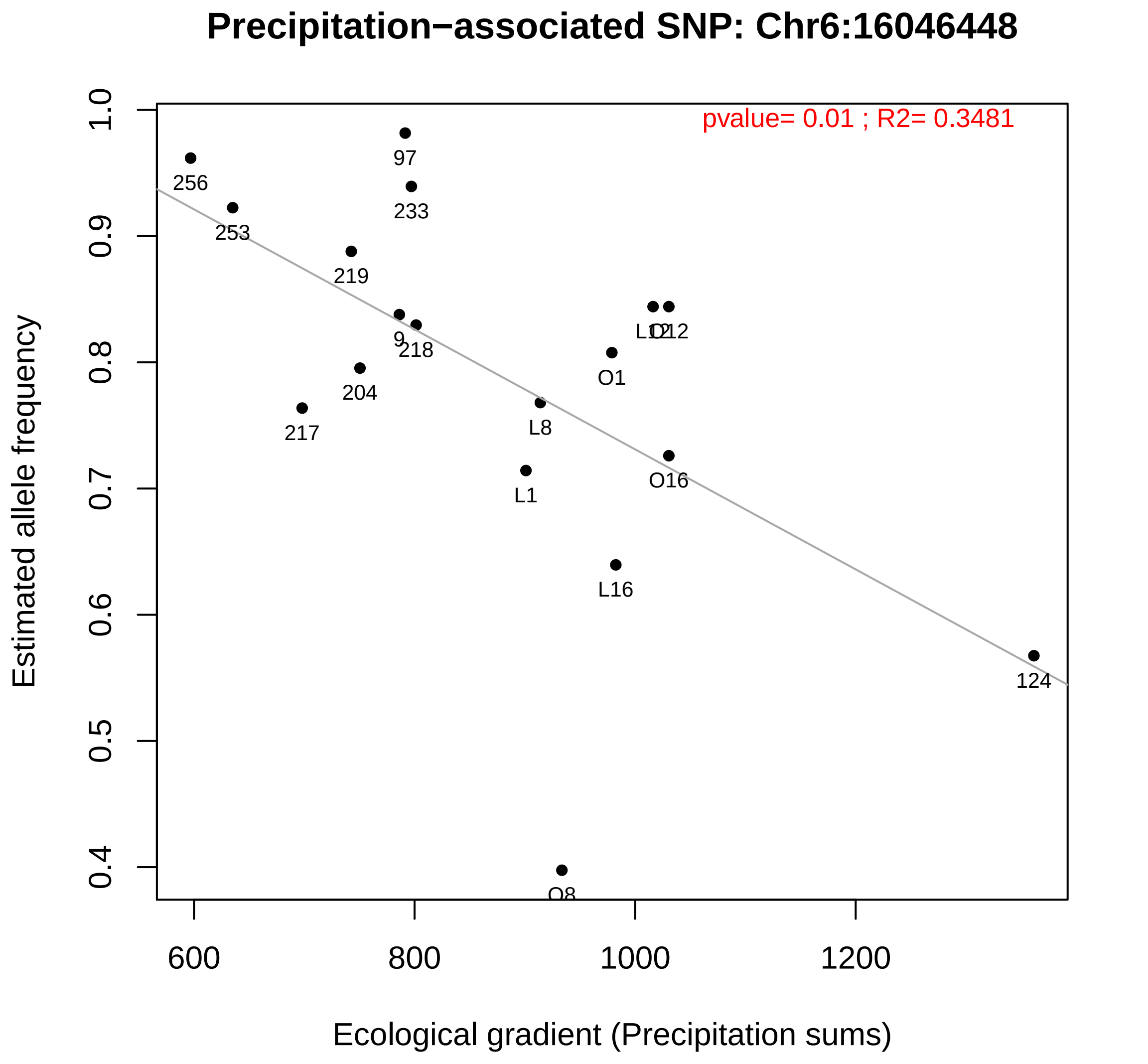

Precipitation−associated SNP: Chr6:16046448
pvalue= 0.01 ; R2= 0.3481
Estimated allele frequency
Ecological gradient (Precipitation sums)
0.4
0.5
0.6
0.7
0.8
0.9
1.0
600
800
1000
1200
256
253
97
233
219
9
218
204
217
L8
L1
O1
L12
O12
O16
L16
O8
124

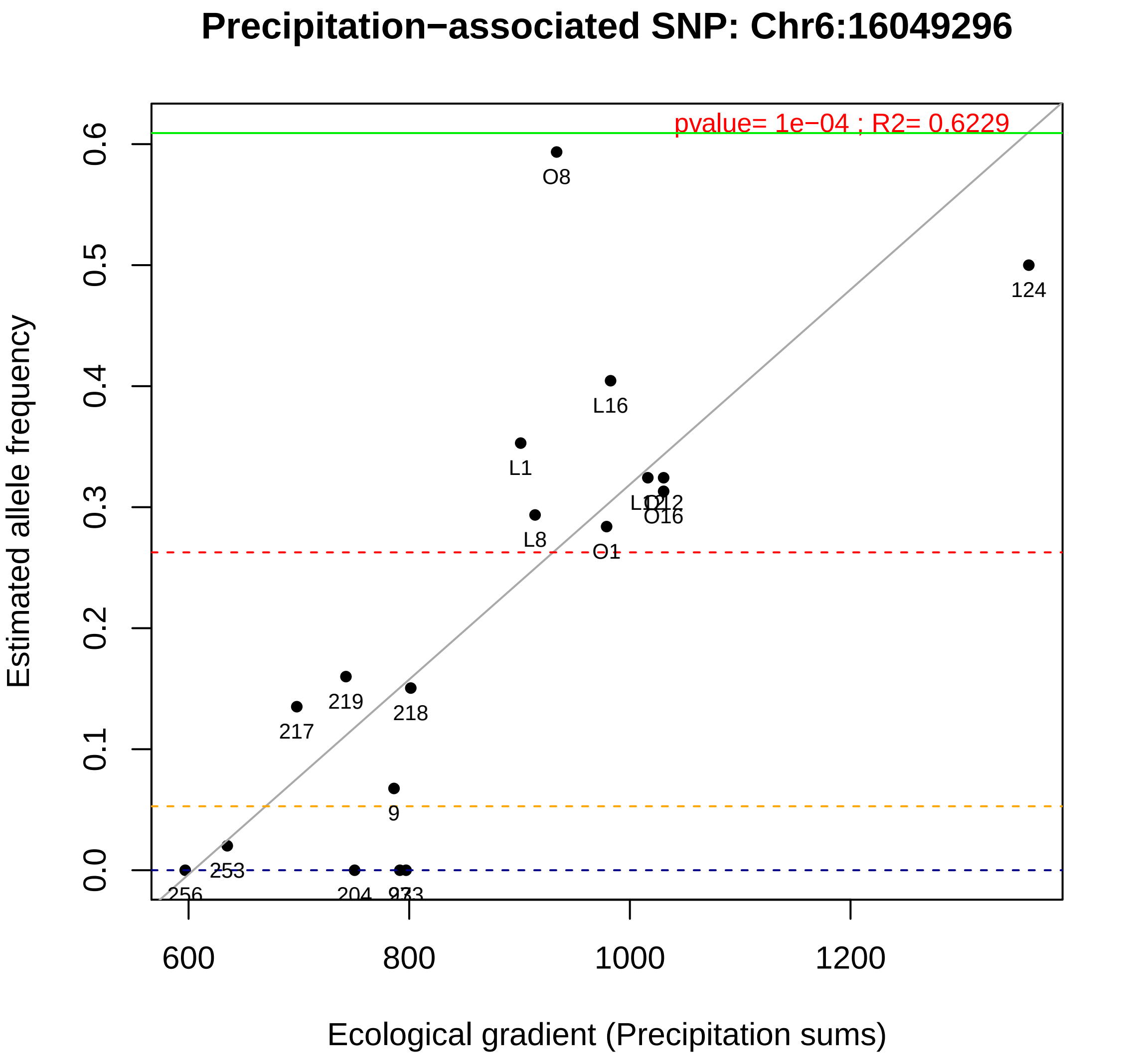
Precipitation−associated SNP: Chr6:16049296
pvalue= 1e−04 ; R2= 0.6229
Estimated allele frequency
Ecological gradient (Precipitation sums)
0.0
0.1
0.2
0.3
0.4
0.5
0.6
600
800
1000
1200
O8
124
L16
L1
O16
L8
O1
219
218
217
9
253
256
204

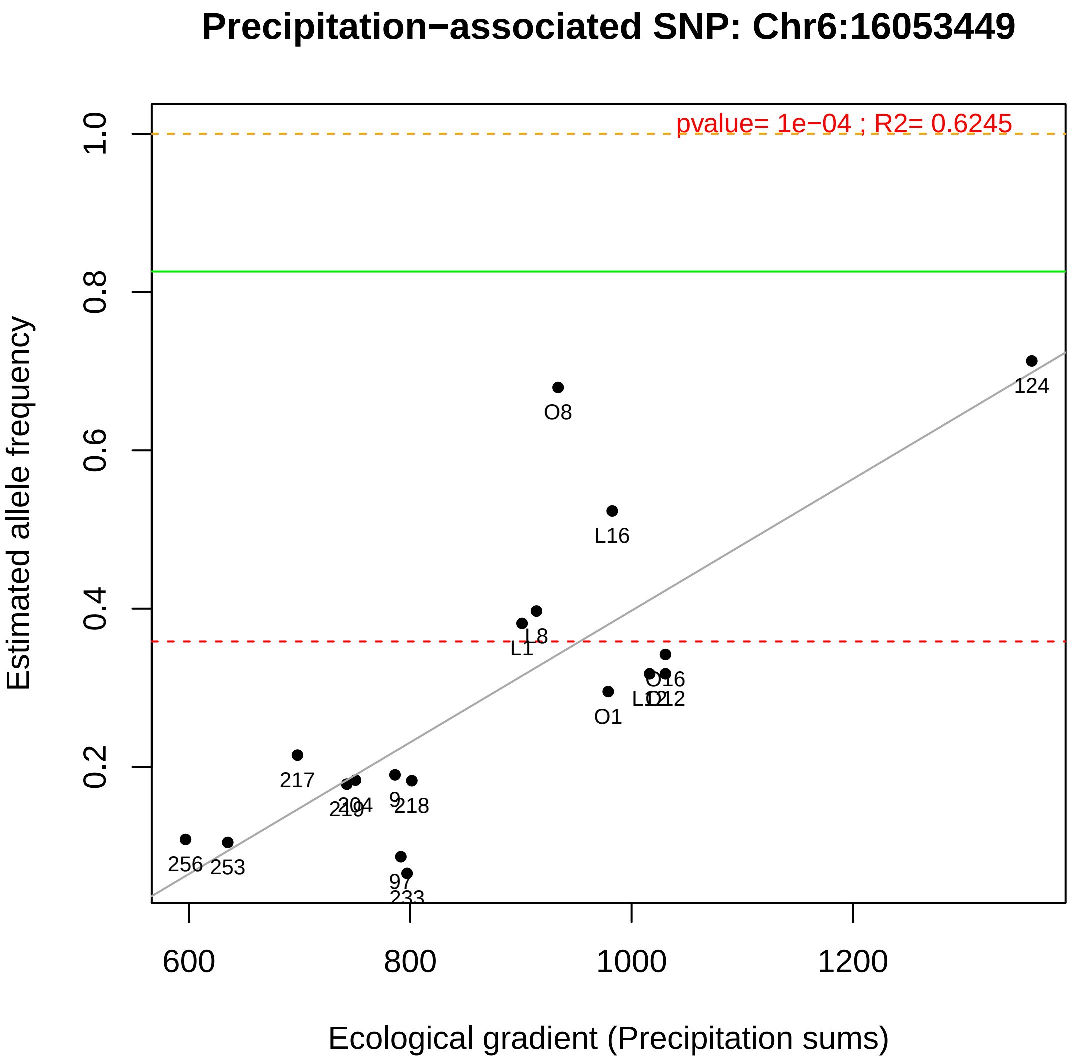

Precipitation−associated SNP: Chr6:16053449
pvalue= 1e−04 ; R2= 0.6245
Estimated allele frequency
Ecological gradient (Precipitation sums)
0.2
0.4
0.6
0.8
1.0
600
800
1000
1200
124
O8
L16
L8
L1
O16
L12
O12
O1
217
204
219
9
218
256
253
97
233

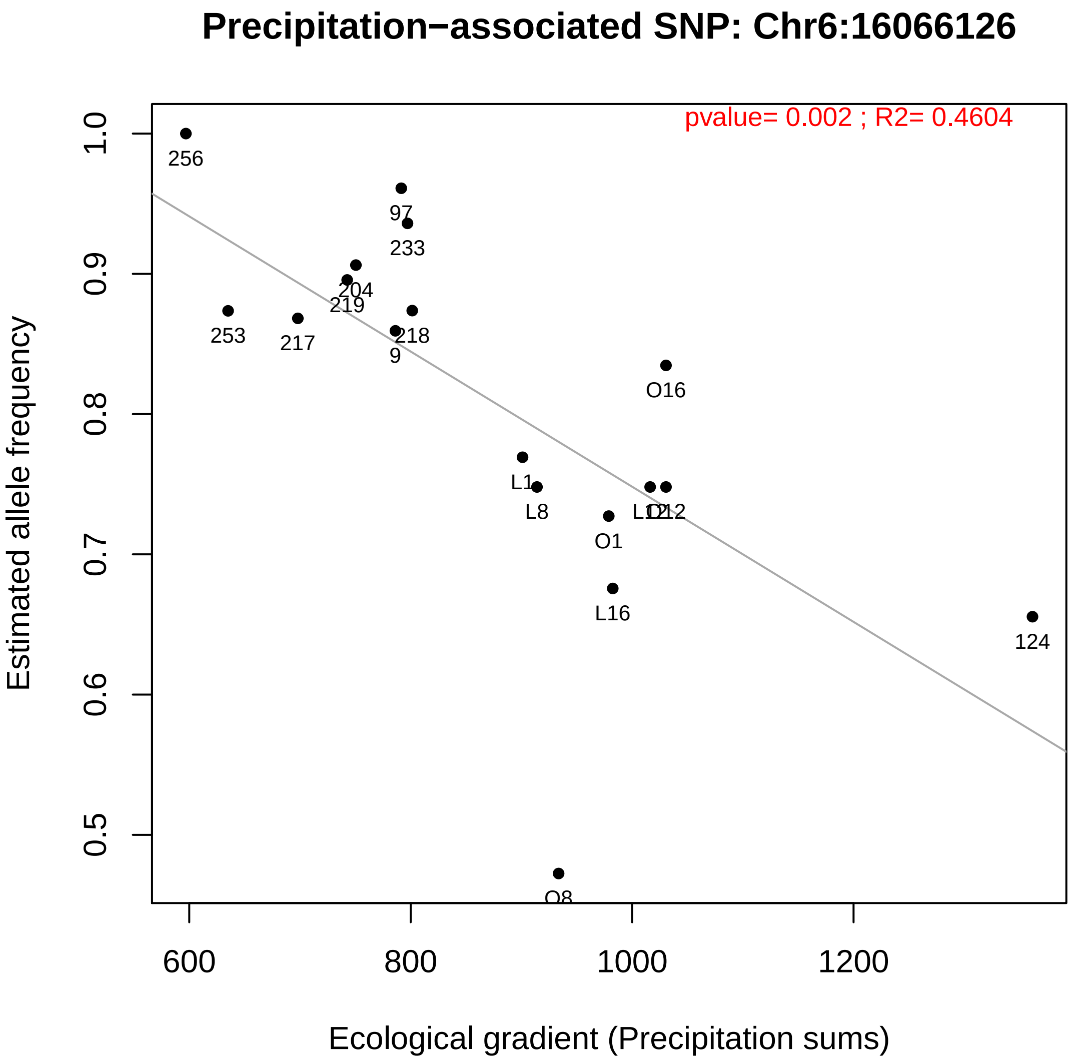

Precipitation−associated SNP: Chr6:16066126
pvalue= 0.002 ; R2= 0.4604
256
97
233
204
219
253
217
218
9
O16
L1
L8
L12
O12
O1
L16
124
O8
1.0
0.9
0.8
0.7
0.6
0.5
Estimated allele frequency
600
800
1000
1200
Ecological gradient (Precipitation sums)

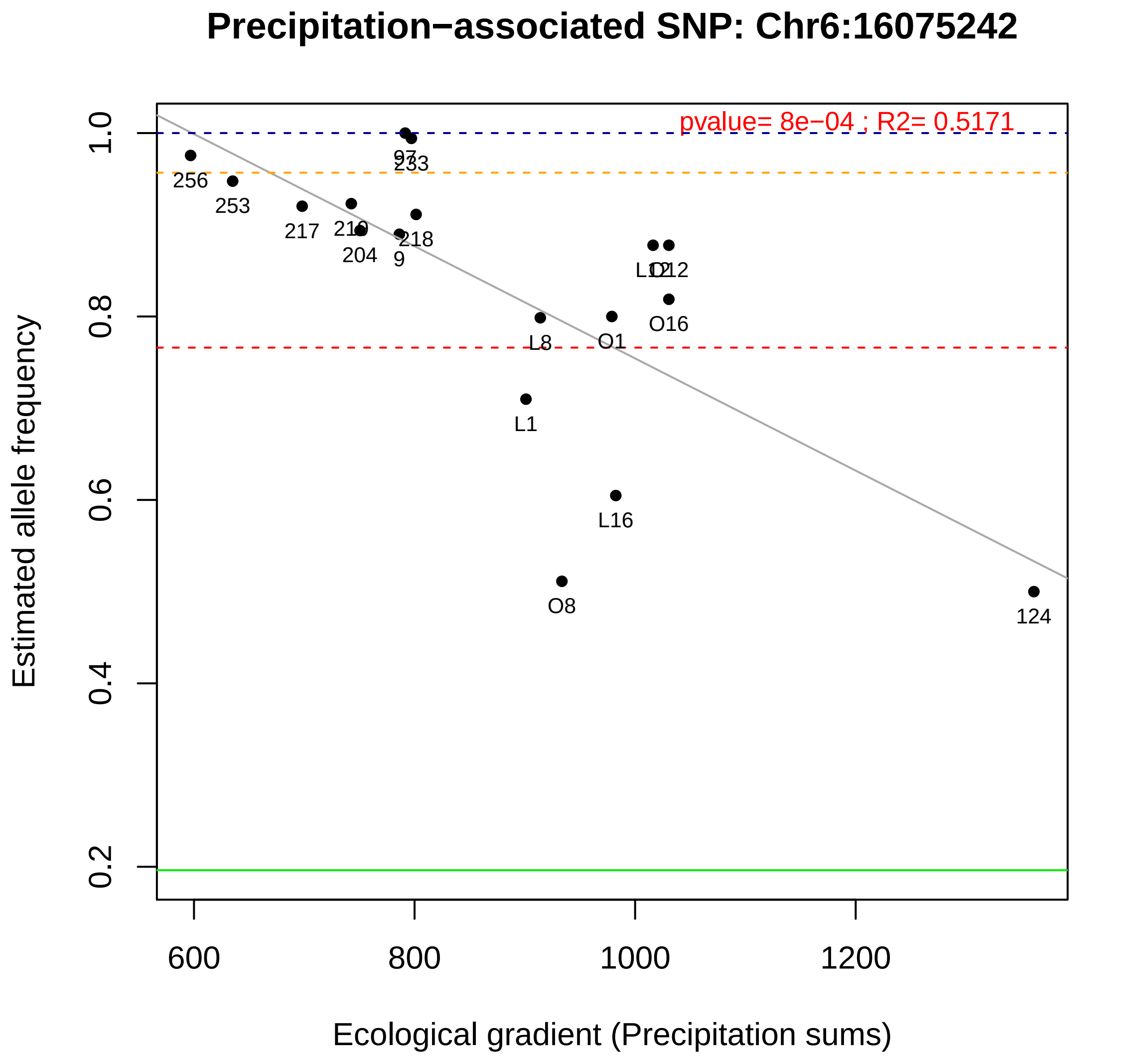

Precipitation−associated SNP: Chr6:16075242
pvalue= 8e−04 ; R2= 0.5171
Estimated allele frequency
Ecological gradient (Precipitation sums)
1.0
0.8
0.6
0.4
0.2
600
800
1000
1200
256
253
217
204
97
233
218
9
L12
O12
O16
L8
O1
L1
L16
O8
124

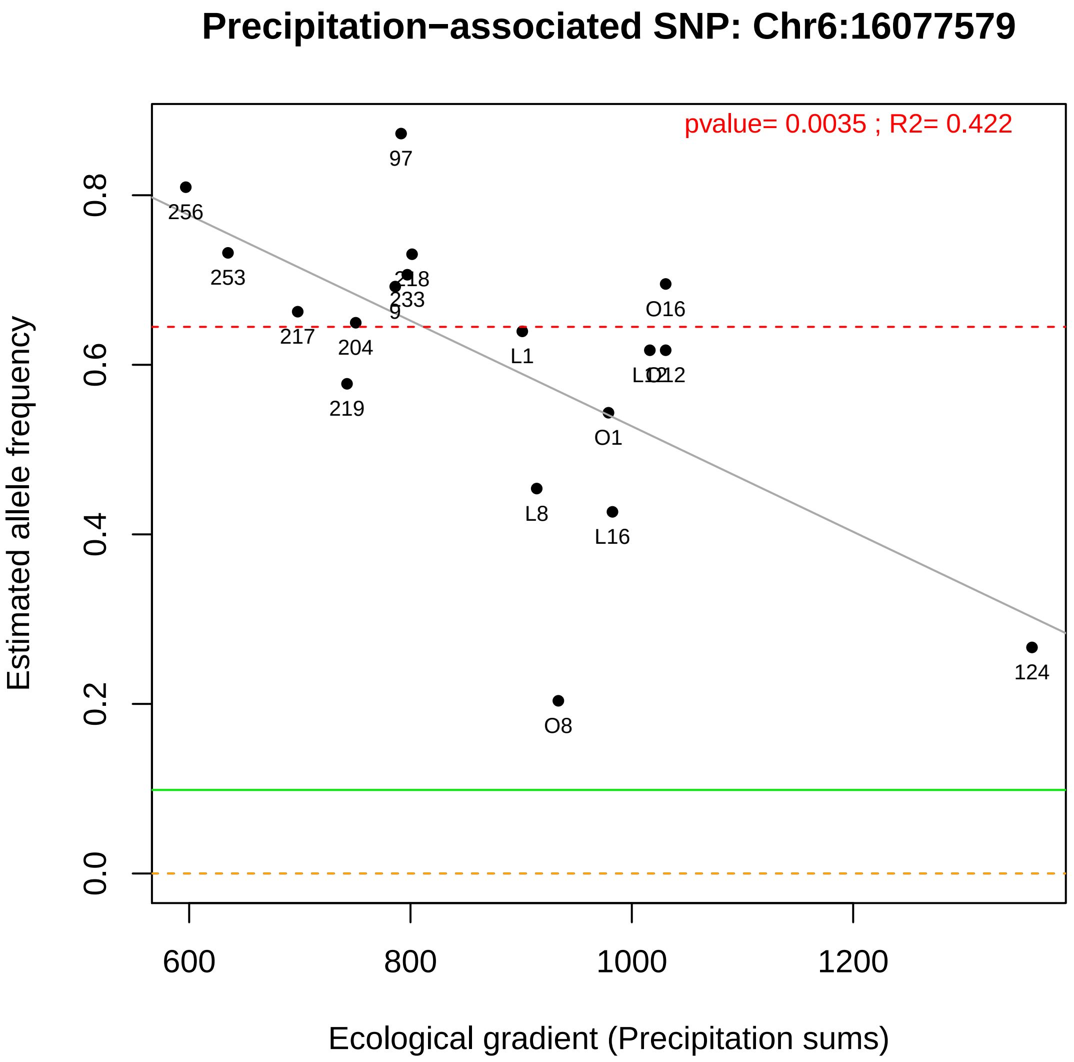

Precipitation−associated SNP: Chr6:16077579
pvalue= 0.0035 ; R2= 0.422
Estimated allele frequency
Ecological gradient (Precipitation sums)
0.0
0.2
0.4
0.6
0.8
600
800
1000
1200
97
256
253
218
233
9
217
204
219
L1
O16
O1
L8
L16
O8
124

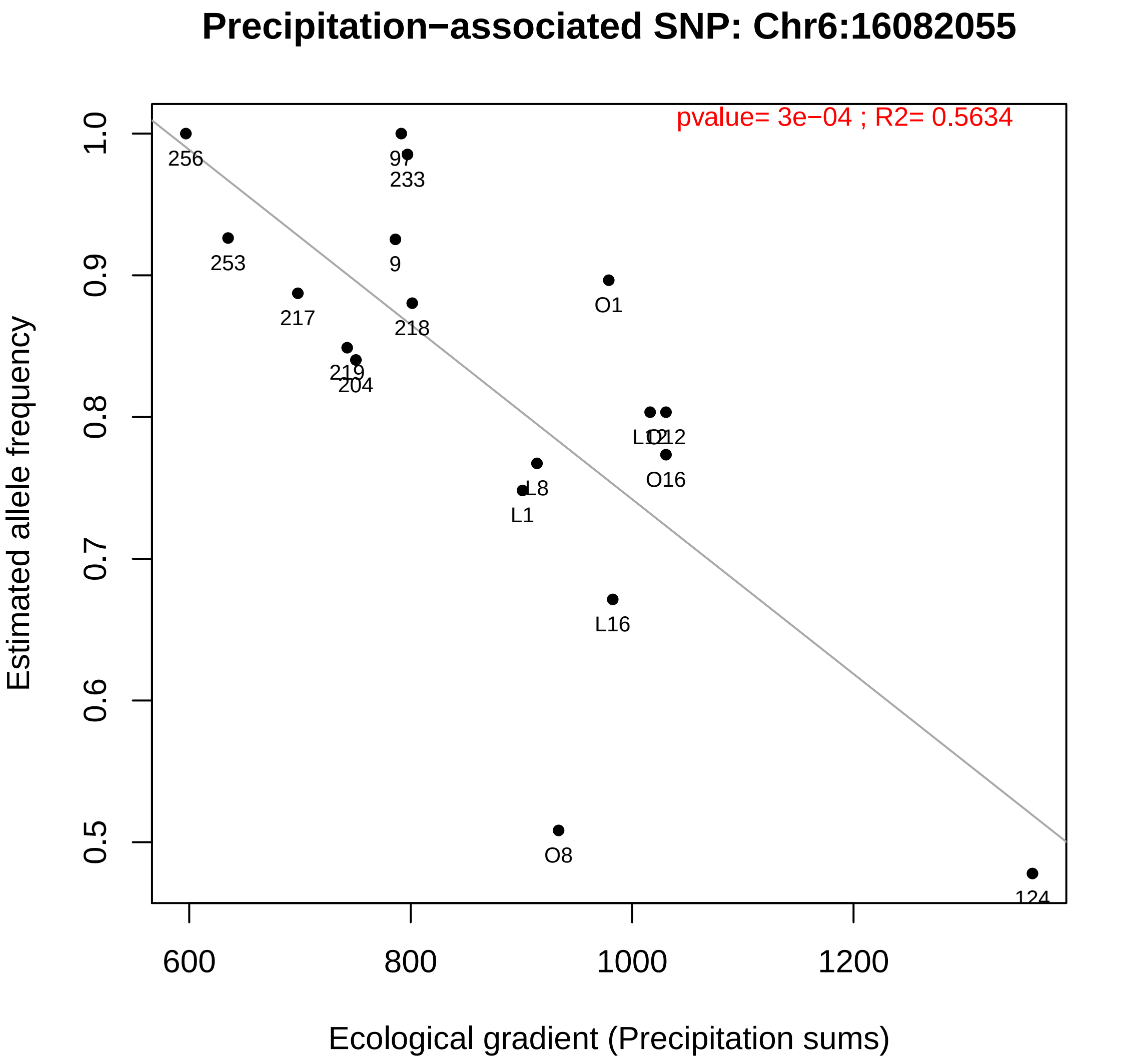

Precipitation−associated SNP: Chr6:16082055
pvalue= 3e−04 ; R2= 0.5634
Estimated allele frequency
Ecological gradient (Precipitation sums)
0.5
0.6
0.7
0.8
0.9
1.0
600
800
1000
1200
256
97
233
253
9
217
218
219
204
O1
L12
O12
O16
L8
L1
L16
O8
124

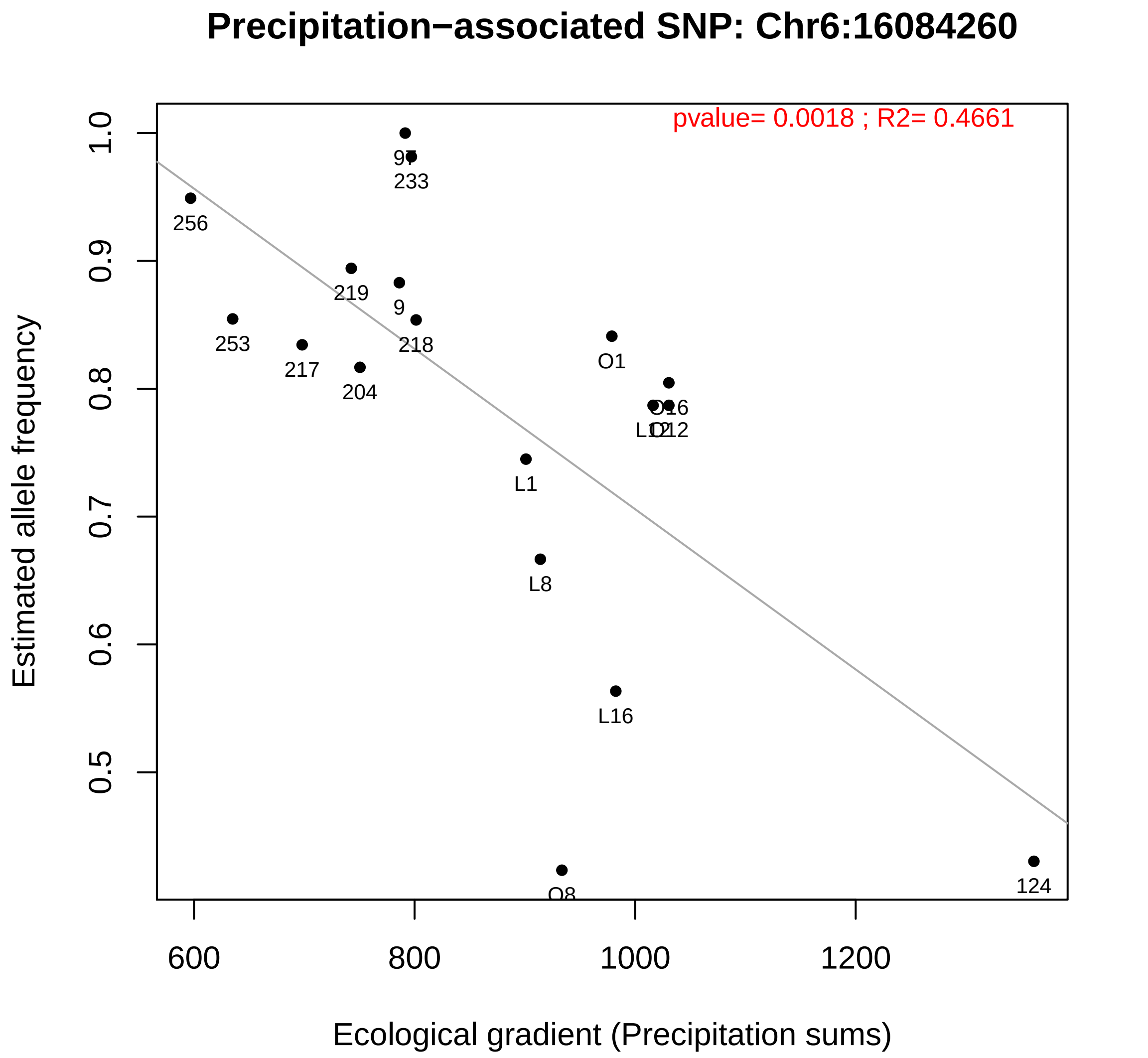
Precipitation−associated SNP: Chr6:16084260
pvalue= 0.0018 ; R2= 0.4661
Estimated allele frequency
1.0
0.9
0.8
0.7
0.6
0.5
600
800
1000
1200
Ecological gradient (Precipitation sums)
233
256
219
9
253
218
217
204
O1
L1
L8
L16
124

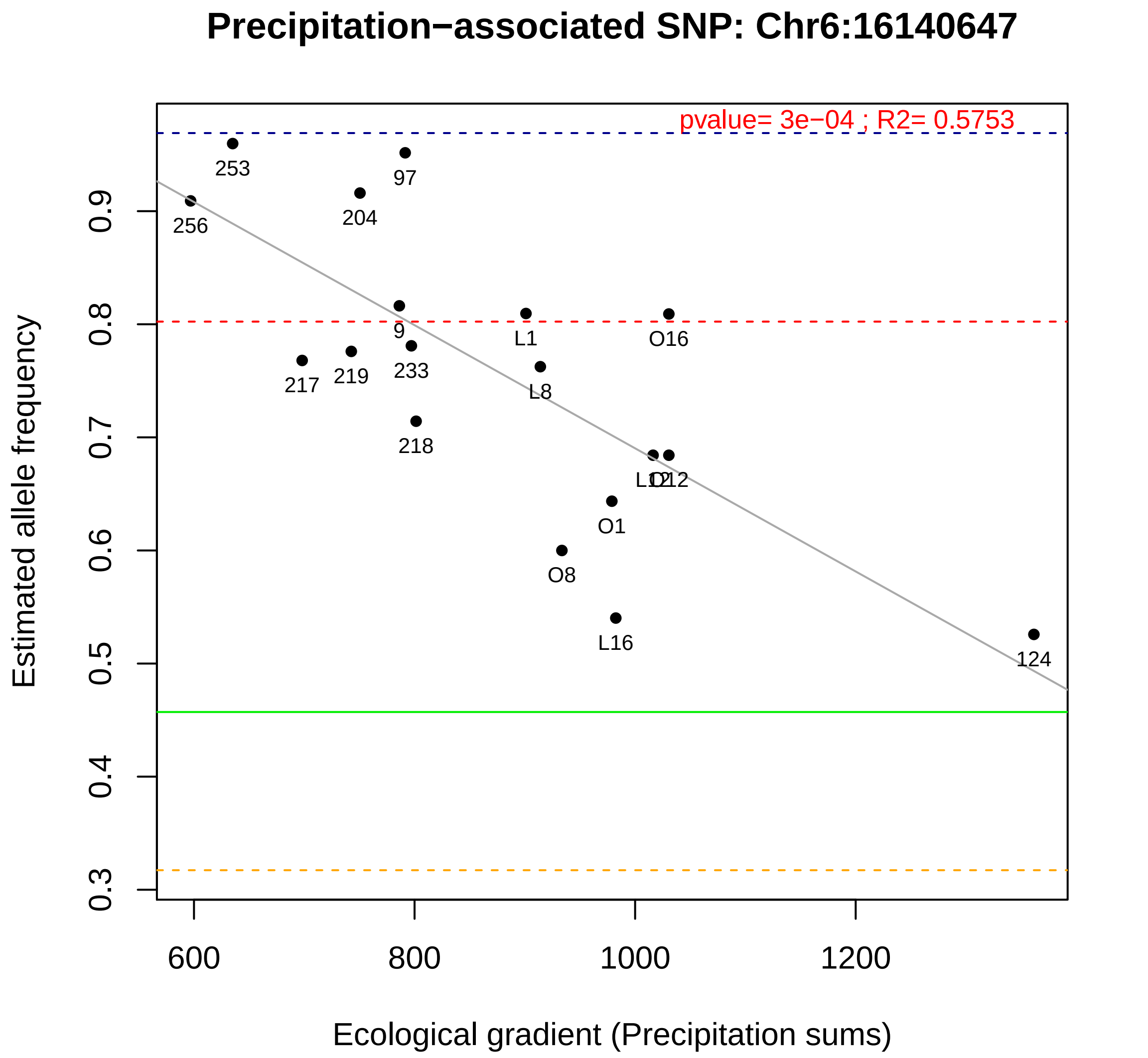

Precipitation−associated SNP: Chr6:16140647
pvalue= 3e−04 ; R2= 0.5753
Estimated allele frequency
0.3
0.4
0.5
0.6
0.7
0.8
0.9
600
800
1000
1200
Ecological gradient (Precipitation sums)
253
97
256
204
9
L1
O16
217
219
233
L8
218
L12
O12
O1
O8
L16
124

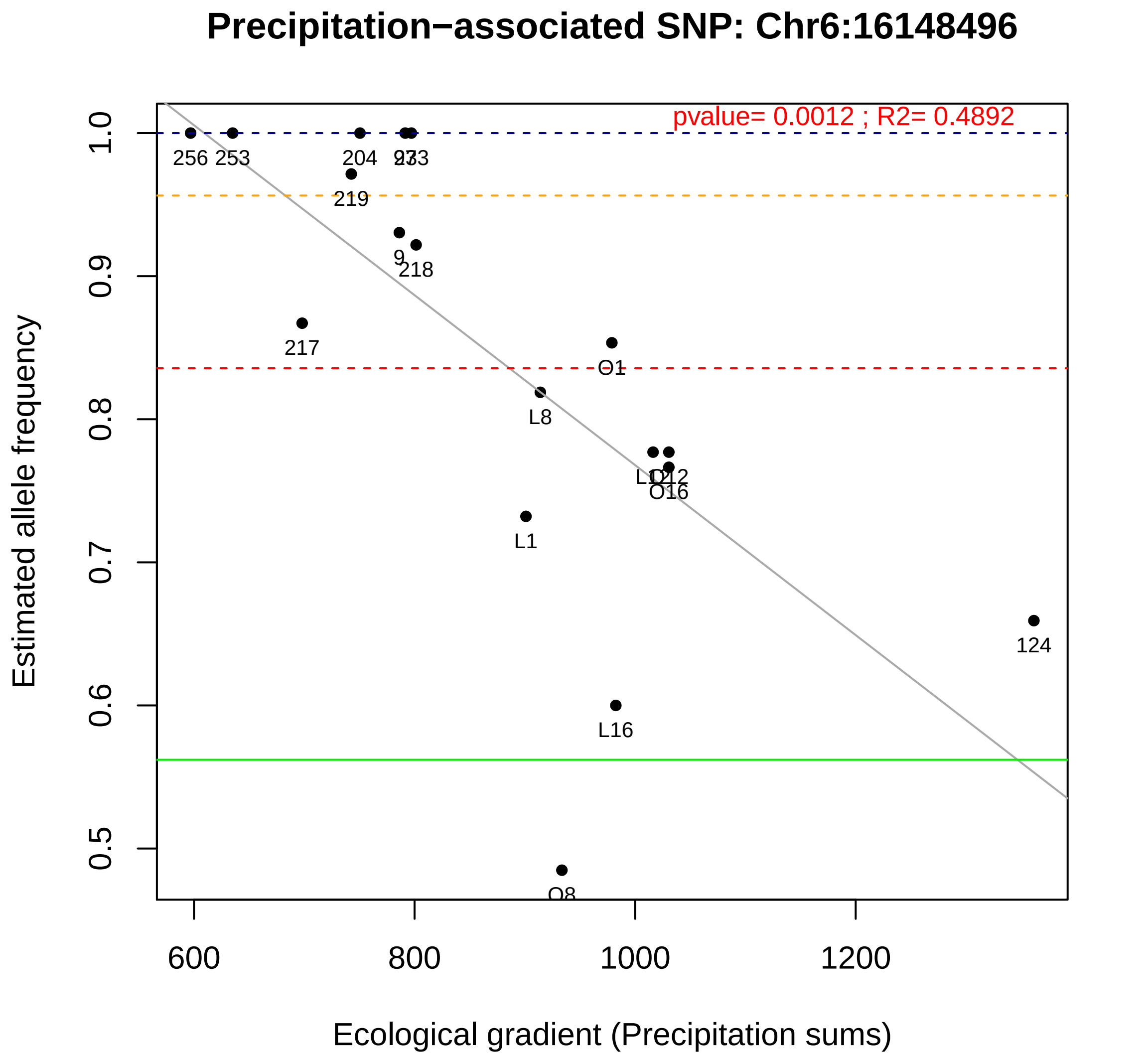

Precipitation−associated SNP: Chr6:16148496
pvalue= 0.0012 ; R2= 0.4892
Estimated allele frequency
Ecological gradient (Precipitation sums)
1.0
0.9
0.8
0.7
0.6
0.5
600
800
1000
1200
256
253
204
219
9
218
217
O1
L8
O16
L1
124
L16
O8

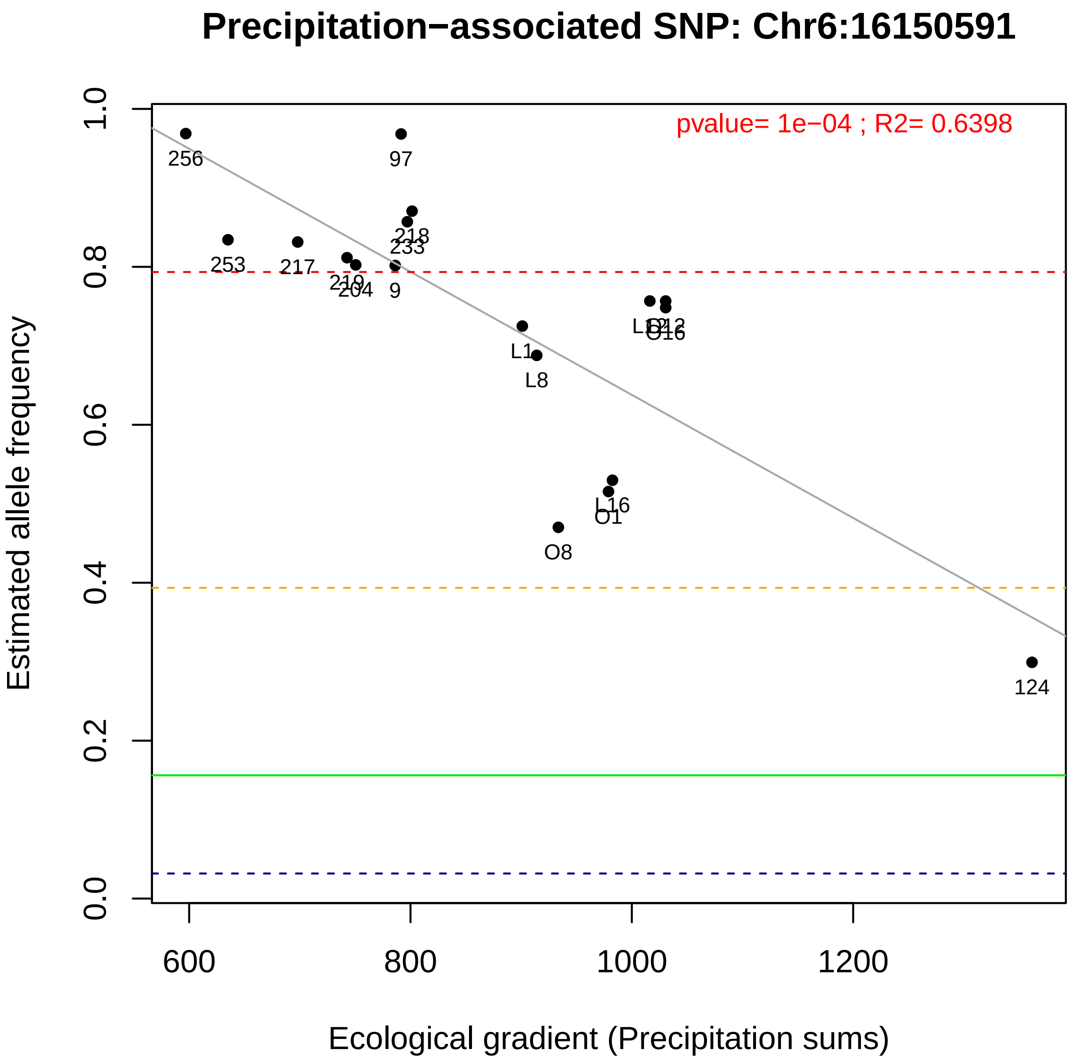

Precipitation−associated SNP: Chr6:16150591
pvalue= 1e−04 ; R2= 0.6398
Estimated allele frequency
Ecological gradient (Precipitation sums)
0.0
0.2
0.4
0.6
0.8
1.0
600
800
1000
1200
256
97
218
233
253
217
204
9
L1
L8
O16
L16
O1
O8
124

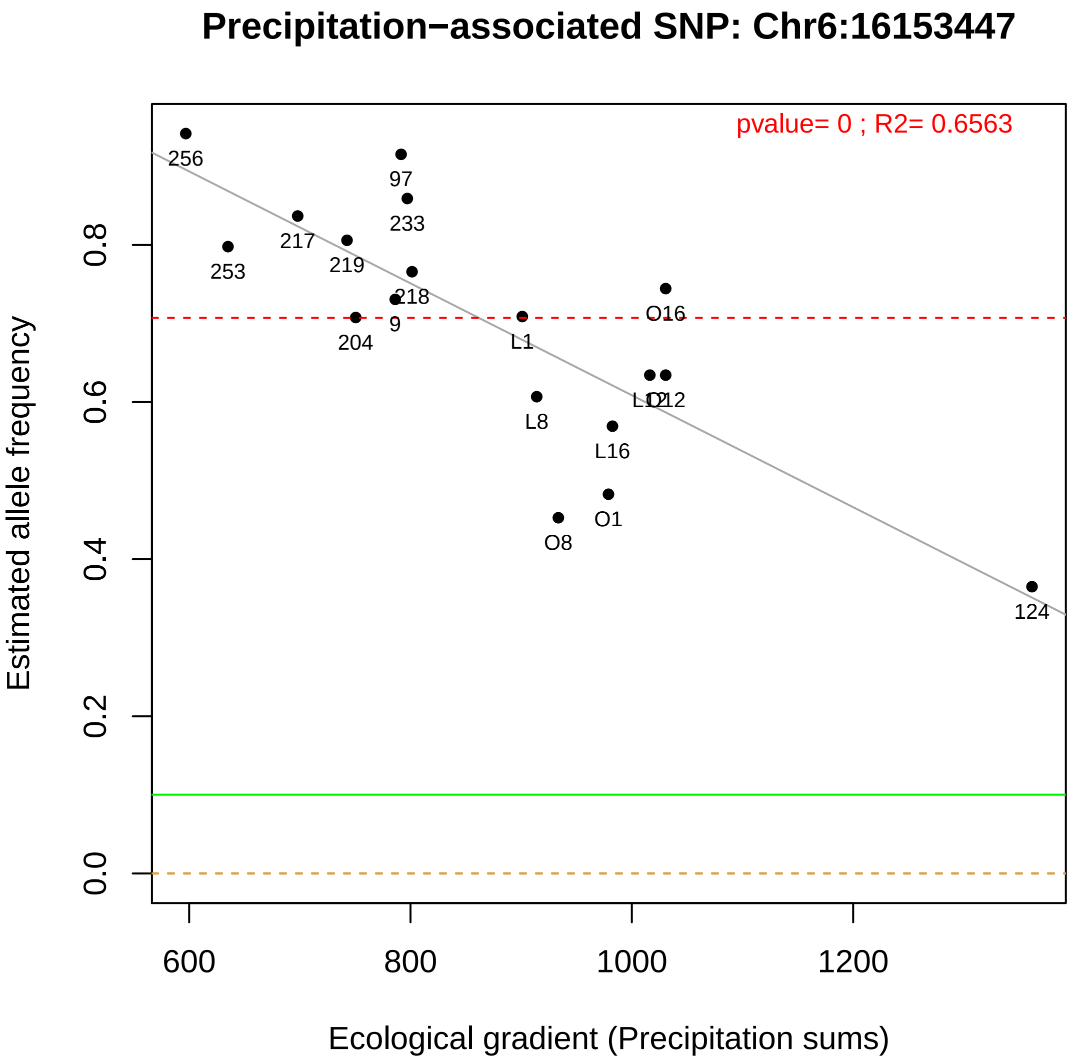

Precipitation−associated SNP: Chr6:16153447
pvalue= 0 ; R2= 0.6563
Estimated allele frequency
Ecological gradient (Precipitation sums)
0.0
0.2
0.4
0.6
0.8
600
800
1000
1200
256
97
233
217
253
219
218
9
204
L1
O16
L12
O12
L8
L16
O1
O8
124

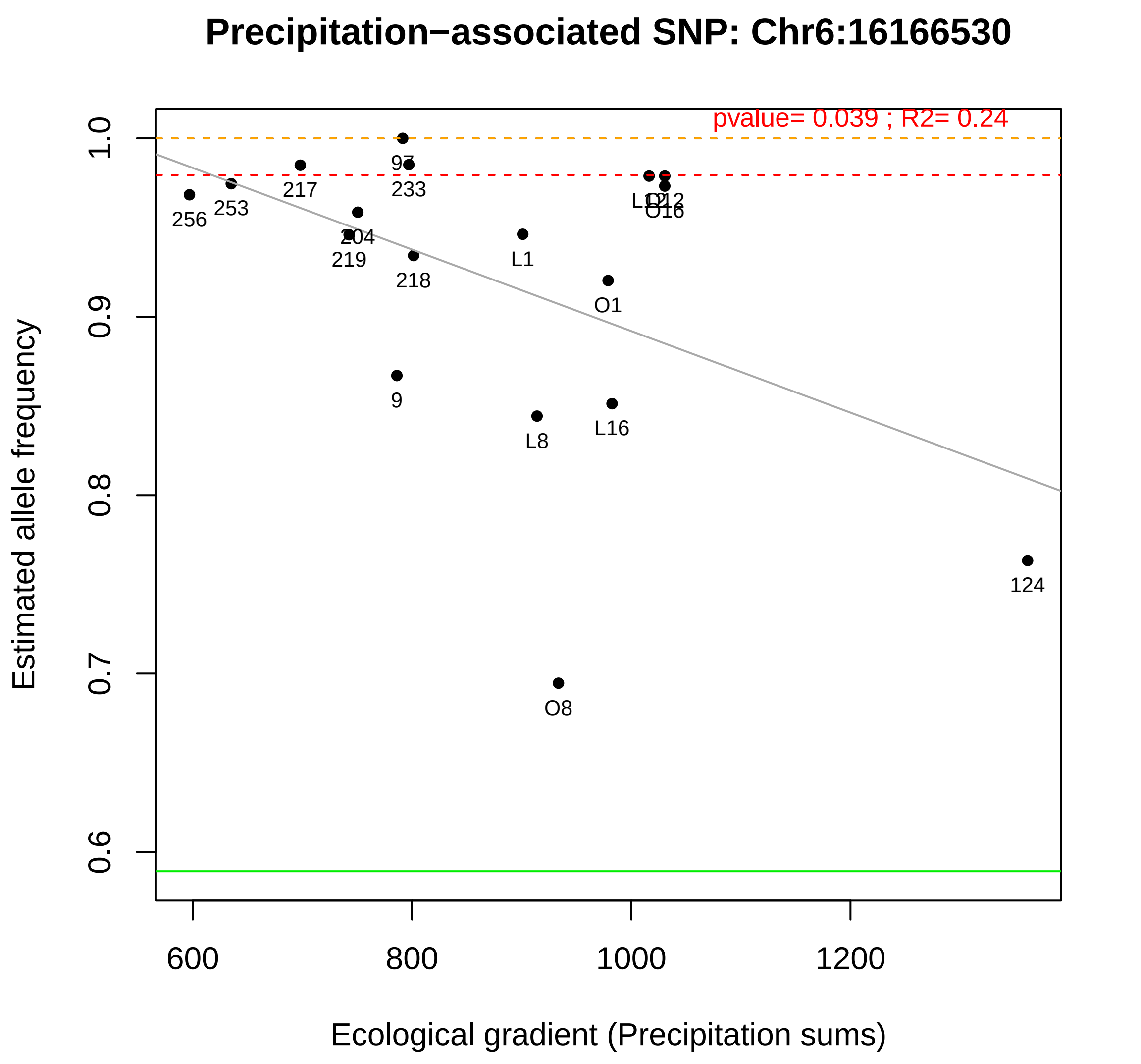
Precipitation−associated SNP: Chr6:16166530
pvalue= 0.039 ; R2= 0.24
Estimated allele frequency
Ecological gradient (Precipitation sums)
0.6
0.7
0.8
0.9
1.0
600
800
1000
1200
256
253
217
97
233
204
219
218
L1
O1
9
L8
L16
O8
124
O16

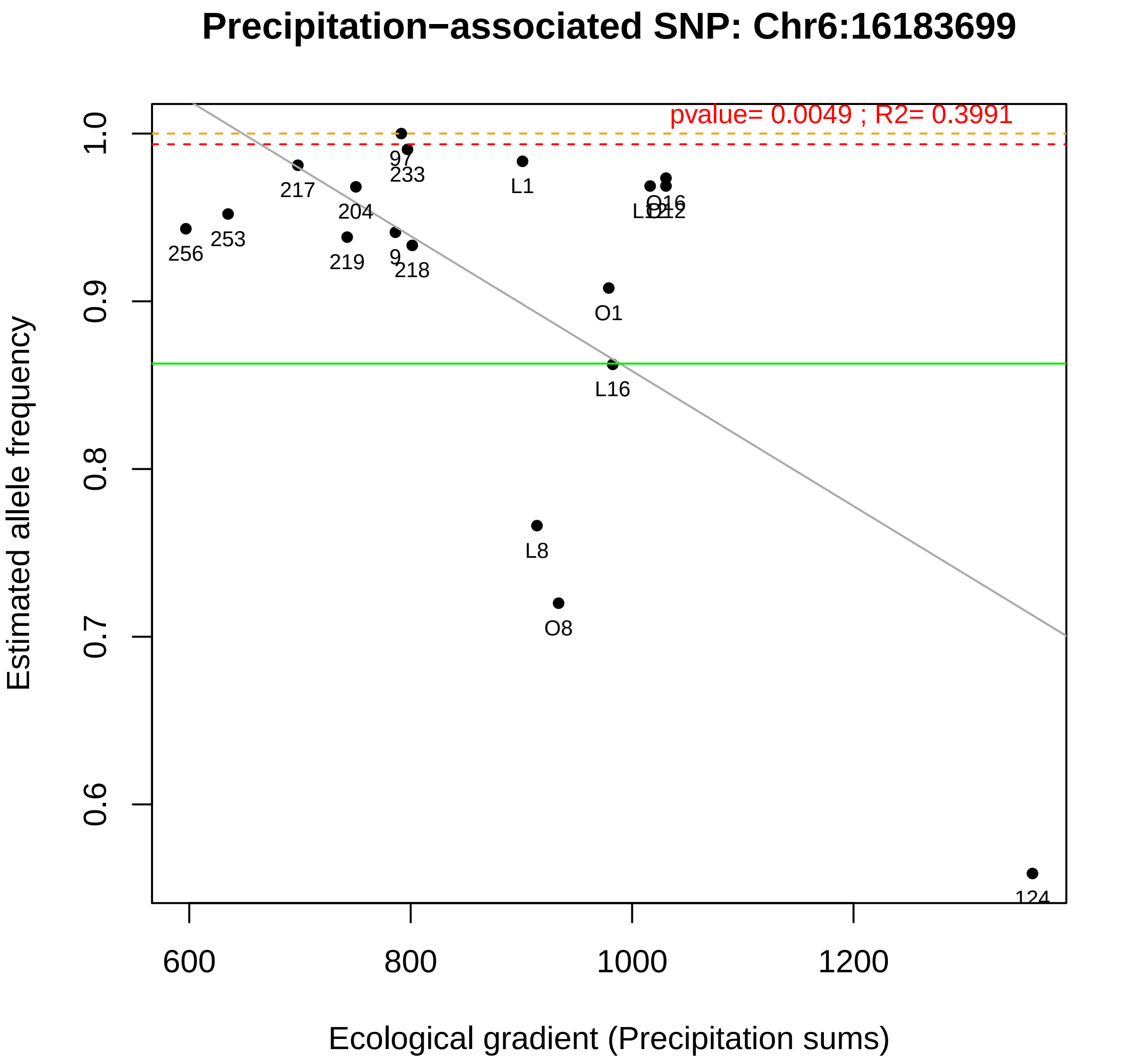
Precipitation−associated SNP: Chr6:16183699
pvalue= 0.0049 ; R2= 0.3991
Estimated allele frequency
Ecological gradient (Precipitation sums)
1.0
0.9
0.8
0.7
0.6
600
800
1000
1200
97
233
217
204
L1
O16
L12
O12
253
256
219
9
218
O1
L16
L8
O8
124

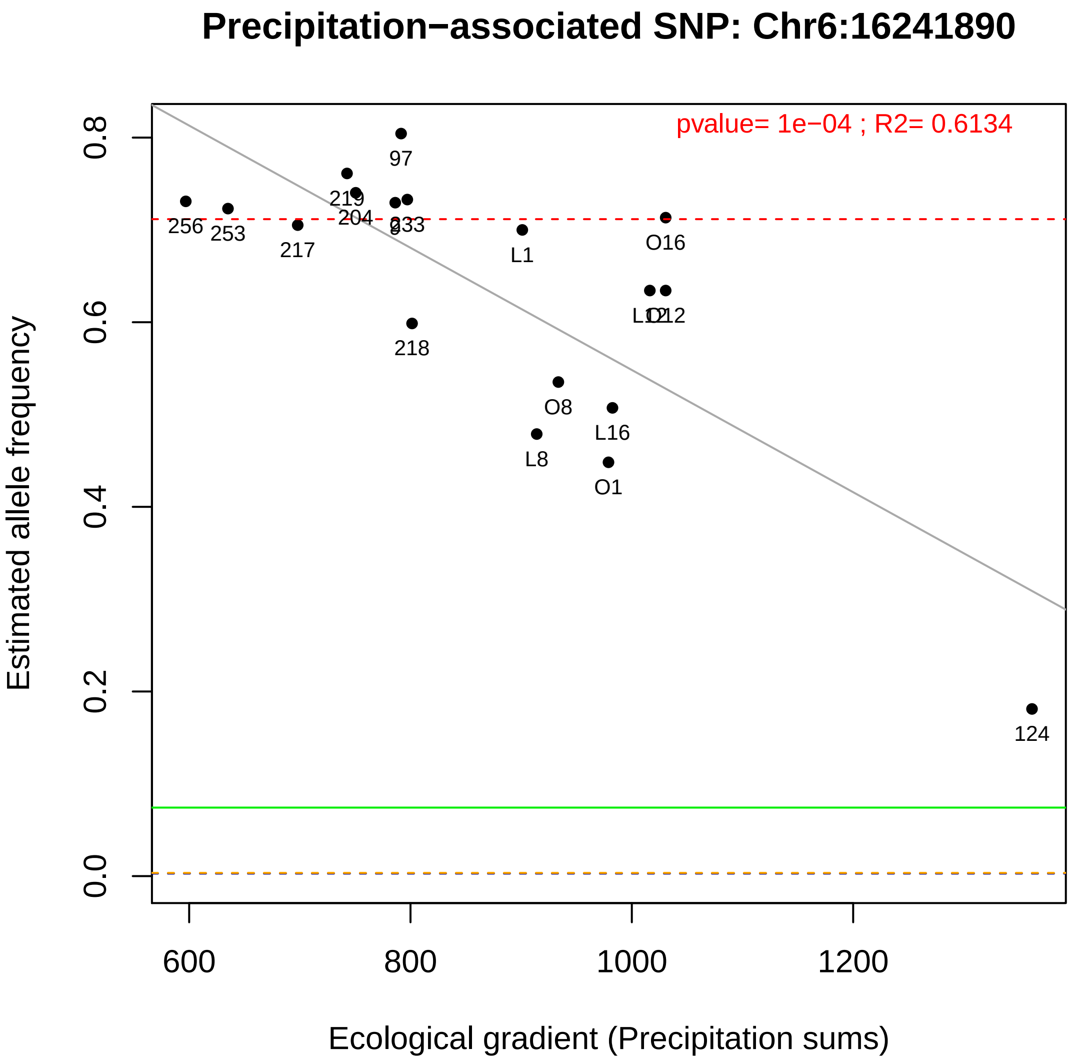

Precipitation−associated SNP: Chr6:16241890
pvalue= 1e−04 ; R2= 0.6134
97
219
204
256
253
217
233
L1
O16
L12
O12
218
O8
L16
L8
O1
124
0.0
0.2
0.4
0.6
0.8
Estimated allele frequency
600
800
1000
1200
Ecological gradient (Precipitation sums)

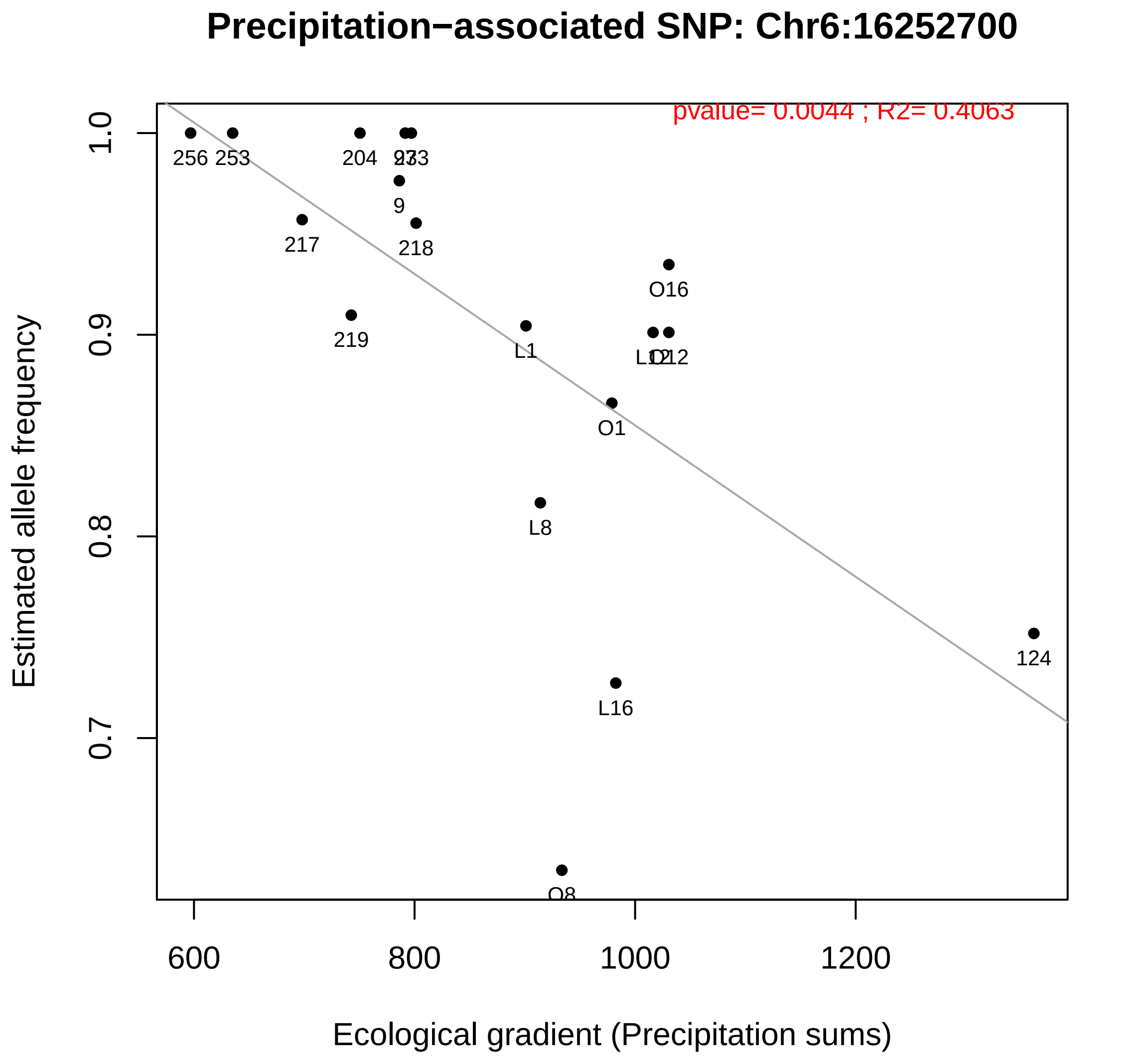
Precipitation−associated SNP: Chr6:16252700
pvalue= 0.0044 ; R2= 0.4063
Estimated allele frequency
1.0
0.9
0.8
0.7
600
800
1000
1200
Ecological gradient (Precipitation sums)
256
253
204
9
217
218
219
O16
L1
O1
L8
124
L16
O8

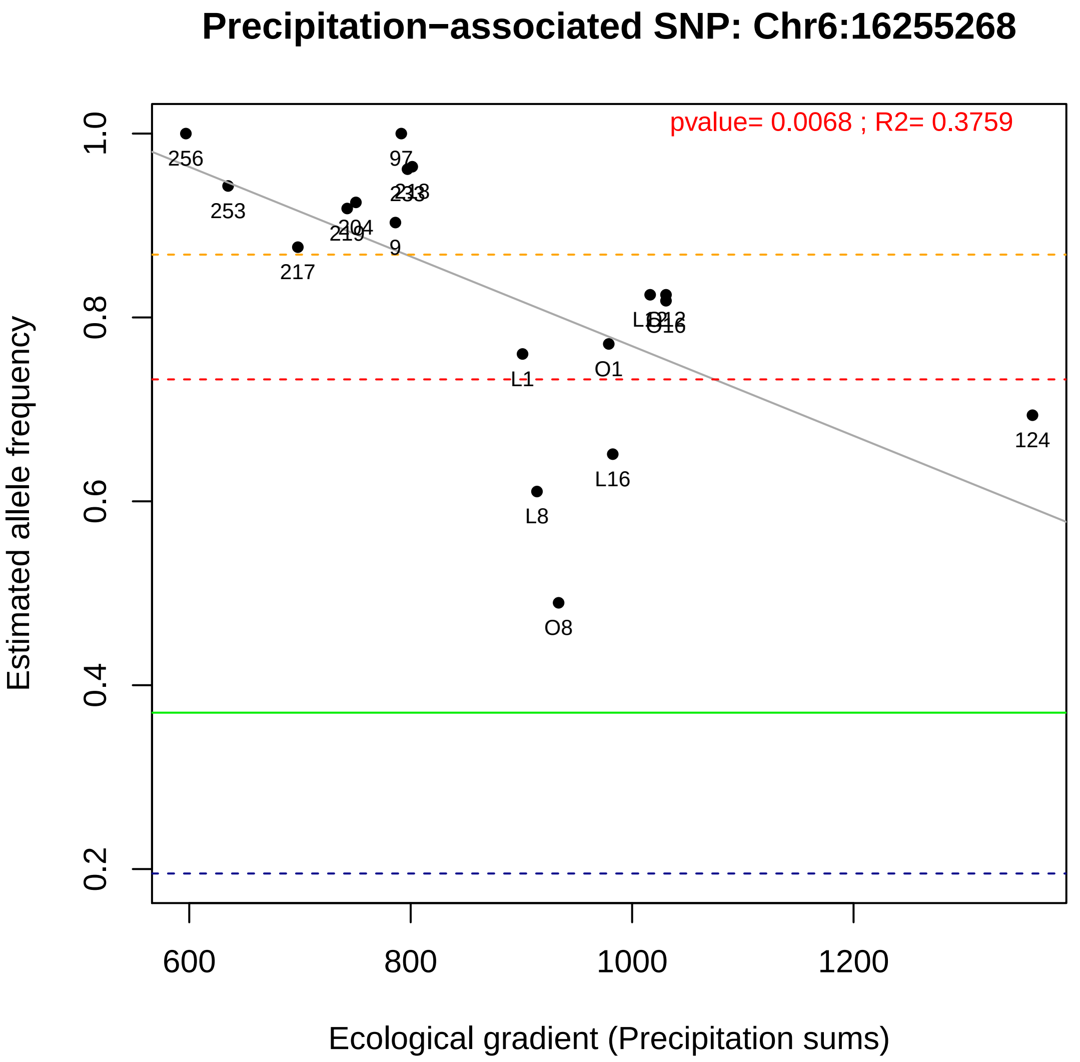

Precipitation−associated SNP: Chr6:16255268
pvalue= 0.0068 ; R2= 0.3759
Estimated allele frequency
Ecological gradient (Precipitation sums)
0.2
0.4
0.6
0.8
1.0
600
800
1000
1200
256
253
217
219
204
97
233
218
9
L1
L8
O8
O1
L16
L12
O12
O16
124

#### Precipitation−associated SNP: Chr6:16260071

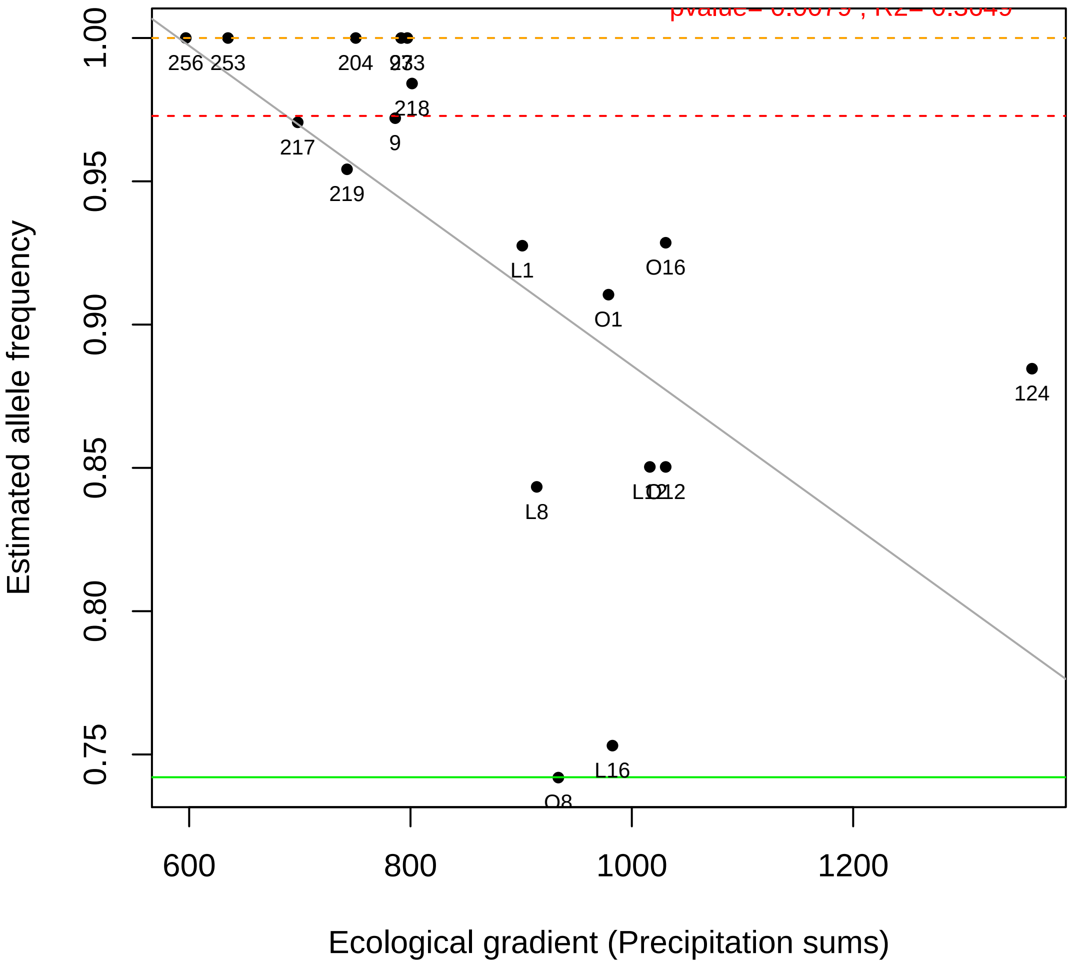

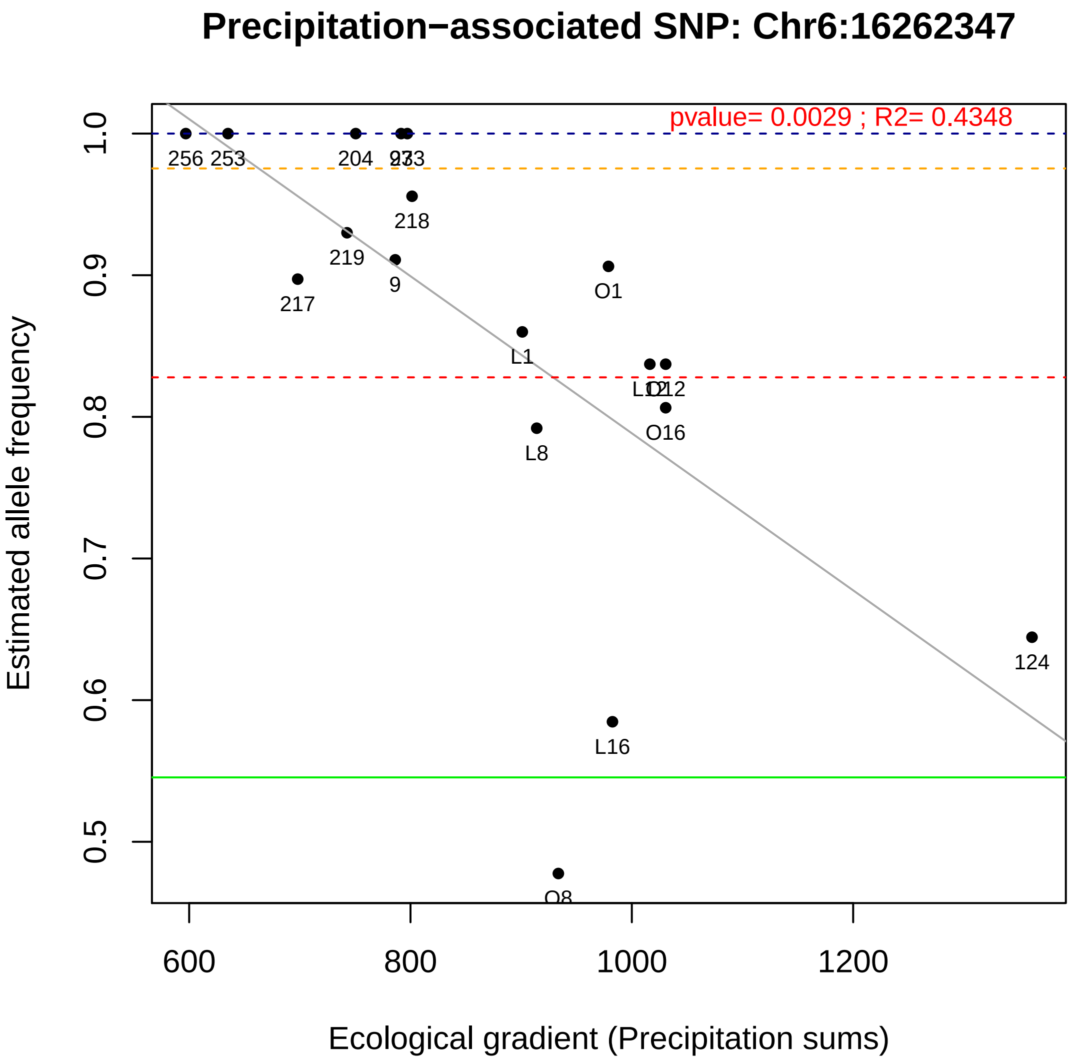
Precipitation−associated SNP: Chr6:16262347
pvalue= 0.0029 ; R2= 0.4348
Estimated allele frequency
Ecological gradient (Precipitation sums)
1.0
0.9
0.8
0.7
0.6
0.5
600
800
1000
1200
256
253
204
218
219
9
217
O1
L1
O12
O16
L8
124
L16

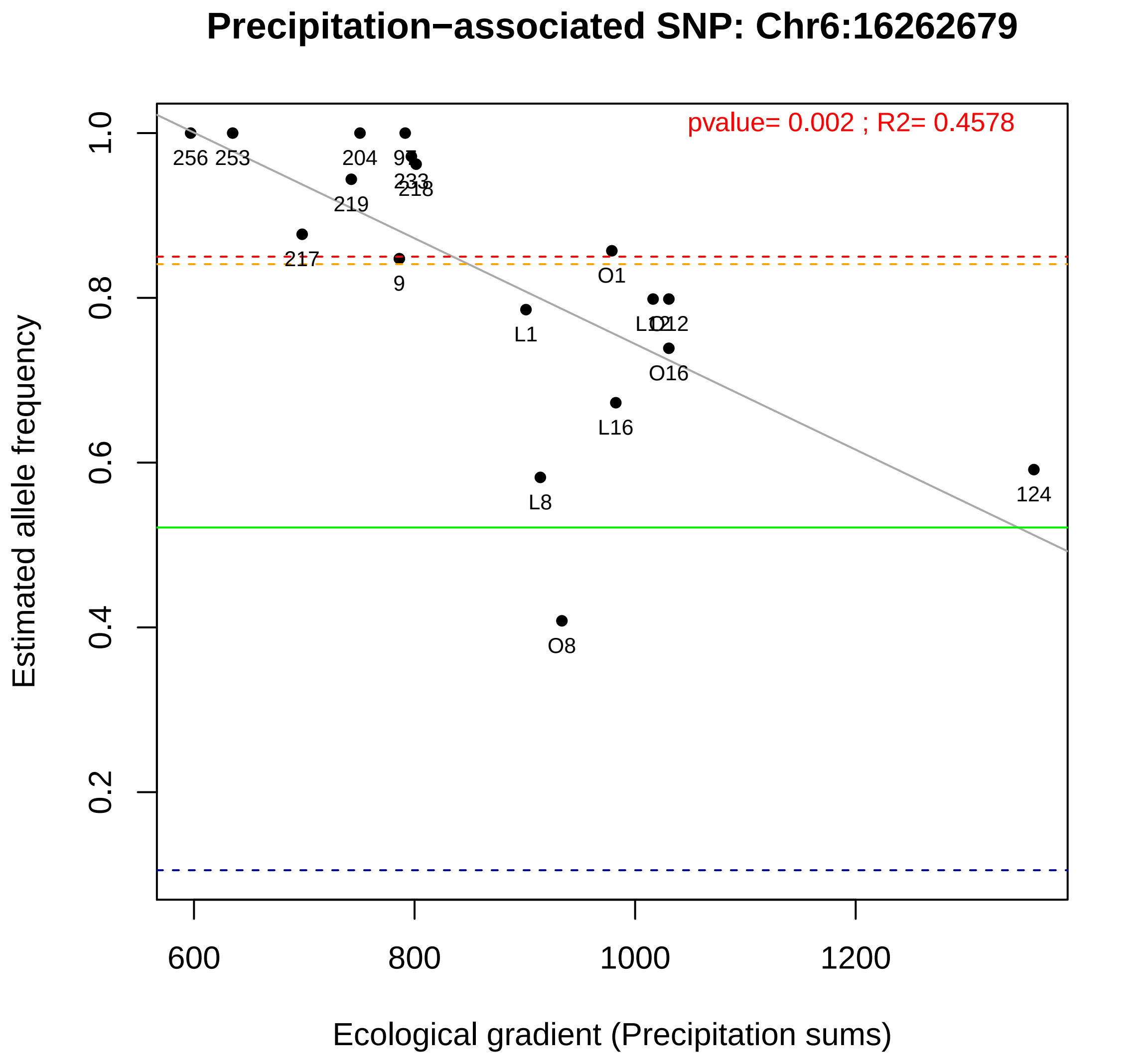

Precipitation−associated SNP: Chr6:16262679
pvalue= 0.002 ; R2= 0.4578
Estimated allele frequency
1.0
0.8
0.6
0.4
0.2
600
800
1000
1200
Ecological gradient (Precipitation sums)
256
253
204
219
217
9
L1
L8
O8
O1
L16
O16
124

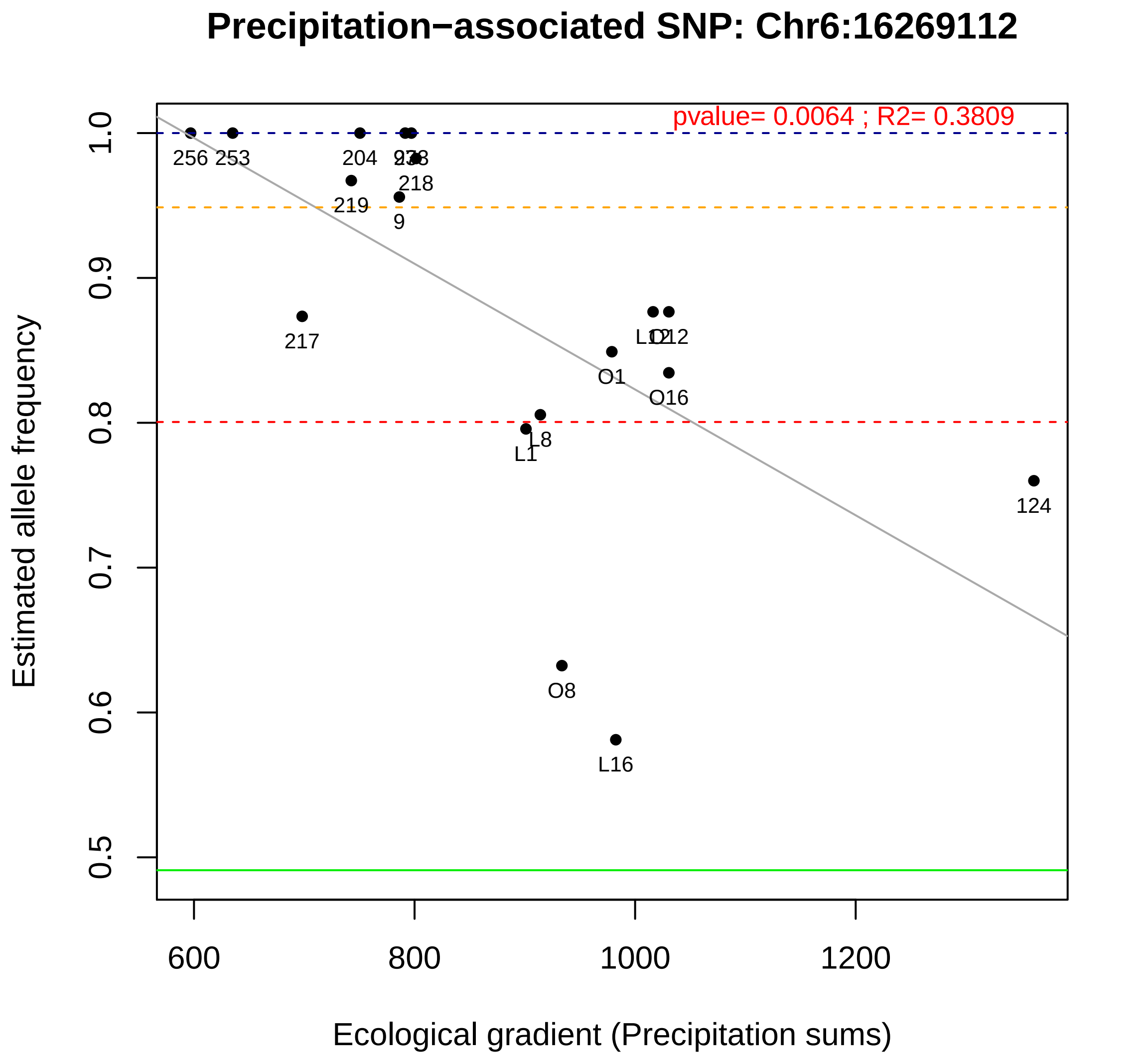
Precipitation−associated SNP: Chr6:16269112
pvalue= 0.0064 ; R2= 0.3809
Estimated allele frequency
Ecological gradient (Precipitation sums)
0.5
0.6
0.7
0.8
0.9
1.0
600
800
1000
1200
256
253
204
218
219
9
217
L12
O12
O1
O16
L8
L1
124
O8
L16

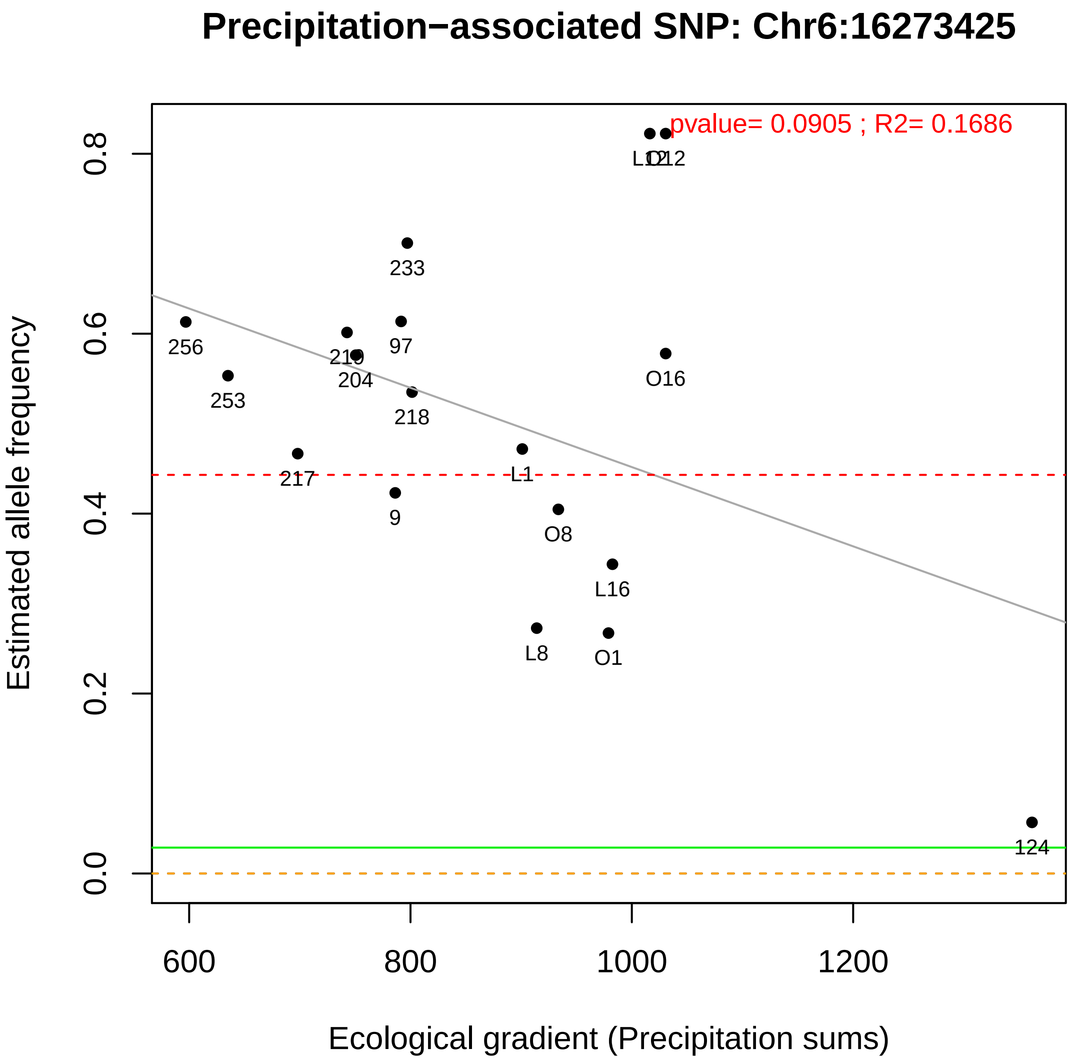
Precipitation−associated SNP: Chr6:16273425
pvalue= 0.0905 ; R2= 0.1686
Estimated allele frequency
Ecological gradient (Precipitation sums)
0.0
0.2
0.4
0.6
0.8
600
800
1000
1200
L12
O12
233
256
219
204
97
O16
253
218
217
L1
9
O8
L16
L8
O1
124

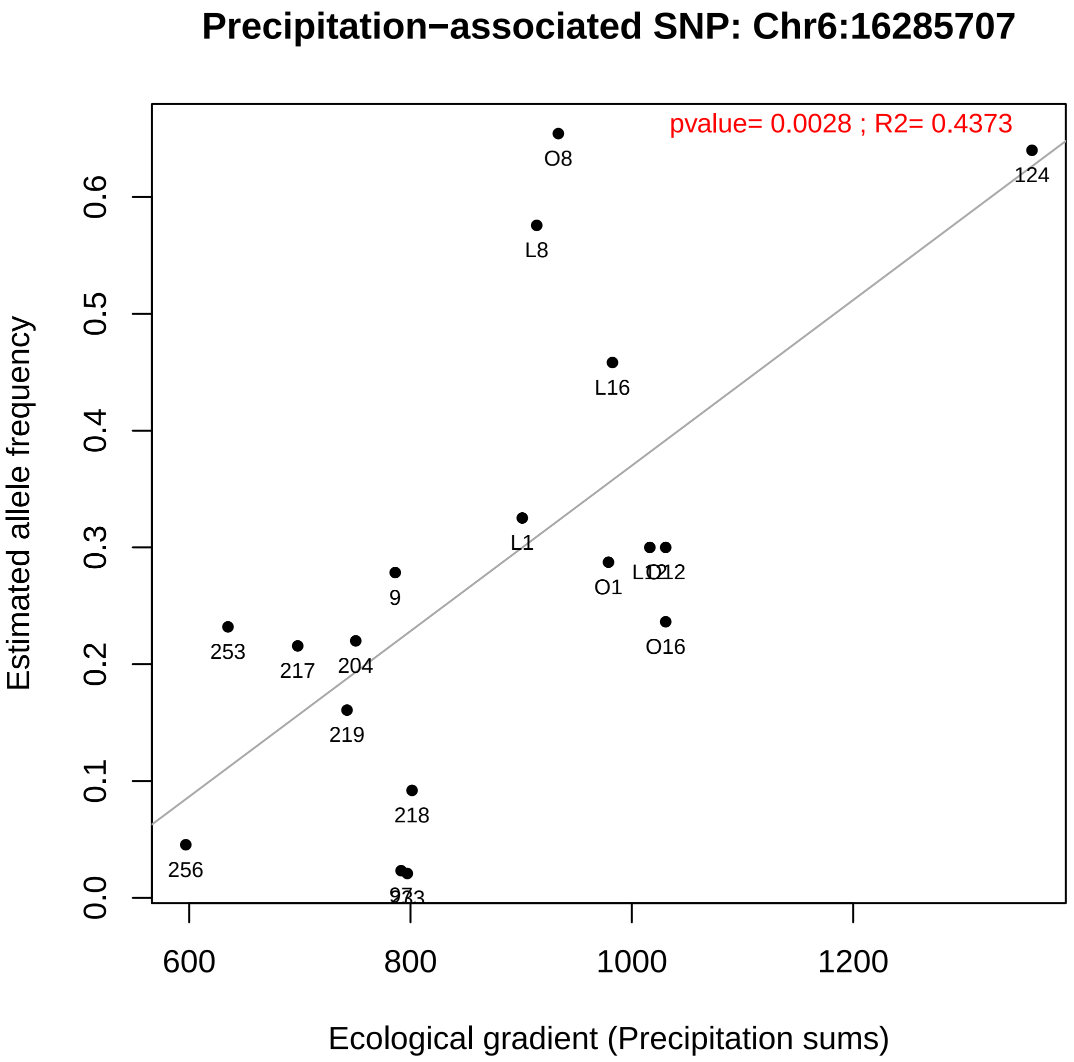

Precipitation−associated SNP: Chr6:16285707
pvalue= 0.0028 ; R2= 0.4373
Estimated allele frequency
Ecological gradient (Precipitation sums)
0.0
0.1
0.2
0.3
0.4
0.5
0.6
600
800
1000
1200
O8
124
L8
L16
L1
O1
O16
9
253
217
204
219
218
256

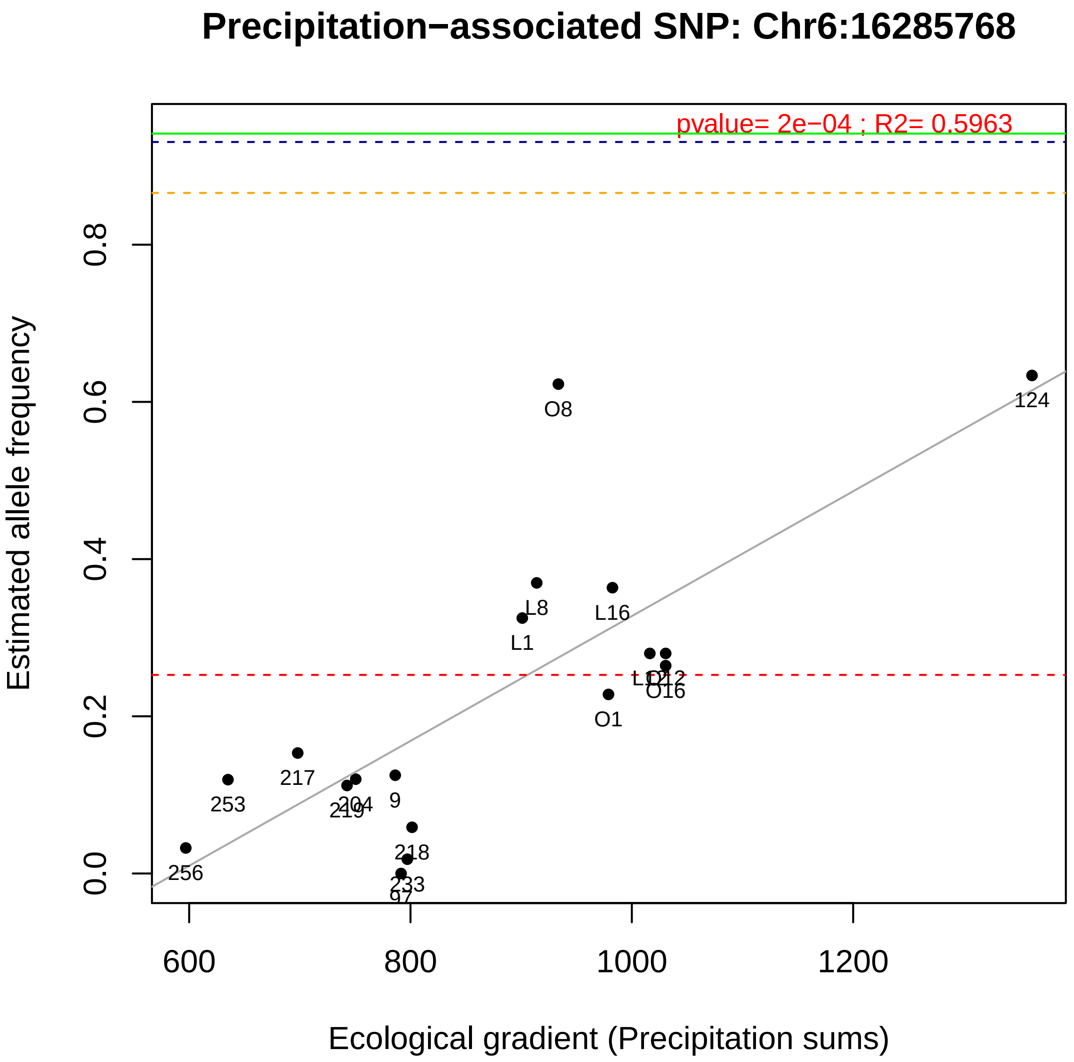
Precipitation−associated SNP: Chr6:16285768
pvalue= 2e−04 ; R2= 0.5963
Estimated allele frequency
0.0
0.2
0.4
0.6
0.8
600
800
1000
1200
Ecological gradient (Precipitation sums)
O8
124
L8
L16
L1
O16
O1
217
253
204
9
218
233
256

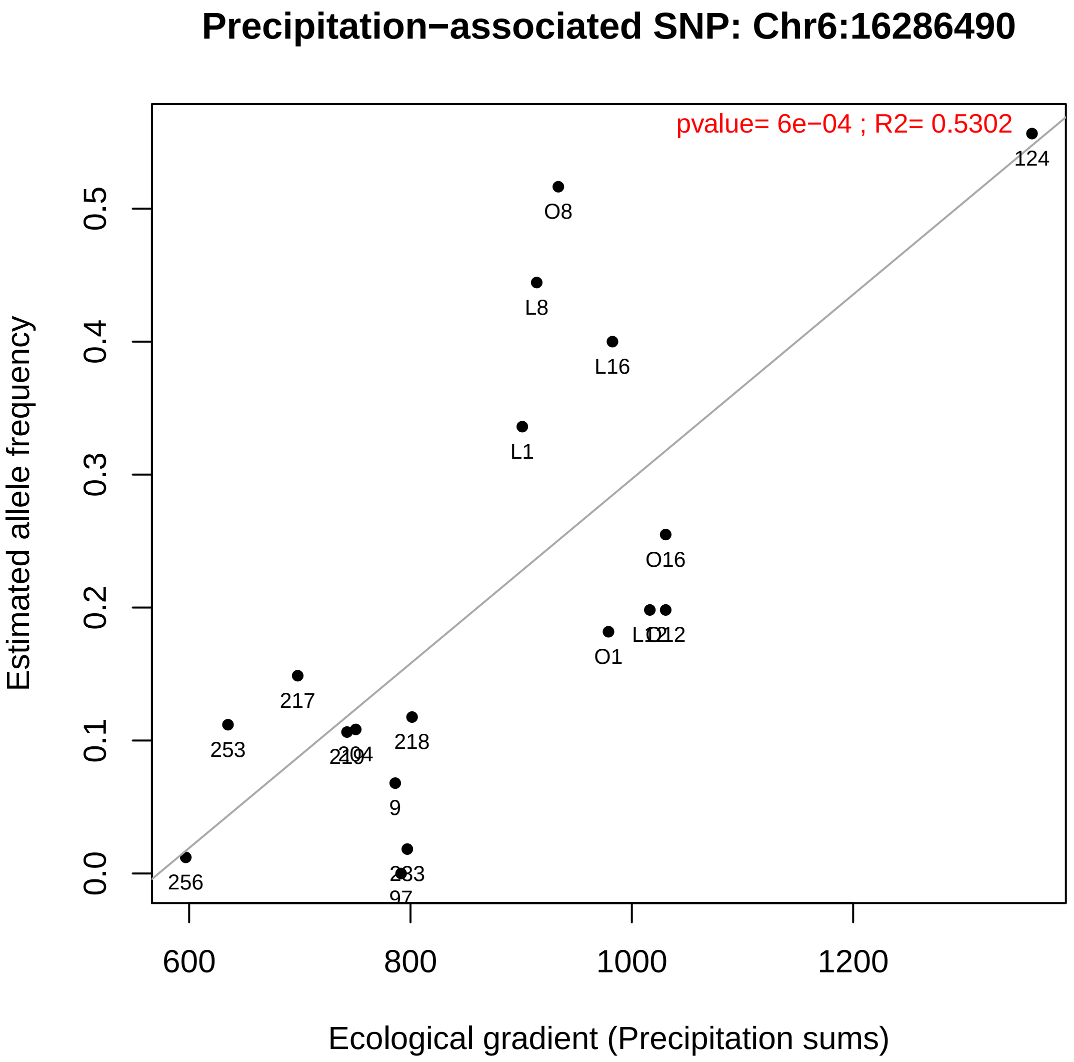
Precipitation−associated SNP: Chr6:16286490
pvalue= 6e−04 ; R2= 0.5302
Estimated allele frequency
Ecological gradient (Precipitation sums)
0.0
0.1
0.2
0.3
0.4
0.5
600
800
1000
1200
124
O8
L8
L16
L1
O16
O1
217
253
218
9
256
97

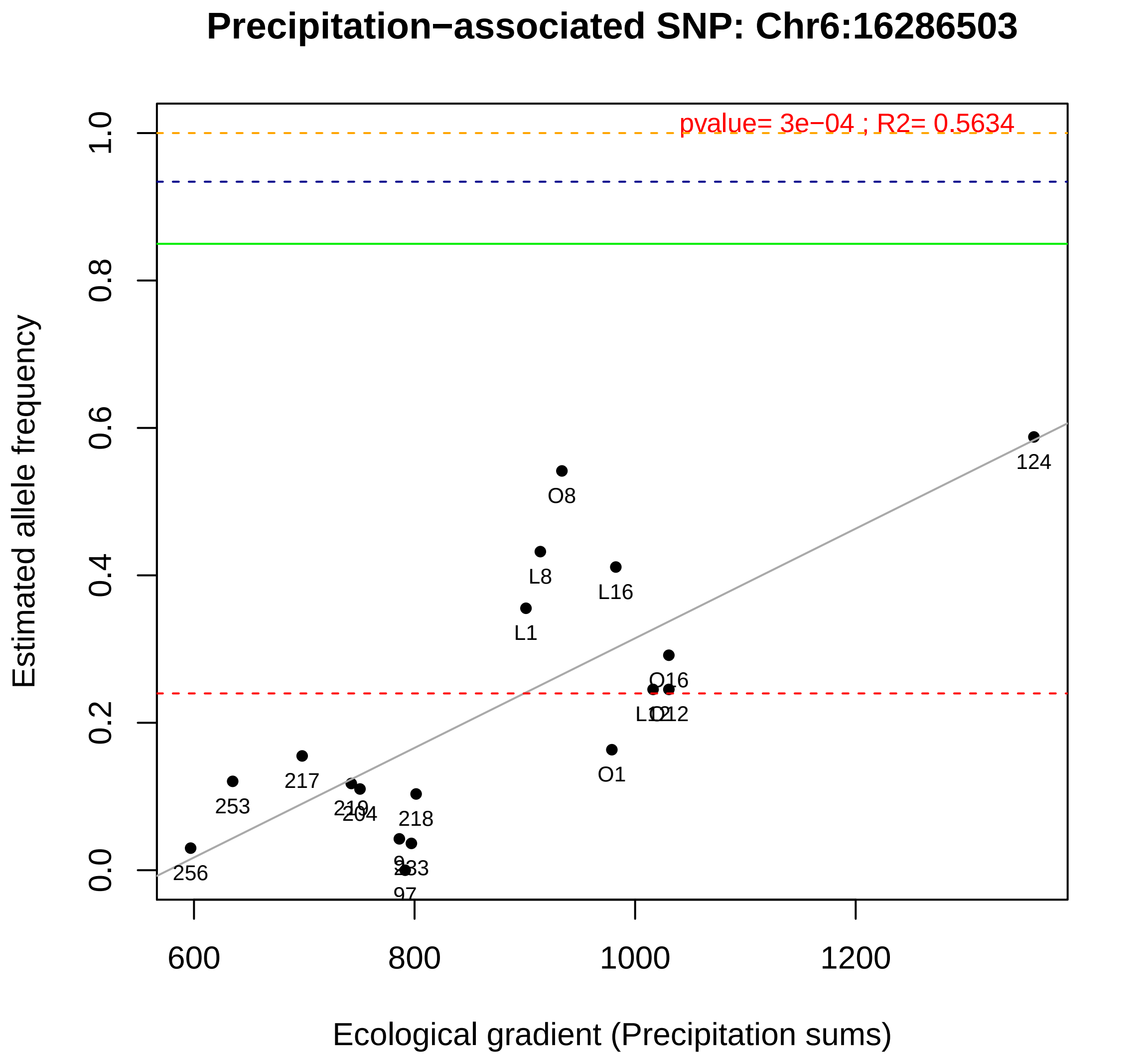
Precipitation−associated SNP: Chr6:16286503
pvalue= 3e−04 ; R2= 0.5634
Estimated allele frequency
Ecological gradient (Precipitation sums)
0.0
0.2
0.4
0.6
0.8
1.0
600
800
1000
1200
124
O8
L8
L16
L1
O16
L12
O12
O1
217
253
219
204
218
256
9
233
97

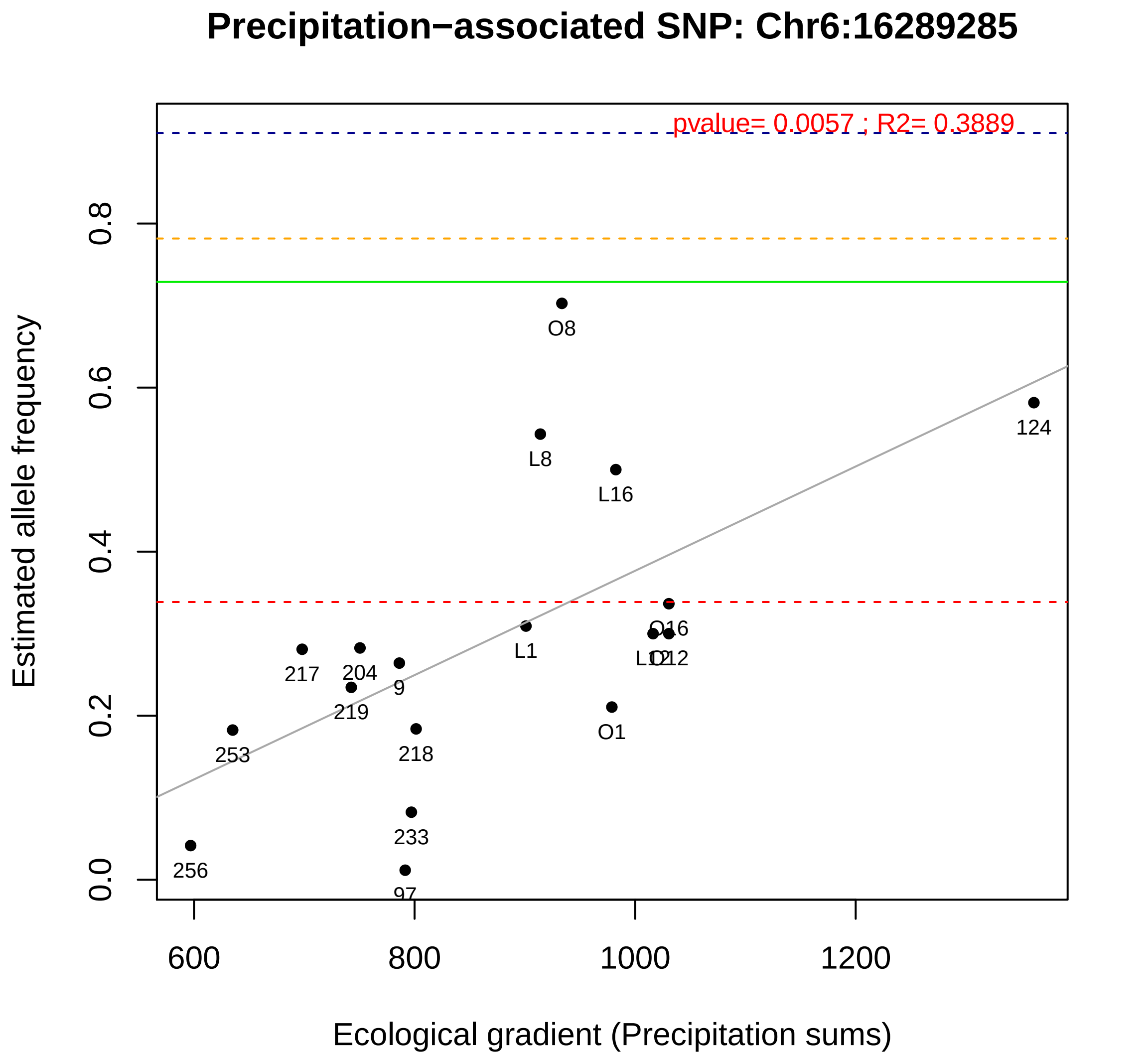

Precipitation−associated SNP: Chr6:16289285
pvalue= 0.0057 ; R2= 0.3889
Estimated allele frequency
0.0
0.2
0.4
0.6
0.8
600
800
1000
1200
Ecological gradient (Precipitation sums)
O8
124
L8
L16
O16
L1
L12
O12
217
204
9
219
O1
253
218
233
256
97

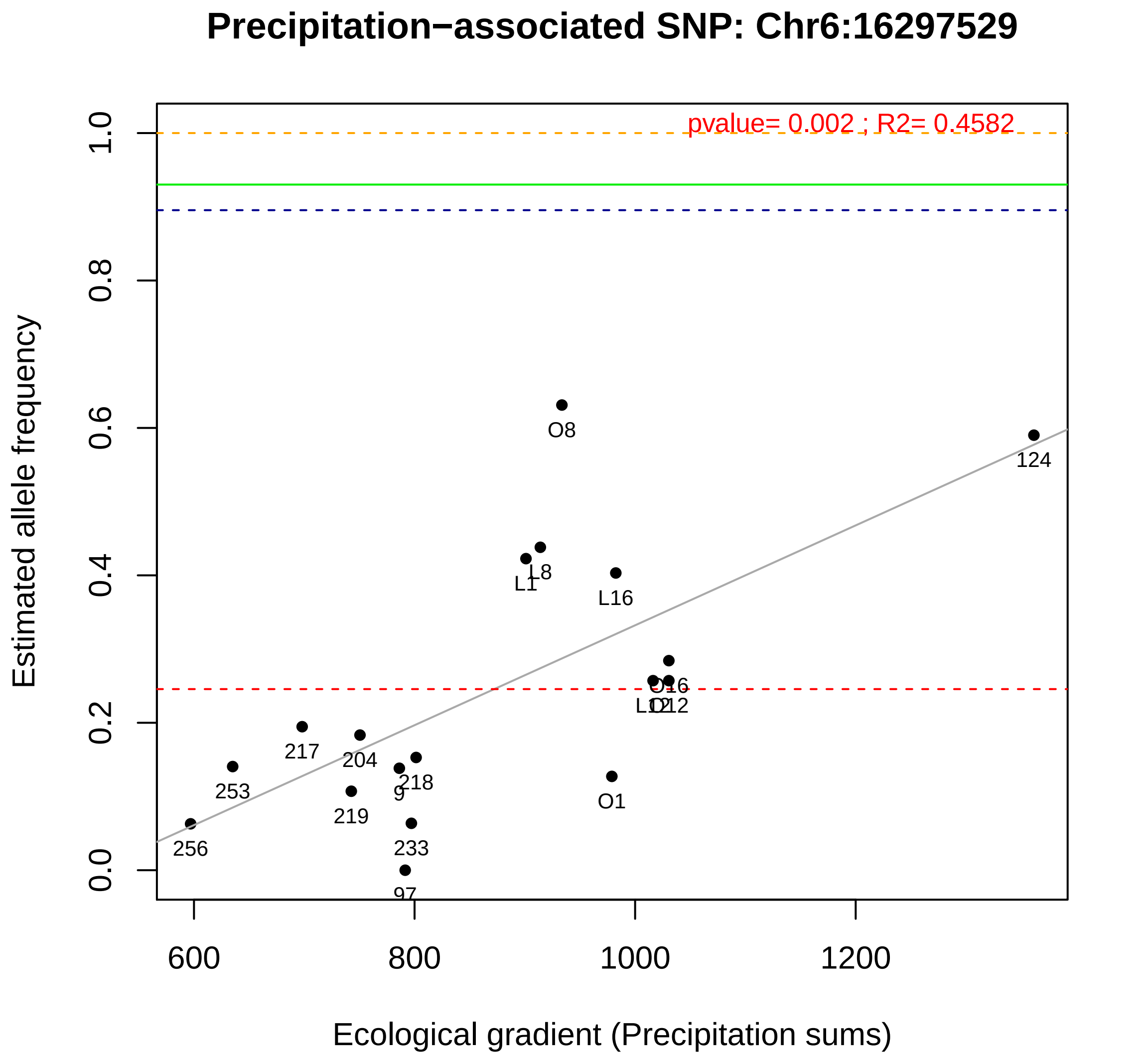

Precipitation−associated SNP: Chr6:16297529
pvalue= 0.002 ; R2= 0.4582
Estimated allele frequency
0.0
0.2
0.4
0.6
0.8
1.0
600
800
1000
1200
Ecological gradient (Precipitation sums)
O8
124
L8
L1
L16
O16
L12
O12
217
204
253
218
9
219
O1
256
233
97

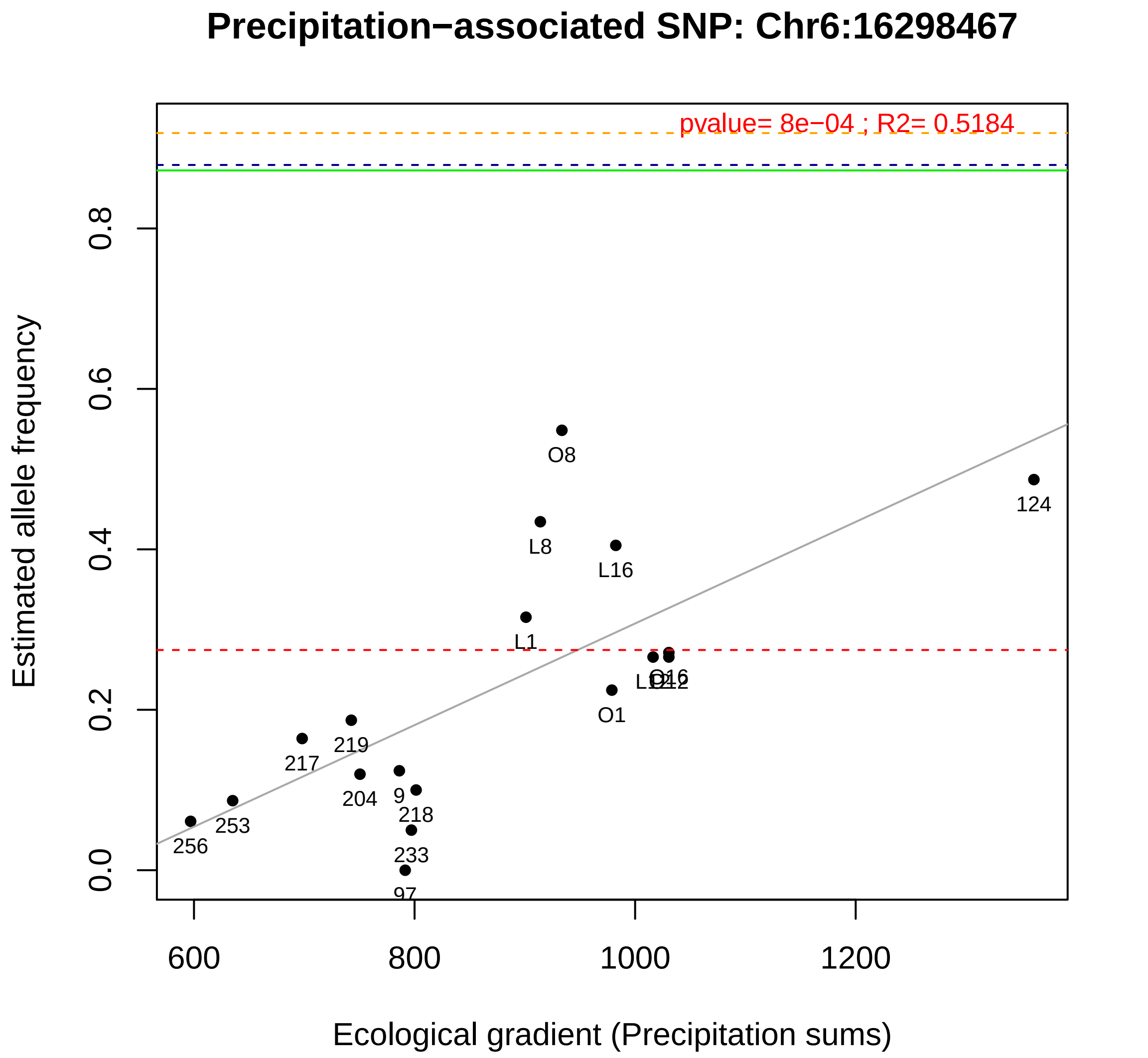
Precipitation−associated SNP: Chr6:16298467
pvalue= 8e−04 ; R2= 0.5184
Estimated allele frequency
0.0
0.2
0.4
0.6
0.8
600
800
1000
1200
Ecological gradient (Precipitation sums)
O8
124
L8
L16
L1
O1
219
217
204
9
218
253
256
233
97

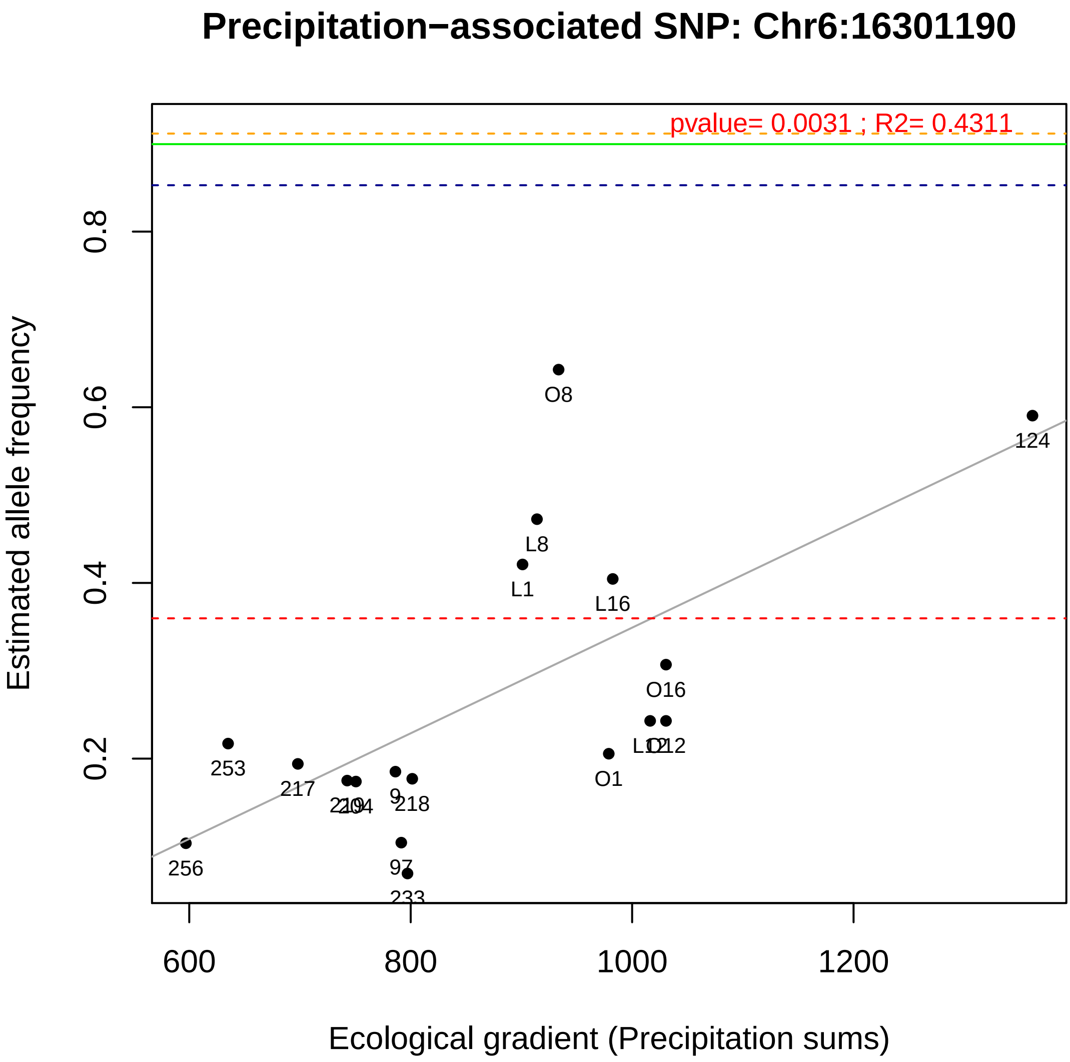

Precipitation−associated SNP: Chr6:16301190
pvalue= 0.0031 ; R2= 0.4311
Estimated allele frequency
0.2
0.4
0.6
0.8
600
800
1000
1200
Ecological gradient (Precipitation sums)
O8
124
L8
L1
L16
O16
O1
253
217
218
9
256
97
233

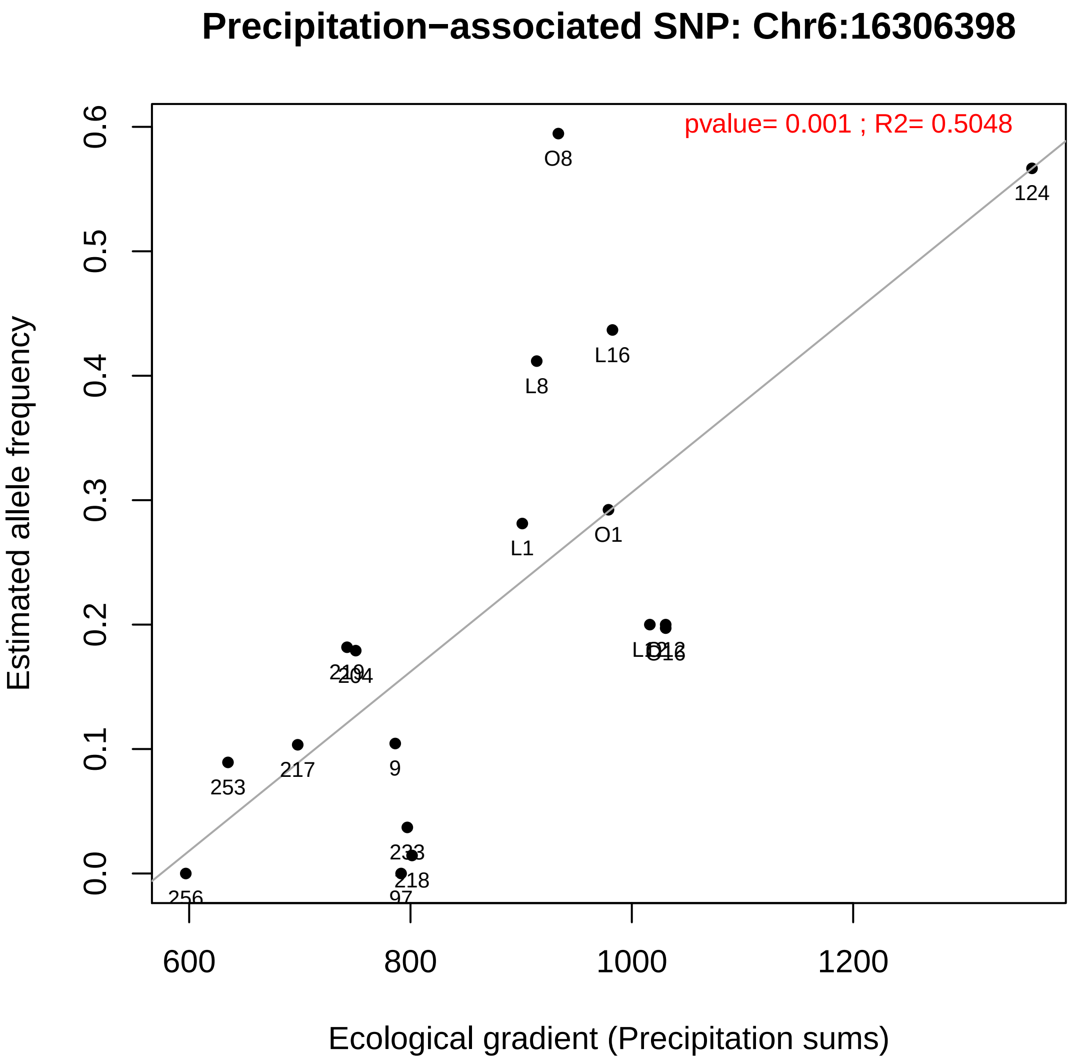

Precipitation−associated SNP: Chr6:16306398
pvalue= 0.001 ; R2= 0.5048
O8
124
L16
L8
O1
L1
219
204
253
217
9
233
218
97
256
0.0
0.1
0.2
0.3
0.4
0.5
0.6
600
800
1000
1200
Estimated allele frequency
Ecological gradient (Precipitation sums)

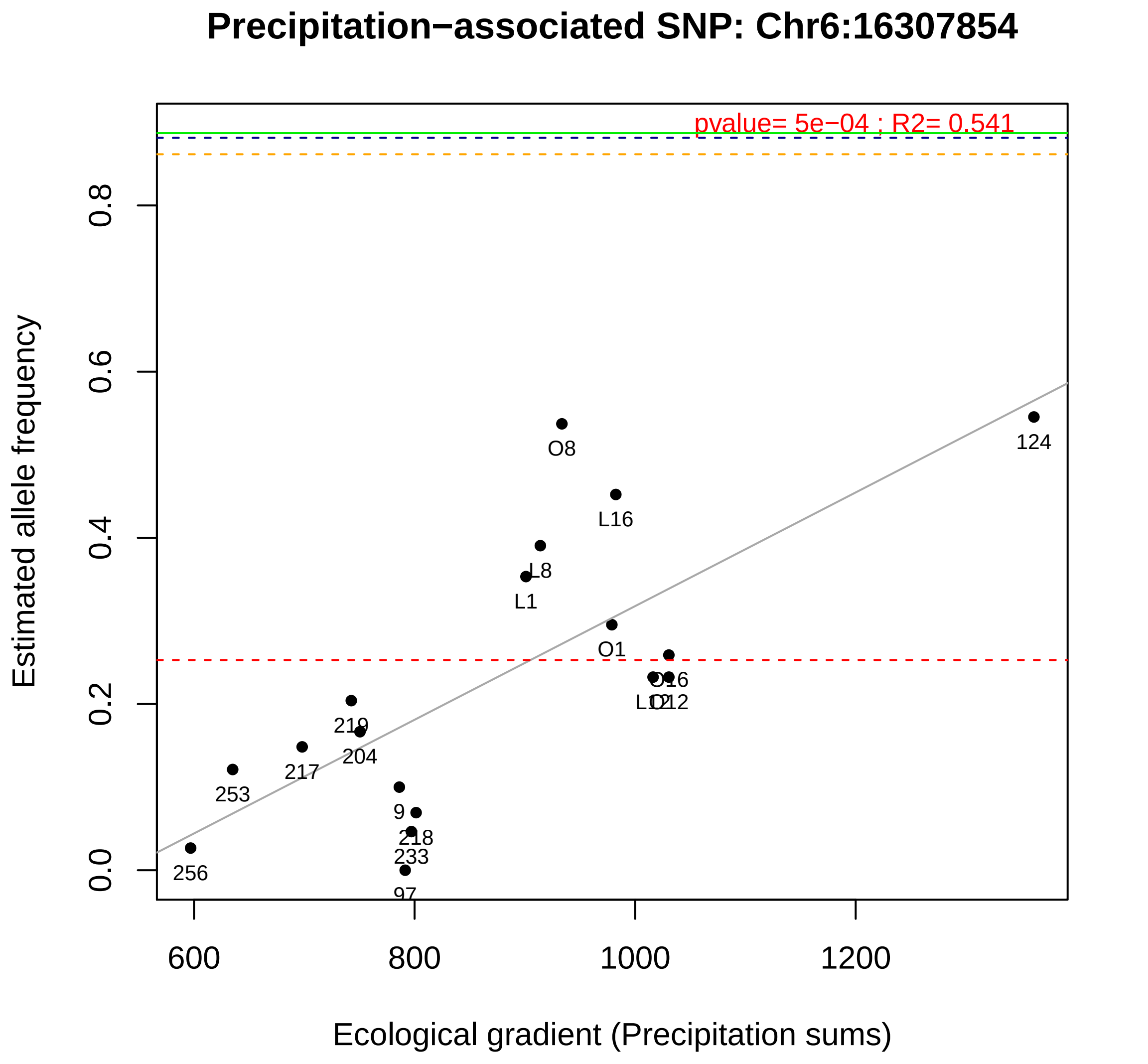

Precipitation−associated SNP: Chr6:16307854
pvalue= 5e−04 ; R2= 0.541
Estimated allele frequency
0.0
0.2
0.4
0.6
0.8
600
800
1000
1200
Ecological gradient (Precipitation sums)
O8
124
L16
L8
L1
O1
O16
L12
O12
219
204
217
253
9
218
233
256
97

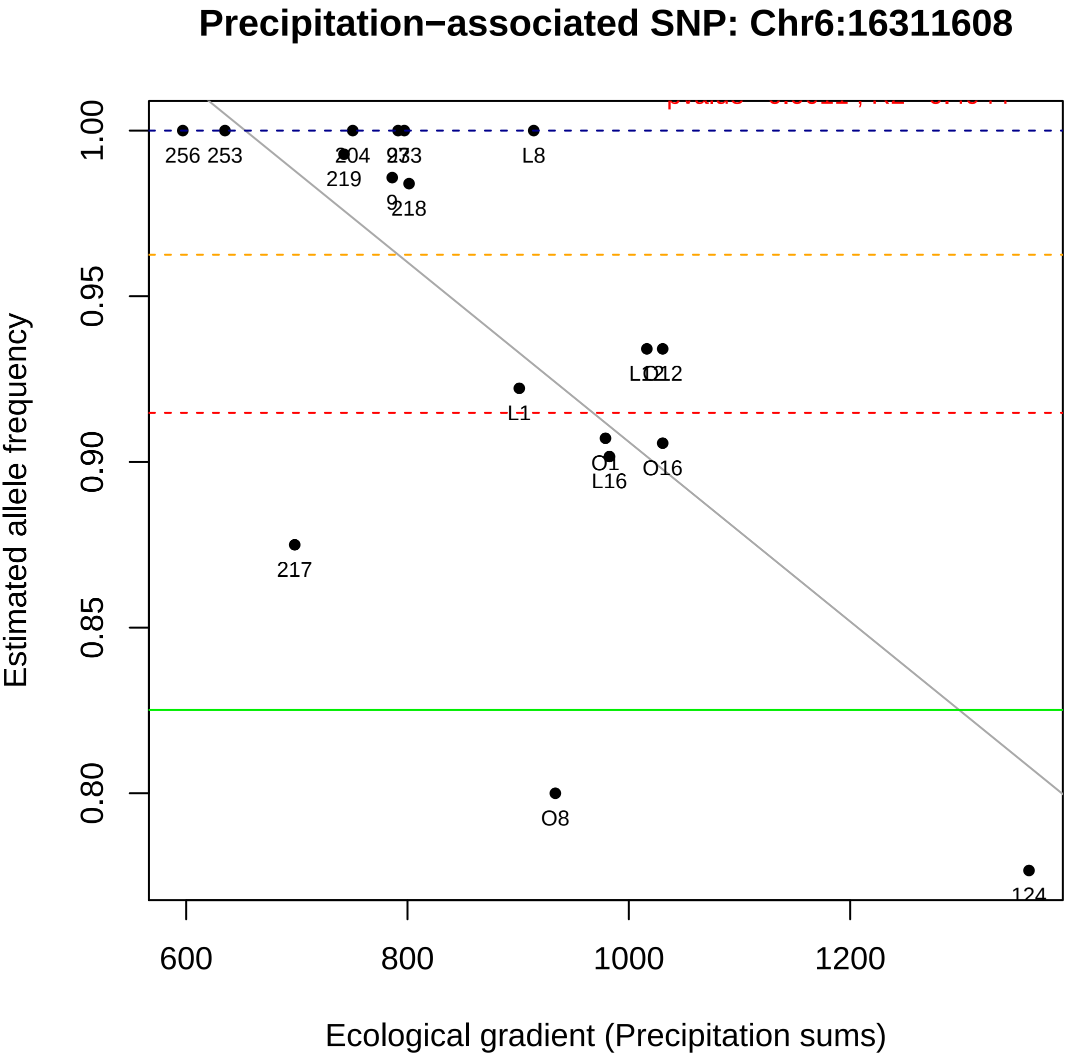

Precipitation−associated SNP: Chr6:16311608
Estimated allele frequency
Ecological gradient (Precipitation sums)
1.00
0.95
0.90
0.85
0.80
600
800
1000
1200
256
253
204
219
27
233
9
218
L8
L1
L12
O12
O1
L16
O16
217
O8
124

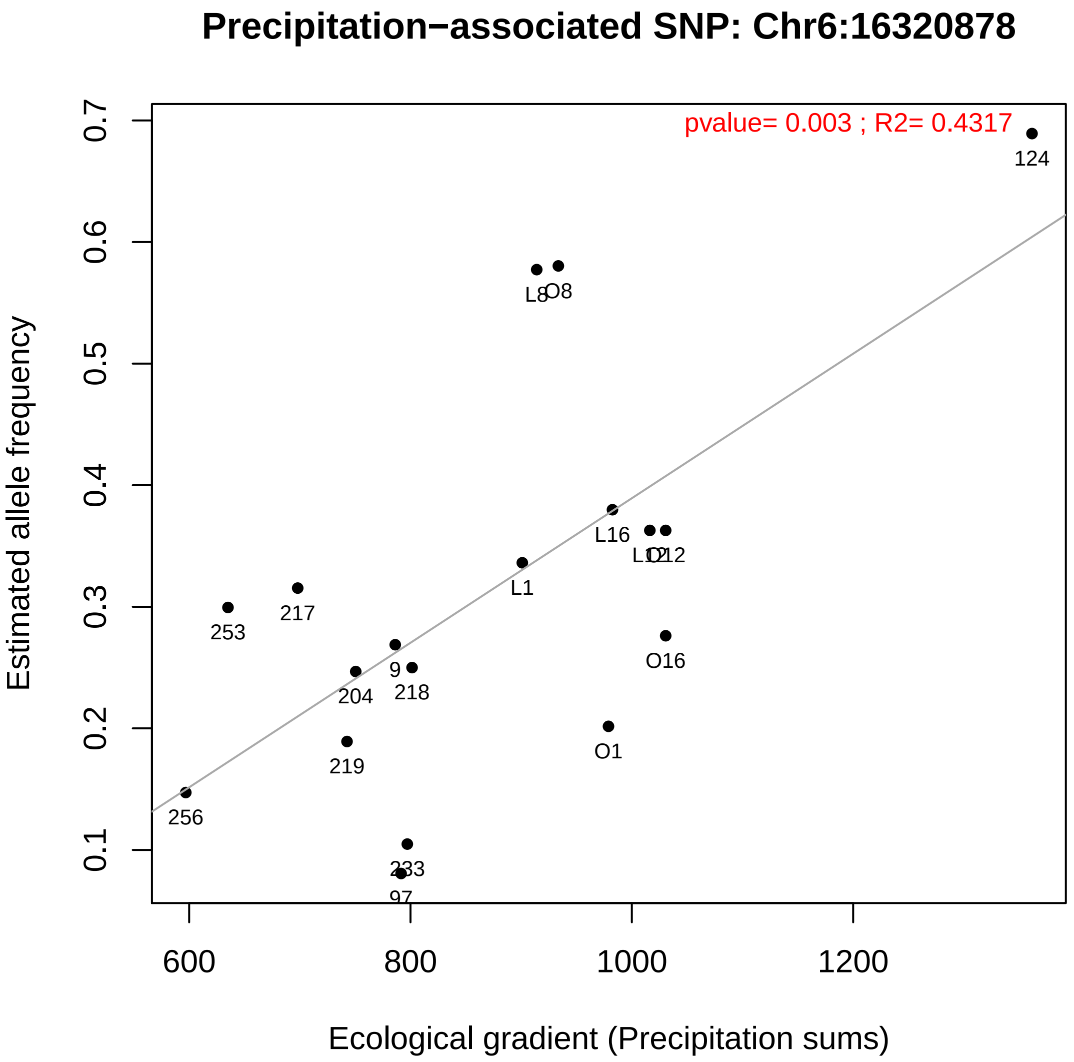

Precipitation−associated SNP: Chr6:16320878
pvalue= 0.003 ; R2= 0.4317
Estimated allele frequency
Ecological gradient (Precipitation sums)
0.1
0.2
0.3
0.4
0.5
0.6
0.7
600
800
1000
1200
124
L8
O8
L16
L12
O12
L1
217
253
9
218
204
O16
O1
219
256
233
97

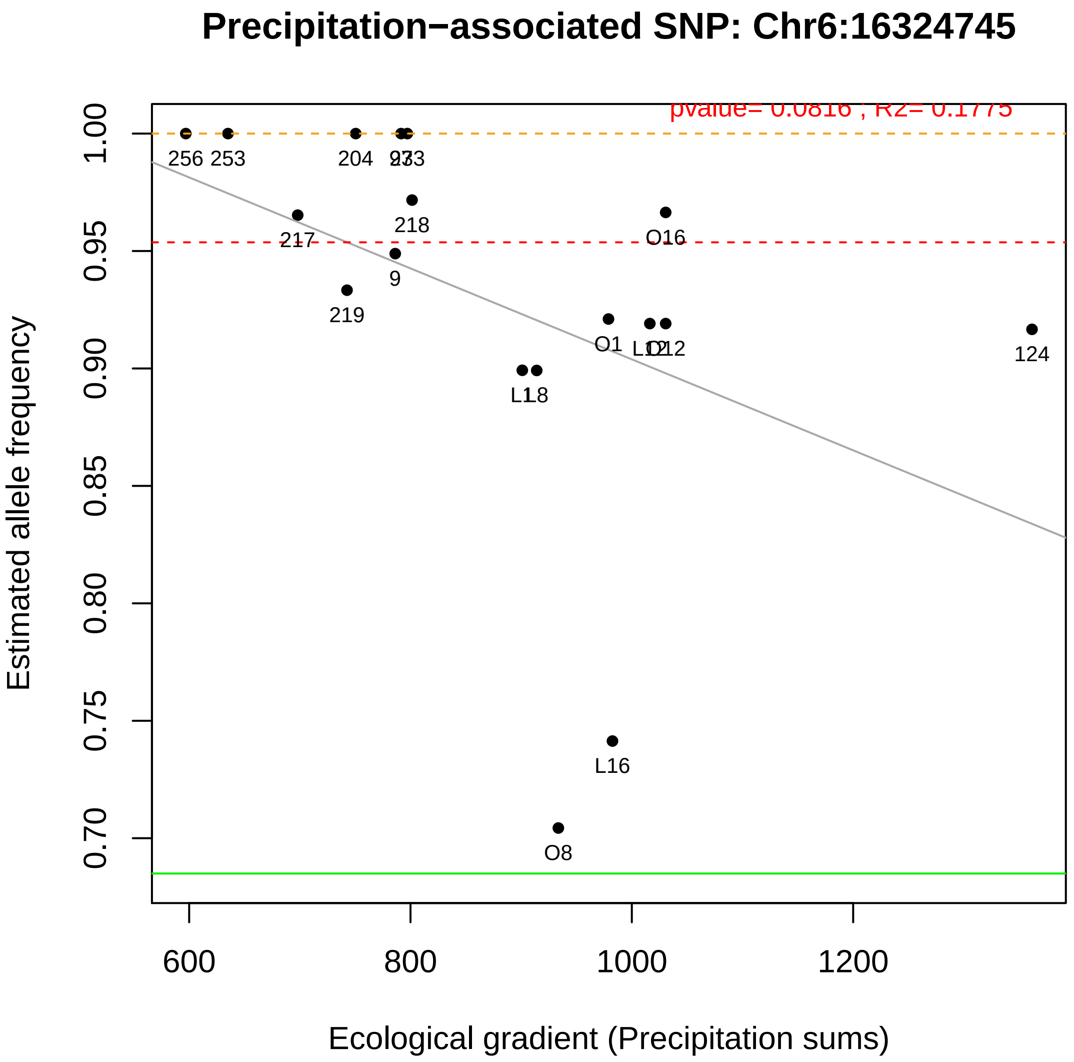

Precipitation−associated SNP: Chr6:16324745
pvalue= 0.0816 ; R2= 0.1775
Estimated allele frequency
Ecological gradient (Precipitation sums)
1.00
0.95
0.90
0.85
0.80
0.75
0.70
600
800
1000
1200
256
253
204
217
218
O16
9
219
O1
124
L16
O8

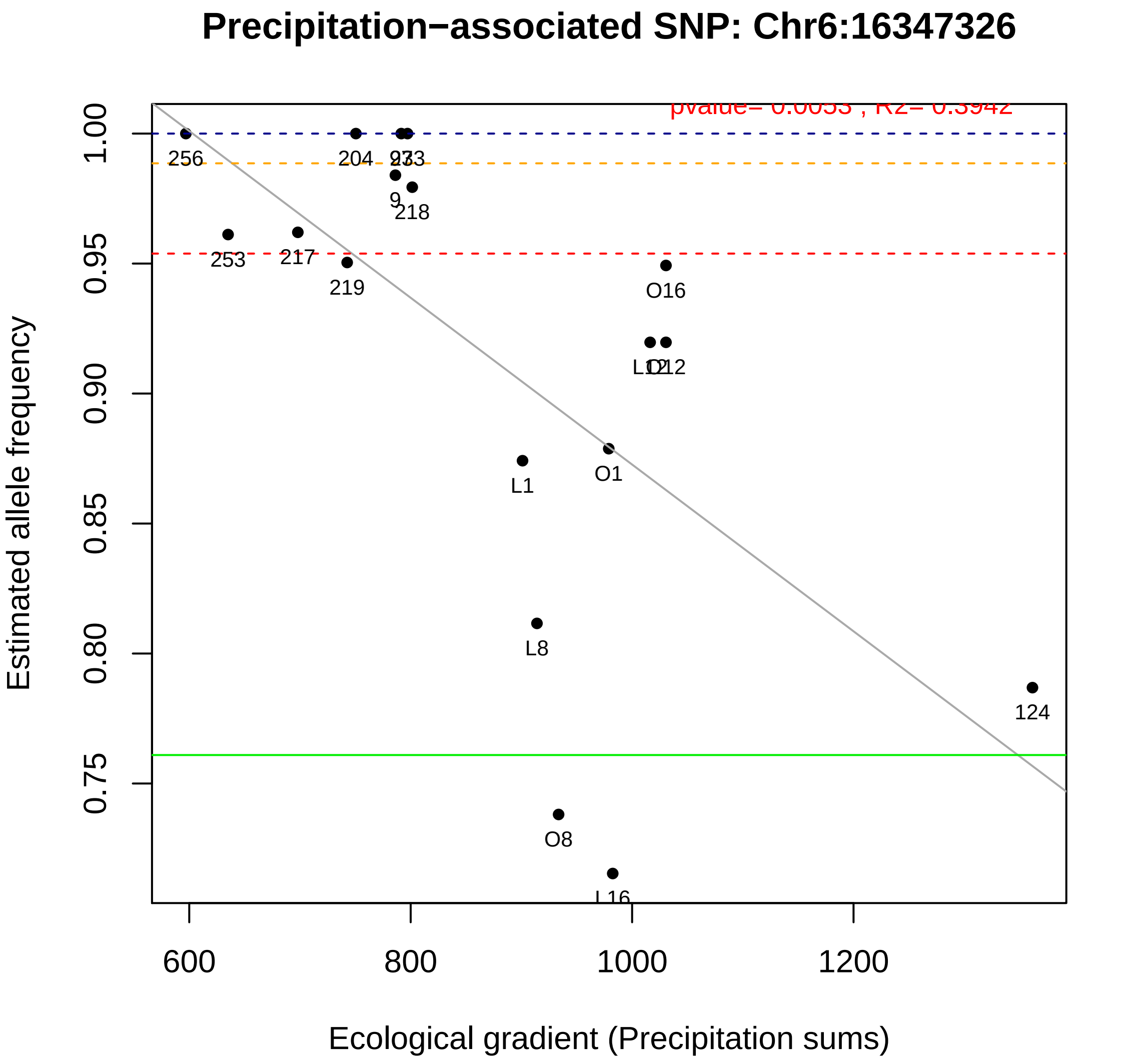

Precipitation−associated SNP: Chr6:16347326
pvalue= 0.0053 , R2= 0.3942
Estimated allele frequency
1.00
0.95
0.90
0.85
0.80
0.75
256
204
97
233
9
218
253
217
219
O16
L12
O12
L1
O1
L8
124
O8
L16
600
800
1000
1200
Ecological gradient (Precipitation sums)

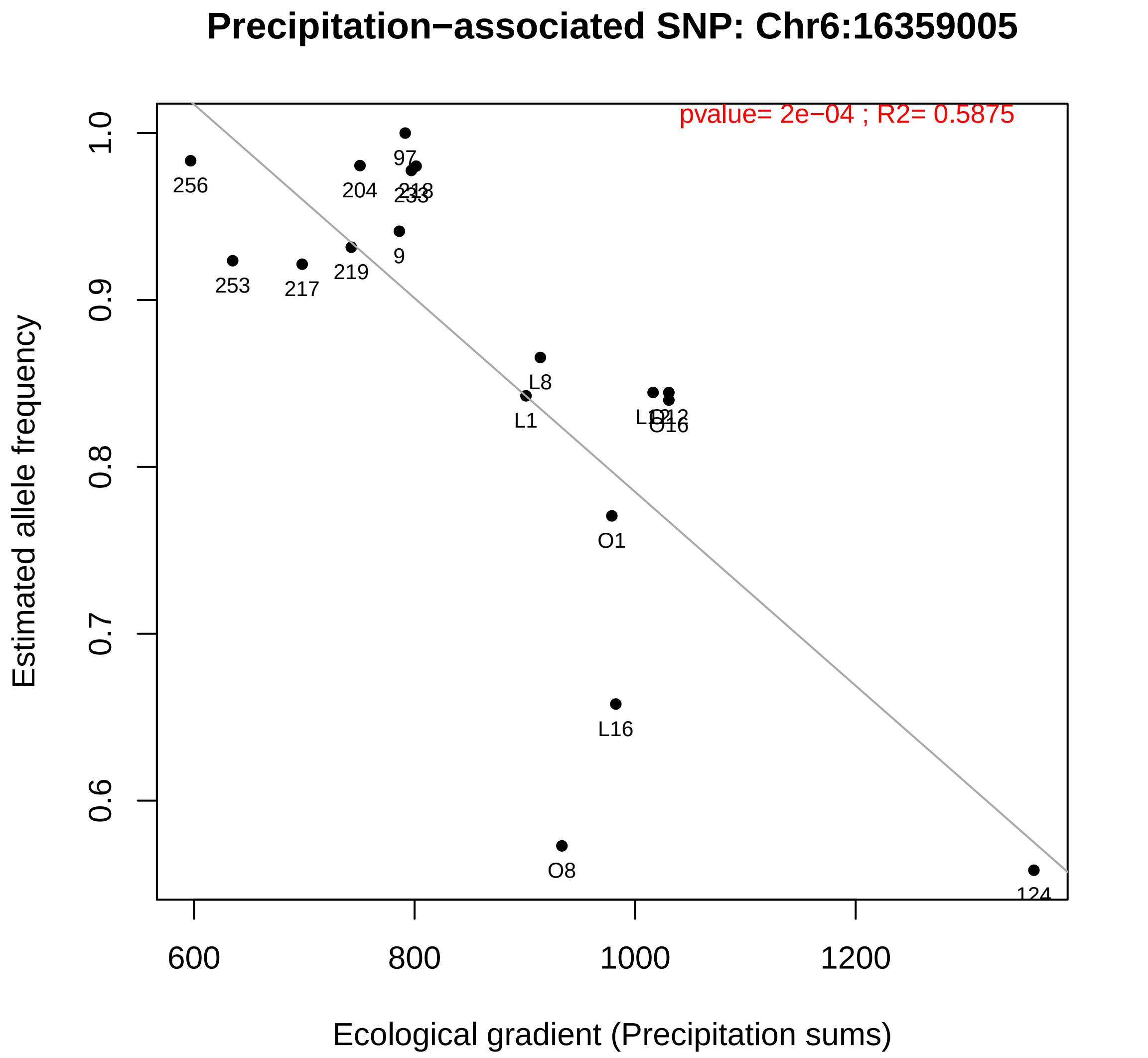
Precipitation−associated SNP: Chr6:16359005
pvalue= 2e−04 ; R2= 0.5875
Estimated allele frequency
Ecological gradient (Precipitation sums)
0.6
0.7
0.8
0.9
1.0
600
800
1000
1200
256
204
97
218
233
9
253
217
219
L8
L1
L12
O12
O16
O1
L16
O8
124

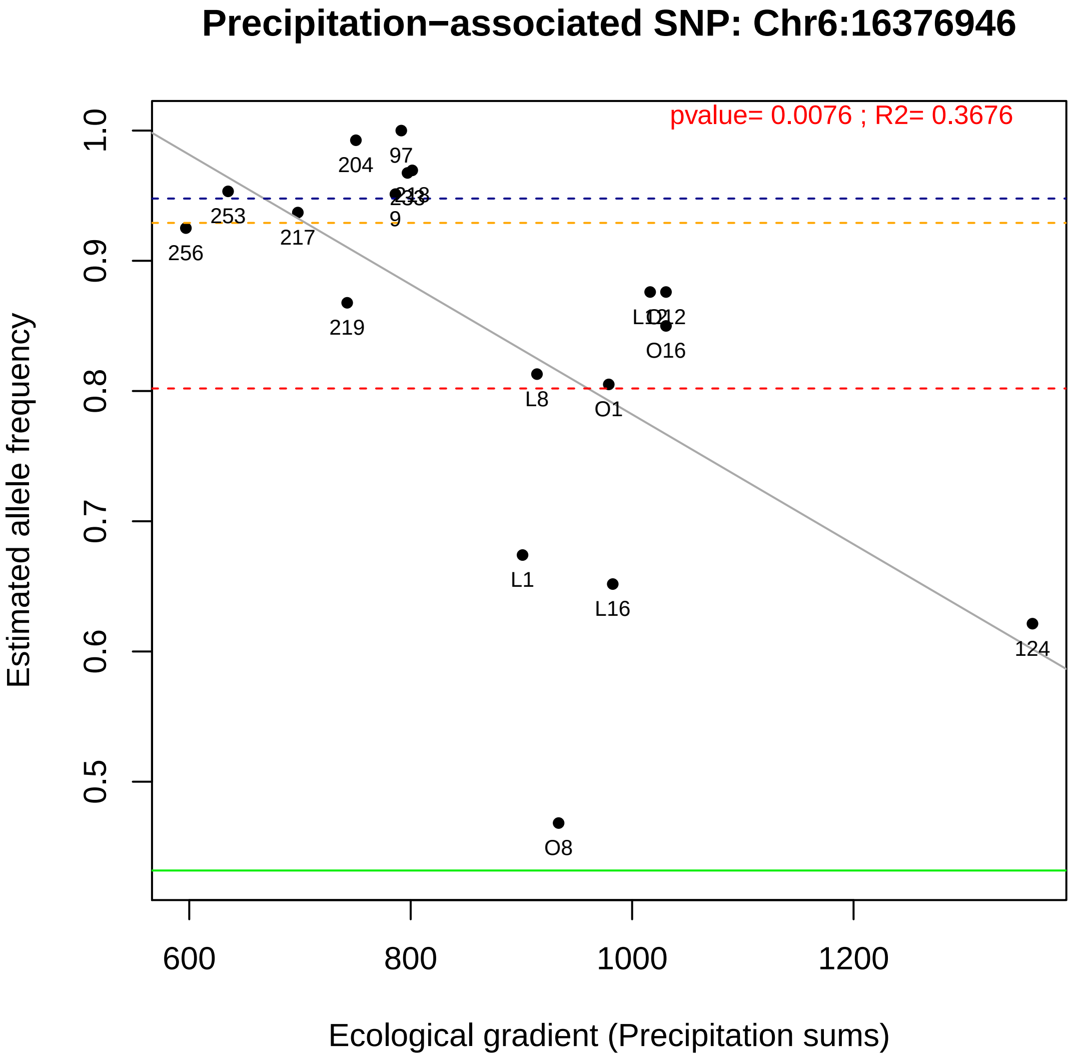
Precipitation−associated SNP: Chr6:16376946
pvalue= 0.0076 ; R2= 0.3676
Estimated allele frequency
0.5
0.6
0.7
0.8
0.9
1.0
600
800
1000
1200
Ecological gradient (Precipitation sums)
204
97
253
256
217
9
219
O16
L8
O1
L1
L16
124
O8

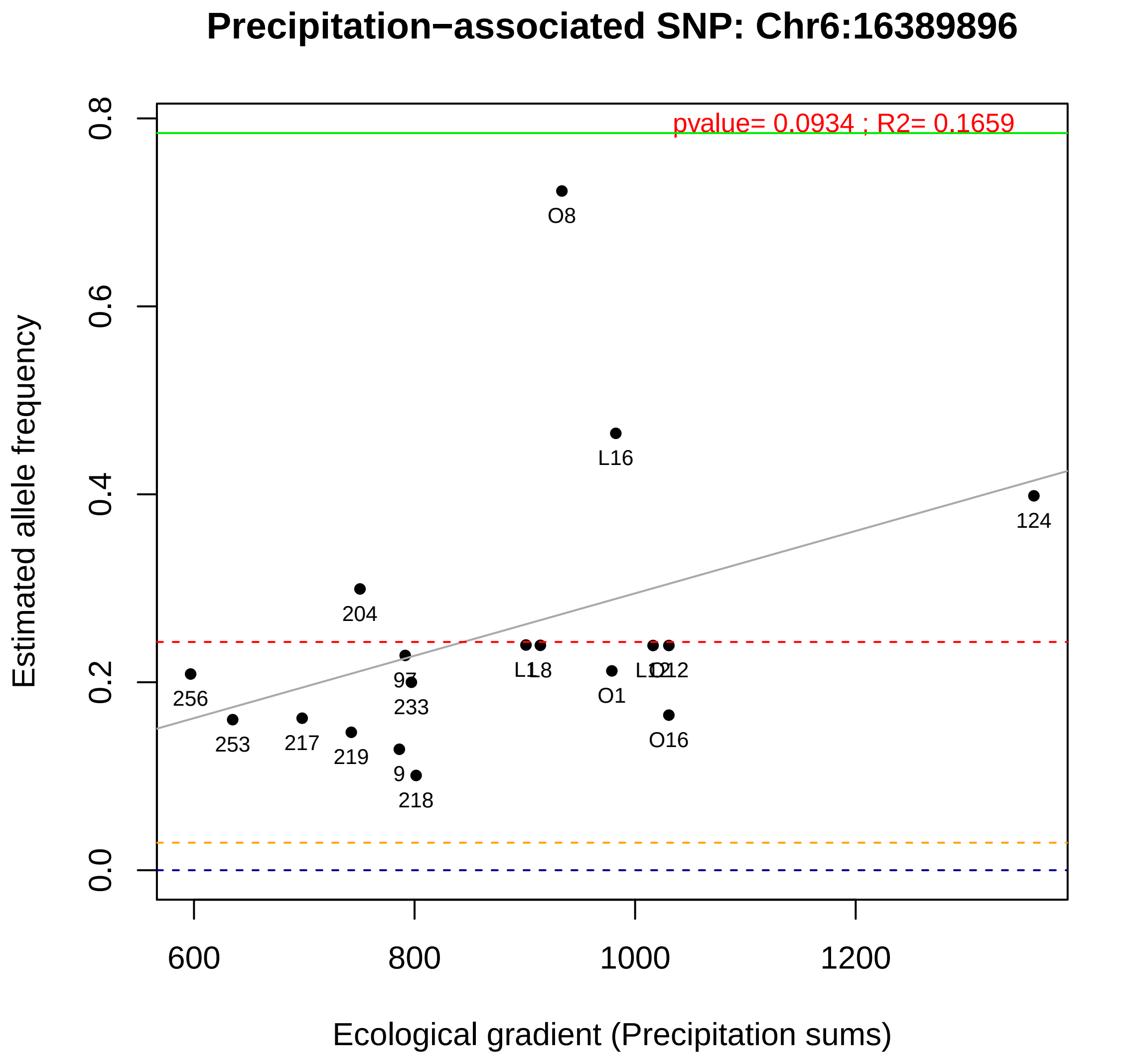

Precipitation−associated SNP: Chr6:16389896
pvalue= 0.0934 ; R2= 0.1659
Estimated allele frequency
0.0
0.2
0.4
0.6
0.8
O8
L16
124
204
256
253
217
219
97
233
9
218
L18
O1
L12
O12
O16
600
800
1000
1200
Ecological gradient (Precipitation sums)

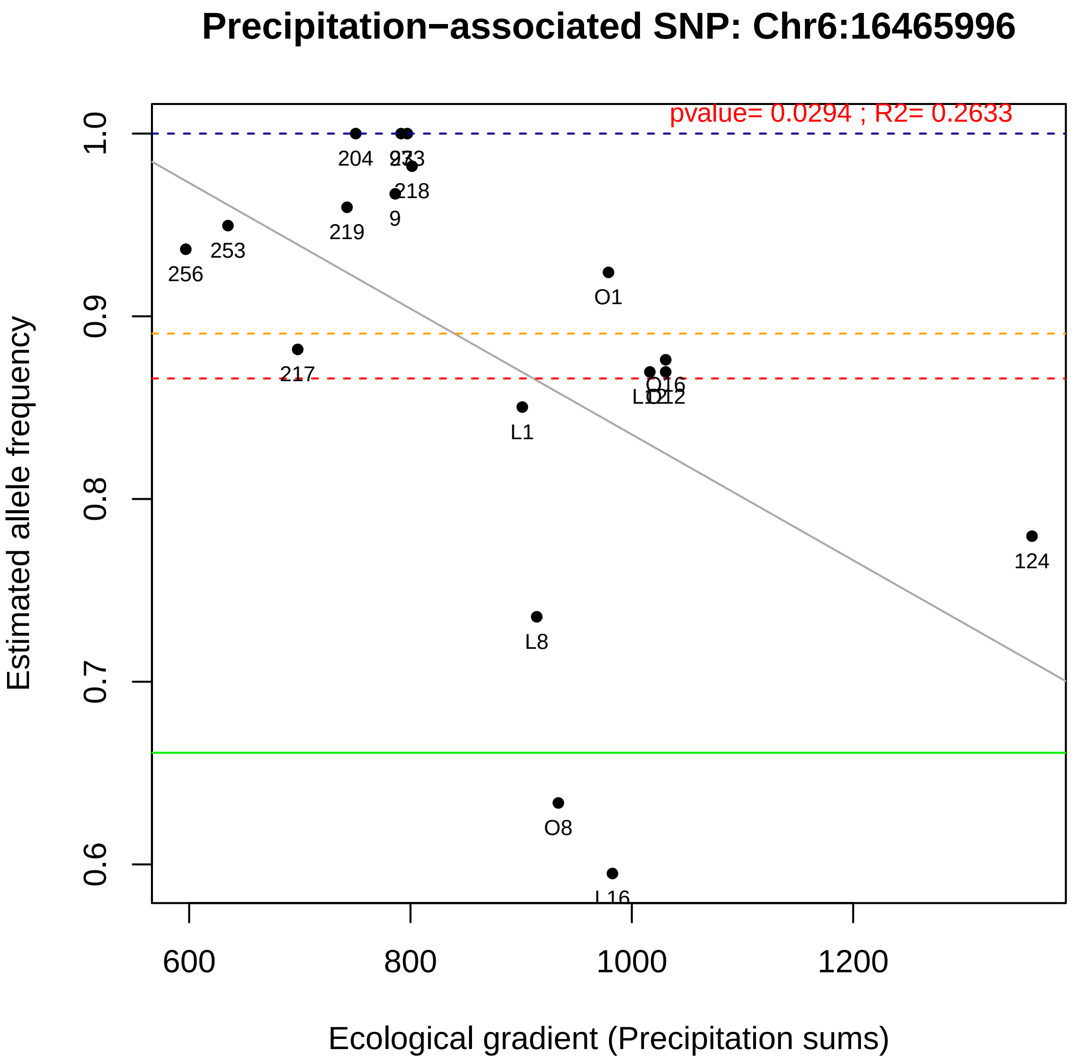

Precipitation−associated SNP: Chr6:16465996
pvalue= 0.0294 ; R2= 0.2633
Estimated allele frequency
Ecological gradient (Precipitation sums)
1.0
0.9
0.8
0.7
0.6
600
800
1000
1200
204
218
219
9
253
256
O1
217
O16
O12
L1
124
L8
O8
L16

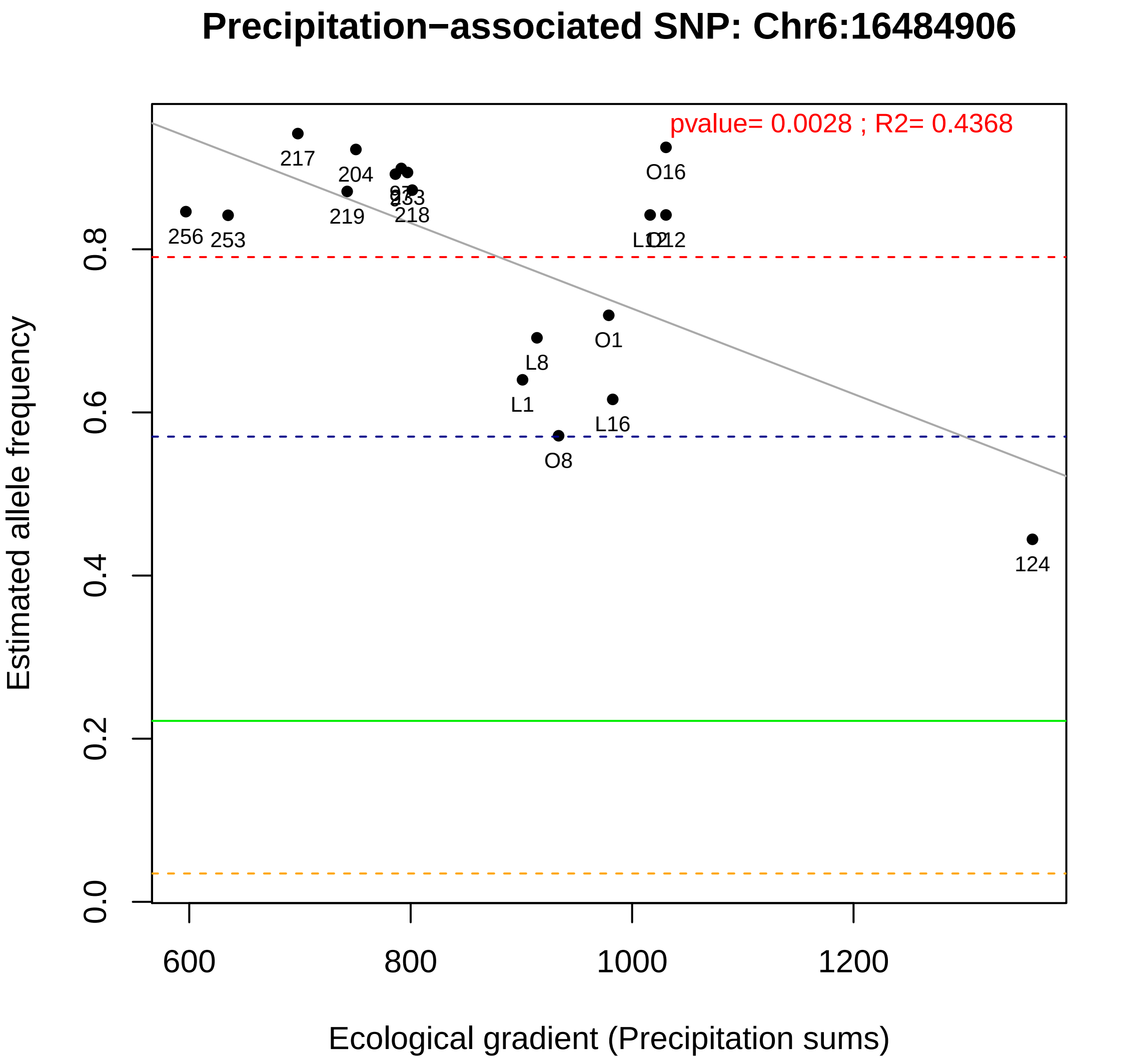
Precipitation−associated SNP: Chr6:16484906
pvalue= 0.0028 ; R2= 0.4368
217
204
233
218
219
256
253
O16
O1
L8
L1
L16
O8
124
Estimated allele frequency
0.0
0.2
0.4
0.6
0.8
600
800
1000
1200
Ecological gradient (Precipitation sums)

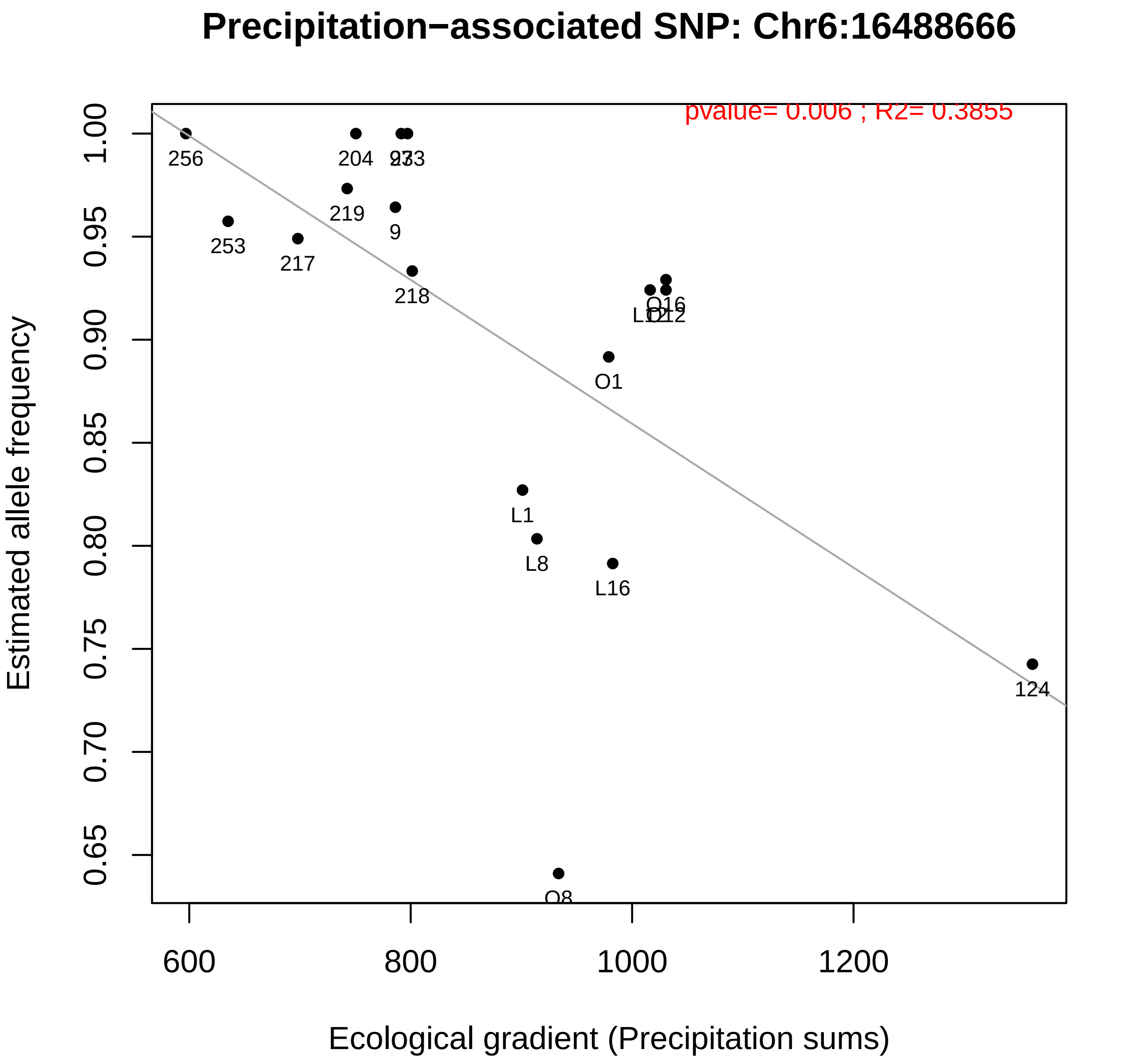

Precipitation−associated SNP: Chr6:16488666
pvalue= 0.006 ; R2= 0.3855
Estimated allele frequency
Ecological gradient (Precipitation sums)
1.00
0.95
0.90
0.85
0.80
0.75
0.70
0.65
600
800
1000
1200
256
204
219
9
253
217
218
O1
L1
L8
L16
O8
124
O16

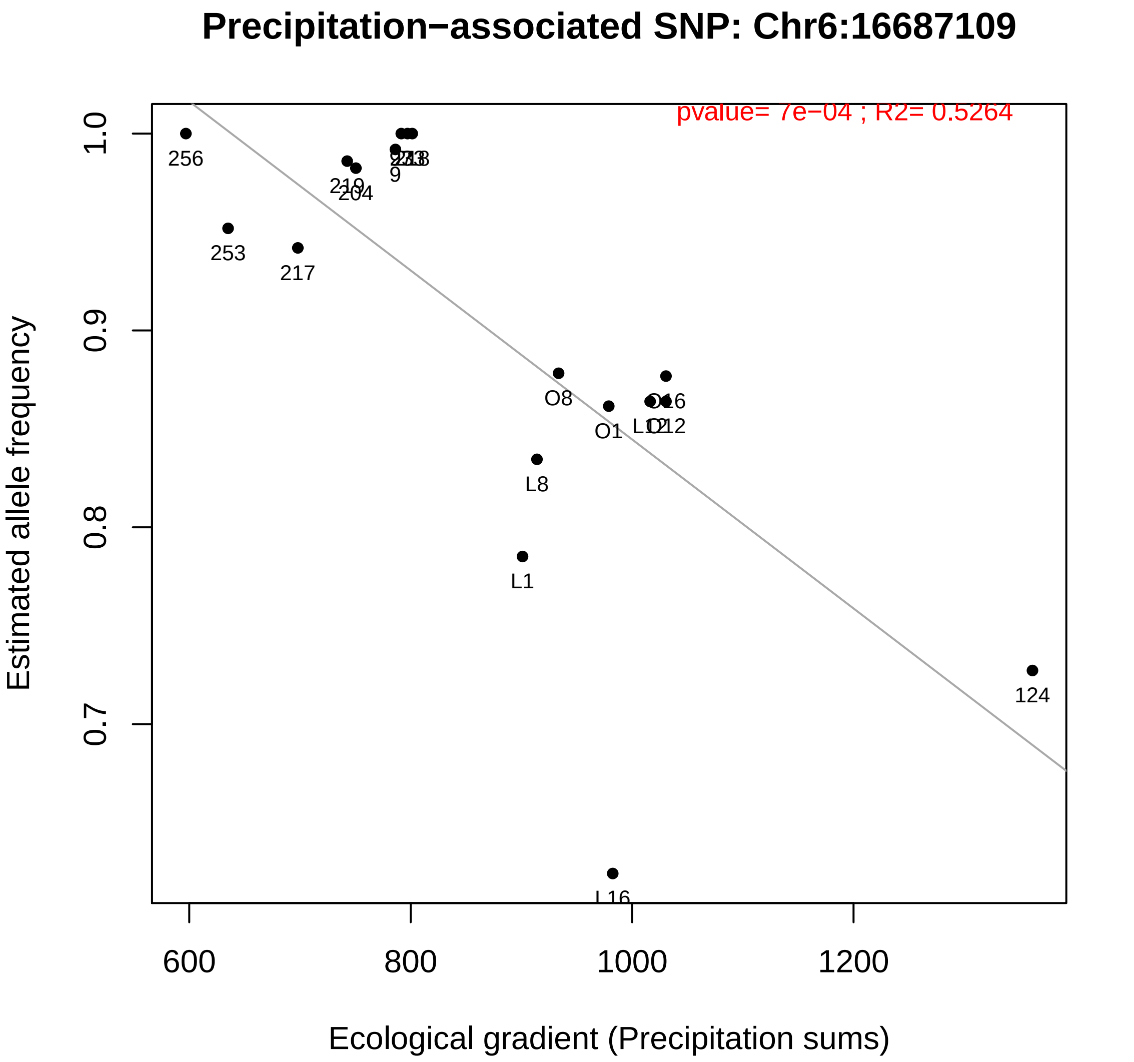
Precipitation−associated SNP: Chr6:16687109
pvalue= 7e−04 ; R2= 0.5264
Estimated allele frequency
1.0
0.9
0.8
0.7
600
800
1000
1200
Ecological gradient (Precipitation sums)
256
253
217
219
204
O8
O1
O16
L8
L1
124

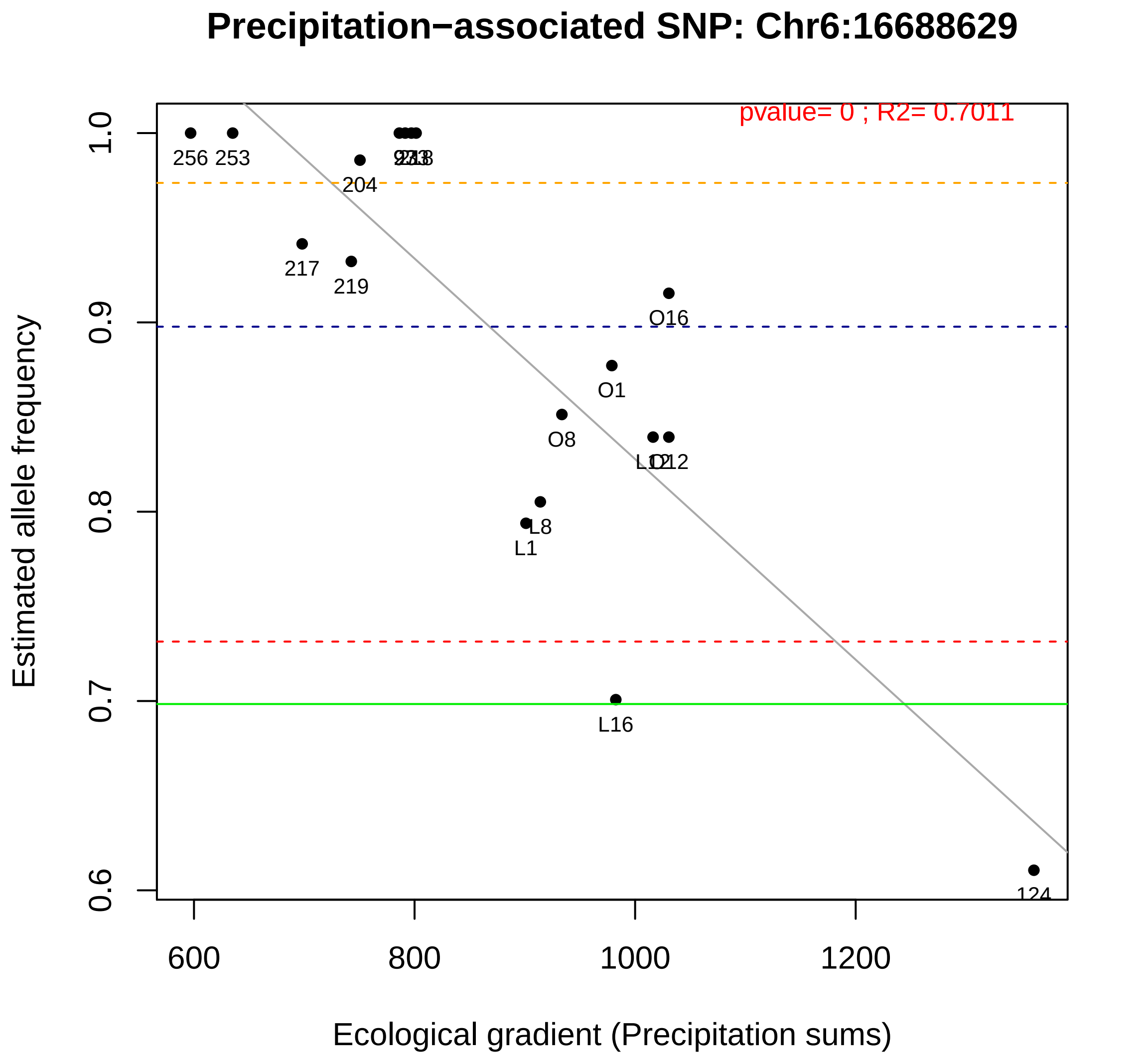

Precipitation−associated SNP: Chr6:16688629
pvalue= 0 ; R2= 0.7011
Estimated allele frequency
Ecological gradient (Precipitation sums)
1.0
0.9
0.8
0.7
0.6
600
800
1000
1200
256
253
204
217
219
O16
O1
O8
L1
L8
L16
124

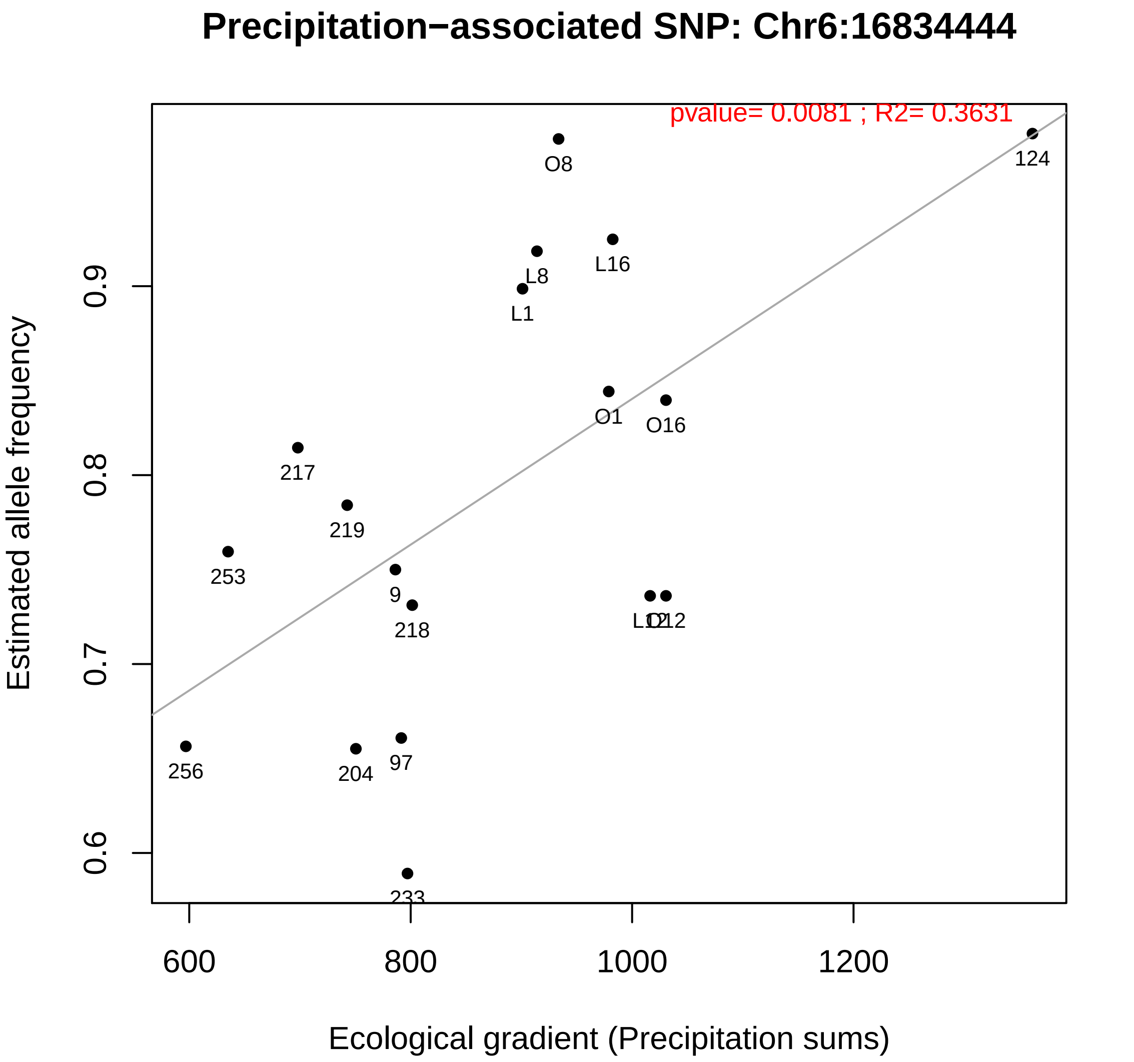

Precipitation−associated SNP: Chr6:16834444
pvalue= 0.0081 ; R2= 0.3631
Estimated allele frequency
Ecological gradient (Precipitation sums)
0.6
0.7
0.8
0.9
600
800
1000
1200
O8
124
L16
L8
L1
O1
O16
217
219
253
9
218
L12
O12
256
204
97
233

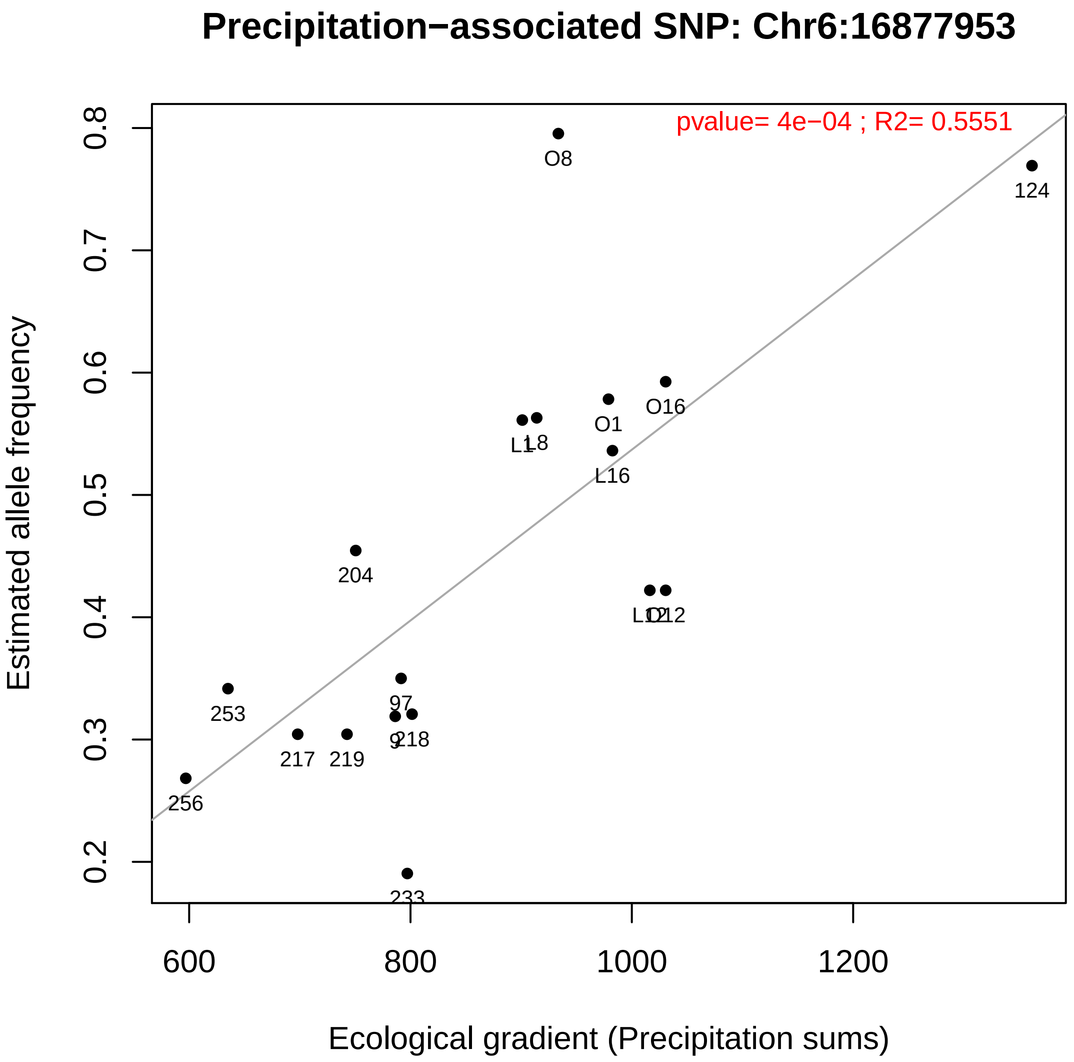

Precipitation−associated SNP: Chr6:16877953
pvalue= 4e−04 ; R2= 0.5551
O8
124
O16
O1
L1
L8
L16
204
L10
O12
253
97
9
218
217
219
256
233
0.8
0.7
0.6
0.5
0.4
0.3
0.2
600
800
1000
1200
Estimated allele frequency
Ecological gradient (Precipitation sums)

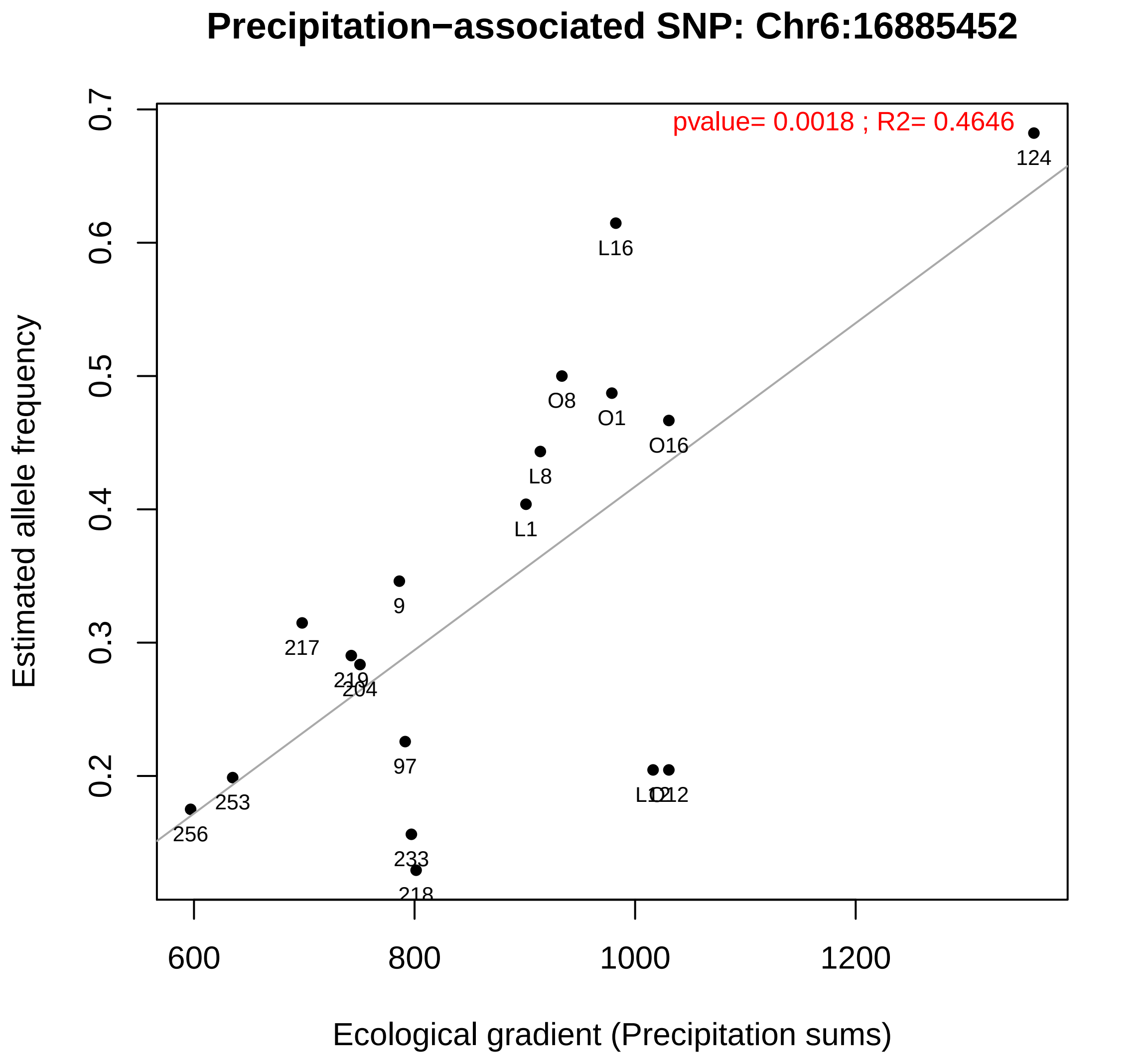
Precipitation−associated SNP: Chr6:16885452
pvalue= 0.0018 ; R2= 0.4646
Estimated allele frequency
Ecological gradient (Precipitation sums)
0.2
0.3
0.4
0.5
0.6
0.7
600
800
1000
1200
124
L16
O8
O1
O16
L8
L1
9
217
219
204
97
253
256
233
218

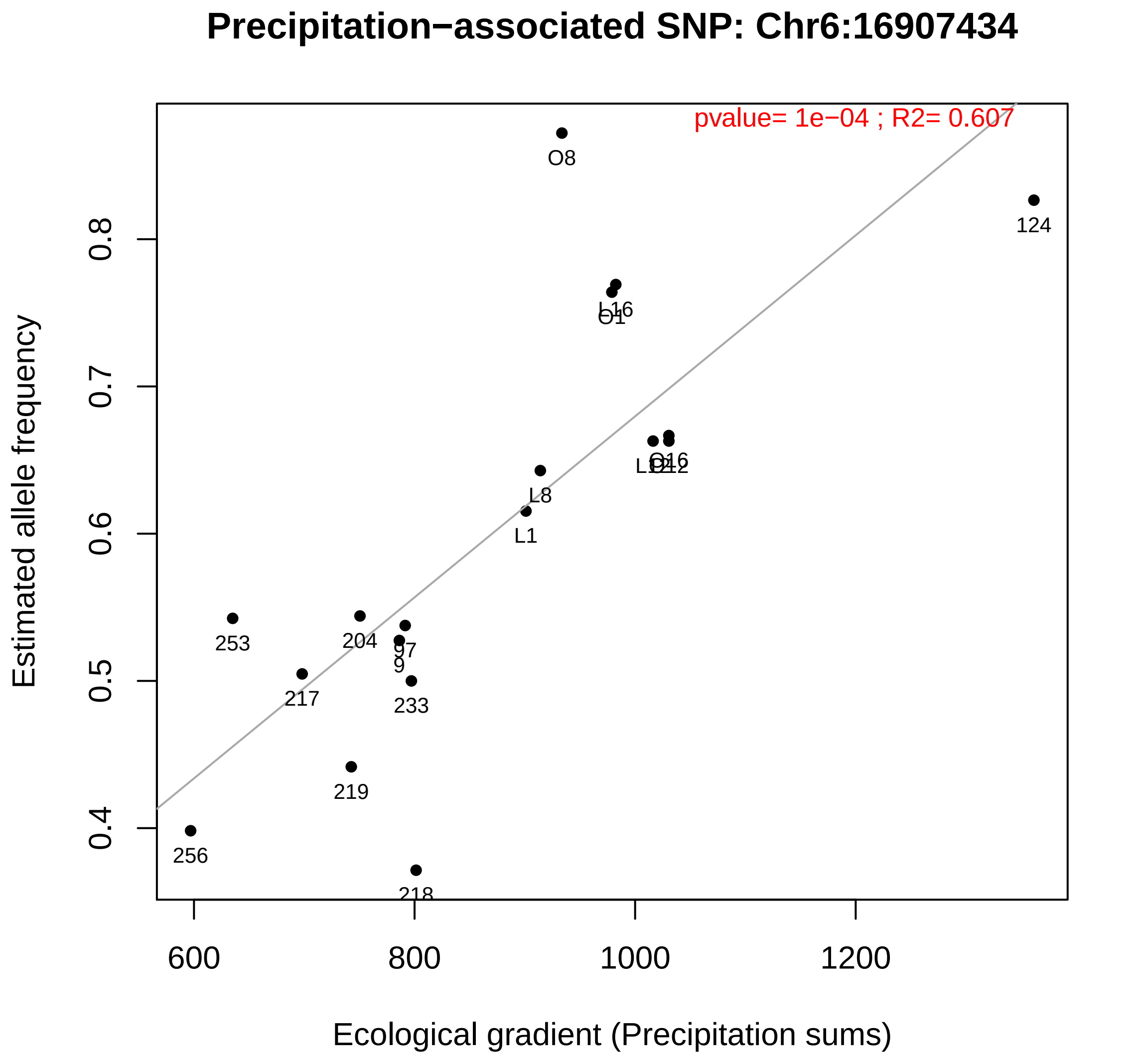

Precipitation−associated SNP: Chr6:16907434
pvalue= 1e−04 ; R2= 0.607
O8
124
L16
O1
O16
L12
L8
L1
253
204
97
9
217
233
219
256
218
0.4
0.5
0.6
0.7
0.8
Estimated allele frequency
600
800
1000
1200
Ecological gradient (Precipitation sums)

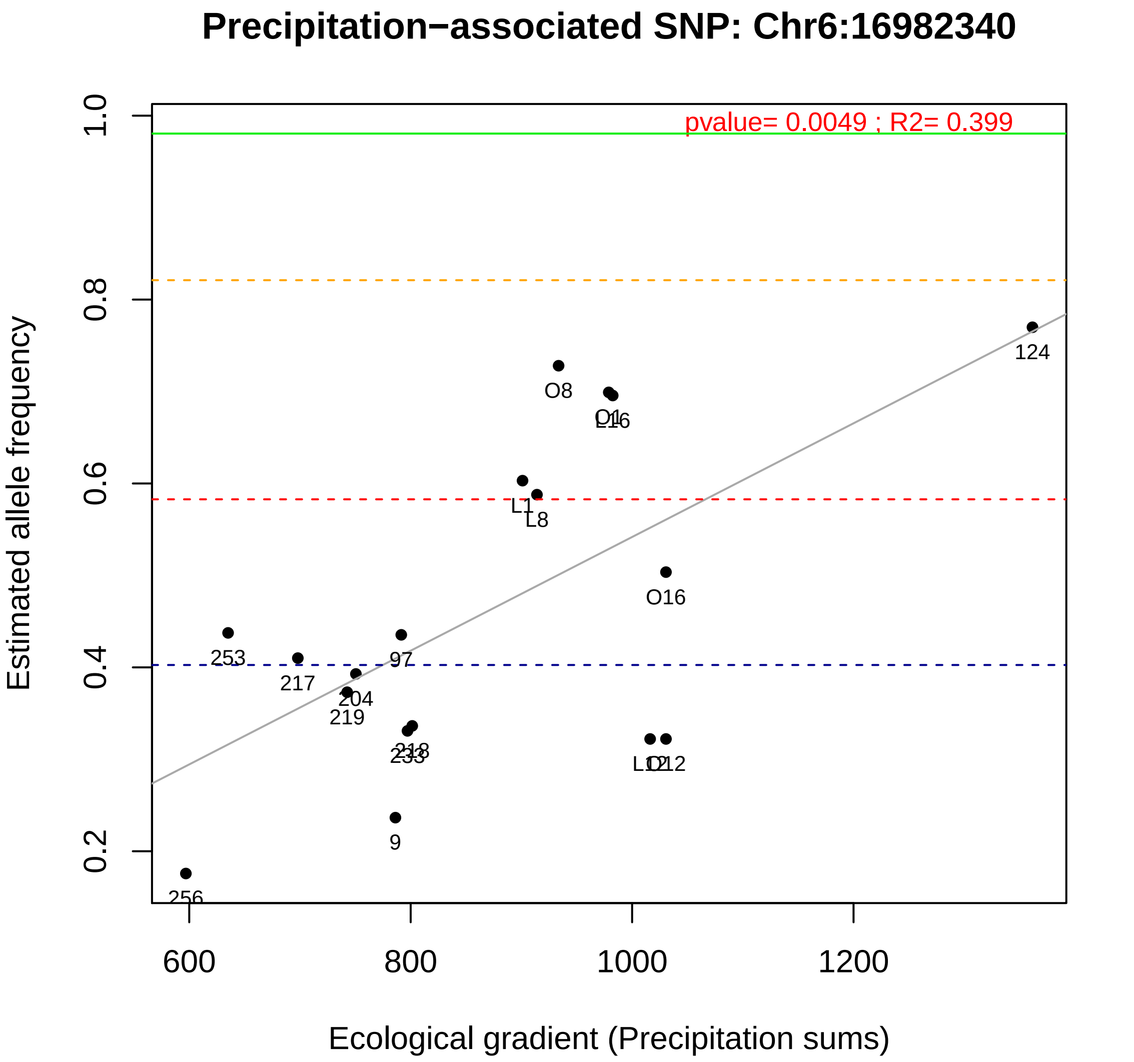

Precipitation−associated SNP: Chr6:16982340
pvalue= 0.0049 ; R2= 0.399
Estimated allele frequency
0.2
0.4
0.6
0.8
1.0
600
800
1000
1200
Ecological gradient (Precipitation sums)
124
O8
L16
L1
L8
O16
253
97
217
204
219
218
233
L12
O12
9
256

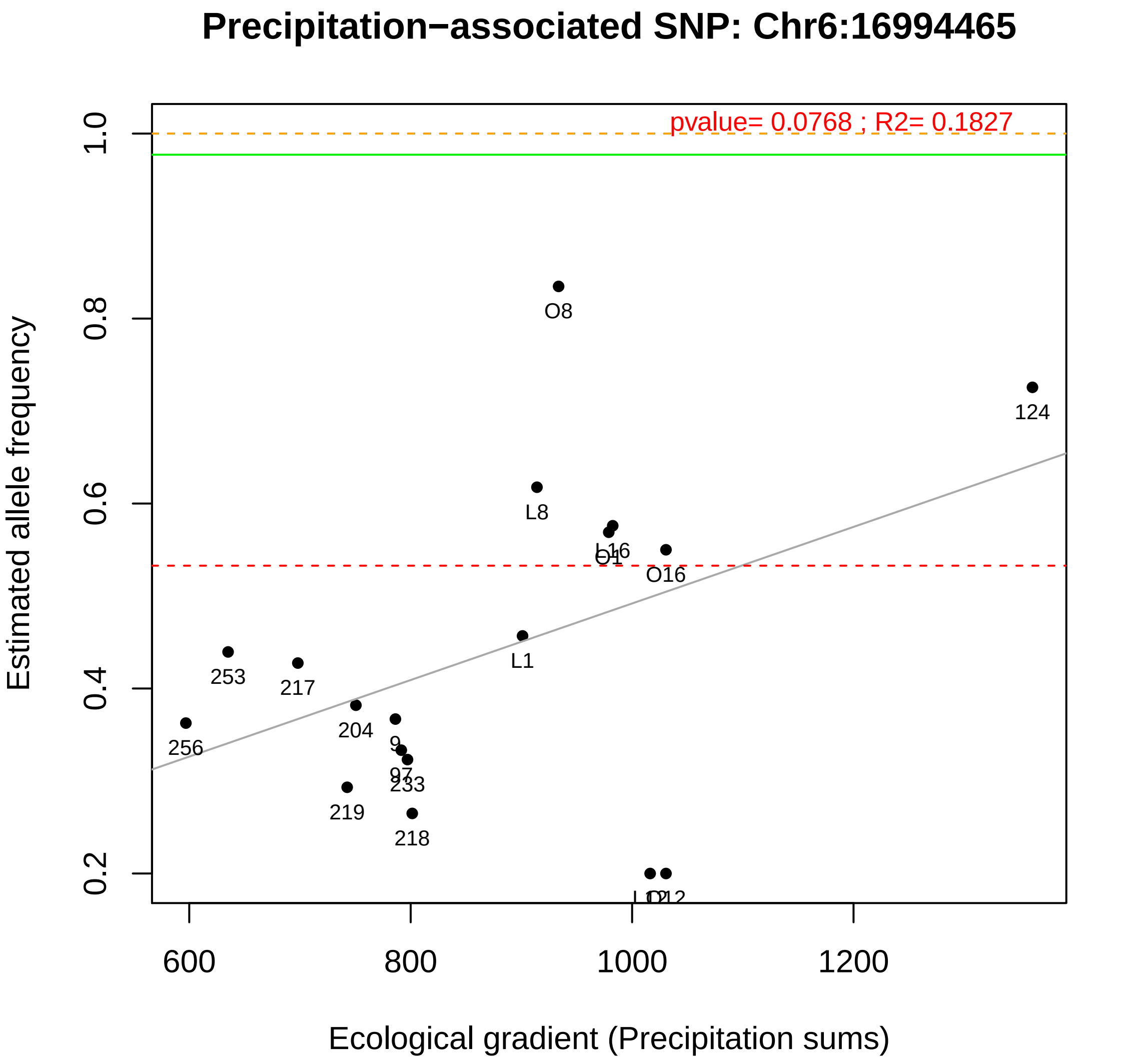
Precipitation−associated SNP: Chr6:16994465
pvalue= 0.0768 ; R2= 0.1827
Estimated allele frequency
Ecological gradient (Precipitation sums)
0.2
0.4
0.6
0.8
1.0
600
800
1000
1200
O8
124
L8
L16
O1
O16
L1
253
217
204
256
9
97
233
219
218
O12

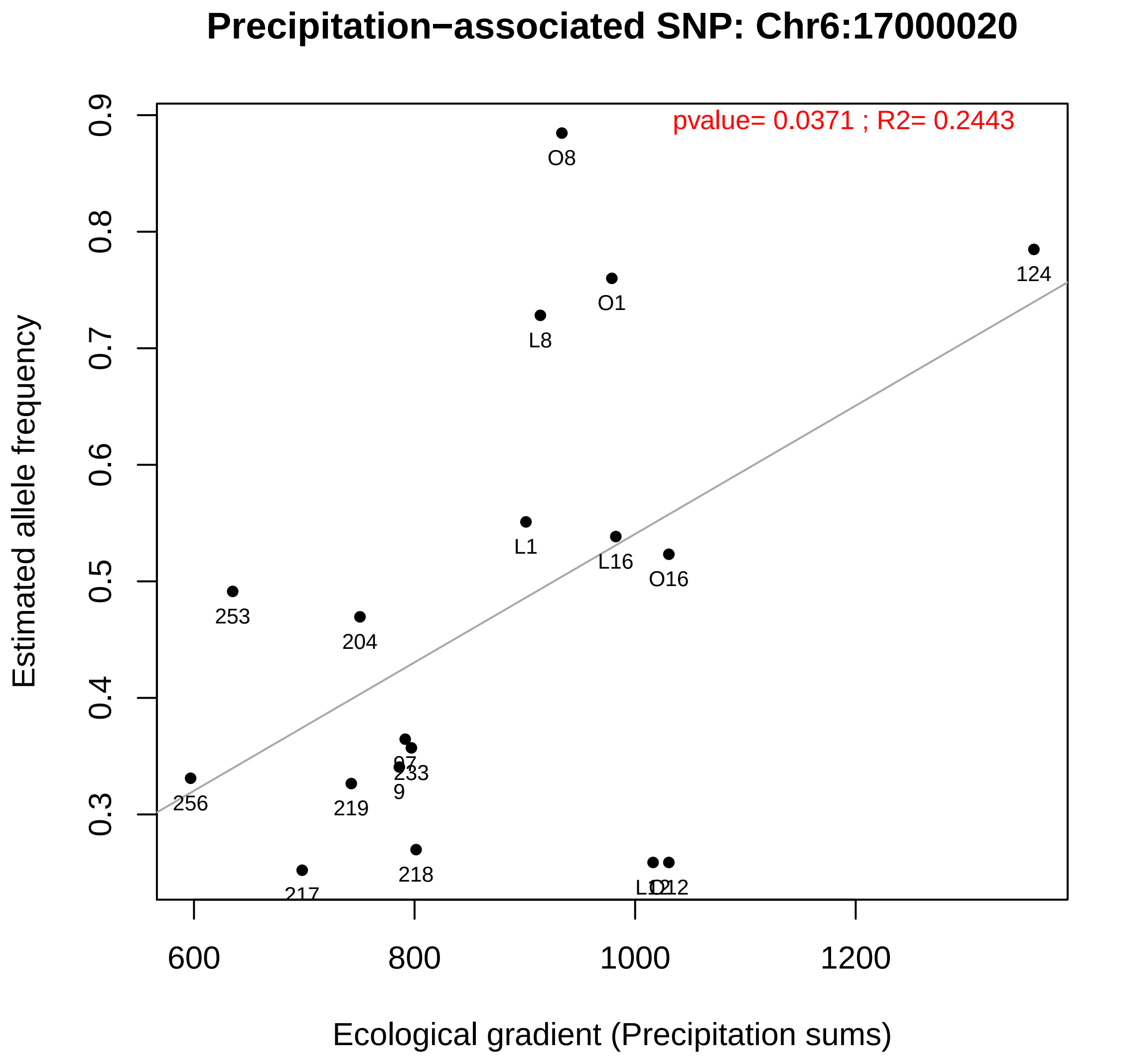

Precipitation−associated SNP: Chr6:17000020
pvalue= 0.0371 ; R2= 0.2443
Estimated allele frequency
Ecological gradient (Precipitation sums)
0.3
0.4
0.5
0.6
0.7
0.8
0.9
600
800
1000
1200
O8
124
O1
L8
L1
L16
O16
253
204
256
219
233
9
218
217

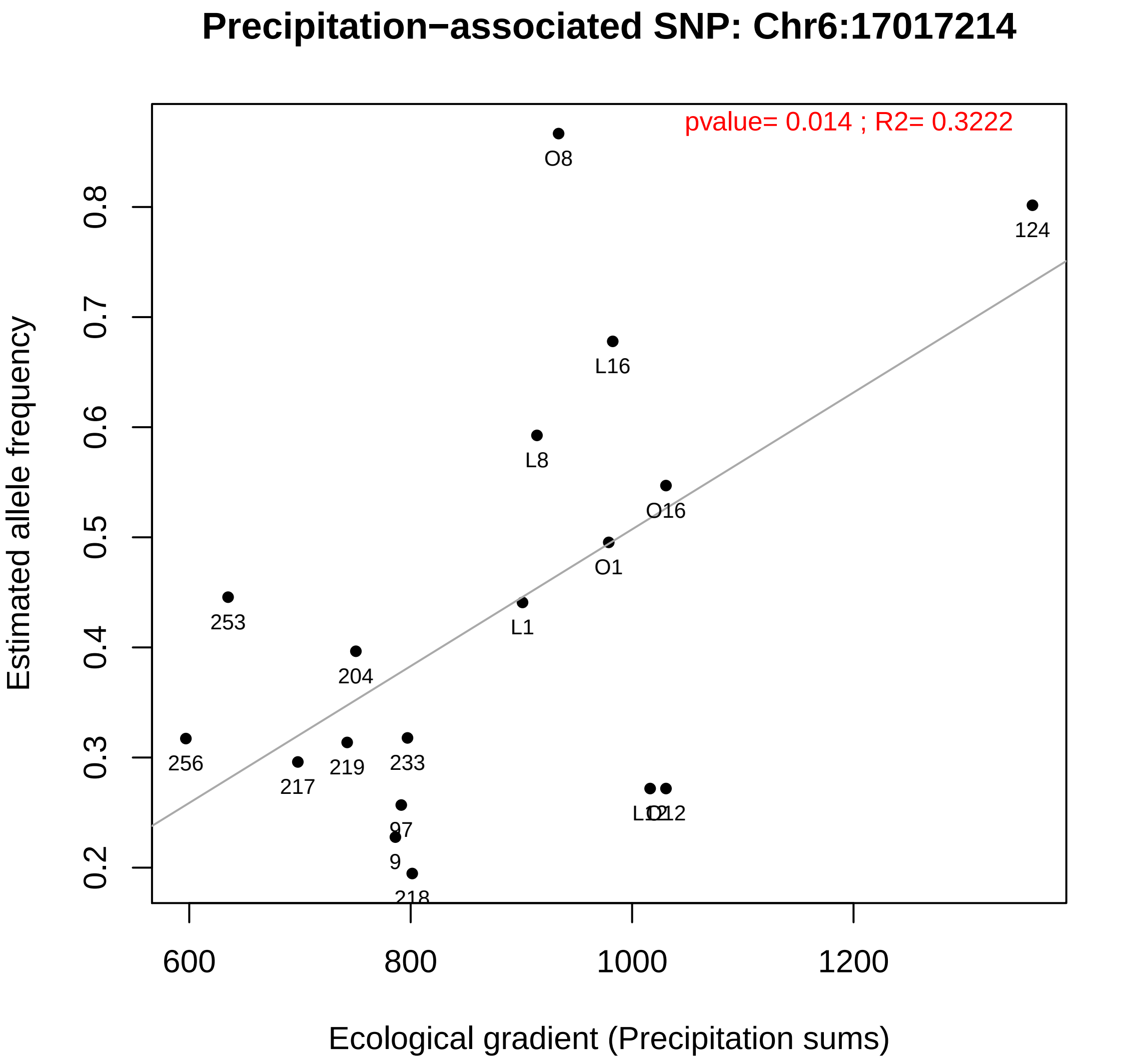

Precipitation−associated SNP: Chr6:17017214
pvalue= 0.014 ; R2= 0.3222
O8
124
L16
L8
O16
O1
L1
253
204
256
217
219
233
97
9
218
L12
O12
0.2
0.3
0.4
0.5
0.6
0.7
0.8
Estimated allele frequency
600
800
1000
1200
Ecological gradient (Precipitation sums)

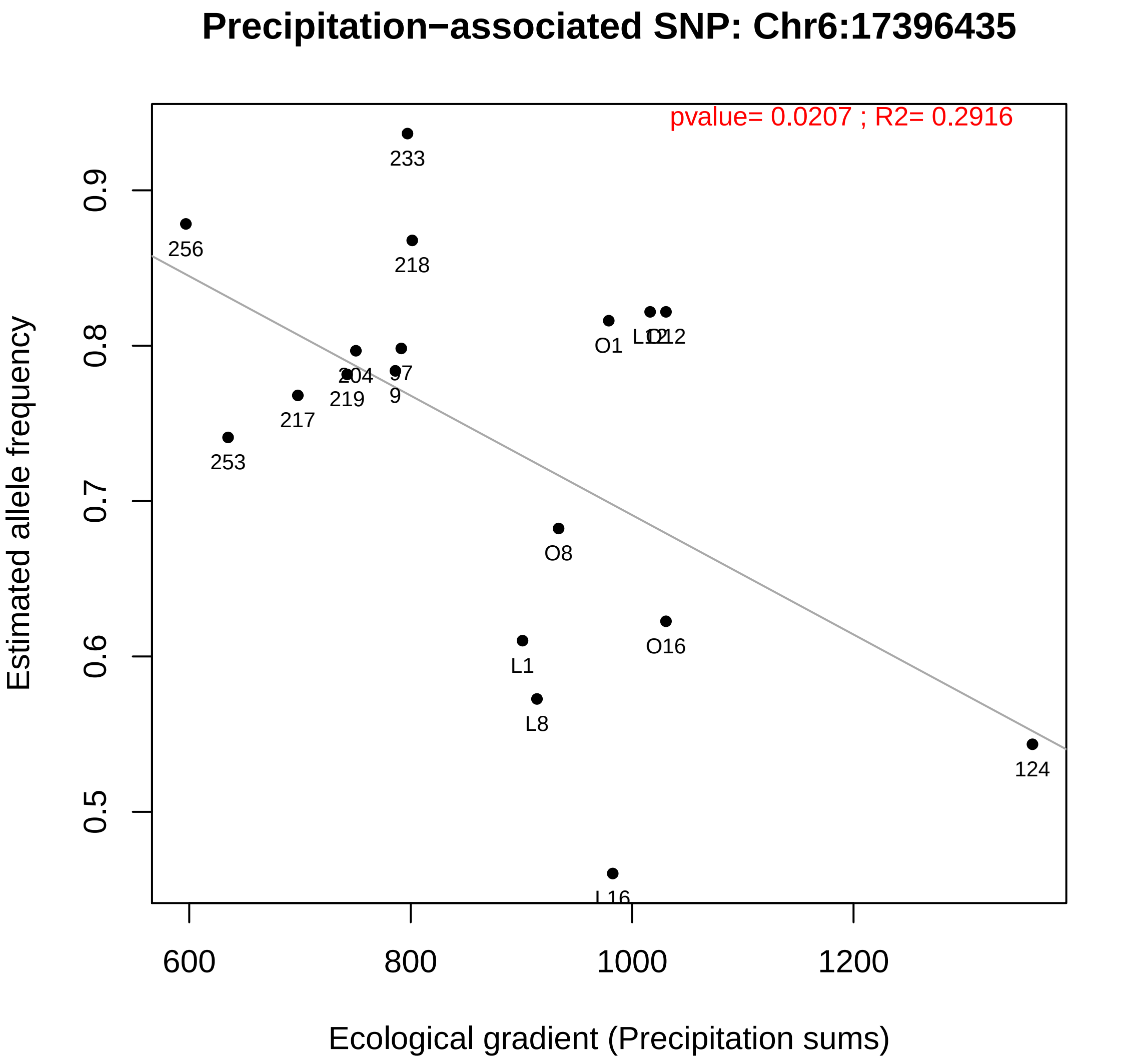
Precipitation−associated SNP: Chr6:17396435
pvalue= 0.0207 ; R2= 0.2916
Estimated allele frequency
Ecological gradient (Precipitation sums)
0.5
0.6
0.7
0.8
0.9
600
800
1000
1200
233
256
218
O1
L12
O12
204
97
219
9
217
253
O8
O16
L1
L8
124
L16

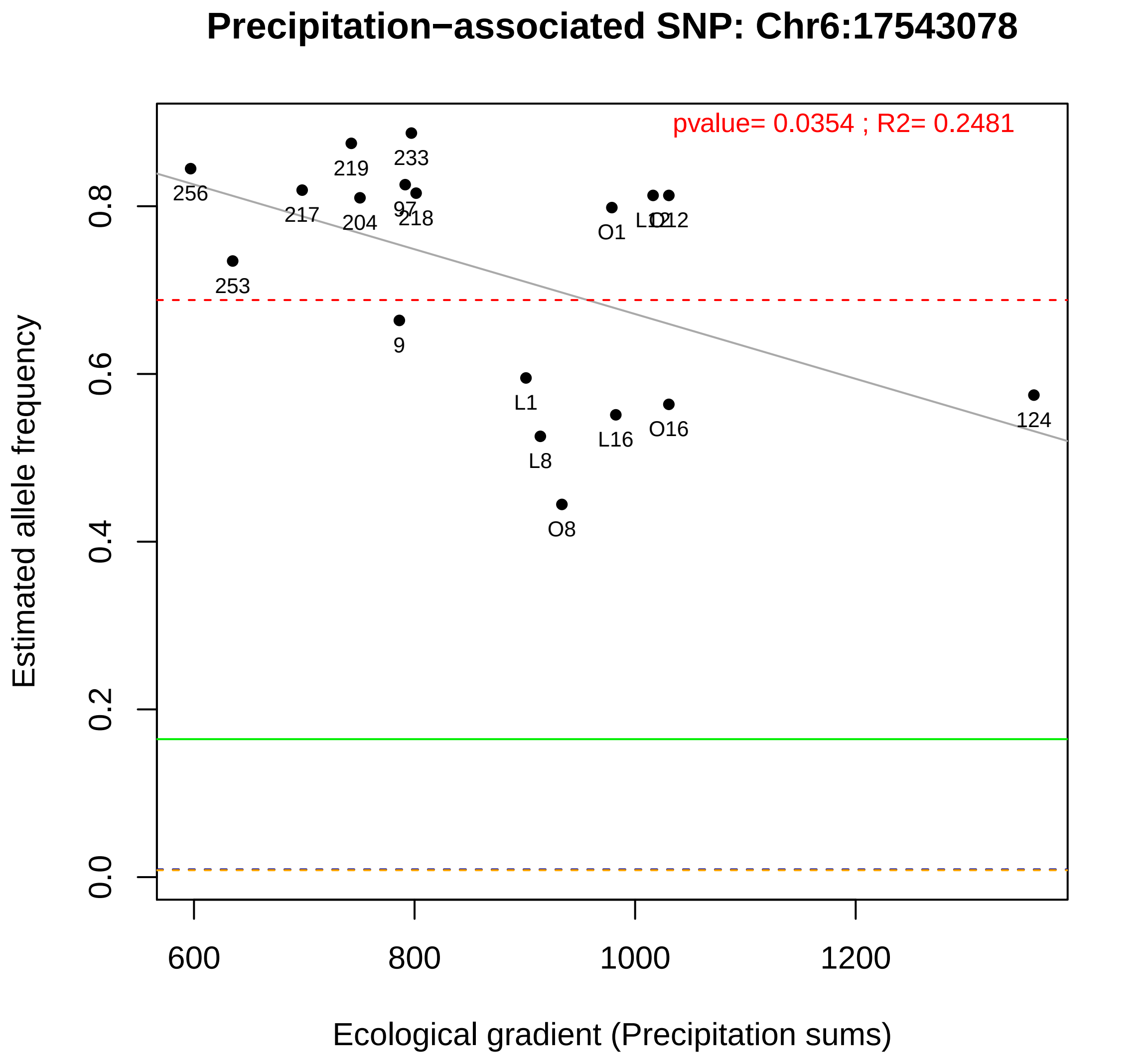
Precipitation−associated SNP: Chr6:17543078
pvalue= 0.0354 ; R2= 0.2481
256
253
217
219
204
233
97
218
9
L1
L8
O8
L16
O1
L12
O12
O16
124
Estimated allele frequency
0.0
0.2
0.4
0.6
0.8
600
800
1000
1200
Ecological gradient (Precipitation sums)

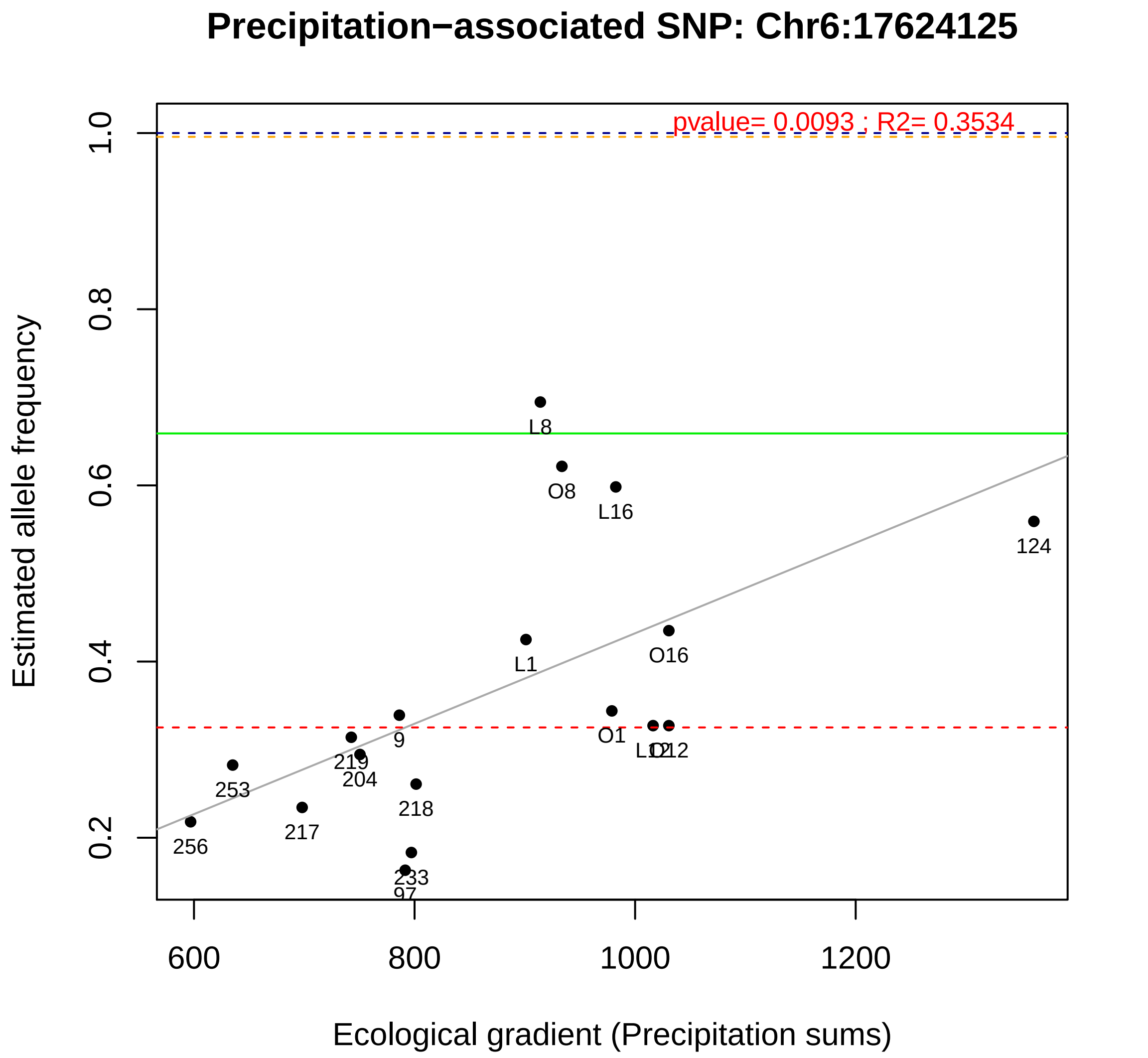
Precipitation−associated SNP: Chr6:17624125
pvalue= 0.0093 ; R2= 0.3534
Estimated allele frequency
0.2
0.4
0.6
0.8
1.0
600
800
1000
1200
Ecological gradient (Precipitation sums)
L8
O8
L16
124
L1
O16
O1
9
219
204
253
218
217
256
233
97

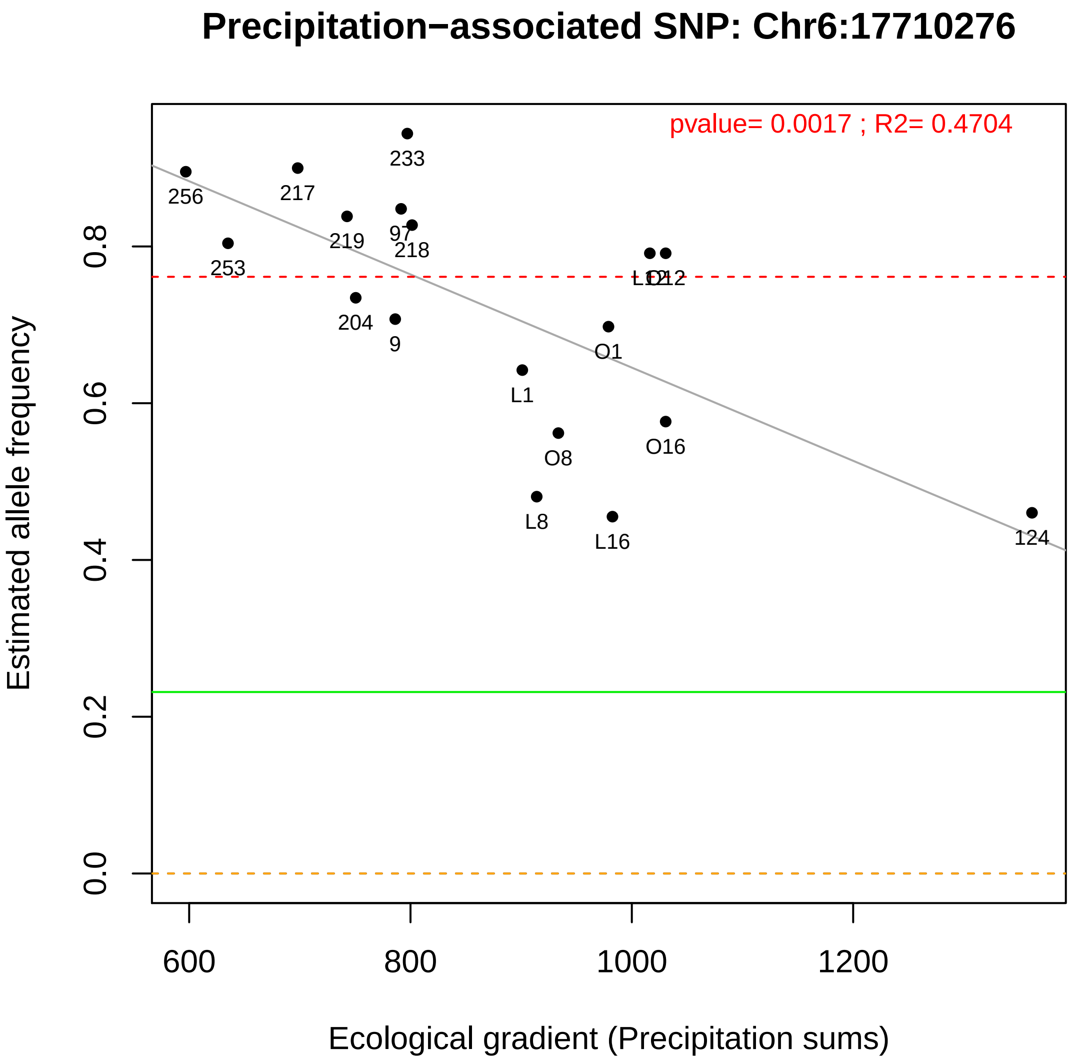

Precipitation−associated SNP: Chr6:17710276
pvalue= 0.0017 ; R2= 0.4704
Estimated allele frequency
Ecological gradient (Precipitation sums)
0.0
0.2
0.4
0.6
0.8
600
800
1000
1200
256
217
233
219
97
218
253
204
9
L1
O1
O8
O16
L8
L16
124
O12

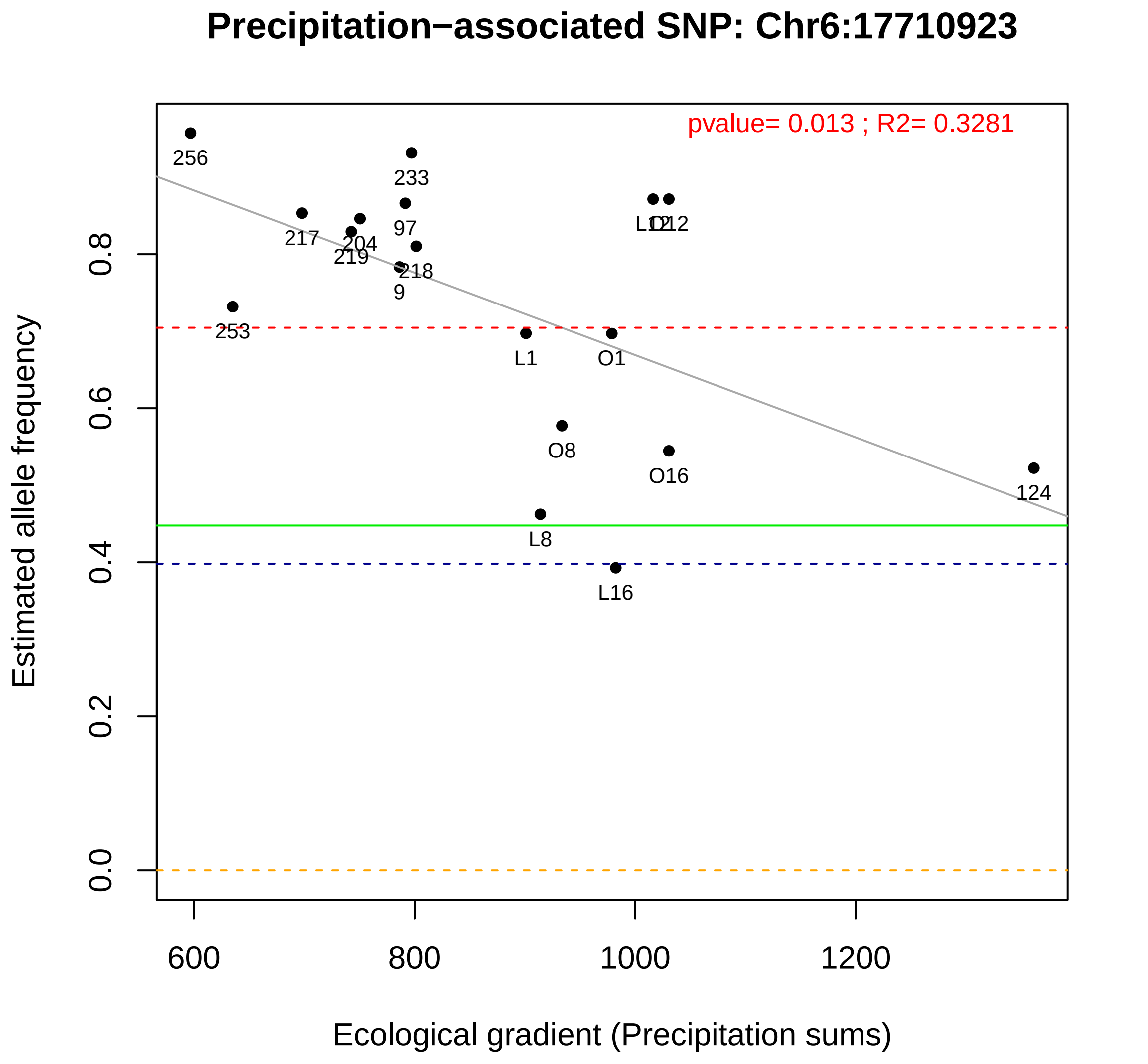

Precipitation−associated SNP: Chr6:17710923
pvalue= 0.013 ; R2= 0.3281
256
233
217
204
219
97
218
9
253
L1
O1
L12
O12
O8
O16
124
L8
L16
0.0
0.2
0.4
0.6
0.8
600
800
1000
1200
Estimated allele frequency
Ecological gradient (Precipitation sums)

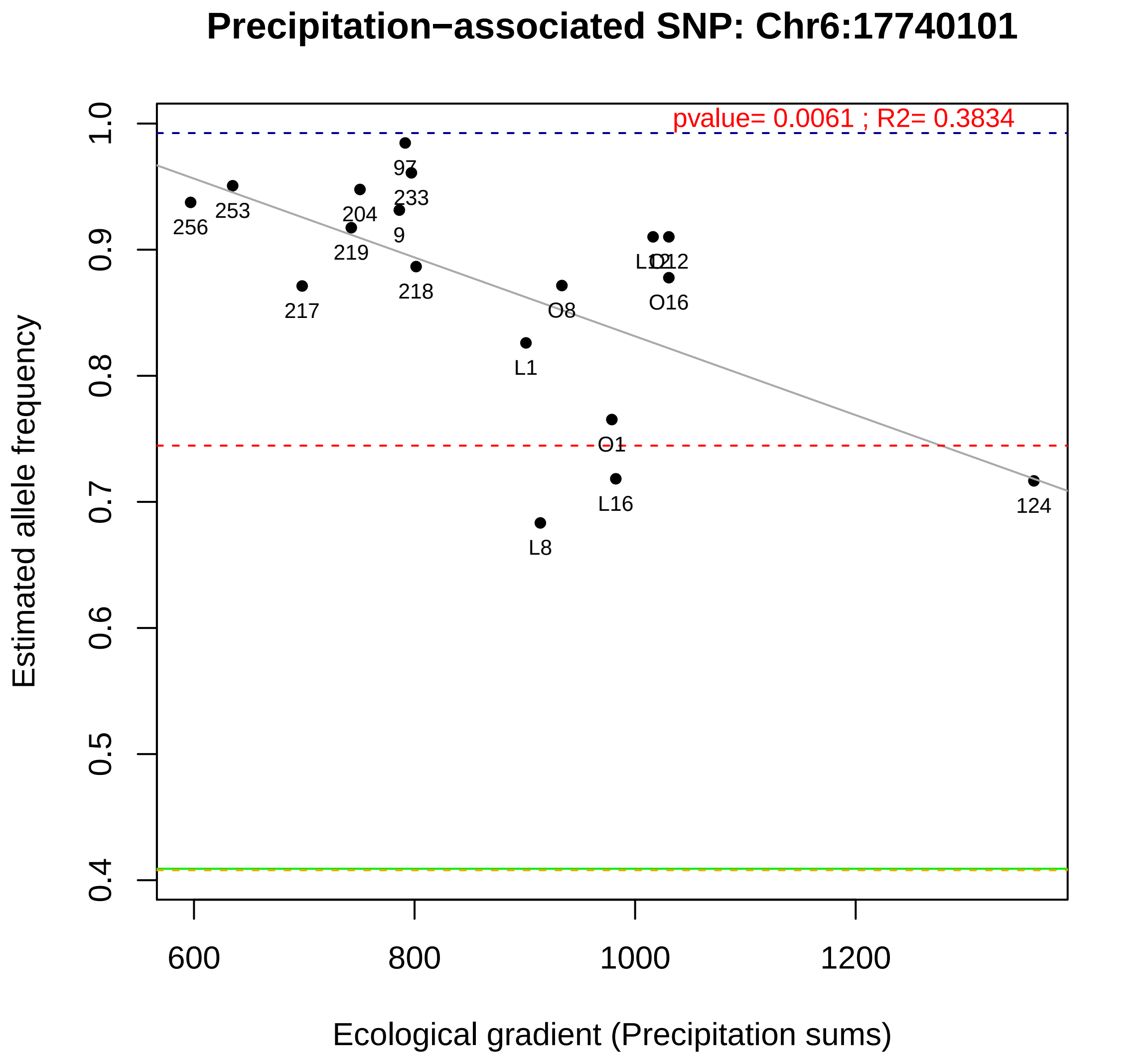
Precipitation−associated SNP: Chr6:17740101
pvalue= 0.0061 ; R2= 0.3834
Estimated allele frequency
1.0
0.9
0.8
0.7
0.6
0.5
0.4
600
800
1000
1200
Ecological gradient (Precipitation sums)
97
233
253
256
204
9
219
218
217
O8
L1
O16
O1
L16
L8
124

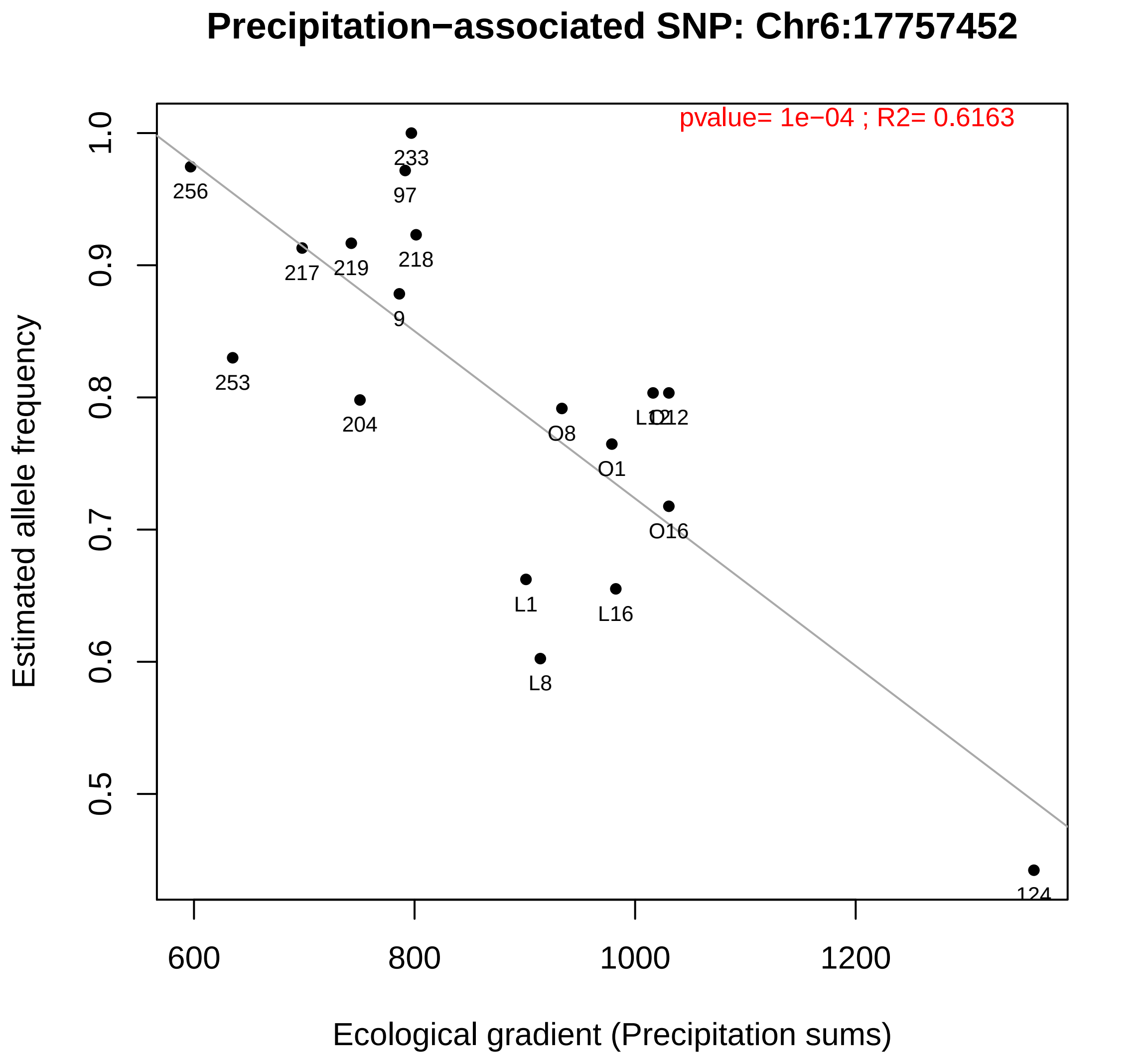
Precipitation−associated SNP: Chr6:17757452
pvalue= 1e−04 ; R2= 0.6163
Estimated allele frequency
Ecological gradient (Precipitation sums)
1.0
0.9
0.8
0.7
0.6
0.5
600
800
1000
1200
233
256
97
217
219
218
9
253
204
O8
L12
O12
O1
O16
L1
L16
L8
124

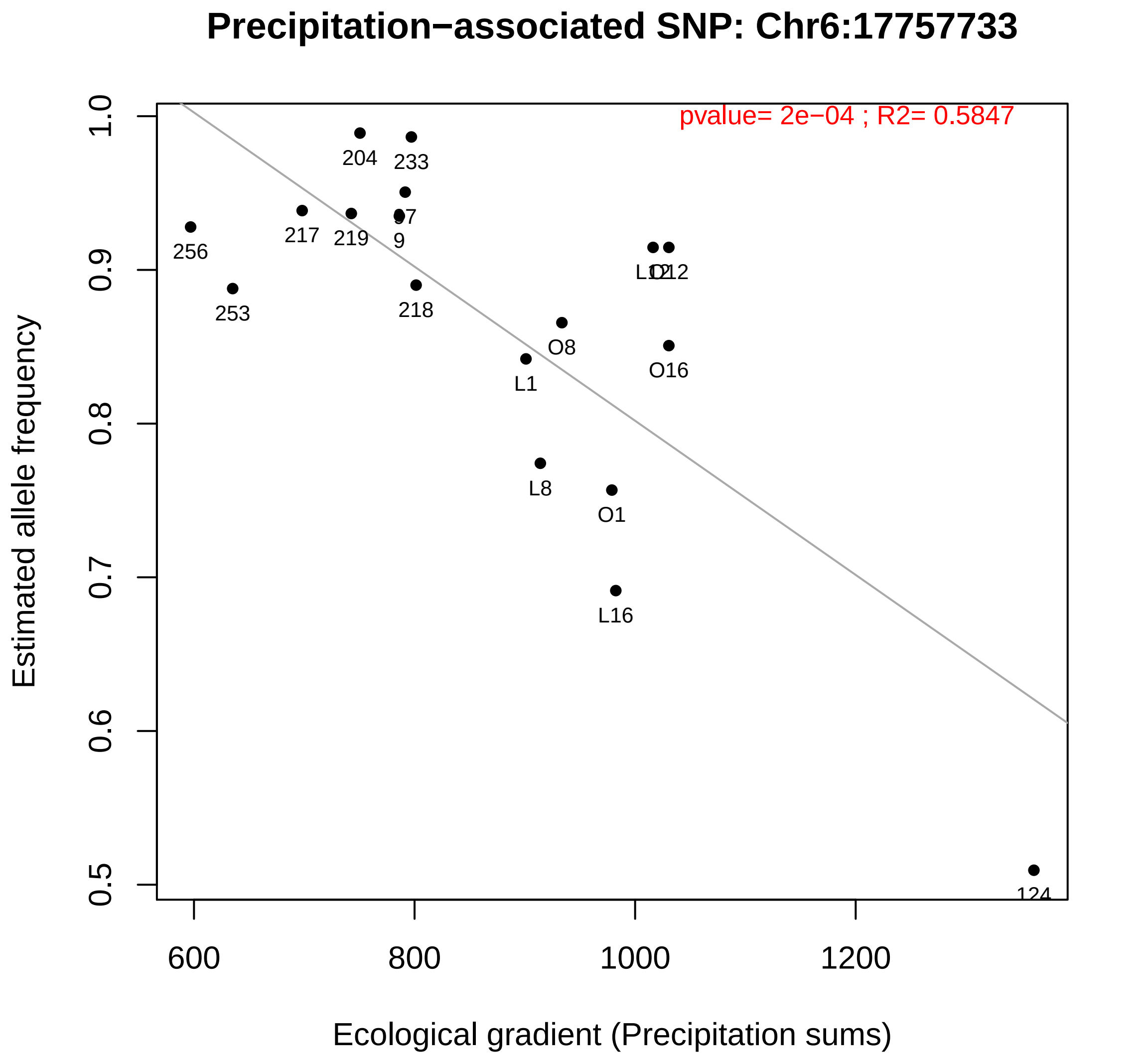

Precipitation−associated SNP: Chr6:17757733
pvalue= 2e−04 ; R2= 0.5847
Estimated allele frequency
Ecological gradient (Precipitation sums)
1.0
0.9
0.8
0.7
0.6
0.5
600
800
1000
1200
204
233
256
217
219
97
9
253
218
L12
O12
O8
L1
O16
L8
O1
L16
124

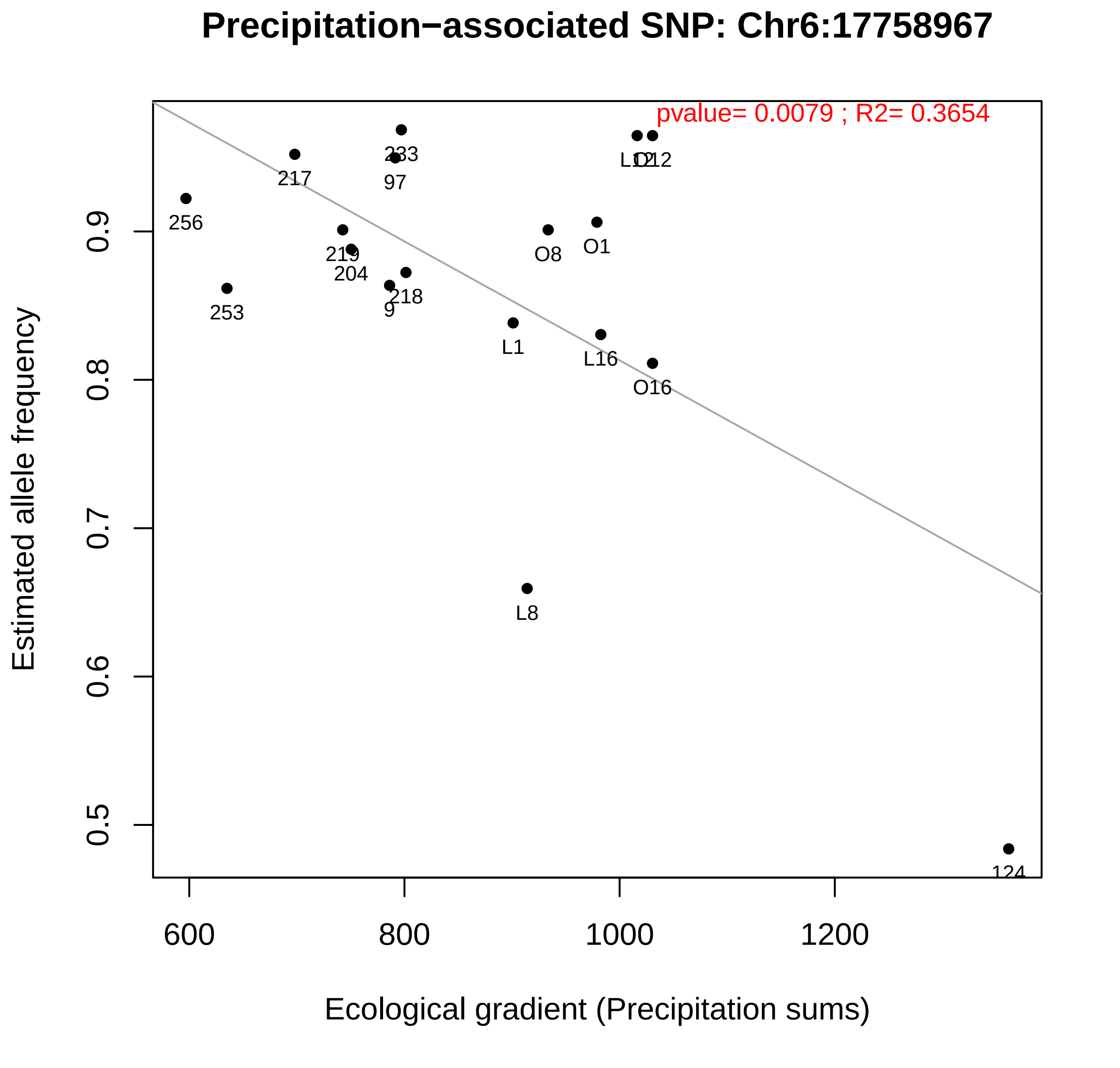
Precipitation−associated SNP: Chr6:17758967
pvalue= 0.0079 ; R2= 0.3654
Estimated allele frequency
Ecological gradient (Precipitation sums)
0.5
0.6
0.7
0.8
0.9
600
800
1000
1200
256
217
233
97
219
204
253
218
9
L1
O8
O1
L16
O16
L8
124

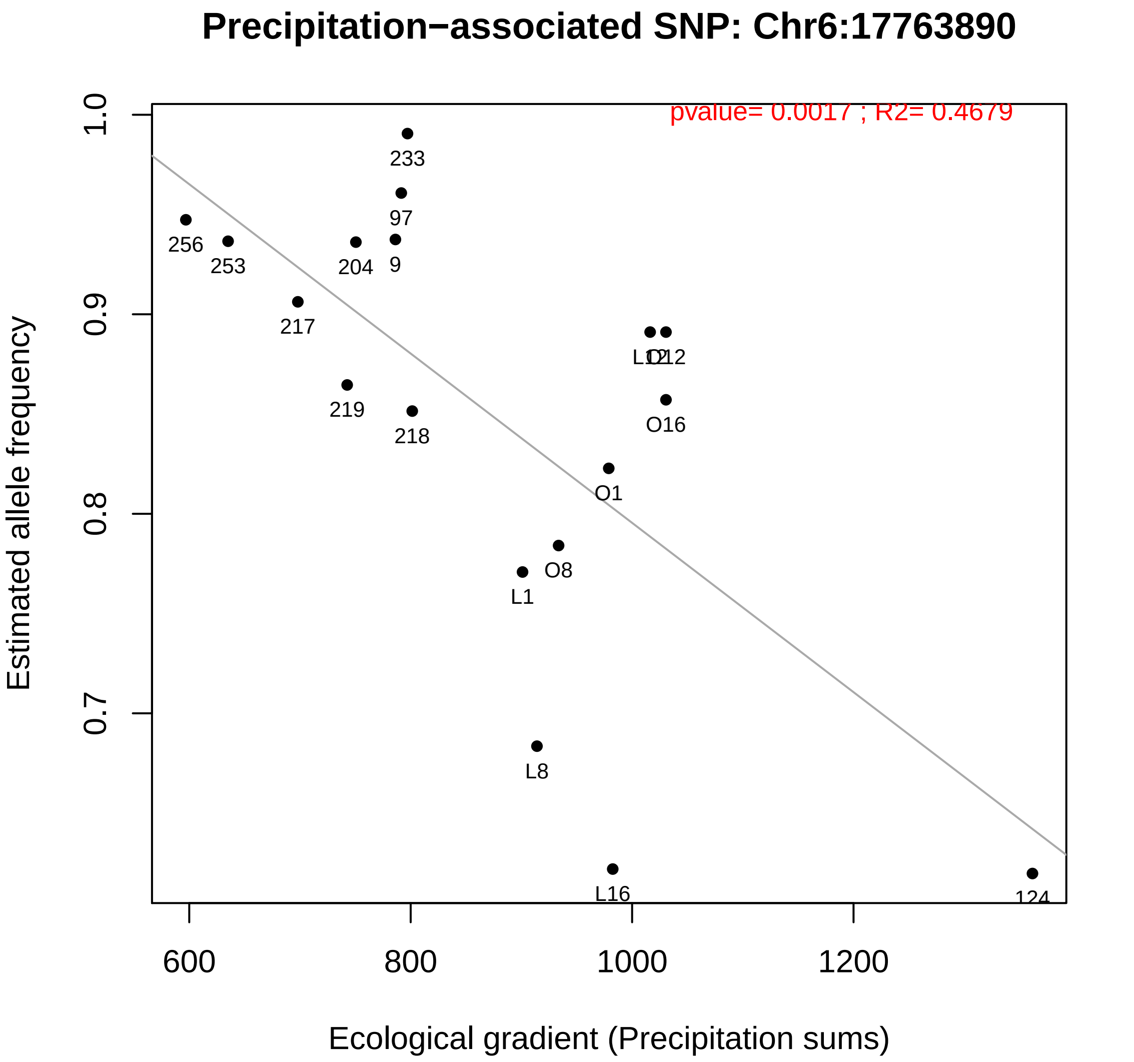

Precipitation−associated SNP: Chr6:17763890
pvalue= 0.0017 ; R2= 0.4679
Estimated allele frequency
Ecological gradient (Precipitation sums)
1.0
0.9
0.8
0.7
600
800
1000
1200
233
97
256
253
204
9
217
219
218
L12
O12
O16
O1
O8
L1
L8
L16
124

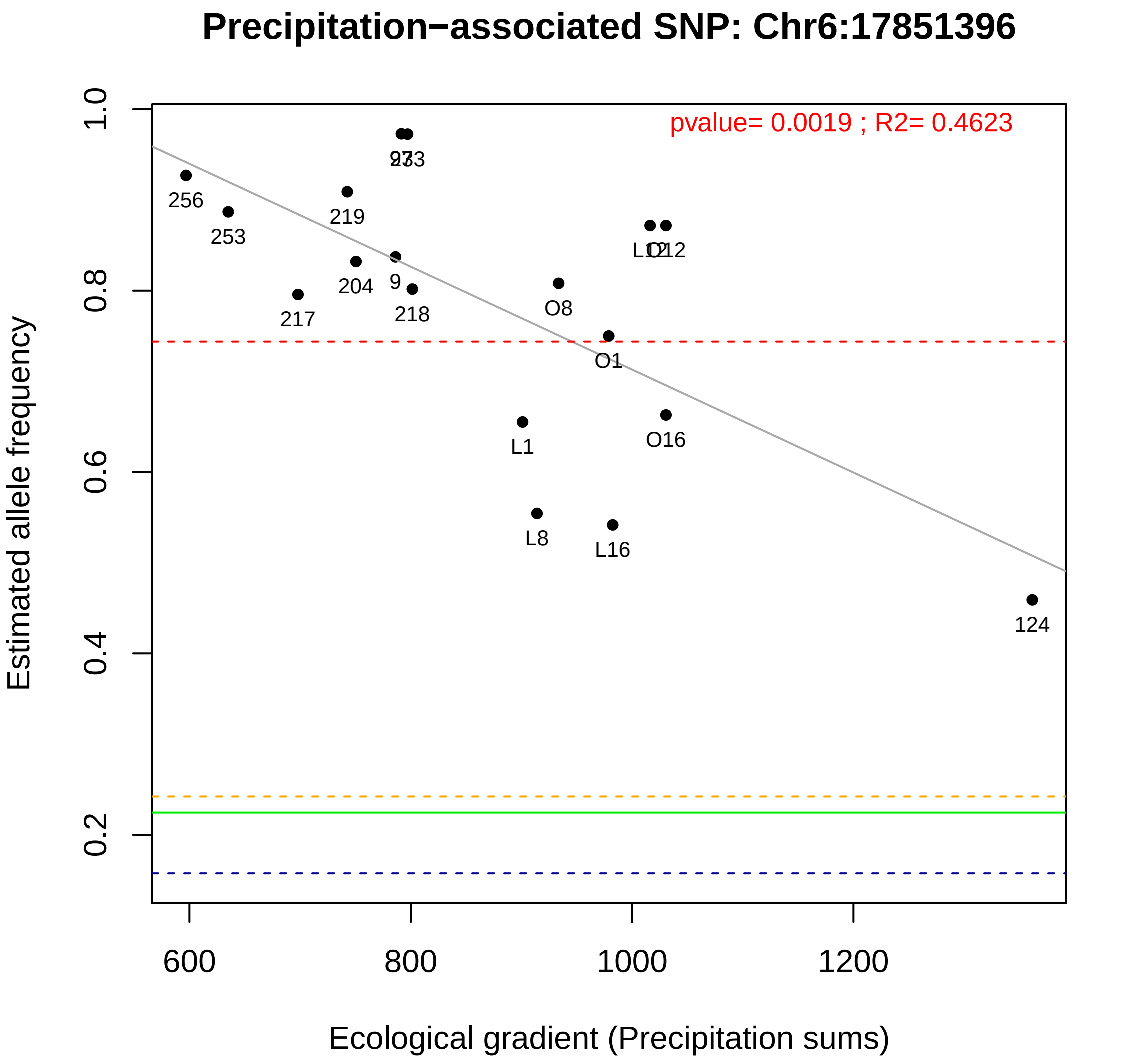

Precipitation−associated SNP: Chr6:17851396
pvalue= 0.0019 ; R2= 0.4623
Estimated allele frequency
Ecological gradient (Precipitation sums)
0.2
0.4
0.6
0.8
1.0
600
800
1000
1200
256
253
219
217
204
9
218
O8
O1
L1
L8
L16
O16
124

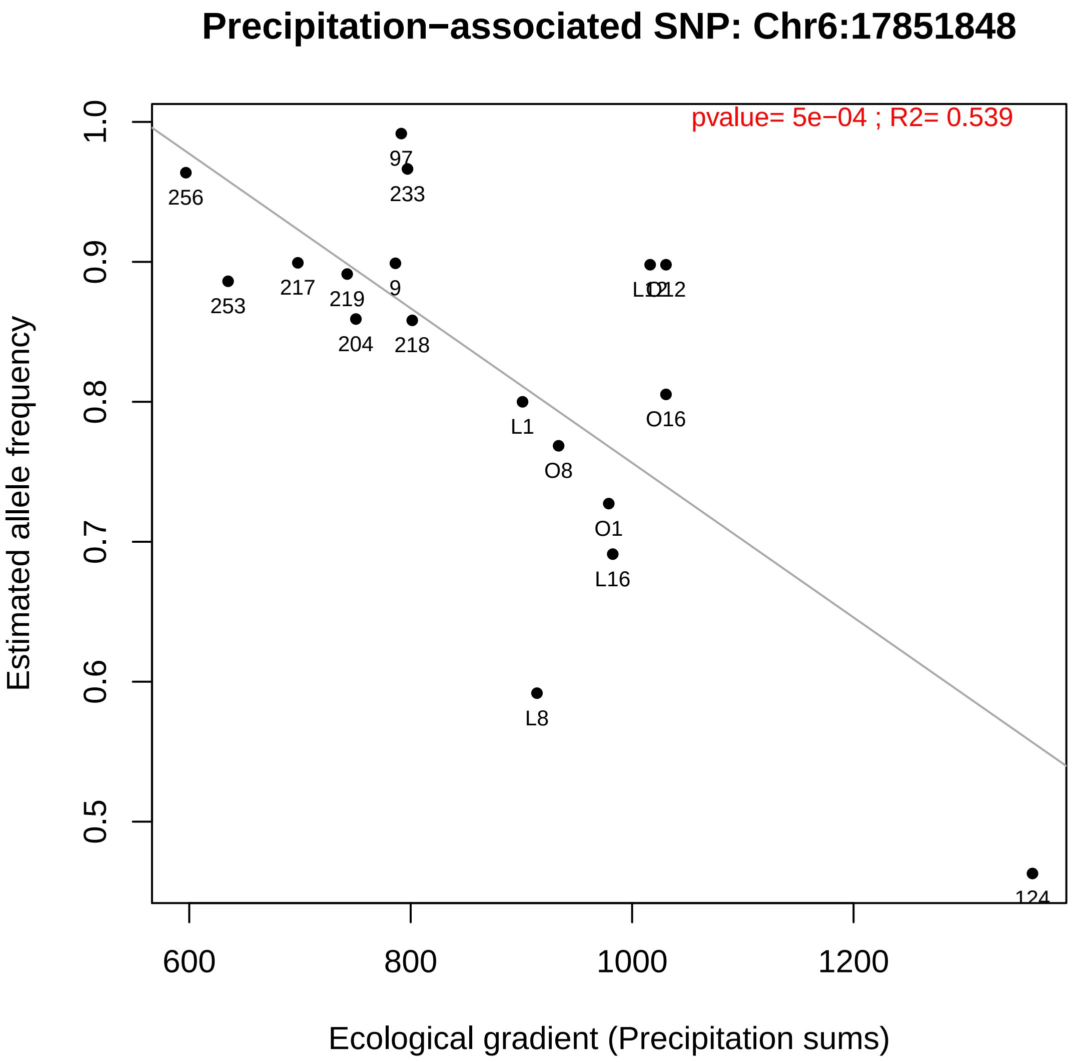
Precipitation−associated SNP: Chr6:17851848
pvalue= 5e−04 ; R2= 0.539
Estimated allele frequency
1.0
0.9
0.8
0.7
0.6
0.5
600
800
1000
1200
Ecological gradient (Precipitation sums)
97
233
256
217
253
219
9
204
218
O12
O16
L1
O8
O1
L16
L8
124

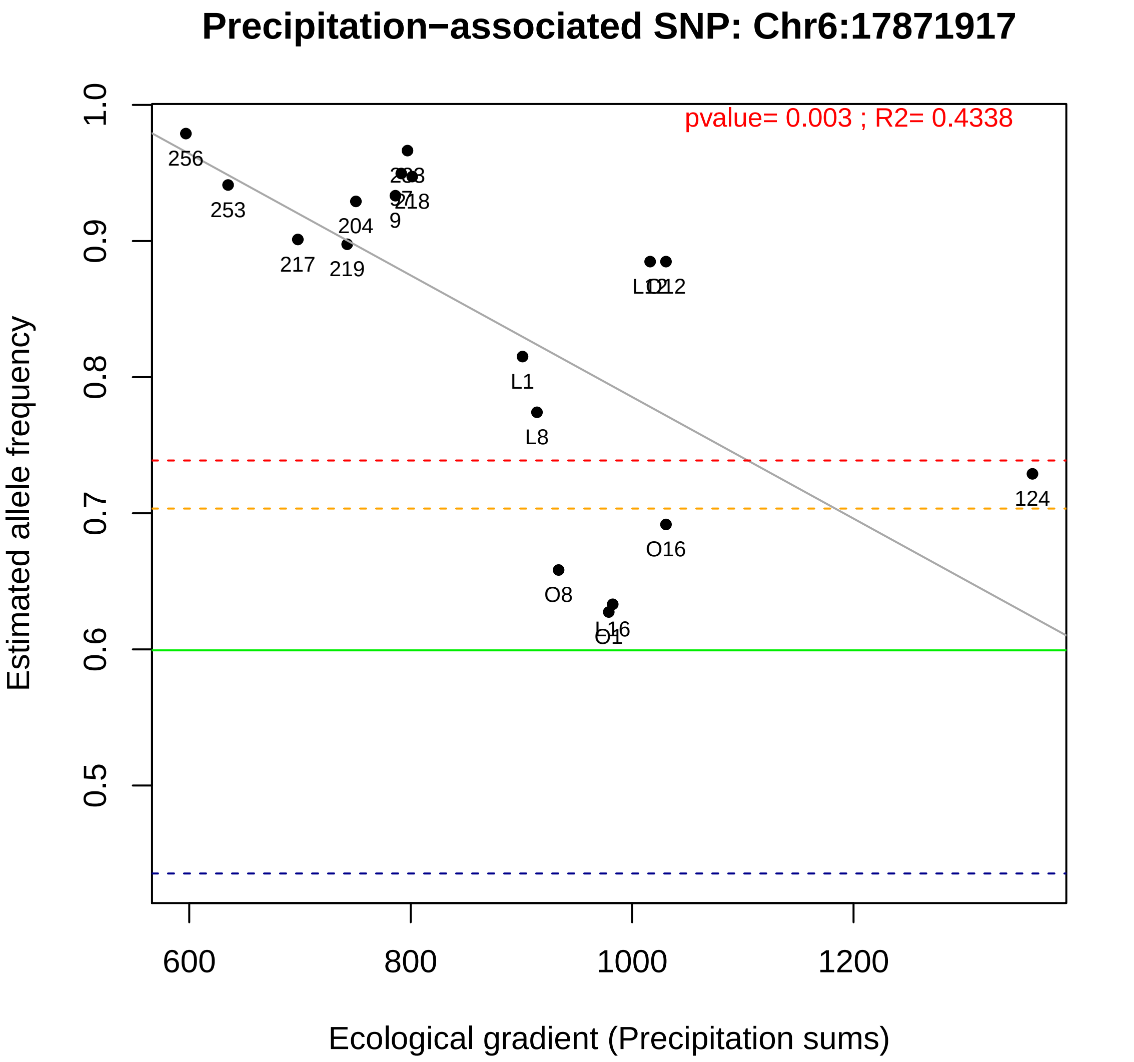

Precipitation−associated SNP: Chr6:17871917
pvalue= 0.003 ; R2= 0.4338
Estimated allele frequency
Ecological gradient (Precipitation sums)
0.5
0.6
0.7
0.8
0.9
1.0
600
800
1000
1200
256
253
217
219
204
218
L1
L8
O16
O8
L16
O1
L12
O12
124

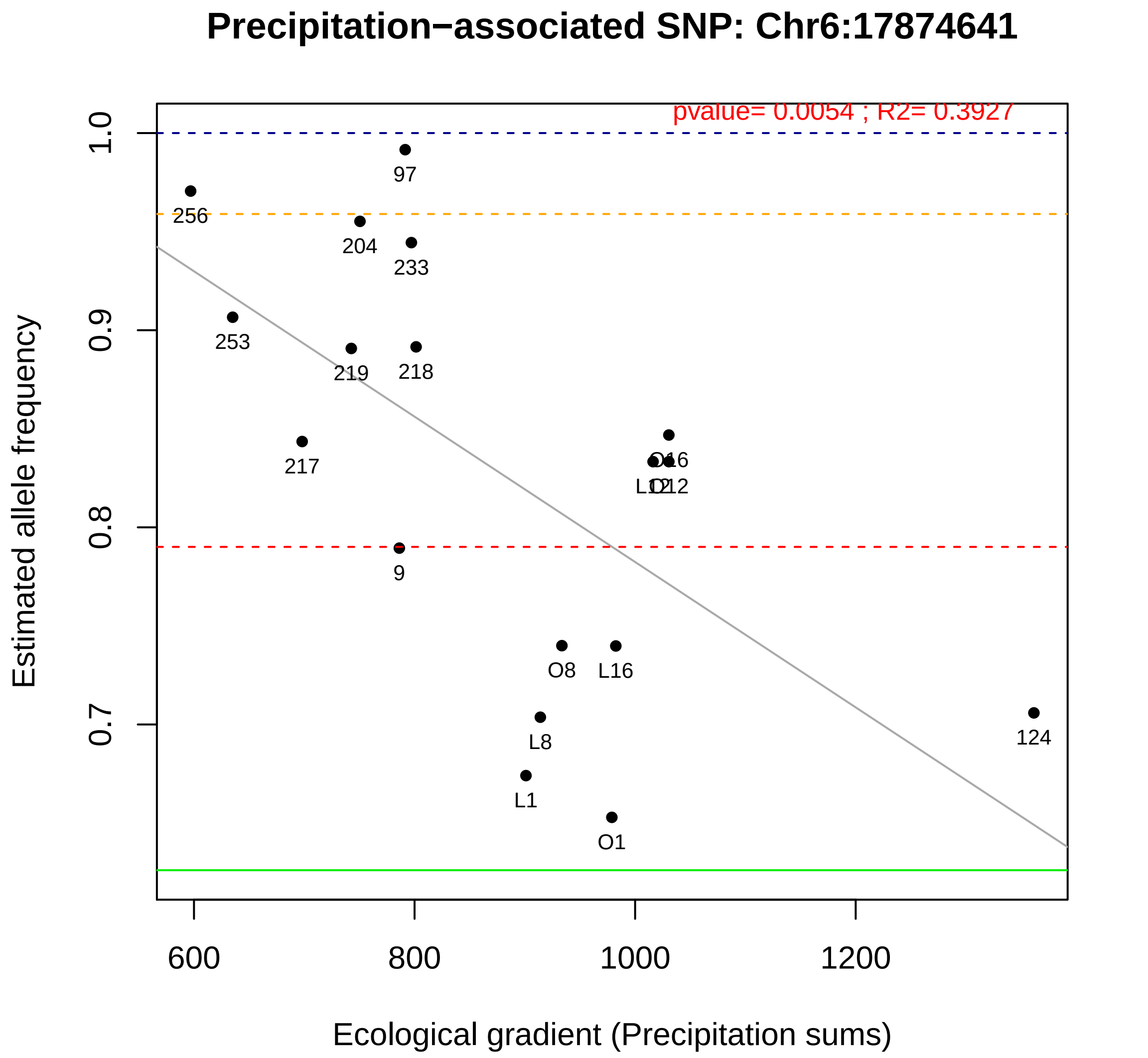
Precipitation−associated SNP: Chr6:17874641
pvalue= 0.0054 ; R2= 0.3927
Estimated allele frequency
1.0
0.9
0.8
0.7
Ecological gradient (Precipitation sums)
600
800
1000
1200
97
256
204
233
253
219
218
217
9
O8
L16
L8
L1
O1
124

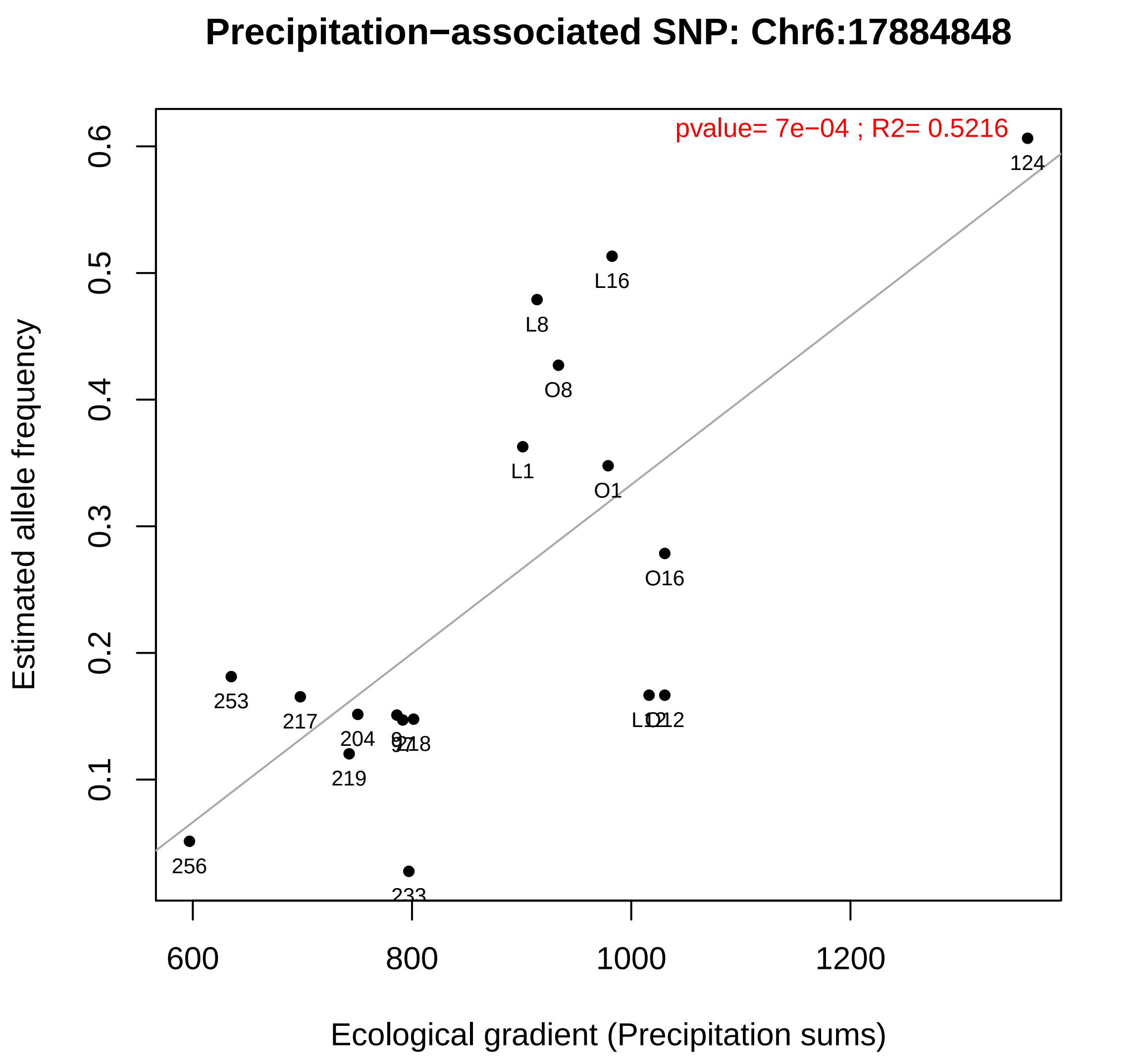

Precipitation−associated SNP: Chr6:17884848
pvalue= 7e−04 ; R2= 0.5216
Estimated allele frequency
Ecological gradient (Precipitation sums)
0.1
0.2
0.3
0.4
0.5
0.6
600
800
1000
1200
124
L16
L8
O8
L1
O1
O16
253
217
204
219
218
256
233
O12

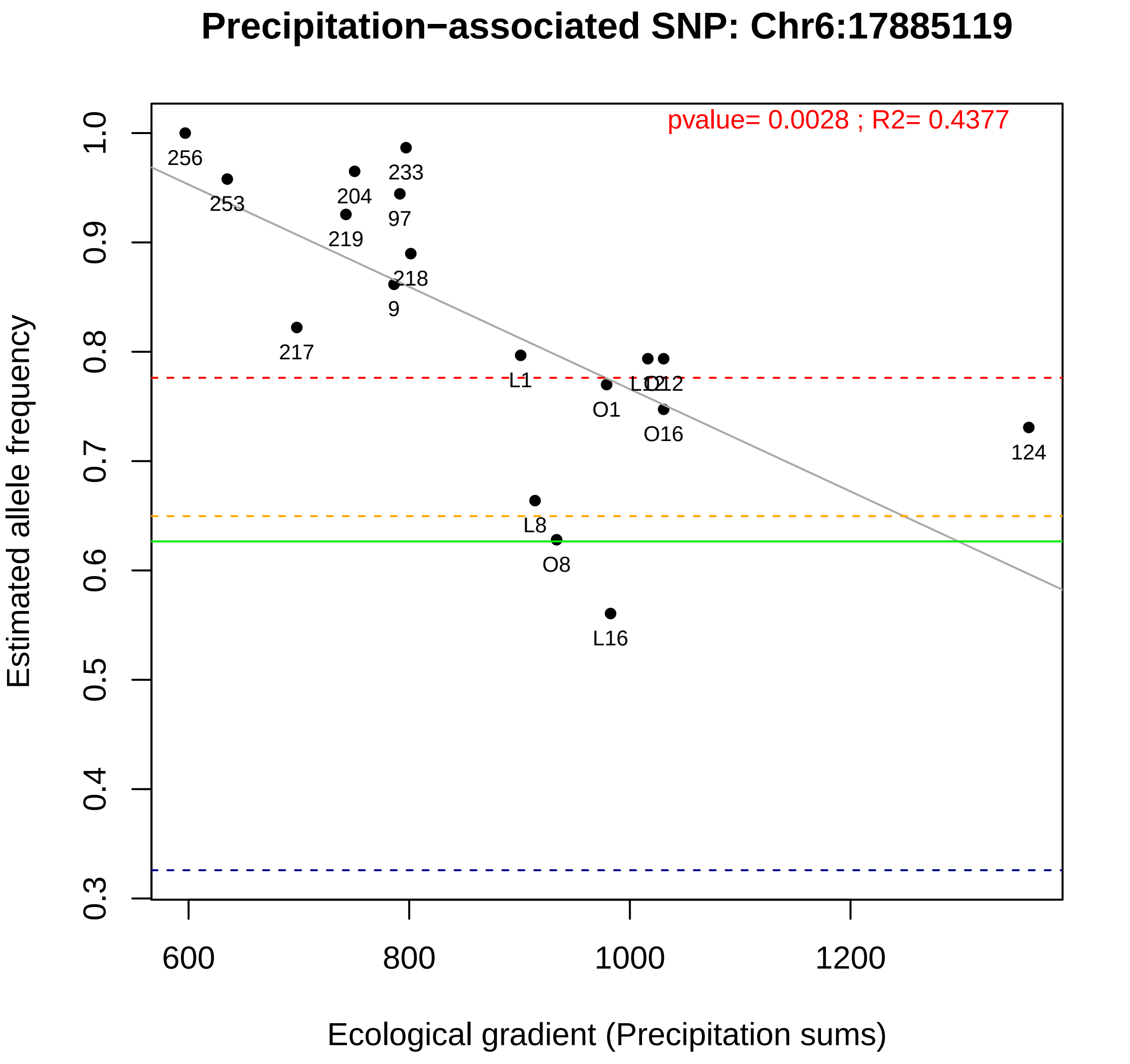
Precipitation−associated SNP: Chr6:17885119
pvalue= 0.0028 ; R2= 0.4377
Estimated allele frequency
Ecological gradient (Precipitation sums)
0.3
0.4
0.5
0.6
0.7
0.8
0.9
1.0
600
800
1000
1200
256
253
204
233
97
219
218
9
217
L1
L12
O12
O1
O16
124
L8
O8
L16

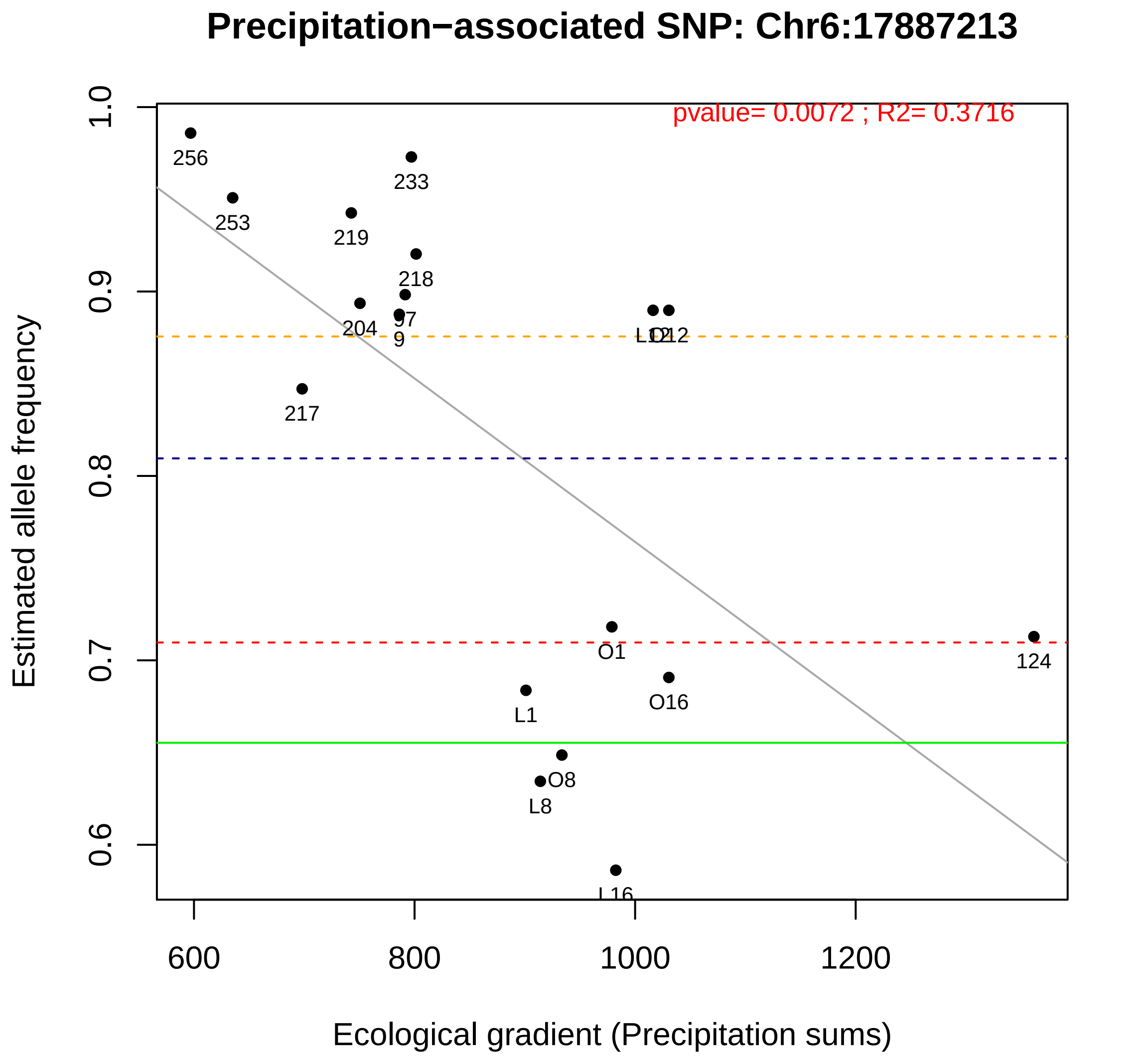
Precipitation−associated SNP: Chr6:17887213
pvalue= 0.0072 ; R2= 0.3716
Estimated allele frequency
Ecological gradient (Precipitation sums)
1.0
0.9
0.8
0.7
0.6
600
800
1000
1200
256
233
253
219
218
204
97
9
L10
O12
217
O1
124
O16
L1
O8
L8
L16

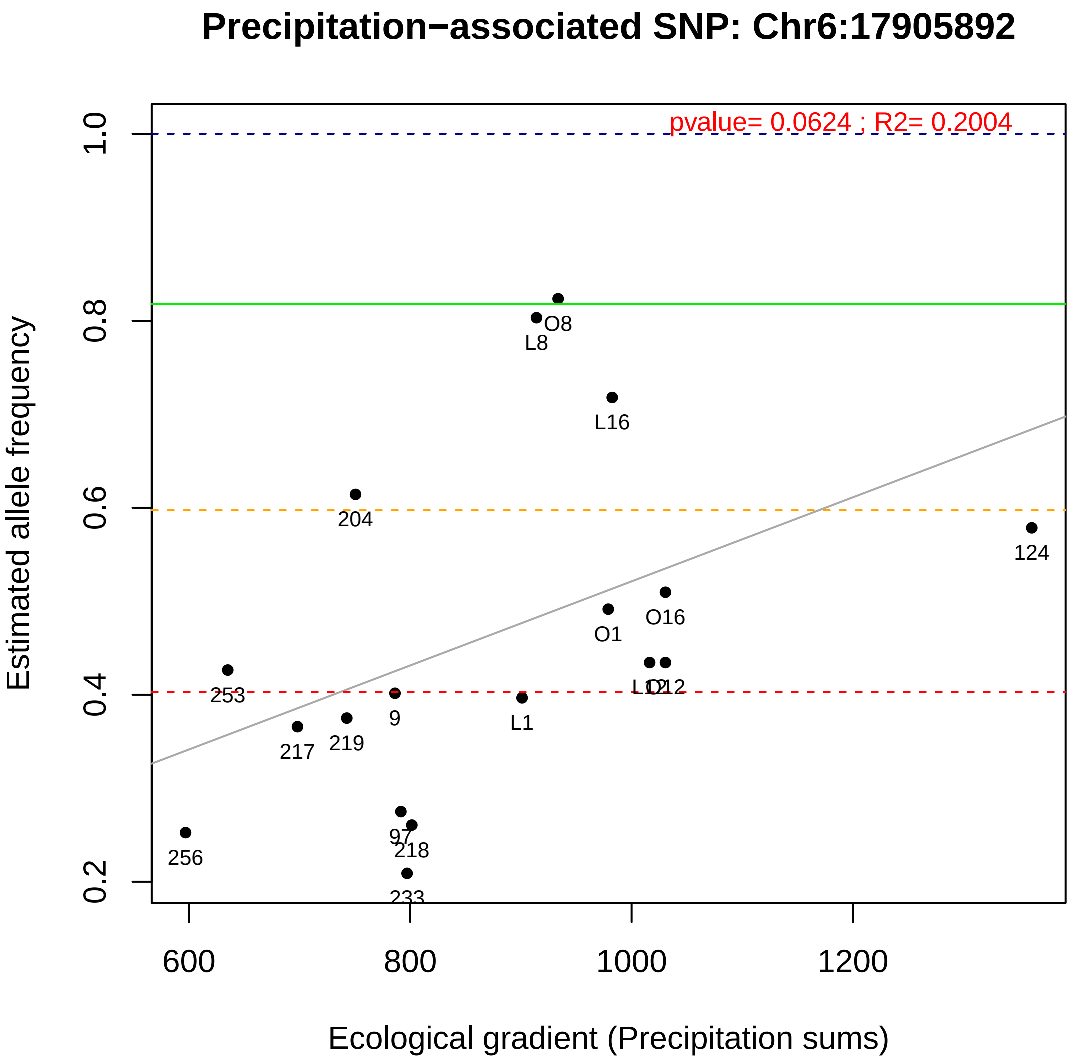
Precipitation−associated SNP: Chr6:17905892
pvalue= 0.0624 ; R2= 0.2004
Estimated allele frequency
Ecological gradient (Precipitation sums)
0.2
0.4
0.6
0.8
1.0
600
800
1000
1200
O8
L8
L16
204
124
O16
O1
253
9
L1
217
219
97
218
256
233

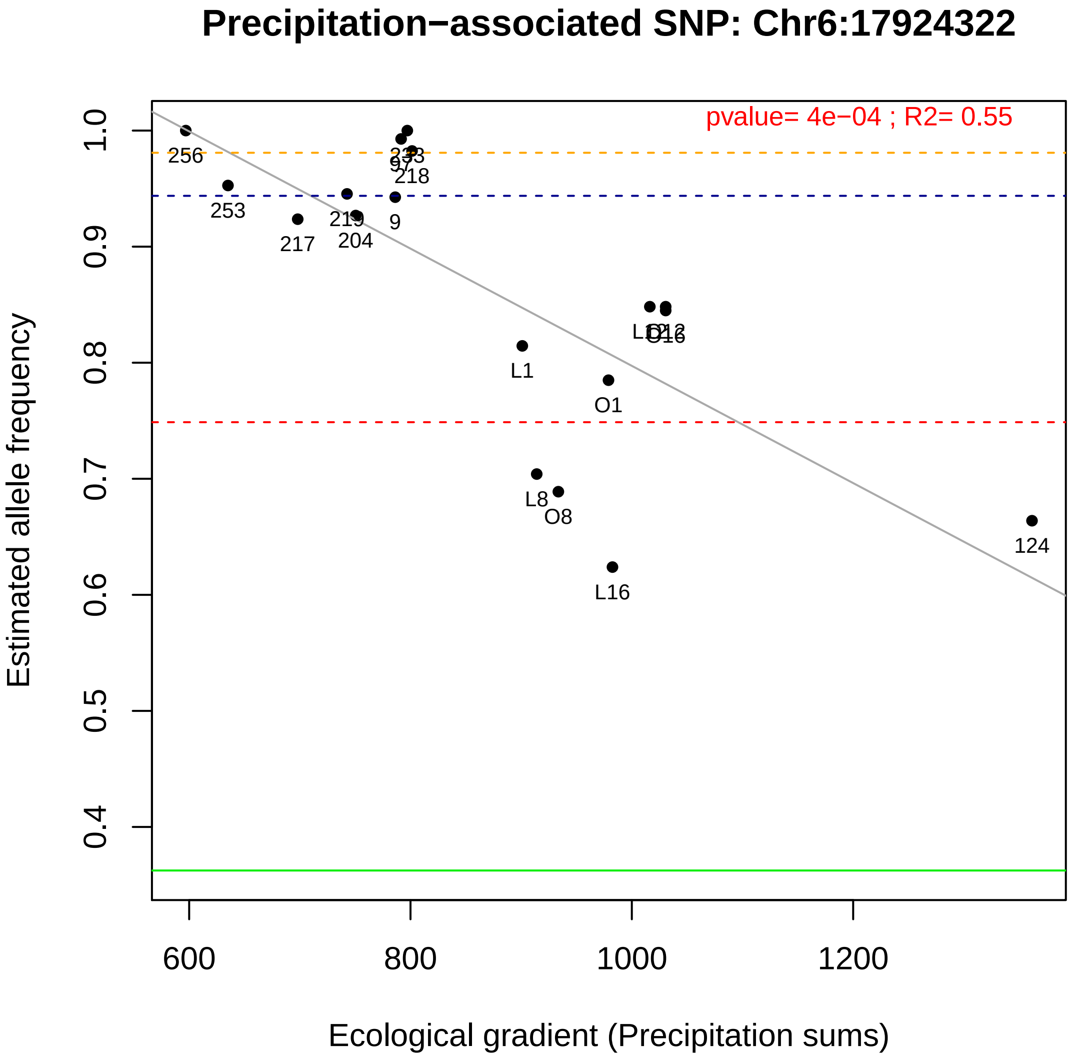

Precipitation−associated SNP: Chr6:17924322
pvalue= 4e−04 ; R2= 0.55
Estimated allele frequency
Ecological gradient (Precipitation sums)
0.4
0.5
0.6
0.7
0.8
0.9
1.0
600
800
1000
1200
256
253
217
219
204
9
218
L1
O1
L8
O8
L16
O16
124

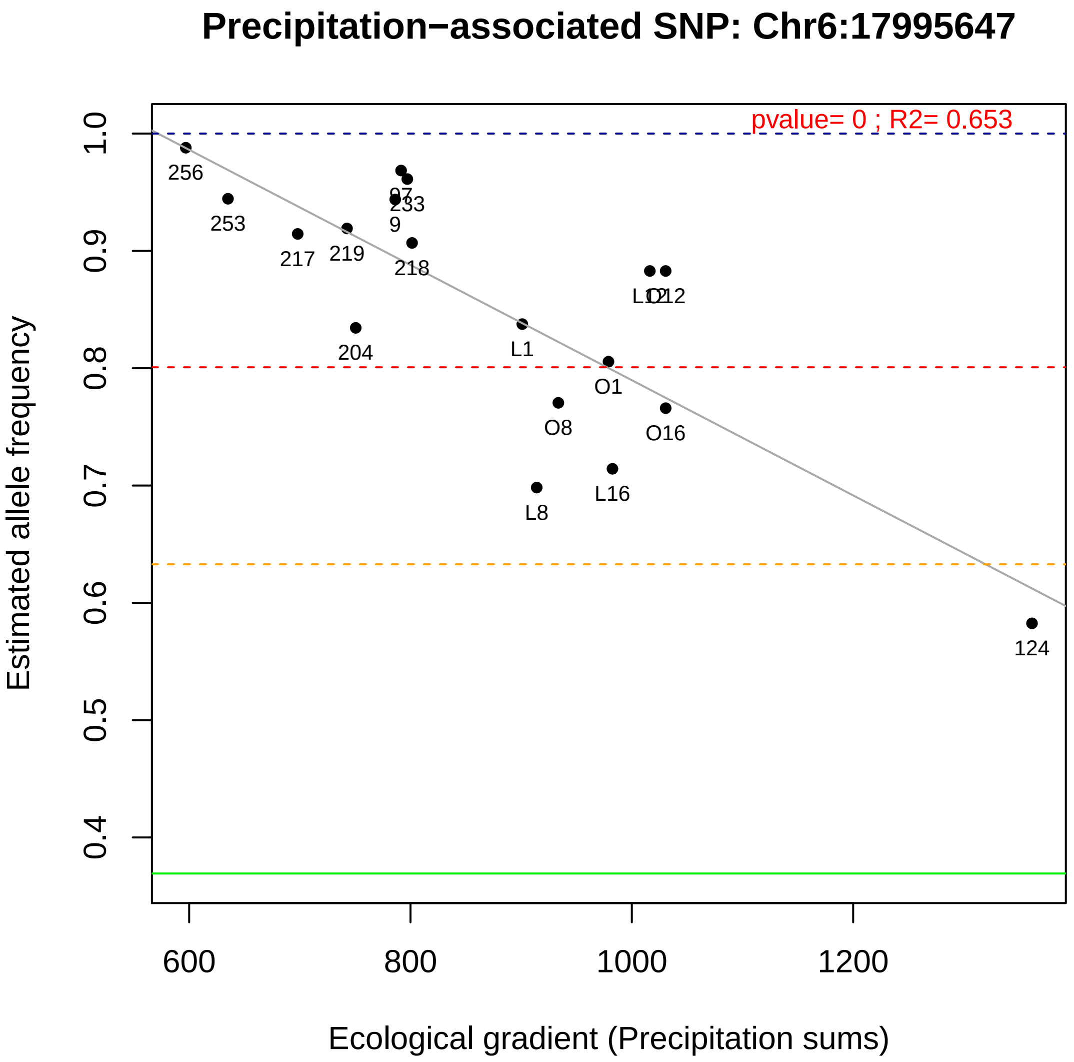

Precipitation−associated SNP: Chr6:17995647
pvalue= 0 ; R2= 0.653
Estimated allele frequency
Ecological gradient (Precipitation sums)
1.0
0.9
0.8
0.7
0.6
0.5
0.4
600
800
1000
1200
256
253
217
219
233
9
218
204
L1
O1
O8
O16
L16
L8
124

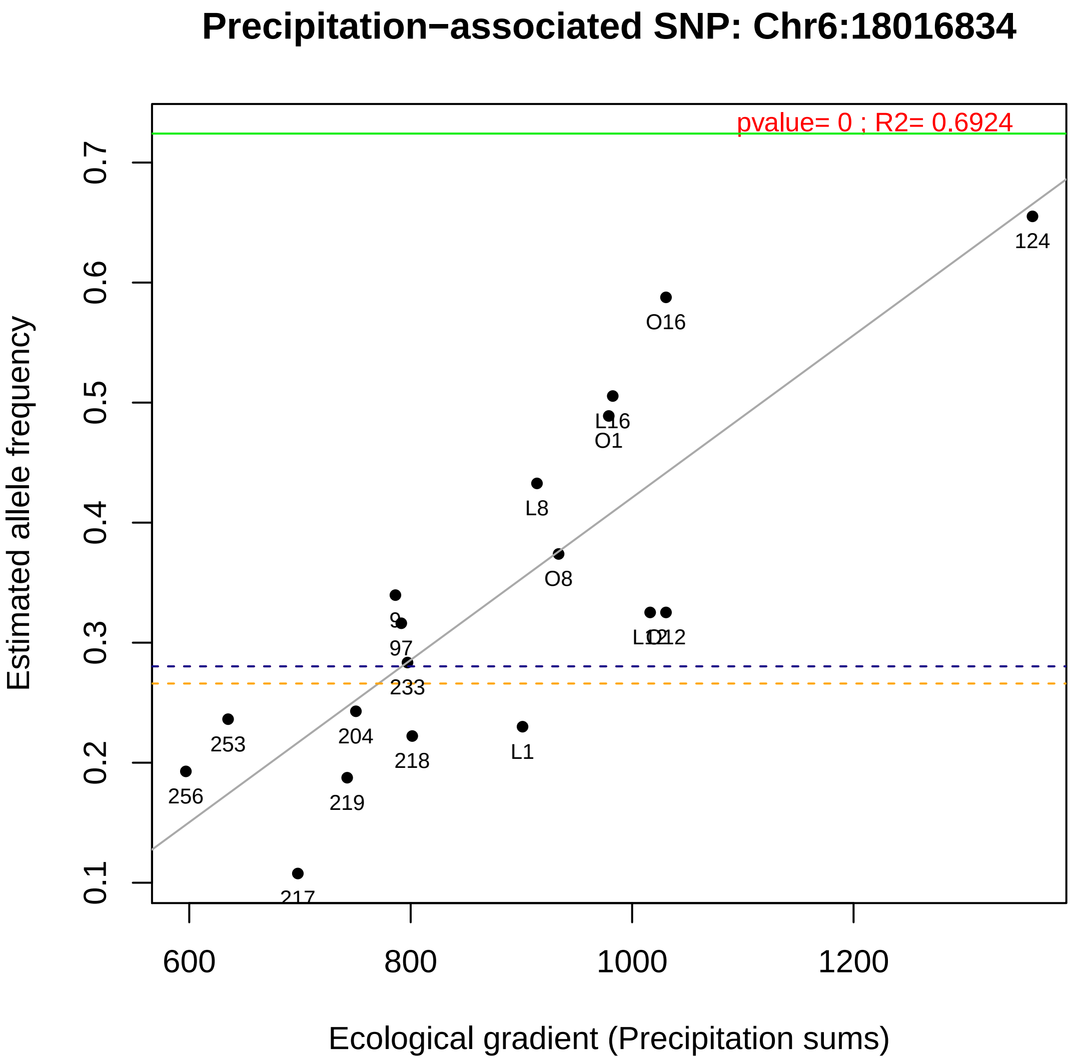

Precipitation−associated SNP: Chr6:18016834
pvalue= 0 ; R2= 0.6924
Estimated allele frequency
Ecological gradient (Precipitation sums)
0.1
0.2
0.3
0.4
0.5
0.6
0.7
600
800
1000
1200
124
O16
L16
O1
L8
O8
L10
O12
97
233
204
253
256
219
218
L1
217

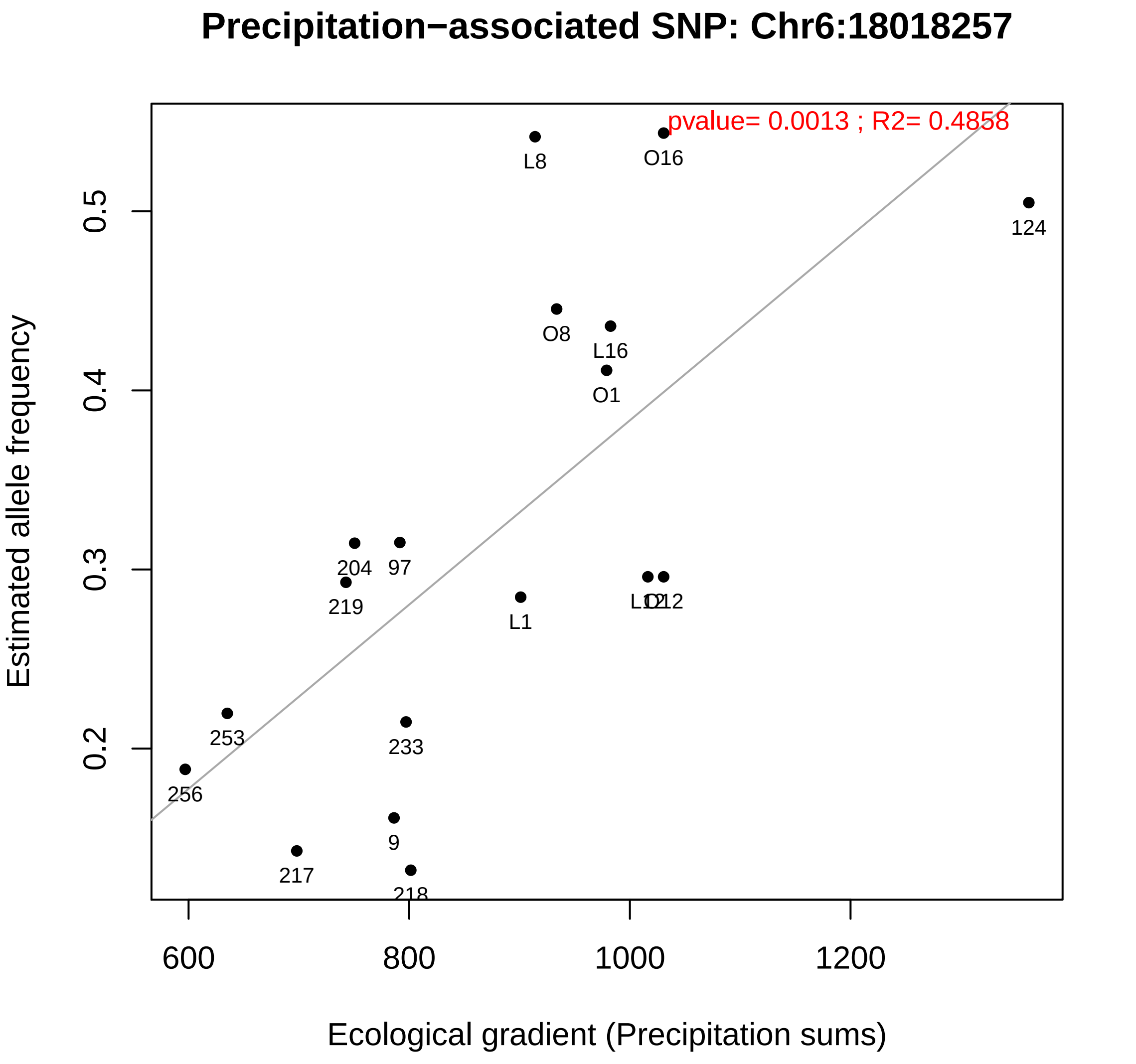
Precipitation−associated SNP: Chr6:18018257
pvalue= 0.0013 ; R2= 0.4858
L8
O16
124
O8
L16
O1
204
97
219
L1
L12
O12
253
233
256
9
217
218
0.2
0.3
0.4
0.5
600
800
1000
1200
Estimated allele frequency
Ecological gradient (Precipitation sums)

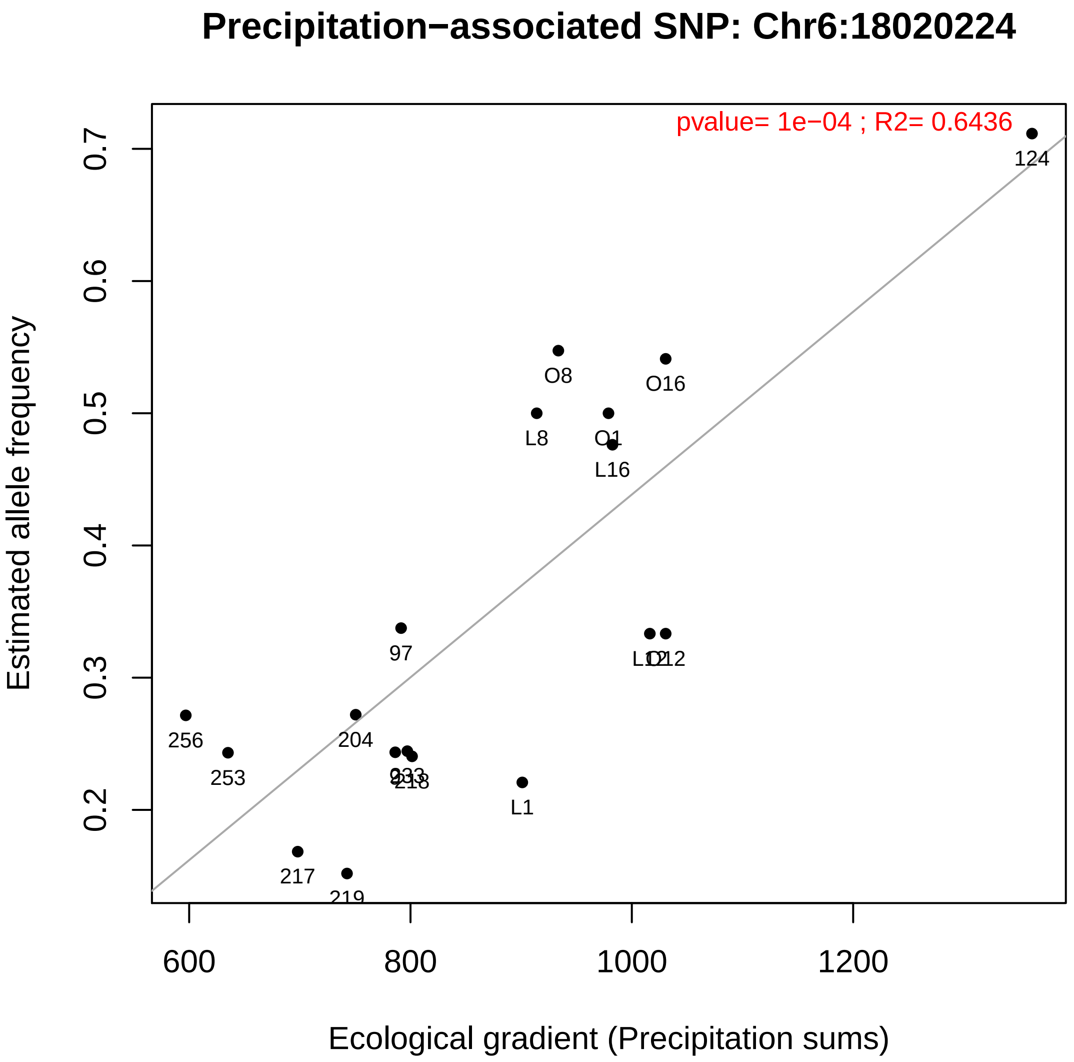

Precipitation−associated SNP: Chr6:18020224
pvalue= 1e−04 ; R2= 0.6436
Estimated allele frequency
Ecological gradient (Precipitation sums)
0.2
0.3
0.4
0.5
0.6
0.7
600
800
1000
1200
124
O8
O16
L8
O1
L16
97
L12
O12
256
204
253
233
218
L1
217
219

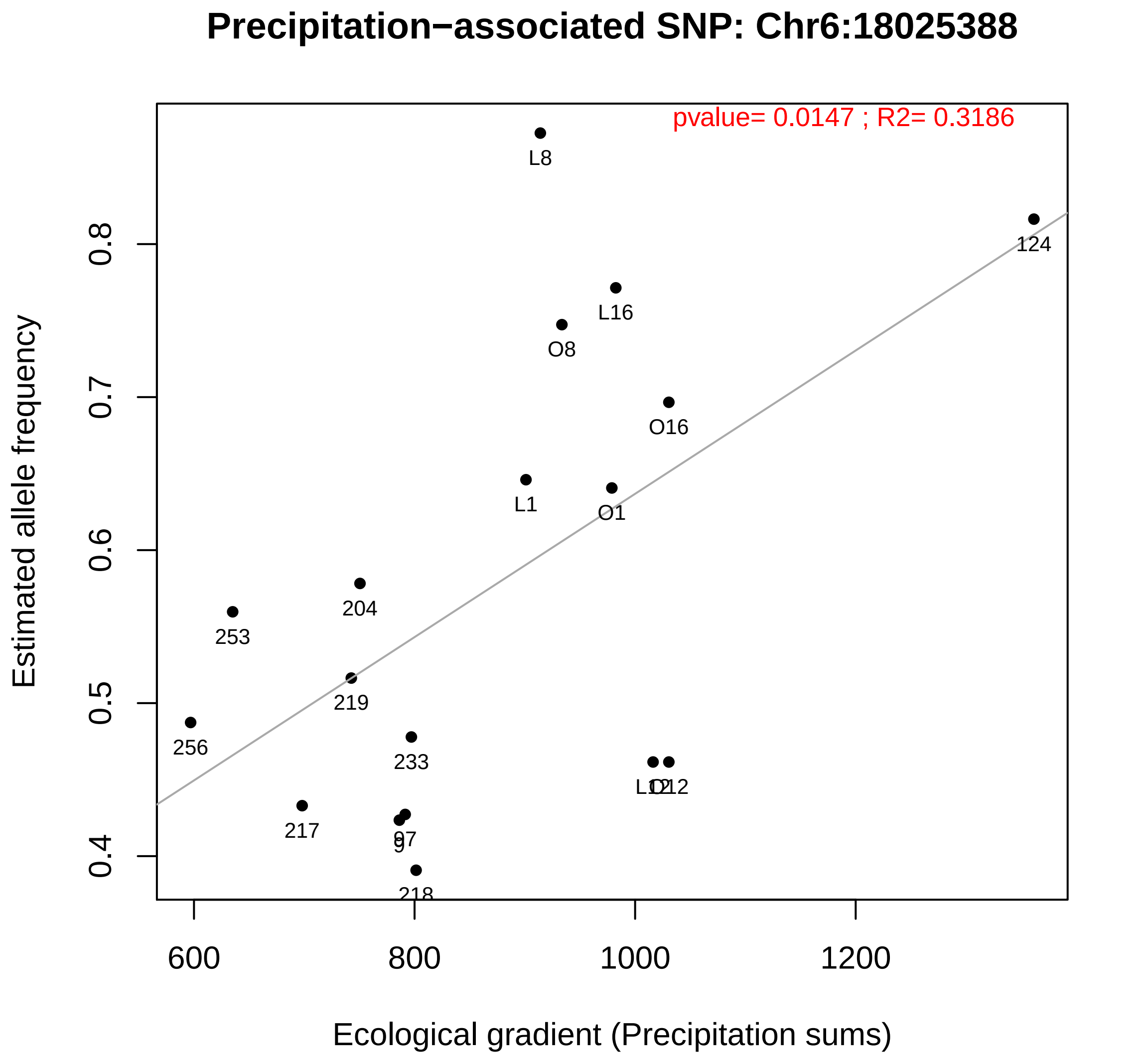

Precipitation−associated SNP: Chr6:18025388
pvalue= 0.0147 ; R2= 0.3186
Estimated allele frequency
0.4
0.5
0.6
0.7
0.8
600
800
1000
1200
Ecological gradient (Precipitation sums)
L8
124
L16
O8
O16
L1
O1
204
253
219
256
233
L12
O12
217
97
9
218

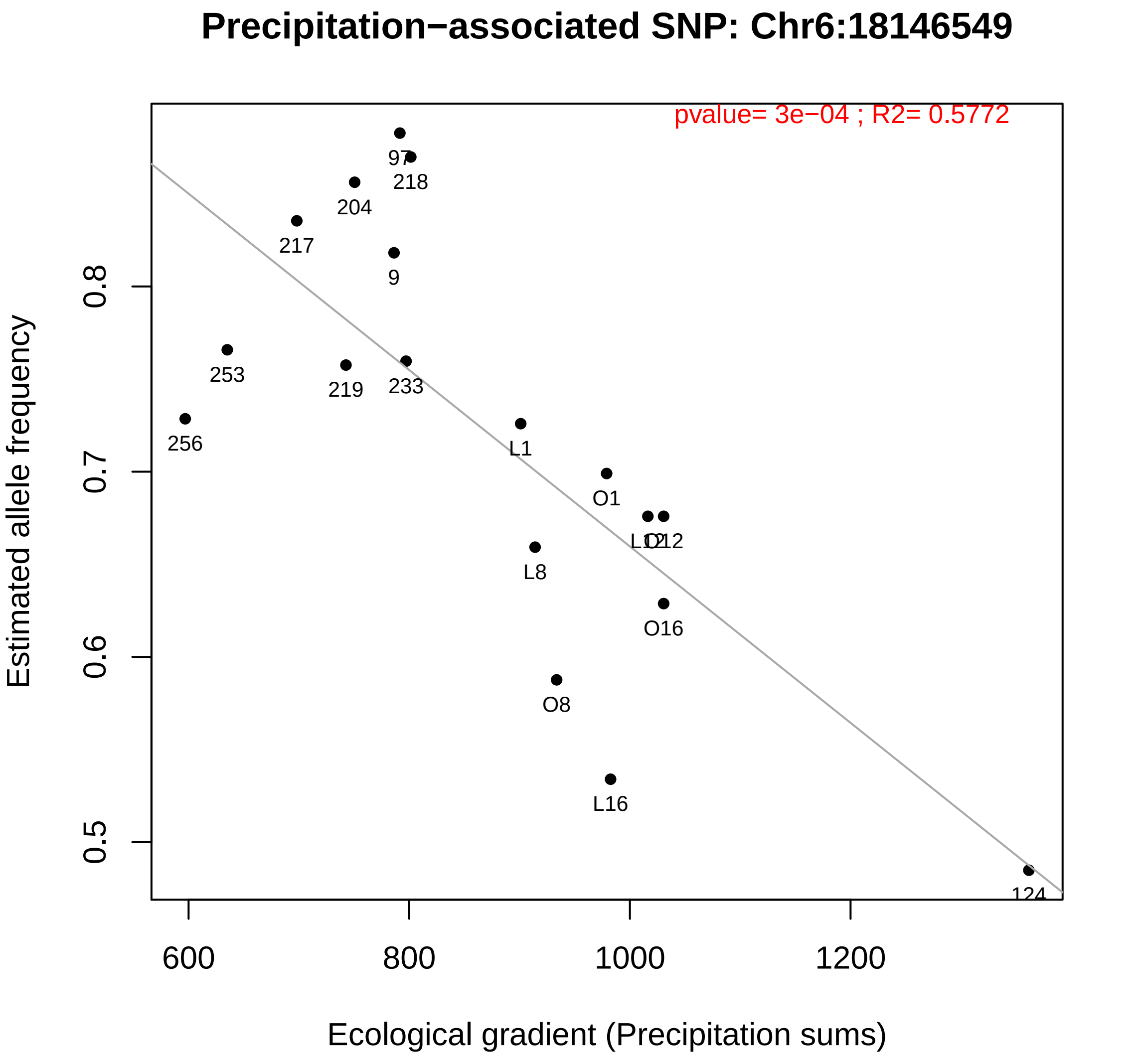

Precipitation−associated SNP: Chr6:18146549
pvalue= 3e−04 ; R2= 0.5772
Estimated allele frequency
0.5
0.6
0.7
0.8
600
800
1000
1200
Ecological gradient (Precipitation sums)
218
204
217
9
253
219
233
256
L1
O1
L8
O16
O8
L16
124

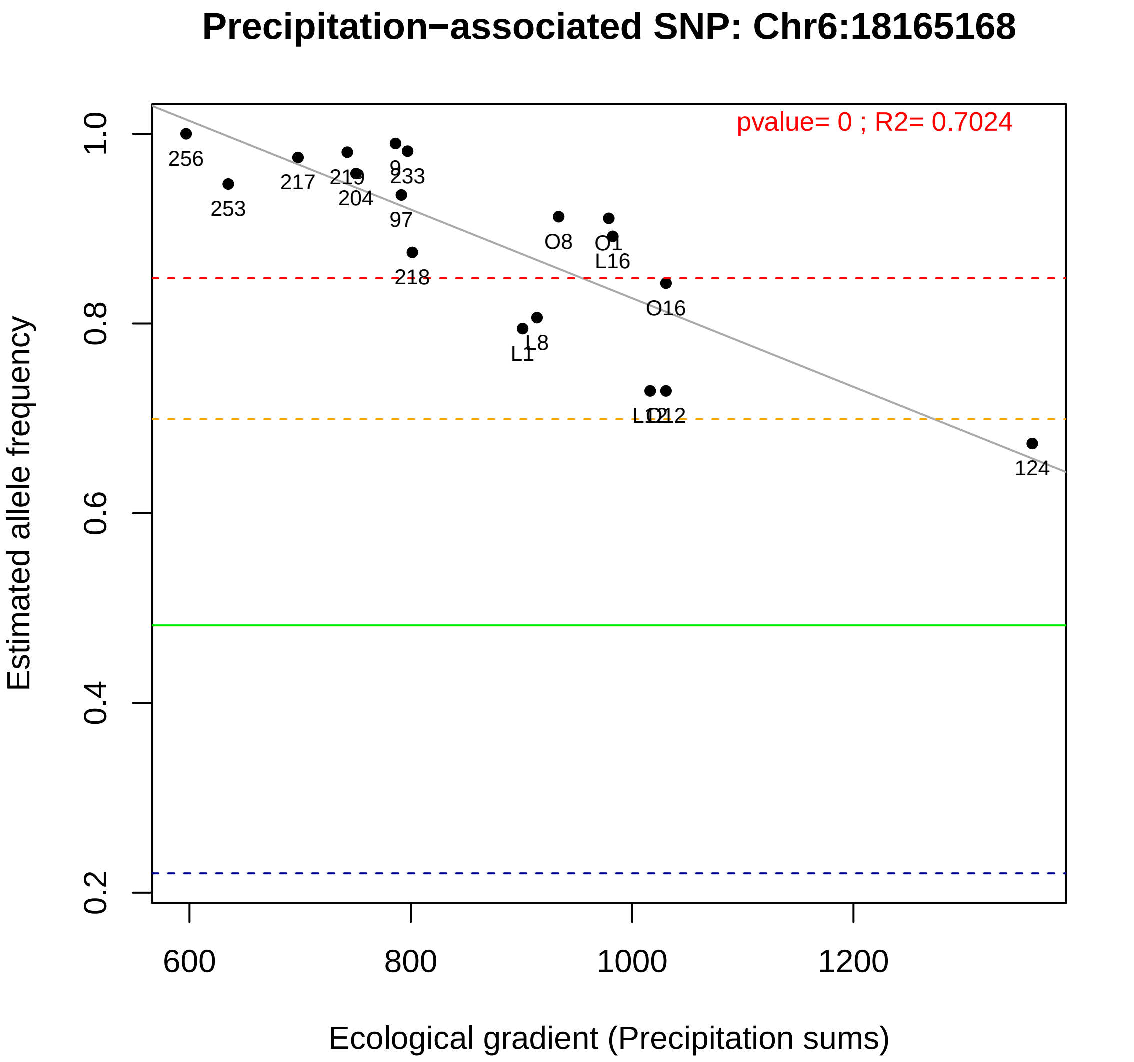

Precipitation−associated SNP: Chr6:18165168
pvalue= 0 ; R2= 0.7024
Estimated allele frequency
Ecological gradient (Precipitation sums)
0.2
0.4
0.6
0.8
1.0
600
800
1000
1200
256
253
217
219
204
9
233
97
218
O8
O1
L16
O16
L8
L1
L12
O12
124

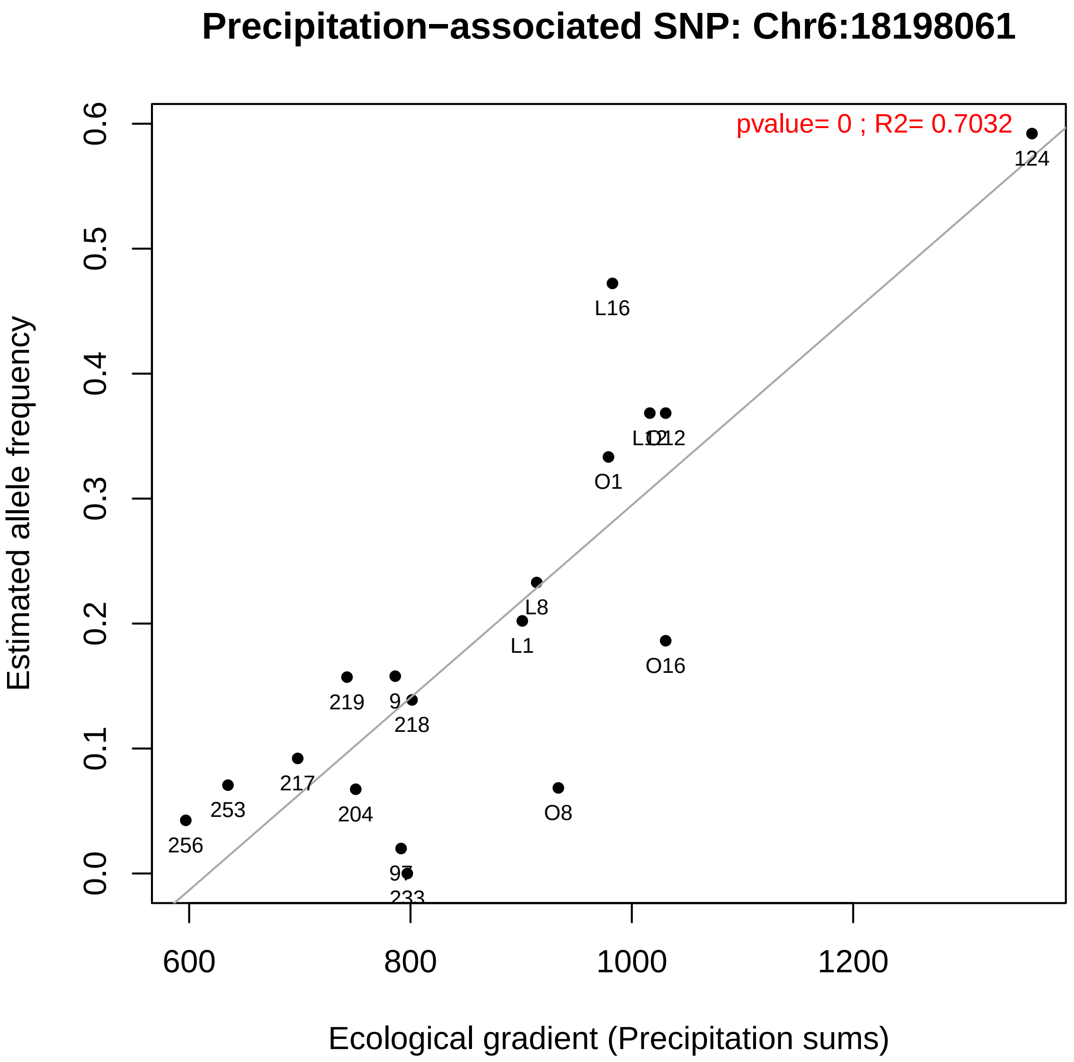

Precipitation−associated SNP: Chr6:18198061
pvalue= 0 ; R2= 0.7032
Estimated allele frequency
Ecological gradient (Precipitation sums)
0.0
0.1
0.2
0.3
0.4
0.5
0.6
600
800
1000
1200
124
L16
L12
O12
O1
L8
L1
O16
219
9
218
217
253
204
O8
256
97
233

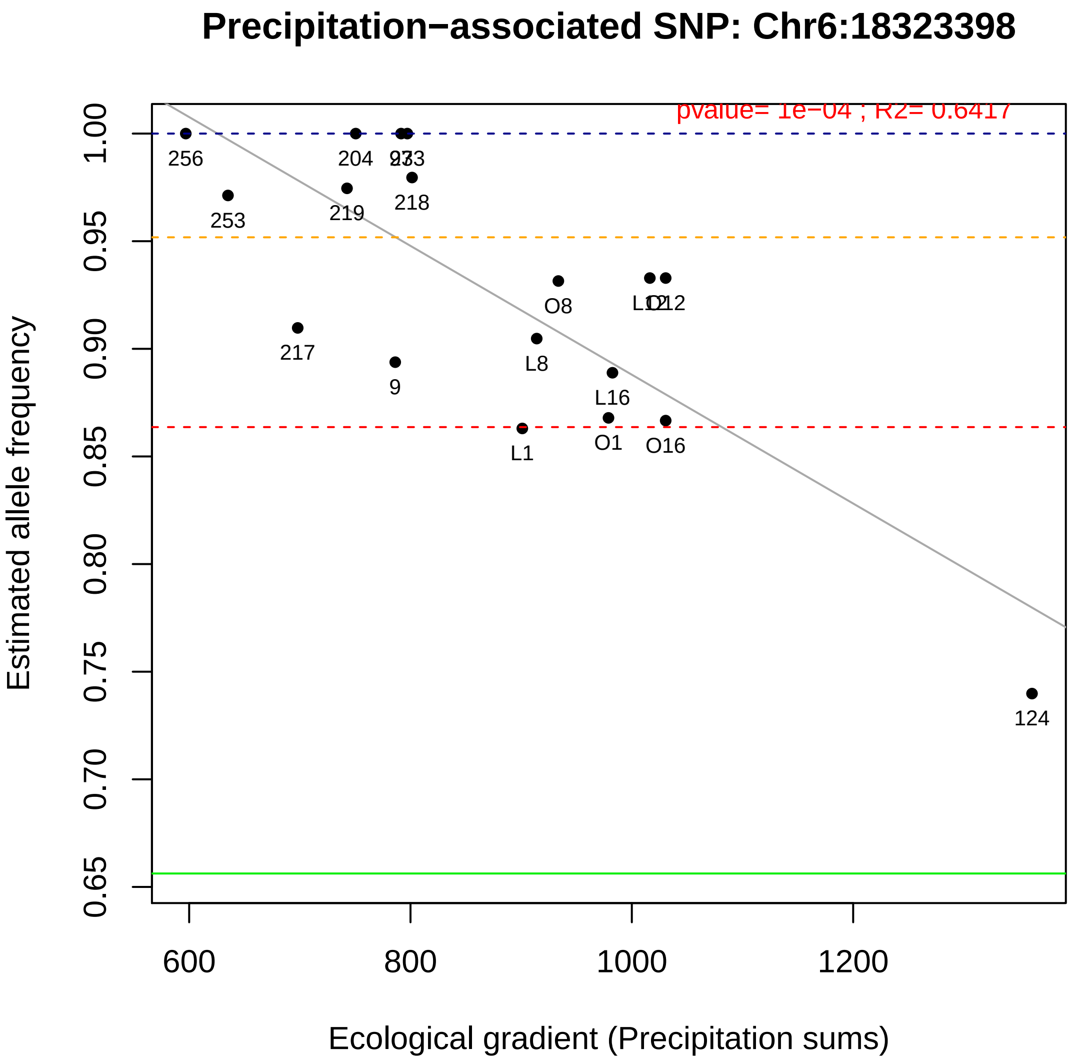

Precipitation−associated SNP: Chr6:18323398
pvalue= 1e−04 ; R2= 0.6417
Estimated allele frequency
Ecological gradient (Precipitation sums)
1.00
0.95
0.90
0.85
0.80
0.75
0.70
0.65
600
800
1000
1200
256
204
253
219
218
217
9
O8
L8
L16
L1
O1
O16
O12
124

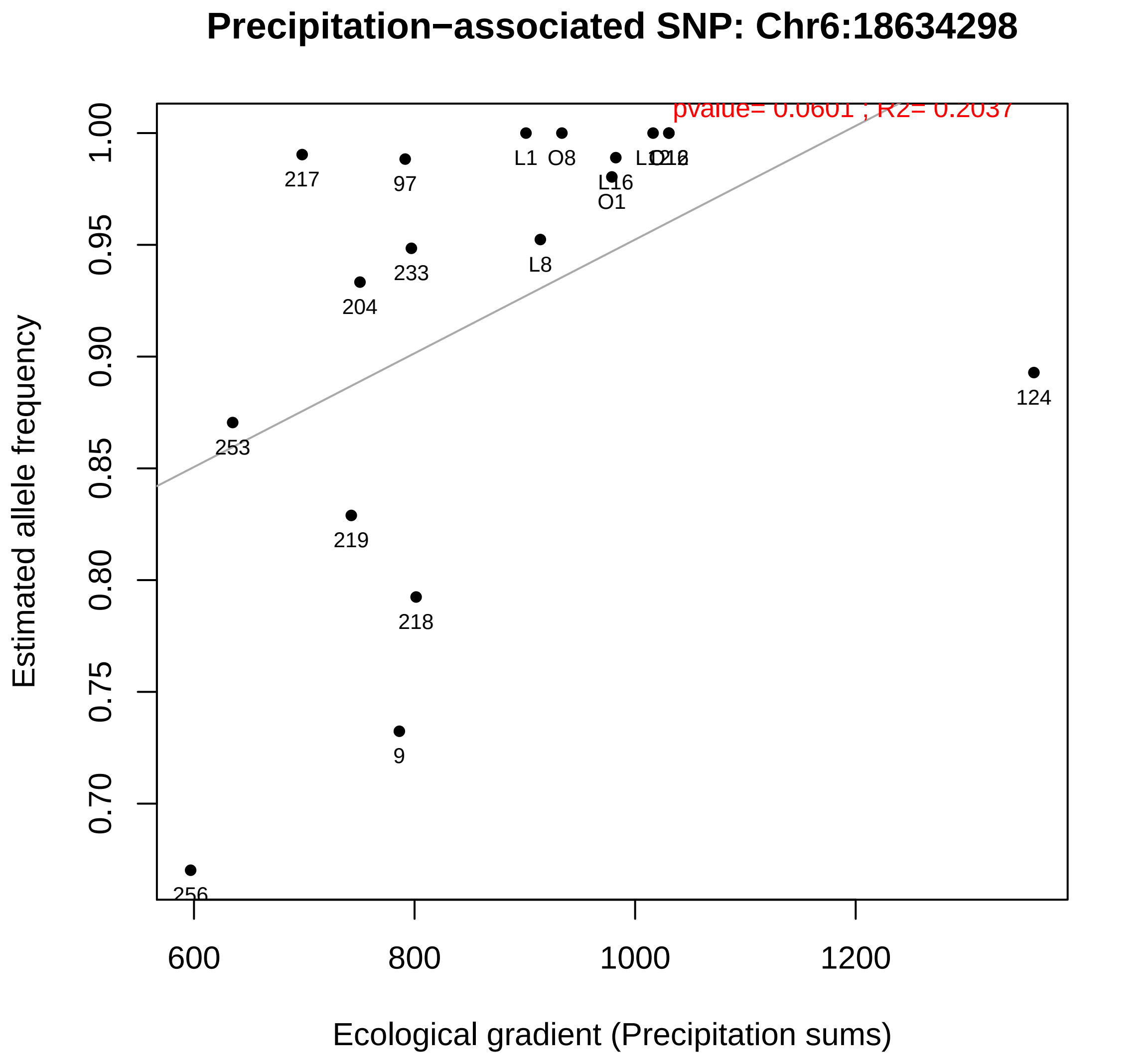

Precipitation−associated SNP: Chr6:18634298
pvalue= 0.0601 ; R2= 0.2037
Estimated allele frequency
Ecological gradient (Precipitation sums)
1.00
0.95
0.90
0.85
0.80
0.75
0.70
600
800
1000
1200
217
97
L1
O8
L16
O1
233
L8
204
253
219
218
9
256
124

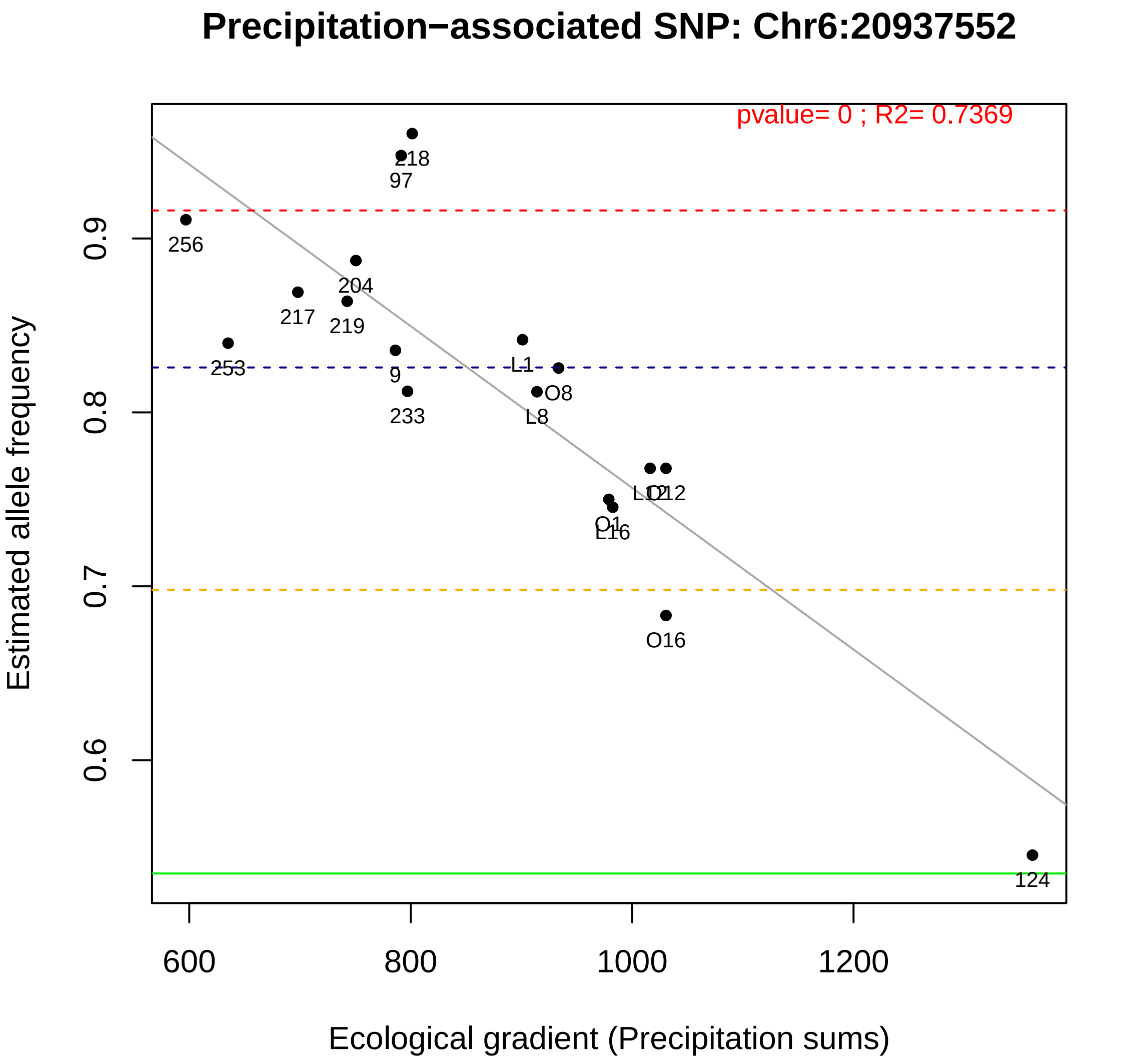

Precipitation−associated SNP: Chr6:20937552
pvalue= 0 ; R2= 0.7369
218
97
256
204
217
219
253
9
233
L1
O8
L8
L12
O12
O1
L16
O16
124
0.9
0.8
0.7
0.6
600
800
1000
1200
Estimated allele frequency
Ecological gradient (Precipitation sums)

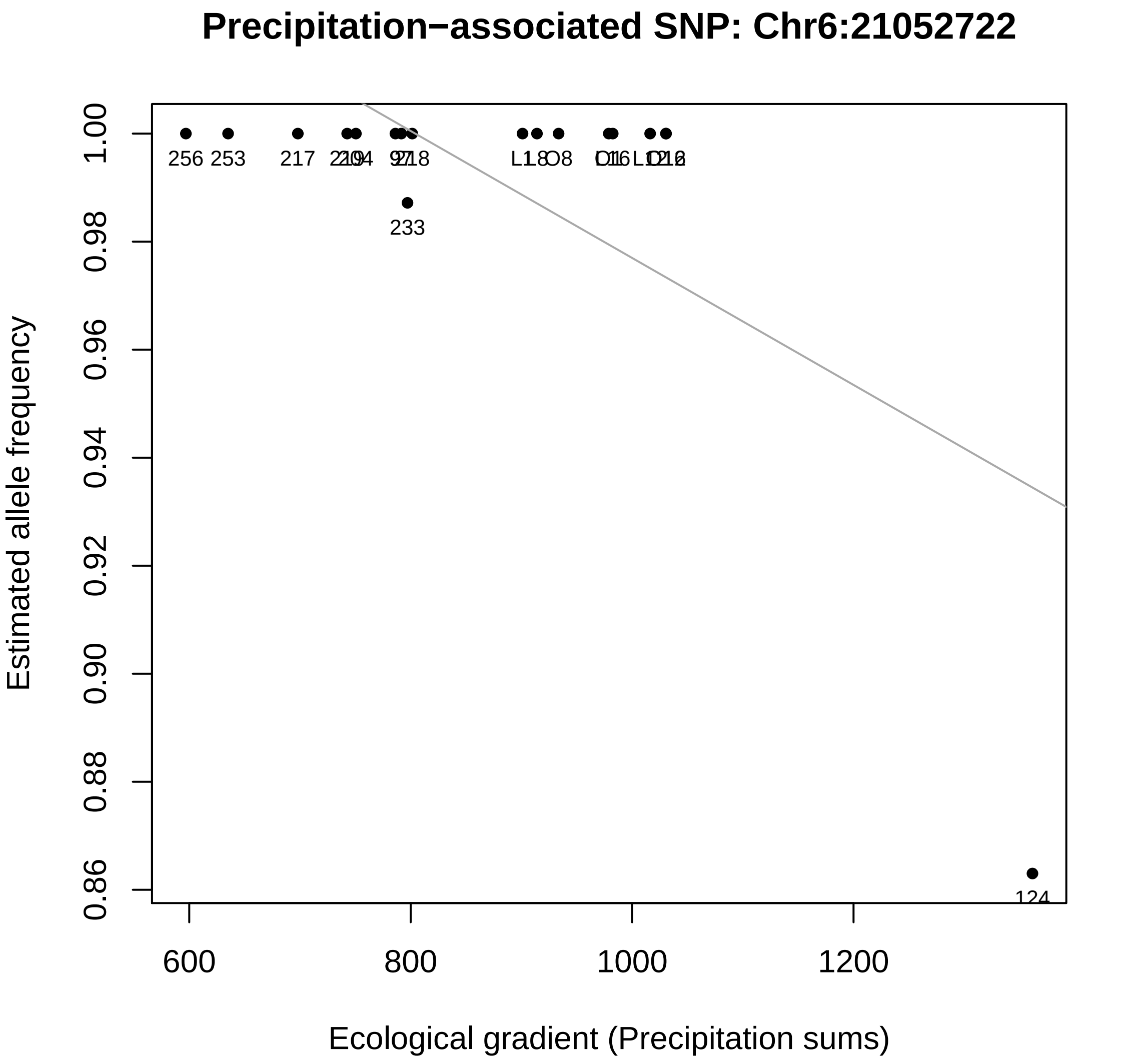

Precipitation−associated SNP: Chr6:21052722
Estimated allele frequency
Ecological gradient (Precipitation sums)
1.00
0.98
0.96
0.94
0.92
0.90
0.88
0.86
600
800
1000
1200
256
253
217
233
124

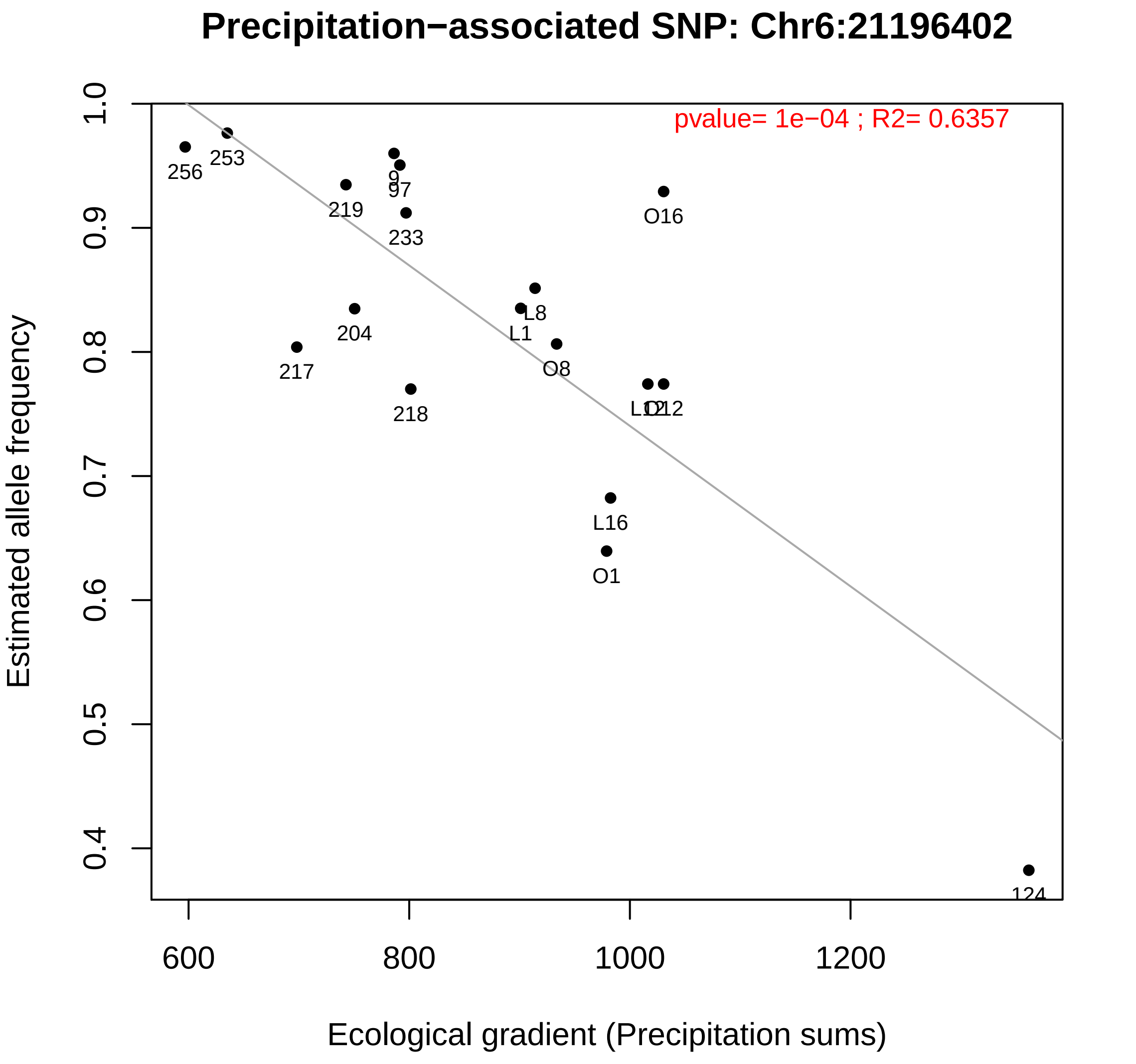

Precipitation−associated SNP: Chr6:21196402
pvalue= 1e−04 ; R2= 0.6357
Estimated allele frequency
Ecological gradient (Precipitation sums)
0.4
0.5
0.6
0.7
0.8
0.9
1.0
600
800
1000
1200
256
253
219
9
97
233
O16
204
L8
L1
217
O8
218
L12
O12
L16
O1
124

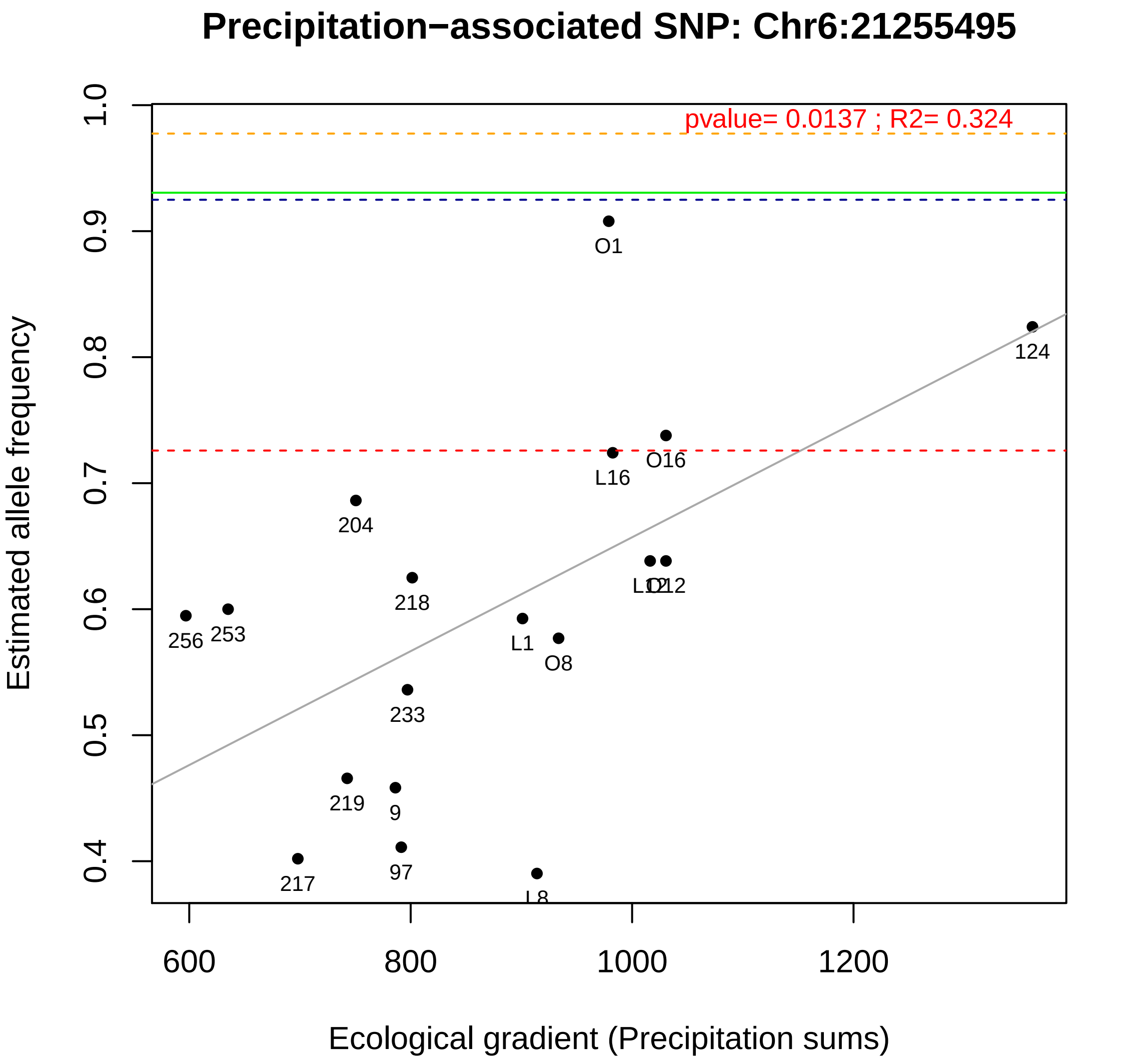
Precipitation−associated SNP: Chr6:21255495
pvalue= 0.0137 ; R2= 0.324
Estimated allele frequency
0.4
0.5
0.6
0.7
0.8
0.9
1.0
600
800
1000
1200
Ecological gradient (Precipitation sums)
O1
124
O16
L16
204
218
L12
O12
256
253
L1
O8
233
219
9
217
97
L8

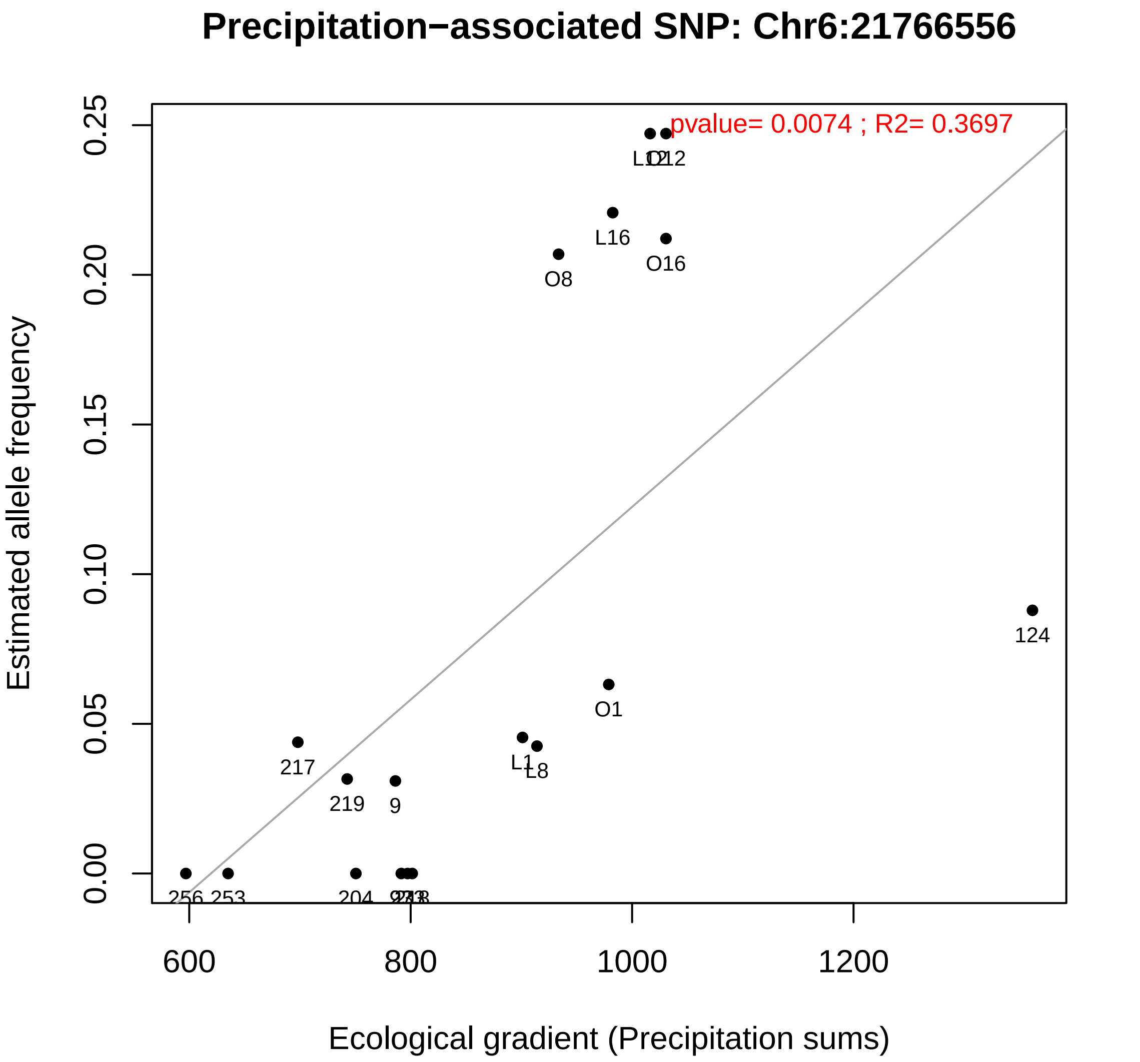
Precipitation−associated SNP: Chr6:21766556
pvalue= 0.0074 ; R2= 0.3697
Estimated allele frequency
0.00
0.05
0.10
0.15
0.20
0.25
Ecological gradient (Precipitation sums)
600
800
1000
1200
O12
L16
O16
O8
124
O1
217
L1
L8
219
9
256
253
204

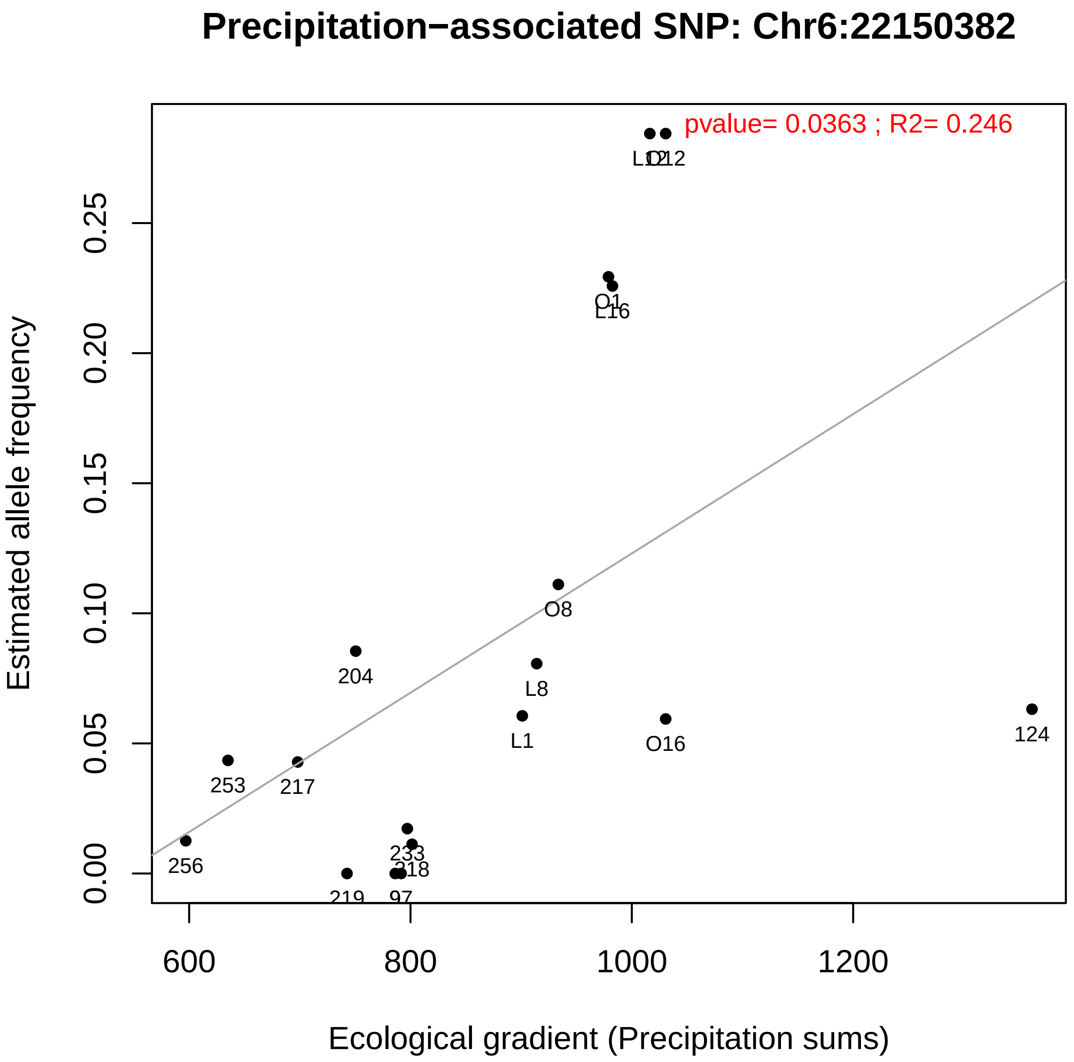

Precipitation−associated SNP: Chr6:22150382
pvalue= 0.0363 ; R2= 0.246
L10
O12
O1
L16
O8
204
L8
L1
O16
124
253
217
256
233
218
219
97
0.00
0.05
0.10
0.15
0.20
0.25
600
800
1000
1200
Estimated allele frequency
Ecological gradient (Precipitation sums)

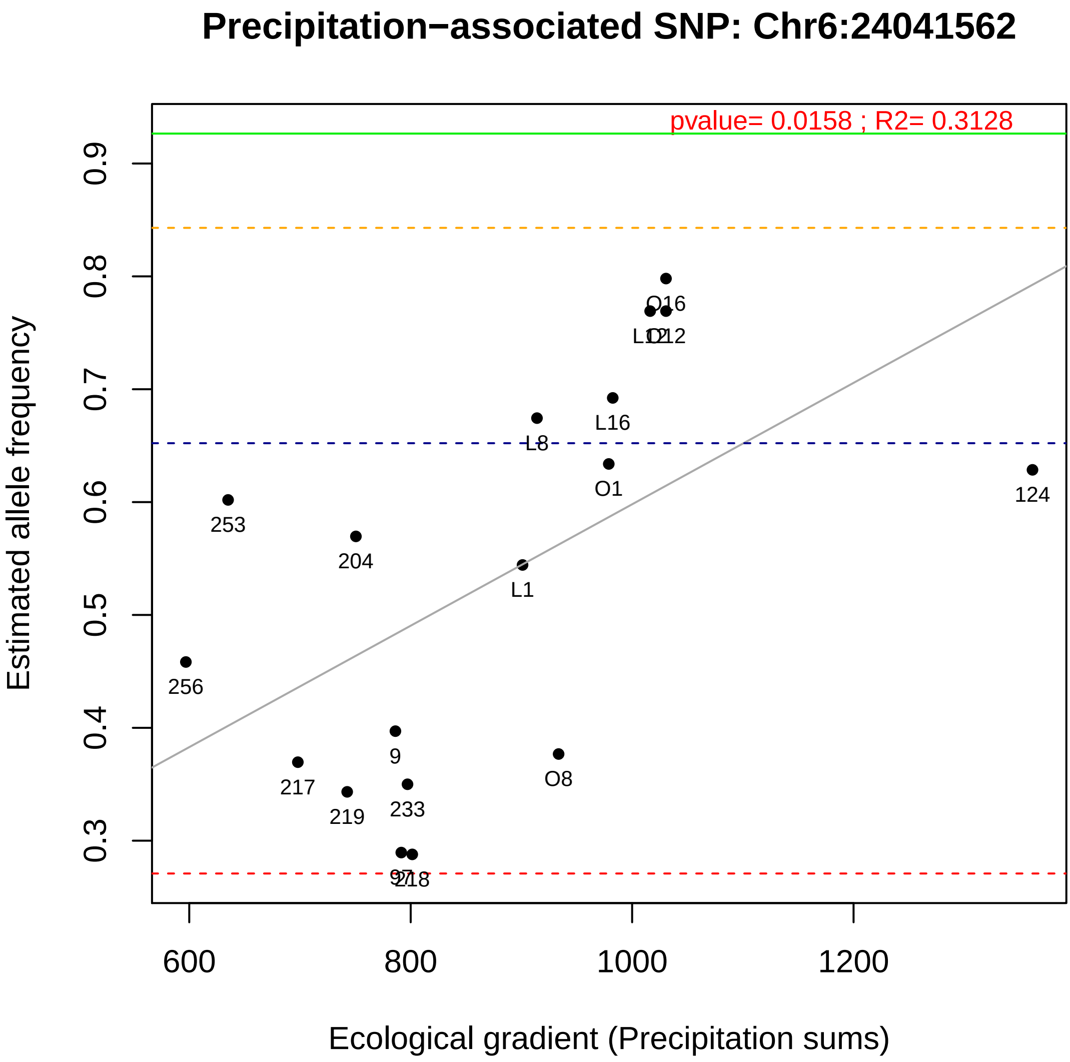

Precipitation−associated SNP: Chr6:24041562
pvalue= 0.0158 ; R2= 0.3128
Estimated allele frequency
0.3
0.4
0.5
0.6
0.7
0.8
0.9
600
800
1000
1200
Ecological gradient (Precipitation sums)
O16
L12
O12
L16
L8
O1
124
253
204
L1
256
9
217
O8
219
233
97
218

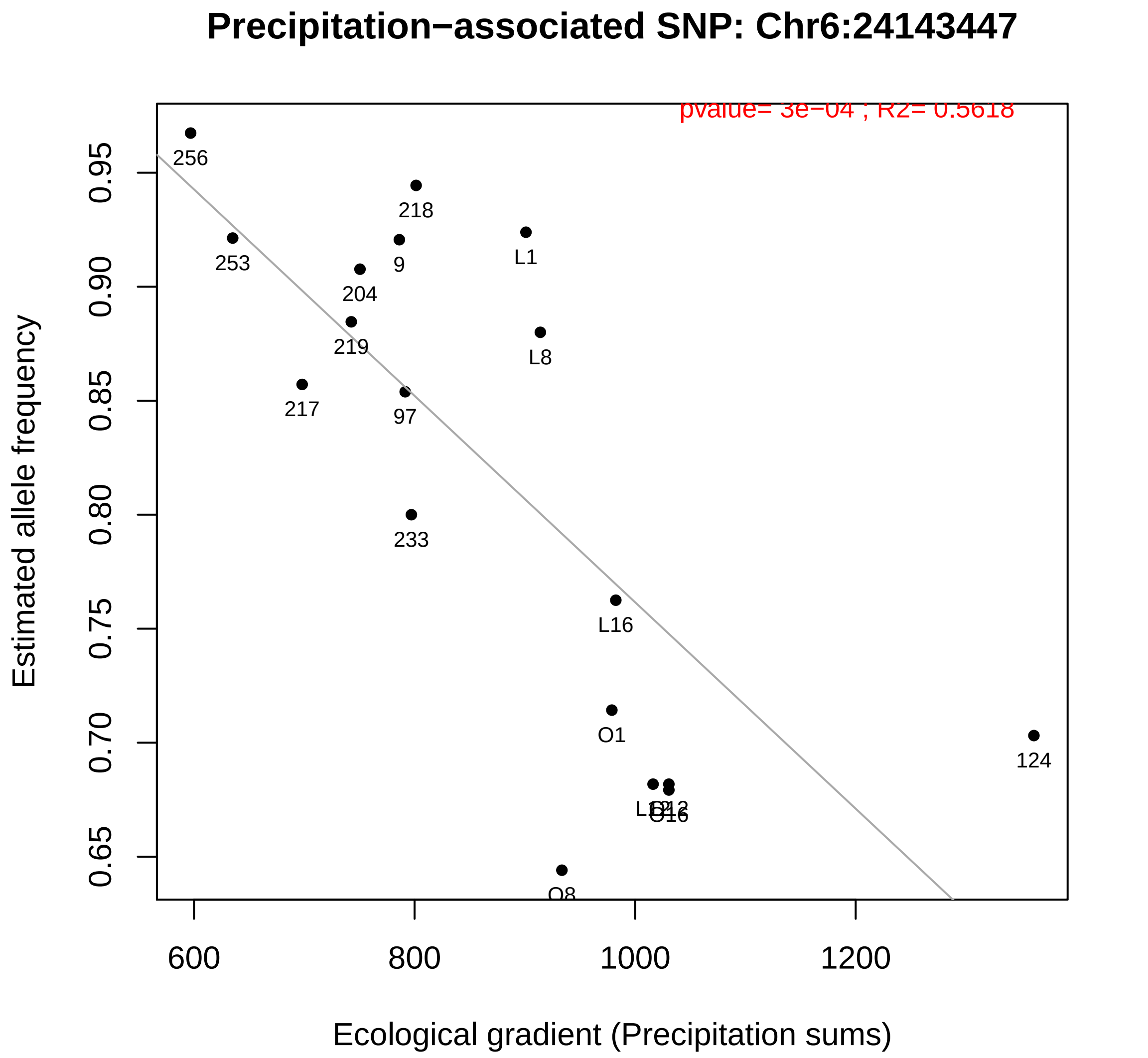

Precipitation−associated SNP: Chr6:24143447
pvalue= 3e−04 ; R2= 0.5618
Estimated allele frequency
0.65
0.70
0.75
0.80
0.85
0.90
0.95
600
800
1000
1200
Ecological gradient (Precipitation sums)
256
218
253
9
L1
204
219
L8
217
97
233
L16
O1
124
O16
O8

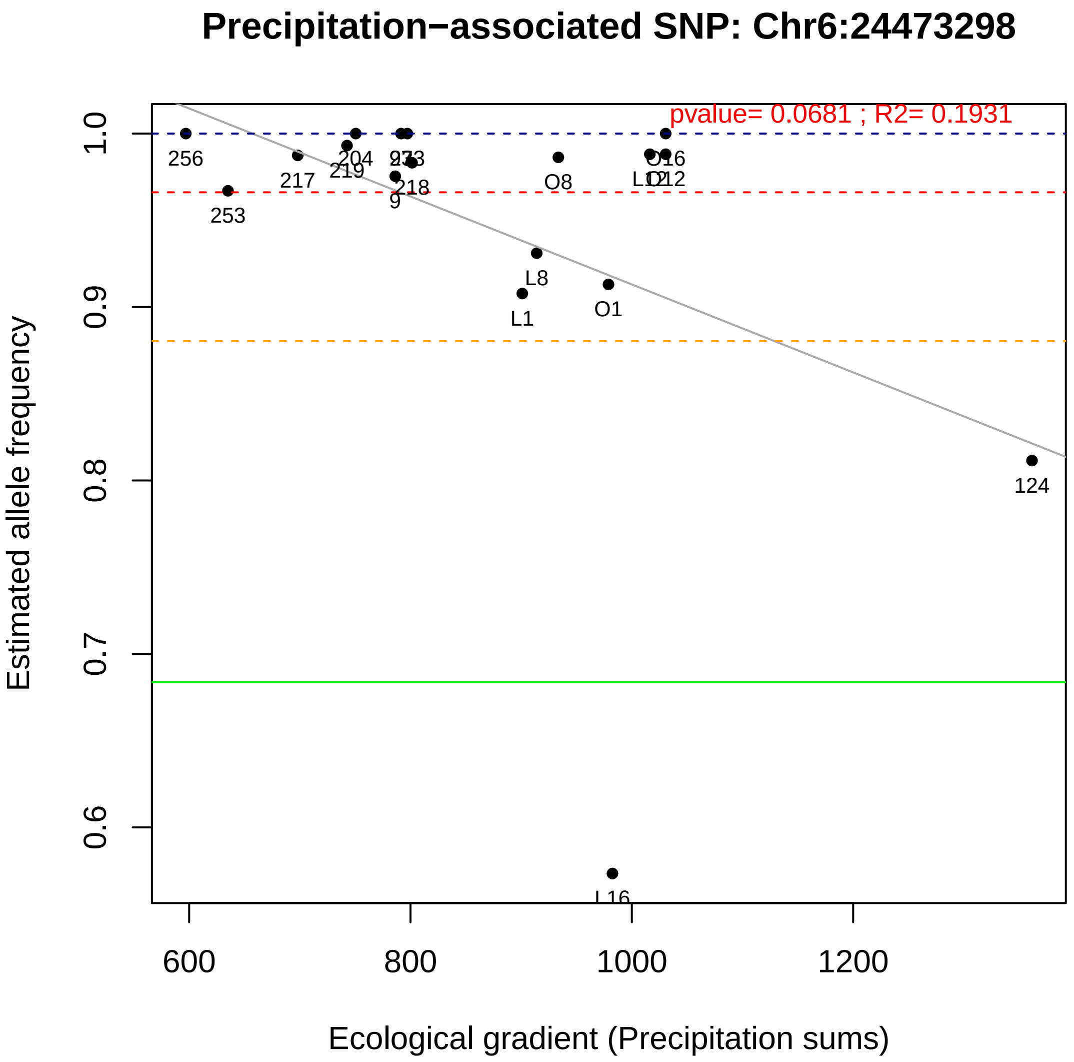
Precipitation−associated SNP: Chr6:24473298
pvalue= 0.0681 ; R2= 0.1931
Estimated allele frequency
Ecological gradient (Precipitation sums)
1.0
0.9
0.8
0.7
0.6
600
800
1000
1200
256
253
217
219
204
218
9
O8
L8
L1
O1
O16
O12
124
L16

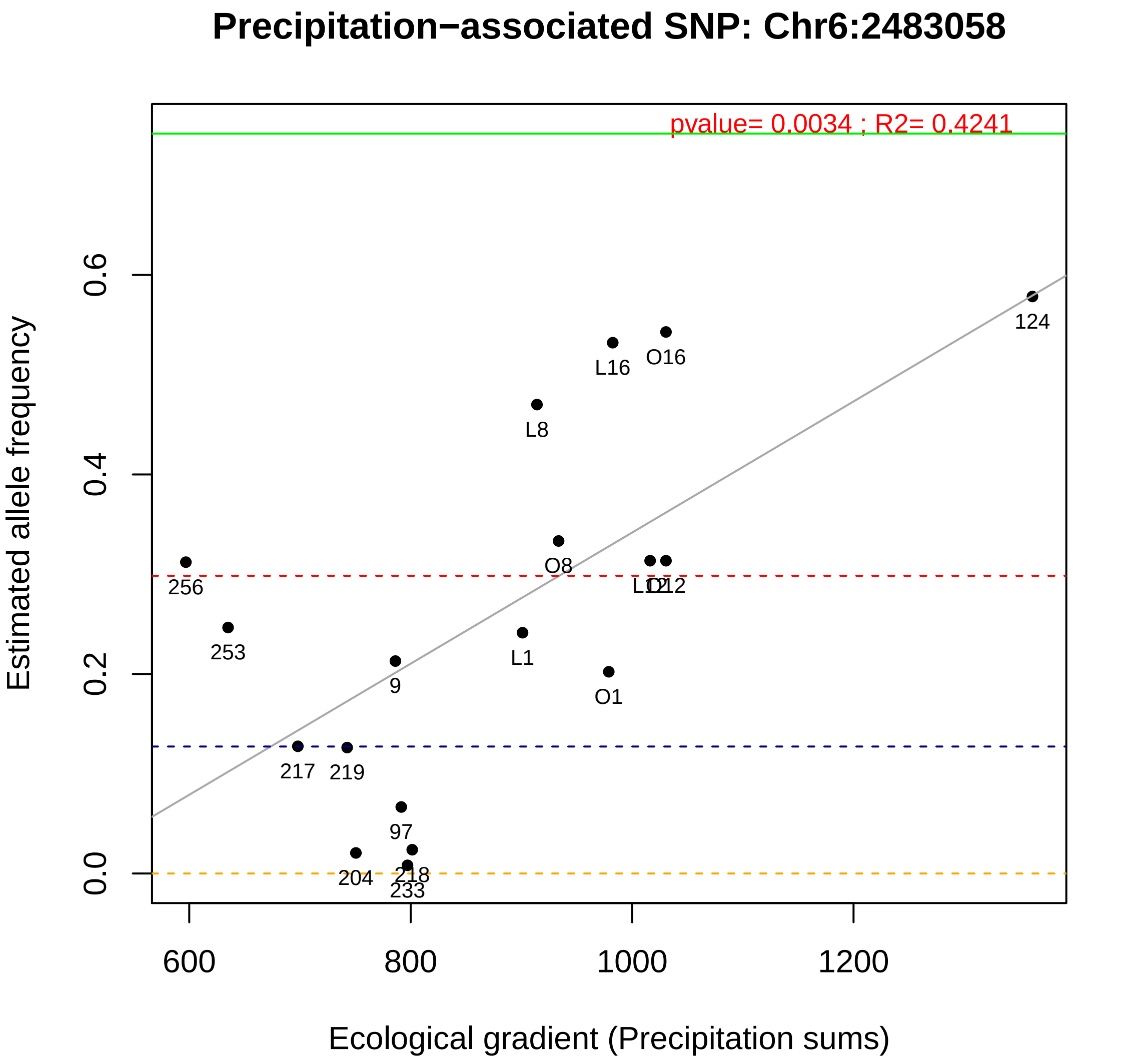
Precipitation−associated SNP: Chr6:2483058
pvalue= 0.0034 ; R2= 0.4241
Estimated allele frequency
0.0
0.2
0.4
0.6
600
800
1000
1200
Ecological gradient (Precipitation sums)
124
O16
L16
L8
O8
256
253
9
L1
O1
217
219
97
204
218
233

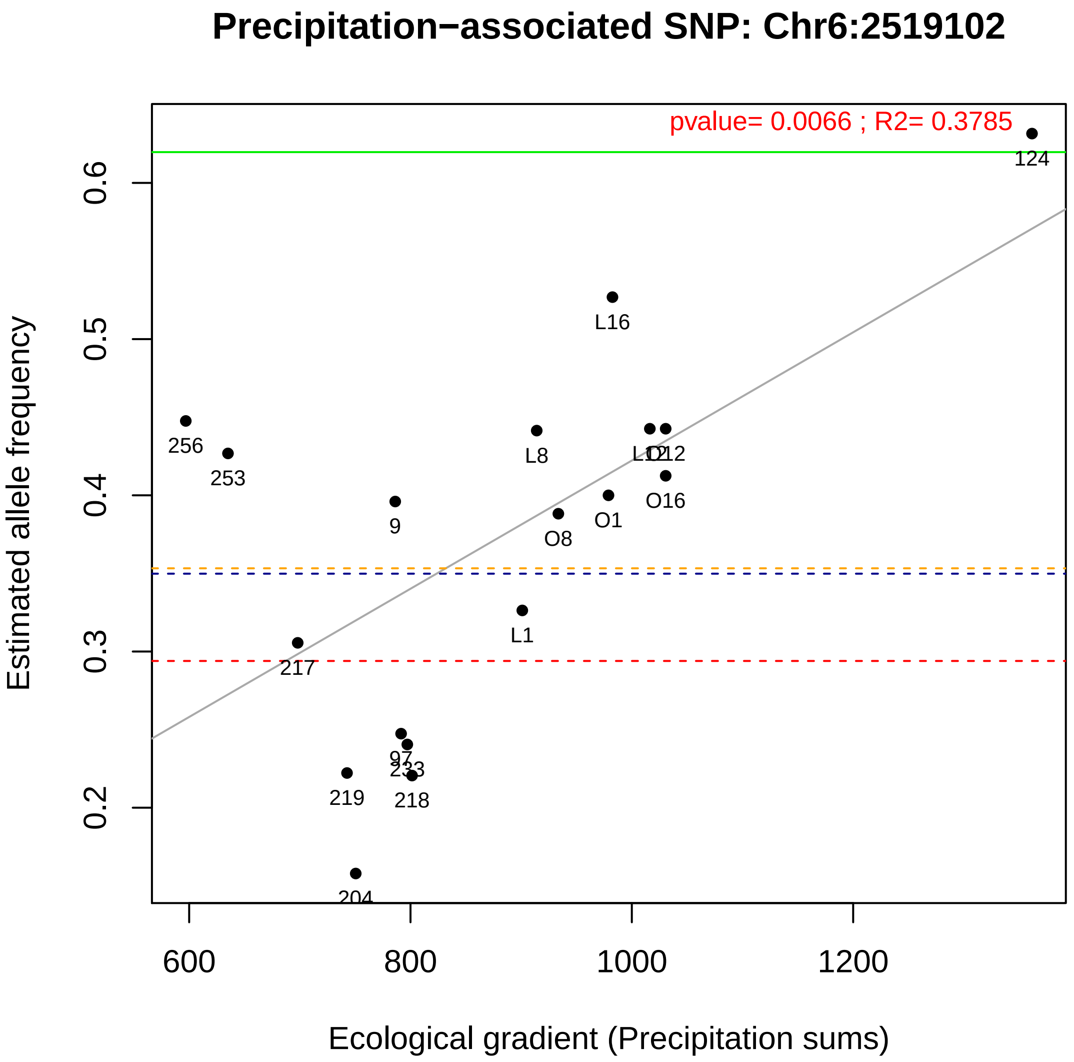
Precipitation−associated SNP: Chr6:2519102
pvalue= 0.0066 ; R2= 0.3785
Estimated allele frequency
0.2
0.3
0.4
0.5
0.6
600
800
1000
1200
Ecological gradient (Precipitation sums)
124
L16
256
253
L8
L10
O12
O16
9
O1
O8
L1
217
97
233
219
218
204

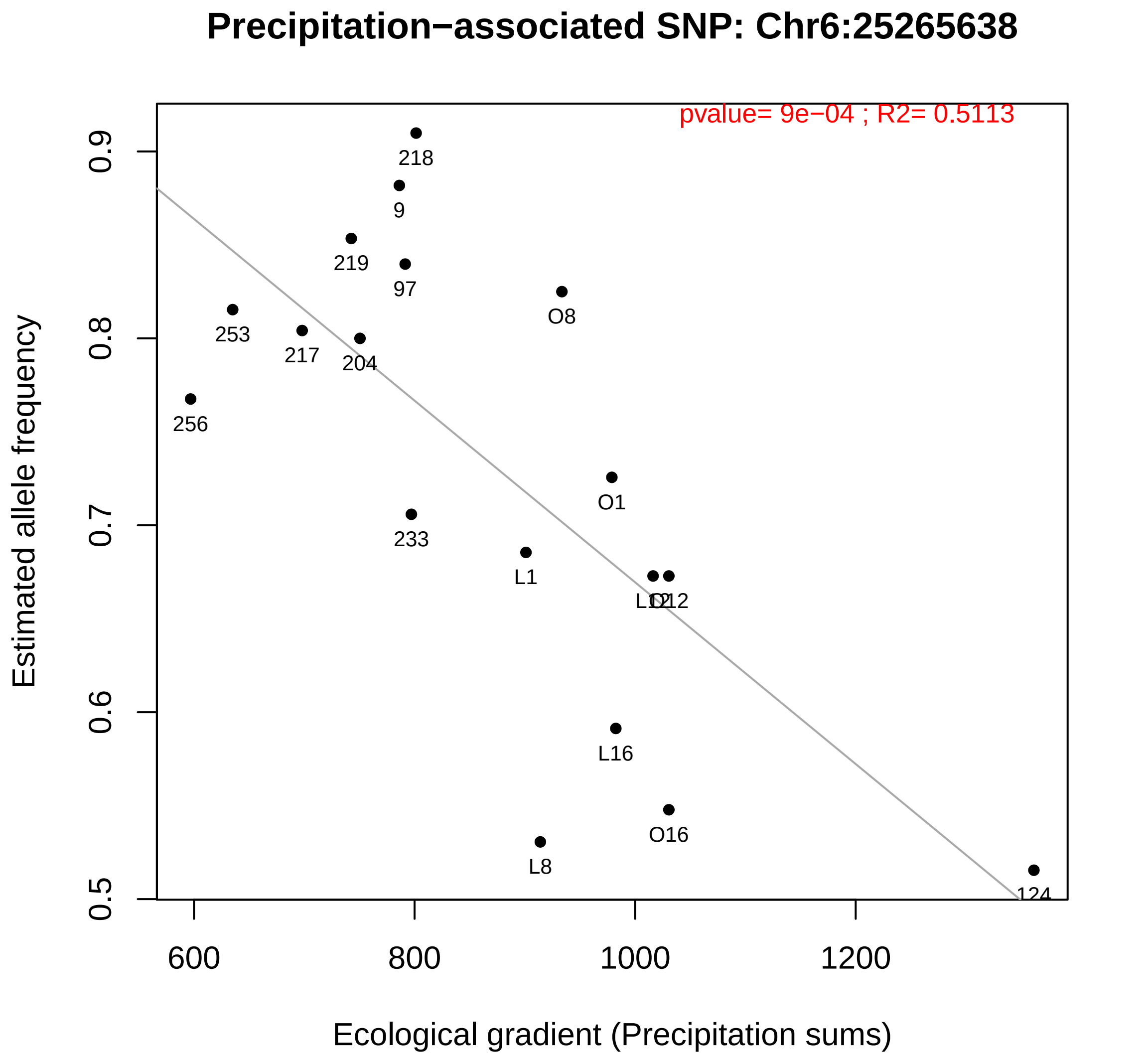
Precipitation−associated SNP: Chr6:25265638
pvalue= 9e−04 ; R2= 0.5113
Estimated allele frequency
0.5
0.6
0.7
0.8
0.9
600
800
1000
1200
Ecological gradient (Precipitation sums)
218
9
219
97
O8
253
217
204
256
O1
233
L1
O12
L16
O16
L8
124

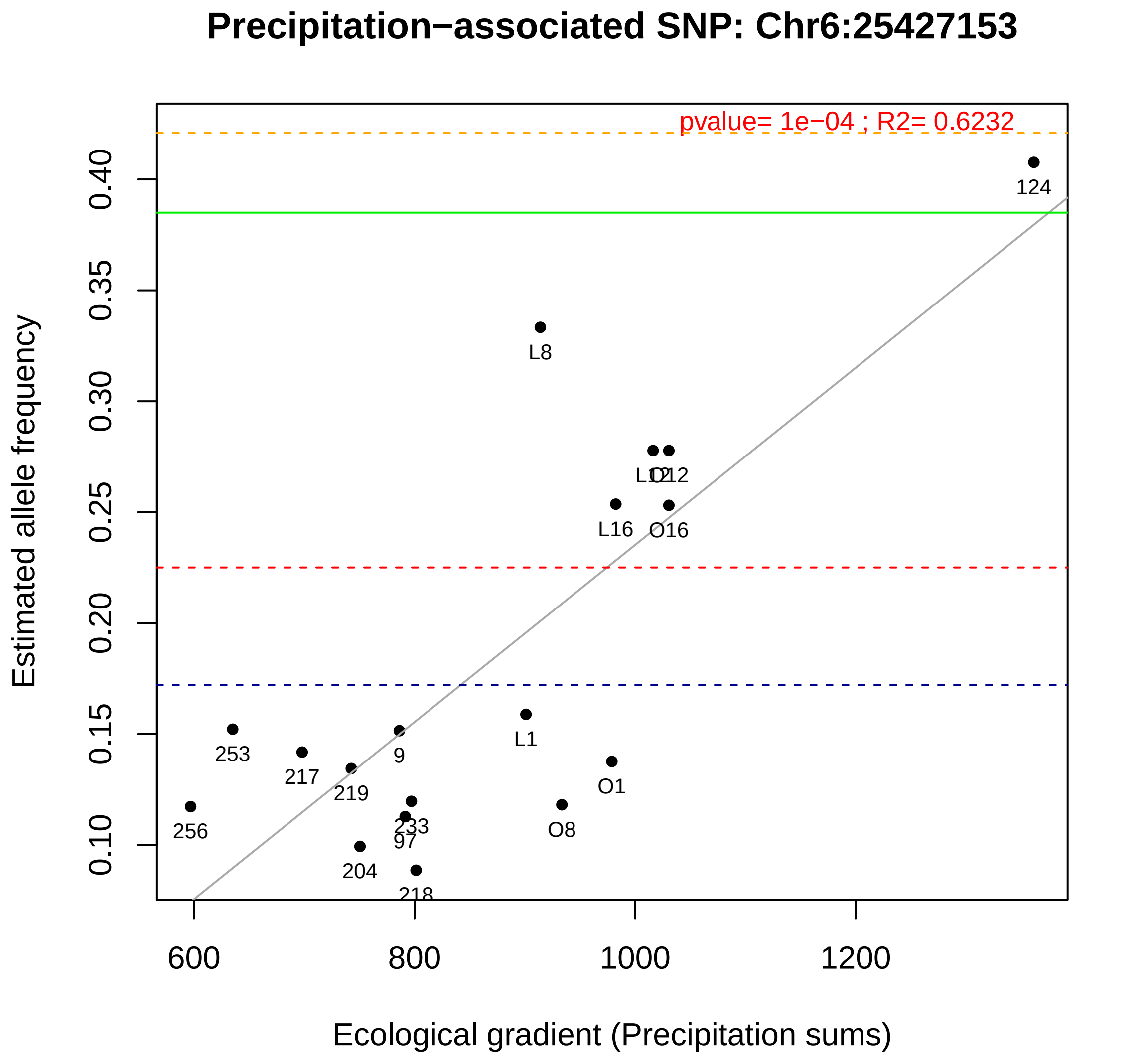
Precipitation−associated SNP: Chr6:25427153
pvalue= 1e−04 ; R2= 0.6232
Estimated allele frequency
0.10
0.15
0.20
0.25
0.30
0.35
0.40
124
L8
L12
O12
L16
O16
L1
O1
O8
253
217
219
9
256
233
97
204
218
600
800
1000
1200
Ecological gradient (Precipitation sums)

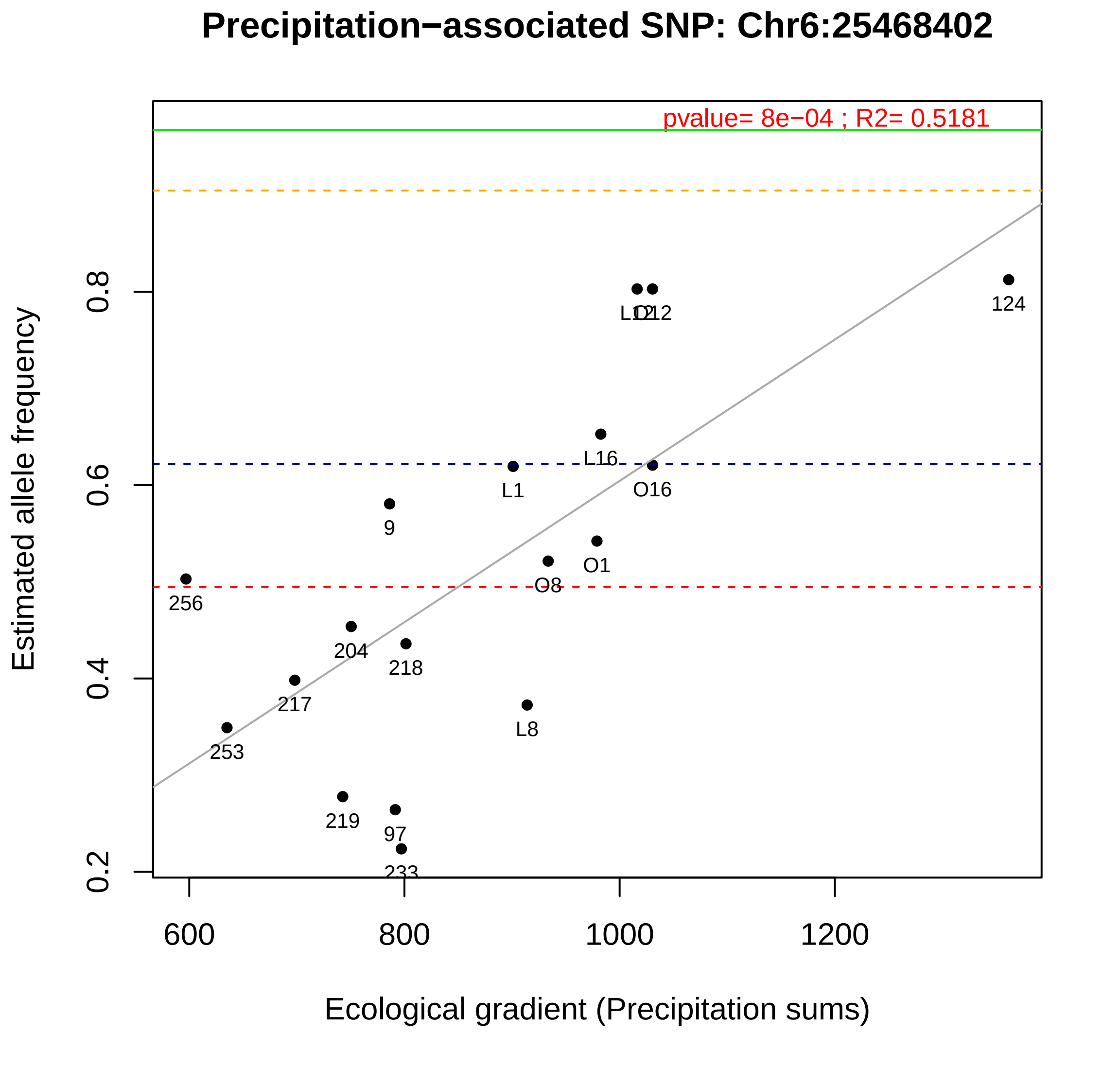

Precipitation−associated SNP: Chr6:25468402
pvalue= 8e−04 ; R2= 0.5181
Estimated allele frequency
0.2
0.4
0.6
0.8
600
800
1000
1200
Ecological gradient (Precipitation sums)
124
L16
L1
O16
9
O1
O8
256
204
218
217
L8
253
219
97
233

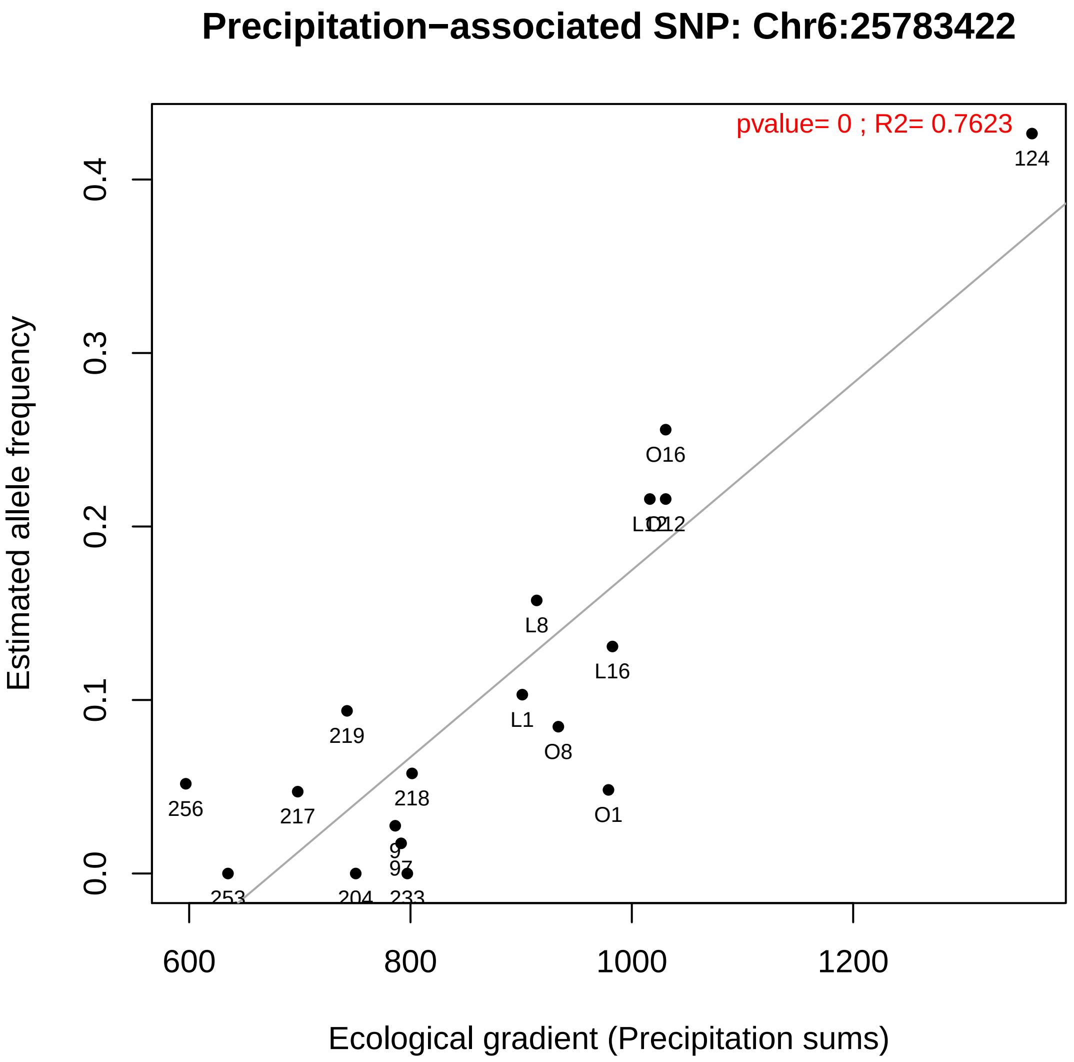
Precipitation−associated SNP: Chr6:25783422
pvalue= 0 ; R2= 0.7623
Estimated allele frequency
0.0
0.1
0.2
0.3
0.4
600
800
1000
1200
Ecological gradient (Precipitation sums)
124
O16
L12
O12
L8
L16
L1
O8
219
256
217
218
O1
9
97
253
204
233

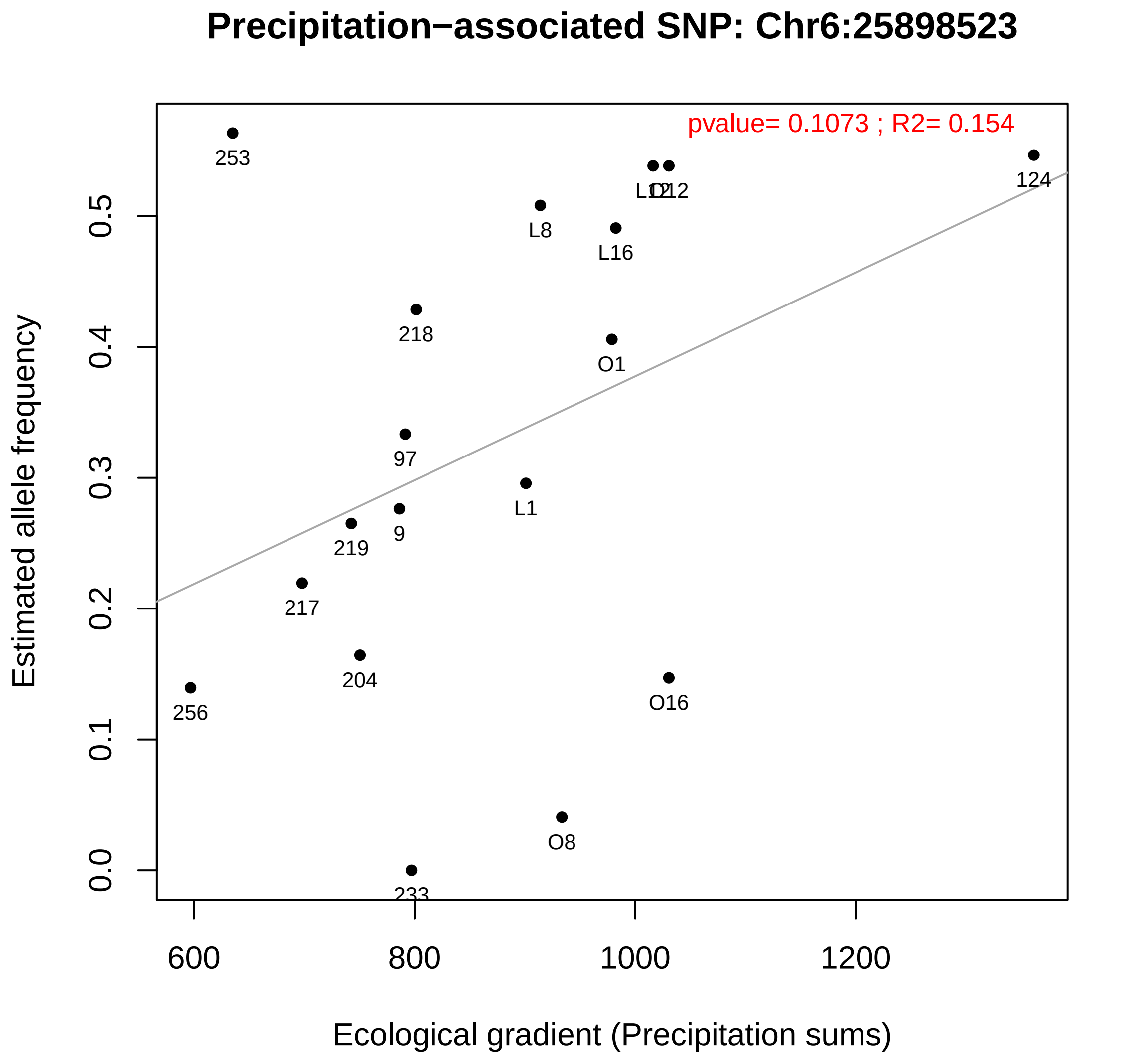
Precipitation−associated SNP: Chr6:25898523
pvalue= 0.1073 ; R2= 0.154
253
124
L8
L16
218
O1
97
L1
9
219
217
204
256
O16
O8
233
0.0
0.1
0.2
0.3
0.4
0.5
600
800
1000
1200
Estimated allele frequency
Ecological gradient (Precipitation sums)

**Precipitation−associated SNP: Chr6:26118176**

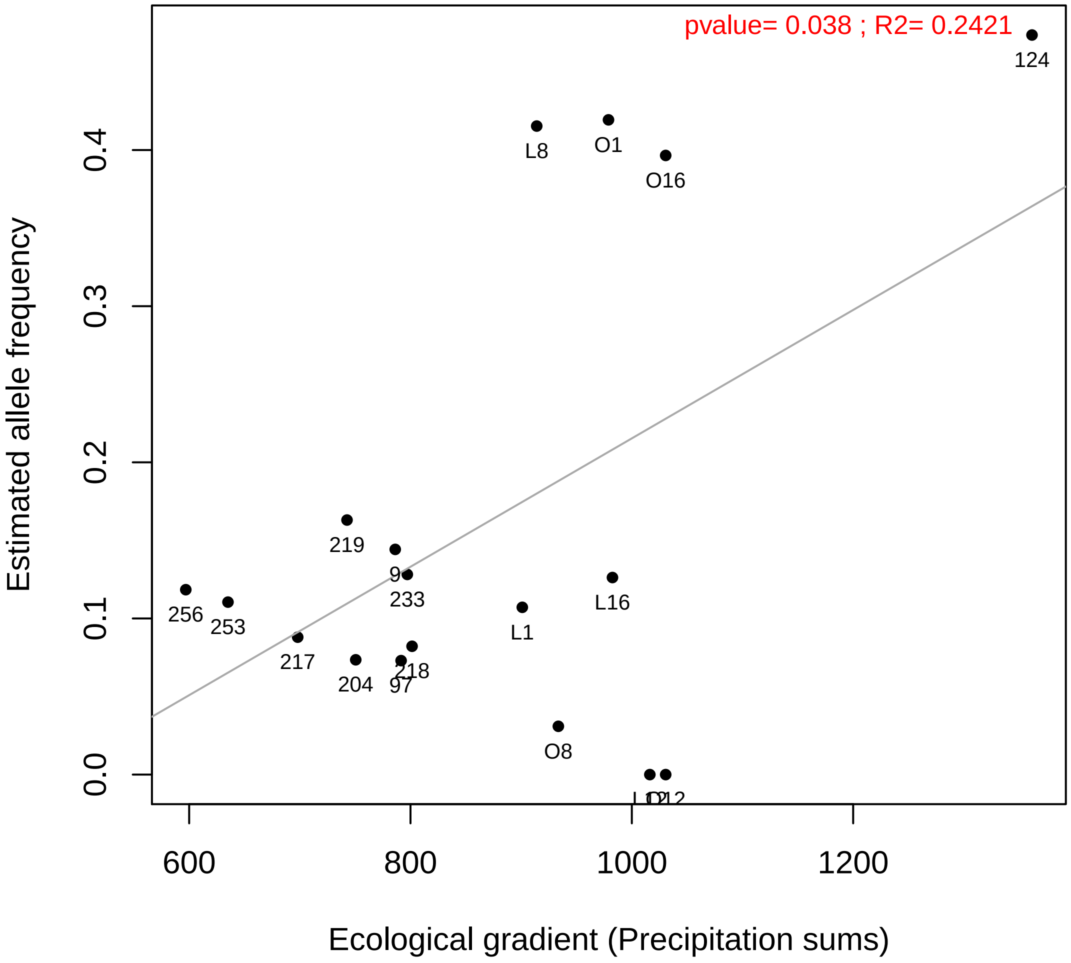

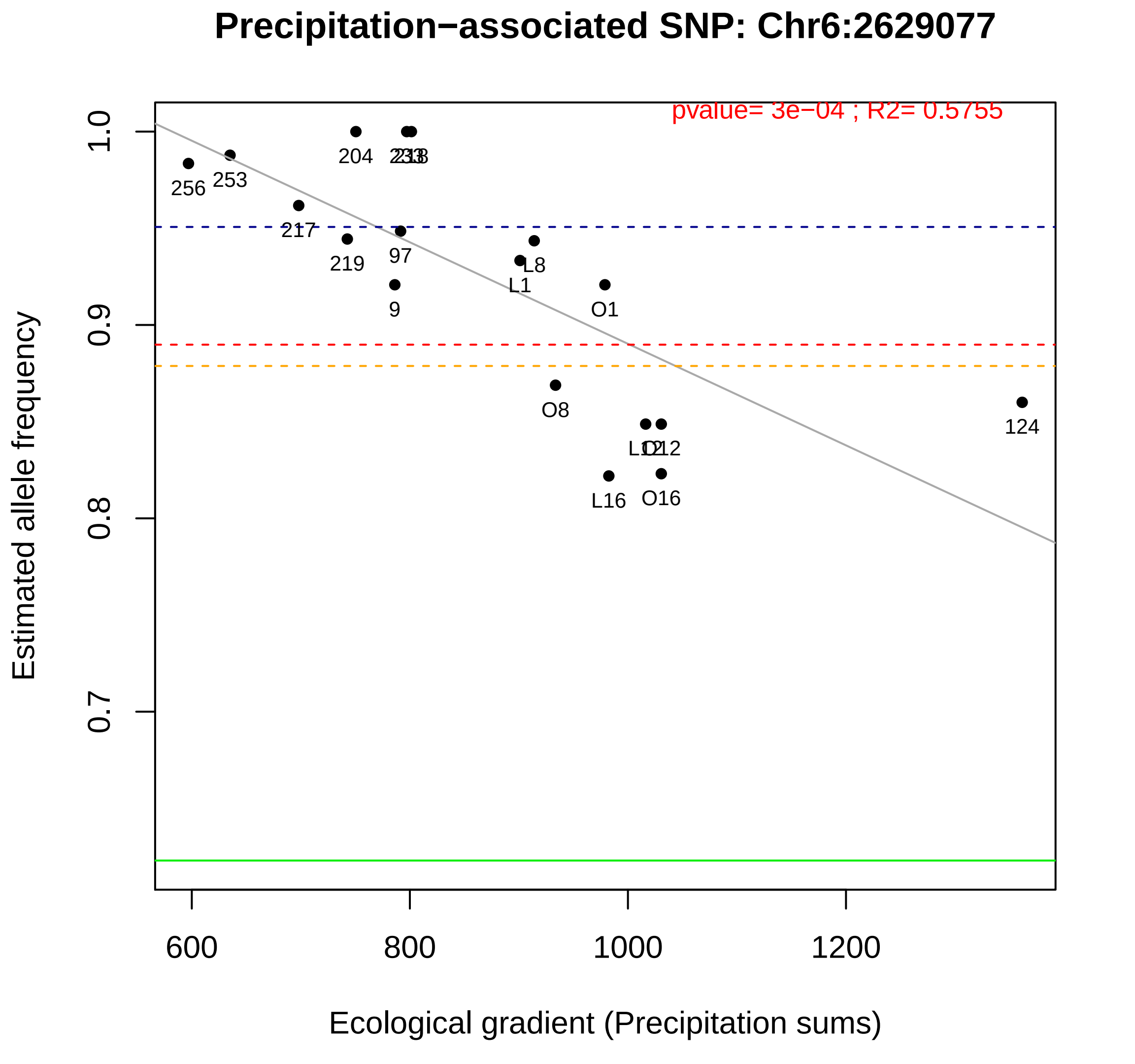

Precipitation−associated SNP: Chr6:2629077
pvalue= 3e−04 ; R2= 0.5755
Estimated allele frequency
1.0
0.9
0.8
0.7
600
800
1000
1200
Ecological gradient (Precipitation sums)
256
253
204
217
219
97
9
L1
L8
O1
O8
L16
O16
124

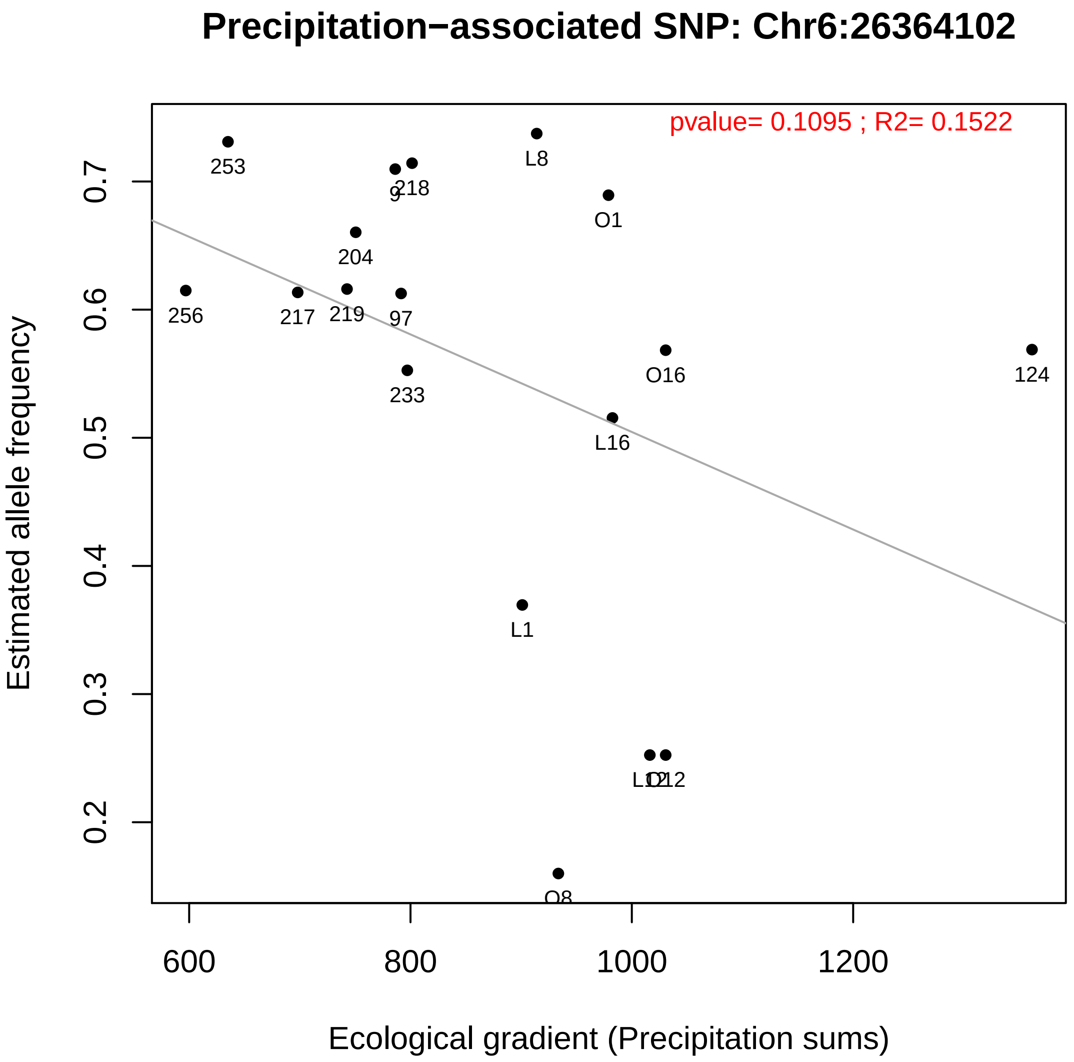
Precipitation−associated SNP: Chr6:26364102
pvalue= 0.1095 ; R2= 0.1522
Estimated allele frequency
Ecological gradient (Precipitation sums)
0.2
0.3
0.4
0.5
0.6
0.7
600
800
1000
1200
253
L8
218
9
O1
204
256
217
219
97
O16
124
233
L16
L1
L12
O12
O8

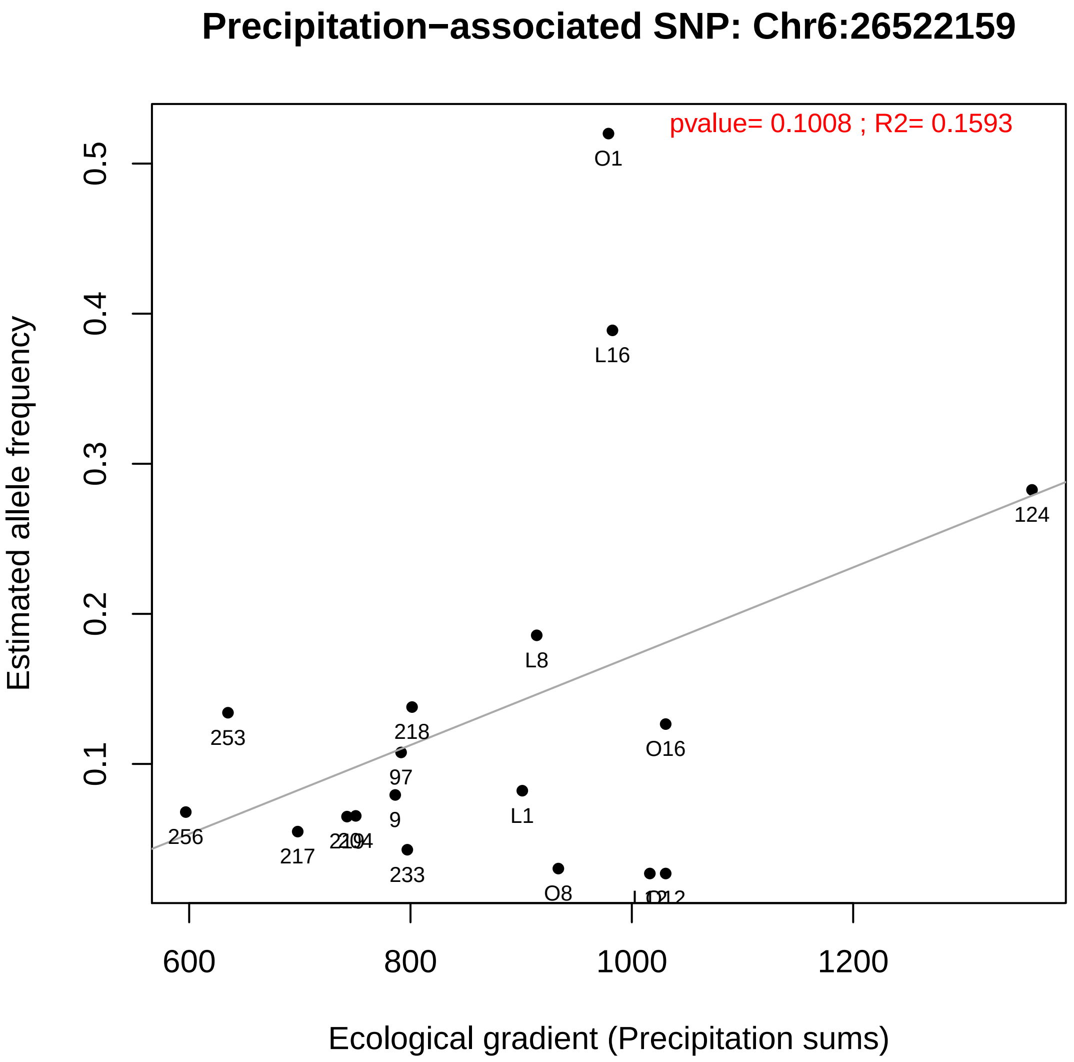
Precipitation−associated SNP: Chr6:26522159
pvalue= 0.1008 ; R2= 0.1593
Estimated allele frequency
Ecological gradient (Precipitation sums)
0.1
0.2
0.3
0.4
0.5
600
800
1000
1200
O1
L16
124
L8
253
218
O16
97
9
L1
256
217
233
O8

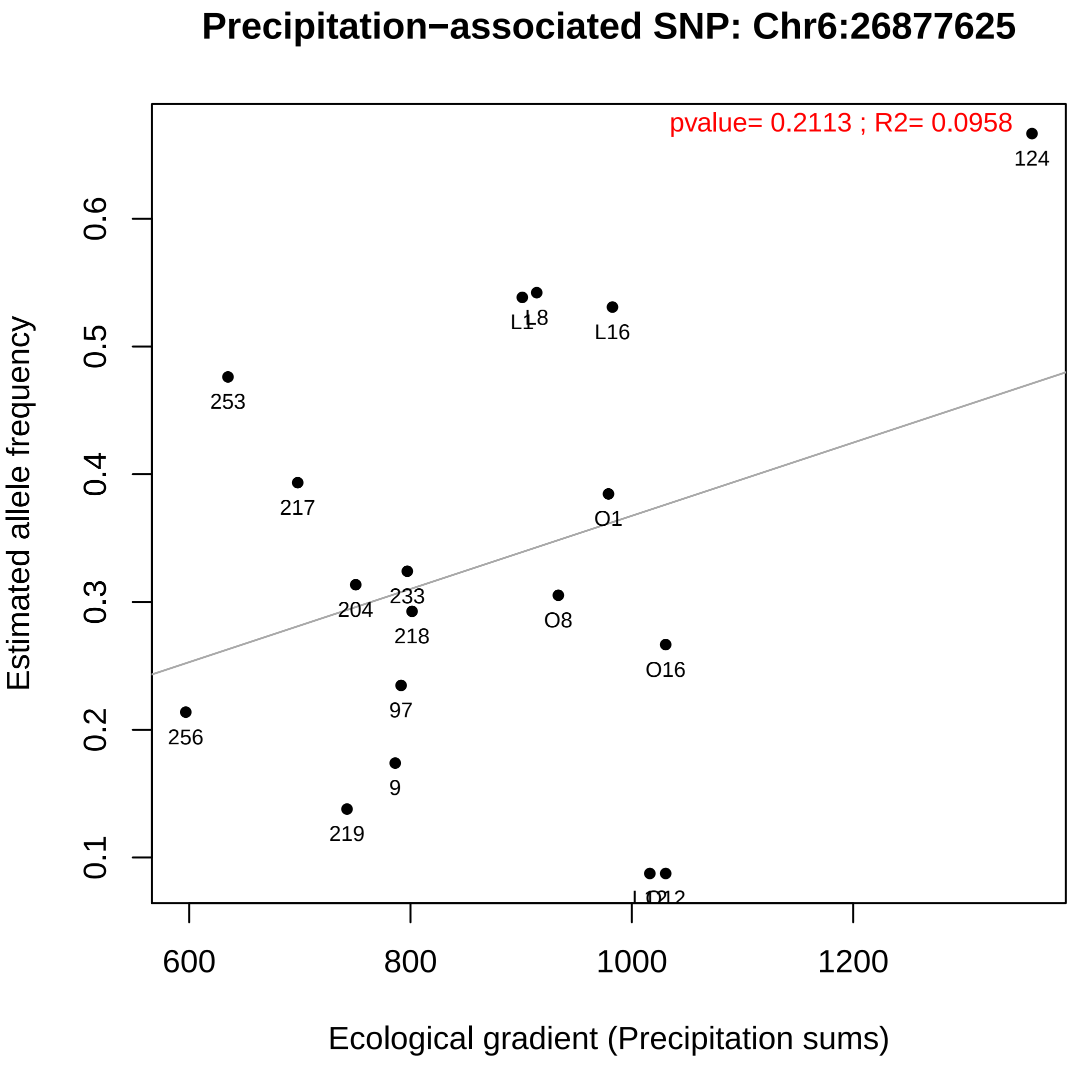

Precipitation−associated SNP: Chr6:26877625
pvalue= 0.2113 ; R2= 0.0958
124
L1
L8
L16
253
217
O1
233
204
218
O8
O16
97
256
9
219
O12
Estimated allele frequency
0.1
0.2
0.3
0.4
0.5
0.6
600
800
1000
1200
Ecological gradient (Precipitation sums)

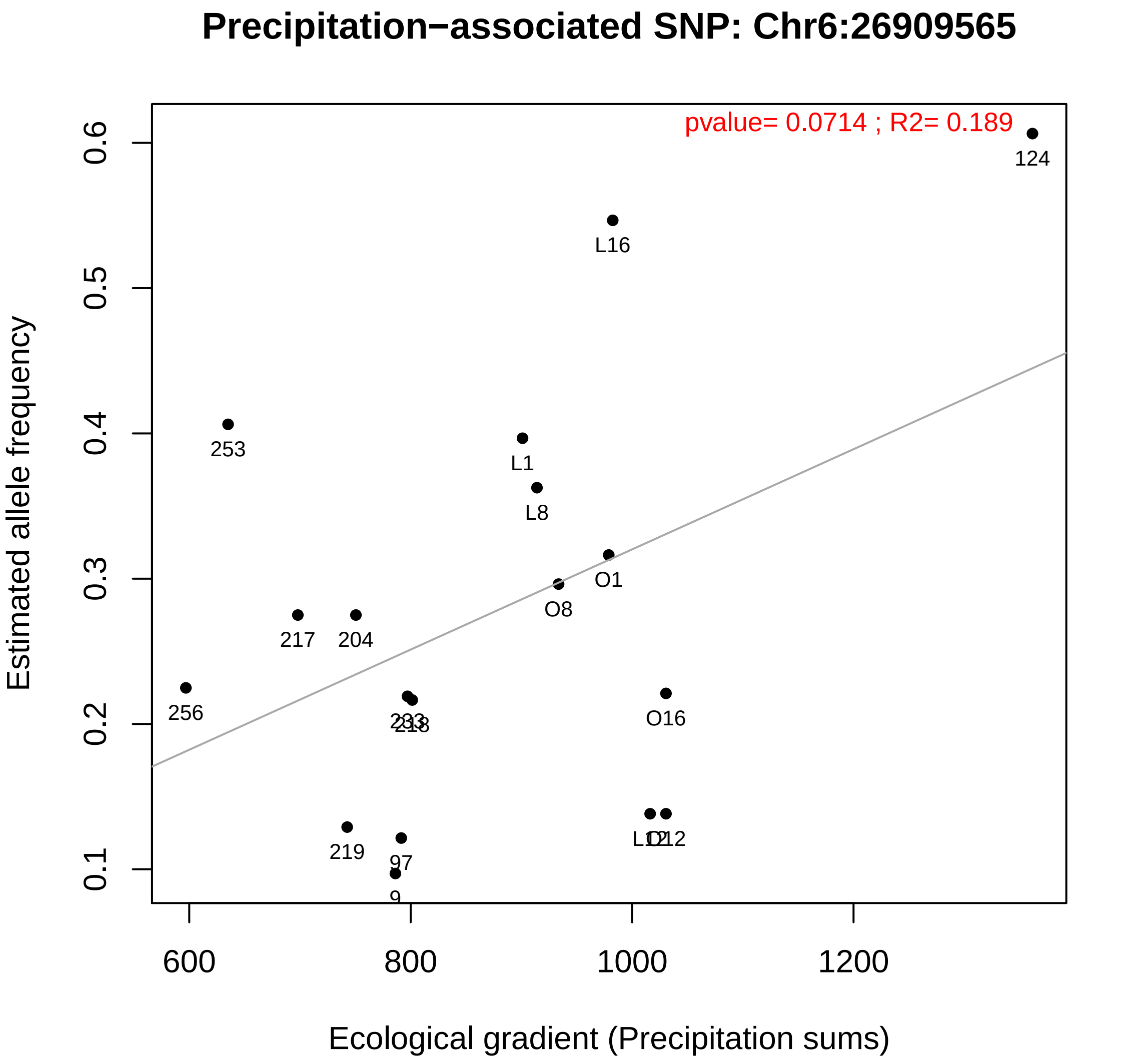
Precipitation−associated SNP: Chr6:26909565
pvalue= 0.0714 ; R2= 0.189
Estimated allele frequency
Ecological gradient (Precipitation sums)
0.1
0.2
0.3
0.4
0.5
0.6
600
800
1000
1200
124
L16
253
L1
L8
O1
O8
217
204
256
O16
219
97
9

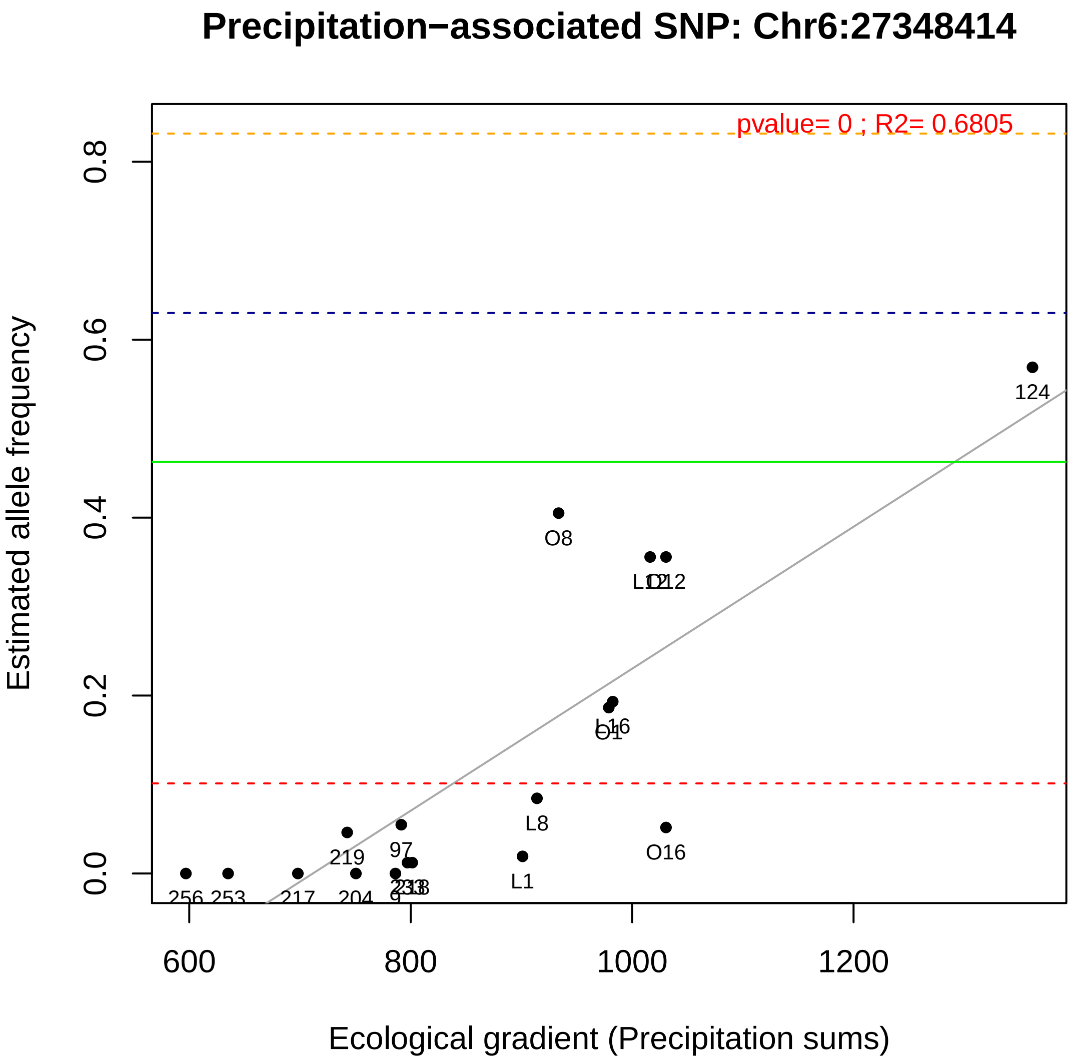

Precipitation−associated SNP: Chr6:27348414
pvalue= 0 ; R2= 0.6805
Estimated allele frequency
Ecological gradient (Precipitation sums)
0.0
0.2
0.4
0.6
0.8
600
800
1000
1200
124
O8
O12
L8
O16
L1
219
97

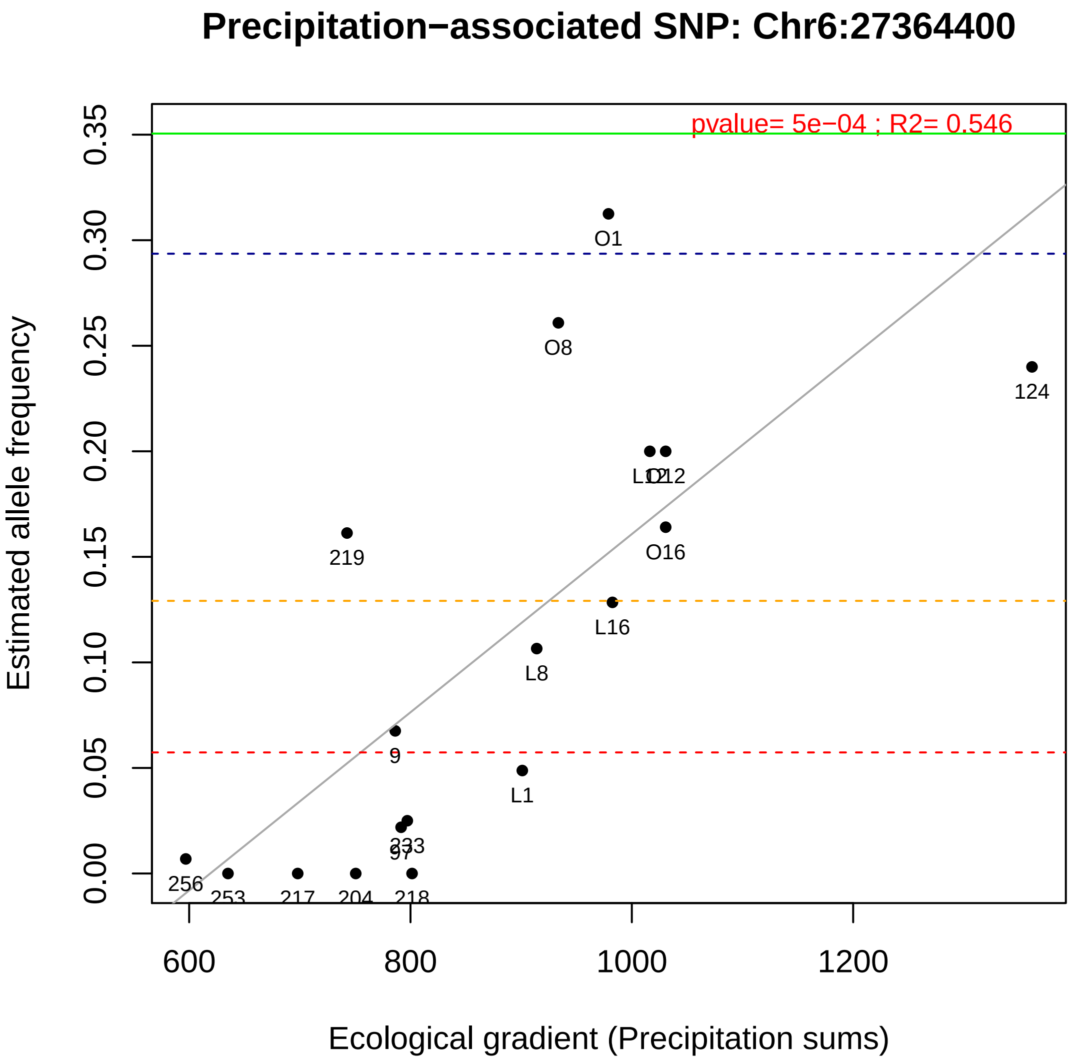

Precipitation−associated SNP: Chr6:27364400
pvalue= 5e−04 ; R2= 0.546
Estimated allele frequency
0.00
0.05
0.10
0.15
0.20
0.25
0.30
0.35
600
800
1000
1200
Ecological gradient (Precipitation sums)
O1
O8
124
O12
O16
219
L16
L8
9
L1
256
253
217
204
218

**Precipitation−associated SNP: Chr6:27470969**

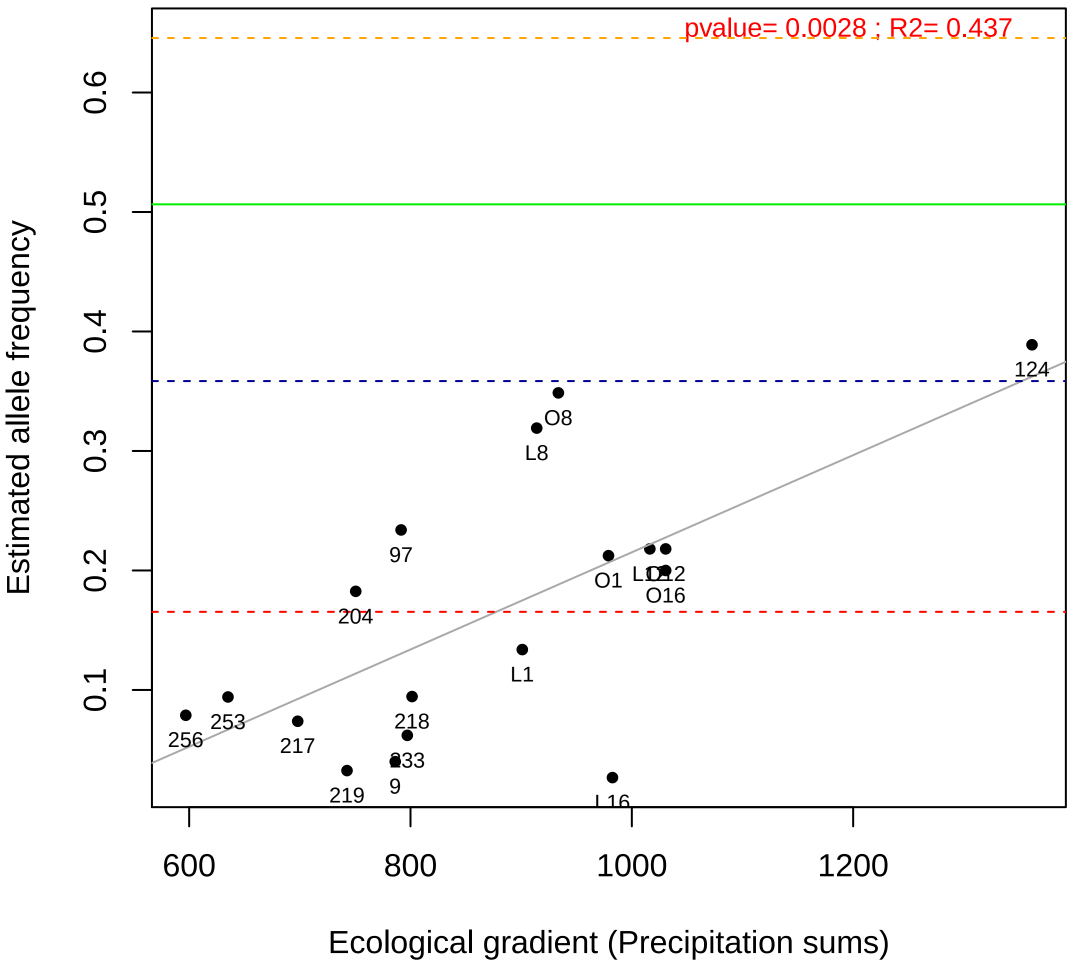

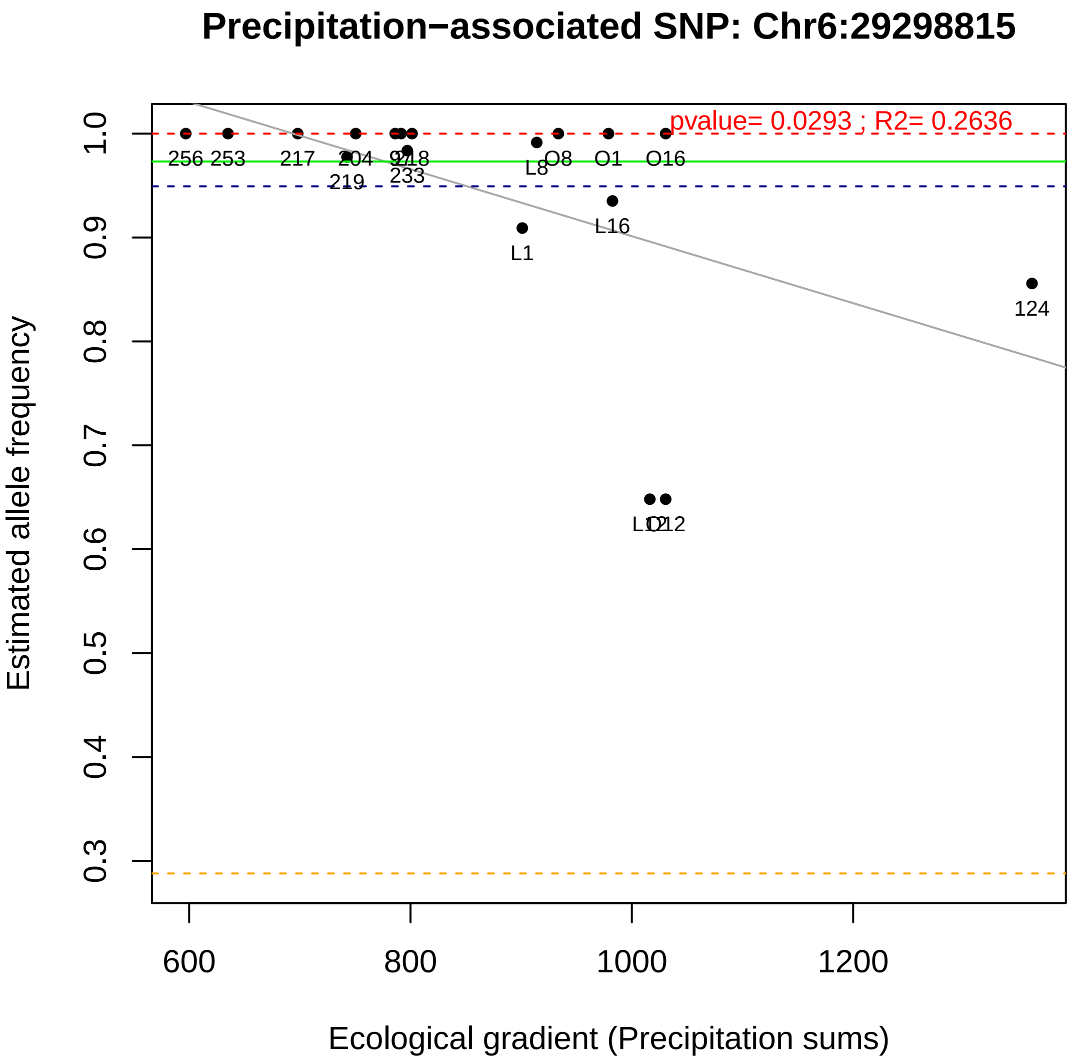

Precipitation−associated SNP: Chr6:29298815
pvalue= 0.0293 ; R2= 0.2636
256
253
217
204
219
92
218
233
L8
O8
O1
O16
L16
L1
124
L12
O12
1.0
0.9
0.8
0.7
0.6
0.5
0.4
0.3
600
800
1000
1200
Estimated allele frequency
Ecological gradient (Precipitation sums)

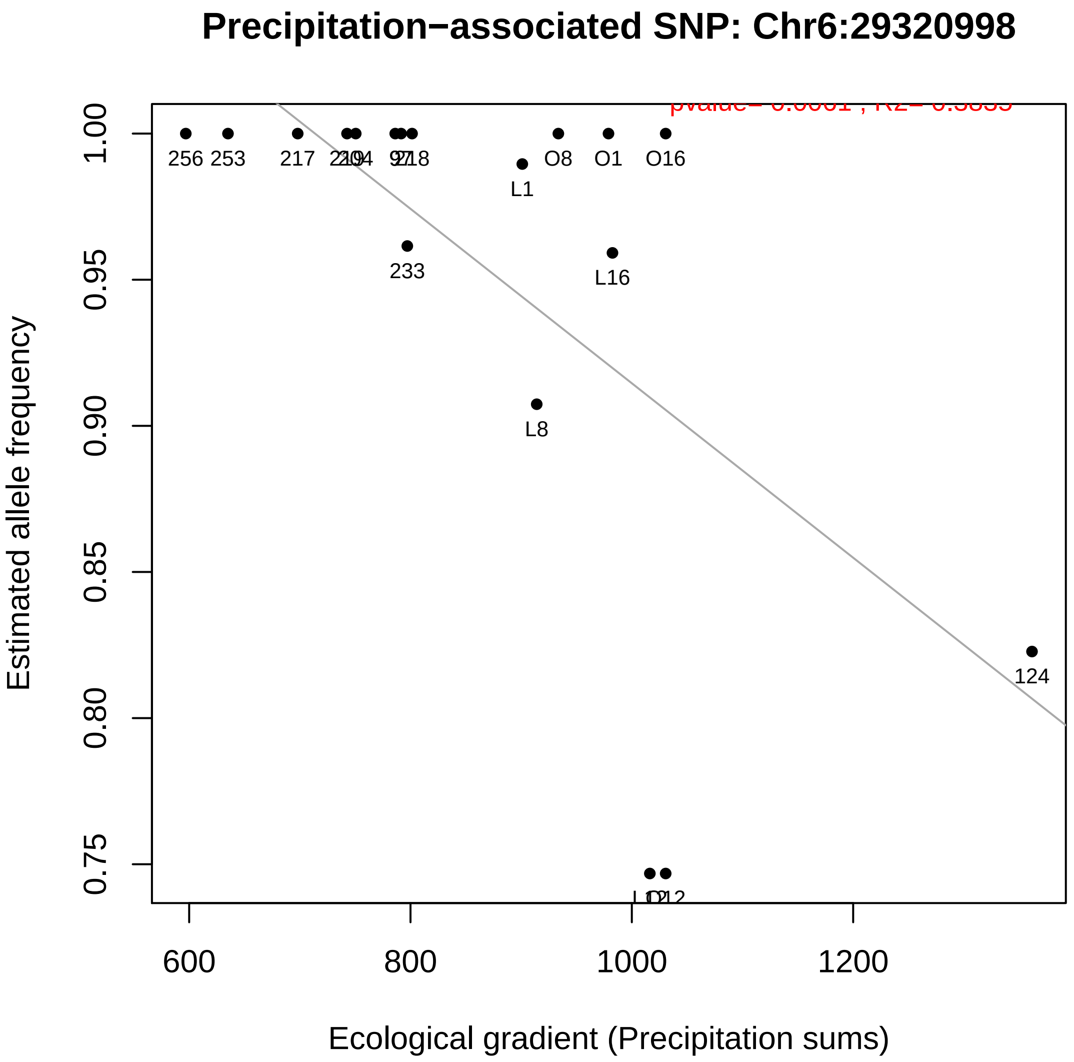

Precipitation−associated SNP: Chr6:29320998
Estimated allele frequency
Ecological gradient (Precipitation sums)
1.00
0.95
0.90
0.85
0.80
0.75
600
800
1000
1200
256
253
217
218
L1
O8
O1
O16
233
L16
L8
124

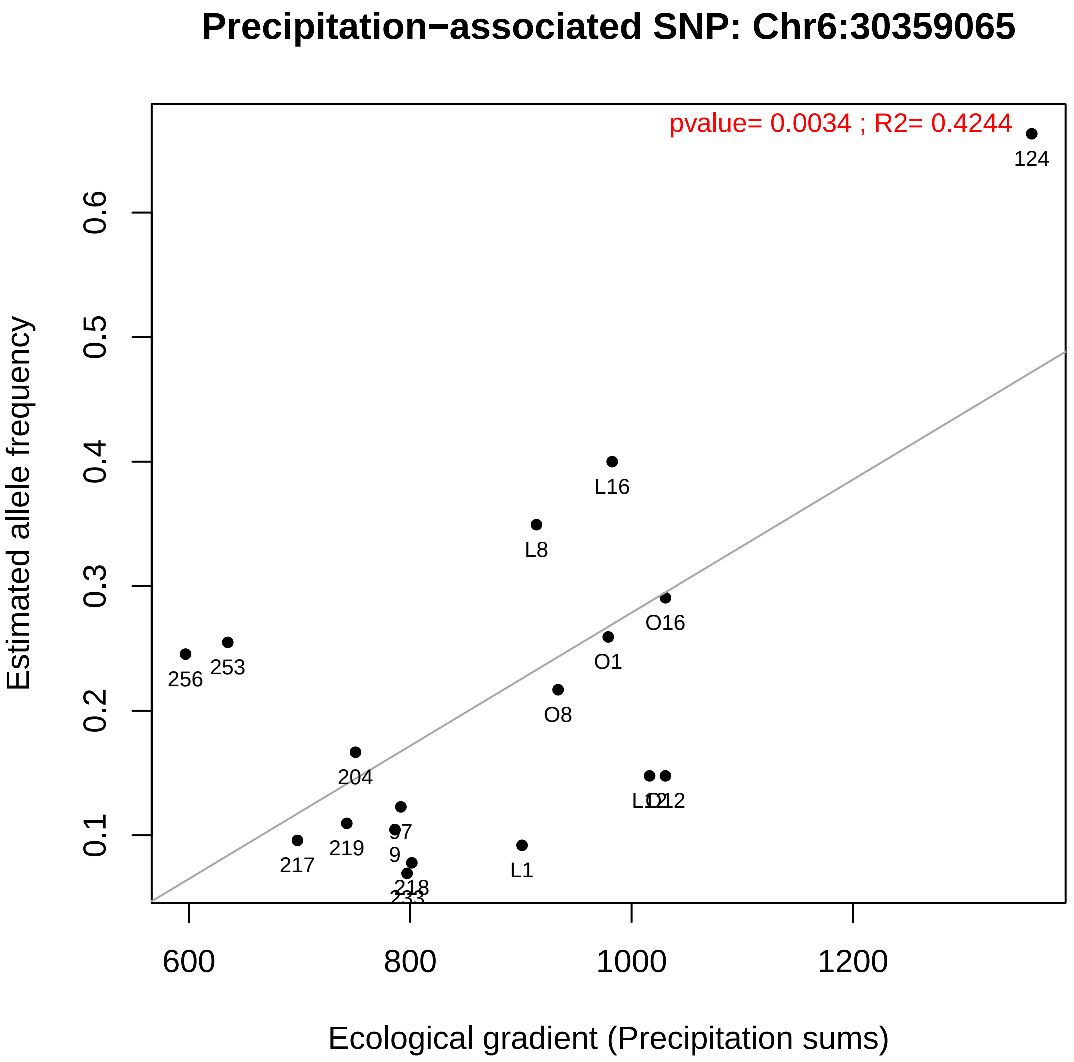
Precipitation−associated SNP: Chr6:30359065
pvalue= 0.0034 ; R2= 0.4244
124
L16
L8
O16
O1
O8
256
253
204
L12
O12
97
9
219
217
218
233
L1
0.1
0.2
0.3
0.4
0.5
0.6
600
800
1000
1200
Estimated allele frequency
Ecological gradient (Precipitation sums)

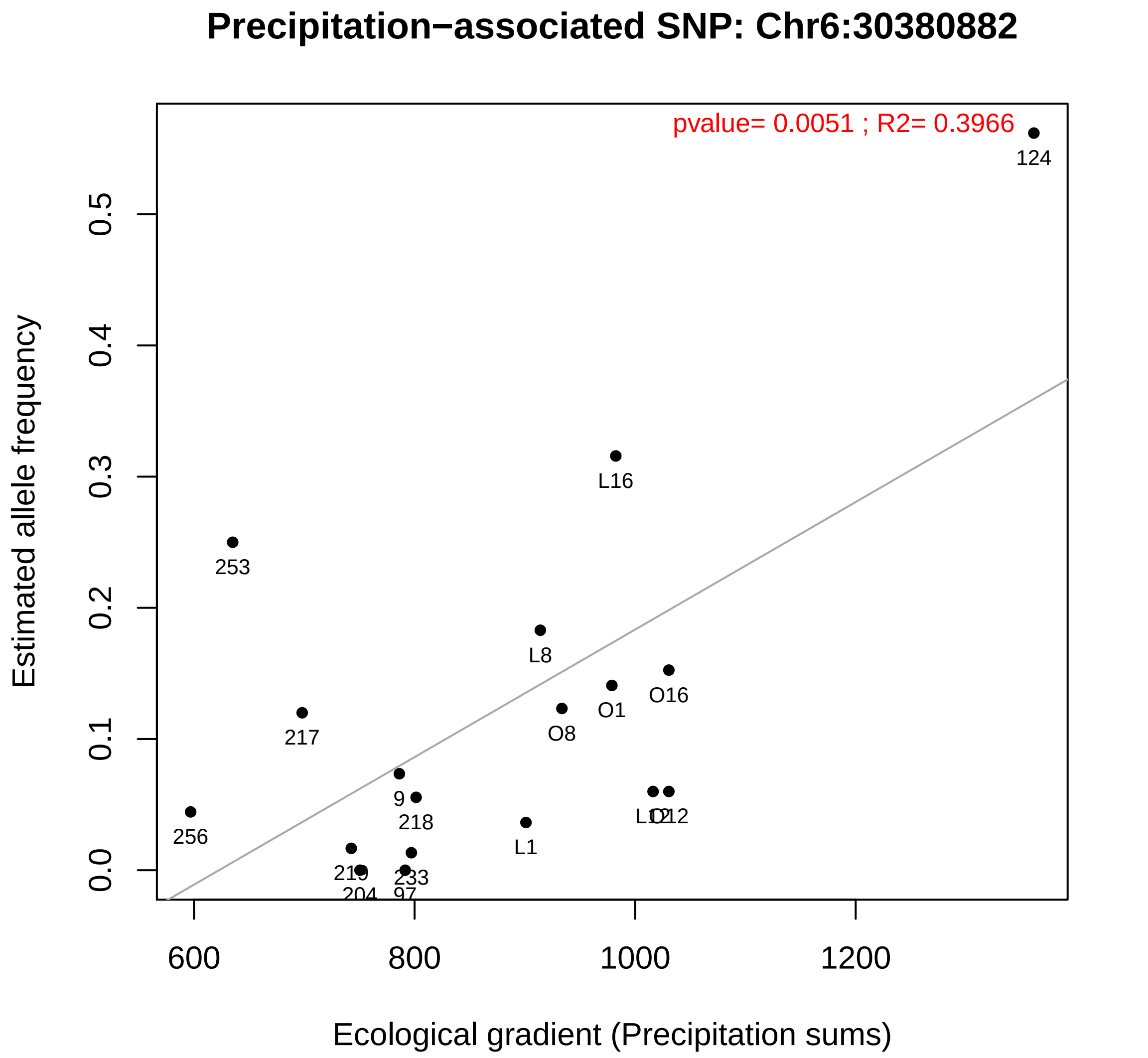
Precipitation−associated SNP: Chr6:30380882
pvalue= 0.0051 ; R2= 0.3966
124
L16
253
L8
O16
O1
O8
217
9
218
256
L1
L12
O12
219
204
233
97
Estimated allele frequency
0.0
0.1
0.2
0.3
0.4
0.5
600
800
1000
1200
Ecological gradient (Precipitation sums)

#### Precipitation−associated SNP: Chr6:30411668

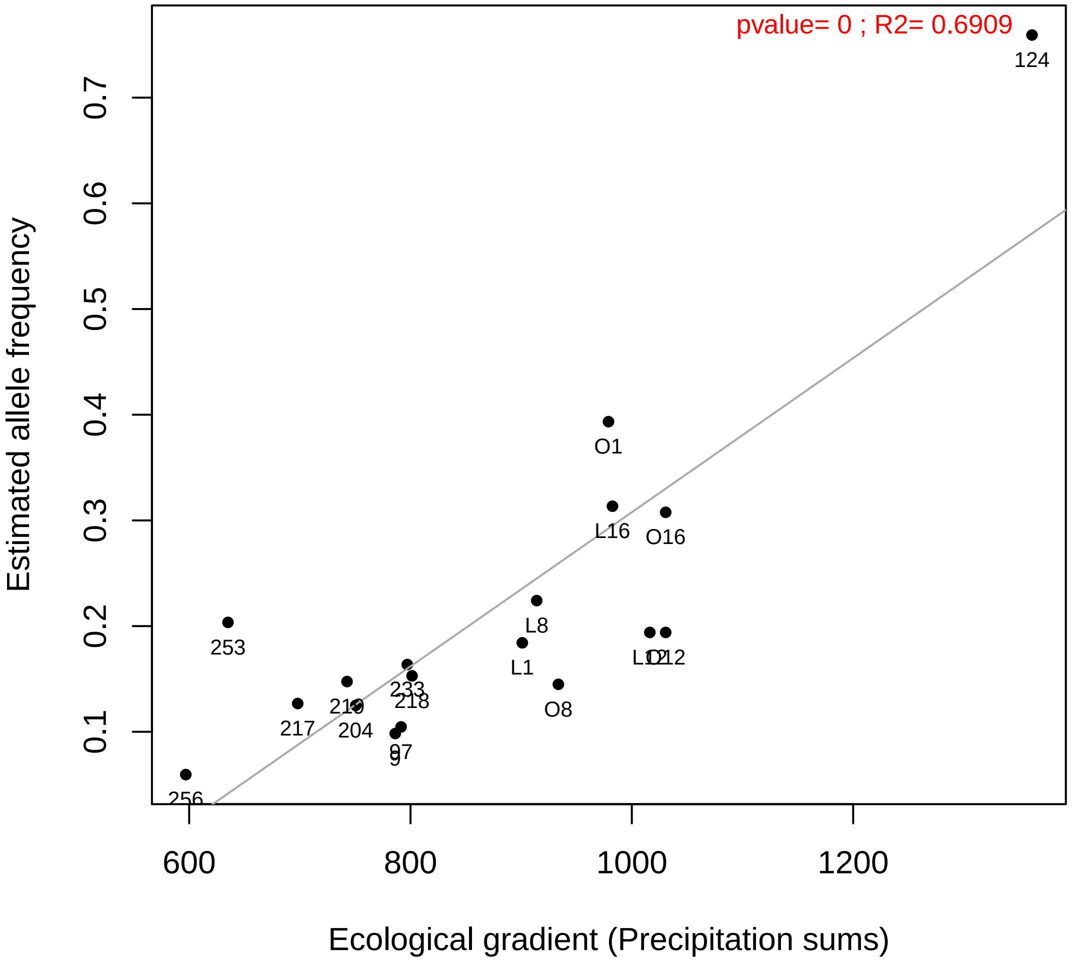

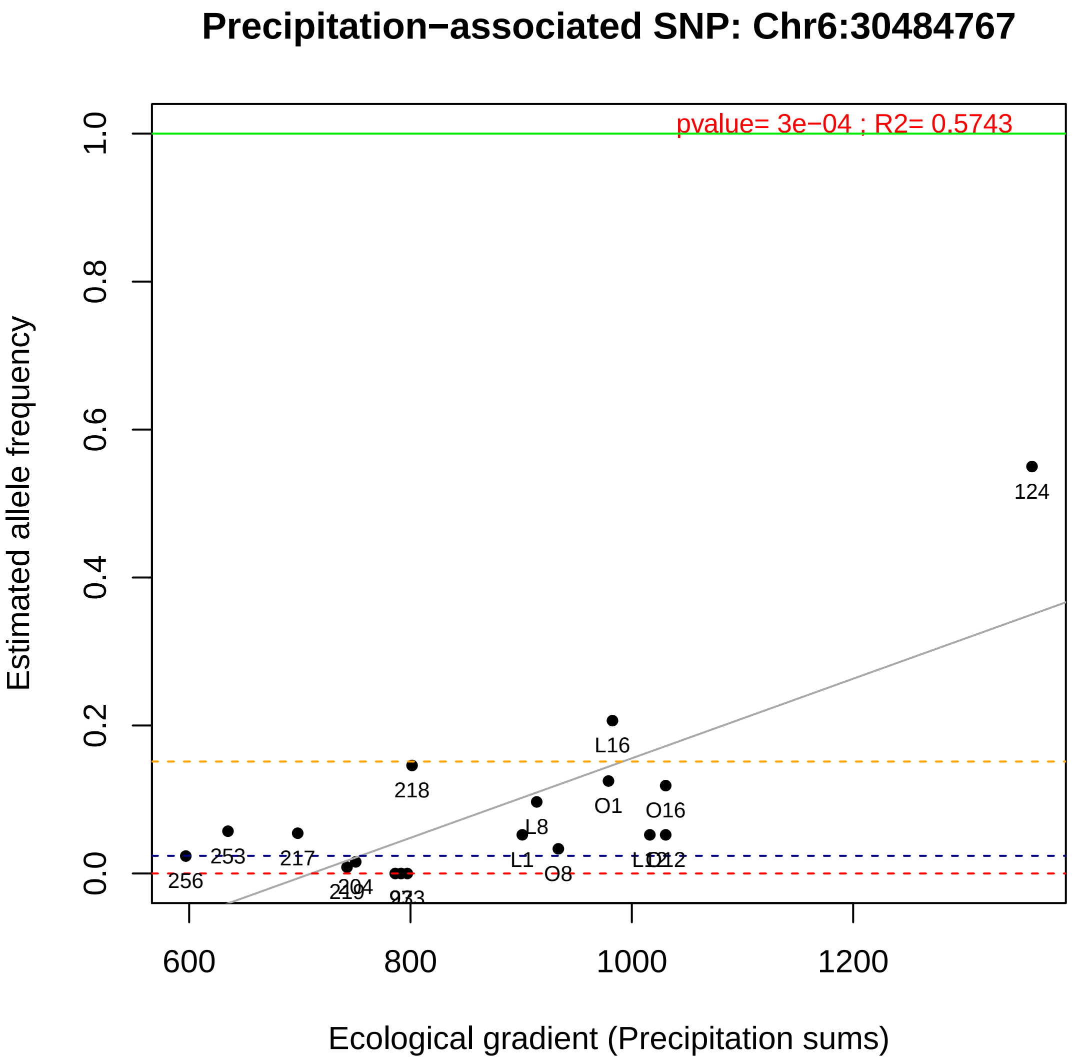
Precipitation−associated SNP: Chr6:30484767
pvalue= 3e−04 ; R2= 0.5743
Estimated allele frequency
0.0
0.2
0.4
0.6
0.8
1.0
Ecological gradient (Precipitation sums)
600
800
1000
1200
124
L16
218
O1
O16
L8
256
253
217
L1
O8

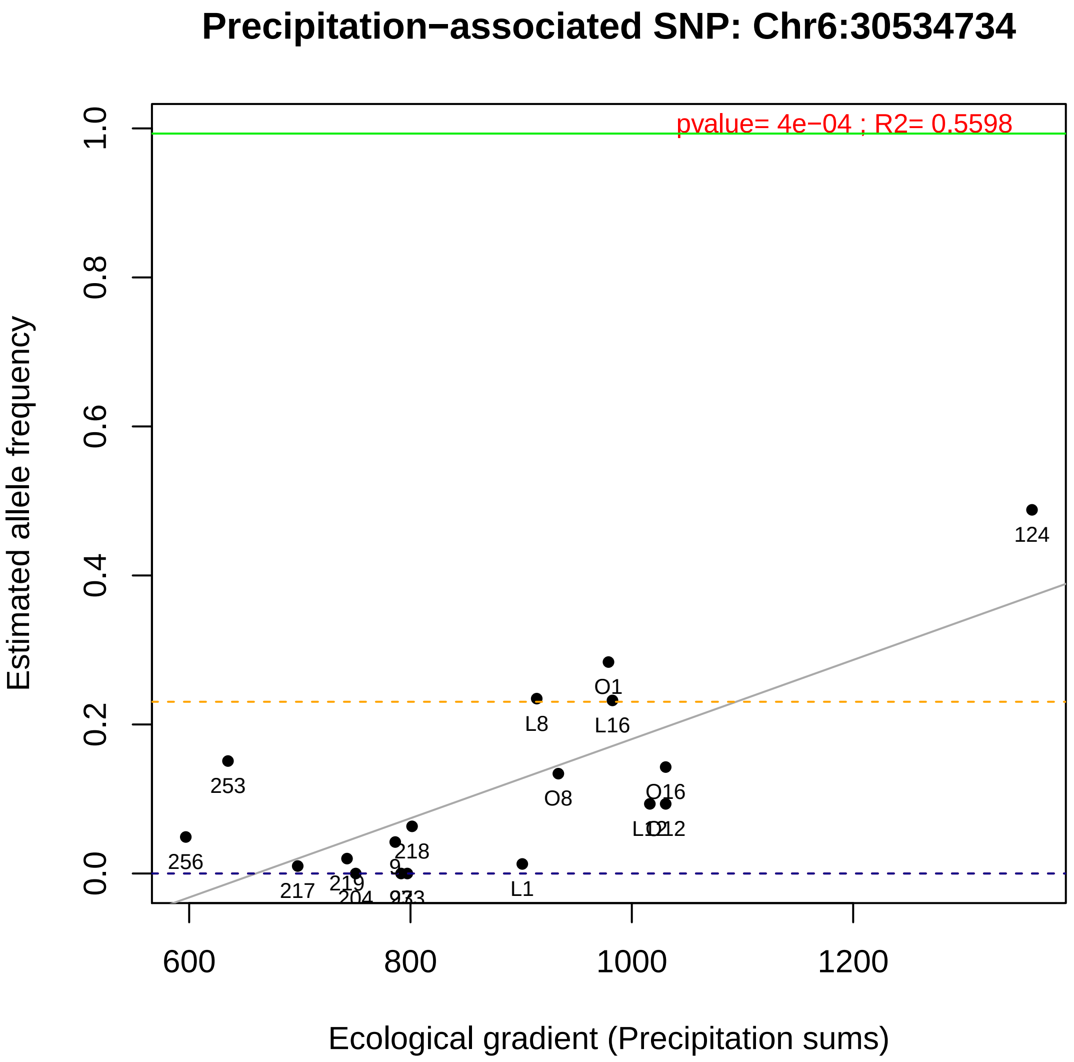
Precipitation−associated SNP: Chr6:30534734
pvalue= 4e−04 ; R2= 0.5598
Estimated allele frequency
0.0
0.2
0.4
0.6
0.8
1.0
600
800
1000
1200
Ecological gradient (Precipitation sums)
124
O1
L8
L16
253
O8
O16
218
256
9
219
217
L1

#### Precipitation−associated SNP: Chr6:30535347

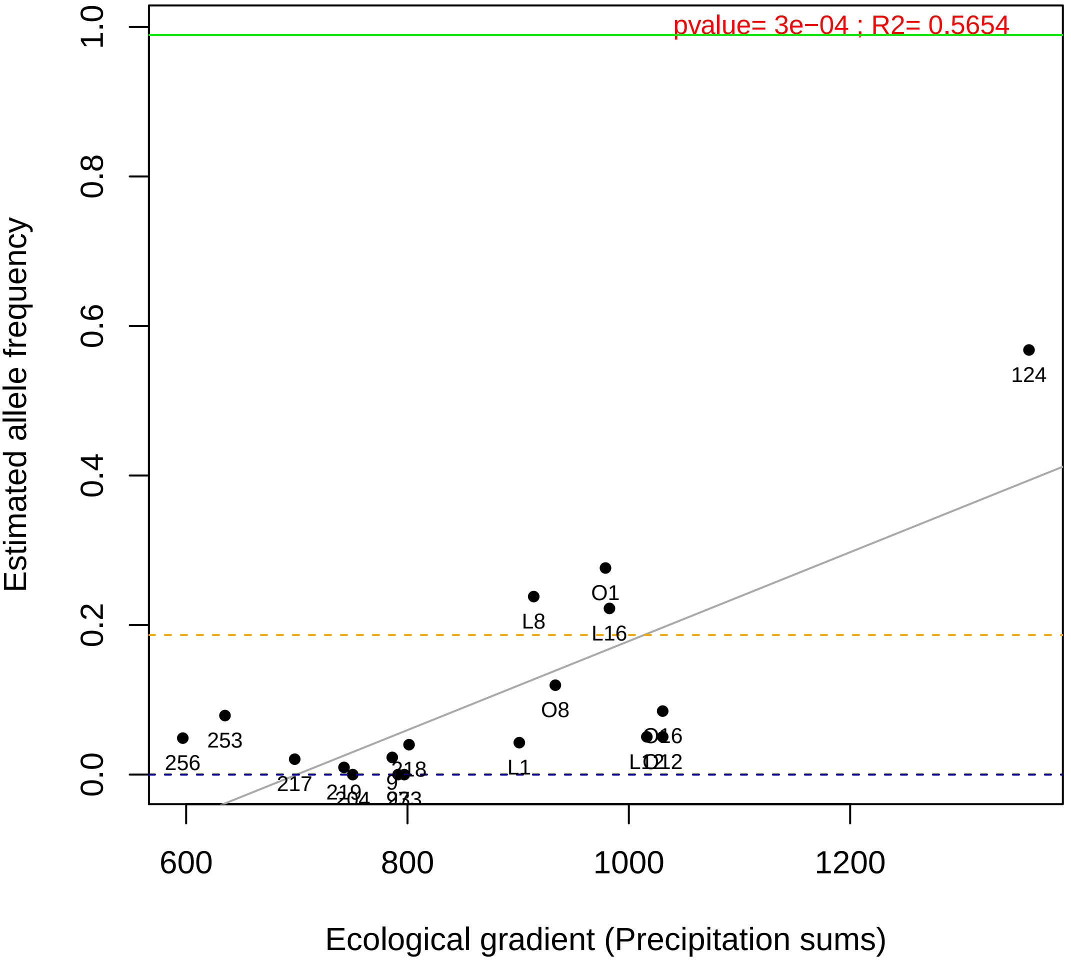

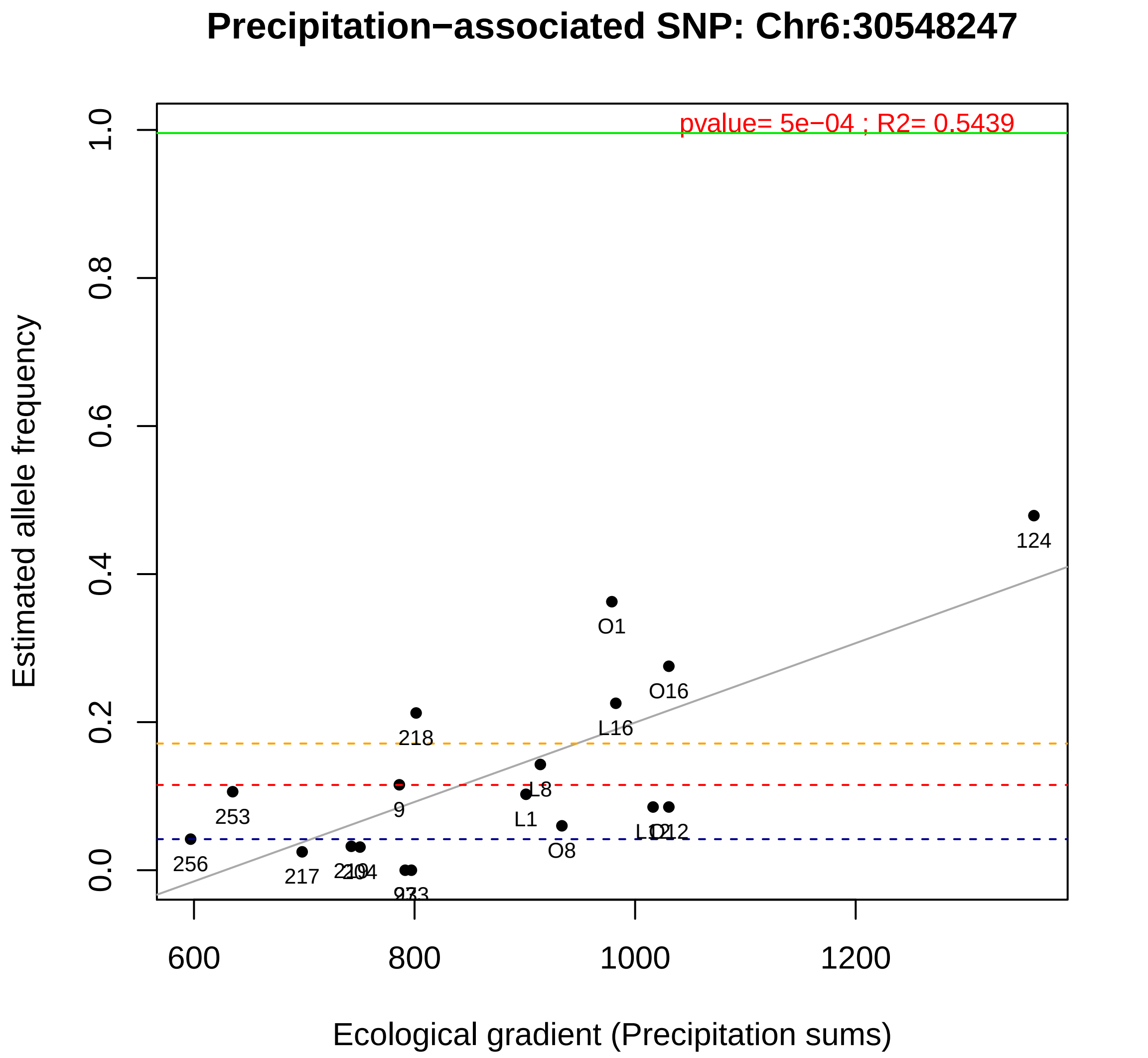

Precipitation−associated SNP: Chr6:30548247
pvalue= 5e−04 ; R2= 0.5439
Estimated allele frequency
0.0
0.2
0.4
0.6
0.8
1.0
600
800
1000
1200
Ecological gradient (Precipitation sums)
124
O1
O16
L16
218
L8
9
L1
253
O8
O12
256
217

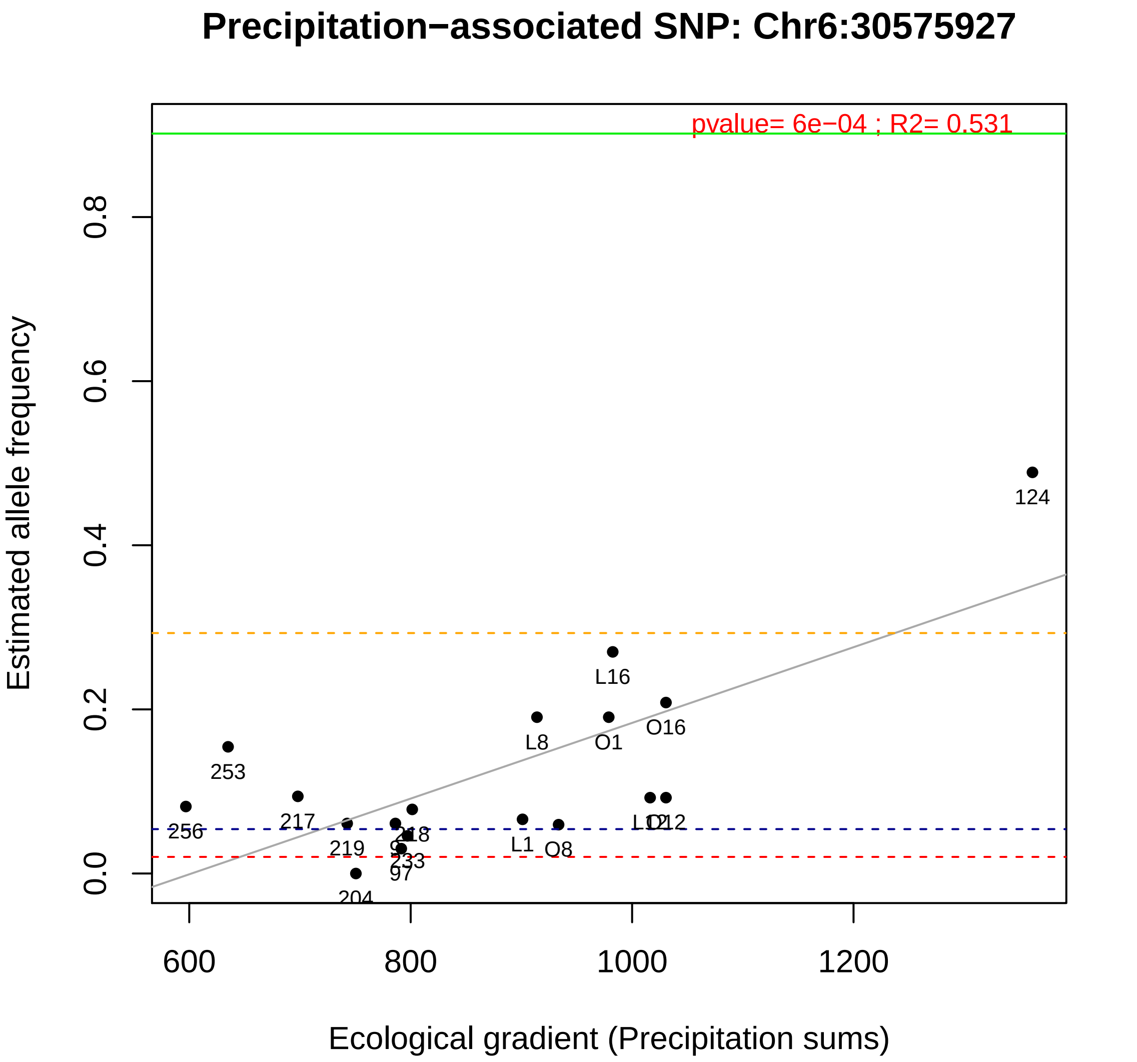

Precipitation−associated SNP: Chr6:30575927
pvalue= 6e−04 ; R2= 0.531
Estimated allele frequency
0.0
0.2
0.4
0.6
0.8
Ecological gradient (Precipitation sums)
600
800
1000
1200
124
L16
O16
L8
O1
253
217
256
219
218
233
97
204
L1
O8
O12

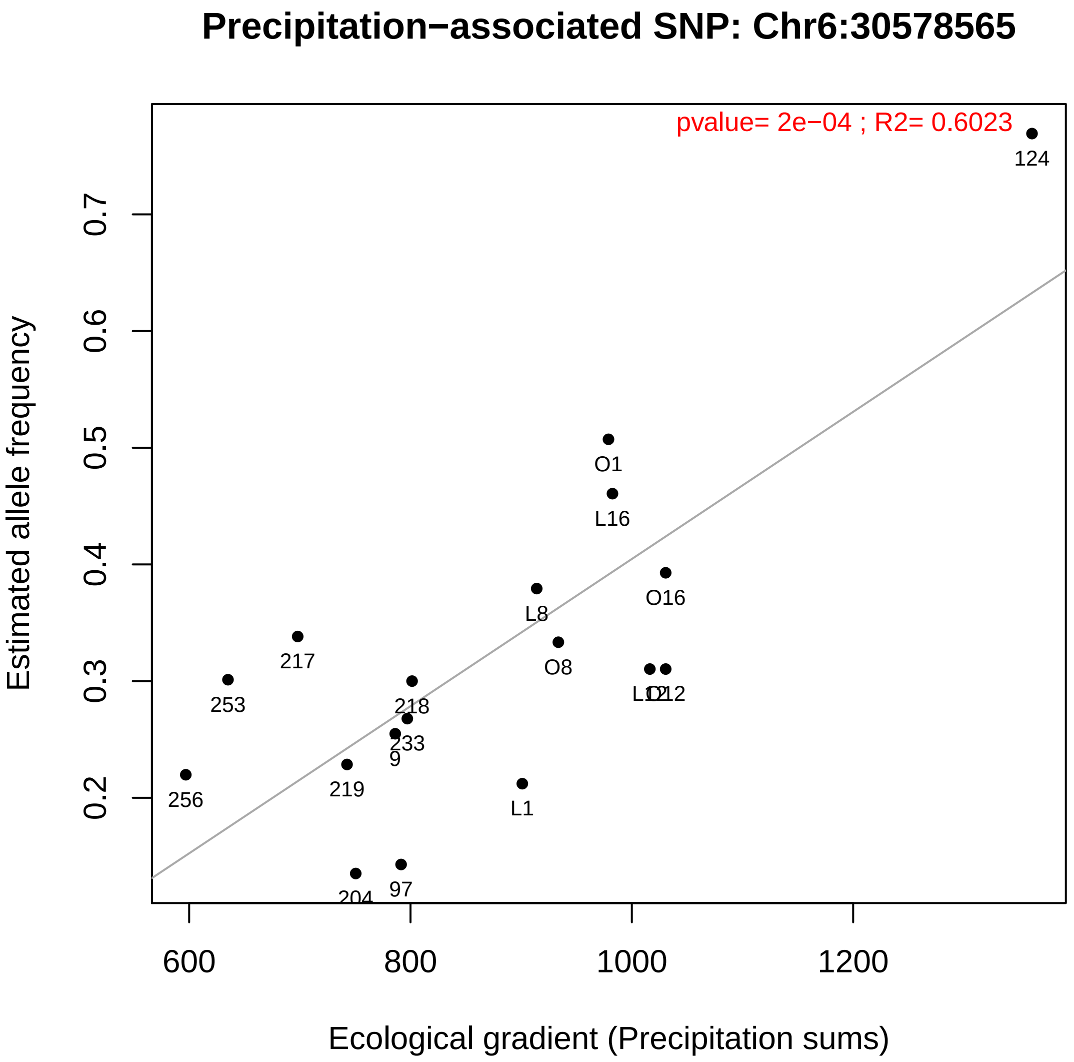
Precipitation−associated SNP: Chr6:30578565
pvalue= 2e−04 ; R2= 0.6023
Estimated allele frequency
Ecological gradient (Precipitation sums)
0.2
0.3
0.4
0.5
0.6
0.7
600
800
1000
1200
124
O1
L16
O16
L8
O8
217
253
218
233
9
219
256
L1
204
97

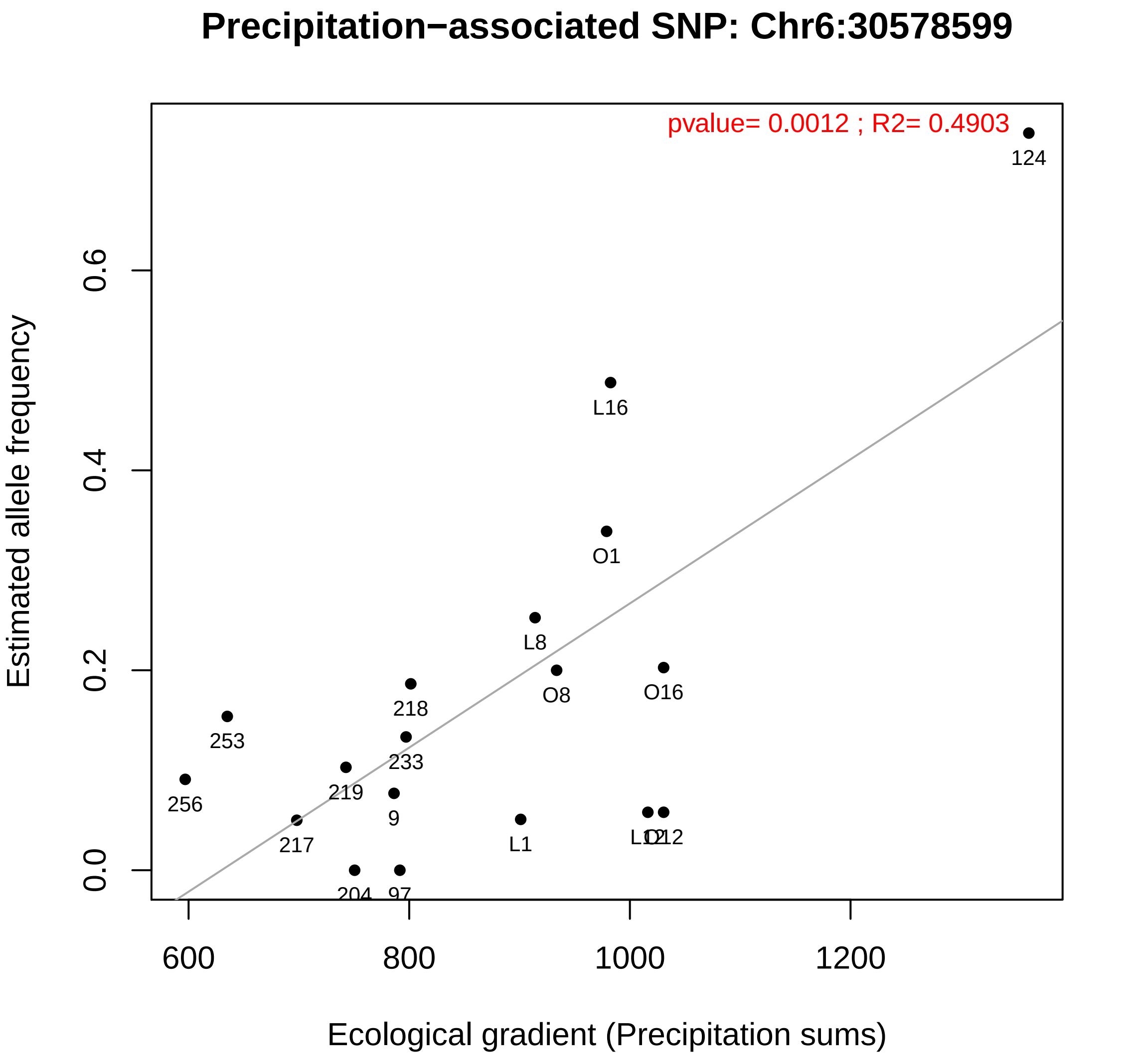

Precipitation−associated SNP: Chr6:30578599
pvalue= 0.0012 ; R2= 0.4903
Estimated allele frequency
0.0
0.2
0.4
0.6
600
800
1000
1200
Ecological gradient (Precipitation sums)
124
L16
O1
L8
O8
O16
218
253
233
219
256
9
217
L1
L12
O12
204
97

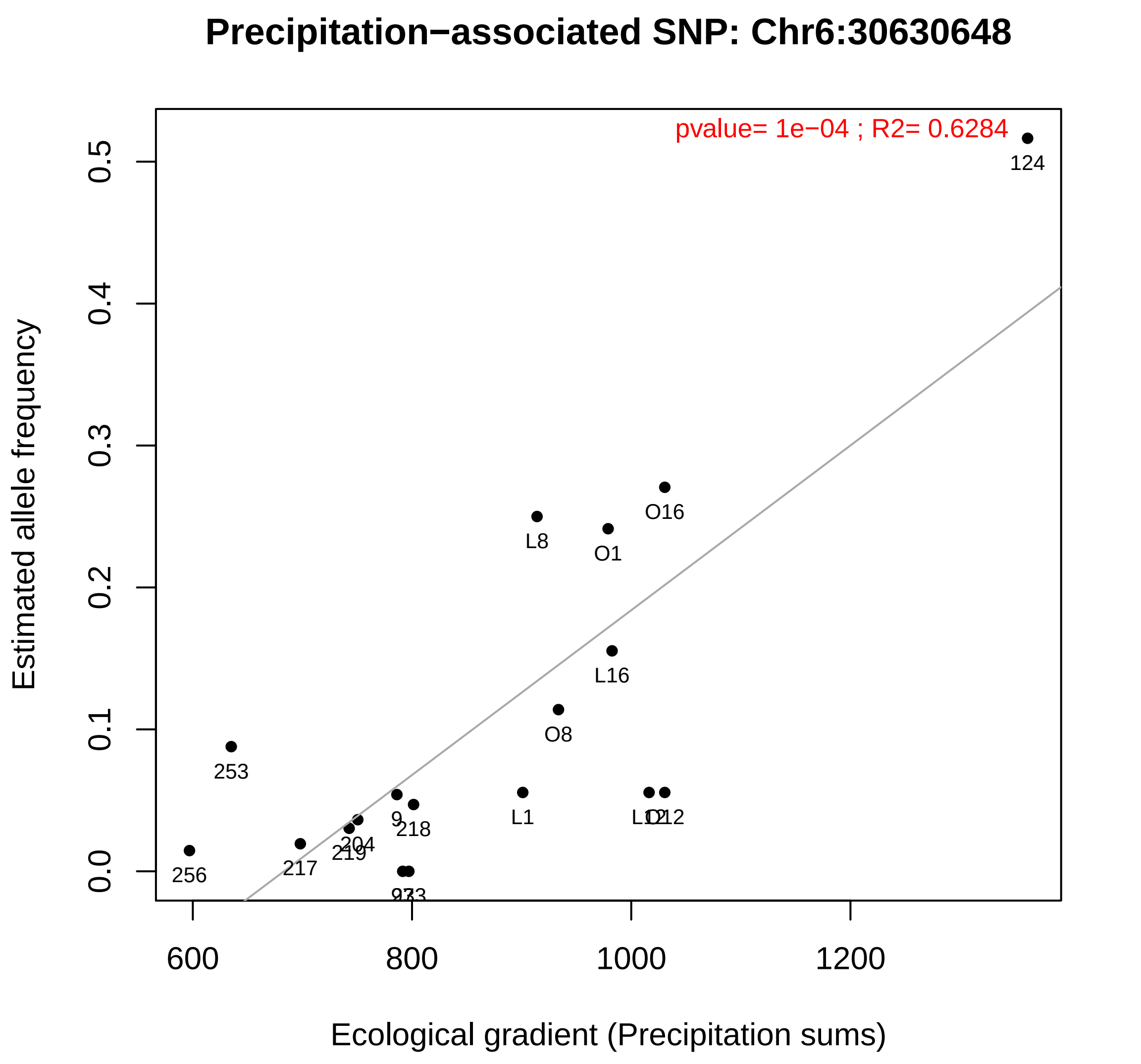
Precipitation−associated SNP: Chr6:30630648
pvalue= 1e−04 ; R2= 0.6284
Estimated allele frequency
0.0
0.1
0.2
0.3
0.4
0.5
600
800
1000
1200
Ecological gradient (Precipitation sums)
124
O16
L8
O1
L16
O8
253
L1
9
218
204
219
217
256
O12

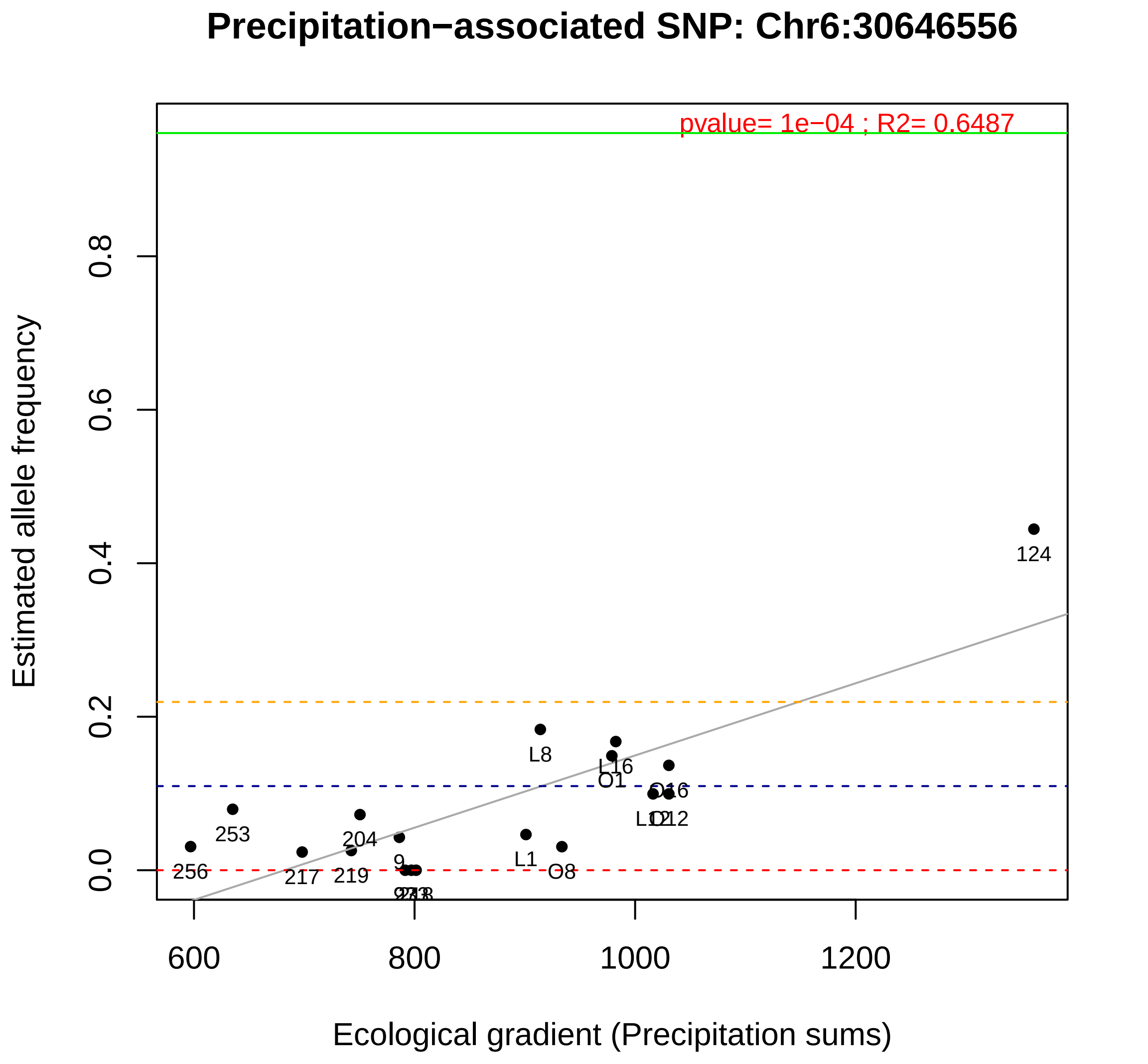
Precipitation−associated SNP: Chr6:30646556
pvalue= 1e−04 ; R2= 0.6487
Estimated allele frequency
0.0
0.2
0.4
0.6
0.8
600
800
1000
1200
Ecological gradient (Precipitation sums)
124
L8
L16
O1
O16
L12
O12
253
204
256
217
219
9
L1
O8

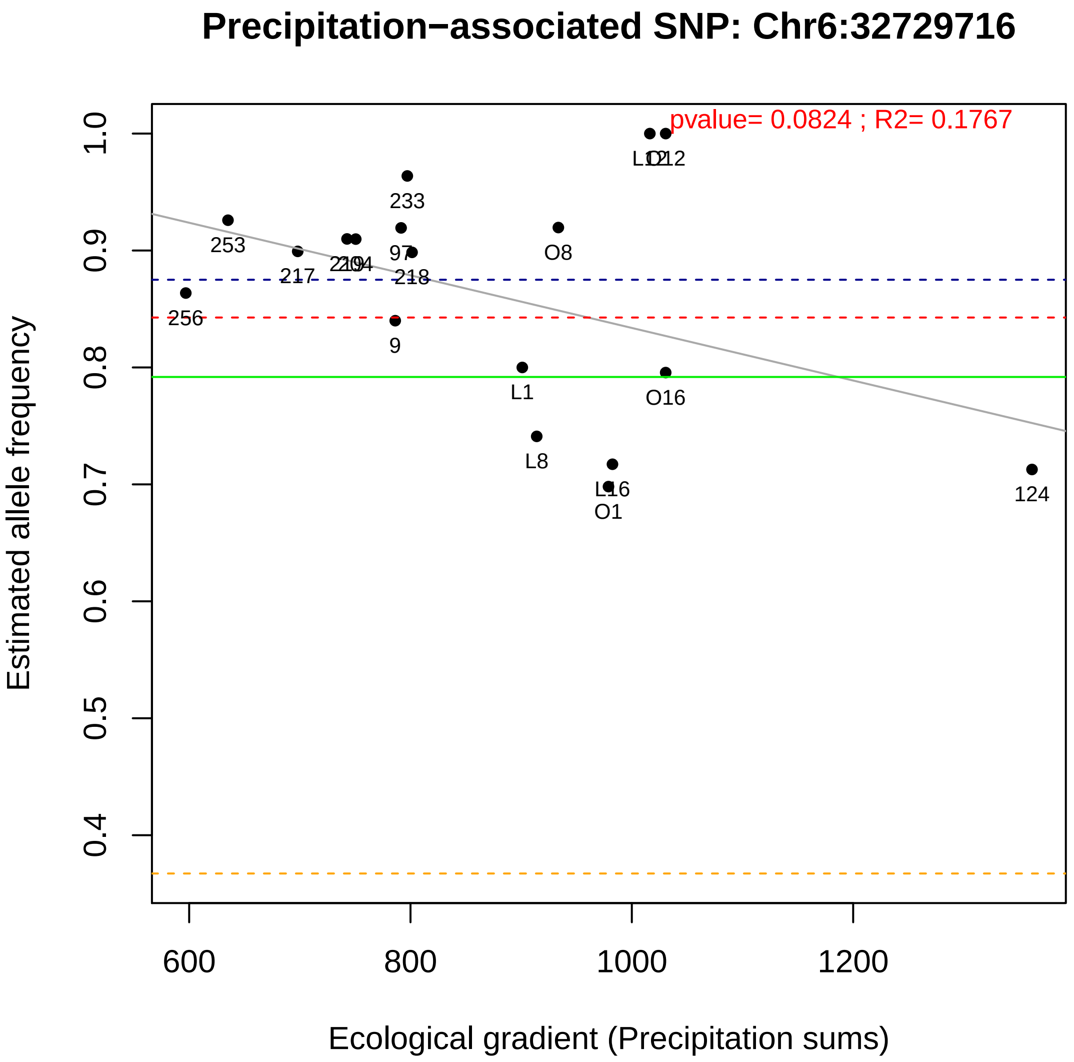

Precipitation−associated SNP: Chr6:32729716
pvalue= 0.0824 ; R2= 0.1767
Estimated allele frequency
Ecological gradient (Precipitation sums)
0.4
0.5
0.6
0.7
0.8
0.9
1.0
600
800
1000
1200
256
253
217
233
97
218
9
O8
L1
L8
O16
L16
O1
124

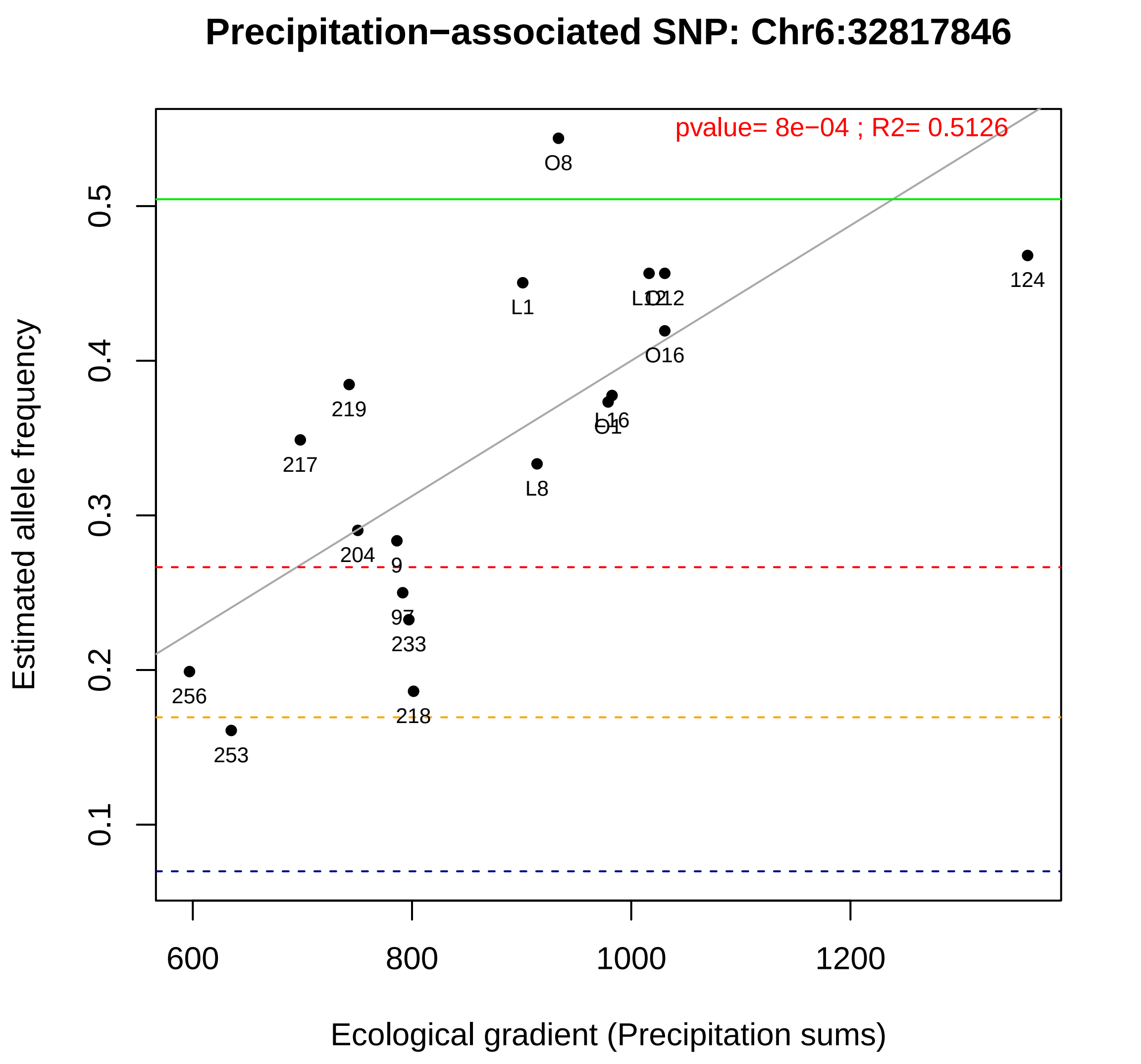
Precipitation−associated SNP: Chr6:32817846
pvalue= 8e−04 ; R2= 0.5126
Estimated allele frequency
0.1
0.2
0.3
0.4
0.5
600
800
1000
1200
Ecological gradient (Precipitation sums)
O8
124
L1
O12
O16
219
217
L8
204
9
233
256
218
253

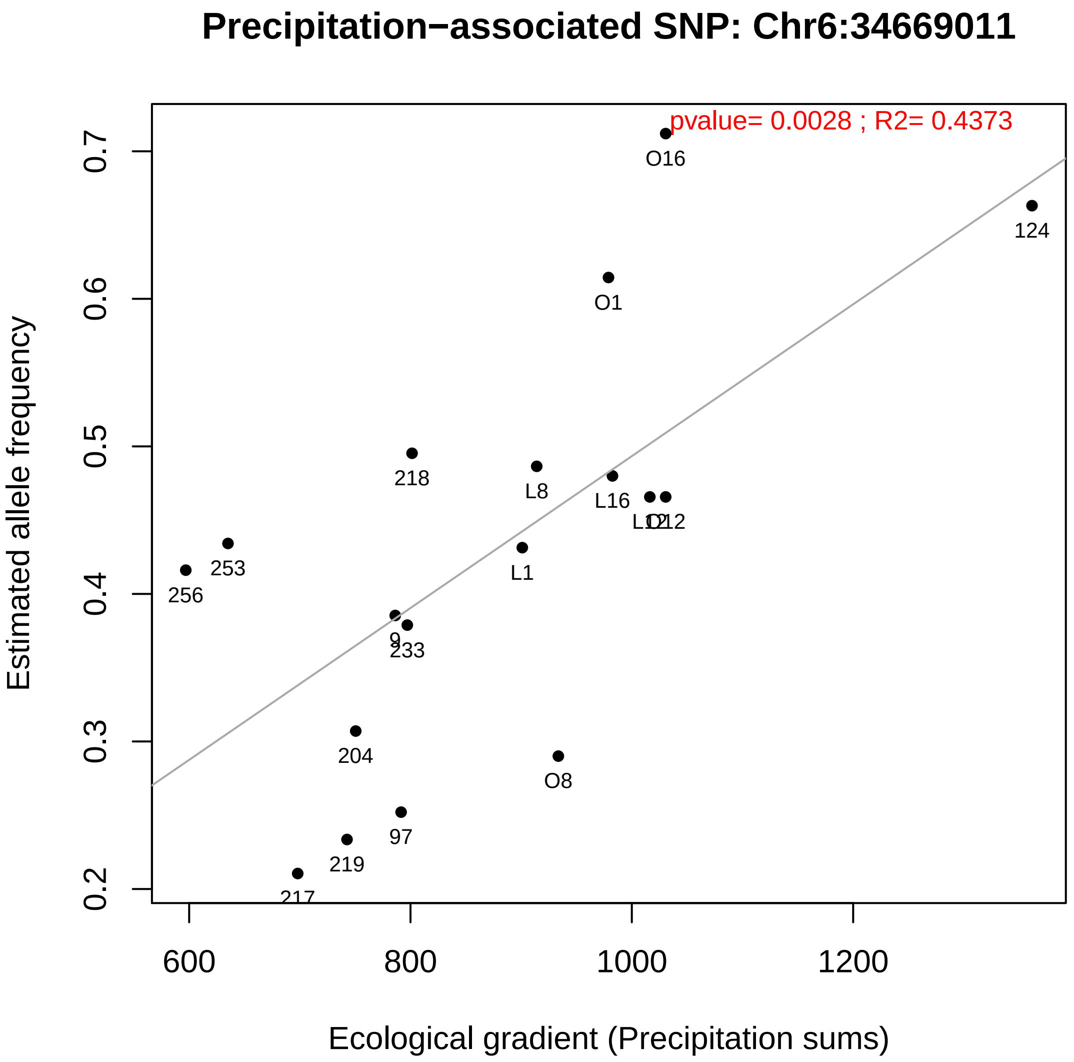

Precipitation−associated SNP: Chr6:34669011
pvalue= 0.0028 ; R2= 0.4373
Estimated allele frequency
Ecological gradient (Precipitation sums)
0.2
0.3
0.4
0.5
0.6
0.7
600
800
1000
1200
O16
124
O1
218
L8
L16
L12
O12
253
256
L1
9
233
204
O8
97
219
217

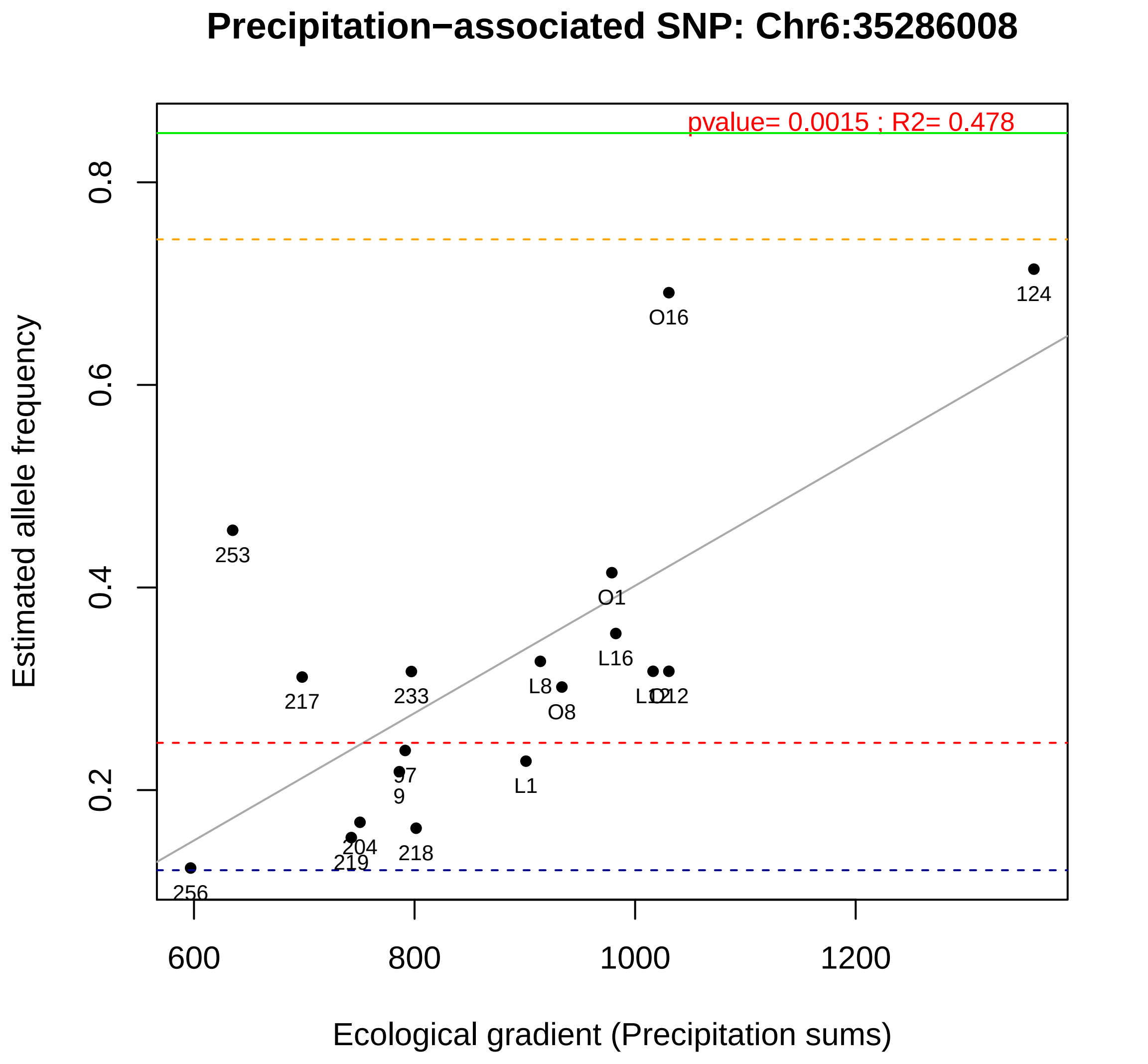

Precipitation−associated SNP: Chr6:35286008
pvalue= 0.0015 ; R2= 0.478
Estimated allele frequency
0.2
0.4
0.6
0.8
600
800
1000
1200
Ecological gradient (Precipitation sums)
124
O16
253
O1
L16
217
233
L8
O8
L1
218
204
219
256

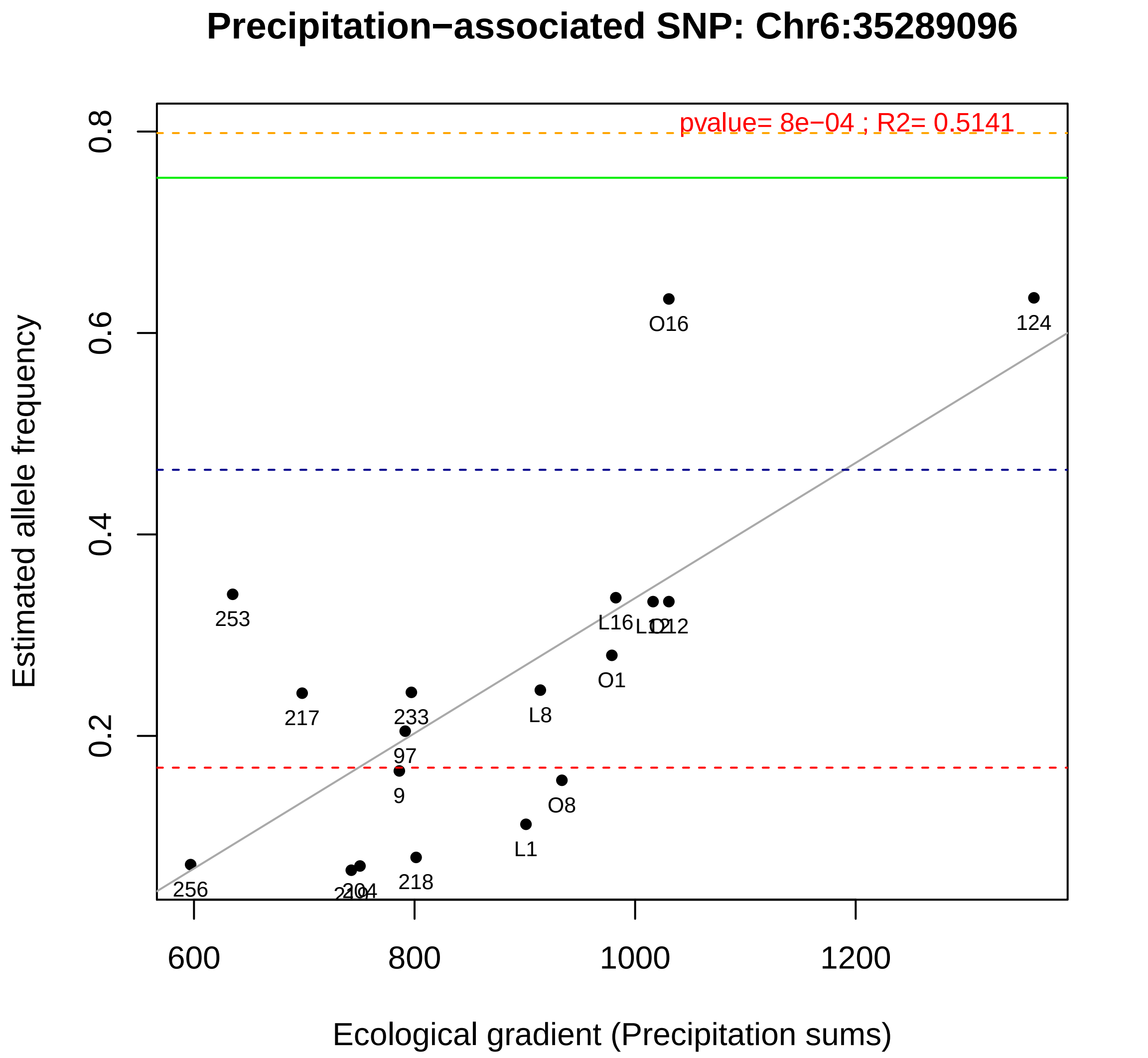
Precipitation−associated SNP: Chr6:35289096
pvalue= 8e−04 ; R2= 0.5141
Estimated allele frequency
0.2
0.4
0.6
0.8
600
800
1000
1200
Ecological gradient (Precipitation sums)
O16
124
253
L16
O12
O1
217
233
L8
97
9
O8
L1
256
204
218

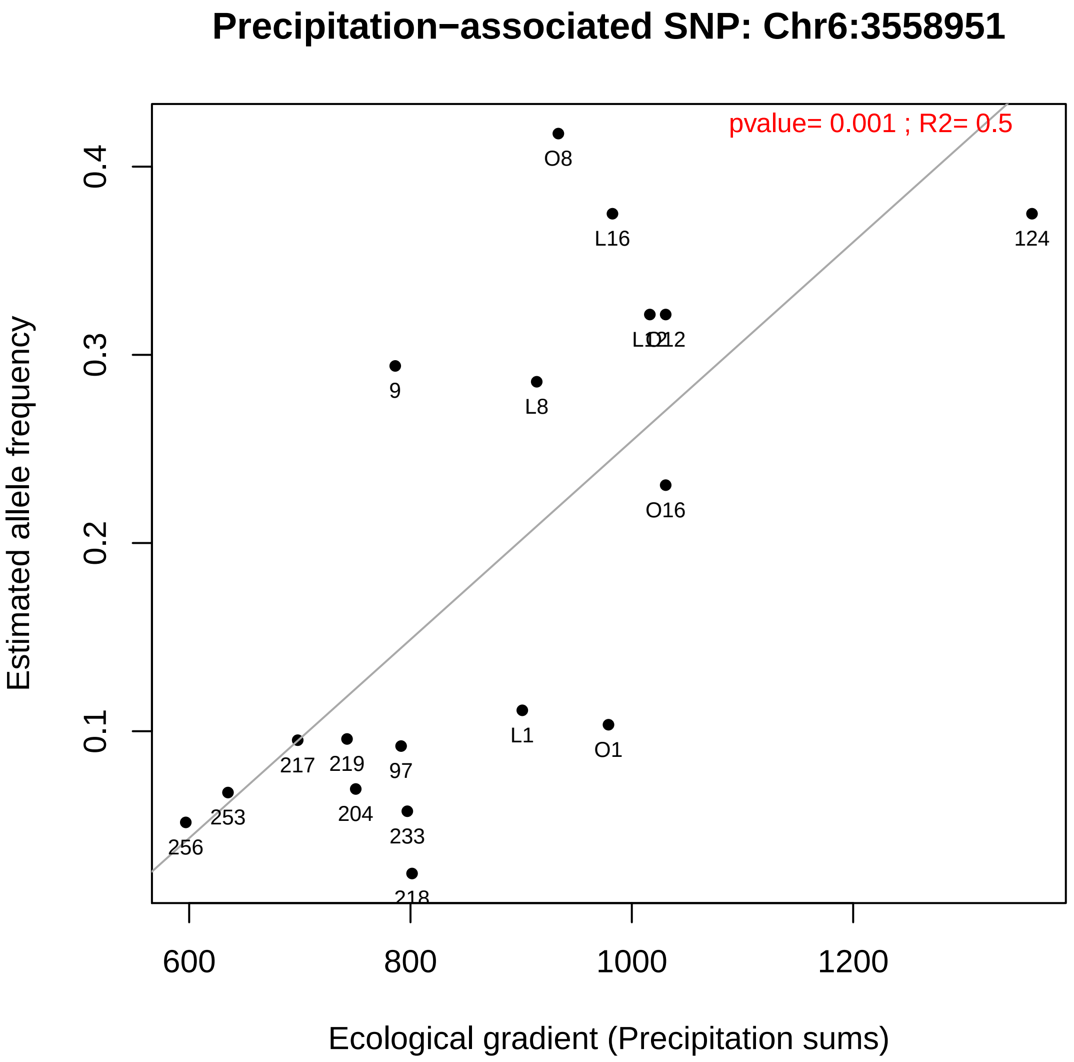
Precipitation−associated SNP: Chr6:3558951
pvalue= 0.001 ; R2= 0.5
Estimated allele frequency
0.1
0.2
0.3
0.4
600
800
1000
1200
Ecological gradient (Precipitation sums)
O8
L16
124
L12
O12
9
L8
O16
L1
O1
217
219
97
253
204
233
256
218

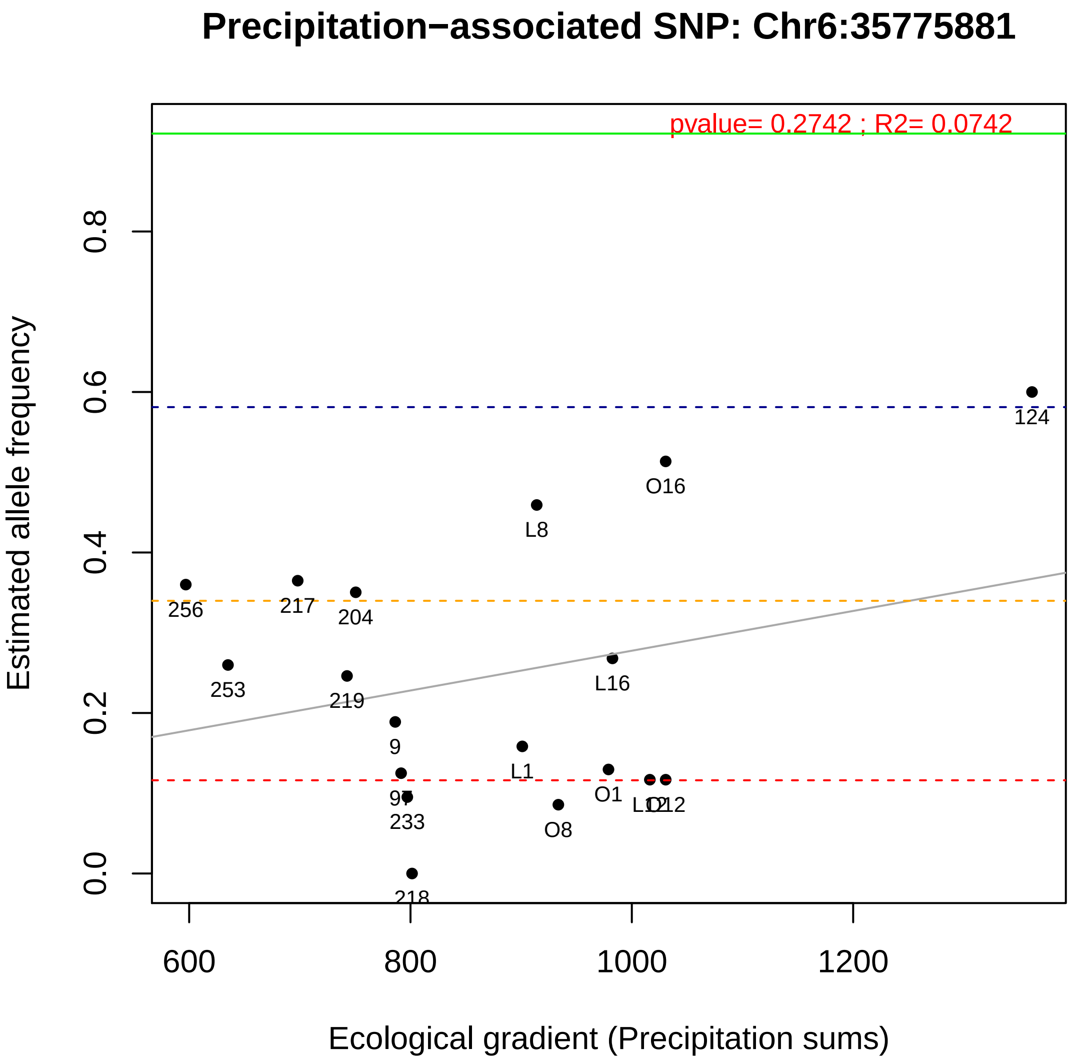

Precipitation−associated SNP: Chr6:35775881
pvalue= 0.2742 ; R2= 0.0742
Estimated allele frequency
Ecological gradient (Precipitation sums)
0.0
0.2
0.4
0.6
0.8
600
800
1000
1200
124
O16
L8
256
217
204
253
219
L16
9
L1
O1
233
O8
218

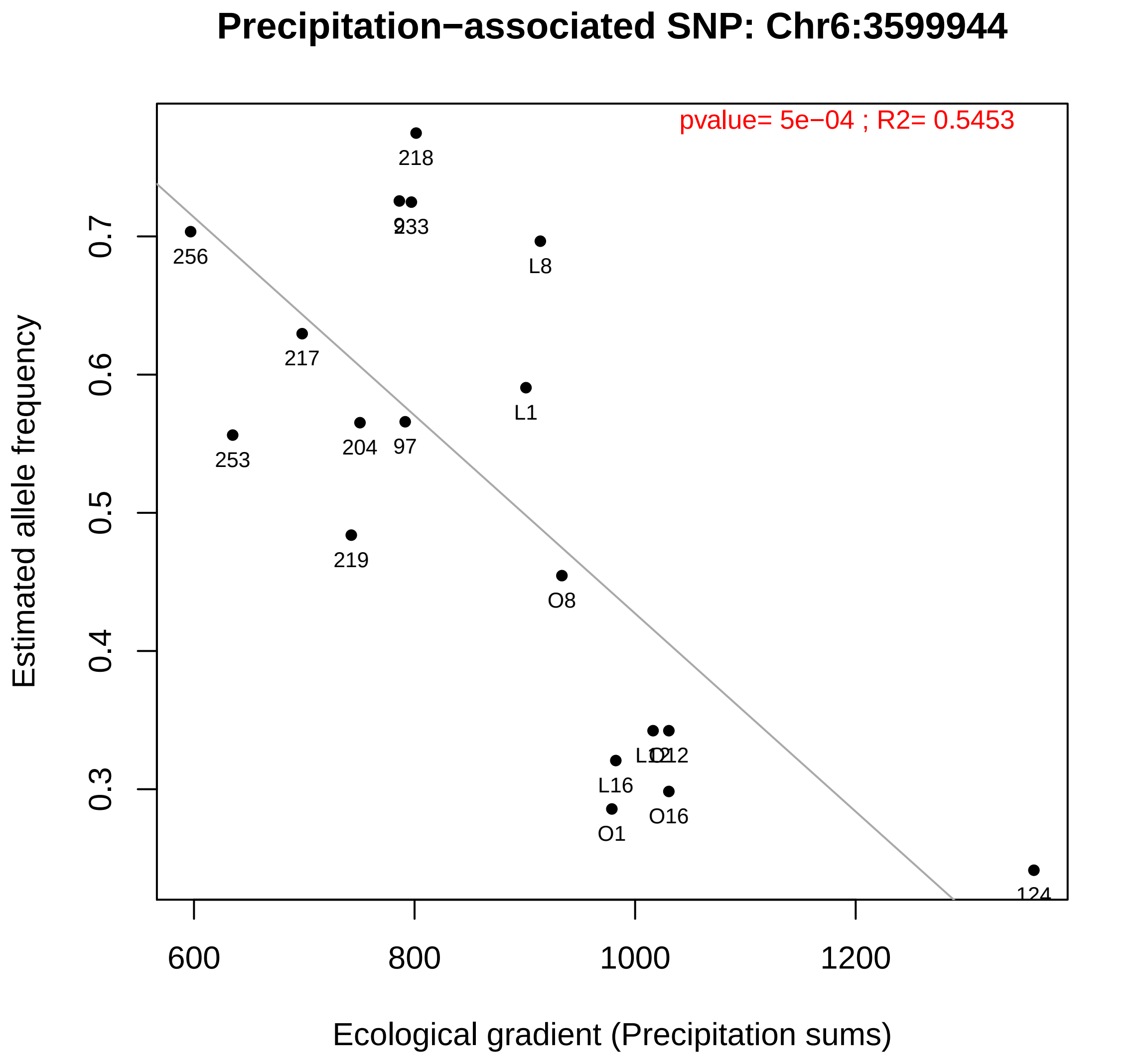

Precipitation−associated SNP: Chr6:3599944
pvalue= 5e−04 ; R2= 0.5453
218
233
256
L8
217
L1
253
204
97
219
O8
L12
O12
L16
O16
O1
124
0.3
0.4
0.5
0.6
0.7
600
800
1000
1200
Estimated allele frequency
Ecological gradient (Precipitation sums)

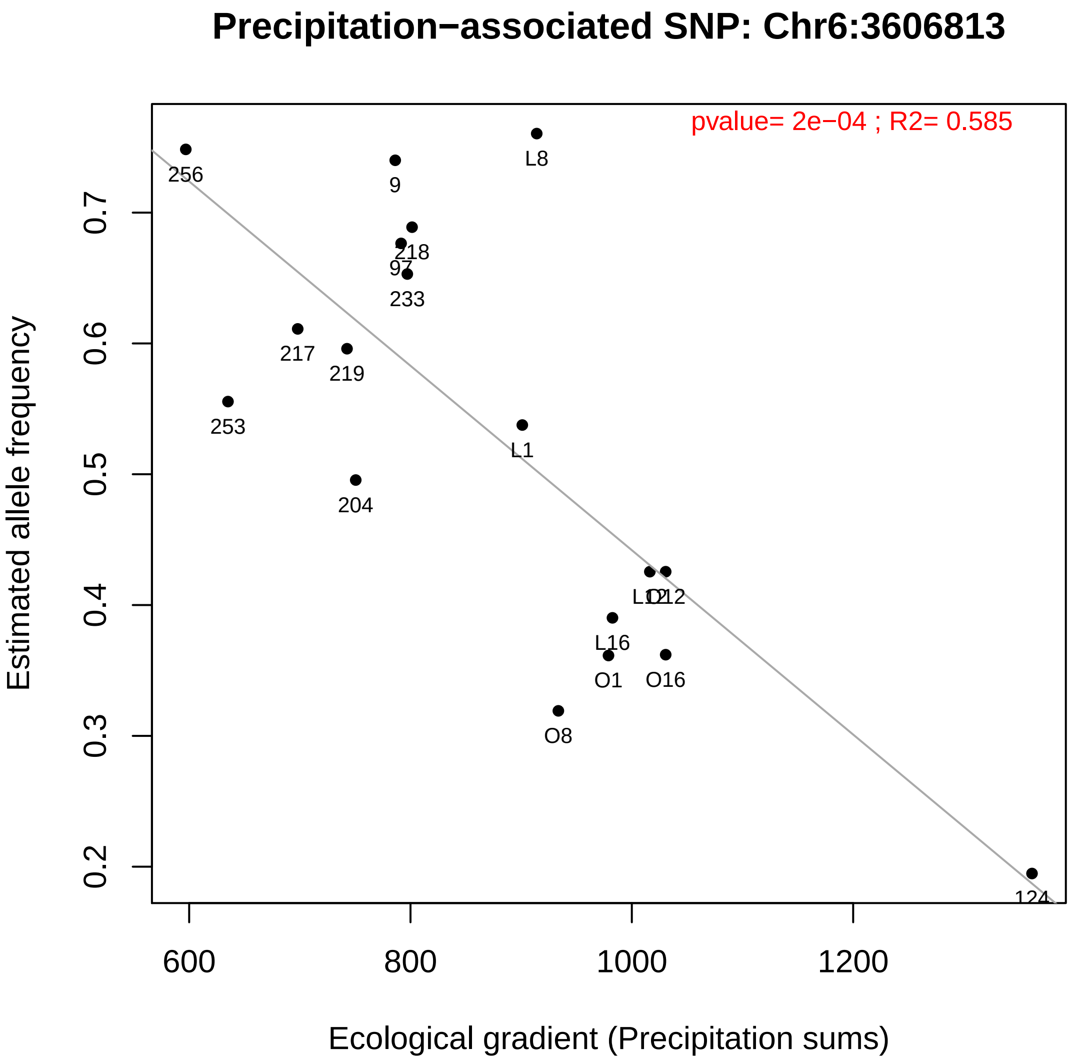
Precipitation−associated SNP: Chr6:3606813
pvalue= 2e−04 ; R2= 0.585
256
9
L8
218
97
233
217
219
253
L1
204
L12
O12
L16
O1
O16
O8
124
0.2
0.3
0.4
0.5
0.6
0.7
600
800
1000
1200
Estimated allele frequency
Ecological gradient (Precipitation sums)

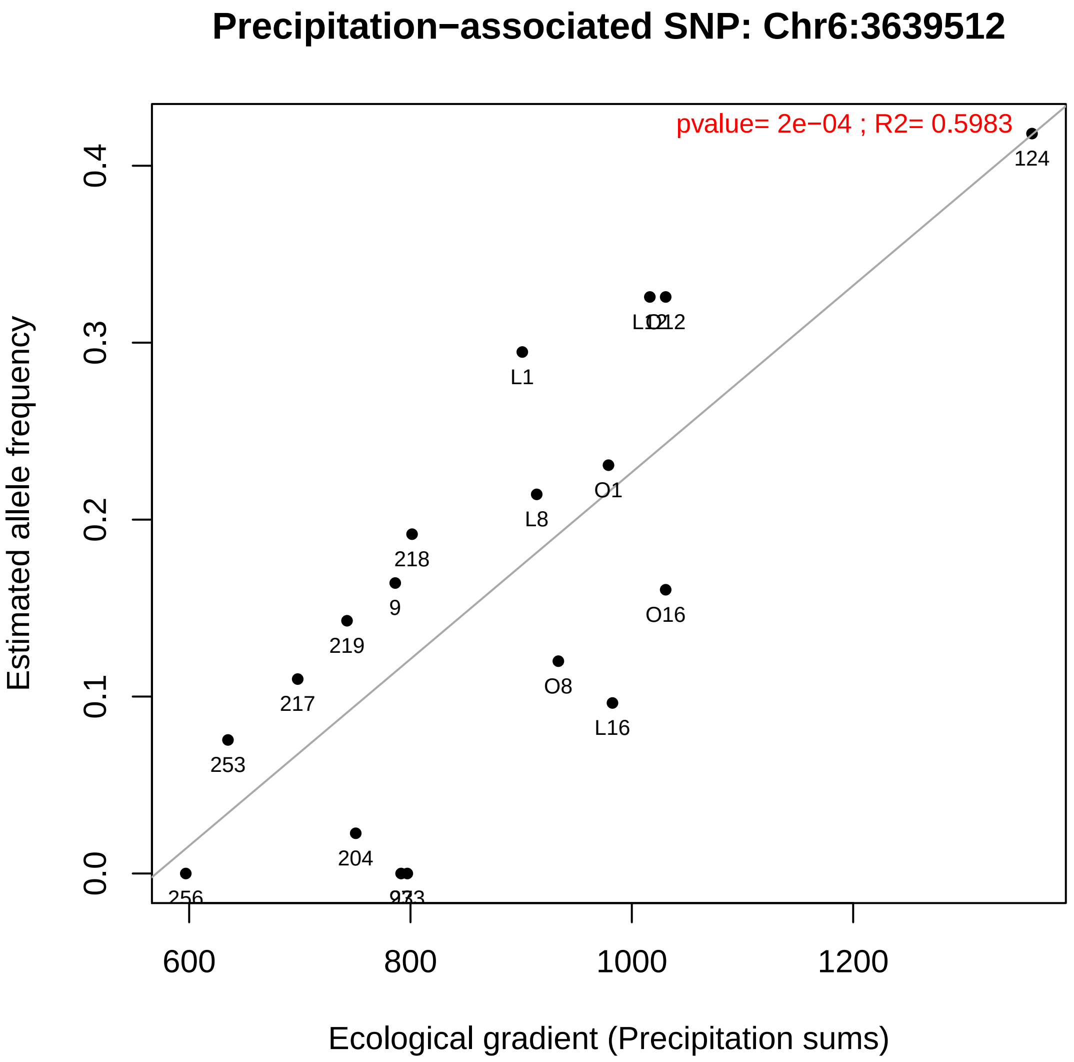
Precipitation−associated SNP: Chr6:3639512
pvalue= 2e−04 ; R2= 0.5983
Estimated allele frequency
0.0
0.1
0.2
0.3
0.4
Ecological gradient (Precipitation sums)
600
800
1000
1200
124
L1
O1
L8
218
9
O16
219
O8
217
L16
253
204
256

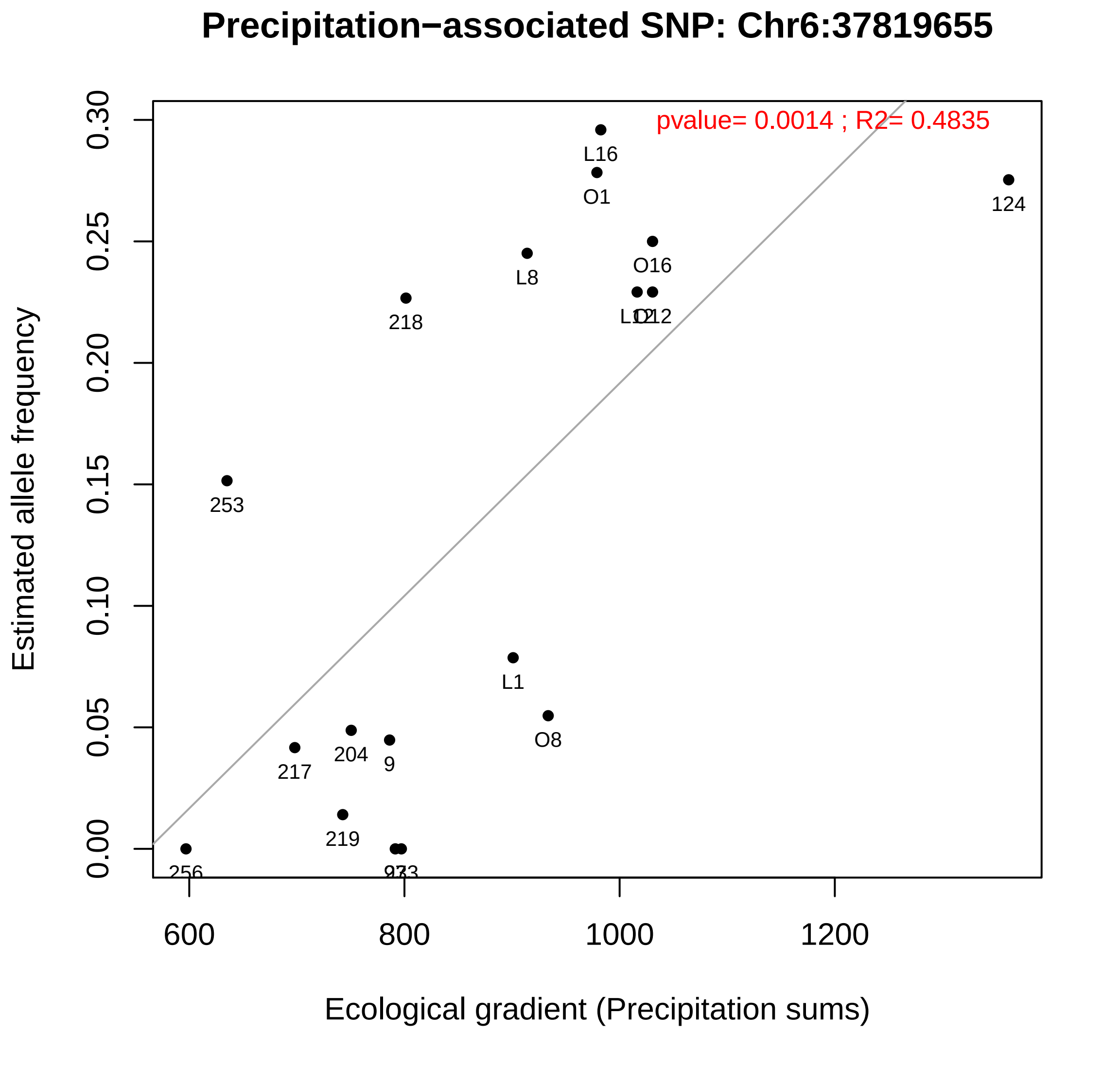
Precipitation−associated SNP: Chr6:37819655
pvalue= 0.0014 ; R2= 0.4835
Estimated allele frequency
Ecological gradient (Precipitation sums)
0.00
0.05
0.10
0.15
0.20
0.25
0.30
600
800
1000
1200
L16
O1
124
O16
L8
218
253
L1
O8
204
217
9
219
256

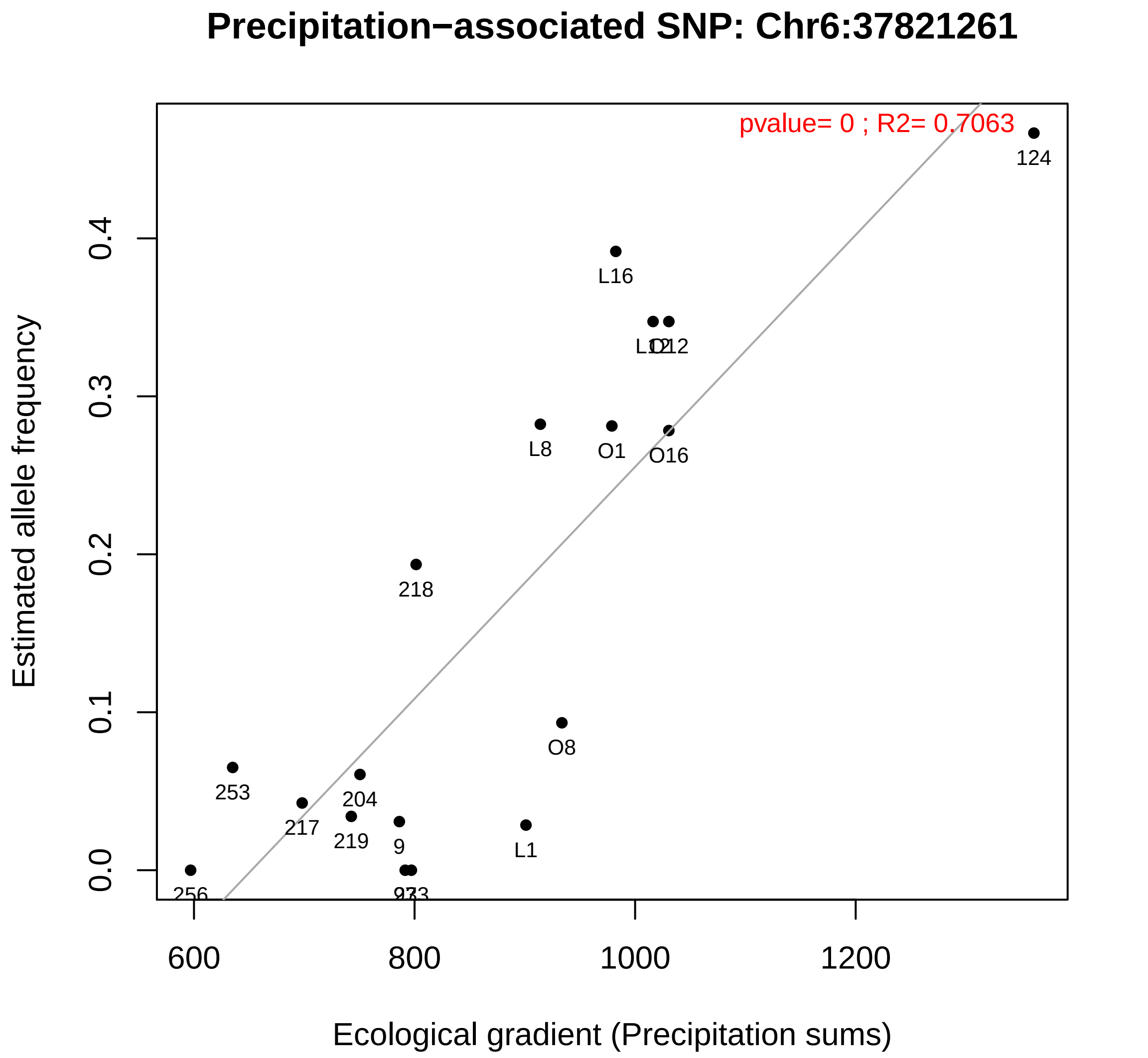

Precipitation−associated SNP: Chr6:37821261
pvalue= 0 ; R2= 0.7063
124
L16
L12
O12
L8
O1
O16
218
O8
253
204
217
219
9
L1
256
97
73
0.0
0.1
0.2
0.3
0.4
Estimated allele frequency
600
800
1000
1200
Ecological gradient (Precipitation sums)

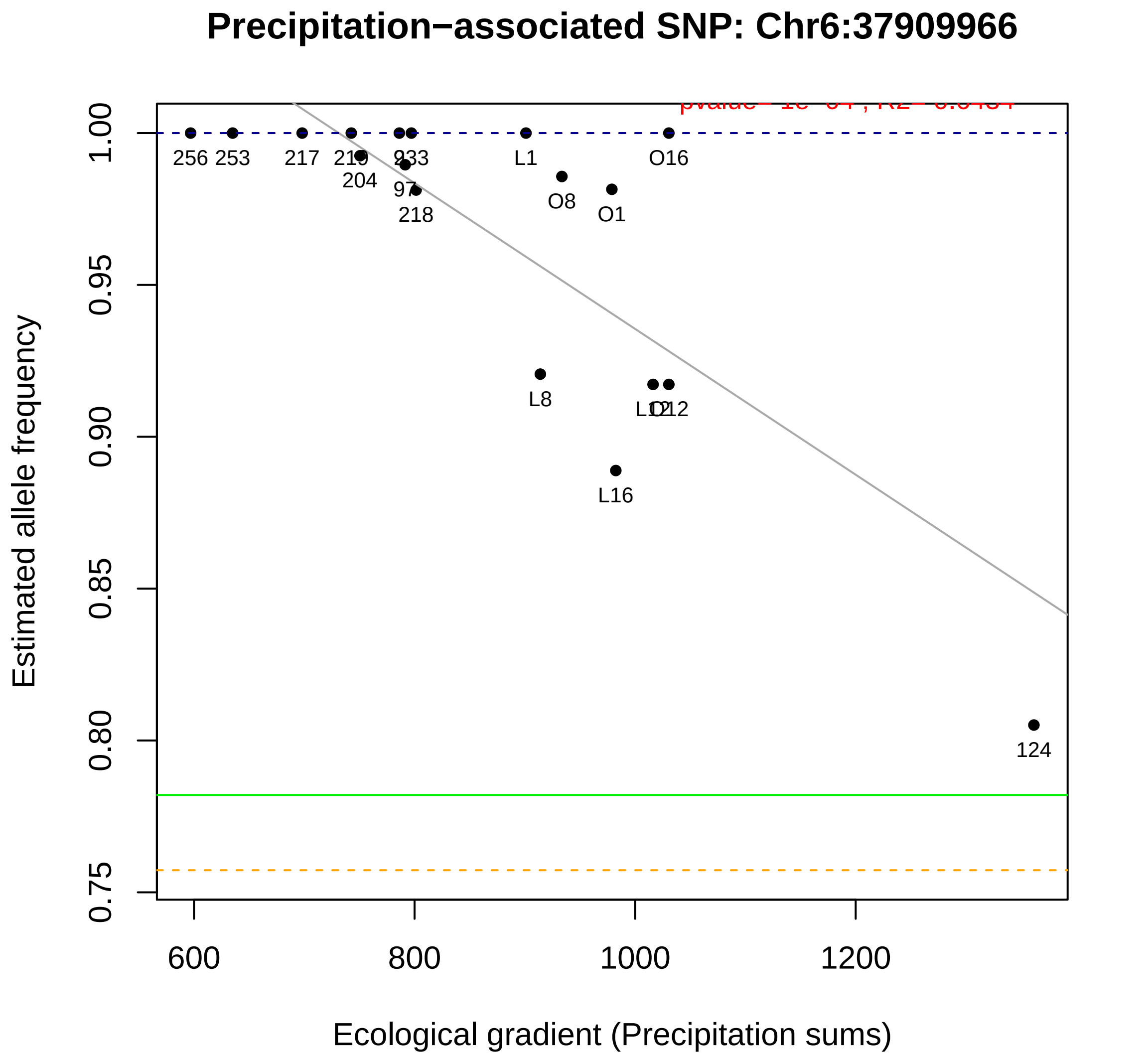

Precipitation−associated SNP: Chr6:37909966
Estimated allele frequency
Ecological gradient (Precipitation sums)
1.00
0.95
0.90
0.85
0.80
0.75
600
800
1000
1200
256
253
217
204
218
L1
O8
O1
O16
L8
L16
124

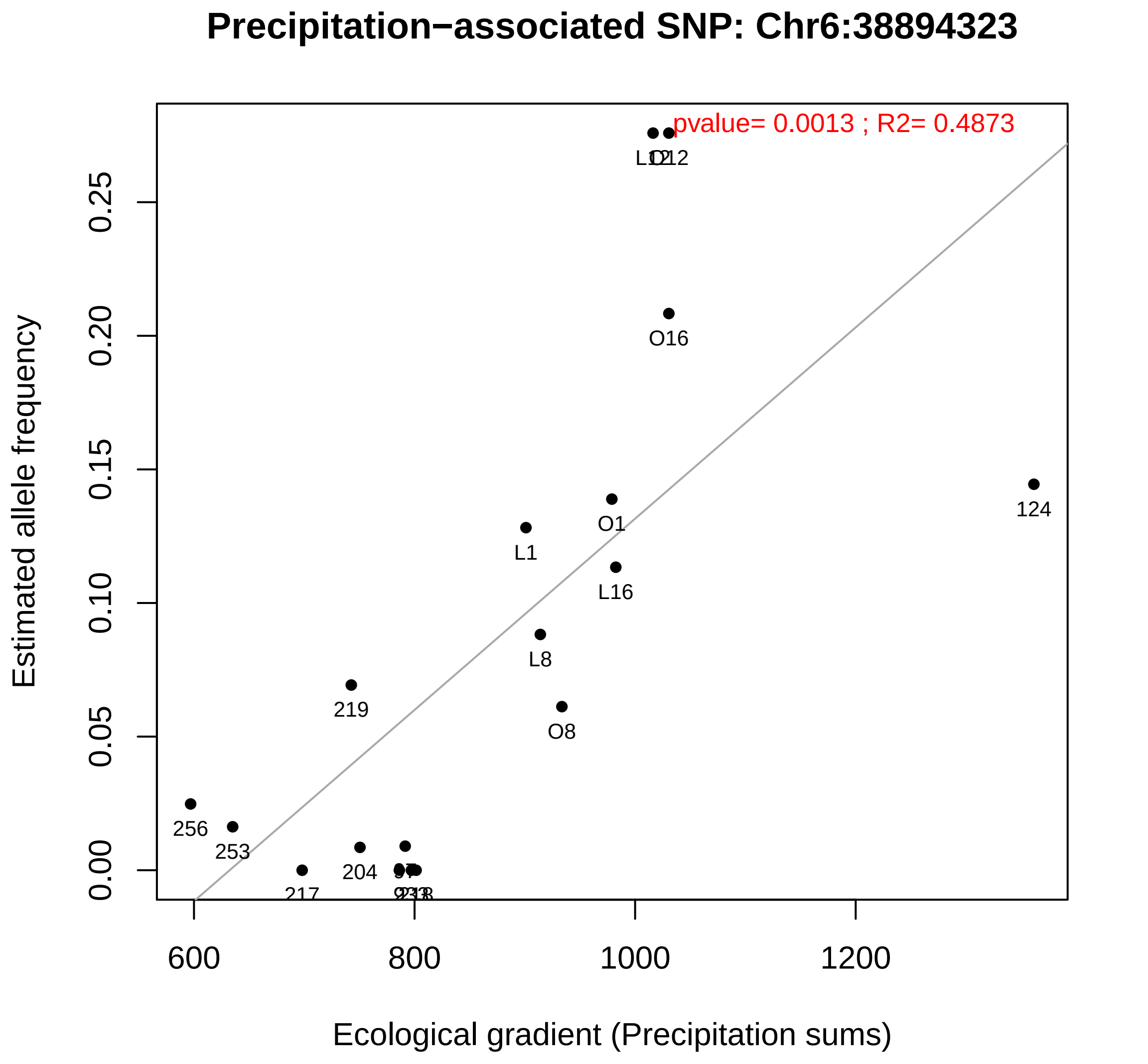

Precipitation−associated SNP: Chr6:38894323
pvalue= 0.0013 ; R2= 0.4873
Estimated allele frequency
Ecological gradient (Precipitation sums)
0.00
0.05
0.10
0.15
0.20
0.25
600
800
1000
1200
O16
124
O1
L1
L16
L8
219
O8
256
253
204
217

**Precipitation−associated SNP: Chr6:38926820**

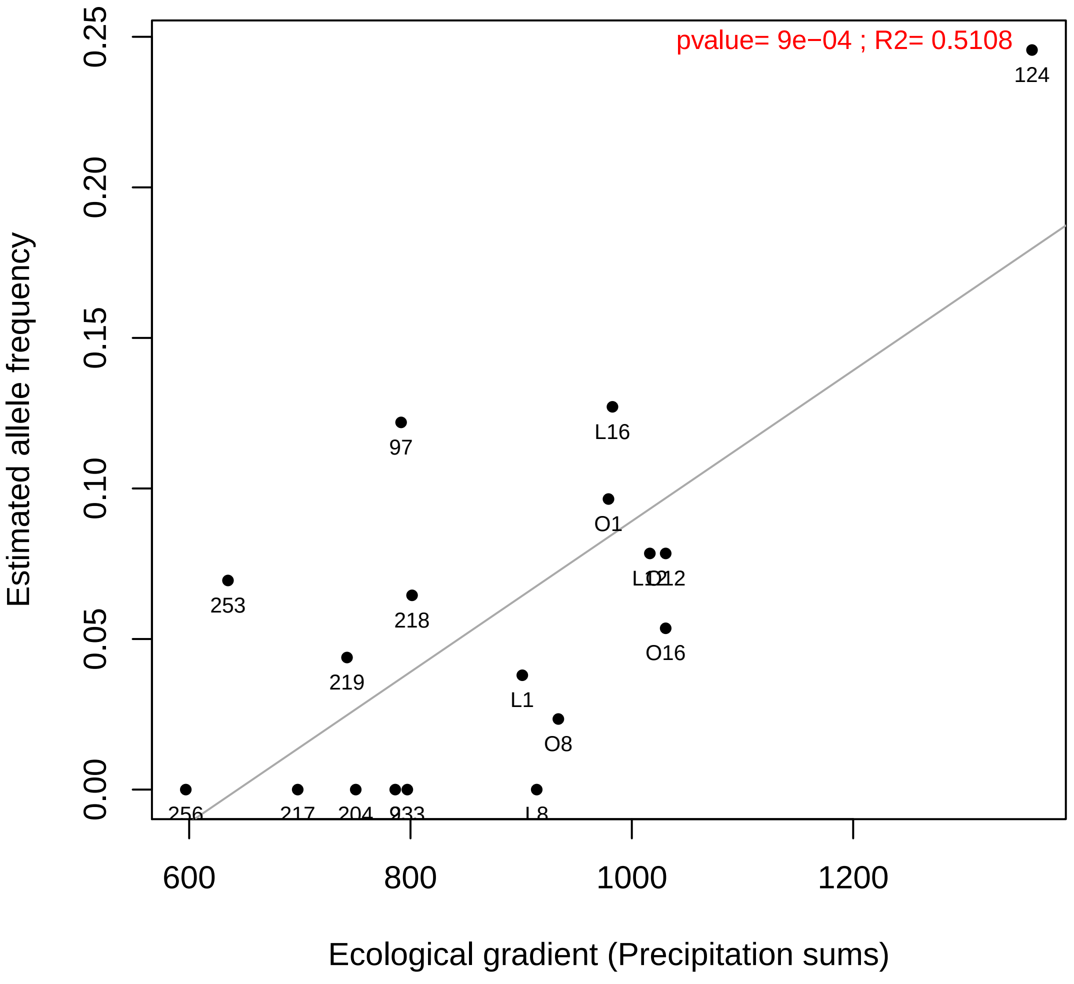

**Precipitation−associated SNP: Chr6:40245701**

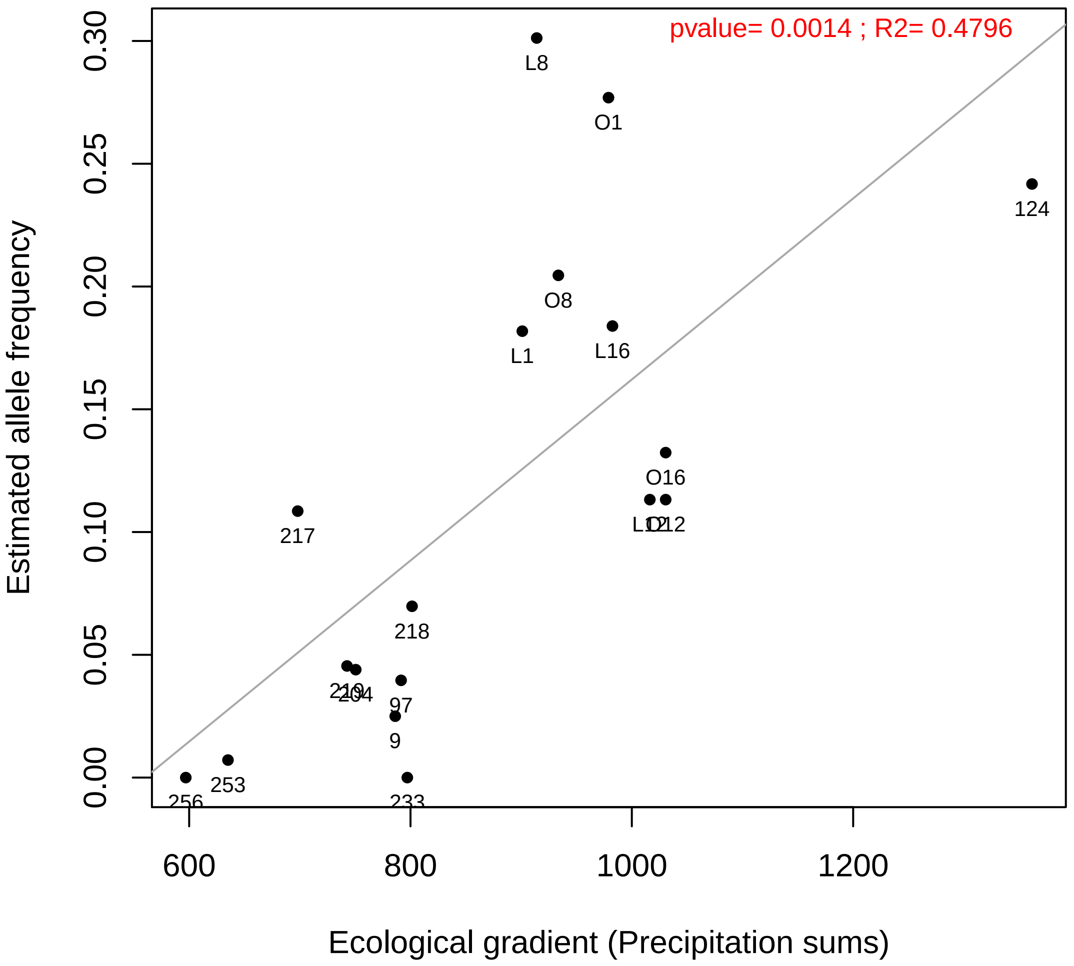

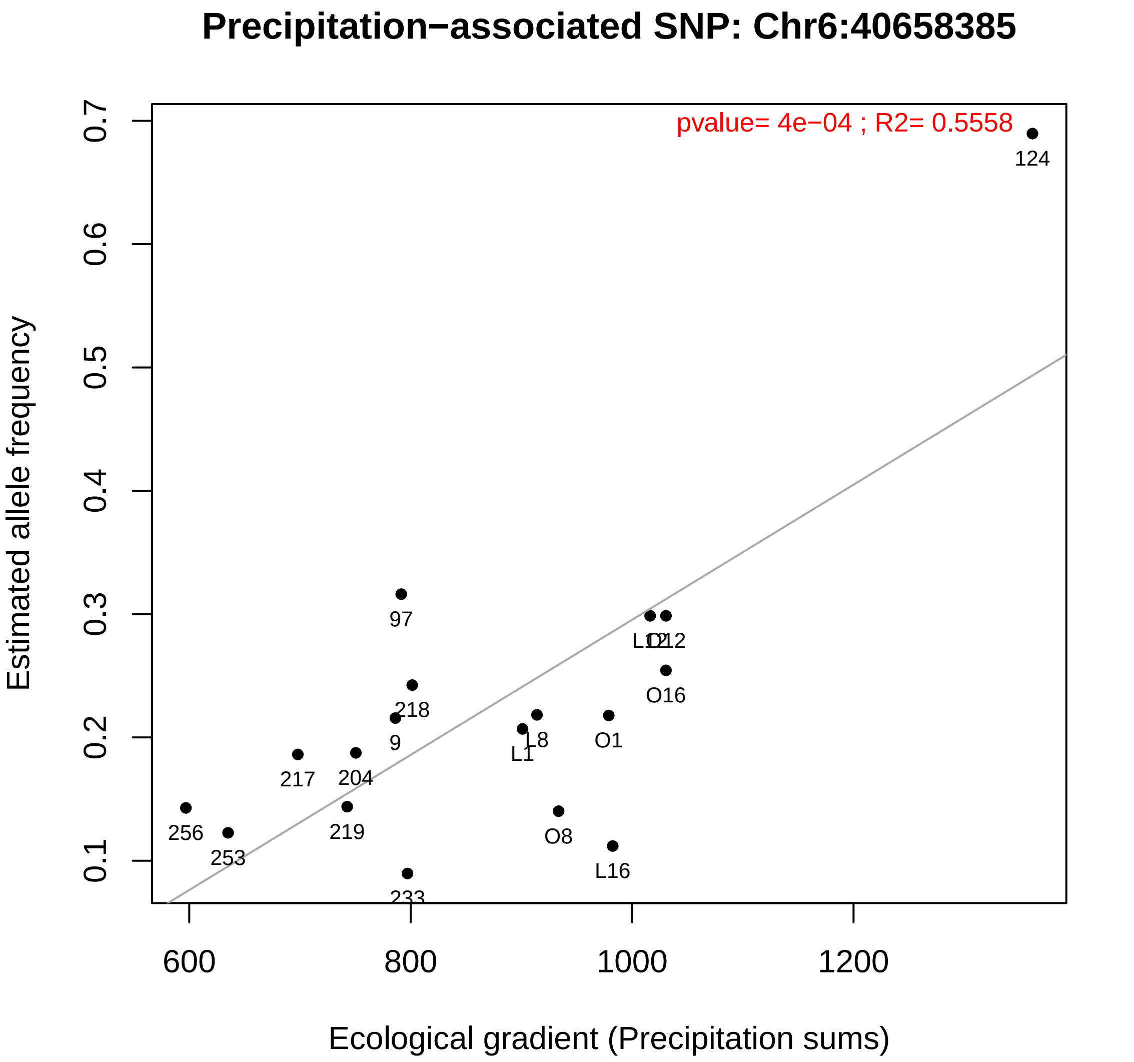
Precipitation−associated SNP: Chr6:40658385
pvalue= 4e−04 ; R2= 0.5558
Estimated allele frequency
Ecological gradient (Precipitation sums)
0.1
0.2
0.3
0.4
0.5
0.6
0.7
600
800
1000
1200
124
97
O12
O16
218
9
L8
L1
O1
217
204
256
219
253
O8
L16
233

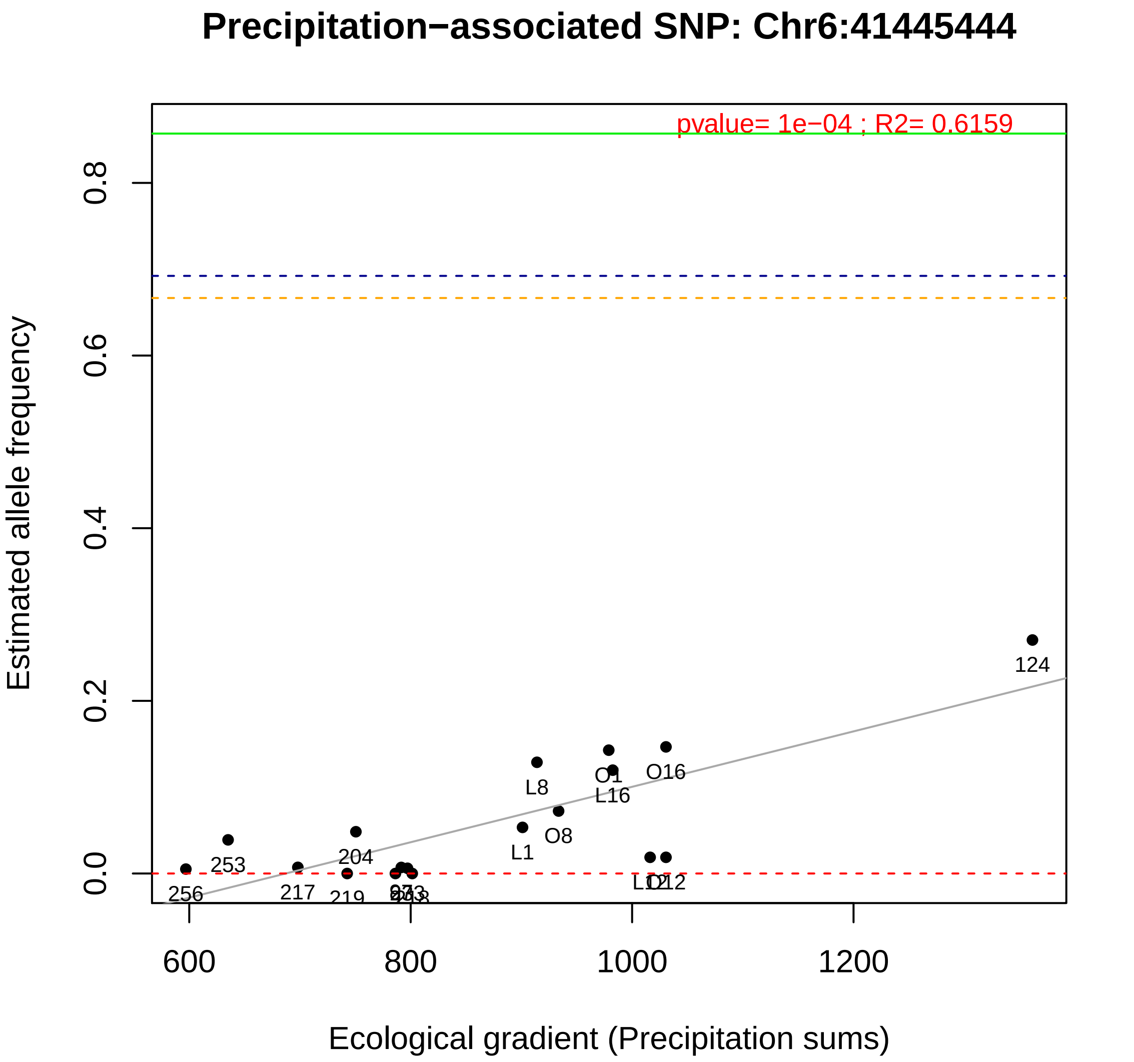
Precipitation−associated SNP: Chr6:41445444
pvalue= 1e−04 ; R2= 0.6159
Estimated allele frequency
0.0
0.2
0.4
0.6
0.8
600
800
1000
1200
Ecological gradient (Precipitation sums)
256
253
217
219
204
L1
L8
O8
O1
L16
O16
124

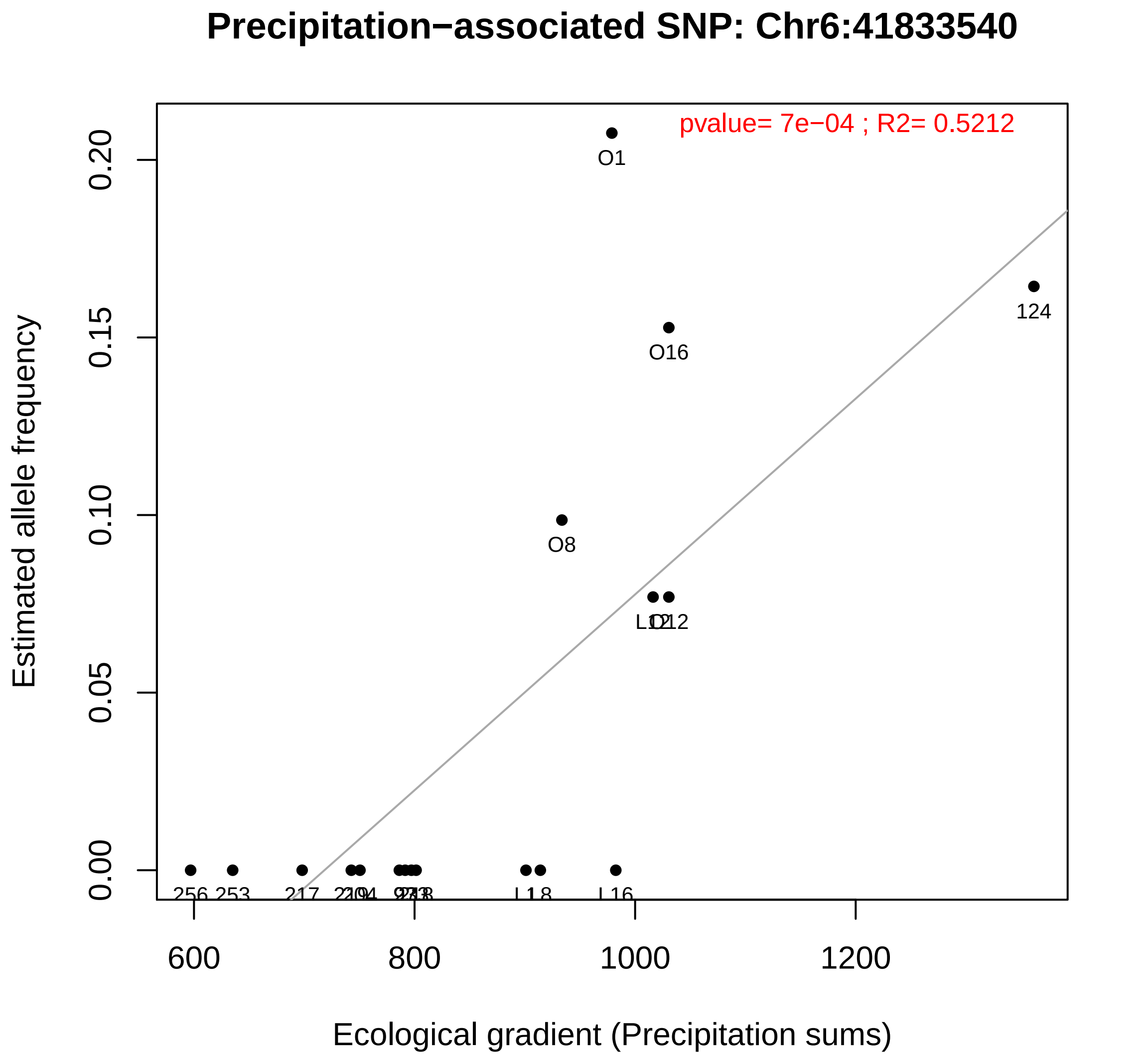

Precipitation−associated SNP: Chr6:41833540
pvalue= 7e−04 ; R2= 0.5212
O1
124
O16
O8
0.20
0.15
0.10
0.05
0.00
600
800
1000
1200
Estimated allele frequency
Ecological gradient (Precipitation sums)

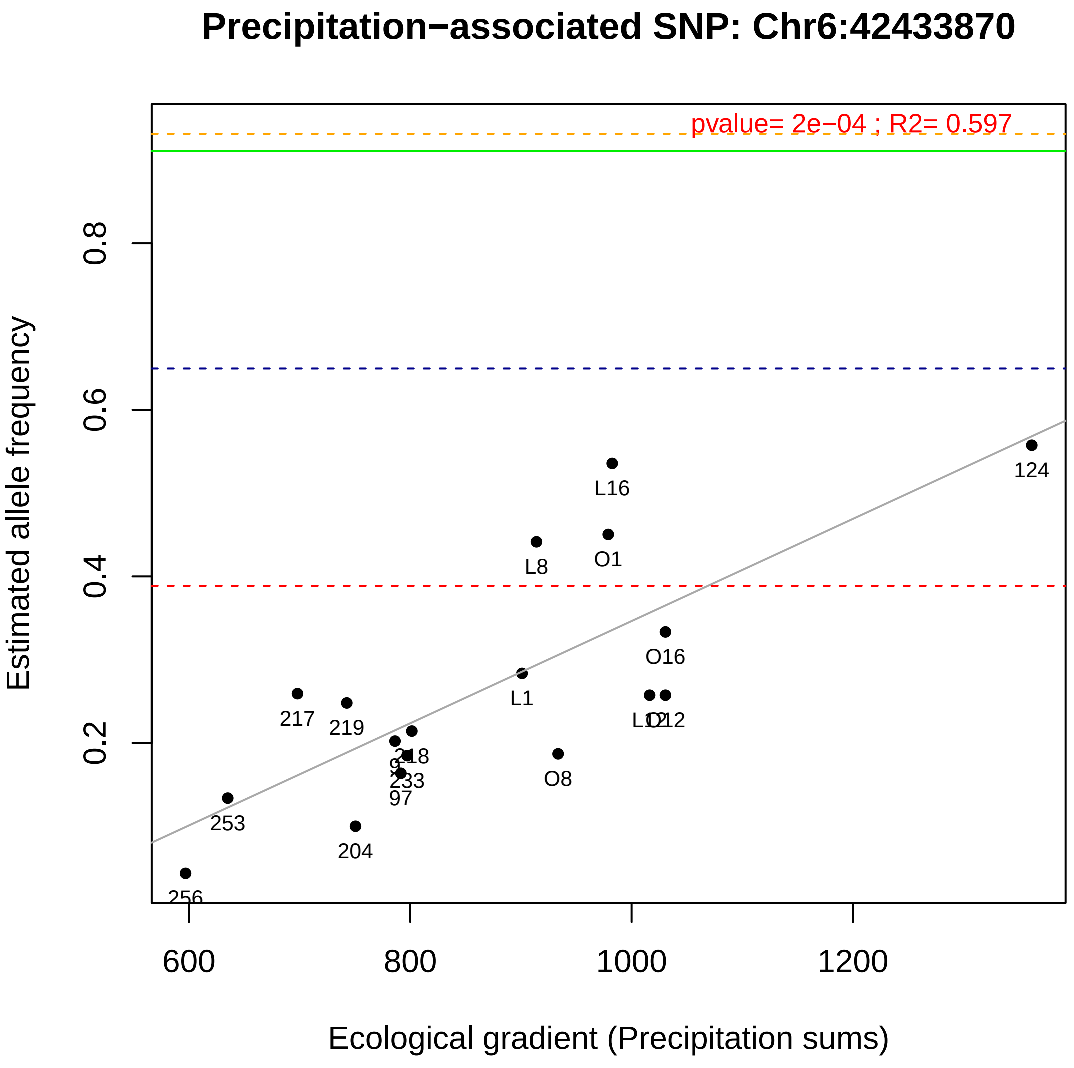

Precipitation−associated SNP: Chr6:42433870
pvalue= 2e−04 ; R2= 0.597
Estimated allele frequency
0.2
0.4
0.6
0.8
600
800
1000
1200
Ecological gradient (Precipitation sums)
124
L16
L8
O1
O16
L1
217
219
218
233
97
O8
253
204
256

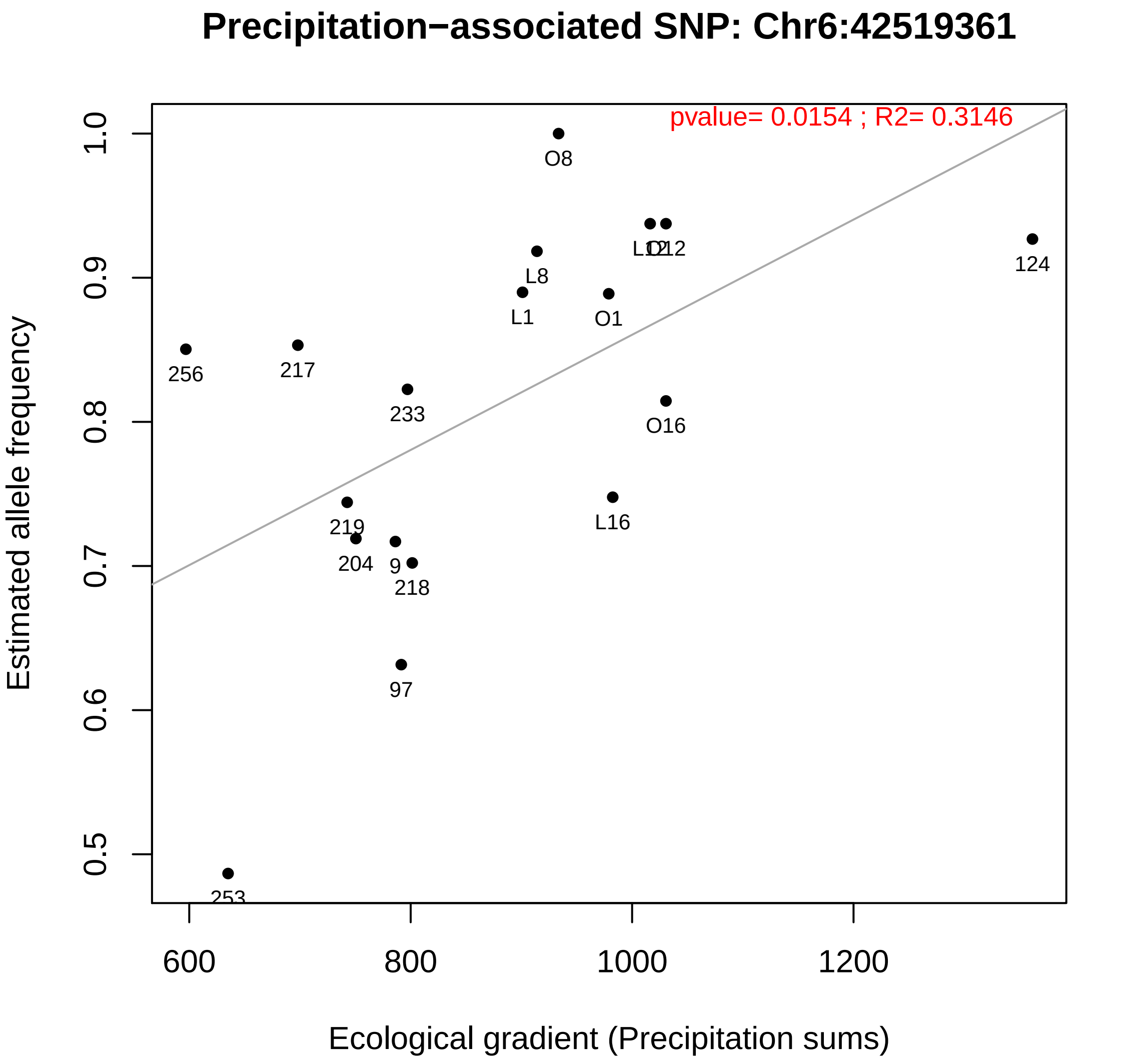

Precipitation−associated SNP: Chr6:42519361
pvalue= 0.0154 ; R2= 0.3146
Estimated allele frequency
1.0
0.9
0.8
0.7
0.6
0.5
600
800
1000
1200
Ecological gradient (Precipitation sums)
O8
L10
O12
124
L8
L1
O1
256
217
233
O16
219
204
9
218
L16
97
253

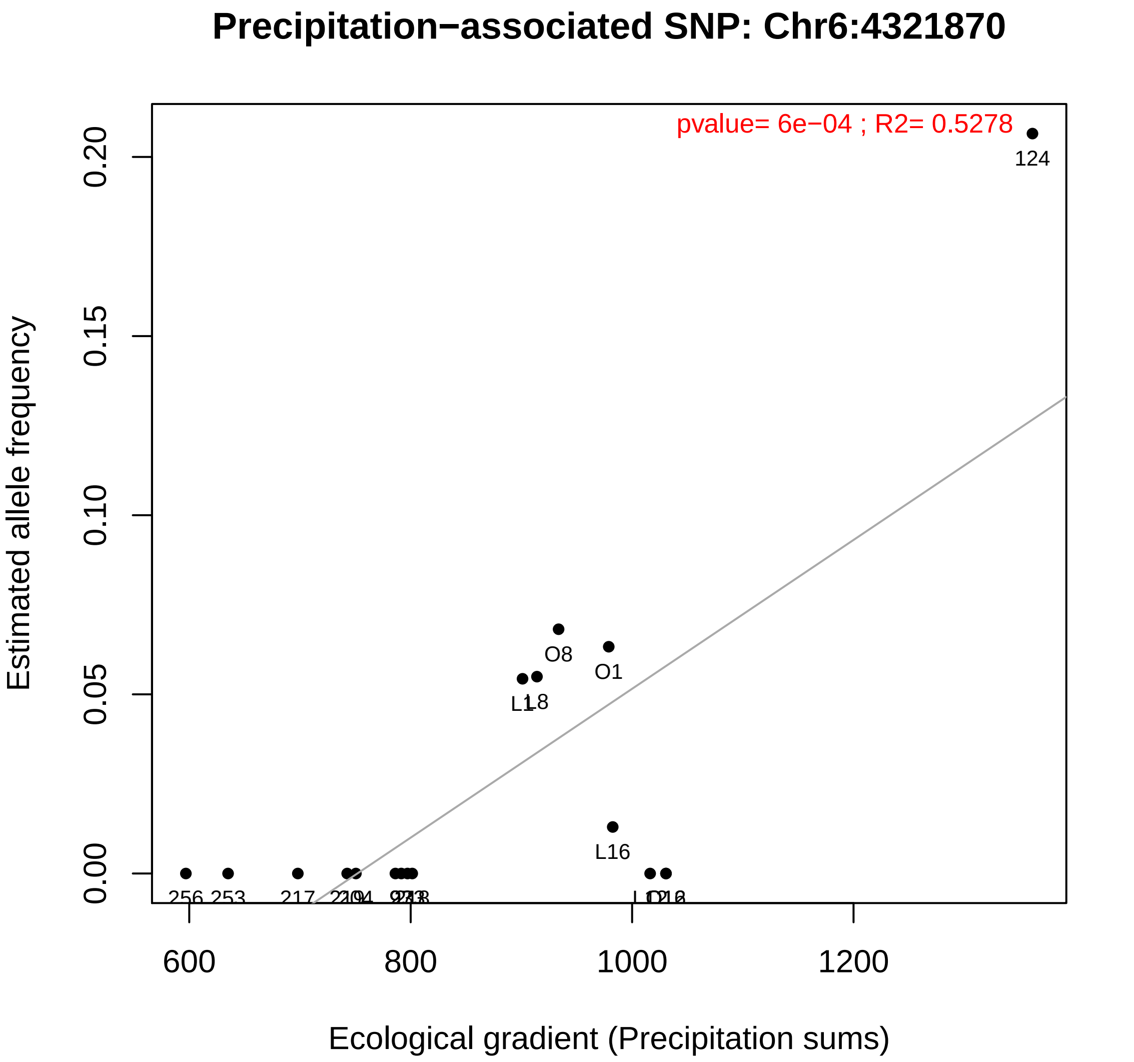
Precipitation−associated SNP: Chr6:4321870
pvalue= 6e−04 ; R2= 0.5278
124
O8
O1
L1
L8
L16
256
253
217
0.00
0.05
0.10
0.15
0.20
600
800
1000
1200
Estimated allele frequency
Ecological gradient (Precipitation sums)

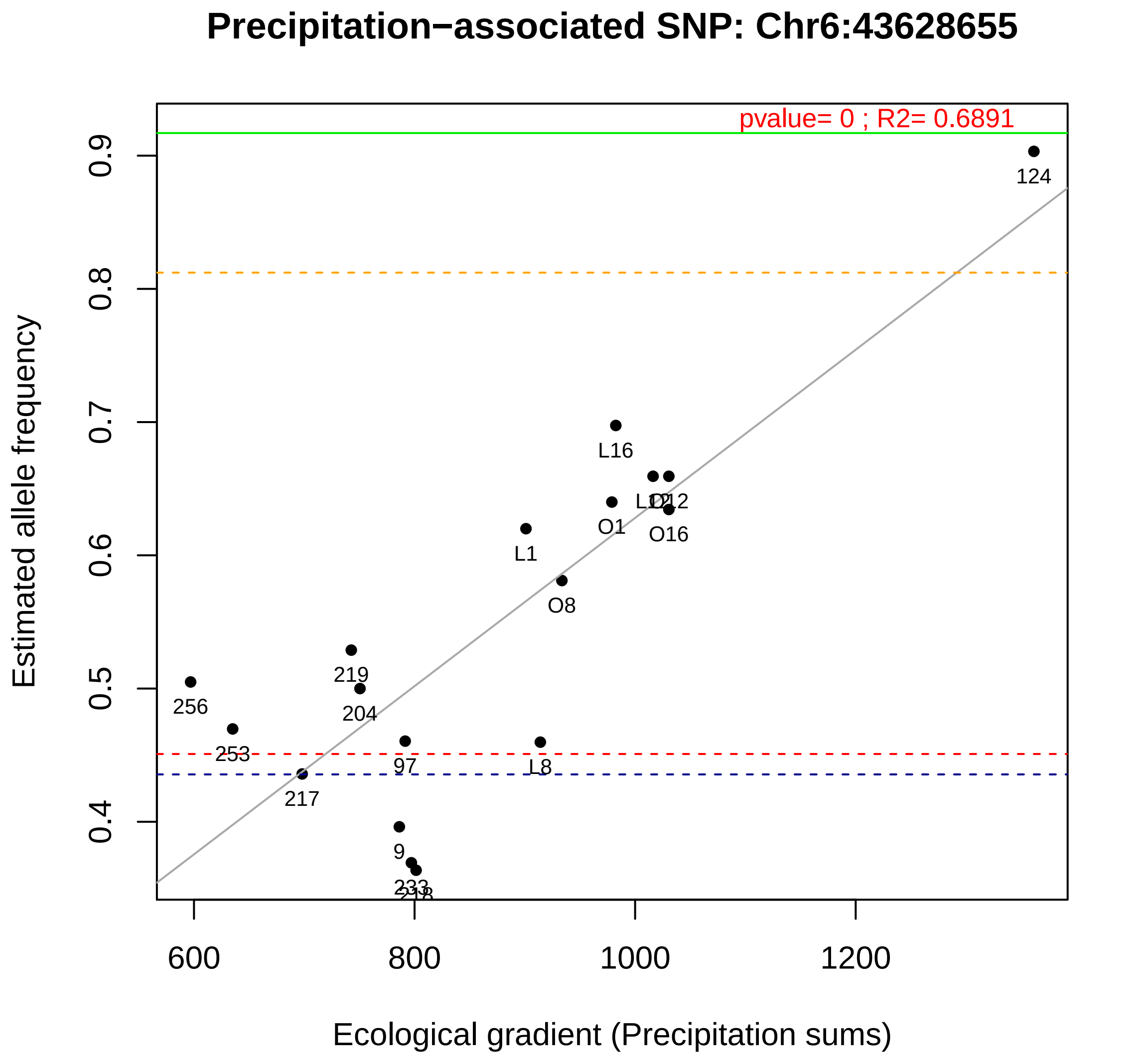

Precipitation−associated SNP: Chr6:43628655
pvalue= 0 ; R2= 0.6891
Estimated allele frequency
Ecological gradient (Precipitation sums)
0.4
0.5
0.6
0.7
0.8
0.9
600
800
1000
1200
124
L16
O1
O16
L1
O8
219
256
204
253
97
L8
217
9

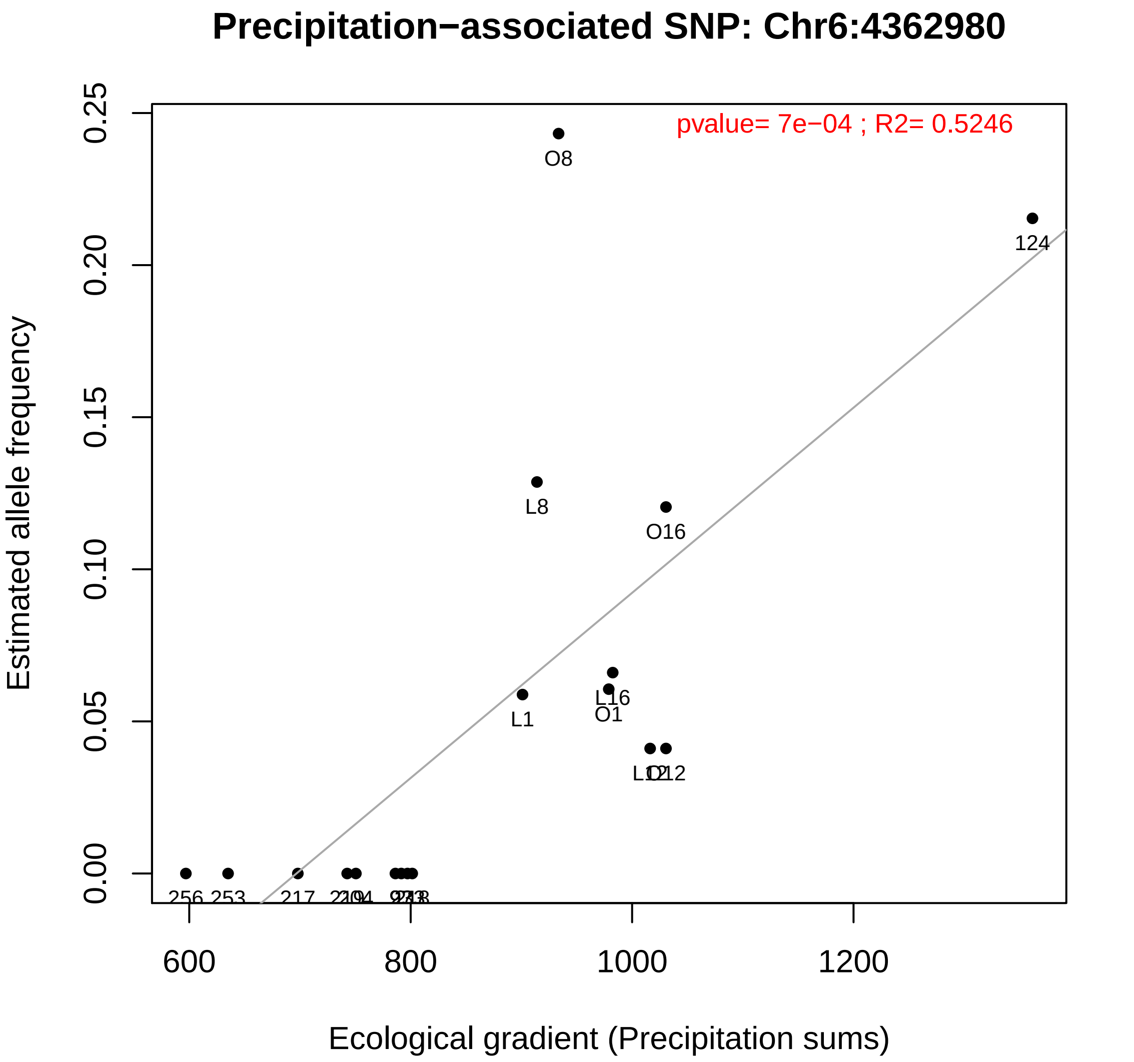
Precipitation−associated SNP: Chr6:4362980
pvalue= 7e−04 ; R2= 0.5246
Estimated allele frequency
Ecological gradient (Precipitation sums)
0.00
0.05
0.10
0.15
0.20
0.25
600
800
1000
1200
O8
124
L8
O16
L1
L16
O1
O12
256
253
217

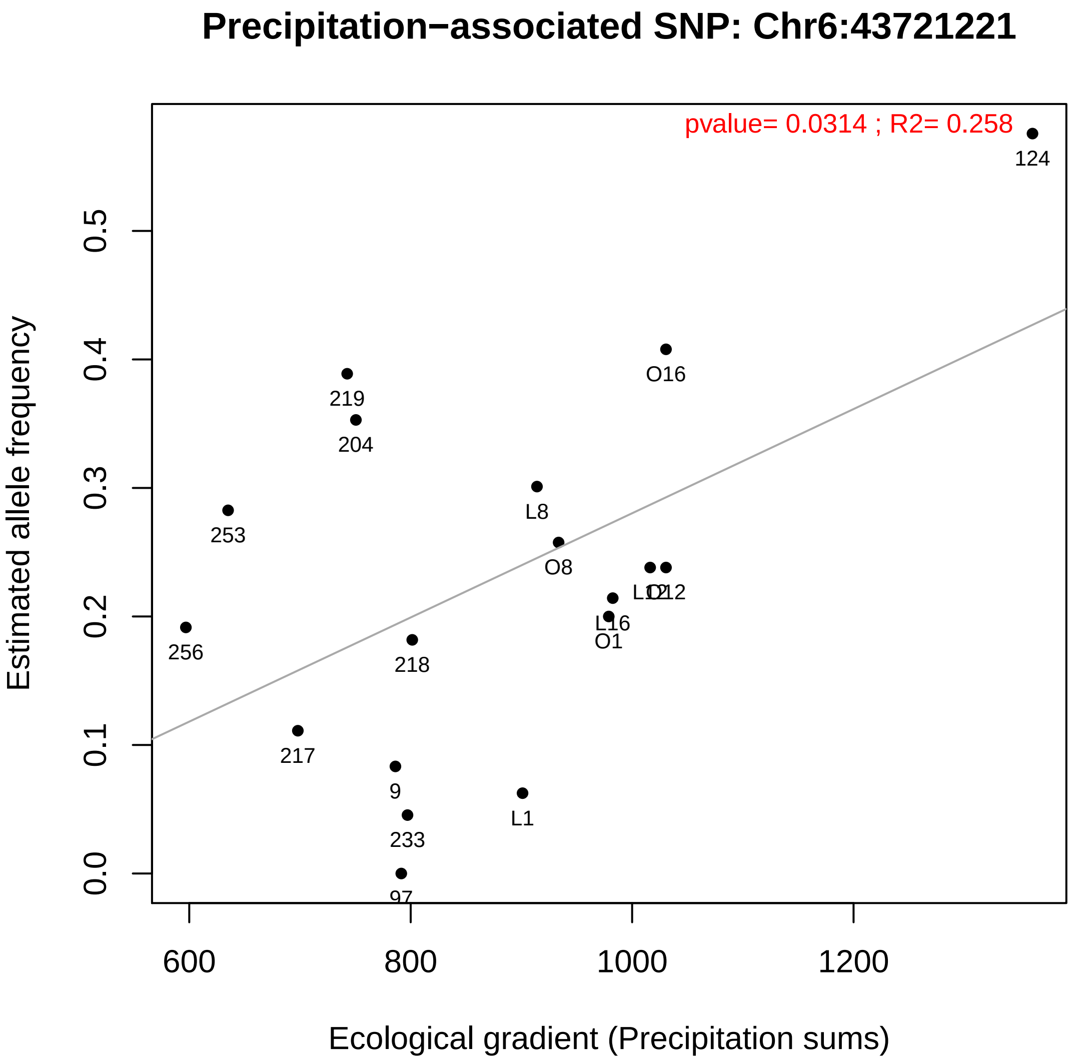

Precipitation−associated SNP: Chr6:43721221
pvalue= 0.0314 ; R2= 0.258
Estimated allele frequency
Ecological gradient (Precipitation sums)
0.0
0.1
0.2
0.3
0.4
0.5
600
800
1000
1200
124
O16
219
204
L8
253
O8
L12
O12
L16
O1
256
218
217
9
L1
233
97

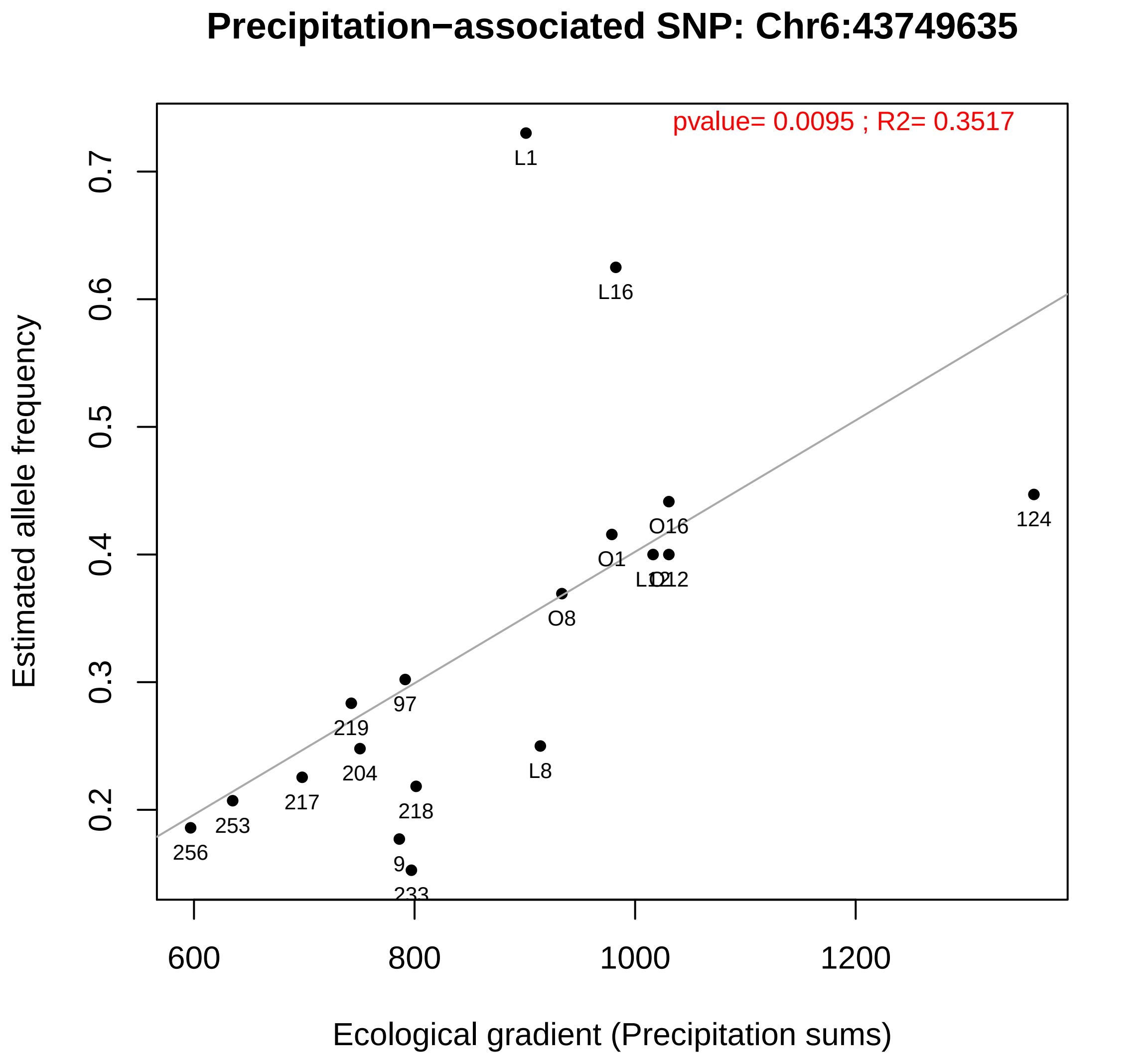
Precipitation−associated SNP: Chr6:43749635
pvalue= 0.0095 ; R2= 0.3517
Estimated allele frequency
Ecological gradient (Precipitation sums)
0.2
0.3
0.4
0.5
0.6
0.7
600
800
1000
1200
L1
L16
O16
124
O1
L12
O12
O8
97
219
204
L8
217
218
253
256
9
233

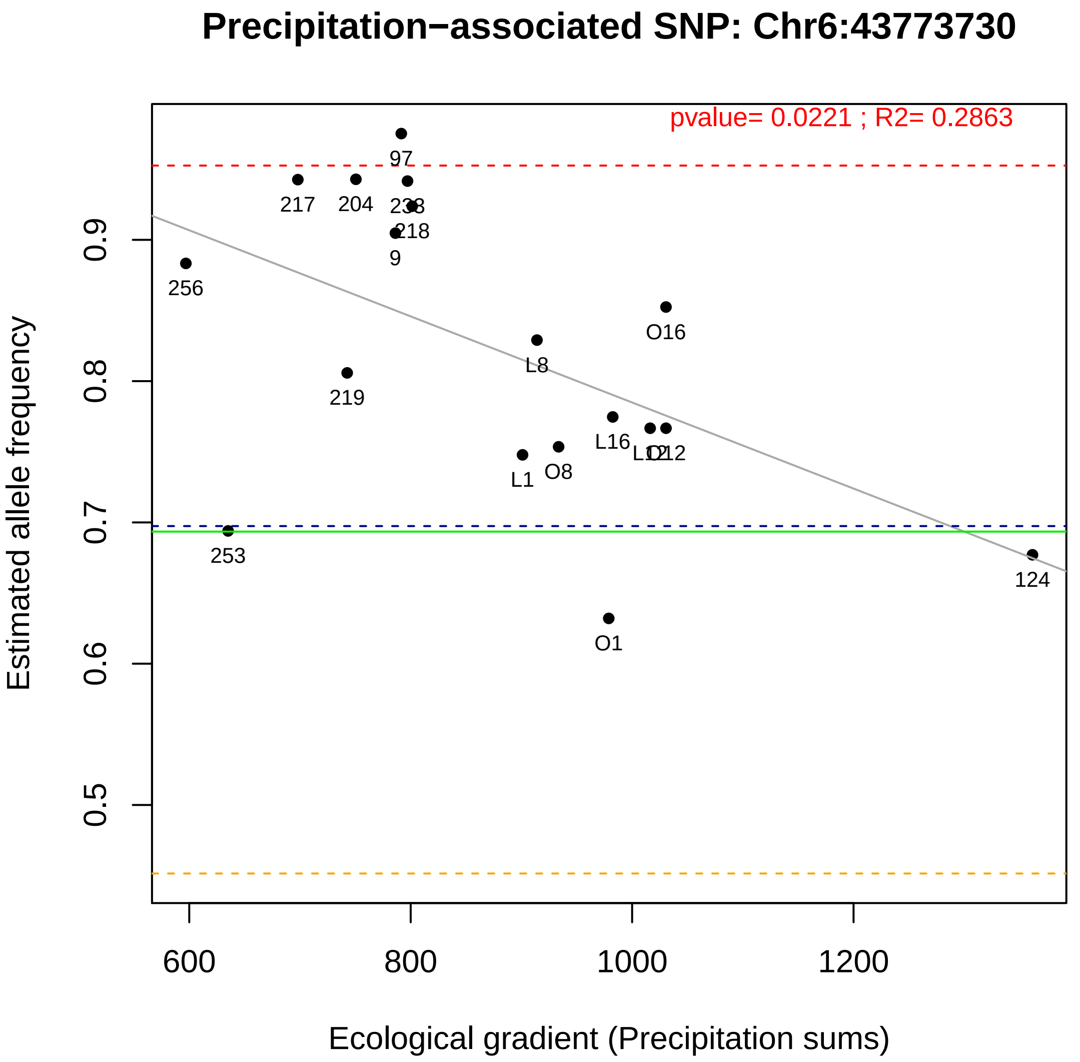
Precipitation−associated SNP: Chr6:43773730
pvalue= 0.0221 ; R2= 0.2863
Estimated allele frequency
0.5
0.6
0.7
0.8
0.9
600
800
1000
1200
Ecological gradient (Precipitation sums)
97
217
204
218
9
256
O16
L8
219
L16
O12
O8
L1
253
124
O1

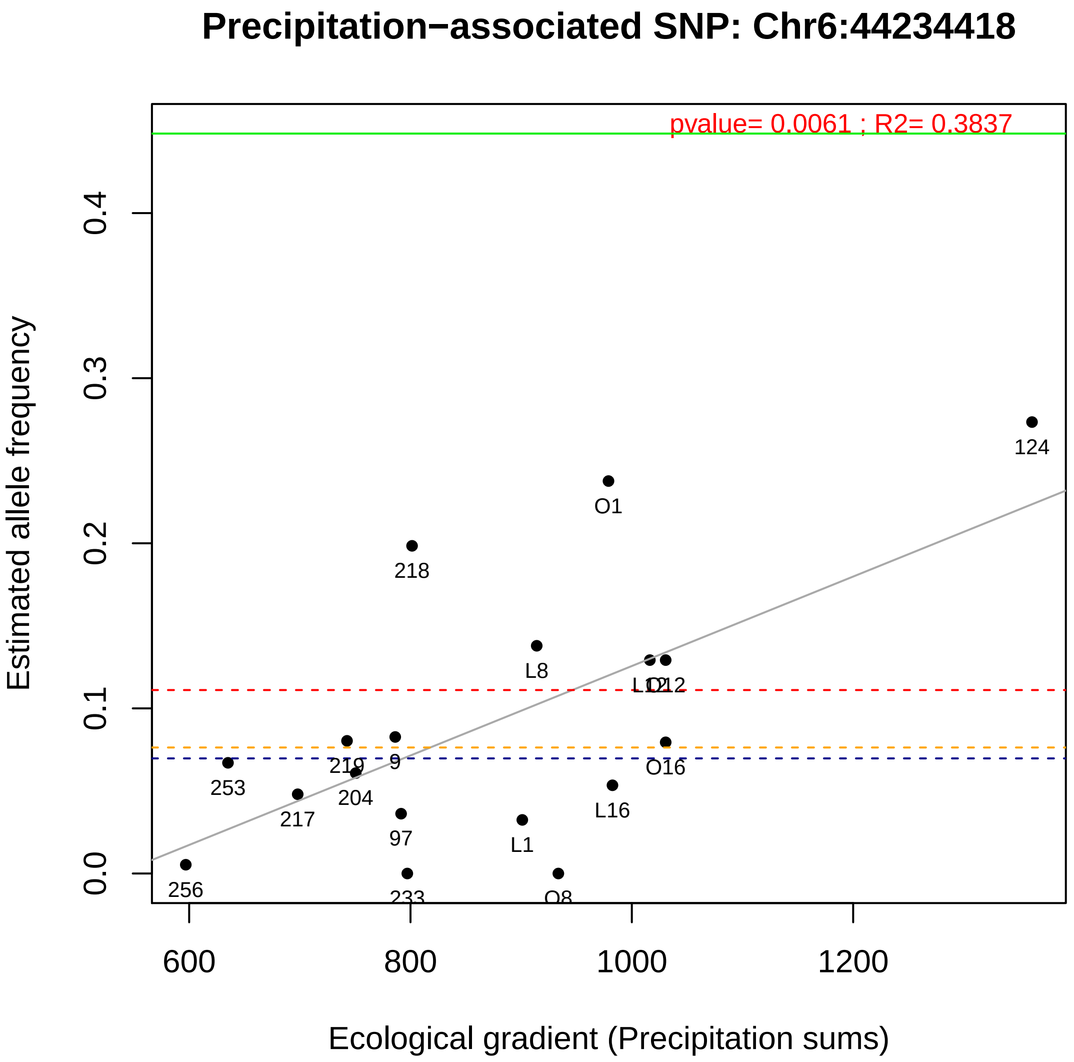
Precipitation−associated SNP: Chr6:44234418
pvalue= 0.0061 ; R2= 0.3837
Estimated allele frequency
0.0
0.1
0.2
0.3
0.4
600
800
1000
1200
Ecological gradient (Precipitation sums)
124
O1
218
L8
O12
O16
L16
219
9
253
204
217
97
L1
256
233
O8

**Precipitation−associated SNP: Chr6:45938151**

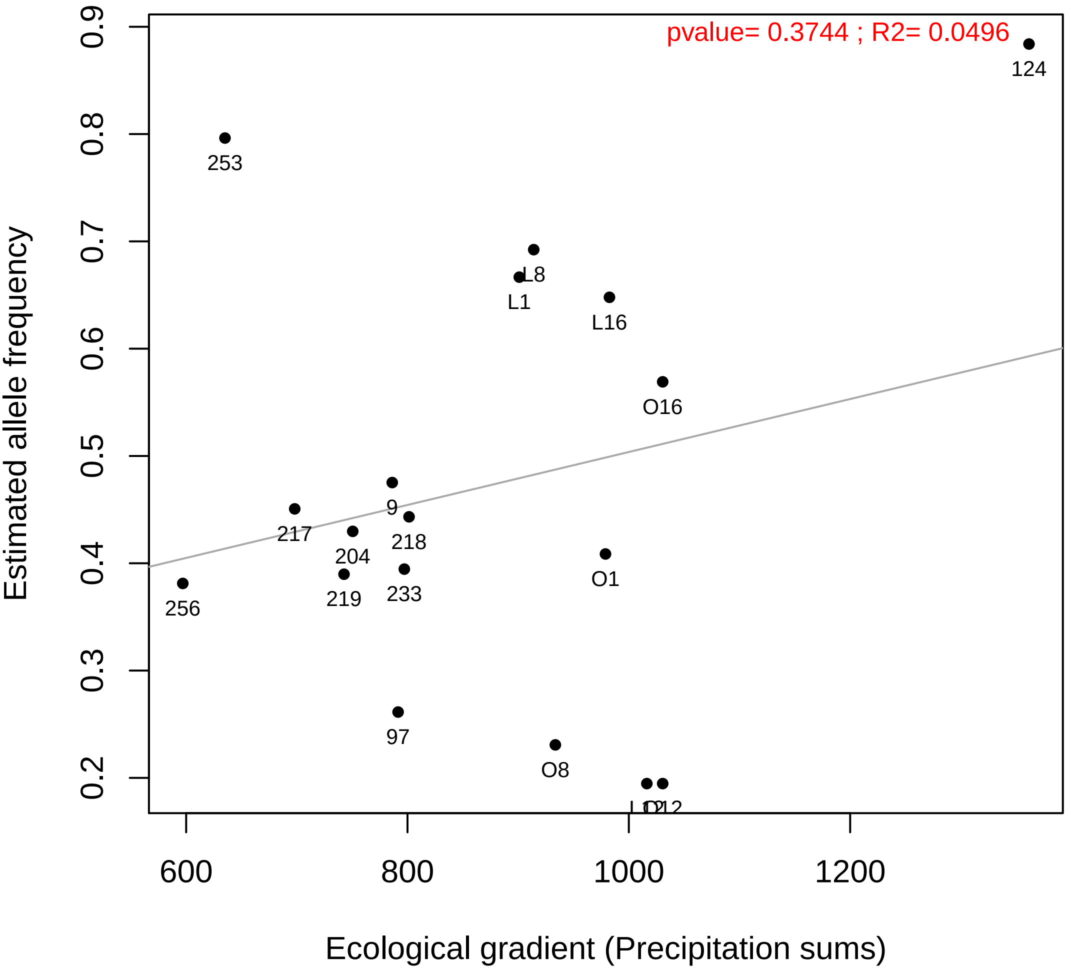

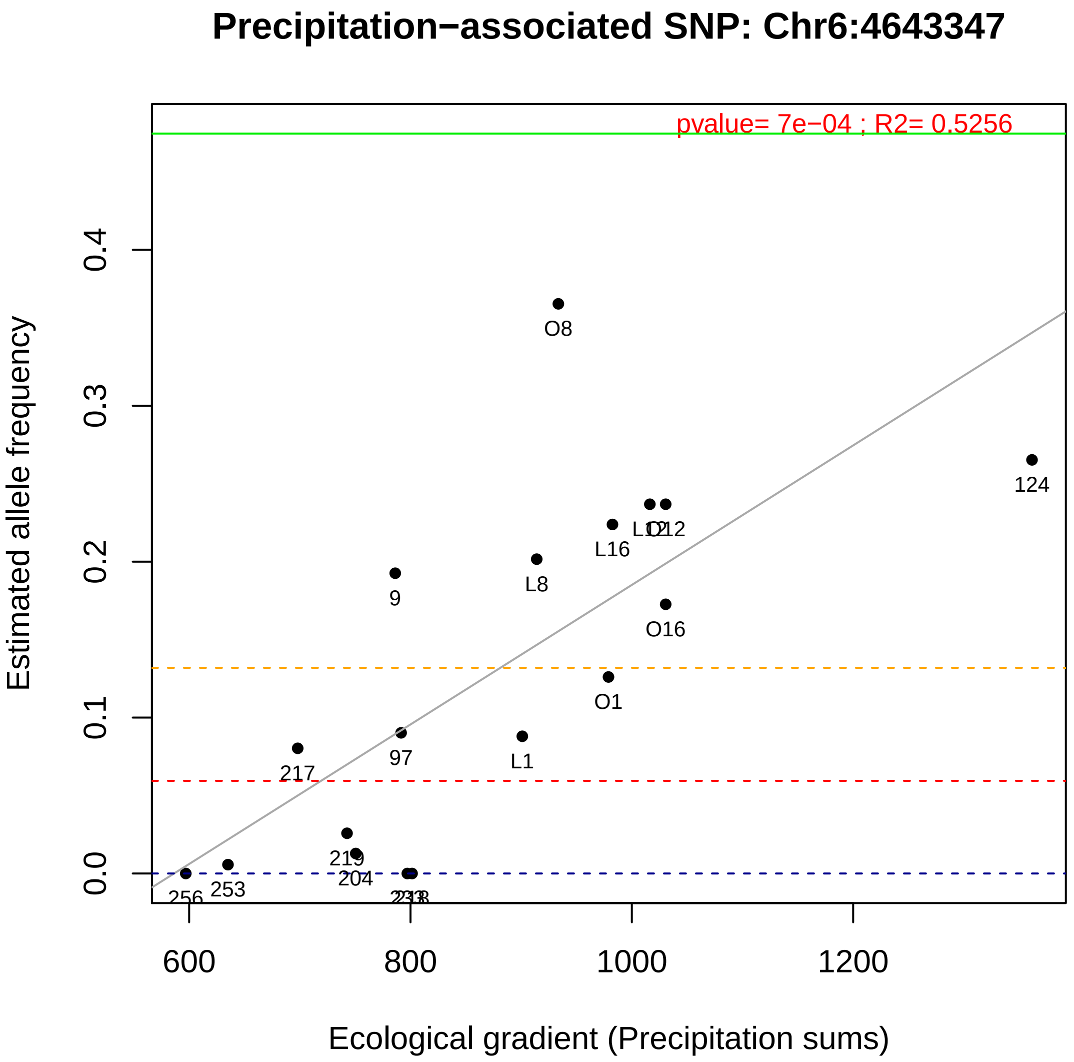

Precipitation−associated SNP: Chr6:4643347
pvalue= 7e−04 ; R2= 0.5256
Estimated allele frequency
0.0
0.1
0.2
0.3
0.4
600
800
1000
1200
Ecological gradient (Precipitation sums)
O8
124
L12
O12
L16
L8
9
O16
O1
217
97
L1
219
204
253
256

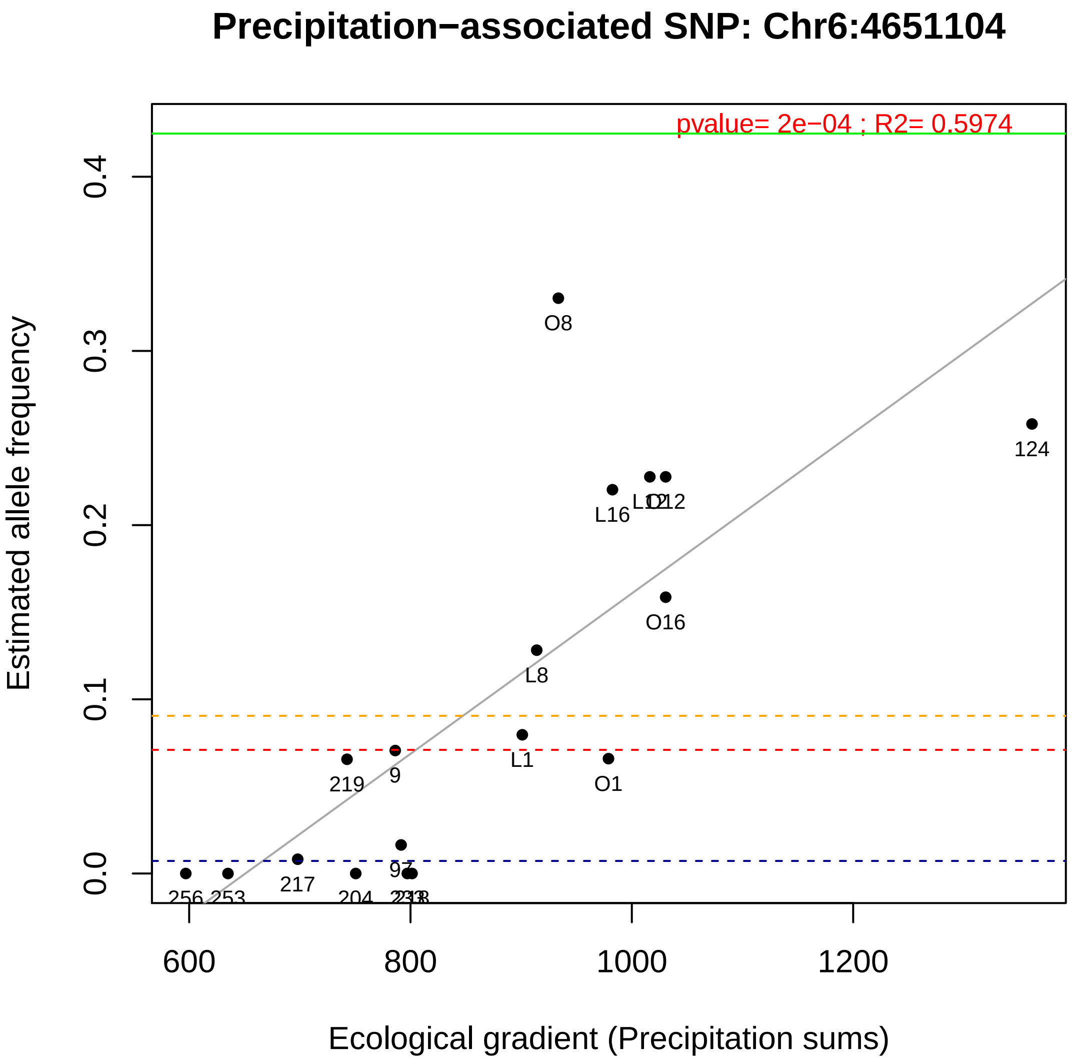
Precipitation−associated SNP: Chr6:4651104
pvalue= 2e−04 ; R2= 0.5974
Estimated allele frequency
0.0
0.1
0.2
0.3
0.4
600
800
1000
1200
Ecological gradient (Precipitation sums)
O8
124
L16
O16
L8
L1
219
9
O1
217
97
256
253
204

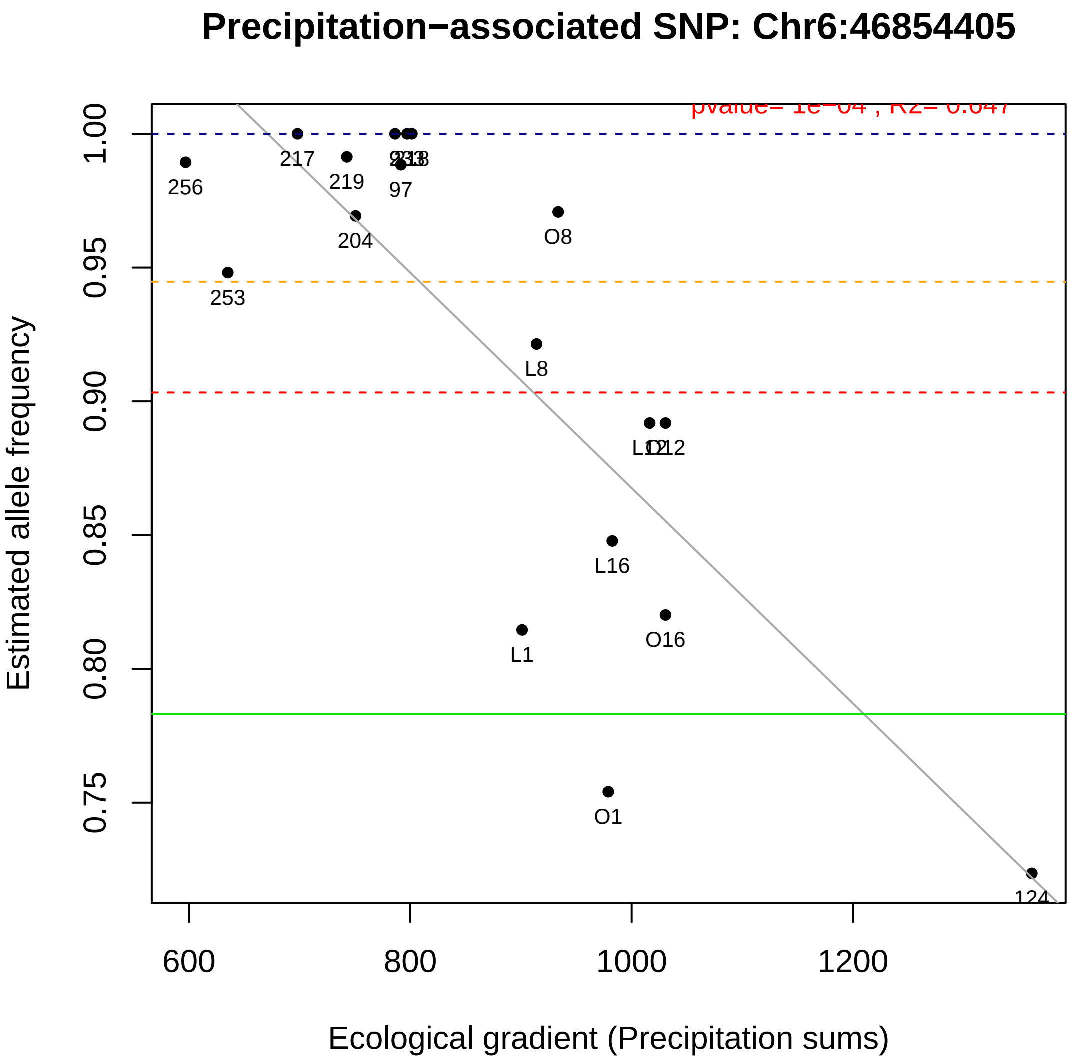
Precipitation−associated SNP: Chr6:46854405
pvalue= 1e−04 , R2= 0.647
Estimated allele frequency
Ecological gradient (Precipitation sums)
0.75
0.80
0.85
0.90
0.95
1.00
600
800
1000
1200
256
217
219
97
204
O8
253
L8
L12
O12
L16
O16
L1
O1
124

**Precipitation−associated SNP: Chr6:47902690**

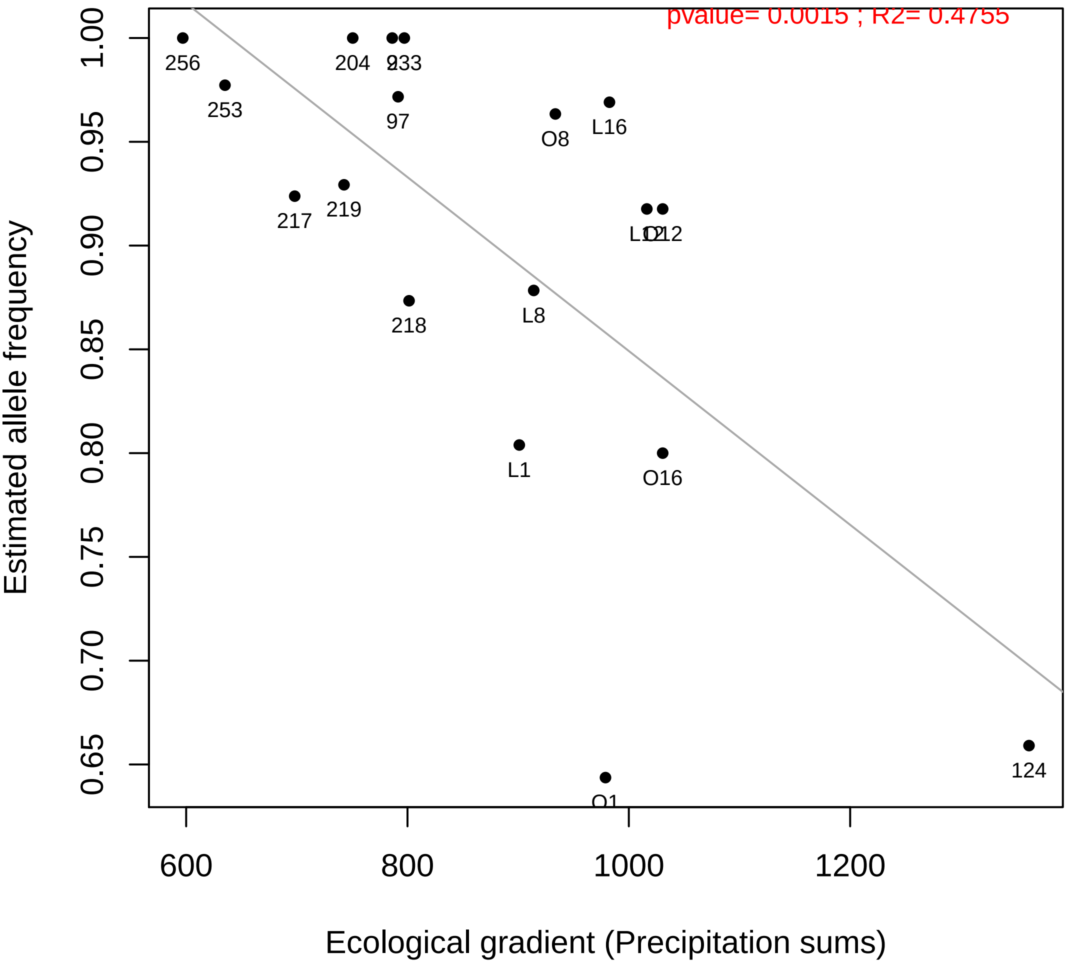

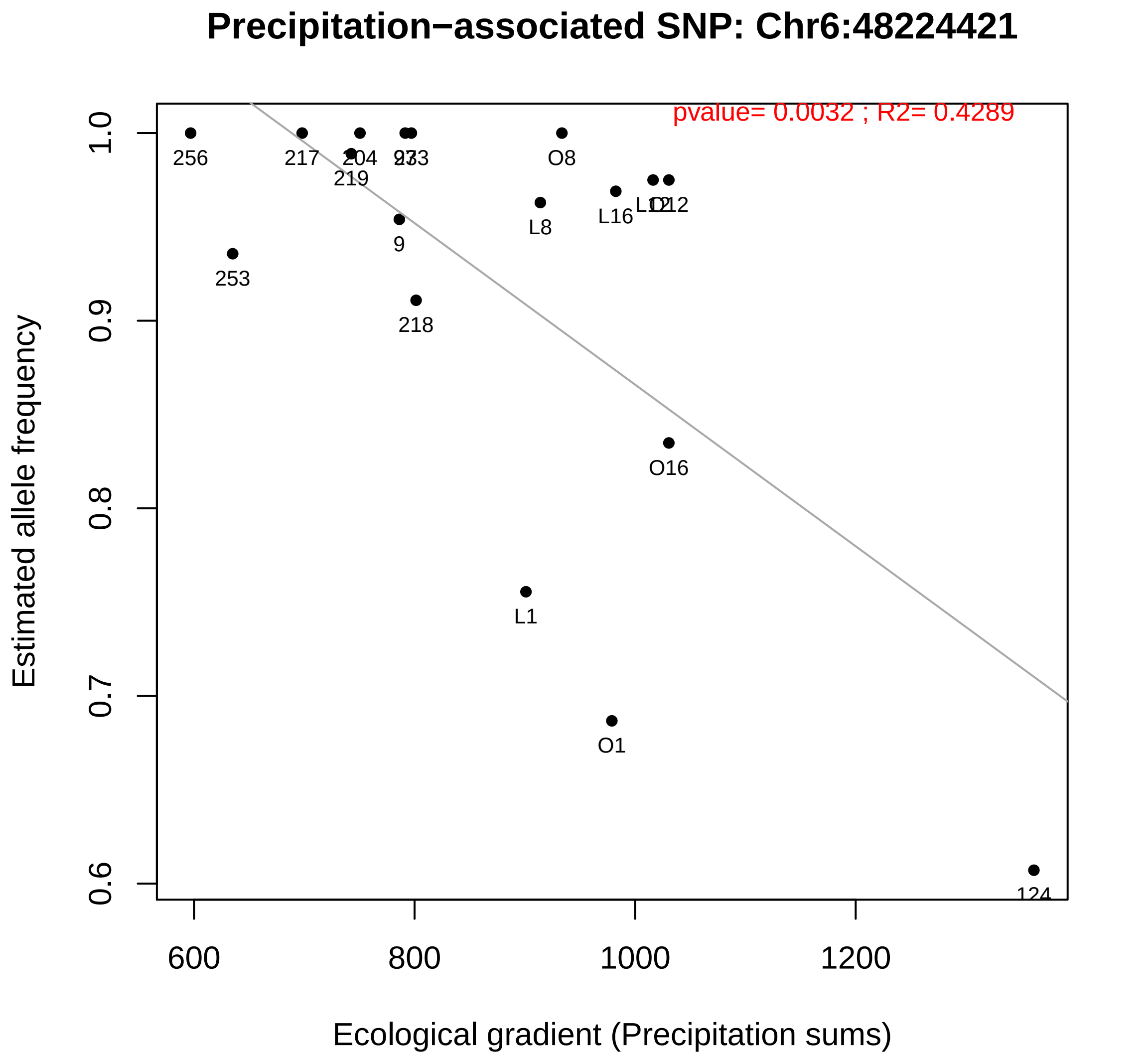
Precipitation−associated SNP: Chr6:48224421
pvalue= 0.0032 ; R2= 0.4289
Estimated allele frequency
Ecological gradient (Precipitation sums)
0.6
0.7
0.8
0.9
1.0
600
800
1000
1200
256
217
204
219
253
9
218
O8
L8
L16
O16
L1
O1
124

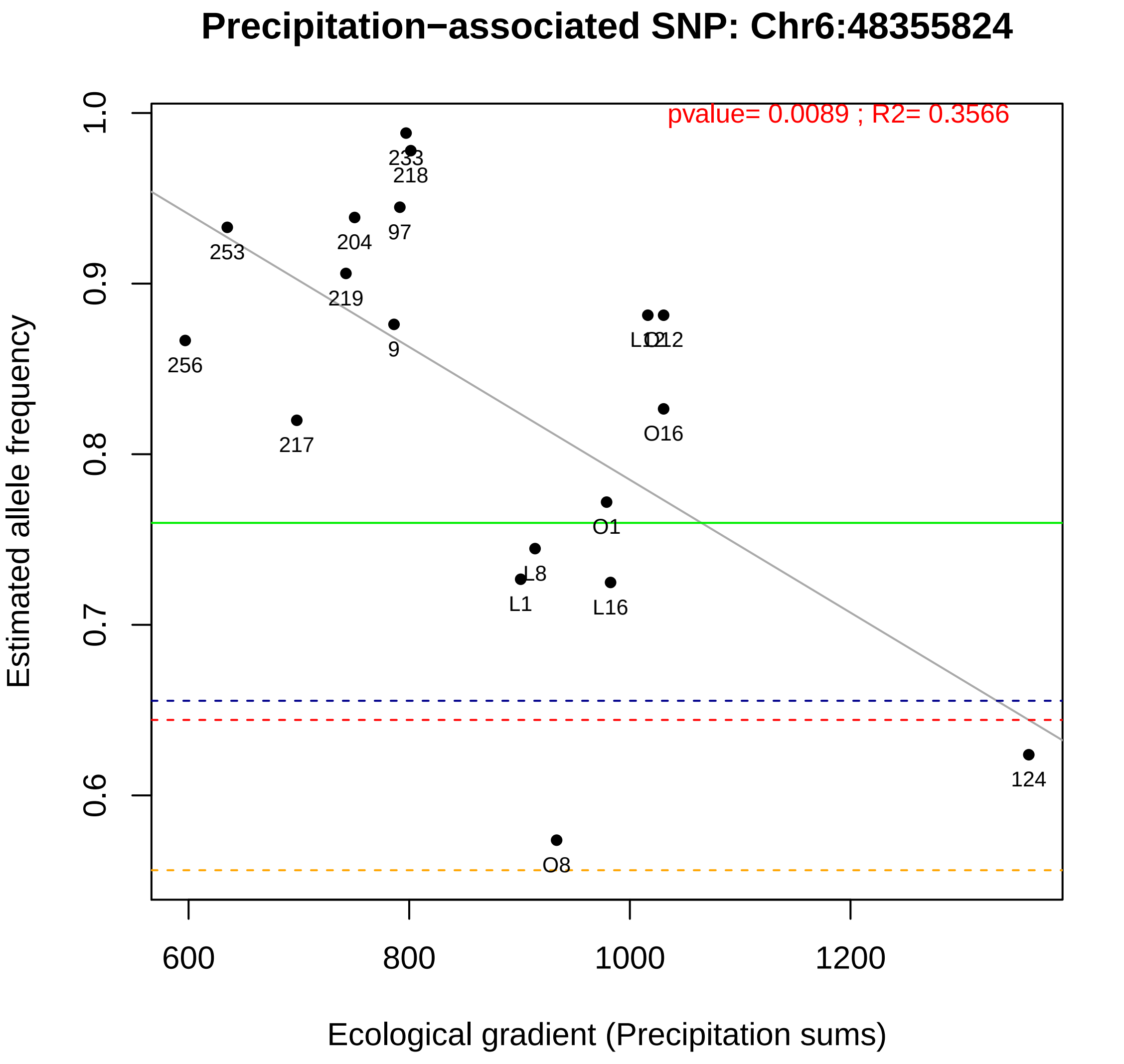

Precipitation−associated SNP: Chr6:48355824
pvalue= 0.0089 ; R2= 0.3566
Estimated allele frequency
Ecological gradient (Precipitation sums)
1.0
0.9
0.8
0.7
0.6
600
800
1000
1200
233
218
97
204
253
219
9
256
217
L12
O12
O16
O1
L8
L1
L16
124
O8

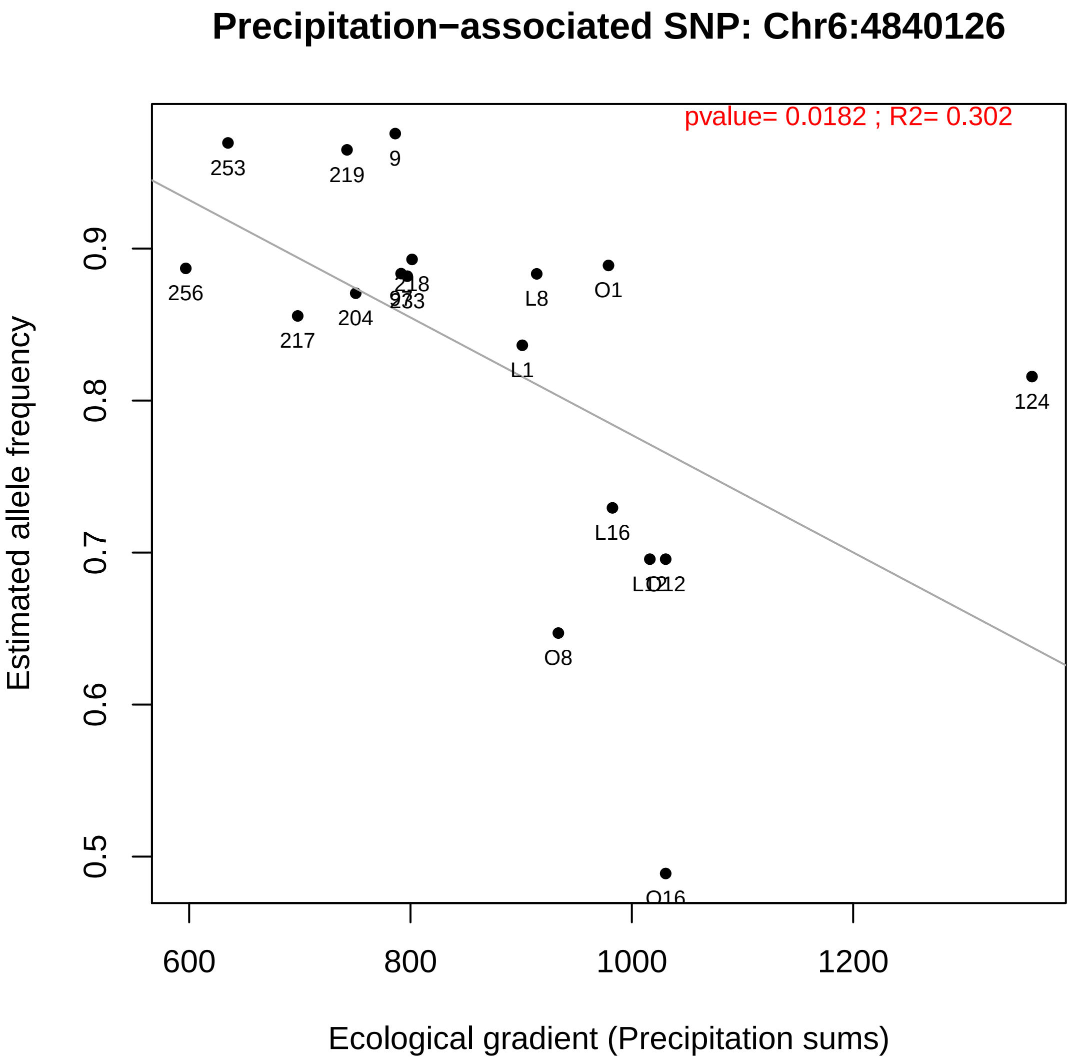
Precipitation−associated SNP: Chr6:4840126
pvalue= 0.0182 ; R2= 0.302
253
219
9
256
218
204
217
L8
O1
L1
124
L16
O12
O8
O16
0.5
0.6
0.7
0.8
0.9
600
800
1000
1200
Estimated allele frequency
Ecological gradient (Precipitation sums)

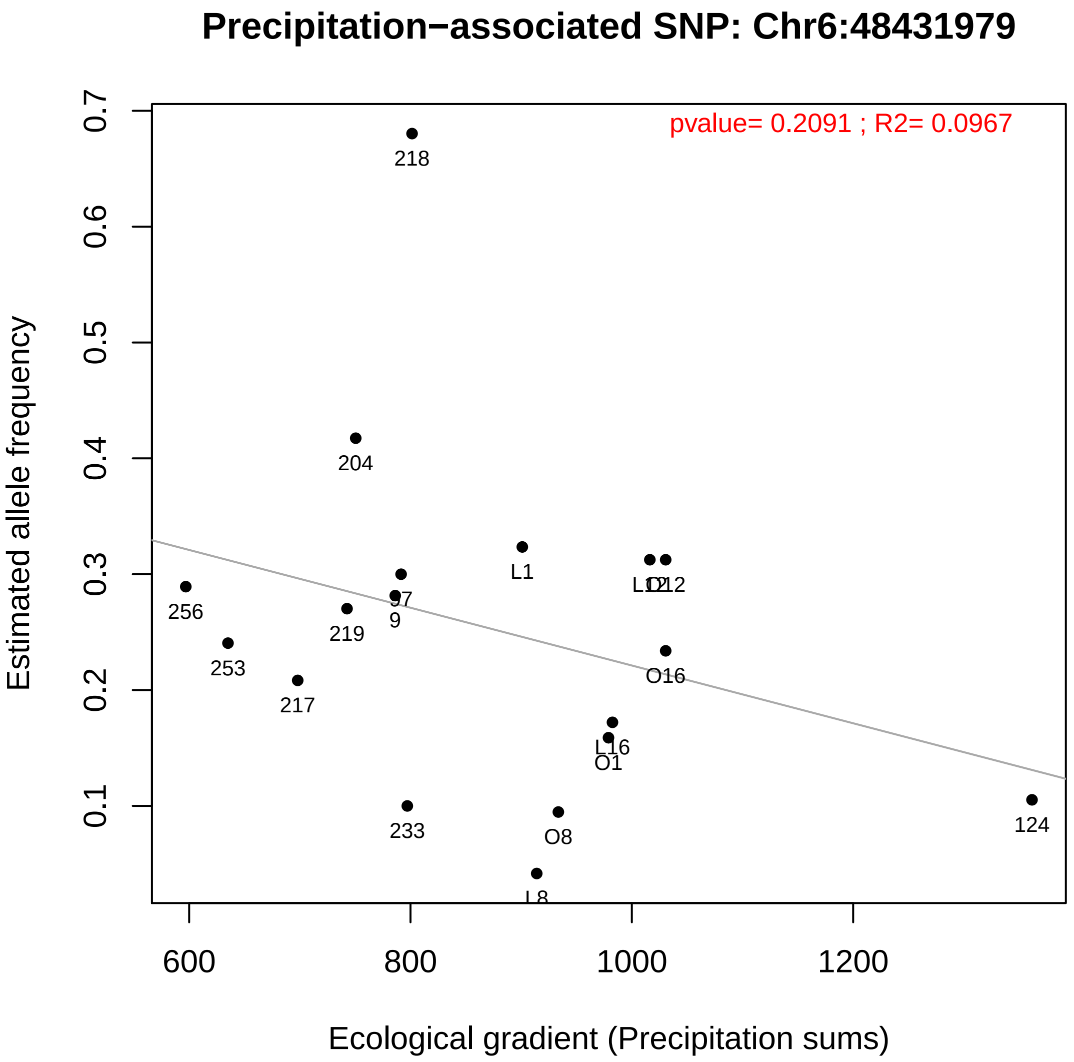

Precipitation−associated SNP: Chr6:48431979
pvalue= 0.2091 ; R2= 0.0967
Estimated allele frequency
0.1
0.2
0.3
0.4
0.5
0.6
0.7
600
800
1000
1200
Ecological gradient (Precipitation sums)
218
204
L1
256
253
217
219
9
233
O8
O16
L16
O1
124

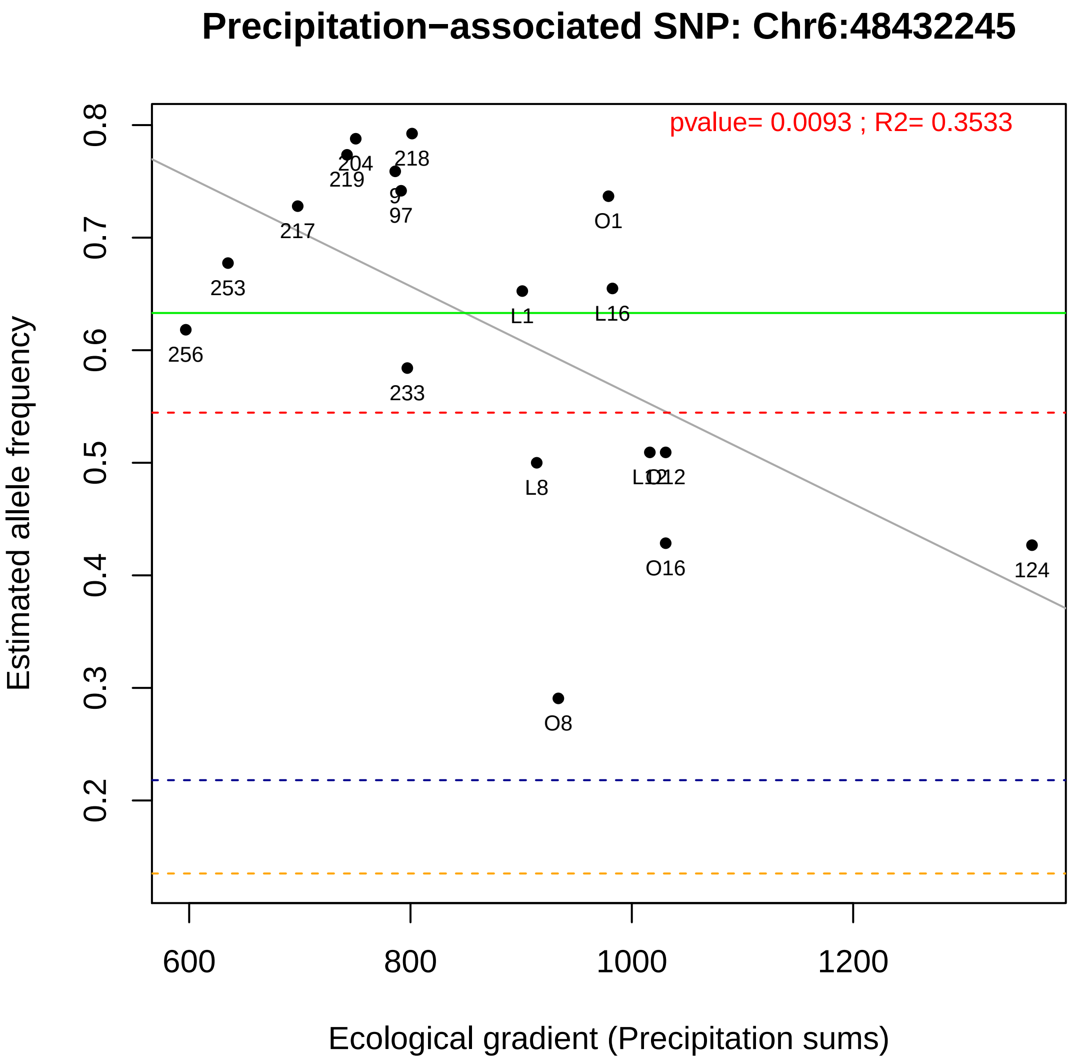

Precipitation−associated SNP: Chr6:48432245
pvalue= 0.0093 ; R2= 0.3533
Estimated allele frequency
Ecological gradient (Precipitation sums)
0.2
0.3
0.4
0.5
0.6
0.7
0.8
600
800
1000
1200
204
219
218
9
97
217
253
256
233
O1
L1
L16
L8
L12
O12
O16
O8
124

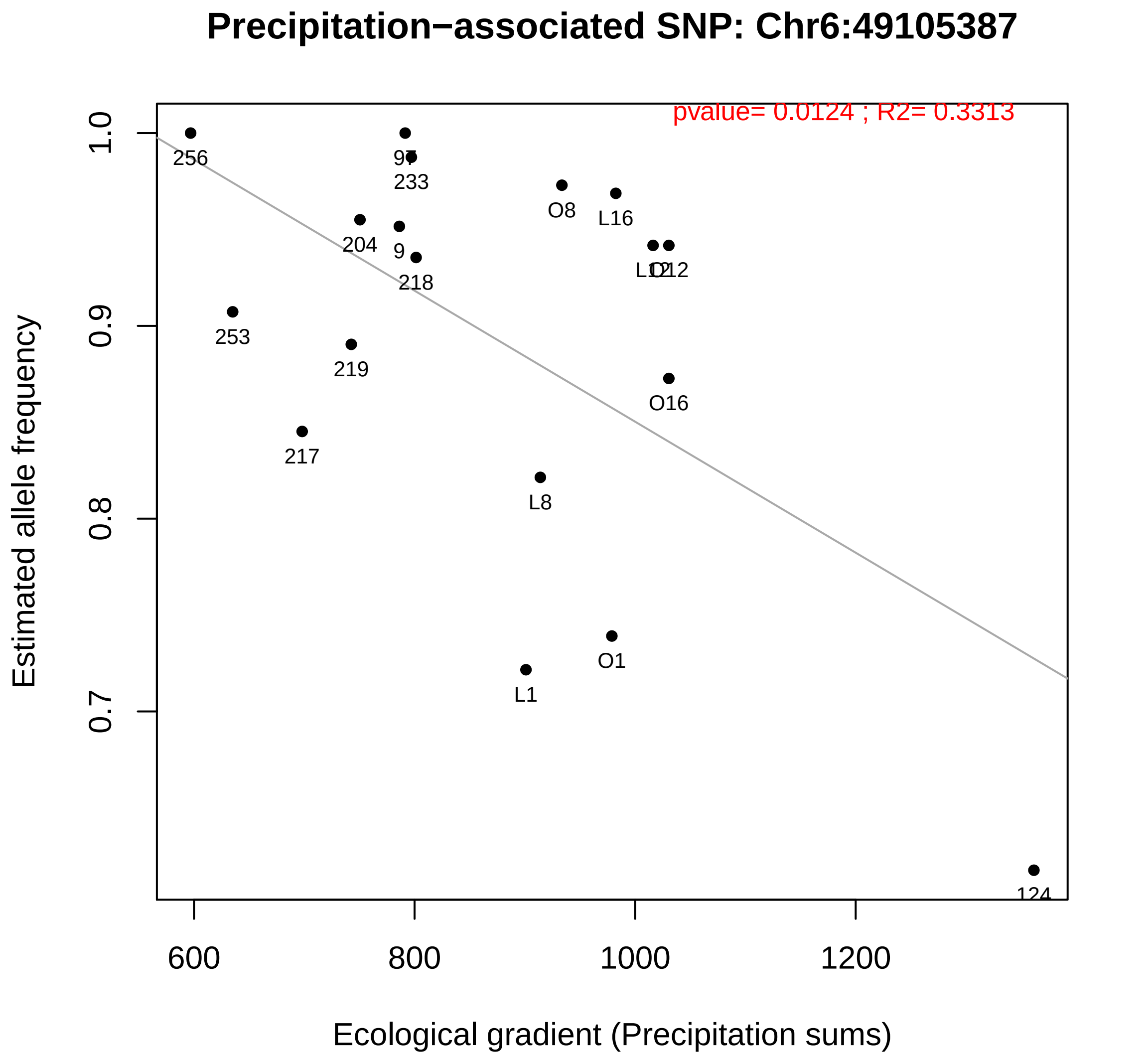
Precipitation−associated SNP: Chr6:49105387
pvalue= 0.0124 ; R2= 0.3313
Estimated allele frequency
Ecological gradient (Precipitation sums)
1.0
0.9
0.8
0.7
600
800
1000
1200
256
97
233
O8
L16
204
9
218
L12
O12
253
219
O16
217
L8
O1
L1
124

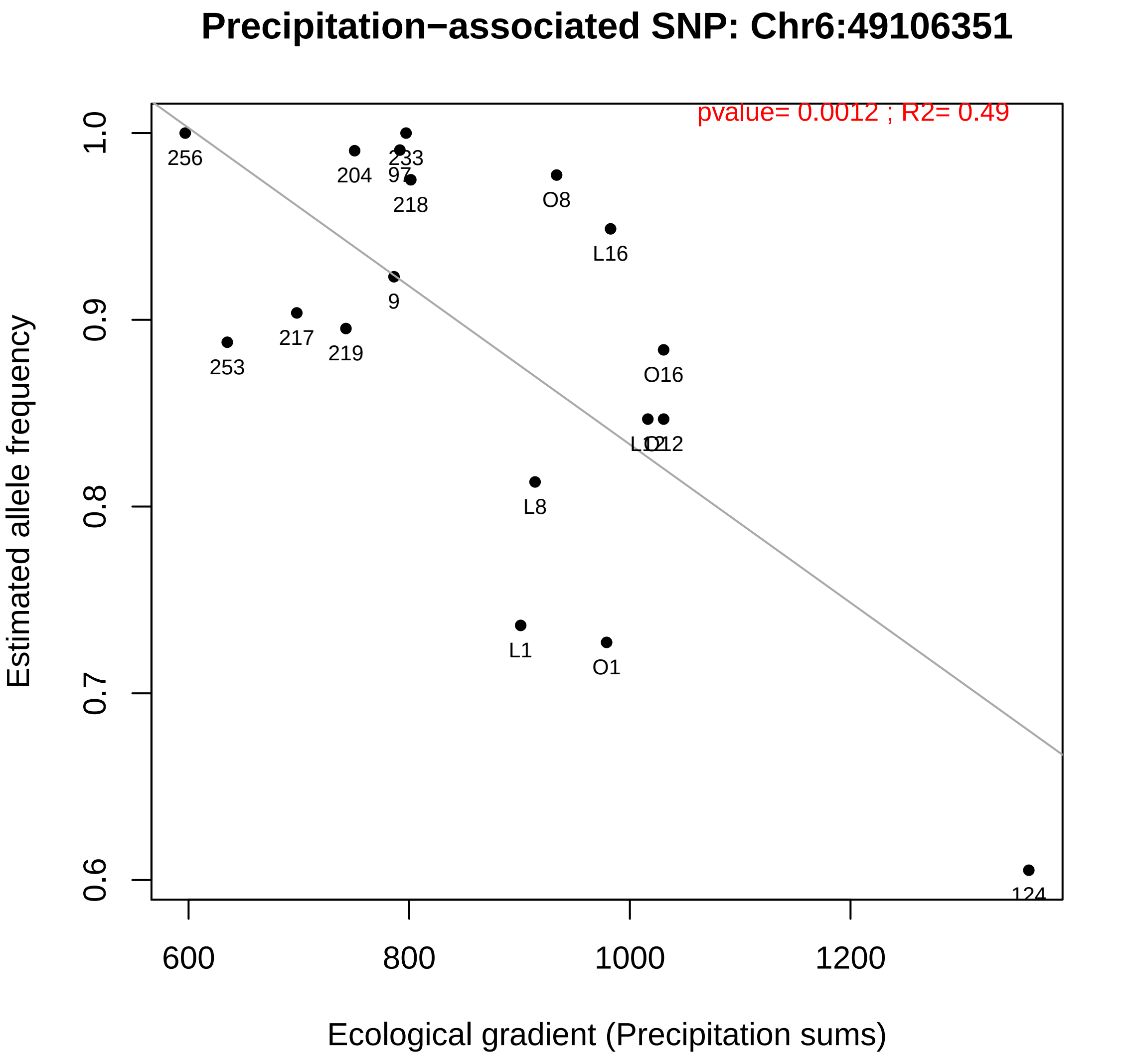
Precipitation−associated SNP: Chr6:49106351
pvalue= 0.0012 ; R2= 0.49
Estimated allele frequency
Ecological gradient (Precipitation sums)
0.6
0.7
0.8
0.9
1.0
600
800
1000
1200
256
204
233
97
218
O8
L16
9
217
219
253
O16
L12
O12
L8
L1
O1
124

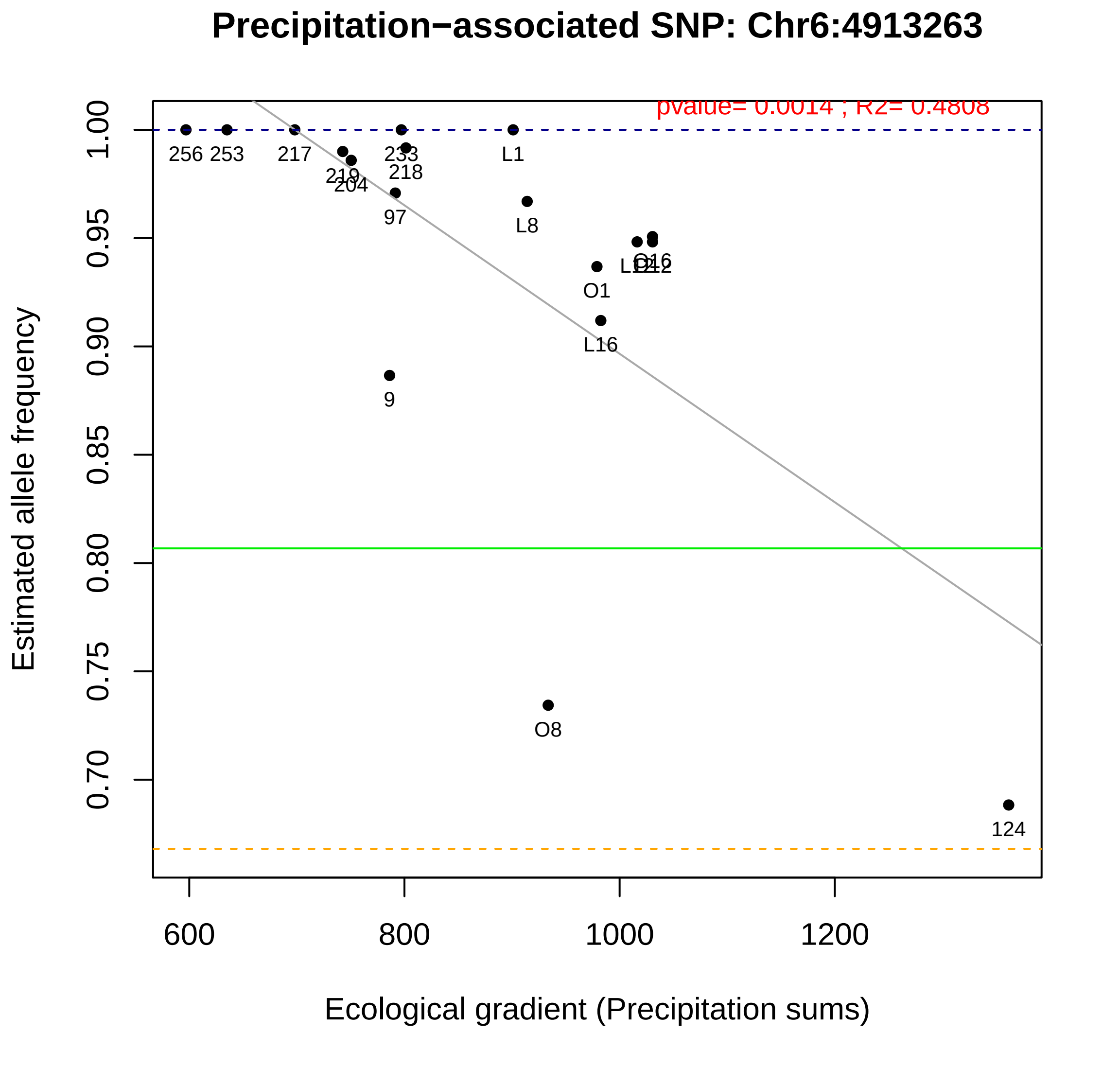

Precipitation−associated SNP: Chr6:4913263
pvalue= 0.0014 ; R2= 0.4808
Estimated allele frequency
1.00
0.95
0.90
0.85
0.80
0.75
0.70
256
253
217
219
204
233
218
97
L1
L8
9
O1
L16
L12
O16
O8
124
600
800
1000
1200
Ecological gradient (Precipitation sums)

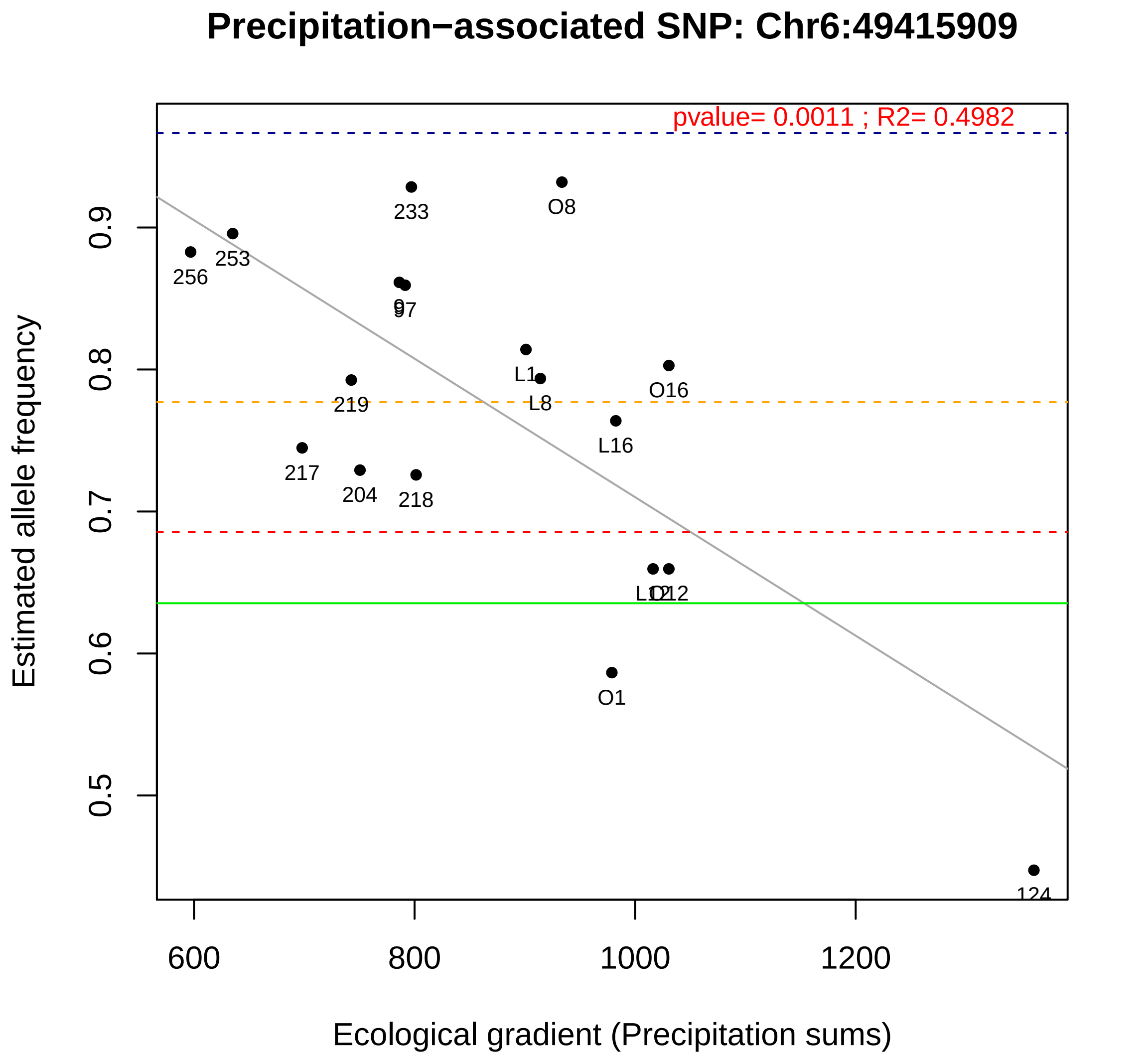
Precipitation−associated SNP: Chr6:49415909
pvalue= 0.0011 ; R2= 0.4982
Estimated allele frequency
0.5
0.6
0.7
0.8
0.9
600
800
1000
1200
Ecological gradient (Precipitation sums)
233
O8
253
256
97
219
L1
L8
O16
L16
217
204
218
O12
O1
124

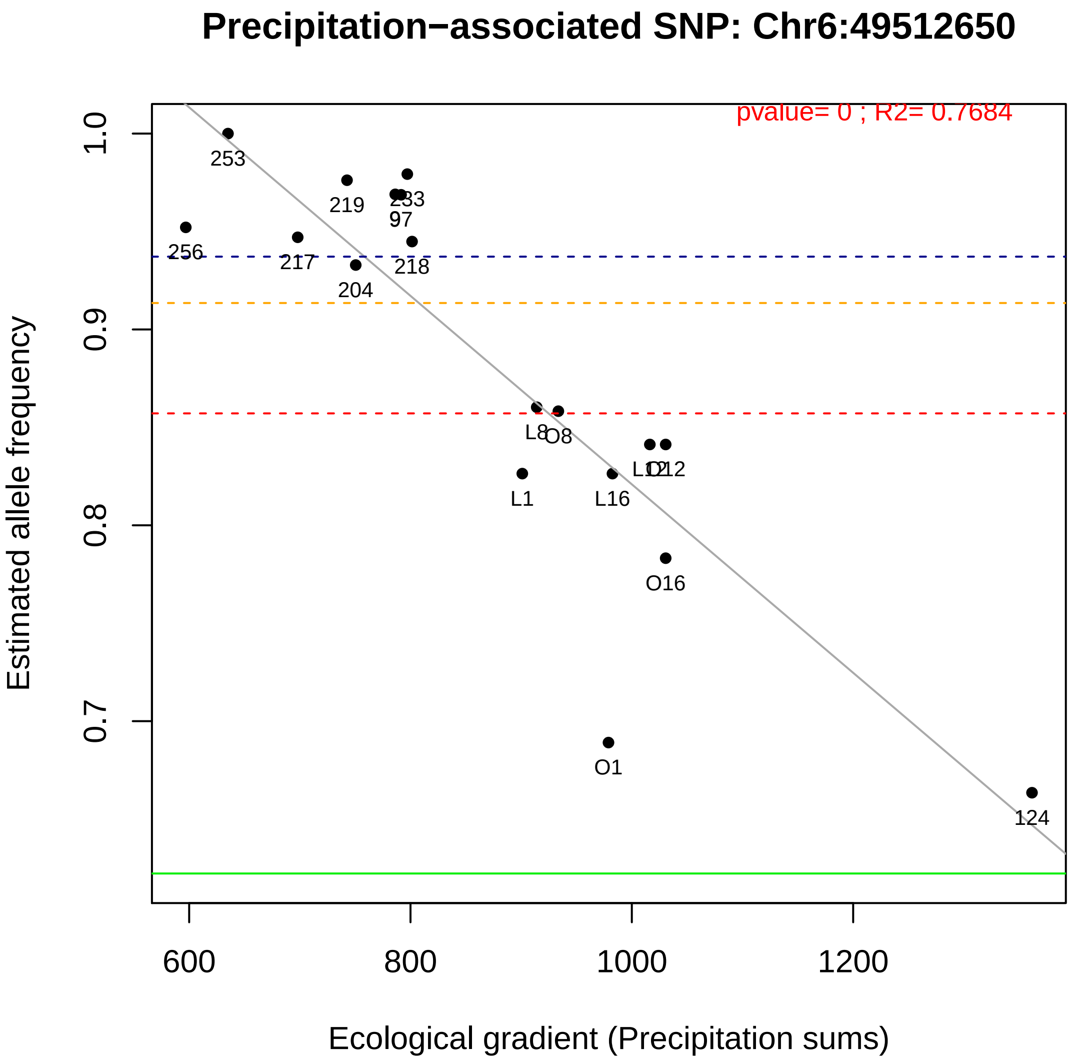

Precipitation−associated SNP: Chr6:49512650
pvalue= 0 ; R2= 0.7684
Estimated allele frequency
Ecological gradient (Precipitation sums)
0.7
0.8
0.9
1.0
600
800
1000
1200
253
219
233
97
256
217
218
204
L8
O8
L1
L16
L12
O12
O16
O1
124

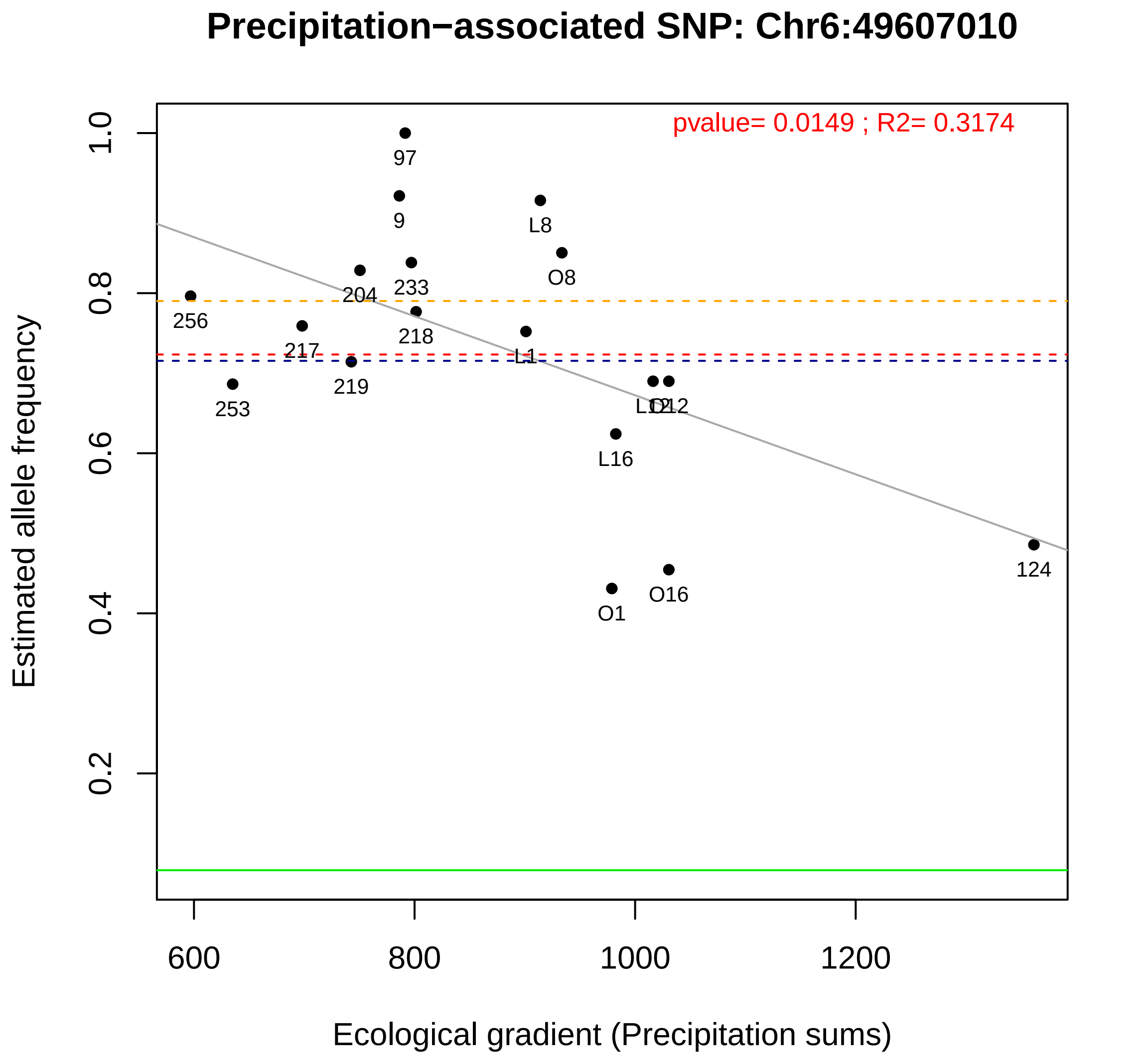

Precipitation−associated SNP: Chr6:49607010
pvalue= 0.0149 ; R2= 0.3174
Estimated allele frequency
Ecological gradient (Precipitation sums)
0.2
0.4
0.6
0.8
1.0
600
800
1000
1200
97
9
L8
O8
233
204
256
218
217
L1
219
253
L12
O12
L16
124
O16
O1

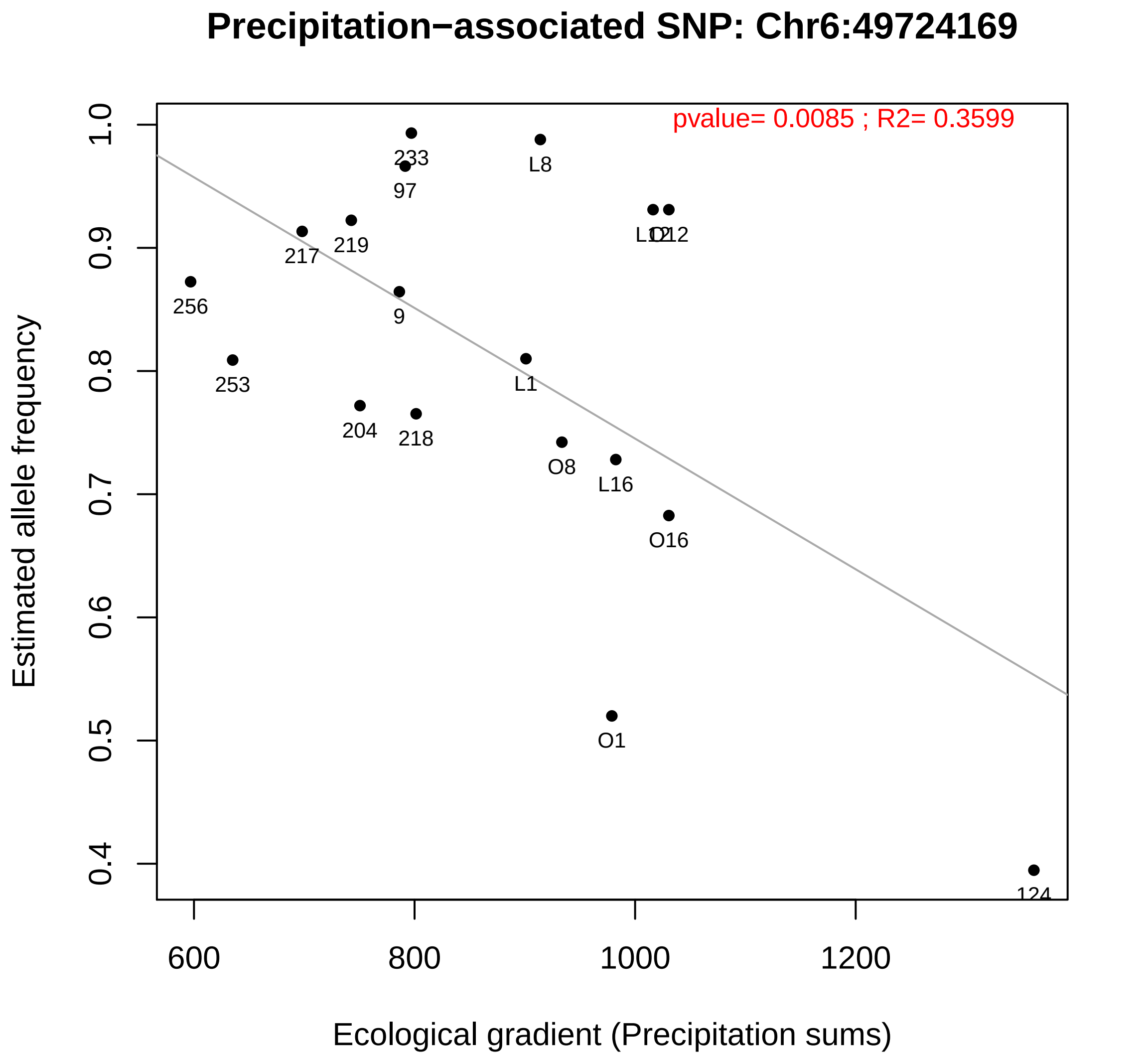

Precipitation−associated SNP: Chr6:49724169
pvalue= 0.0085 ; R2= 0.3599
Estimated allele frequency
Ecological gradient (Precipitation sums)
1.0
0.9
0.8
0.7
0.6
0.5
0.4
600
800
1000
1200
233
L8
97
217
219
256
9
253
L1
204
218
O8
L16
O16
O1
124

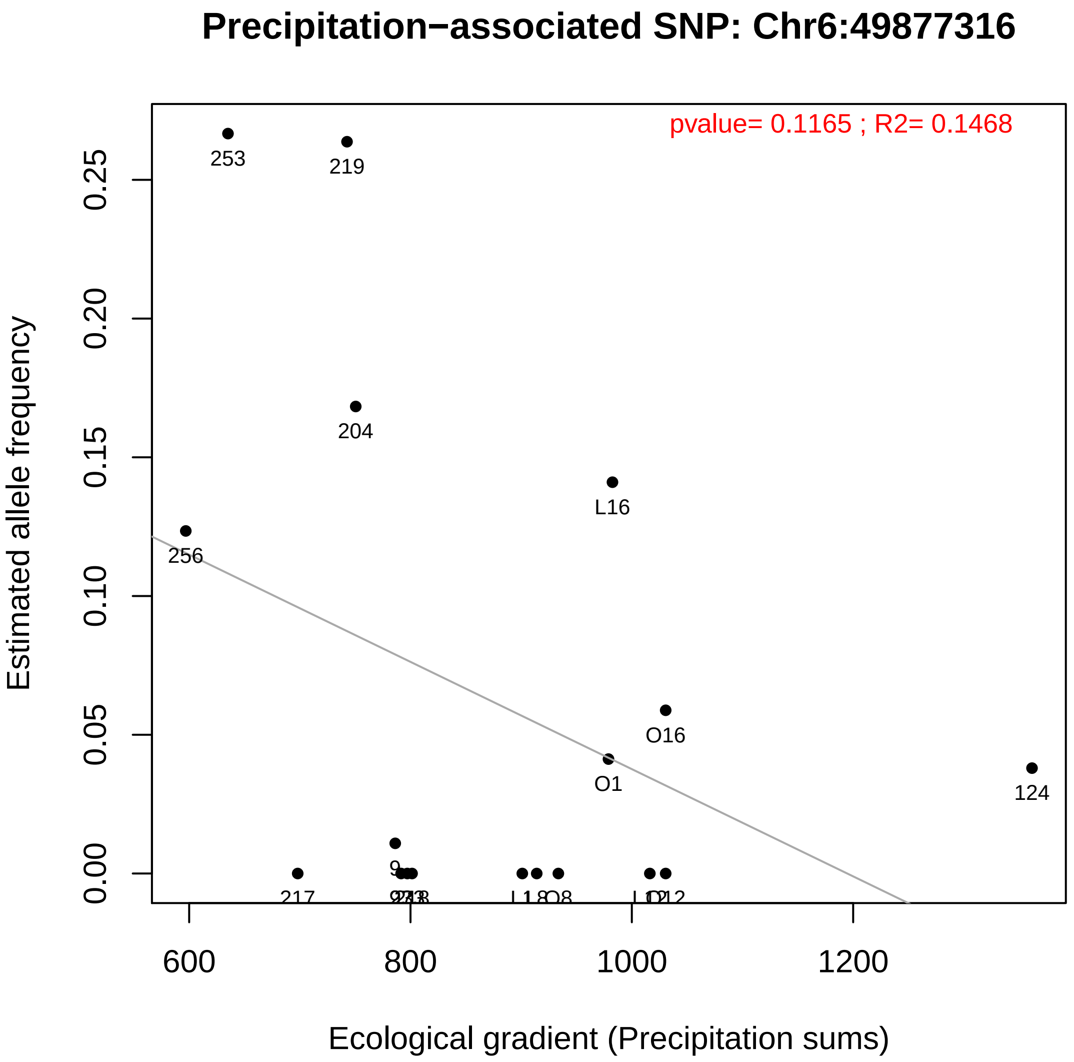
Precipitation−associated SNP: Chr6:49877316
pvalue= 0.1165 ; R2= 0.1468
253
219
204
L16
256
O16
O1
124
9
217
0.00
0.05
0.10
0.15
0.20
0.25
600
800
1000
1200
Estimated allele frequency
Ecological gradient (Precipitation sums)

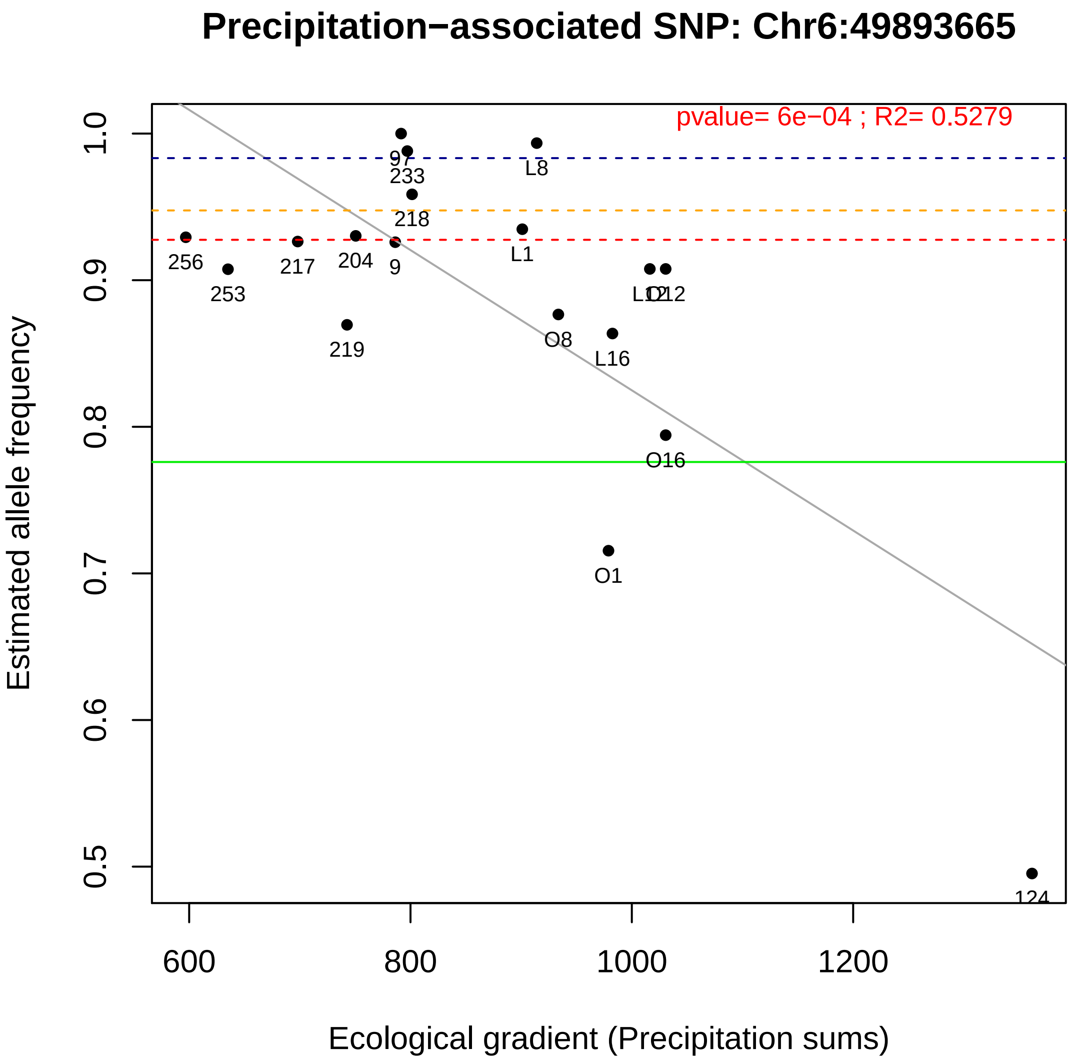

Precipitation−associated SNP: Chr6:49893665
pvalue= 6e−04 ; R2= 0.5279
Estimated allele frequency
Ecological gradient (Precipitation sums)
0.5
0.6
0.7
0.8
0.9
1.0
600
800
1000
1200
256
253
217
204
219
9
97
233
218
L8
L1
O8
L16
L12
O12
O16
O1
124

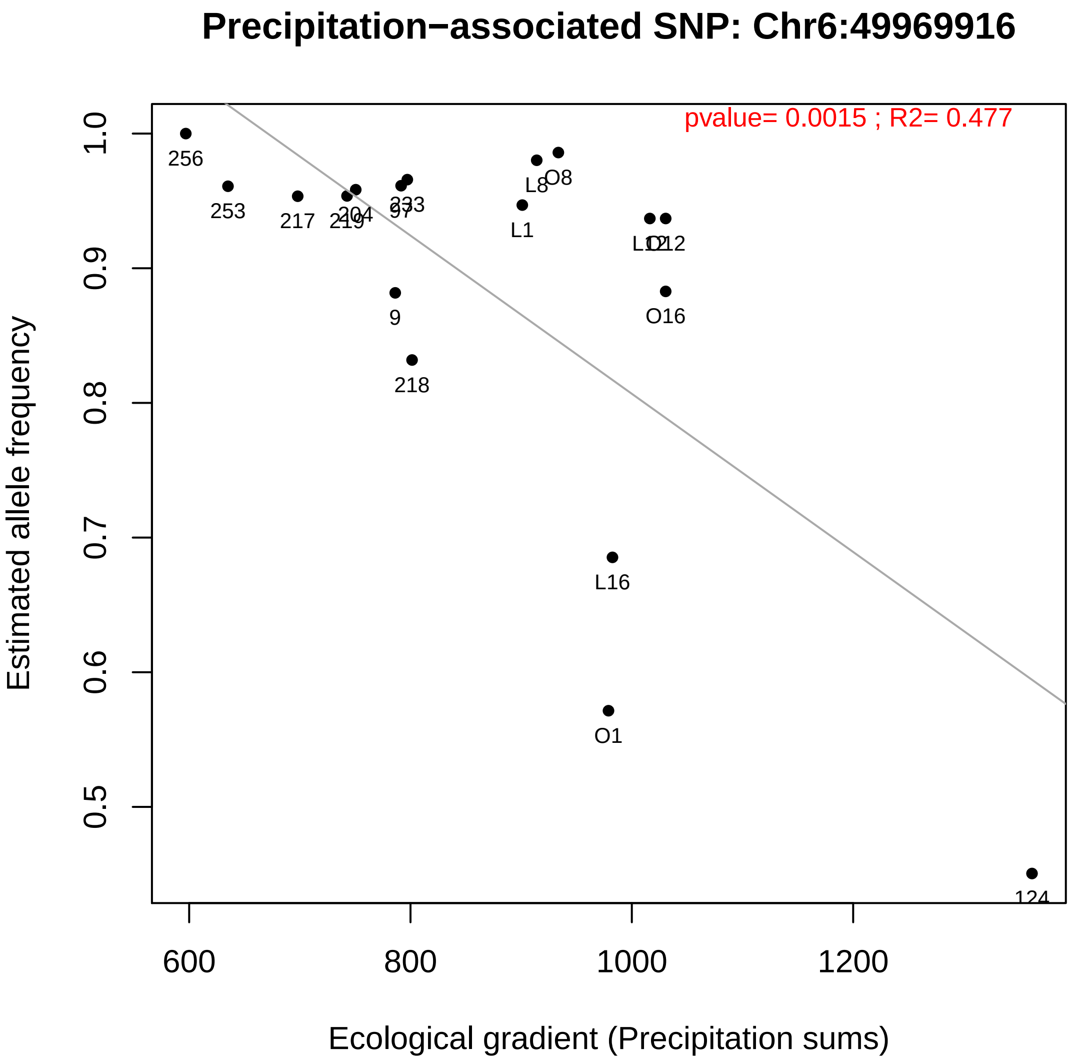

Precipitation−associated SNP: Chr6:49969916
pvalue= 0.0015 ; R2= 0.477
Estimated allele frequency
Ecological gradient (Precipitation sums)
0.5
0.6
0.7
0.8
0.9
1.0
600
800
1000
1200
256
253
217
219
204
233
97
9
218
L1
L8
O8
L12
O12
O16
L16
O1
124

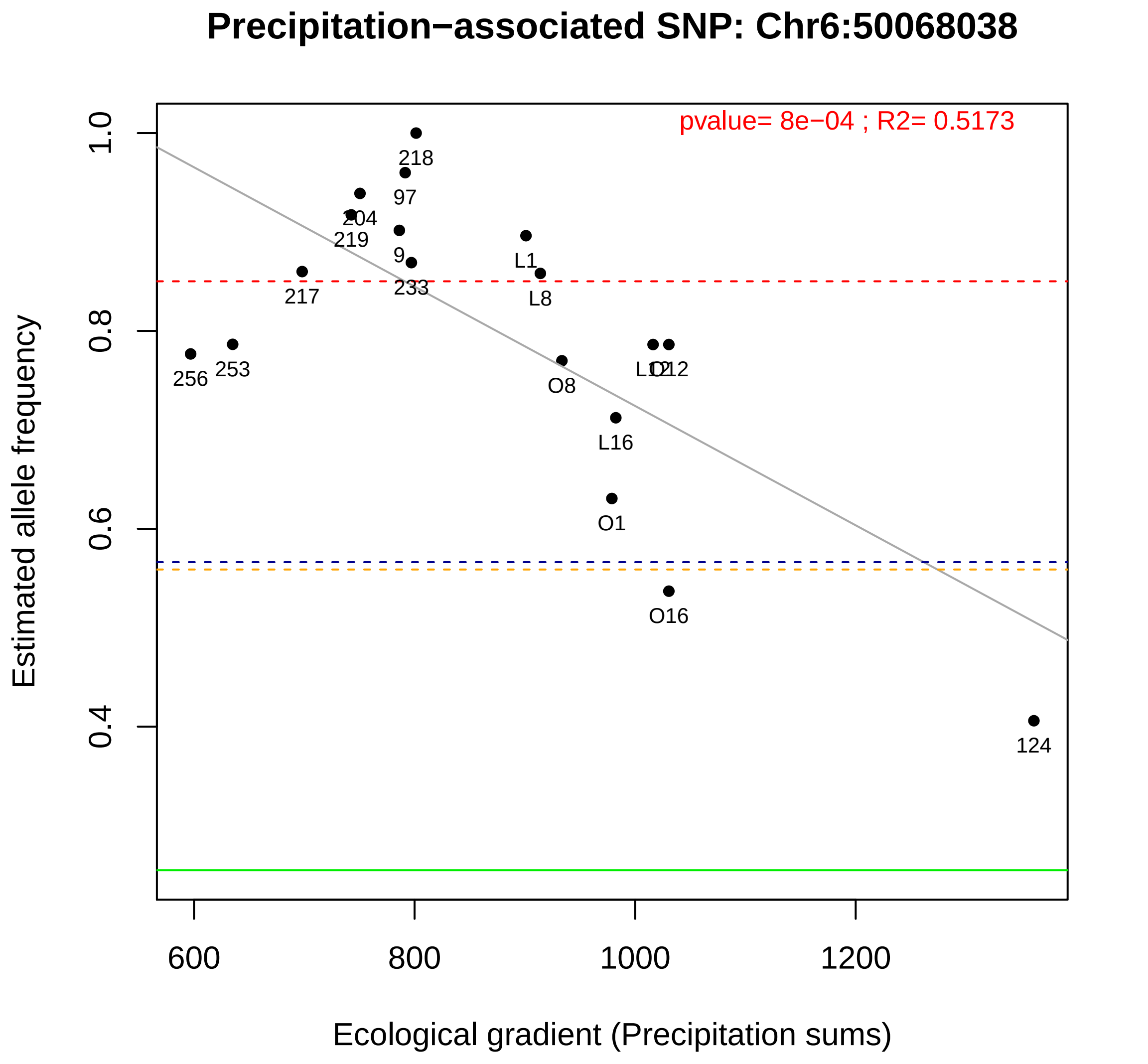

Precipitation−associated SNP: Chr6:50068038
pvalue= 8e−04 ; R2= 0.5173
218
97
204
219
9
233
217
L1
L8
256
253
O8
L12
O12
L16
O1
O16
124
Estimated allele frequency
0.4
0.6
0.8
1.0
600
800
1000
1200
Ecological gradient (Precipitation sums)

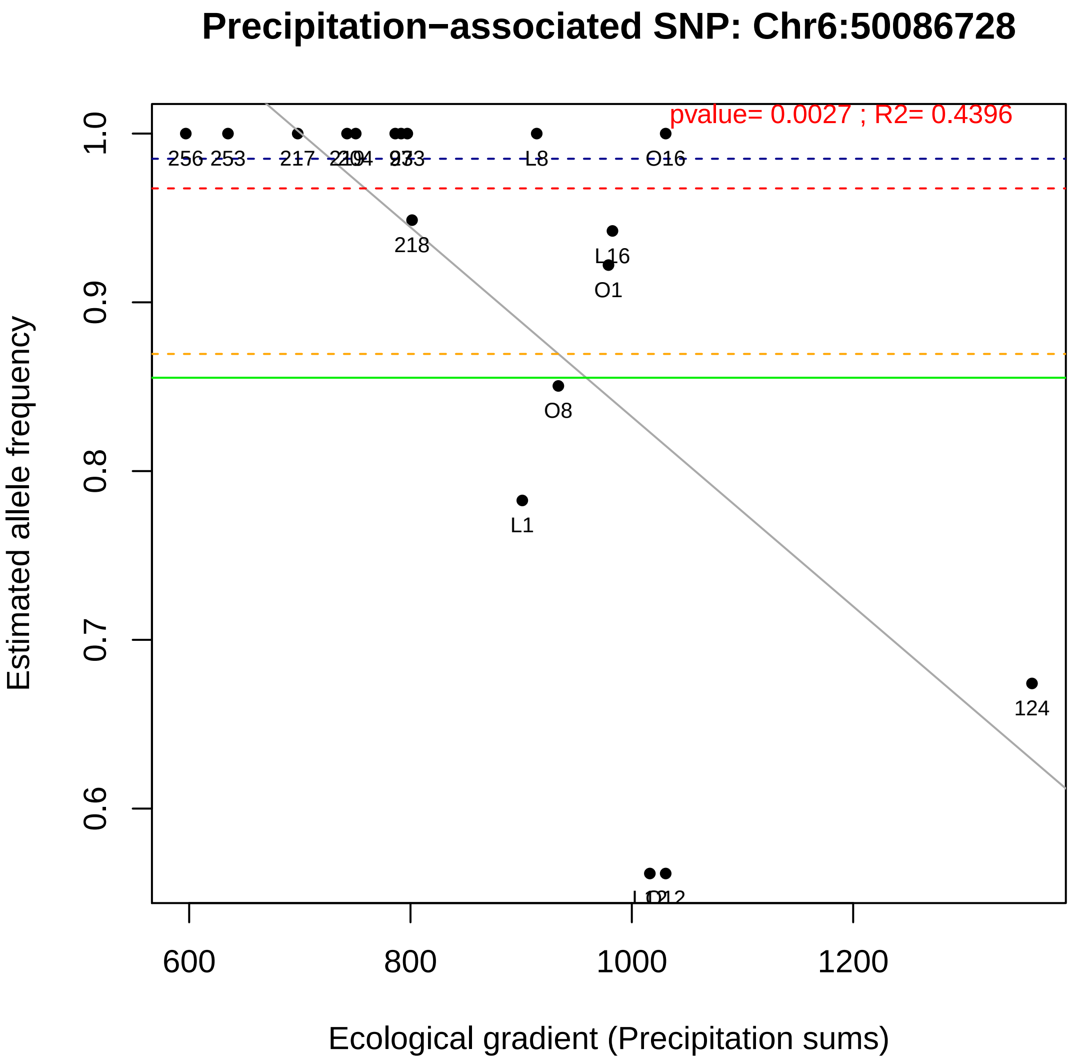

Precipitation−associated SNP: Chr6:50086728
pvalue= 0.0027 ; R2= 0.4396
256
253
217
218
L8
O16
L16
O1
O8
L1
124
1.0
0.9
0.8
0.7
0.6
600
800
1000
1200
Estimated allele frequency
Ecological gradient (Precipitation sums)

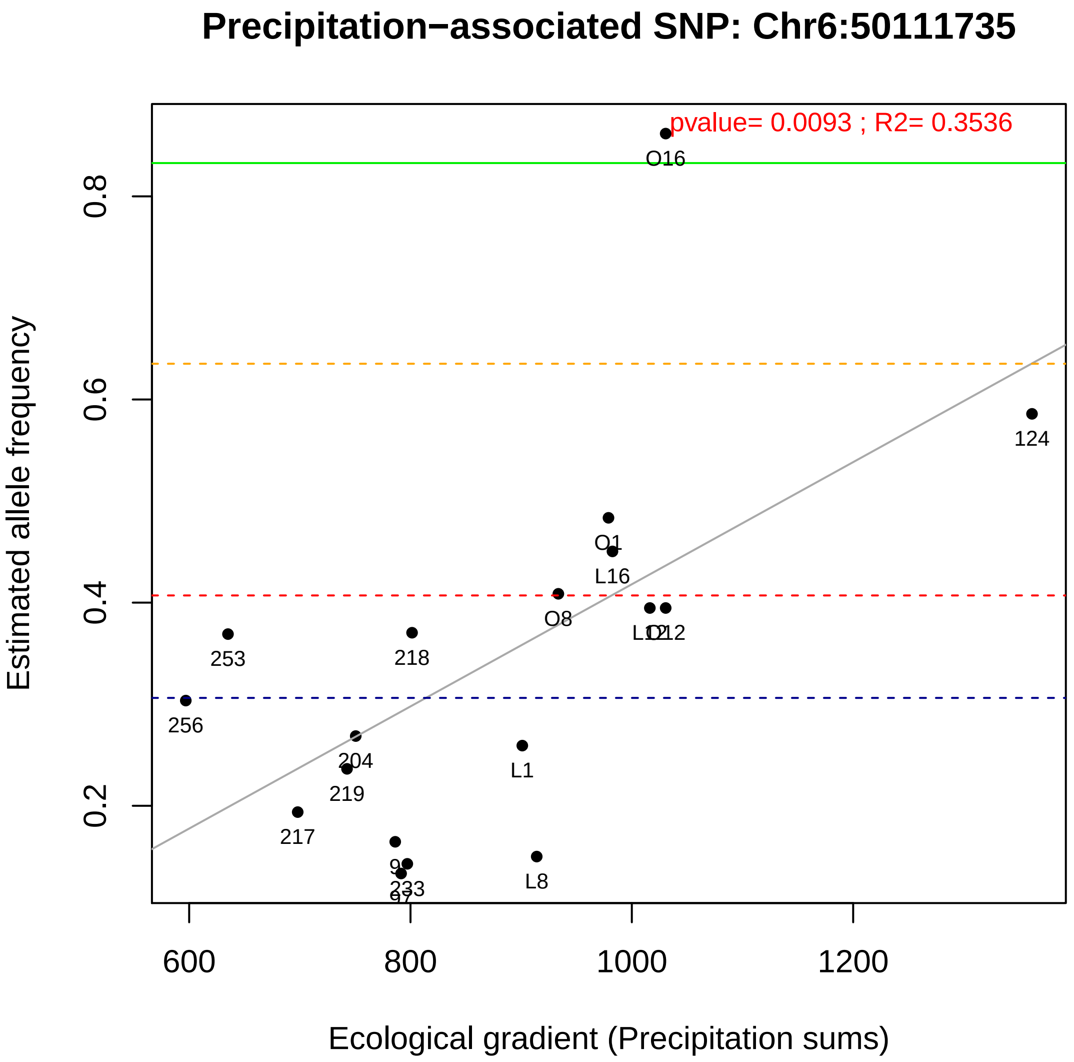
Precipitation−associated SNP: Chr6:50111735
pvalue= 0.0093 ; R2= 0.3536
Estimated allele frequency
0.2
0.4
0.6
0.8
600
800
1000
1200
Ecological gradient (Precipitation sums)
O16
124
O1
L16
O8
L12
O12
253
218
256
204
219
L1
217
233
L8

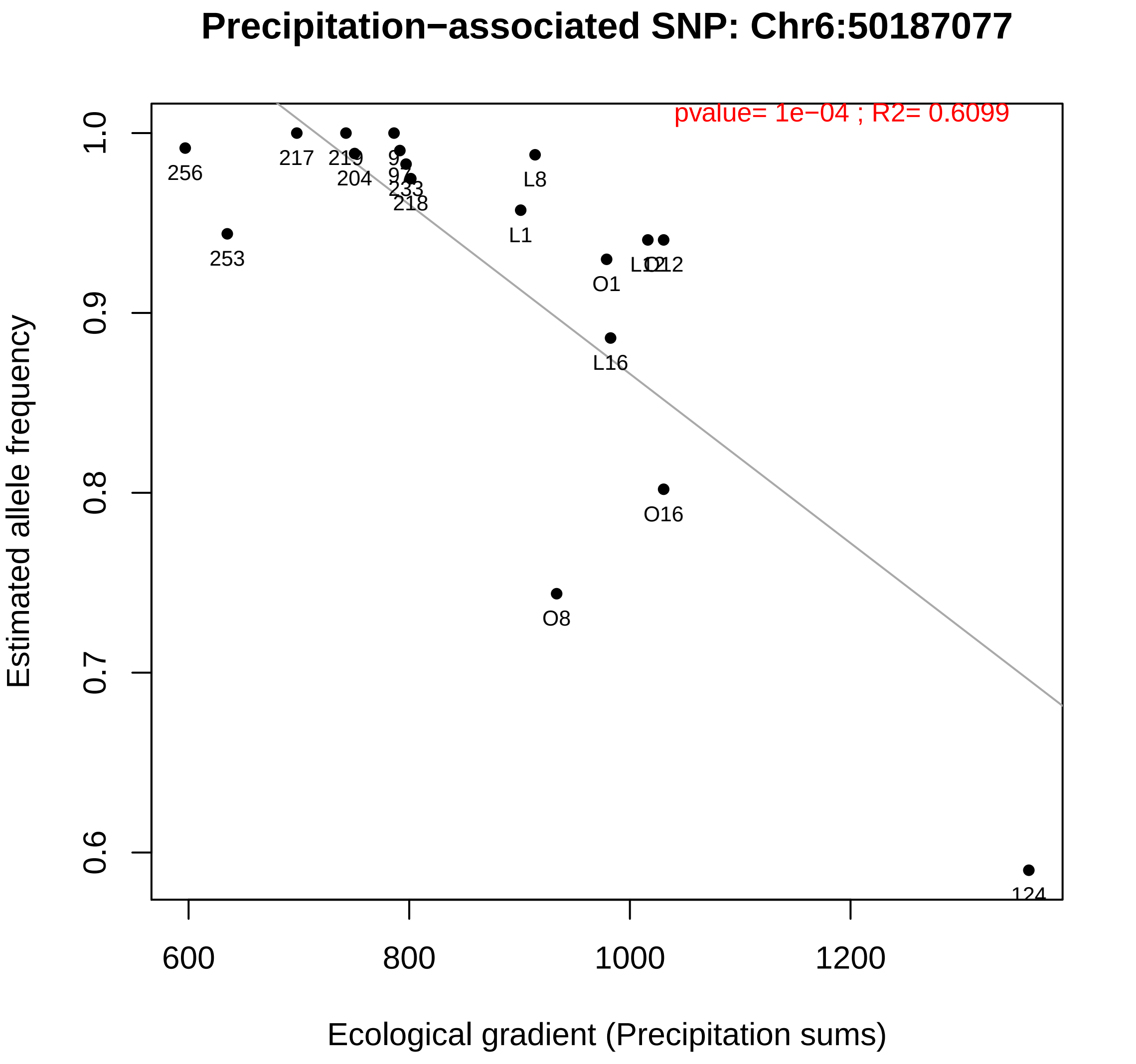
Precipitation−associated SNP: Chr6:50187077
pvalue= 1e−04 ; R2= 0.6099
Estimated allele frequency
Ecological gradient (Precipitation sums)
1.0
0.9
0.8
0.7
0.6
600
800
1000
1200
256
253
217
219
204
9
97
233
218
L8
L1
L12
O12
O1
L16
O16
O8
124

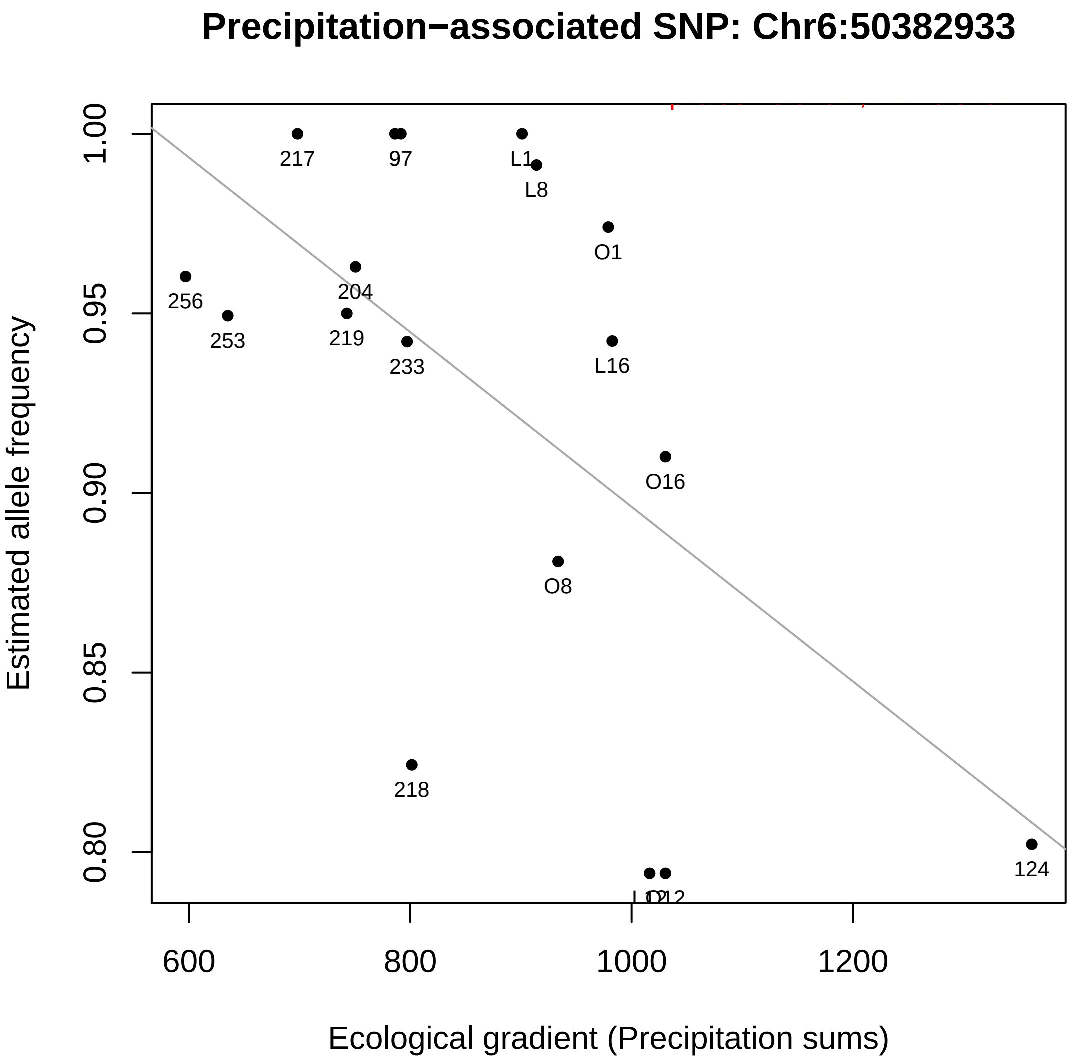

Precipitation−associated SNP: Chr6:50382933
Estimated allele frequency
Ecological gradient (Precipitation sums)
1.00
0.95
0.90
0.85
0.80
600
800
1000
1200
217
97
L1
L8
O1
256
204
253
219
233
L16
O16
O8
218
124

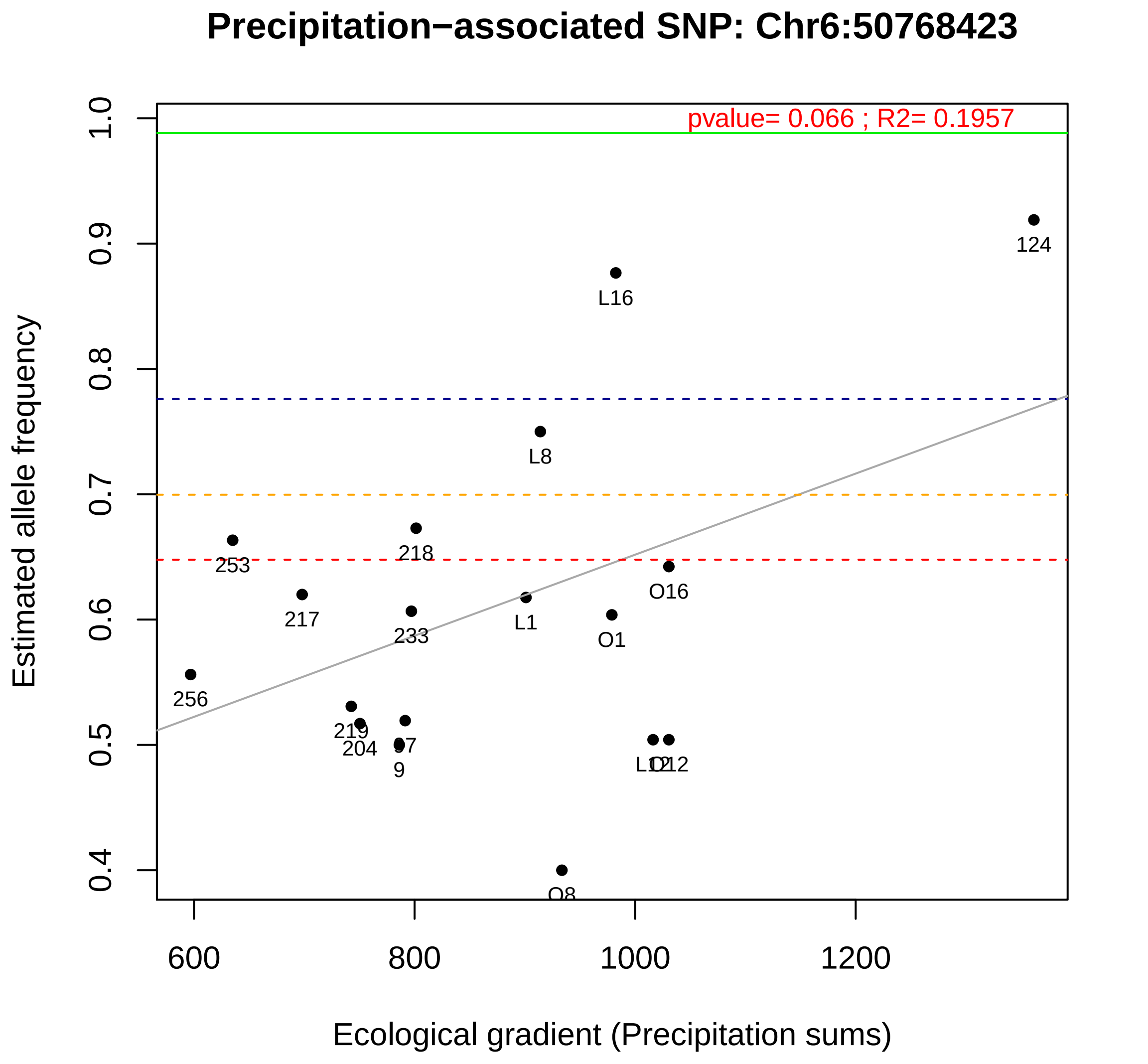
Precipitation−associated SNP: Chr6:50768423
pvalue= 0.066 ; R2= 0.1957
Estimated allele frequency
Ecological gradient (Precipitation sums)
0.4
0.5
0.6
0.7
0.8
0.9
1.0
600
800
1000
1200
124
L16
L8
218
253
O16
217
L1
233
O1
256
219
204
9
O8

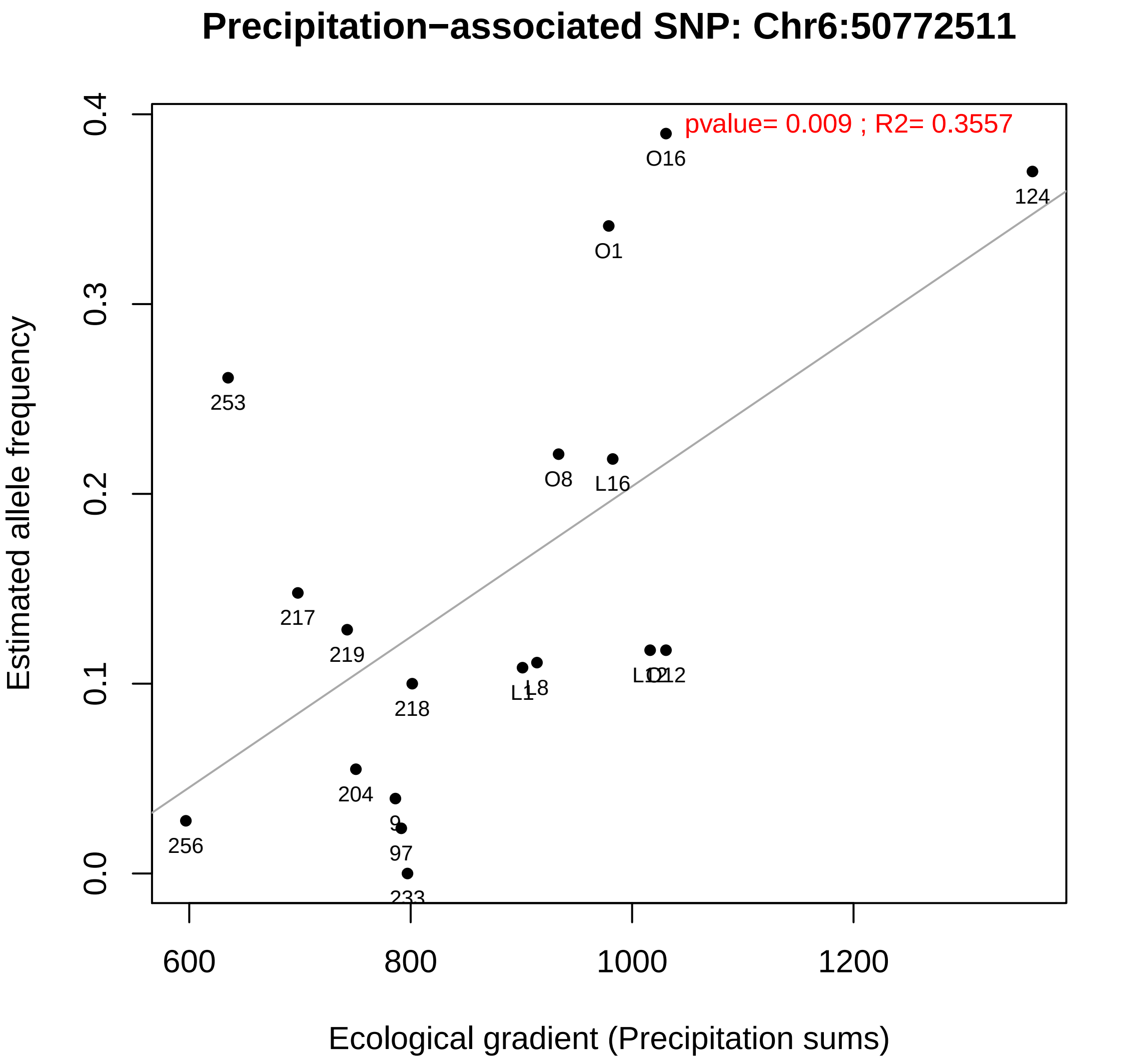
Precipitation−associated SNP: Chr6:50772511
pvalue= 0.009 ; R2= 0.3557
O16
124
O1
253
O8
L16
217
219
218
L1
L8
L12
O12
204
9
256
97
233
0.0
0.1
0.2
0.3
0.4
600
800
1000
1200
Estimated allele frequency
Ecological gradient (Precipitation sums)

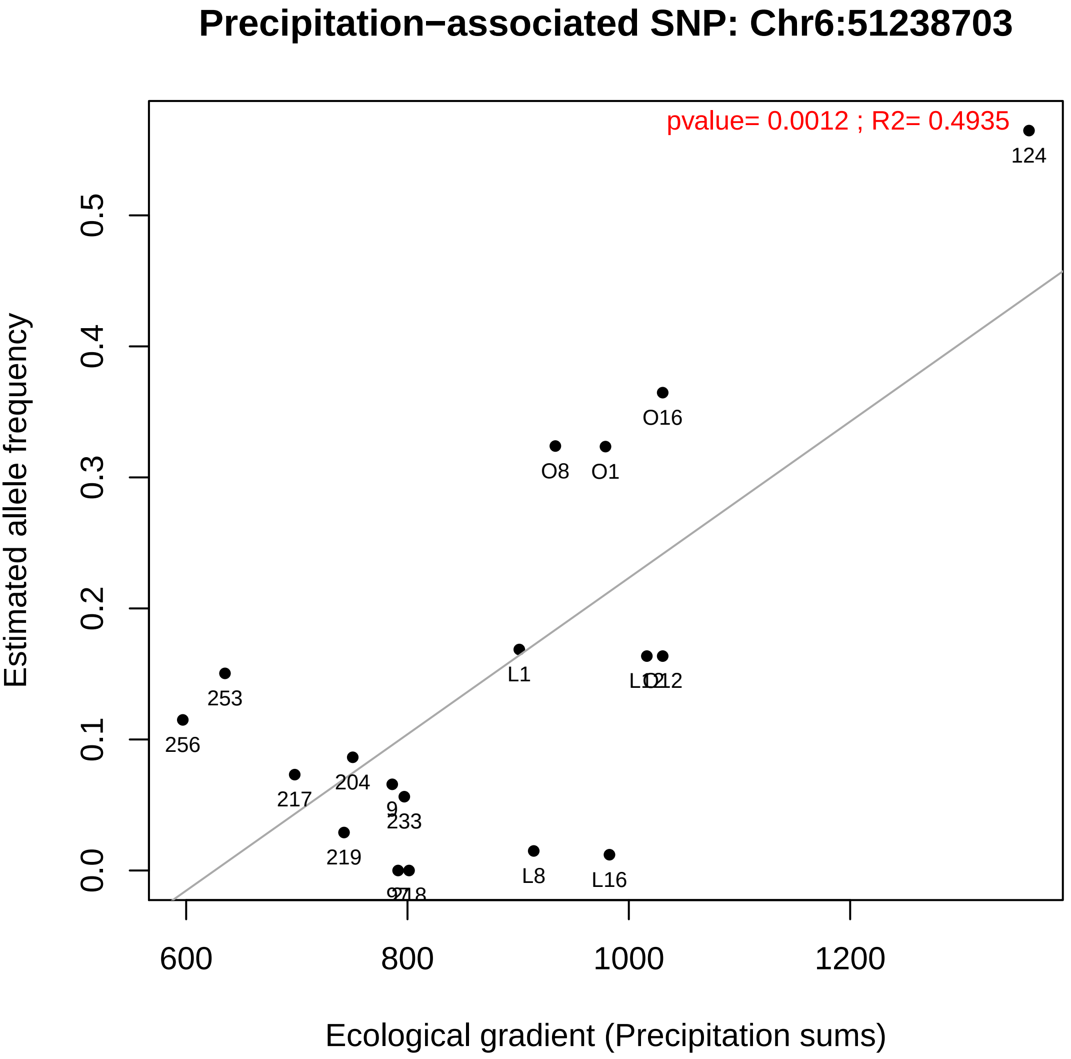
Precipitation−associated SNP: Chr6:51238703
pvalue= 0.0012 ; R2= 0.4935
Estimated allele frequency
0.0
0.1
0.2
0.3
0.4
0.5
Ecological gradient (Precipitation sums)
600
800
1000
1200
124
O16
O8
O1
L1
253
256
204
217
233
219
L8
L16

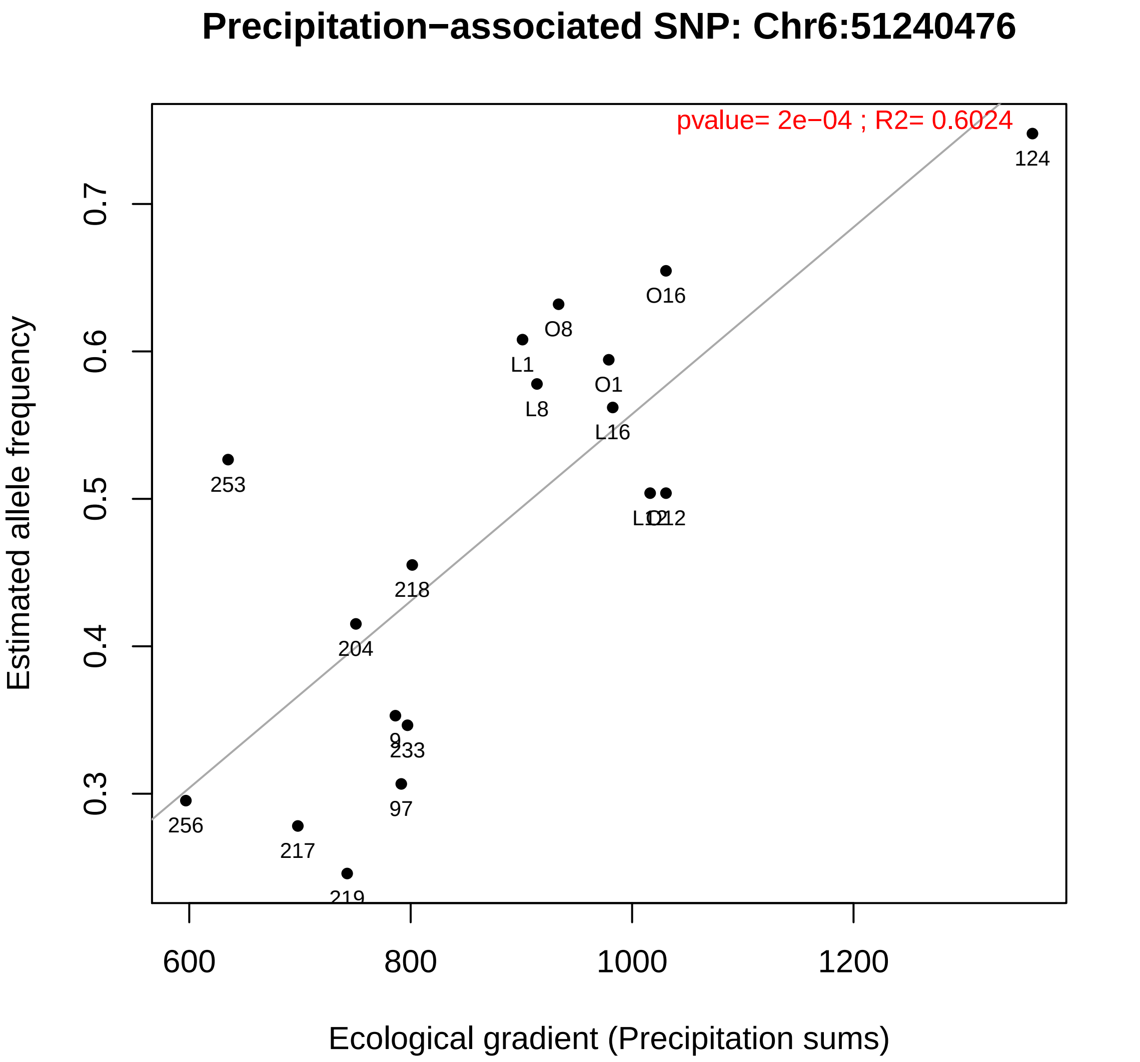

Precipitation−associated SNP: Chr6:51240476
pvalue= 2e−04 ; R2= 0.6024
124
O16
O8
L1
O1
L8
L16
253
L12
O12
218
204
9
233
97
256
217
219
0.3
0.4
0.5
0.6
0.7
600
800
1000
1200
Estimated allele frequency
Ecological gradient (Precipitation sums)

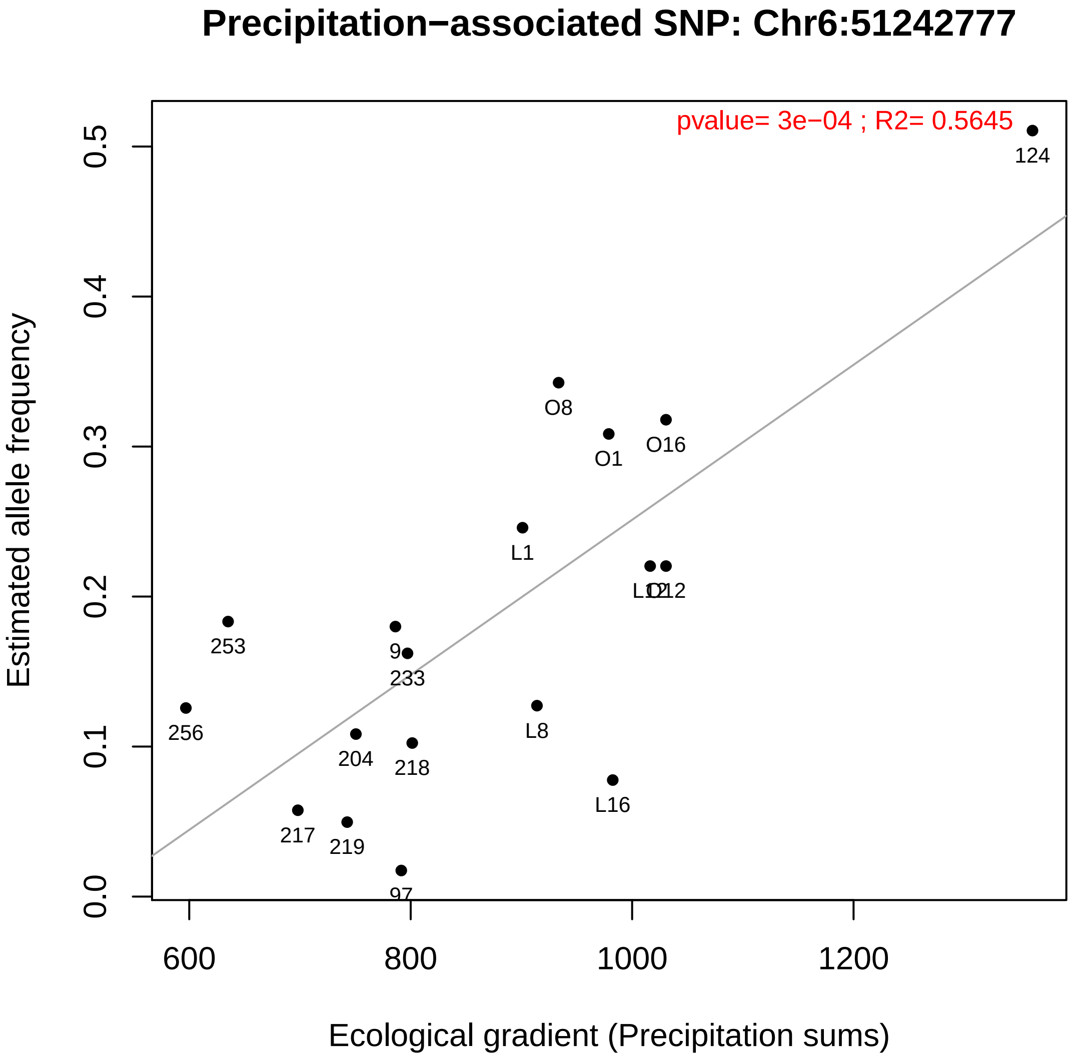
Precipitation−associated SNP: Chr6:51242777
pvalue= 3e−04 ; R2= 0.5645
Estimated allele frequency
Ecological gradient (Precipitation sums)
0.0
0.1
0.2
0.3
0.4
0.5
600
800
1000
1200
124
O8
O1
O16
L1
O12
253
9
233
256
L8
204
218
L16
217
219
97

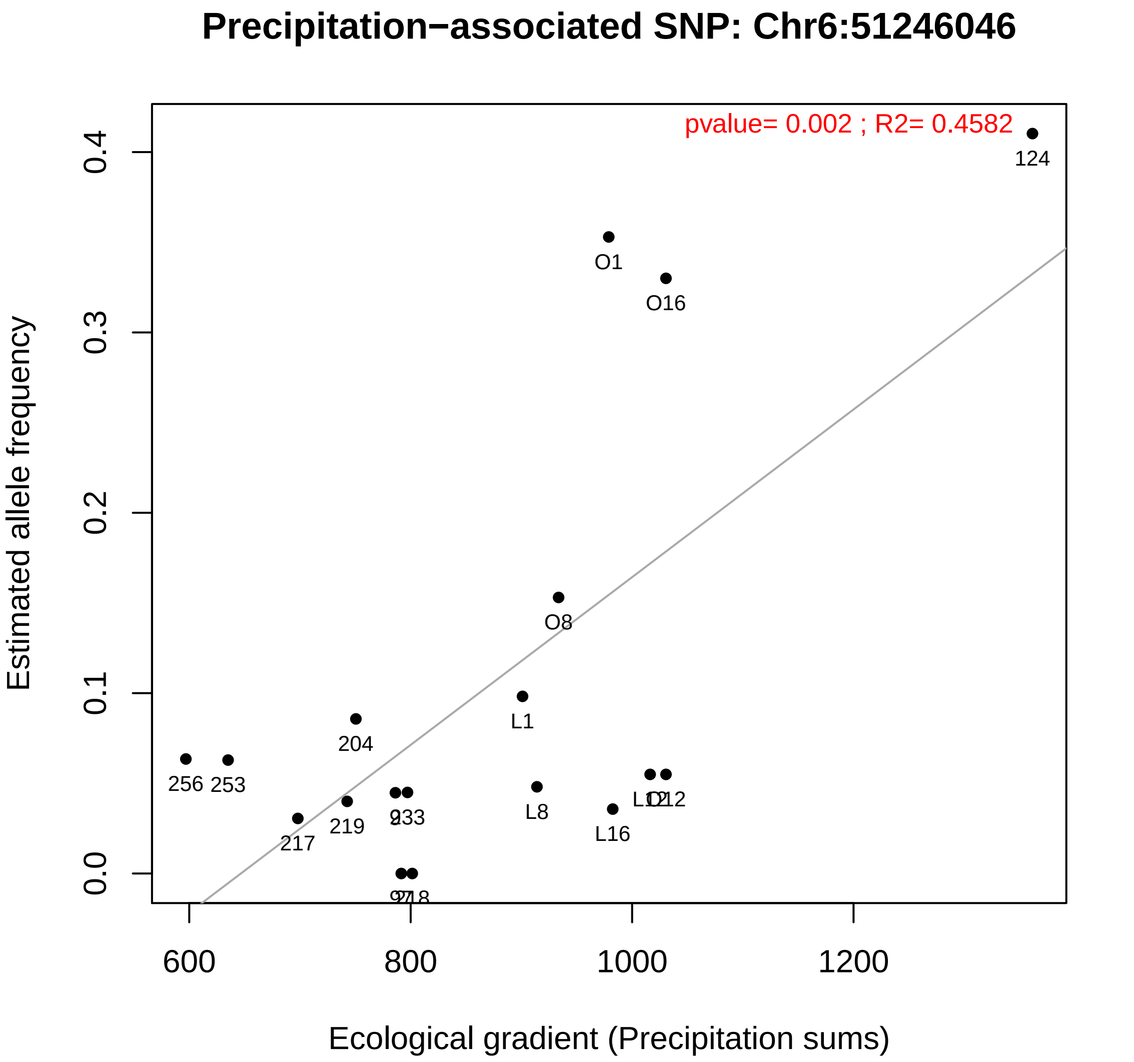
Precipitation−associated SNP: Chr6:51246046
pvalue= 0.002 ; R2= 0.4582
Estimated allele frequency
Ecological gradient (Precipitation sums)
0.0
0.1
0.2
0.3
0.4
600
800
1000
1200
124
O1
O16
O8
L1
204
256
253
217
219
L8
L16

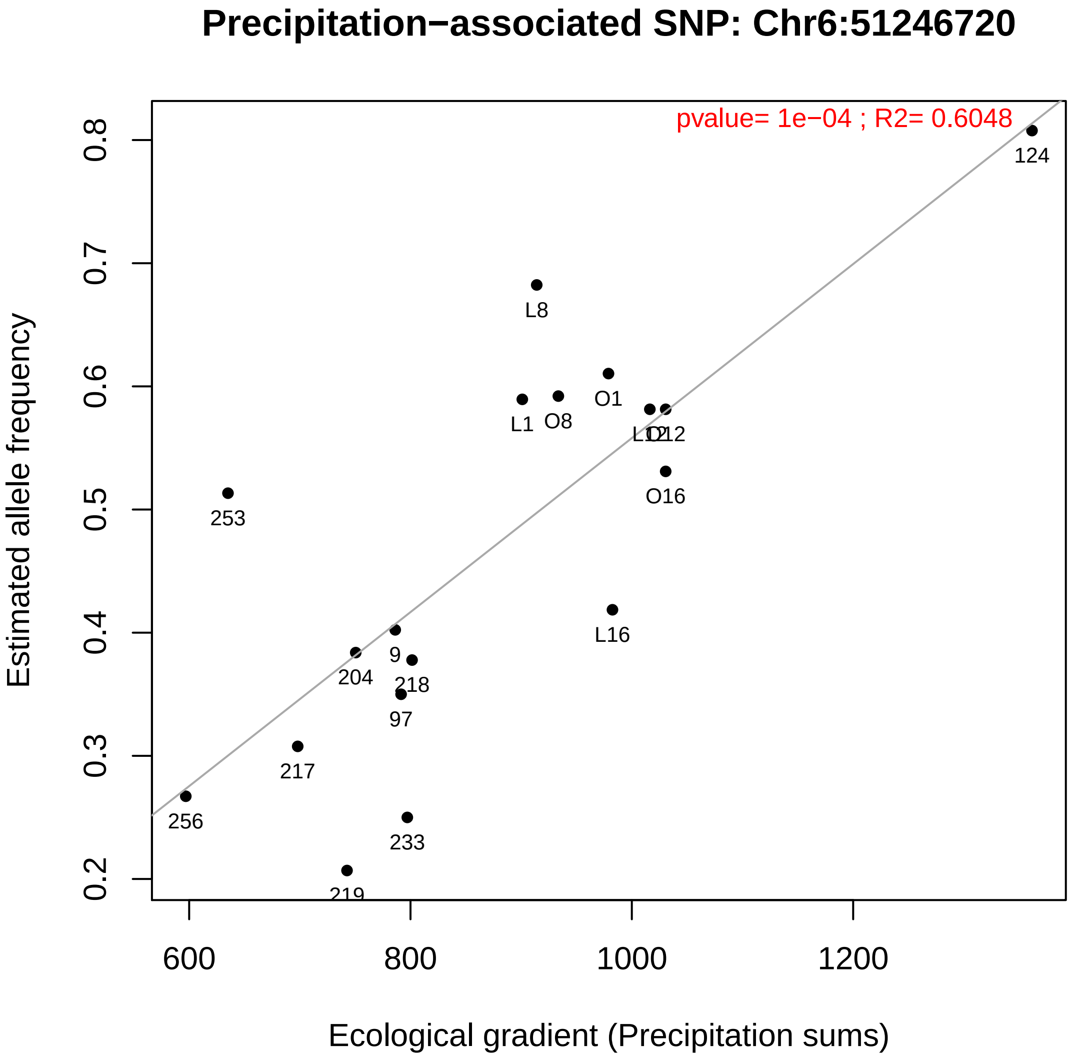
Precipitation−associated SNP: Chr6:51246720
pvalue= 1e−04 ; R2= 0.6048
Estimated allele frequency
Ecological gradient (Precipitation sums)
0.2
0.3
0.4
0.5
0.6
0.7
0.8
600
800
1000
1200
124
L8
O1
L1
O8
L12
O12
O16
253
L16
9
204
218
97
217
256
233
219

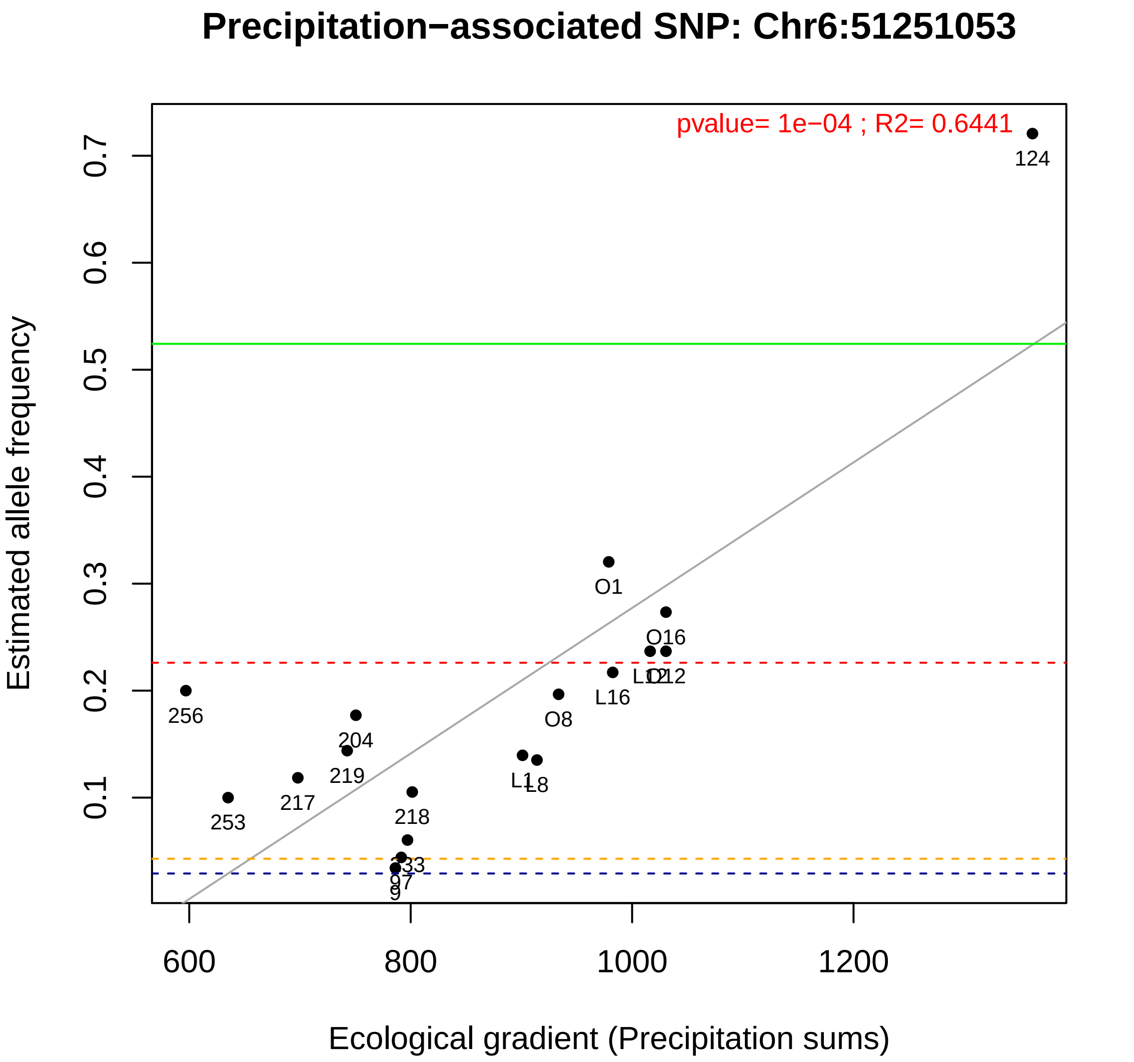

Precipitation−associated SNP: Chr6:51251053
pvalue= 1e−04 ; R2= 0.6441
Estimated allele frequency
0.1
0.2
0.3
0.4
0.5
0.6
0.7
Ecological gradient (Precipitation sums)
600
800
1000
1200
124
O1
O16
L10
O12
L16
O8
256
204
219
217
253
218
L1
L8
233
97
9

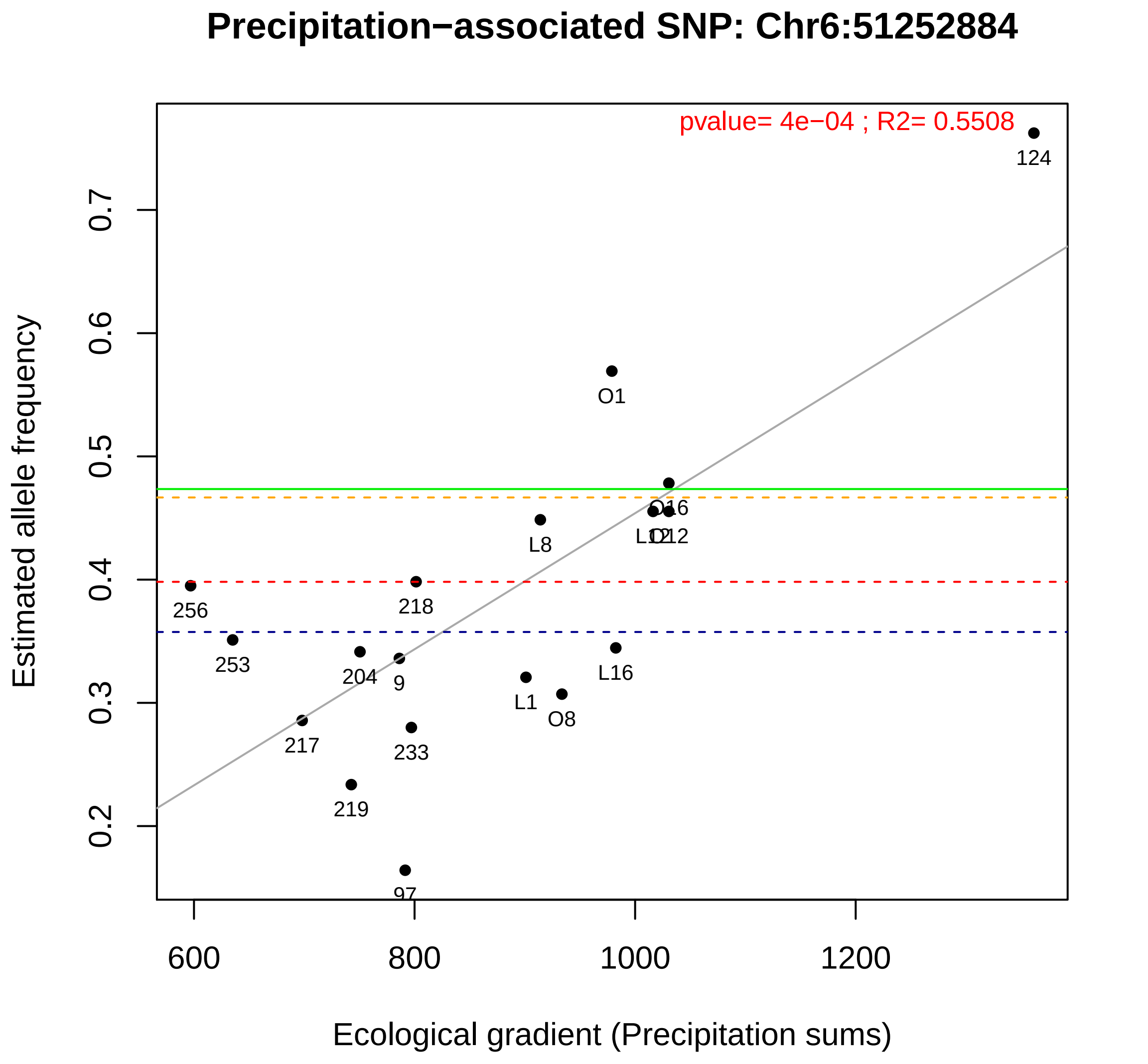

Precipitation−associated SNP: Chr6:51252884
pvalue= 4e−04 ; R2= 0.5508
Estimated allele frequency
Ecological gradient (Precipitation sums)
0.2
0.3
0.4
0.5
0.6
0.7
600
800
1000
1200
124
O1
O16
L8
L12
O12
256
218
253
204
9
L16
L1
O8
217
233
219
97

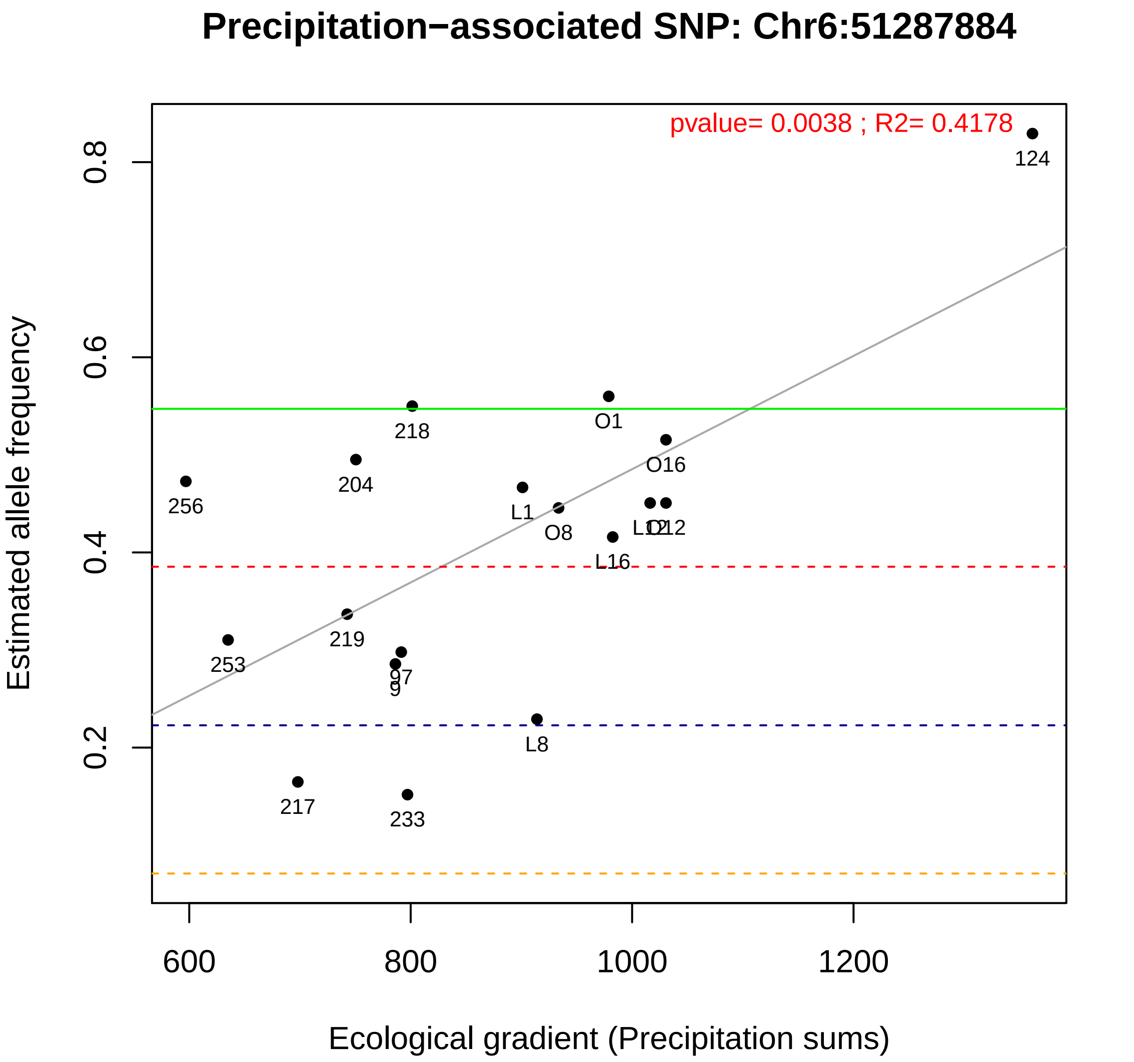

Precipitation−associated SNP: Chr6:51287884
pvalue= 0.0038 ; R2= 0.4178
Estimated allele frequency
Ecological gradient (Precipitation sums)
0.2
0.4
0.6
0.8
600
800
1000
1200
124
218
O1
O16
204
256
L1
O8
O12
L16
219
253
97
9
L8
217
233

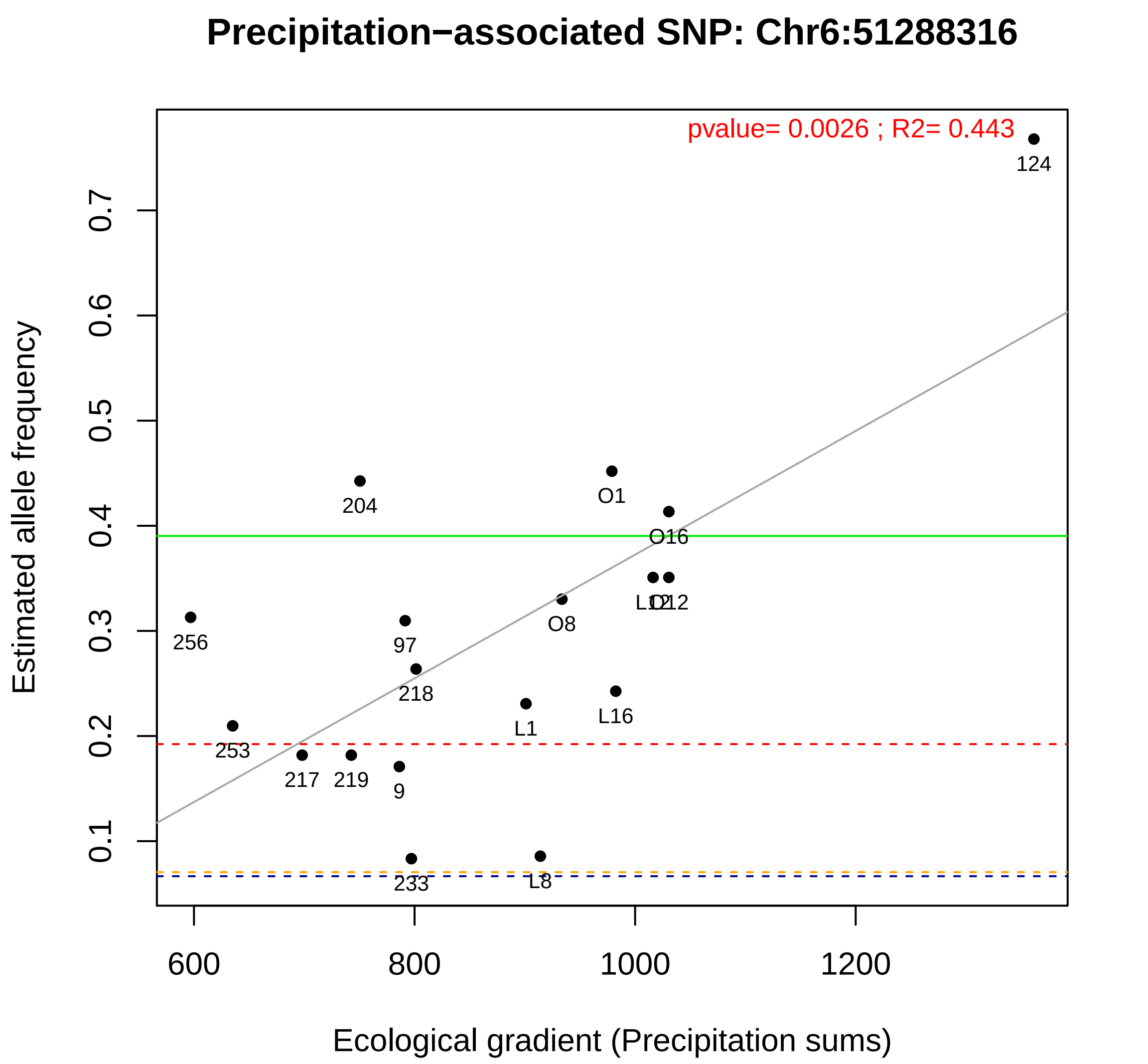

Precipitation−associated SNP: Chr6:51288316
pvalue= 0.0026 ; R2= 0.443
Estimated allele frequency
0.1
0.2
0.3
0.4
0.5
0.6
0.7
600
800
1000
1200
Ecological gradient (Precipitation sums)
124
204
O1
O16
L12
O12
256
97
O8
218
L1
L16
253
217
219
9
233
L8

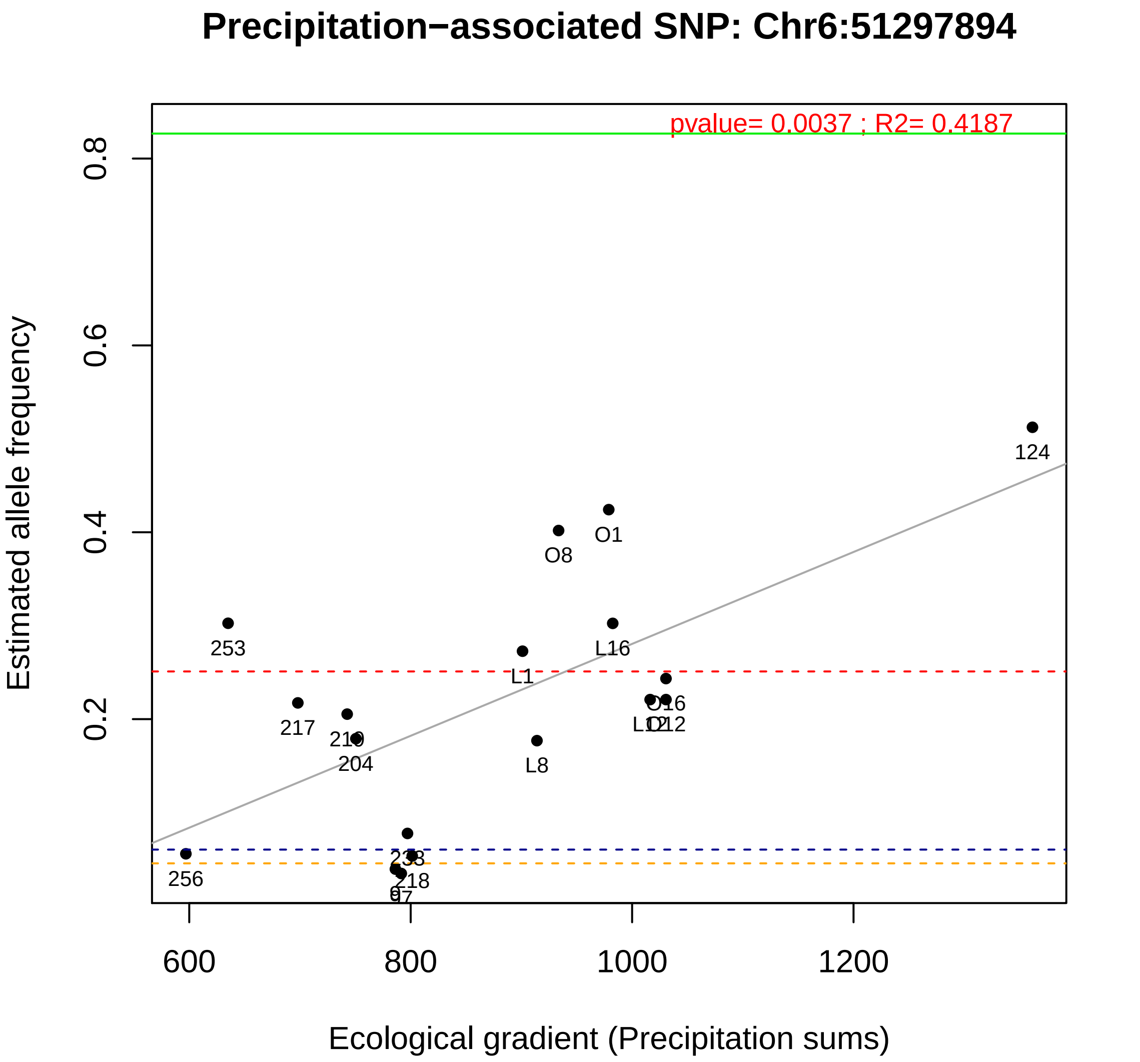

Precipitation−associated SNP: Chr6:51297894
pvalue= 0.0037 ; R2= 0.4187
Estimated allele frequency
0.2
0.4
0.6
0.8
600
800
1000
1200
Ecological gradient (Precipitation sums)
124
O1
O8
253
L16
L1
O16
L12
O12
217
210
204
L8
233
218
256
9
97

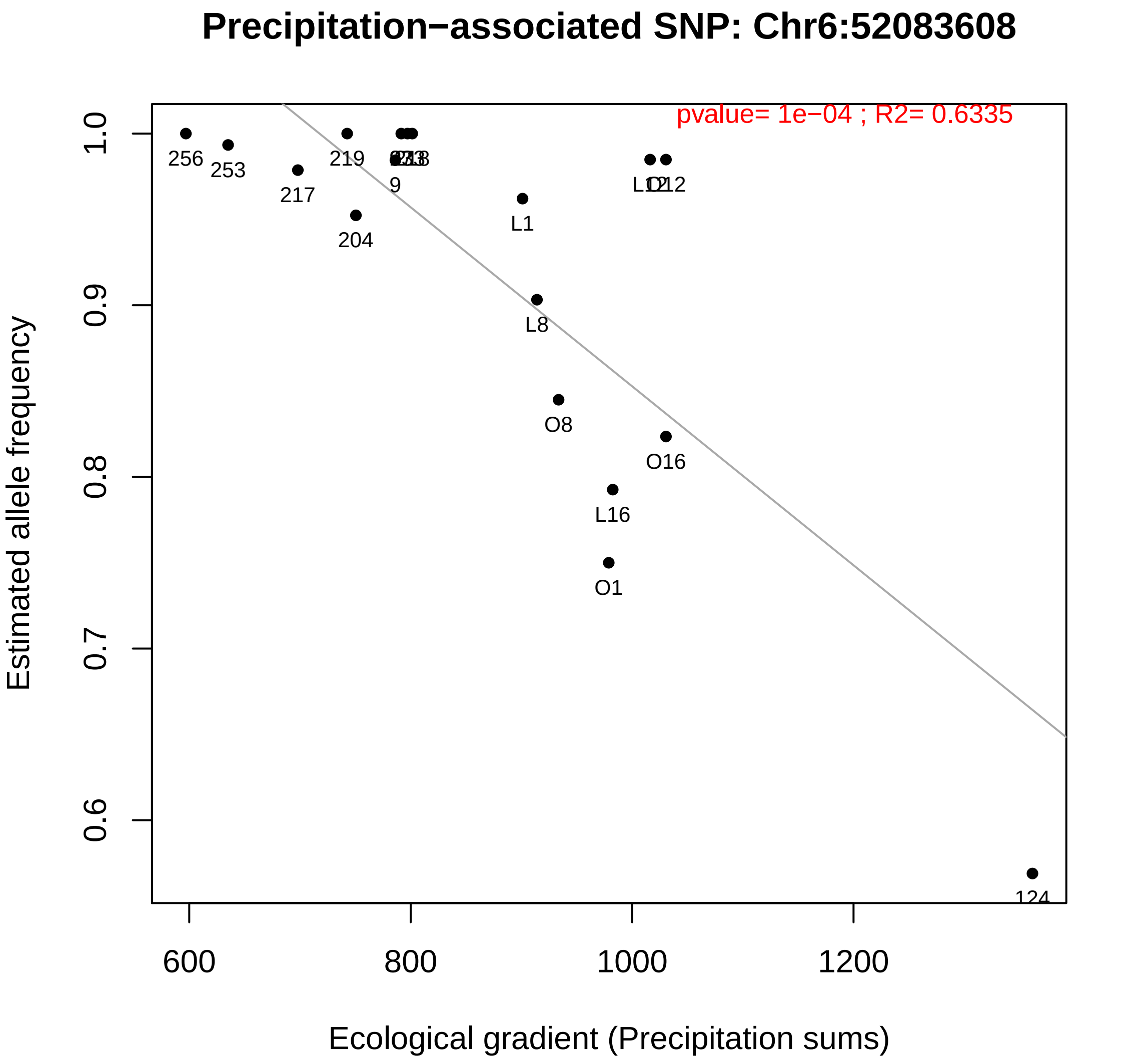
Precipitation−associated SNP: Chr6:52083608
pvalue= 1e−04 ; R2= 0.6335
Estimated allele frequency
Ecological gradient (Precipitation sums)
1.0
0.9
0.8
0.7
0.6
600
800
1000
1200
256
253
217
219
204
9
L1
L8
O8
O16
L16
O1
124

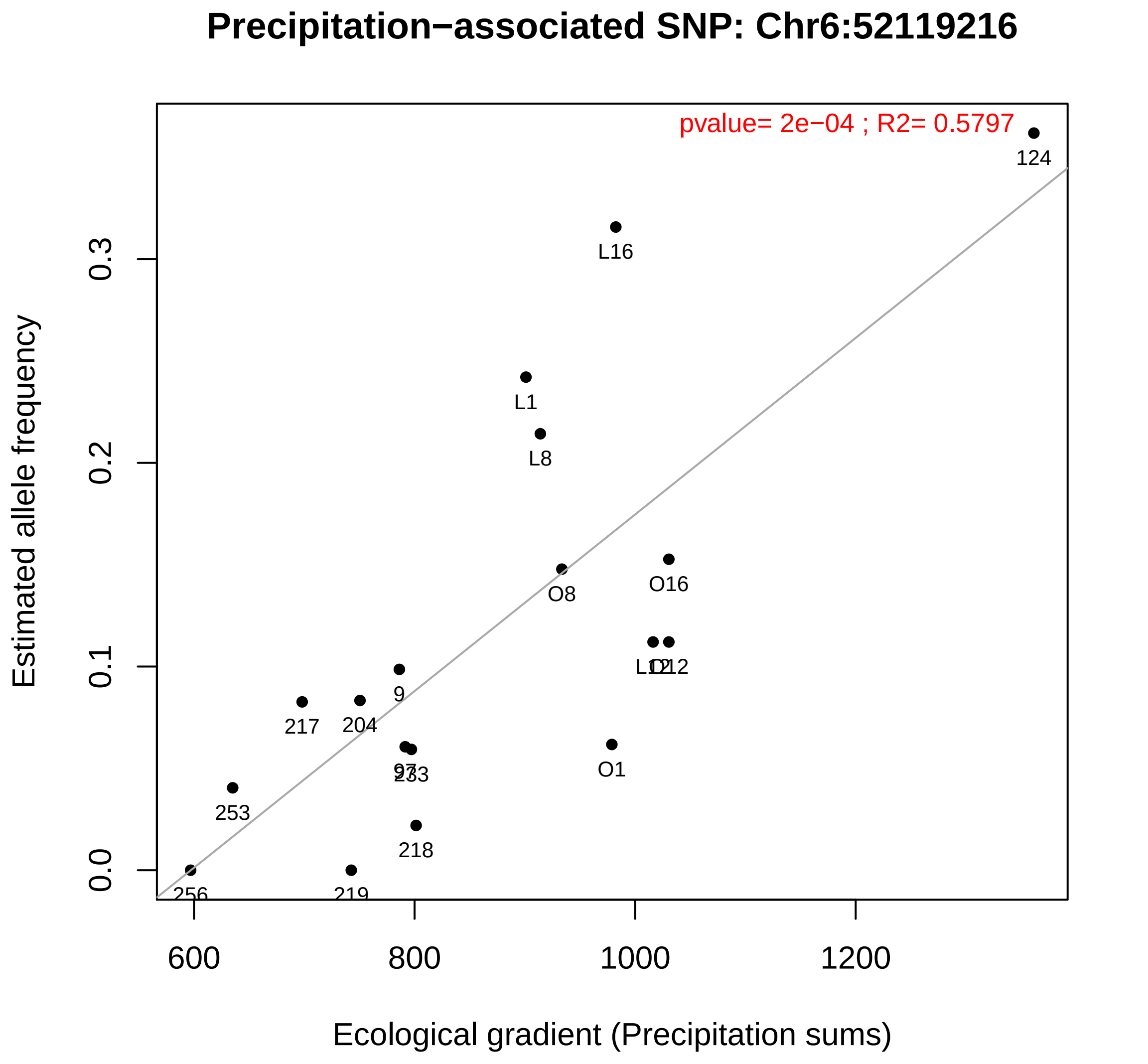

Precipitation−associated SNP: Chr6:52119216
pvalue= 2e−04 ; R2= 0.5797
124
L16
L1
L8
O8
O16
L10
O12
9
217
204
97
233
O1
253
218
256
219
0.0
0.1
0.2
0.3
600
800
1000
1200
Estimated allele frequency
Ecological gradient (Precipitation sums)

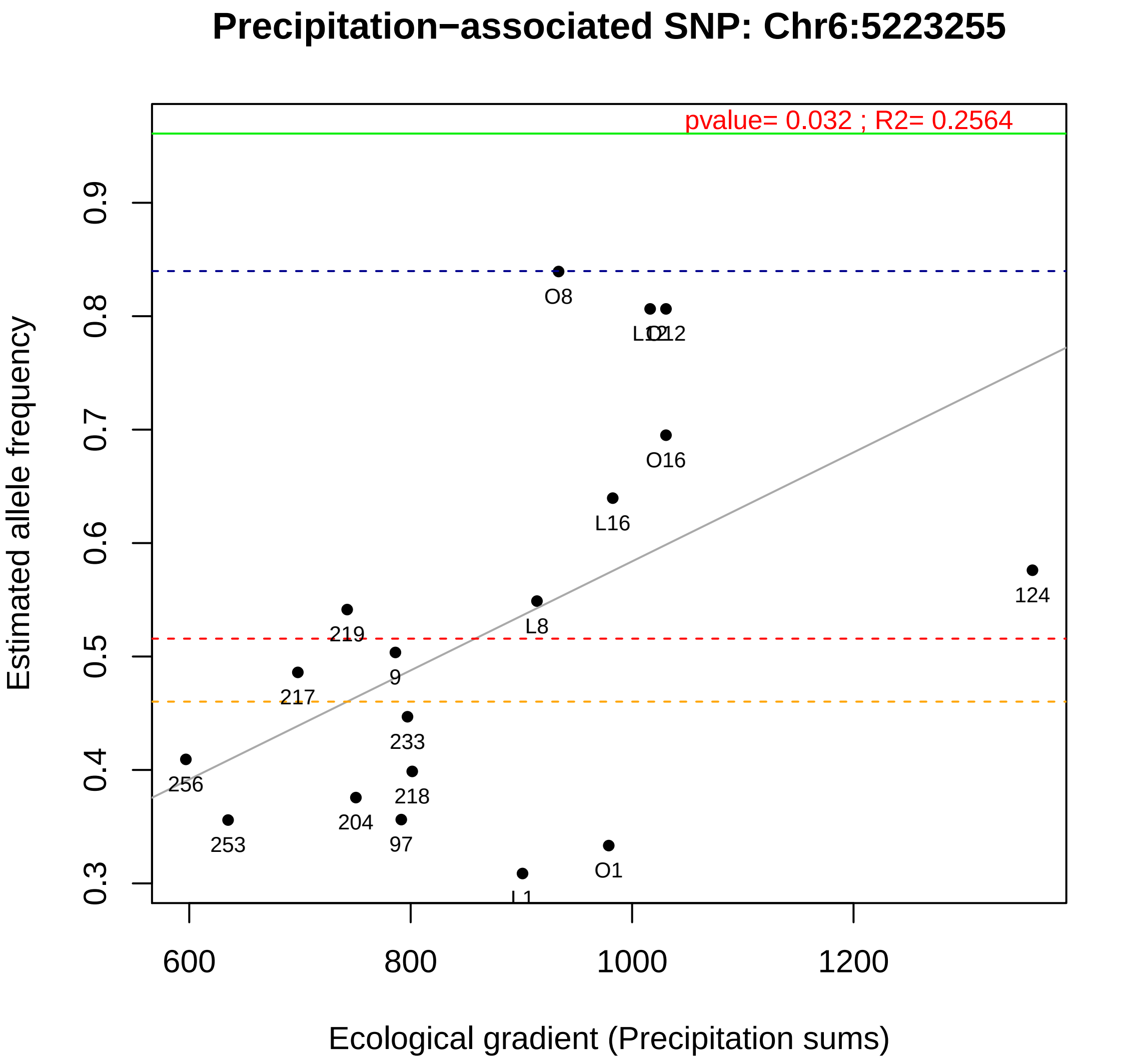
Precipitation−associated SNP: Chr6:5223255
pvalue= 0.032 ; R2= 0.2564
Estimated allele frequency
0.3
0.4
0.5
0.6
0.7
0.8
0.9
600
800
1000
1200
Ecological gradient (Precipitation sums)
O8
L12
O12
O16
L16
124
L8
219
9
217
233
256
218
204
253
97
O1
L1

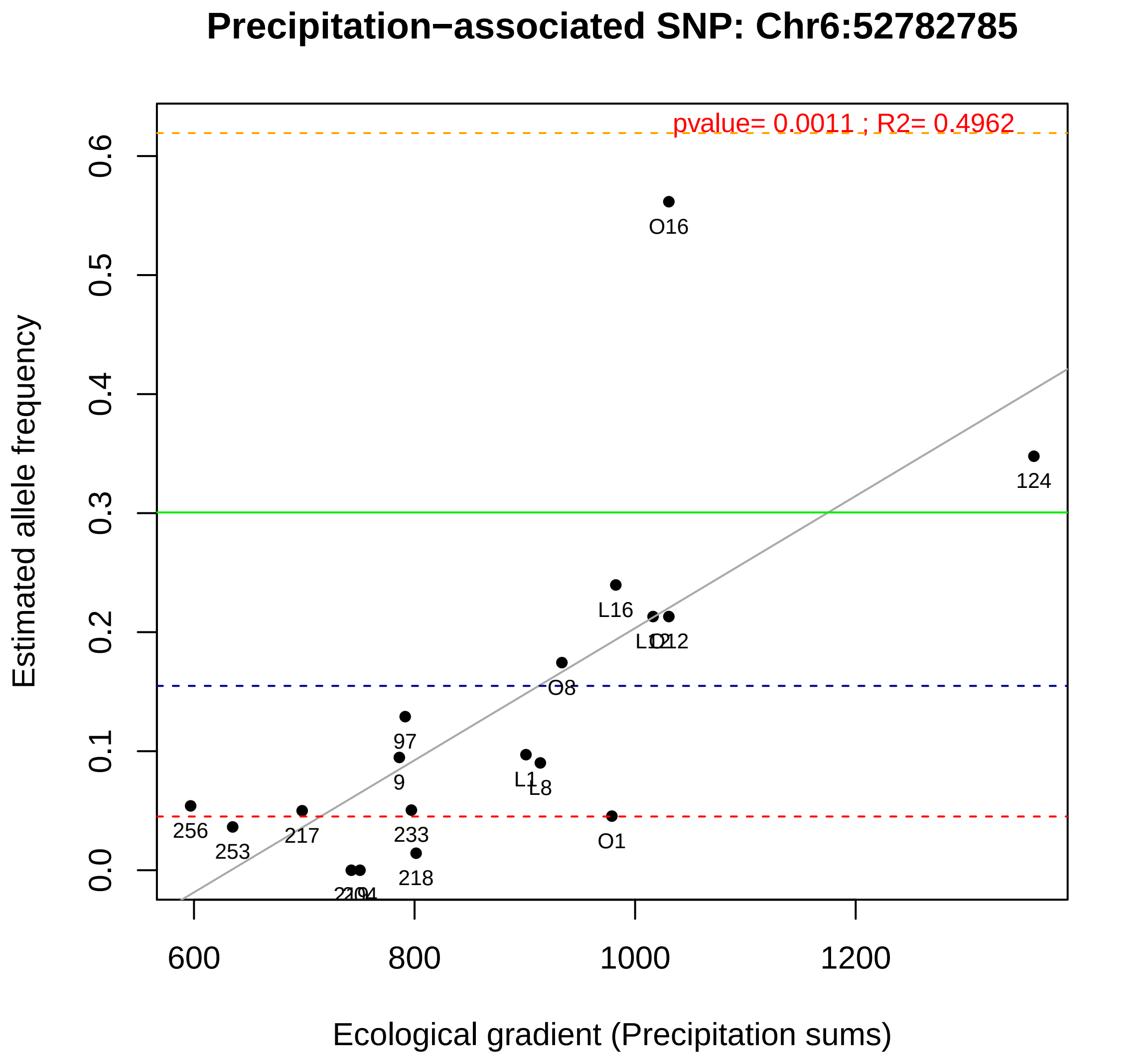

Precipitation−associated SNP: Chr6:52782785
pvalue= 0.0011 ; R2= 0.4962
Estimated allele frequency
0.0
0.1
0.2
0.3
0.4
0.5
0.6
O16
124
L16
L12
O12
O8
97
9
L1
L8
256
253
217
233
O1
218
210
214
600
800
1000
1200
Ecological gradient (Precipitation sums)

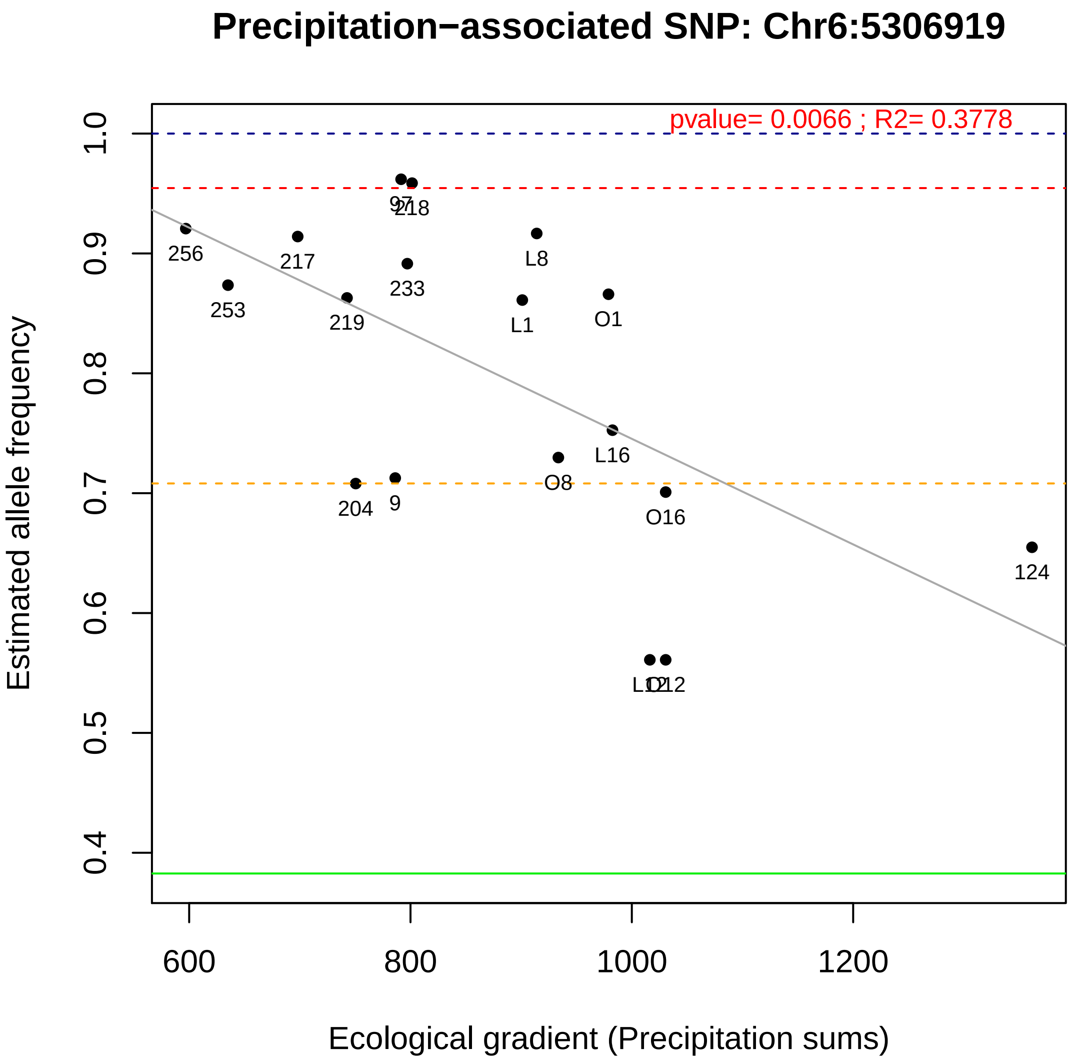

Precipitation−associated SNP: Chr6:5306919
pvalue= 0.0066 ; R2= 0.3778
Estimated allele frequency
Ecological gradient (Precipitation sums)
0.4
0.5
0.6
0.7
0.8
0.9
1.0
600
800
1000
1200
256
253
217
219
97
218
233
204
9
L8
L1
O1
O8
L16
O16
L12
O12
124

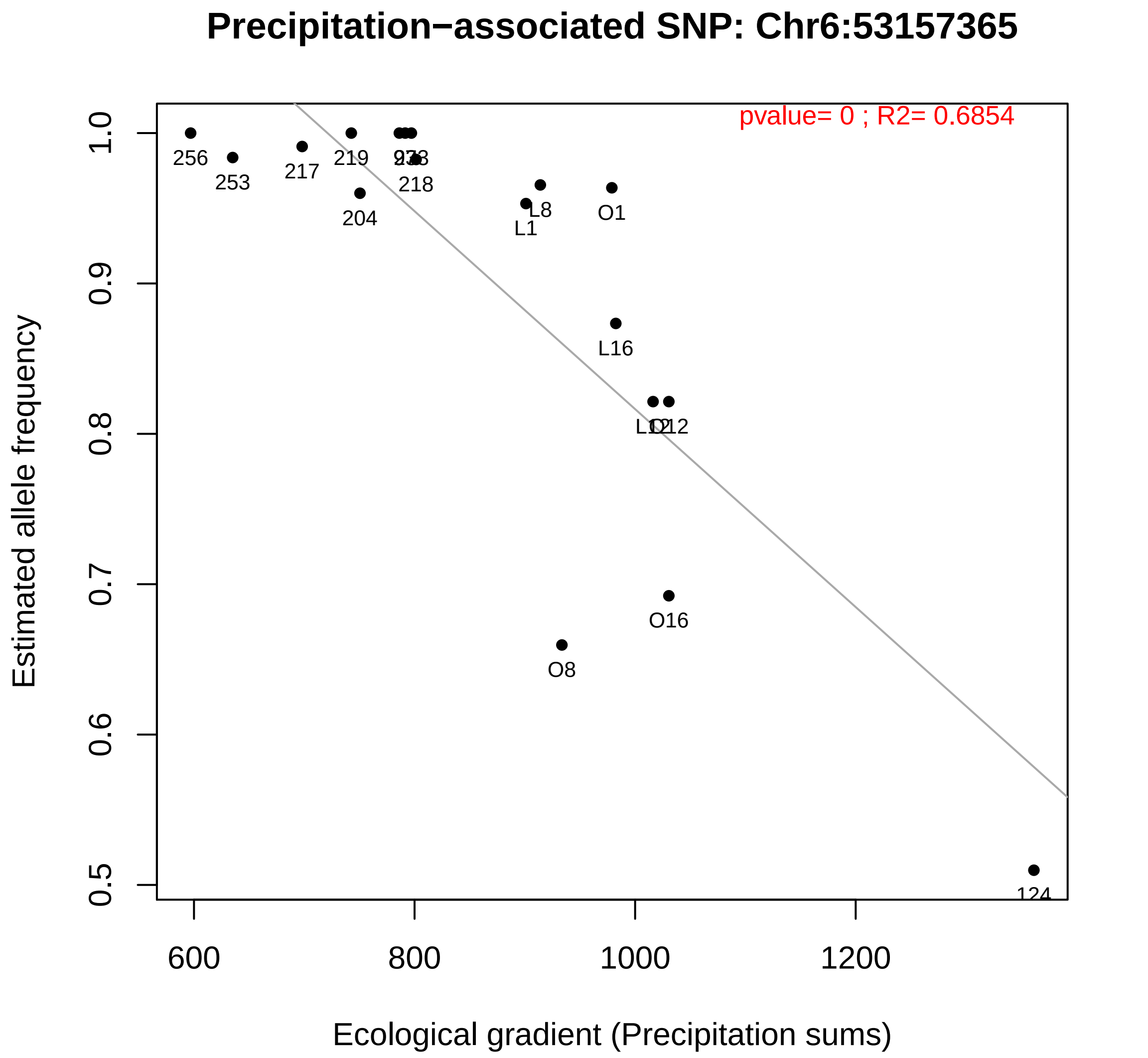

Precipitation−associated SNP: Chr6:53157365
pvalue= 0 ; R2= 0.6854
Estimated allele frequency
Ecological gradient (Precipitation sums)
0.5
0.6
0.7
0.8
0.9
1.0
600
800
1000
1200
256
253
217
219
204
218
L8
L1
O1
L16
O12
O16
O8
124

**Precipitation−associated SNP: Chr6:53272764**

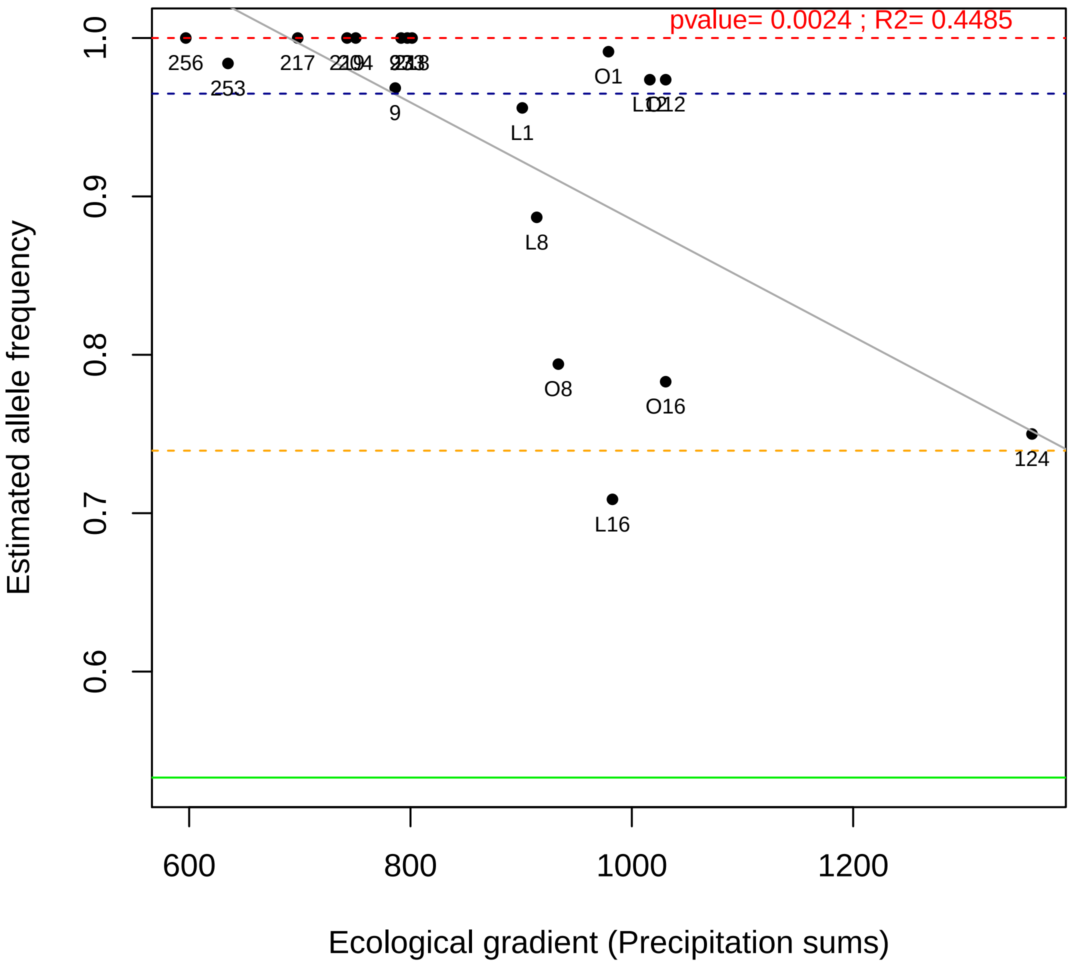

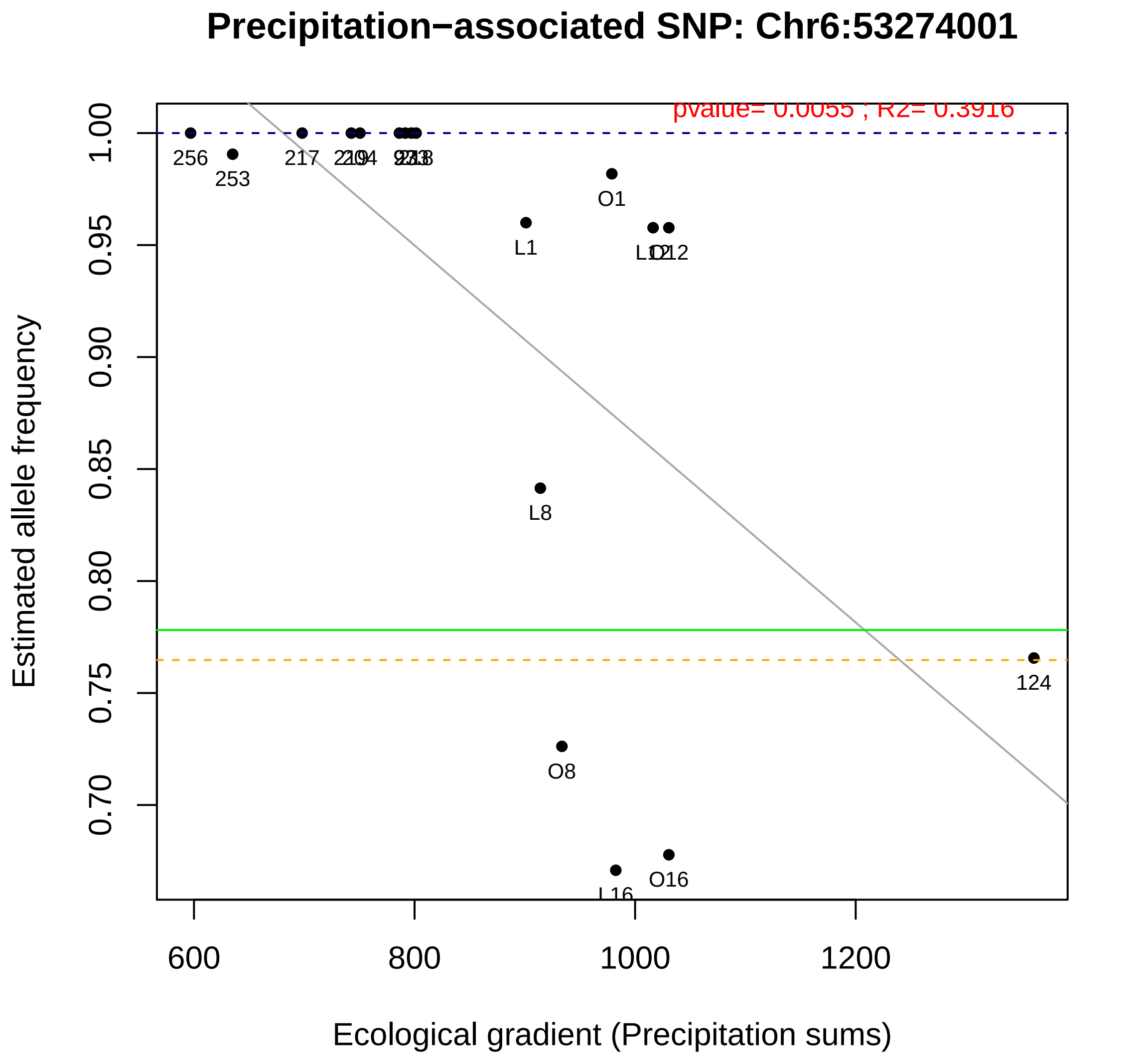
Precipitation−associated SNP: Chr6:53274001
pvalue= 0.0055 ; R2= 0.3916
Estimated allele frequency
Ecological gradient (Precipitation sums)
1.00
0.95
0.90
0.85
0.80
0.75
0.70
600
800
1000
1200
256
253
217
O1
L1
L8
124
O8
O16
L16

**Precipitation−associated SNP: Chr6:53276034**

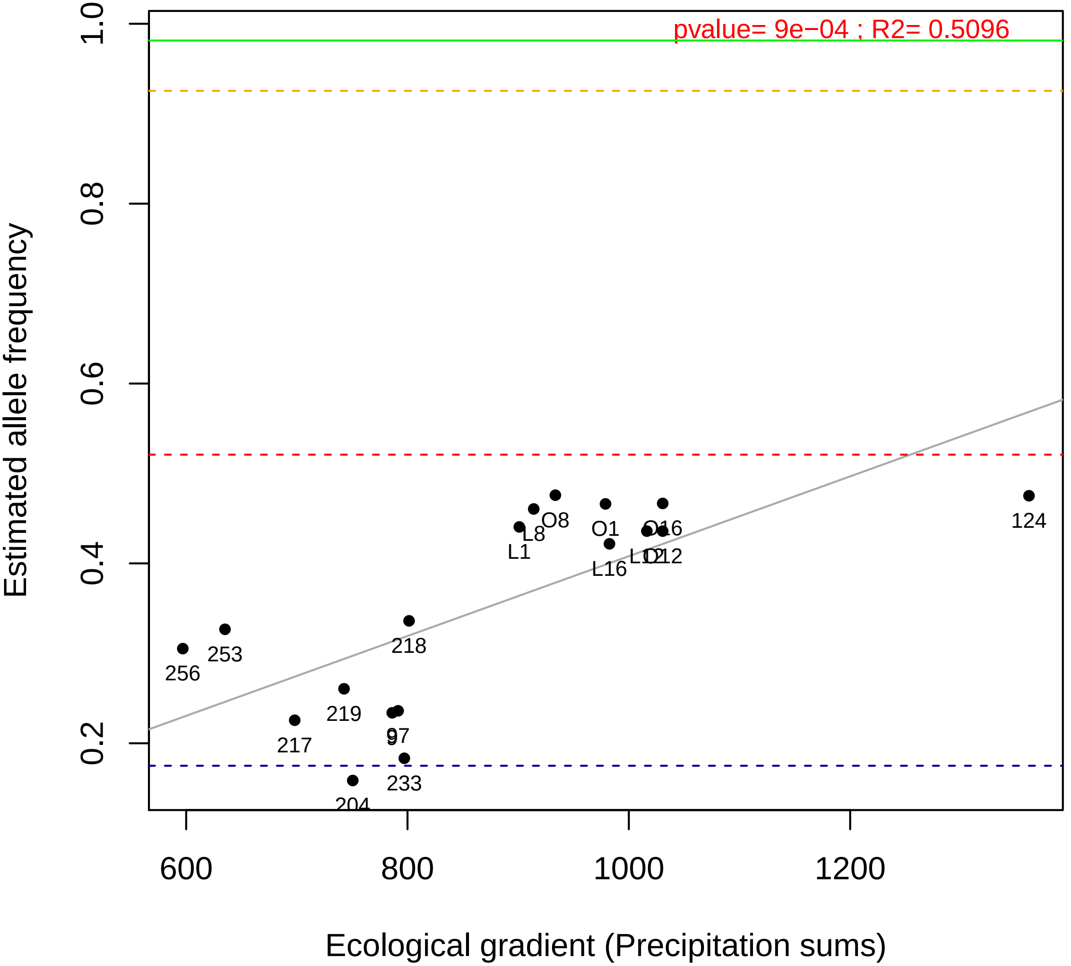

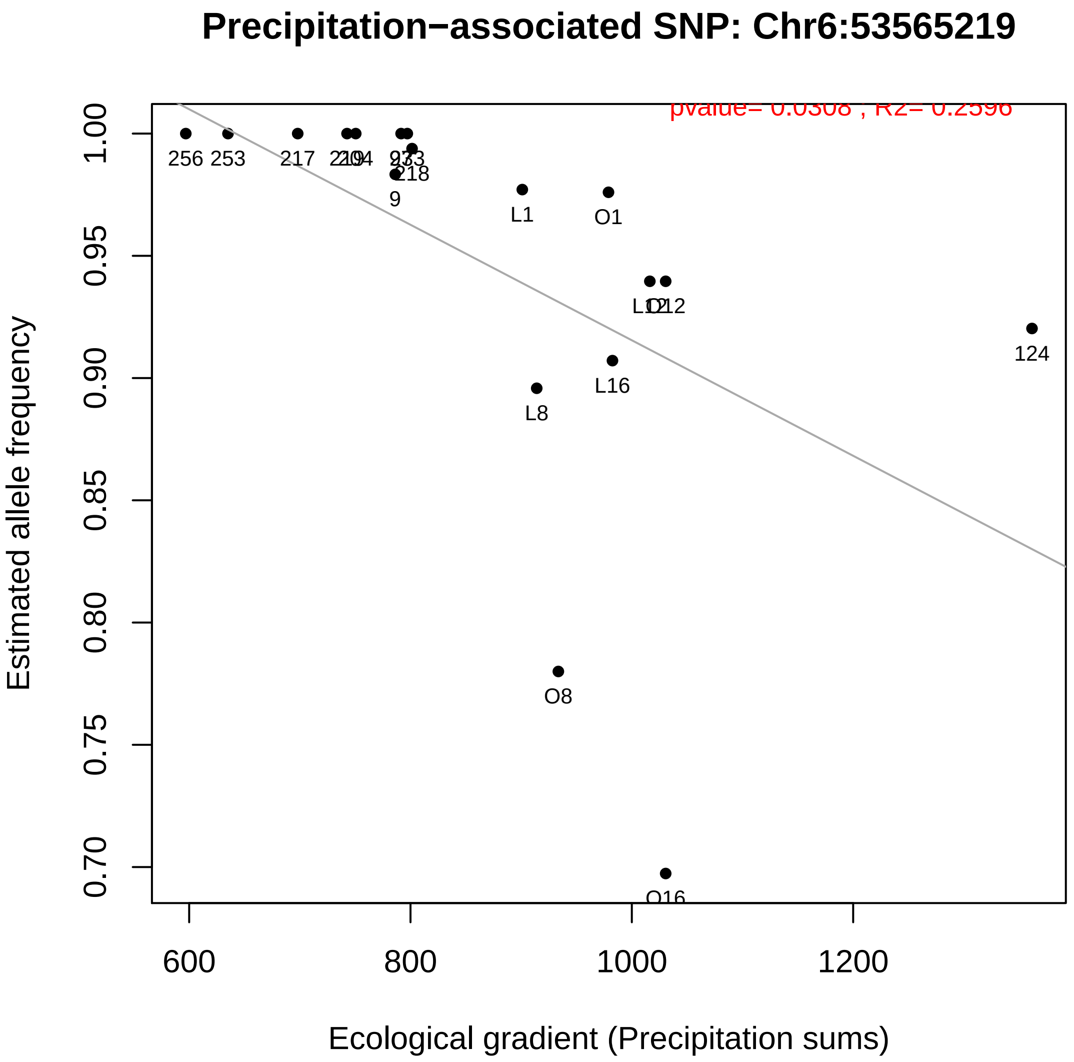
Precipitation−associated SNP: Chr6:53565219
pvalue= 0.0308 , R2= 0.2596
Estimated allele frequency
Ecological gradient (Precipitation sums)
1.00
0.95
0.90
0.85
0.80
0.75
0.70
600
800
1000
1200
256
253
217
218
9
L1
O1
124
L16
L8
O8

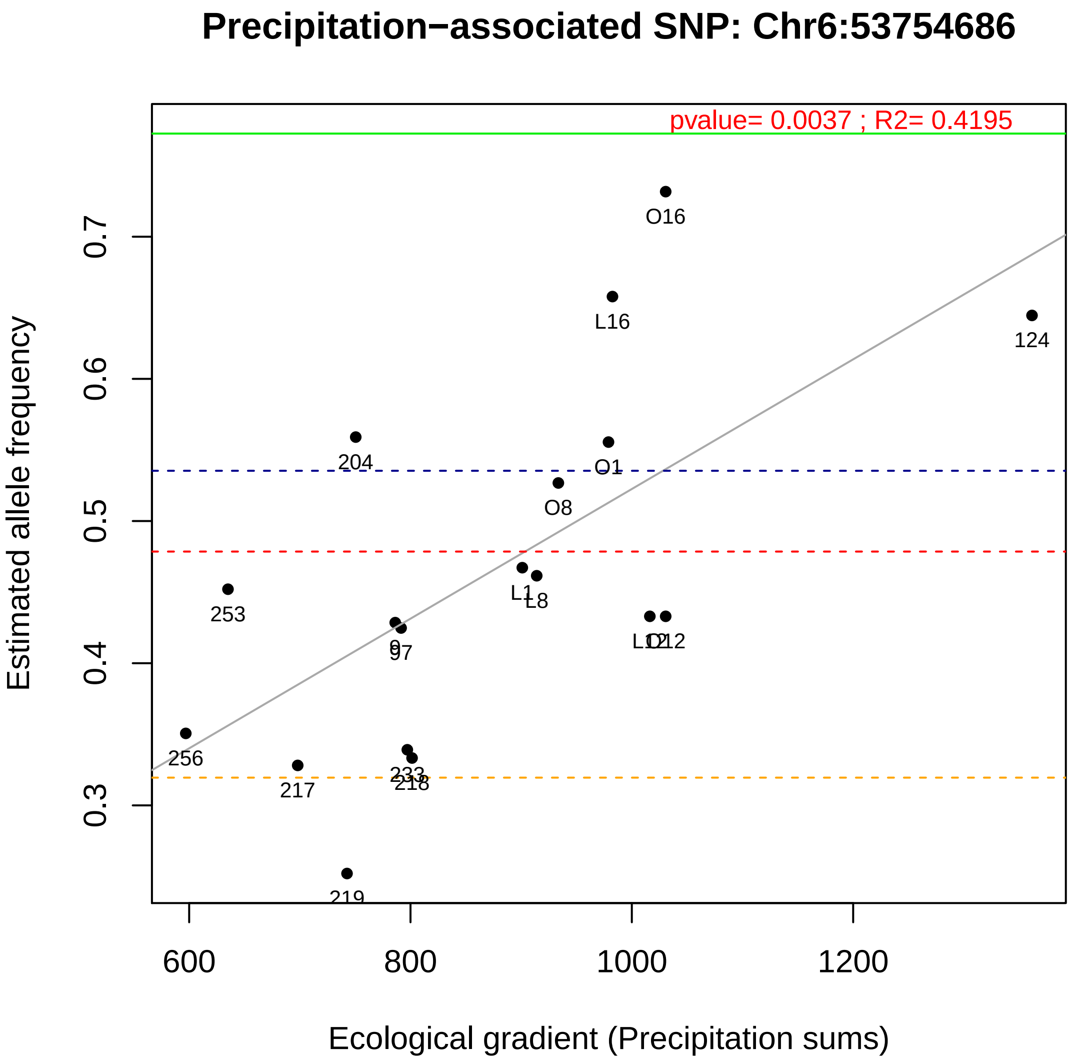

Precipitation−associated SNP: Chr6:53754686
pvalue= 0.0037 ; R2= 0.4195
Estimated allele frequency
Ecological gradient (Precipitation sums)
0.3
0.4
0.5
0.6
0.7
600
800
1000
1200
O16
L16
124
204
O1
O8
L1
L8
253
9
97
L12
O12
256
217
233
218
219

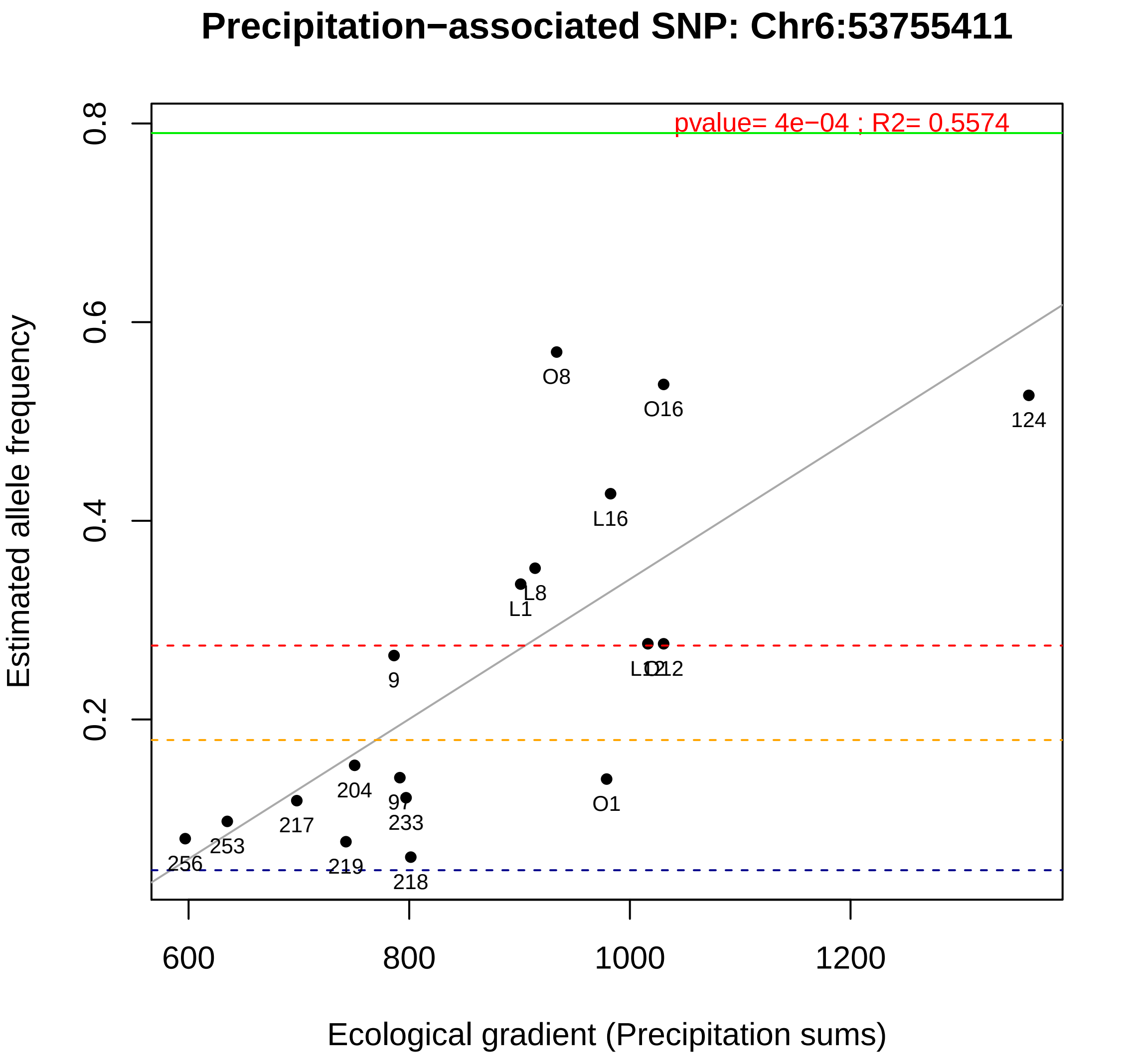
Precipitation−associated SNP: Chr6:53755411
pvalue= 4e−04 ; R2= 0.5574
Estimated allele frequency
0.2
0.4
0.6
0.8
Ecological gradient (Precipitation sums)
600
800
1000
1200
O8
O16
124
L16
L8
L1
9
L10
O12
204
97
233
O1
217
253
256
219
218

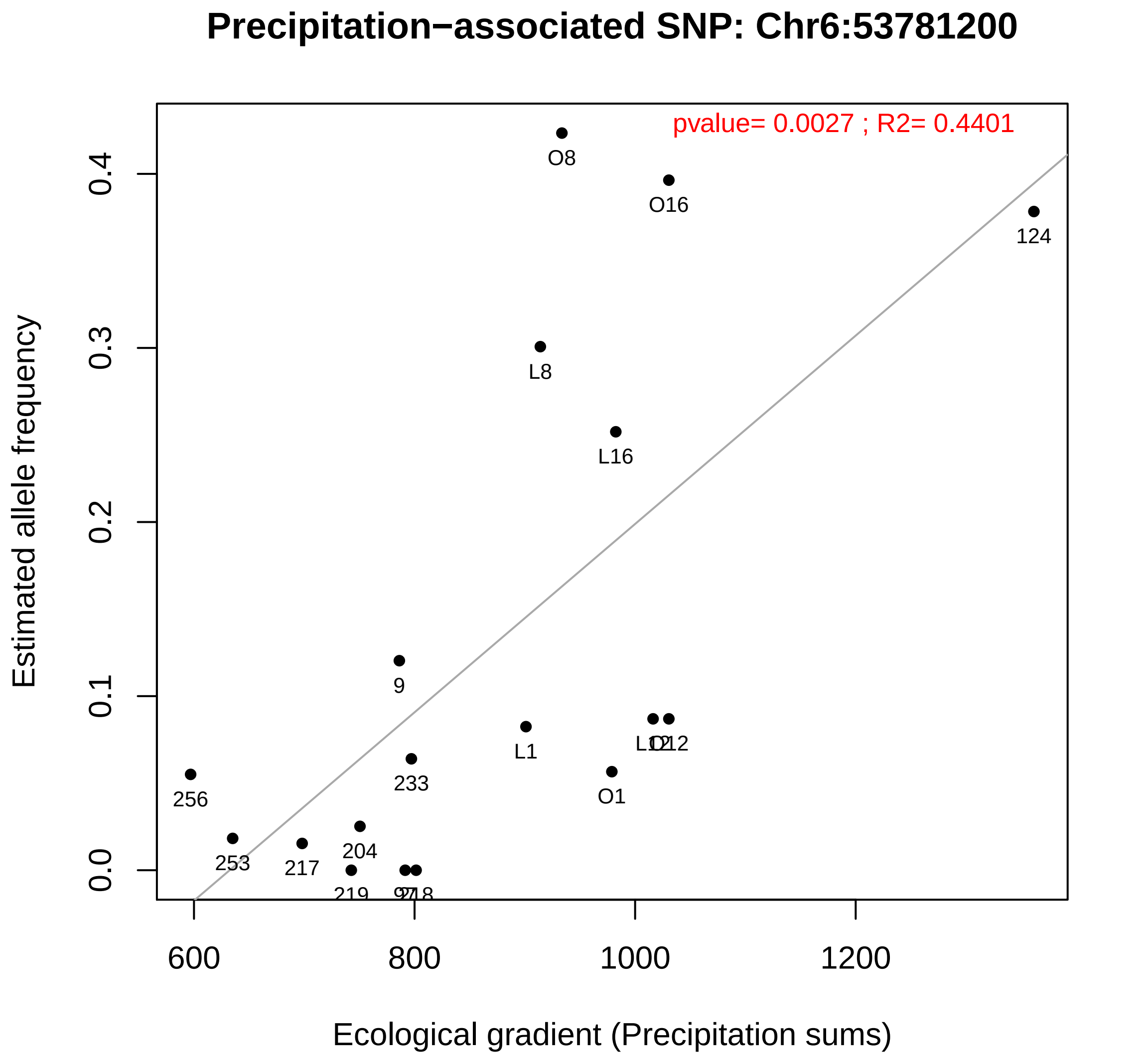
Precipitation−associated SNP: Chr6:53781200
pvalue= 0.0027 ; R2= 0.4401
Estimated allele frequency
Ecological gradient (Precipitation sums)
0.0
0.1
0.2
0.3
0.4
600
800
1000
1200
O8
O16
124
L8
L16
9
L1
O12
233
O1
256
253
217
204
219

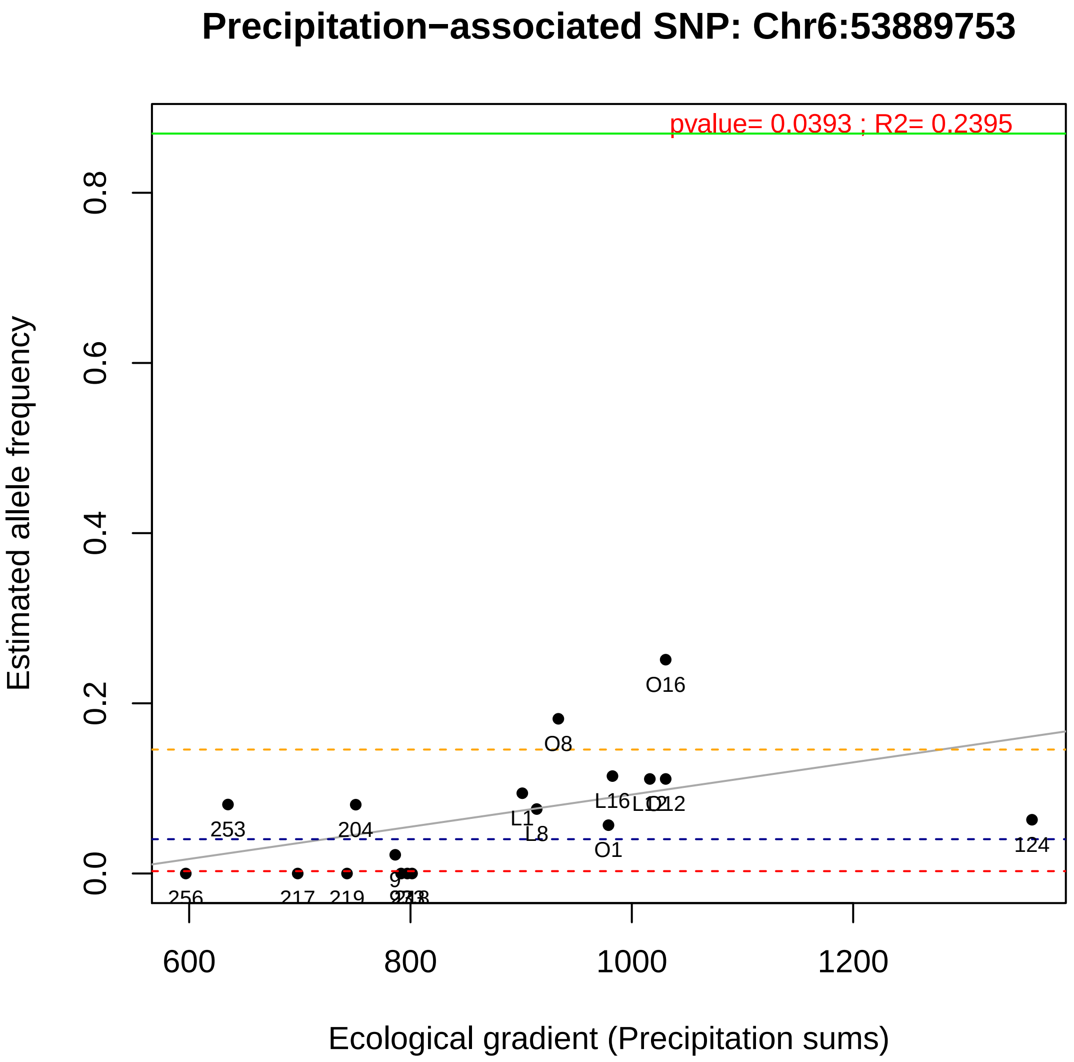
Precipitation−associated SNP: Chr6:53889753
pvalue= 0.0393 ; R2= 0.2395
Estimated allele frequency
0.0
0.2
0.4
0.6
0.8
600
800
1000
1200
Ecological gradient (Precipitation sums)
256
253
217
219
204
9
L1
L8
O8
O16
L16
O1
O12
124

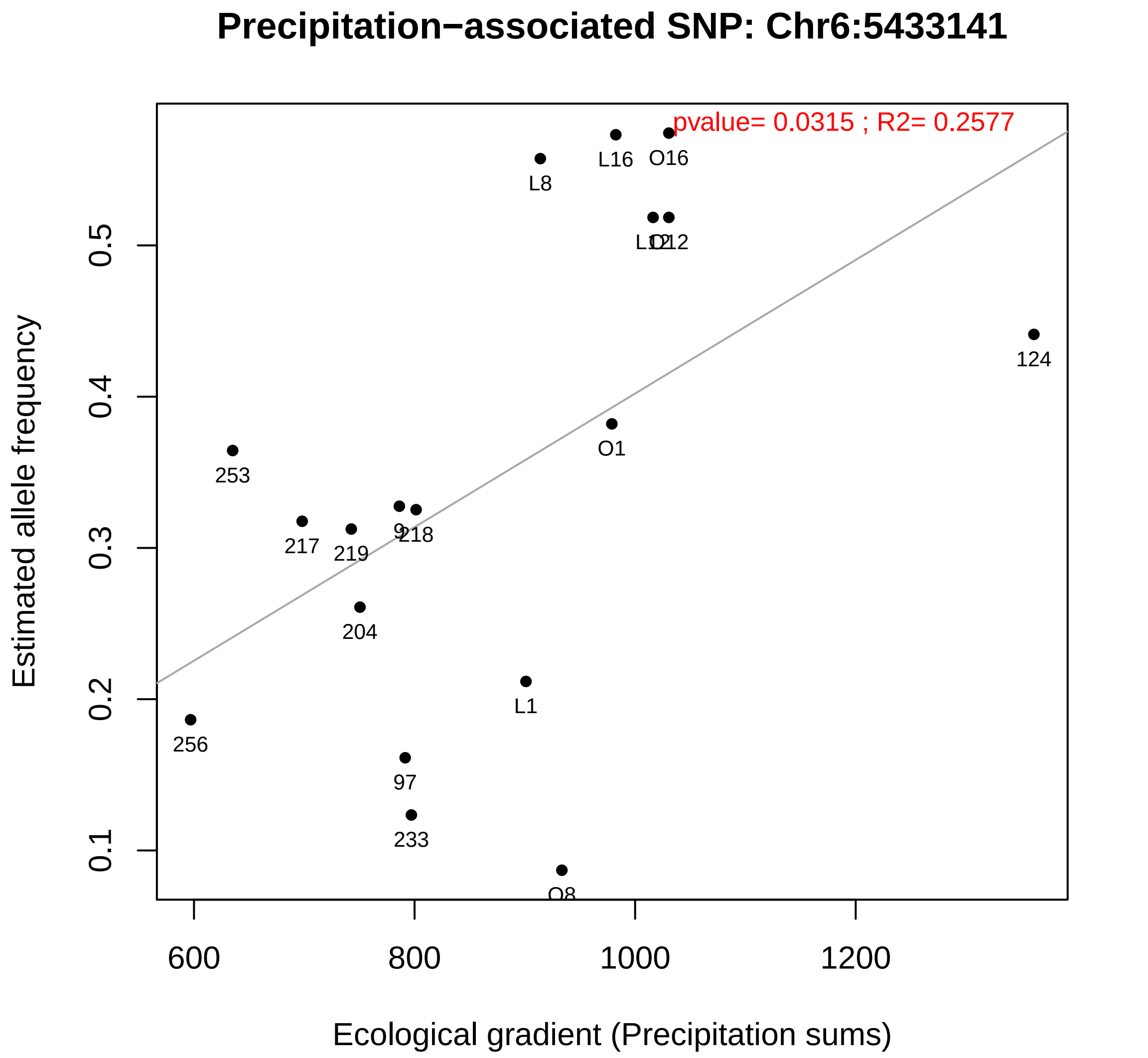

Precipitation−associated SNP: Chr6:5433141
pvalue= 0.0315 ; R2= 0.2577
L16
O16
L8
L12
O12
124
O1
253
217
219
9
218
204
L1
256
97
233
O8
0.1
0.2
0.3
0.4
0.5
600
800
1000
1200
Estimated allele frequency
Ecological gradient (Precipitation sums)

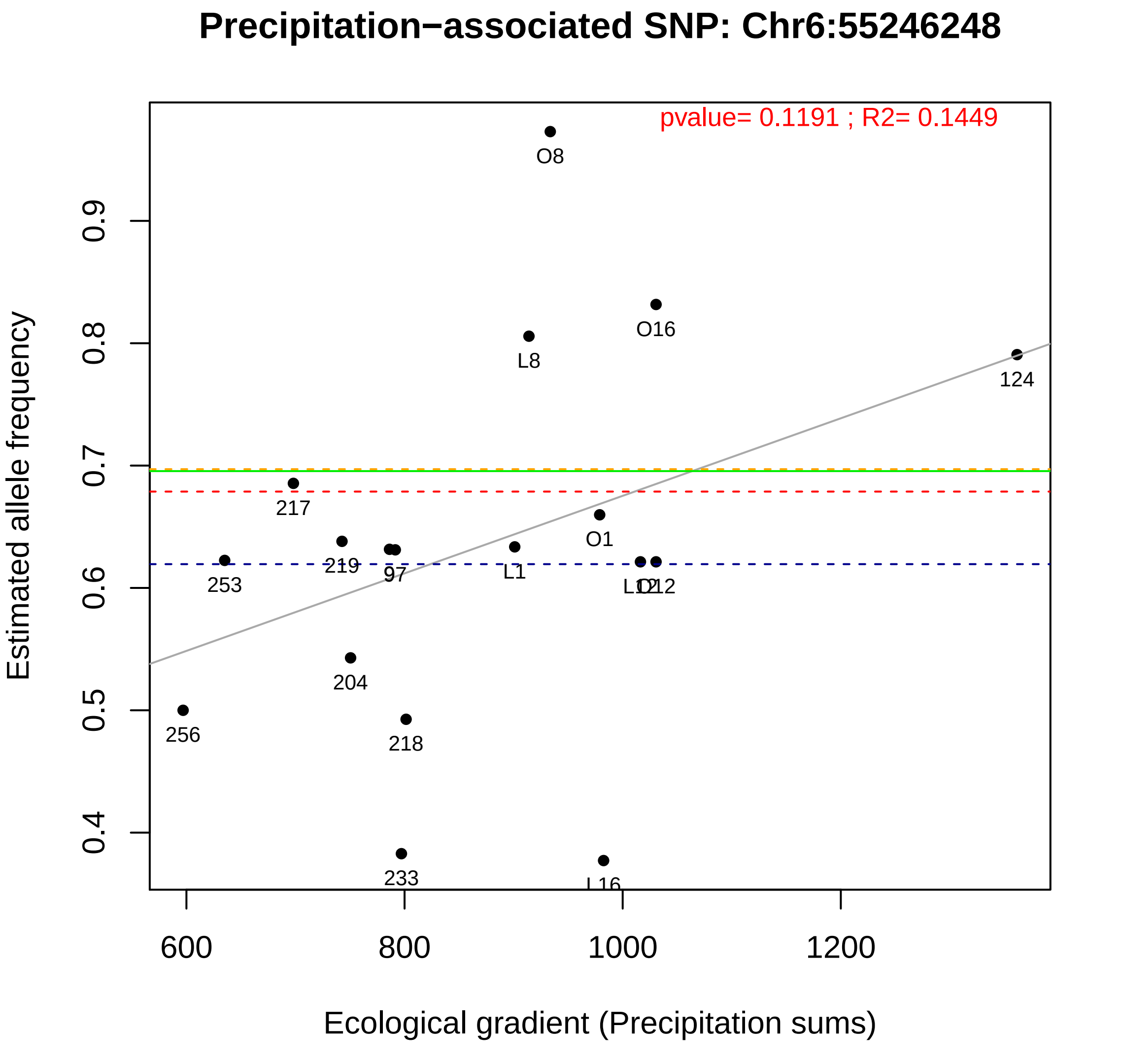
Precipitation−associated SNP: Chr6:55246248
pvalue= 0.1191 ; R2= 0.1449
Estimated allele frequency
0.4
0.5
0.6
0.7
0.8
0.9
600
800
1000
1200
Ecological gradient (Precipitation sums)
O8
O16
L8
124
217
O1
219
97
253
L1
O12
204
256
218
233
L16

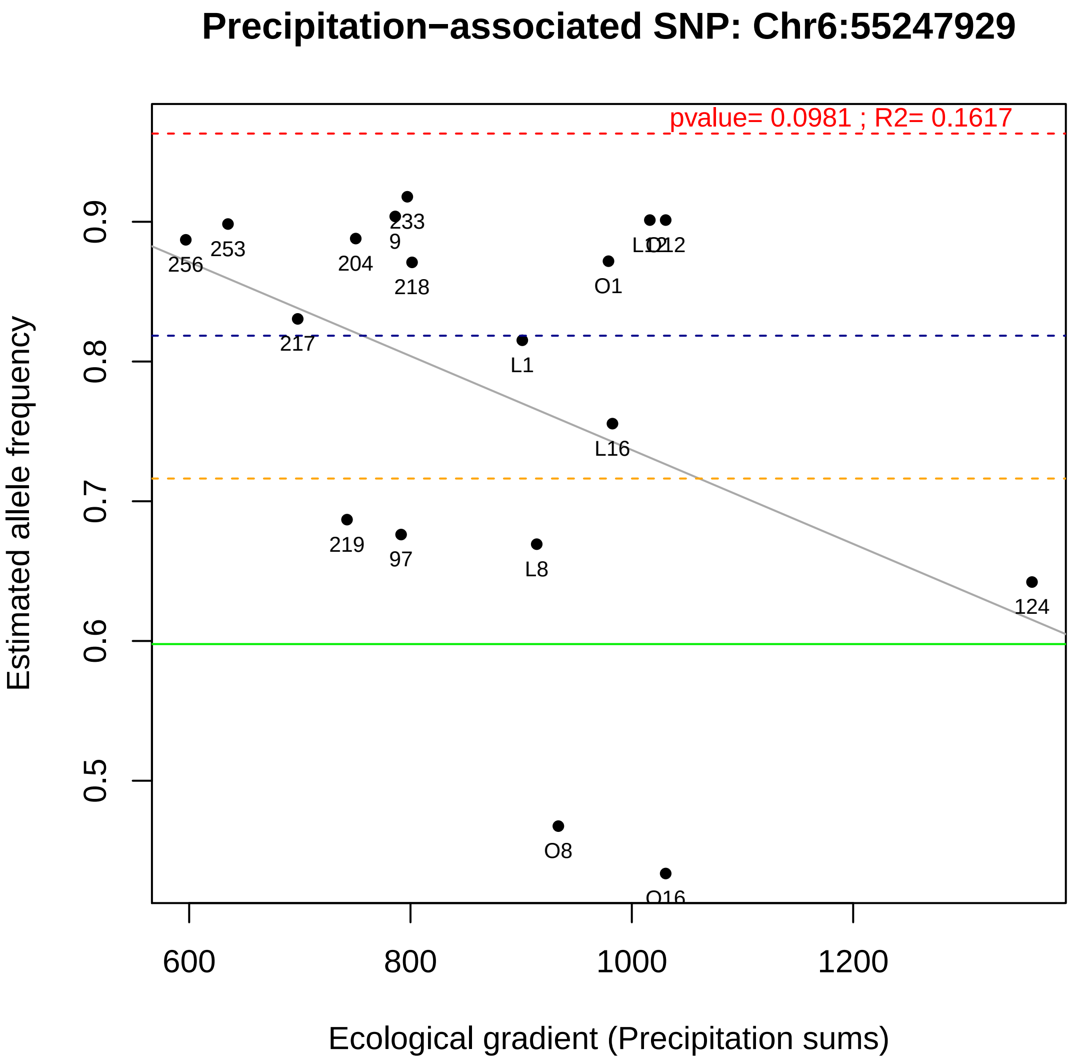

Precipitation−associated SNP: Chr6:55247929
pvalue= 0.0981 ; R2= 0.1617
Estimated allele frequency
0.5
0.6
0.7
0.8
0.9
600
800
1000
1200
Ecological gradient (Precipitation sums)
256
253
204
233
9
218
217
L1
O1
L16
219
97
L8
124
O8
O16

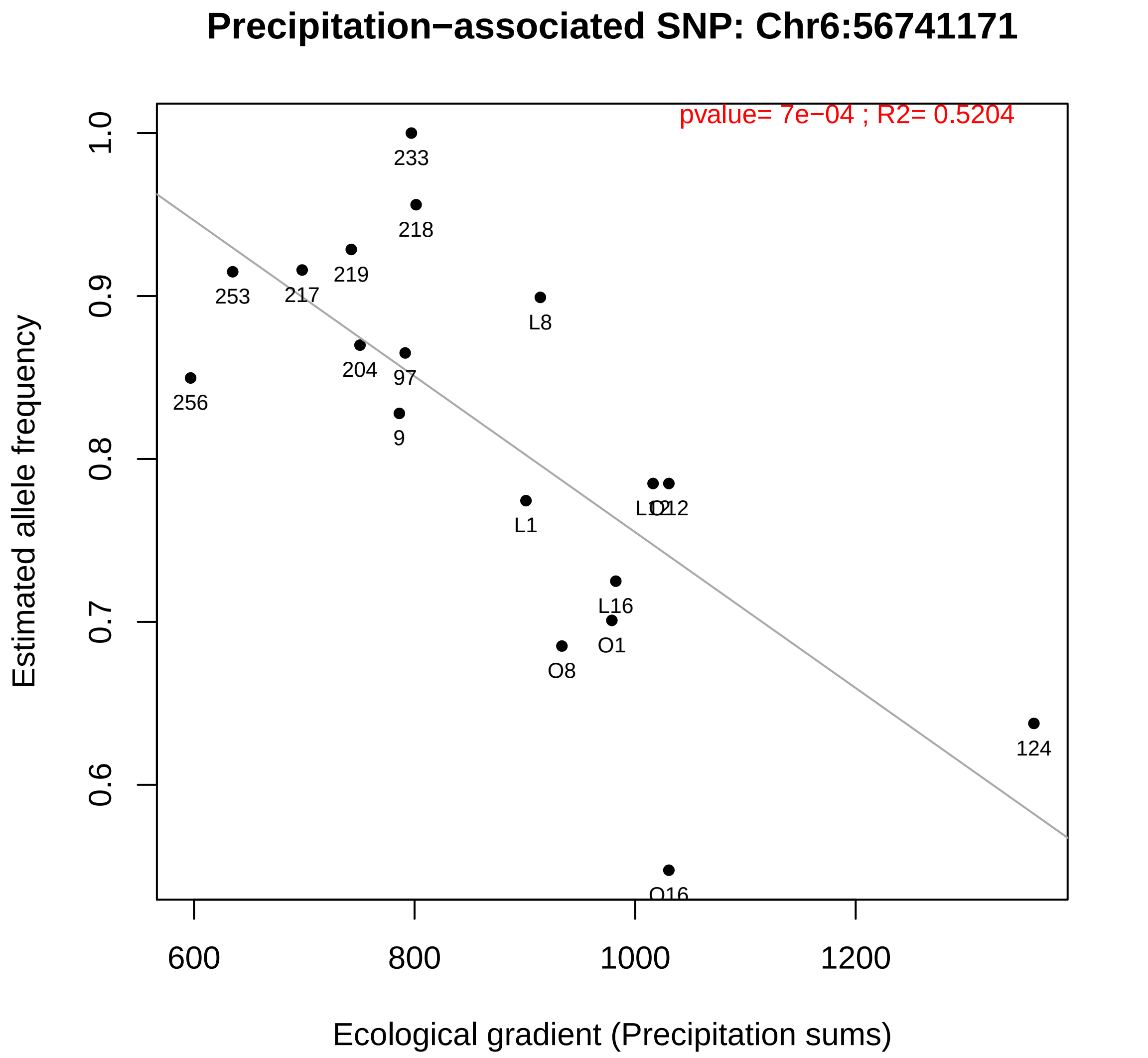
Precipitation−associated SNP: Chr6:56741171
pvalue= 7e−04 ; R2= 0.5204
Estimated allele frequency
Ecological gradient (Precipitation sums)
1.0
0.9
0.8
0.7
0.6
600
800
1000
1200
233
218
219
253
217
L8
204
97
256
9
L1
L12
O12
L16
O1
O8
124
O16

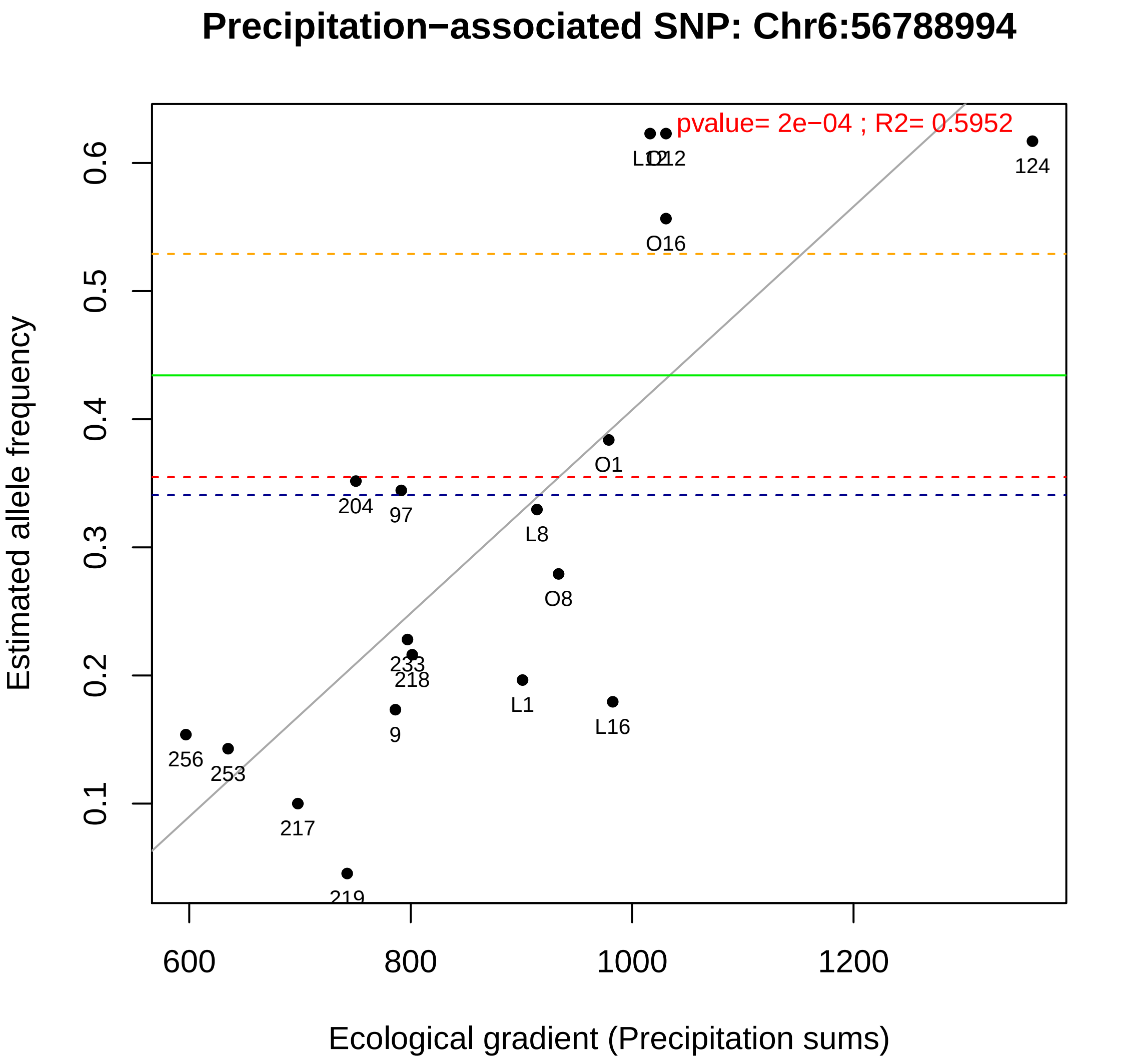
Precipitation−associated SNP: Chr6:56788994
pvalue= 2e−04 ; R2= 0.5952
Estimated allele frequency
Ecological gradient (Precipitation sums)
0.1
0.2
0.3
0.4
0.5
0.6
600
800
1000
1200
L10
O12
124
O16
O1
204
97
L8
O8
233
218
L1
L16
9
256
253
217
219

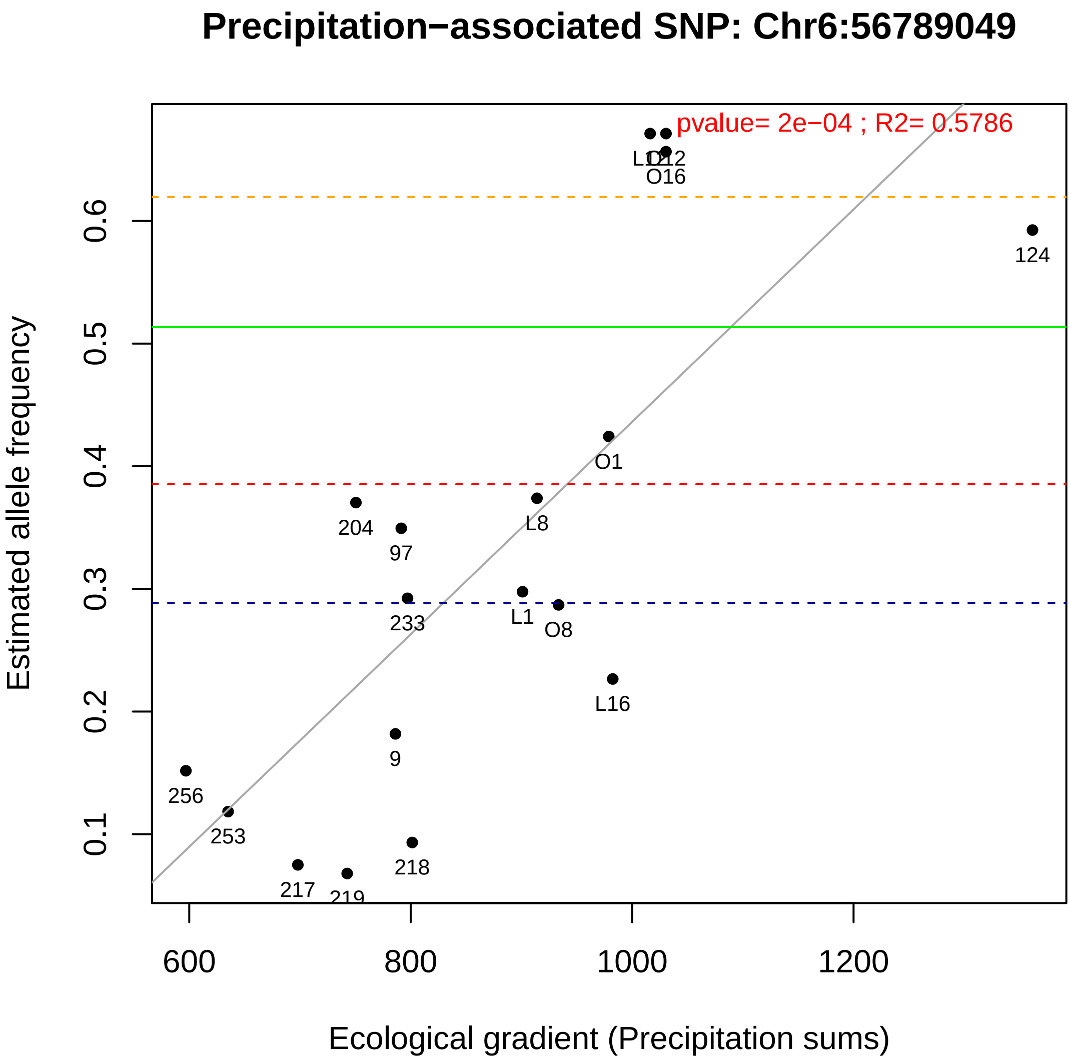

Precipitation−associated SNP: Chr6:56789049
pvalue= 2e−04 ; R2= 0.5786
Estimated allele frequency
Ecological gradient (Precipitation sums)
0.1
0.2
0.3
0.4
0.5
0.6
600
800
1000
1200
L12
O12
O16
124
O1
204
97
L8
233
L1
O8
L16
9
256
253
217
219
218

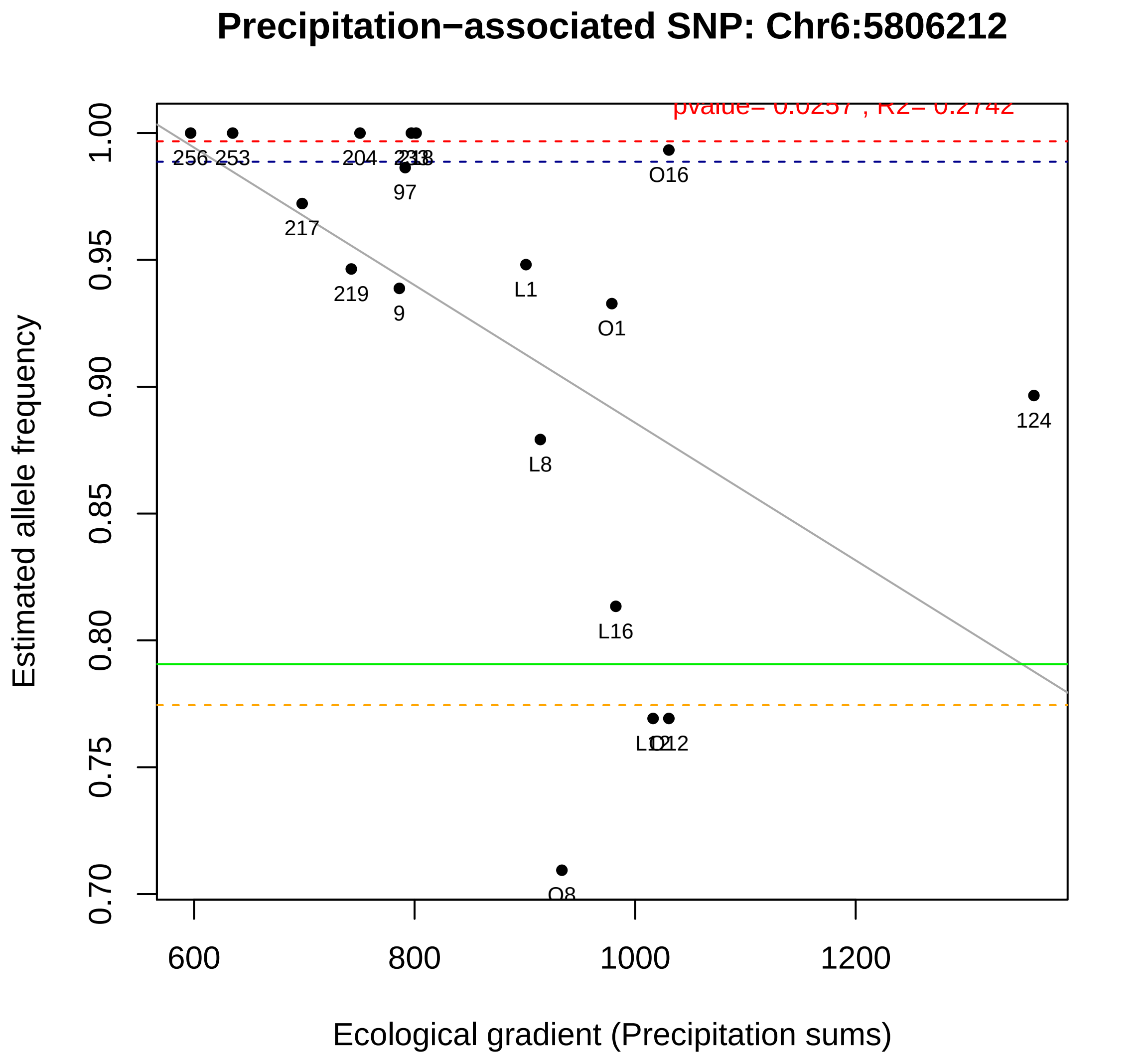
Precipitation−associated SNP: Chr6:5806212
pvalue= 0.0257 , R2= 0.2742
Estimated allele frequency
Ecological gradient (Precipitation sums)
0.70
0.75
0.80
0.85
0.90
0.95
1.00
600
800
1000
1200
256
253
204
97
217
219
9
L1
O1
O16
124
L8
L16
O8

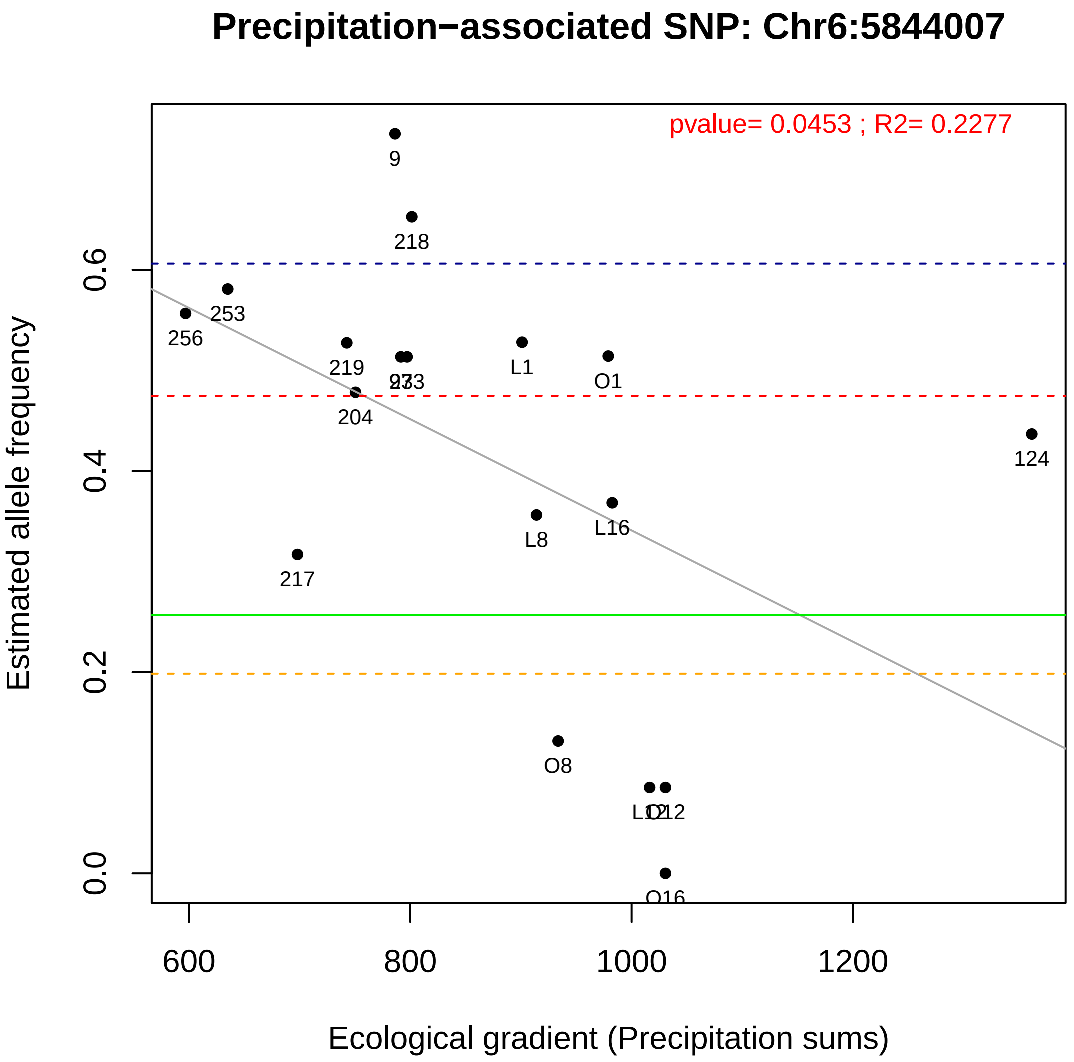

Precipitation−associated SNP: Chr6:5844007
pvalue= 0.0453 ; R2= 0.2277
Estimated allele frequency
Ecological gradient (Precipitation sums)
0.0
0.2
0.4
0.6
600
800
1000
1200
9
218
253
256
219
204
L1
O1
124
L16
L8
217
O8
O16

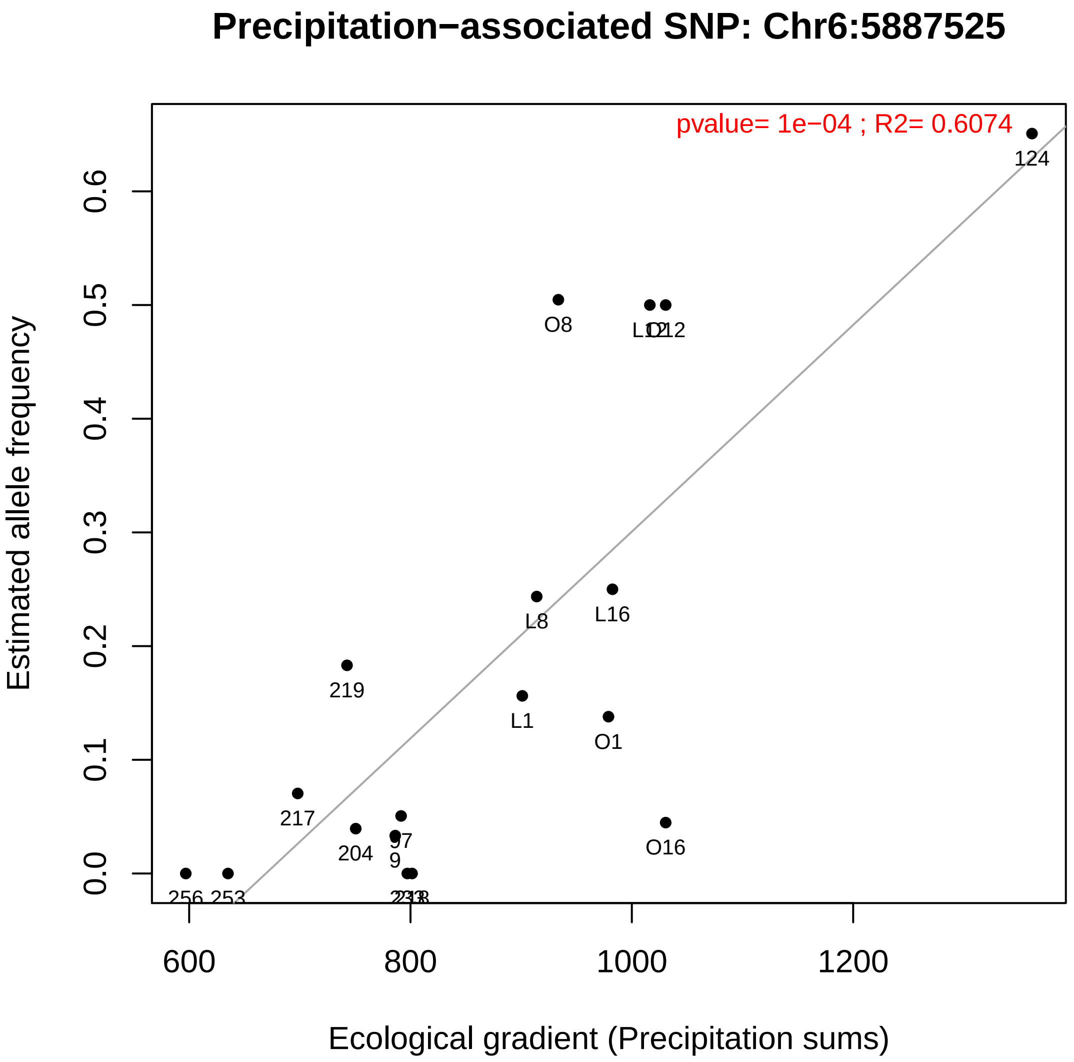
Precipitation−associated SNP: Chr6:5887525
pvalue= 1e−04 ; R2= 0.6074
Estimated allele frequency
0.0
0.1
0.2
0.3
0.4
0.5
0.6
600
800
1000
1200
Ecological gradient (Precipitation sums)
124
O8
219
L8
L16
L1
O1
217
204
O16
9
256
253

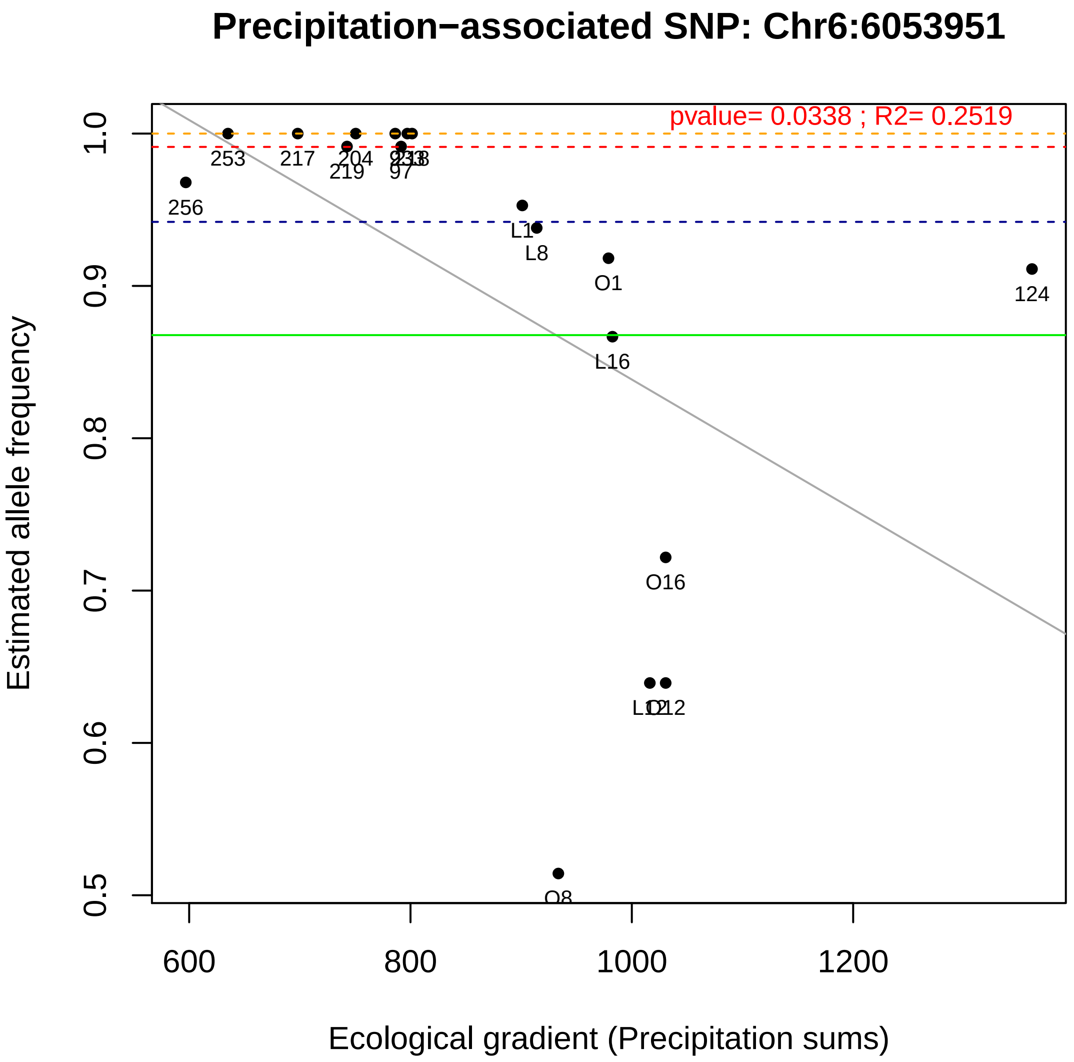

Precipitation−associated SNP: Chr6:6053951
pvalue= 0.0338 ; R2= 0.2519
Estimated allele frequency
Ecological gradient (Precipitation sums)
0.5
0.6
0.7
0.8
0.9
1.0
600
800
1000
1200
253
217
204
219
97
256
L1
L8
O1
124
L16
O16
O12
O8

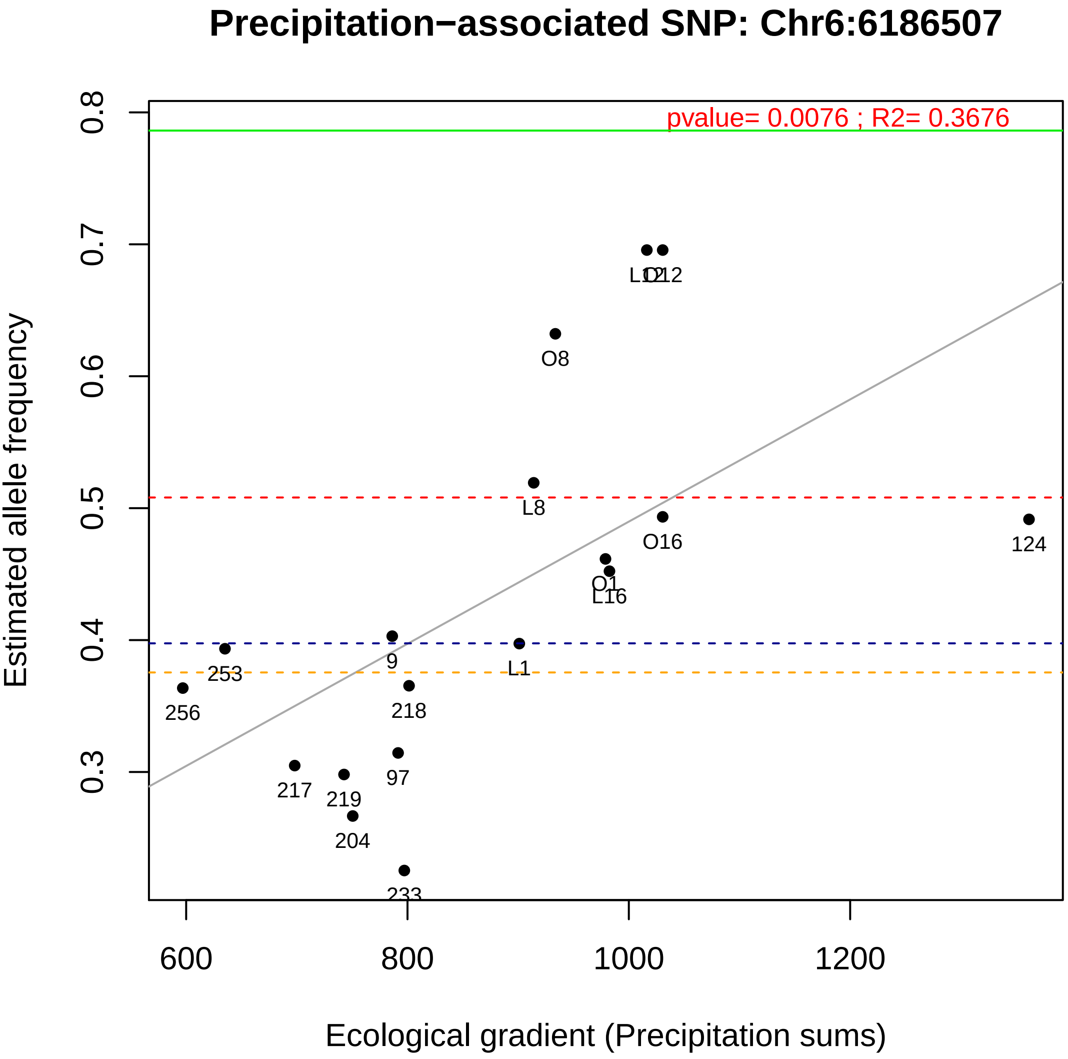
Precipitation−associated SNP: Chr6:6186507
pvalue= 0.0076 ; R2= 0.3676
Estimated allele frequency
0.8
0.7
0.6
0.5
0.4
0.3
600
800
1000
1200
Ecological gradient (Precipitation sums)
O8
L8
O16
124
O1
L16
9
L1
253
256
218
97
217
219
204
233

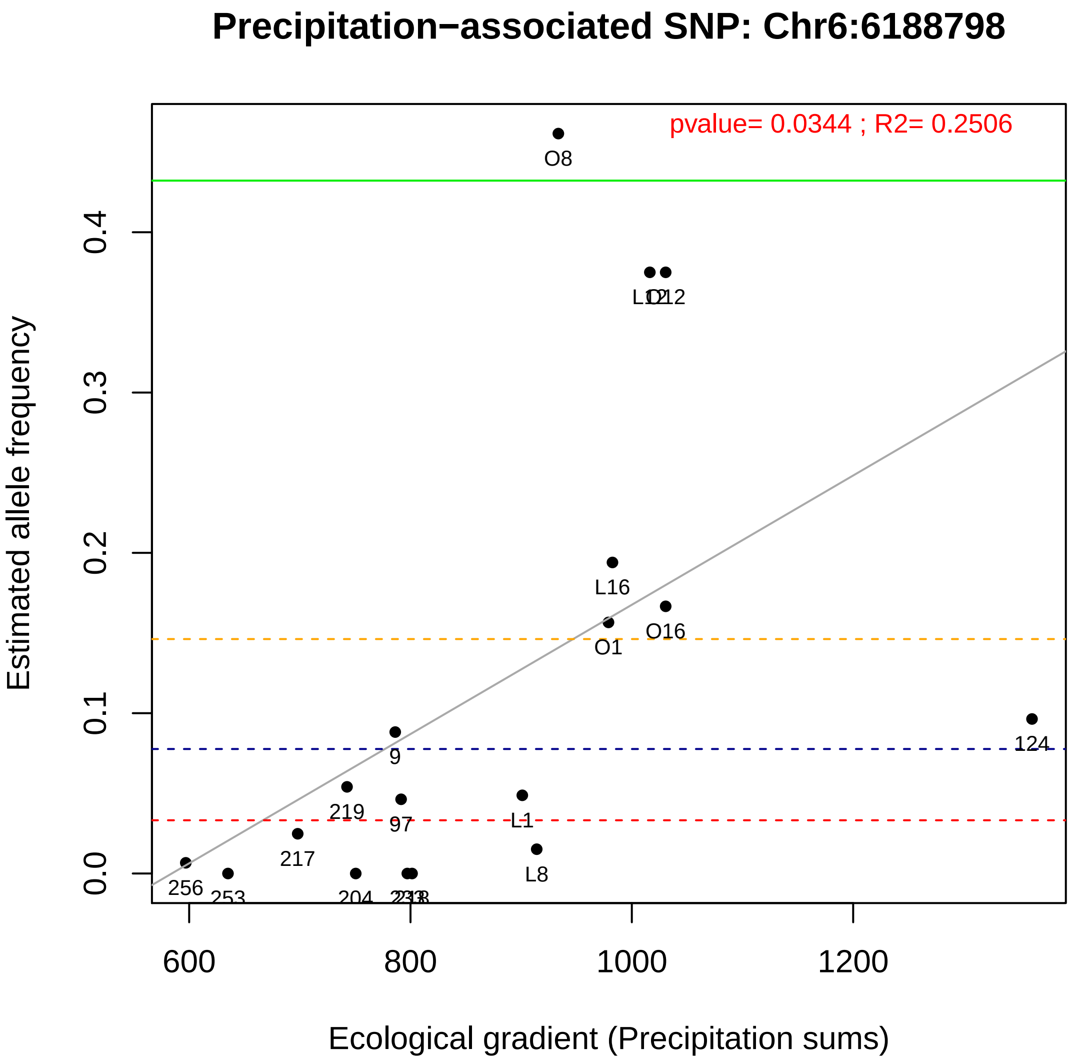

Precipitation−associated SNP: Chr6:6188798
pvalue= 0.0344 ; R2= 0.2506
O8
L12
O12
L16
O16
O1
124
9
219
97
L1
217
L8
256
253
204
0.0
0.1
0.2
0.3
0.4
600
800
1000
1200
Estimated allele frequency
Ecological gradient (Precipitation sums)

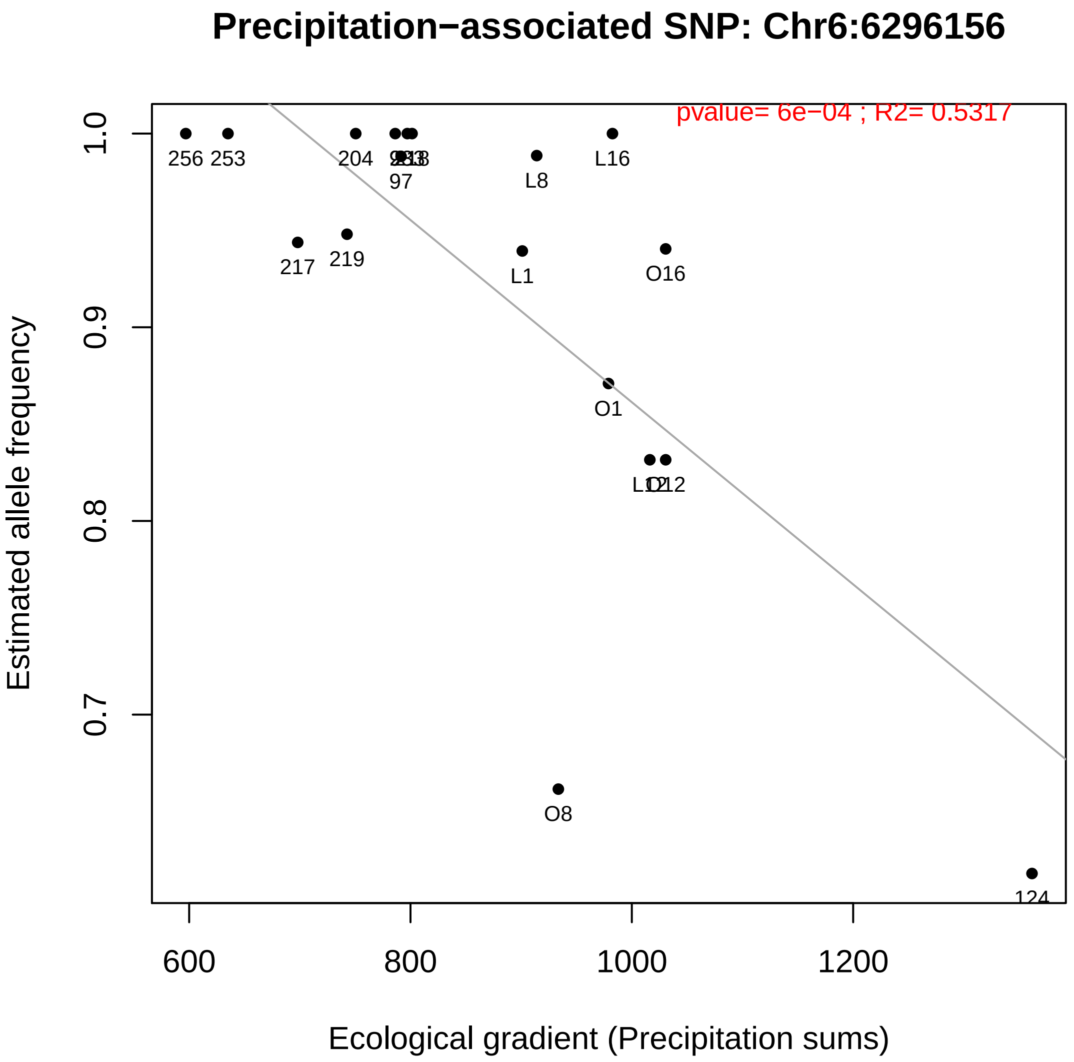

Precipitation−associated SNP: Chr6:6296156
pvalue= 6e−04 ; R2= 0.5317
Estimated allele frequency
Ecological gradient (Precipitation sums)
0.7
0.8
0.9
1.0
600
800
1000
1200
256
253
204
97
L8
L16
217
219
L1
O16
O1
O12
O8
124

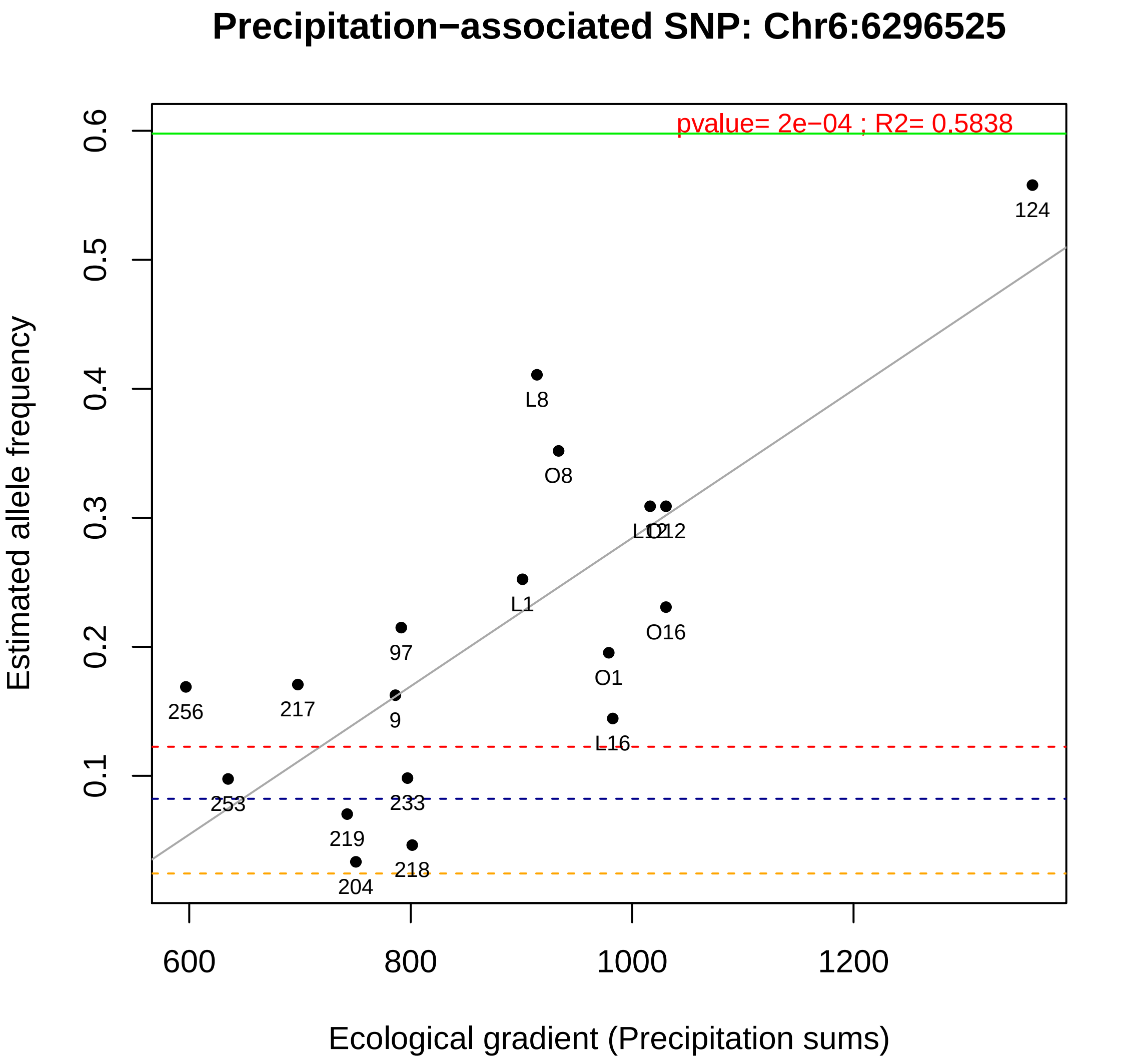

Precipitation−associated SNP: Chr6:6296525
pvalue= 2e−04 ; R2= 0.5838
Estimated allele frequency
Ecological gradient (Precipitation sums)
0.1
0.2
0.3
0.4
0.5
0.6
600
800
1000
1200
124
L8
O8
L1
97
O16
O1
256
217
9
L16
253
233
219
218
204

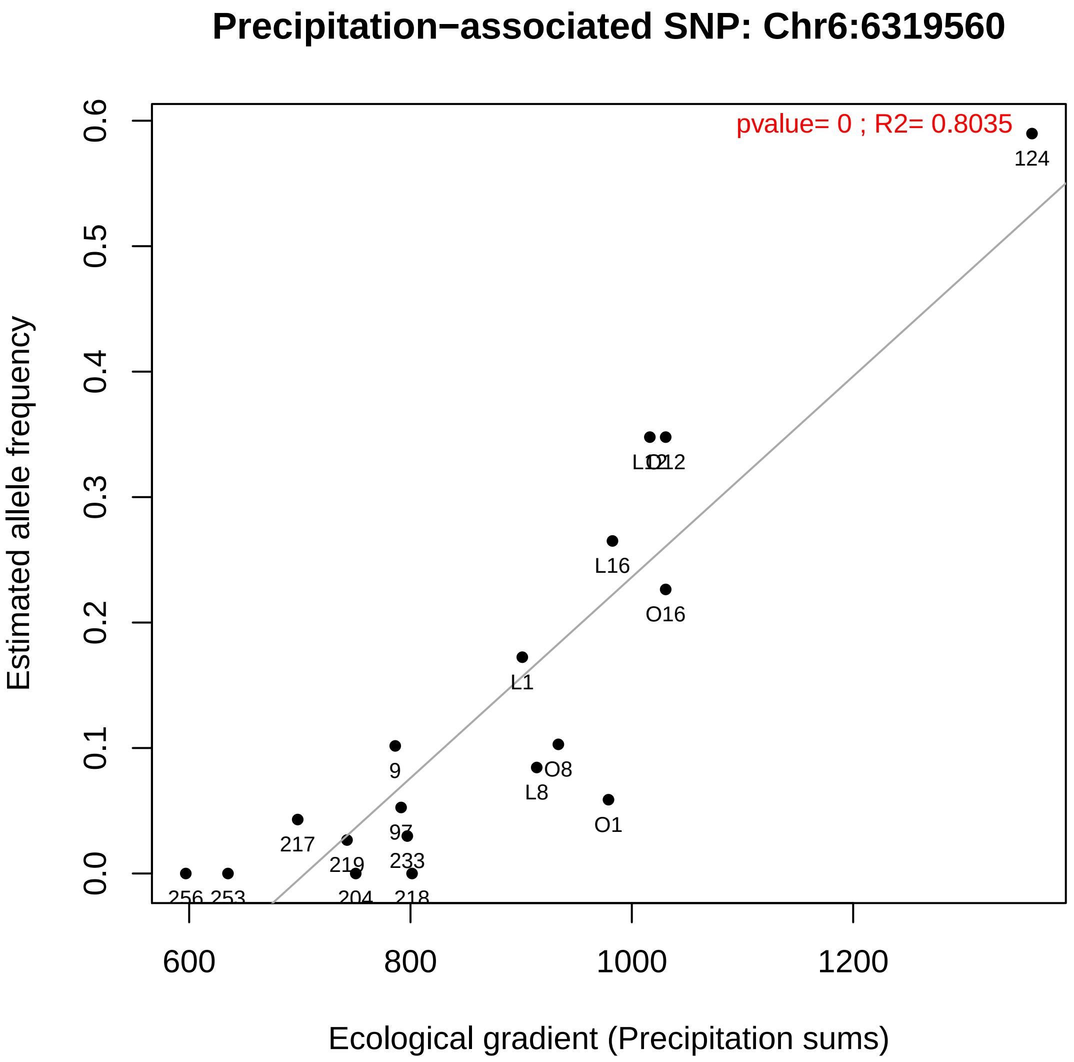
Precipitation−associated SNP: Chr6:6319560
pvalue= 0 ; R2= 0.8035
Estimated allele frequency
Ecological gradient (Precipitation sums)
0.0
0.1
0.2
0.3
0.4
0.5
0.6
600
800
1000
1200
124
L12
O12
L16
O16
L1
9
O8
L8
O1
217
97
233
219
204
218
256
253

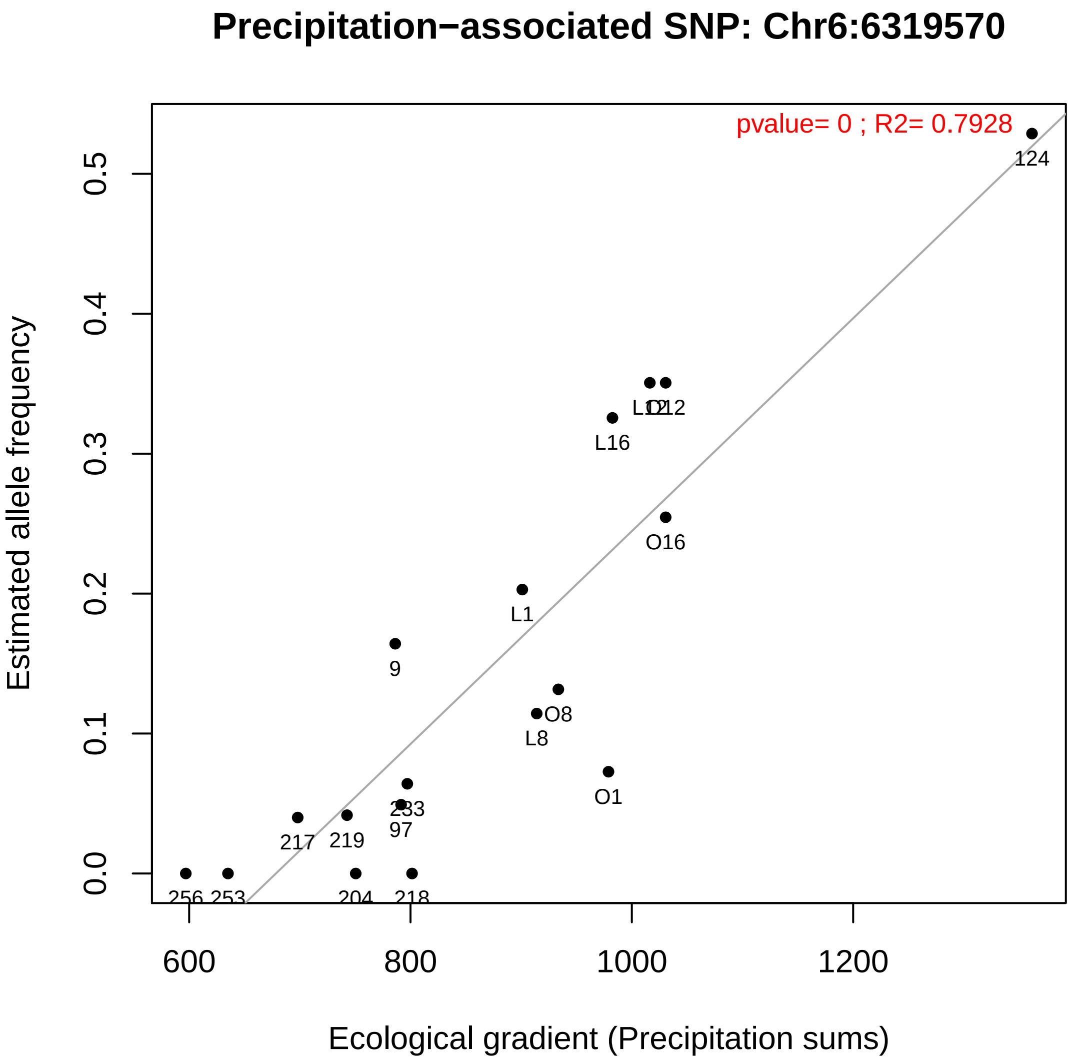

Precipitation−associated SNP: Chr6:6319570
pvalue= 0 ; R2= 0.7928
124
L12
O12
L16
O16
L1
9
O8
L8
O1
233
97
217
219
256
253
204
218
0.0
0.1
0.2
0.3
0.4
0.5
600
800
1000
1200
Estimated allele frequency
Ecological gradient (Precipitation sums)

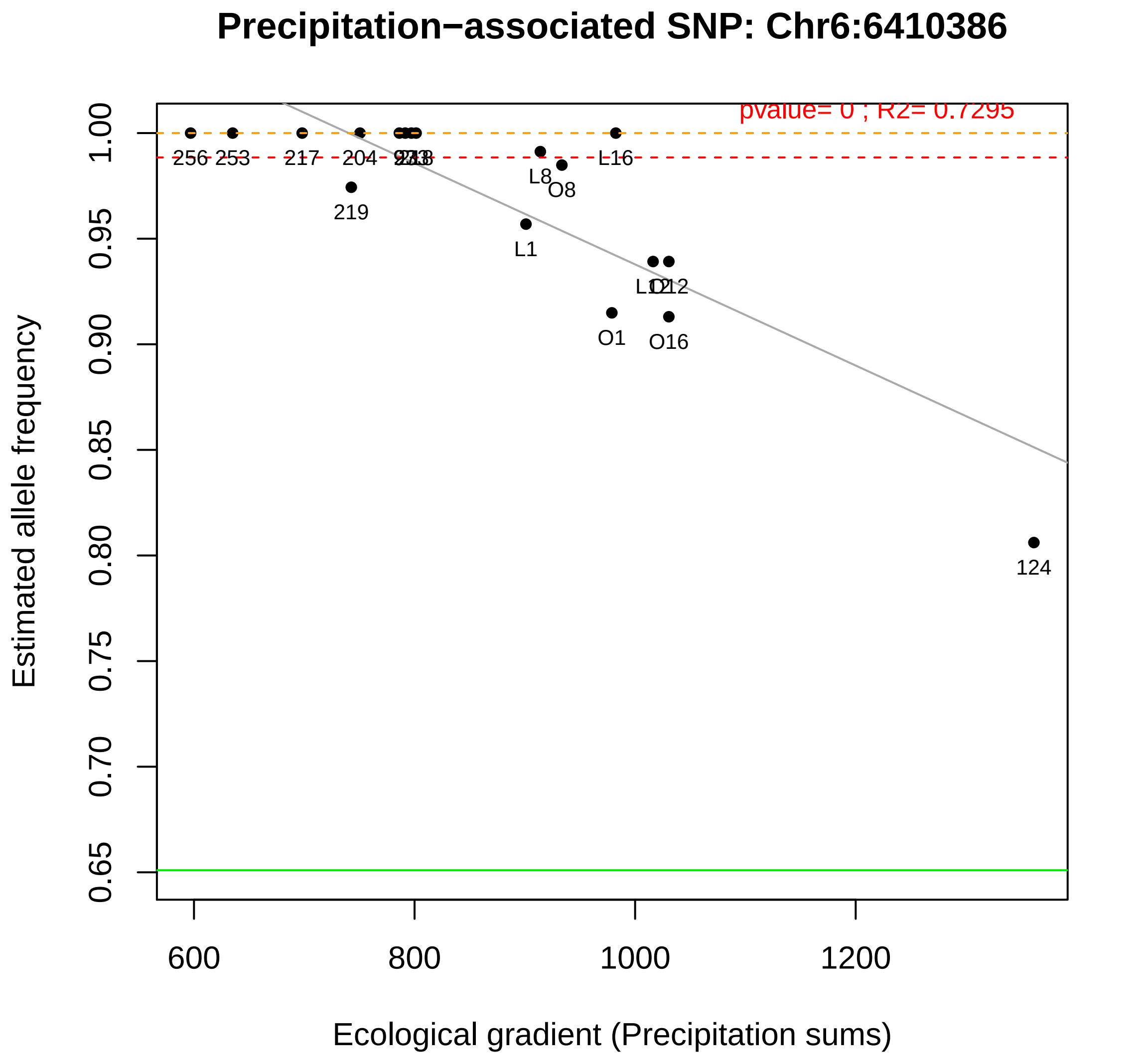

Precipitation−associated SNP: Chr6:6410386
pvalue= 0 ; R2= 0.7295
Estimated allele frequency
1.00
0.95
0.90
0.85
0.80
0.75
0.70
0.65
600
800
1000
1200
Ecological gradient (Precipitation sums)
256
253
217
204
219
L8
O8
L16
L1
O1
O16
124

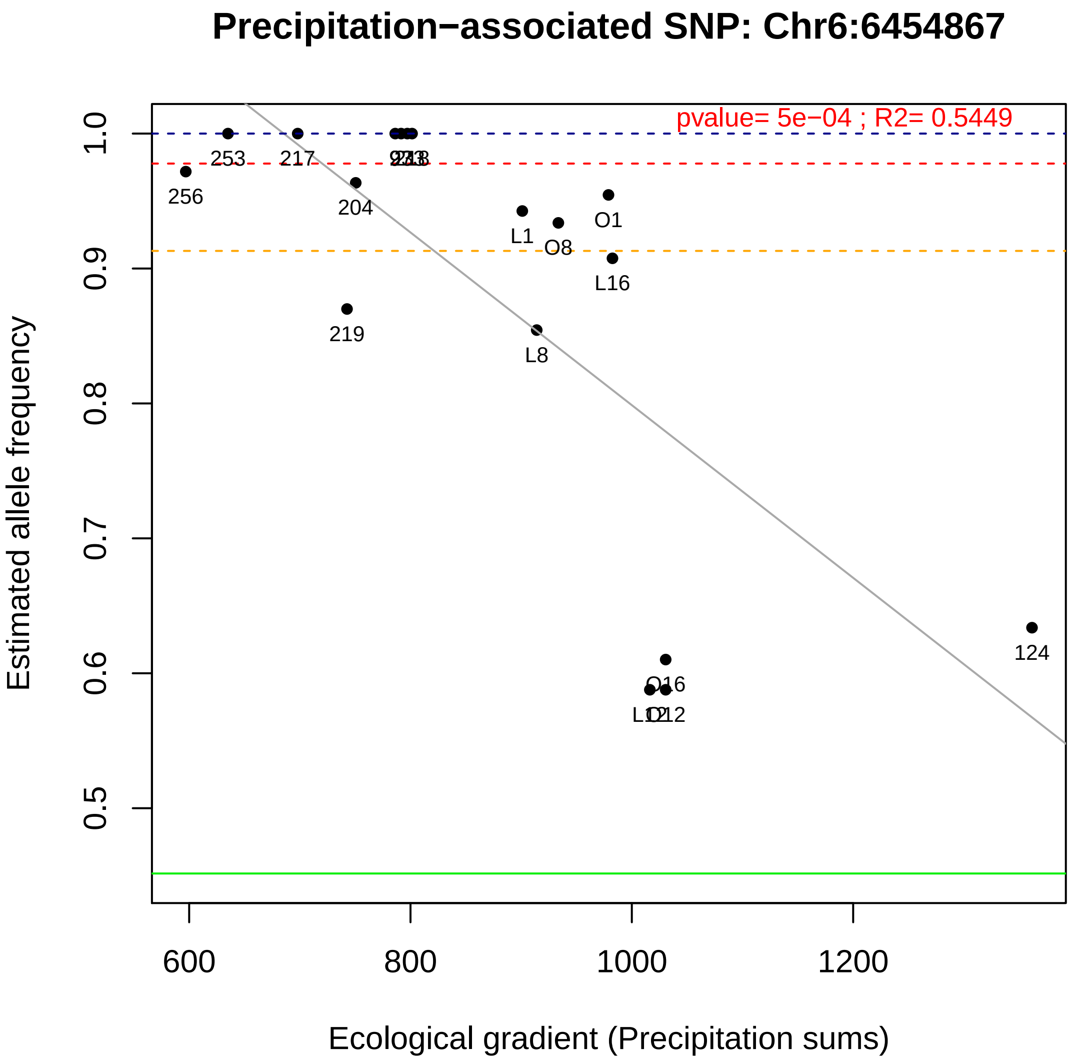
Precipitation−associated SNP: Chr6:6454867
pvalue= 5e−04 ; R2= 0.5449
Estimated allele frequency
Ecological gradient (Precipitation sums)
0.5
0.6
0.7
0.8
0.9
1.0
600
800
1000
1200
253
217
256
204
L1
O8
O1
L16
219
L8
124
O16

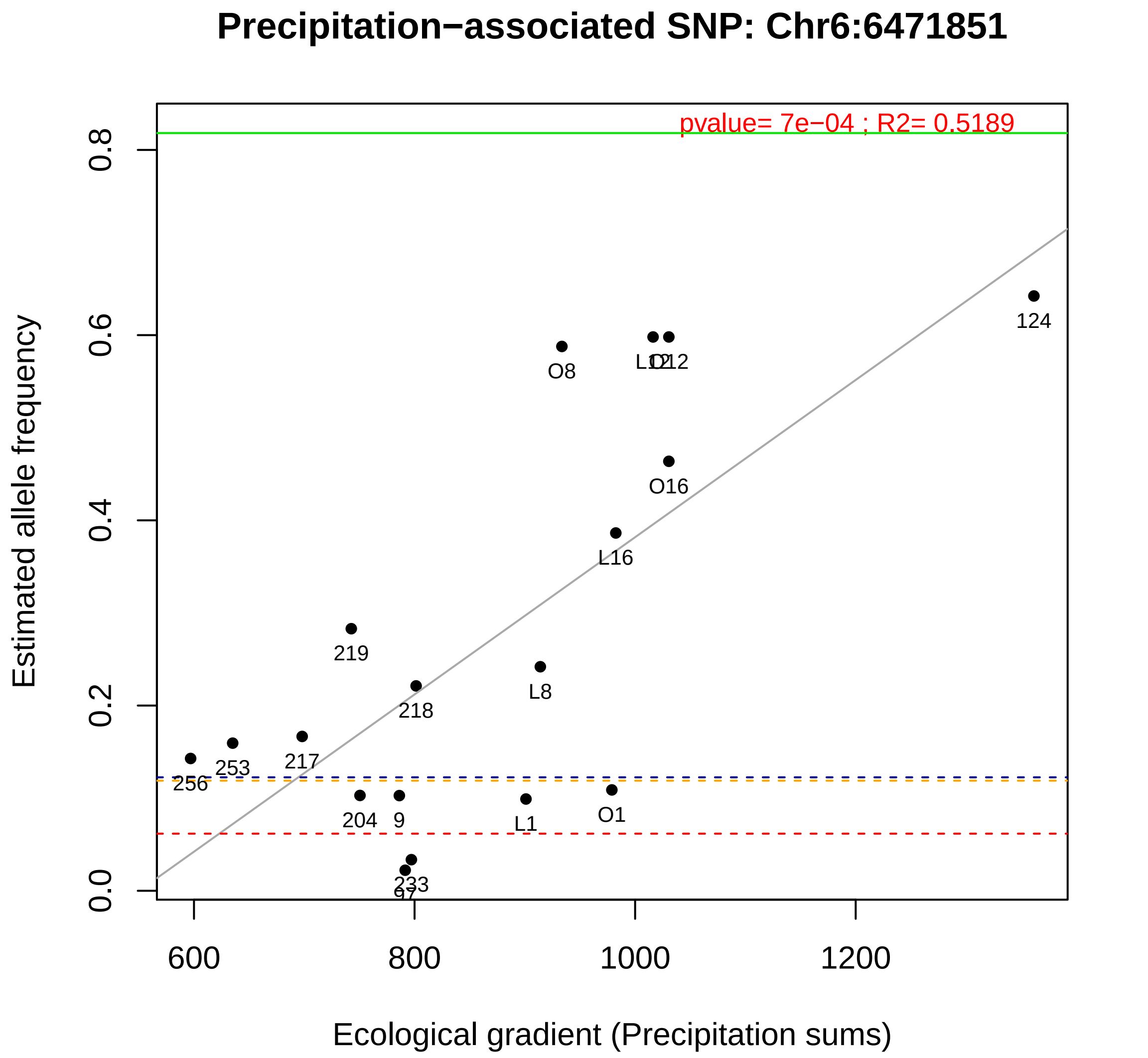
Precipitation−associated SNP: Chr6:6471851
pvalue= 7e−04 ; R2= 0.5189
Estimated allele frequency
0.0
0.2
0.4
0.6
0.8
600
800
1000
1200
Ecological gradient (Precipitation sums)
124
O8
O12
O16
L16
219
218
L8
256
253
217
204
9
L1
O1
233

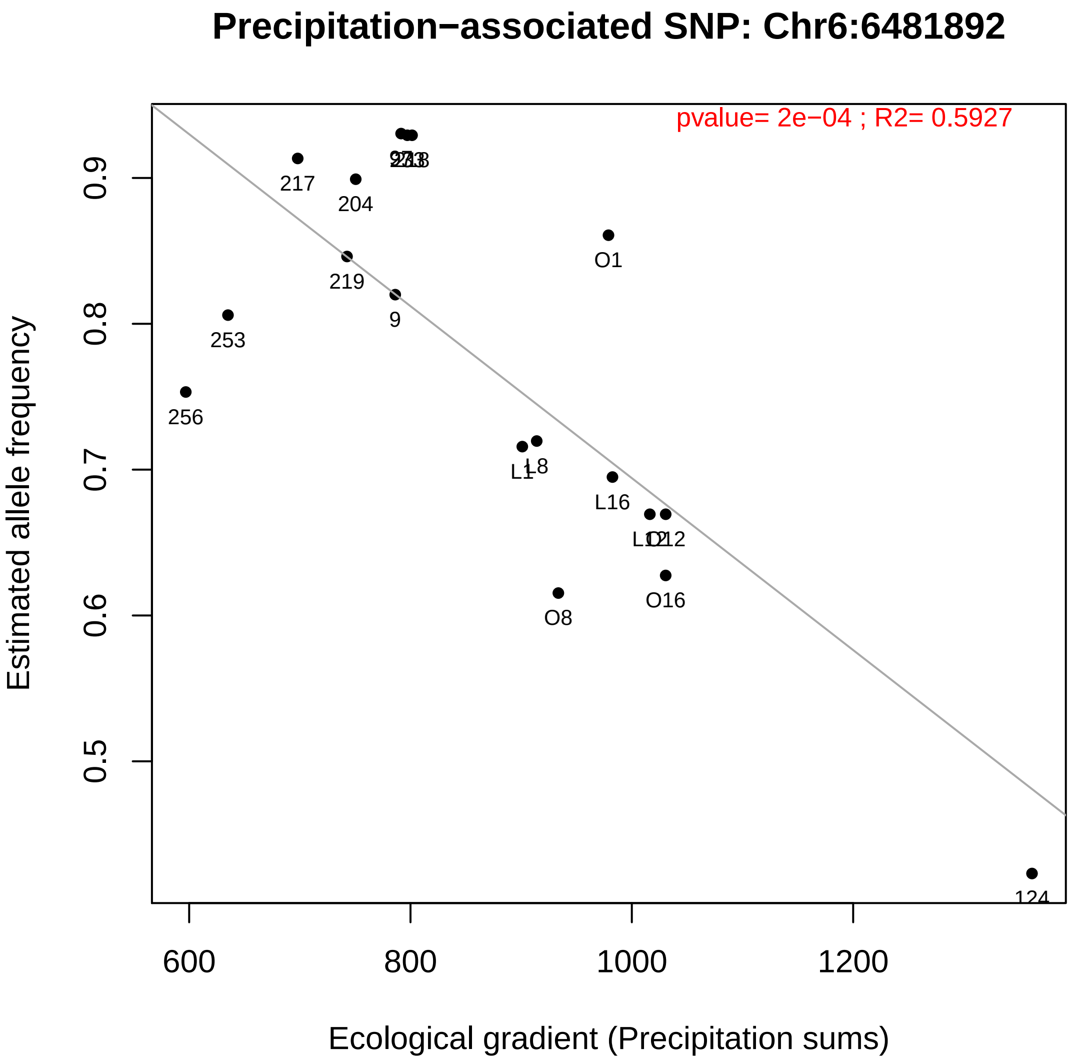

Precipitation−associated SNP: Chr6:6481892
pvalue= 2e−04 ; R2= 0.5927
Estimated allele frequency
Ecological gradient (Precipitation sums)
0.5
0.6
0.7
0.8
0.9
600
800
1000
1200
217
204
219
9
253
256
O1
L1
L8
L16
O16
O8
124

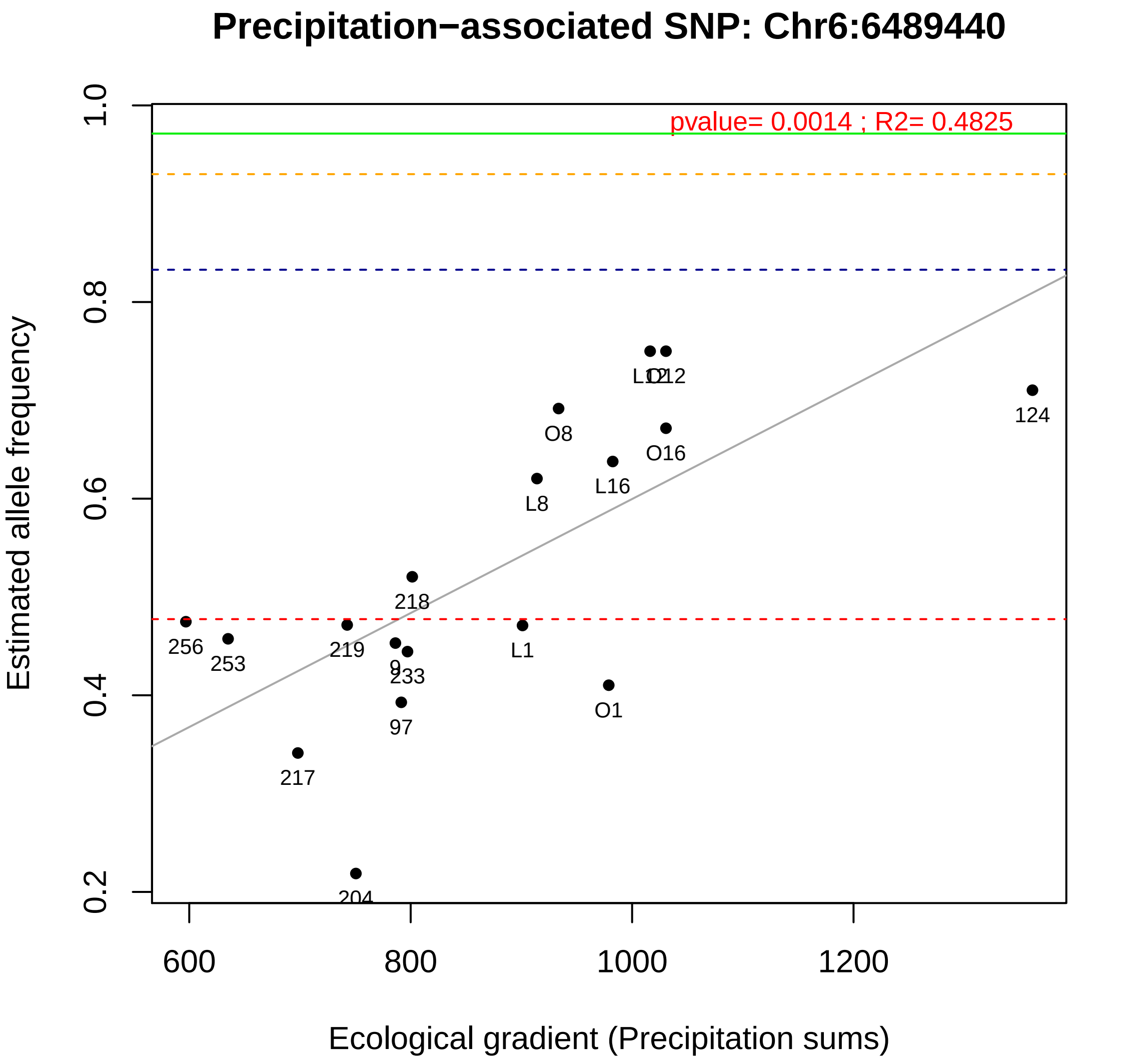

Precipitation−associated SNP: Chr6:6489440
pvalue= 0.0014 ; R2= 0.4825
Estimated allele frequency
0.2
0.4
0.6
0.8
1.0
600
800
1000
1200
Ecological gradient (Precipitation sums)
L12
O12
124
O8
O16
L16
L8
218
256
253
219
9
233
L1
97
O1
217
204

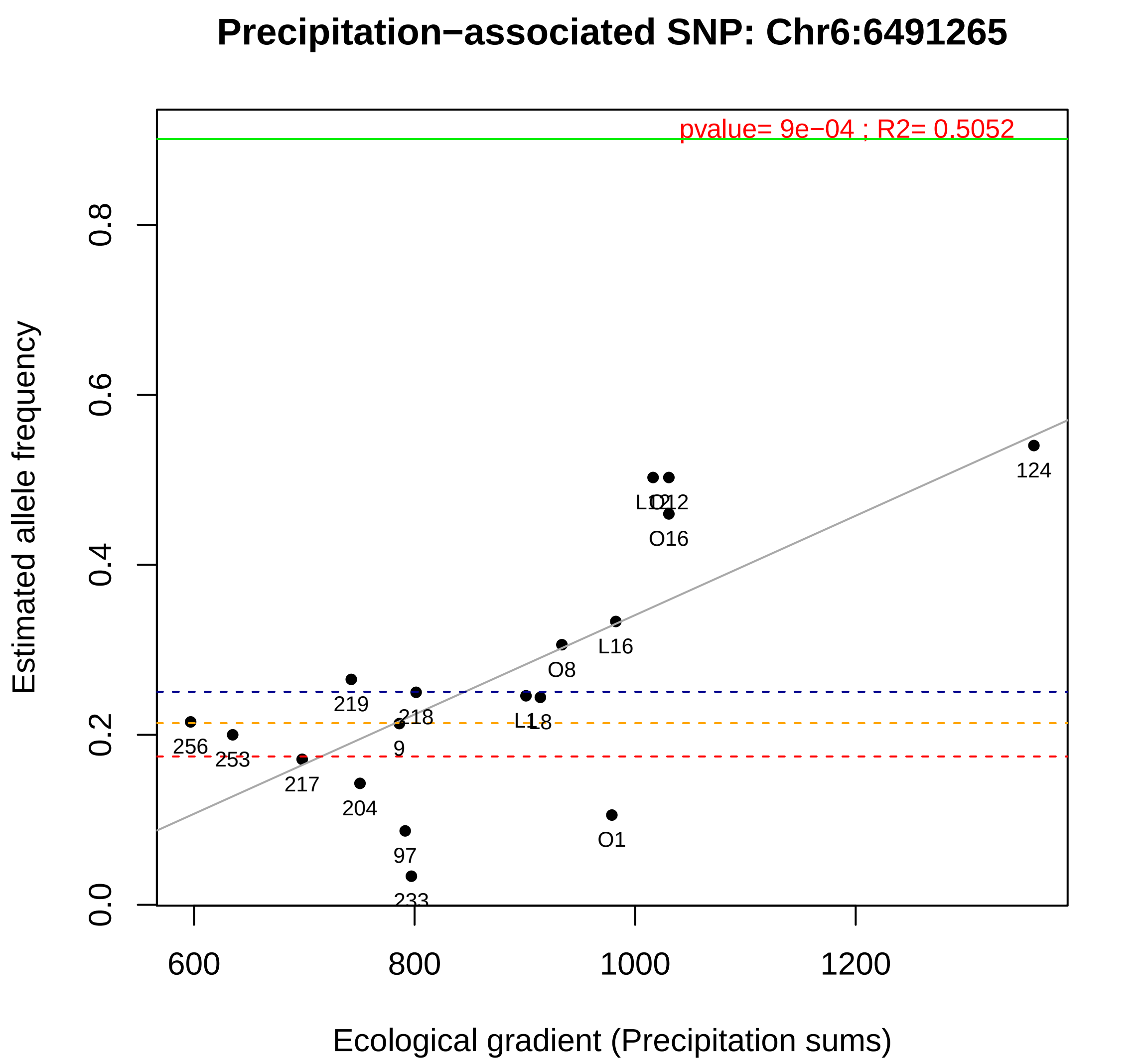
Precipitation−associated SNP: Chr6:6491265
pvalue= 9e−04 ; R2= 0.5052
Estimated allele frequency
0.0
0.2
0.4
0.6
0.8
600
800
1000
1200
Ecological gradient (Precipitation sums)
124
O16
L16
O8
219
218
256
253
9
217
204
97
233
O1

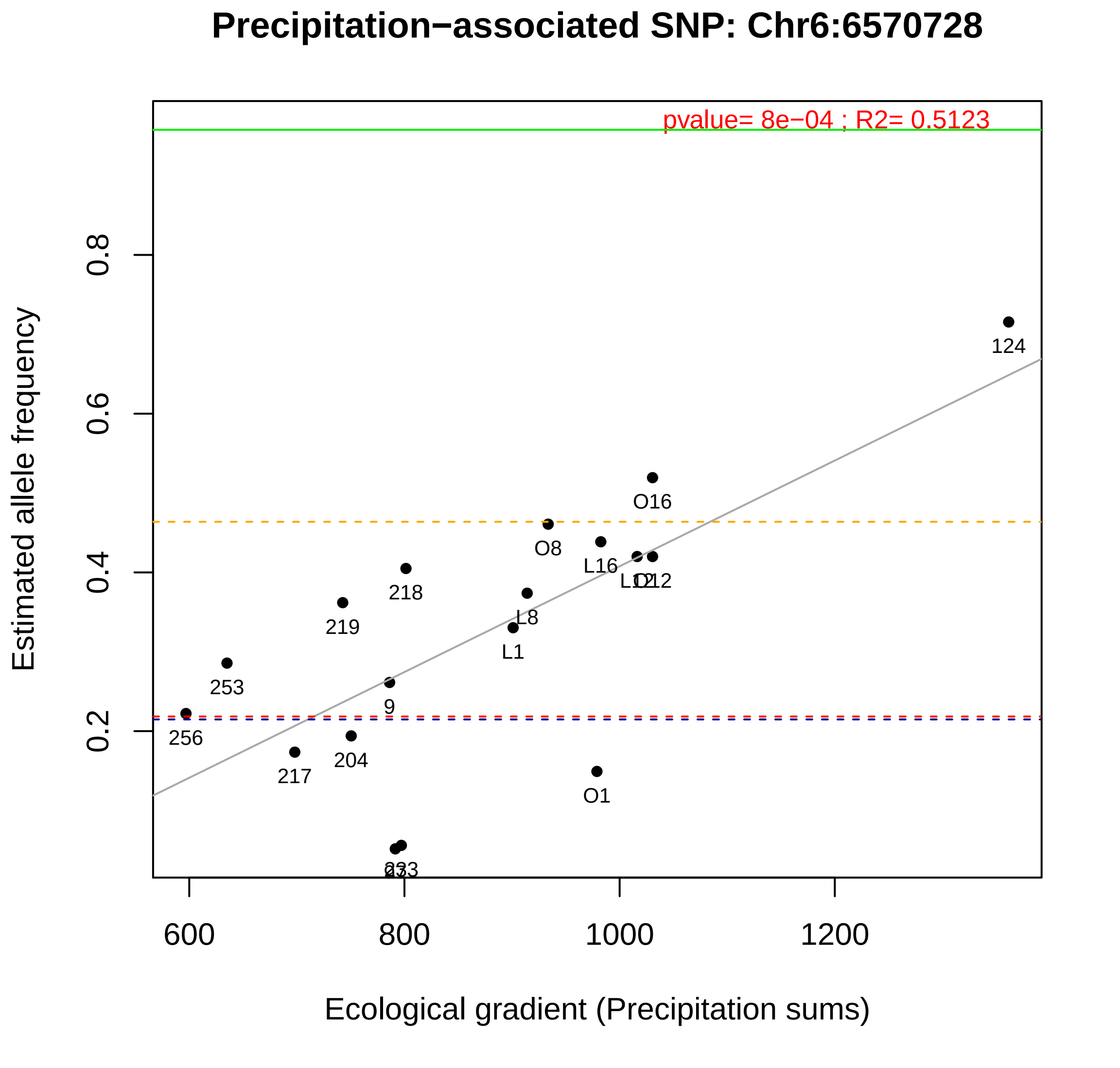
Precipitation−associated SNP: Chr6:6570728
pvalue= 8e−04 ; R2= 0.5123
Estimated allele frequency
0.2
0.4
0.6
0.8
600
800
1000
1200
Ecological gradient (Precipitation sums)
124
O16
O8
L16
218
219
L8
L1
253
9
256
204
217
O1

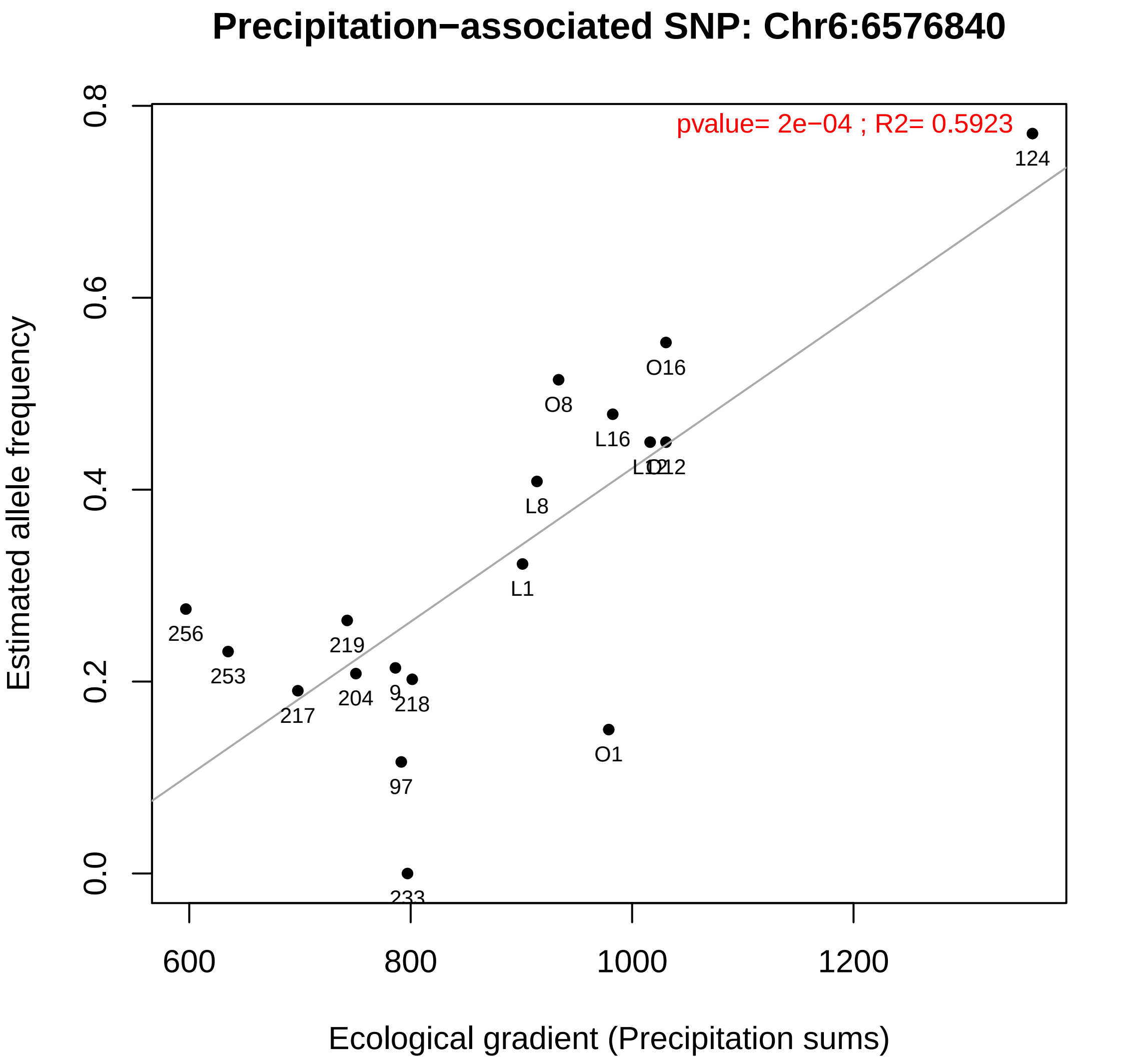

Precipitation−associated SNP: Chr6:6576840
pvalue= 2e−04 ; R2= 0.5923
Estimated allele frequency
Ecological gradient (Precipitation sums)
0.0
0.2
0.4
0.6
0.8
600
800
1000
1200
124
O16
O8
L16
L10
O12
L8
L1
256
219
253
217
204
9
218
O1
97
233

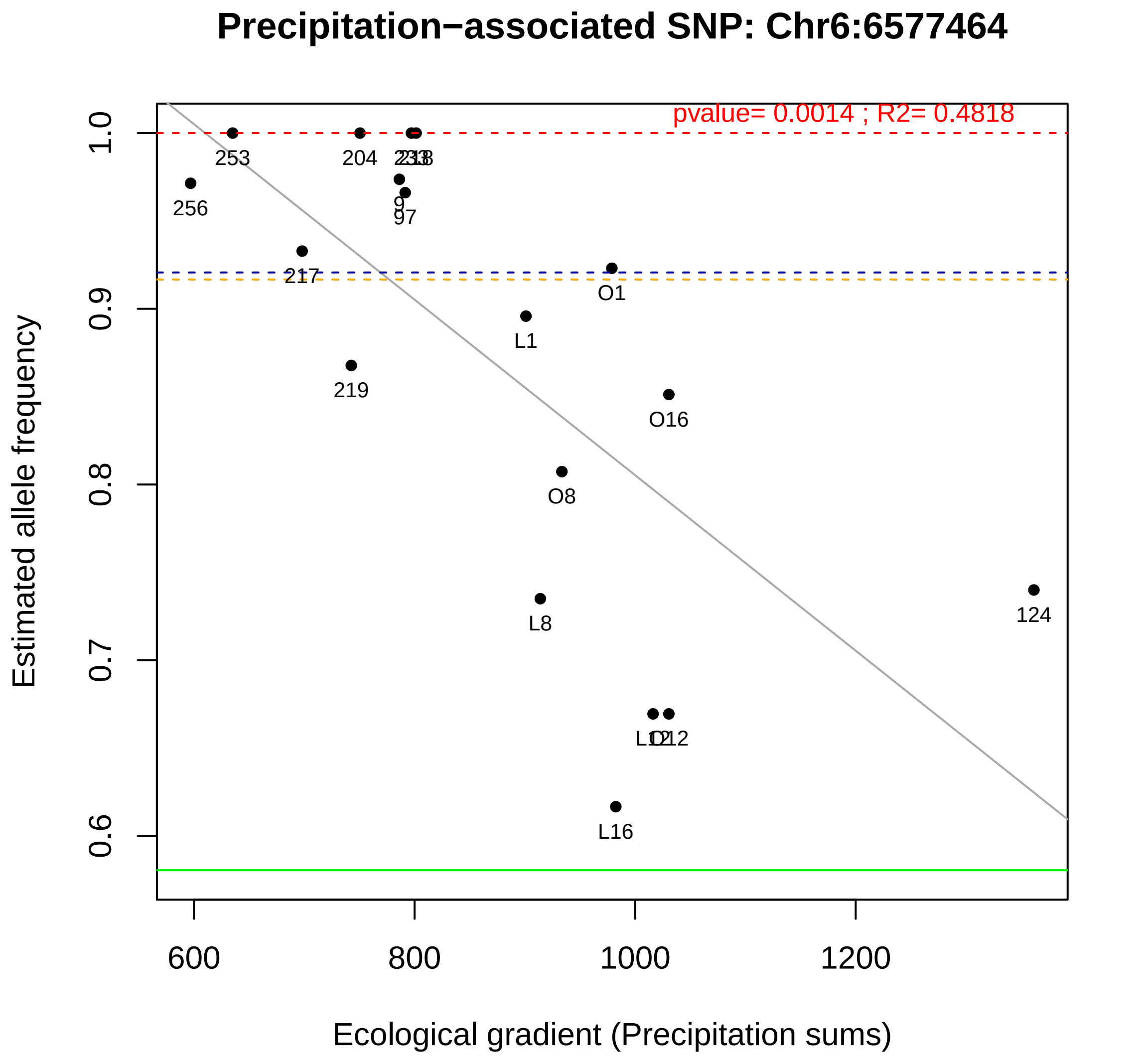
Precipitation−associated SNP: Chr6:6577464
pvalue= 0.0014 ; R2= 0.4818
Estimated allele frequency
Ecological gradient (Precipitation sums)
0.6
0.7
0.8
0.9
1.0
600
800
1000
1200
253
204
256
217
219
O1
L1
O16
O8
L8
124
L16
97
9

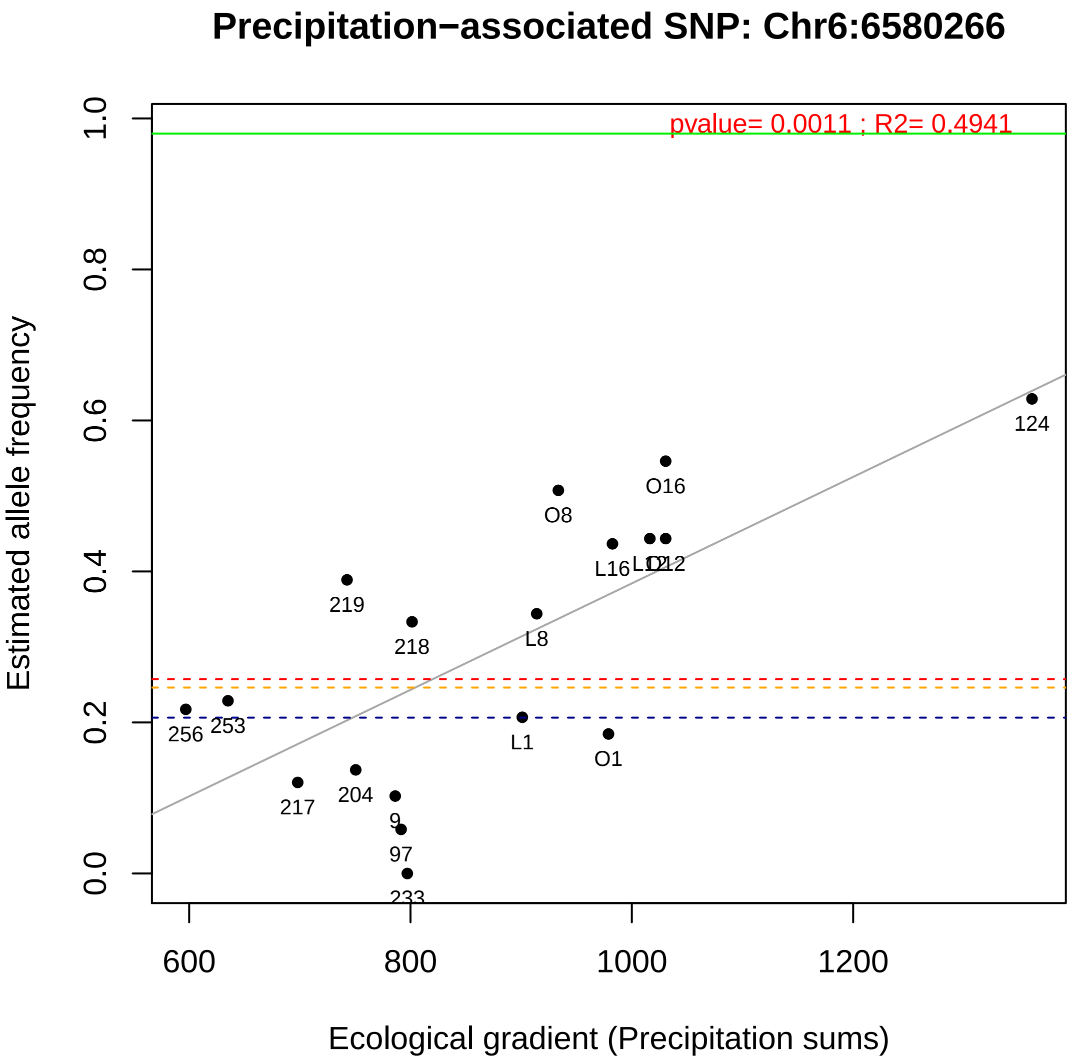

Precipitation−associated SNP: Chr6:6580266
pvalue= 0.0011 ; R2= 0.4941
Estimated allele frequency
0.0
0.2
0.4
0.6
0.8
1.0
600
800
1000
1200
Ecological gradient (Precipitation sums)
124
O16
O8
L16
L12
O12
219
218
L8
256
253
L1
O1
217
204
9
97
233

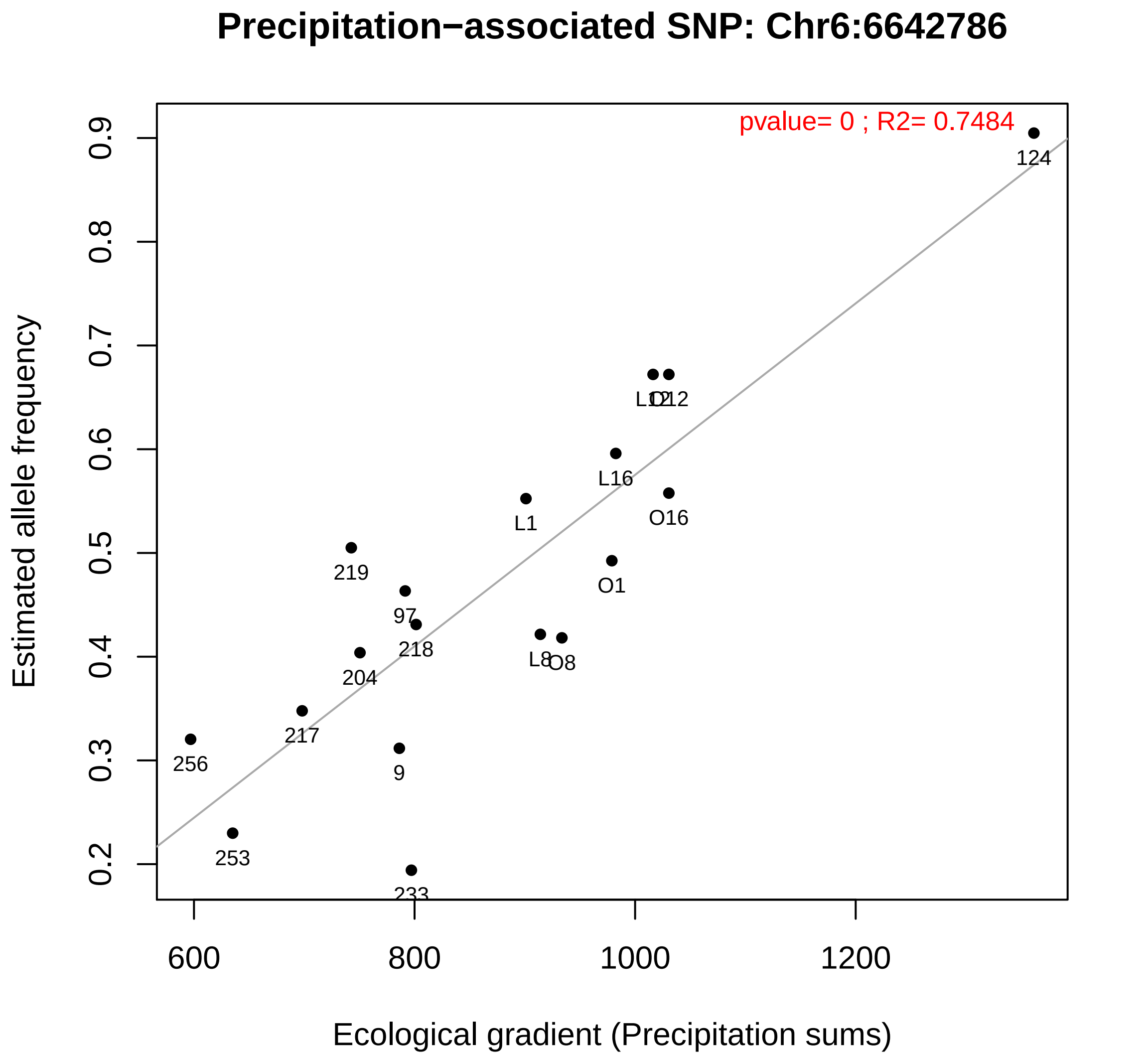

Precipitation−associated SNP: Chr6:6642786
pvalue= 0 ; R2= 0.7484
Estimated allele frequency
Ecological gradient (Precipitation sums)
0.2
0.3
0.4
0.5
0.6
0.7
0.8
0.9
600
800
1000
1200
124
L12
O12
L16
O16
L1
O1
219
97
218
204
L8
O8
217
256
9
253
233

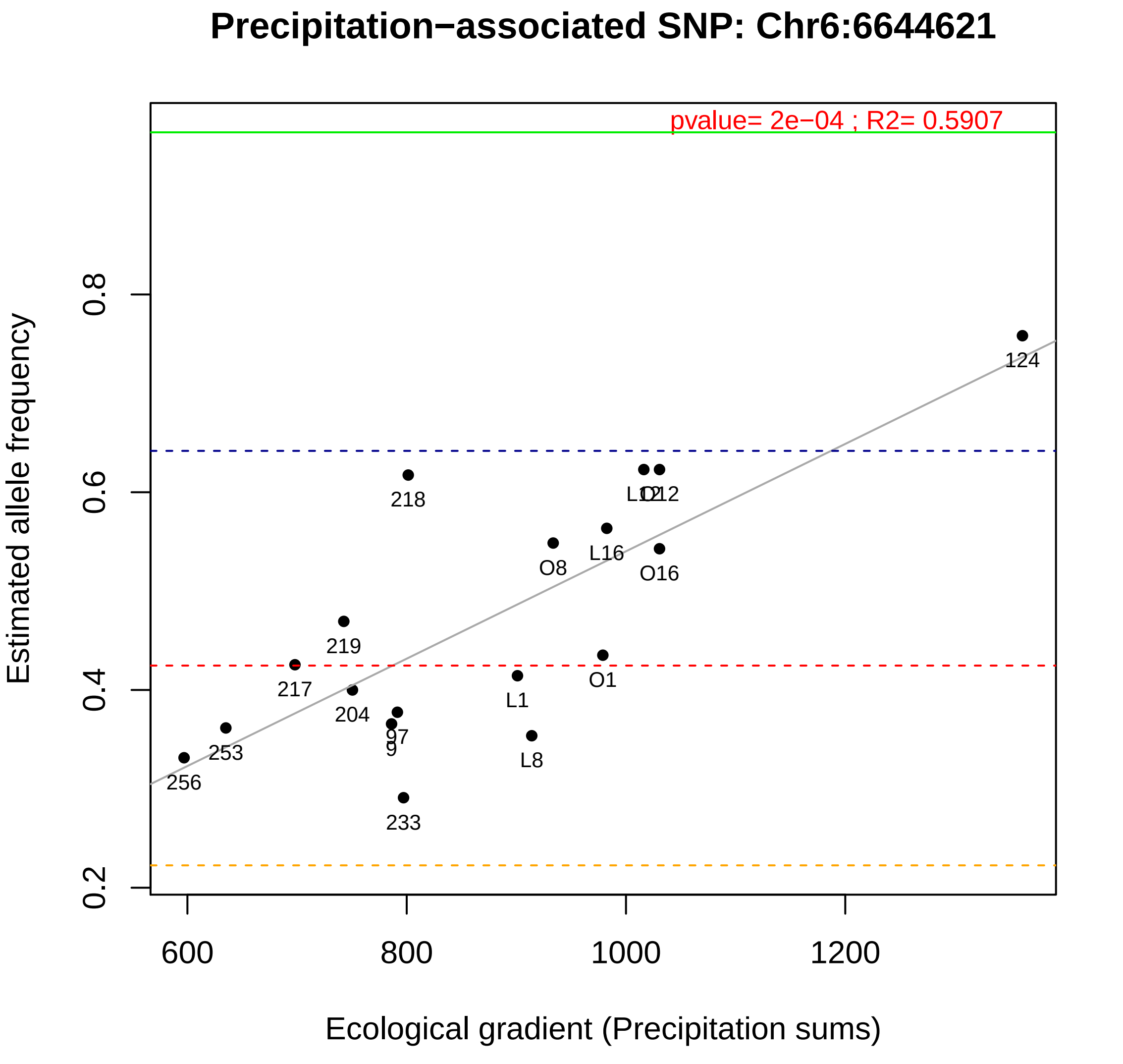

Precipitation−associated SNP: Chr6:6644621
pvalue= 2e−04 ; R2= 0.5907
Estimated allele frequency
0.2
0.4
0.6
0.8
600
800
1000
1200
Ecological gradient (Precipitation sums)
124
218
L12
O12
L16
O8
O16
219
217
204
O1
L1
97
9
253
256
L8
233

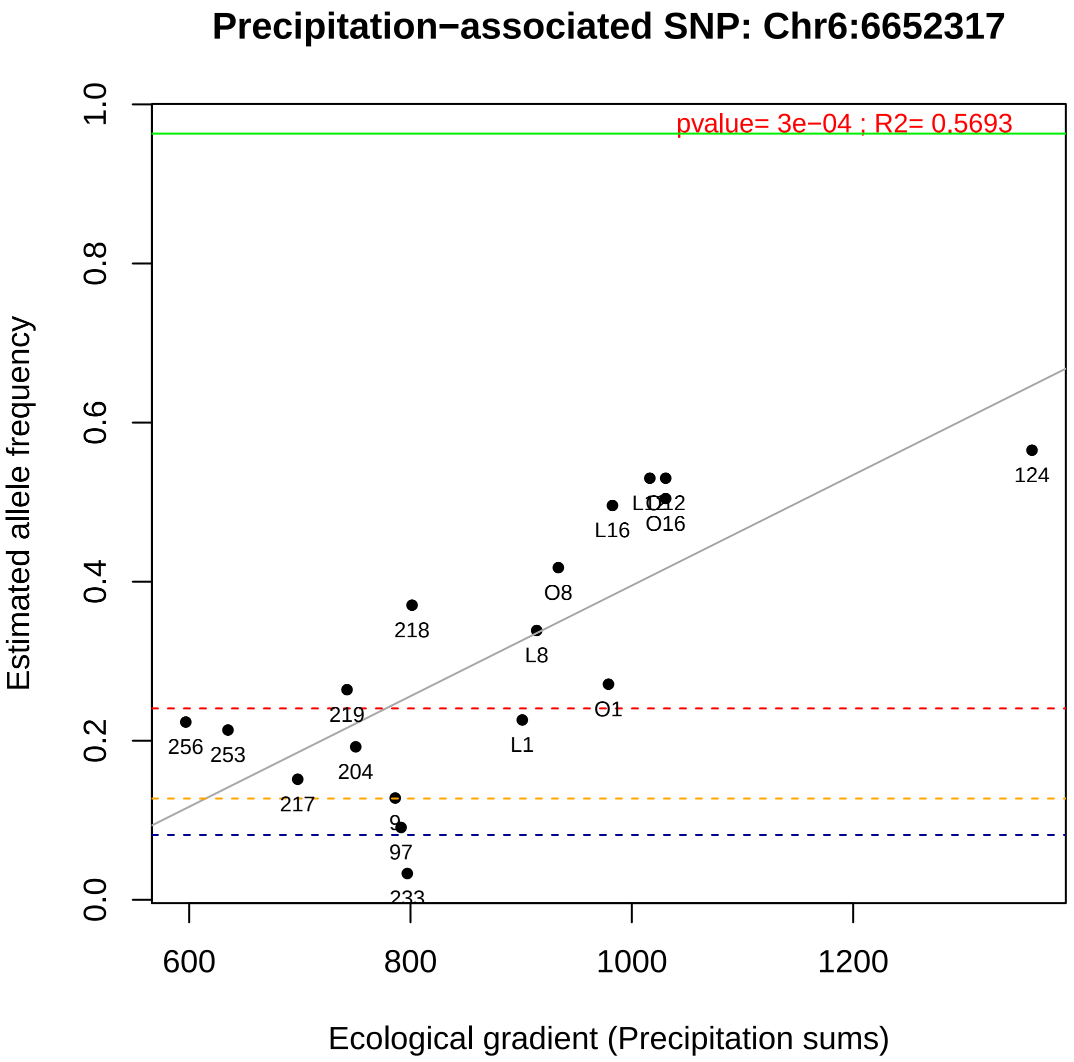
Precipitation−associated SNP: Chr6:6652317
pvalue= 3e−04 ; R2= 0.5693
Estimated allele frequency
0.0
0.2
0.4
0.6
0.8
1.0
600
800
1000
1200
Ecological gradient (Precipitation sums)
124
L12
O12
O16
L16
O8
218
L8
O1
219
256
253
204
L1
217
9
97
233

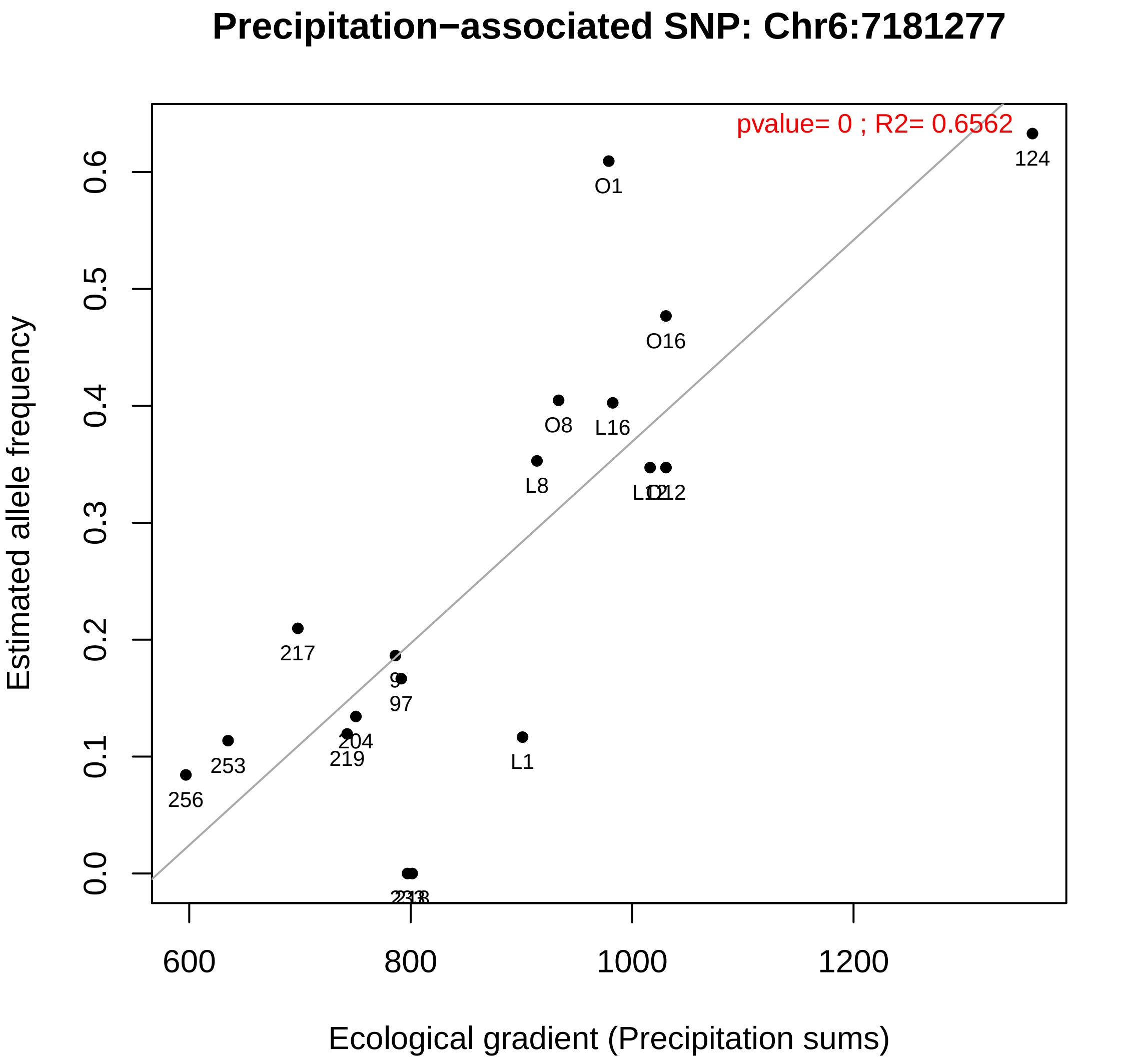

Precipitation−associated SNP: Chr6:7181277
pvalue= 0 ; R2= 0.6562
124
O1
O16
O8
L16
L8
L12
O12
217
9
97
204
219
253
256
L1
0.0
0.1
0.2
0.3
0.4
0.5
0.6
600
800
1000
1200
Estimated allele frequency
Ecological gradient (Precipitation sums)

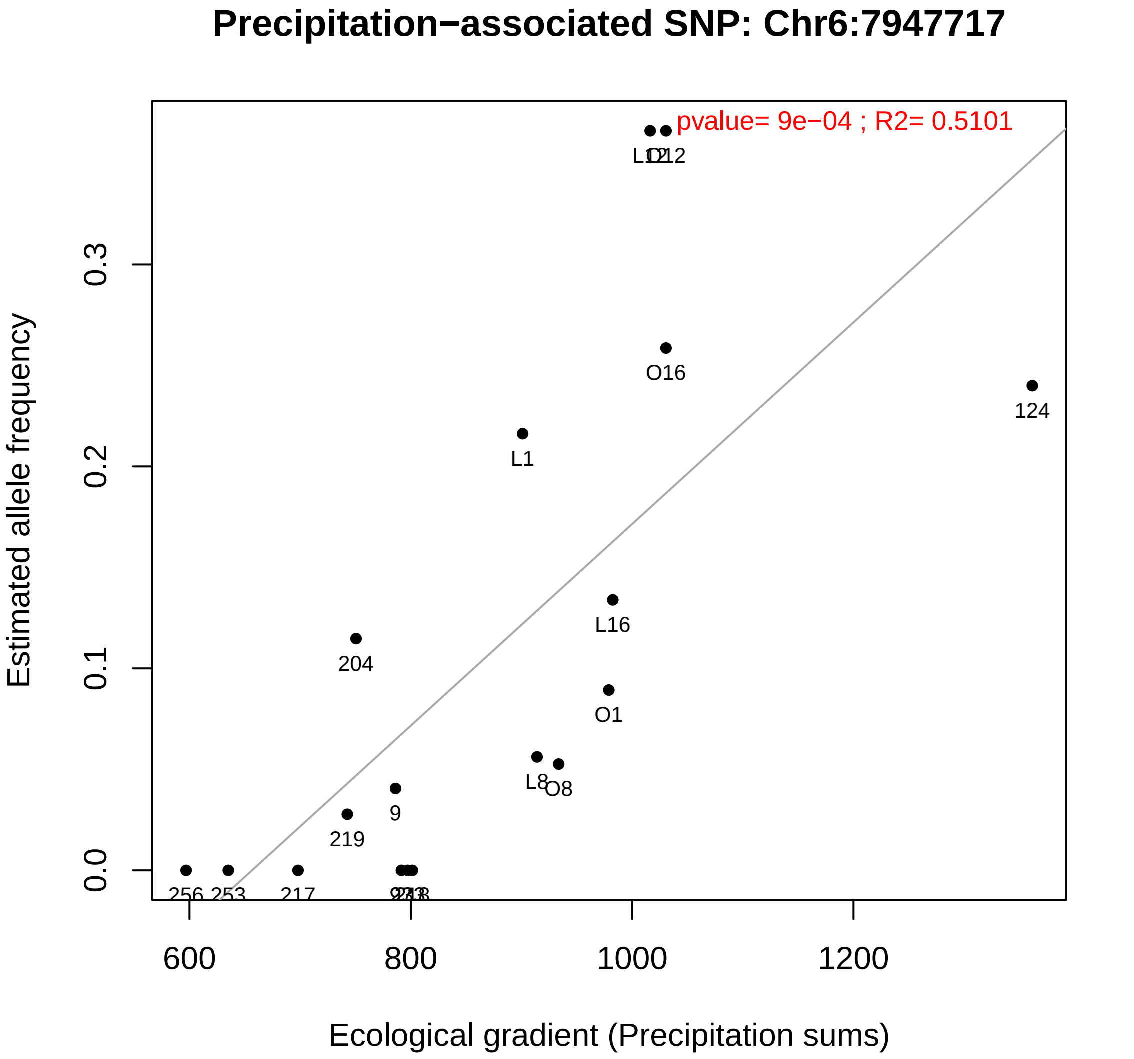
Precipitation−associated SNP: Chr6:7947717
pvalue= 9e−04 ; R2= 0.5101
O16
124
L1
L16
204
O1
L8
O8
9
219
256
253
217
Estimated allele frequency
0.0
0.1
0.2
0.3
600
800
1000
1200
Ecological gradient (Precipitation sums)

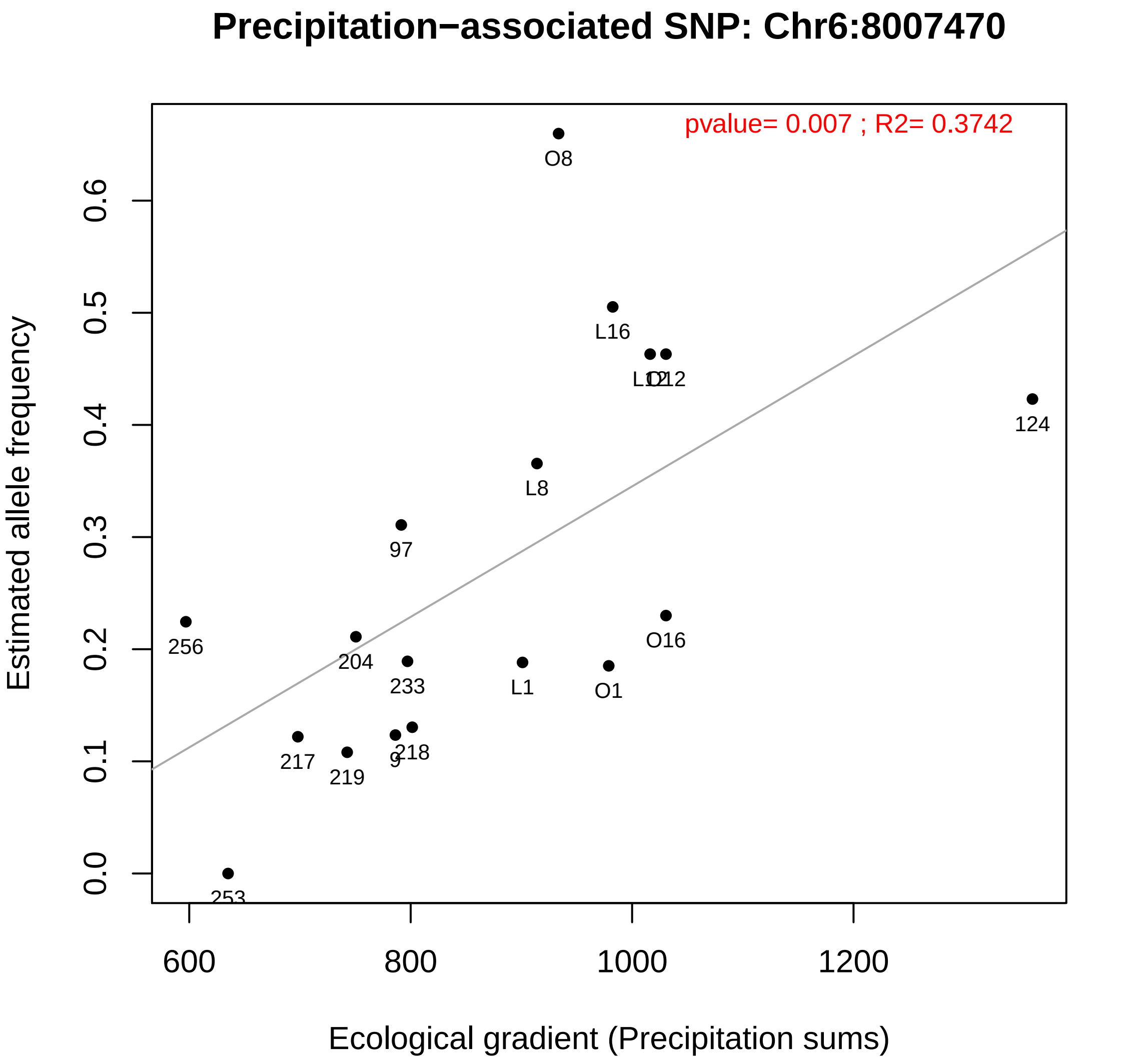
Precipitation−associated SNP: Chr6:8007470
pvalue= 0.007 ; R2= 0.3742
O8
L16
L12
O12
124
L8
97
256
204
233
L1
O1
O16
217
219
9
218
253
0.0
0.1
0.2
0.3
0.4
0.5
0.6
600
800
1000
1200
Estimated allele frequency
Ecological gradient (Precipitation sums)

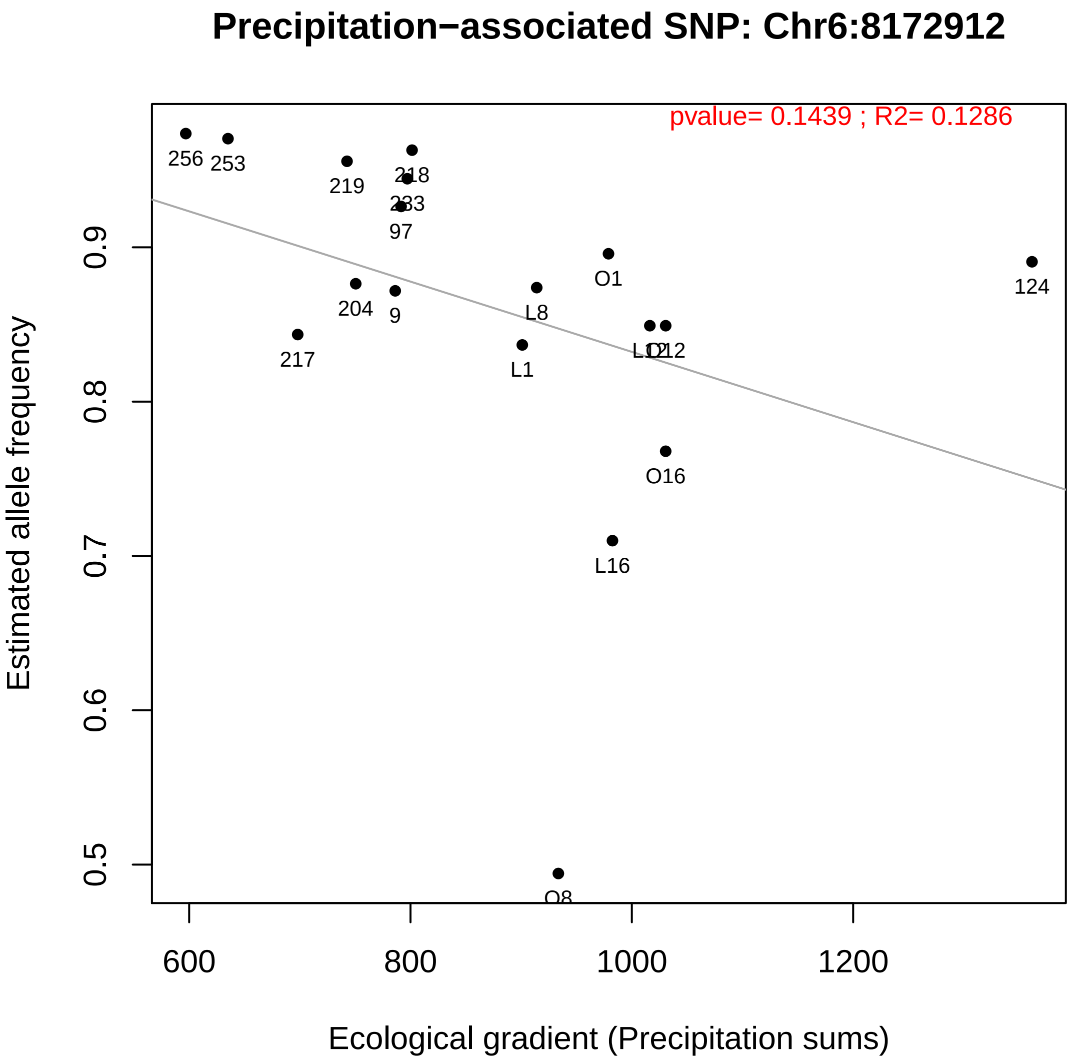
Precipitation−associated SNP: Chr6:8172912
pvalue= 0.1439 ; R2= 0.1286
256
253
219
218
233
97
204
9
217
L8
L1
O1
L12
O12
O16
L16
124
O8
0.5
0.6
0.7
0.8
0.9
600
800
1000
1200
Estimated allele frequency
Ecological gradient (Precipitation sums)

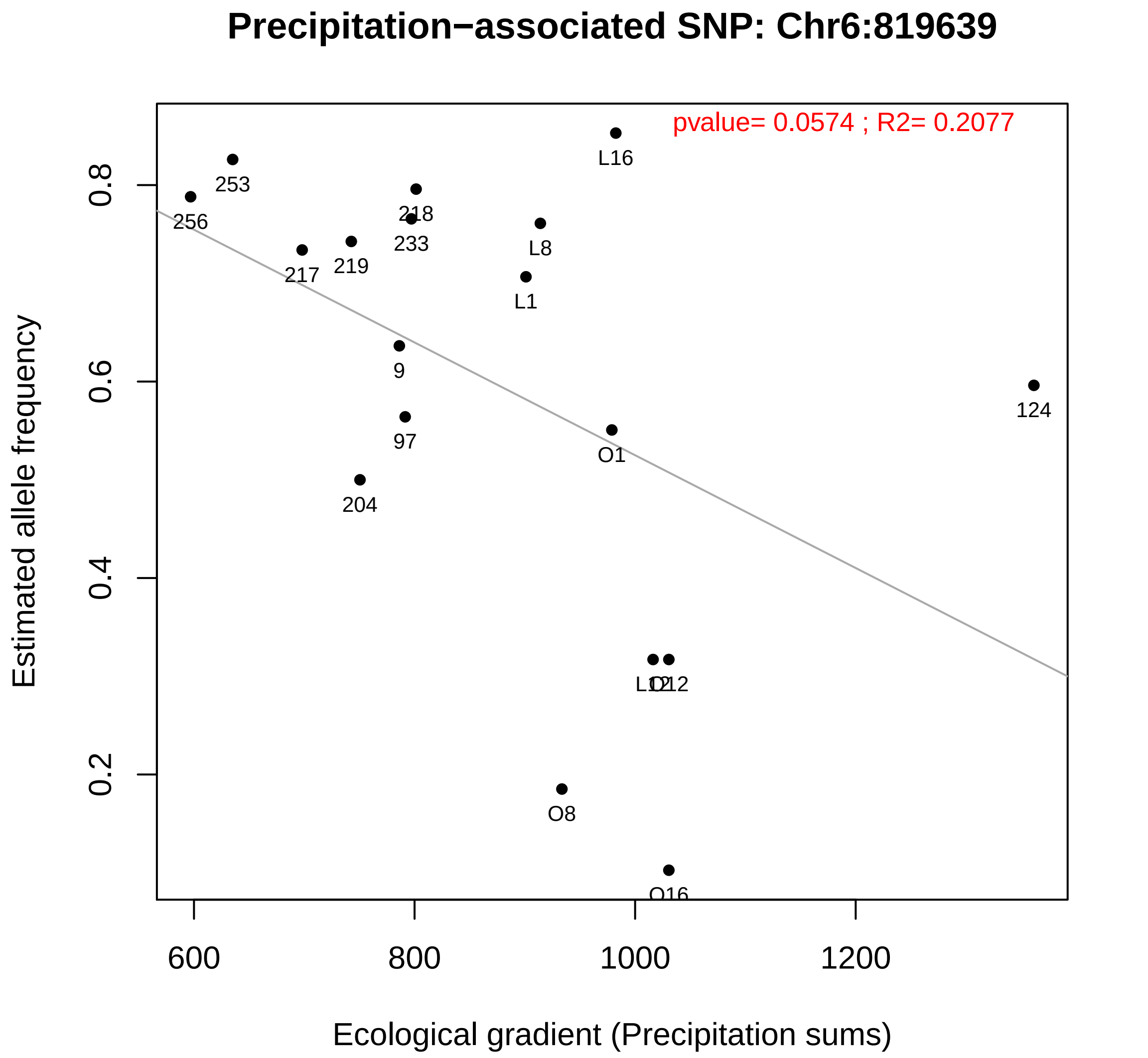
Precipitation−associated SNP: Chr6:819639
pvalue= 0.0574 ; R2= 0.2077
Estimated allele frequency
0.2
0.4
0.6
0.8
600
800
1000
1200
Ecological gradient (Precipitation sums)
256
253
218
233
217
219
L8
L1
L16
9
97
204
O1
124
L12
O12
O8
O16

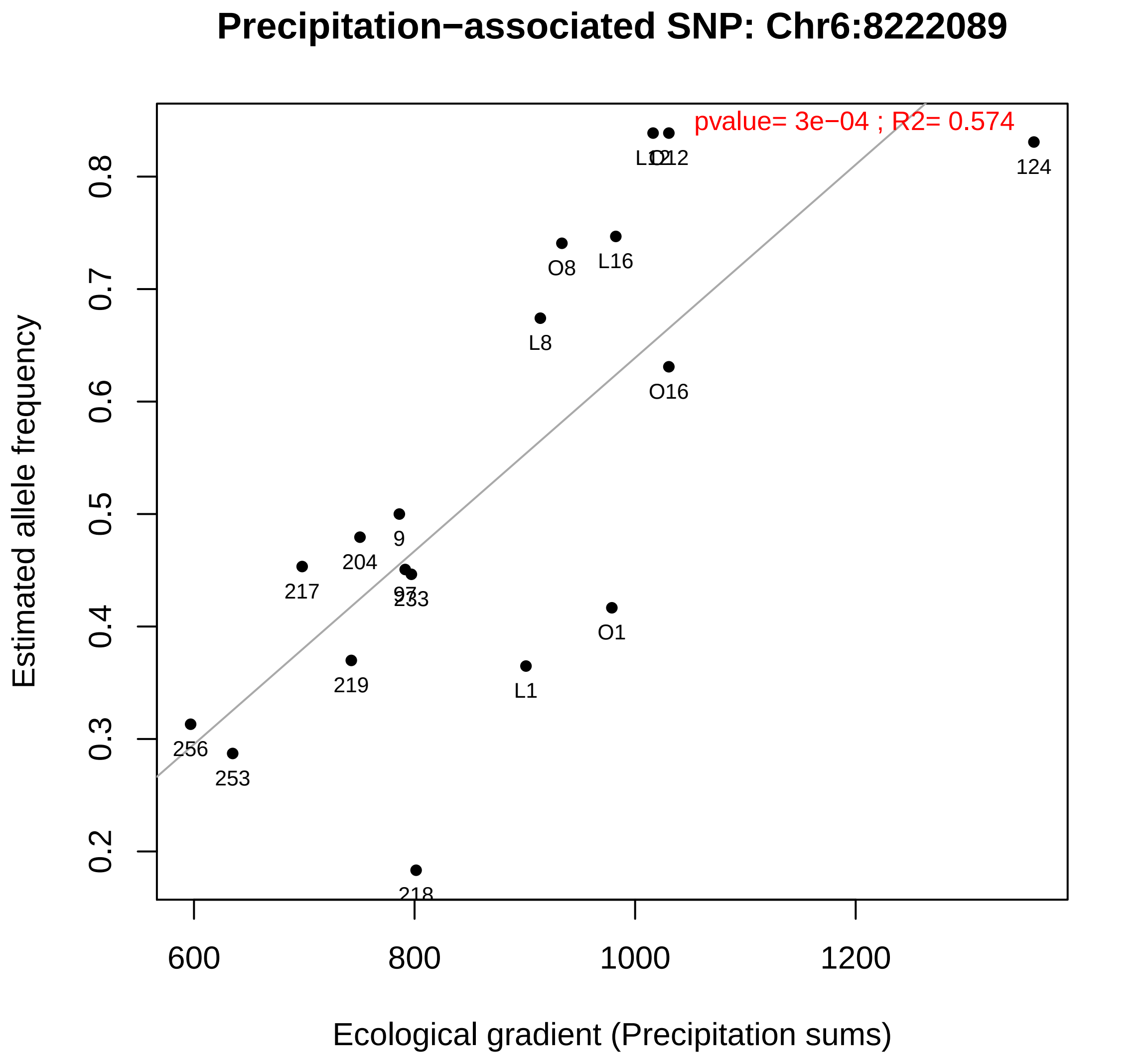
Precipitation−associated SNP: Chr6:8222089
pvalue= 3e−04 ; R2= 0.574
Estimated allele frequency
0.2
0.3
0.4
0.5
0.6
0.7
0.8
600
800
1000
1200
Ecological gradient (Precipitation sums)
124
L16
O8
L8
O16
9
204
217
97
233
O1
219
L1
256
253
218

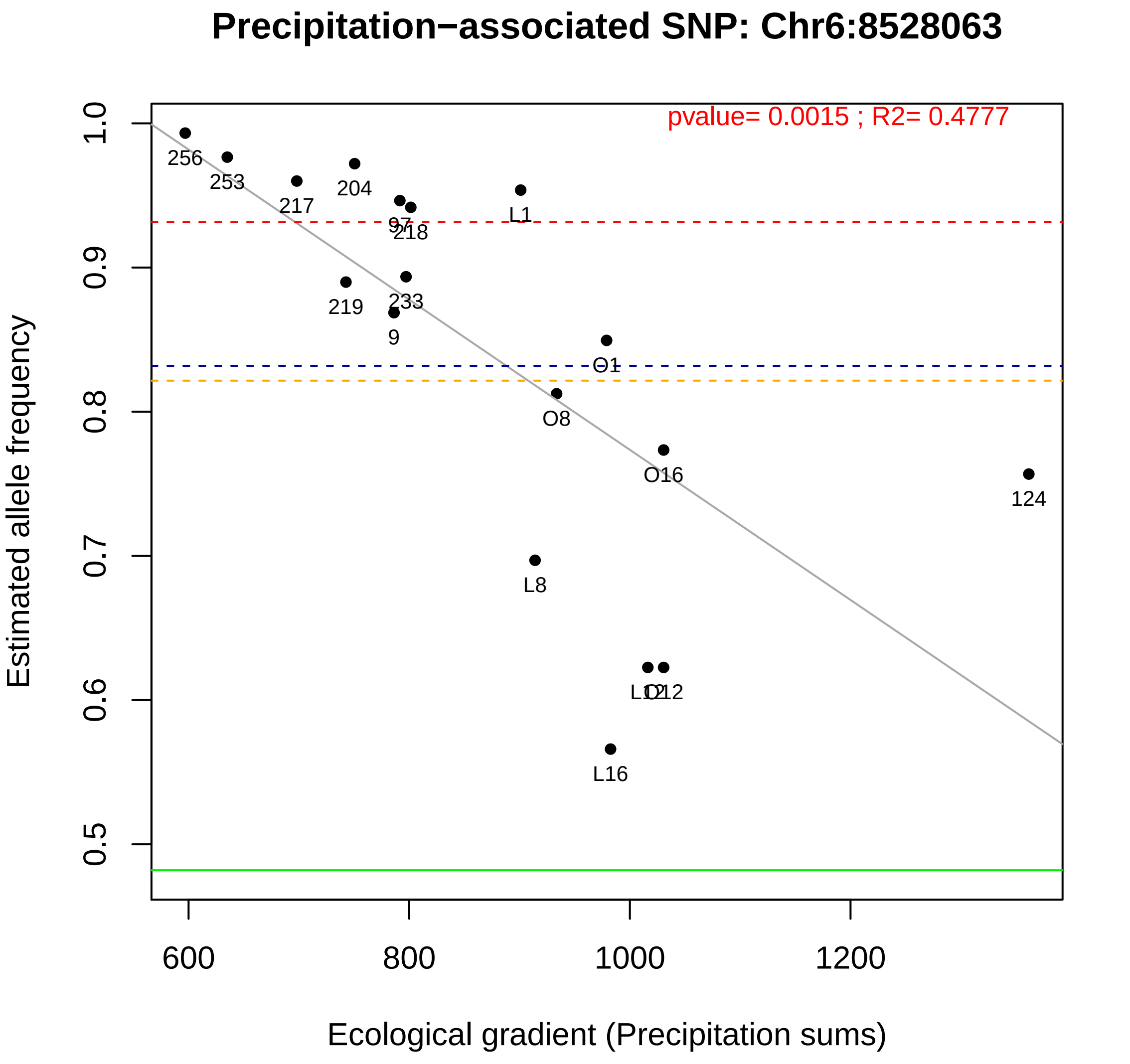
Precipitation−associated SNP: Chr6:8528063
pvalue= 0.0015 ; R2= 0.4777
Estimated allele frequency
Ecological gradient (Precipitation sums)
1.0
0.9
0.8
0.7
0.6
0.5
600
800
1000
1200
256
253
217
204
97
218
L1
219
233
9
O1
O8
O16
124
L8
L12
O12
L16

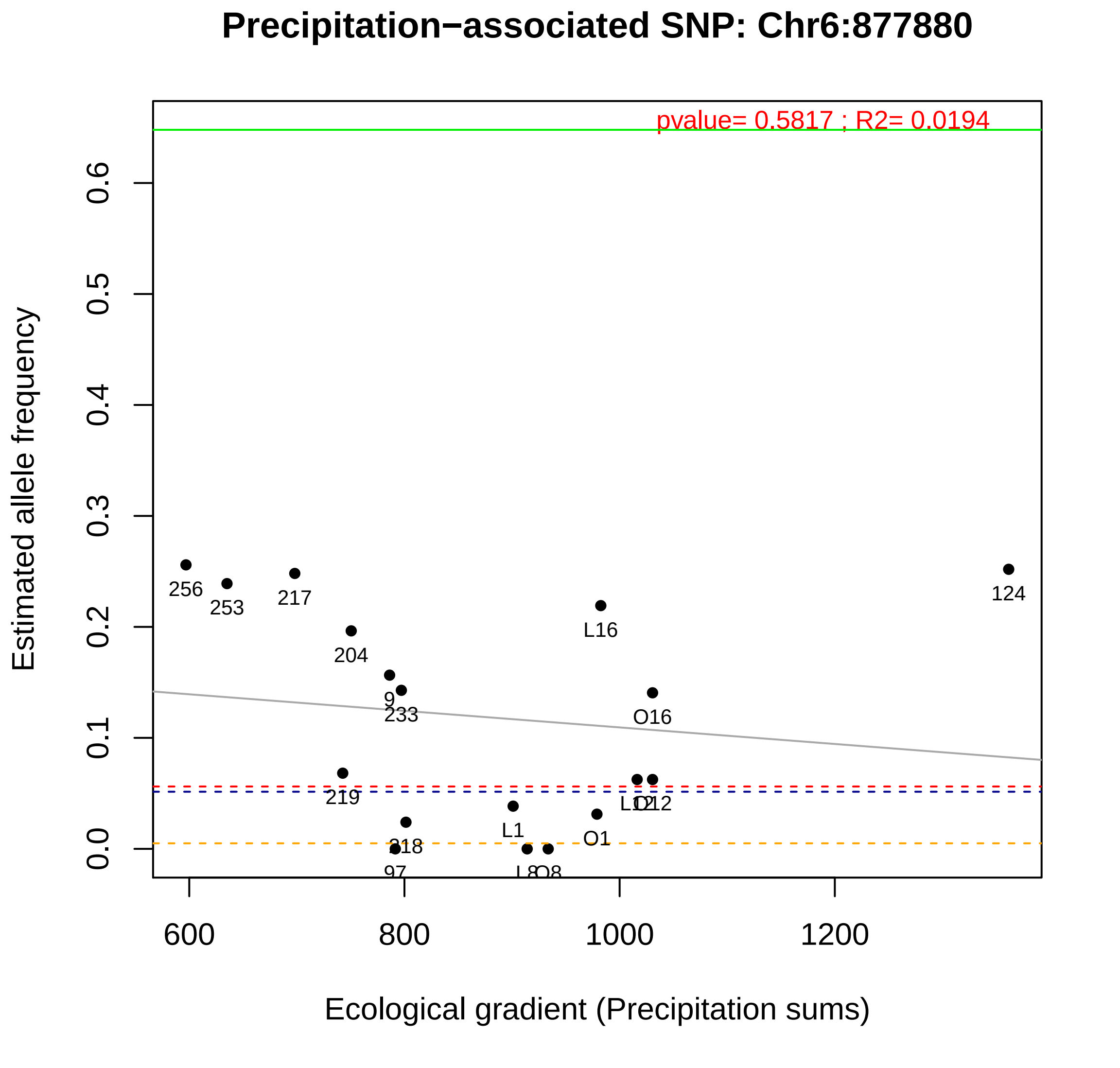

Precipitation−associated SNP: Chr6:877880
pvalue= 0.5817 ; R2= 0.0194
Estimated allele frequency
0.0
0.1
0.2
0.3
0.4
0.5
0.6
600
800
1000
1200
Ecological gradient (Precipitation sums)
256
253
217
204
9
233
219
218
97
L1
L8
O8
O1
L16
O16
L12
O12
124

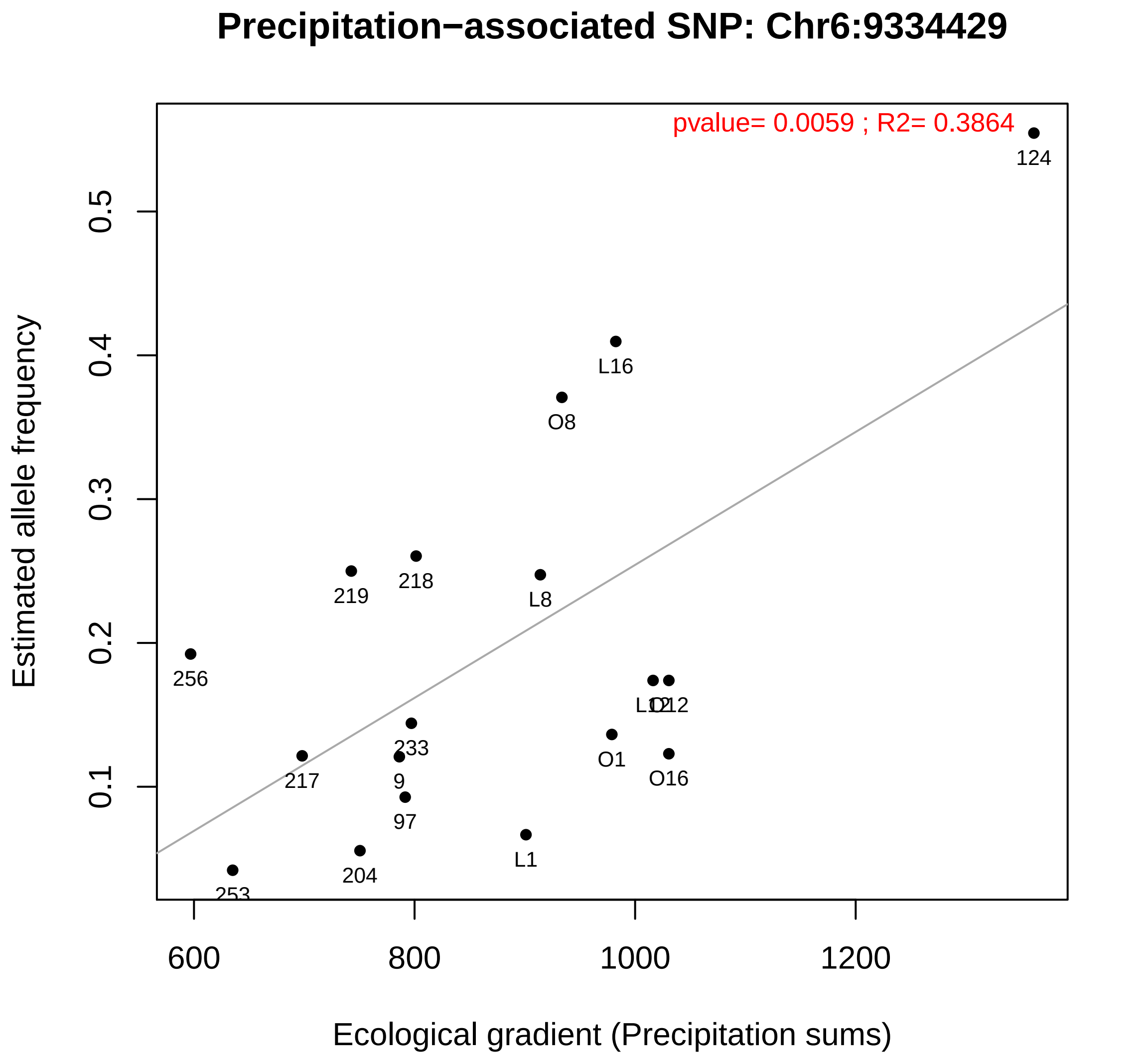
Precipitation−associated SNP: Chr6:9334429
pvalue= 0.0059 ; R2= 0.3864
124
L16
O8
219
218
L8
256
L12
O12
233
O1
217
9
O16
97
L1
204
253
0.1
0.2
0.3
0.4
0.5
600
800
1000
1200
Estimated allele frequency
Ecological gradient (Precipitation sums)

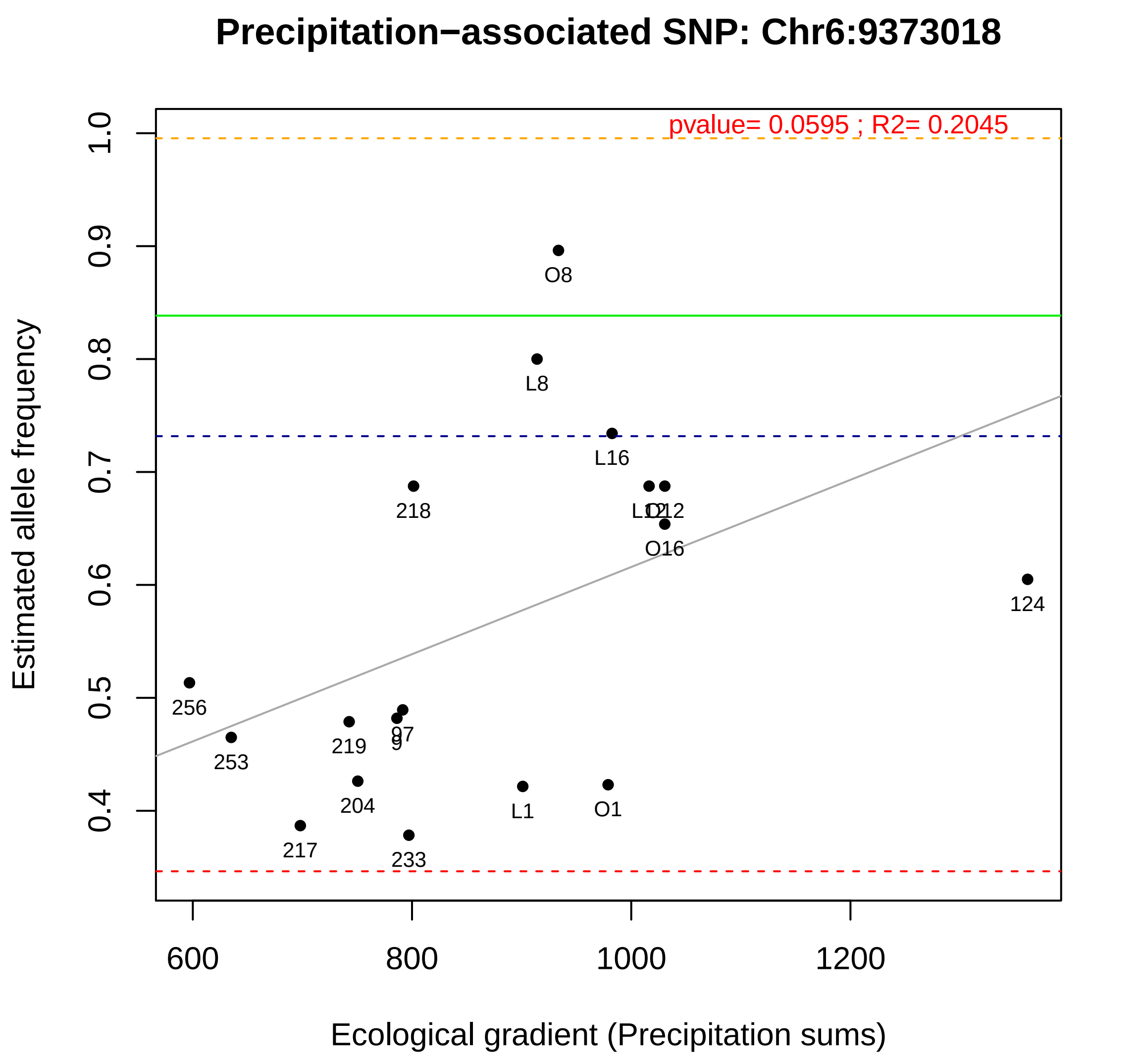
Precipitation−associated SNP: Chr6:9373018
pvalue= 0.0595 ; R2= 0.2045
Estimated allele frequency
0.4
0.5
0.6
0.7
0.8
0.9
1.0
600
800
1000
1200
Ecological gradient (Precipitation sums)
O8
L8
L16
218
O12
O16
124
256
253
219
97
9
204
217
233
L1
O1

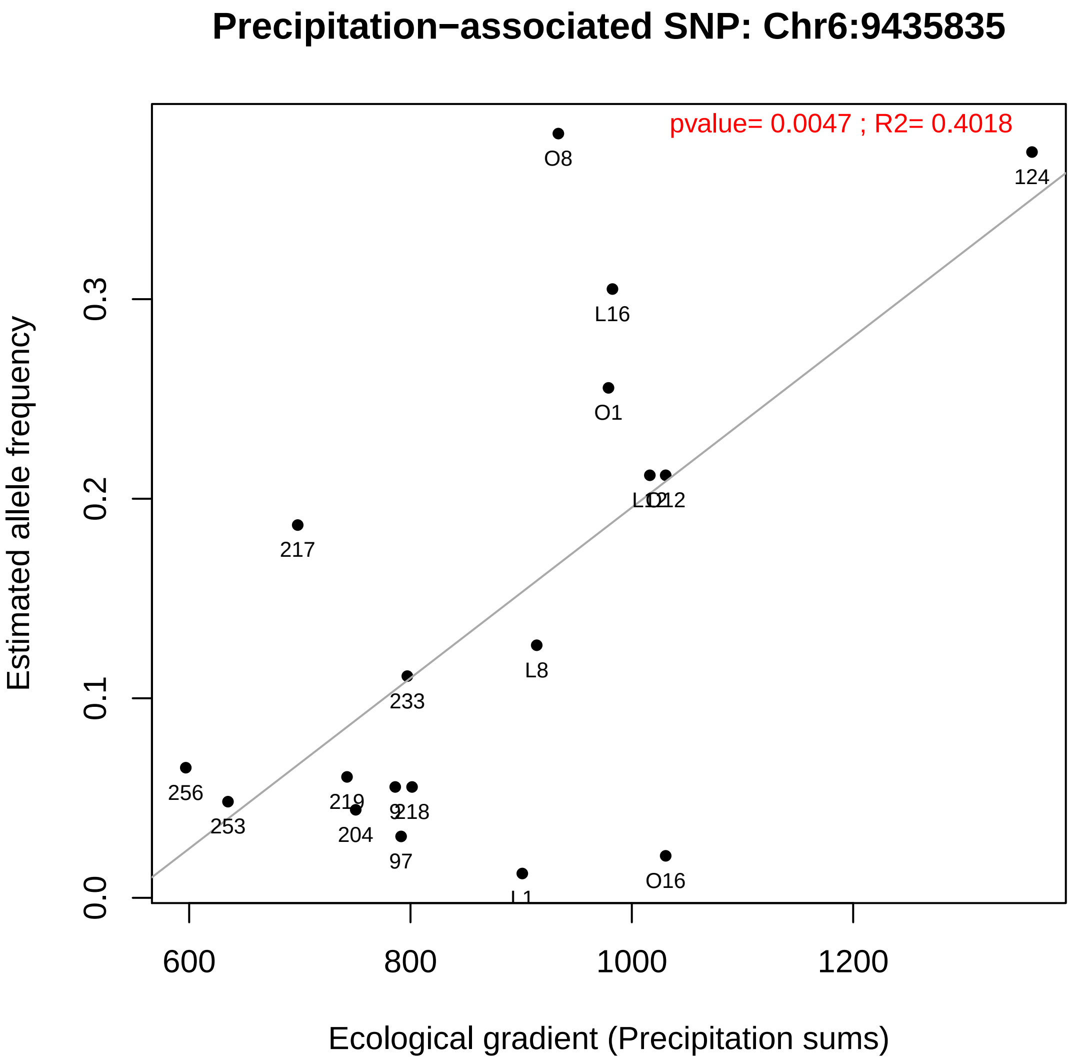

Precipitation−associated SNP: Chr6:9435835
pvalue= 0.0047 ; R2= 0.4018
Estimated allele frequency
Ecological gradient (Precipitation sums)
0.0
0.1
0.2
0.3
600
800
1000
1200
O8
124
L16
O1
L10
O12
217
L8
233
256
253
219
204
9
218
97
L1
O16

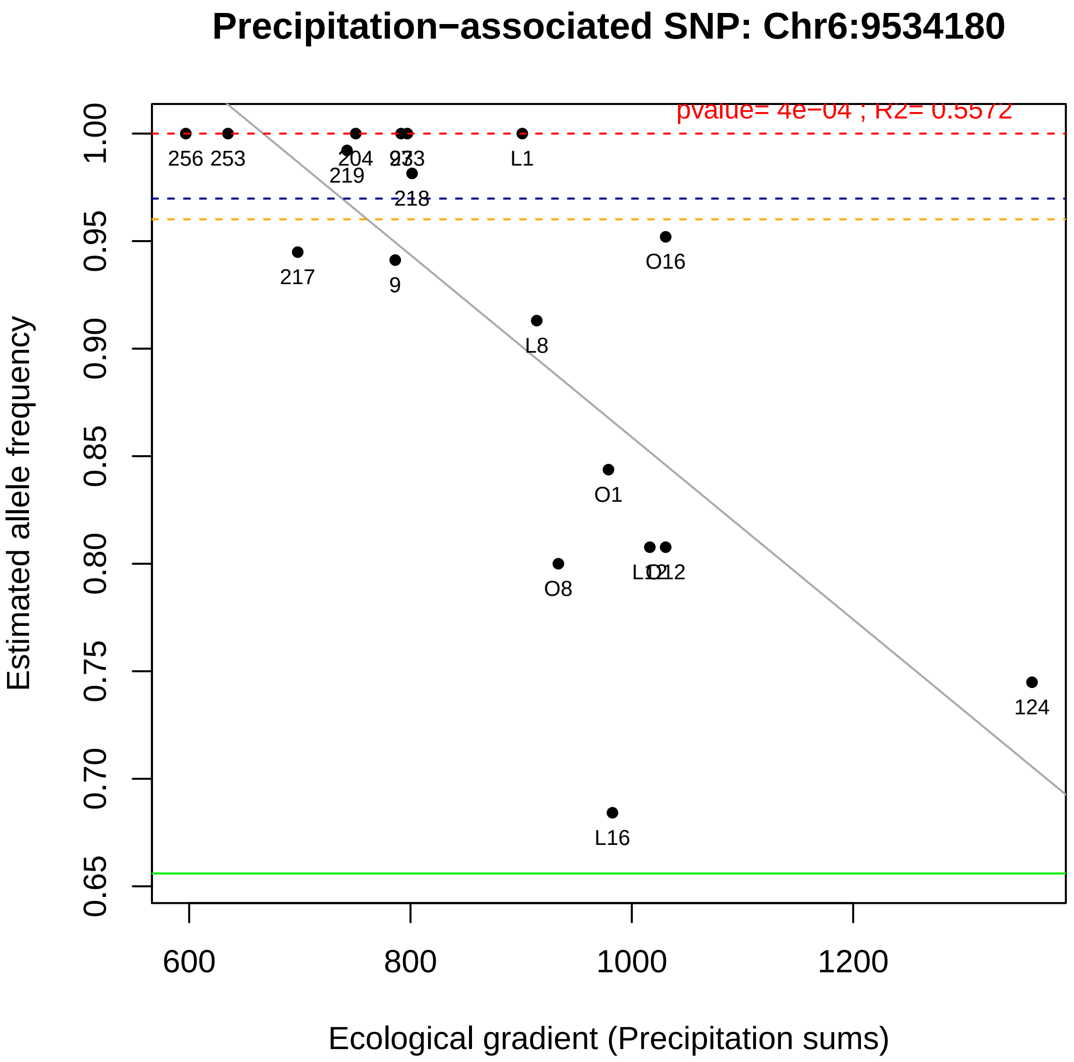

Precipitation−associated SNP: Chr6:9534180
pvalue= 4e−04 ; R2= 0.5572
Estimated allele frequency
Ecological gradient (Precipitation sums)
0.65
0.70
0.75
0.80
0.85
0.90
0.95
1.00
600
800
1000
1200
256
253
204
219
L1
218
O16
217
9
L8
O1
O8
124
L16

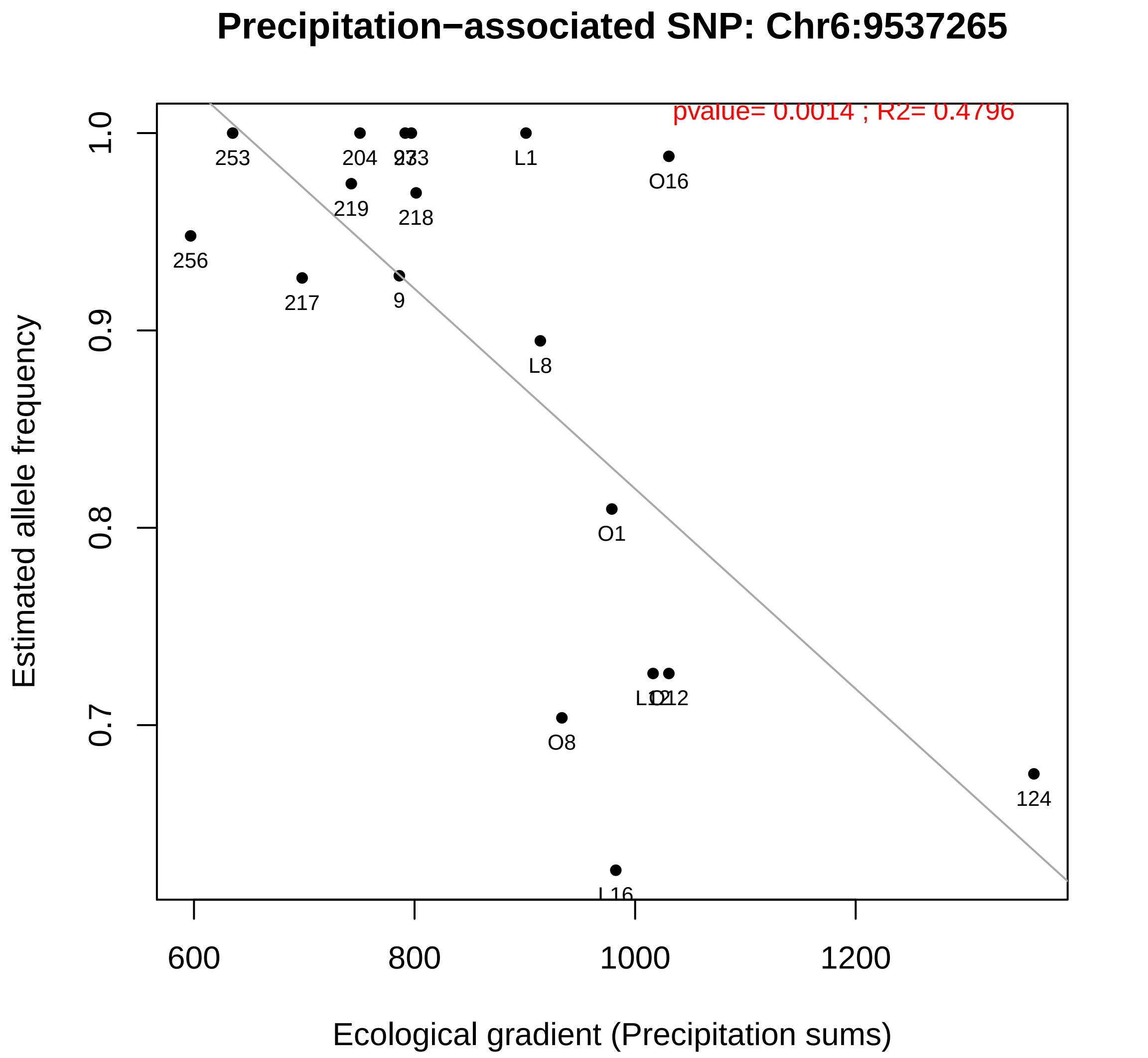

Precipitation−associated SNP: Chr6:9537265
pvalue= 0.0014 ; R2= 0.4796
253
204
97
233
L1
O16
219
218
256
217
9
L8
O1
L12
O12
O8
124
L16
1.0
0.9
0.8
0.7
600
800
1000
1200
Estimated allele frequency
Ecological gradient (Precipitation sums)

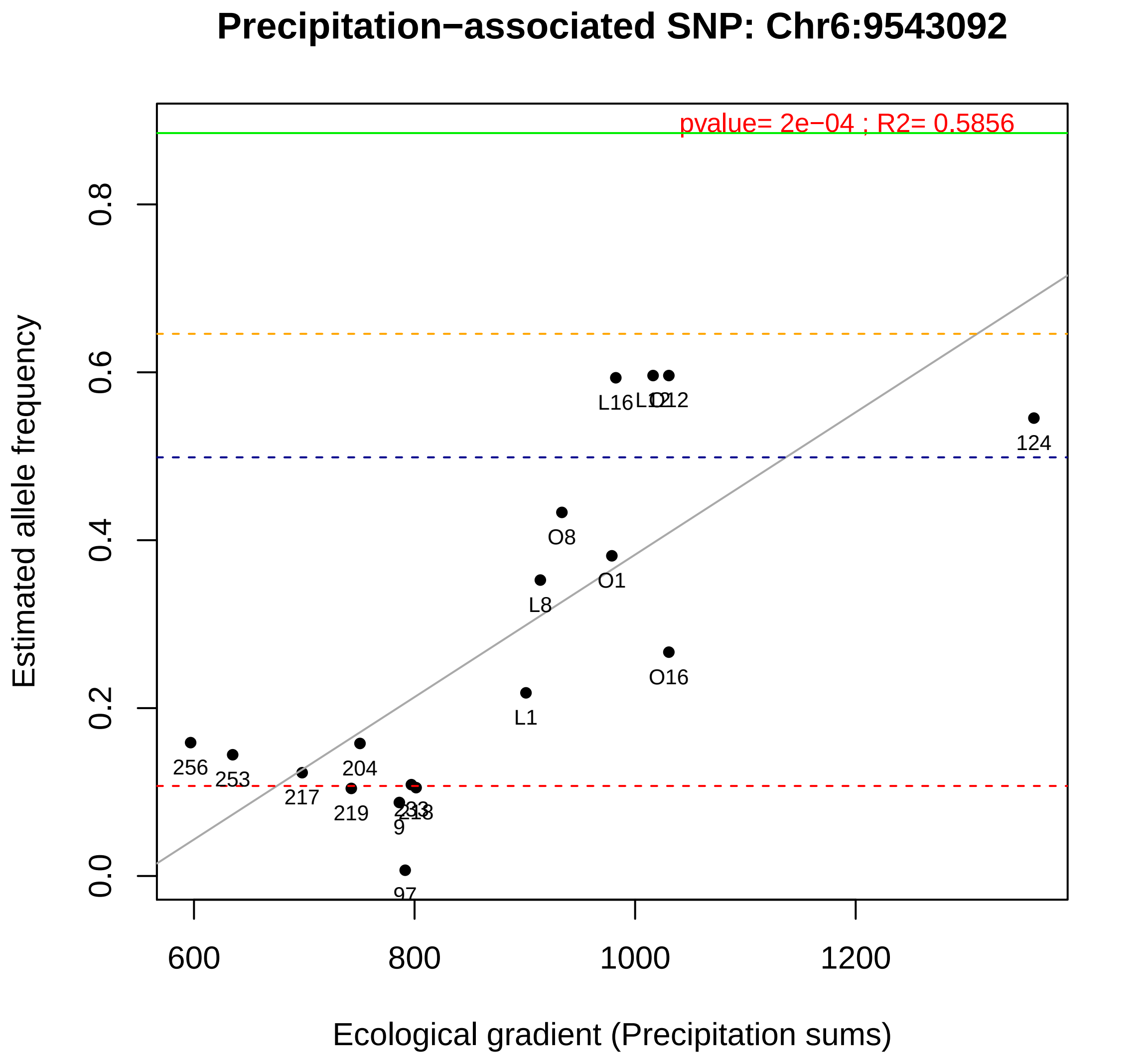
Precipitation−associated SNP: Chr6:9543092
pvalue= 2e−04 ; R2= 0.5856
Estimated allele frequency
0.0
0.2
0.4
0.6
0.8
600
800
1000
1200
Ecological gradient (Precipitation sums)
256
253
217
204
219
9
97
L1
L8
O8
O1
O16
L16
O12
124

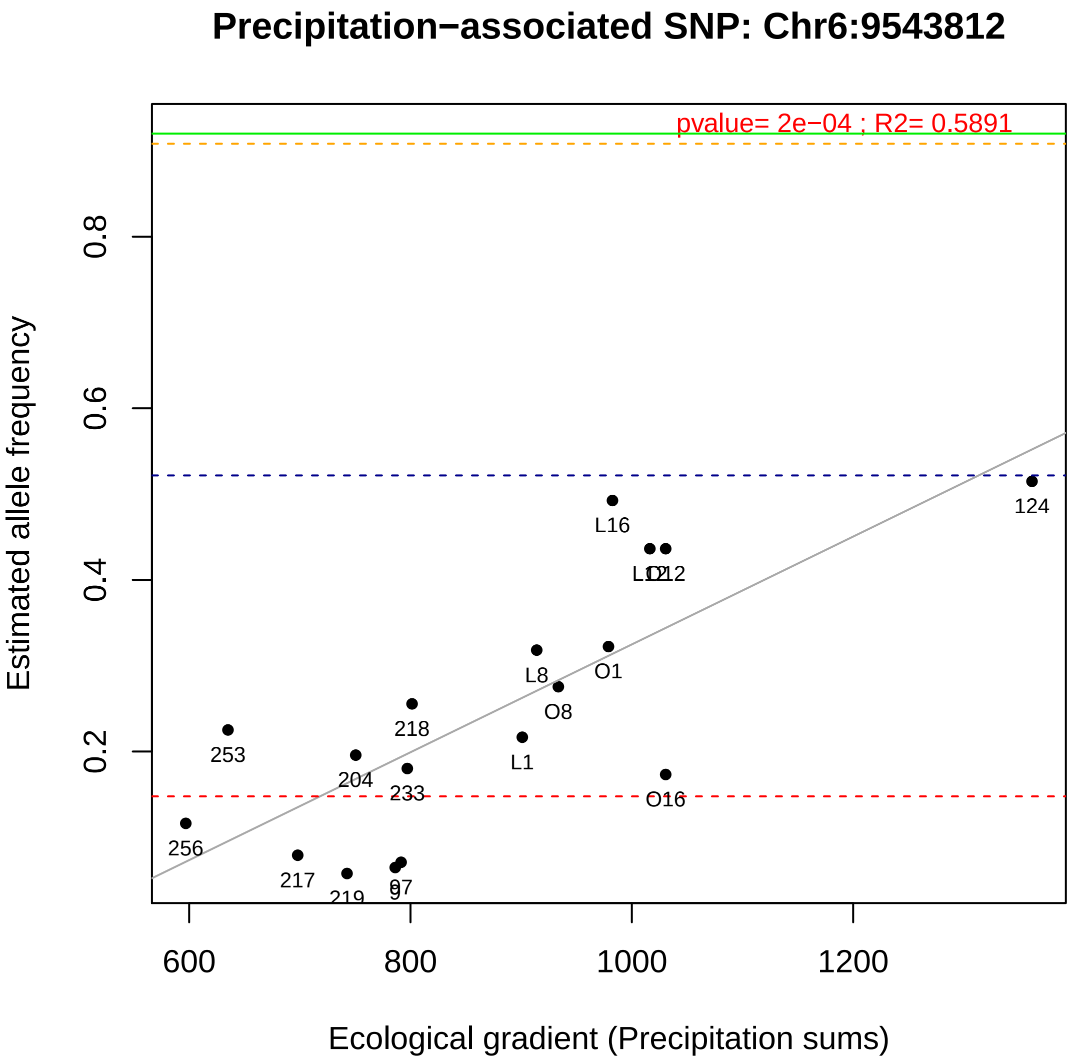
Precipitation−associated SNP: Chr6:9543812
pvalue= 2e−04 ; R2= 0.5891
Estimated allele frequency
0.2
0.4
0.6
0.8
600
800
1000
1200
Ecological gradient (Precipitation sums)
124
L16
O1
L8
O8
218
253
204
233
L1
O16
256
217
219
97

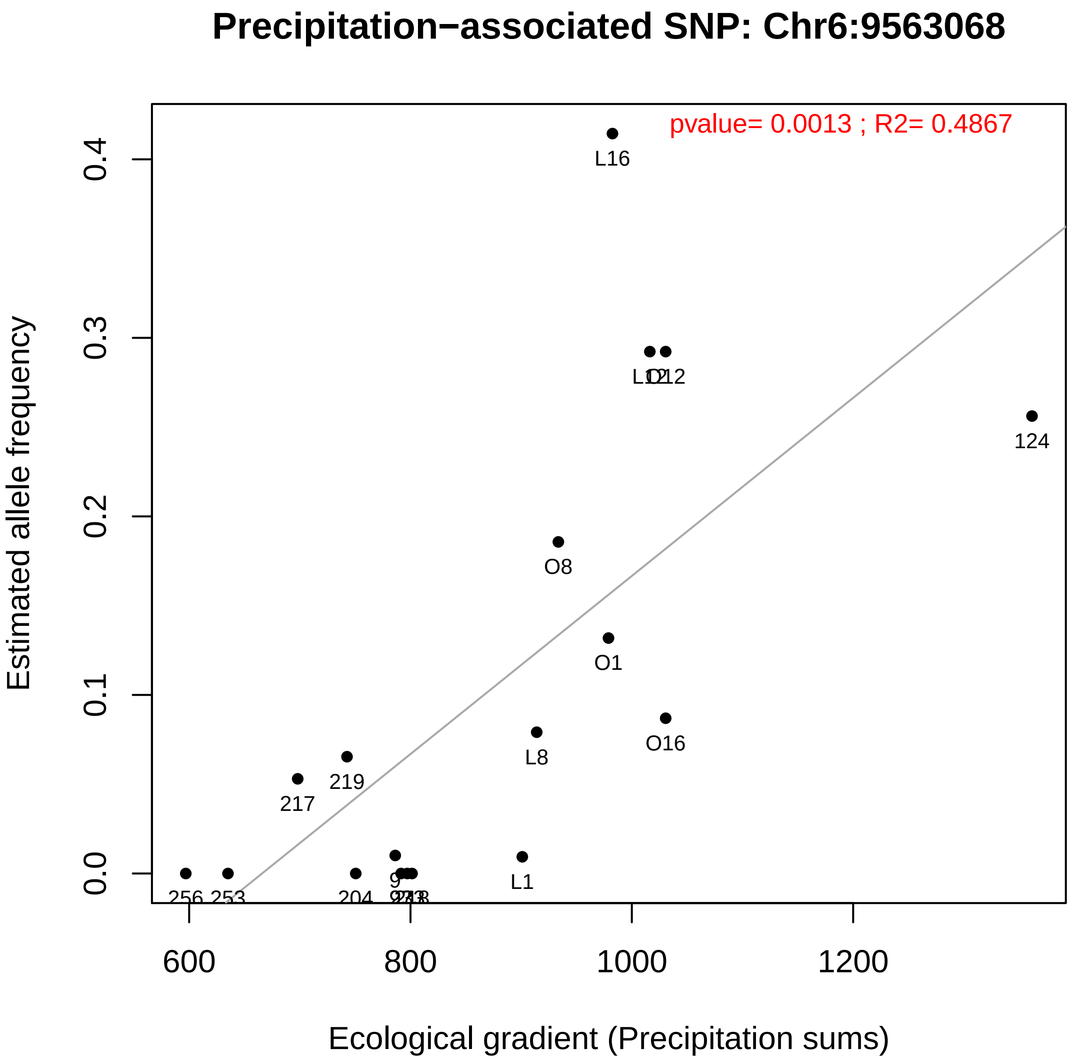
Precipitation−associated SNP: Chr6:9563068
pvalue= 0.0013 ; R2= 0.4867
Estimated allele frequency
Ecological gradient (Precipitation sums)
0.0
0.1
0.2
0.3
0.4
600
800
1000
1200
L16
124
O8
O1
O16
L8
219
217
L1
204
256
253

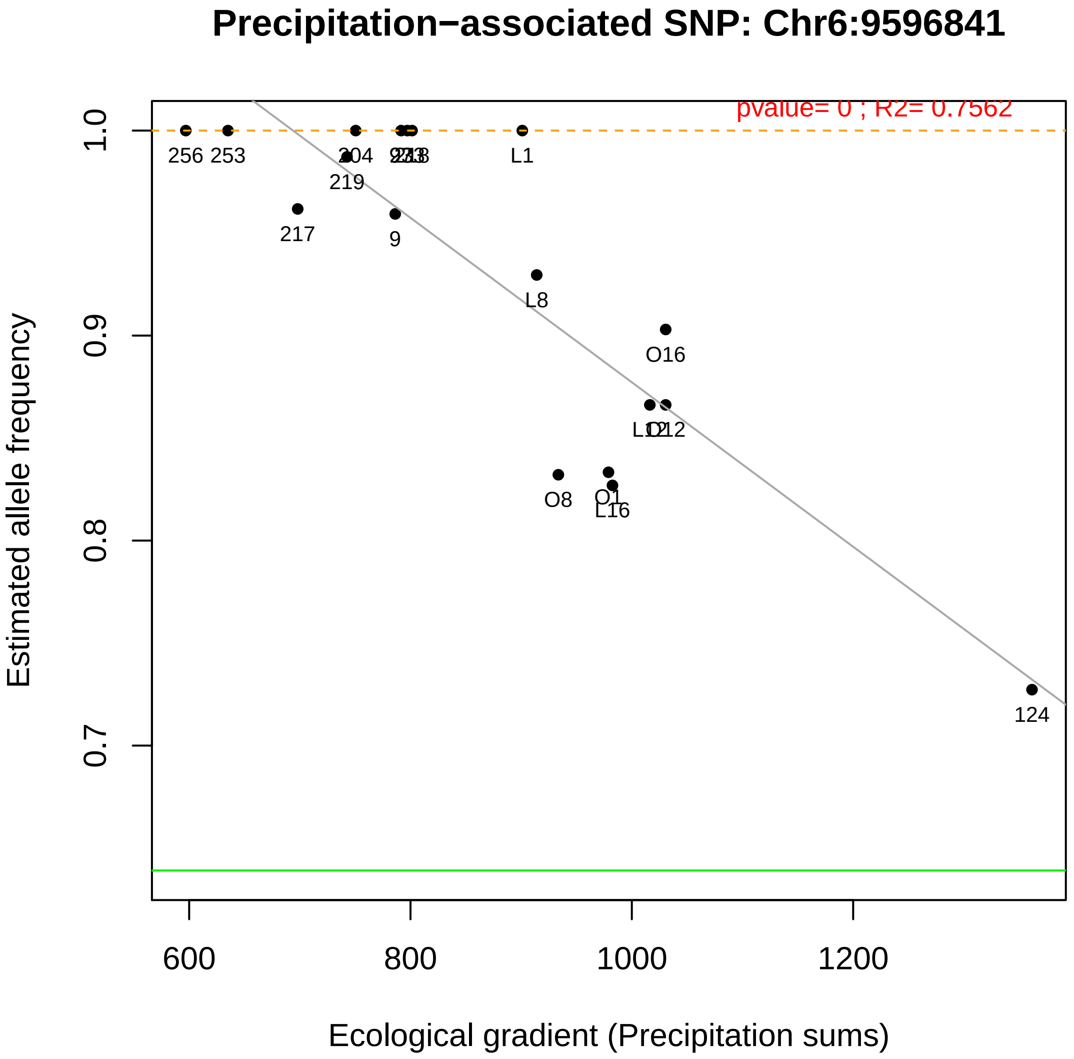

Precipitation−associated SNP: Chr6:9596841
pvalue= 0 ; R2= 0.7562
Estimated allele frequency
Ecological gradient (Precipitation sums)
1.0
0.9
0.8
0.7
600
800
1000
1200
256
253
204
219
217
9
L1
L8
O16
O12
O8
O1
L16
124

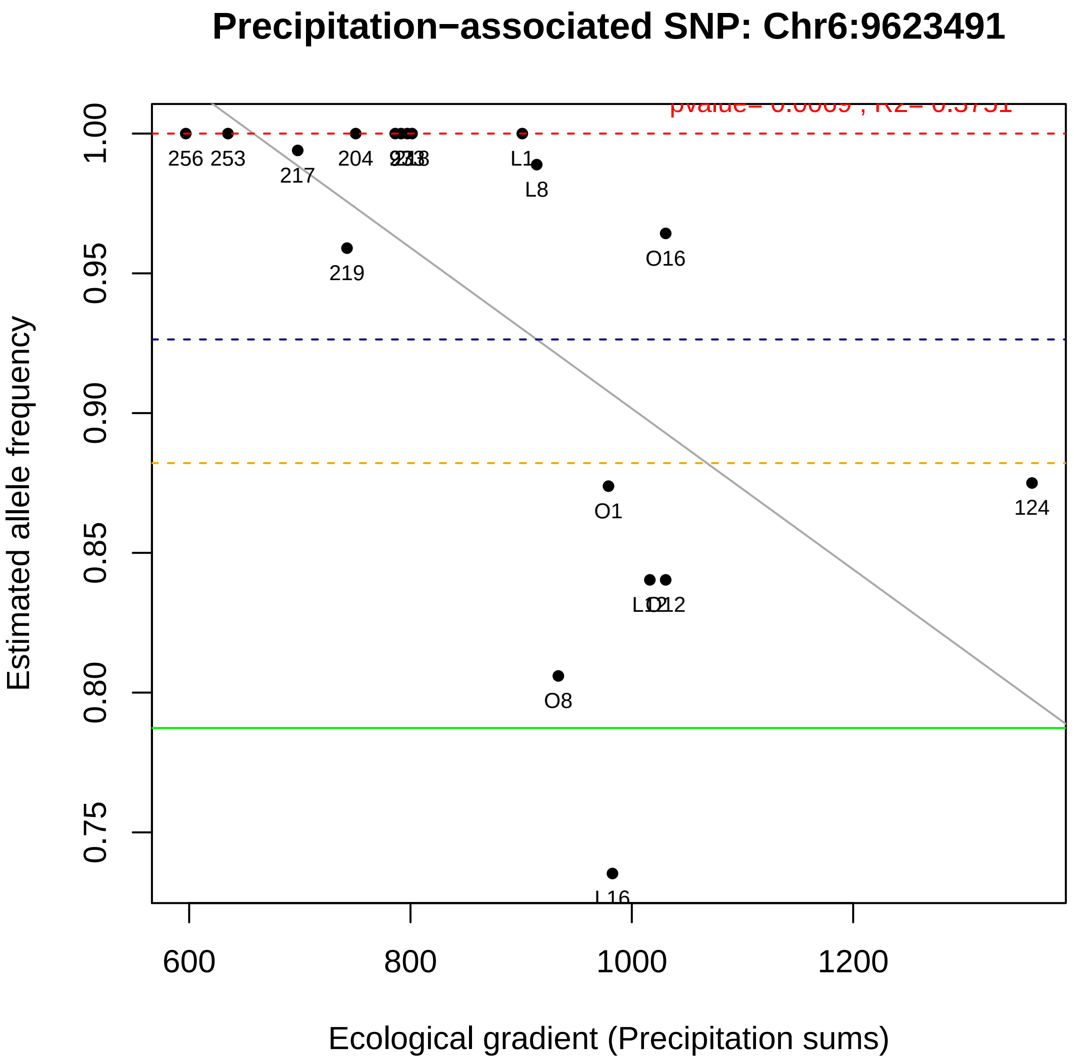
Precipitation−associated SNP: Chr6:9623491
Estimated allele frequency
Ecological gradient (Precipitation sums)
0.75
0.80
0.85
0.90
0.95
1.00
600
800
1000
1200
256
253
217
204
219
L1
L8
O16
O1
124
O8

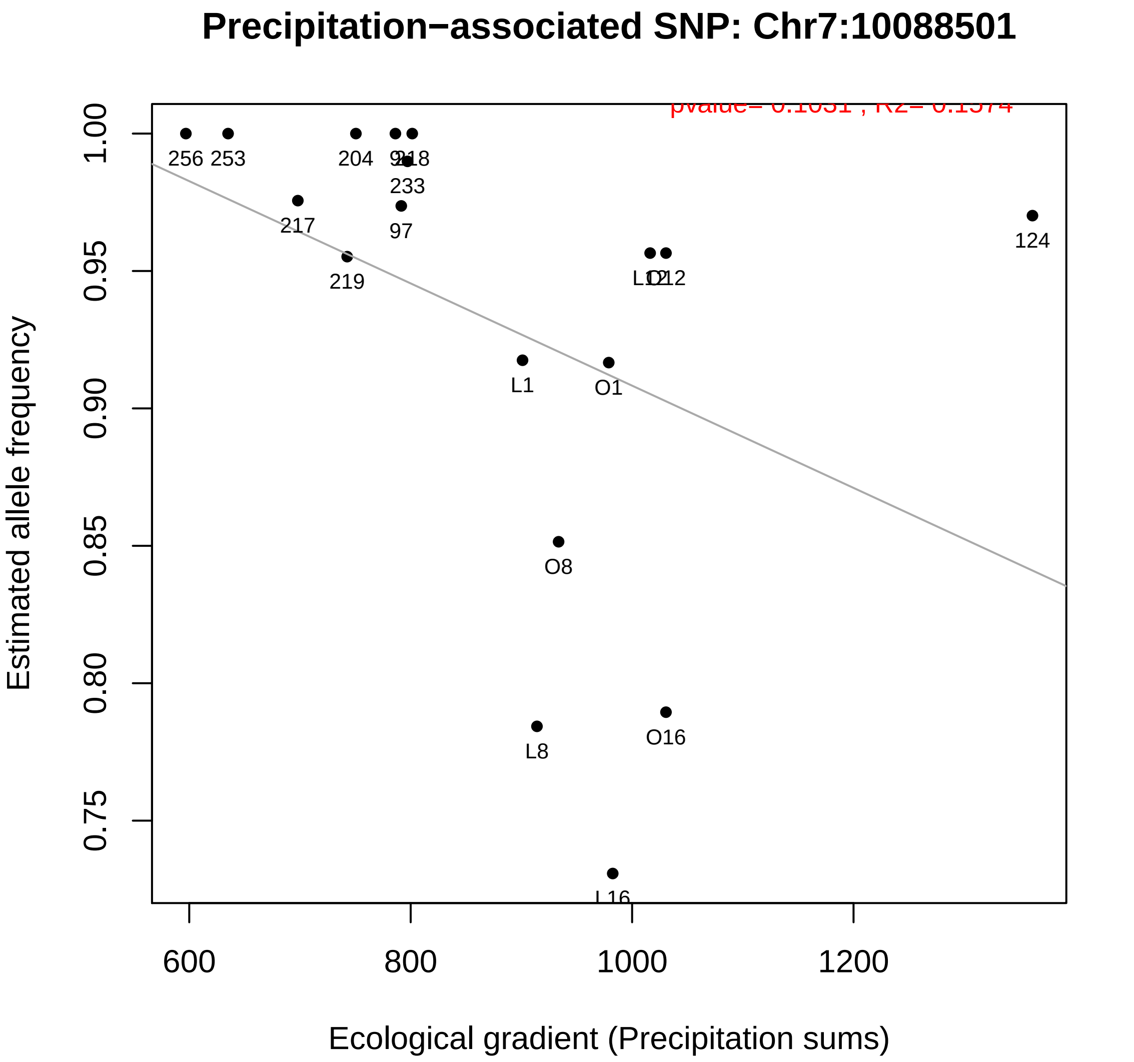

Precipitation−associated SNP: Chr7:10088501
pvalue= 0.1031 ; R2= 0.1574
Estimated allele frequency
Ecological gradient (Precipitation sums)
1.00
0.95
0.90
0.85
0.80
0.75
600
800
1000
1200
256
253
204
218
233
217
97
219
124
L12
O12
L1
O1
O8
L8
O16
L16

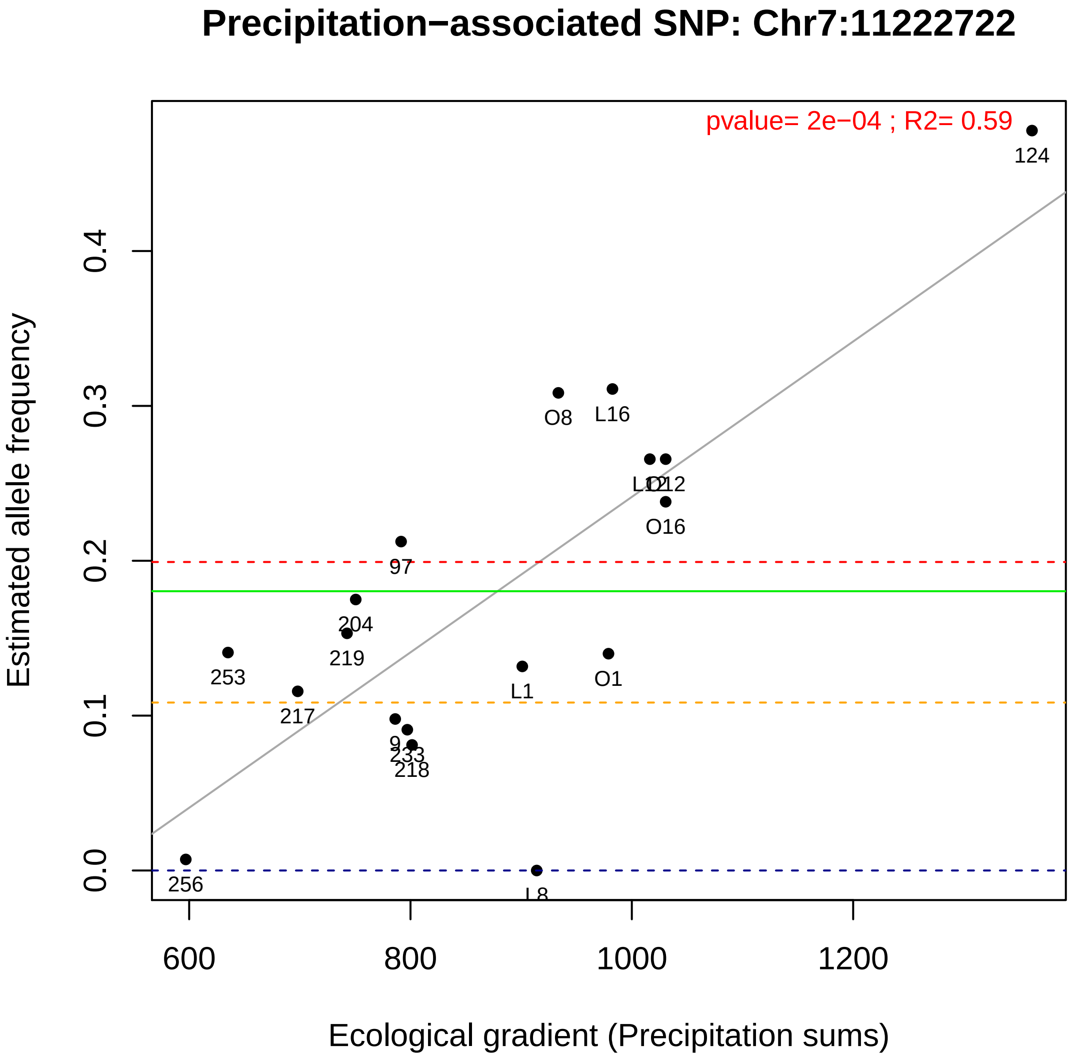

Precipitation−associated SNP: Chr7:11222722
pvalue= 2e−04 ; R2= 0.59
124
O8
L16
L12
O12
O16
97
204
219
253
217
L1
O1
9
233
218
256
L8
0.0
0.1
0.2
0.3
0.4
600
800
1000
1200
Estimated allele frequency
Ecological gradient (Precipitation sums)

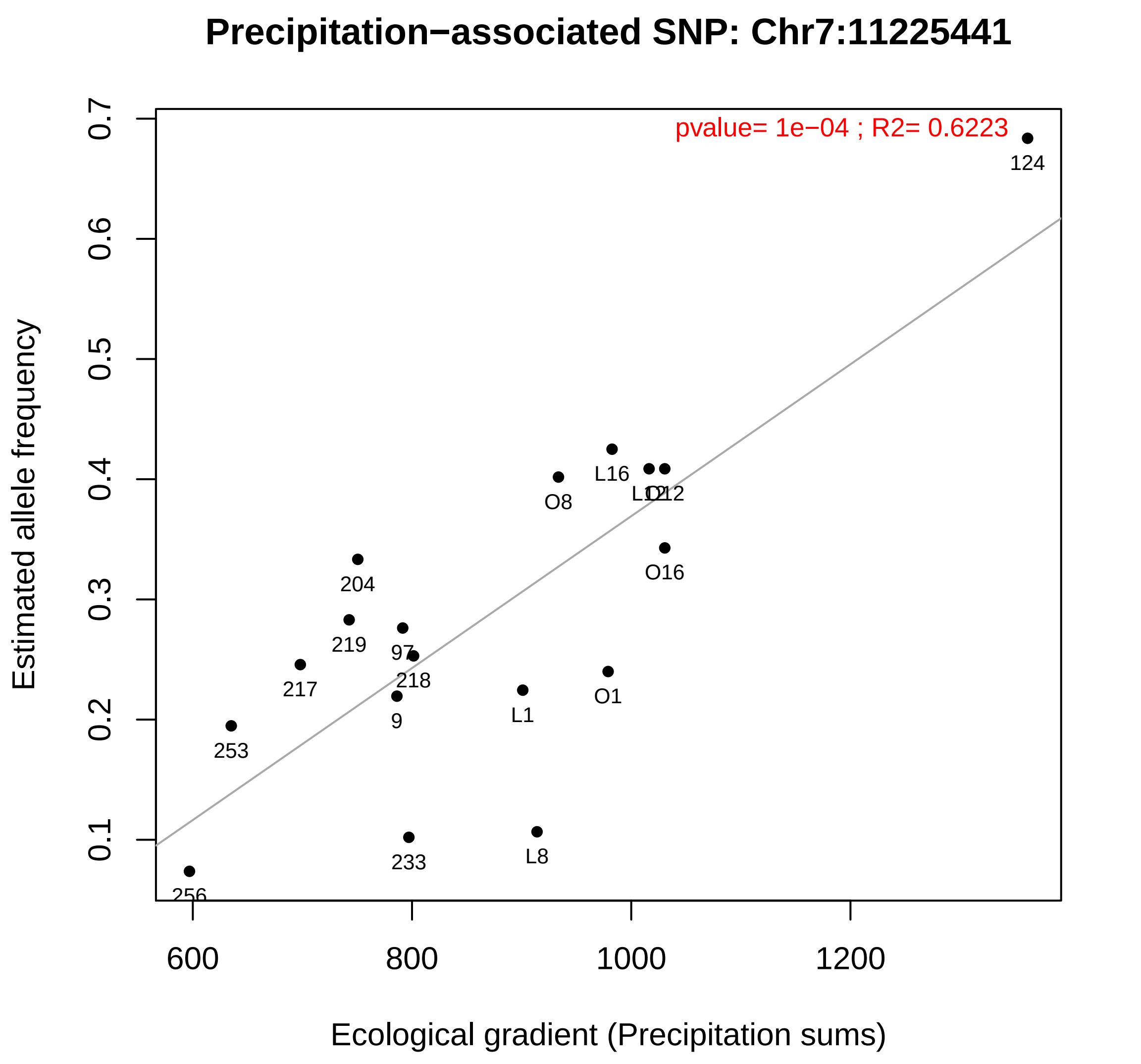

Precipitation−associated SNP: Chr7:11225441
pvalue= 1e−04 ; R2= 0.6223
Estimated allele frequency
Ecological gradient (Precipitation sums)
0.1
0.2
0.3
0.4
0.5
0.6
0.7
600
800
1000
1200
124
L16
O8
O12
O16
204
219
97
218
217
9
L1
O1
253
233
L8
256

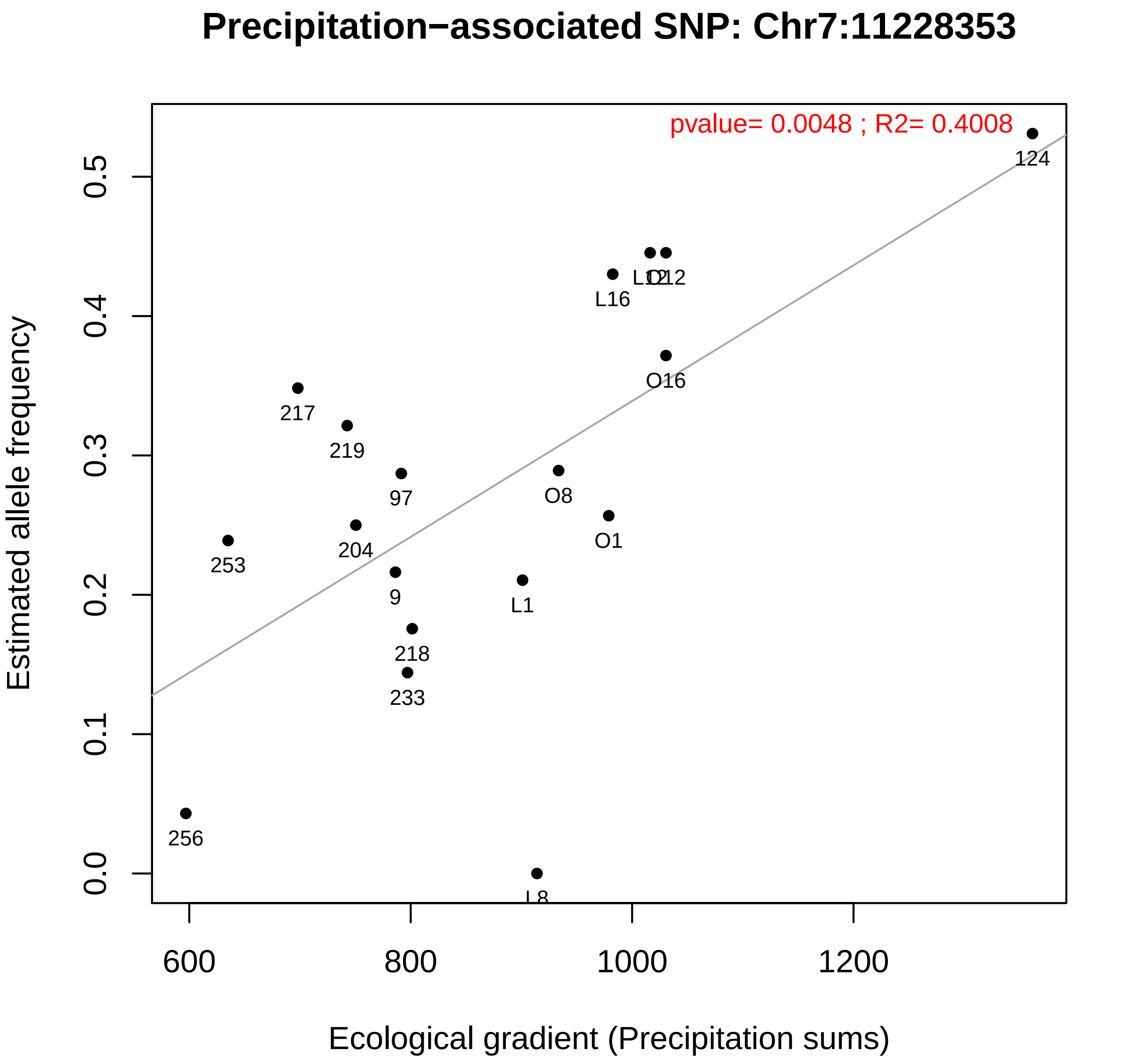

Precipitation−associated SNP: Chr7:11228353
pvalue= 0.0048 ; R2= 0.4008
Estimated allele frequency
Ecological gradient (Precipitation sums)
0.0
0.1
0.2
0.3
0.4
0.5
600
800
1000
1200
124
L16
O16
217
219
97
O8
O1
253
204
9
L1
218
233
256

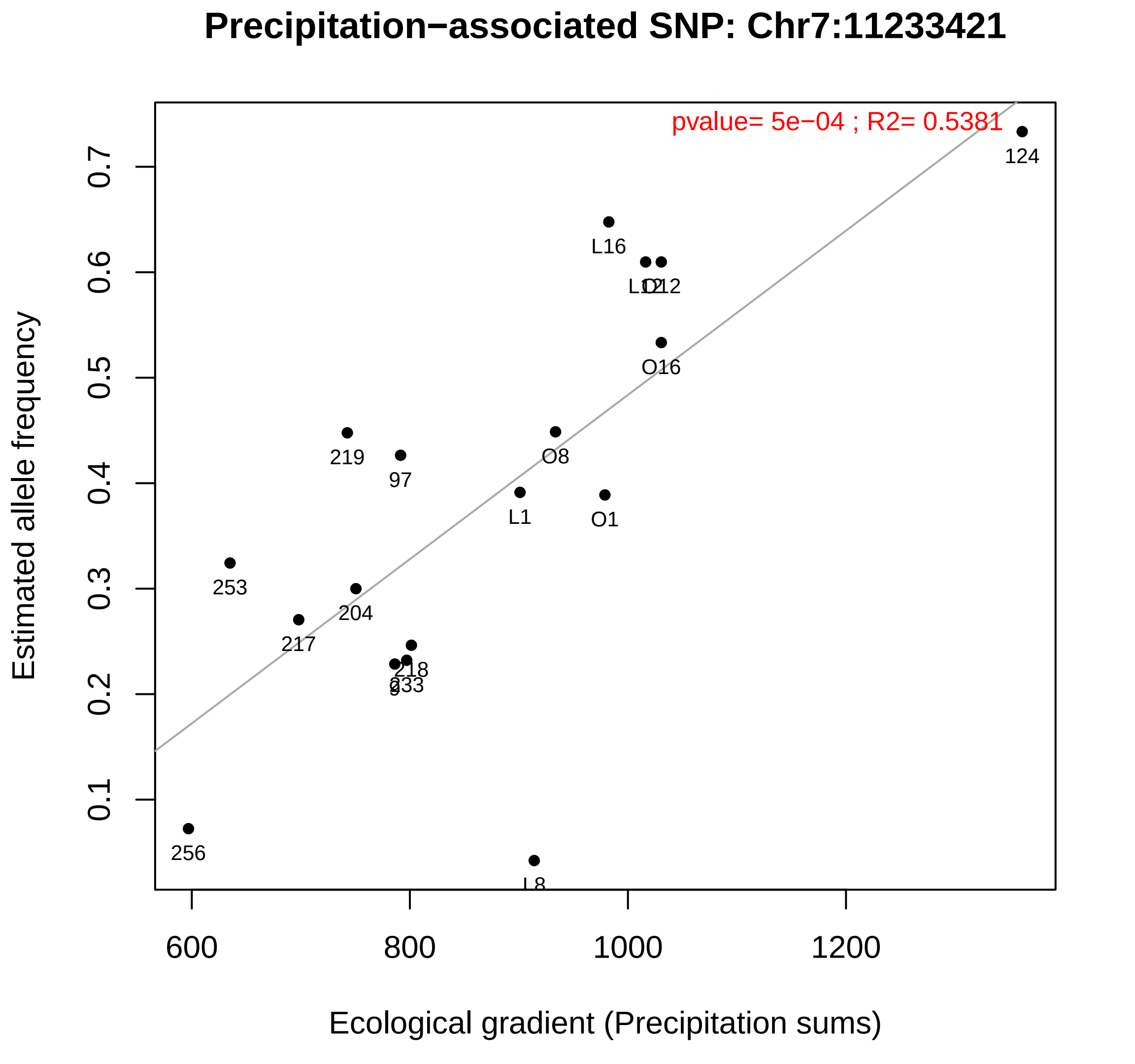
Precipitation−associated SNP: Chr7:11233421
pvalue= 5e−04 ; R2= 0.5381
124
L16
L12
O12
O16
219
97
O8
L1
O1
253
204
217
218
233
256
L8
0.1
0.2
0.3
0.4
0.5
0.6
0.7
600
800
1000
1200
Estimated allele frequency
Ecological gradient (Precipitation sums)

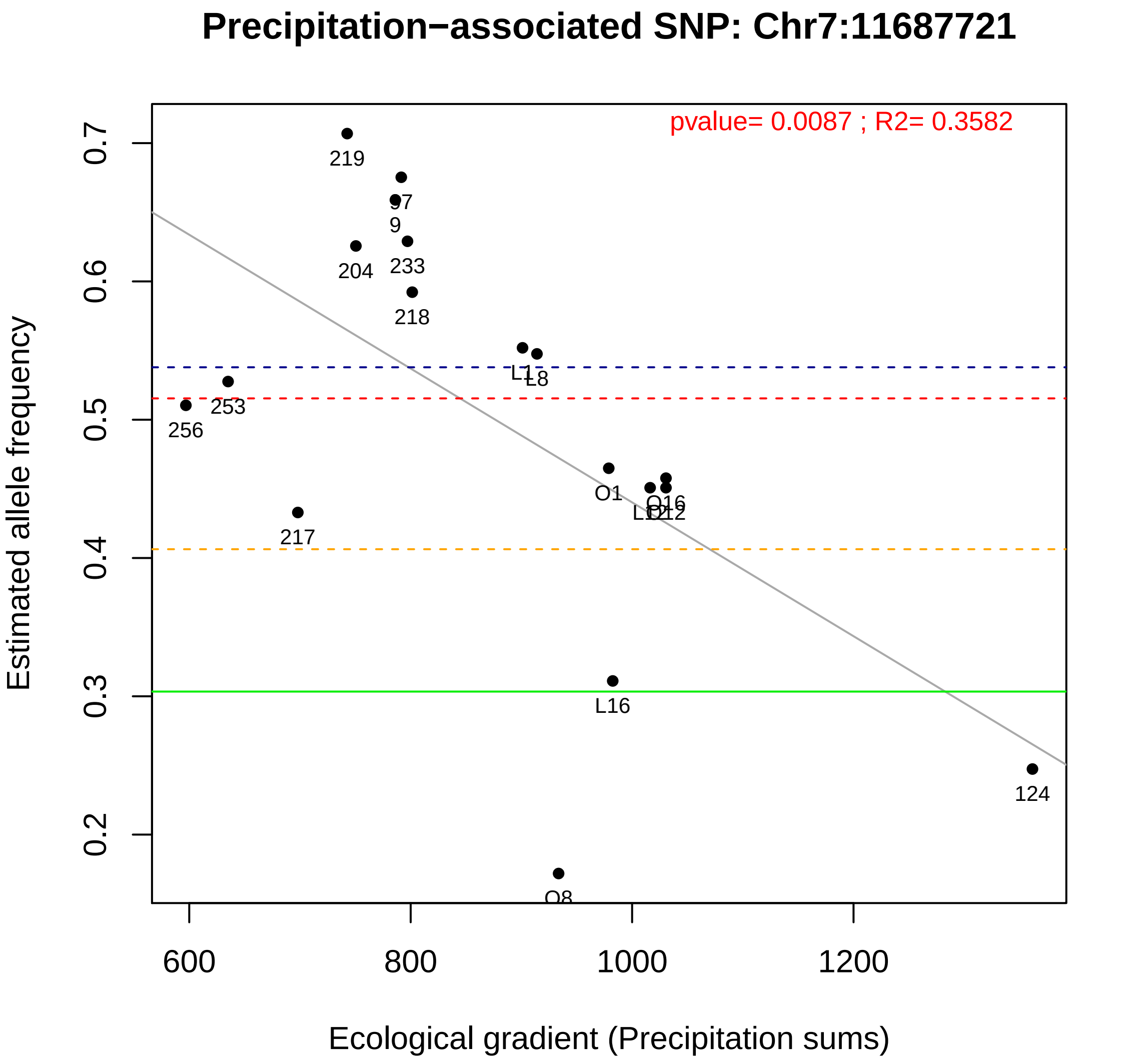

Precipitation−associated SNP: Chr7:11687721
pvalue= 0.0087 ; R2= 0.3582
Estimated allele frequency
Ecological gradient (Precipitation sums)
0.2
0.3
0.4
0.5
0.6
0.7
600
800
1000
1200
219
97
9
204
233
218
L1
L8
253
256
217
O1
O16
L10
O12
L16
124
O8

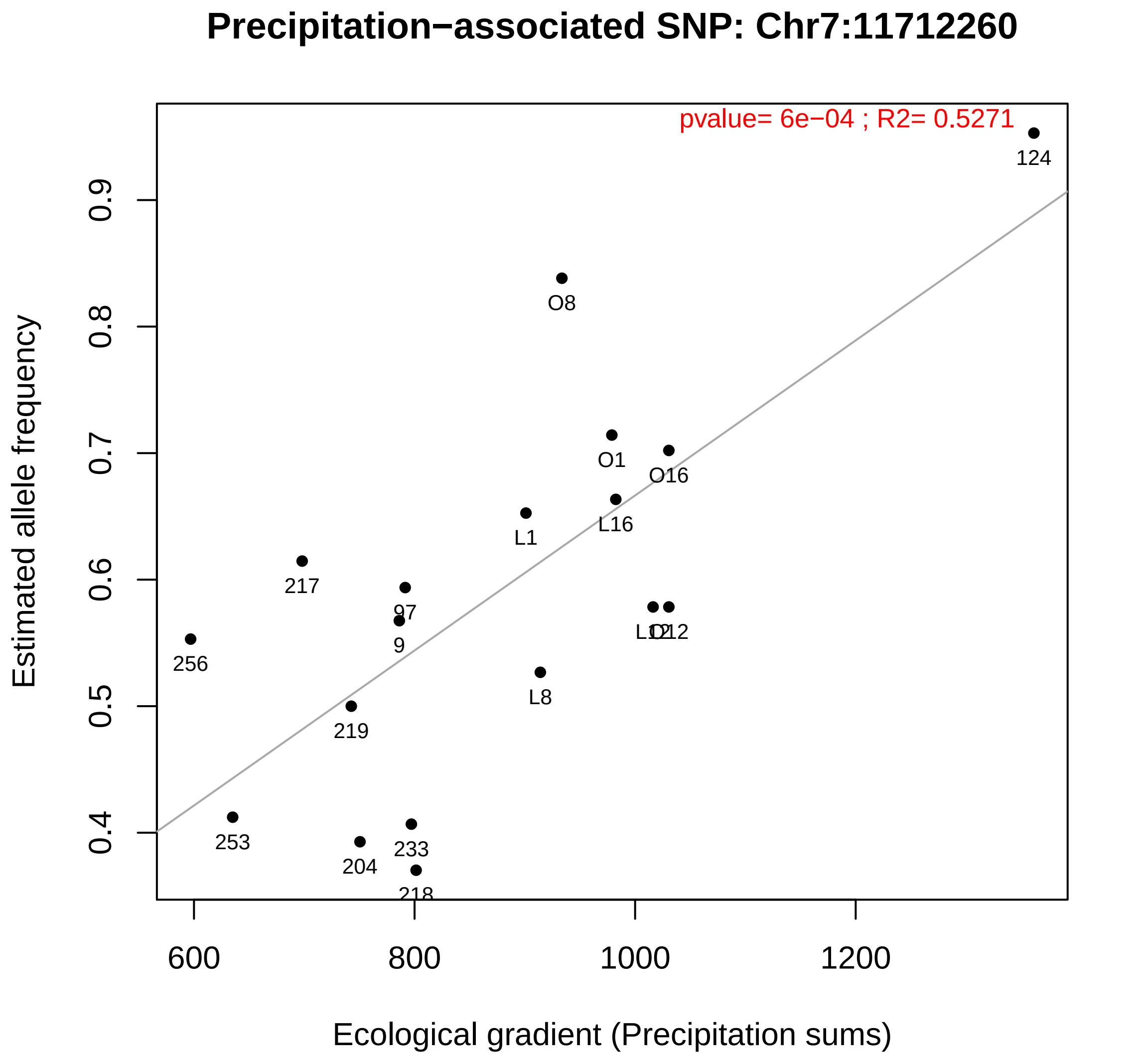

Precipitation−associated SNP: Chr7:11712260
pvalue= 6e−04 ; R2= 0.5271
Estimated allele frequency
0.4
0.5
0.6
0.7
0.8
0.9
600
800
1000
1200
Ecological gradient (Precipitation sums)
124
O8
O1
O16
L16
L1
217
97
9
256
L8
219
253
204
233
218

#### Precipitation−associated SNP: Chr7:11770392

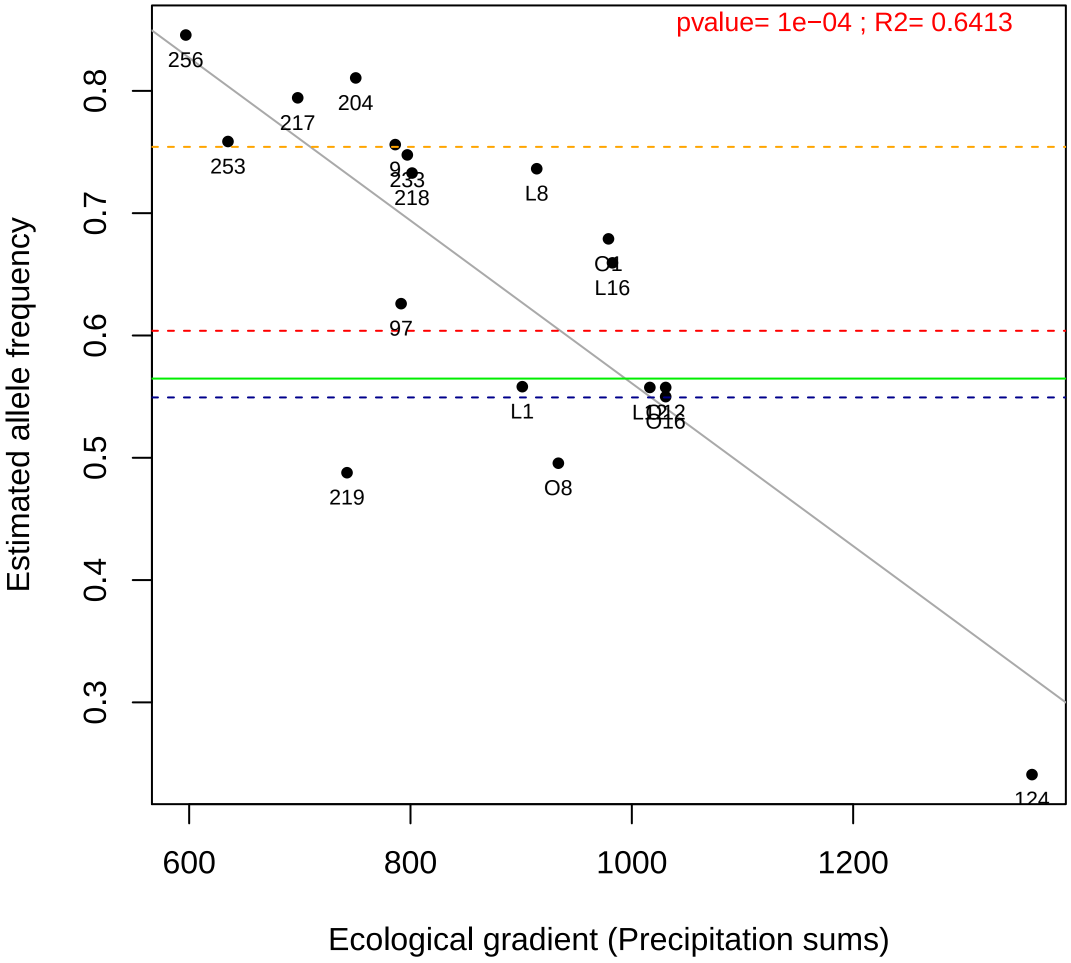

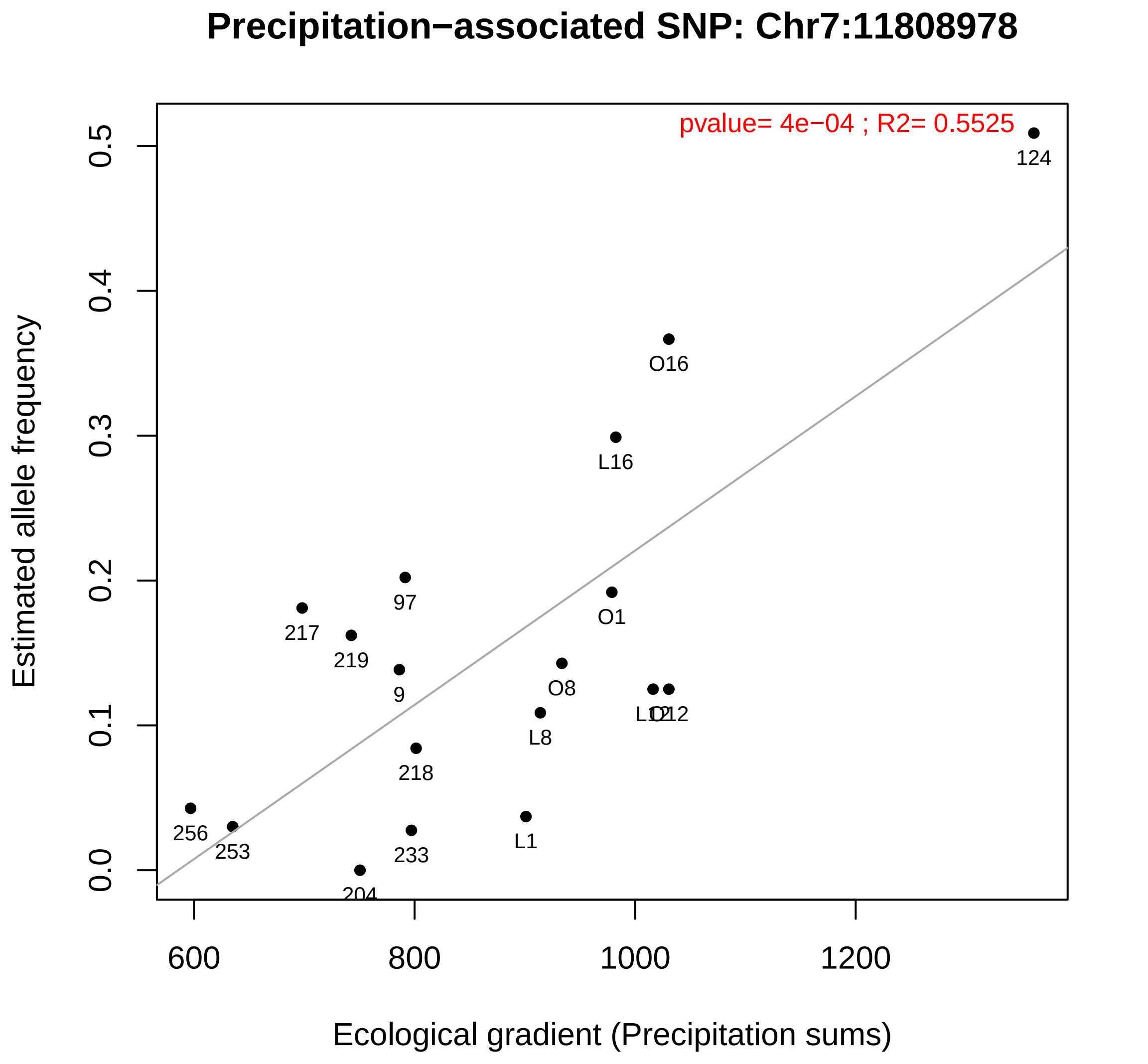
Precipitation−associated SNP: Chr7:11808978
pvalue= 4e−04 ; R2= 0.5525
Estimated allele frequency
Ecological gradient (Precipitation sums)
0.0
0.1
0.2
0.3
0.4
0.5
600
800
1000
1200
124
O16
L16
97
217
219
O1
9
O8
L12
O12
L8
218
256
253
233
L1
204

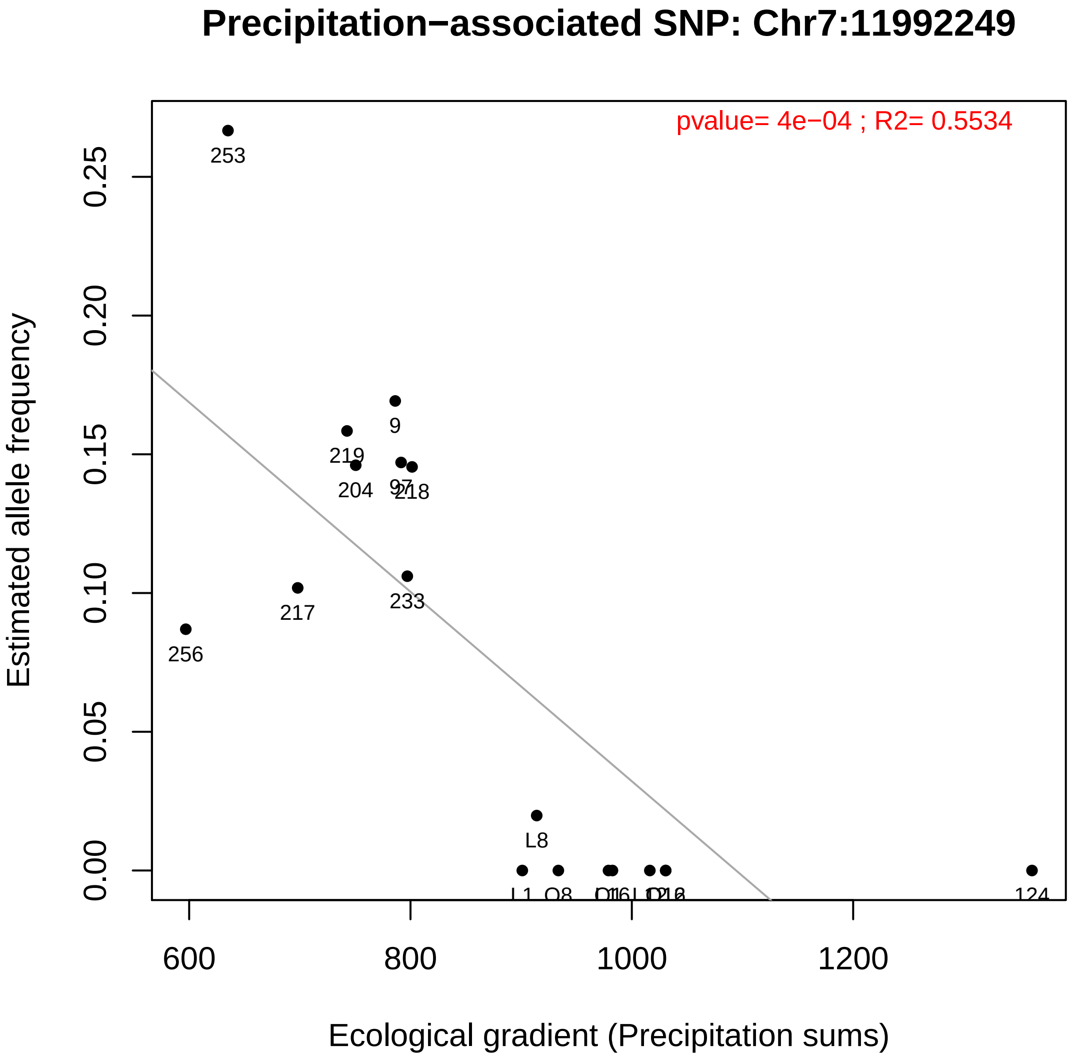
Precipitation−associated SNP: Chr7:11992249
pvalue= 4e−04 ; R2= 0.5534
Estimated allele frequency
Ecological gradient (Precipitation sums)
0.00
0.05
0.10
0.15
0.20
0.25
600
800
1000
1200
253
9
219
204
218
233
217
256
L8
124

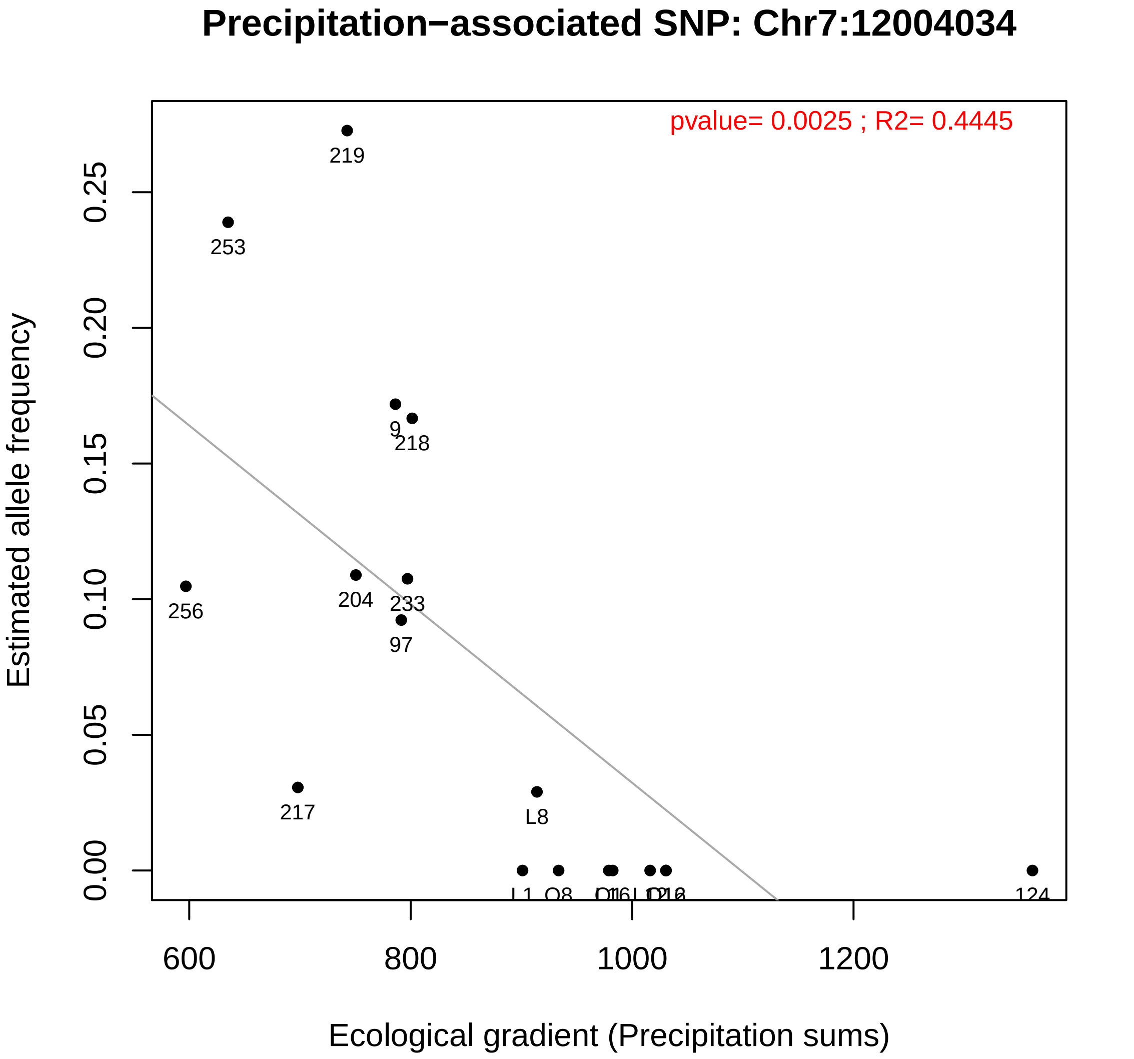
Precipitation−associated SNP: Chr7:12004034
pvalue= 0.0025 ; R2= 0.4445
Estimated allele frequency
0.00
0.05
0.10
0.15
0.20
0.25
600
800
1000
1200
Ecological gradient (Precipitation sums)
219
253
9
218
256
204
233
97
217
L8
L1
O8
124

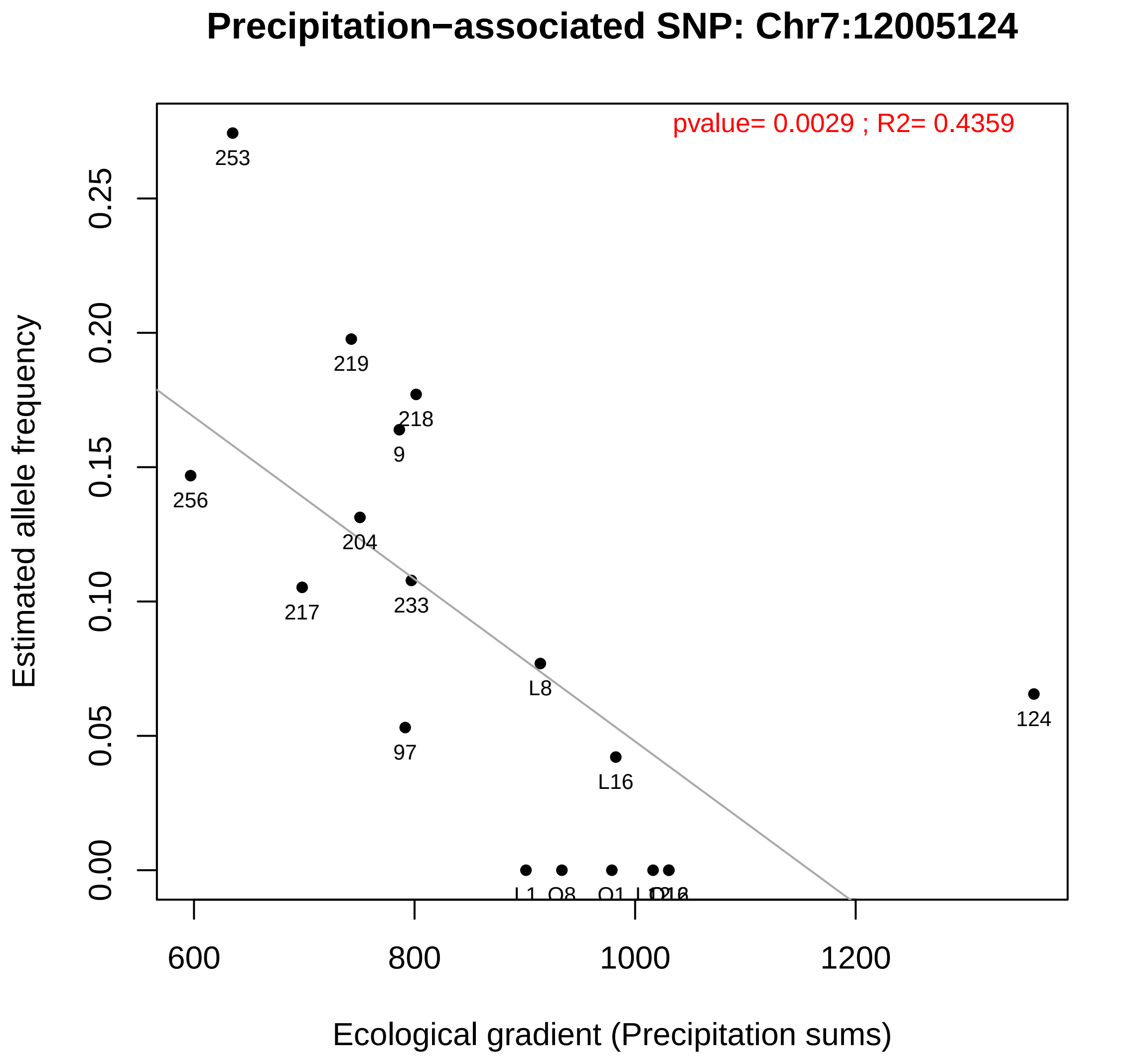
Precipitation−associated SNP: Chr7:12005124
pvalue= 0.0029 ; R2= 0.4359
Estimated allele frequency
Ecological gradient (Precipitation sums)
0.00
0.05
0.10
0.15
0.20
0.25
600
800
1000
1200
253
219
218
9
256
204
217
233
L8
124
97
L16

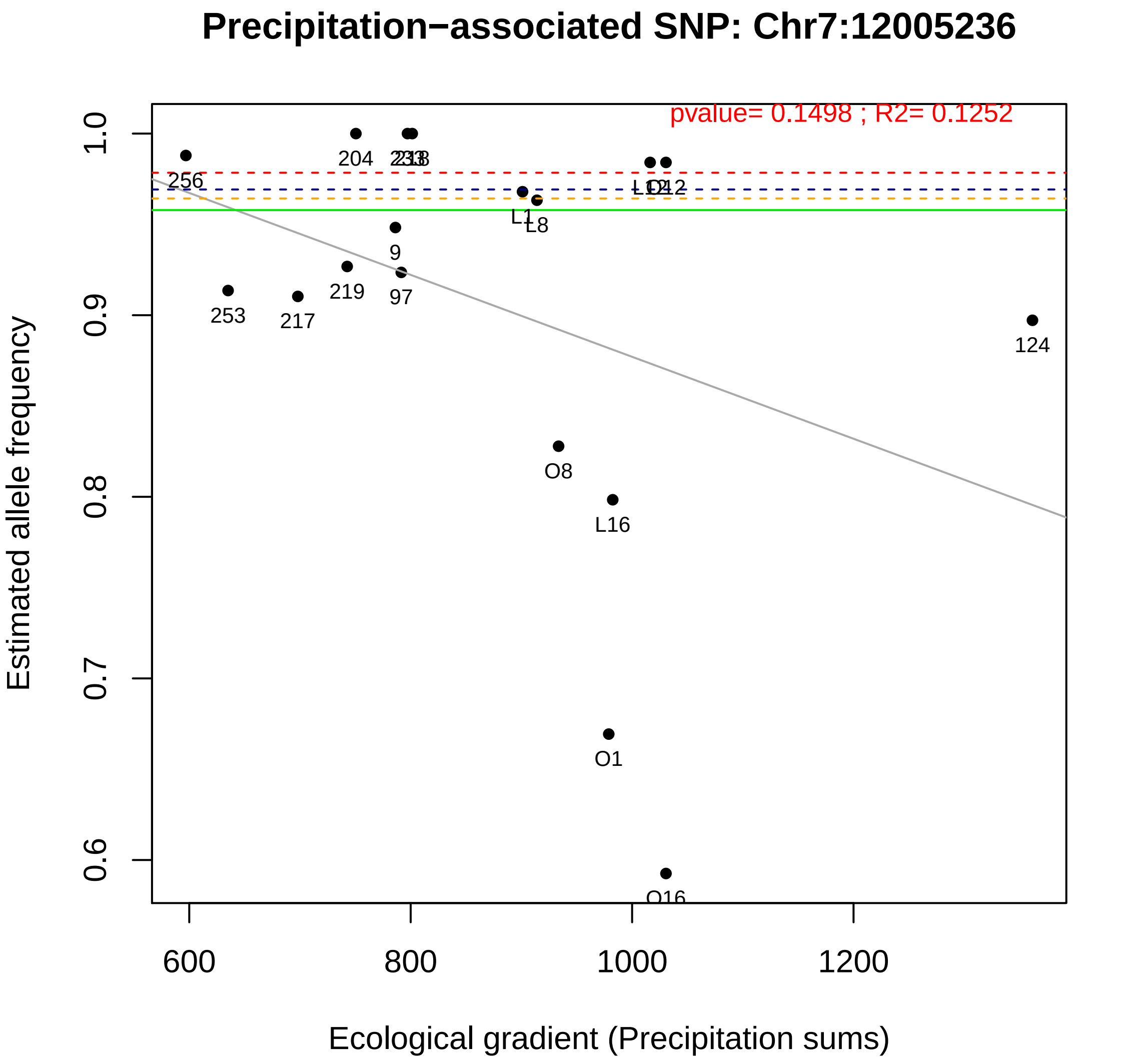
Precipitation−associated SNP: Chr7:12005236
pvalue= 0.1498 ; R2= 0.1252
Estimated allele frequency
Ecological gradient (Precipitation sums)
1.0
0.9
0.8
0.7
0.6
600
800
1000
1200
256
204
9
219
97
253
217
L1
L8
O8
L16
O1
O16
124

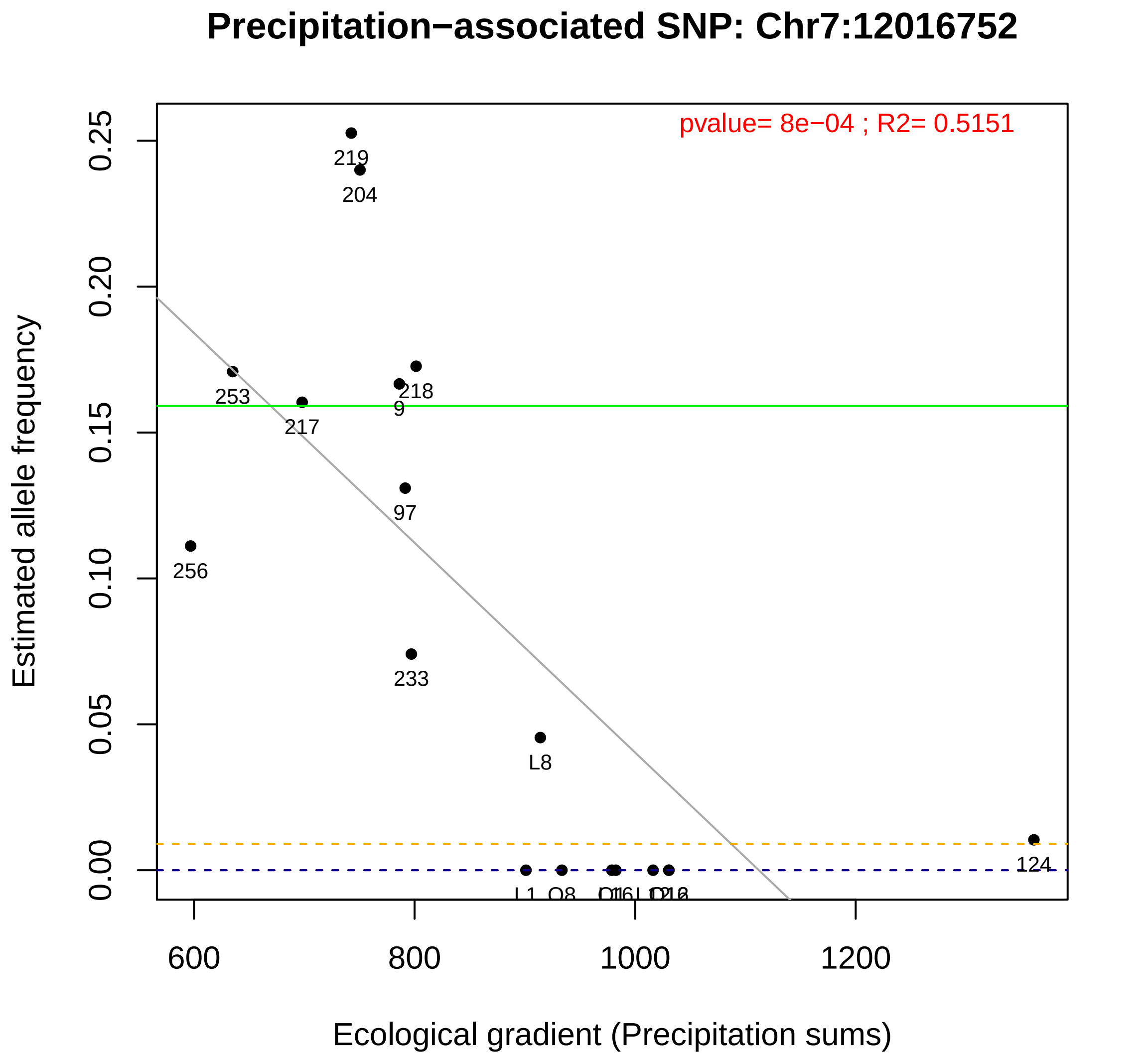

Precipitation−associated SNP: Chr7:12016752
pvalue= 8e−04 ; R2= 0.5151
Estimated allele frequency
Ecological gradient (Precipitation sums)
0.00
0.05
0.10
0.15
0.20
0.25
600
800
1000
1200
219
204
253
217
218
9
97
256
233
L8
124

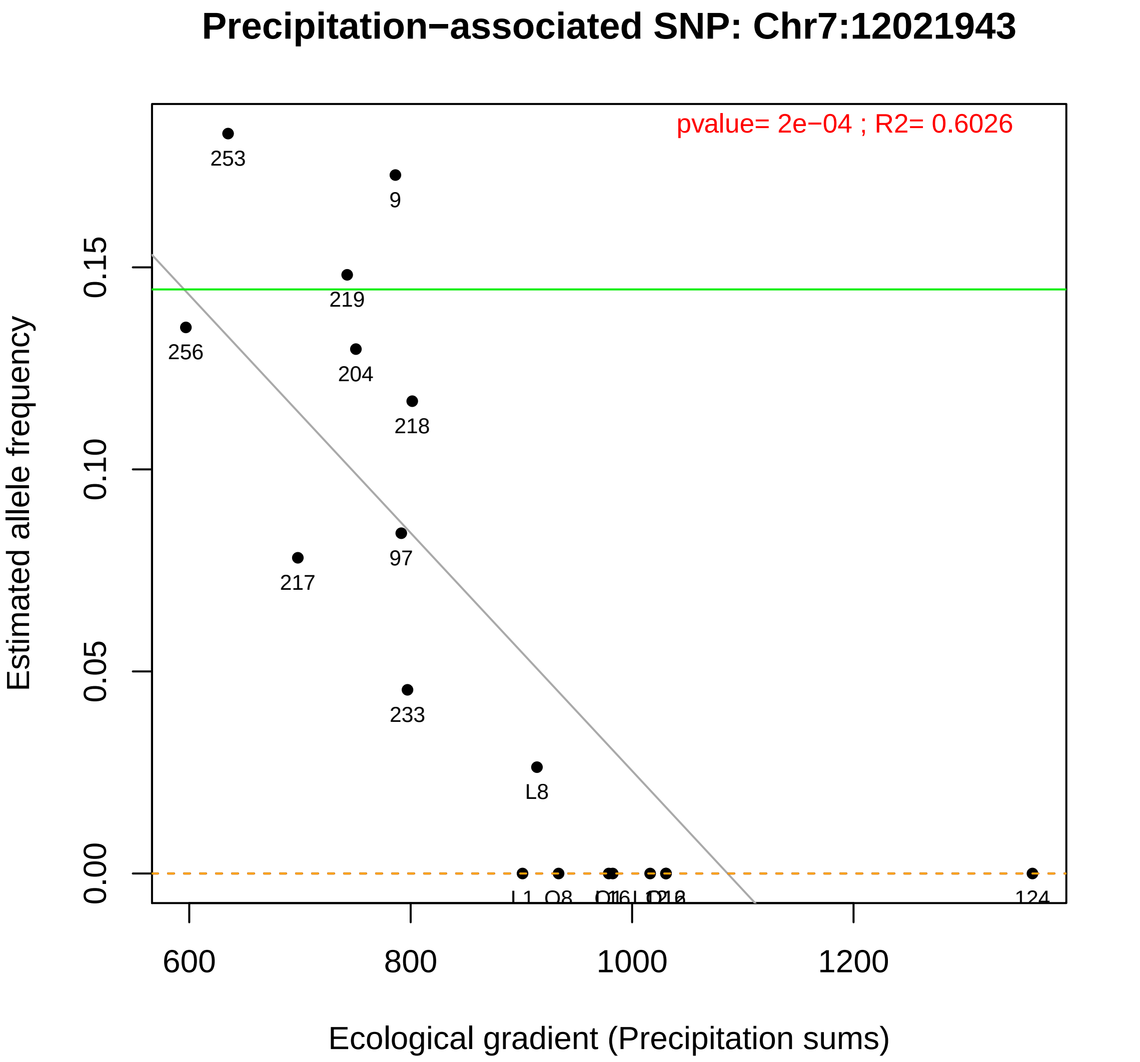
Precipitation−associated SNP: Chr7:12021943
pvalue= 2e−04 ; R2= 0.6026
Estimated allele frequency
0.00
0.05
0.10
0.15
600
800
1000
1200
Ecological gradient (Precipitation sums)
253
9
219
256
204
218
97
217
233
L8
L1
O8
124

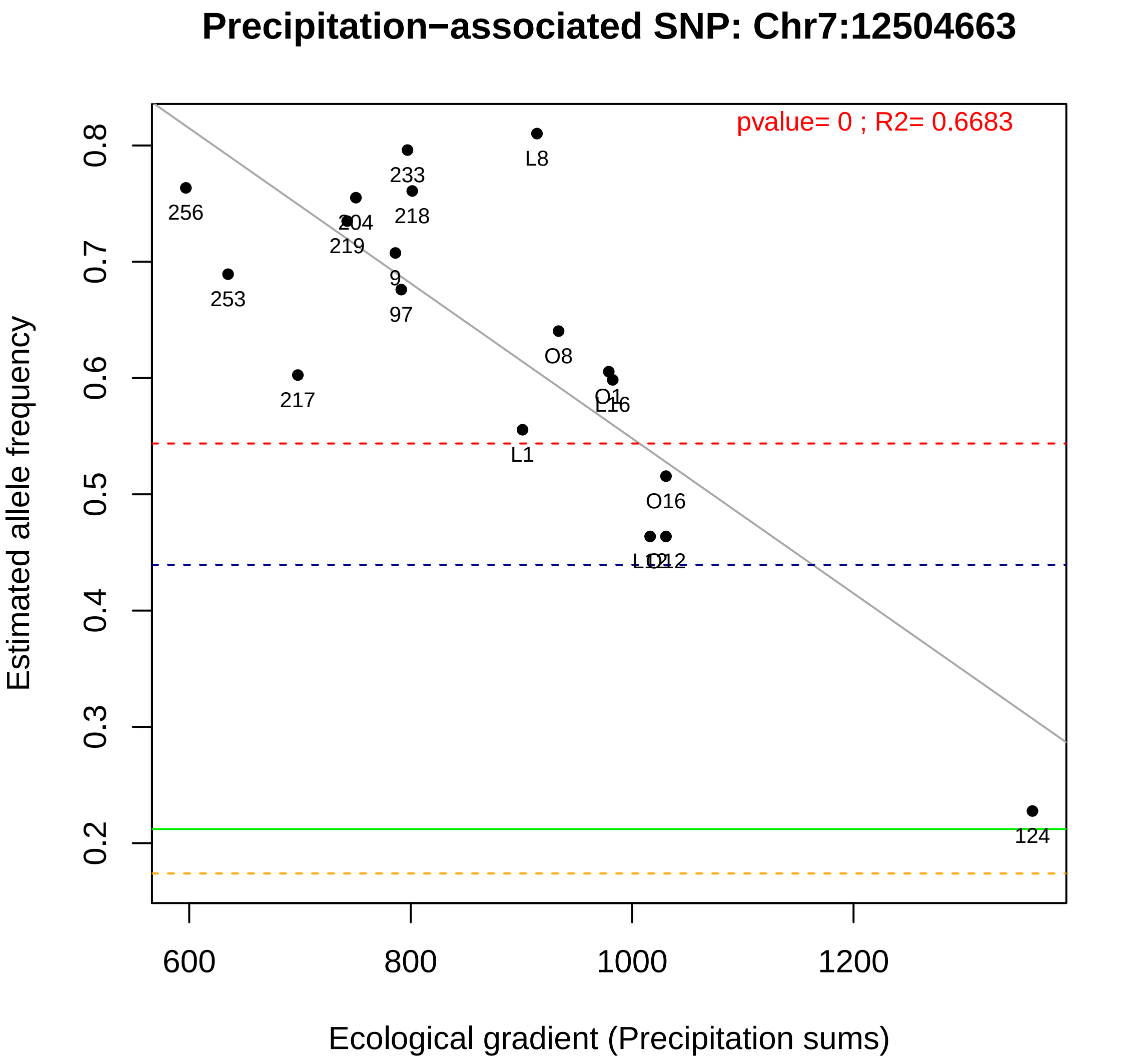

Precipitation−associated SNP: Chr7:12504663
pvalue= 0 ; R2= 0.6683
Estimated allele frequency
Ecological gradient (Precipitation sums)
0.2
0.3
0.4
0.5
0.6
0.7
0.8
600
800
1000
1200
256
253
217
219
204
233
218
9
97
L8
L1
O8
O1
L16
O16
L10
O12
124

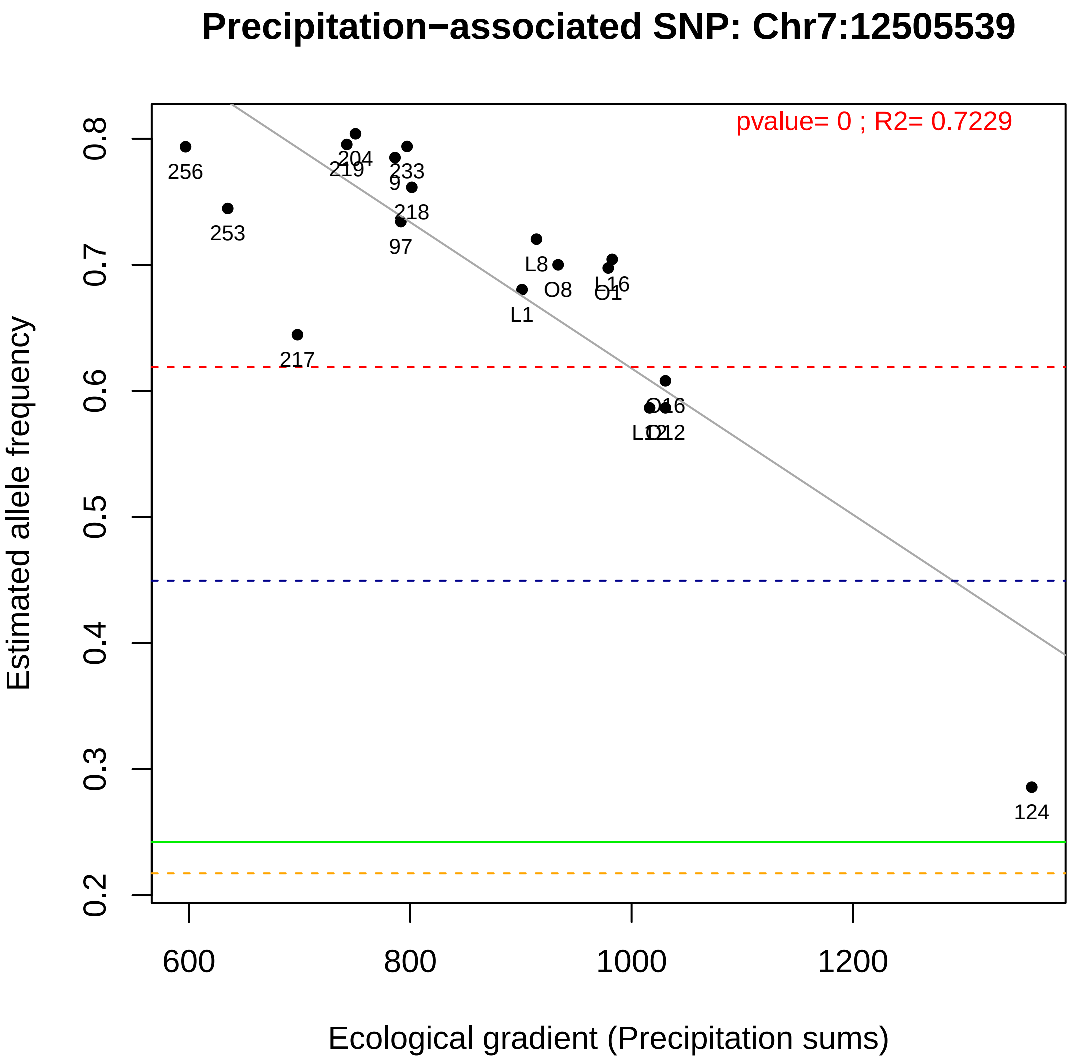

Precipitation−associated SNP: Chr7:12505539
pvalue= 0 ; R2= 0.7229
Estimated allele frequency
Ecological gradient (Precipitation sums)
0.2
0.3
0.4
0.5
0.6
0.7
0.8
600
800
1000
1200
256
253
217
204
219
233
9
218
97
L8
O8
L1
L16
O1
O16
L12
O12
124

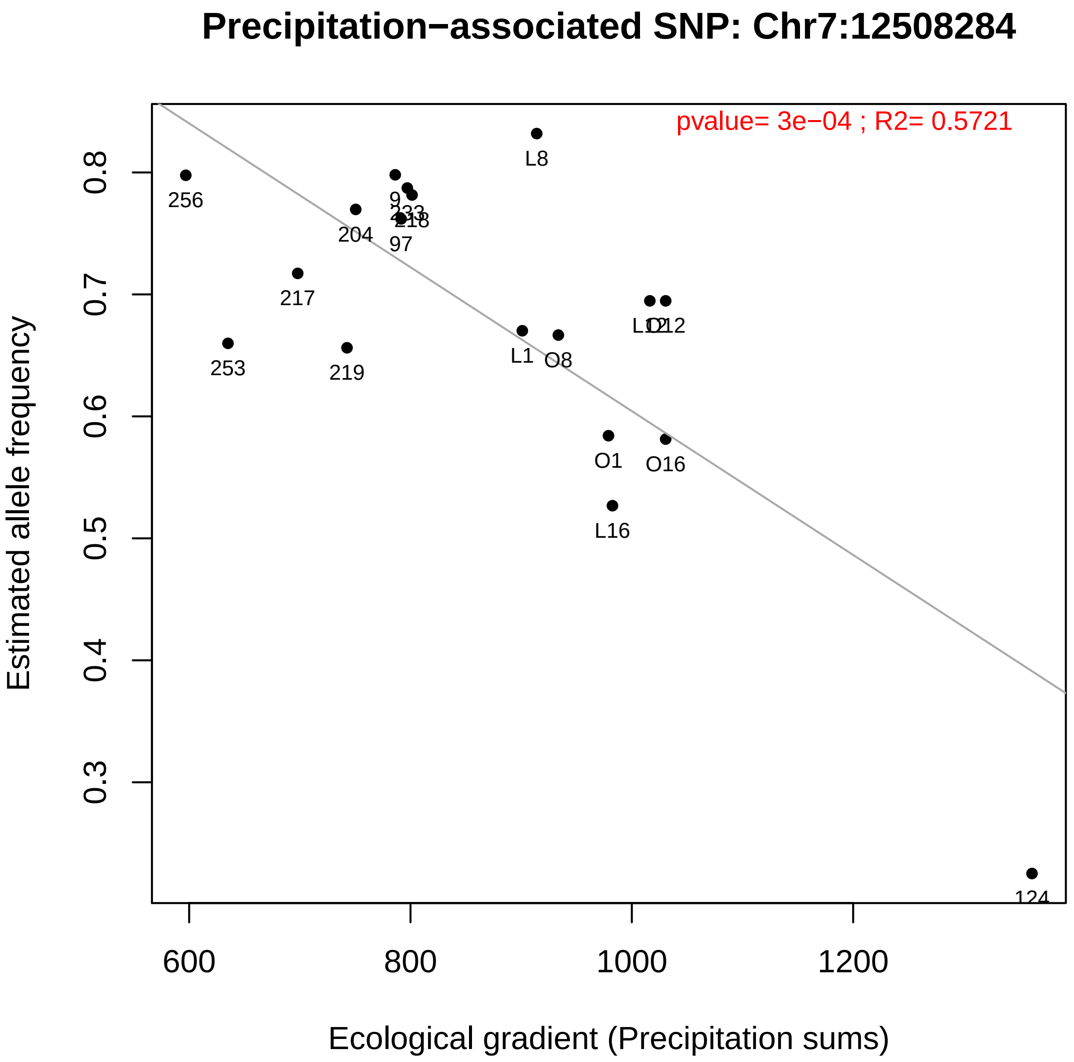
Precipitation−associated SNP: Chr7:12508284
pvalue= 3e−04 ; R2= 0.5721
Estimated allele frequency
Ecological gradient (Precipitation sums)
0.3
0.4
0.5
0.6
0.7
0.8
600
800
1000
1200
256
253
217
219
204
L8
L1
O8
O1
O16
L16
O12
124

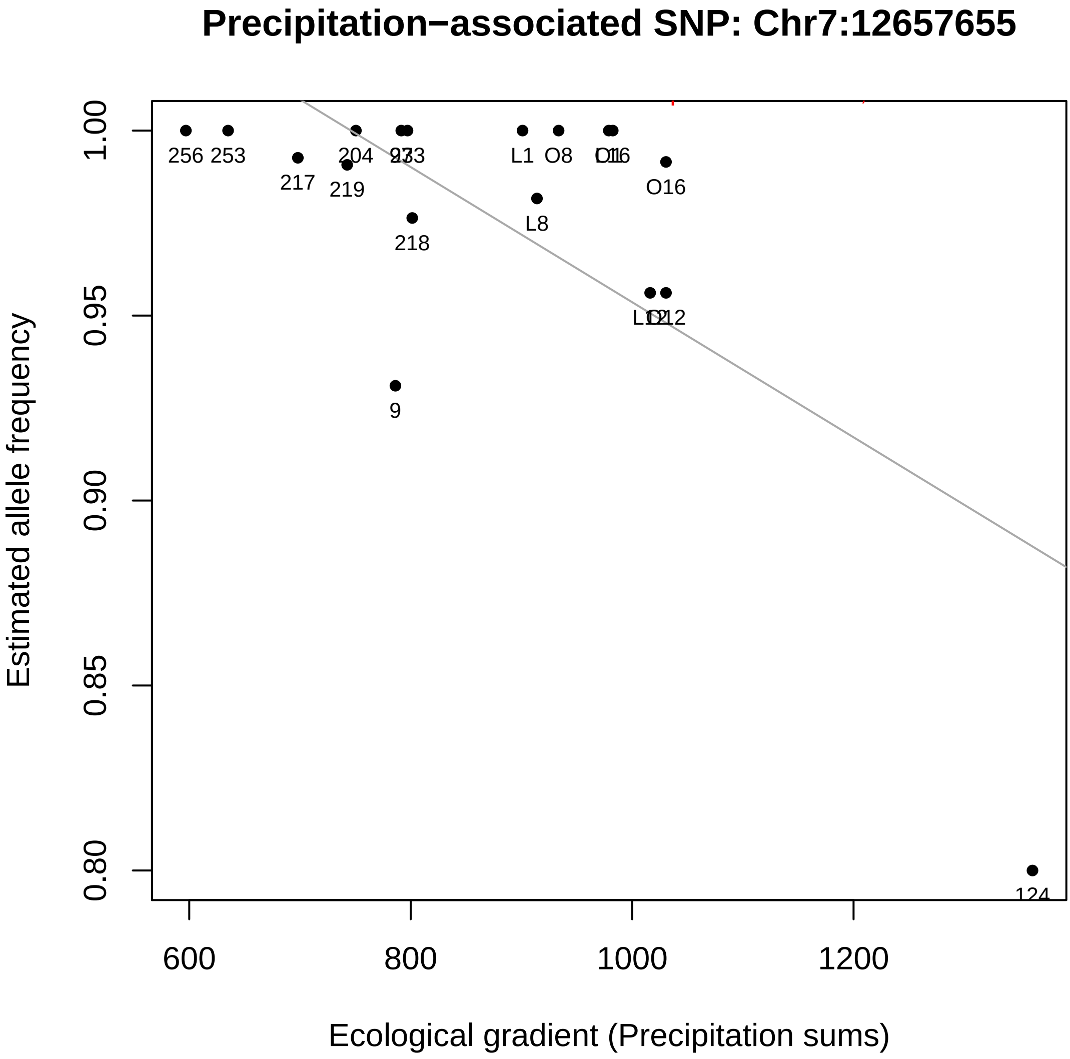

Precipitation−associated SNP: Chr7:12657655
Estimated allele frequency
Ecological gradient (Precipitation sums)
0.80
0.85
0.90
0.95
1.00
600
800
1000
1200
256
253
217
219
204
218
9
L1
O8
L8
O16
124

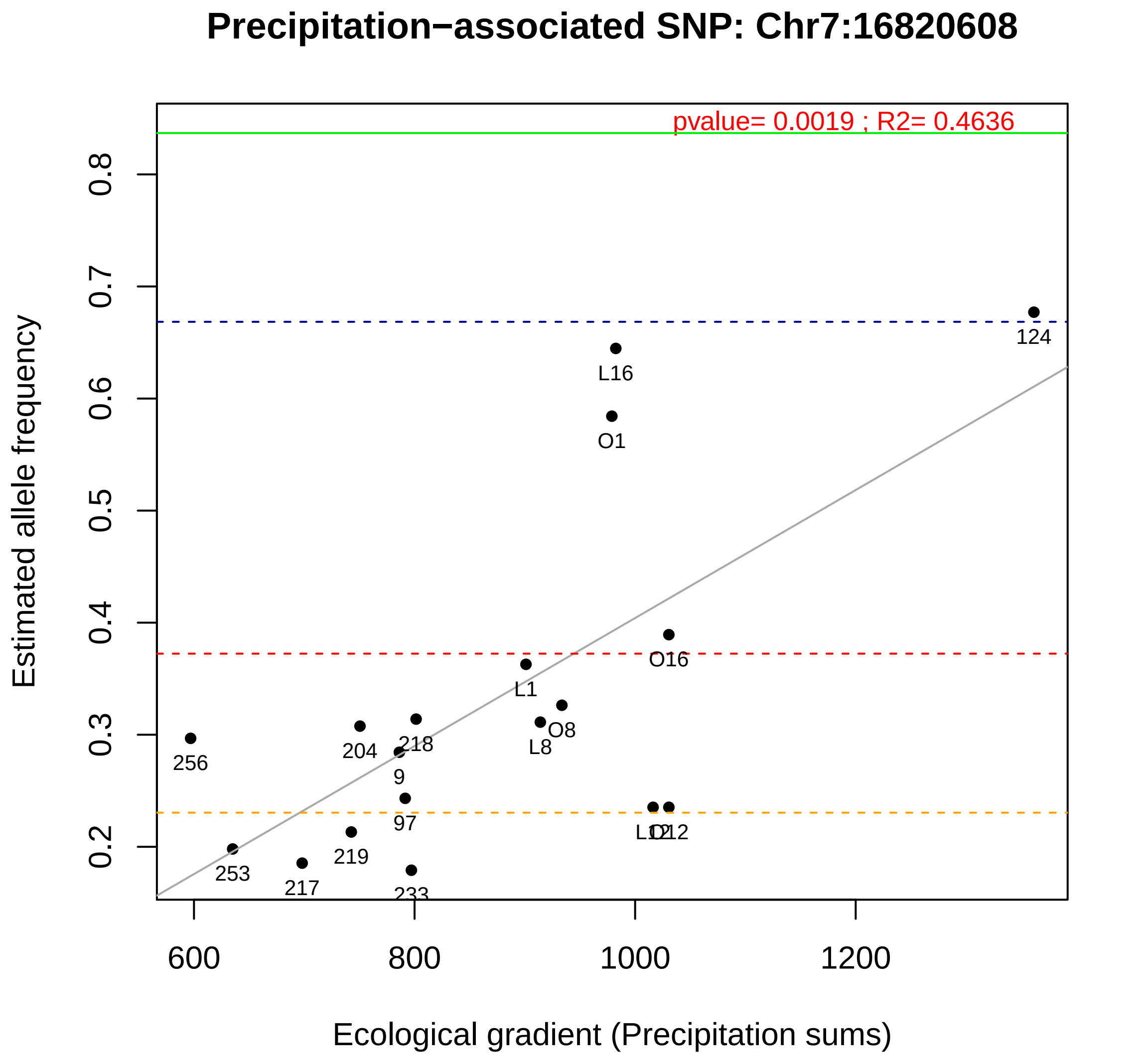

Precipitation−associated SNP: Chr7:16820608
pvalue= 0.0019 ; R2= 0.4636
Estimated allele frequency
0.2
0.3
0.4
0.5
0.6
0.7
0.8
600
800
1000
1200
Ecological gradient (Precipitation sums)
124
L16
O1
O16
L1
O8
L8
218
204
256
9
97
219
253
217
233

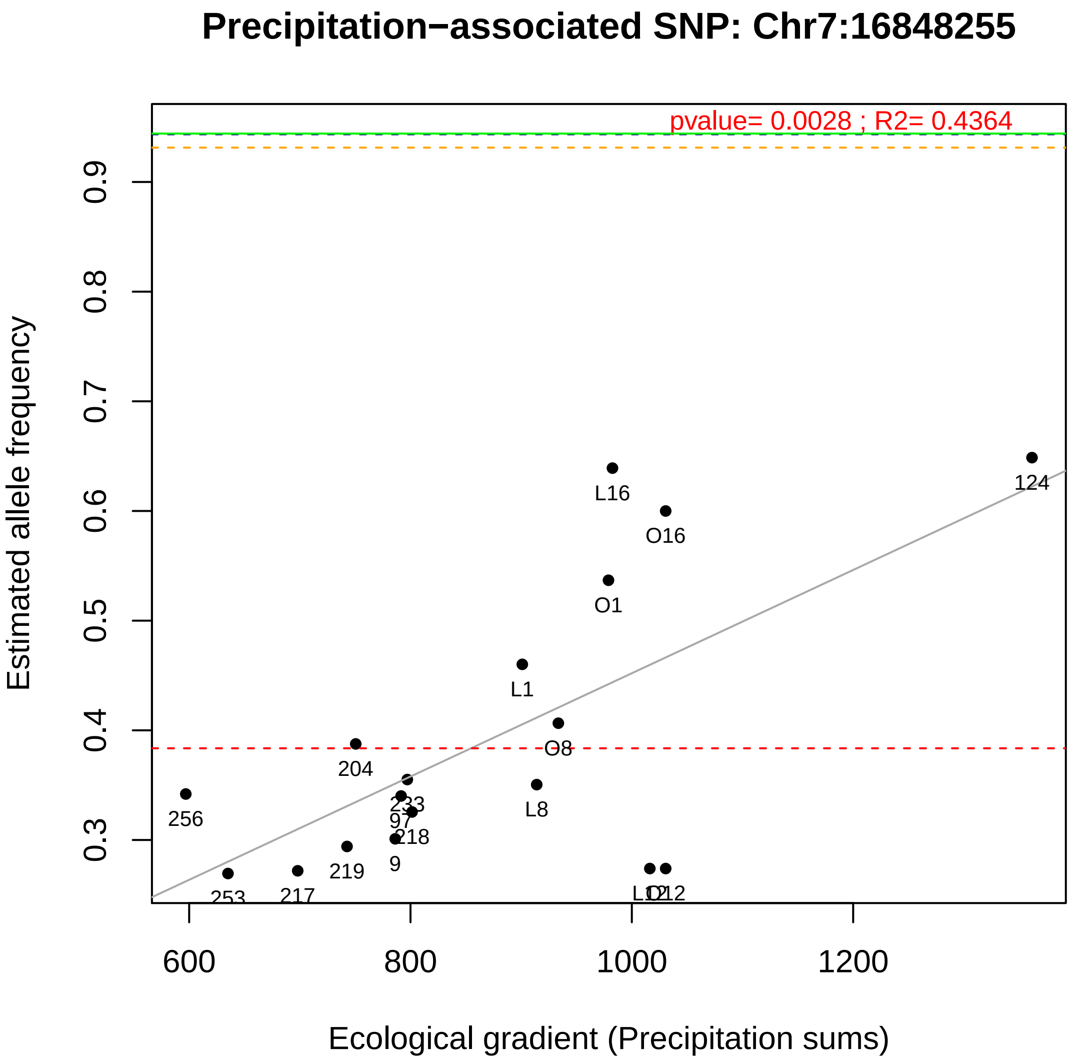

Precipitation−associated SNP: Chr7:16848255
pvalue= 0.0028 ; R2= 0.4364
Estimated allele frequency
0.3
0.4
0.5
0.6
0.7
0.8
0.9
600
800
1000
1200
Ecological gradient (Precipitation sums)
L16
124
O16
O1
L1
O8
204
256
233
97
218
L8
219
9
253
217

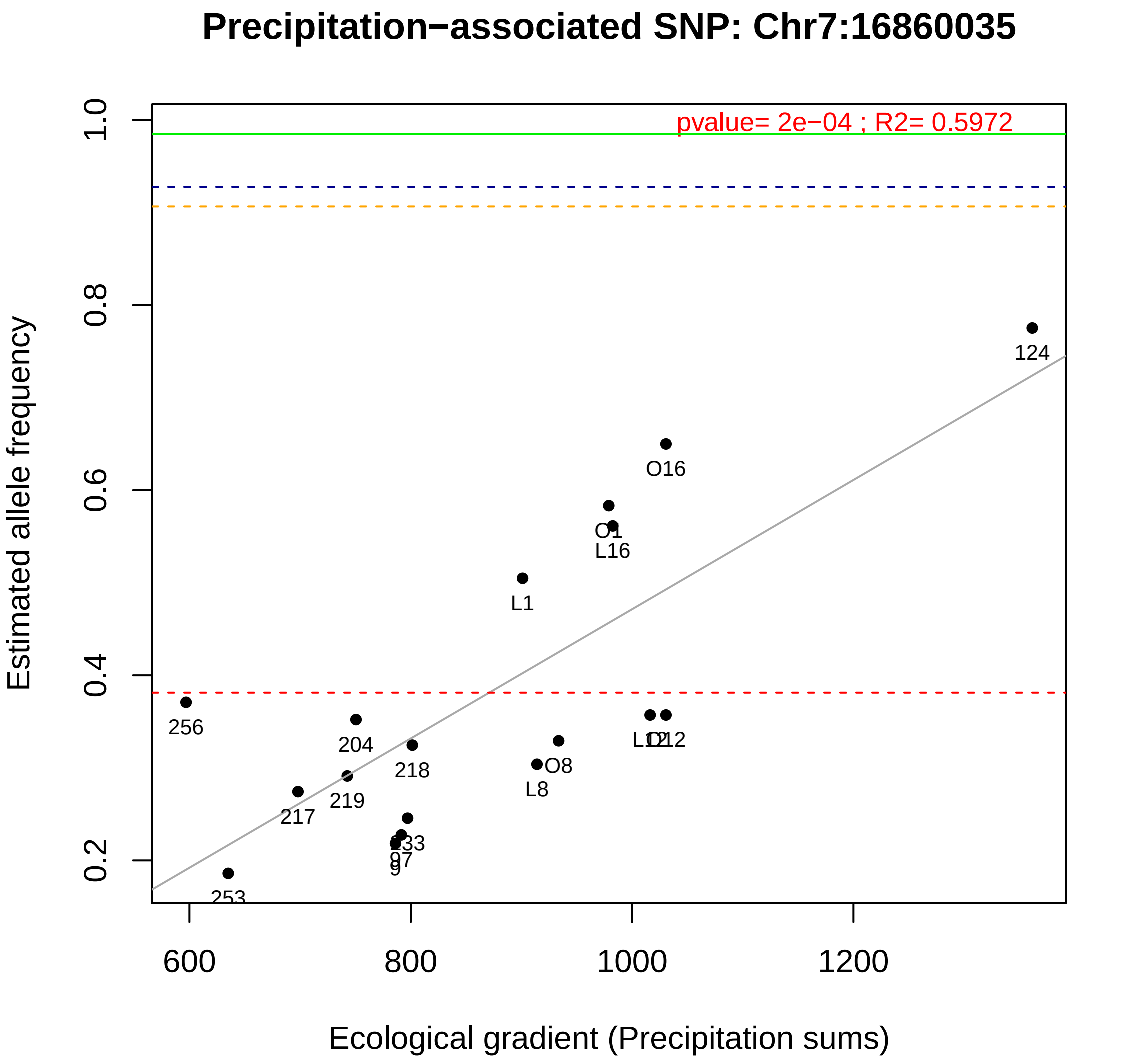
Precipitation−associated SNP: Chr7:16860035
pvalue= 2e−04 ; R2= 0.5972
Estimated allele frequency
0.2
0.4
0.6
0.8
1.0
Ecological gradient (Precipitation sums)
600
800
1000
1200
124
O16
O1
L16
L1
256
204
218
219
217
O8
L8
L12
O12
233
97
9
253

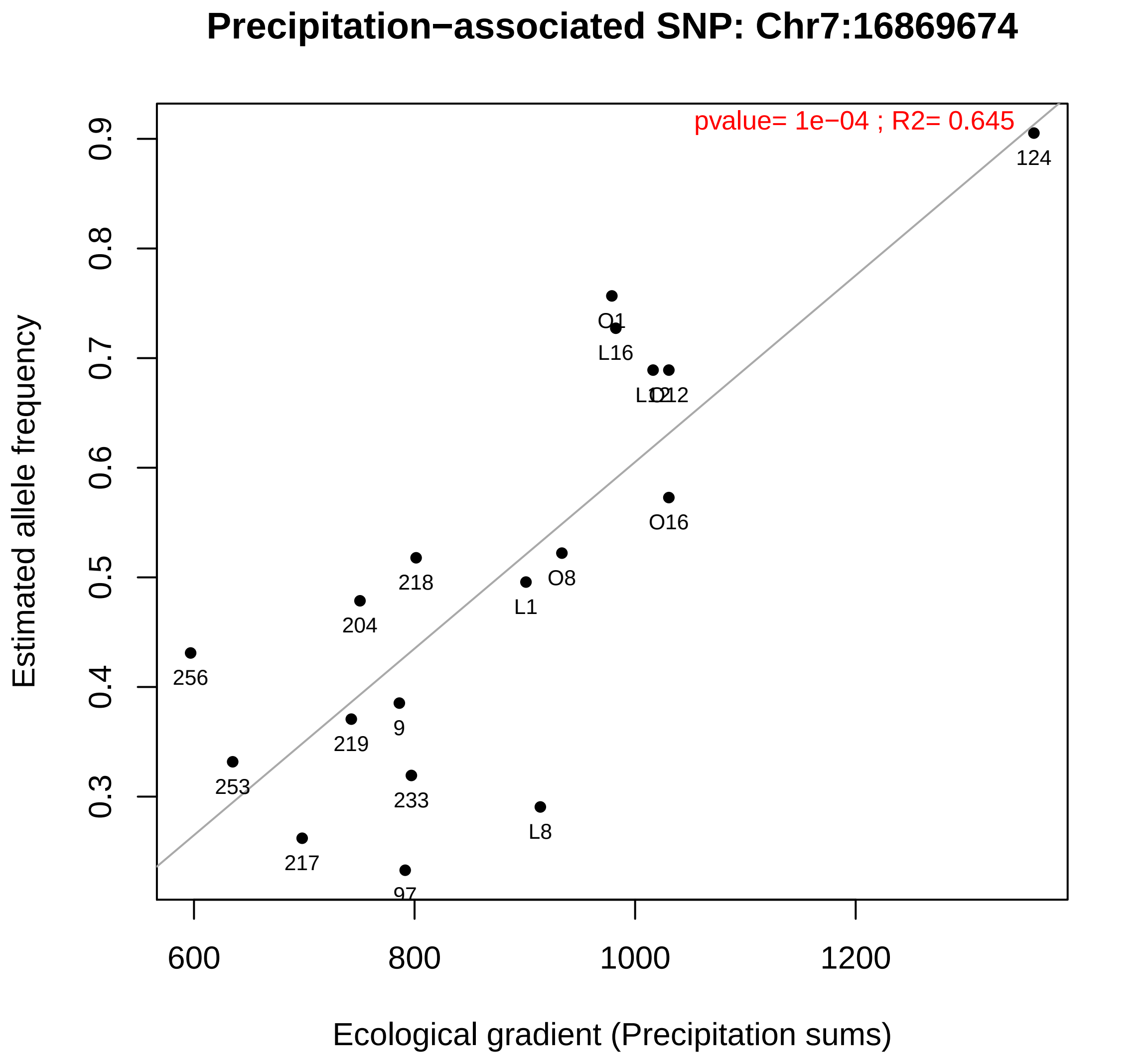
Precipitation−associated SNP: Chr7:16869674
pvalue= 1e−04 ; R2= 0.645
124
O1
L16
L12
O12
O16
O8
218
L1
204
256
9
219
253
233
L8
217
97
0.3
0.4
0.5
0.6
0.7
0.8
0.9
Estimated allele frequency
600
800
1000
1200
Ecological gradient (Precipitation sums)

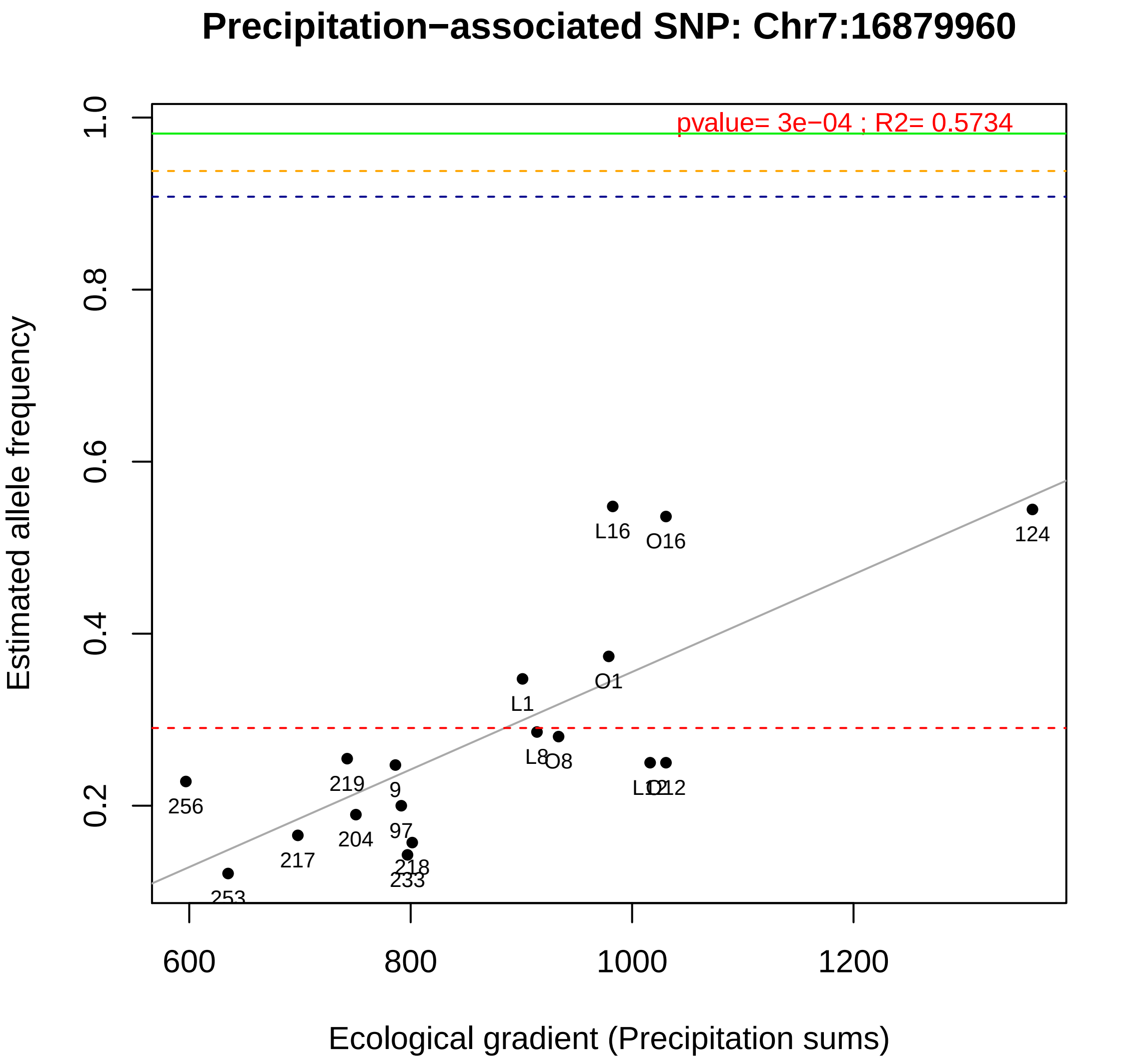
Precipitation−associated SNP: Chr7:16879960
pvalue= 3e−04 ; R2= 0.5734
Estimated allele frequency
0.2
0.4
0.6
0.8
1.0
600
800
1000
1200
Ecological gradient (Precipitation sums)
L16
O16
124
O1
L1
L8
O8
L12
O12
256
219
9
97
204
217
218
233
253

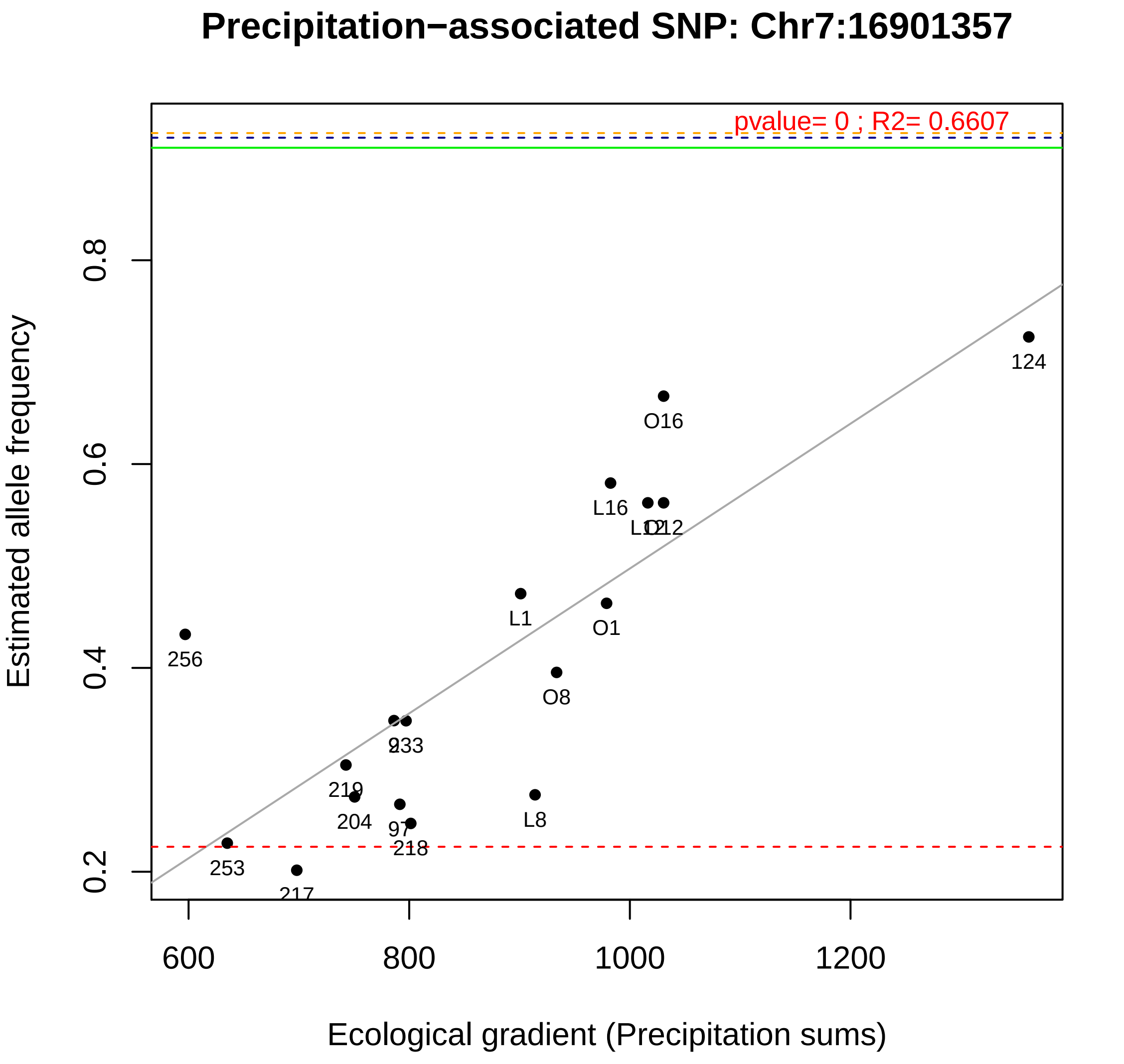

Precipitation−associated SNP: Chr7:16901357
pvalue= 0 ; R2= 0.6607
Estimated allele frequency
0.2
0.4
0.6
0.8
600
800
1000
1200
Ecological gradient (Precipitation sums)
124
O16
L16
L12
O12
L1
O1
256
O8
9
233
219
204
97
218
L8
253
217

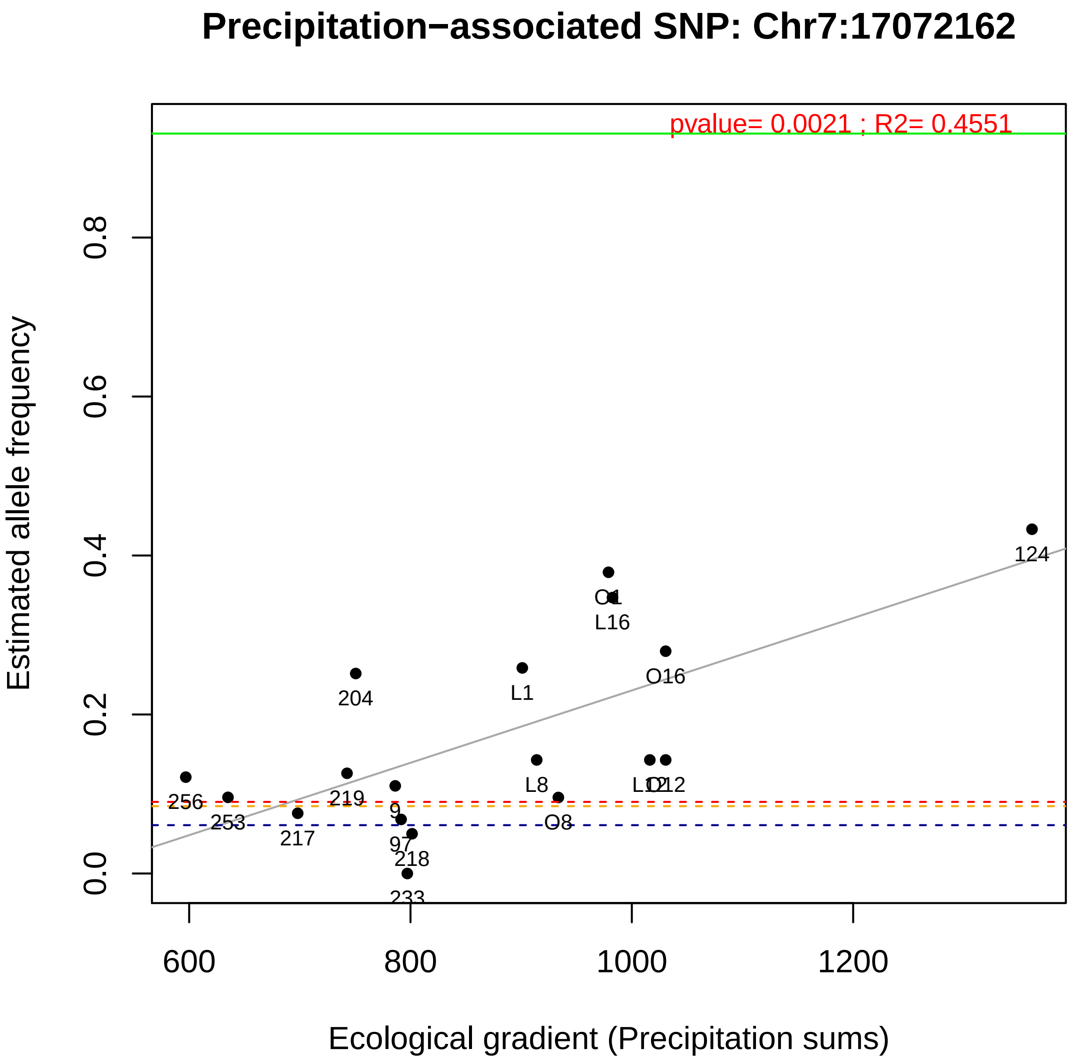
Precipitation−associated SNP: Chr7:17072162
pvalue= 0.0021 ; R2= 0.4551
Estimated allele frequency
0.0
0.2
0.4
0.6
0.8
600
800
1000
1200
Ecological gradient (Precipitation sums)
124
L16
O16
L1
204
L8
O8
256
253
217
219
97
218
233

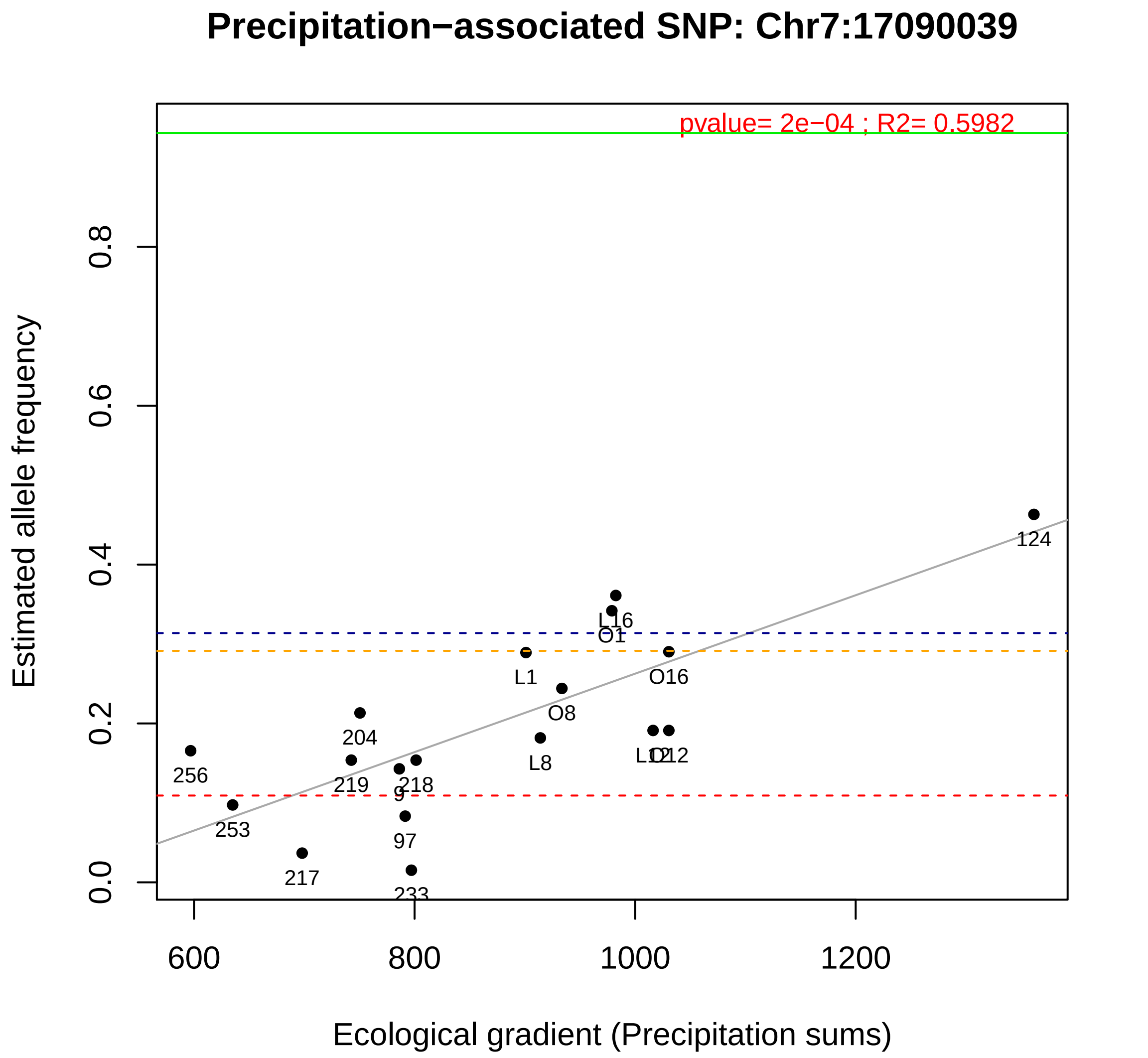
Precipitation−associated SNP: Chr7:17090039
pvalue= 2e−04 ; R2= 0.5982
Estimated allele frequency
0.0
0.2
0.4
0.6
0.8
600
800
1000
1200
Ecological gradient (Precipitation sums)
124
L16
O1
L1
O16
O8
204
L8
L12
O12
256
219
218
9
253
97
217
233

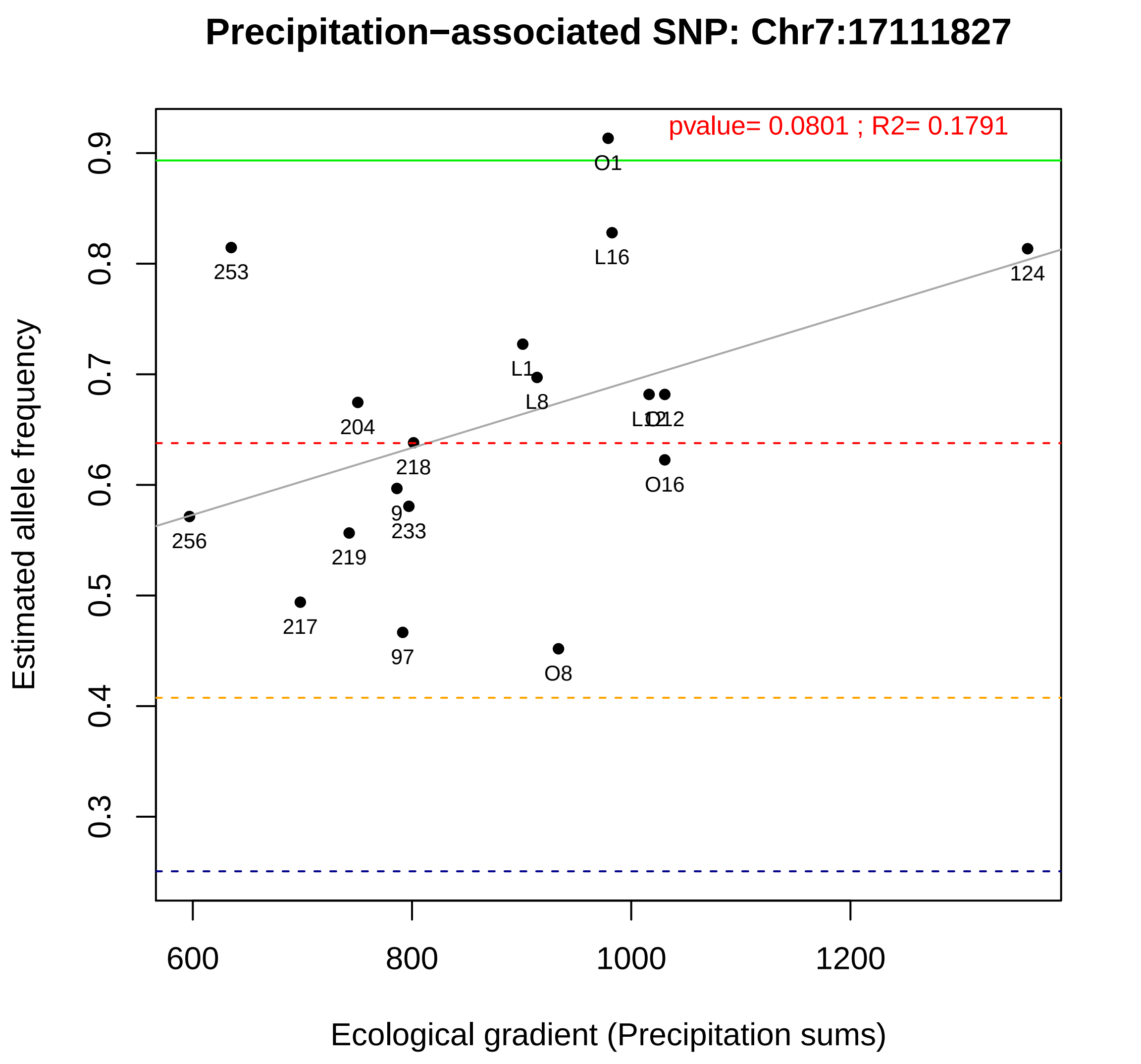

Precipitation−associated SNP: Chr7:17111827
pvalue= 0.0801 ; R2= 0.1791
Estimated allele frequency
Ecological gradient (Precipitation sums)
0.3
0.4
0.5
0.6
0.7
0.8
0.9
600
800
1000
1200
O1
L16
253
124
L1
L8
204
L12
O12
218
O16
9
233
256
219
217
97
O8

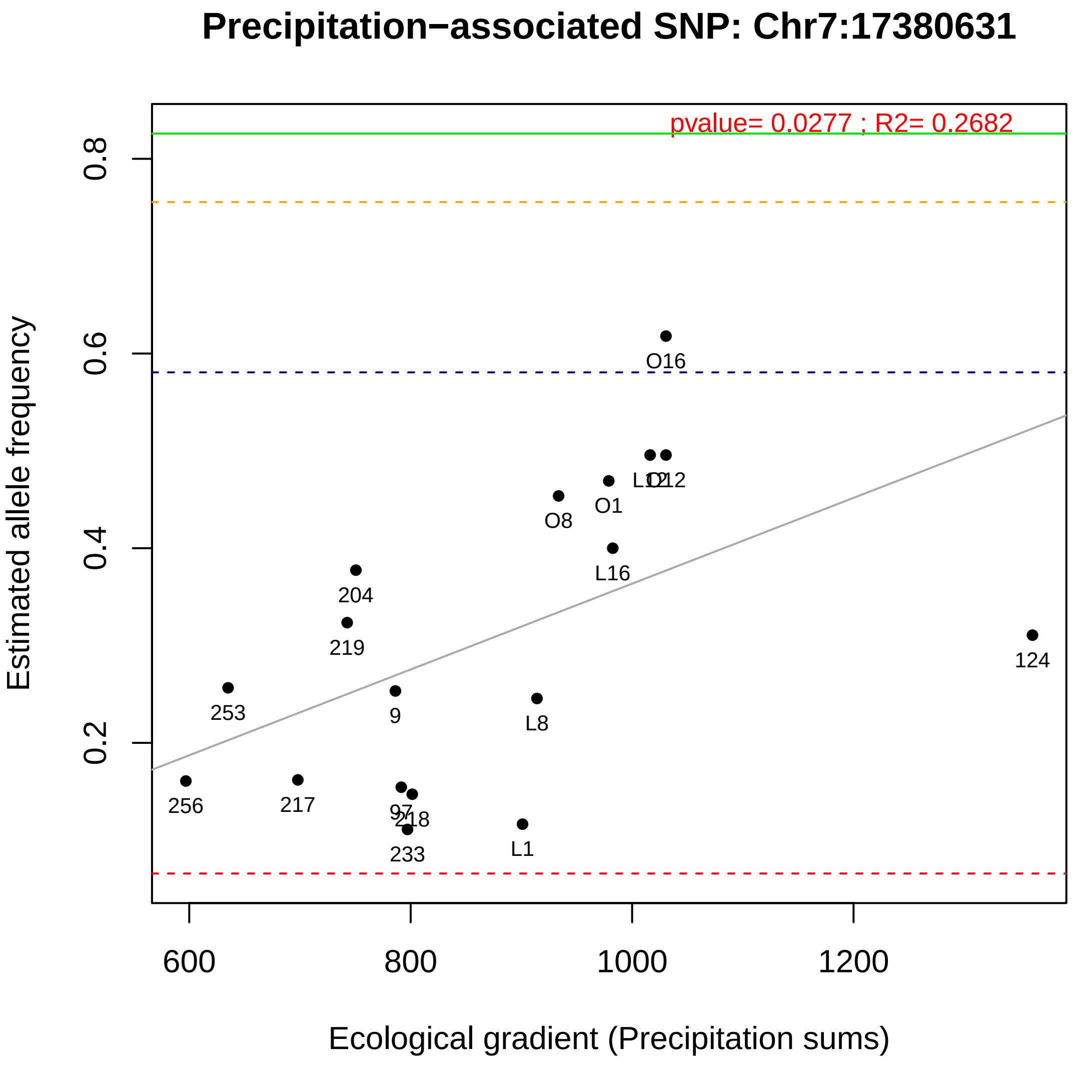
Precipitation−associated SNP: Chr7:17380631
pvalue= 0.0277 ; R2= 0.2682
Estimated allele frequency
0.2
0.4
0.6
0.8
600
800
1000
1200
Ecological gradient (Precipitation sums)
O16
L12
O12
O1
O8
L16
204
219
124
253
9
L8
256
217
97
218
233
L1

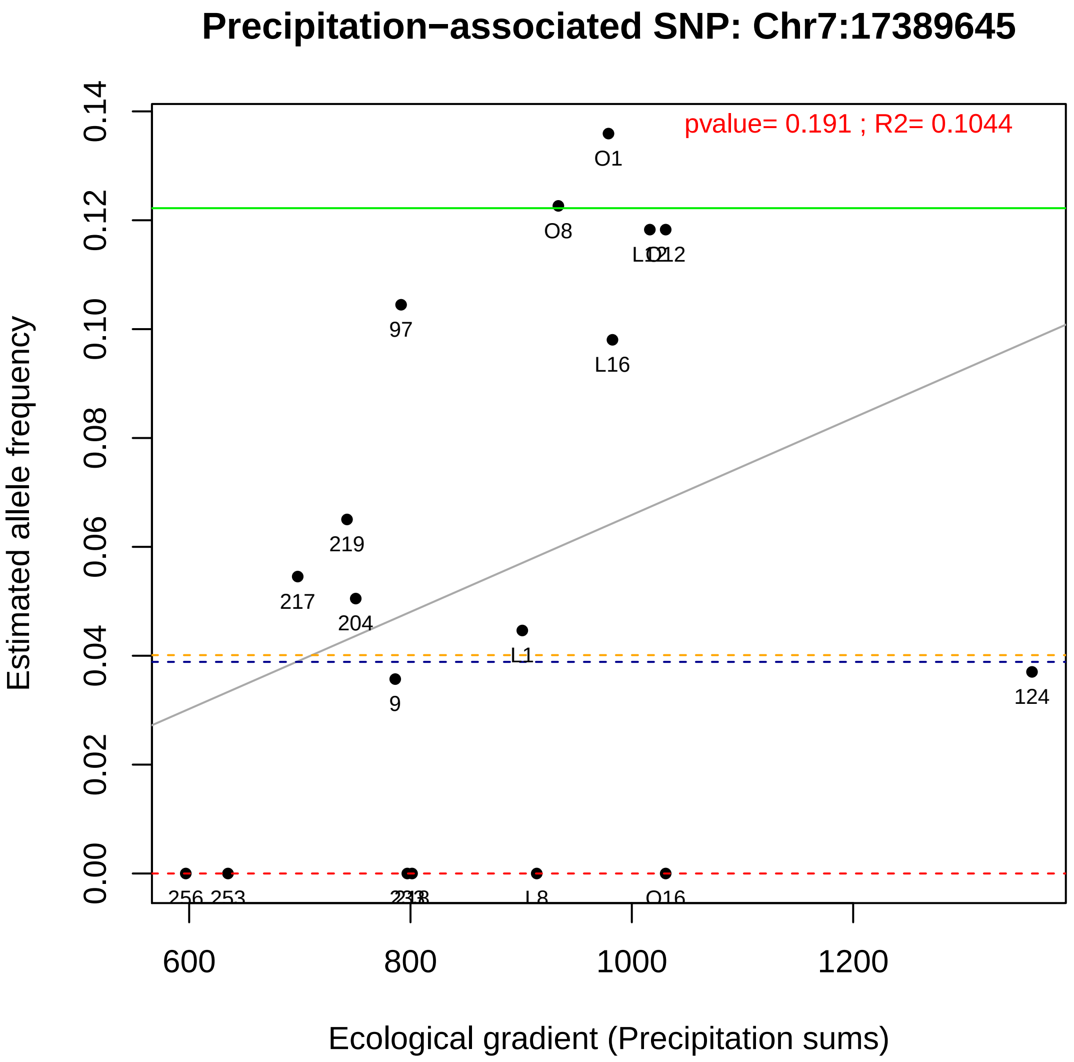
Precipitation−associated SNP: Chr7:17389645
pvalue= 0.191 ; R2= 0.1044
Estimated allele frequency
Ecological gradient (Precipitation sums)
0.00
0.02
0.04
0.06
0.08
0.10
0.12
0.14
600
800
1000
1200
O1
O8
97
L16
219
217
204
L1
9
124

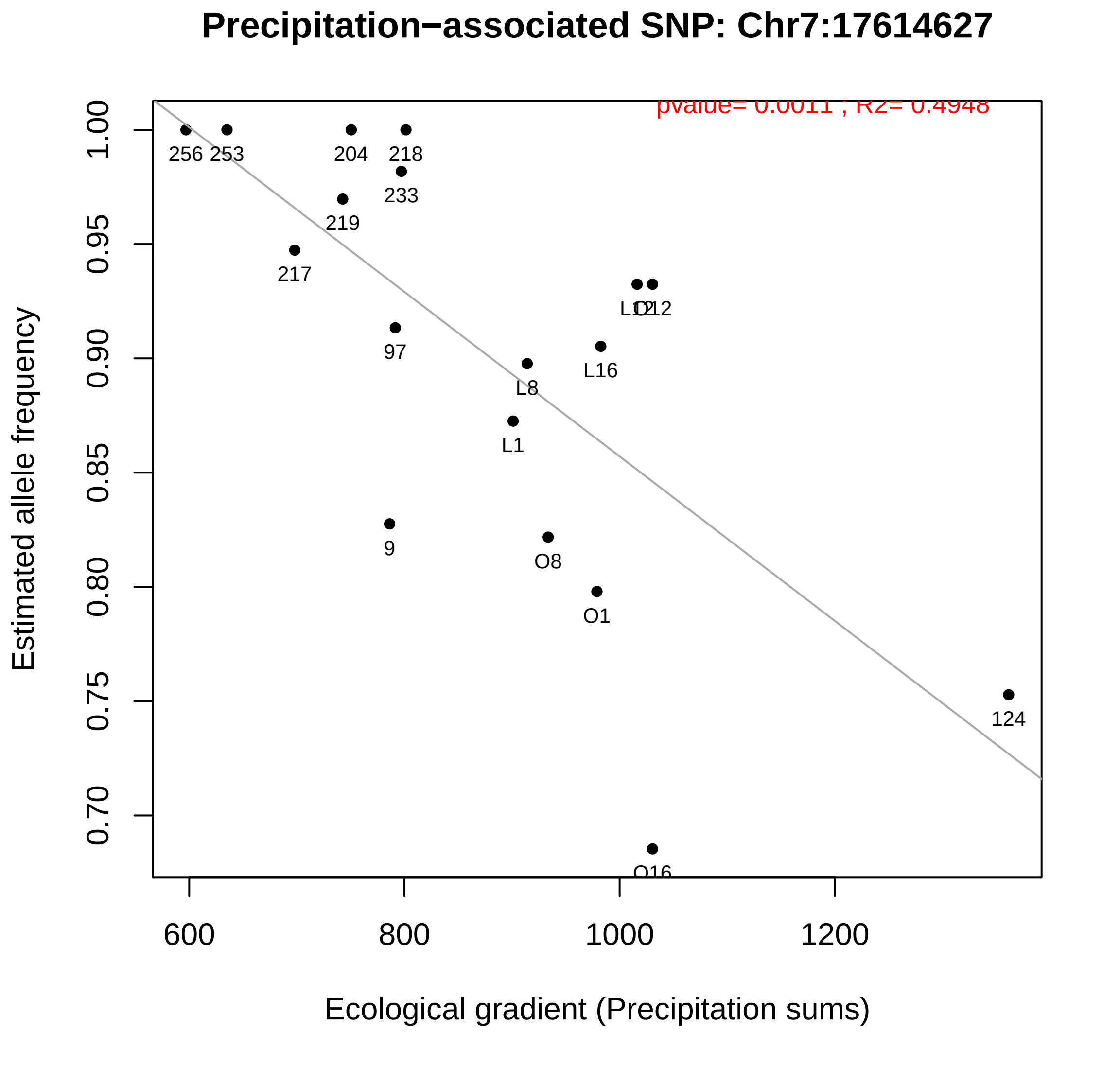

Precipitation−associated SNP: Chr7:17614627
pvalue= 0.0011 ; R2= 0.4948
Estimated allele frequency
1.00
0.95
0.90
0.85
0.80
0.75
0.70
256
253
204
218
233
219
217
L12
O12
97
L16
L8
L1
9
O8
O1
124
O16
600
800
1000
1200
Ecological gradient (Precipitation sums)

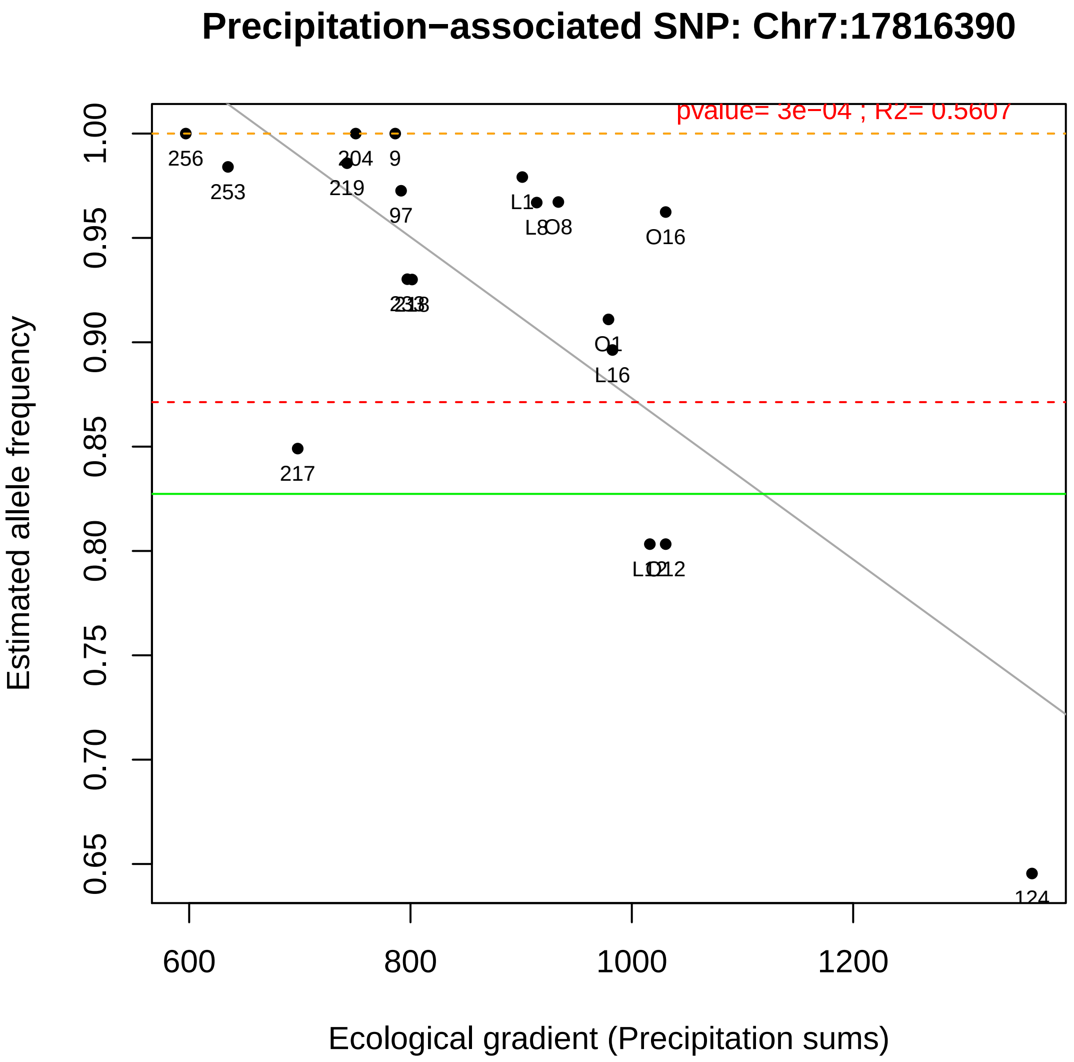

Precipitation−associated SNP: Chr7:17816390
pvalue= 3e−04 ; R2= 0.5607
Estimated allele frequency
Ecological gradient (Precipitation sums)
1.00
0.95
0.90
0.85
0.80
0.75
0.70
0.65
600
800
1000
1200
256
253
204
219
9
97
L1
L8
O8
O16
O1
L16
217
L12
O12
124

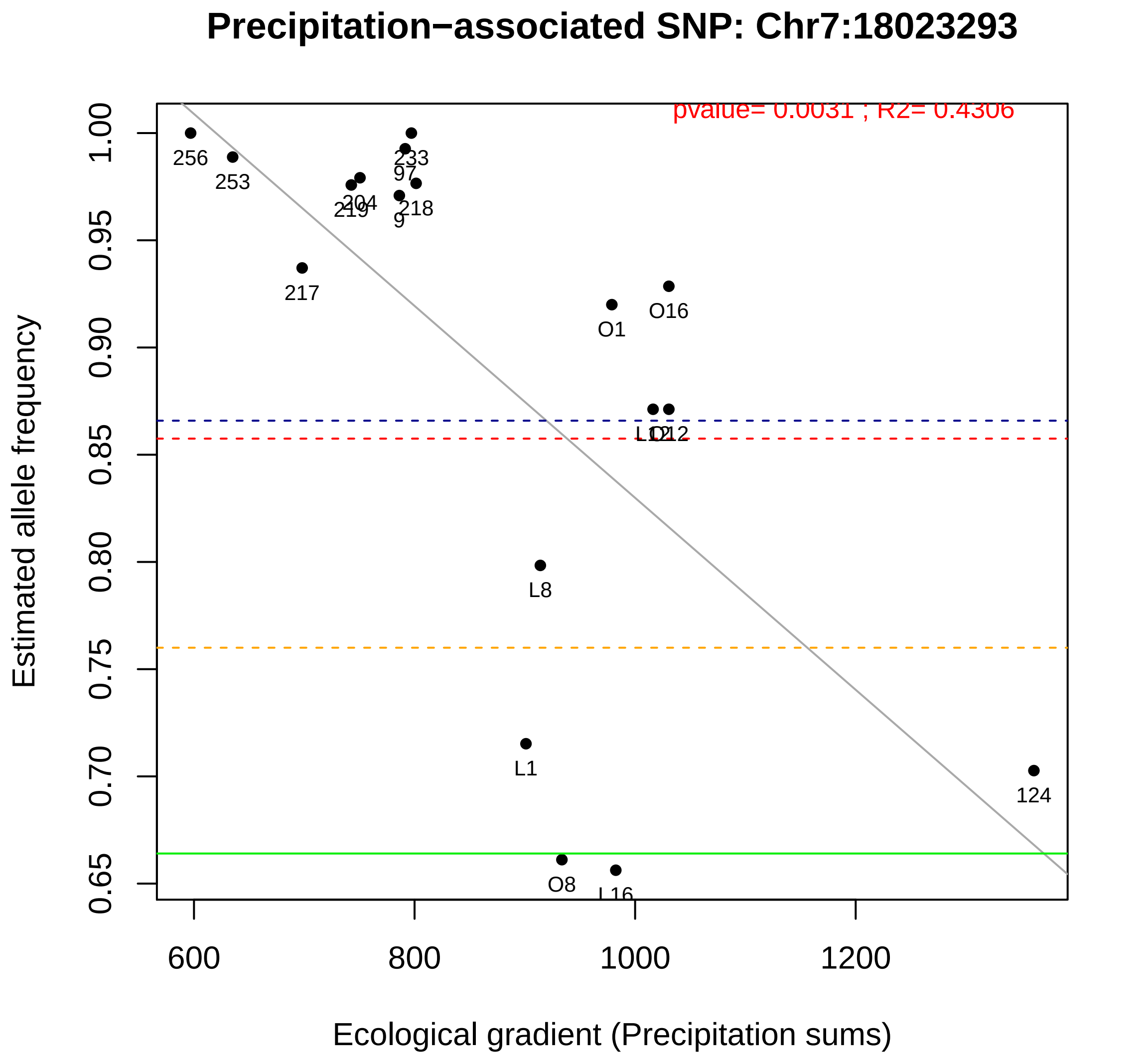

Precipitation−associated SNP: Chr7:18023293
pvalue= 0.0031 ; R2= 0.4306
Estimated allele frequency
Ecological gradient (Precipitation sums)
0.65
0.70
0.75
0.80
0.85
0.90
0.95
1.00
600
800
1000
1200
256
253
233
97
204
219
218
9
217
O1
O16
L8
L1
124
O8

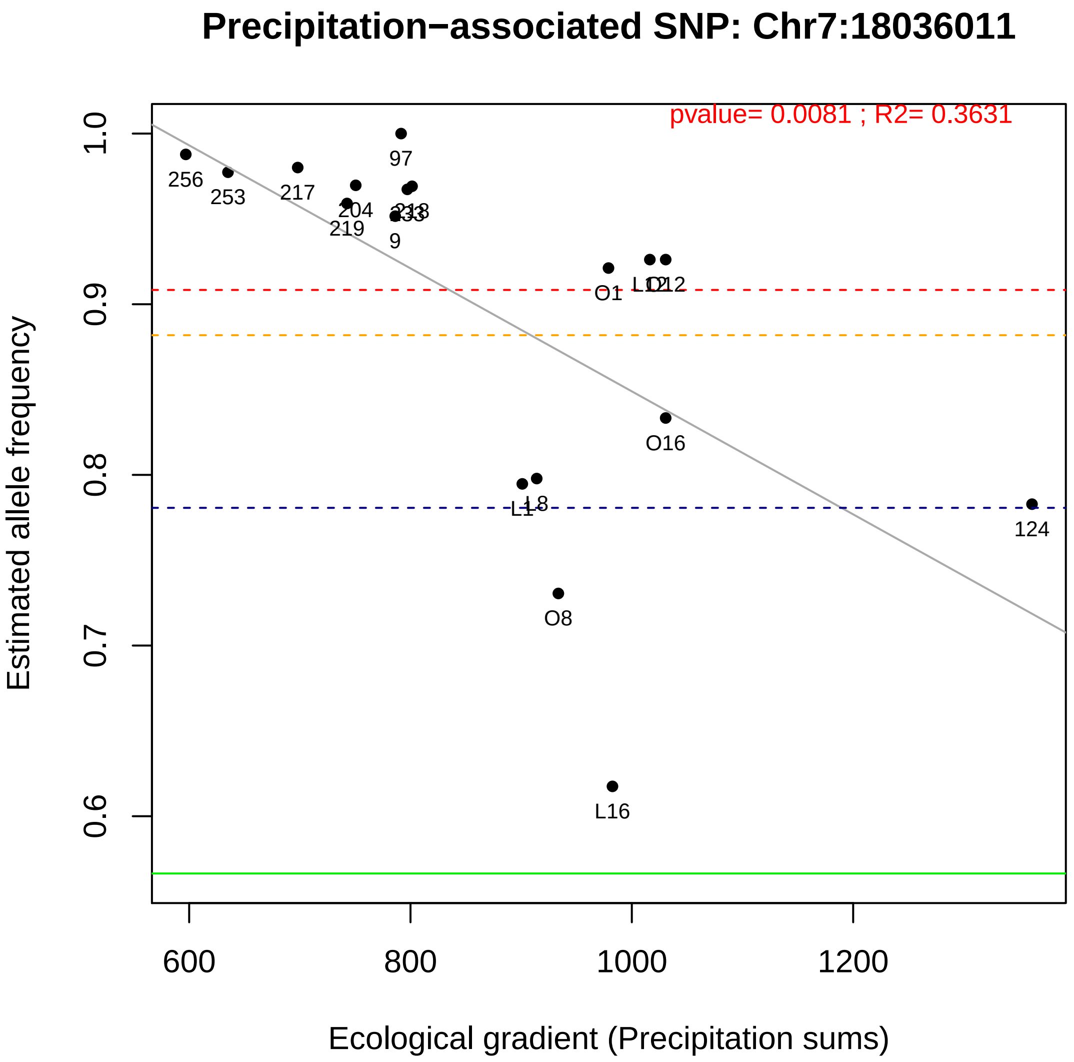
Precipitation−associated SNP: Chr7:18036011
pvalue= 0.0081 ; R2= 0.3631
Estimated allele frequency
Ecological gradient (Precipitation sums)
0.6
0.7
0.8
0.9
1.0
600
800
1000
1200
256
253
217
204
219
97
9
O1
O16
L8
O8
L16
124

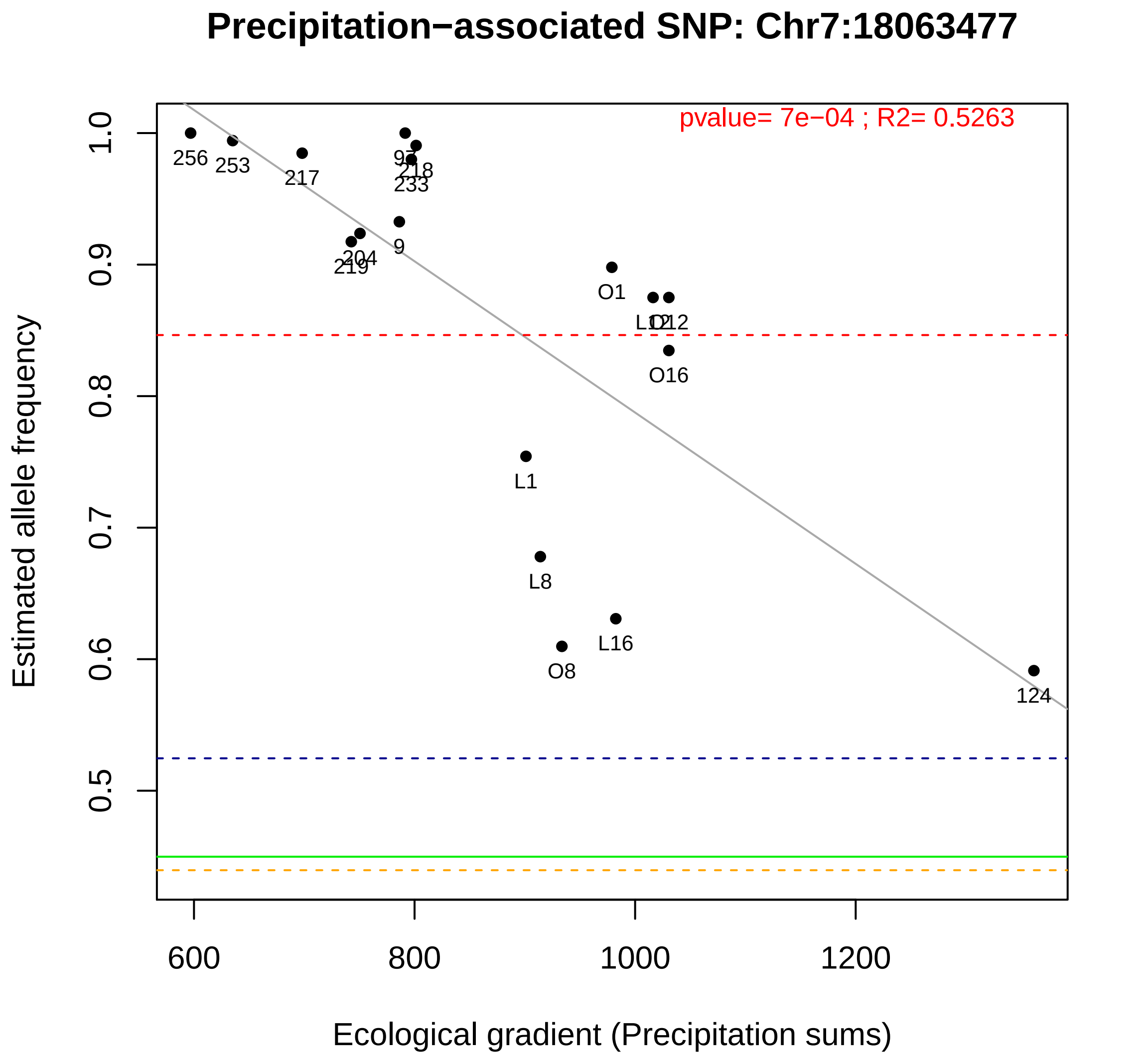

Precipitation−associated SNP: Chr7:18063477
pvalue= 7e−04 ; R2= 0.5263
Estimated allele frequency
Ecological gradient (Precipitation sums)
0.5
0.6
0.7
0.8
0.9
1.0
600
800
1000
1200
256
253
217
97
218
233
204
219
9
O1
L10
O12
O16
L1
L8
L16
O8
124

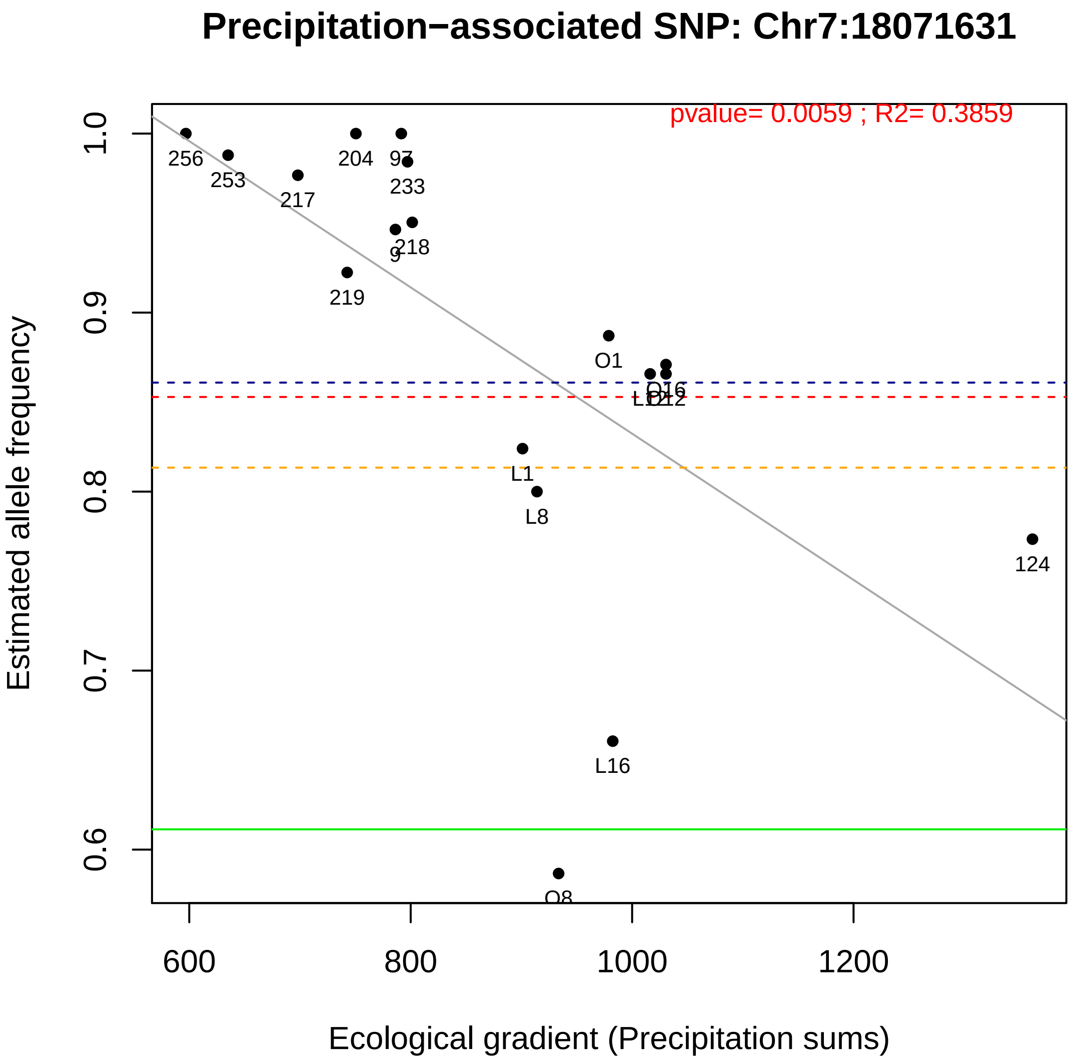

Precipitation−associated SNP: Chr7:18071631
pvalue= 0.0059 ; R2= 0.3859
Estimated allele frequency
Ecological gradient (Precipitation sums)
1.0
0.9
0.8
0.7
0.6
600
800
1000
1200
256
253
217
204
97
233
218
9
219
O1
O16
L12
L1
L8
124
L16
O8

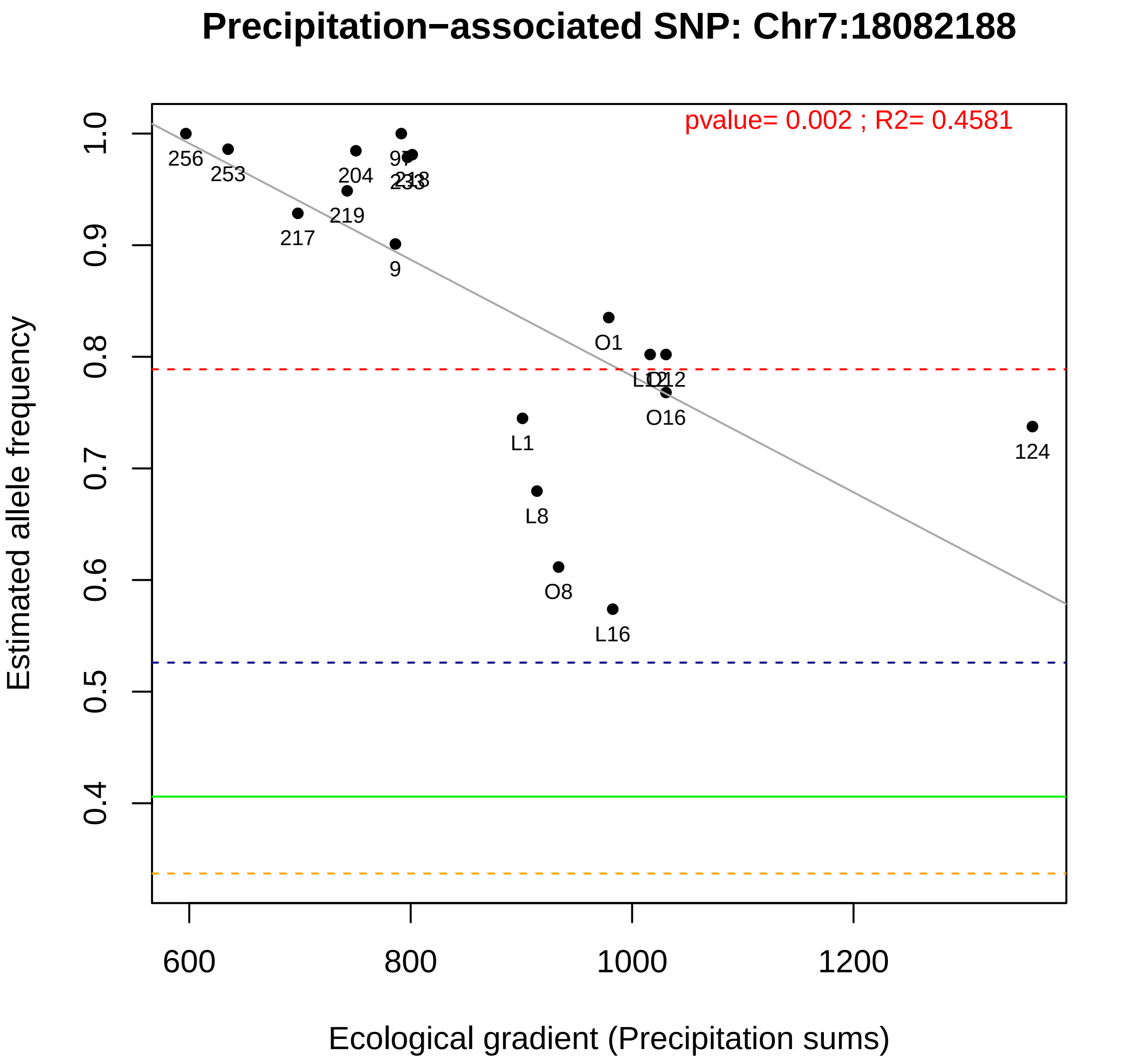
Precipitation−associated SNP: Chr7:18082188
pvalue= 0.002 ; R2= 0.4581
Estimated allele frequency
1.0
0.9
0.8
0.7
0.6
0.5
0.4
600
800
1000
1200
Ecological gradient (Precipitation sums)
256
253
204
219
217
9
O1
O16
L1
L8
O8
L16
124

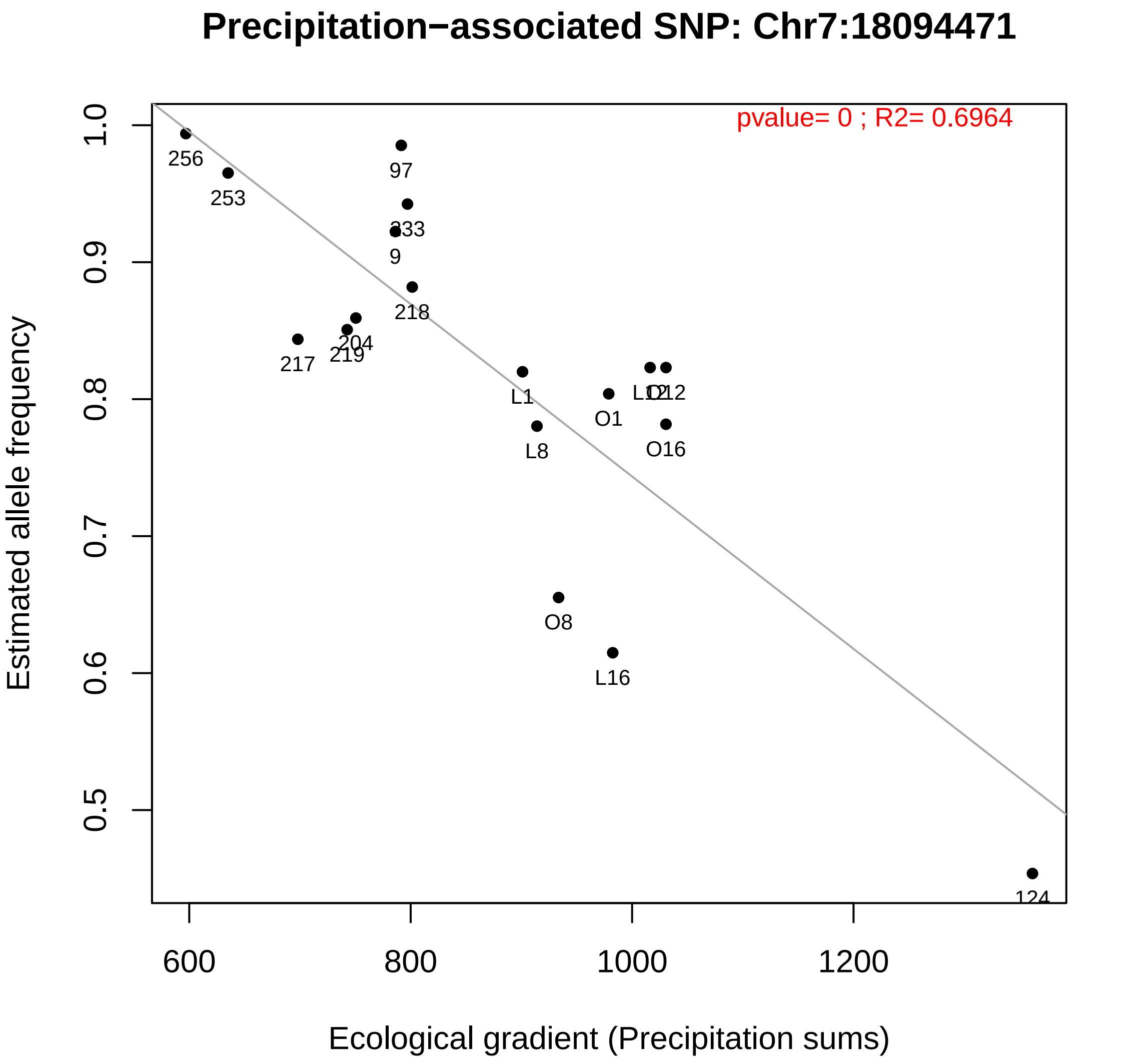
Precipitation−associated SNP: Chr7:18094471
pvalue= 0 ; R2= 0.6964
Estimated allele frequency
Ecological gradient (Precipitation sums)
1.0
0.9
0.8
0.7
0.6
0.5
600
800
1000
1200
256
253
97
333
9
218
204
219
217
L1
L8
O1
L12
O12
O16
O8
L16
124

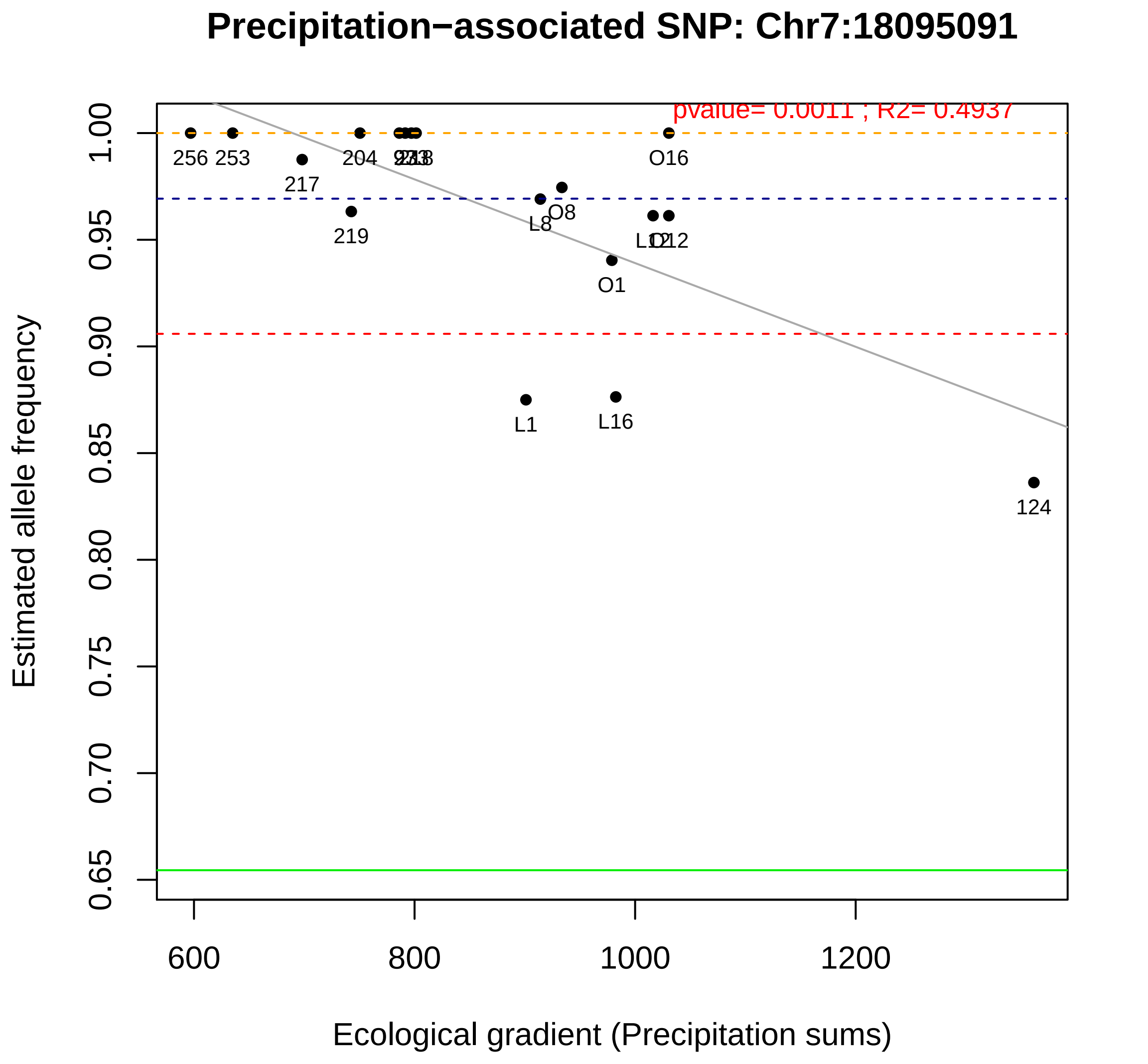
Precipitation−associated SNP: Chr7:18095091
pvalue= 0.0011 ; R2= 0.4937
Estimated allele frequency
Ecological gradient (Precipitation sums)
0.65
0.70
0.75
0.80
0.85
0.90
0.95
1.00
600
800
1000
1200
256
253
217
204
219
L8
O8
O16
O1
L1
L16
124

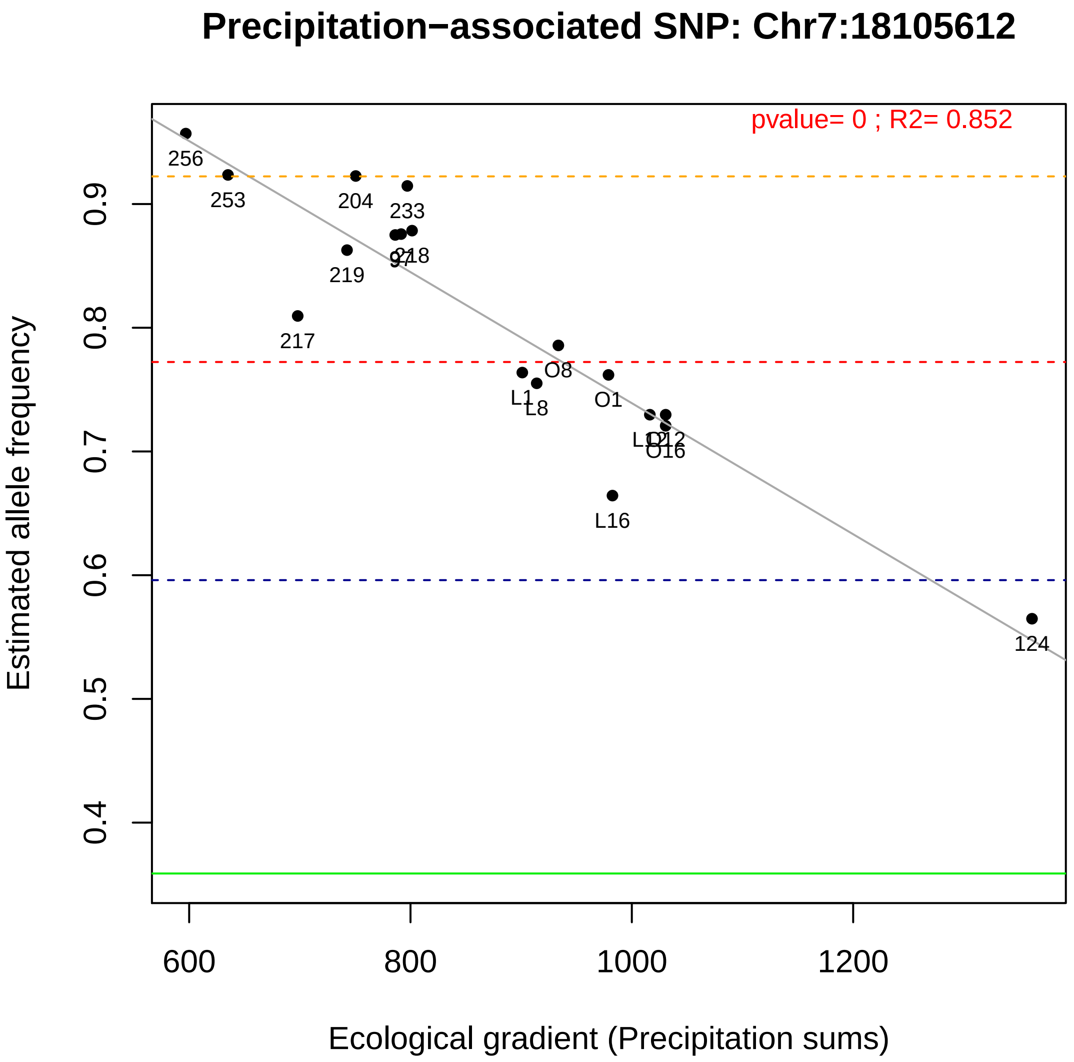

Precipitation−associated SNP: Chr7:18105612
pvalue= 0 ; R2= 0.852
Estimated allele frequency
Ecological gradient (Precipitation sums)
0.4
0.5
0.6
0.7
0.8
0.9
600
800
1000
1200
256
253
204
233
218
97
219
217
O8
L1
L8
O1
L12
O12
O16
L16
124

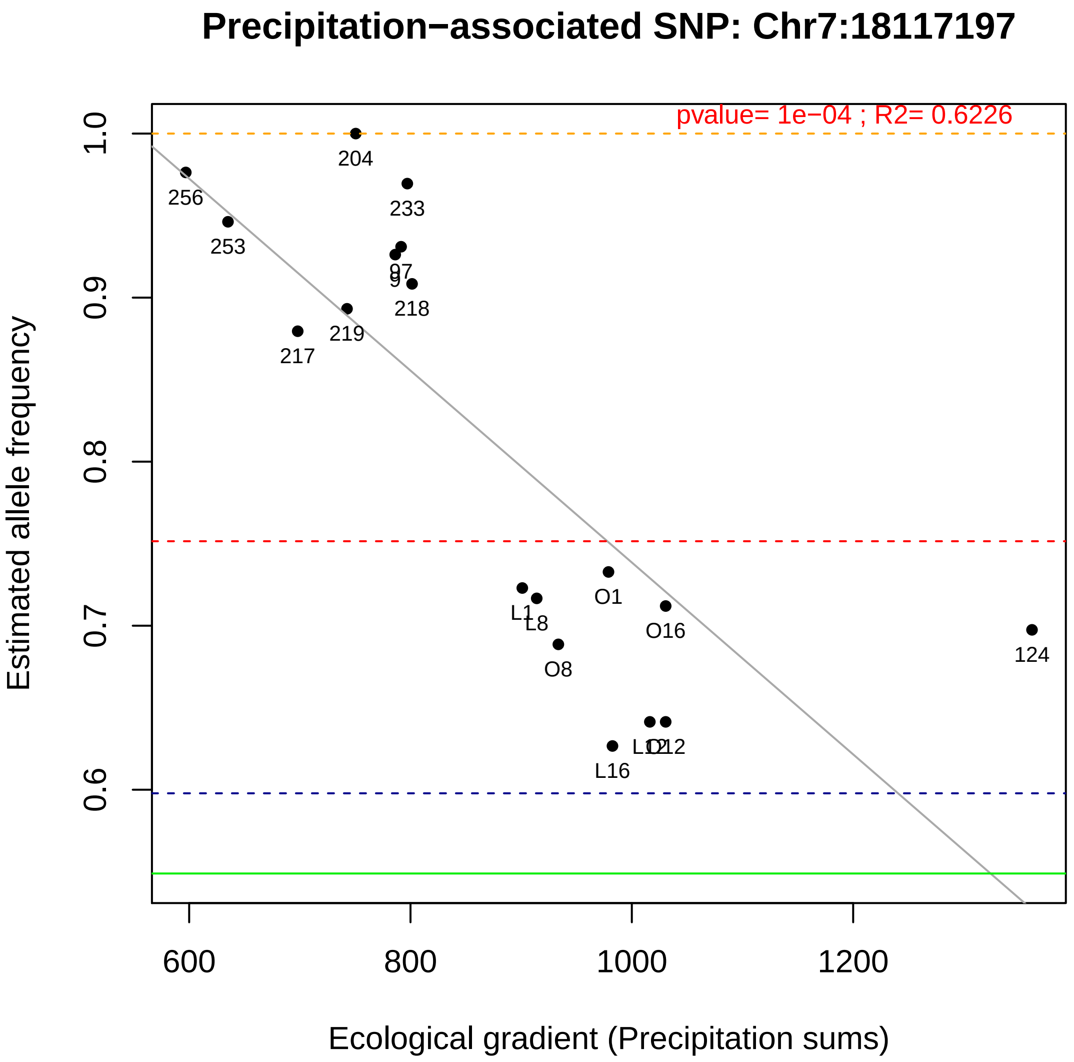
Precipitation−associated SNP: Chr7:18117197
pvalue= 1e−04 ; R2= 0.6226
Estimated allele frequency
Ecological gradient (Precipitation sums)
0.6
0.7
0.8
0.9
1.0
600
800
1000
1200
204
256
233
253
97
9
218
219
217
O1
L1
L8
O16
124
O8
L12
O12
L16

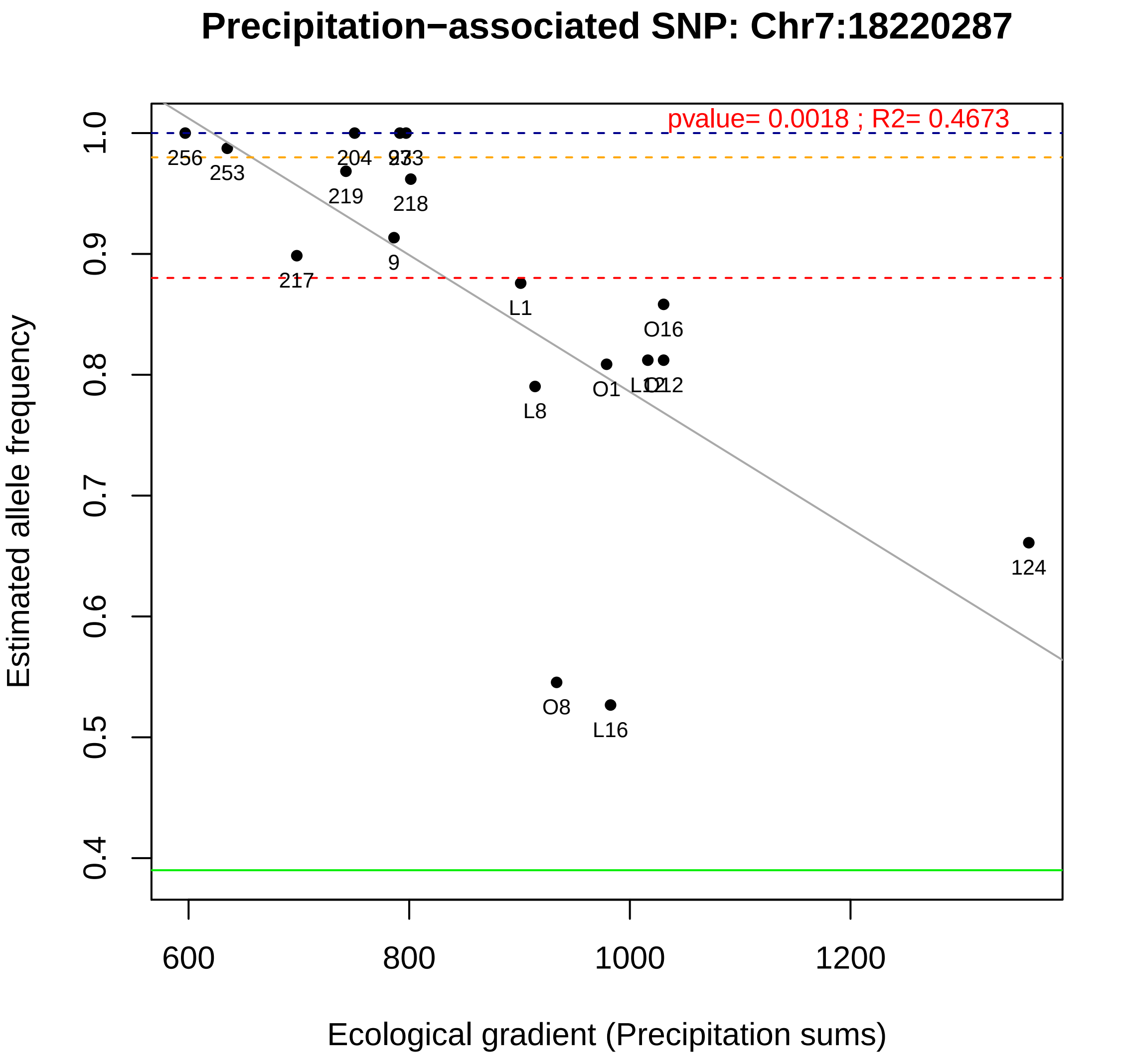
Precipitation−associated SNP: Chr7:18220287
pvalue= 0.0018 ; R2= 0.4673
Estimated allele frequency
Ecological gradient (Precipitation sums)
1.0
0.9
0.8
0.7
0.6
0.5
0.4
600
800
1000
1200
256
253
204
219
218
217
9
L1
O16
O1
L8
124
O8
L16

#### Precipitation−associated SNP: Chr7:18226898

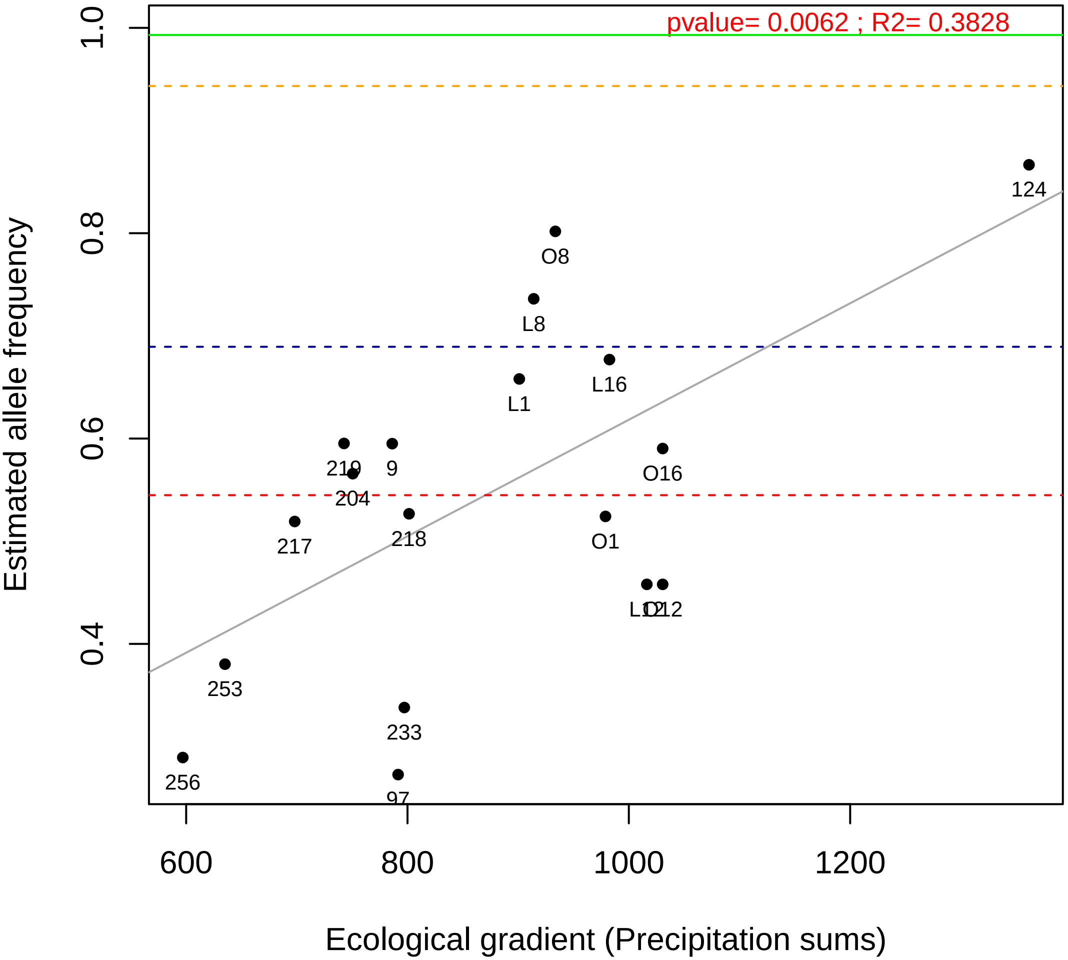

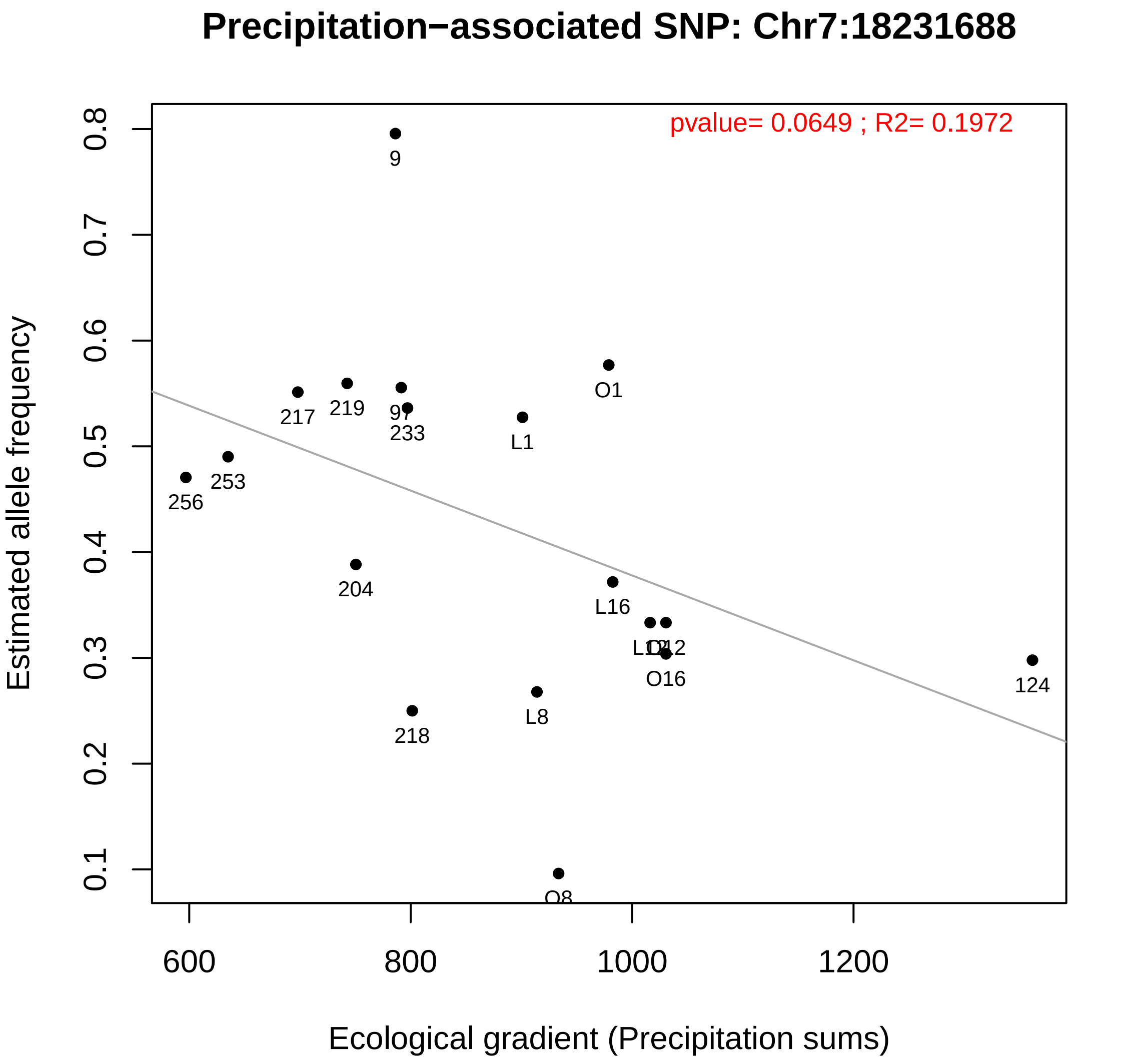

Precipitation−associated SNP: Chr7:18231688
pvalue= 0.0649 ; R2= 0.1972
Estimated allele frequency
Ecological gradient (Precipitation sums)
0.1
0.2
0.3
0.4
0.5
0.6
0.7
0.8
600
800
1000
1200
9
217
219
97
233
L1
O1
256
253
204
L16
L12
O12
O16
124
218
L8
O8

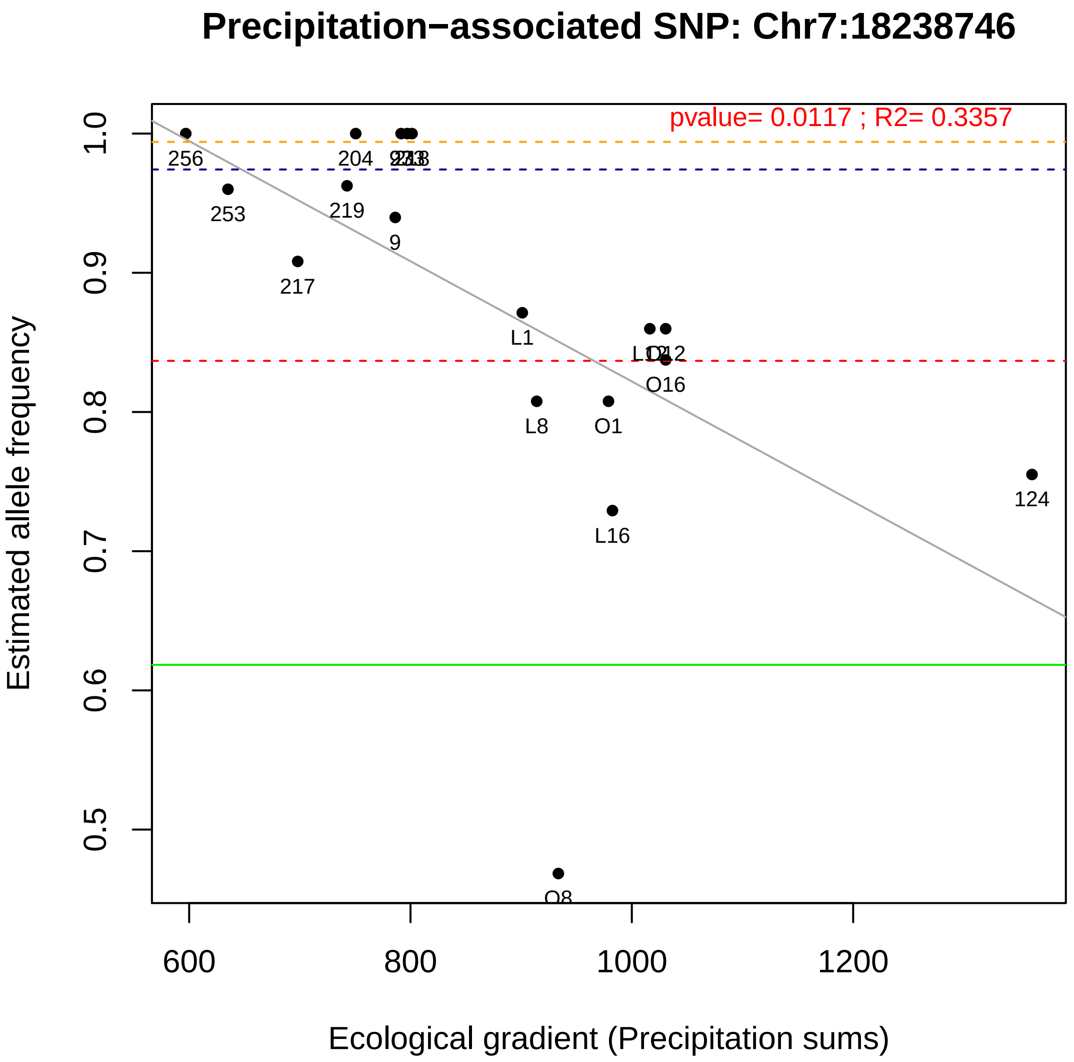
Precipitation−associated SNP: Chr7:18238746
pvalue= 0.0117 ; R2= 0.3357
Estimated allele frequency
Ecological gradient (Precipitation sums)
0.5
0.6
0.7
0.8
0.9
1.0
600
800
1000
1200
256
204
253
219
9
217
L1
O16
L8
O1
124
L16
O8

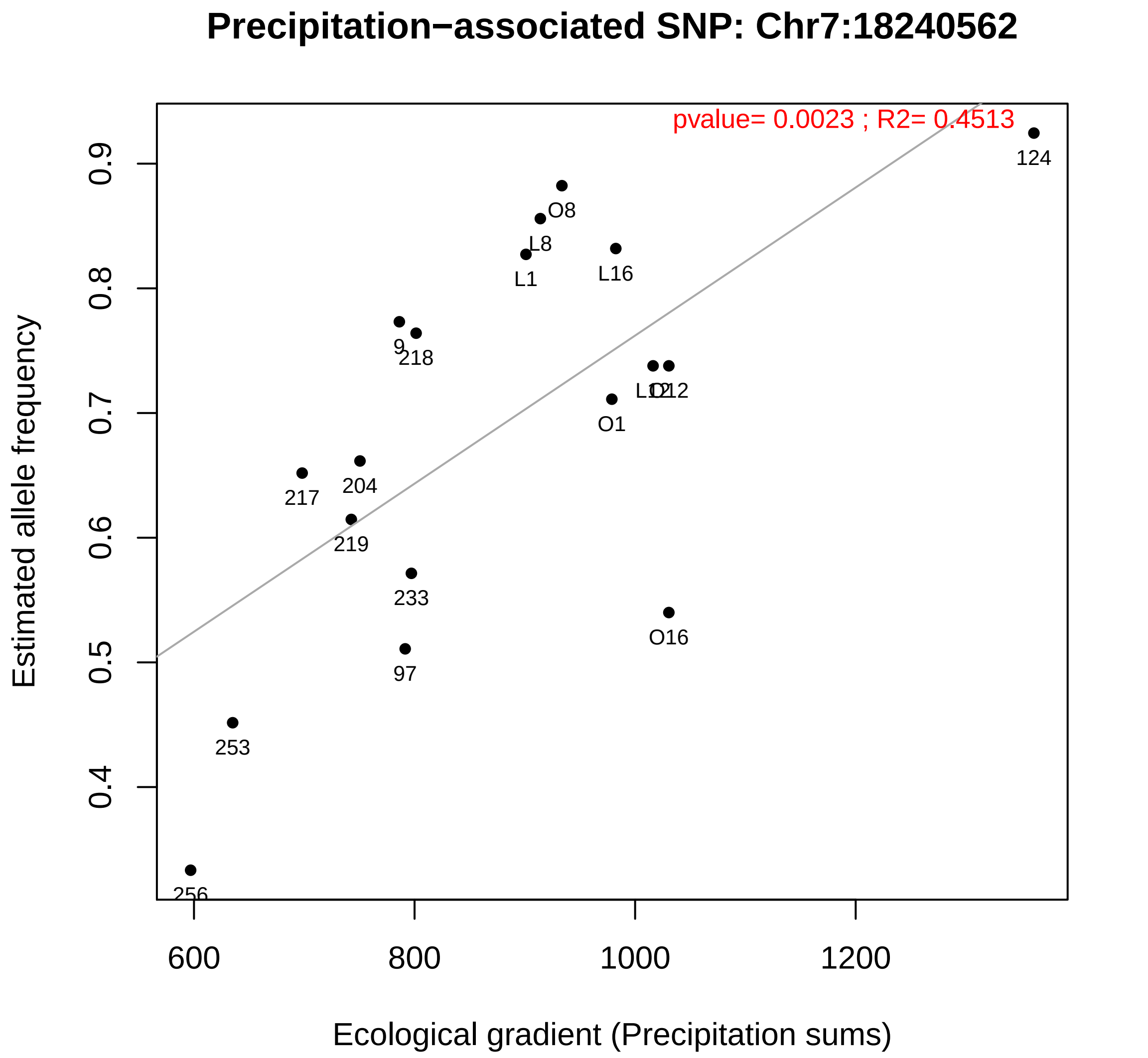
Precipitation−associated SNP: Chr7:18240562
pvalue= 0.0023 ; R2= 0.4513
Estimated allele frequency
0.4
0.5
0.6
0.7
0.8
0.9
600
800
1000
1200
Ecological gradient (Precipitation sums)
124
O8
L8
L1
L16
9
218
O1
O12
O16
217
204
219
233
97
253
256

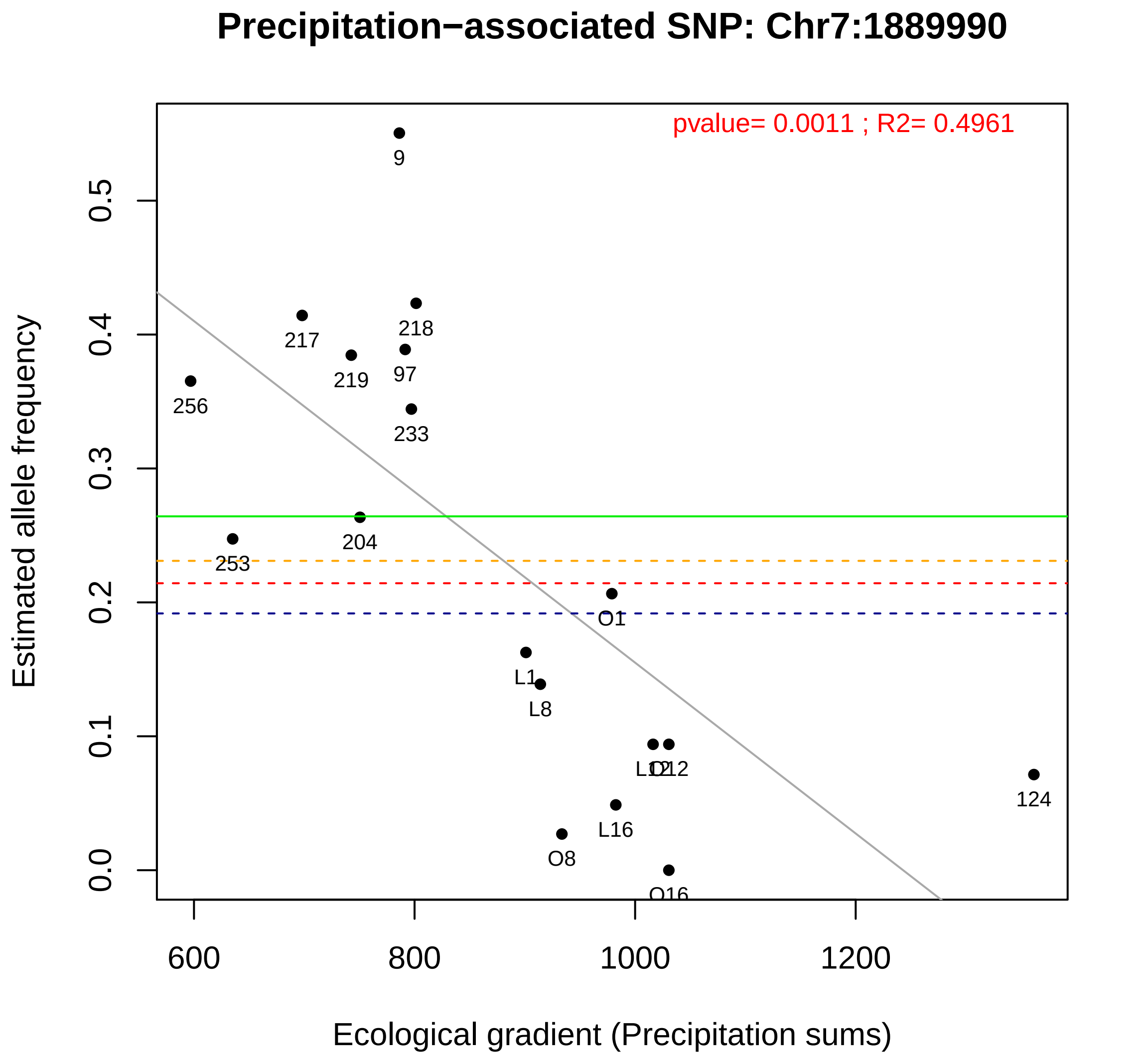

Precipitation−associated SNP: Chr7:1889990
pvalue= 0.0011 ; R2= 0.4961
Estimated allele frequency
Ecological gradient (Precipitation sums)
0.0
0.1
0.2
0.3
0.4
0.5
600
800
1000
1200
9
218
217
219
97
256
233
204
253
O1
L1
L8
L12
O12
124
L16
O8
O16

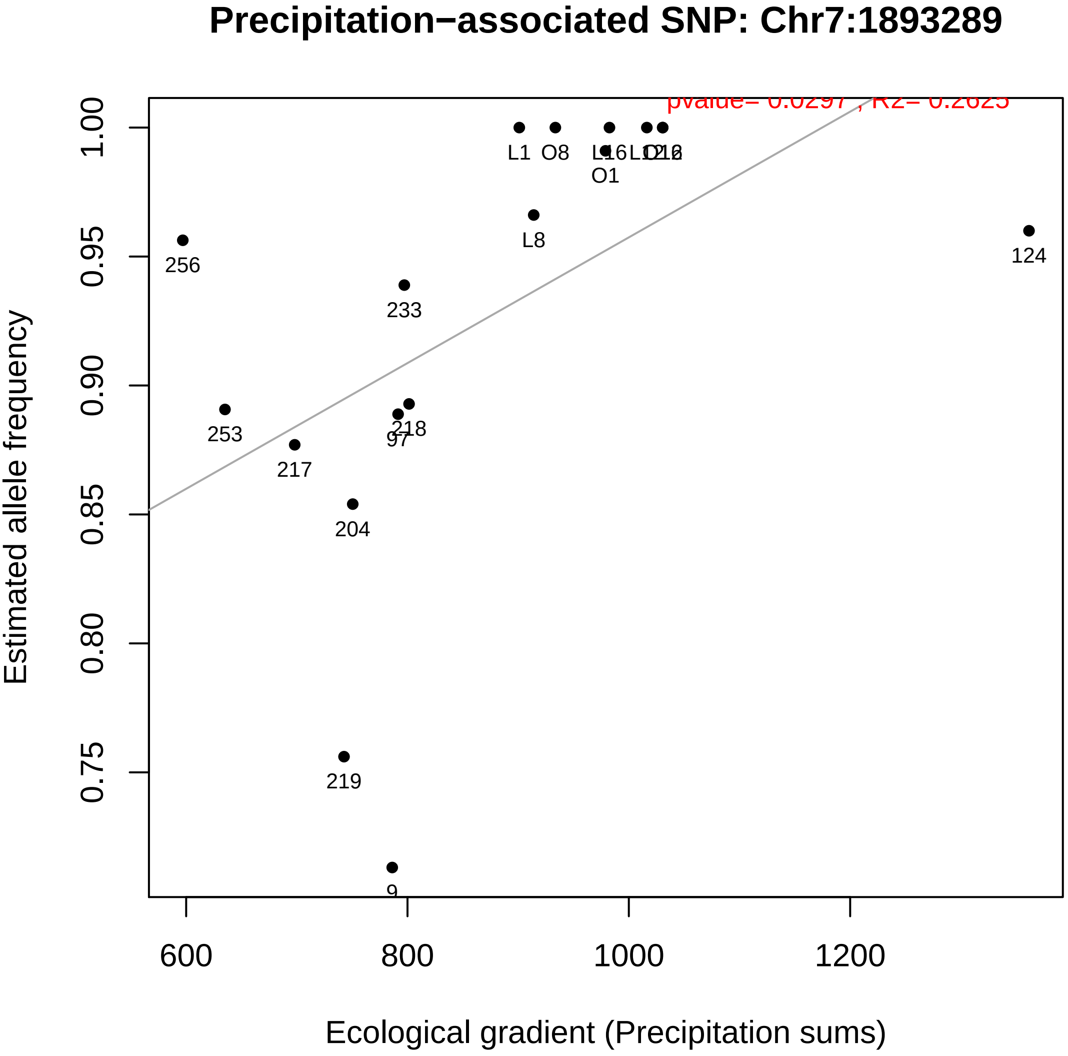
Precipitation−associated SNP: Chr7:1893289
pvalue= 0.0297 , R2= 0.2625
Estimated allele frequency
1.00
0.95
0.90
0.85
0.80
0.75
600
800
1000
1200
Ecological gradient (Precipitation sums)
L1
O8
L16
L12
O2
O1
L8
256
124
233
253
218
97
217
204
219
9

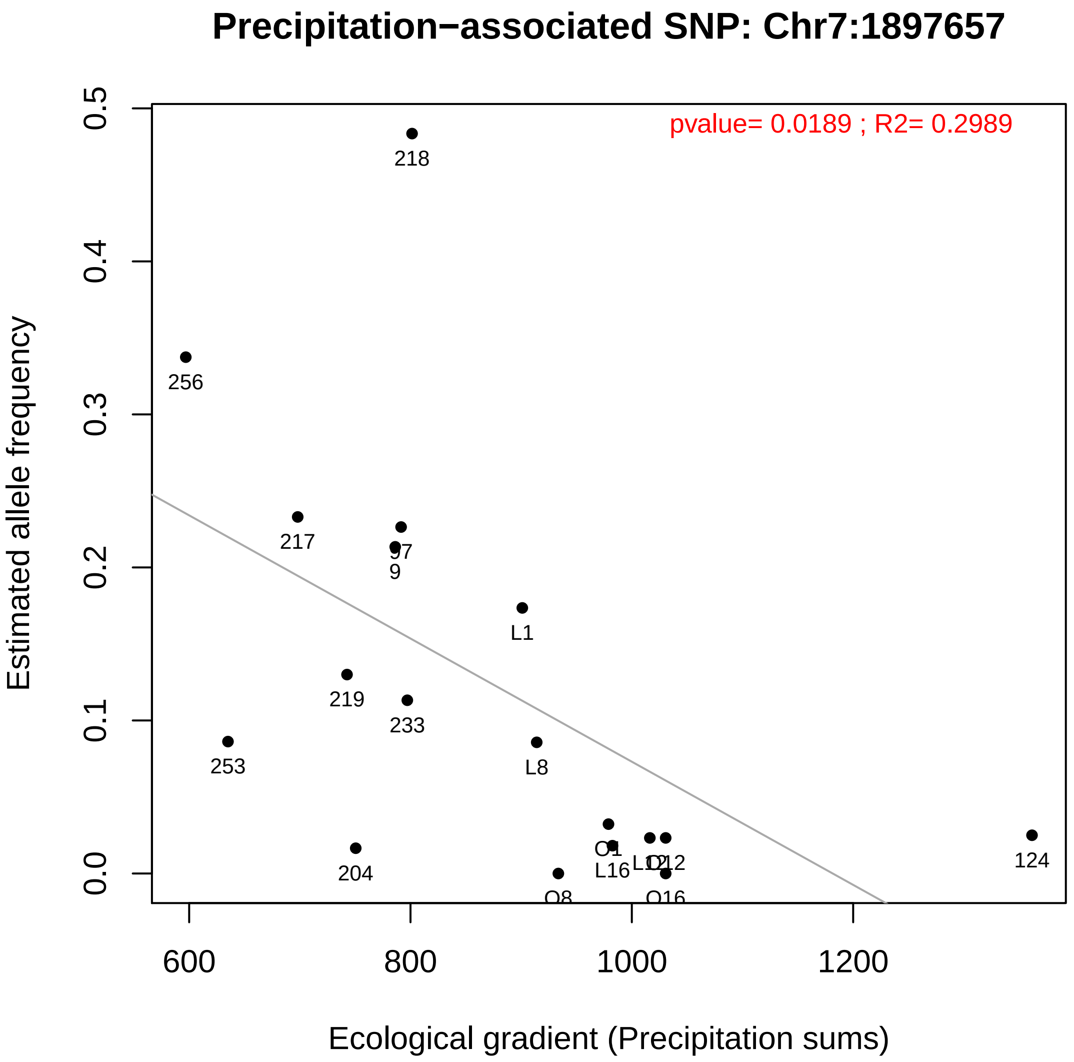

Precipitation−associated SNP: Chr7:1897657
pvalue= 0.0189 ; R2= 0.2989
Estimated allele frequency
0.0
0.1
0.2
0.3
0.4
0.5
Ecological gradient (Precipitation sums)
600
800
1000
1200
218
256
217
97
9
L1
219
233
253
L8
204
O1
L16
O8
O16
124

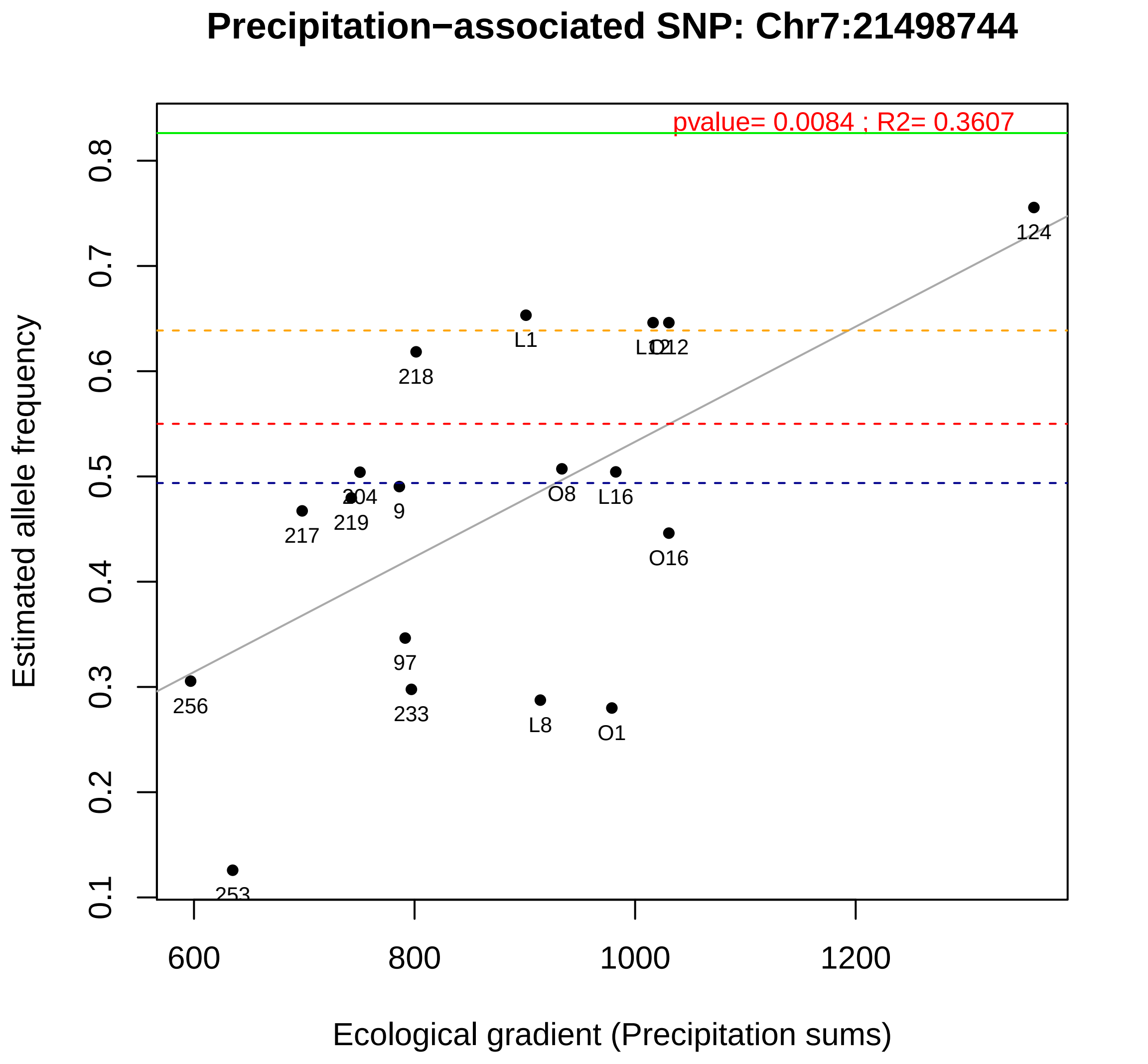

Precipitation−associated SNP: Chr7:21498744
pvalue= 0.0084 ; R2= 0.3607
Estimated allele frequency
0.1
0.2
0.3
0.4
0.5
0.6
0.7
0.8
600
800
1000
1200
Ecological gradient (Precipitation sums)
124
L1
L12
O12
218
204
219
9
217
O8
L16
O16
97
256
233
L8
O1
253

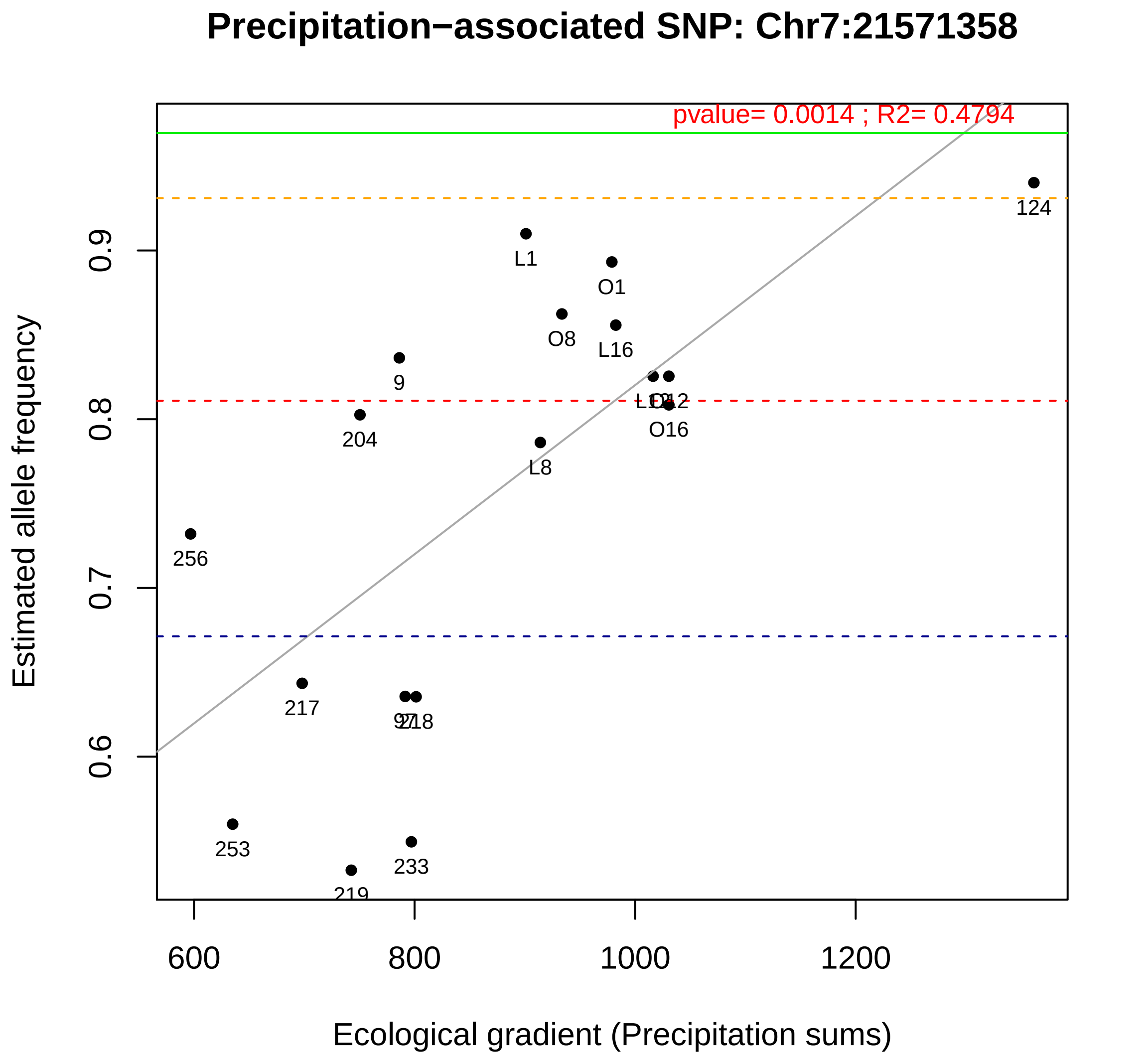
Precipitation−associated SNP: Chr7:21571358
pvalue= 0.0014 ; R2= 0.4794
Estimated allele frequency
Ecological gradient (Precipitation sums)
0.6
0.7
0.8
0.9
600
800
1000
1200
124
L1
O1
O8
L16
9
204
L8
O16
256
217
218
253
233
219

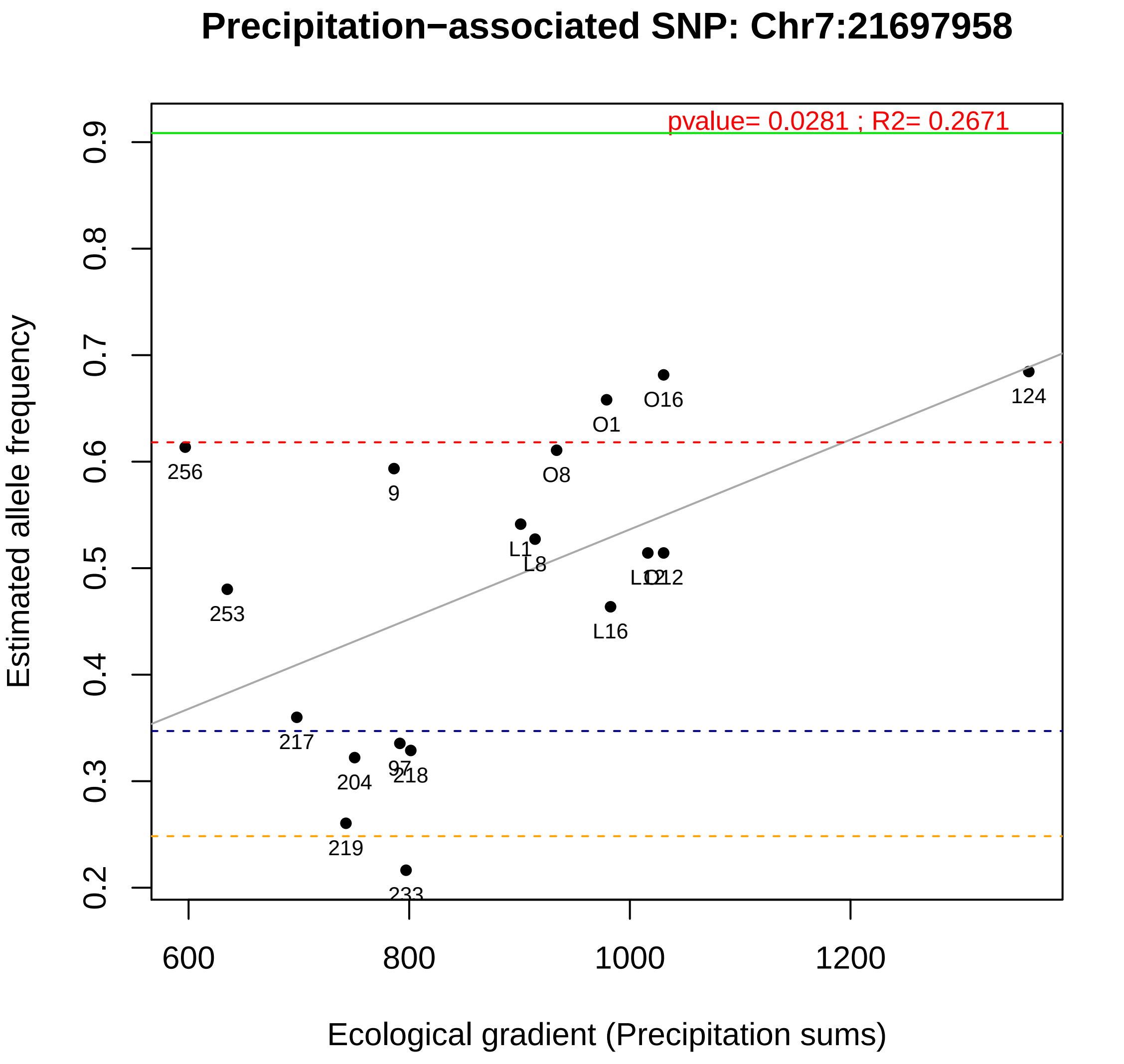

Precipitation−associated SNP: Chr7:21697958
pvalue= 0.0281 ; R2= 0.2671
Estimated allele frequency
0.2
0.3
0.4
0.5
0.6
0.7
0.8
0.9
600
800
1000
1200
Ecological gradient (Precipitation sums)
256
253
217
204
219
9
97
218
233
L1
L8
O8
O1
O16
L16
124

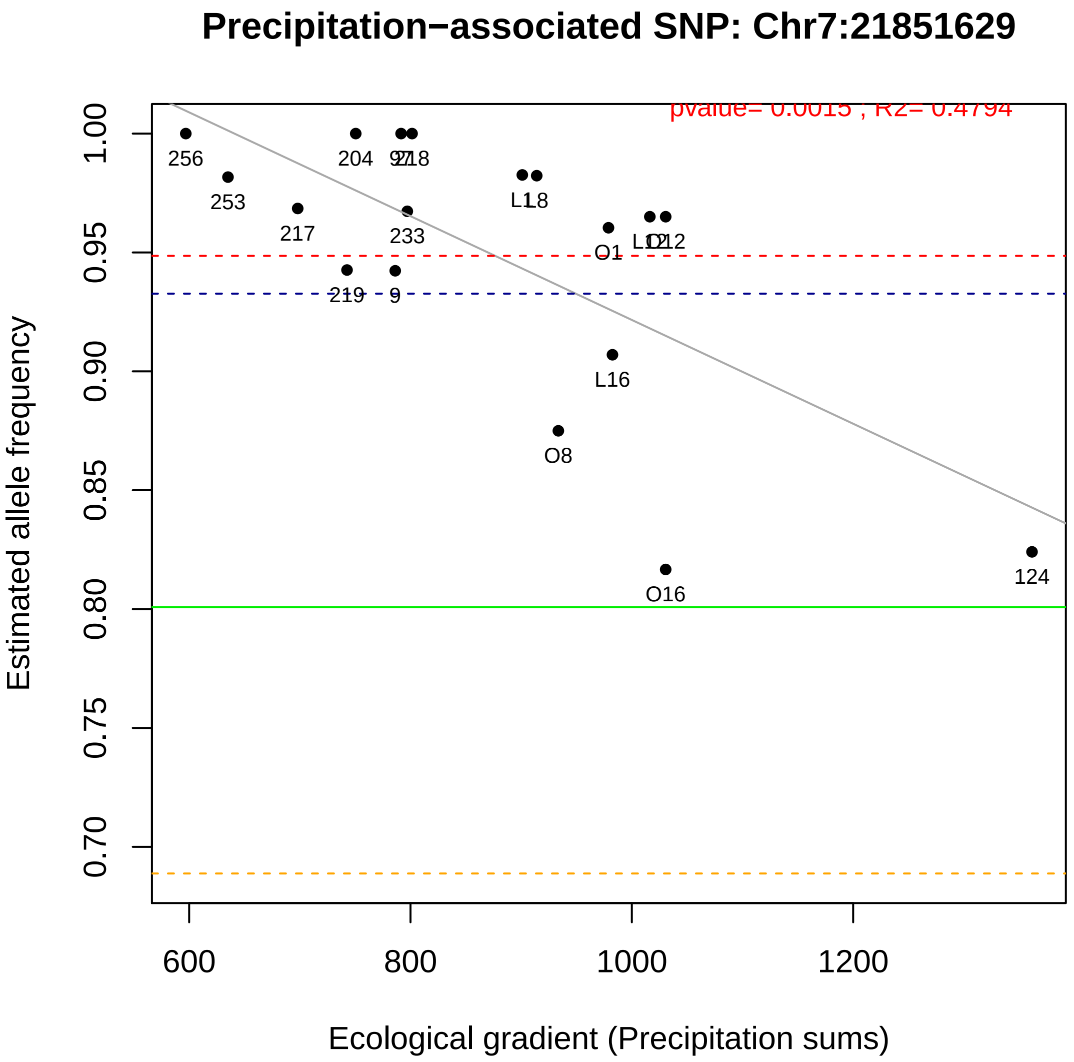

Precipitation−associated SNP: Chr7:21851629
pvalue= 0.0015 ; R2= 0.4794
Estimated allele frequency
Ecological gradient (Precipitation sums)
1.00
0.95
0.90
0.85
0.80
0.75
0.70
600
800
1000
1200
256
253
204
217
233
219
9
O1
L16
O8
O16
124

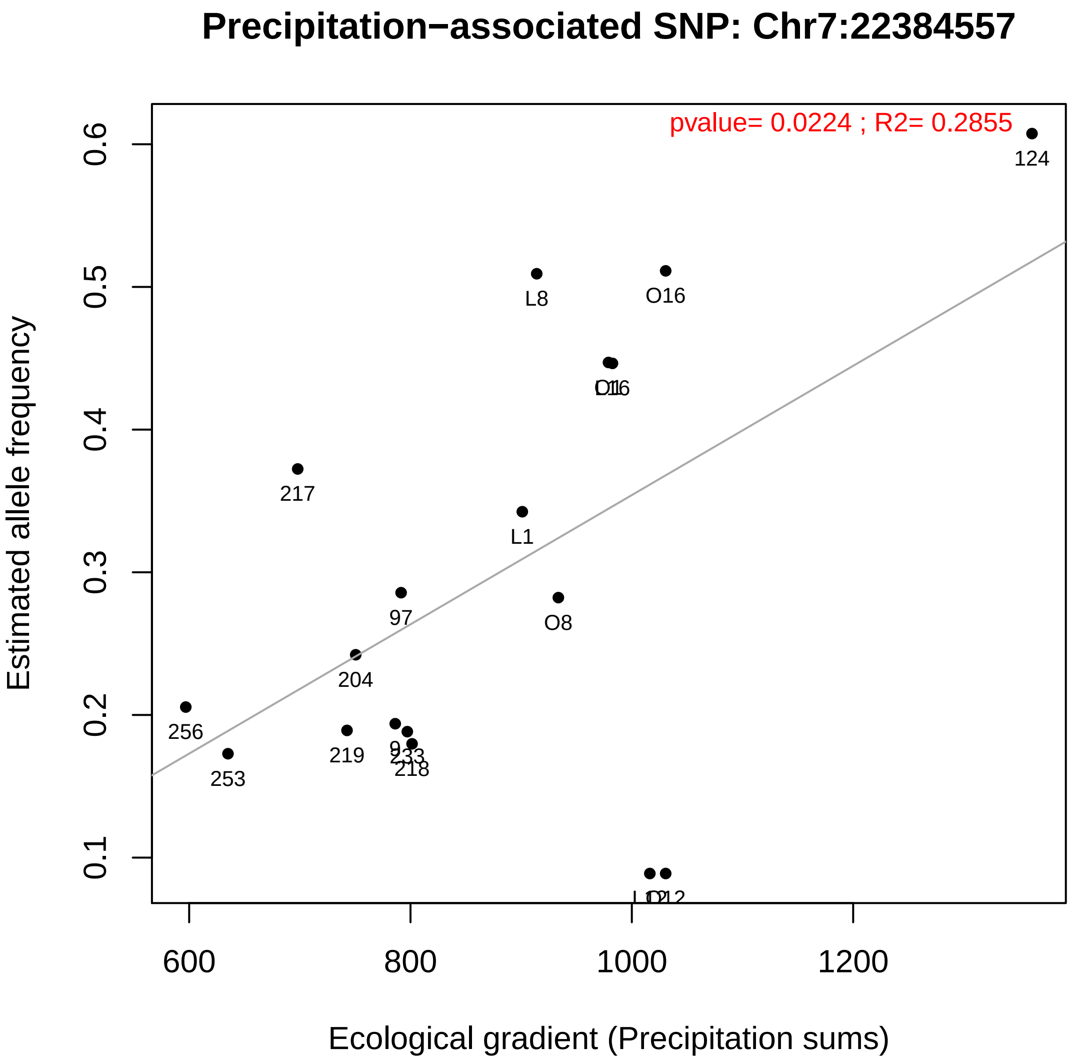

Precipitation−associated SNP: Chr7:22384557
pvalue= 0.0224 ; R2= 0.2855
124
L8
O16
217
L1
97
O8
204
256
253
219
233
218
0.6
0.5
0.4
0.3
0.2
0.1
600
800
1000
1200
Estimated allele frequency
Ecological gradient (Precipitation sums)

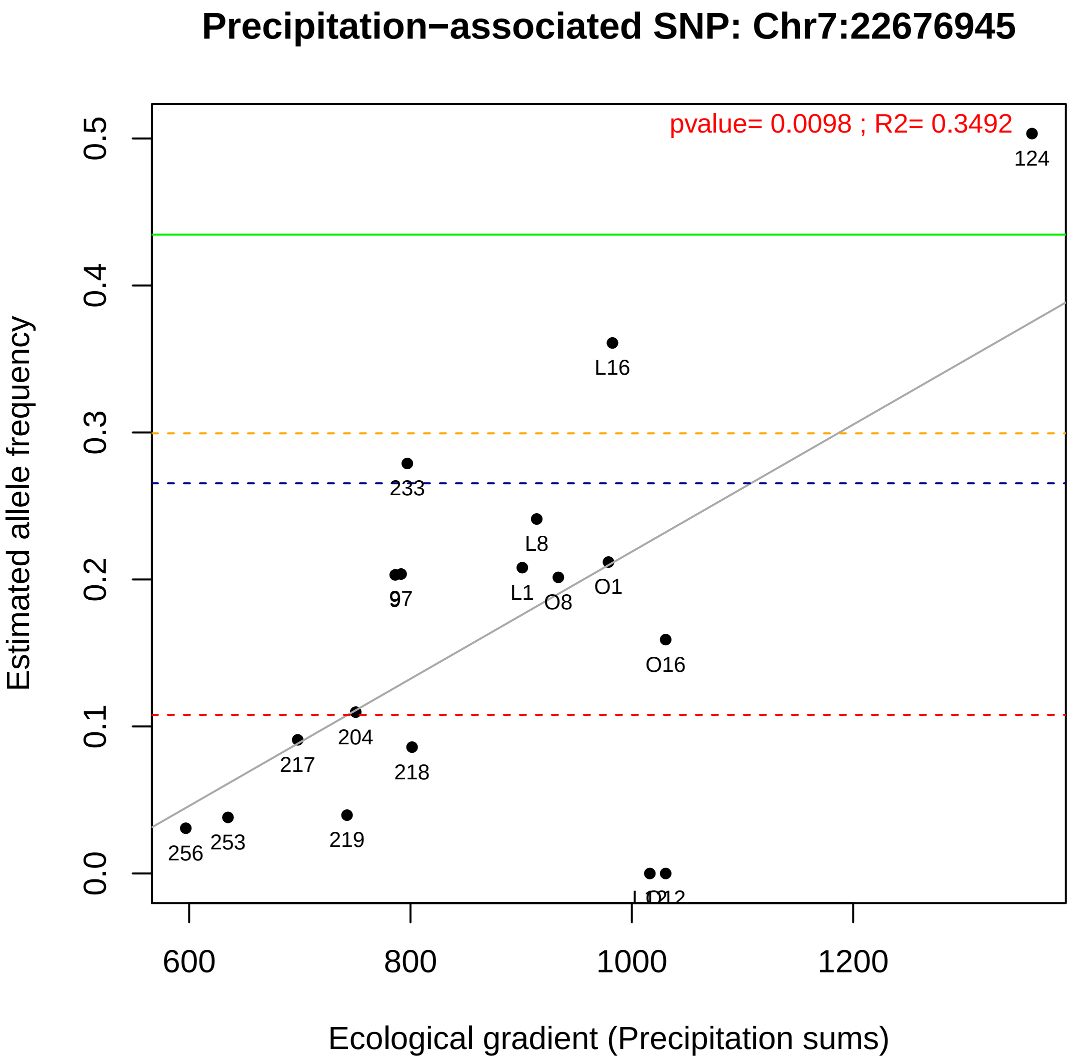

Precipitation−associated SNP: Chr7:22676945
pvalue= 0.0098 ; R2= 0.3492
Estimated allele frequency
Ecological gradient (Precipitation sums)
0.0
0.1
0.2
0.3
0.4
0.5
600
800
1000
1200
124
L16
233
L8
L1
O8
O1
O16
204
217
218
256
253
219

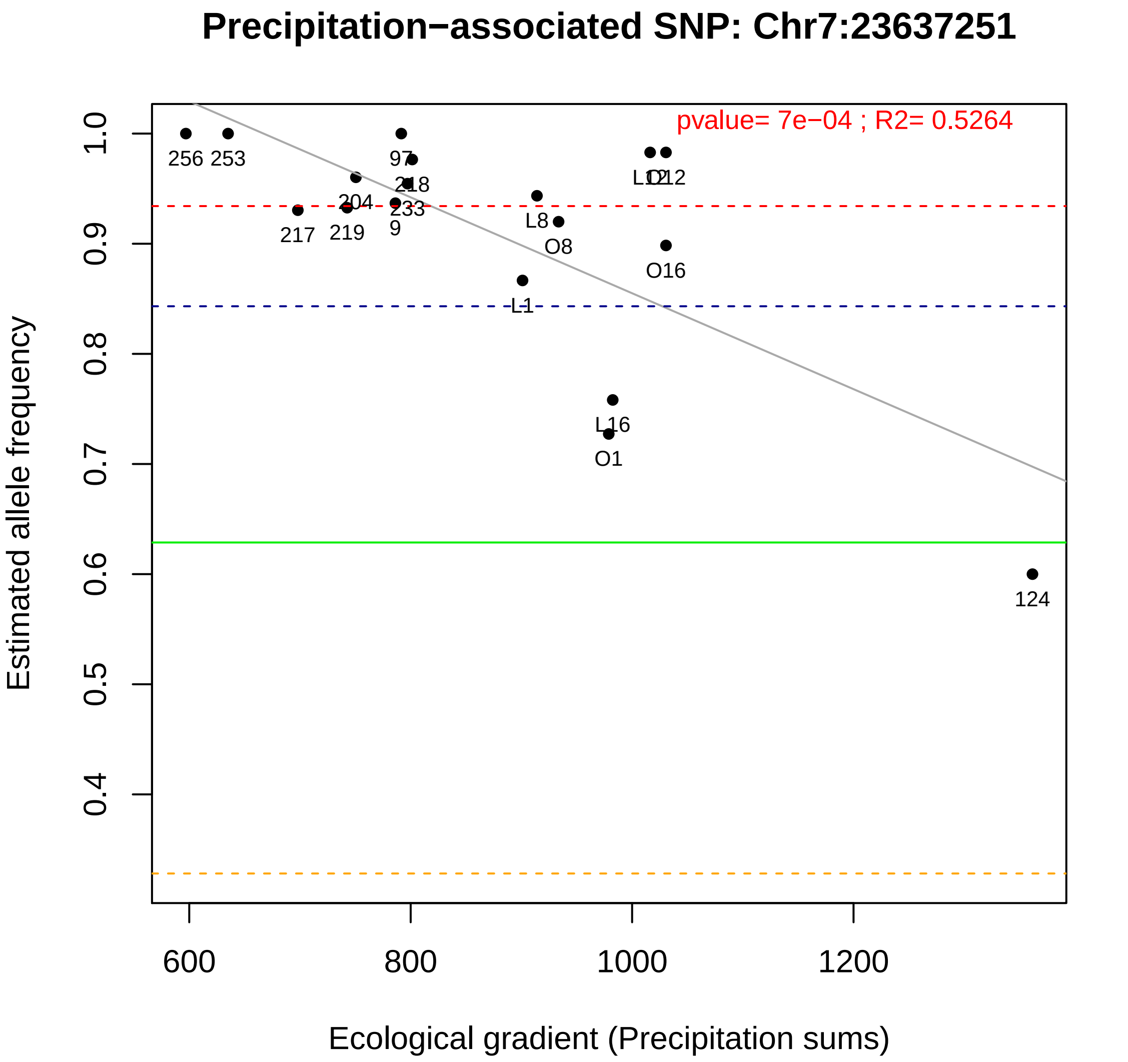
Precipitation−associated SNP: Chr7:23637251
pvalue= 7e−04 ; R2= 0.5264
Estimated allele frequency
Ecological gradient (Precipitation sums)
0.4
0.5
0.6
0.7
0.8
0.9
1.0
600
800
1000
1200
256
253
97
218
204
233
9
217
219
L8
O8
L12
O12
O16
L1
L16
O1
124

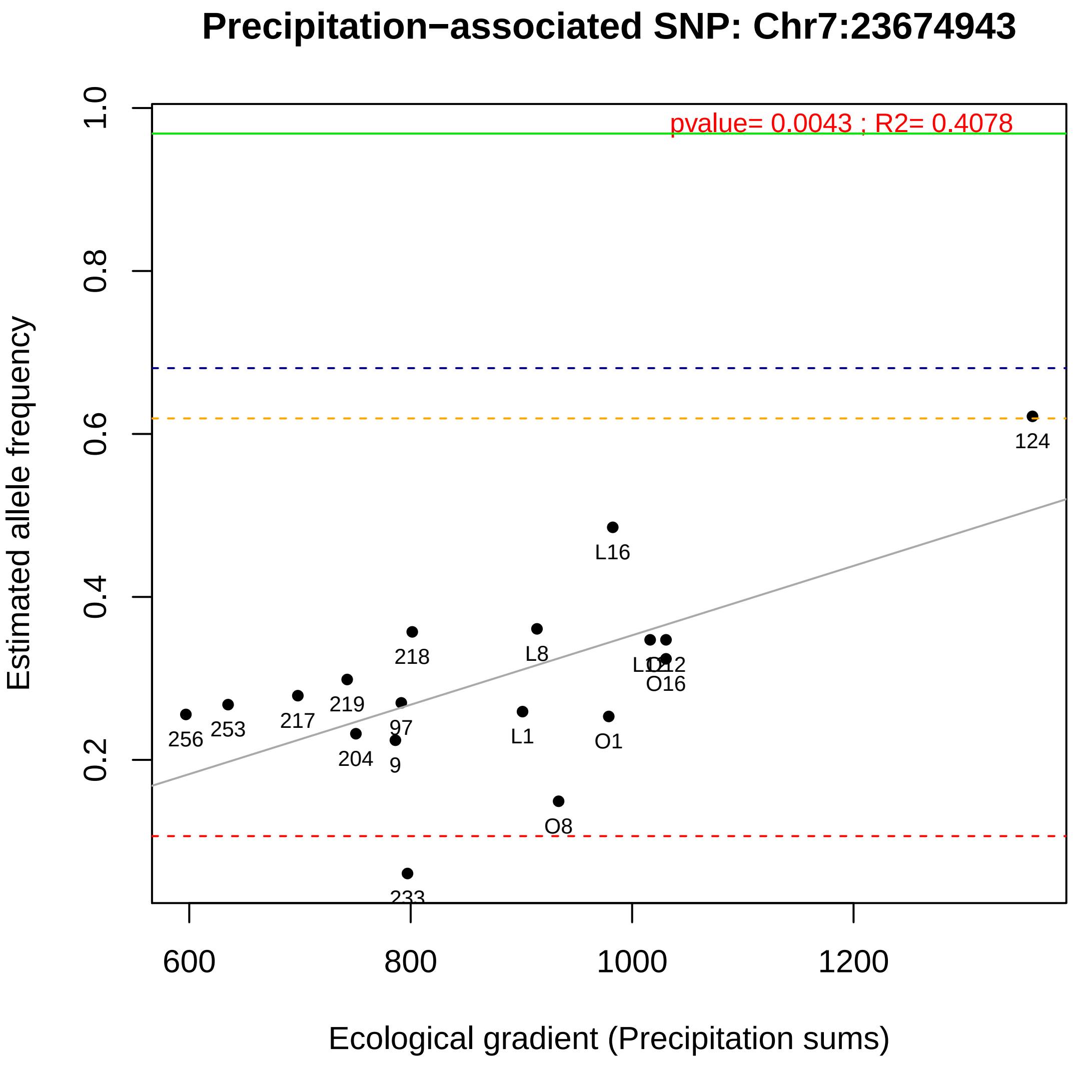

Precipitation−associated SNP: Chr7:23674943
pvalue= 0.0043 ; R2= 0.4078
Estimated allele frequency
1.0
0.8
0.6
0.4
0.2
124
L16
218
L8
O16
219
217
97
253
256
L1
O1
204
9
O8
233
600
800
1000
1200
Ecological gradient (Precipitation sums)

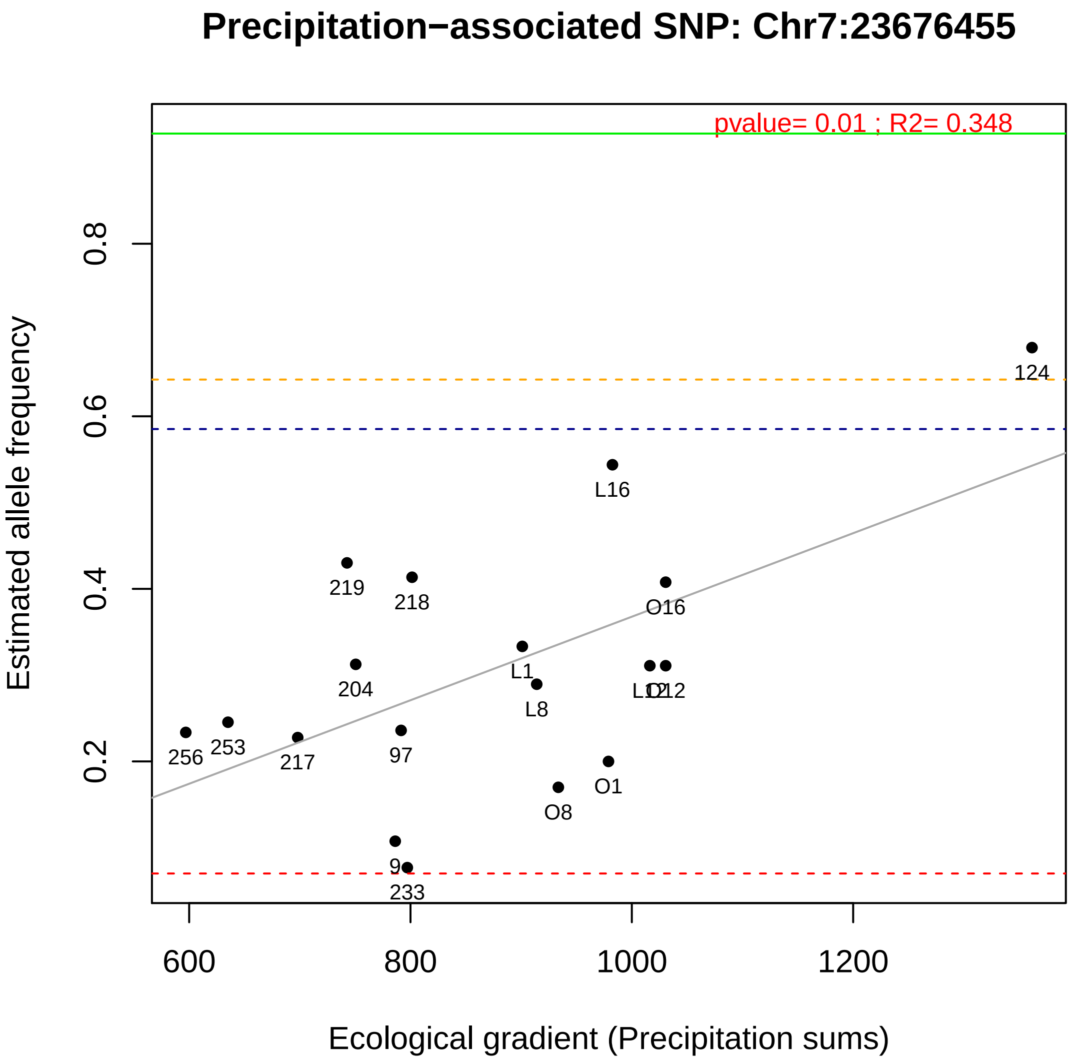
Precipitation−associated SNP: Chr7:23676455
pvalue= 0.01 ; R2= 0.348
Estimated allele frequency
0.2
0.4
0.6
0.8
Ecological gradient (Precipitation sums)
600
800
1000
1200
124
L16
219
218
O16
204
L1
L8
L1O12
256
253
217
97
O1
O8
9
233

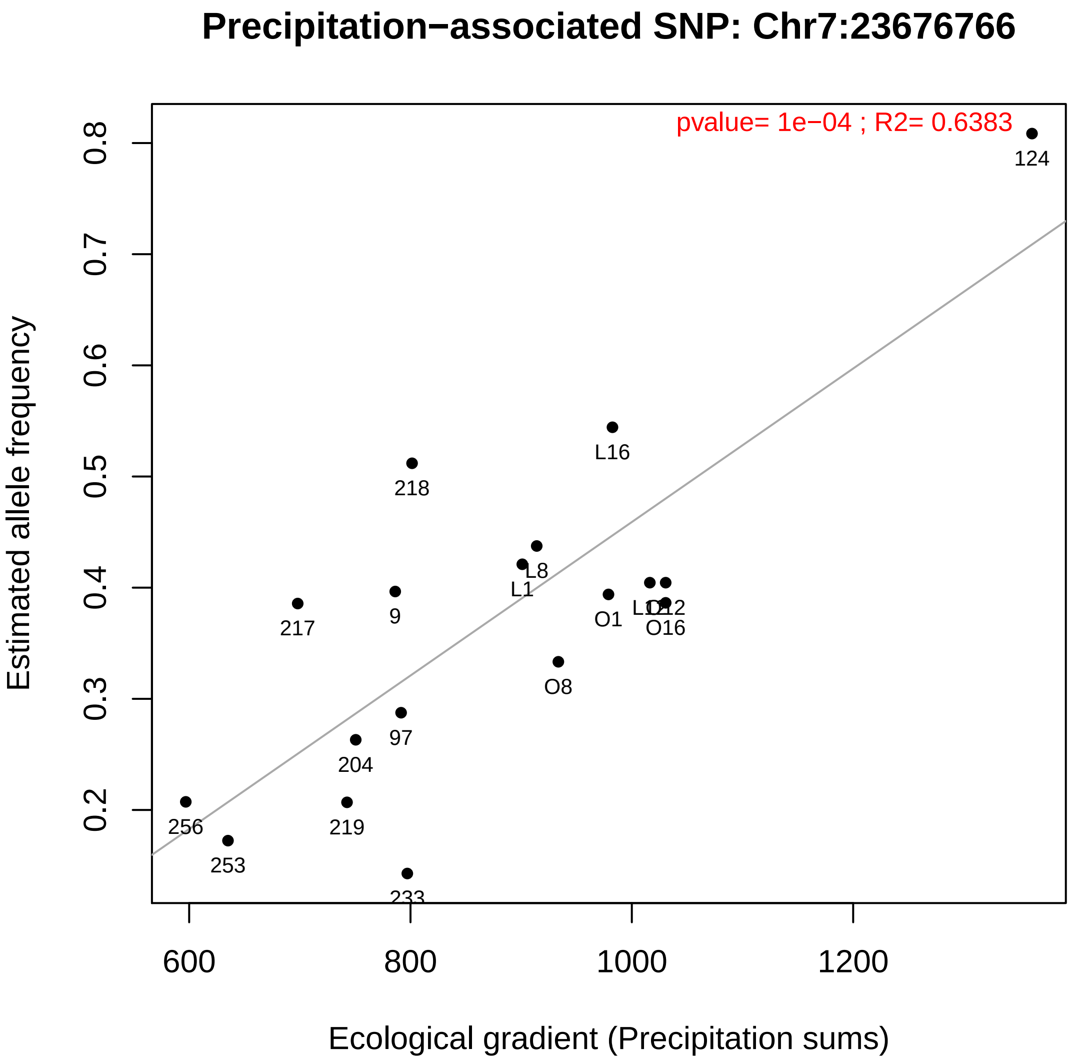
Precipitation−associated SNP: Chr7:23676766
pvalue= 1e−04 ; R2= 0.6383
Estimated allele frequency
Ecological gradient (Precipitation sums)
0.2
0.3
0.4
0.5
0.6
0.7
0.8
600
800
1000
1200
124
L16
218
L8
L1
9
217
O1
O16
O8
97
204
256
219
253
233

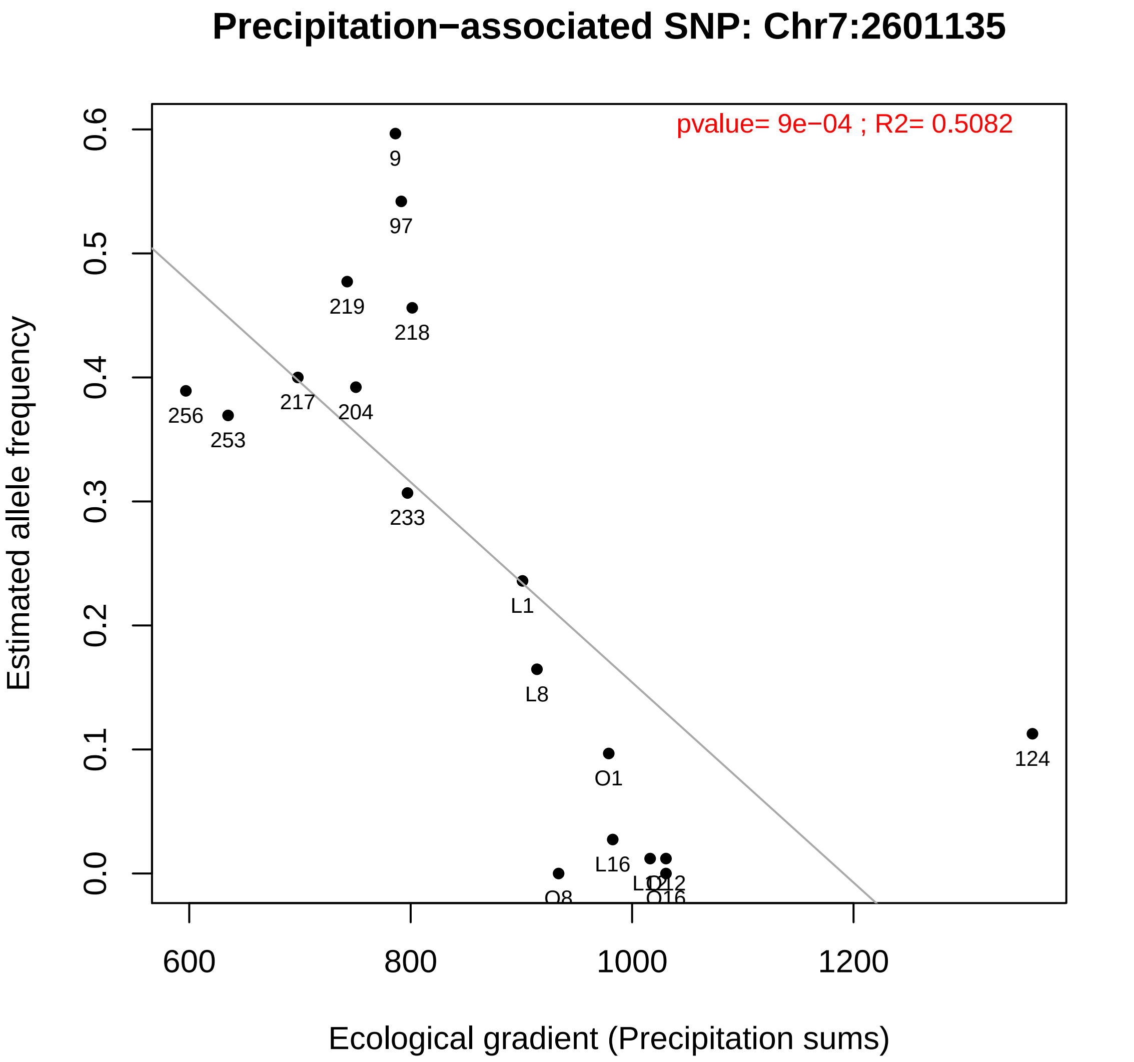
Precipitation−associated SNP: Chr7:2601135
pvalue= 9e−04 ; R2= 0.5082
Estimated allele frequency
0.0
0.1
0.2
0.3
0.4
0.5
0.6
600
800
1000
1200
Ecological gradient (Precipitation sums)
9
97
219
218
256
253
217
204
233
L1
L8
O1
124
L16
O8
O12
O16

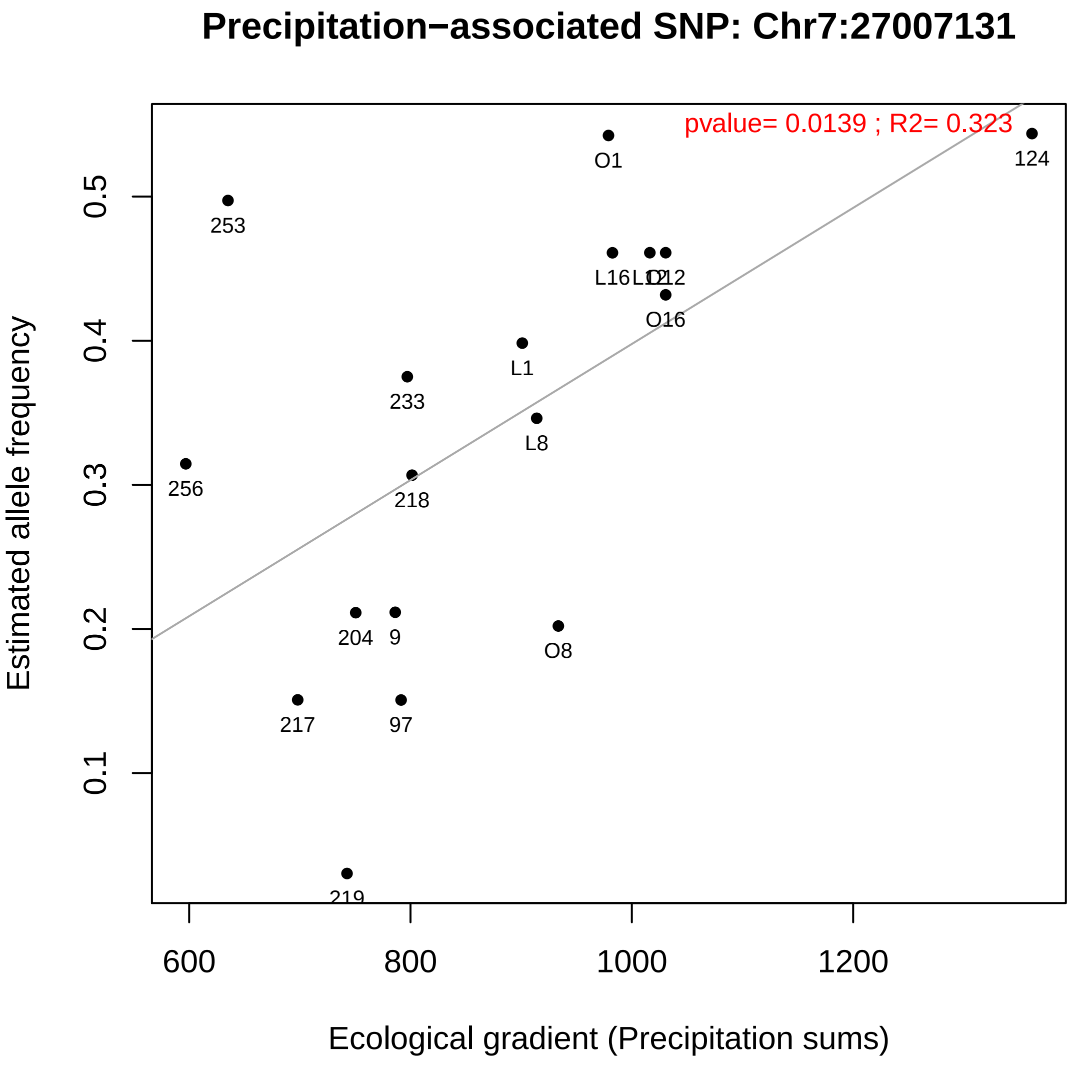

Precipitation−associated SNP: Chr7:27007131
pvalue= 0.0139 ; R2= 0.323
O1
124
253
L16
L12
O12
O16
L1
233
L8
256
218
204
9
O8
217
97
219
0.1
0.2
0.3
0.4
0.5
600
800
1000
1200
Estimated allele frequency
Ecological gradient (Precipitation sums)

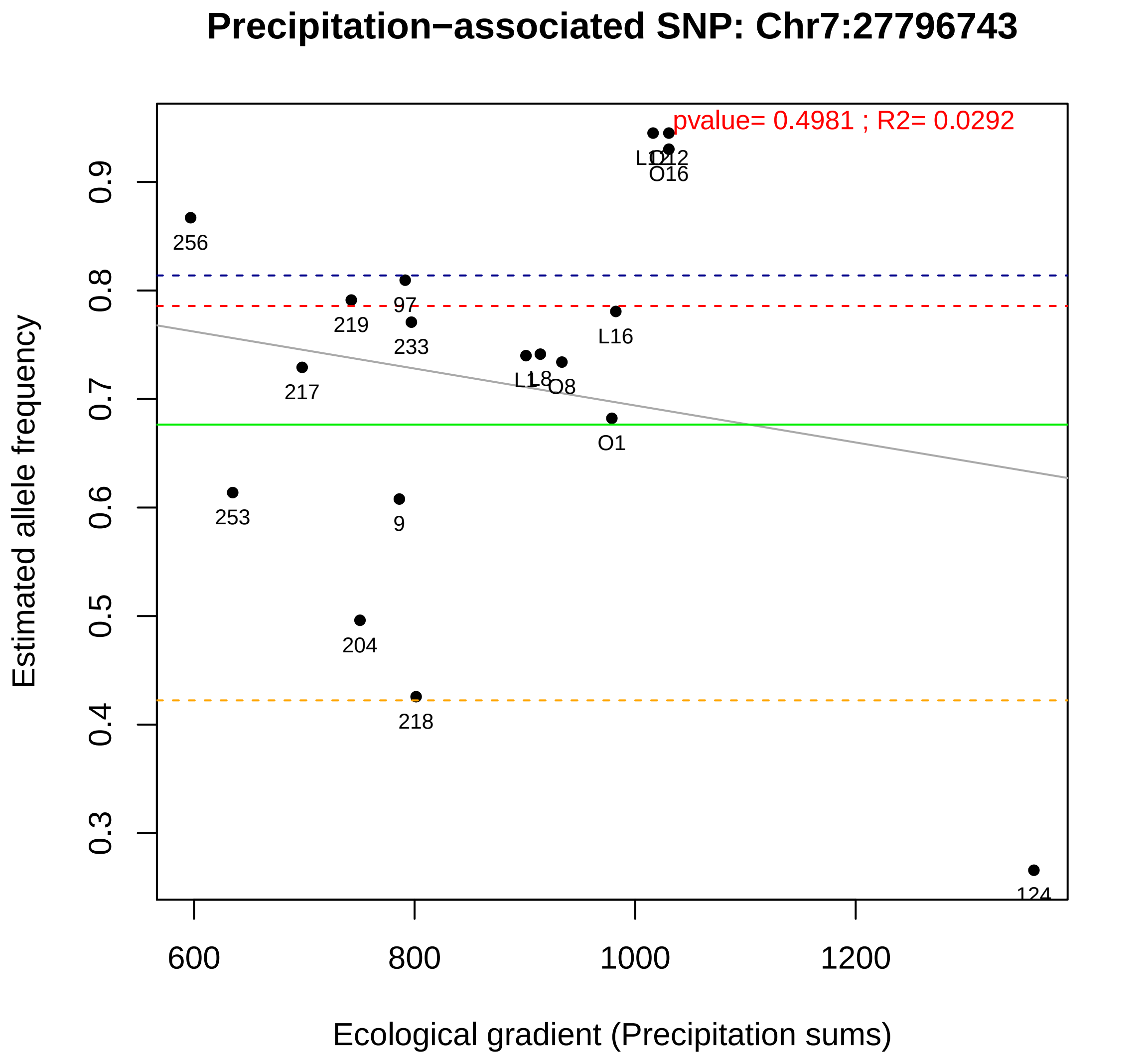
Precipitation−associated SNP: Chr7:27796743
pvalue= 0.4981 ; R2= 0.0292
Estimated allele frequency
0.3
0.4
0.5
0.6
0.7
0.8
0.9
600
800
1000
1200
Ecological gradient (Precipitation sums)
256
219
97
233
217
253
9
204
218
L16
O8
O1
O16
124

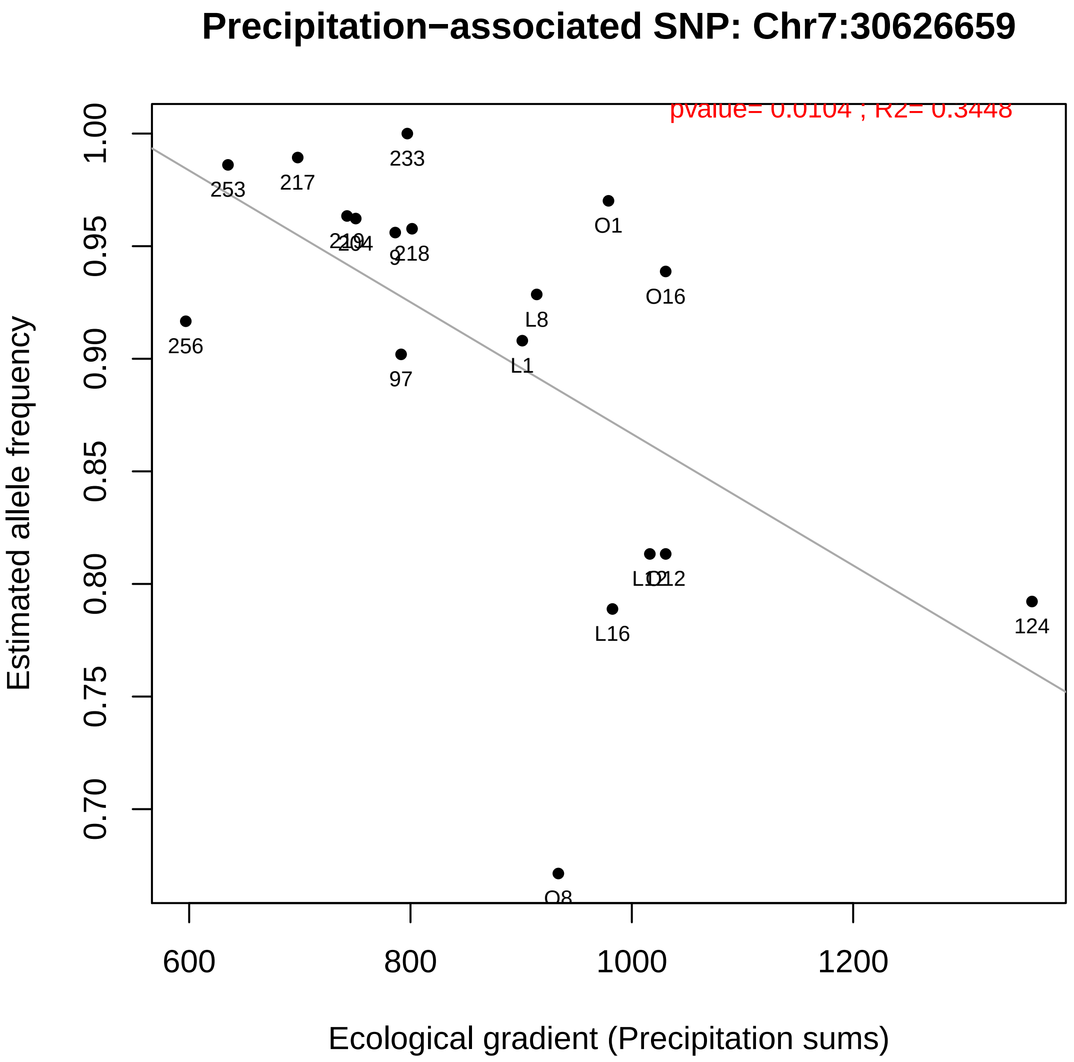

Precipitation−associated SNP: Chr7:30626659
pvalue= 0.0104 ; R2= 0.3448
Estimated allele frequency
Ecological gradient (Precipitation sums)
1.00
0.95
0.90
0.85
0.80
0.75
0.70
600
800
1000
1200
233
253
217
O1
9
218
O16
L8
256
97
L1
L16
124

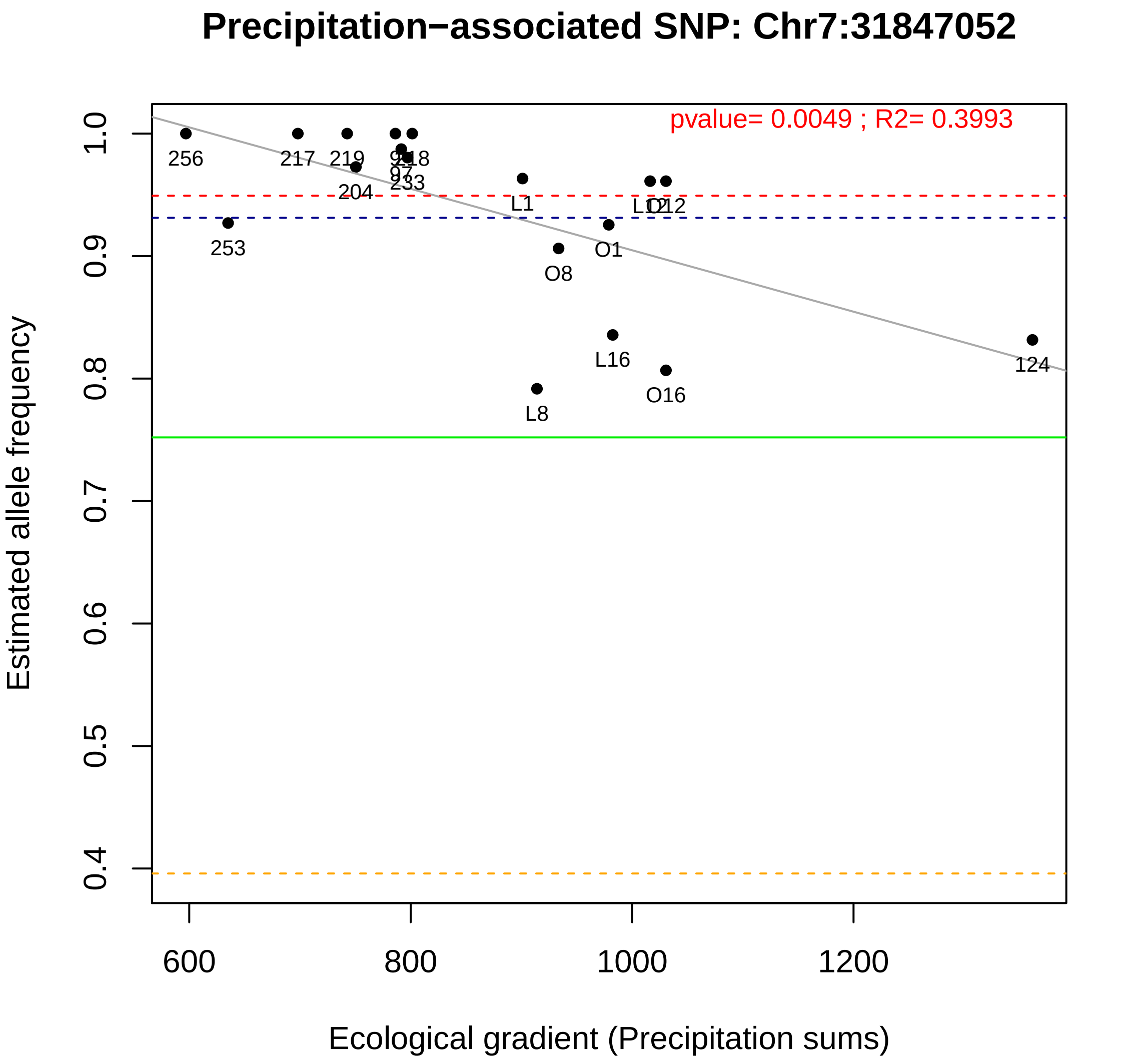

Precipitation−associated SNP: Chr7:31847052
pvalue= 0.0049 ; R2= 0.3993
256
217
219
204
233
218
97
253
L1
L12
O12
O1
O8
L16
O16
L8
124
1.0
0.9
0.8
0.7
0.6
0.5
0.4
Estimated allele frequency
600
800
1000
1200
Ecological gradient (Precipitation sums)

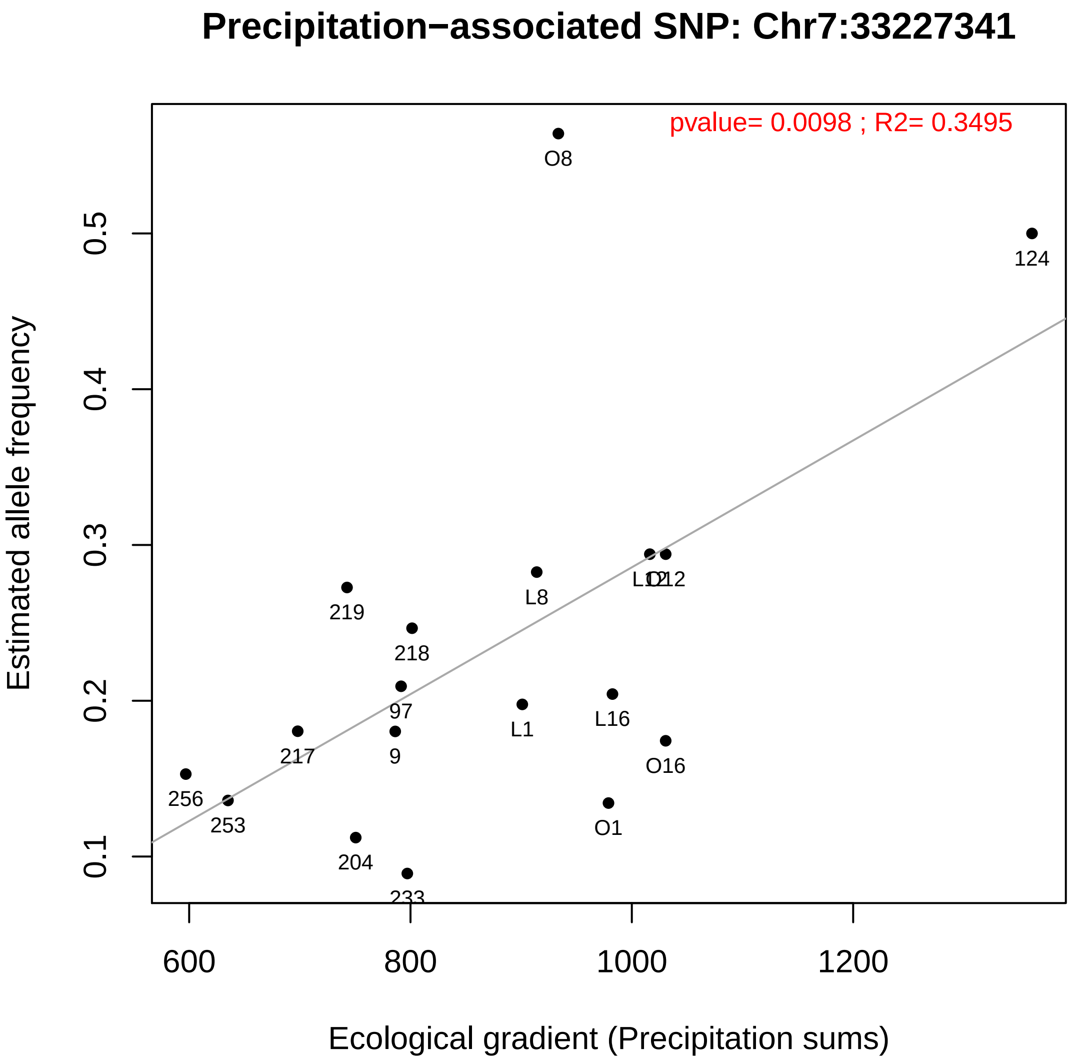

Precipitation−associated SNP: Chr7:33227341
pvalue= 0.0098 ; R2= 0.3495
Estimated allele frequency
0.1
0.2
0.3
0.4
0.5
600
800
1000
1200
Ecological gradient (Precipitation sums)
O8
124
L8
219
218
97
9
217
256
253
204
233
L1
L16
O16
O1

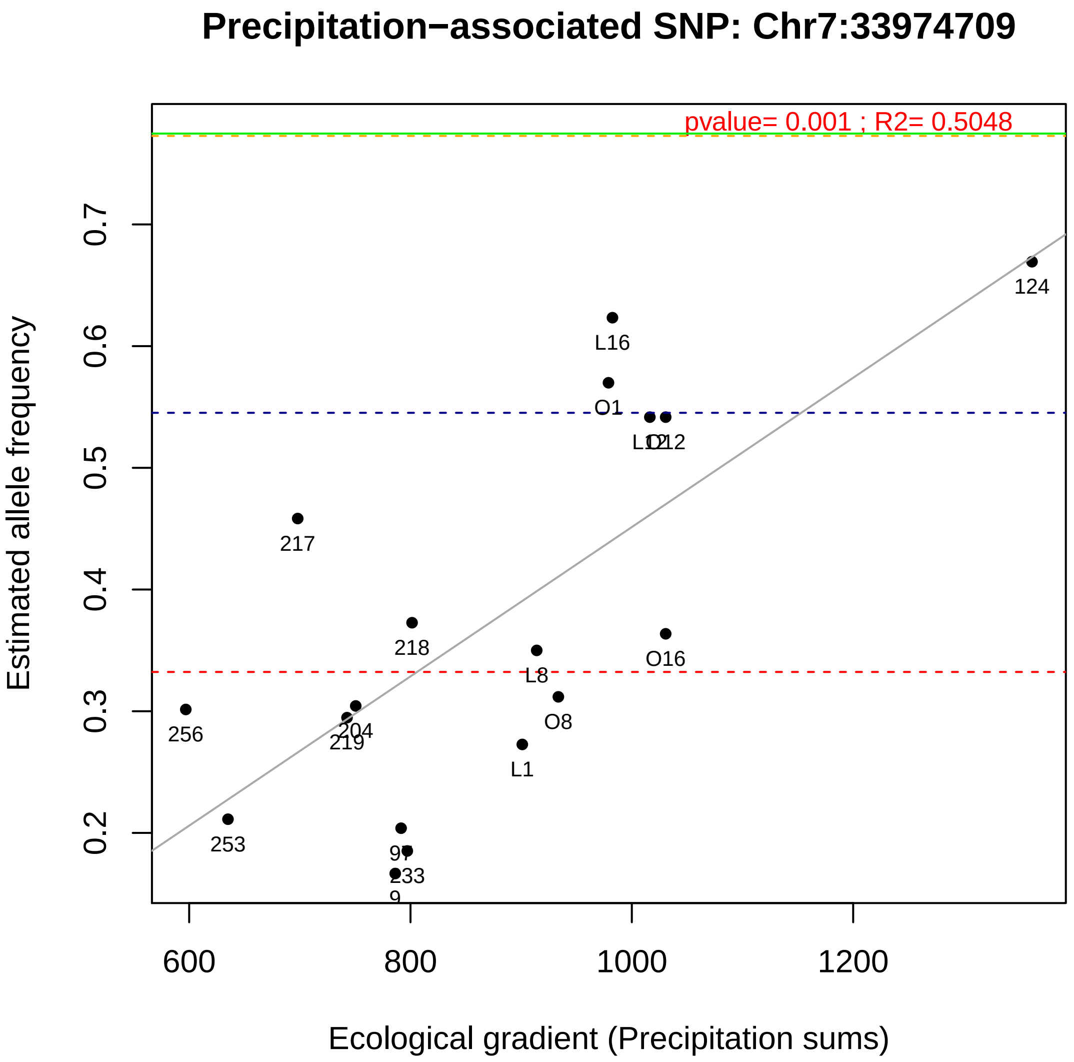

Precipitation−associated SNP: Chr7:33974709
pvalue= 0.001 ; R2= 0.5048
Estimated allele frequency
Ecological gradient (Precipitation sums)
0.2
0.3
0.4
0.5
0.6
0.7
600
800
1000
1200
124
L16
O1
L12
O12
217
218
O16
L8
O8
256
204
219
L1
253
97
233
9

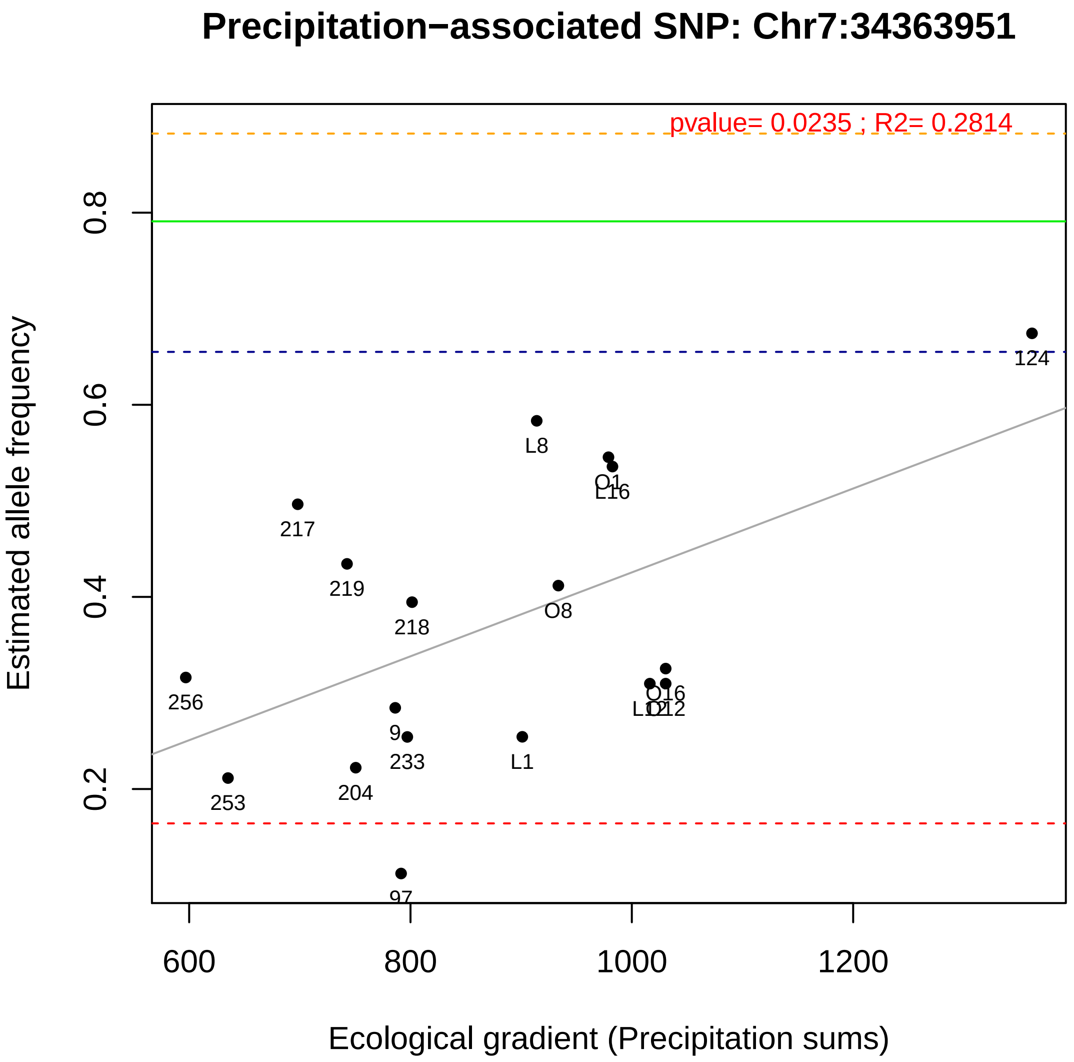
Precipitation−associated SNP: Chr7:34363951
pvalue= 0.0235 ; R2= 0.2814
Estimated allele frequency
0.2
0.4
0.6
0.8
600
800
1000
1200
Ecological gradient (Precipitation sums)
124
L8
217
219
218
O8
256
253
204
9
233
L1
97

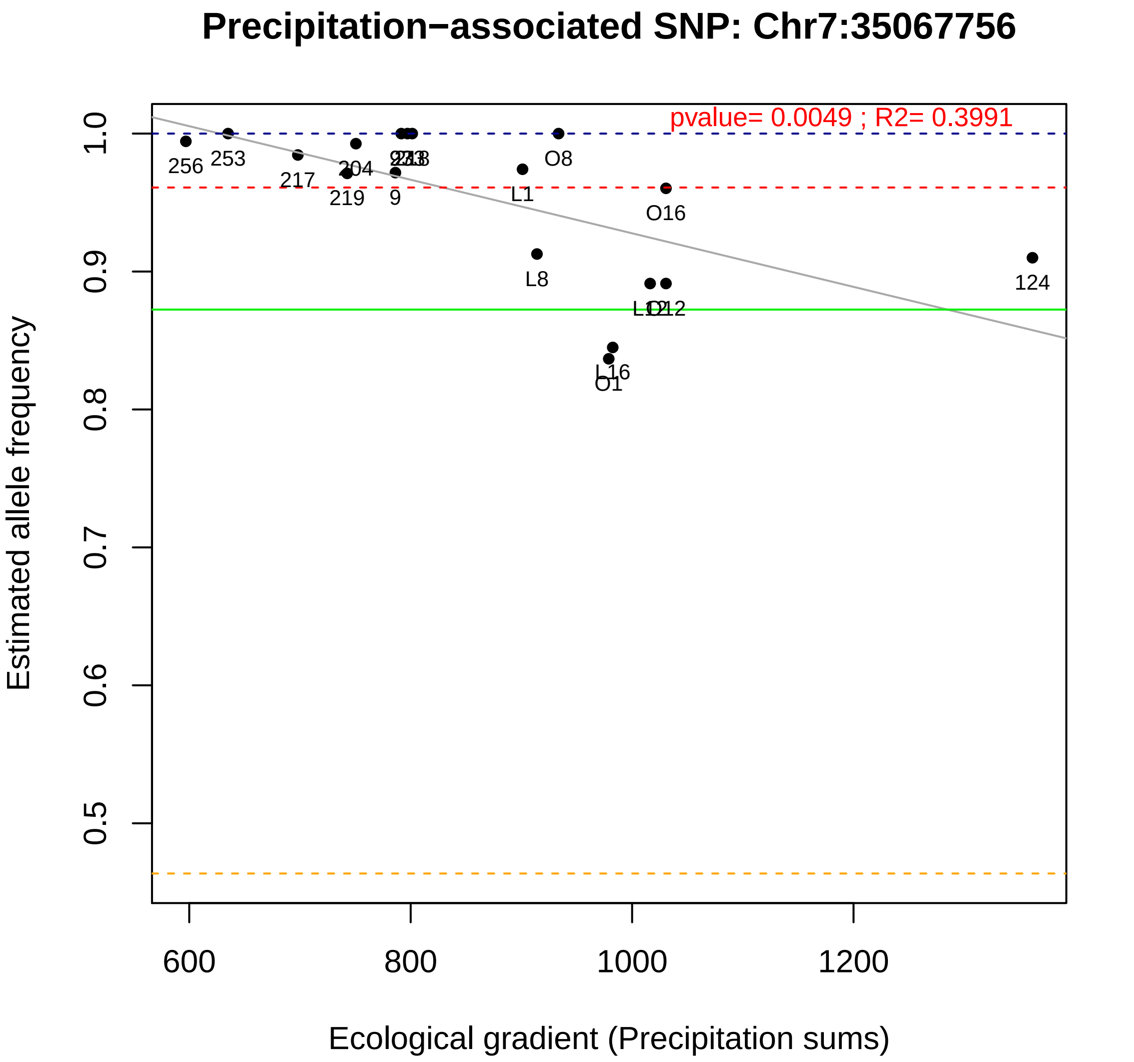

Precipitation−associated SNP: Chr7:35067756
pvalue= 0.0049 ; R2= 0.3991
Estimated allele frequency
Ecological gradient (Precipitation sums)
1.0
0.9
0.8
0.7
0.6
0.5
600
800
1000
1200
256
253
217
204
219
9
L1
O8
O16
L8
124
L16
O1

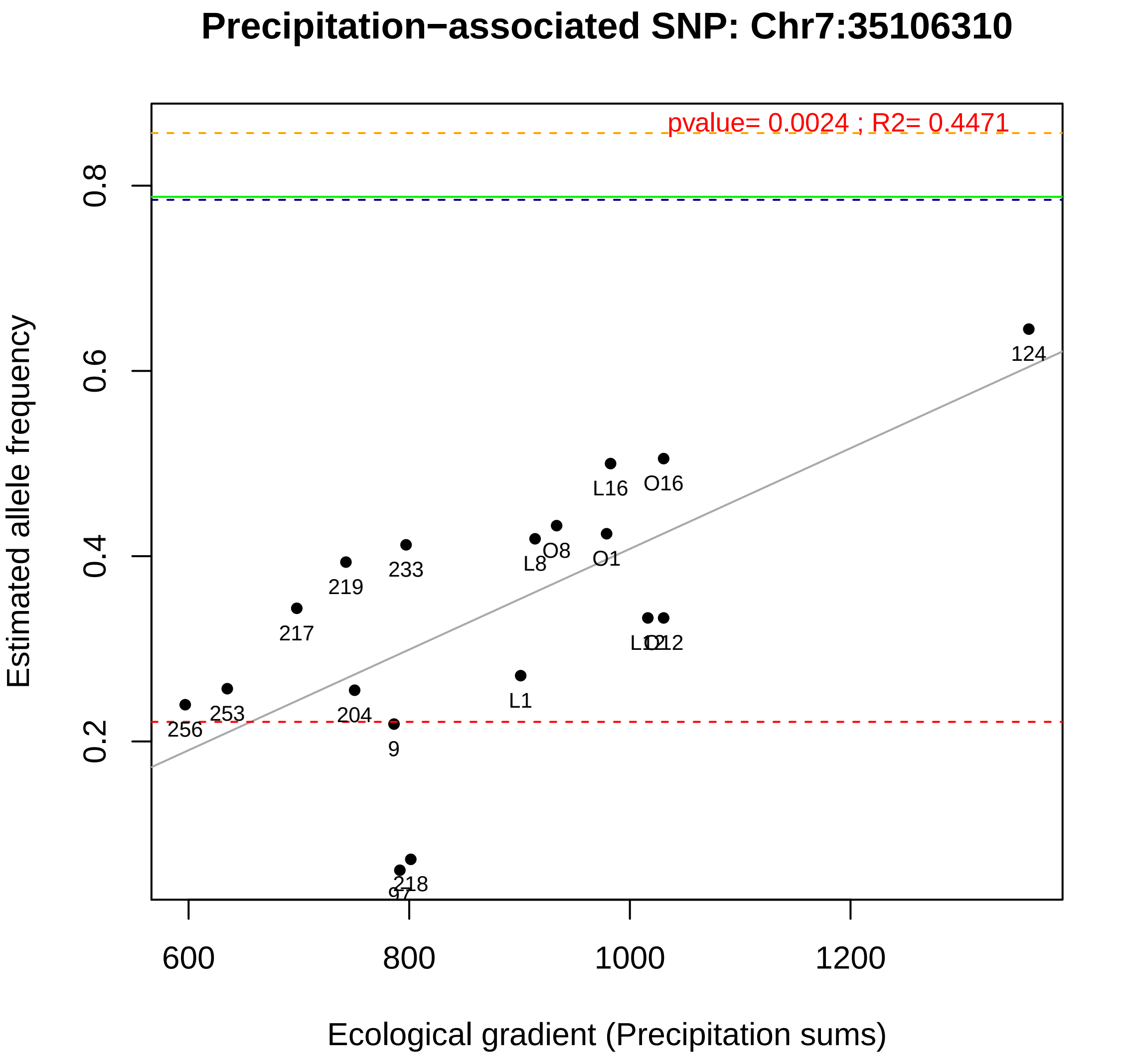
Precipitation−associated SNP: Chr7:35106310
pvalue= 0.0024 ; R2= 0.4471
Estimated allele frequency
0.2
0.4
0.6
0.8
600
800
1000
1200
Ecological gradient (Precipitation sums)
124
L16
O16
O8
L8
O1
233
219
217
L1
256
253
204
9
218
O12

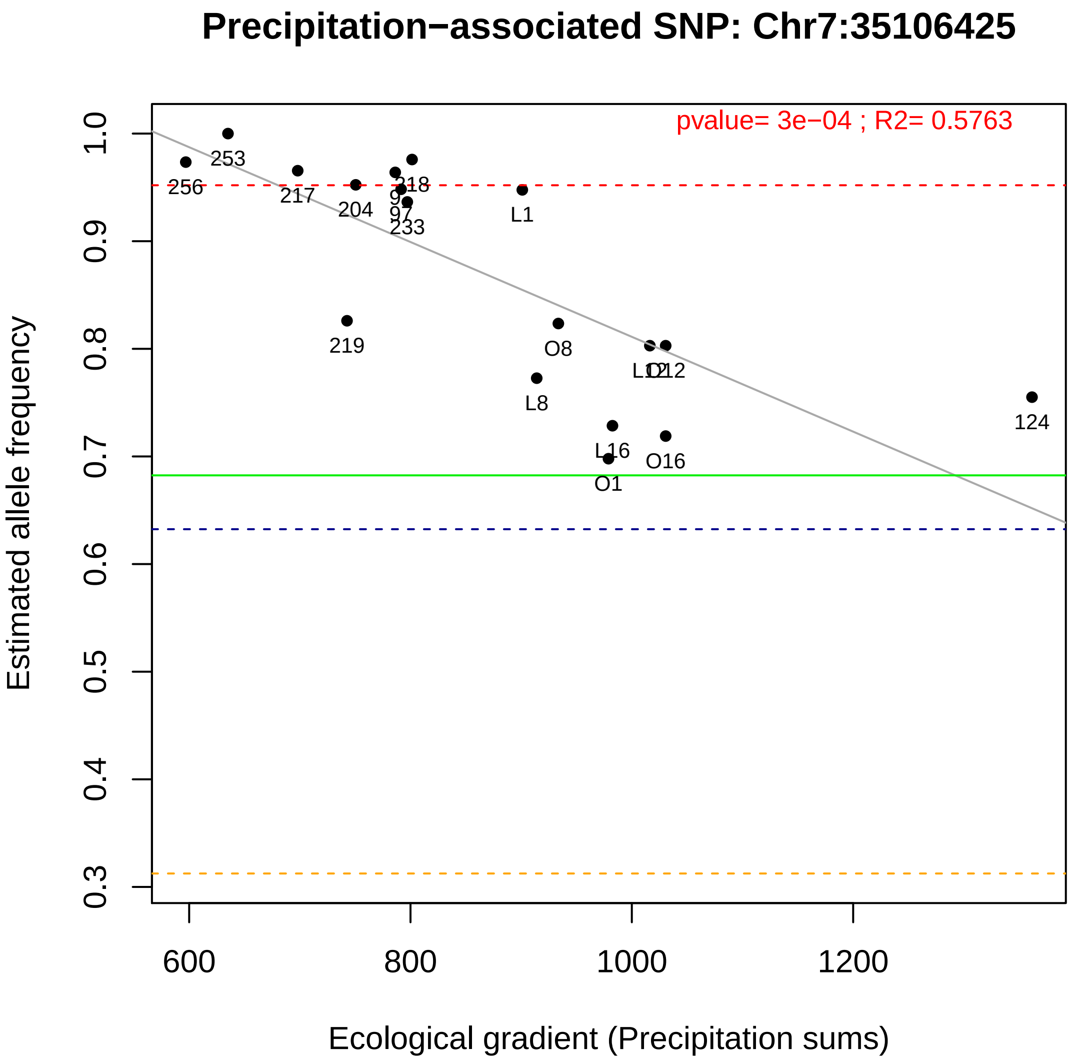
Precipitation−associated SNP: Chr7:35106425
pvalue= 3e−04 ; R2= 0.5763
Estimated allele frequency
Ecological gradient (Precipitation sums)
0.3
0.4
0.5
0.6
0.7
0.8
0.9
1.0
600
800
1000
1200
256
253
217
204
218
233
L1
219
O8
L8
O12
L16
O16
O1
124

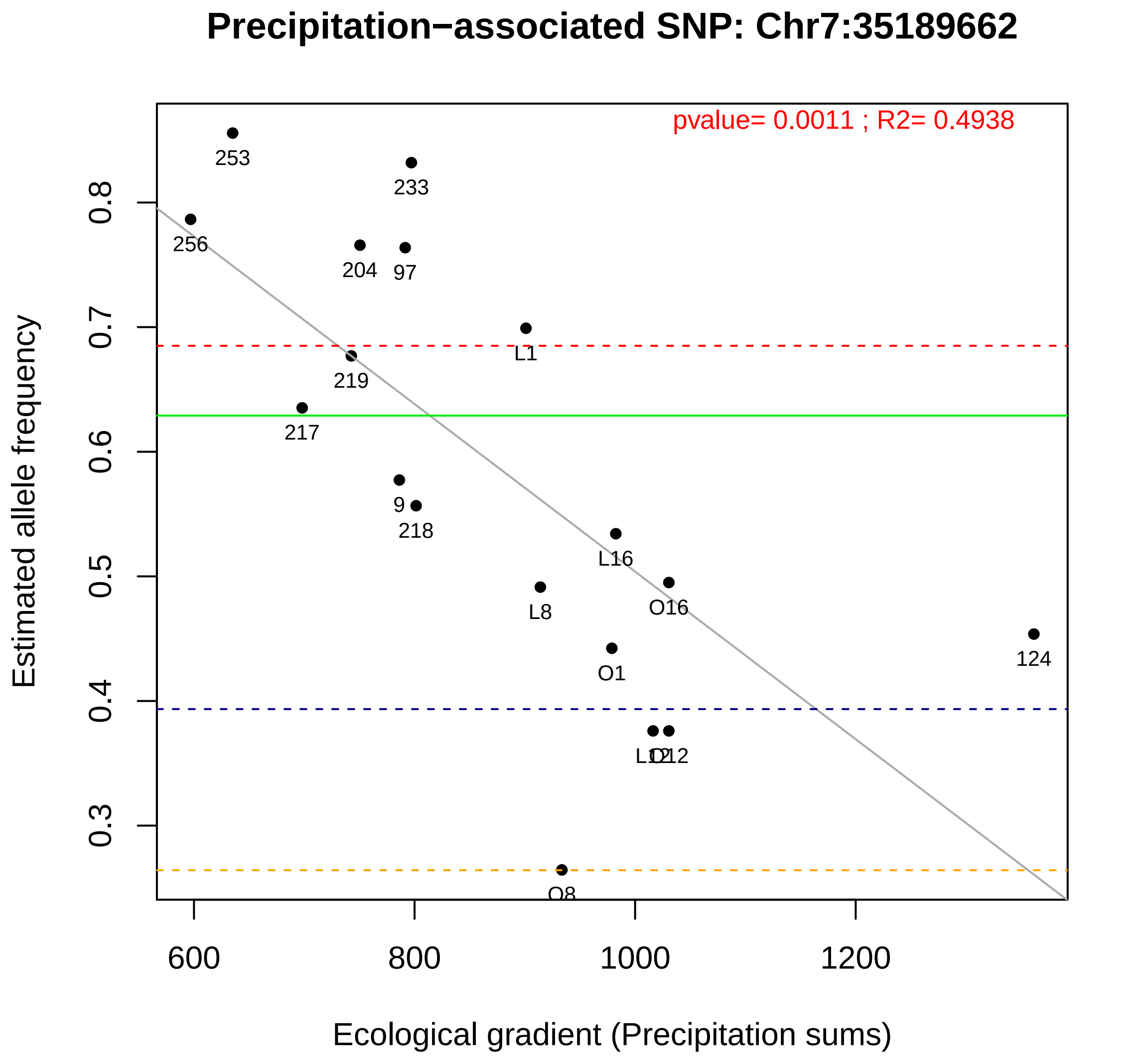
Precipitation−associated SNP: Chr7:35189662
pvalue= 0.0011 ; R2= 0.4938
Estimated allele frequency
Ecological gradient (Precipitation sums)
0.3
0.4
0.5
0.6
0.7
0.8
600
800
1000
1200
253
233
256
204
97
L1
219
217
9
218
L16
L8
O16
O1
124
O8

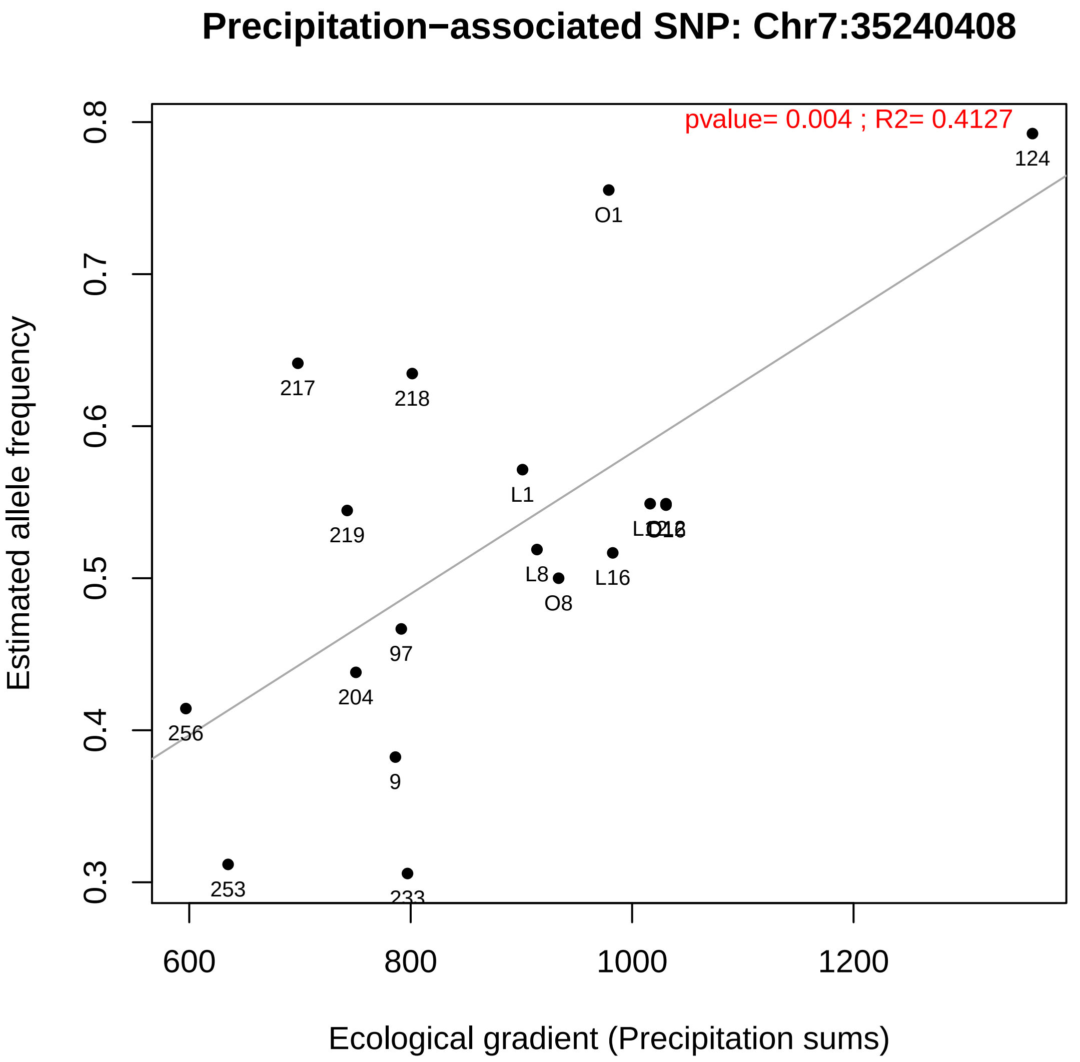
Precipitation−associated SNP: Chr7:35240408
pvalue= 0.004 ; R2= 0.4127
Estimated allele frequency
Ecological gradient (Precipitation sums)
0.3
0.4
0.5
0.6
0.7
0.8
600
800
1000
1200
124
O1
217
218
L1
219
L8
O8
L16
97
204
256
9
253
233

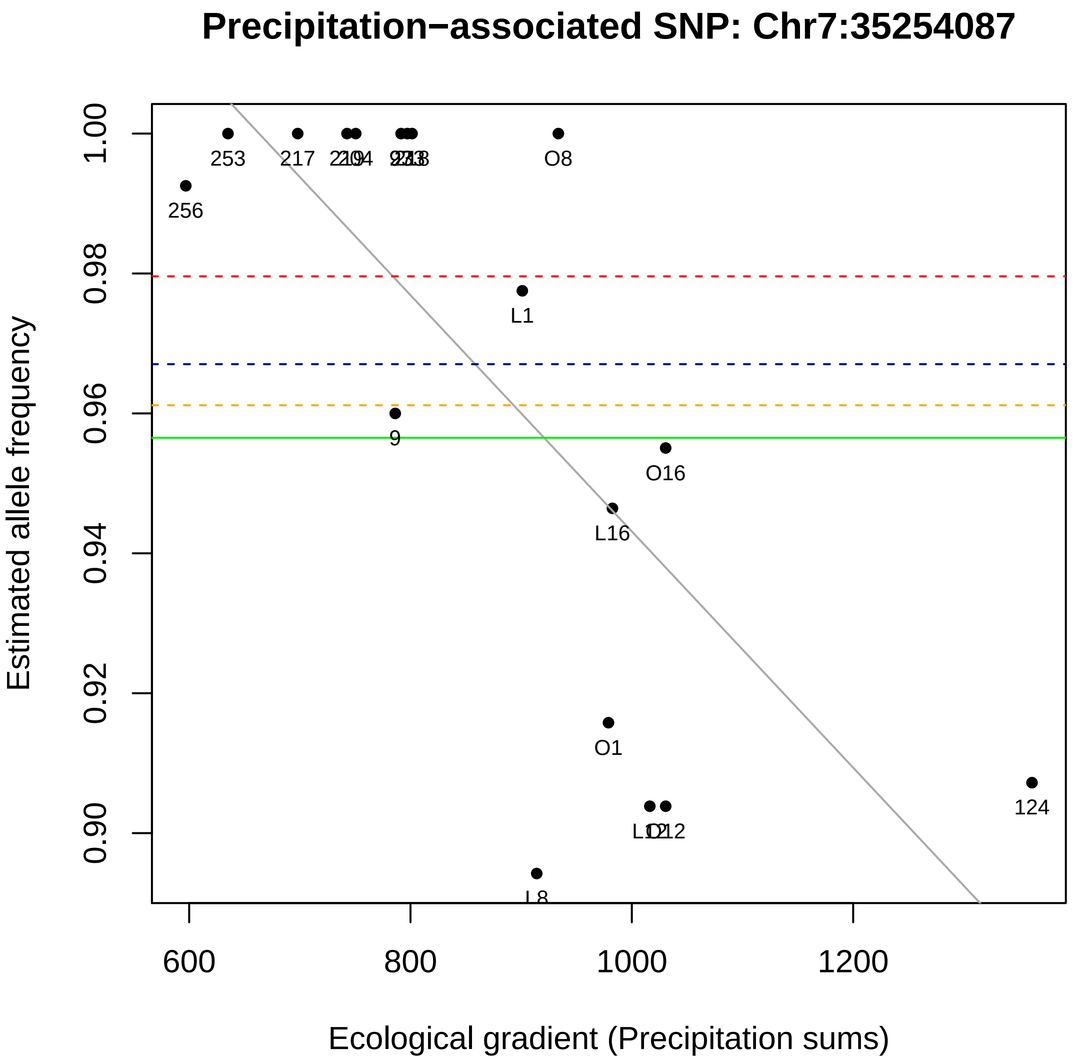
Precipitation−associated SNP: Chr7:35254087
Estimated allele frequency
Ecological gradient (Precipitation sums)
1.00
0.98
0.96
0.94
0.92
0.90
600
800
1000
1200
253
217
256
O8
L1
9
O16
L16
O1
124
L8

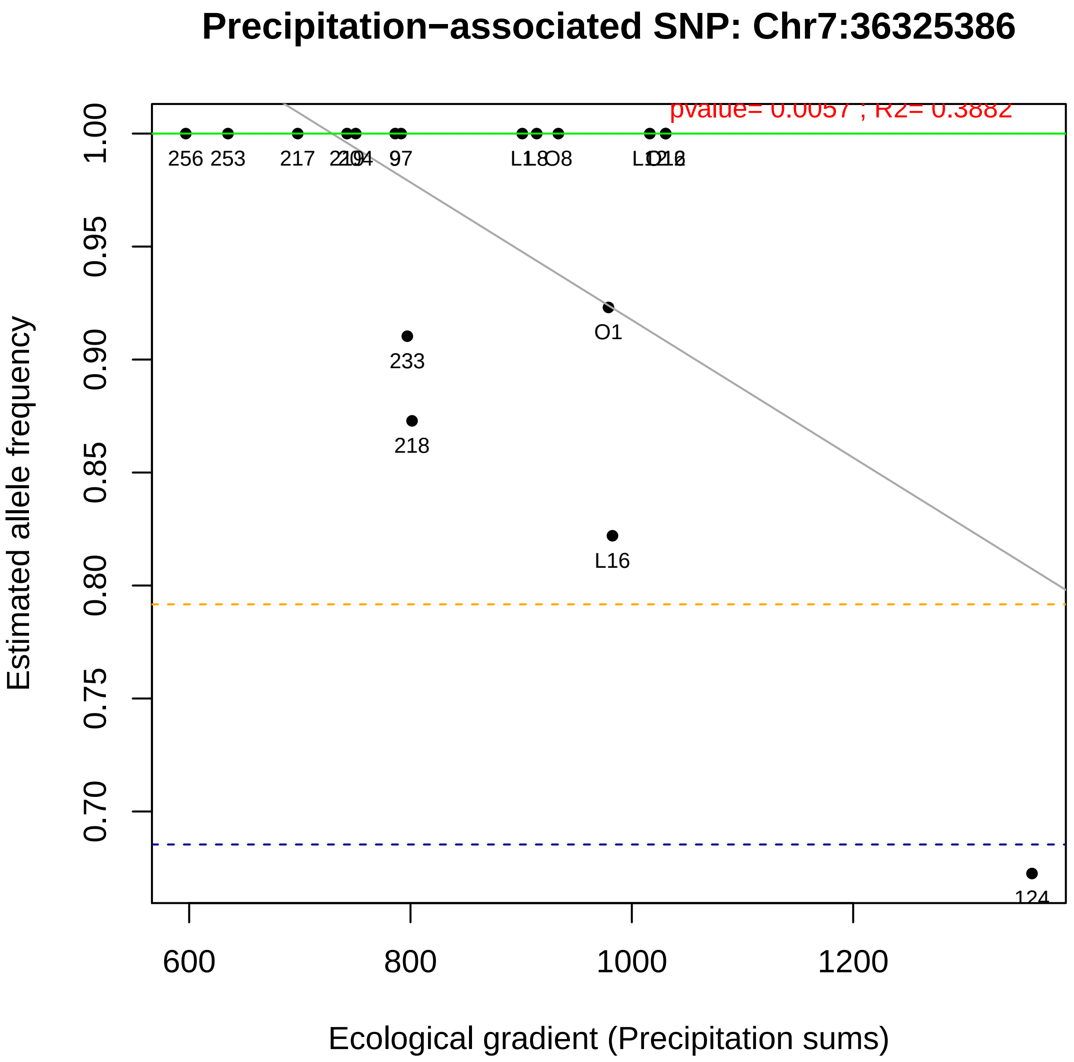

Precipitation−associated SNP: Chr7:36325386
pvalue= 0.0057 ; R2= 0.3882
Estimated allele frequency
1.00
0.95
0.90
0.85
0.80
0.75
0.70
256
253
217
97
O8
233
218
O1
L16
124
600
800
1000
1200
Ecological gradient (Precipitation sums)

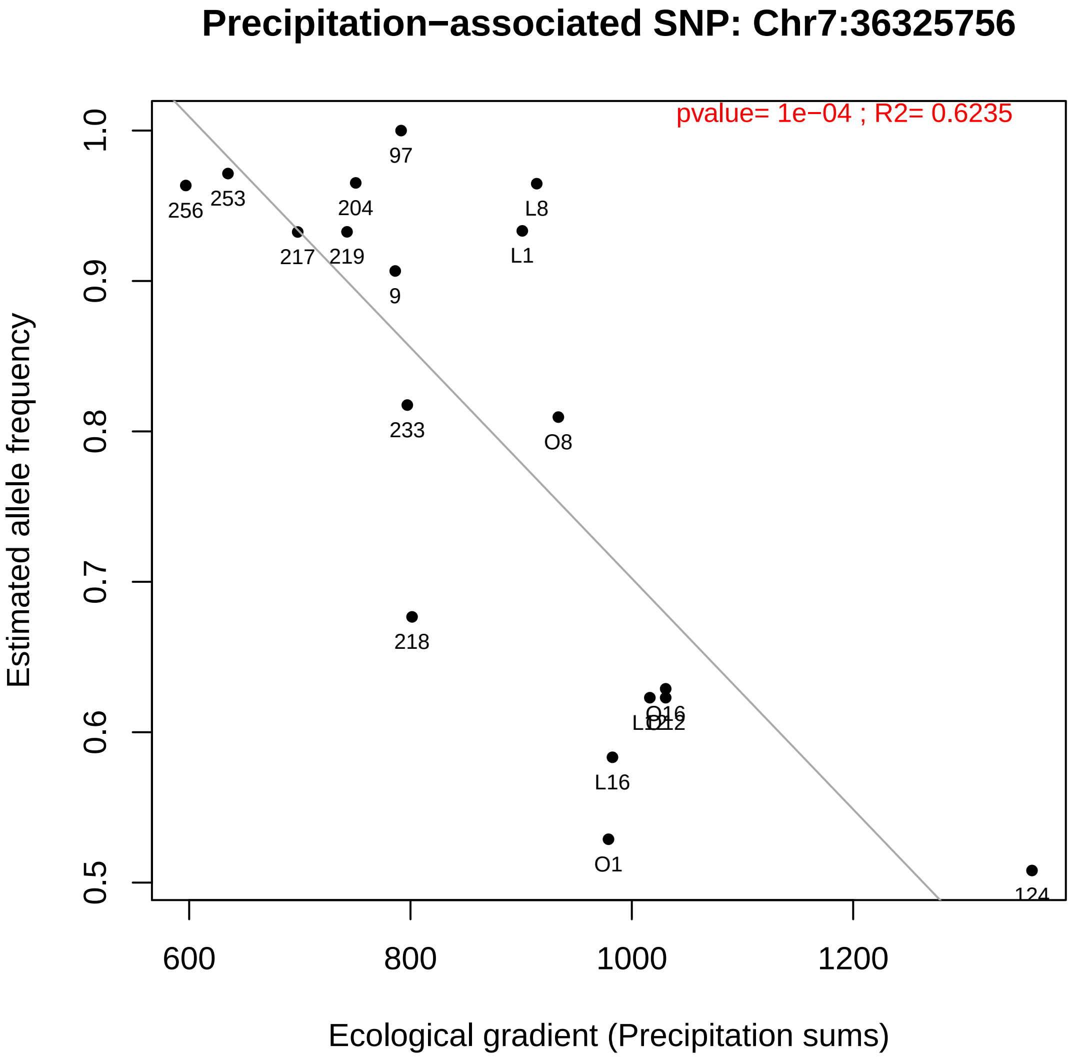
Precipitation−associated SNP: Chr7:36325756
pvalue= 1e−04 ; R2= 0.6235
Estimated allele frequency
0.5
0.6
0.7
0.8
0.9
1.0
Ecological gradient (Precipitation sums)
600
800
1000
1200
97
256
253
204
L8
217
219
L1
9
233
O8
218
O16
L16
O1
124

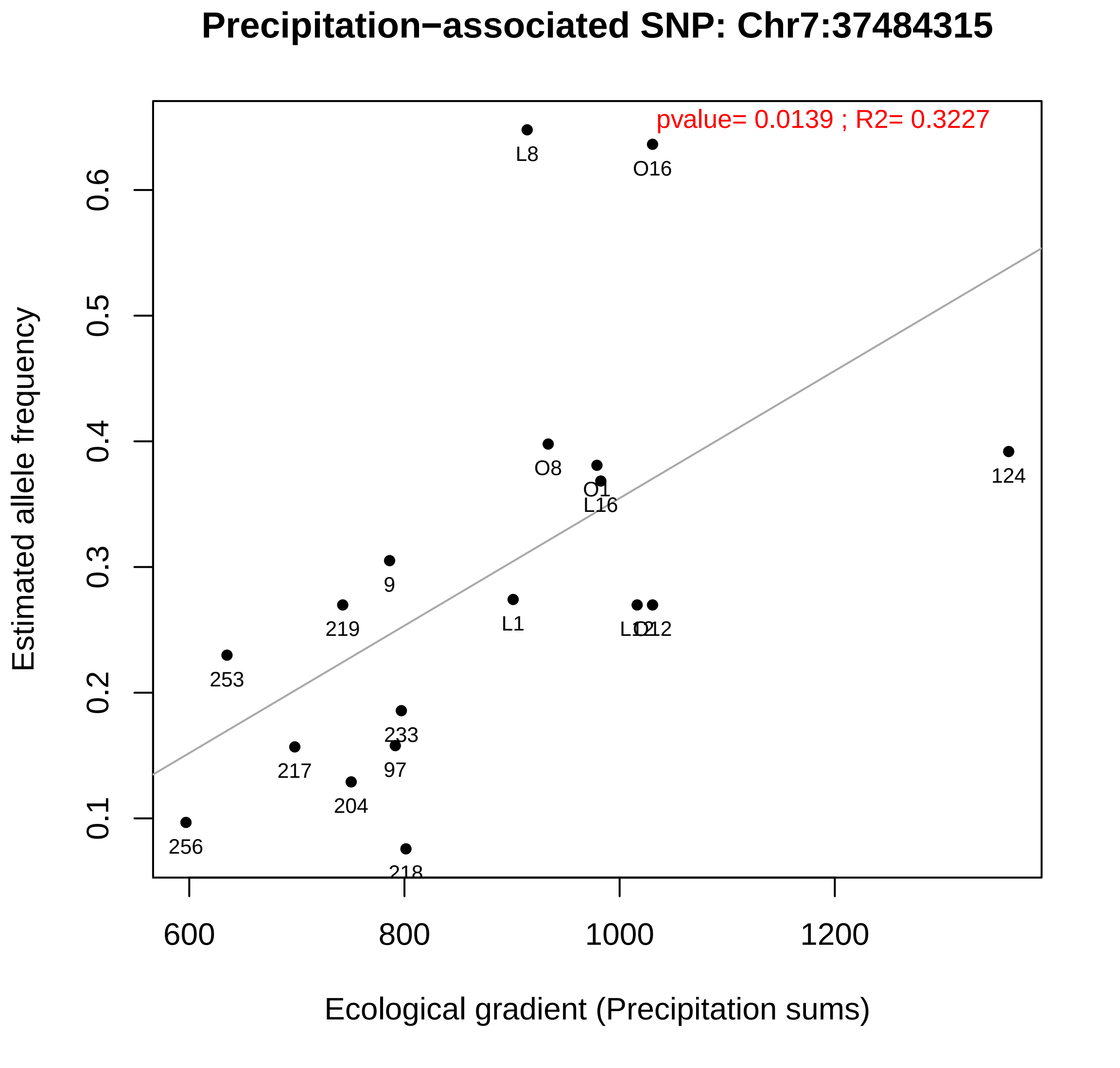
Precipitation−associated SNP: Chr7:37484315
pvalue= 0.0139 ; R2= 0.3227
L8
O16
O8
O1
L16
124
9
219
L1
L12
O12
253
233
217
97
204
256
218
0.1
0.2
0.3
0.4
0.5
0.6
600
800
1000
1200
Estimated allele frequency
Ecological gradient (Precipitation sums)

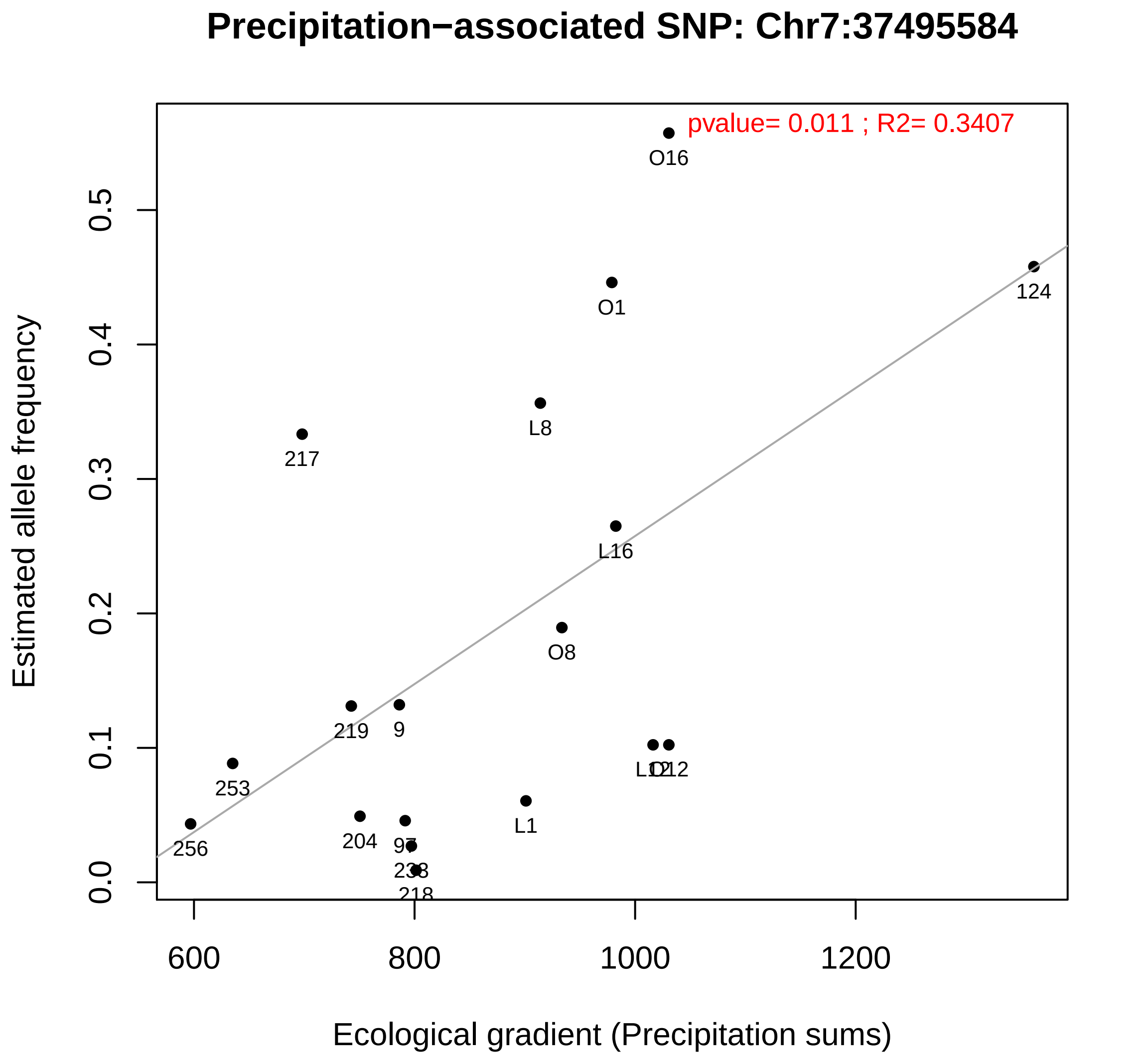
Precipitation−associated SNP: Chr7:37495584
pvalue= 0.011 ; R2= 0.3407
O16
O1
124
L8
217
L16
O8
219
9
253
L12
O12
256
204
L1
97
238
218
0.0
0.1
0.2
0.3
0.4
0.5
600
800
1000
1200
Estimated allele frequency
Ecological gradient (Precipitation sums)

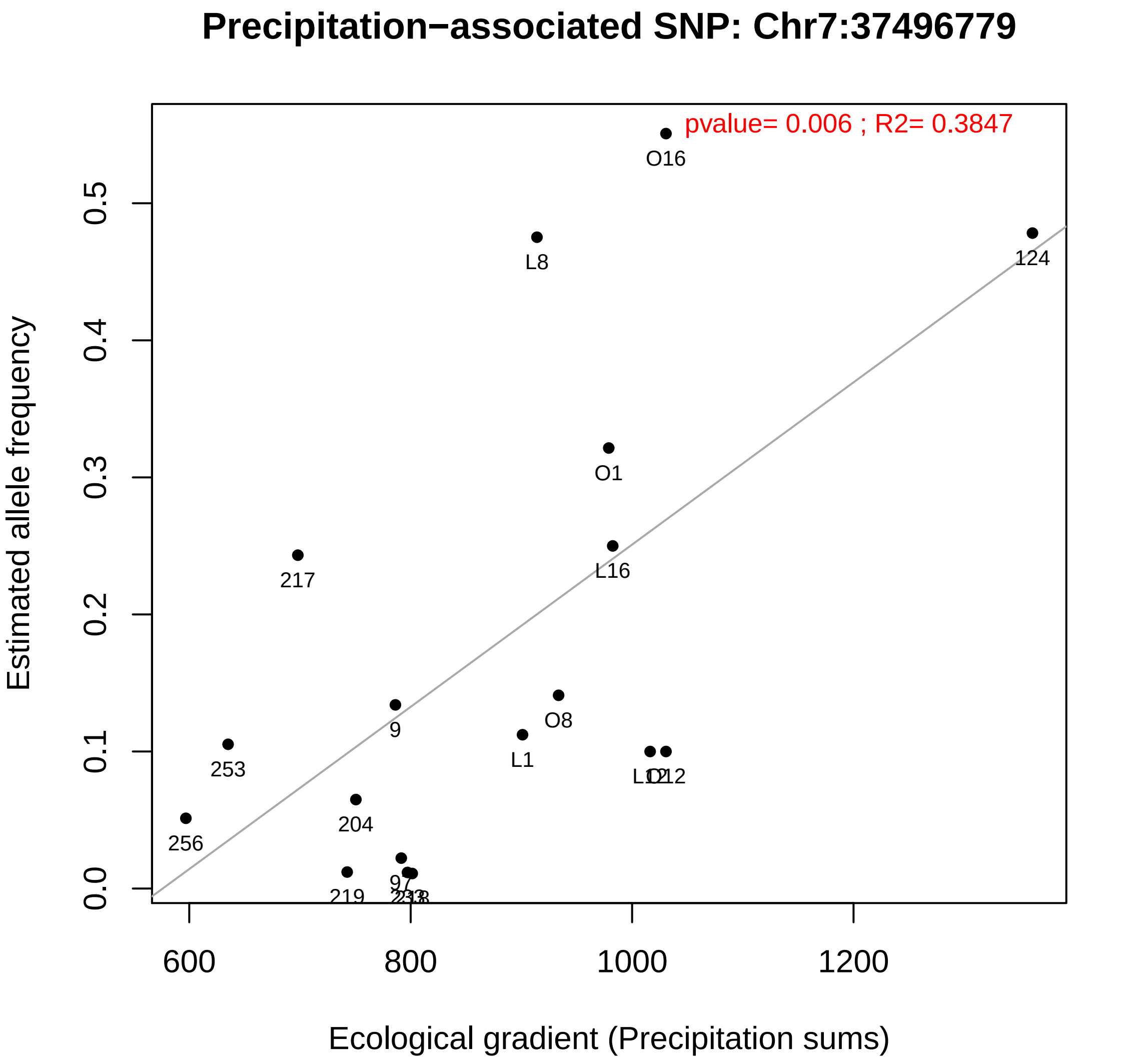

Precipitation−associated SNP: Chr7:37496779
pvalue= 0.006 ; R2= 0.3847
O16
L8
124
O1
L16
217
O8
9
L1
253
204
256
219
Estimated allele frequency
Ecological gradient (Precipitation sums)
0.0
0.1
0.2
0.3
0.4
0.5
600
800
1000
1200

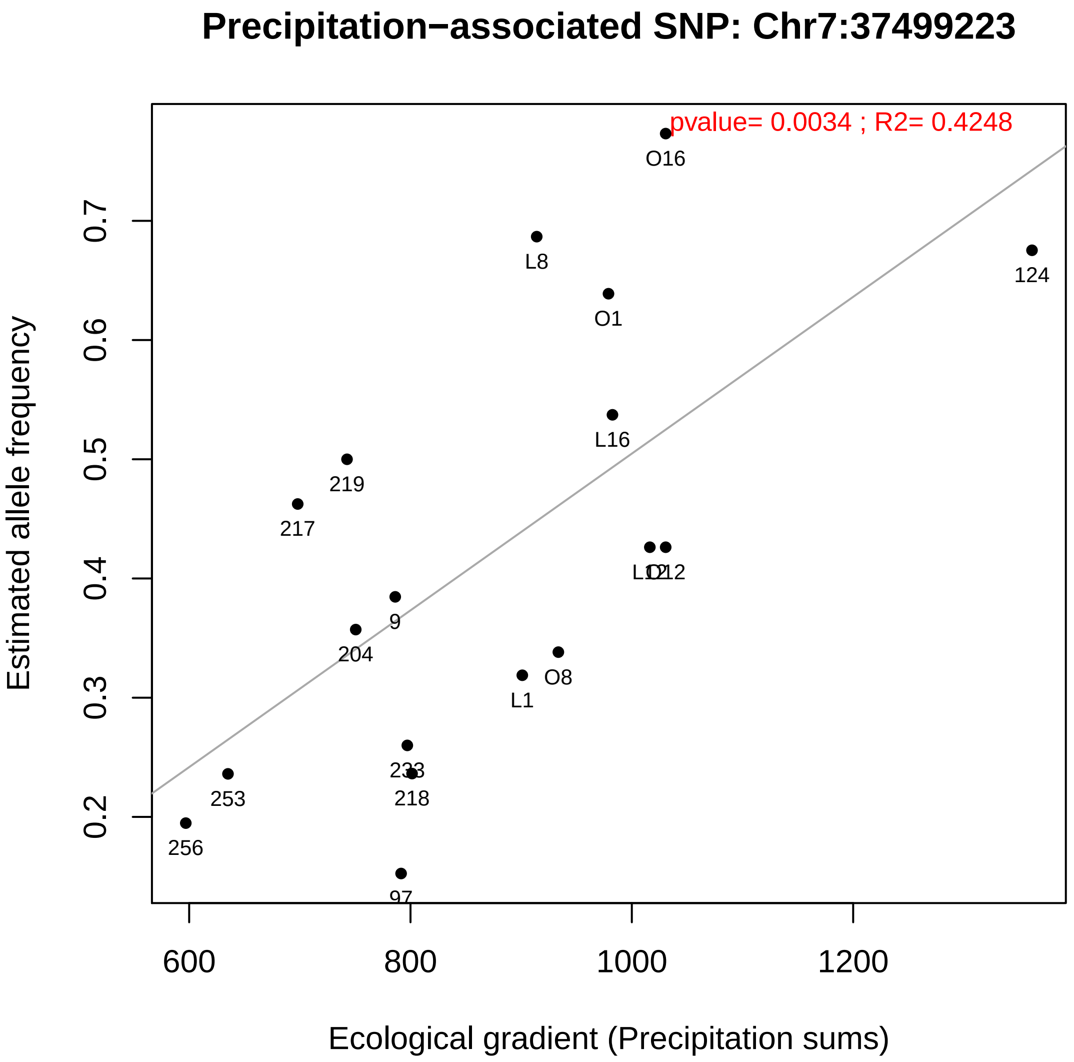
Precipitation−associated SNP: Chr7:37499223
pvalue= 0.0034 ; R2= 0.4248
O16
L8
124
O1
L16
219
217
L12
O12
9
204
O8
L1
233
253
218
256
97
0.2
0.3
0.4
0.5
0.6
0.7
600
800
1000
1200
Estimated allele frequency
Ecological gradient (Precipitation sums)

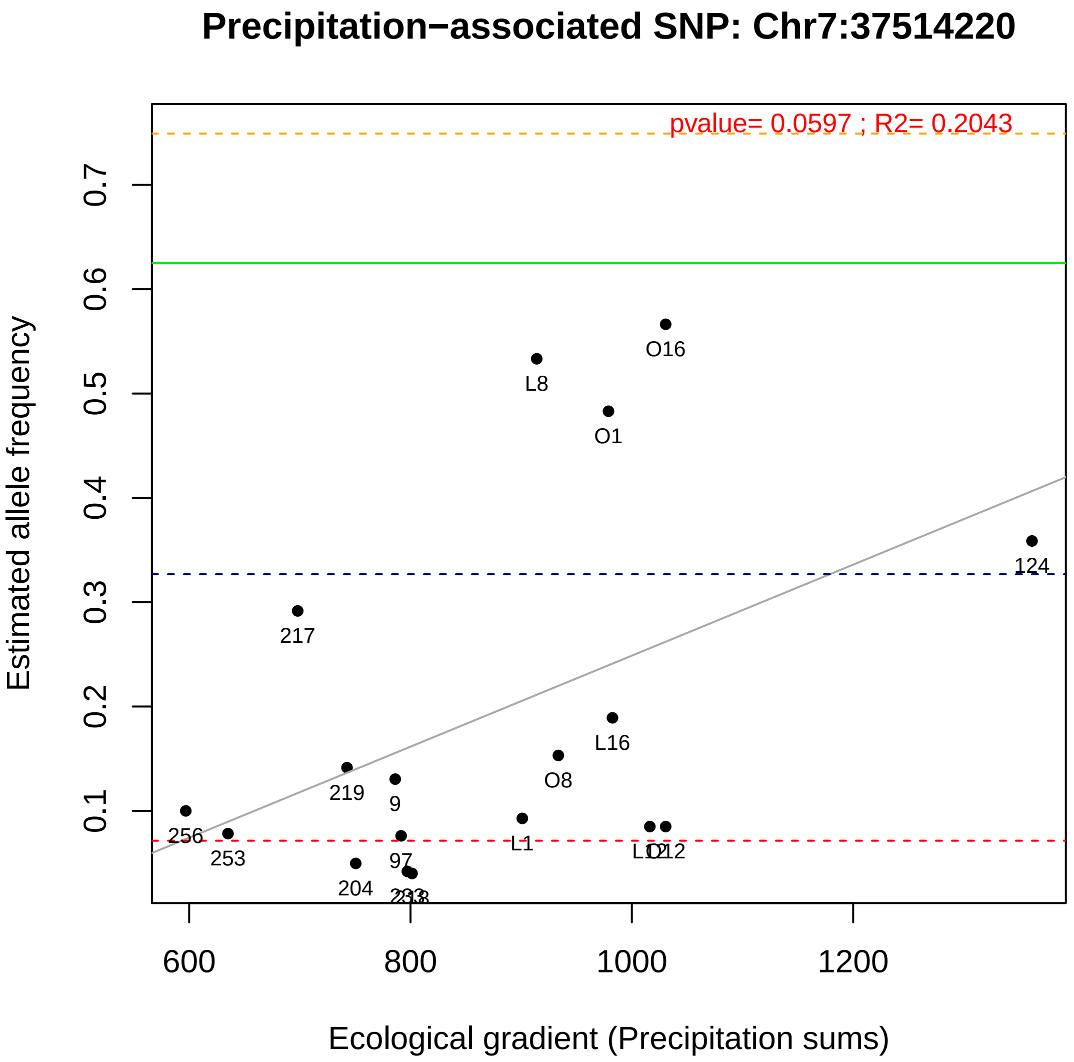

Precipitation−associated SNP: Chr7:37514220
pvalue= 0.0597 ; R2= 0.2043
Estimated allele frequency
0.1
0.2
0.3
0.4
0.5
0.6
0.7
600
800
1000
1200
Ecological gradient (Precipitation sums)
O16
L8
O1
124
217
L16
O8
219
9
256
253
L1
97
204

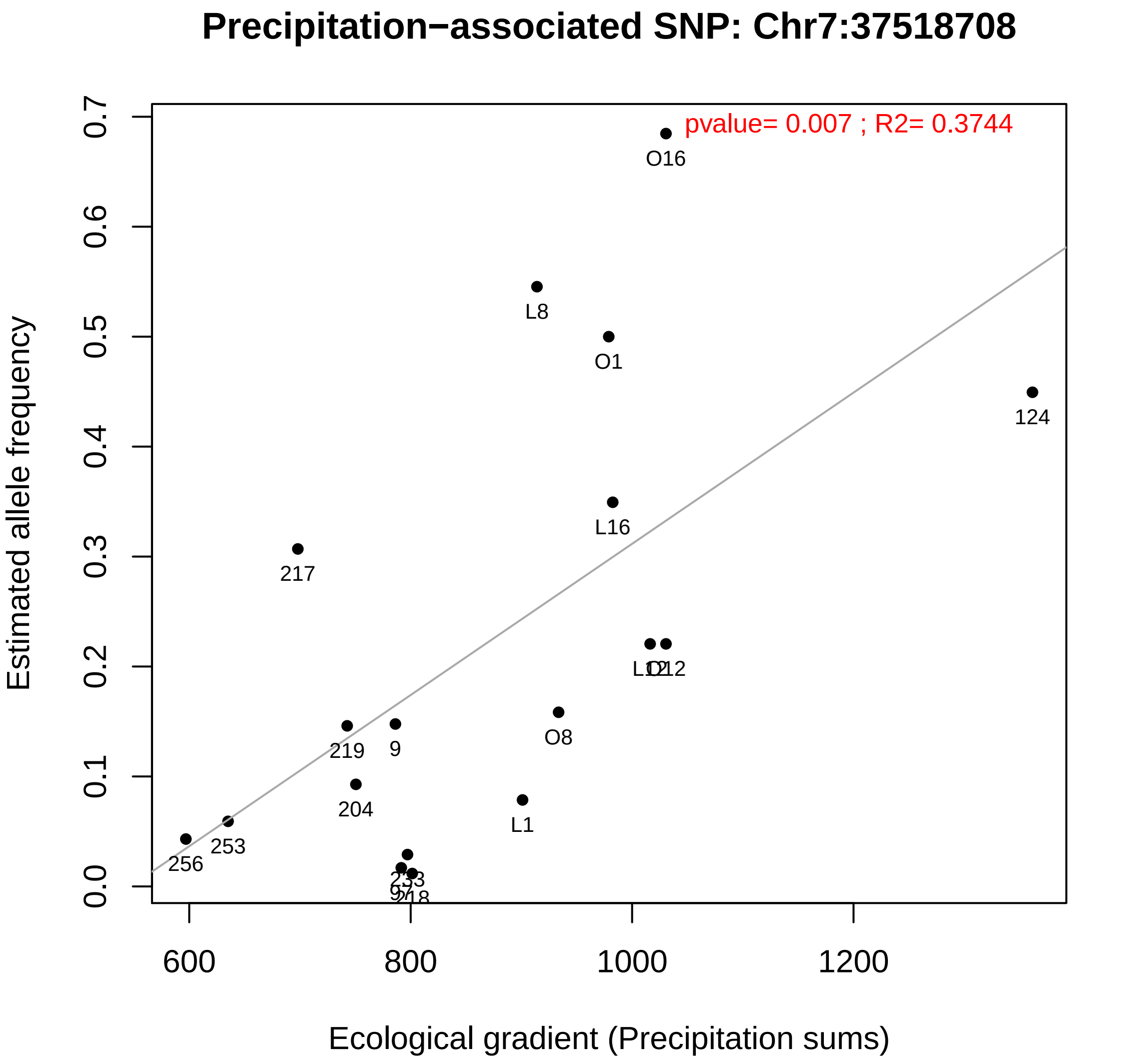
Precipitation−associated SNP: Chr7:37518708
pvalue= 0.007 ; R2= 0.3744
Estimated allele frequency
Ecological gradient (Precipitation sums)
0.0
0.1
0.2
0.3
0.4
0.5
0.6
0.7
600
800
1000
1200
O16
L8
O1
124
L16
217
O8
219
9
204
L1
253
256

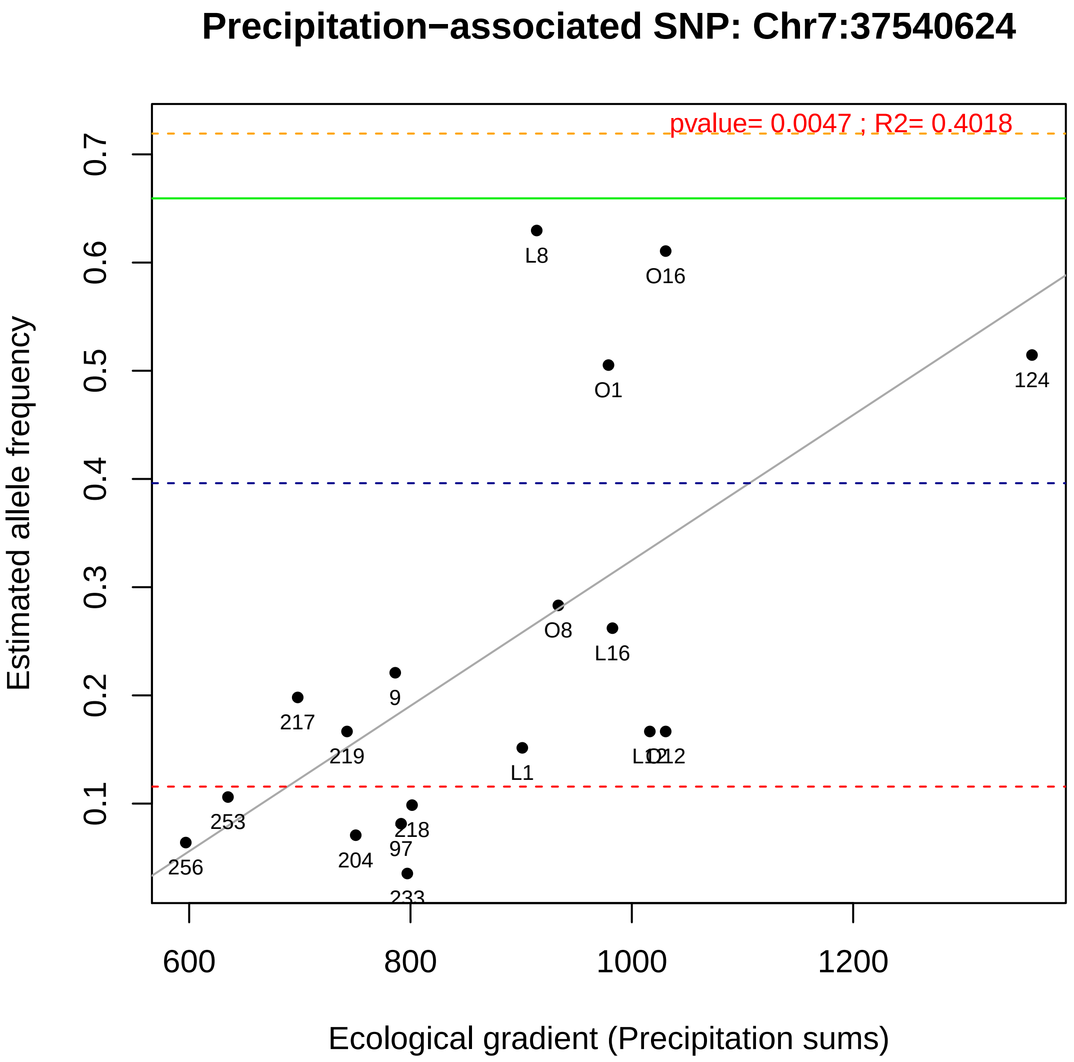

Precipitation−associated SNP: Chr7:37540624
pvalue= 0.0047 ; R2= 0.4018
Estimated allele frequency
0.1
0.2
0.3
0.4
0.5
0.6
0.7
600
800
1000
1200
Ecological gradient (Precipitation sums)
L8
O16
O1
124
O8
L16
9
217
219
L1
253
218
97
256
204
233

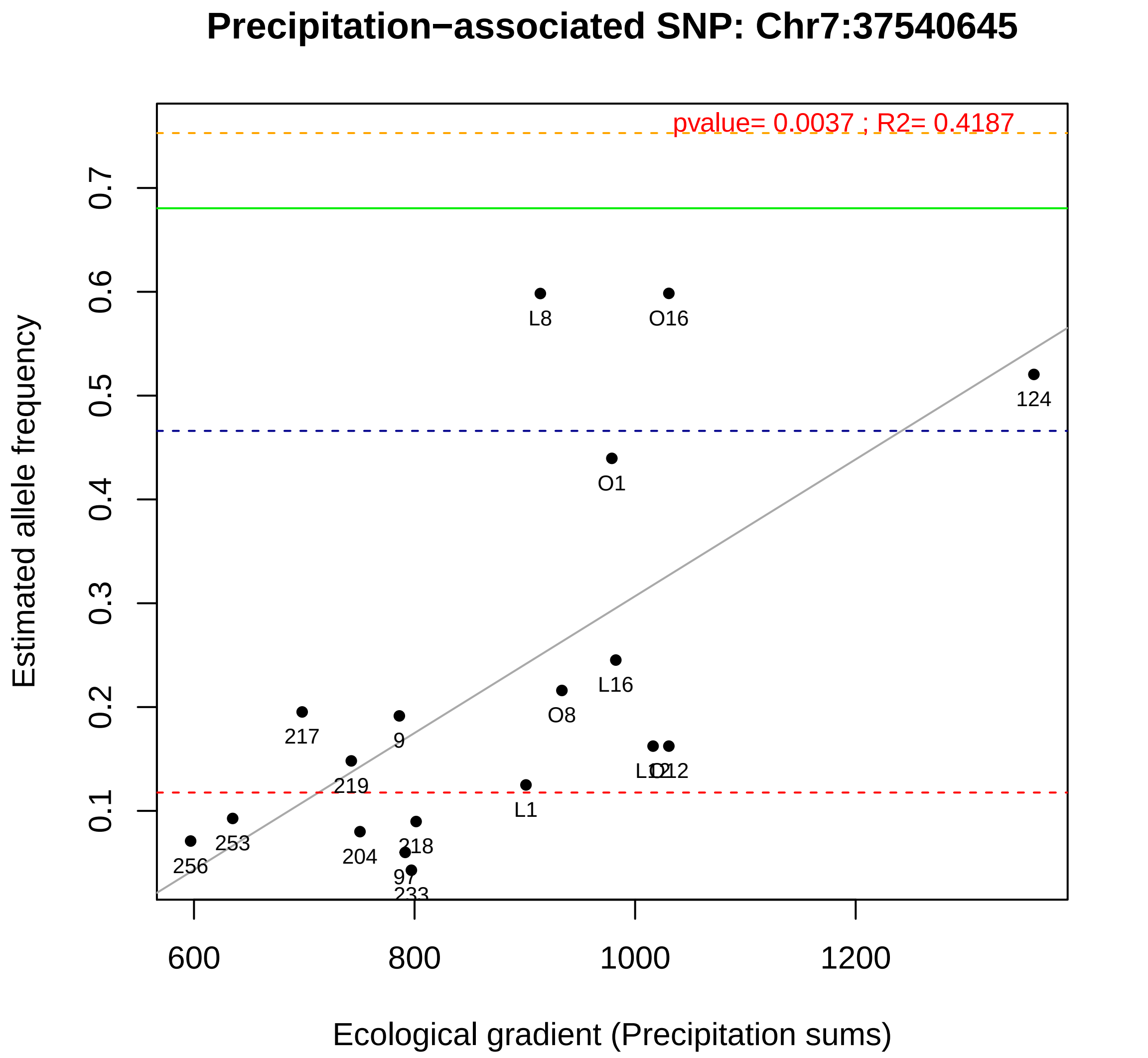

Precipitation−associated SNP: Chr7:37540645
pvalue= 0.0037 ; R2= 0.4187
Estimated allele frequency
0.1
0.2
0.3
0.4
0.5
0.6
0.7
600
800
1000
1200
Ecological gradient (Precipitation sums)
L8
O16
124
O1
L16
O8
217
9
219
L1
253
256
204
218
97
233

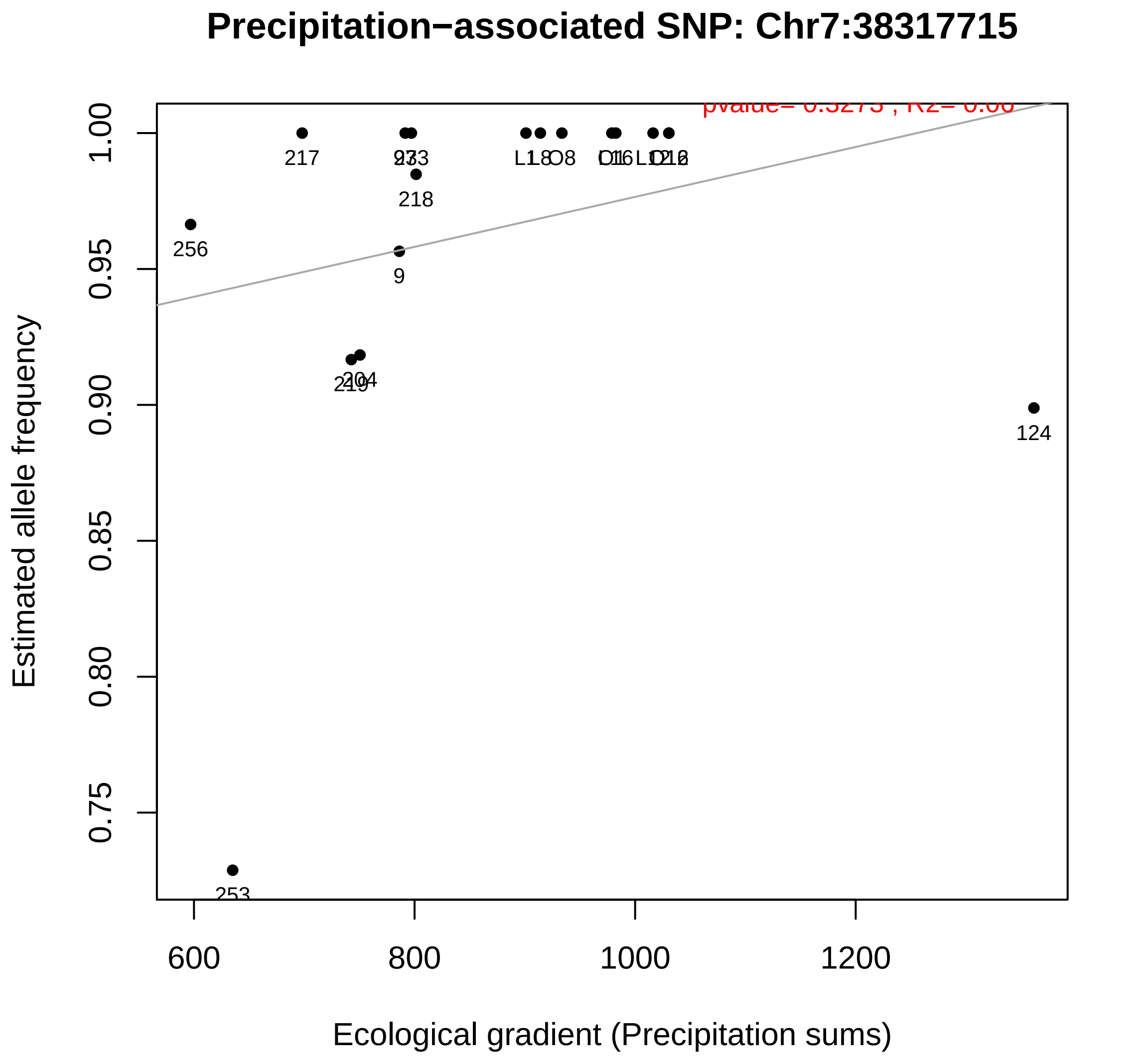

Precipitation−associated SNP: Chr7:38317715
pvalue= 0.3273 ; R2= 0.06
Estimated allele frequency
Ecological gradient (Precipitation sums)
0.75
0.80
0.85
0.90
0.95
1.00
600
800
1000
1200
217
256
9
218
124
253

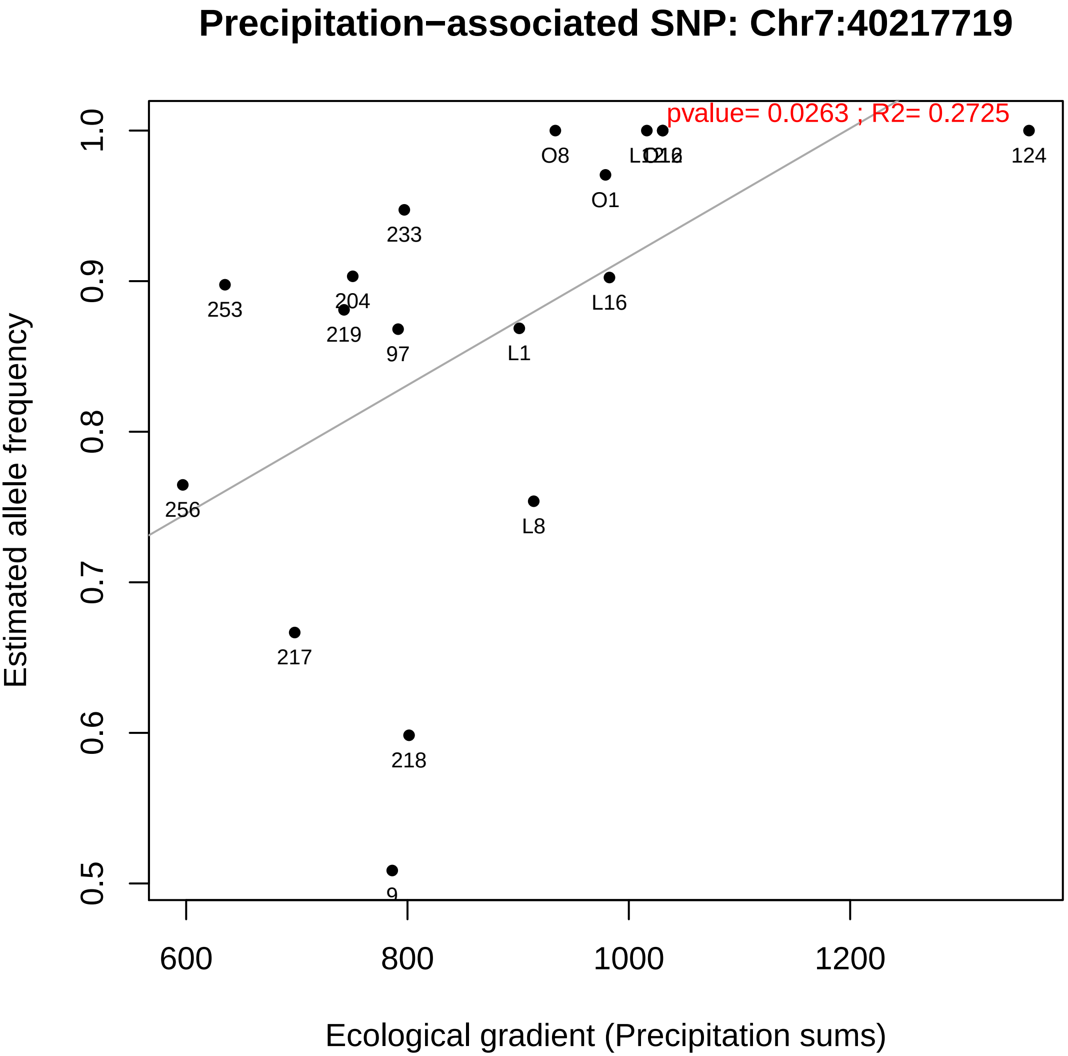
Precipitation−associated SNP: Chr7:40217719
pvalue= 0.0263 ; R2= 0.2725
Estimated allele frequency
0.5
0.6
0.7
0.8
0.9
1.0
600
800
1000
1200
Ecological gradient (Precipitation sums)
O8
124
O1
233
204
253
219
L16
97
L1
256
L8
217
218
9

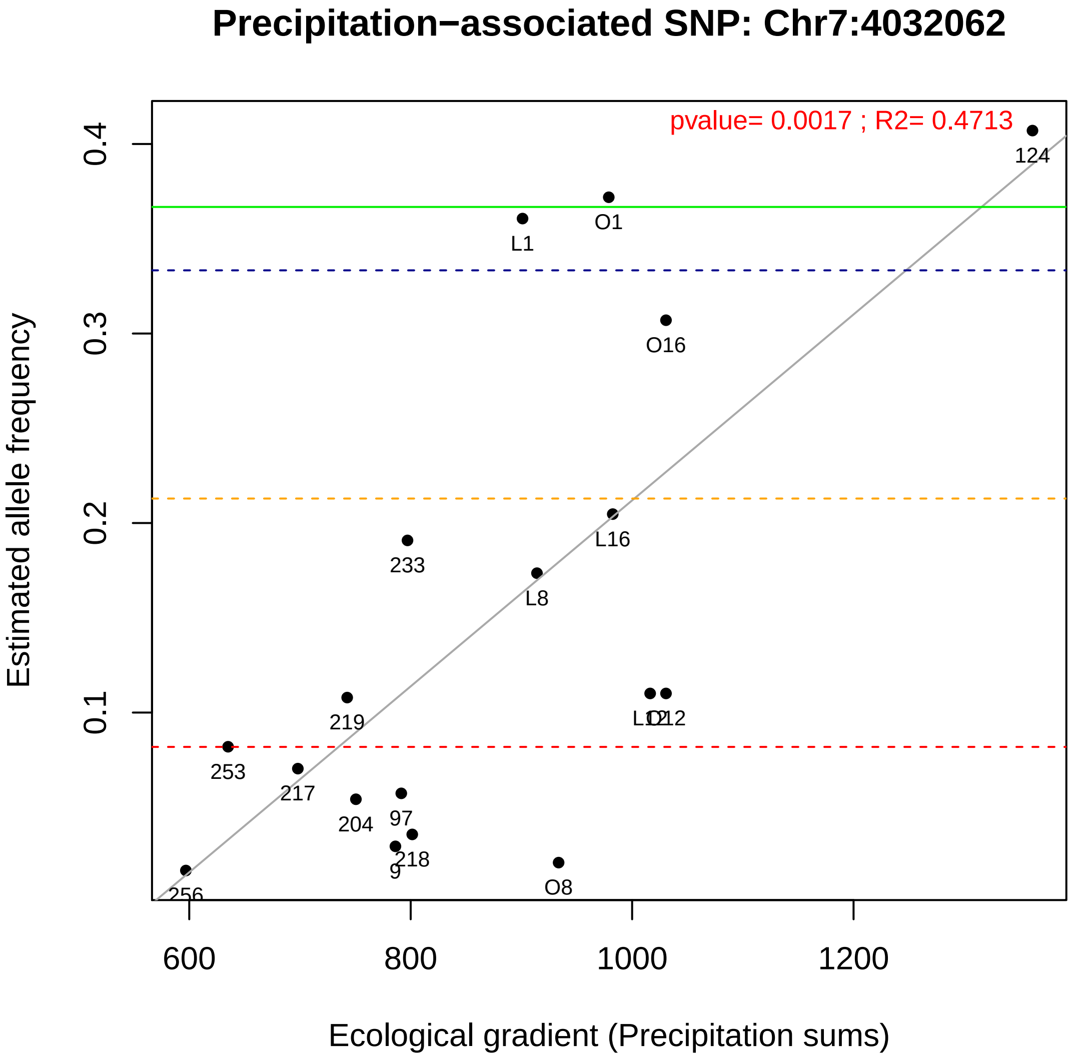
Precipitation−associated SNP: Chr7:4032062
pvalue= 0.0017 ; R2= 0.4713
Estimated allele frequency
0.1
0.2
0.3
0.4
600
800
1000
1200
Ecological gradient (Precipitation sums)
124
O1
L1
O16
L16
233
L8
219
L12
O12
253
217
204
97
218
9
O8
256

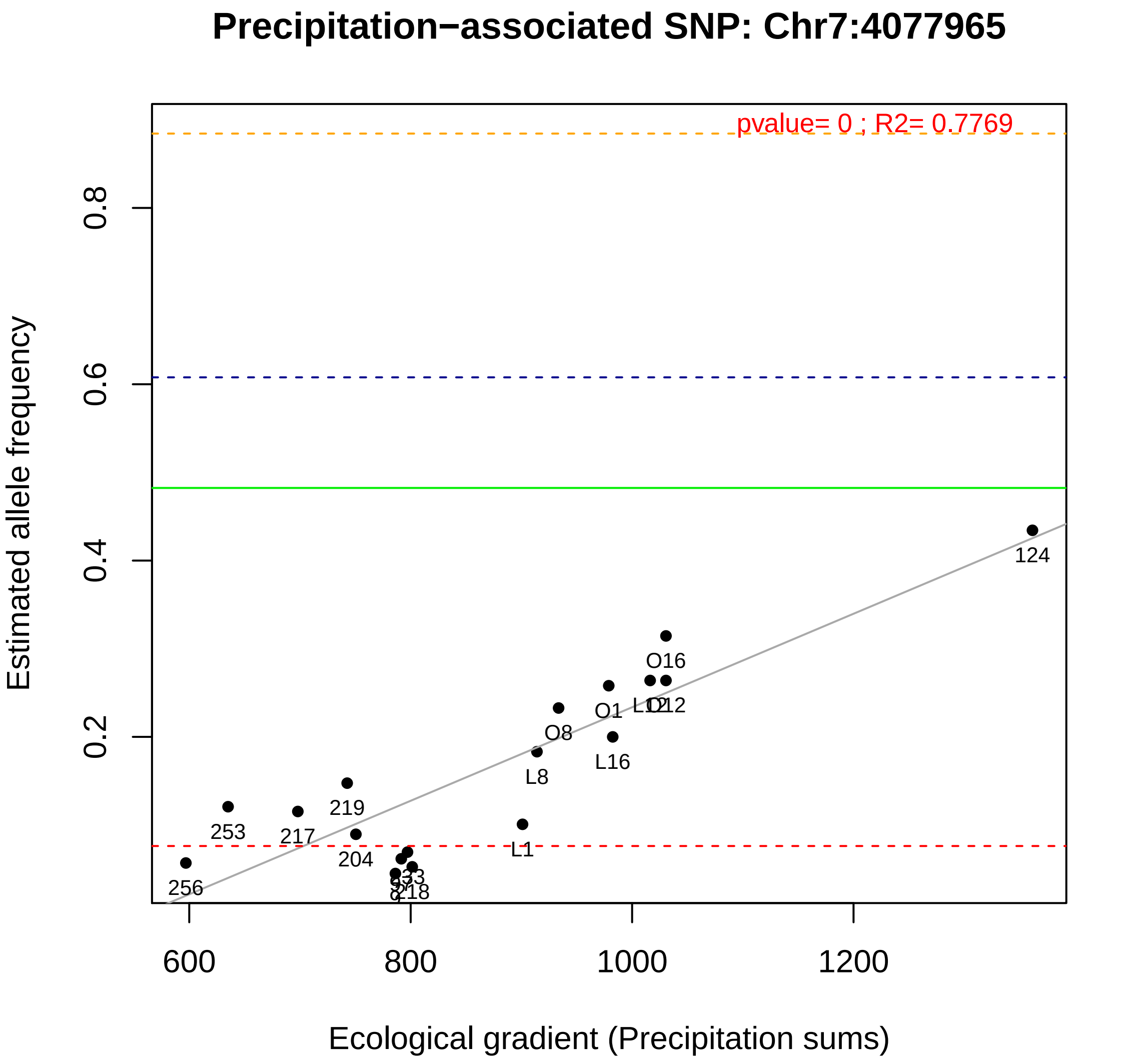

Precipitation−associated SNP: Chr7:4077965
pvalue= 0 ; R2= 0.7769
Estimated allele frequency
0.2
0.4
0.6
0.8
600
800
1000
1200
Ecological gradient (Precipitation sums)
124
O16
O1
O12
O8
L16
L8
L1
219
253
217
204
256
218

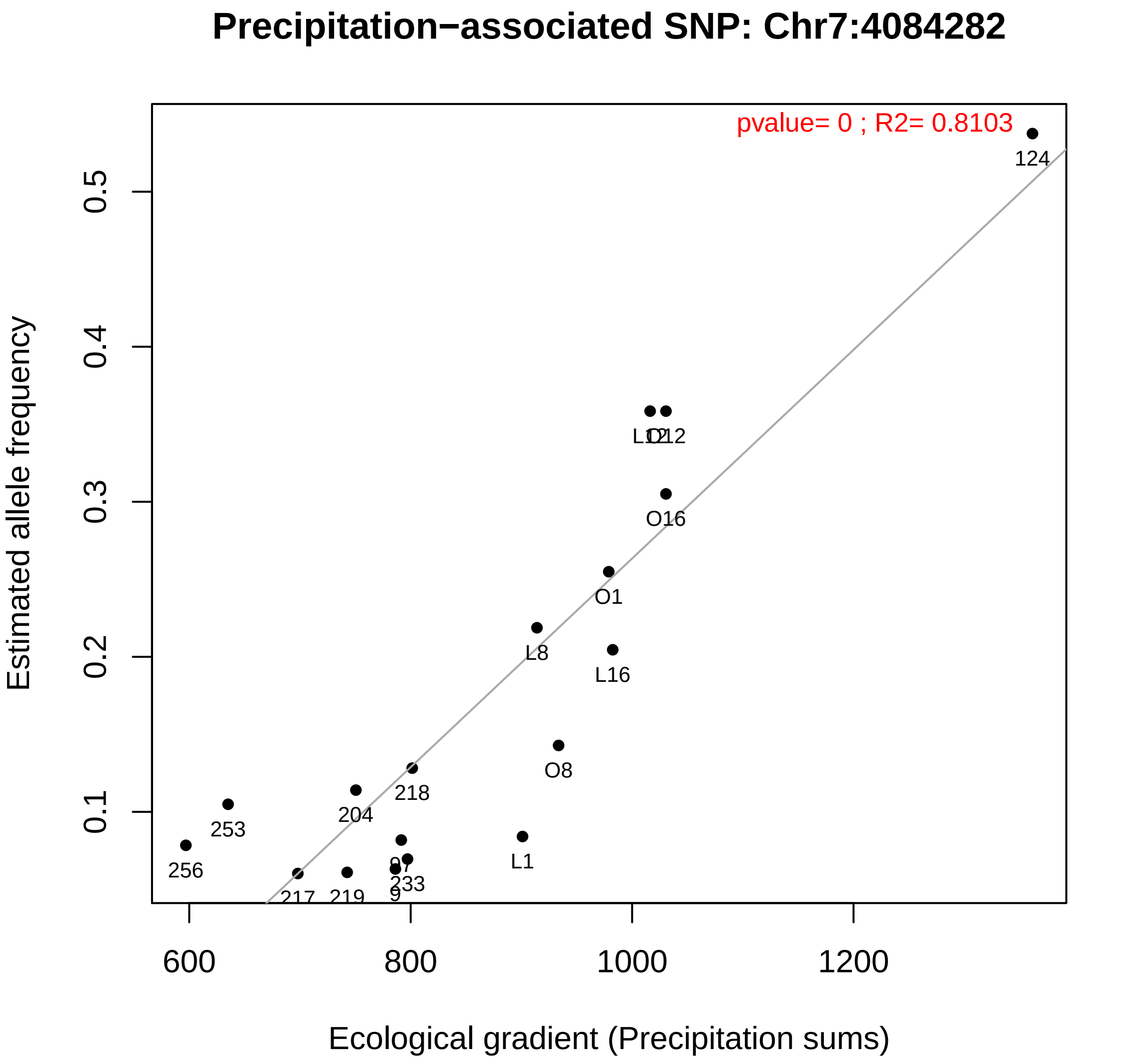

Precipitation−associated SNP: Chr7:4084282
pvalue= 0 ; R2= 0.8103
124
L12
O12
O16
O1
L8
L16
O8
218
204
253
256
97
233
9
217
219
L1
0.1
0.2
0.3
0.4
0.5
Estimated allele frequency
600
800
1000
1200
Ecological gradient (Precipitation sums)

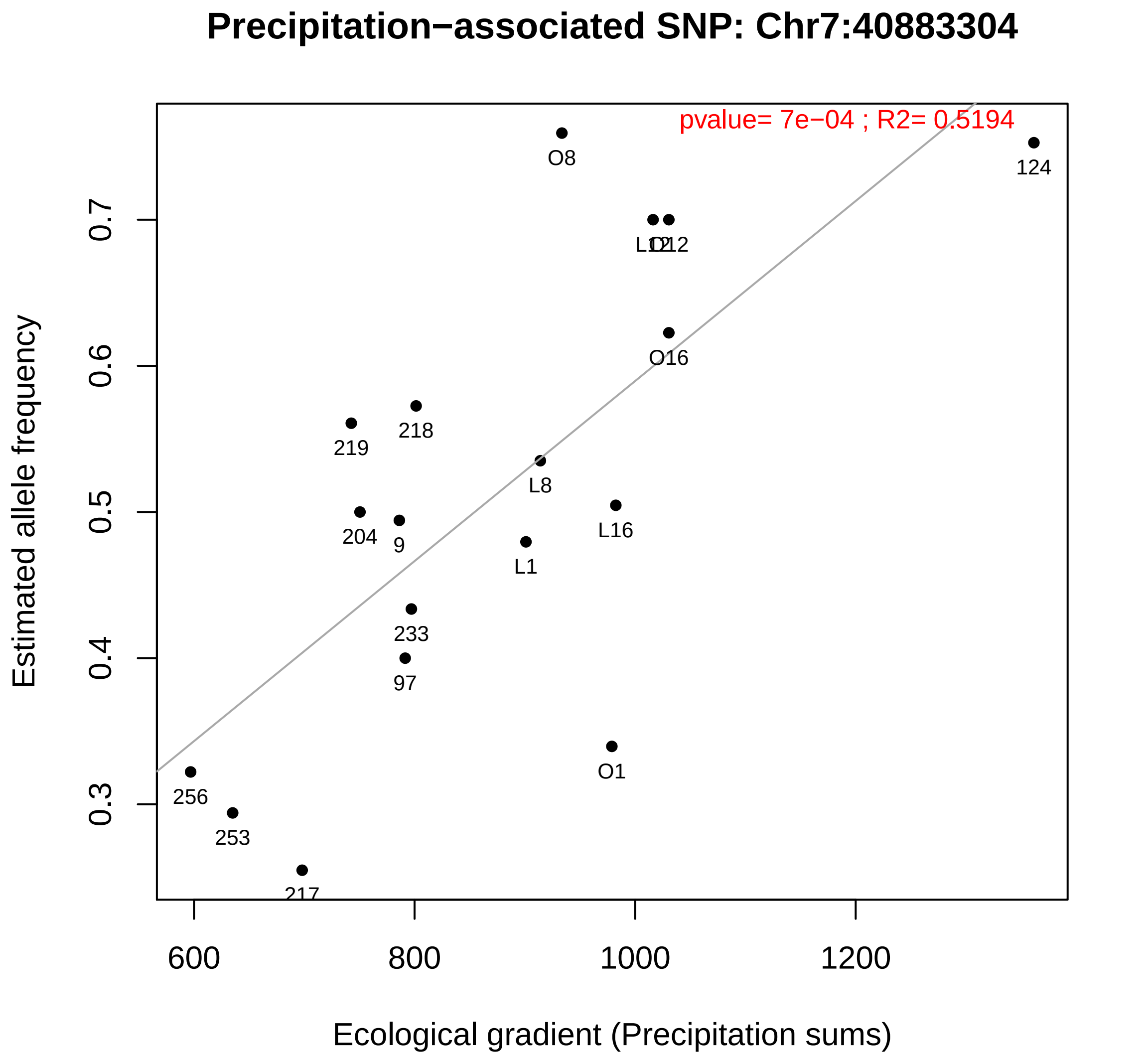

Precipitation−associated SNP: Chr7:40883304
pvalue= 7e−04 ; R2= 0.5194
Estimated allele frequency
0.3
0.4
0.5
0.6
0.7
600
800
1000
1200
Ecological gradient (Precipitation sums)
O8
124
L12
O12
O16
218
219
L8
204
9
L16
L1
233
97
O1
256
253
217

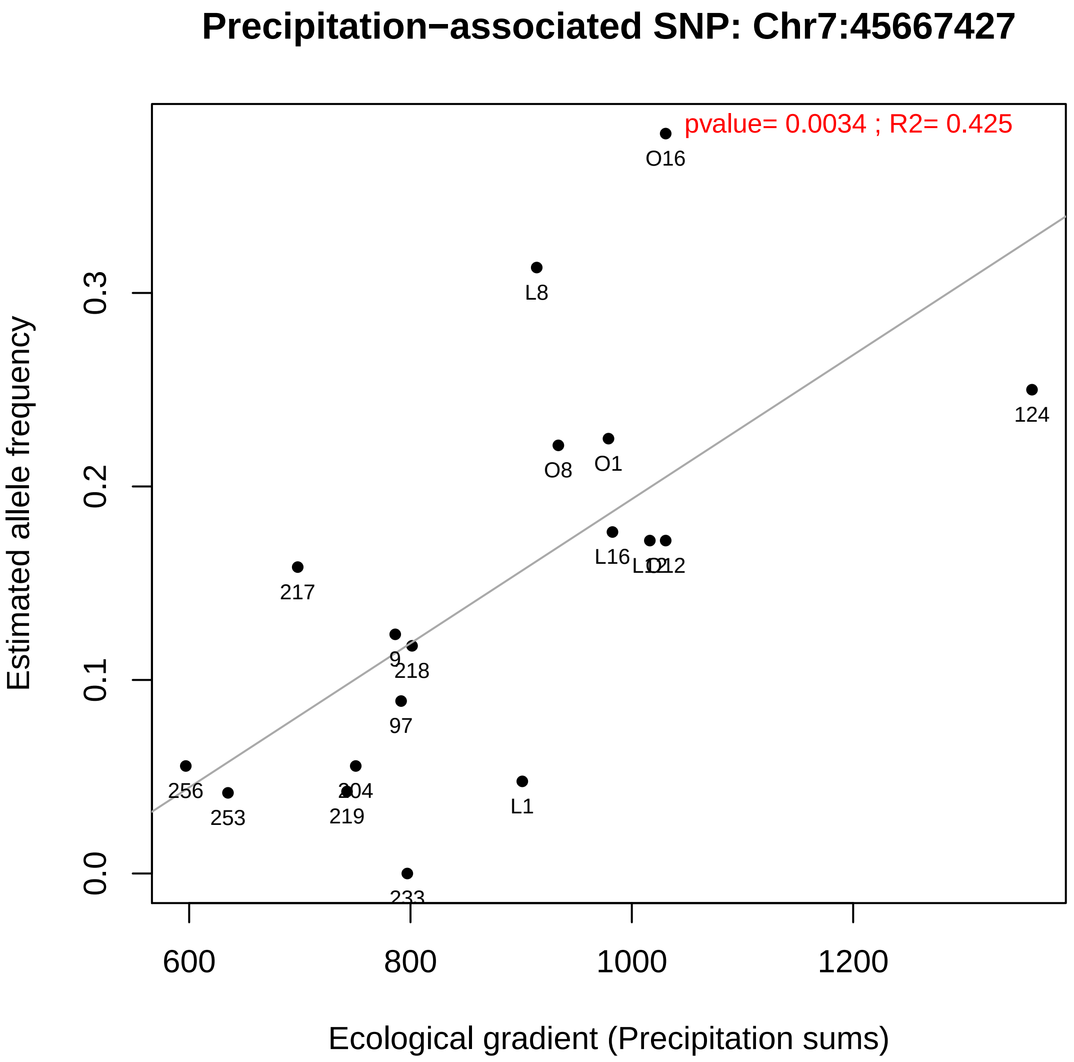
Precipitation−associated SNP: Chr7:45667427
pvalue= 0.0034 ; R2= 0.425
O16
L8
124
O8
O1
L16
L12
O12
217
9
218
97
256
253
204
219
L1
233
0.0
0.1
0.2
0.3
600
800
1000
1200
Estimated allele frequency
Ecological gradient (Precipitation sums)

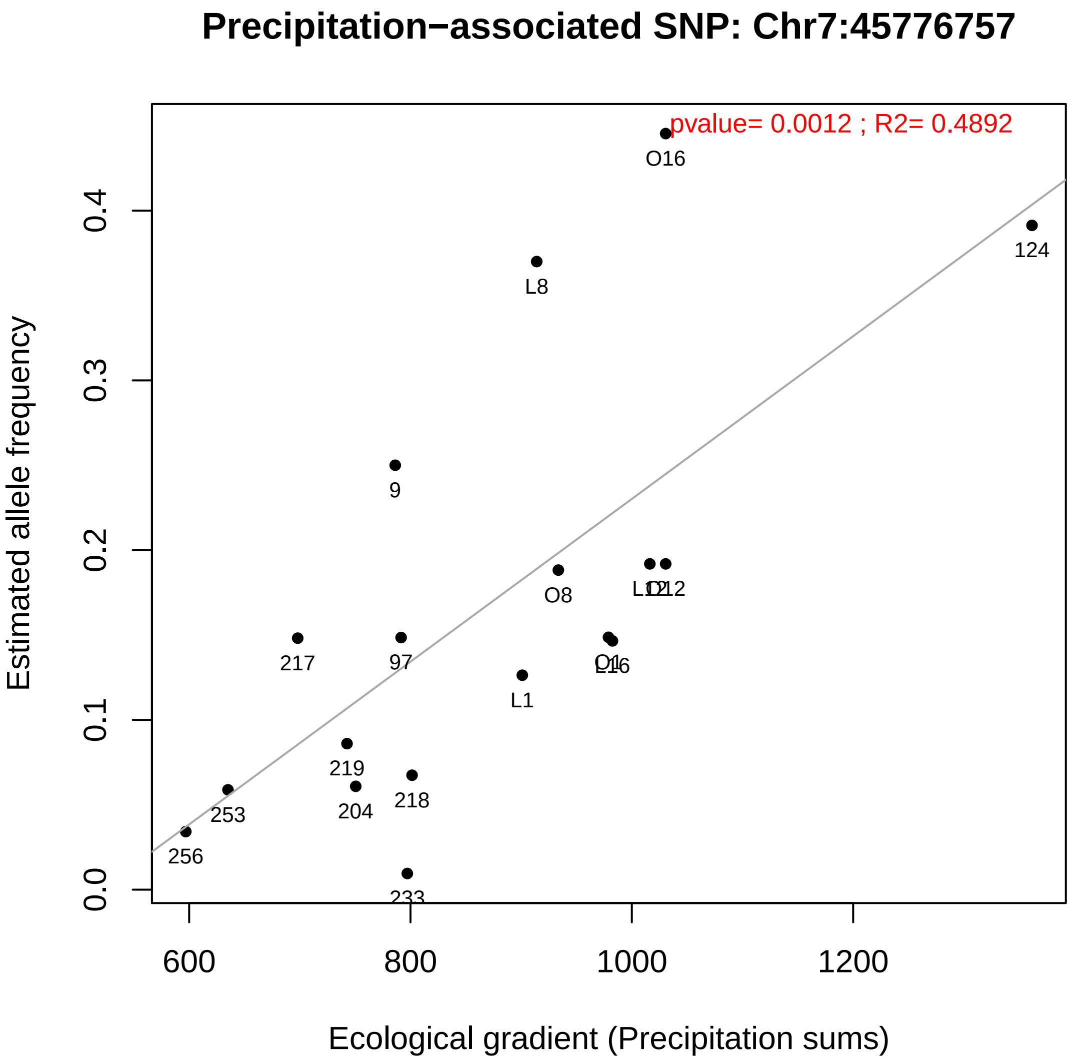

Precipitation−associated SNP: Chr7:45776757
pvalue= 0.0012 ; R2= 0.4892
O16
124
L8
9
O8
L12
O12
217
97
O1
L16
L1
219
218
253
204
256
233
0.0
0.1
0.2
0.3
0.4
600
800
1000
1200
Estimated allele frequency
Ecological gradient (Precipitation sums)

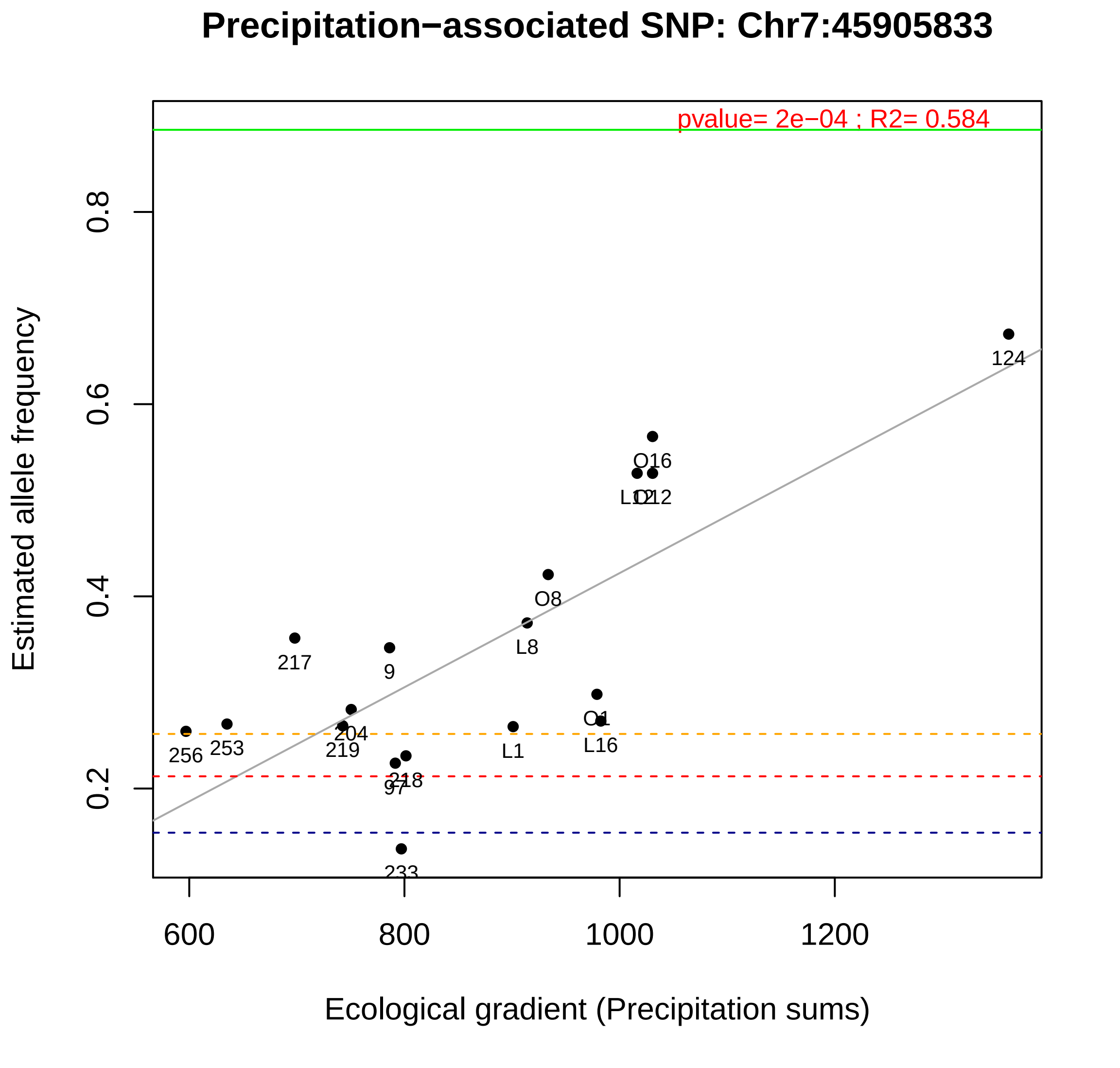

Precipitation−associated SNP: Chr7:45905833
pvalue= 2e−04 ; R2= 0.584
Estimated allele frequency
0.2
0.4
0.6
0.8
Ecological gradient (Precipitation sums)
600
800
1000
1200
124
O16
L12
O12
O8
L8
217
9
O1
L16
L1
204
219
256
253
218
97
233

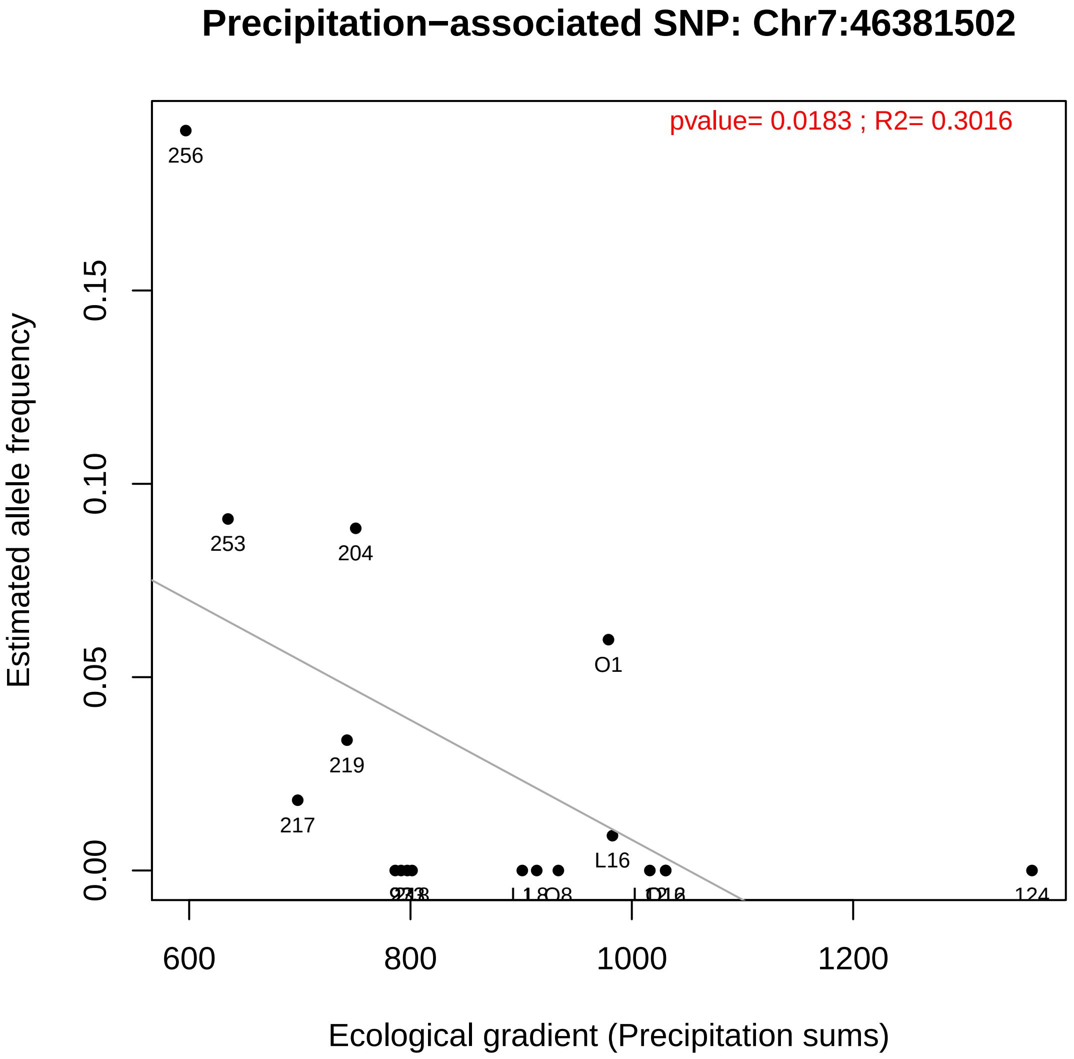
Precipitation−associated SNP: Chr7:46381502
pvalue= 0.0183 ; R2= 0.3016
Estimated allele frequency
0.00
0.05
0.10
0.15
Ecological gradient (Precipitation sums)
600
800
1000
1200
256
253
204
O1
219
217
L16
124

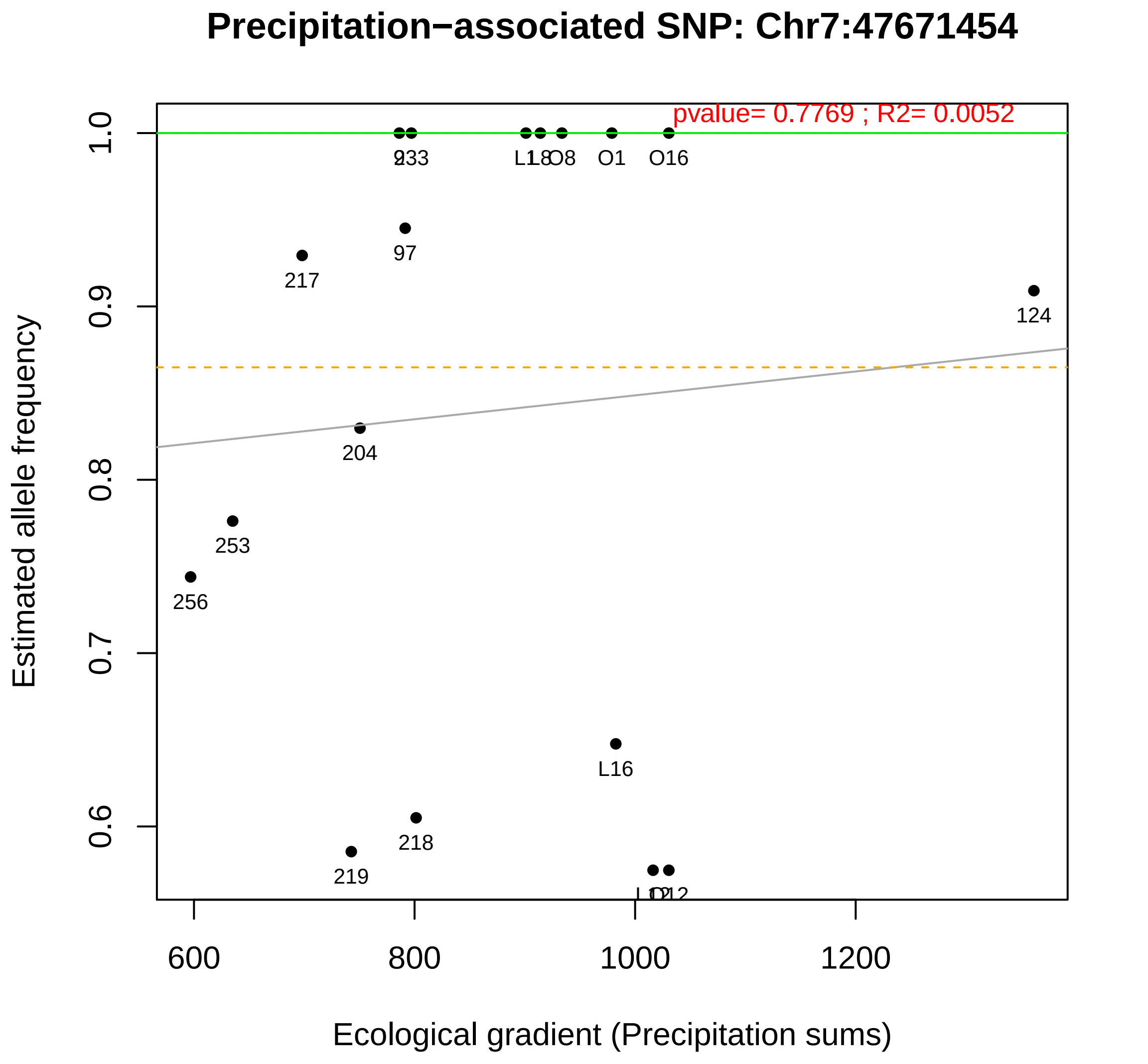
Precipitation−associated SNP: Chr7:47671454
pvalue= 0.7769 ; R2= 0.0052
Estimated allele frequency
Ecological gradient (Precipitation sums)
0.6
0.7
0.8
0.9
1.0
600
800
1000
1200
97
217
124
204
253
256
L16
218
219
O1
O16

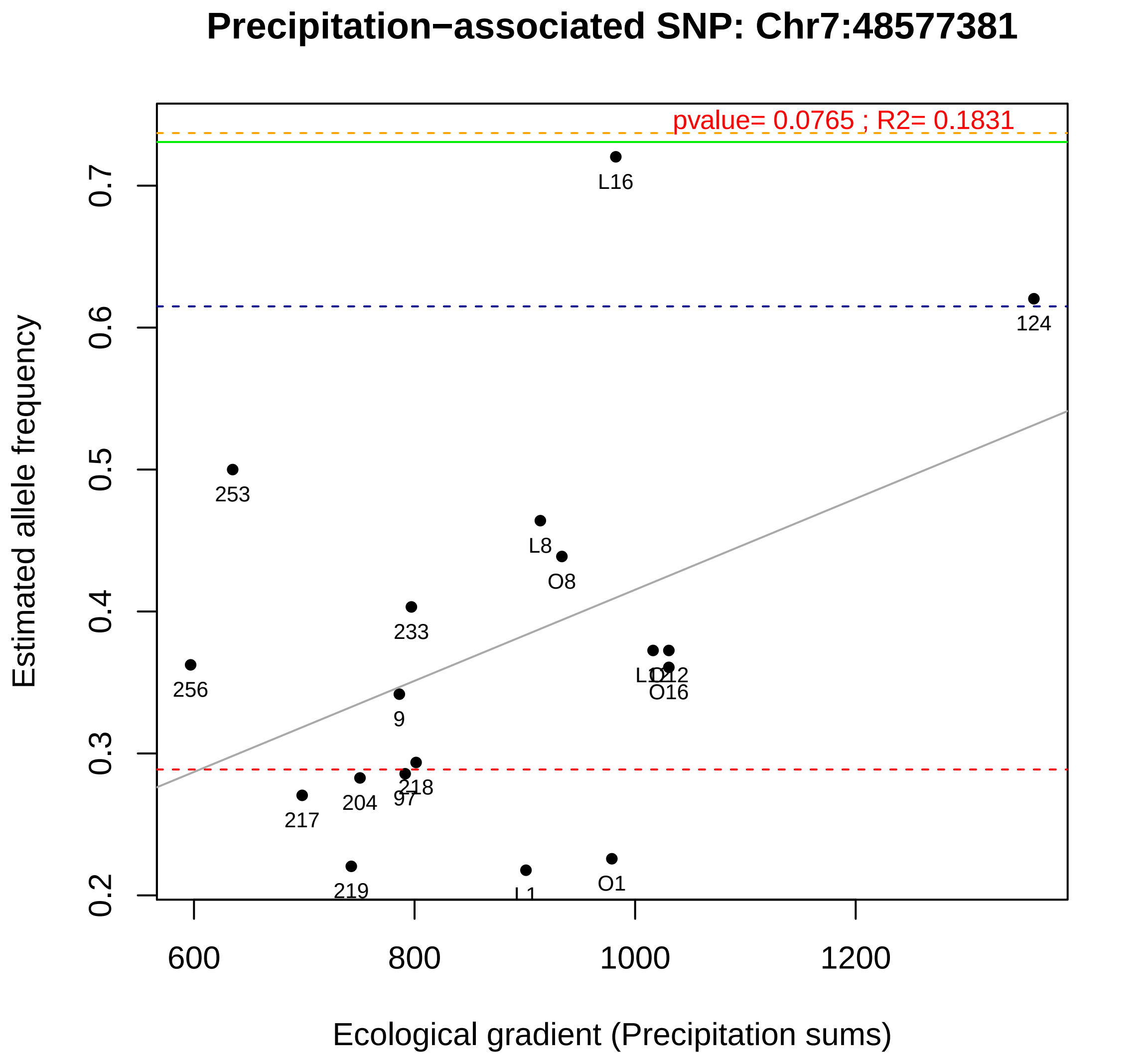

Precipitation−associated SNP: Chr7:48577381
pvalue= 0.0765 ; R2= 0.1831
Estimated allele frequency
0.2
0.3
0.4
0.5
0.6
0.7
600
800
1000
1200
Ecological gradient (Precipitation sums)
L16
124
253
L8
O8
233
256
9
O12
O16
218
204
97
217
219
O1

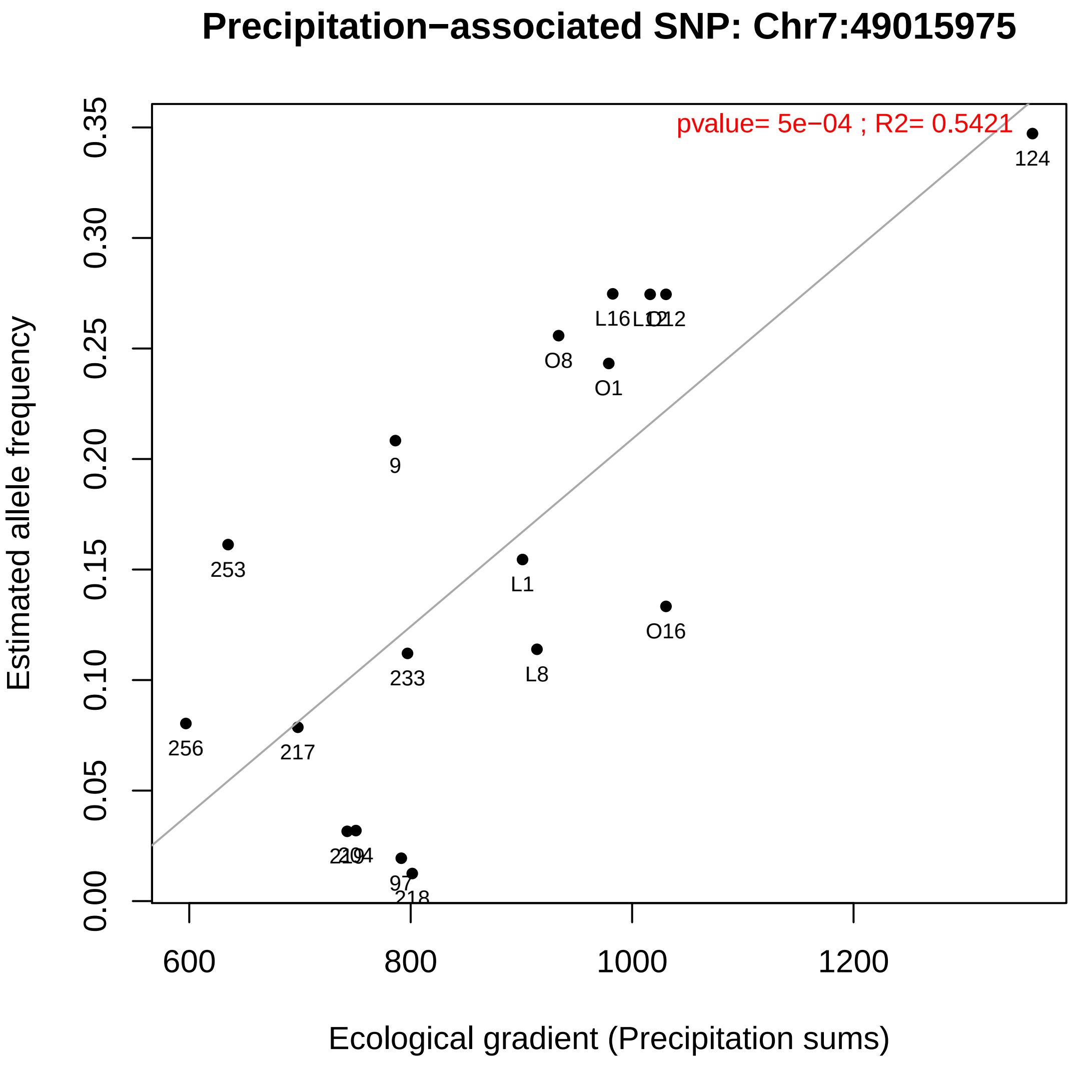

Precipitation−associated SNP: Chr7:49015975
pvalue= 5e−04 ; R2= 0.5421
Estimated allele frequency
Ecological gradient (Precipitation sums)
0.00
0.05
0.10
0.15
0.20
0.25
0.30
0.35
600
800
1000
1200
124
L16
O12
O8
O1
9
253
L1
O16
233
L8
256
217
97
218

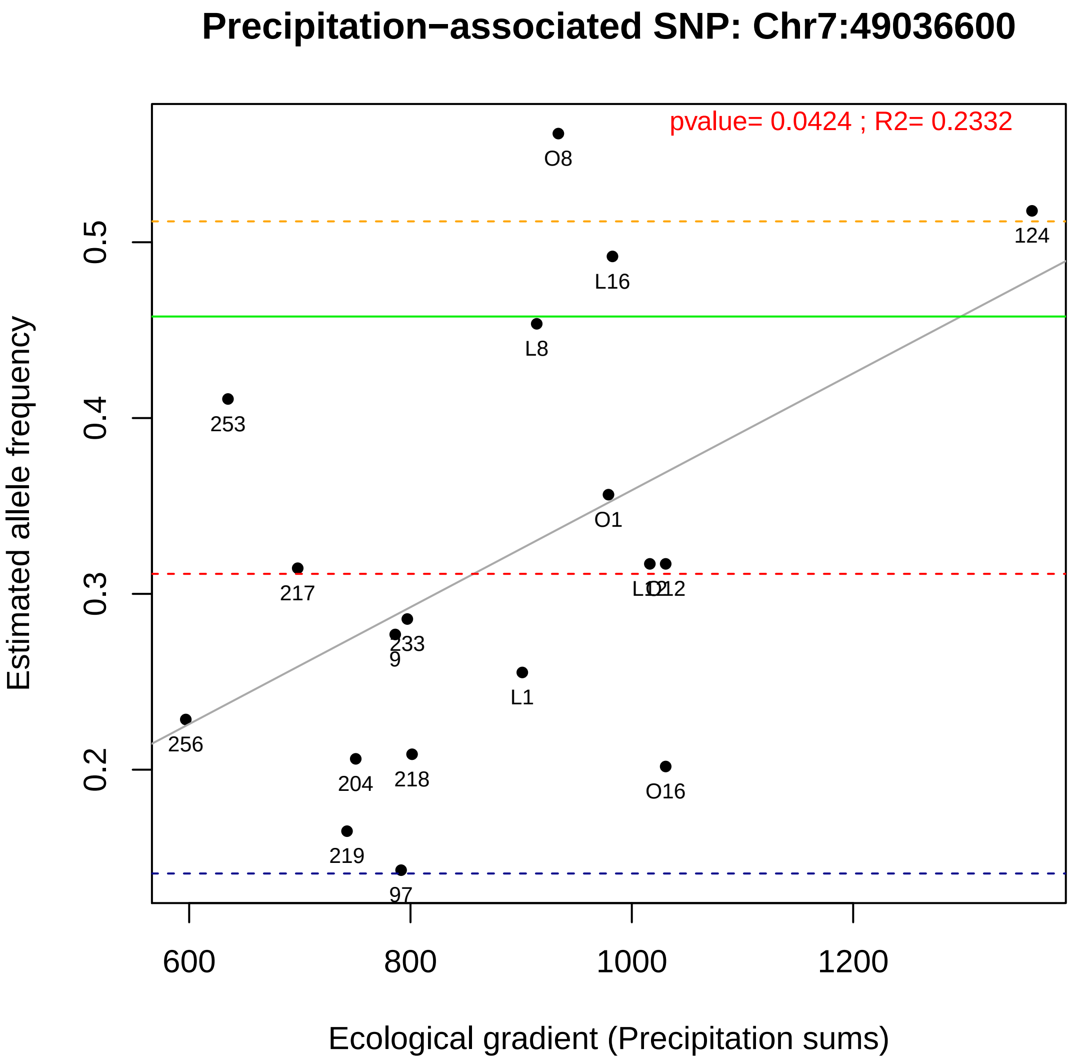
Precipitation−associated SNP: Chr7:49036600
pvalue= 0.0424 ; R2= 0.2332
Estimated allele frequency
0.2
0.3
0.4
0.5
600
800
1000
1200
Ecological gradient (Precipitation sums)
O8
124
L16
L8
253
O1
217
L12
O12
233
9
L1
256
204
218
O16
219
97

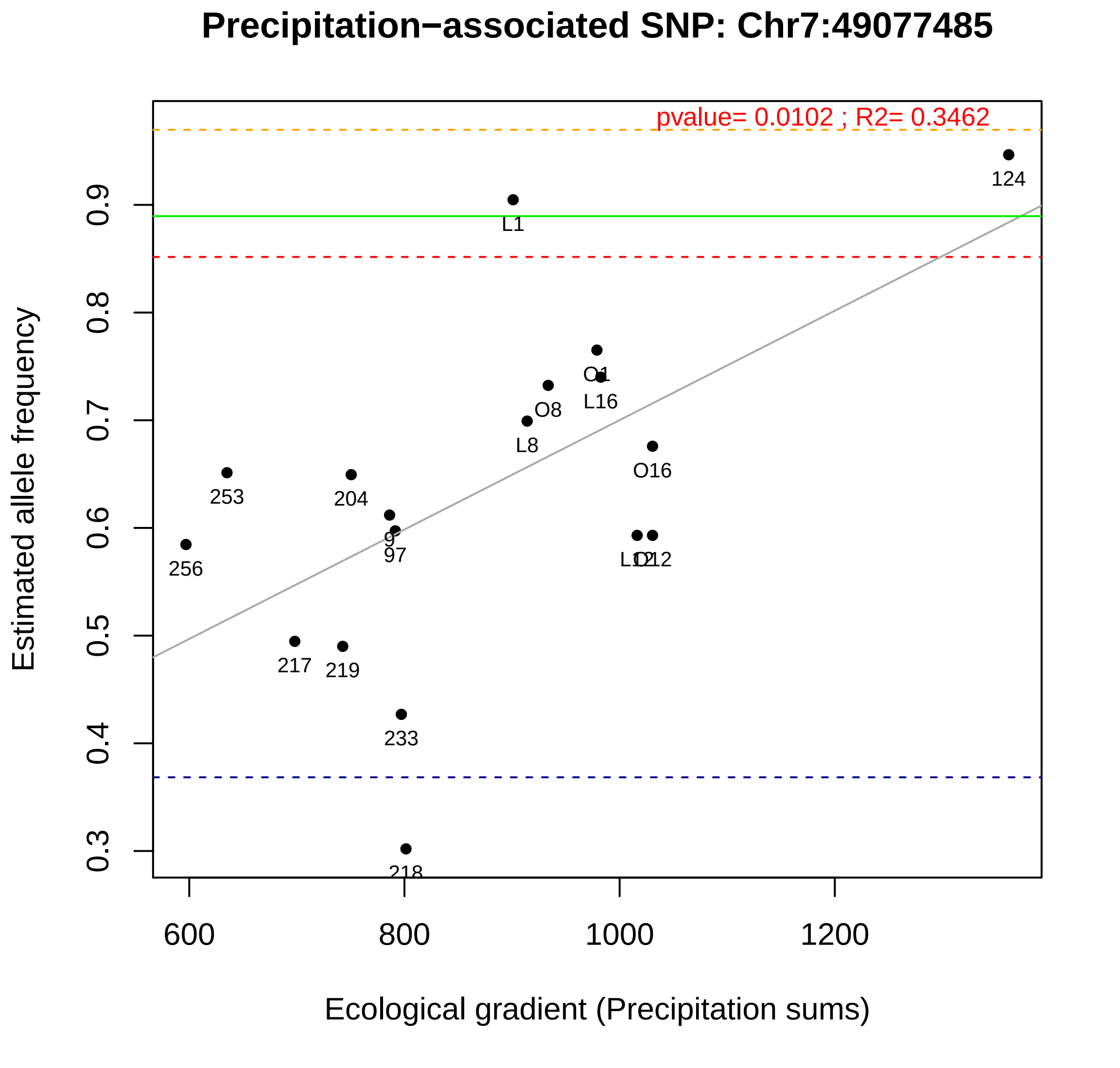

Precipitation−associated SNP: Chr7:49077485
pvalue= 0.0102 ; R2= 0.3462
Estimated allele frequency
0.3
0.4
0.5
0.6
0.7
0.8
0.9
600
800
1000
1200
Ecological gradient (Precipitation sums)
124
L1
O1
L16
O8
L8
O16
253
204
256
9
97
L12
O12
217
219
233
218

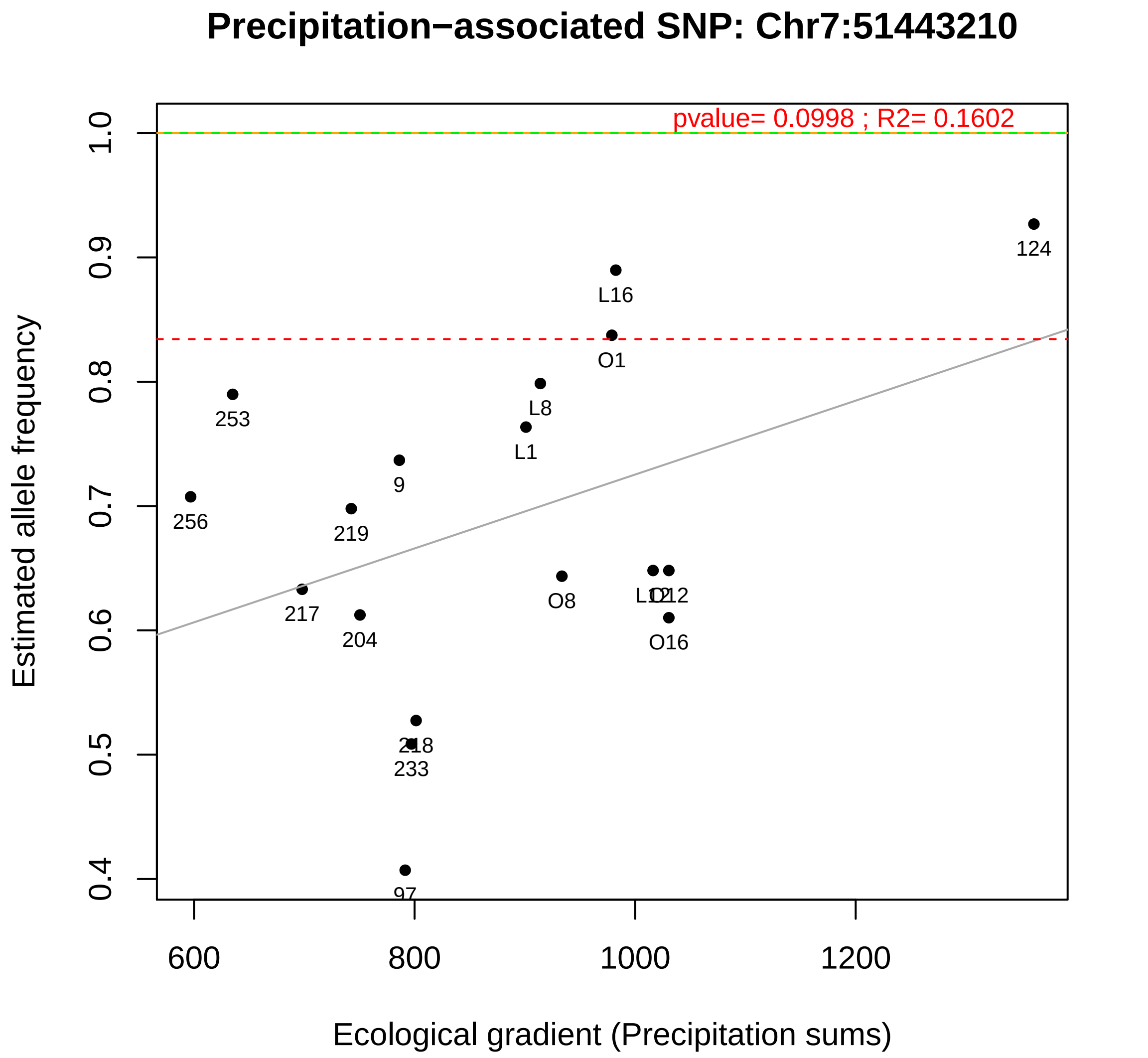

Precipitation−associated SNP: Chr7:51443210
pvalue= 0.0998 ; R2= 0.1602
Estimated allele frequency
Ecological gradient (Precipitation sums)
0.4
0.5
0.6
0.7
0.8
0.9
1.0
600
800
1000
1200
124
L16
O1
L8
L1
253
9
256
219
217
204
O8
L12
O12
O16
218
233
97

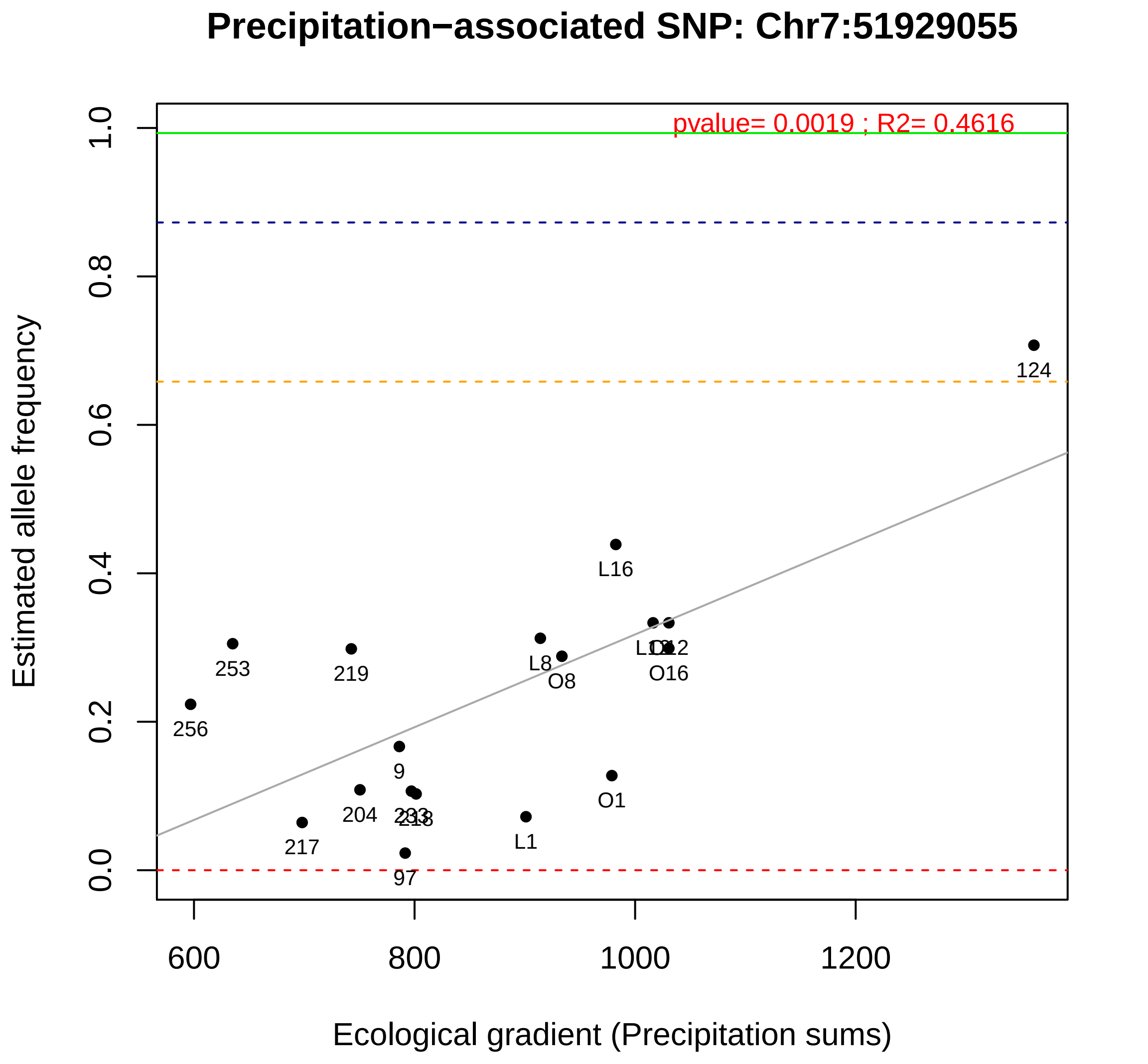

Precipitation−associated SNP: Chr7:51929055
pvalue= 0.0019 ; R2= 0.4616
Estimated allele frequency
0.0
0.2
0.4
0.6
0.8
1.0
600
800
1000
1200
Ecological gradient (Precipitation sums)
124
L16
L12
O12
O16
L8
O8
253
219
256
9
204
233
218
217
97
L1
O1

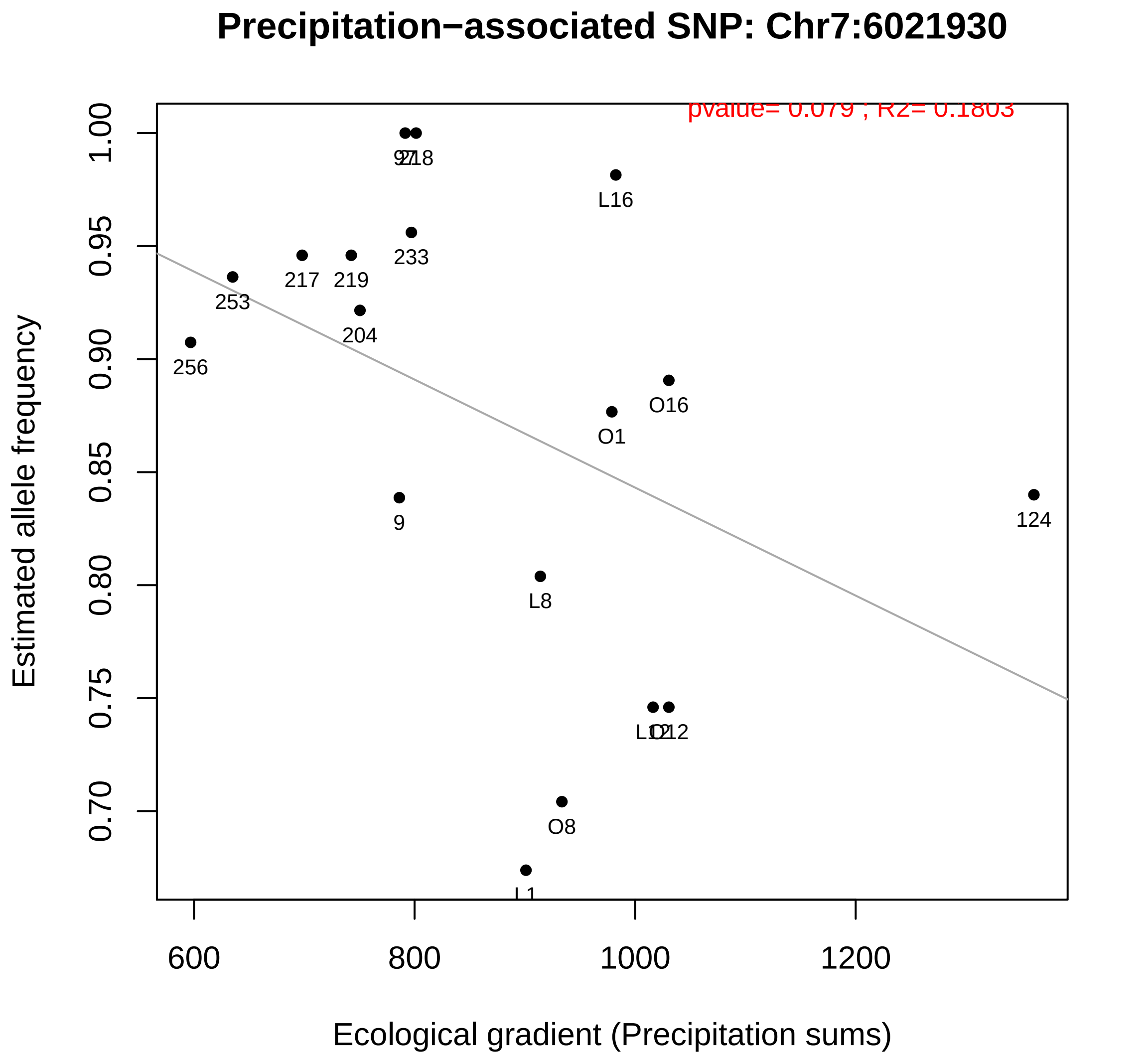

Precipitation−associated SNP: Chr7:6021930
pvalue= 0.079 ; R2= 0.1803
Estimated allele frequency
Ecological gradient (Precipitation sums)
1.00
0.95
0.90
0.85
0.80
0.75
0.70
600
800
1000
1200
218
L16
233
217
219
253
204
256
O16
O1
9
124
L8
O12
O8

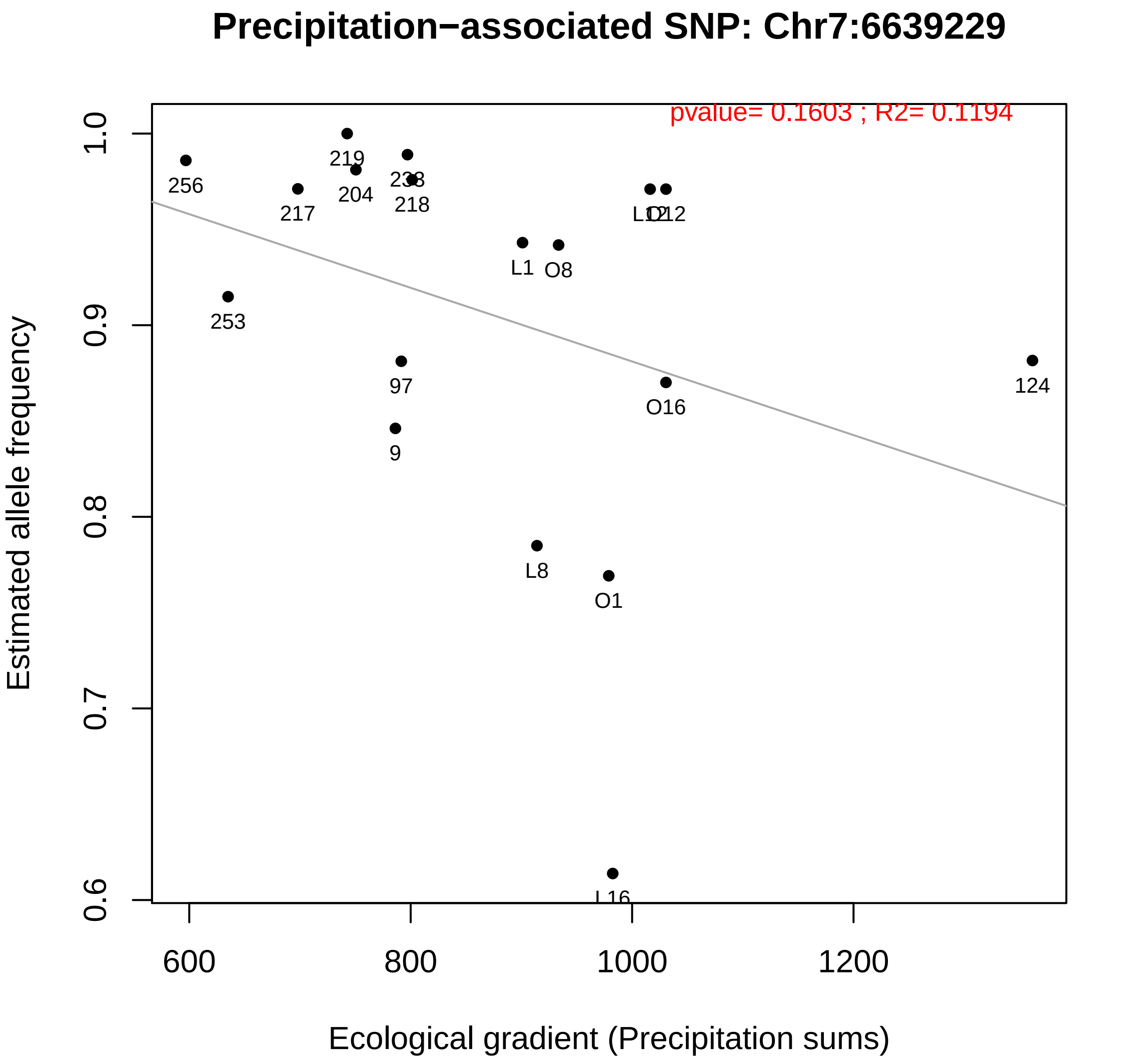
Precipitation−associated SNP: Chr7:6639229
pvalue= 0.1603 ; R2= 0.1194
256
253
217
219
204
238
218
97
9
L1
O8
L12
O12
L8
O1
L16
O16
124
Estimated allele frequency
0.6
0.7
0.8
0.9
1.0
600
800
1000
1200
Ecological gradient (Precipitation sums)

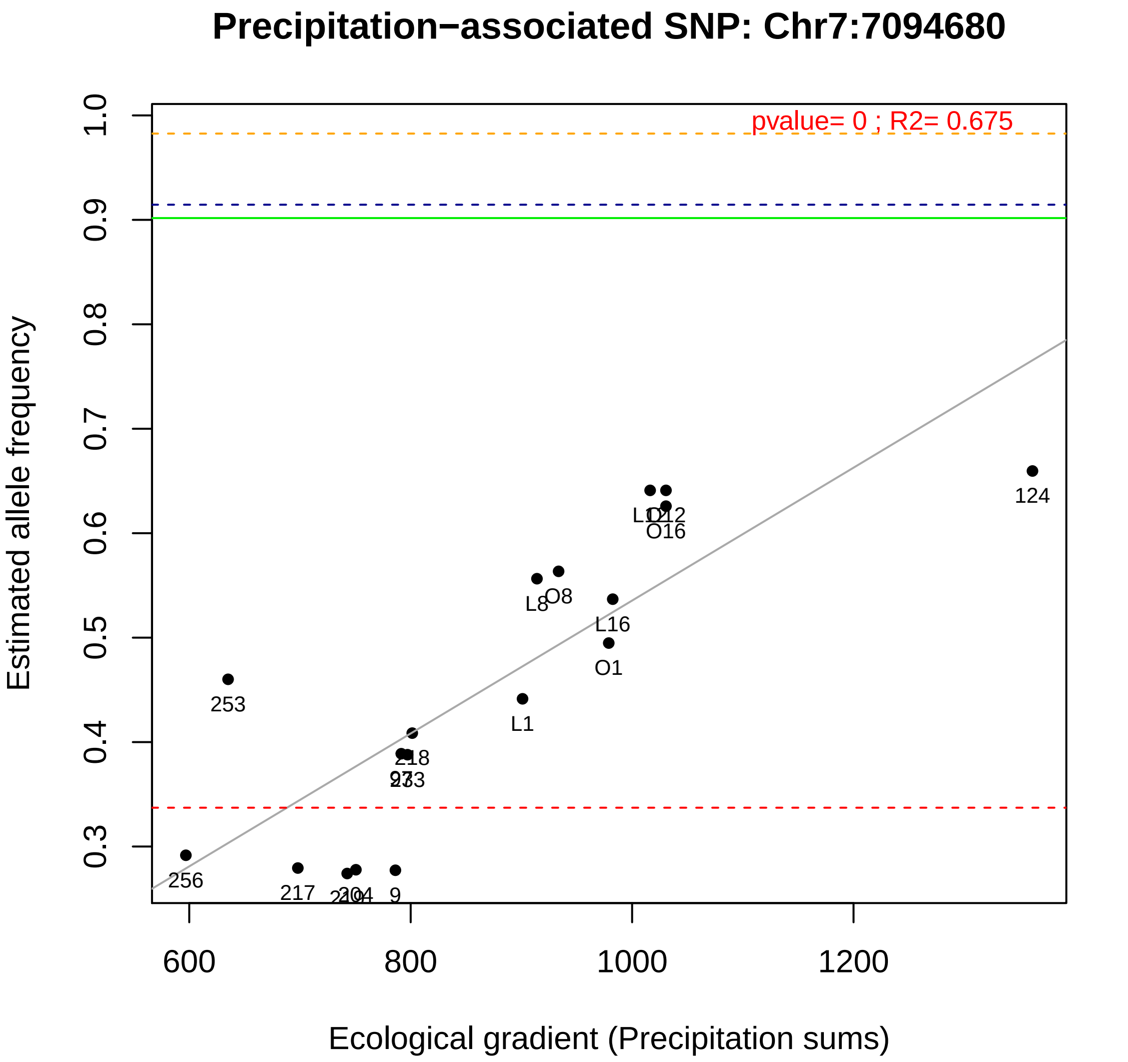
Precipitation−associated SNP: Chr7:7094680
pvalue= 0 ; R2= 0.675
Estimated allele frequency
Ecological gradient (Precipitation sums)
0.3
0.4
0.5
0.6
0.7
0.8
0.9
1.0
600
800
1000
1200
124
L12
O12
O16
O8
L8
L16
O1
L1
253
218
97
233
256
217
204
9

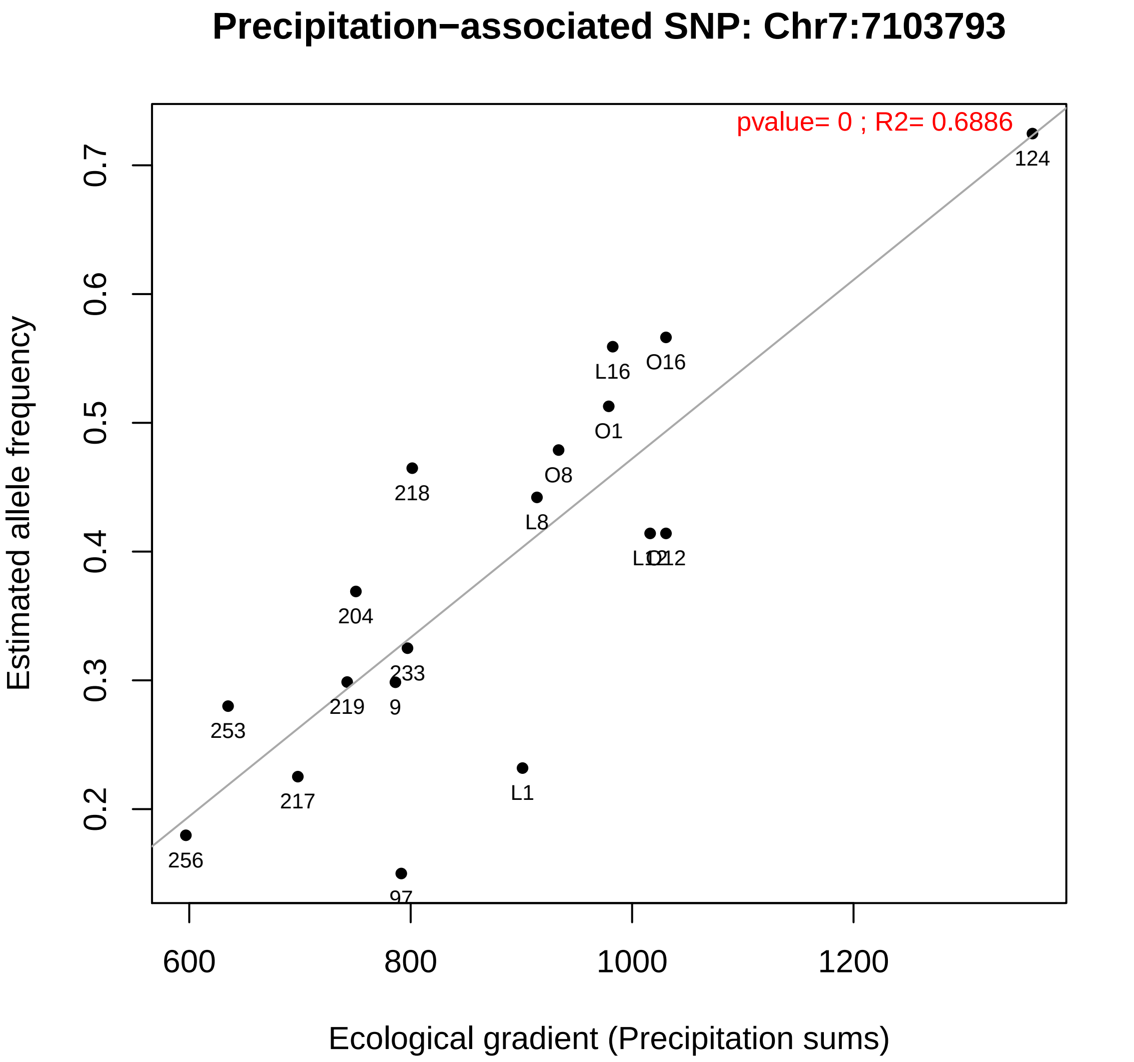

Precipitation−associated SNP: Chr7:7103793
pvalue= 0 ; R2= 0.6886
124
O16
L16
O1
O8
218
L8
L12
O12
204
233
219
9
253
217
L1
256
97
0.2
0.3
0.4
0.5
0.6
0.7
600
800
1000
1200
Estimated allele frequency
Ecological gradient (Precipitation sums)

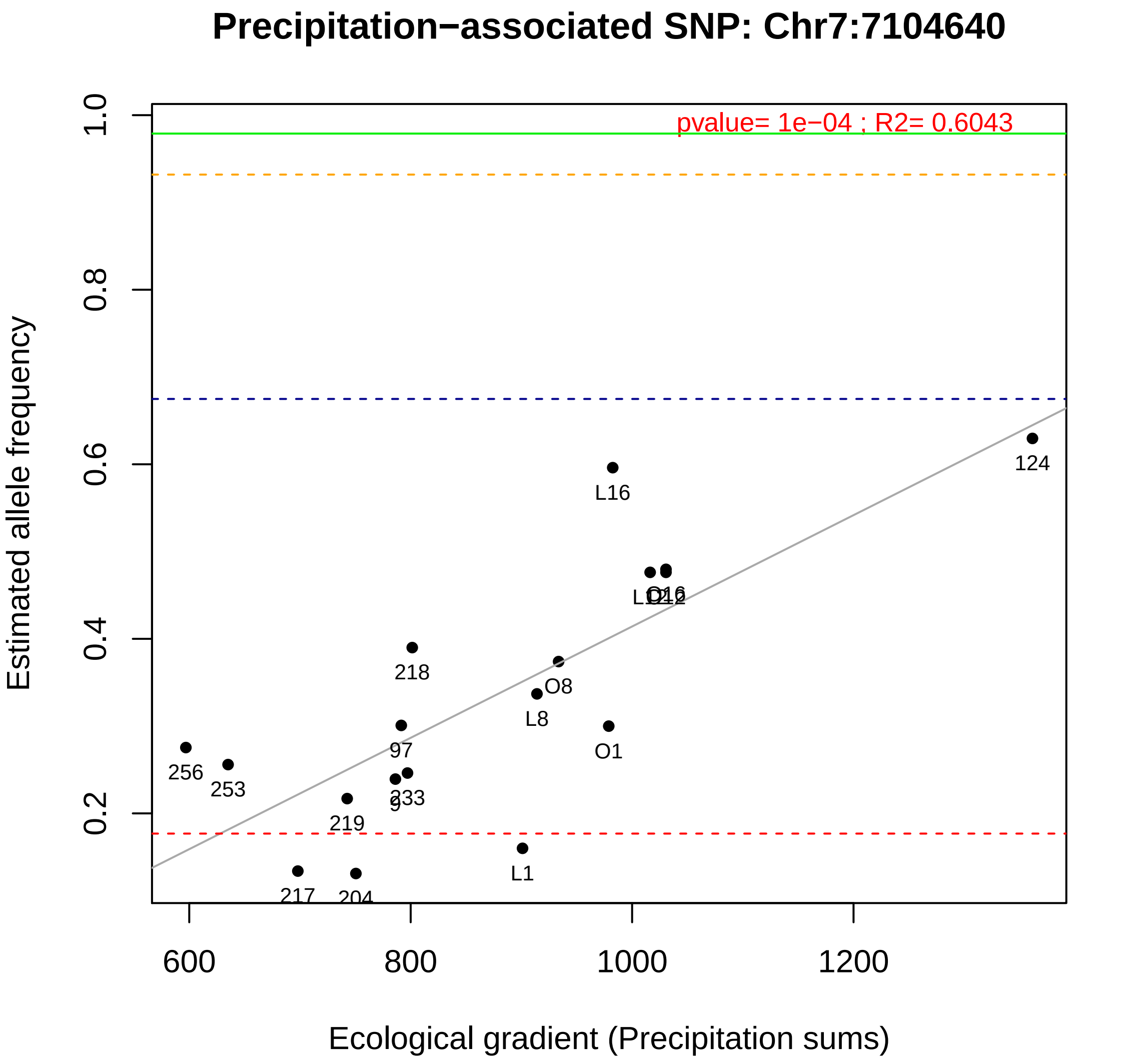

Precipitation−associated SNP: Chr7:7104640
pvalue= 1e−04 ; R2= 0.6043
Estimated allele frequency
Ecological gradient (Precipitation sums)
0.2
0.4
0.6
0.8
1.0
600
800
1000
1200
124
L16
218
O8
L8
O1
97
256
253
219
217
204
L1

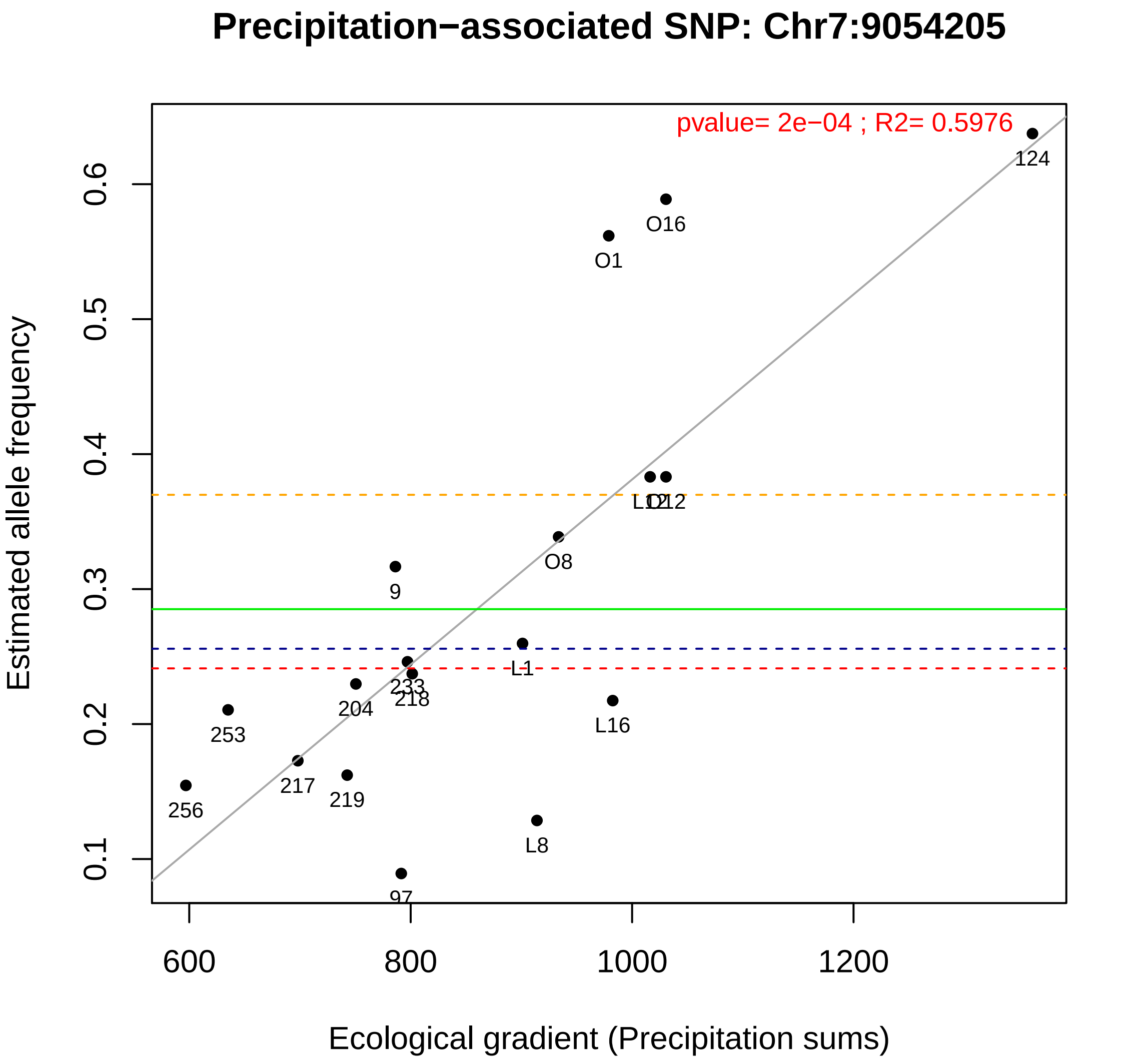

Precipitation−associated SNP: Chr7:9054205
pvalue= 2e−04 ; R2= 0.5976
Estimated allele frequency
Ecological gradient (Precipitation sums)
0.1
0.2
0.3
0.4
0.5
0.6
600
800
1000
1200
124
O16
O1
O8
9
L1
233
218
204
253
L16
217
219
256
L8
97

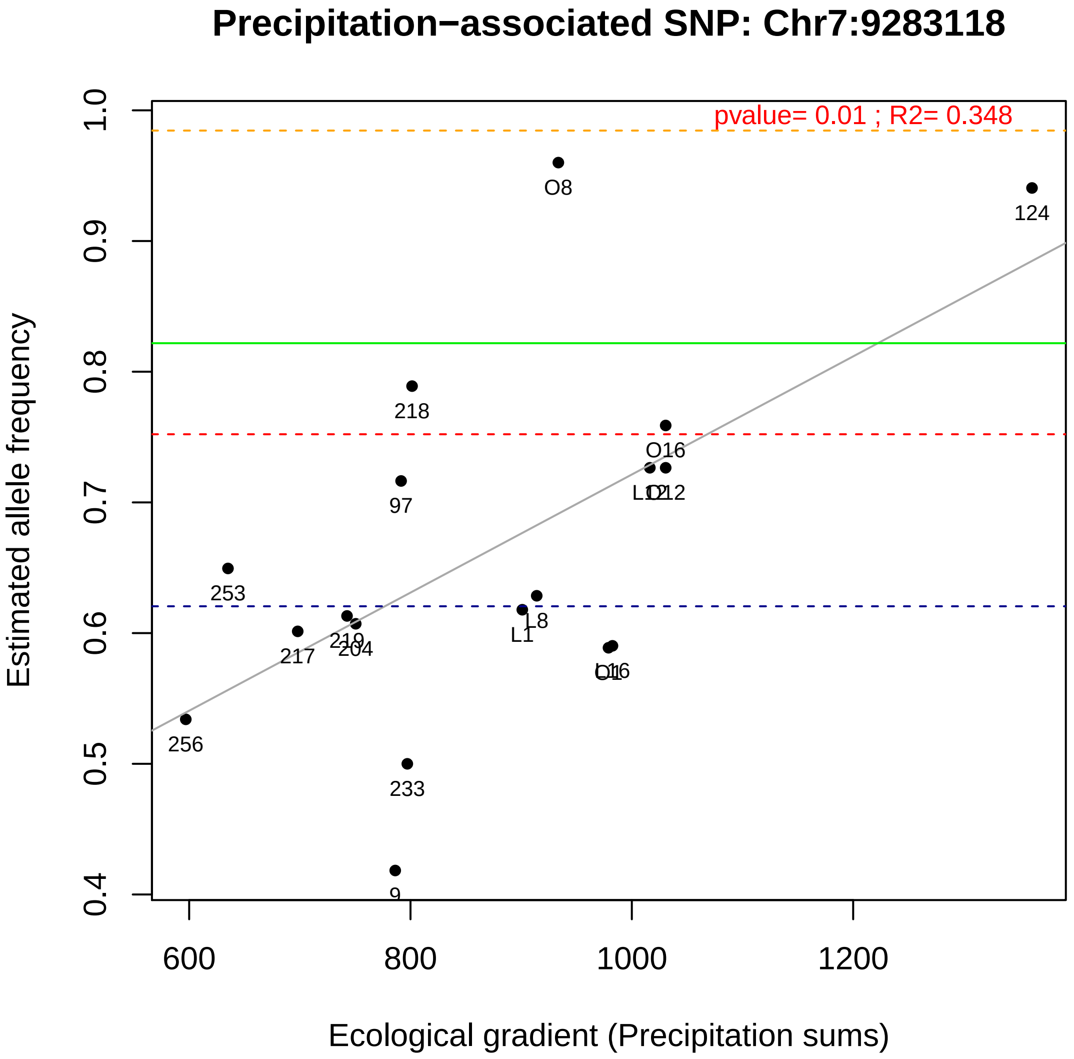
Precipitation−associated SNP: Chr7:9283118
pvalue= 0.01 ; R2= 0.348
Estimated allele frequency
Ecological gradient (Precipitation sums)
0.4
0.5
0.6
0.7
0.8
0.9
1.0
600
800
1000
1200
O8
124
218
O16
L12
O12
97
253
219
204
217
L8
L1
L16
O1
256
233
9

**Precipitation−associated SNP: Chr8:11501602**

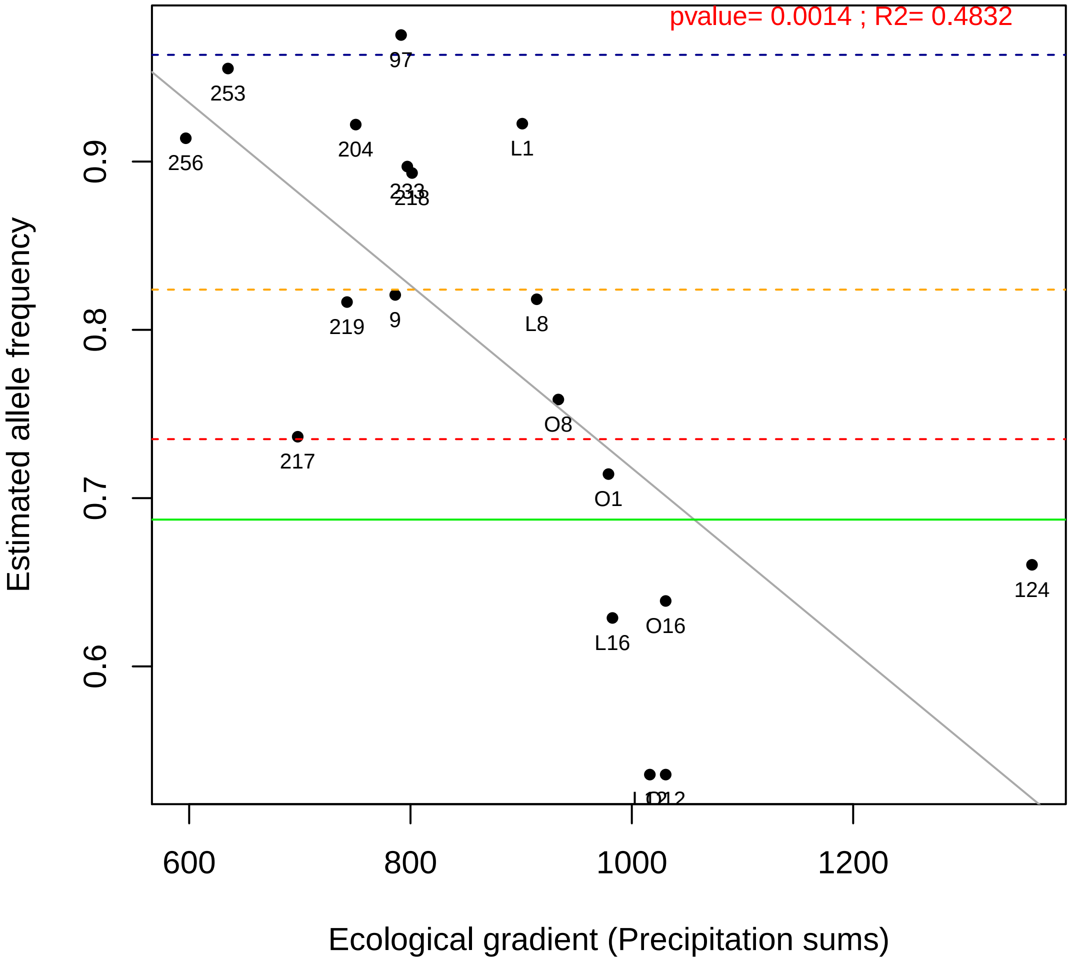

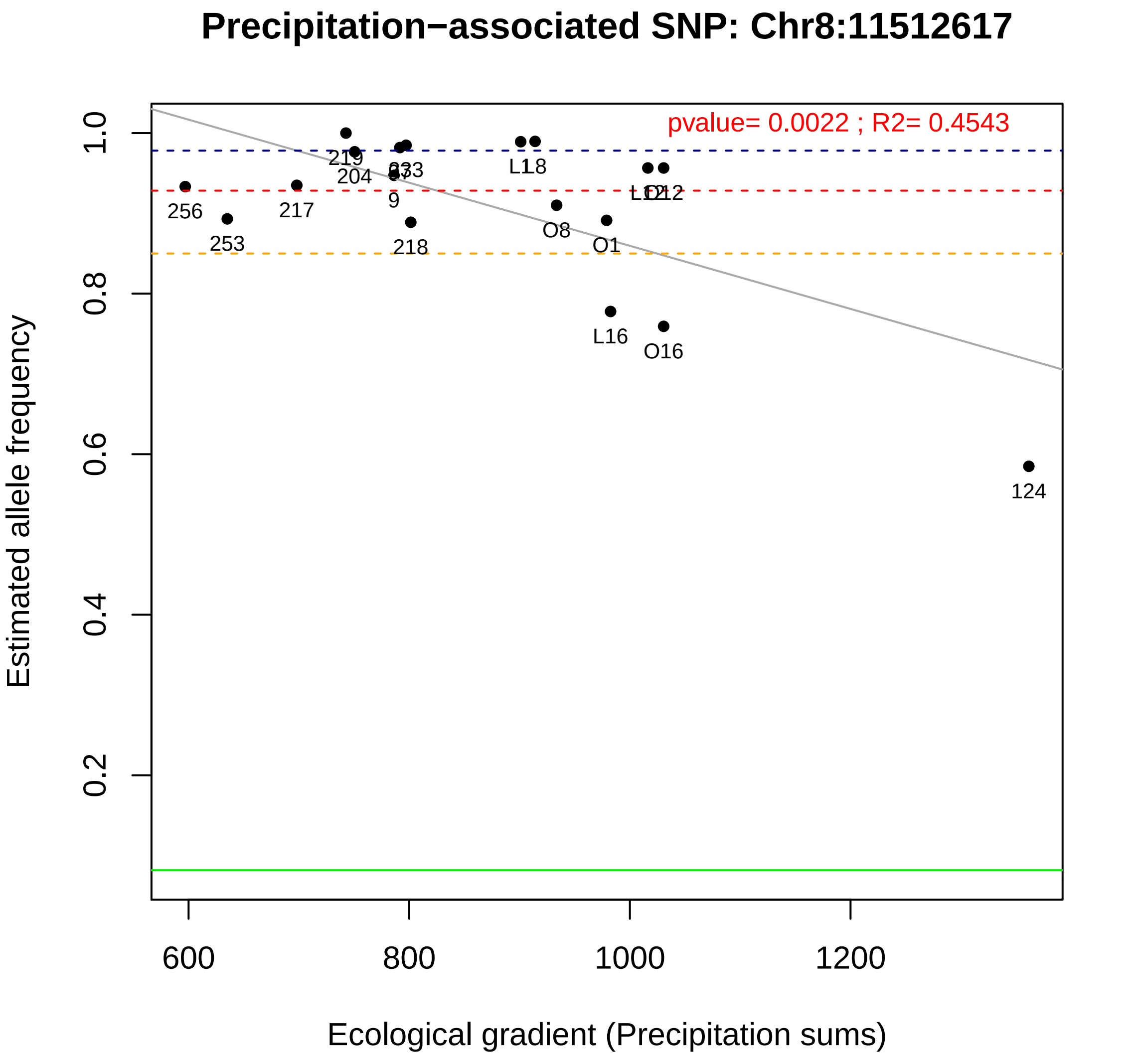

Precipitation−associated SNP: Chr8:11512617
pvalue= 0.0022 ; R2= 0.4543
Estimated allele frequency
Ecological gradient (Precipitation sums)
0.2
0.4
0.6
0.8
1.0
600
800
1000
1200
256
253
217
219
204
218
O8
O1
L16
O16
124

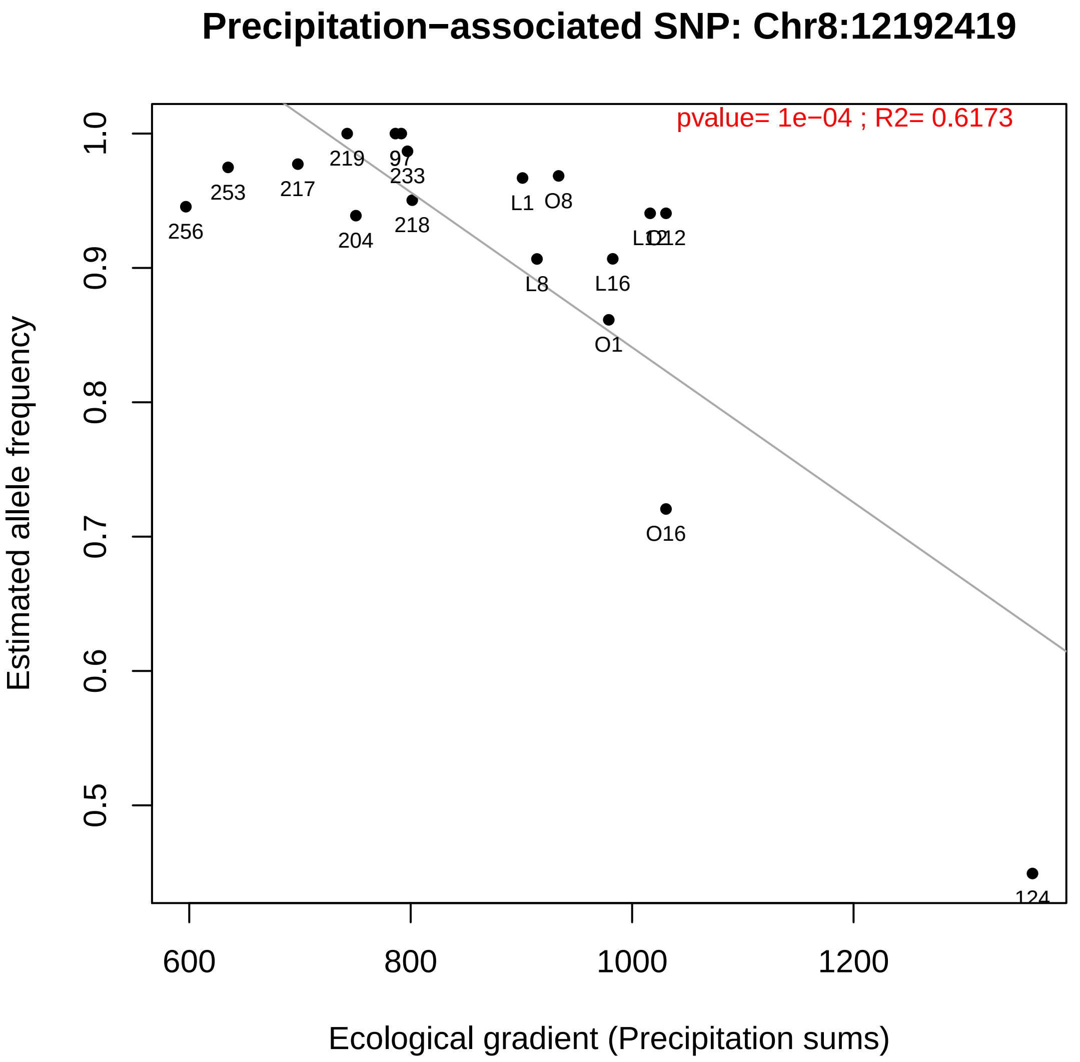
Precipitation−associated SNP: Chr8:12192419
pvalue= 1e−04 ; R2= 0.6173
Estimated allele frequency
Ecological gradient (Precipitation sums)
1.0
0.9
0.8
0.7
0.6
0.5
600
800
1000
1200
219
97
233
253
217
256
204
218
L1
O8
L8
L16
O1
O16
124

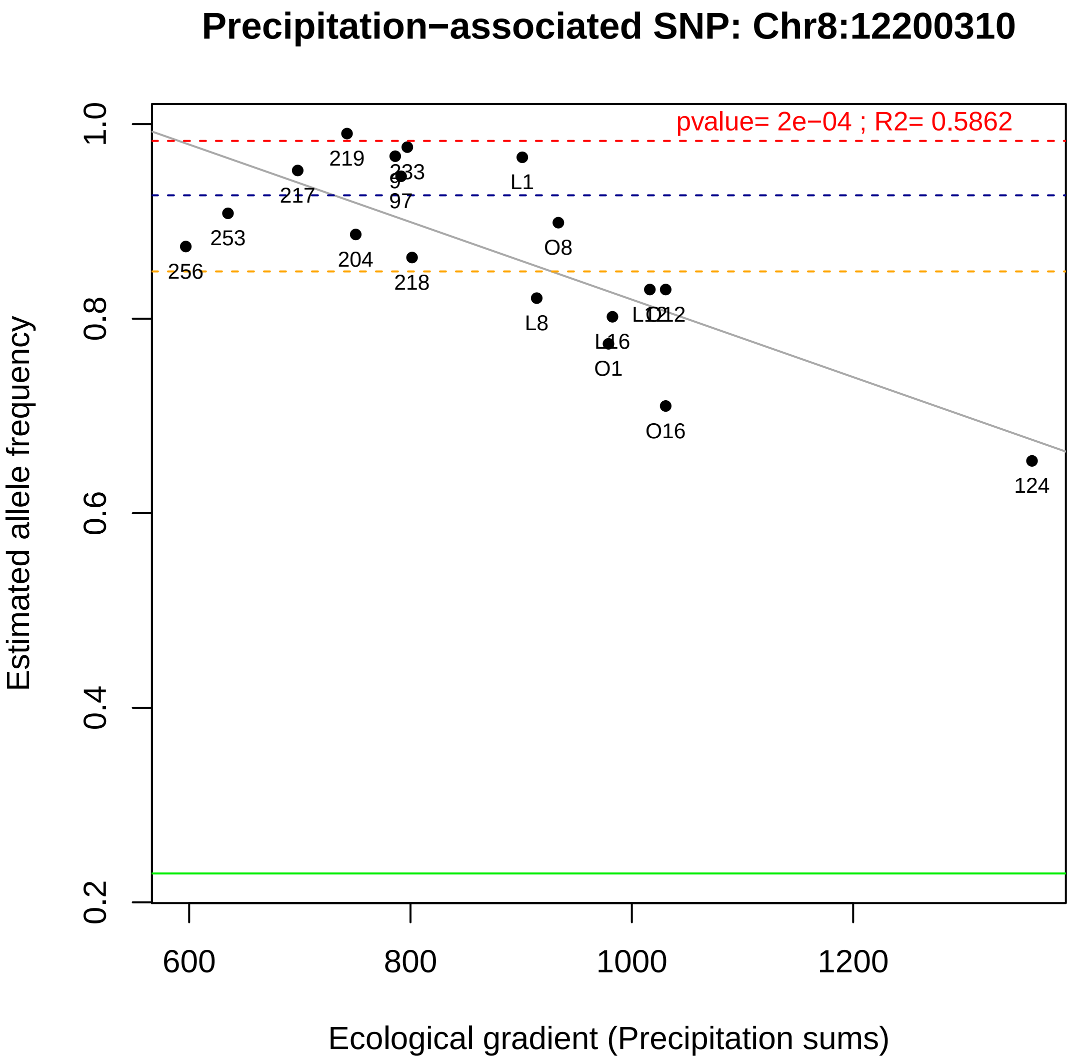

Precipitation−associated SNP: Chr8:12200310
pvalue= 2e−04 ; R2= 0.5862
Estimated allele frequency
Ecological gradient (Precipitation sums)
1.0
0.8
0.6
0.4
0.2
600
800
1000
1200
219
233
9
97
217
253
256
204
218
L1
O8
L8
L12
O12
L16
O1
O16
124

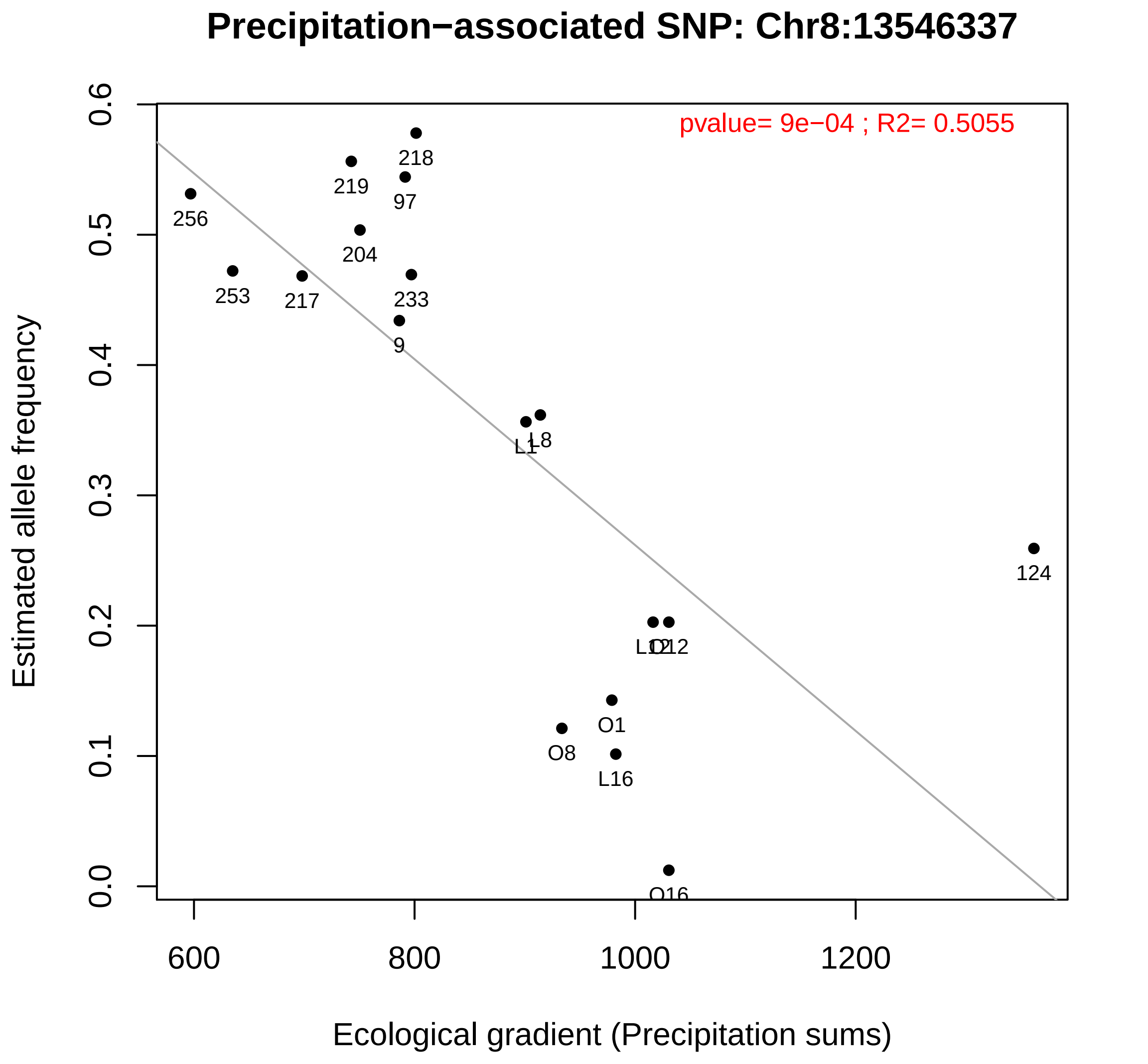

Precipitation−associated SNP: Chr8:13546337
pvalue= 9e−04 ; R2= 0.5055
Estimated allele frequency
Ecological gradient (Precipitation sums)
0.0
0.1
0.2
0.3
0.4
0.5
0.6
600
800
1000
1200
218
219
97
256
204
253
217
233
9
L1
L8
124
L12
O12
O1
O8
L16
O16

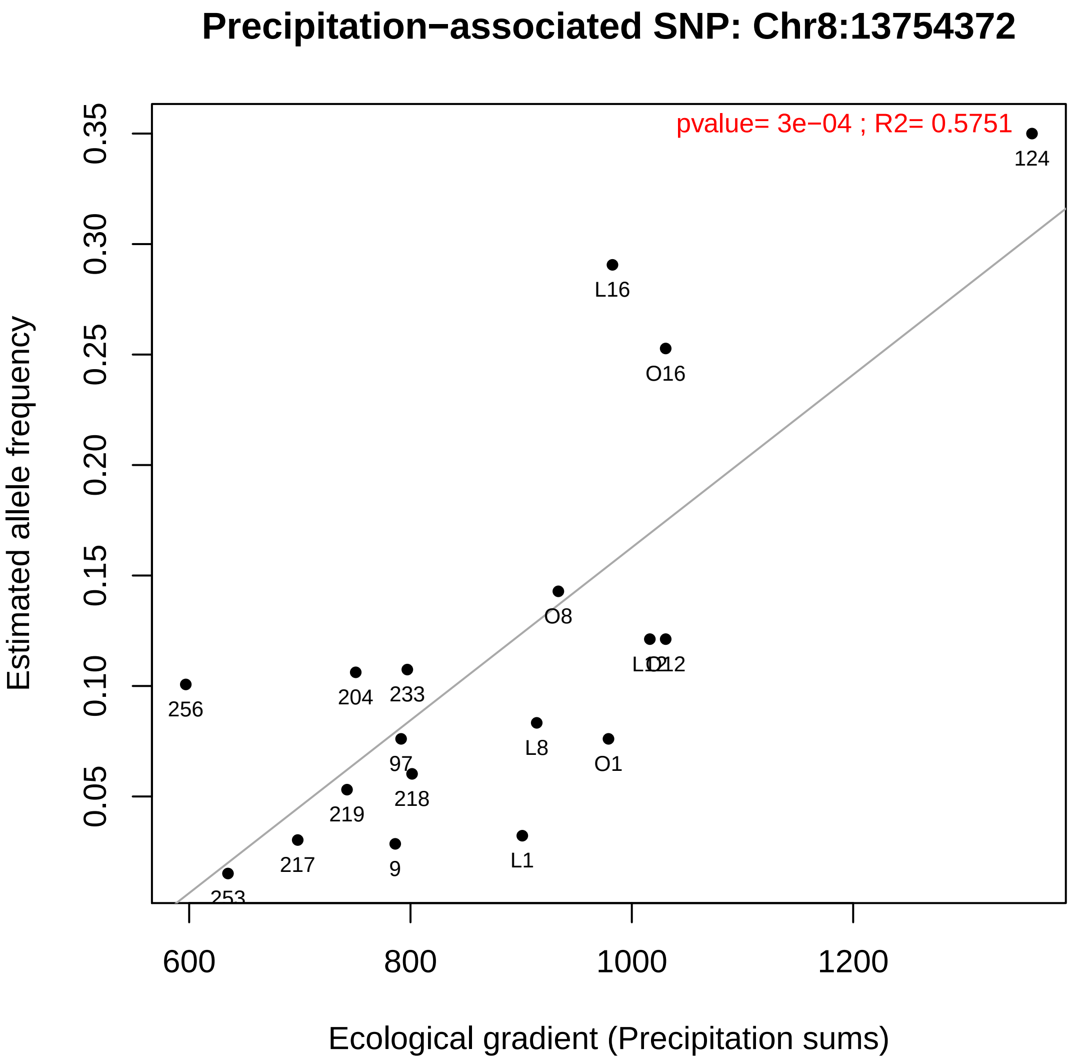

Precipitation−associated SNP: Chr8:13754372
pvalue= 3e−04 ; R2= 0.5751
Estimated allele frequency
0.05
0.10
0.15
0.20
0.25
0.30
0.35
600
800
1000
1200
Ecological gradient (Precipitation sums)
124
L16
O16
O8
L12
O12
256
204
233
97
218
L8
O1
219
217
9
L1
253

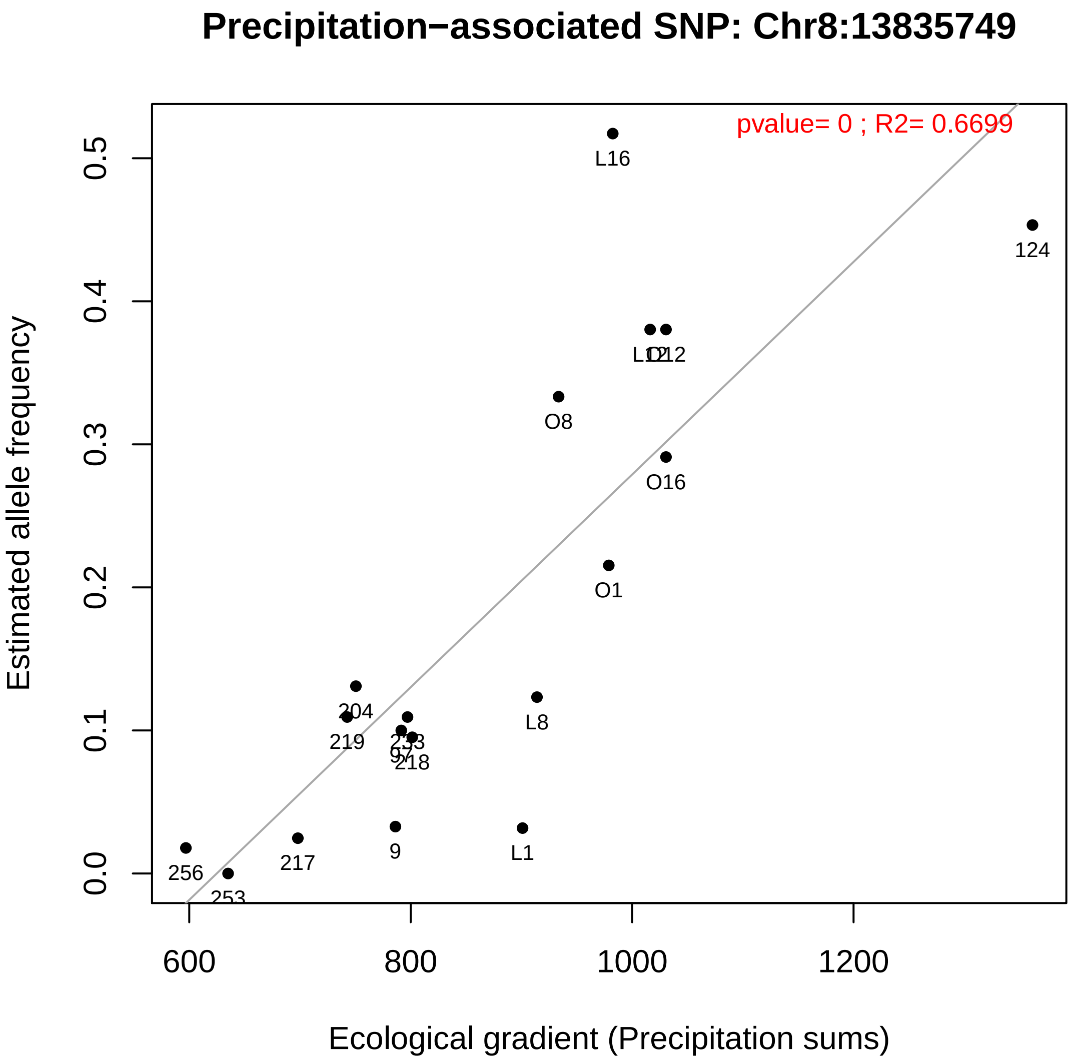

Precipitation−associated SNP: Chr8:13835749
pvalue= 0 ; R2= 0.6699
Estimated allele frequency
Ecological gradient (Precipitation sums)
0.0
0.1
0.2
0.3
0.4
0.5
600
800
1000
1200
L16
124
O8
O16
O1
L8
204
219
218
9
L1
256
253
217

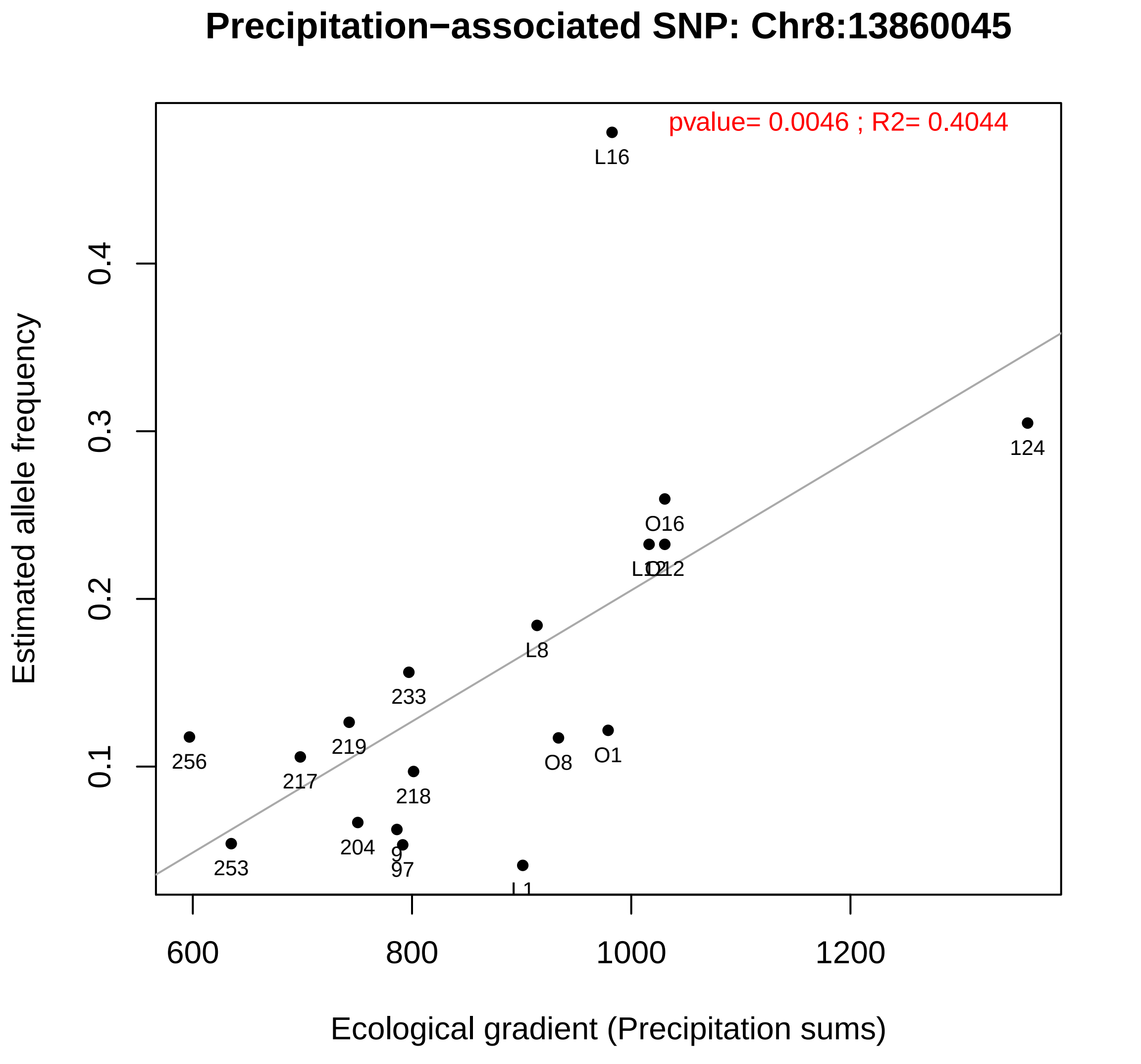

Precipitation−associated SNP: Chr8:13860045
pvalue= 0.0046 ; R2= 0.4044
Estimated allele frequency
Ecological gradient (Precipitation sums)
0.1
0.2
0.3
0.4
600
800
1000
1200
L16
124
O16
L10
O12
L8
233
219
256
217
218
O8
O1
204
253
9
97
L1

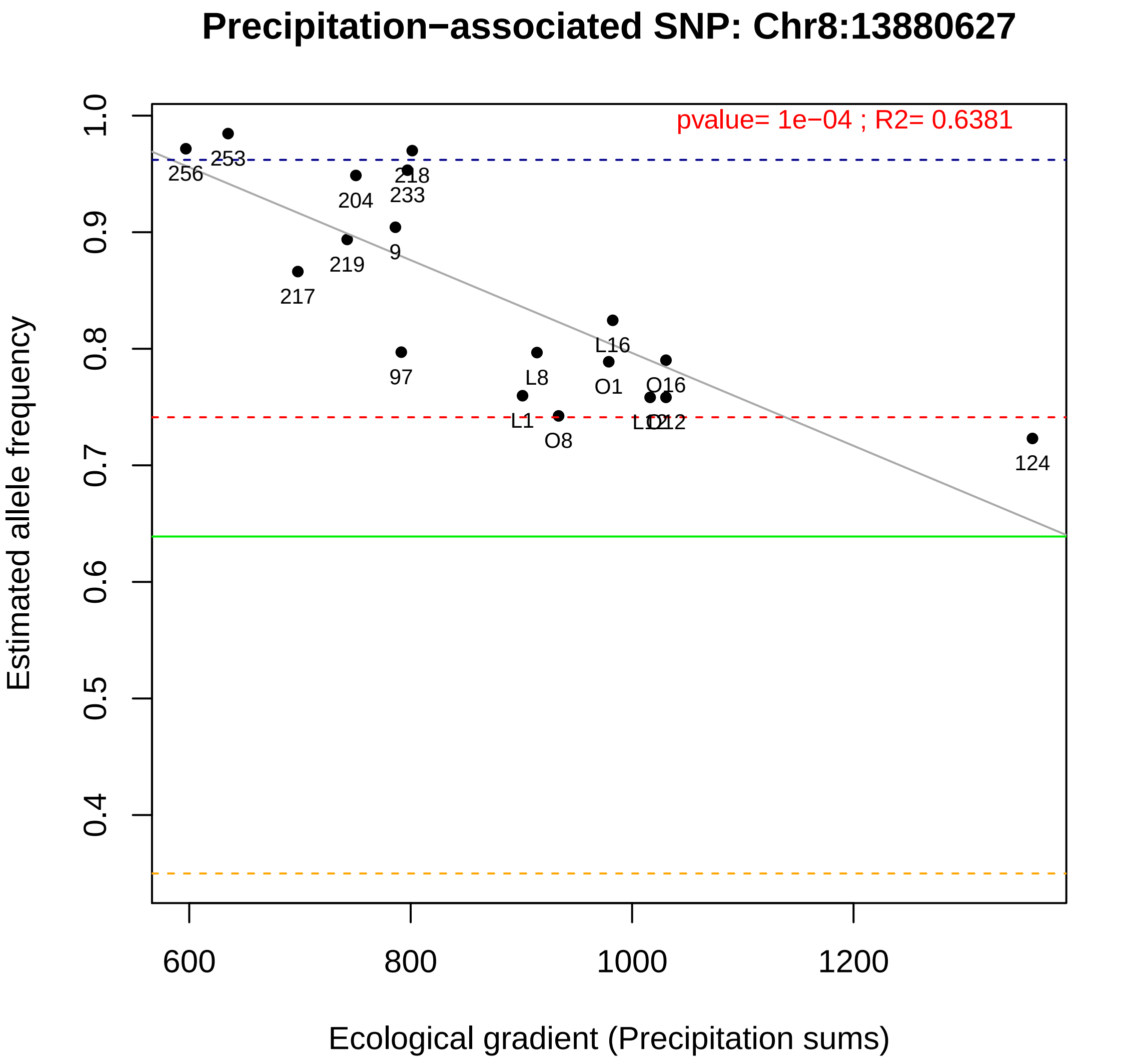
Precipitation−associated SNP: Chr8:13880627
pvalue= 1e−04 ; R2= 0.6381
Estimated allele frequency
Ecological gradient (Precipitation sums)
1.0
0.9
0.8
0.7
0.6
0.5
0.4
600
800
1000
1200
256
253
218
233
204
219
9
217
97
L8
L16
O1
O16
L1
O8
124

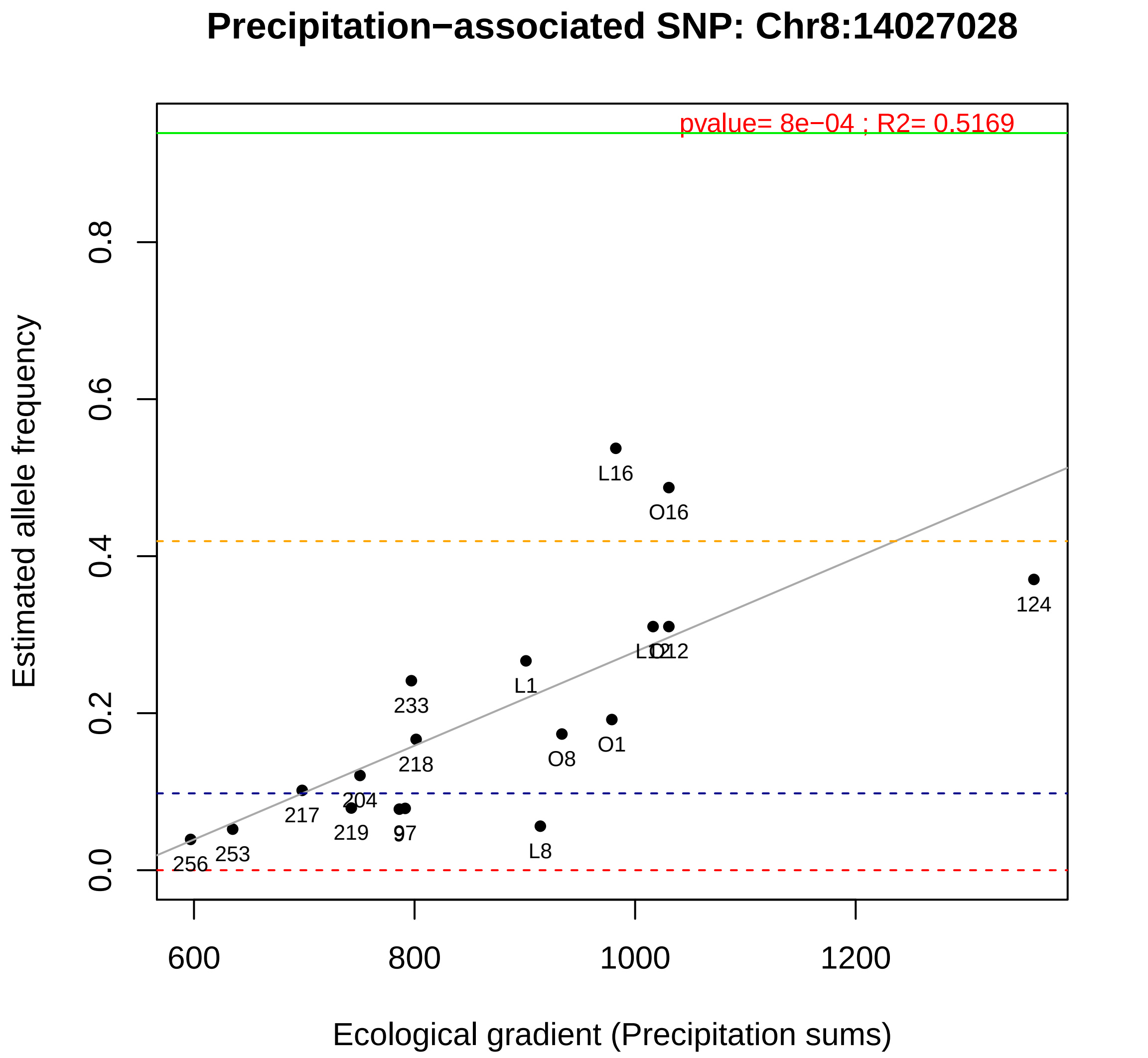

Precipitation−associated SNP: Chr8:14027028
pvalue= 8e−04 ; R2= 0.5169
Estimated allele frequency
0.0
0.2
0.4
0.6
0.8
600
800
1000
1200
Ecological gradient (Precipitation sums)
L16
O16
124
L12
O12
L1
233
218
O8
O1
204
217
219
97
L8
256
253

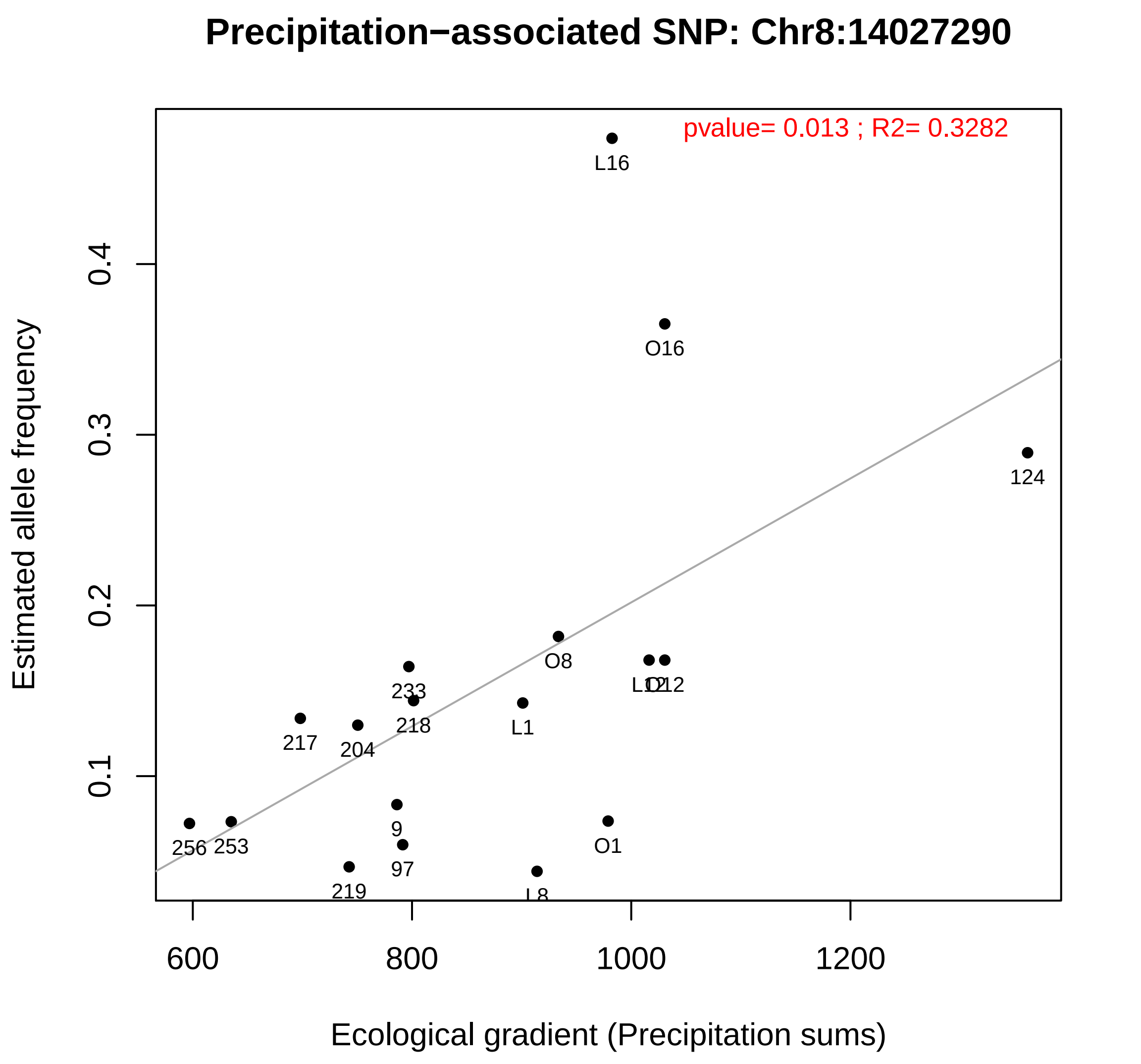

Precipitation−associated SNP: Chr8:14027290
pvalue= 0.013 ; R2= 0.3282
L16
O16
124
O8
L12
O12
233
218
L1
217
204
9
O1
256
253
97
219
L8
0.1
0.2
0.3
0.4
600
800
1000
1200
Estimated allele frequency
Ecological gradient (Precipitation sums)

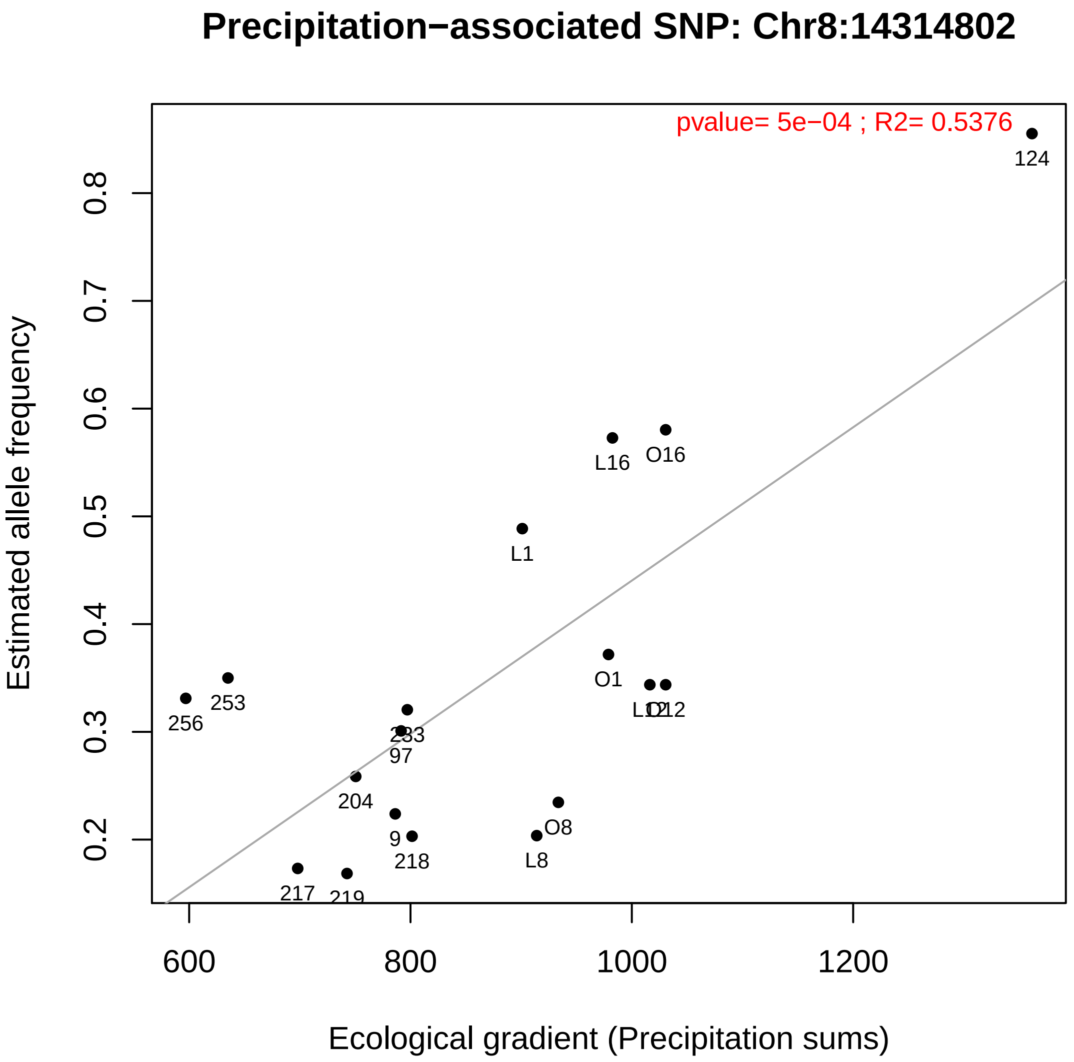

Precipitation−associated SNP: Chr8:14314802
pvalue= 5e−04 ; R2= 0.5376
Estimated allele frequency
0.2
0.3
0.4
0.5
0.6
0.7
0.8
600
800
1000
1200
Ecological gradient (Precipitation sums)
124
L16
O16
L1
O1
L12
O12
256
253
233
97
204
9
218
L8
O8
217
219

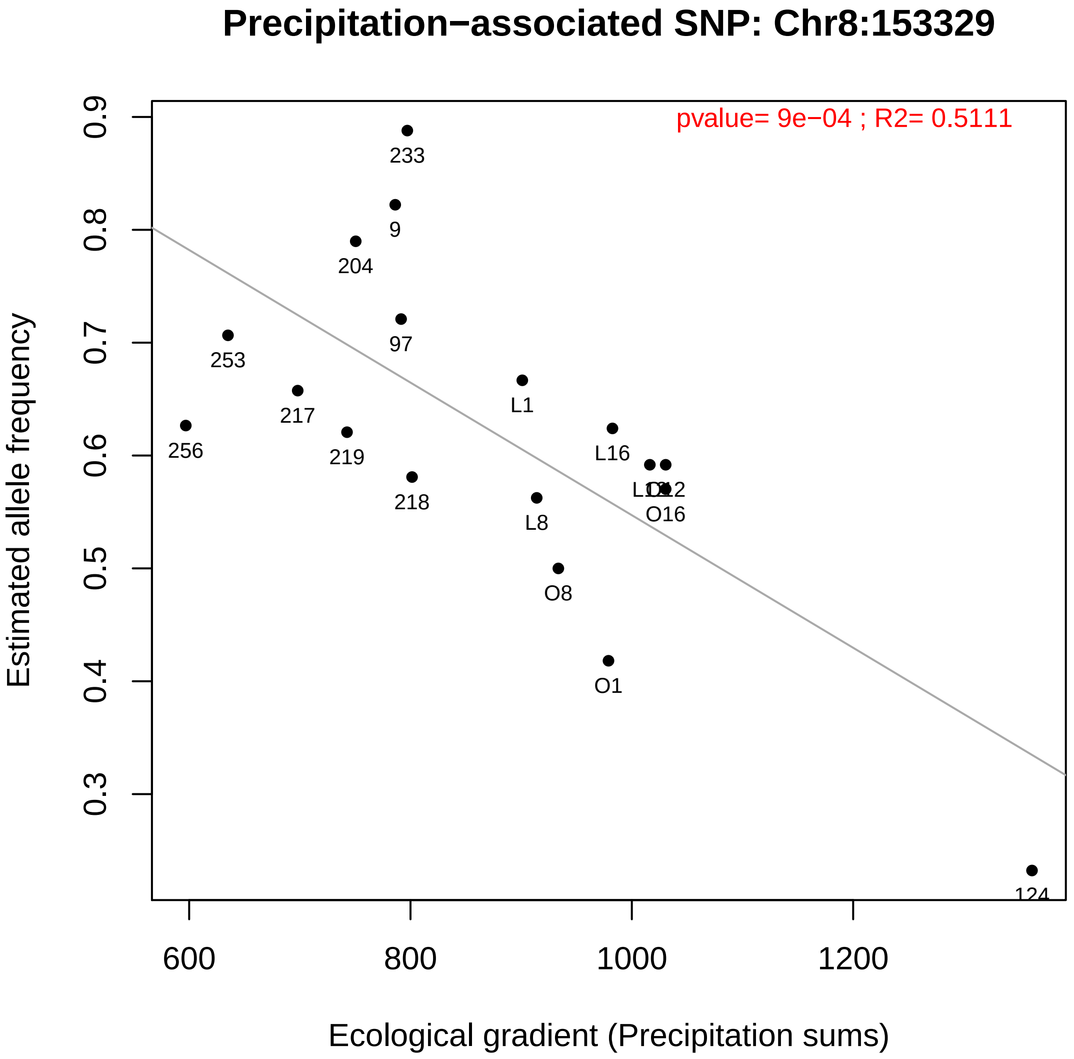
Precipitation−associated SNP: Chr8:153329
pvalue= 9e−04 ; R2= 0.5111
Estimated allele frequency
0.3
0.4
0.5
0.6
0.7
0.8
0.9
600
800
1000
1200
Ecological gradient (Precipitation sums)
233
9
204
97
253
217
256
219
218
L1
L16
L8
O16
O8
O1
124

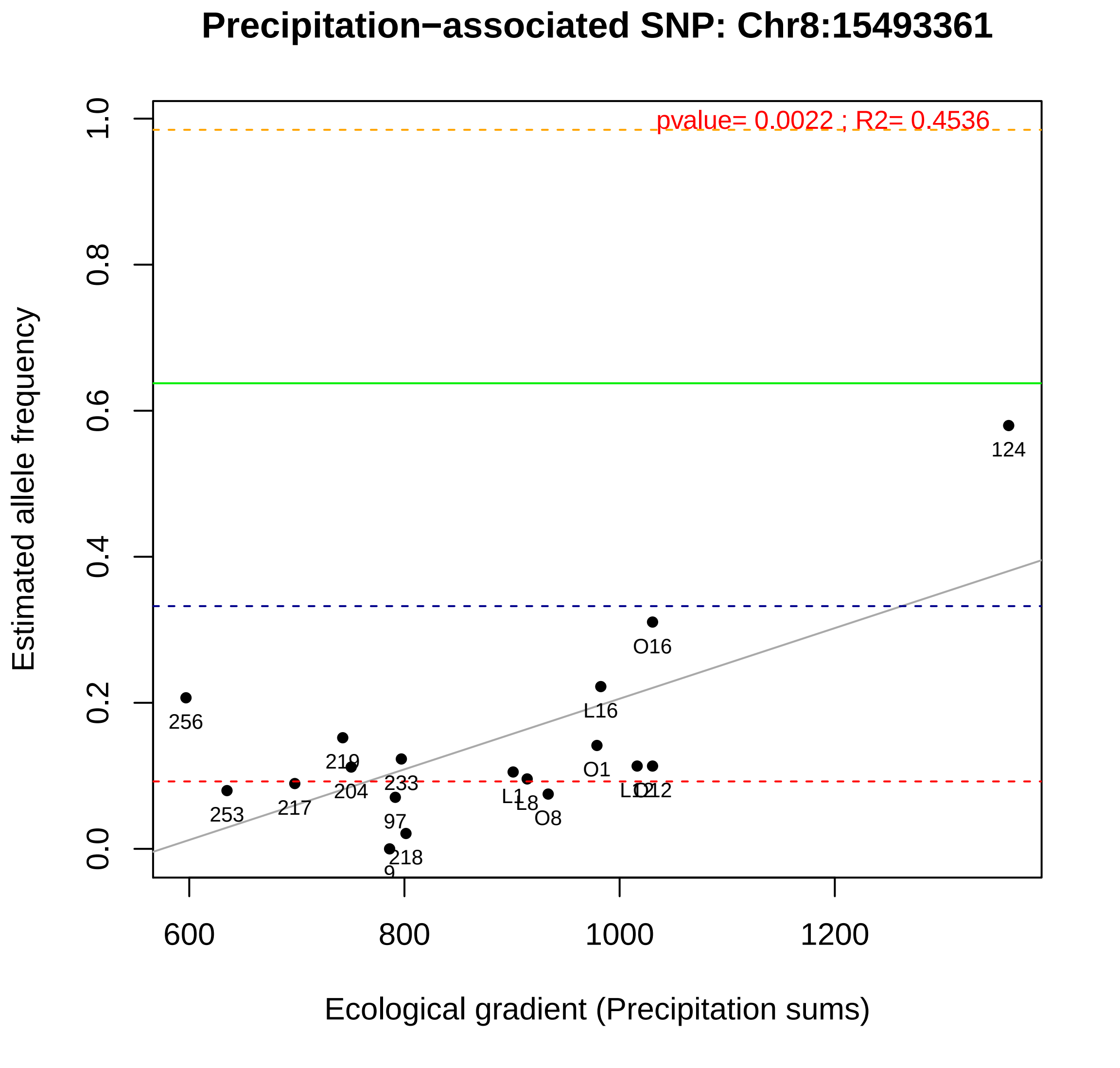
Precipitation−associated SNP: Chr8:15493361
pvalue= 0.0022 ; R2= 0.4536
Estimated allele frequency
0.0
0.2
0.4
0.6
0.8
1.0
600
800
1000
1200
Ecological gradient (Precipitation sums)
124
O16
L16
256
O1
219
204
233
253
217
97
L1
L8
O8
218
9

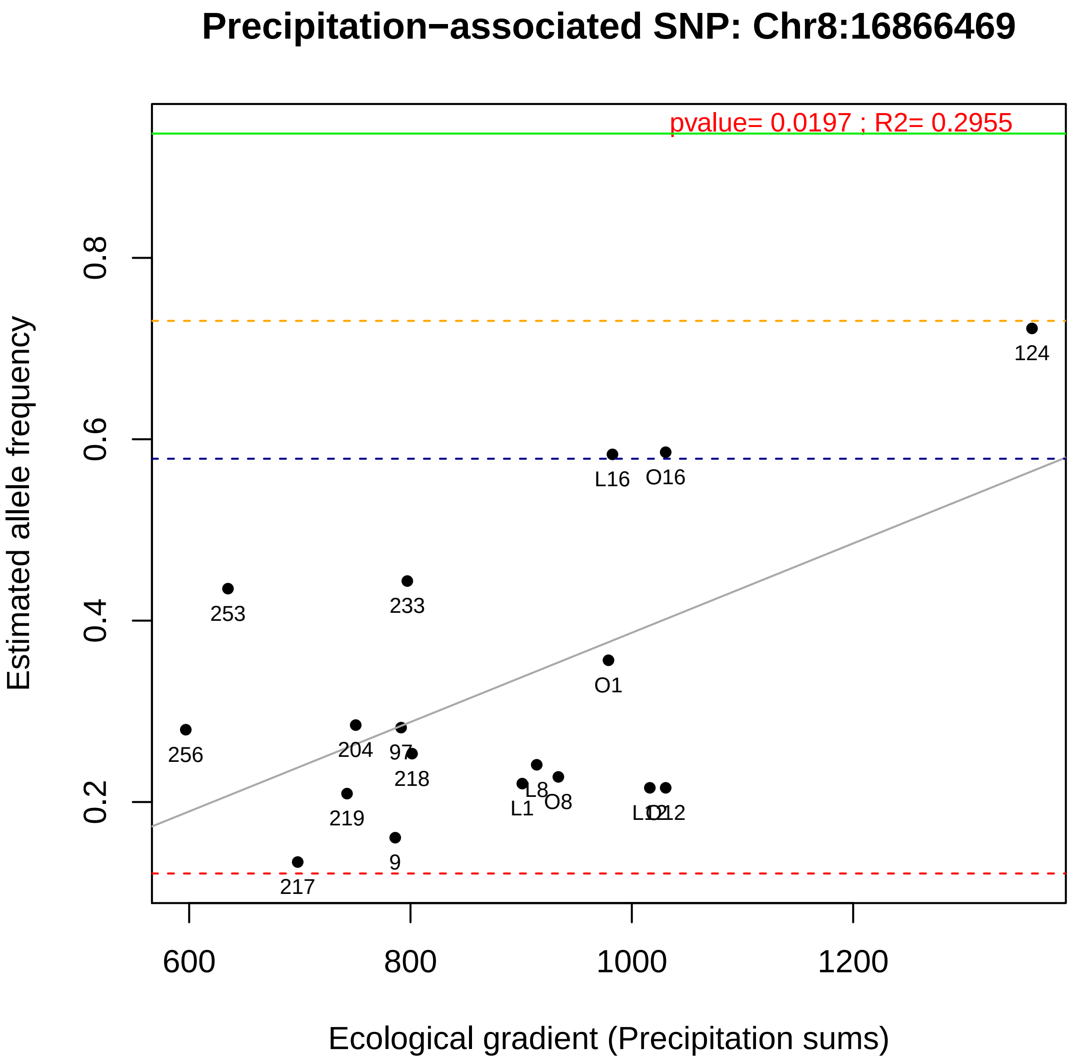
Precipitation−associated SNP: Chr8:16866469
pvalue= 0.0197 ; R2= 0.2955
Estimated allele frequency
0.2
0.4
0.6
0.8
600
800
1000
1200
Ecological gradient (Precipitation sums)
124
L16
O16
253
233
O1
256
204
97
218
219
L8
O8
L1
L12
O12
9
217

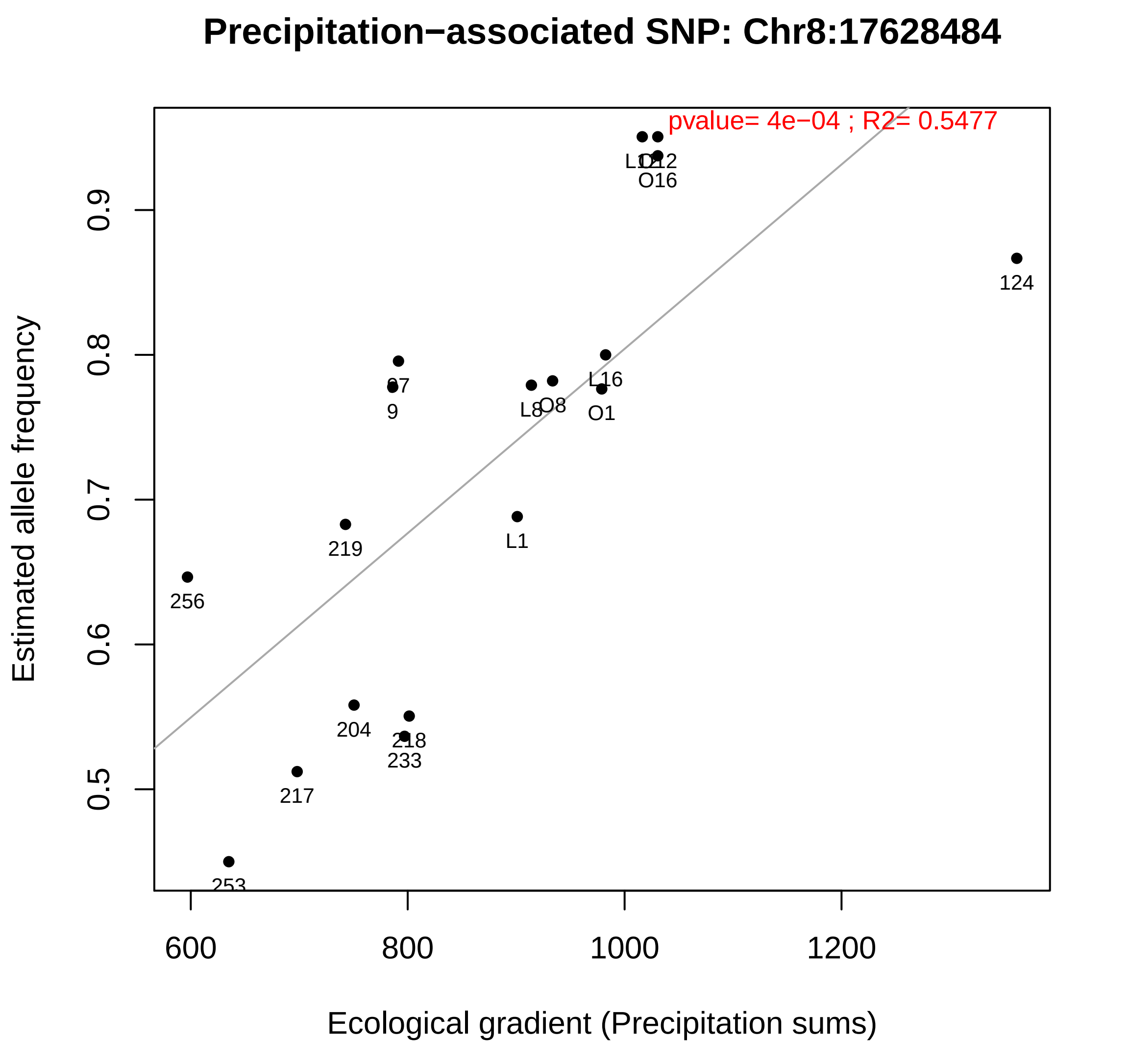

Precipitation−associated SNP: Chr8:17628484
pvalue= 4e−04 ; R2= 0.5477
Estimated allele frequency
Ecological gradient (Precipitation sums)
0.5
0.6
0.7
0.8
0.9
600
800
1000
1200
O16
124
9
L8
O8
L16
O1
219
L1
256
204
218
233
217

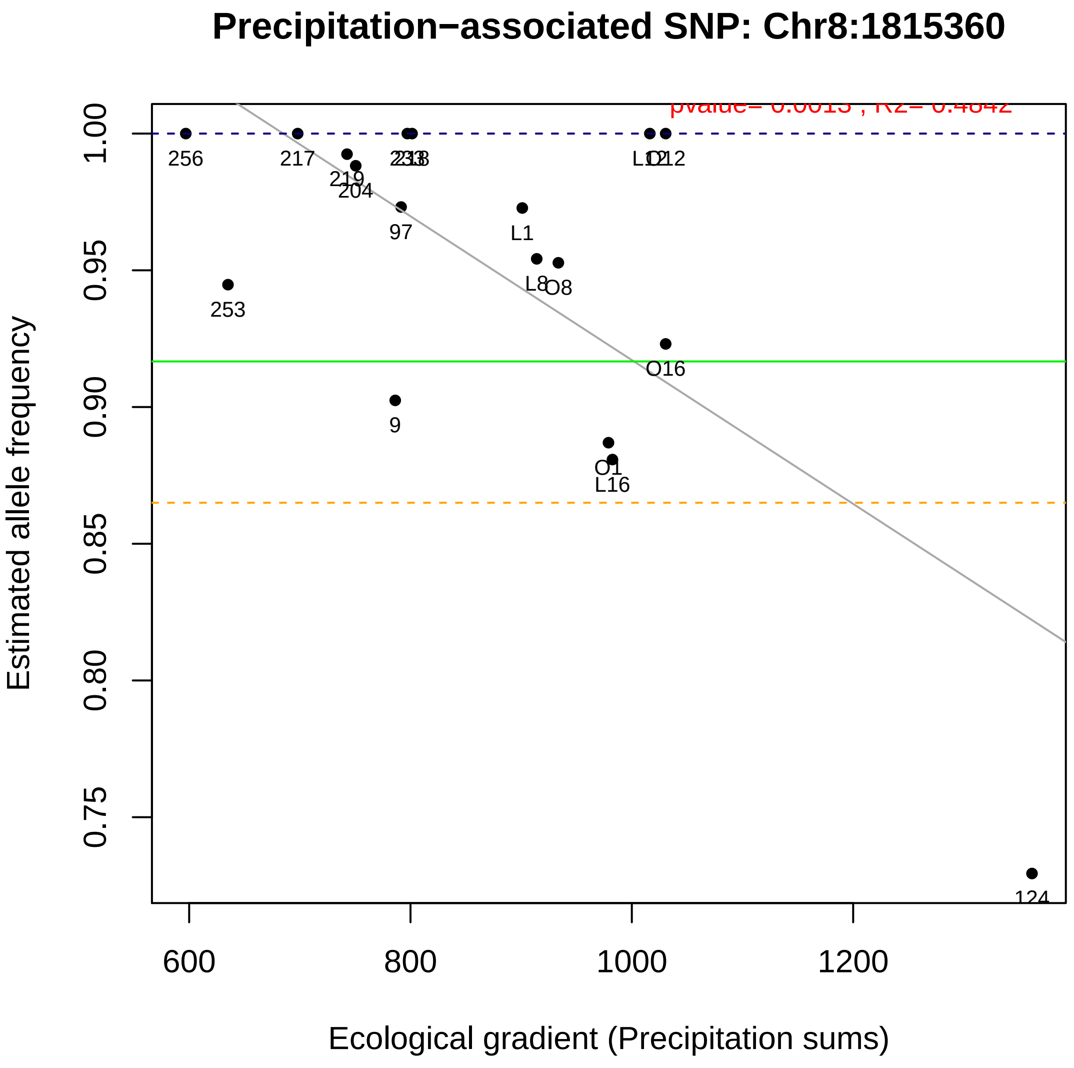
Precipitation−associated SNP: Chr8:1815360
pvalue= 0.0013 , R2= 0.4842
Estimated allele frequency
Ecological gradient (Precipitation sums)
0.75
0.80
0.85
0.90
0.95
1.00
600
800
1000
1200
256
217
219
204
97
253
9
L1
L8
O8
O16
O1
L16
124

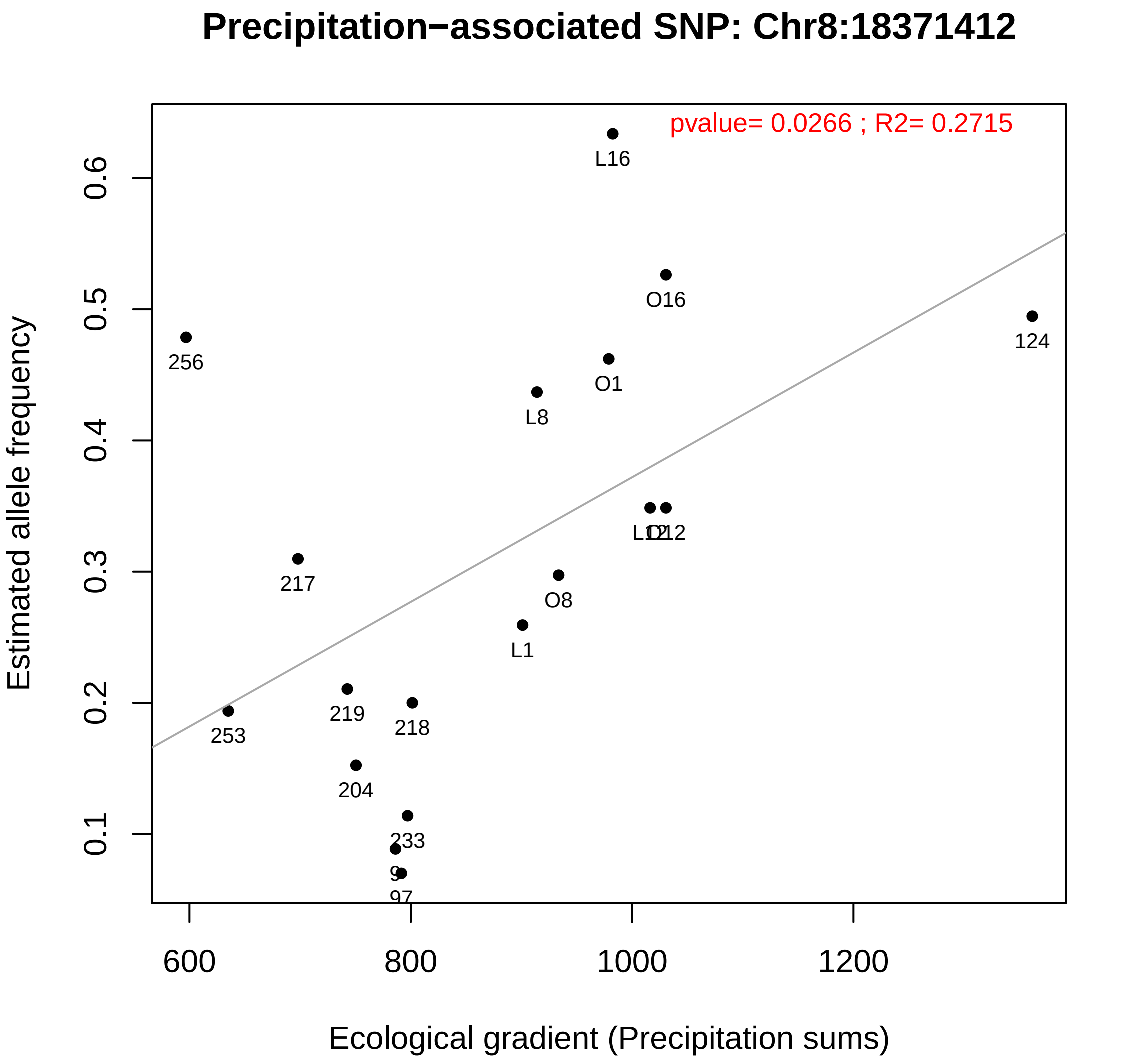
Precipitation−associated SNP: Chr8:18371412
pvalue= 0.0266 ; R2= 0.2715
Estimated allele frequency
0.1
0.2
0.3
0.4
0.5
0.6
600
800
1000
1200
Ecological gradient (Precipitation sums)
L16
O16
124
256
O1
L8
L12
O12
217
O8
L1
219
218
253
204
233
97

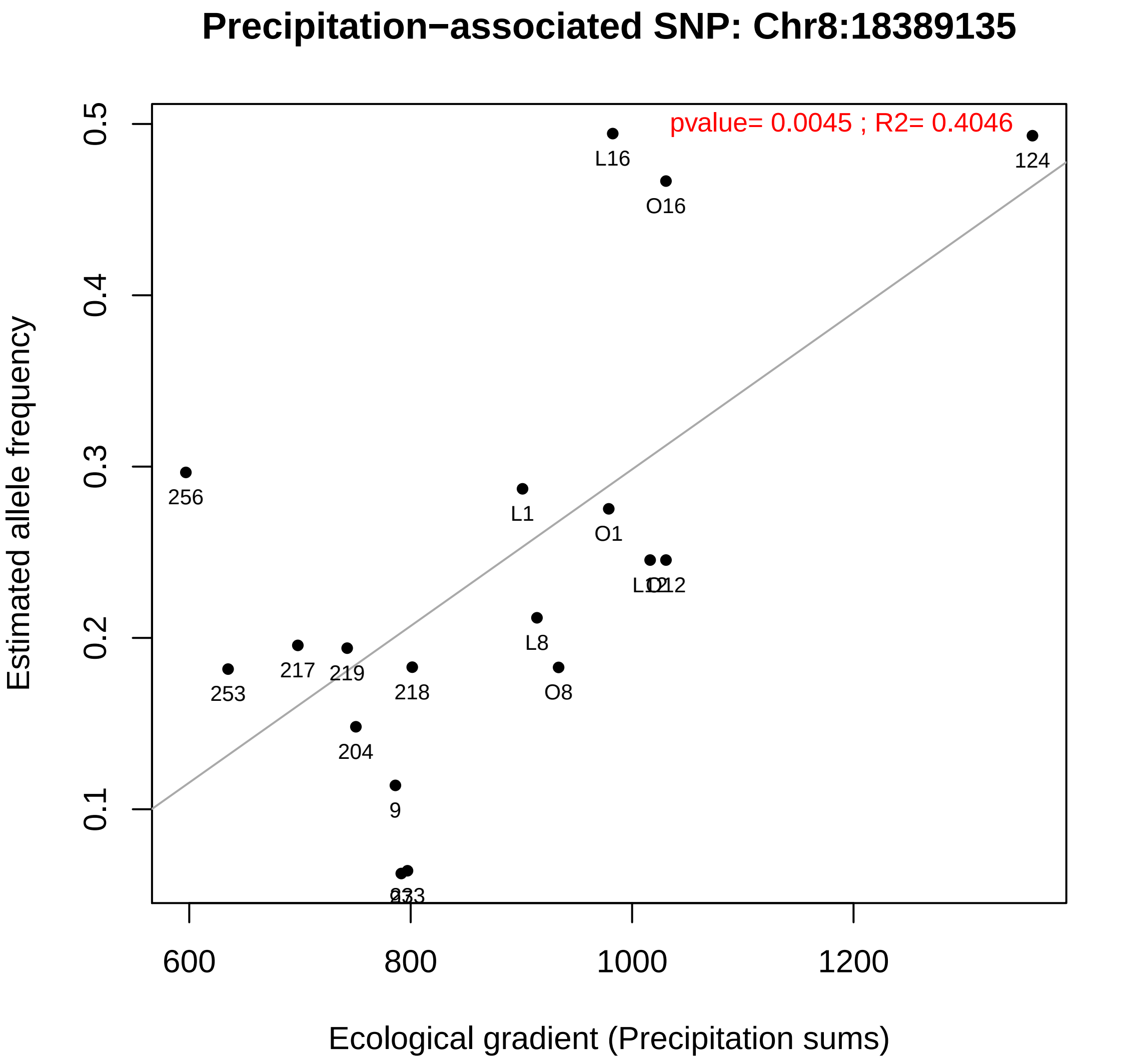

Precipitation−associated SNP: Chr8:18389135
pvalue= 0.0045 ; R2= 0.4046
Estimated allele frequency
0.1
0.2
0.3
0.4
0.5
600
800
1000
1200
Ecological gradient (Precipitation sums)
L16
124
O16
256
L1
O1
L8
217
219
253
218
O8
204
9

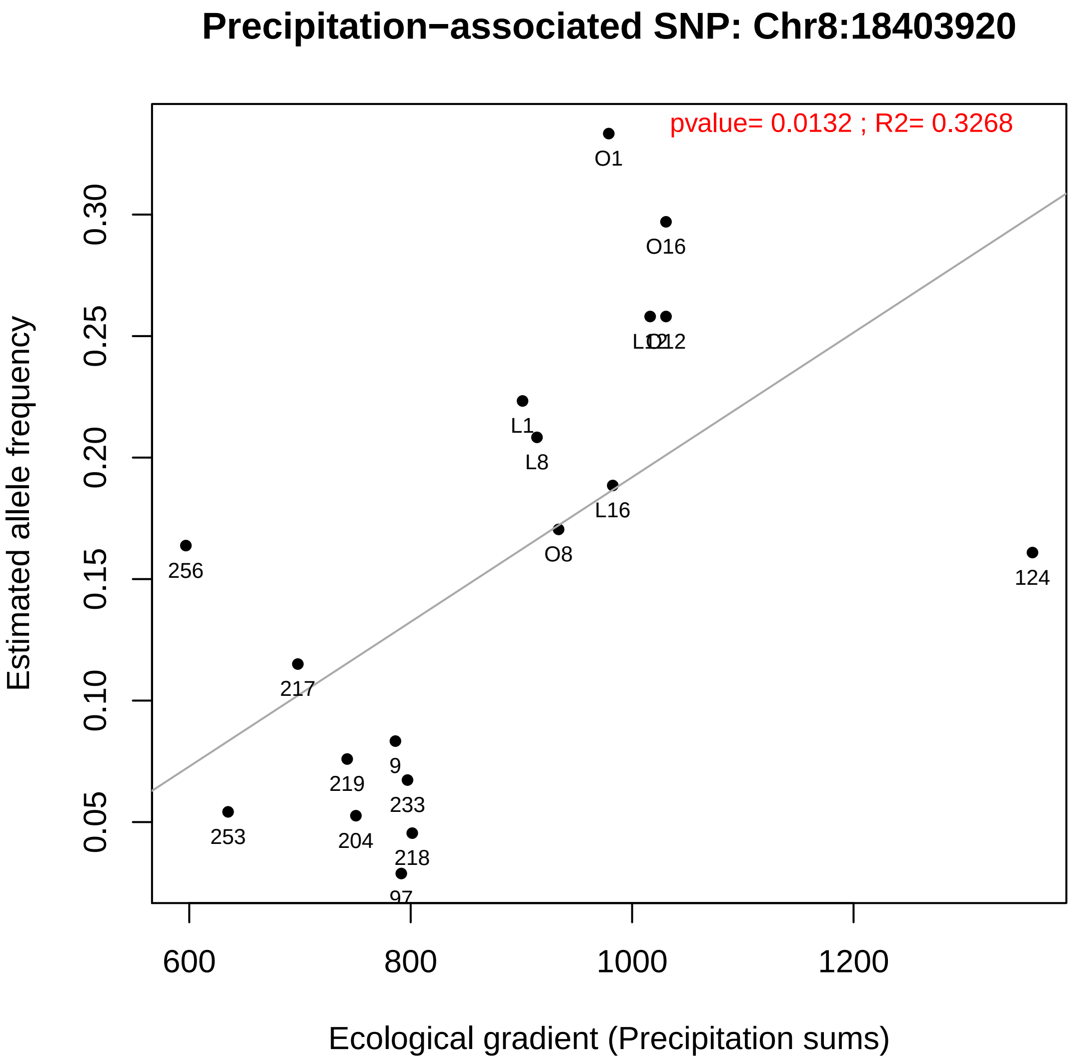
Precipitation−associated SNP: Chr8:18403920
pvalue= 0.0132 ; R2= 0.3268
Estimated allele frequency
0.05
0.10
0.15
0.20
0.25
0.30
600
800
1000
1200
Ecological gradient (Precipitation sums)
O1
O16
L12
O12
L1
L8
L16
O8
256
124
217
9
219
233
253
204
218
97

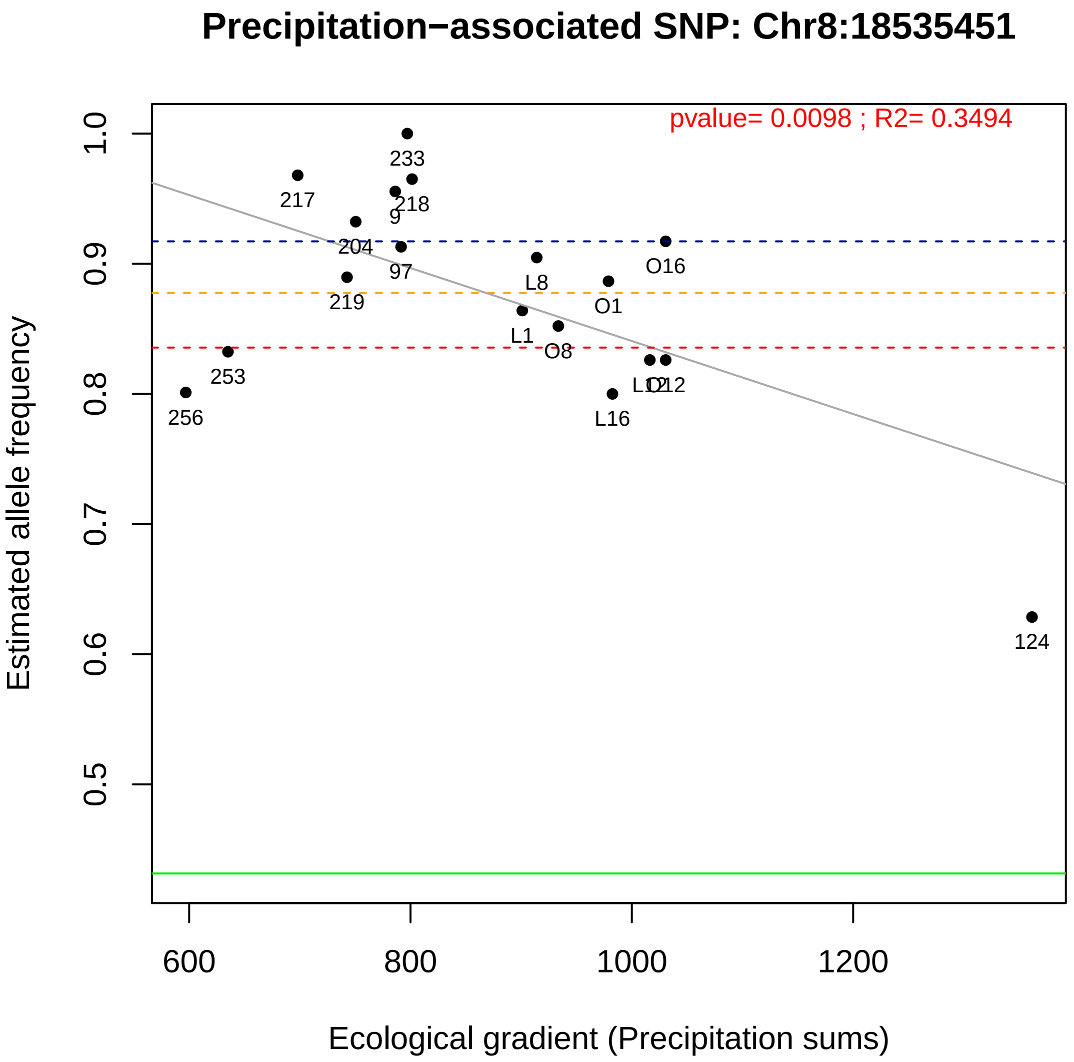
Precipitation−associated SNP: Chr8:18535451
pvalue= 0.0098 ; R2= 0.3494
Estimated allele frequency
0.5
0.6
0.7
0.8
0.9
1.0
600
800
1000
1200
Ecological gradient (Precipitation sums)
233
217
218
9
204
97
219
L8
O16
O1
L1
O8
253
256
L12
O12
L16
124

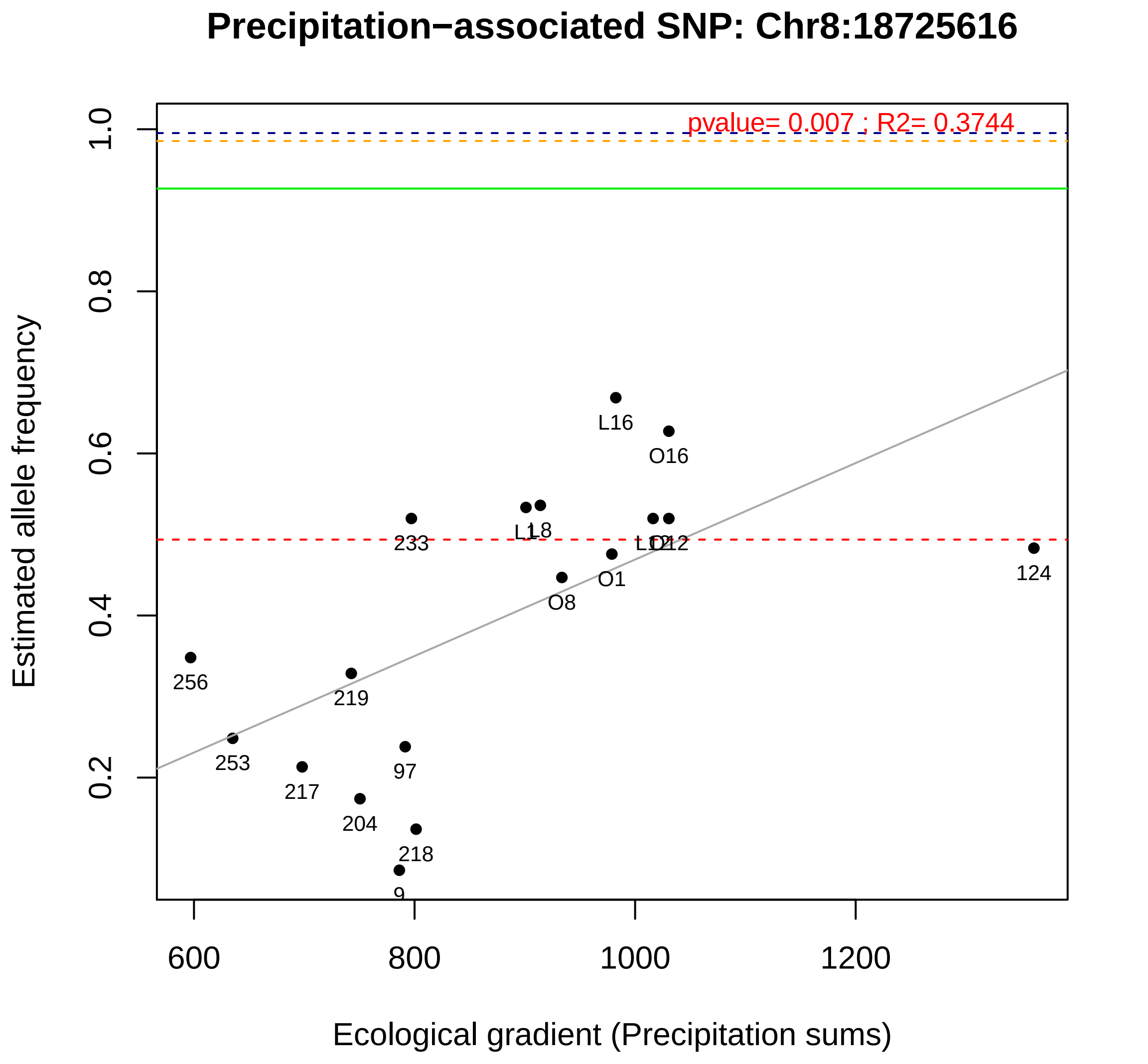

Precipitation−associated SNP: Chr8:18725616
pvalue= 0.007 ; R2= 0.3744
Estimated allele frequency
Ecological gradient (Precipitation sums)
0.2
0.4
0.6
0.8
1.0
600
800
1000
1200
L16
O16
233
O8
O1
124
256
219
253
217
97
204
218
9

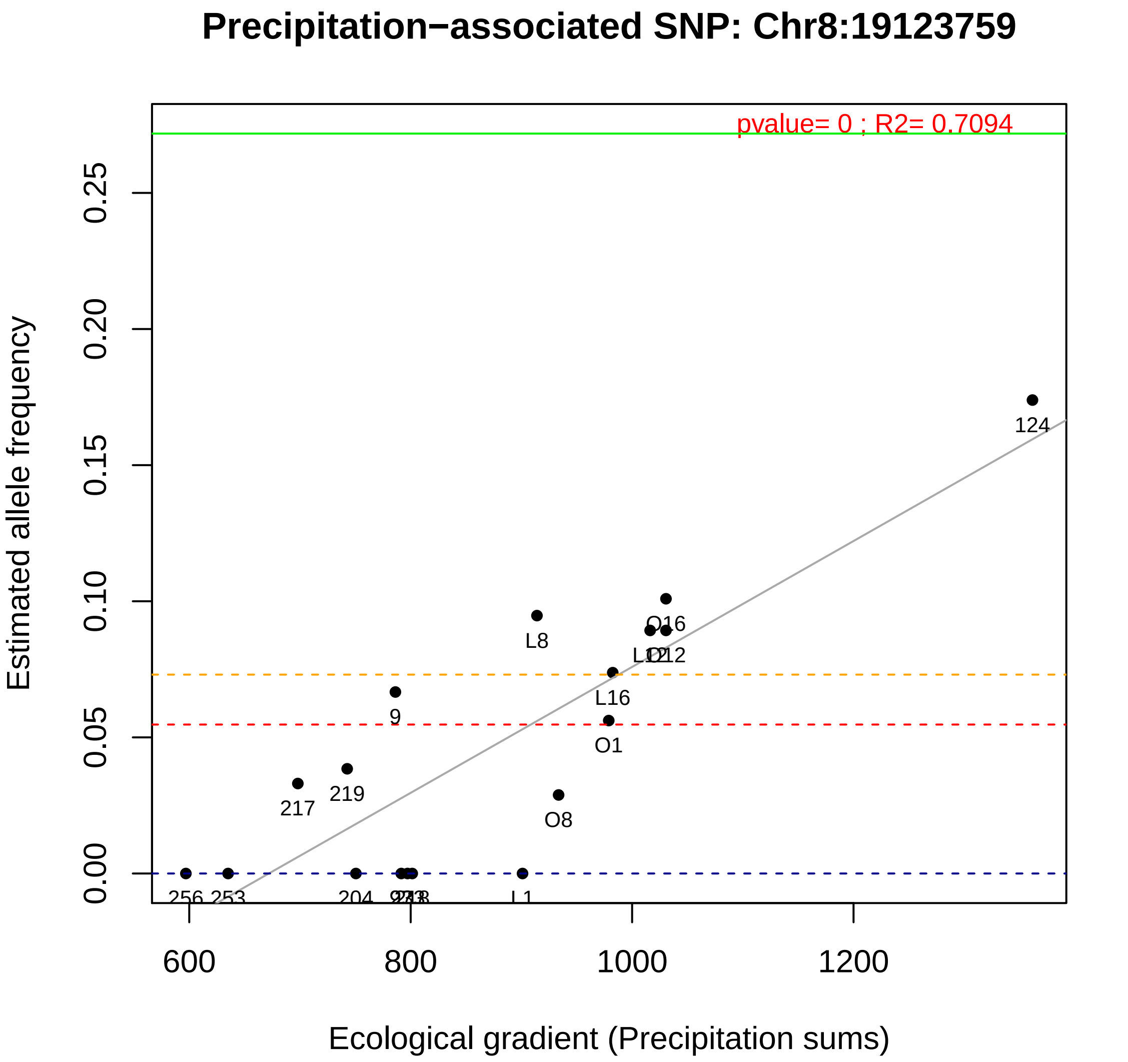

Precipitation−associated SNP: Chr8:19123759
pvalue= 0 ; R2= 0.7094
Estimated allele frequency
0.00
0.05
0.10
0.15
0.20
0.25
600
800
1000
1200
Ecological gradient (Precipitation sums)
124
L8
O16
L16
O1
9
219
217
O8
L1
204
253

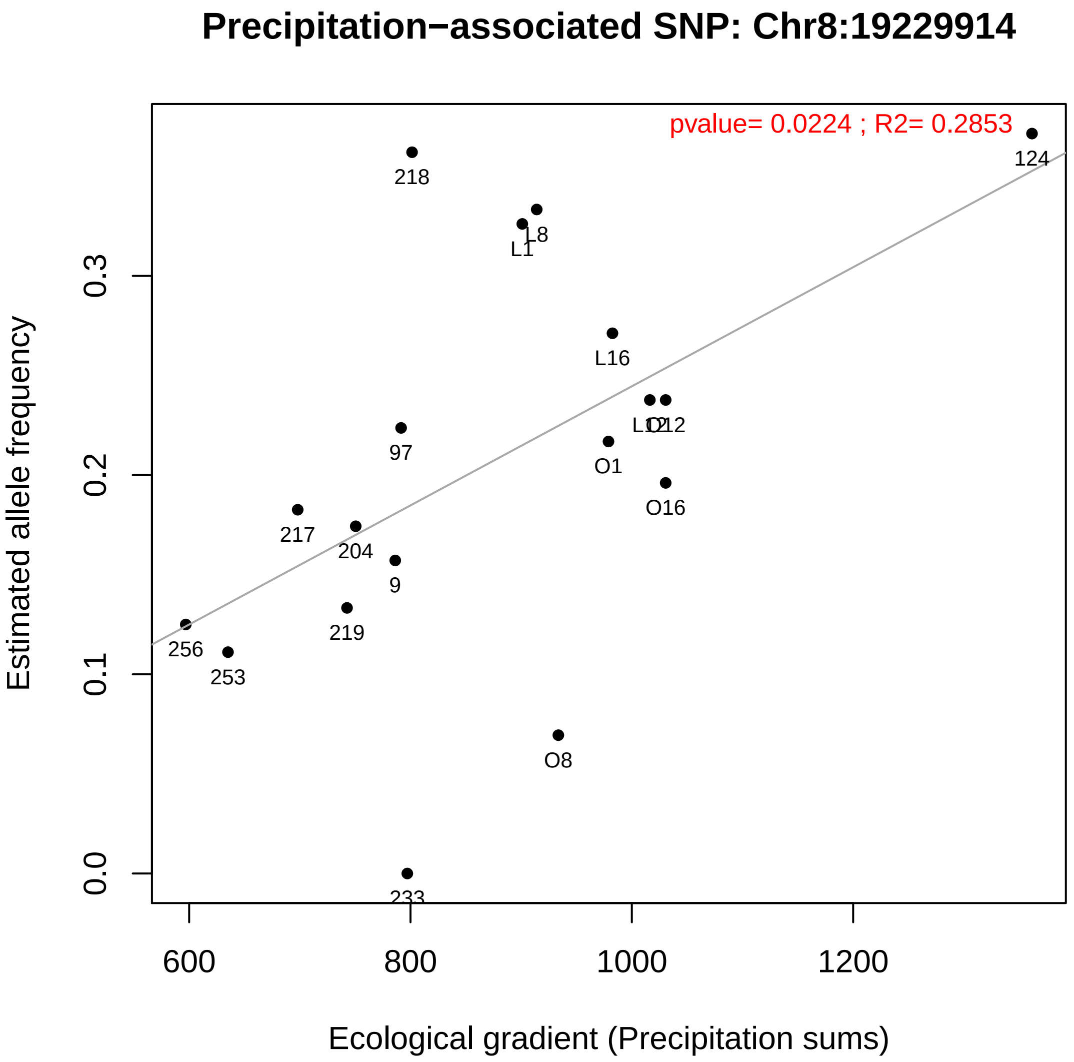
Precipitation−associated SNP: Chr8:19229914
pvalue= 0.0224 ; R2= 0.2853
Estimated allele frequency
Ecological gradient (Precipitation sums)
0.0
0.1
0.2
0.3
600
800
1000
1200
218
124
L8
L1
L16
L12
O12
97
O1
O16
217
204
9
219
256
253
O8
233

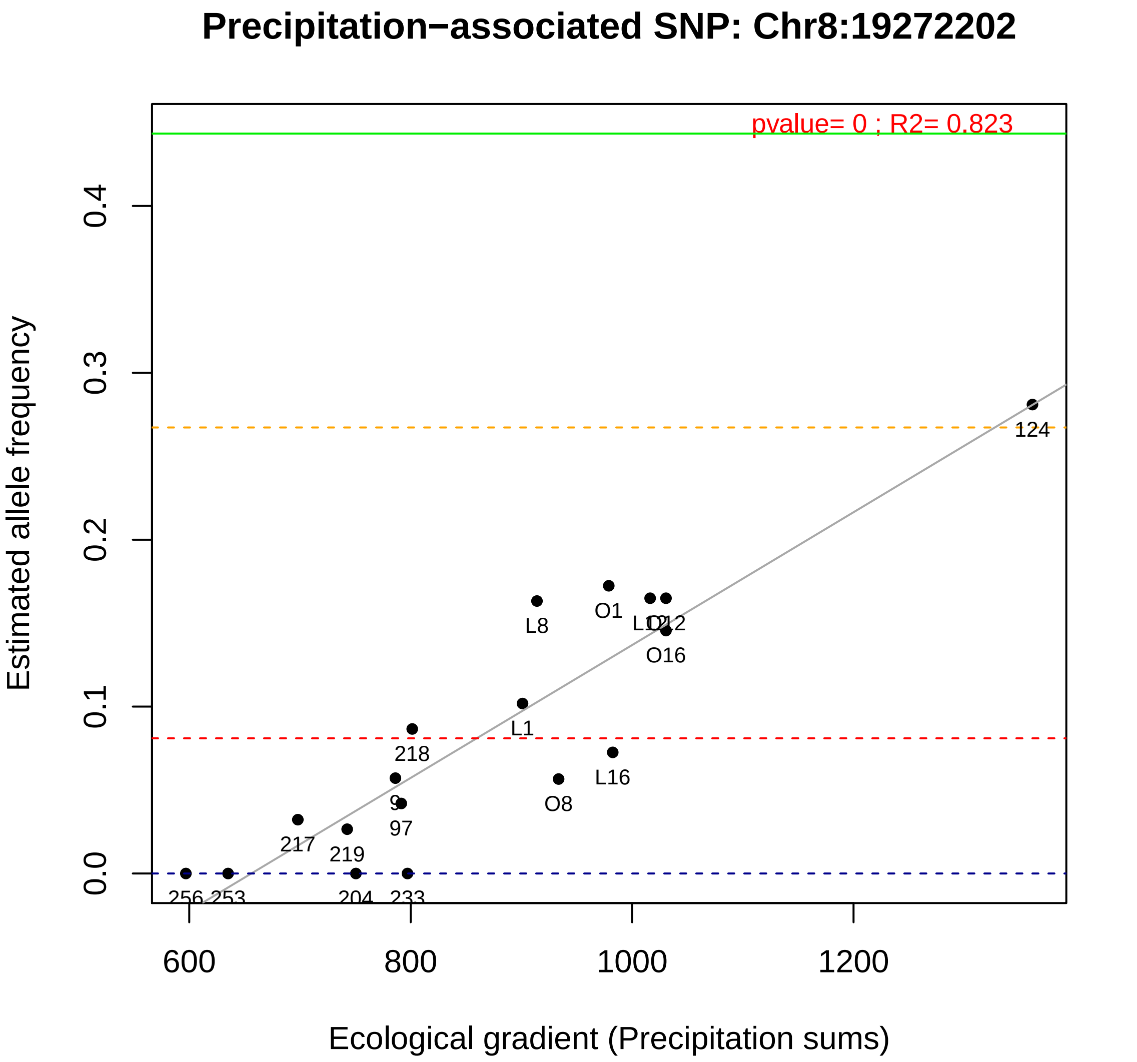
Precipitation−associated SNP: Chr8:19272202
pvalue= 0 ; R2= 0.823
Estimated allele frequency
0.0
0.1
0.2
0.3
0.4
600
800
1000
1200
Ecological gradient (Precipitation sums)
124
O1
L8
L12
O12
O16
L1
218
L16
O8
9
97
217
219
256
253
204
233

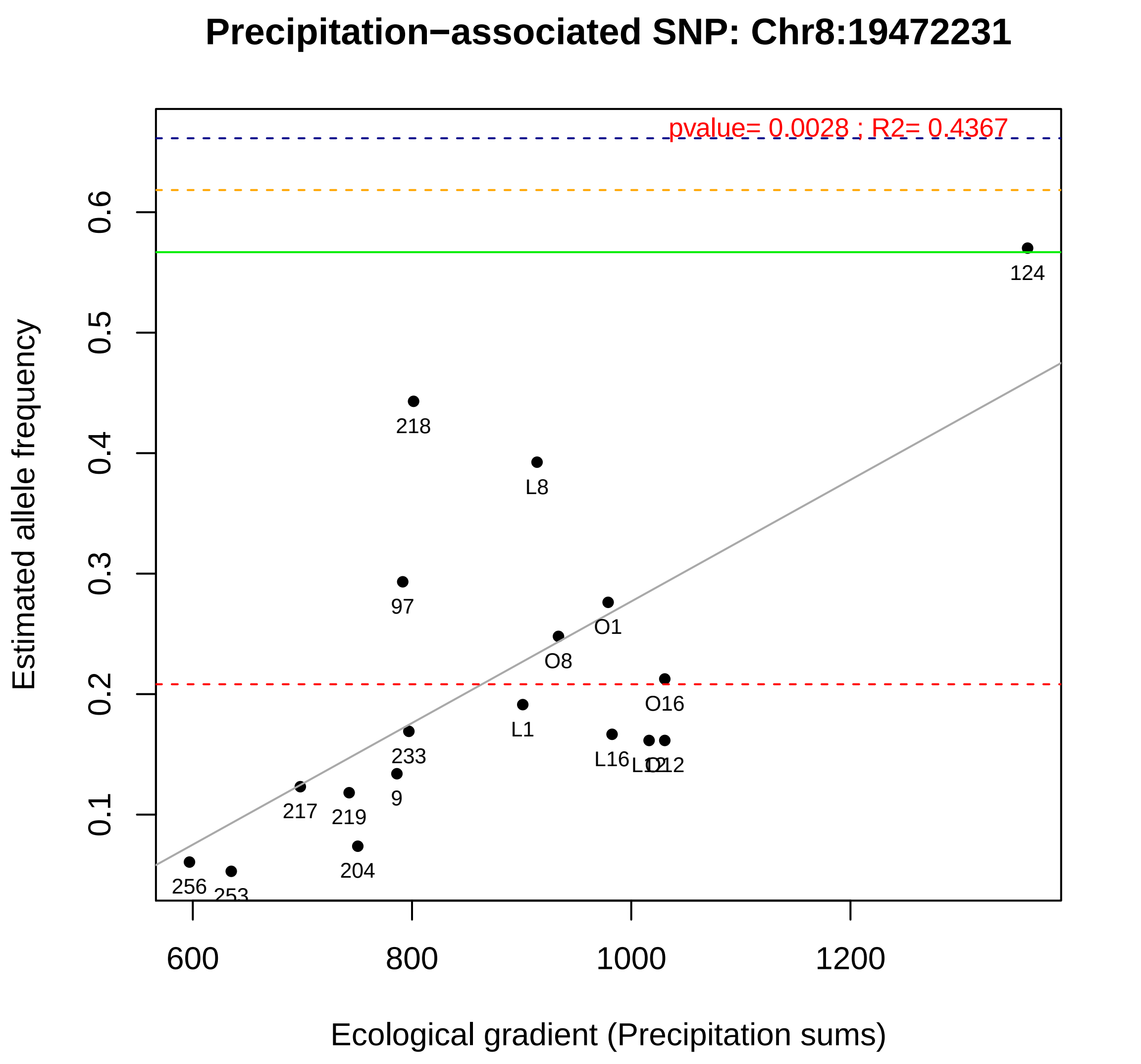
Precipitation−associated SNP: Chr8:19472231
pvalue= 0.0028 ; R2= 0.4367
Estimated allele frequency
0.1
0.2
0.3
0.4
0.5
0.6
600
800
1000
1200
Ecological gradient (Precipitation sums)
124
218
L8
97
O1
O8
O16
L1
233
L16
L12
O12
217
219
9
204
256
253

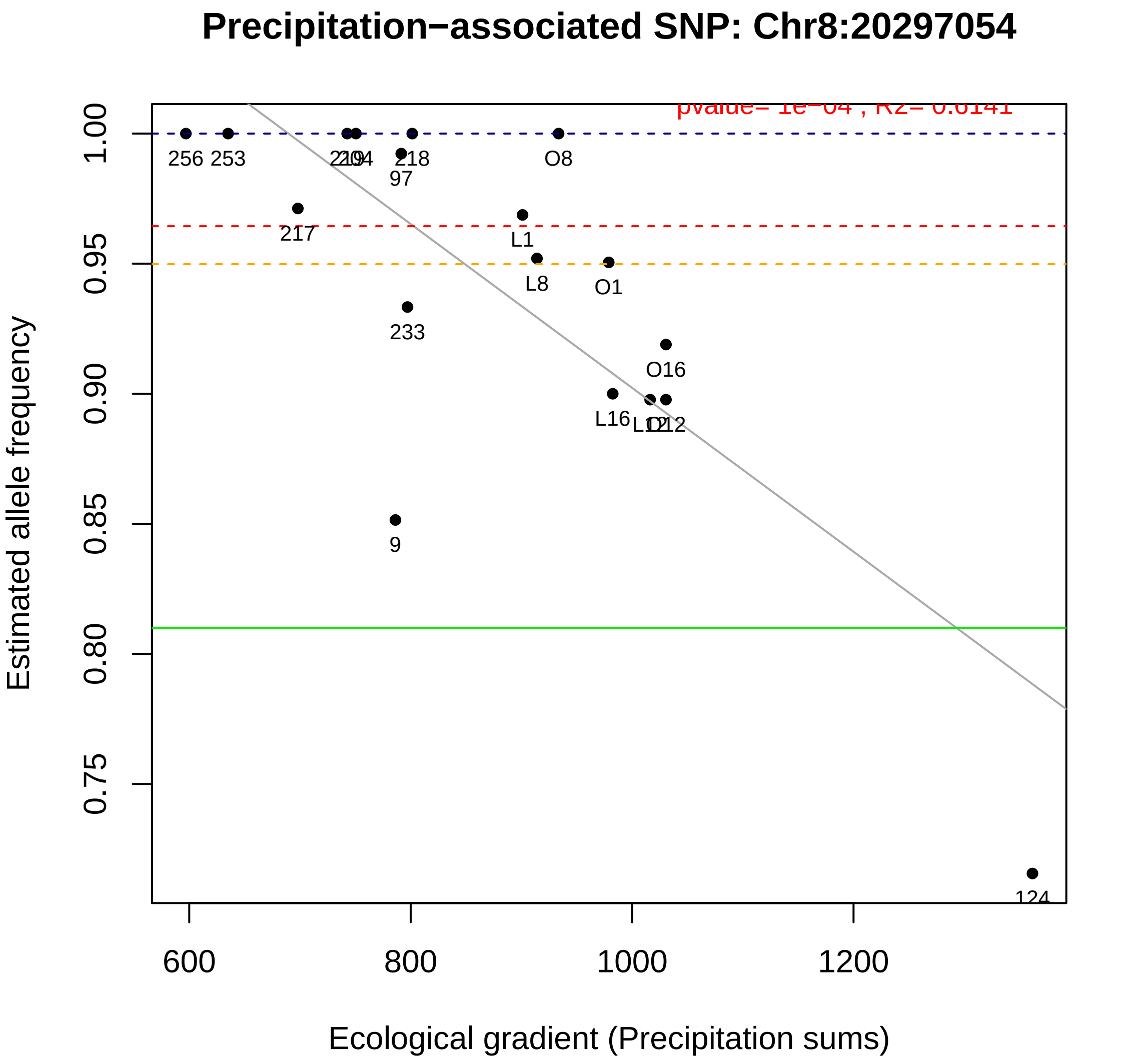
Precipitation−associated SNP: Chr8:20297054
pvalue= 1e−04 , R2= 0.6141
Estimated allele frequency
1.00
0.95
0.90
0.85
0.80
0.75
600
800
1000
1200
Ecological gradient (Precipitation sums)
256
253
218
97
O8
217
L1
L8
O1
233
O16
L16
9
124

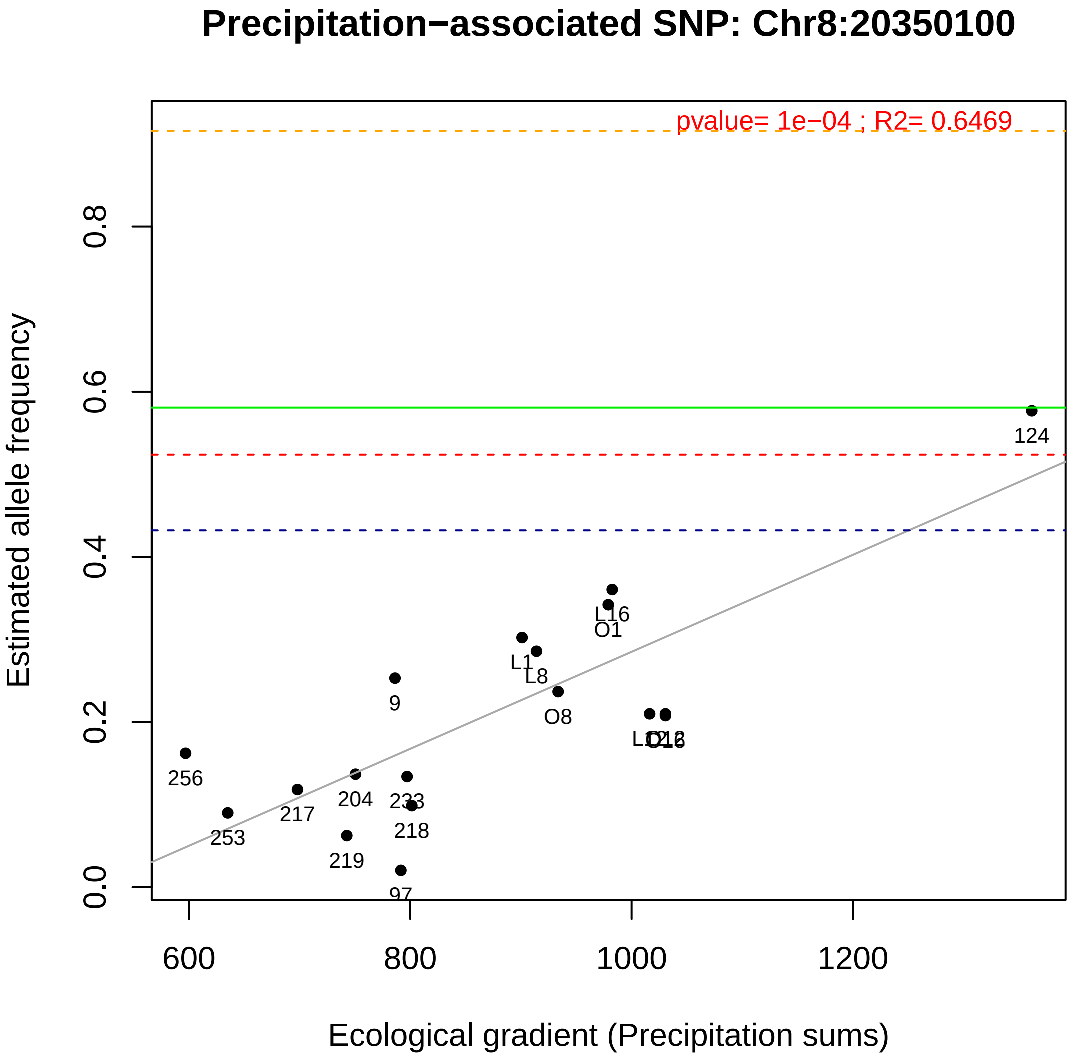

Precipitation−associated SNP: Chr8:20350100
pvalue= 1e−04 ; R2= 0.6469
Estimated allele frequency
0.0
0.2
0.4
0.6
0.8
600
800
1000
1200
Ecological gradient (Precipitation sums)
124
L16
O1
L1
L8
O8
9
256
204
233
217
218
253
219
97

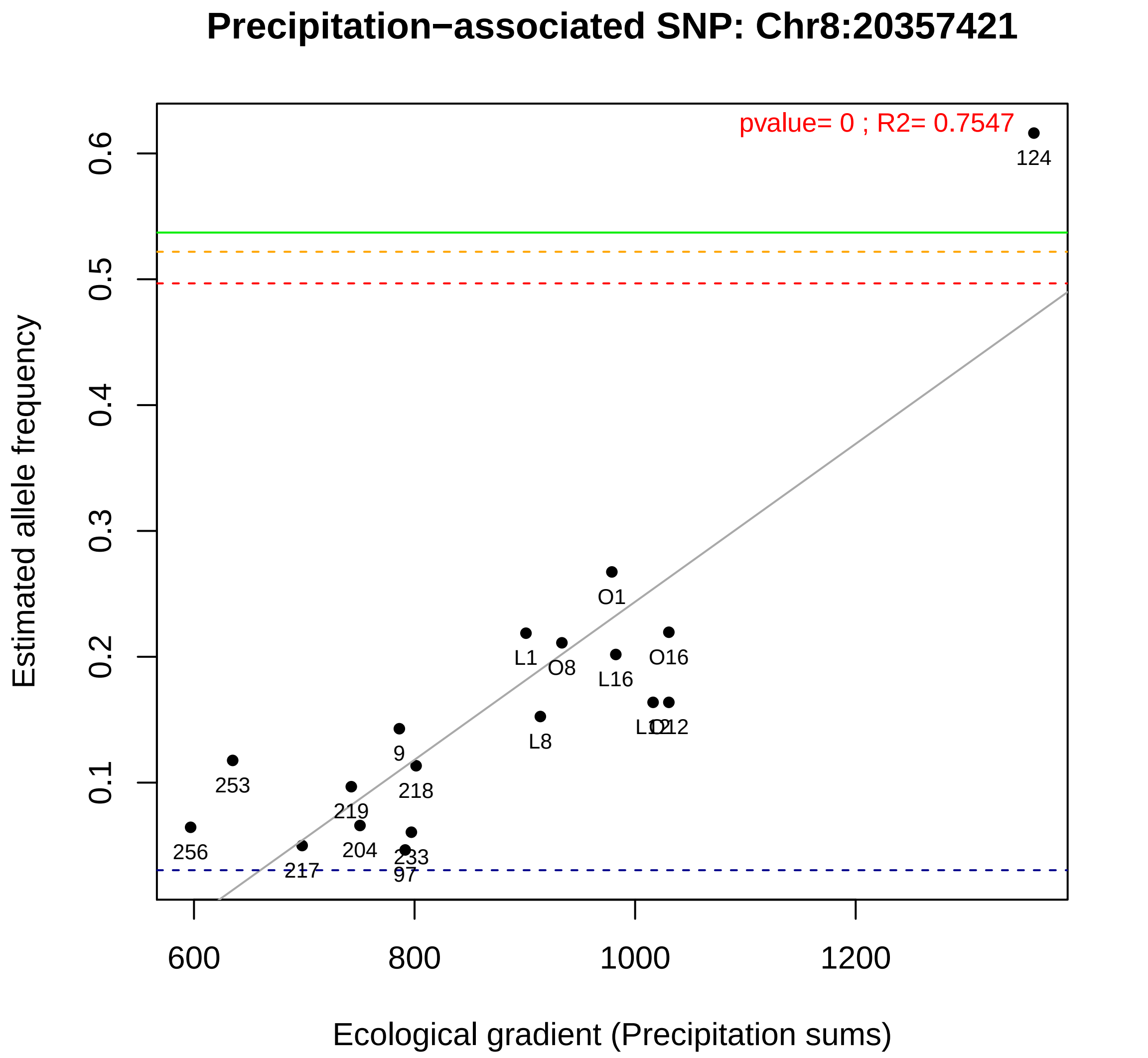

Precipitation−associated SNP: Chr8:20357421
pvalue= 0 ; R2= 0.7547
124
O1
L1
O8
L16
O16
L12
O12
L8
9
253
218
219
256
204
217
233
97
0.1
0.2
0.3
0.4
0.5
0.6
600
800
1000
1200
Estimated allele frequency
Ecological gradient (Precipitation sums)

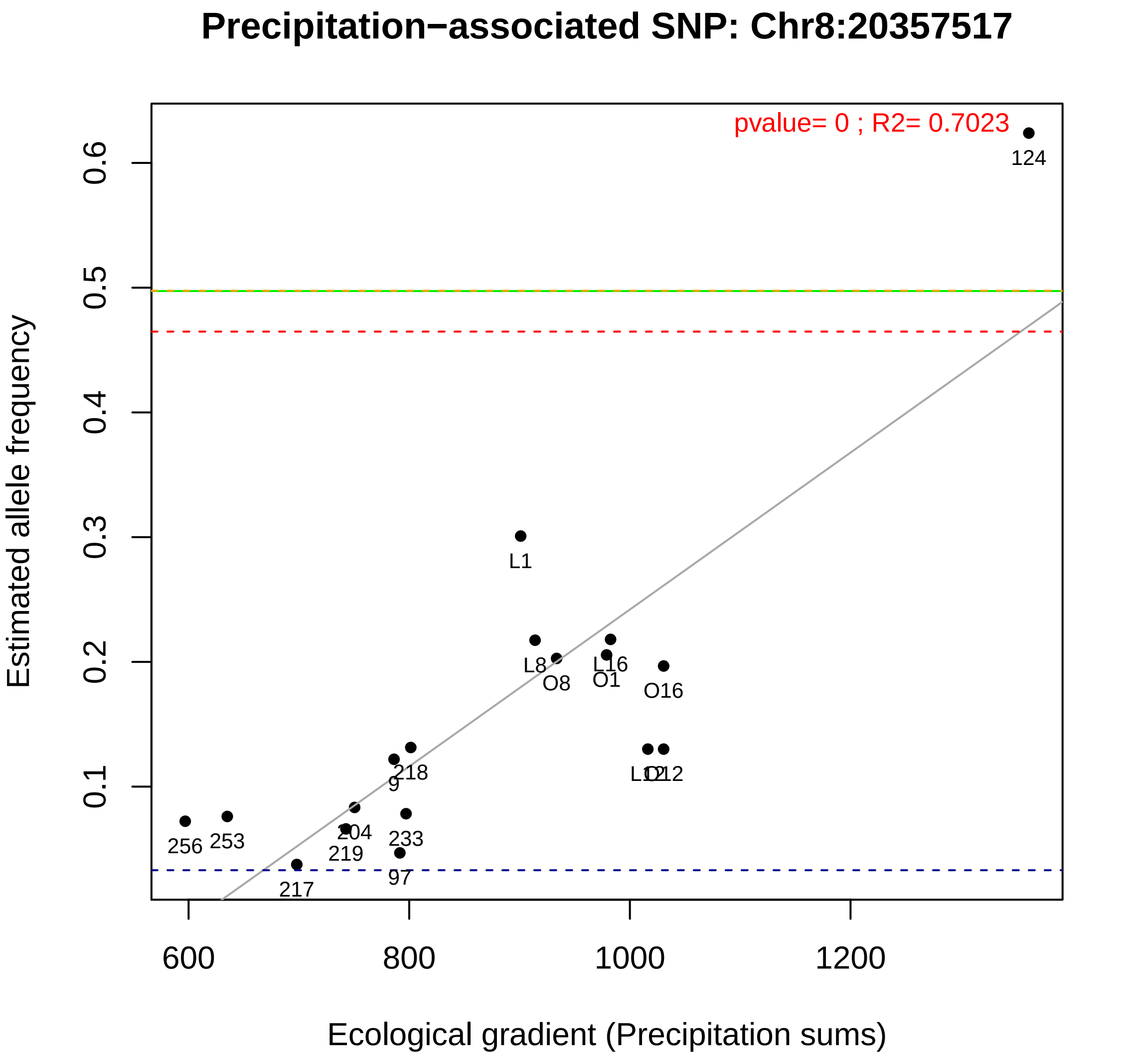
Precipitation−associated SNP: Chr8:20357517
pvalue= 0 ; R2= 0.7023
Estimated allele frequency
Ecological gradient (Precipitation sums)
0.1
0.2
0.3
0.4
0.5
0.6
600
800
1000
1200
124
L1
L8
O8
L16
O1
O16
L12
O12
218
9
204
219
233
97
256
253
217

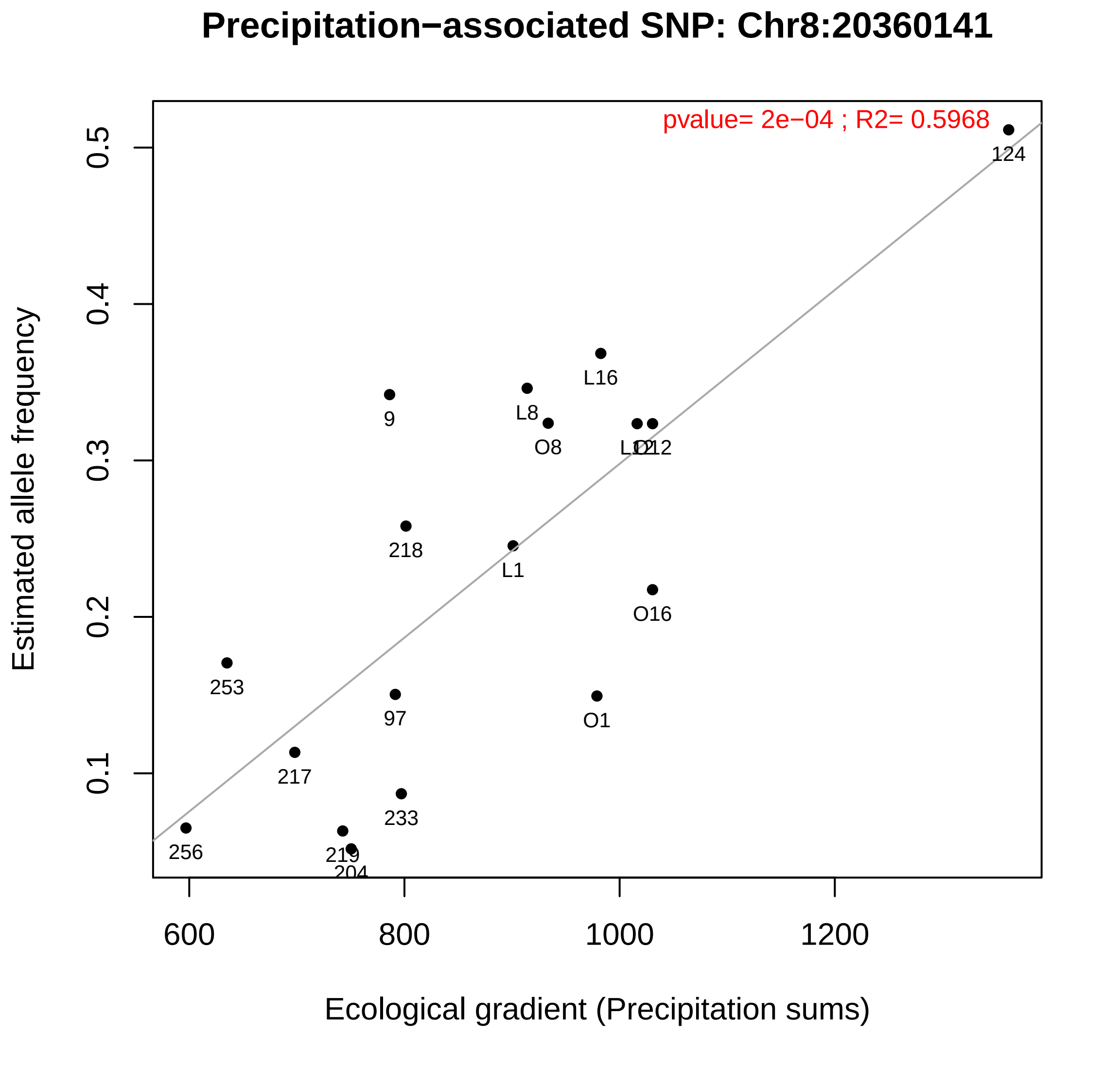
Precipitation−associated SNP: Chr8:20360141
pvalue= 2e−04 ; R2= 0.5968
Estimated allele frequency
Ecological gradient (Precipitation sums)
0.1
0.2
0.3
0.4
0.5
600
800
1000
1200
124
L16
L8
9
O8
O12
218
L1
O16
253
97
O1
217
233
256
219
204

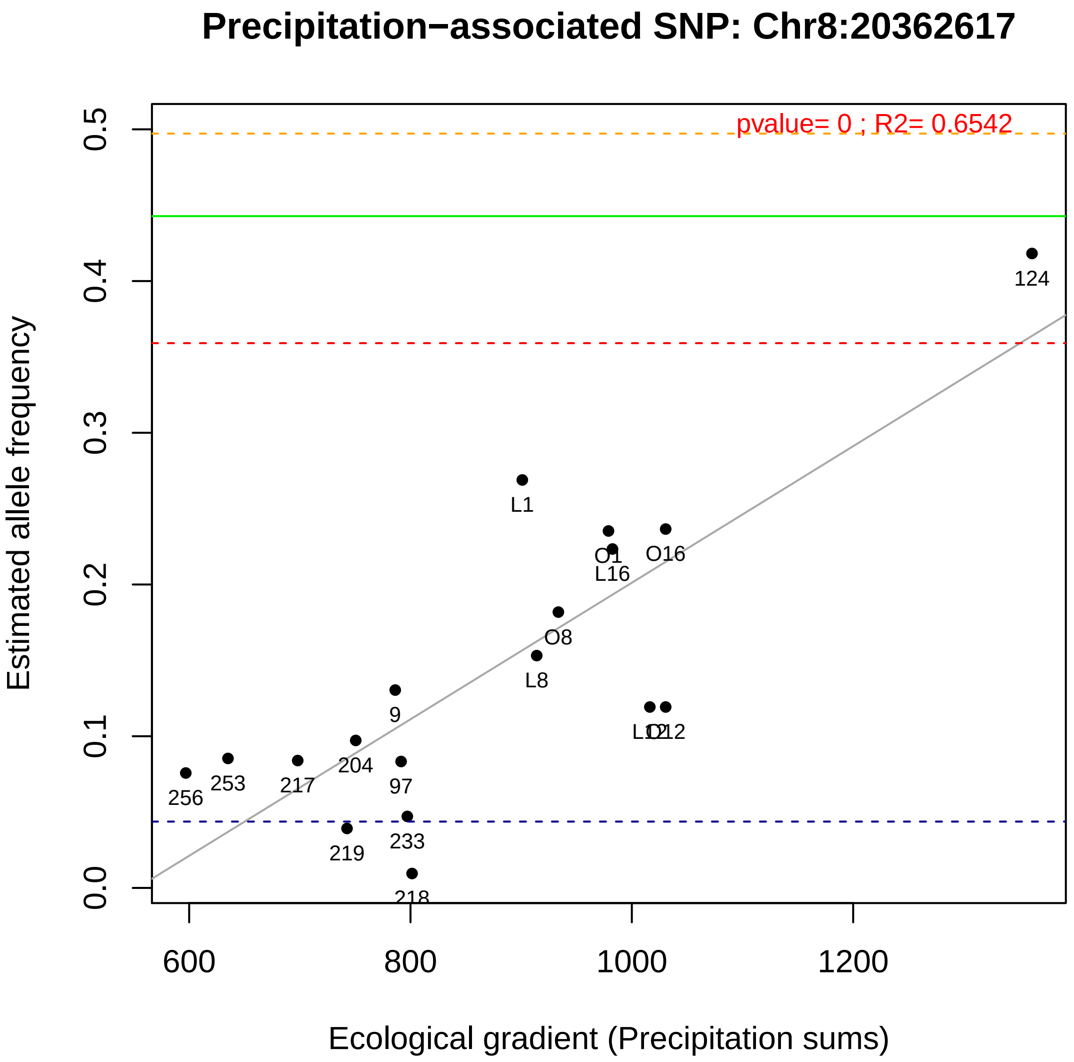
Precipitation−associated SNP: Chr8:20362617
pvalue= 0 ; R2= 0.6542
Estimated allele frequency
0.0
0.1
0.2
0.3
0.4
0.5
Ecological gradient (Precipitation sums)
600
800
1000
1200
124
L1
O1
L16
O16
O8
L8
9
L12
O12
204
97
253
217
256
233
219
218

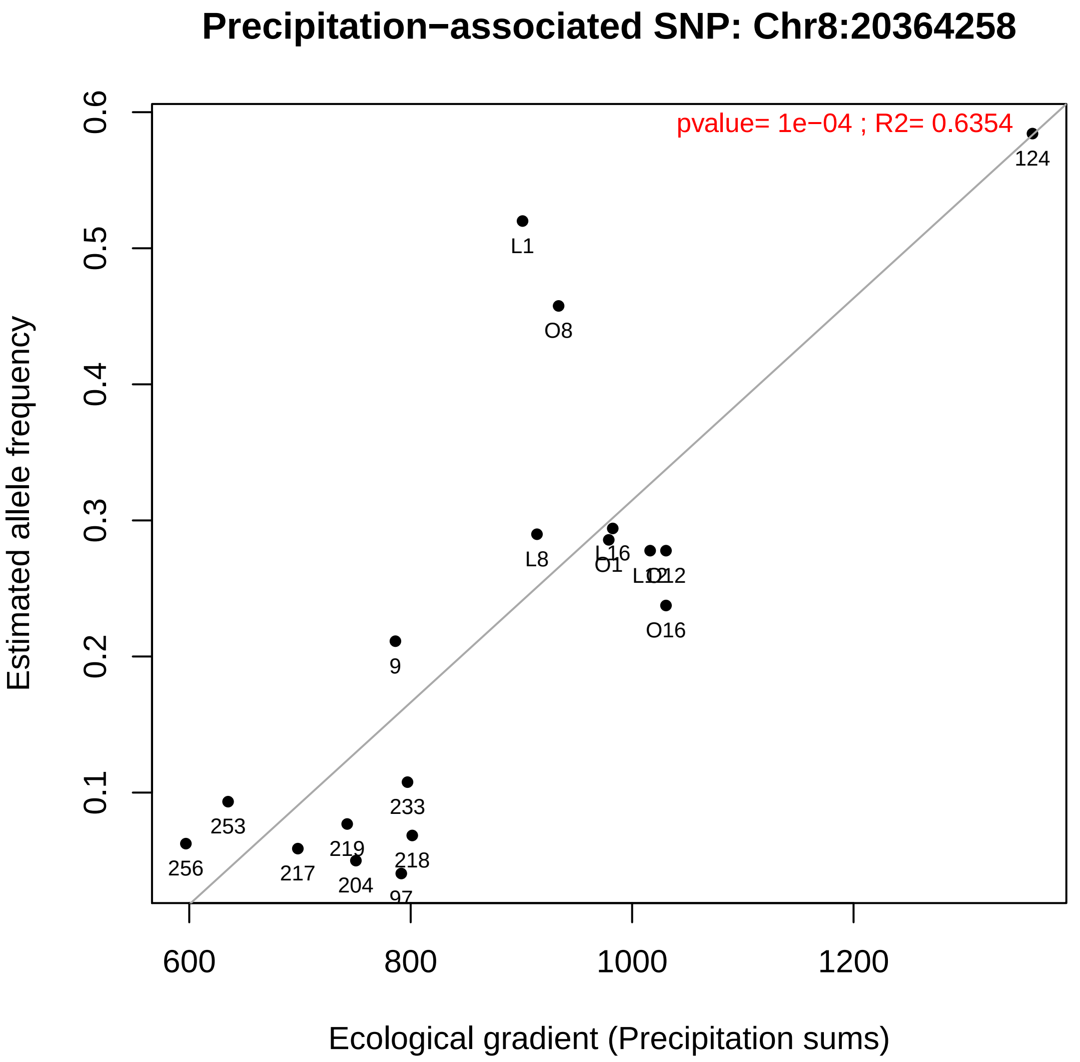
Precipitation−associated SNP: Chr8:20364258
pvalue= 1e−04 ; R2= 0.6354
124
L1
O8
L8
L16
O1
O12
O16
9
233
253
219
218
256
217
204
97
0.1
0.2
0.3
0.4
0.5
0.6
600
800
1000
1200
Estimated allele frequency
Ecological gradient (Precipitation sums)

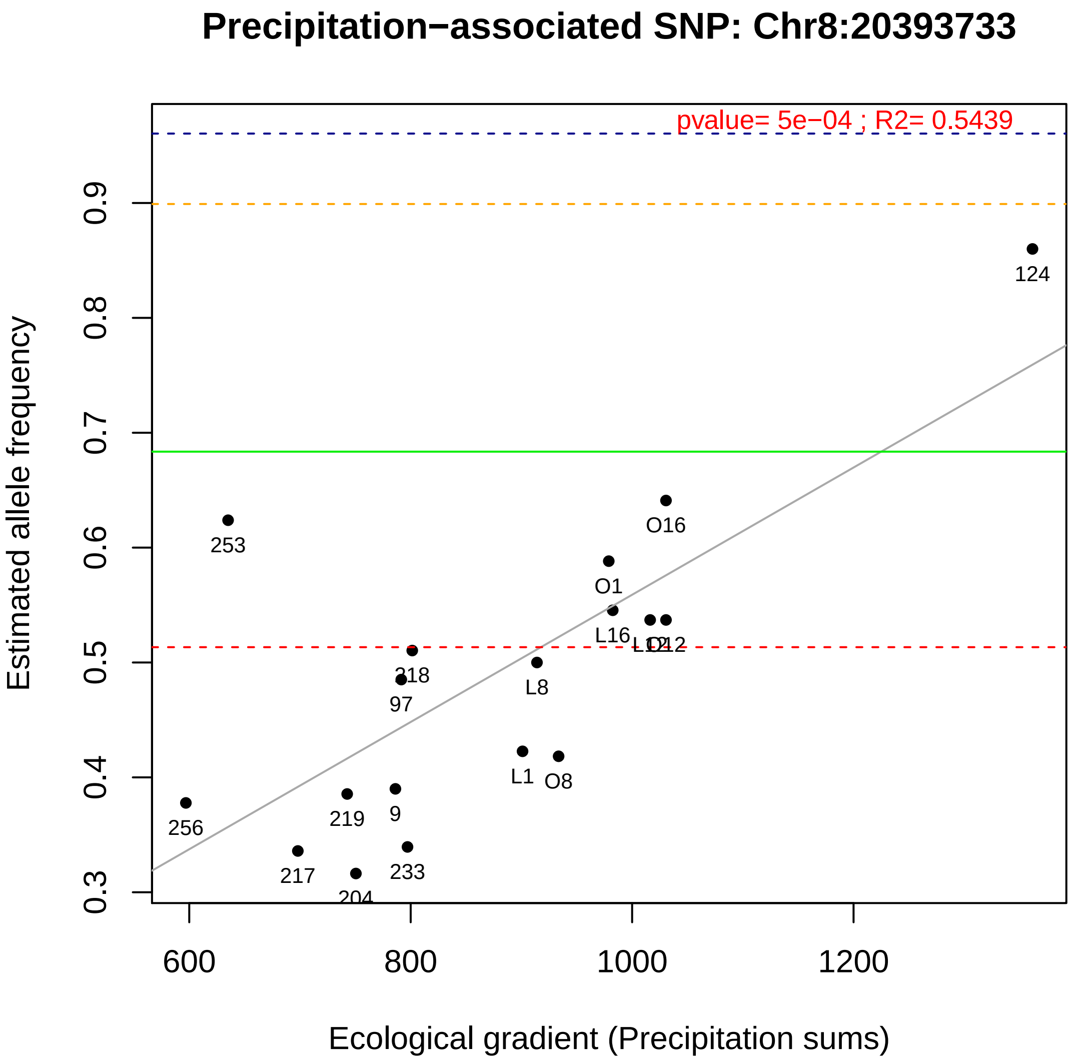
Precipitation−associated SNP: Chr8:20393733
pvalue= 5e−04 ; R2= 0.5439
Estimated allele frequency
0.3
0.4
0.5
0.6
0.7
0.8
0.9
600
800
1000
1200
Ecological gradient (Precipitation sums)
124
O16
253
O1
L16
L12
O12
218
97
L8
L1
O8
256
219
9
217
204
233

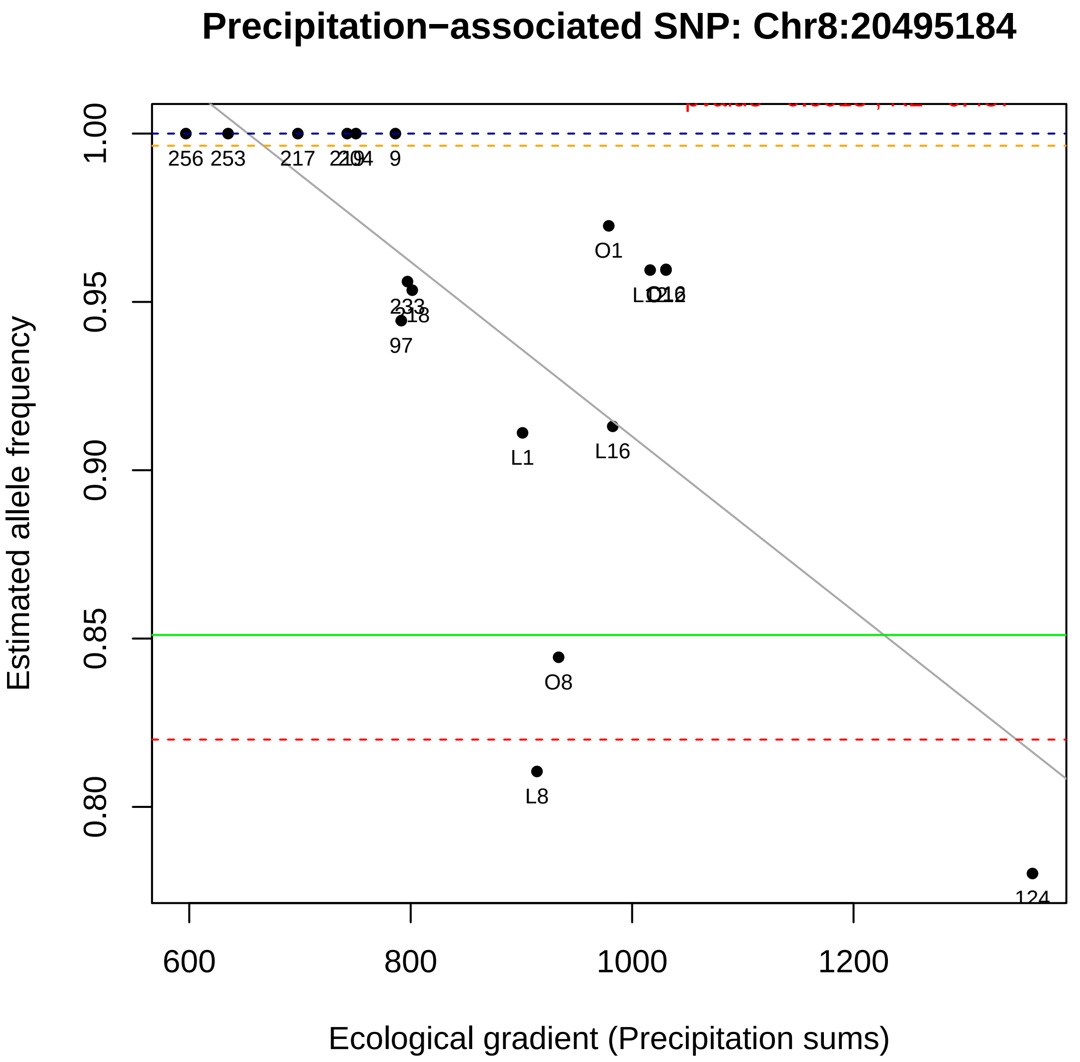
Precipitation−associated SNP: Chr8:20495184
Estimated allele frequency
1.00
0.95
0.90
0.85
0.80
600
800
1000
1200
Ecological gradient (Precipitation sums)
256
253
217
9
233
97
O1
L1
L16
O8
L8
124

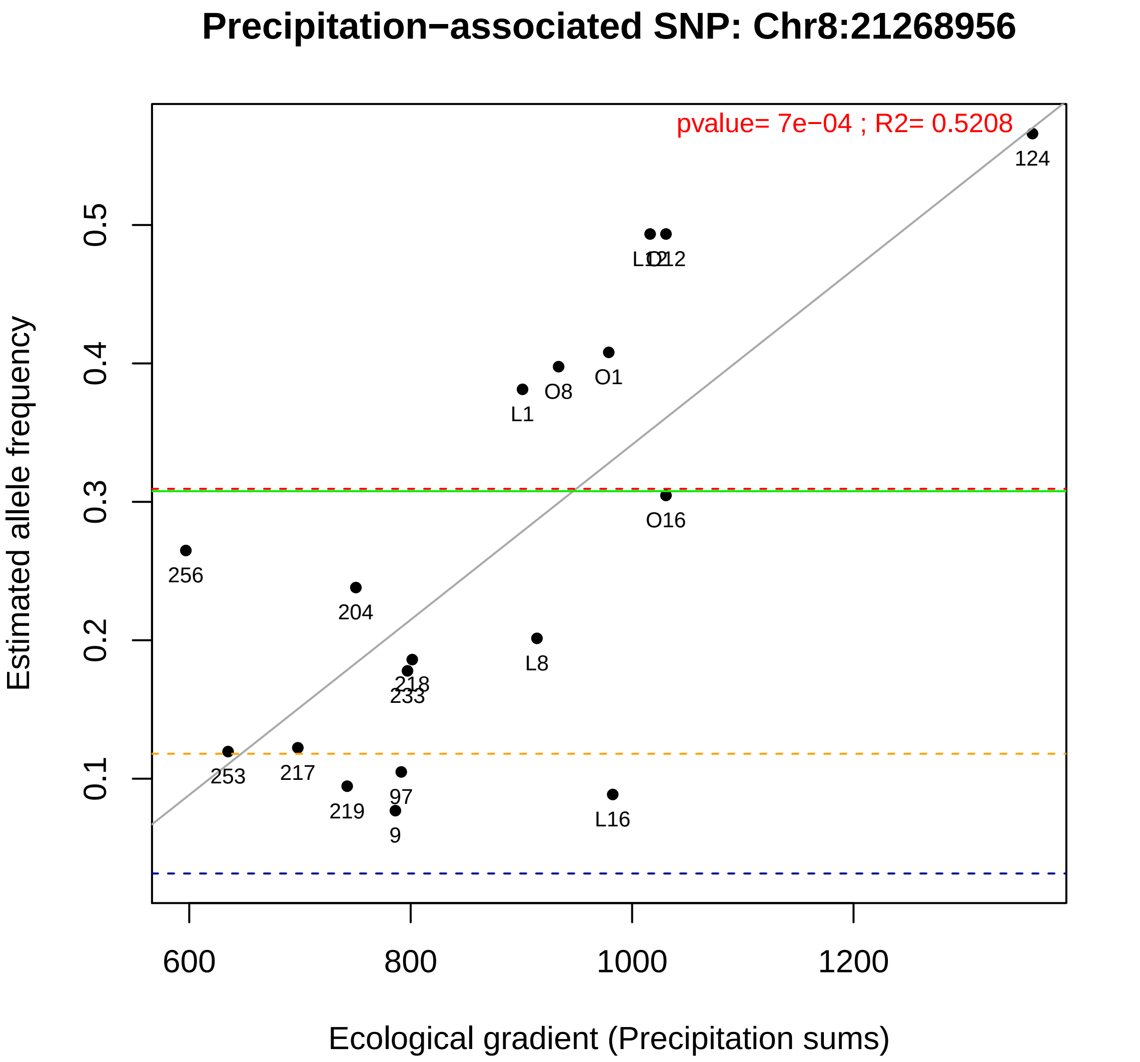
Precipitation−associated SNP: Chr8:21268956
pvalue= 7e−04 ; R2= 0.5208
124
L12
O12
O1
O8
L1
O16
256
204
L8
218
233
253
217
97
219
9
L16
0.1
0.2
0.3
0.4
0.5
600
800
1000
1200
Estimated allele frequency
Ecological gradient (Precipitation sums)

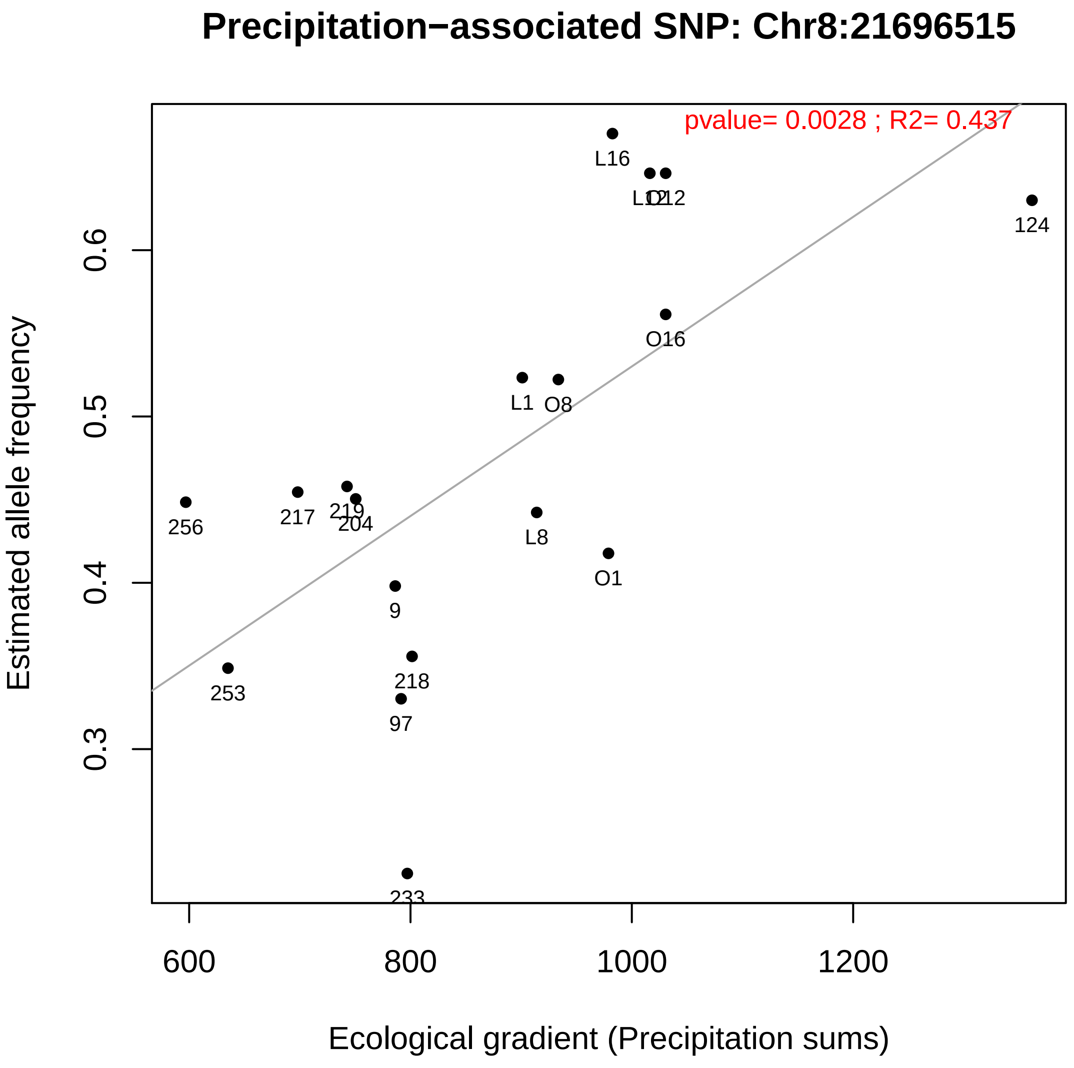

Precipitation−associated SNP: Chr8:21696515
pvalue= 0.0028 ; R2= 0.437
L16
L12
O12
124
O16
L1
O8
256
217
219
204
L8
O1
9
253
218
97
233
0.3
0.4
0.5
0.6
Estimated allele frequency
600
800
1000
1200
Ecological gradient (Precipitation sums)

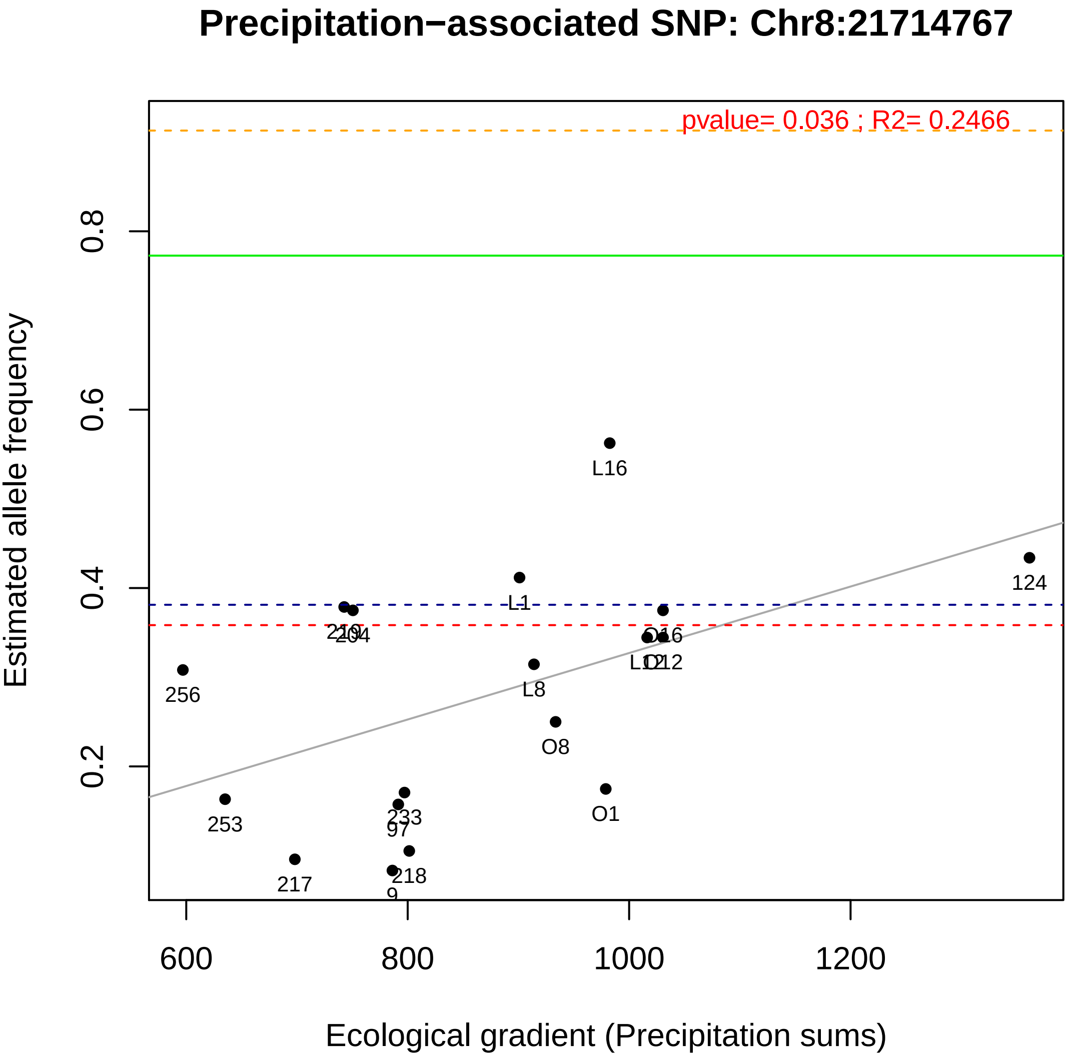
Precipitation−associated SNP: Chr8:21714767
pvalue= 0.036 ; R2= 0.2466
Estimated allele frequency
0.2
0.4
0.6
0.8
Ecological gradient (Precipitation sums)
600
800
1000
1200
L16
124
L1
204
256
L8
O8
253
233
97
O1
217
218
9

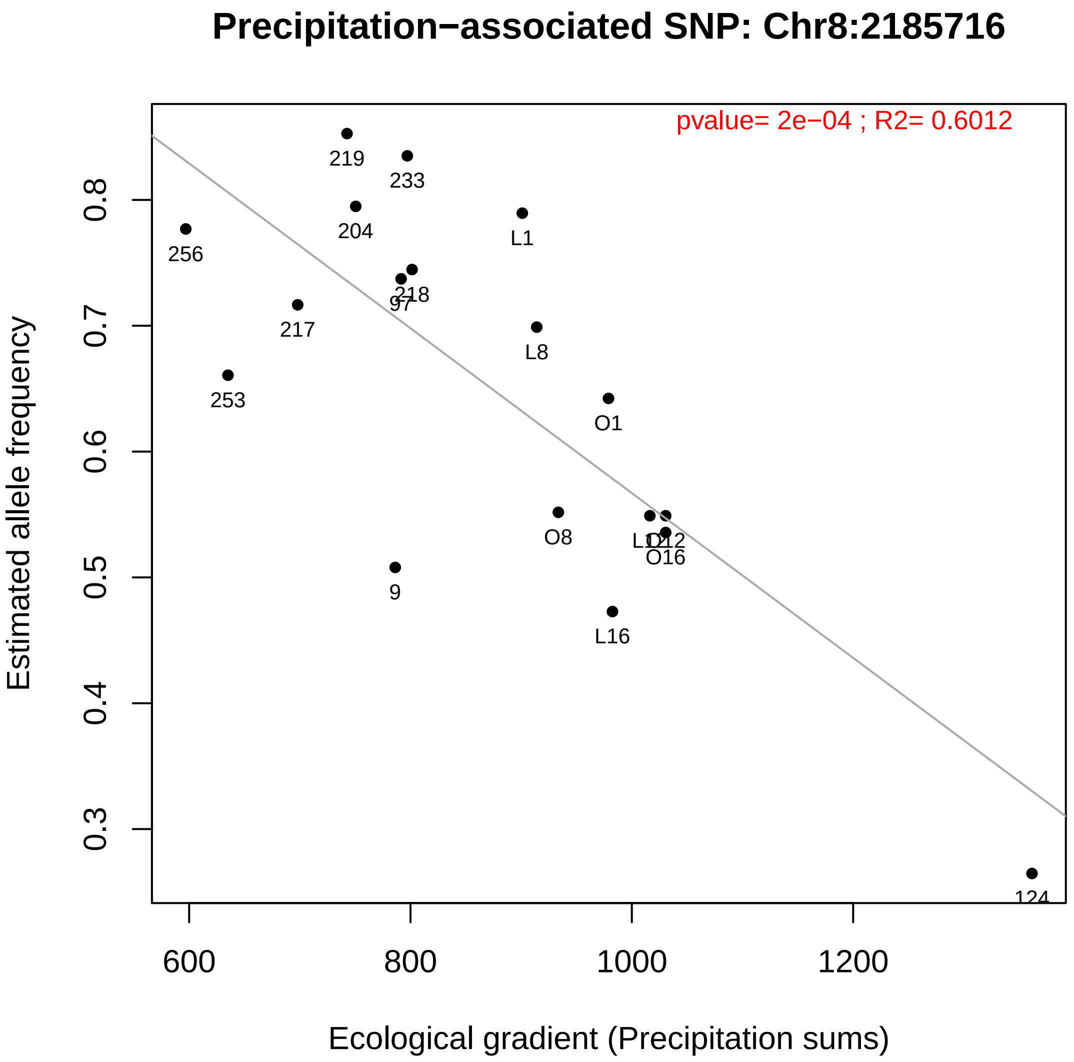
Precipitation−associated SNP: Chr8:2185716
pvalue= 2e−04 ; R2= 0.6012
Estimated allele frequency
0.3
0.4
0.5
0.6
0.7
0.8
600
800
1000
1200
Ecological gradient (Precipitation sums)
219
233
204
256
L1
218
97
217
L8
253
O1
O8
O16
9
L16
124

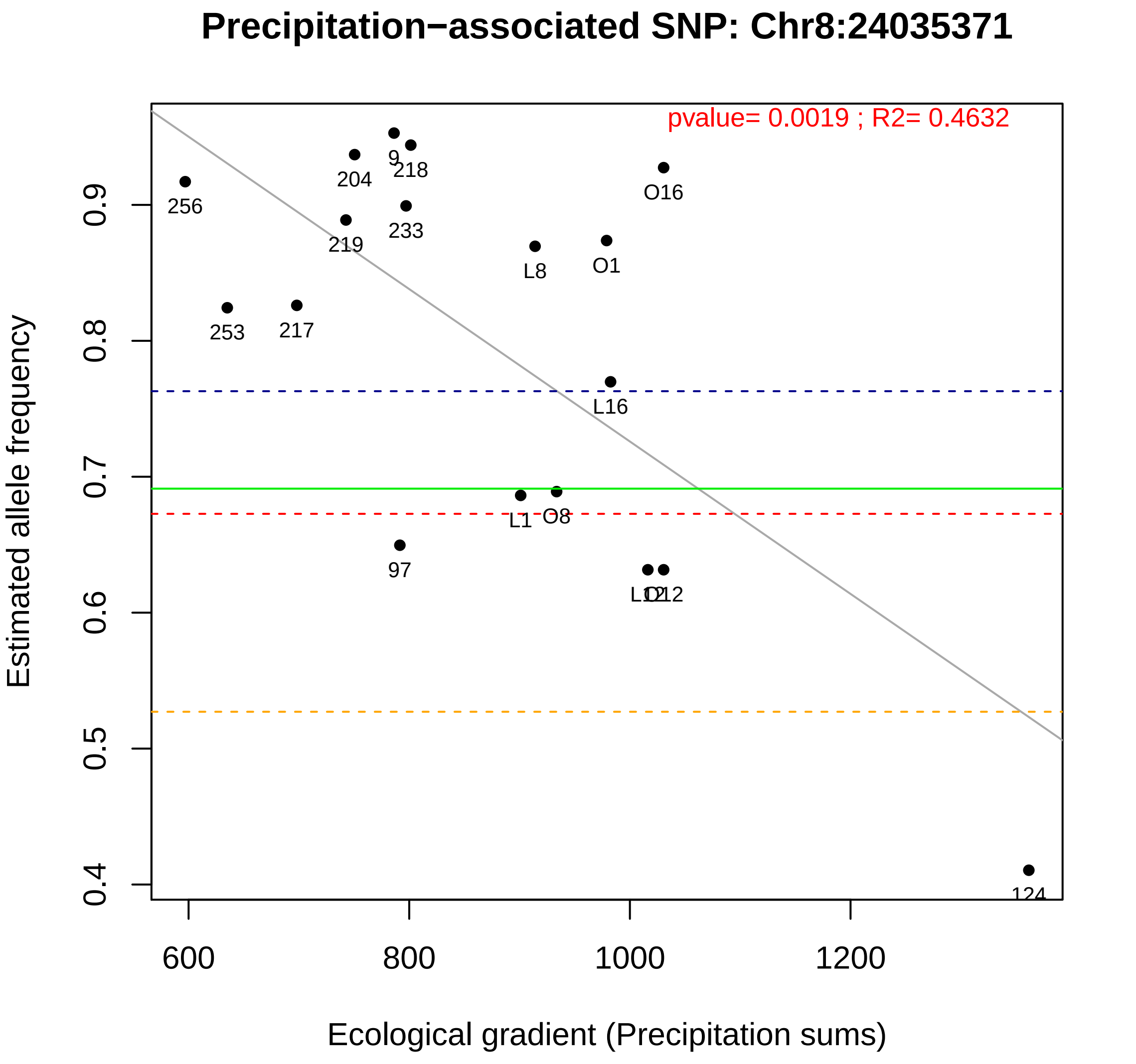
Precipitation−associated SNP: Chr8:24035371
pvalue= 0.0019 ; R2= 0.4632
Estimated allele frequency
0.4
0.5
0.6
0.7
0.8
0.9
600
800
1000
1200
Ecological gradient (Precipitation sums)
256
204
9
218
O16
219
233
L8
O1
253
217
L16
L1
O8
97
L12
O12
124

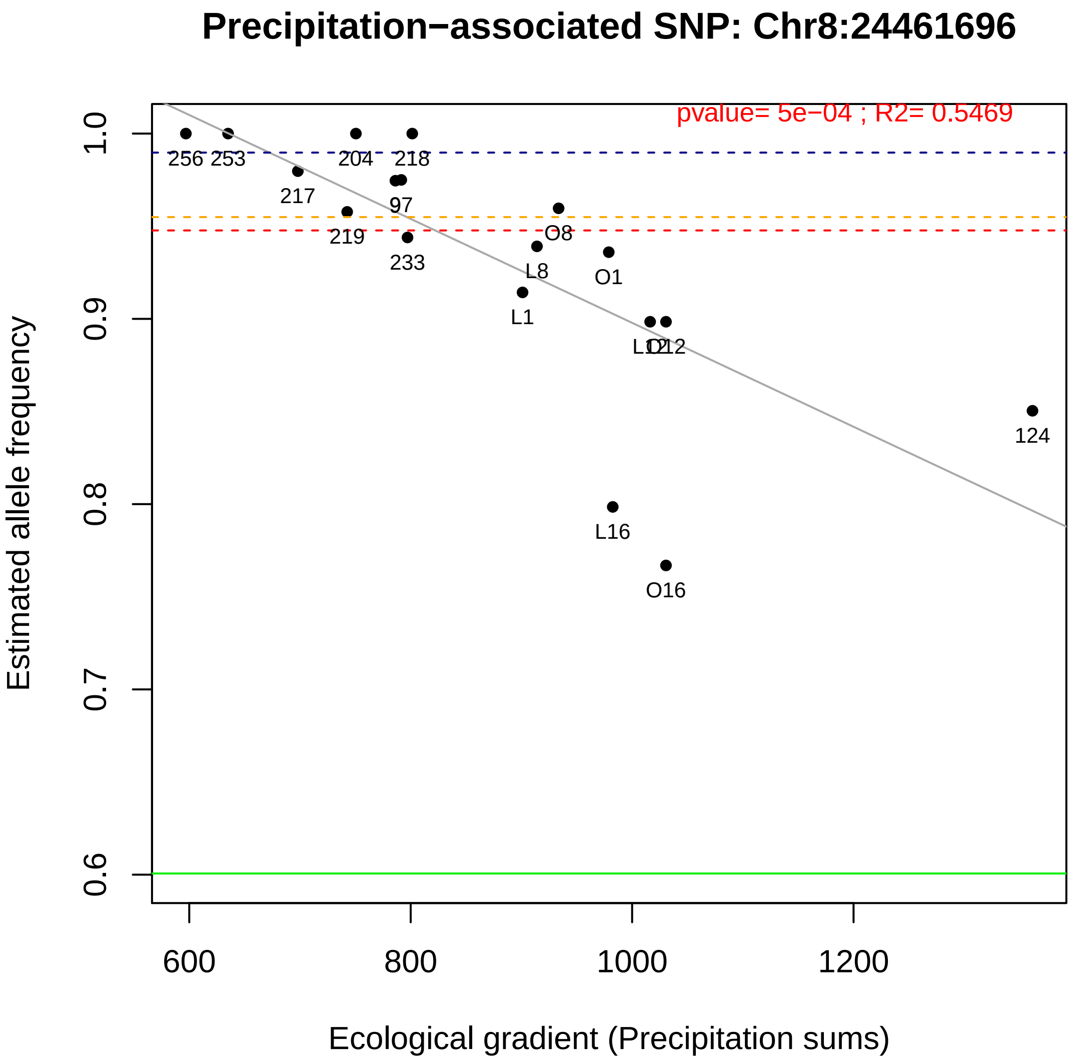

Precipitation−associated SNP: Chr8:24461696
pvalue= 5e−04 ; R2= 0.5469
256
253
204
218
217
97
219
233
O8
L8
O1
L1
L12
O12
124
L16
O16
1.0
0.9
0.8
0.7
0.6
600
800
1000
1200
Estimated allele frequency
Ecological gradient (Precipitation sums)

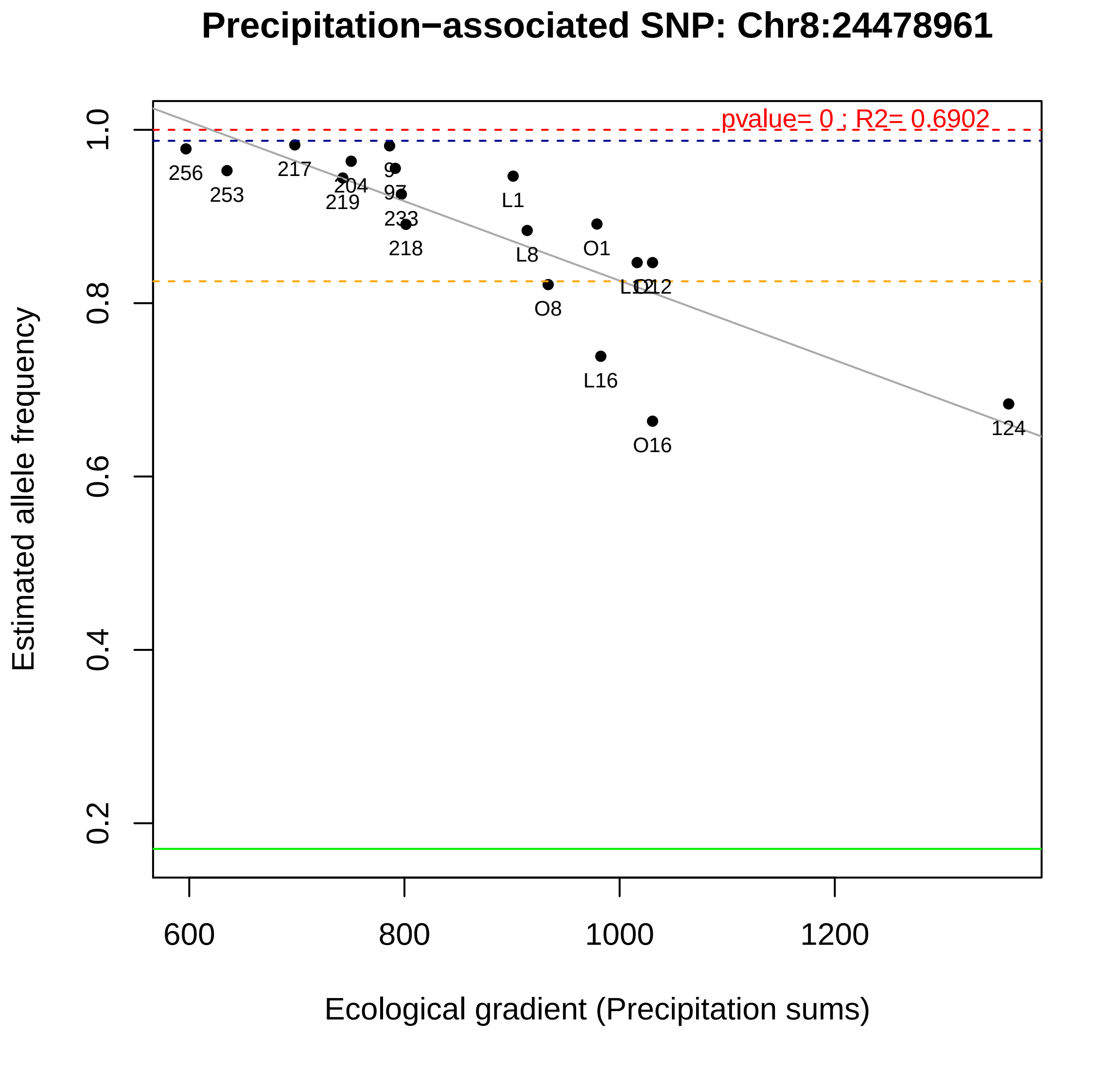
Precipitation−associated SNP: Chr8:24478961
pvalue= 0 ; R2= 0.6902
Estimated allele frequency
0.2
0.4
0.6
0.8
1.0
600
800
1000
1200
Ecological gradient (Precipitation sums)
256
253
217
219
204
9
97
233
218
L1
L8
O8
O1
L10
O12
L16
O16
124

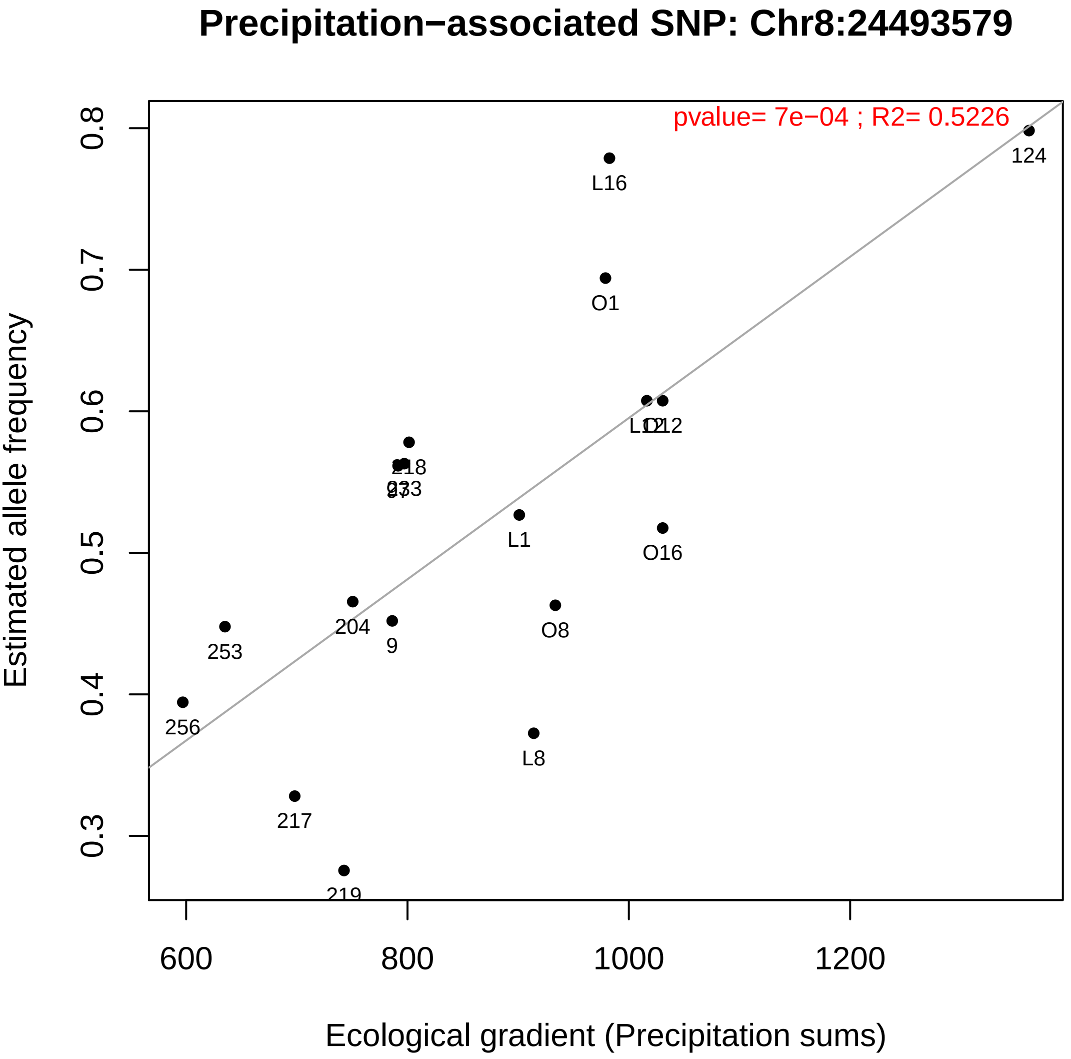
Precipitation−associated SNP: Chr8:24493579
pvalue= 7e−04 ; R2= 0.5226
Estimated allele frequency
Ecological gradient (Precipitation sums)
0.3
0.4
0.5
0.6
0.7
0.8
600
800
1000
1200
124
L16
O1
218
L1
O16
204
9
O8
253
256
L8
217
219

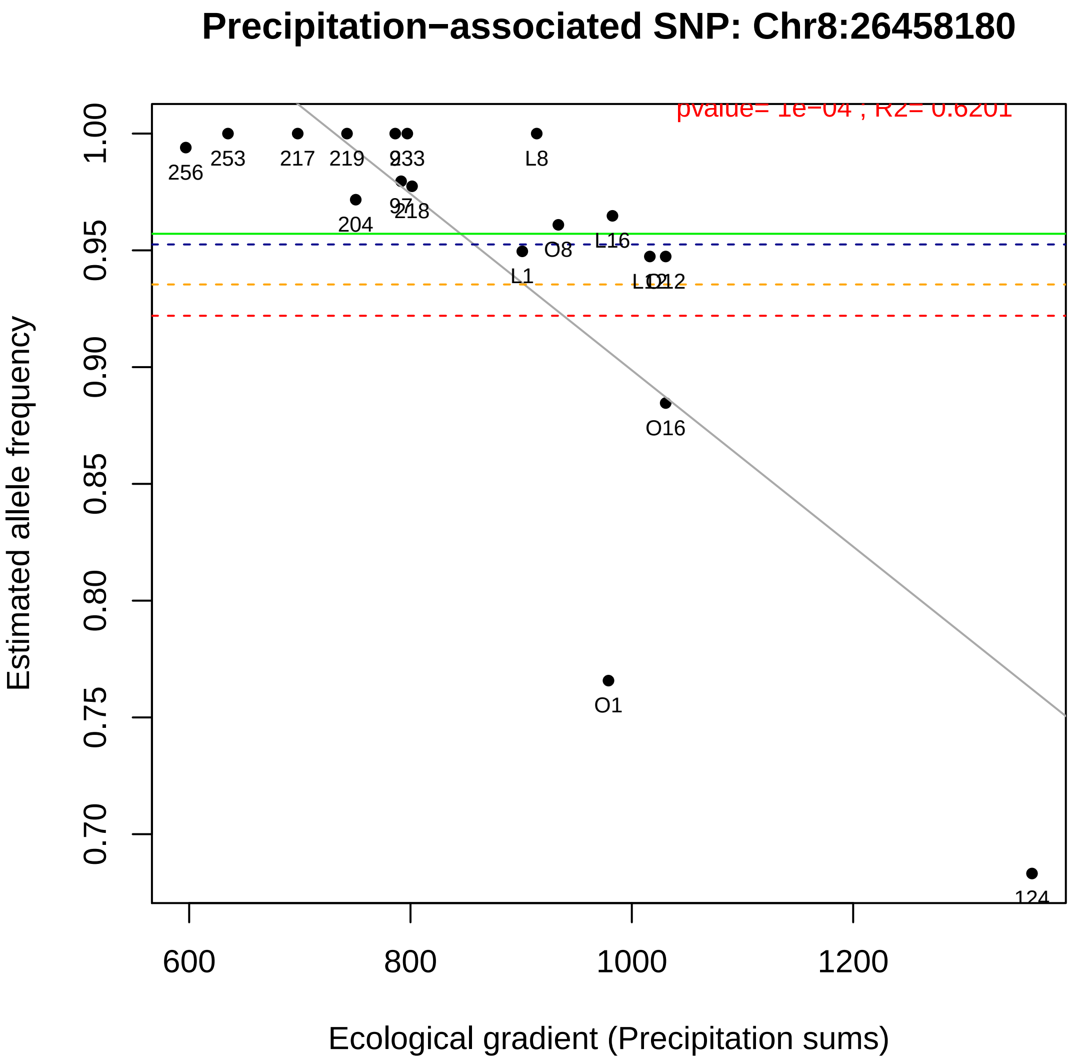
Precipitation−associated SNP: Chr8:26458180
pvalue= 1e−04 ; R2= 0.6201
Estimated allele frequency
Ecological gradient (Precipitation sums)
1.00
0.95
0.90
0.85
0.80
0.75
0.70
600
800
1000
1200
256
253
217
219
233
L8
97
218
204
L16
O8
L1
O12
O16
O1
124

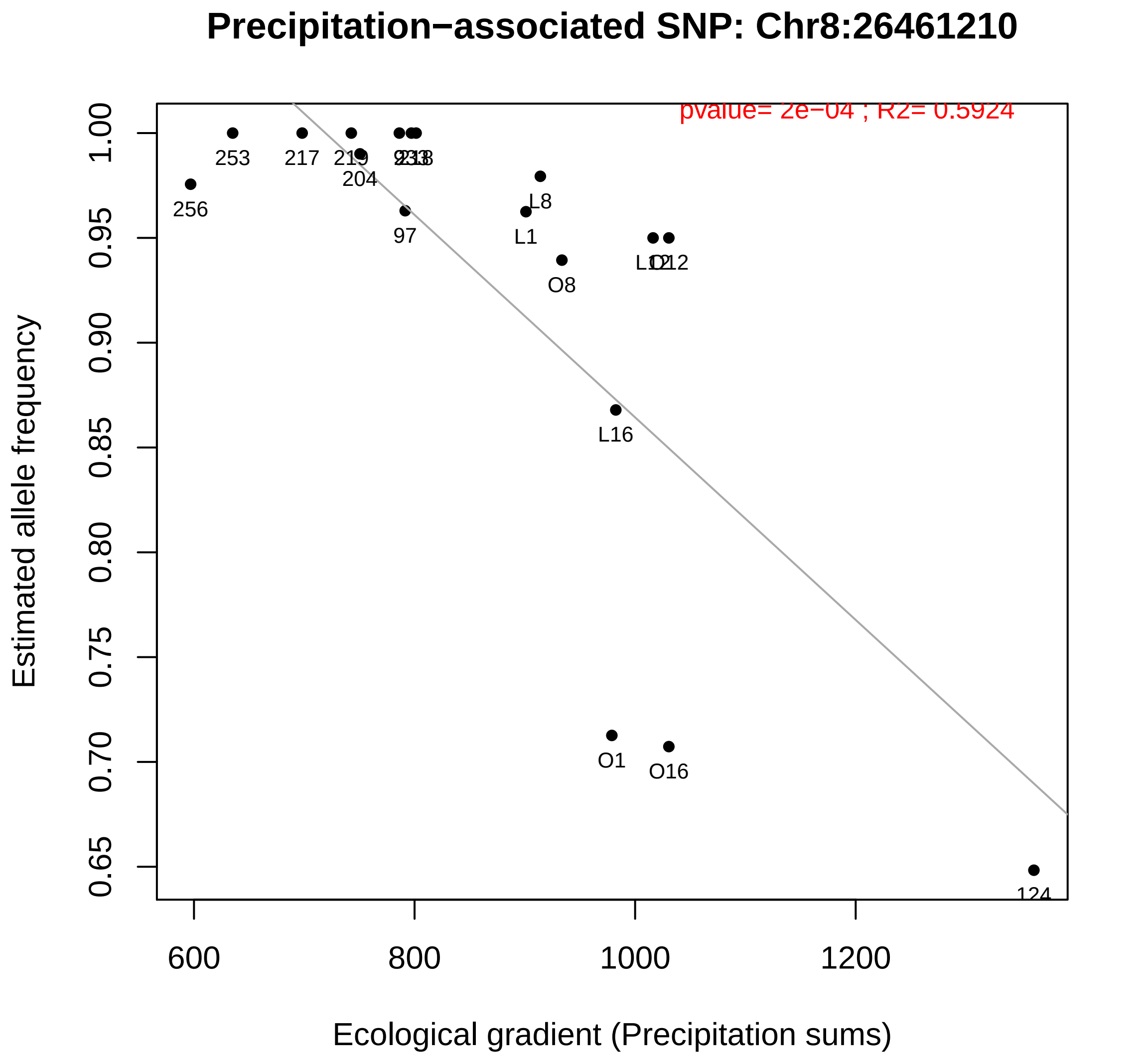

Precipitation−associated SNP: Chr8:26461210
pvalue= 2e−04 ; R2= 0.5924
Estimated allele frequency
Ecological gradient (Precipitation sums)
1.00
0.95
0.90
0.85
0.80
0.75
0.70
0.65
600
800
1000
1200
253
217
219
204
256
97
L8
L1
O8
L12
O12
L16
O1
O16
124

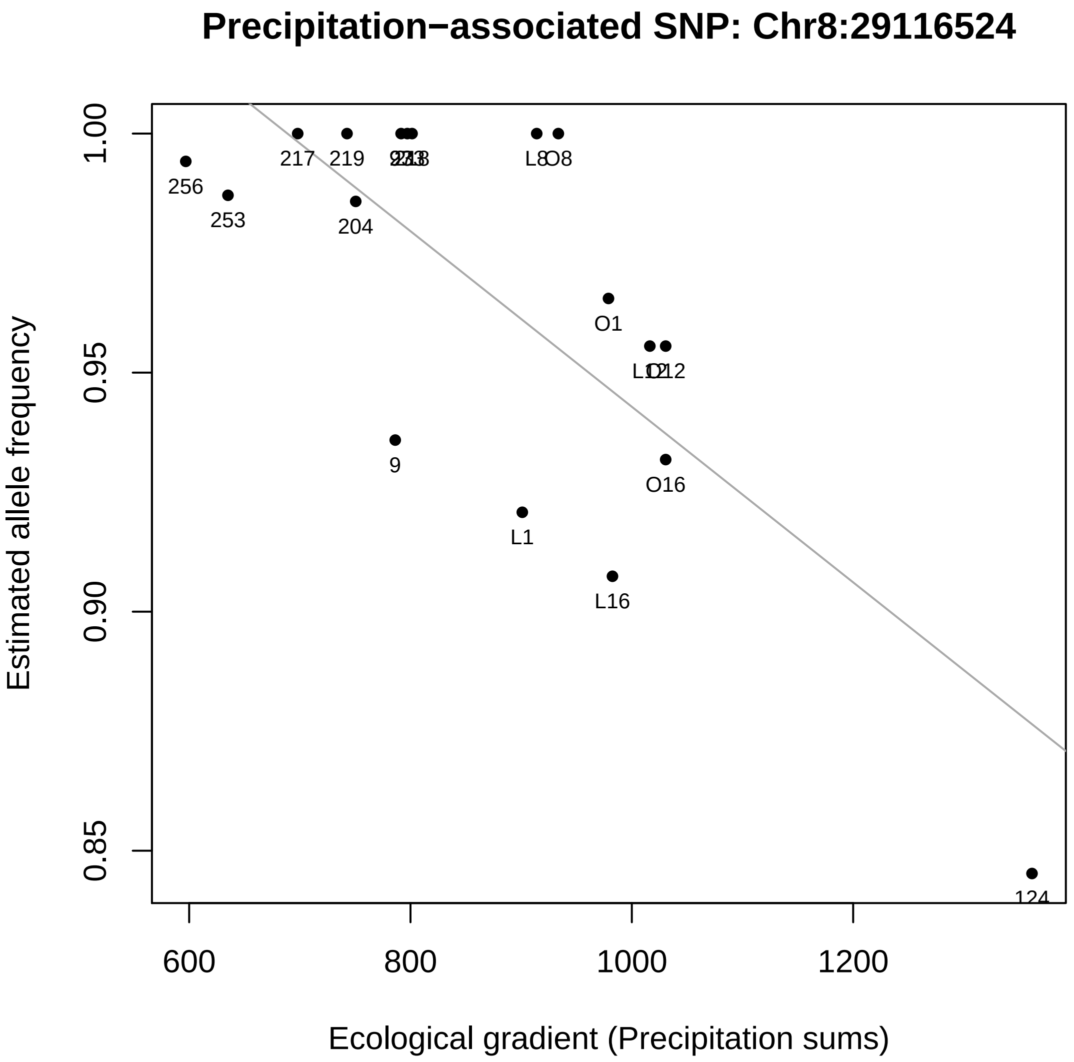
Precipitation−associated SNP: Chr8:29116524
Estimated allele frequency
Ecological gradient (Precipitation sums)
1.00
0.95
0.90
0.85
600
800
1000
1200
256
253
217
219
204
L8
O8
O1
O12
O16
9
L1
L16
124

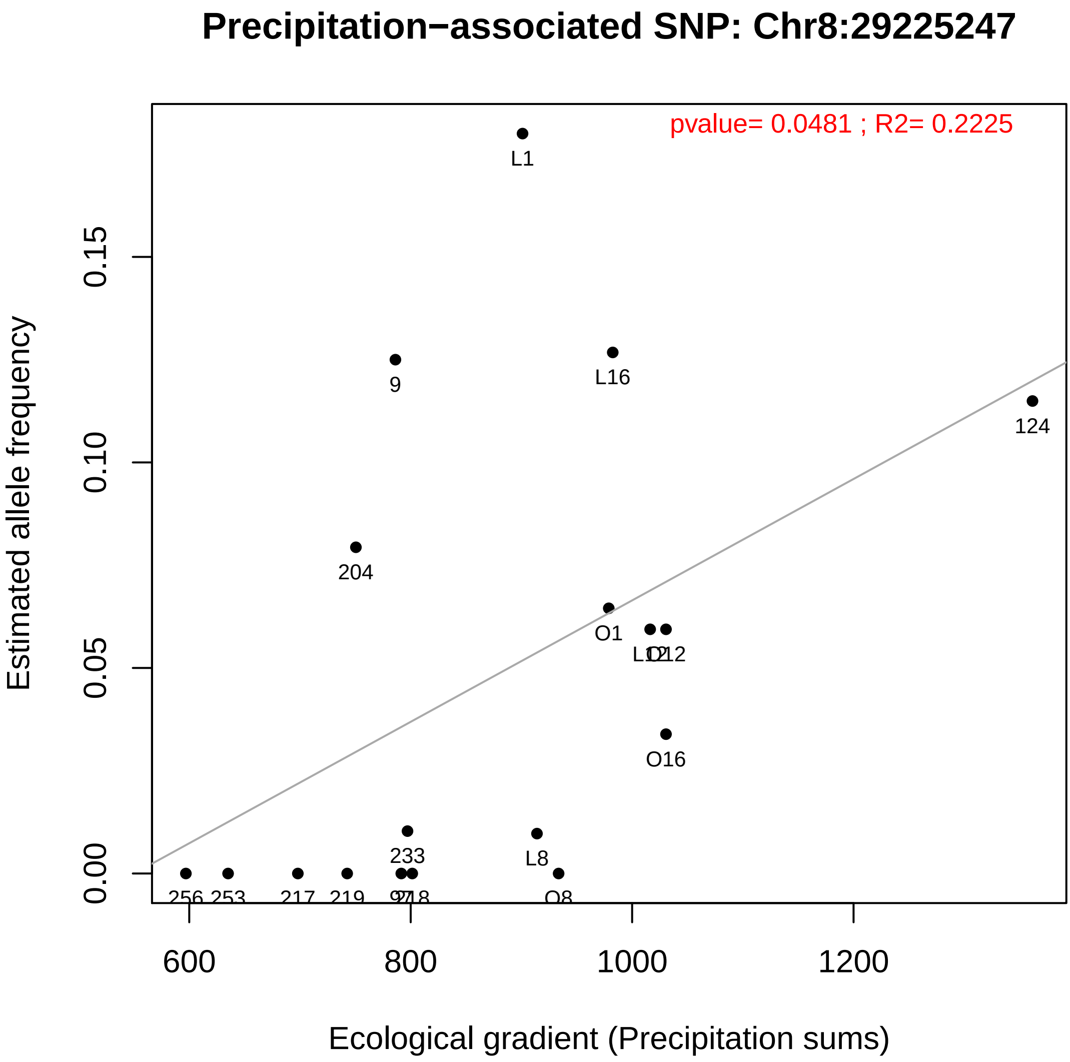

Precipitation−associated SNP: Chr8:29225247
pvalue= 0.0481 ; R2= 0.2225
Estimated allele frequency
0.00
0.05
0.10
0.15
600
800
1000
1200
Ecological gradient (Precipitation sums)
L1
9
L16
124
204
O1
O12
O16
233
L8
O8
256
253
217
219

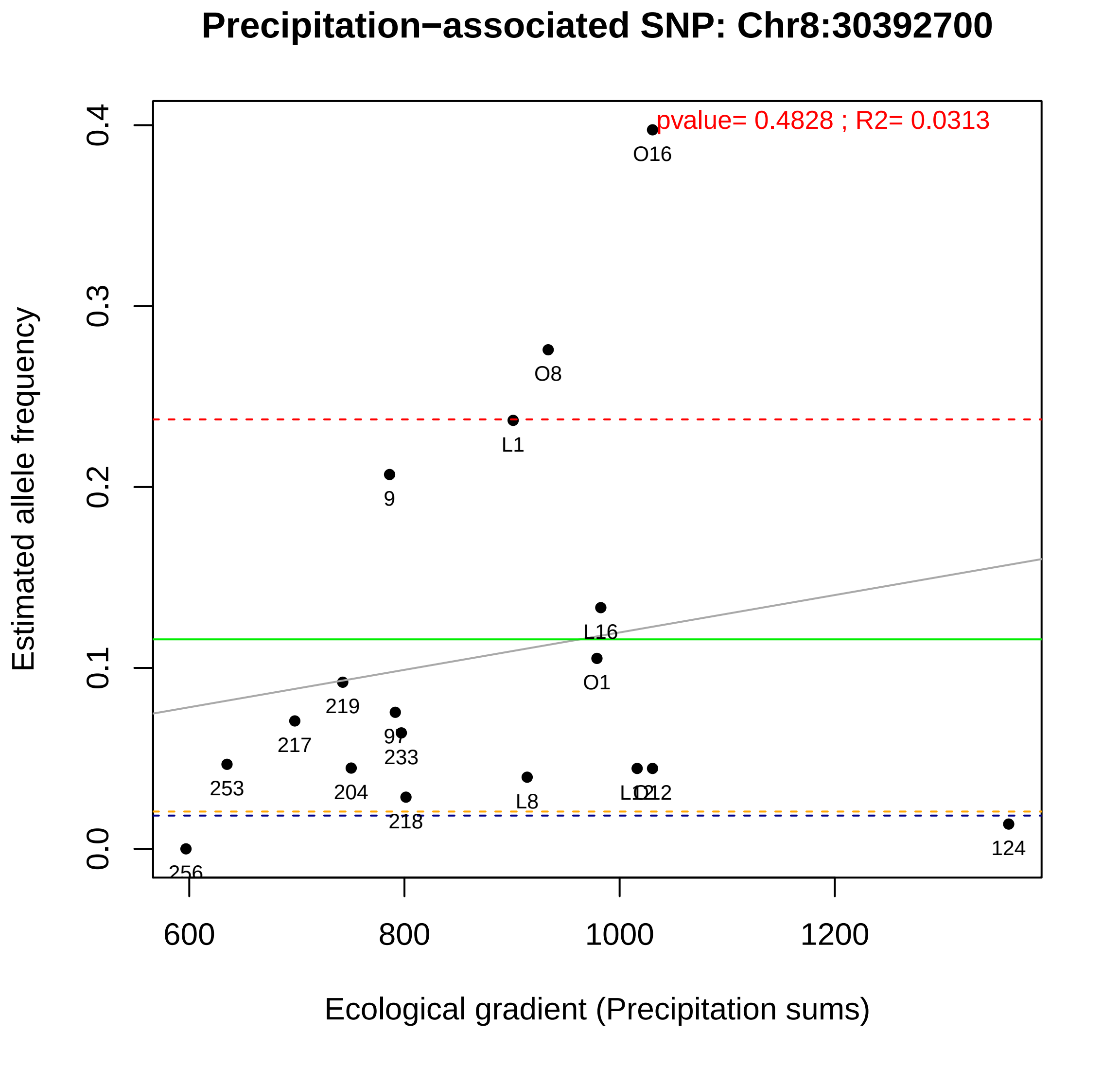

Precipitation−associated SNP: Chr8:30392700
pvalue= 0.4828 ; R2= 0.0313
Estimated allele frequency
0.0
0.1
0.2
0.3
0.4
600
800
1000
1200
Ecological gradient (Precipitation sums)
O16
O8
L1
9
L16
O1
219
217
97
233
253
204
218
L8
L12
O12
124
256

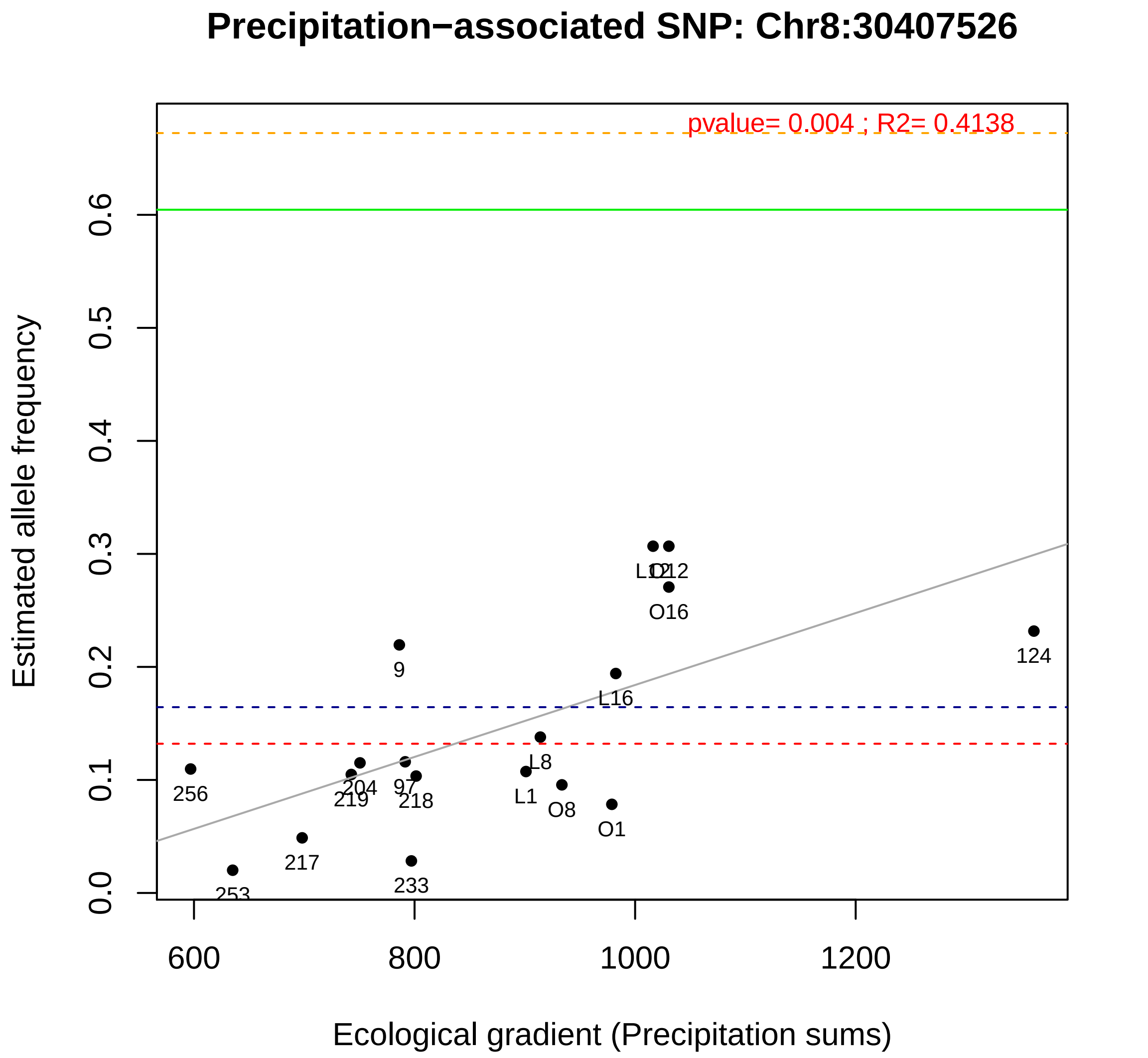

Precipitation−associated SNP: Chr8:30407526
pvalue= 0.004 ; R2= 0.4138
Estimated allele frequency
0.0
0.1
0.2
0.3
0.4
0.5
0.6
600
800
1000
1200
Ecological gradient (Precipitation sums)
256
253
217
219
204
9
97
218
233
L1
L8
O8
O1
L16
L12
O12
O16
124

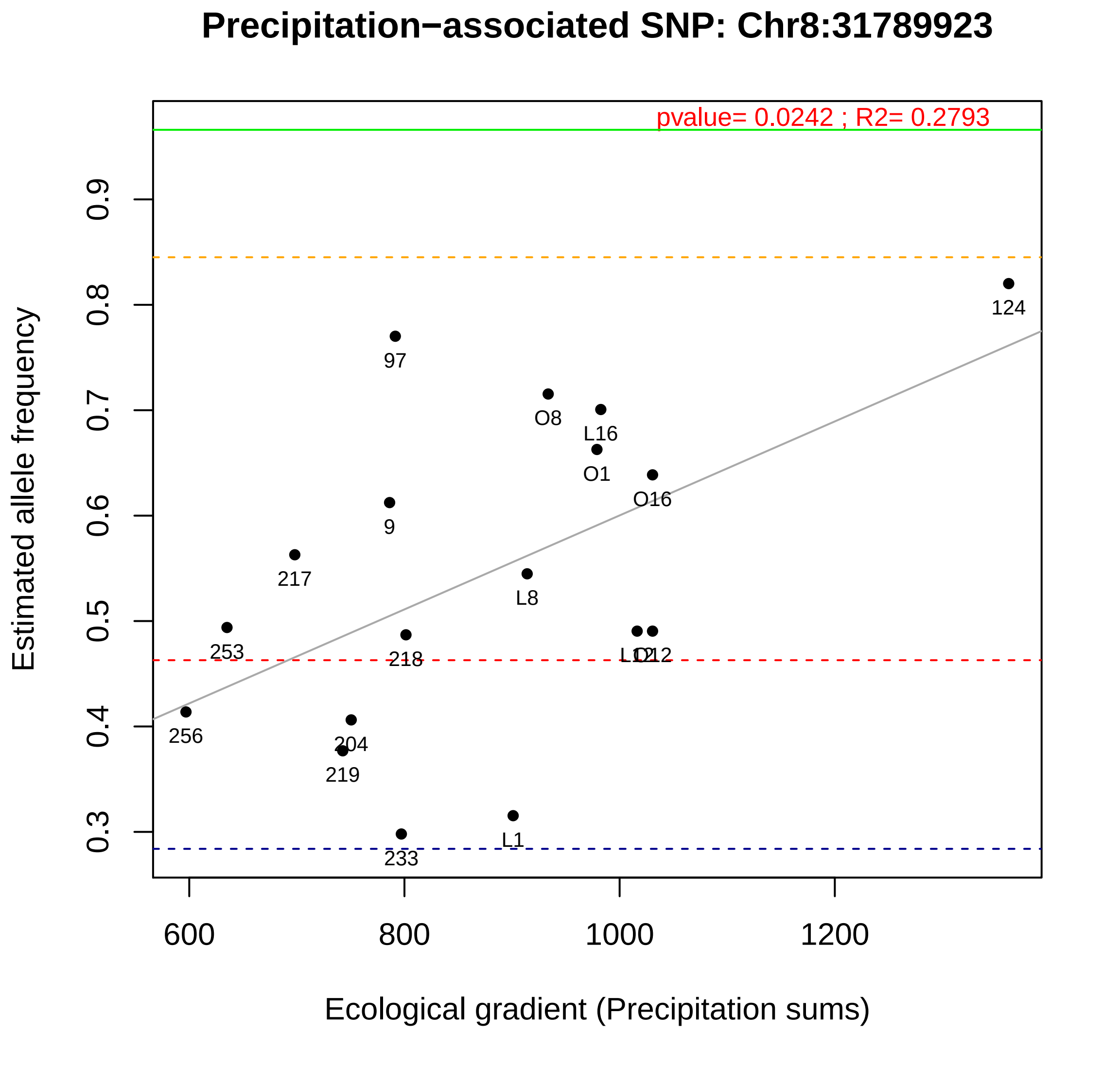

Precipitation−associated SNP: Chr8:31789923
pvalue= 0.0242 ; R2= 0.2793
Estimated allele frequency
0.3
0.4
0.5
0.6
0.7
0.8
0.9
600
800
1000
1200
Ecological gradient (Precipitation sums)
124
97
O8
L16
O1
O16
9
217
L8
253
218
L12
O12
256
204
219
233
L1

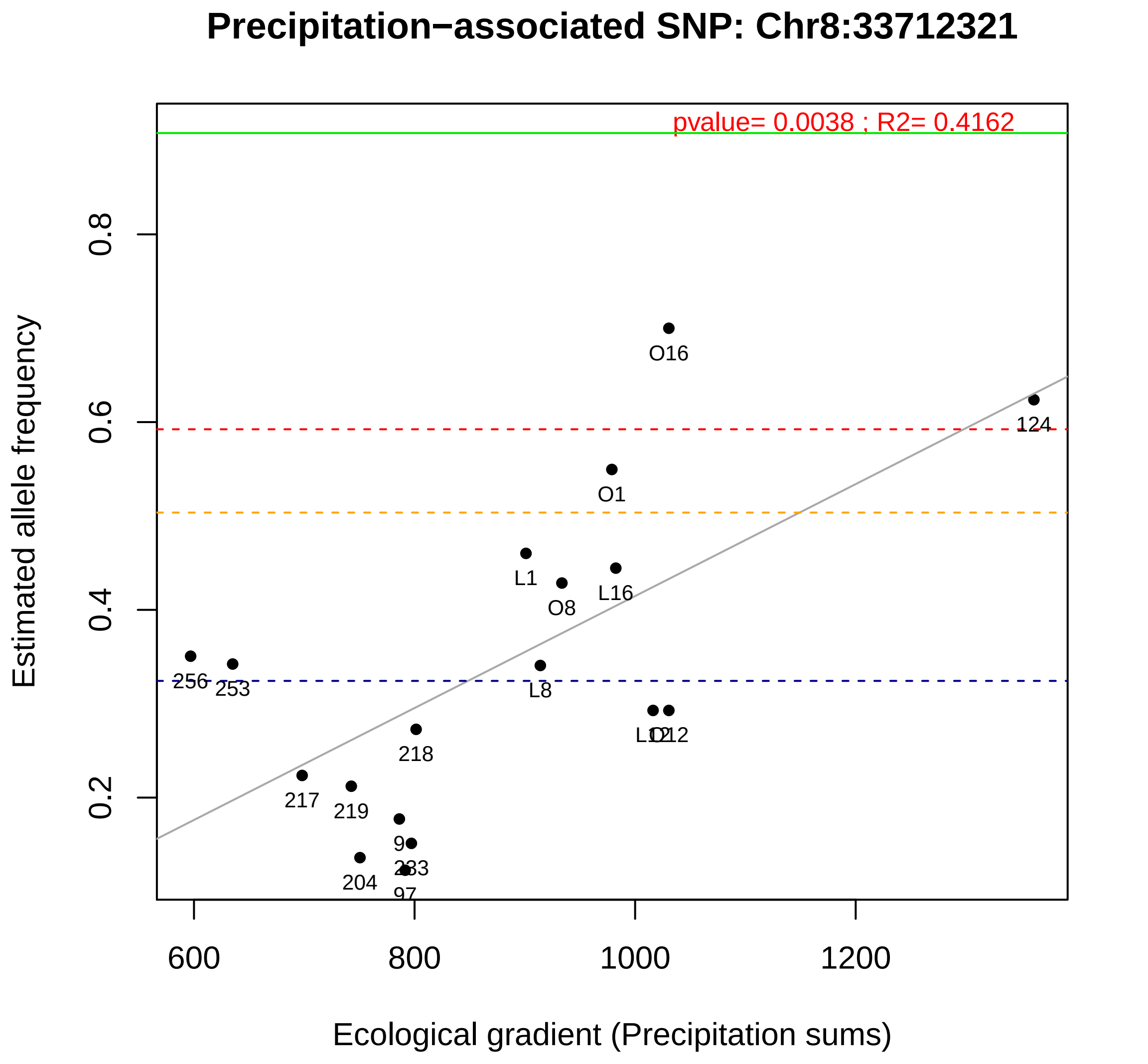
Precipitation−associated SNP: Chr8:33712321
pvalue= 0.0038 ; R2= 0.4162
Estimated allele frequency
0.2
0.4
0.6
0.8
600
800
1000
1200
Ecological gradient (Precipitation sums)
O16
124
O1
L1
O8
L16
256
253
L8
L12
O12
218
217
219
204
97
233
9

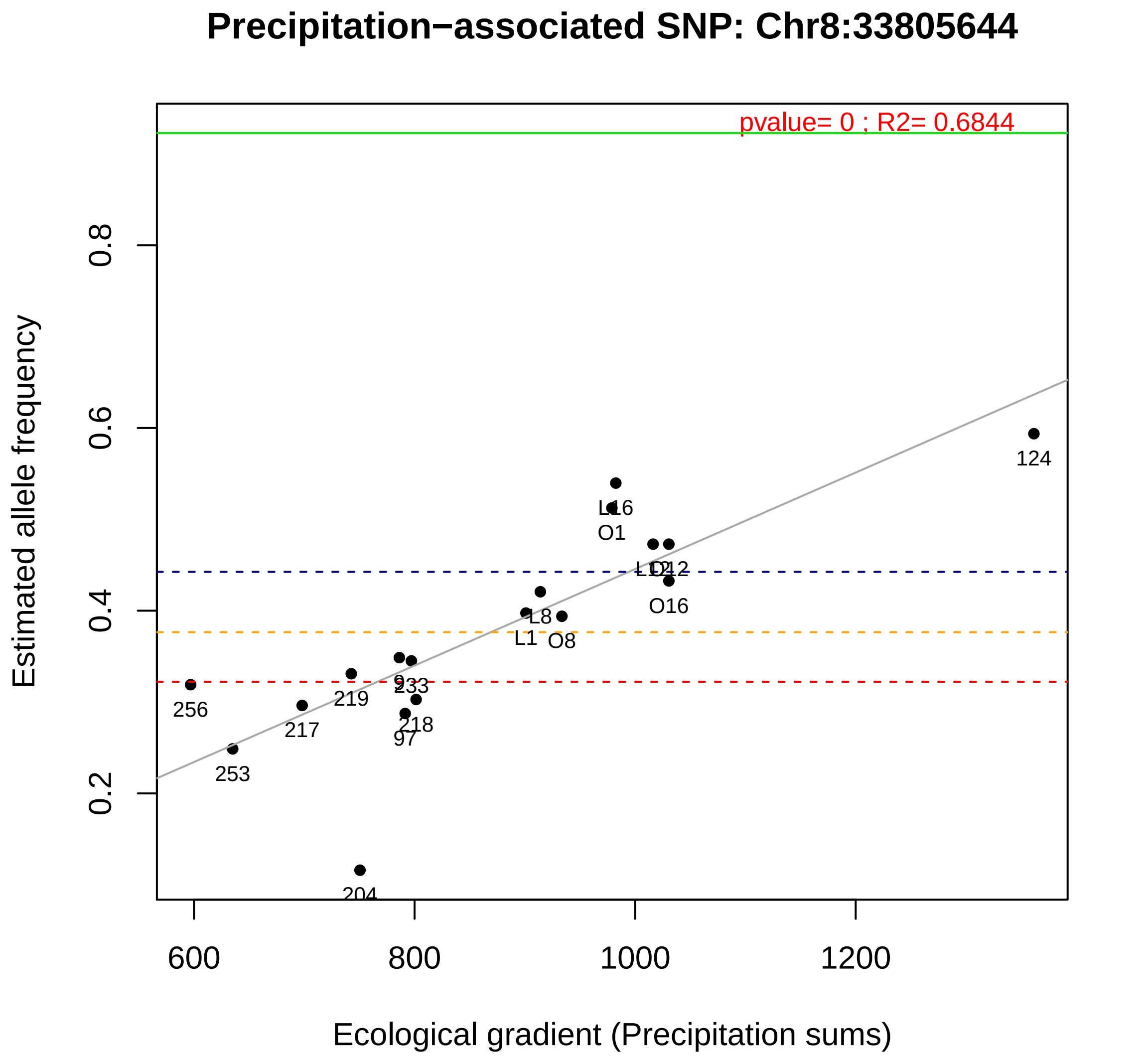
Precipitation−associated SNP: Chr8:33805644
pvalue= 0 ; R2= 0.6844
Estimated allele frequency
0.2
0.4
0.6
0.8
600
800
1000
1200
Ecological gradient (Precipitation sums)
124
L16
O1
L12
O12
O16
L8
L1
O8
256
219
233
217
218
97
253
204

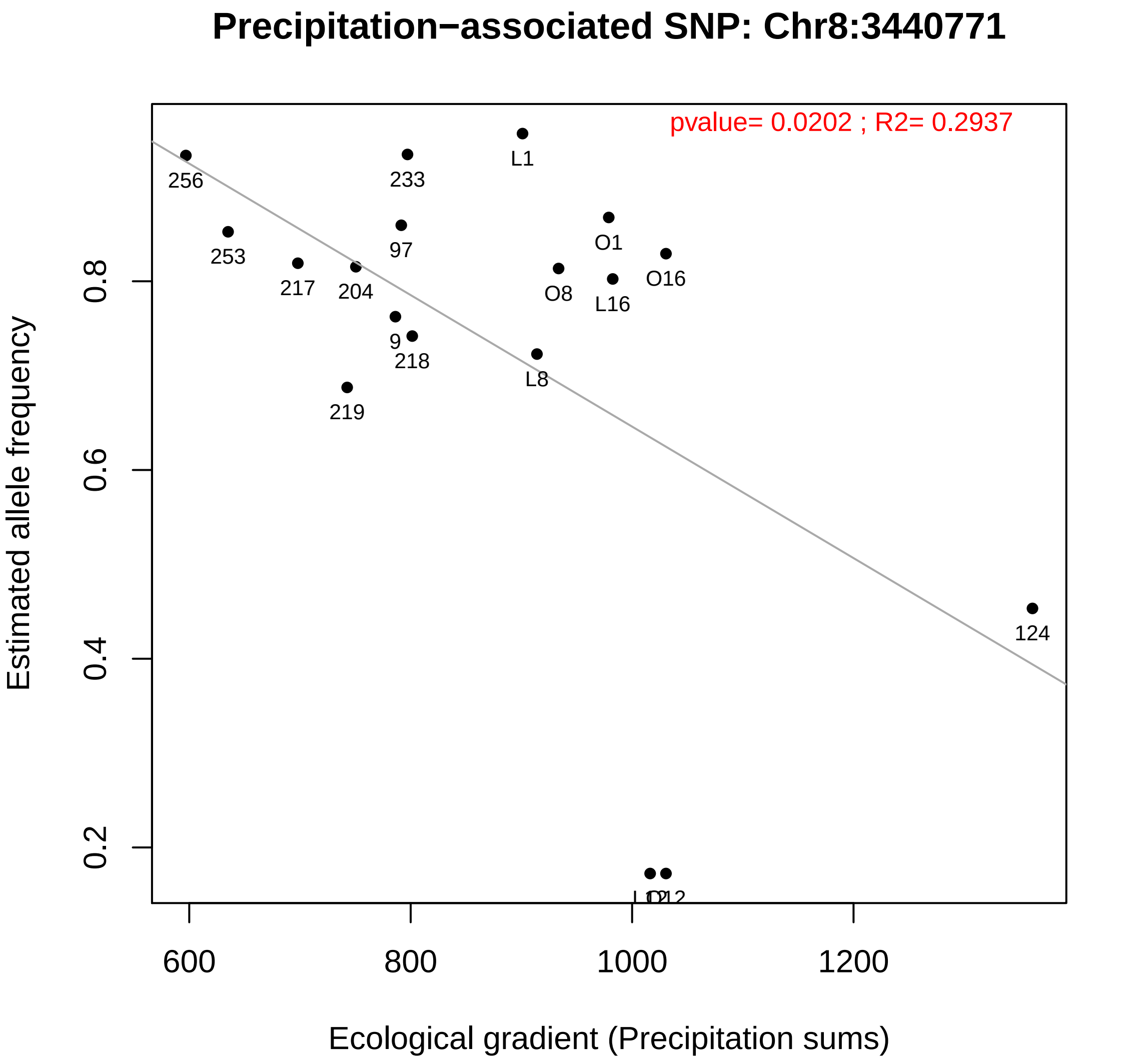

Precipitation−associated SNP: Chr8:3440771
pvalue= 0.0202 ; R2= 0.2937
Estimated allele frequency
0.2
0.4
0.6
0.8
600
800
1000
1200
Ecological gradient (Precipitation sums)
256
233
L1
253
97
O1
217
204
O8
O16
L16
9
218
L8
219
124
O12

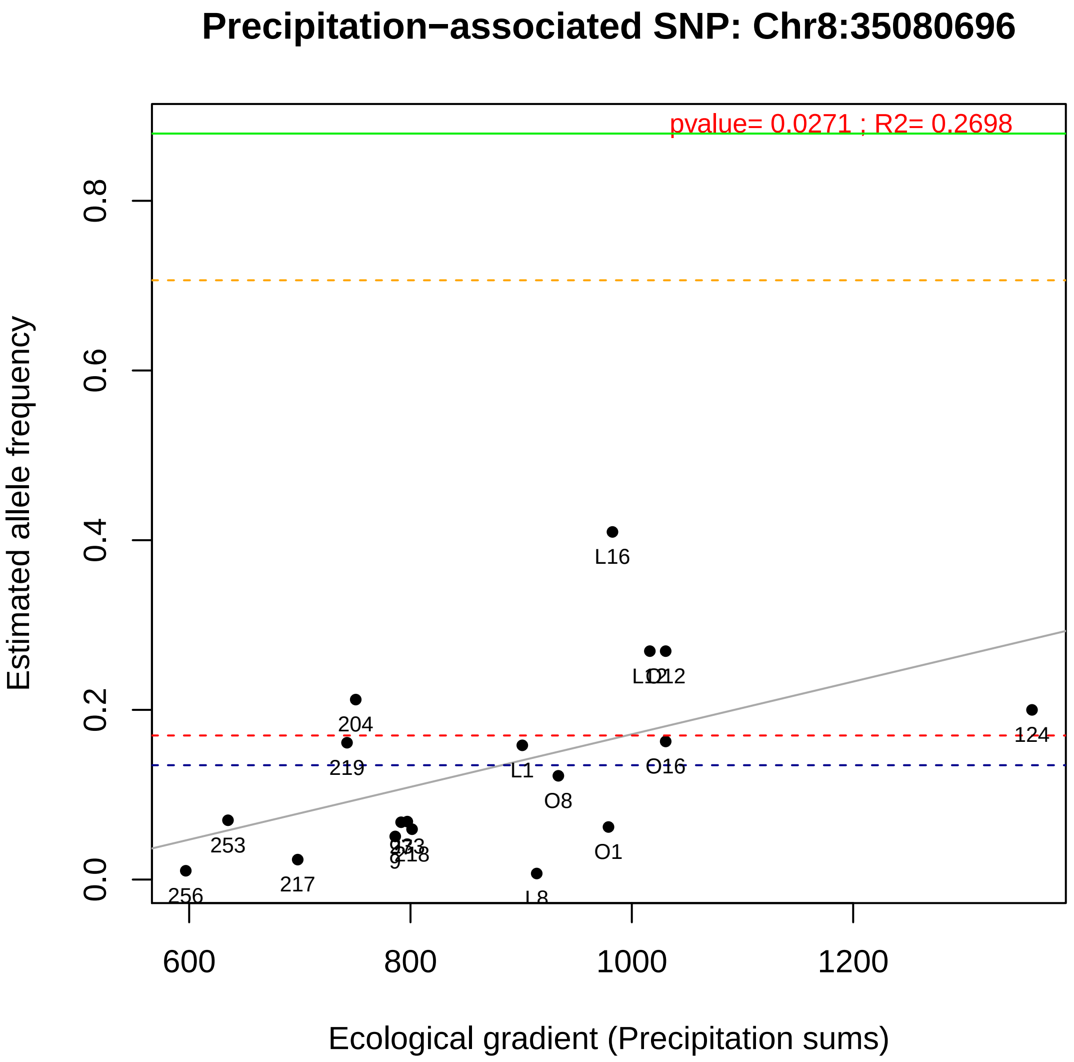
Precipitation−associated SNP: Chr8:35080696
pvalue= 0.0271 ; R2= 0.2698
Estimated allele frequency
0.0
0.2
0.4
0.6
0.8
Ecological gradient (Precipitation sums)
600
800
1000
1200
L16
204
219
L1
O8
124
O16
253
217
256
218
O1

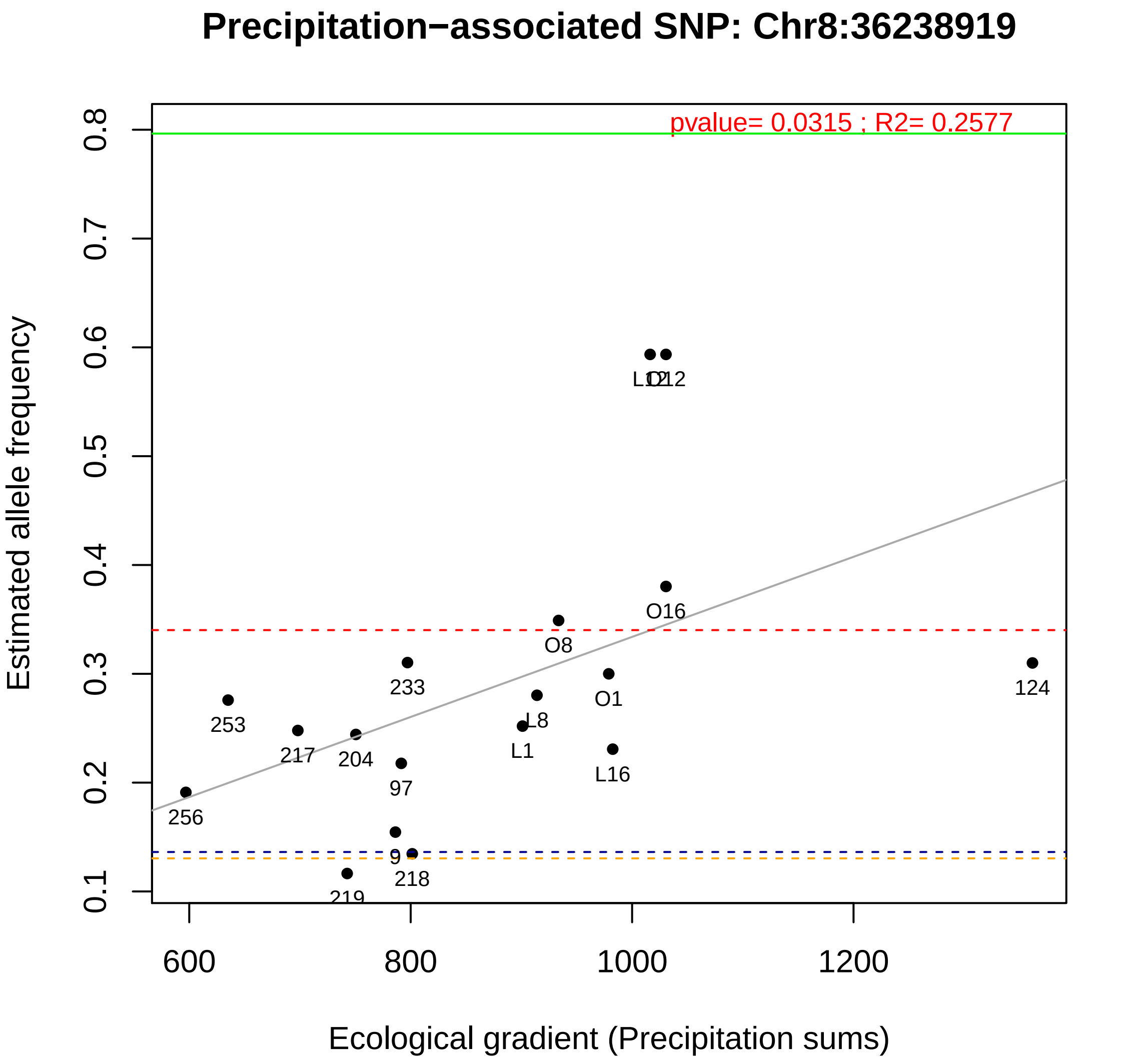
Precipitation−associated SNP: Chr8:36238919
pvalue= 0.0315 ; R2= 0.2577
Estimated allele frequency
0.1
0.2
0.3
0.4
0.5
0.6
0.7
0.8
600
800
1000
1200
Ecological gradient (Precipitation sums)
256
253
217
204
97
233
219
218
L1
L8
O8
O1
L16
O16
124
O12

**Precipitation−associated SNP: Chr8:36282269**

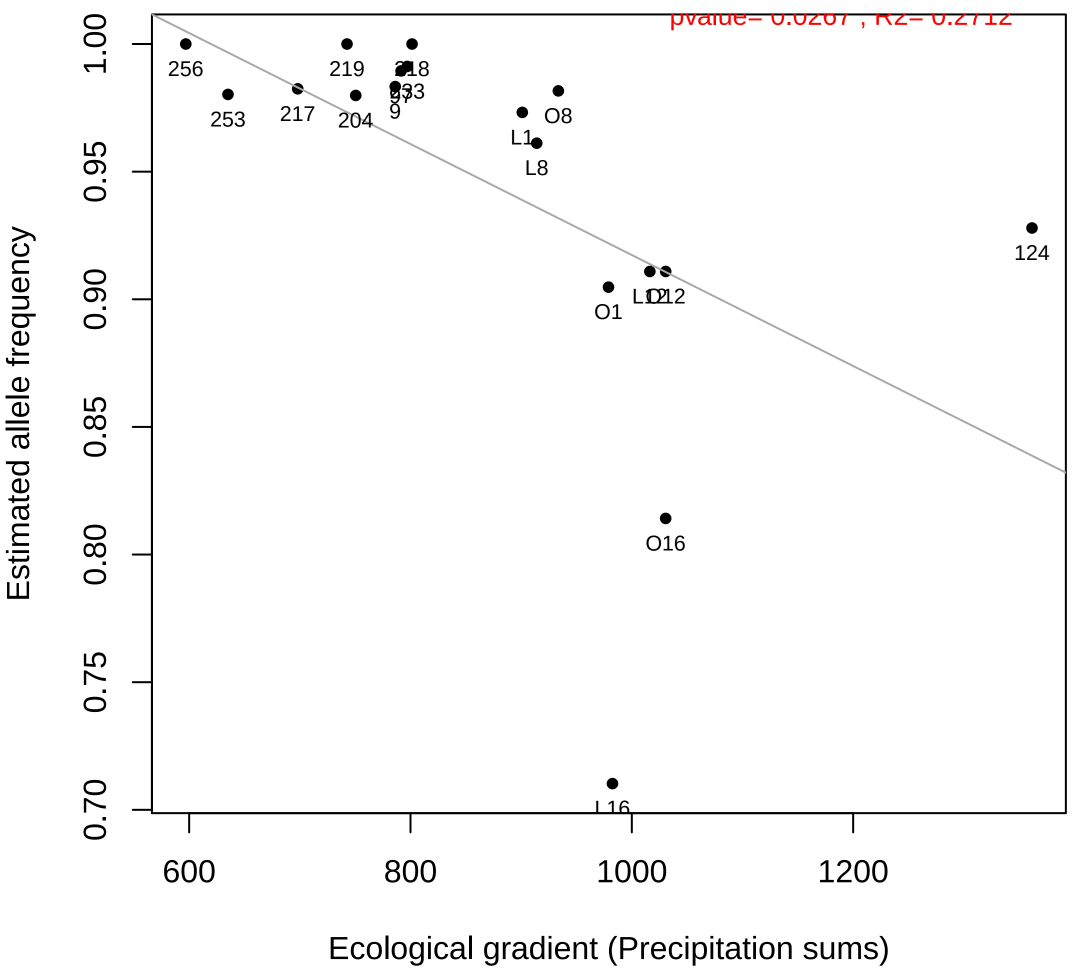

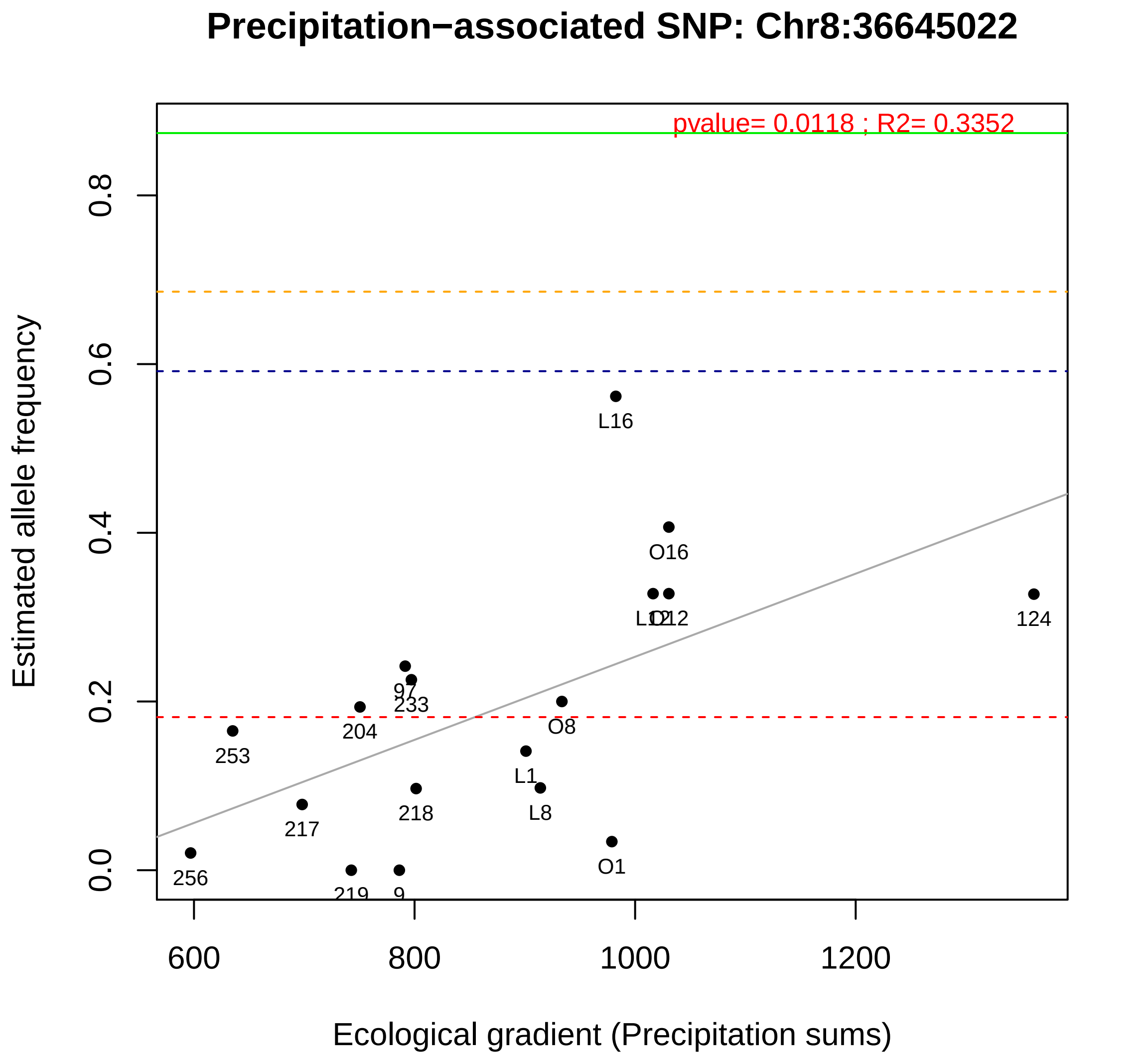

Precipitation−associated SNP: Chr8:36645022
pvalue= 0.0118 ; R2= 0.3352
Estimated allele frequency
0.0
0.2
0.4
0.6
0.8
600
800
1000
1200
Ecological gradient (Precipitation sums)
L16
O16
124
204
253
233
97
O8
L1
L8
217
218
256
219
9
O1

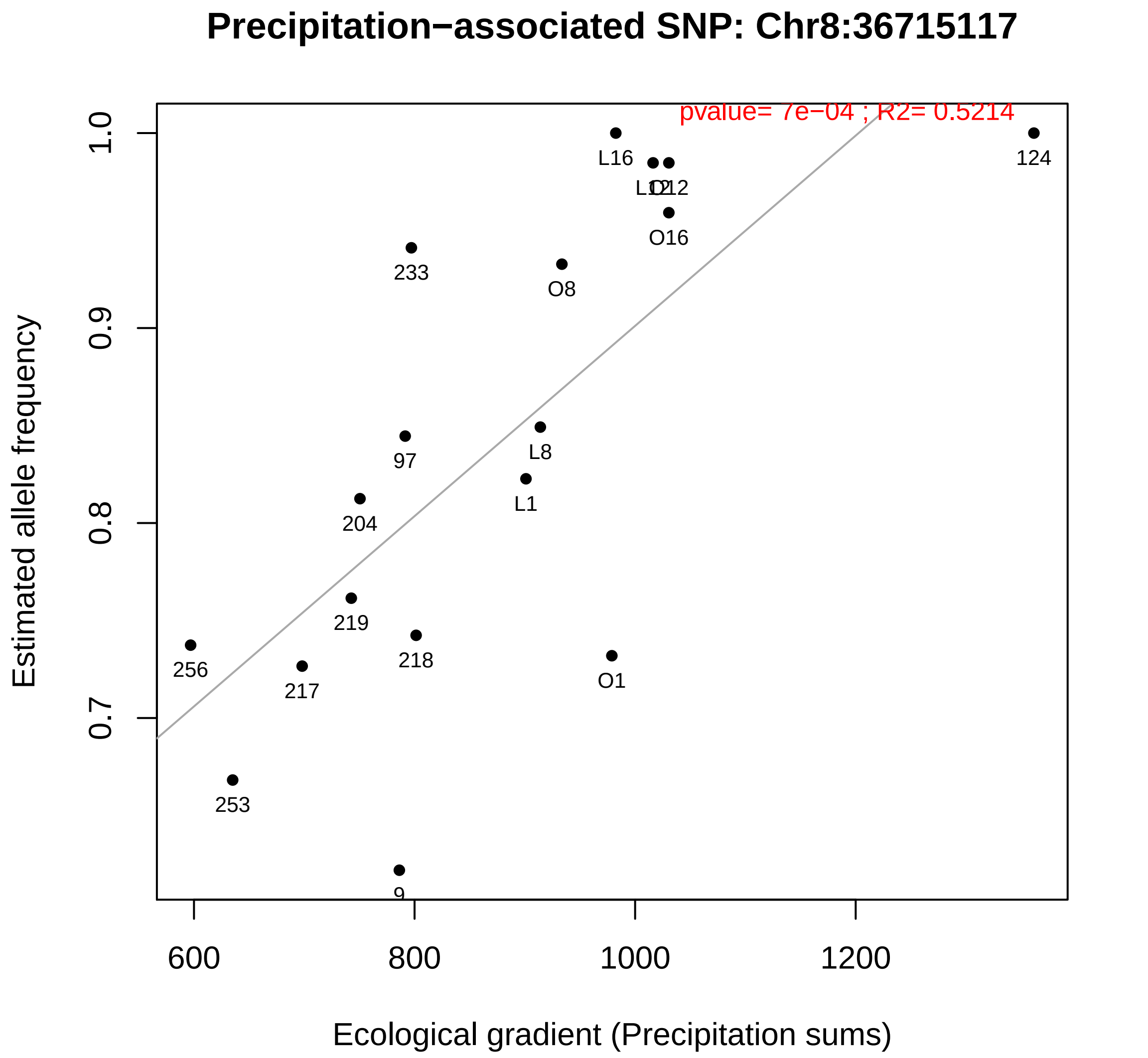
Precipitation−associated SNP: Chr8:36715117
pvalue= 7e−04 ; R2= 0.5214
Estimated allele frequency
Ecological gradient (Precipitation sums)
0.7
0.8
0.9
1.0
600
800
1000
1200
L16
124
L12
O12
O16
233
O8
97
L8
L1
204
219
218
256
217
O1
253
9

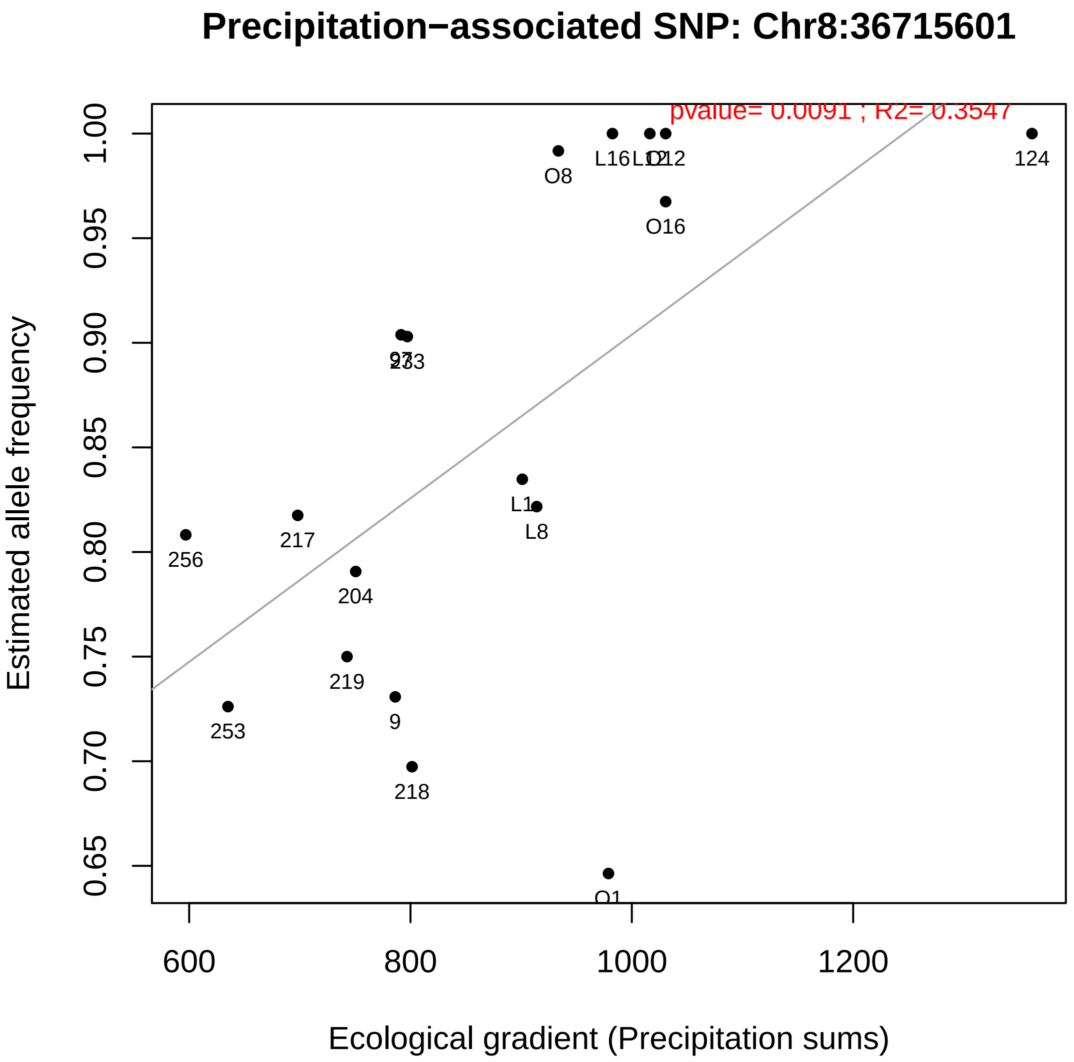

Precipitation−associated SNP: Chr8:36715601
pvalue= 0.0091 ; R2= 0.3547
Estimated allele frequency
Ecological gradient (Precipitation sums)
0.65
0.70
0.75
0.80
0.85
0.90
0.95
1.00
600
800
1000
1200
L16
O12
124
O8
O16
L1
L8
217
256
204
219
253
9
218
O1

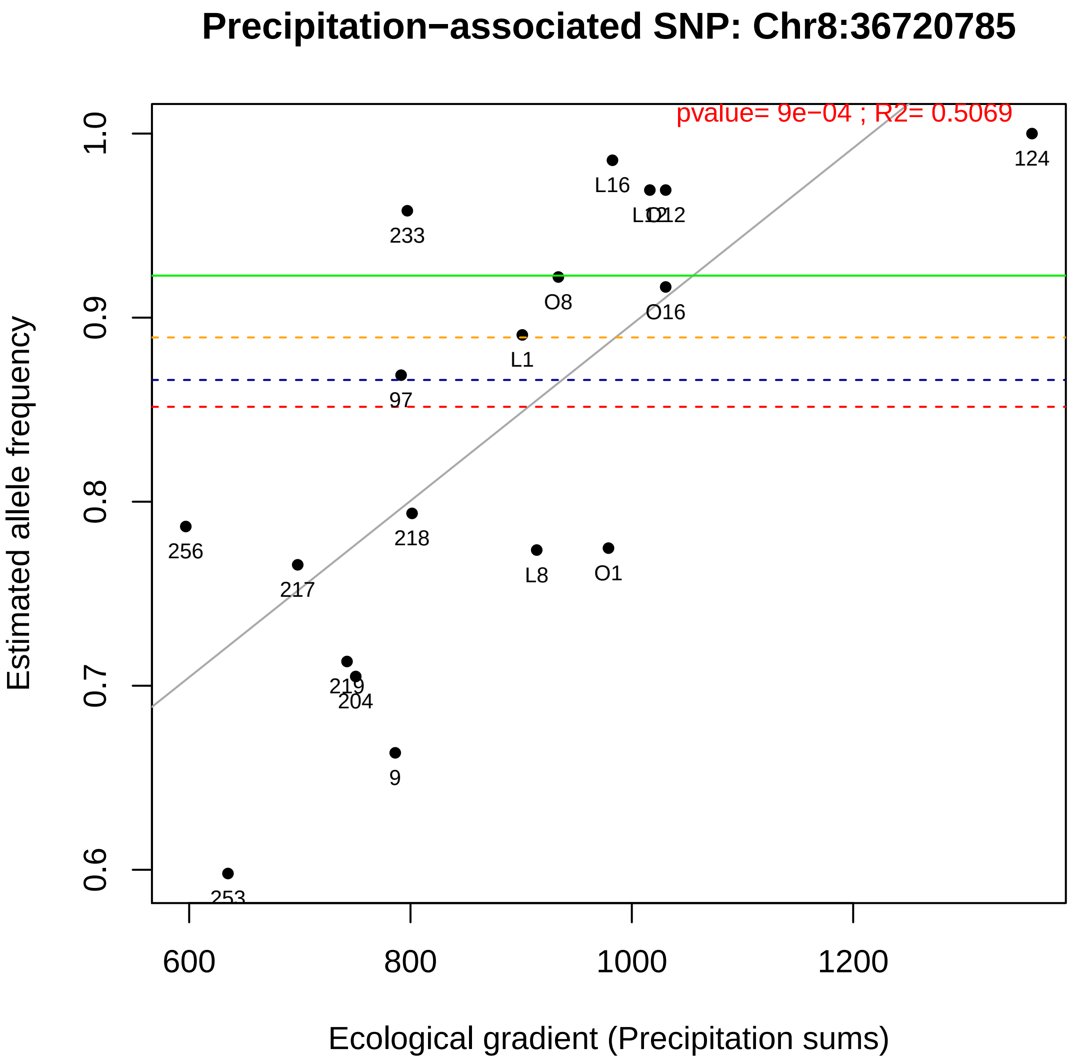

Precipitation−associated SNP: Chr8:36720785
pvalue= 9e−04 ; R2= 0.5069
Estimated allele frequency
Ecological gradient (Precipitation sums)
0.6
0.7
0.8
0.9
1.0
600
800
1000
1200
124
L16
233
O8
O16
L1
97
218
256
217
L8
O1
219
204
9
253

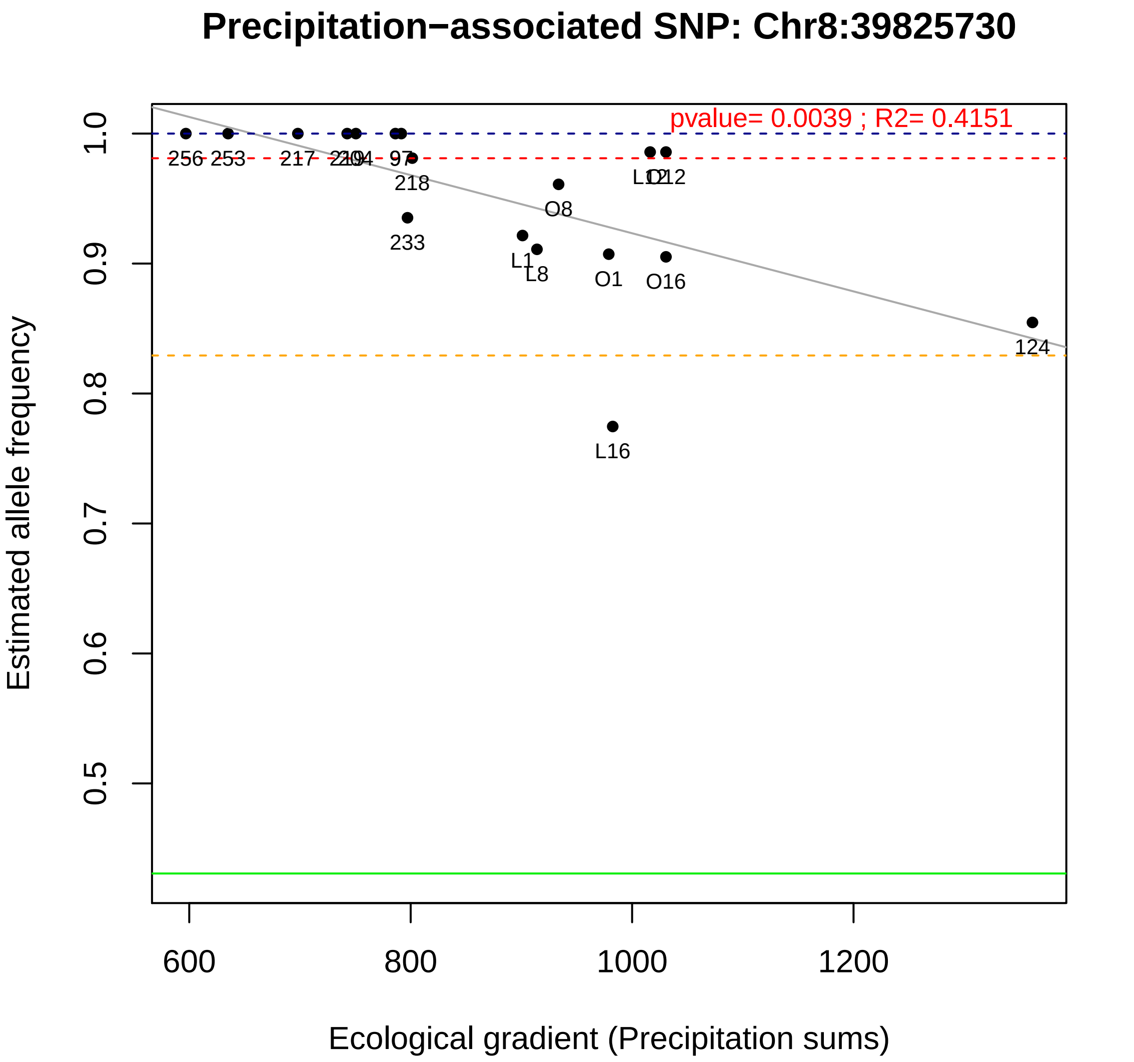

Precipitation−associated SNP: Chr8:39825730
pvalue= 0.0039 ; R2= 0.4151
Estimated allele frequency
Ecological gradient (Precipitation sums)
1.0
0.9
0.8
0.7
0.6
0.5
600
800
1000
1200
256
253
217
97
218
233
O8
L1
L8
O1
O16
L16
124

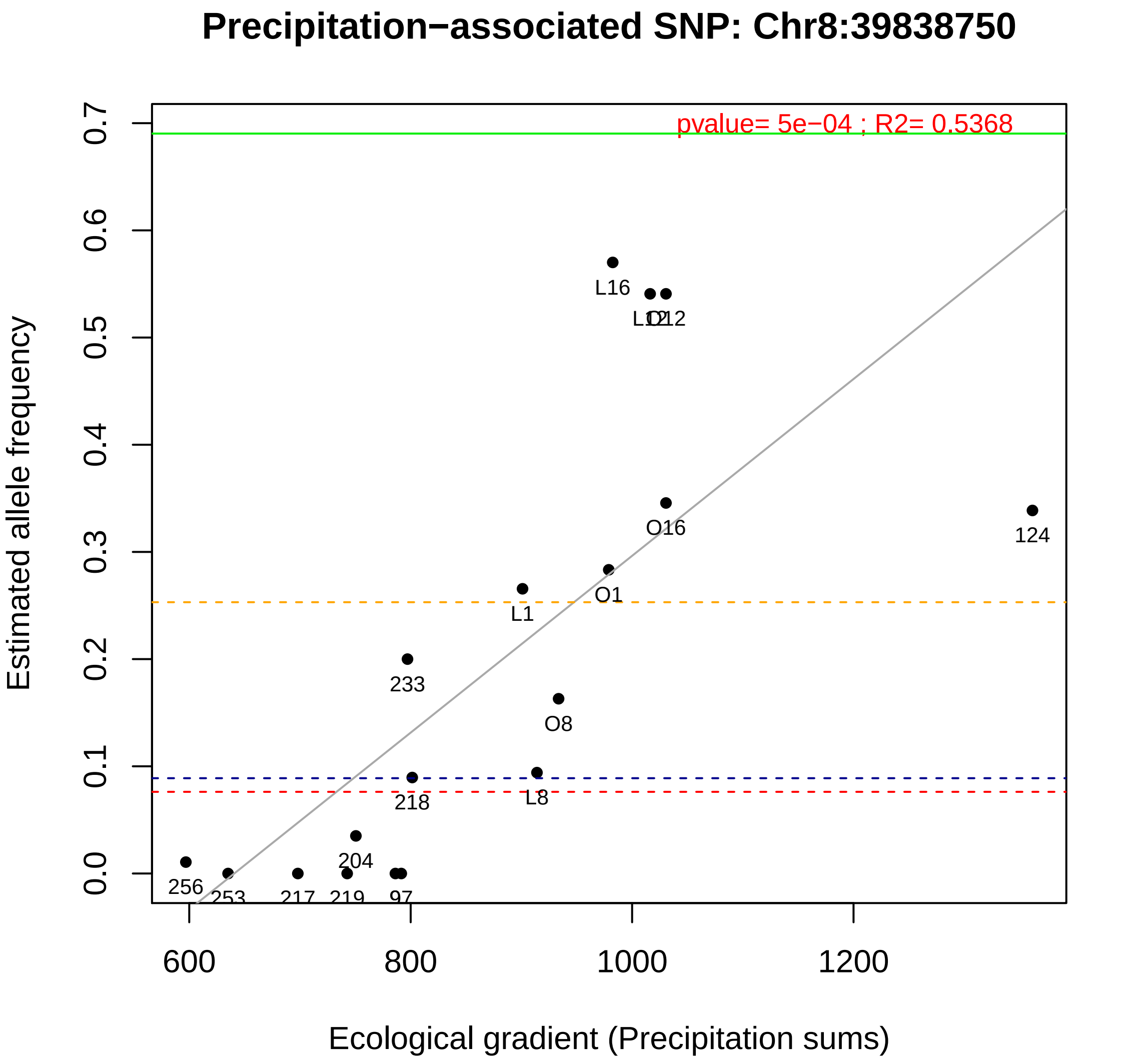

Precipitation−associated SNP: Chr8:39838750
pvalue= 5e−04 ; R2= 0.5368
Estimated allele frequency
0.0
0.1
0.2
0.3
0.4
0.5
0.6
0.7
L16
L12
O12
O16
124
O1
L1
233
O8
218
L8
204
256
253
217
219
97
600
800
1000
1200
Ecological gradient (Precipitation sums)

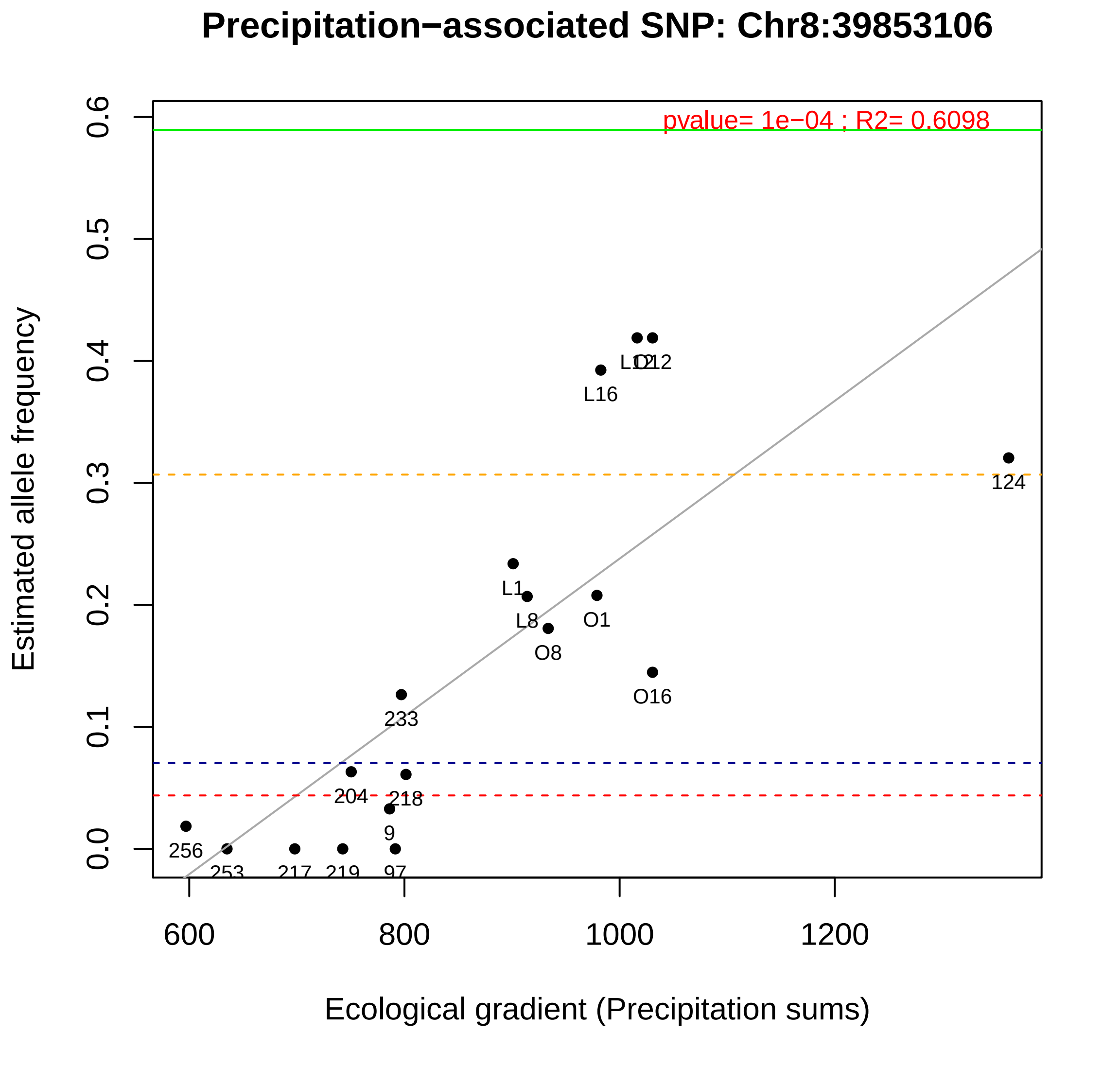

Precipitation−associated SNP: Chr8:39853106
pvalue= 1e−04 ; R2= 0.6098
Estimated allele frequency
0.0
0.1
0.2
0.3
0.4
0.5
0.6
600
800
1000
1200
Ecological gradient (Precipitation sums)
L12
O12
L16
124
L1
L8
O1
O8
O16
233
204
218
9
256
253
217
219
97

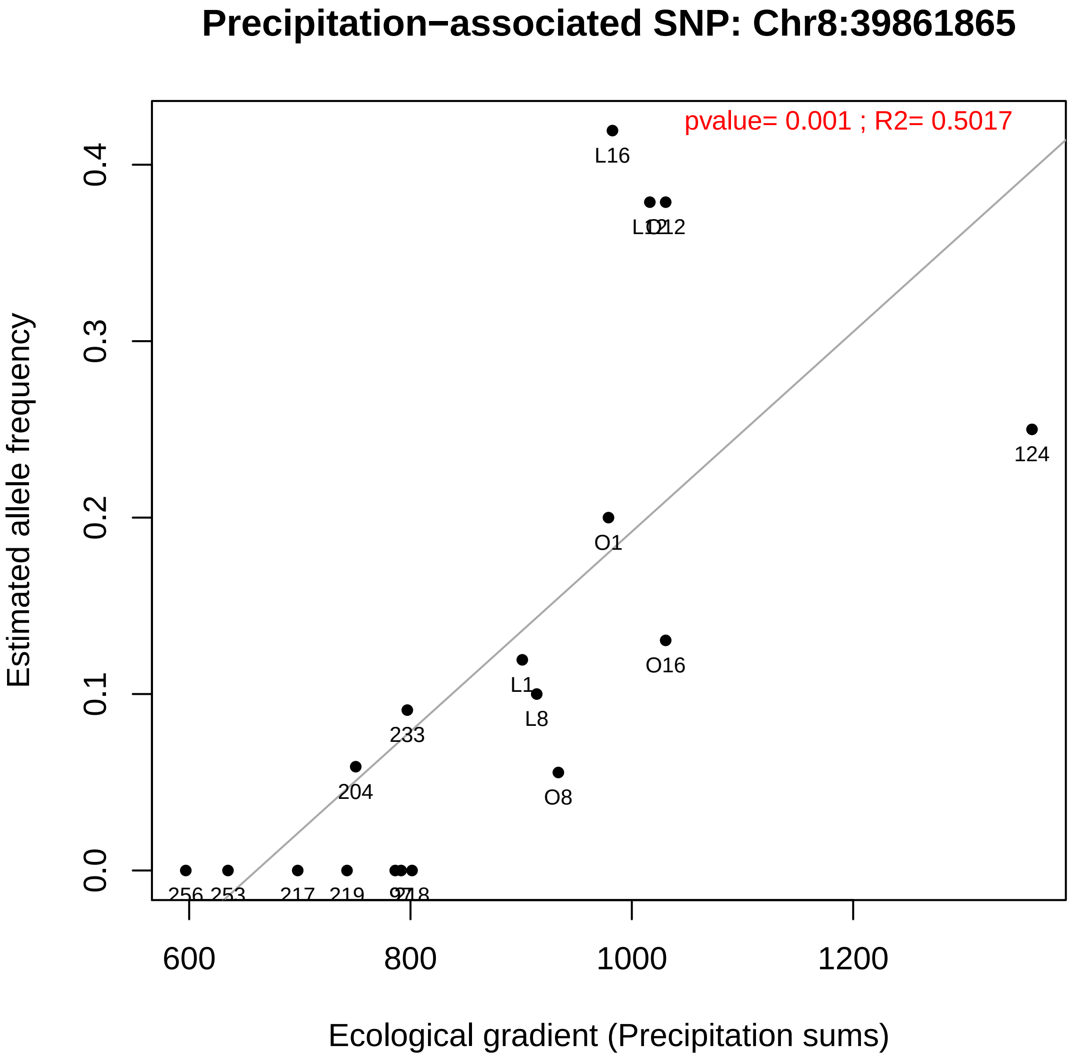
Precipitation−associated SNP: Chr8:39861865
pvalue= 0.001 ; R2= 0.5017
Estimated allele frequency
0.0
0.1
0.2
0.3
0.4
Ecological gradient (Precipitation sums)
600
800
1000
1200
L16
124
O1
O16
L1
L8
233
204
O8
217
219

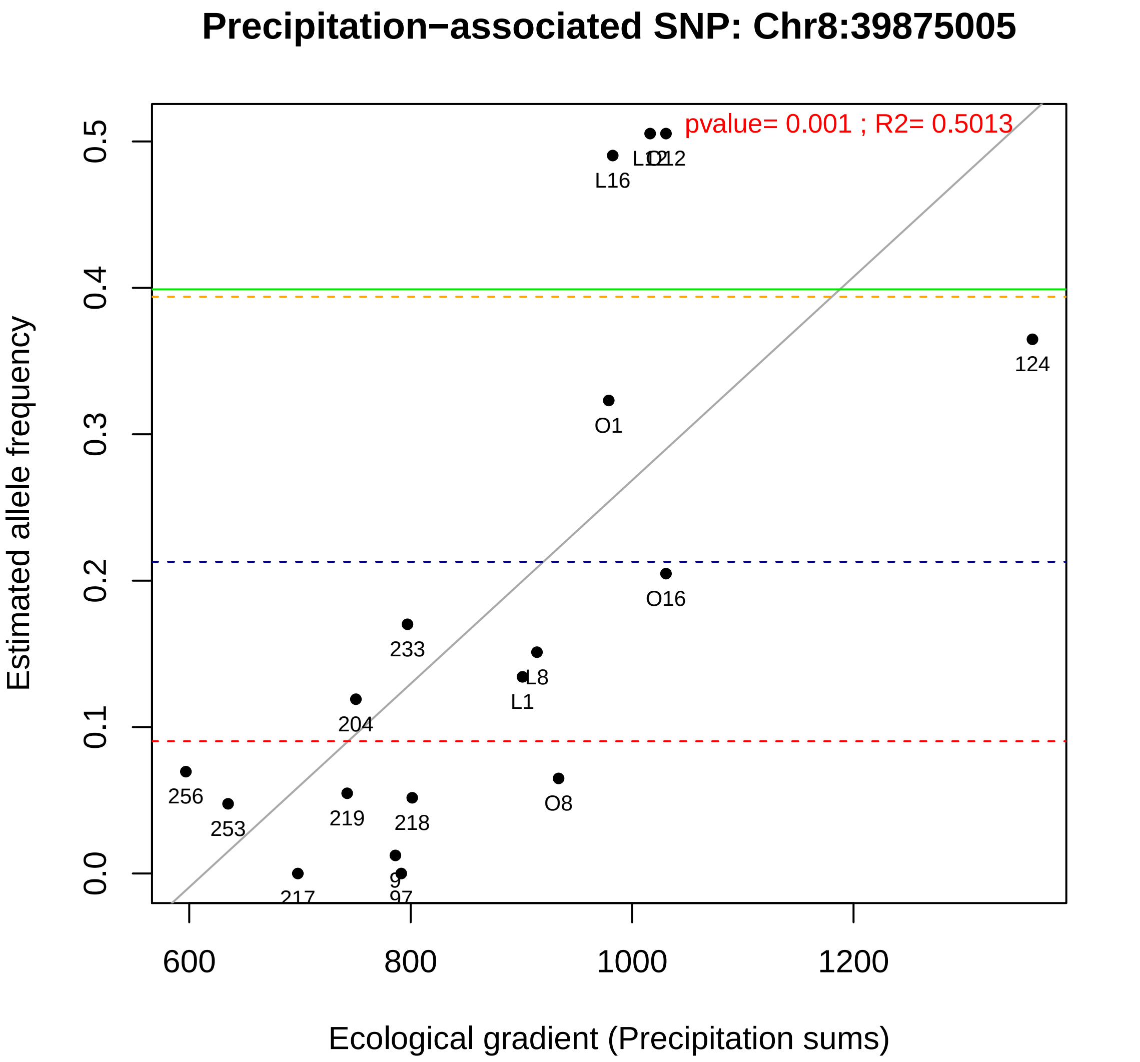
Precipitation−associated SNP: Chr8:39875005
pvalue= 0.001 ; R2= 0.5013
Estimated allele frequency
Ecological gradient (Precipitation sums)
0.0
0.1
0.2
0.3
0.4
0.5
600
800
1000
1200
L16
L12
O12
124
O1
O16
233
L8
L1
204
256
253
219
218
O8
217
9
97

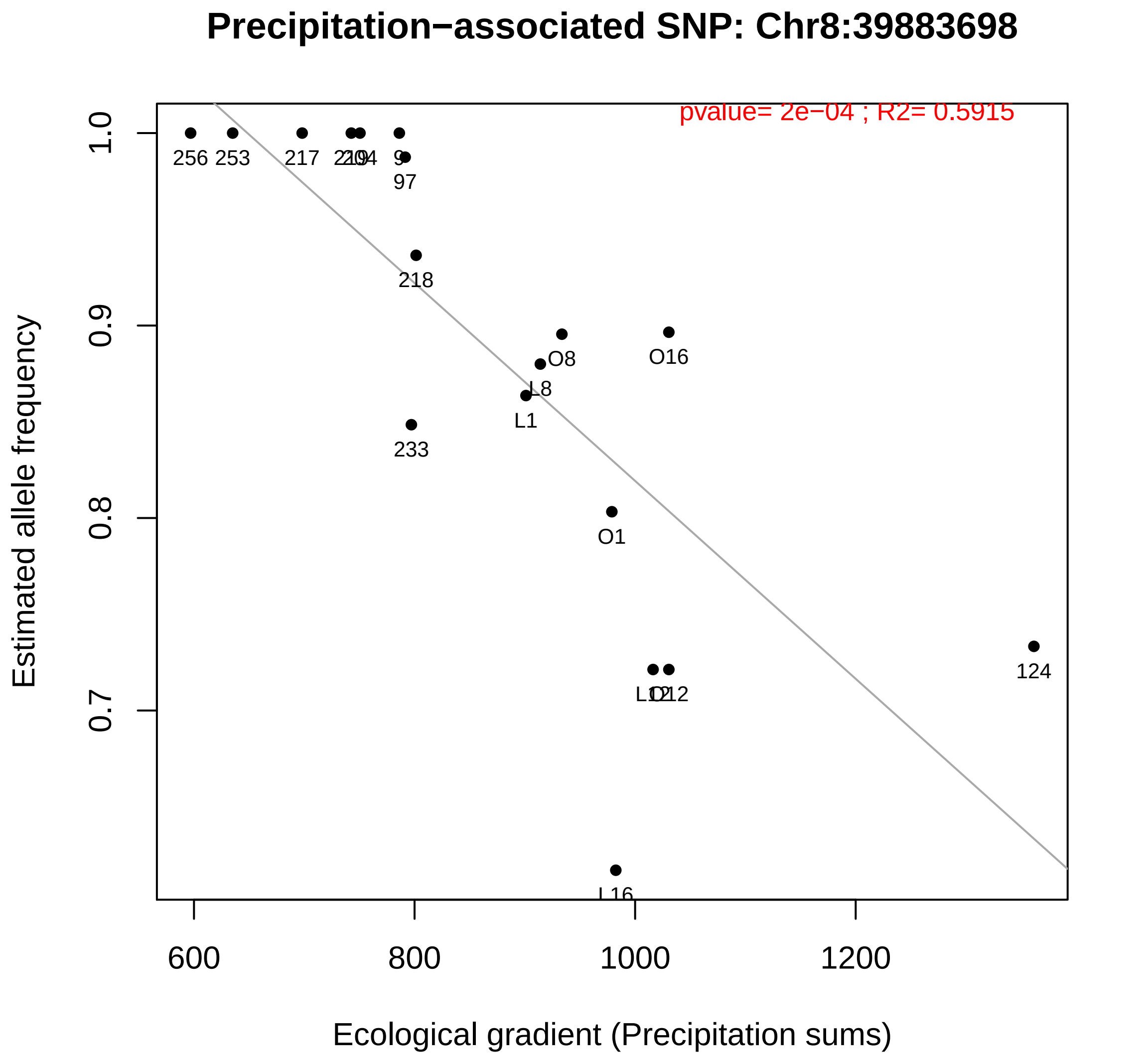

Precipitation−associated SNP: Chr8:39883698
pvalue= 2e−04 ; R2= 0.5915
256
253
217
210
204
9
97
218
O8
O16
L8
L1
233
O1
124
L12
O12
L16
0.7
0.8
0.9
1.0
600
800
1000
1200
Estimated allele frequency
Ecological gradient (Precipitation sums)

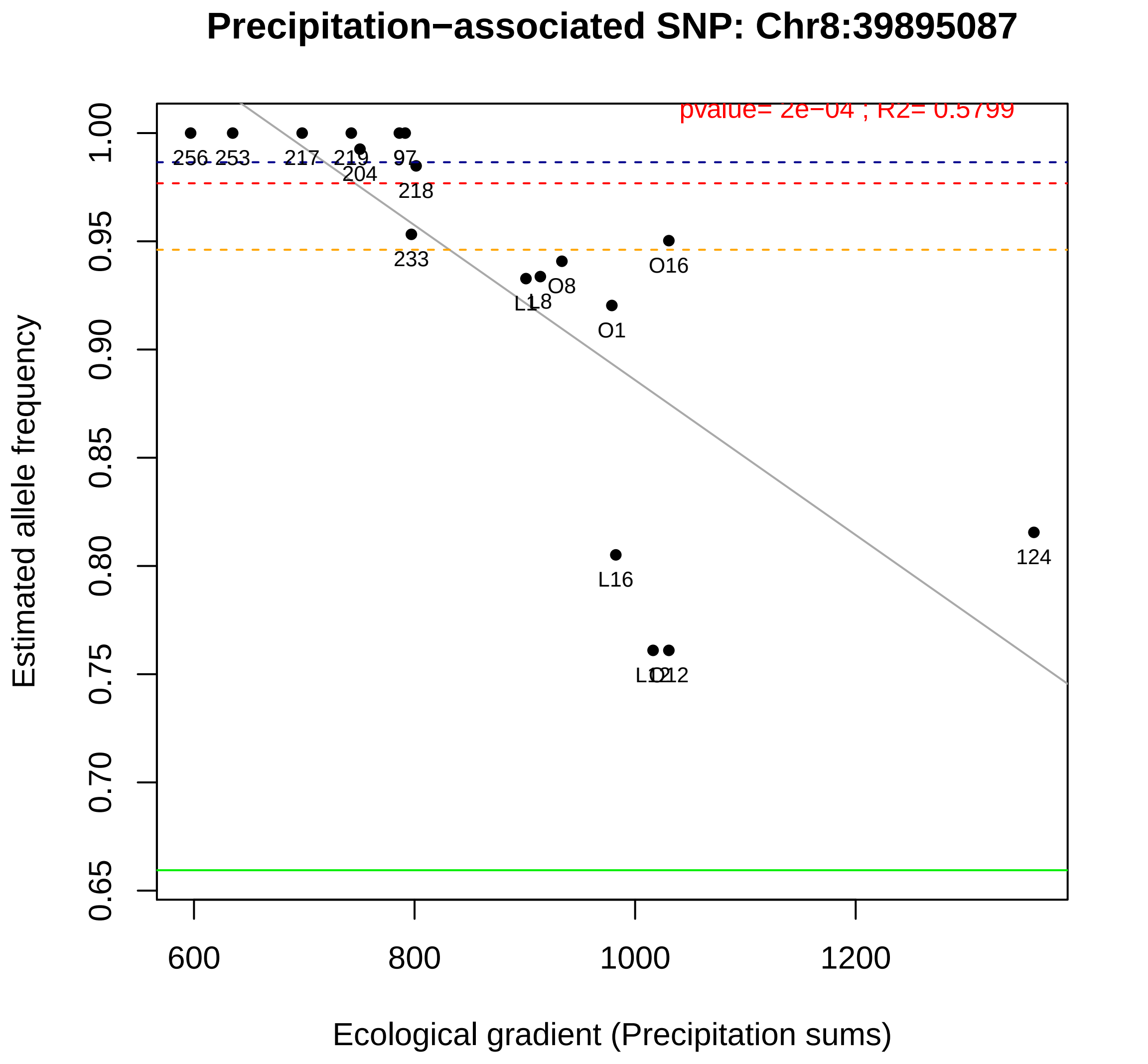

Precipitation−associated SNP: Chr8:39895087
pvalue= 2e−04 ; R2= 0.5799
Estimated allele frequency
1.00
0.95
0.90
0.85
0.80
0.75
0.70
0.65
600
800
1000
1200
Ecological gradient (Precipitation sums)
256
253
217
219
204
97
218
233
L1
L8
O8
O16
O1
L16
124
L12
O12

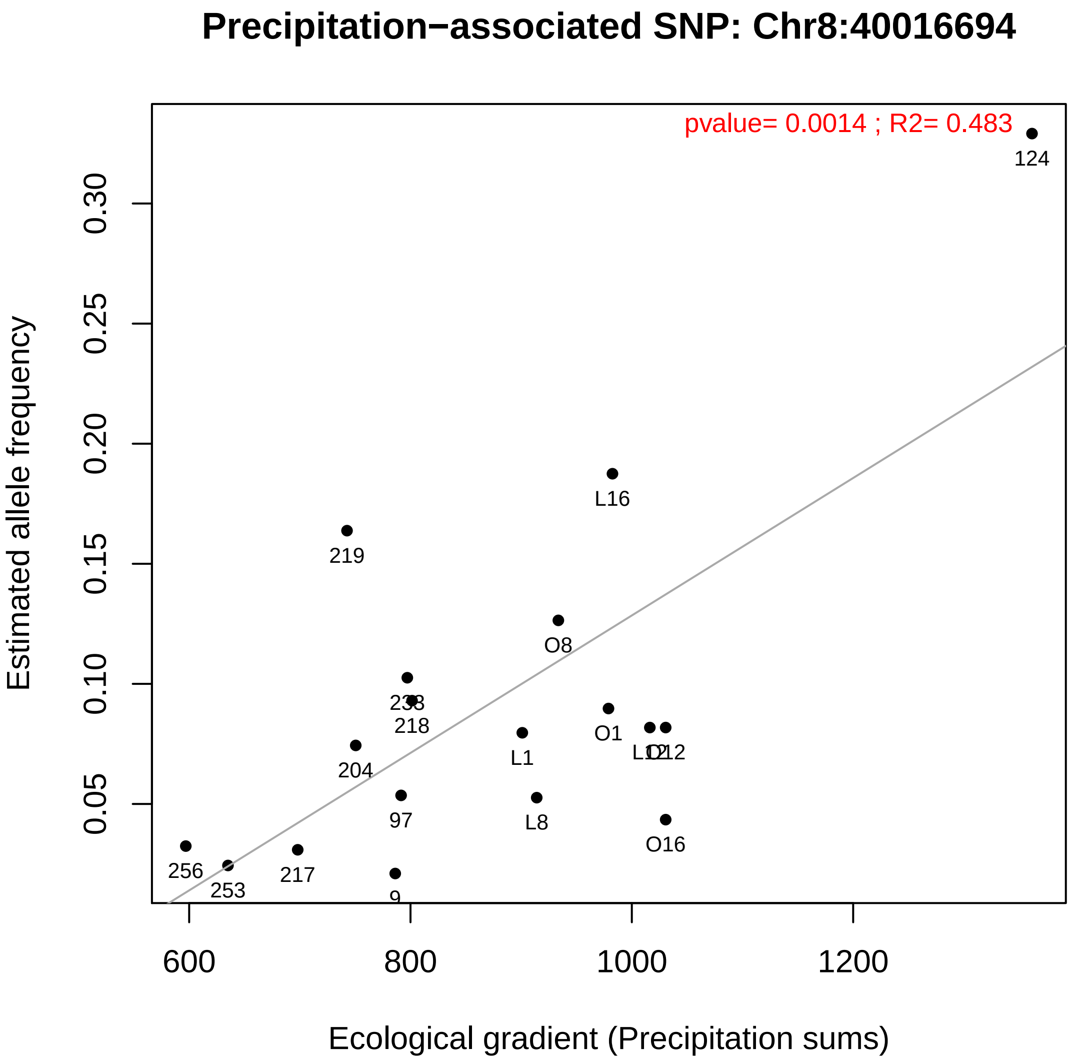
Precipitation−associated SNP: Chr8:40016694
pvalue= 0.0014 ; R2= 0.483
Estimated allele frequency
0.05
0.10
0.15
0.20
0.25
0.30
600
800
1000
1200
Ecological gradient (Precipitation sums)
124
L16
219
O8
238
218
O1
L10
O12
204
L1
97
L8
O16
256
253
217
9

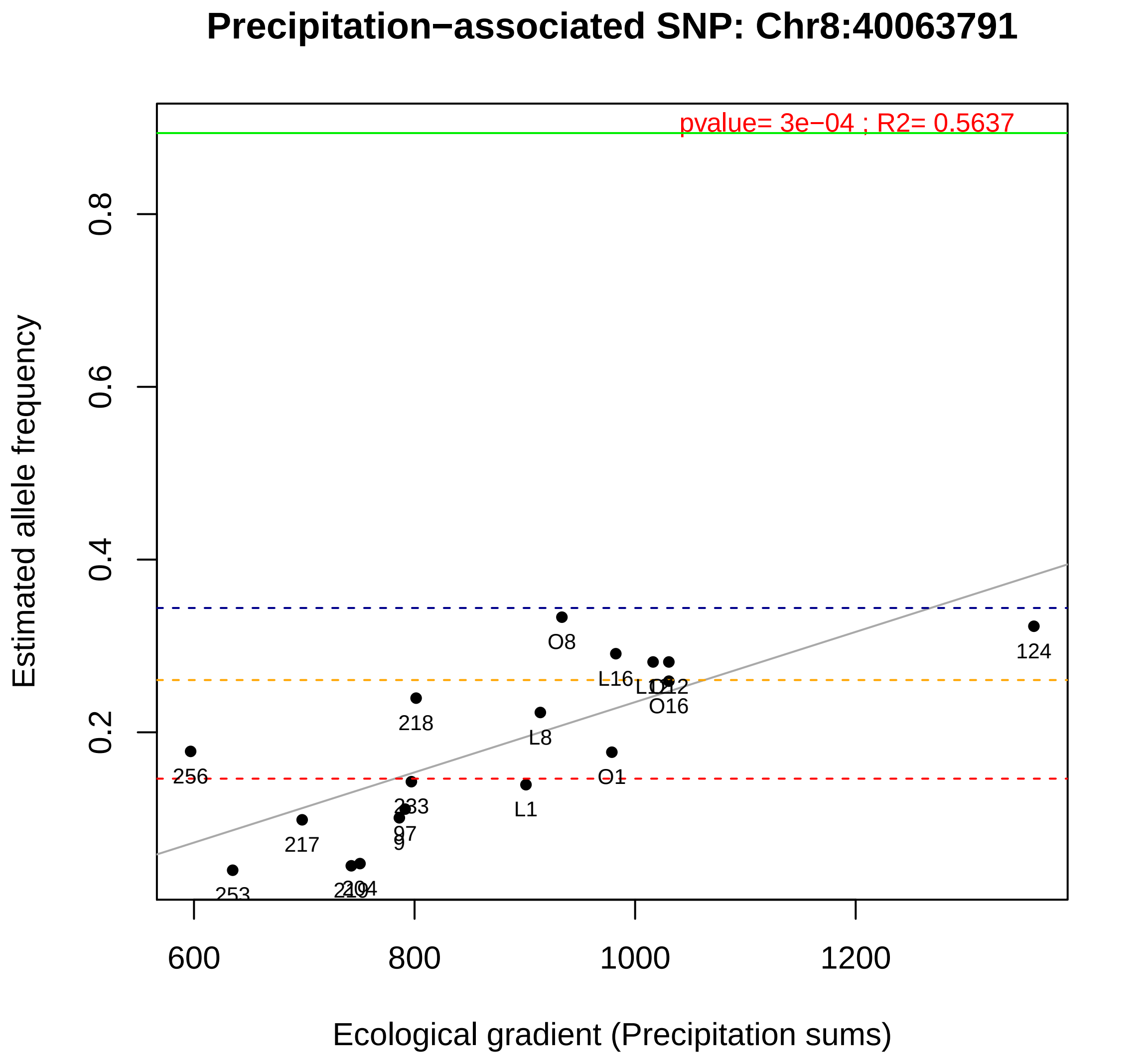

Precipitation−associated SNP: Chr8:40063791
pvalue= 3e−04 ; R2= 0.5637
Estimated allele frequency
0.2
0.4
0.6
0.8
600
800
1000
1200
Ecological gradient (Precipitation sums)
O8
124
L16
O16
218
L8
256
O1
233
L1
217
97
9
253

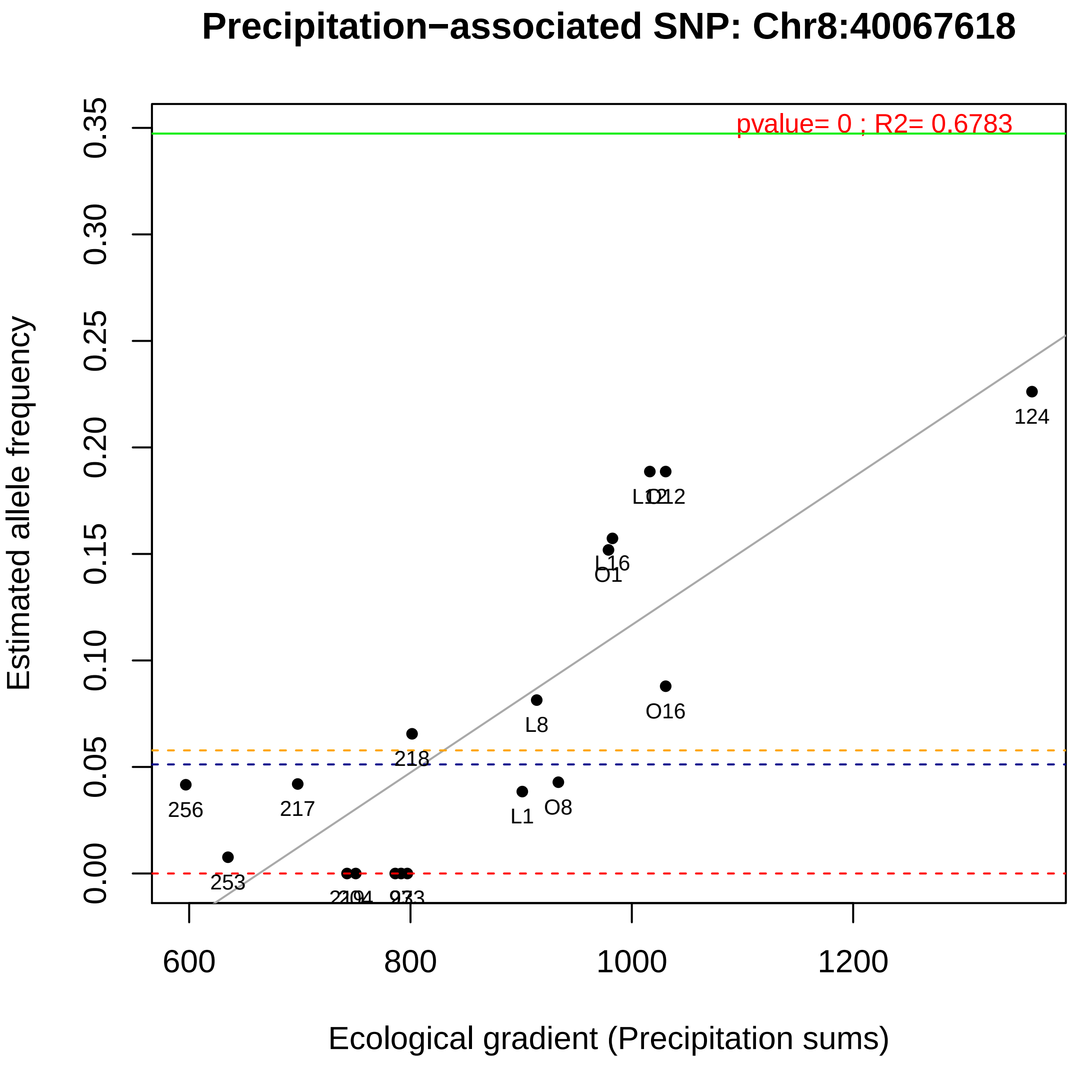

Precipitation−associated SNP: Chr8:40067618
pvalue= 0 ; R2= 0.6783
Estimated allele frequency
0.00
0.05
0.10
0.15
0.20
0.25
0.30
0.35
600
800
1000
1200
Ecological gradient (Precipitation sums)
124
O12
L16
O1
L8
O16
218
256
217
L1
O8
253

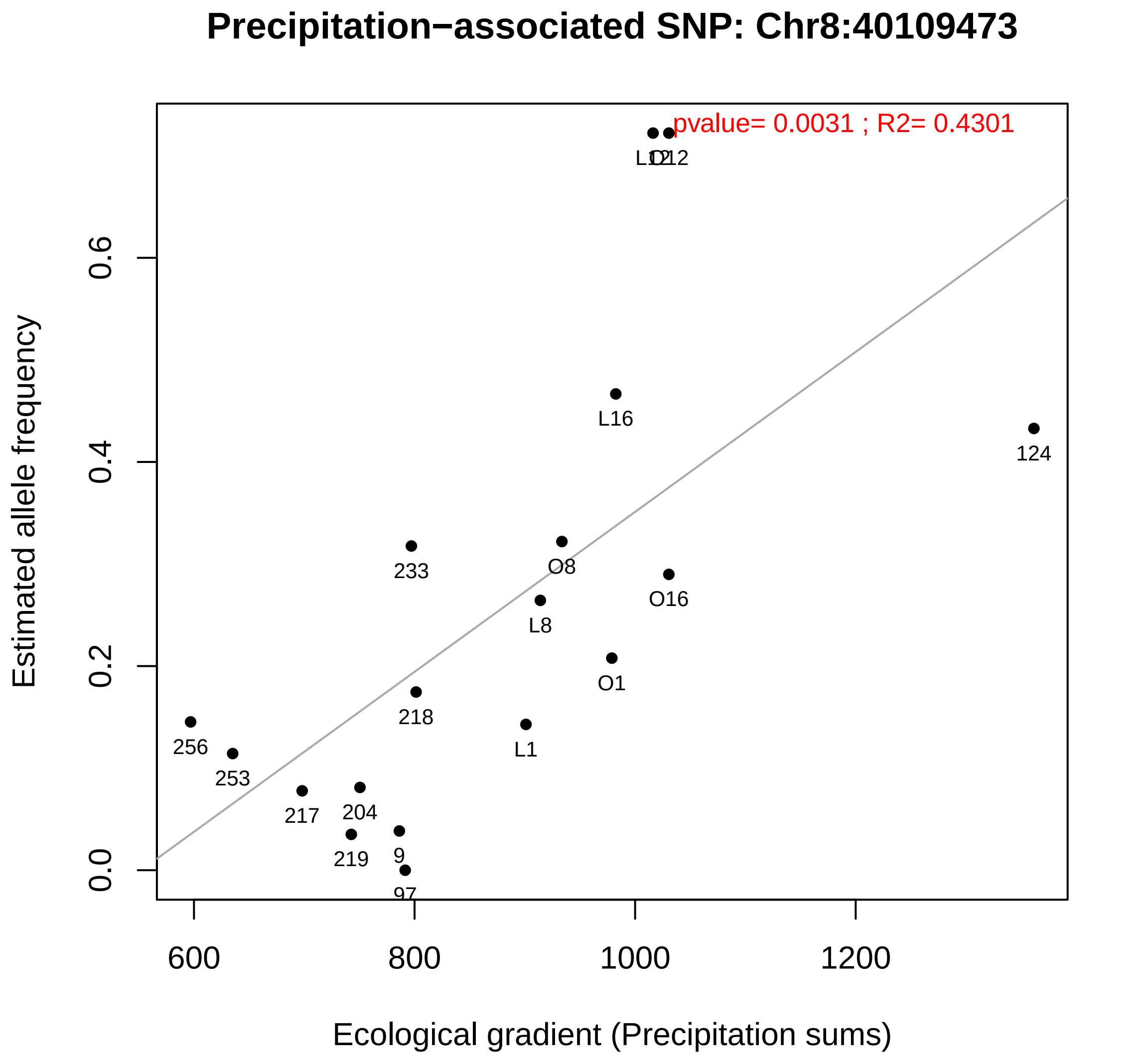

Precipitation−associated SNP: Chr8:40109473
pvalue= 0.0031 ; R2= 0.4301
Estimated allele frequency
0.0
0.2
0.4
0.6
600
800
1000
1200
Ecological gradient (Precipitation sums)
O12
L16
124
233
O8
O16
L8
O1
218
L1
256
253
217
204
219
9
97

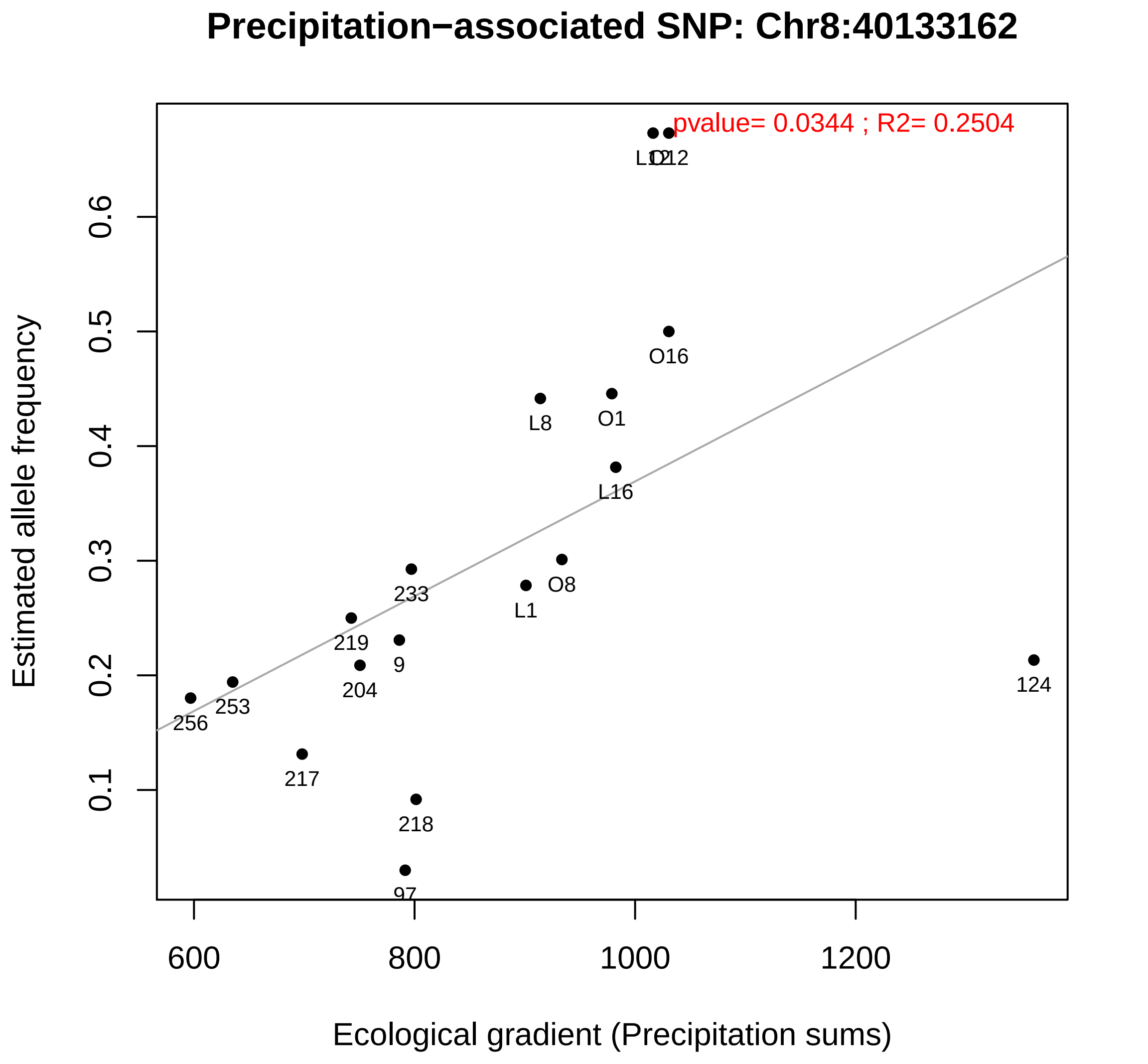

Precipitation−associated SNP: Chr8:40133162
pvalue= 0.0344 ; R2= 0.2504
Estimated allele frequency
0.1
0.2
0.3
0.4
0.5
0.6
600
800
1000
1200
Ecological gradient (Precipitation sums)
L12
O12
O16
L8
O1
L16
O8
L1
233
219
9
204
253
256
217
218
97
124

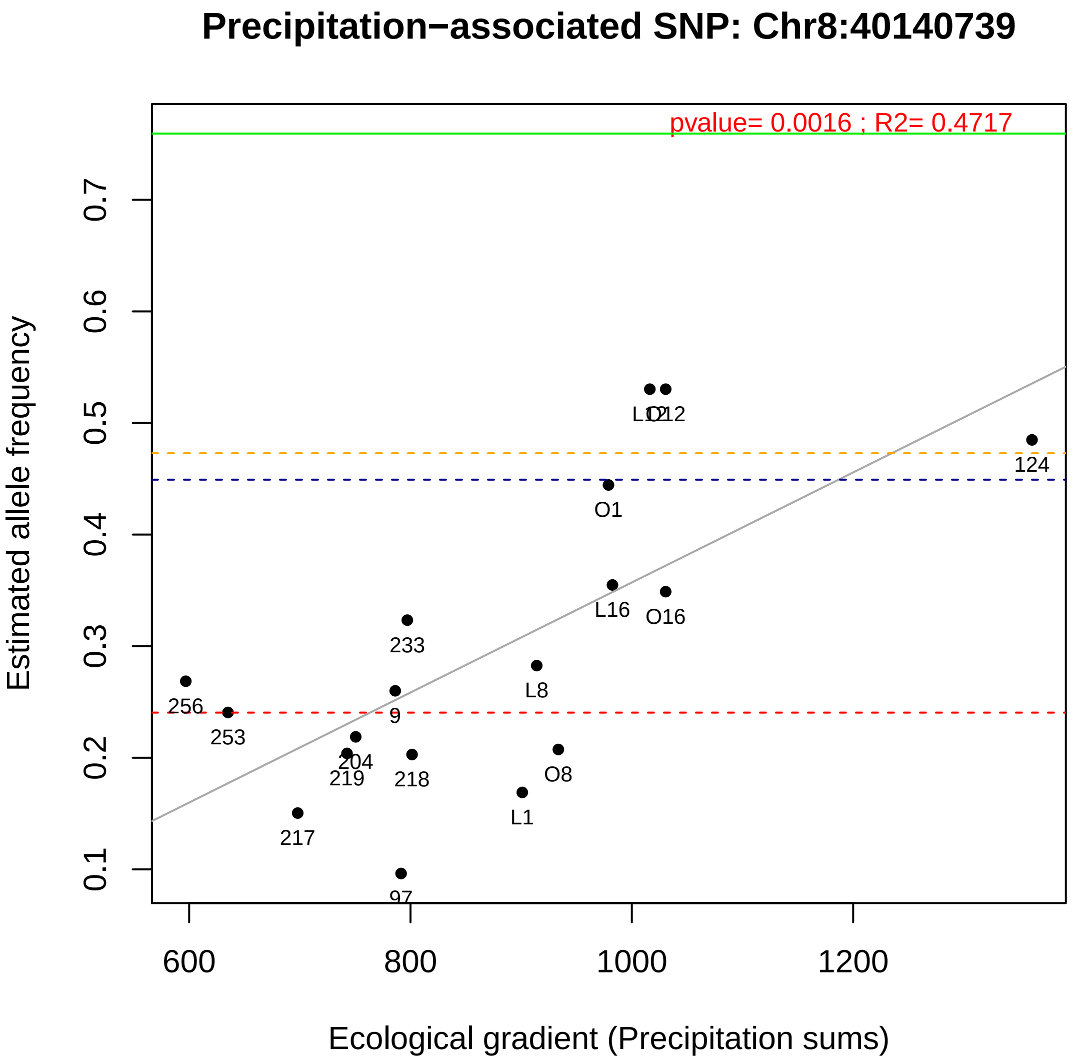

Precipitation−associated SNP: Chr8:40140739
pvalue= 0.0016 ; R2= 0.4717
Estimated allele frequency
0.1
0.2
0.3
0.4
0.5
0.6
0.7
600
800
1000
1200
Ecological gradient (Precipitation sums)
L12
O12
124
O1
L16
O16
233
L8
256
253
9
204
219
218
O8
L1
217
97

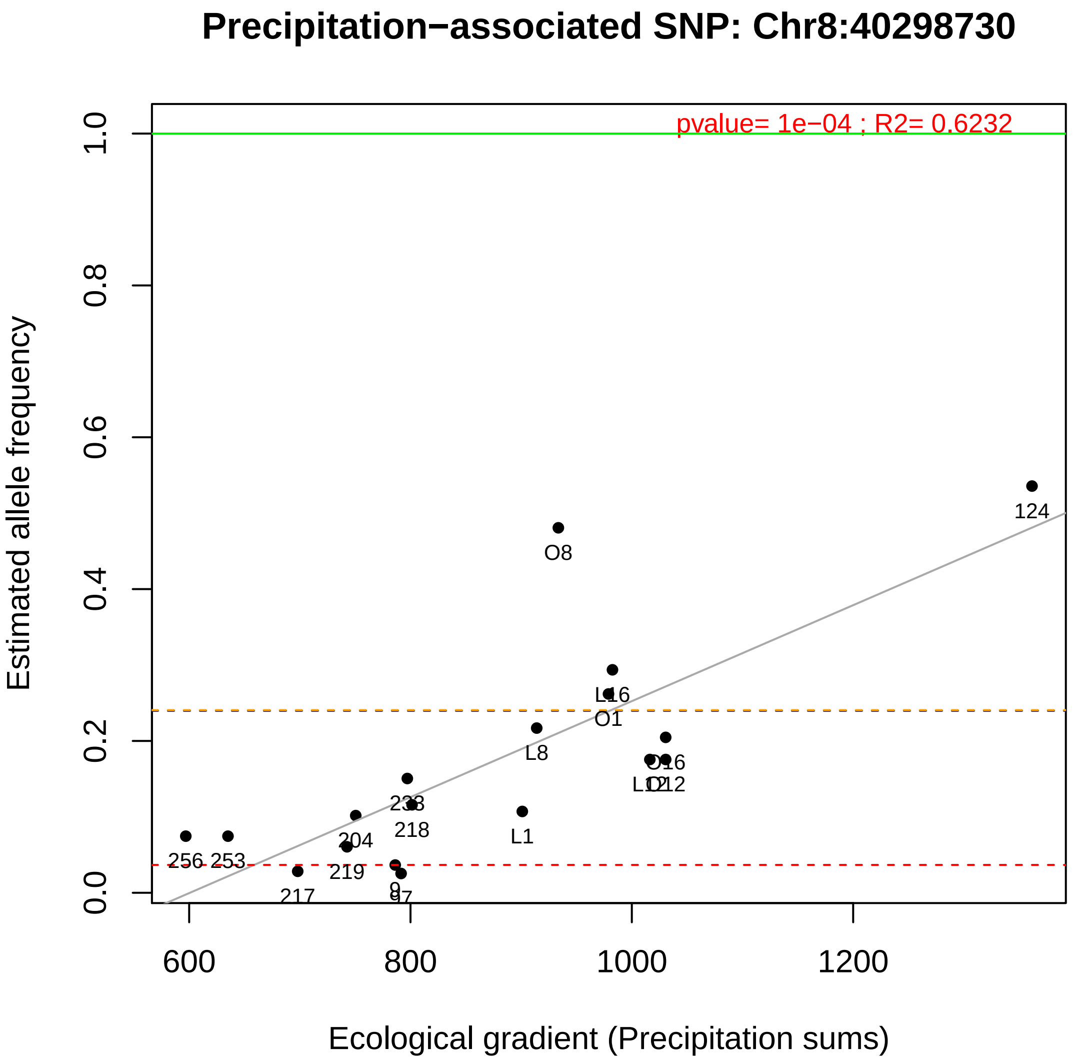
Precipitation−associated SNP: Chr8:40298730
pvalue= 1e−04 ; R2= 0.6232
Estimated allele frequency
Ecological gradient (Precipitation sums)
0.0
0.2
0.4
0.6
0.8
1.0
600
800
1000
1200
124
O8
L16
O1
L8
O16
L12
O12
233
218
L1
204
256
253
219
217
9
97

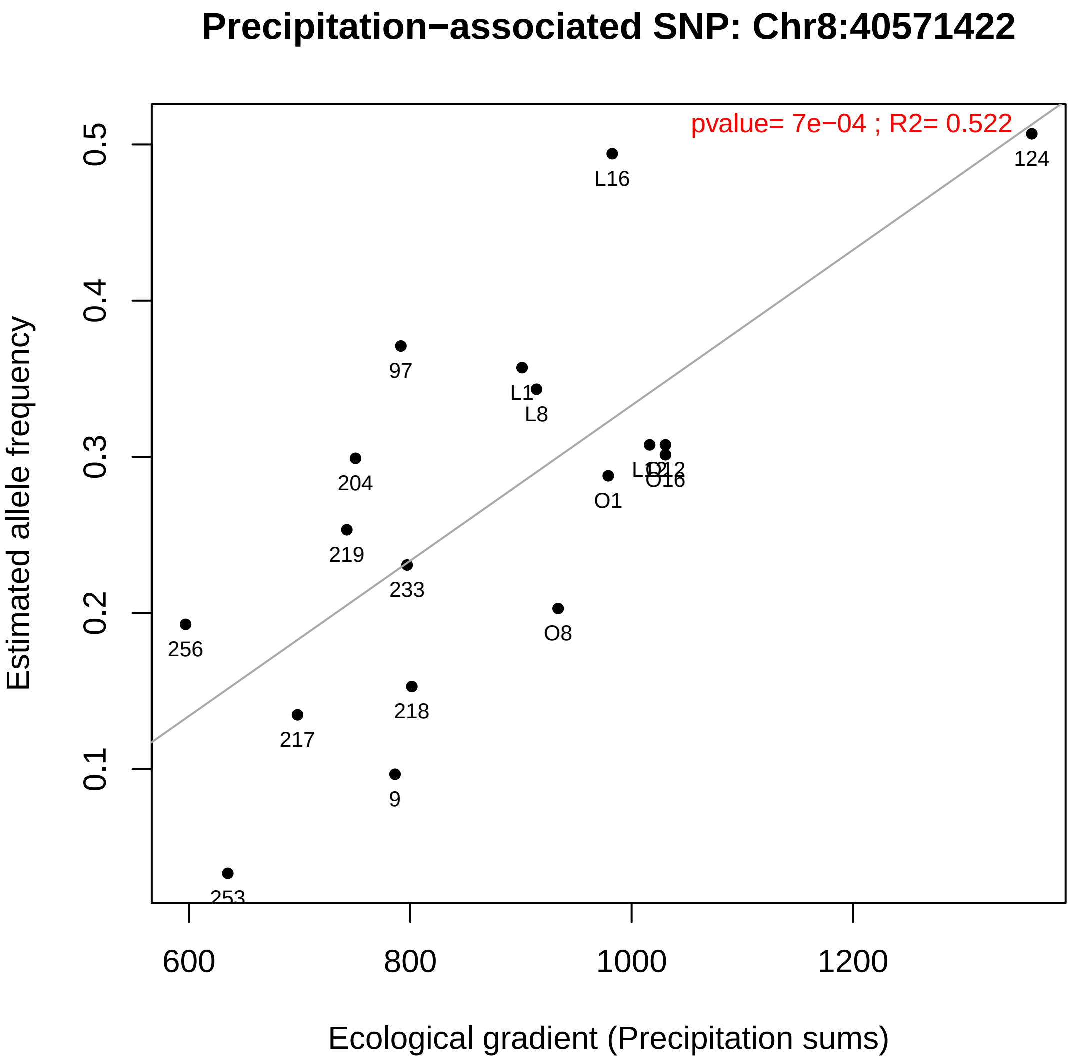

Precipitation−associated SNP: Chr8:40571422
pvalue= 7e−04 ; R2= 0.522
Estimated allele frequency
Ecological gradient (Precipitation sums)
0.1
0.2
0.3
0.4
0.5
600
800
1000
1200
124
L16
97
L1
L8
204
O1
O16
219
233
O8
256
218
217
9
253

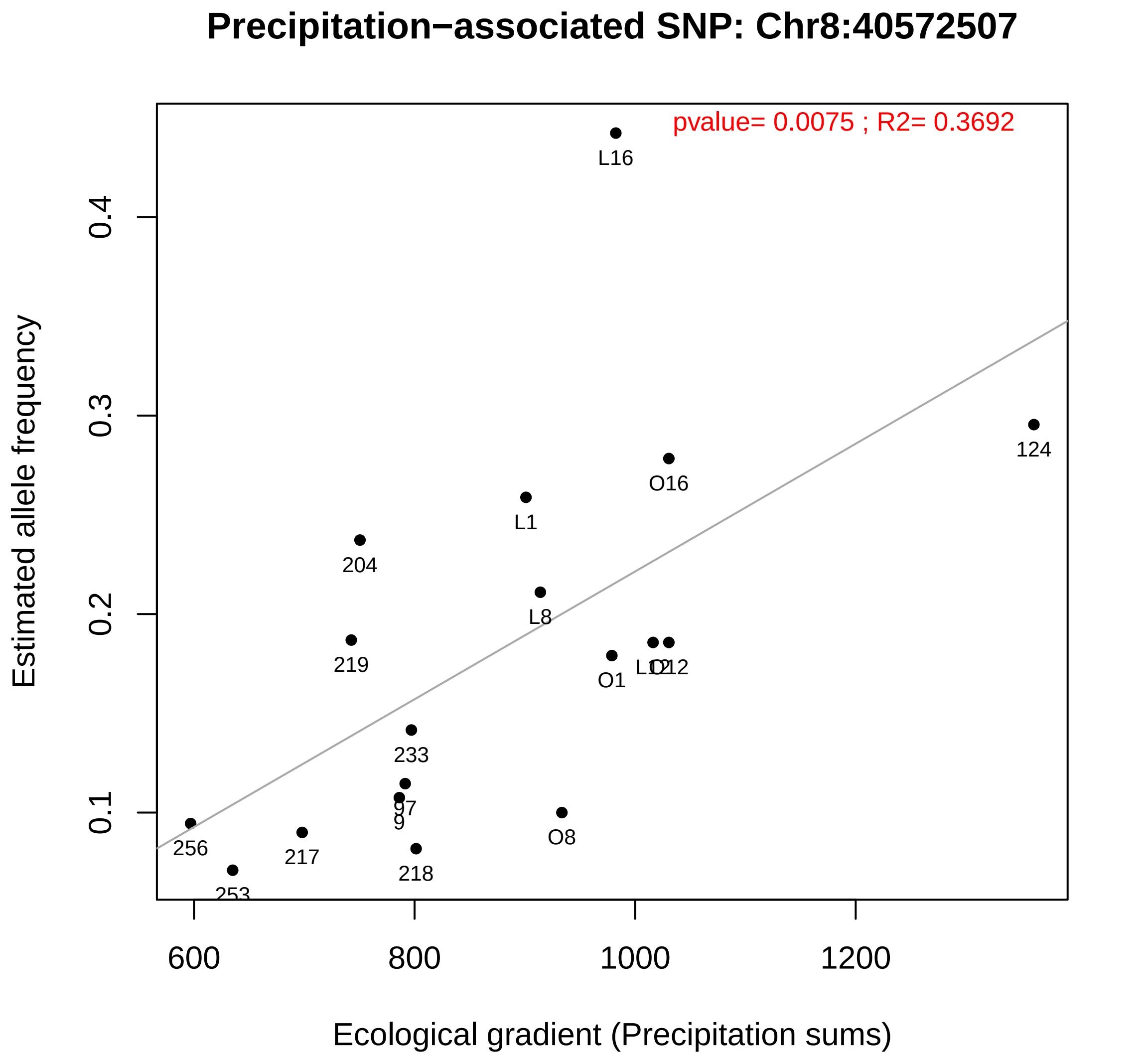
Precipitation−associated SNP: Chr8:40572507
pvalue= 0.0075 ; R2= 0.3692
Estimated allele frequency
Ecological gradient (Precipitation sums)
0.1
0.2
0.3
0.4
600
800
1000
1200
L16
124
O16
L1
204
L8
219
O1
L10
O12
233
97
9
O8
256
217
218
253

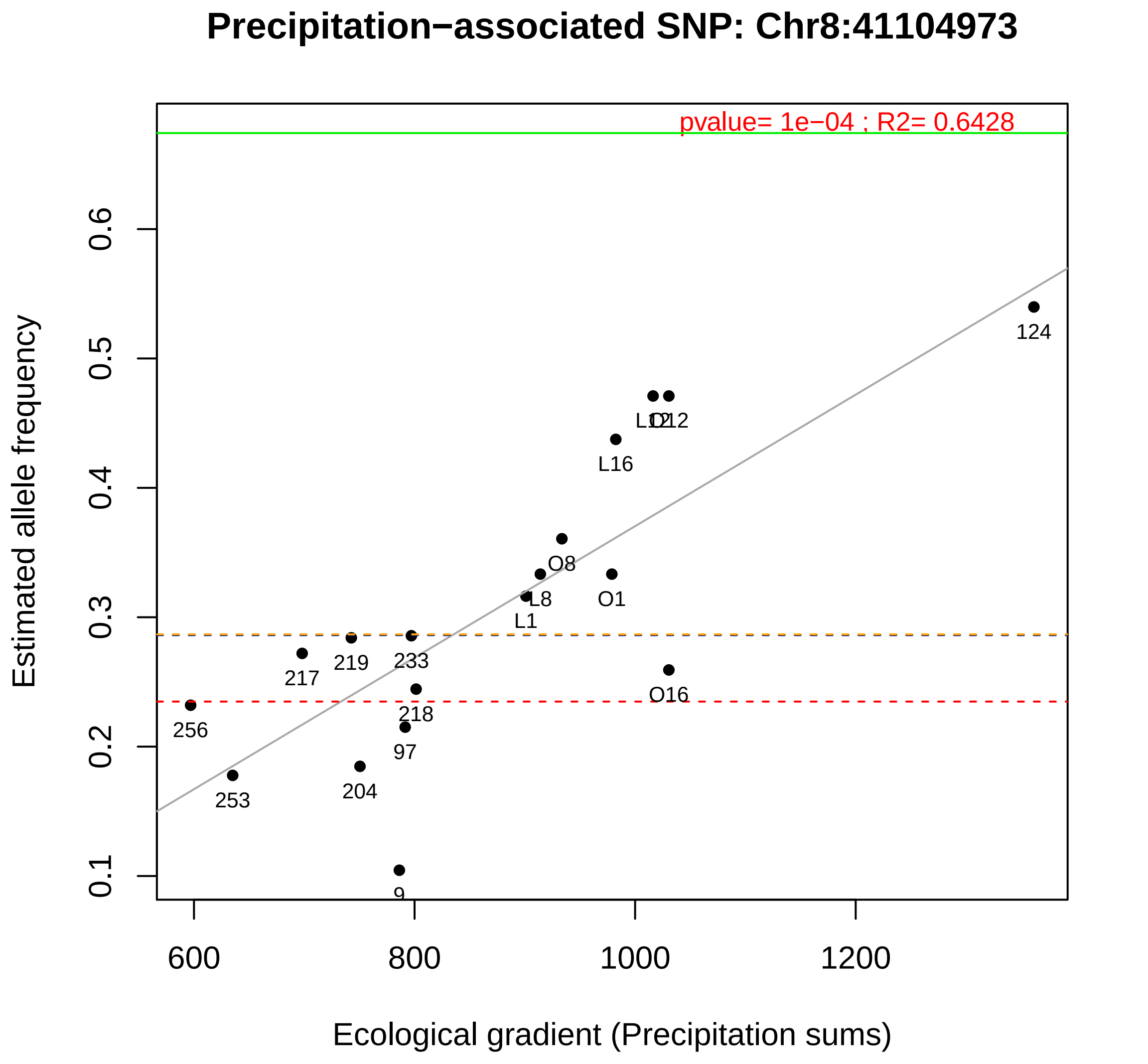

Precipitation−associated SNP: Chr8:41104973
pvalue= 1e−04 ; R2= 0.6428
Estimated allele frequency
0.1
0.2
0.3
0.4
0.5
0.6
Ecological gradient (Precipitation sums)
600
800
1000
1200
124
L12
O12
L16
O8
L8
O1
L1
219
233
217
O16
218
256
97
253
204
9

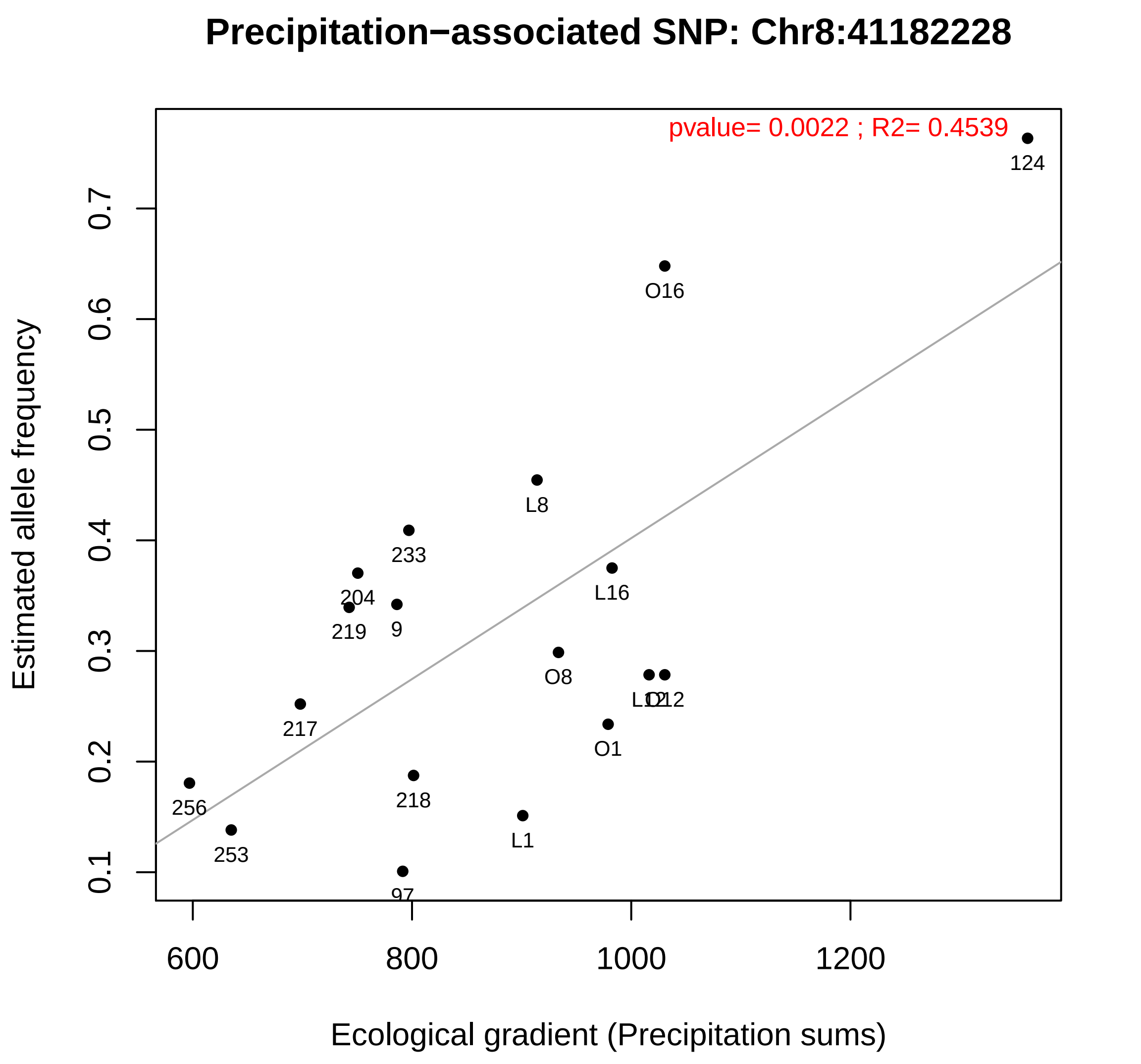
Precipitation−associated SNP: Chr8:41182228
pvalue= 0.0022 ; R2= 0.4539
Estimated allele frequency
Ecological gradient (Precipitation sums)
0.1
0.2
0.3
0.4
0.5
0.6
0.7
600
800
1000
1200
124
O16
L8
233
204
219
9
L16
O8
L10
O12
217
O1
256
218
L1
253
97

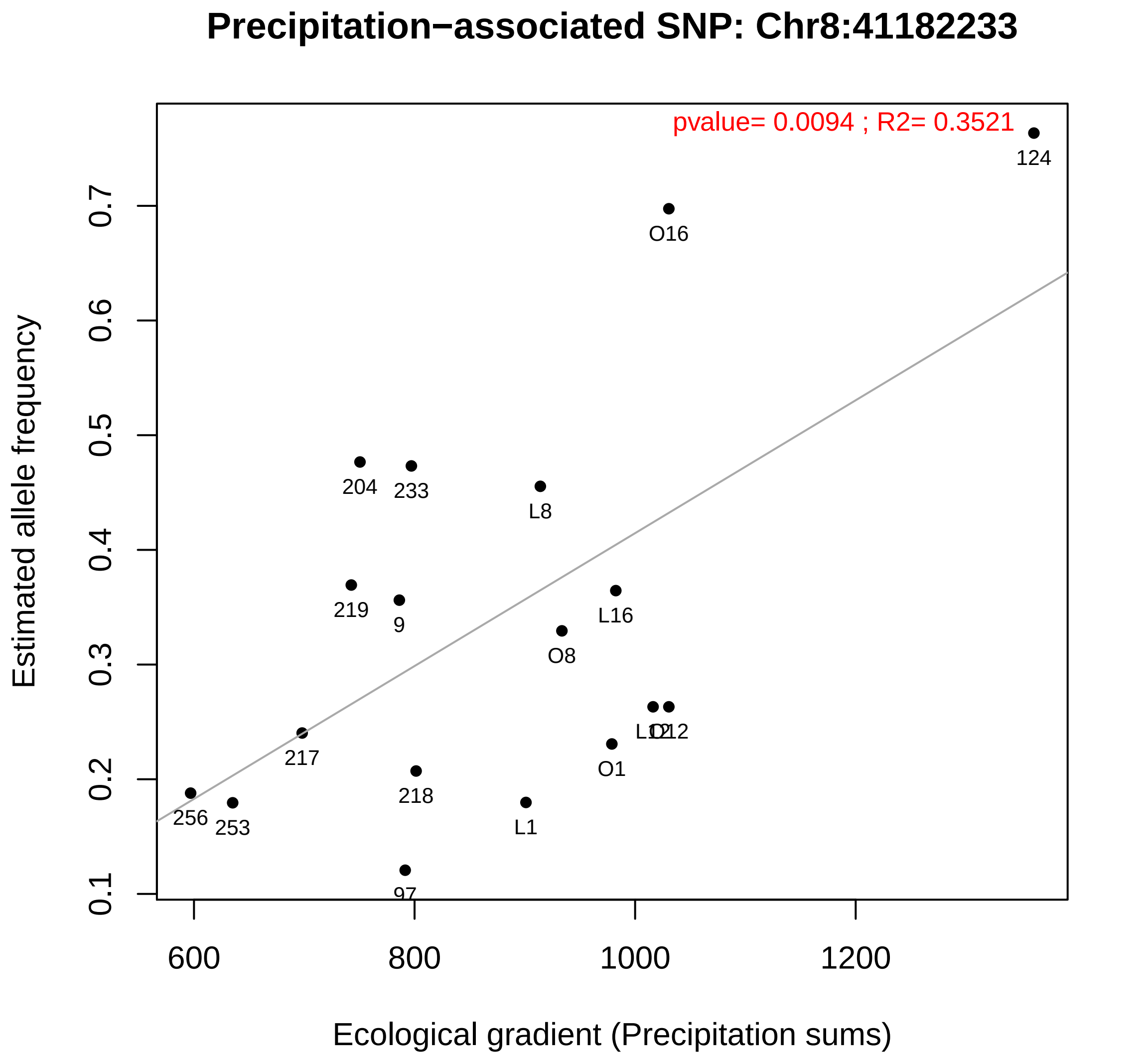

Precipitation−associated SNP: Chr8:41182233
pvalue= 0.0094 ; R2= 0.3521
124
O16
204
233
L8
219
9
L16
O8
L12
O12
O1
217
218
256
253
L1
97
0.1
0.2
0.3
0.4
0.5
0.6
0.7
600
800
1000
1200
Estimated allele frequency
Ecological gradient (Precipitation sums)

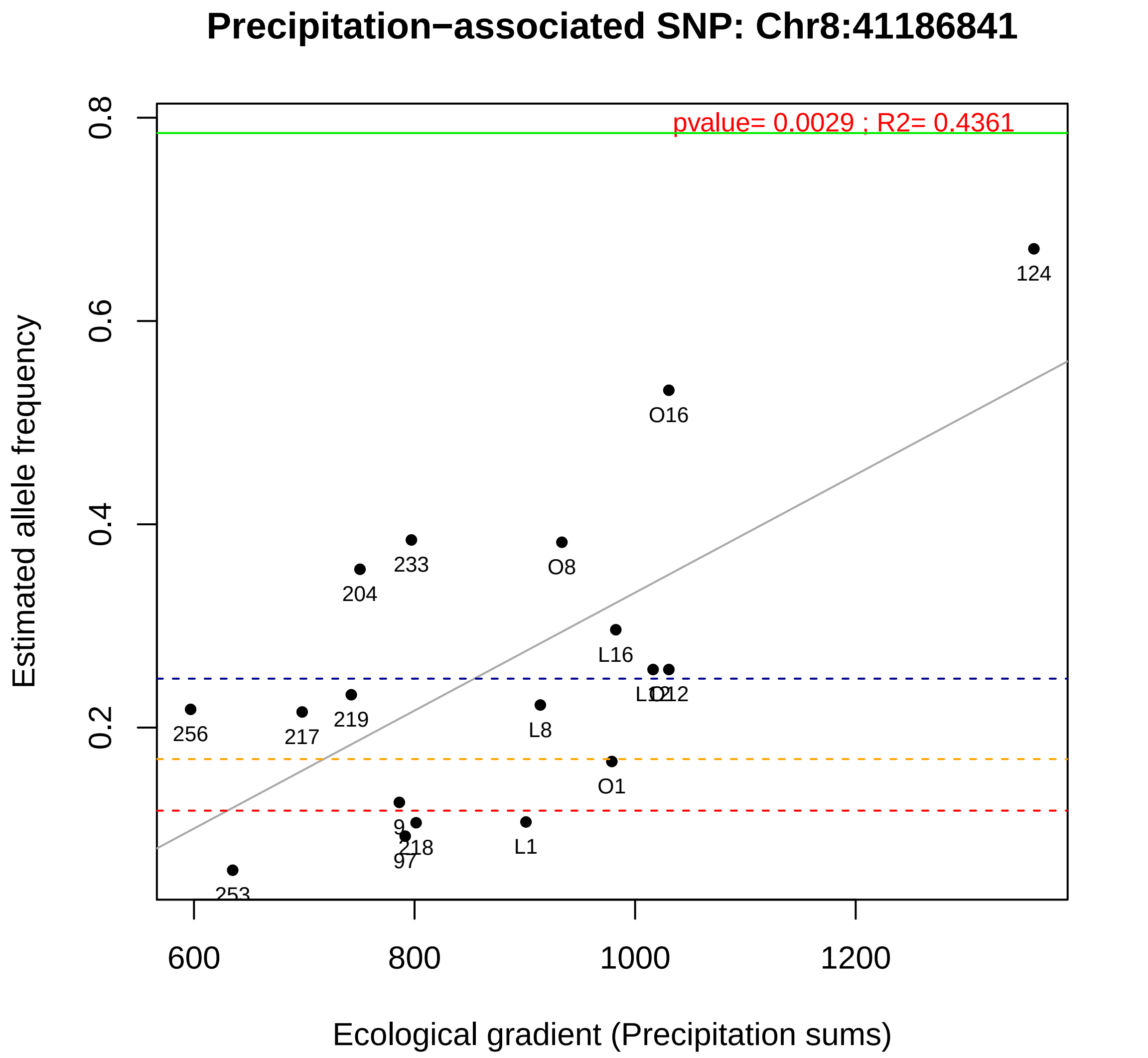
Precipitation−associated SNP: Chr8:41186841
pvalue= 0.0029 ; R2= 0.4361
124
O16
233
O8
204
L16
L12
O12
256
217
219
L8
O1
9
218
97
L1
253
0.8
0.6
0.4
0.2
600
800
1000
1200
Estimated allele frequency
Ecological gradient (Precipitation sums)

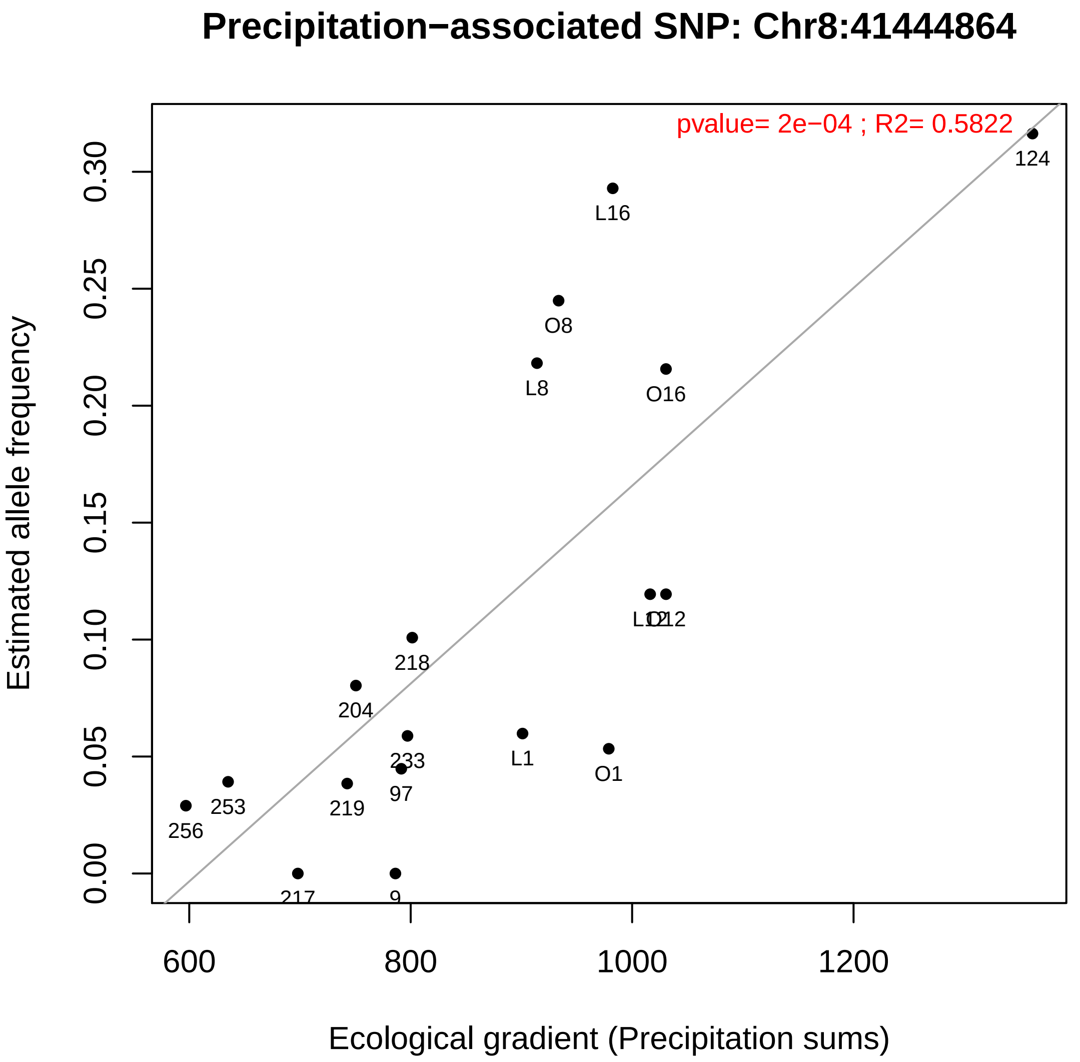

Precipitation−associated SNP: Chr8:41444864
pvalue= 2e−04 ; R2= 0.5822
124
L16
O8
L8
O16
L12
O12
218
204
233
L1
O1
97
253
219
256
217
9
0.00
0.05
0.10
0.15
0.20
0.25
0.30
600
800
1000
1200
Estimated allele frequency
Ecological gradient (Precipitation sums)

**Precipitation−associated SNP: Chr8:41688375**

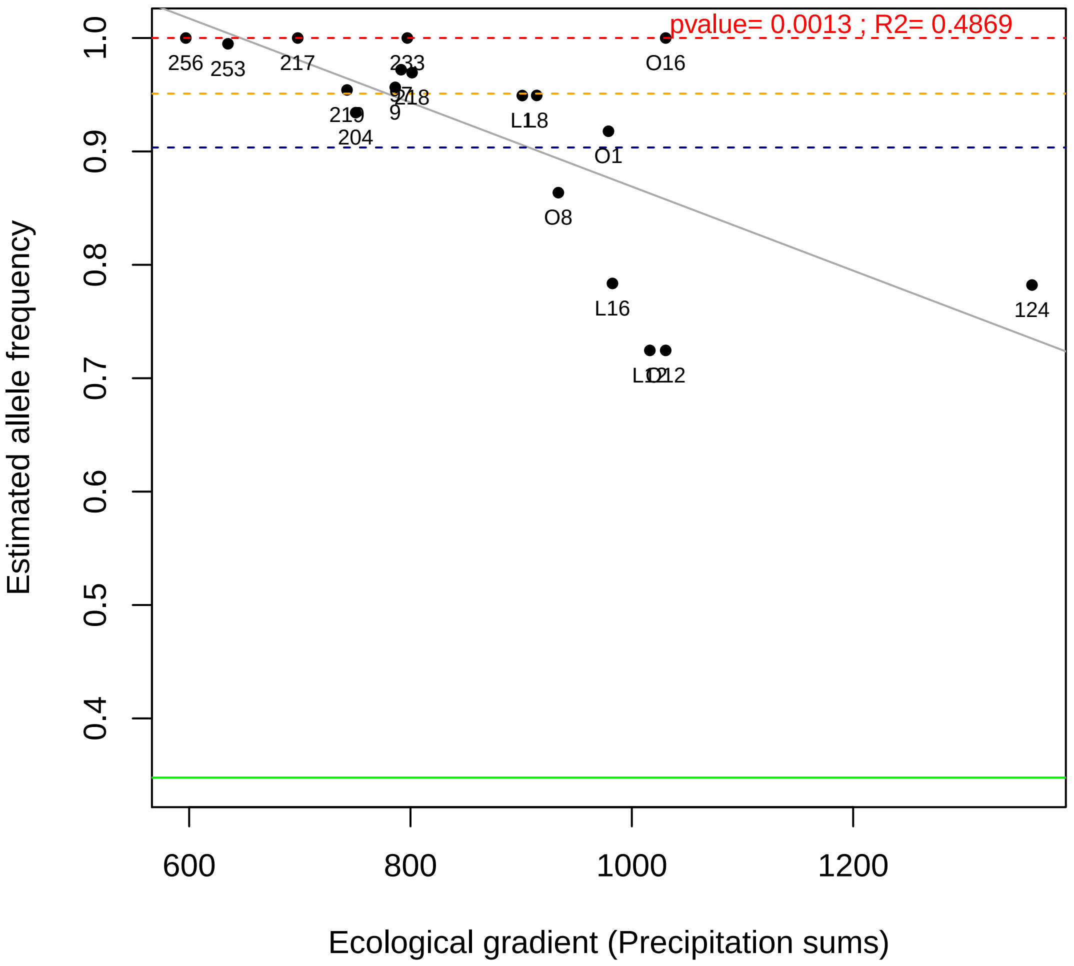

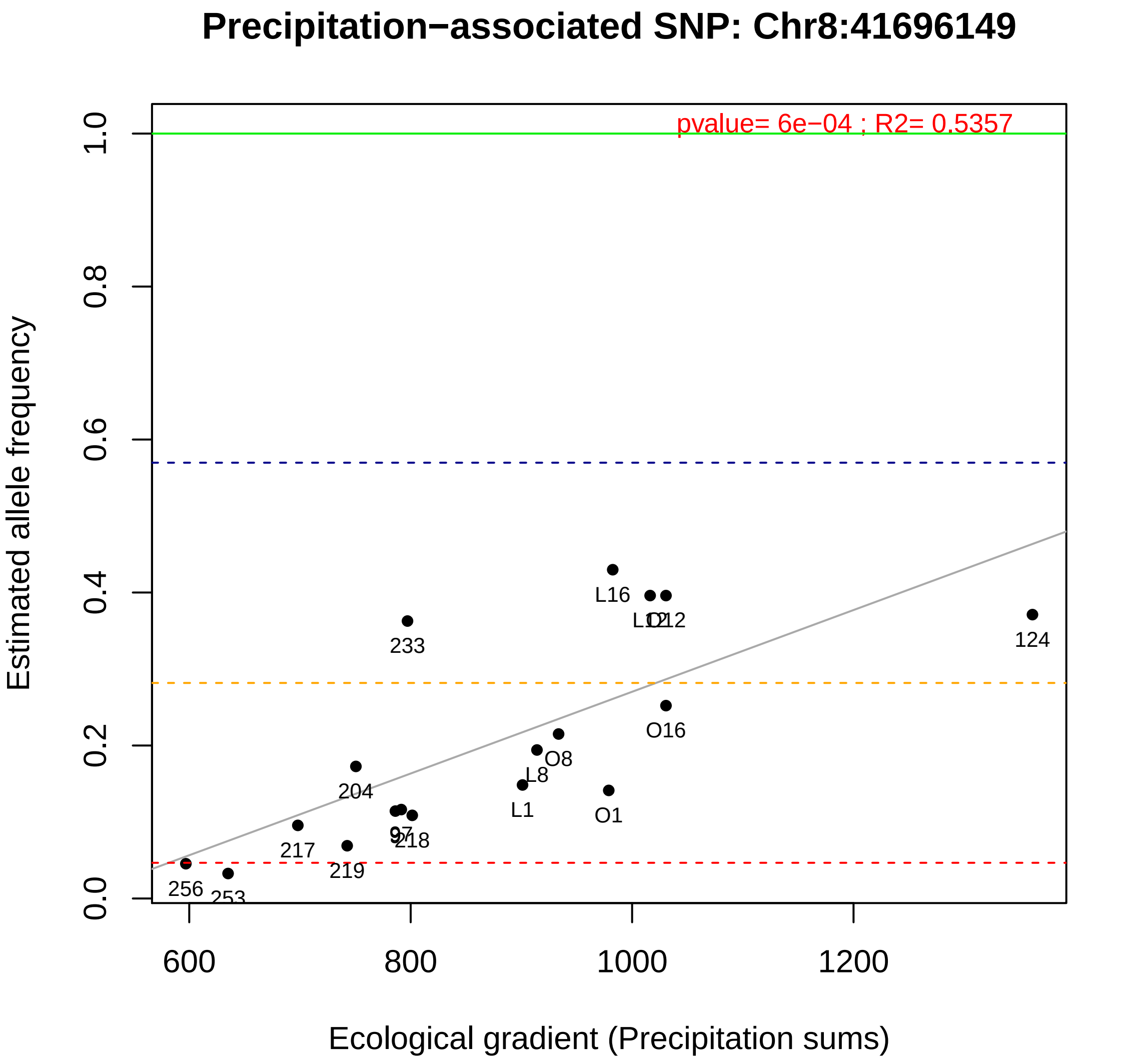

Precipitation−associated SNP: Chr8:41696149
pvalue= 6e−04 ; R2= 0.5357
Estimated allele frequency
0.0
0.2
0.4
0.6
0.8
1.0
600
800
1000
1200
Ecological gradient (Precipitation sums)
256
253
217
219
204
233
218
L1
L8
O8
O1
L16
O12
O16
124

### Precipitation−associated SNP: Chr8:41720129

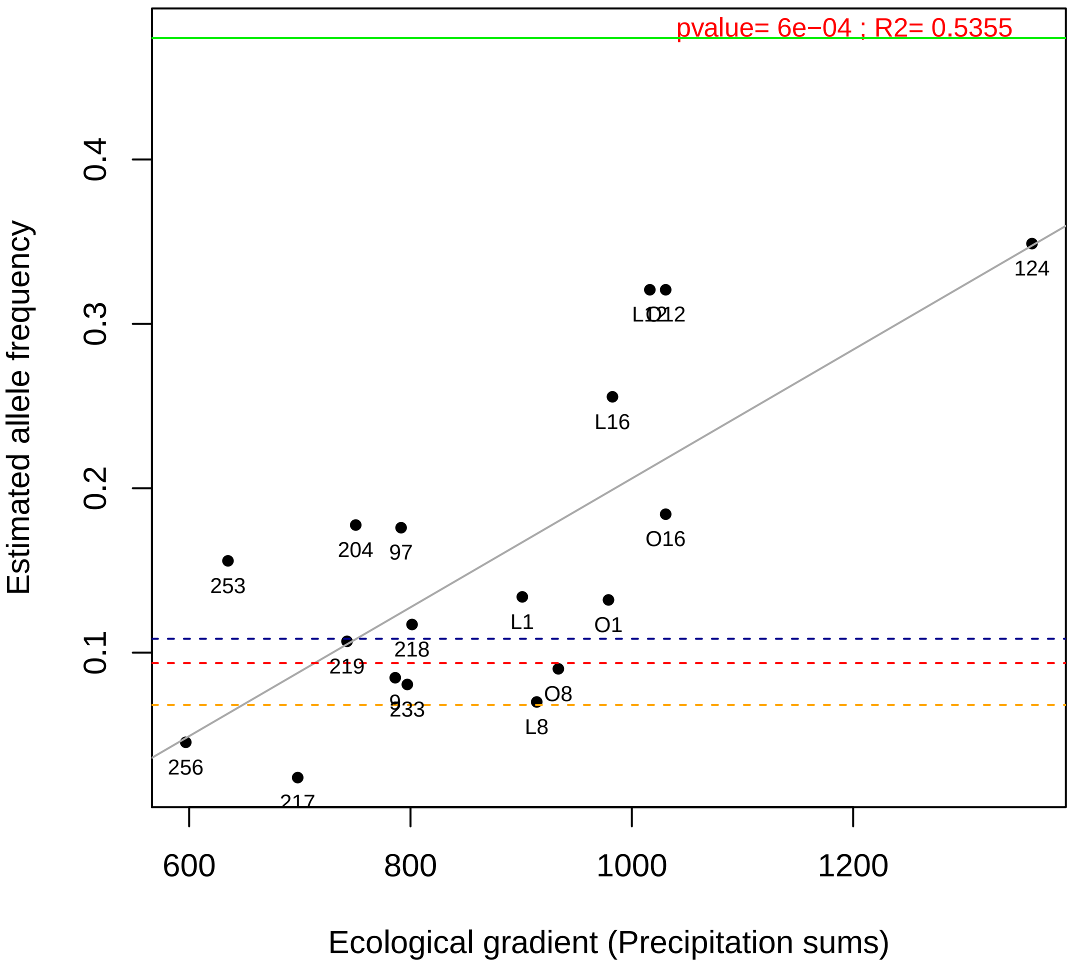

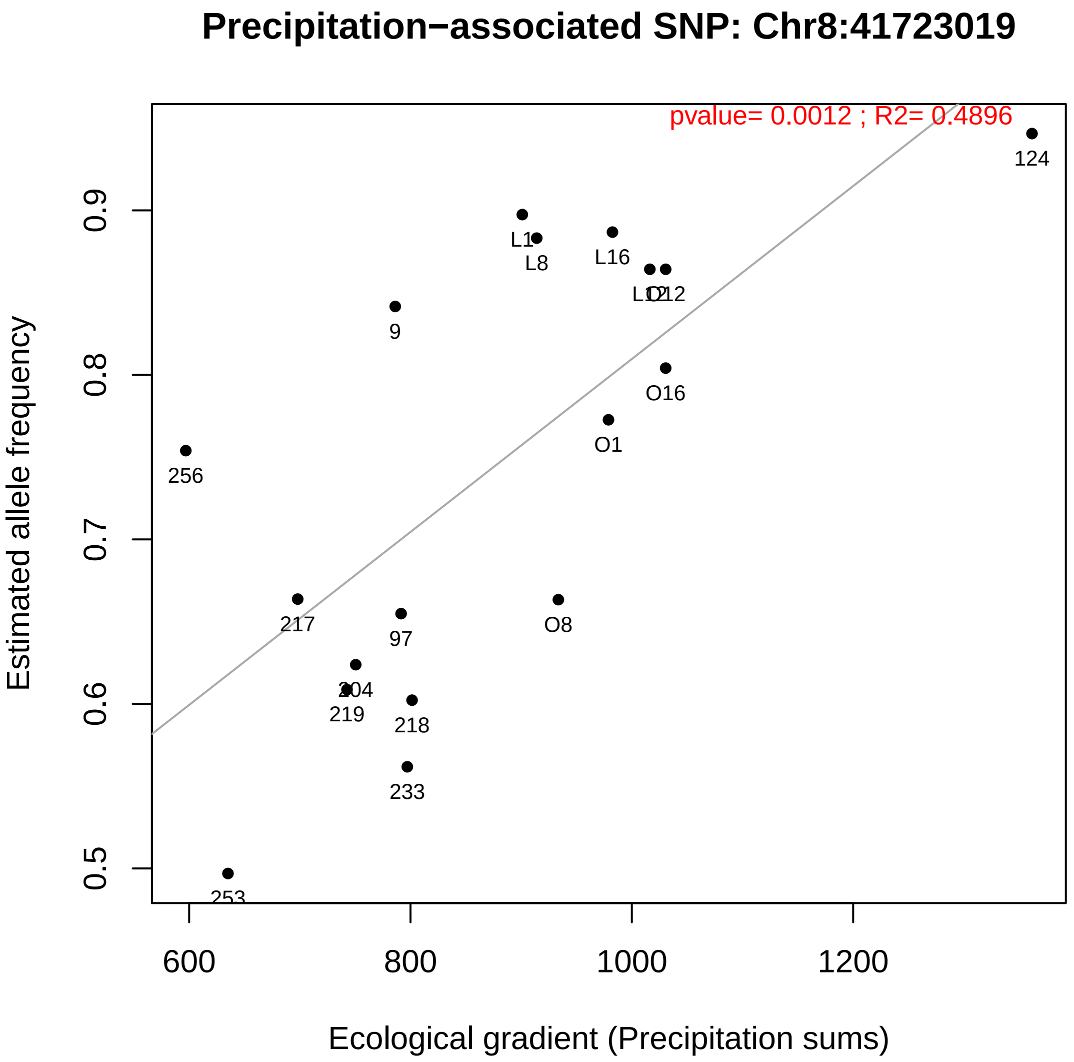
Precipitation−associated SNP: Chr8:41723019
pvalue= 0.0012 ; R2= 0.4896
Estimated allele frequency
Ecological gradient (Precipitation sums)
0.5
0.6
0.7
0.8
0.9
600
800
1000
1200
124
L1
L8
L16
L12
O12
9
O16
O1
256
217
97
O8
204
219
218
233
253

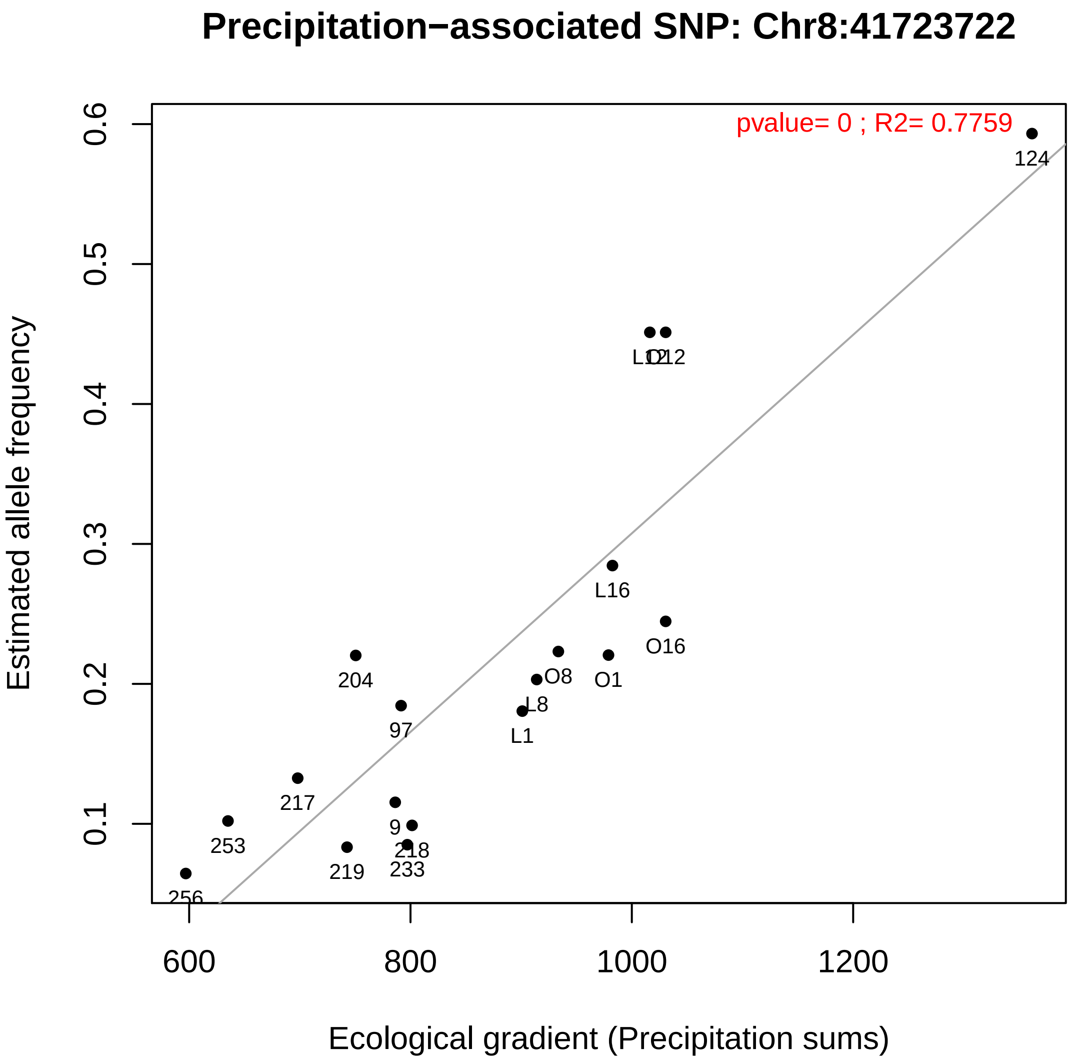

Precipitation−associated SNP: Chr8:41723722
pvalue= 0 ; R2= 0.7759
124
L12
O12
L16
O16
204
O8
O1
L8
97
L1
217
9
253
218
219
233
256
0.1
0.2
0.3
0.4
0.5
0.6
600
800
1000
1200
Estimated allele frequency
Ecological gradient (Precipitation sums)

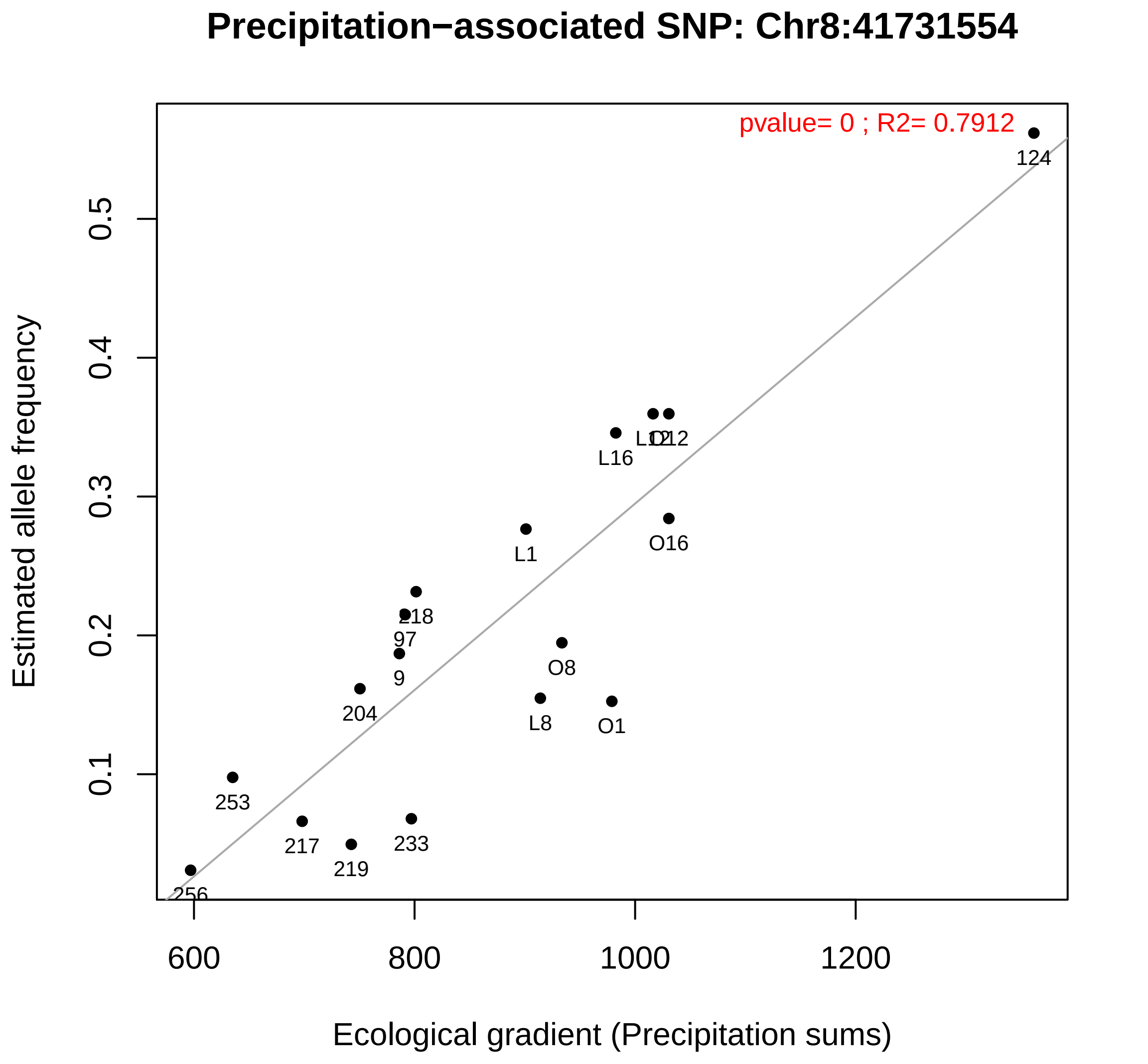

Precipitation−associated SNP: Chr8:41731554
pvalue= 0 ; R2= 0.7912
124
L12
O12
L16
O16
L1
218
97
9
204
O8
L8
O1
253
217
233
219
256
0.1
0.2
0.3
0.4
0.5
600
800
1000
1200
Estimated allele frequency
Ecological gradient (Precipitation sums)

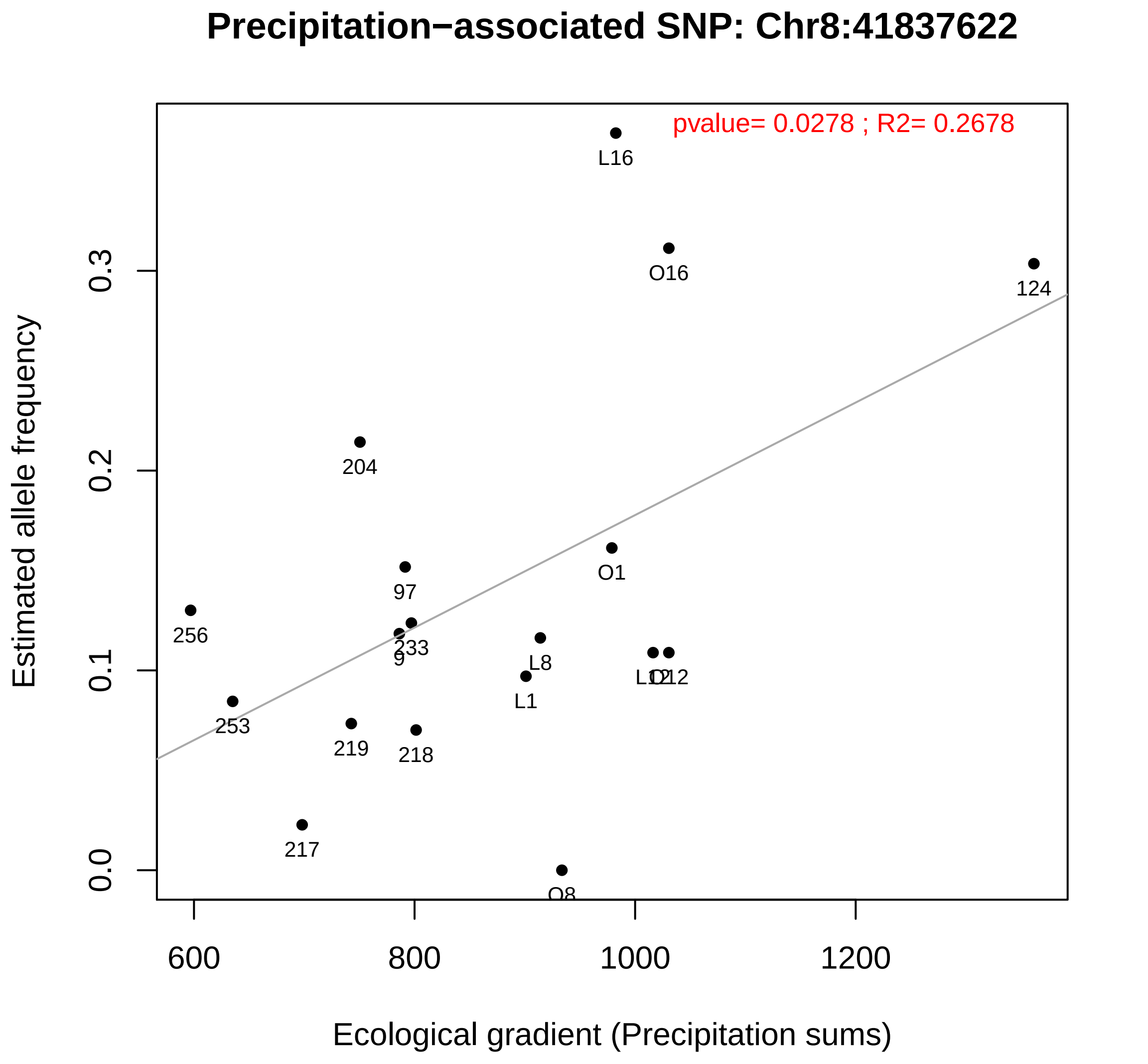

Precipitation−associated SNP: Chr8:41837622
pvalue= 0.0278 ; R2= 0.2678
L16
O16
124
204
O1
97
256
233
9
L8
L1
L12
O12
253
219
218
217
O8
0.0
0.1
0.2
0.3
Estimated allele frequency
600
800
1000
1200
Ecological gradient (Precipitation sums)

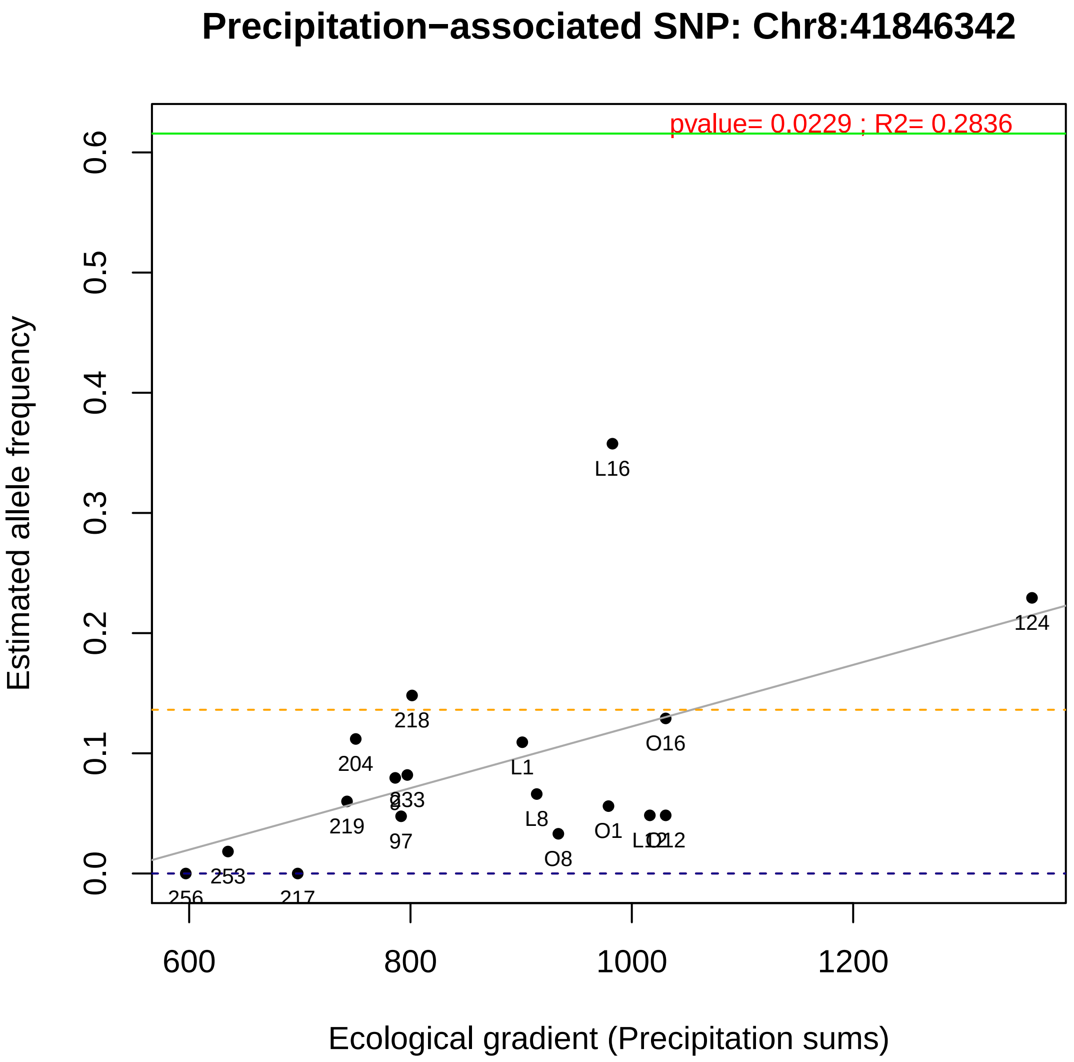
Precipitation−associated SNP: Chr8:41846342
pvalue= 0.0229 ; R2= 0.2836
Estimated allele frequency
0.0
0.1
0.2
0.3
0.4
0.5
0.6
600
800
1000
1200
Ecological gradient (Precipitation sums)
L16
124
218
O16
204
L1
233
219
L8
97
O1
O8
253
256
217

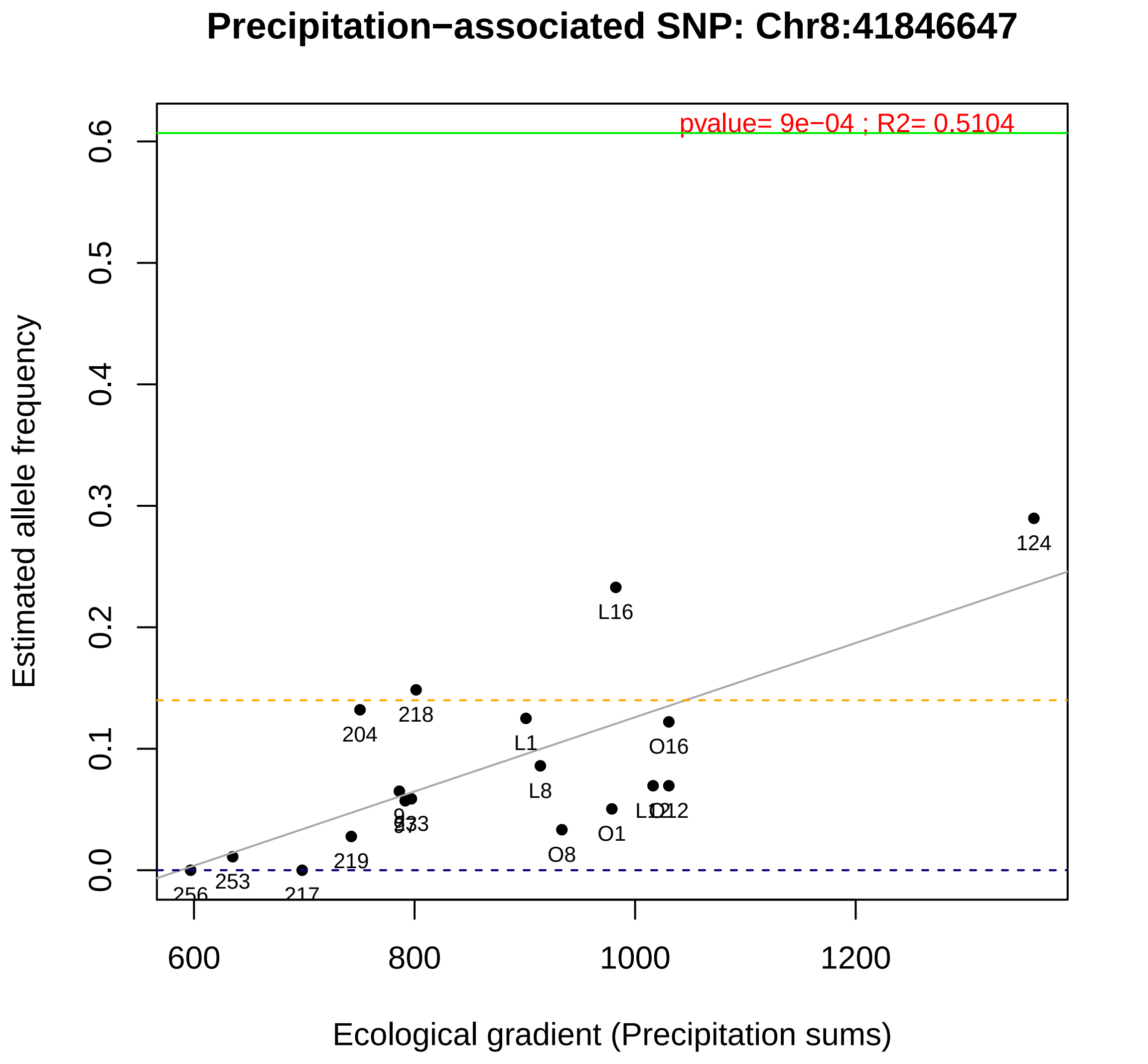

Precipitation−associated SNP: Chr8:41846647
pvalue= 9e−04 ; R2= 0.5104
Estimated allele frequency
0.0
0.1
0.2
0.3
0.4
0.5
0.6
124
L16
218
204
L1
O16
L8
O8
O1
219
253
256
217
600
800
1000
1200
Ecological gradient (Precipitation sums)

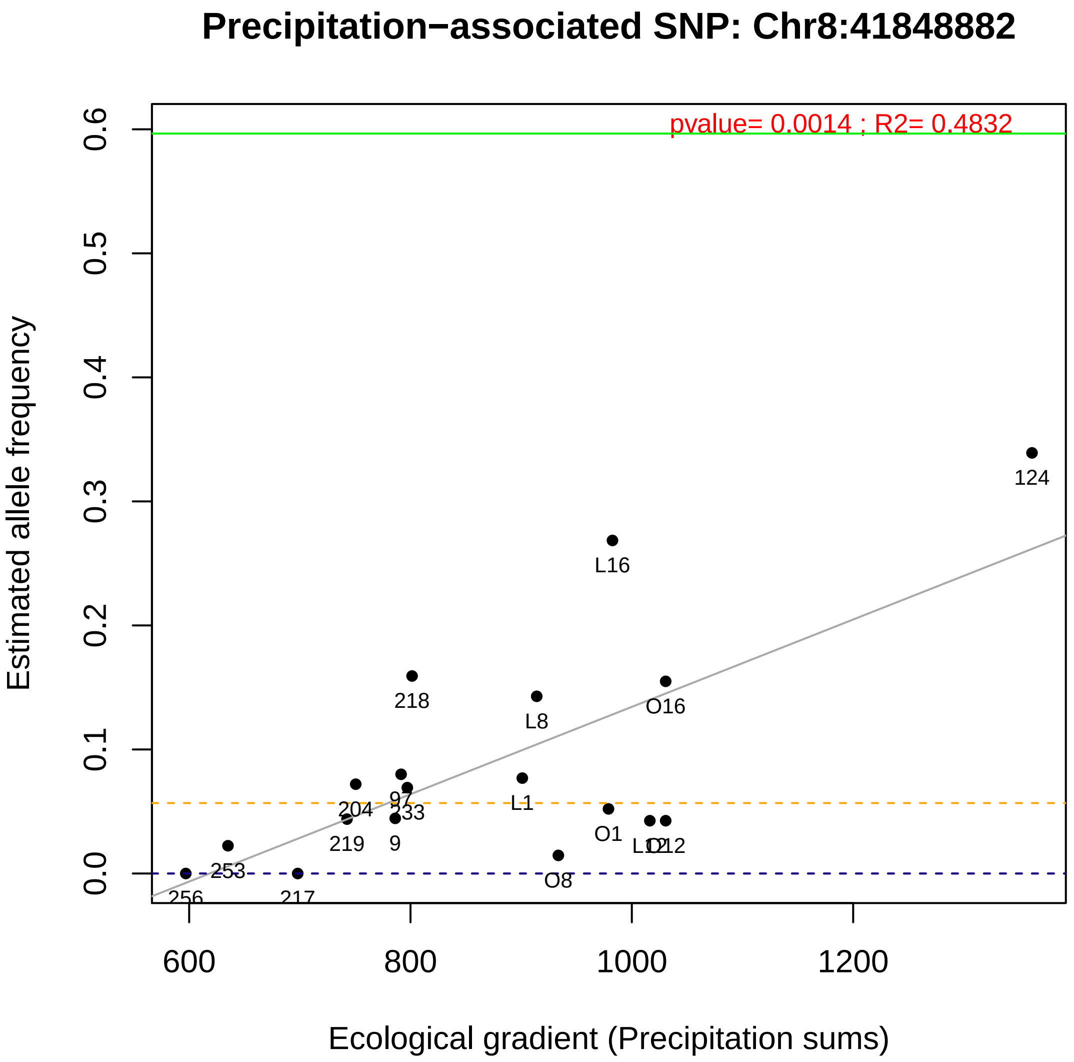
Precipitation−associated SNP: Chr8:41848882
pvalue= 0.0014 ; R2= 0.4832
Estimated allele frequency
0.0
0.1
0.2
0.3
0.4
0.5
0.6
600
800
1000
1200
Ecological gradient (Precipitation sums)
124
L16
218
L8
O16
97
204
233
L1
219
9
O1
L12
O12
253
O8
256
217

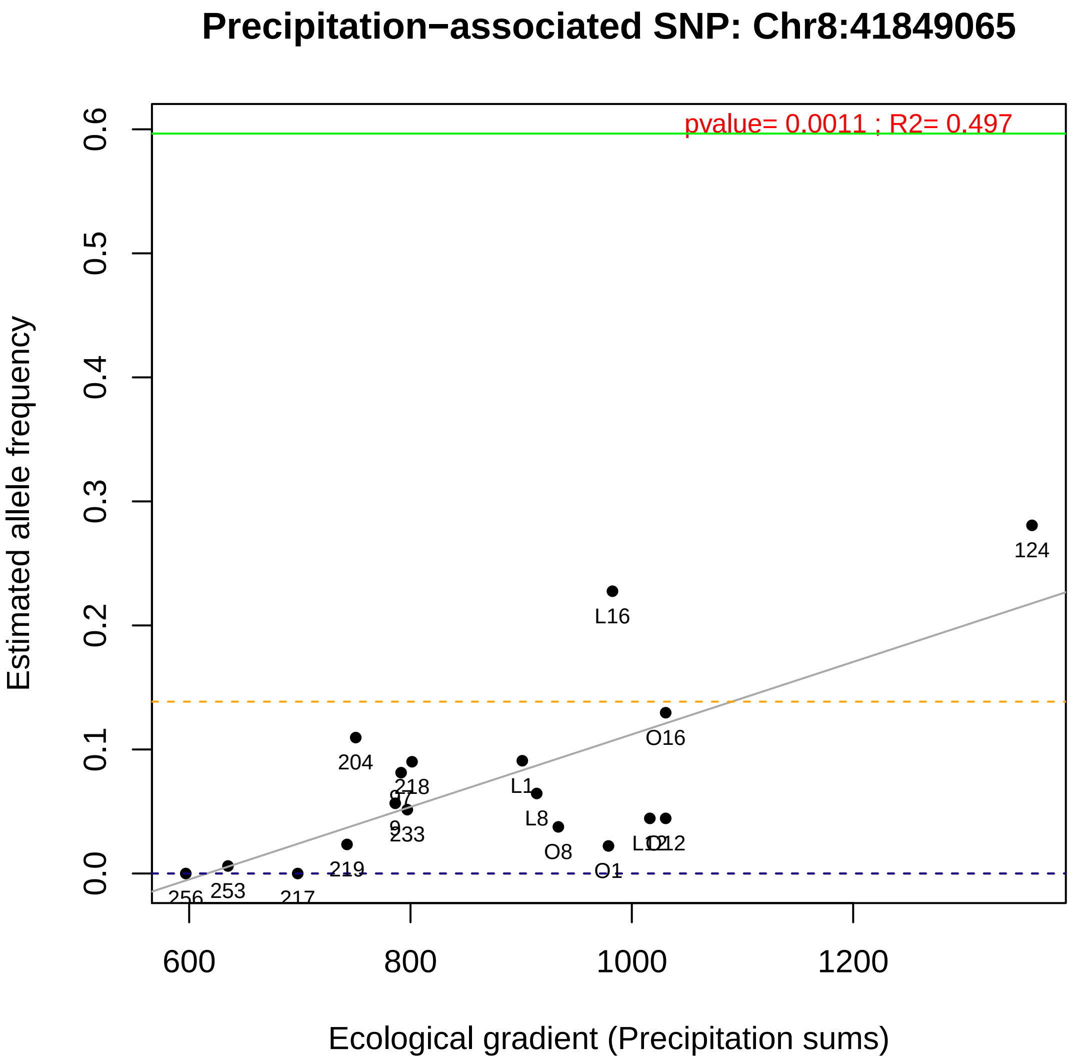

Precipitation−associated SNP: Chr8:41849065
pvalue= 0.0011 ; R2= 0.497
Estimated allele frequency
0.0
0.1
0.2
0.3
0.4
0.5
0.6
600
800
1000
1200
Ecological gradient (Precipitation sums)
124
L16
O16
204
218
97
9
233
L1
L8
O8
O1
L12
O12
219
256
253
217

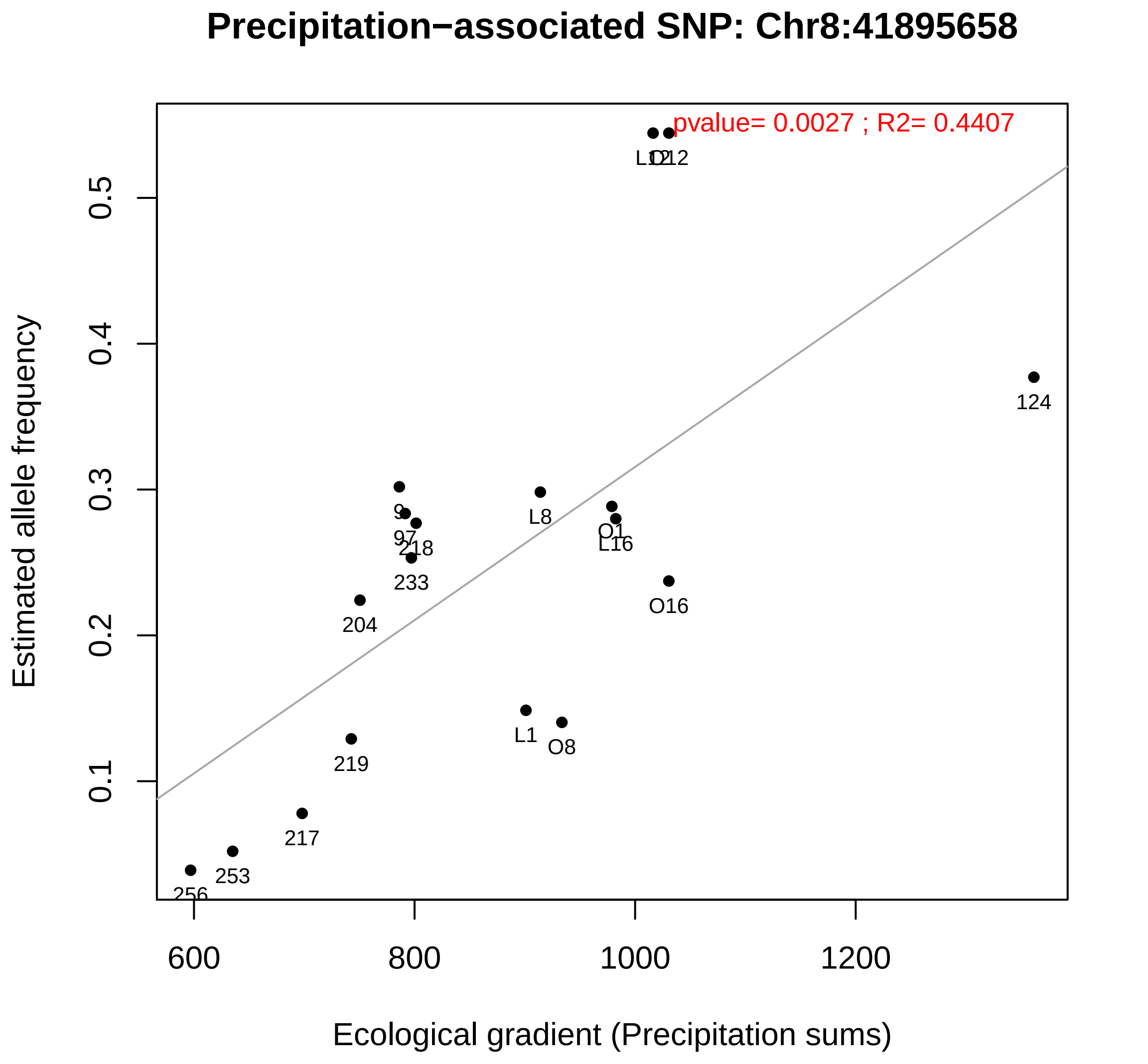
Precipitation−associated SNP: Chr8:41895658
pvalue= 0.0027 ; R2= 0.4407
Estimated allele frequency
Ecological gradient (Precipitation sums)
0.1
0.2
0.3
0.4
0.5
600
800
1000
1200
124
L8
O1
L16
O16
204
233
218
97
219
L1
O8
217
253
256

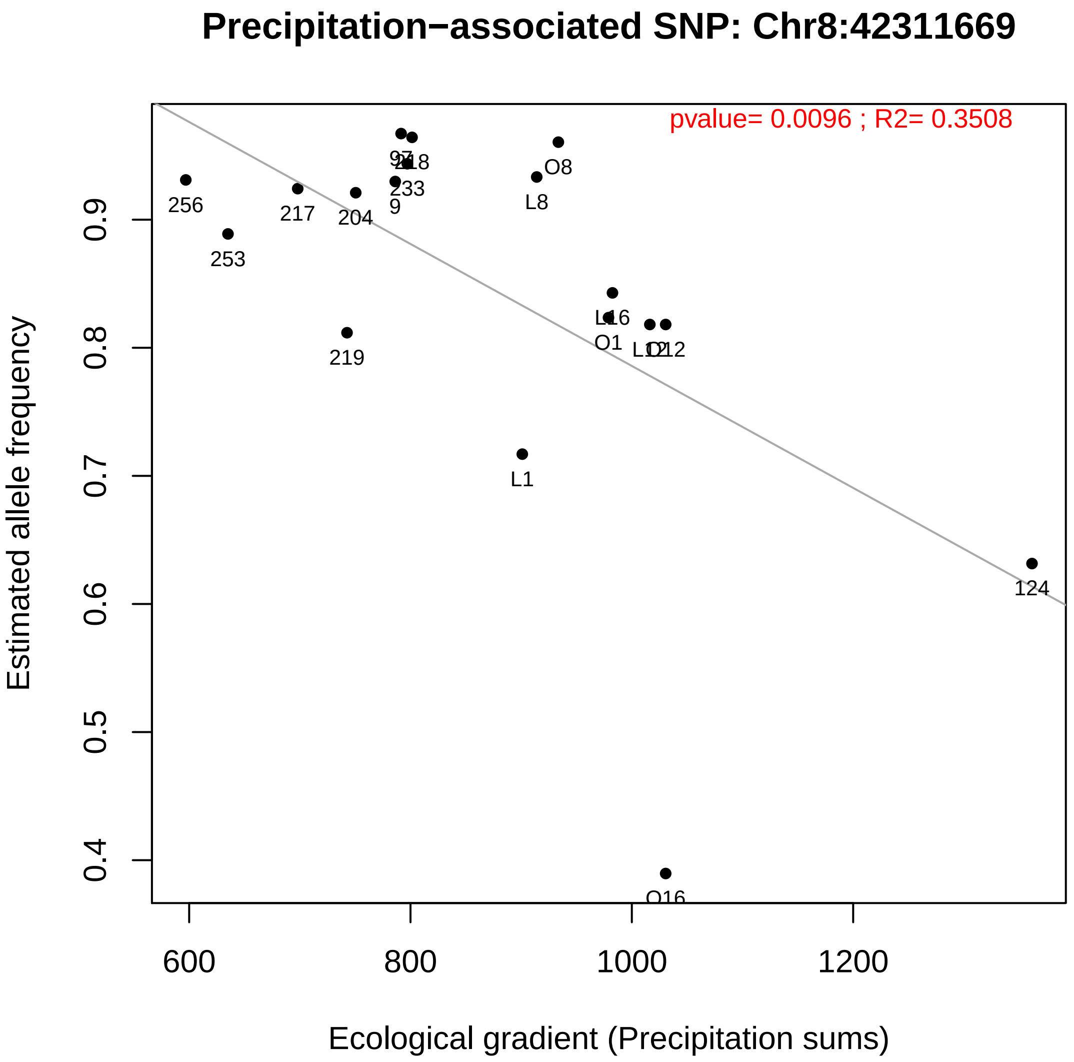
Precipitation−associated SNP: Chr8:42311669
pvalue= 0.0096 ; R2= 0.3508
Estimated allele frequency
0.4
0.5
0.6
0.7
0.8
0.9
600
800
1000
1200
Ecological gradient (Precipitation sums)
256
253
217
204
219
233
9
O8
L8
L1
O1
124
O16

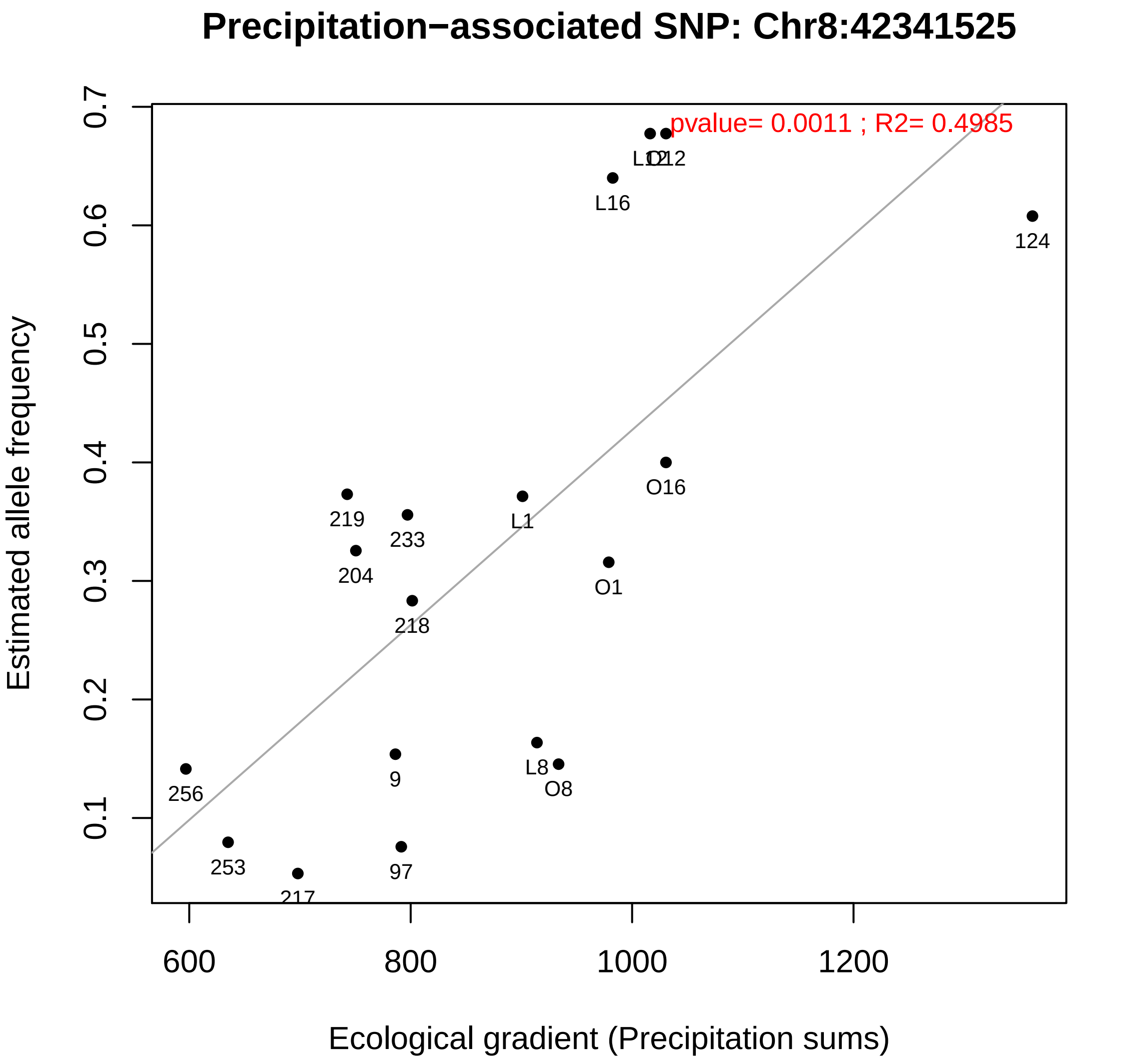

Precipitation−associated SNP: Chr8:42341525
pvalue= 0.0011 ; R2= 0.4985
Estimated allele frequency
0.1
0.2
0.3
0.4
0.5
0.6
0.7
600
800
1000
1200
Ecological gradient (Precipitation sums)
L16
124
O16
219
233
L1
204
O1
218
256
9
L8
O8
253
97
217

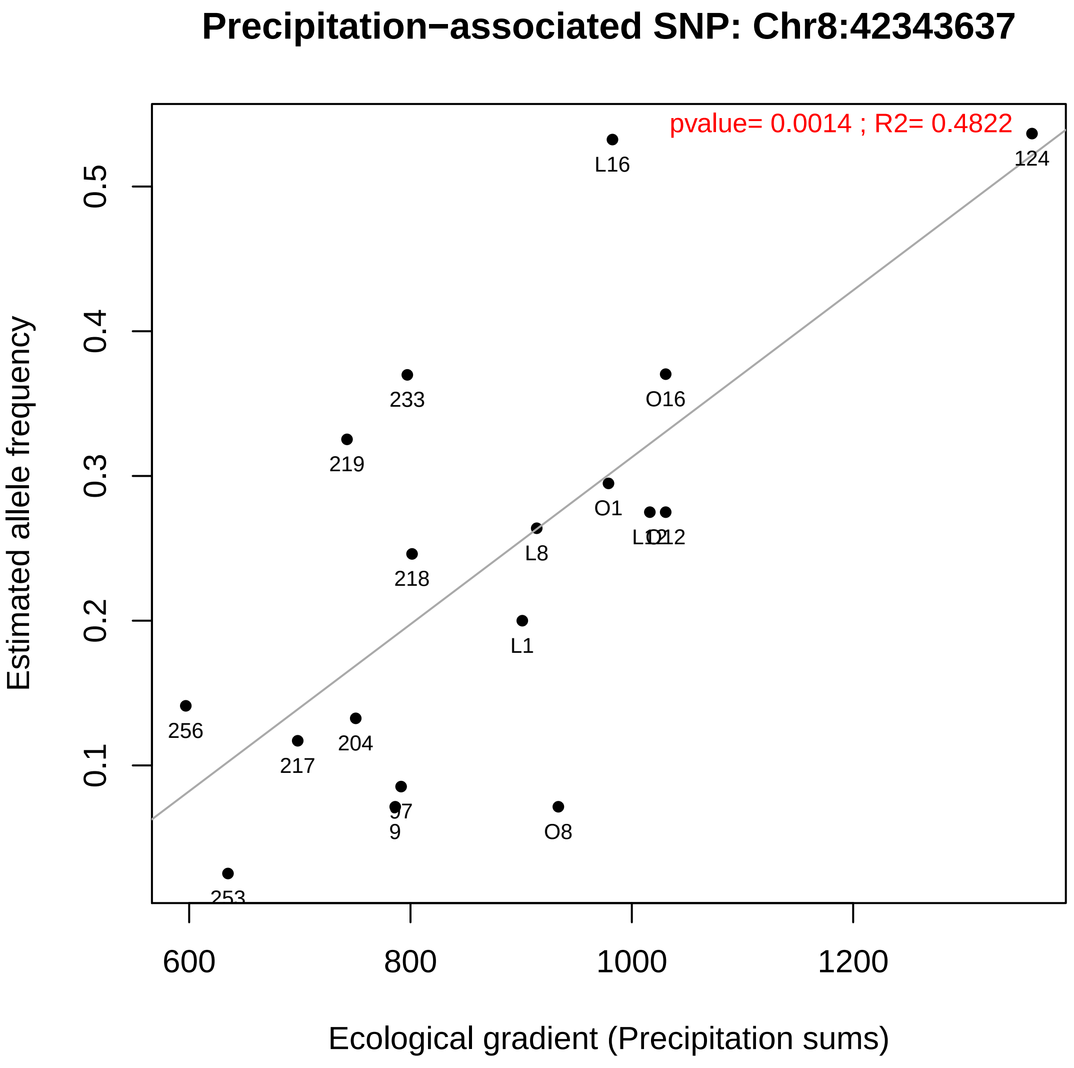
Precipitation−associated SNP: Chr8:42343637
pvalue= 0.0014 ; R2= 0.4822
Estimated allele frequency
Ecological gradient (Precipitation sums)
0.1
0.2
0.3
0.4
0.5
600
800
1000
1200
L16
124
233
O16
219
O1
L8
218
L1
256
204
217
97
9
O8
253

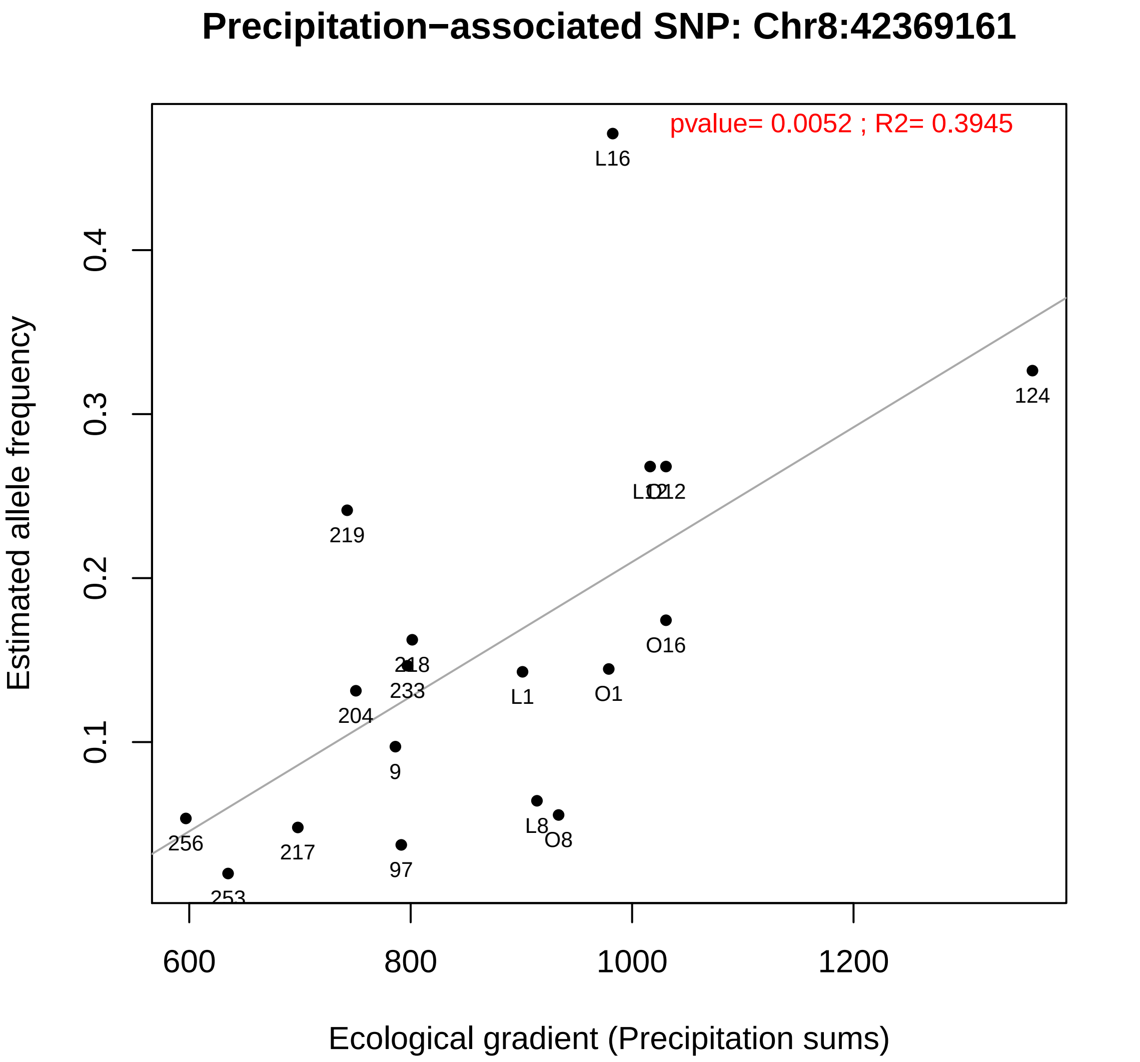

Precipitation−associated SNP: Chr8:42369161
pvalue= 0.0052 ; R2= 0.3945
L16
124
L12
O12
219
O16
218
233
L1
O1
204
9
L8
O8
256
217
97
253
0.1
0.2
0.3
0.4
600
800
1000
1200
Estimated allele frequency
Ecological gradient (Precipitation sums)

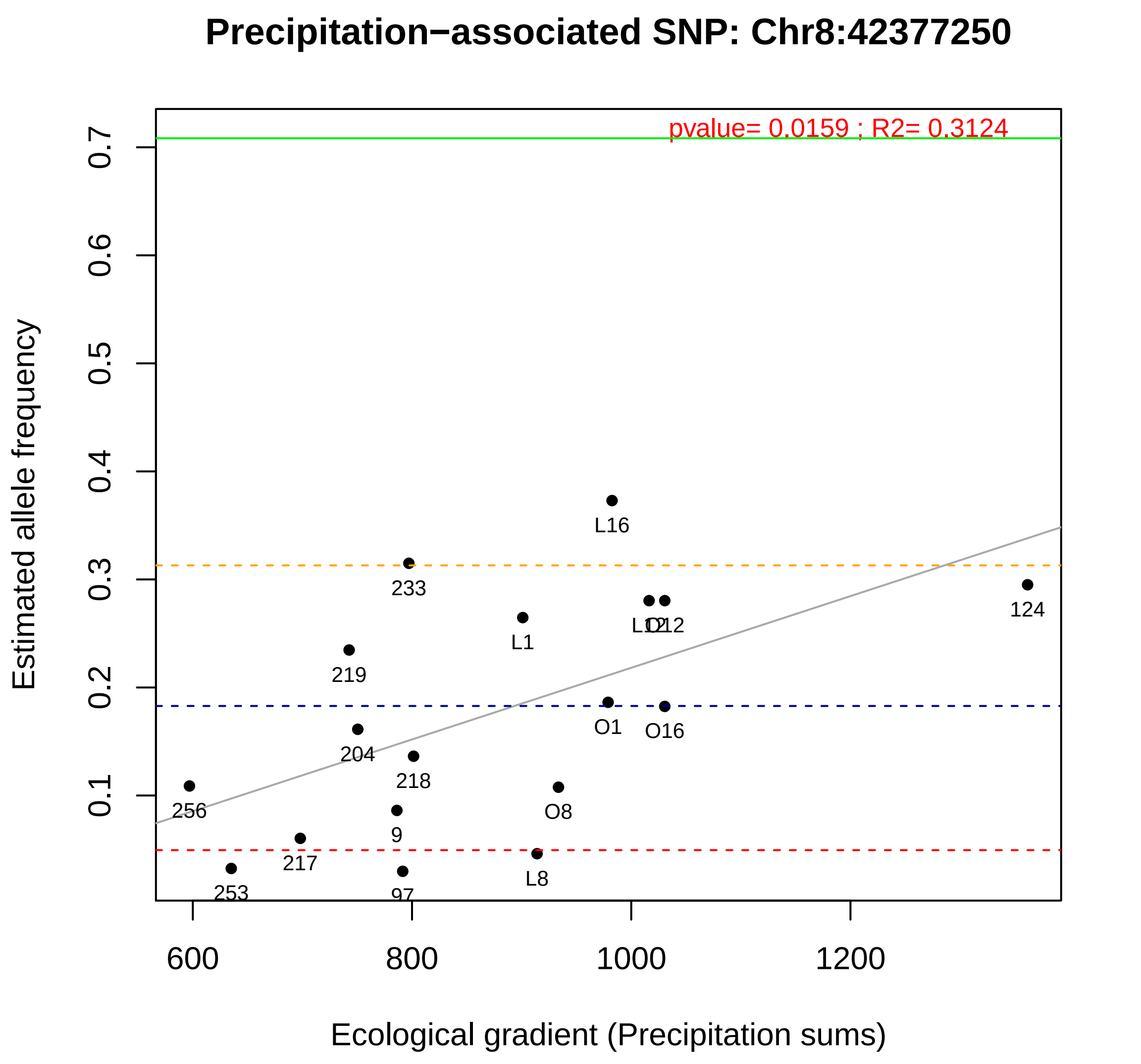

Precipitation−associated SNP: Chr8:42377250
pvalue= 0.0159 ; R2= 0.3124
Estimated allele frequency
0.1
0.2
0.3
0.4
0.5
0.6
0.7
L16
233
124
L1
L12
O12
219
O1
O16
204
218
256
O8
9
217
L8
253
97
600
800
1000
1200
Ecological gradient (Precipitation sums)

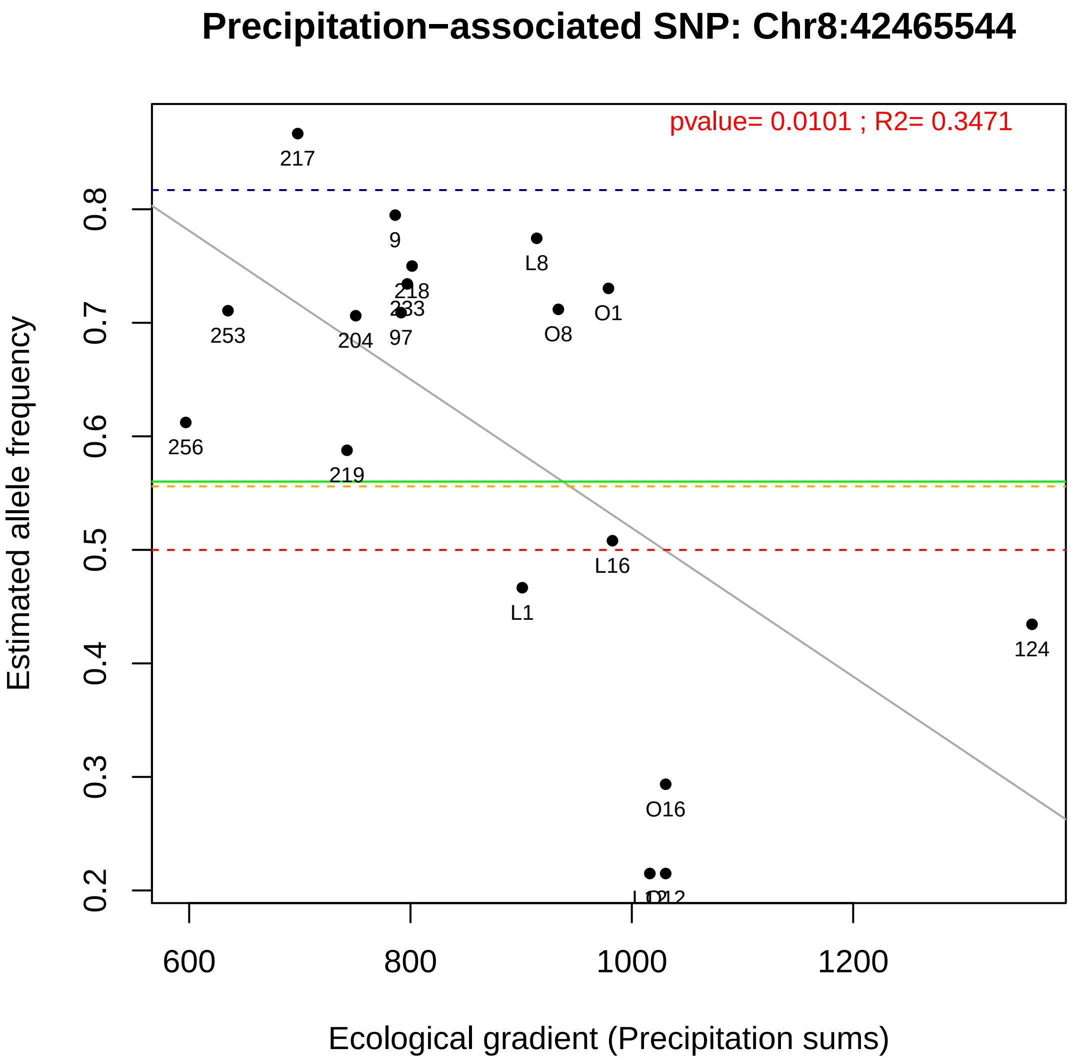
Precipitation−associated SNP: Chr8:42465544
pvalue= 0.0101 ; R2= 0.3471
217
9
L8
218
233
O1
253
204
97
O8
256
219
L16
L1
124
O16
0.2
0.3
0.4
0.5
0.6
0.7
0.8
600
800
1000
1200
Estimated allele frequency
Ecological gradient (Precipitation sums)

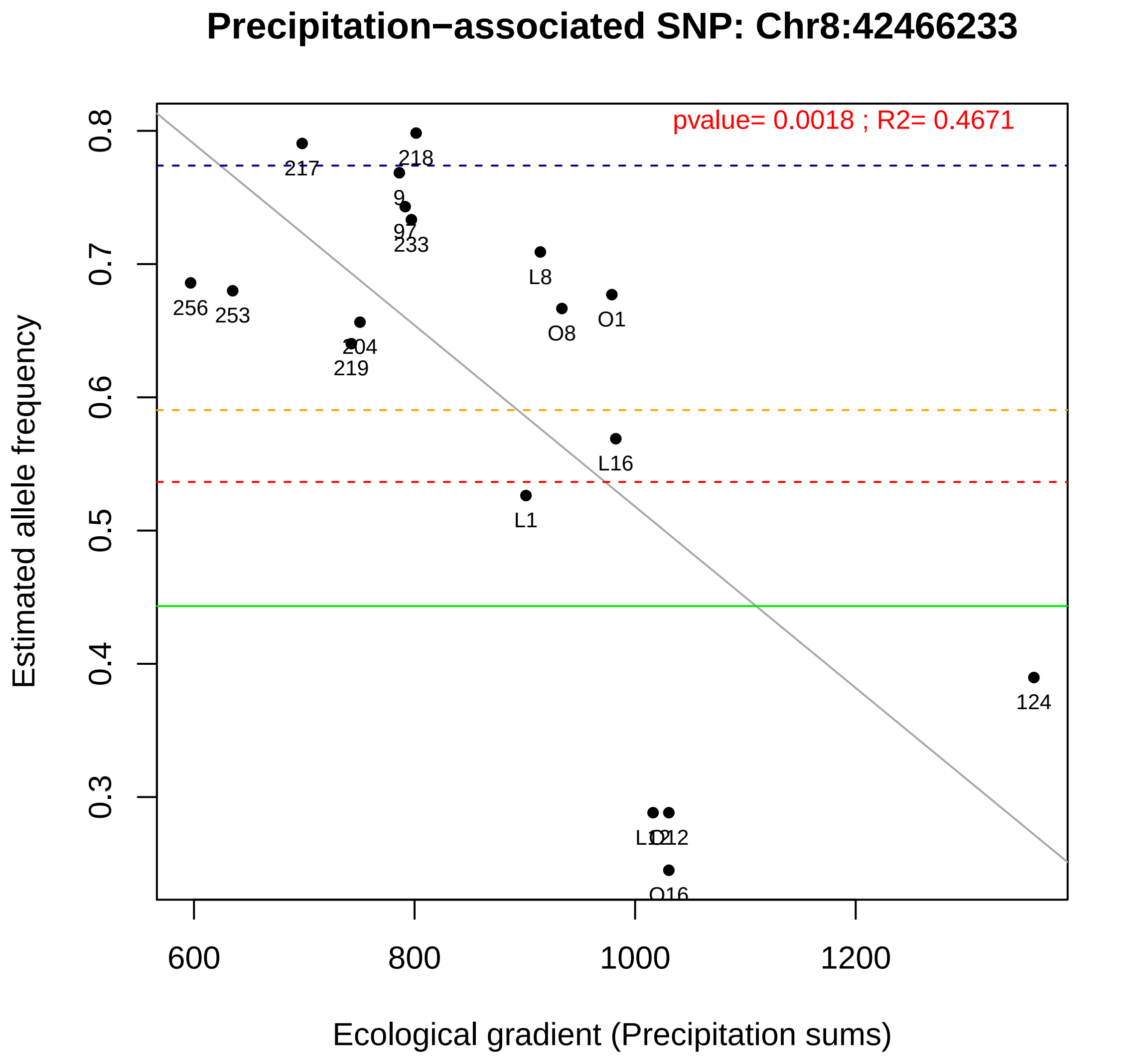

Precipitation−associated SNP: Chr8:42466233
pvalue= 0.0018 ; R2= 0.4671
Estimated allele frequency
Ecological gradient (Precipitation sums)
0.3
0.4
0.5
0.6
0.7
0.8
600
800
1000
1200
217
218
9
97
233
256
253
204
219
L8
O1
O8
L16
L1
124
O12
O16

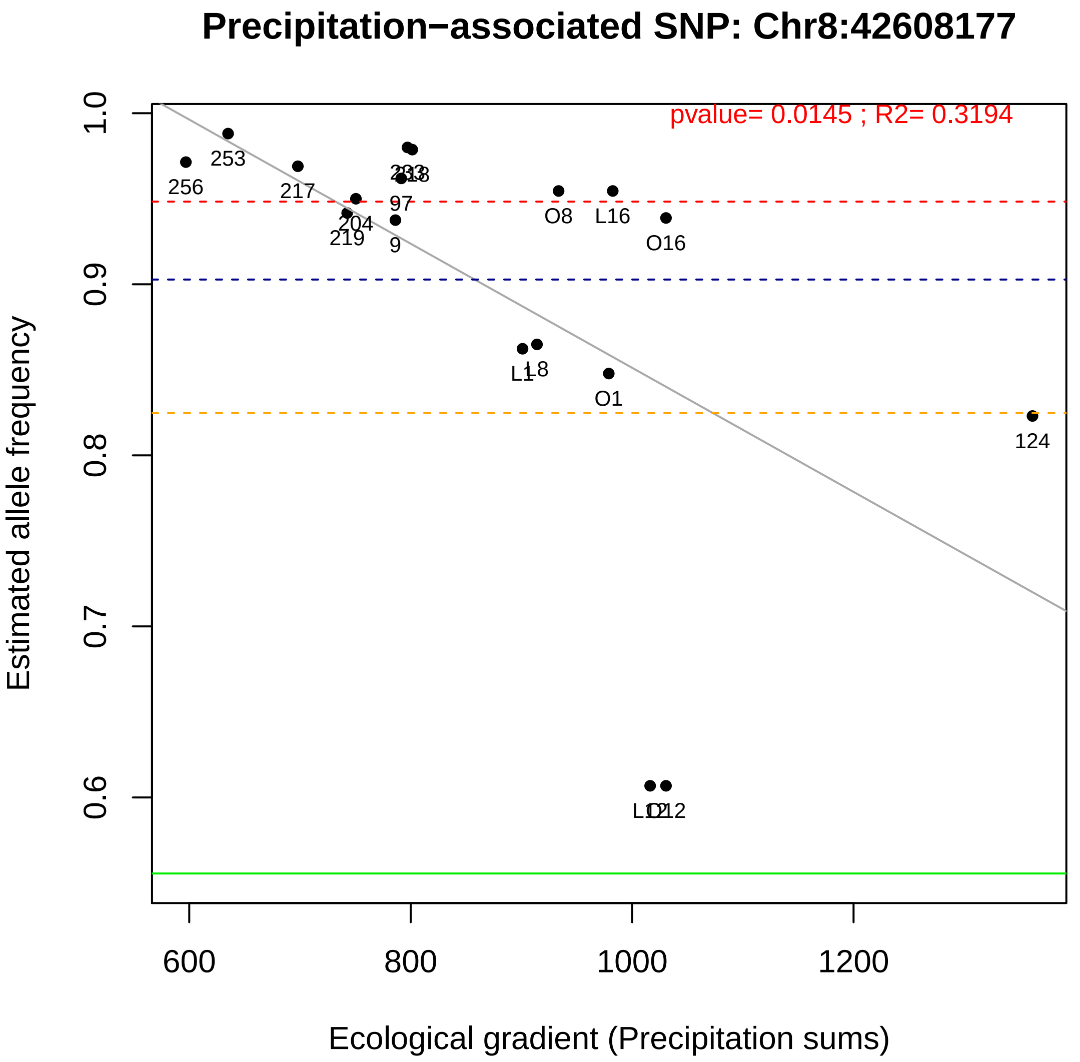
Precipitation−associated SNP: Chr8:42608177
pvalue= 0.0145 ; R2= 0.3194
Estimated allele frequency
Ecological gradient (Precipitation sums)
1.0
0.9
0.8
0.7
0.6
600
800
1000
1200
256
253
217
204
219
97
9
O8
L16
O16
L1
L8
O1
124
O12

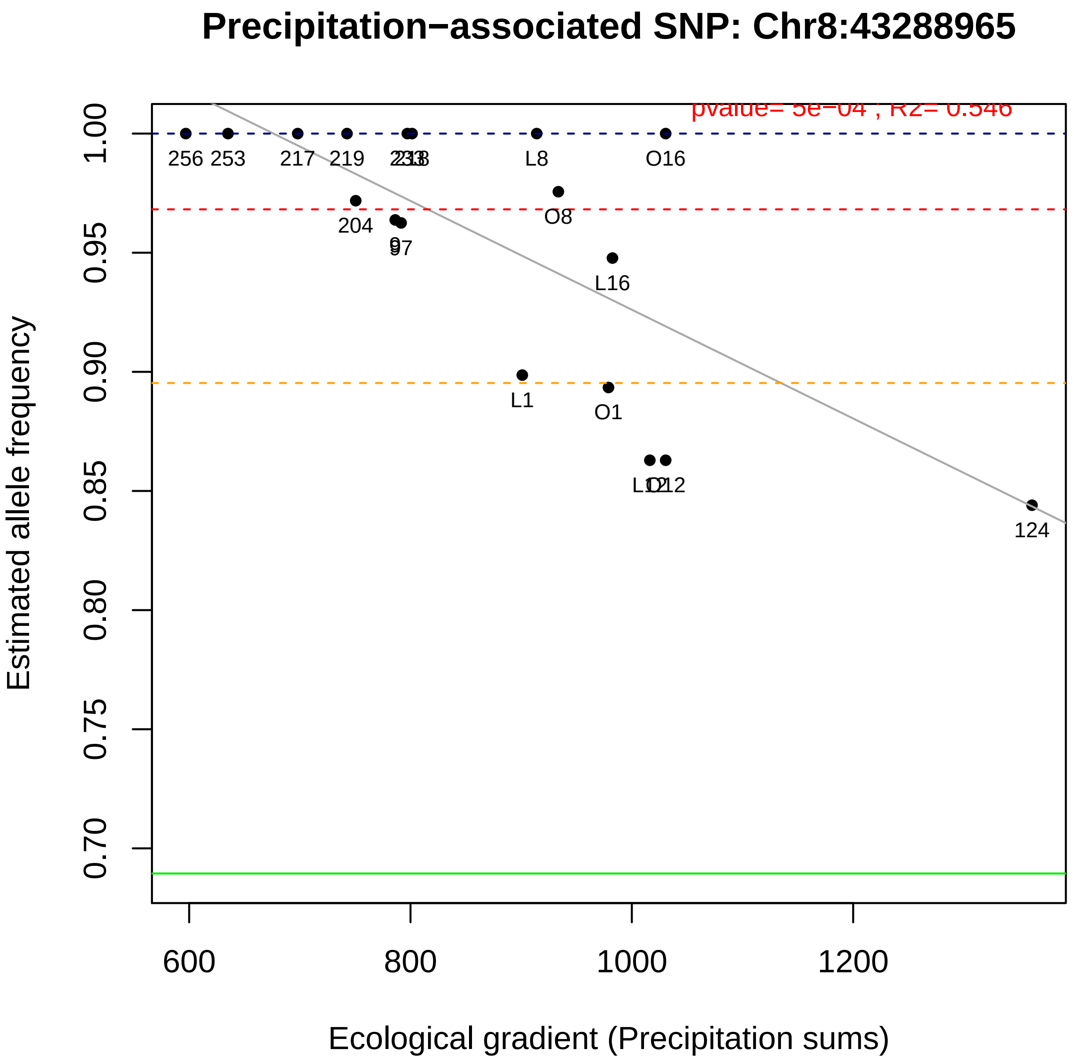
Precipitation−associated SNP: Chr8:43288965
pvalue= 5e−04 ; R2= 0.546
Estimated allele frequency
Ecological gradient (Precipitation sums)
1.00
0.95
0.90
0.85
0.80
0.75
0.70
600
800
1000
1200
256
253
217
219
L8
O16
204
O8
L16
L1
O1
124

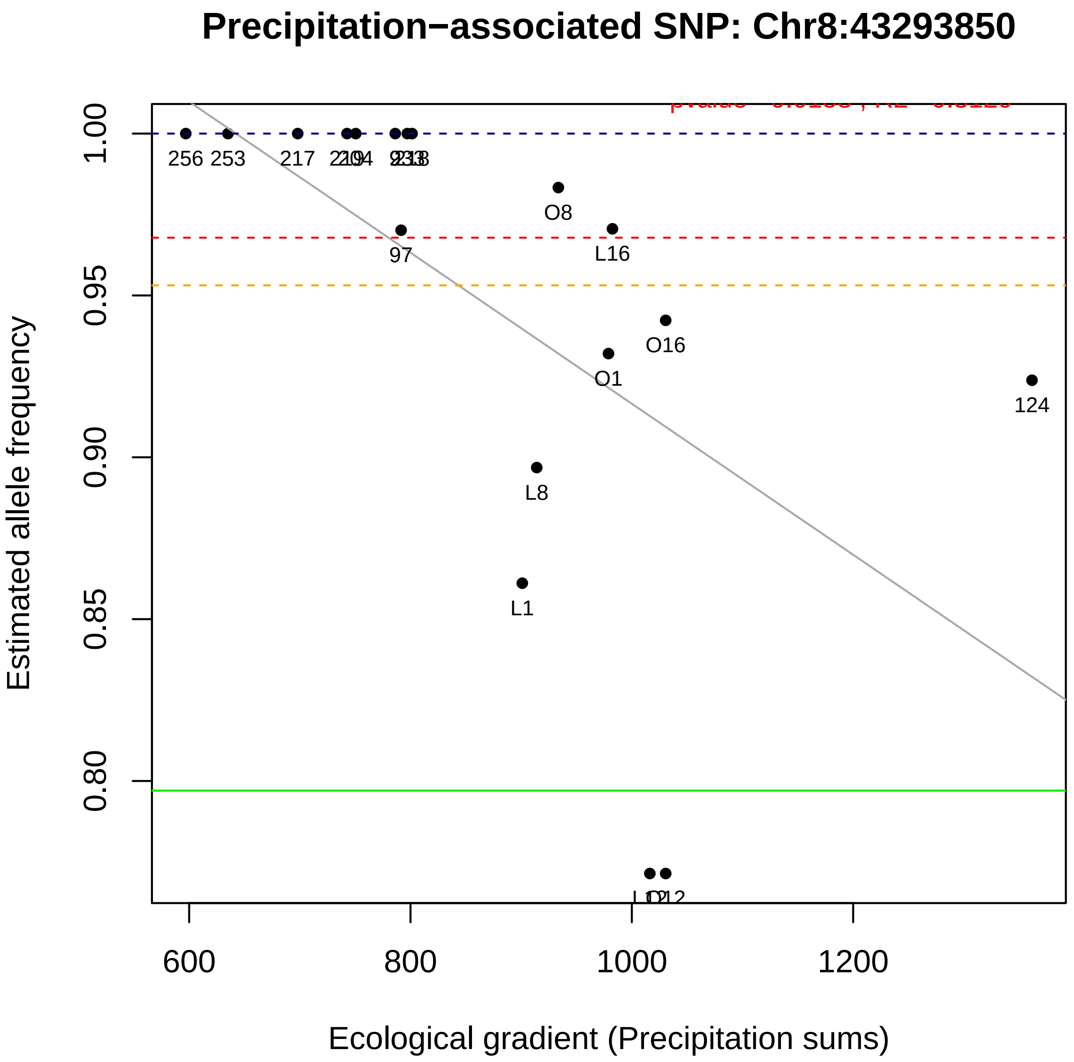
Precipitation−associated SNP: Chr8:43293850
Estimated allele frequency
Ecological gradient (Precipitation sums)
1.00
0.95
0.90
0.85
0.80
600
800
1000
1200
256
253
217
97
O8
L16
O16
O1
124
L8
L1

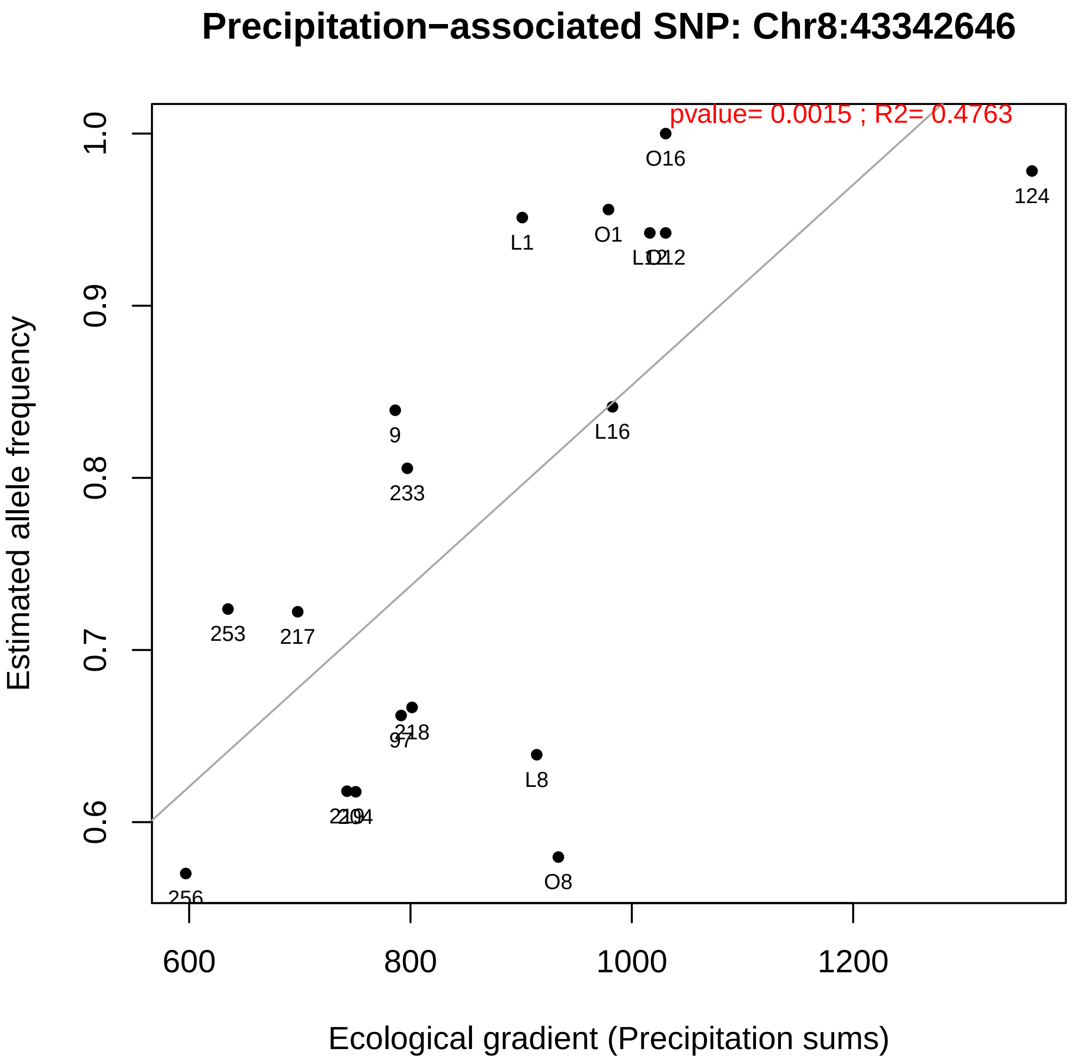

Precipitation−associated SNP: Chr8:43342646
pvalue= 0.0015 ; R2= 0.4763
Estimated allele frequency
Ecological gradient (Precipitation sums)
0.6
0.7
0.8
0.9
1.0
600
800
1000
1200
O16
124
L1
O1
L16
9
233
253
217
218
97
L8
O8
256

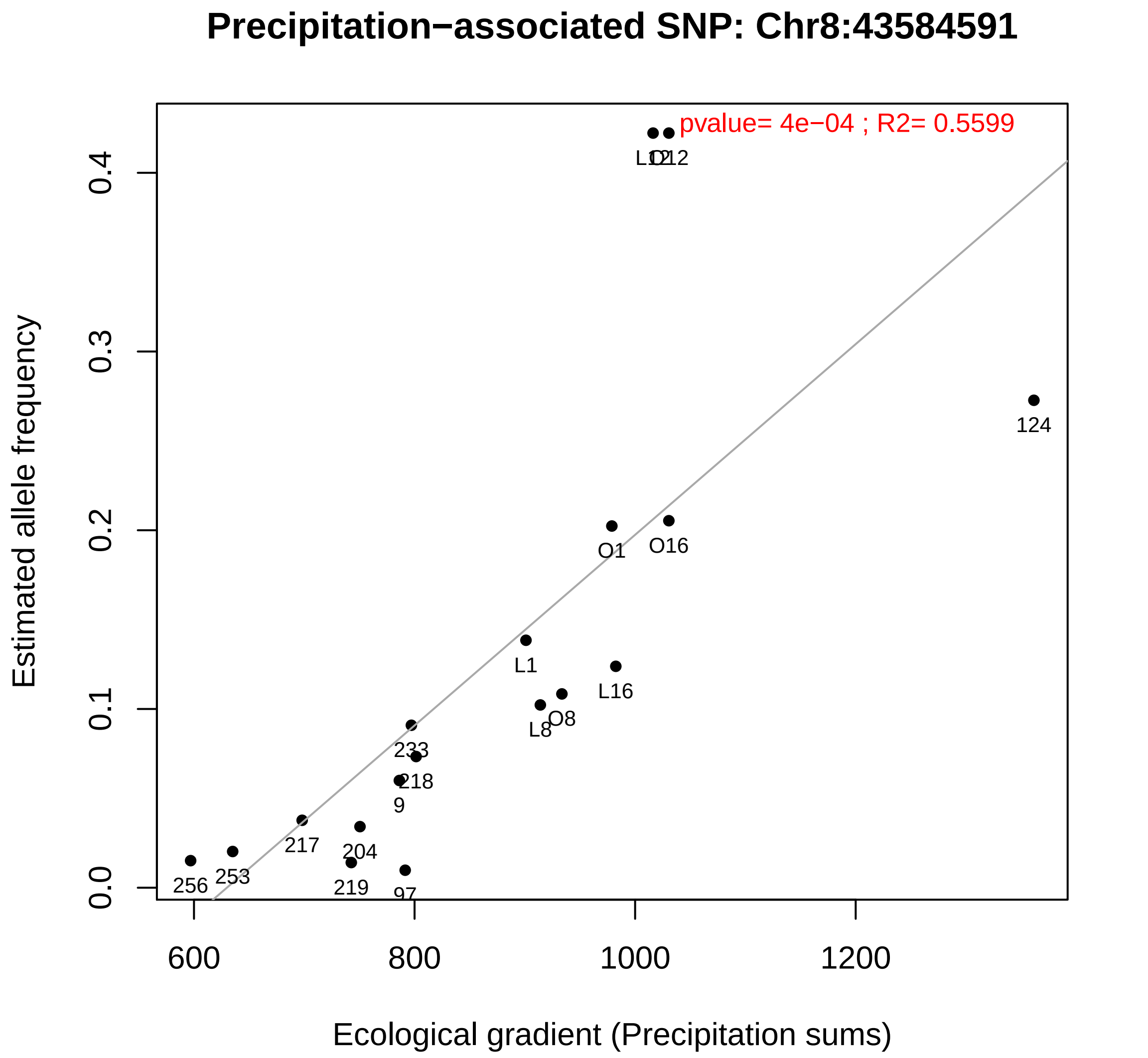

Precipitation−associated SNP: Chr8:43584591
pvalue= 4e−04 ; R2= 0.5599
Estimated allele frequency
Ecological gradient (Precipitation sums)
0.0
0.1
0.2
0.3
0.4
600
800
1000
1200
L12
O12
124
O1
O16
L1
L16
O8
L8
233
218
9
217
204
219
97
256
253

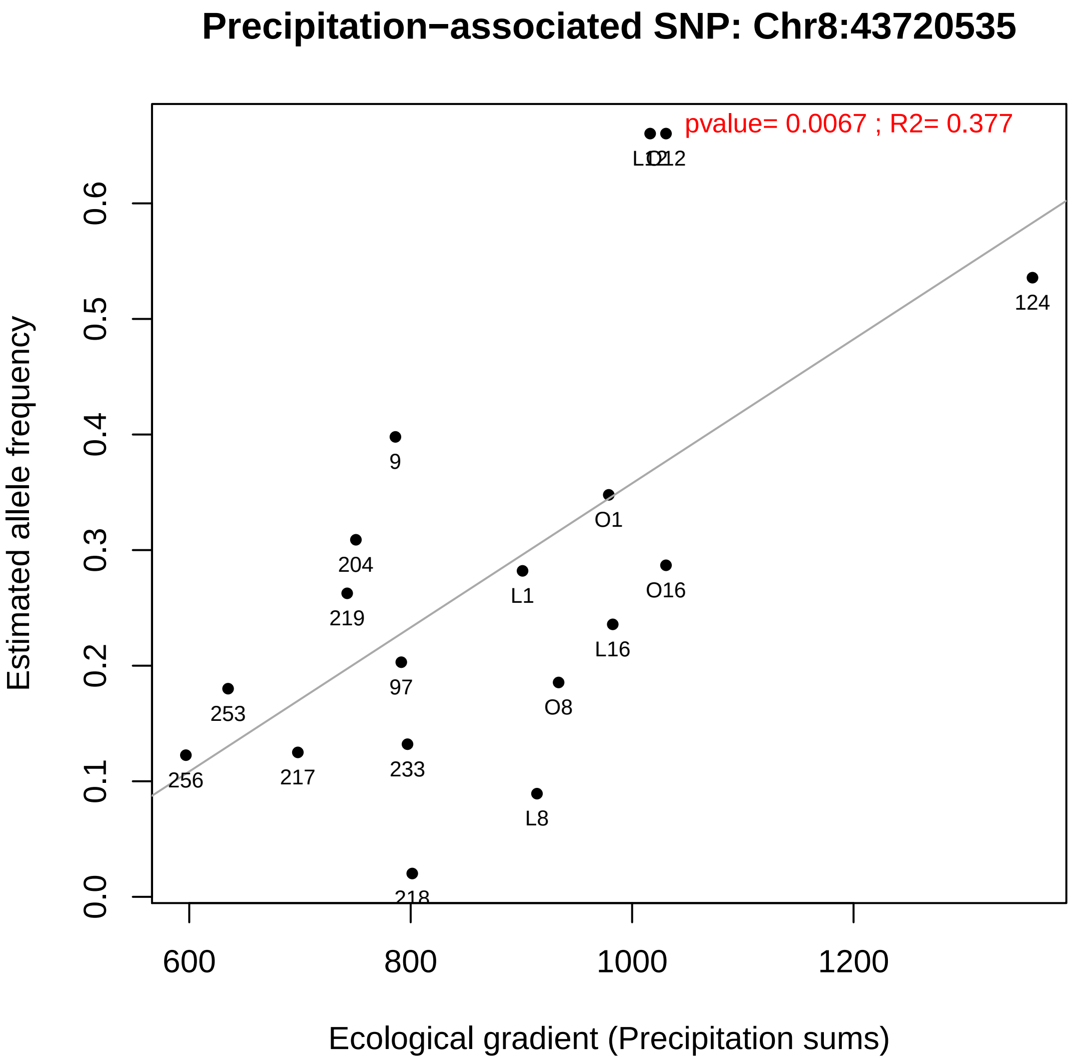
Precipitation−associated SNP: Chr8:43720535
pvalue= 0.0067 ; R2= 0.377
Estimated allele frequency
0.0
0.1
0.2
0.3
0.4
0.5
0.6
600
800
1000
1200
Ecological gradient (Precipitation sums)
L12
O12
124
9
O1
204
219
L1
O16
L16
97
O8
253
256
217
233
L8
218

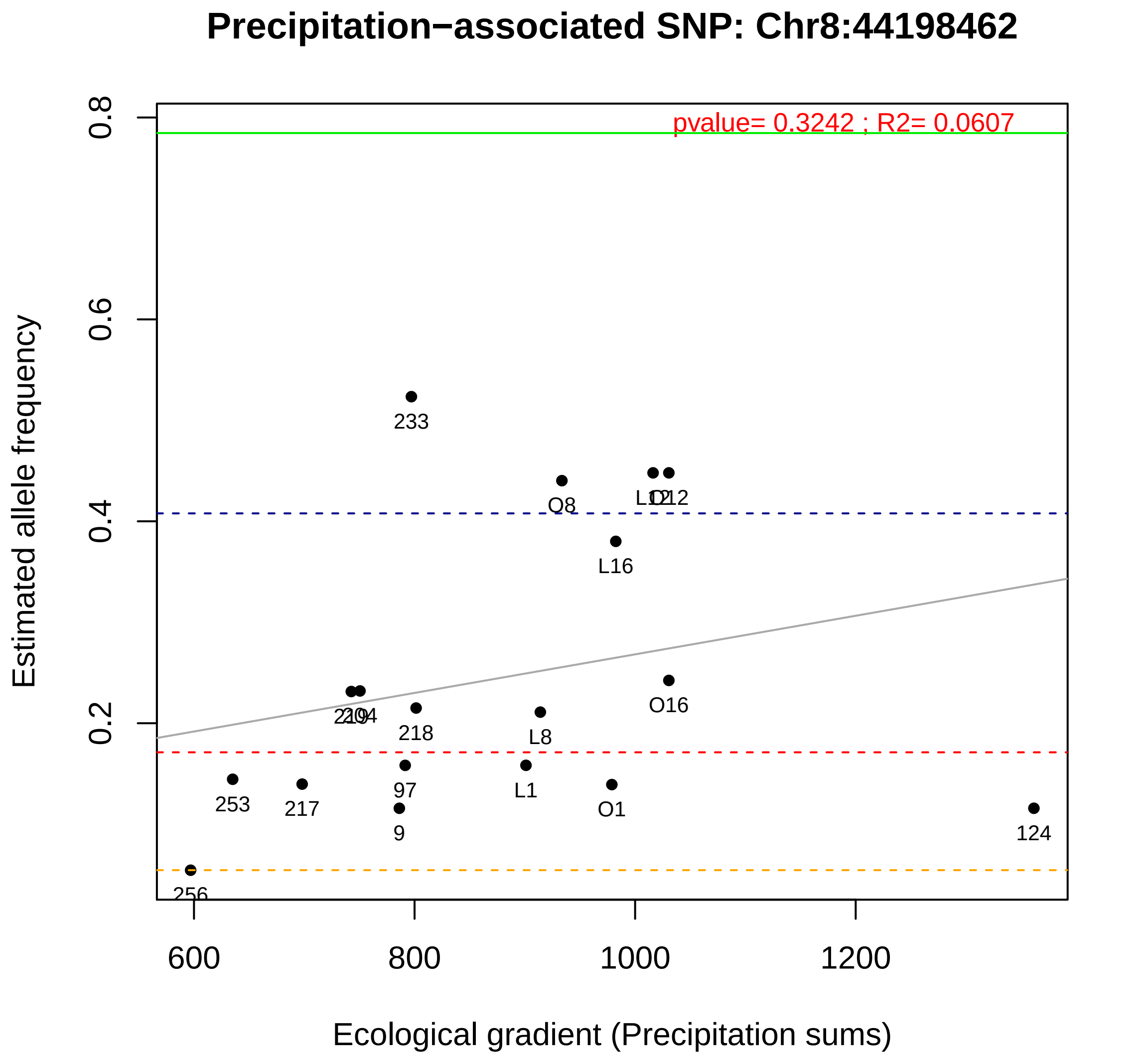

Precipitation−associated SNP: Chr8:44198462
pvalue= 0.3242 ; R2= 0.0607
Estimated allele frequency
0.2
0.4
0.6
0.8
600
800
1000
1200
Ecological gradient (Precipitation sums)
233
Q8
L16
O16
218
L8
97
L1
O1
253
217
9
124
256

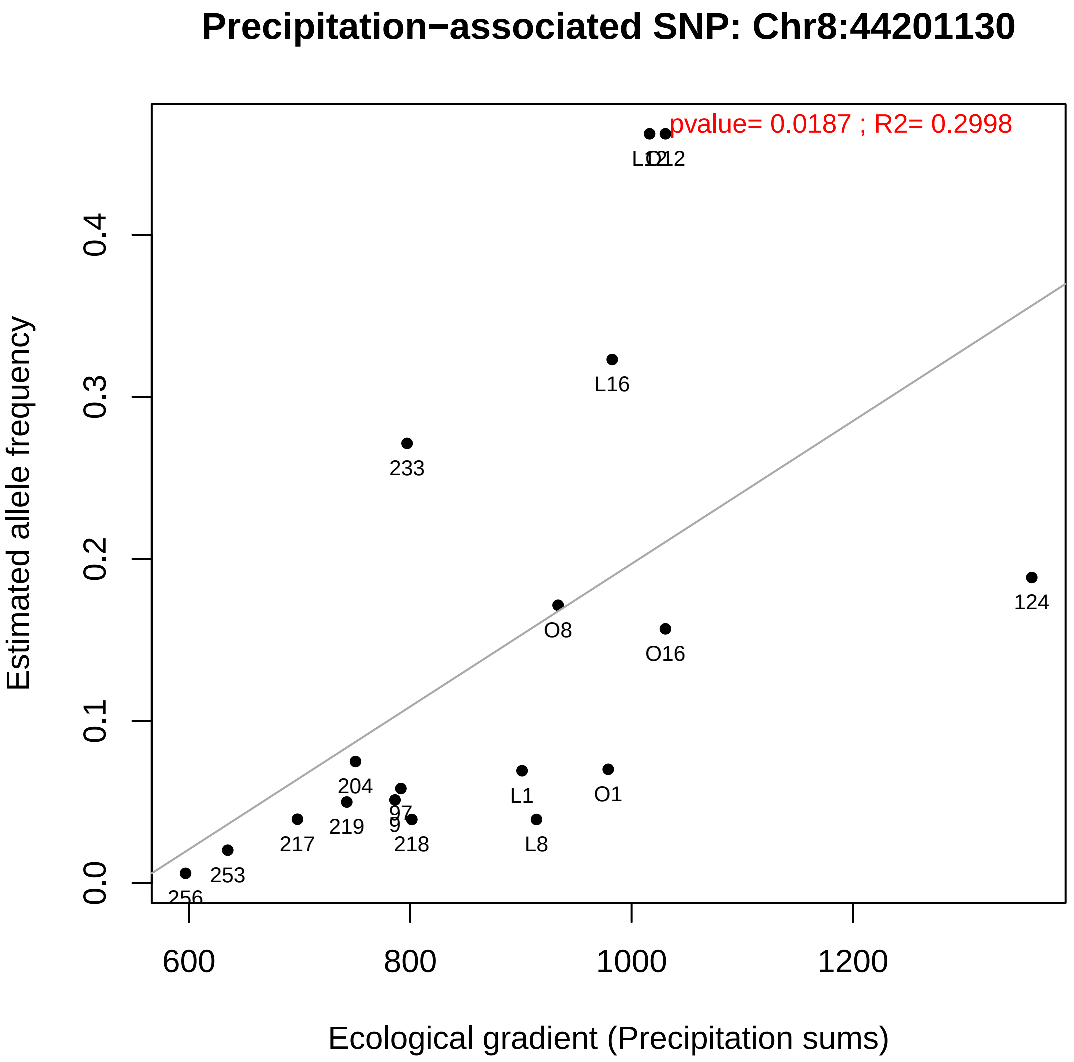

Precipitation−associated SNP: Chr8:44201130
pvalue= 0.0187 ; R2= 0.2998
L12
O12
L16
233
O8
124
O16
204
L1
O1
97
9
219
217
218
L8
253
256
0.0
0.1
0.2
0.3
0.4
600
800
1000
1200
Estimated allele frequency
Ecological gradient (Precipitation sums)

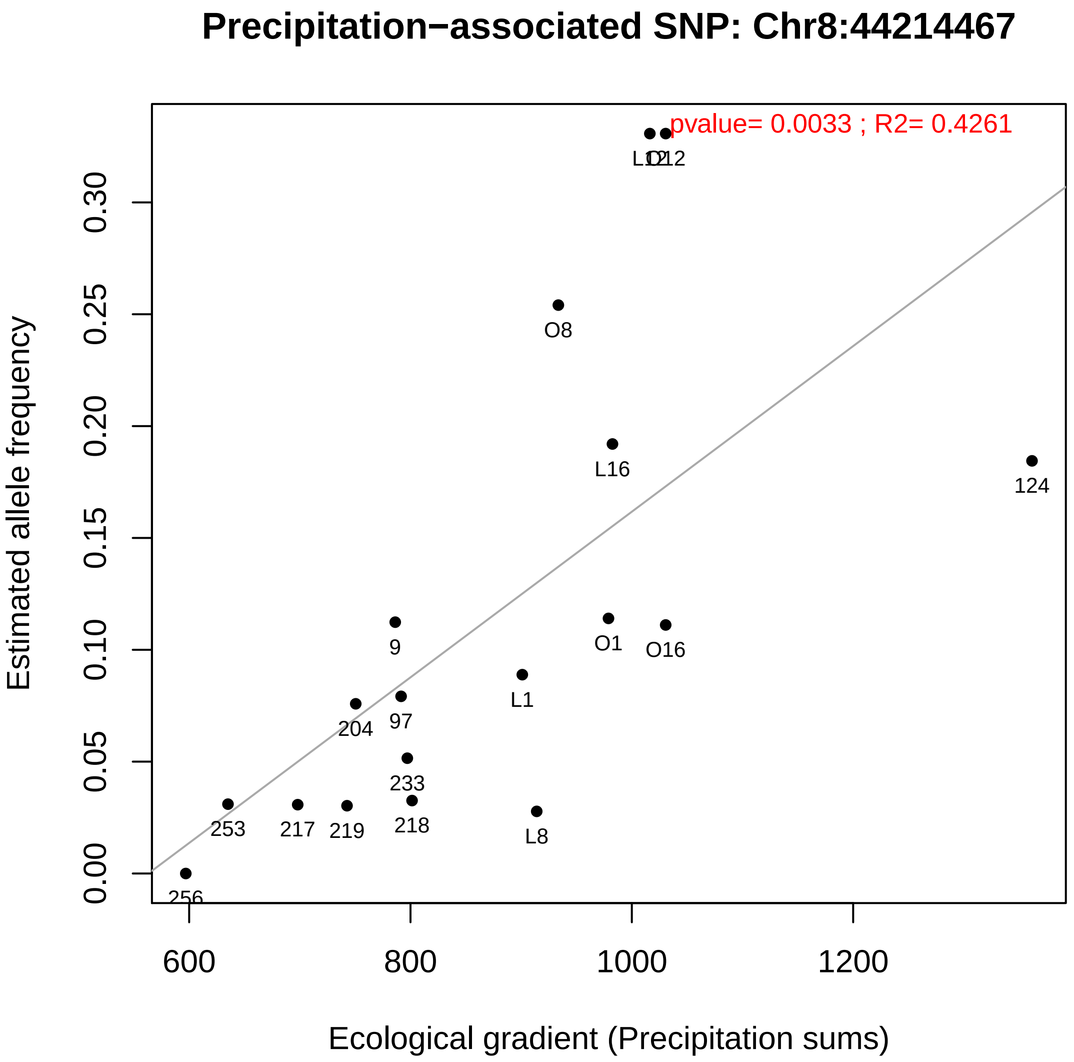

Precipitation−associated SNP: Chr8:44214467
pvalue= 0.0033 ; R2= 0.4261
Estimated allele frequency
0.00
0.05
0.10
0.15
0.20
0.25
0.30
600
800
1000
1200
Ecological gradient (Precipitation sums)
O8
L16
124
9
O1
O16
L1
97
204
233
253
217
219
218
L8
256

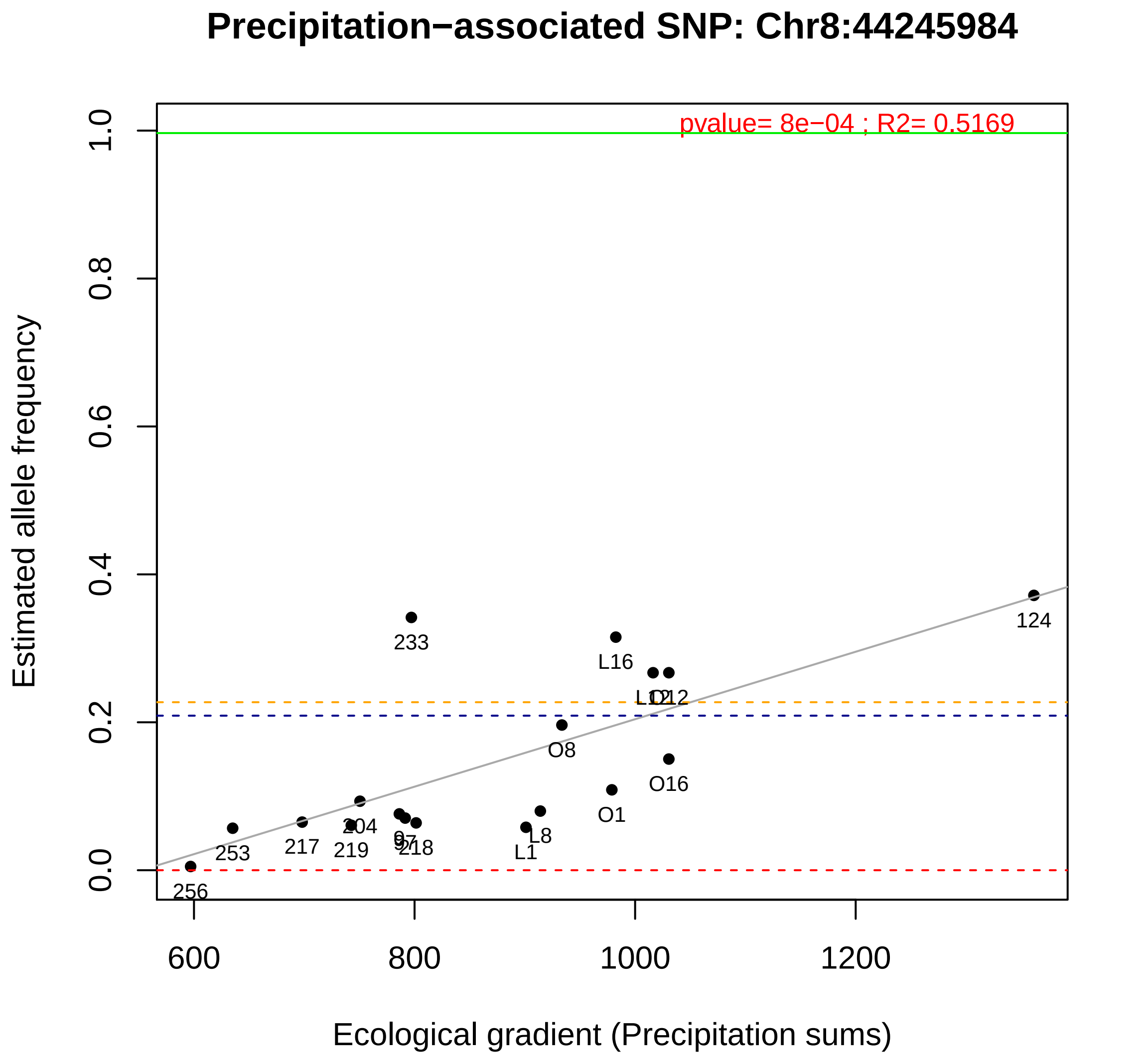
Precipitation−associated SNP: Chr8:44245984
pvalue= 8e−04 ; R2= 0.5169
Estimated allele frequency
Ecological gradient (Precipitation sums)
0.0
0.2
0.4
0.6
0.8
1.0
600
800
1000
1200
233
L16
124
L10
O12
O8
O16
O1
L8
L1
204
97
218
219
217
253
256

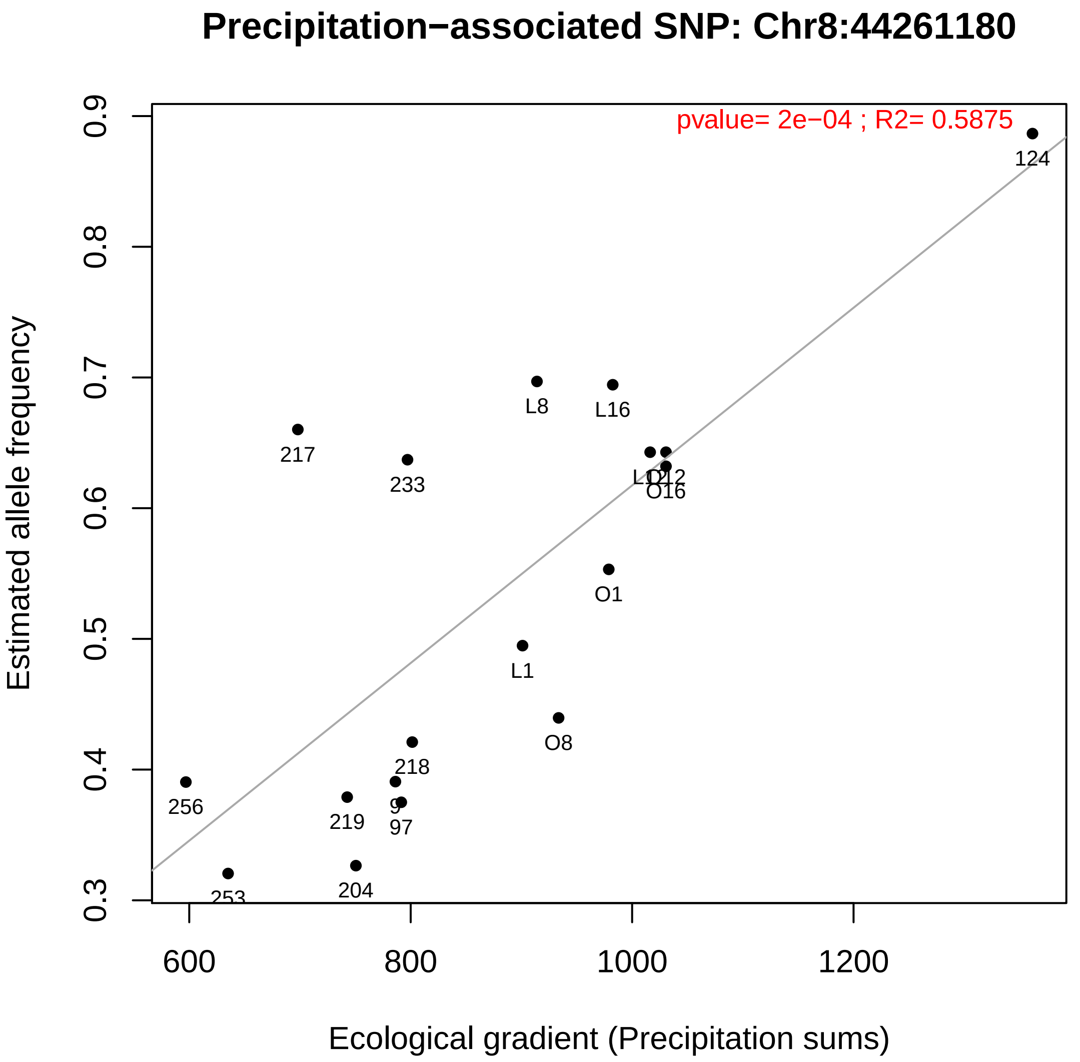

Precipitation−associated SNP: Chr8:44261180
pvalue= 2e−04 ; R2= 0.5875
124
L8
L16
217
233
O16
O1
L1
O8
218
256
219
97
204
253
0.3
0.4
0.5
0.6
0.7
0.8
0.9
600
800
1000
1200
Estimated allele frequency
Ecological gradient (Precipitation sums)

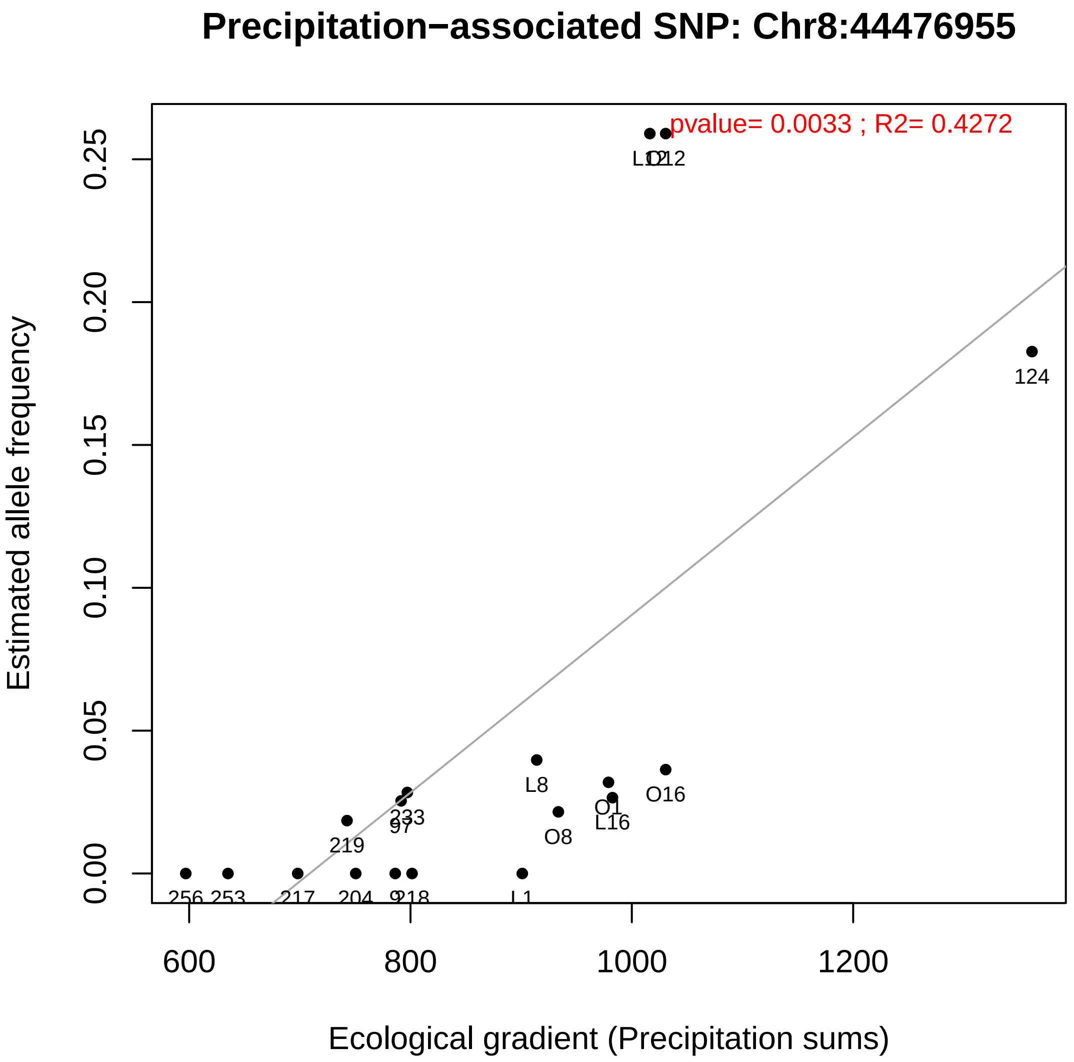

Precipitation−associated SNP: Chr8:44476955
pvalue= 0.0033 ; R2= 0.4272
Estimated allele frequency
0.00
0.05
0.10
0.15
0.20
0.25
600
800
1000
1200
Ecological gradient (Precipitation sums)
124
L8
O16
O8
L16
219
233
97

**Precipitation−associated SNP: Chr8:44481380**

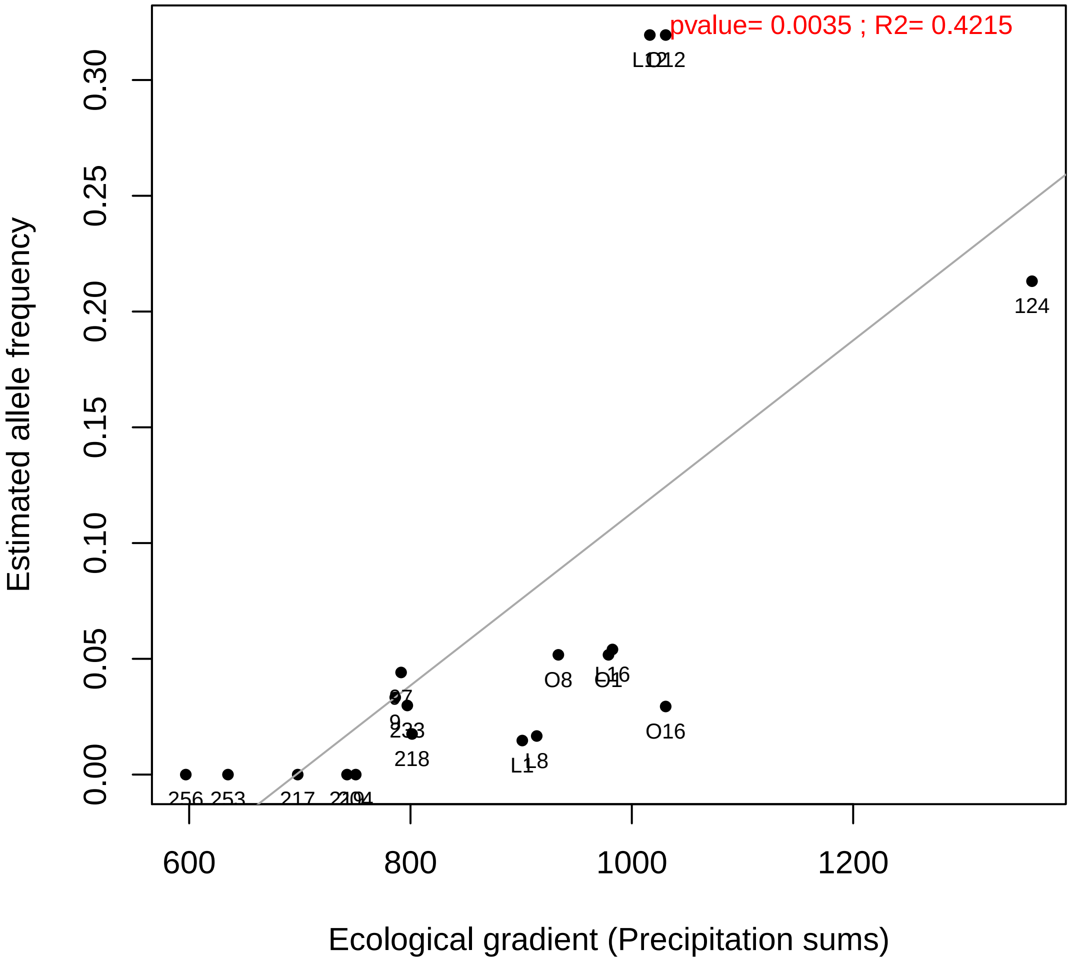

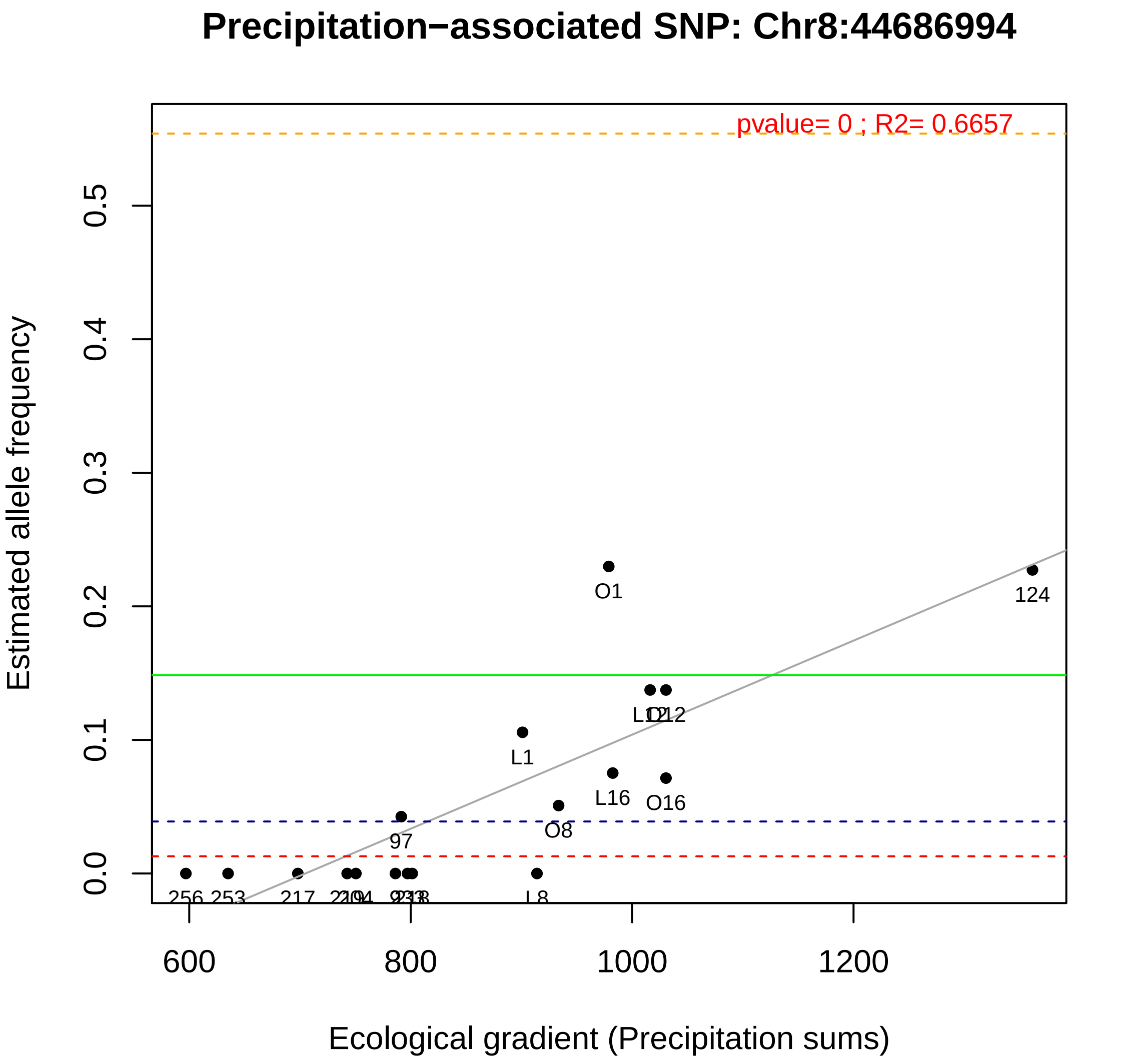
Precipitation−associated SNP: Chr8:44686994
pvalue= 0 ; R2= 0.6657
Estimated allele frequency
0.0
0.1
0.2
0.3
0.4
0.5
600
800
1000
1200
Ecological gradient (Precipitation sums)
O1
124
L1
L16
O16
O8
97
256
253
217
L8

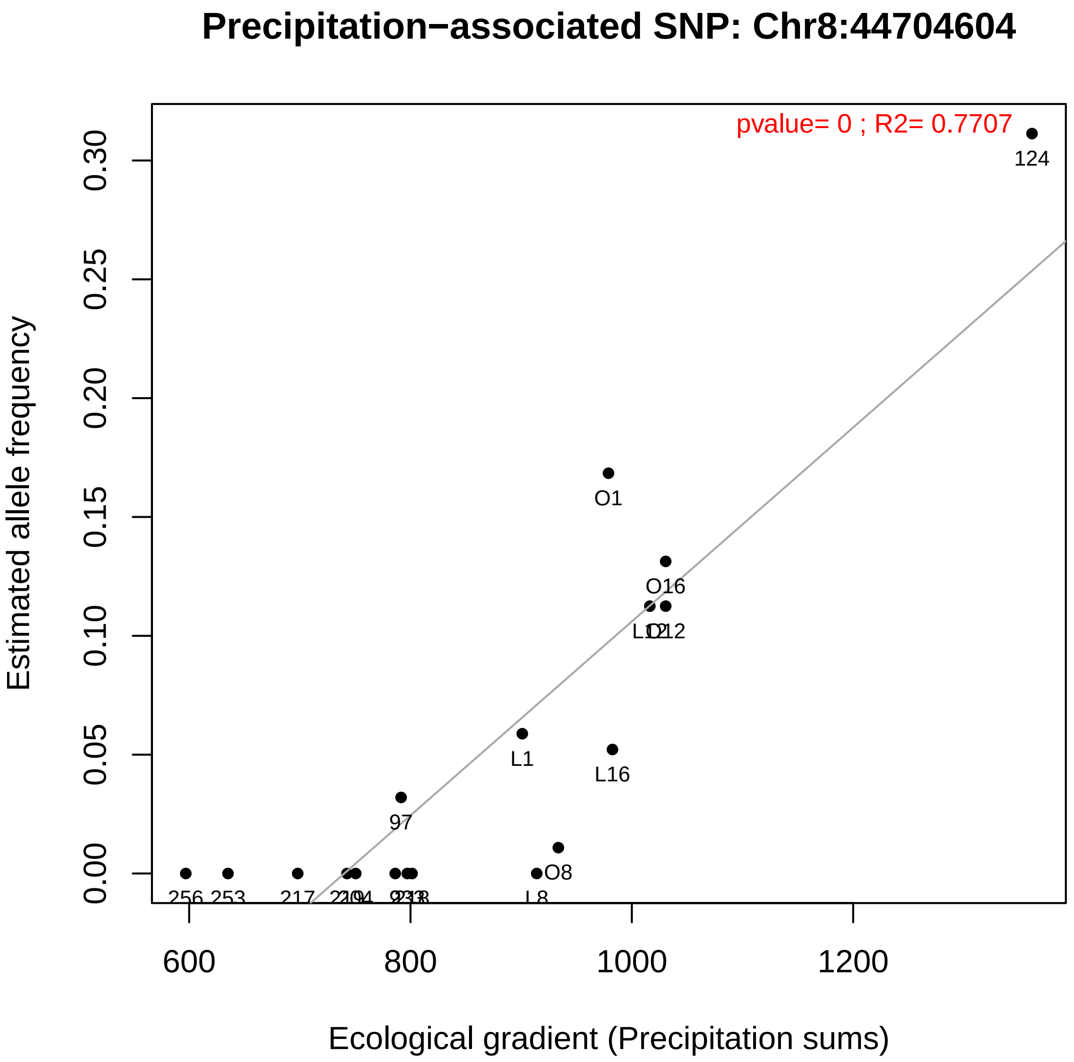

Precipitation−associated SNP: Chr8:44704604
pvalue= 0 ; R2= 0.7707
124
O1
O16
L12
O12
L1
L16
97
O8
L8
256
253
217
0.00
0.05
0.10
0.15
0.20
0.25
0.30
Estimated allele frequency
600
800
1000
1200
Ecological gradient (Precipitation sums)

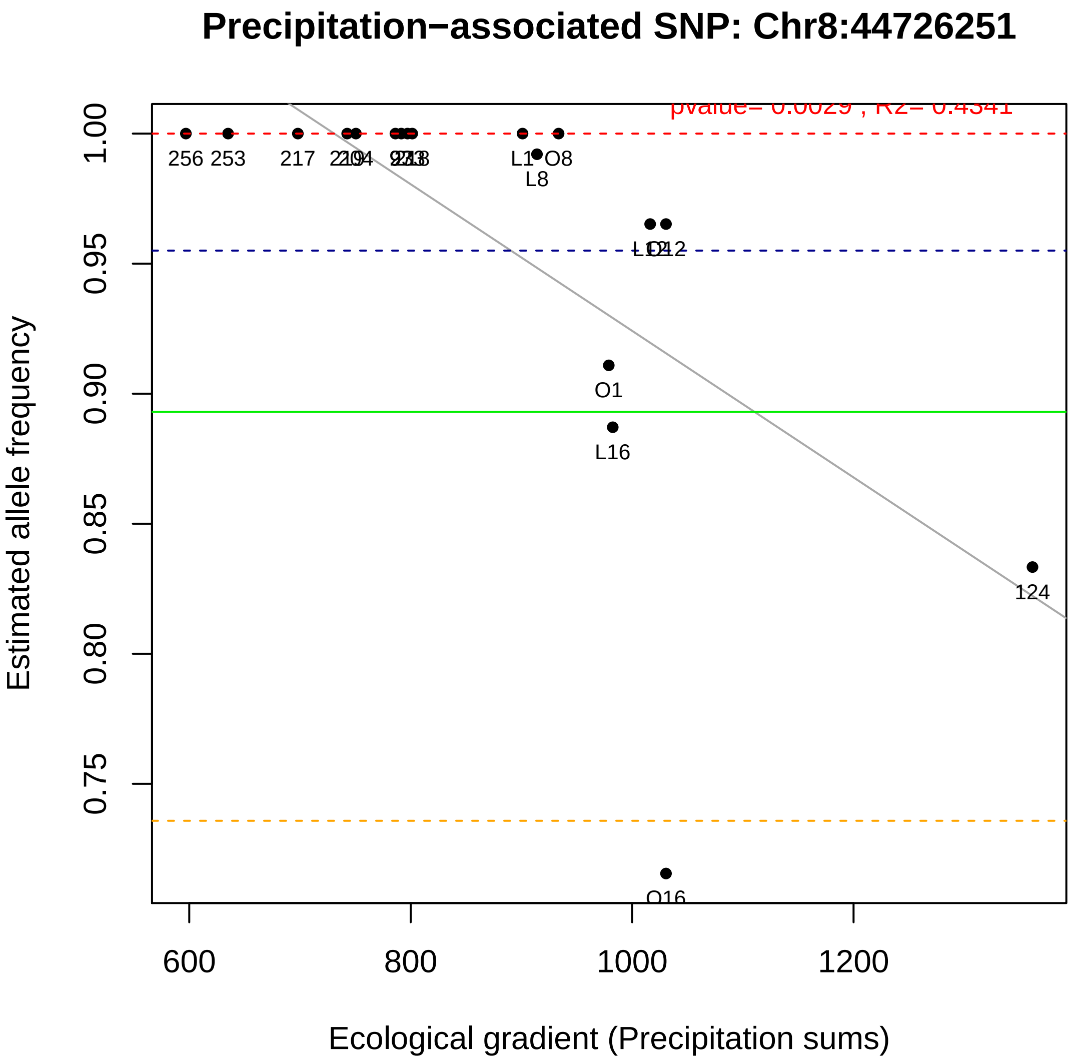

Precipitation−associated SNP: Chr8:44726251
pvalue= 0.0029 , R2= 0.4341
Estimated allele frequency
Ecological gradient (Precipitation sums)
1.00
0.95
0.90
0.85
0.80
0.75
600
800
1000
1200
256
253
217
L1
O8
L8
O1
L16
124
O16

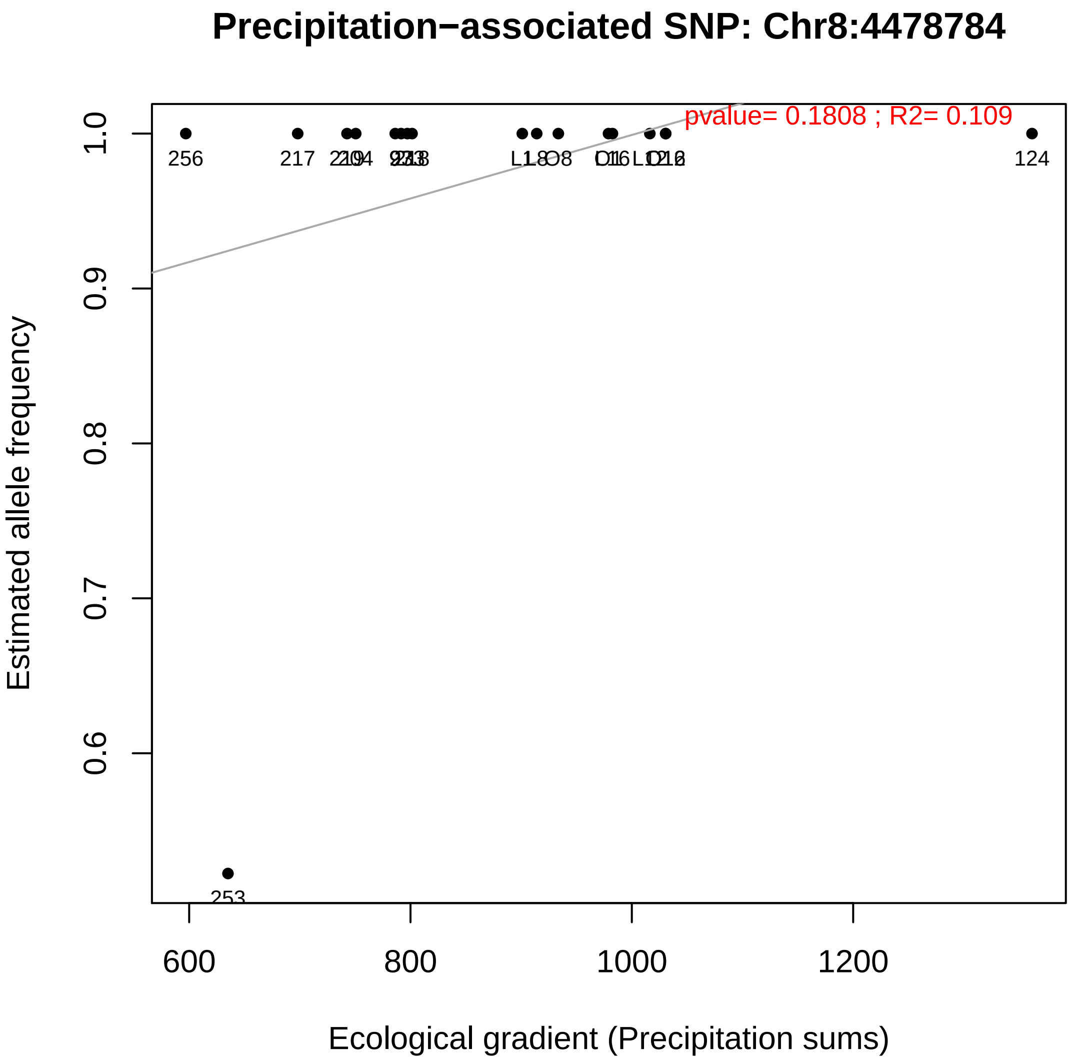

Precipitation−associated SNP: Chr8:4478784
pvalue= 0.1808 ; R2= 0.109
Estimated allele frequency
Ecological gradient (Precipitation sums)
1.0
0.9
0.8
0.7
0.6
600
800
1000
1200
256
217
124
253

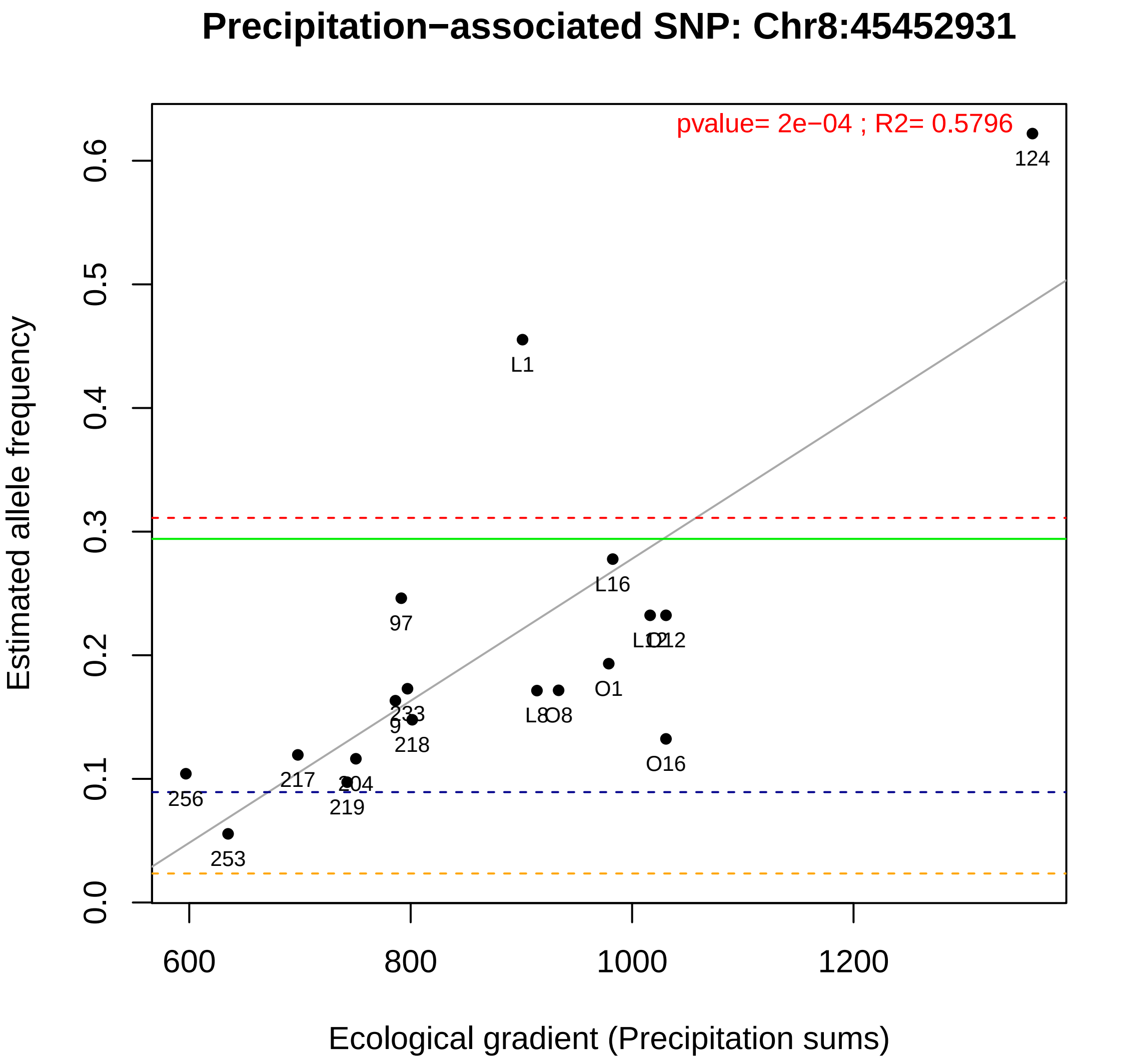

Precipitation−associated SNP: Chr8:45452931
pvalue= 2e−04 ; R2= 0.5796
Estimated allele frequency
Ecological gradient (Precipitation sums)
0.0
0.1
0.2
0.3
0.4
0.5
0.6
600
800
1000
1200
124
L1
L16
97
L12
O12
O1
233
9
218
L8
O8
O16
217
204
219
256
253

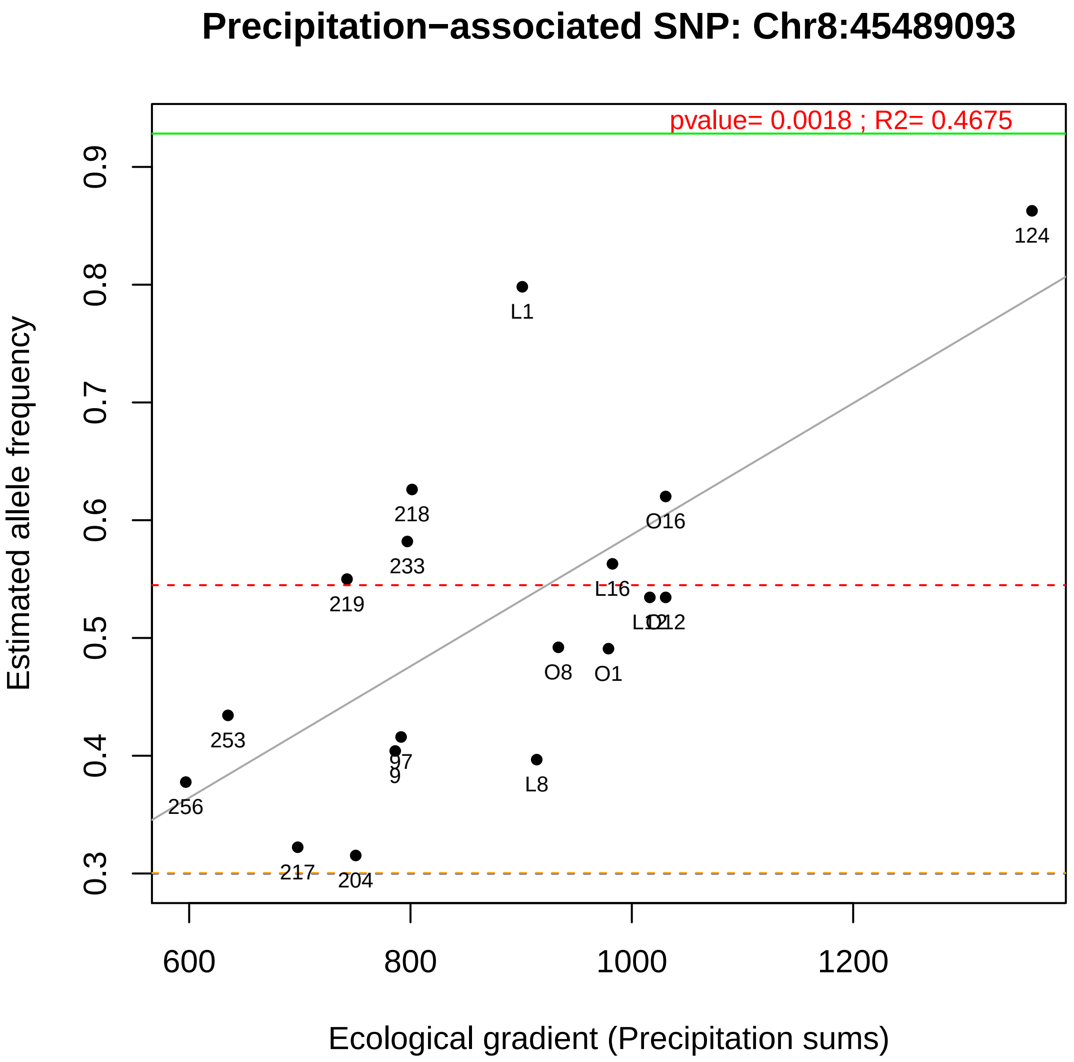
Precipitation−associated SNP: Chr8:45489093
pvalue= 0.0018 ; R2= 0.4675
Estimated allele frequency
0.3
0.4
0.5
0.6
0.7
0.8
0.9
600
800
1000
1200
Ecological gradient (Precipitation sums)
124
L1
218
233
219
O16
L16
L12
O12
O8
O1
253
97
9
L8
256
217
204

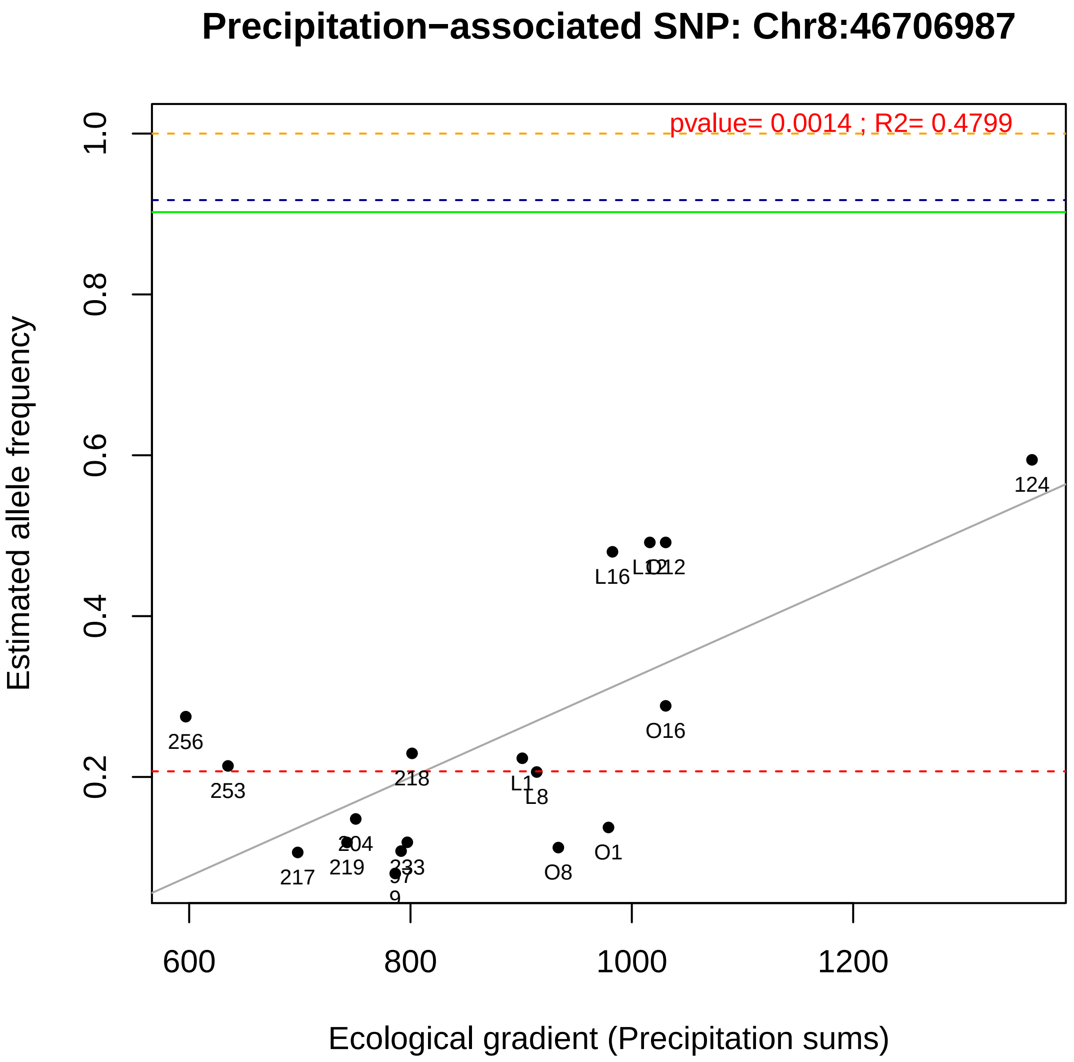

Precipitation−associated SNP: Chr8:46706987
pvalue= 0.0014 ; R2= 0.4799
Estimated allele frequency
Ecological gradient (Precipitation sums)
0.2
0.4
0.6
0.8
1.0
600
800
1000
1200
124
L16
O16
256
253
218
L1
L8
204
219
233
217
O8
O1

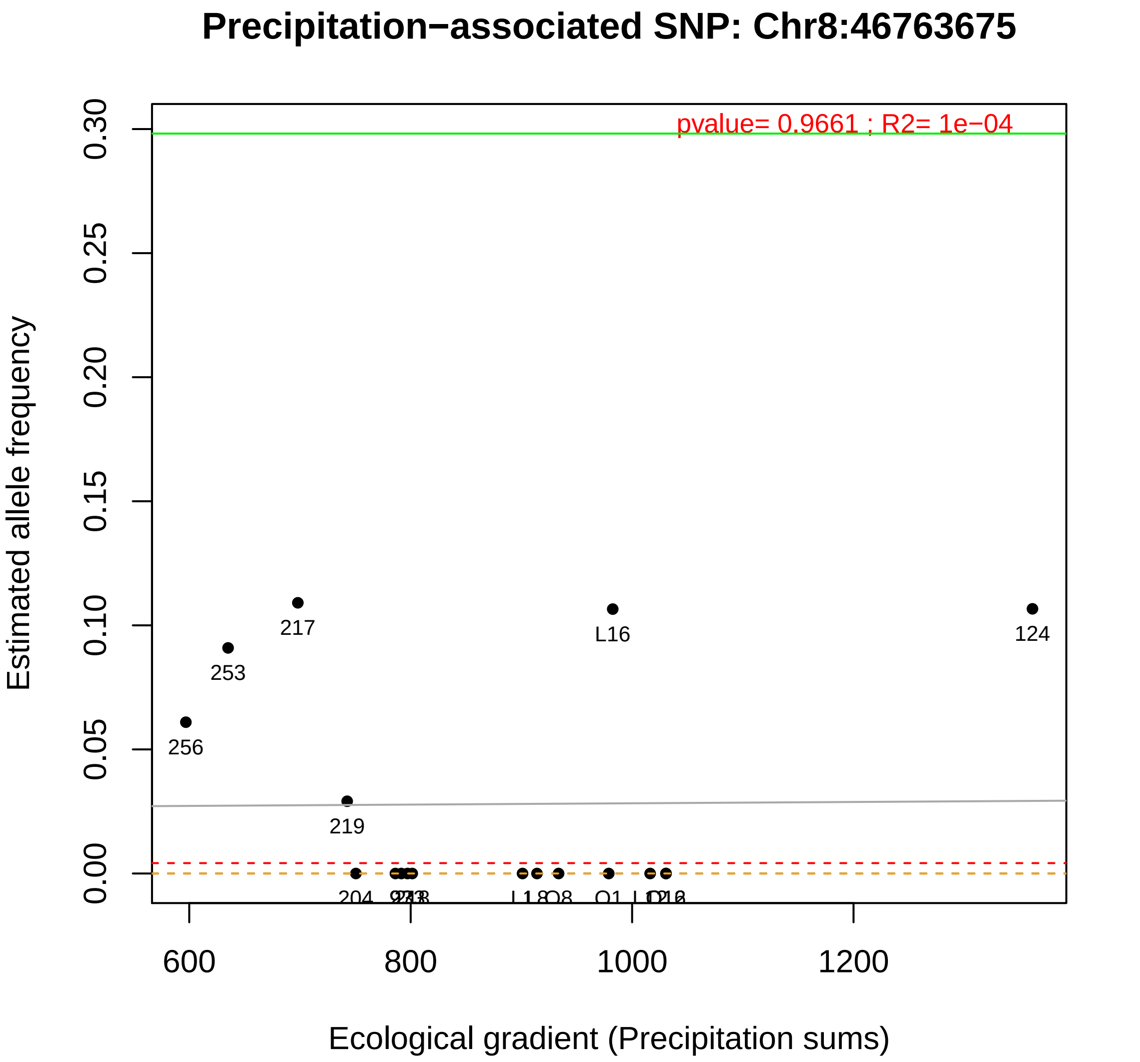

Precipitation−associated SNP: Chr8:46763675
pvalue= 0.9661 ; R2= 1e−04
Estimated allele frequency
0.00
0.05
0.10
0.15
0.20
0.25
0.30
256
253
217
219
L16
124
600
800
1000
1200
Ecological gradient (Precipitation sums)

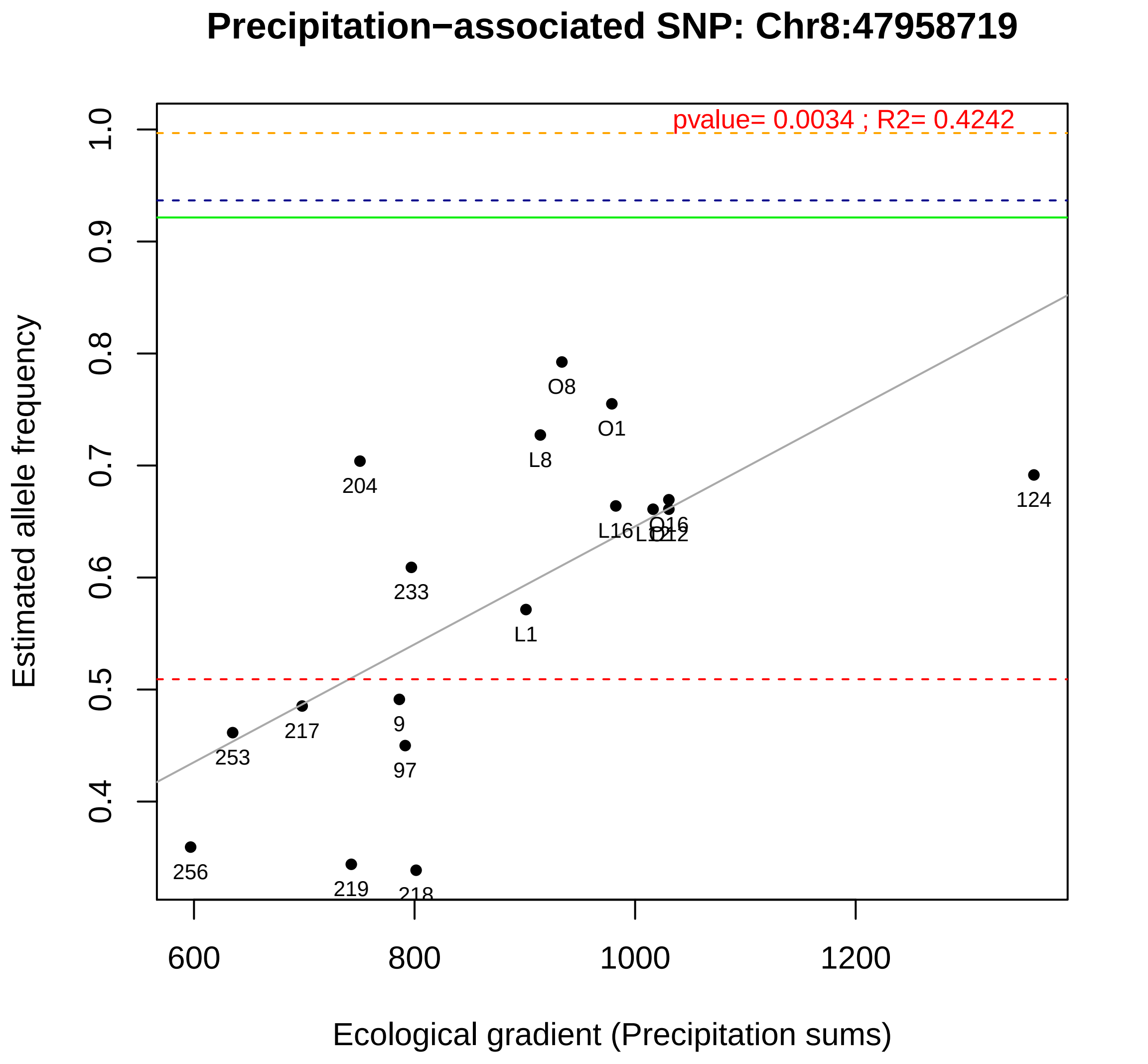

Precipitation−associated SNP: Chr8:47958719
pvalue= 0.0034 ; R2= 0.4242
Estimated allele frequency
Ecological gradient (Precipitation sums)
0.4
0.5
0.6
0.7
0.8
0.9
1.0
600
800
1000
1200
O8
O1
L8
204
124
L16
O16
L12
233
L1
217
9
253
97
256
219
218

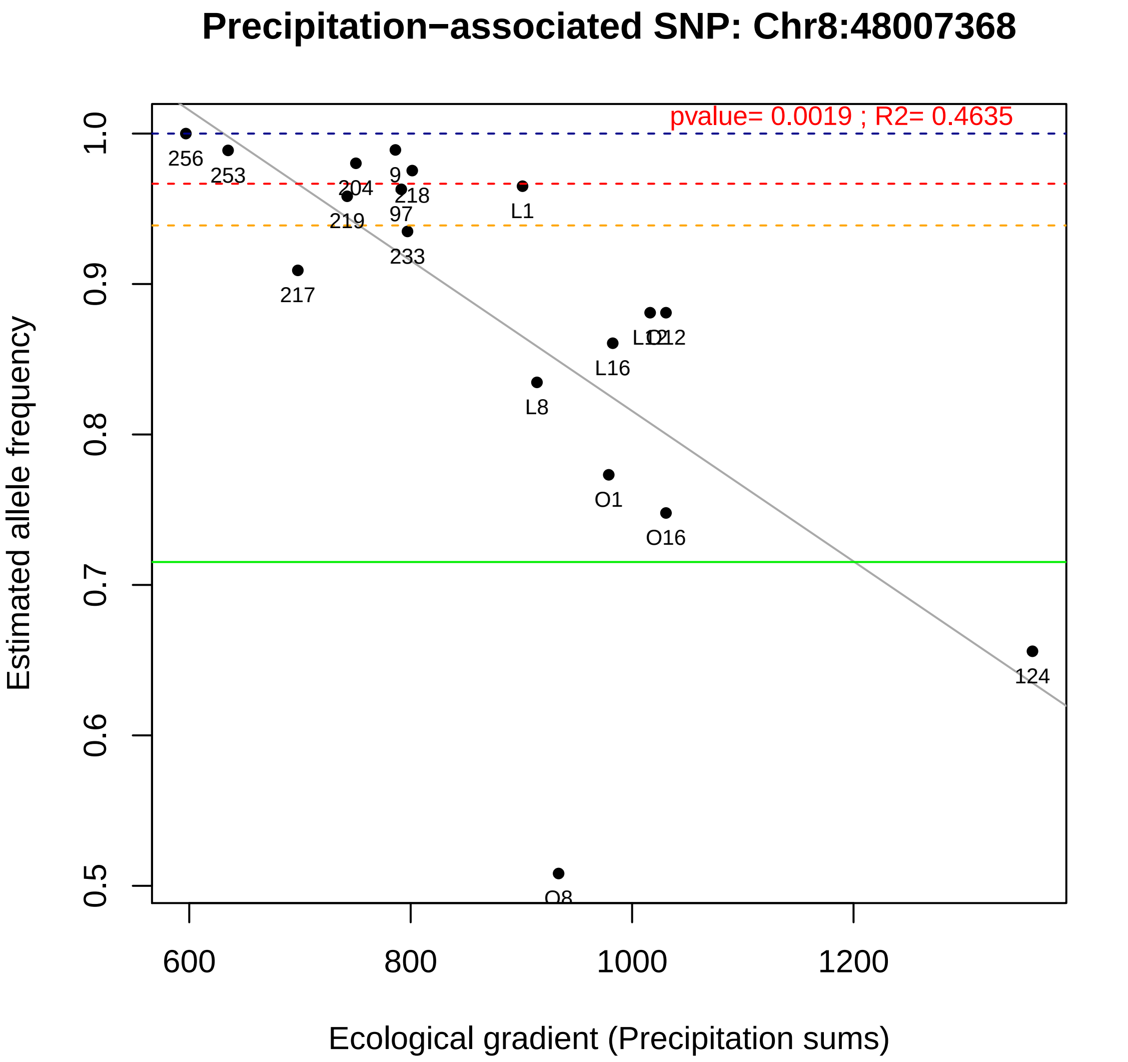

Precipitation−associated SNP: Chr8:48007368
pvalue= 0.0019 ; R2= 0.4635
Estimated allele frequency
Ecological gradient (Precipitation sums)
1.0
0.9
0.8
0.7
0.6
0.5
600
800
1000
1200
256
253
204
9
218
97
219
233
217
L1
L12
O12
L16
L8
O1
O16
O8
124

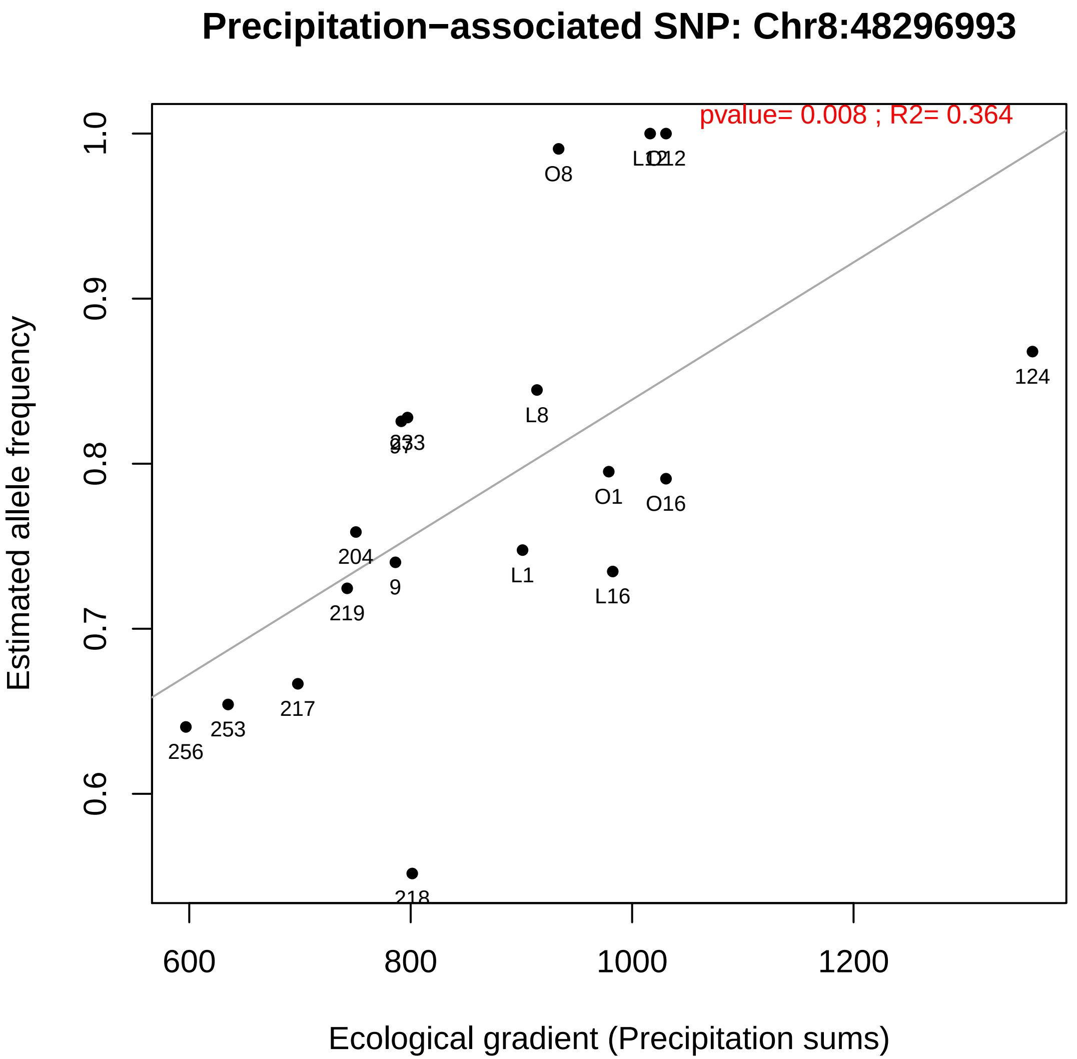

Precipitation−associated SNP: Chr8:48296993
pvalue= 0.008 ; R2= 0.364
Estimated allele frequency
Ecological gradient (Precipitation sums)
0.6
0.7
0.8
0.9
1.0
600
800
1000
1200
O8
124
L8
O1
O16
204
9
L1
L16
219
217
253
256
218

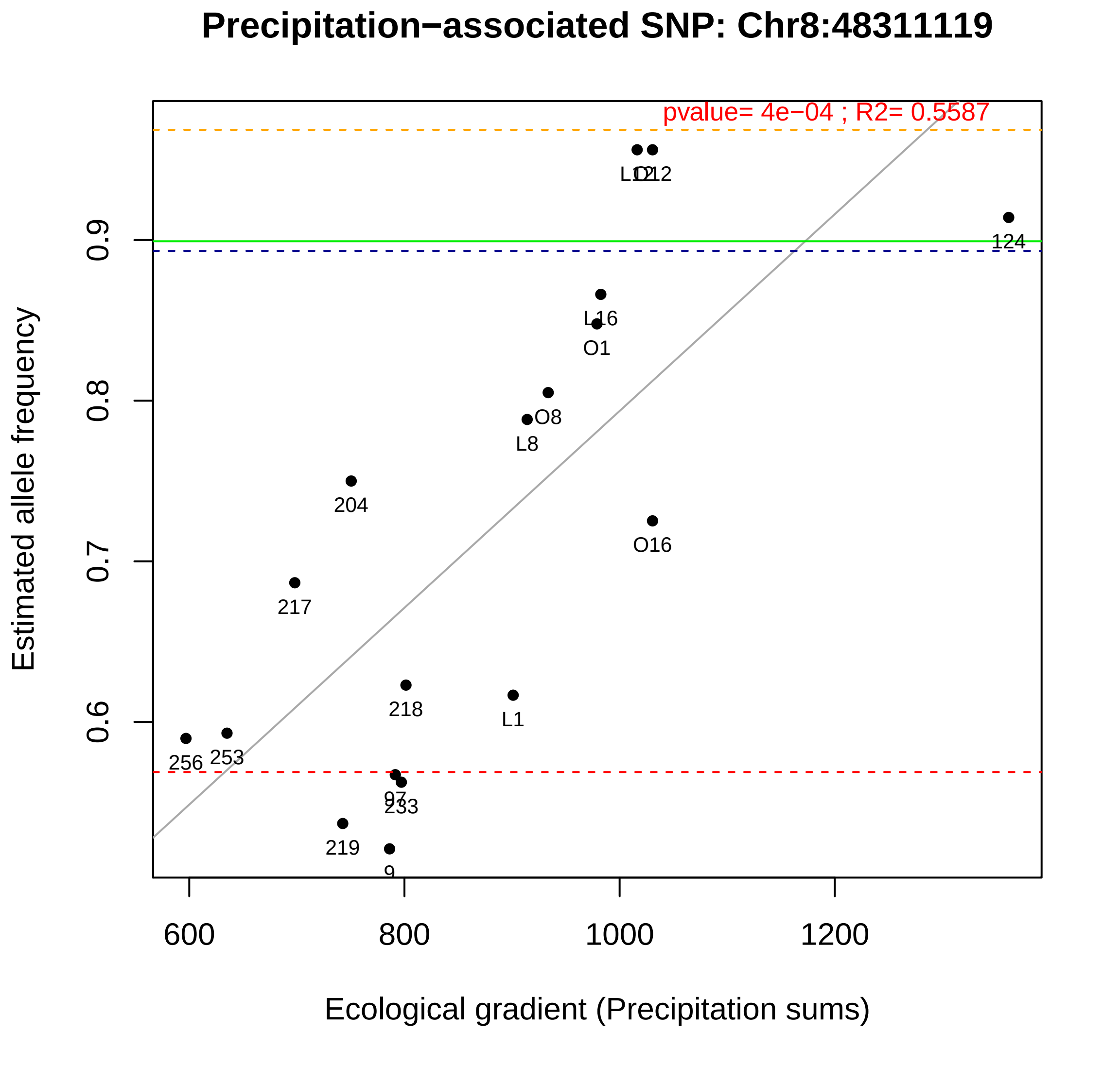
Precipitation−associated SNP: Chr8:48311119
pvalue= 4e−04 ; R2= 0.5587
Estimated allele frequency
0.6
0.7
0.8
0.9
600
800
1000
1200
Ecological gradient (Precipitation sums)
L12
O12
124
L16
O1
O8
L8
204
O16
217
218
L1
256
253
97
233
219
9

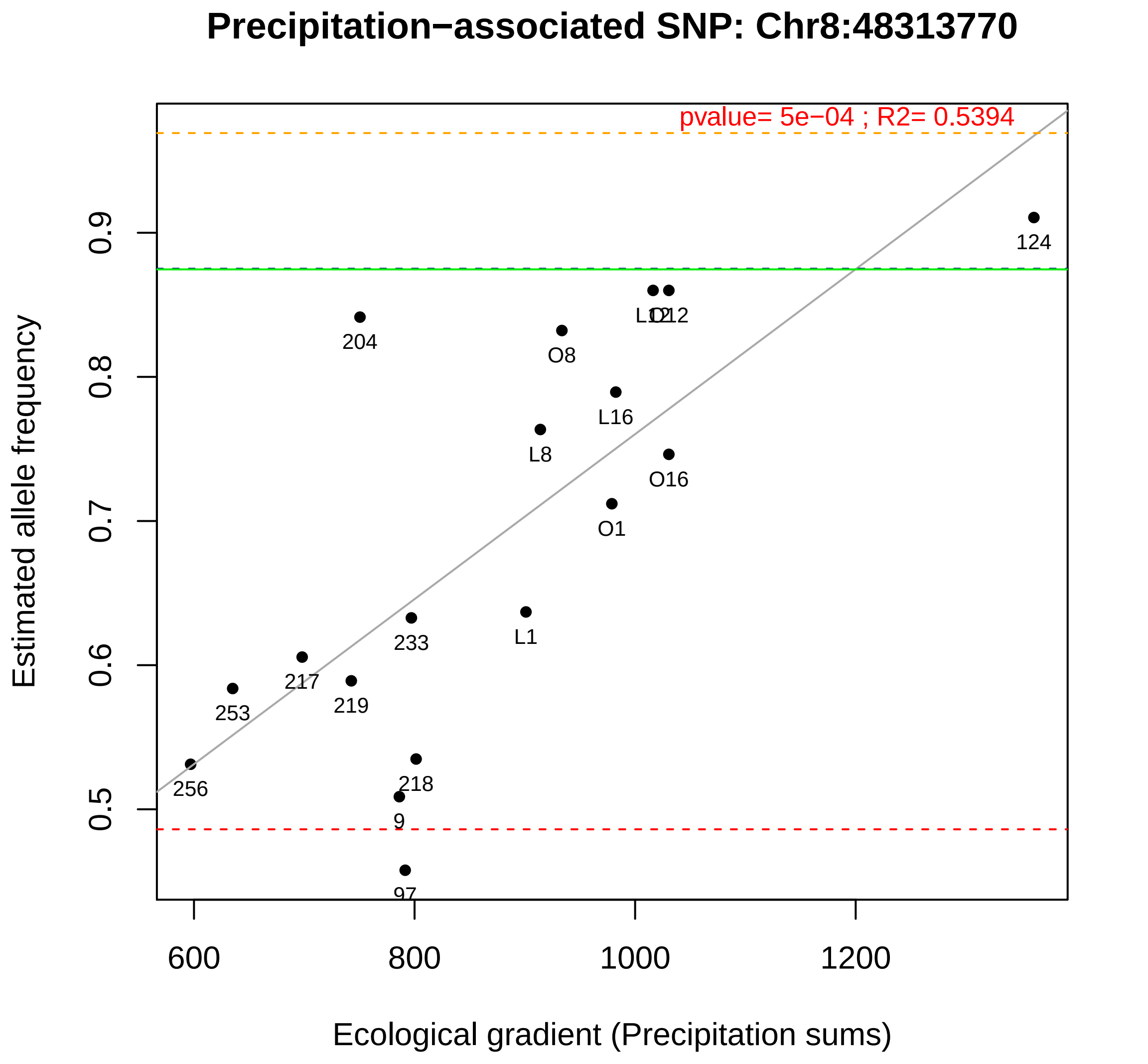

Precipitation−associated SNP: Chr8:48313770
pvalue= 5e−04 ; R2= 0.5394
Estimated allele frequency
0.5
0.6
0.7
0.8
0.9
600
800
1000
1200
Ecological gradient (Precipitation sums)
124
204
O8
L16
L8
O16
O1
233
L1
217
253
219
256
218
9
97

**Precipitation−associated SNP: Chr8:48375828**

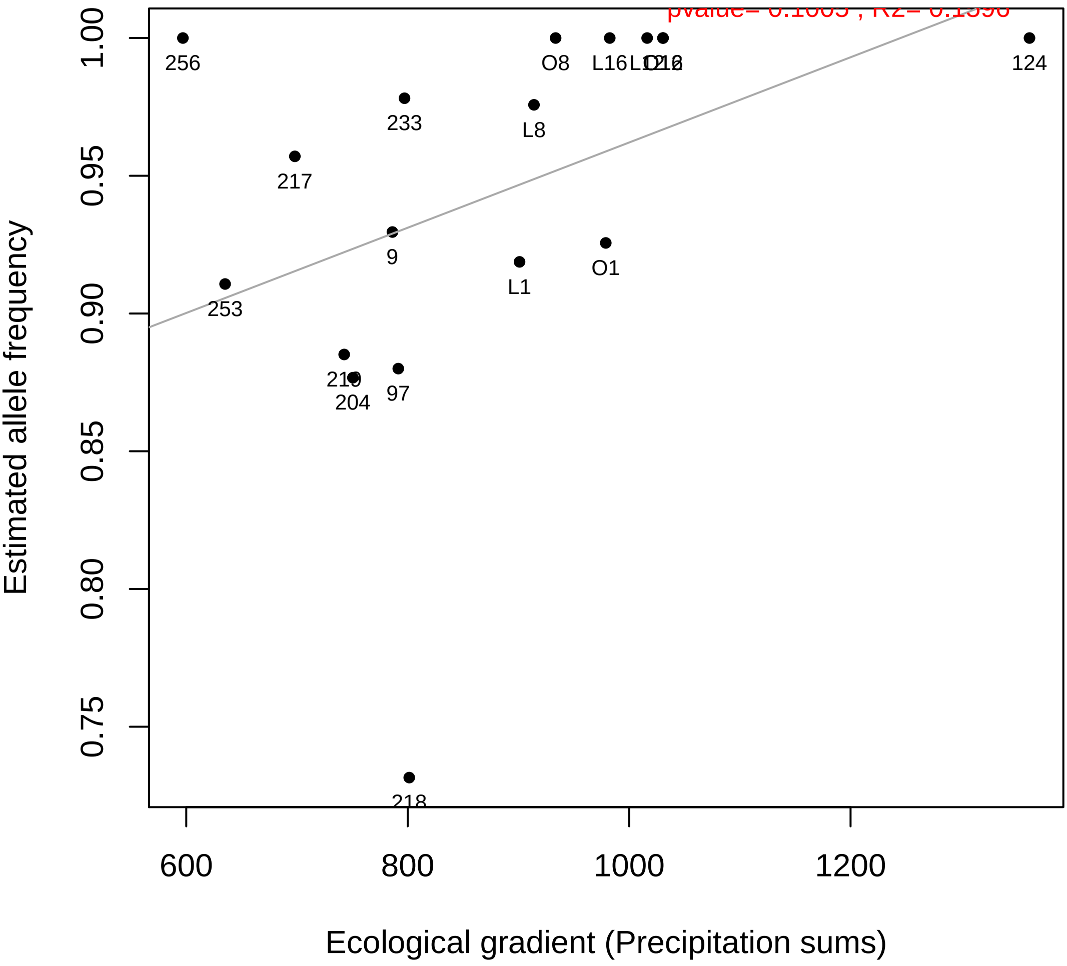

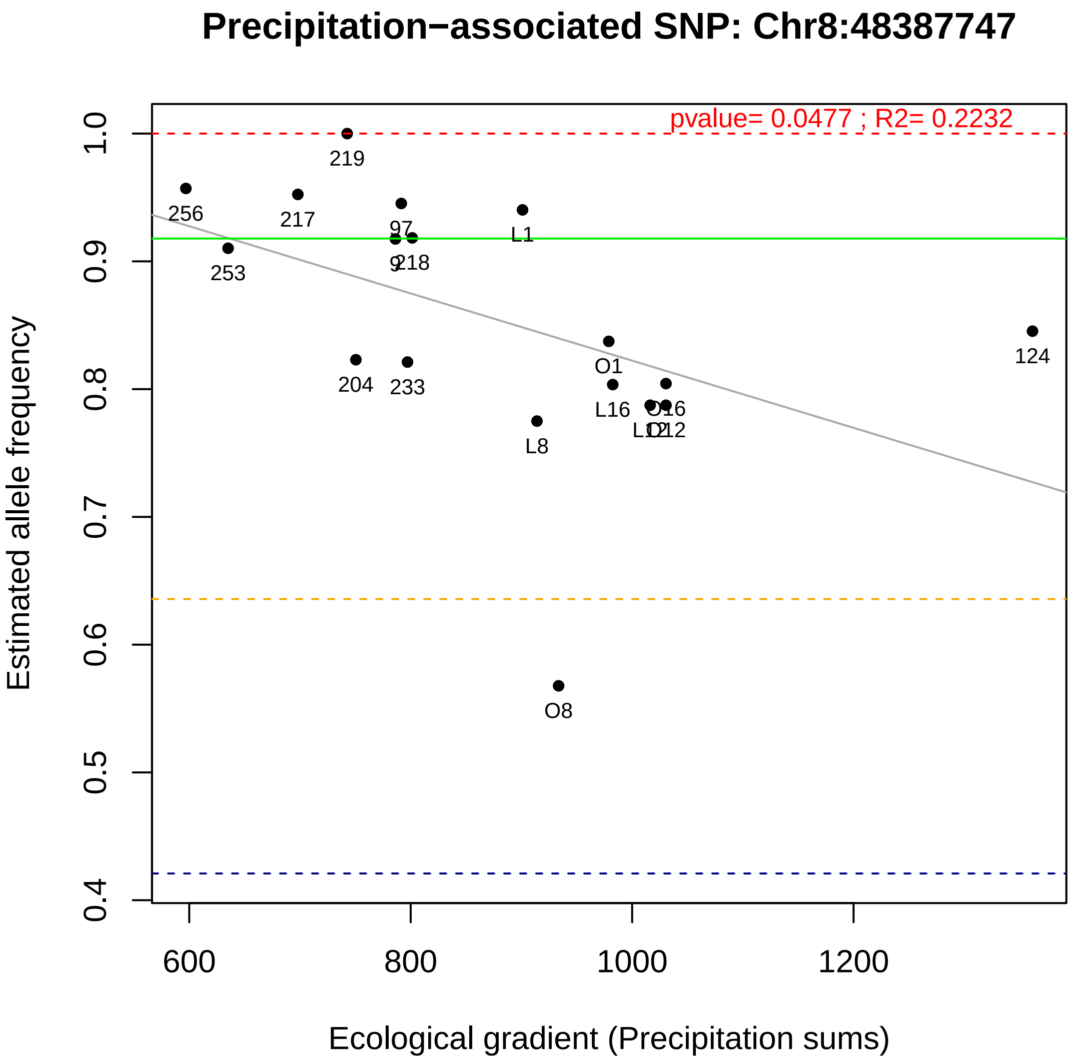

Precipitation−associated SNP: Chr8:48387747
pvalue= 0.0477 ; R2= 0.2232
Estimated allele frequency
Ecological gradient (Precipitation sums)
0.4
0.5
0.6
0.7
0.8
0.9
1.0
600
800
1000
1200
219
256
217
97
L1
253
218
204
233
O1
124
L16
O16
L12
O12
L8
O8

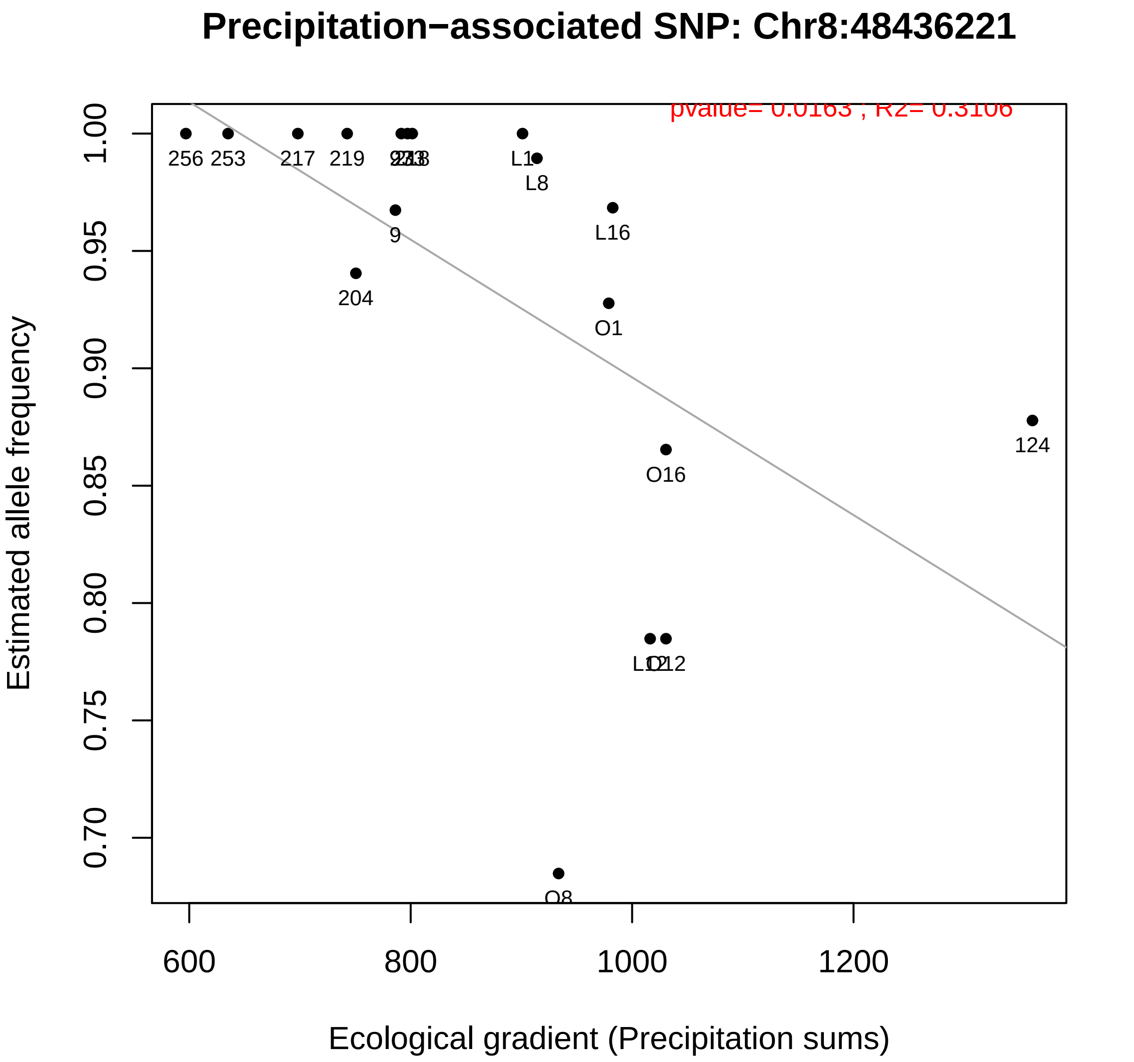
Precipitation−associated SNP: Chr8:48436221
pvalue= 0.0163 ; R2= 0.3106
Estimated allele frequency
Ecological gradient (Precipitation sums)
1.00
0.95
0.90
0.85
0.80
0.75
0.70
600
800
1000
1200
256
253
217
219
L1
L8
9
L16
204
O1
124
O16
O8

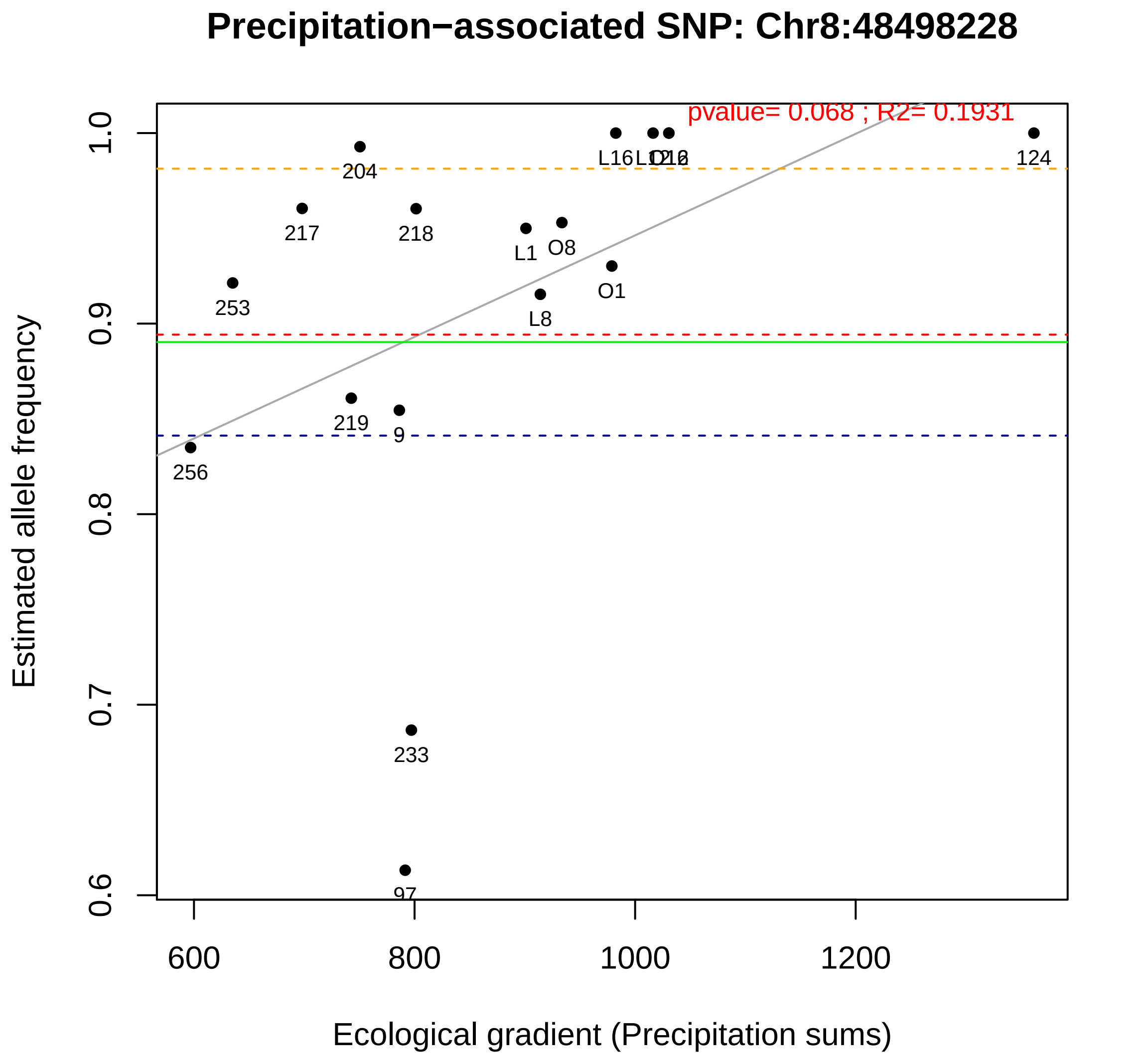

Precipitation−associated SNP: Chr8:48498228
pvalue= 0.068 ; R2= 0.1931
Estimated allele frequency
Ecological gradient (Precipitation sums)
0.6
0.7
0.8
0.9
1.0
600
800
1000
1200
204
L16
124
217
218
L1
O8
253
O1
L8
219
9
256
233
97

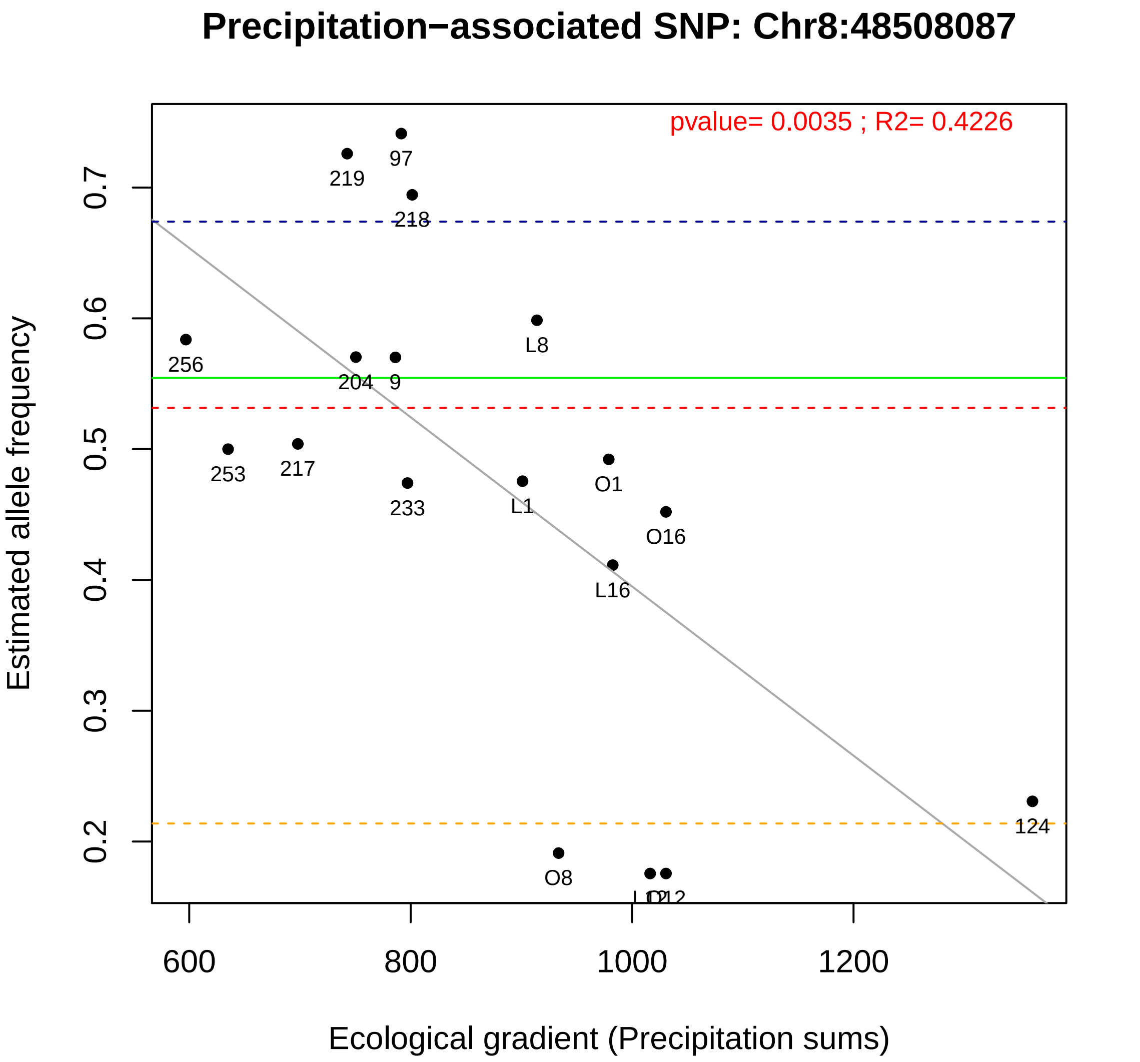
Precipitation−associated SNP: Chr8:48508087
pvalue= 0.0035 ; R2= 0.4226
Estimated allele frequency
0.2
0.3
0.4
0.5
0.6
0.7
600
800
1000
1200
Ecological gradient (Precipitation sums)
97
219
218
L8
256
204
9
253
217
233
L1
O1
O16
L16
124
O8

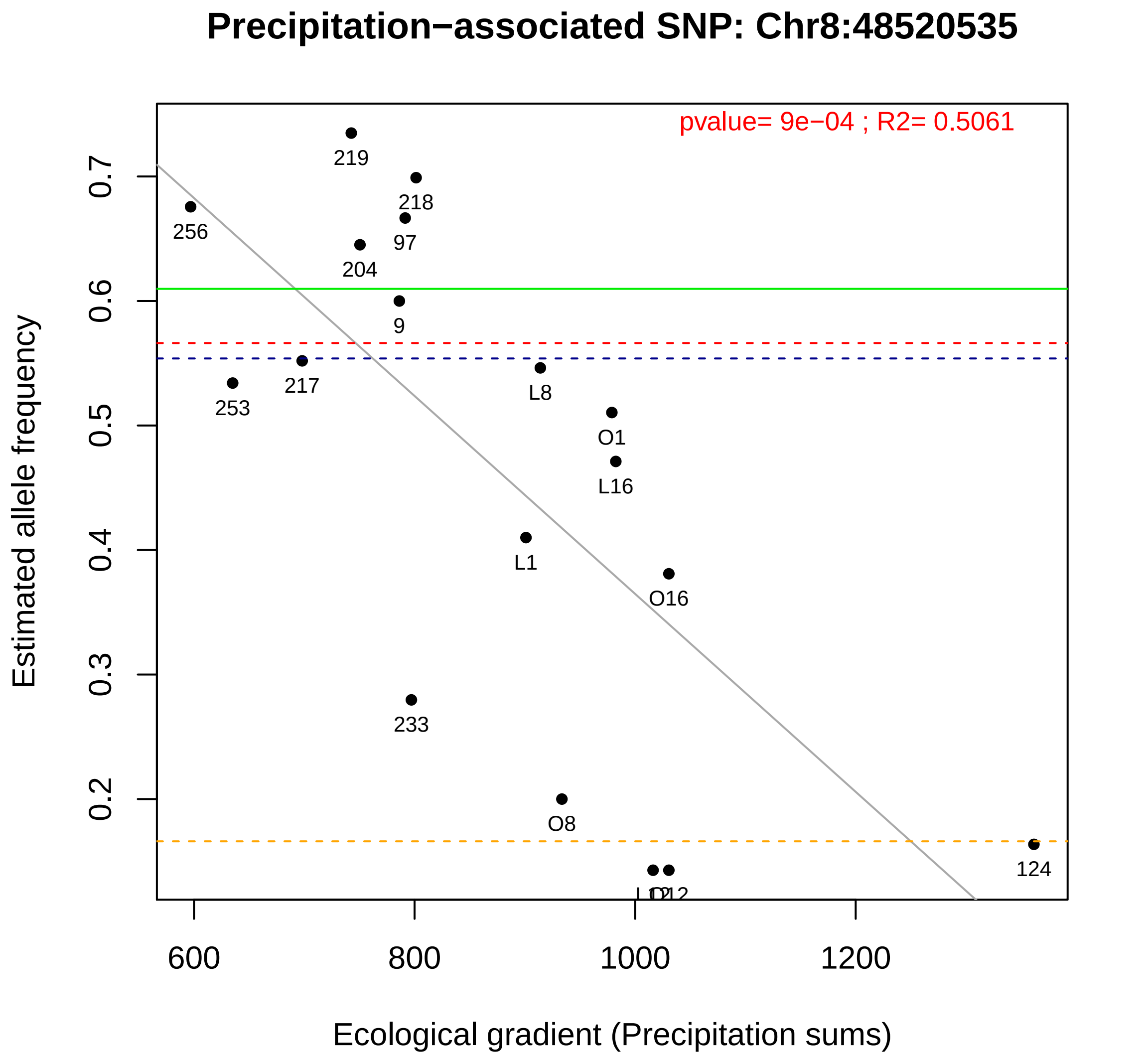

Precipitation−associated SNP: Chr8:48520535
pvalue= 9e−04 ; R2= 0.5061
Estimated allele frequency
Ecological gradient (Precipitation sums)
0.2
0.3
0.4
0.5
0.6
0.7
600
800
1000
1200
219
218
256
97
204
9
217
253
L8
O1
L16
L1
O16
233
O8
124

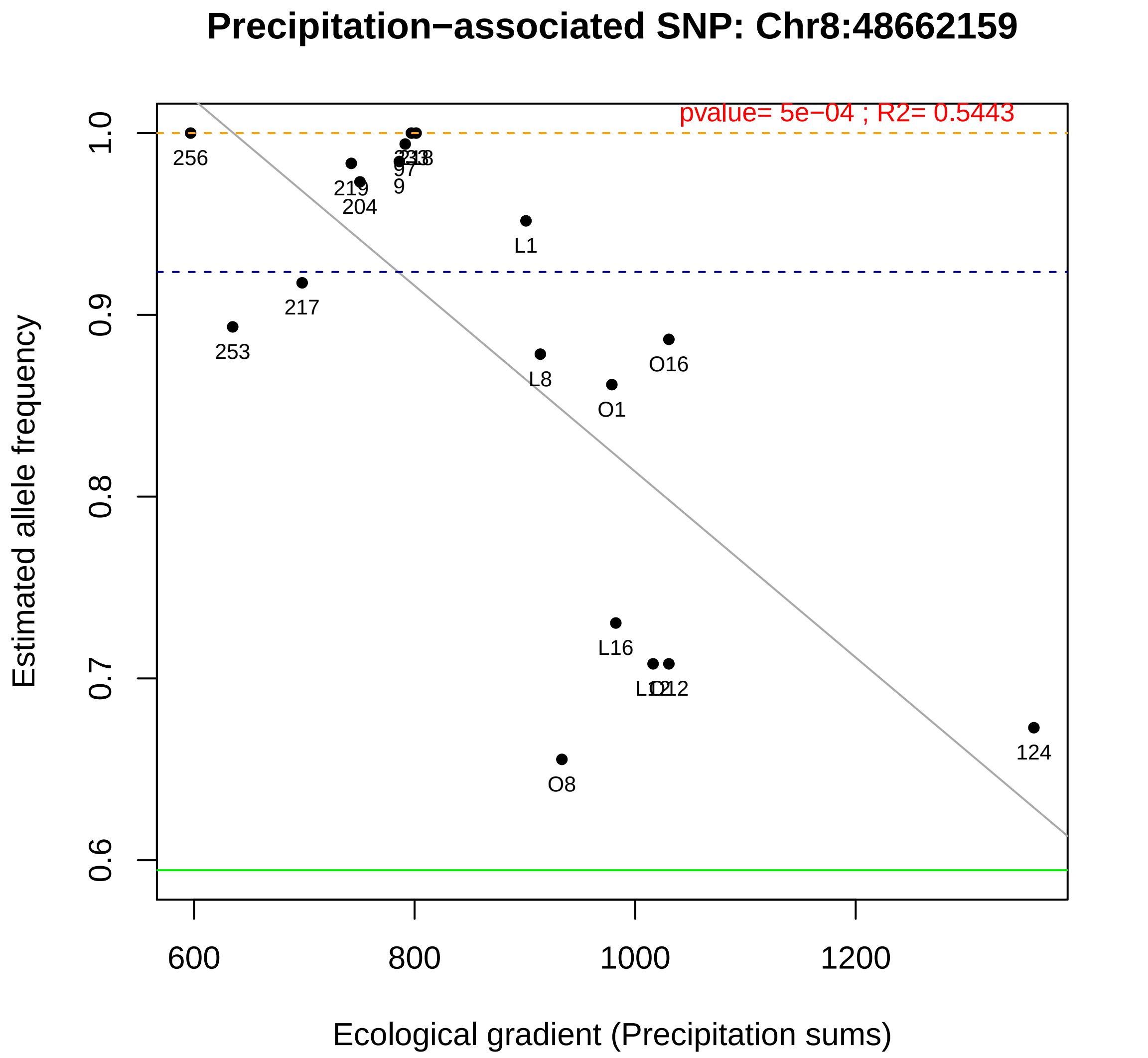

Precipitation−associated SNP: Chr8:48662159
pvalue= 5e−04 ; R2= 0.5443
Estimated allele frequency
Ecological gradient (Precipitation sums)
0.6
0.7
0.8
0.9
1.0
600
800
1000
1200
256
219
204
97
9
L1
217
253
L8
O1
O16
L16
O12
124
O8

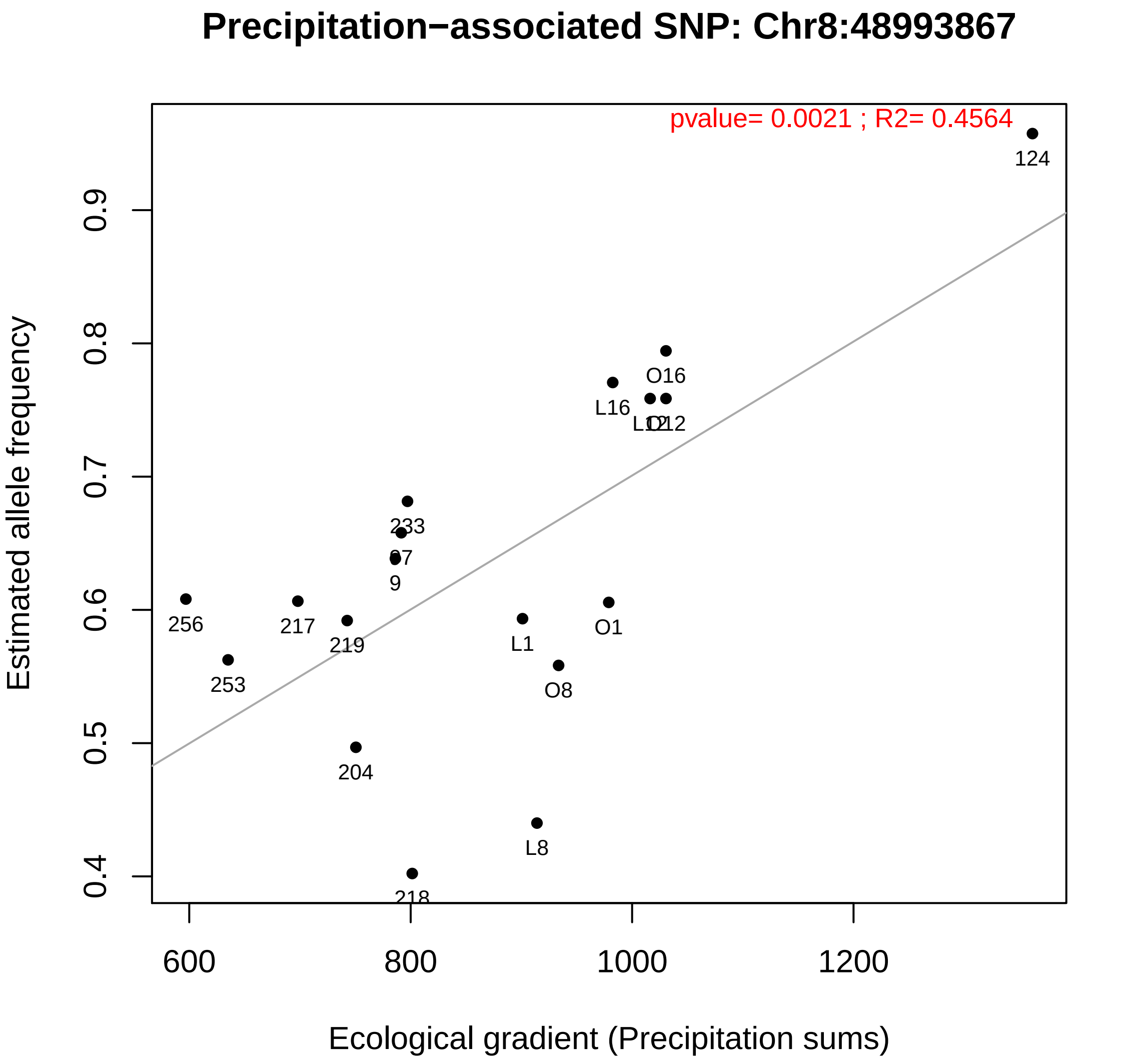

Precipitation−associated SNP: Chr8:48993867
pvalue= 0.0021 ; R2= 0.4564
Estimated allele frequency
Ecological gradient (Precipitation sums)
0.4
0.5
0.6
0.7
0.8
0.9
600
800
1000
1200
124
O16
L16
L12
O12
233
97
9
256
217
219
L1
O1
253
O8
204
L8
218

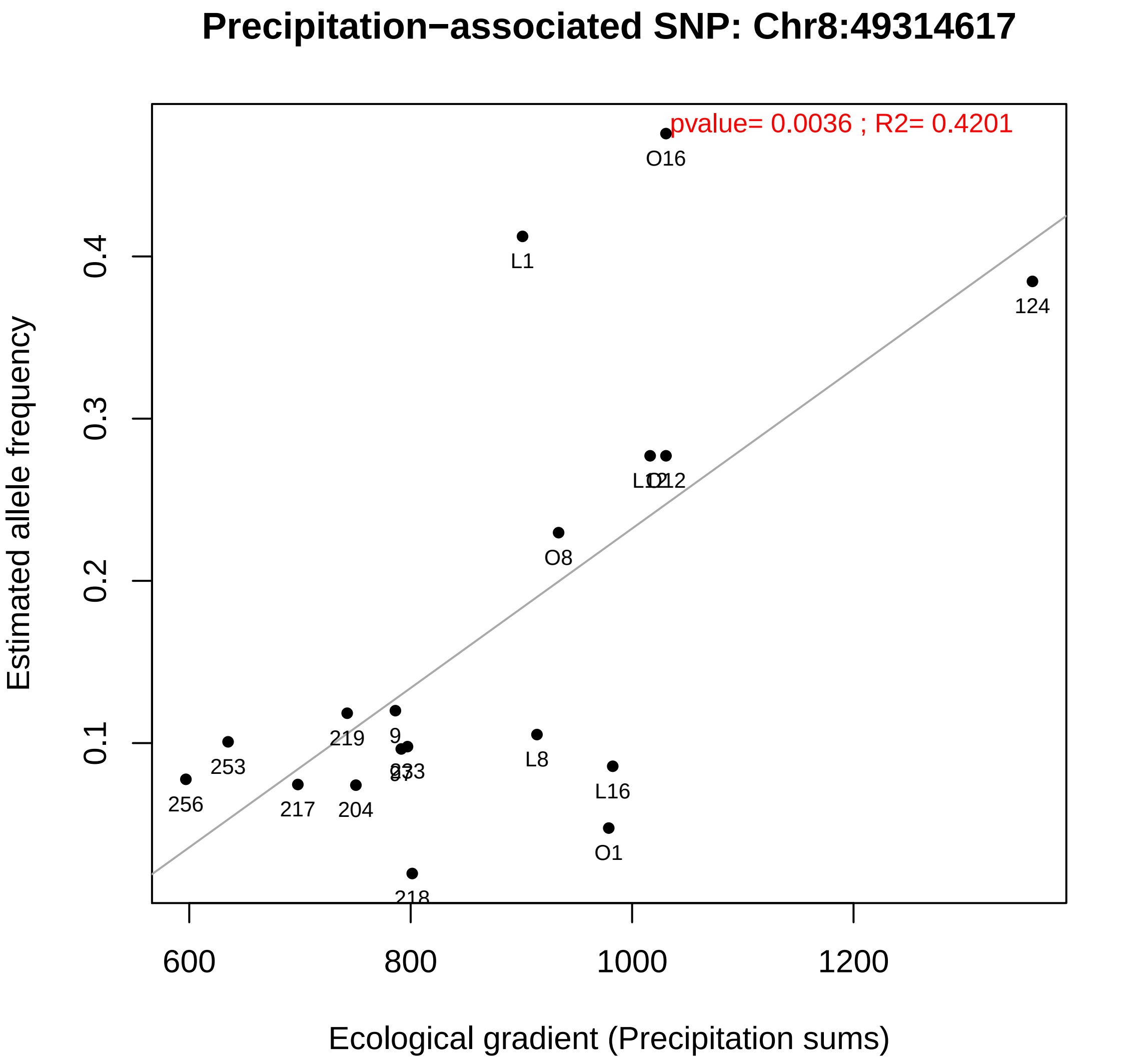

Precipitation−associated SNP: Chr8:49314617
pvalue= 0.0036 ; R2= 0.4201
Estimated allele frequency
Ecological gradient (Precipitation sums)
0.1
0.2
0.3
0.4
600
800
1000
1200
O16
L1
124
O8
219
9
253
256
217
204
L8
L16
O1
218

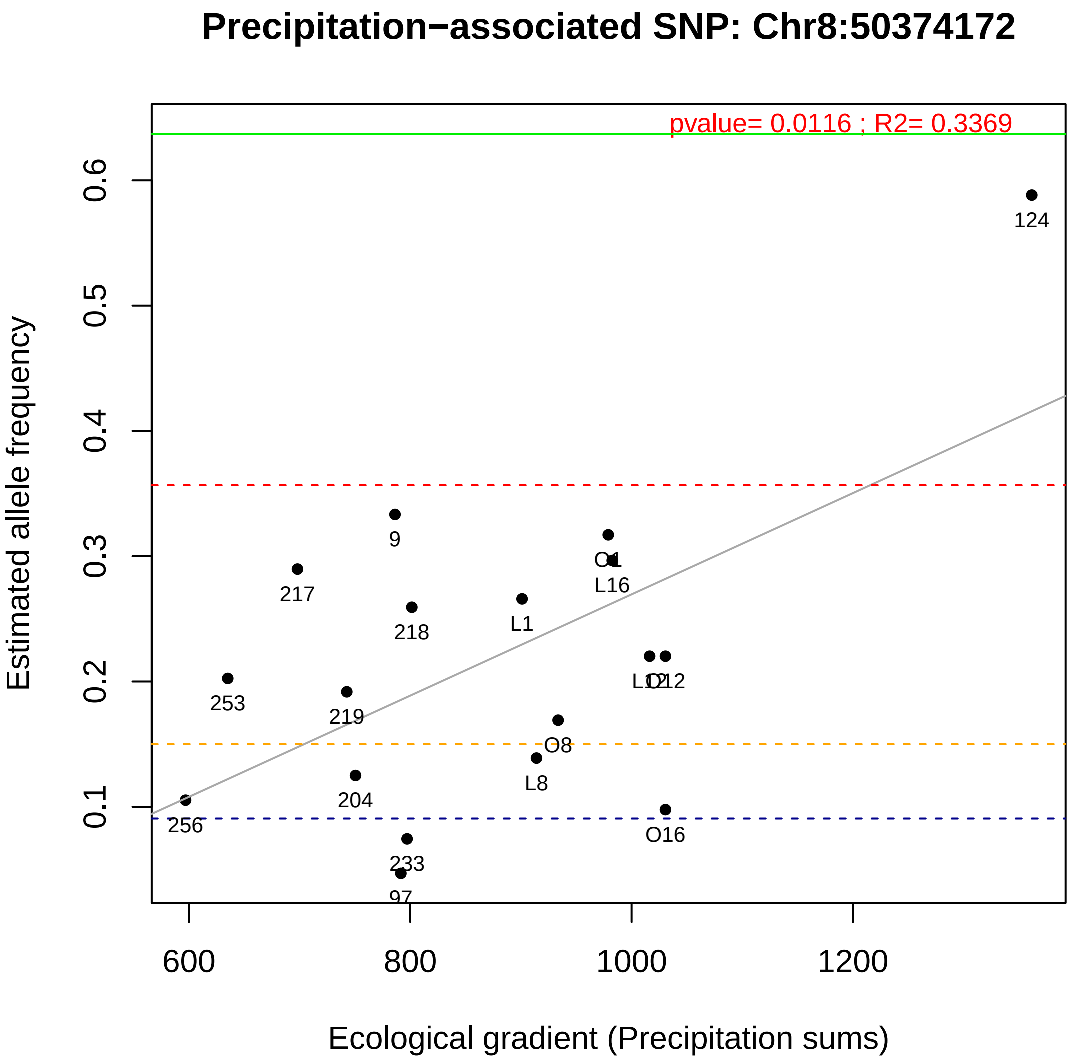

Precipitation−associated SNP: Chr8:50374172
pvalue= 0.0116 ; R2= 0.3369
Estimated allele frequency
0.1
0.2
0.3
0.4
0.5
0.6
124
9
O1
L16
217
218
L1
L12
O12
253
219
O8
L8
204
256
O16
233
97
600
800
1000
1200
Ecological gradient (Precipitation sums)

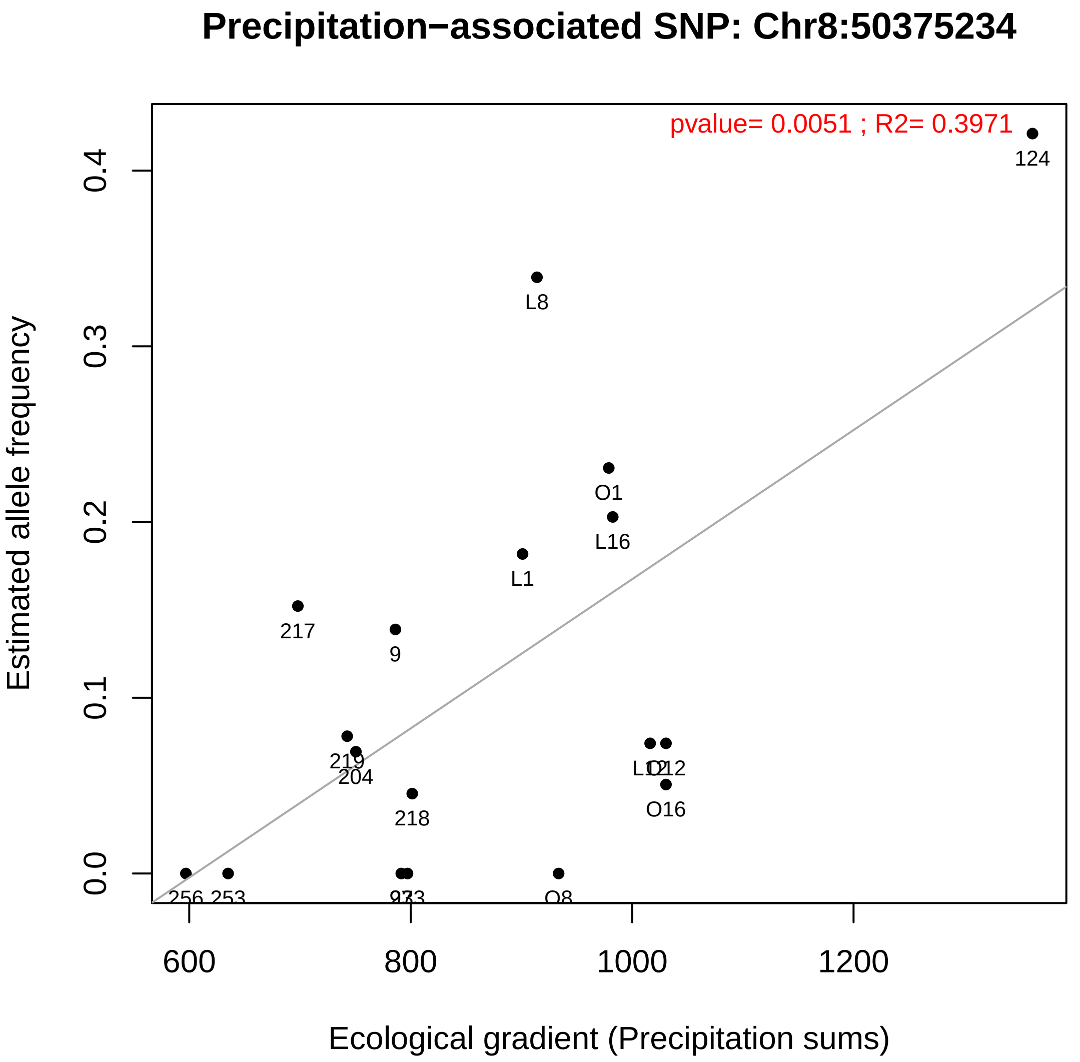
Precipitation−associated SNP: Chr8:50375234
pvalue= 0.0051 ; R2= 0.3971
Estimated allele frequency
Ecological gradient (Precipitation sums)
0.0
0.1
0.2
0.3
0.4
600
800
1000
1200
124
L8
O1
L16
L1
217
9
219
204
218
O16
O8
256
253

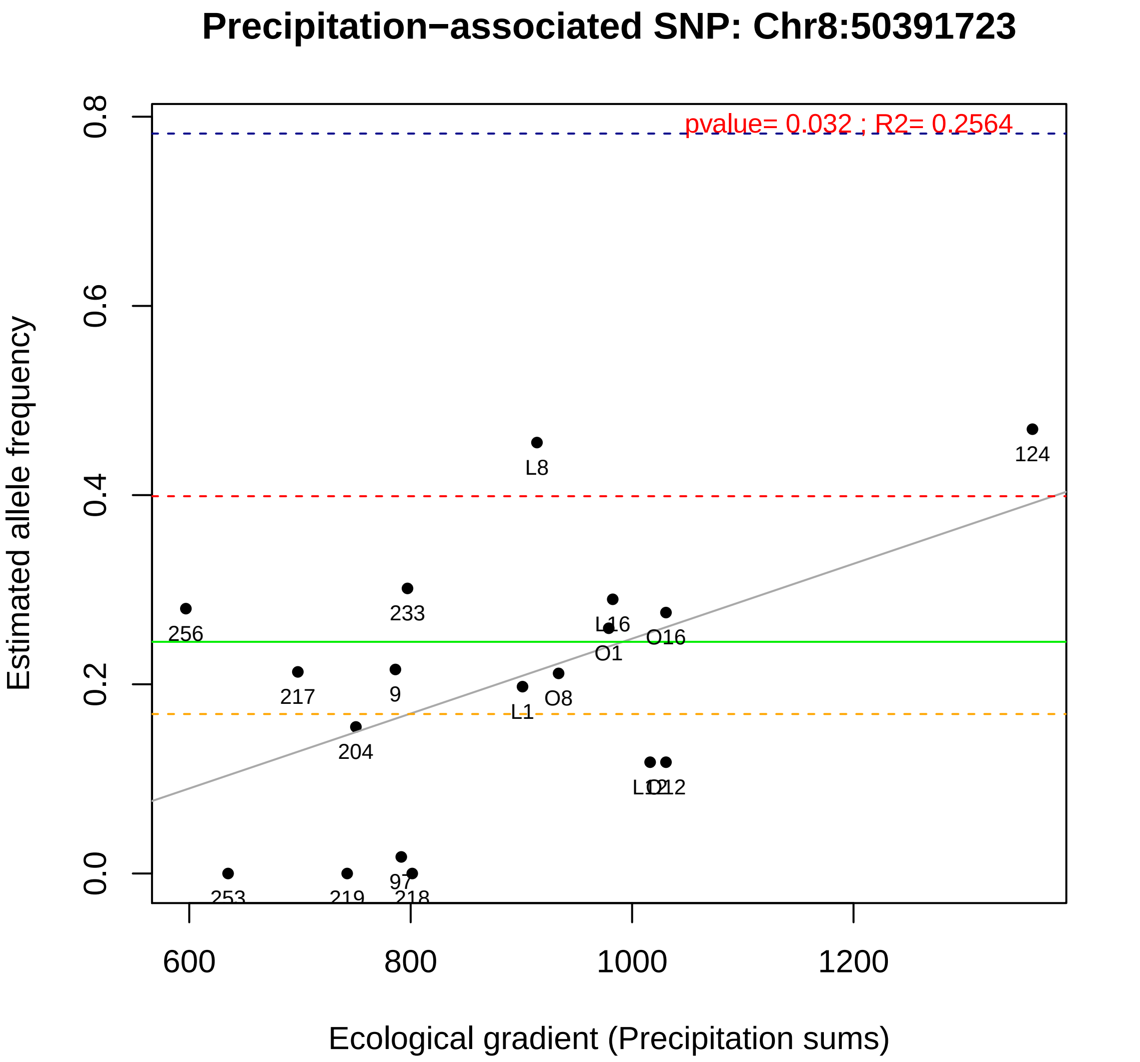
Precipitation−associated SNP: Chr8:50391723
pvalue= 0.032 ; R2= 0.2564
Estimated allele frequency
0.0
0.2
0.4
0.6
0.8
600
800
1000
1200
Ecological gradient (Precipitation sums)
L8
124
233
256
L16
O16
O1
217
9
O8
L1
204
97
253
219
218

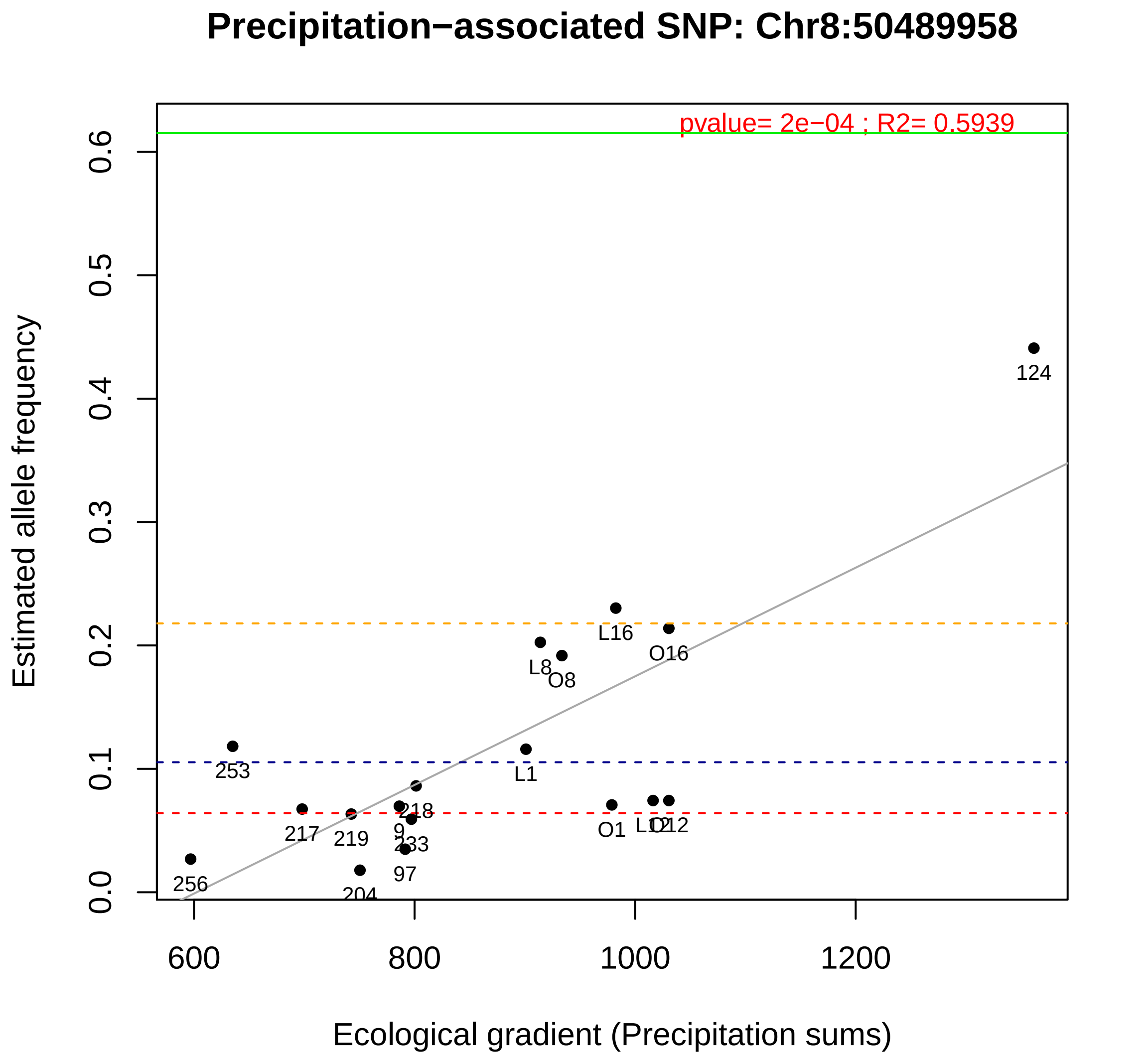

Precipitation−associated SNP: Chr8:50489958
pvalue= 2e−04 ; R2= 0.5939
Estimated allele frequency
0.0
0.1
0.2
0.3
0.4
0.5
0.6
600
800
1000
1200
Ecological gradient (Precipitation sums)
124
L16
O16
L8
O8
L1
253
217
219
218
9
233
97
256
204
O1
L12
O12

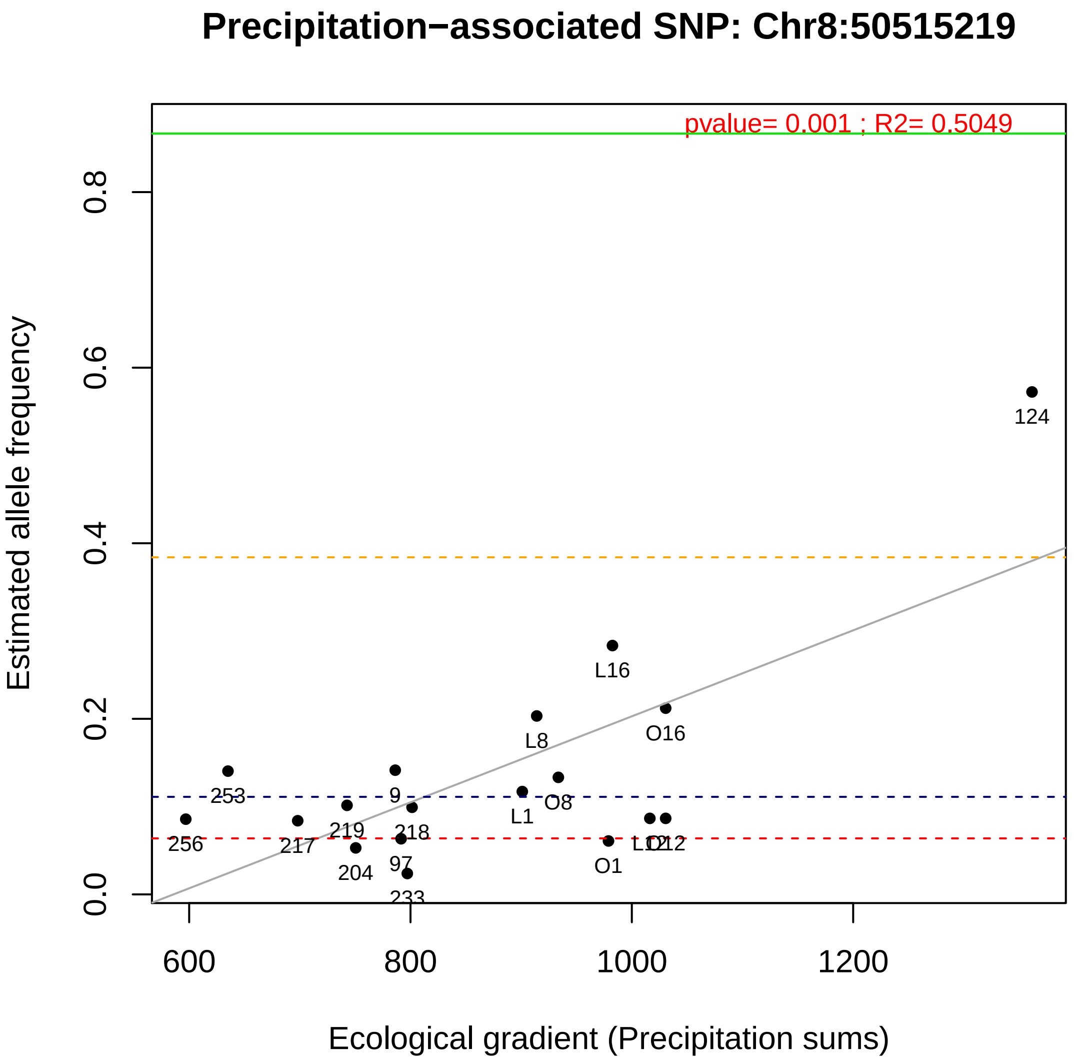

Precipitation−associated SNP: Chr8:50515219
pvalue= 0.001 ; R2= 0.5049
Estimated allele frequency
0.0
0.2
0.4
0.6
0.8
600
800
1000
1200
Ecological gradient (Precipitation sums)
124
L16
O16
L8
253
9
O8
L1
219
218
256
217
204
97
233
O1
O12

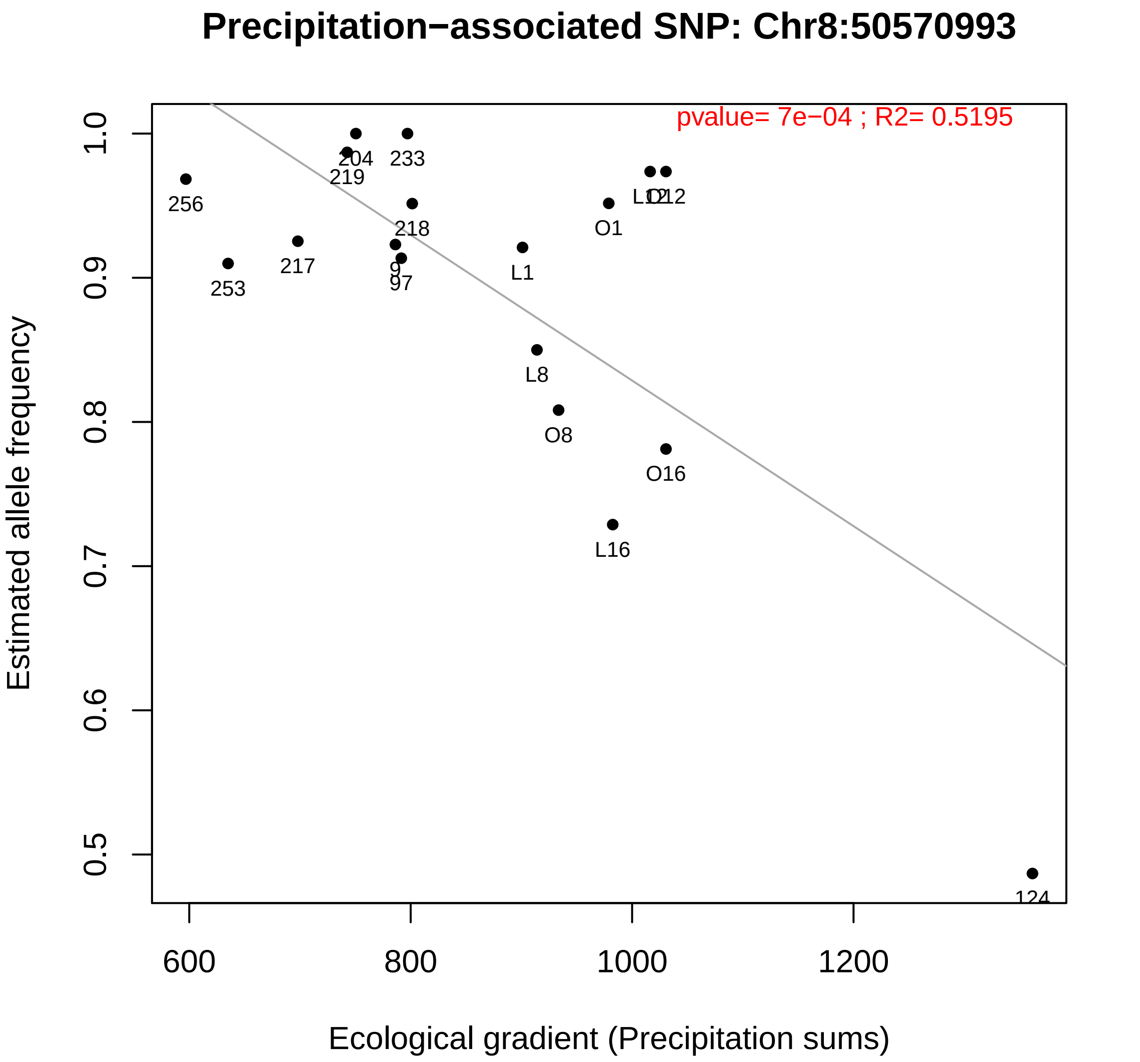
Precipitation−associated SNP: Chr8:50570993
pvalue= 7e−04 ; R2= 0.5195
Estimated allele frequency
Ecological gradient (Precipitation sums)
0.5
0.6
0.7
0.8
0.9
1.0
600
800
1000
1200
256
253
217
219
204
233
218
9
97
L1
L8
O8
O1
L12
O12
O16
L16
124

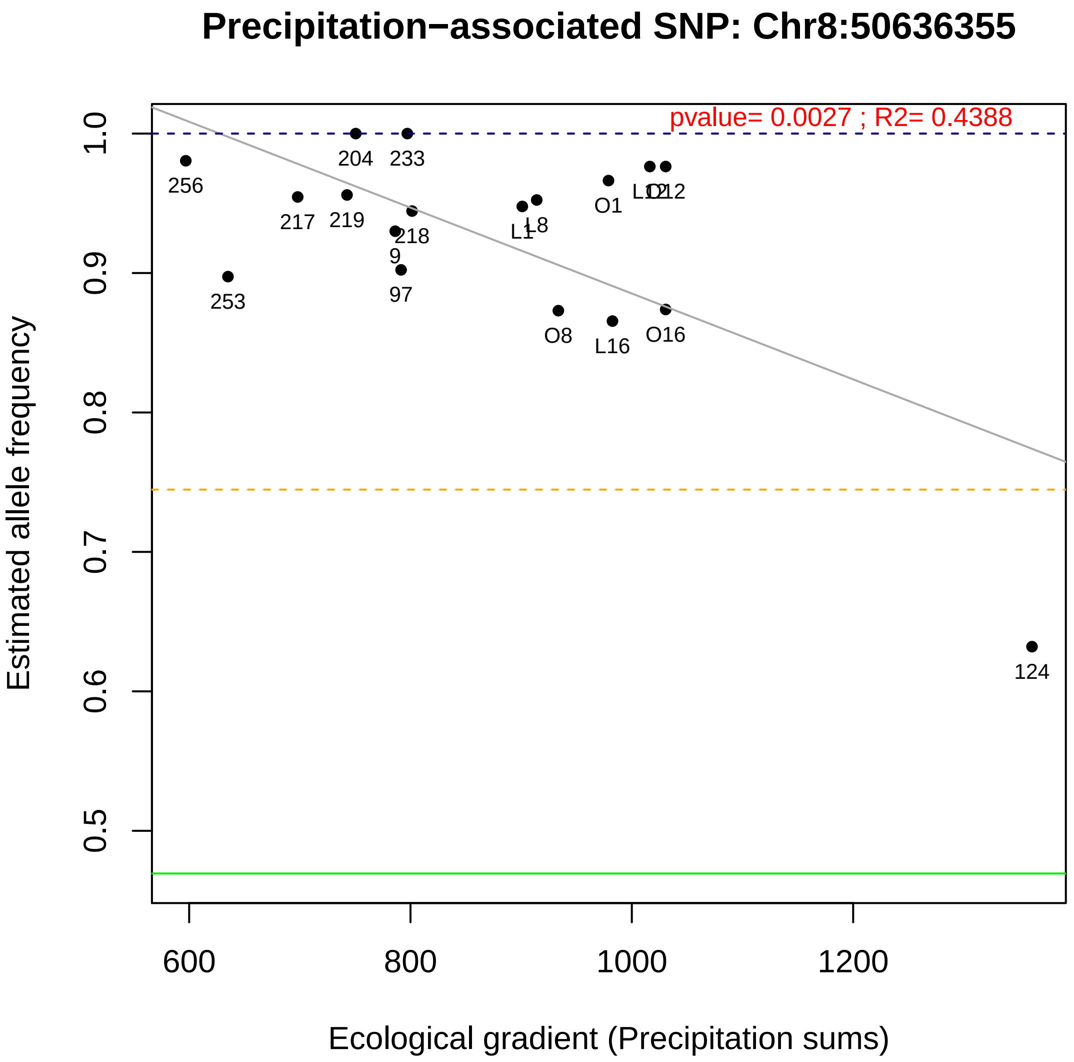

Precipitation−associated SNP: Chr8:50636355
pvalue= 0.0027 ; R2= 0.4388
Estimated allele frequency
Ecological gradient (Precipitation sums)
0.5
0.6
0.7
0.8
0.9
1.0
600
800
1000
1200
204
233
256
217
219
218
9
97
253
L1
L8
O1
L12
O12
O8
L16
O16
124

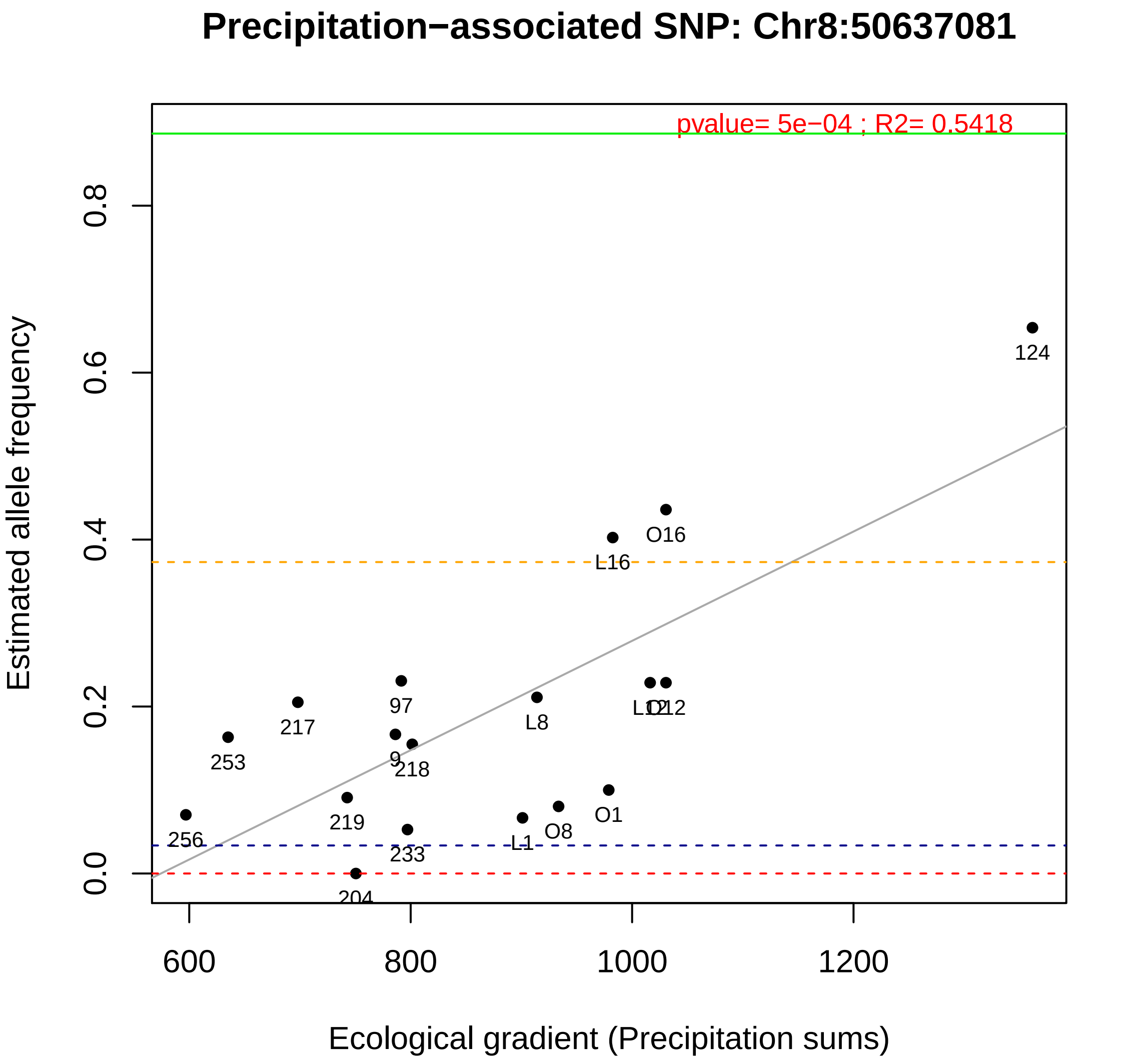

Precipitation−associated SNP: Chr8:50637081
pvalue= 5e−04 ; R2= 0.5418
Estimated allele frequency
0.0
0.2
0.4
0.6
0.8
600
800
1000
1200
Ecological gradient (Precipitation sums)
124
O16
L16
97
L8
L12
O12
217
253
9
218
219
O1
256
O8
L1
233
204

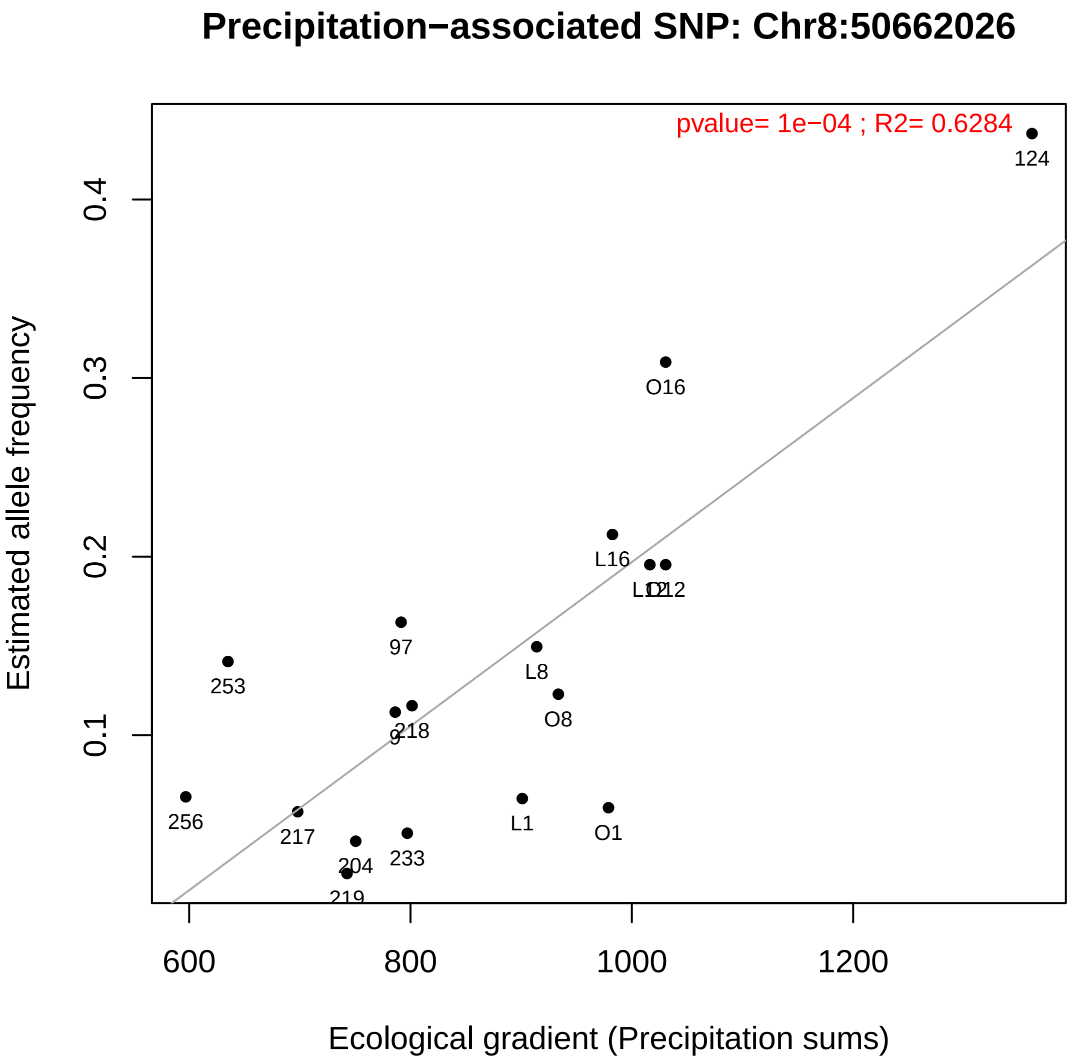

Precipitation−associated SNP: Chr8:50662026
pvalue= 1e−04 ; R2= 0.6284
Estimated allele frequency
0.1
0.2
0.3
0.4
Ecological gradient (Precipitation sums)
600
800
1000
1200
124
O16
L16
L12
O12
97
L8
253
O8
218
9
256
217
L1
O1
204
233
219

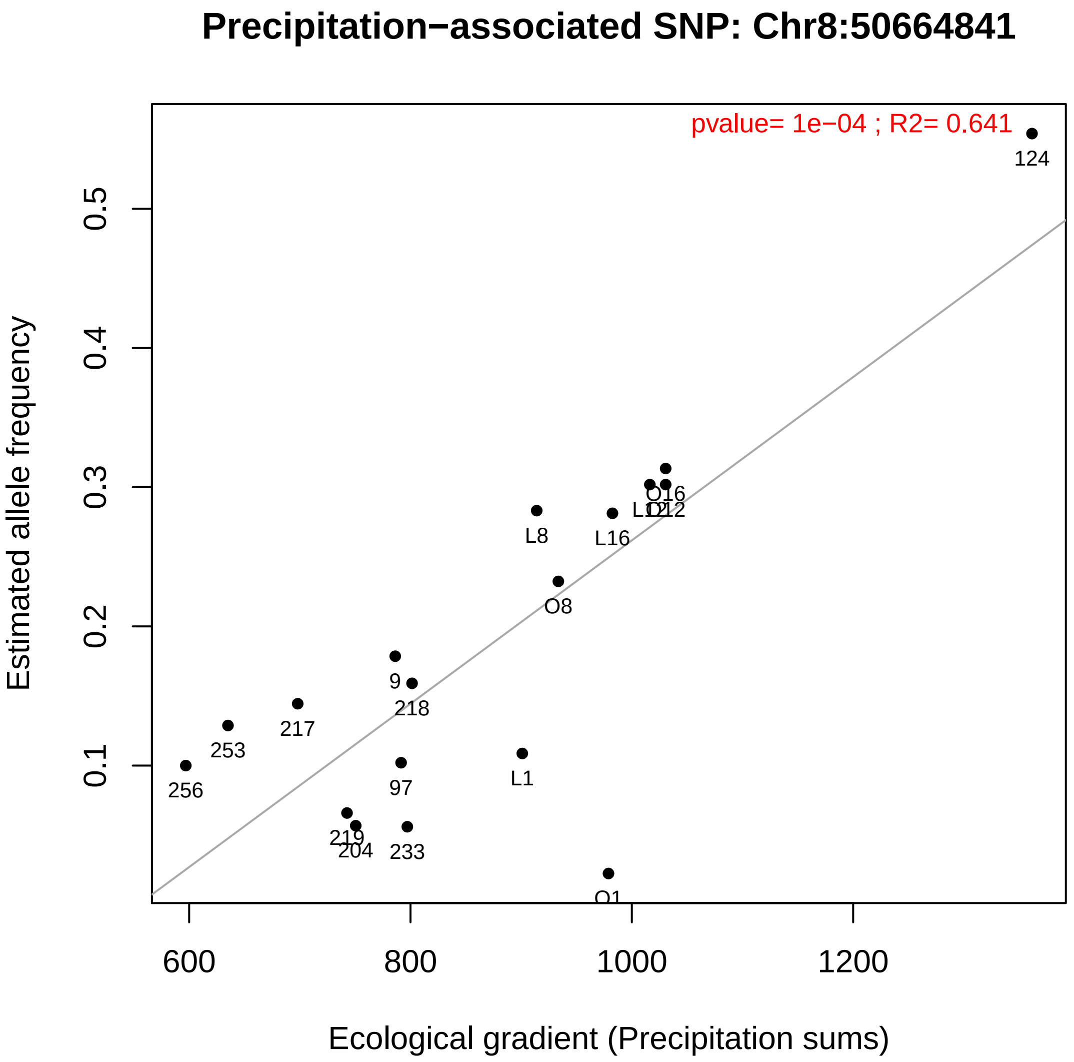
Precipitation−associated SNP: Chr8:50664841
pvalue= 1e−04 ; R2= 0.641
Estimated allele frequency
0.1
0.2
0.3
0.4
0.5
600
800
1000
1200
Ecological gradient (Precipitation sums)
124
O16
L12
O12
L8
L16
O8
9
218
217
253
256
97
L1
219
204
233

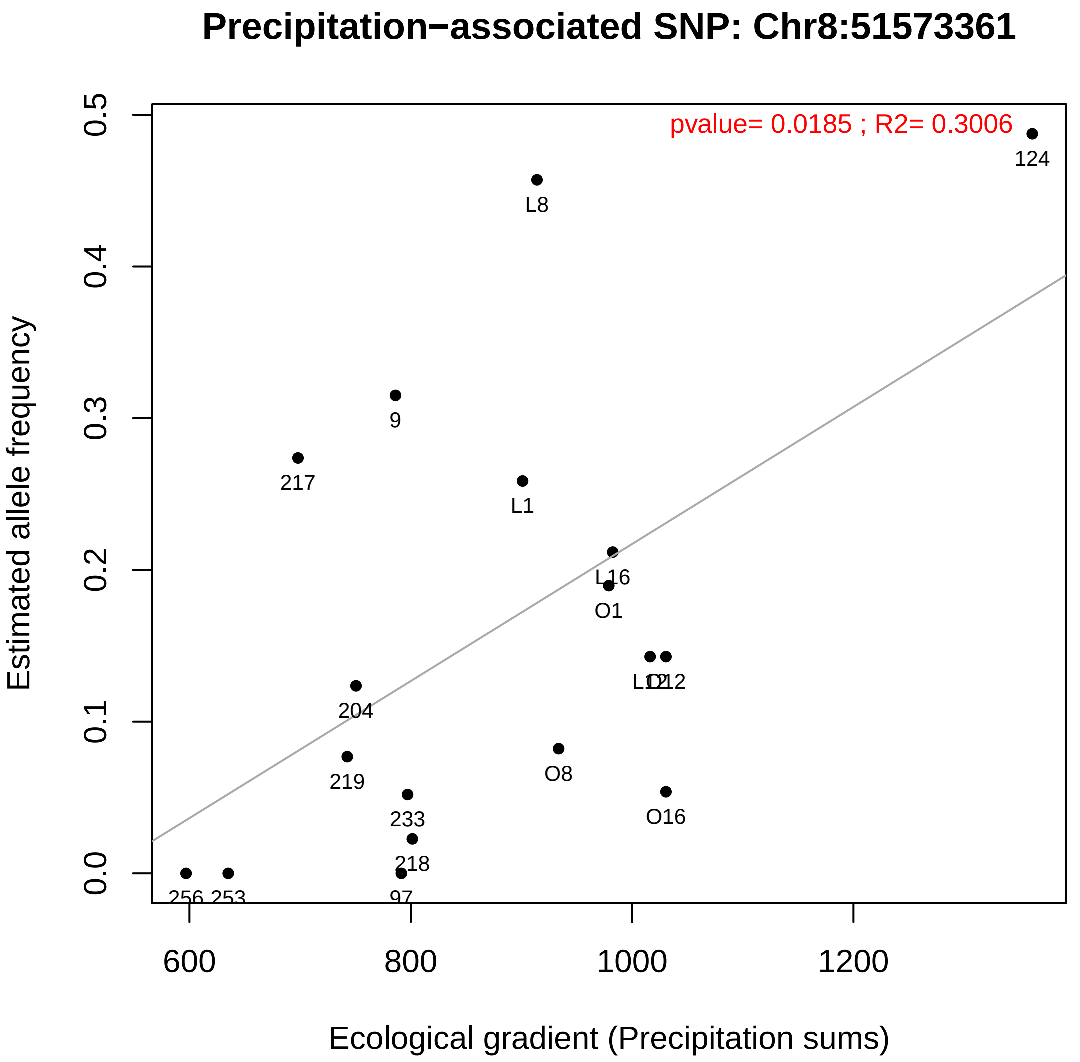

Precipitation−associated SNP: Chr8:51573361
pvalue= 0.0185 ; R2= 0.3006
Estimated allele frequency
Ecological gradient (Precipitation sums)
0.0
0.1
0.2
0.3
0.4
0.5
600
800
1000
1200
124
L8
9
217
L1
L16
O1
L12
O12
204
219
O8
233
O16
218
256
253
97

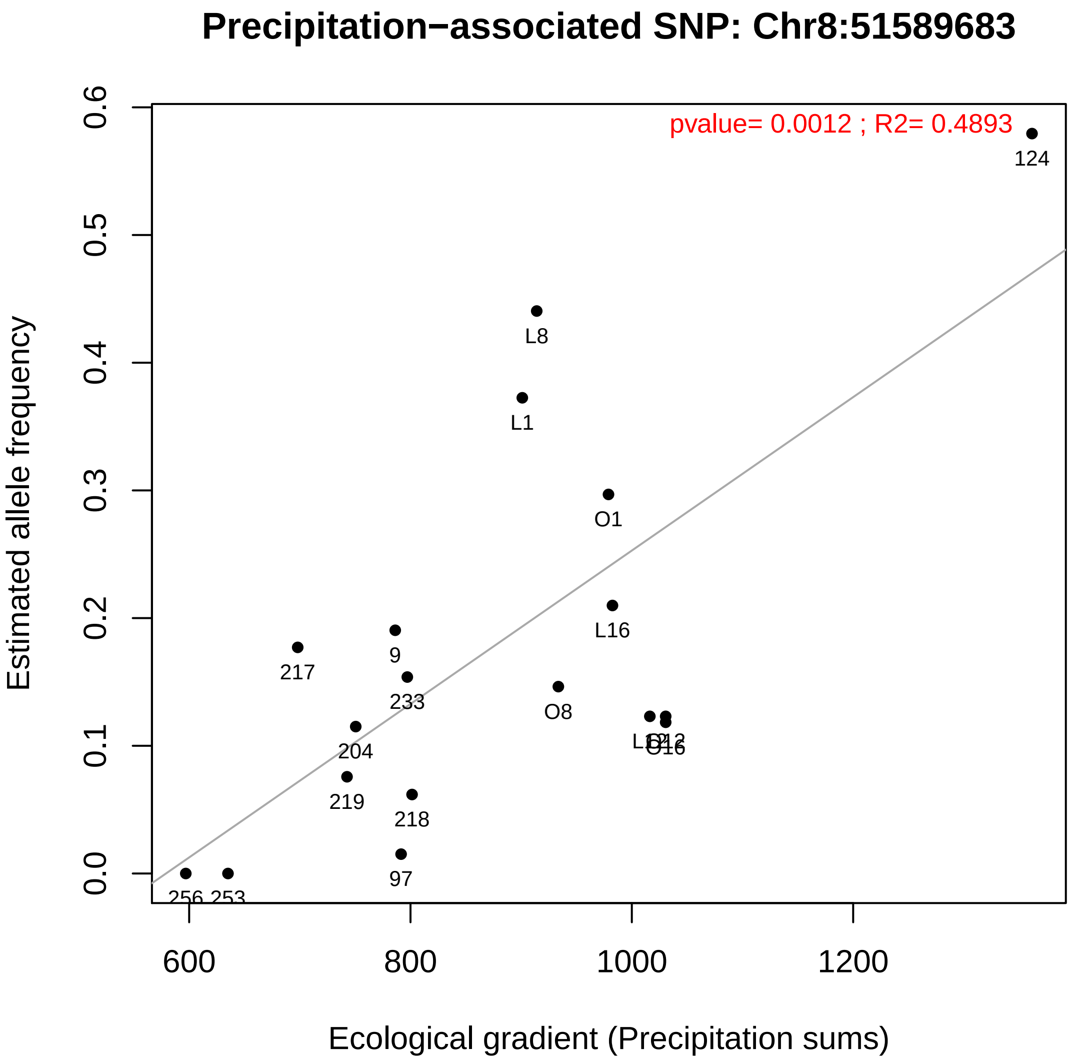
Precipitation−associated SNP: Chr8:51589683
pvalue= 0.0012 ; R2= 0.4893
Estimated allele frequency
0.0
0.1
0.2
0.3
0.4
0.5
0.6
600
800
1000
1200
Ecological gradient (Precipitation sums)
124
L8
L1
O1
L16
217
9
233
O8
204
219
218
97
256
253

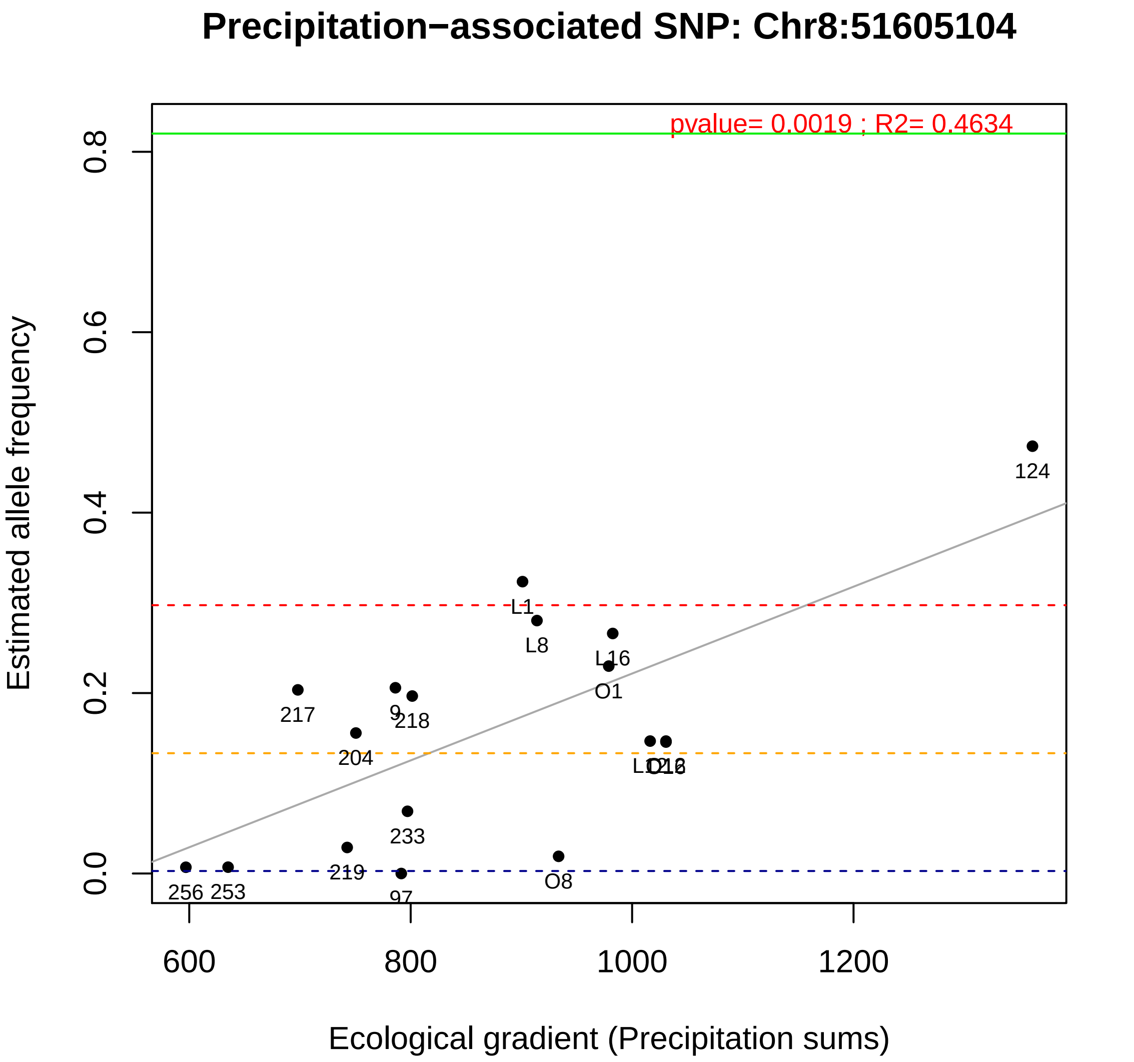

Precipitation−associated SNP: Chr8:51605104
pvalue= 0.0019 ; R2= 0.4634
Estimated allele frequency
0.0
0.2
0.4
0.6
0.8
124
L1
L8
L16
O1
217
9
218
204
L12
O16
233
219
256
253
97
O8
600
800
1000
1200
Ecological gradient (Precipitation sums)

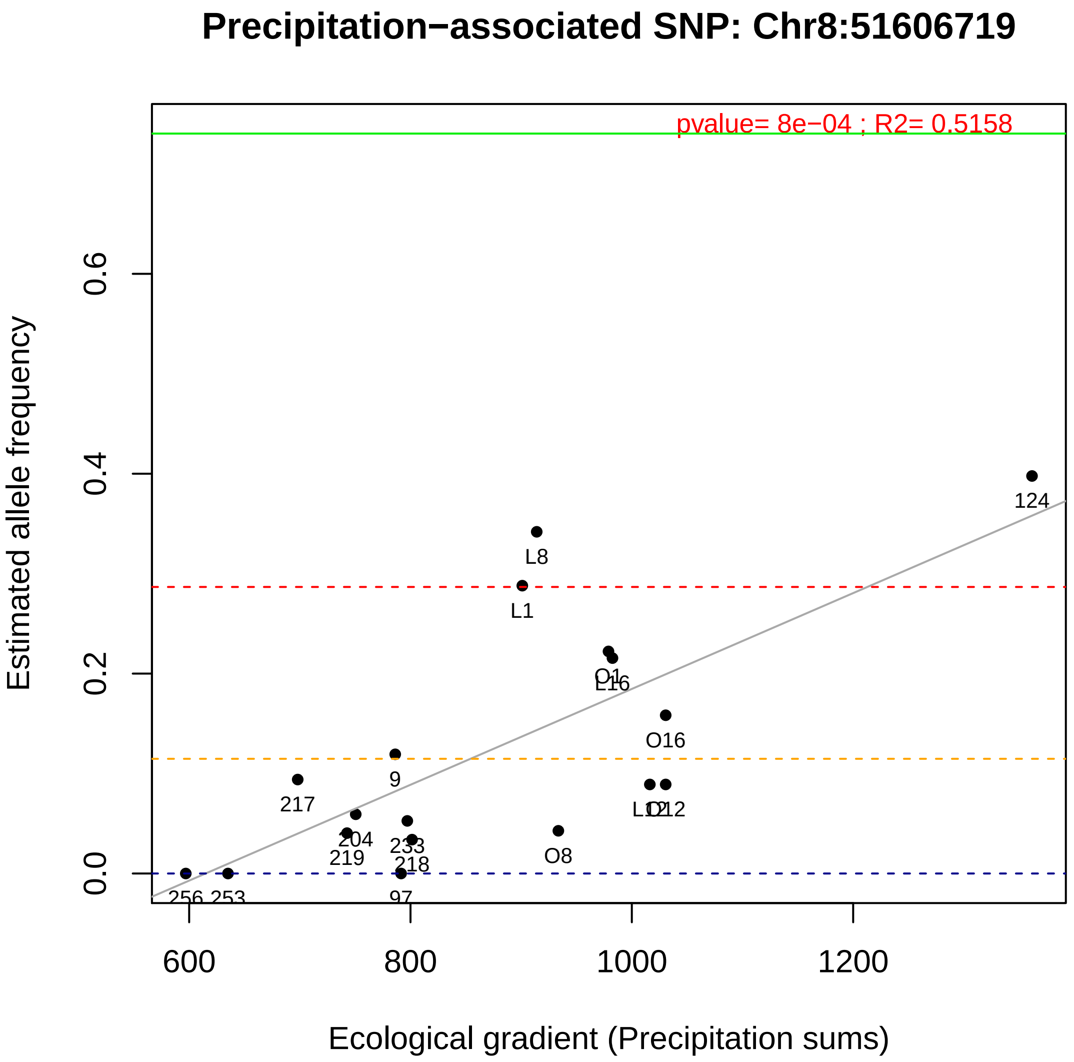

Precipitation−associated SNP: Chr8:51606719
pvalue= 8e−04 ; R2= 0.5158
Estimated allele frequency
0.0
0.2
0.4
0.6
600
800
1000
1200
Ecological gradient (Precipitation sums)
124
L8
L1
O1
L16
O16
9
217
L12
O12
204
233
219
218
O8
97
256
253

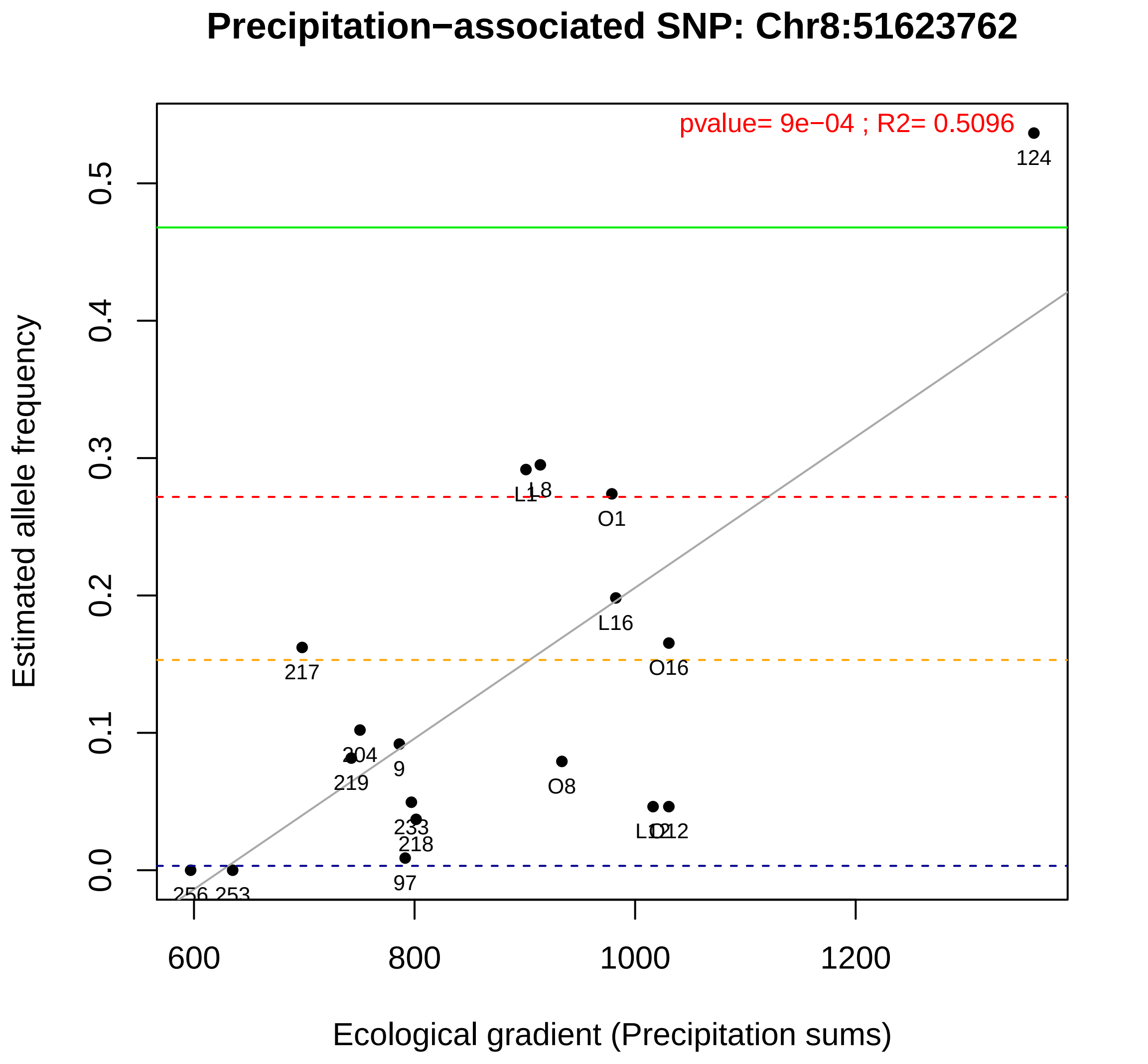

Precipitation−associated SNP: Chr8:51623762
pvalue= 9e−04 ; R2= 0.5096
124
L1
L8
O1
L16
O16
217
204
219
9
O8
233
218
L12
O12
97
256
253
0.0
0.1
0.2
0.3
0.4
0.5
600
800
1000
1200
Estimated allele frequency
Ecological gradient (Precipitation sums)

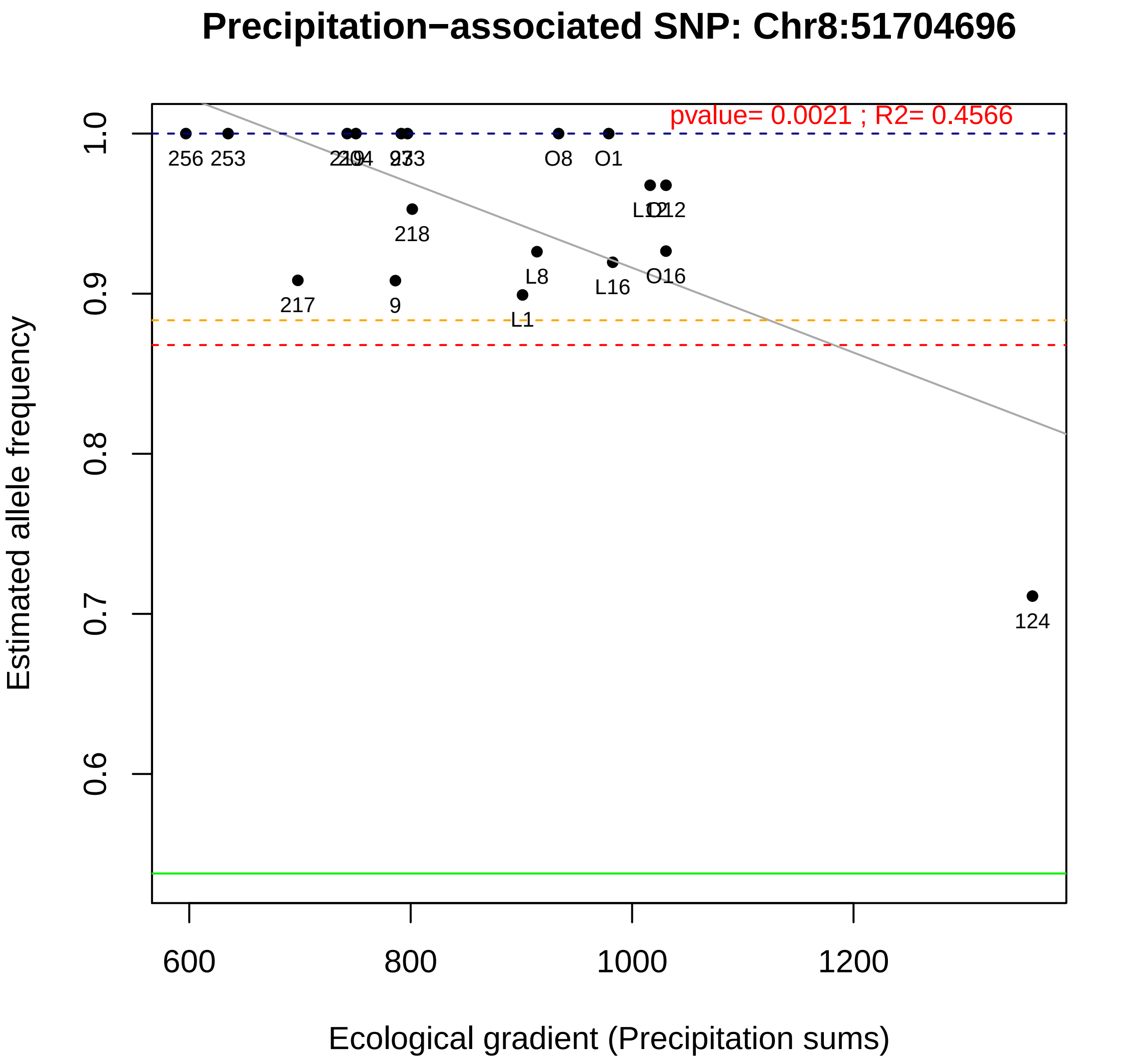
Precipitation−associated SNP: Chr8:51704696
pvalue= 0.0021 ; R2= 0.4566
Estimated allele frequency
Ecological gradient (Precipitation sums)
0.6
0.7
0.8
0.9
1.0
600
800
1000
1200
256
253
O8
O1
218
217
9
L8
L1
L16
O16
124

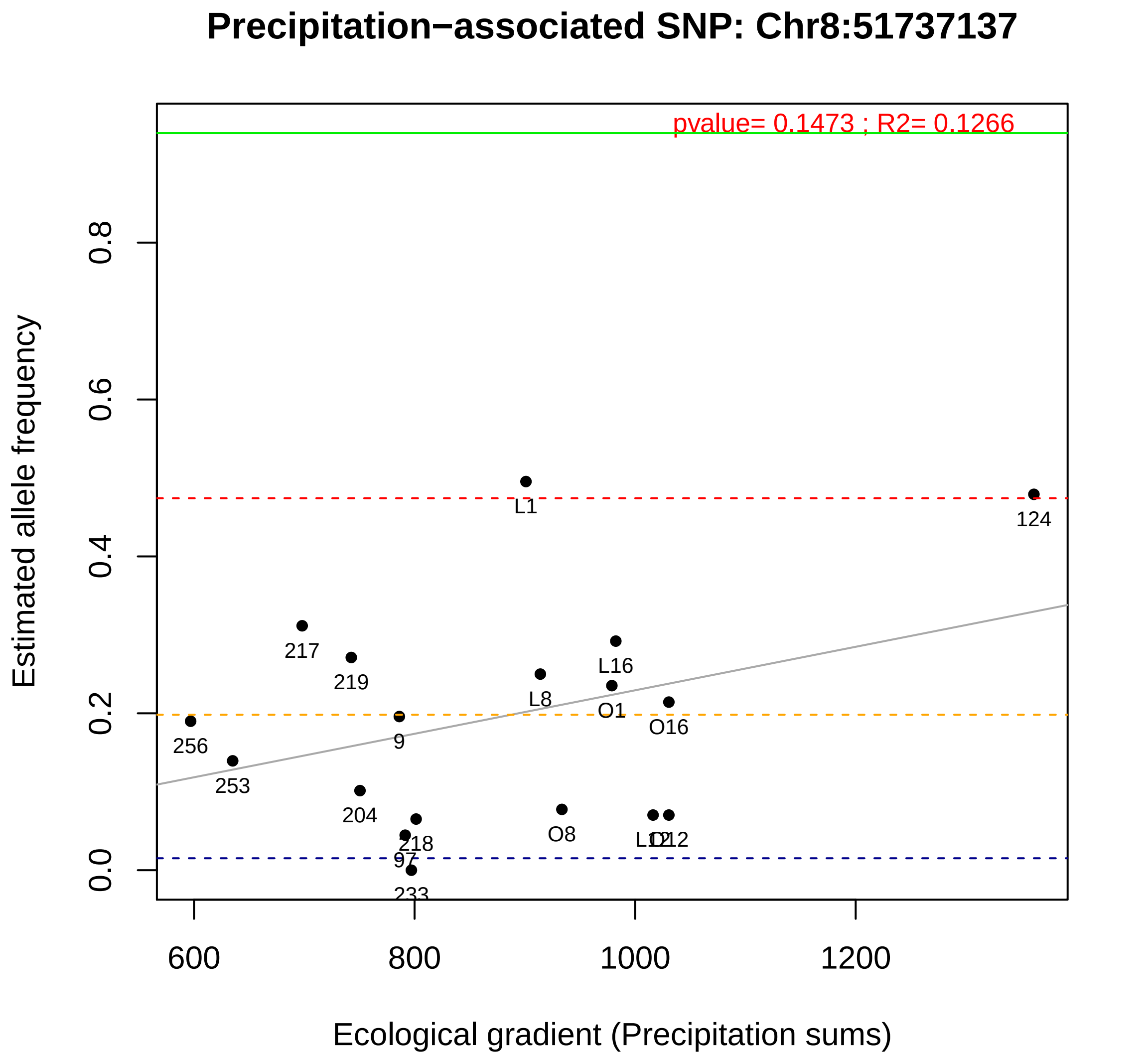
Precipitation−associated SNP: Chr8:51737137
pvalue= 0.1473 ; R2= 0.1266
Estimated allele frequency
0.0
0.2
0.4
0.6
0.8
600
800
1000
1200
Ecological gradient (Precipitation sums)
L1
124
217
219
L16
L8
O1
O16
256
9
253
204
O8
218
97
233

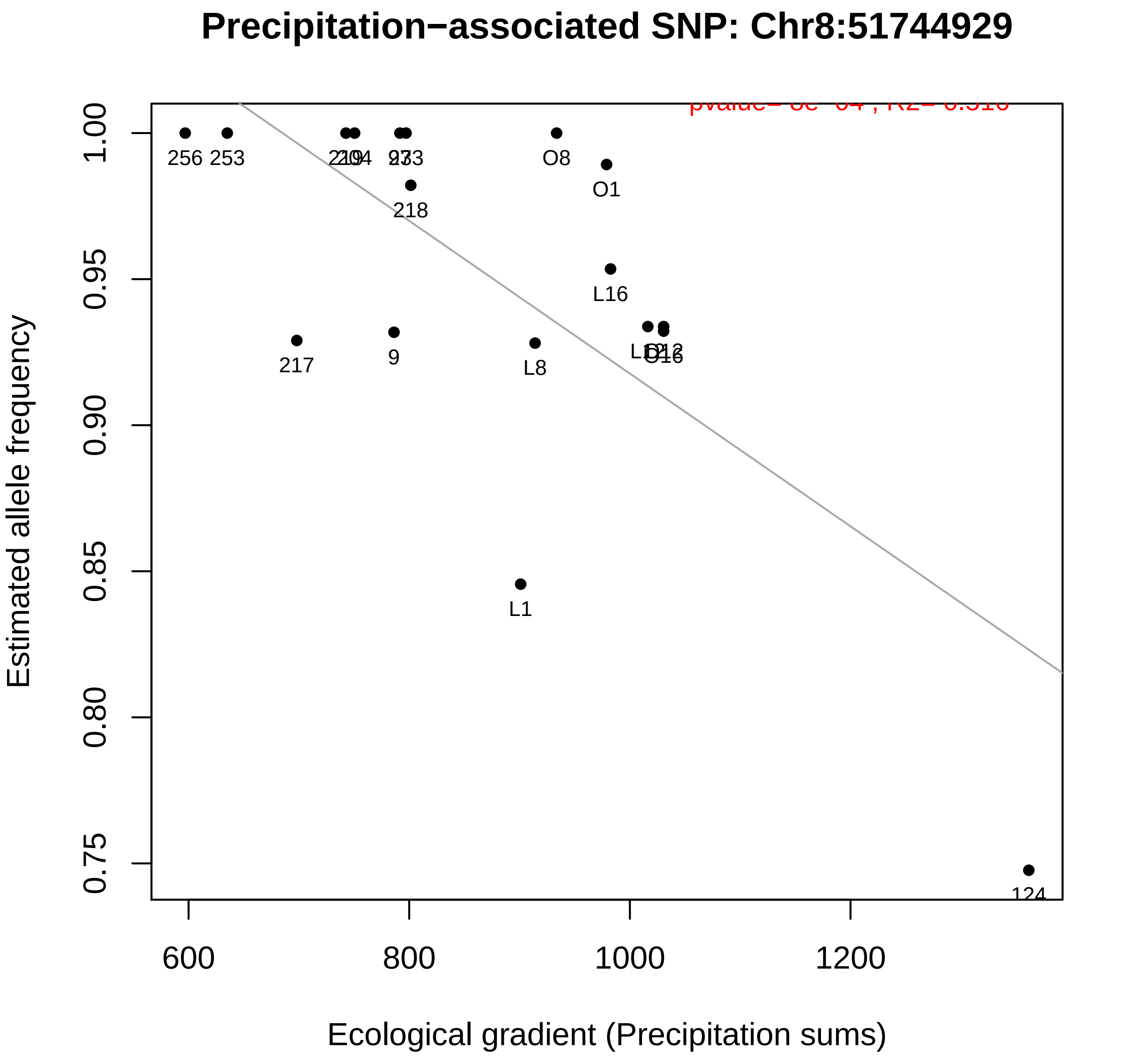
Precipitation−associated SNP: Chr8:51744929
Estimated allele frequency
Ecological gradient (Precipitation sums)
1.00
0.95
0.90
0.85
0.80
0.75
600
800
1000
1200
256
253
O8
O1
218
L16
217
9
L8
L1
124

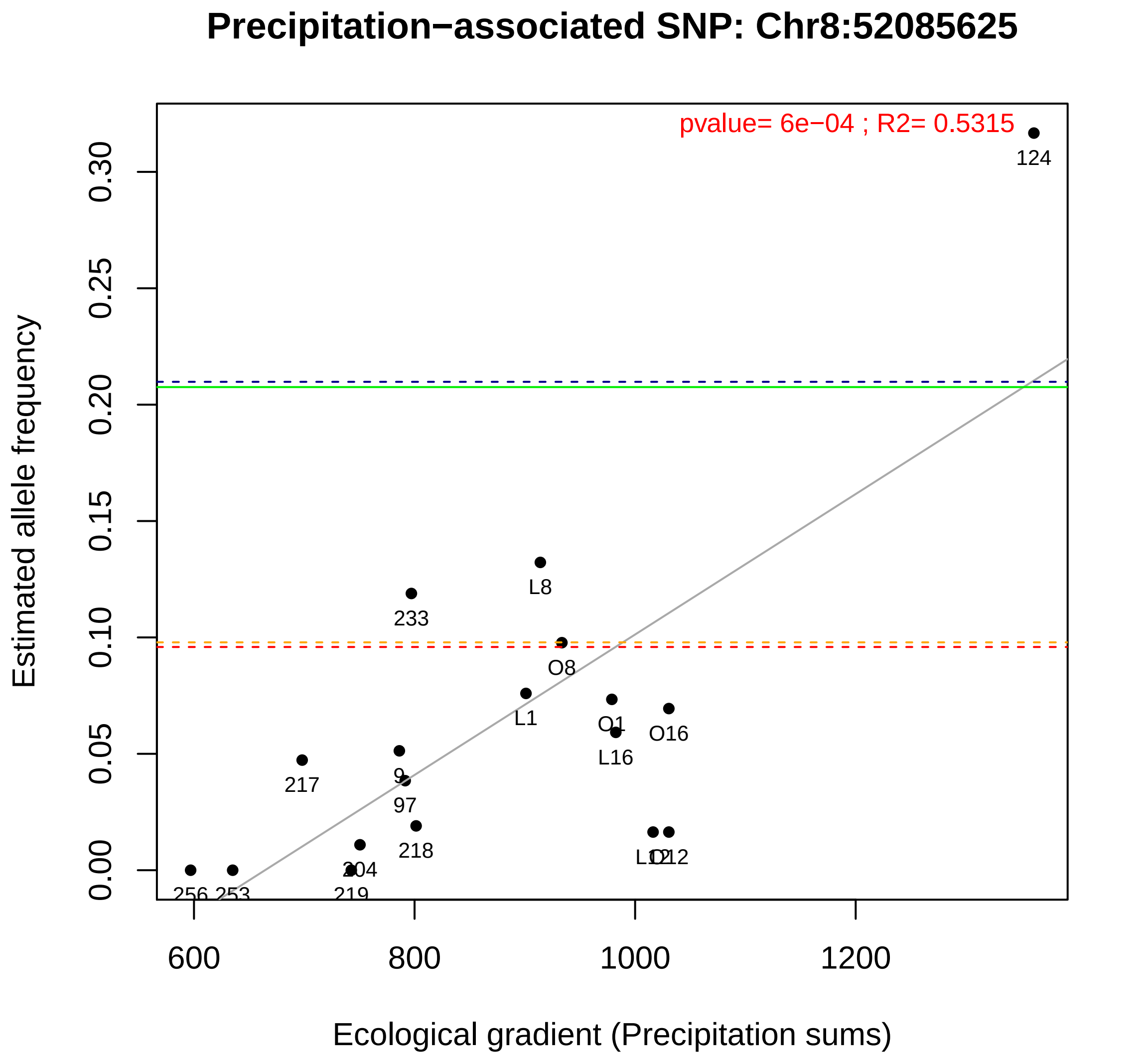

Precipitation−associated SNP: Chr8:52085625
pvalue= 6e−04 ; R2= 0.5315
Estimated allele frequency
Ecological gradient (Precipitation sums)
0.00
0.05
0.10
0.15
0.20
0.25
0.30
600
800
1000
1200
124
L8
233
O8
L1
O1
O16
L16
217
9
97
218
204
256
253
219

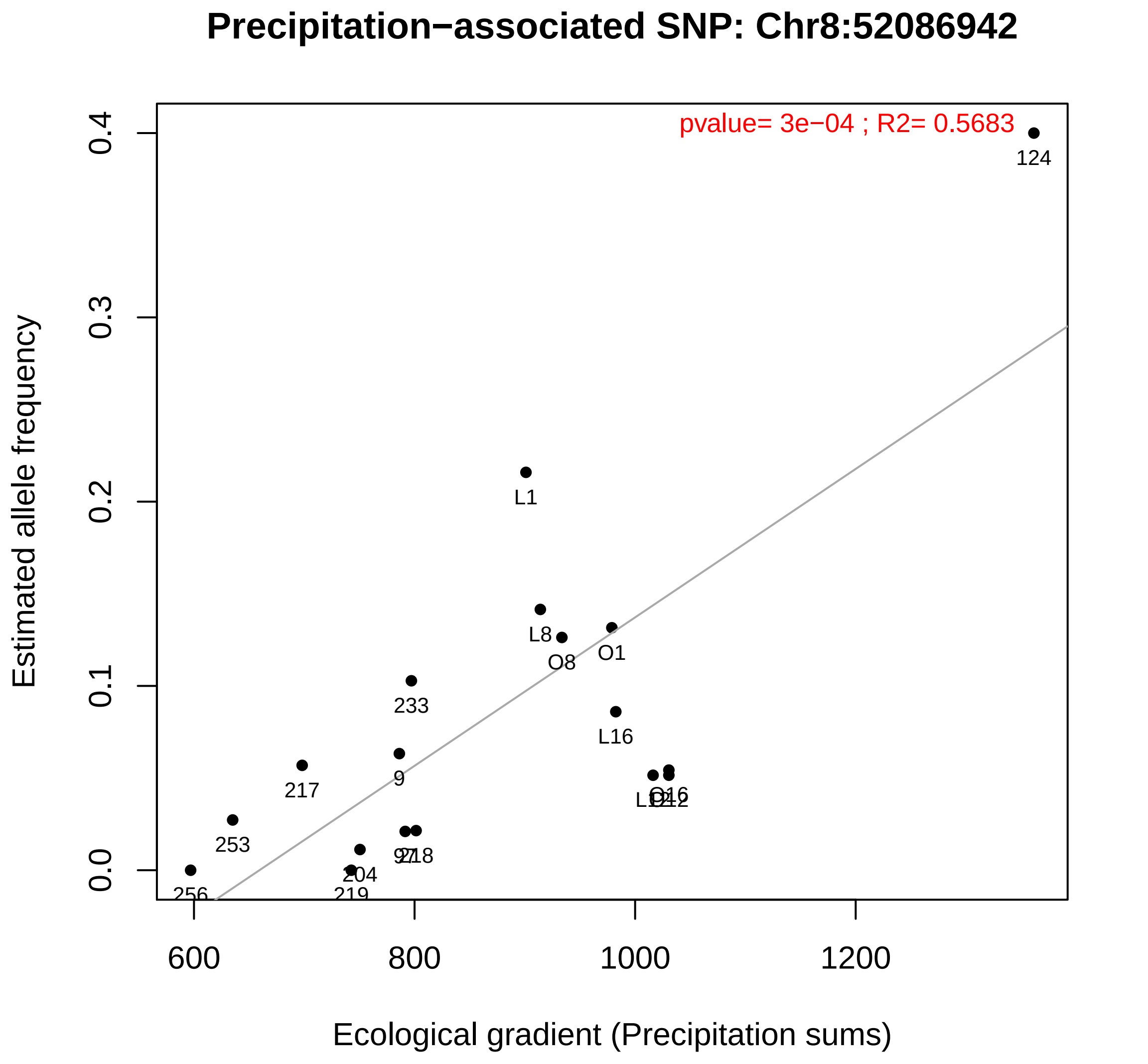

Precipitation−associated SNP: Chr8:52086942
pvalue= 3e−04 ; R2= 0.5683
Estimated allele frequency
Ecological gradient (Precipitation sums)
0.0
0.1
0.2
0.3
0.4
600
800
1000
1200
124
L1
L8
O8
O1
233
L16
217
9
253
204
218
256
219

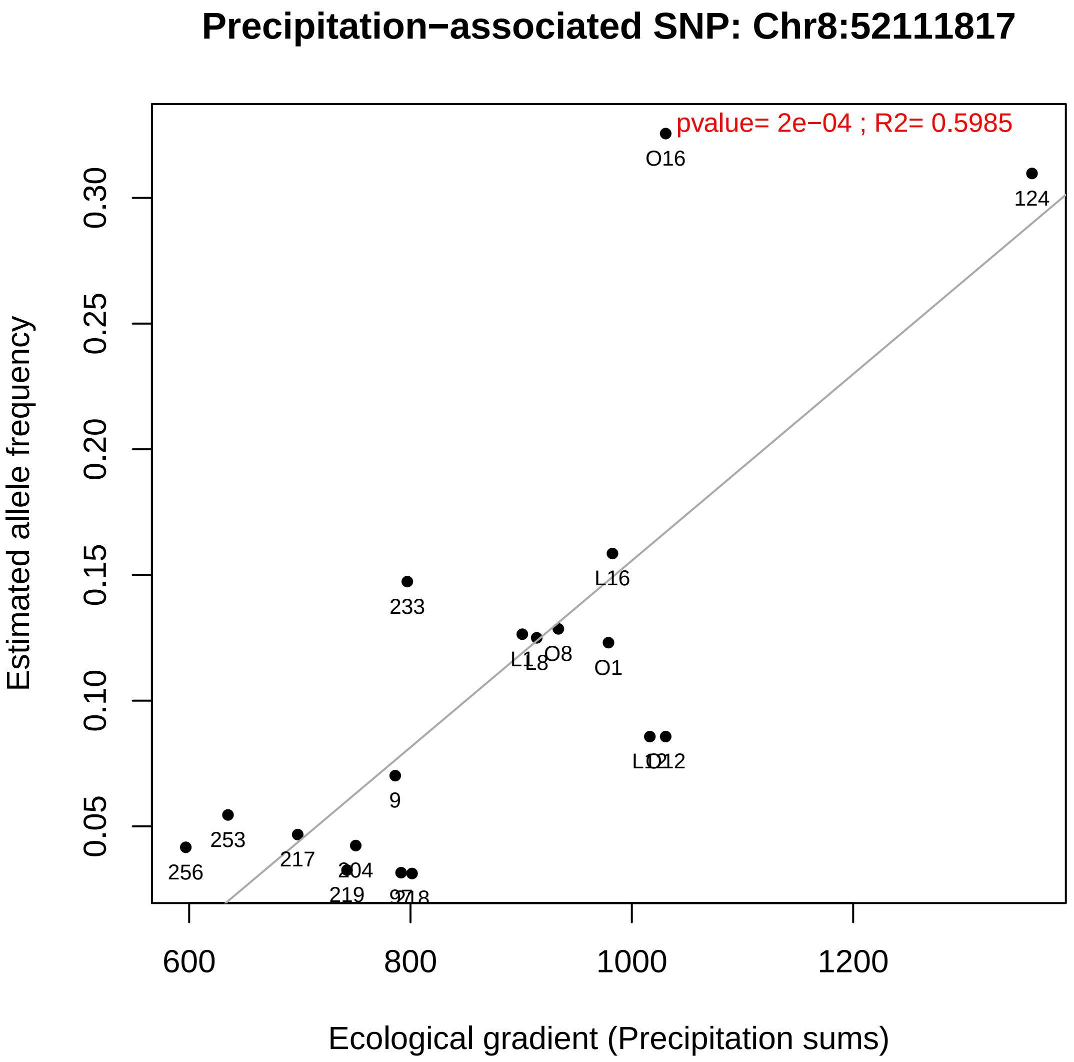
Precipitation−associated SNP: Chr8:52111817
pvalue= 2e−04 ; R2= 0.5985
Estimated allele frequency
0.05
0.10
0.15
0.20
0.25
0.30
600
800
1000
1200
Ecological gradient (Precipitation sums)
O16
124
L16
233
O8
L8
O1
L12
O12
9
253
256
217
204
219
218

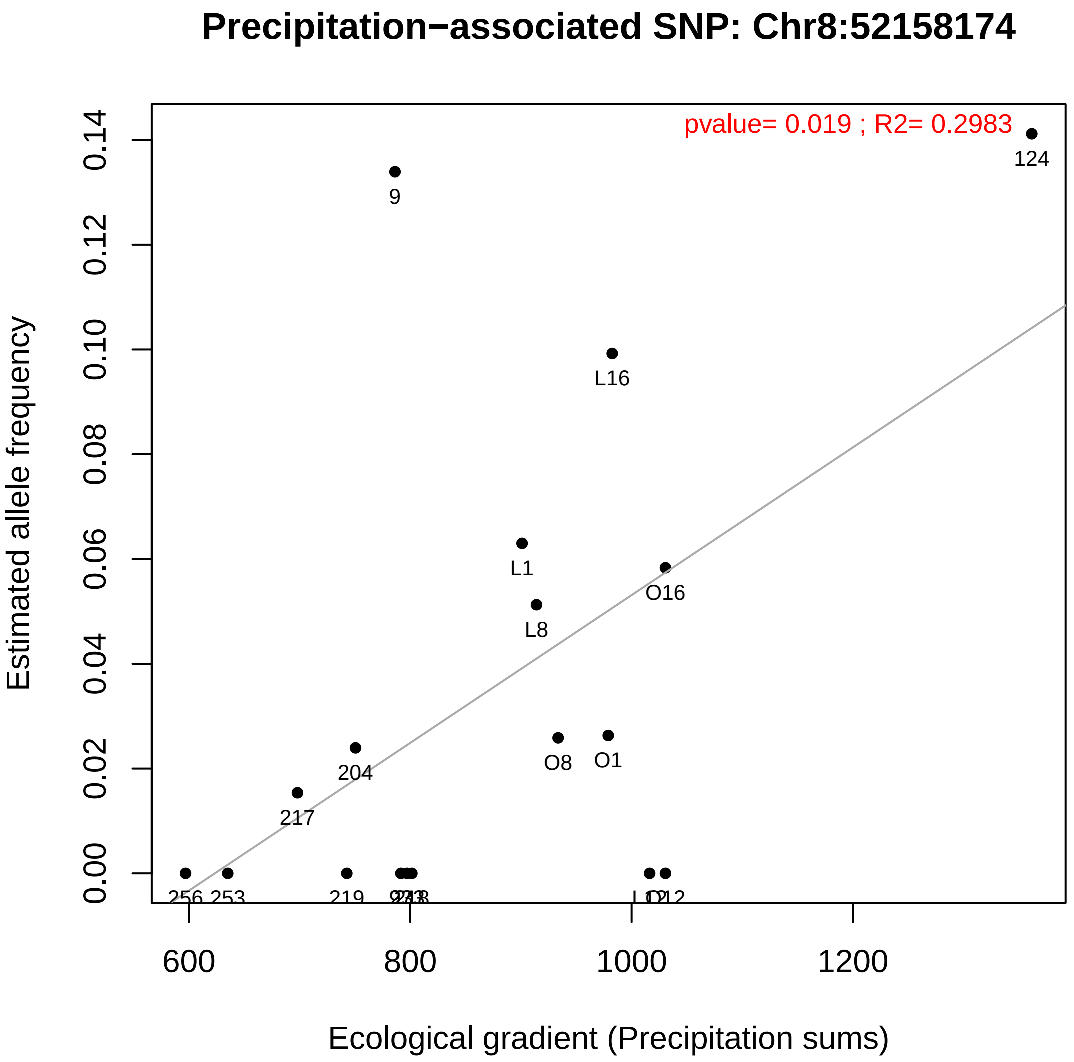

Precipitation−associated SNP: Chr8:52158174
pvalue= 0.019 ; R2= 0.2983
Estimated allele frequency
0.00
0.02
0.04
0.06
0.08
0.10
0.12
0.14
Ecological gradient (Precipitation sums)
600
800
1000
1200
124
9
L16
L1
O16
L8
O8
O1
204
217
256
253
219

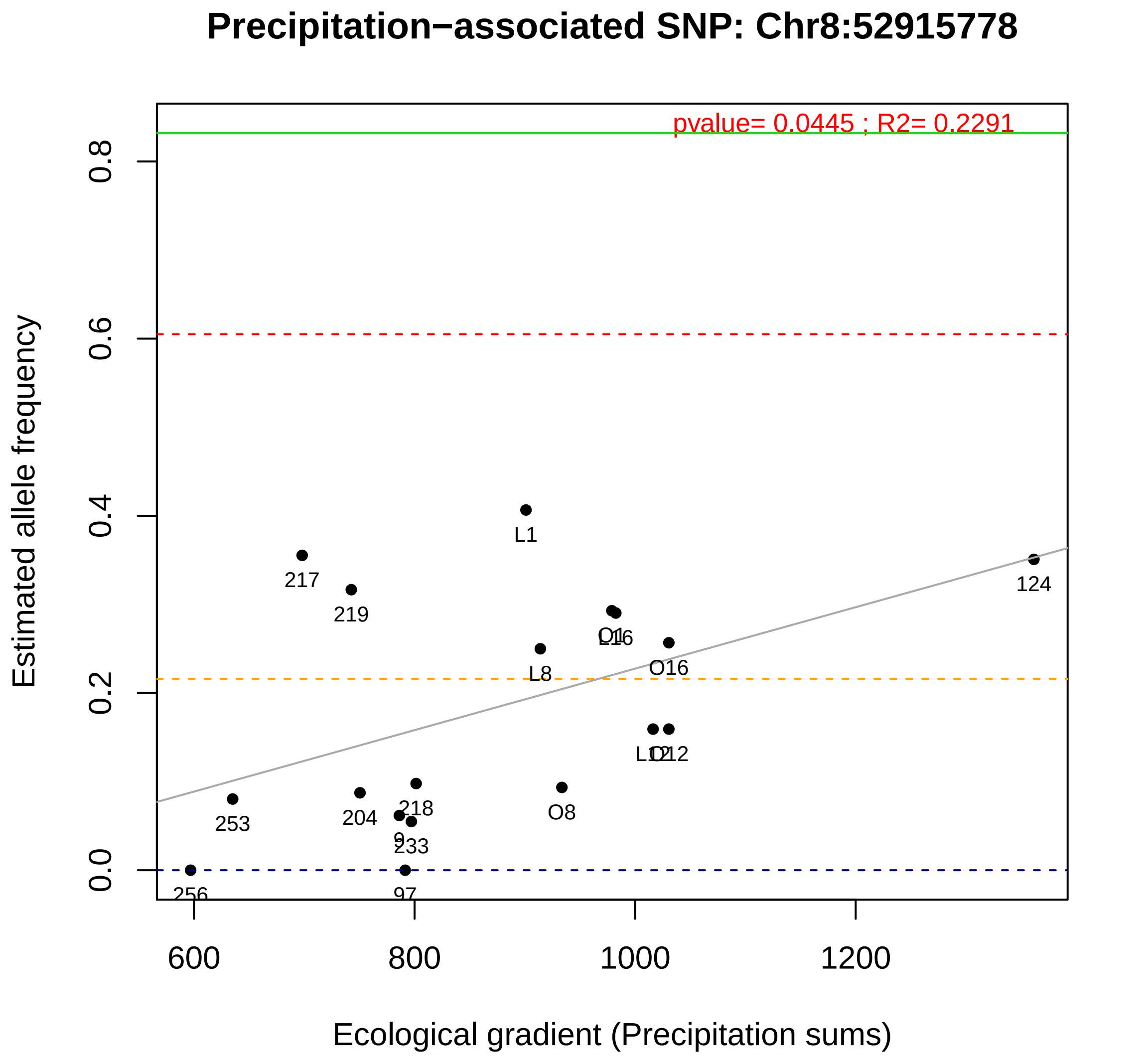

Precipitation−associated SNP: Chr8:52915778
pvalue= 0.0445 ; R2= 0.2291
Estimated allele frequency
0.0
0.2
0.4
0.6
0.8
600
800
1000
1200
Ecological gradient (Precipitation sums)
L1
217
219
124
O1
L16
O16
L8
L12
O12
253
204
218
O8
9
233
256
97

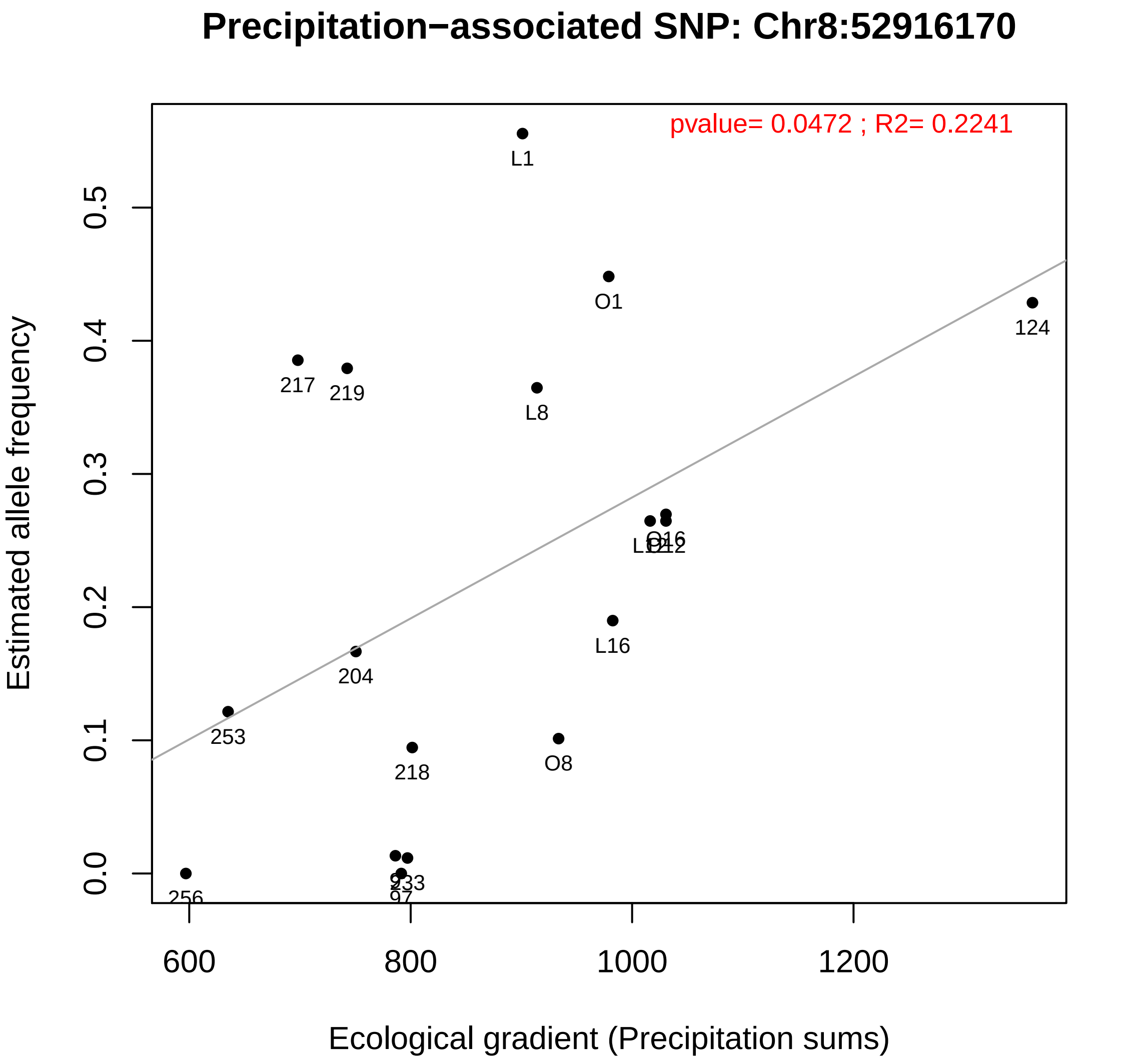

Precipitation−associated SNP: Chr8:52916170
pvalue= 0.0472 ; R2= 0.2241
Estimated allele frequency
0.0
0.1
0.2
0.3
0.4
0.5
600
800
1000
1200
Ecological gradient (Precipitation sums)
L1
O1
124
217
219
L8
O16
L16
204
253
218
O8
233
97
256

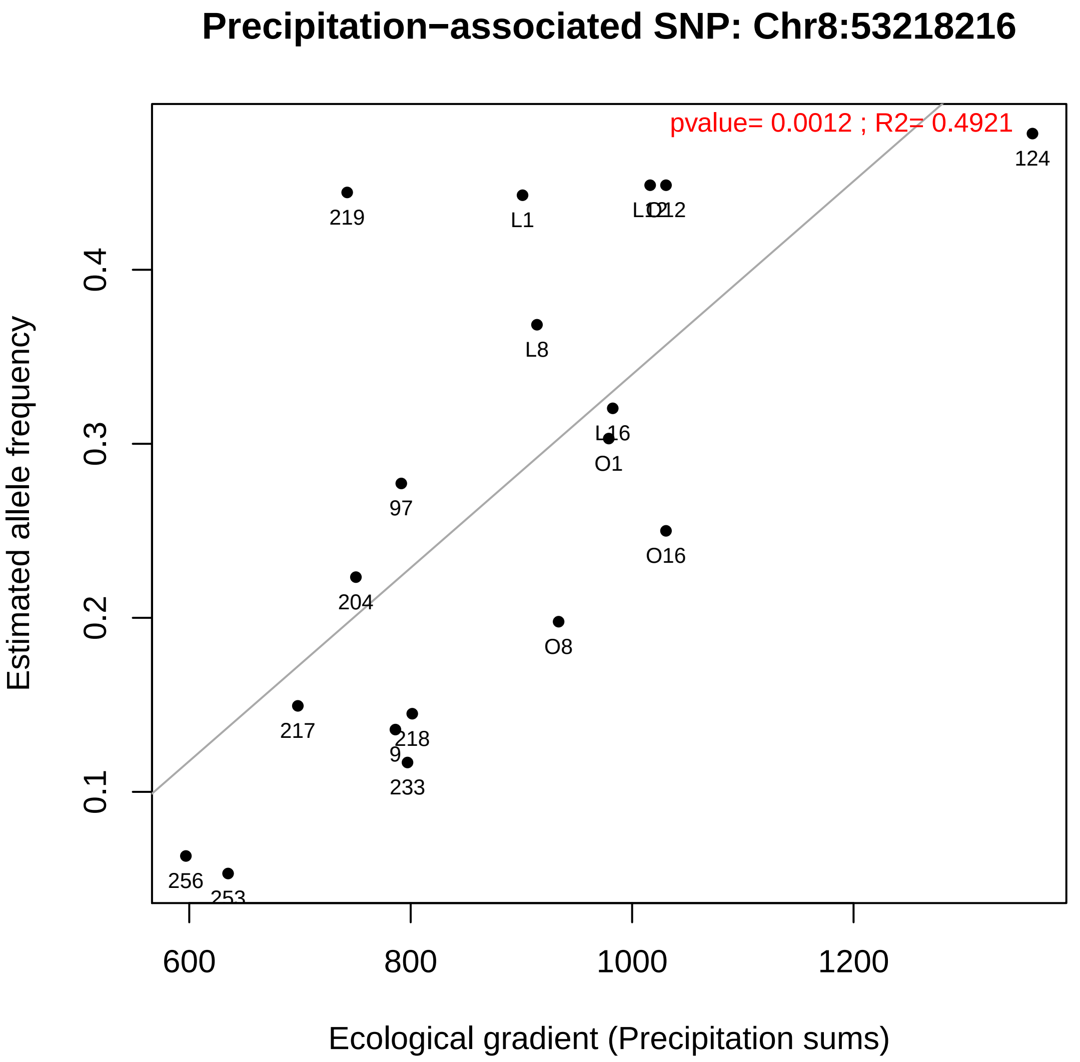
Precipitation−associated SNP: Chr8:53218216
pvalue= 0.0012 ; R2= 0.4921
Estimated allele frequency
0.1
0.2
0.3
0.4
600
800
1000
1200
Ecological gradient (Precipitation sums)
124
219
L1
L12
O12
L8
L16
O1
97
O16
204
O8
217
218
9
233
256
253

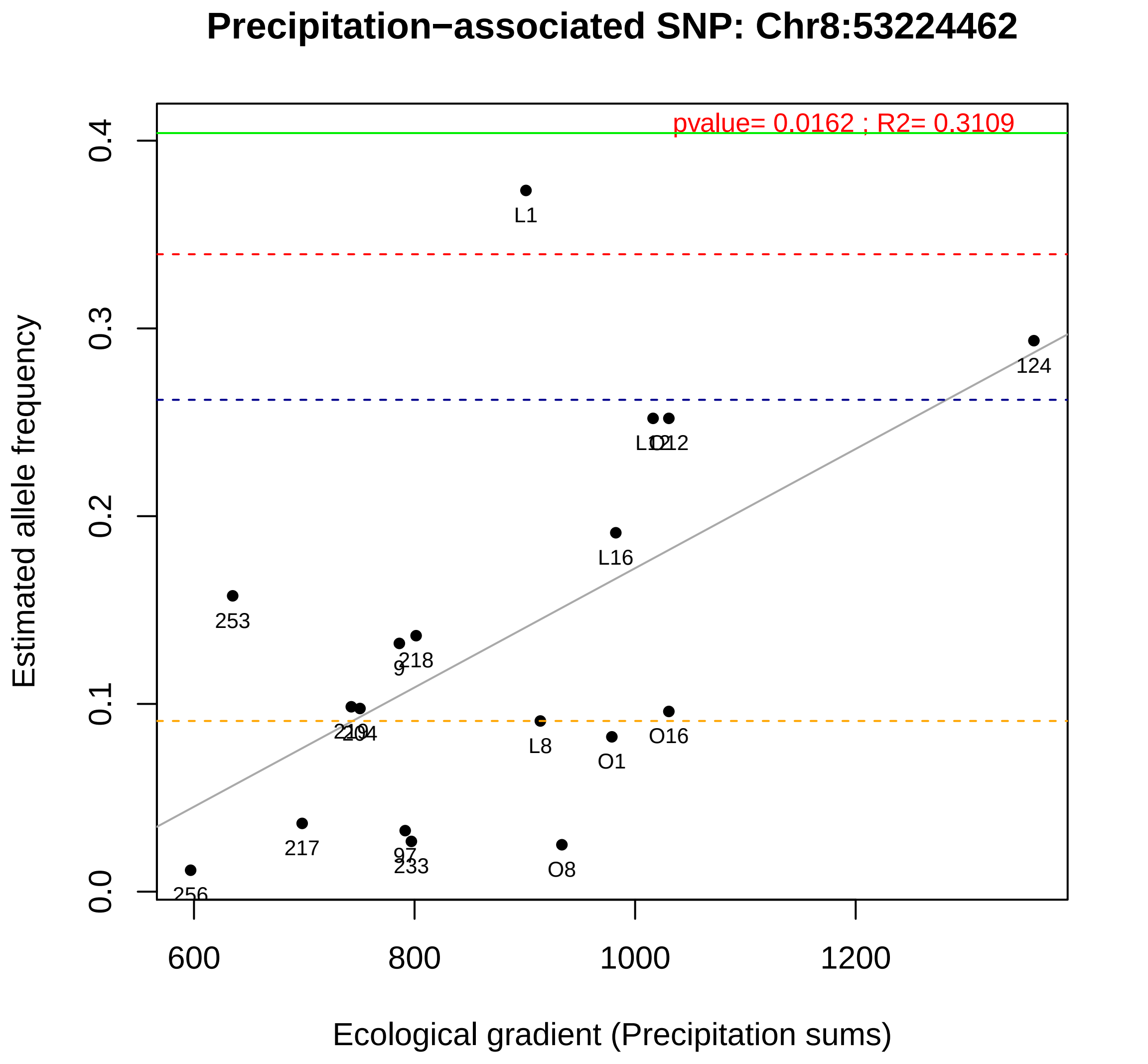
Precipitation−associated SNP: Chr8:53224462
pvalue= 0.0162 ; R2= 0.3109
Estimated allele frequency
Ecological gradient (Precipitation sums)
0.0
0.1
0.2
0.3
0.4
600
800
1000
1200
L1
124
L16
O12
253
218
9
L8
O1
O16
217
97
233
O8
256

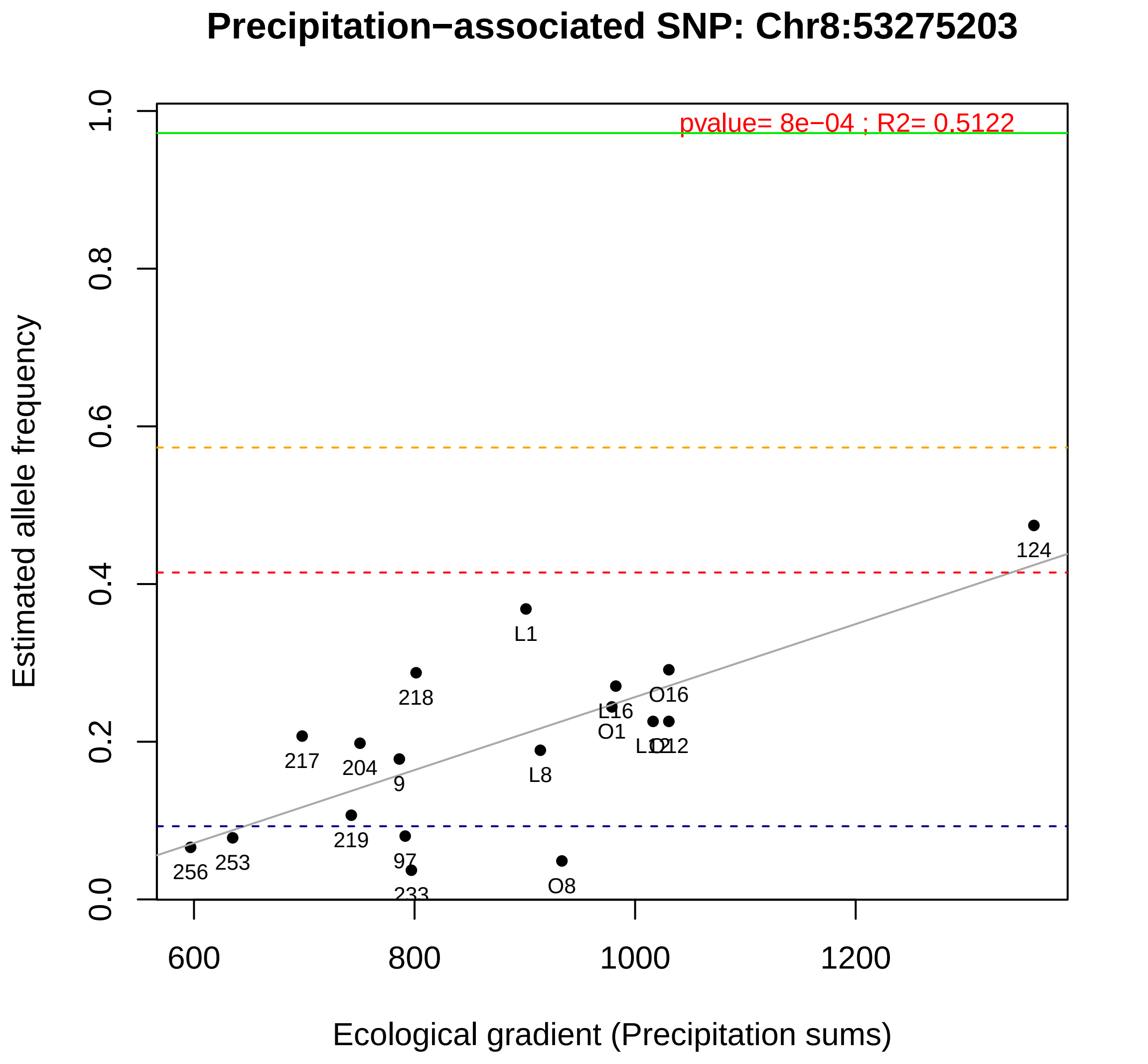

Precipitation−associated SNP: Chr8:53275203
pvalue= 8e−04 ; R2= 0.5122
Estimated allele frequency
0.0
0.2
0.4
0.6
0.8
1.0
600
800
1000
1200
Ecological gradient (Precipitation sums)
124
L1
218
O16
L16
O1
L12
O12
217
204
9
L8
219
253
256
97
233
O8

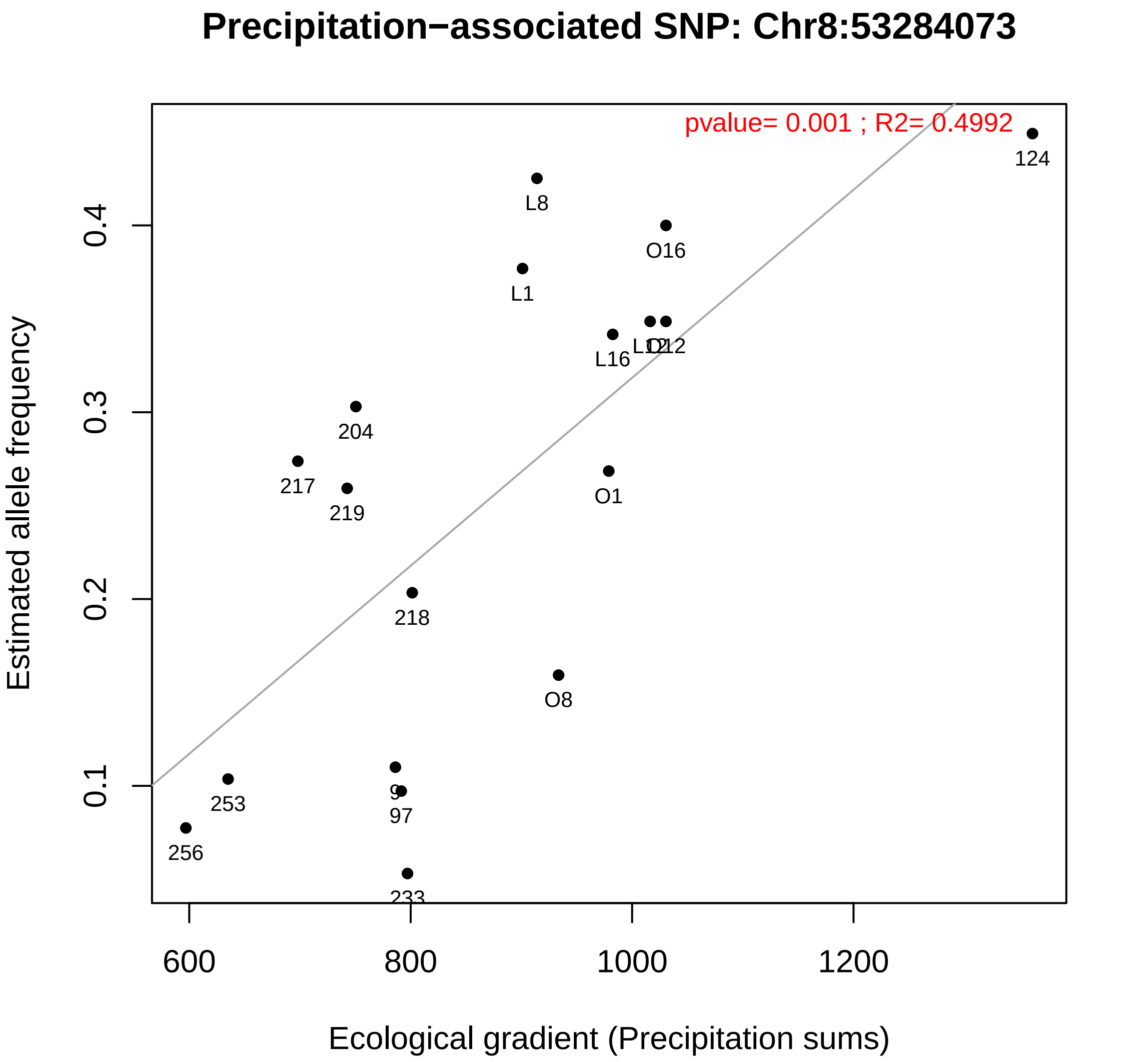
Precipitation−associated SNP: Chr8:53284073
pvalue= 0.001 ; R2= 0.4992
Estimated allele frequency
0.1
0.2
0.3
0.4
600
800
1000
1200
Ecological gradient (Precipitation sums)
124
L8
O16
L1
L16
O12
204
217
219
O1
218
O8
253
97
256
233

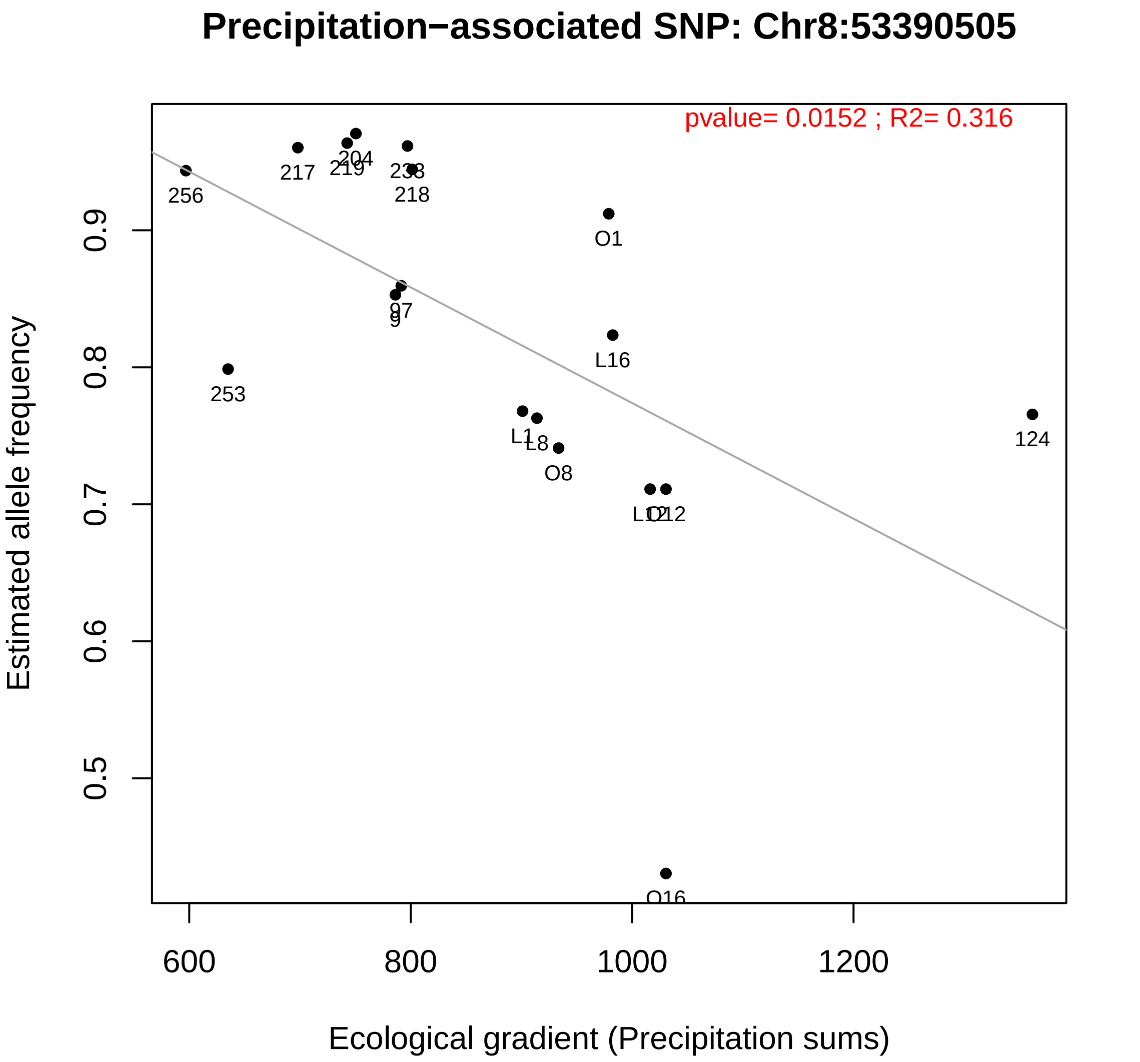
Precipitation−associated SNP: Chr8:53390505
pvalue= 0.0152 ; R2= 0.316
Estimated allele frequency
0.5
0.6
0.7
0.8
0.9
600
800
1000
1200
Ecological gradient (Precipitation sums)
256
217
219
204
218
O1
253
L16
O8
124
O16

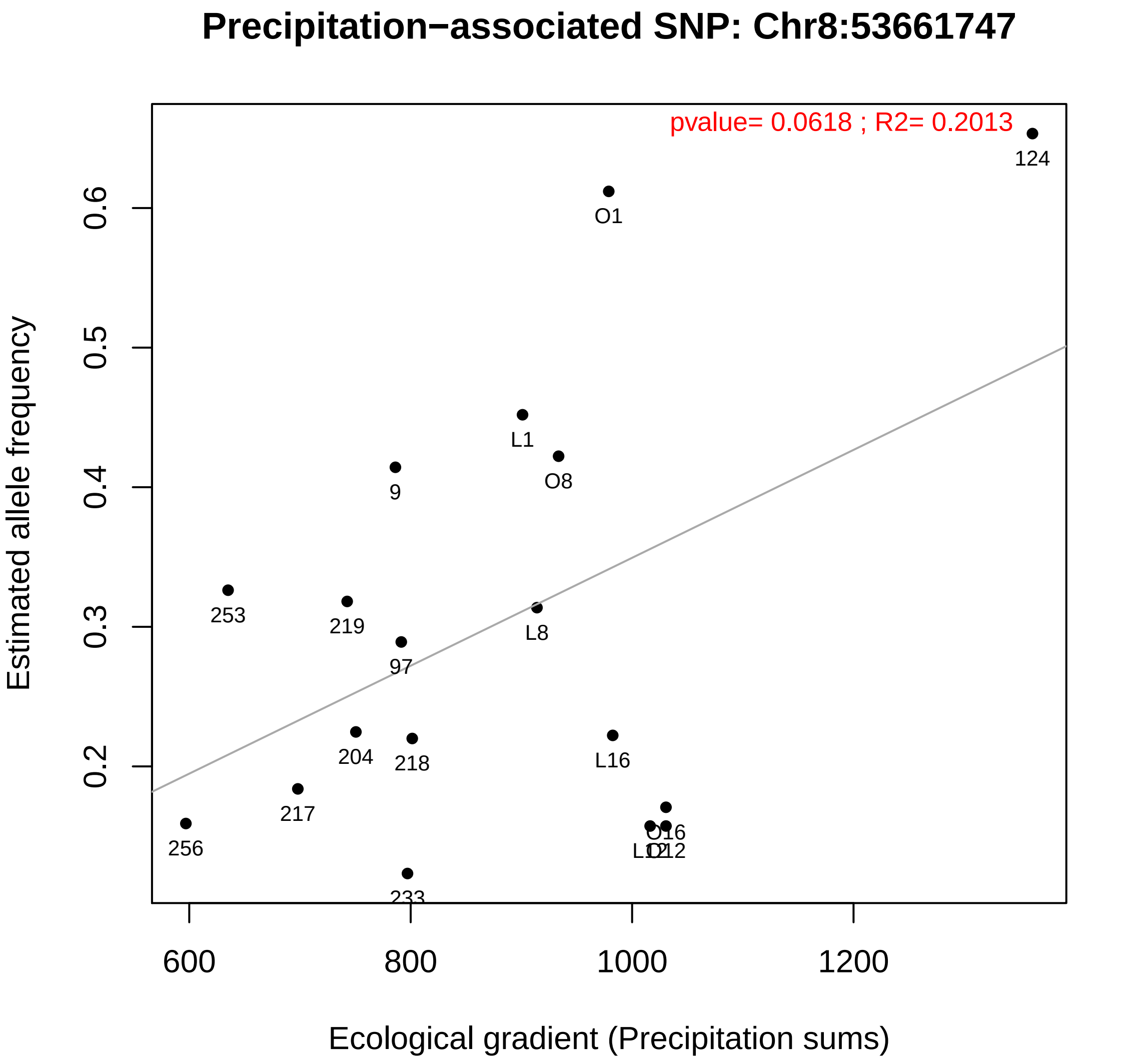
Precipitation−associated SNP: Chr8:53661747
pvalue= 0.0618 ; R2= 0.2013
Estimated allele frequency
0.2
0.3
0.4
0.5
0.6
600
800
1000
1200
Ecological gradient (Precipitation sums)
124
O1
L1
O8
9
253
219
L8
97
204
218
L16
217
256
O16
L12
O12
233

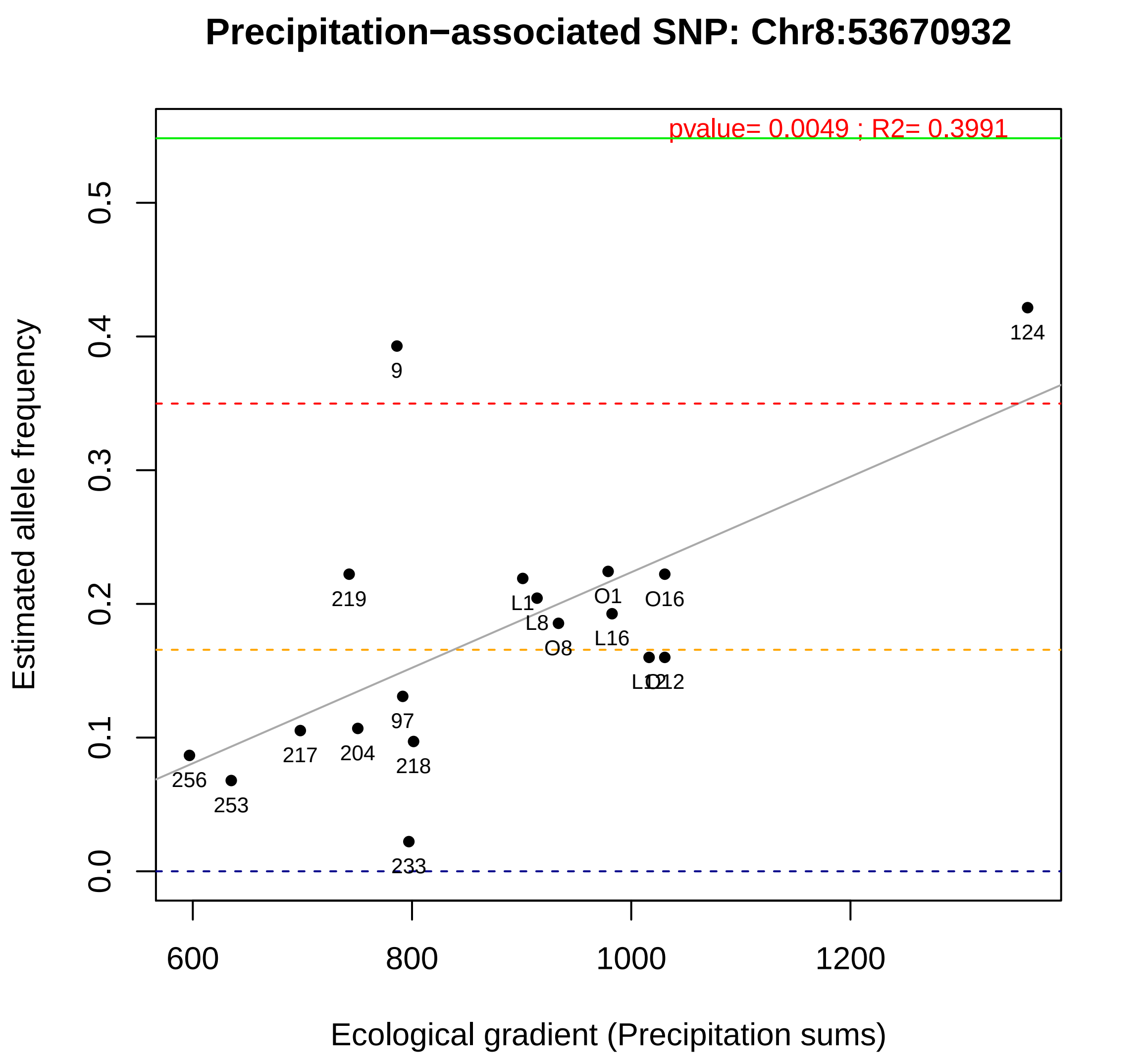

Precipitation−associated SNP: Chr8:53670932
pvalue= 0.0049 ; R2= 0.3991
Estimated allele frequency
0.0
0.1
0.2
0.3
0.4
0.5
600
800
1000
1200
Ecological gradient (Precipitation sums)
124
9
219
L1
L8
O8
O1
L16
O16
L12
O12
97
217
204
218
256
253
233

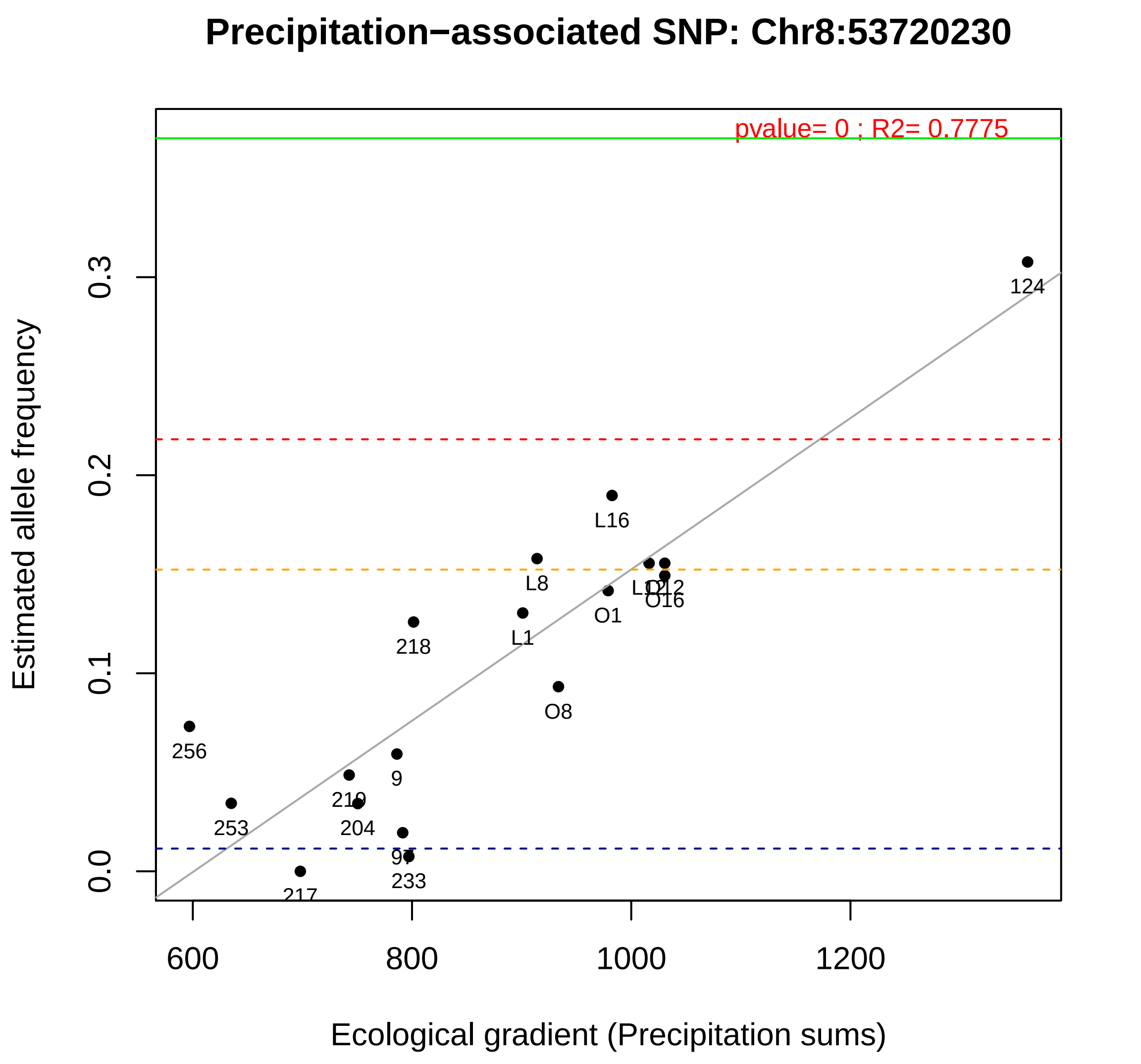

Precipitation−associated SNP: Chr8:53720230
pvalue= 0 ; R2= 0.7775
Estimated allele frequency
0.0
0.1
0.2
0.3
600
800
1000
1200
Ecological gradient (Precipitation sums)
124
L16
L8
O1
O16
L1
218
O8
256
9
253
204
233
217

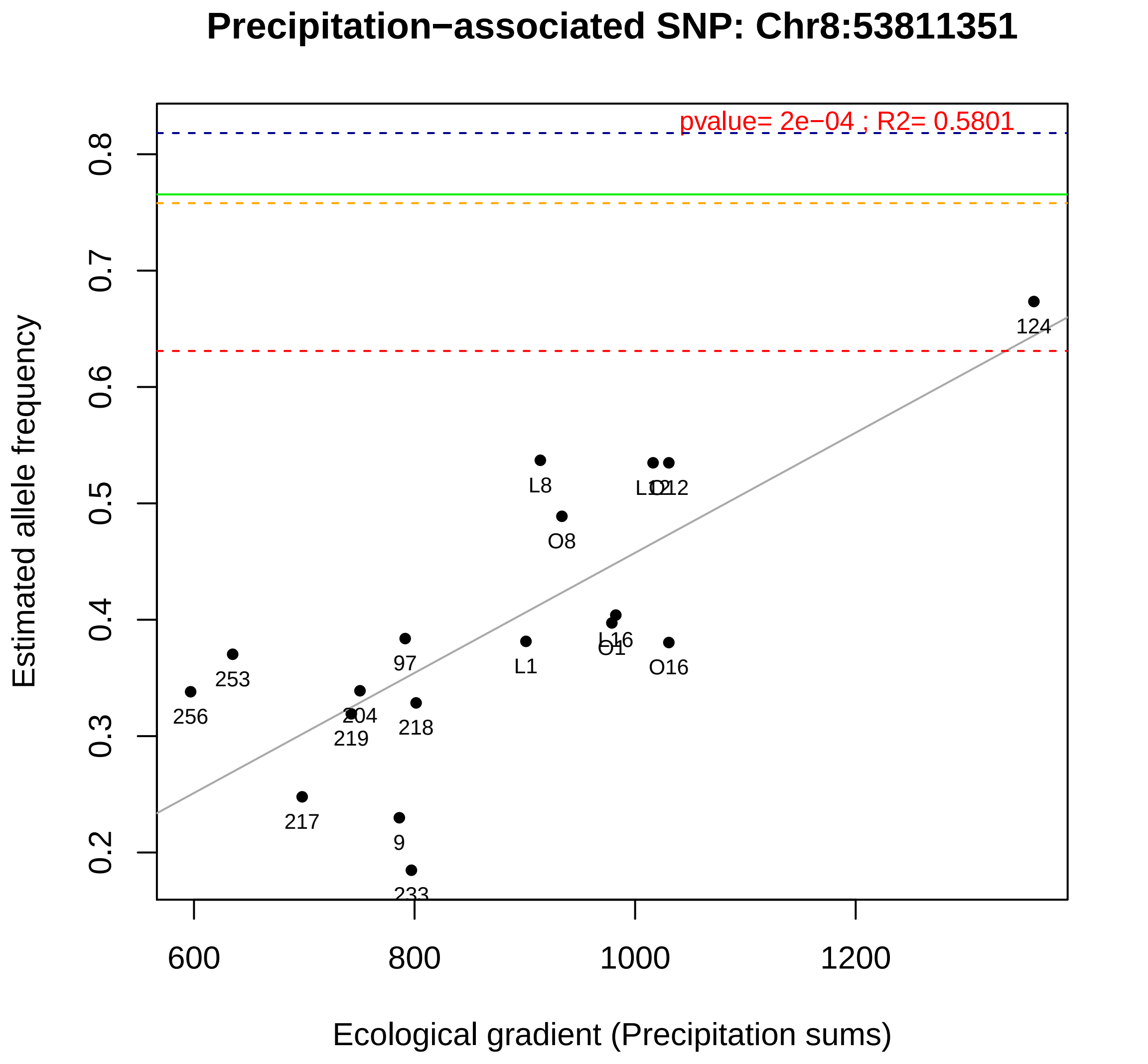

Precipitation−associated SNP: Chr8:53811351
pvalue= 2e−04 ; R2= 0.5801
Estimated allele frequency
0.2
0.3
0.4
0.5
0.6
0.7
0.8
600
800
1000
1200
Ecological gradient (Precipitation sums)
124
L8
L12
O12
O8
L16
O1
L1
O16
97
253
256
204
219
218
217
9
233

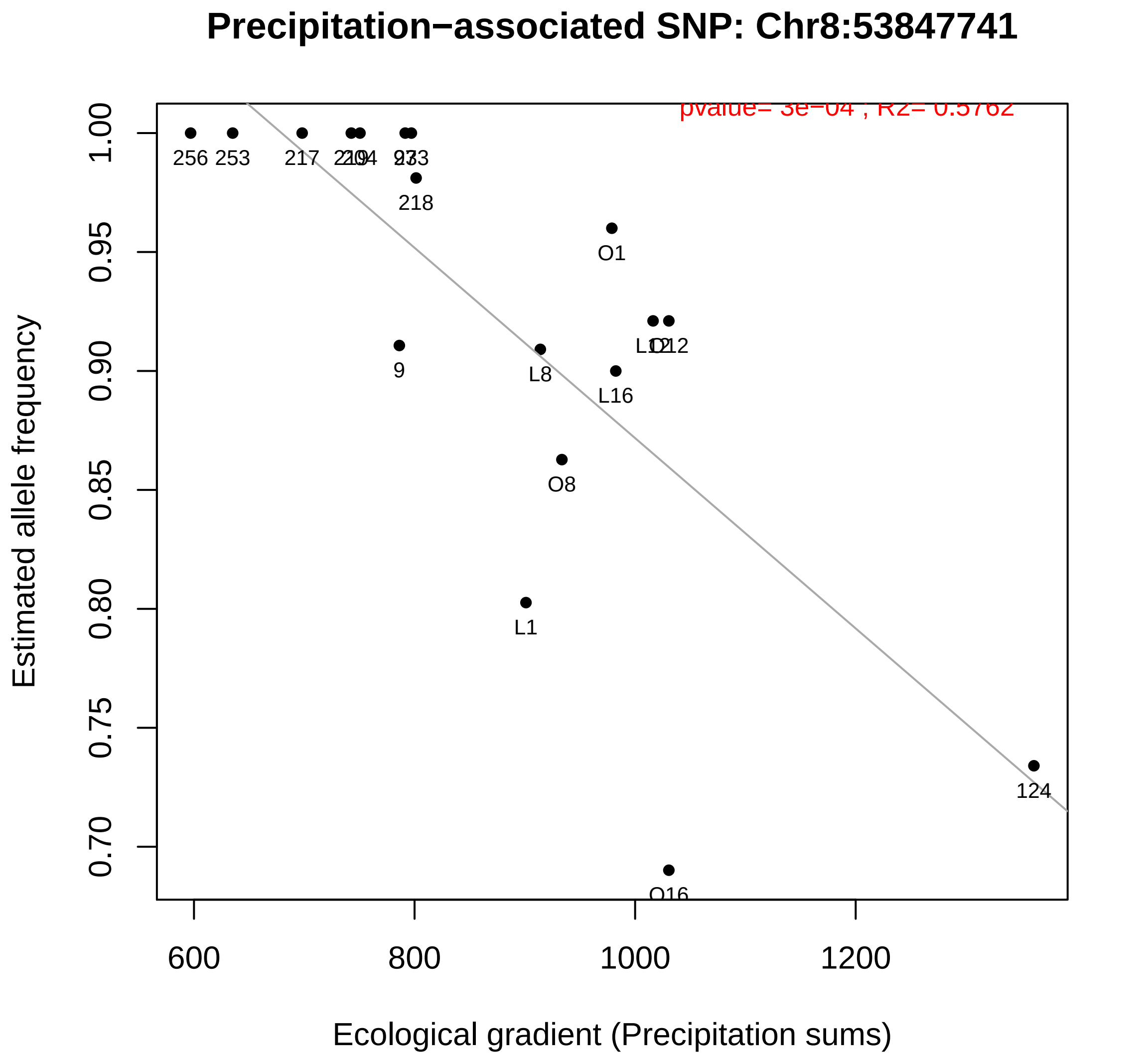
Precipitation−associated SNP: Chr8:53847741
pvalue= 3e−04 ; R2= 0.5762
Estimated allele frequency
Ecological gradient (Precipitation sums)
1.00
0.95
0.90
0.85
0.80
0.75
0.70
600
800
1000
1200
256
253
217
218
O1
9
L8
L16
O8
L1
124
O16

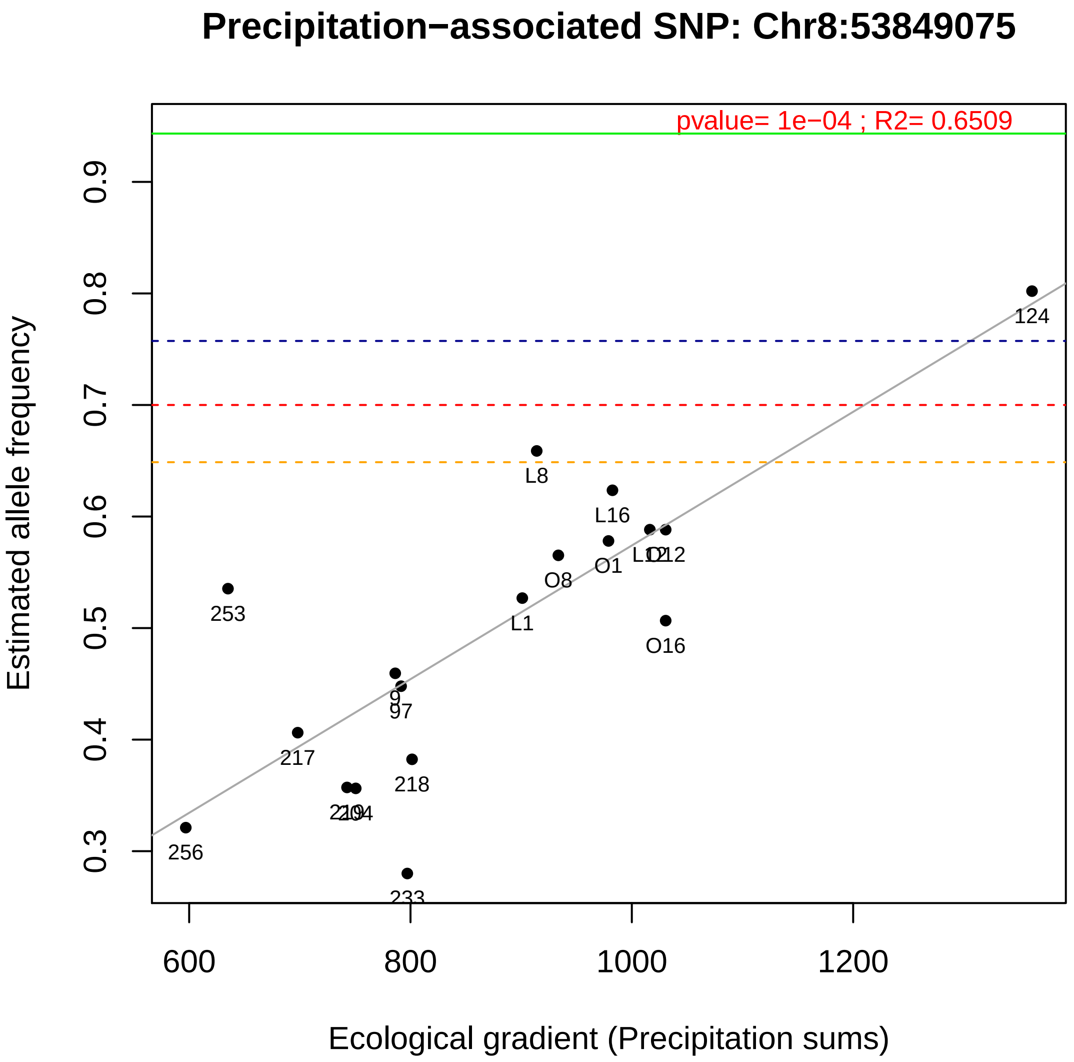
Precipitation−associated SNP: Chr8:53849075
pvalue= 1e−04 ; R2= 0.6509
Estimated allele frequency
0.3
0.4
0.5
0.6
0.7
0.8
0.9
600
800
1000
1200
Ecological gradient (Precipitation sums)
124
L8
L16
L12
O12
O1
O8
253
L1
O16
9
97
217
218
210
204
256
233

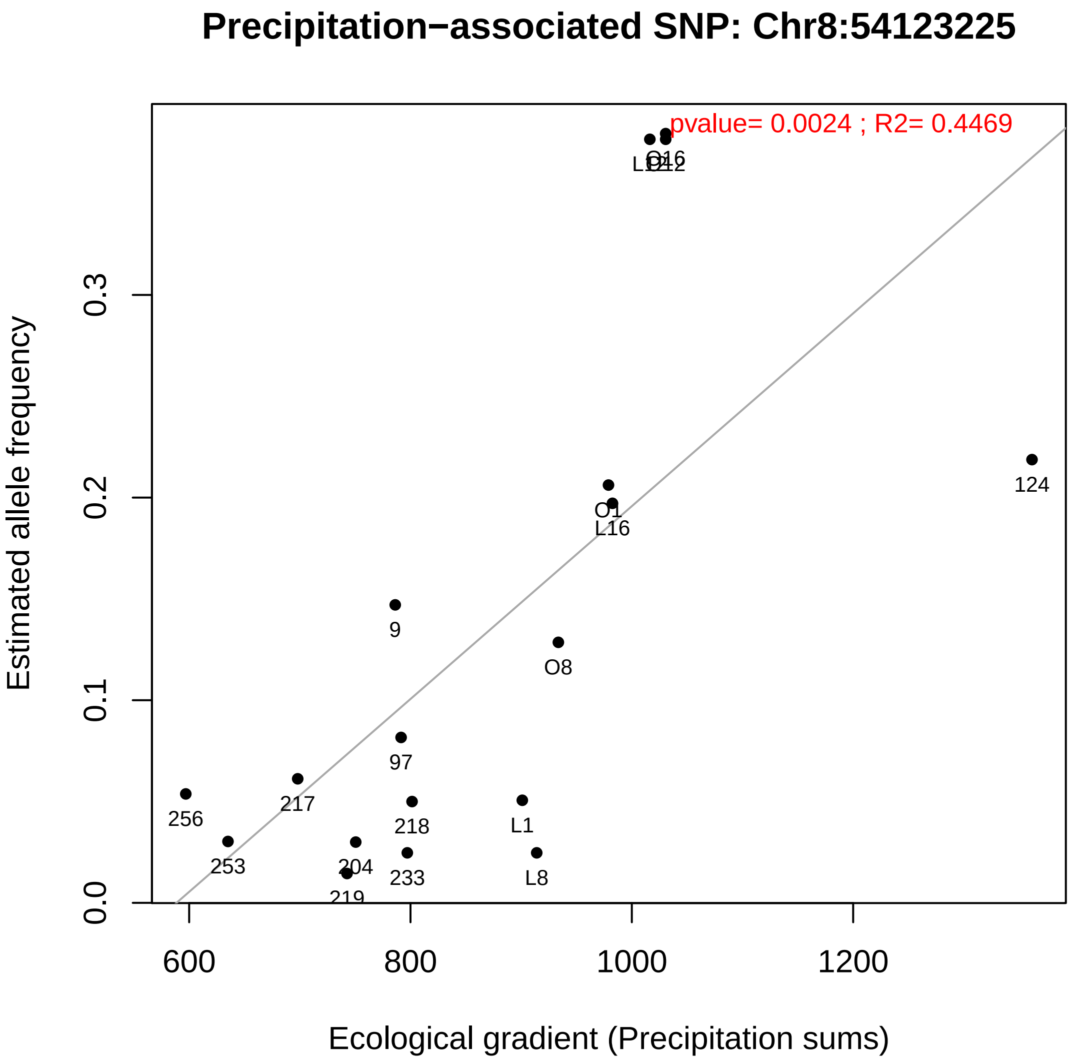

Precipitation−associated SNP: Chr8:54123225
pvalue= 0.0024 ; R2= 0.4469
Estimated allele frequency
Ecological gradient (Precipitation sums)
0.0
0.1
0.2
0.3
600
800
1000
1200
O16
L12
124
O1
L16
9
O8
97
217
256
218
L1
253
204
233
L8
219

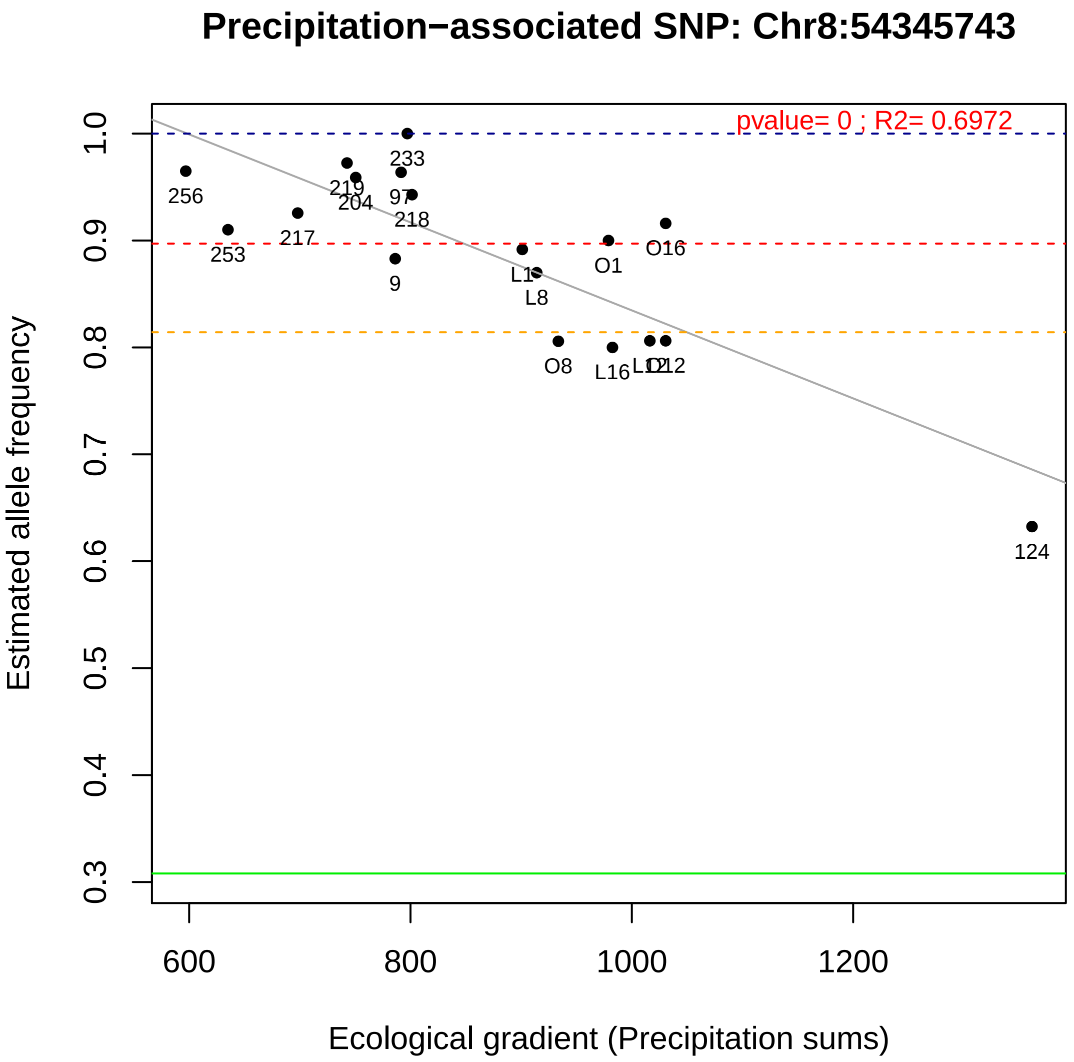

Precipitation−associated SNP: Chr8:54345743
pvalue= 0 ; R2= 0.6972
Estimated allele frequency
1.0
0.9
0.8
0.7
0.6
0.5
0.4
0.3
600
800
1000
1200
Ecological gradient (Precipitation sums)
256
253
217
219
204
233
97
218
9
L1
L8
O1
O16
O8
L16
124

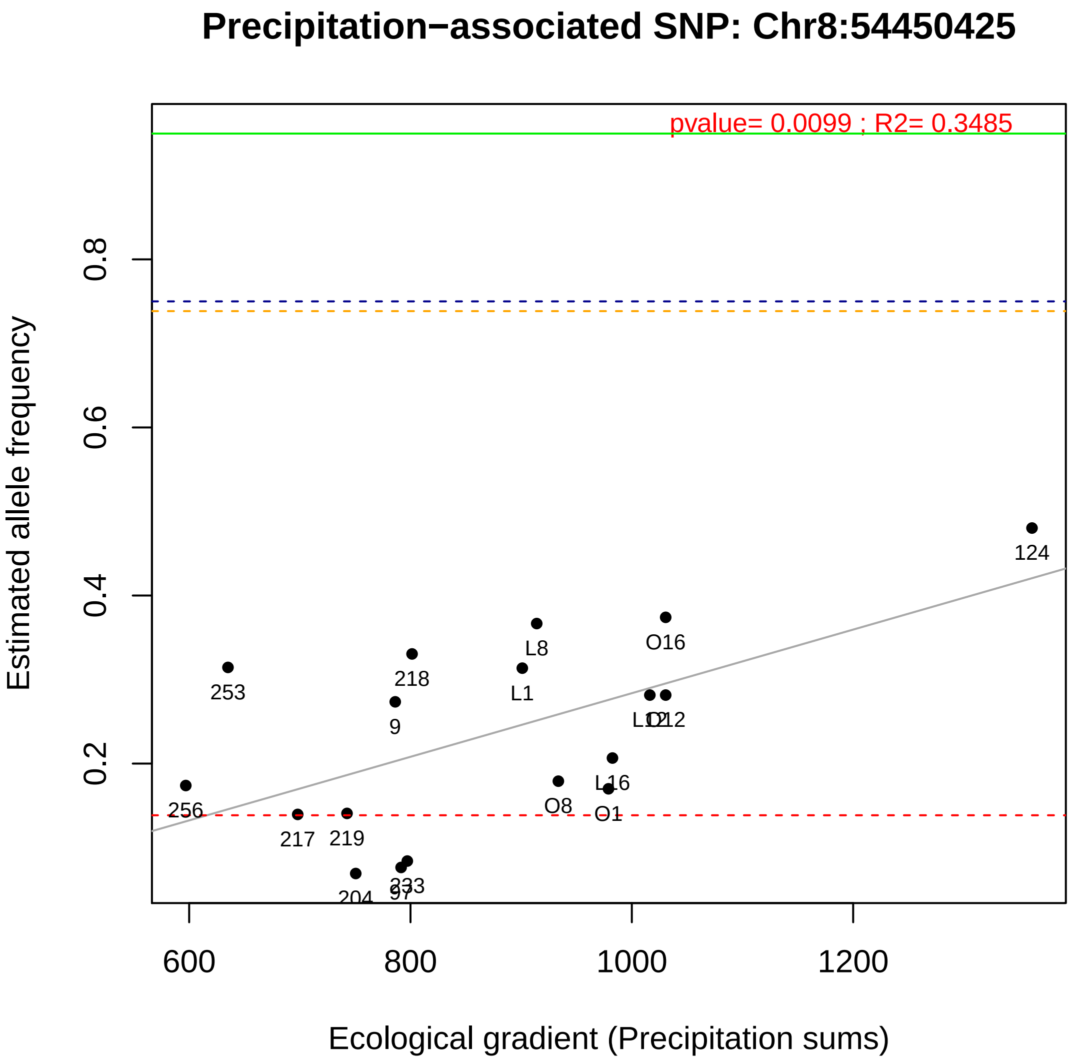
Precipitation−associated SNP: Chr8:54450425
pvalue= 0.0099 ; R2= 0.3485
Estimated allele frequency
0.2
0.4
0.6
0.8
600
800
1000
1200
Ecological gradient (Precipitation sums)
124
O16
L8
218
253
L1
9
L12
O12
L16
O8
O1
256
217
219
233
97
204

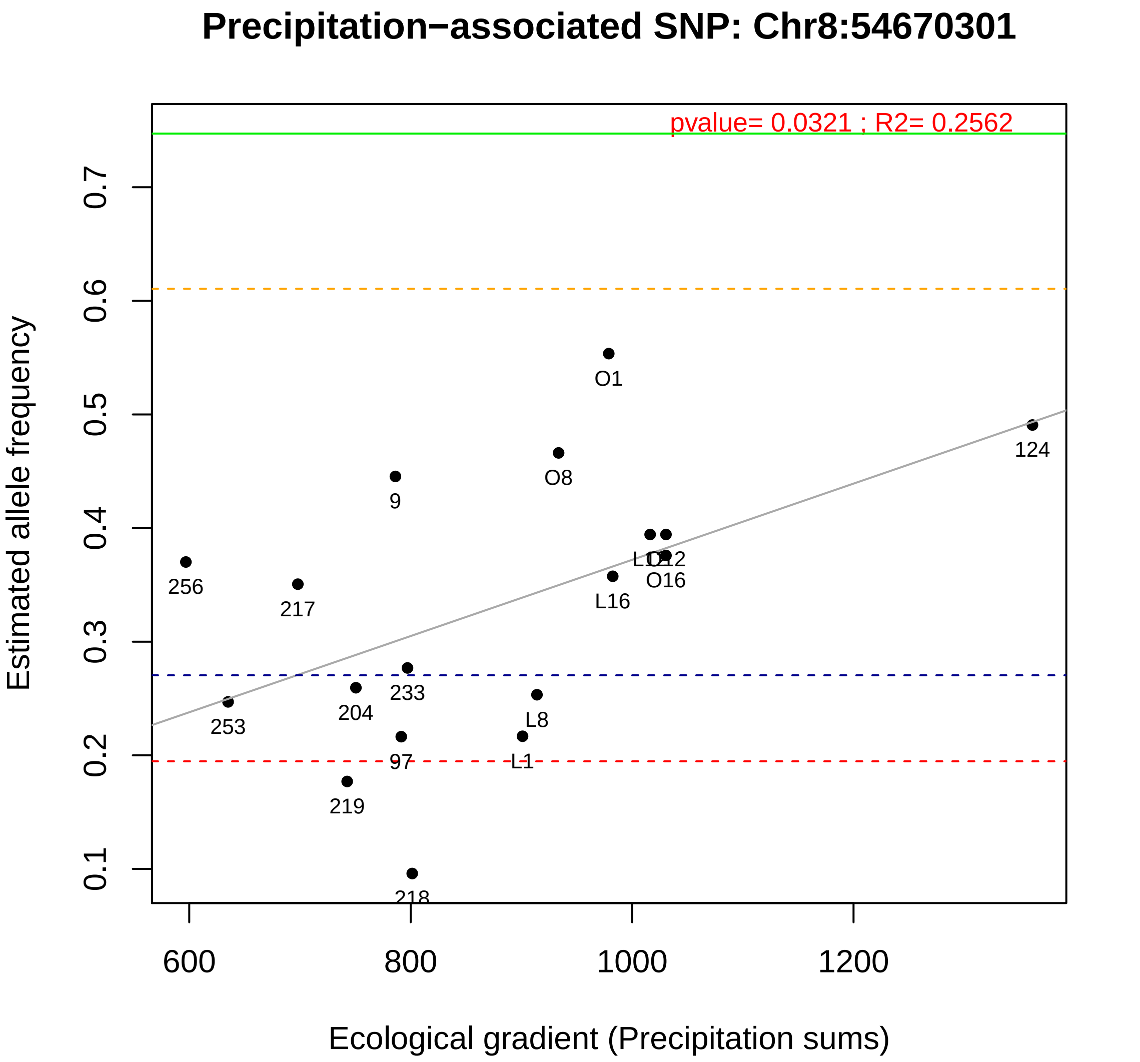
Precipitation−associated SNP: Chr8:54670301
pvalue= 0.0321 ; R2= 0.2562
Estimated allele frequency
0.1
0.2
0.3
0.4
0.5
0.6
0.7
600
800
1000
1200
Ecological gradient (Precipitation sums)
O1
124
O8
9
256
217
L16
O16
233
204
253
L8
97
L1
219
218

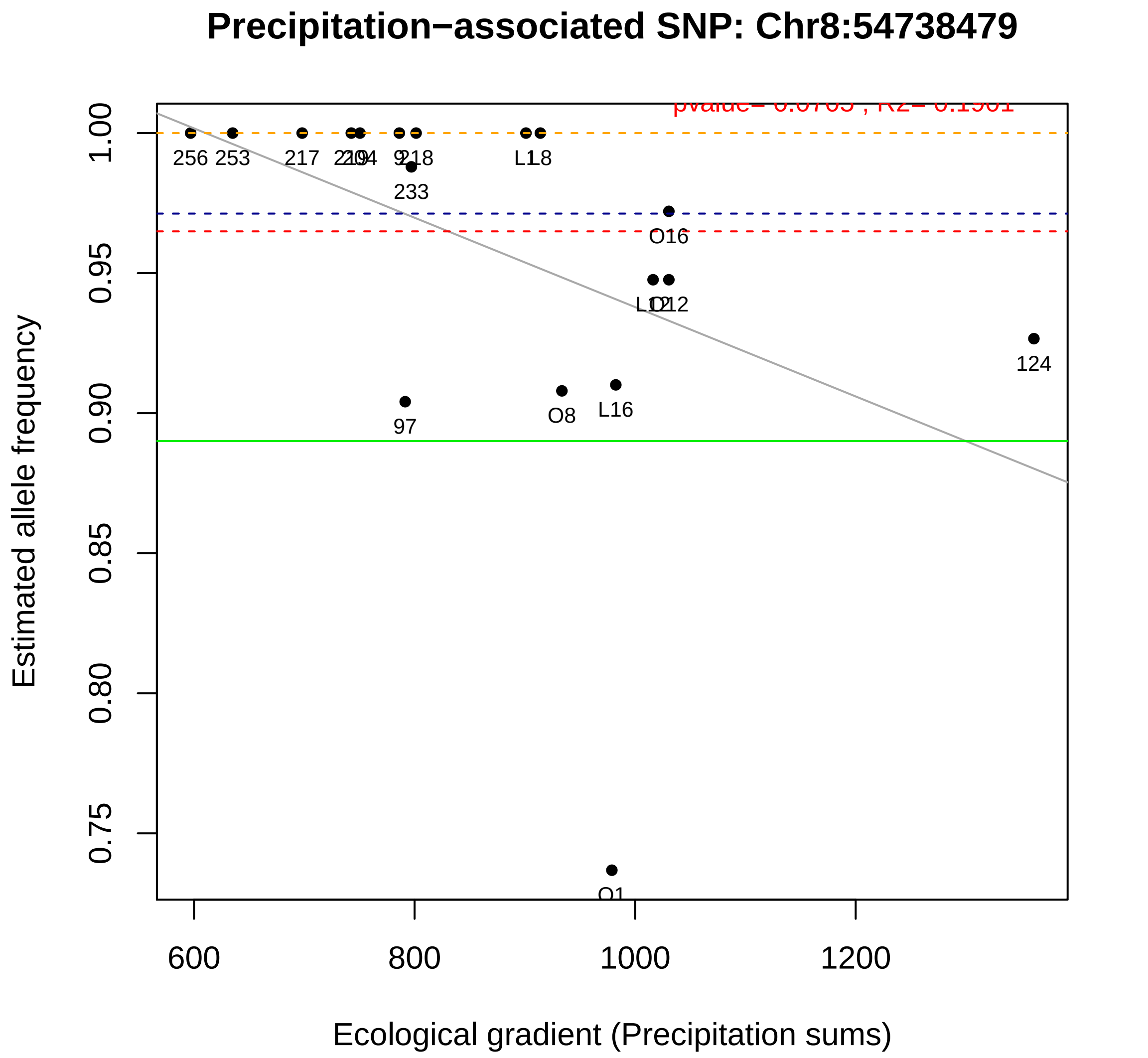
Precipitation−associated SNP: Chr8:54738479
Estimated allele frequency
Ecological gradient (Precipitation sums)
1.00
0.95
0.90
0.85
0.80
0.75
600
800
1000
1200
256
253
217
233
O16
124
97
O8
L16
O1

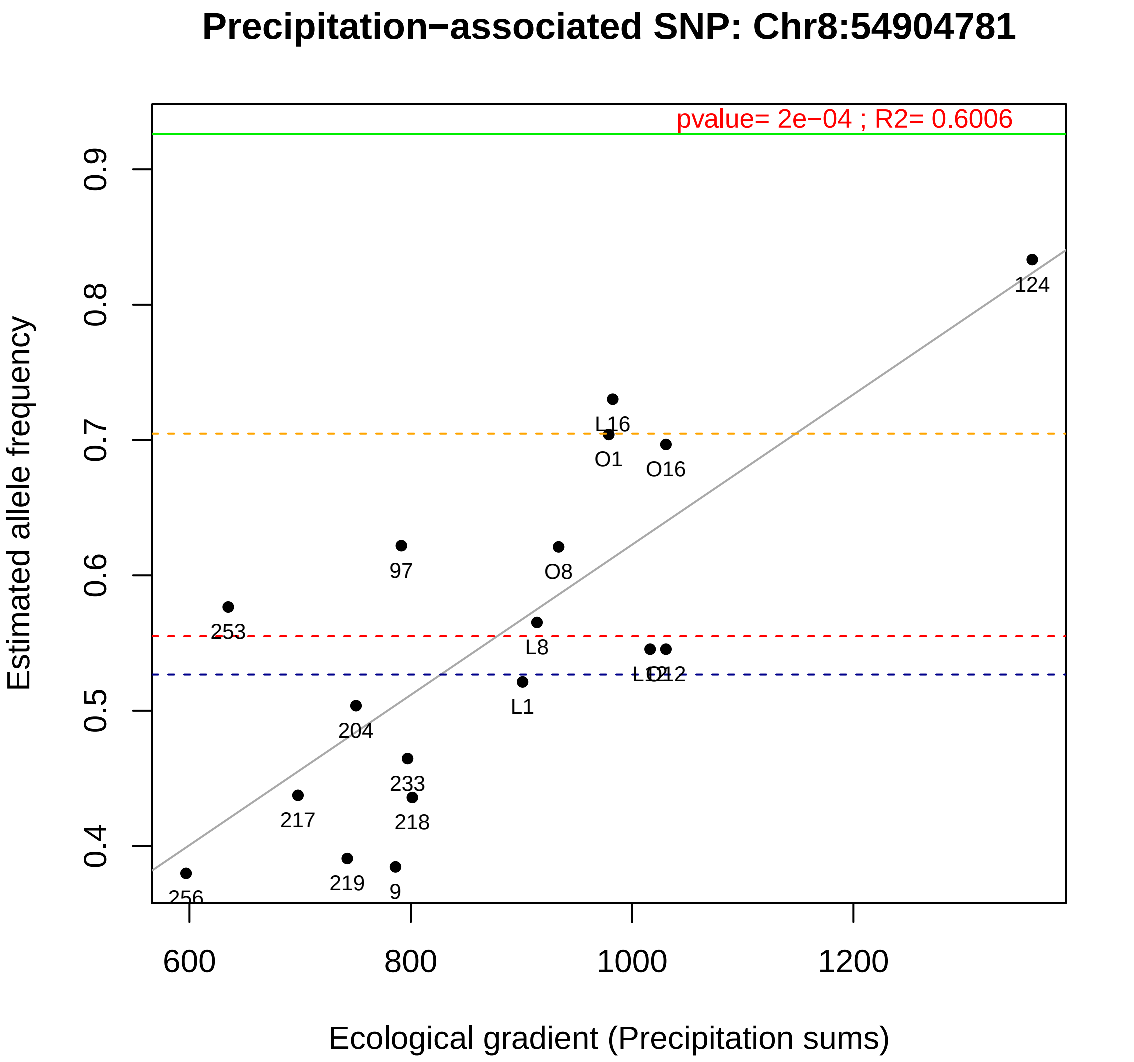
Precipitation−associated SNP: Chr8:54904781
pvalue= 2e−04 ; R2= 0.6006
Estimated allele frequency
0.4
0.5
0.6
0.7
0.8
0.9
600
800
1000
1200
Ecological gradient (Precipitation sums)
124
L16
O1
O16
97
O8
253
L8
L1
204
233
217
218
219
9
256

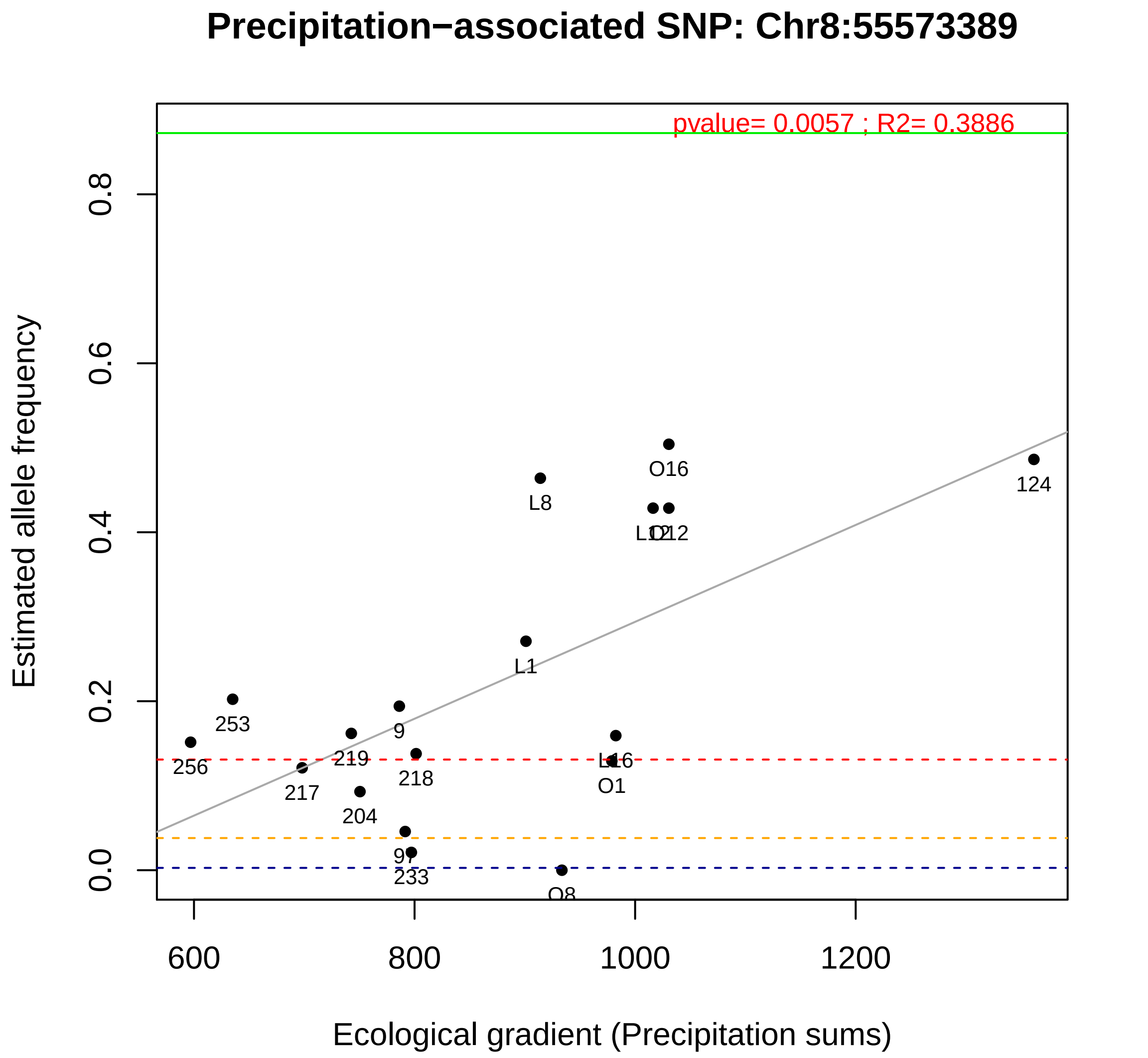

Precipitation−associated SNP: Chr8:55573389
pvalue= 0.0057 ; R2= 0.3886
Estimated allele frequency
0.0
0.2
0.4
0.6
0.8
600
800
1000
1200
Ecological gradient (Precipitation sums)
O16
124
L8
L12
O12
L1
253
9
256
219
218
L16
O1
217
204
97
233
O8

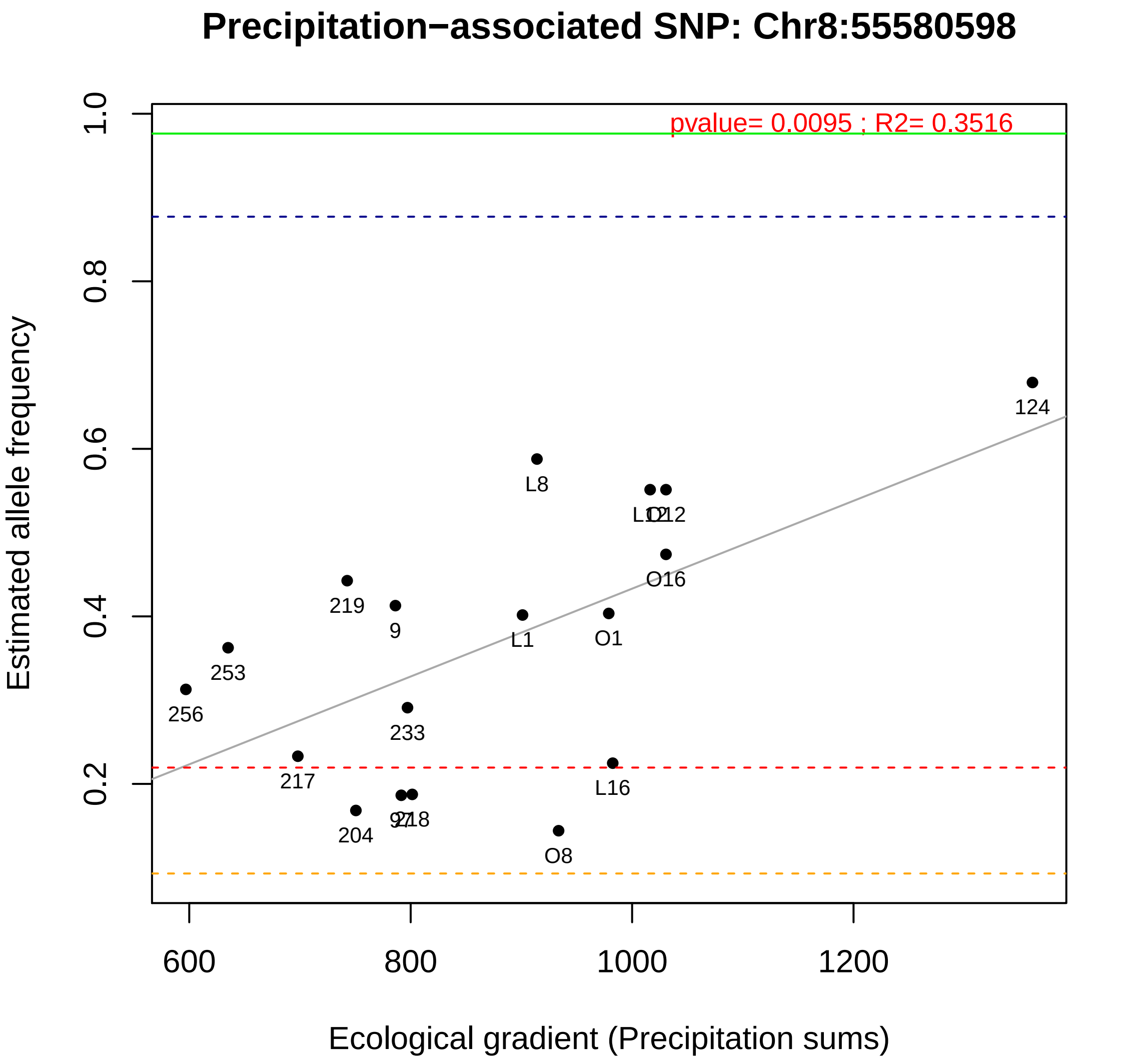

Precipitation−associated SNP: Chr8:55580598
pvalue= 0.0095 ; R2= 0.3516
Estimated allele frequency
0.2
0.4
0.6
0.8
1.0
600
800
1000
1200
Ecological gradient (Precipitation sums)
124
L8
O16
219
9
L1
O1
253
256
233
217
L16
204
218
O8

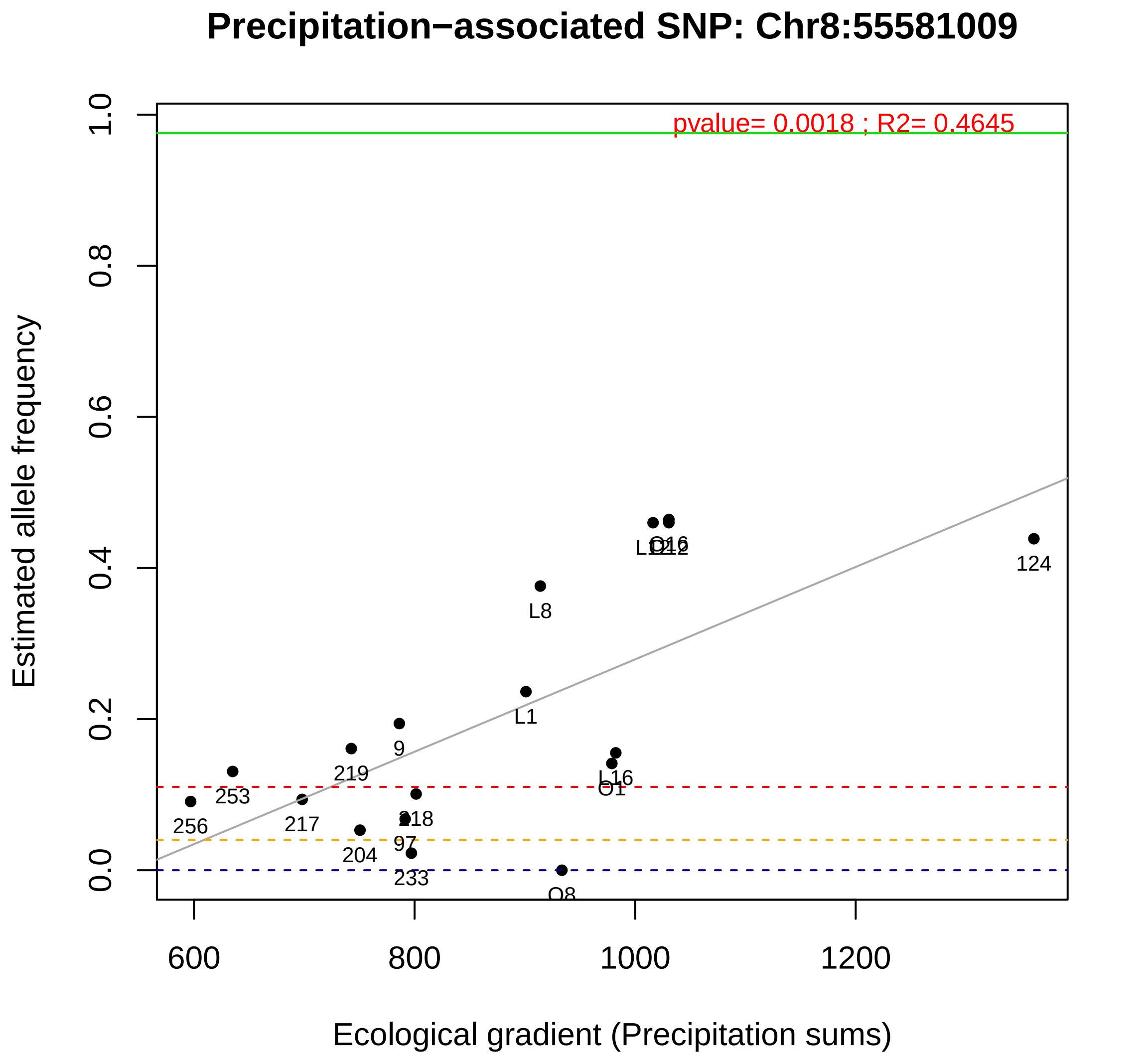

Precipitation−associated SNP: Chr8:55581009
pvalue= 0.0018 ; R2= 0.4645
Estimated allele frequency
Ecological gradient (Precipitation sums)
0.0
0.2
0.4
0.6
0.8
1.0
600
800
1000
1200
256
253
217
219
204
9
218
97
233
L1
L8
O8
L16
O1
124

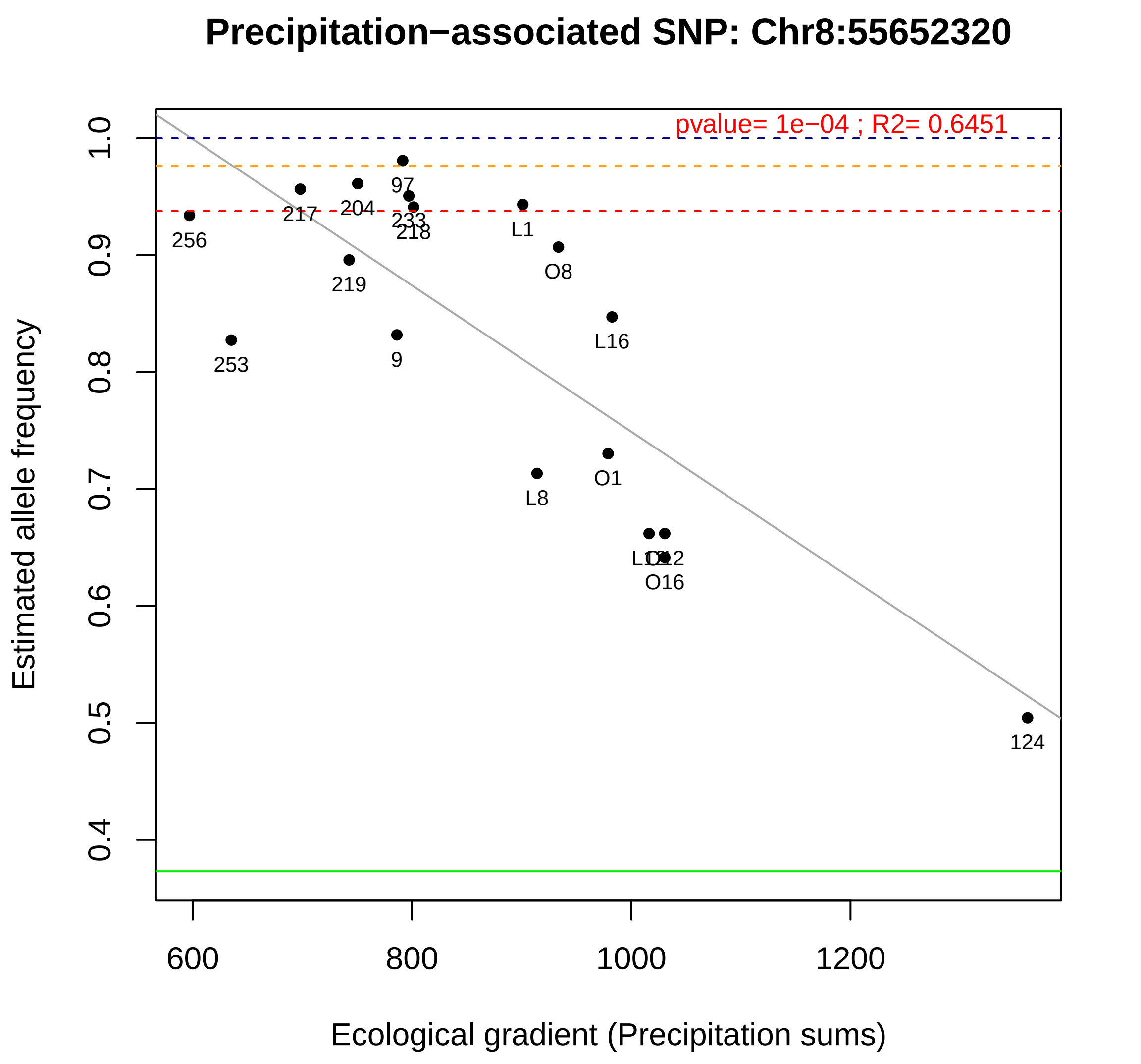
Precipitation−associated SNP: Chr8:55652320
pvalue= 1e−04 ; R2= 0.6451
Estimated allele frequency
Ecological gradient (Precipitation sums)
0.4
0.5
0.6
0.7
0.8
0.9
1.0
600
800
1000
1200
256
217
204
97
233
218
L1
O8
219
253
9
L16
O1
L8
O16
124

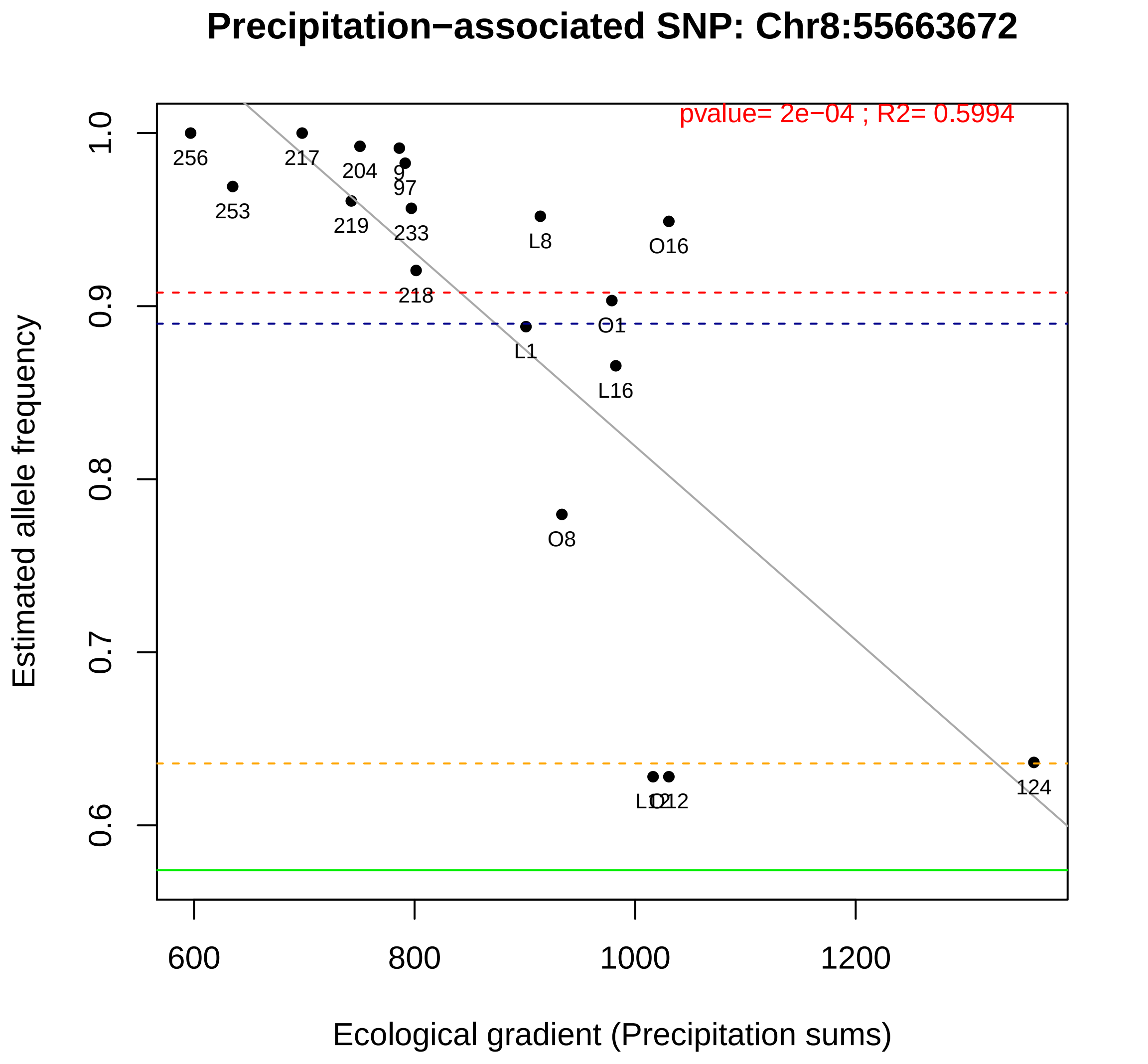

Precipitation−associated SNP: Chr8:55663672
pvalue= 2e−04 ; R2= 0.5994
256
217
204
9
97
253
219
233
L8
O16
218
O1
L1
L16
O8
L12
O12
124
1.0
0.9
0.8
0.7
0.6
600
800
1000
1200
Estimated allele frequency
Ecological gradient (Precipitation sums)

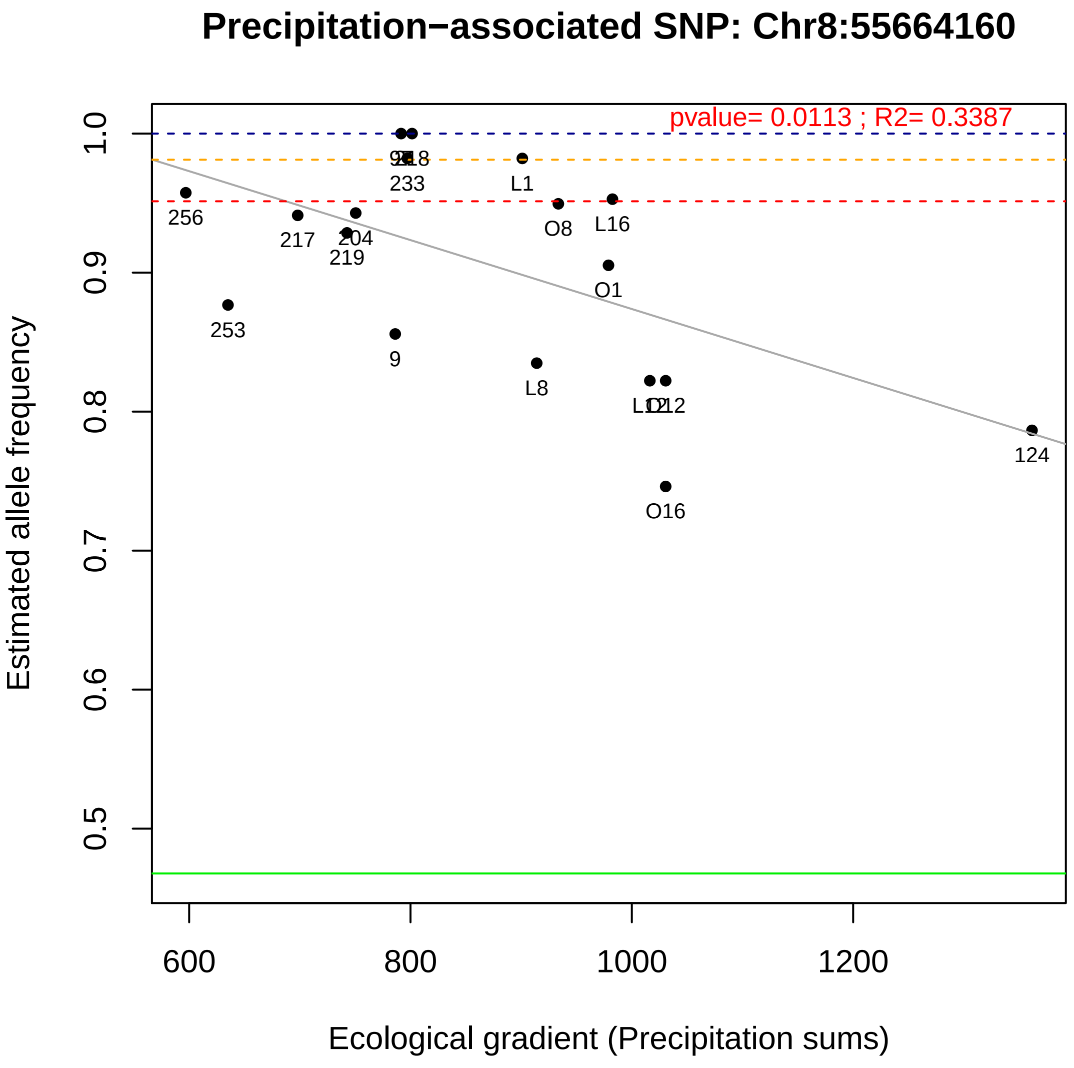

Precipitation−associated SNP: Chr8:55664160
pvalue= 0.0113 ; R2= 0.3387
Estimated allele frequency
0.5
0.6
0.7
0.8
0.9
1.0
600
800
1000
1200
Ecological gradient (Precipitation sums)
256
253
217
219
204
9
233
L1
O8
L16
O1
L8
O12
124
O16

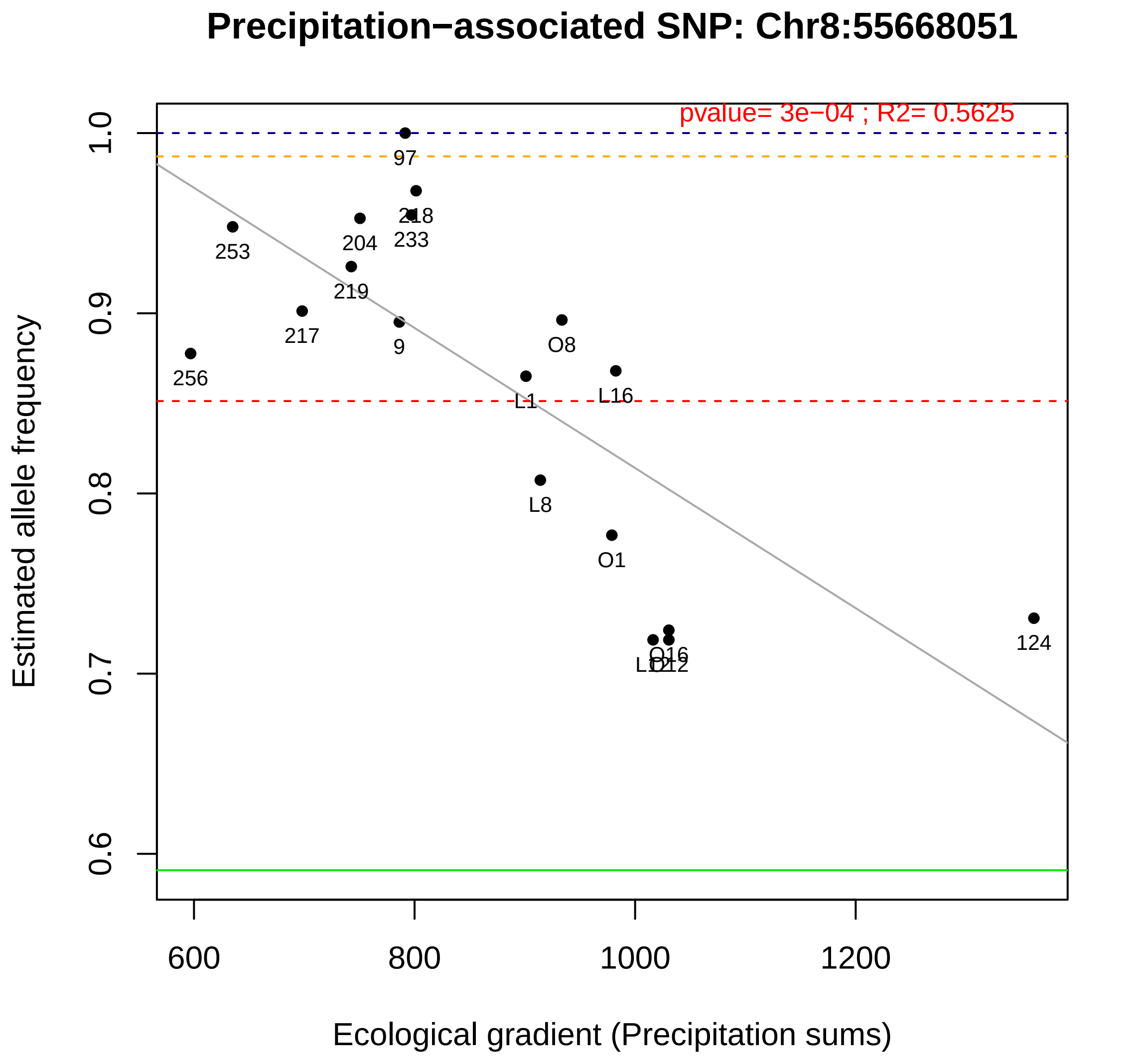

Precipitation−associated SNP: Chr8:55668051
pvalue= 3e−04 ; R2= 0.5625
Estimated allele frequency
Ecological gradient (Precipitation sums)
0.6
0.7
0.8
0.9
1.0
600
800
1000
1200
97
218
233
204
253
219
217
9
256
O8
L1
L16
L8
O1
O16
O12
124

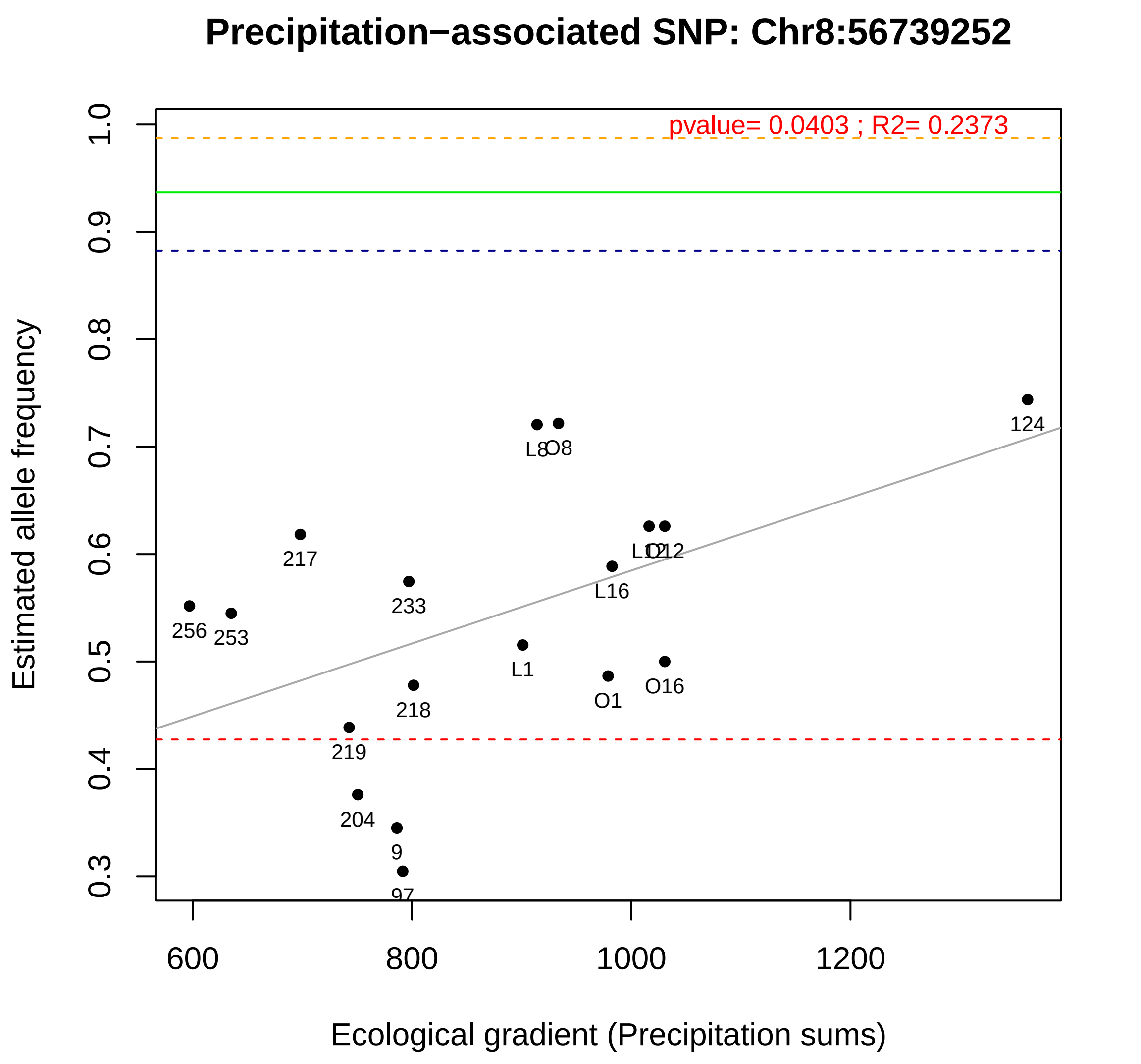

Precipitation−associated SNP: Chr8:56739252
pvalue= 0.0403 ; R2= 0.2373
Estimated allele frequency
Ecological gradient (Precipitation sums)
1.0
0.9
0.8
0.7
0.6
0.5
0.4
0.3
600
800
1000
1200
124
L8
O8
217
L12
O12
L16
233
256
253
L1
O16
O1
218
219
204
9
97

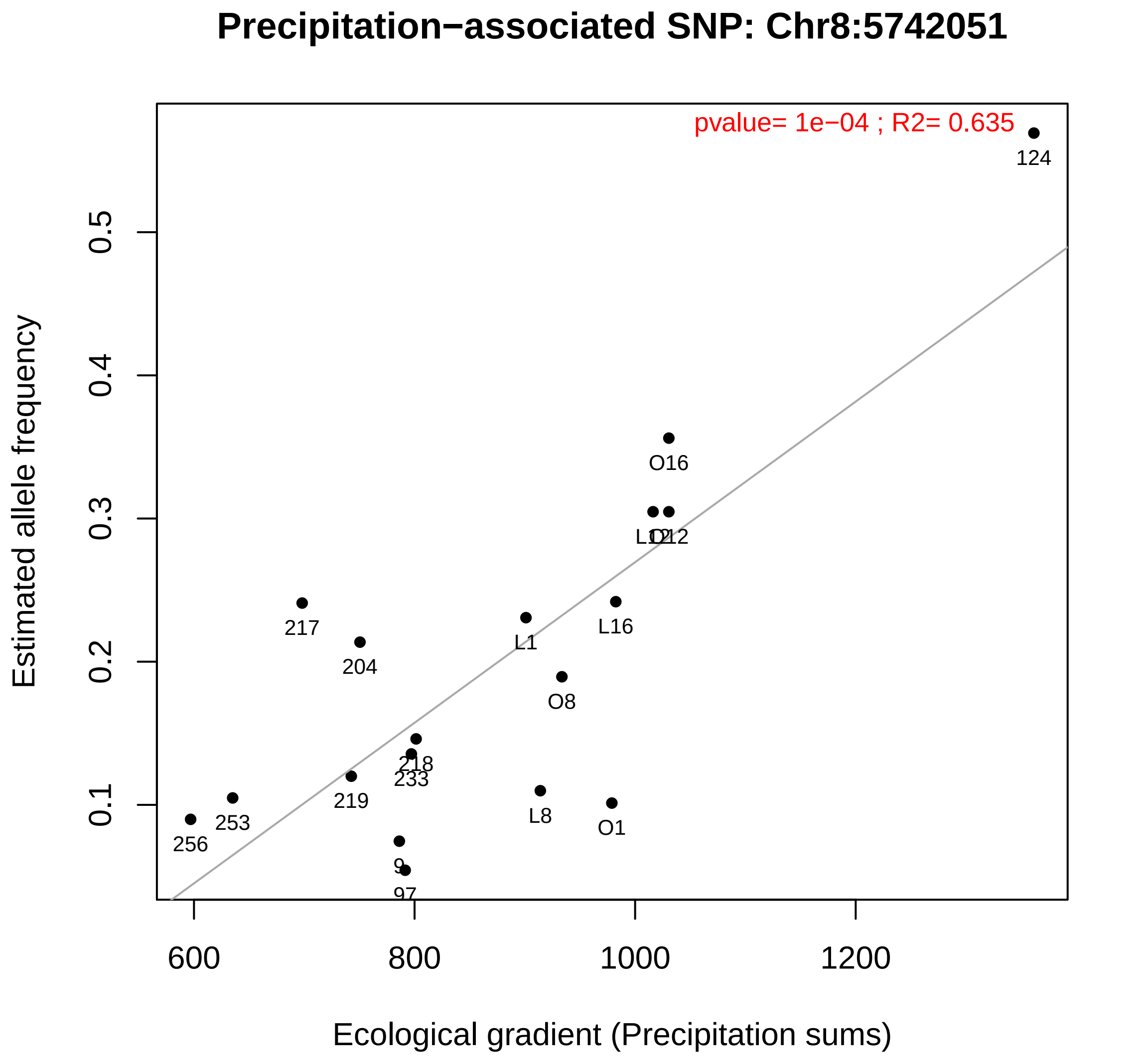

Precipitation−associated SNP: Chr8:5742051
pvalue= 1e−04 ; R2= 0.635
Estimated allele frequency
Ecological gradient (Precipitation sums)
0.1
0.2
0.3
0.4
0.5
600
800
1000
1200
124
O16
L12
O12
217
204
L1
L16
O8
218
233
219
253
256
L8
O1
9
97

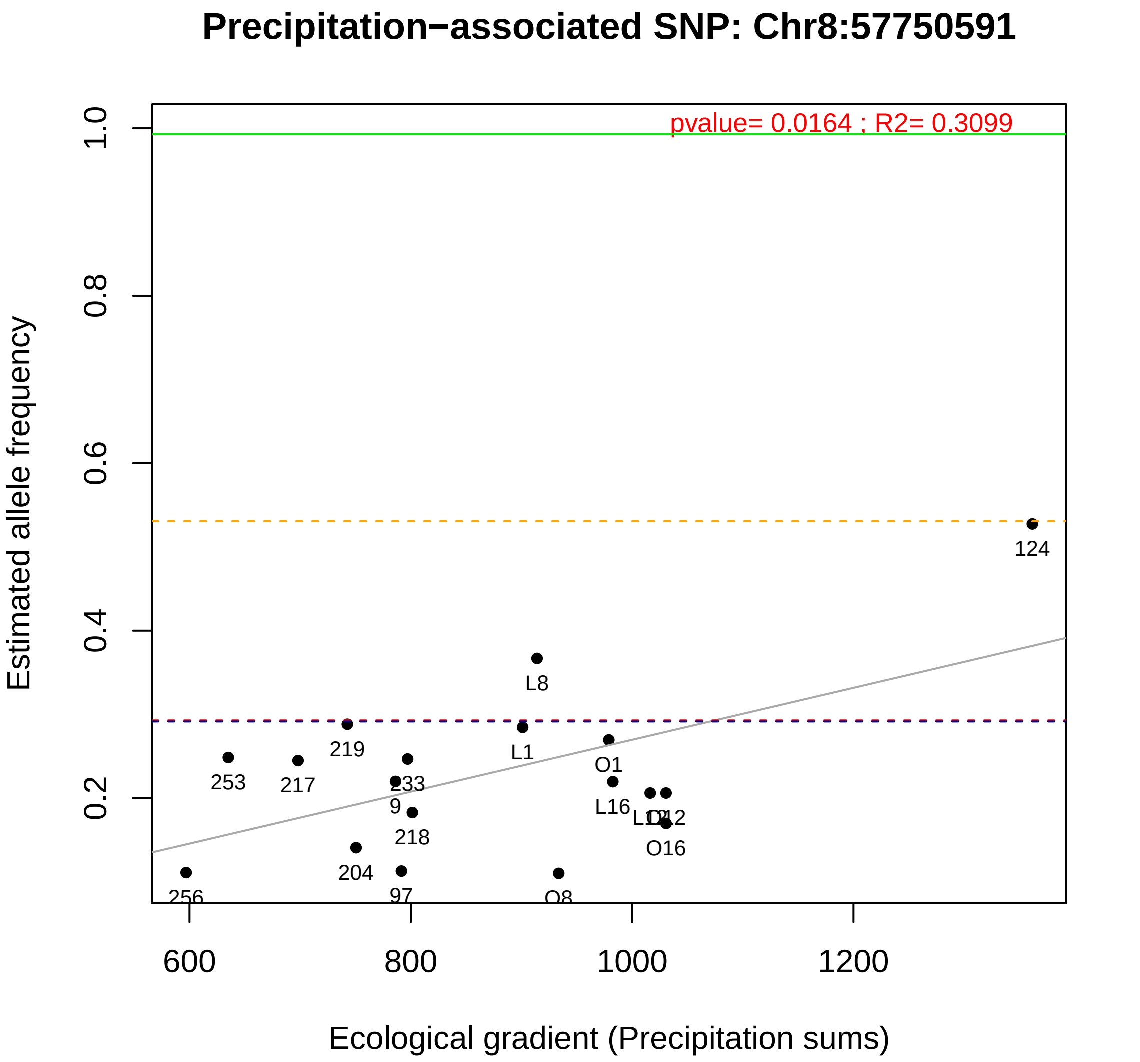

Precipitation−associated SNP: Chr8:57750591
pvalue= 0.0164 ; R2= 0.3099
Estimated allele frequency
Ecological gradient (Precipitation sums)
0.2
0.4
0.6
0.8
1.0
600
800
1000
1200
124
L8
219
L1
253
217
O1
233
9
L16
218
O16
204
256
97
O8

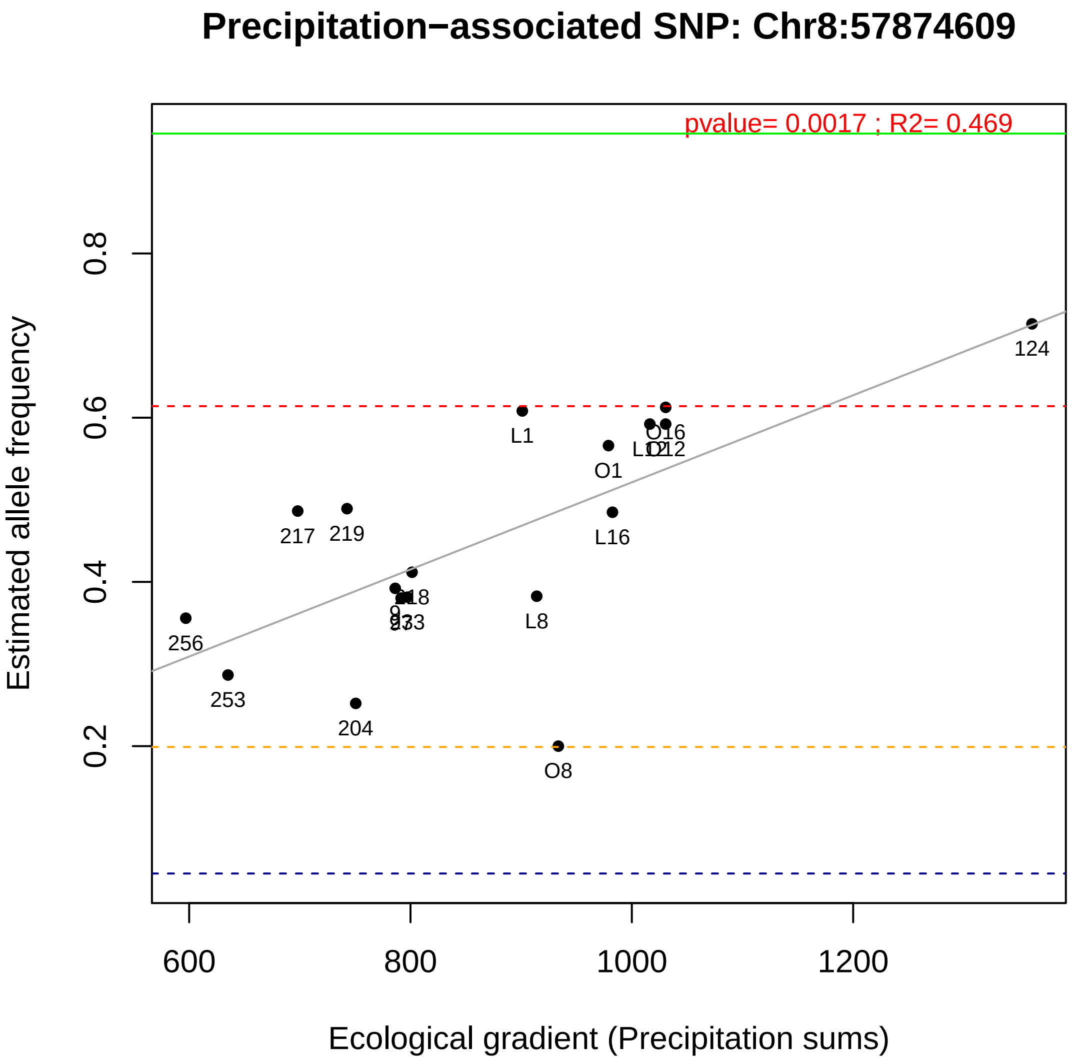

Precipitation−associated SNP: Chr8:57874609
pvalue= 0.0017 ; R2= 0.469
Estimated allele frequency
0.2
0.4
0.6
0.8
600
800
1000
1200
Ecological gradient (Precipitation sums)
124
L1
O16
L12
O12
O1
L16
217
219
L8
256
253
204
O8

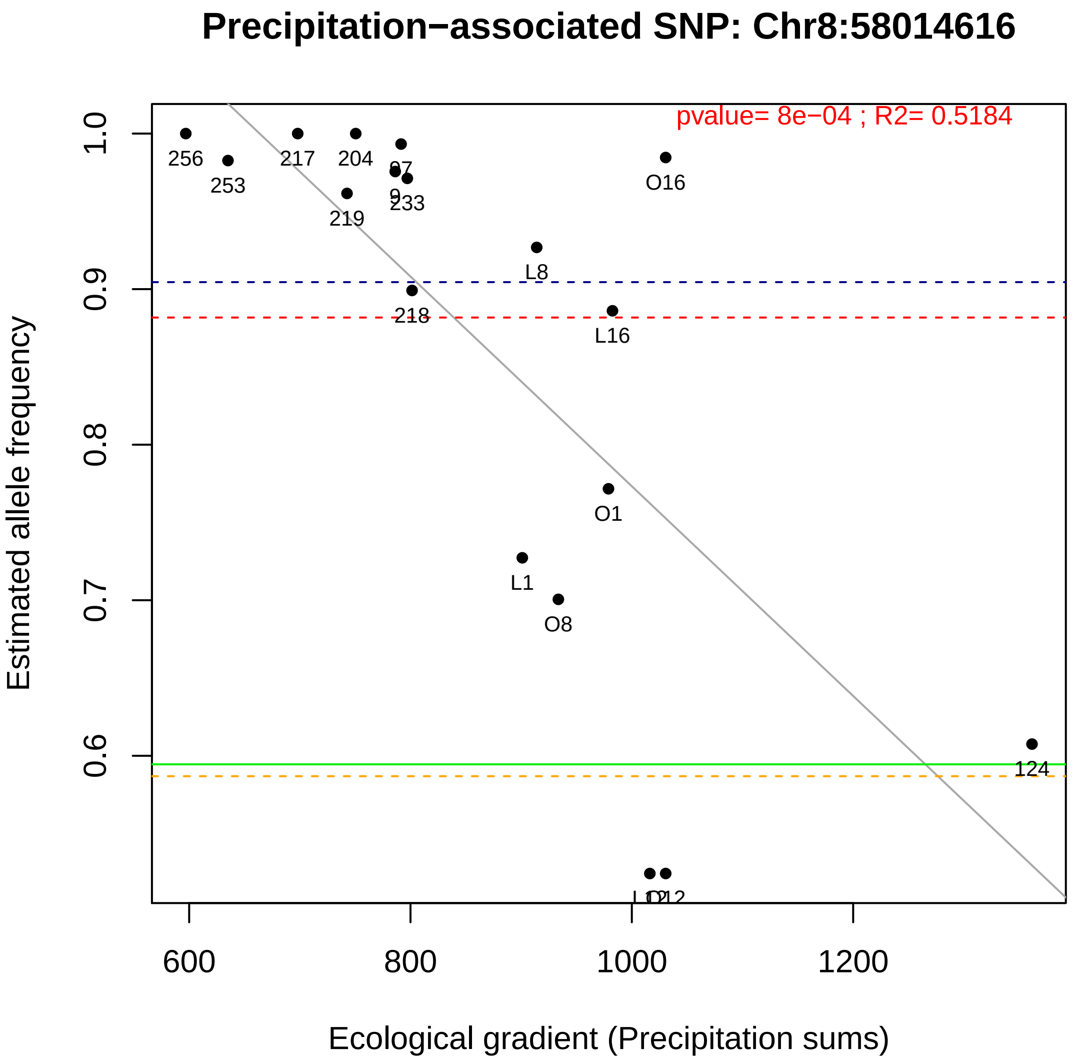
Precipitation−associated SNP: Chr8:58014616
pvalue= 8e−04 ; R2= 0.5184
Estimated allele frequency
Ecological gradient (Precipitation sums)
1.0
0.9
0.8
0.7
0.6
600
800
1000
1200
256
253
217
204
219
233
O16
L8
218
L16
O1
L1
O8
124

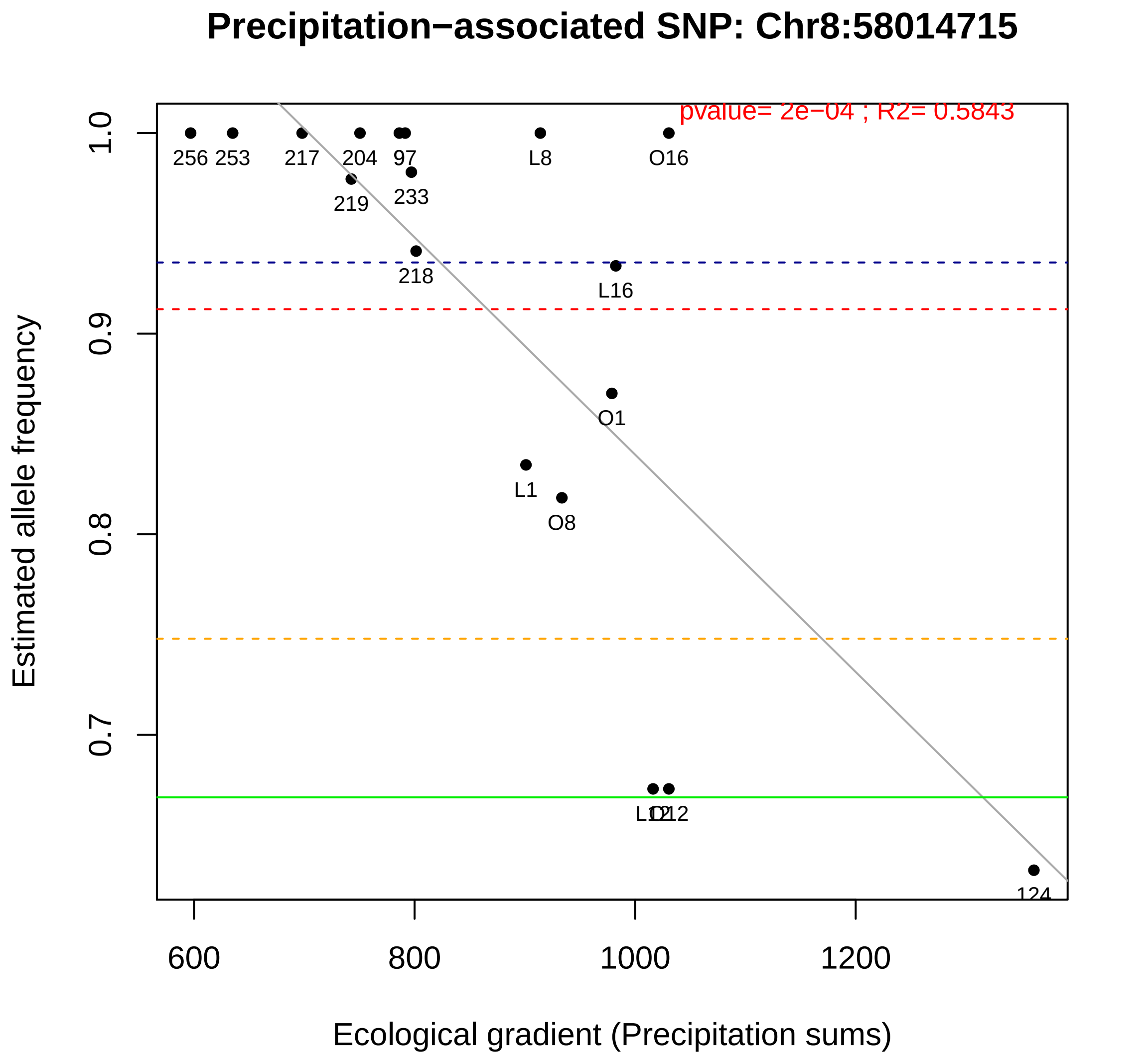
Precipitation−associated SNP: Chr8:58014715
pvalue= 2e−04 ; R2= 0.5843
Estimated allele frequency
1.0
0.9
0.8
0.7
256
253
217
204
97
L8
O16
219
233
218
L16
O1
L1
O8
L12
O12
124
600
800
1000
1200
Ecological gradient (Precipitation sums)

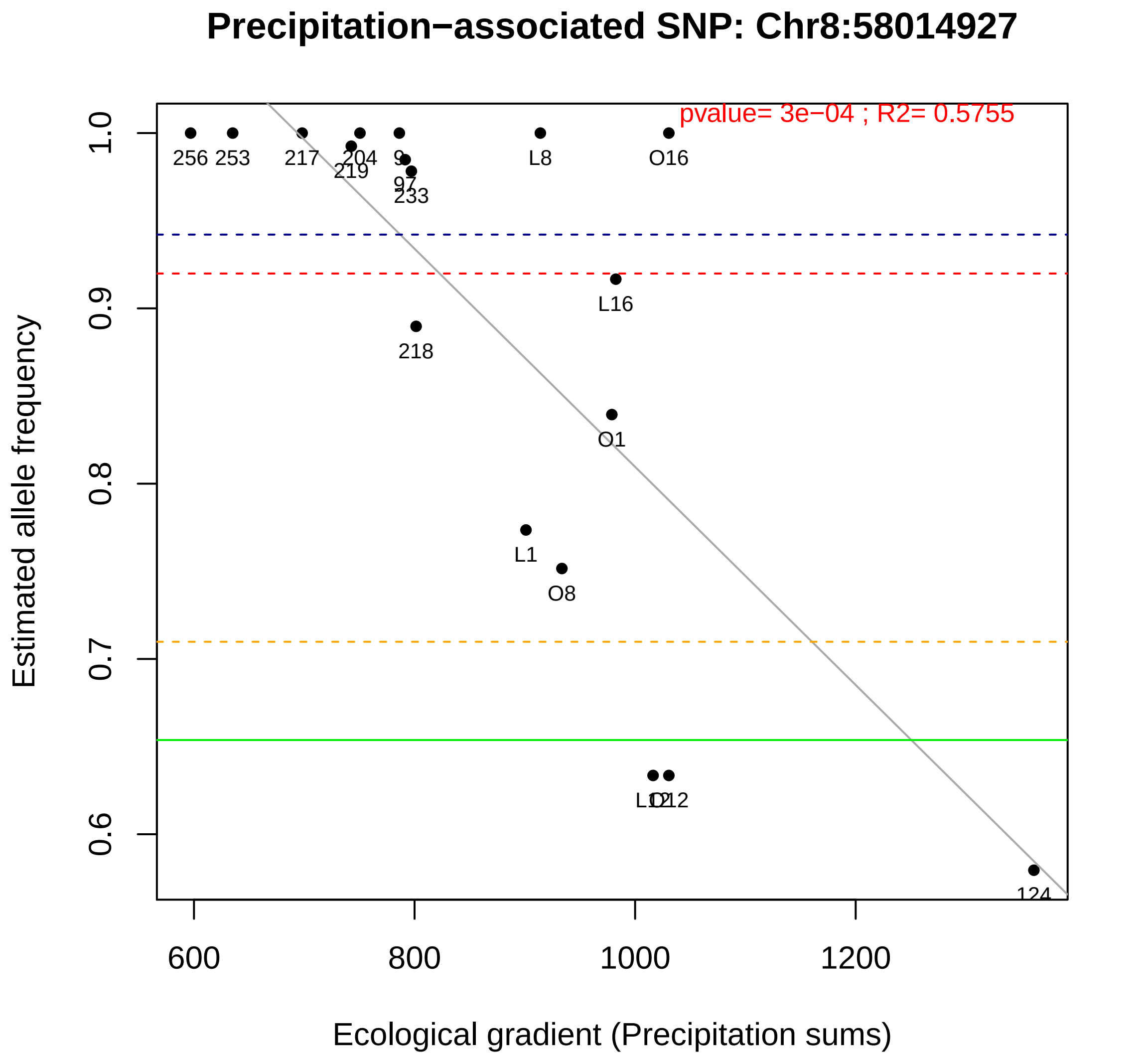

Precipitation−associated SNP: Chr8:58014927
pvalue= 3e−04 ; R2= 0.5755
Estimated allele frequency
0.6
0.7
0.8
0.9
1.0
600
800
1000
1200
Ecological gradient (Precipitation sums)
256
253
217
204
219
9
97
233
L8
O16
L16
218
O1
L1
O8
L12
O12
124

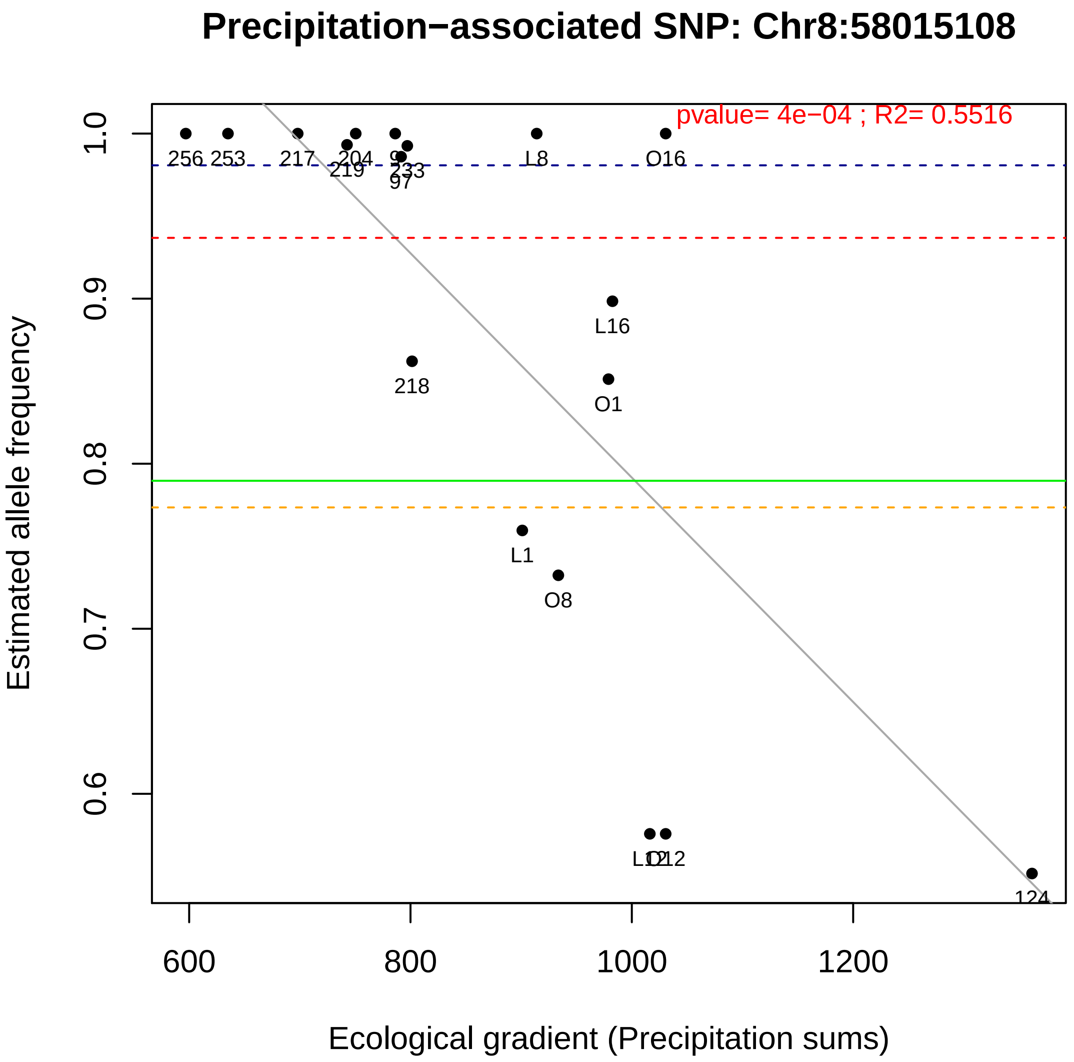
Precipitation−associated SNP: Chr8:58015108
pvalue= 4e−04 ; R2= 0.5516
Estimated allele frequency
Ecological gradient (Precipitation sums)
1.0
0.9
0.8
0.7
0.6
600
800
1000
1200
256
253
217
204
219
233
97
L8
O16
L16
218
O1
L1
O8
O12
124

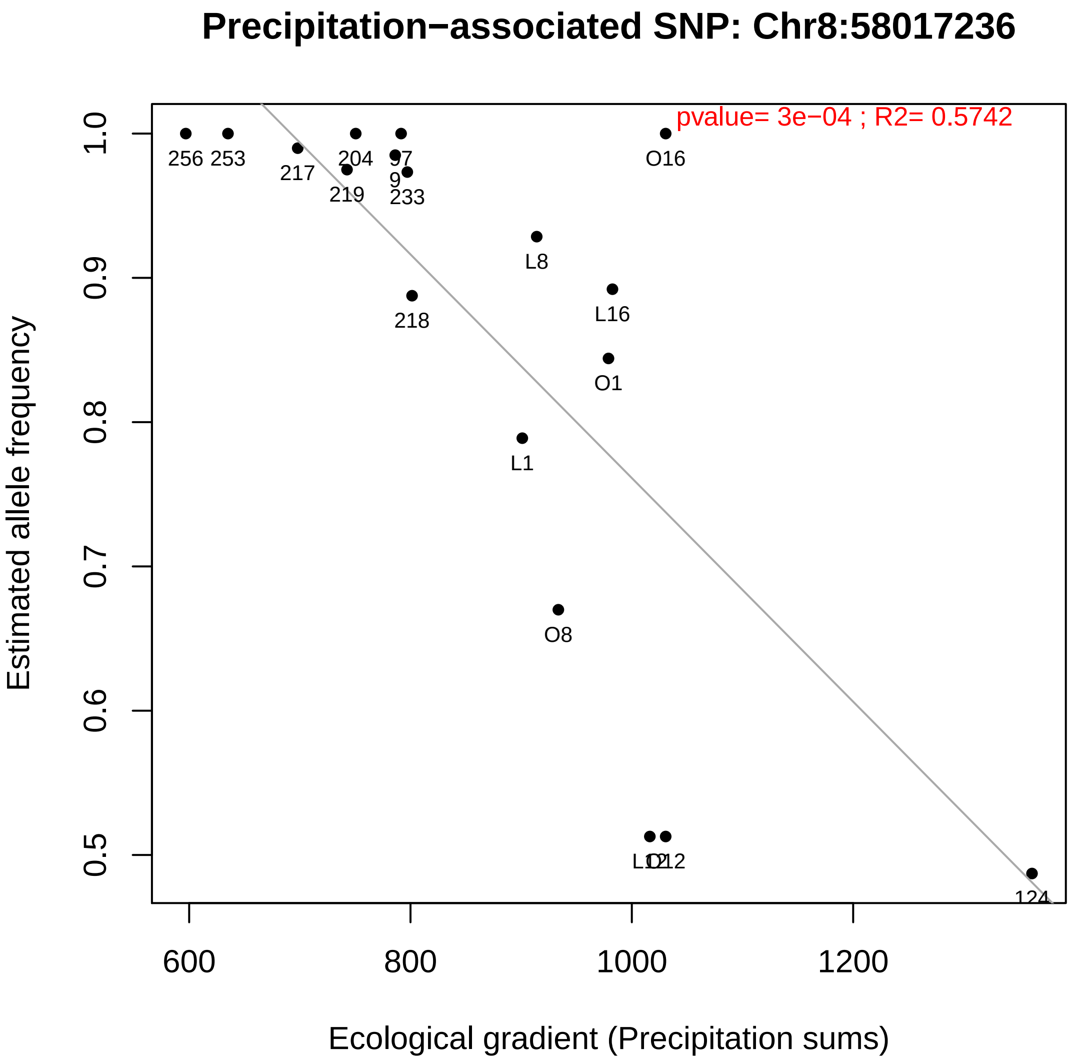
Precipitation−associated SNP: Chr8:58017236
pvalue= 3e−04 ; R2= 0.5742
256
253
217
219
204
97
9
233
O16
L8
218
L16
O1
L1
O8
L12
O12
124
1.0
0.9
0.8
0.7
0.6
0.5
600
800
1000
1200
Estimated allele frequency
Ecological gradient (Precipitation sums)

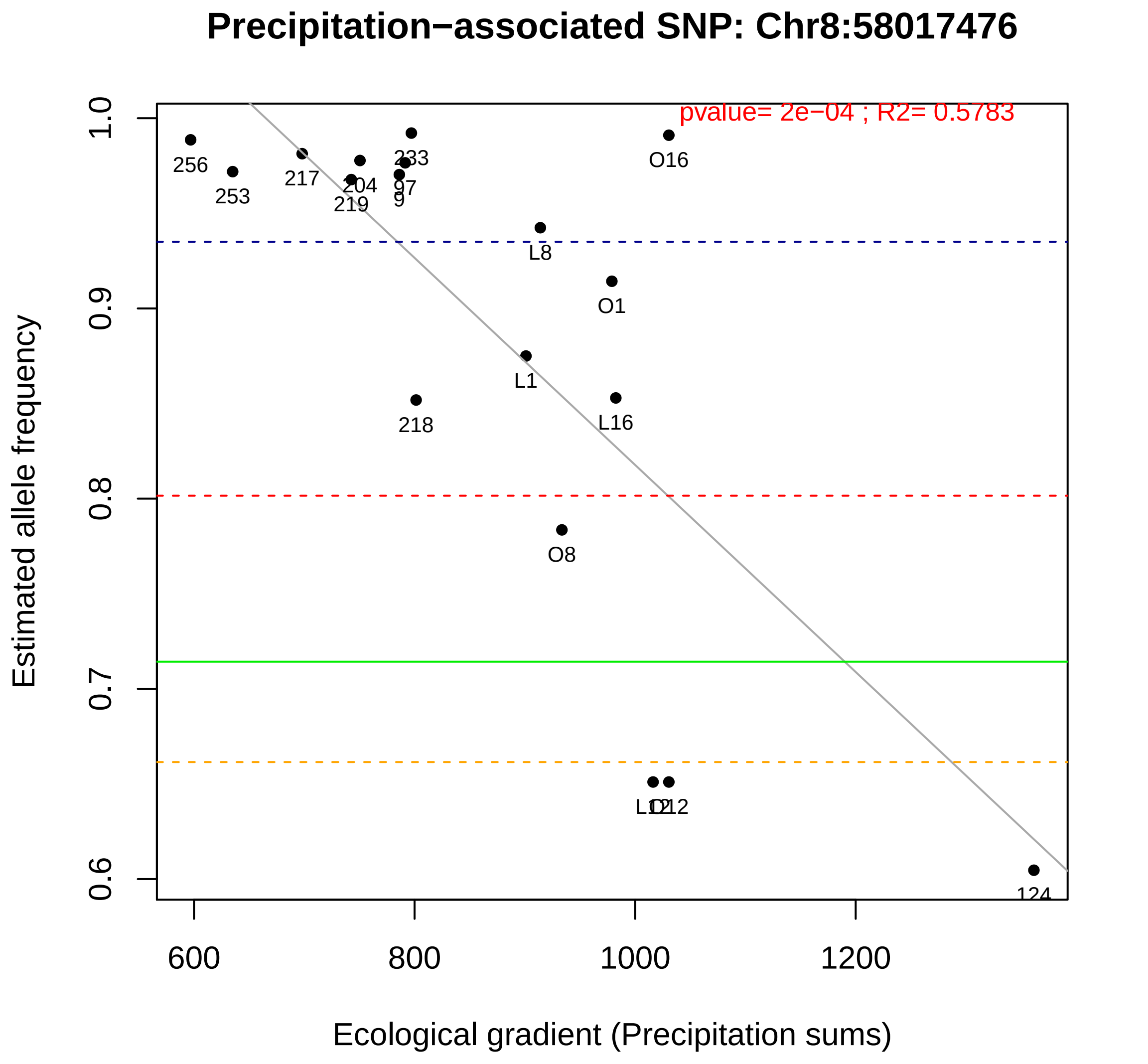
Precipitation−associated SNP: Chr8:58017476
pvalue= 2e−04 ; R2= 0.5783
Estimated allele frequency
Ecological gradient (Precipitation sums)
0.6
0.7
0.8
0.9
1.0
600
800
1000
1200
256
253
217
204
219
233
97
9
O16
L8
O1
L1
218
L16
O8
L12
O12
124

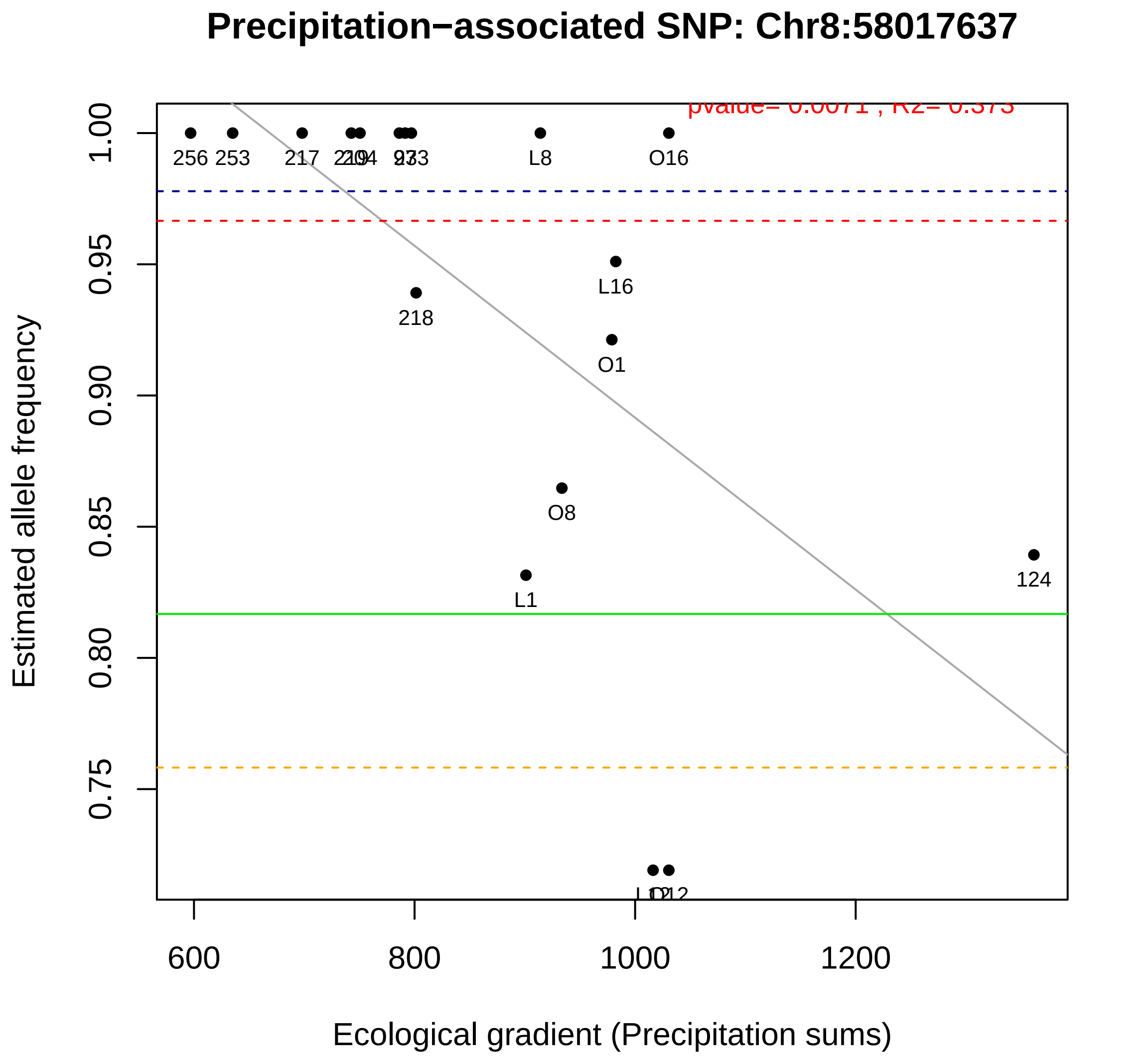

Precipitation−associated SNP: Chr8:58017637
pvalue= 0.0071 , R2= 0.373
Estimated allele frequency
Ecological gradient (Precipitation sums)
0.75
0.80
0.85
0.90
0.95
1.00
600
800
1000
1200
256
253
217
L8
O16
L16
218
O1
O8
L1
124

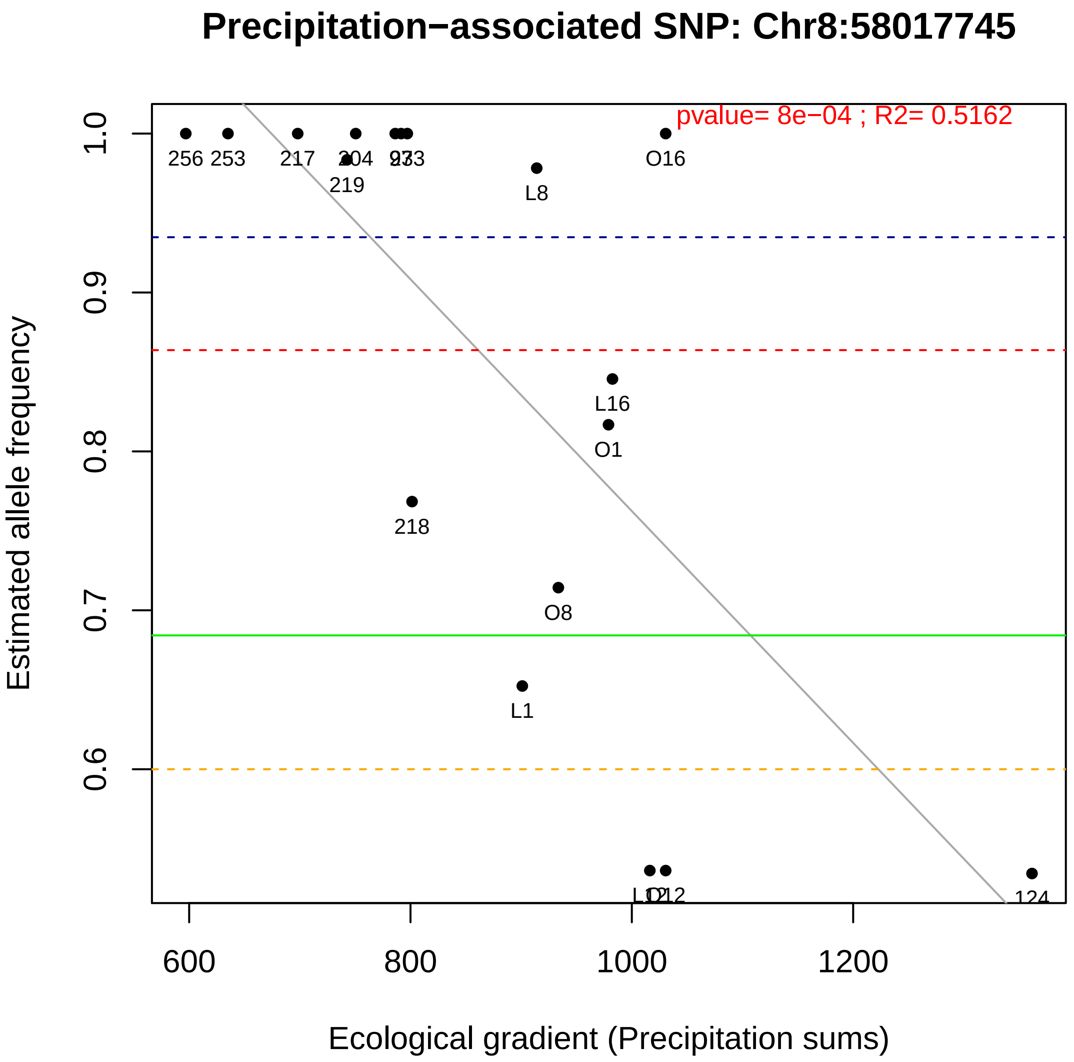

Precipitation−associated SNP: Chr8:58017745
pvalue= 8e−04 ; R2= 0.5162
Estimated allele frequency
Ecological gradient (Precipitation sums)
0.6
0.7
0.8
0.9
1.0
600
800
1000
1200
256
253
217
204
219
233
L8
O16
L16
O1
218
O8
L1
124

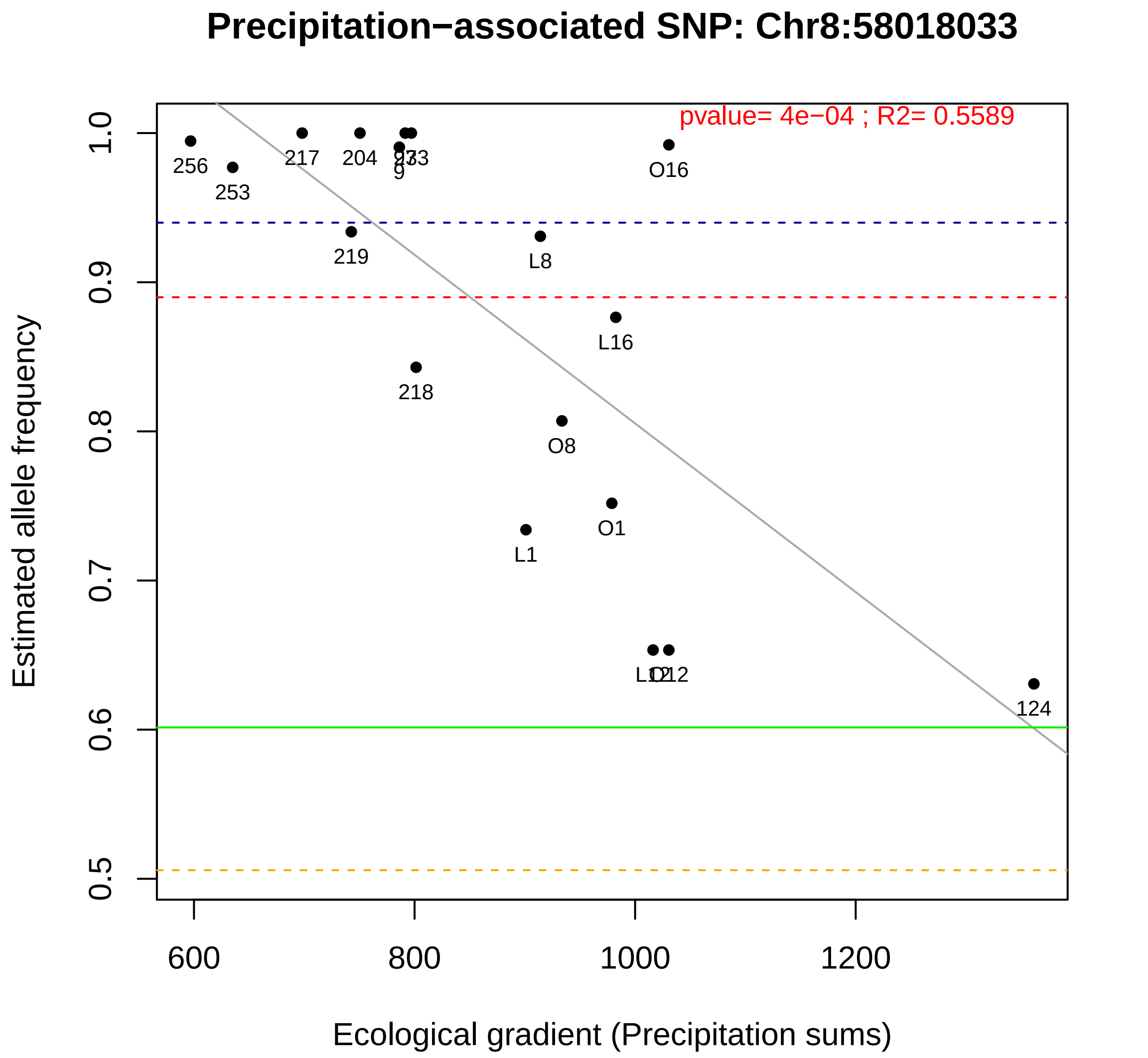

Precipitation−associated SNP: Chr8:58018033
pvalue= 4e−04 ; R2= 0.5589
256
253
217
204
219
218
L8
O16
L16
O8
O1
L1
124
0.5
0.6
0.7
0.8
0.9
1.0
600
800
1000
1200
Estimated allele frequency
Ecological gradient (Precipitation sums)

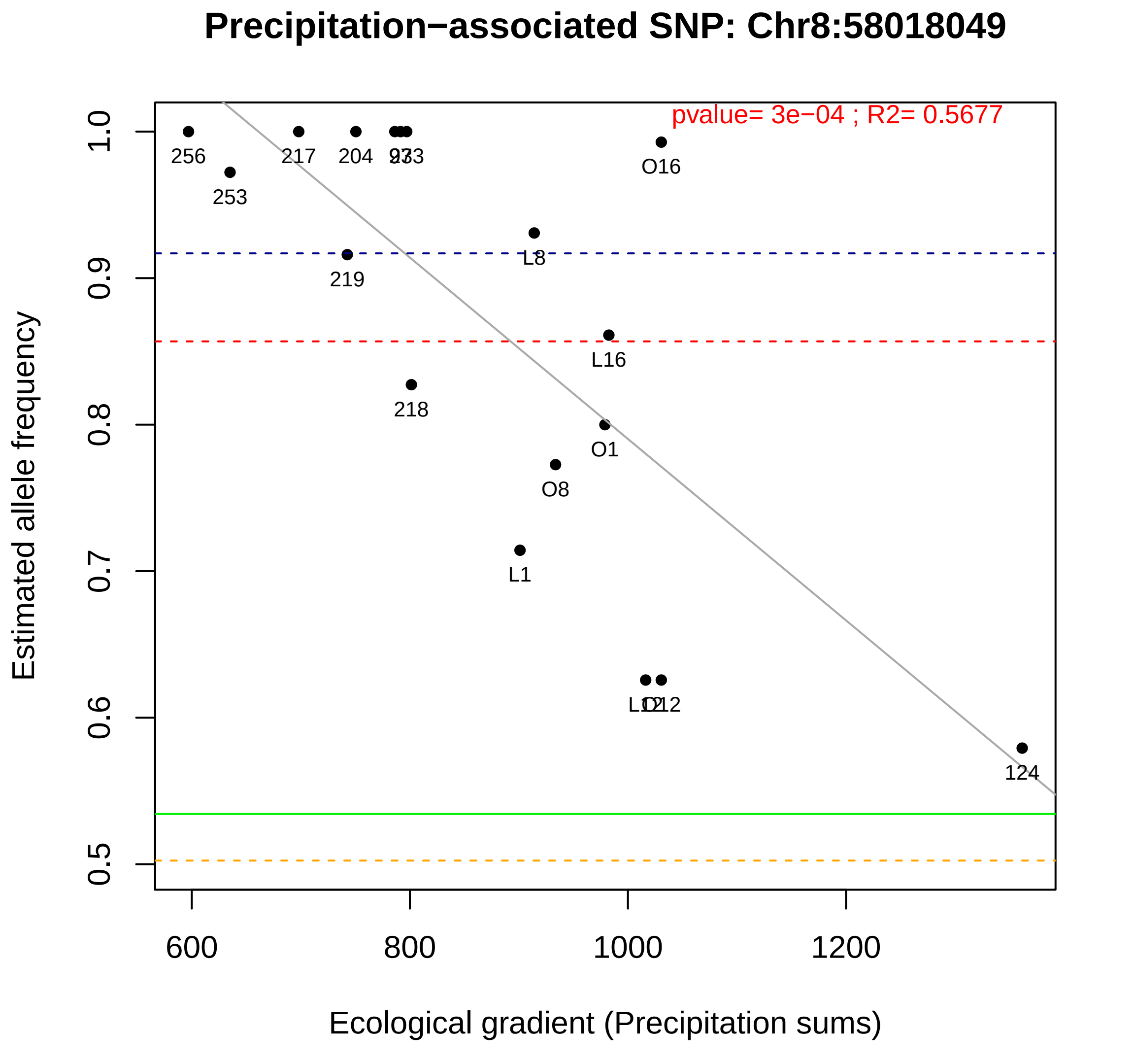

Precipitation−associated SNP: Chr8:58018049
pvalue= 3e−04 ; R2= 0.5677
256
253
217
204
97
233
219
218
L8
O16
L16
O1
O8
L1
L12
O12
124
Estimated allele frequency
0.5
0.6
0.7
0.8
0.9
1.0
600
800
1000
1200
Ecological gradient (Precipitation sums)

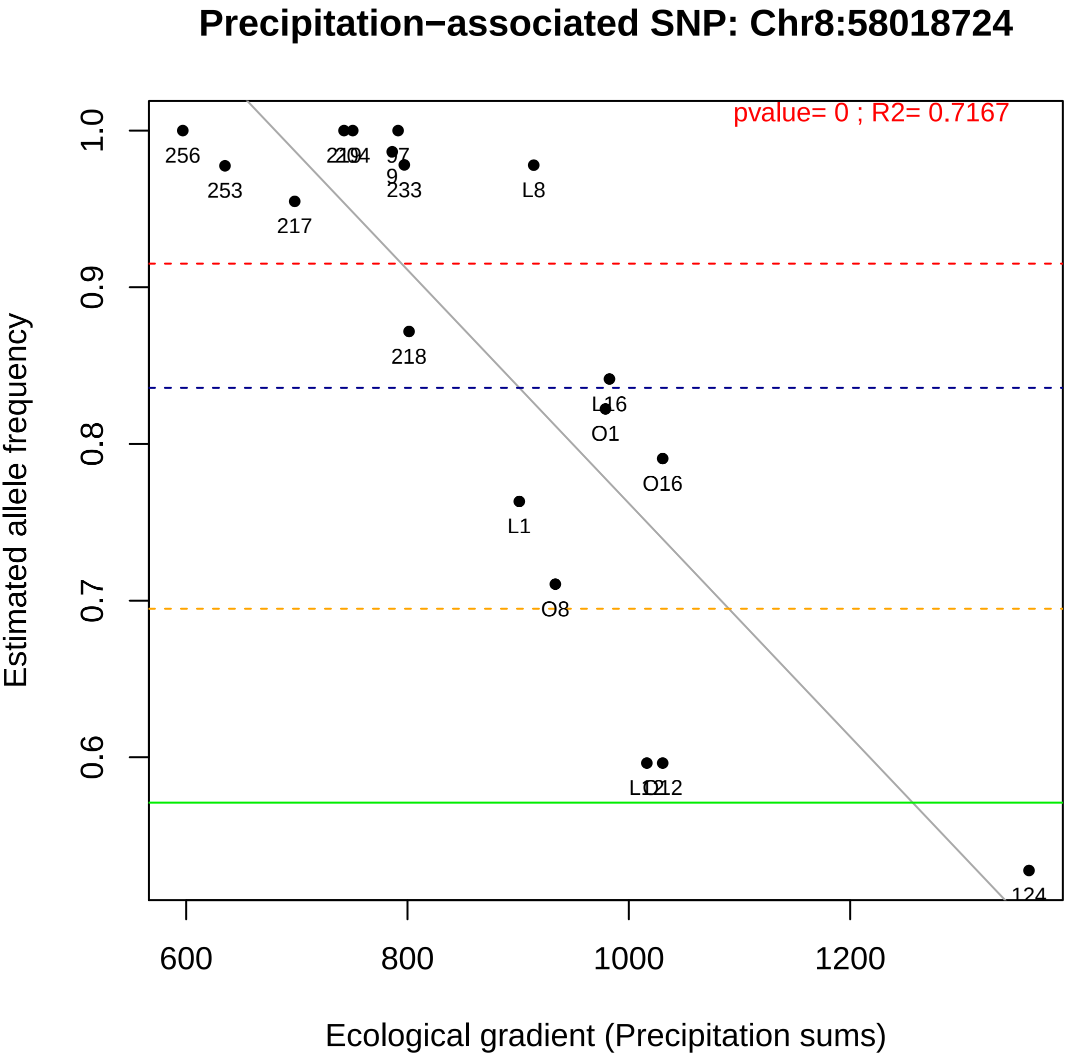

Precipitation−associated SNP: Chr8:58018724
pvalue= 0 ; R2= 0.7167
Estimated allele frequency
Ecological gradient (Precipitation sums)
1.0
0.9
0.8
0.7
0.6
600
800
1000
1200
256
253
217
233
L8
218
L16
O1
O16
L1
O8
124

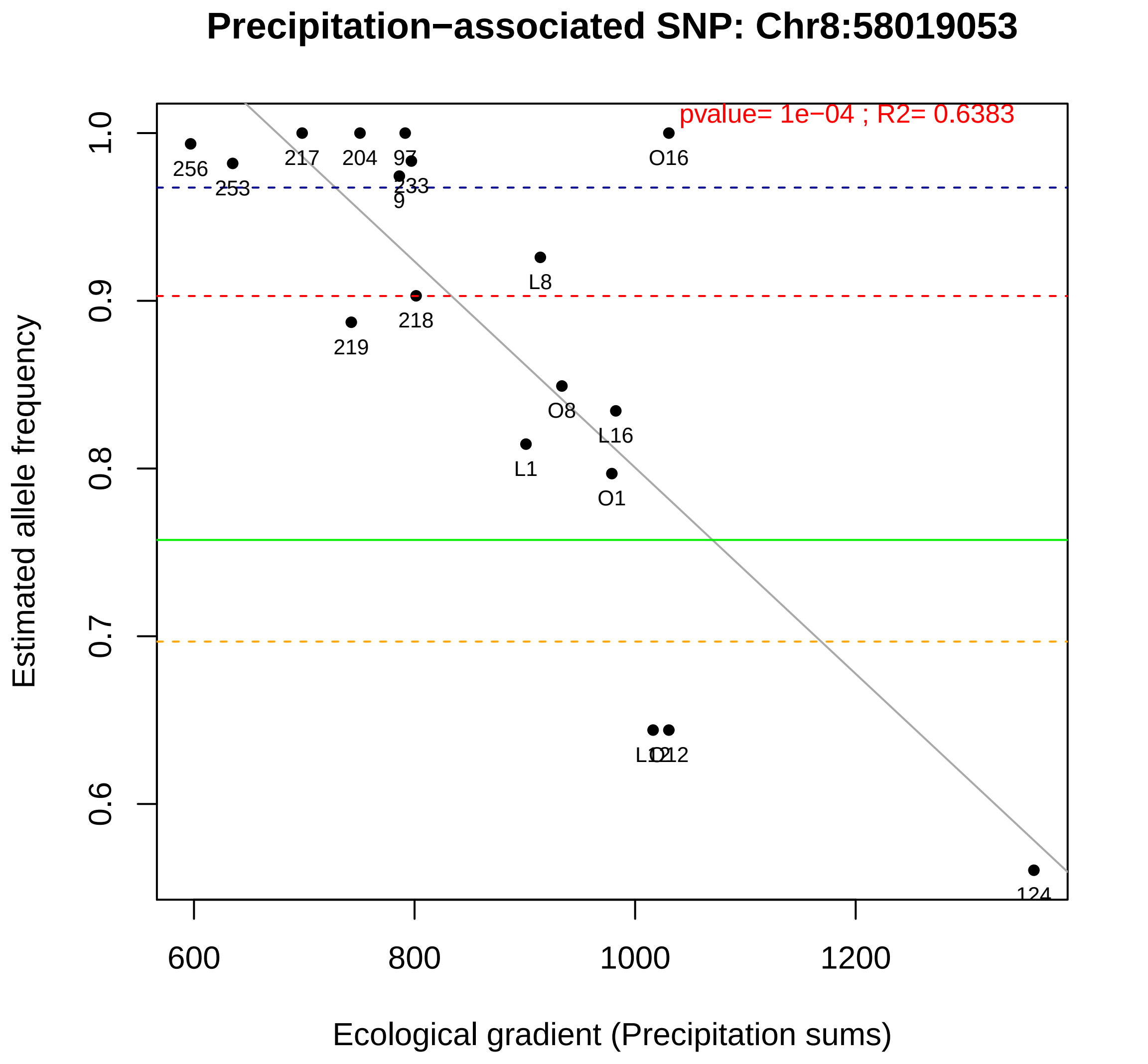

Precipitation−associated SNP: Chr8:58019053
pvalue= 1e−04 ; R2= 0.6383
Estimated allele frequency
Ecological gradient (Precipitation sums)
0.6
0.7
0.8
0.9
1.0
600
800
1000
1200
256
253
217
204
97
233
9
O16
L8
218
219
O8
L16
L1
O1
L12
O12
124

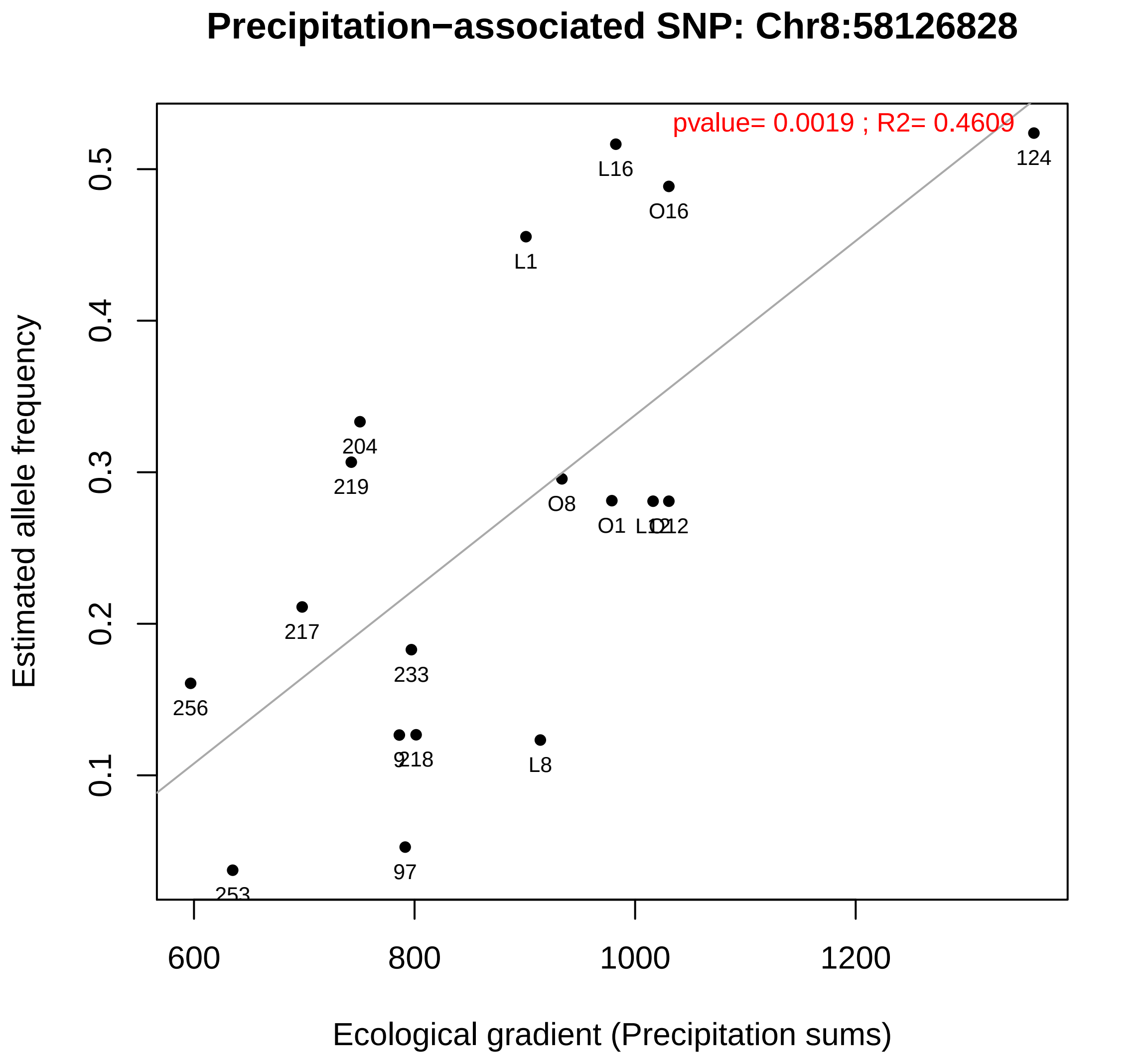
Precipitation−associated SNP: Chr8:58126828
pvalue= 0.0019 ; R2= 0.4609
Estimated allele frequency
0.1
0.2
0.3
0.4
0.5
600
800
1000
1200
Ecological gradient (Precipitation sums)
124
L16
O16
L1
204
219
O8
O1
217
233
256
218
L8
97
253

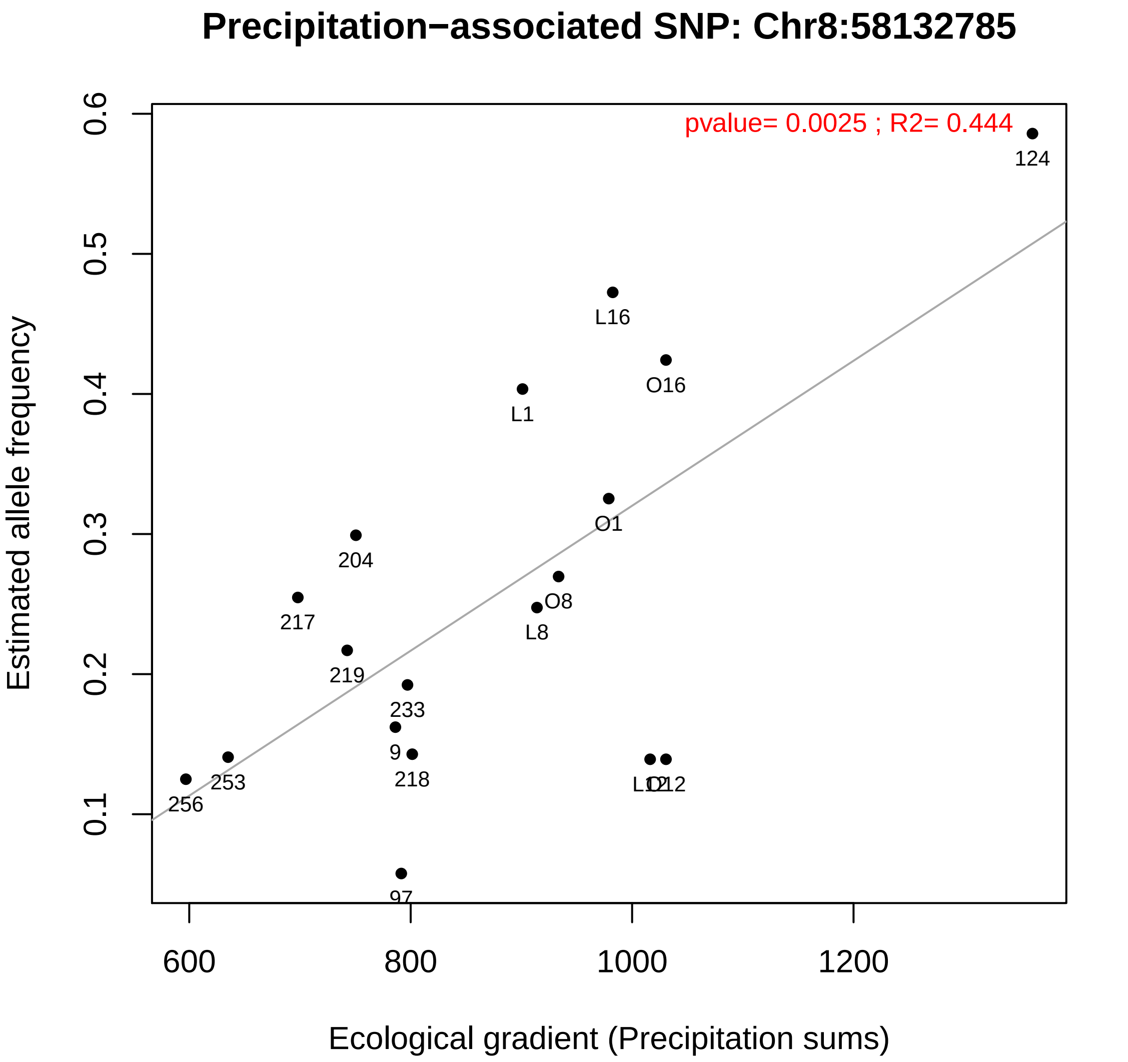
Precipitation−associated SNP: Chr8:58132785
pvalue= 0.0025 ; R2= 0.444
Estimated allele frequency
0.1
0.2
0.3
0.4
0.5
0.6
600
800
1000
1200
Ecological gradient (Precipitation sums)
124
L16
O16
L1
O1
204
217
O8
L8
219
233
9
218
253
256
97

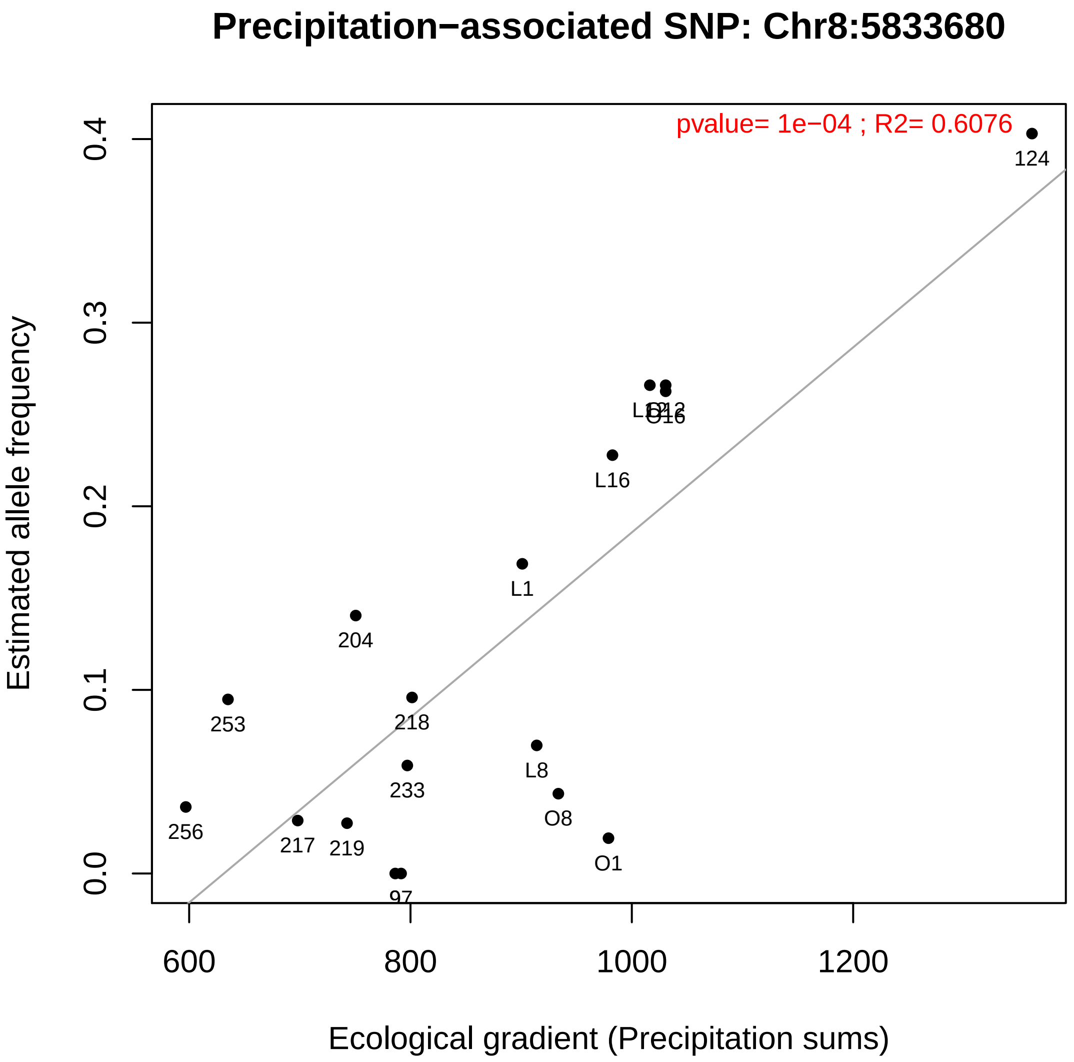

Precipitation−associated SNP: Chr8:5833680
pvalue= 1e−04 ; R2= 0.6076
Estimated allele frequency
0.0
0.1
0.2
0.3
0.4
Ecological gradient (Precipitation sums)
600
800
1000
1200
124
L16
L1
204
253
218
L8
233
O8
256
217
219
O1
97

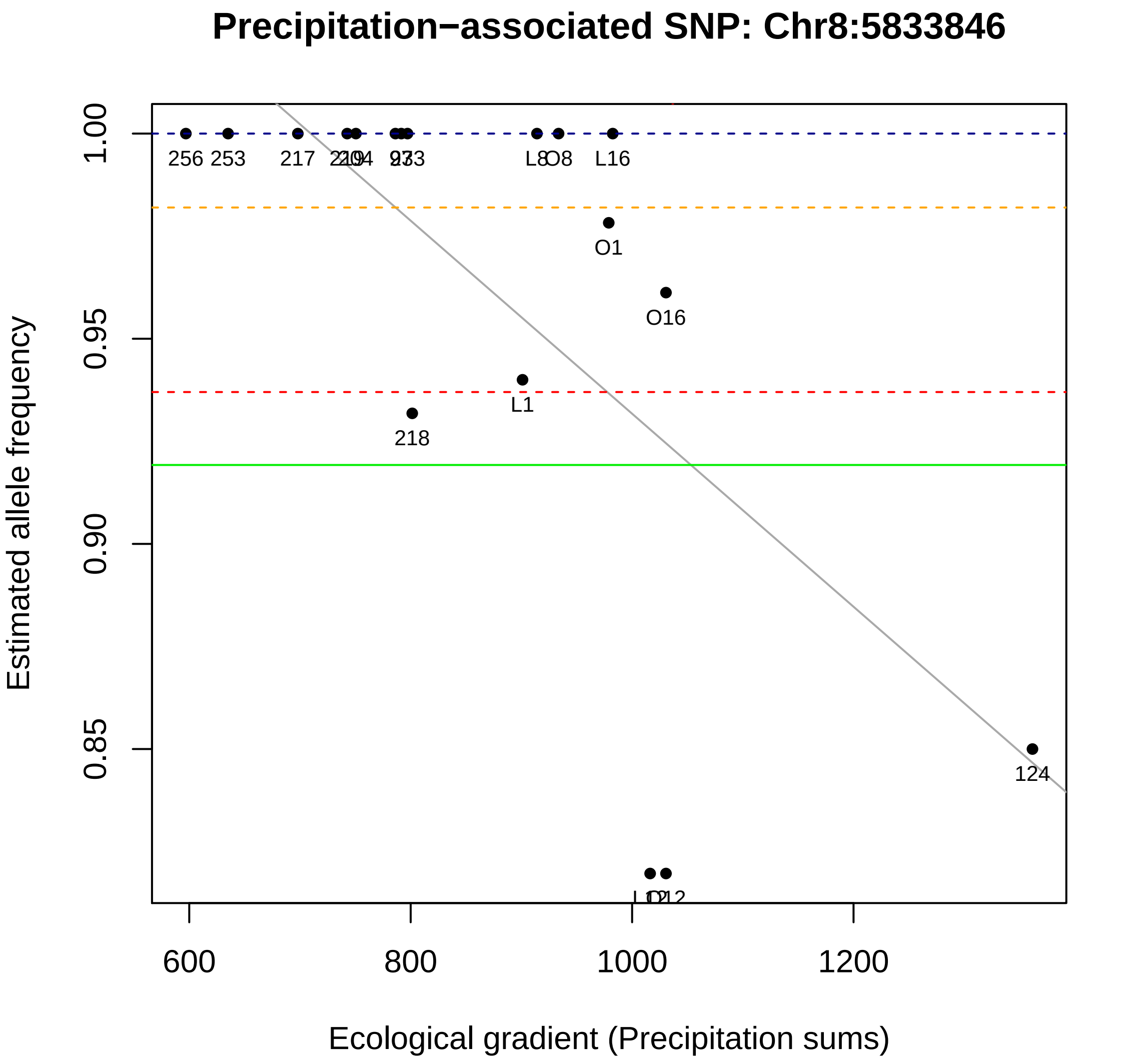

Precipitation−associated SNP: Chr8:5833846
Estimated allele frequency
Ecological gradient (Precipitation sums)
1.00
0.95
0.90
0.85
600
800
1000
1200
256
253
217
L8
O8
L16
O1
O16
L1
218
124

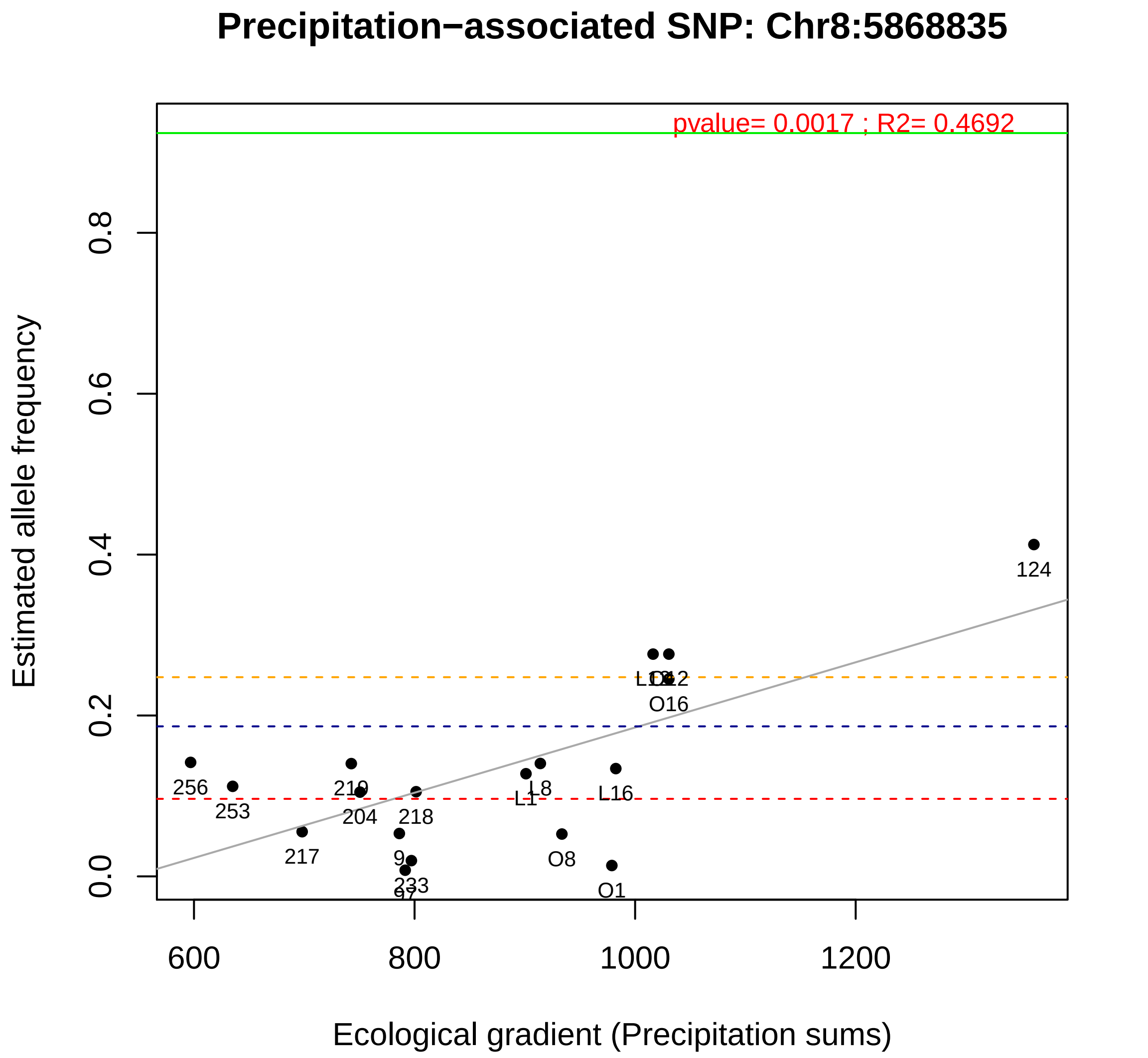

Precipitation−associated SNP: Chr8:5868835
pvalue= 0.0017 ; R2= 0.4692
Estimated allele frequency
0.0
0.2
0.4
0.6
0.8
600
800
1000
1200
Ecological gradient (Precipitation sums)
124
O16
256
253
217
204
218
L8
L1
L16
O8
O1
233

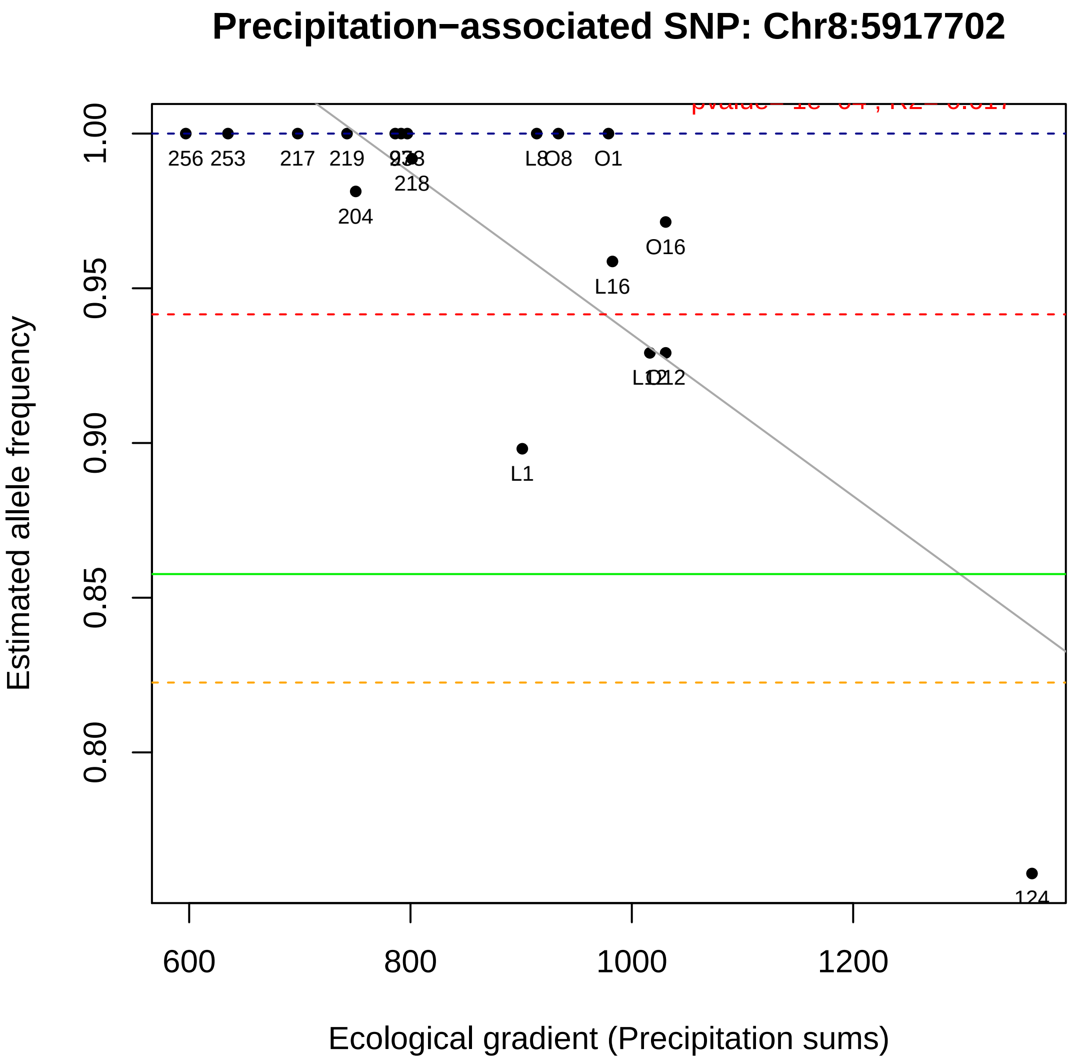
Precipitation−associated SNP: Chr8:5917702
Estimated allele frequency
Ecological gradient (Precipitation sums)
1.00
0.95
0.90
0.85
0.80
600
800
1000
1200
256
253
217
219
218
204
L8
O8
O1
O16
L16
L1
124

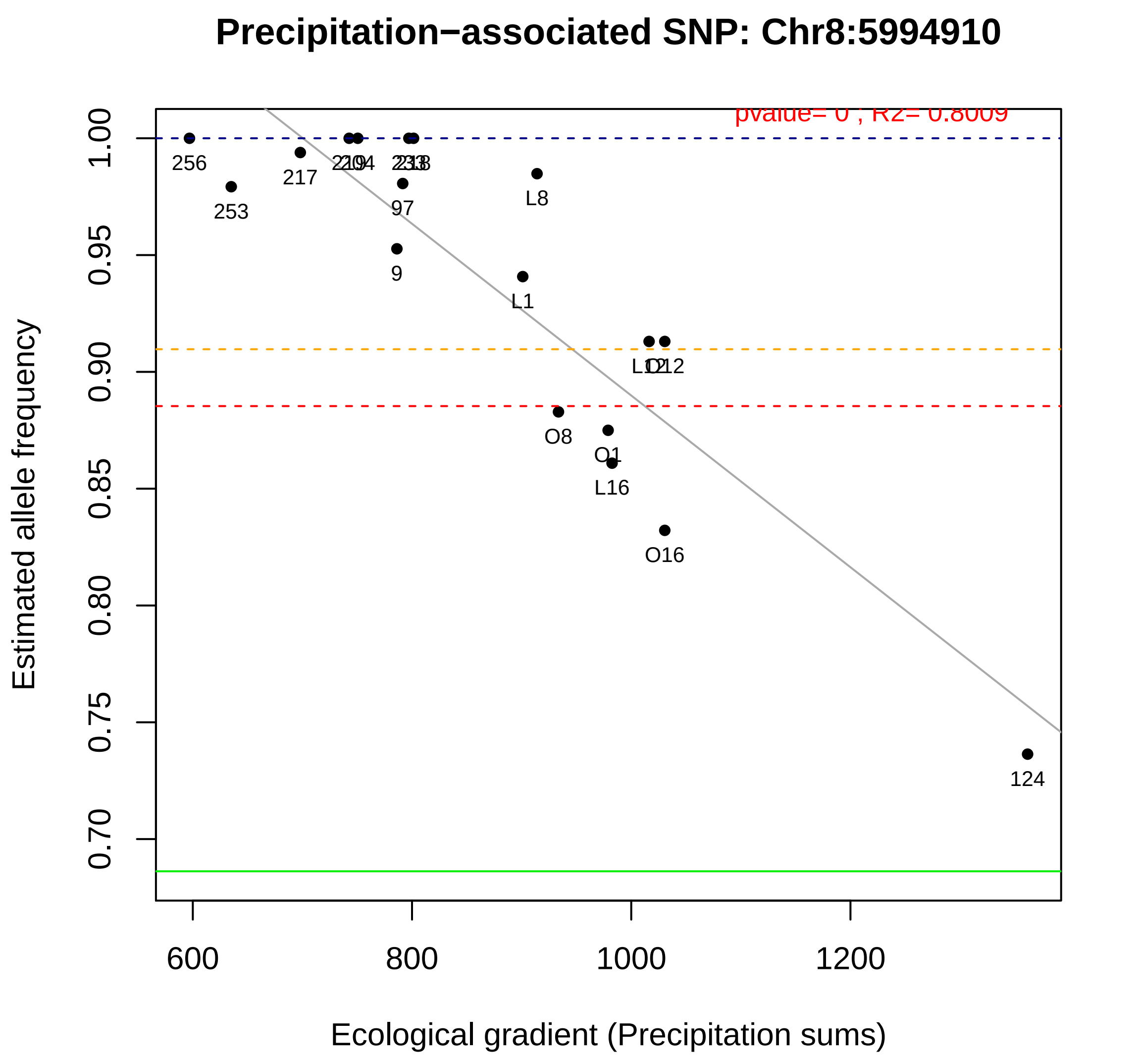
Precipitation−associated SNP: Chr8:5994910
pvalue= 0 ; R2= 0.8009
Estimated allele frequency
Ecological gradient (Precipitation sums)
1.00
0.95
0.90
0.85
0.80
0.75
0.70
600
800
1000
1200
256
253
217
97
9
L8
L1
O8
O1
L16
O16
124

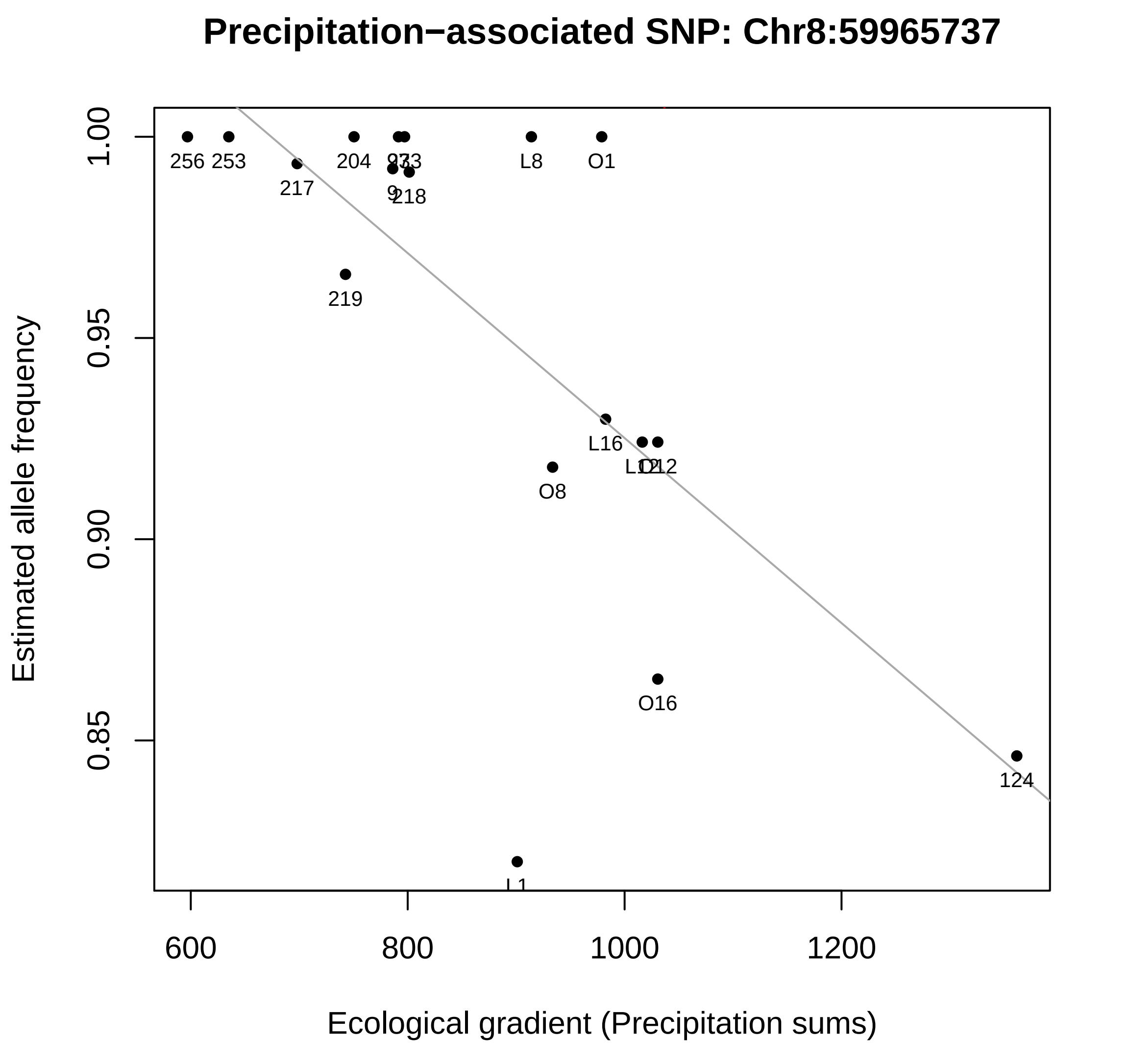

Precipitation−associated SNP: Chr8:59965737
Estimated allele frequency
1.00
0.95
0.90
0.85
256
253
217
204
9
233
218
L8
O1
219
L16
L12
O12
O8
O16
124
L1
600
800
1000
1200
Ecological gradient (Precipitation sums)

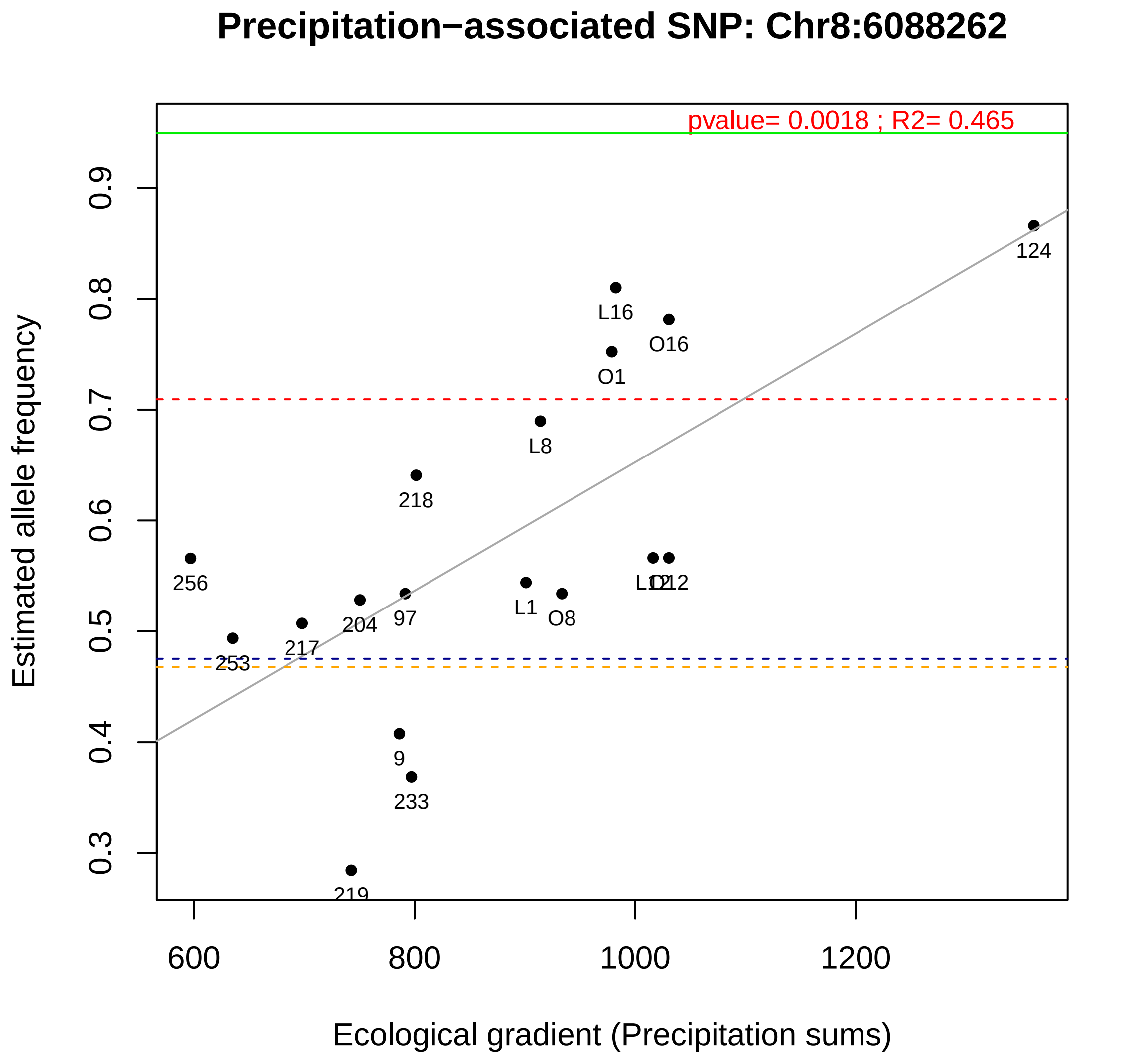
Precipitation−associated SNP: Chr8:6088262
pvalue= 0.0018 ; R2= 0.465
Estimated allele frequency
0.3
0.4
0.5
0.6
0.7
0.8
0.9
600
800
1000
1200
Ecological gradient (Precipitation sums)
124
L16
O16
O1
L8
218
256
253
217
204
97
L1
O8
L12
O12
9
233
219

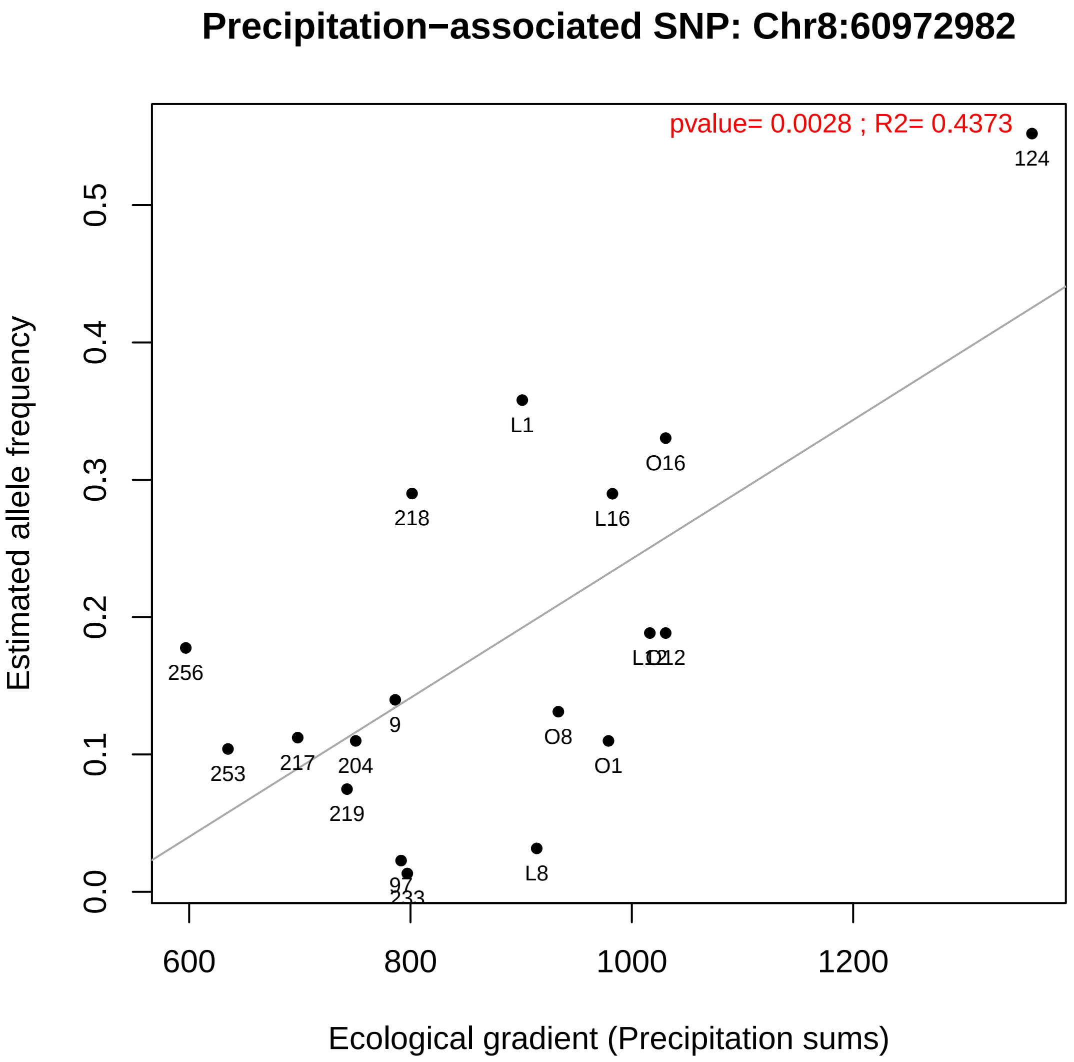
Precipitation−associated SNP: Chr8:60972982
pvalue= 0.0028 ; R2= 0.4373
Estimated allele frequency
Ecological gradient (Precipitation sums)
0.0
0.1
0.2
0.3
0.4
0.5
600
800
1000
1200
124
L1
O16
218
L16
256
9
O8
253
217
204
O1
219
L8
97
233

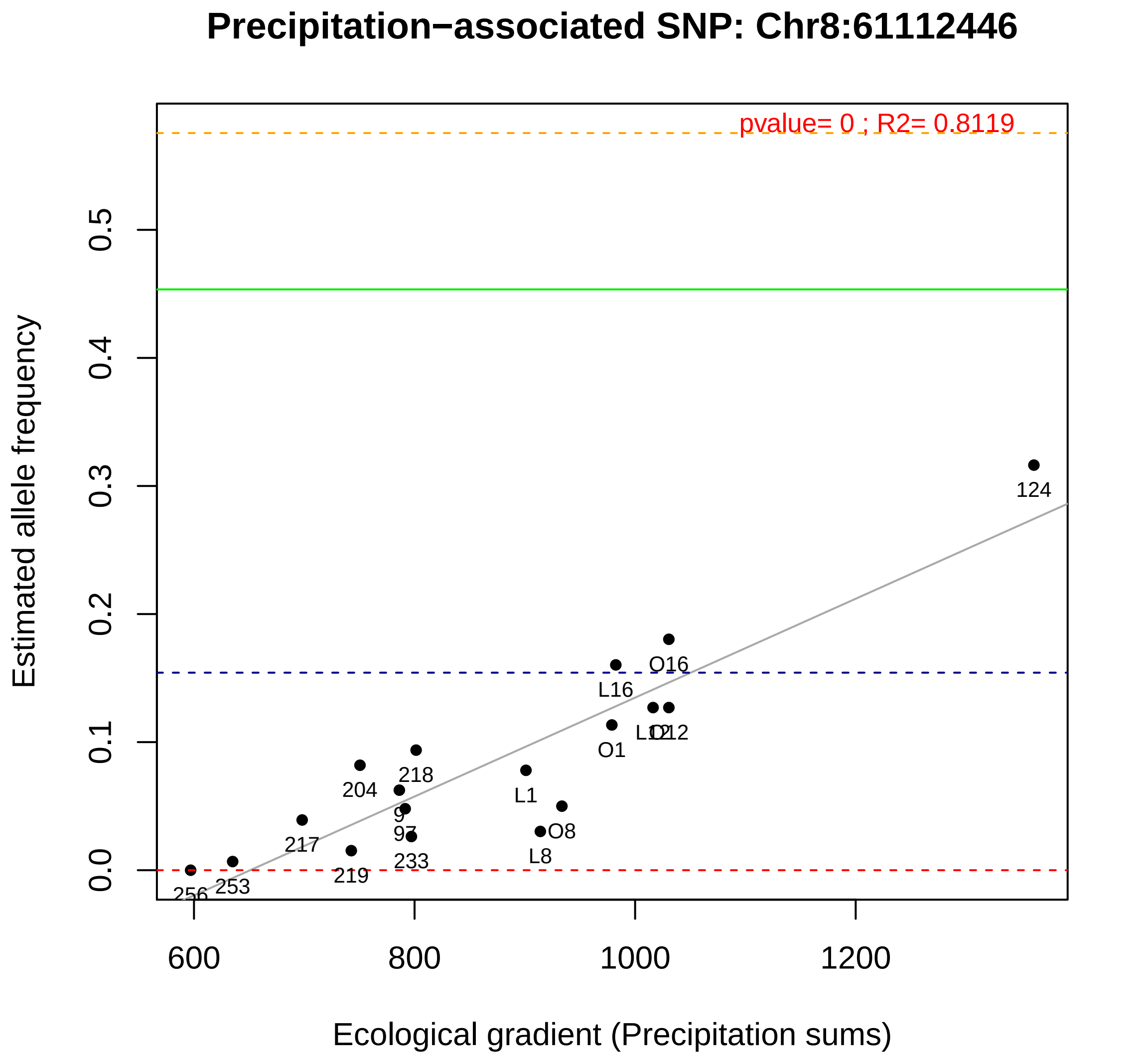

Precipitation−associated SNP: Chr8:61112446
pvalue= 0 ; R2= 0.8119
Estimated allele frequency
0.0
0.1
0.2
0.3
0.4
0.5
600
800
1000
1200
Ecological gradient (Precipitation sums)
124
O16
L16
O1
O12
L1
O8
L8
218
204
217
219
233
253
256

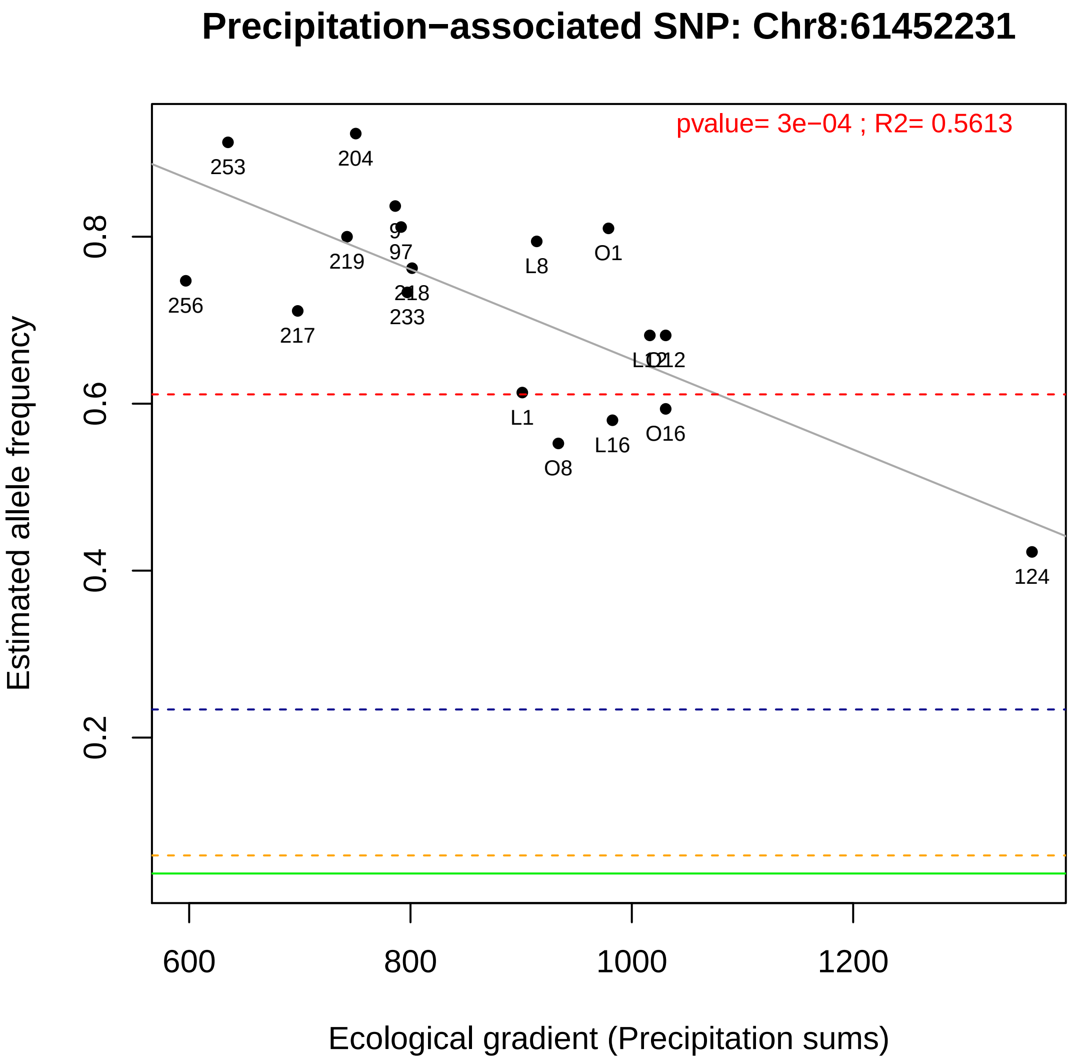
Precipitation−associated SNP: Chr8:61452231
pvalue= 3e−04 ; R2= 0.5613
253
204
219
97
256
217
233
L8
O1
L1
O8
L16
O16
124
0.2
0.4
0.6
0.8
600
800
1000
1200
Estimated allele frequency
Ecological gradient (Precipitation sums)

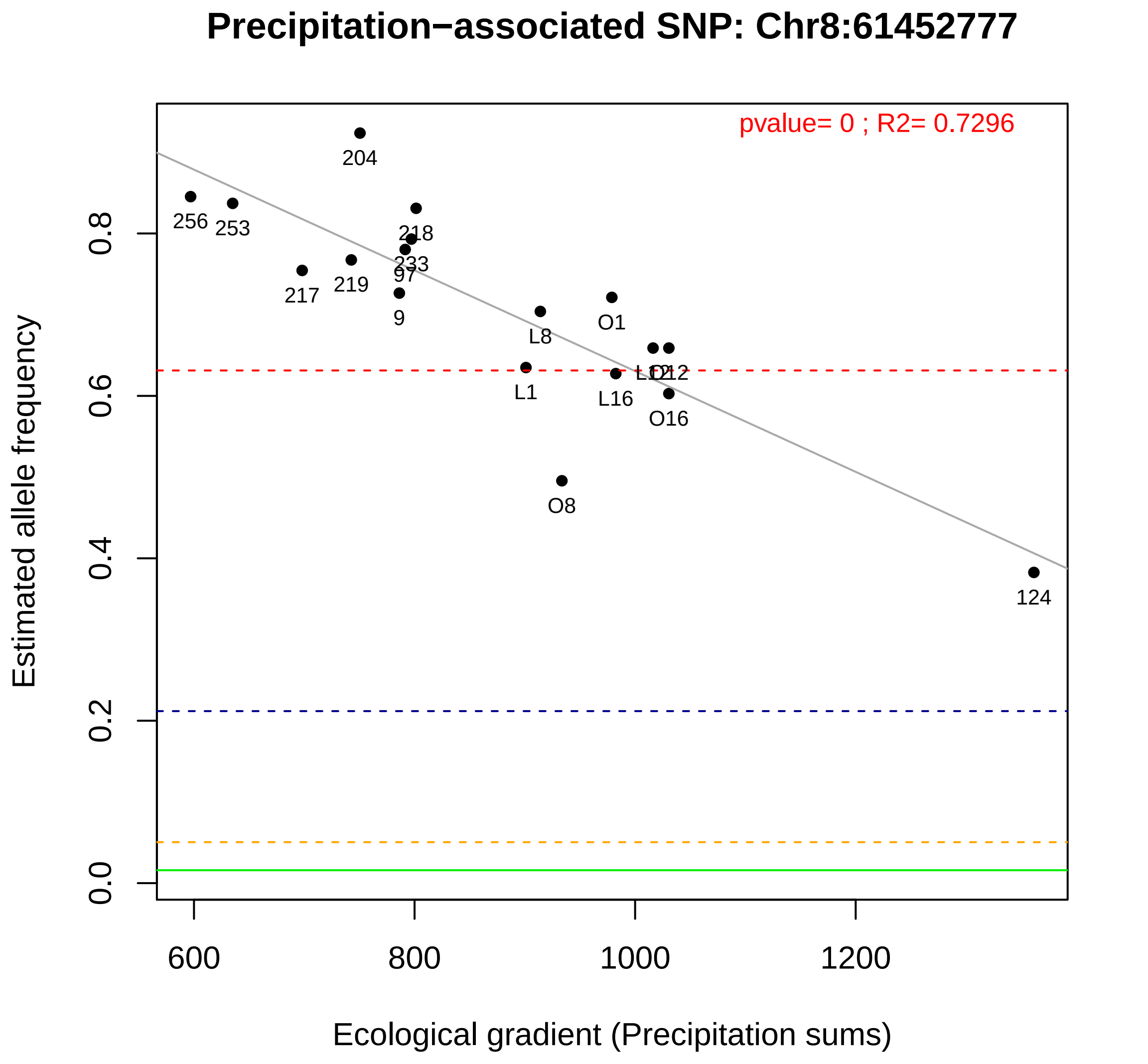

Precipitation−associated SNP: Chr8:61452777
pvalue= 0 ; R2= 0.7296
204
256
253
218
233
97
217
219
9
L8
O1
L1
L16
O16
O8
124
Estimated allele frequency
0.0
0.2
0.4
0.6
0.8
600
800
1000
1200
Ecological gradient (Precipitation sums)

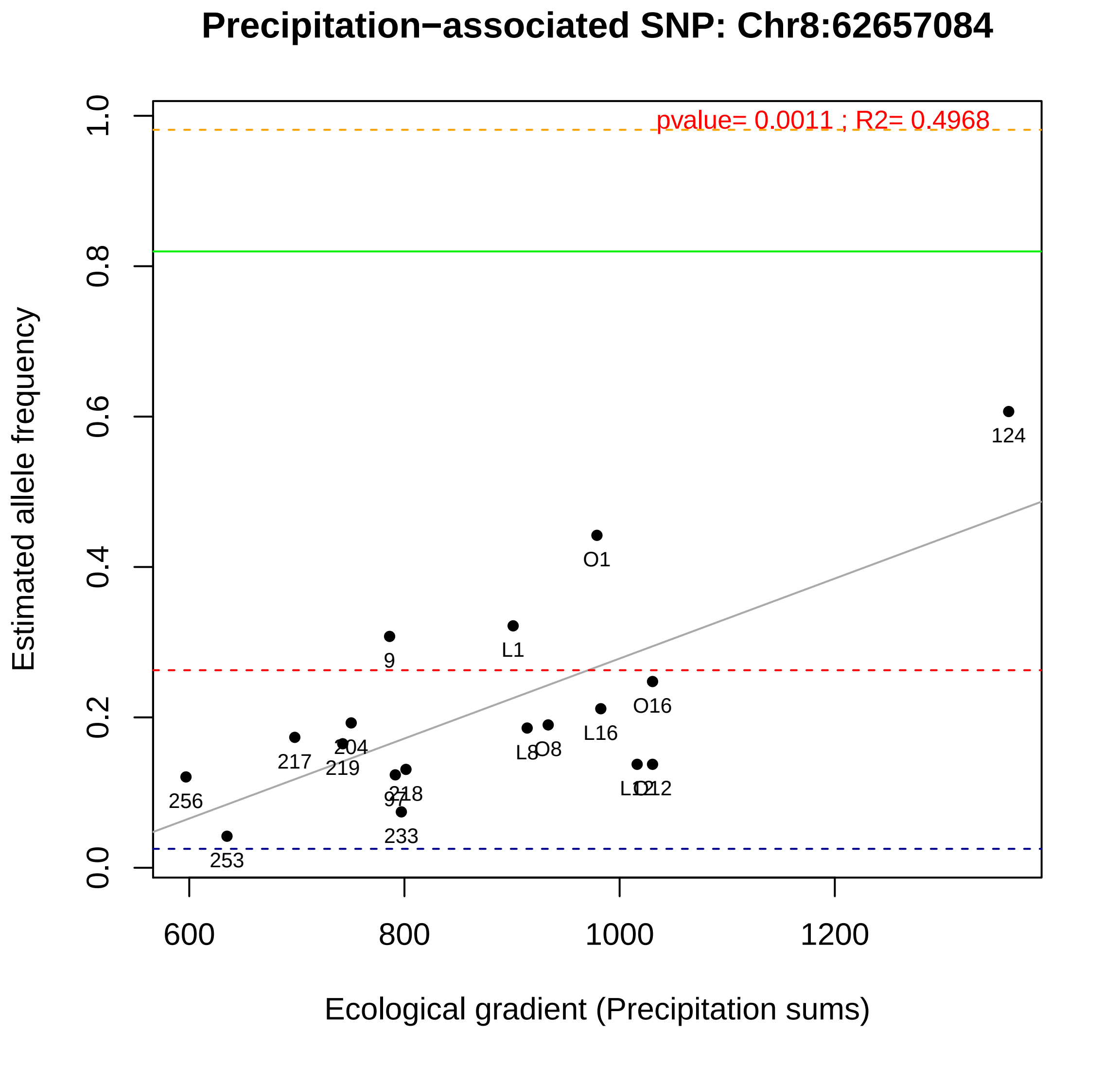
Precipitation−associated SNP: Chr8:62657084
pvalue= 0.0011 ; R2= 0.4968
Estimated allele frequency
0.0
0.2
0.4
0.6
0.8
1.0
Ecological gradient (Precipitation sums)
600
800
1000
1200
124
O1
L1
9
O16
L16
L8
O8
204
219
217
256
218
97
233
253
L12
O12

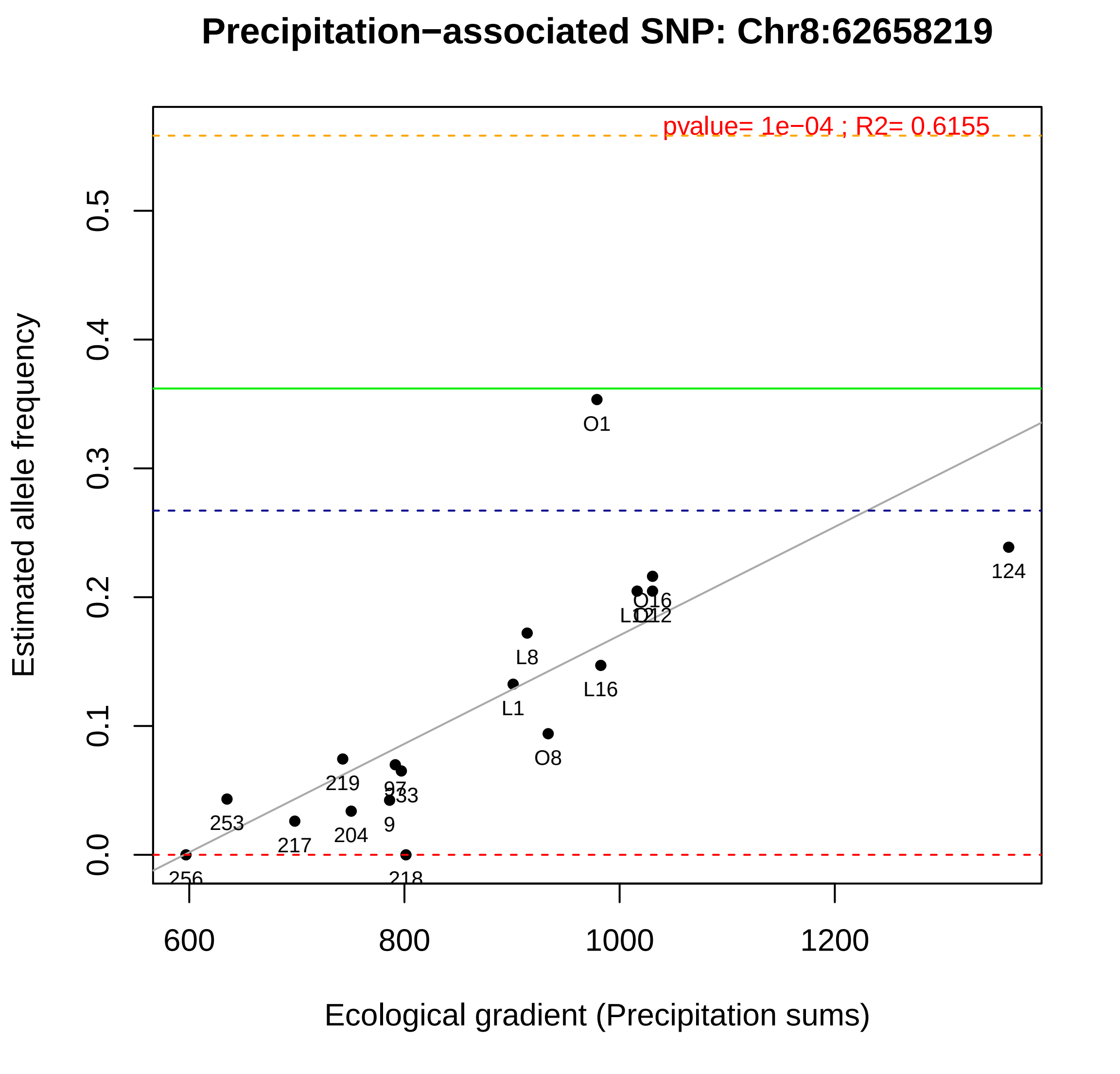

Precipitation−associated SNP: Chr8:62658219
pvalue= 1e−04 ; R2= 0.6155
Estimated allele frequency
0.0
0.1
0.2
0.3
0.4
0.5
600
800
1000
1200
Ecological gradient (Precipitation sums)
O1
124
O16
L12
O12
L8
L16
L1
O8
219
97
233
9
253
204
217
256
218

#### Precipitation−associated SNP: Chr8:62746254

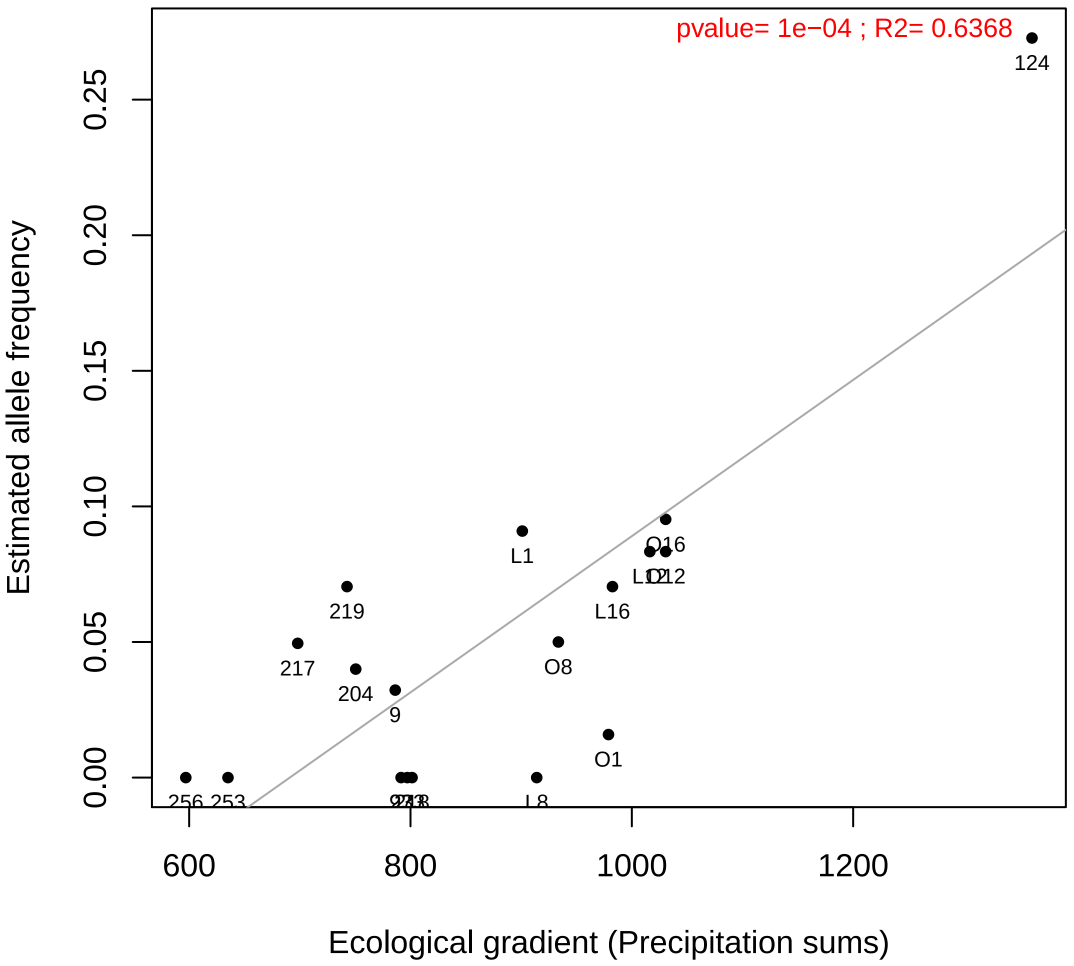

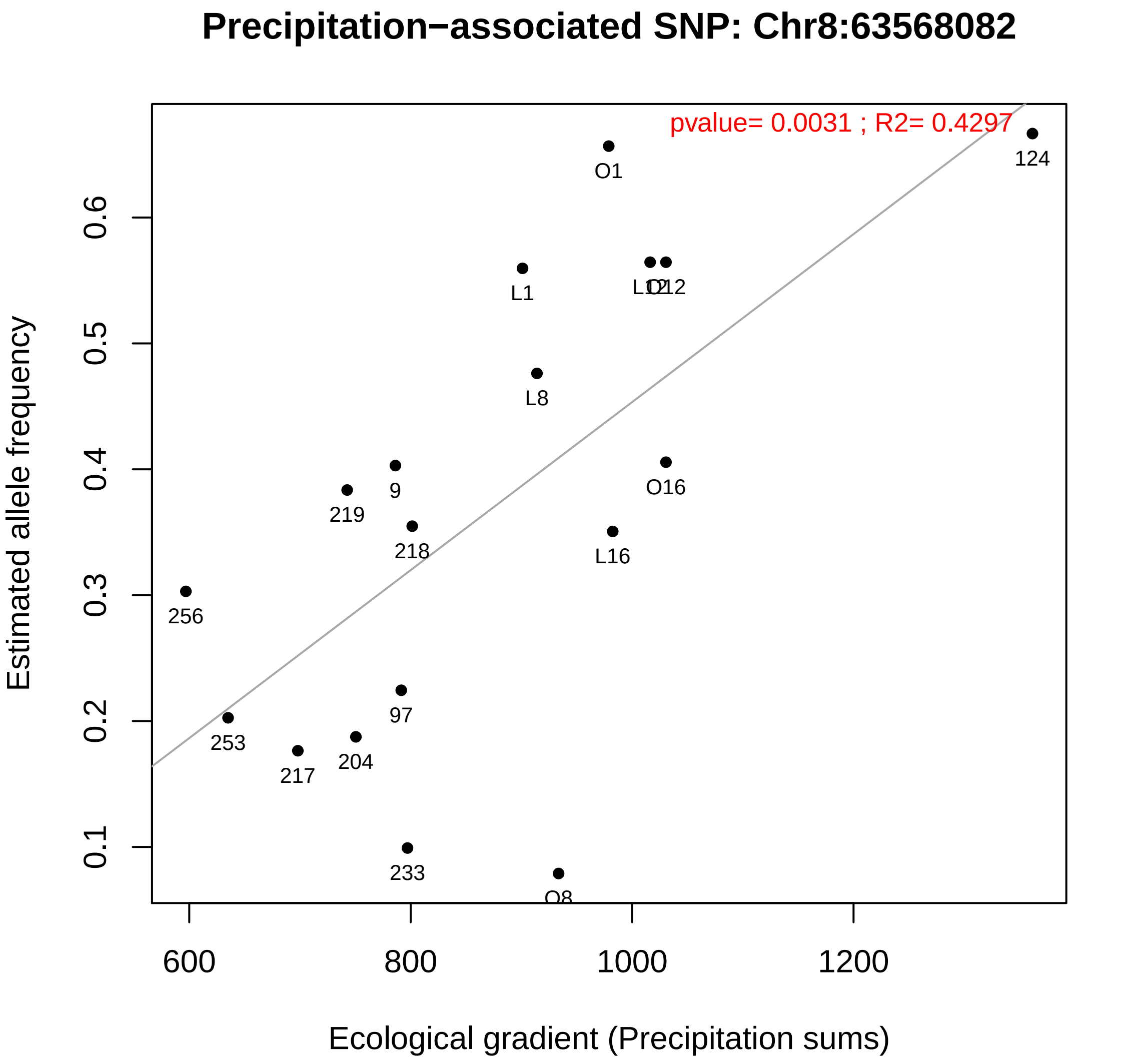

Precipitation−associated SNP: Chr8:63568082
pvalue= 0.0031 ; R2= 0.4297
Estimated allele frequency
Ecological gradient (Precipitation sums)
0.1
0.2
0.3
0.4
0.5
0.6
600
800
1000
1200
124
O1
L1
L8
9
219
218
O16
L16
256
253
217
204
97
233
O8

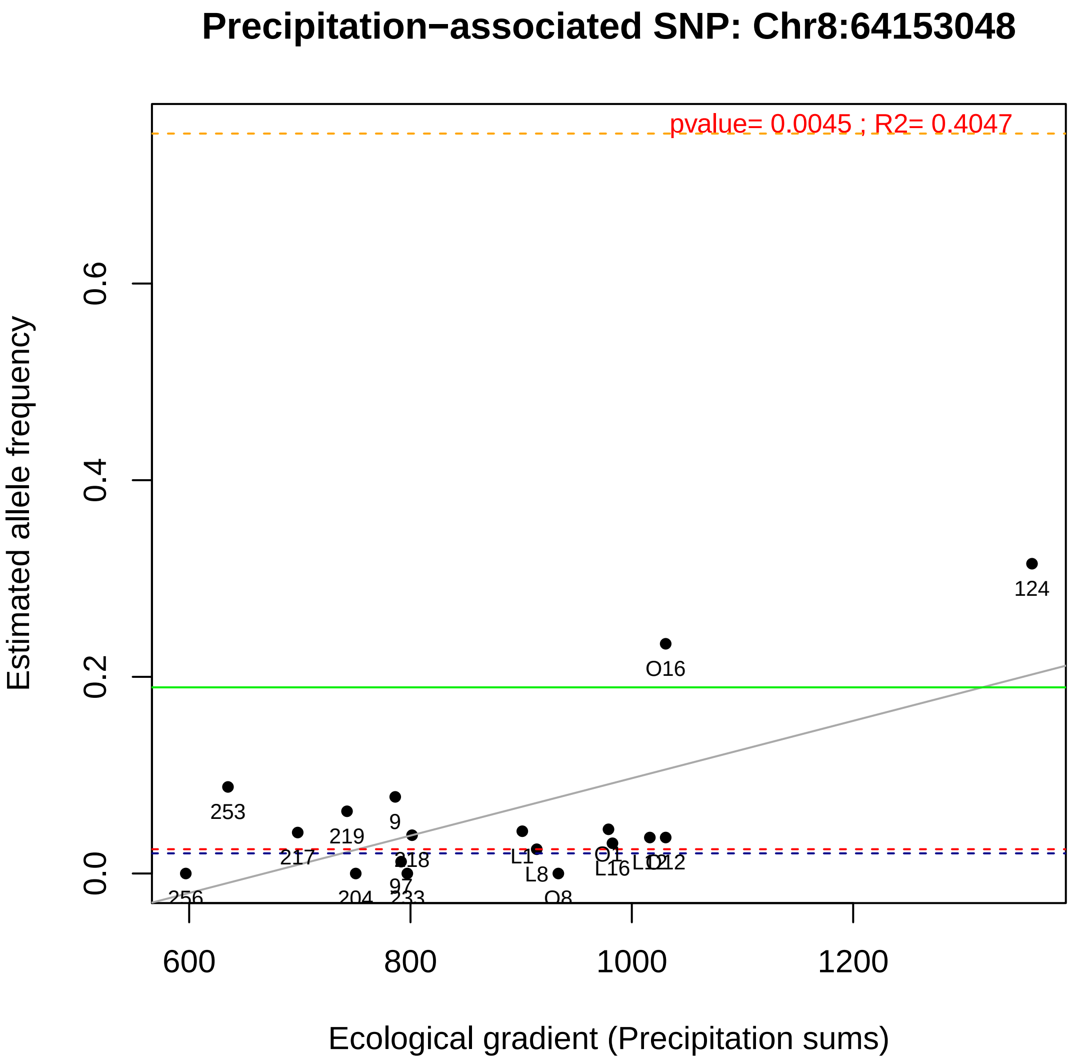

Precipitation−associated SNP: Chr8:64153048
pvalue= 0.0045 ; R2= 0.4047
Estimated allele frequency
0.0
0.2
0.4
0.6
600
800
1000
1200
Ecological gradient (Precipitation sums)
124
O16
253
219
9
217
218
L1
L8
O8
L16
L12
O12
97
233
204
256

**Precipitation−associated SNP: Chr8:64282196**

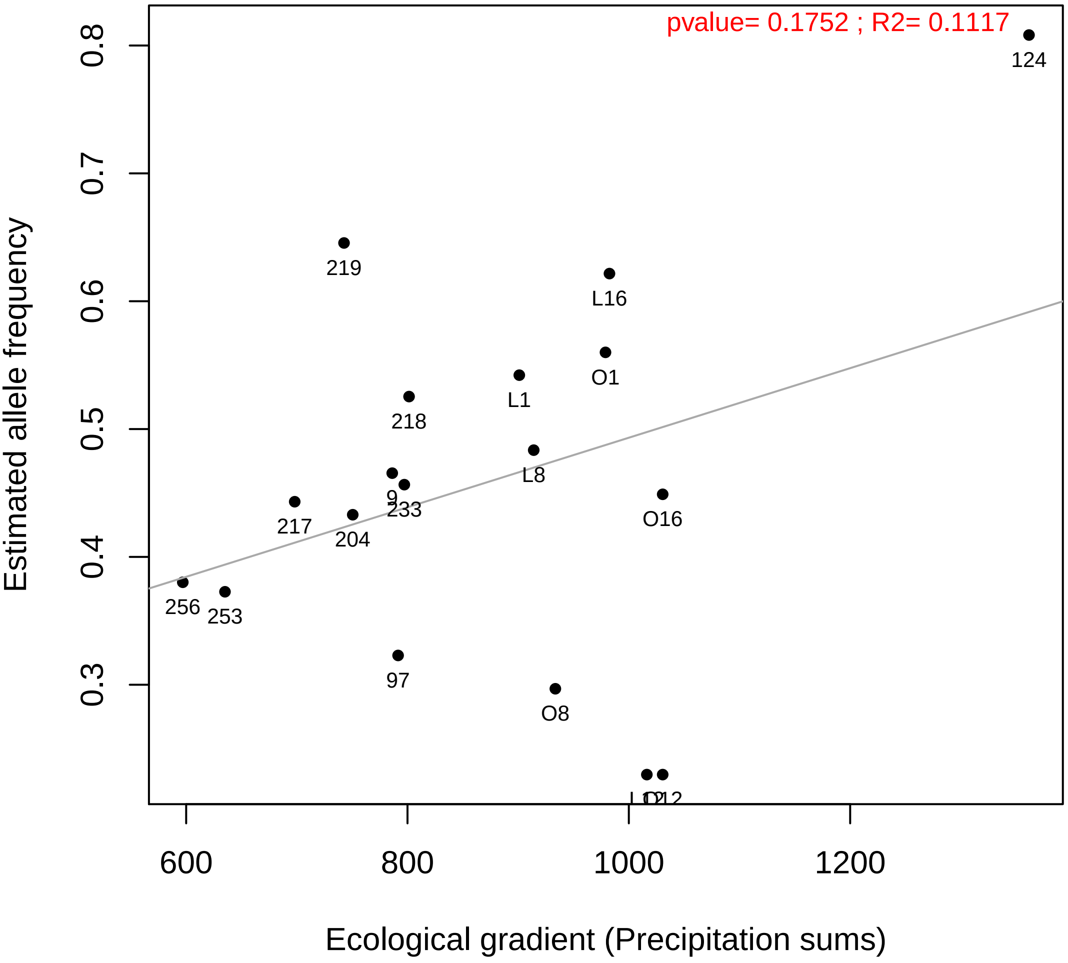

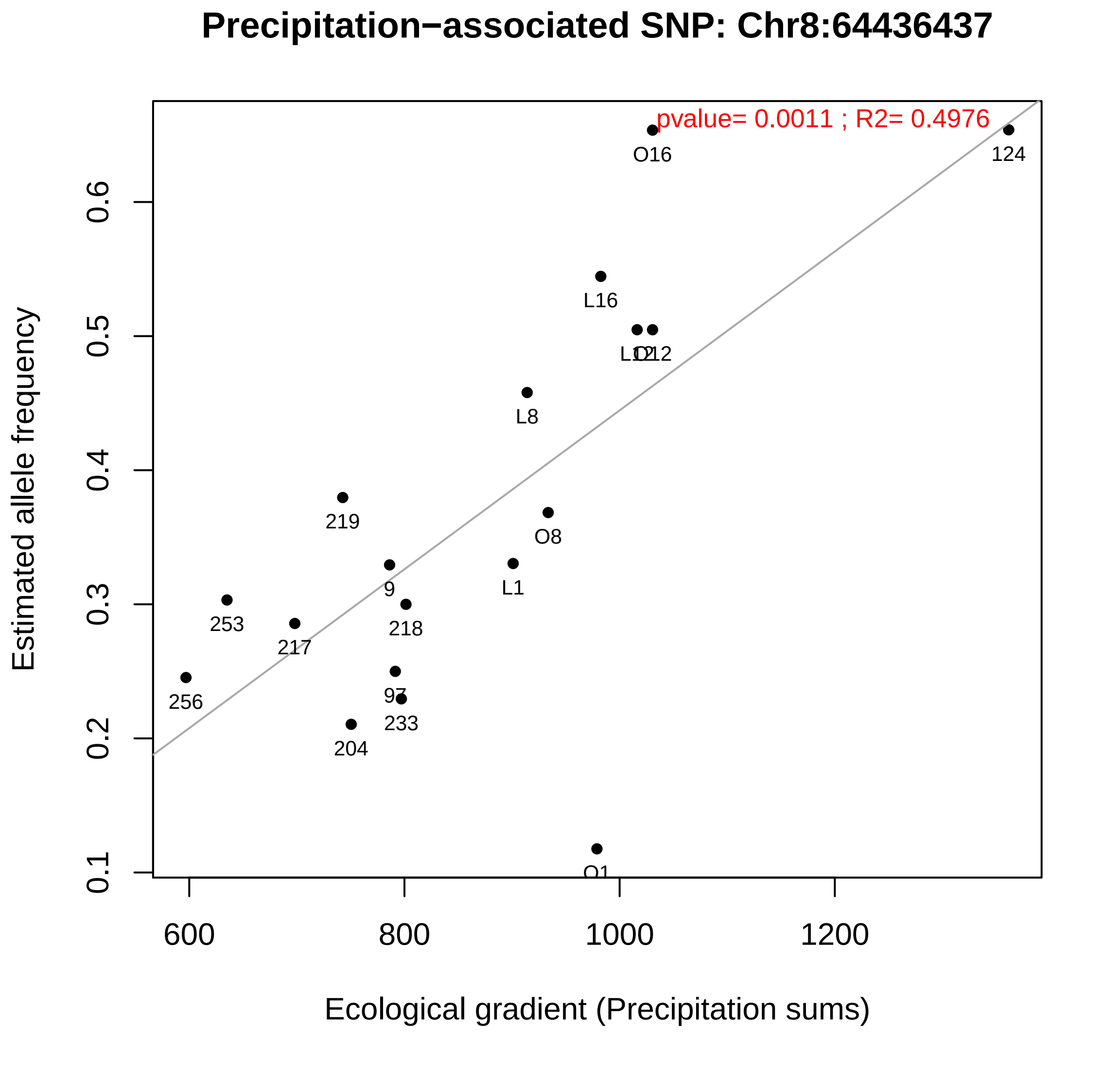

Precipitation−associated SNP: Chr8:64436437
pvalue= 0.0011 ; R2= 0.4976
Estimated allele frequency
Ecological gradient (Precipitation sums)
0.1
0.2
0.3
0.4
0.5
0.6
600
800
1000
1200
O16
124
L16
L12
O12
L8
219
O8
9
L1
253
218
217
256
97
233
204
O1

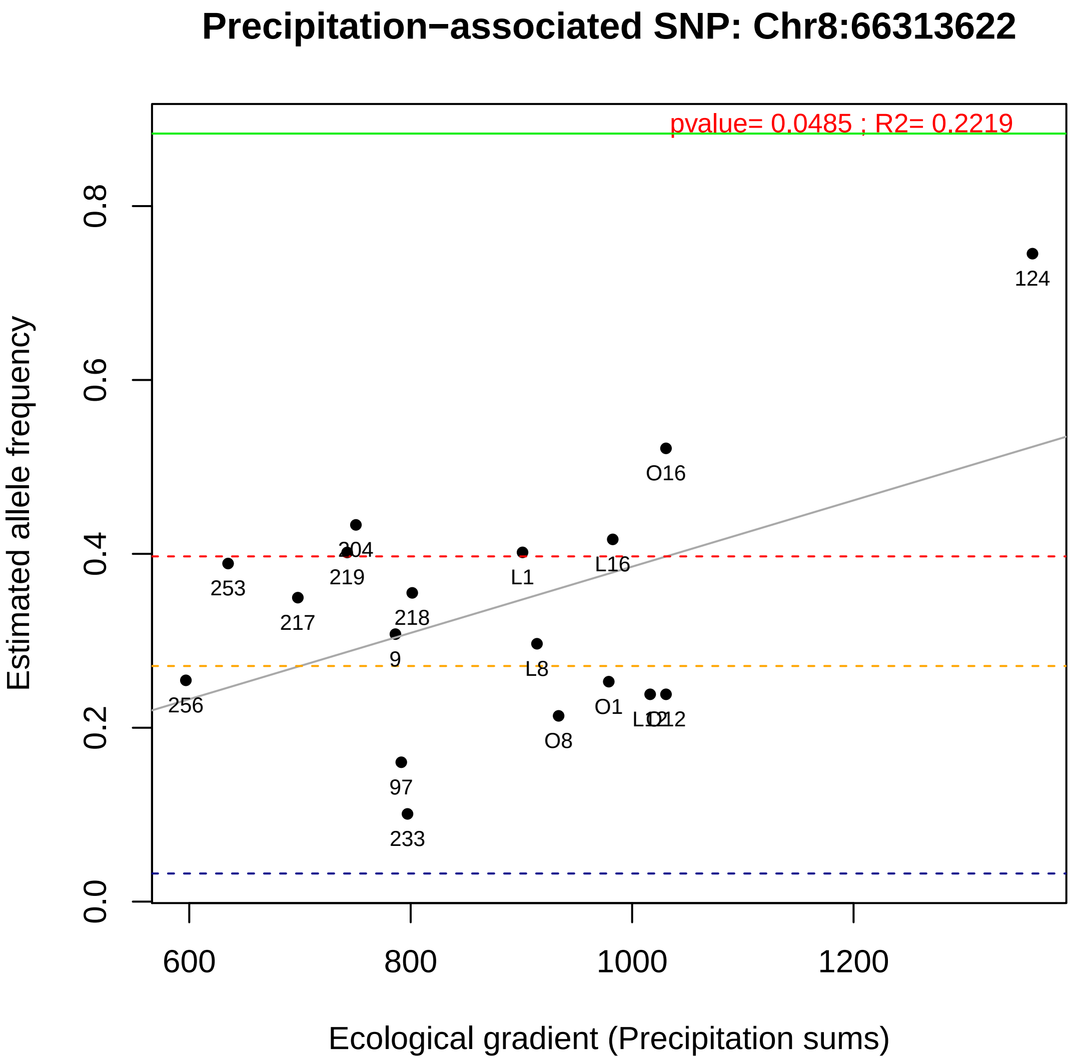
Precipitation−associated SNP: Chr8:66313622
pvalue= 0.0485 ; R2= 0.2219
Estimated allele frequency
Ecological gradient (Precipitation sums)
0.0
0.2
0.4
0.6
0.8
600
800
1000
1200
124
O16
L16
L1
204
219
253
217
218
9
L8
256
O1
L12
O12
O8
97
233

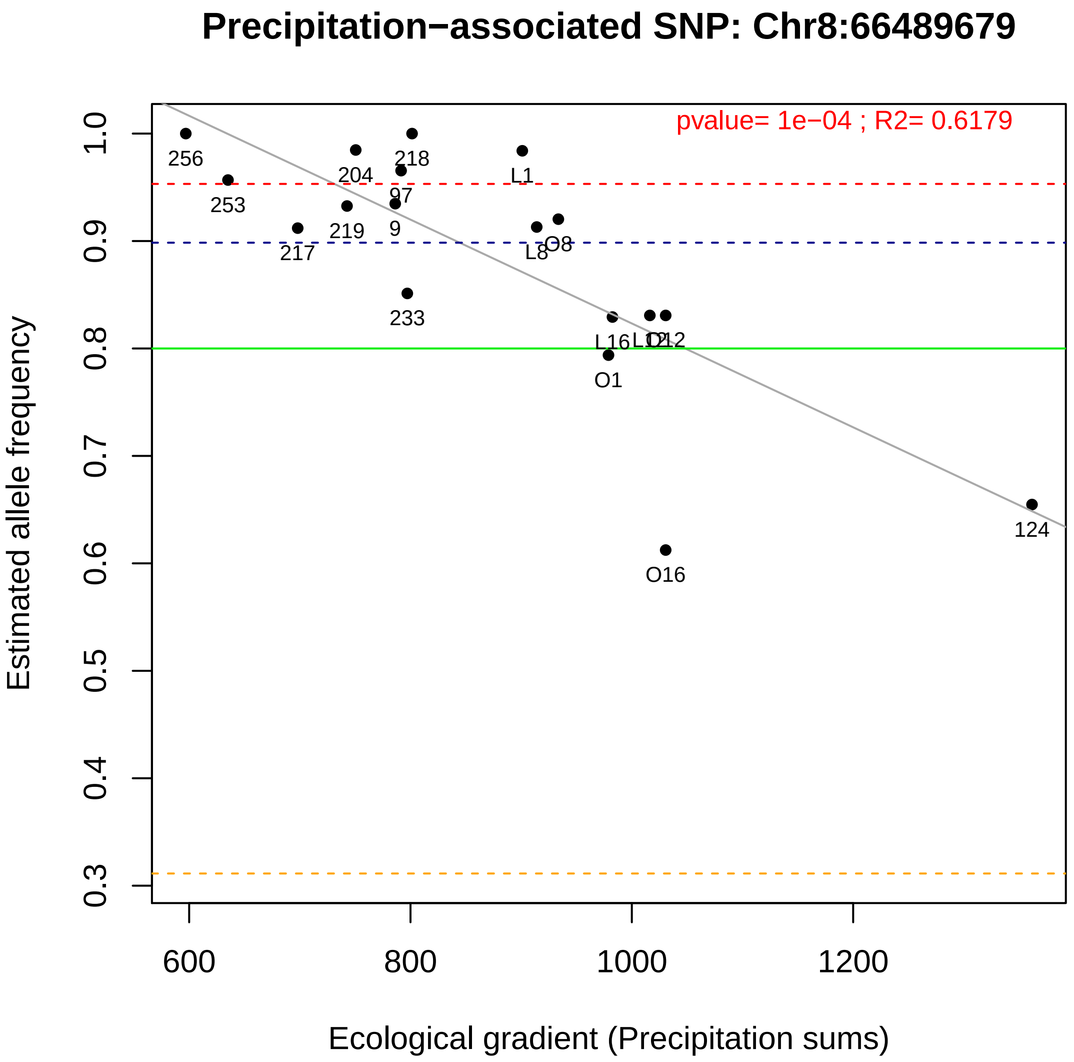
Precipitation−associated SNP: Chr8:66489679
pvalue= 1e−04 ; R2= 0.6179
Estimated allele frequency
Ecological gradient (Precipitation sums)
0.3
0.4
0.5
0.6
0.7
0.8
0.9
1.0
600
800
1000
1200
256
253
204
218
219
217
97
9
233
L1
L8
O8
L16
O1
O12
O16
124

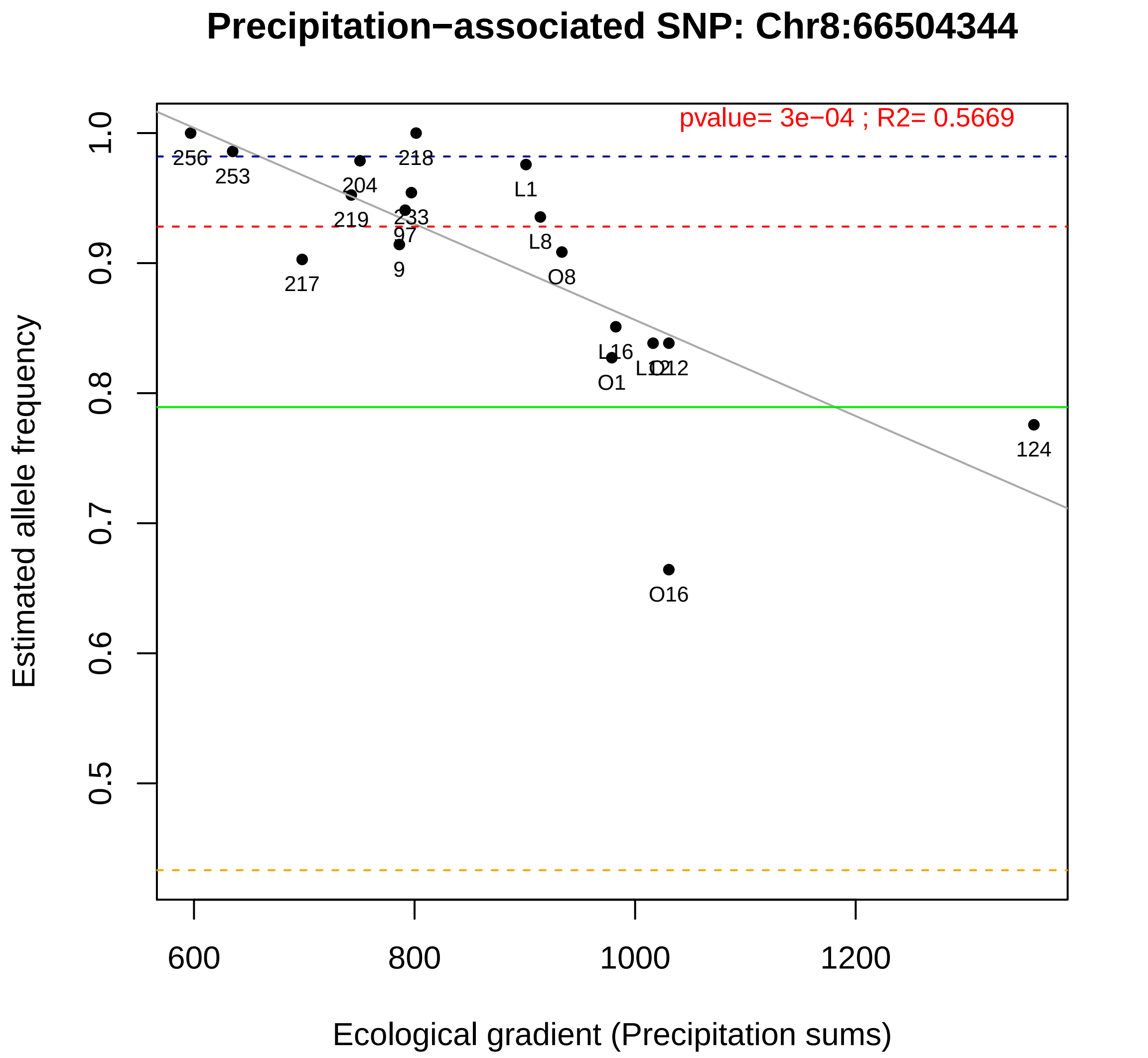
Precipitation−associated SNP: Chr8:66504344
pvalue= 3e−04 ; R2= 0.5669
Estimated allele frequency
Ecological gradient (Precipitation sums)
1.0
0.9
0.8
0.7
0.6
0.5
600
800
1000
1200
256
253
204
218
219
233
97
9
217
L1
L8
O8
L16
O1
O12
O16
124

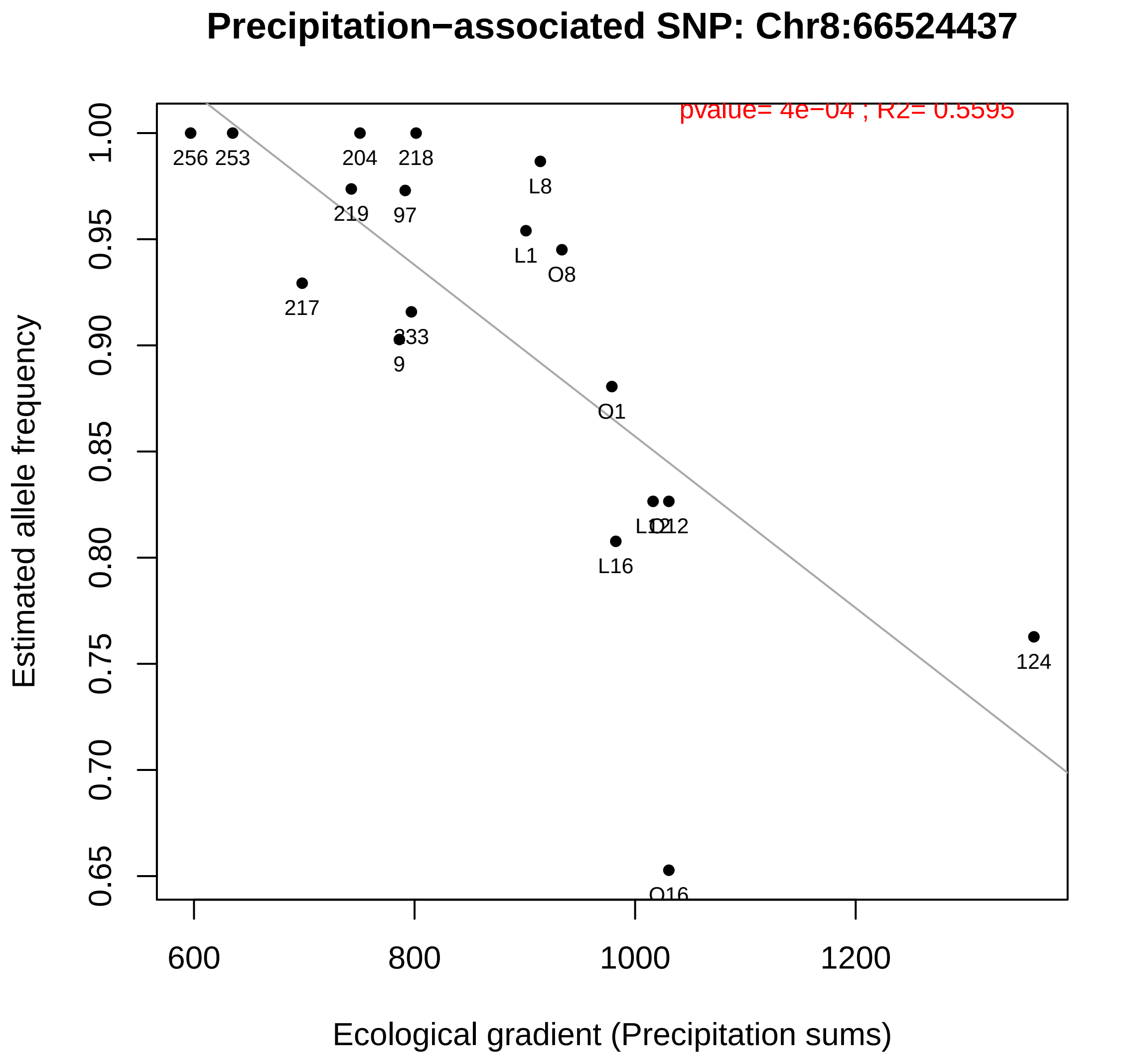
Precipitation−associated SNP: Chr8:66524437
pvalue= 4e−04 ; R2= 0.5595
Estimated allele frequency
Ecological gradient (Precipitation sums)
0.65
0.70
0.75
0.80
0.85
0.90
0.95
1.00
600
800
1000
1200
256
253
204
218
L8
219
97
L1
O8
217
233
9
O1
L12
O12
L16
124
O16

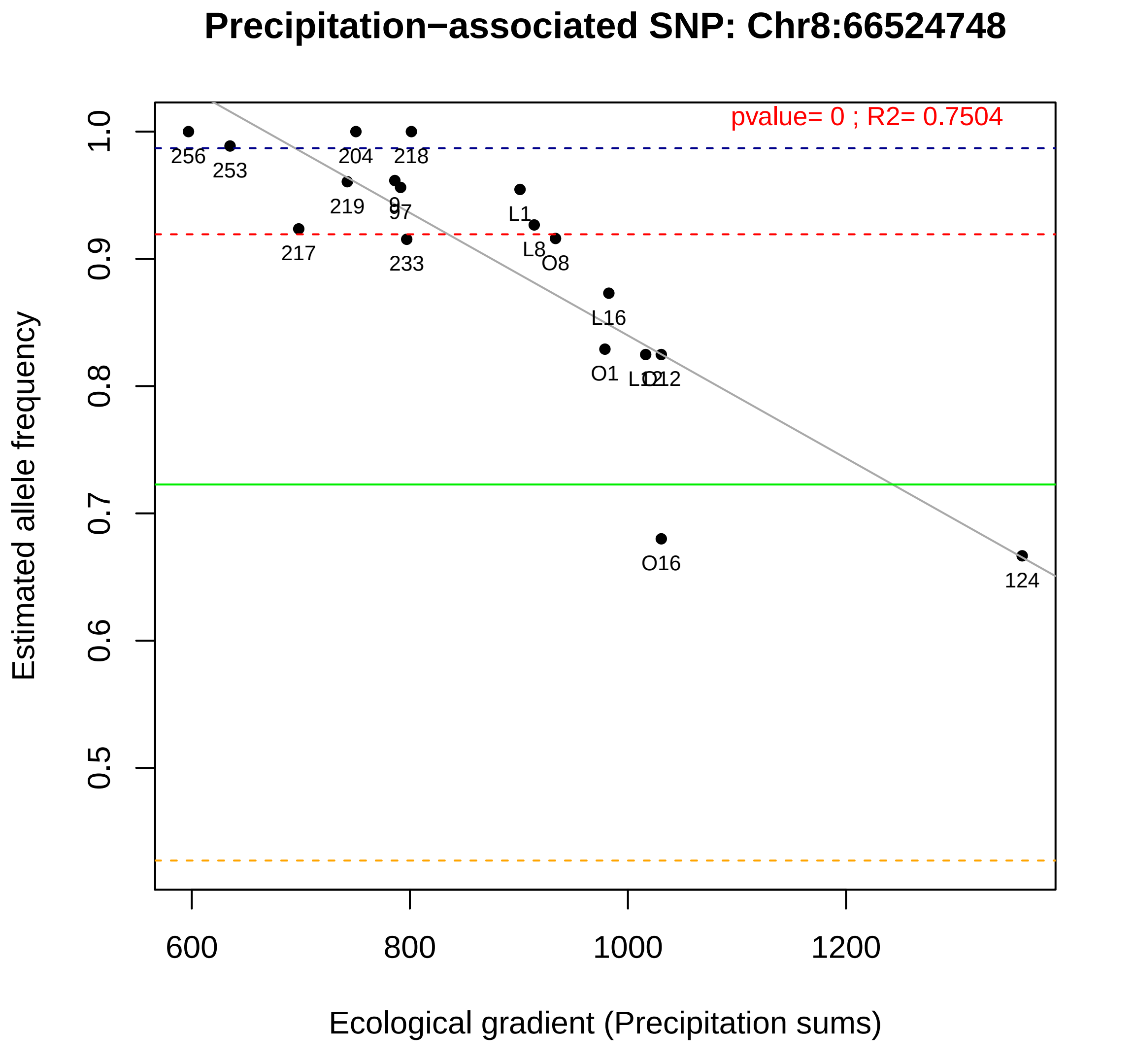

Precipitation−associated SNP: Chr8:66524748
pvalue= 0 ; R2= 0.7504
256
253
204
218
219
9
97
217
233
L1
L8
O8
L16
O1
L12
O12
O16
124
Estimated allele frequency
0.5
0.6
0.7
0.8
0.9
1.0
600
800
1000
1200
Ecological gradient (Precipitation sums)

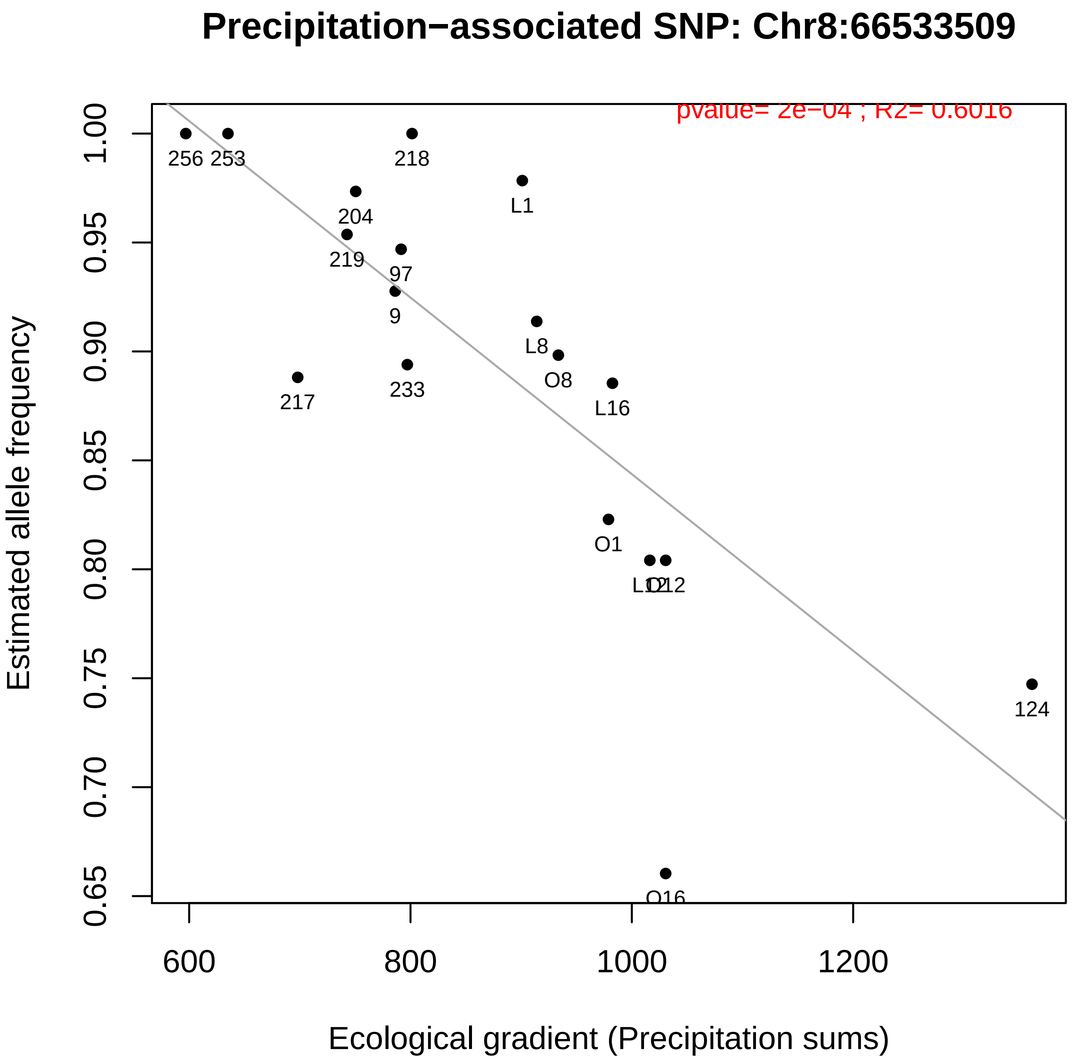

Precipitation−associated SNP: Chr8:66533509
pvalue= 2e−04 ; R2= 0.6016
Estimated allele frequency
Ecological gradient (Precipitation sums)
0.65
0.70
0.75
0.80
0.85
0.90
0.95
1.00
600
800
1000
1200
256
253
218
204
L1
219
97
9
L8
O8
217
233
L16
O1
L12
O12
124
O16

**Precipitation−associated SNP: Chr8:66562283**

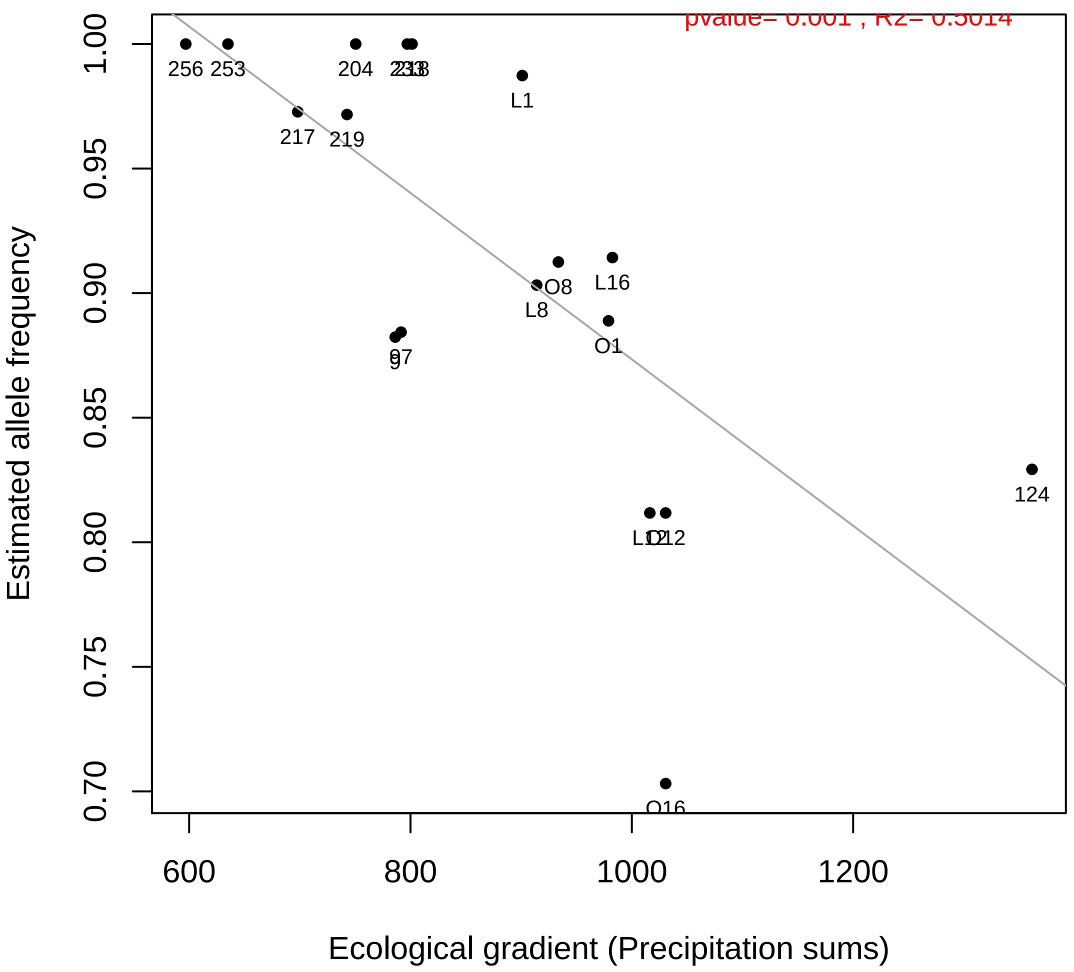

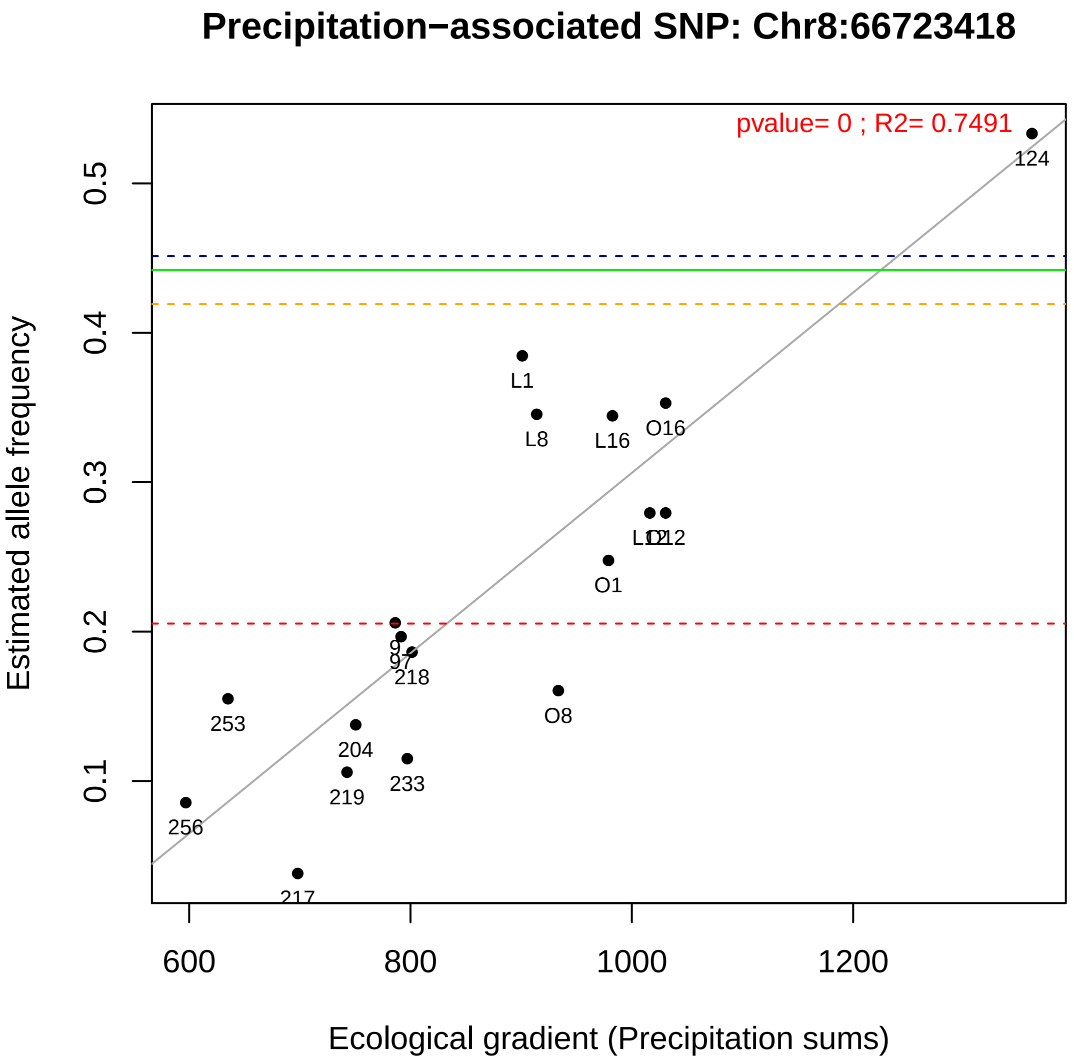

Precipitation−associated SNP: Chr8:66723418
pvalue= 0 ; R2= 0.7491
Estimated allele frequency
Ecological gradient (Precipitation sums)
0.1
0.2
0.3
0.4
0.5
600
800
1000
1200
124
L1
L8
L16
O16
O12
O1
O8
218
253
204
233
219
256
217

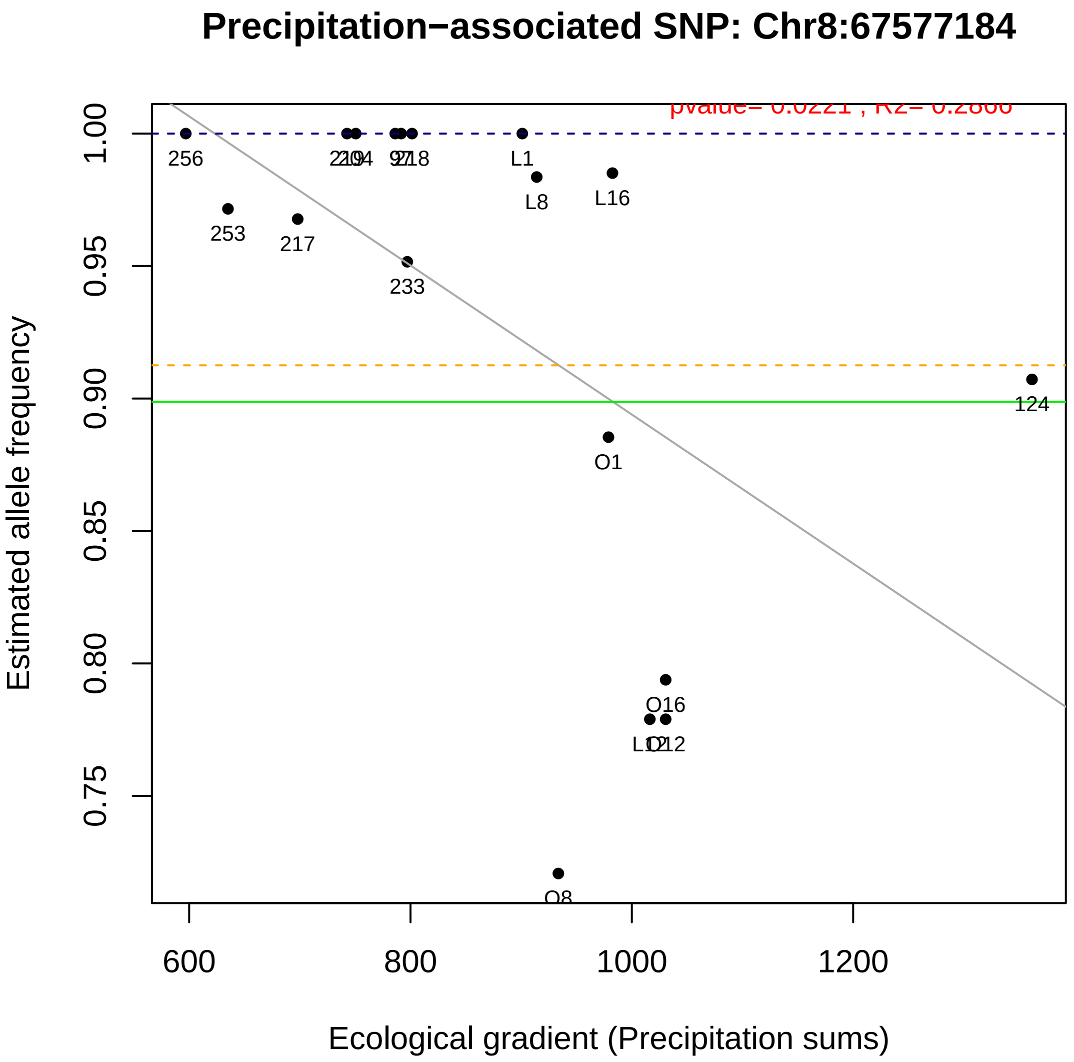

Precipitation−associated SNP: Chr8:67577184
pvalue= 0.0221 , R2= 0.2866
Estimated allele frequency
Ecological gradient (Precipitation sums)
0.75
0.80
0.85
0.90
0.95
1.00
600
800
1000
1200
256
253
217
233
L1
L8
L16
124
O1
O16
O12
O8

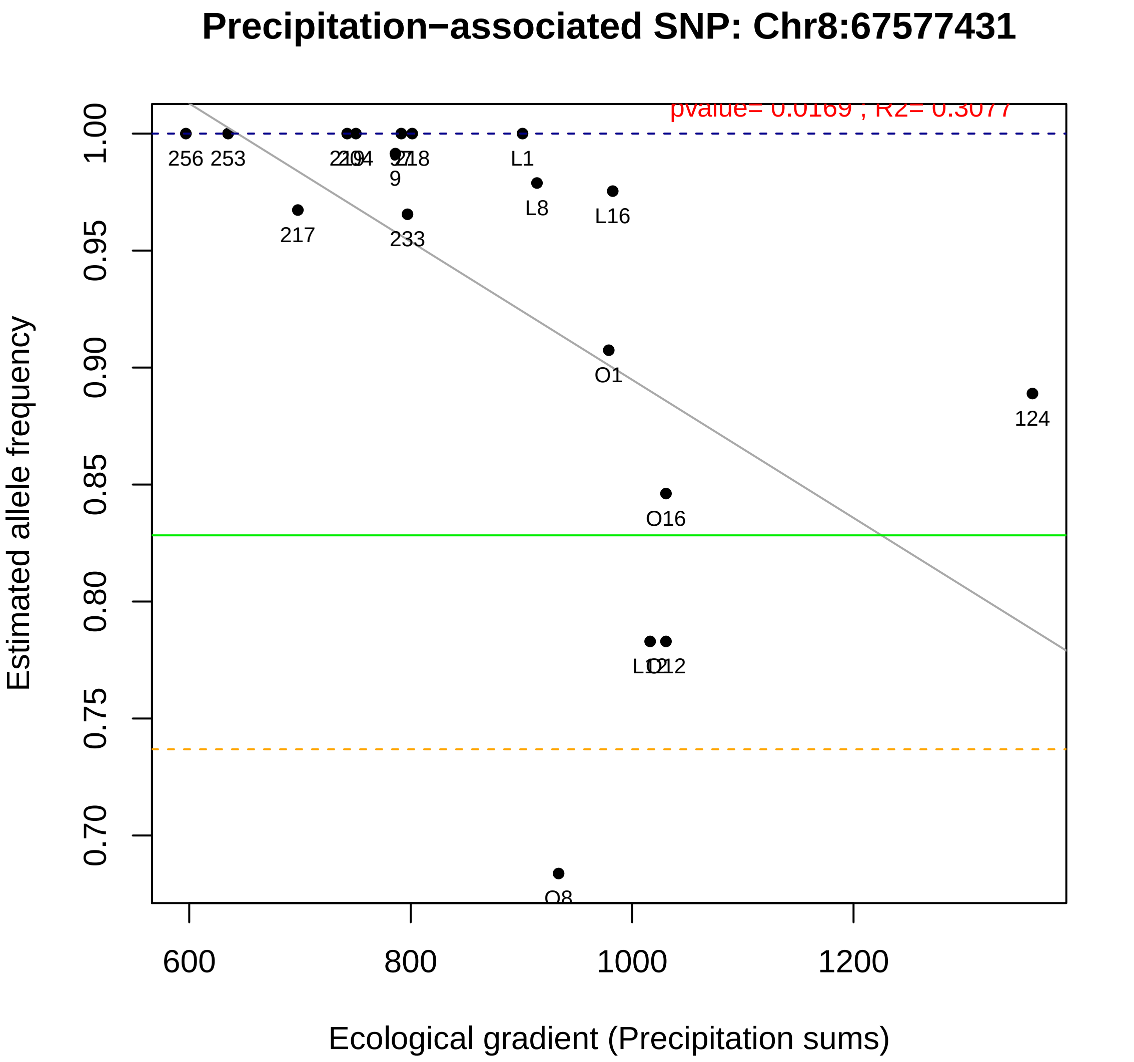
Precipitation−associated SNP: Chr8:67577431
pvalue= 0.0169 ; R2= 0.3077
Estimated allele frequency
Ecological gradient (Precipitation sums)
1.00
0.95
0.90
0.85
0.80
0.75
0.70
600
800
1000
1200
256
253
217
233
L1
L8
L16
O1
124
O16
O8

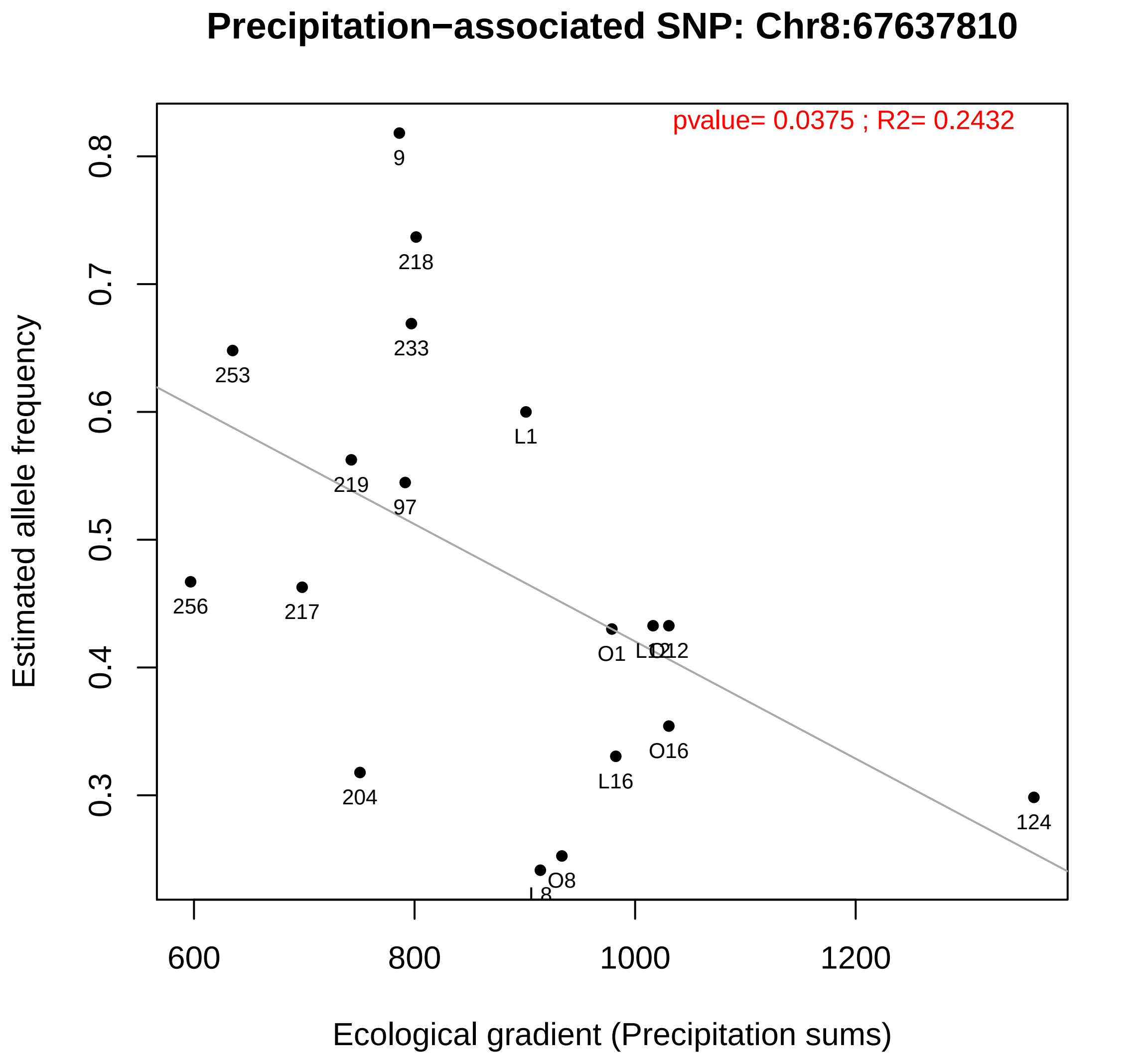
Precipitation−associated SNP: Chr8:67637810
pvalue= 0.0375 ; R2= 0.2432
Estimated allele frequency
0.3
0.4
0.5
0.6
0.7
0.8
600
800
1000
1200
Ecological gradient (Precipitation sums)
9
218
233
253
L1
219
97
256
217
O1
O12
O16
L16
204
124
O8

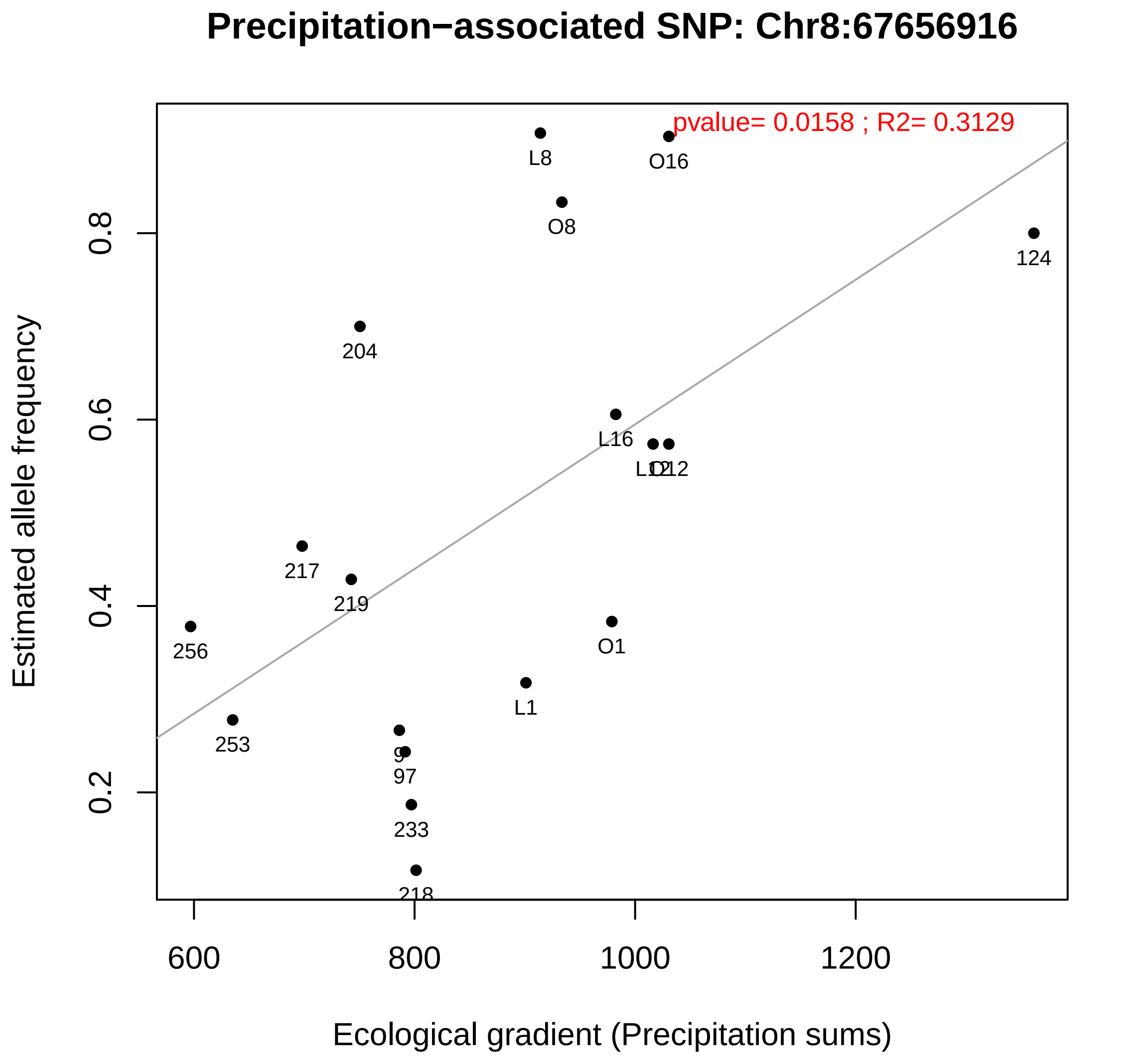

Precipitation−associated SNP: Chr8:67656916
pvalue= 0.0158 ; R2= 0.3129
Estimated allele frequency
Ecological gradient (Precipitation sums)
0.2
0.4
0.6
0.8
600
800
1000
1200
L8
O16
O8
124
204
L16
L12
O12
217
219
256
O1
L1
253
9
97
233
218

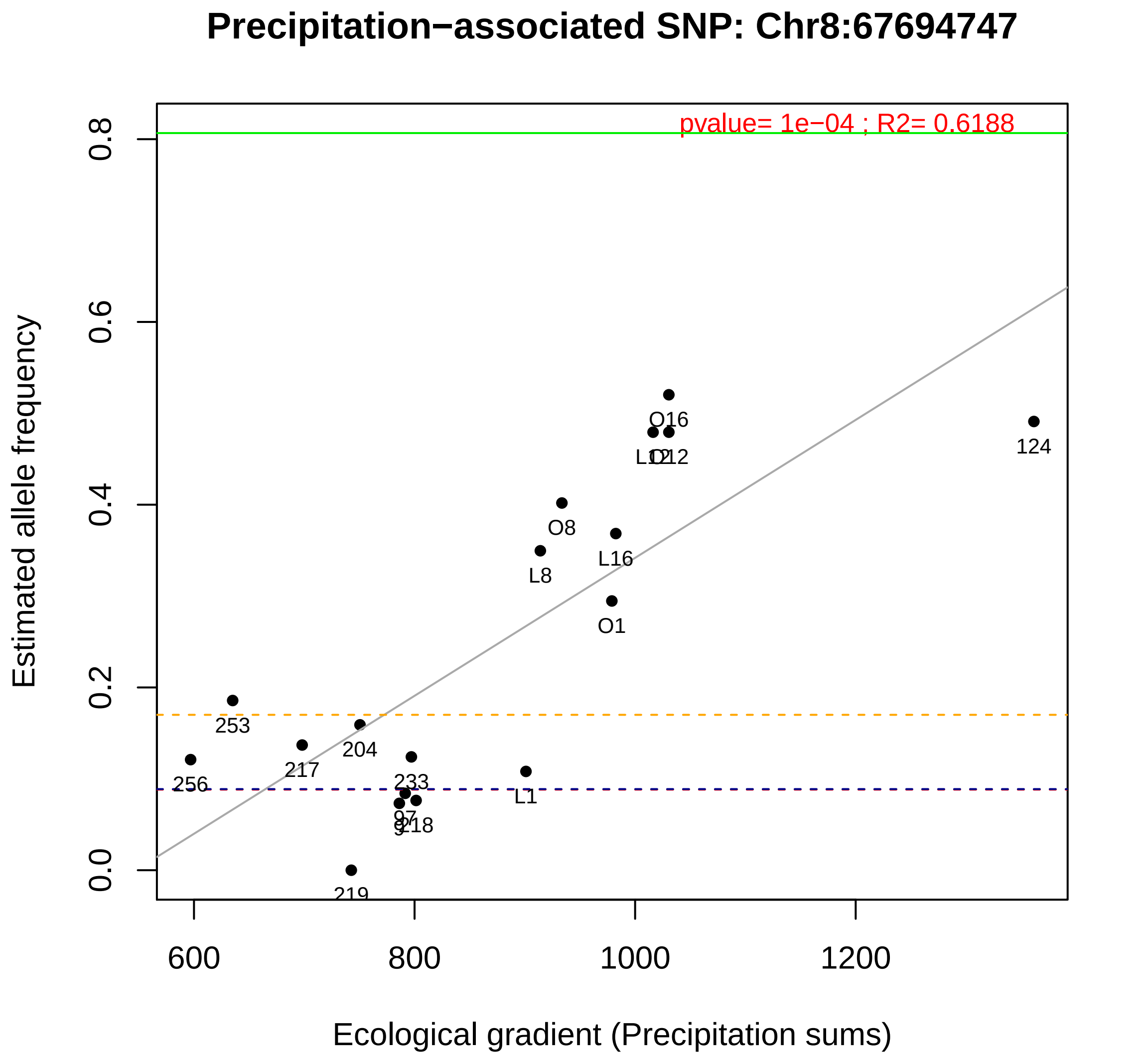
Precipitation−associated SNP: Chr8:67694747
pvalue= 1e−04 ; R2= 0.6188
Estimated allele frequency
Ecological gradient (Precipitation sums)
0.0
0.2
0.4
0.6
0.8
600
800
1000
1200
O16
L12
O12
124
O8
L16
L8
O1
253
204
217
256
233
L1
97
9
218
219

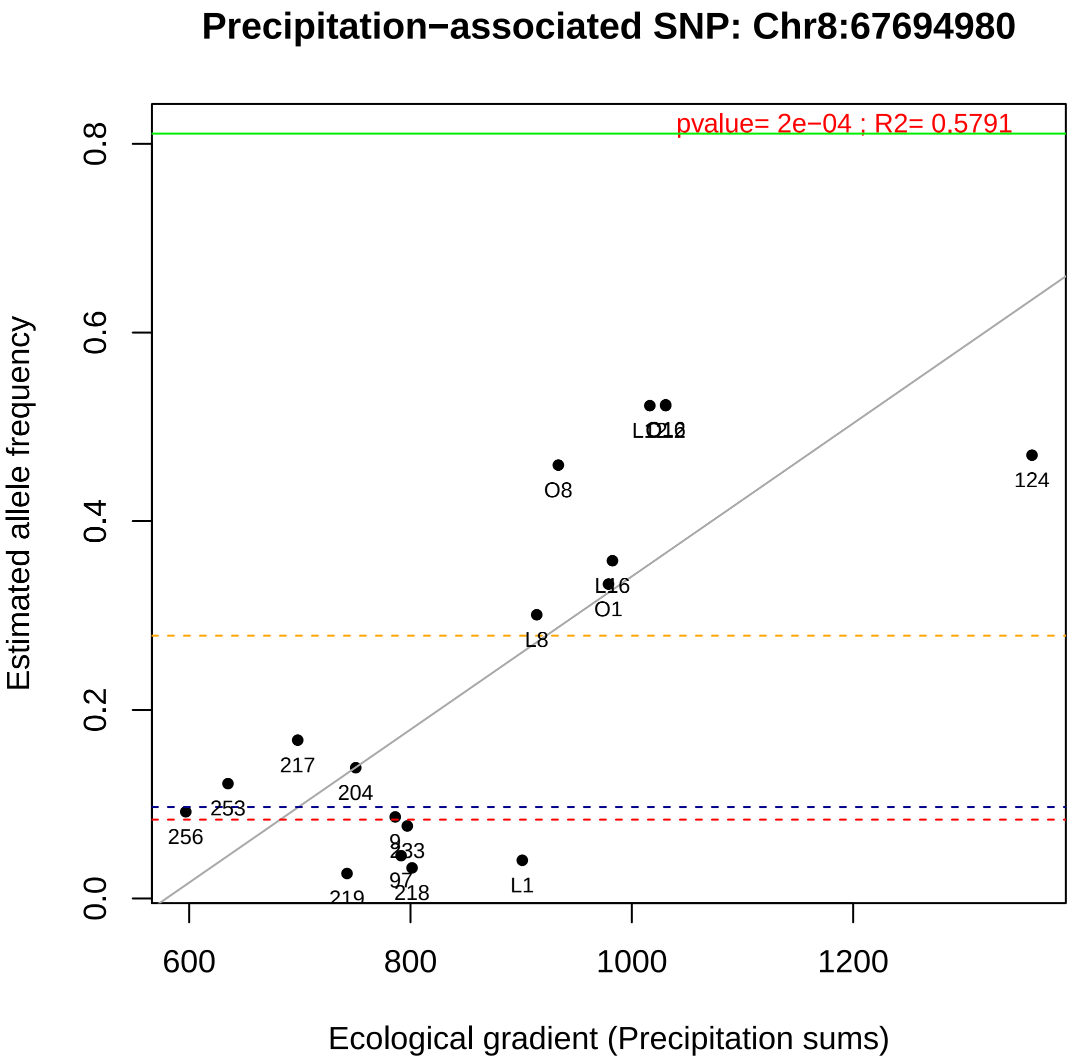

Precipitation−associated SNP: Chr8:67694980
pvalue= 2e−04 ; R2= 0.5791
Estimated allele frequency
0.0
0.2
0.4
0.6
0.8
600
800
1000
1200
Ecological gradient (Precipitation sums)
124
O8
L16
O1
L8
217
204
253
256
9
233
97
218
219
L1

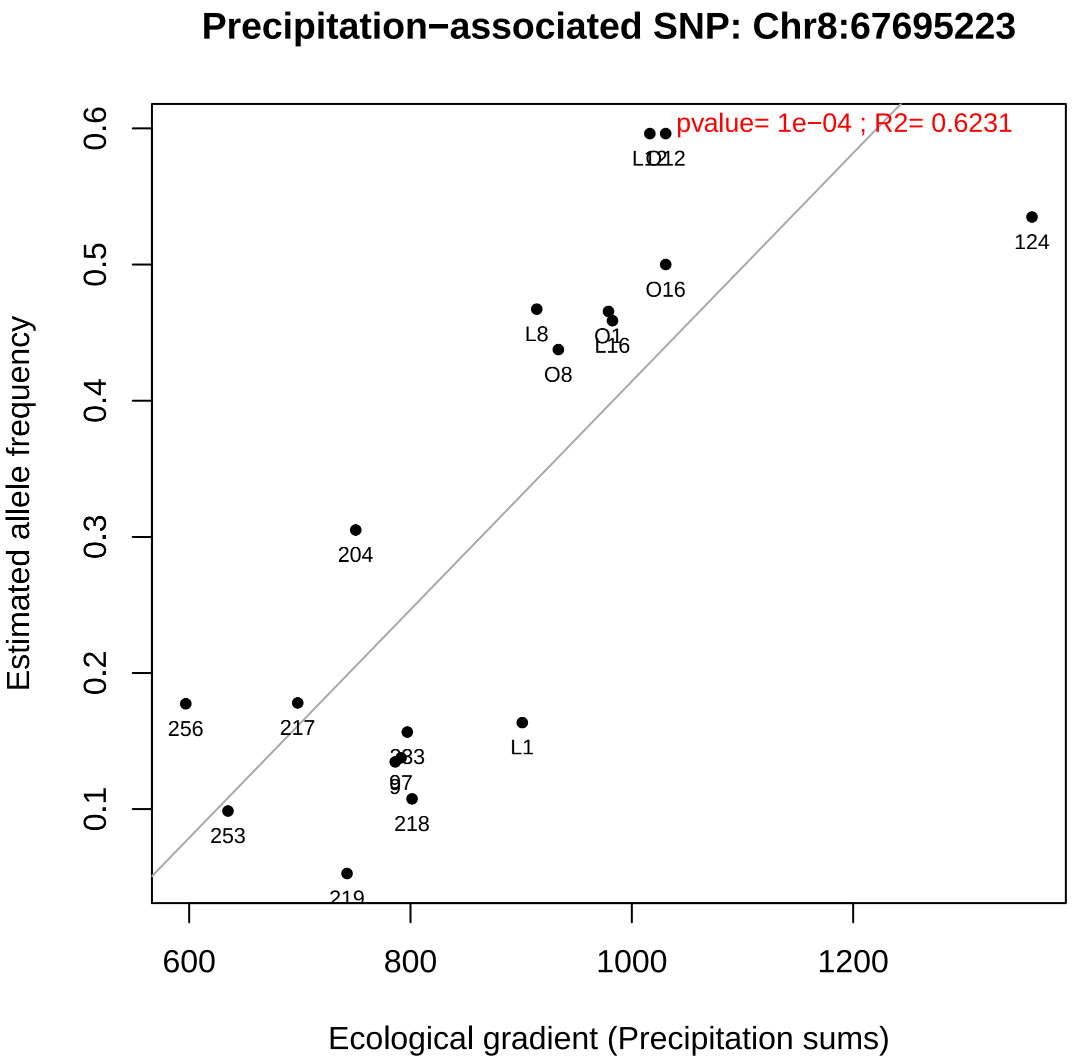
Precipitation−associated SNP: Chr8:67695223
pvalue= 1e−04 ; R2= 0.6231
Estimated allele frequency
Ecological gradient (Precipitation sums)
0.1
0.2
0.3
0.4
0.5
0.6
600
800
1000
1200
124
O16
L8
O1
L16
O8
204
256
217
233
L1
97
218
253
219

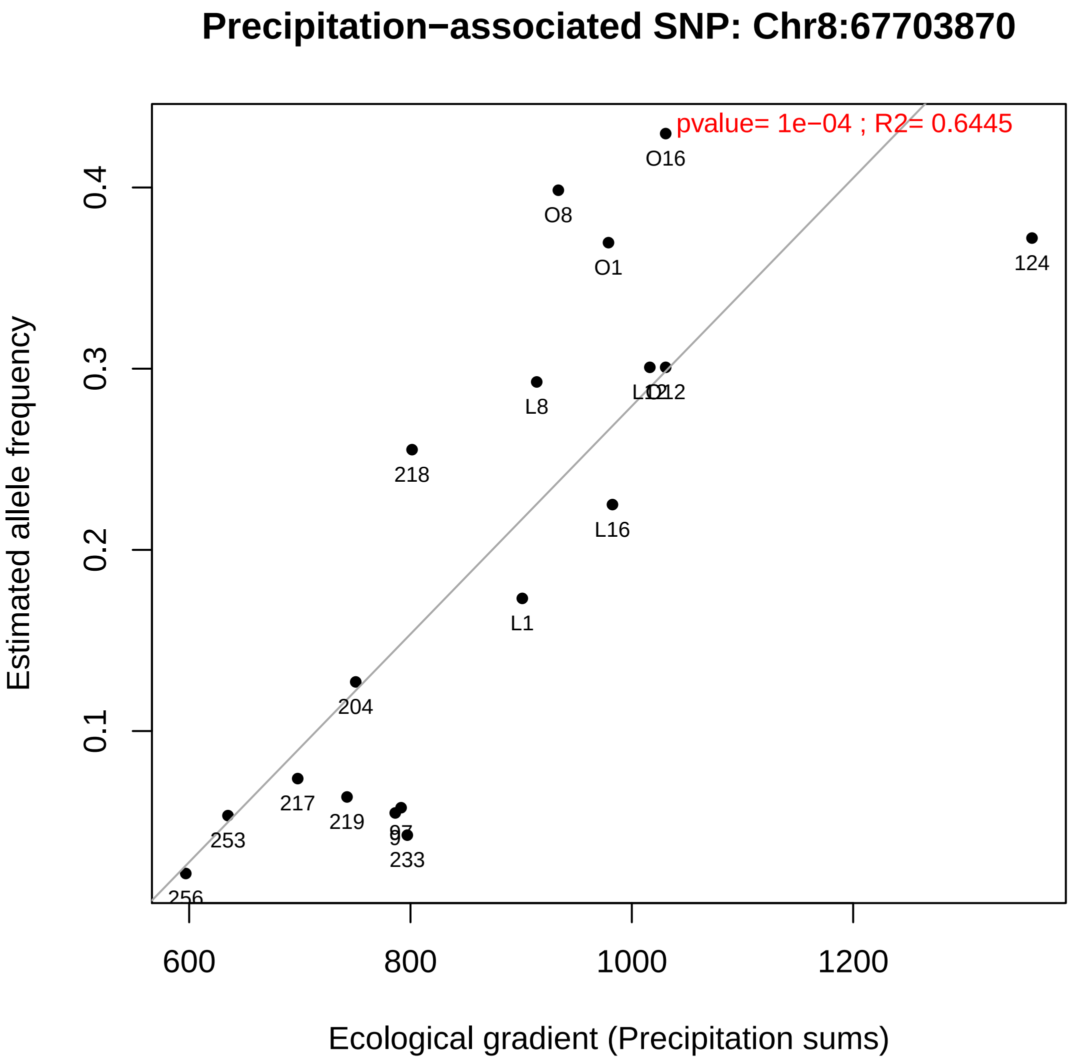
Precipitation−associated SNP: Chr8:67703870
pvalue= 1e−04 ; R2= 0.6445
O16
O8
O1
124
L12
O12
L8
218
L16
L1
204
217
219
253
233
256
0.1
0.2
0.3
0.4
Estimated allele frequency
600
800
1000
1200
Ecological gradient (Precipitation sums)

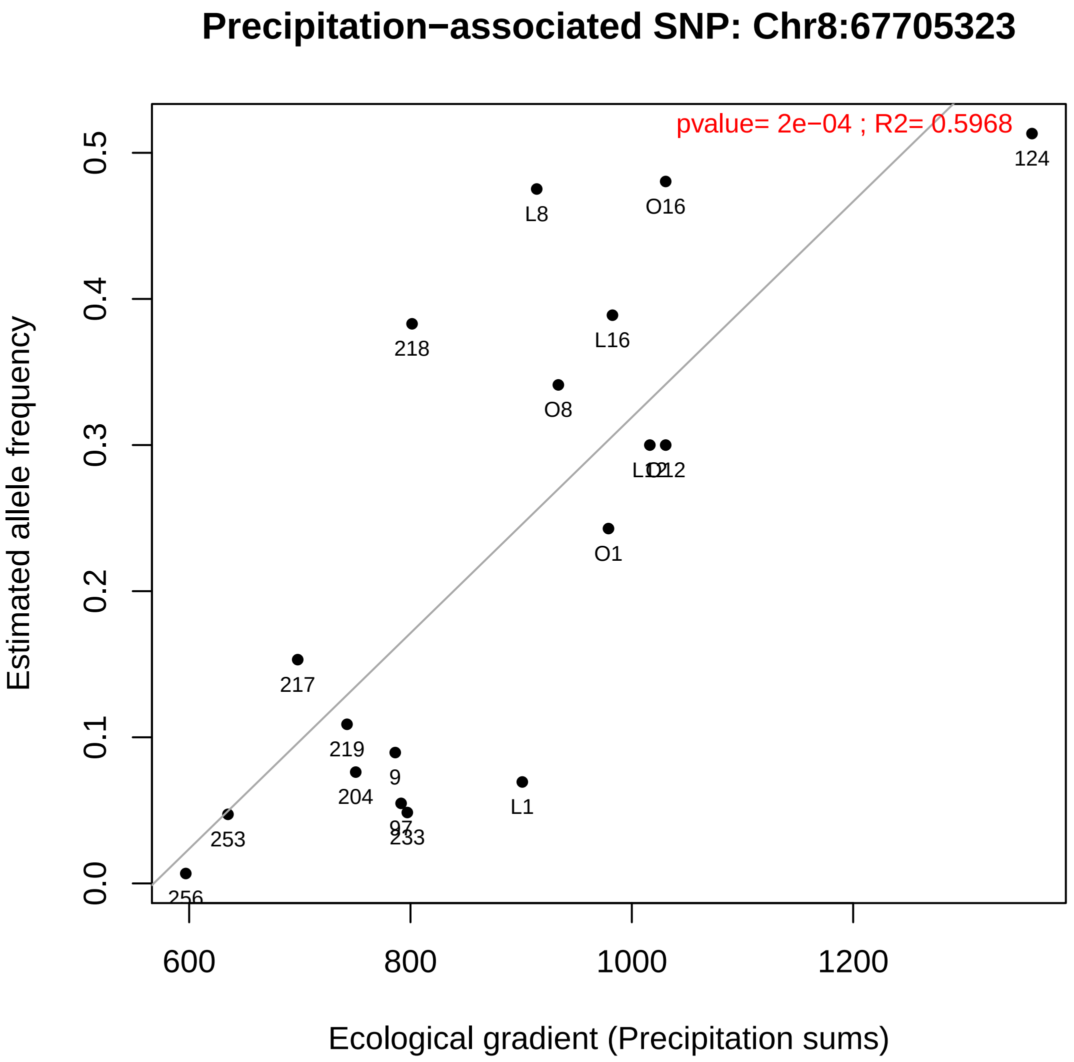

Precipitation−associated SNP: Chr8:67705323
pvalue= 2e−04 ; R2= 0.5968
124
L8
O16
218
L16
O8
L10
O12
O1
217
219
9
204
L1
253
97
233
256
0.0
0.1
0.2
0.3
0.4
0.5
Estimated allele frequency
600
800
1000
1200
Ecological gradient (Precipitation sums)

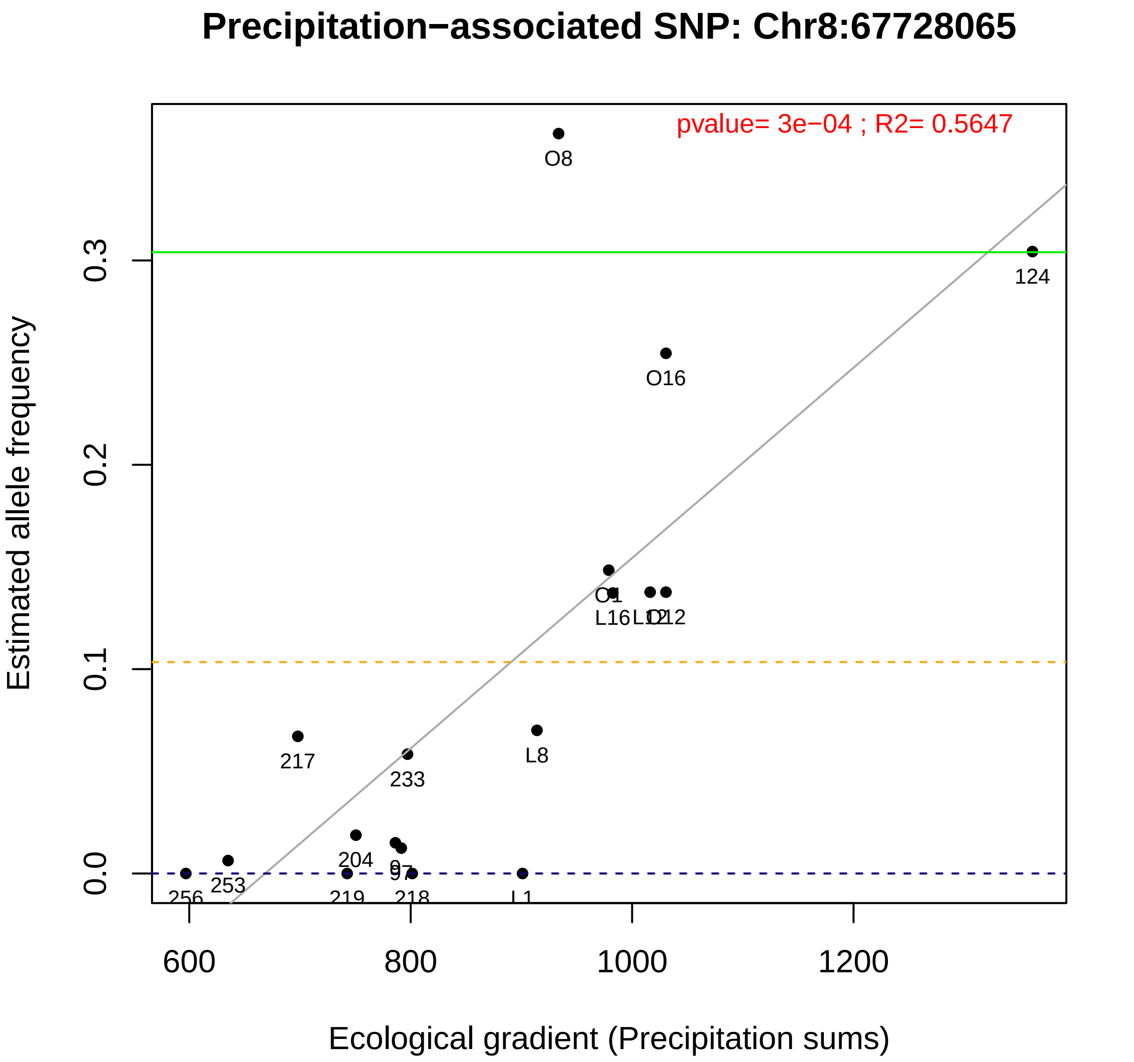
Precipitation−associated SNP: Chr8:67728065
pvalue= 3e−04 ; R2= 0.5647
O8
124
O16
O1
L16
L12
O12
217
233
L8
204
253
256
219
218
L1
0.0
0.1
0.2
0.3
600
800
1000
1200
Estimated allele frequency
Ecological gradient (Precipitation sums)

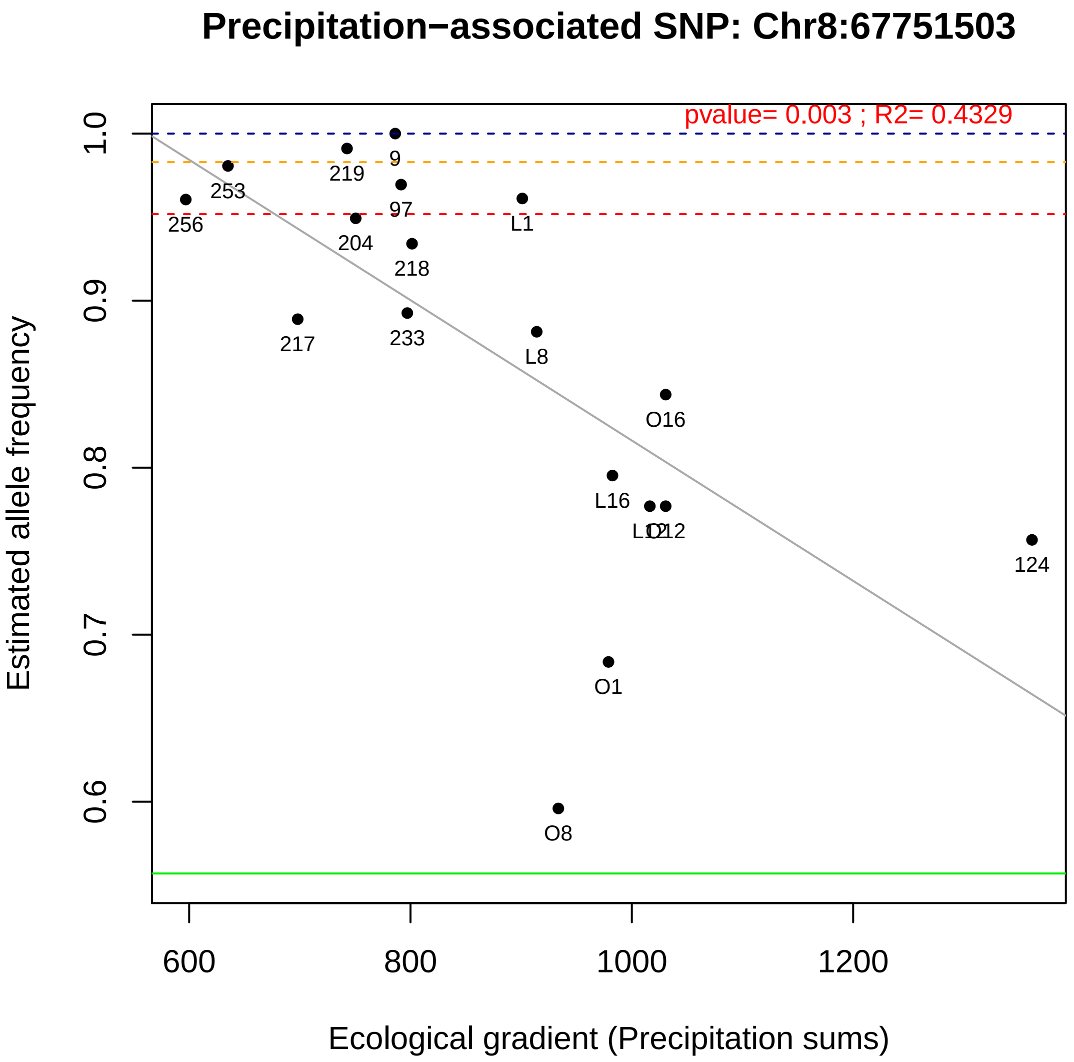

Precipitation−associated SNP: Chr8:67751503
pvalue= 0.003 ; R2= 0.4329
Estimated allele frequency
Ecological gradient (Precipitation sums)
0.6
0.7
0.8
0.9
1.0
600
800
1000
1200
256
253
219
9
97
204
218
L1
217
233
L8
O16
L16
L12
O12
124
O1
O8

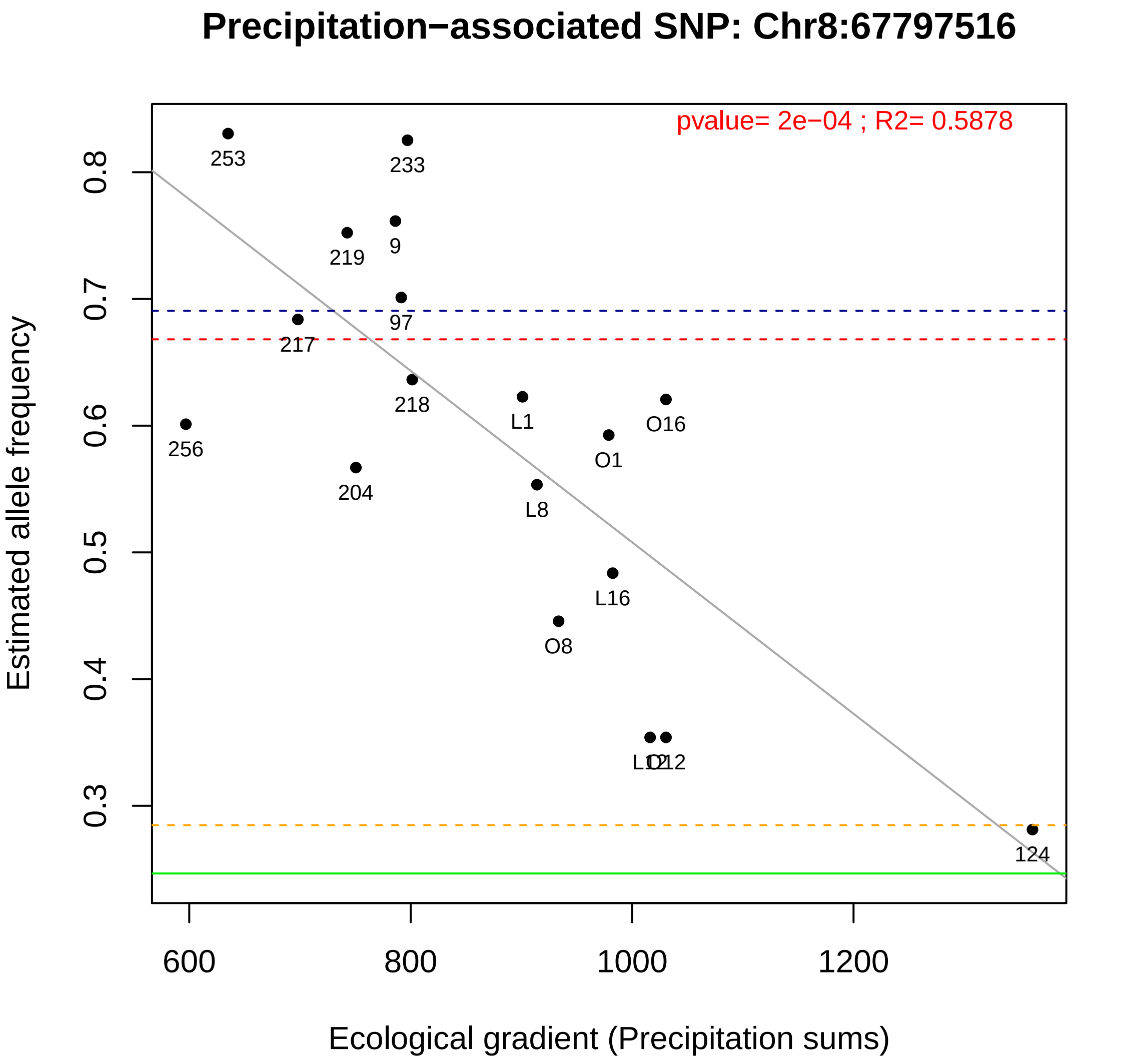

Precipitation−associated SNP: Chr8:67797516
pvalue= 2e−04 ; R2= 0.5878
Estimated allele frequency
0.3
0.4
0.5
0.6
0.7
0.8
253
233
219
9
97
217
218
256
L1
O16
O1
204
L8
L16
O8
L12
O12
124
600
800
1000
1200
Ecological gradient (Precipitation sums)

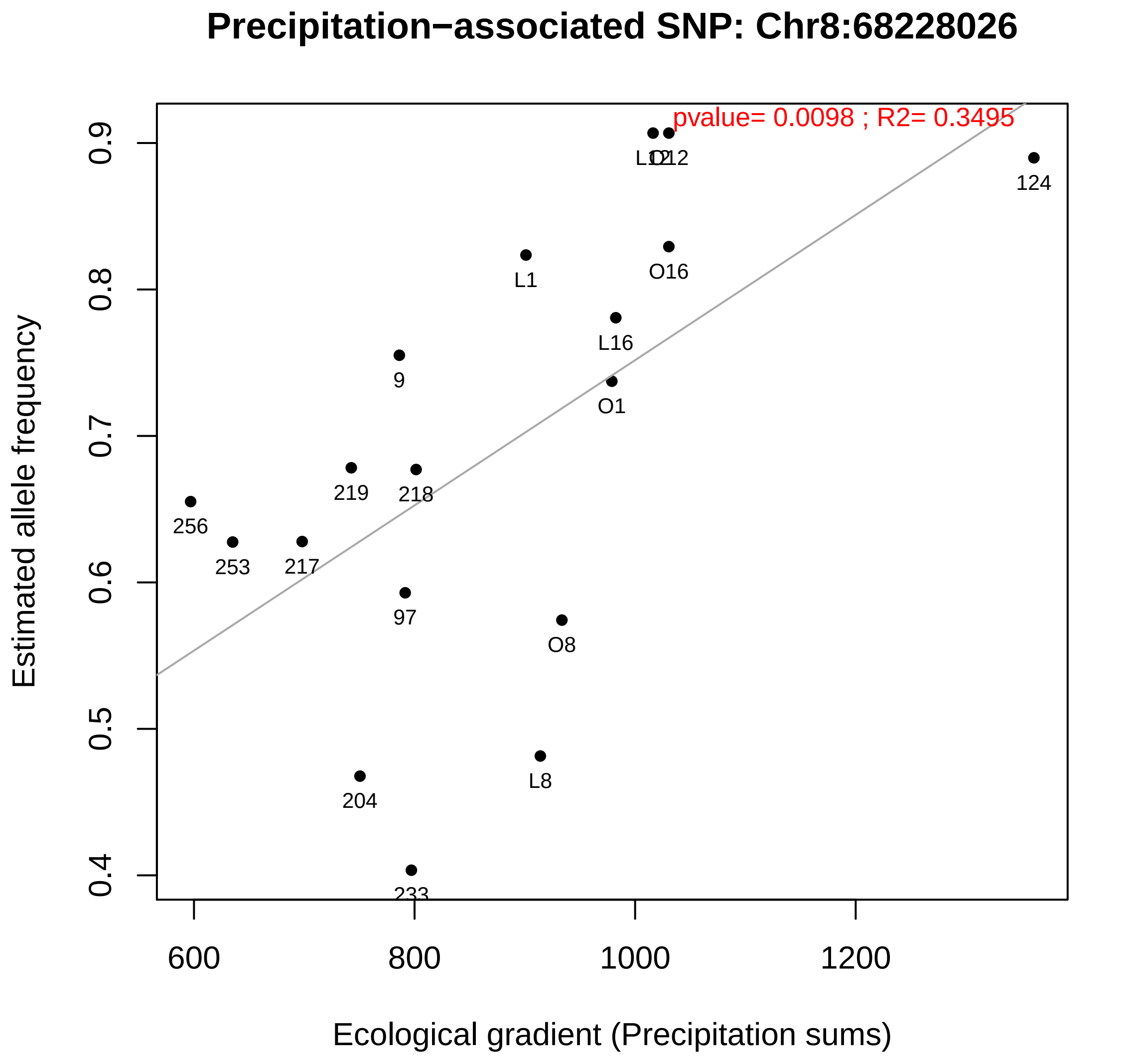
Precipitation−associated SNP: Chr8:68228026
pvalue= 0.0098 ; R2= 0.3495
Estimated allele frequency
0.4
0.5
0.6
0.7
0.8
0.9
600
800
1000
1200
Ecological gradient (Precipitation sums)
L10
O12
124
L1
O16
L16
9
O1
219
218
256
253
217
97
O8
L8
204
233

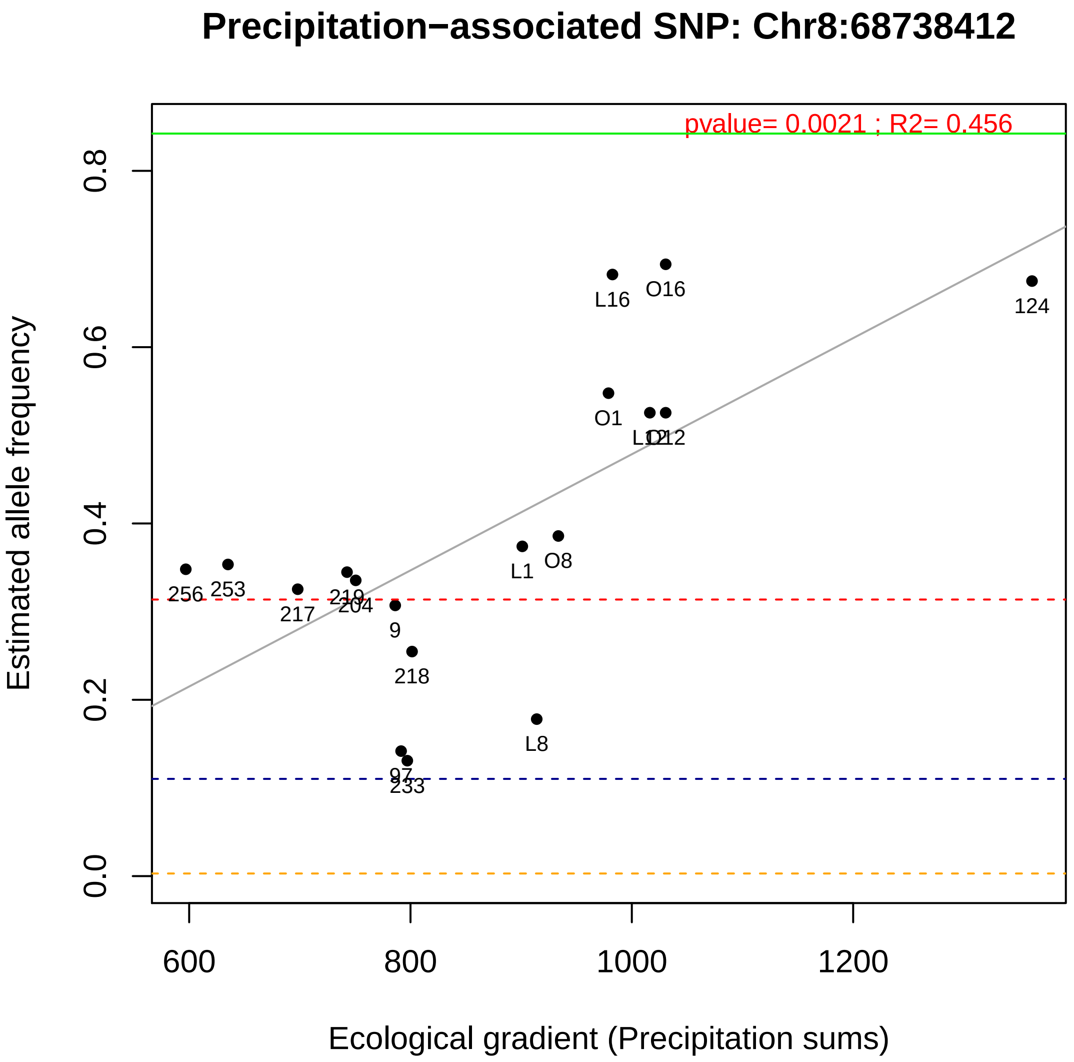

Precipitation−associated SNP: Chr8:68738412
pvalue= 0.0021 ; R2= 0.456
Estimated allele frequency
0.0
0.2
0.4
0.6
0.8
600
800
1000
1200
Ecological gradient (Precipitation sums)
L16
O16
124
O1
L12
O12
L1
O8
256
253
217
219
204
9
218
L8
97
233

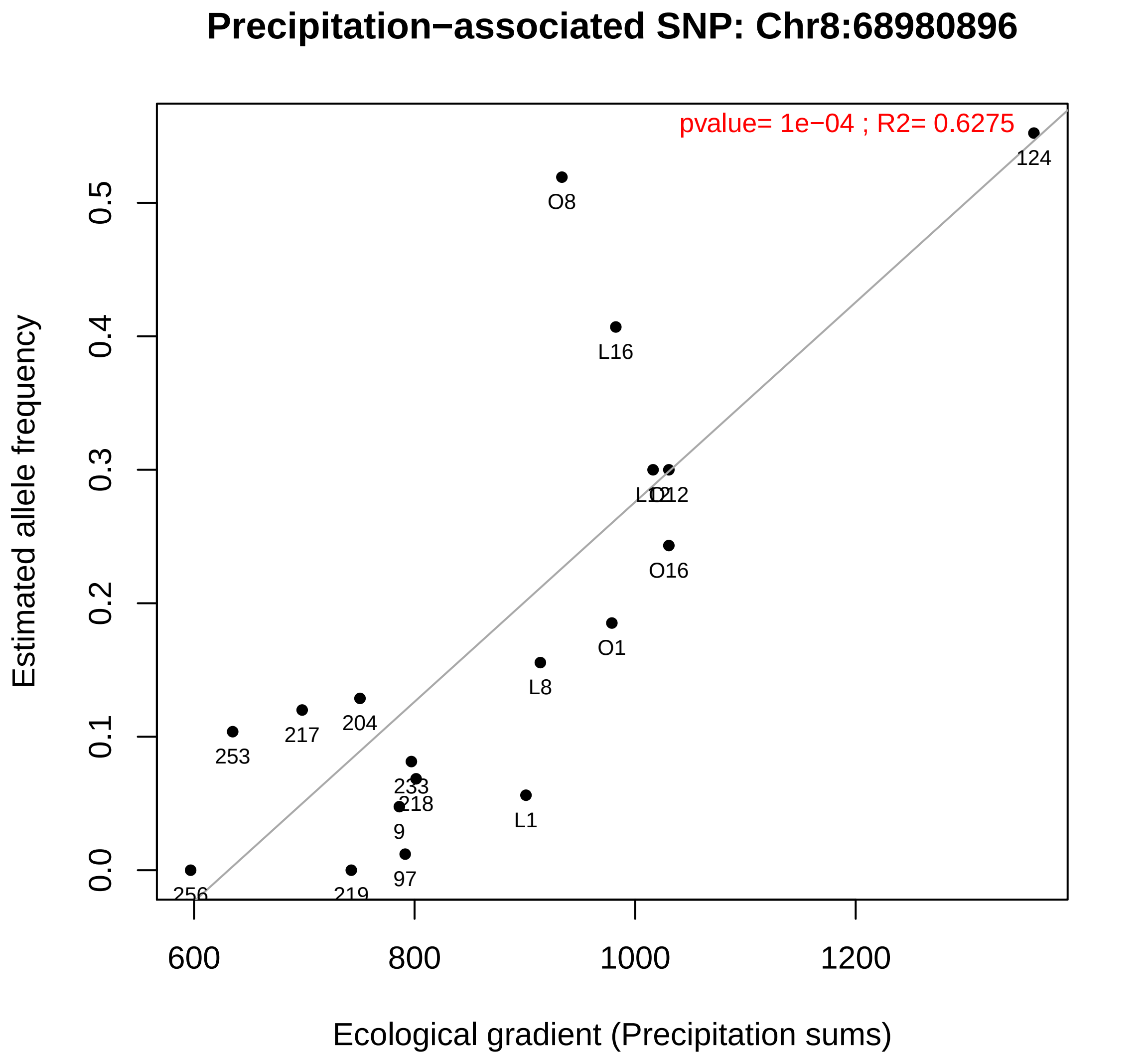

Precipitation−associated SNP: Chr8:68980896
pvalue= 1e−04 ; R2= 0.6275
Estimated allele frequency
Ecological gradient (Precipitation sums)
0.0
0.1
0.2
0.3
0.4
0.5
600
800
1000
1200
124
O8
L16
L12
O12
O16
O1
L8
204
217
253
233
218
9
L1
97
256
219

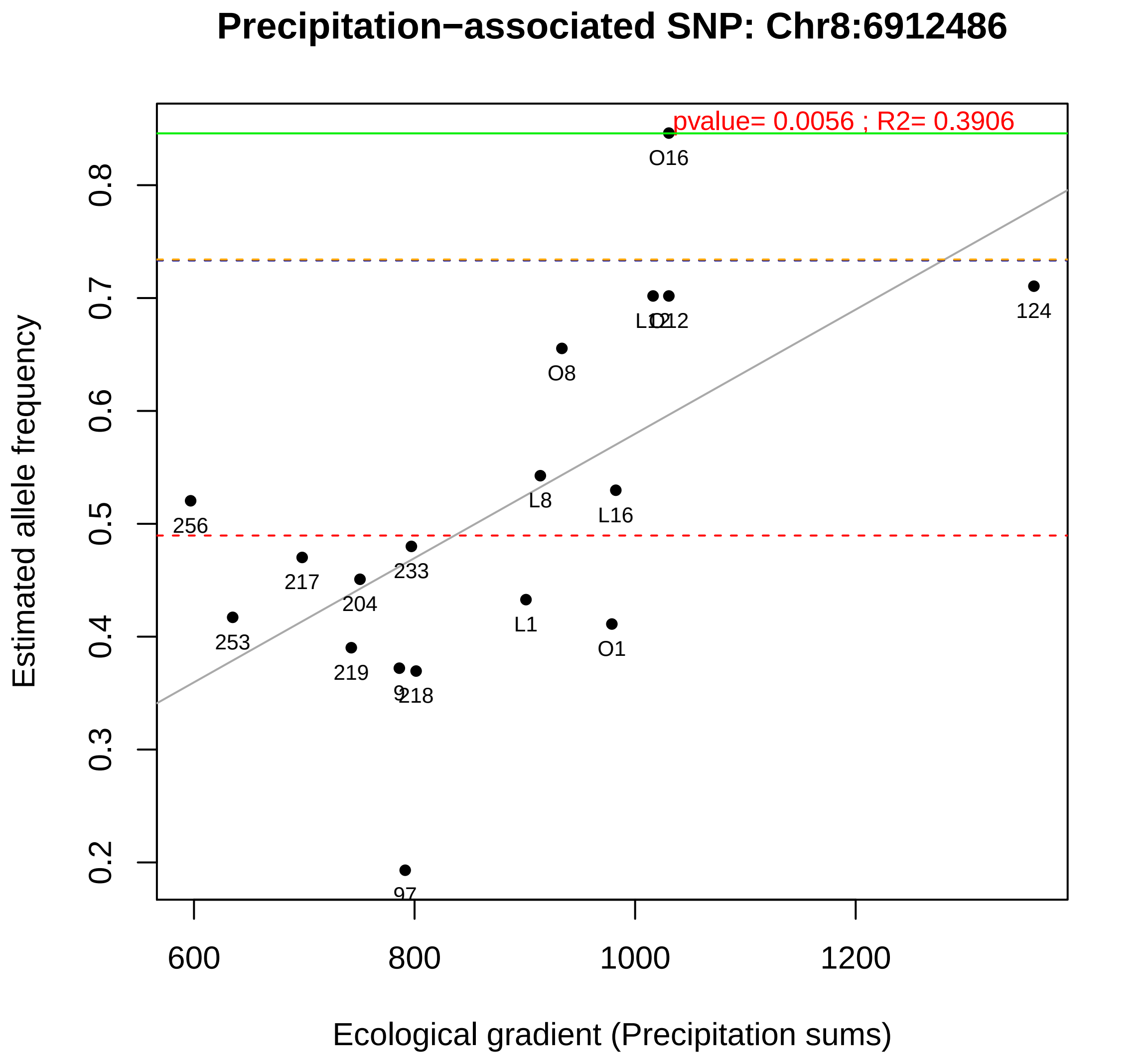
Precipitation−associated SNP: Chr8:6912486
pvalue= 0.0056 ; R2= 0.3906
O16
L12
O12
124
O8
L8
L16
256
217
233
204
253
L1
O1
219
9
218
97
0.2
0.3
0.4
0.5
0.6
0.7
0.8
600
800
1000
1200
Estimated allele frequency
Ecological gradient (Precipitation sums)

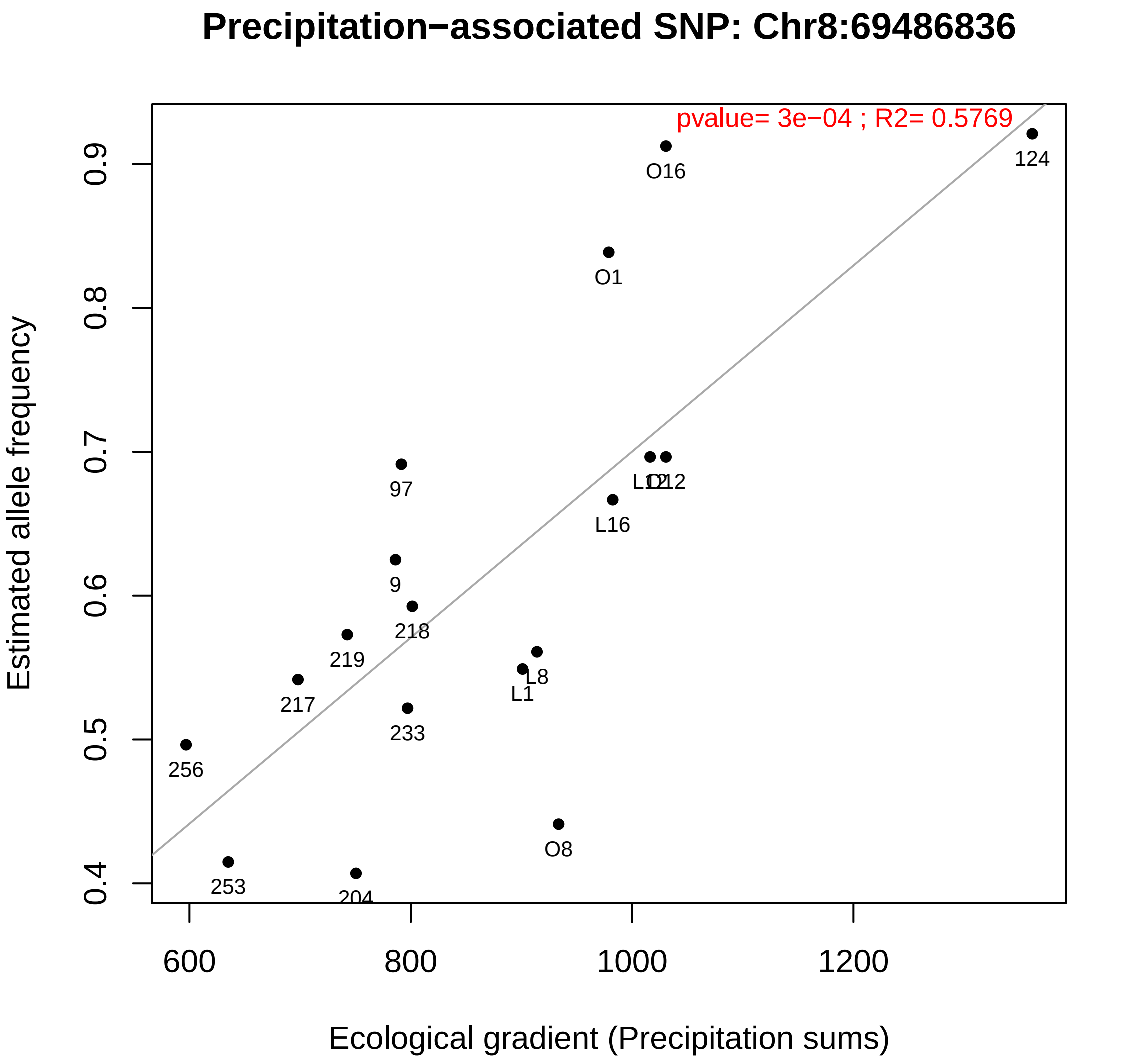
Precipitation−associated SNP: Chr8:69486836
pvalue= 3e−04 ; R2= 0.5769
Estimated allele frequency
Ecological gradient (Precipitation sums)
0.4
0.5
0.6
0.7
0.8
0.9
600
800
1000
1200
O16
124
O1
97
L12
O12
L16
9
218
219
L8
217
L1
233
256
O8
253
204

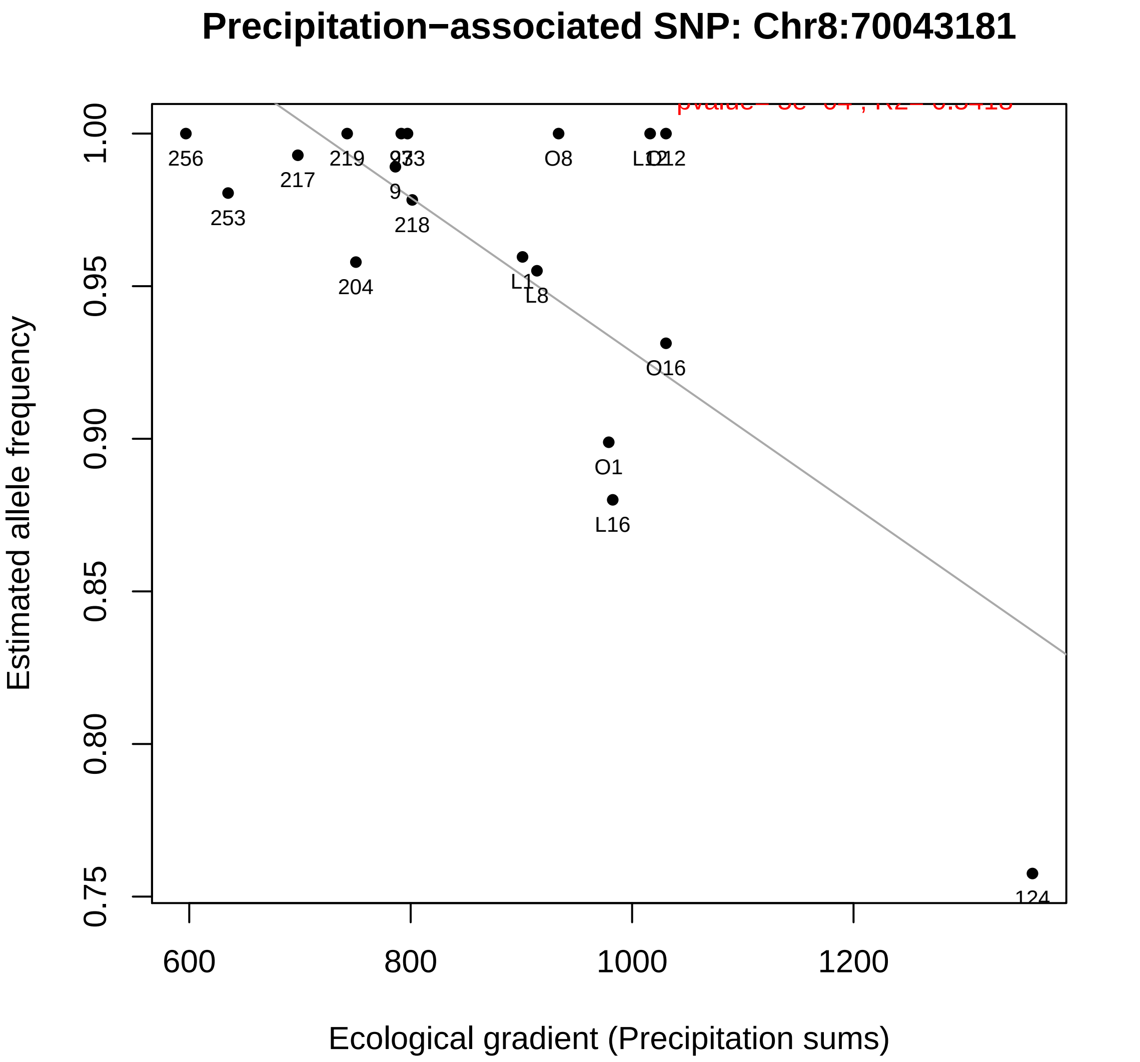

Precipitation−associated SNP: Chr8:70043181
Estimated allele frequency
0.75
0.80
0.85
0.90
0.95
1.00
600
800
1000
1200
Ecological gradient (Precipitation sums)
256
253
217
219
204
9
218
O8
L1
L8
O16
O1
L16
124

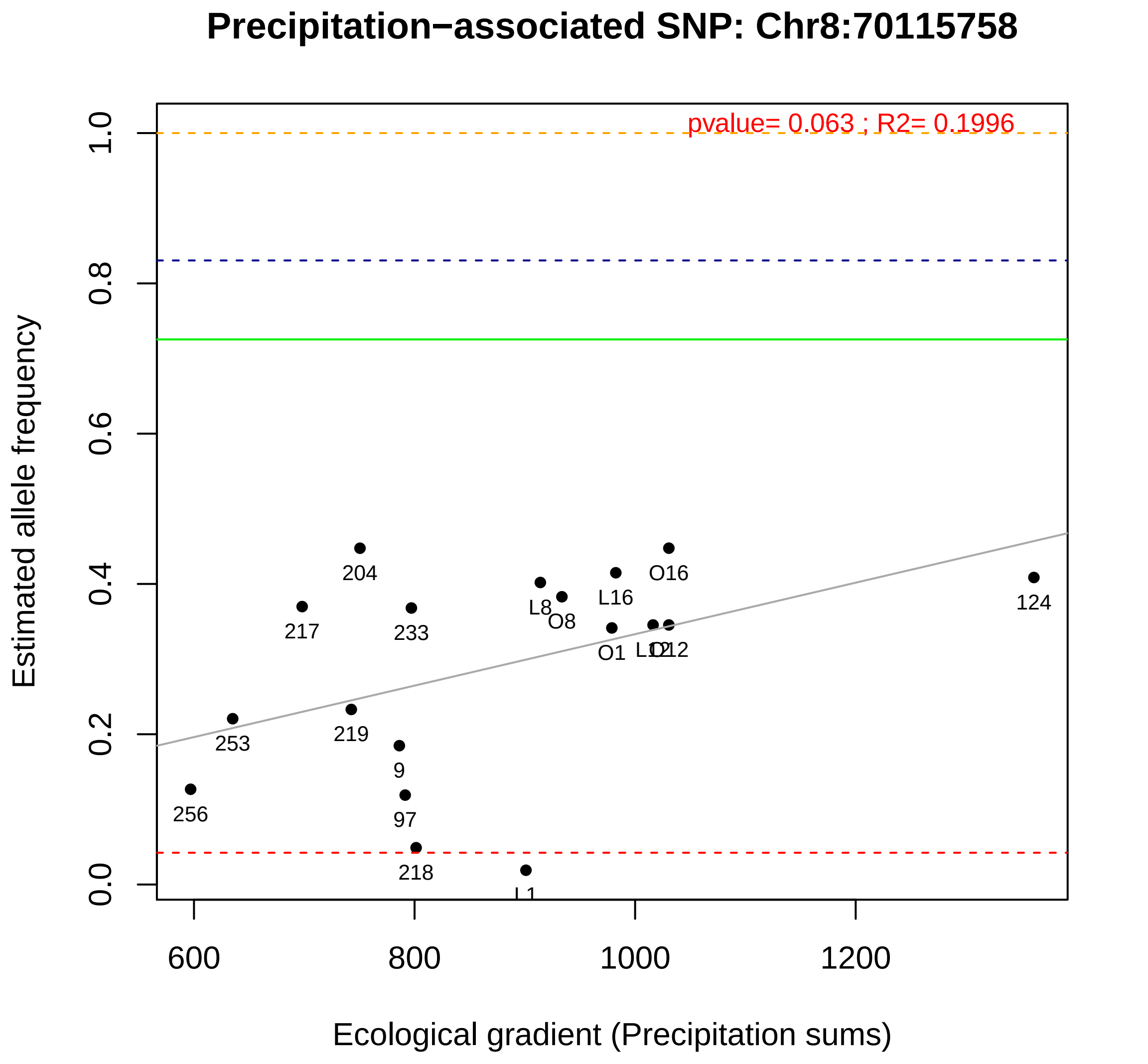

Precipitation−associated SNP: Chr8:70115758
pvalue= 0.063 ; R2= 0.1996
Estimated allele frequency
0.0
0.2
0.4
0.6
0.8
1.0
204
217
233
L8
O8
L16
O16
124
O1
L12
O12
253
219
9
256
97
218
L1
600
800
1000
1200
Ecological gradient (Precipitation sums)

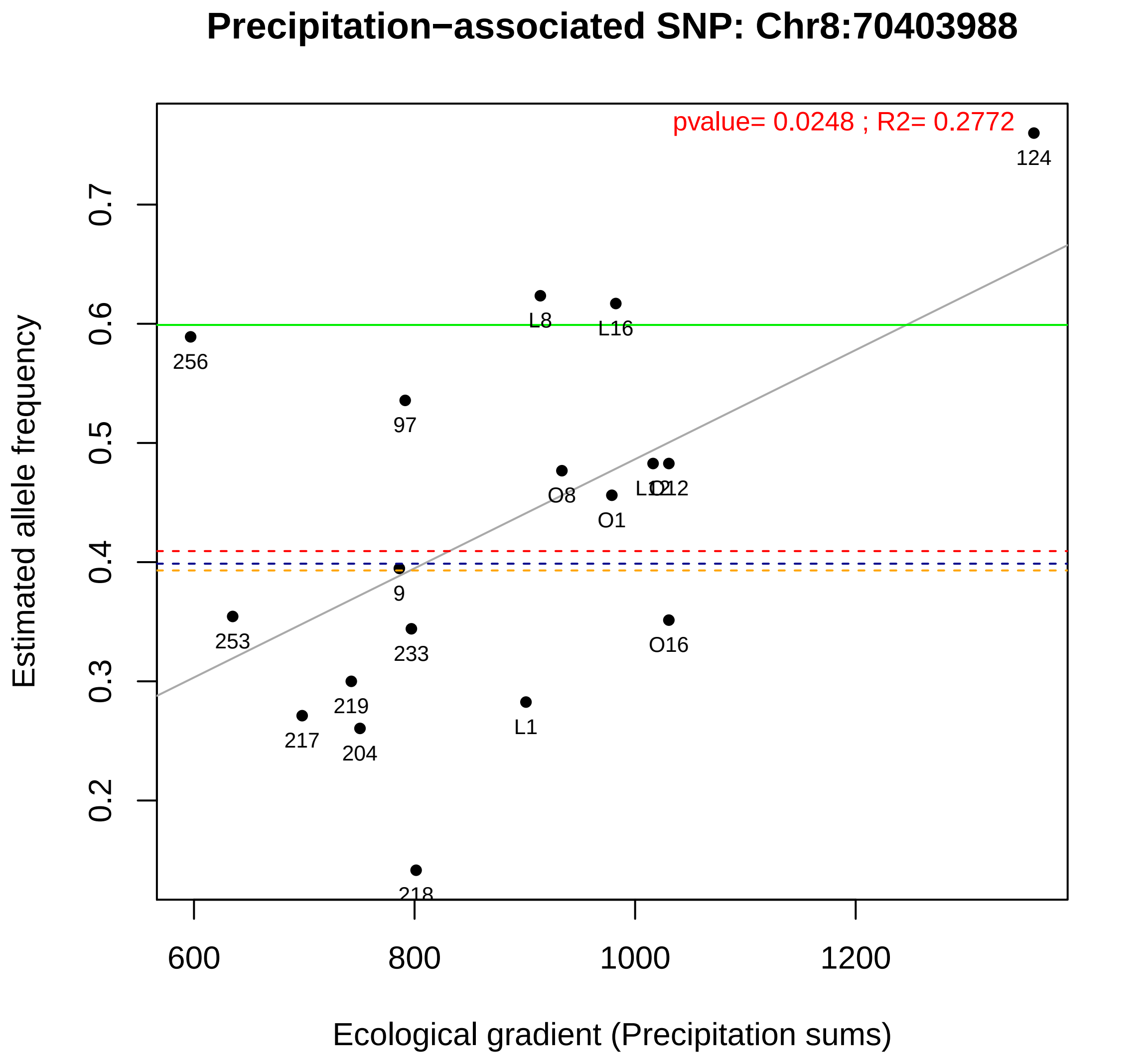
Precipitation−associated SNP: Chr8:70403988
pvalue= 0.0248 ; R2= 0.2772
124
L8
L16
256
97
O8
L12
O12
O1
9
253
233
O16
219
217
204
L1
218
0.2
0.3
0.4
0.5
0.6
0.7
600
800
1000
1200
Estimated allele frequency
Ecological gradient (Precipitation sums)

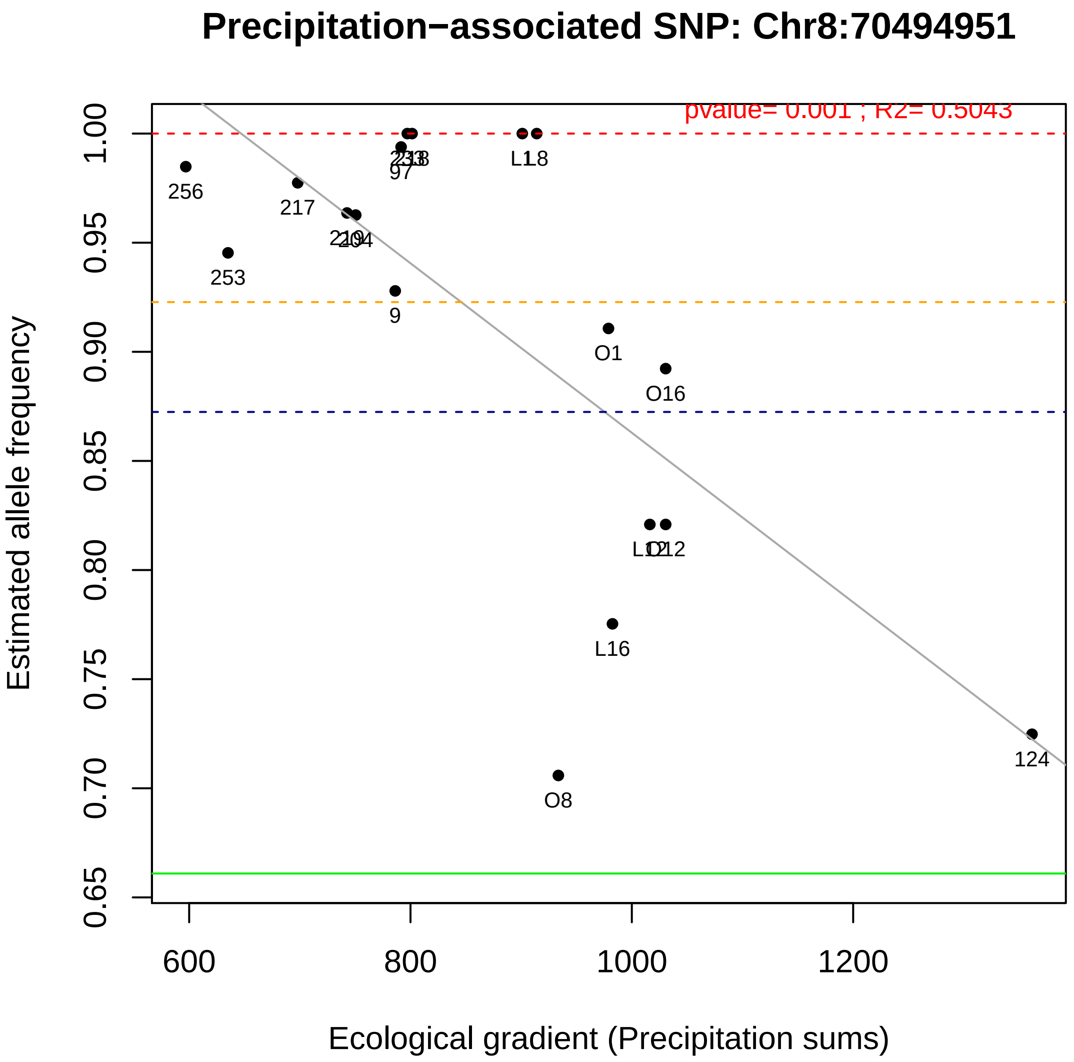

Precipitation−associated SNP: Chr8:70494951
pvalue= 0.001 ; R2= 0.5043
Estimated allele frequency
Ecological gradient (Precipitation sums)
0.65
0.70
0.75
0.80
0.85
0.90
0.95
1.00
600
800
1000
1200
256
217
253
9
97
O1
O16
L16
O8
124

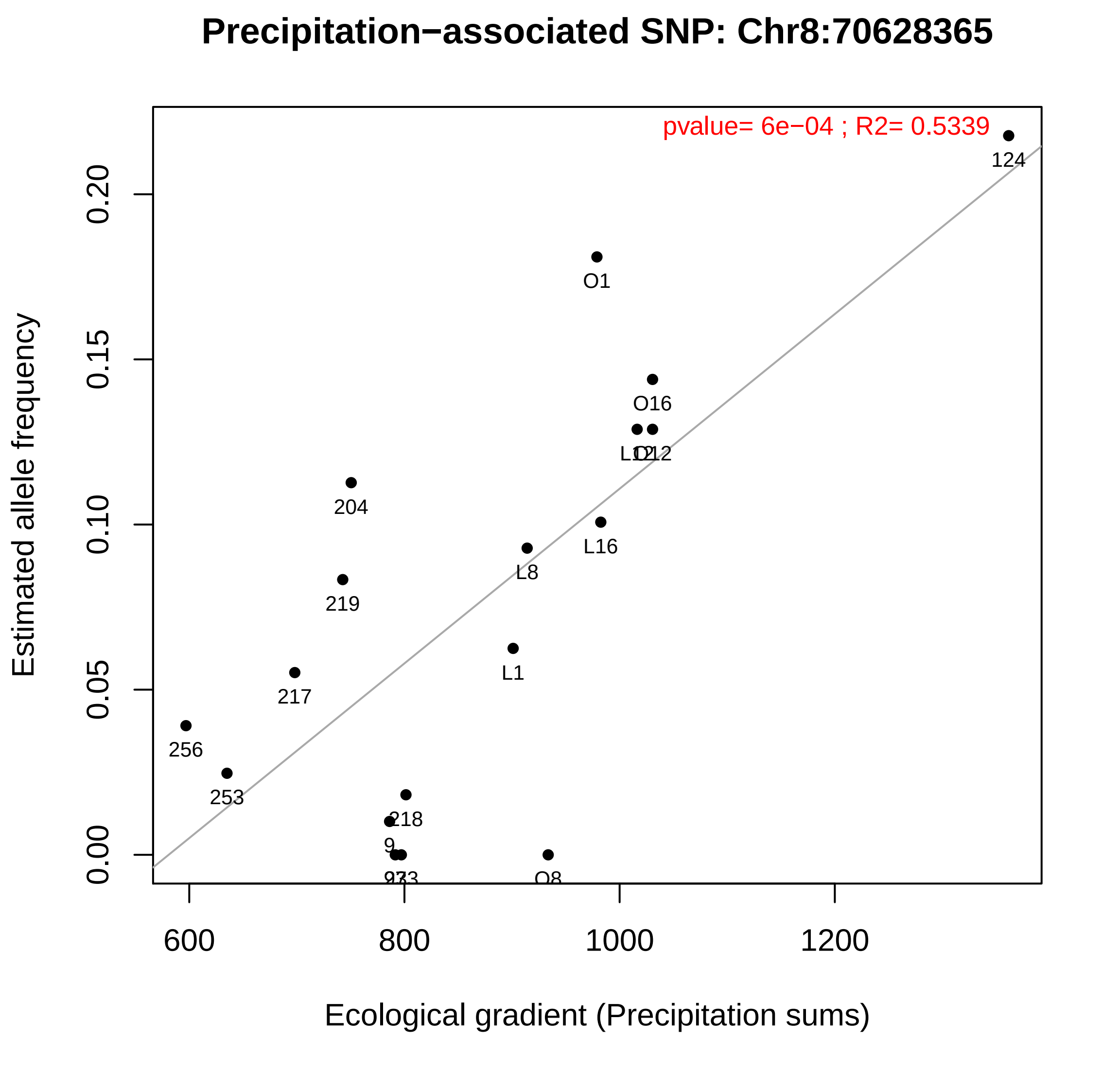

Precipitation−associated SNP: Chr8:70628365
pvalue= 6e−04 ; R2= 0.5339
Estimated allele frequency
0.00
0.05
0.10
0.15
0.20
600
800
1000
1200
Ecological gradient (Precipitation sums)
124
O1
O16
L16
L8
219
204
217
L1
256
253
218
9
O8

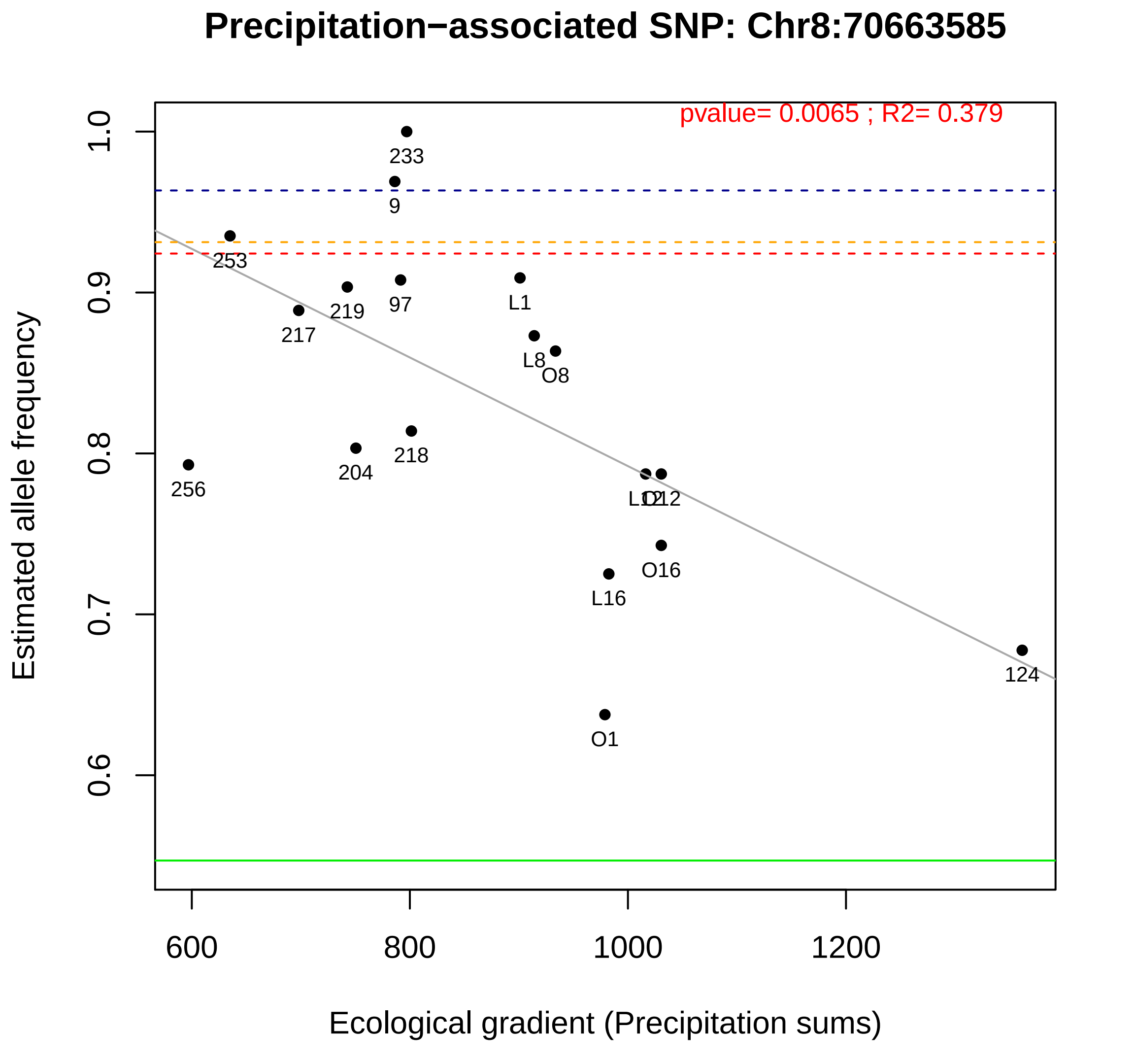
Precipitation−associated SNP: Chr8:70663585
pvalue= 0.0065 ; R2= 0.379
Estimated allele frequency
Ecological gradient (Precipitation sums)
1.0
0.9
0.8
0.7
0.6
600
800
1000
1200
233
9
253
219
97
L1
217
L8
O8
218
204
256
L12
O12
O16
L16
124
O1

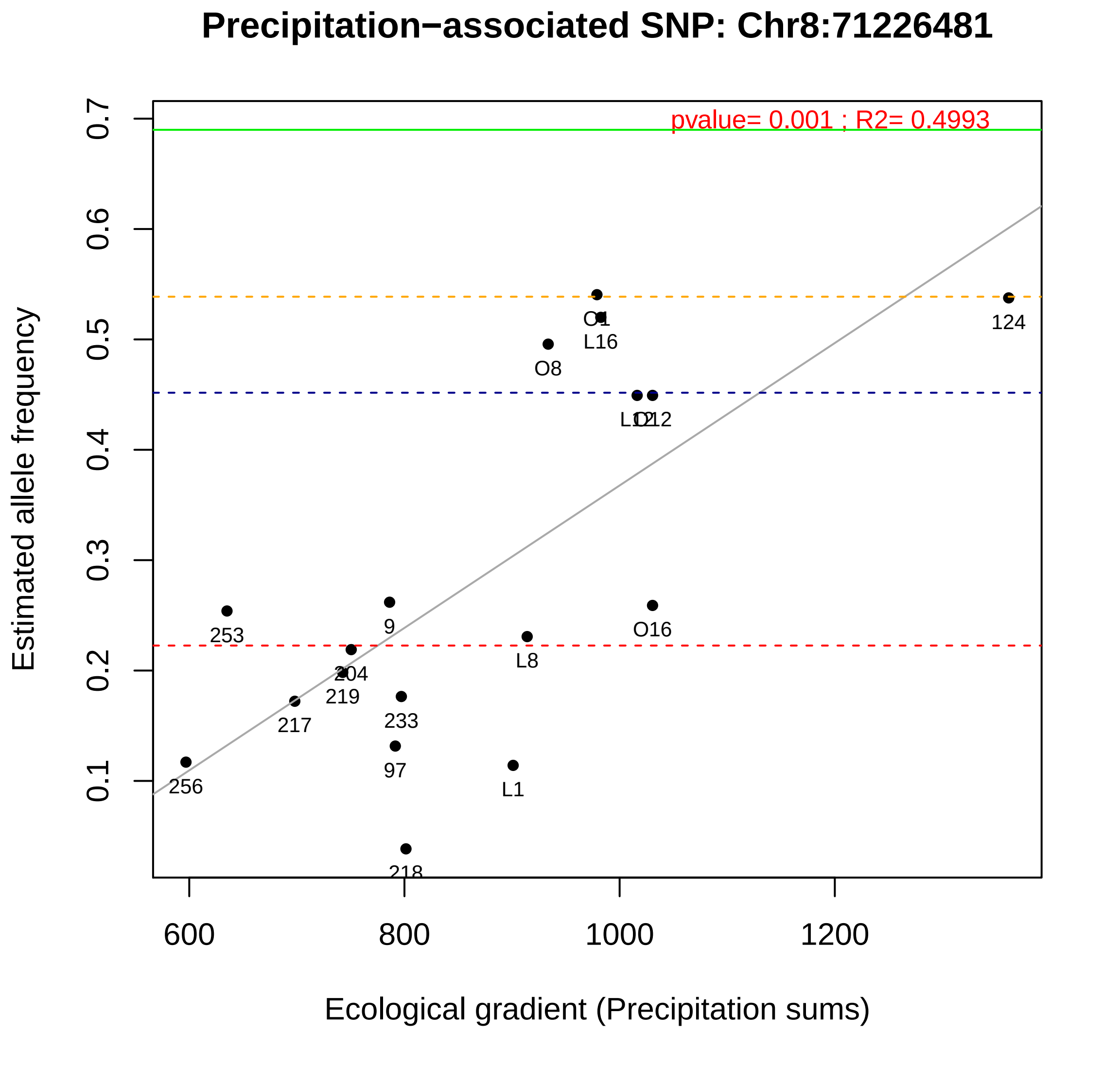
Precipitation−associated SNP: Chr8:71226481
pvalue= 0.001 ; R2= 0.4993
Estimated allele frequency
0.1
0.2
0.3
0.4
0.5
0.6
0.7
Ecological gradient (Precipitation sums)
600
800
1000
1200
256
253
217
219
204
9
233
97
218
L1
L8
O8
L16
L12
O12
O16
124

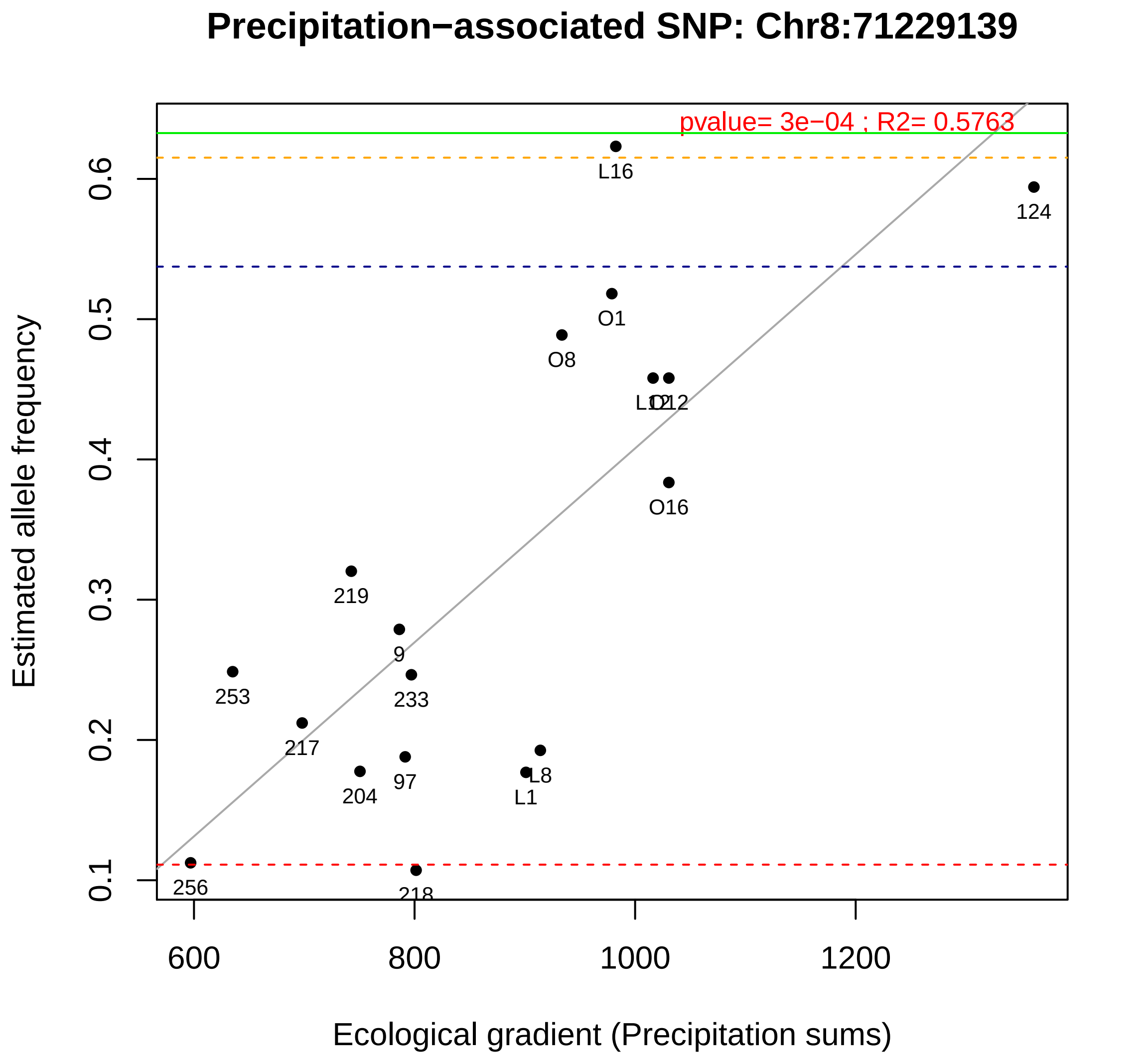

Precipitation−associated SNP: Chr8:71229139
pvalue= 3e−04 ; R2= 0.5763
Estimated allele frequency
0.1
0.2
0.3
0.4
0.5
0.6
600
800
1000
1200
Ecological gradient (Precipitation sums)
L16
124
O1
O8
O16
219
9
253
233
217
204
97
L8
L1
256
218

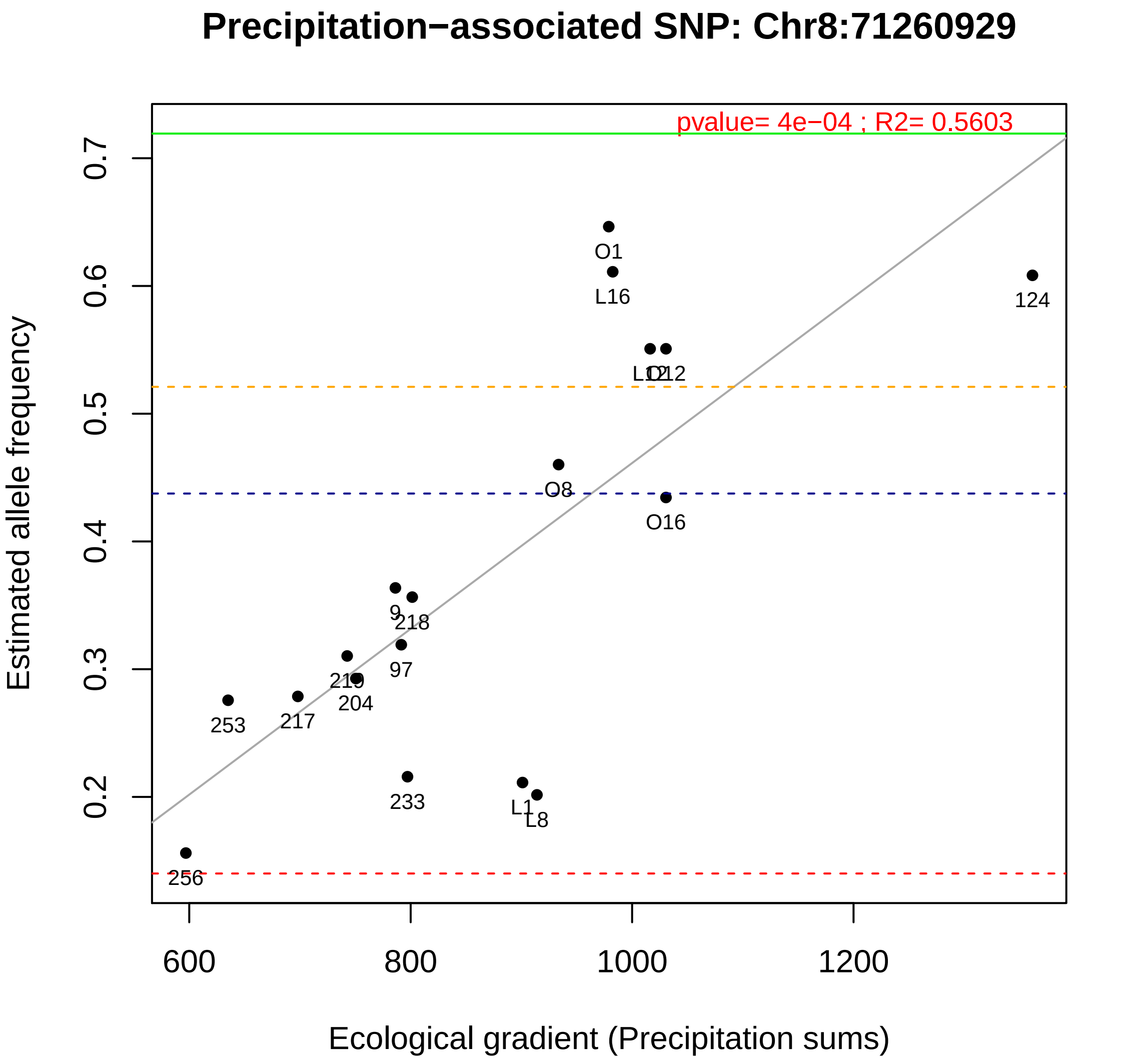

Precipitation−associated SNP: Chr8:71260929
pvalue= 4e−04 ; R2= 0.5603
Estimated allele frequency
0.2
0.3
0.4
0.5
0.6
0.7
600
800
1000
1200
Ecological gradient (Precipitation sums)
O1
L16
124
L12
O12
O8
O16
9
218
97
219
204
253
217
233
L1
L8
256

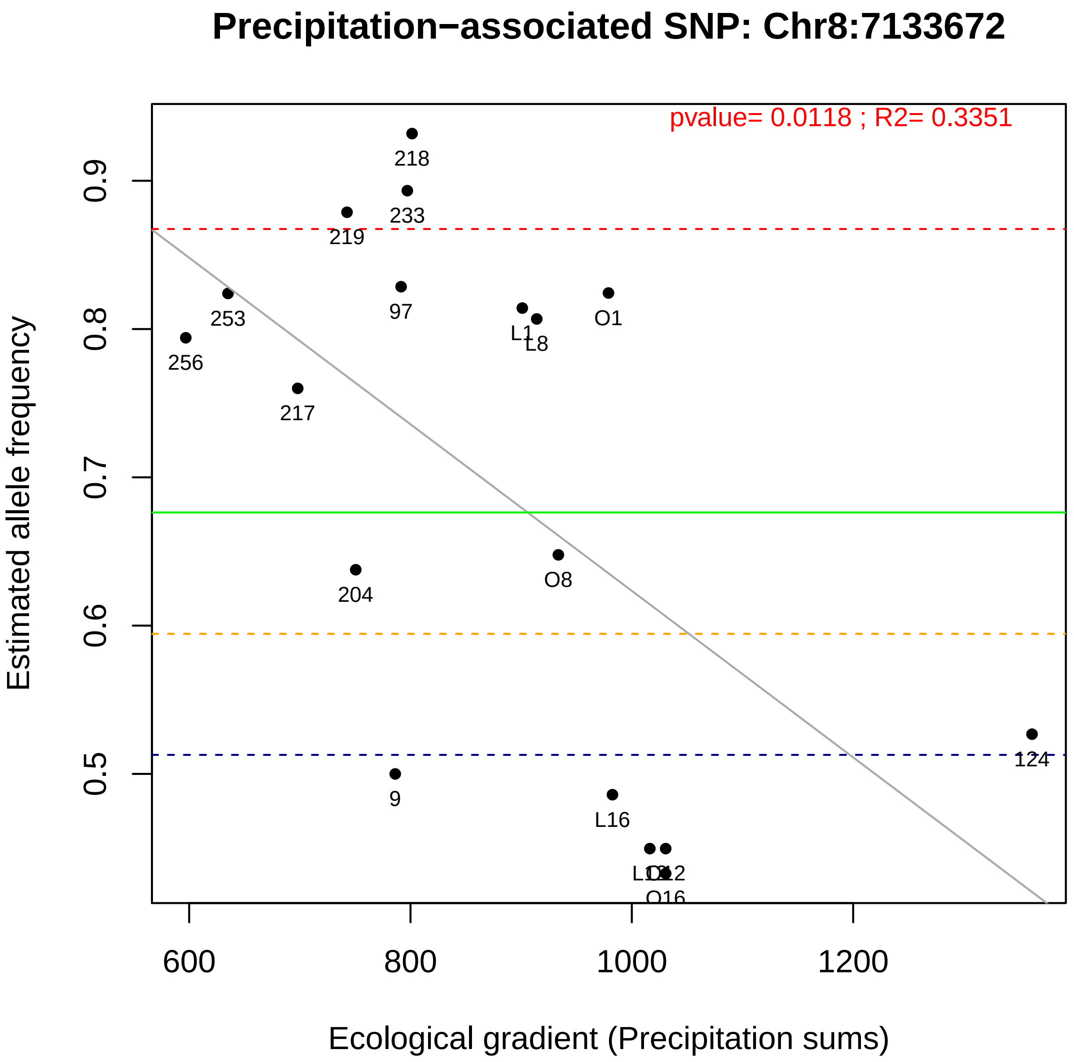
Precipitation−associated SNP: Chr8:7133672
pvalue= 0.0118 ; R2= 0.3351
Estimated allele frequency
Ecological gradient (Precipitation sums)
0.5
0.6
0.7
0.8
0.9
600
800
1000
1200
218
233
219
253
97
L1
L8
O1
256
217
204
O8
9
L16
124
O16

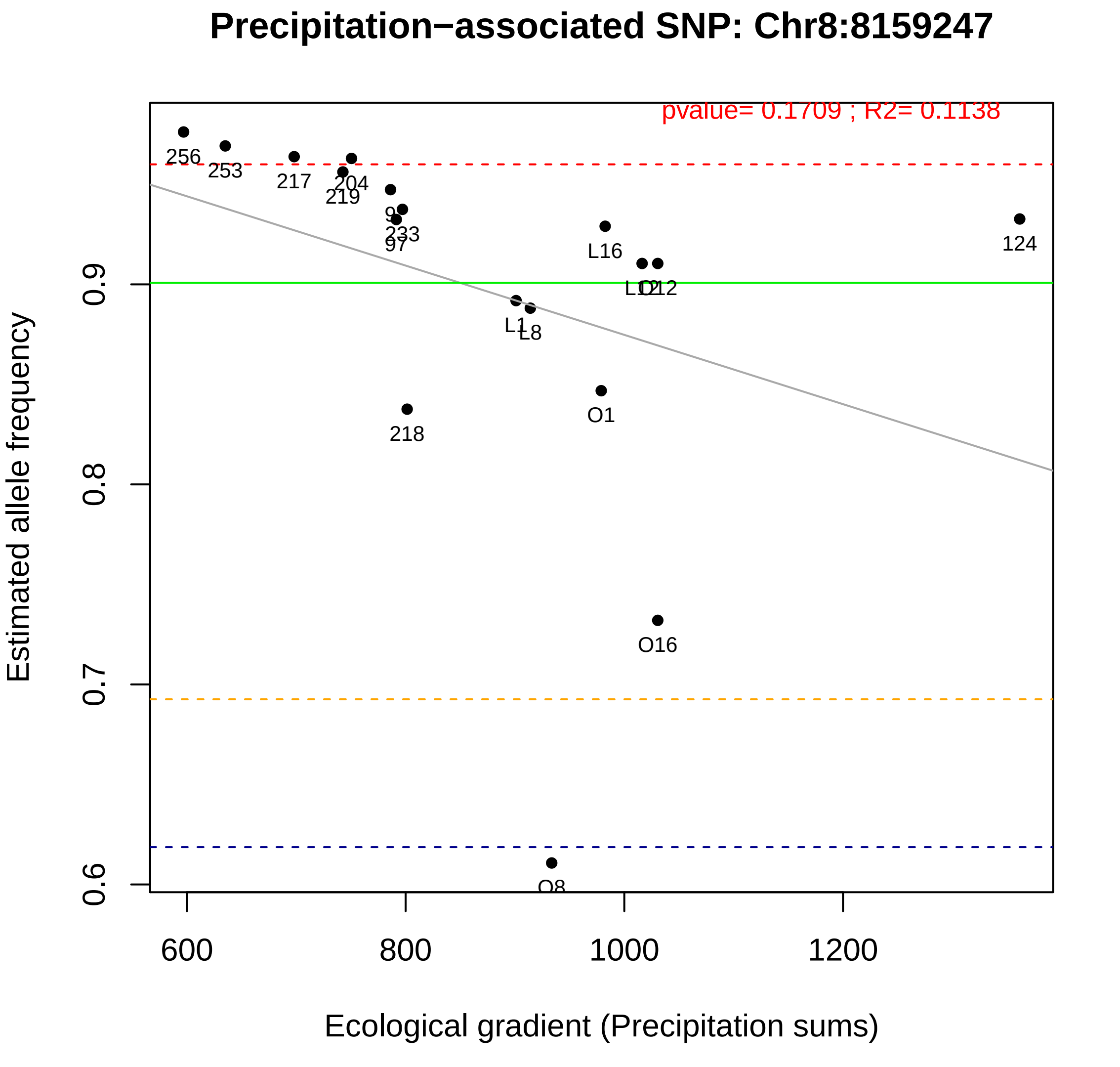
Precipitation−associated SNP: Chr8:8159247
pvalue= 0.1709 ; R2= 0.1138
256
253
217
204
219
233
97
L16
124
L1
L8
O1
218
O16
O8
0.6
0.7
0.8
0.9
600
800
1000
1200
Estimated allele frequency
Ecological gradient (Precipitation sums)

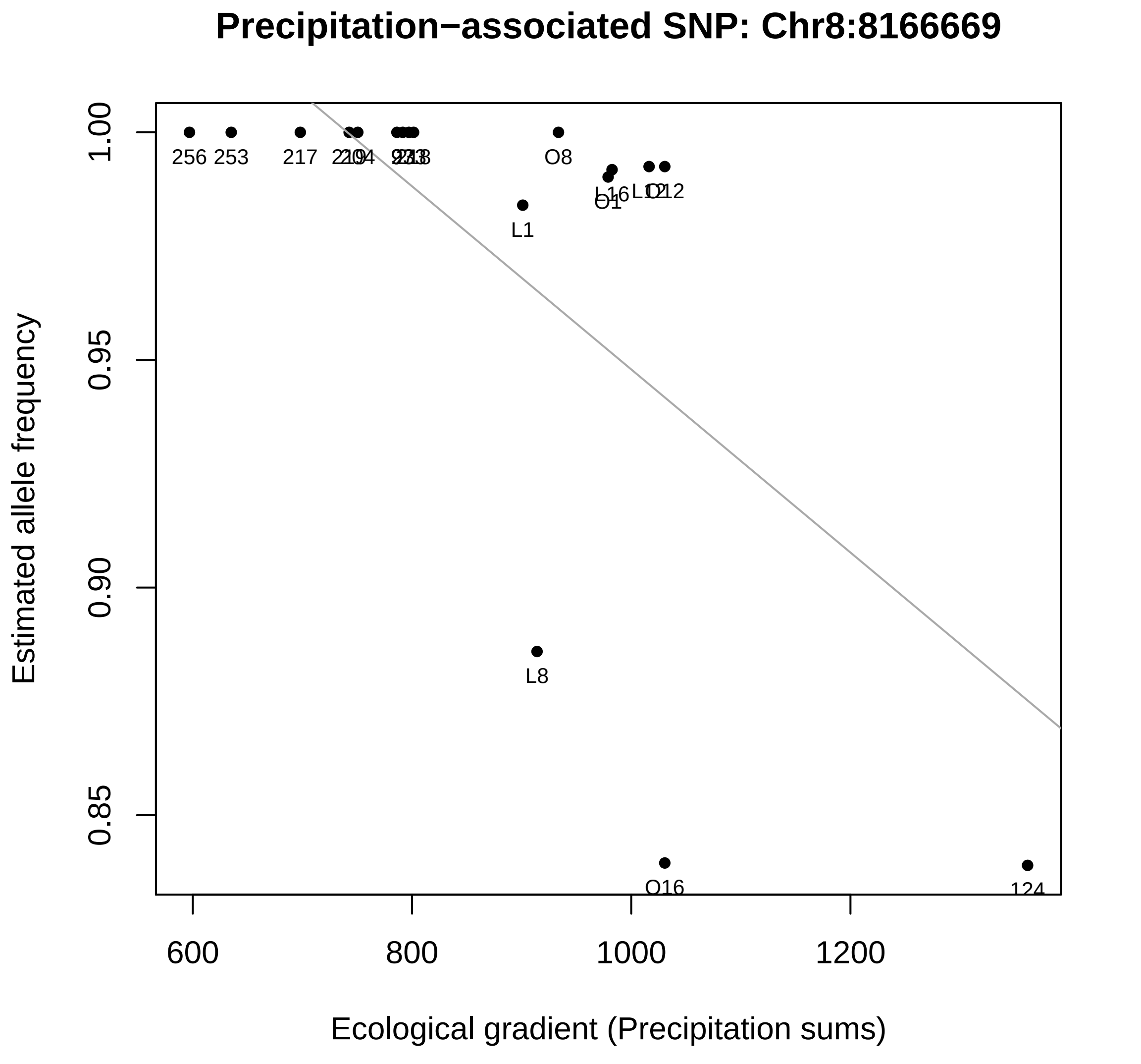

Precipitation−associated SNP: Chr8:8166669
Estimated allele frequency
Ecological gradient (Precipitation sums)
1.00
0.95
0.90
0.85
600
800
1000
1200
256
253
217
O8
L1
L8
O16
124

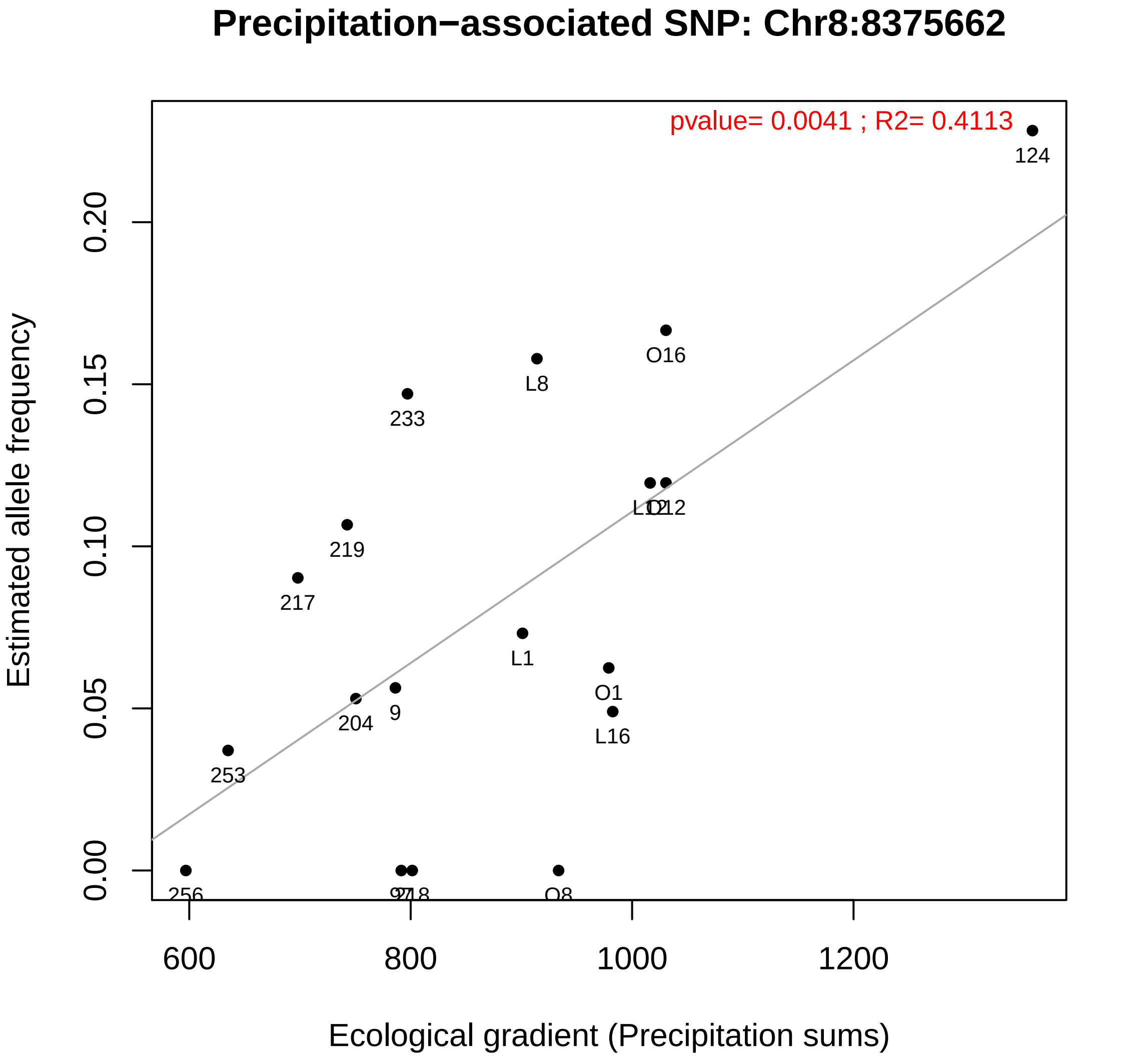
Precipitation−associated SNP: Chr8:8375662
pvalue= 0.0041 ; R2= 0.4113
124
O16
L8
233
219
217
L1
O1
L16
9
204
253
O8
Estimated allele frequency
0.00
0.05
0.10
0.15
0.20
600
800
1000
1200
Ecological gradient (Precipitation sums)

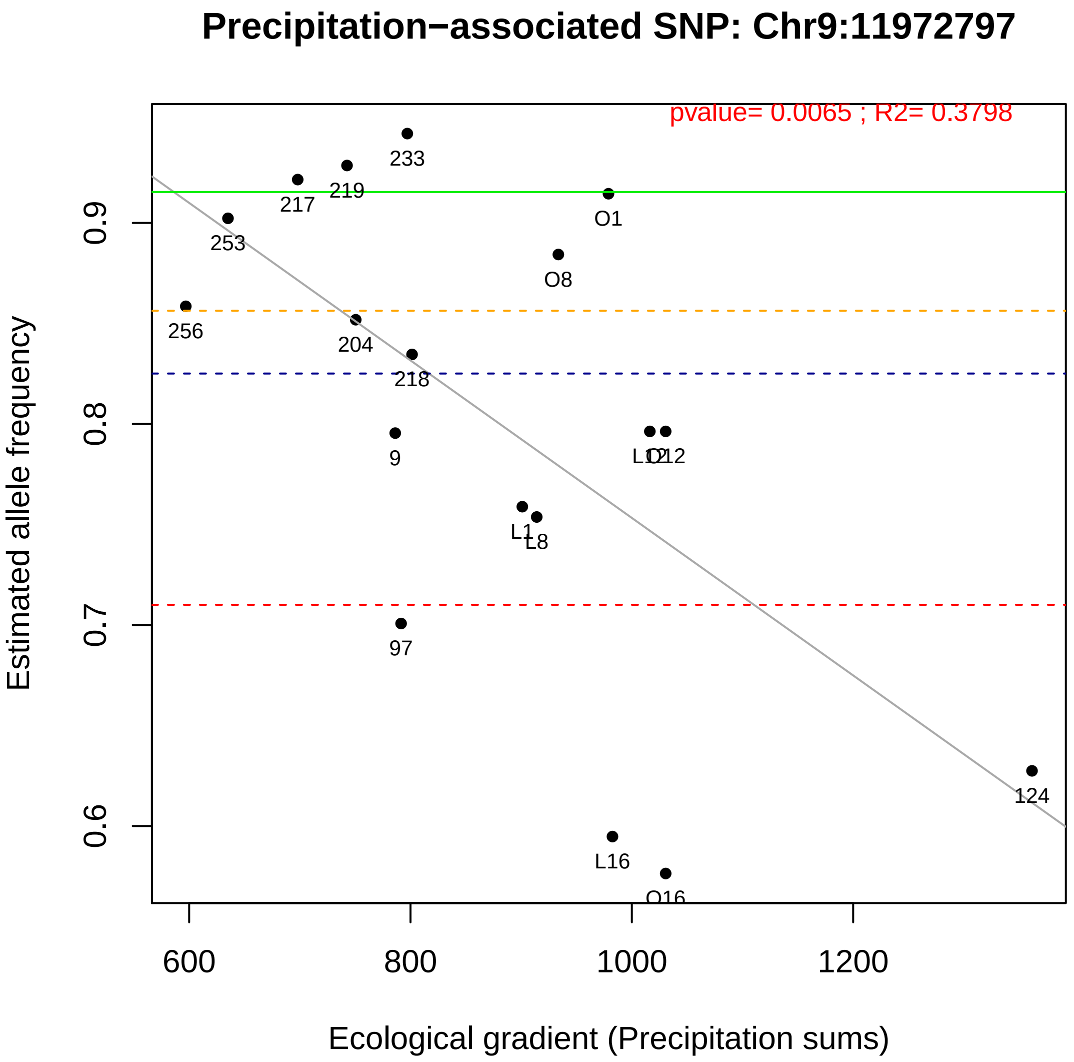

Precipitation−associated SNP: Chr9:11972797
pvalue= 0.0065 ; R2= 0.3798
Estimated allele frequency
Ecological gradient (Precipitation sums)
0.6
0.7
0.8
0.9
600
800
1000
1200
233
219
217
253
O1
O8
256
204
218
9
L12
O12
L1
L8
97
124
L16
O16

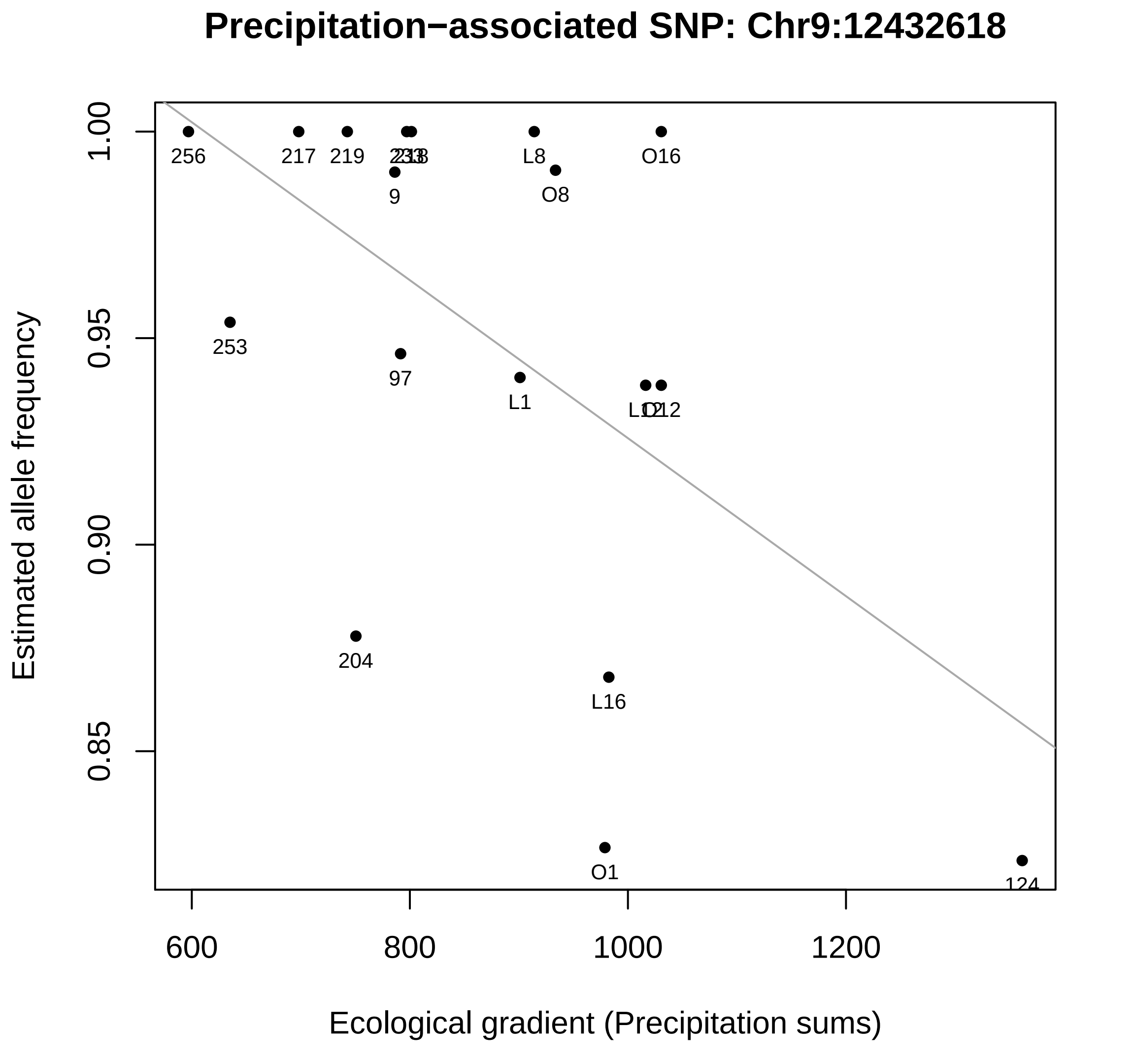
Precipitation−associated SNP: Chr9:12432618
Estimated allele frequency
1.00
0.95
0.90
0.85
Ecological gradient (Precipitation sums)
600
800
1000
1200
256
217
219
L8
O16
9
O8
253
97
L1
O12
204
L16
O1
124

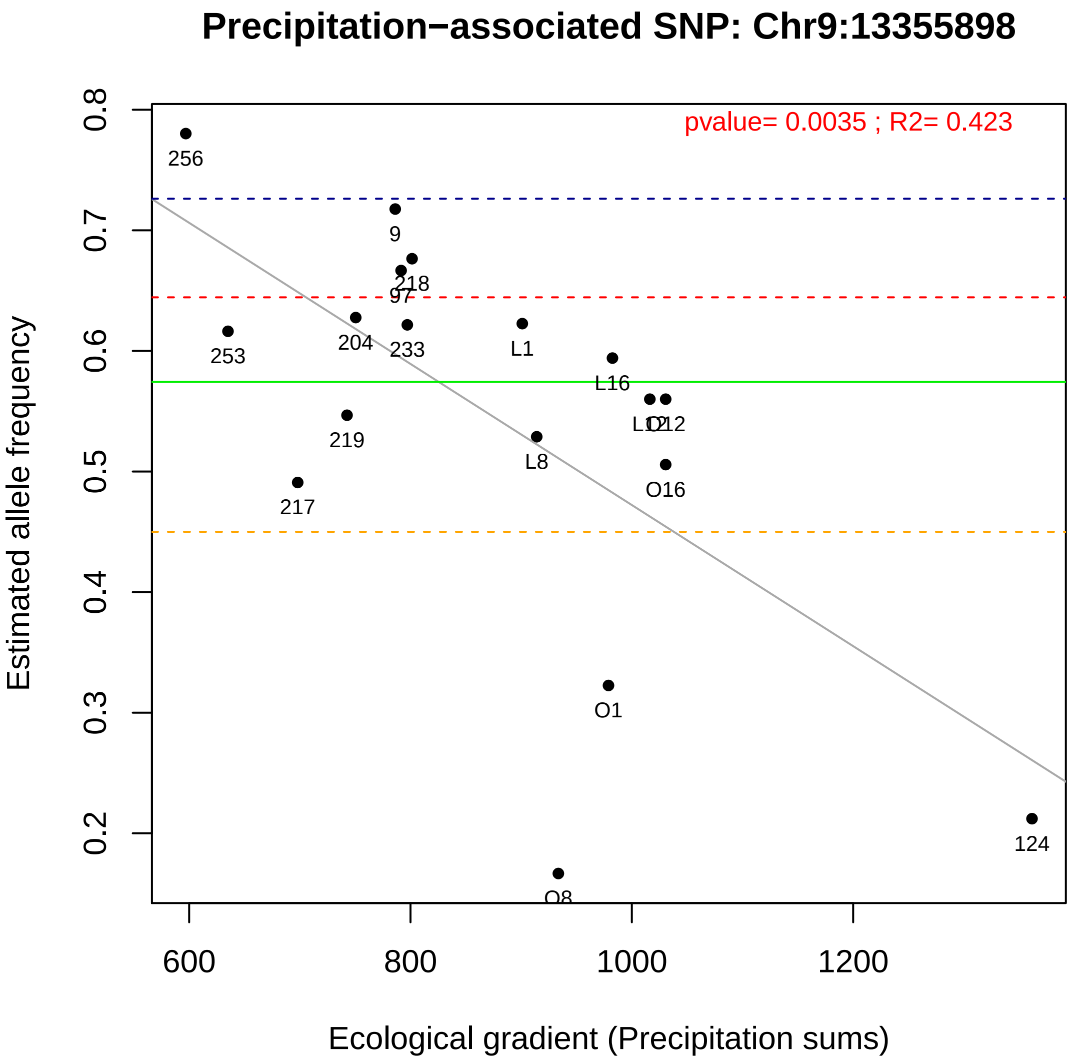
Precipitation−associated SNP: Chr9:13355898
pvalue= 0.0035 ; R2= 0.423
Estimated allele frequency
0.8
0.7
0.6
0.5
0.4
0.3
0.2
256
9
218
97
253
204
233
L1
L16
L12
O12
219
L8
O16
217
O1
124
O8
600
800
1000
1200
Ecological gradient (Precipitation sums)

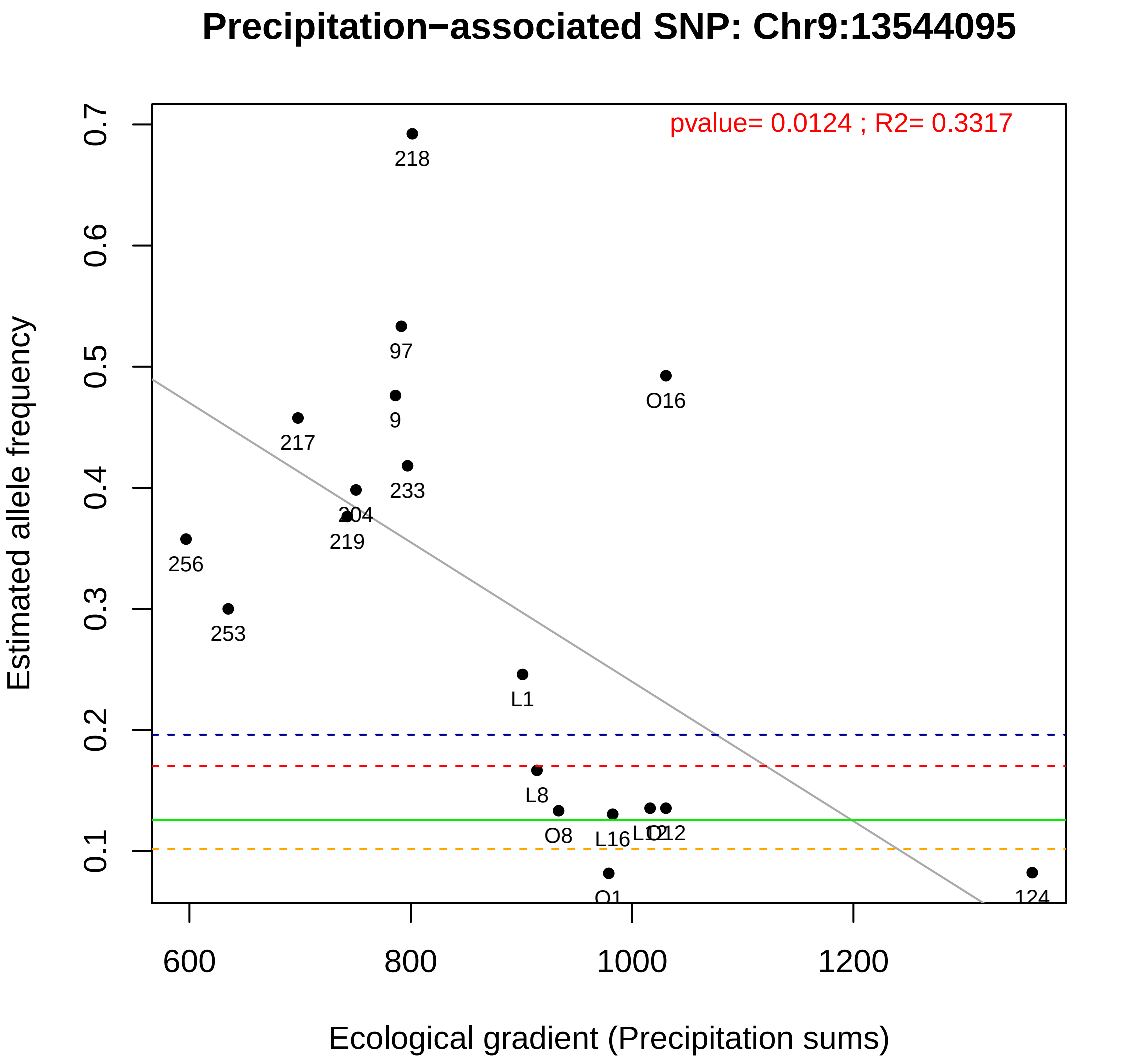

Precipitation−associated SNP: Chr9:13544095
pvalue= 0.0124 ; R2= 0.3317
Estimated allele frequency
Ecological gradient (Precipitation sums)
0.1
0.2
0.3
0.4
0.5
0.6
0.7
600
800
1000
1200
218
97
O16
9
217
233
204
219
256
253
L1
L8
O8
L16
O12
O1
124

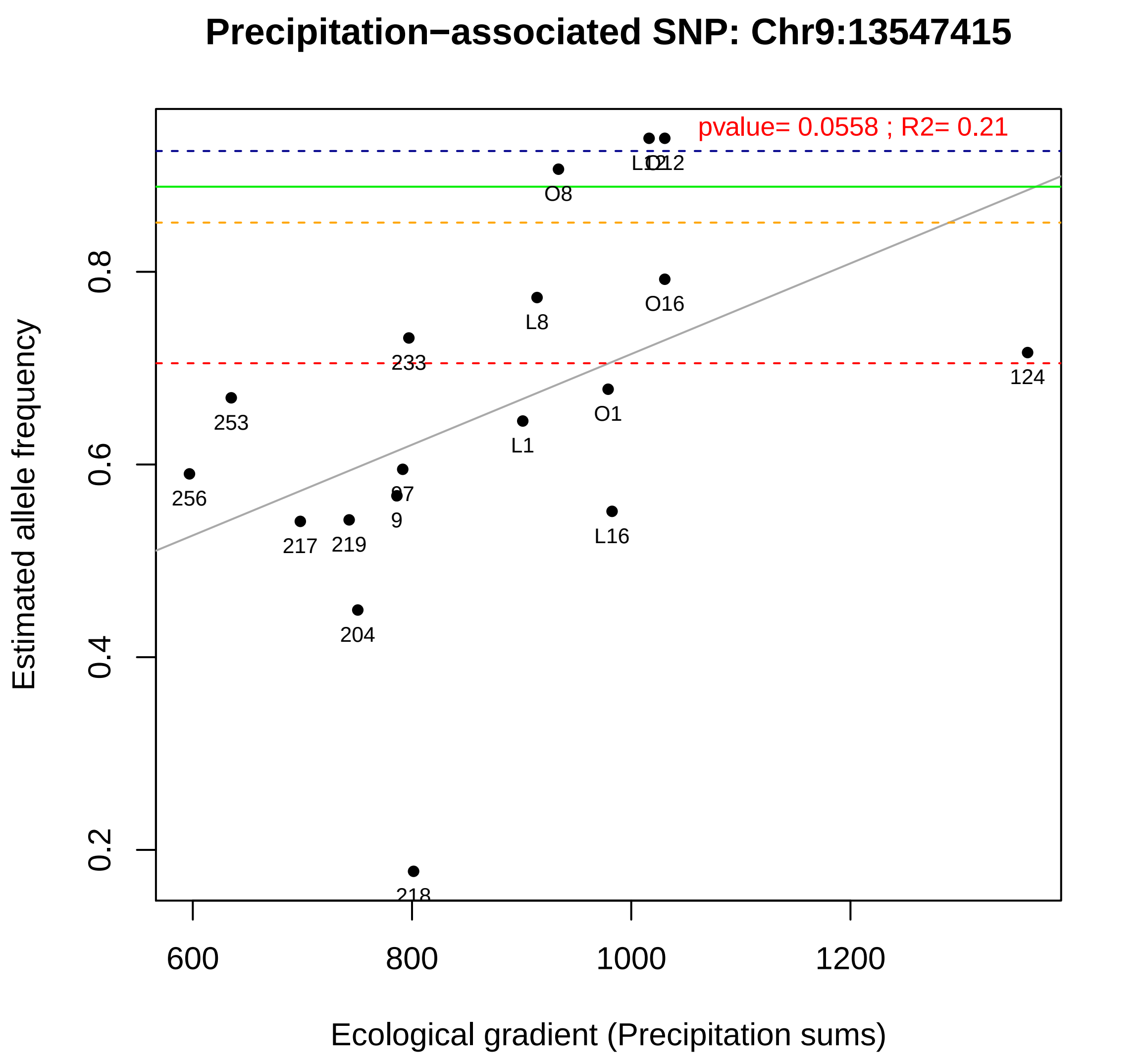
Precipitation−associated SNP: Chr9:13547415
pvalue= 0.0558 ; R2= 0.21
Estimated allele frequency
0.2
0.4
0.6
0.8
600
800
1000
1200
Ecological gradient (Precipitation sums)
O8
O16
L8
233
124
253
O1
L1
256
217
219
9
L16
204
218

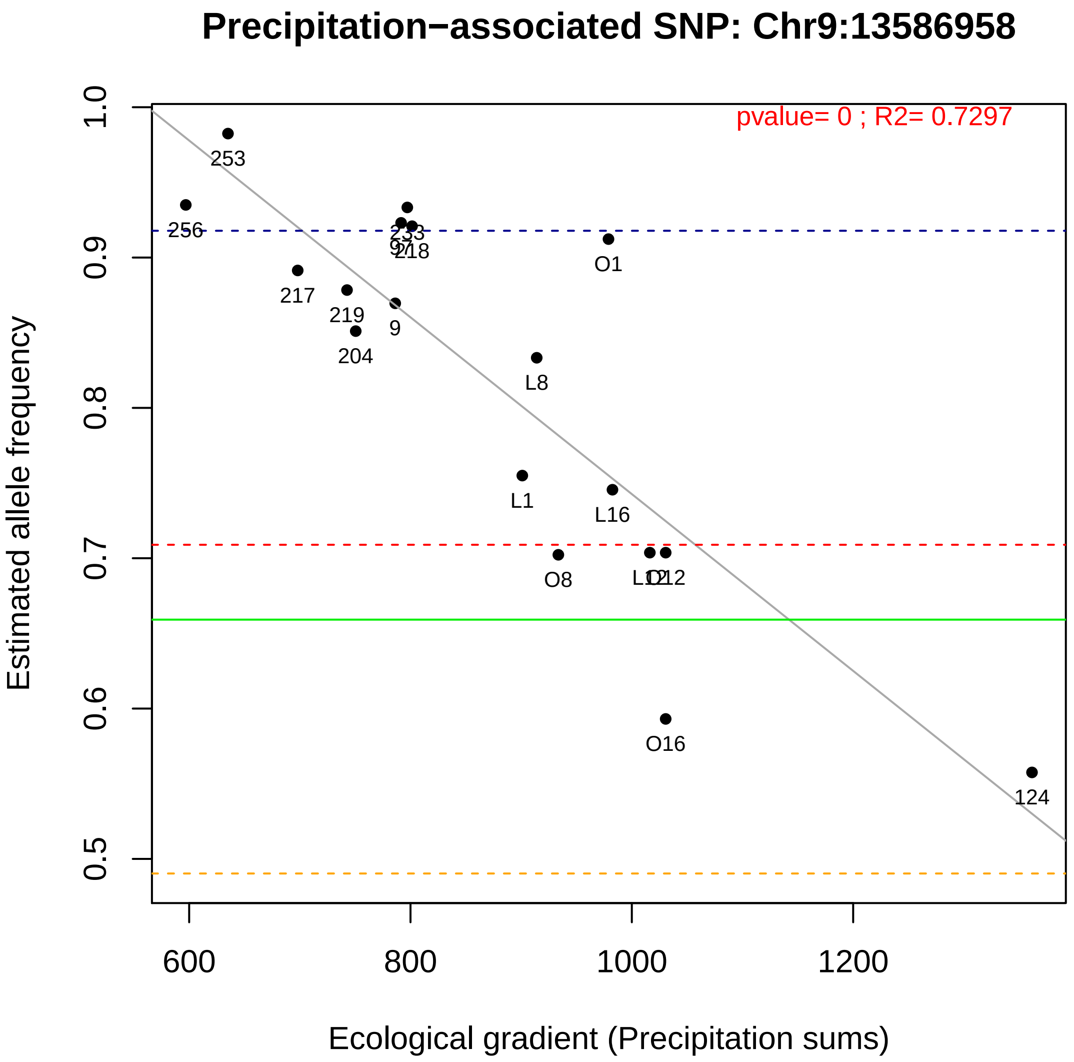

Precipitation−associated SNP: Chr9:13586958
pvalue= 0 ; R2= 0.7297
Estimated allele frequency
Ecological gradient (Precipitation sums)
1.0
0.9
0.8
0.7
0.6
0.5
600
800
1000
1200
253
256
233
97
218
O1
217
219
9
204
L8
L1
L16
O8
L12
O12
O16
124

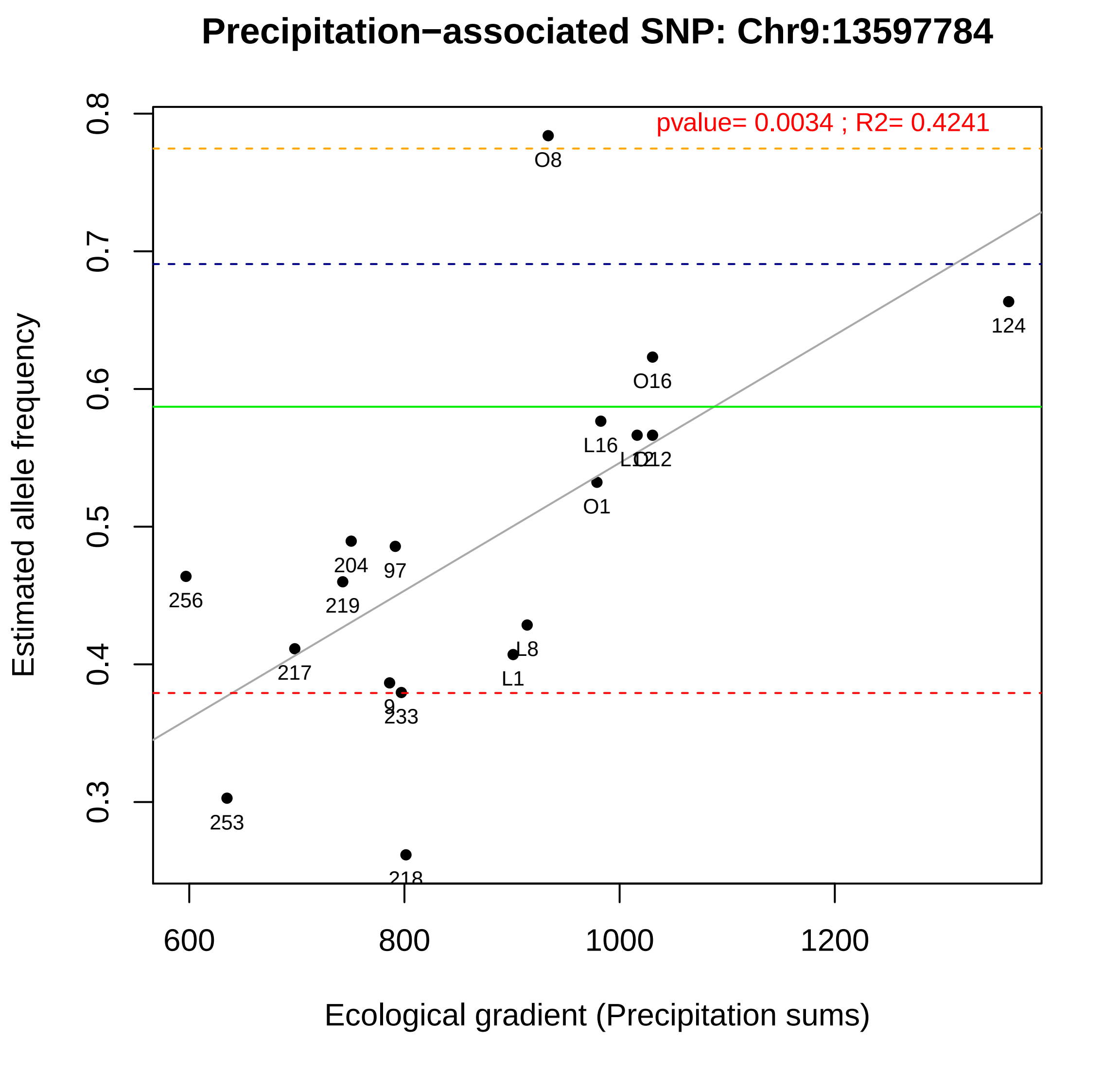
Precipitation−associated SNP: Chr9:13597784
pvalue= 0.0034 ; R2= 0.4241
Estimated allele frequency
0.8
0.7
0.6
0.5
0.4
0.3
O8
124
O16
L16
L12
O12
O1
204
97
256
219
L8
217
L1
9
233
253
218
600
800
1000
1200
Ecological gradient (Precipitation sums)

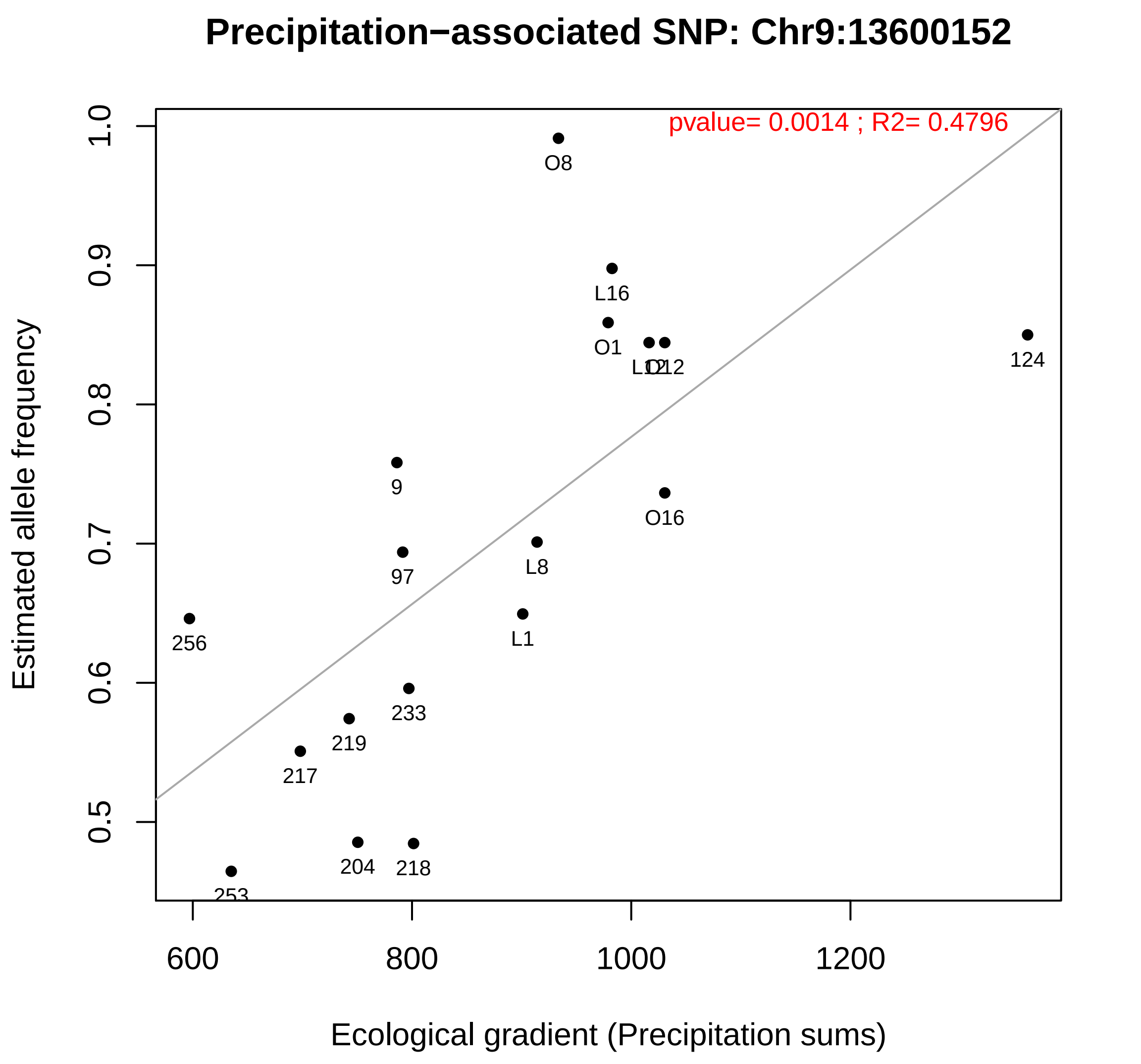
Precipitation−associated SNP: Chr9:13600152
pvalue= 0.0014 ; R2= 0.4796
Estimated allele frequency
Ecological gradient (Precipitation sums)
1.0
0.9
0.8
0.7
0.6
0.5
600
800
1000
1200
O8
L16
O1
O12
124
9
O16
97
L8
256
L1
233
219
217
204
218
253

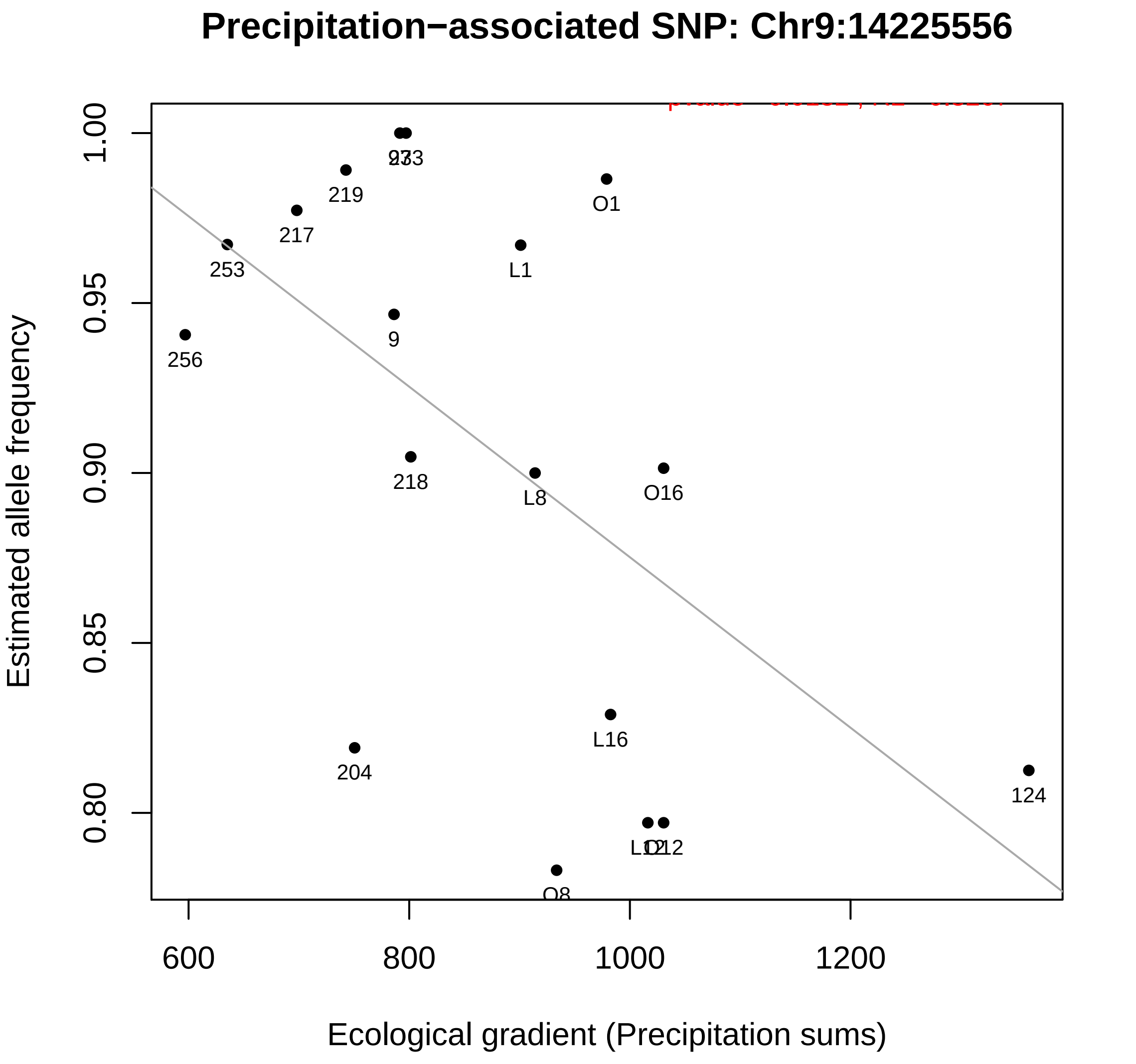
Precipitation−associated SNP: Chr9:14225556
Estimated allele frequency
Ecological gradient (Precipitation sums)
1.00
0.95
0.90
0.85
0.80
600
800
1000
1200
219
O1
217
253
L1
9
256
218
L8
O16
L16
204
124
O8

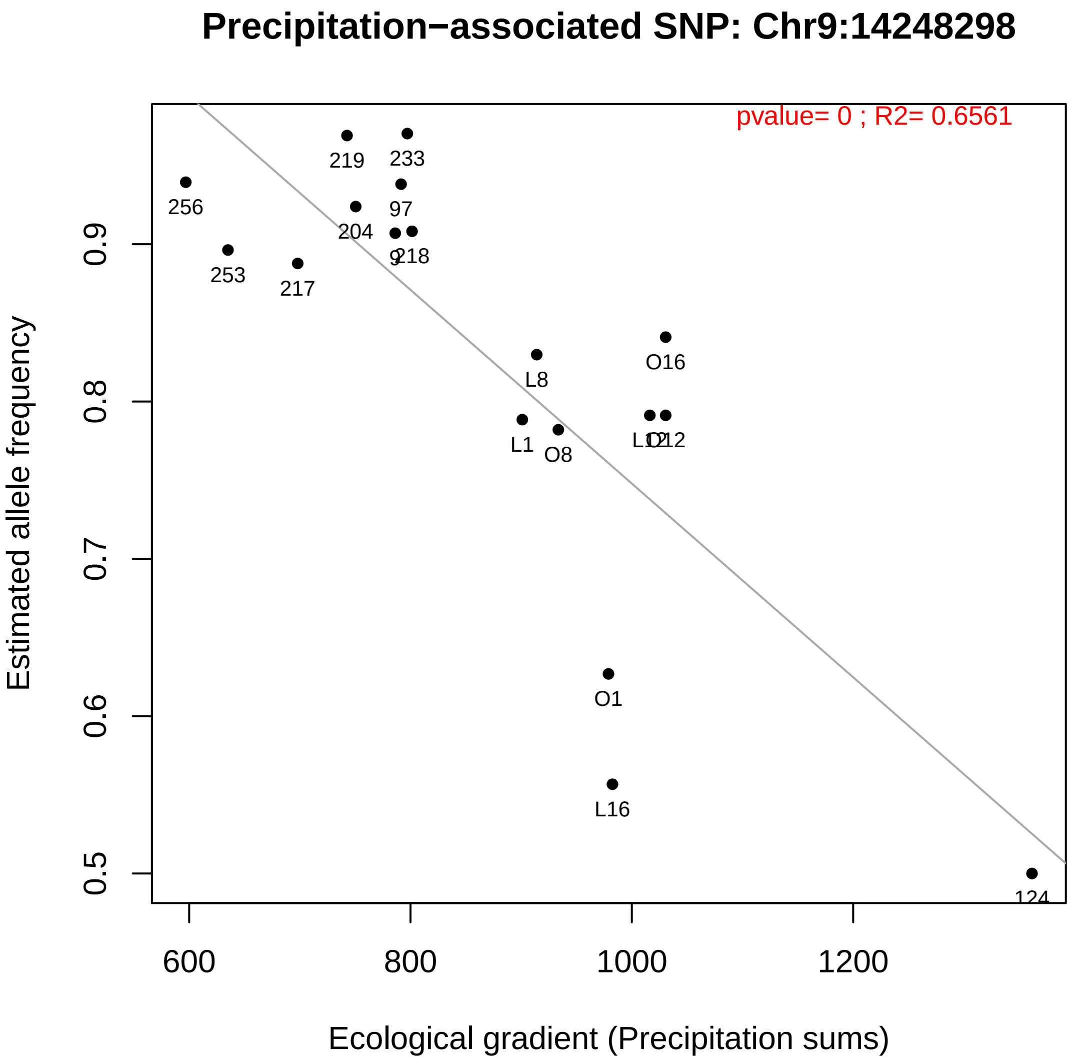

Precipitation−associated SNP: Chr9:14248298
pvalue= 0 ; R2= 0.6561
Estimated allele frequency
Ecological gradient (Precipitation sums)
0.5
0.6
0.7
0.8
0.9
600
800
1000
1200
256
219
233
97
204
253
217
218
L8
O16
L1
O8
O12
O1
L16
124

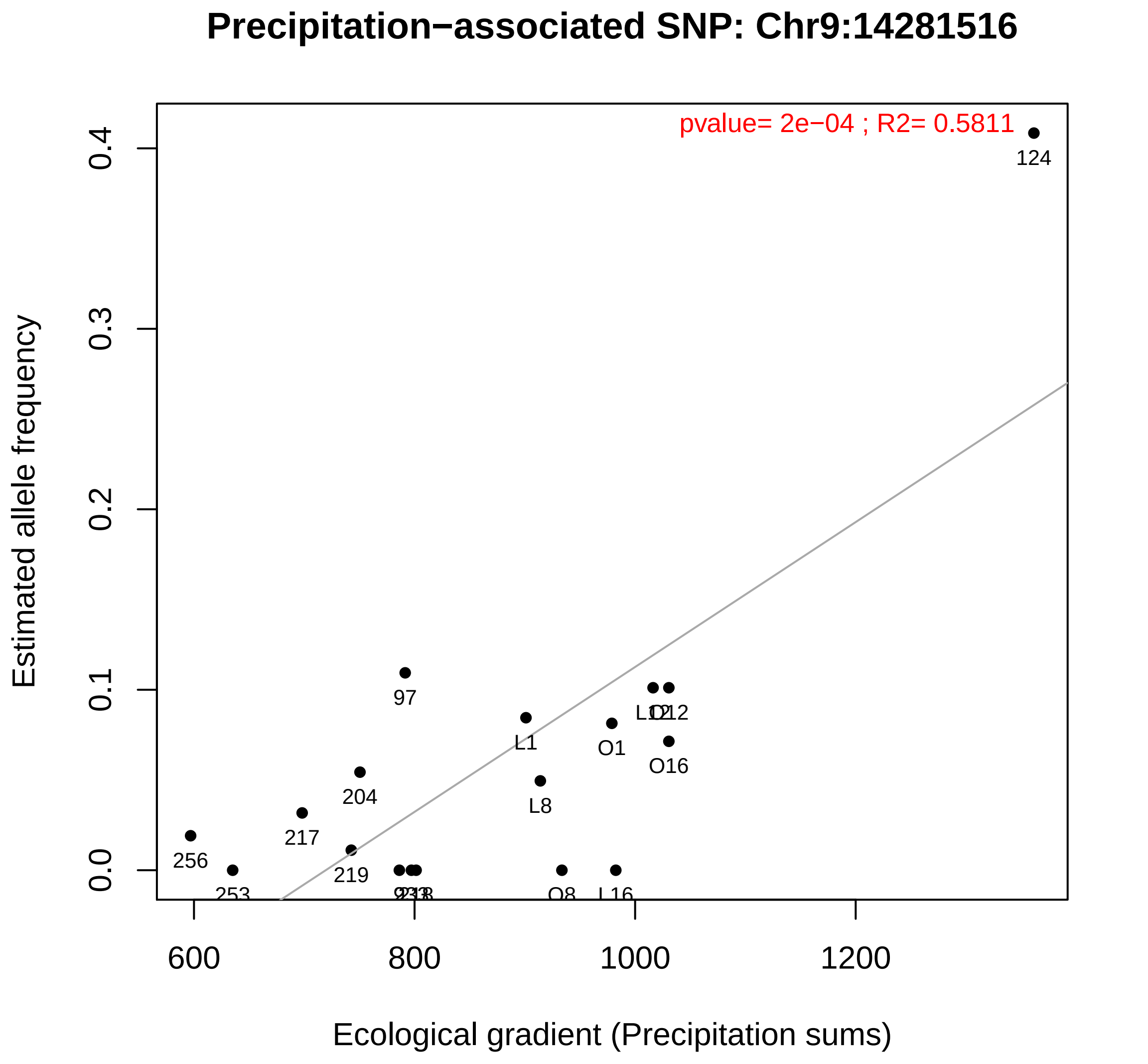

Precipitation−associated SNP: Chr9:14281516
pvalue= 2e−04 ; R2= 0.5811
Estimated allele frequency
0.0
0.1
0.2
0.3
0.4
600
800
1000
1200
Ecological gradient (Precipitation sums)
124
97
L1
O1
O16
L8
204
217
256
219
253
O8
L16

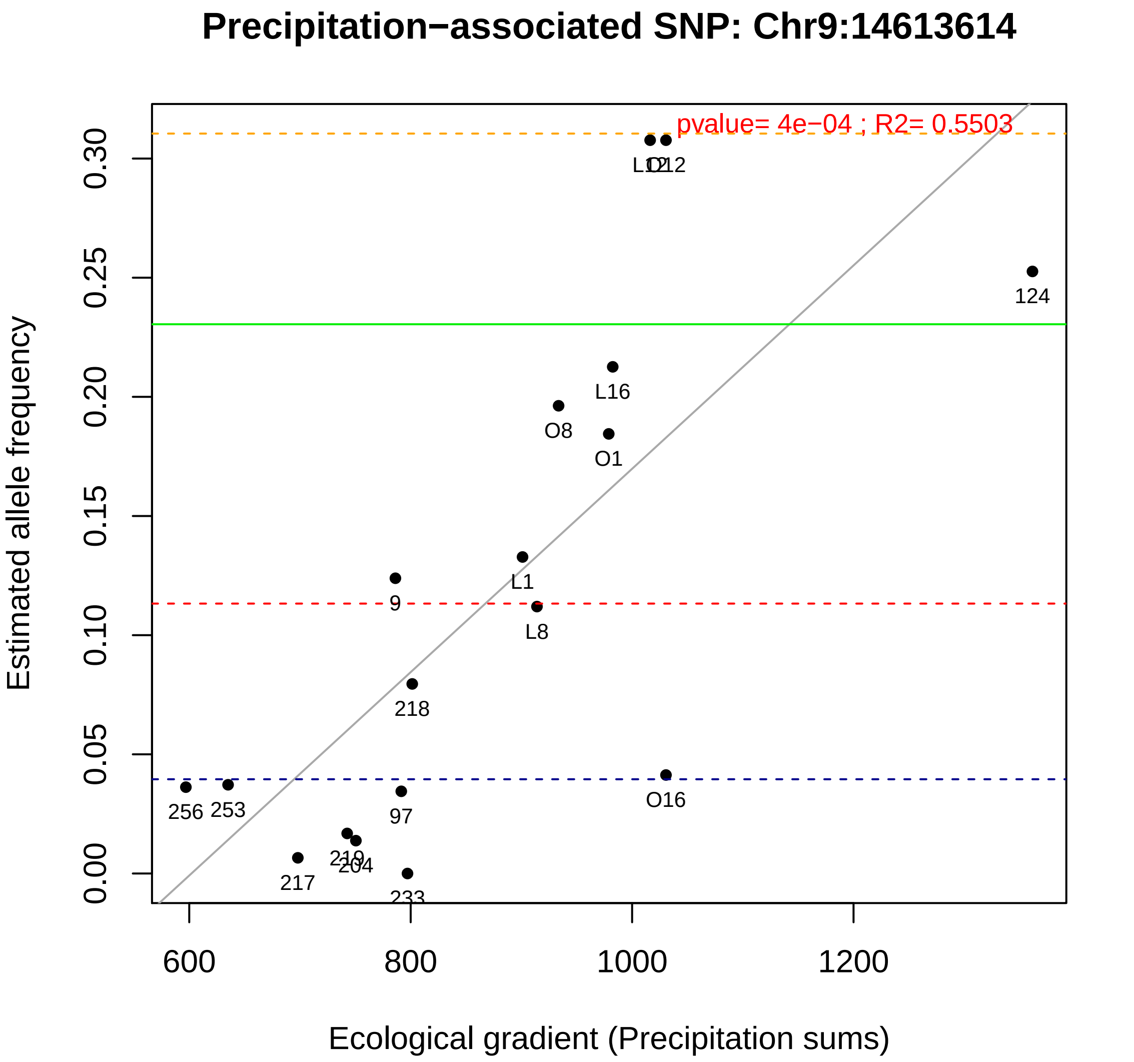

Precipitation−associated SNP: Chr9:14613614
pvalue= 4e−04 ; R2= 0.5503
Estimated allele frequency
Ecological gradient (Precipitation sums)
0.00
0.05
0.10
0.15
0.20
0.25
0.30
600
800
1000
1200
L16
O8
O1
124
L1
9
L8
218
O16
256
253
97
217
204
233

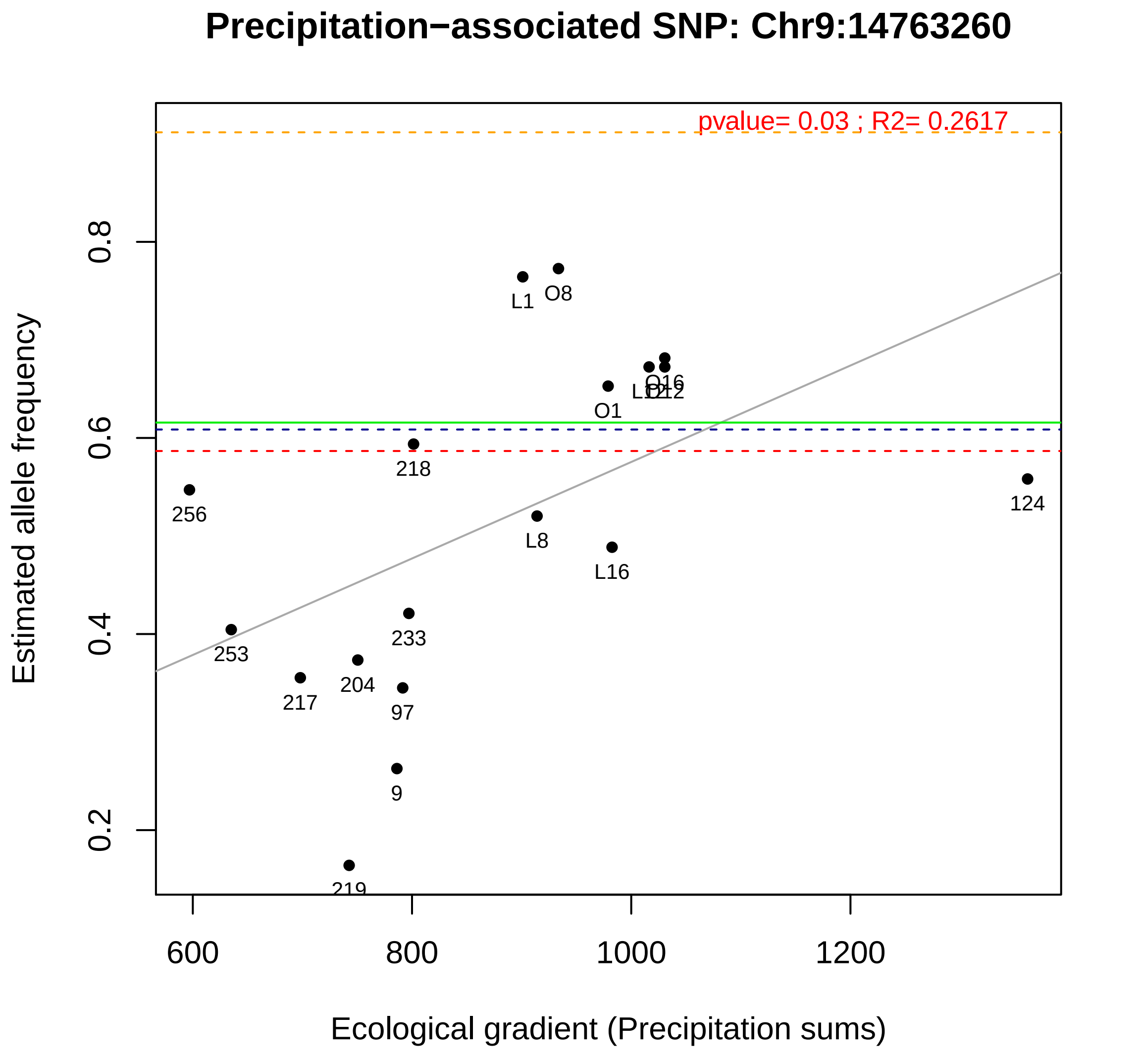
Precipitation−associated SNP: Chr9:14763260
pvalue= 0.03 ; R2= 0.2617
Estimated allele frequency
0.2
0.4
0.6
0.8
Ecological gradient (Precipitation sums)
600
800
1000
1200
L1
O8
O16
O1
218
256
124
L8
L16
233
253
204
217
97
9
219

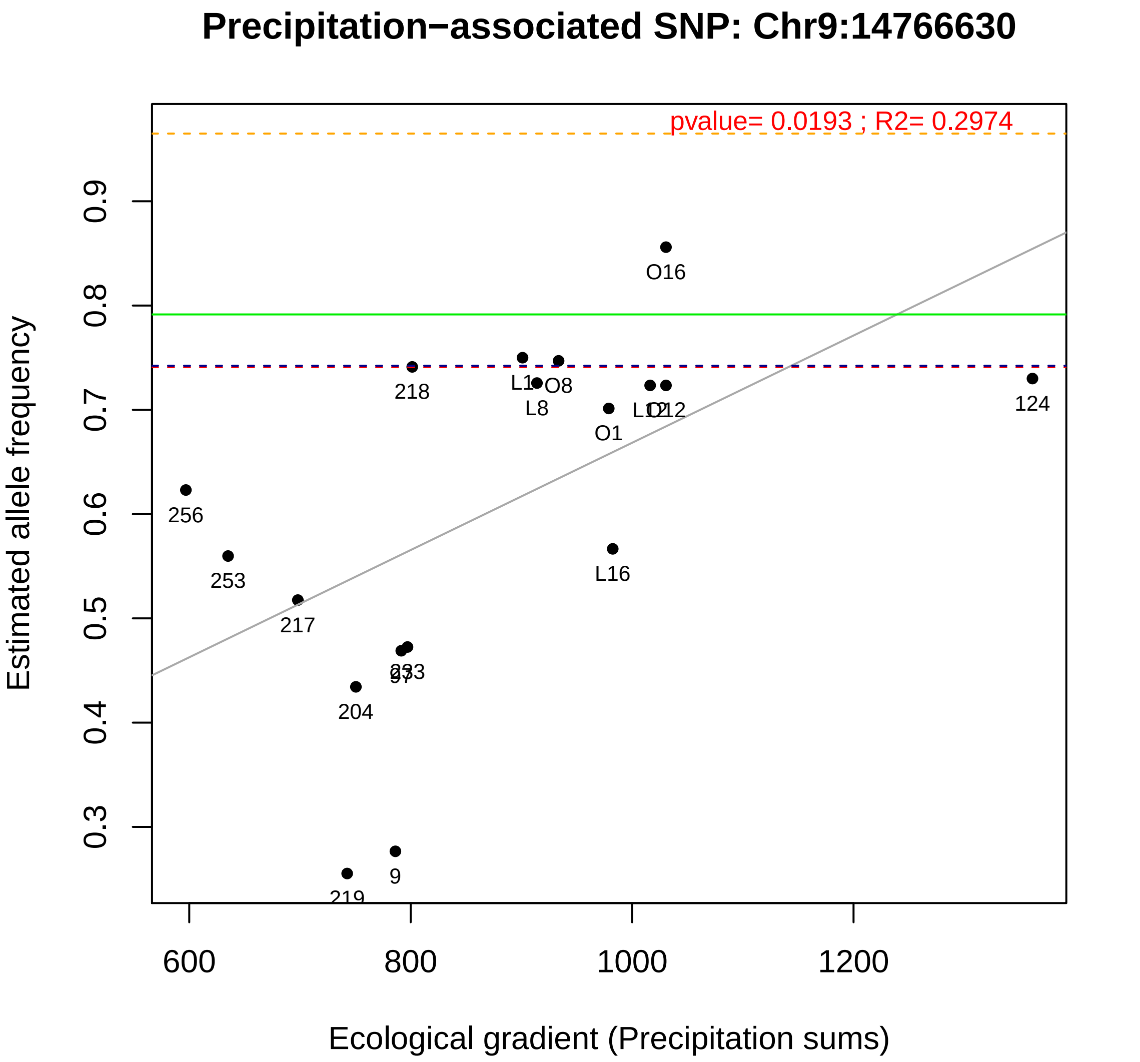

Precipitation−associated SNP: Chr9:14766630
pvalue= 0.0193 ; R2= 0.2974
Estimated allele frequency
0.3
0.4
0.5
0.6
0.7
0.8
0.9
600
800
1000
1200
Ecological gradient (Precipitation sums)
O16
218
L1
O8
L8
O1
124
256
253
217
L16
204
219
9

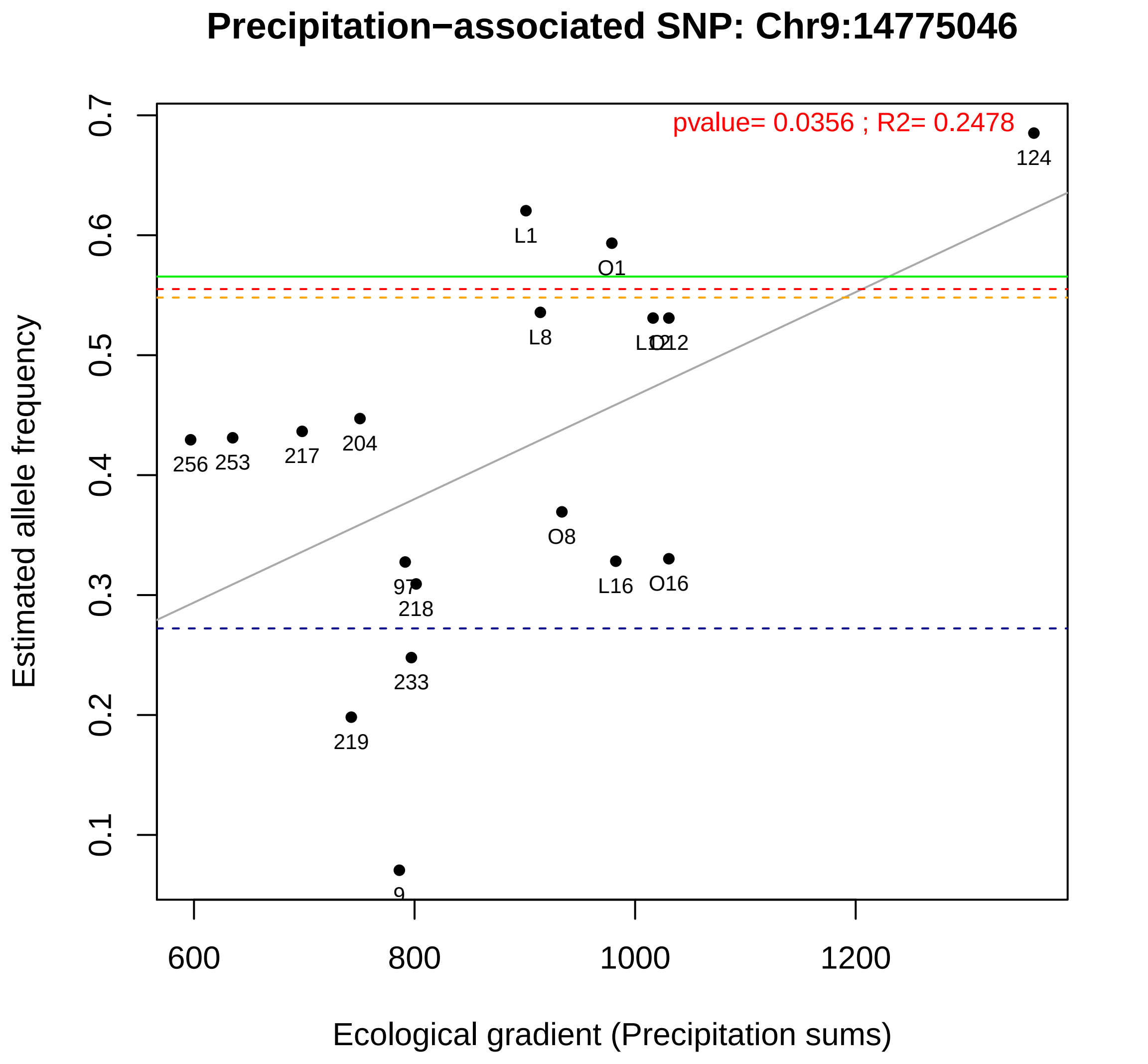

Precipitation−associated SNP: Chr9:14775046
pvalue= 0.0356 ; R2= 0.2478
124
L1
O1
L8
L12
O12
204
256
253
217
O8
97
218
L16
O16
233
219
9
0.1
0.2
0.3
0.4
0.5
0.6
0.7
600
800
1000
1200
Estimated allele frequency
Ecological gradient (Precipitation sums)

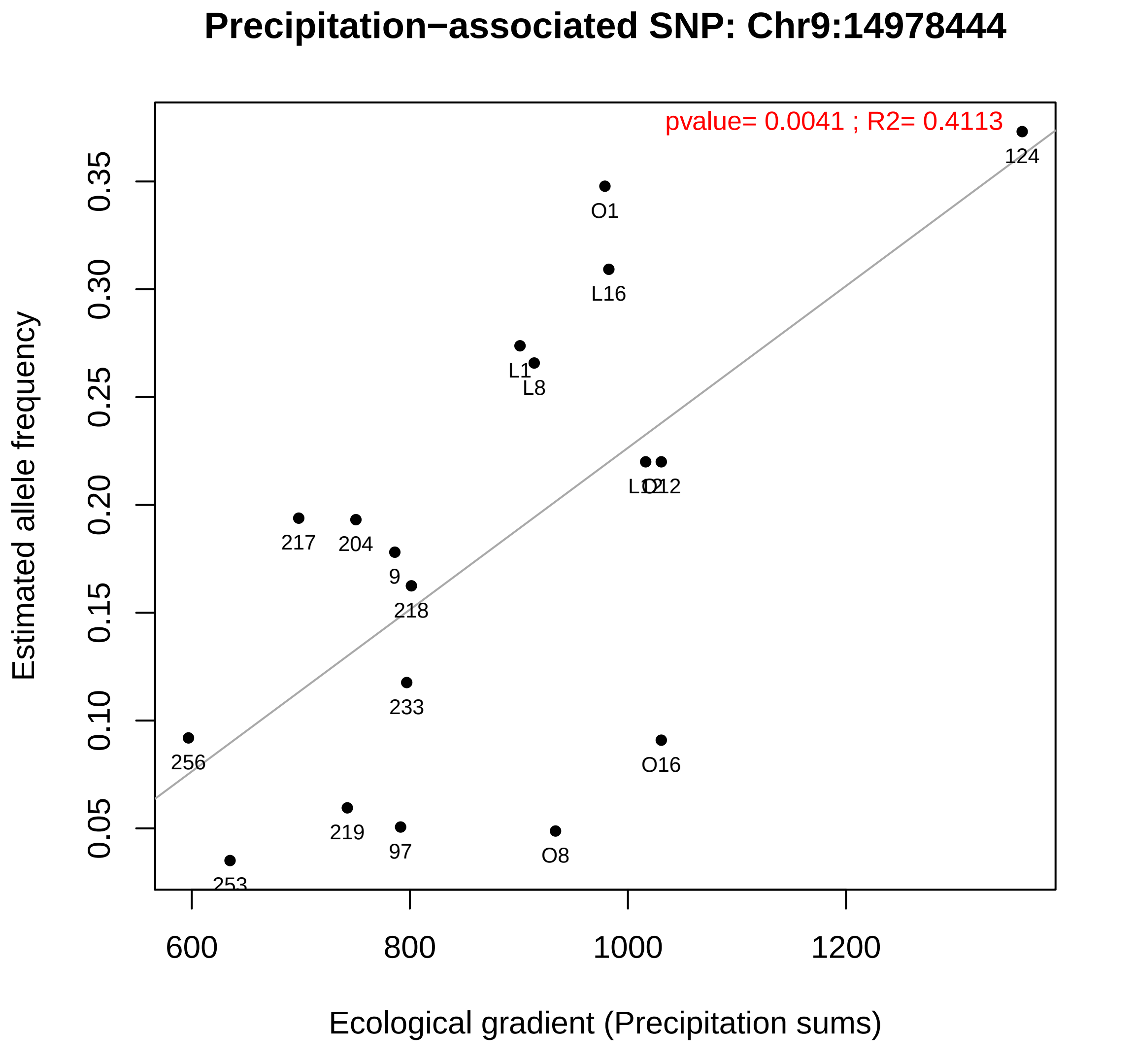

Precipitation−associated SNP: Chr9:14978444
pvalue= 0.0041 ; R2= 0.4113
Estimated allele frequency
0.05
0.10
0.15
0.20
0.25
0.30
0.35
600
800
1000
1200
Ecological gradient (Precipitation sums)
124
O1
L16
L1
L8
O12
217
204
9
218
233
256
O16
219
97
O8
253

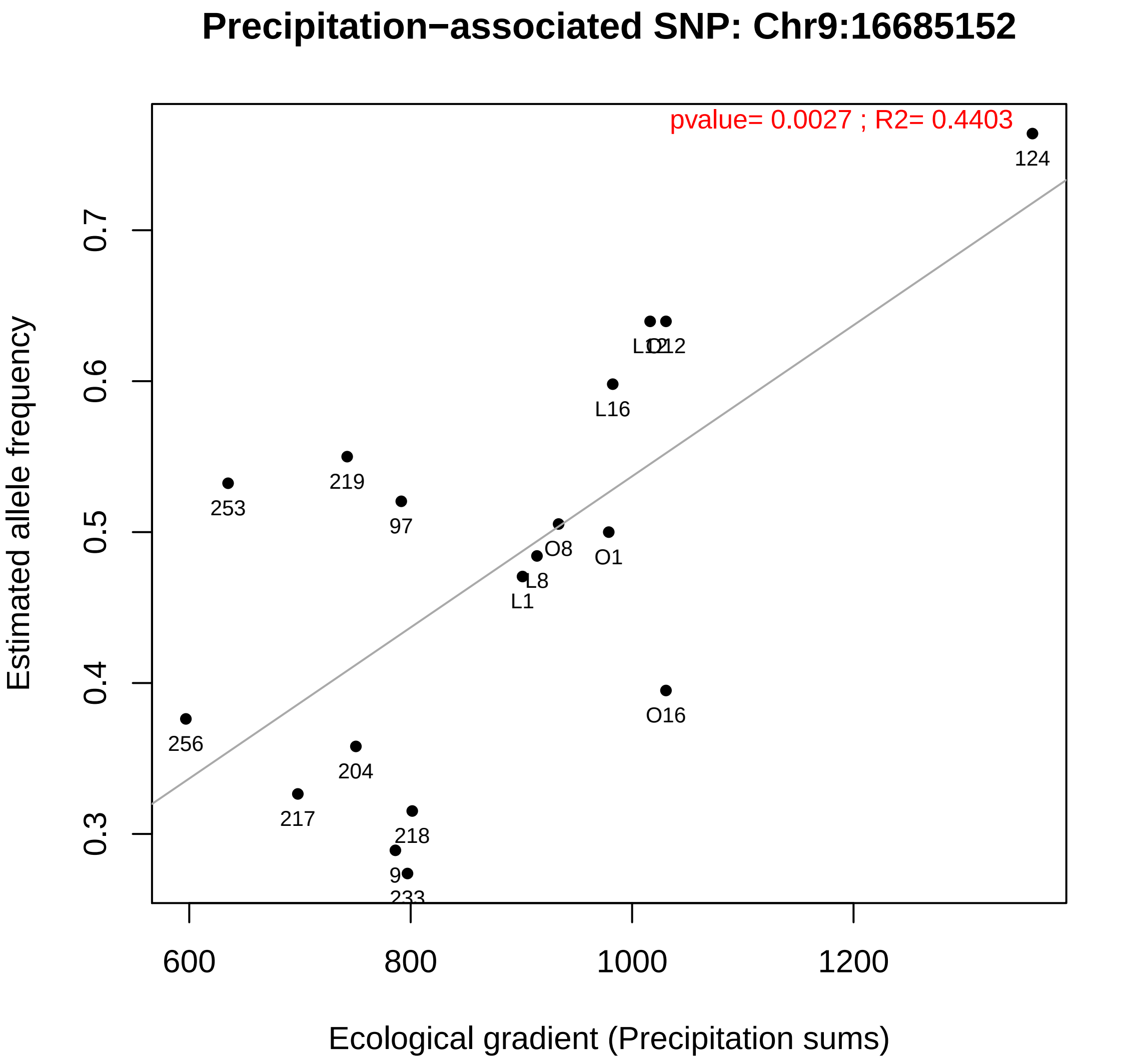

Precipitation−associated SNP: Chr9:16685152
pvalue= 0.0027 ; R2= 0.4403
124
L10
O12
L16
219
253
97
O8
O1
L8
L1
O16
256
204
217
218
9
233
0.3
0.4
0.5
0.6
0.7
600
800
1000
1200
Estimated allele frequency
Ecological gradient (Precipitation sums)

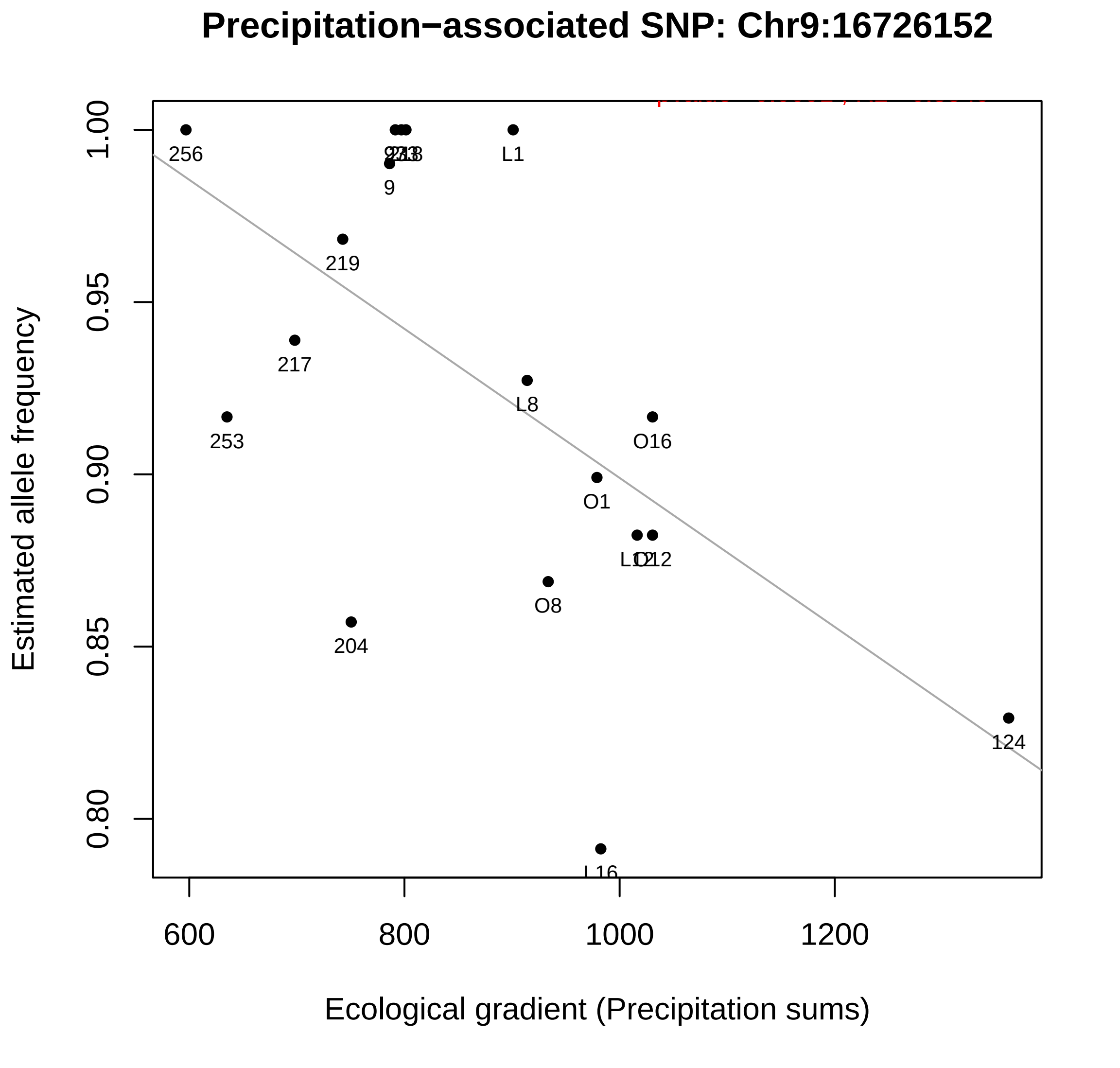
Precipitation−associated SNP: Chr9:16726152
Estimated allele frequency
Ecological gradient (Precipitation sums)
1.00
0.95
0.90
0.85
0.80
600
800
1000
1200
256
9
L1
219
217
L8
253
O16
O1
O8
204
124

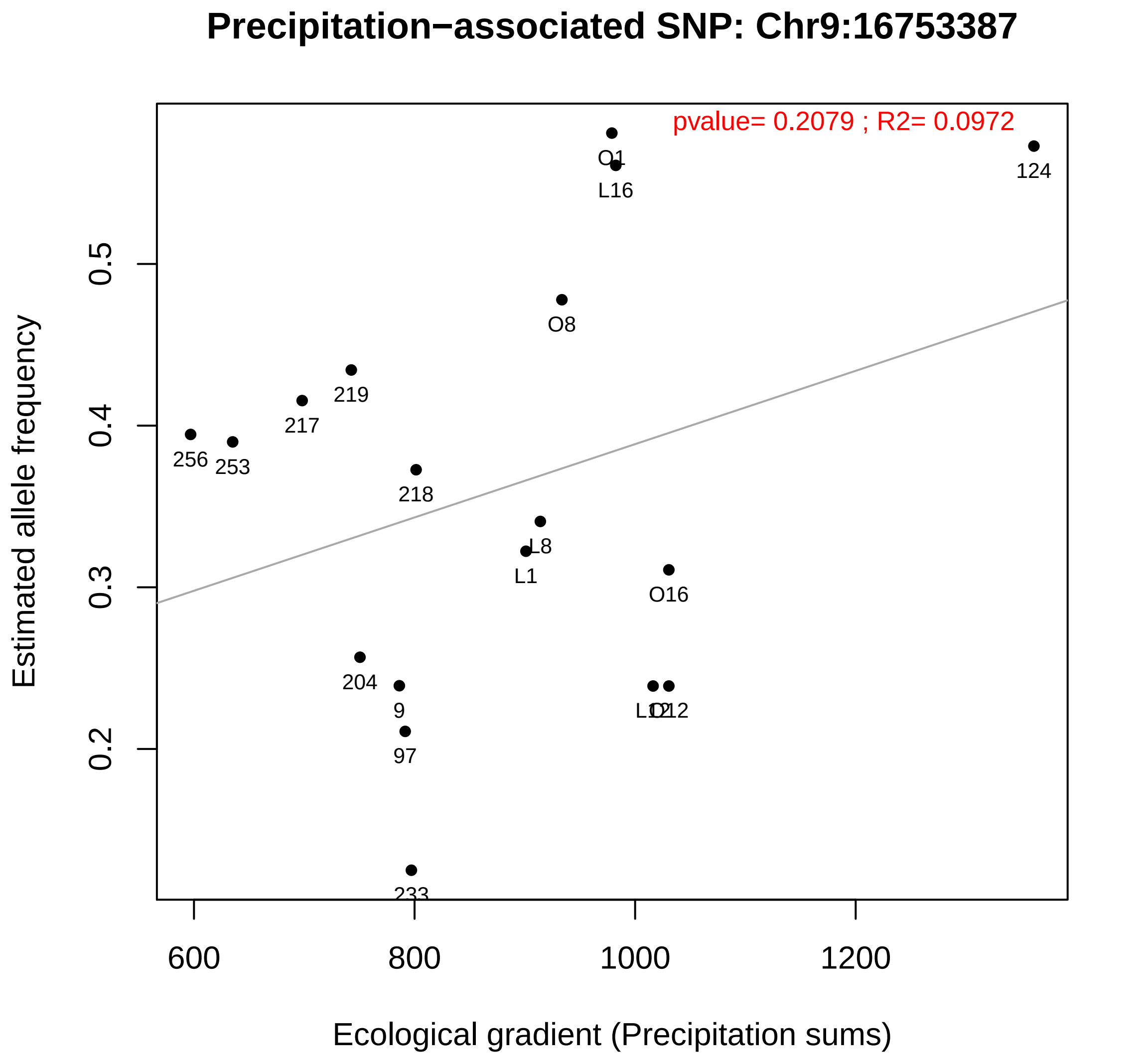

Precipitation−associated SNP: Chr9:16753387
pvalue= 0.2079 ; R2= 0.0972
Estimated allele frequency
0.2
0.3
0.4
0.5
600
800
1000
1200
Ecological gradient (Precipitation sums)
O1
L16
124
O8
219
217
256
253
218
L8
L1
O16
204
9
L12
O12
97
233

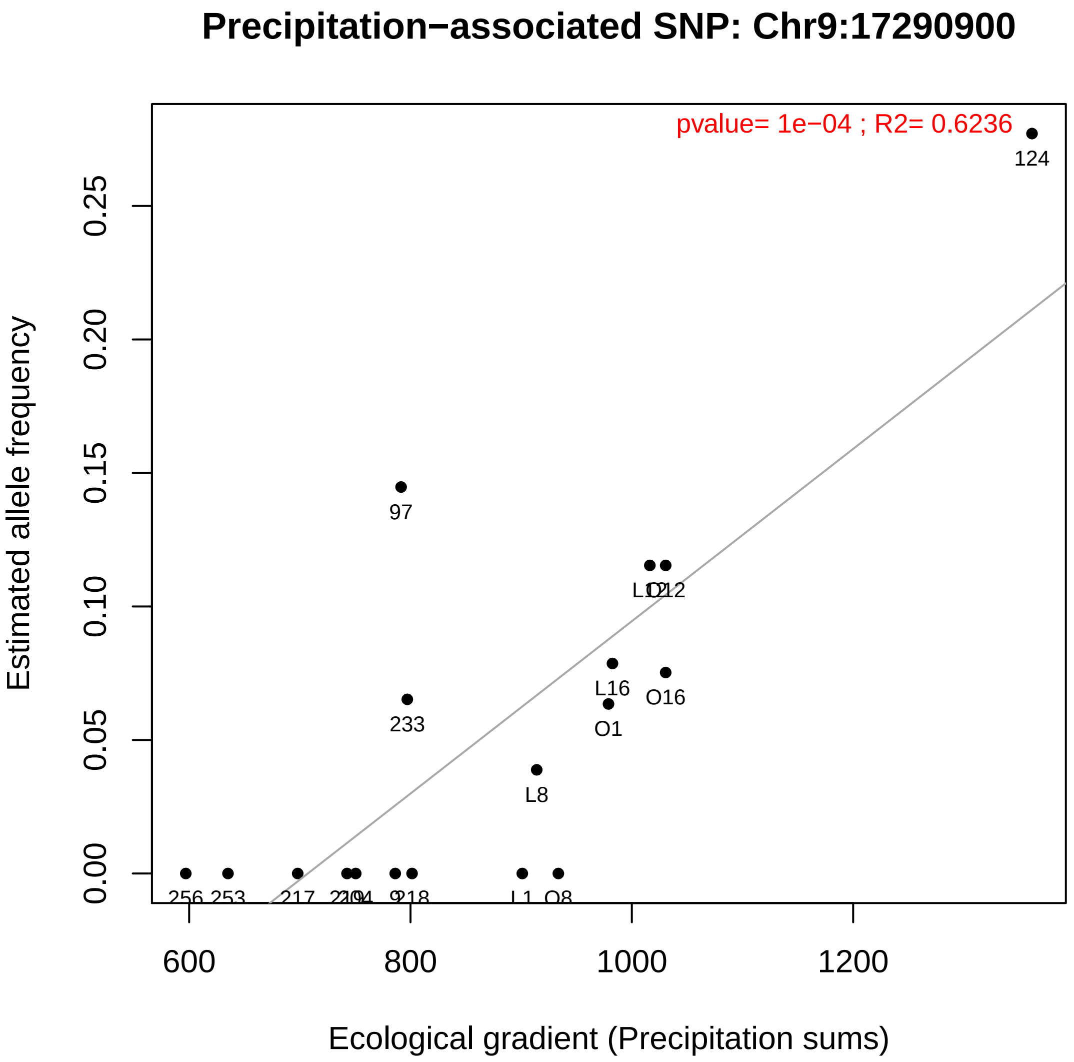
Precipitation−associated SNP: Chr9:17290900
pvalue= 1e−04 ; R2= 0.6236
Estimated allele frequency
0.00
0.05
0.10
0.15
0.20
0.25
600
800
1000
1200
Ecological gradient (Precipitation sums)
124
97
L16
O16
233
O1
L8

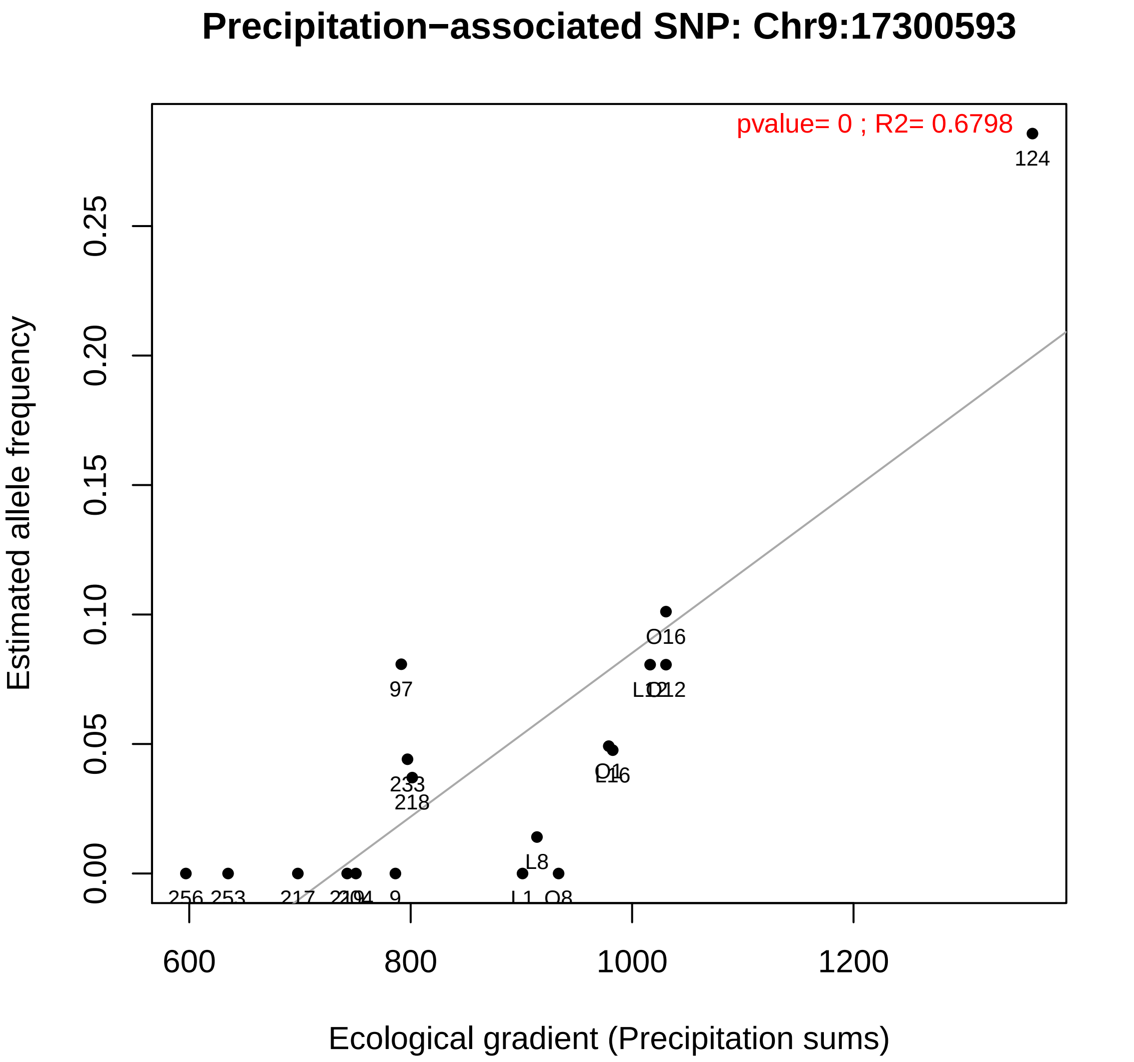

Precipitation−associated SNP: Chr9:17300593
pvalue= 0 ; R2= 0.6798
124
O16
97
L10
O12
233
218
O1
L16
L8
256
253
217
9
L1
O8
0.00
0.05
0.10
0.15
0.20
0.25
600
800
1000
1200
Estimated allele frequency
Ecological gradient (Precipitation sums)

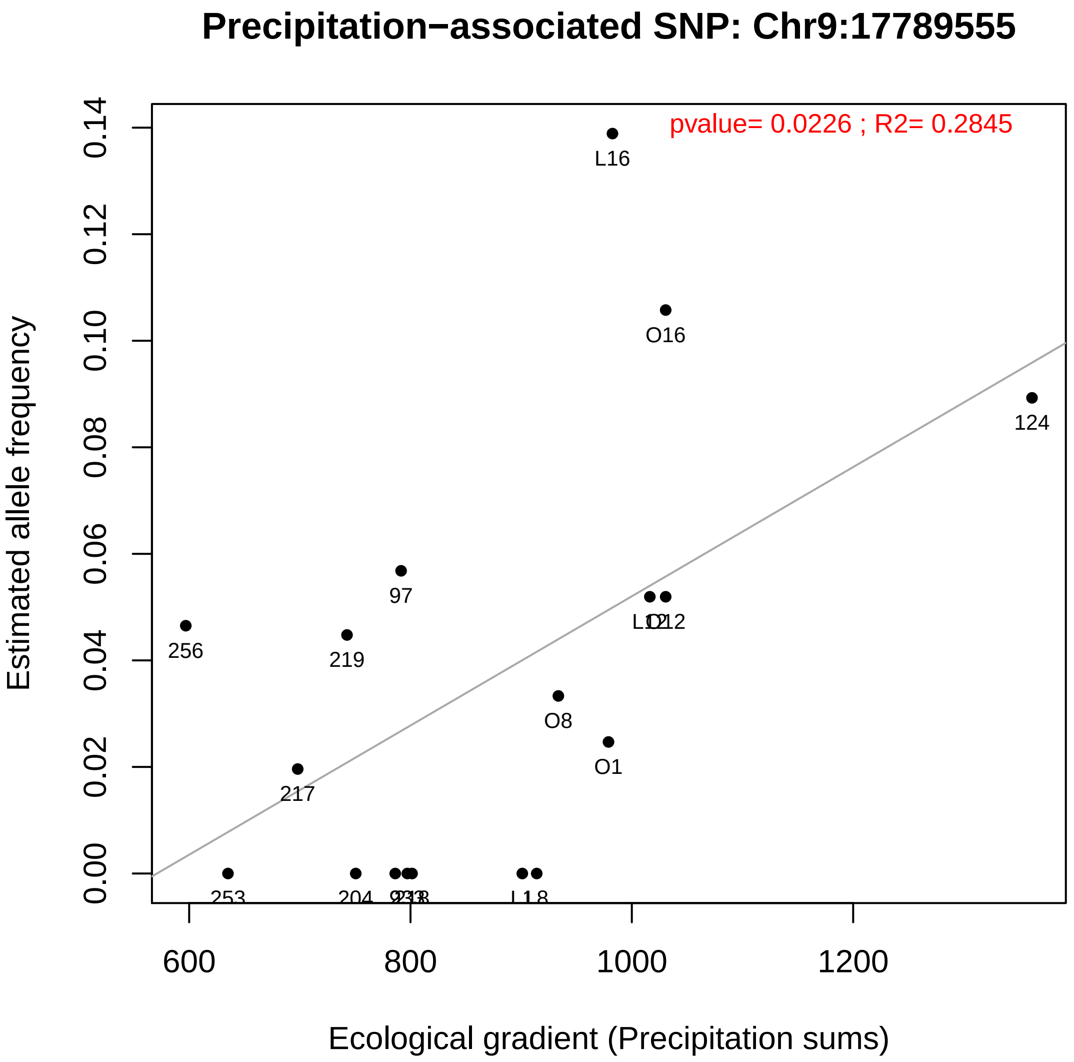

Precipitation−associated SNP: Chr9:17789555
pvalue= 0.0226 ; R2= 0.2845
Estimated allele frequency
Ecological gradient (Precipitation sums)
0.00
0.02
0.04
0.06
0.08
0.10
0.12
0.14
600
800
1000
1200
L16
O16
124
97
256
219
O8
O1
217
253
204

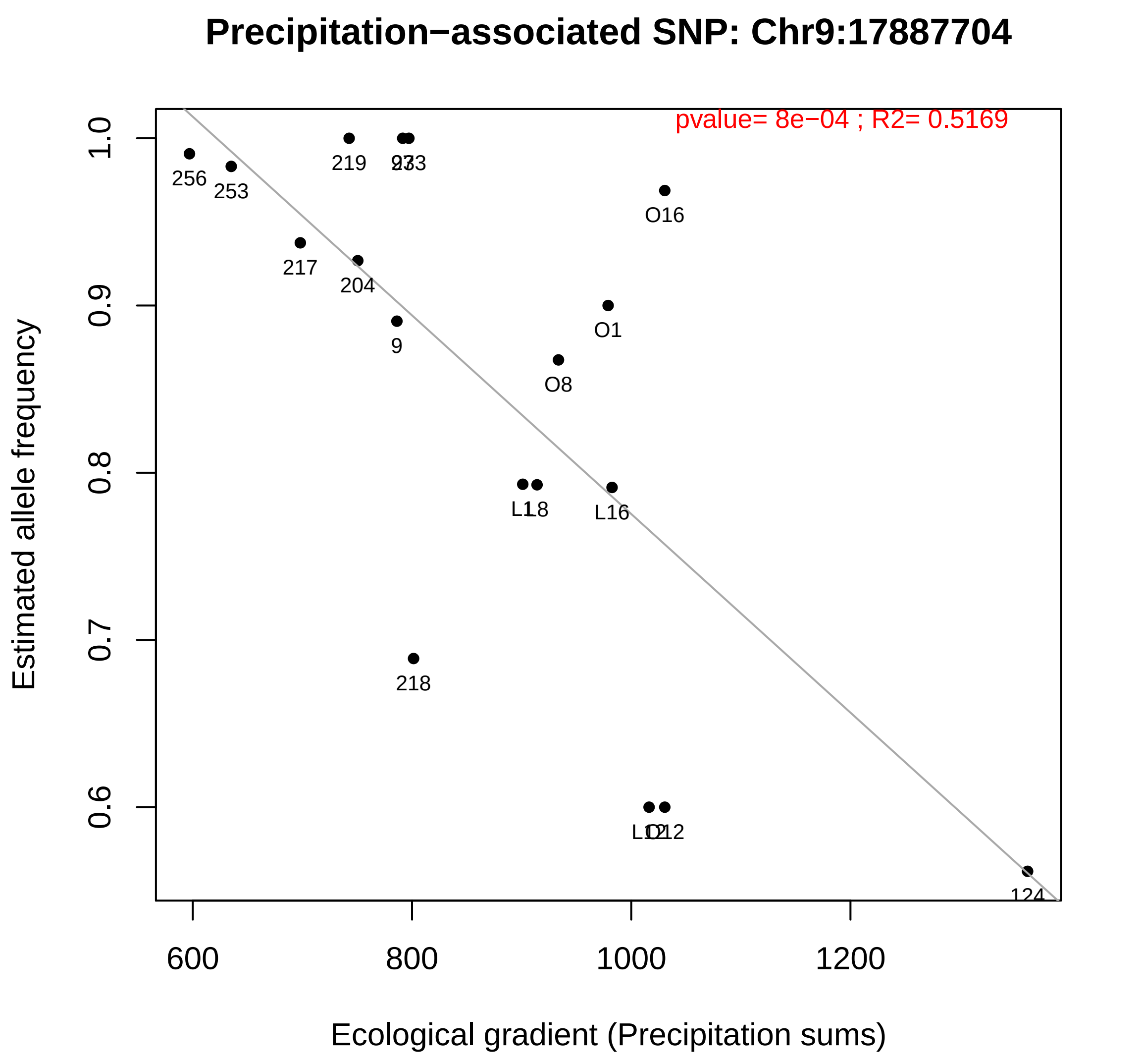

Precipitation−associated SNP: Chr9:17887704
pvalue= 8e−04 ; R2= 0.5169
Estimated allele frequency
Ecological gradient (Precipitation sums)
0.6
0.7
0.8
0.9
1.0
600
800
1000
1200
256
253
219
217
204
9
O16
O1
O8
L16
218
124

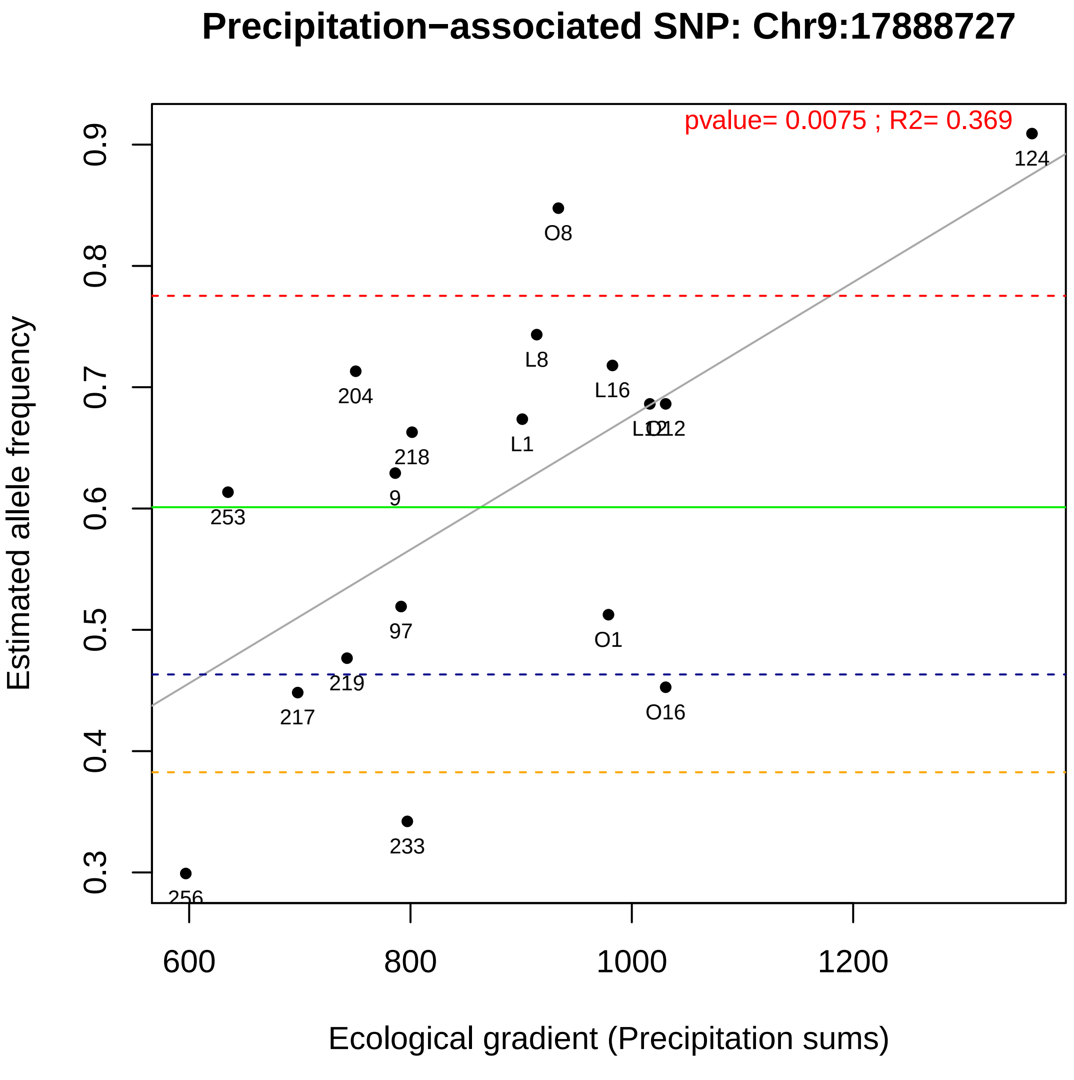

Precipitation−associated SNP: Chr9:17888727
pvalue= 0.0075 ; R2= 0.369
124
O8
L8
L16
204
L10
O12
L1
218
9
253
97
O1
219
217
O16
233
256
0.3
0.4
0.5
0.6
0.7
0.8
0.9
600
800
1000
1200
Estimated allele frequency
Ecological gradient (Precipitation sums)

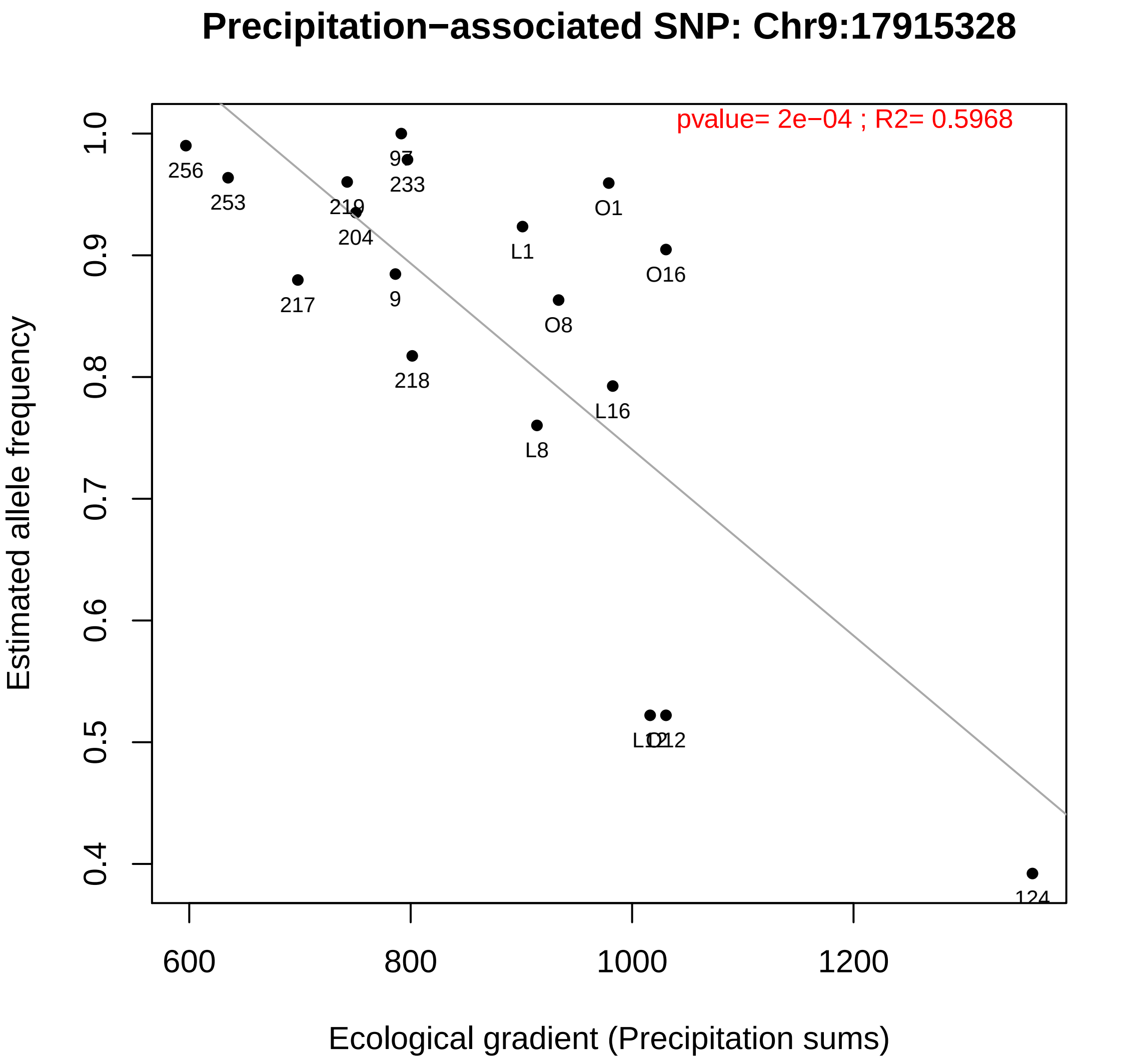
Precipitation−associated SNP: Chr9:17915328
pvalue= 2e−04 ; R2= 0.5968
Estimated allele frequency
Ecological gradient (Precipitation sums)
0.4
0.5
0.6
0.7
0.8
0.9
1.0
600
800
1000
1200
256
253
219
204
97
233
217
9
218
L1
O1
O16
O8
L16
L8
L12
O12
124

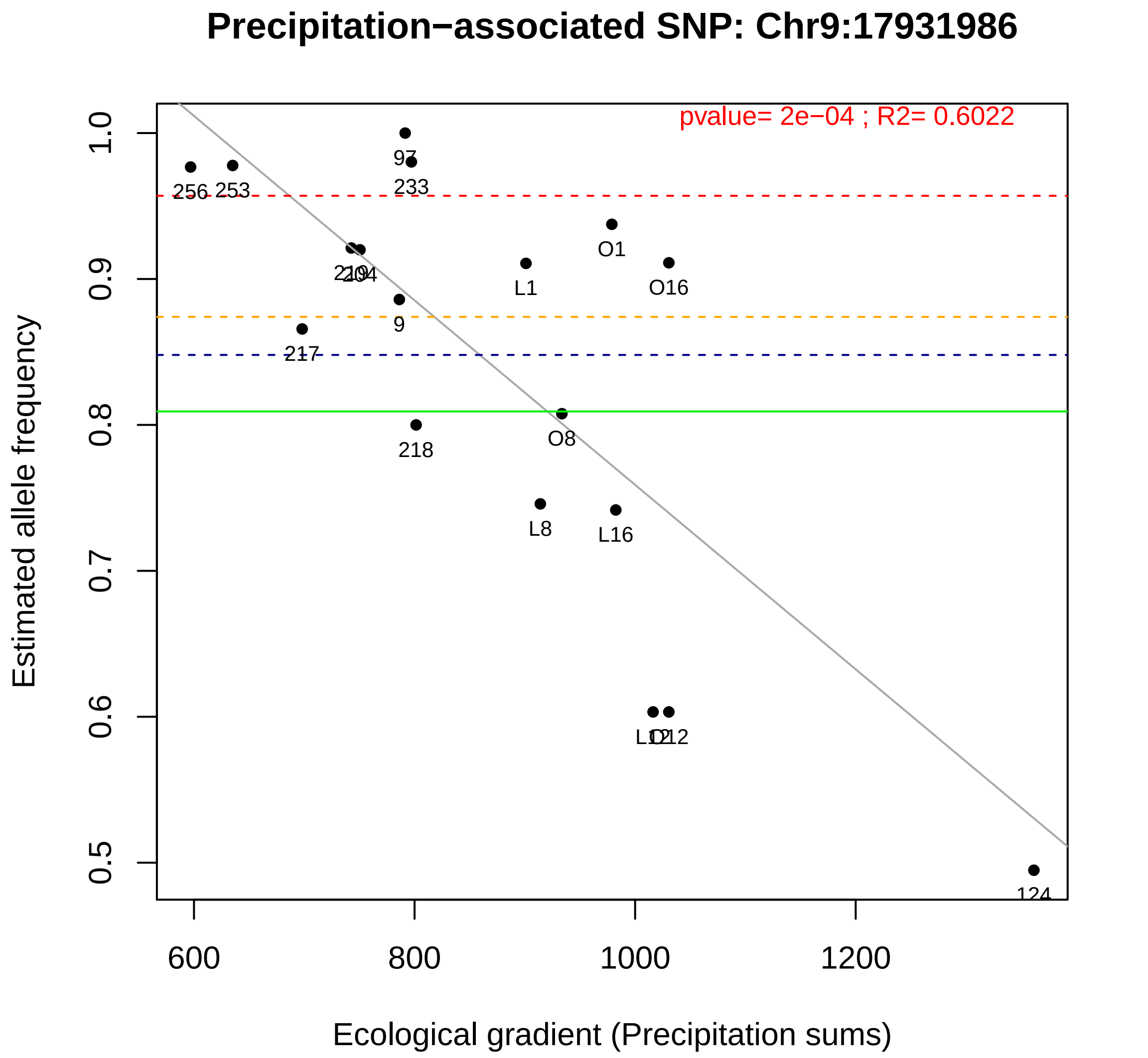
Precipitation−associated SNP: Chr9:17931986
pvalue= 2e−04 ; R2= 0.6022
Estimated allele frequency
1.0
0.9
0.8
0.7
0.6
0.5
600
800
1000
1200
Ecological gradient (Precipitation sums)
256
253
97
233
210
204
217
9
L1
O1
O16
218
O8
L8
L16
L12
O12
124

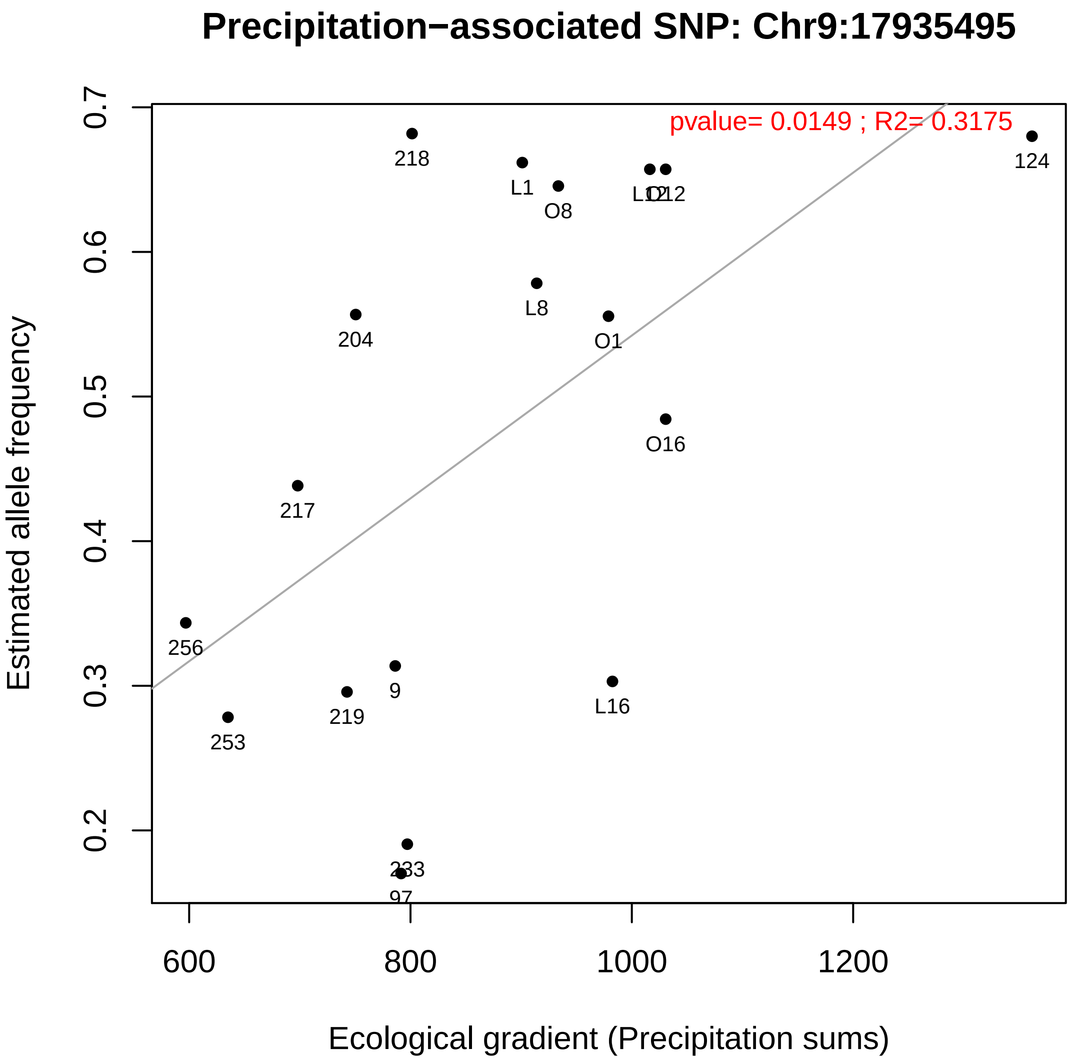
Precipitation−associated SNP: Chr9:17935495
pvalue= 0.0149 ; R2= 0.3175
Estimated allele frequency
Ecological gradient (Precipitation sums)
0.2
0.3
0.4
0.5
0.6
0.7
600
800
1000
1200
218
L1
O8
L12
O12
124
L8
204
O1
O16
217
256
9
219
L16
253
233
97

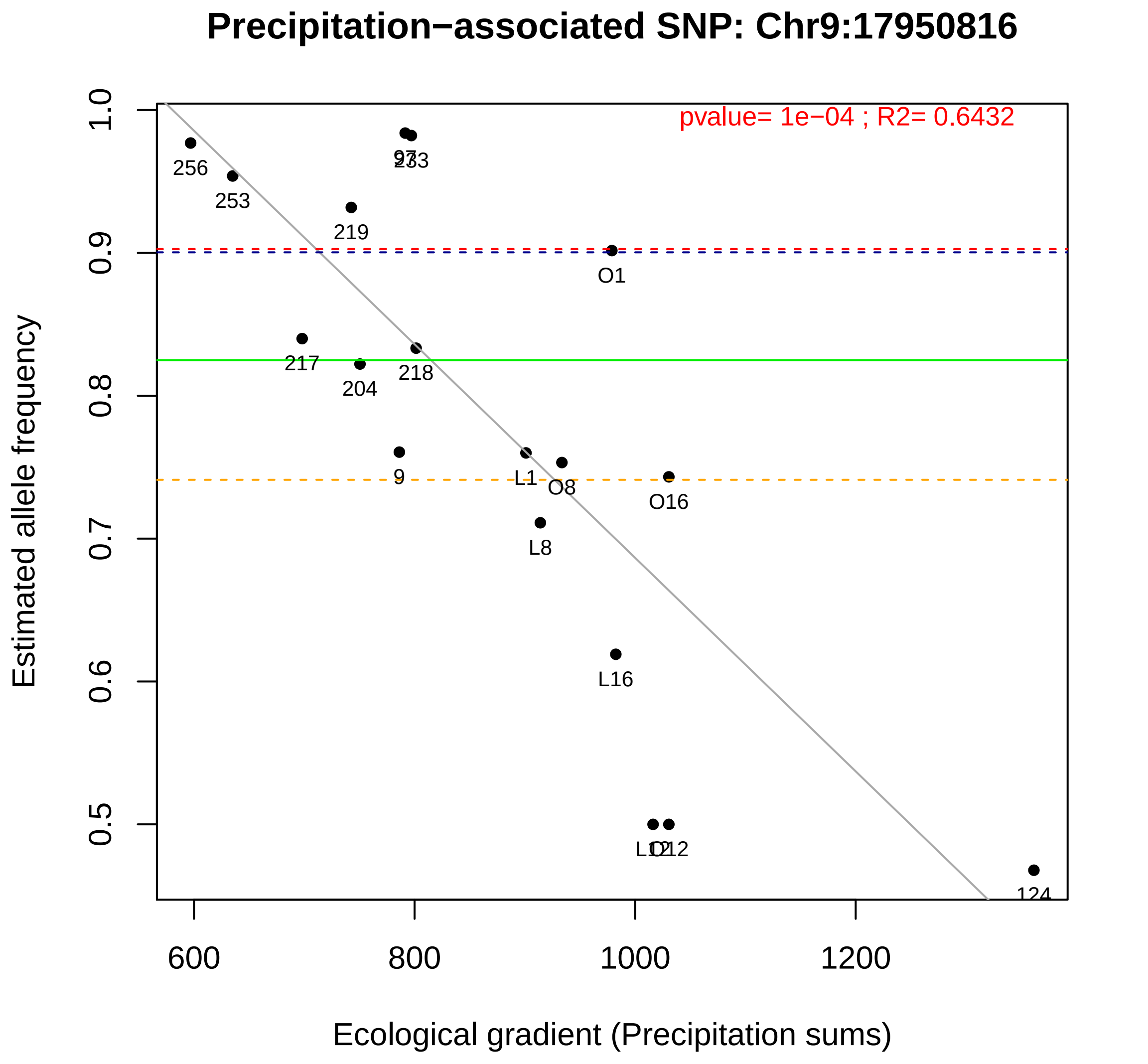

Precipitation−associated SNP: Chr9:17950816
pvalue= 1e−04 ; R2= 0.6432
Estimated allele frequency
Ecological gradient (Precipitation sums)
0.5
0.6
0.7
0.8
0.9
1.0
600
800
1000
1200
256
253
219
217
204
218
9
L1
O8
O1
O16
L8
L16
124

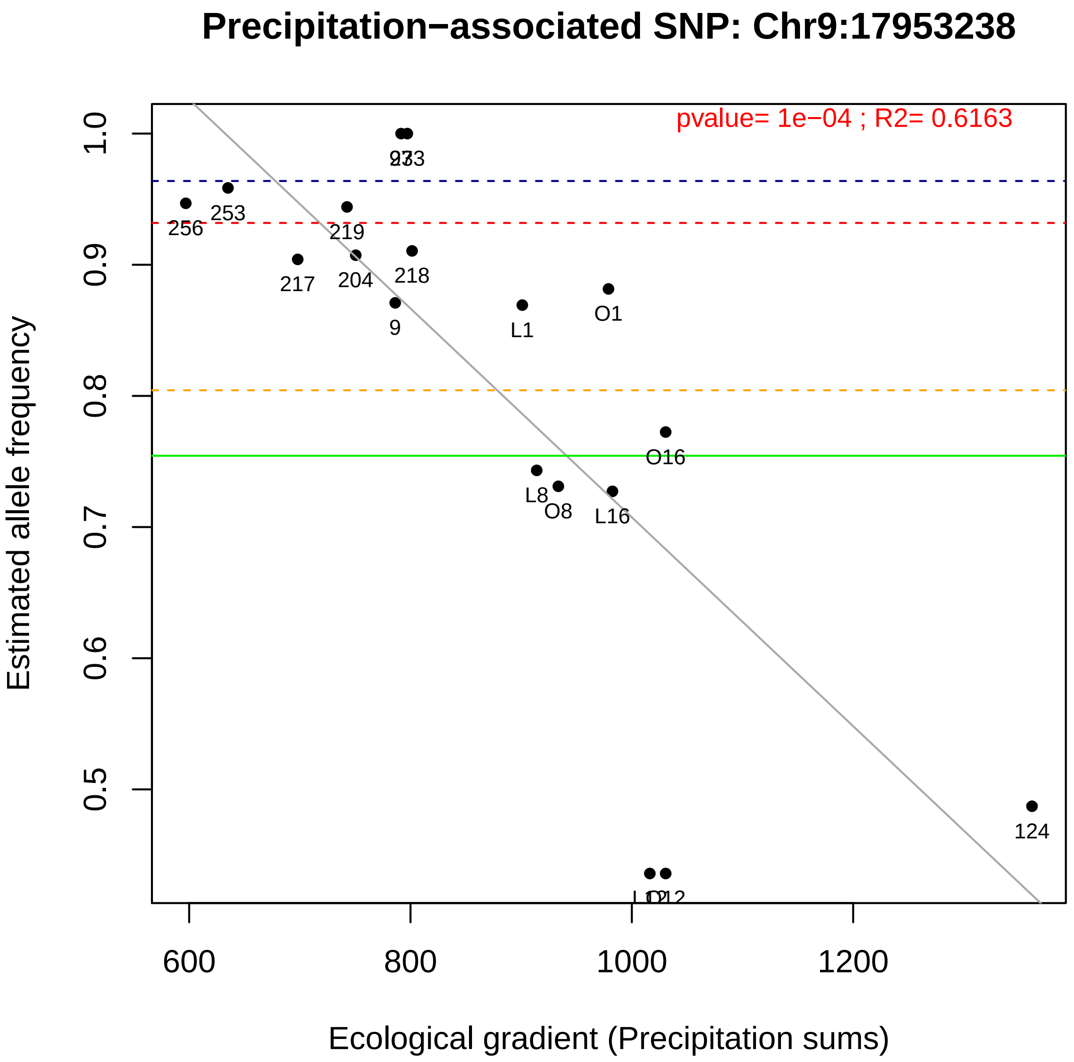
Precipitation−associated SNP: Chr9:17953238
pvalue= 1e−04 ; R2= 0.6163
Estimated allele frequency
0.5
0.6
0.7
0.8
0.9
1.0
256
253
219
217
204
218
9
L1
O1
O16
L8
O8
L16
124
600
800
1000
1200
Ecological gradient (Precipitation sums)

**Precipitation−associated SNP: Chr9:17959227**

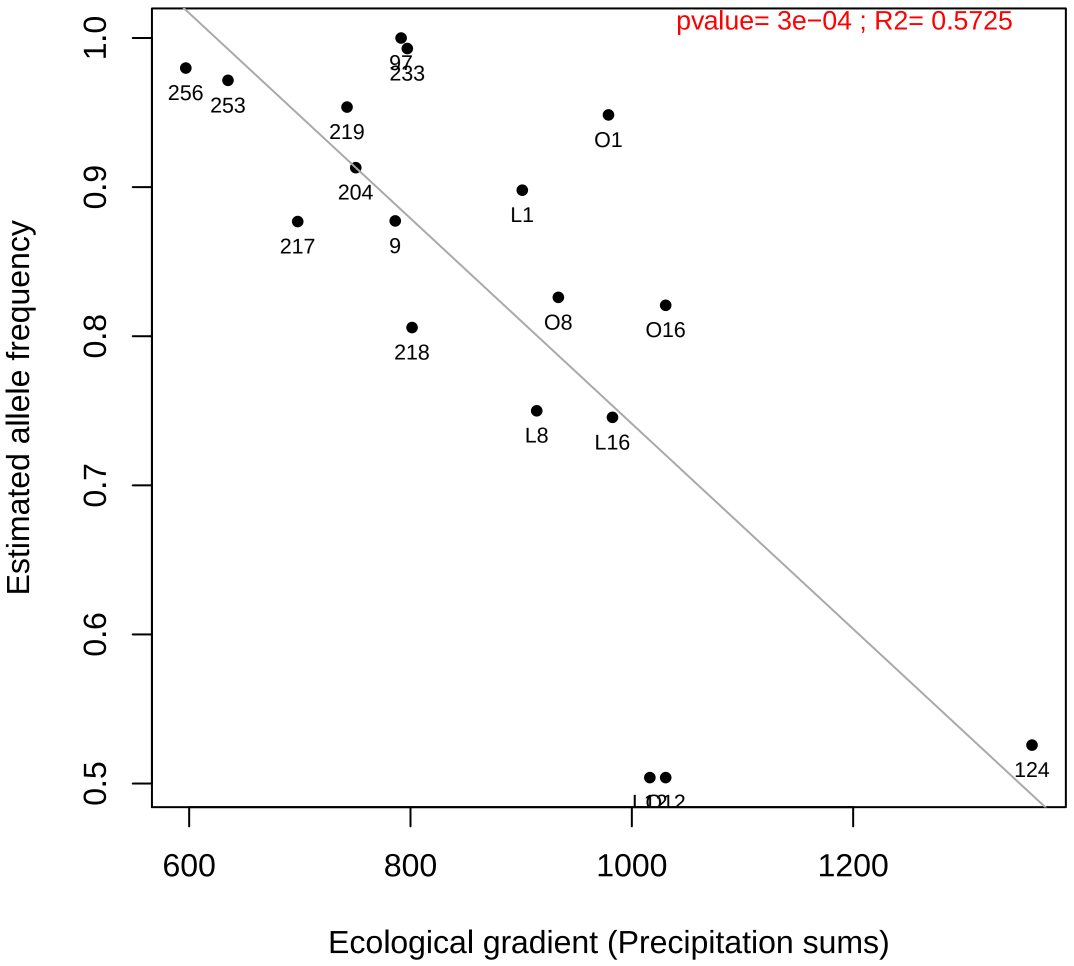

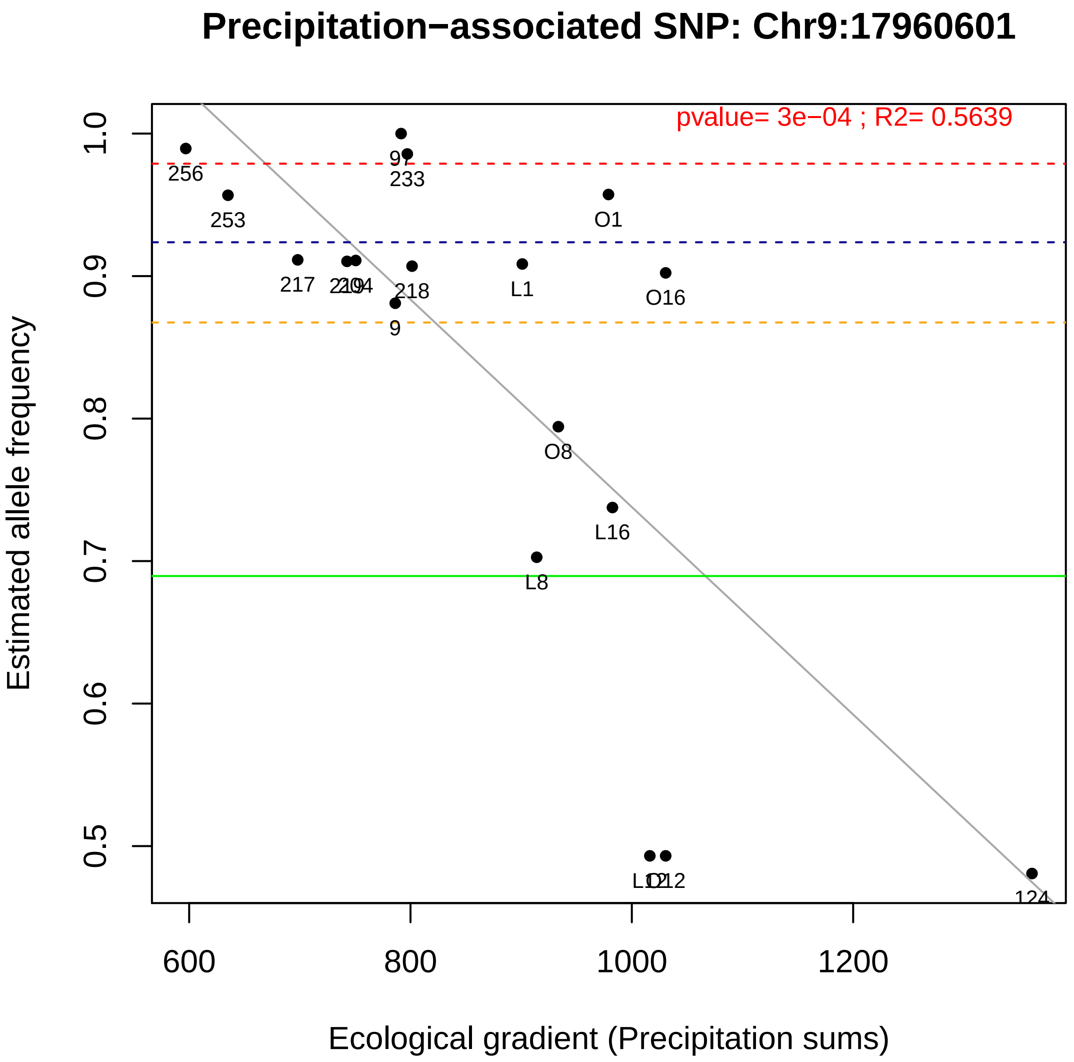
Precipitation−associated SNP: Chr9:17960601
pvalue= 3e−04 ; R2= 0.5639
Estimated allele frequency
Ecological gradient (Precipitation sums)
1.0
0.9
0.8
0.7
0.6
0.5
600
800
1000
1200
256
97
233
253
O1
217
219
204
218
L1
O16
9
O8
L16
L8
L12
O12
124

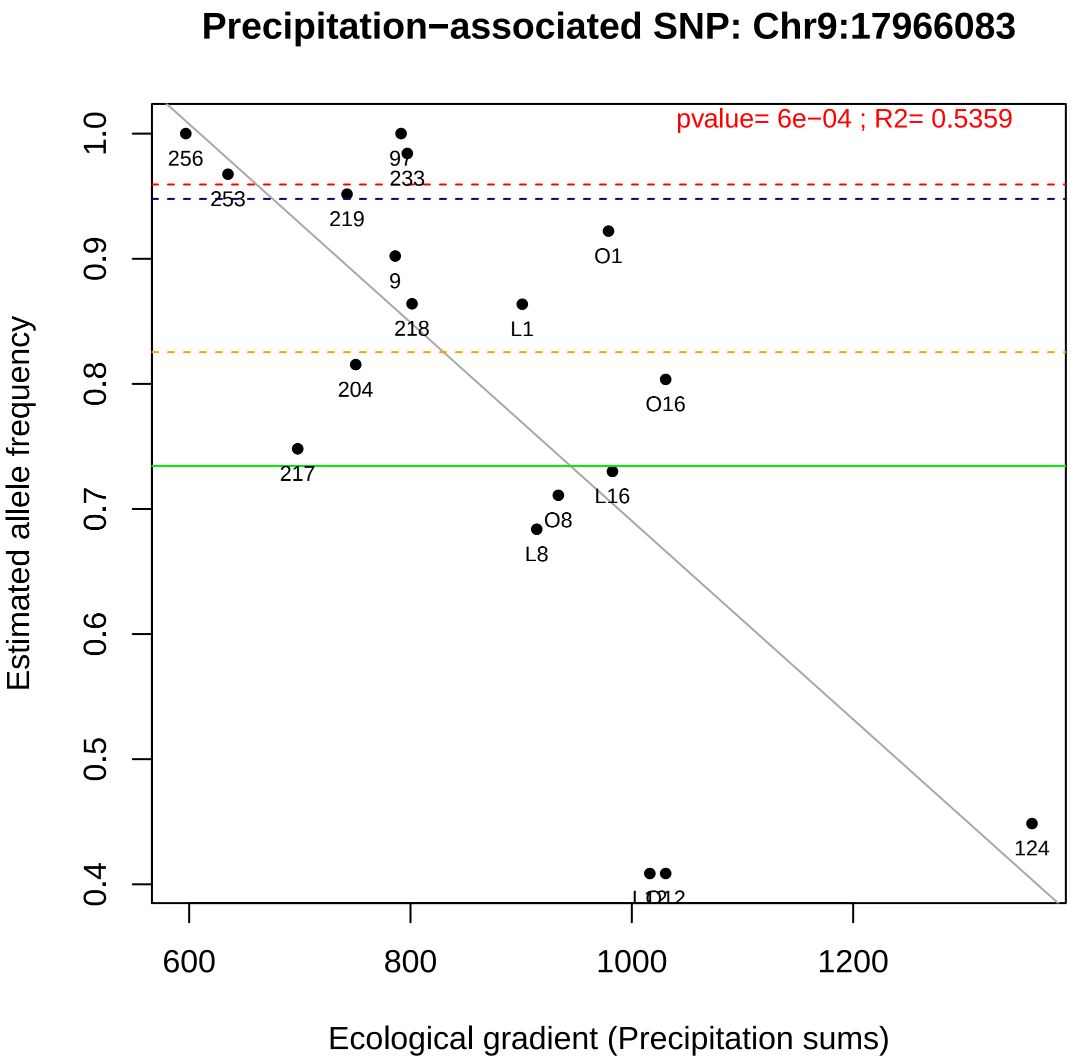

Precipitation−associated SNP: Chr9:17966083
pvalue= 6e−04 ; R2= 0.5359
Estimated allele frequency
Ecological gradient (Precipitation sums)
1.0
0.9
0.8
0.7
0.6
0.5
0.4
600
800
1000
1200
256
253
97
233
219
9
218
L1
O1
204
O16
217
L16
O8
L8
124

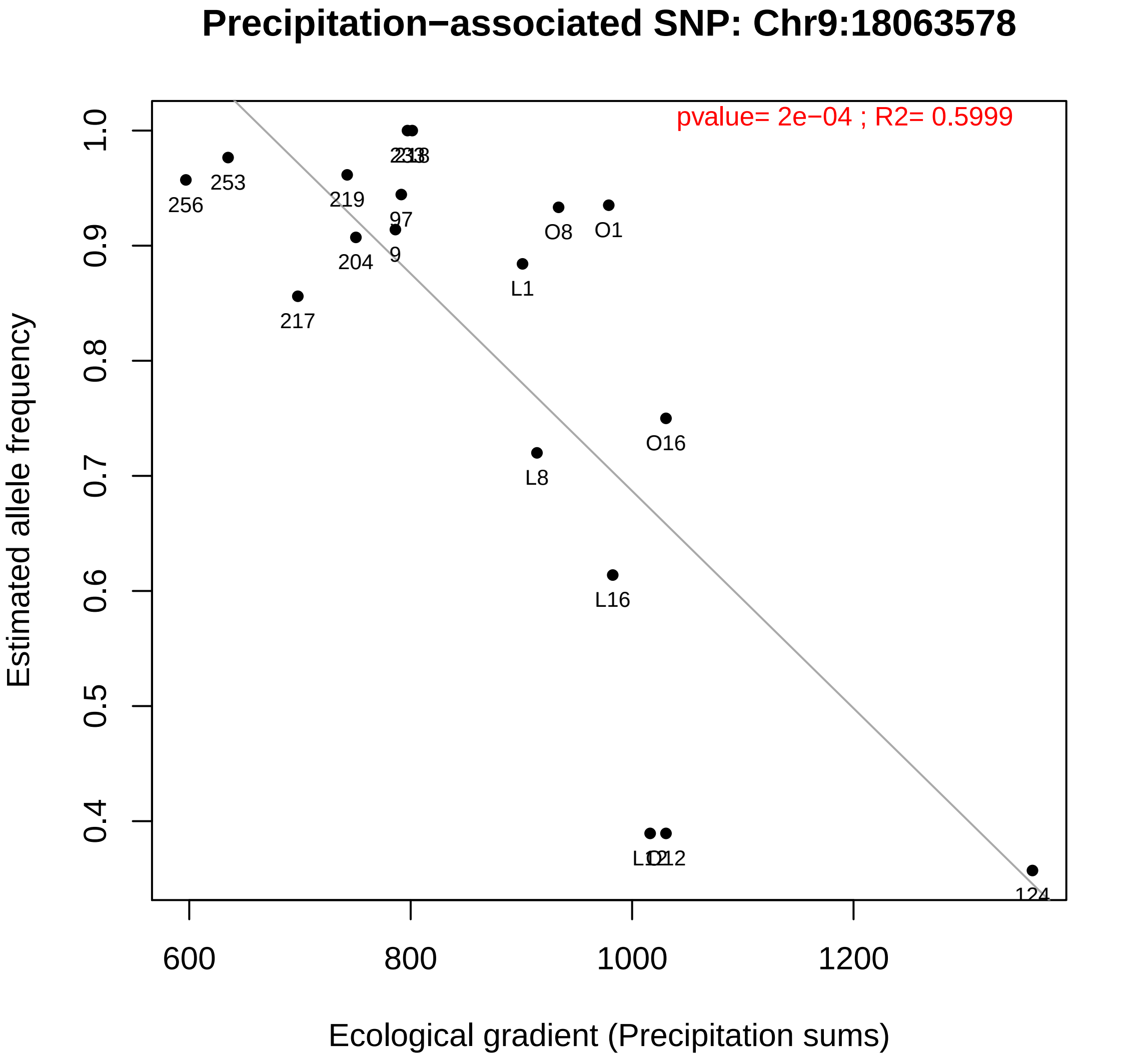
Precipitation−associated SNP: Chr9:18063578
pvalue= 2e−04 ; R2= 0.5999
Estimated allele frequency
Ecological gradient (Precipitation sums)
0.4
0.5
0.6
0.7
0.8
0.9
1.0
600
800
1000
1200
256
253
219
97
204
9
217
O8
O1
L1
O16
L8
L16
124

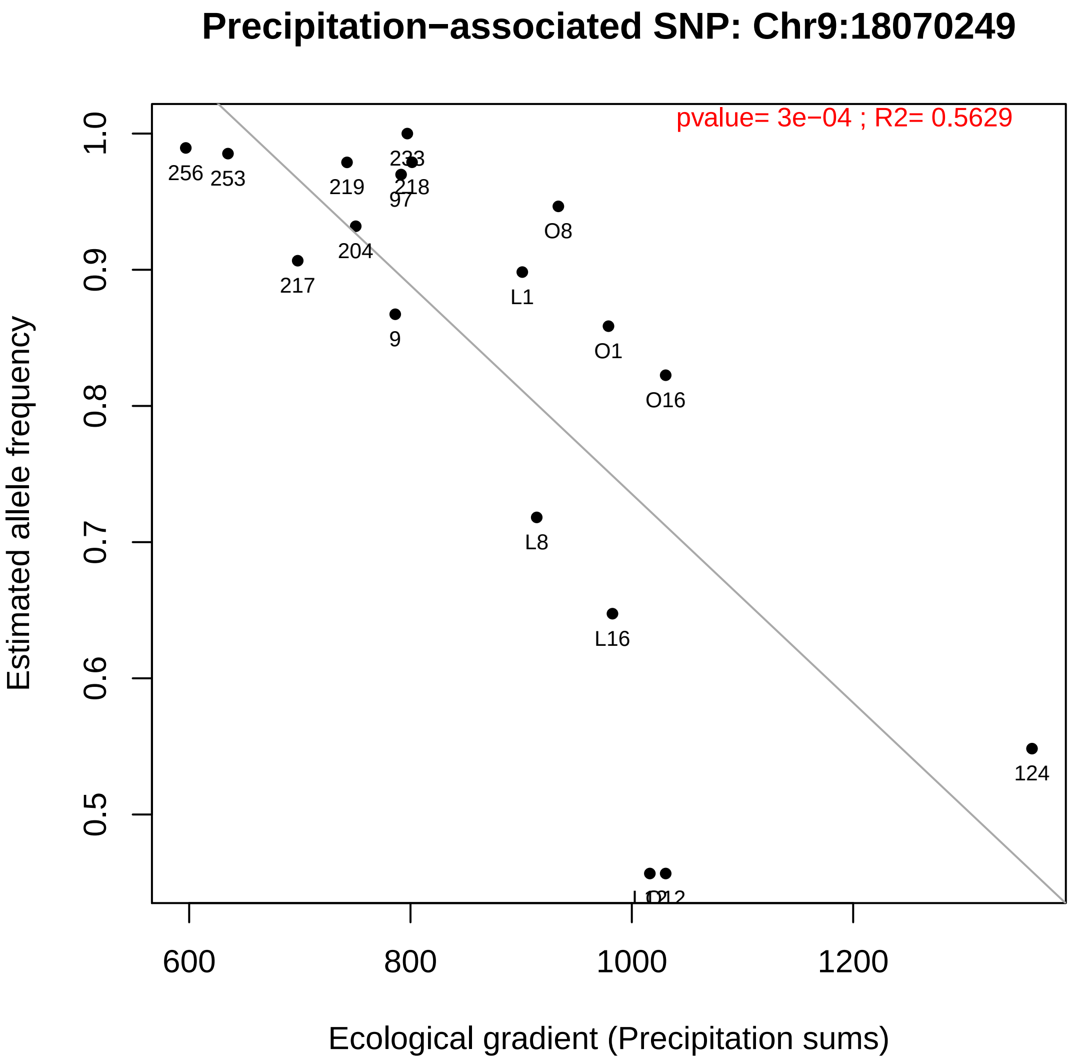

Precipitation−associated SNP: Chr9:18070249
pvalue= 3e−04 ; R2= 0.5629
Estimated allele frequency
Ecological gradient (Precipitation sums)
1.0
0.9
0.8
0.7
0.6
0.5
600
800
1000
1200
256
253
219
233
218
97
204
217
O8
L1
9
O1
O16
L8
L16
124

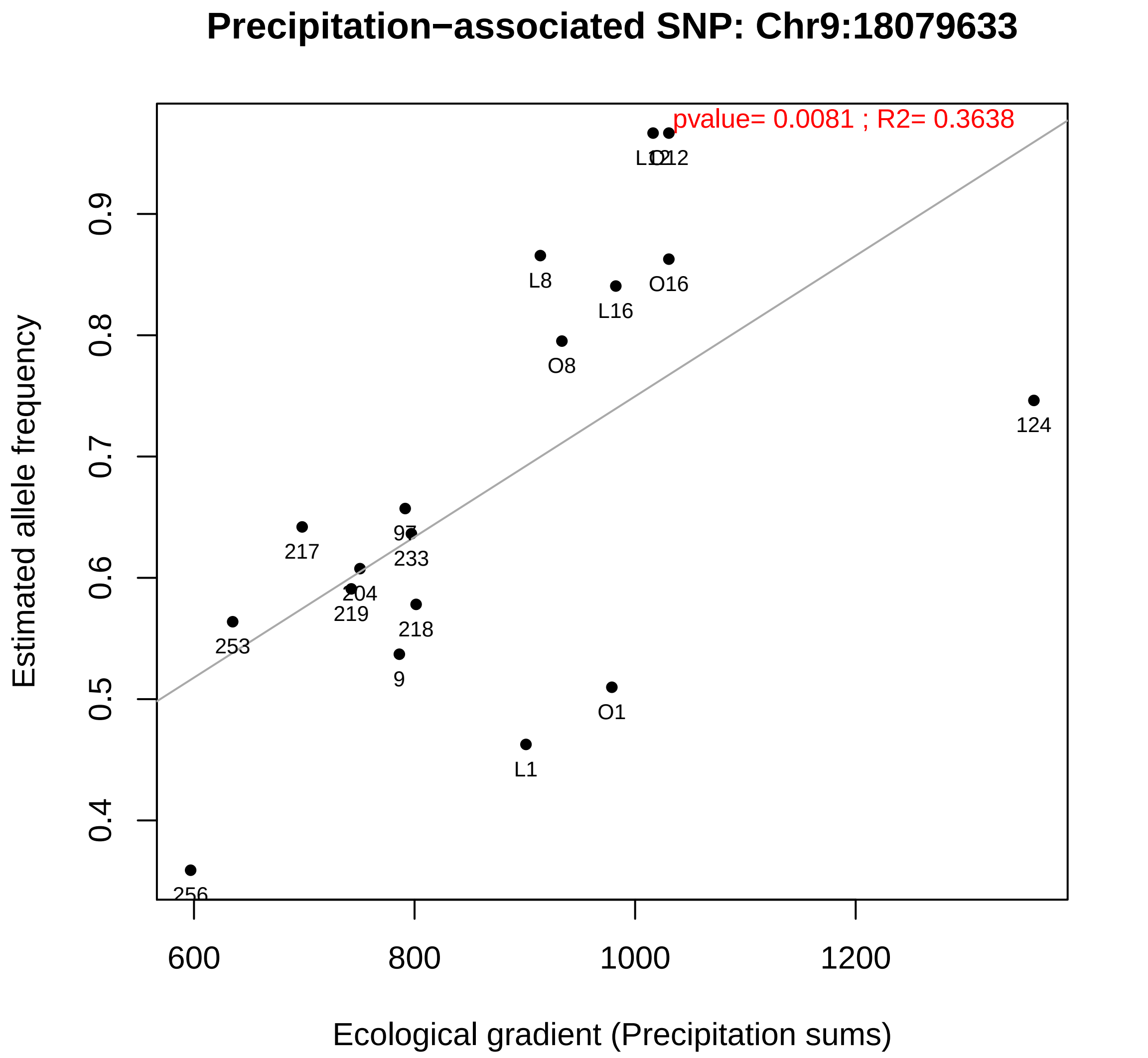

Precipitation−associated SNP: Chr9:18079633
pvalue= 0.0081 ; R2= 0.3638
L12
O12
L8
O16
L16
O8
124
97
217
233
204
219
218
253
9
O1
L1
256
0.4
0.5
0.6
0.7
0.8
0.9
600
800
1000
1200
Estimated allele frequency
Ecological gradient (Precipitation sums)

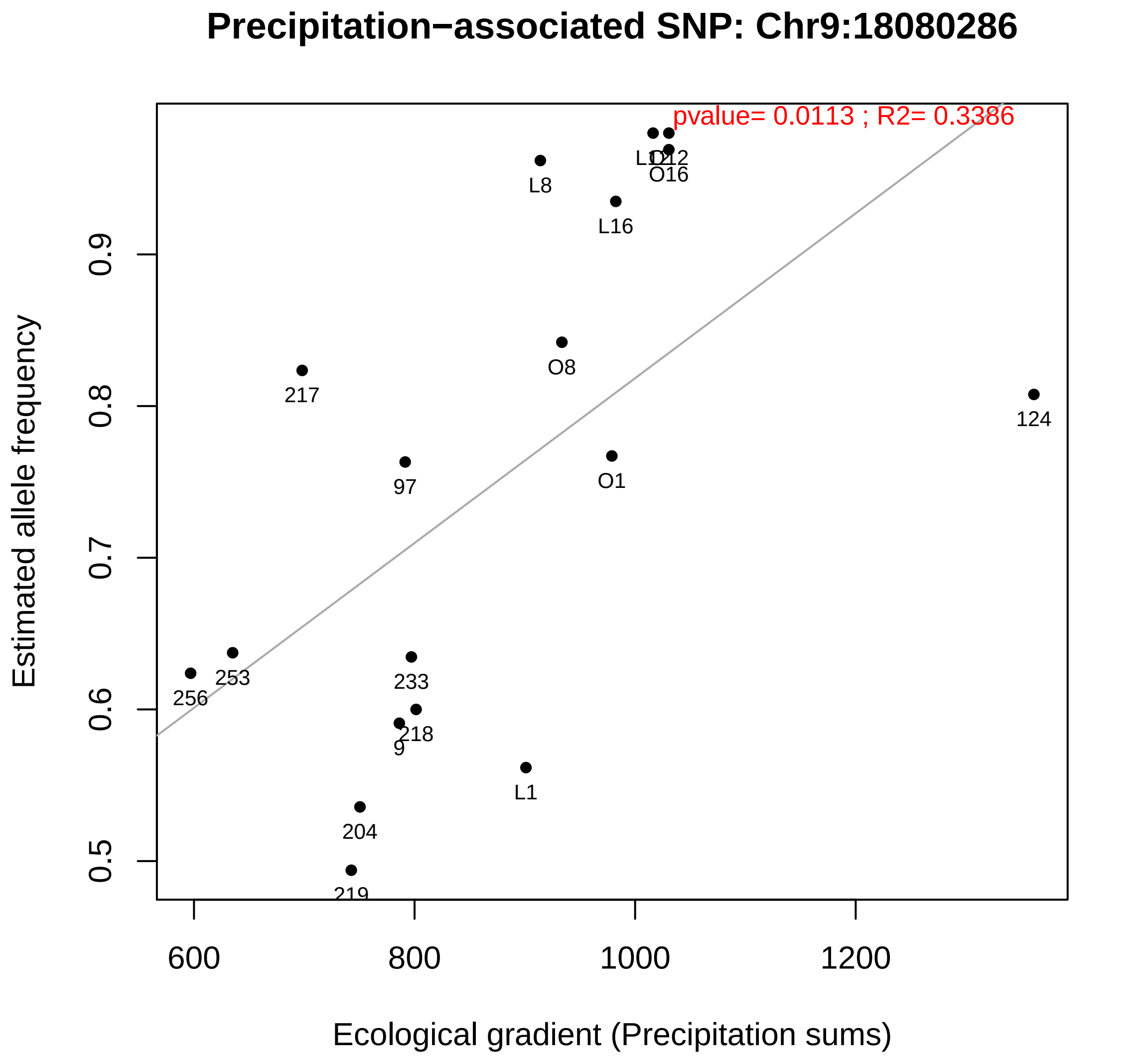
Precipitation−associated SNP: Chr9:18080286
pvalue= 0.0113 ; R2= 0.3386
Estimated allele frequency
Ecological gradient (Precipitation sums)
0.5
0.6
0.7
0.8
0.9
600
800
1000
1200
O16
L8
L16
O8
217
124
97
O1
253
256
233
218
9
L1
204
219

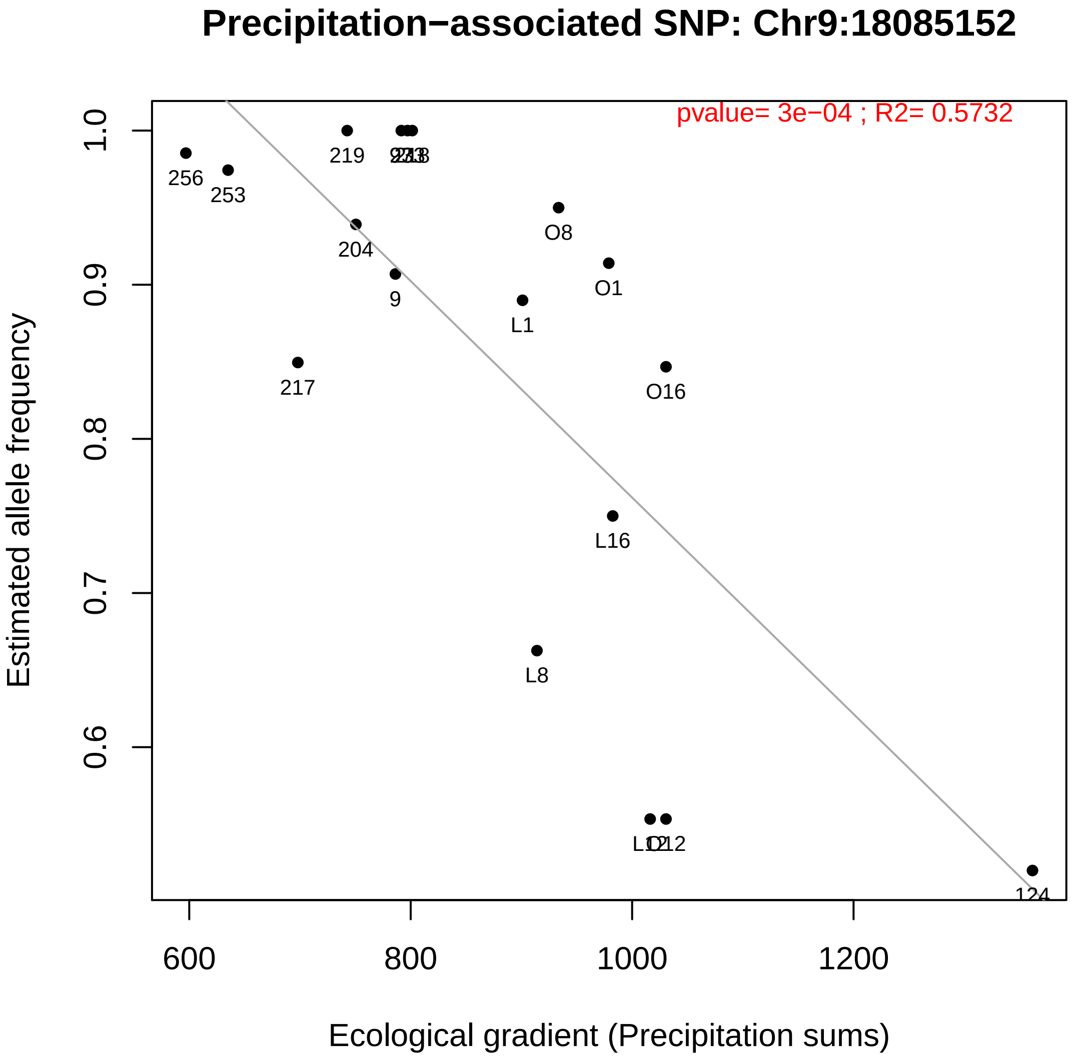
Precipitation−associated SNP: Chr9:18085152
pvalue= 3e−04 ; R2= 0.5732
Estimated allele frequency
Ecological gradient (Precipitation sums)
0.6
0.7
0.8
0.9
1.0
600
800
1000
1200
256
253
219
204
9
217
O8
O1
L1
O16
L16
L8
124

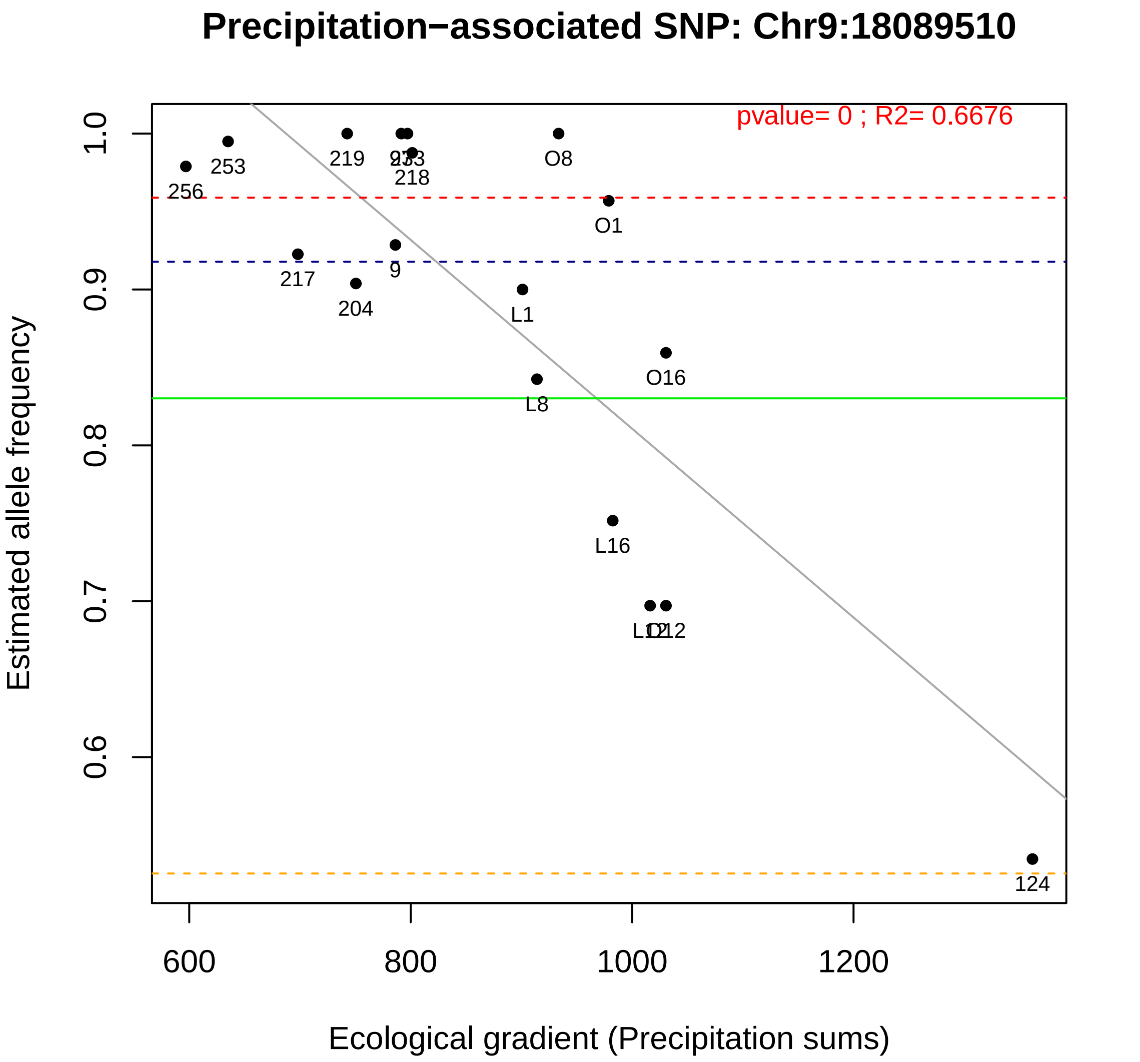

Precipitation−associated SNP: Chr9:18089510
pvalue= 0 ; R2= 0.6676
Estimated allele frequency
Ecological gradient (Precipitation sums)
1.0
0.9
0.8
0.7
0.6
600
800
1000
1200
256
253
219
218
O8
O1
217
9
204
L1
O16
L8
L16
124

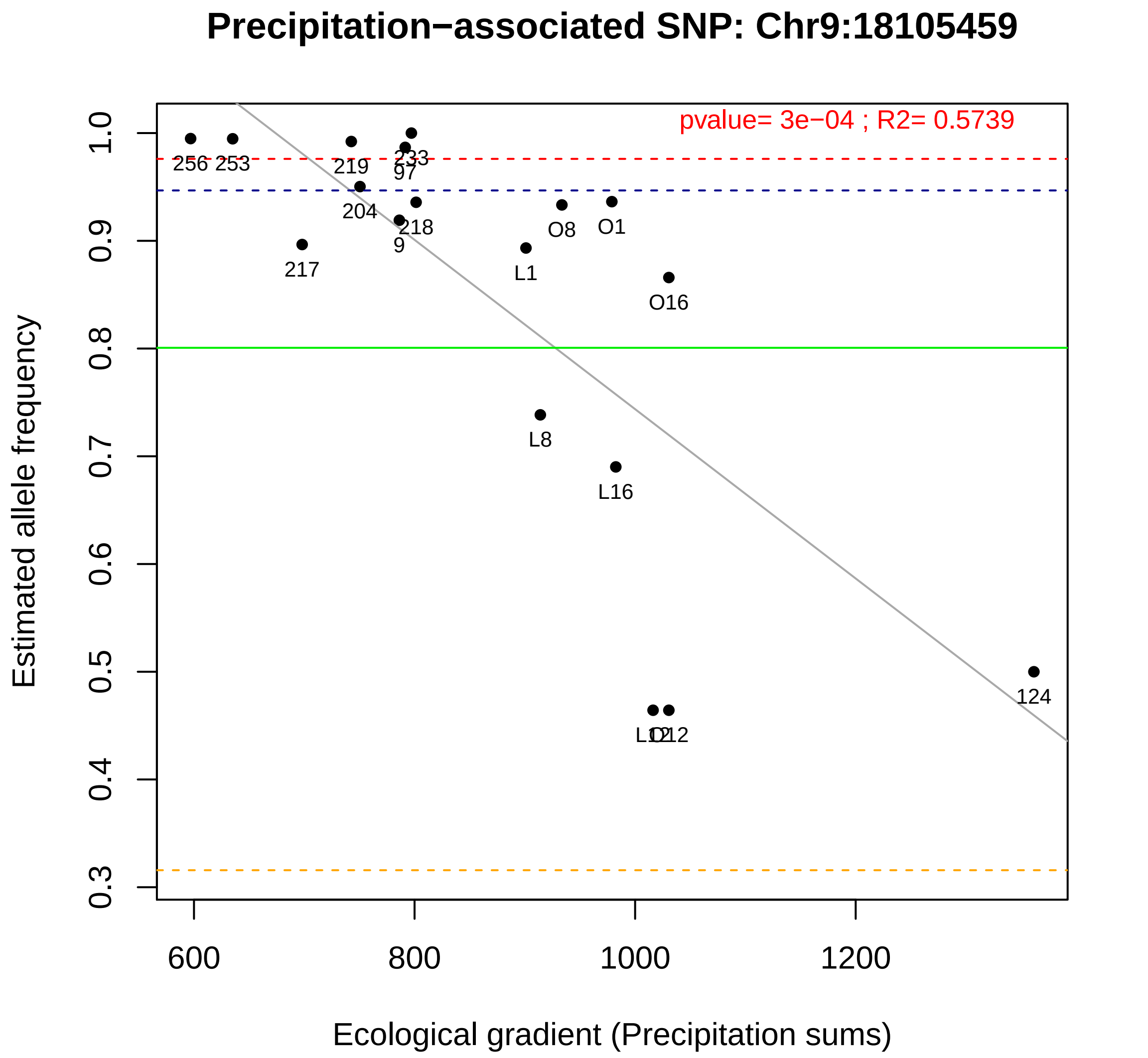

Precipitation−associated SNP: Chr9:18105459
pvalue= 3e−04 ; R2= 0.5739
Estimated allele frequency
0.3
0.4
0.5
0.6
0.7
0.8
0.9
1.0
600
800
1000
1200
Ecological gradient (Precipitation sums)
256
253
219
233
97
204
218
9
217
O8
O1
L1
O16
L8
L16
124
L12
O12

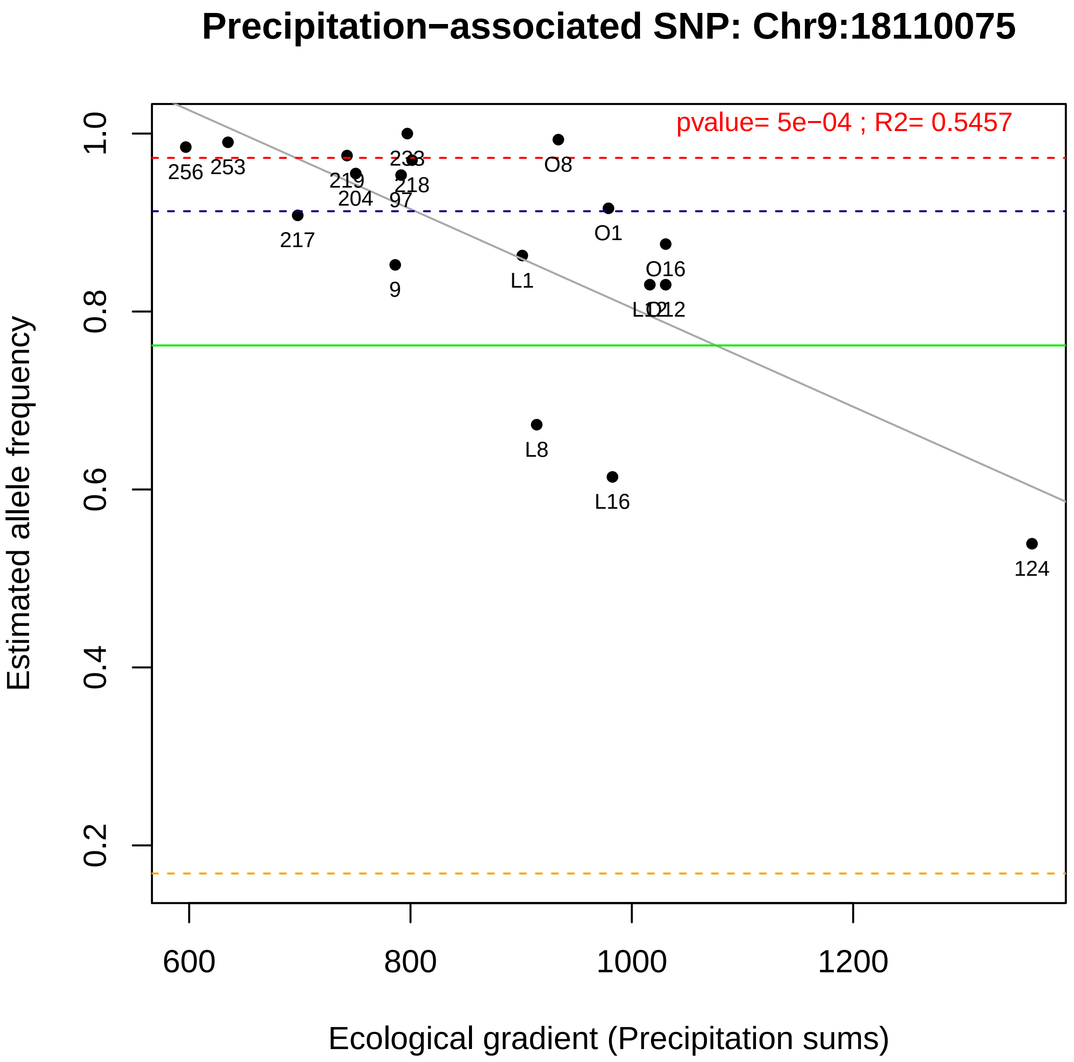
Precipitation−associated SNP: Chr9:18110075
pvalue= 5e−04 ; R2= 0.5457
Estimated allele frequency
Ecological gradient (Precipitation sums)
0.2
0.4
0.6
0.8
1.0
600
800
1000
1200
256
253
219
204
217
9
97
218
L1
O8
O1
O16
L12
O12
L8
L16
124

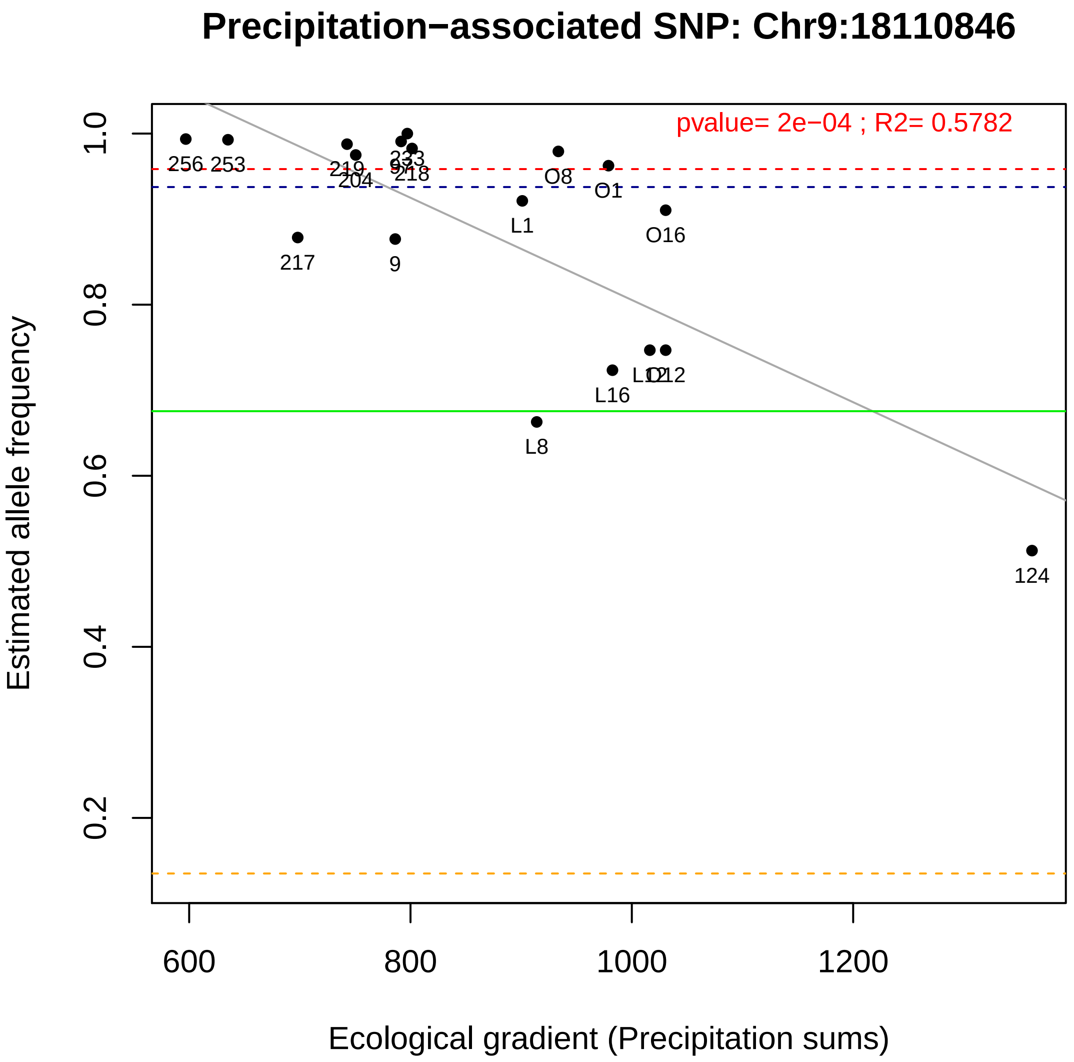
Precipitation−associated SNP: Chr9:18110846
pvalue= 2e−04 ; R2= 0.5782
Estimated allele frequency
Ecological gradient (Precipitation sums)
1.0
0.8
0.6
0.4
0.2
600
800
1000
1200
256
253
219
204
233
97
218
217
9
O8
O1
L1
O16
L10
O12
L16
L8
124

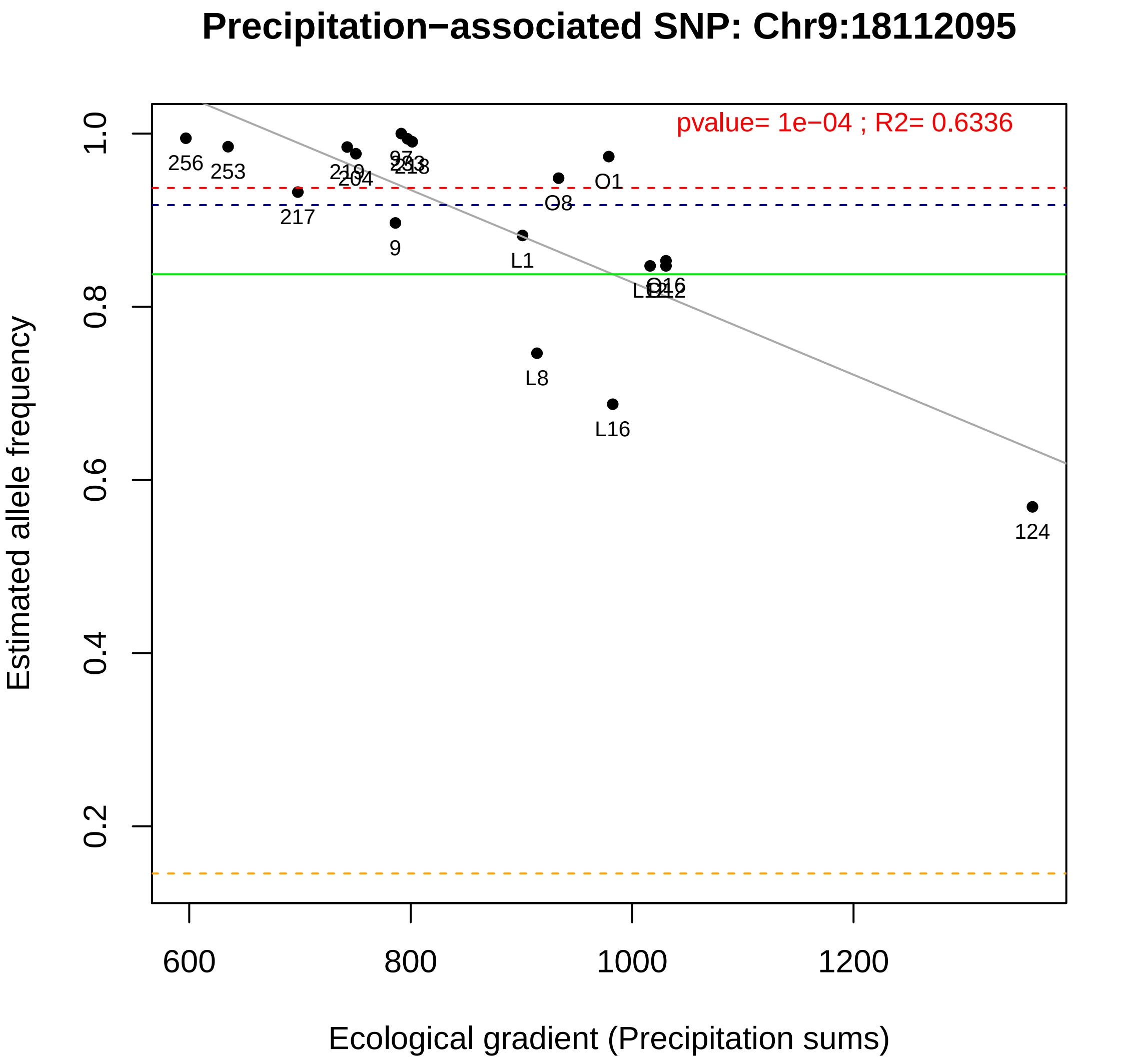

Precipitation−associated SNP: Chr9:18112095
pvalue= 1e−04 ; R2= 0.6336
Estimated allele frequency
Ecological gradient (Precipitation sums)
0.2
0.4
0.6
0.8
1.0
600
800
1000
1200
256
253
217
9
L1
O8
O1
L8
L16
124

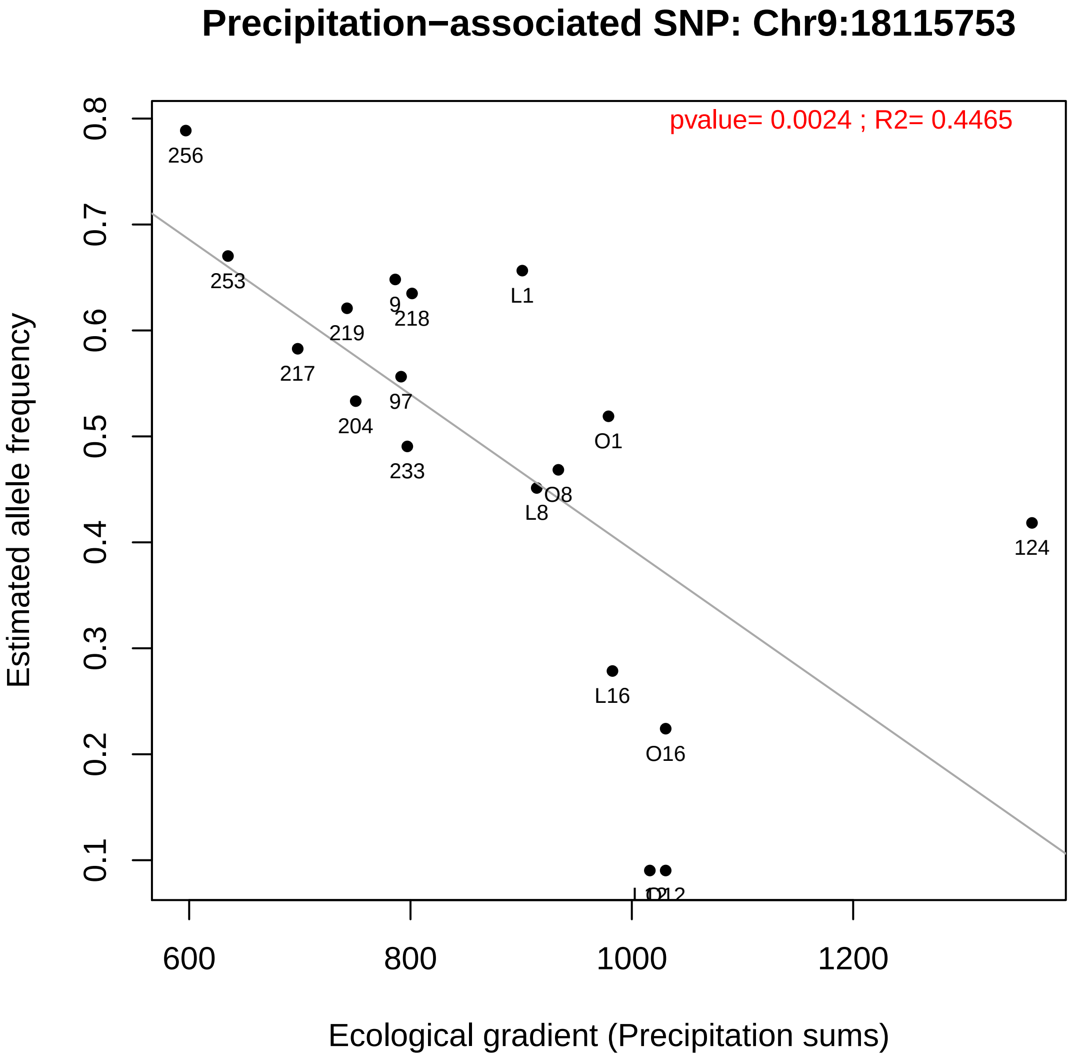
Precipitation−associated SNP: Chr9:18115753
pvalue= 0.0024 ; R2= 0.4465
Estimated allele frequency
0.1
0.2
0.3
0.4
0.5
0.6
0.7
0.8
600
800
1000
1200
Ecological gradient (Precipitation sums)
256
253
219
217
204
9
218
97
233
L1
O1
O8
L8
124
L16
O16

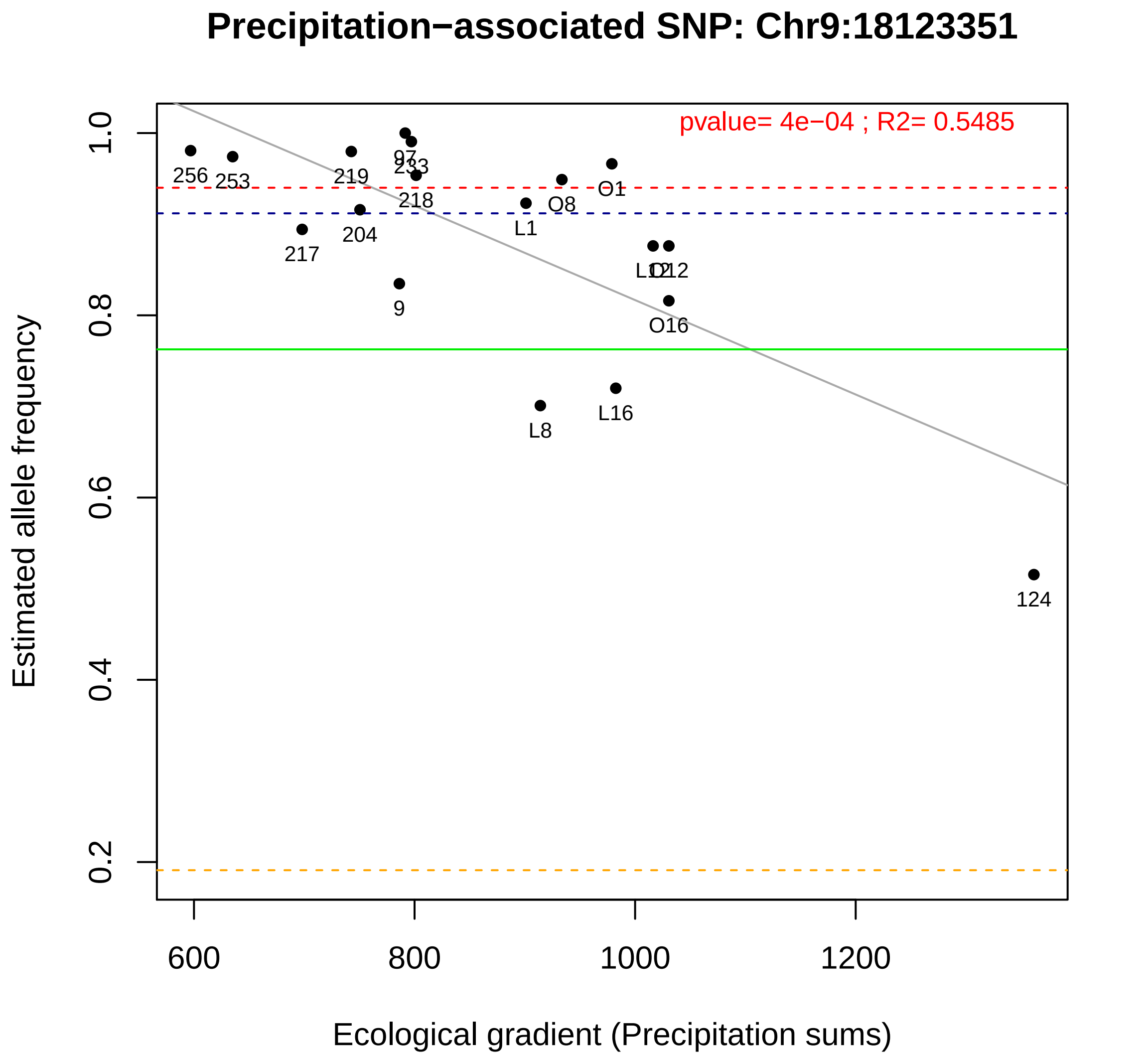

Precipitation−associated SNP: Chr9:18123351
pvalue= 4e−04 ; R2= 0.5485
Estimated allele frequency
Ecological gradient (Precipitation sums)
1.0
0.8
0.6
0.4
0.2
600
800
1000
1200
256
253
219
97
233
218
204
217
9
L1
O8
O1
L12
O12
O16
L16
L8
124

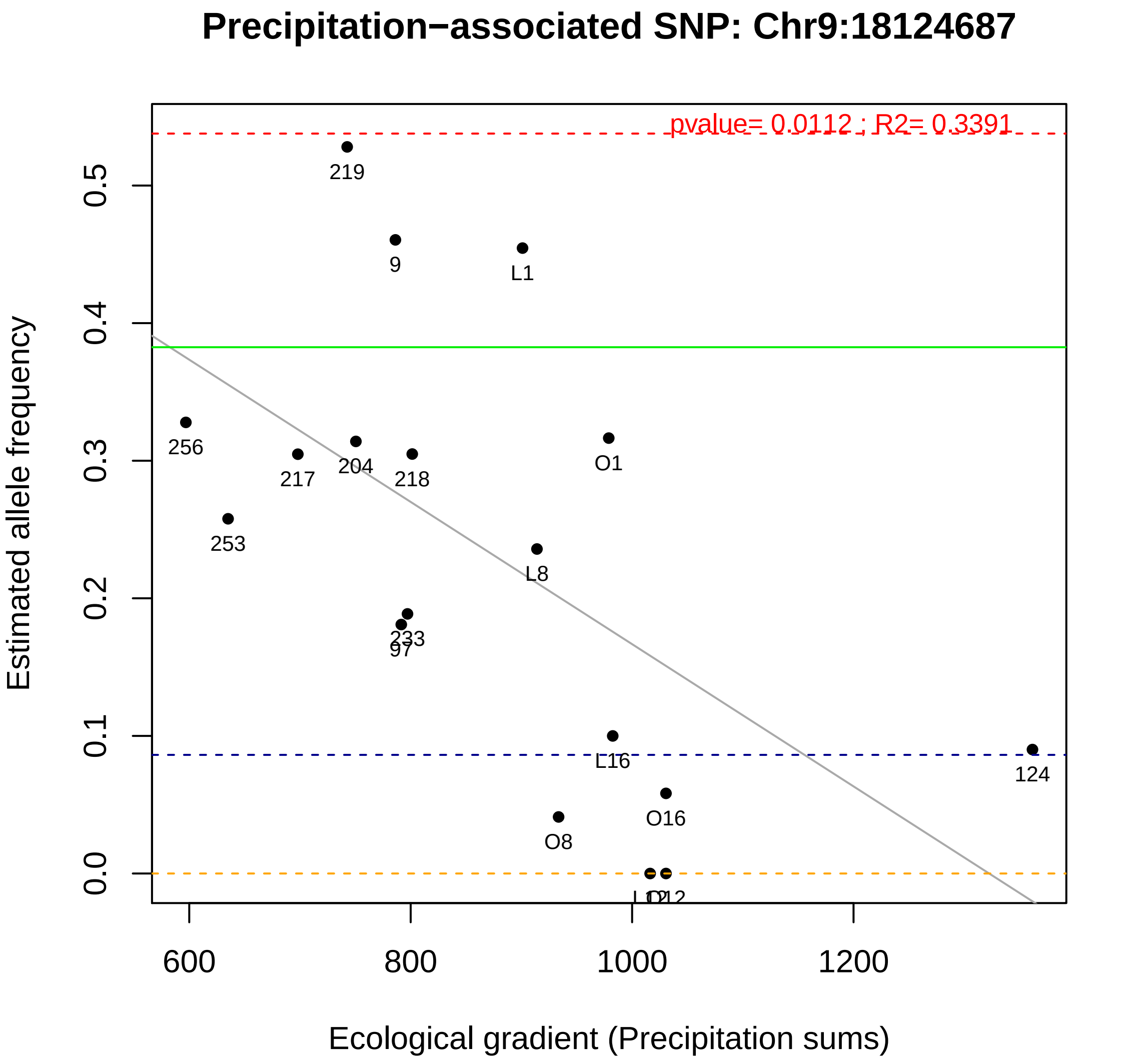
Precipitation−associated SNP: Chr9:18124687
pvalue= 0.0112 ; R2= 0.3391
Estimated allele frequency
0.0
0.1
0.2
0.3
0.4
0.5
600
800
1000
1200
Ecological gradient (Precipitation sums)
219
9
L1
256
217
204
218
O1
253
L8
233
97
L16
124
O16
O8

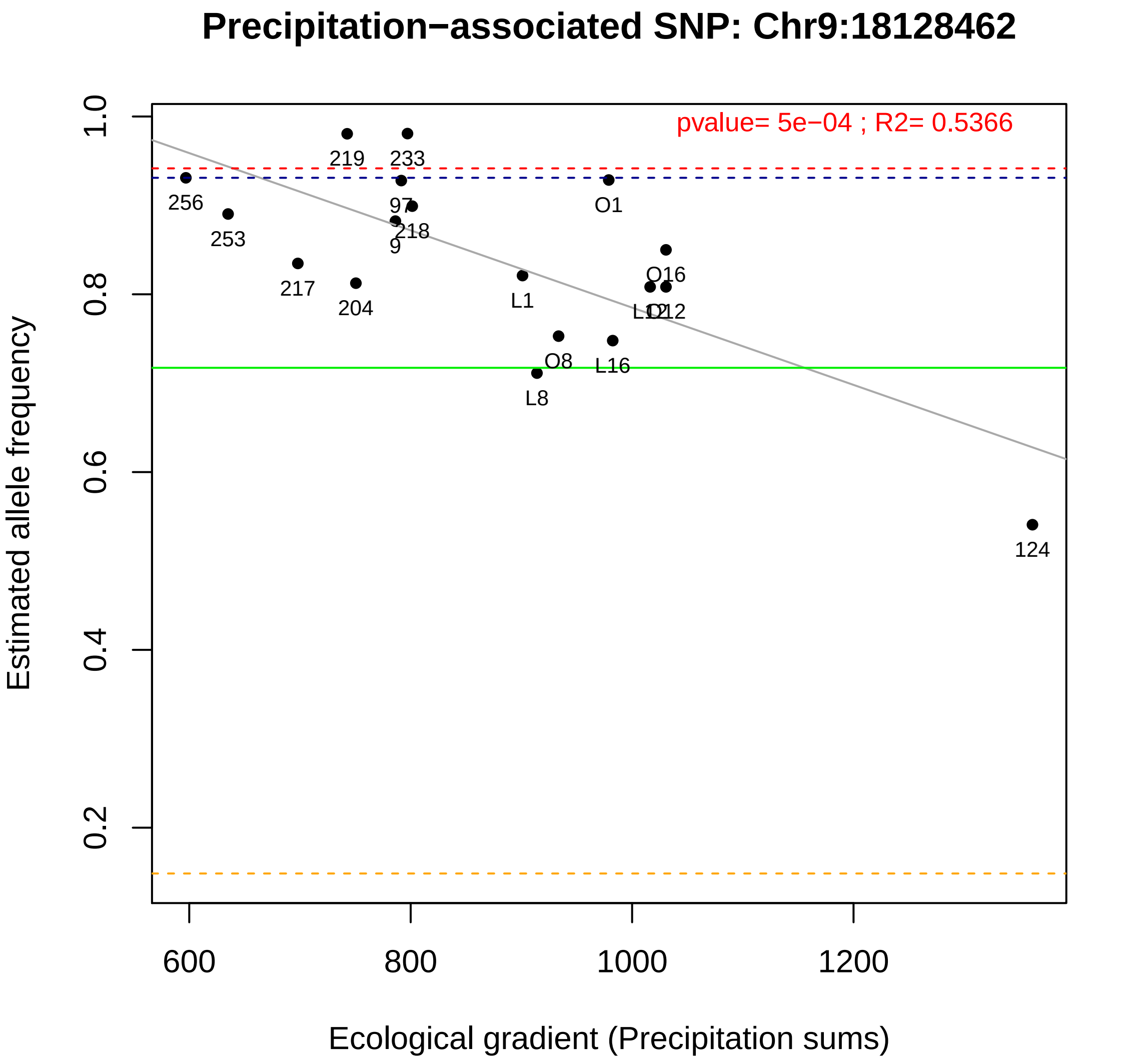

Precipitation−associated SNP: Chr9:18128462
pvalue= 5e−04 ; R2= 0.5366
Estimated allele frequency
Ecological gradient (Precipitation sums)
0.2
0.4
0.6
0.8
1.0
600
800
1000
1200
219
233
256
97
218
9
253
217
204
O1
O16
L1
L12
O12
O8
L16
L8
124

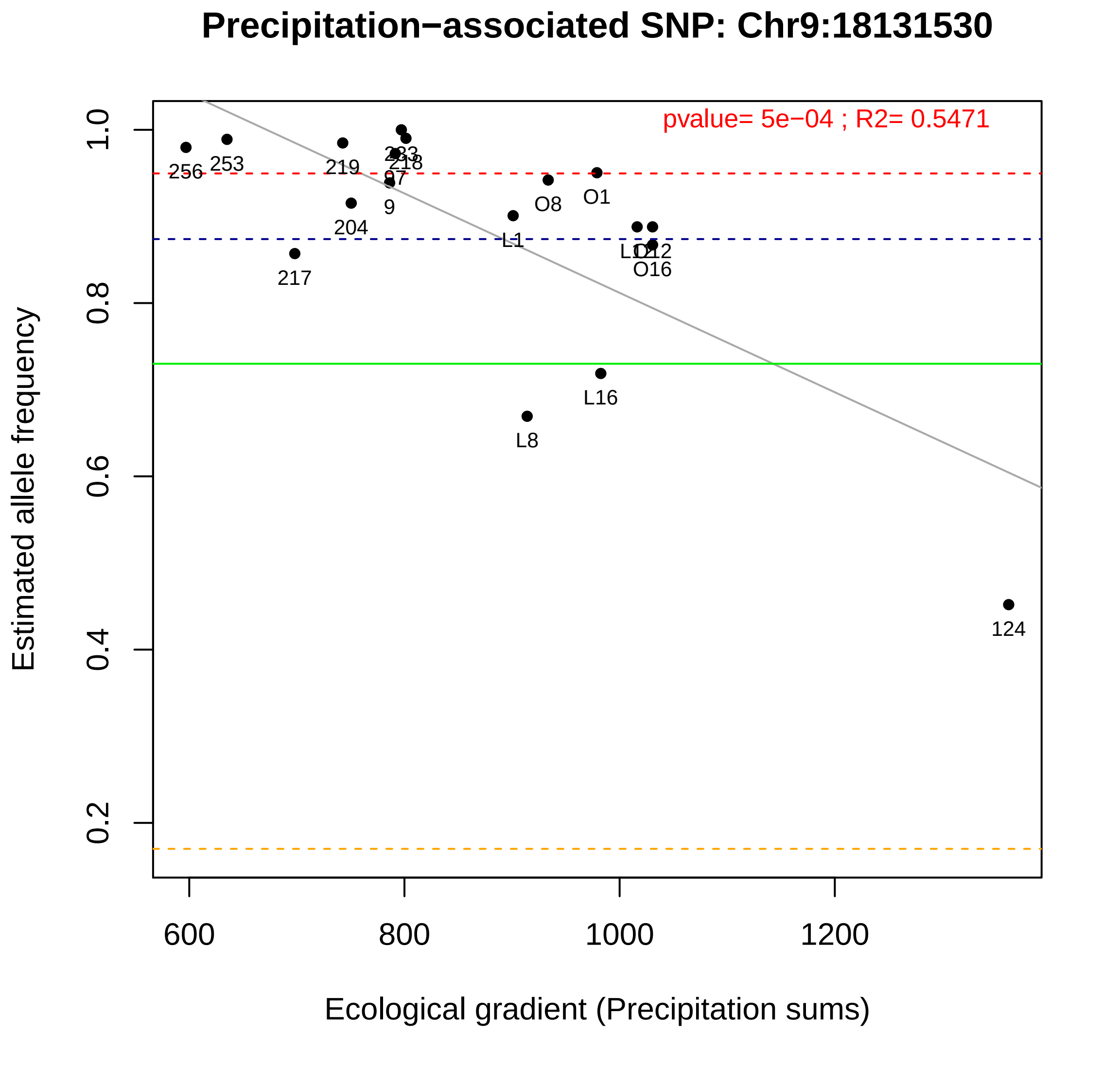
Precipitation−associated SNP: Chr9:18131530
pvalue= 5e−04 ; R2= 0.5471
Estimated allele frequency
Ecological gradient (Precipitation sums)
0.2
0.4
0.6
0.8
1.0
600
800
1000
1200
256
253
219
217
204
218
9
L1
O8
O1
O16
L16
L8
124

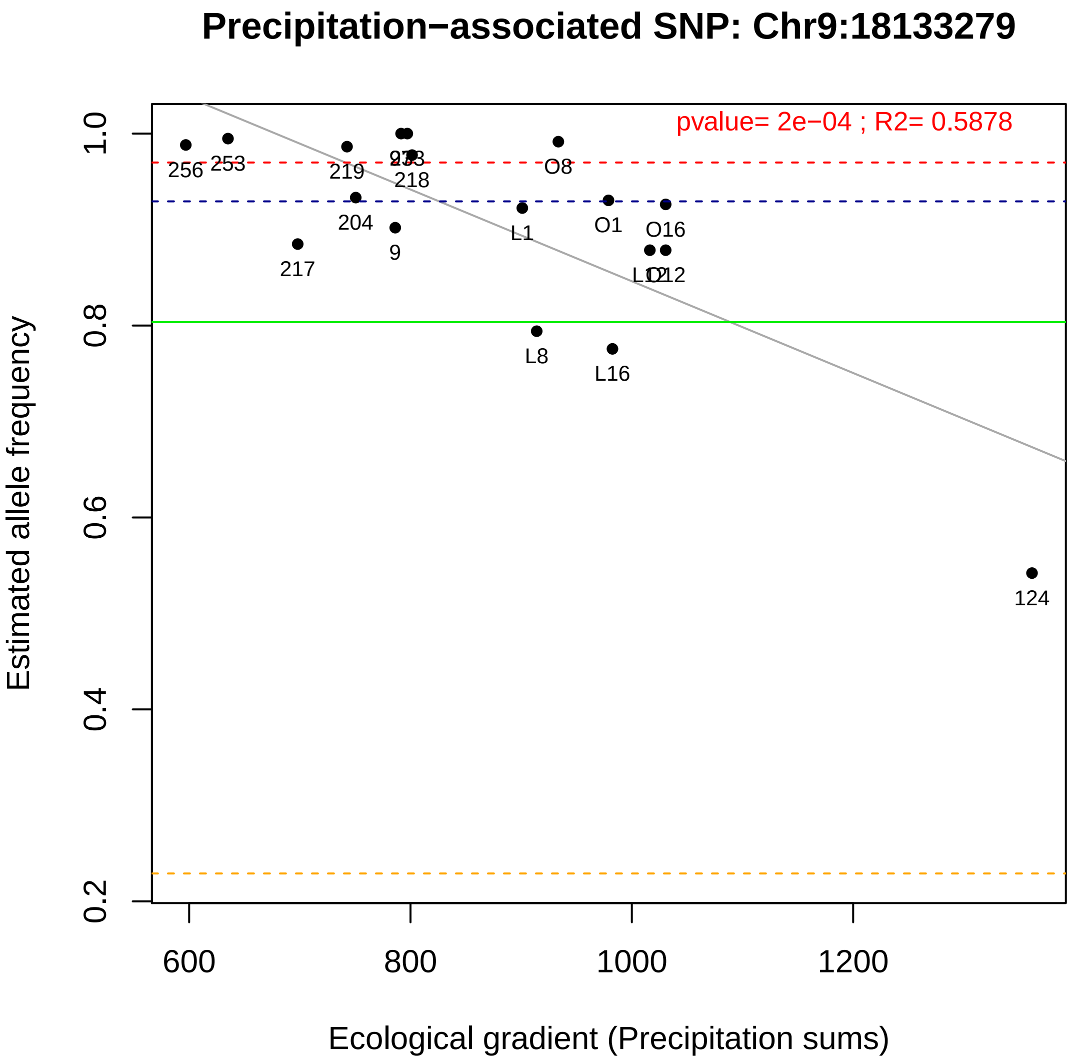

Precipitation−associated SNP: Chr9:18133279
pvalue= 2e−04 ; R2= 0.5878
Estimated allele frequency
Ecological gradient (Precipitation sums)
1.0
0.8
0.6
0.4
0.2
600
800
1000
1200
256
253
219
218
204
217
9
L1
O8
O1
O16
O12
L8
L16
124

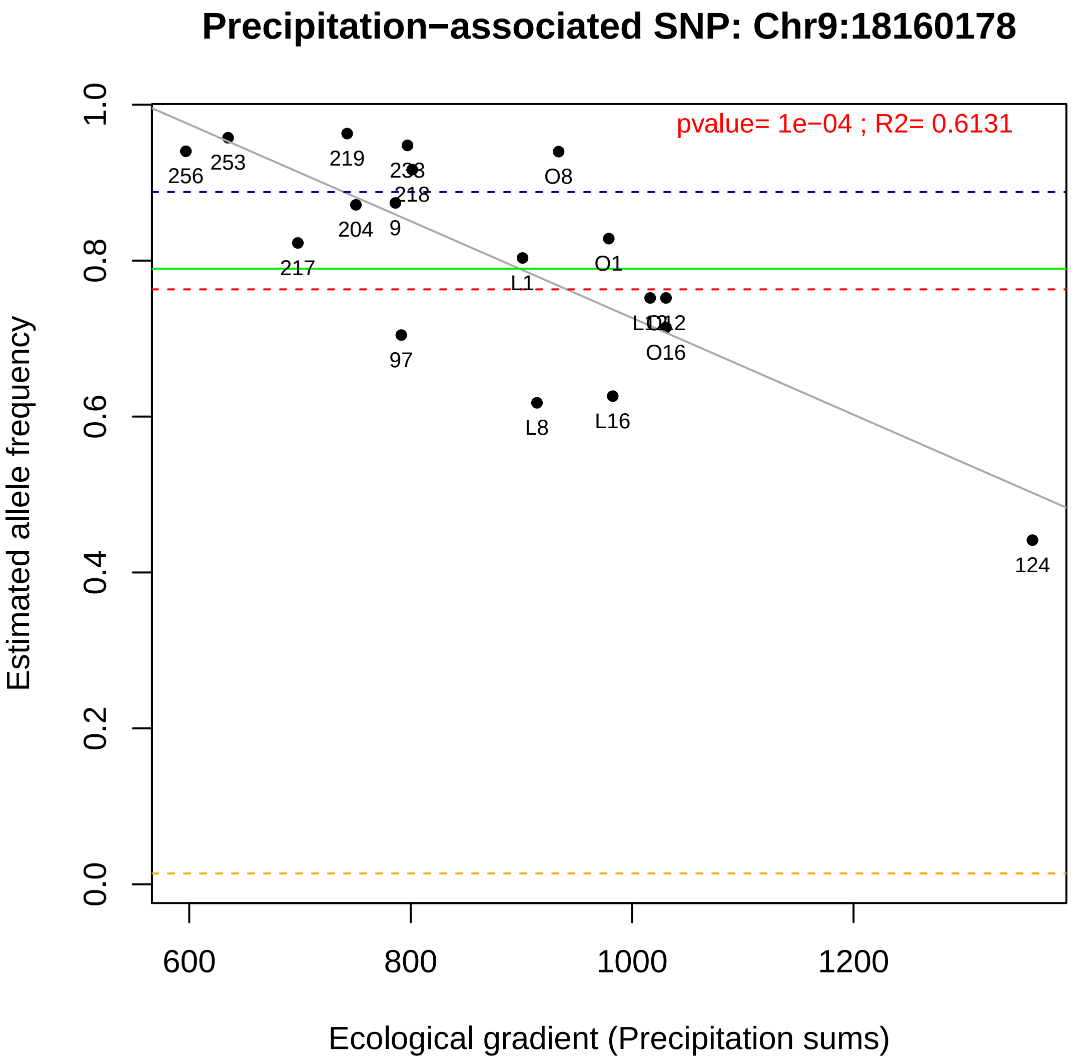
Precipitation−associated SNP: Chr9:18160178
pvalue= 1e−04 ; R2= 0.6131
Estimated allele frequency
Ecological gradient (Precipitation sums)
0.0
0.2
0.4
0.6
0.8
1.0
600
800
1000
1200
256
253
219
218
204
9
217
97
L1
O8
O1
L8
L16
O16
124

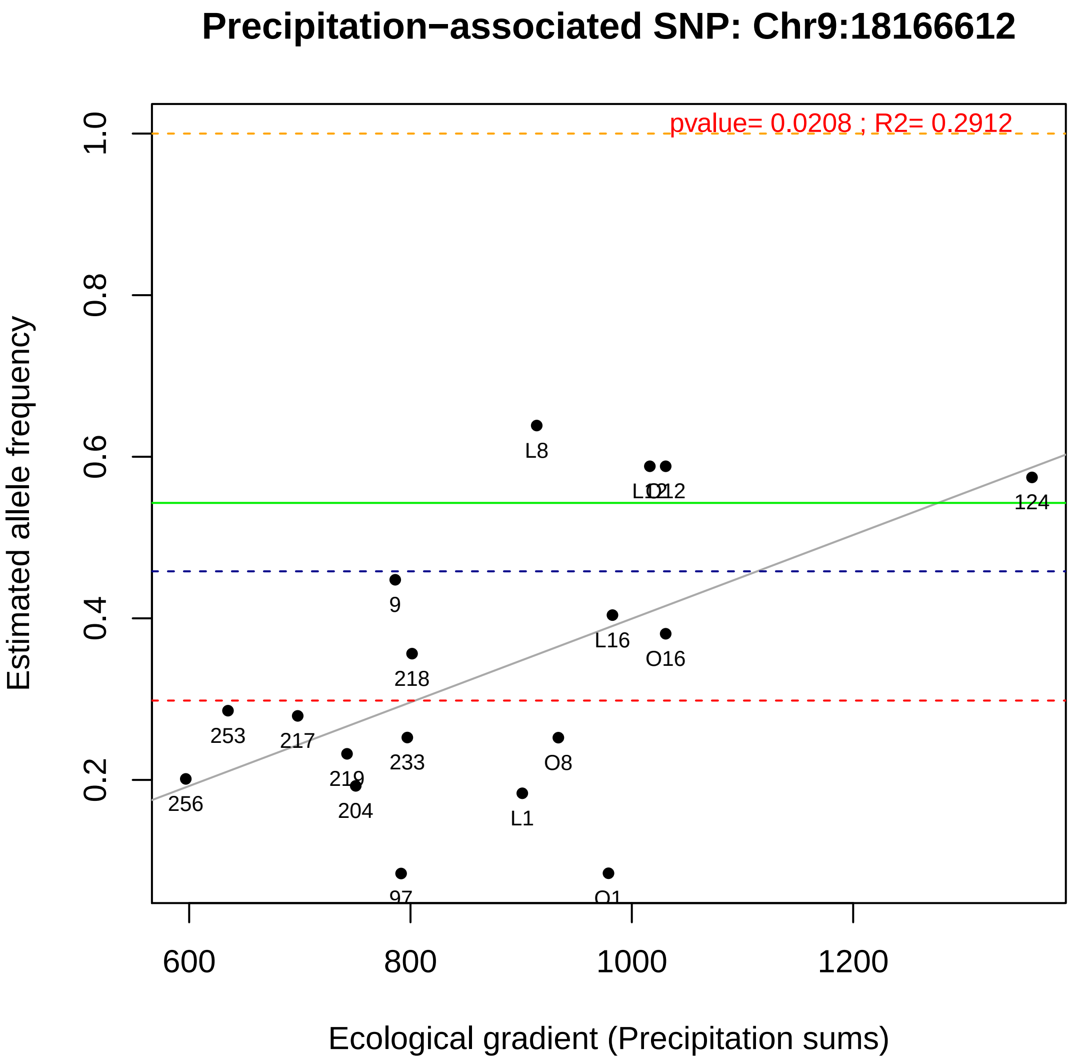
Precipitation−associated SNP: Chr9:18166612
pvalue= 0.0208 ; R2= 0.2912
Estimated allele frequency
Ecological gradient (Precipitation sums)
0.2
0.4
0.6
0.8
1.0
600
800
1000
1200
L8
L12
O12
124
9
L16
O16
218
253
217
233
O8
219
204
256
L1
97
O1

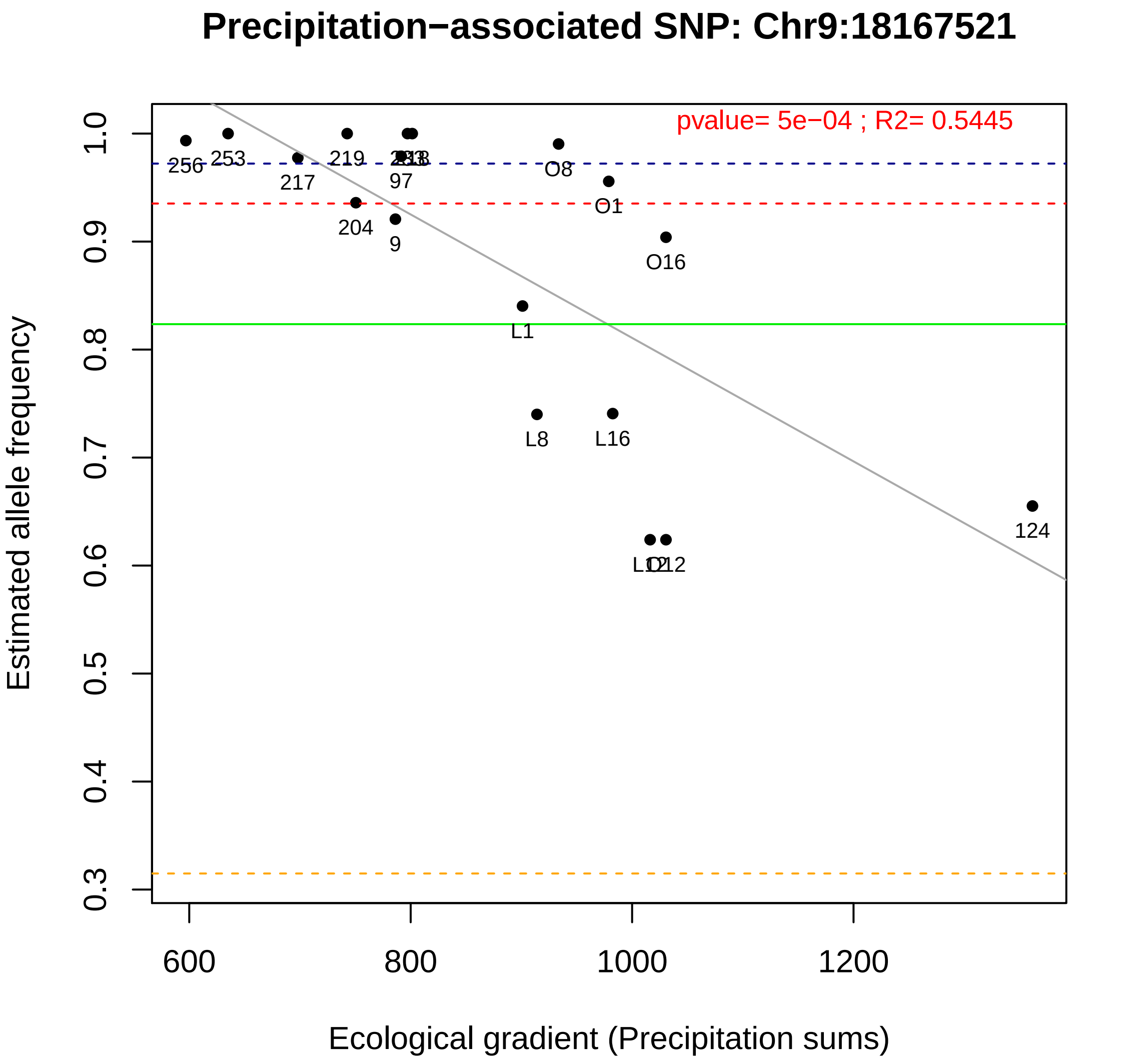

Precipitation−associated SNP: Chr9:18167521
pvalue= 5e−04 ; R2= 0.5445
Estimated allele frequency
Ecological gradient (Precipitation sums)
0.3
0.4
0.5
0.6
0.7
0.8
0.9
1.0
600
800
1000
1200
256
253
217
219
204
97
9
O8
O1
O16
L1
L8
L16
124

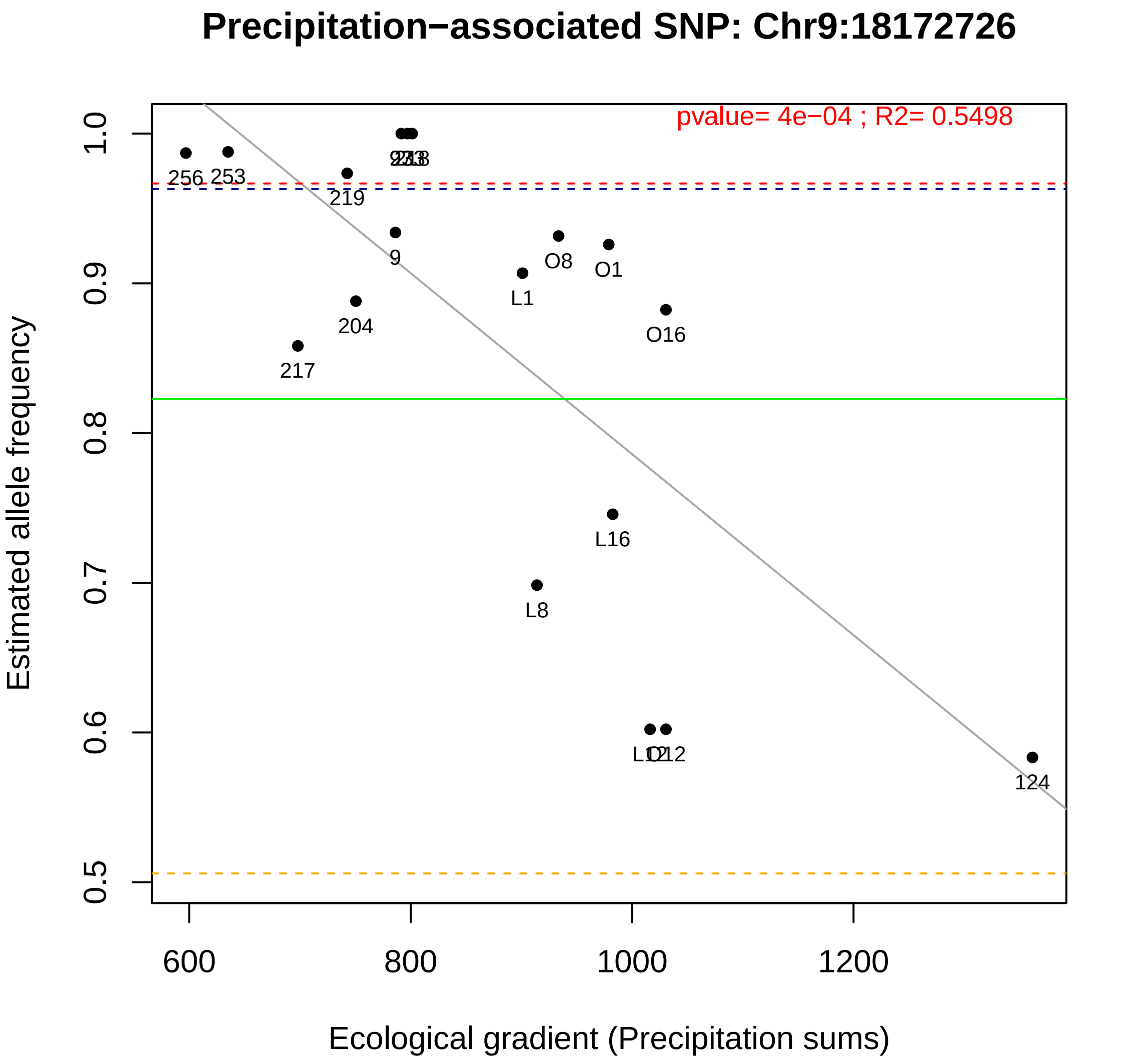
Precipitation−associated SNP: Chr9:18172726
pvalue= 4e−04 ; R2= 0.5498
Estimated allele frequency
Ecological gradient (Precipitation sums)
0.5
0.6
0.7
0.8
0.9
1.0
600
800
1000
1200
256
253
219
9
204
217
L1
O8
O1
O16
L16
L8
124

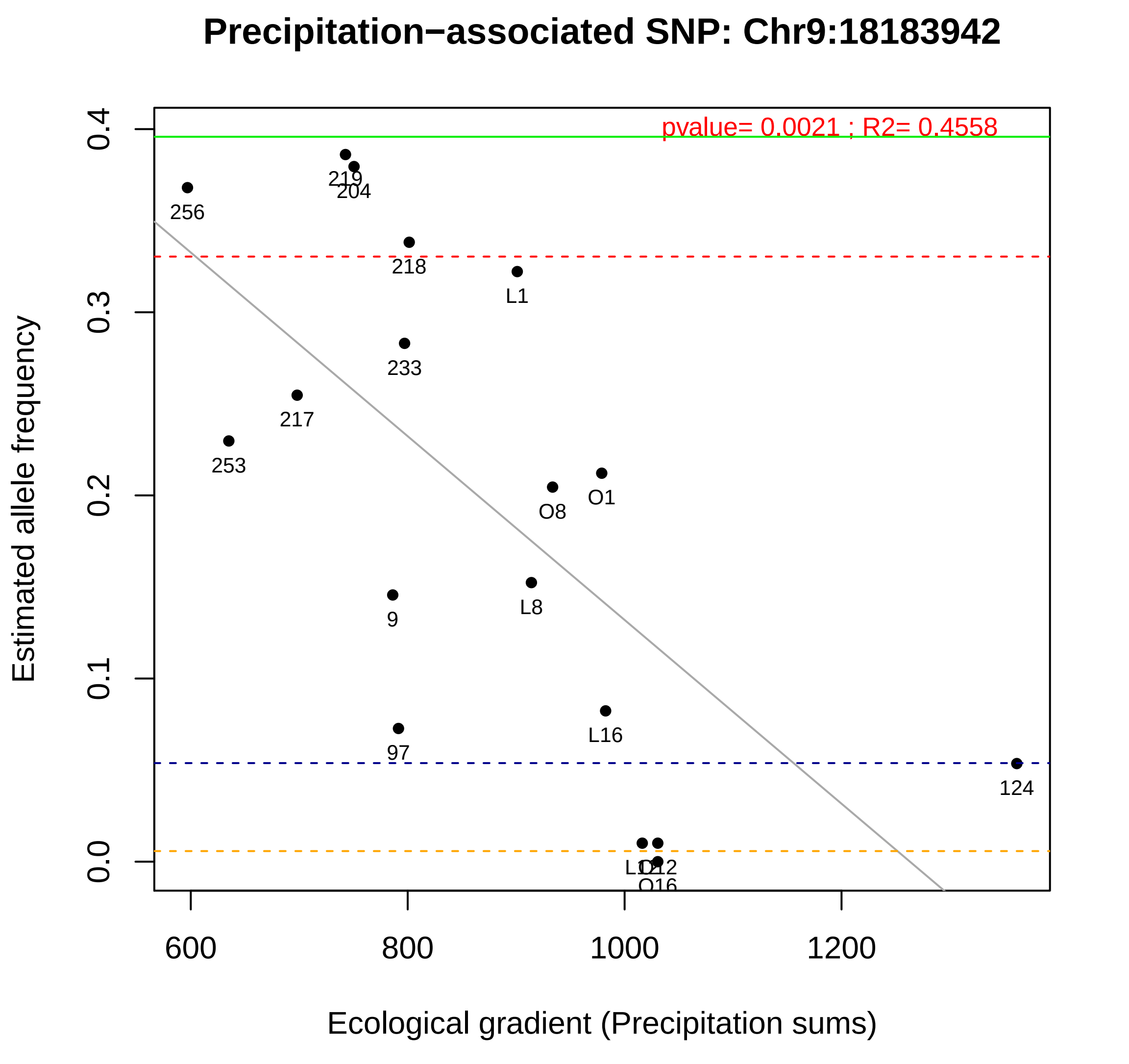
Precipitation−associated SNP: Chr9:18183942
pvalue= 0.0021 ; R2= 0.4558
Estimated allele frequency
Ecological gradient (Precipitation sums)
0.0
0.1
0.2
0.3
0.4
600
800
1000
1200
256
219
204
218
L1
233
217
253
O8
O1
9
L8
L16
97
124
O16

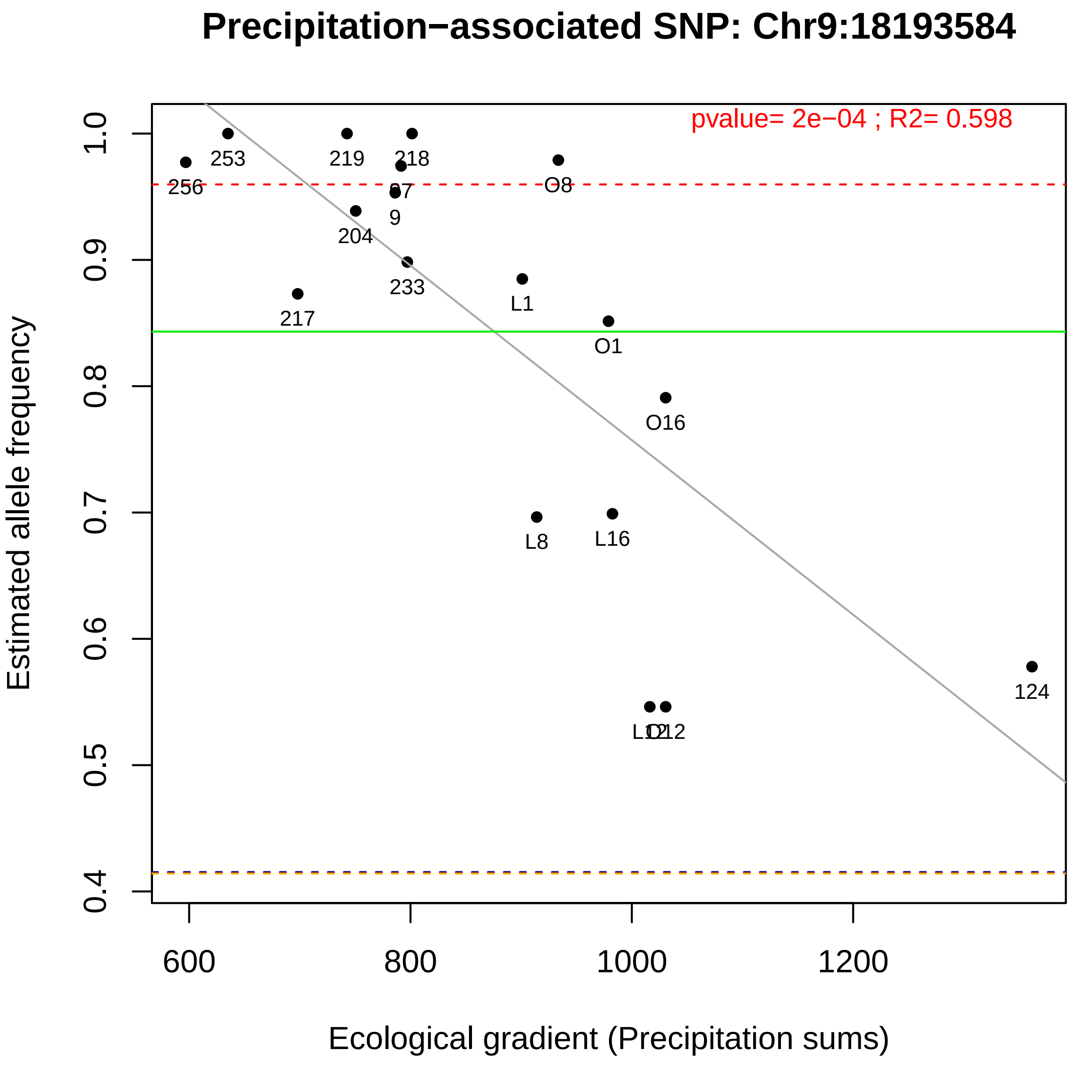
Precipitation−associated SNP: Chr9:18193584
pvalue= 2e−04 ; R2= 0.598
Estimated allele frequency
Ecological gradient (Precipitation sums)
1.0
0.9
0.8
0.7
0.6
0.5
0.4
600
800
1000
1200
256
253
219
218
O8
204
9
233
217
L1
O1
O16
L8
L16
124

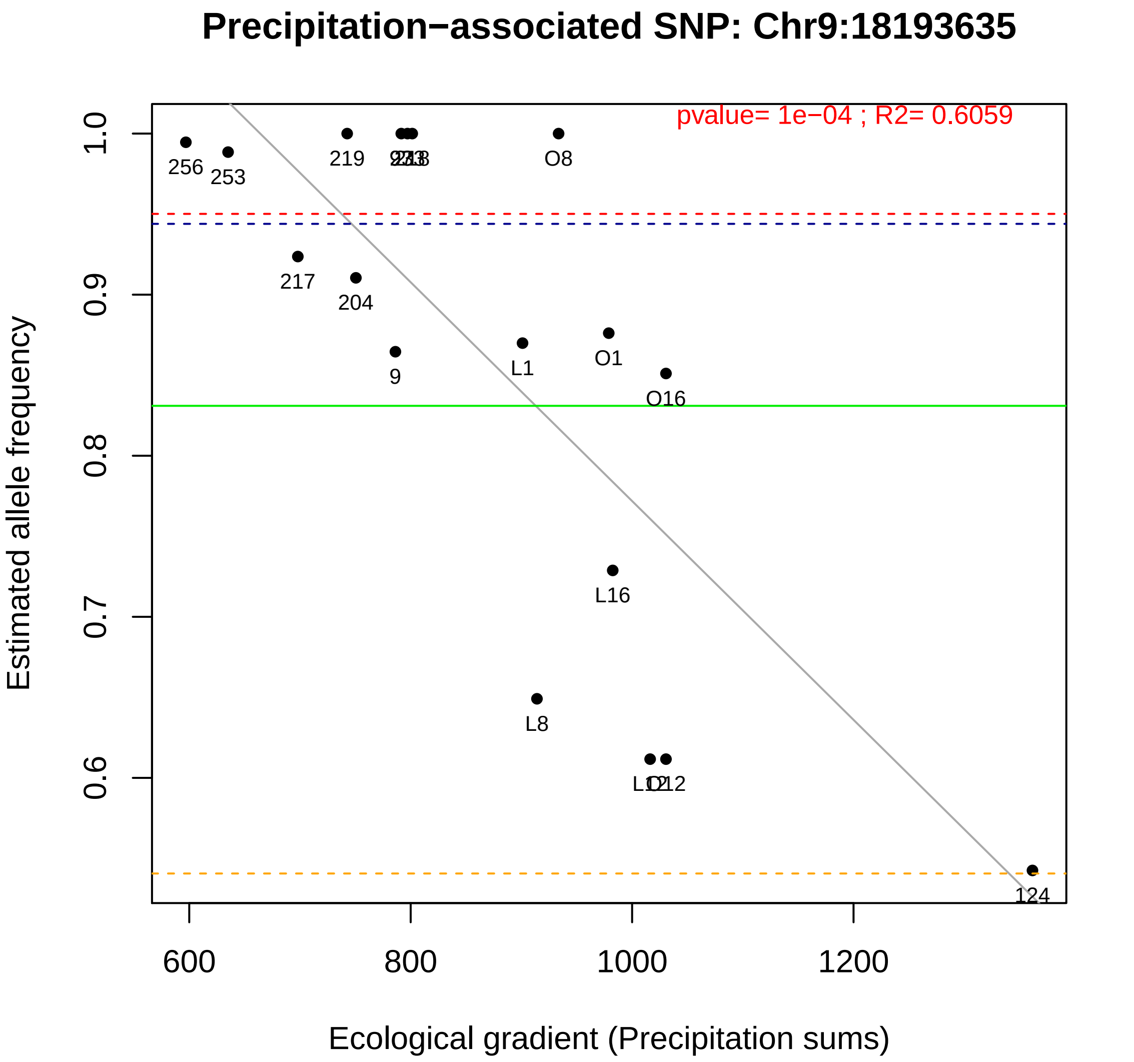

Precipitation−associated SNP: Chr9:18193635
pvalue= 1e−04 ; R2= 0.6059
Estimated allele frequency
1.0
0.9
0.8
0.7
0.6
600
800
1000
1200
Ecological gradient (Precipitation sums)
256
253
219
O8
217
204
9
L1
O1
O16
L16
L8
O12
124

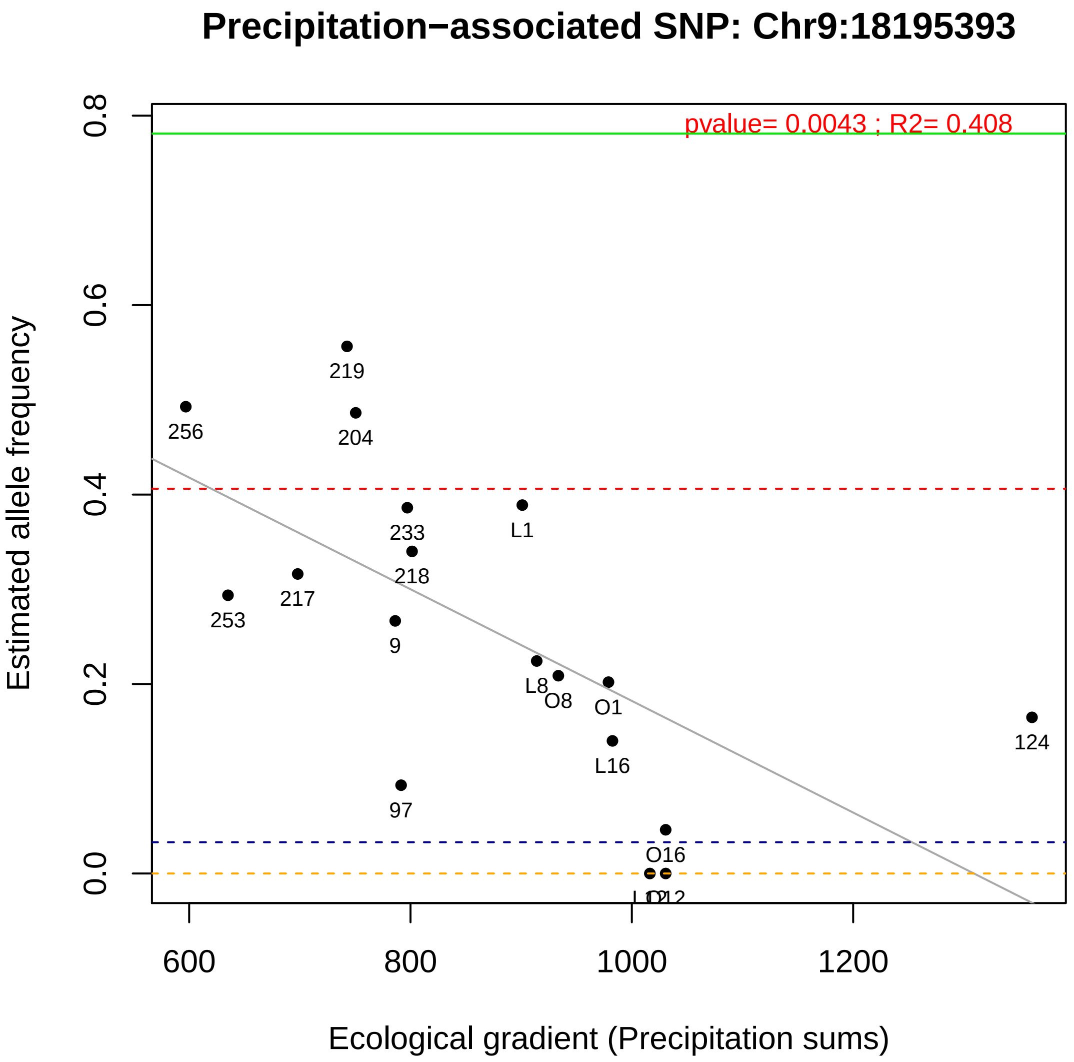

Precipitation−associated SNP: Chr9:18195393
pvalue= 0.0043 ; R2= 0.408
Estimated allele frequency
0.0
0.2
0.4
0.6
0.8
600
800
1000
1200
Ecological gradient (Precipitation sums)
219
256
204
233
L1
218
217
253
9
L8
O8
O1
124
L16
97
O16

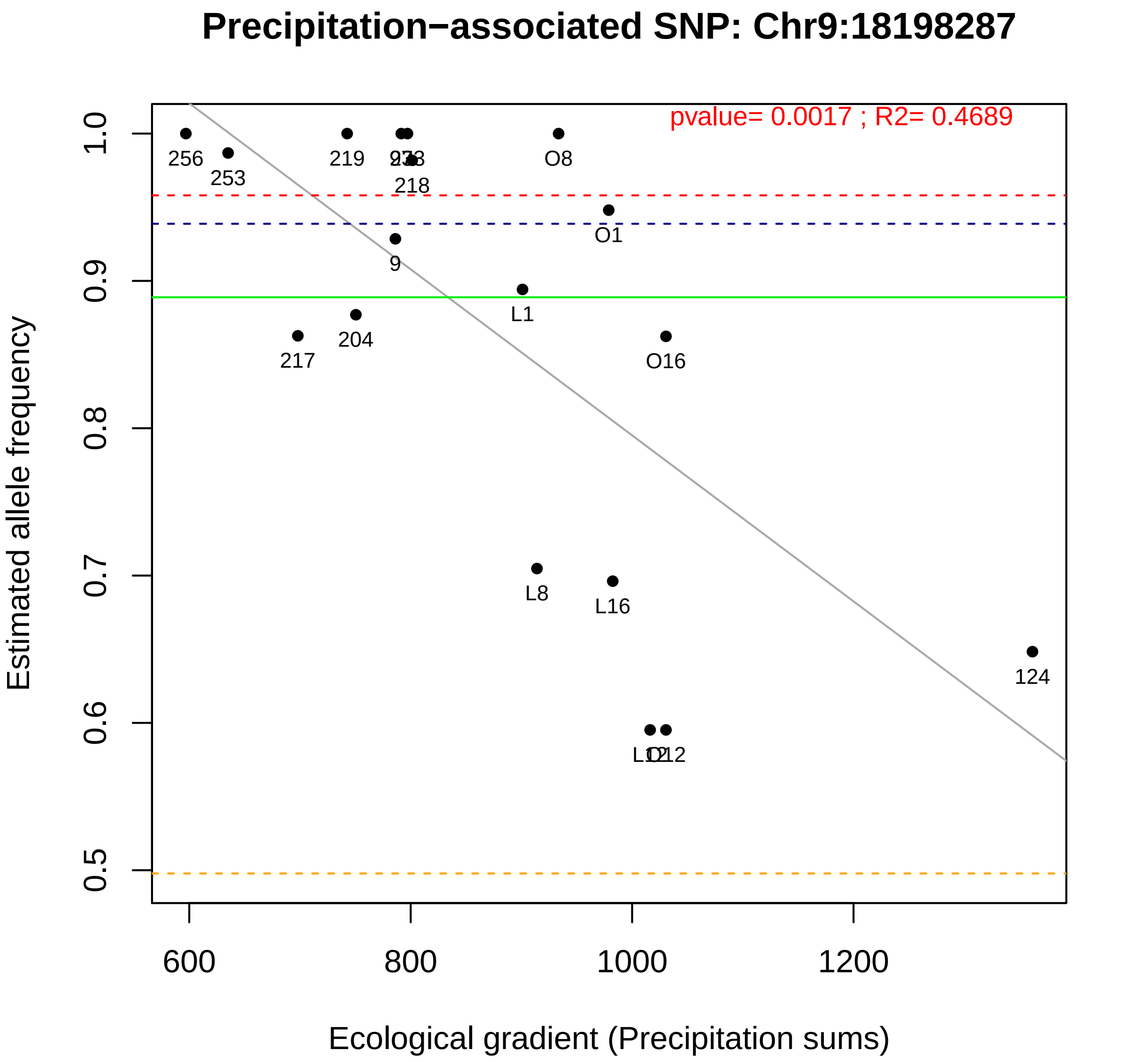
Precipitation−associated SNP: Chr9:18198287
pvalue= 0.0017 ; R2= 0.4689
Estimated allele frequency
Ecological gradient (Precipitation sums)
1.0
0.9
0.8
0.7
0.6
0.5
600
800
1000
1200
256
253
219
218
O8
O1
9
L1
204
217
O16
L8
L16
124

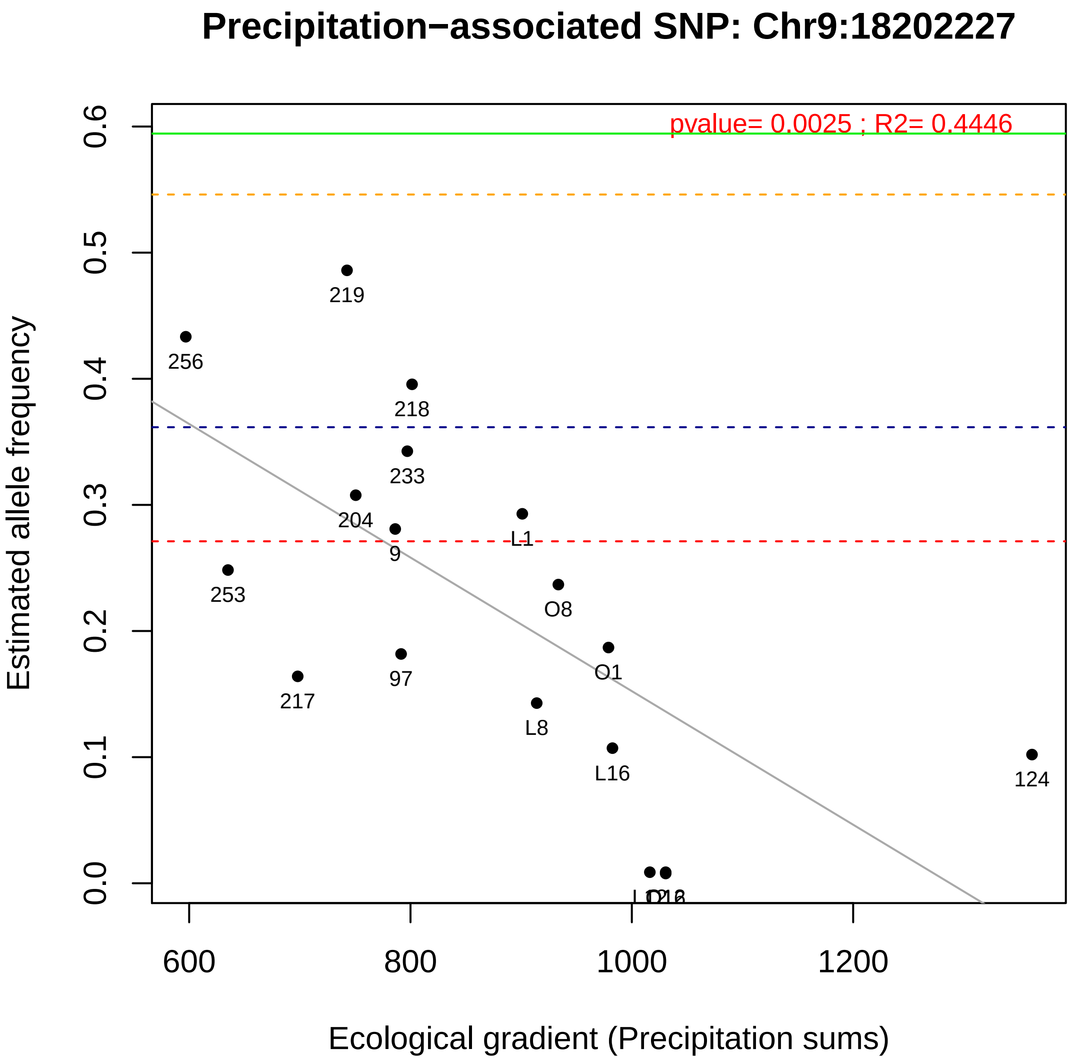
Precipitation−associated SNP: Chr9:18202227
pvalue= 0.0025 ; R2= 0.4446
Estimated allele frequency
0.0
0.1
0.2
0.3
0.4
0.5
0.6
600
800
1000
1200
Ecological gradient (Precipitation sums)
219
256
218
233
204
9
L1
253
O8
O1
97
217
L8
L16
124

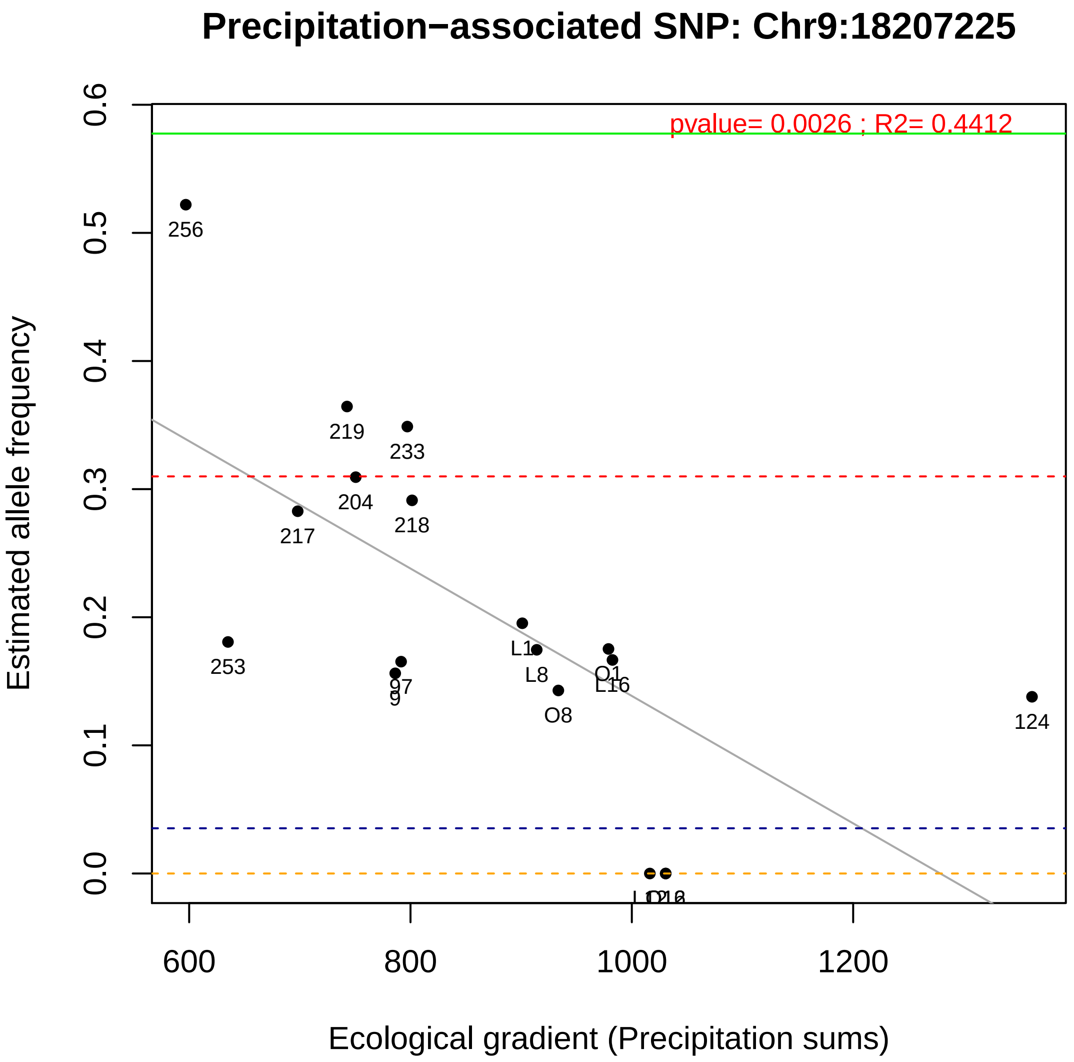

Precipitation−associated SNP: Chr9:18207225
pvalue= 0.0026 ; R2= 0.4412
Estimated allele frequency
Ecological gradient (Precipitation sums)
0.0
0.1
0.2
0.3
0.4
0.5
0.6
600
800
1000
1200
256
219
233
204
218
217
253
97
9
L8
O8
O1
L16
124

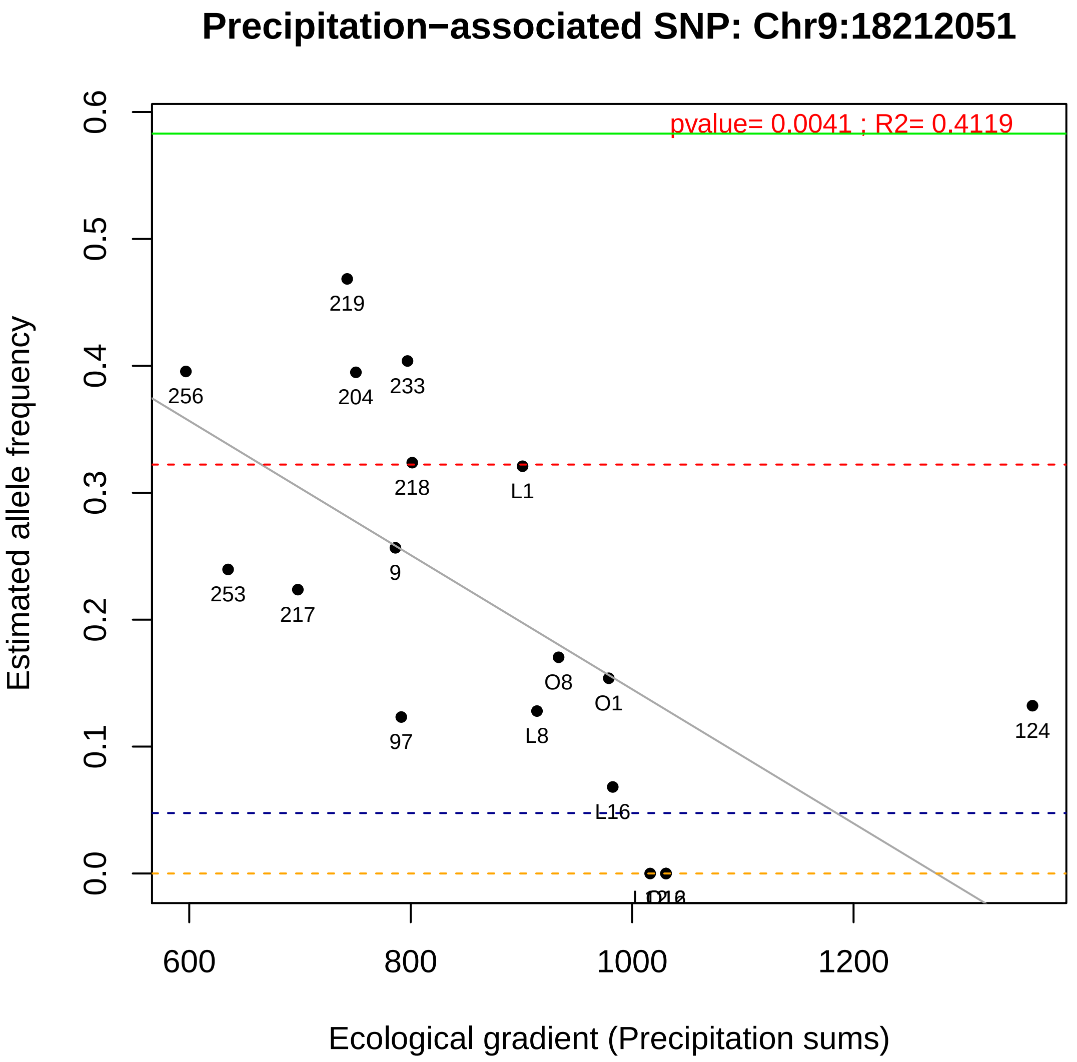
Precipitation−associated SNP: Chr9:18212051
pvalue= 0.0041 ; R2= 0.4119
Estimated allele frequency
0.0
0.1
0.2
0.3
0.4
0.5
0.6
Ecological gradient (Precipitation sums)
600
800
1000
1200
219
256
204
233
218
L1
9
253
217
O8
O1
97
L8
124
L16

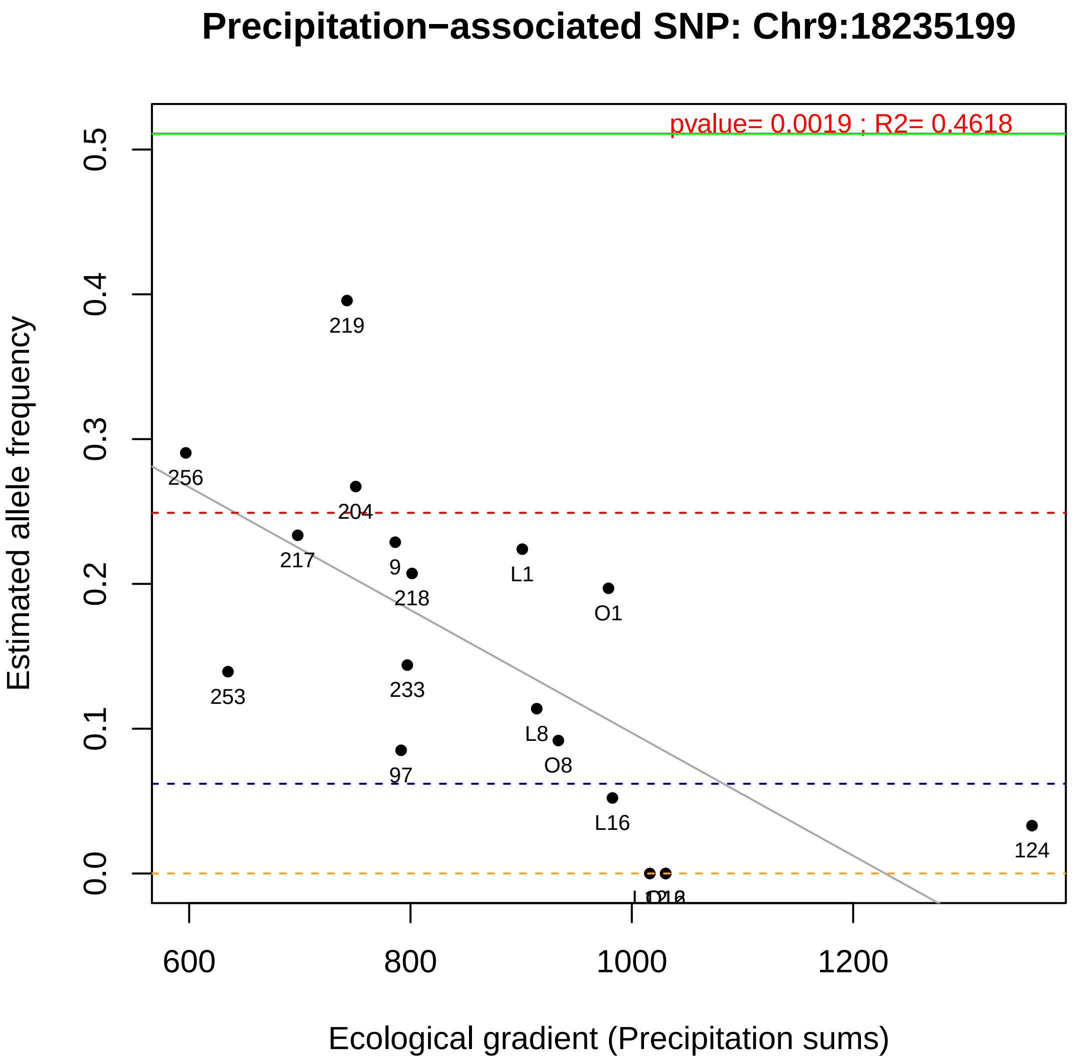
Precipitation−associated SNP: Chr9:18235199
pvalue= 0.0019 ; R2= 0.4618
Estimated allele frequency
0.0
0.1
0.2
0.3
0.4
0.5
600
800
1000
1200
Ecological gradient (Precipitation sums)
219
256
204
217
9
218
L1
O1
253
233
L8
O8
97
L16
124

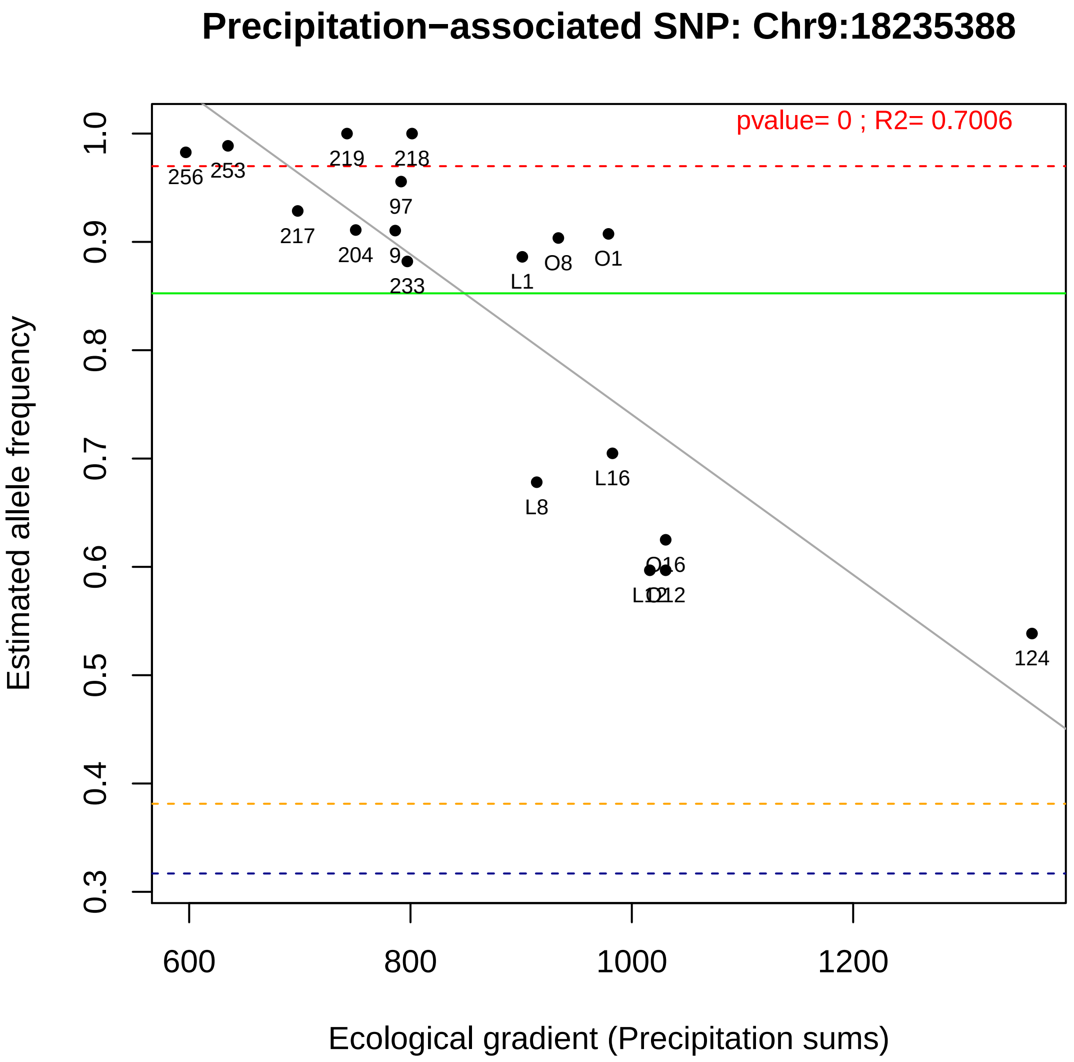

Precipitation−associated SNP: Chr9:18235388
pvalue= 0 ; R2= 0.7006
Estimated allele frequency
Ecological gradient (Precipitation sums)
0.3
0.4
0.5
0.6
0.7
0.8
0.9
1.0
600
800
1000
1200
256
253
219
218
97
217
204
9
233
L1
O8
O1
L8
L16
O16
L10
O12
124

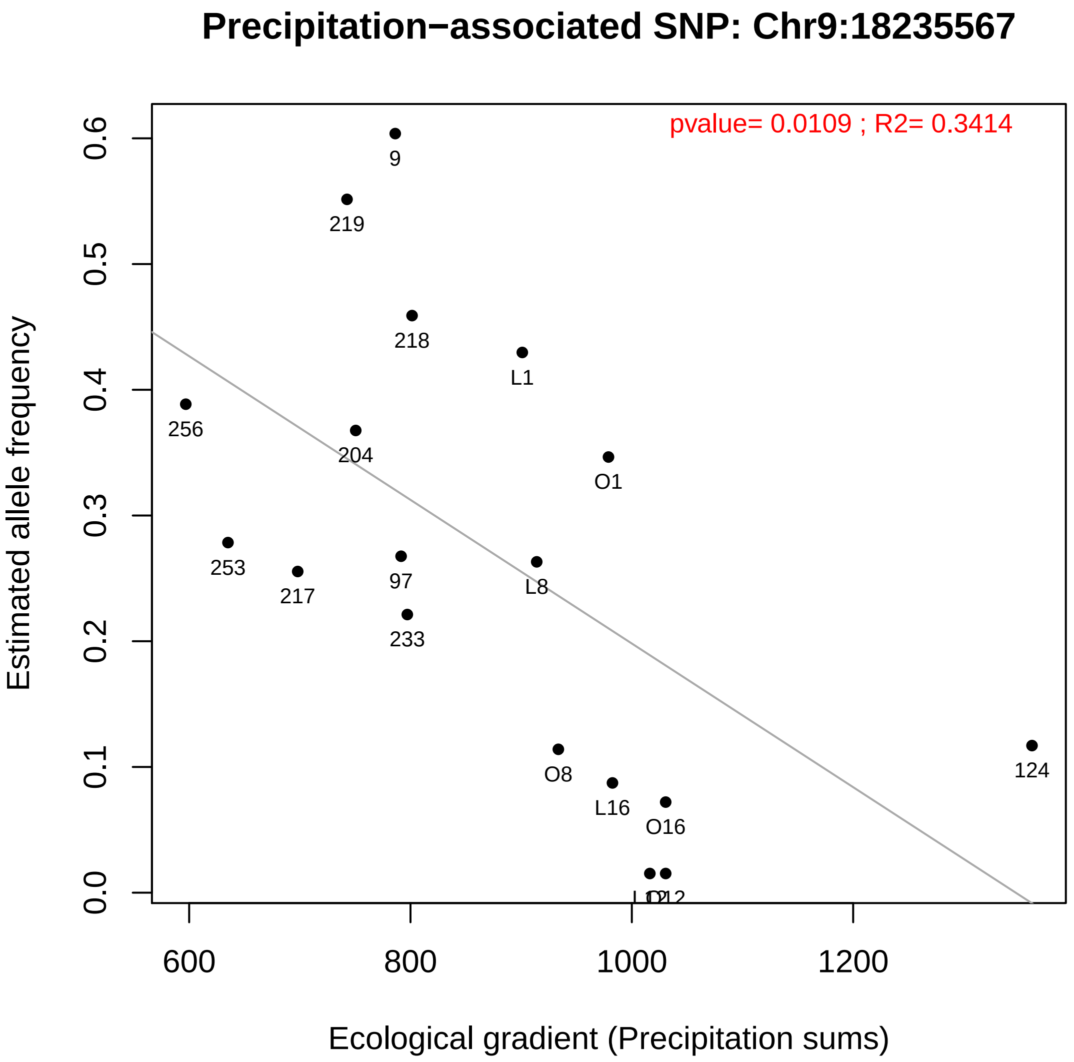
Precipitation−associated SNP: Chr9:18235567
pvalue= 0.0109 ; R2= 0.3414
Estimated allele frequency
Ecological gradient (Precipitation sums)
0.0
0.1
0.2
0.3
0.4
0.5
0.6
600
800
1000
1200
9
219
218
L1
256
204
O1
253
217
97
L8
233
O8
124
L16
O16

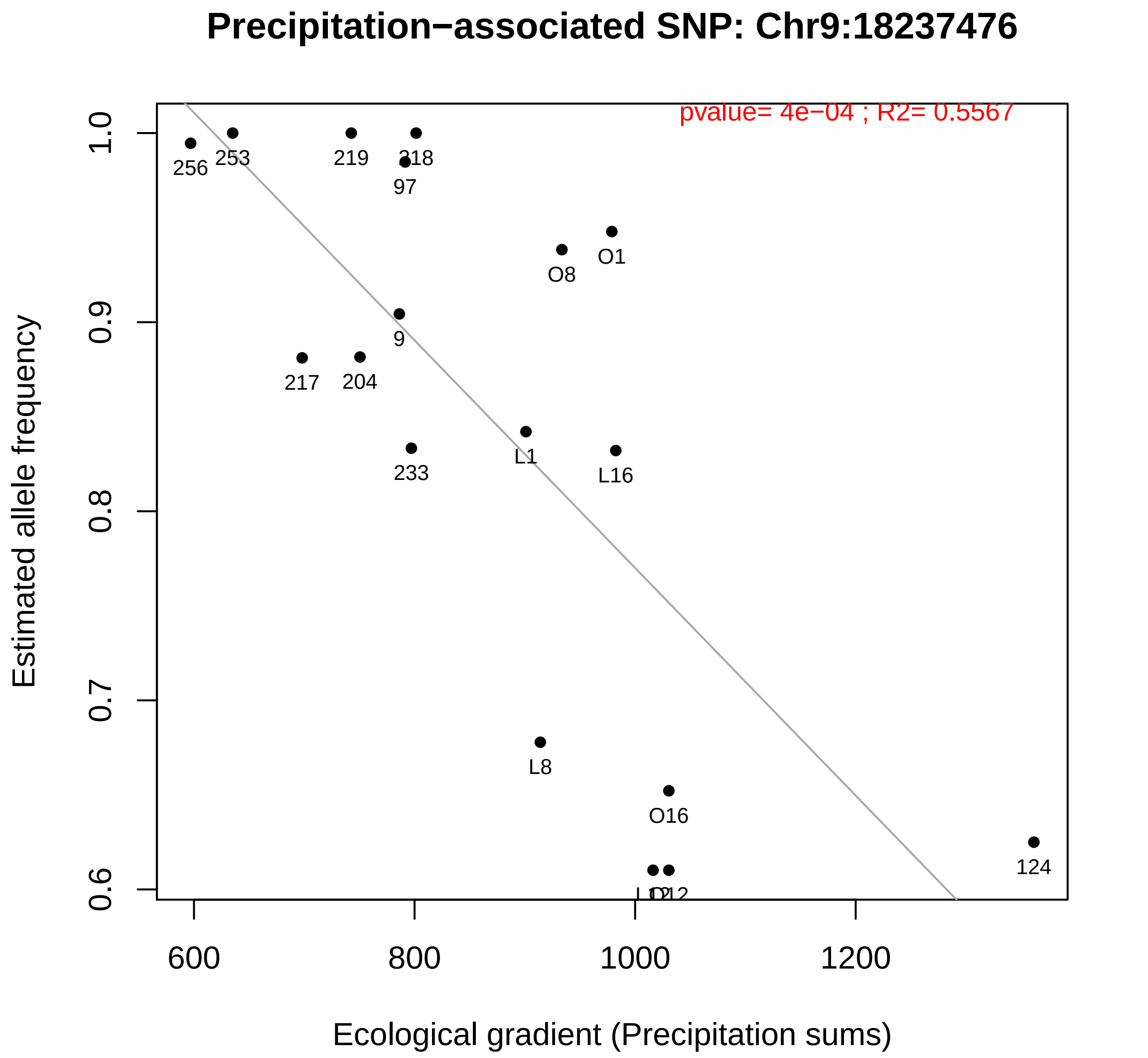

Precipitation−associated SNP: Chr9:18237476
pvalue= 4e−04 ; R2= 0.5567
Estimated allele frequency
Ecological gradient (Precipitation sums)
0.6
0.7
0.8
0.9
1.0
600
800
1000
1200
256
253
219
218
97
O1
O8
9
217
204
233
L1
L16
L8
O16
124

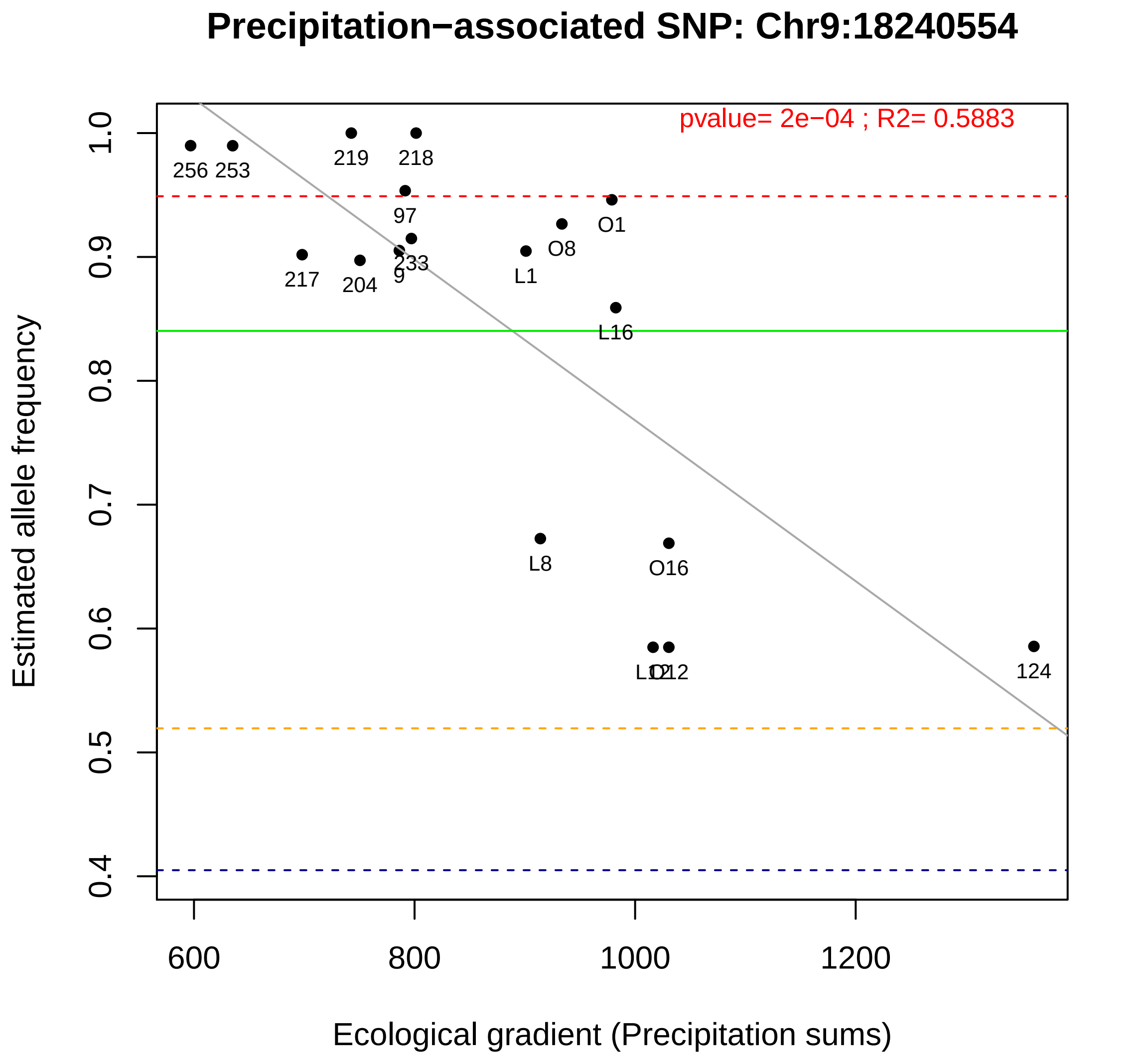
Precipitation−associated SNP: Chr9:18240554
pvalue= 2e−04 ; R2= 0.5883
Estimated allele frequency
Ecological gradient (Precipitation sums)
0.4
0.5
0.6
0.7
0.8
0.9
1.0
600
800
1000
1200
256
253
219
218
97
O1
O8
233
9
217
204
L1
L16
L8
O16
L12
O12
124

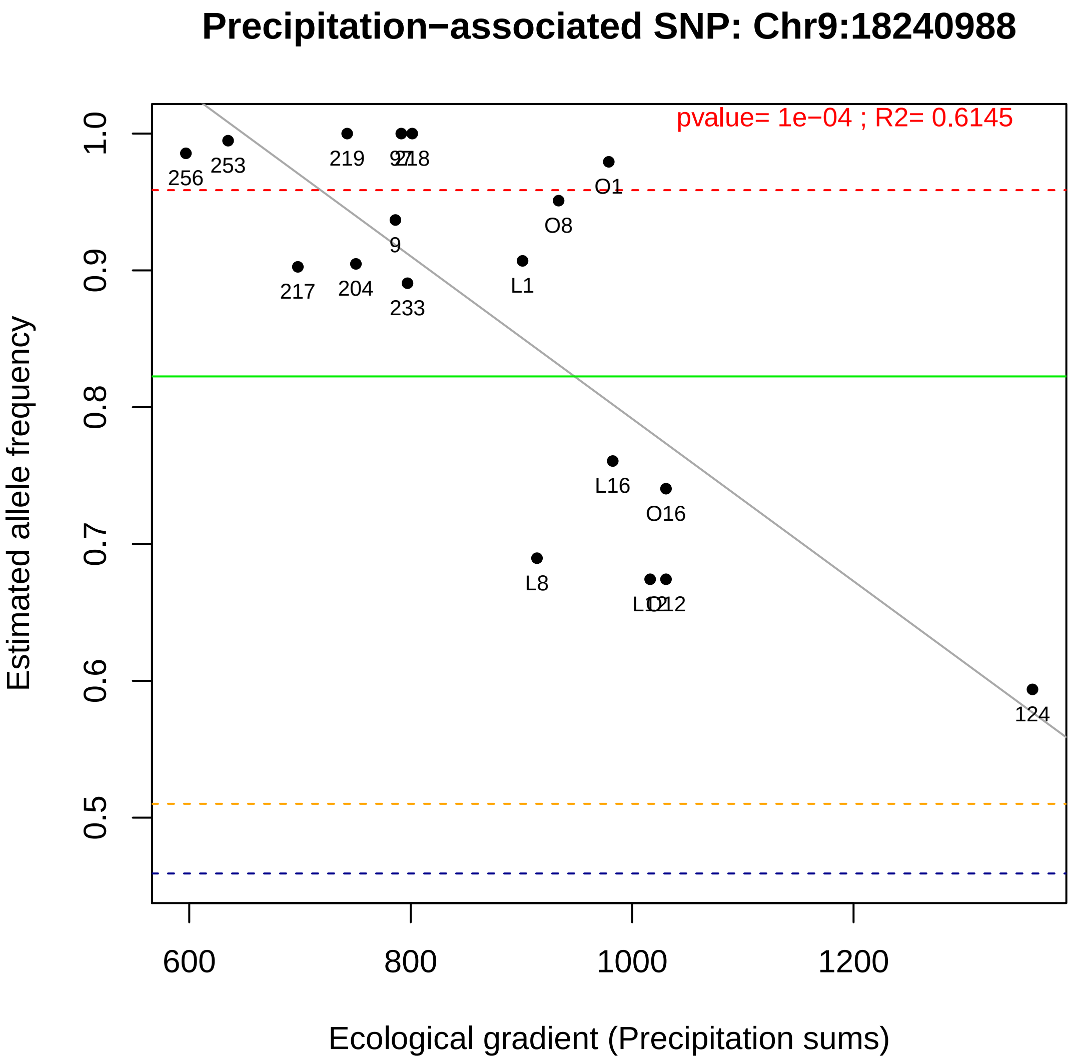
Precipitation−associated SNP: Chr9:18240988
pvalue= 1e−04 ; R2= 0.6145
Estimated allele frequency
Ecological gradient (Precipitation sums)
0.5
0.6
0.7
0.8
0.9
1.0
600
800
1000
1200
256
253
219
97
218
O1
O8
9
217
204
233
L1
L16
O16
L8
L12
O12
124

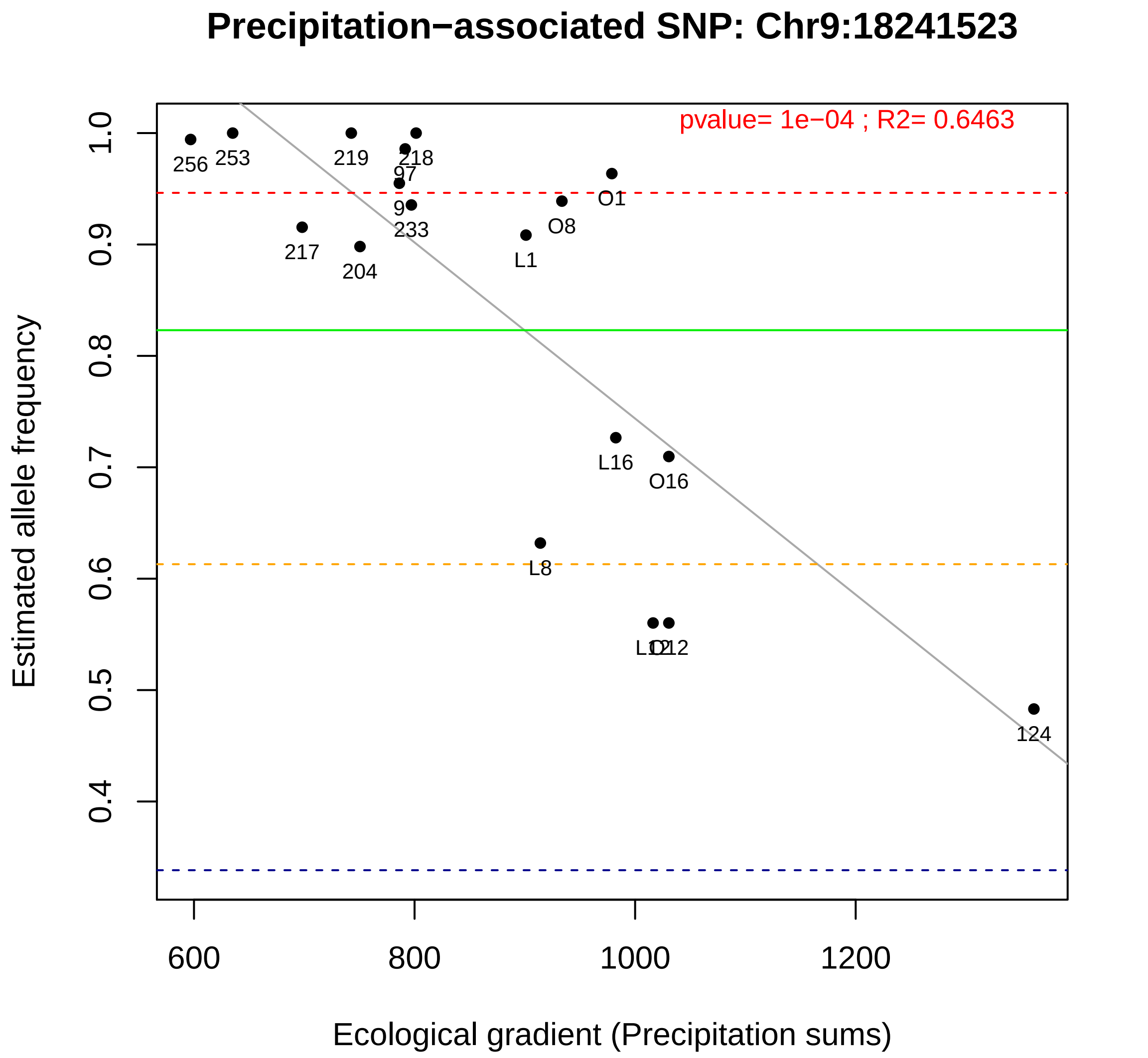
Precipitation−associated SNP: Chr9:18241523
pvalue= 1e−04 ; R2= 0.6463
Estimated allele frequency
Ecological gradient (Precipitation sums)
1.0
0.9
0.8
0.7
0.6
0.5
0.4
600
800
1000
1200
256
253
219
218
97
9
233
217
204
L1
O8
O1
L16
O16
L8
L12
O12
124

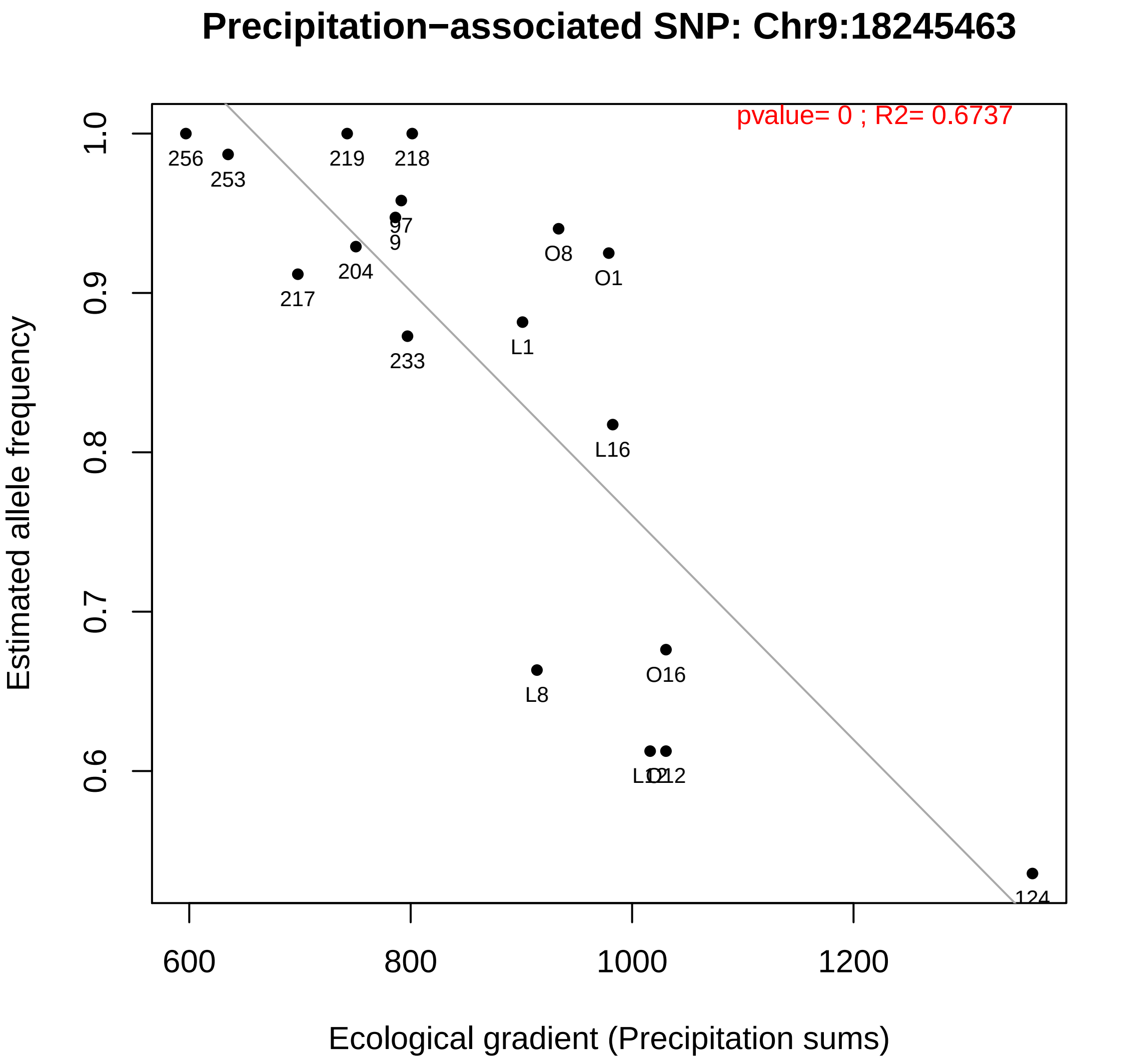
Precipitation−associated SNP: Chr9:18245463
pvalue= 0 ; R2= 0.6737
256
253
219
218
97
9
204
217
O8
O1
L1
233
L16
L8
O16
L12
O12
124
1.0
0.9
0.8
0.7
0.6
600
800
1000
1200
Estimated allele frequency
Ecological gradient (Precipitation sums)

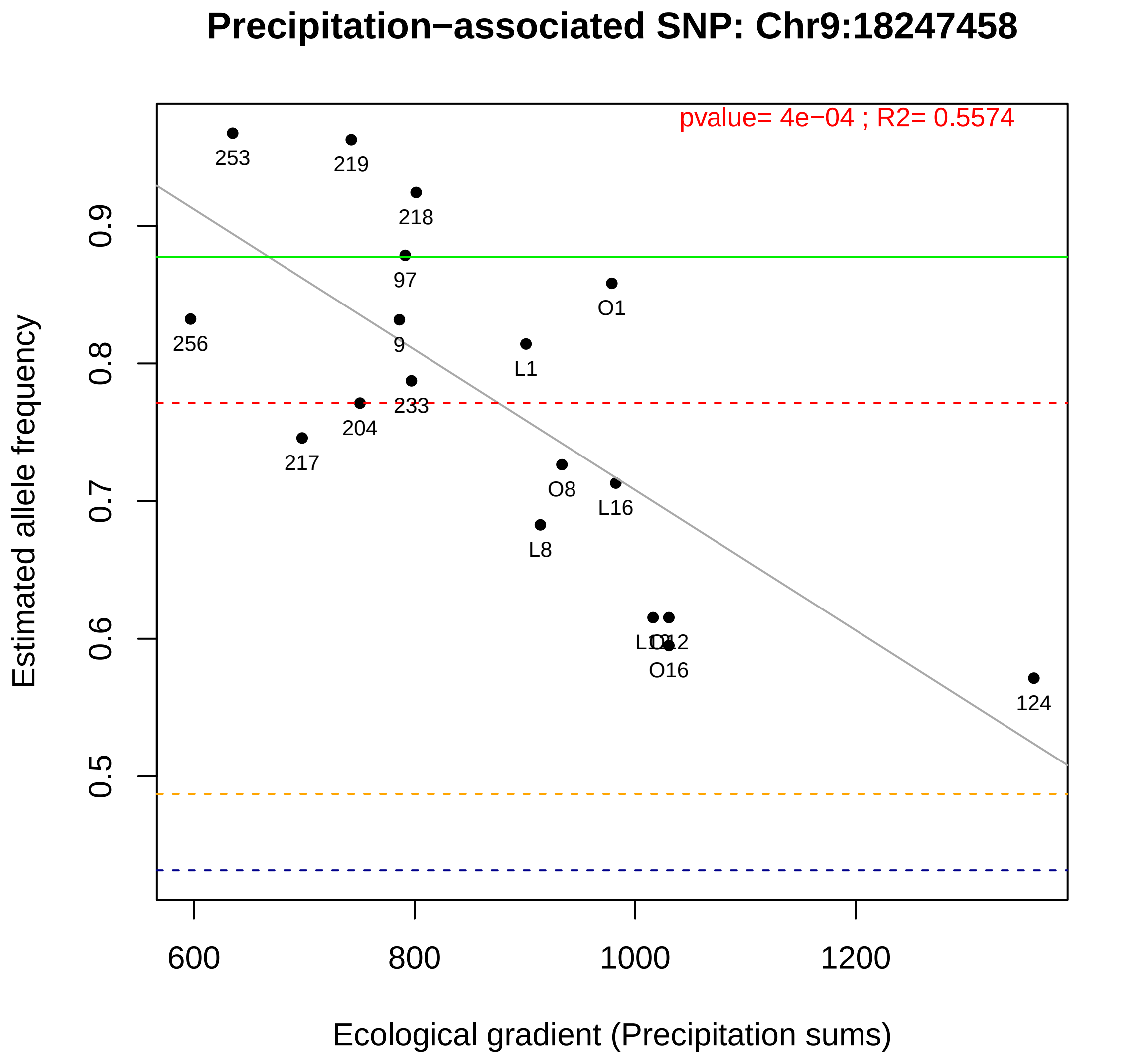

Precipitation−associated SNP: Chr9:18247458
pvalue= 4e−04 ; R2= 0.5574
253
219
218
97
O1
256
9
L1
233
204
217
O8
L16
L8
L12
O12
O16
124
0.5
0.6
0.7
0.8
0.9
600
800
1000
1200
Estimated allele frequency
Ecological gradient (Precipitation sums)

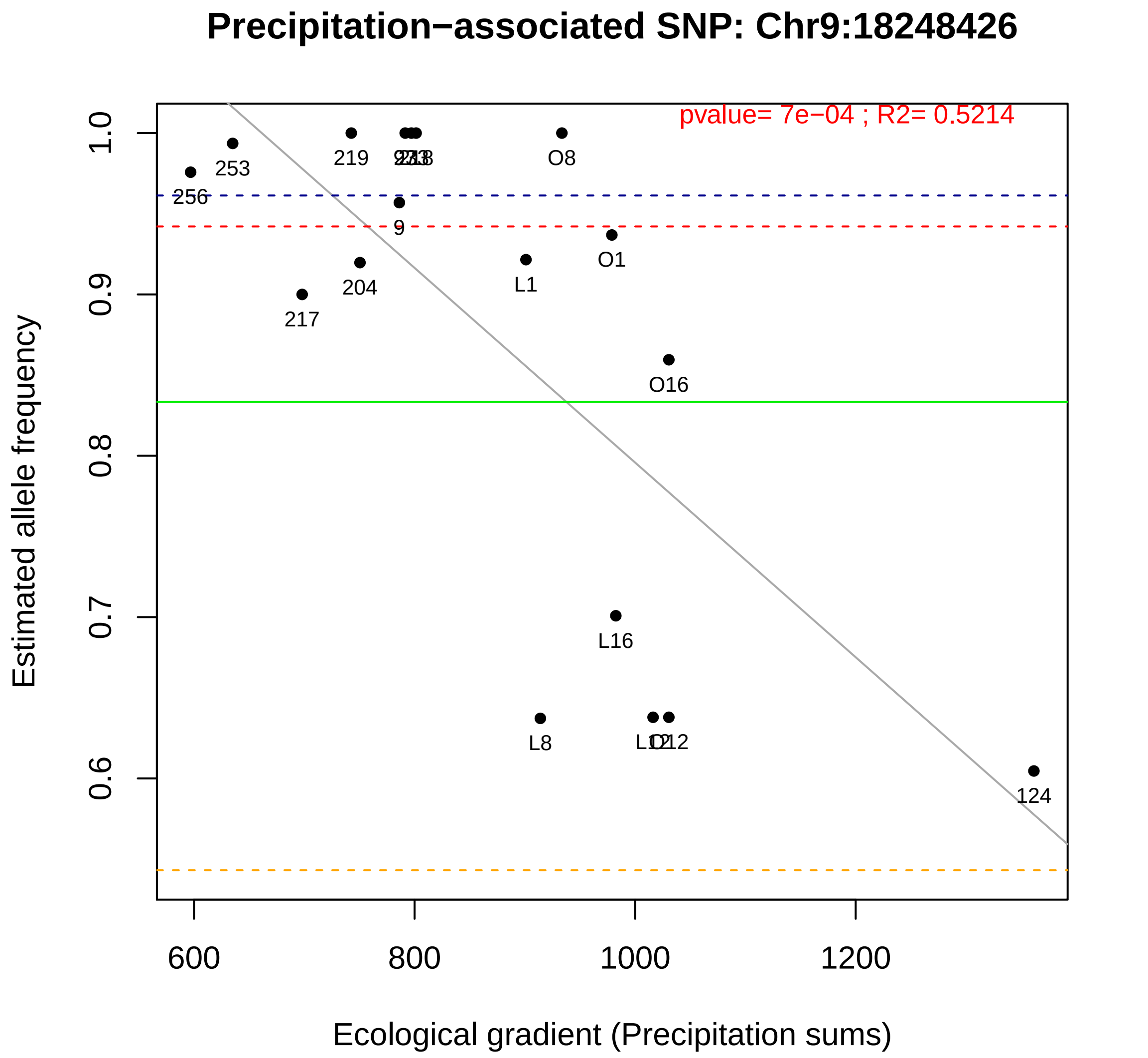
Precipitation−associated SNP: Chr9:18248426
pvalue= 7e−04 ; R2= 0.5214
Estimated allele frequency
Ecological gradient (Precipitation sums)
0.6
0.7
0.8
0.9
1.0
600
800
1000
1200
256
253
219
217
204
9
L1
O8
O1
O16
L16
L8
124

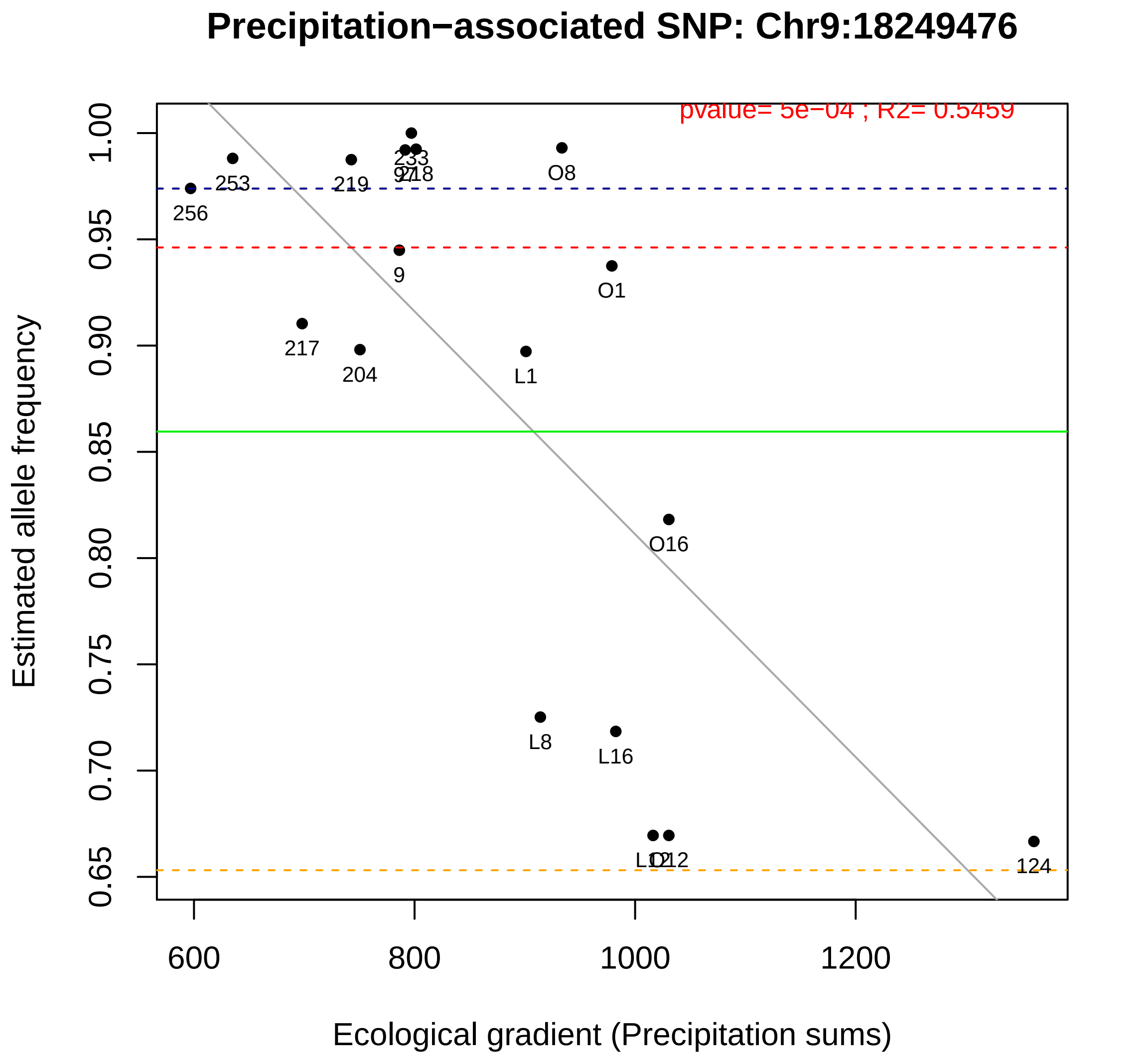

Precipitation−associated SNP: Chr9:18249476
pvalue= 5e−04 ; R2= 0.5459
Estimated allele frequency
1.00
0.95
0.90
0.85
0.80
0.75
0.70
0.65
600
800
1000
1200
Ecological gradient (Precipitation sums)
256
253
219
233
97
218
O8
9
O1
217
204
L1
O16
L8
L16
L12
O12
124

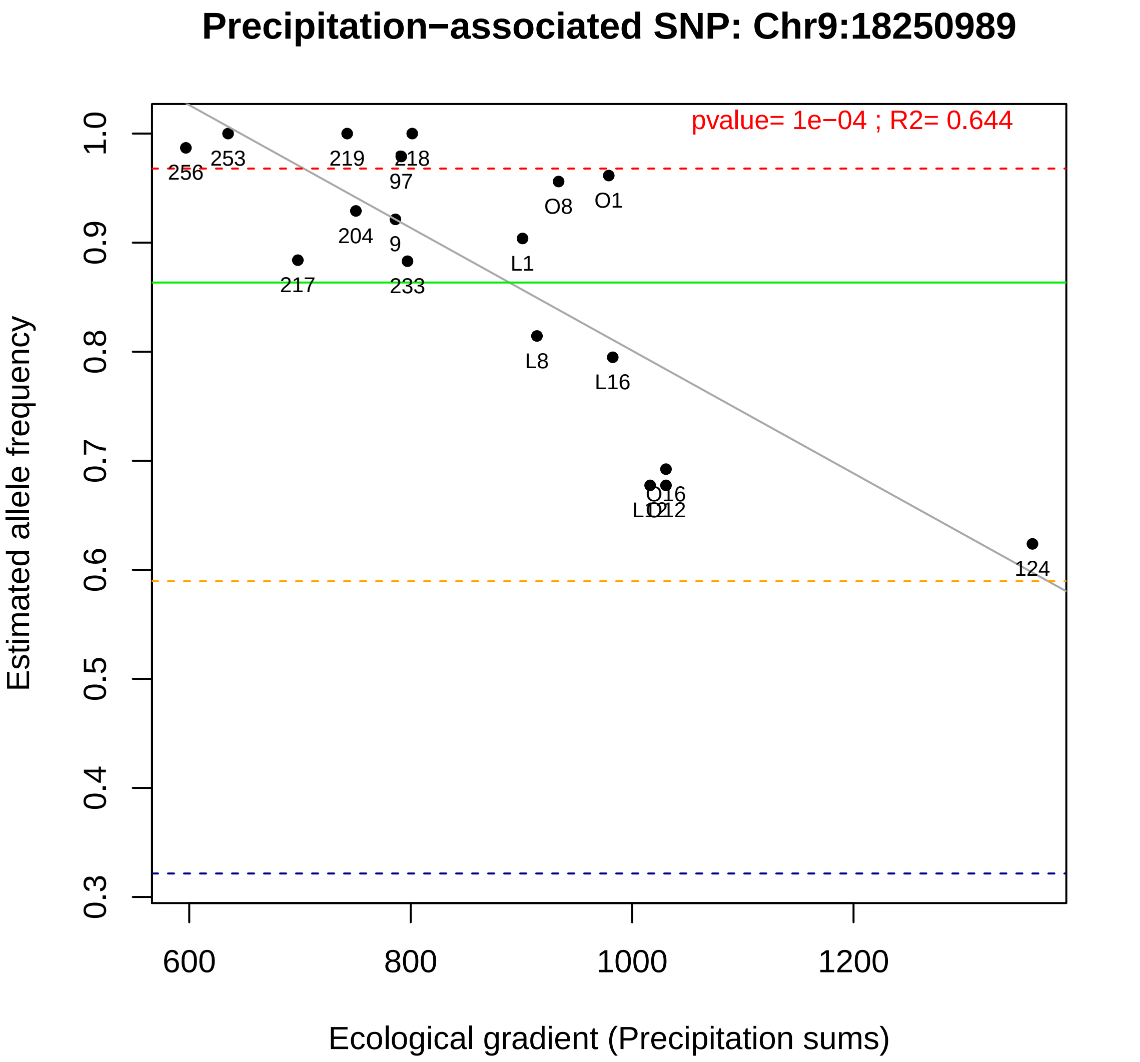
Precipitation−associated SNP: Chr9:18250989
pvalue= 1e−04 ; R2= 0.644
Estimated allele frequency
Ecological gradient (Precipitation sums)
1.0
0.9
0.8
0.7
0.6
0.5
0.4
0.3
600
800
1000
1200
256
253
219
218
97
204
9
217
233
O8
O1
L1
L8
L16
O16
O12
124

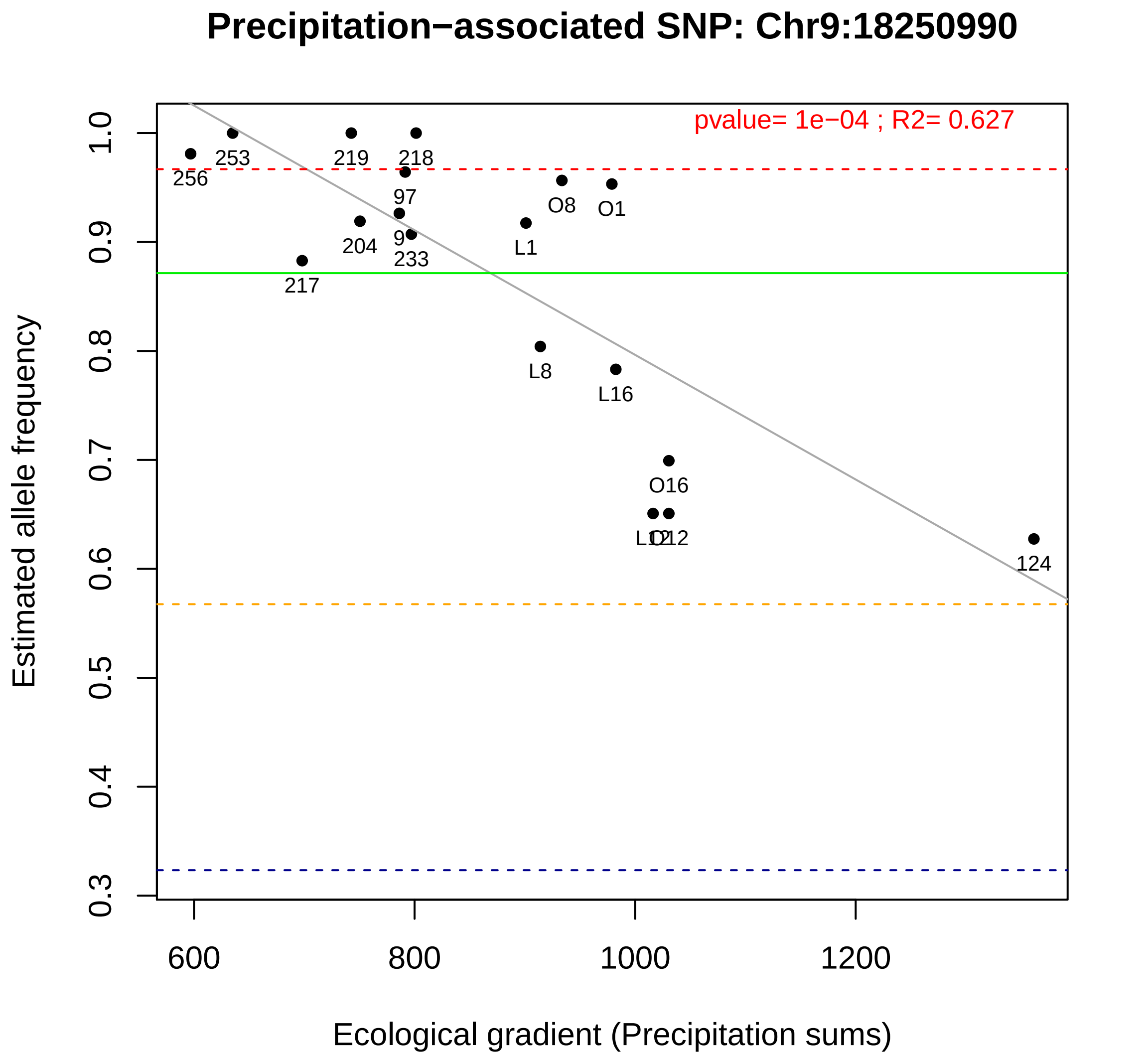
Precipitation−associated SNP: Chr9:18250990
pvalue= 1e−04 ; R2= 0.627
Estimated allele frequency
Ecological gradient (Precipitation sums)
1.0
0.9
0.8
0.7
0.6
0.5
0.4
0.3
600
800
1000
1200
256
253
219
218
97
204
9
233
217
O8
O1
L1
L8
L16
O16
L12
O12
124

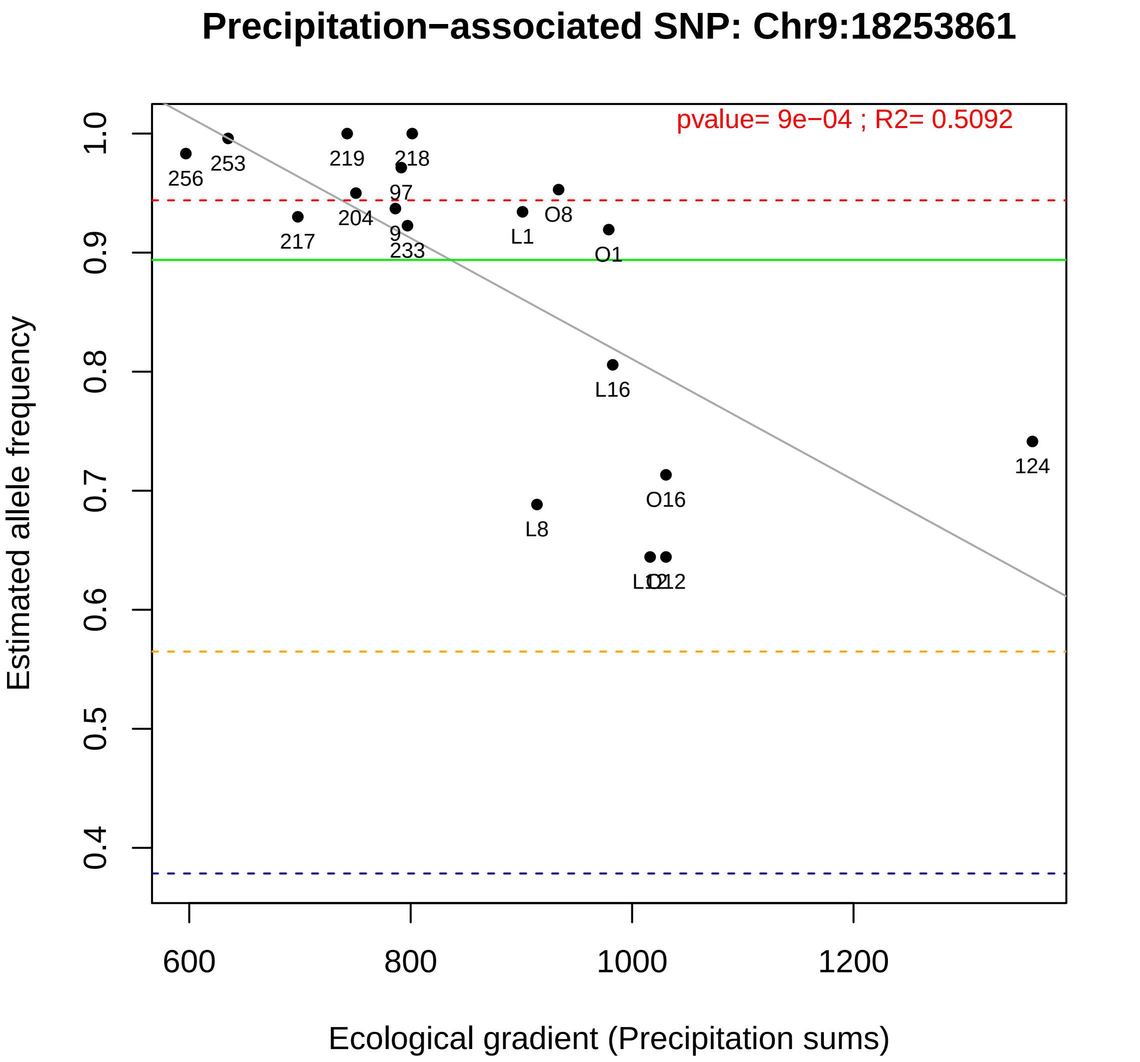

Precipitation−associated SNP: Chr9:18253861
pvalue= 9e−04 ; R2= 0.5092
Estimated allele frequency
Ecological gradient (Precipitation sums)
0.4
0.5
0.6
0.7
0.8
0.9
1.0
600
800
1000
1200
256
253
219
218
97
204
9
233
217
L1
O8
O1
L16
124
O16
L8
L12
O12

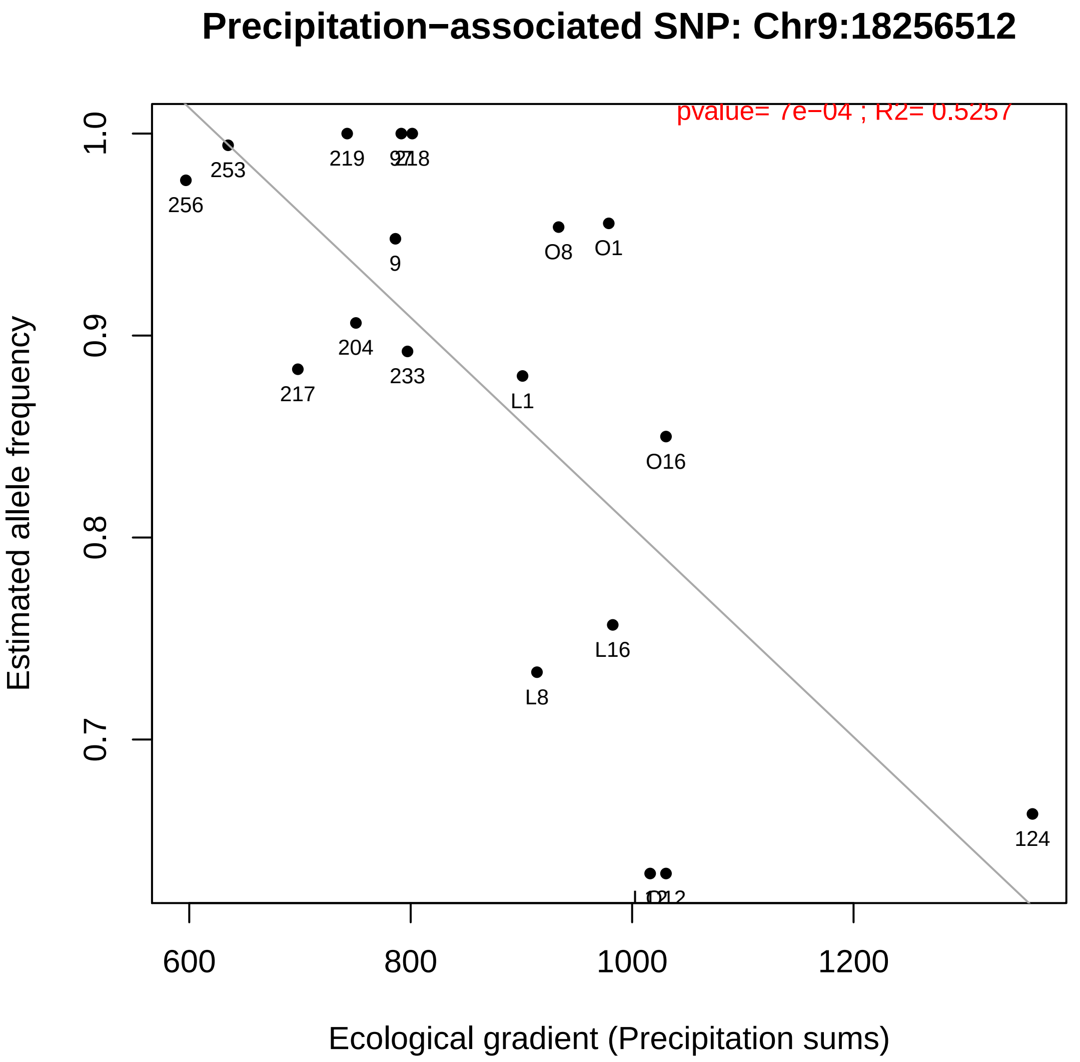
Precipitation−associated SNP: Chr9:18256512
pvalue= 7e−04 ; R2= 0.5257
Estimated allele frequency
Ecological gradient (Precipitation sums)
1.0
0.9
0.8
0.7
600
800
1000
1200
219
253
256
9
O8
O1
204
233
217
L1
O16
L16
L8
124

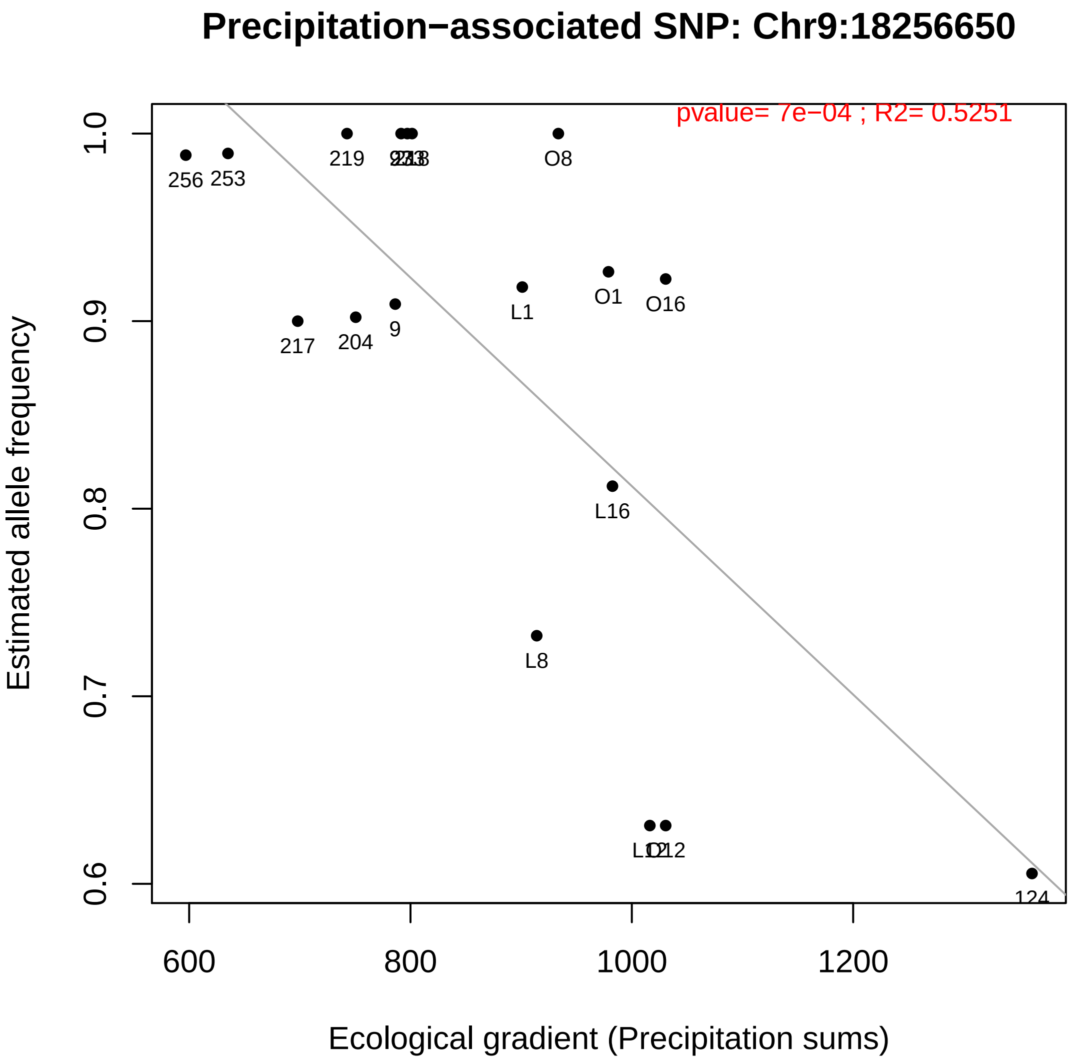
Precipitation−associated SNP: Chr9:18256650
pvalue= 7e−04 ; R2= 0.5251
Estimated allele frequency
Ecological gradient (Precipitation sums)
0.6
0.7
0.8
0.9
1.0
600
800
1000
1200
256
253
219
O8
217
204
9
L1
O1
O16
L16
L8
124

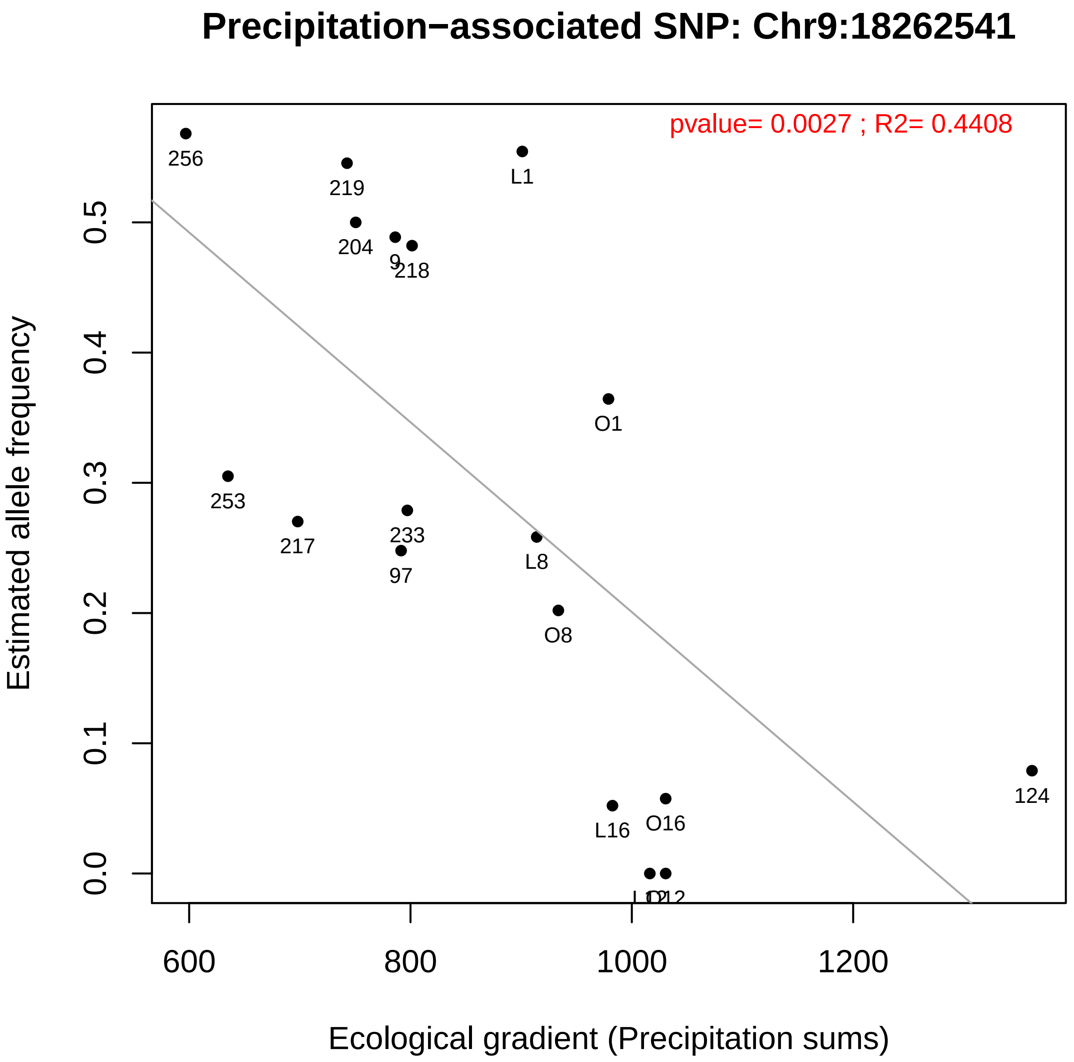
Precipitation−associated SNP: Chr9:18262541
pvalue= 0.0027 ; R2= 0.4408
256
219
L1
204
9
218
O1
253
217
233
97
L8
O8
124
L16
O16
0.0
0.1
0.2
0.3
0.4
0.5
600
800
1000
1200
Estimated allele frequency
Ecological gradient (Precipitation sums)

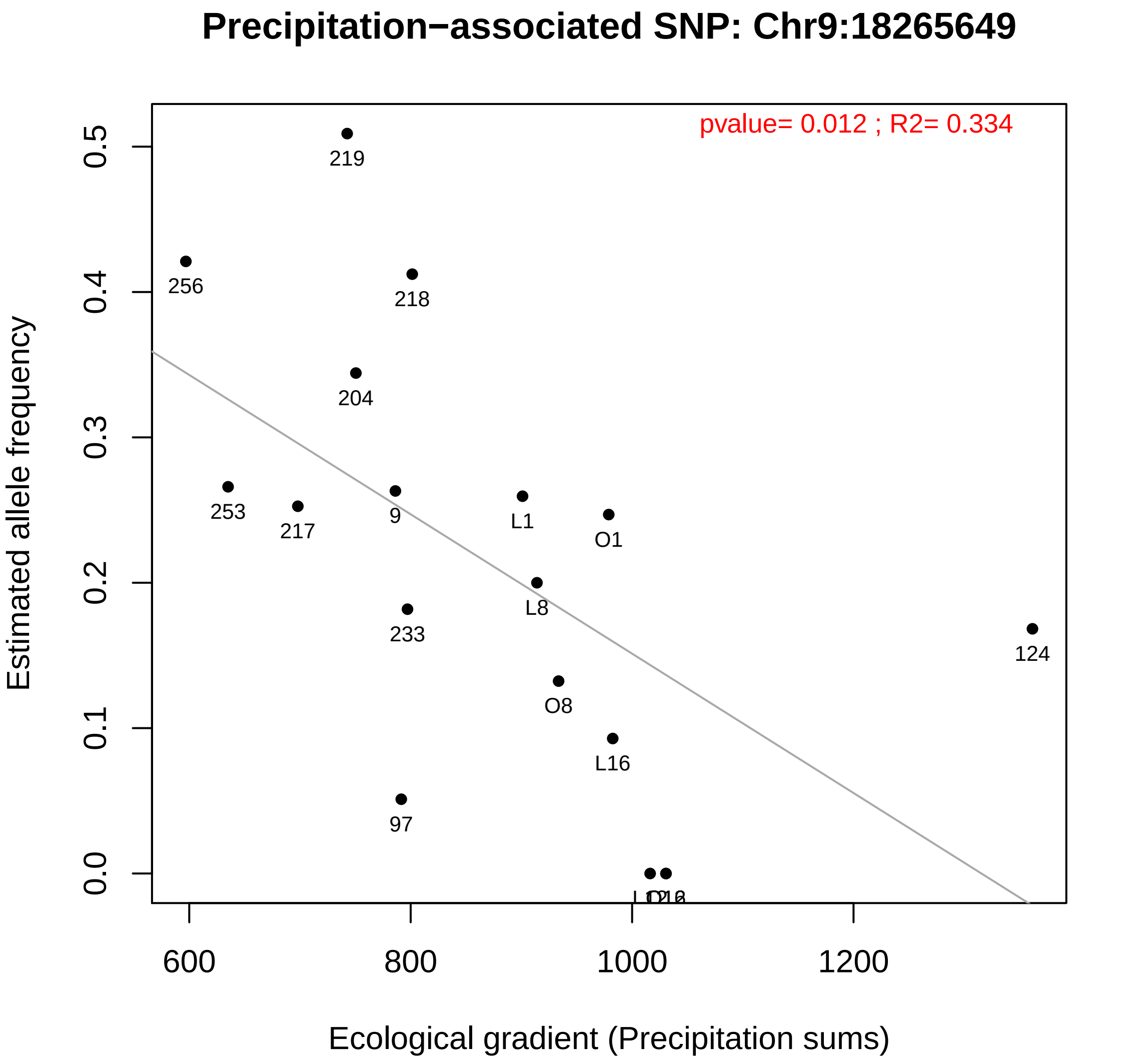
Precipitation−associated SNP: Chr9:18265649
pvalue= 0.012 ; R2= 0.334
Estimated allele frequency
0.0
0.1
0.2
0.3
0.4
0.5
Ecological gradient (Precipitation sums)
600
800
1000
1200
219
256
218
204
253
217
9
L1
O1
L8
233
124
O8
L16
97

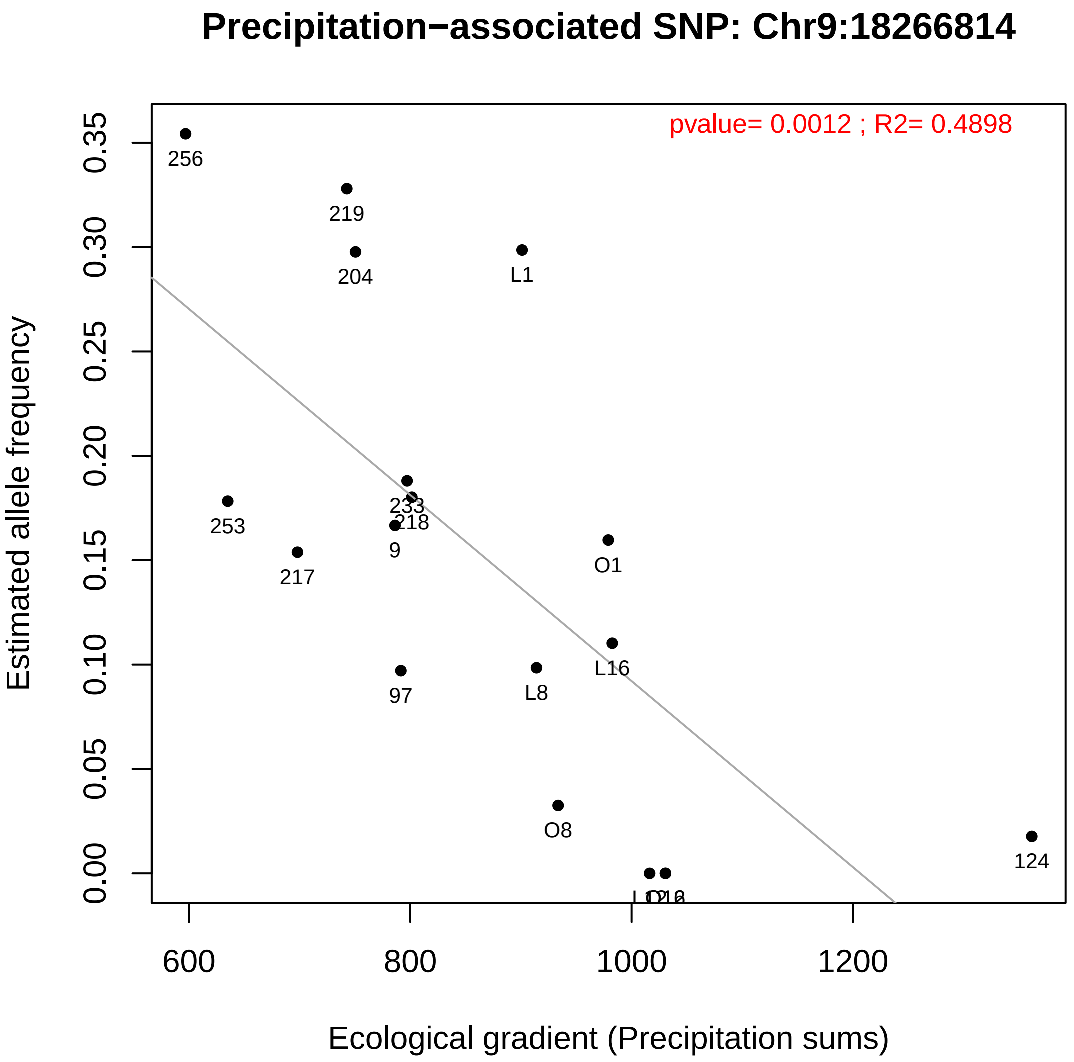
Precipitation−associated SNP: Chr9:18266814
pvalue= 0.0012 ; R2= 0.4898
Estimated allele frequency
0.00
0.05
0.10
0.15
0.20
0.25
0.30
0.35
Ecological gradient (Precipitation sums)
600
800
1000
1200
256
219
204
L1
253
217
233
218
9
O1
L16
L8
97
O8
124

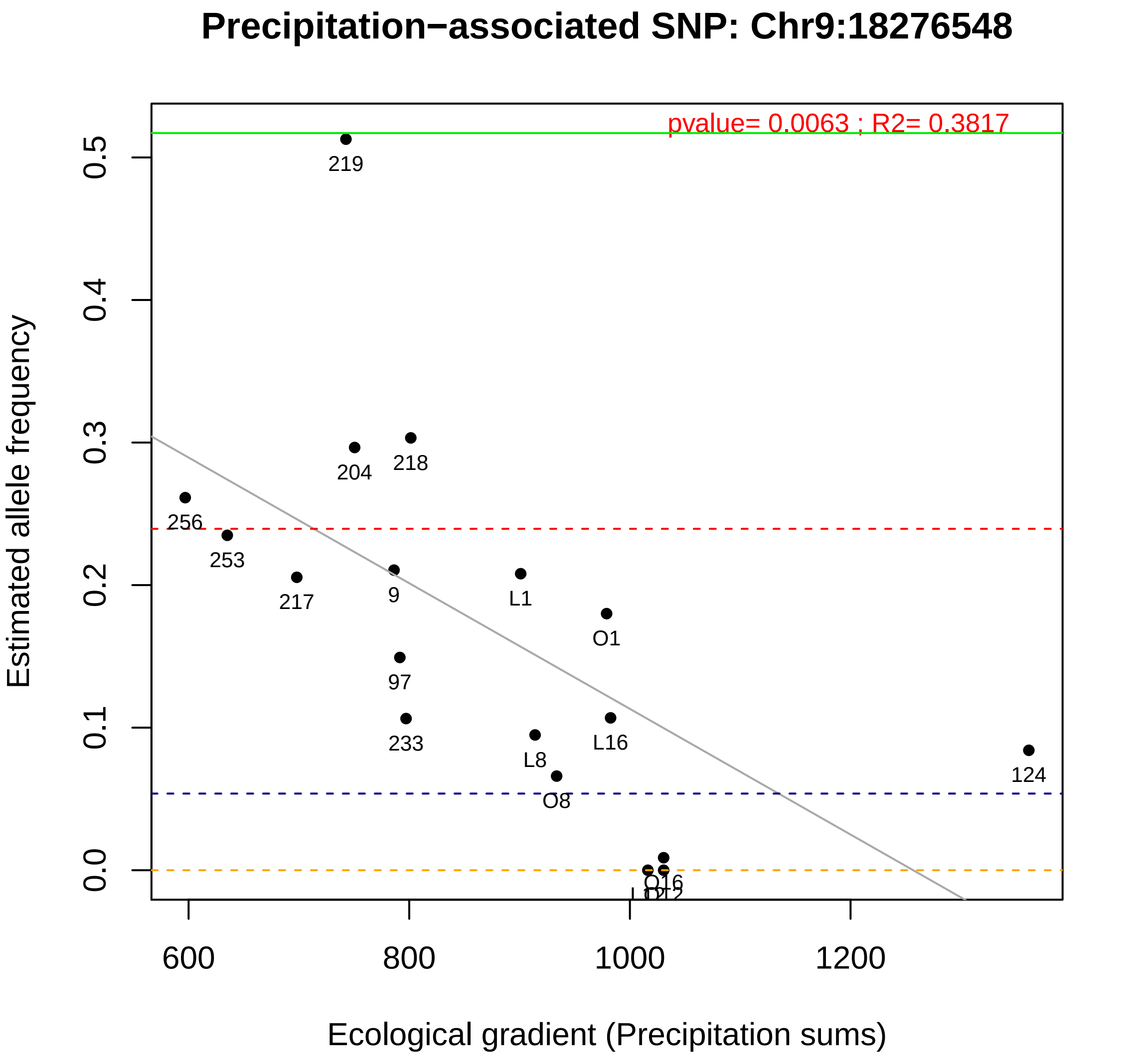
Precipitation−associated SNP: Chr9:18276548
pvalue= 0.0063 ; R2= 0.3817
Estimated allele frequency
0.0
0.1
0.2
0.3
0.4
0.5
Ecological gradient (Precipitation sums)
600
800
1000
1200
219
204
218
256
253
217
9
L1
O1
97
233
L16
L8
O8
124

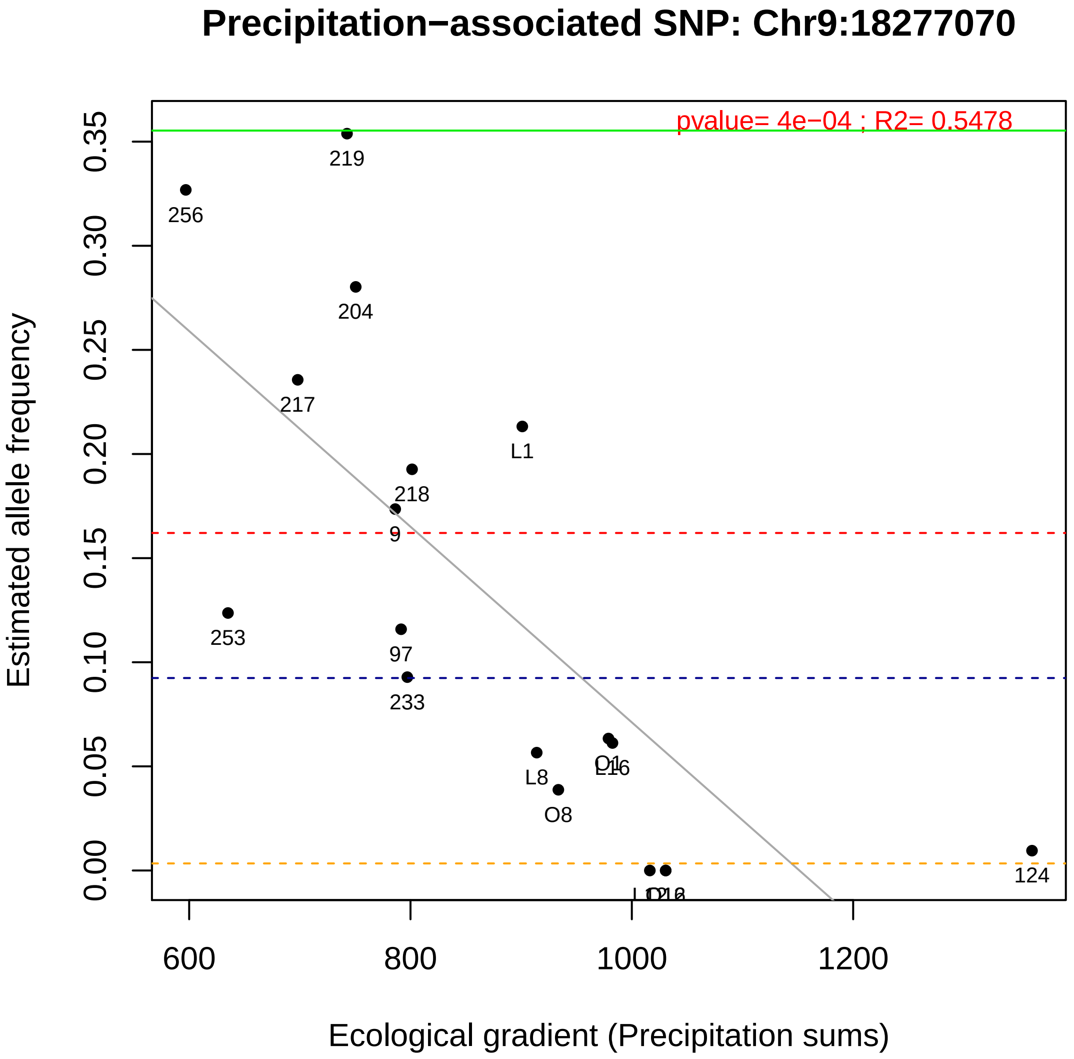
Precipitation−associated SNP: Chr9:18277070
pvalue= 4e−04 ; R2= 0.5478
Estimated allele frequency
0.00
0.05
0.10
0.15
0.20
0.25
0.30
0.35
Ecological gradient (Precipitation sums)
600
800
1000
1200
219
256
204
217
L1
218
9
253
97
233
L8
O8
124

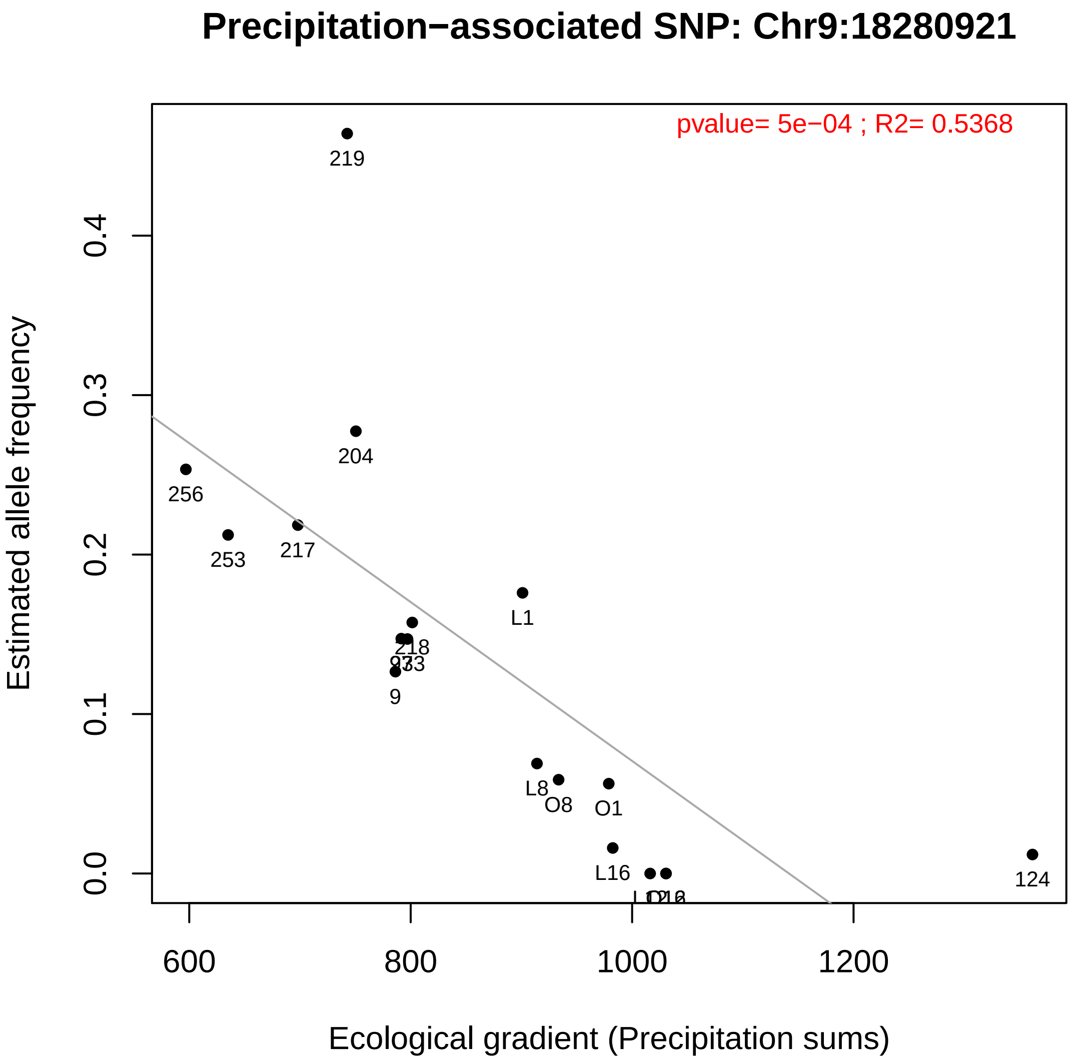

Precipitation−associated SNP: Chr9:18280921
pvalue= 5e−04 ; R2= 0.5368
Estimated allele frequency
Ecological gradient (Precipitation sums)
0.0
0.1
0.2
0.3
0.4
600
800
1000
1200
219
204
256
253
217
L1
218
9
L8
O8
O1
L16
124

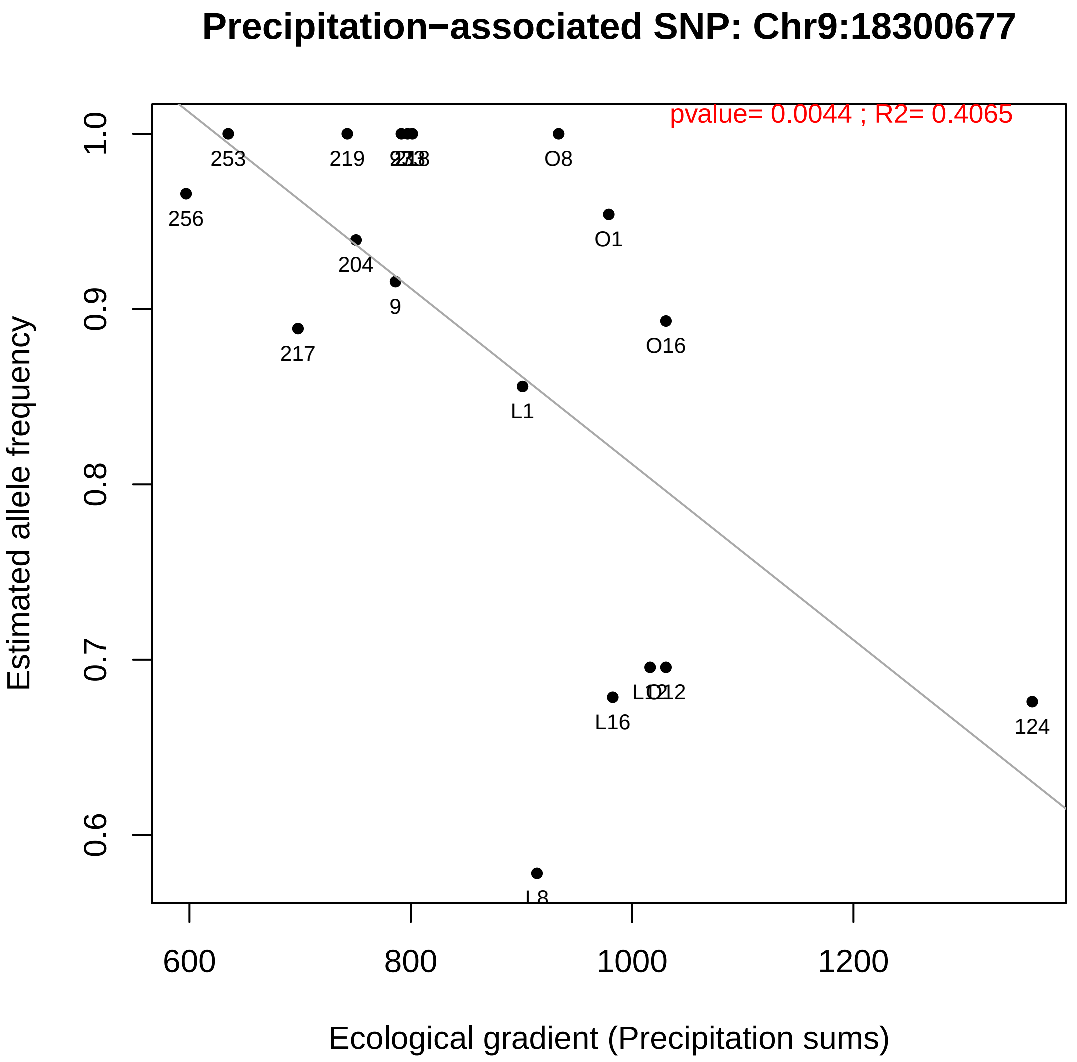
Precipitation−associated SNP: Chr9:18300677
pvalue= 0.0044 ; R2= 0.4065
Estimated allele frequency
Ecological gradient (Precipitation sums)
1.0
0.9
0.8
0.7
0.6
600
800
1000
1200
253
219
O8
256
204
O1
9
217
O16
L1
O12
L16
124

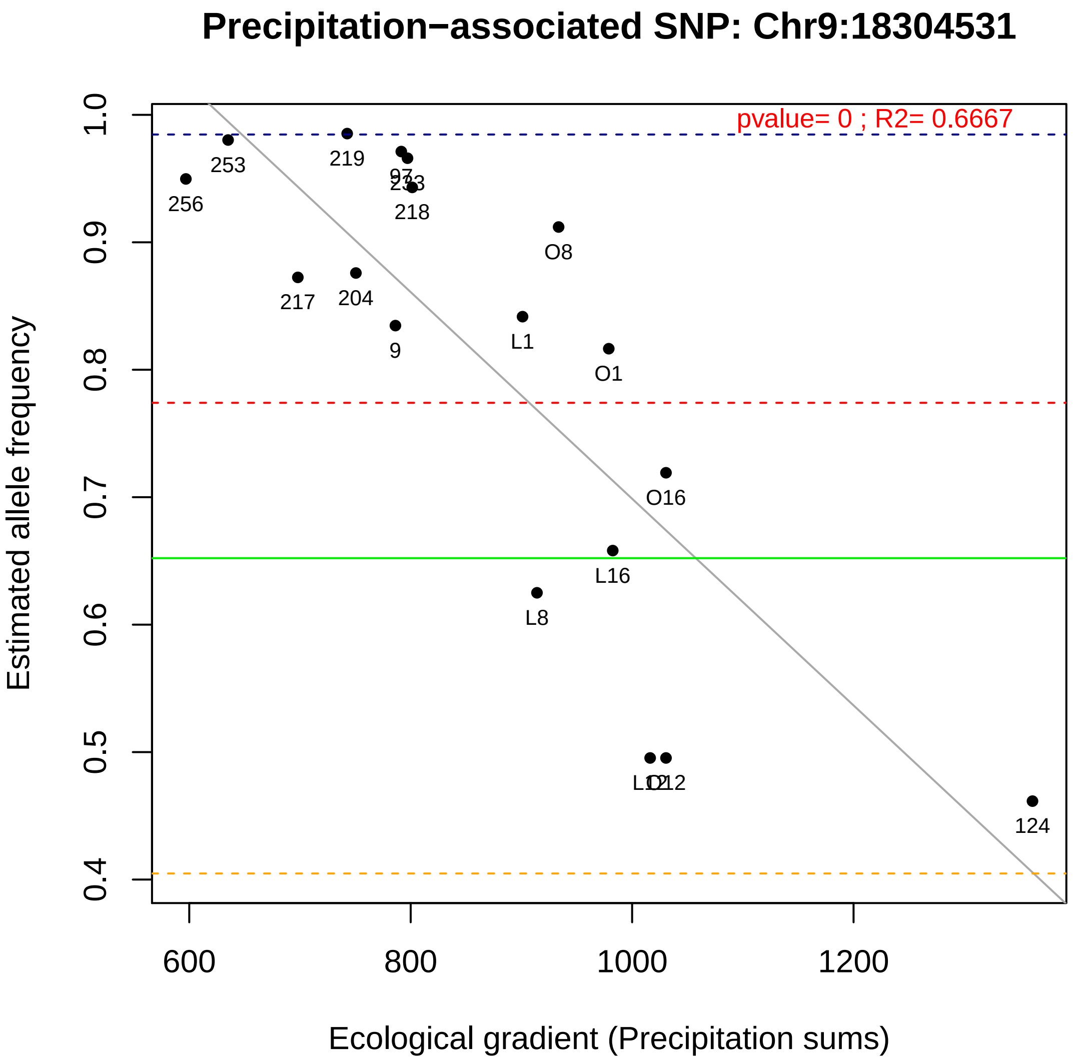

Precipitation−associated SNP: Chr9:18304531
pvalue= 0 ; R2= 0.6667
Estimated allele frequency
Ecological gradient (Precipitation sums)
1.0
0.9
0.8
0.7
0.6
0.5
0.4
600
800
1000
1200
256
253
219
217
204
9
218
O8
L1
O1
O16
L16
L8
124

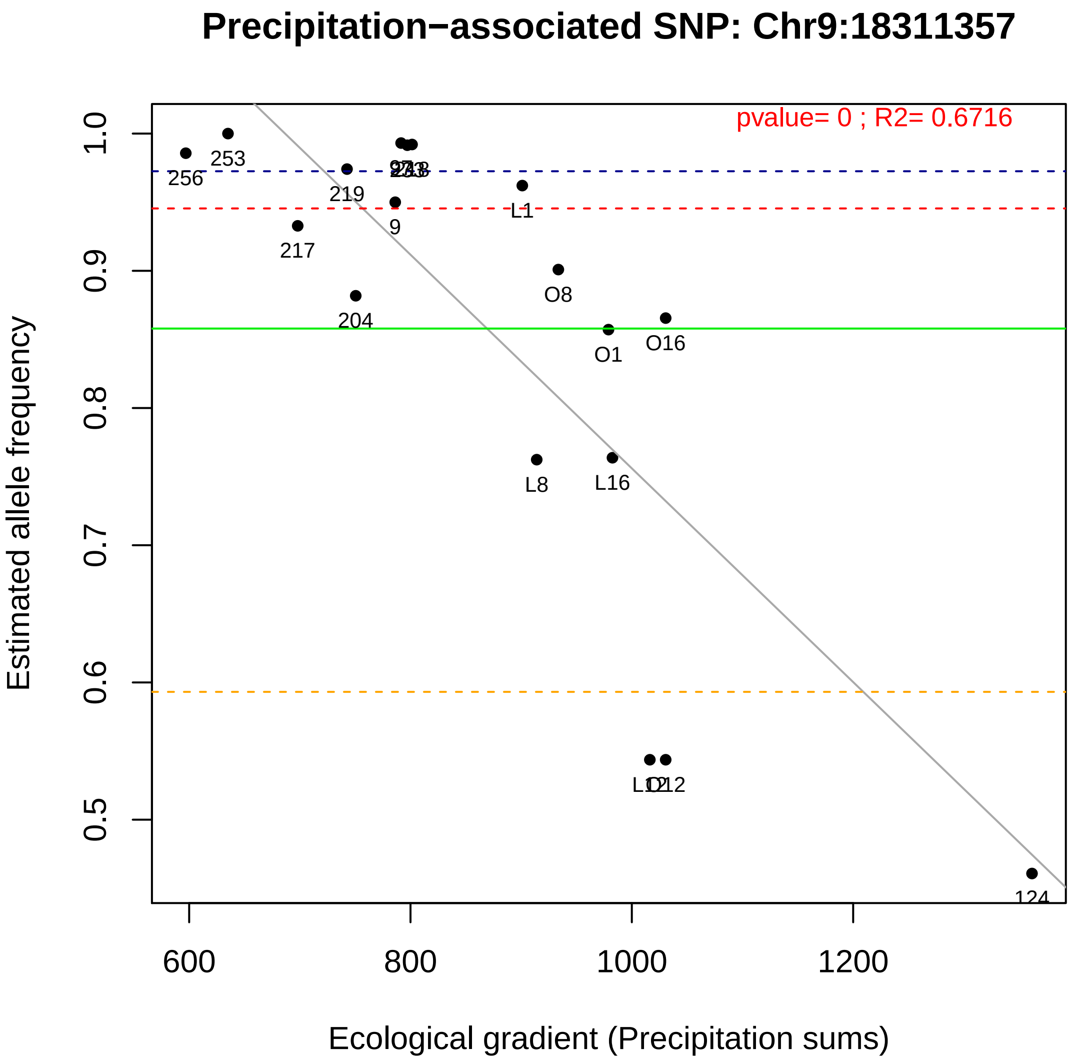
Precipitation−associated SNP: Chr9:18311357
pvalue= 0 ; R2= 0.6716
Estimated allele frequency
1.0
0.9
0.8
0.7
0.6
0.5
600
800
1000
1200
Ecological gradient (Precipitation sums)
256
253
219
217
204
9
L1
O8
O1
O16
L8
L16
124

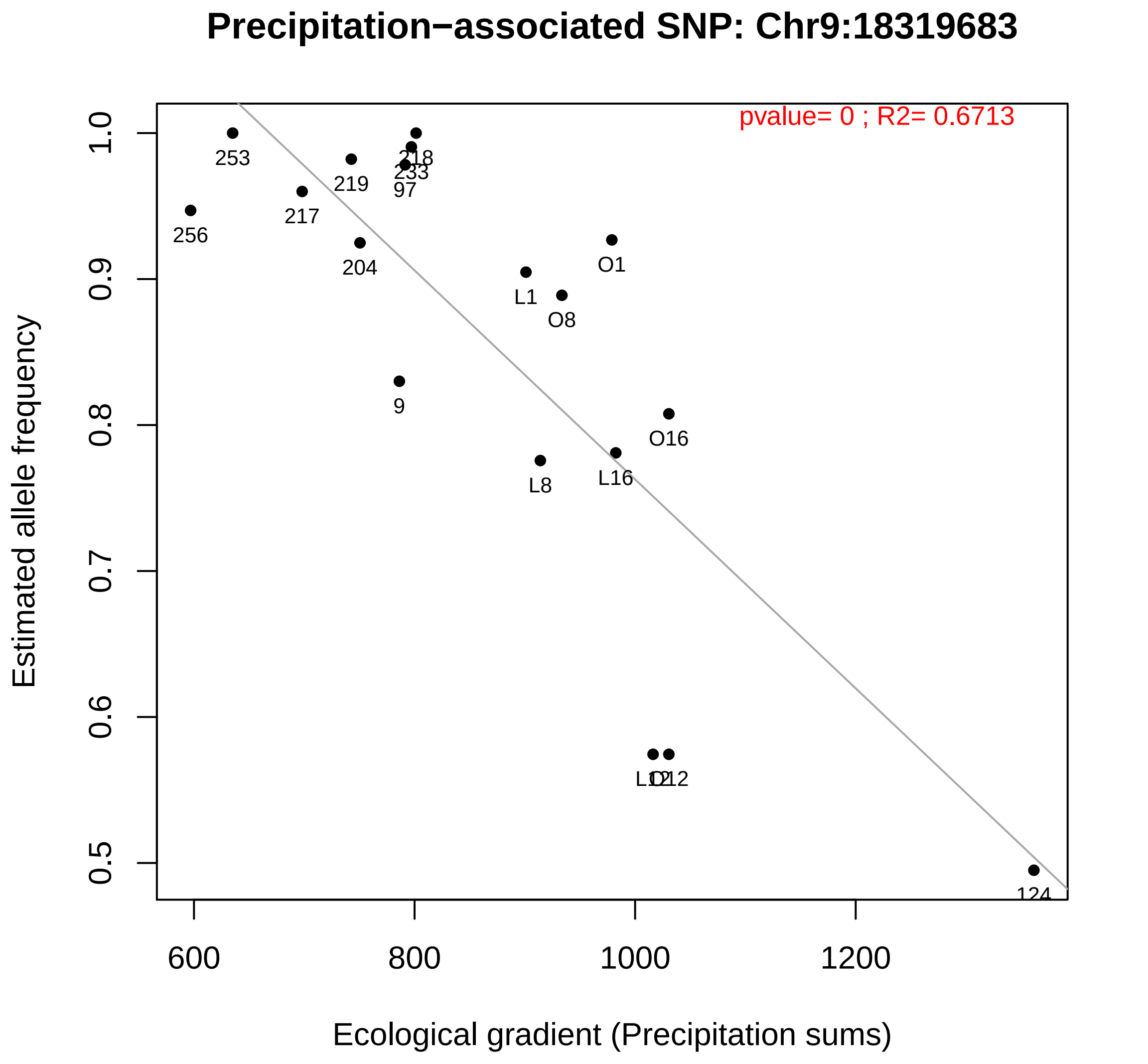

Precipitation−associated SNP: Chr9:18319683
pvalue= 0 ; R2= 0.6713
Estimated allele frequency
Ecological gradient (Precipitation sums)
0.5
0.6
0.7
0.8
0.9
1.0
600
800
1000
1200
253
218
233
97
219
217
256
204
O1
L1
O8
9
O16
L8
L16
L12
O12
124

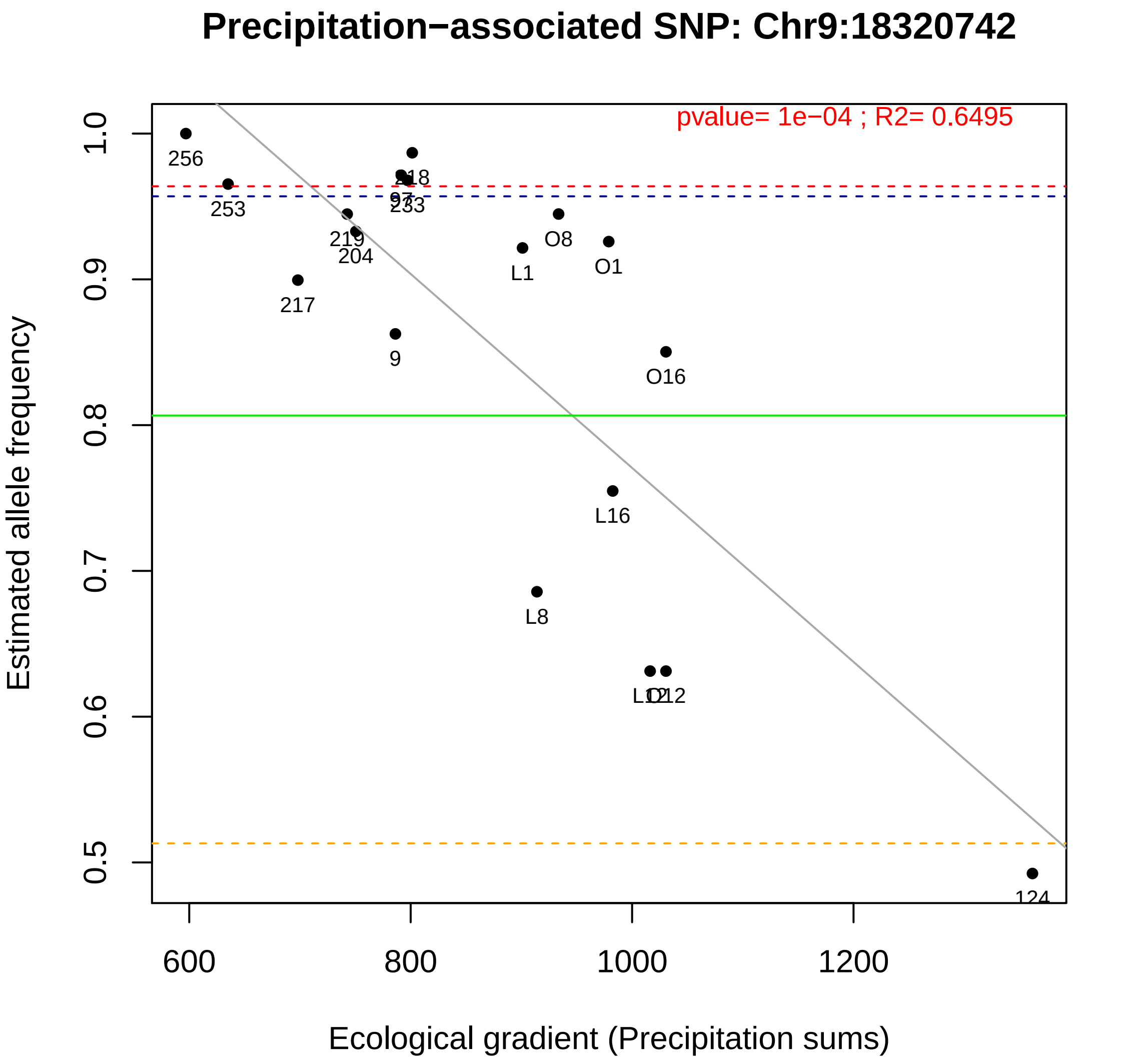

Precipitation−associated SNP: Chr9:18320742
pvalue= 1e−04 ; R2= 0.6495
Estimated allele frequency
Ecological gradient (Precipitation sums)
1.0
0.9
0.8
0.7
0.6
0.5
600
800
1000
1200
256
253
218
97
233
219
204
217
9
L1
O8
O1
O16
L16
L8
L12
O12
124

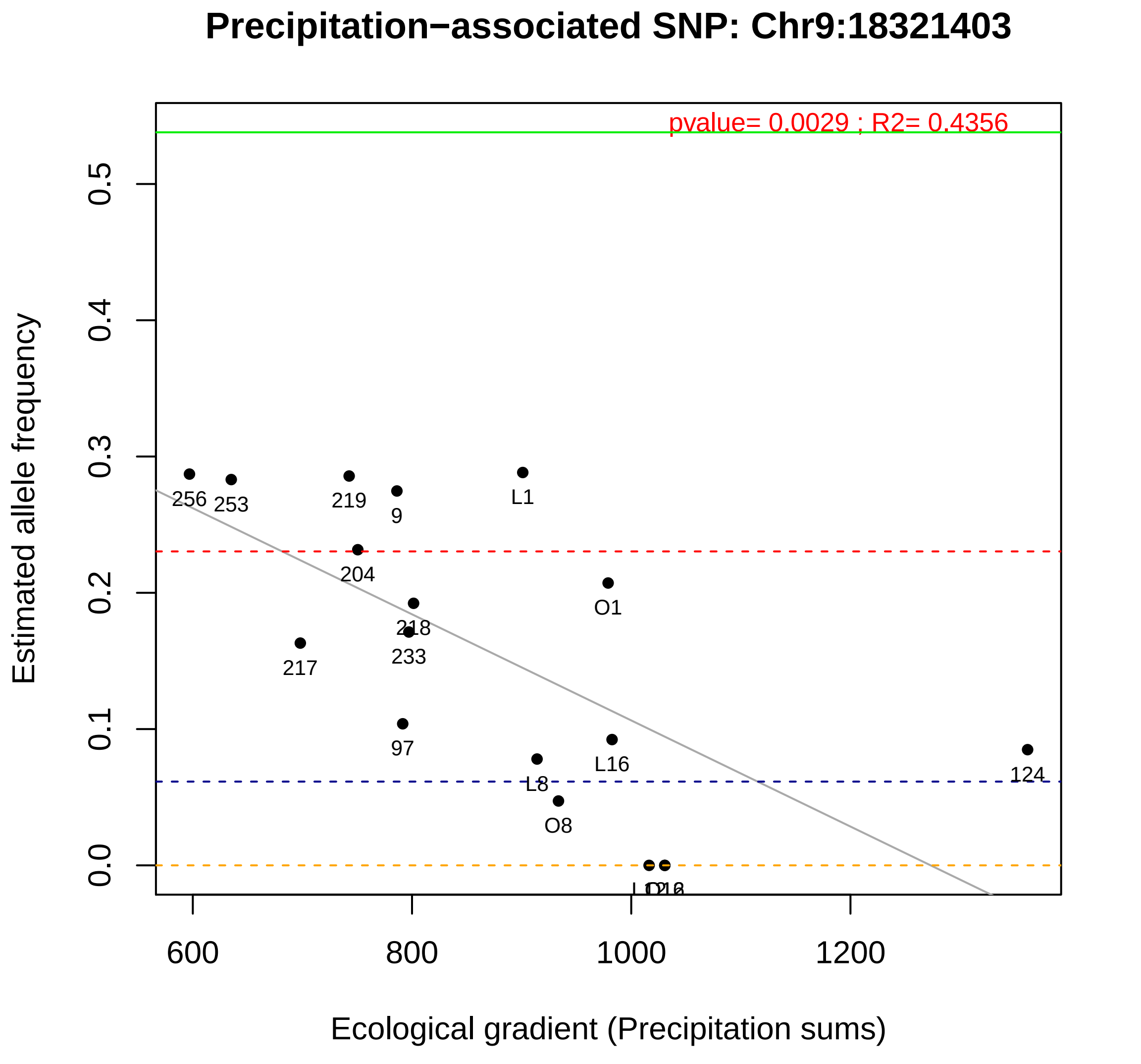
Precipitation−associated SNP: Chr9:18321403
pvalue= 0.0029 ; R2= 0.4356
Estimated allele frequency
0.0
0.1
0.2
0.3
0.4
0.5
600
800
1000
1200
Ecological gradient (Precipitation sums)
256
253
219
9
L1
204
O1
218
233
217
97
L16
124
L8
O8

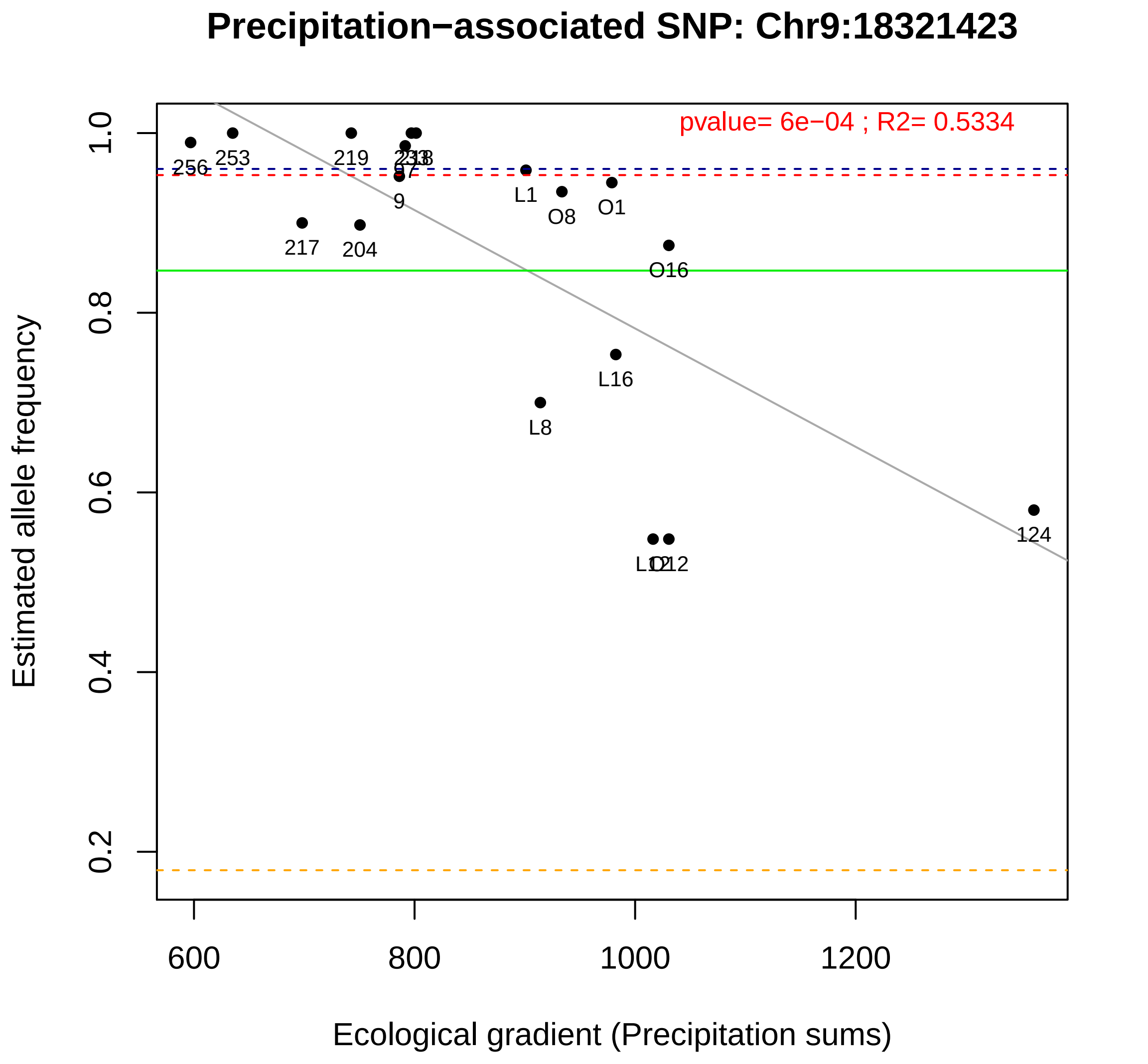

Precipitation−associated SNP: Chr9:18321423
pvalue= 6e−04 ; R2= 0.5334
Estimated allele frequency
Ecological gradient (Precipitation sums)
0.2
0.4
0.6
0.8
1.0
600
800
1000
1200
256
253
219
217
204
9
L1
O8
O1
O16
L16
L8
124

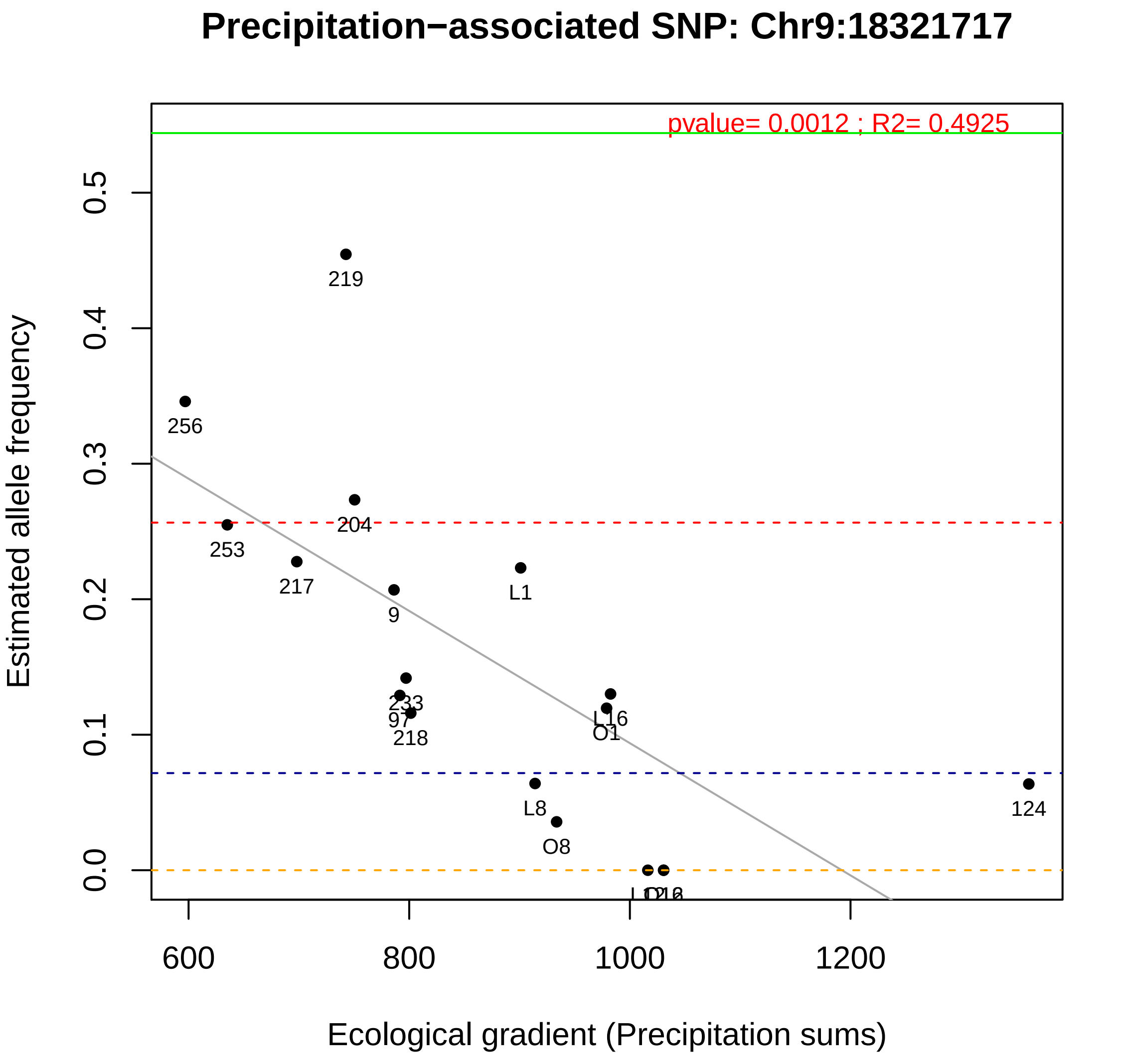

Precipitation−associated SNP: Chr9:18321717
pvalue= 0.0012 ; R2= 0.4925
Estimated allele frequency
Ecological gradient (Precipitation sums)
0.0
0.1
0.2
0.3
0.4
0.5
600
800
1000
1200
219
256
204
253
217
9
L1
233
97
218
L16
O1
L8
O8
124

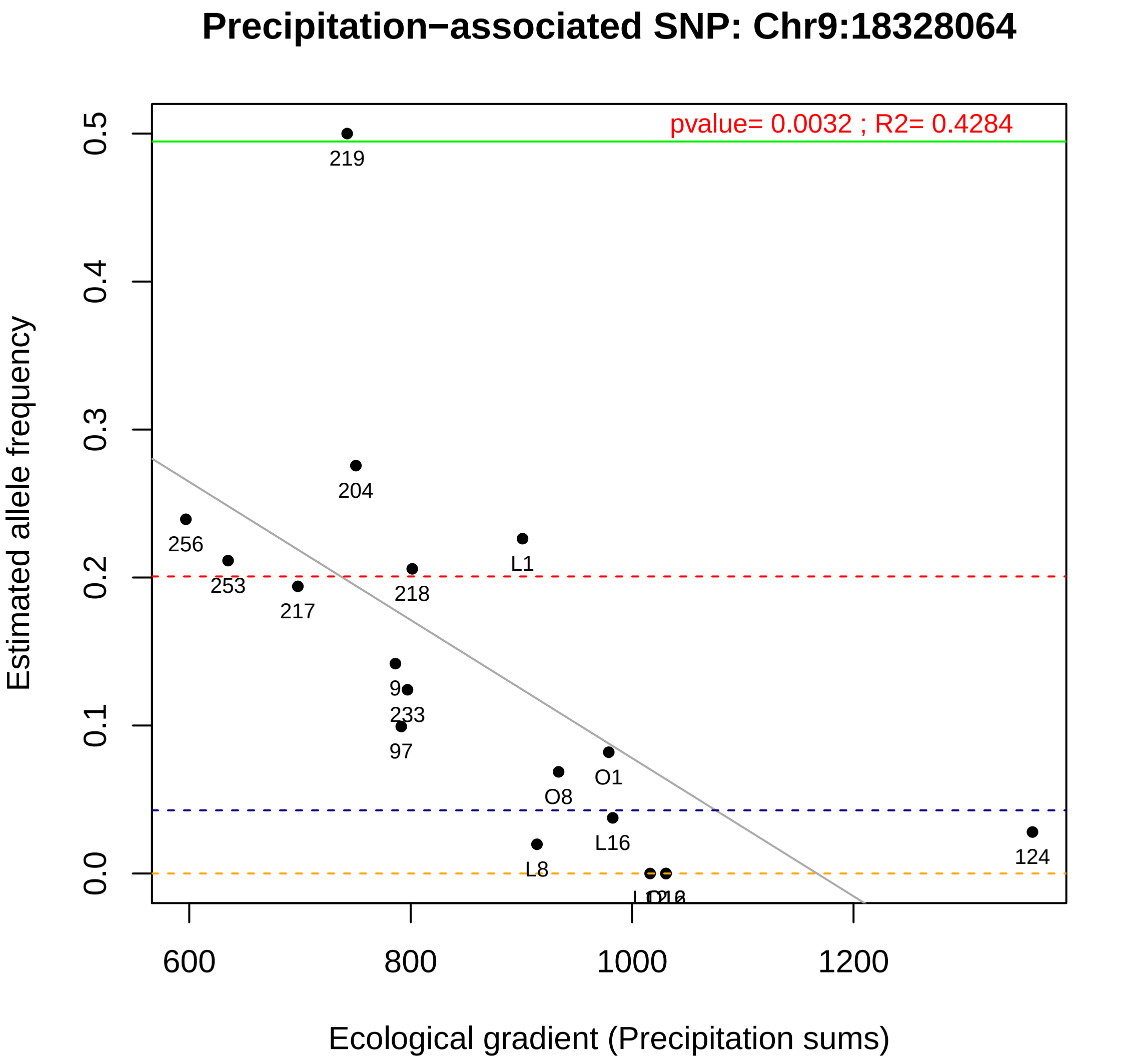
Precipitation−associated SNP: Chr9:18328064
pvalue= 0.0032 ; R2= 0.4284
Estimated allele frequency
0.0
0.1
0.2
0.3
0.4
0.5
Ecological gradient (Precipitation sums)
600
800
1000
1200
219
204
256
253
217
218
L1
9
233
97
O1
O8
L16
L8
124

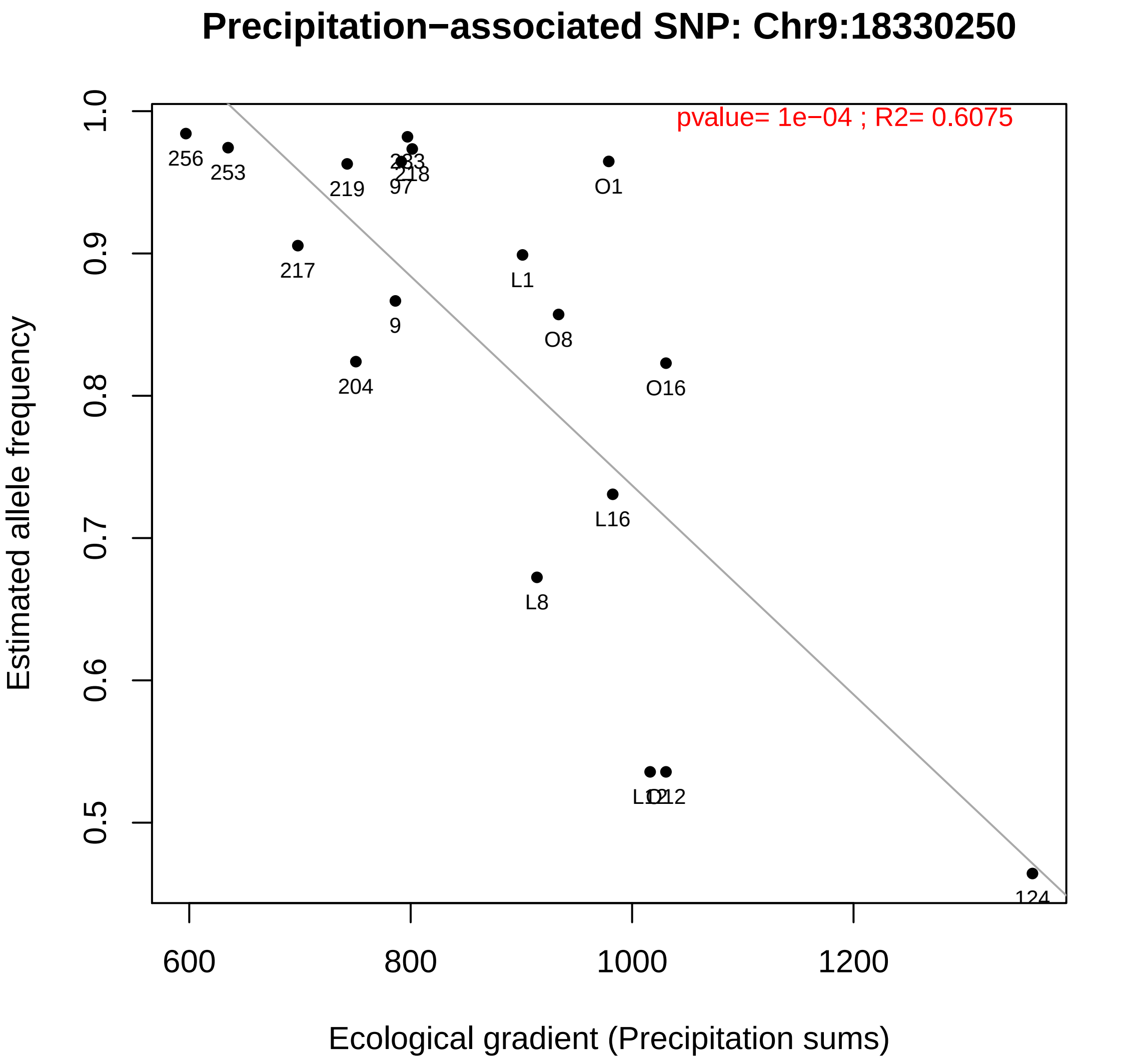
Precipitation−associated SNP: Chr9:18330250
pvalue= 1e−04 ; R2= 0.6075
Estimated allele frequency
Ecological gradient (Precipitation sums)
0.5
0.6
0.7
0.8
0.9
1.0
600
800
1000
1200
256
253
219
233
218
97
O1
217
L1
9
O8
204
O16
L16
L8
L12
O12
124

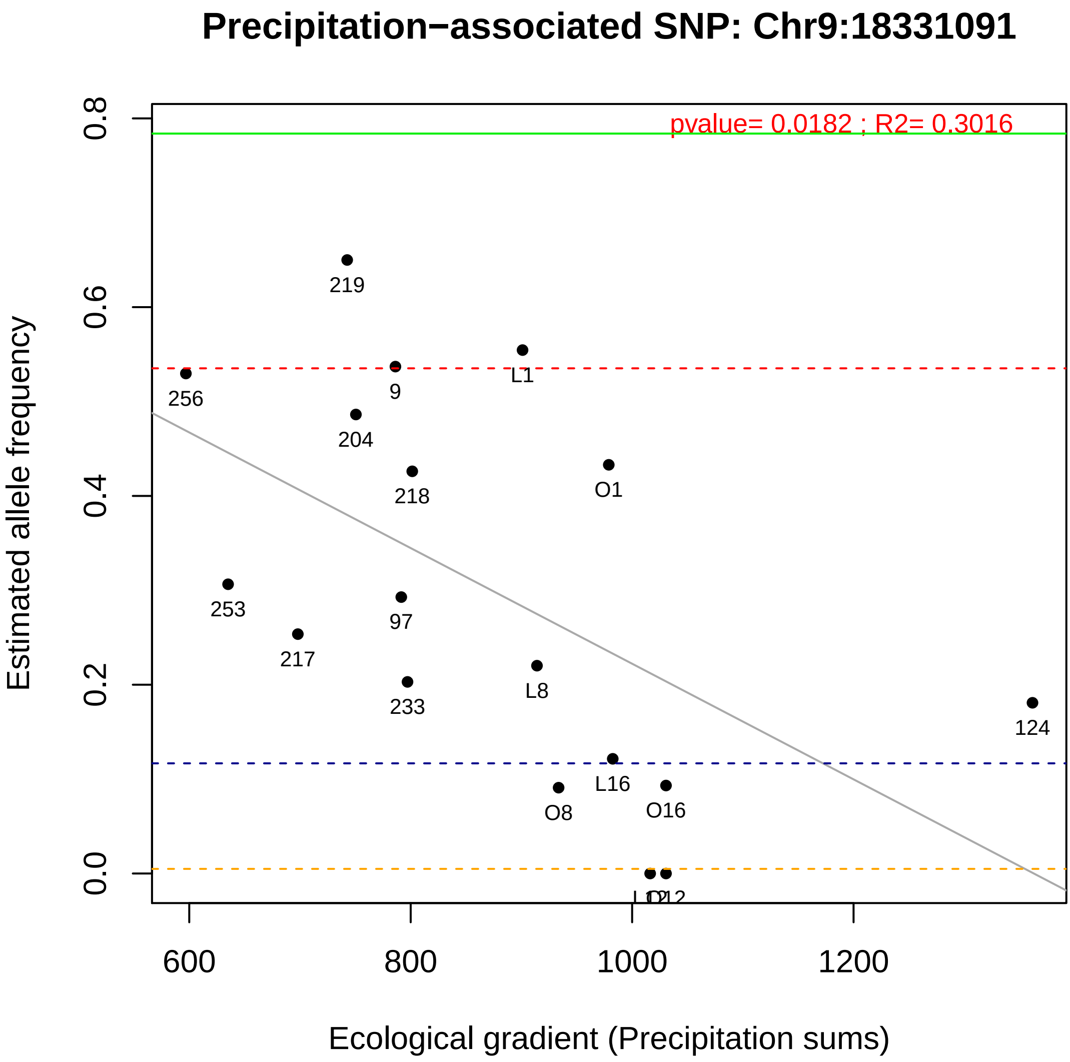
Precipitation−associated SNP: Chr9:18331091
pvalue= 0.0182 ; R2= 0.3016
Estimated allele frequency
0.0
0.2
0.4
0.6
0.8
Ecological gradient (Precipitation sums)
600
800
1000
1200
219
9
L1
256
204
218
O1
253
97
217
233
L8
124
L16
O16
O8

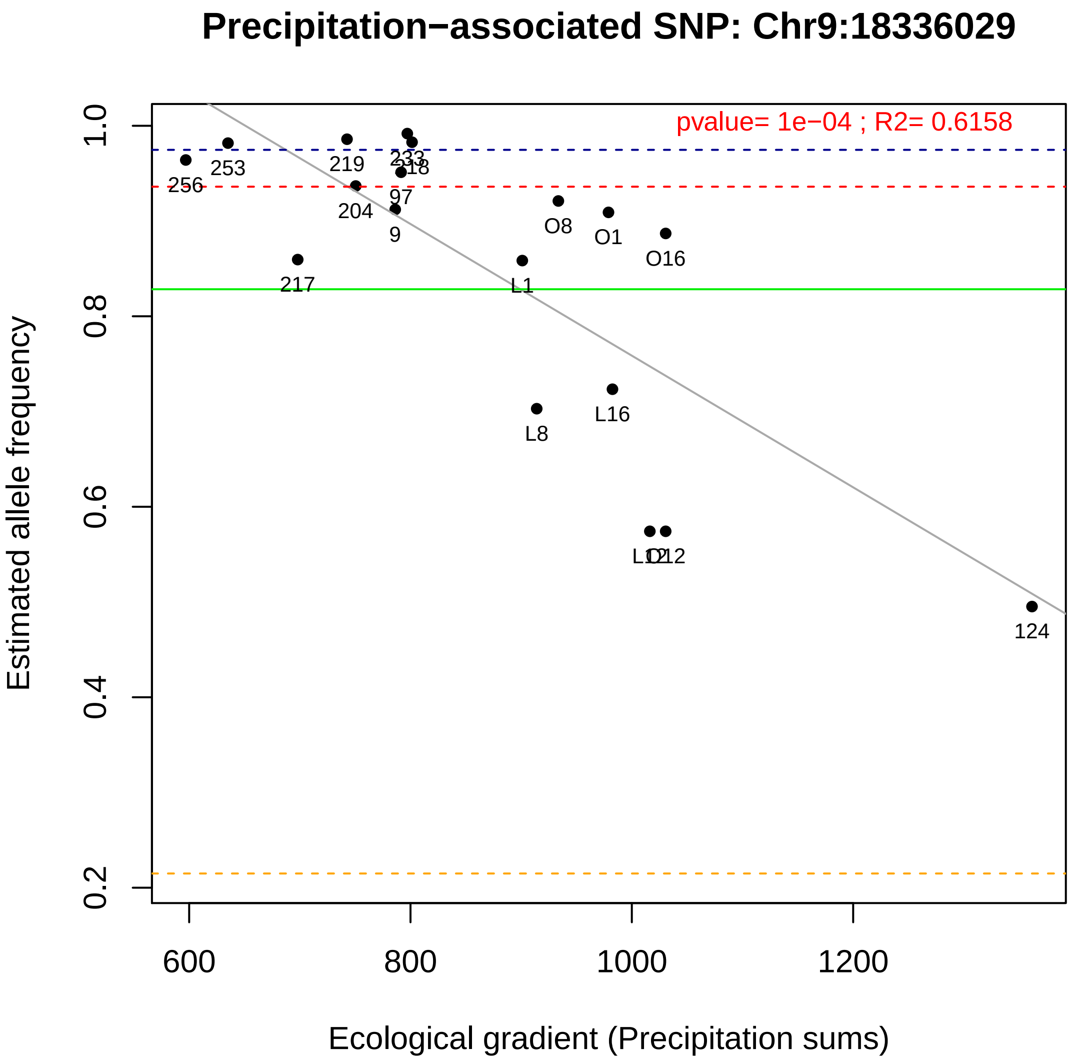

Precipitation−associated SNP: Chr9:18336029
pvalue= 1e−04 ; R2= 0.6158
Estimated allele frequency
Ecological gradient (Precipitation sums)
0.2
0.4
0.6
0.8
1.0
600
800
1000
1200
256
253
219
233
218
97
204
9
217
L1
O8
O1
O16
L8
L16
L12
O12
124

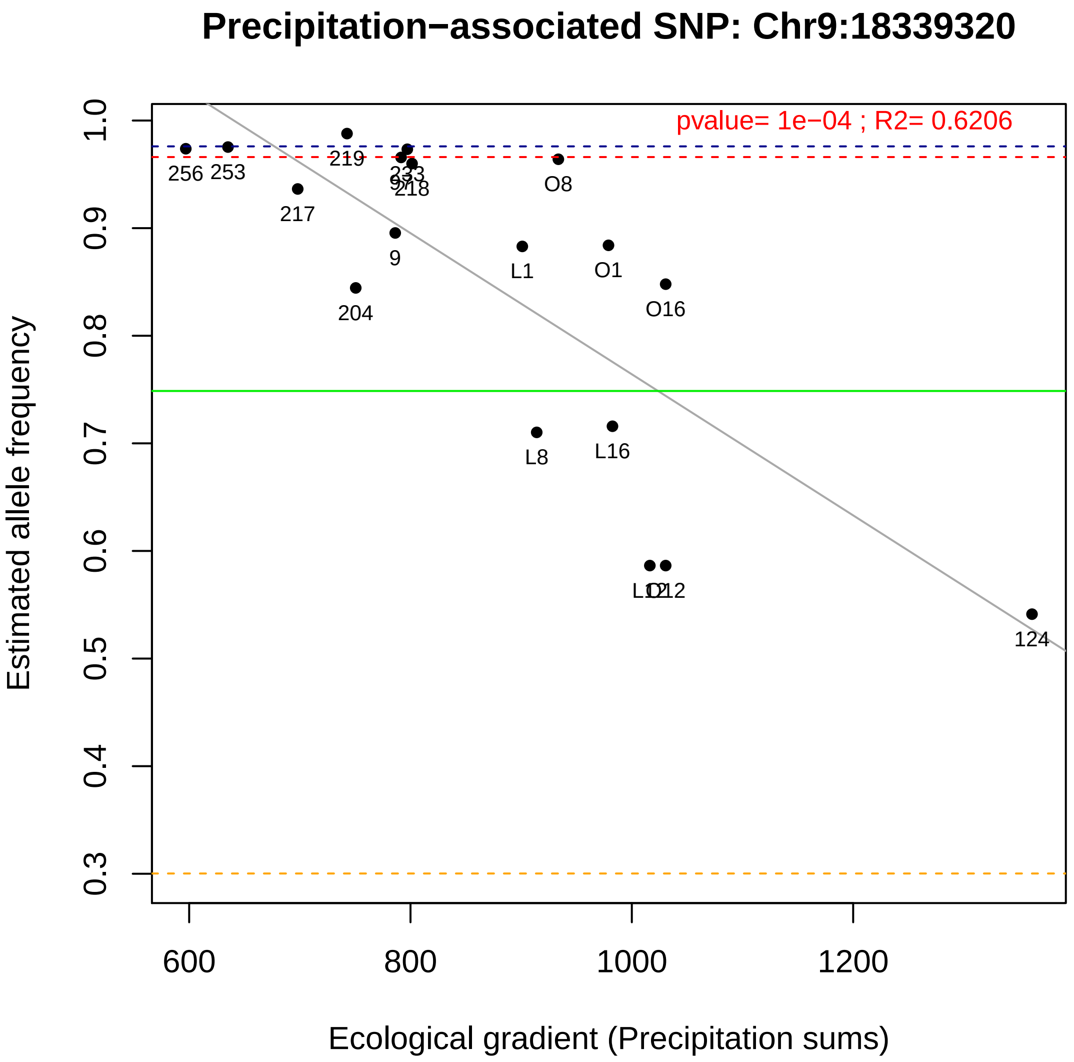
Precipitation−associated SNP: Chr9:18339320
pvalue= 1e−04 ; R2= 0.6206
Estimated allele frequency
0.3
0.4
0.5
0.6
0.7
0.8
0.9
1.0
Ecological gradient (Precipitation sums)
600
800
1000
1200
256
253
219
217
233
218
O8
9
204
L1
O1
O16
L8
L16
124

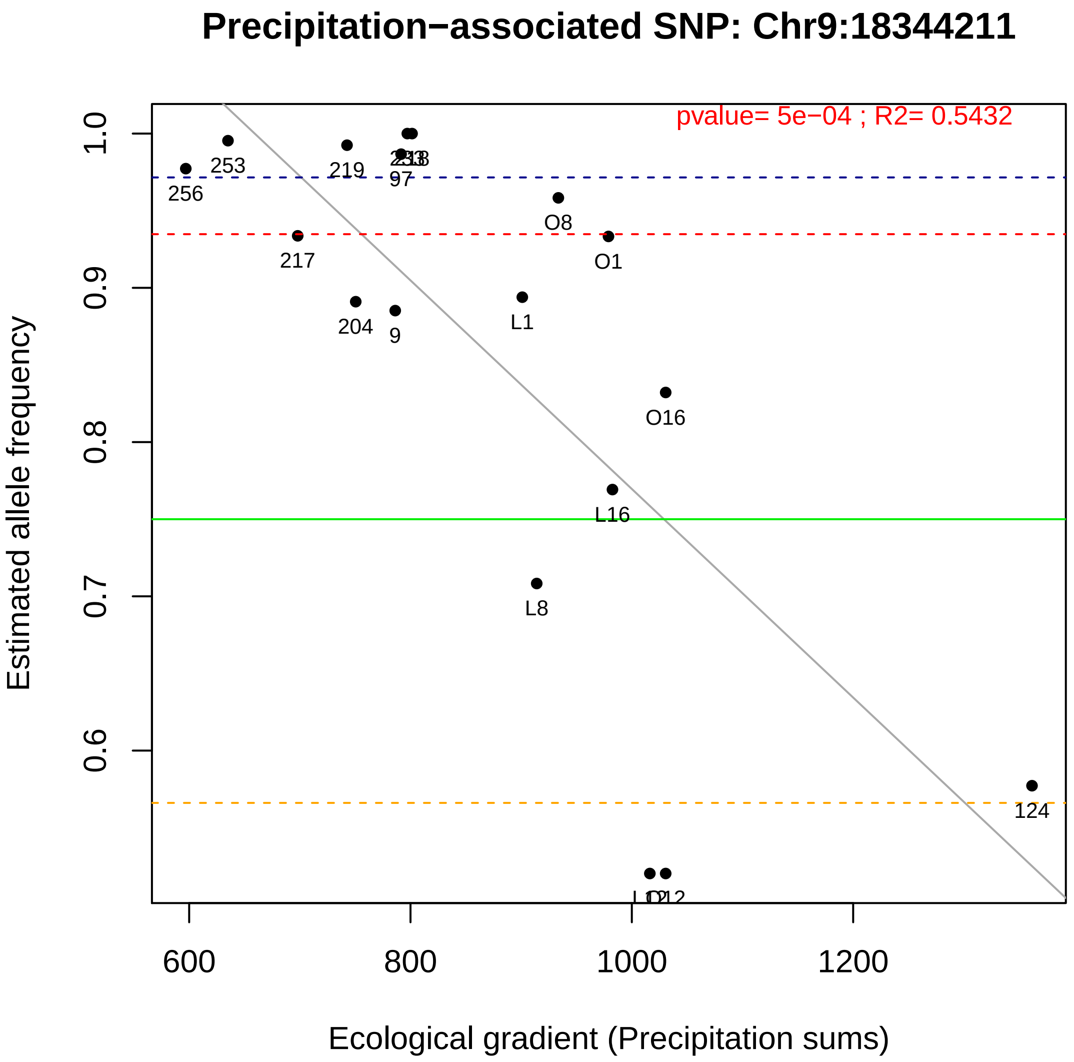
Precipitation−associated SNP: Chr9:18344211
pvalue= 5e−04 ; R2= 0.5432
Estimated allele frequency
1.0
0.9
0.8
0.7
0.6
600
800
1000
1200
Ecological gradient (Precipitation sums)
256
253
219
97
217
204
9
O8
O1
L1
O16
L16
L8
124

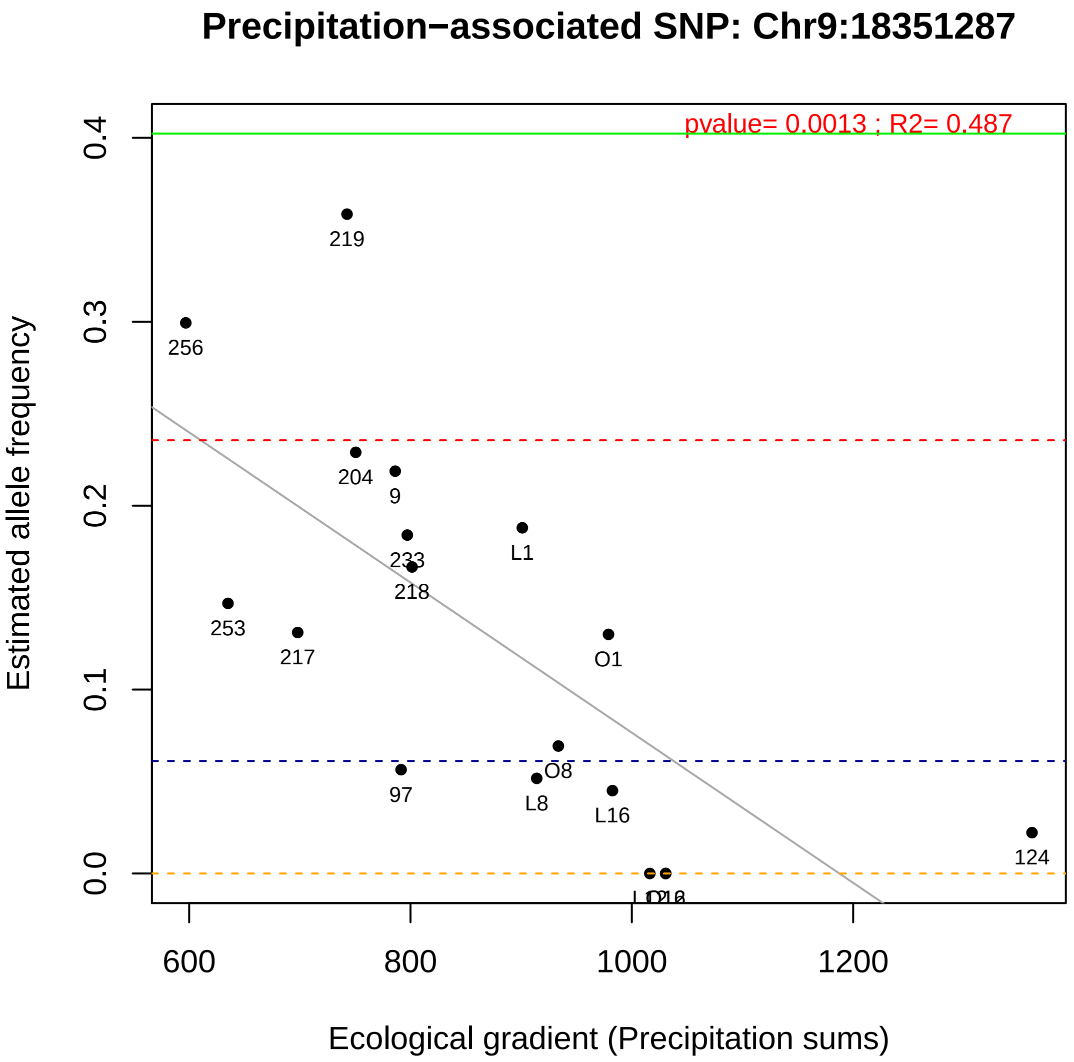
Precipitation−associated SNP: Chr9:18351287
pvalue= 0.0013 ; R2= 0.487
Estimated allele frequency
0.0
0.1
0.2
0.3
0.4
600
800
1000
1200
Ecological gradient (Precipitation sums)
219
256
204
9
233
218
L1
253
217
O1
O8
97
L8
L16
124

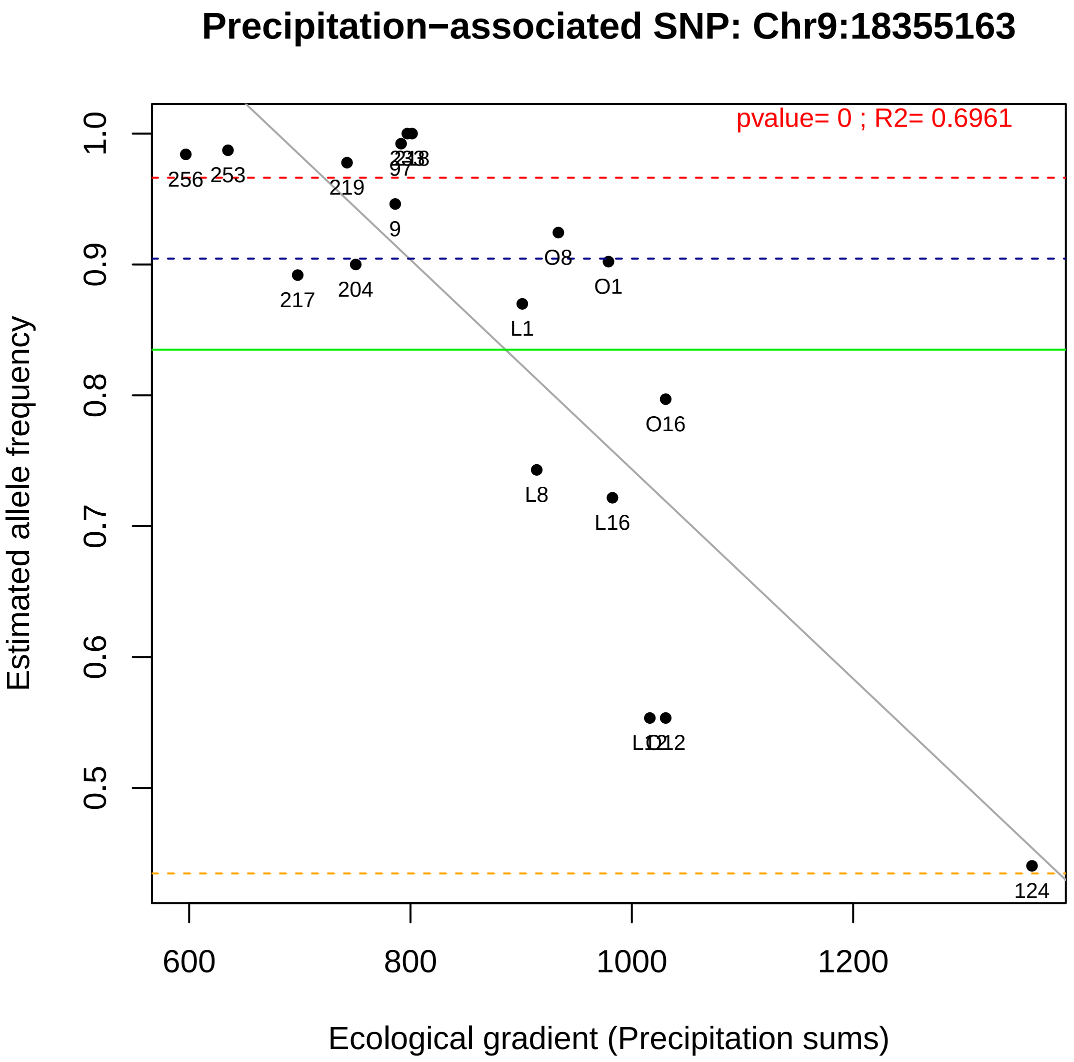

Precipitation−associated SNP: Chr9:18355163
pvalue= 0 ; R2= 0.6961
Estimated allele frequency
Ecological gradient (Precipitation sums)
1.0
0.9
0.8
0.7
0.6
0.5
600
800
1000
1200
256
253
219
97
9
O8
O1
217
204
L1
O16
L8
L16
O12
124

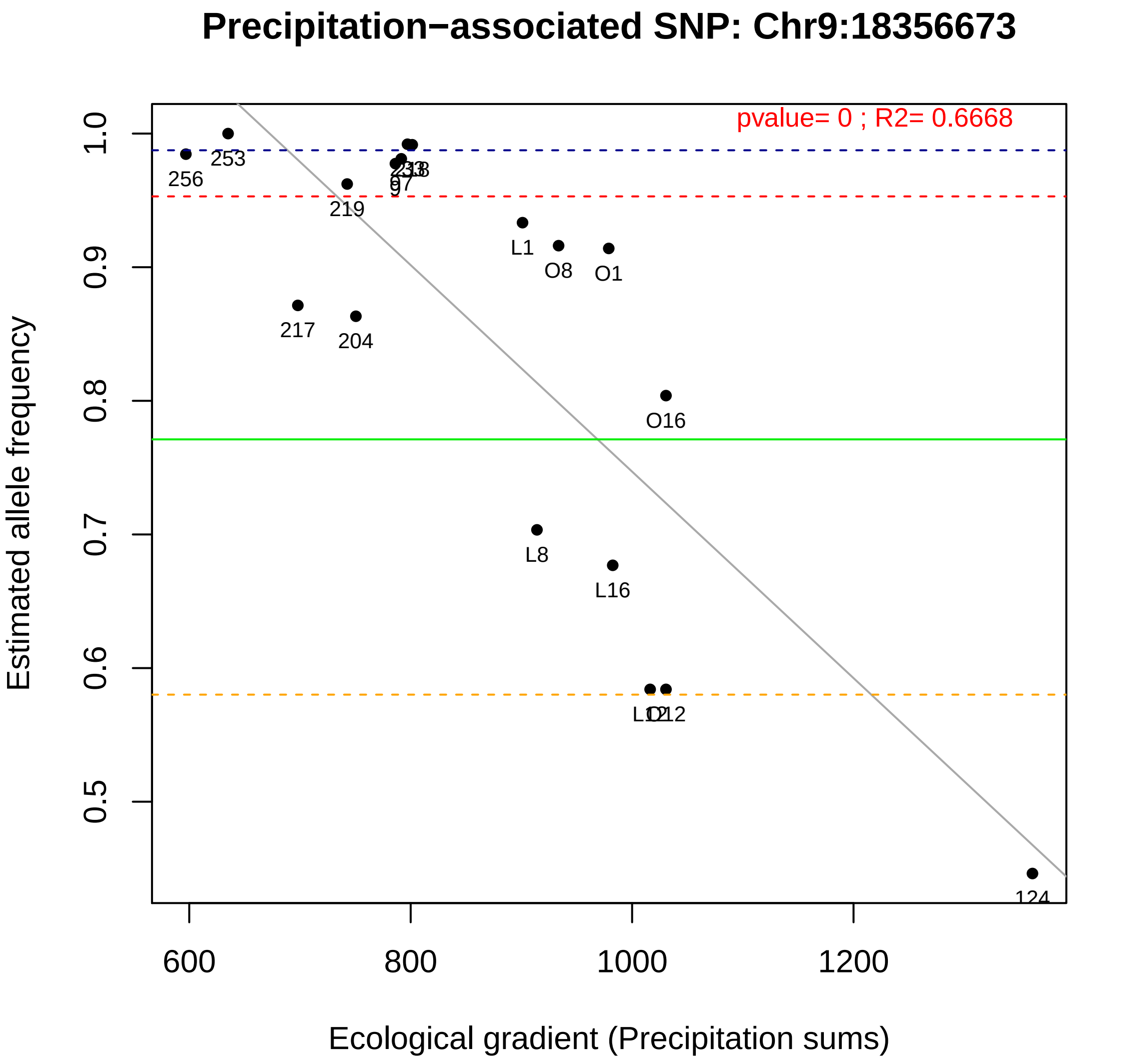
Precipitation−associated SNP: Chr9:18356673
pvalue= 0 ; R2= 0.6668
Estimated allele frequency
Ecological gradient (Precipitation sums)
1.0
0.9
0.8
0.7
0.6
0.5
600
800
1000
1200
256
253
219
217
204
L1
O8
O1
O16
L8
L16
O12
124

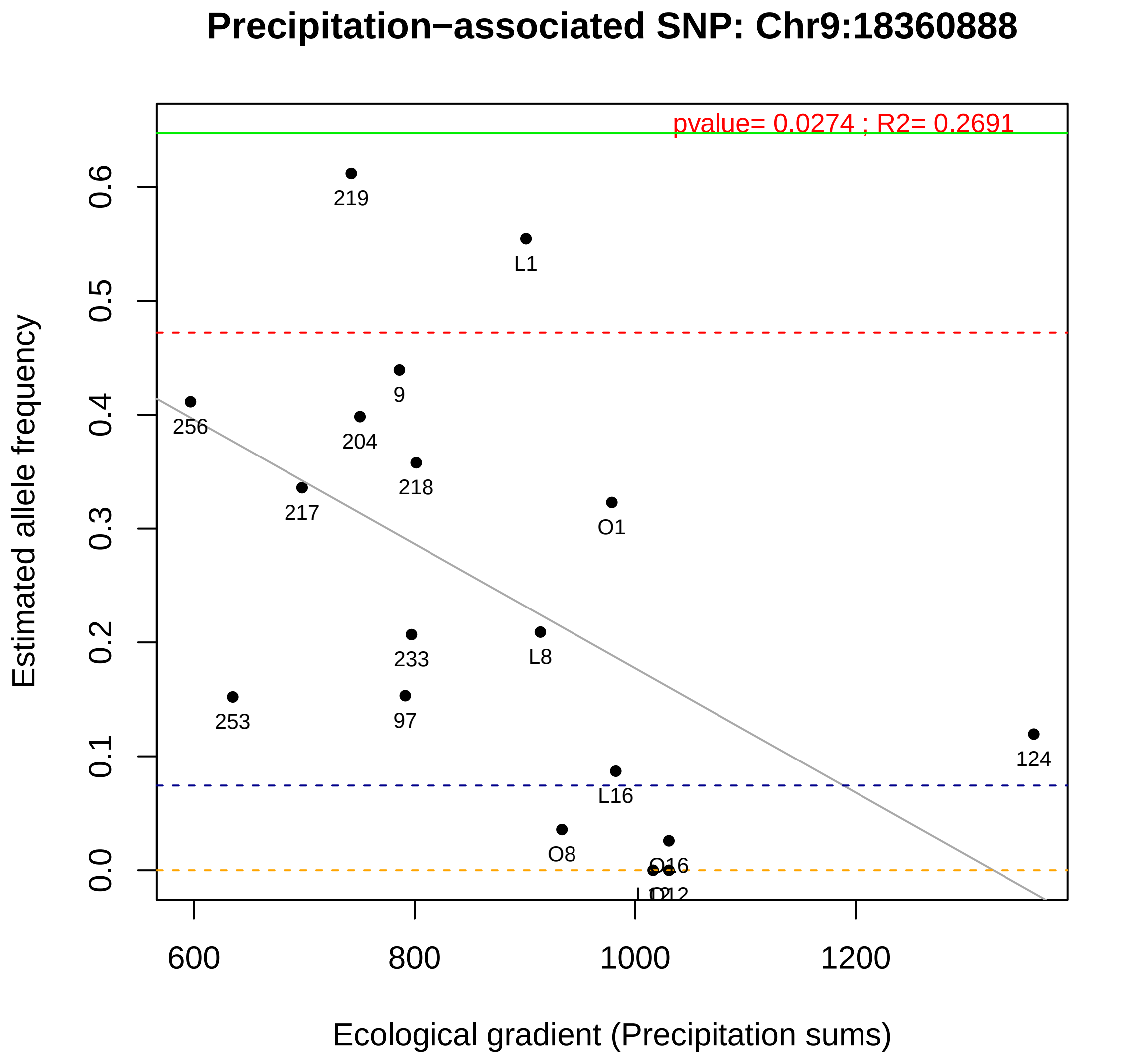

Precipitation−associated SNP: Chr9:18360888
pvalue= 0.0274 ; R2= 0.2691
Estimated allele frequency
Ecological gradient (Precipitation sums)
0.0
0.1
0.2
0.3
0.4
0.5
0.6
600
800
1000
1200
219
L1
9
256
204
218
217
O1
233
L8
253
97
124
L16
O8
O16

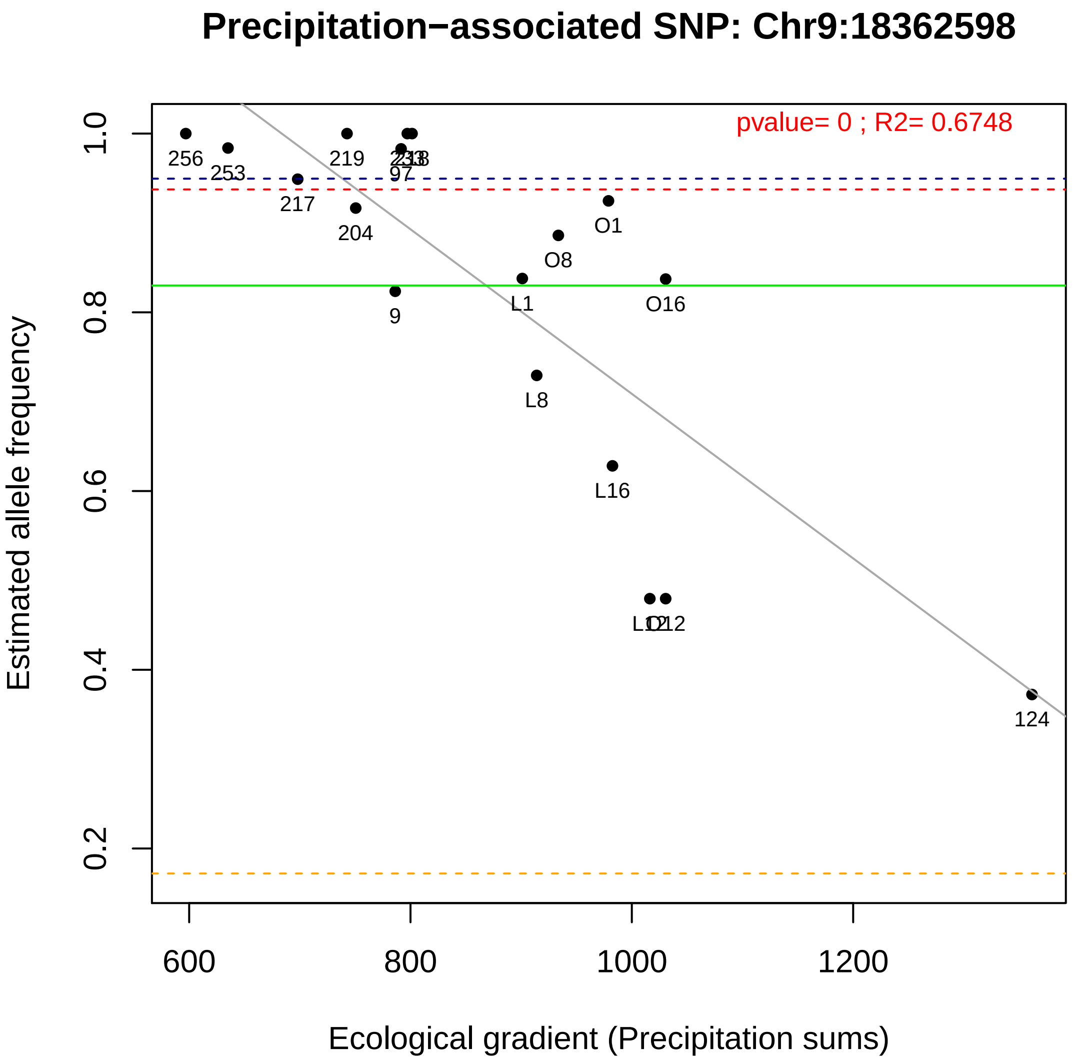
Precipitation−associated SNP: Chr9:18362598
pvalue= 0 ; R2= 0.6748
Estimated allele frequency
Ecological gradient (Precipitation sums)
0.2
0.4
0.6
0.8
1.0
600
800
1000
1200
256
253
219
217
204
97
9
L1
O8
O1
O16
L8
L16
124

**Precipitation−associated SNP: Chr9:18362798**

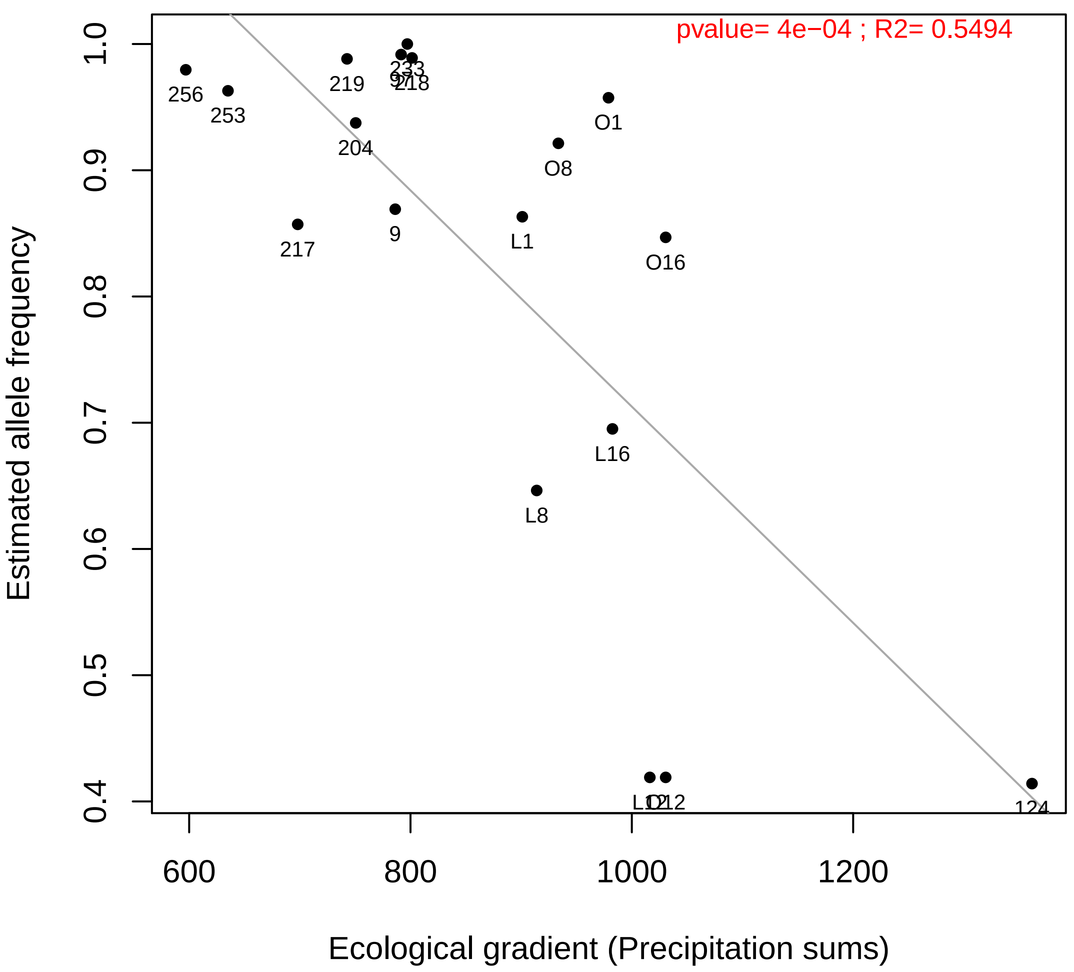

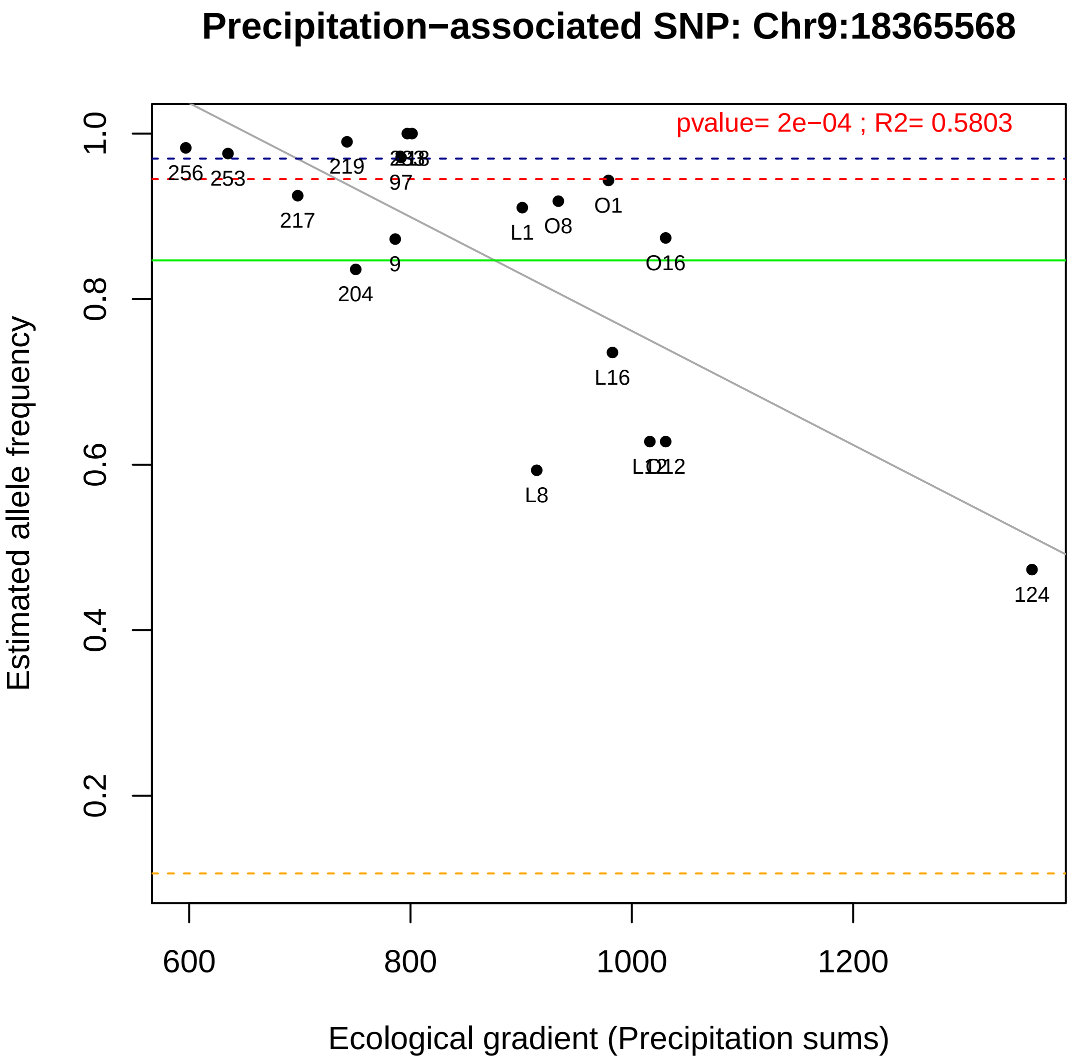
Precipitation−associated SNP: Chr9:18365568
pvalue= 2e−04 ; R2= 0.5803
Estimated allele frequency
Ecological gradient (Precipitation sums)
0.2
0.4
0.6
0.8
1.0
600
800
1000
1200
256
253
219
217
97
9
204
L1
O8
O1
O16
L16
L8
124

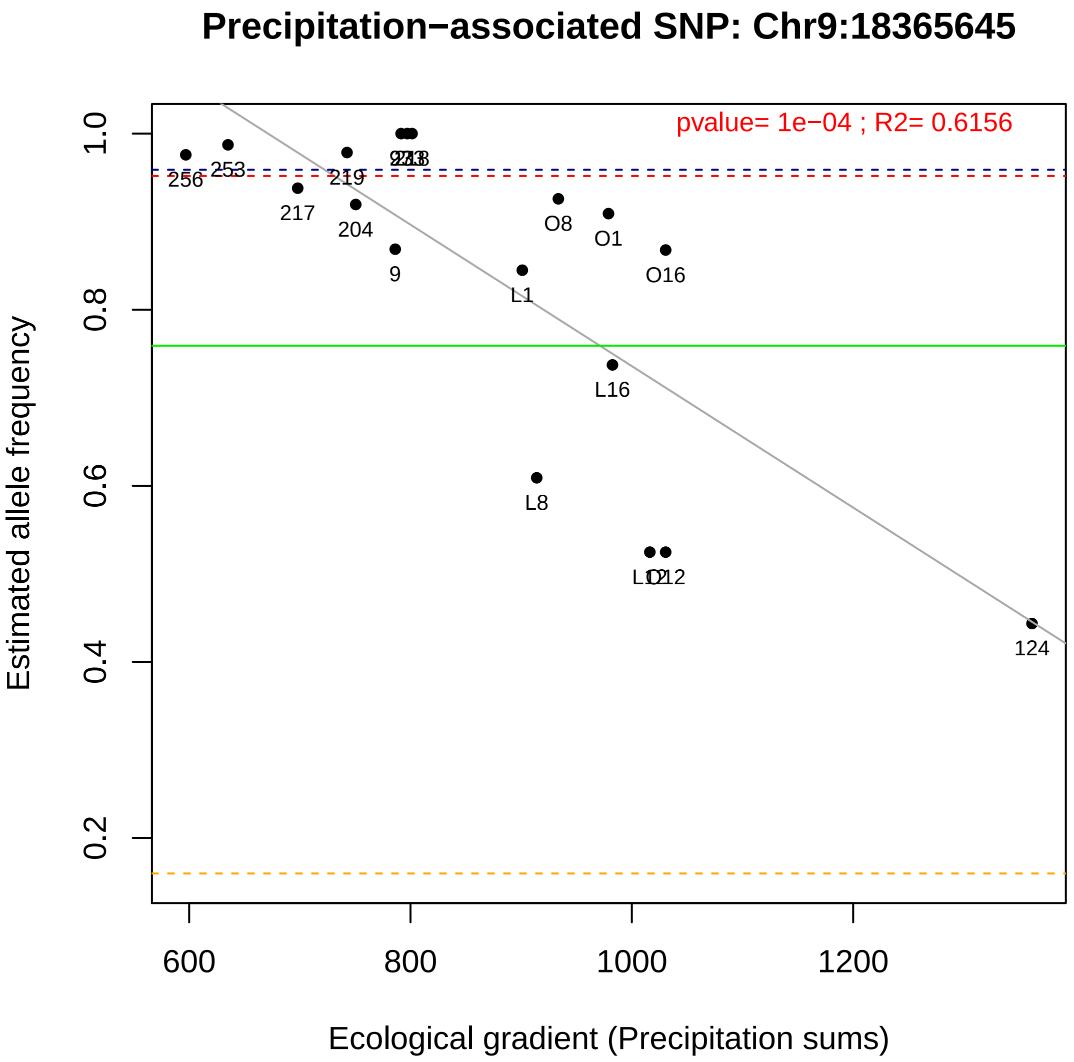
Precipitation−associated SNP: Chr9:18365645
pvalue= 1e−04 ; R2= 0.6156
Estimated allele frequency
Ecological gradient (Precipitation sums)
0.2
0.4
0.6
0.8
1.0
600
800
1000
1200
256
253
219
217
204
9
O8
O1
O16
L1
L16
L8
L12
O12
124

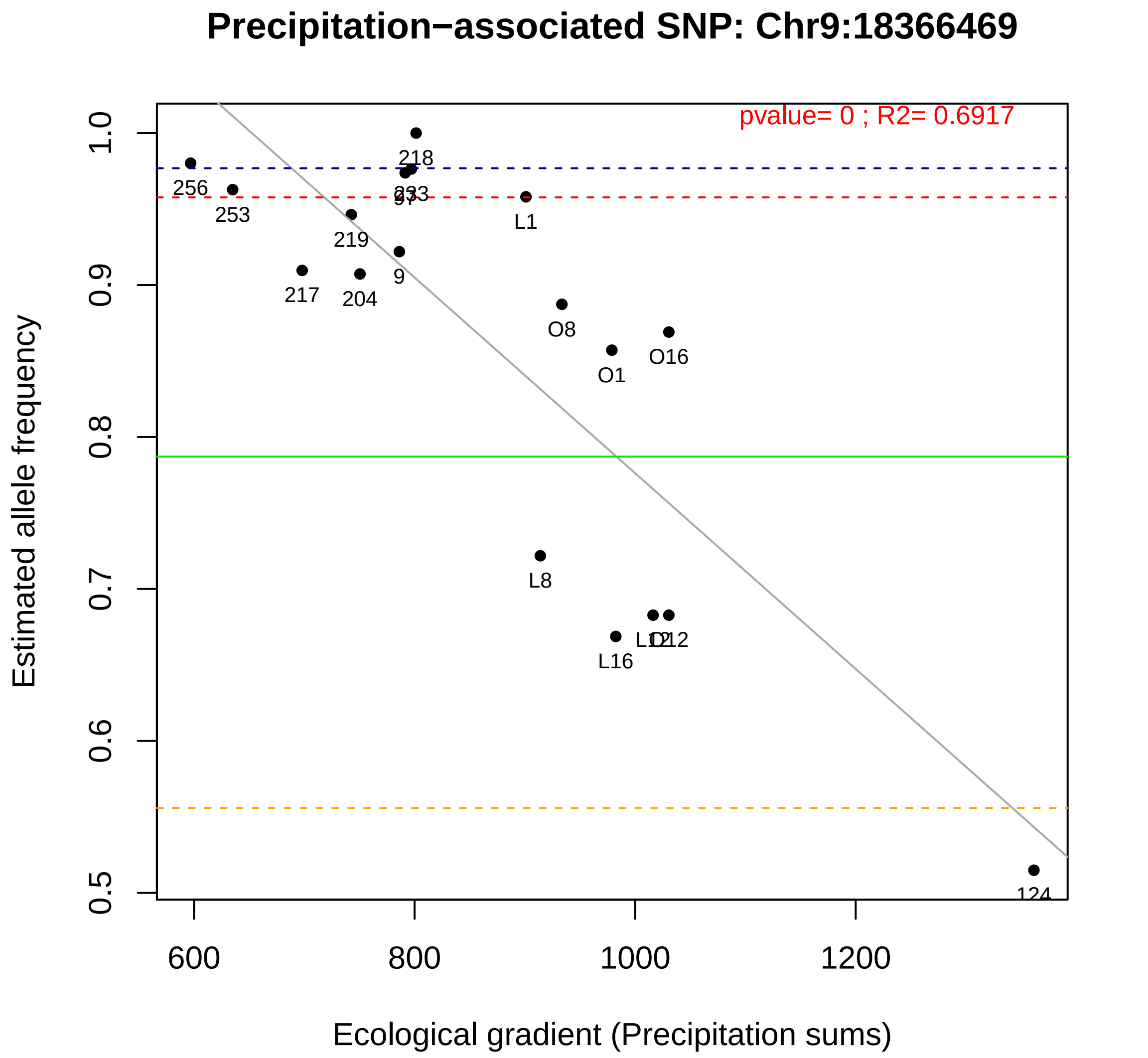

Precipitation−associated SNP: Chr9:18366469
pvalue= 0 ; R2= 0.6917
Estimated allele frequency
Ecological gradient (Precipitation sums)
0.5
0.6
0.7
0.8
0.9
1.0
600
800
1000
1200
256
253
218
219
217
204
9
L1
O8
O1
O16
L8
L16
124

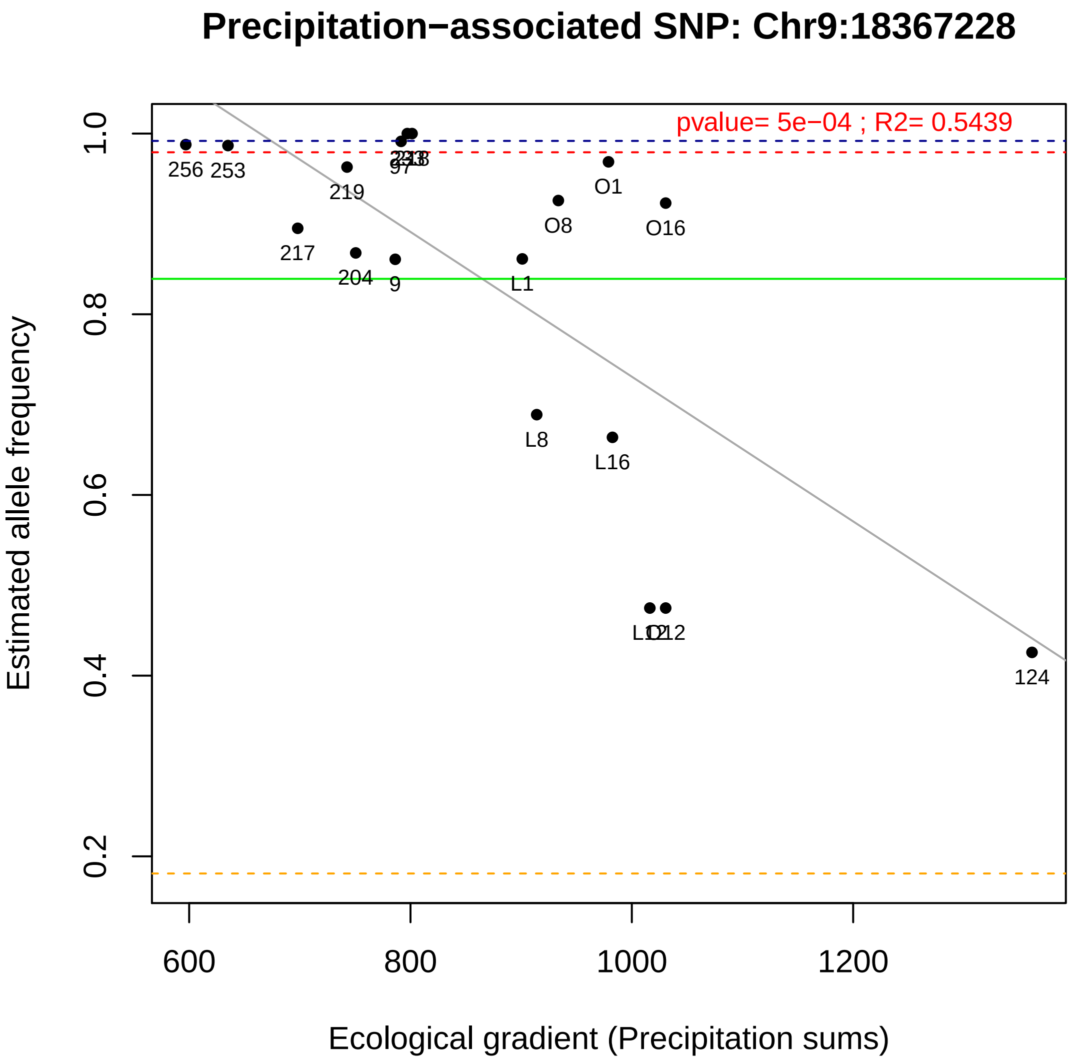

Precipitation−associated SNP: Chr9:18367228
pvalue= 5e−04 ; R2= 0.5439
Estimated allele frequency
Ecological gradient (Precipitation sums)
0.2
0.4
0.6
0.8
1.0
600
800
1000
1200
256
253
219
217
204
9
L1
O8
O1
O16
L8
L16
124

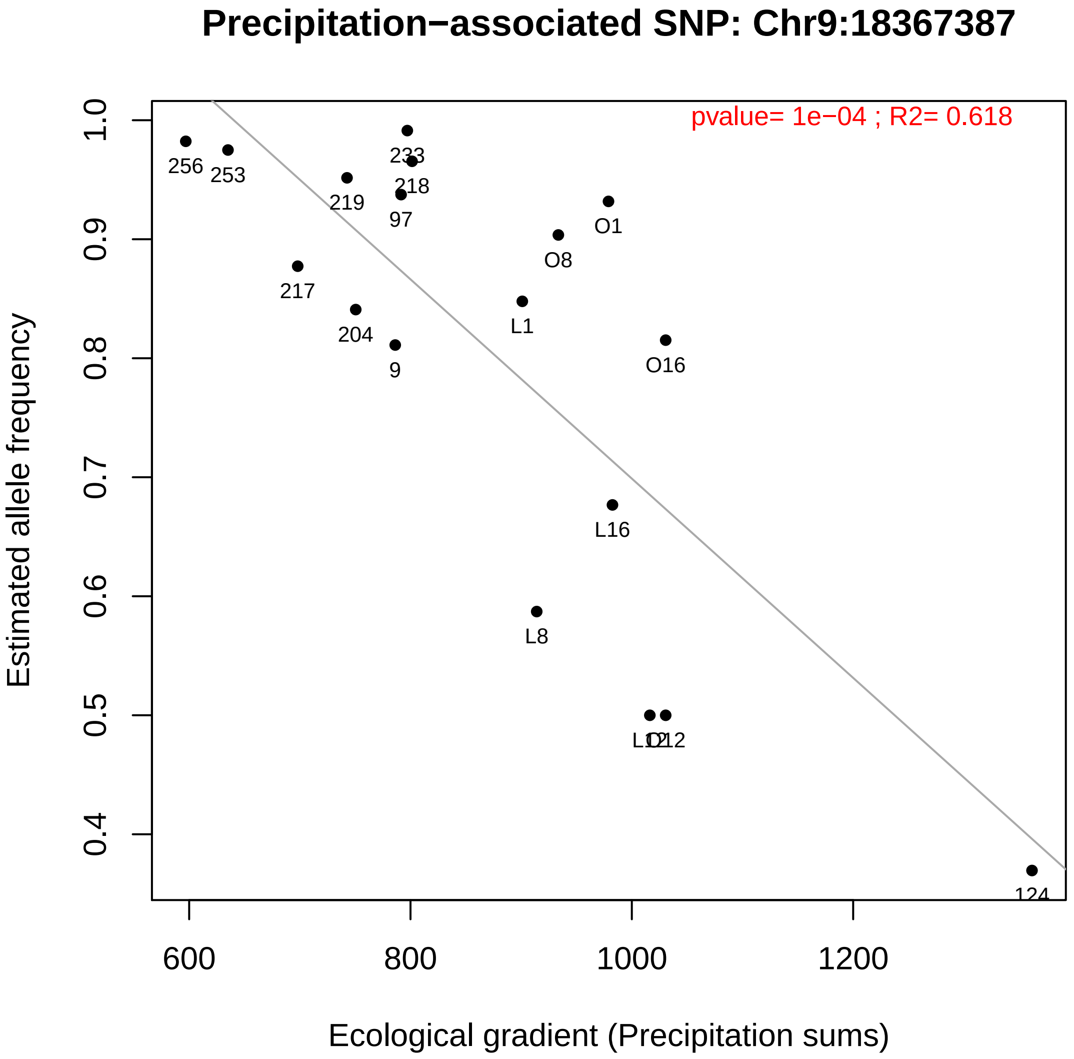
Precipitation−associated SNP: Chr9:18367387
pvalue= 1e−04 ; R2= 0.618
Estimated allele frequency
Ecological gradient (Precipitation sums)
0.4
0.5
0.6
0.7
0.8
0.9
1.0
600
800
1000
1200
256
253
233
218
219
97
O1
O8
217
204
L1
9
O16
L16
L8
L10
O12
124

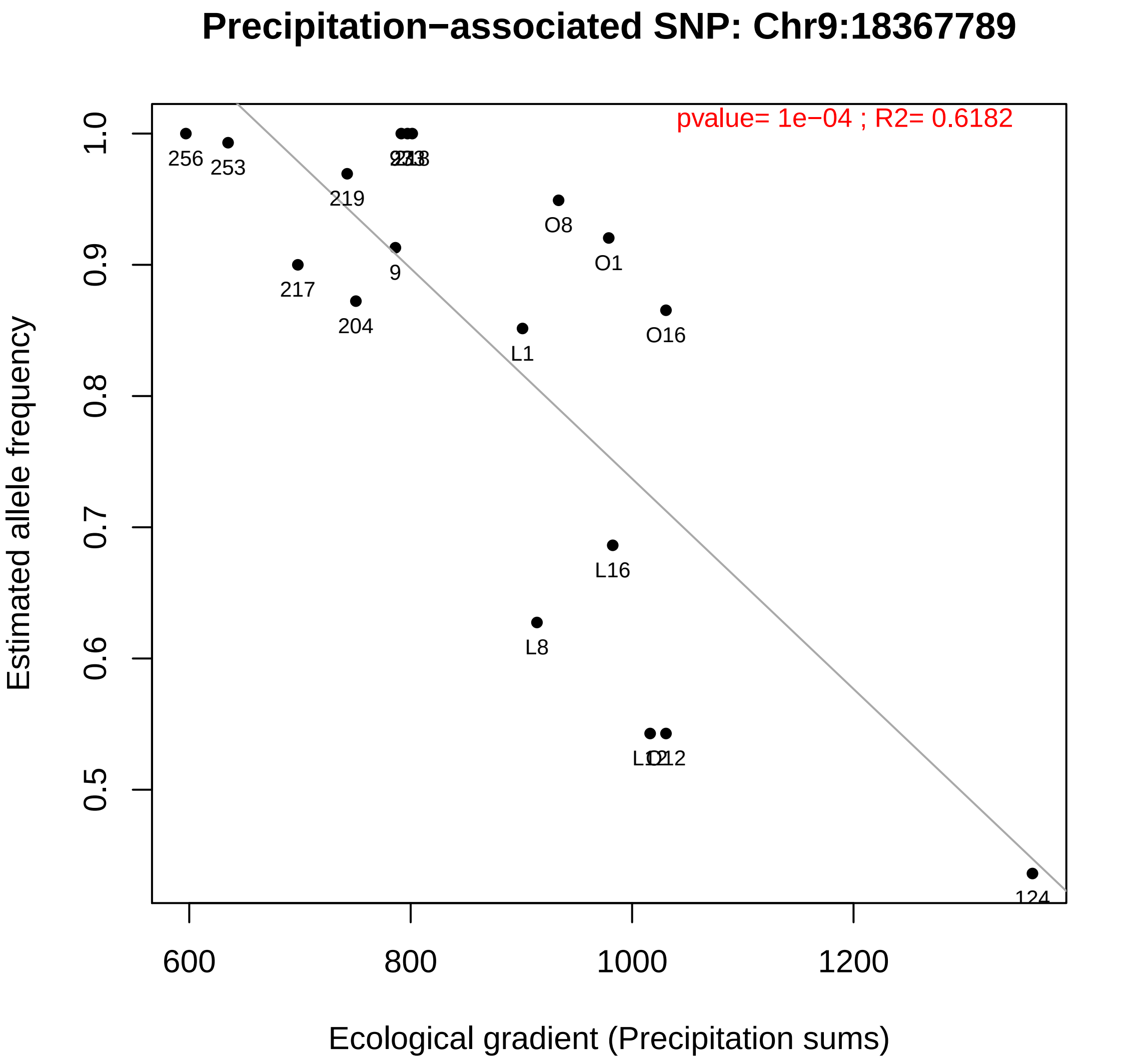

Precipitation−associated SNP: Chr9:18367789
pvalue= 1e−04 ; R2= 0.6182
Estimated allele frequency
Ecological gradient (Precipitation sums)
1.0
0.9
0.8
0.7
0.6
0.5
600
800
1000
1200
256
253
219
217
204
9
O8
O1
O16
L1
L16
L8
124

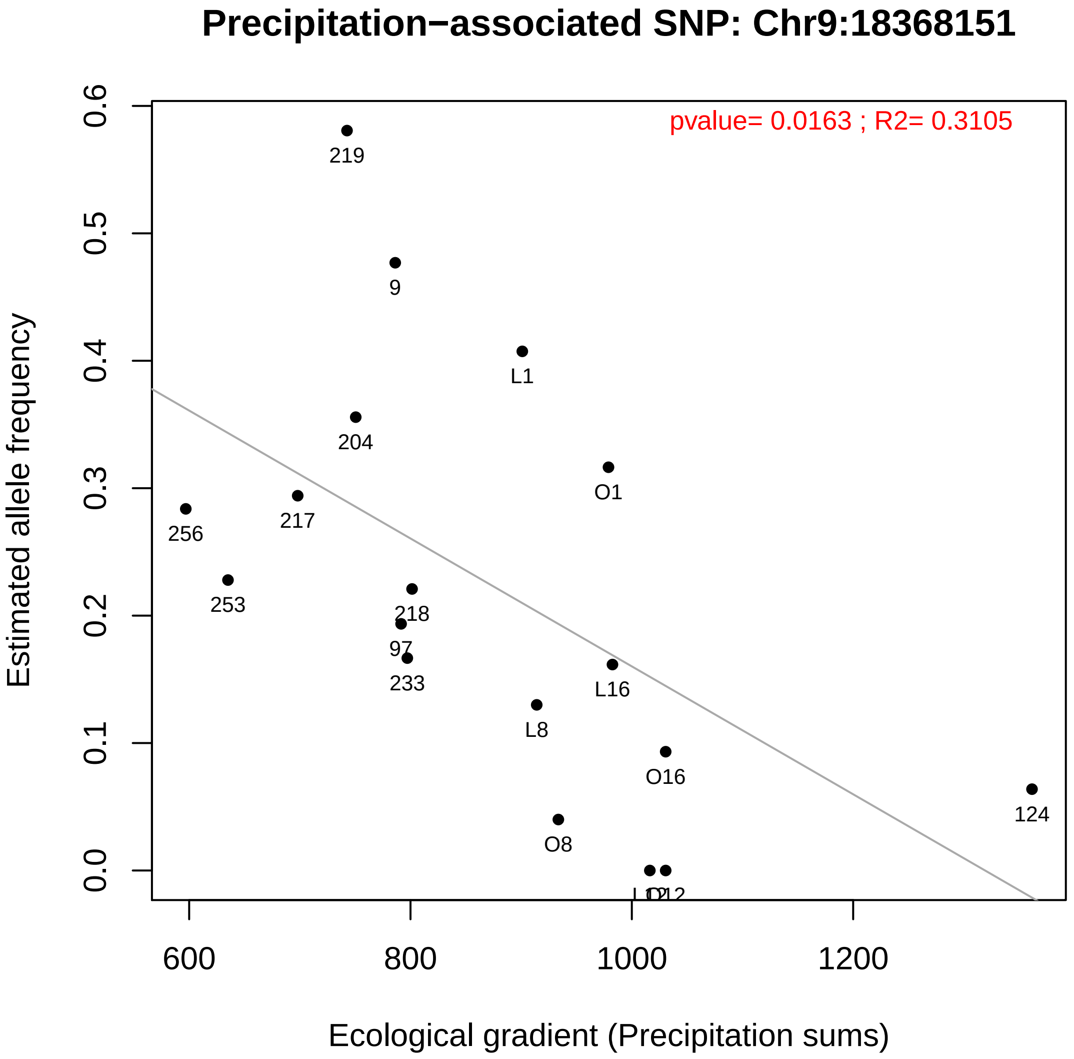
Precipitation−associated SNP: Chr9:18368151
pvalue= 0.0163 ; R2= 0.3105
Estimated allele frequency
0.0
0.1
0.2
0.3
0.4
0.5
0.6
Ecological gradient (Precipitation sums)
600
800
1000
1200
219
9
L1
204
O1
256
217
253
218
97
233
L16
L8
O16
124
O8

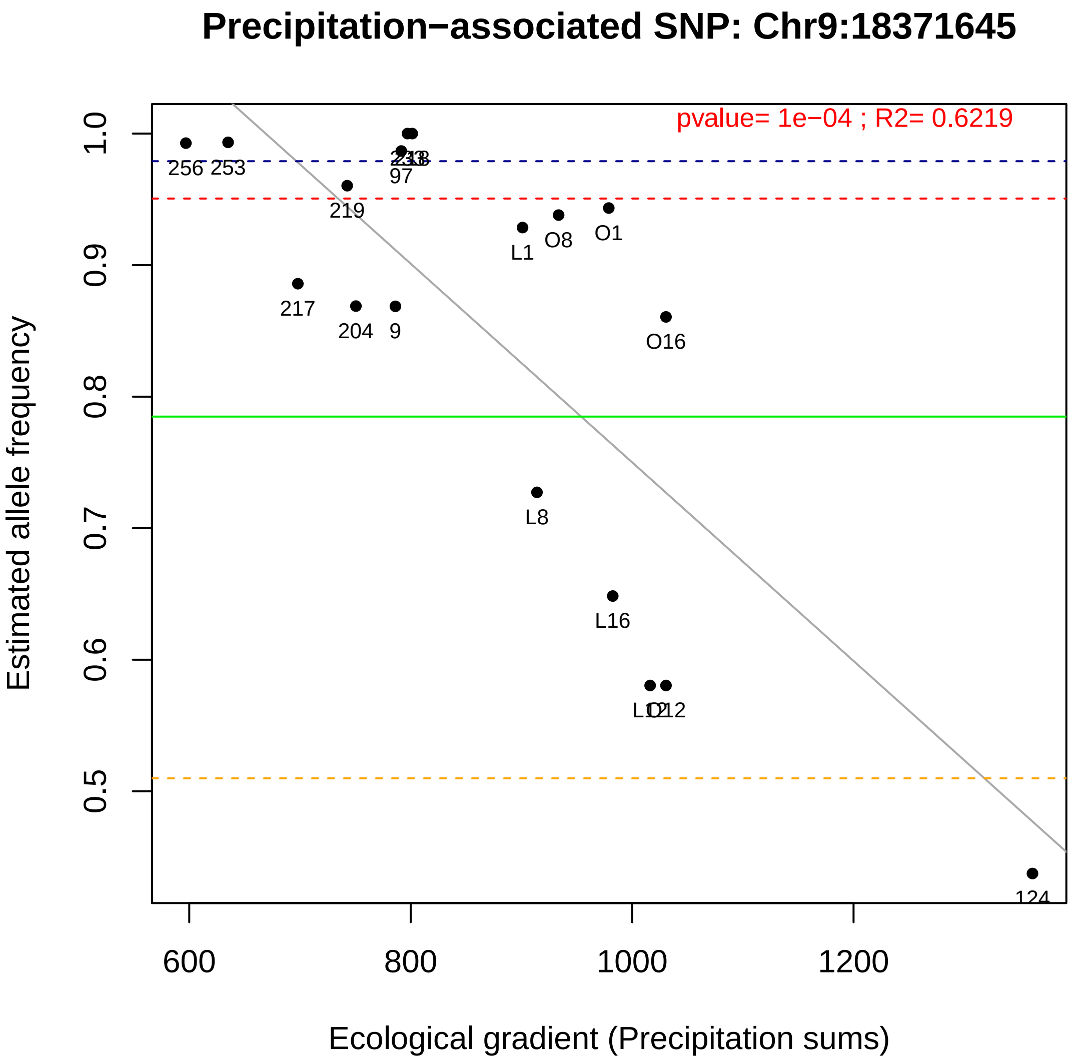
Precipitation−associated SNP: Chr9:18371645
pvalue= 1e−04 ; R2= 0.6219
Estimated allele frequency
Ecological gradient (Precipitation sums)
0.5
0.6
0.7
0.8
0.9
1.0
600
800
1000
1200
256
253
97
219
L1
O8
O1
217
204
9
O16
L8
L16
124

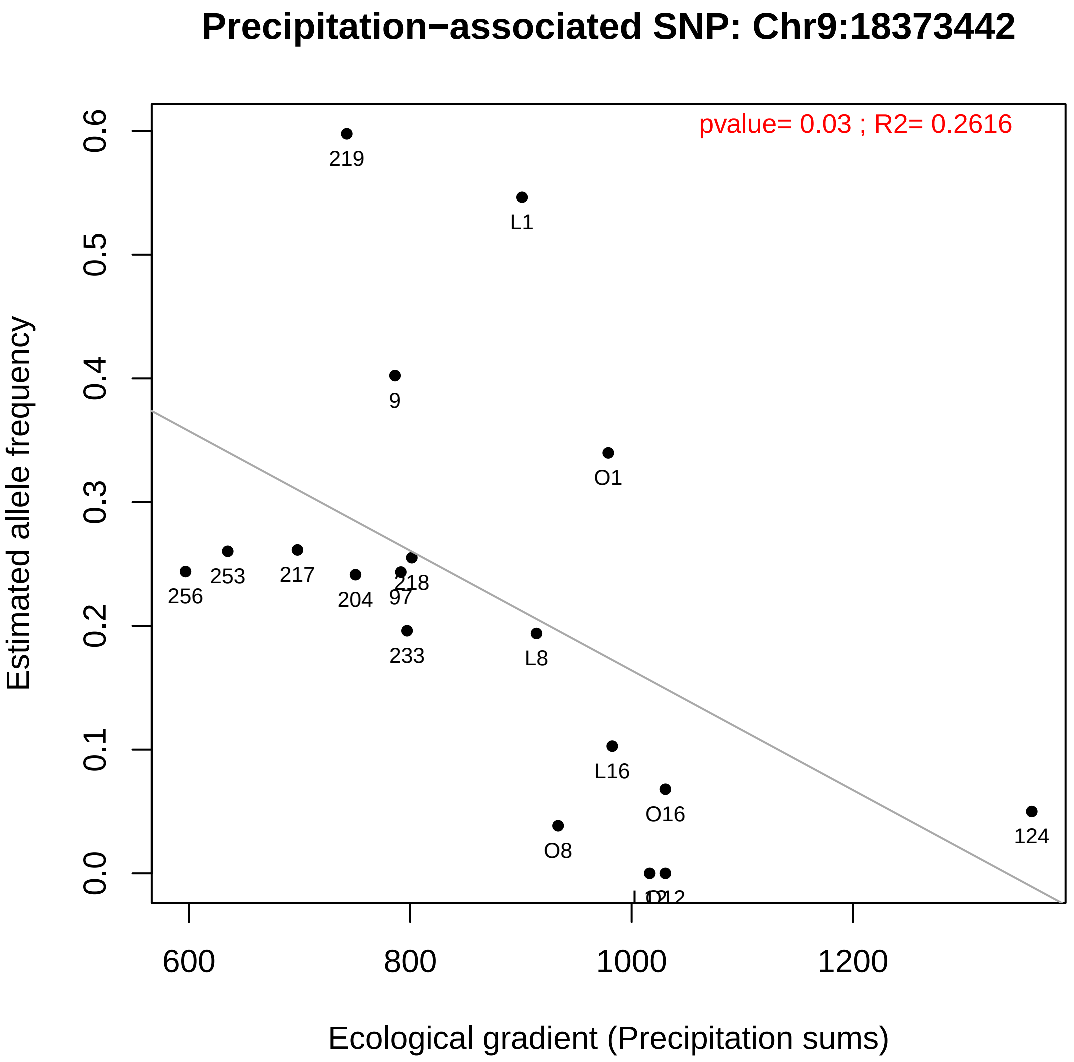
Precipitation−associated SNP: Chr9:18373442
pvalue= 0.03 ; R2= 0.2616
Estimated allele frequency
0.0
0.1
0.2
0.3
0.4
0.5
0.6
Ecological gradient (Precipitation sums)
600
800
1000
1200
219
L1
9
O1
256
253
217
204
97
218
233
L8
L16
O16
O8
124

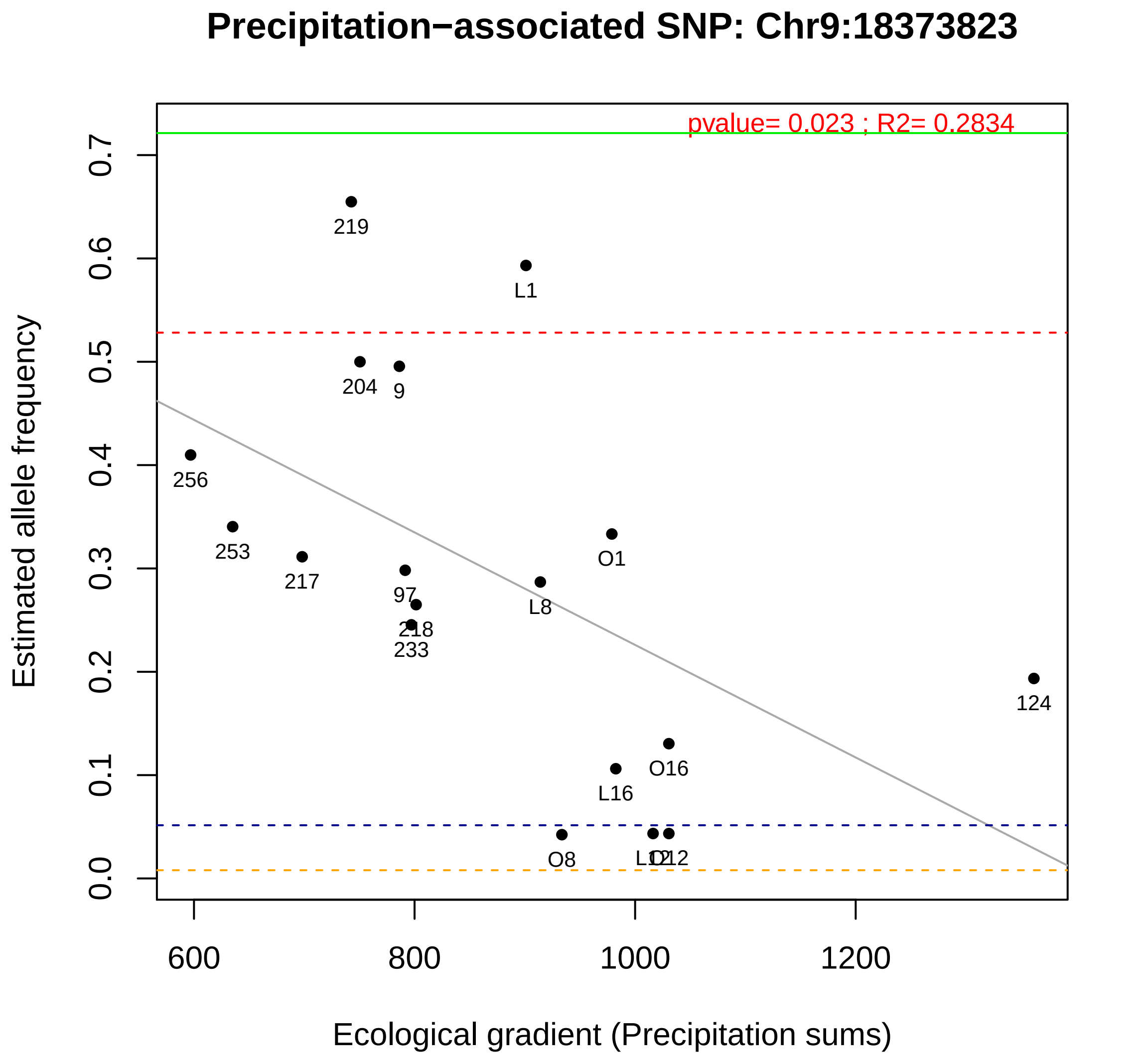

Precipitation−associated SNP: Chr9:18373823
pvalue= 0.023 ; R2= 0.2834
Estimated allele frequency
0.0
0.1
0.2
0.3
0.4
0.5
0.6
0.7
600
800
1000
1200
Ecological gradient (Precipitation sums)
219
L1
204
9
256
253
217
97
218
233
L8
O1
124
O16
L16
O8
L12
O12

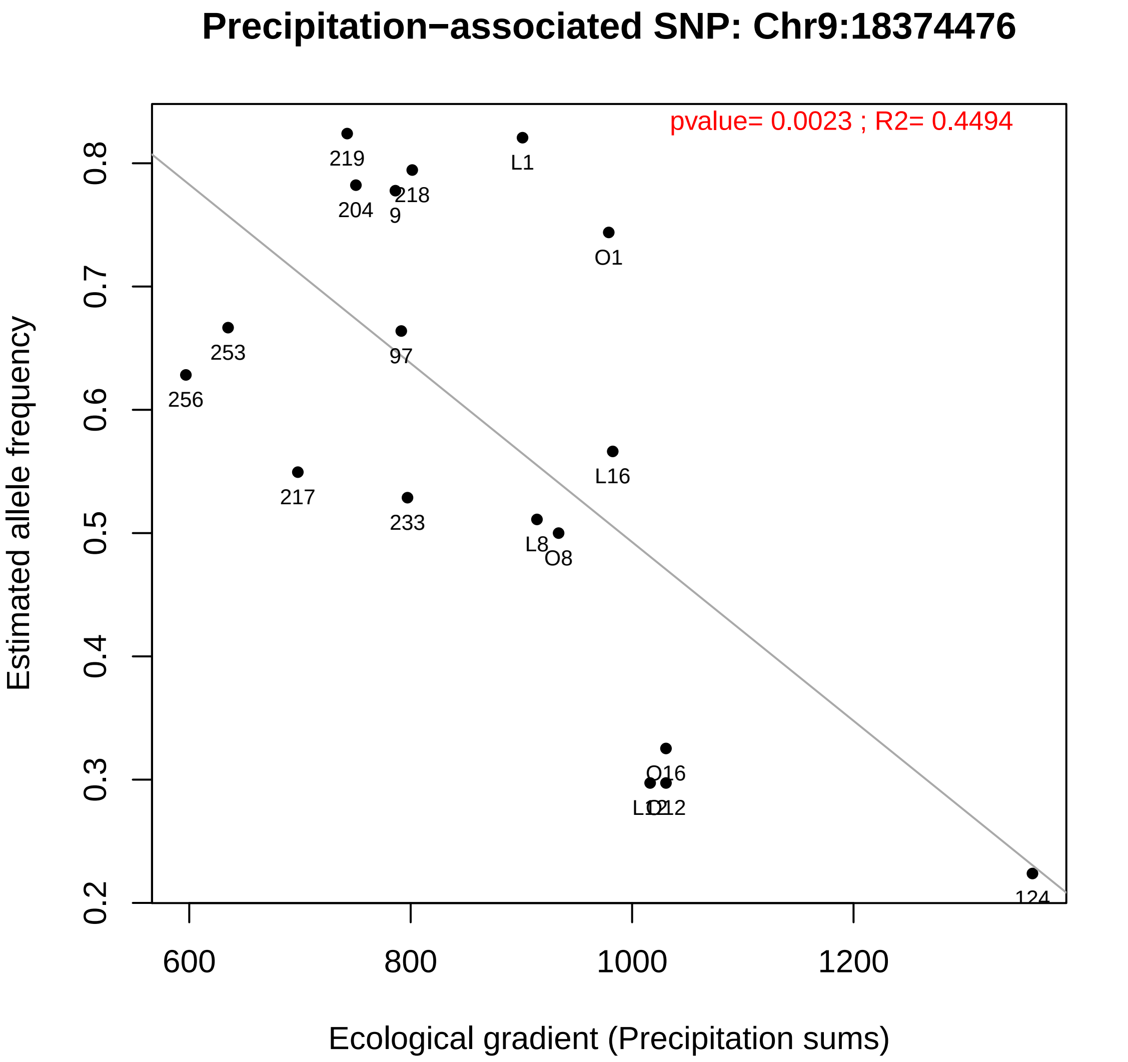
Precipitation−associated SNP: Chr9:18374476
pvalue= 0.0023 ; R2= 0.4494
Estimated allele frequency
Ecological gradient (Precipitation sums)
0.2
0.3
0.4
0.5
0.6
0.7
0.8
600
800
1000
1200
219
L1
218
204
9
O1
253
97
256
L16
217
233
L8
O8
O16
L12
O12
124

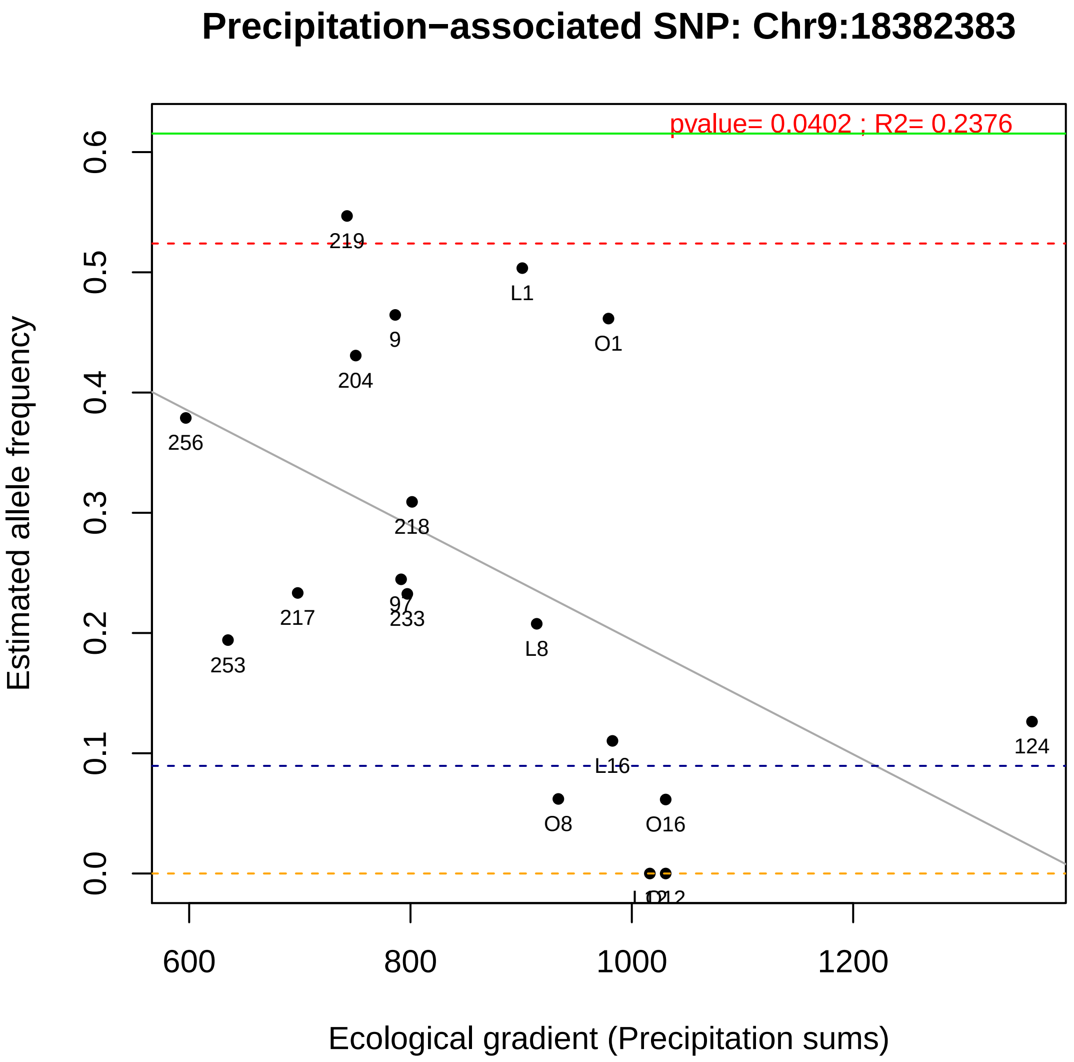
Precipitation−associated SNP: Chr9:18382383
pvalue= 0.0402 ; R2= 0.2376
Estimated allele frequency
0.0
0.1
0.2
0.3
0.4
0.5
0.6
Ecological gradient (Precipitation sums)
600
800
1000
1200
219
L1
9
O1
204
256
218
97
233
217
L8
253
124
L16
O8
O16

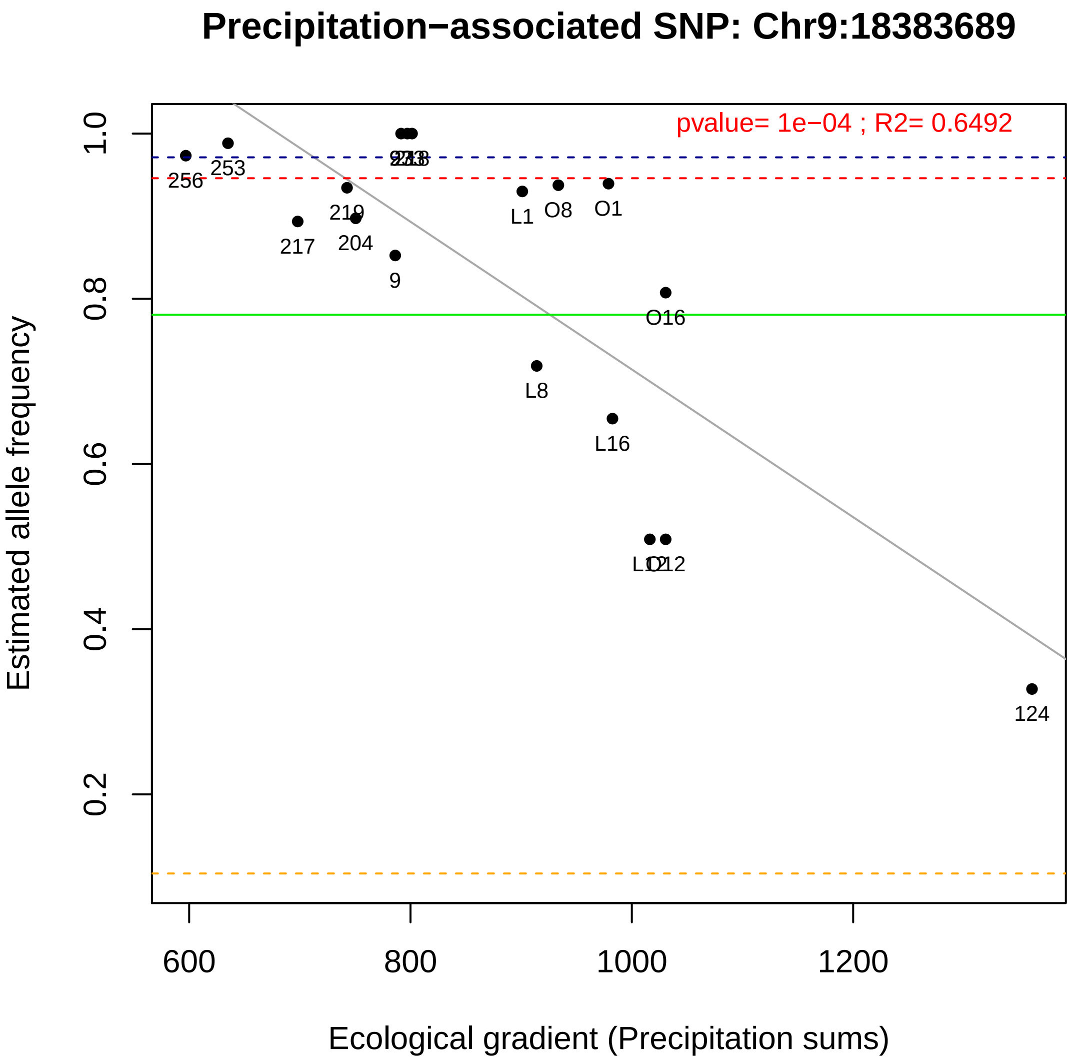

Precipitation−associated SNP: Chr9:18383689
pvalue= 1e−04 ; R2= 0.6492
Estimated allele frequency
Ecological gradient (Precipitation sums)
0.2
0.4
0.6
0.8
1.0
600
800
1000
1200
256
253
219
217
204
9
L1
O8
O1
O16
L8
L16
124

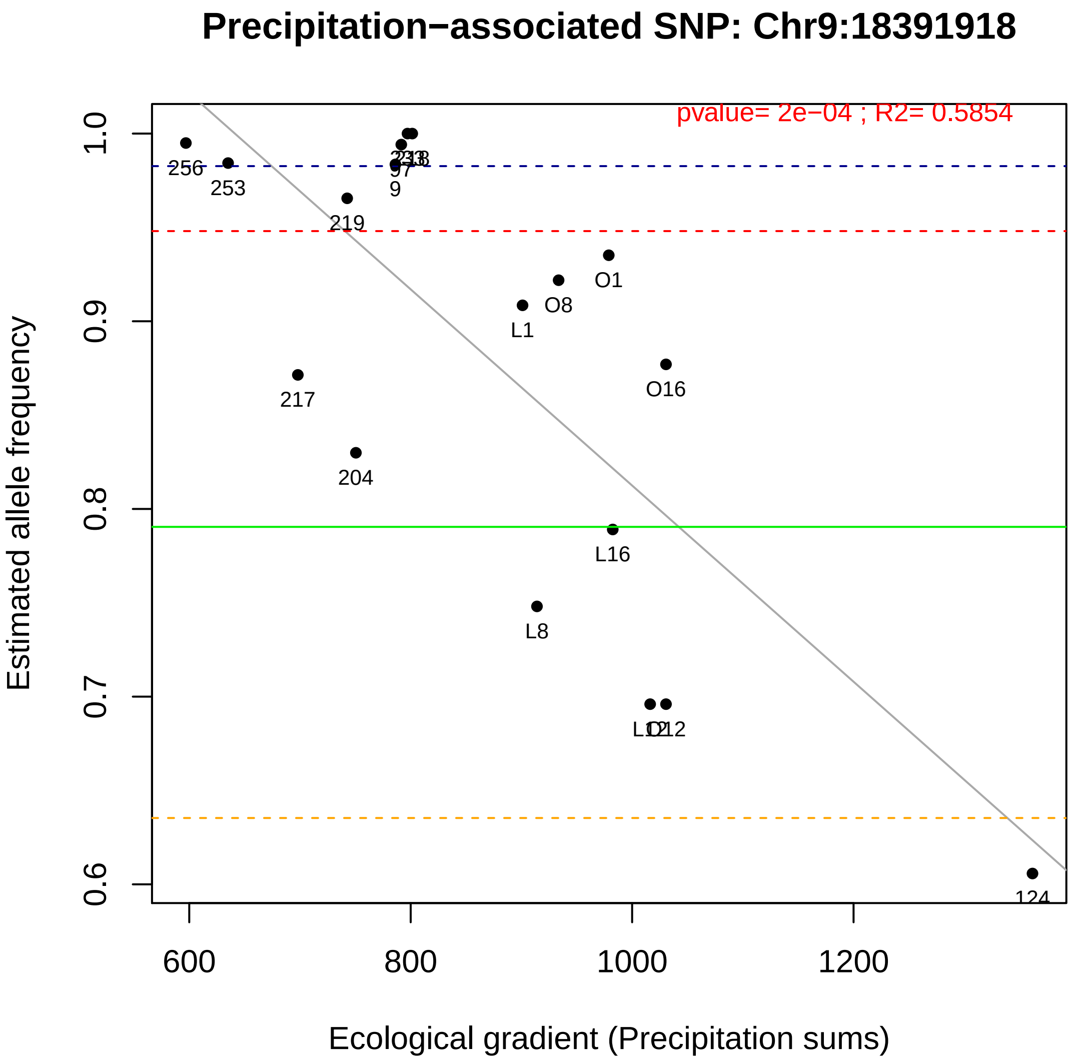
Precipitation−associated SNP: Chr9:18391918
pvalue= 2e−04 ; R2= 0.5854
Estimated allele frequency
1.0
0.9
0.8
0.7
0.6
600
800
1000
1200
Ecological gradient (Precipitation sums)
256
253
97
9
219
O1
O8
L1
217
O16
204
L16
L8
124

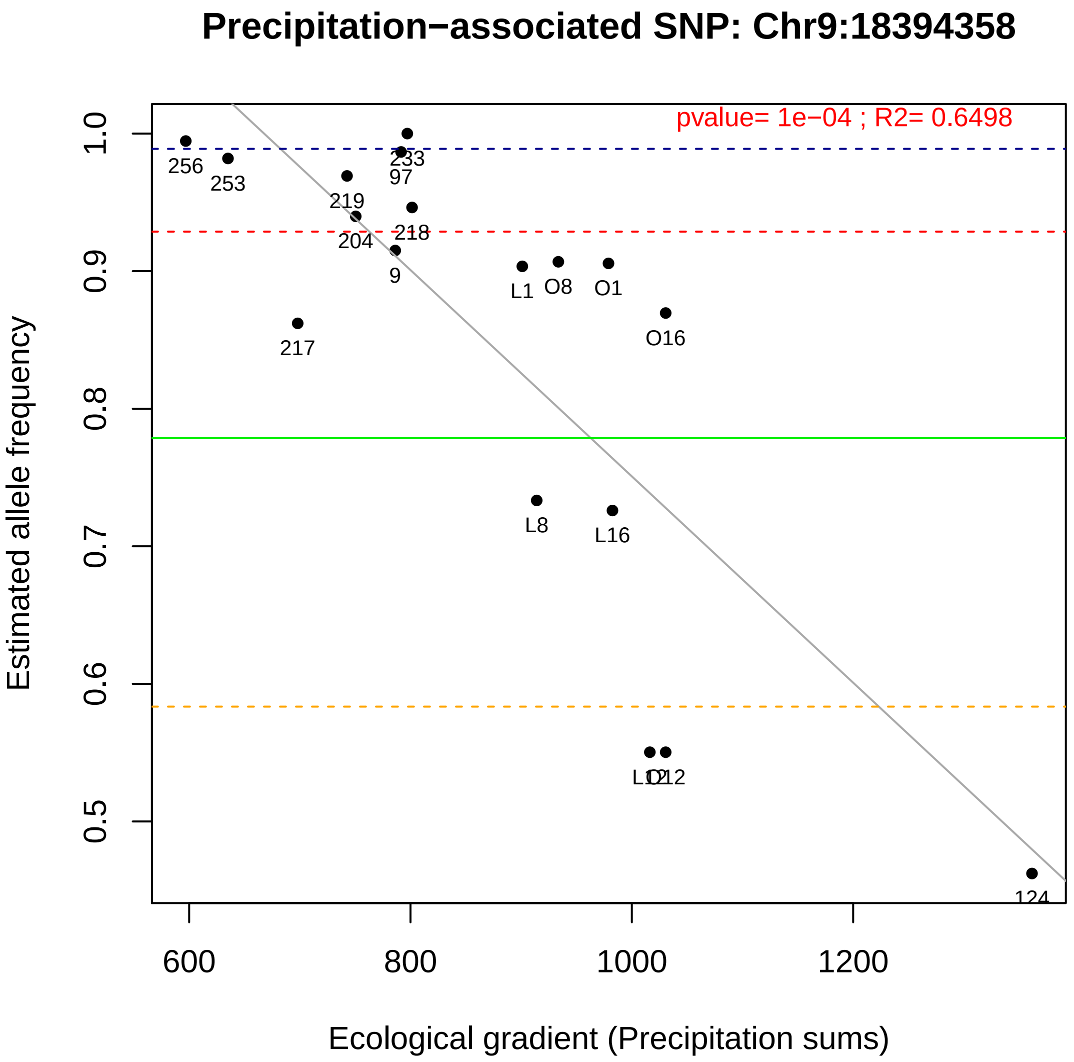
Precipitation−associated SNP: Chr9:18394358
pvalue= 1e−04 ; R2= 0.6498
Estimated allele frequency
Ecological gradient (Precipitation sums)
0.5
0.6
0.7
0.8
0.9
1.0
600
800
1000
1200
256
253
219
204
233
97
218
9
217
L1
O8
O1
O16
L8
L16
124

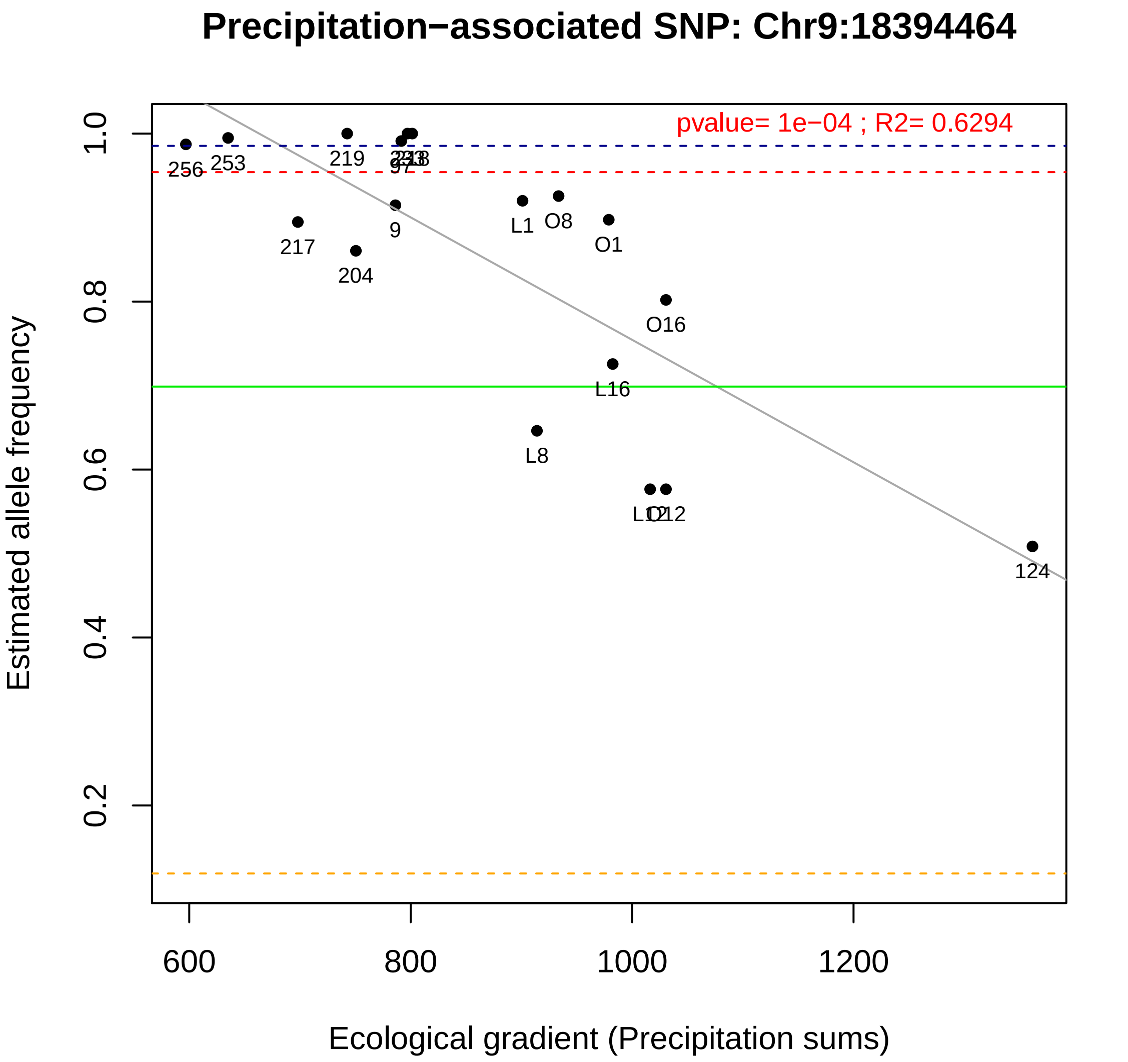

Precipitation−associated SNP: Chr9:18394464
pvalue= 1e−04 ; R2= 0.6294
Estimated allele frequency
Ecological gradient (Precipitation sums)
0.2
0.4
0.6
0.8
1.0
600
800
1000
1200
256
253
219
217
204
9
L1
O8
O1
O16
L16
L8
124

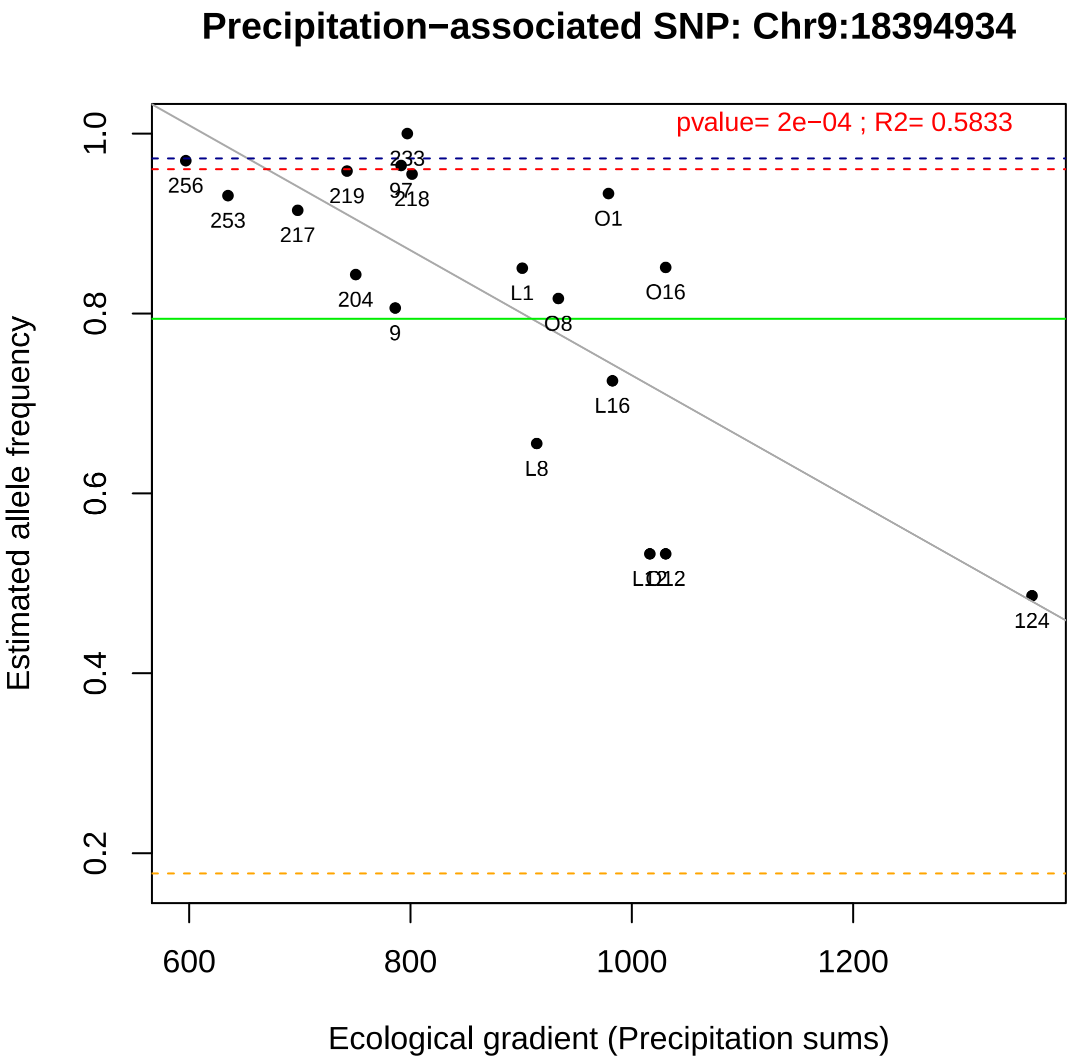

Precipitation−associated SNP: Chr9:18394934
pvalue= 2e−04 ; R2= 0.5833
Estimated allele frequency
0.2
0.4
0.6
0.8
1.0
600
800
1000
1200
Ecological gradient (Precipitation sums)
256
253
217
219
204
233
97
218
9
L1
O8
O1
O16
L16
L8
L12
O12
124

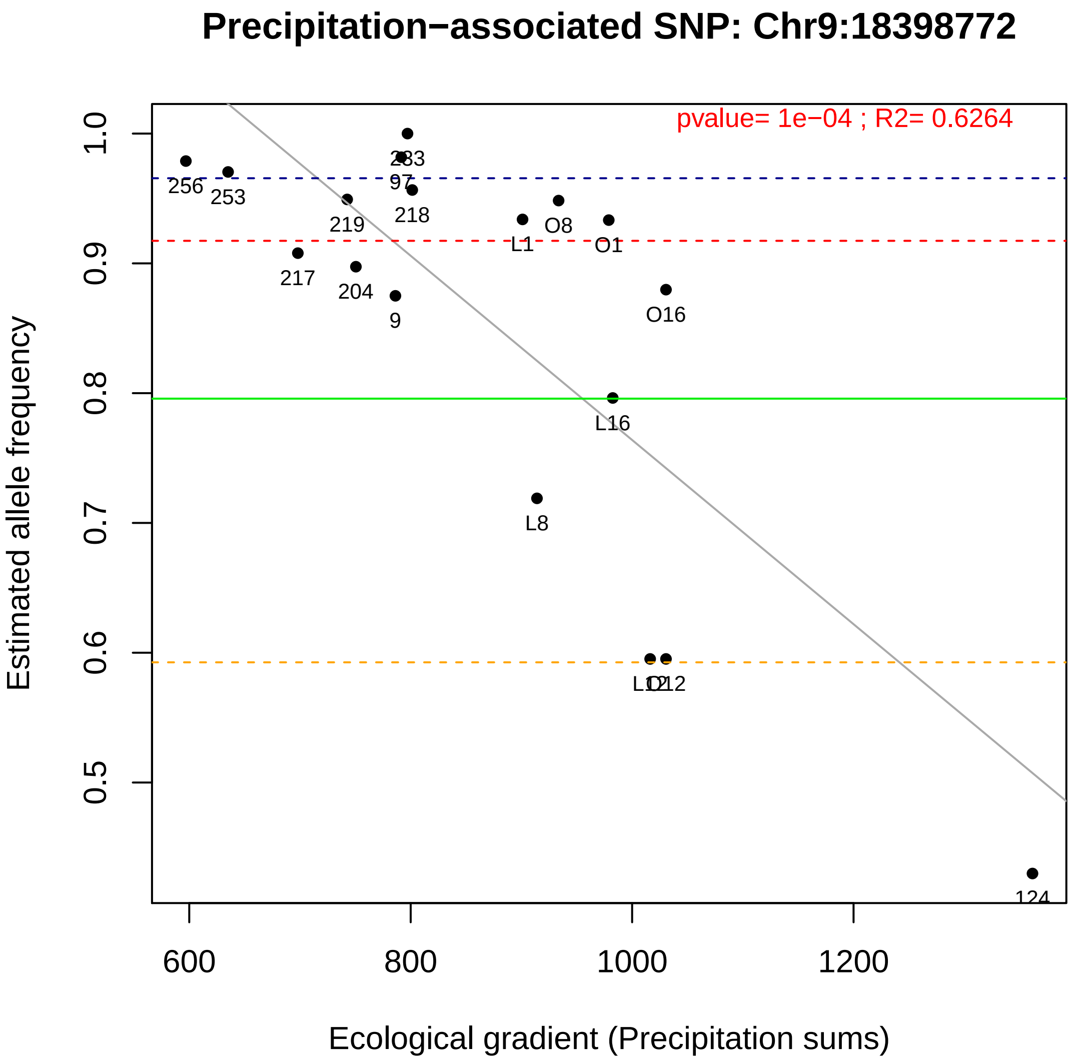
Precipitation−associated SNP: Chr9:18398772
pvalue= 1e−04 ; R2= 0.6264
Estimated allele frequency
1.0
0.9
0.8
0.7
0.6
0.5
600
800
1000
1200
Ecological gradient (Precipitation sums)
256
253
233
97
219
218
217
204
9
L1
O8
O1
O16
L16
L8
L12
O12
124

**Precipitation−associated SNP: Chr9:18402305**

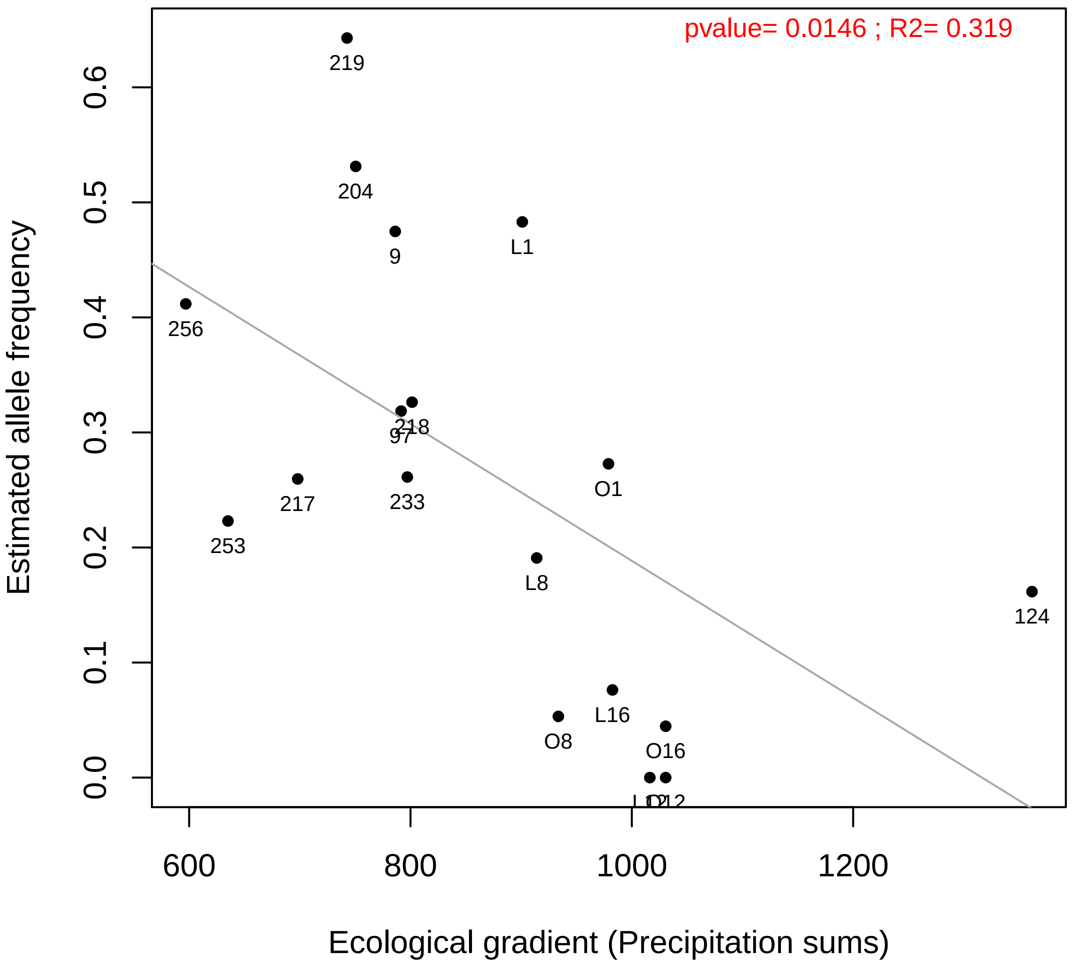

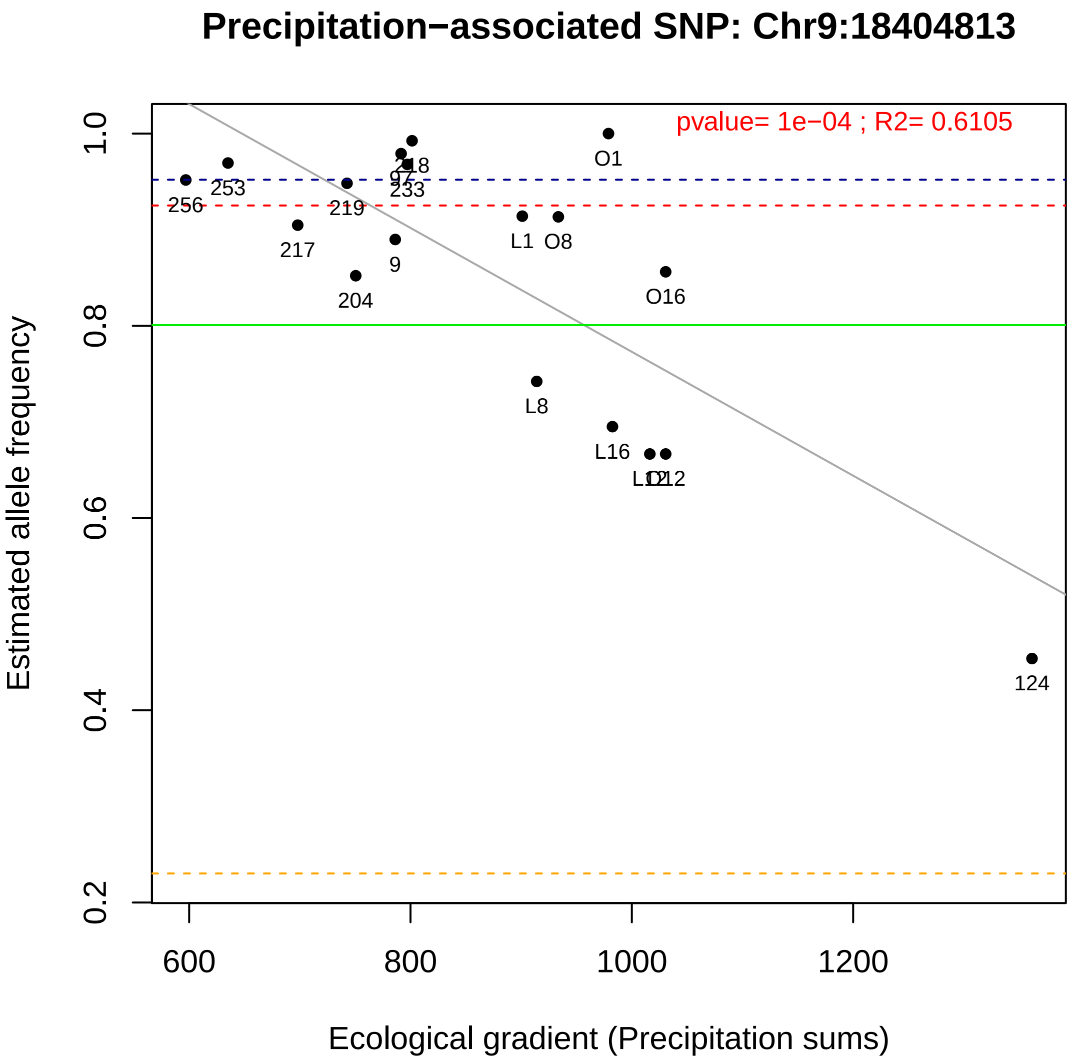
Precipitation−associated SNP: Chr9:18404813
pvalue= 1e−04 ; R2= 0.6105
Estimated allele frequency
Ecological gradient (Precipitation sums)
0.2
0.4
0.6
0.8
1.0
600
800
1000
1200
256
253
219
217
204
9
97
218
233
L1
O8
O1
O16
L8
L16
L12
O12
124

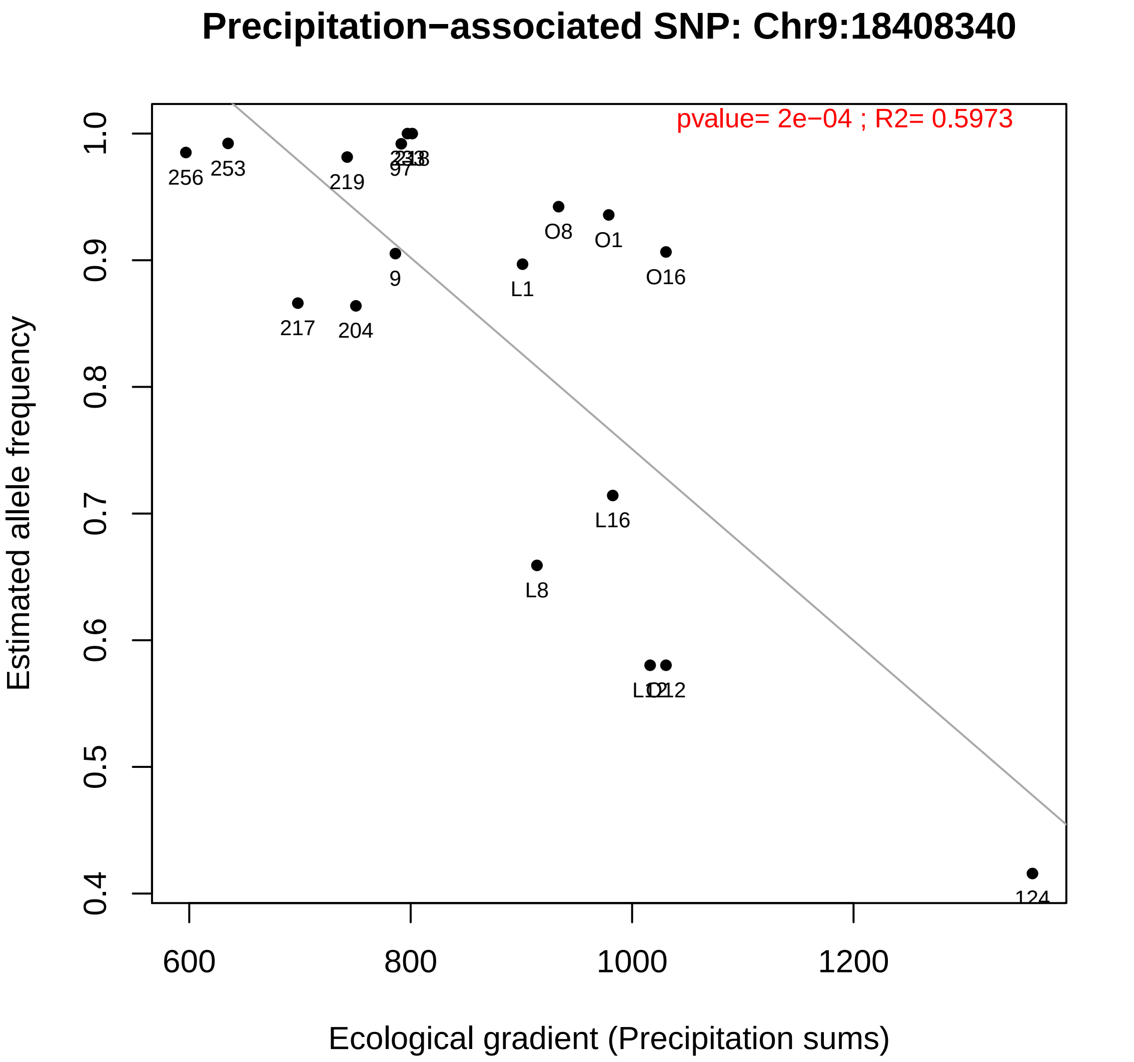
Precipitation−associated SNP: Chr9:18408340
pvalue= 2e−04 ; R2= 0.5973
Estimated allele frequency
Ecological gradient (Precipitation sums)
1.0
0.9
0.8
0.7
0.6
0.5
0.4
600
800
1000
1200
256
253
219
97
9
217
204
O8
O1
O16
L1
L16
L8
124

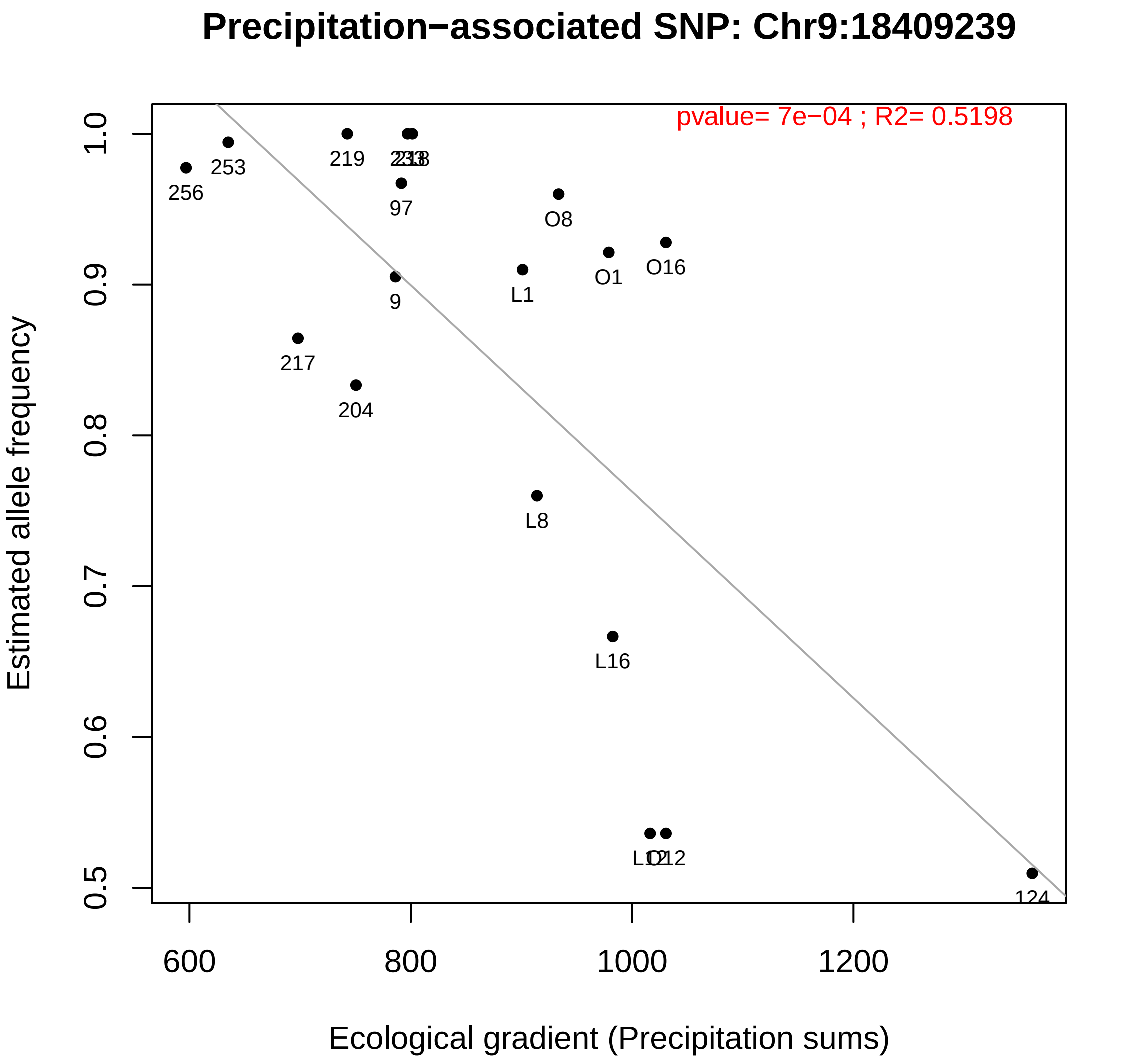
Precipitation−associated SNP: Chr9:18409239
pvalue= 7e−04 ; R2= 0.5198
Estimated allele frequency
0.5
0.6
0.7
0.8
0.9
1.0
600
800
1000
1200
Ecological gradient (Precipitation sums)
256
253
219
97
9
217
204
O8
L1
O1
O16
L8
L16
124

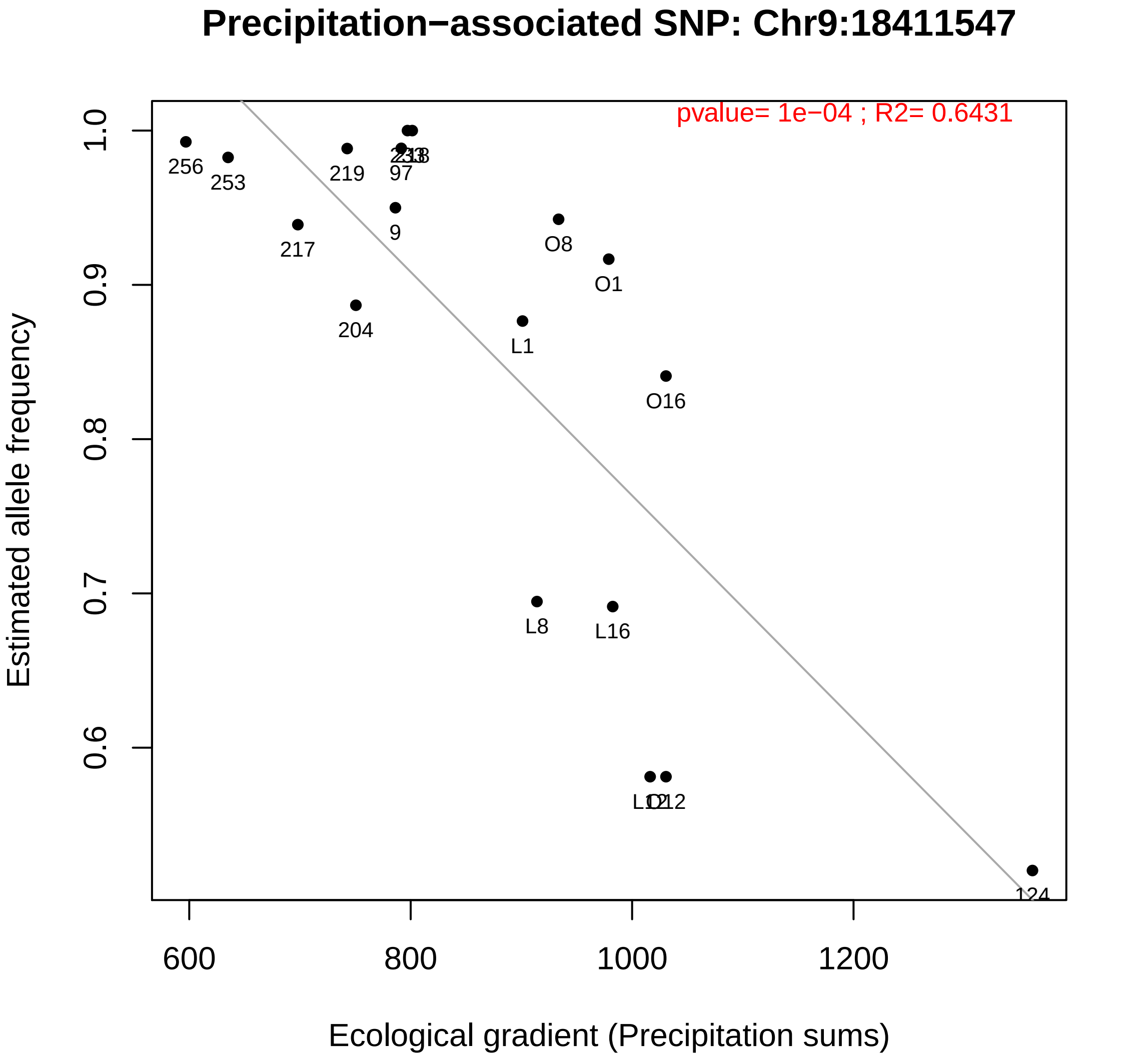
Precipitation−associated SNP: Chr9:18411547
pvalue= 1e−04 ; R2= 0.6431
Estimated allele frequency
Ecological gradient (Precipitation sums)
0.6
0.7
0.8
0.9
1.0
600
800
1000
1200
256
253
219
97
9
217
204
O8
O1
L1
O16
L8
L16
124

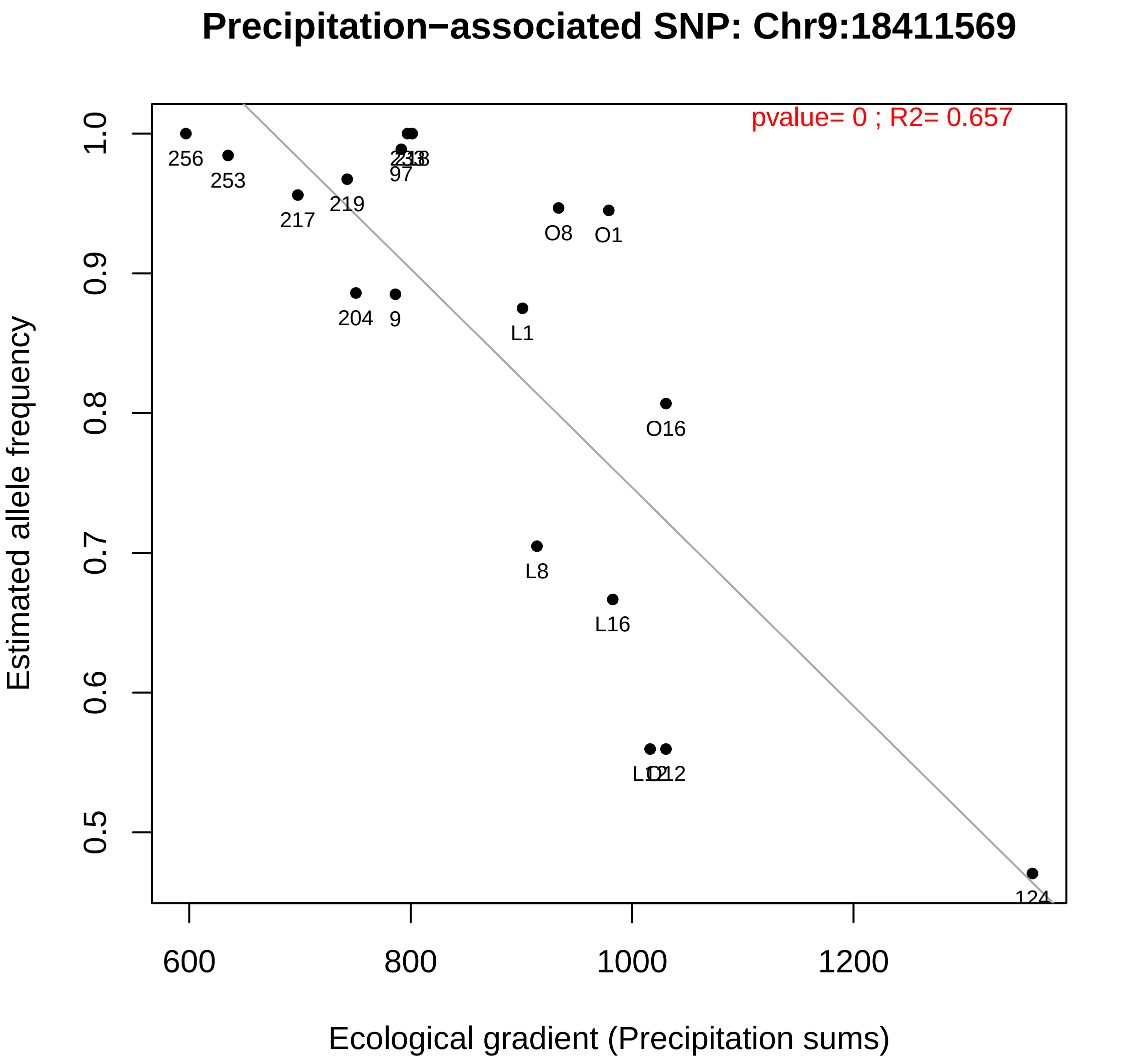
Precipitation−associated SNP: Chr9:18411569
pvalue= 0 ; R2= 0.657
Estimated allele frequency
Ecological gradient (Precipitation sums)
0.5
0.6
0.7
0.8
0.9
1.0
600
800
1000
1200
256
253
217
219
97
204
9
O8
O1
L1
O16
L8
L16
124

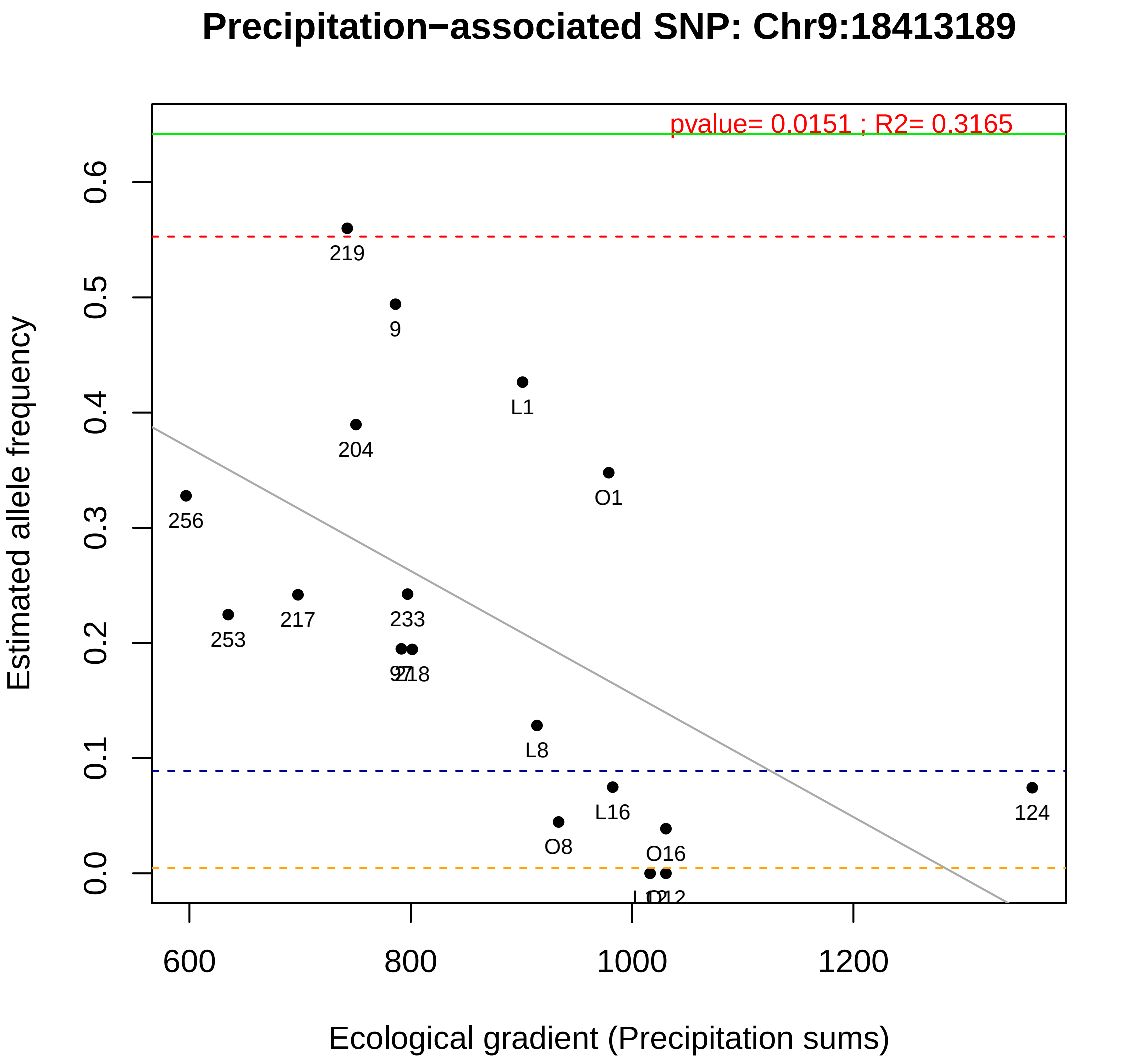
Precipitation−associated SNP: Chr9:18413189
pvalue= 0.0151 ; R2= 0.3165
Estimated allele frequency
0.0
0.1
0.2
0.3
0.4
0.5
0.6
219
9
L1
204
O1
256
217
233
253
97
218
L8
L16
124
O8
O16
O12
600
800
1000
1200
Ecological gradient (Precipitation sums)

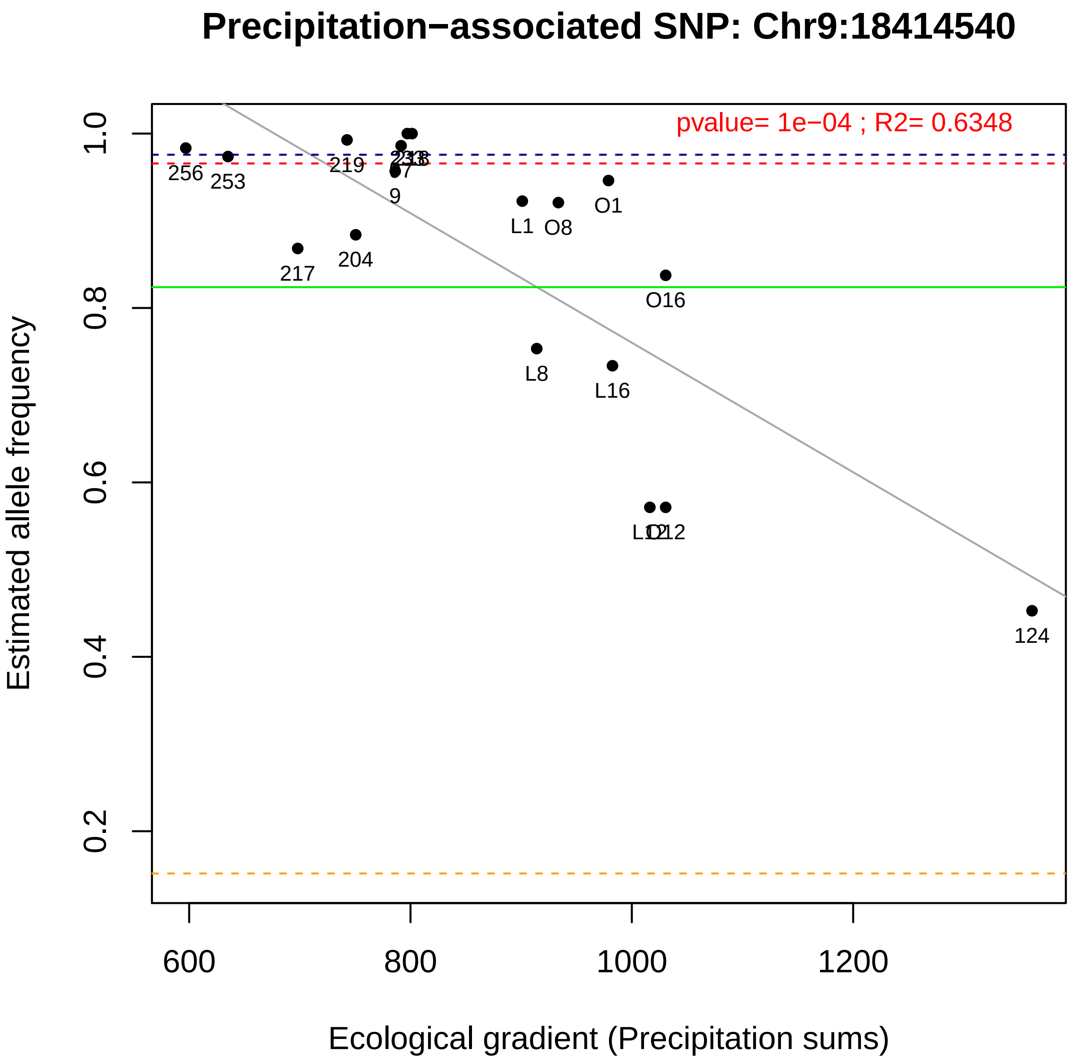

Precipitation−associated SNP: Chr9:18414540
pvalue= 1e−04 ; R2= 0.6348
Estimated allele frequency
Ecological gradient (Precipitation sums)
0.2
0.4
0.6
0.8
1.0
600
800
1000
1200
256
253
219
9
217
204
L1
O8
O1
O16
L8
L16
124

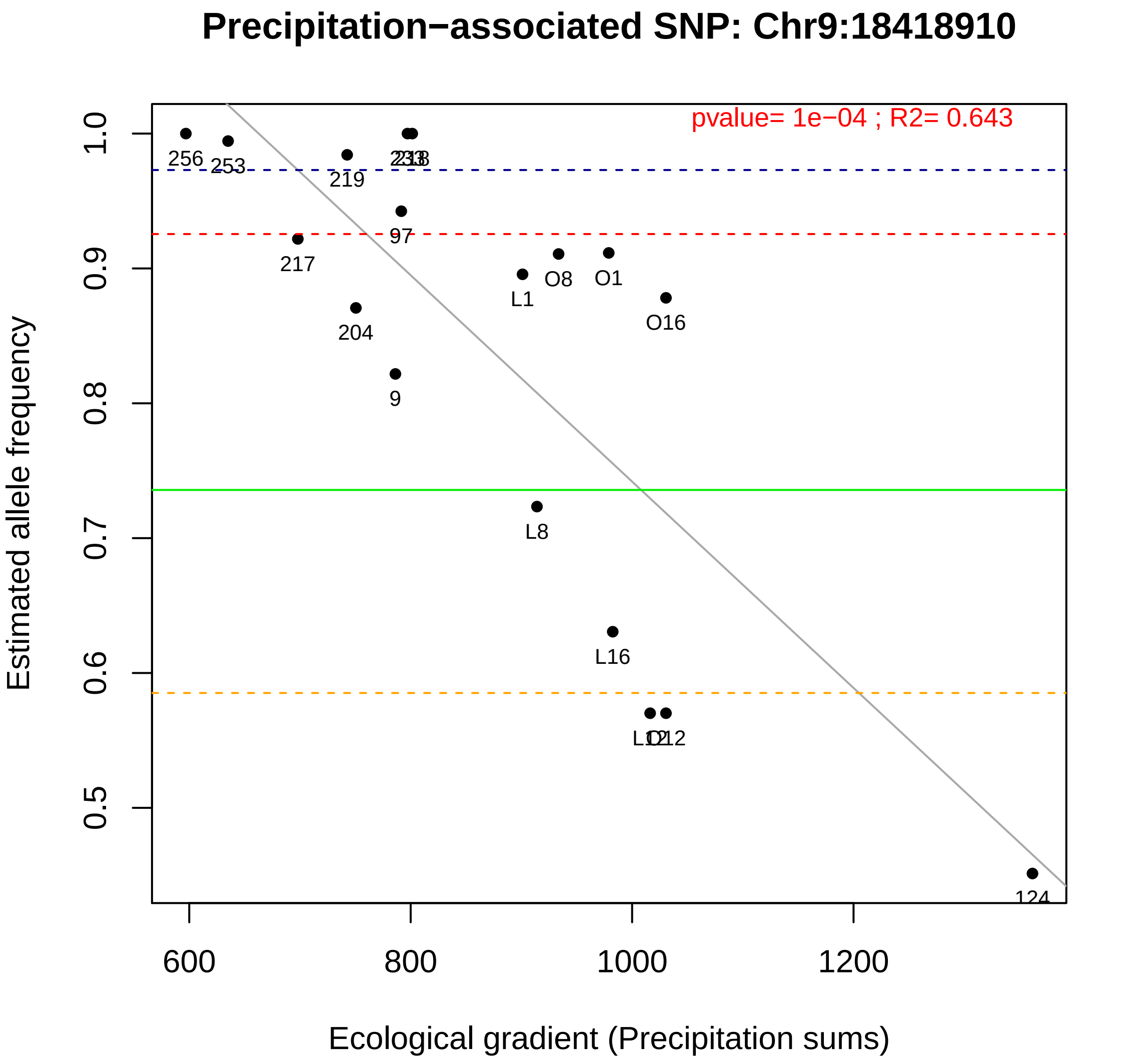

Precipitation−associated SNP: Chr9:18418910
pvalue= 1e−04 ; R2= 0.643
Estimated allele frequency
1.0
0.9
0.8
0.7
0.6
0.5
256
253
219
217
204
97
9
L1
O8
O1
O16
L8
L16
124
600
800
1000
1200
Ecological gradient (Precipitation sums)

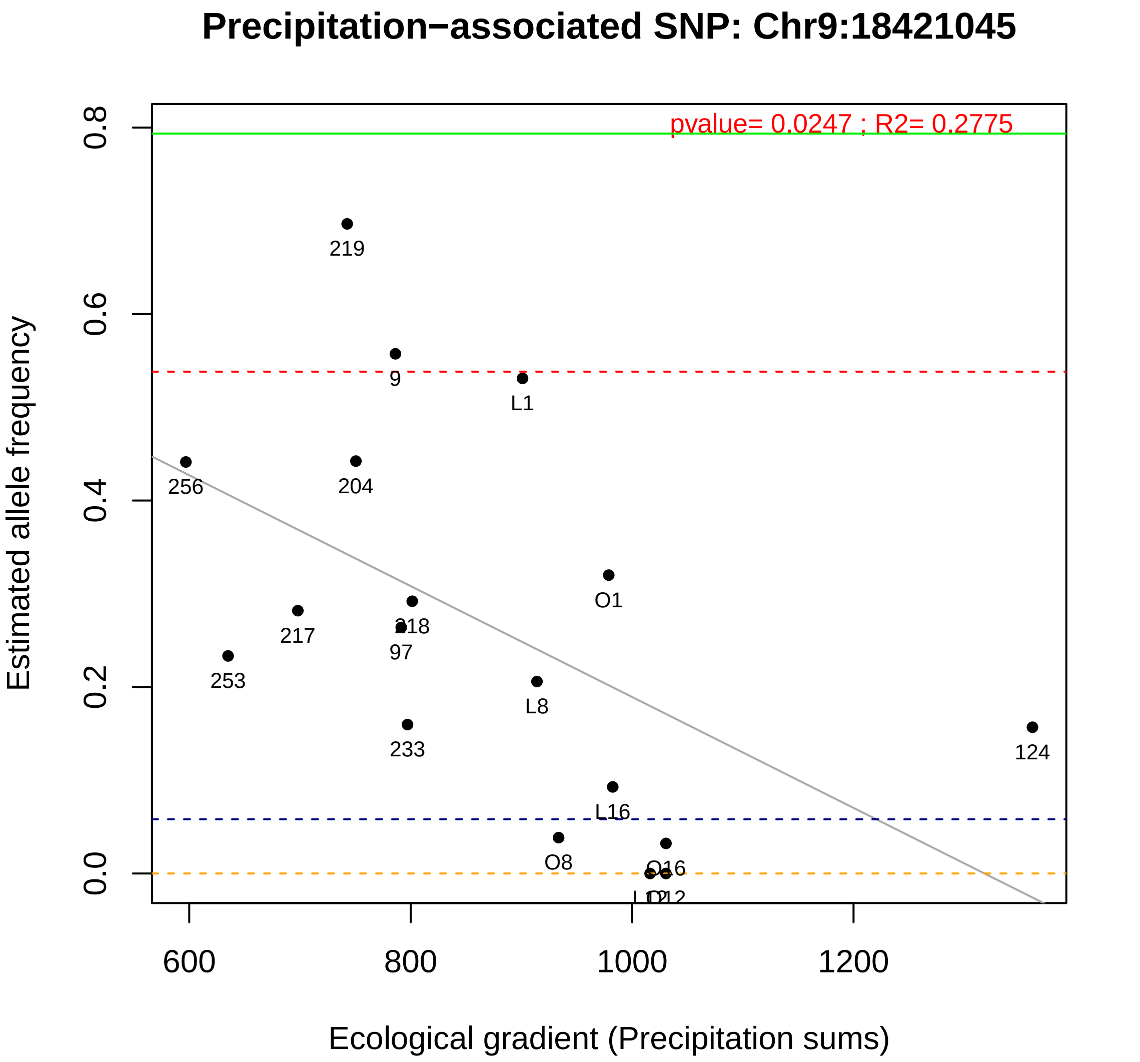

Precipitation−associated SNP: Chr9:18421045
pvalue= 0.0247 ; R2= 0.2775
Estimated allele frequency
Ecological gradient (Precipitation sums)
0.0
0.2
0.4
0.6
0.8
600
800
1000
1200
219
9
L1
256
204
O1
217
218
97
253
L8
233
124
L16
O8
O16
L12
O12

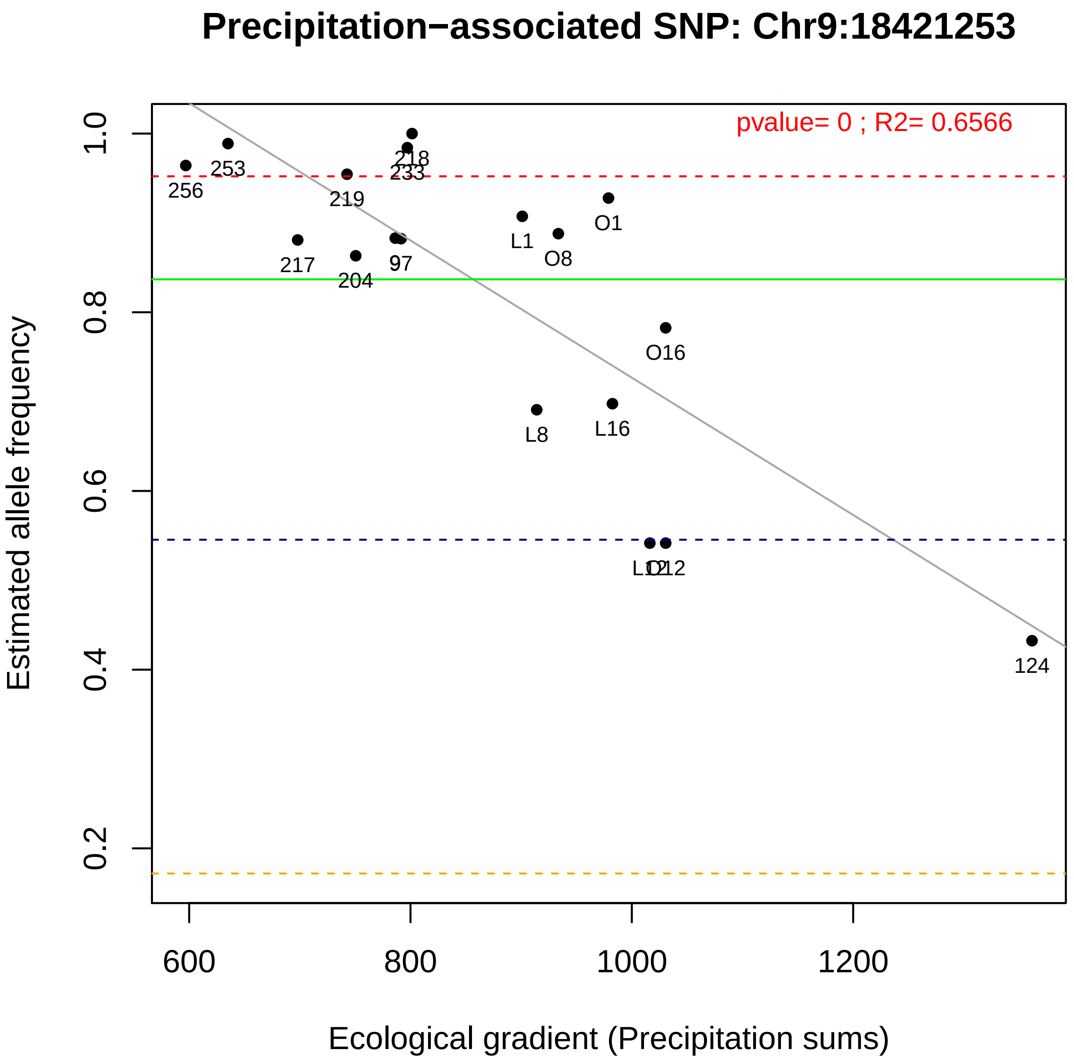

Precipitation−associated SNP: Chr9:18421253
pvalue= 0 ; R2= 0.6566
Estimated allele frequency
Ecological gradient (Precipitation sums)
1.0
0.8
0.6
0.4
0.2
600
800
1000
1200
256
253
219
218
233
217
204
97
L1
O8
O1
O16
L8
L16
L12
O12
124

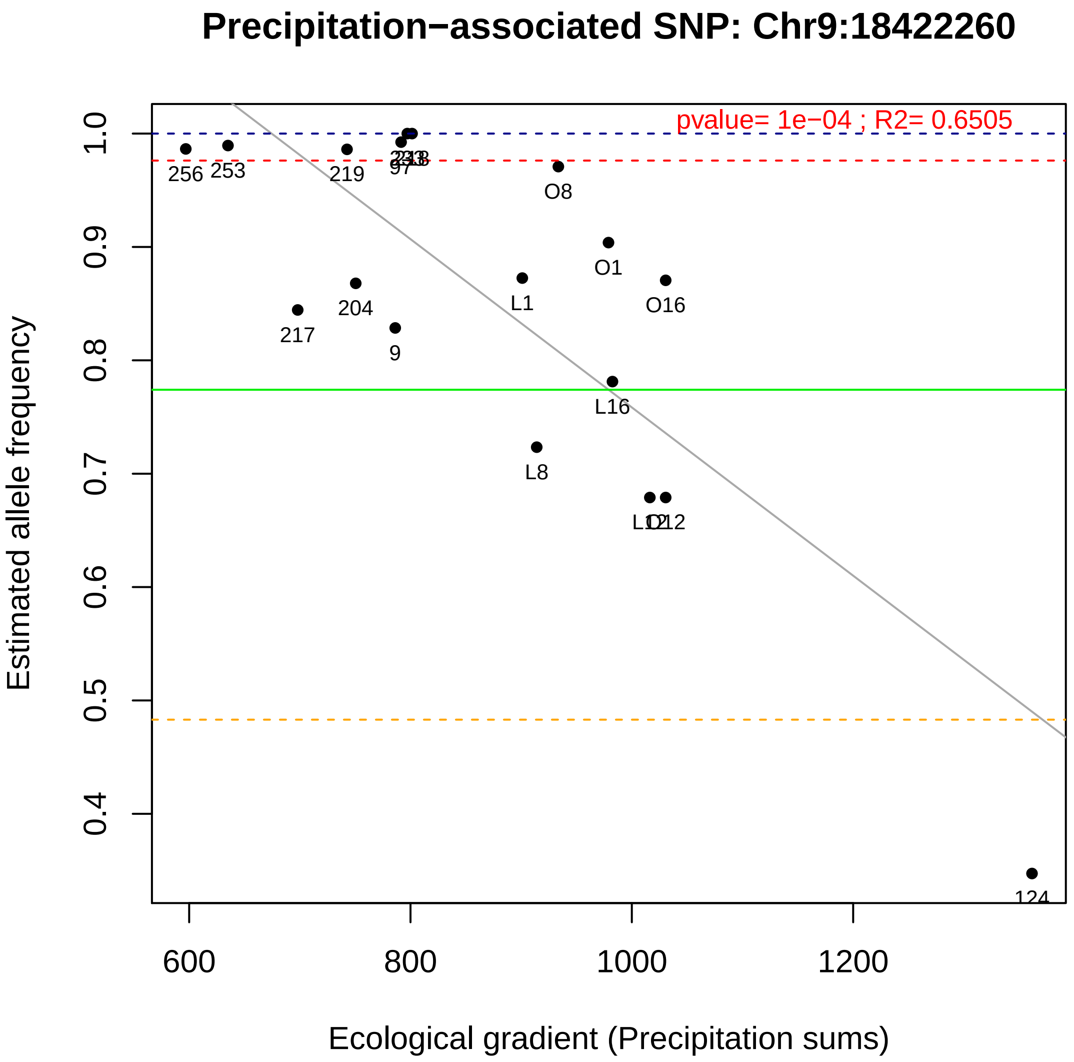

Precipitation−associated SNP: Chr9:18422260
pvalue= 1e−04 ; R2= 0.6505
Estimated allele frequency
Ecological gradient (Precipitation sums)
1.0
0.9
0.8
0.7
0.6
0.5
0.4
600
800
1000
1200
256
253
219
97
O8
O1
L1
O16
204
217
9
L16
L8
124

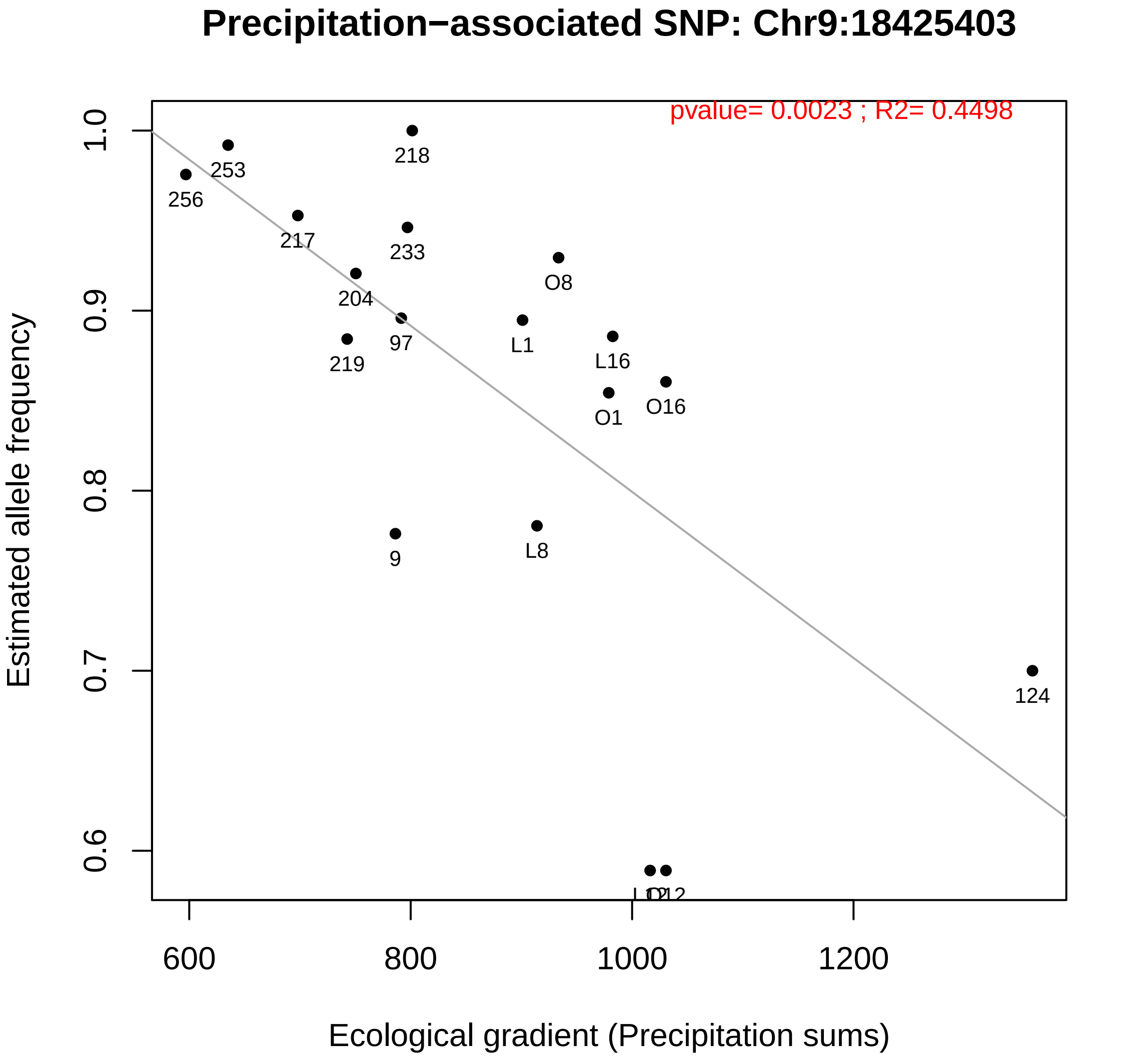
Precipitation−associated SNP: Chr9:18425403
pvalue= 0.0023 ; R2= 0.4498
Estimated allele frequency
1.0
0.9
0.8
0.7
0.6
600
800
1000
1200
Ecological gradient (Precipitation sums)
218
253
256
217
233
204
97
219
O8
L1
L16
O16
O1
9
L8
124

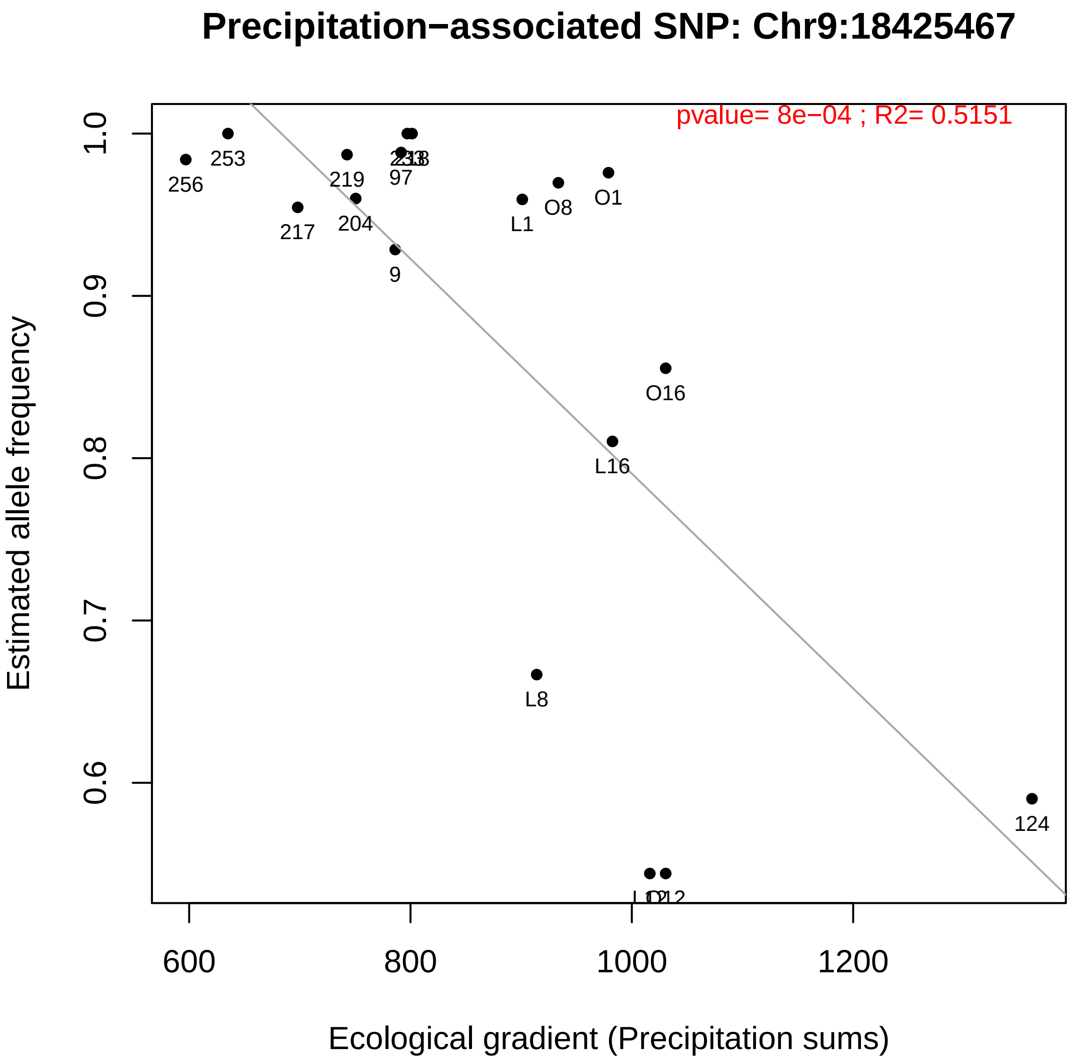
Precipitation−associated SNP: Chr9:18425467
pvalue= 8e−04 ; R2= 0.5151
Estimated allele frequency
1.0
0.9
0.8
0.7
0.6
600
800
1000
1200
Ecological gradient (Precipitation sums)
256
253
219
204
217
97
9
L1
O8
O1
O16
L16
L8
124

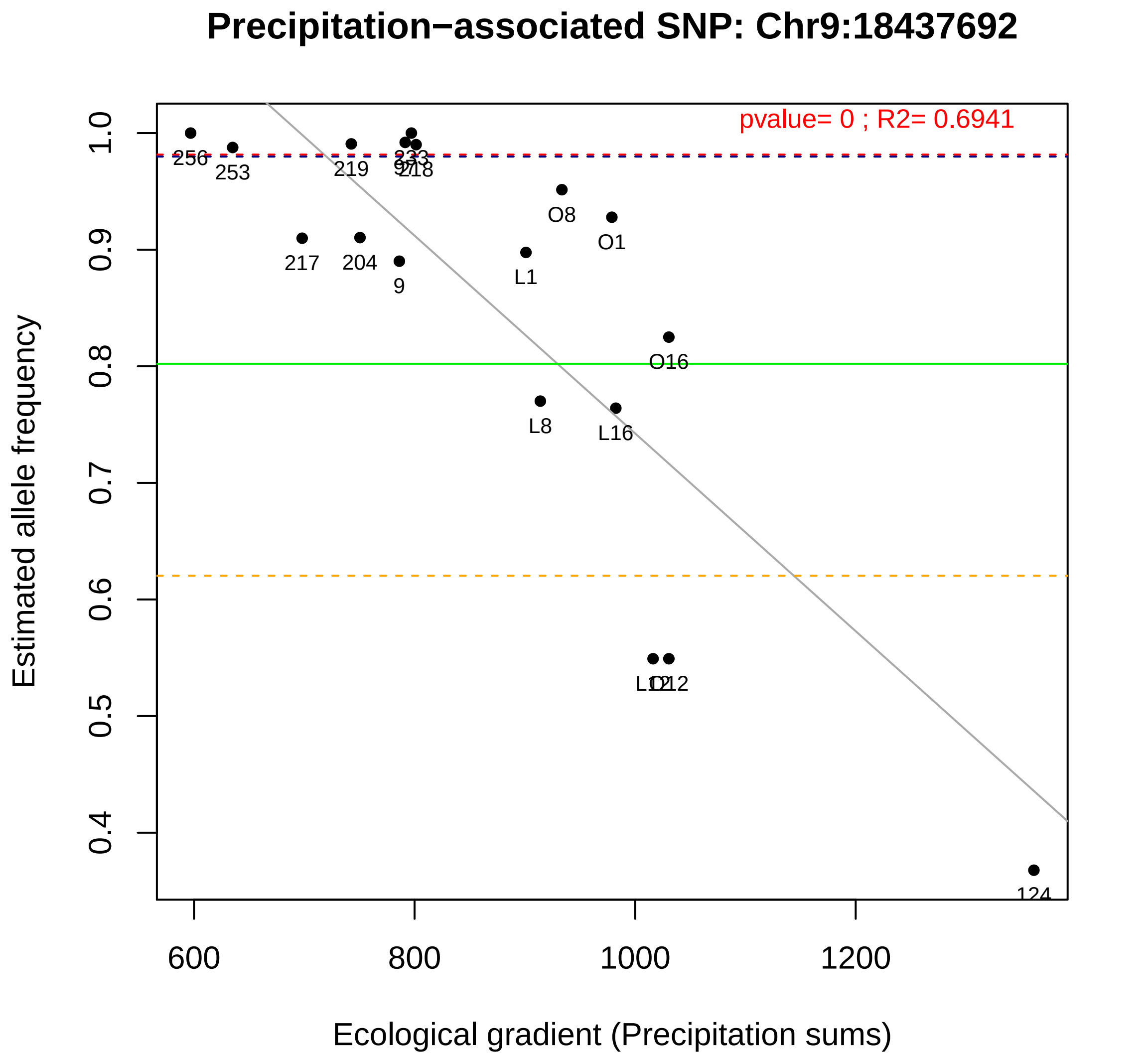

Precipitation−associated SNP: Chr9:18437692
pvalue= 0 ; R2= 0.6941
256
253
219
233
97
218
217
204
9
O8
O1
L1
O16
L8
L16
L12
O12
124
1.0
0.9
0.8
0.7
0.6
0.5
0.4
Estimated allele frequency
600
800
1000
1200
Ecological gradient (Precipitation sums)

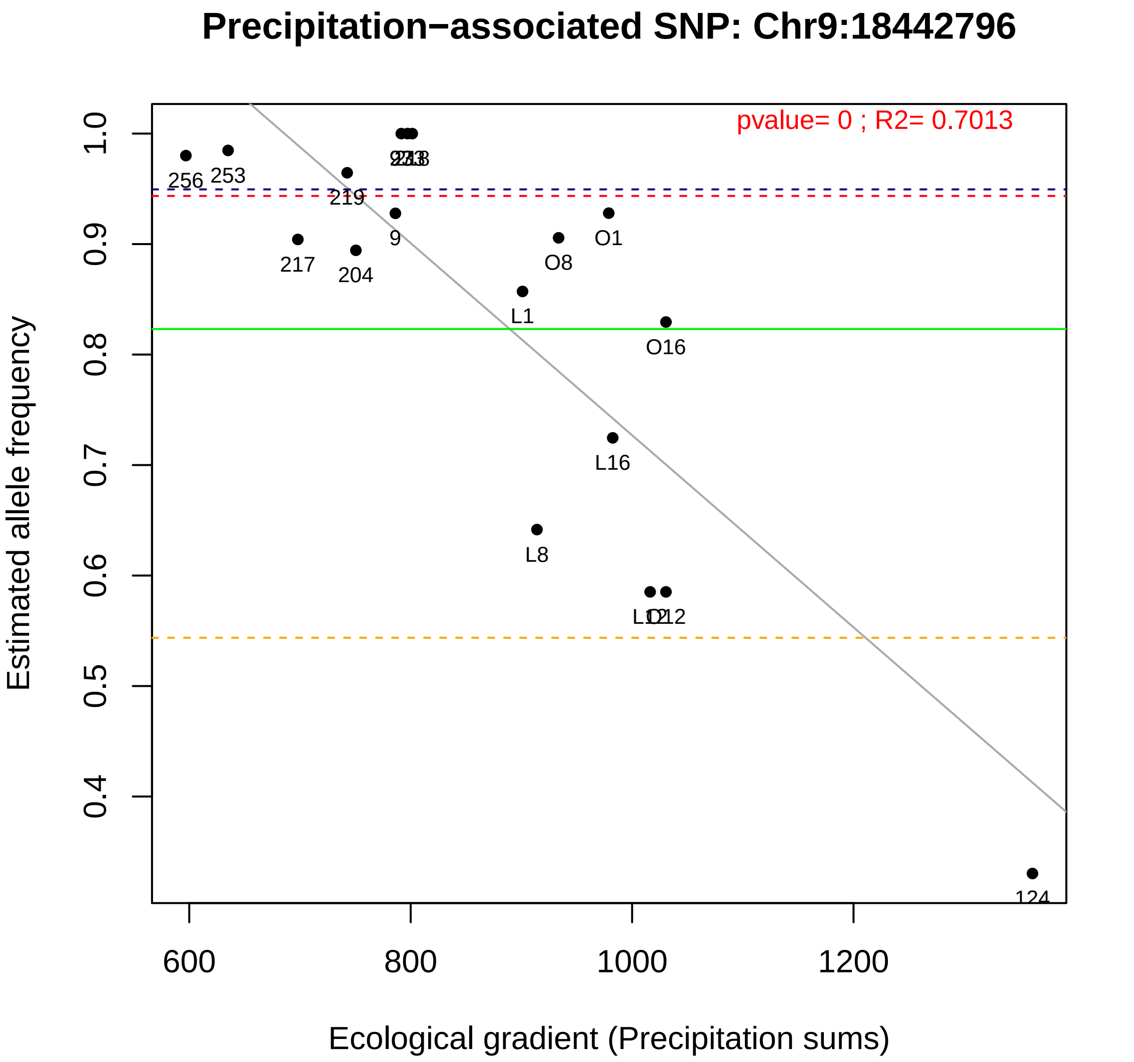
Precipitation−associated SNP: Chr9:18442796
pvalue= 0 ; R2= 0.7013
Estimated allele frequency
Ecological gradient (Precipitation sums)
1.0
0.9
0.8
0.7
0.6
0.5
0.4
600
800
1000
1200
256
253
219
217
204
9
O1
O8
L1
O16
L16
L8
124

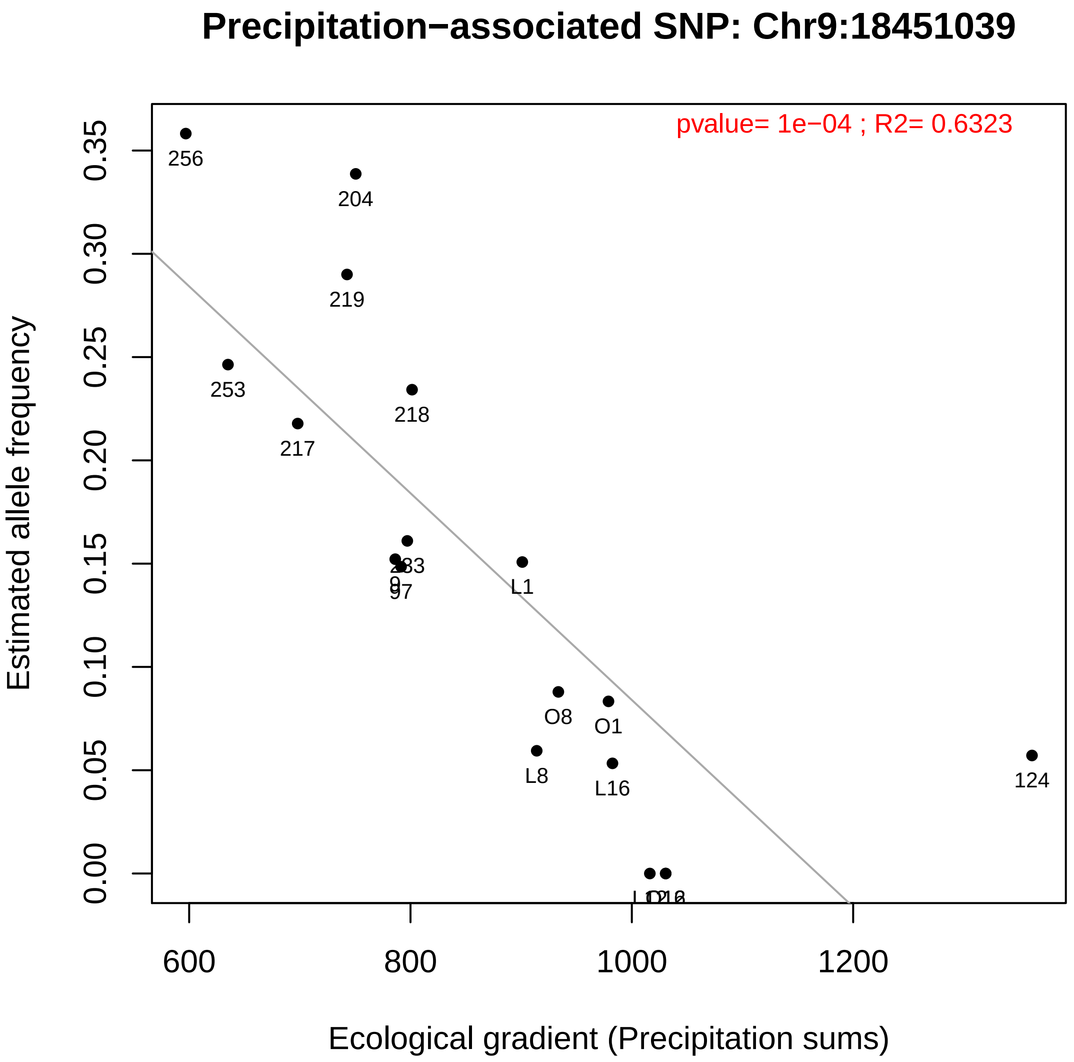
Precipitation−associated SNP: Chr9:18451039
pvalue= 1e−04 ; R2= 0.6323
Estimated allele frequency
Ecological gradient (Precipitation sums)
0.00
0.05
0.10
0.15
0.20
0.25
0.30
0.35
600
800
1000
1200
256
204
219
253
218
217
97
L1
O8
O1
L8
L16
124

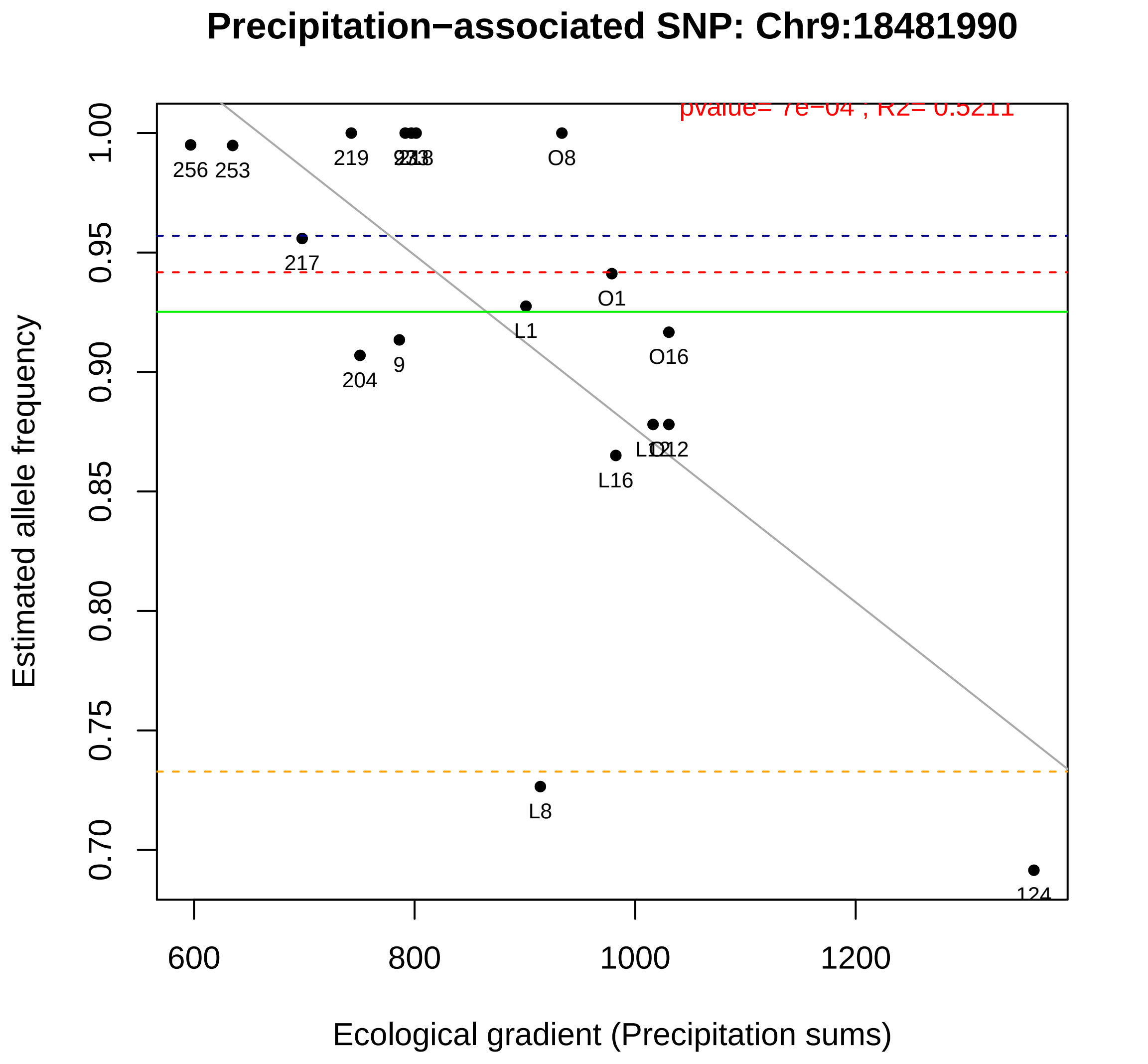

Precipitation−associated SNP: Chr9:18481990
pvalue= 7e−04 ; R2= 0.5211
Estimated allele frequency
Ecological gradient (Precipitation sums)
1.00
0.95
0.90
0.85
0.80
0.75
0.70
600
800
1000
1200
256
253
219
O8
217
O1
L1
O16
204
9
L16
L8
124

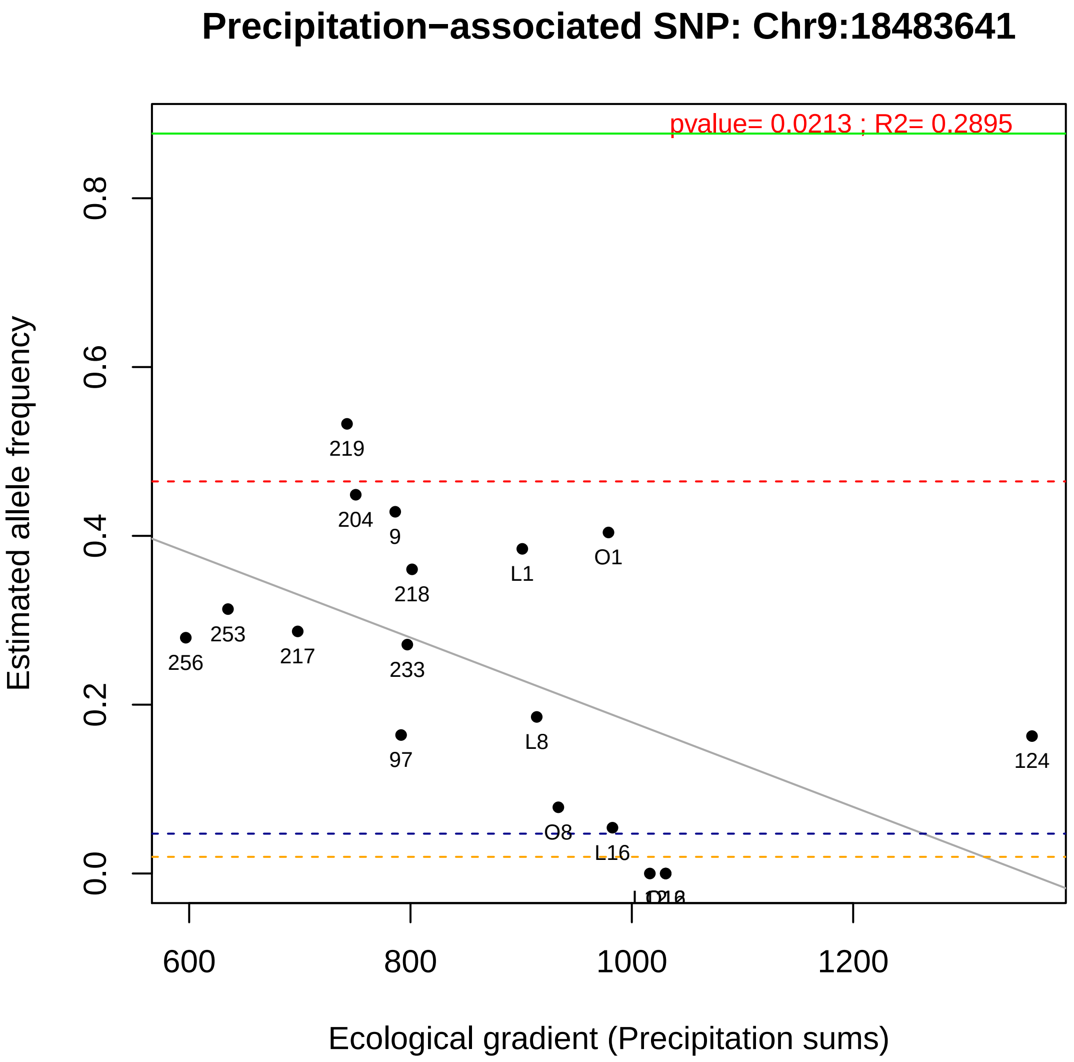
Precipitation−associated SNP: Chr9:18483641
pvalue= 0.0213 ; R2= 0.2895
Estimated allele frequency
0.0
0.2
0.4
0.6
0.8
Ecological gradient (Precipitation sums)
600
800
1000
1200
219
204
9
218
L1
O1
253
256
217
233
97
L8
124
O8
L16

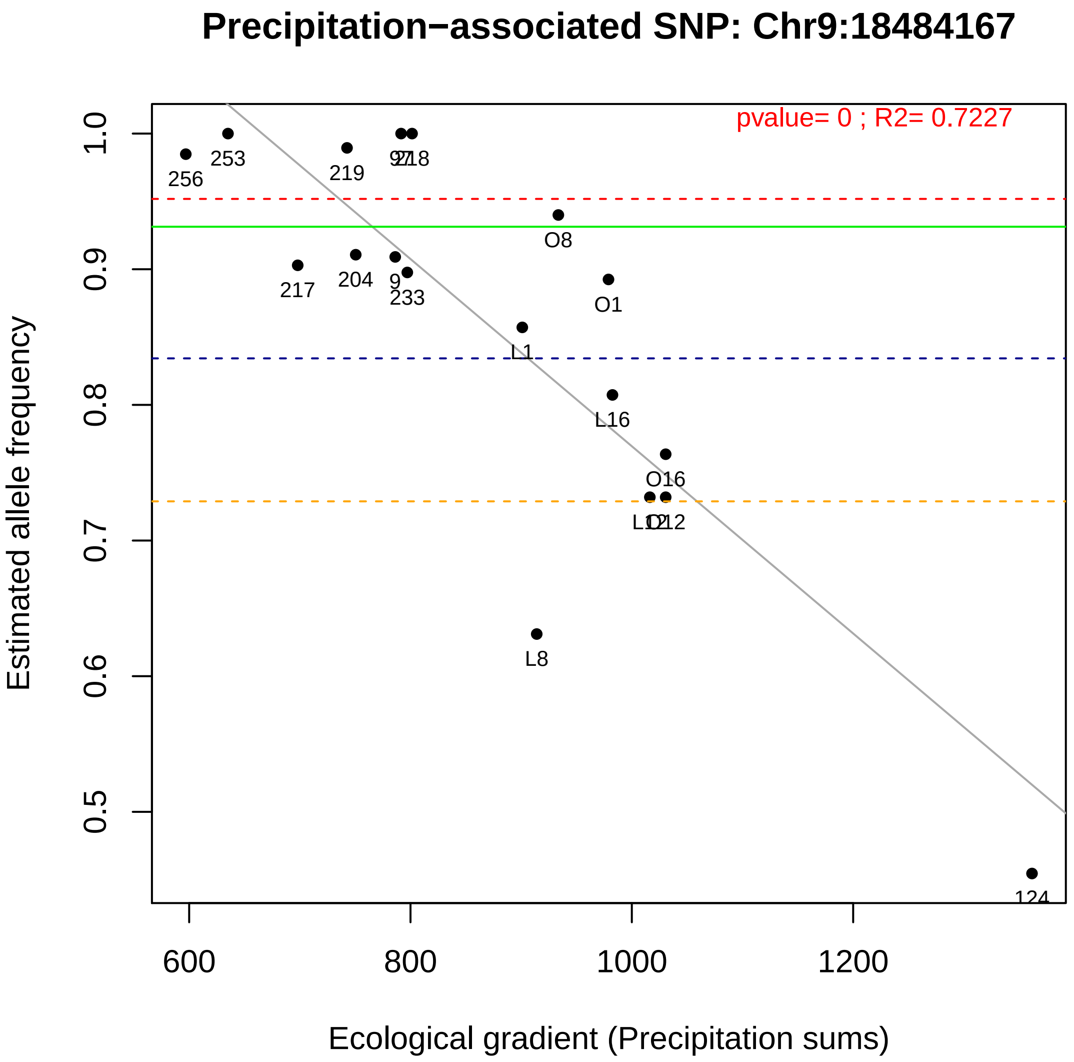

Precipitation−associated SNP: Chr9:18484167
pvalue= 0 ; R2= 0.7227
Estimated allele frequency
Ecological gradient (Precipitation sums)
1.0
0.9
0.8
0.7
0.6
0.5
600
800
1000
1200
256
253
219
97
218
O8
217
204
9
233
O1
L1
L16
O16
L10
O12
L8
124

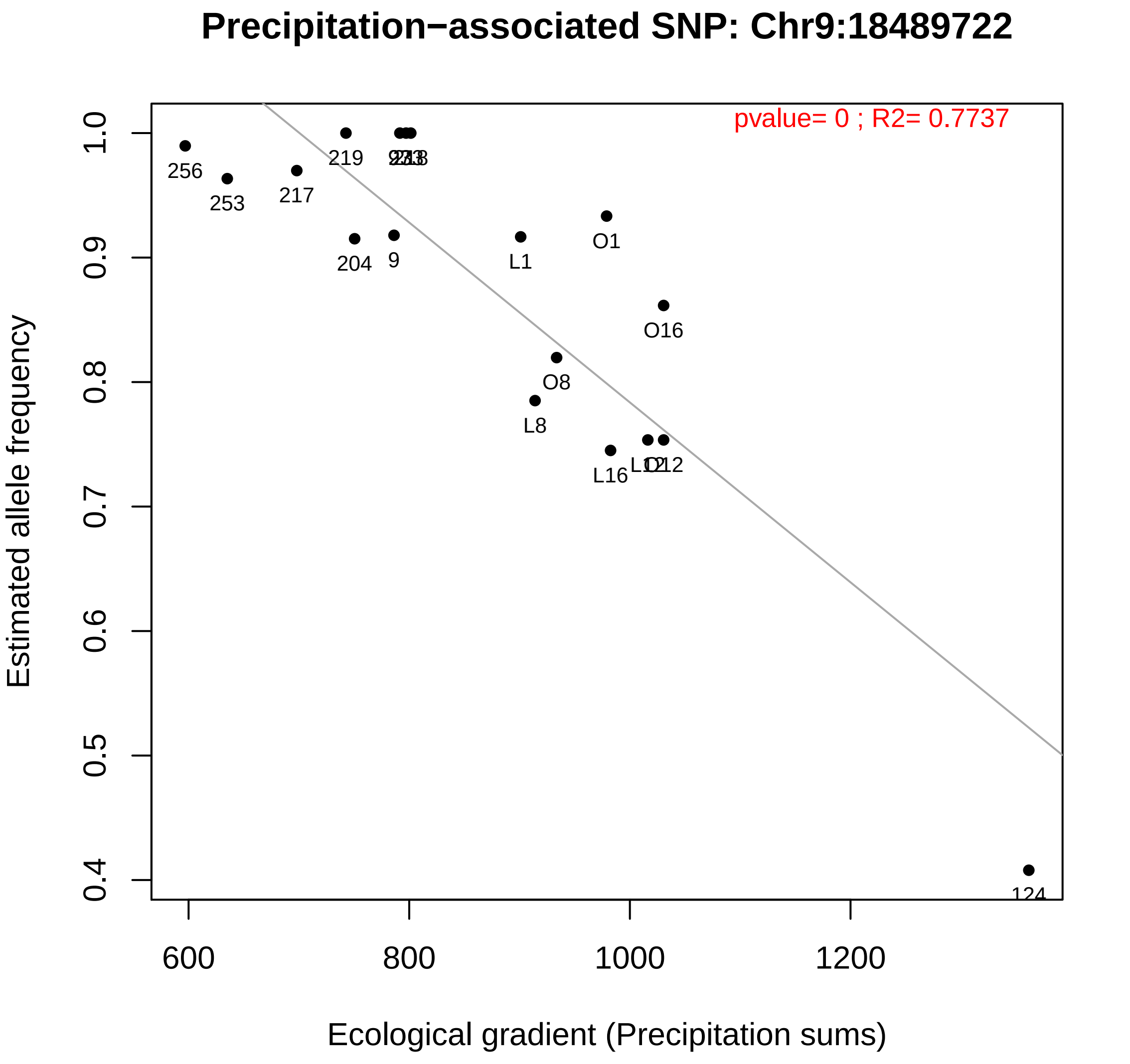
Precipitation−associated SNP: Chr9:18489722
pvalue= 0 ; R2= 0.7737
Estimated allele frequency
Ecological gradient (Precipitation sums)
1.0
0.9
0.8
0.7
0.6
0.5
0.4
600
800
1000
1200
256
253
217
219
204
9
L1
O1
O16
O8
L8
L16
O12
124

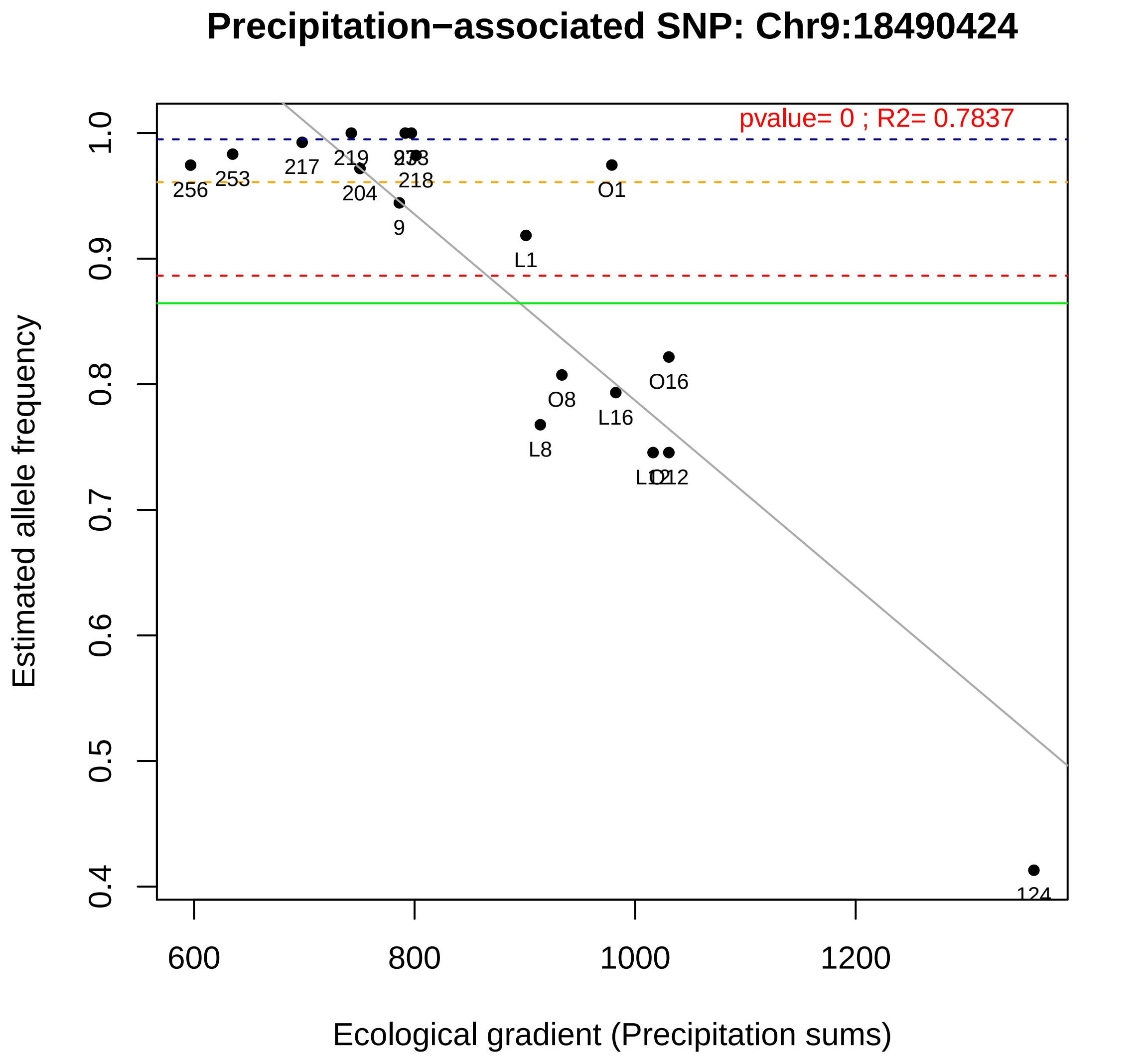

Precipitation−associated SNP: Chr9:18490424
pvalue= 0 ; R2= 0.7837
Estimated allele frequency
Ecological gradient (Precipitation sums)
0.4
0.5
0.6
0.7
0.8
0.9
1.0
600
800
1000
1200
256
253
217
219
204
218
9
L1
O1
O16
O8
L16
L8
124

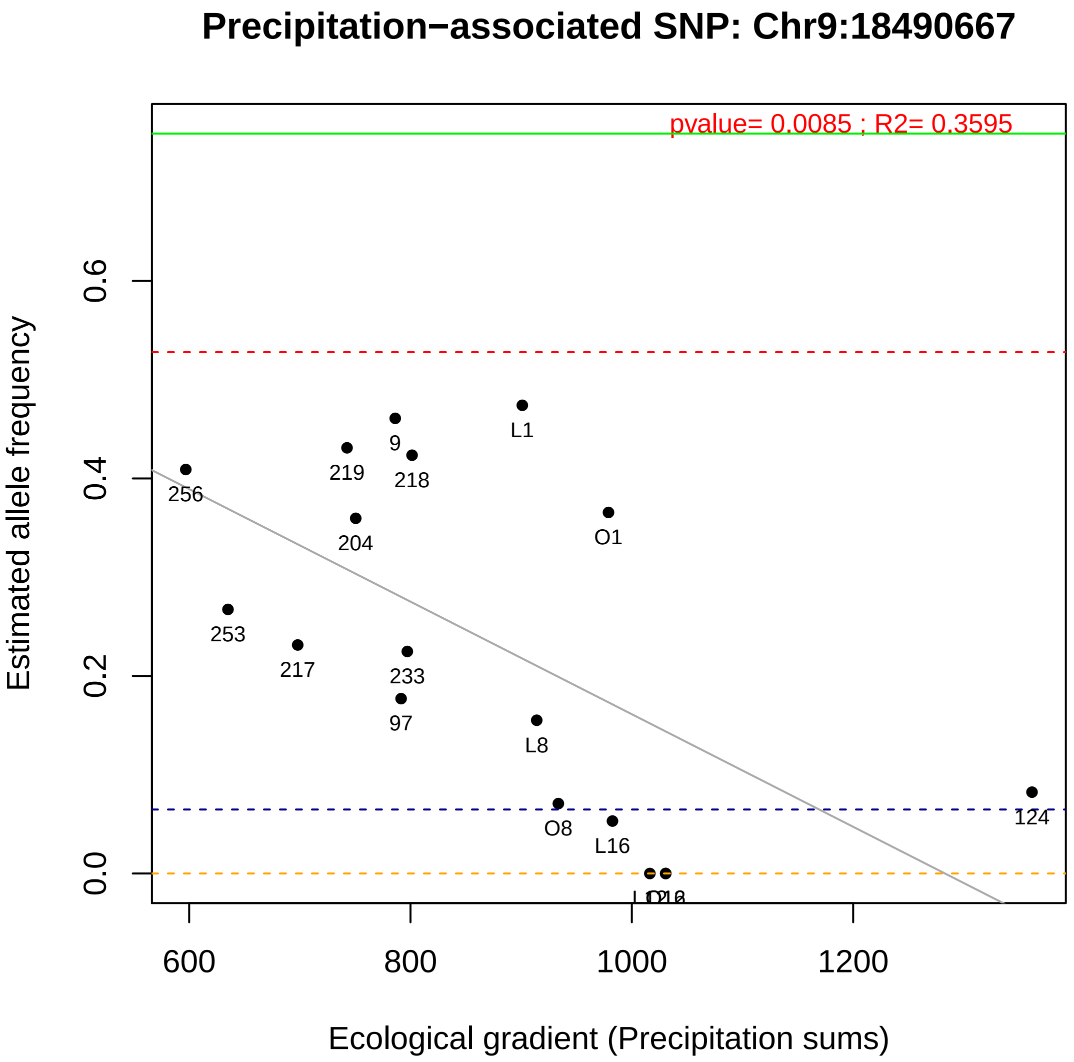
Precipitation−associated SNP: Chr9:18490667
pvalue= 0.0085 ; R2= 0.3595
Estimated allele frequency
0.0
0.2
0.4
0.6
Ecological gradient (Precipitation sums)
600
800
1000
1200
256
253
217
219
204
9
218
233
97
L1
O1
L8
O8
L16
124

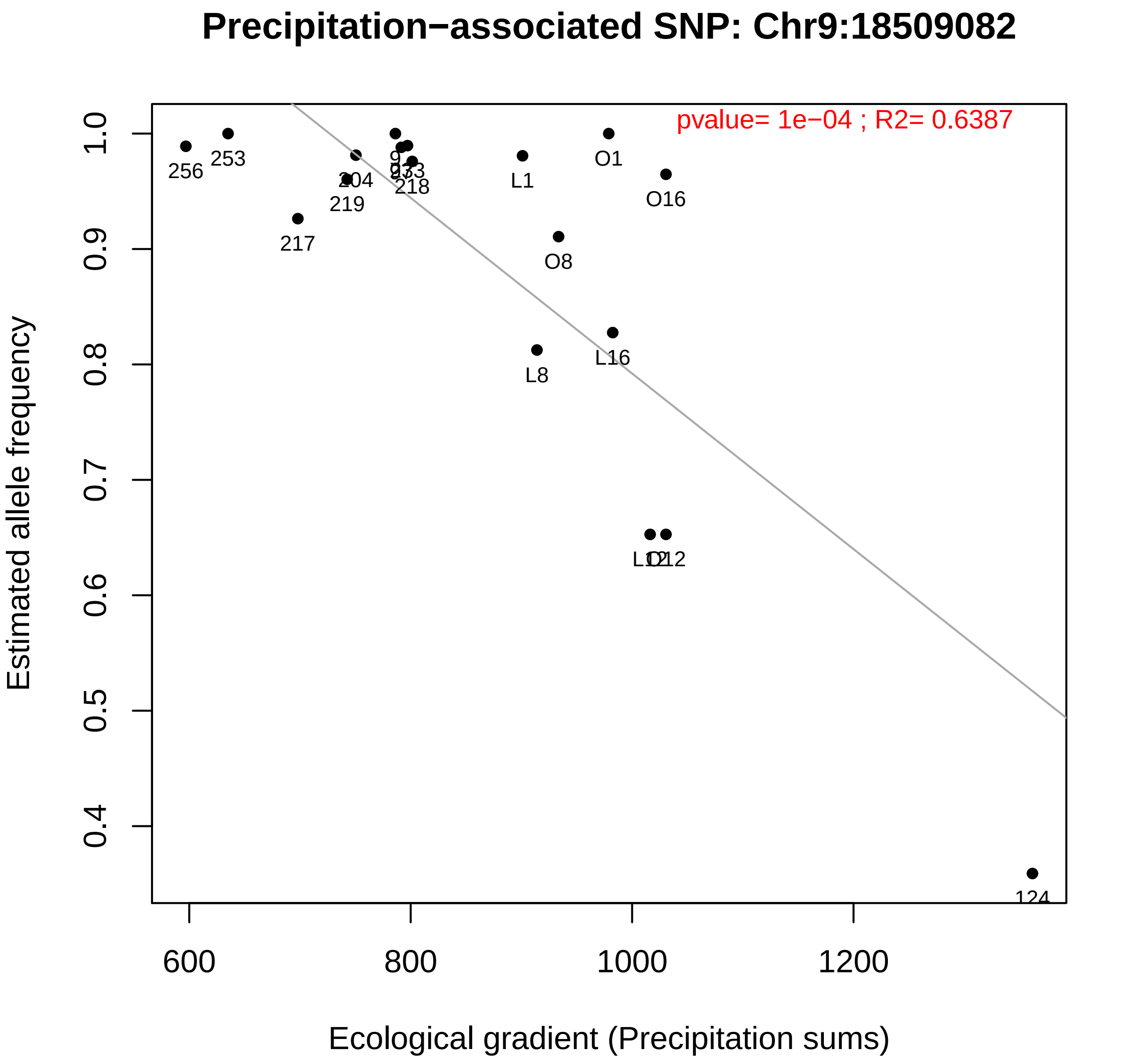

Precipitation−associated SNP: Chr9:18509082
pvalue= 1e−04 ; R2= 0.6387
Estimated allele frequency
Ecological gradient (Precipitation sums)
0.4
0.5
0.6
0.7
0.8
0.9
1.0
600
800
1000
1200
256
253
204
219
217
218
L1
O1
O16
O8
L16
L8
124

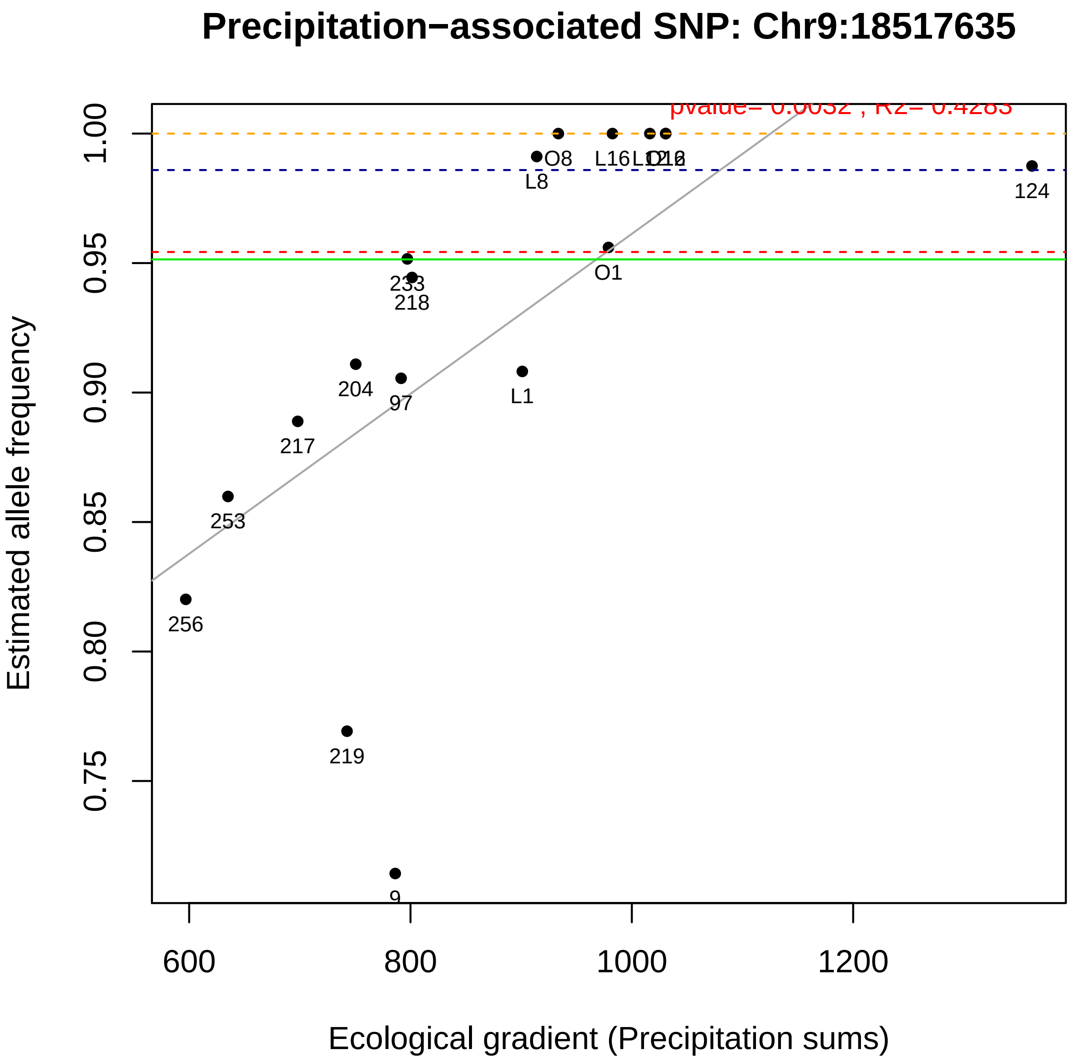
Precipitation−associated SNP: Chr9:18517635
pvalue= 0.0032 , R2= 0.4283
Estimated allele frequency
Ecological gradient (Precipitation sums)
1.00
0.95
0.90
0.85
0.80
0.75
600
800
1000
1200
O8
L16
L8
124
O1
233
218
204
97
L1
217
253
256
219
9

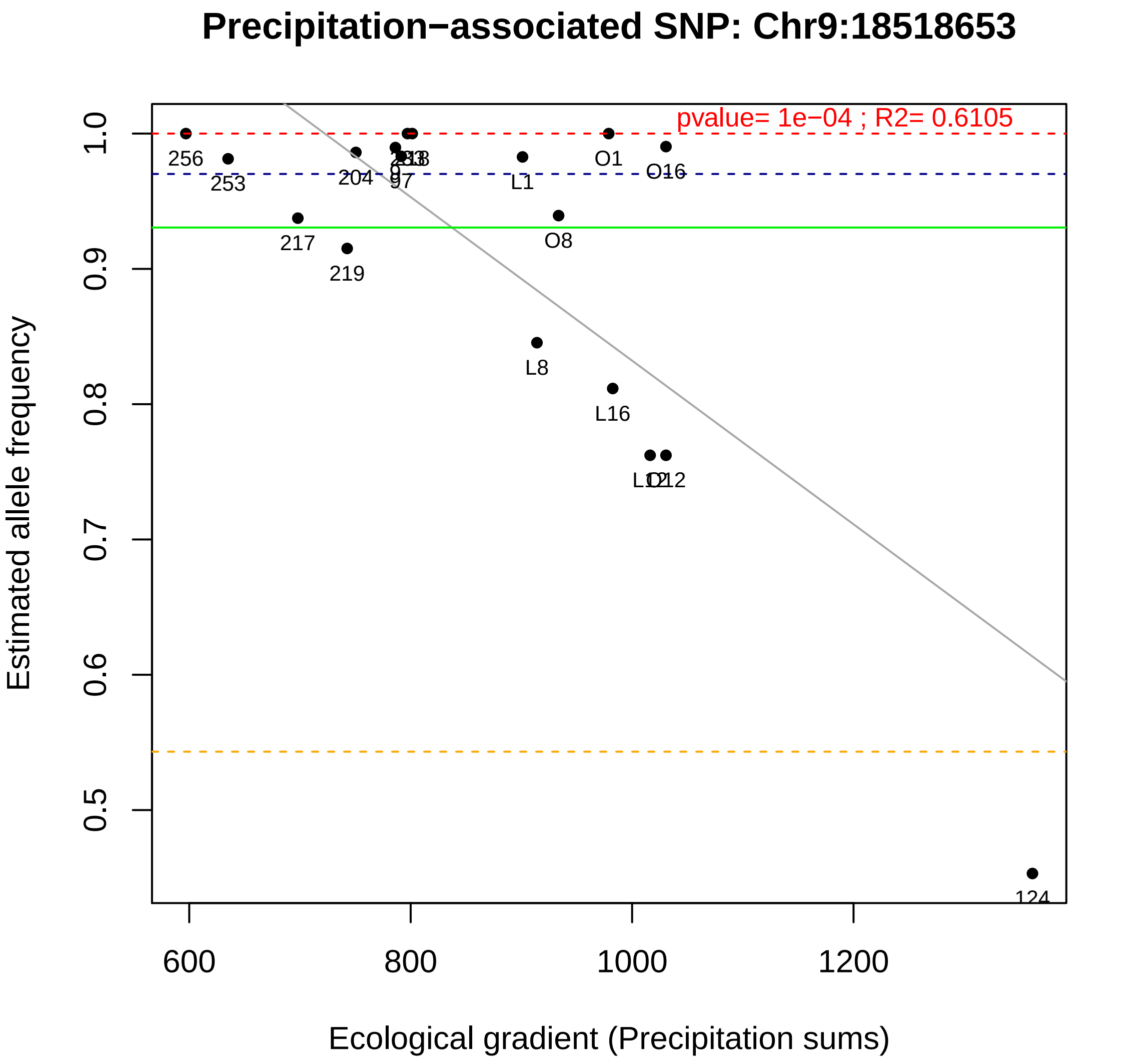
Precipitation−associated SNP: Chr9:18518653
pvalue= 1e−04 ; R2= 0.6105
Estimated allele frequency
0.5
0.6
0.7
0.8
0.9
1.0
600
800
1000
1200
Ecological gradient (Precipitation sums)
256
253
204
217
219
97
L1
O1
O16
O8
L8
L16
124

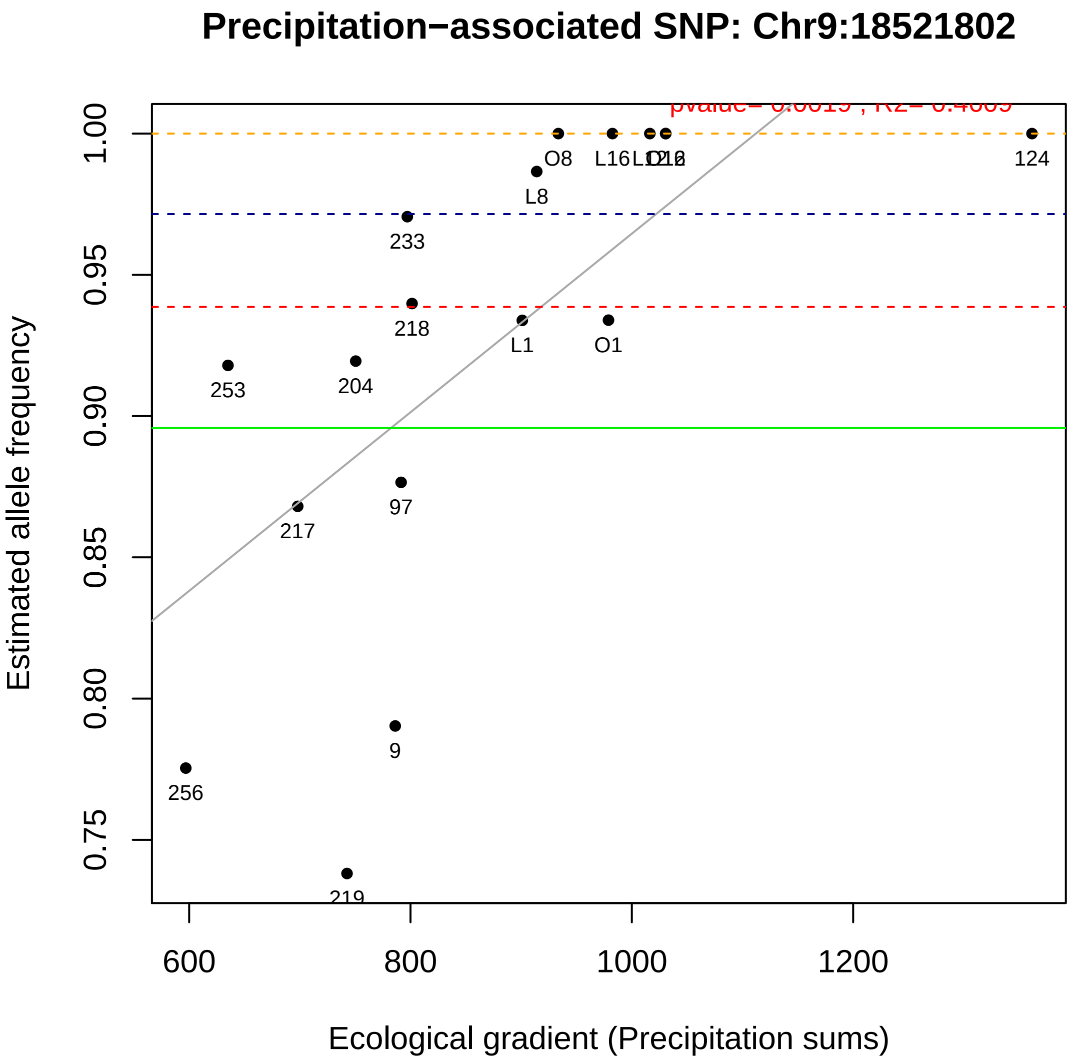

Precipitation−associated SNP: Chr9:18521802
pvalue=
Estimated allele frequency
1.00
0.95
0.90
0.85
0.80
0.75
600
800
1000
1200
Ecological gradient (Precipitation sums)
O8
L16
124
L8
233
218
L1
O1
253
204
97
217
9
256
219

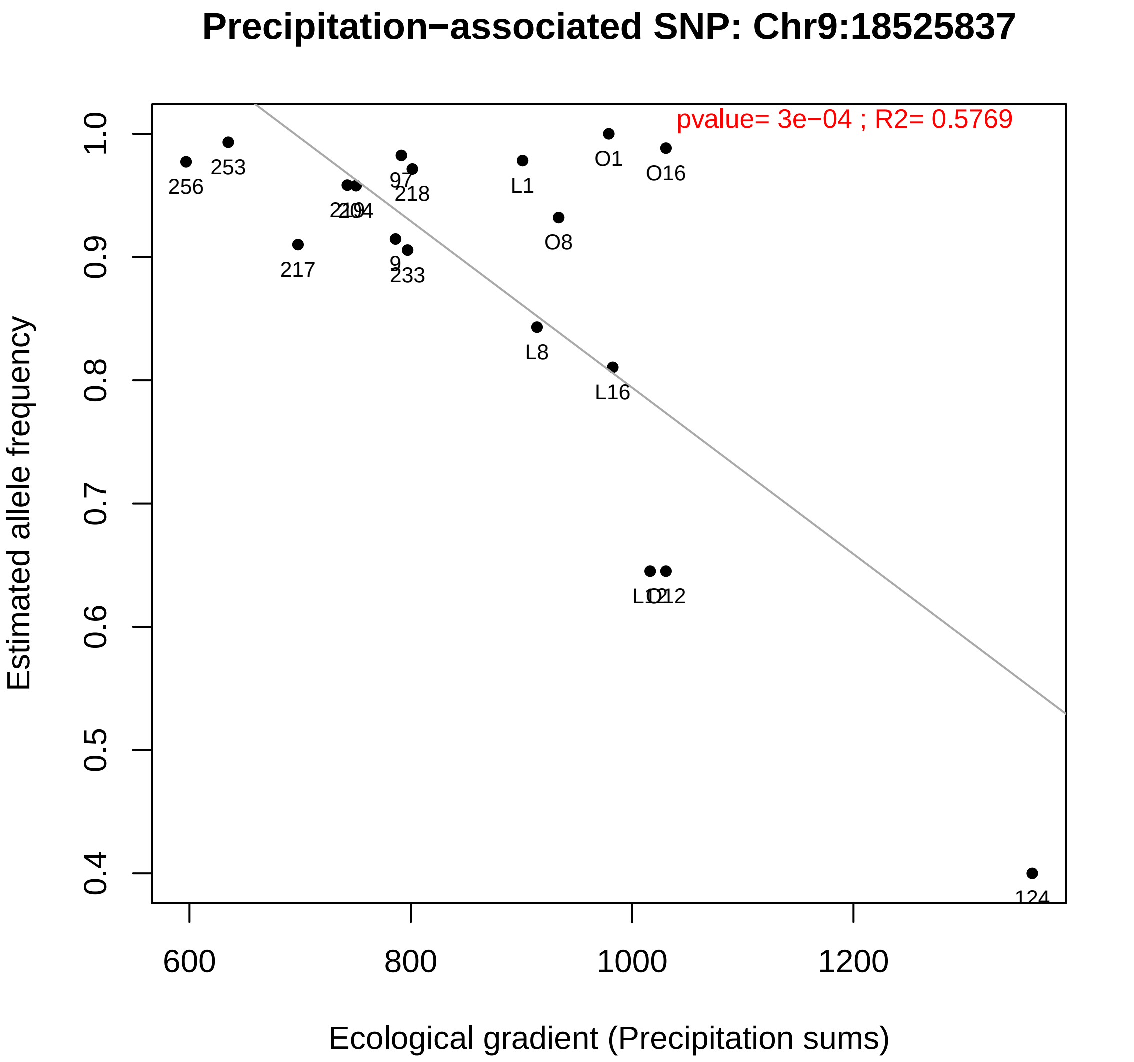
Precipitation−associated SNP: Chr9:18525837
pvalue= 3e−04 ; R2= 0.5769
Estimated allele frequency
Ecological gradient (Precipitation sums)
0.4
0.5
0.6
0.7
0.8
0.9
1.0
600
800
1000
1200
256
253
97
218
L1
O1
O16
O8
217
9
233
L8
L16
124

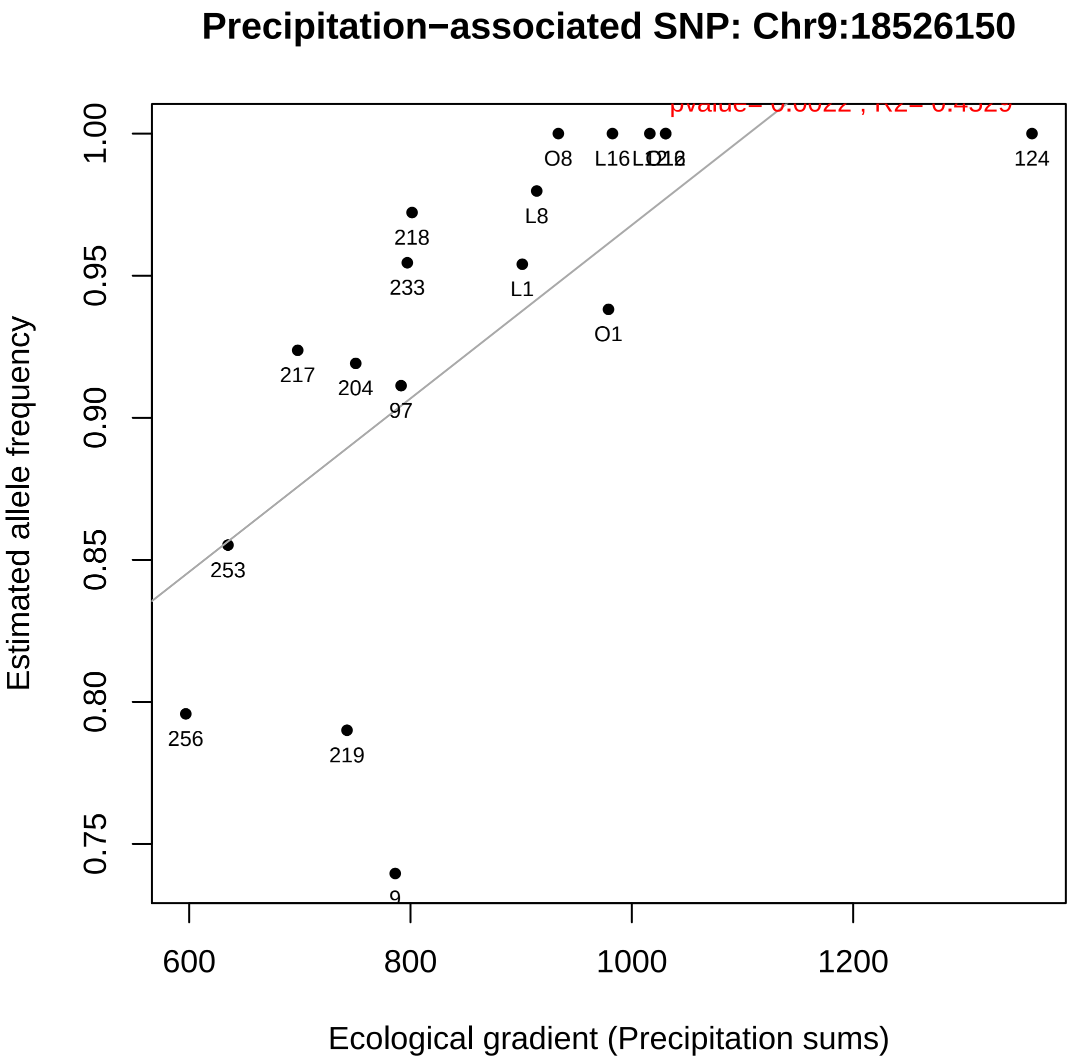

Precipitation−associated SNP: Chr9:18526150
Estimated allele frequency
Ecological gradient (Precipitation sums)
0.75
0.80
0.85
0.90
0.95
1.00
600
800
1000
1200
O8
L16
124
L8
218
233
L1
O1
217
204
97
253
256
219
9

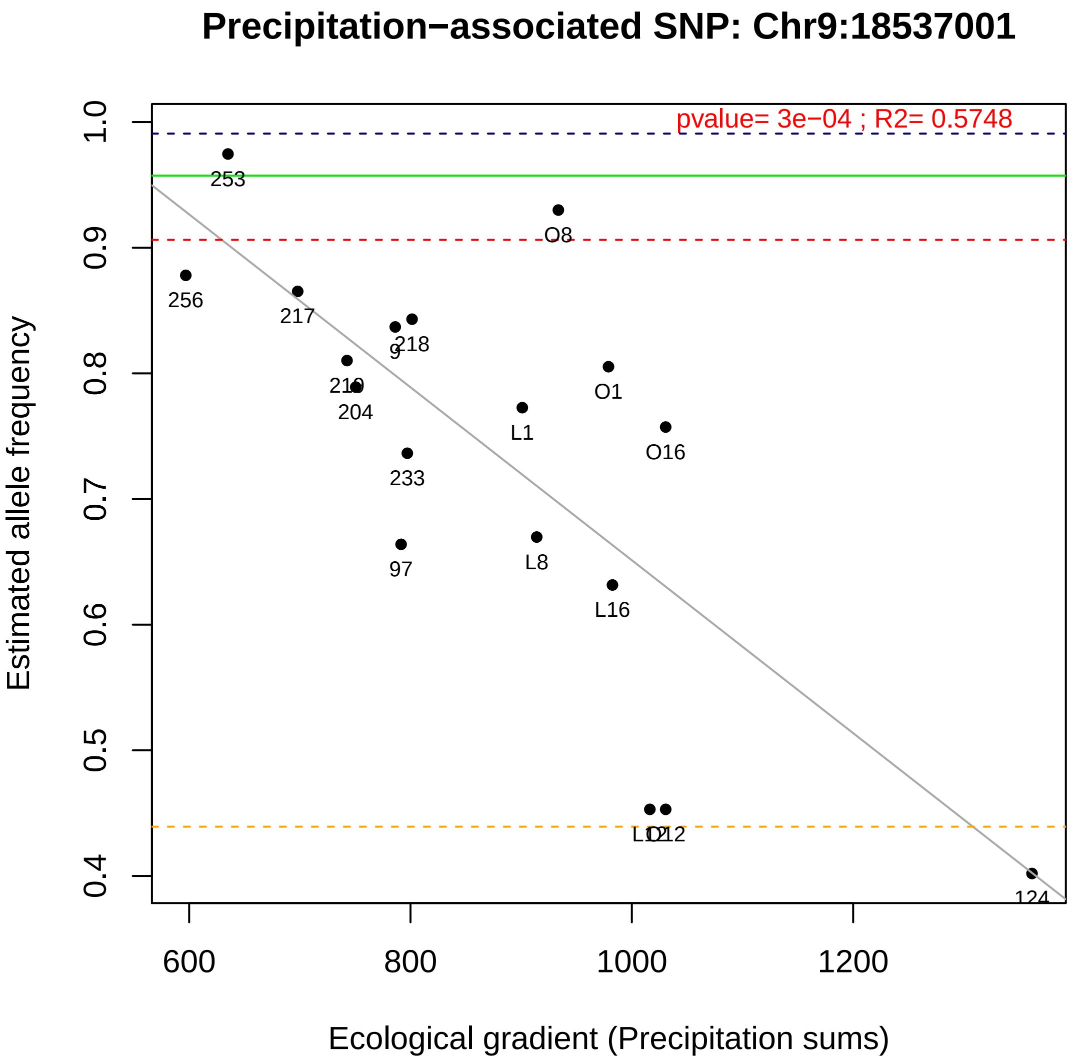

Precipitation−associated SNP: Chr9:18537001
pvalue= 3e−04 ; R2= 0.5748
Estimated allele frequency
Ecological gradient (Precipitation sums)
1.0
0.9
0.8
0.7
0.6
0.5
0.4
600
800
1000
1200
253
O8
256
217
218
9
210
204
O1
L1
O16
233
97
L8
L16
L12
O12
124

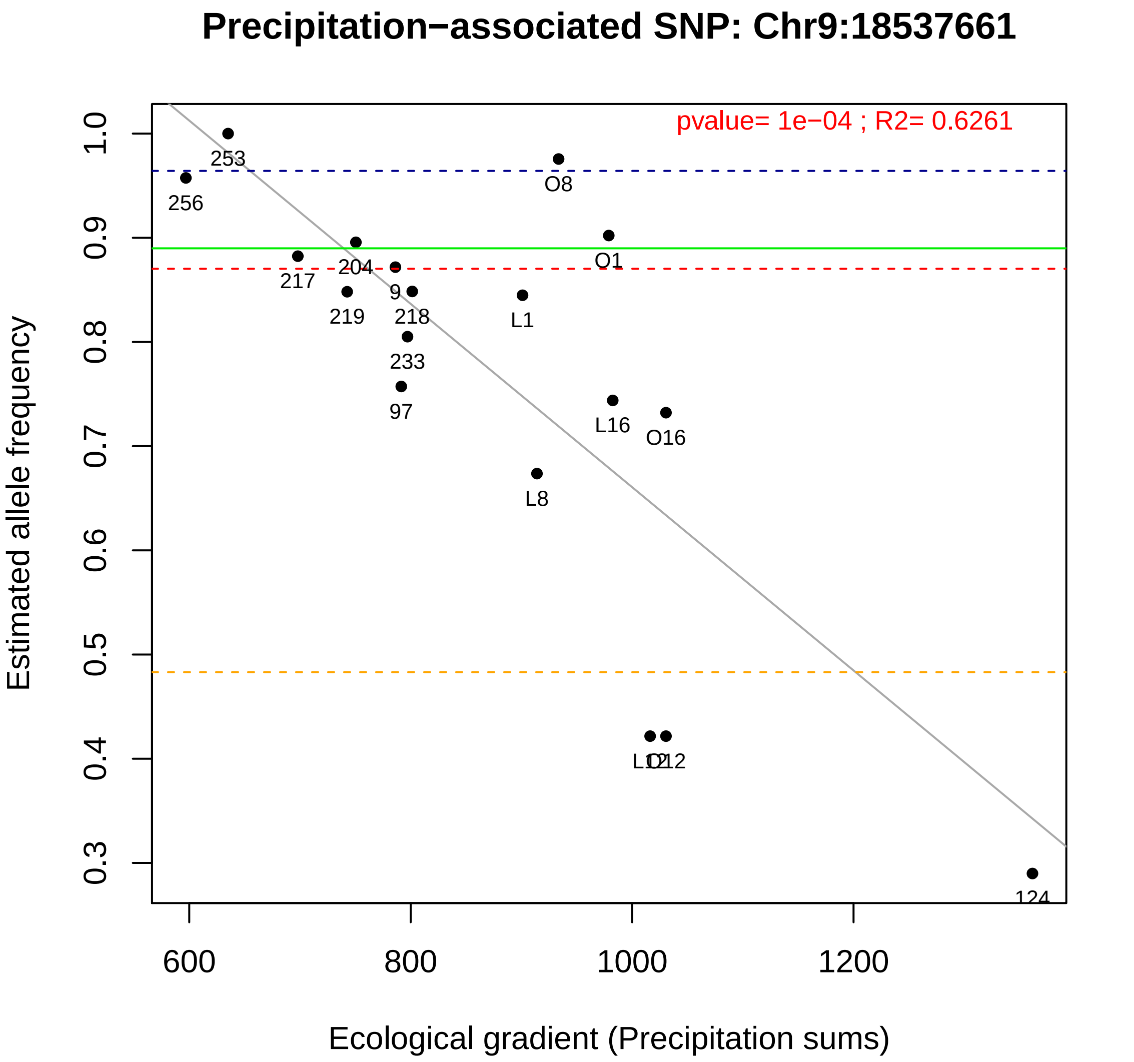
Precipitation−associated SNP: Chr9:18537661
pvalue= 1e−04 ; R2= 0.6261
Estimated allele frequency
Ecological gradient (Precipitation sums)
1.0
0.9
0.8
0.7
0.6
0.5
0.4
0.3
600
800
1000
1200
253
256
O8
217
204
9
218
219
O1
L1
233
97
L16
O16
L8
L12
O12
124

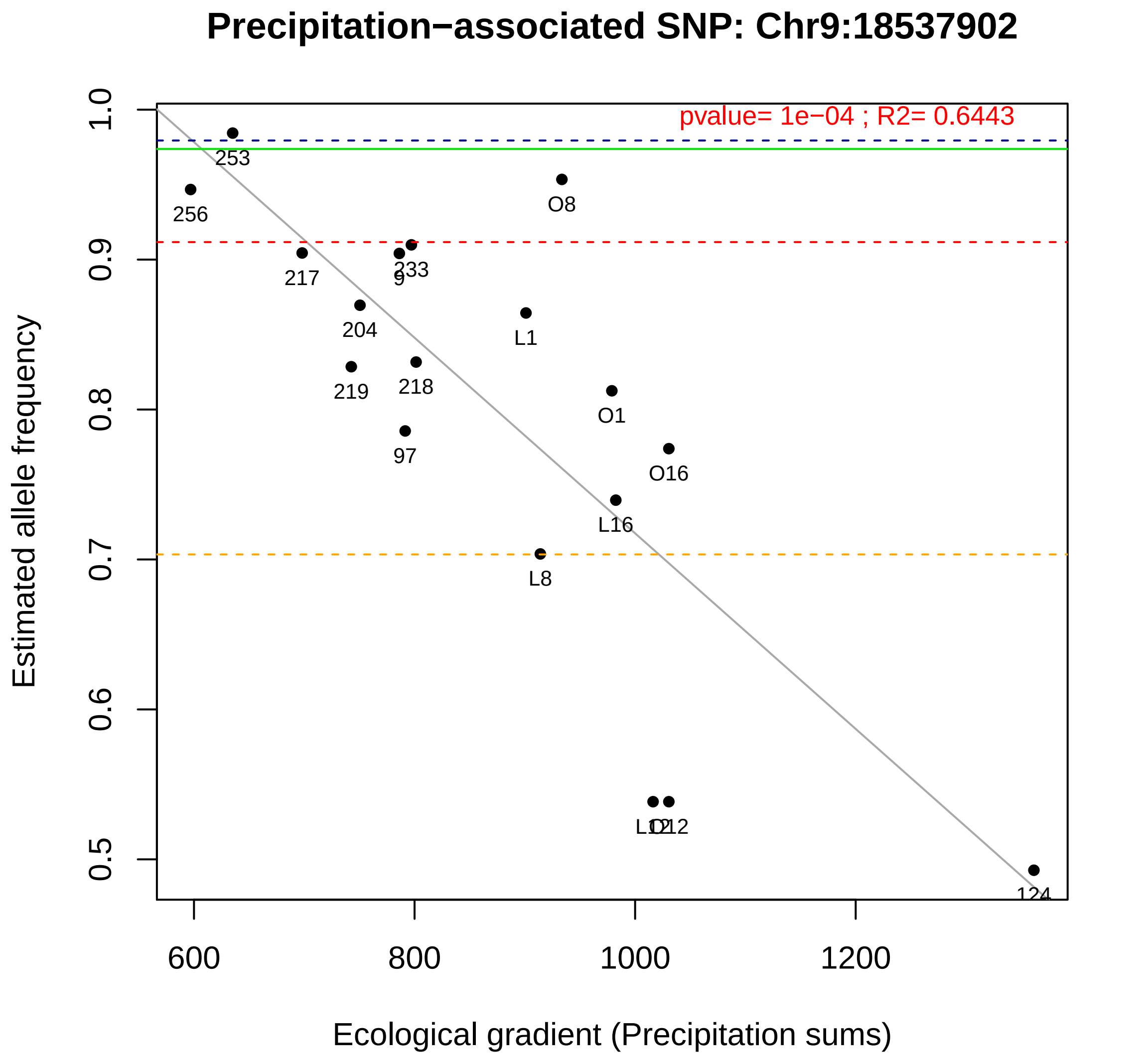

Precipitation−associated SNP: Chr9:18537902
pvalue= 1e−04 ; R2= 0.6443
Estimated allele frequency
Ecological gradient (Precipitation sums)
0.5
0.6
0.7
0.8
0.9
1.0
600
800
1000
1200
253
256
O8
217
233
9
204
L1
219
218
O1
97
O16
L16
L8
124

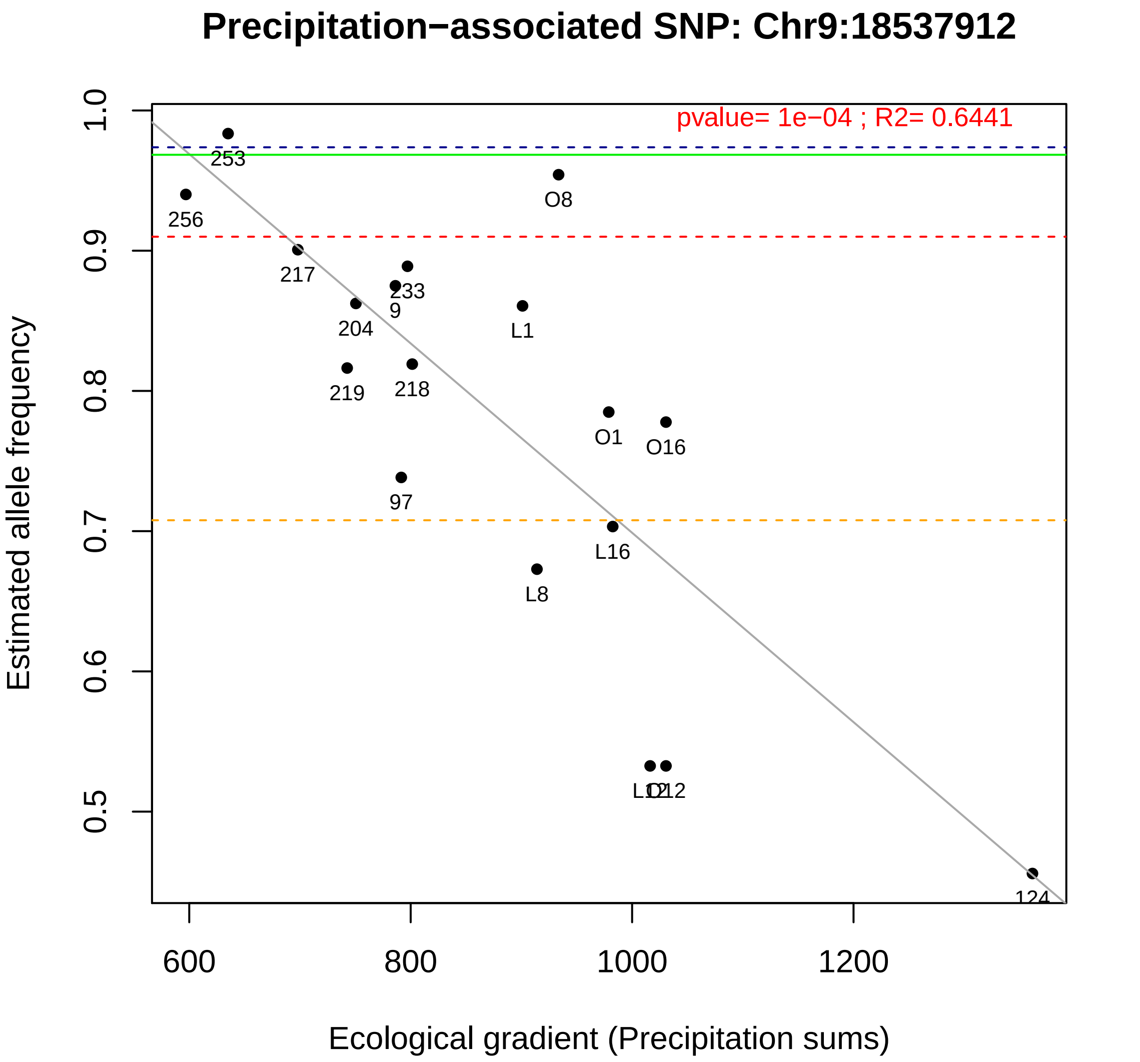
Precipitation−associated SNP: Chr9:18537912
pvalue= 1e−04 ; R2= 0.6441
Estimated allele frequency
Ecological gradient (Precipitation sums)
1.0
0.9
0.8
0.7
0.6
0.5
600
800
1000
1200
253
256
O8
217
233
9
204
L1
219
218
O1
O16
97
L16
L8
L12
O12
124

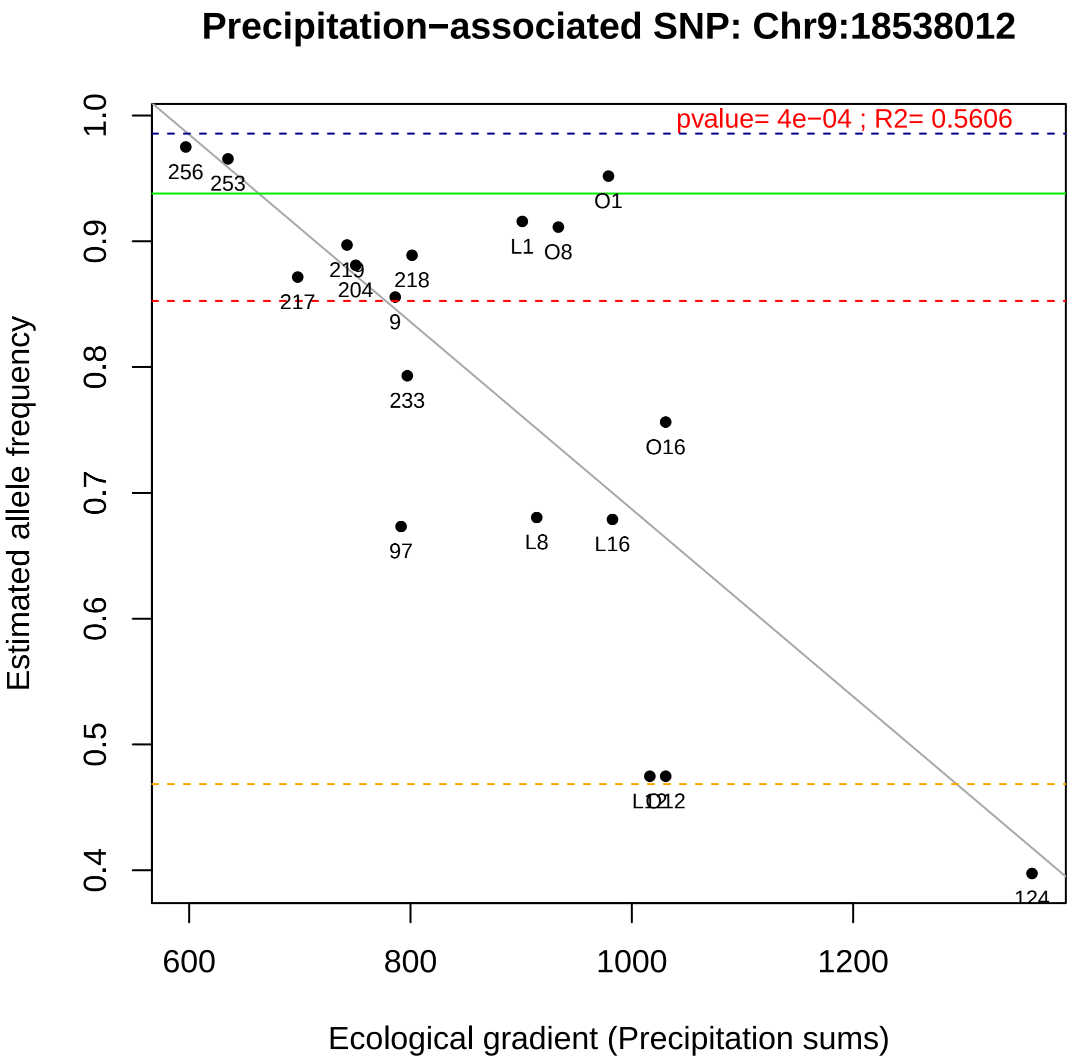

Precipitation−associated SNP: Chr9:18538012
pvalue= 4e−04 ; R2= 0.5606
Estimated allele frequency
Ecological gradient (Precipitation sums)
1.0
0.9
0.8
0.7
0.6
0.5
0.4
600
800
1000
1200
256
253
O1
L1
O8
219
204
218
217
9
233
O16
97
L8
L16
L12
O12
124

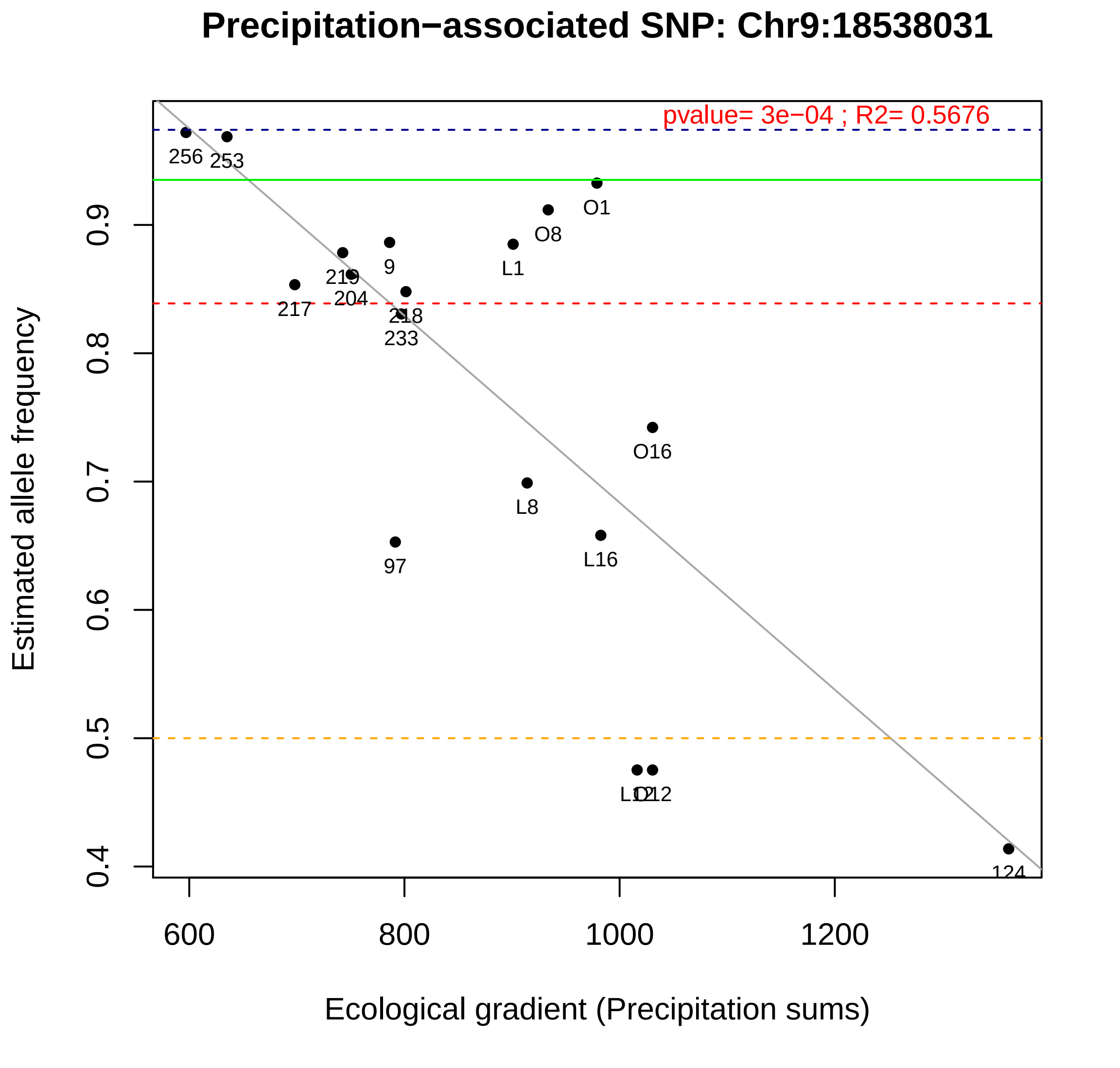
Precipitation−associated SNP: Chr9:18538031
pvalue= 3e−04 ; R2= 0.5676
Estimated allele frequency
0.4
0.5
0.6
0.7
0.8
0.9
600
800
1000
1200
Ecological gradient (Precipitation sums)
256
253
O1
O8
9
L1
219
204
217
218
233
O16
L8
97
L16
L12
O12
124

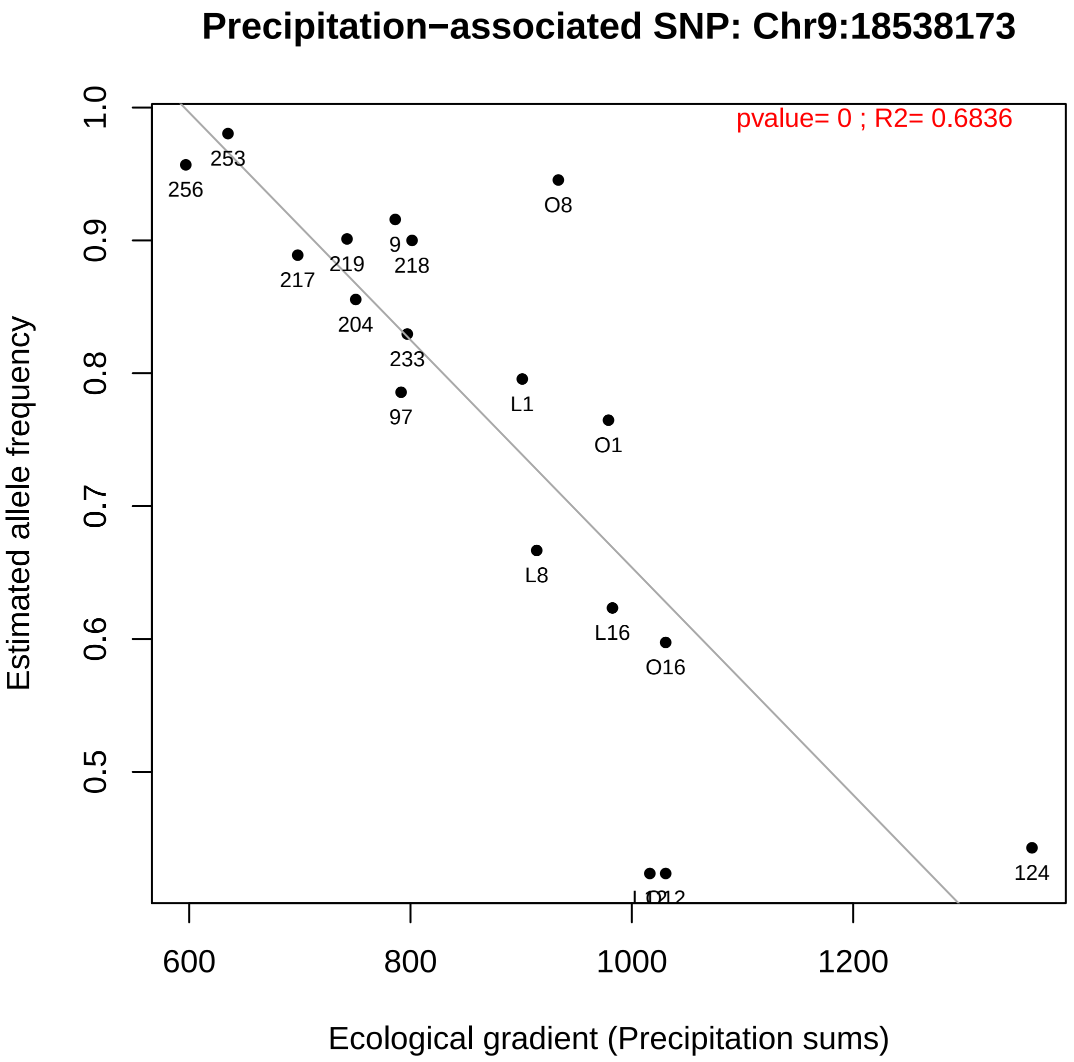

Precipitation−associated SNP: Chr9:18538173
pvalue= 0 ; R2= 0.6836
Estimated allele frequency
Ecological gradient (Precipitation sums)
1.0
0.9
0.8
0.7
0.6
0.5
600
800
1000
1200
253
256
O8
9
219
218
217
204
233
L1
97
O1
L8
L16
O16
124
O12

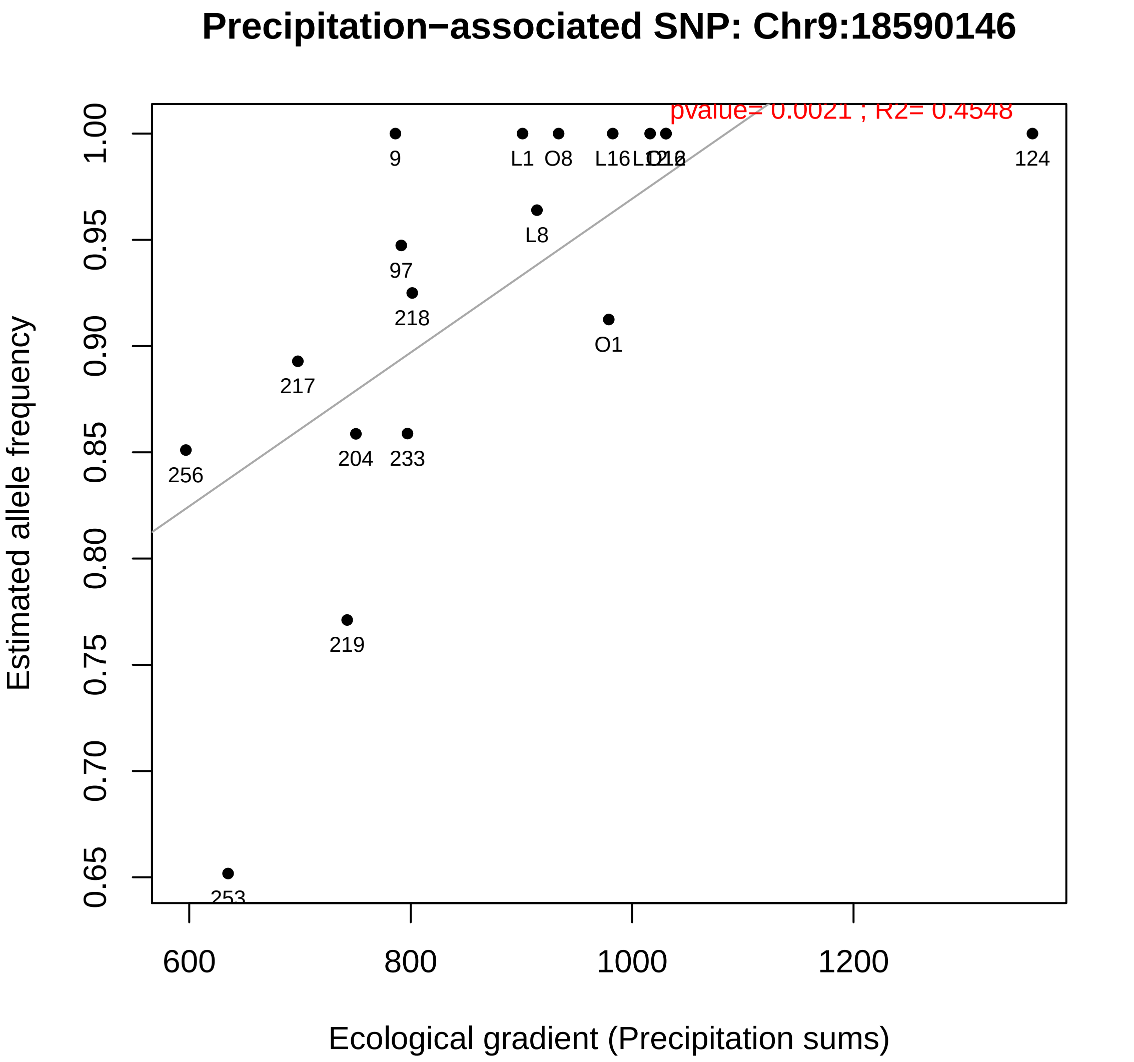

Precipitation−associated SNP: Chr9:18590146
pvalue= 0.0021 ; R2= 0.4548
Estimated allele frequency
Ecological gradient (Precipitation sums)
0.65
0.70
0.75
0.80
0.85
0.90
0.95
1.00
600
800
1000
1200
9
L1
O8
L16
124
L8
97
218
O1
217
256
204
233
219
253

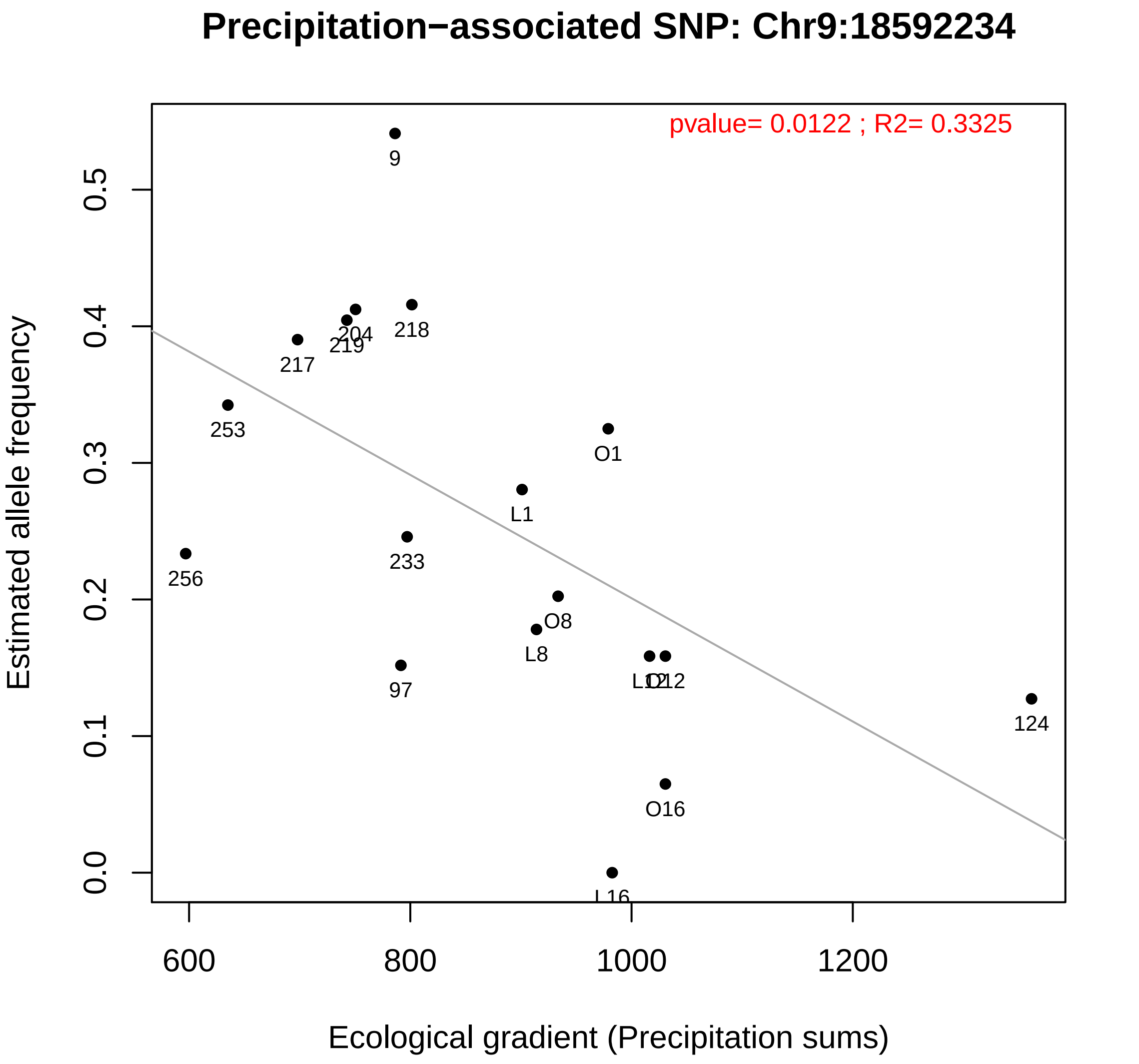
Precipitation−associated SNP: Chr9:18592234
pvalue= 0.0122 ; R2= 0.3325
Estimated allele frequency
0.0
0.1
0.2
0.3
0.4
0.5
Ecological gradient (Precipitation sums)
600
800
1000
1200
9
204
219
218
217
253
O1
L1
256
233
O8
L8
97
124
O16
L16

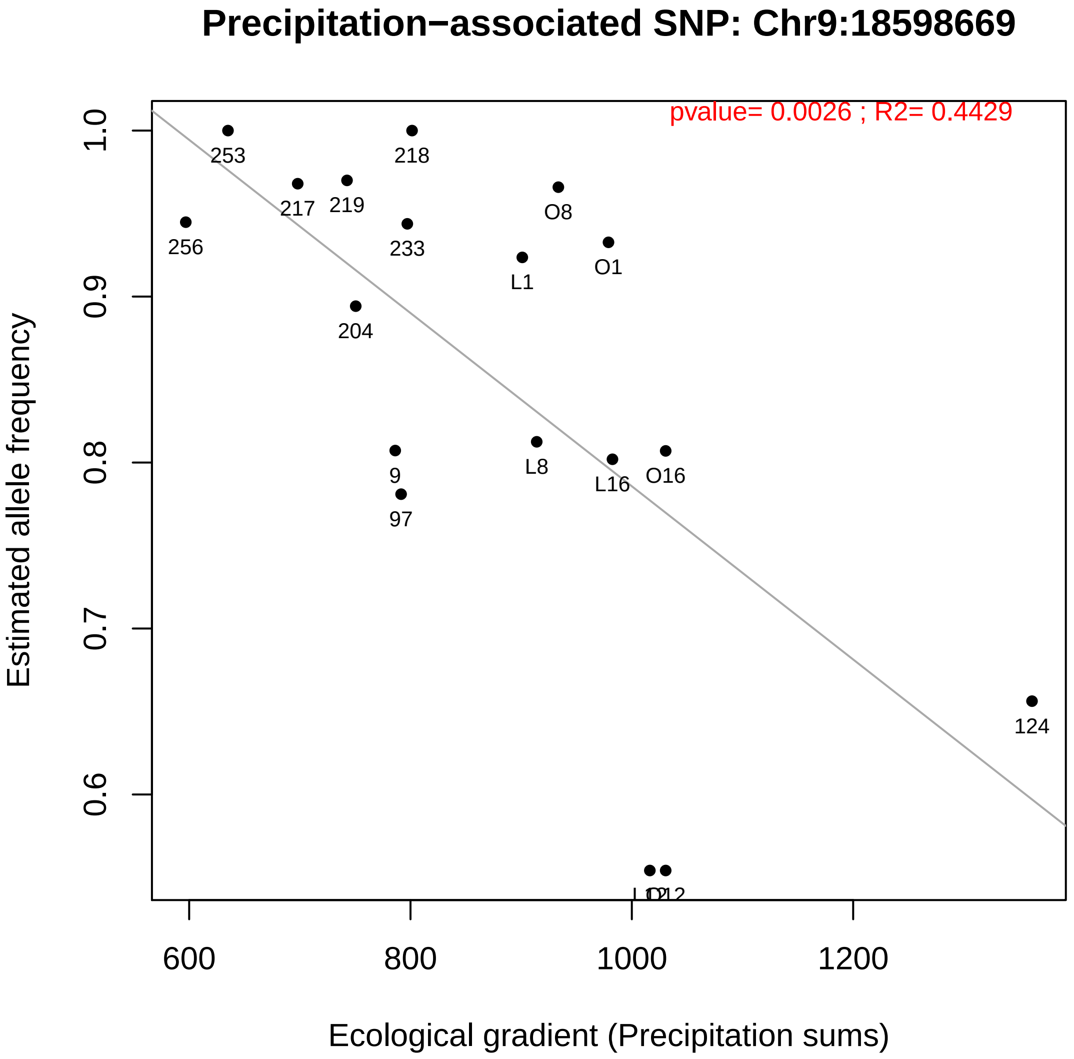
Precipitation−associated SNP: Chr9:18598669
pvalue= 0.0026 ; R2= 0.4429
Estimated allele frequency
1.0
0.9
0.8
0.7
0.6
600
800
1000
1200
Ecological gradient (Precipitation sums)
253
218
217
219
256
233
O8
O1
L1
204
9
97
L8
L16
O16
124
O12

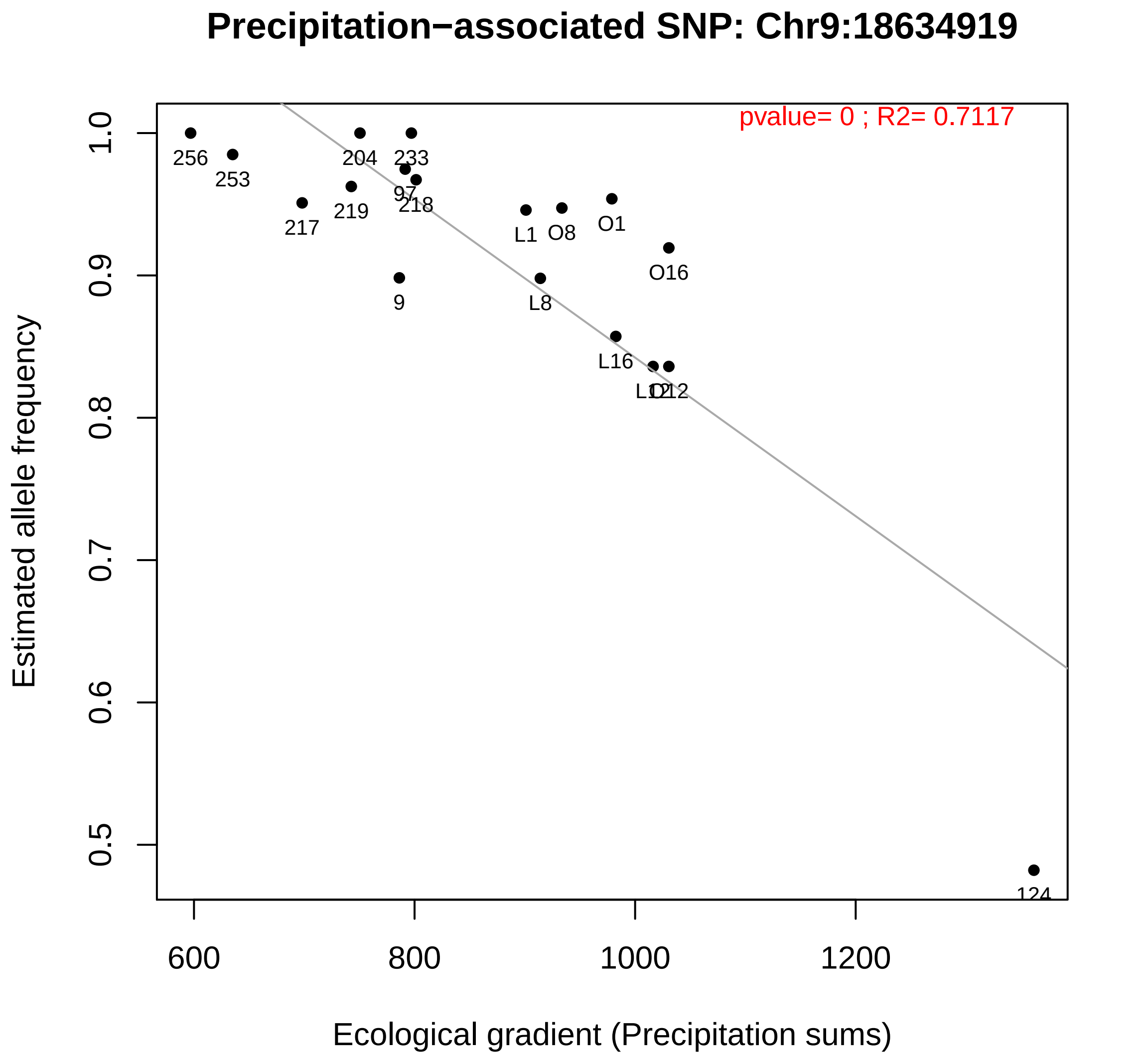

Precipitation−associated SNP: Chr9:18634919
pvalue= 0 ; R2= 0.7117
Estimated allele frequency
Ecological gradient (Precipitation sums)
0.5
0.6
0.7
0.8
0.9
1.0
600
800
1000
1200
256
253
204
233
97
218
219
217
9
L1
O8
O1
O16
L8
L16
124

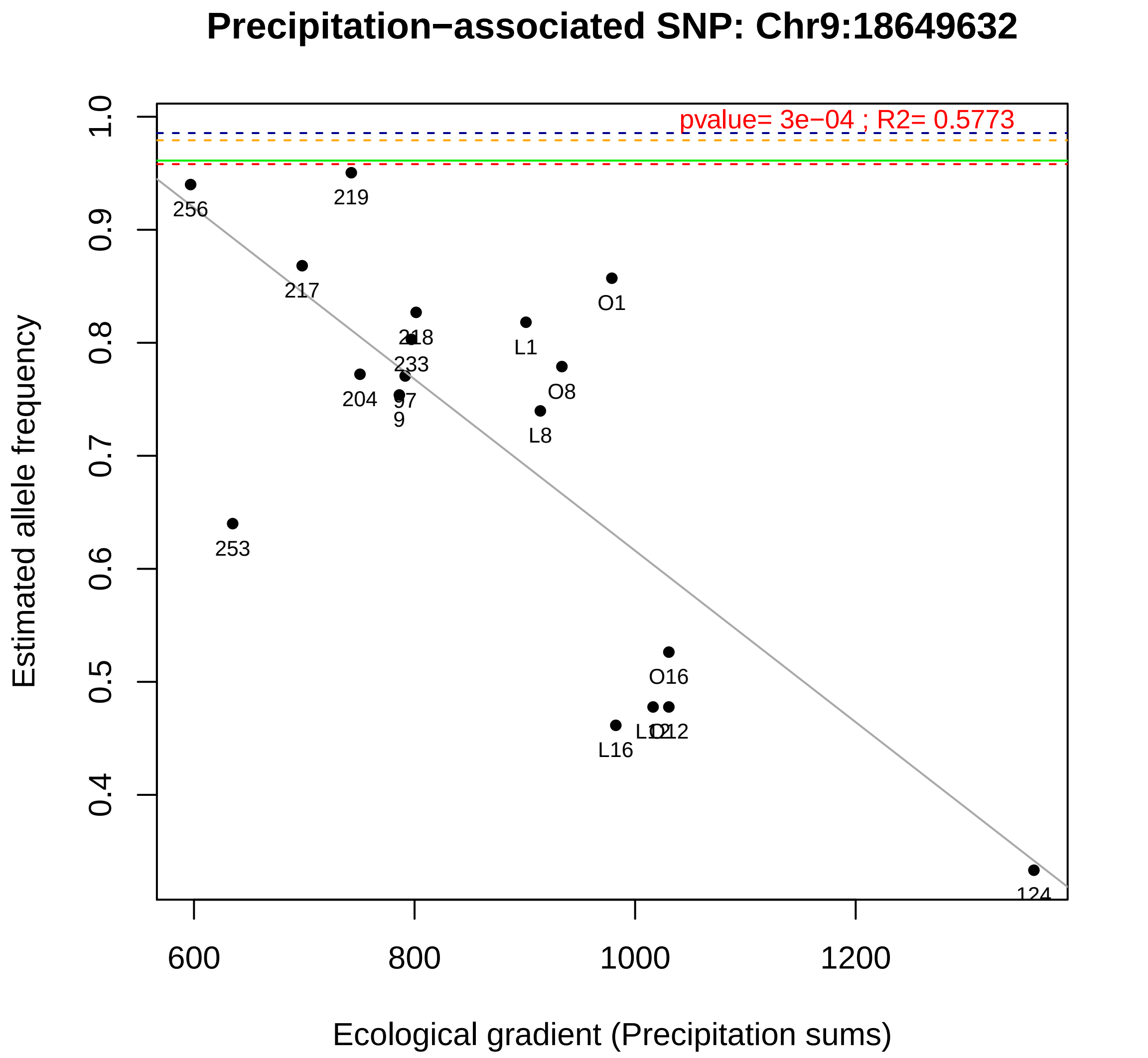

Precipitation−associated SNP: Chr9:18649632
pvalue= 3e−04 ; R2= 0.5773
Estimated allele frequency
Ecological gradient (Precipitation sums)
0.4
0.5
0.6
0.7
0.8
0.9
1.0
600
800
1000
1200
256
219
217
218
233
204
97
9
L1
O1
O8
L8
253
O16
L16
L12
O12
124

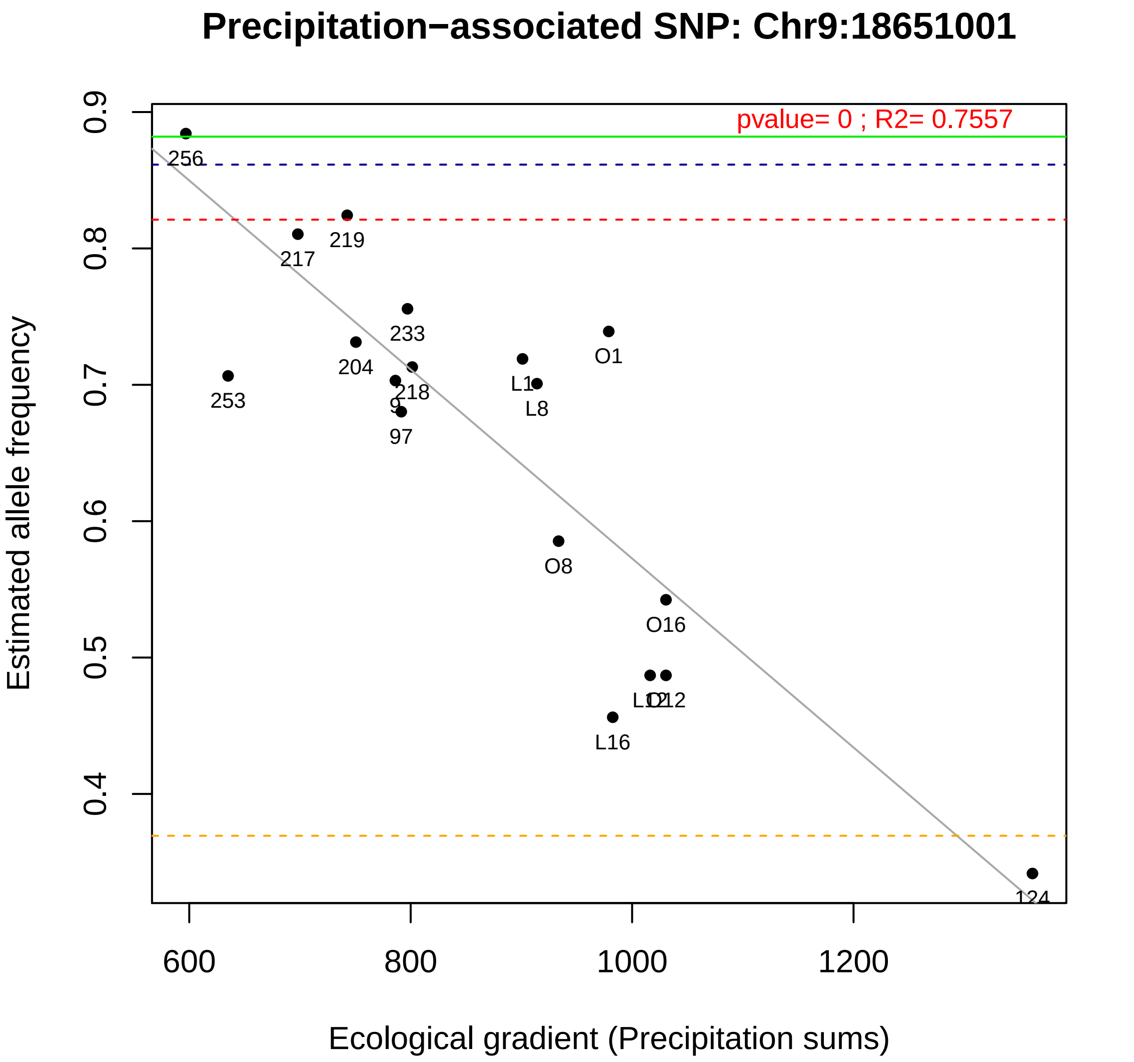
Precipitation−associated SNP: Chr9:18651001
pvalue= 0 ; R2= 0.7557
Estimated allele frequency
0.9
0.8
0.7
0.6
0.5
0.4
600
800
1000
1200
Ecological gradient (Precipitation sums)
256
219
217
233
204
253
218
97
L1
L8
O1
O8
O16
L12
O12
L16
124

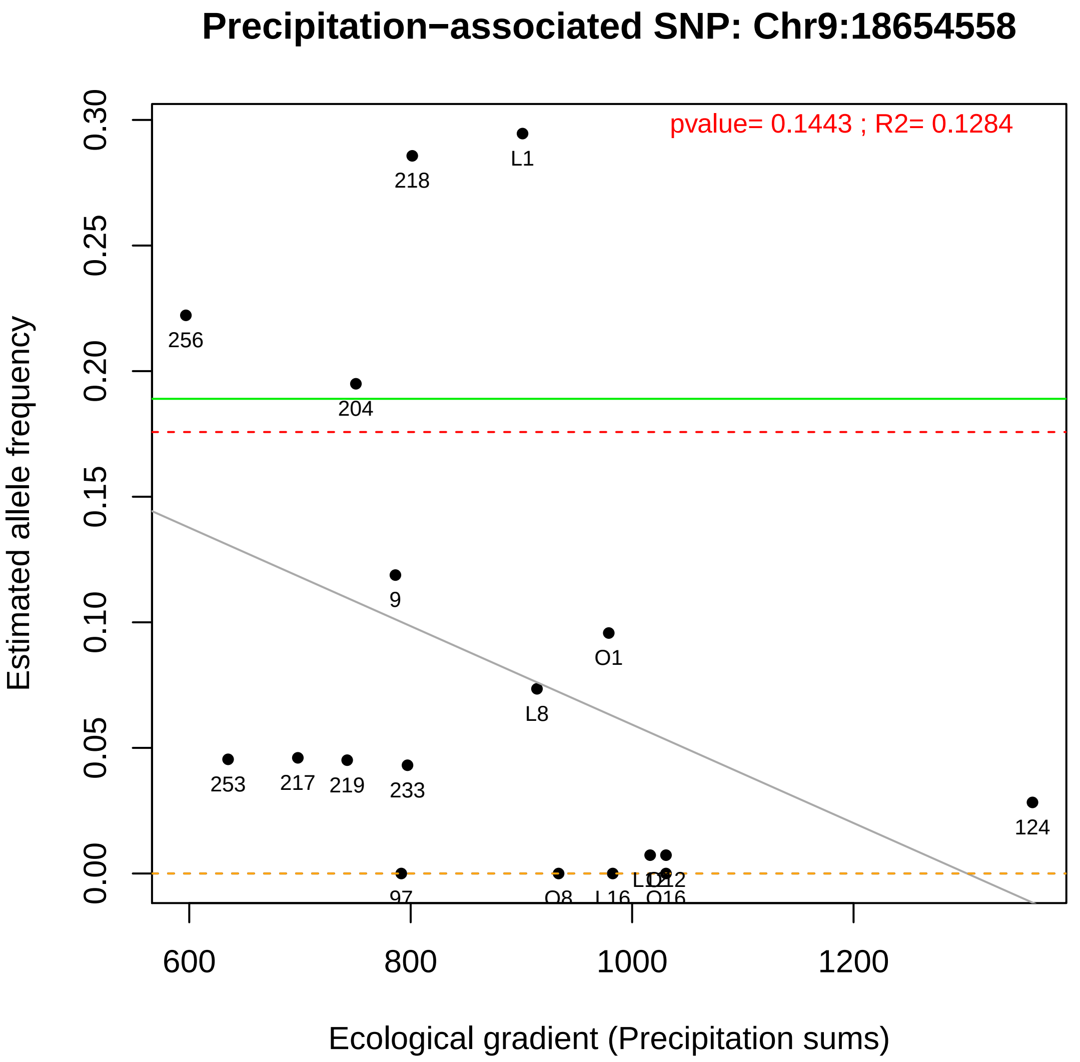

Precipitation−associated SNP: Chr9:18654558
pvalue= 0.1443 ; R2= 0.1284
Estimated allele frequency
Ecological gradient (Precipitation sums)
0.00
0.05
0.10
0.15
0.20
0.25
0.30
600
800
1000
1200
218
L1
256
204
9
O1
L8
253
217
219
233
124
97
O8
L16
O16

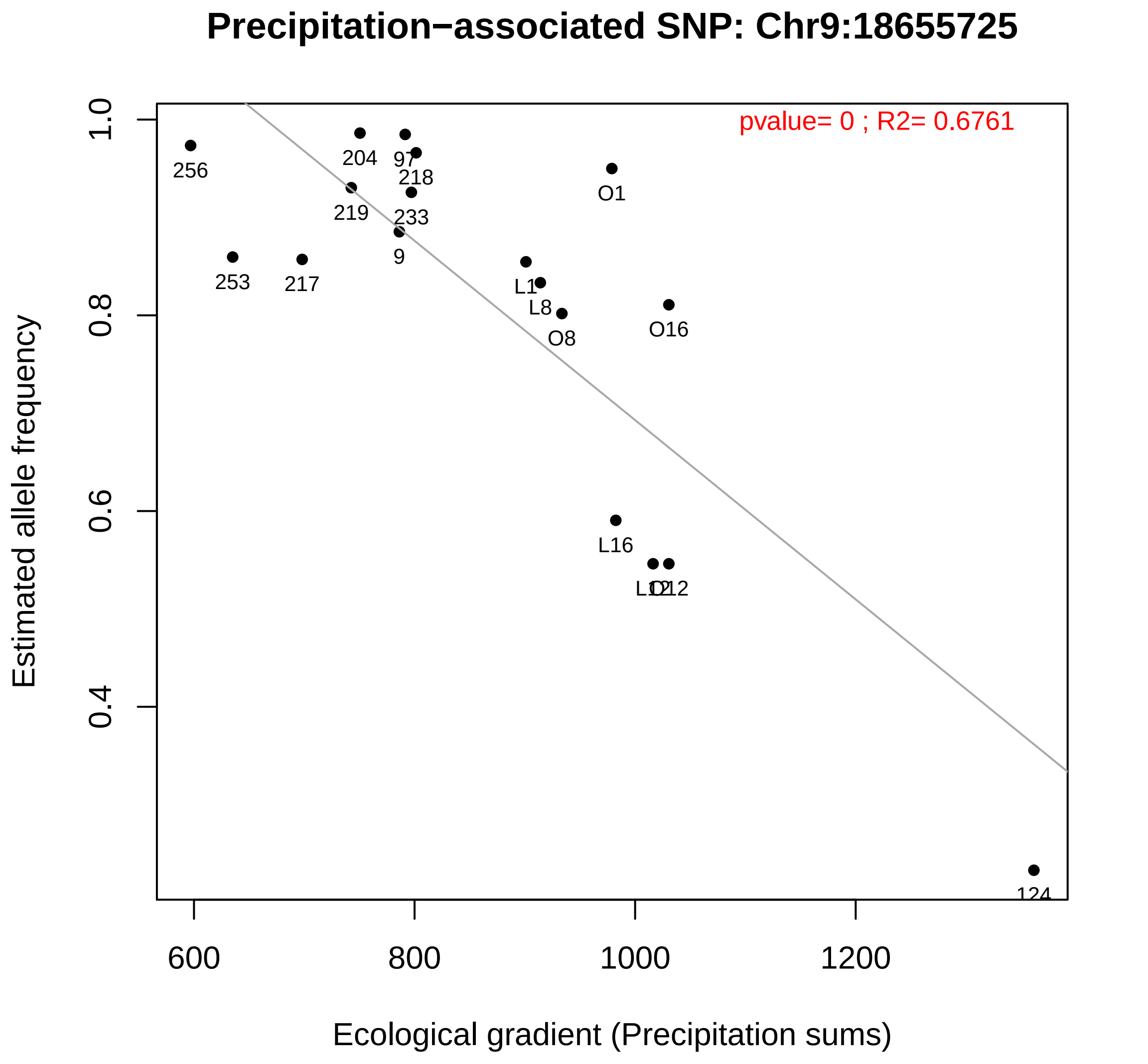
Precipitation−associated SNP: Chr9:18655725
pvalue= 0 ; R2= 0.6761
Estimated allele frequency
0.4
0.6
0.8
1.0
600
800
1000
1200
Ecological gradient (Precipitation sums)
256
204
97
218
219
233
9
253
217
O1
L1
L8
O8
O16
L16
L12
O12
124

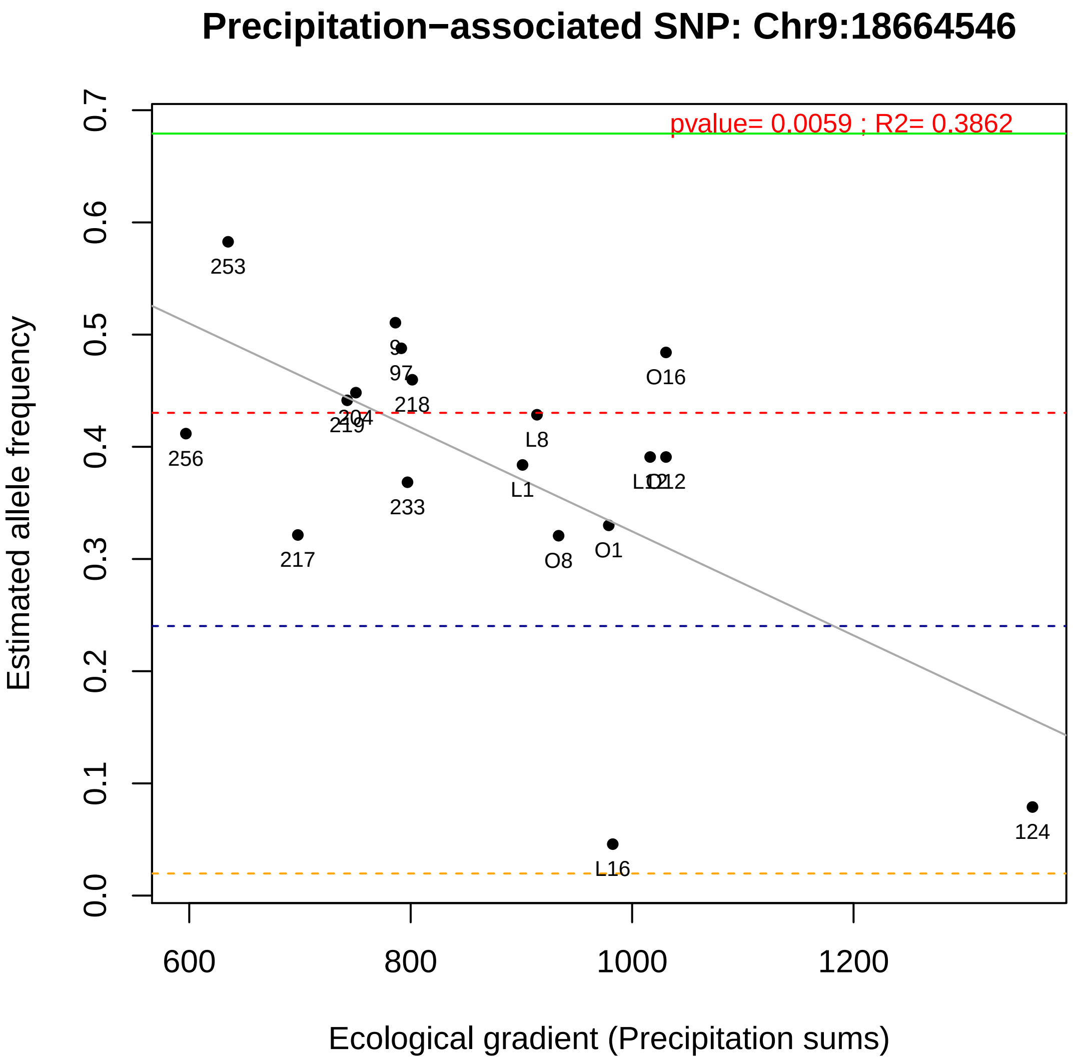

Precipitation−associated SNP: Chr9:18664546
pvalue= 0.0059 ; R2= 0.3862
Estimated allele frequency
Ecological gradient (Precipitation sums)
0.0
0.1
0.2
0.3
0.4
0.5
0.6
0.7
600
800
1000
1200
253
256
217
219
204
9
97
218
233
L1
L8
O8
O1
O16
L12
O12
L16
124

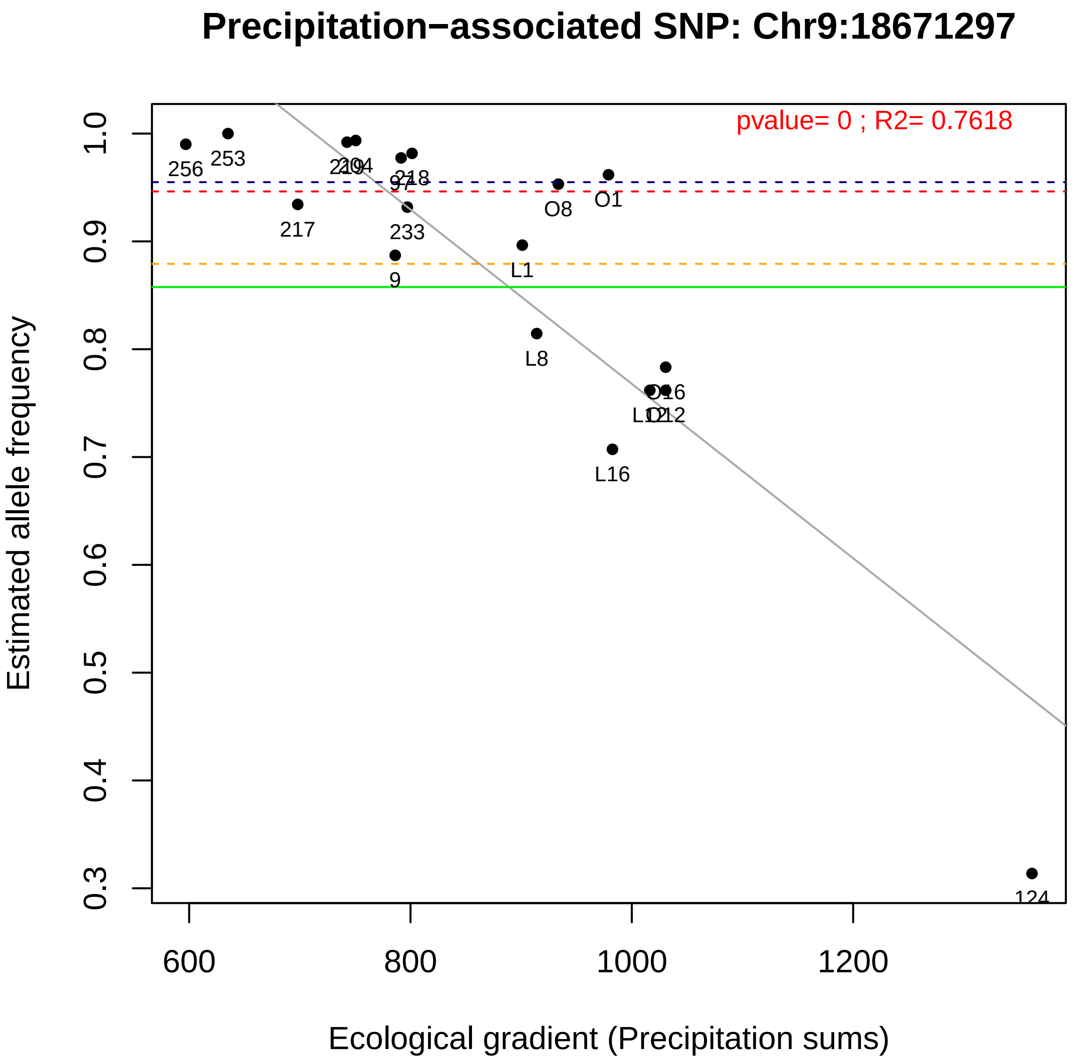

Precipitation−associated SNP: Chr9:18671297
pvalue= 0 ; R2= 0.7618
Estimated allele frequency
Ecological gradient (Precipitation sums)
1.0
0.9
0.8
0.7
0.6
0.5
0.4
0.3
600
800
1000
1200
256
253
217
233
9
L1
O8
O1
L8
L16
124

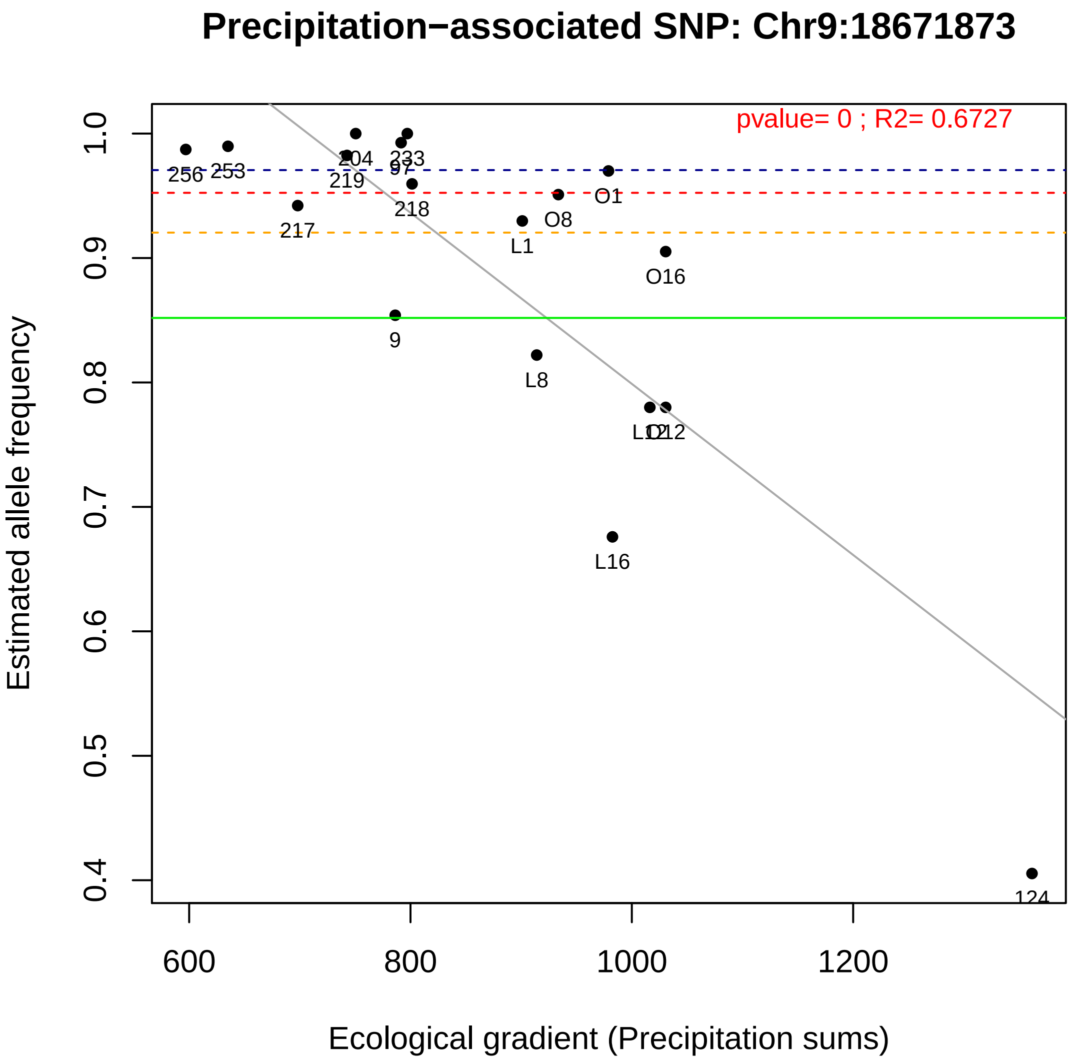

Precipitation−associated SNP: Chr9:18671873
pvalue= 0 ; R2= 0.6727
Estimated allele frequency
Ecological gradient (Precipitation sums)
1.0
0.9
0.8
0.7
0.6
0.5
0.4
600
800
1000
1200
256
253
204
219
233
97
218
217
O1
O8
L1
O16
9
L8
L12
O12
L16
124

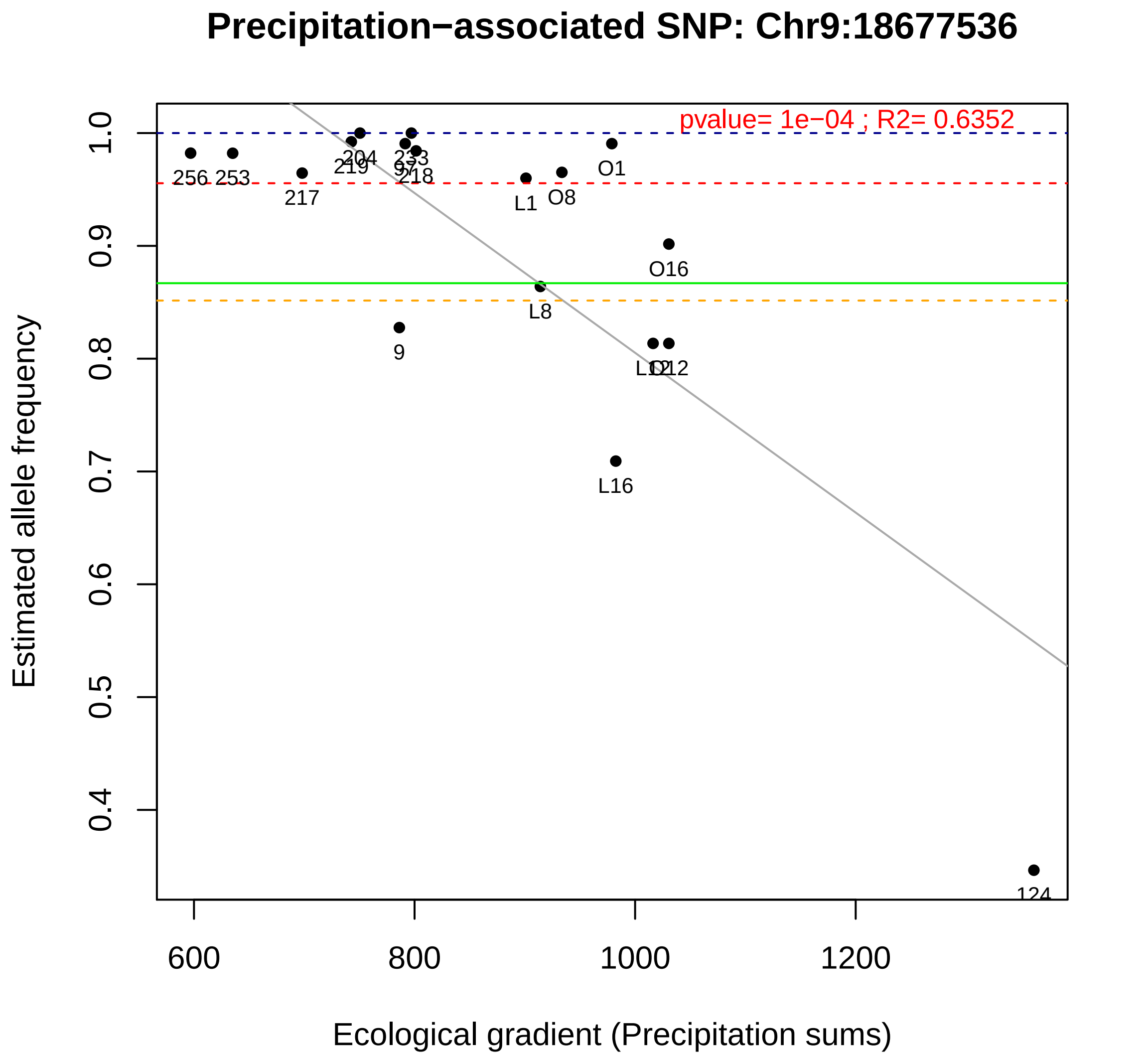

Precipitation−associated SNP: Chr9:18677536
pvalue= 1e−04 ; R2= 0.6352
Estimated allele frequency
Ecological gradient (Precipitation sums)
1.0
0.9
0.8
0.7
0.6
0.5
0.4
600
800
1000
1200
256
253
217
219
204
233
97
218
L1
O8
O1
O16
L8
9
L10
O12
L16
124

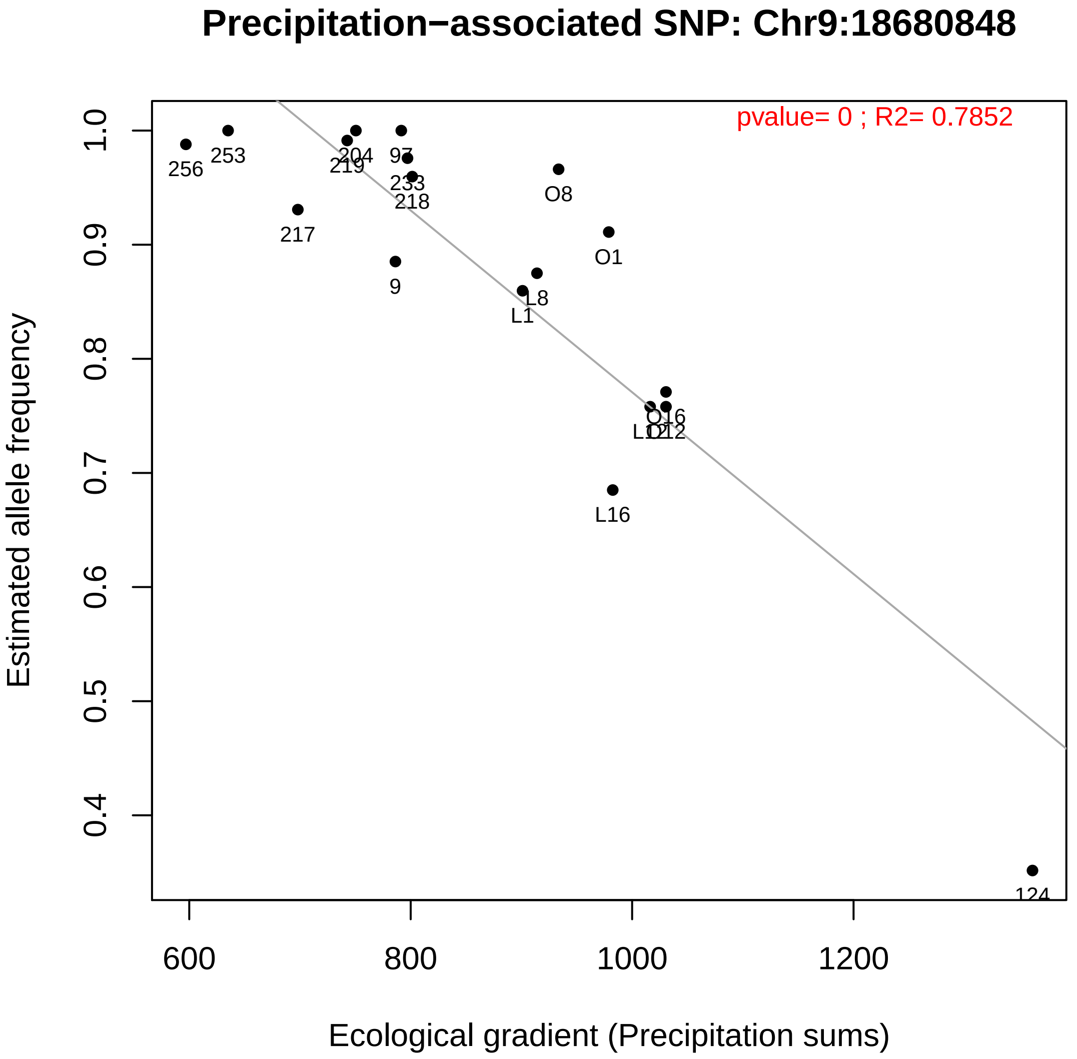
Precipitation−associated SNP: Chr9:18680848
pvalue= 0 ; R2= 0.7852
Estimated allele frequency
Ecological gradient (Precipitation sums)
1.0
0.9
0.8
0.7
0.6
0.5
0.4
600
800
1000
1200
256
253
204
219
97
233
218
217
9
O8
O1
L8
L1
O16
O12
L16
124

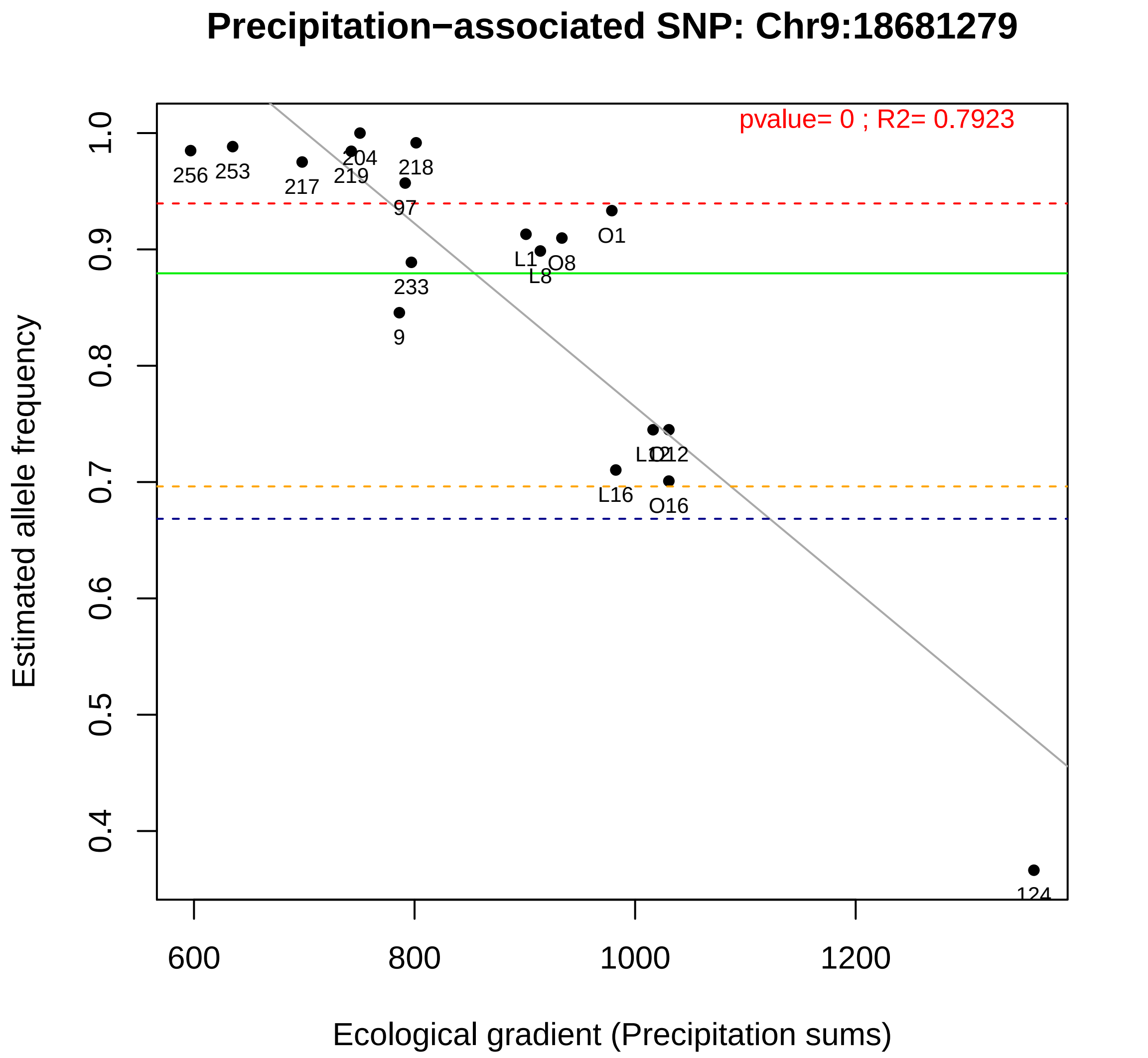

Precipitation−associated SNP: Chr9:18681279
pvalue= 0 ; R2= 0.7923
Estimated allele frequency
Ecological gradient (Precipitation sums)
1.0
0.9
0.8
0.7
0.6
0.5
0.4
600
800
1000
1200
256
253
217
219
204
218
97
233
9
L1
L8
O8
O1
L12
O12
L16
O16
124

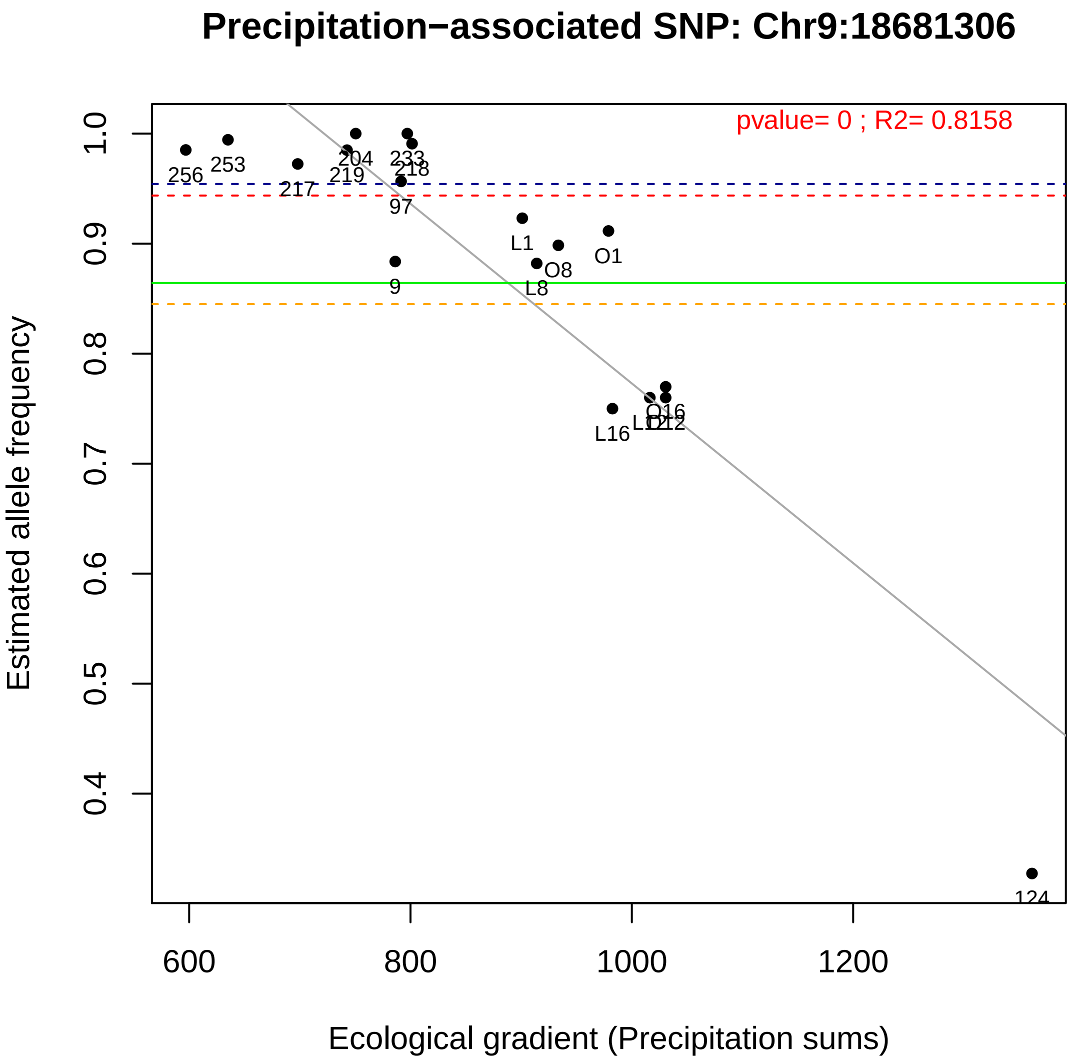

Precipitation−associated SNP: Chr9:18681306
pvalue= 0 ; R2= 0.8158
Estimated allele frequency
Ecological gradient (Precipitation sums)
1.0
0.9
0.8
0.7
0.6
0.5
0.4
600
800
1000
1200
256
253
204
219
217
233
218
97
9
L1
L8
O8
O1
L16
O16
O12
124

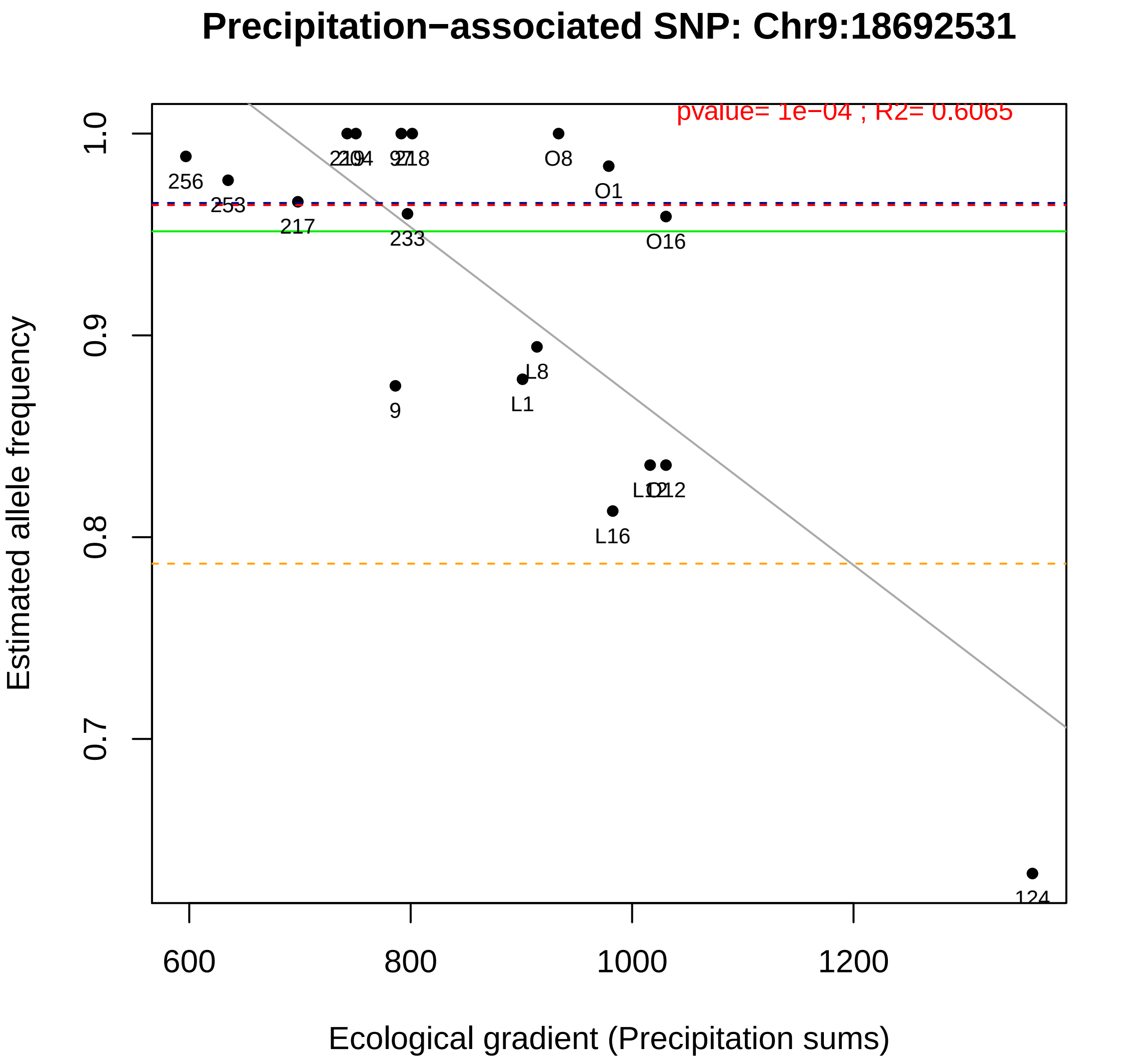
Precipitation−associated SNP: Chr9:18692531
pvalue= 1e−04 ; R2= 0.6065
Estimated allele frequency
Ecological gradient (Precipitation sums)
1.0
0.9
0.8
0.7
600
800
1000
1200
256
253
217
233
9
L1
L8
O8
O1
O16
L16
124

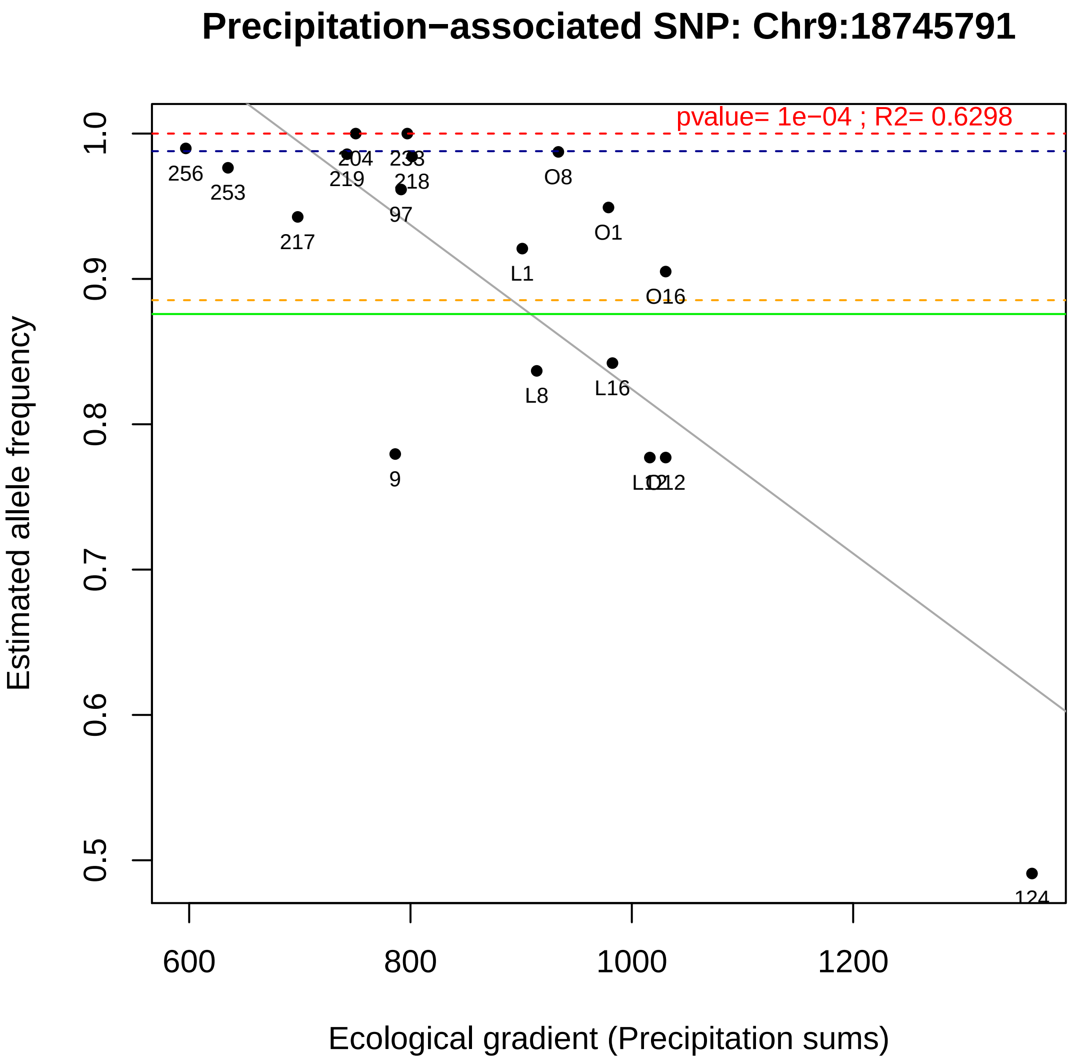
Precipitation−associated SNP: Chr9:18745791
pvalue= 1e−04 ; R2= 0.6298
Estimated allele frequency
Ecological gradient (Precipitation sums)
1.0
0.9
0.8
0.7
0.6
0.5
600
800
1000
1200
256
253
219
204
238
218
97
217
9
L1
O8
O1
O16
L8
L16
L12
O12
124

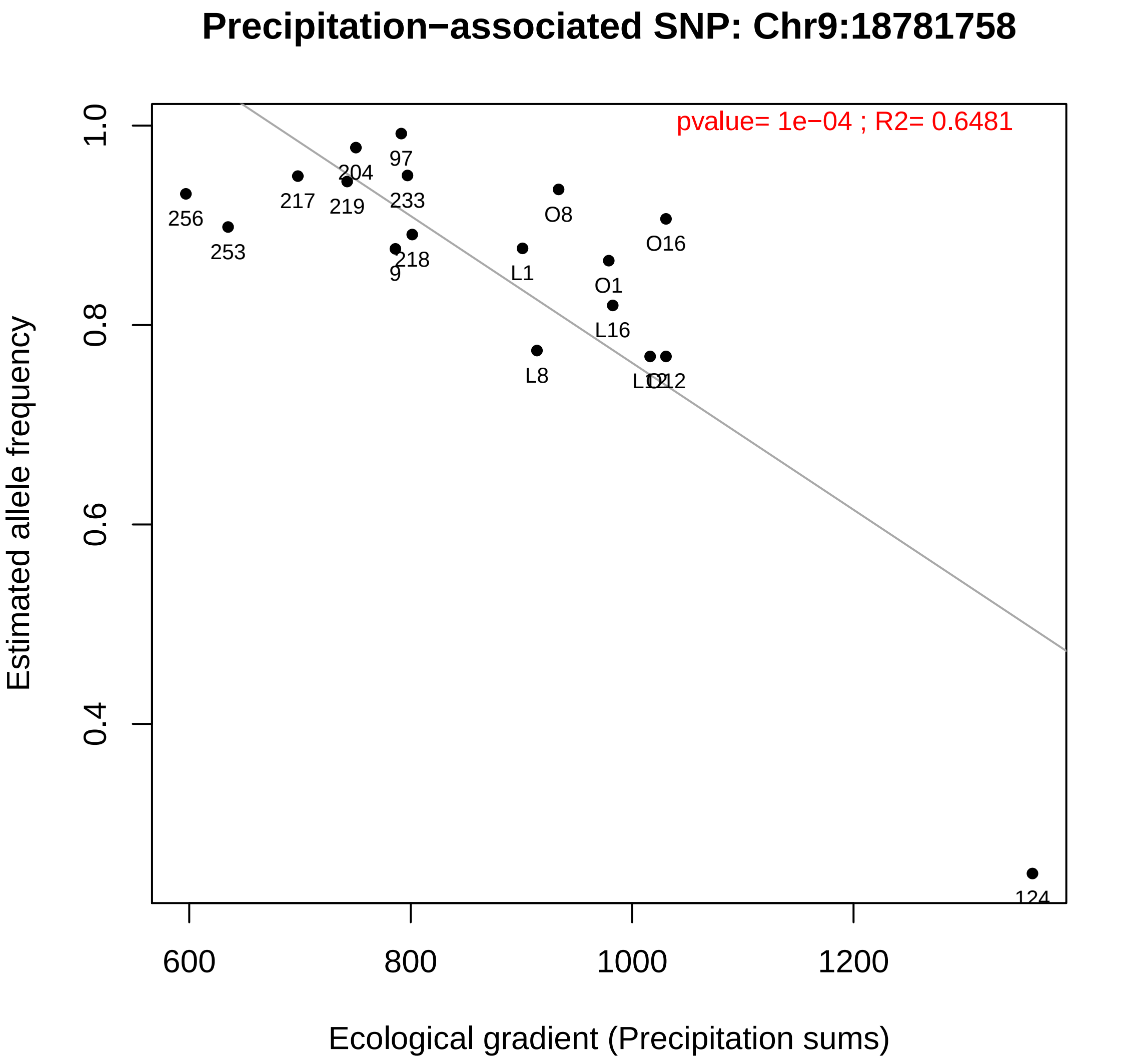
Precipitation−associated SNP: Chr9:18781758
pvalue= 1e−04 ; R2= 0.6481
Estimated allele frequency
Ecological gradient (Precipitation sums)
1.0
0.8
0.6
0.4
600
800
1000
1200
256
253
217
219
204
97
233
218
9
L1
L8
O8
O1
L16
O16
L12
O12
124

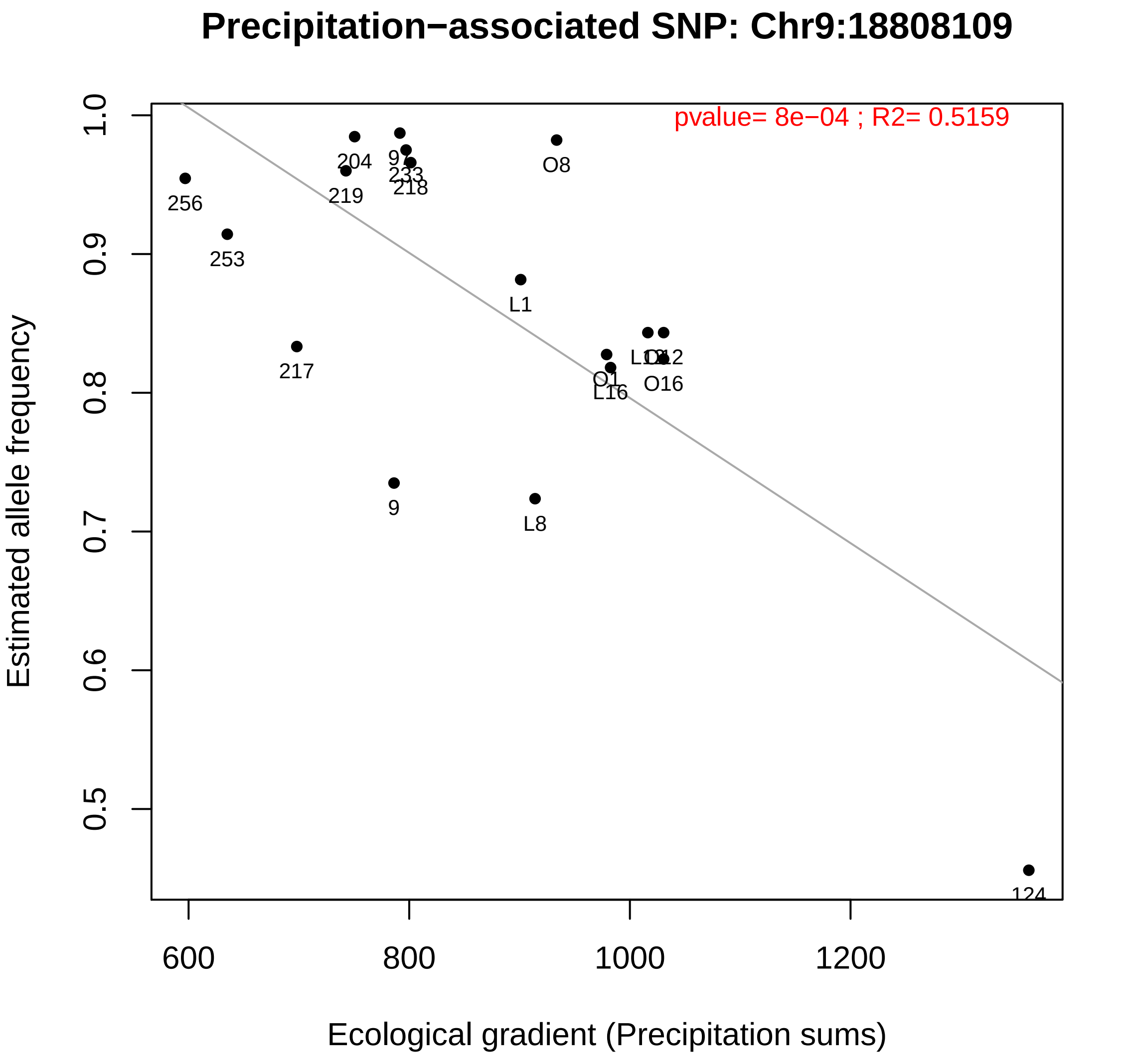
Precipitation−associated SNP: Chr9:18808109
pvalue= 8e−04 ; R2= 0.5159
Estimated allele frequency
Ecological gradient (Precipitation sums)
0.5
0.6
0.7
0.8
0.9
1.0
600
800
1000
1200
256
253
217
204
219
97
233
218
9
L1
L8
O8
O1
L16
O16
124

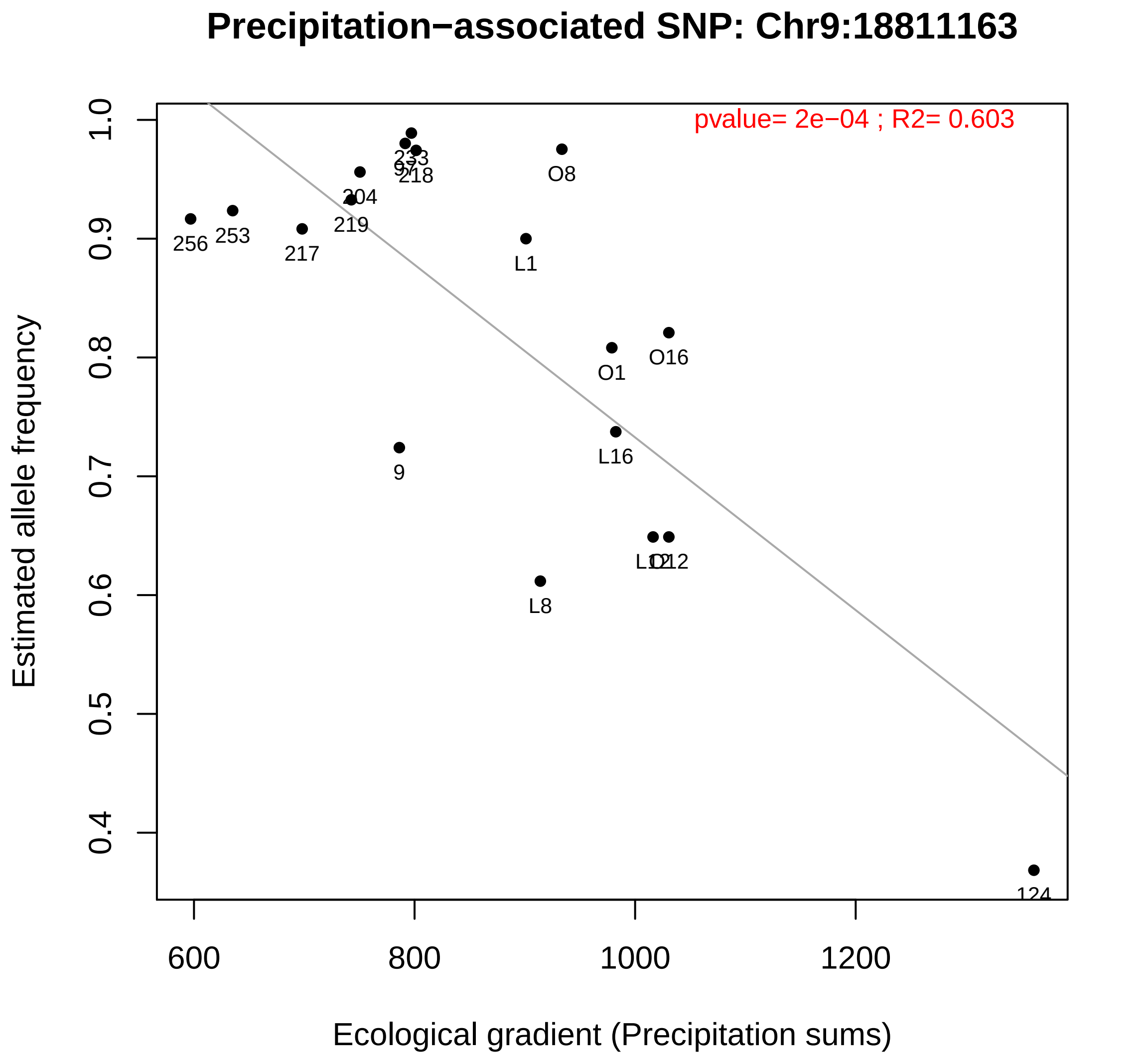
Precipitation−associated SNP: Chr9:18811163
pvalue= 2e−04 ; R2= 0.603
Estimated allele frequency
0.4
0.5
0.6
0.7
0.8
0.9
1.0
600
800
1000
1200
Ecological gradient (Precipitation sums)
256
253
217
219
204
233
97
218
O8
L1
O1
O16
L16
9
L8
L12
O12
124

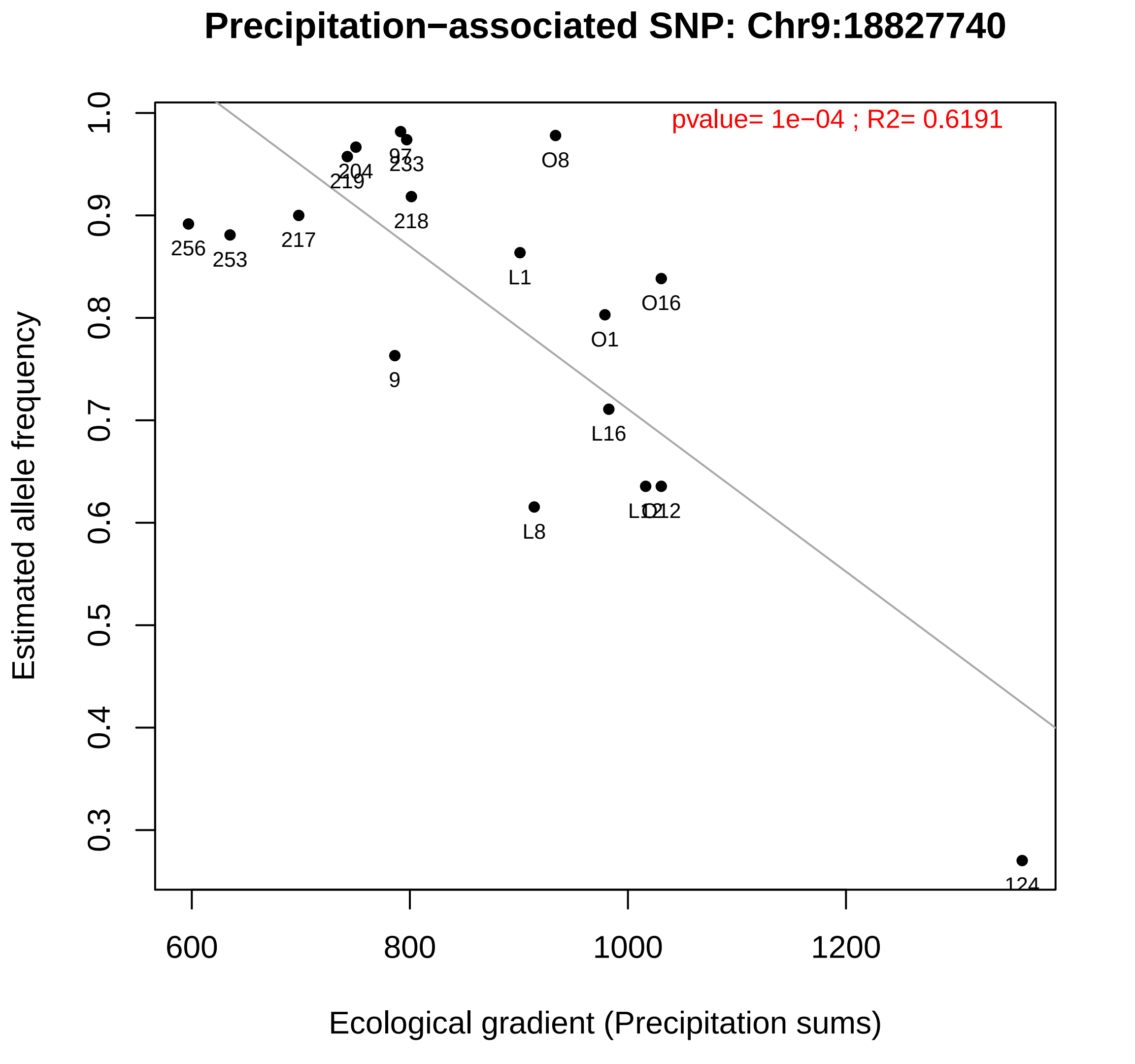

Precipitation−associated SNP: Chr9:18827740
pvalue= 1e−04 ; R2= 0.6191
Estimated allele frequency
Ecological gradient (Precipitation sums)
0.3
0.4
0.5
0.6
0.7
0.8
0.9
1.0
600
800
1000
1200
256
253
217
219
204
97
233
218
9
L1
O8
O1
O16
L16
L8
L12
O12
124

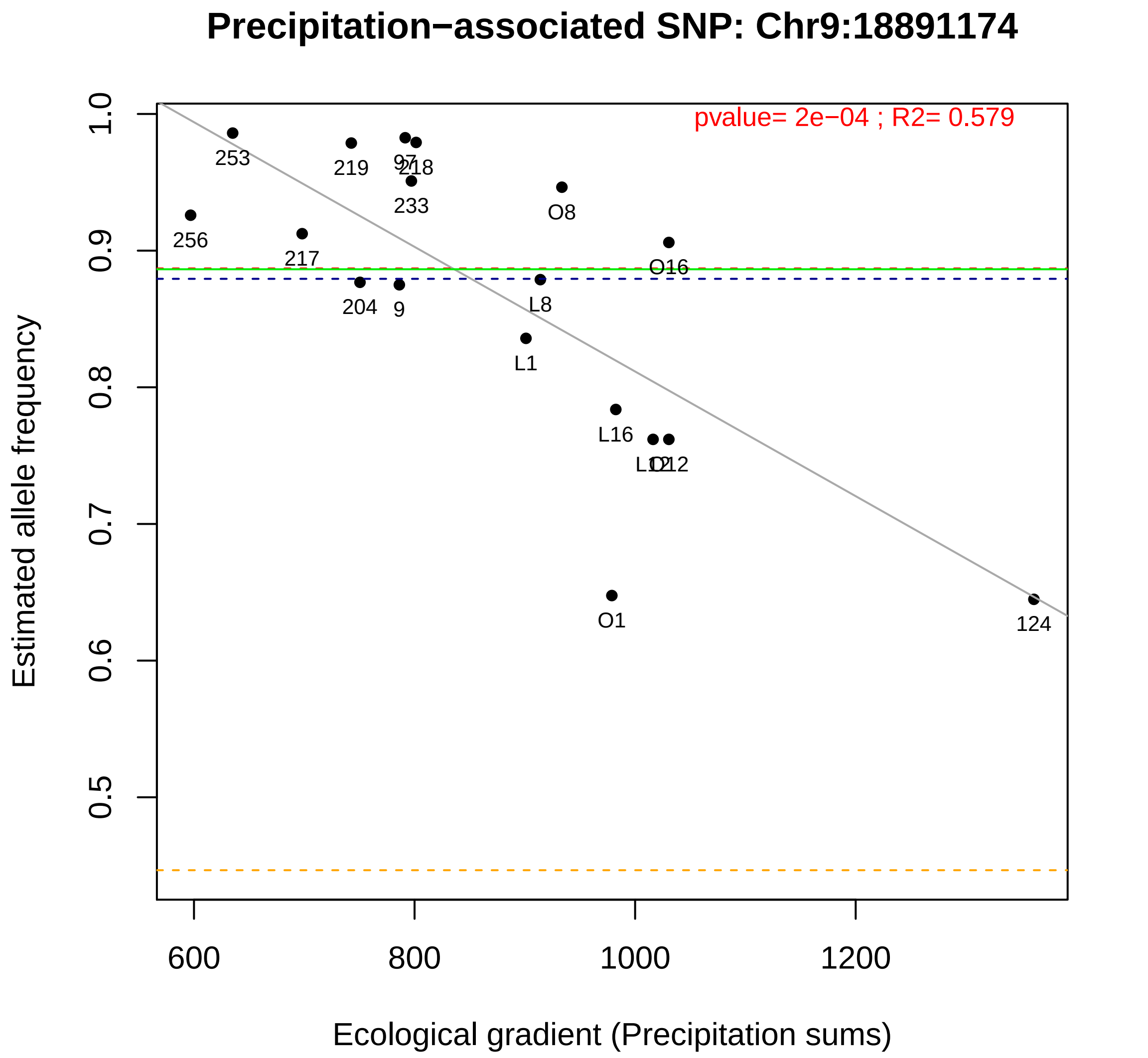

Precipitation−associated SNP: Chr9:18891174
pvalue= 2e−04 ; R2= 0.579
Estimated allele frequency
Ecological gradient (Precipitation sums)
0.5
0.6
0.7
0.8
0.9
1.0
600
800
1000
1200
253
219
97
218
233
O8
256
217
O16
204
9
L8
L1
L16
L12
O12
O1
124

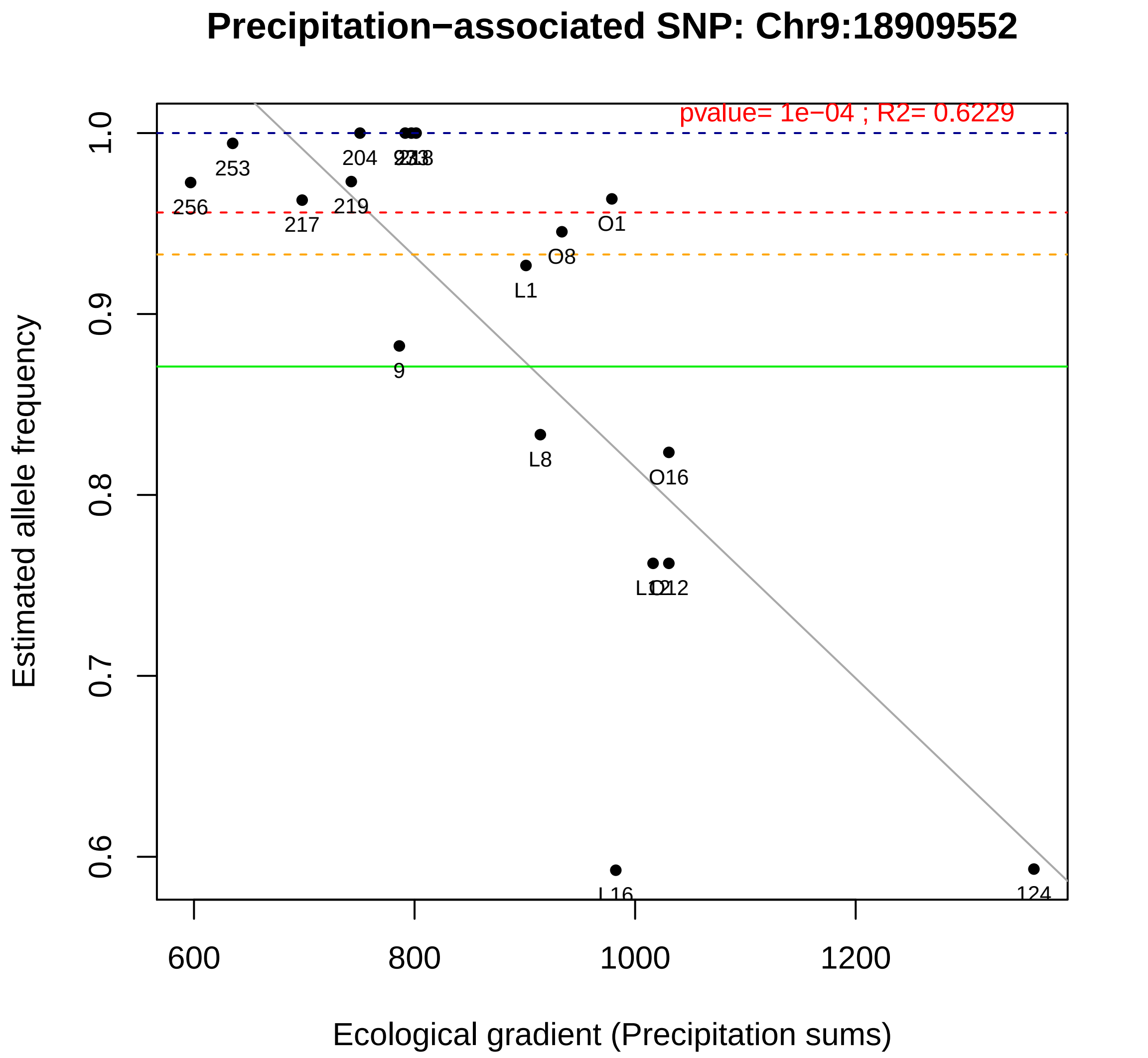
Precipitation−associated SNP: Chr9:18909552
pvalue= 1e−04 ; R2= 0.6229
Estimated allele frequency
Ecological gradient (Precipitation sums)
1.0
0.9
0.8
0.7
0.6
600
800
1000
1200
253
204
256
219
217
O1
O8
L1
9
L8
O16
L16
124

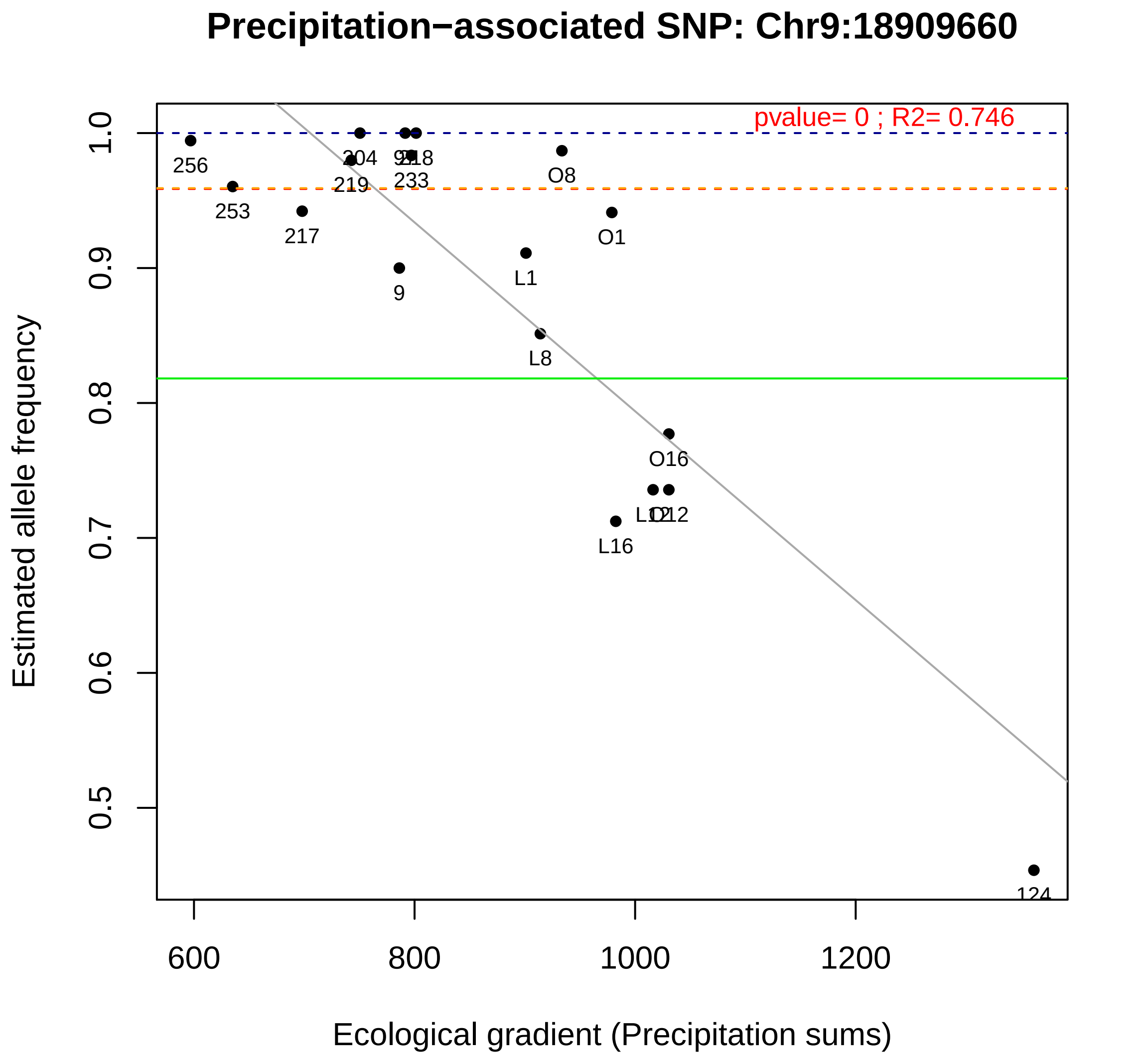

Precipitation−associated SNP: Chr9:18909660
pvalue= 0 ; R2= 0.746
Estimated allele frequency
Ecological gradient (Precipitation sums)
1.0
0.9
0.8
0.7
0.6
0.5
600
800
1000
1200
256
253
217
204
219
9
218
233
O8
O1
L1
L8
O16
O12
L16
124

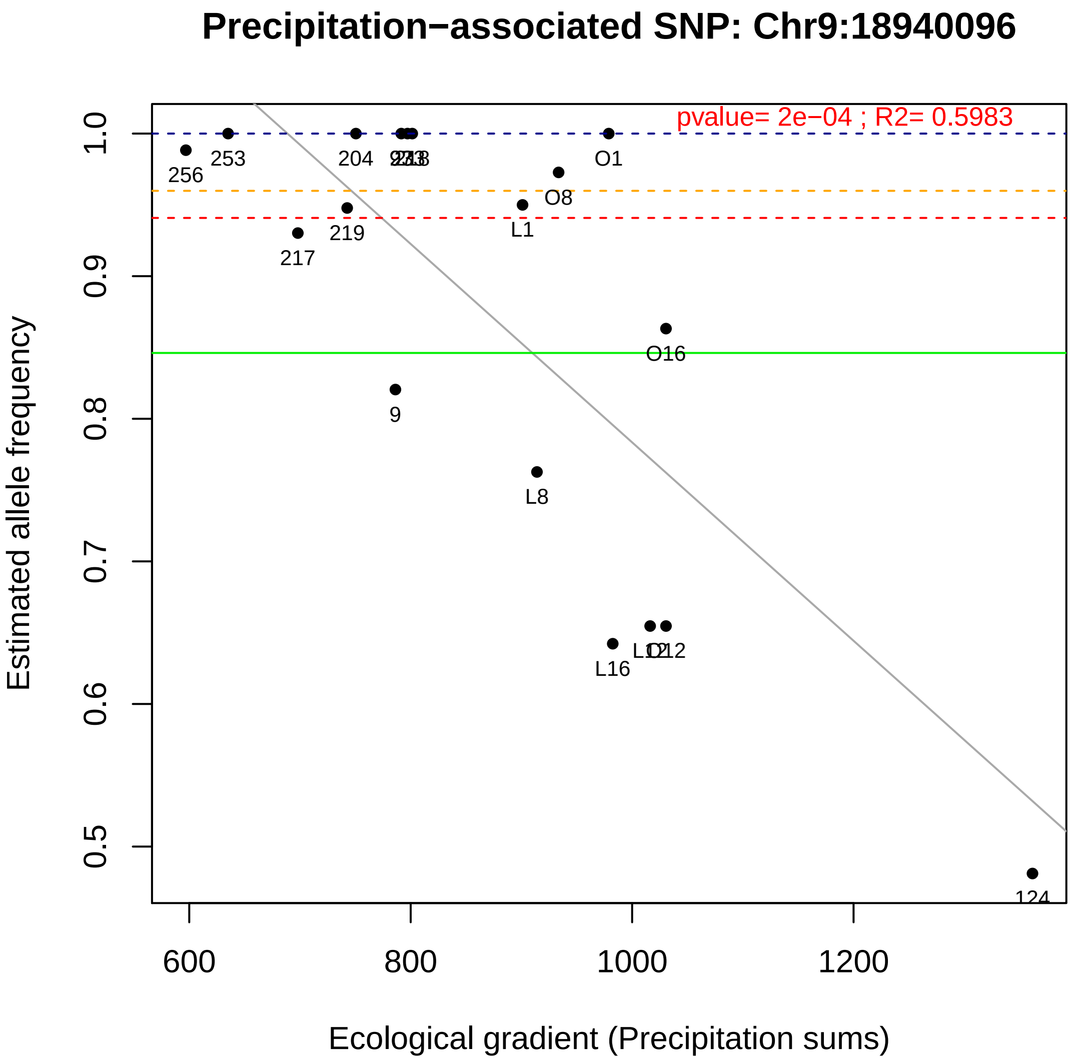
Precipitation−associated SNP: Chr9:18940096
pvalue= 2e−04 ; R2= 0.5983
Estimated allele frequency
Ecological gradient (Precipitation sums)
1.0
0.9
0.8
0.7
0.6
0.5
600
800
1000
1200
256
253
204
O1
O8
219
217
L1
O16
9
L8
L16
O12
124

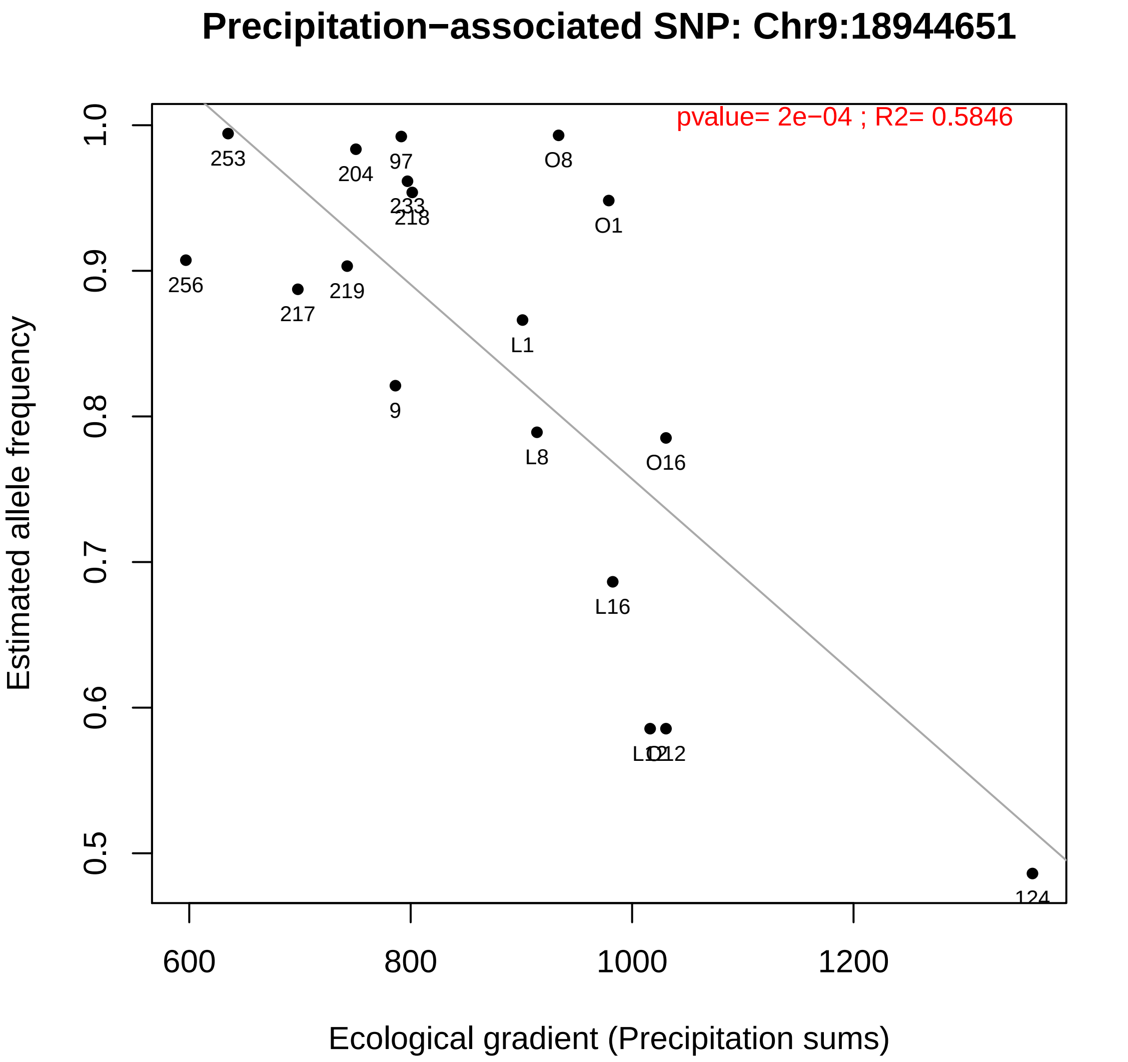
Precipitation−associated SNP: Chr9:18944651
pvalue= 2e−04 ; R2= 0.5846
Estimated allele frequency
Ecological gradient (Precipitation sums)
1.0
0.9
0.8
0.7
0.6
0.5
600
800
1000
1200
253
204
97
O8
233
218
O1
256
219
217
L1
9
L8
O16
L16
L12
O12
124

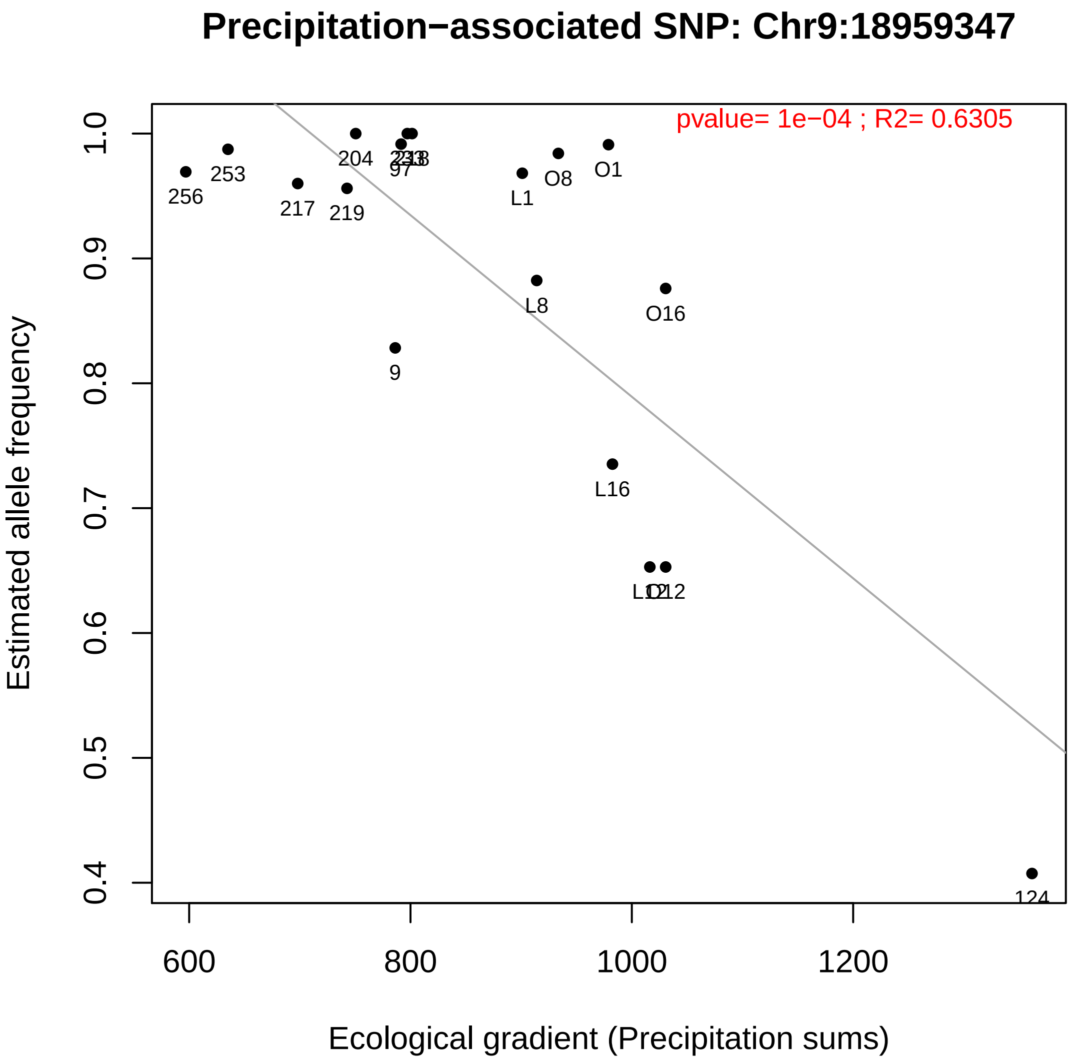

Precipitation−associated SNP: Chr9:18959347
pvalue= 1e−04 ; R2= 0.6305
Estimated allele frequency
Ecological gradient (Precipitation sums)
0.4
0.5
0.6
0.7
0.8
0.9
1.0
600
800
1000
1200
256
253
217
219
204
97
9
L1
O8
O1
L8
O16
L16
124

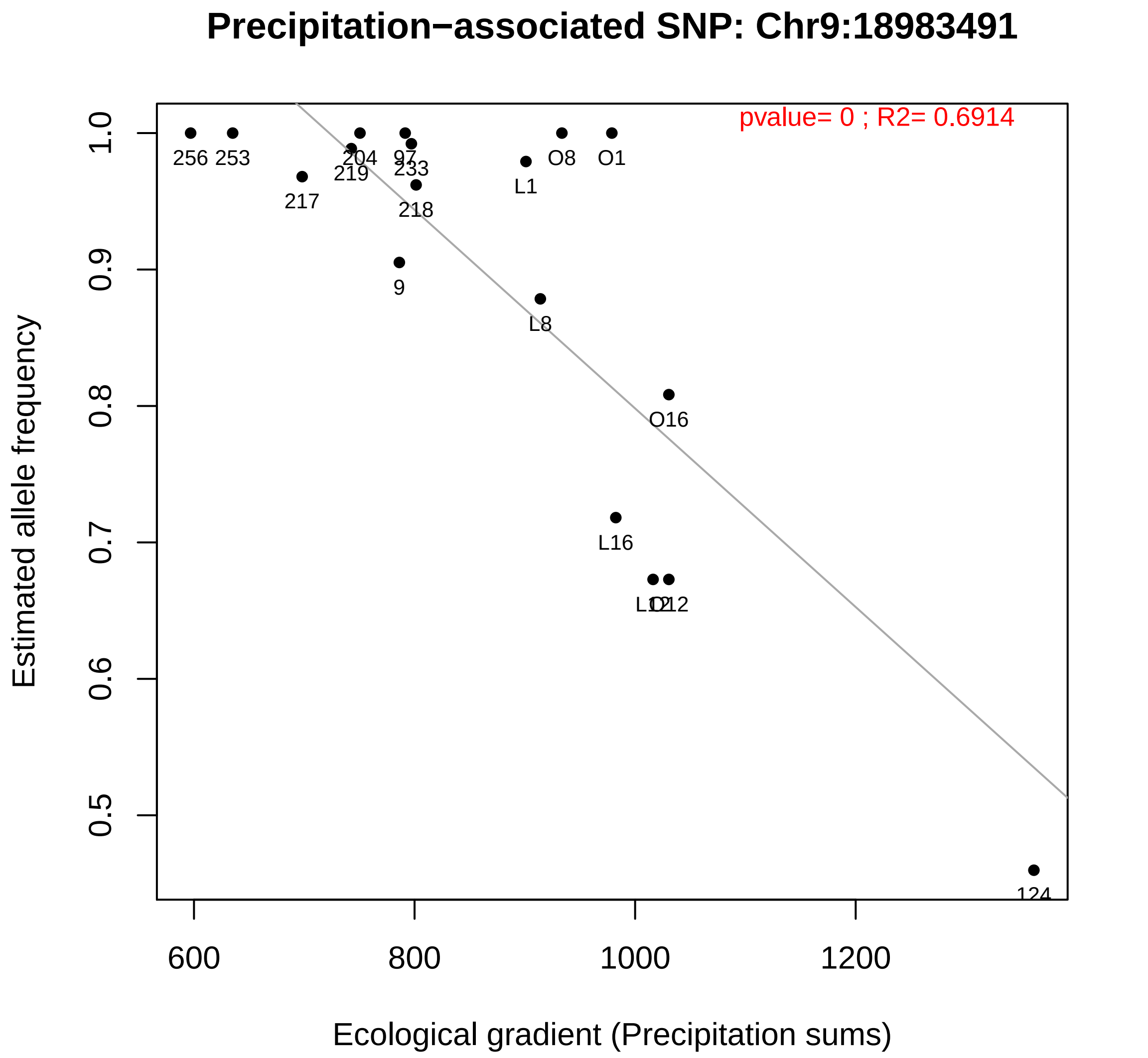
Precipitation−associated SNP: Chr9:18983491
pvalue= 0 ; R2= 0.6914
Estimated allele frequency
Ecological gradient (Precipitation sums)
1.0
0.9
0.8
0.7
0.6
0.5
600
800
1000
1200
256
253
204
219
97
233
217
218
9
L1
O8
O1
L8
O16
L16
L12
O12
124

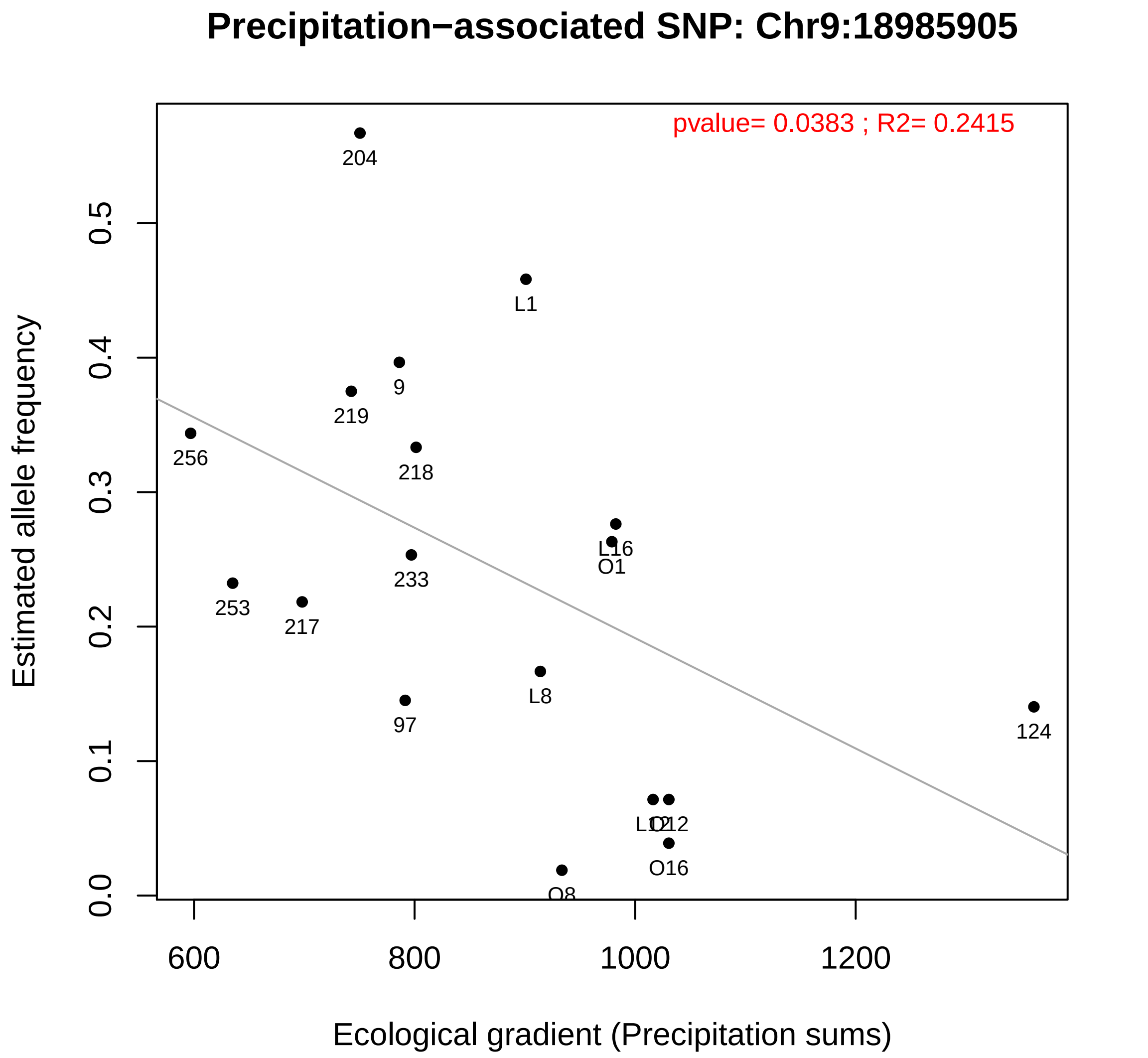

Precipitation−associated SNP: Chr9:18985905
pvalue= 0.0383 ; R2= 0.2415
Estimated allele frequency
Ecological gradient (Precipitation sums)
0.0
0.1
0.2
0.3
0.4
0.5
600
800
1000
1200
204
L1
9
219
256
218
L16
O1
233
253
217
L8
97
124
L10
O12
O16
O8

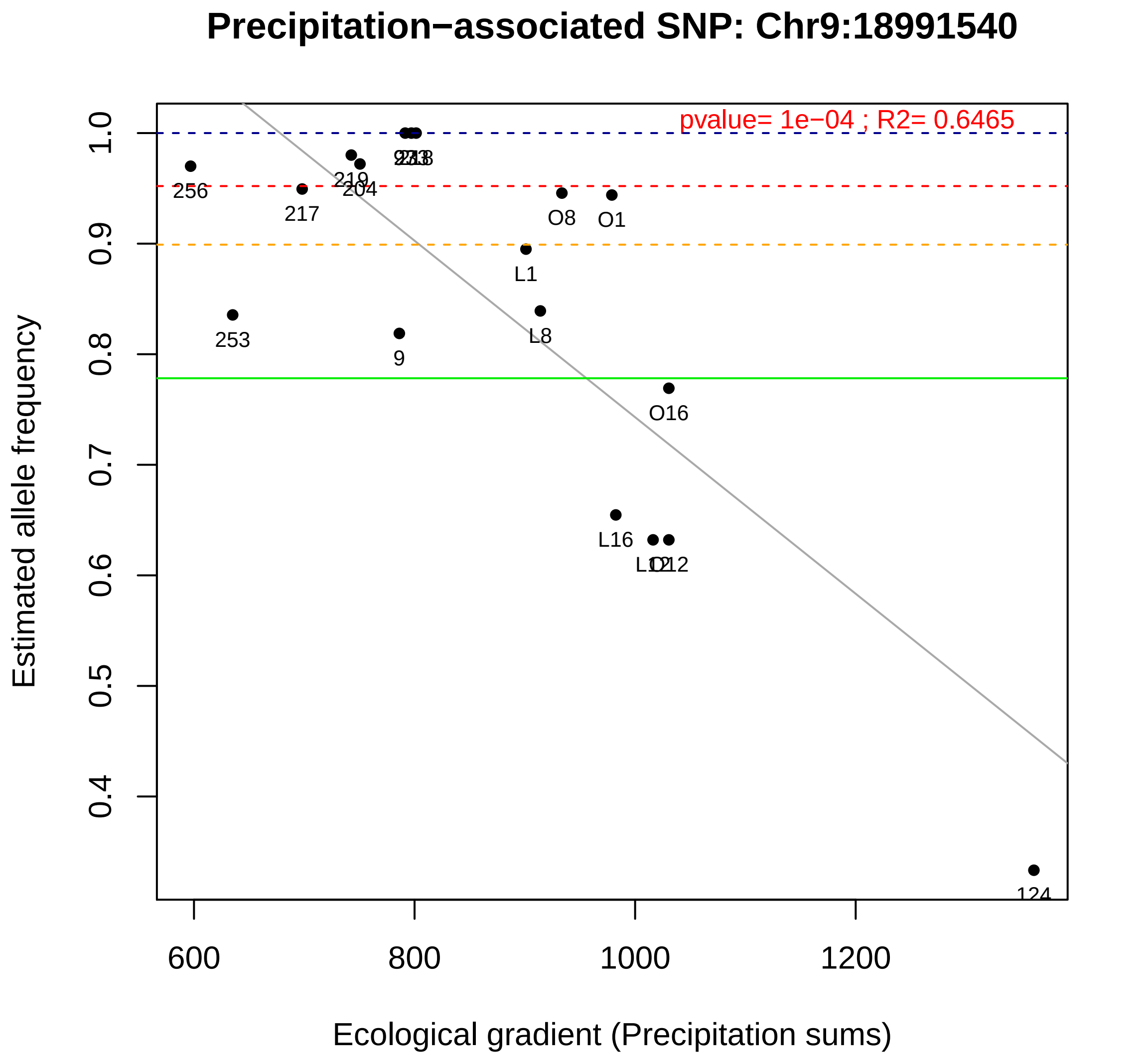

Precipitation−associated SNP: Chr9:18991540
pvalue= 1e−04 ; R2= 0.6465
Estimated allele frequency
Ecological gradient (Precipitation sums)
0.4
0.5
0.6
0.7
0.8
0.9
1.0
600
800
1000
1200
256
253
217
219
204
9
L1
L8
O8
O1
O16
L16
124

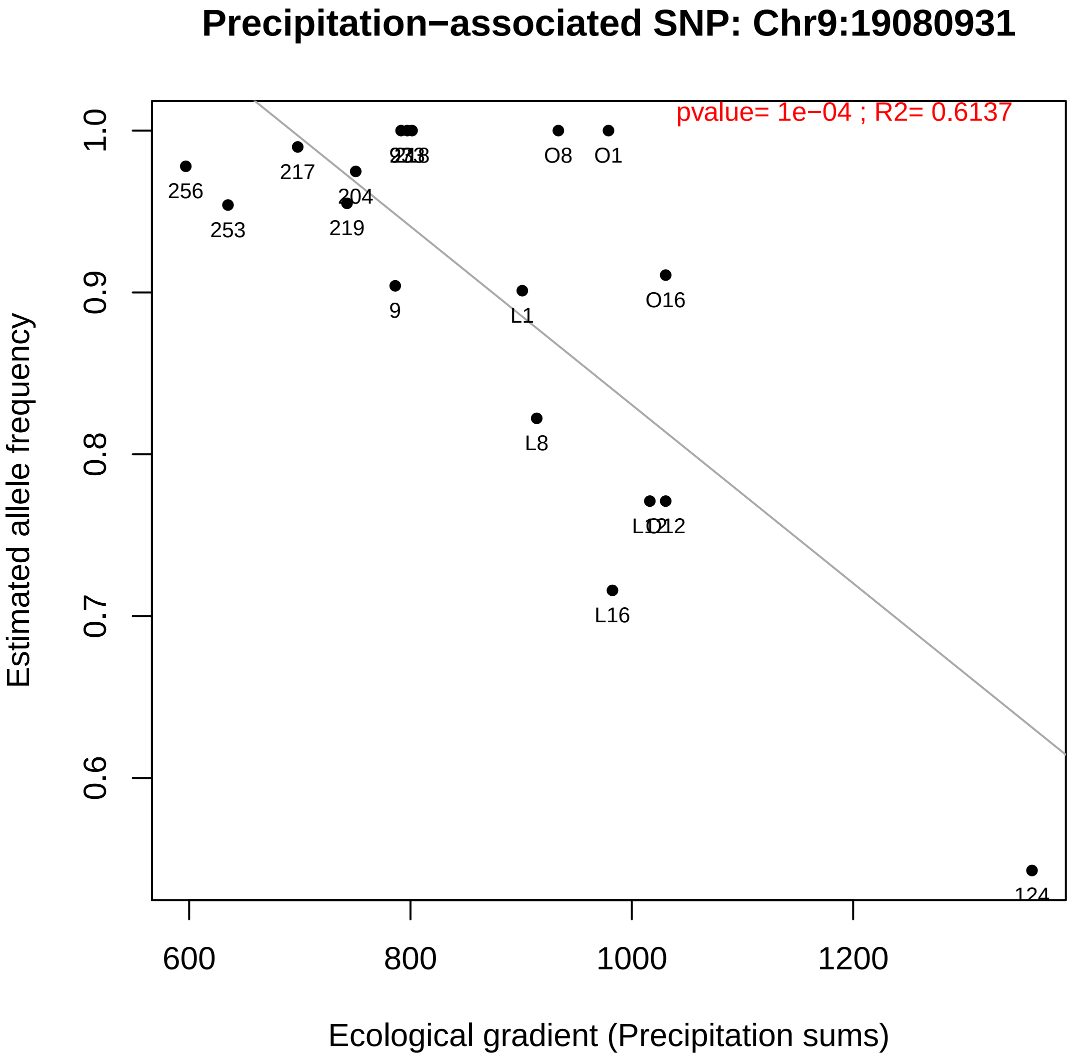
Precipitation−associated SNP: Chr9:19080931
pvalue= 1e−04 ; R2= 0.6137
Estimated allele frequency
Ecological gradient (Precipitation sums)
0.6
0.7
0.8
0.9
1.0
600
800
1000
1200
256
253
217
204
219
9
O8
O1
L1
O16
L8
L16
124

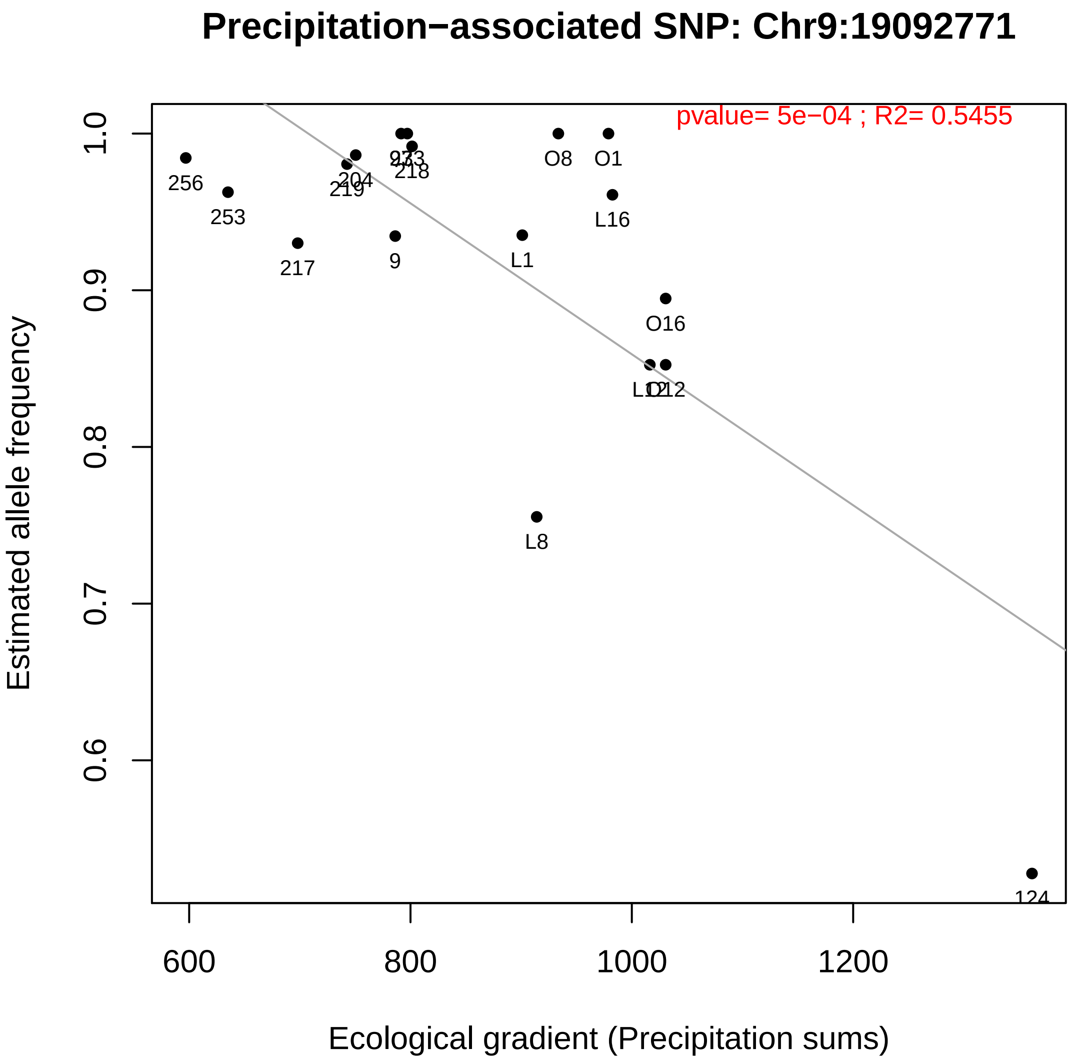

Precipitation−associated SNP: Chr9:19092771
pvalue= 5e−04 ; R2= 0.5455
Estimated allele frequency
Ecological gradient (Precipitation sums)
1.0
0.9
0.8
0.7
0.6
600
800
1000
1200
256
253
217
204
219
9
218
O8
O1
L16
L1
O16
L8
124

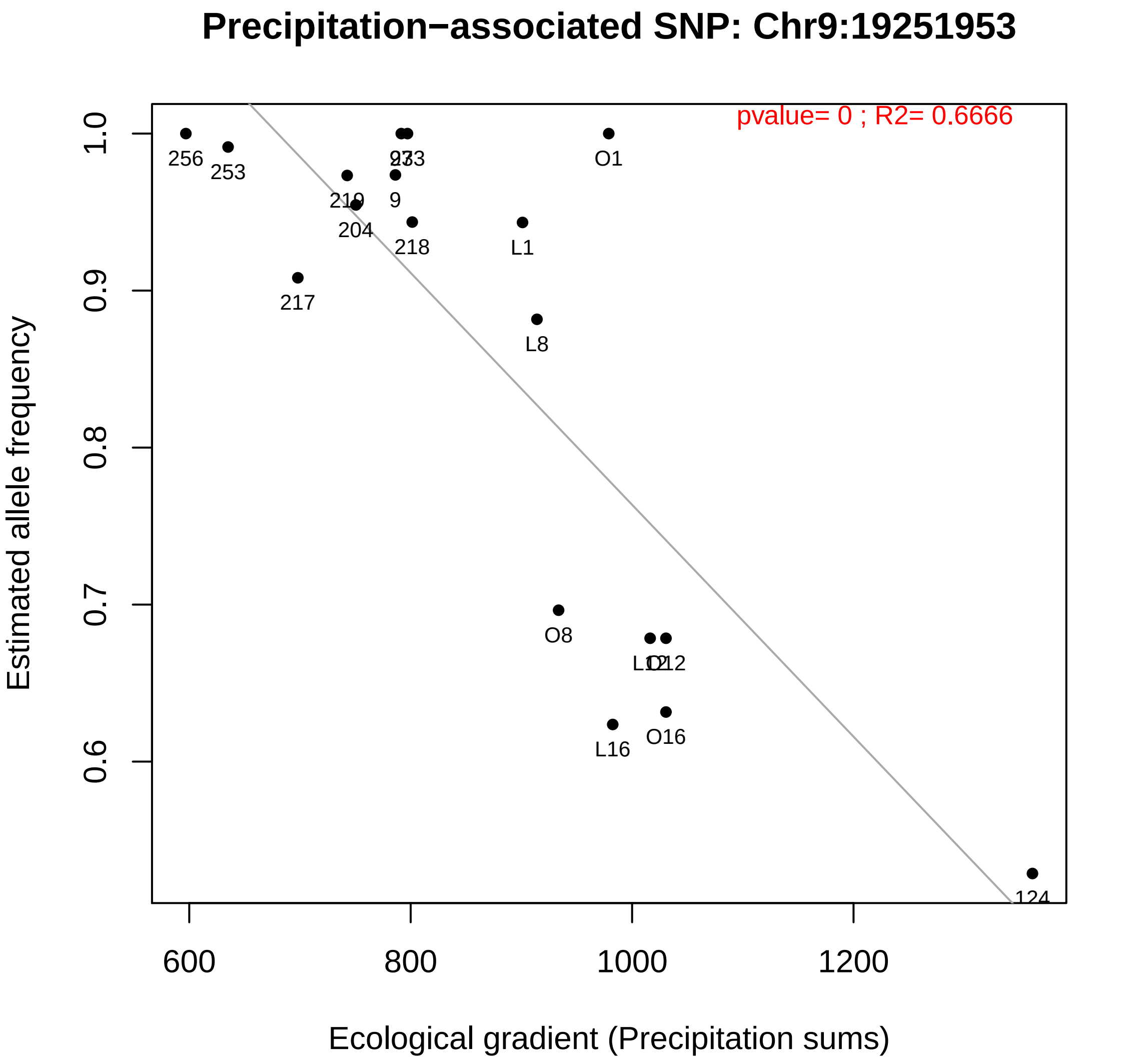
Precipitation−associated SNP: Chr9:19251953
pvalue= 0 ; R2= 0.6666
Estimated allele frequency
Ecological gradient (Precipitation sums)
1.0
0.9
0.8
0.7
0.6
600
800
1000
1200
256
253
219
204
217
9
218
L1
O1
L8
O8
O12
O16
L16
124

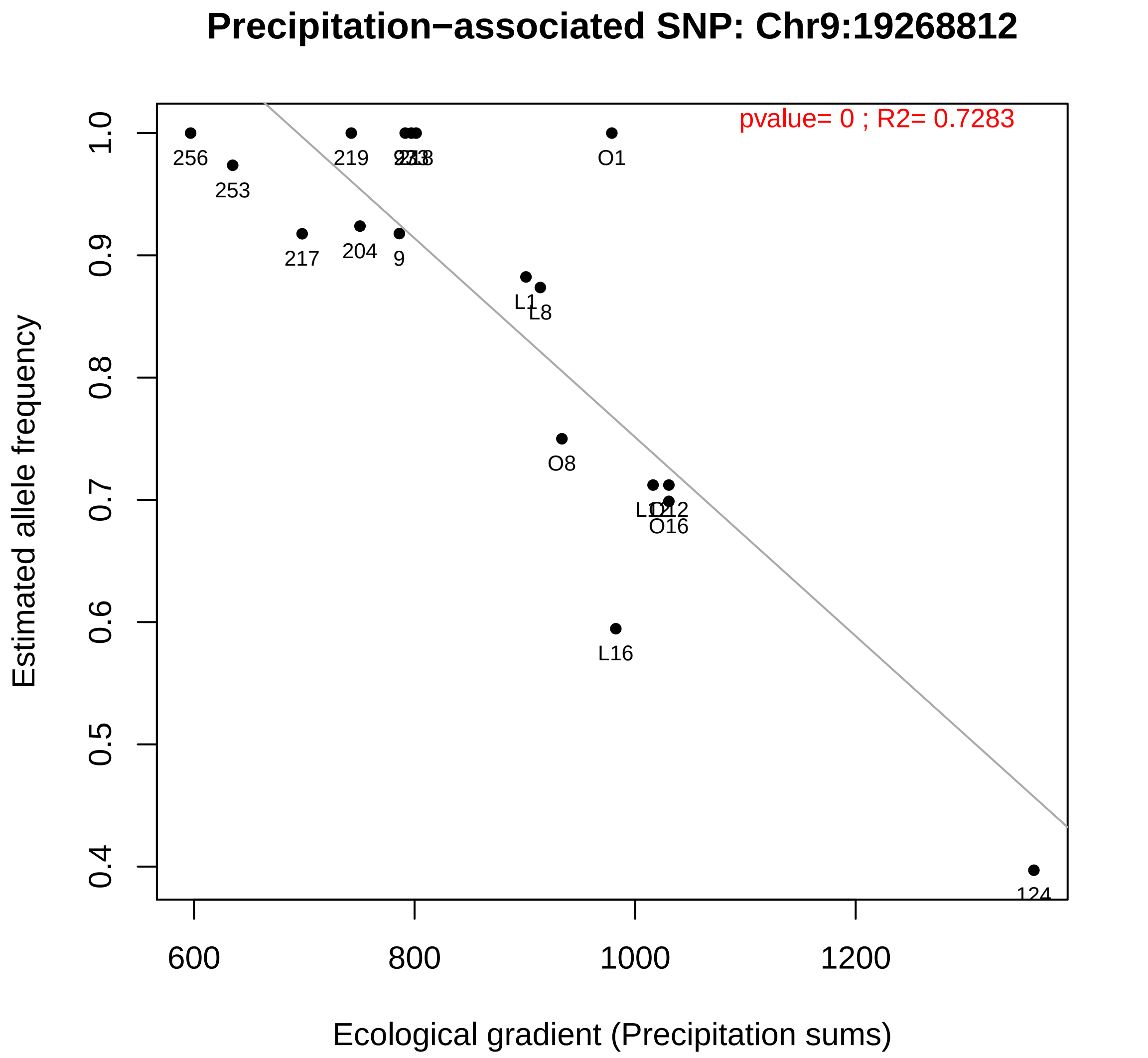

Precipitation−associated SNP: Chr9:19268812
pvalue= 0 ; R2= 0.7283
Estimated allele frequency
Ecological gradient (Precipitation sums)
1.0
0.9
0.8
0.7
0.6
0.5
0.4
600
800
1000
1200
256
253
219
217
204
9
O1
L1
L8
O8
O16
L16
124

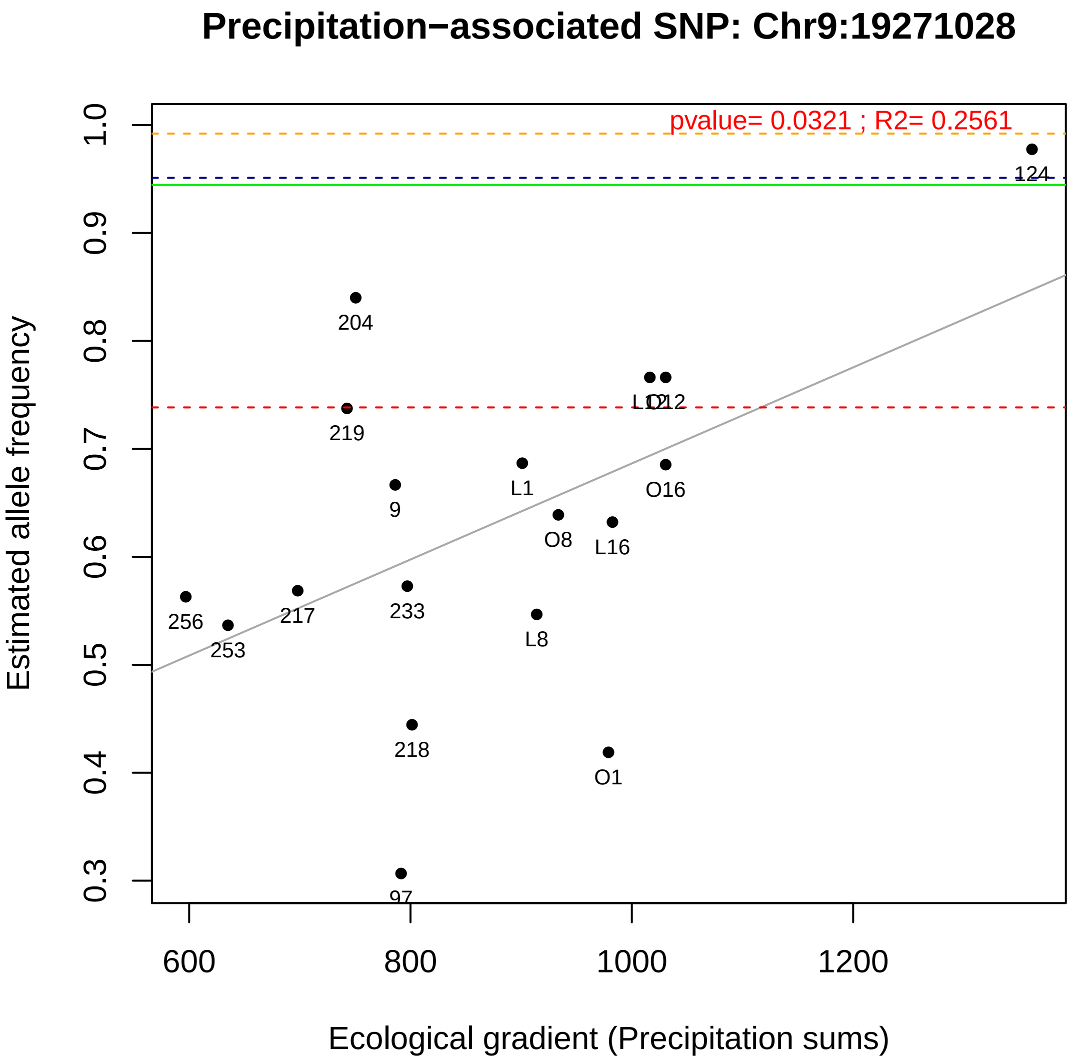

Precipitation−associated SNP: Chr9:19271028
pvalue= 0.0321 ; R2= 0.2561
Estimated allele frequency
Ecological gradient (Precipitation sums)
0.3
0.4
0.5
0.6
0.7
0.8
0.9
1.0
600
800
1000
1200
124
204
219
9
L1
O8
L16
O16
O12
256
253
217
233
L8
218
O1
97

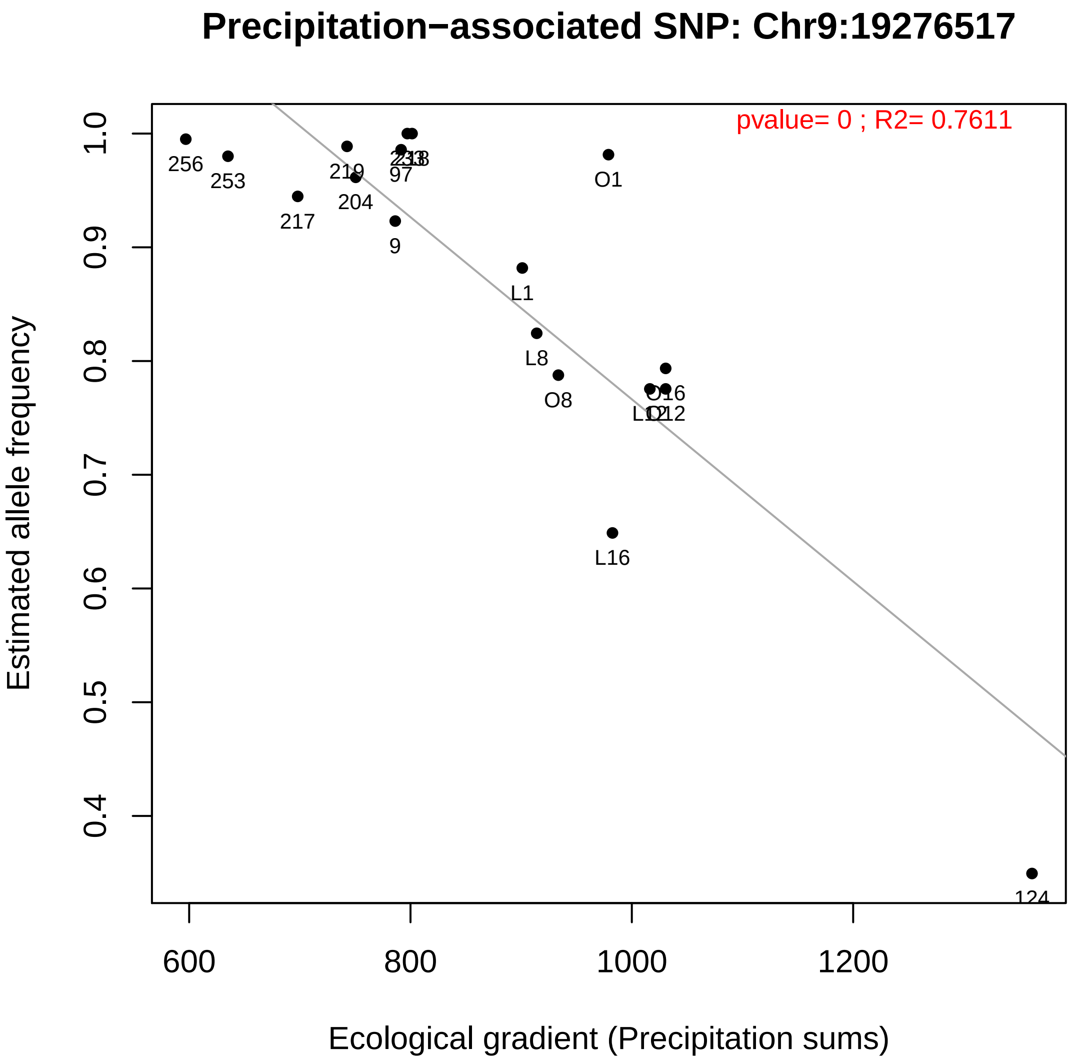

Precipitation−associated SNP: Chr9:19276517
pvalue= 0 ; R2= 0.7611
Estimated allele frequency
Ecological gradient (Precipitation sums)
0.4
0.5
0.6
0.7
0.8
0.9
1.0
600
800
1000
1200
256
253
219
204
217
97
9
O1
L1
L8
O8
O16
O12
L16
124

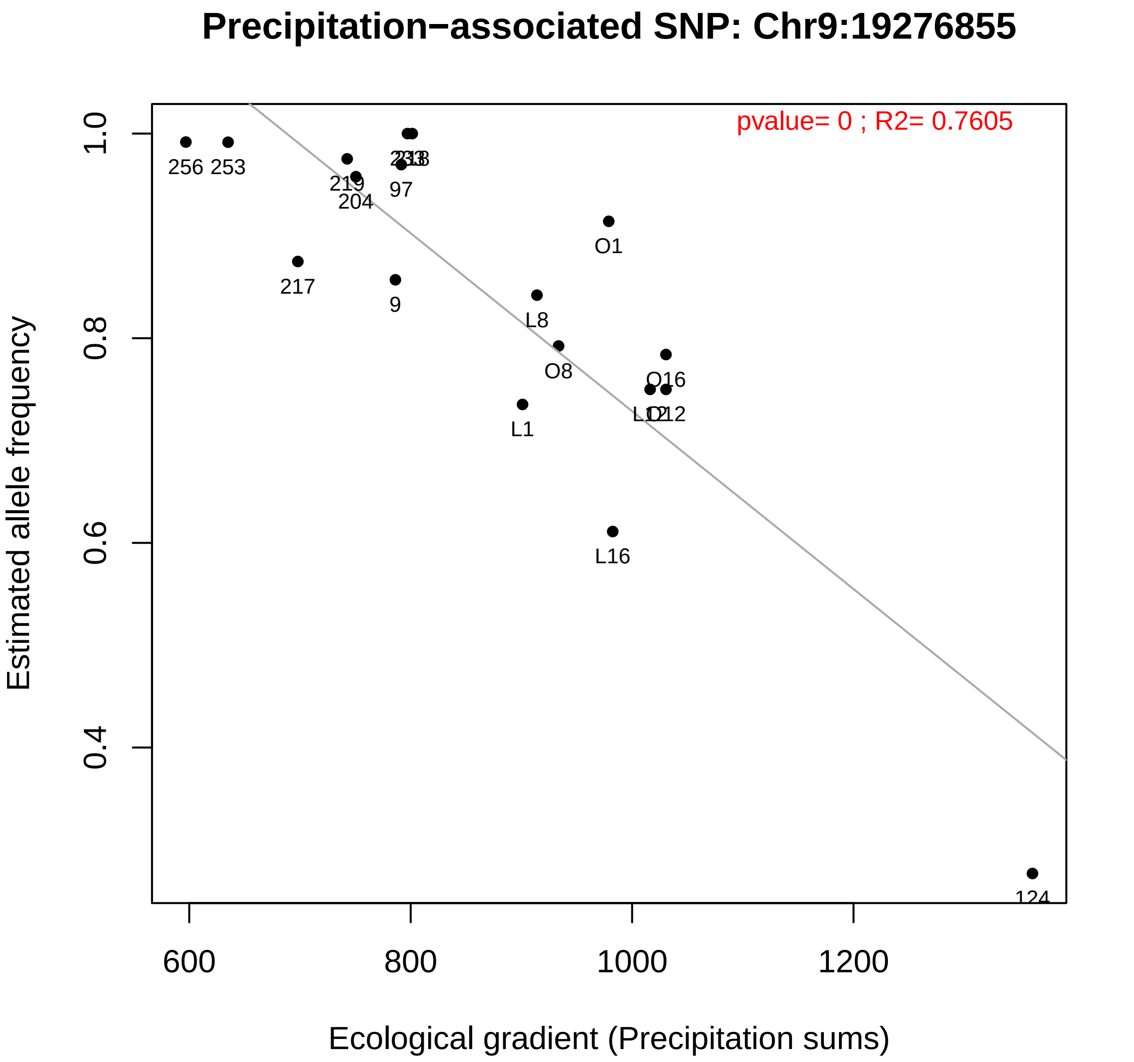
Precipitation−associated SNP: Chr9:19276855
pvalue= 0 ; R2= 0.7605
Estimated allele frequency
Ecological gradient (Precipitation sums)
1.0
0.8
0.6
0.4
600
800
1000
1200
256
253
219
204
97
217
9
O1
L8
O8
O16
L1
L12
O12
L16
124

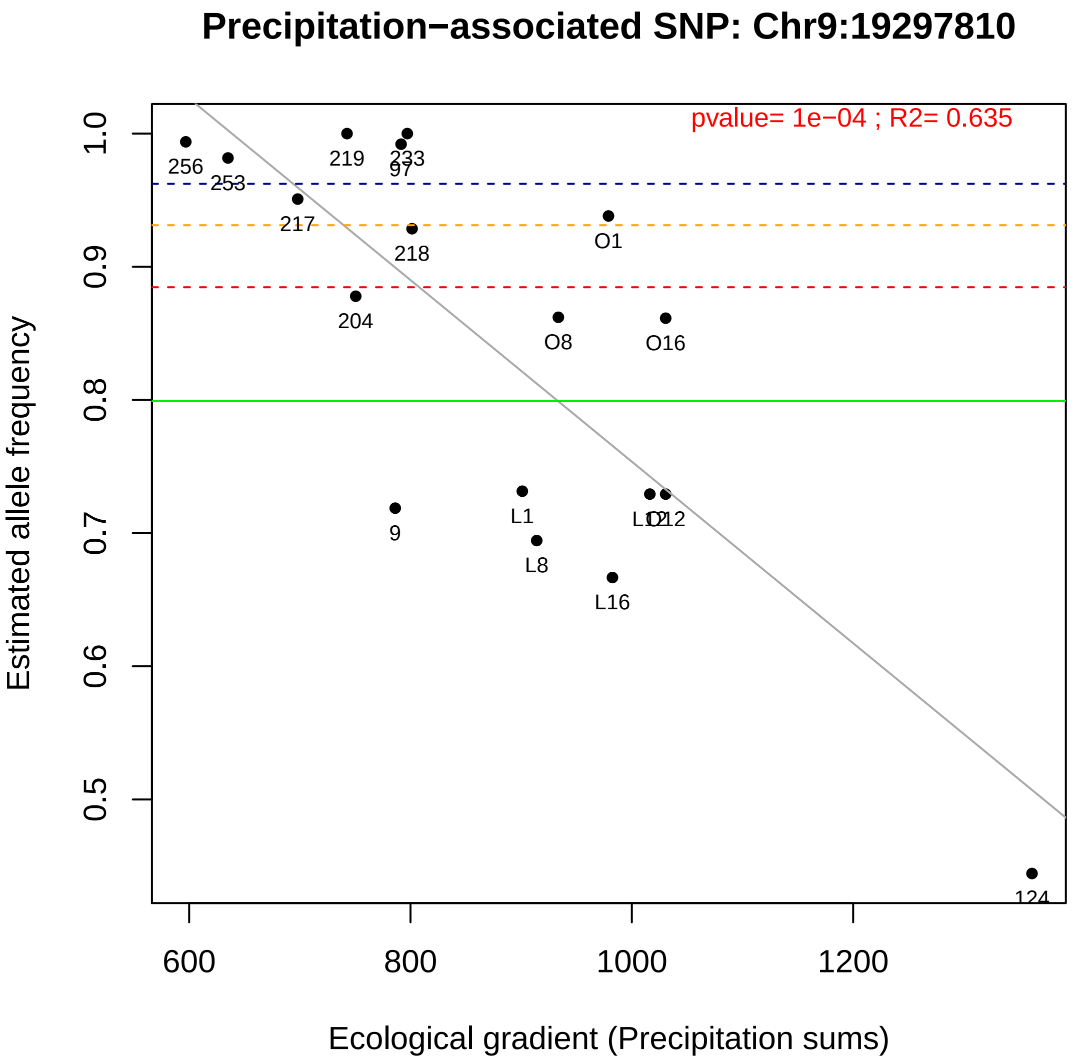

Precipitation−associated SNP: Chr9:19297810
pvalue= 1e−04 ; R2= 0.635
Estimated allele frequency
Ecological gradient (Precipitation sums)
0.5
0.6
0.7
0.8
0.9
1.0
600
800
1000
1200
256
253
219
233
97
217
218
O1
204
O8
O16
9
L1
L8
L16
L12
O12
124

**Precipitation−associated SNP: Chr9:19299379**

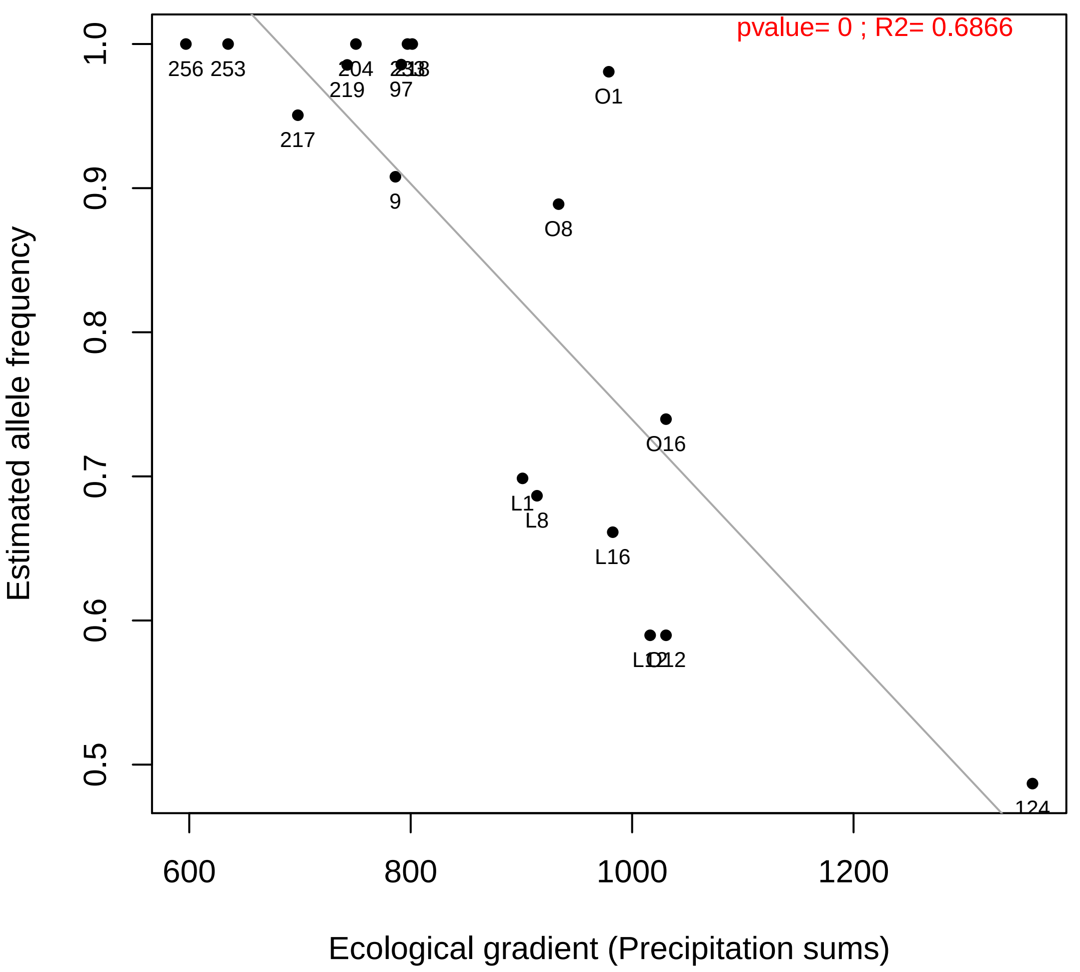

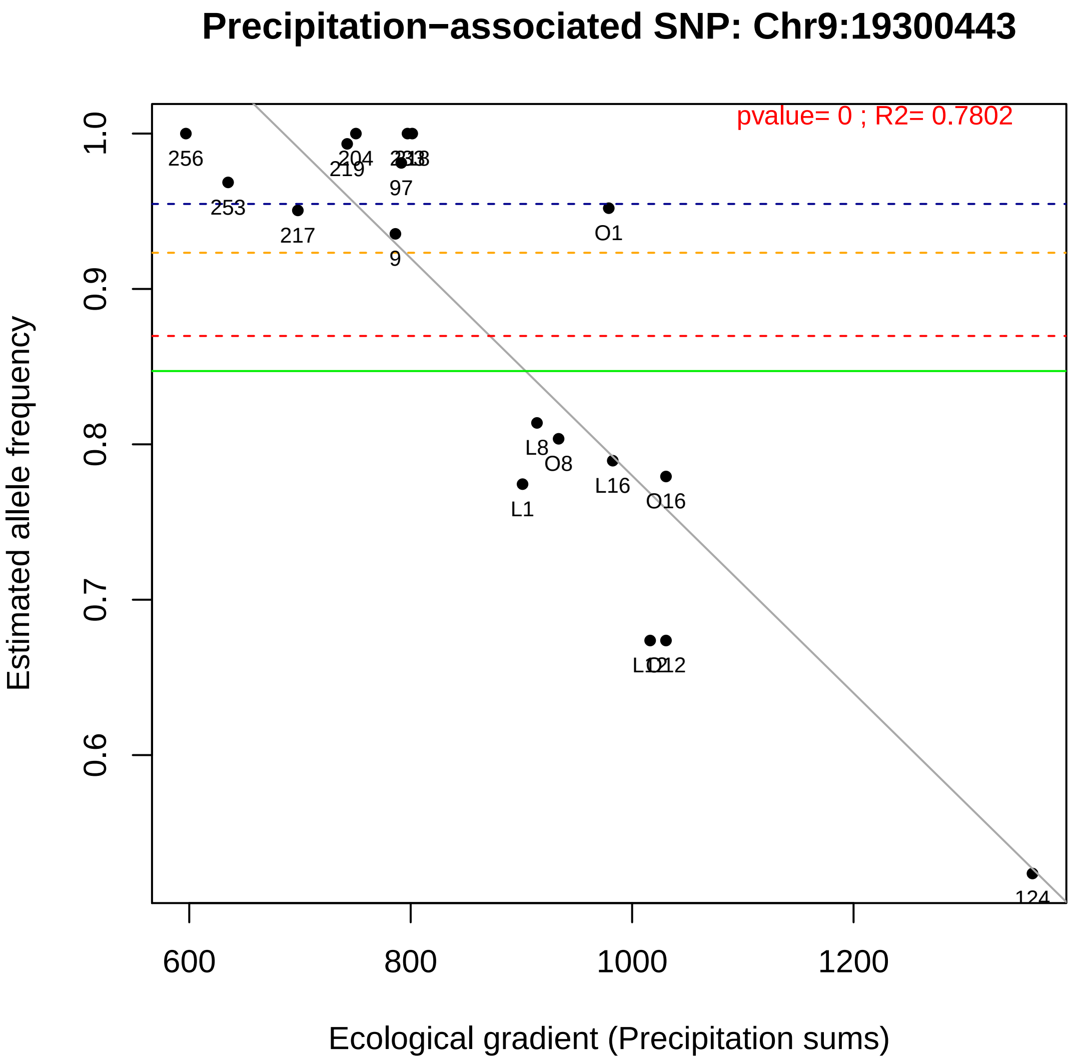
Precipitation−associated SNP: Chr9:19300443
pvalue= 0 ; R2= 0.7802
Estimated allele frequency
1.0
0.9
0.8
0.7
0.6
600
800
1000
1200
Ecological gradient (Precipitation sums)
256
253
217
204
219
97
9
O1
L8
O8
L16
O16
L1
124

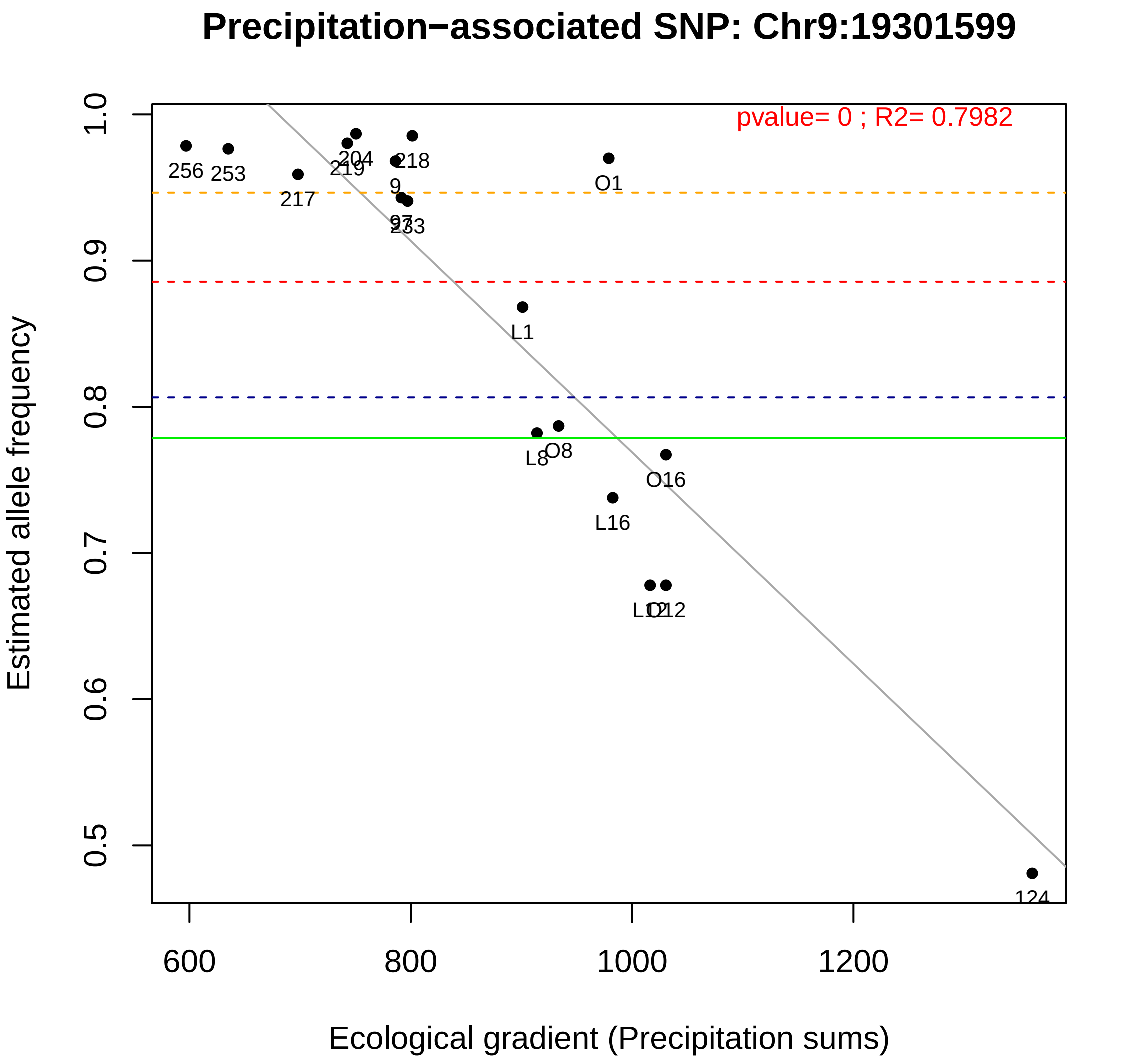

Precipitation−associated SNP: Chr9:19301599
pvalue= 0 ; R2= 0.7982
Estimated allele frequency
0.5
0.6
0.7
0.8
0.9
1.0
600
800
1000
1200
Ecological gradient (Precipitation sums)
256
253
217
204
219
218
9
O1
L1
L8
O8
O16
L16
O12
124

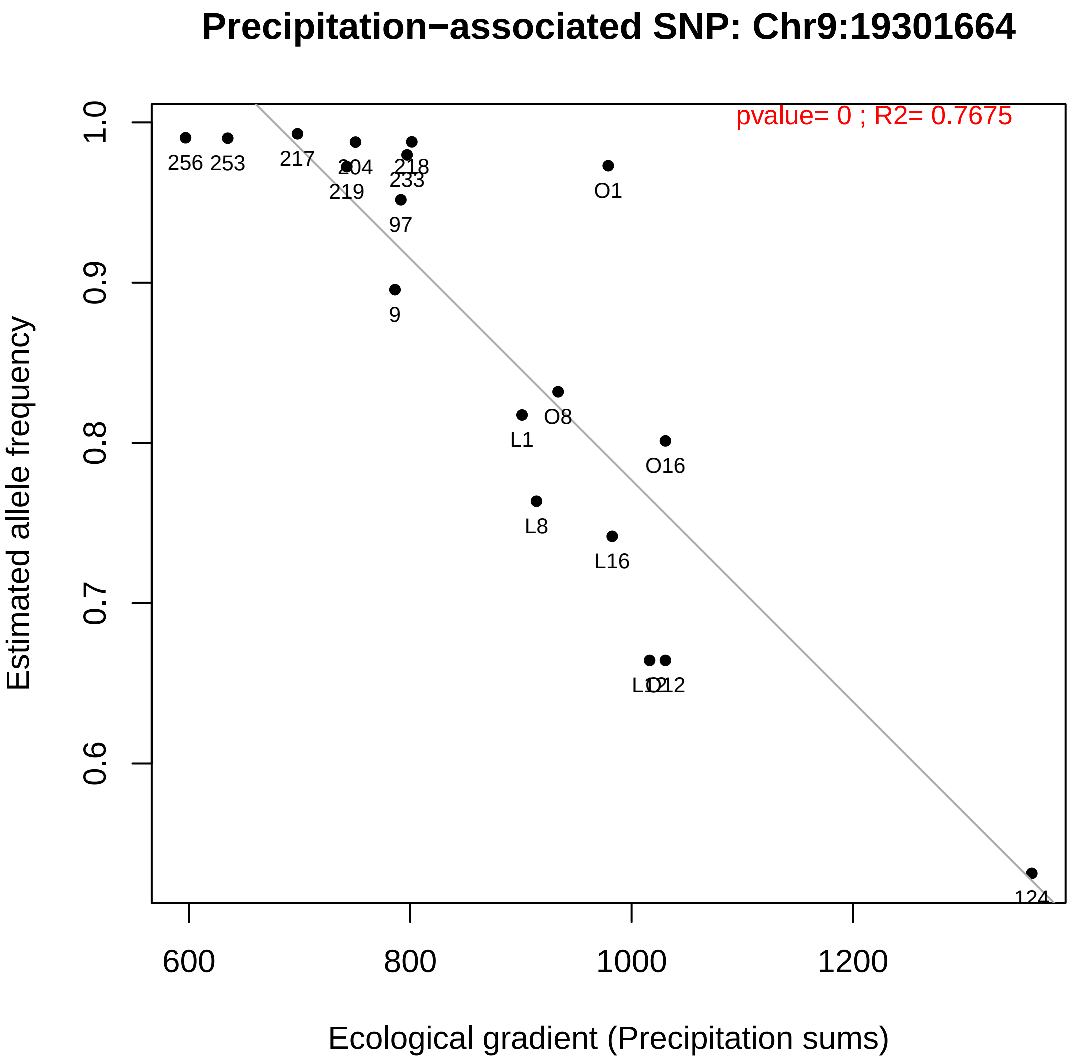
Precipitation−associated SNP: Chr9:19301664
pvalue= 0 ; R2= 0.7675
Estimated allele frequency
Ecological gradient (Precipitation sums)
1.0
0.9
0.8
0.7
0.6
600
800
1000
1200
256
253
217
204
219
218
233
97
9
O1
O8
L1
O16
L8
L16
L12
O12
124

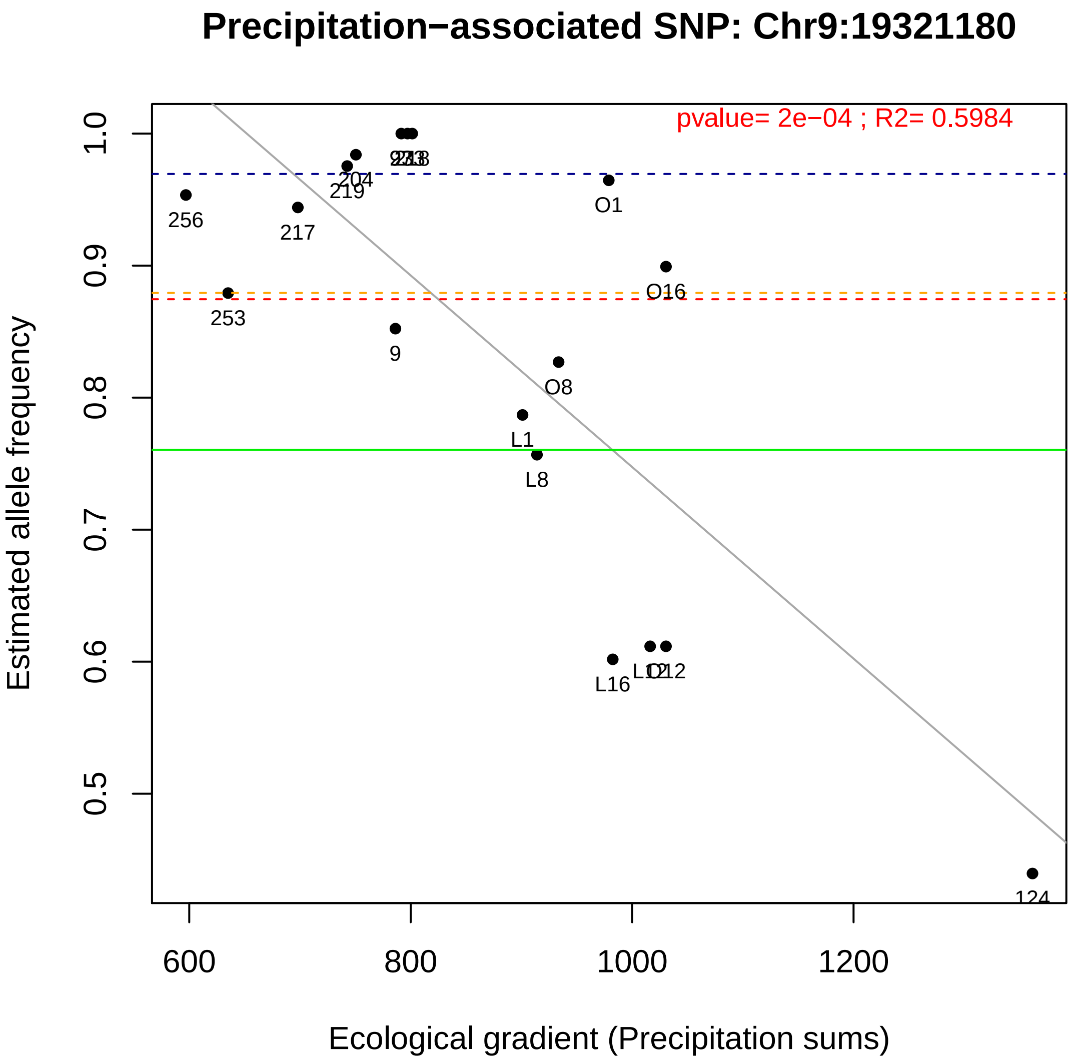

Precipitation−associated SNP: Chr9:19321180
pvalue= 2e−04 ; R2= 0.5984
Estimated allele frequency
Ecological gradient (Precipitation sums)
0.5
0.6
0.7
0.8
0.9
1.0
600
800
1000
1200
256
253
217
219
204
9
O1
O16
O8
L1
L8
L16
124

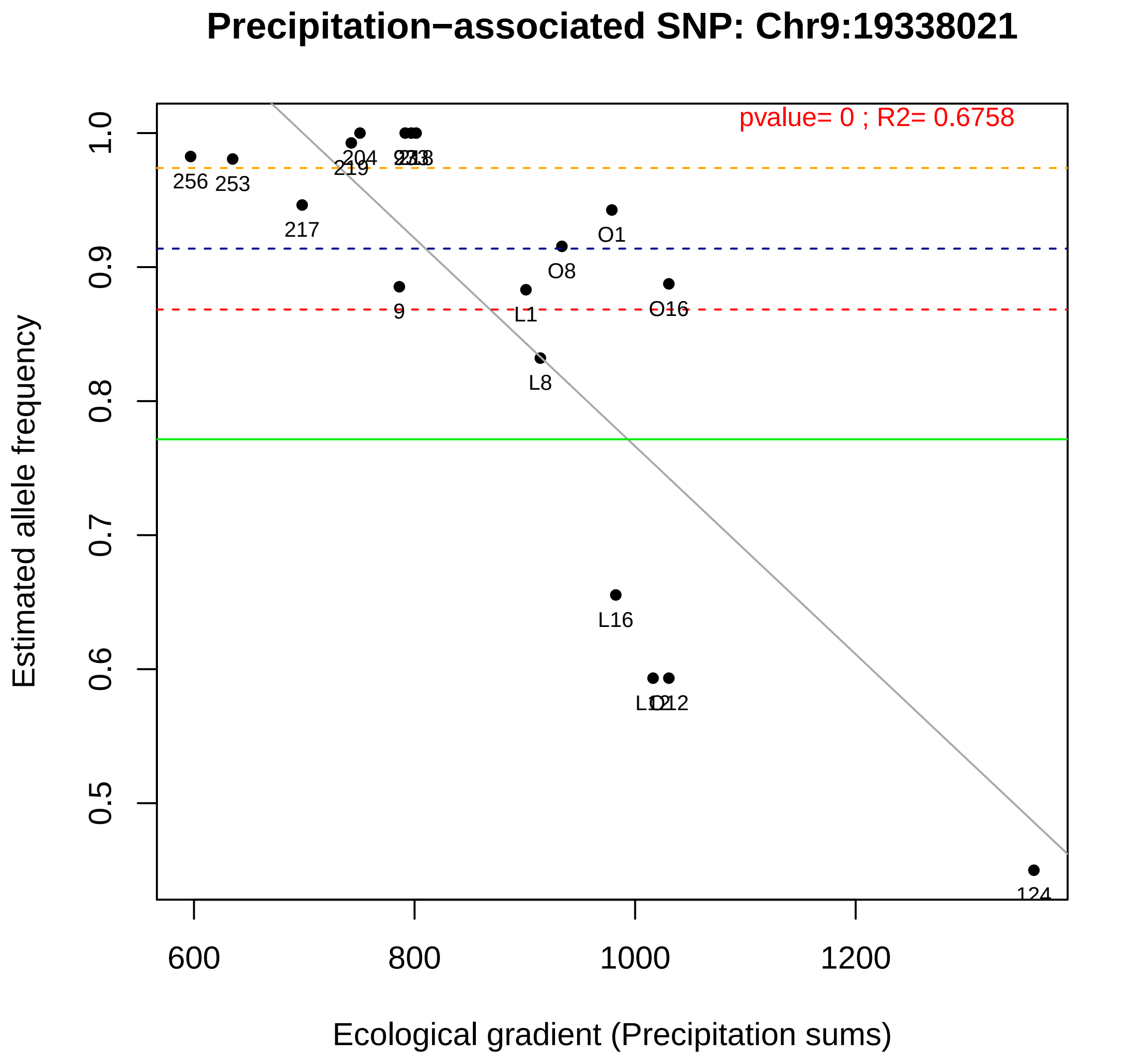

Precipitation−associated SNP: Chr9:19338021
pvalue= 0 ; R2= 0.6758
Estimated allele frequency
1.0
0.9
0.8
0.7
0.6
0.5
600
800
1000
1200
Ecological gradient (Precipitation sums)
256
253
217
204
219
9
L1
L8
O8
O1
O16
L16
O12
124

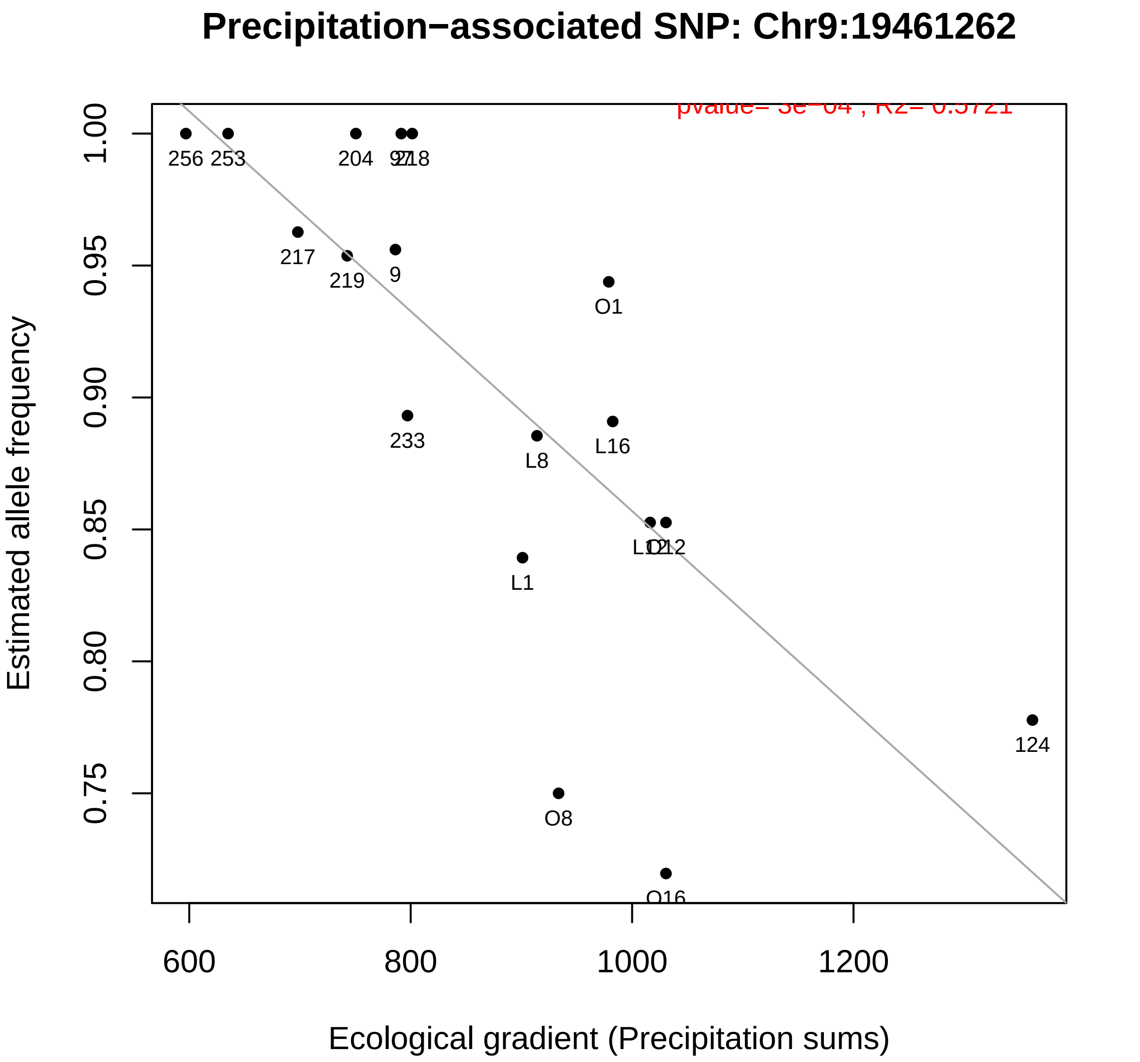

Precipitation−associated SNP: Chr9:19461262
pvalue= 3e−04 , R2= 0.5721
Estimated allele frequency
1.00
0.95
0.90
0.85
0.80
0.75
256
253
204
97
218
217
219
9
O1
233
L8
L16
L10
O12
L1
124
O8
O16
600
800
1000
1200
Ecological gradient (Precipitation sums)

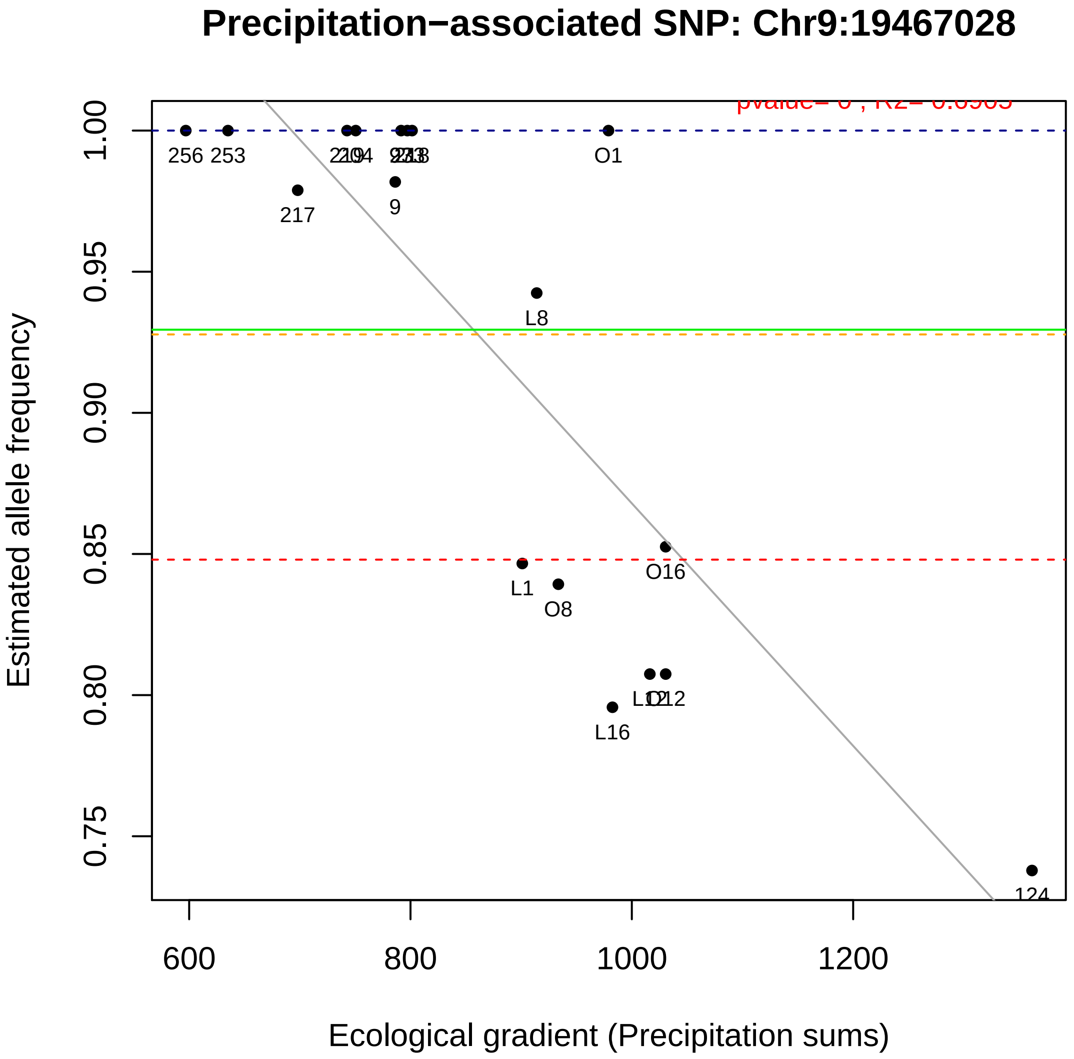

Precipitation−associated SNP: Chr9:19467028
Estimated allele frequency
1.00
0.95
0.90
0.85
0.80
0.75
600
800
1000
1200
Ecological gradient (Precipitation sums)
256
253
217
9
O1
L8
L1
O8
O16
L16
124

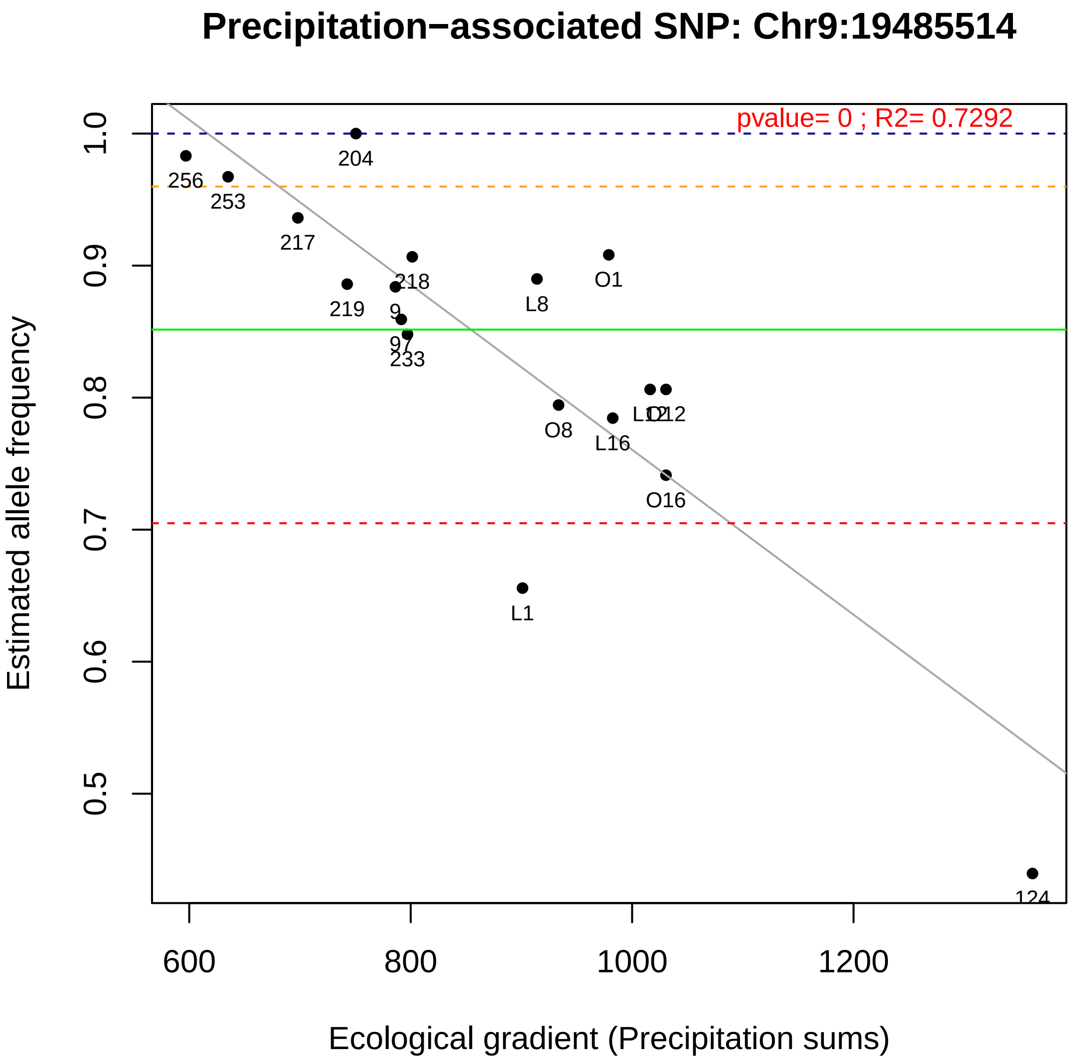

Precipitation−associated SNP: Chr9:19485514
pvalue= 0 ; R2= 0.7292
Estimated allele frequency
Ecological gradient (Precipitation sums)
1.0
0.9
0.8
0.7
0.6
0.5
600
800
1000
1200
256
253
204
217
219
218
9
97
233
L8
O1
O8
L16
L12
O12
O16
L1
124

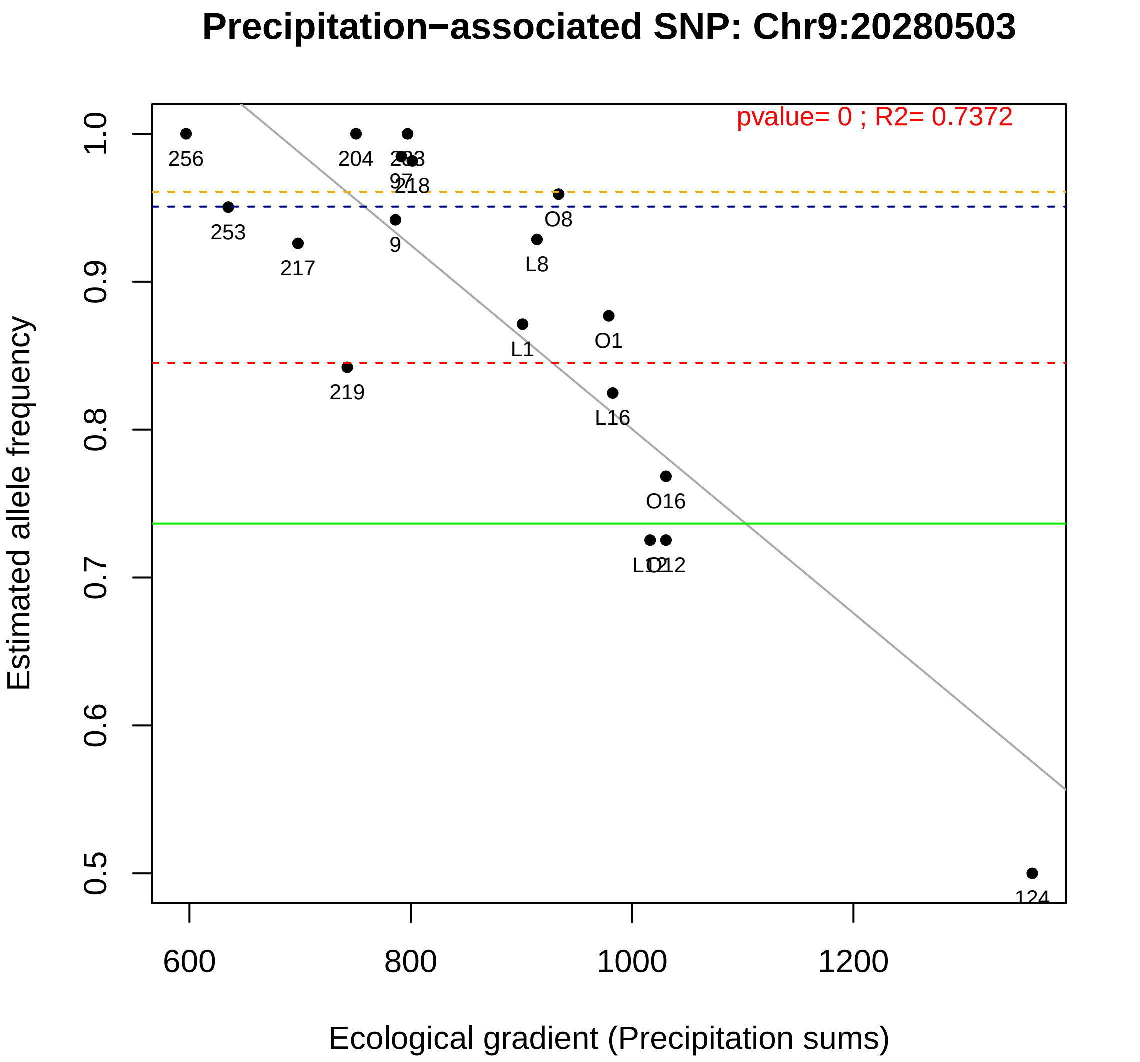
Precipitation−associated SNP: Chr9:20280503
pvalue= 0 ; R2= 0.7372
Estimated allele frequency
Ecological gradient (Precipitation sums)
0.5
0.6
0.7
0.8
0.9
1.0
600
800
1000
1200
256
204
253
217
219
9
L8
O8
L1
O1
L16
O16
124

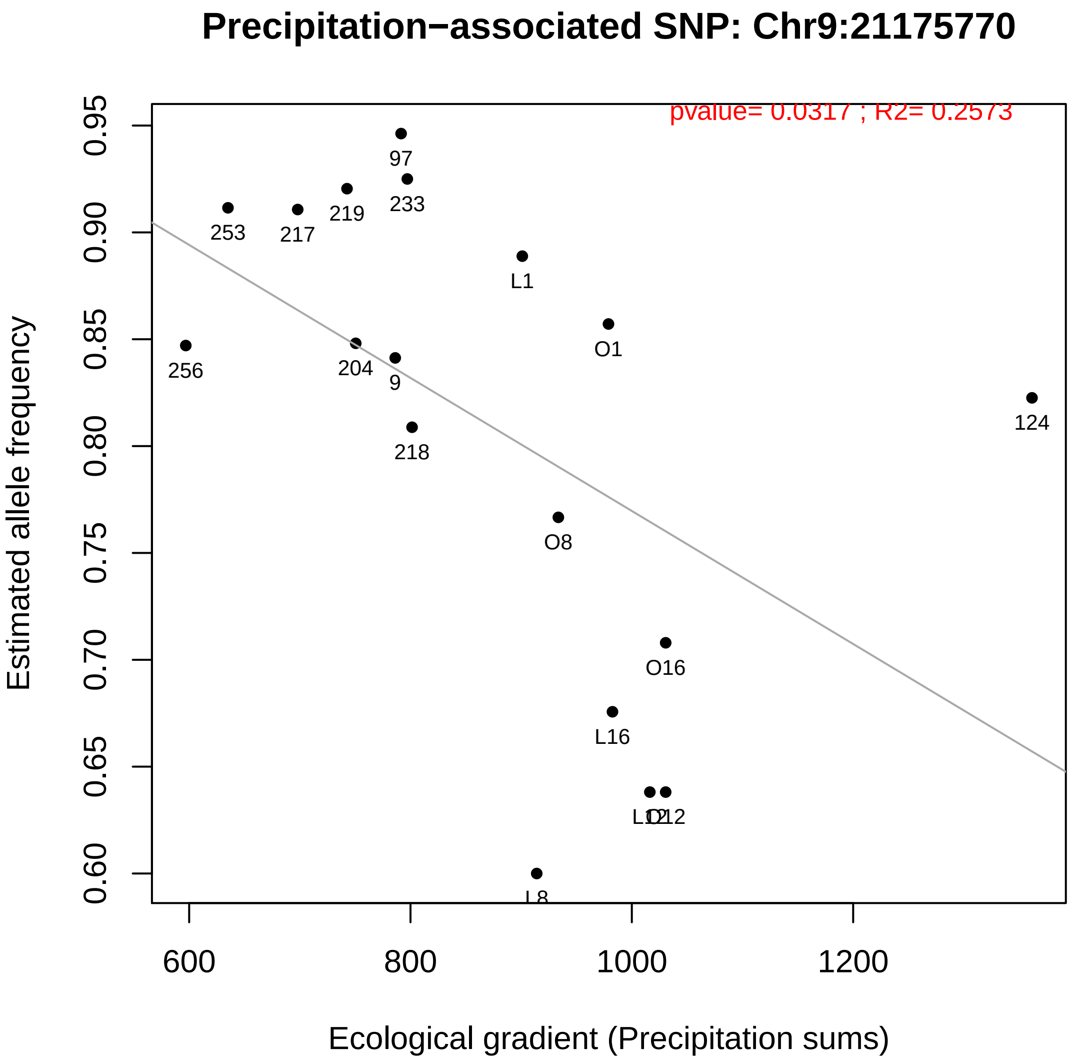

Precipitation−associated SNP: Chr9:21175770
pvalue= 0.0317 ; R2= 0.2573
Estimated allele frequency
0.60
0.65
0.70
0.75
0.80
0.85
0.90
0.95
600
800
1000
1200
Ecological gradient (Precipitation sums)
97
233
219
253
217
L1
O1
256
204
9
124
218
O8
O16
L16

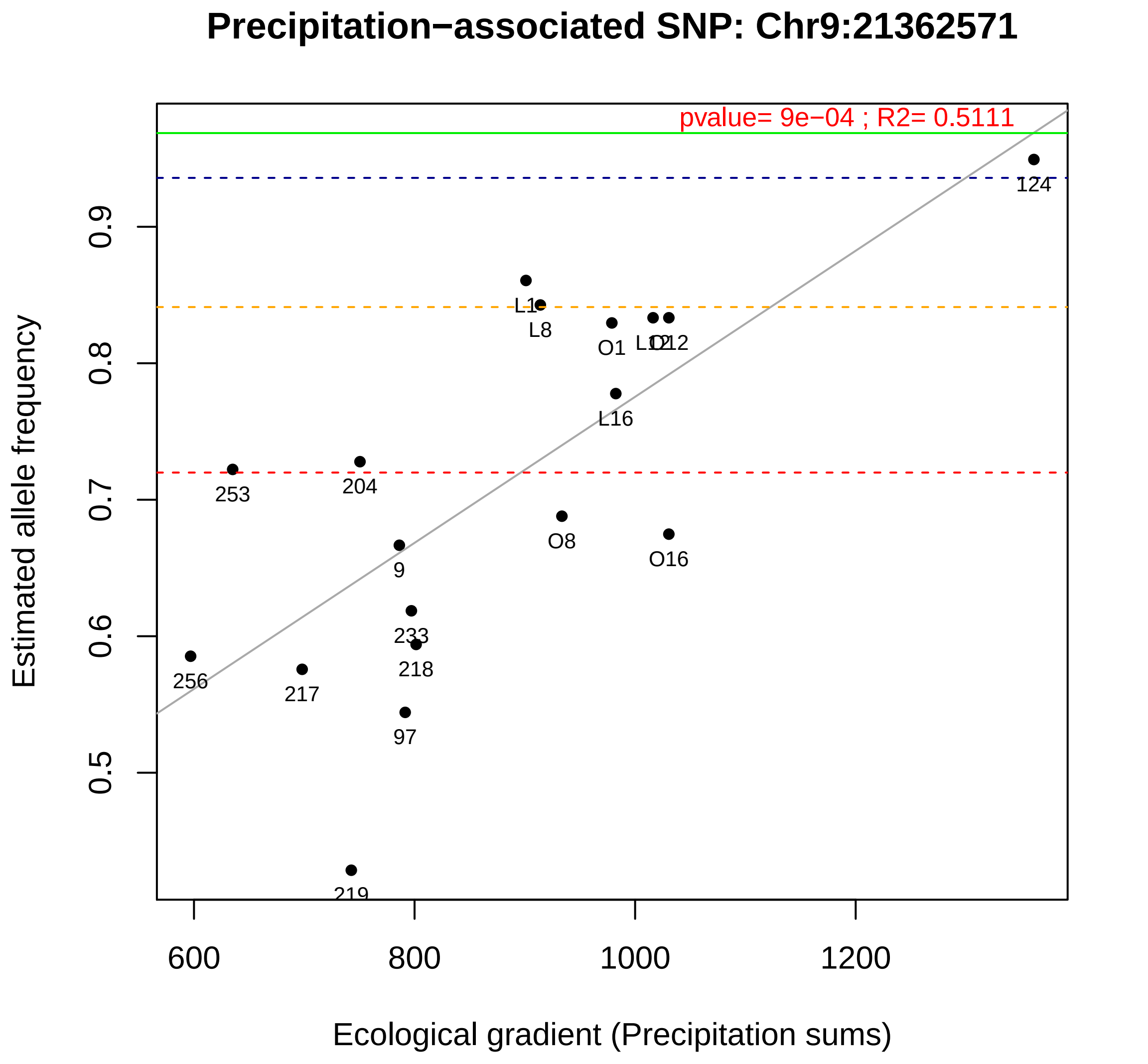

Precipitation−associated SNP: Chr9:21362571
pvalue= 9e−04 ; R2= 0.5111
Estimated allele frequency
0.5
0.6
0.7
0.8
0.9
124
L1
L8
O1
L12
O12
L16
253
204
O8
O16
9
233
218
256
217
97
219
600
800
1000
1200
Ecological gradient (Precipitation sums)

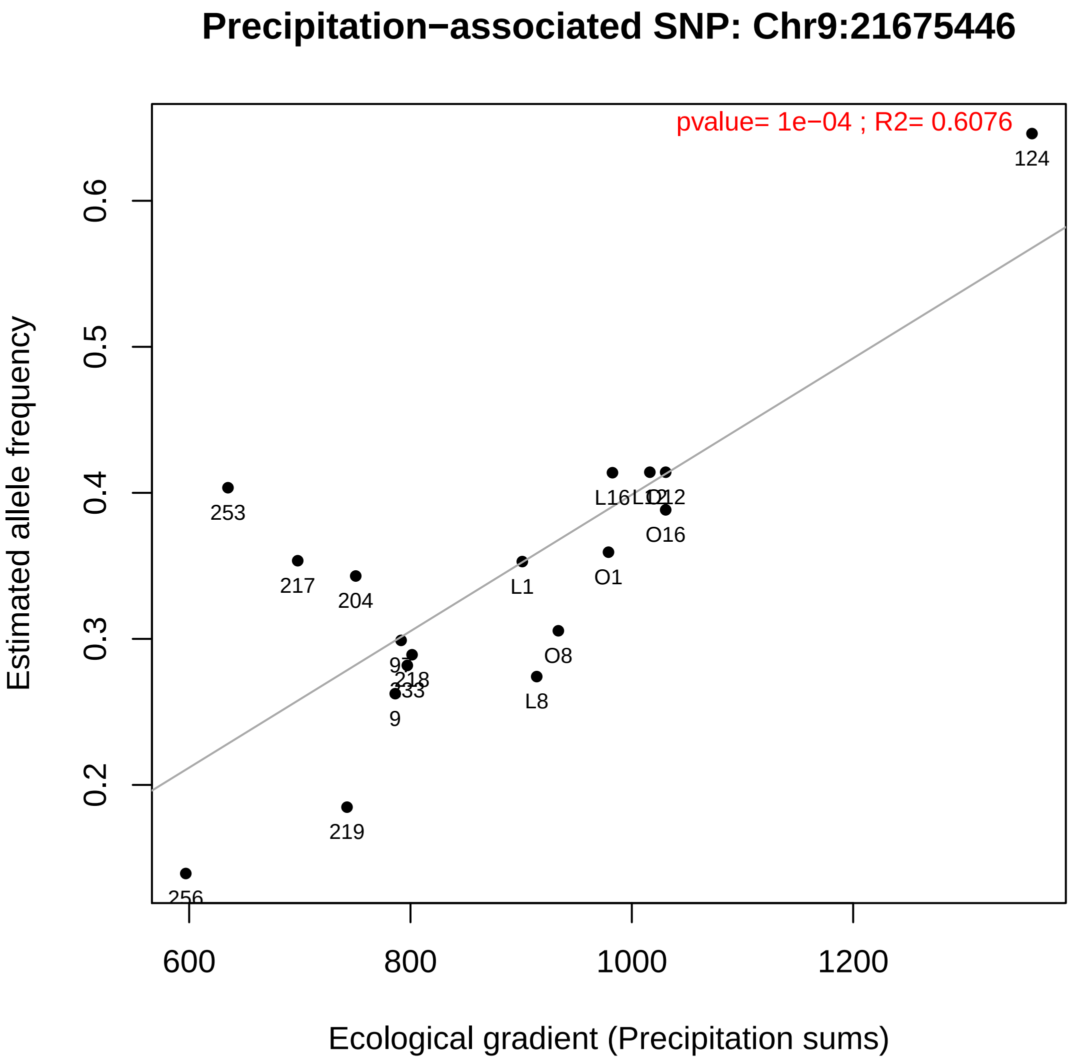
Precipitation−associated SNP: Chr9:21675446
pvalue= 1e−04 ; R2= 0.6076
Estimated allele frequency
0.2
0.3
0.4
0.5
0.6
600
800
1000
1200
Ecological gradient (Precipitation sums)
124
253
217
204
L1
O1
O16
L16
O8
L8
218
9
219
256

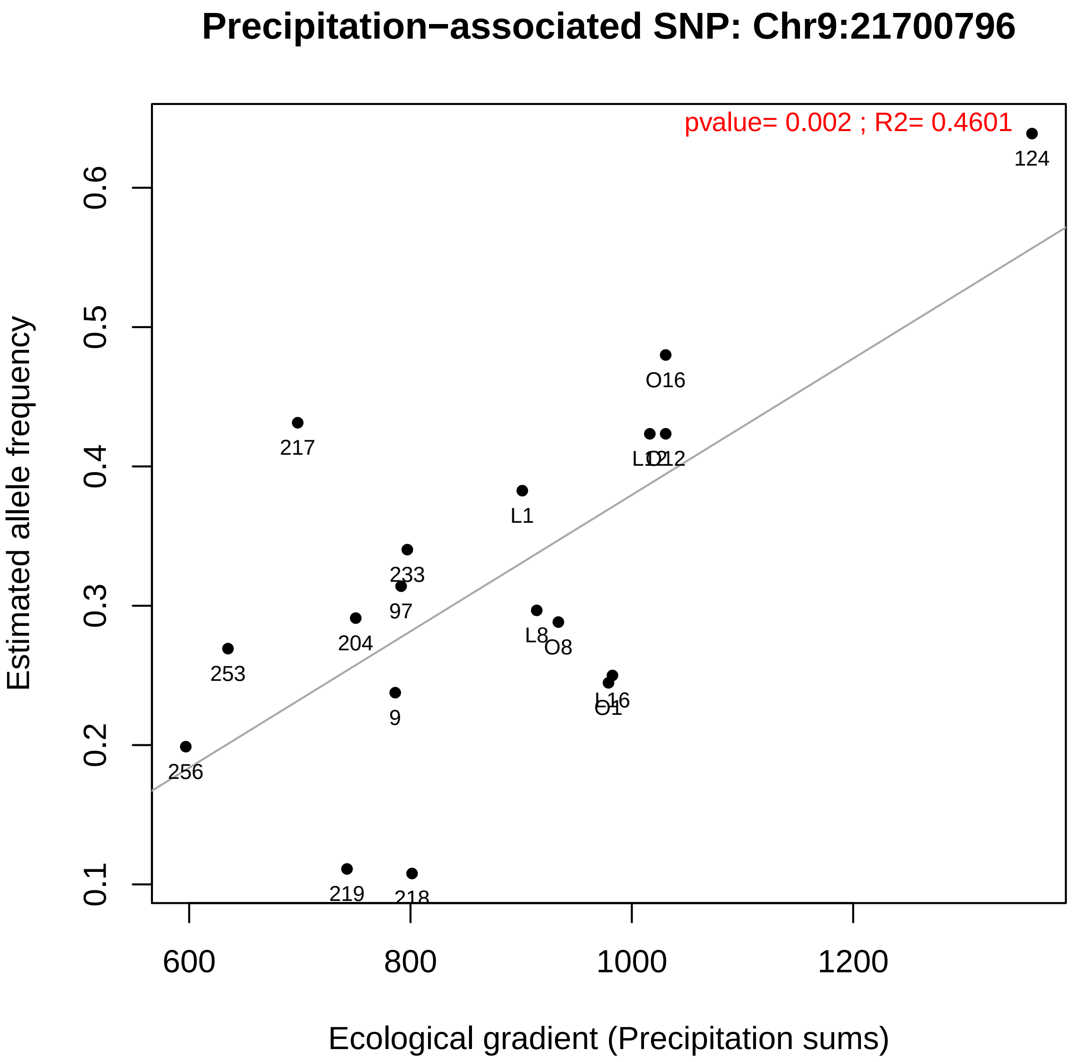
Precipitation−associated SNP: Chr9:21700796
pvalue= 0.002 ; R2= 0.4601
Estimated allele frequency
Ecological gradient (Precipitation sums)
0.1
0.2
0.3
0.4
0.5
0.6
600
800
1000
1200
124
O16
217
L1
233
97
204
L8
O8
253
9
L16
O1
256
219
218

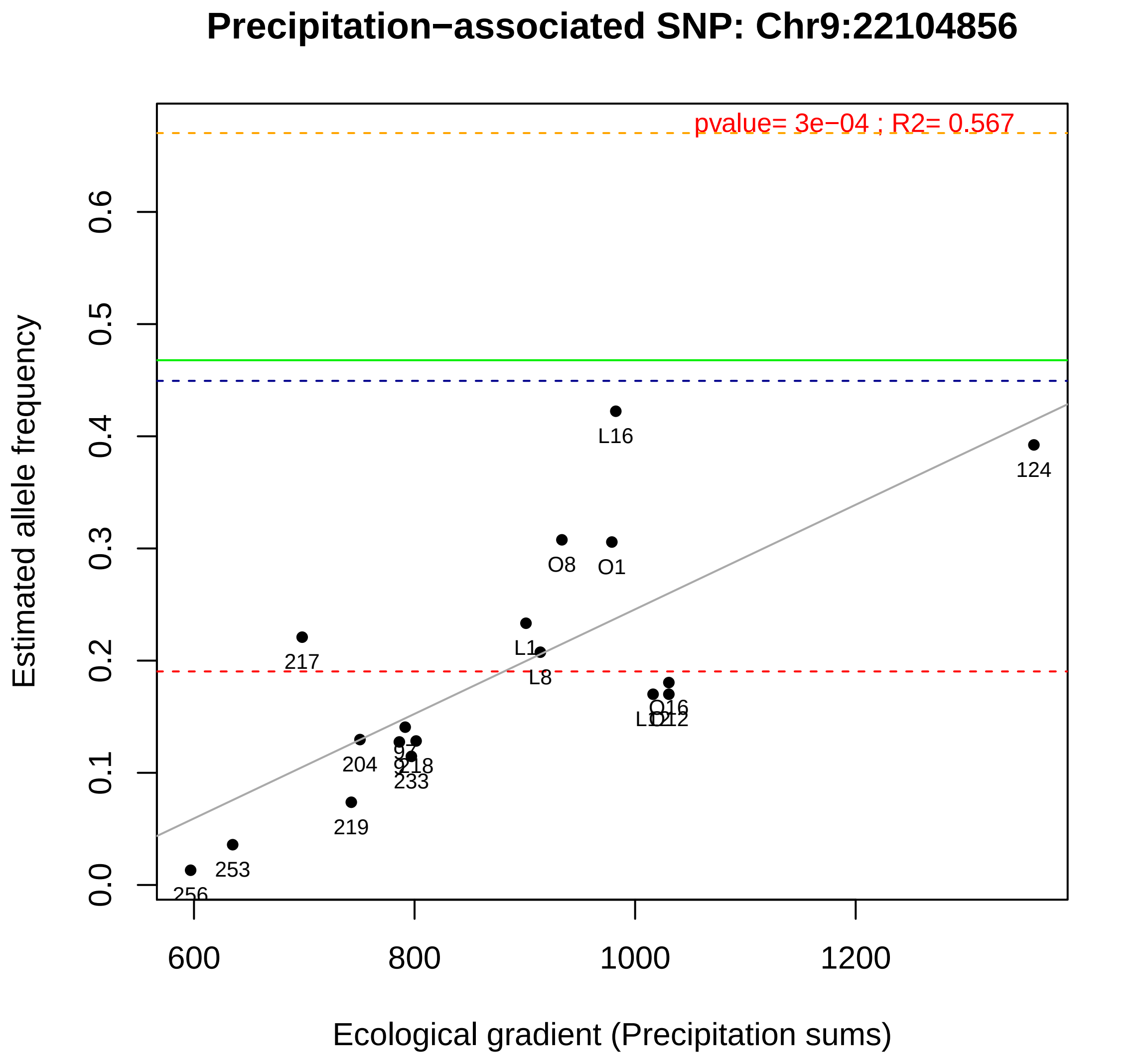

Precipitation−associated SNP: Chr9:22104856
pvalue= 3e−04 ; R2= 0.567
Estimated allele frequency
0.0
0.1
0.2
0.3
0.4
0.5
0.6
600
800
1000
1200
Ecological gradient (Precipitation sums)
L16
124
O8
O1
217
L1
L8
O16
L12
O12
204
97
218
233
219
253
256

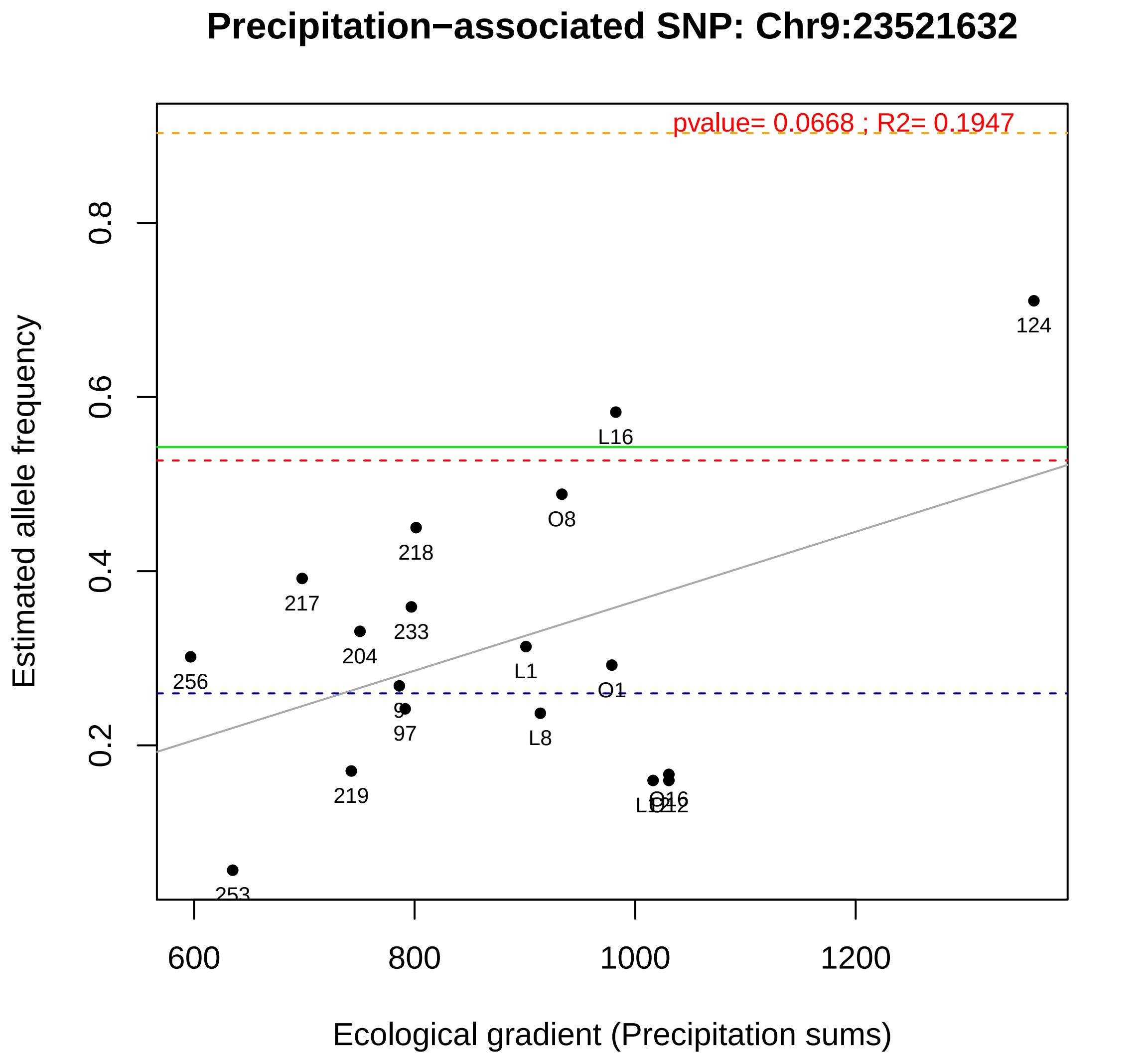

Precipitation−associated SNP: Chr9:23521632
pvalue= 0.0668 ; R2= 0.1947
Estimated allele frequency
0.2
0.4
0.6
0.8
600
800
1000
1200
Ecological gradient (Precipitation sums)
124
L16
O8
218
217
233
204
256
L1
O1
97
L8
219
253

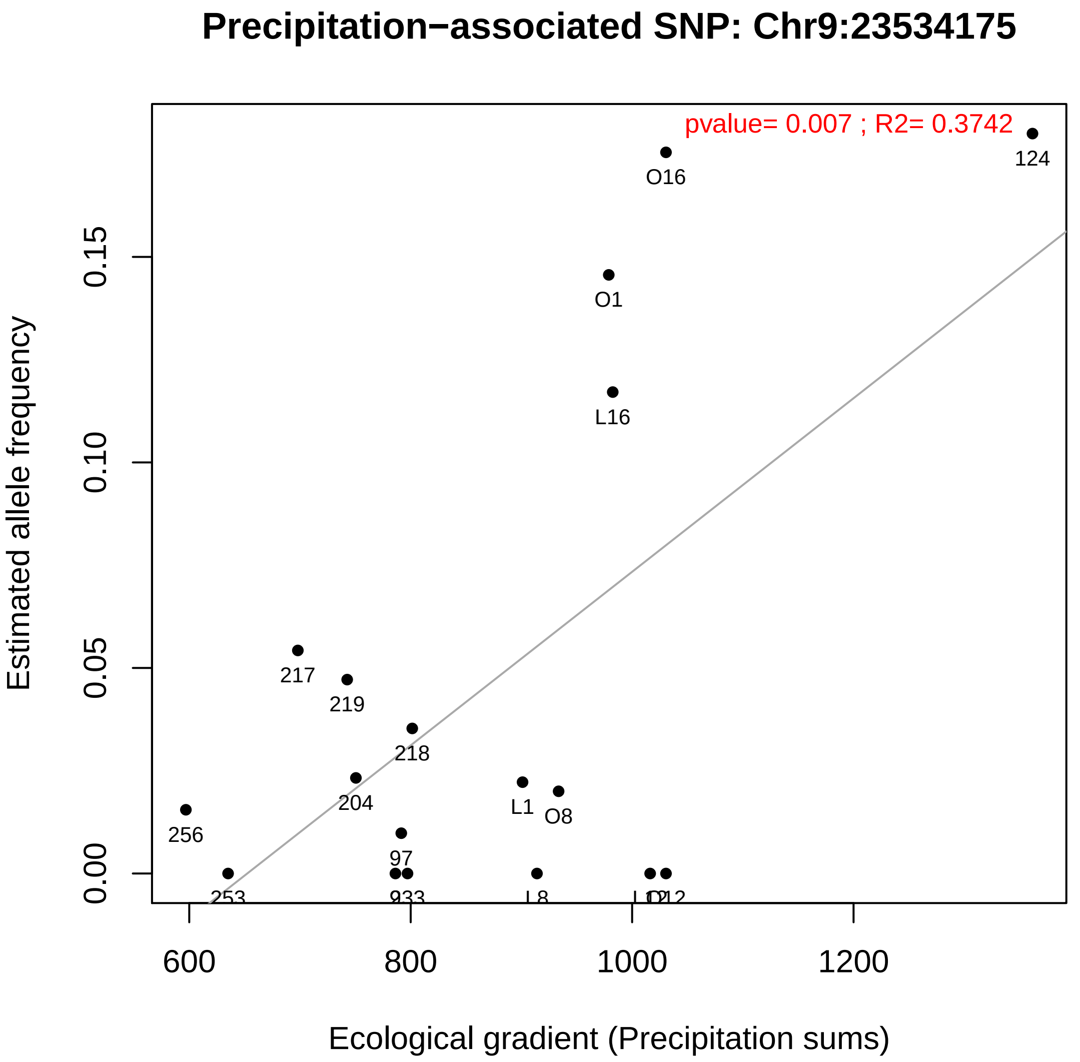
Precipitation−associated SNP: Chr9:23534175
pvalue= 0.007 ; R2= 0.3742
Estimated allele frequency
Ecological gradient (Precipitation sums)
0.00
0.05
0.10
0.15
600
800
1000
1200
124
O16
O1
L16
217
219
218
204
L1
O8
256
97
253
L8

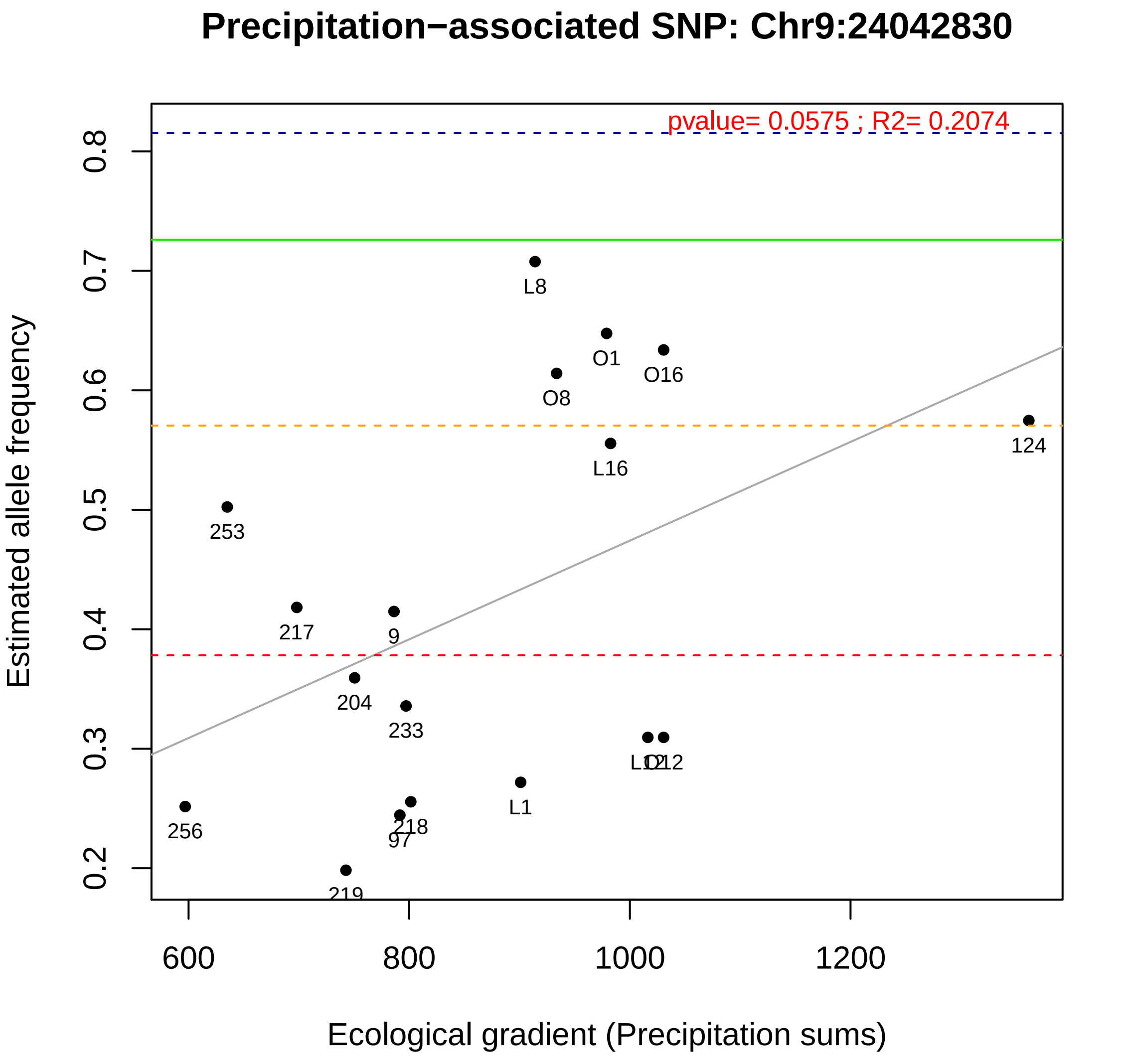
Precipitation−associated SNP: Chr9:24042830
pvalue= 0.0575 ; R2= 0.2074
Estimated allele frequency
0.2
0.3
0.4
0.5
0.6
0.7
0.8
600
800
1000
1200
Ecological gradient (Precipitation sums)
L8
O1
O16
O8
124
L16
253
217
9
204
233
L12
O12
L1
256
218
97
219

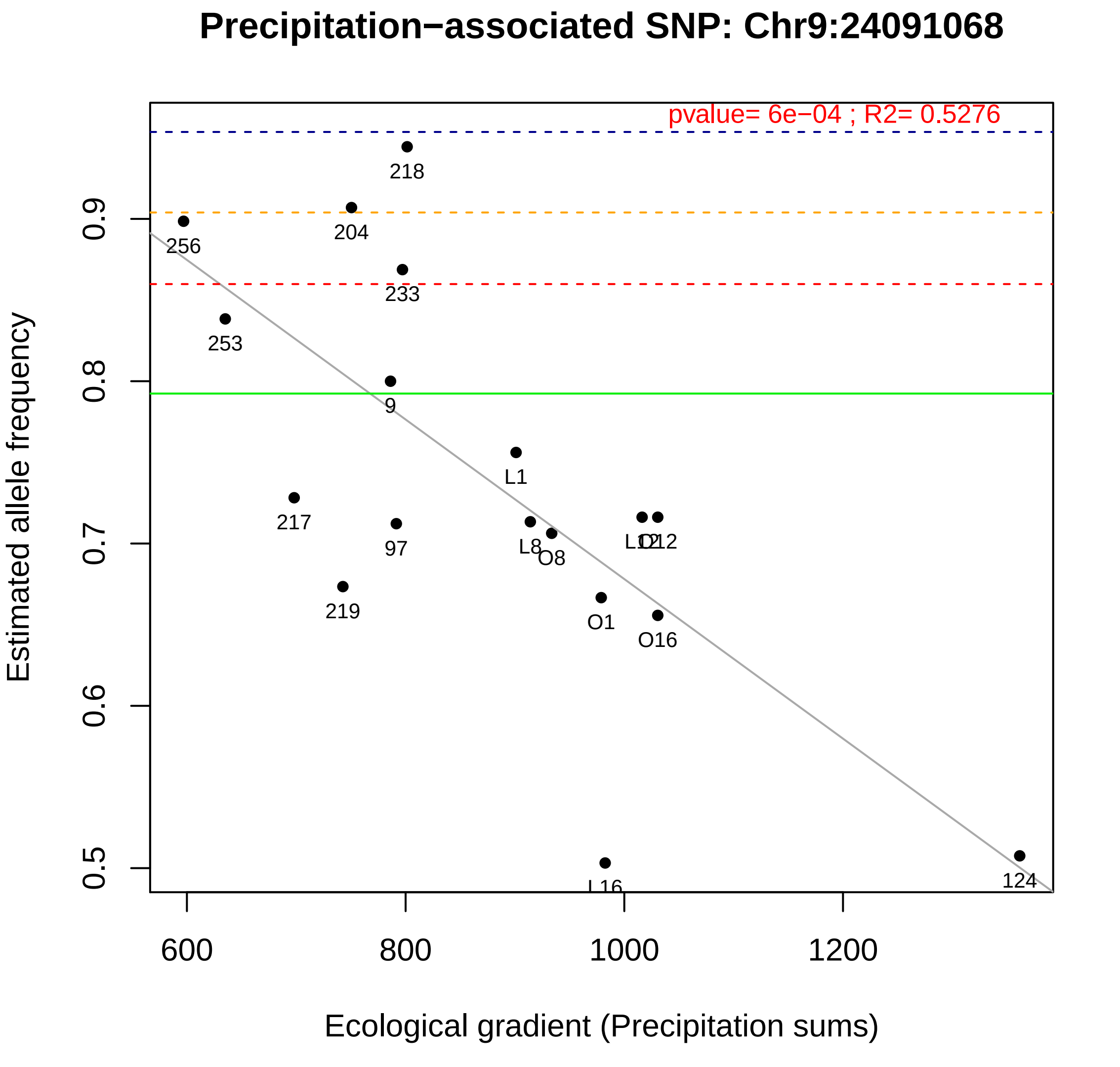
Precipitation−associated SNP: Chr9:24091068
pvalue= 6e−04 ; R2= 0.5276
Estimated allele frequency
0.5
0.6
0.7
0.8
0.9
600
800
1000
1200
Ecological gradient (Precipitation sums)
218
204
256
233
253
9
L1
217
97
L8
O8
L12
O12
219
O1
O16
L16
124

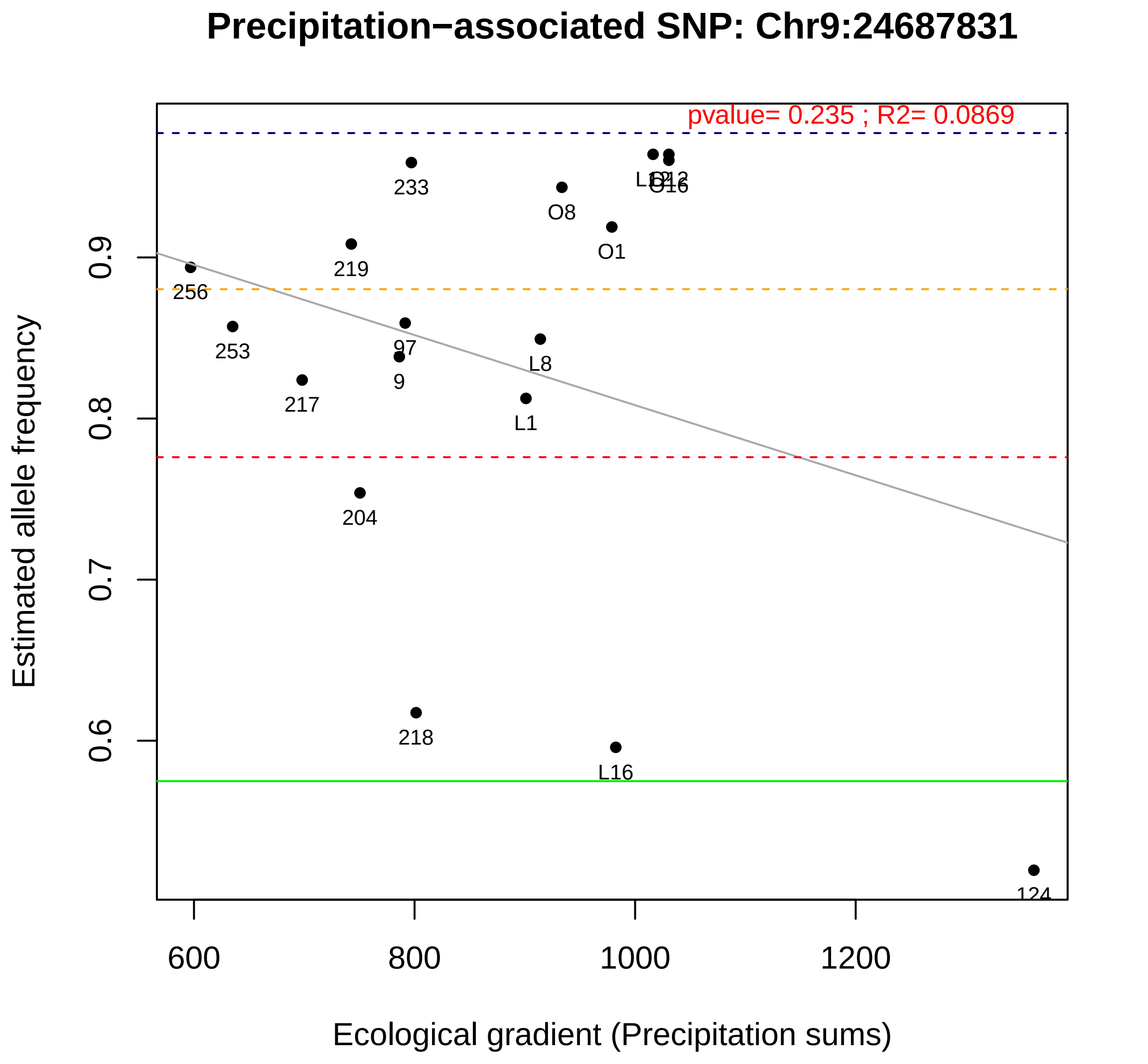

Precipitation−associated SNP: Chr9:24687831
pvalue= 0.235 ; R2= 0.0869
Estimated allele frequency
0.6
0.7
0.8
0.9
600
800
1000
1200
Ecological gradient (Precipitation sums)
233
O8
O1
219
256
253
97
9
L8
217
L1
204
218
L16
124

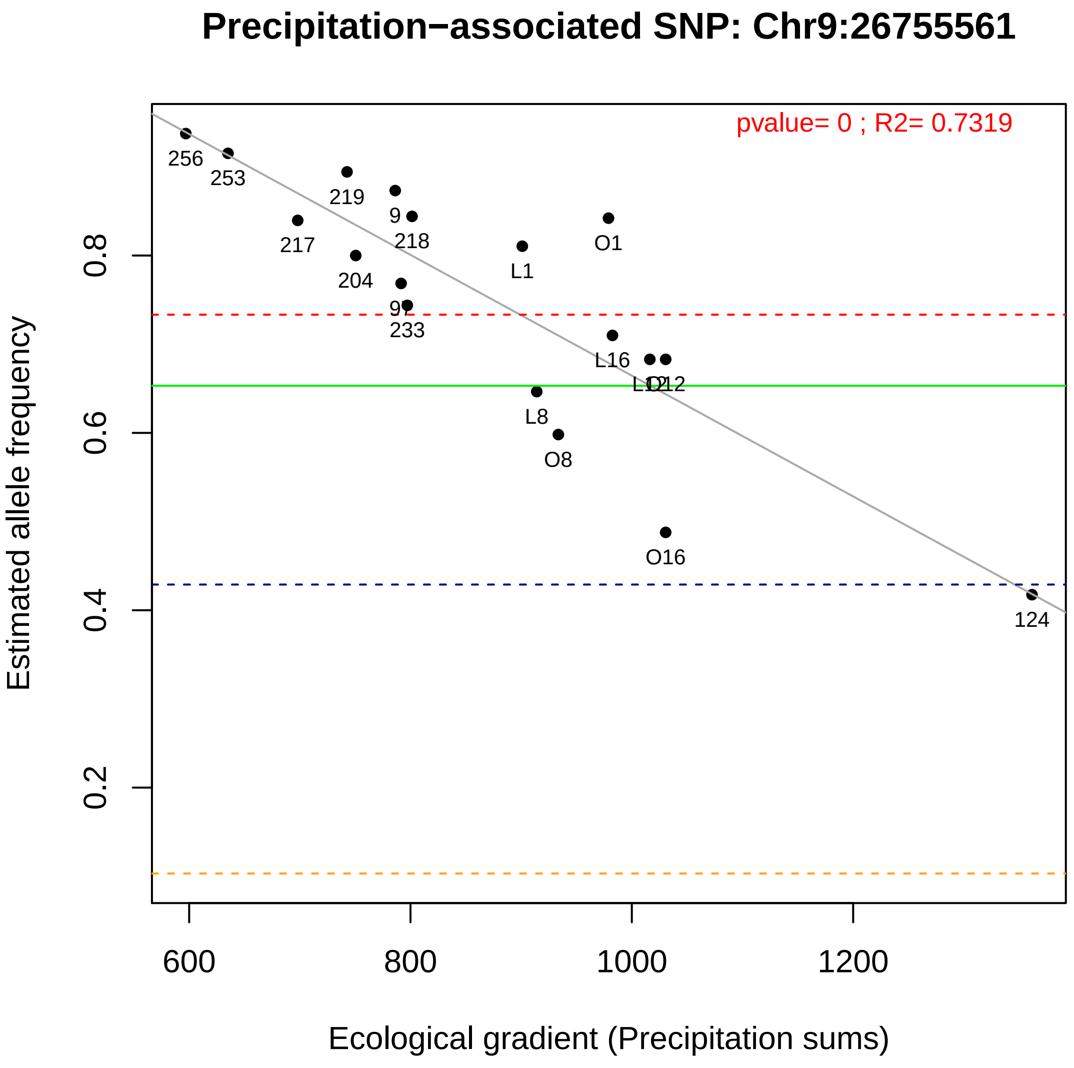

Precipitation−associated SNP: Chr9:26755561
pvalue= 0 ; R2= 0.7319
Estimated allele frequency
Ecological gradient (Precipitation sums)
0.2
0.4
0.6
0.8
600
800
1000
1200
256
253
219
9
218
217
204
L1
O1
97
233
L16
L8
O8
O16
124

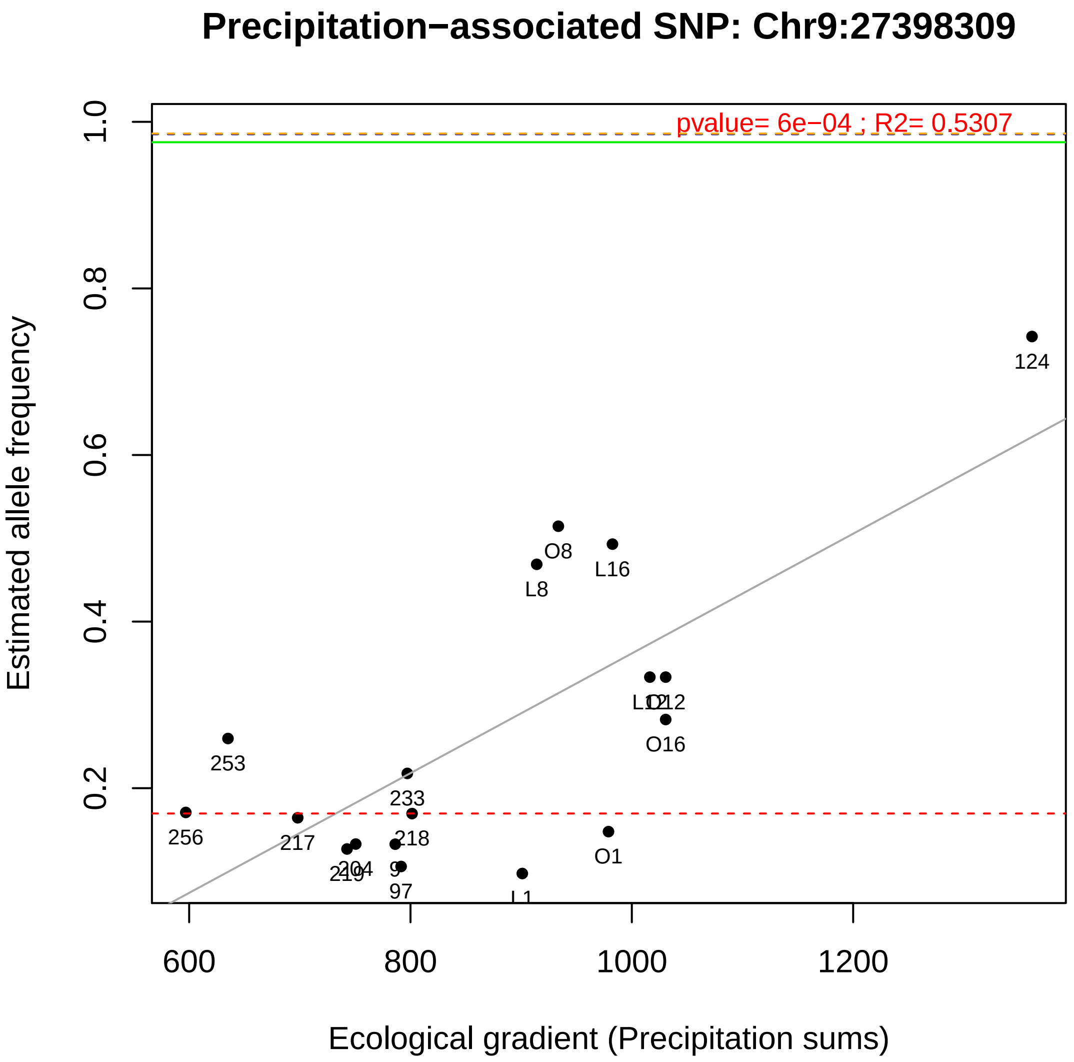

Precipitation−associated SNP: Chr9:27398309
pvalue= 6e−04 ; R2= 0.5307
Estimated allele frequency
Ecological gradient (Precipitation sums)
0.2
0.4
0.6
0.8
1.0
600
800
1000
1200
124
O8
L16
L8
O12
O16
253
233
256
217
218
204
219
97
O1

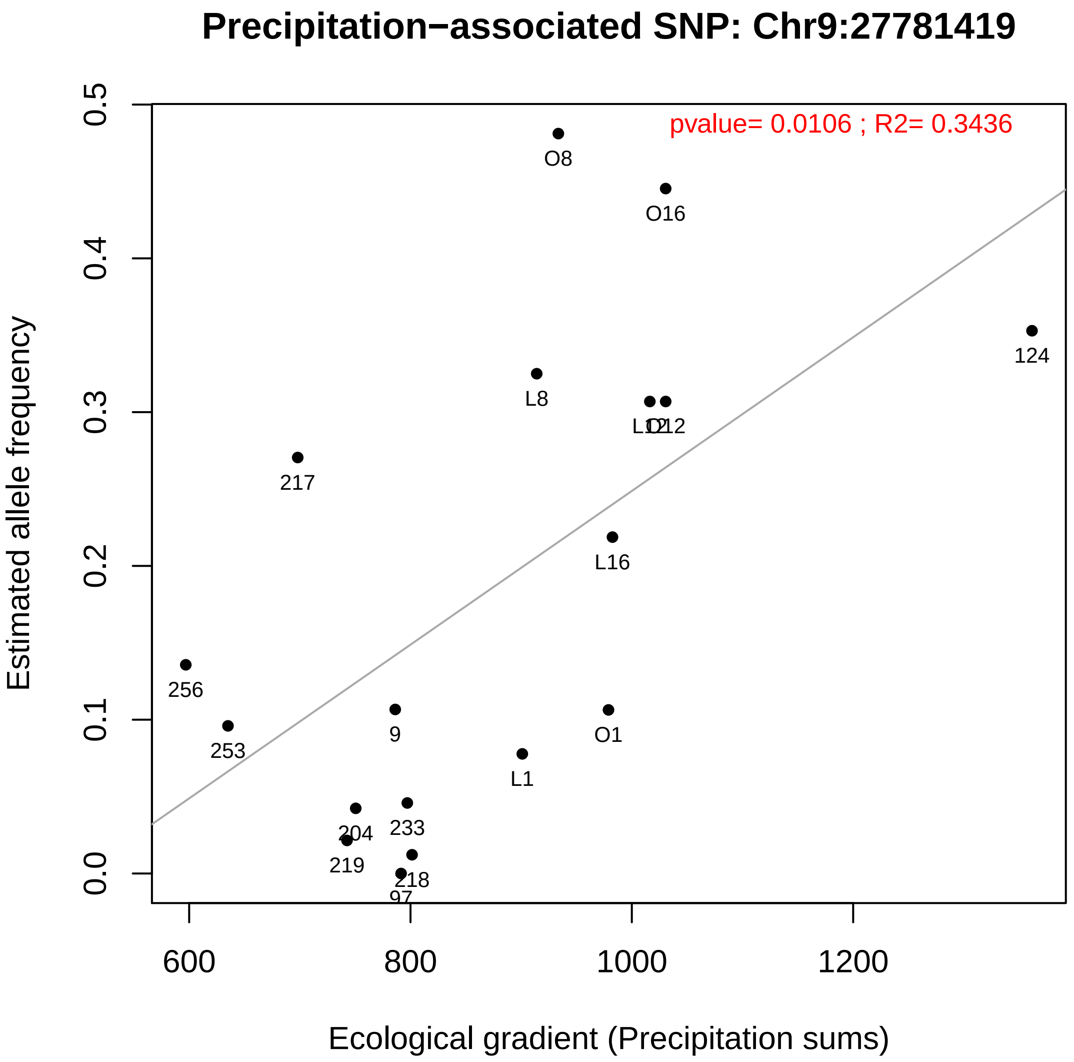
Precipitation−associated SNP: Chr9:27781419
pvalue= 0.0106 ; R2= 0.3436
O8
O16
124
L8
L10
O12
217
L16
256
253
9
O1
L1
204
233
219
218
97
0.0
0.1
0.2
0.3
0.4
0.5
600
800
1000
1200
Estimated allele frequency
Ecological gradient (Precipitation sums)

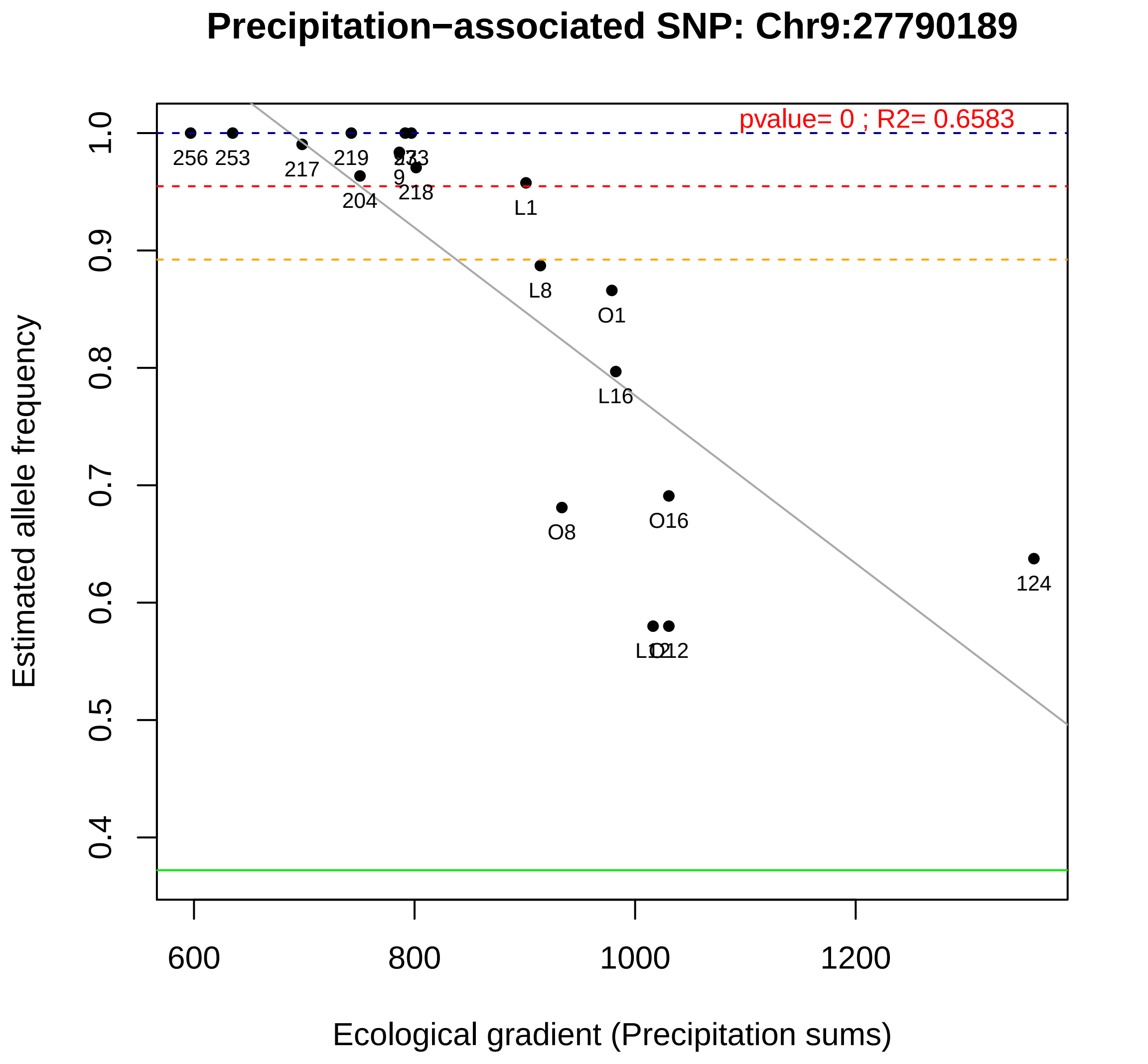

Precipitation−associated SNP: Chr9:27790189
pvalue= 0 ; R2= 0.6583
256
253
217
219
204
9
218
L1
L8
O1
L16
O8
O16
124
1.0
0.9
0.8
0.7
0.6
0.5
0.4
600
800
1000
1200
Estimated allele frequency
Ecological gradient (Precipitation sums)

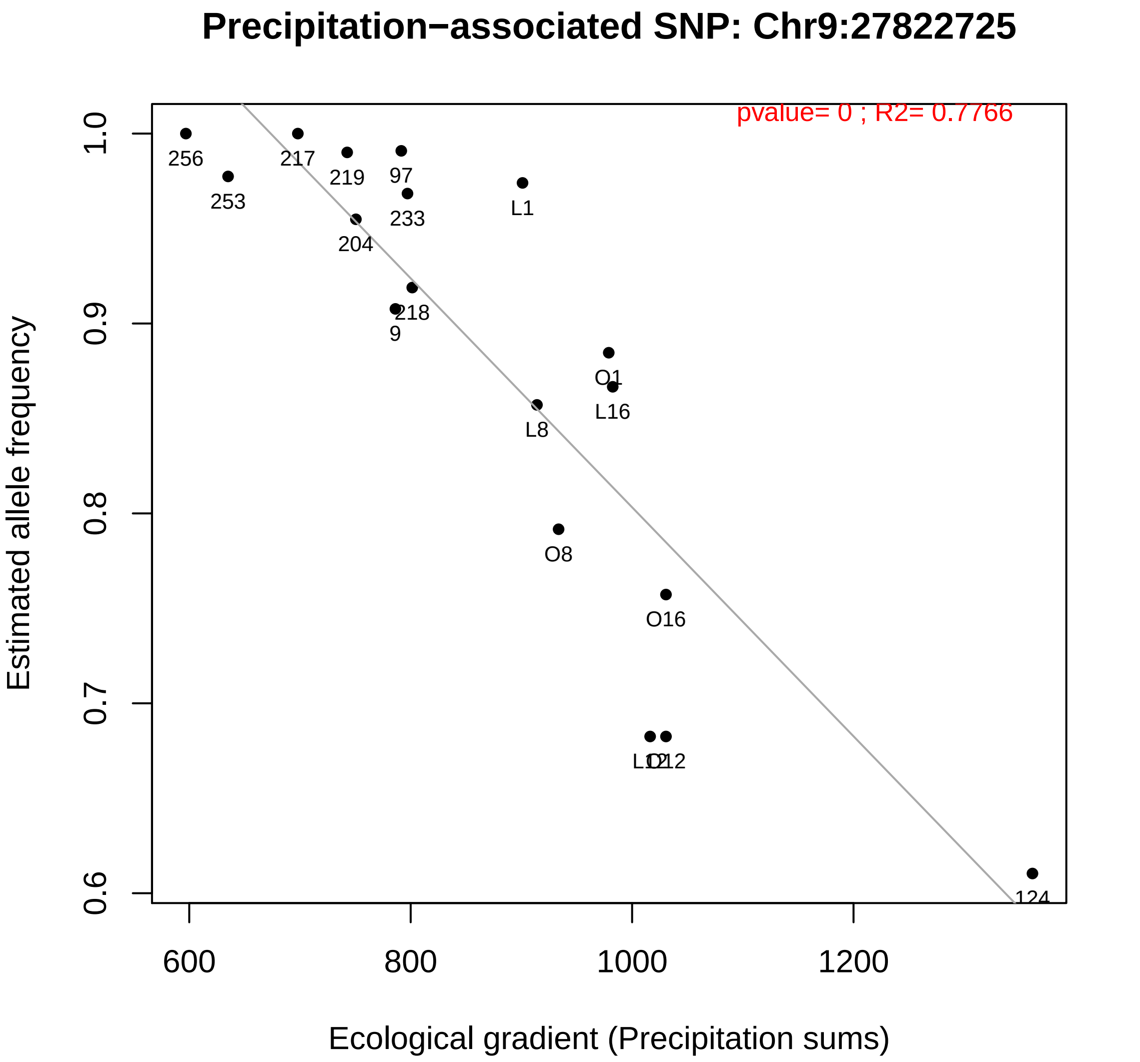
Precipitation−associated SNP: Chr9:27822725
pvalue= 0 ; R2= 0.7766
Estimated allele frequency
1.0
0.9
0.8
0.7
0.6
600
800
1000
1200
Ecological gradient (Precipitation sums)
256
253
217
219
97
233
204
L1
218
9
O1
L16
L8
O8
O16
L12
O12
124

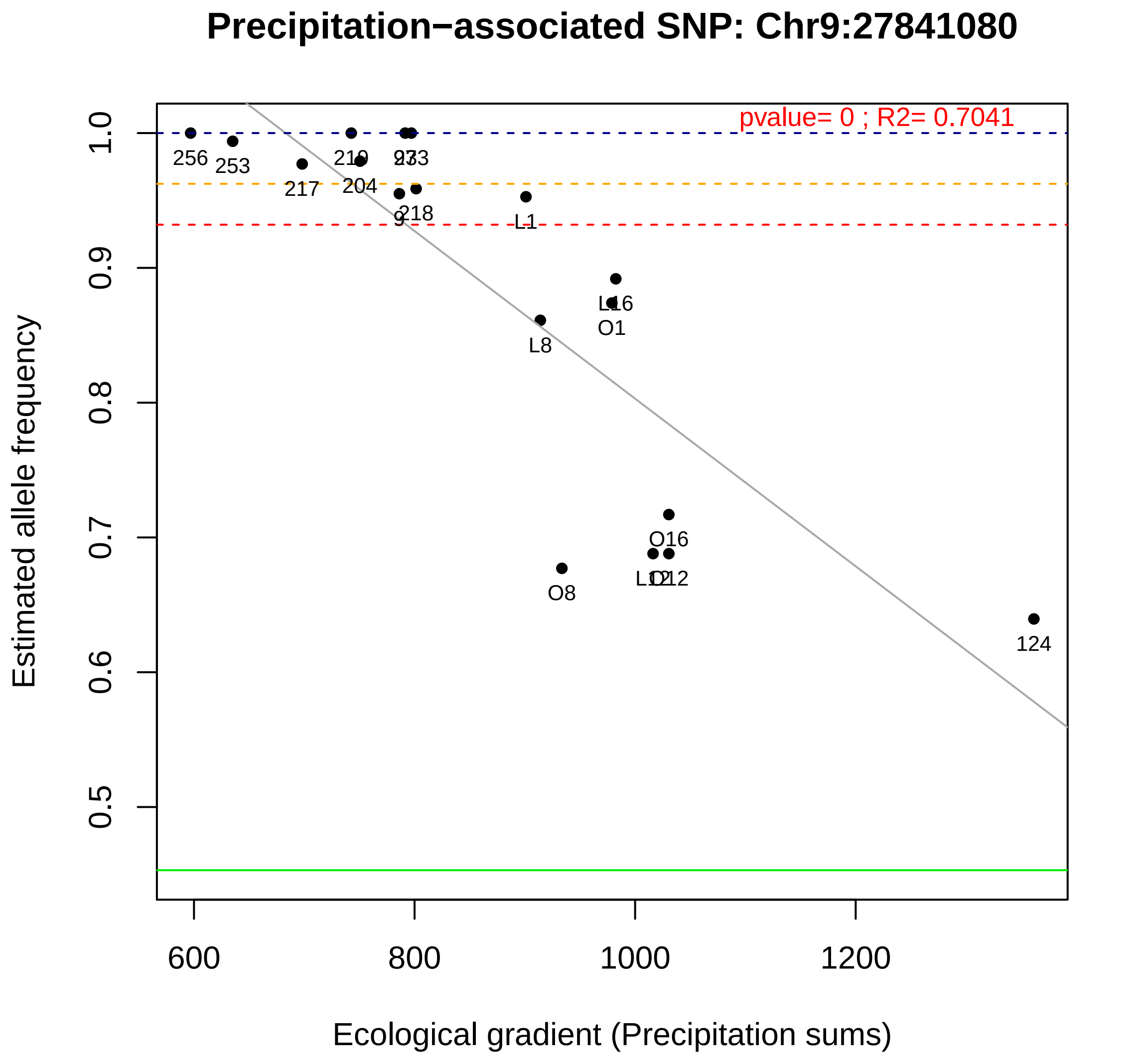
Precipitation−associated SNP: Chr9:27841080
pvalue= 0 ; R2= 0.7041
Estimated allele frequency
0.5
0.6
0.7
0.8
0.9
1.0
600
800
1000
1200
Ecological gradient (Precipitation sums)
256
253
217
210
204
97
233
218
9
L1
L16
O1
L8
O16
O8
L12
O12
124

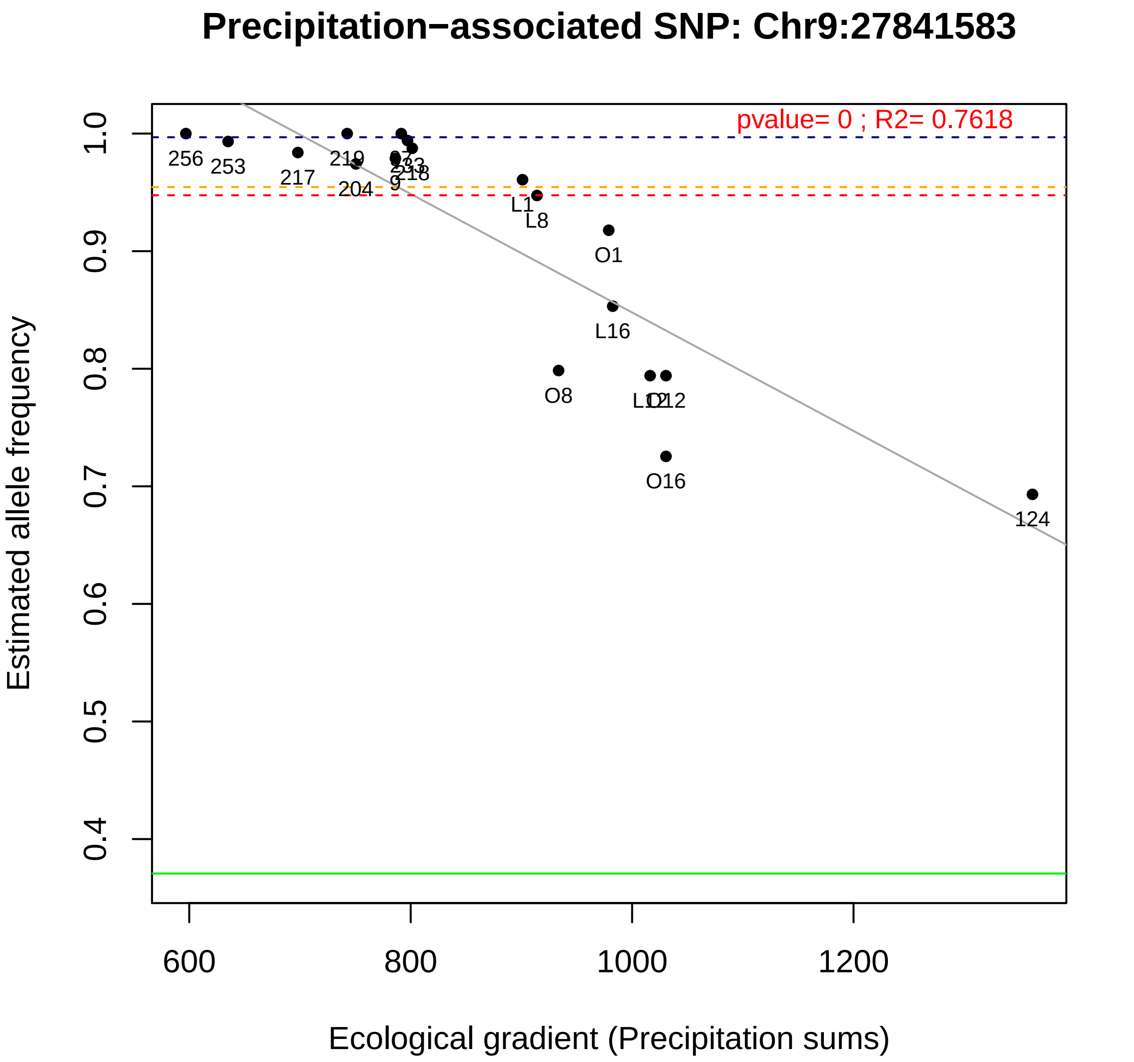
Precipitation−associated SNP: Chr9:27841583
pvalue= 0 ; R2= 0.7618
Estimated allele frequency
Ecological gradient (Precipitation sums)
0.4
0.5
0.6
0.7
0.8
0.9
1.0
600
800
1000
1200
256
253
217
219
204
9
218
L1
L8
O1
L16
O8
O12
O16
124

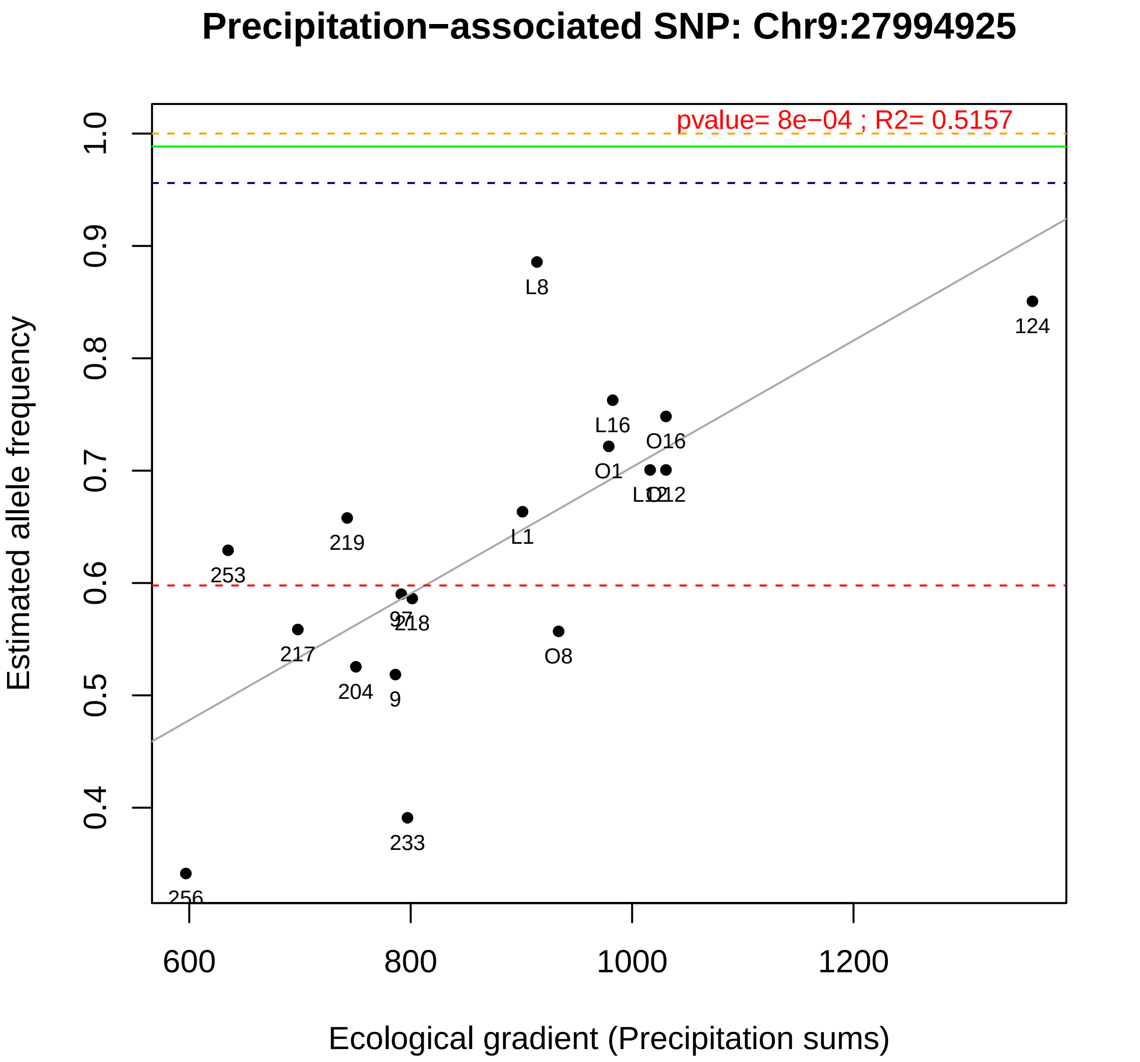

Precipitation−associated SNP: Chr9:27994925
pvalue= 8e−04 ; R2= 0.5157
Estimated allele frequency
Ecological gradient (Precipitation sums)
0.4
0.5
0.6
0.7
0.8
0.9
1.0
600
800
1000
1200
L8
124
L16
O16
O1
L10
O12
L1
219
253
97
218
217
O8
204
9
233
256

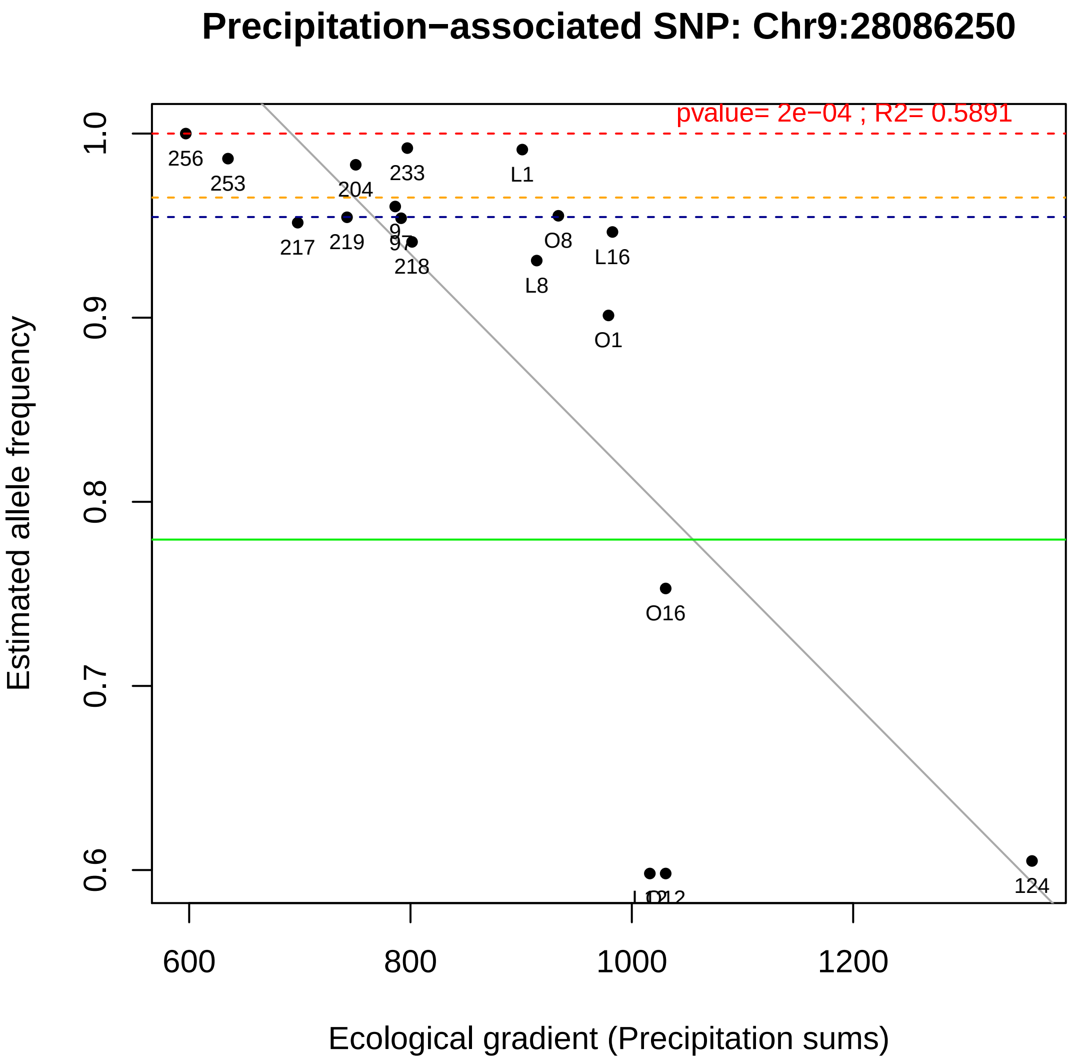

Precipitation−associated SNP: Chr9:28086250
pvalue= 2e−04 ; R2= 0.5891
Estimated allele frequency
Ecological gradient (Precipitation sums)
1.0
0.9
0.8
0.7
0.6
600
800
1000
1200
256
253
204
233
L1
217
219
9
97
218
O8
L16
L8
O1
O16
124

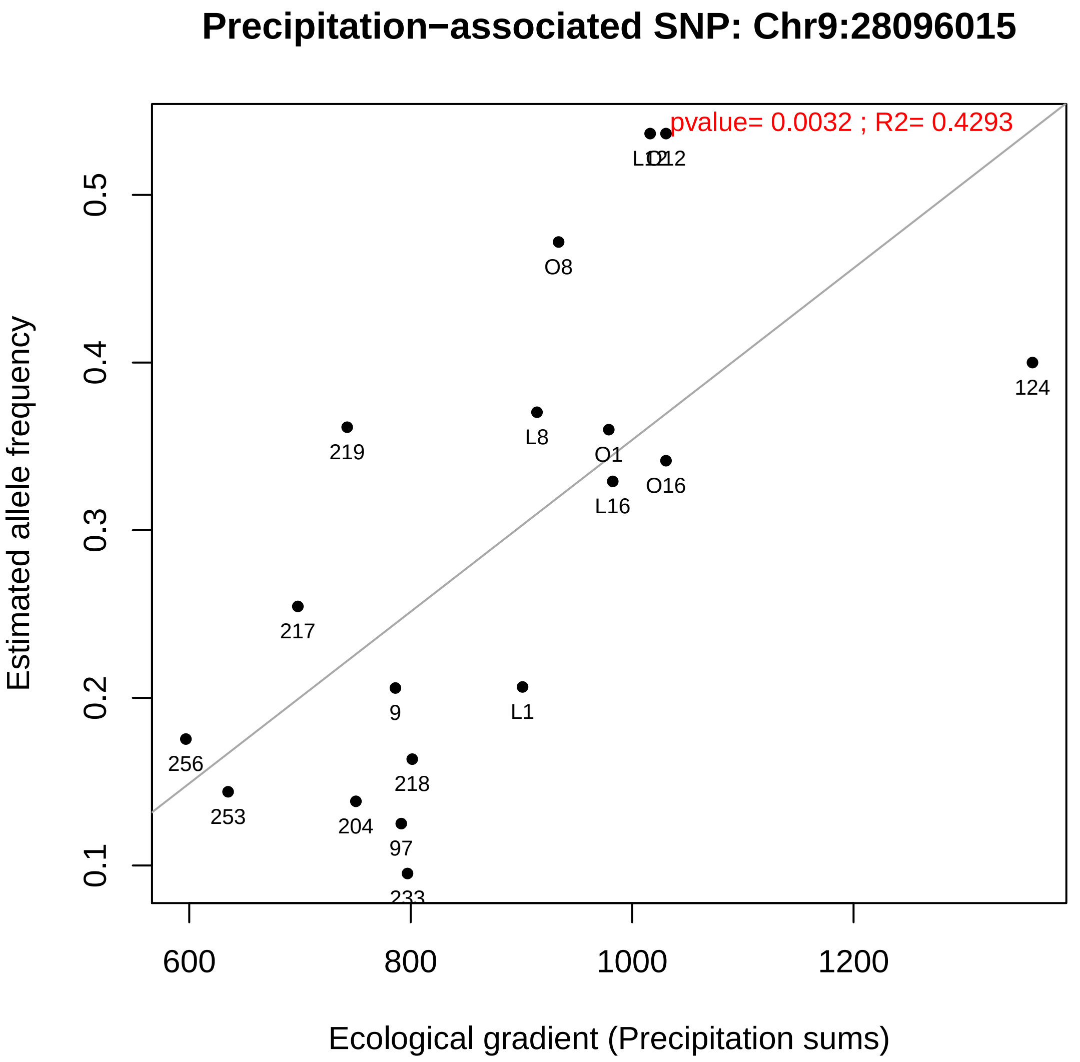

Precipitation−associated SNP: Chr9:28096015
pvalue= 0.0032 ; R2= 0.4293
Estimated allele frequency
0.1
0.2
0.3
0.4
0.5
600
800
1000
1200
Ecological gradient (Precipitation sums)
O8
124
219
L8
O1
O16
L16
217
9
L1
256
218
253
204
97
233

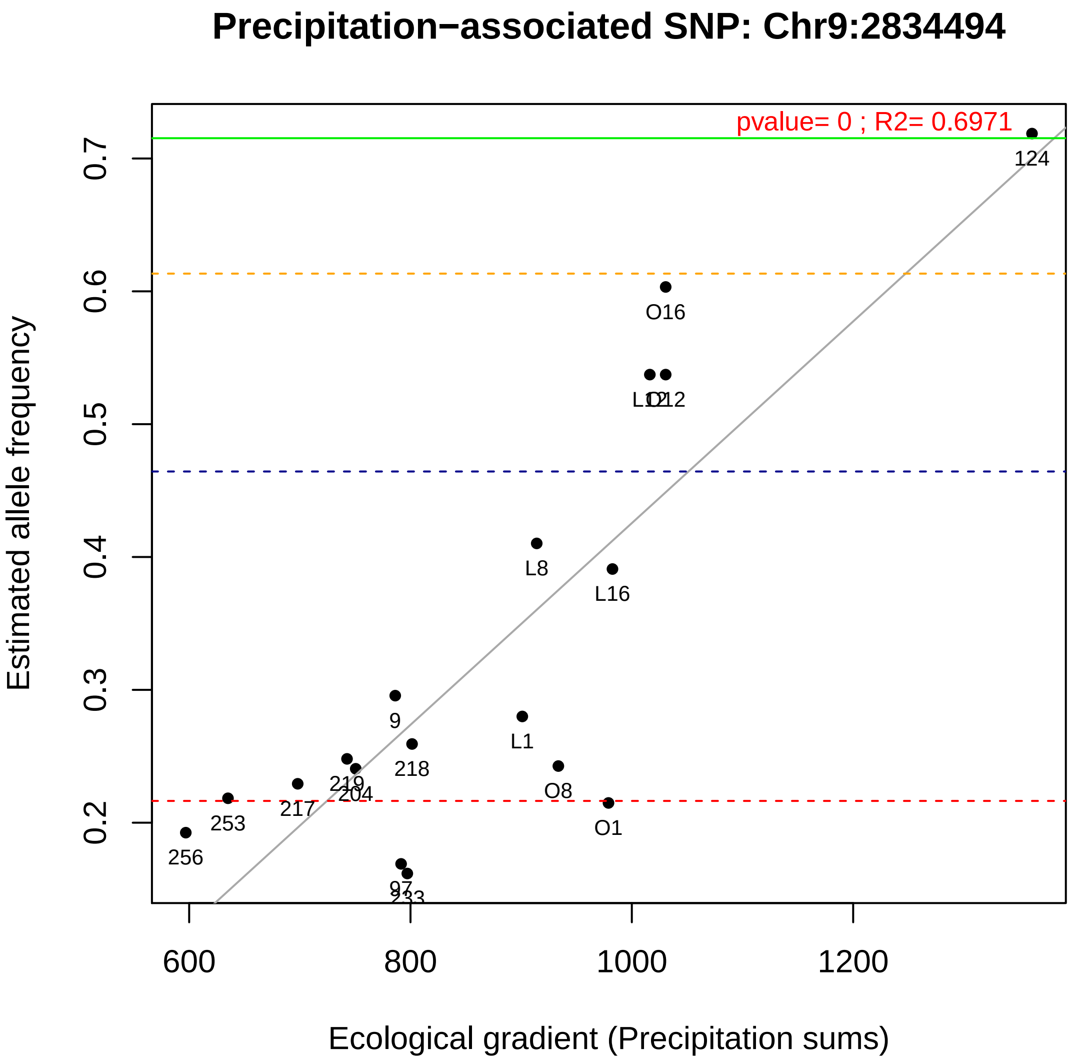

Precipitation−associated SNP: Chr9:2834494
pvalue= 0 ; R2= 0.6971
Estimated allele frequency
Ecological gradient (Precipitation sums)
0.2
0.3
0.4
0.5
0.6
0.7
600
800
1000
1200
124
O16
L12
O12
L8
L16
9
L1
218
219
204
O8
217
253
O1
256
97
233

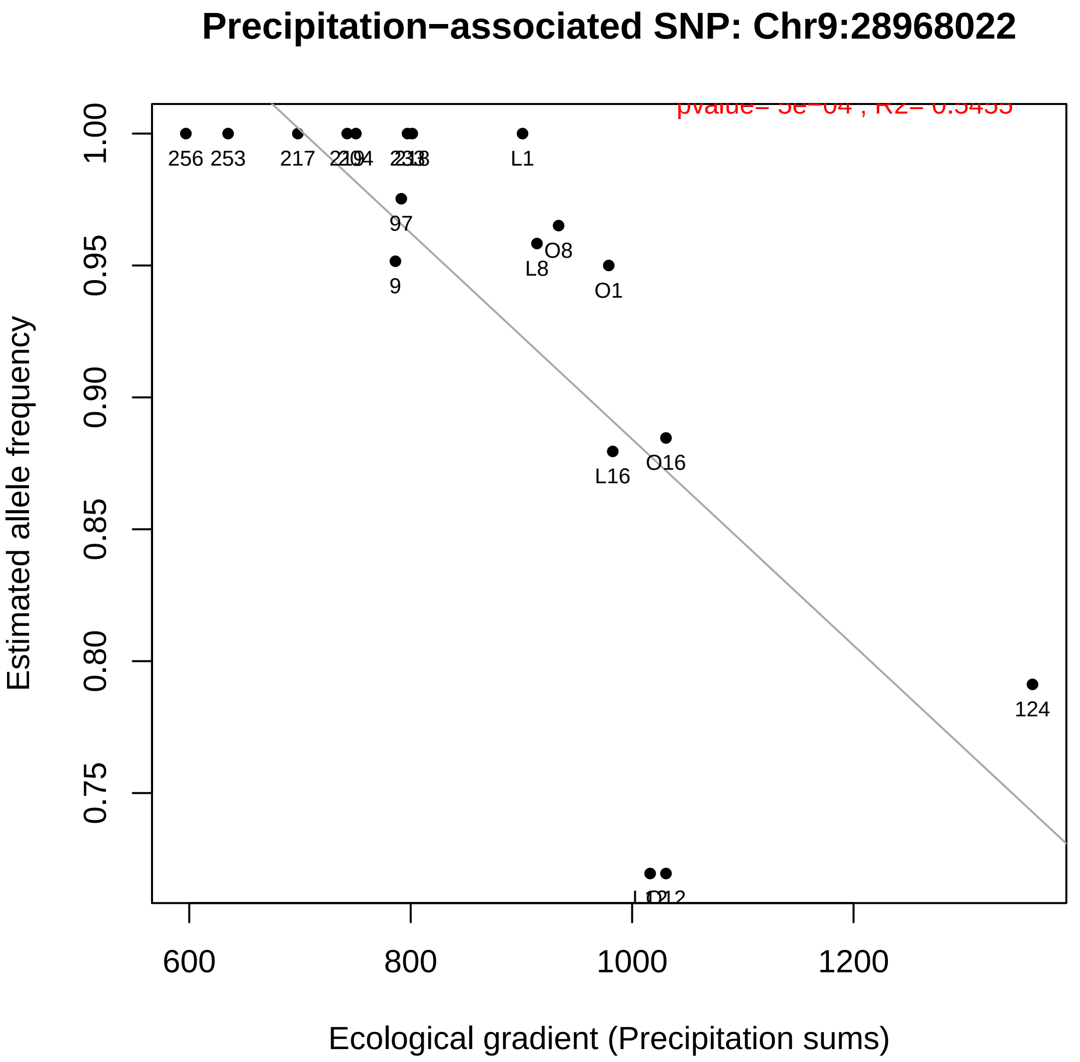
Precipitation−associated SNP: Chr9:28968022
pvalue= 5e−04 , R2= 0.5455
Estimated allele frequency
Ecological gradient (Precipitation sums)
1.00
0.95
0.90
0.85
0.80
0.75
600
800
1000
1200
256
253
217
L1
97
9
O8
L8
O1
L16
O16
124

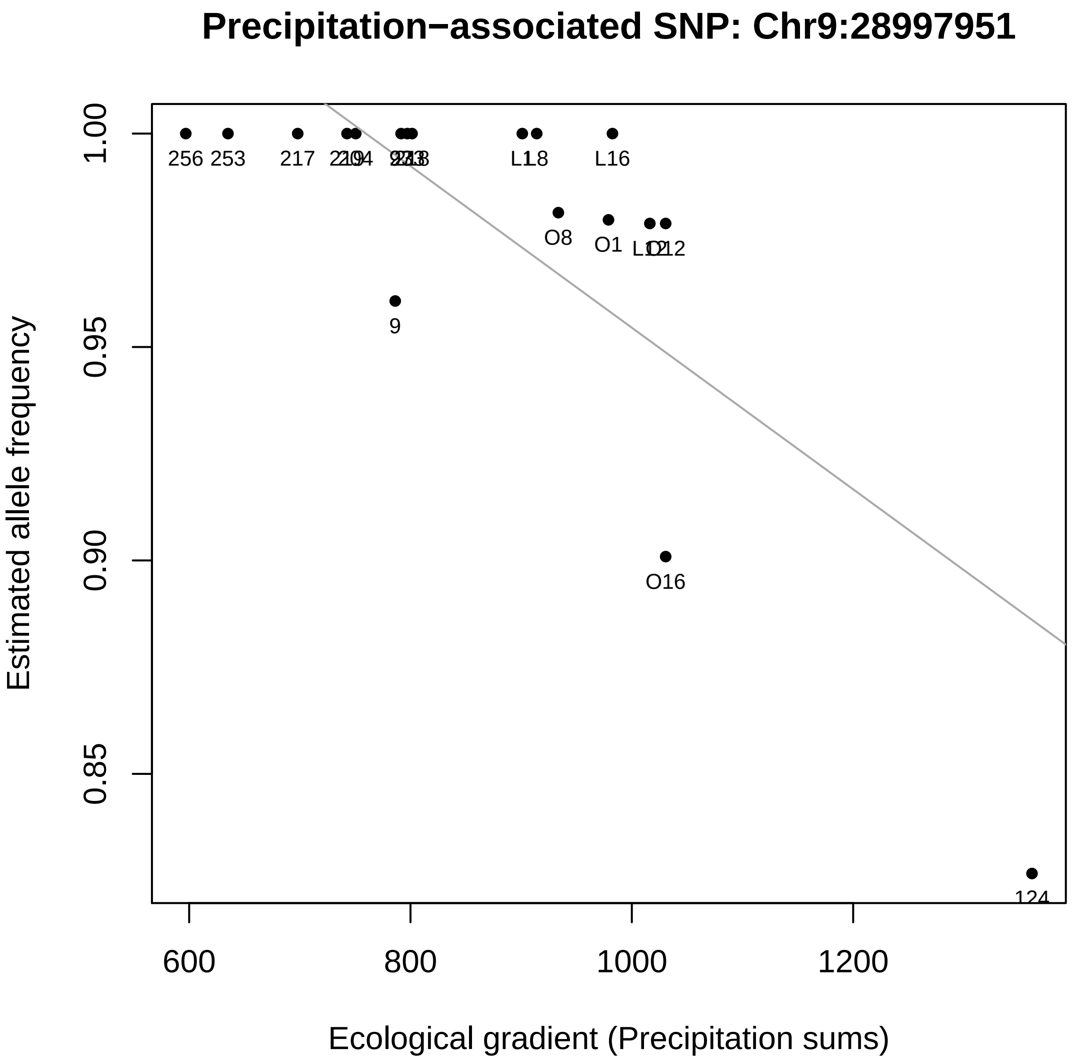
Precipitation−associated SNP: Chr9:28997951
Estimated allele frequency
Ecological gradient (Precipitation sums)
1.00
0.95
0.90
0.85
600
800
1000
1200
256
253
217
L16
O8
O1
9
O16
124

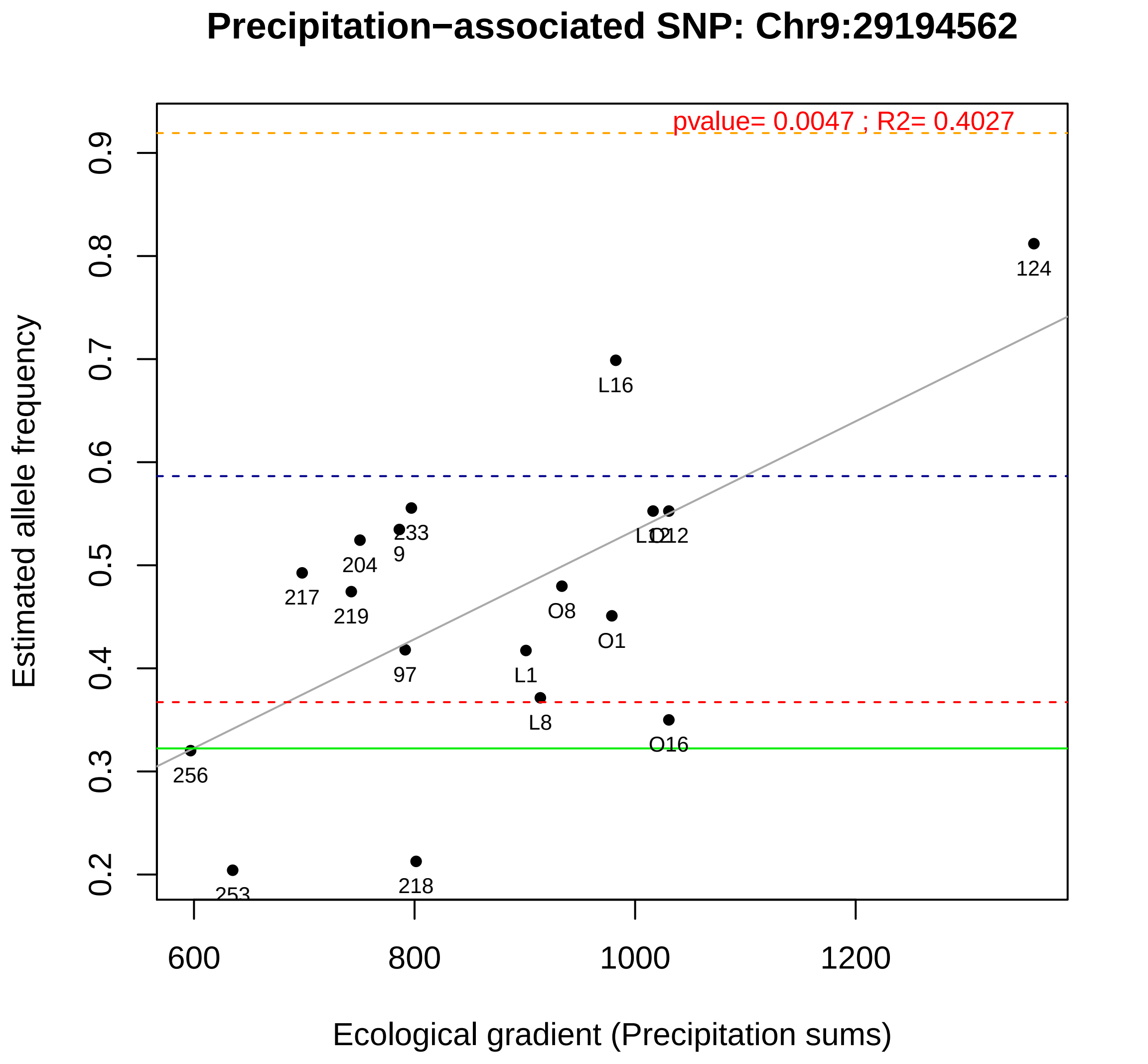

Precipitation−associated SNP: Chr9:29194562
pvalue= 0.0047 ; R2= 0.4027
Estimated allele frequency
0.2
0.3
0.4
0.5
0.6
0.7
0.8
0.9
600
800
1000
1200
Ecological gradient (Precipitation sums)
124
L16
233
9
204
217
219
O8
O1
97
L1
L8
O16
256
253
218
L12
O12

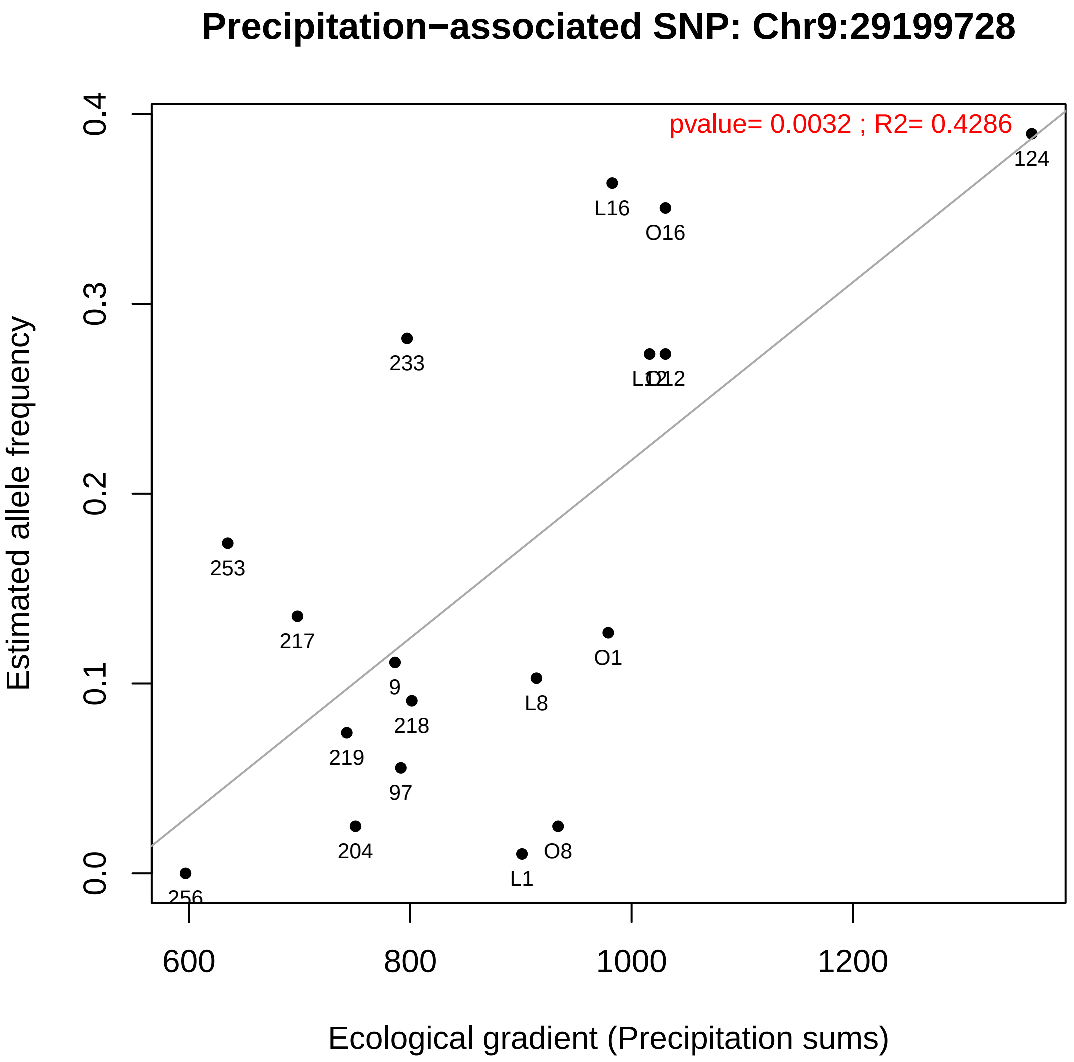

Precipitation−associated SNP: Chr9:29199728
pvalue= 0.0032 ; R2= 0.4286
124
L16
O16
233
L12
O12
253
217
O1
9
L8
218
219
97
204
O8
L1
256
0.0
0.1
0.2
0.3
0.4
600
800
1000
1200
Estimated allele frequency
Ecological gradient (Precipitation sums)

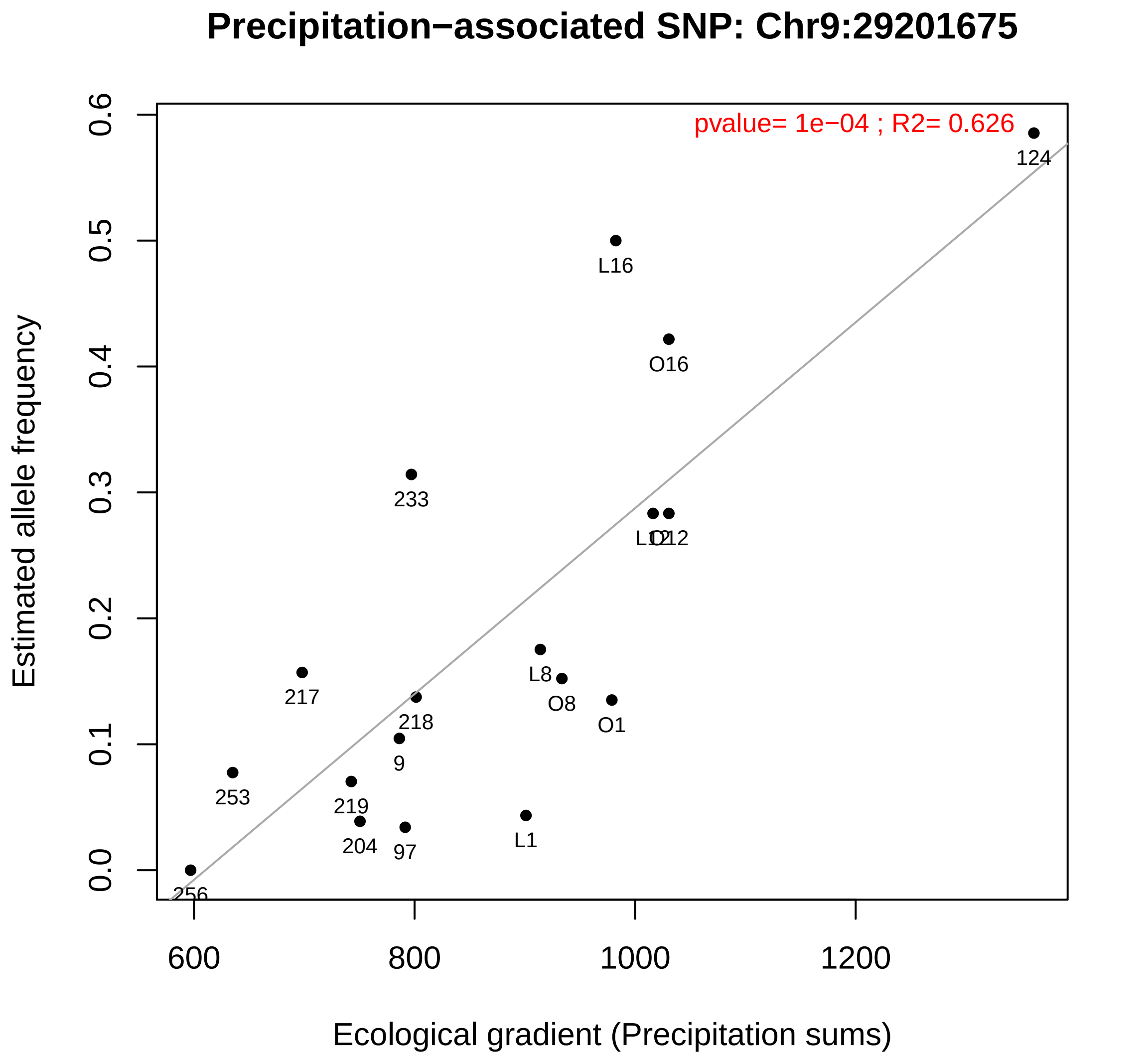
Precipitation−associated SNP: Chr9:29201675
pvalue= 1e−04 ; R2= 0.626
Estimated allele frequency
Ecological gradient (Precipitation sums)
0.0
0.1
0.2
0.3
0.4
0.5
0.6
600
800
1000
1200
124
L16
O16
233
L12
O12
217
218
L8
O8
O1
9
253
219
204
97
L1
256

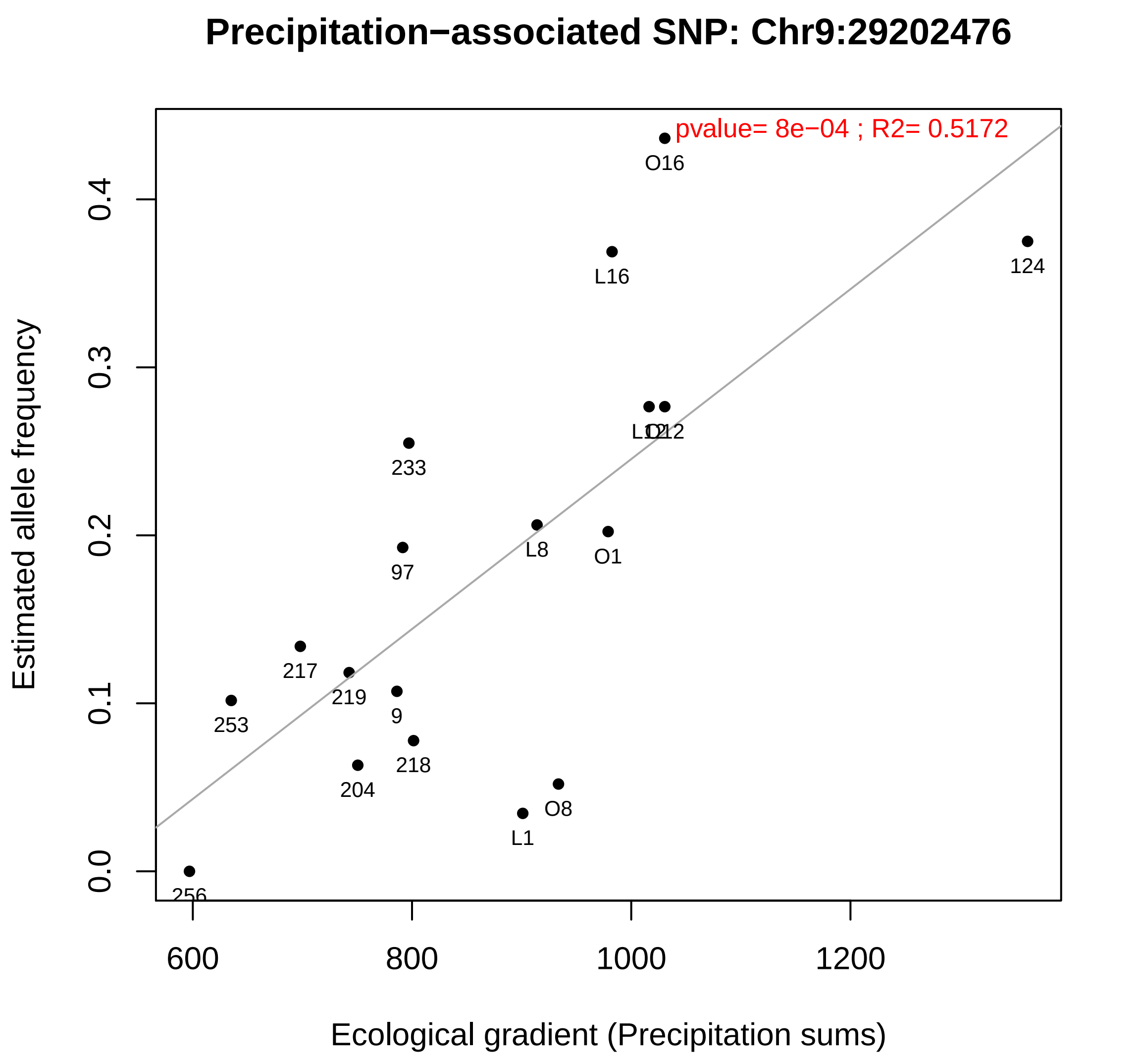
Precipitation−associated SNP: Chr9:29202476
pvalue= 8e−04 ; R2= 0.5172
Estimated allele frequency
Ecological gradient (Precipitation sums)
0.0
0.1
0.2
0.3
0.4
600
800
1000
1200
O16
L16
124
233
97
L8
O1
217
219
9
253
218
204
L1
O8
256

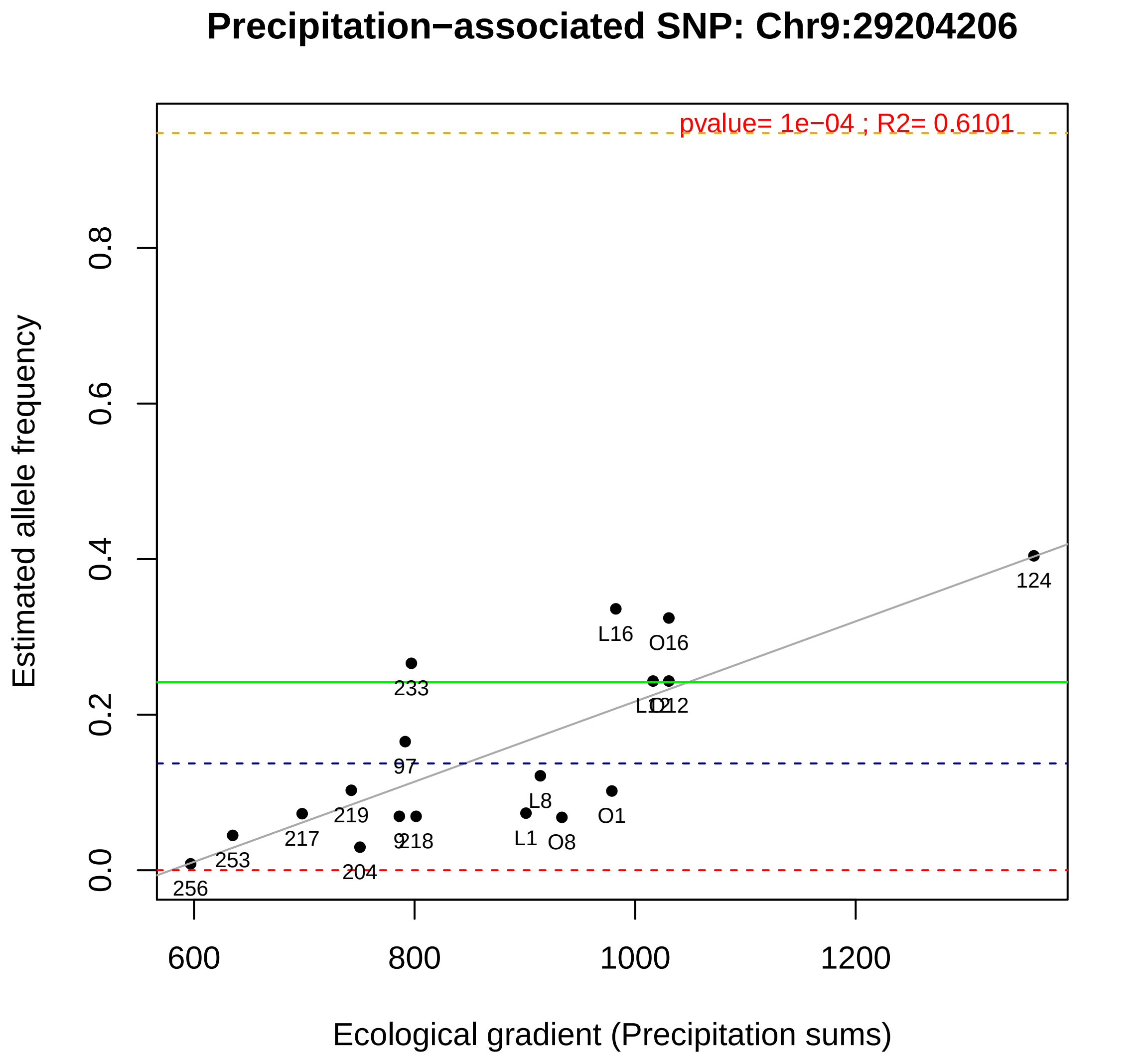

Precipitation−associated SNP: Chr9:29204206
pvalue= 1e−04 ; R2= 0.6101
Estimated allele frequency
0.0
0.2
0.4
0.6
0.8
600
800
1000
1200
Ecological gradient (Precipitation sums)
256
253
204
217
219
97
233
L1
L8
O8
O1
L16
O16
124

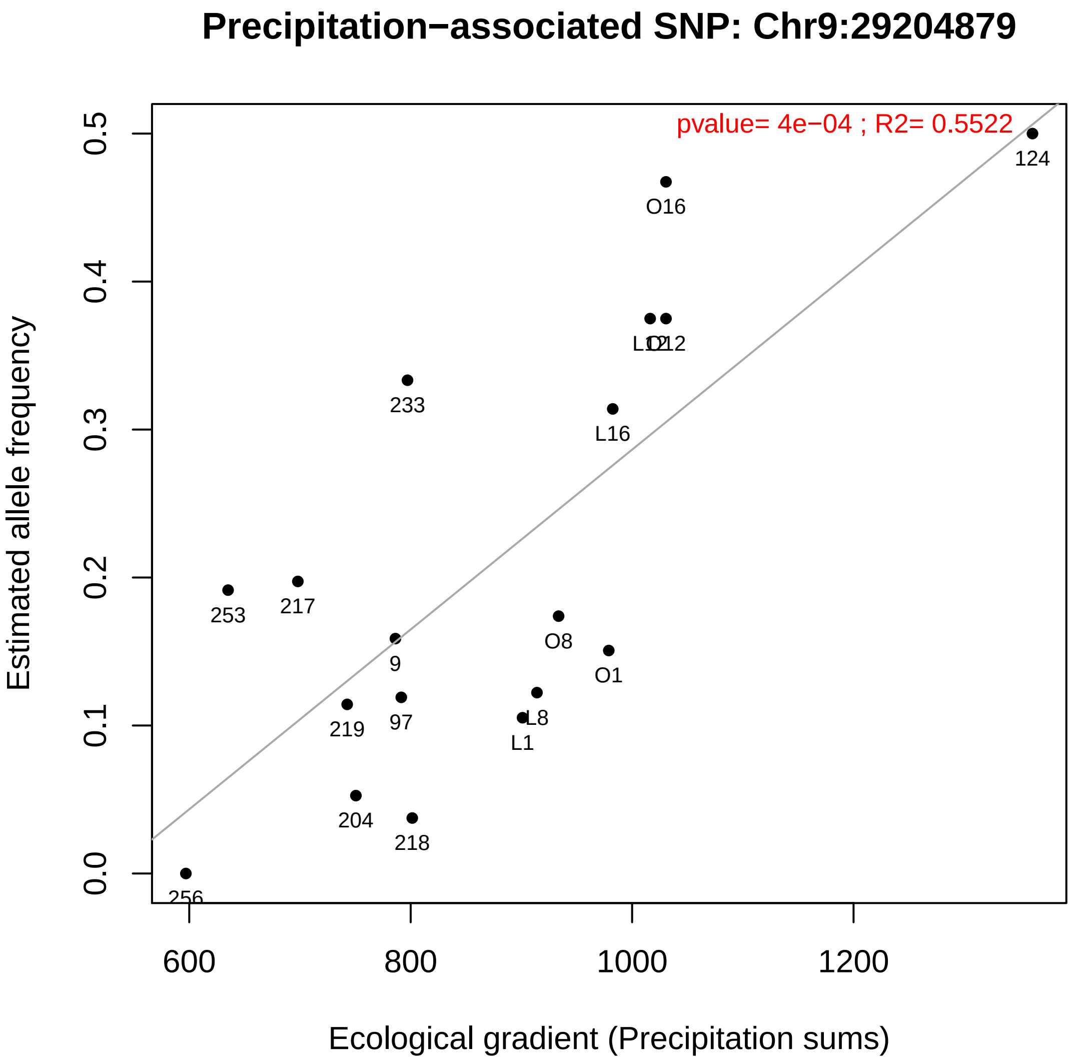
Precipitation−associated SNP: Chr9:29204879
pvalue= 4e−04 ; R2= 0.5522
Estimated allele frequency
0.0
0.1
0.2
0.3
0.4
0.5
600
800
1000
1200
Ecological gradient (Precipitation sums)
124
O16
L12
O12
233
L16
253
217
O8
9
O1
219
97
L8
L1
204
218
256

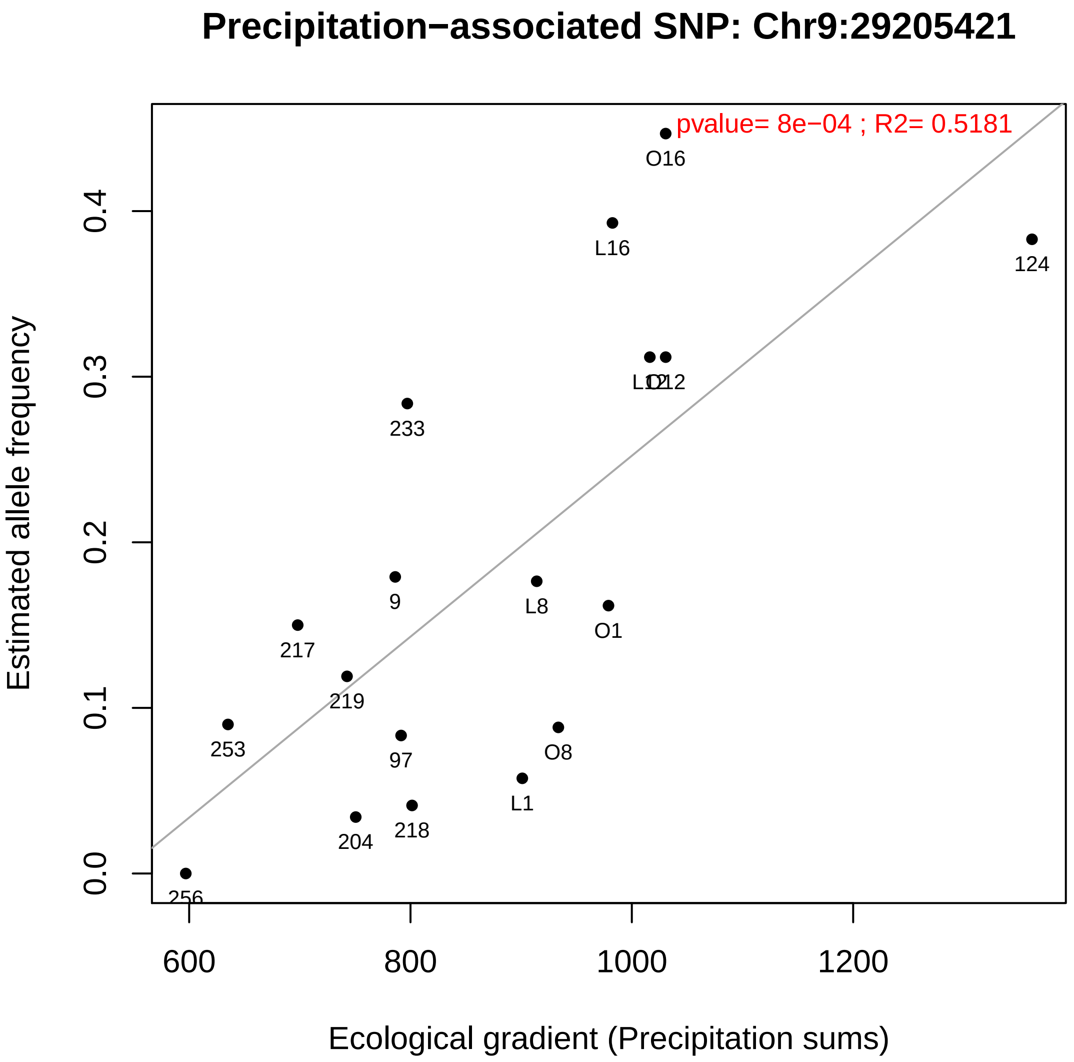

Precipitation−associated SNP: Chr9:29205421
pvalue= 8e−04 ; R2= 0.5181
O16
L16
124
L12
O12
233
9
L8
O1
217
219
253
97
O8
L1
204
218
256
0.0
0.1
0.2
0.3
0.4
Estimated allele frequency
600
800
1000
1200
Ecological gradient (Precipitation sums)

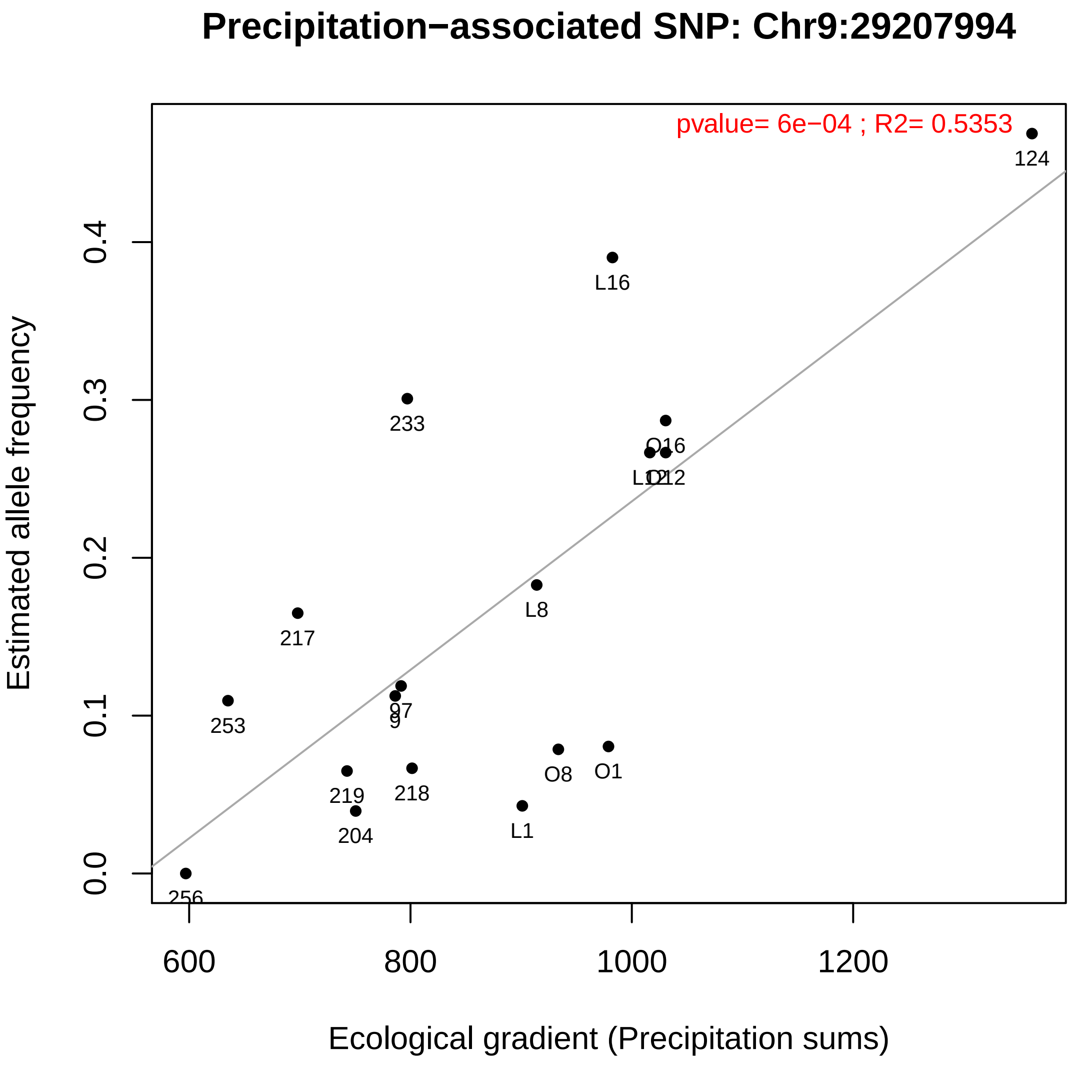

Precipitation−associated SNP: Chr9:29207994
pvalue= 6e−04 ; R2= 0.5353
Estimated allele frequency
Ecological gradient (Precipitation sums)
0.0
0.1
0.2
0.3
0.4
600
800
1000
1200
124
L16
233
O16
L10
O12
L8
217
253
97
9
O8
O1
219
218
204
L1
256

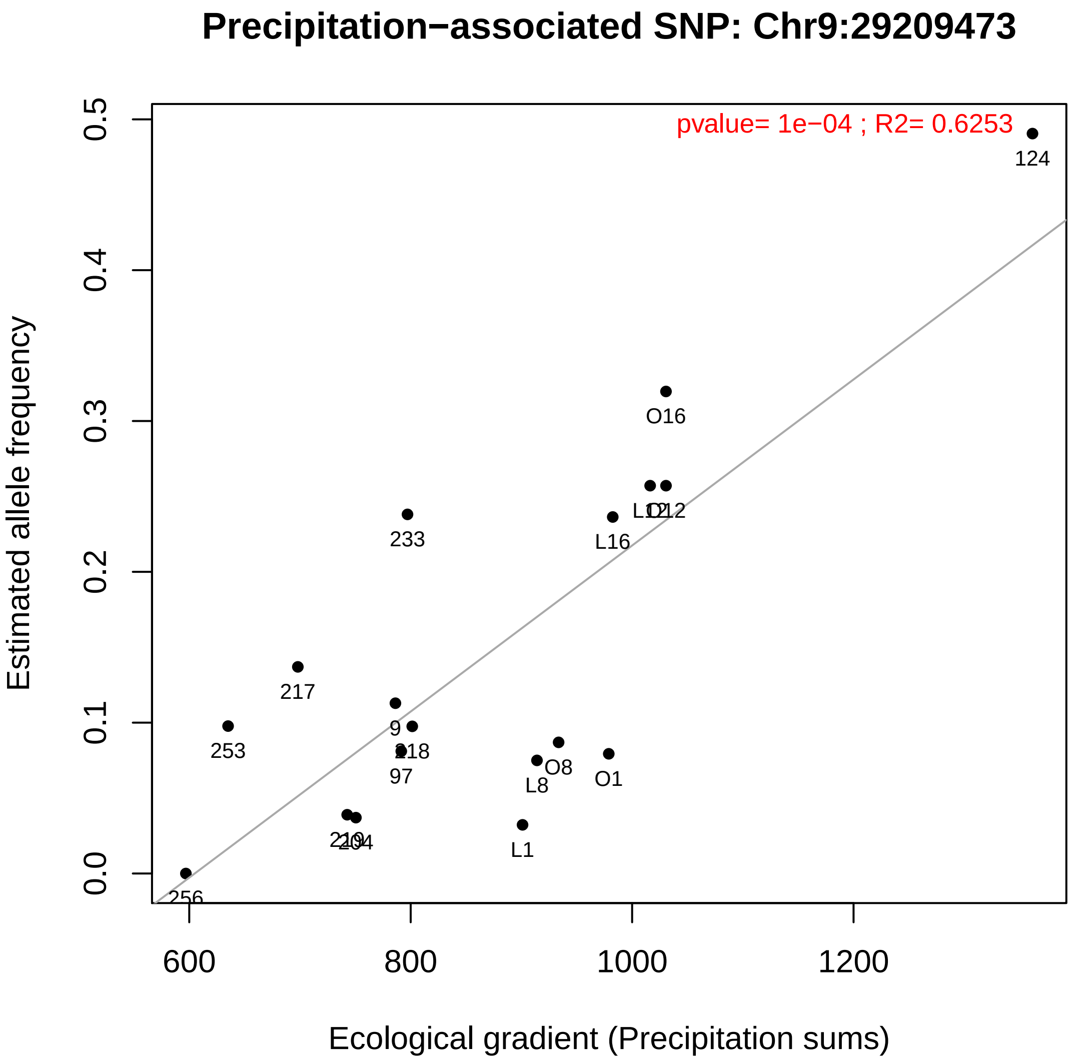
Precipitation−associated SNP: Chr9:29209473
pvalue= 1e−04 ; R2= 0.6253
Estimated allele frequency
0.0
0.1
0.2
0.3
0.4
0.5
600
800
1000
1200
Ecological gradient (Precipitation sums)
124
O16
L12
O12
L16
233
217
253
9
218
97
O8
L8
O1
210
204
L1
256

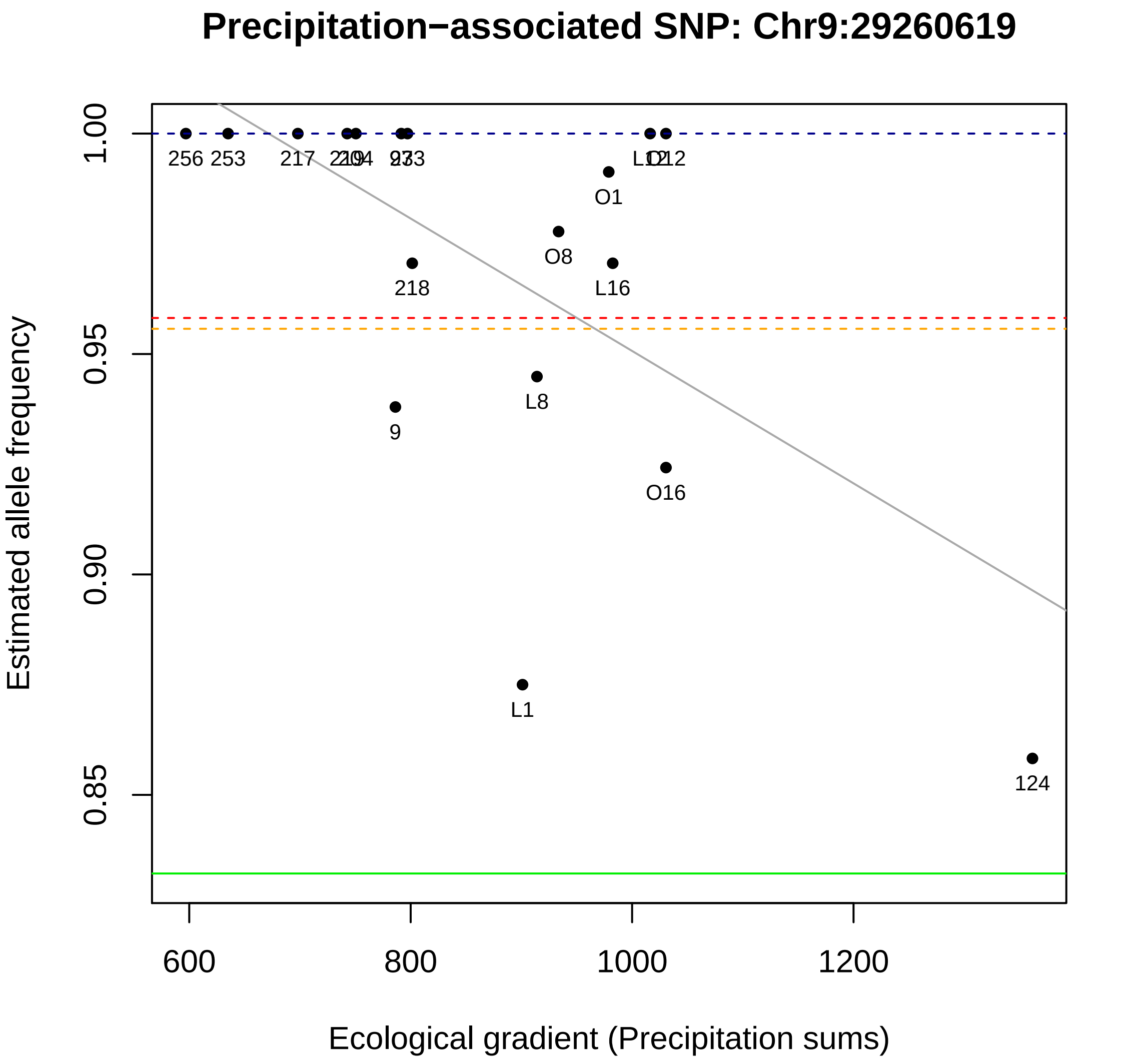
Precipitation−associated SNP: Chr9:29260619
Estimated allele frequency
Ecological gradient (Precipitation sums)
1.00
0.95
0.90
0.85
600
800
1000
1200
256
253
217
O1
O8
218
L16
L8
9
O16
L1
124

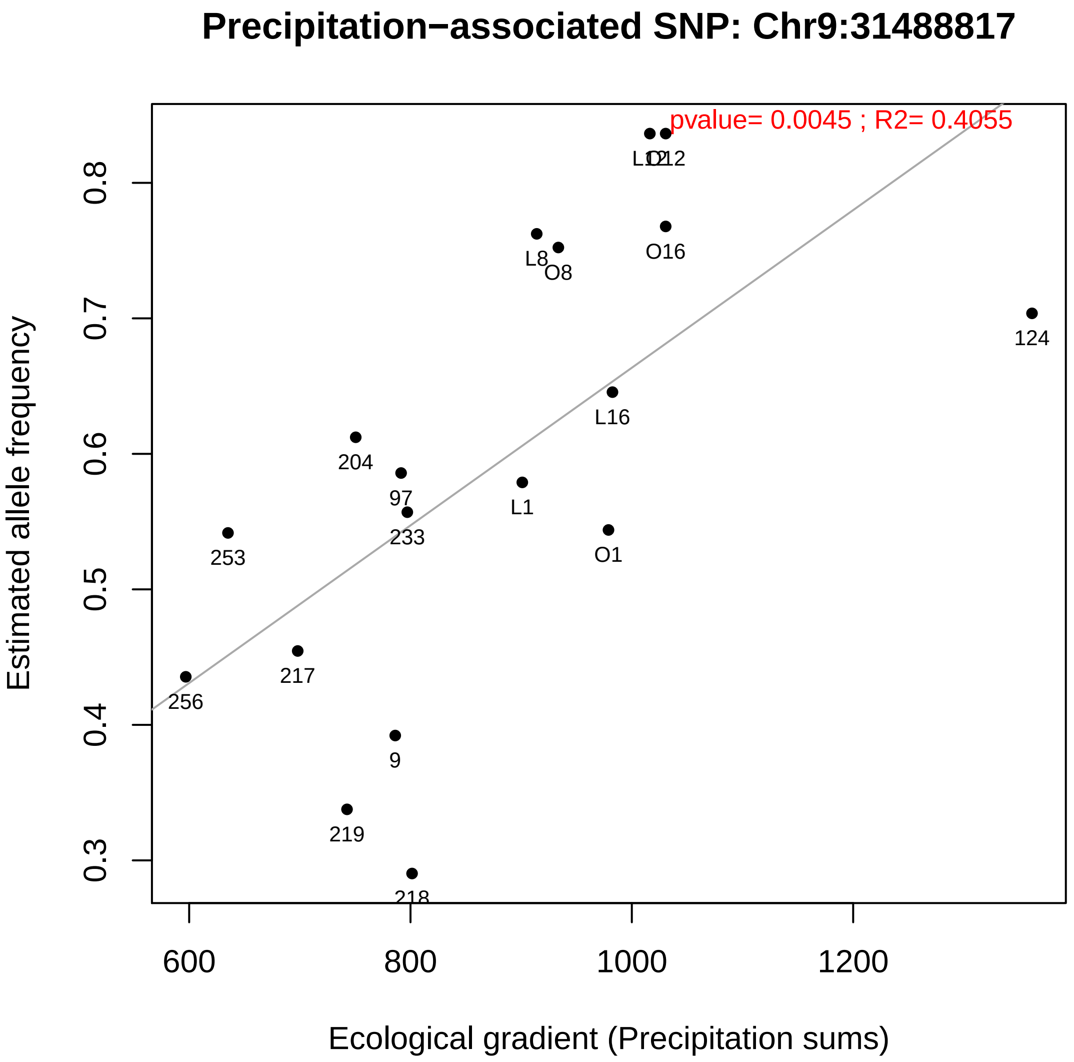
Precipitation−associated SNP: Chr9:31488817
pvalue= 0.0045 ; R2= 0.4055
Estimated allele frequency
0.3
0.4
0.5
0.6
0.7
0.8
600
800
1000
1200
Ecological gradient (Precipitation sums)
O12
O16
L8
O8
124
L16
204
97
233
L1
O1
253
217
256
9
219
218

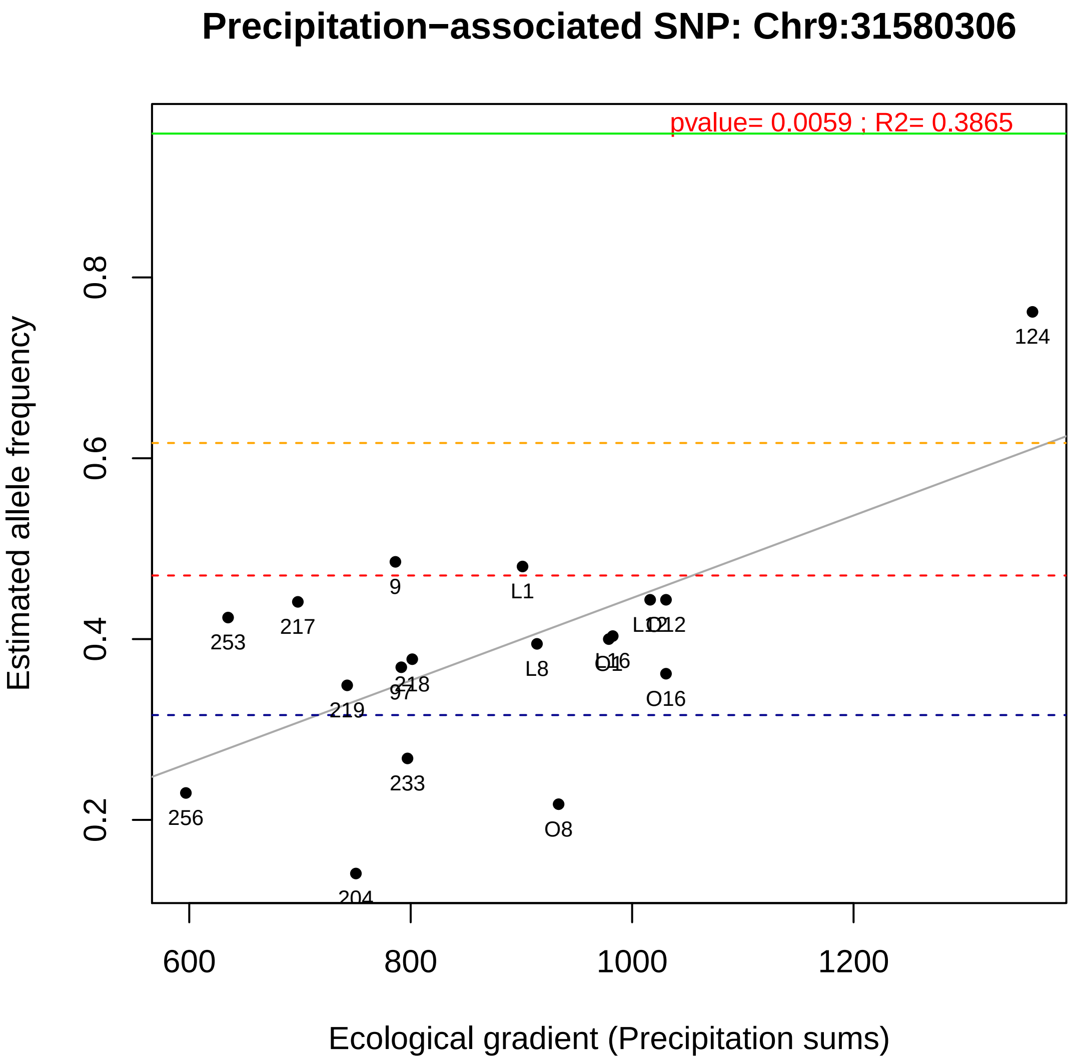

Precipitation−associated SNP: Chr9:31580306
pvalue= 0.0059 ; R2= 0.3865
Estimated allele frequency
0.2
0.4
0.6
0.8
600
800
1000
1200
Ecological gradient (Precipitation sums)
124
9
L1
217
253
L12
O12
L16
O1
L8
218
97
219
O16
233
256
O8
204

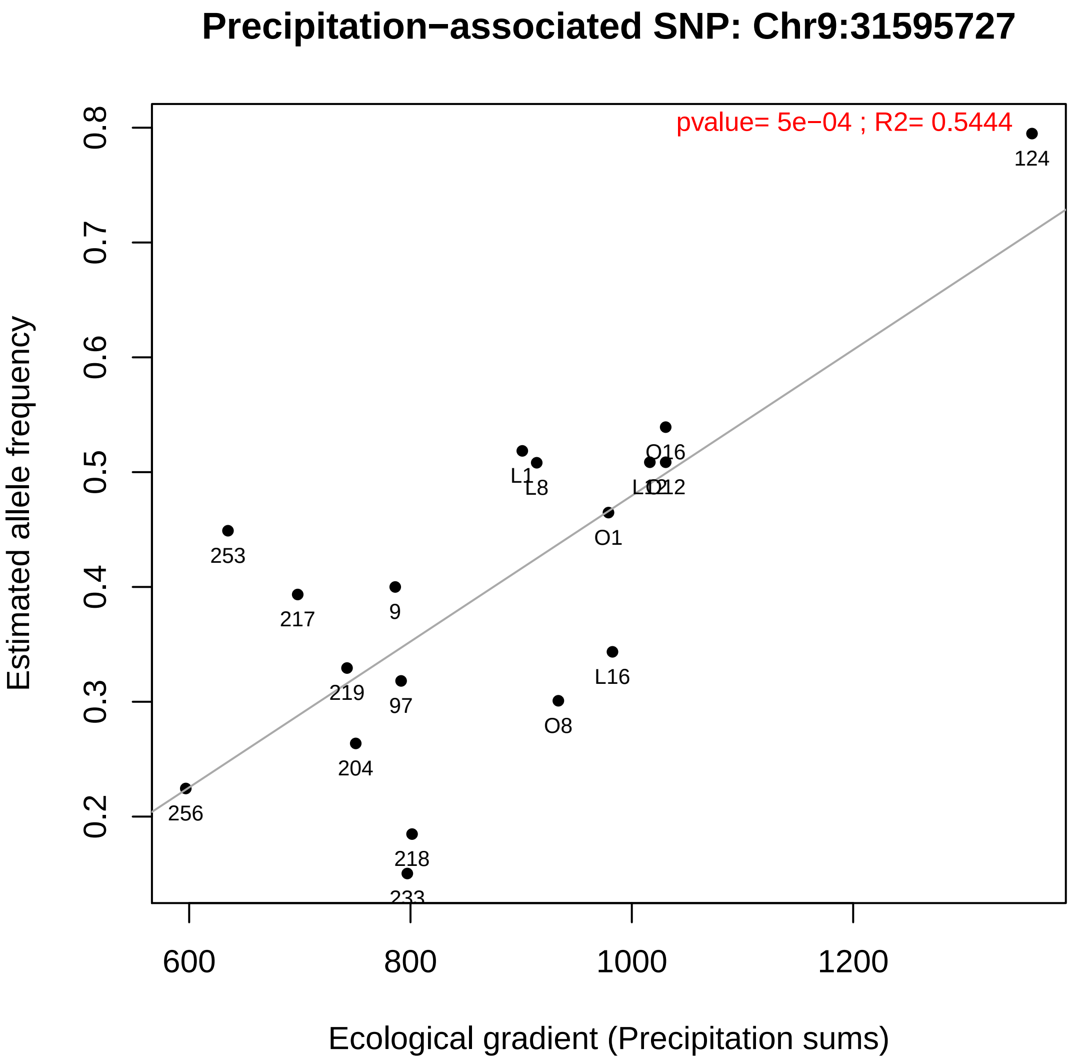

Precipitation−associated SNP: Chr9:31595727
pvalue= 5e−04 ; R2= 0.5444
Estimated allele frequency
0.2
0.3
0.4
0.5
0.6
0.7
0.8
600
800
1000
1200
Ecological gradient (Precipitation sums)
124
O16
L1
L8
L12
O12
O1
253
217
9
L16
219
97
O8
204
256
218
233

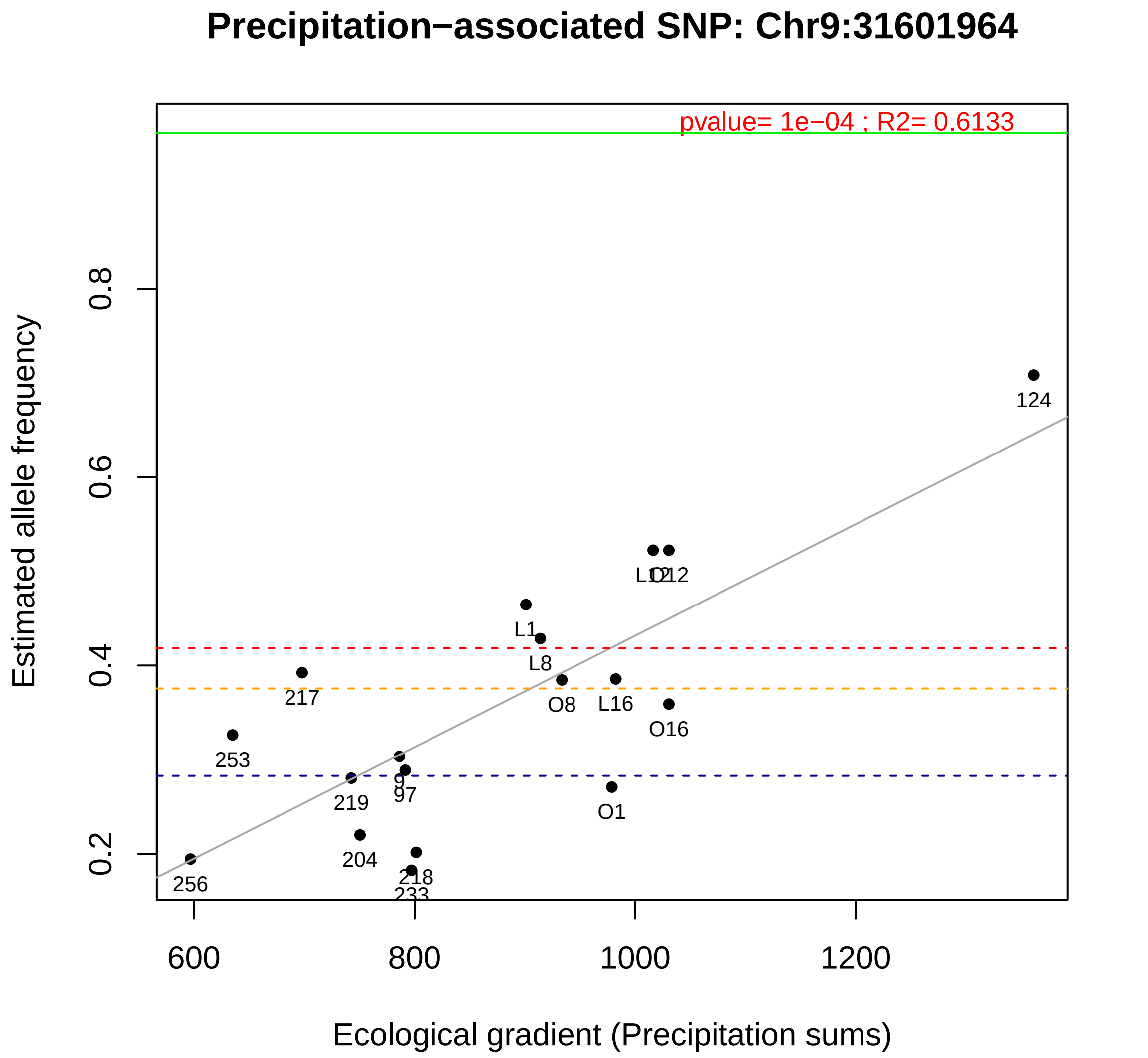
Precipitation−associated SNP: Chr9:31601964
pvalue= 1e−04 ; R2= 0.6133
Estimated allele frequency
0.2
0.4
0.6
0.8
600
800
1000
1200
Ecological gradient (Precipitation sums)
124
L12
O12
L1
L8
O8
L16
O16
O1
217
253
219
9
97
204
218
233
256

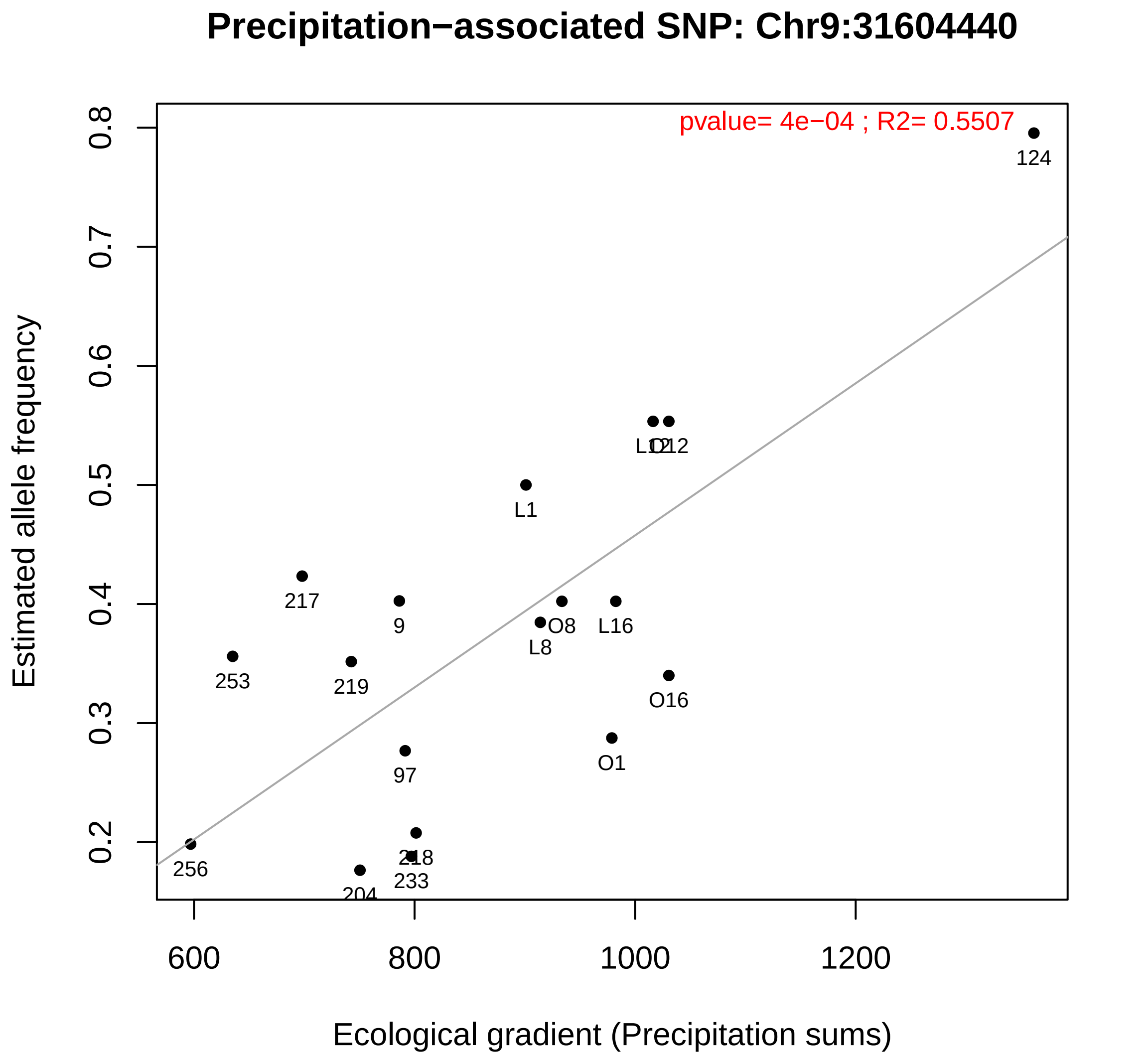

Precipitation−associated SNP: Chr9:31604440
pvalue= 4e−04 ; R2= 0.5507
124
L12
O12
L1
217
9
O8
L16
L8
253
219
O16
97
O1
256
218
233
204
0.2
0.3
0.4
0.5
0.6
0.7
0.8
Estimated allele frequency
600
800
1000
1200
Ecological gradient (Precipitation sums)

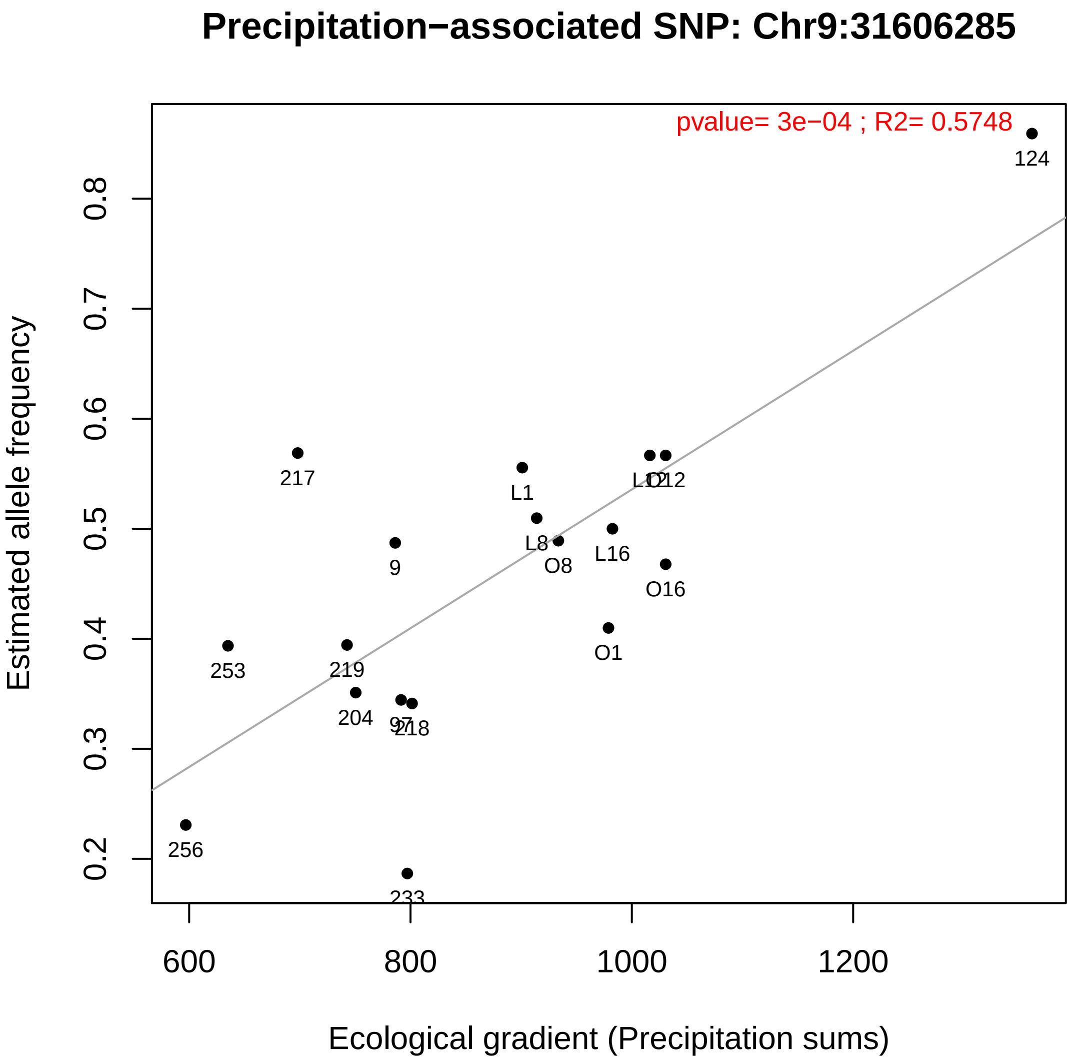
Precipitation−associated SNP: Chr9:31606285
pvalue= 3e−04 ; R2= 0.5748
Estimated allele frequency
0.2
0.3
0.4
0.5
0.6
0.7
0.8
600
800
1000
1200
Ecological gradient (Precipitation sums)
124
217
L1
O12
L8
O8
L16
9
O16
O1
253
219
204
218
256
233

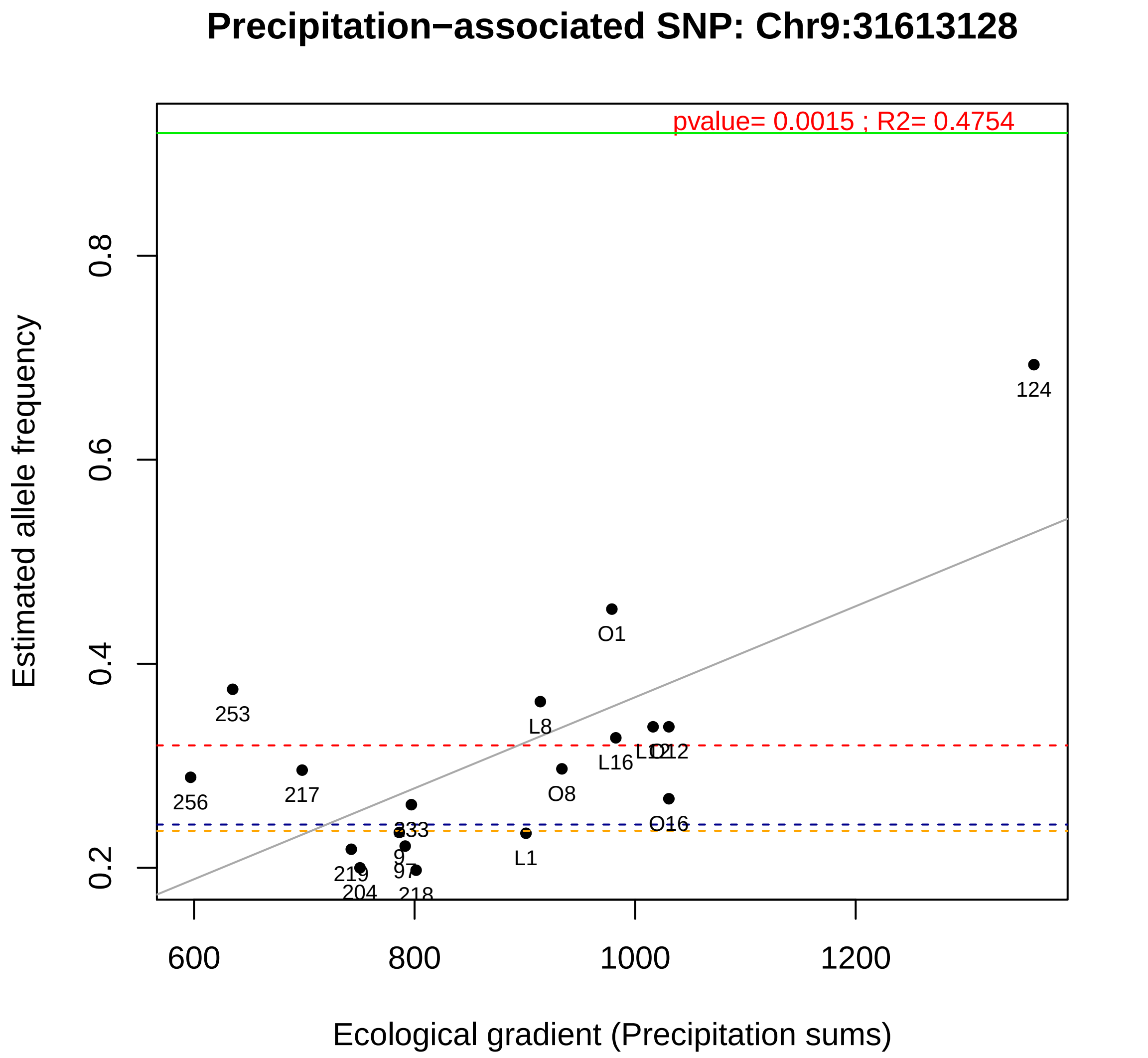

Precipitation−associated SNP: Chr9:31613128
pvalue= 0.0015 ; R2= 0.4754
Estimated allele frequency
0.2
0.4
0.6
0.8
600
800
1000
1200
Ecological gradient (Precipitation sums)
124
O1
253
L8
L16
O12
O8
256
217
O16
233
L1
219
204
218

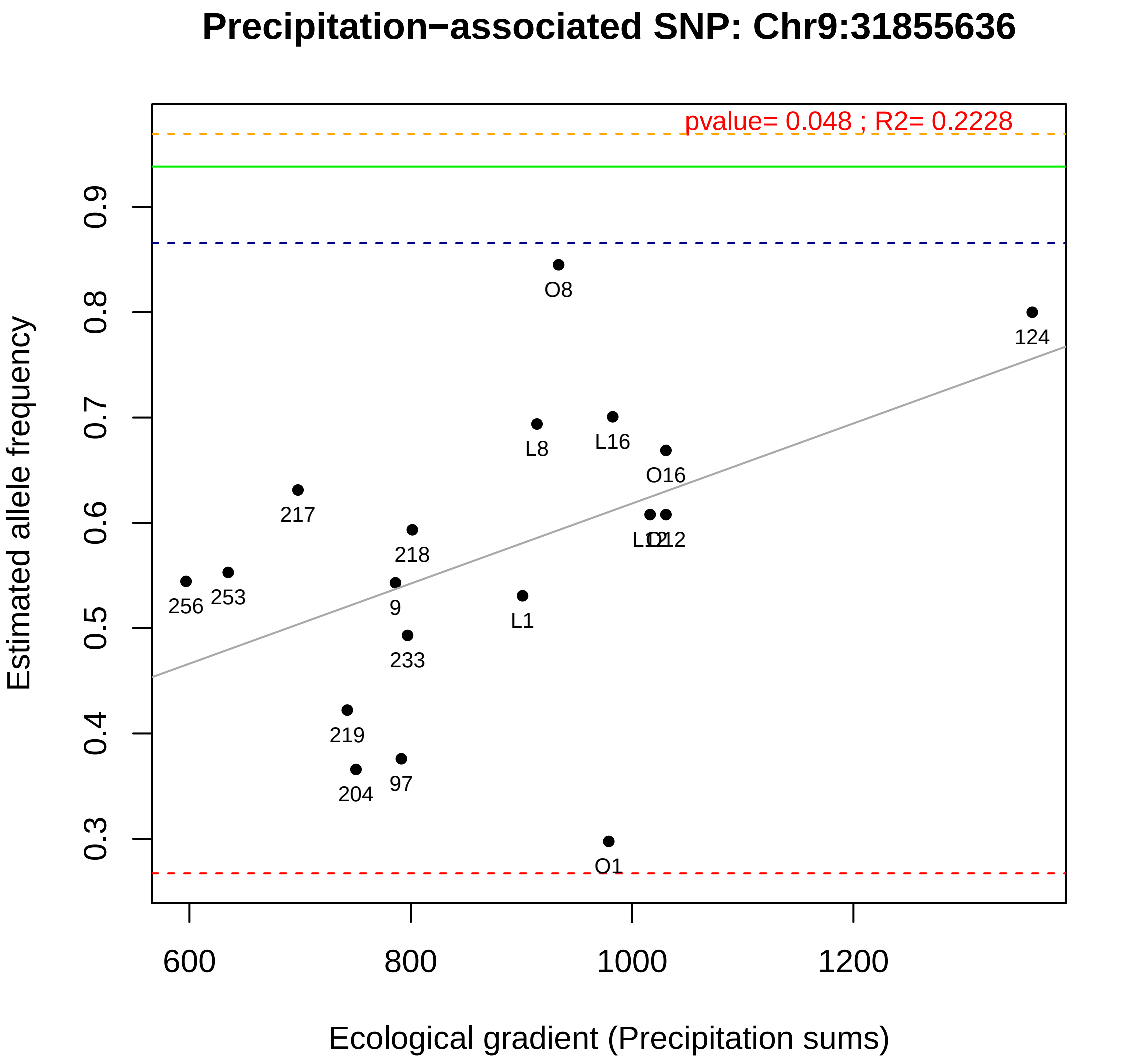
Precipitation−associated SNP: Chr9:31855636
pvalue= 0.048 ; R2= 0.2228
Estimated allele frequency
0.3
0.4
0.5
0.6
0.7
0.8
0.9
600
800
1000
1200
Ecological gradient (Precipitation sums)
O8
124
L8
L16
O16
217
218
256
253
9
L1
233
219
204
97
O1

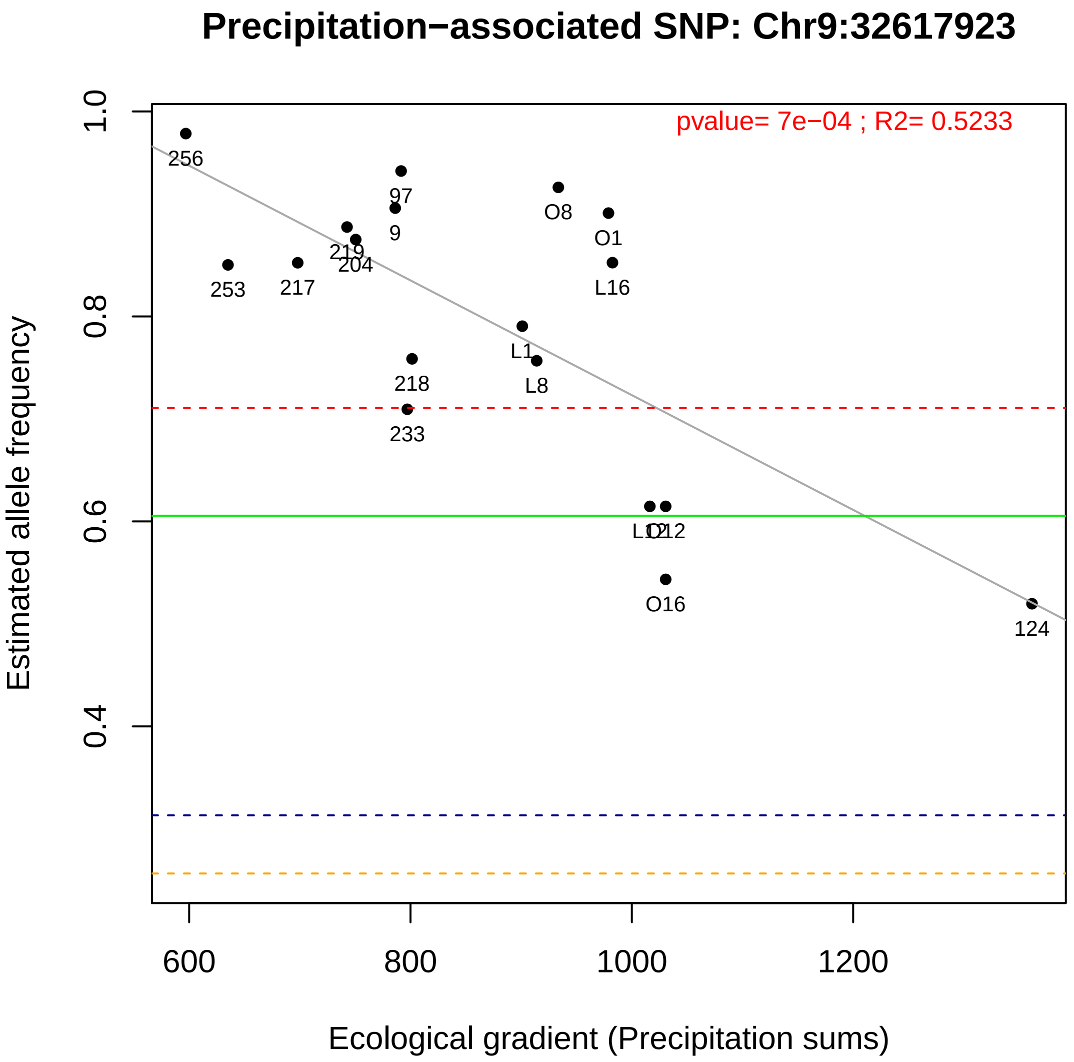

Precipitation−associated SNP: Chr9:32617923
pvalue= 7e−04 ; R2= 0.5233
Estimated allele frequency
Ecological gradient (Precipitation sums)
1.0
0.8
0.6
0.4
600
800
1000
1200
256
97
9
O8
O1
219
204
253
217
L16
L1
L8
218
233
L12
O12
O16
124

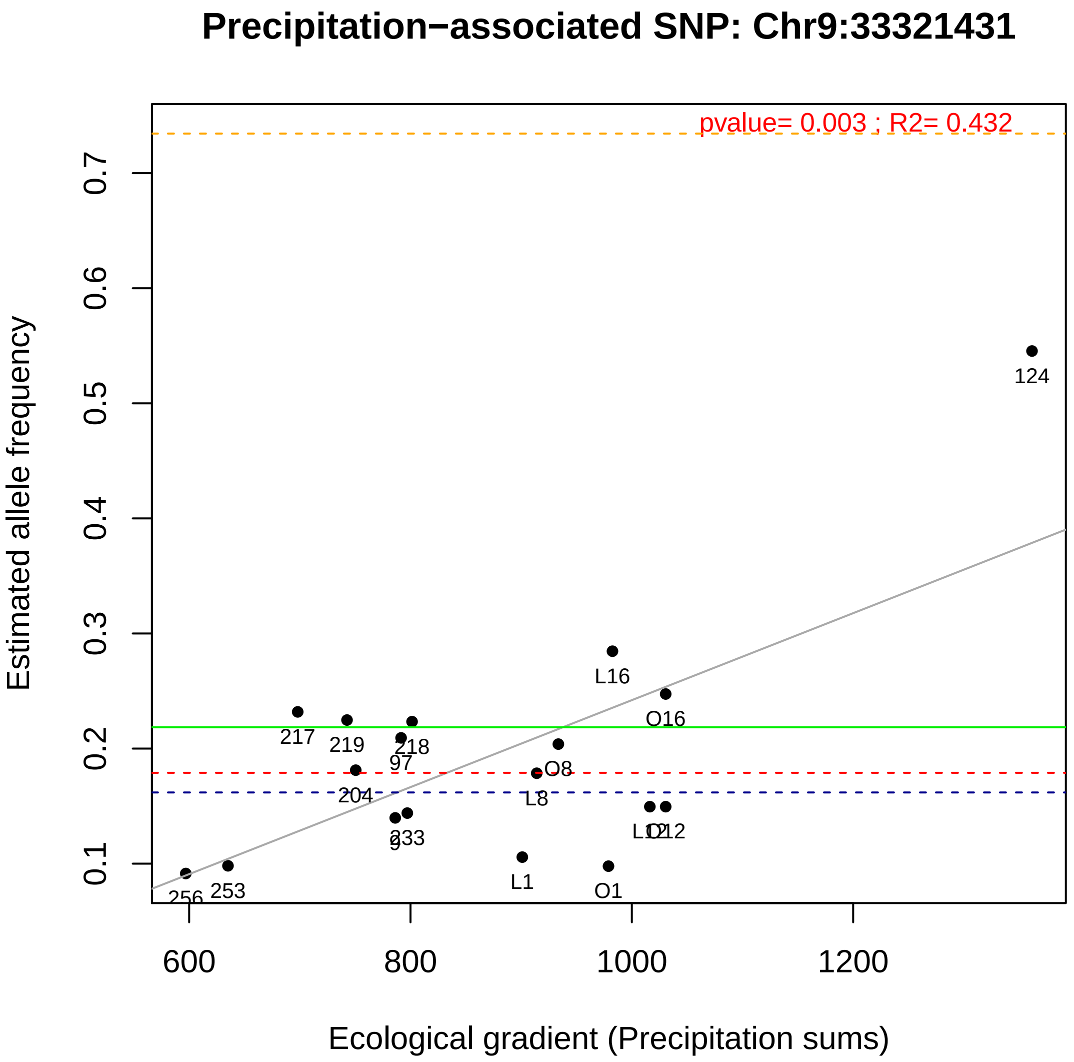

Precipitation−associated SNP: Chr9:33321431
pvalue= 0.003 ; R2= 0.432
Estimated allele frequency
0.1
0.2
0.3
0.4
0.5
0.6
0.7
600
800
1000
1200
Ecological gradient (Precipitation sums)
124
L16
O16
217
219
218
97
204
O8
L8
233
L1
O1
O12
253
256

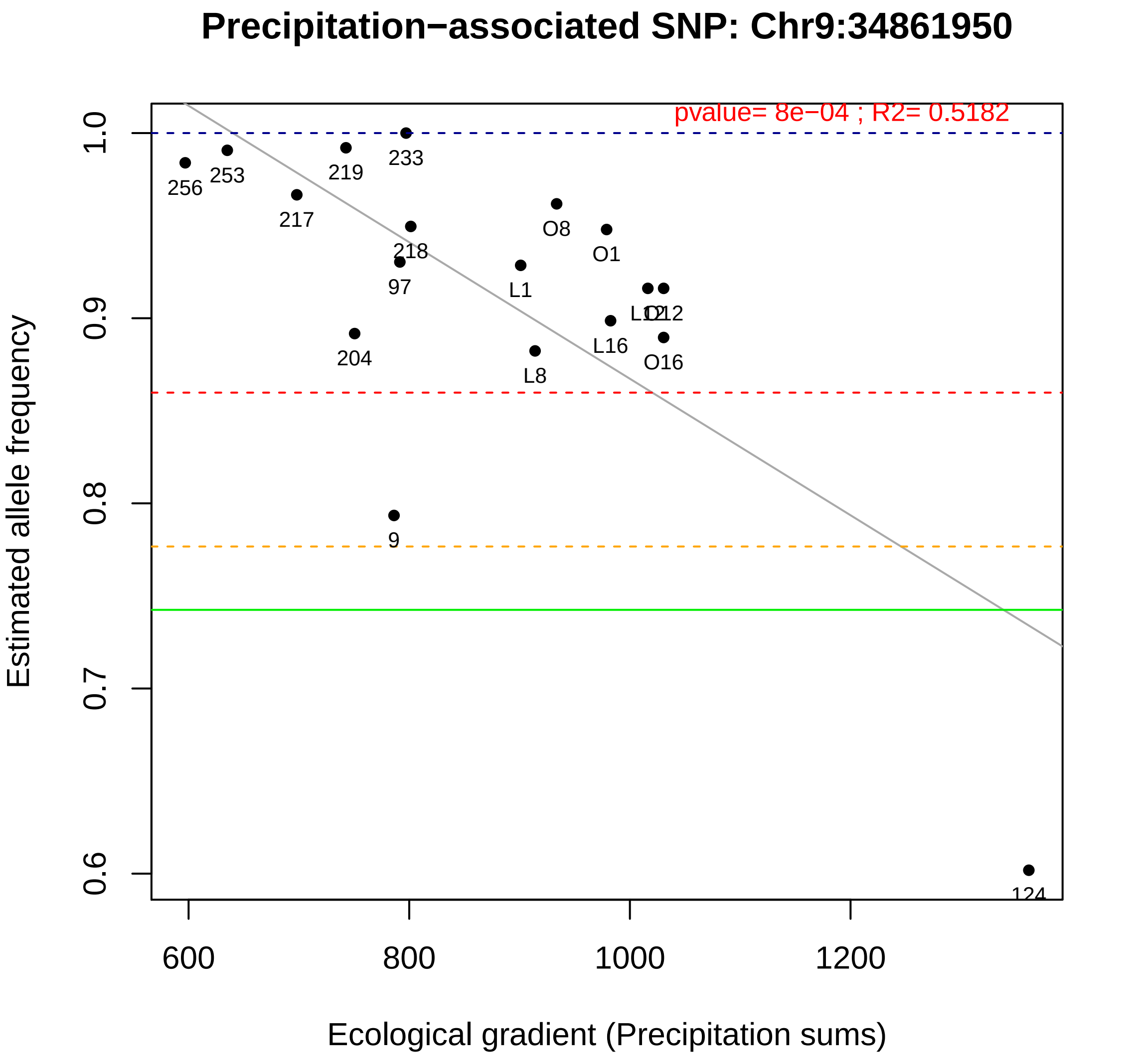
Precipitation−associated SNP: Chr9:34861950
pvalue= 8e−04 ; R2= 0.5182
Estimated allele frequency
0.6
0.7
0.8
0.9
1.0
600
800
1000
1200
Ecological gradient (Precipitation sums)
256
253
219
233
217
218
97
O8
O1
L1
L12
O12
L16
O16
204
L8
9
124

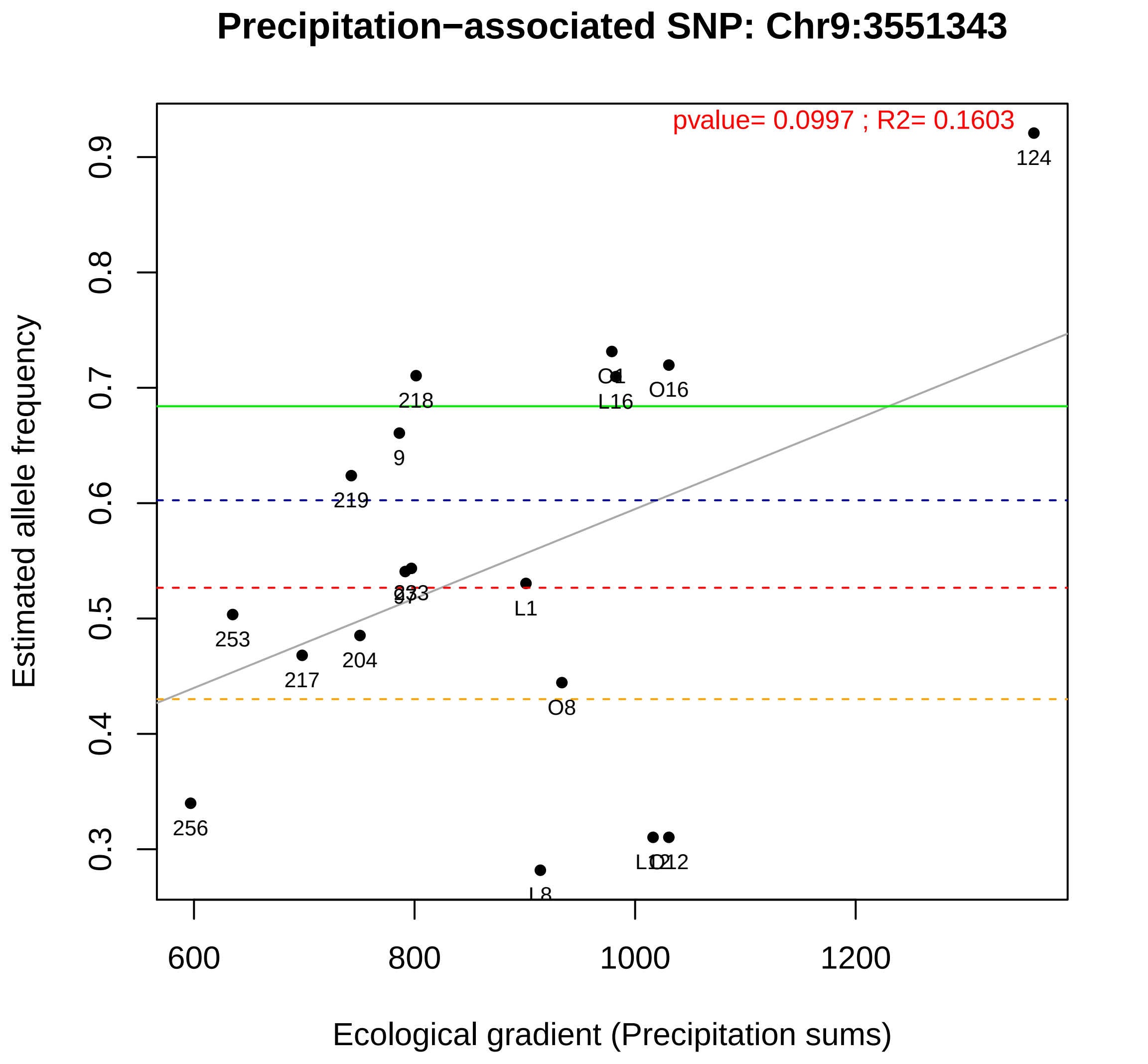

Precipitation−associated SNP: Chr9:3551343
pvalue= 0.0997 ; R2= 0.1603
Estimated allele frequency
Ecological gradient (Precipitation sums)
0.3
0.4
0.5
0.6
0.7
0.8
0.9
600
800
1000
1200
124
218
O1
L16
O16
9
219
L1
253
204
217
O8
256
L8

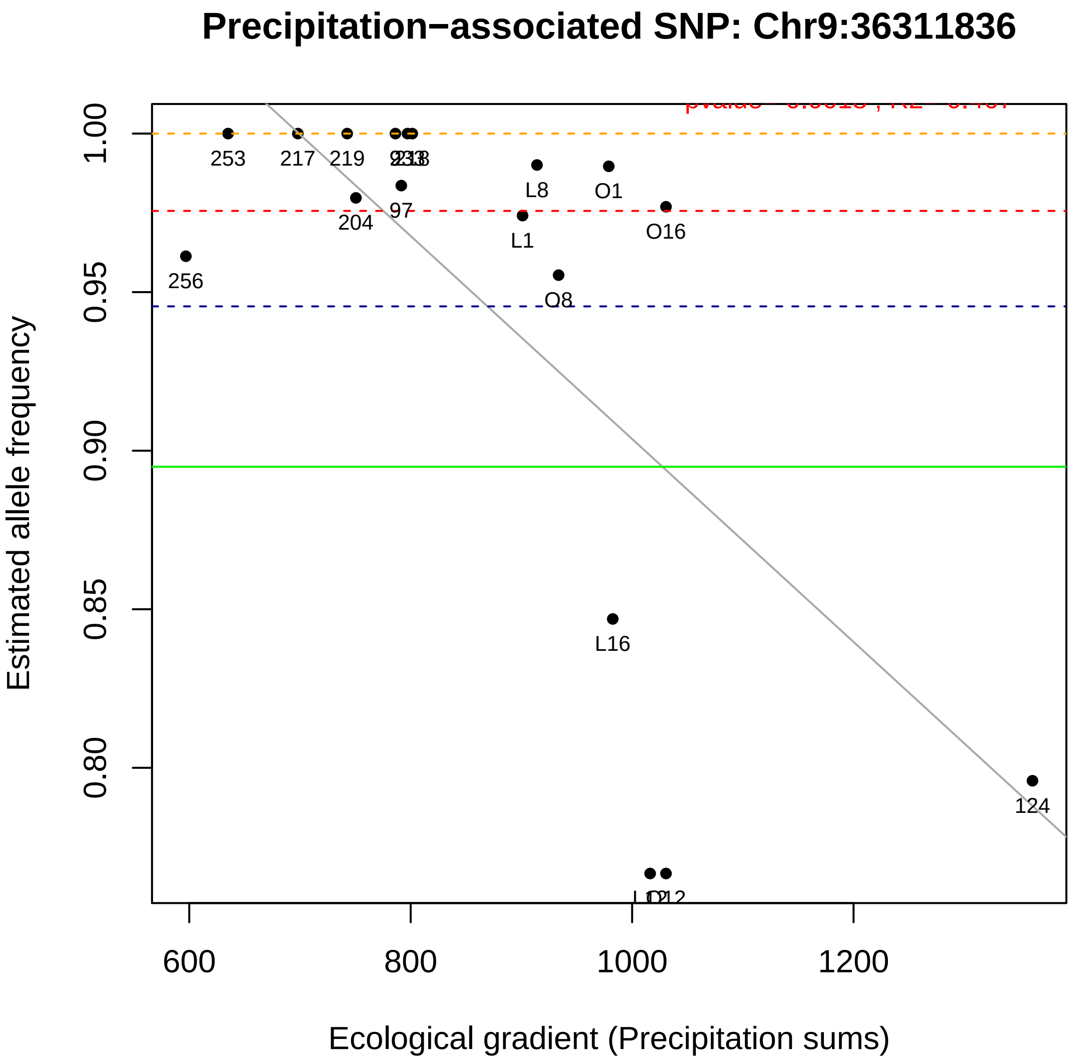
Precipitation−associated SNP: Chr9:36311836
Estimated allele frequency
Ecological gradient (Precipitation sums)
0.80
0.85
0.90
0.95
1.00
600
800
1000
1200
253
217
219
204
97
L8
O1
O16
L1
256
O8
L16
124

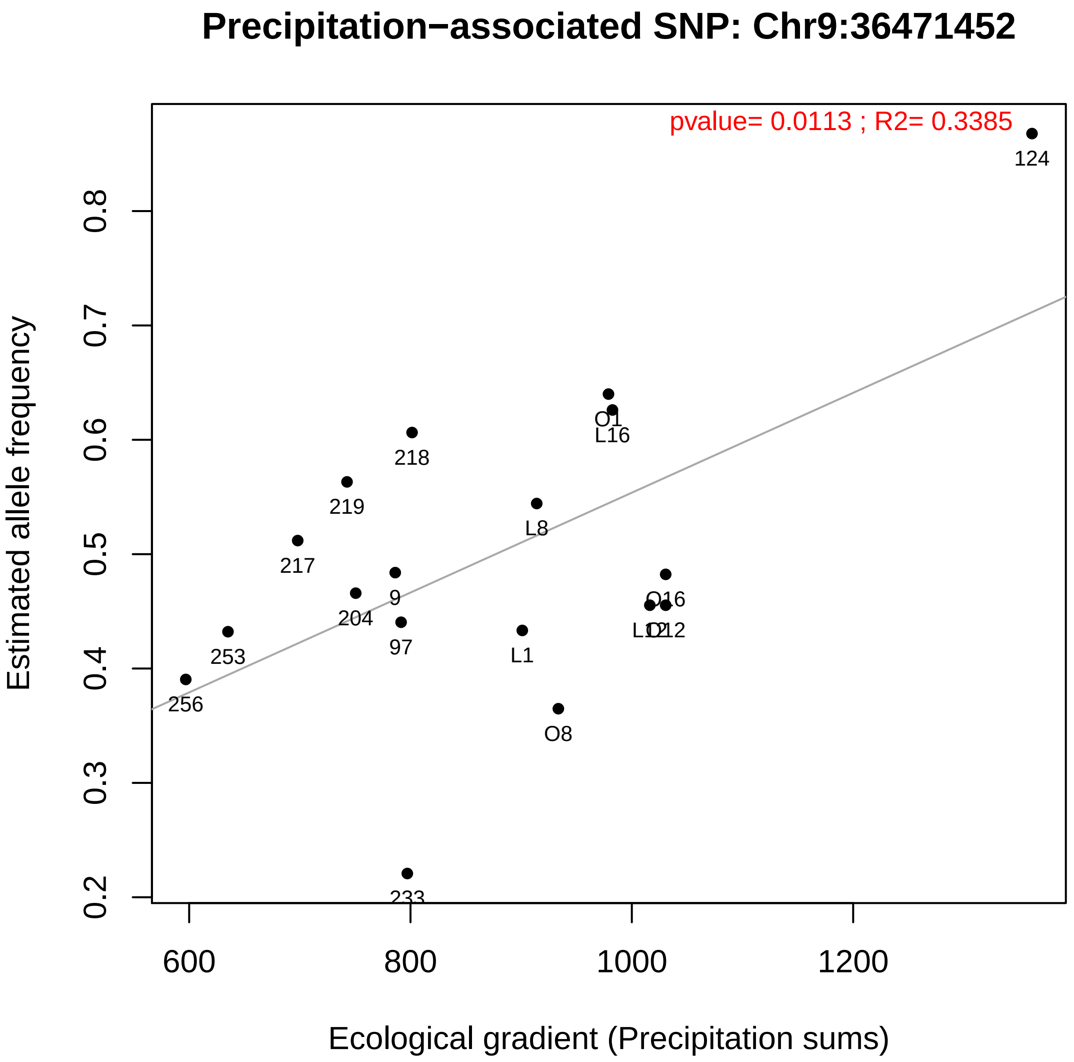
Precipitation−associated SNP: Chr9:36471452
pvalue= 0.0113 ; R2= 0.3385
Estimated allele frequency
Ecological gradient (Precipitation sums)
0.2
0.3
0.4
0.5
0.6
0.7
0.8
600
800
1000
1200
124
O1
L16
218
219
L8
217
9
204
97
L1
253
256
O8
O16
L12
O12
233

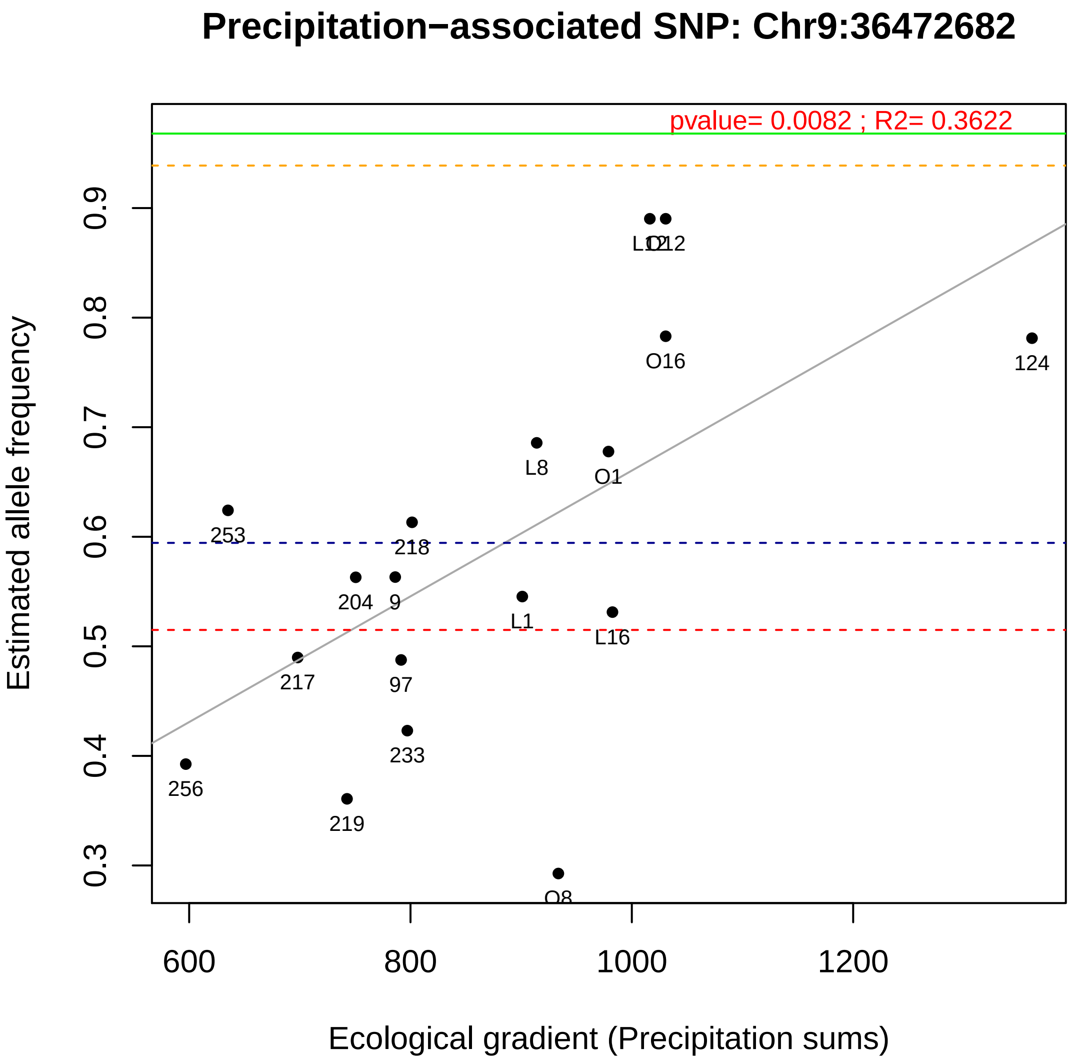

Precipitation−associated SNP: Chr9:36472682
pvalue= 0.0082 ; R2= 0.3622
Estimated allele frequency
0.3
0.4
0.5
0.6
0.7
0.8
0.9
600
800
1000
1200
Ecological gradient (Precipitation sums)
L12
O12
O16
124
L8
O1
253
218
204
9
L1
L16
217
97
233
256
219
O8

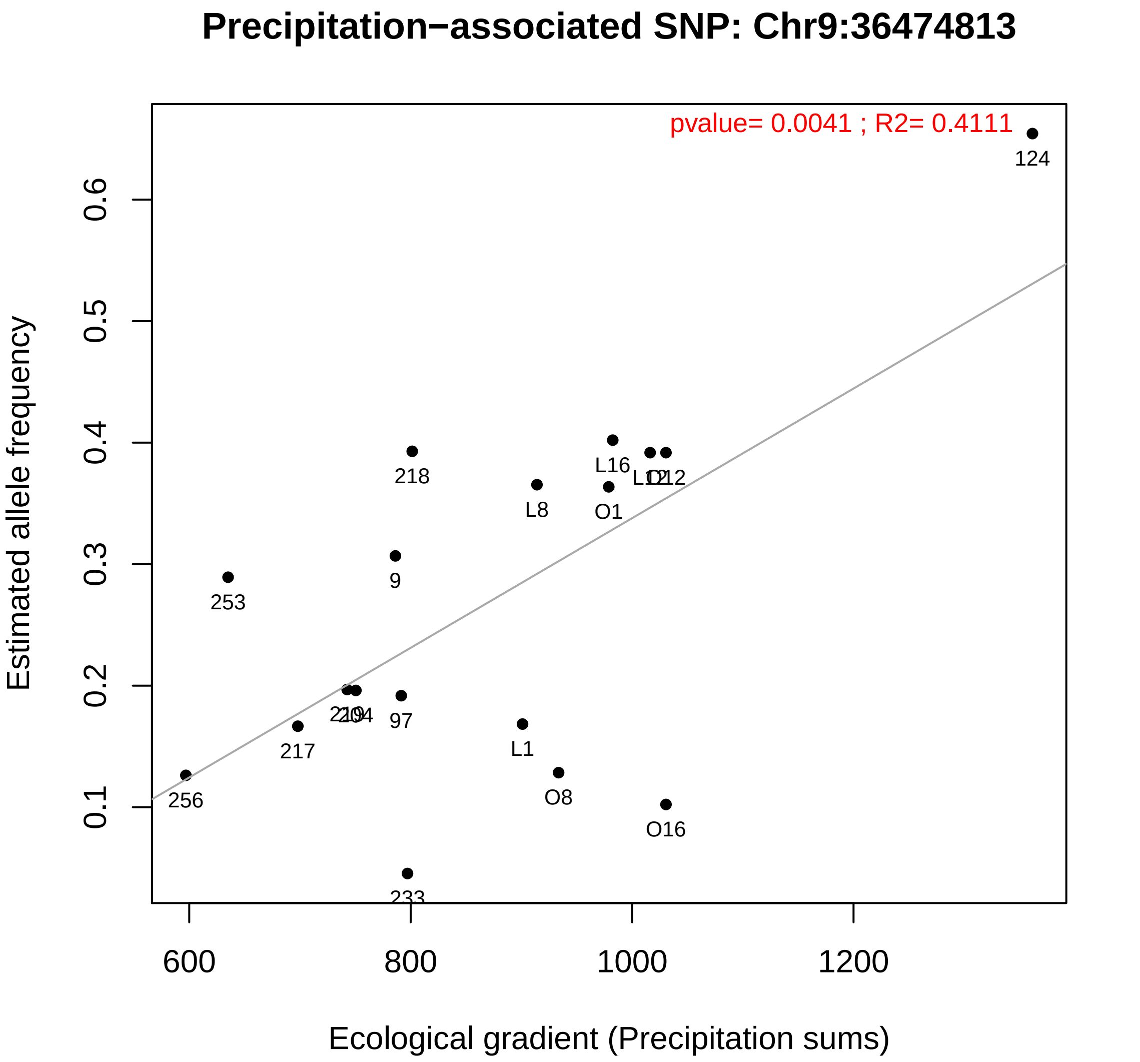

Precipitation−associated SNP: Chr9:36474813
pvalue= 0.0041 ; R2= 0.4111
Estimated allele frequency
0.1
0.2
0.3
0.4
0.5
0.6
600
800
1000
1200
Ecological gradient (Precipitation sums)
124
218
L16
L8
O1
O12
9
253
97
217
L1
256
O8
O16
233

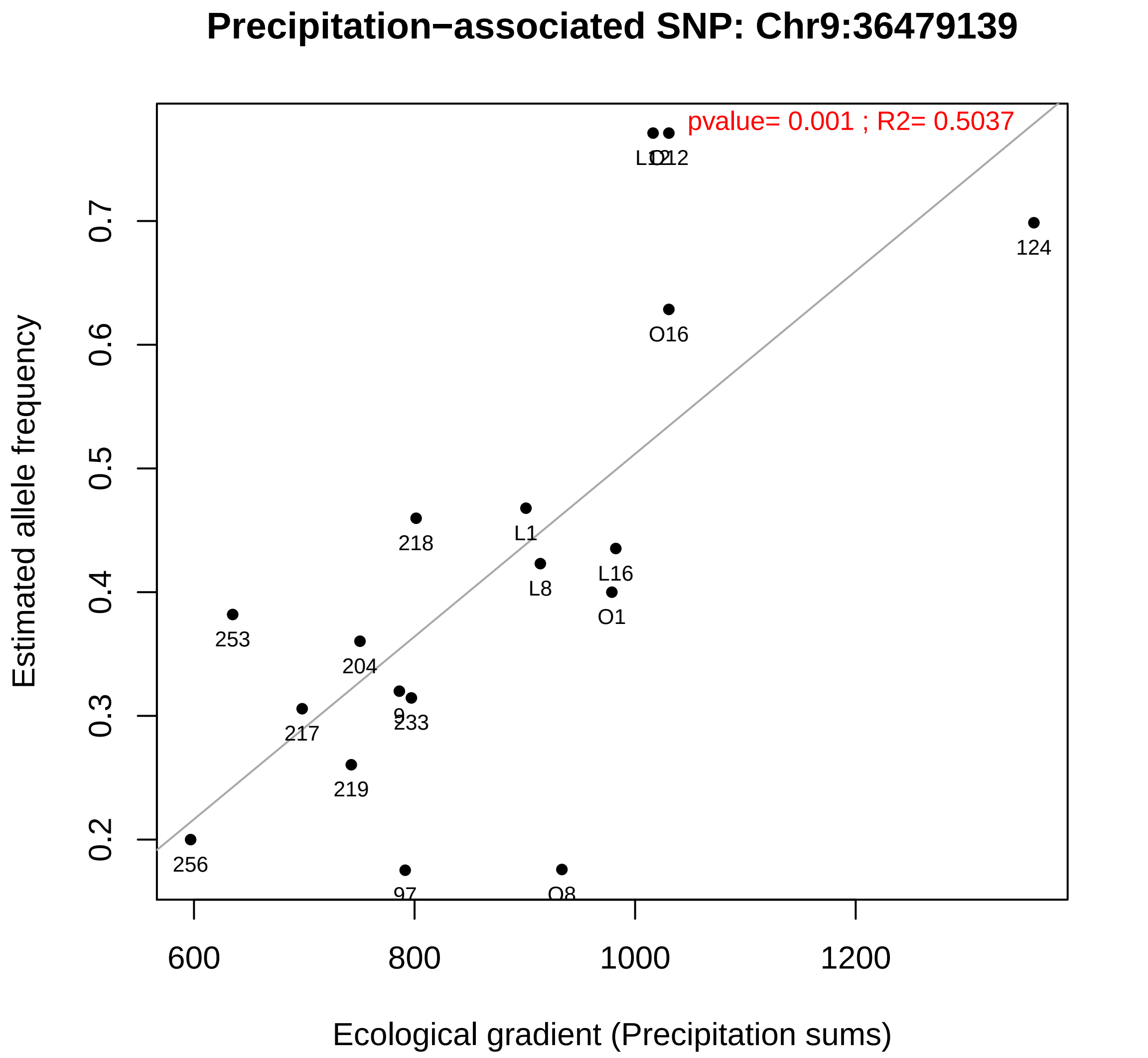
Precipitation−associated SNP: Chr9:36479139
pvalue= 0.001 ; R2= 0.5037
L12
O12
124
O16
218
L1
L8
L16
O1
253
204
217
9
233
219
256
97
O8
0.2
0.3
0.4
0.5
0.6
0.7
Estimated allele frequency
600
800
1000
1200
Ecological gradient (Precipitation sums)

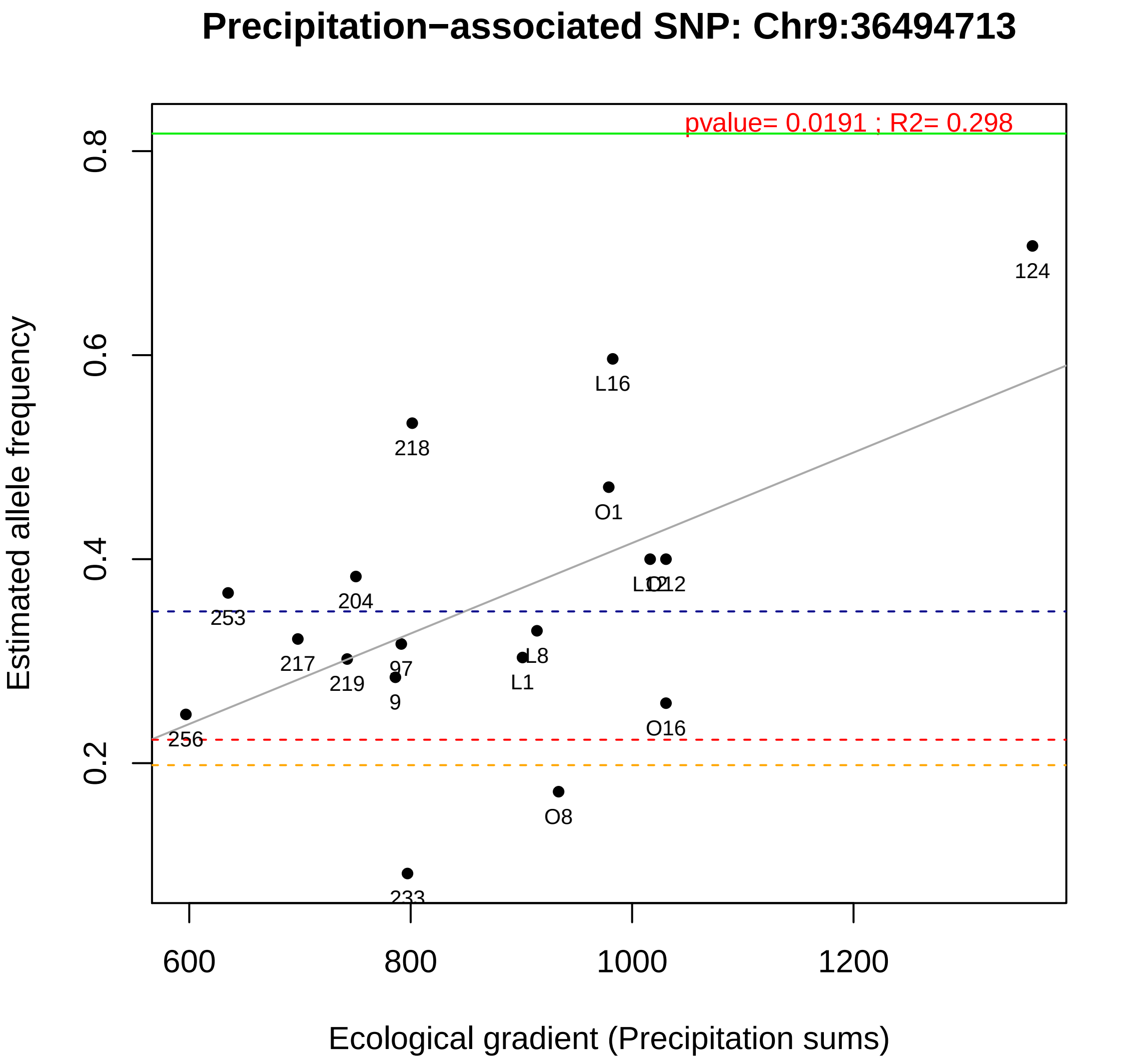

Precipitation−associated SNP: Chr9:36494713
pvalue= 0.0191 ; R2= 0.298
Estimated allele frequency
0.2
0.4
0.6
0.8
124
L16
218
O1
L12
O12
204
253
217
219
97
9
L8
L1
O16
256
O8
233
600
800
1000
1200
Ecological gradient (Precipitation sums)

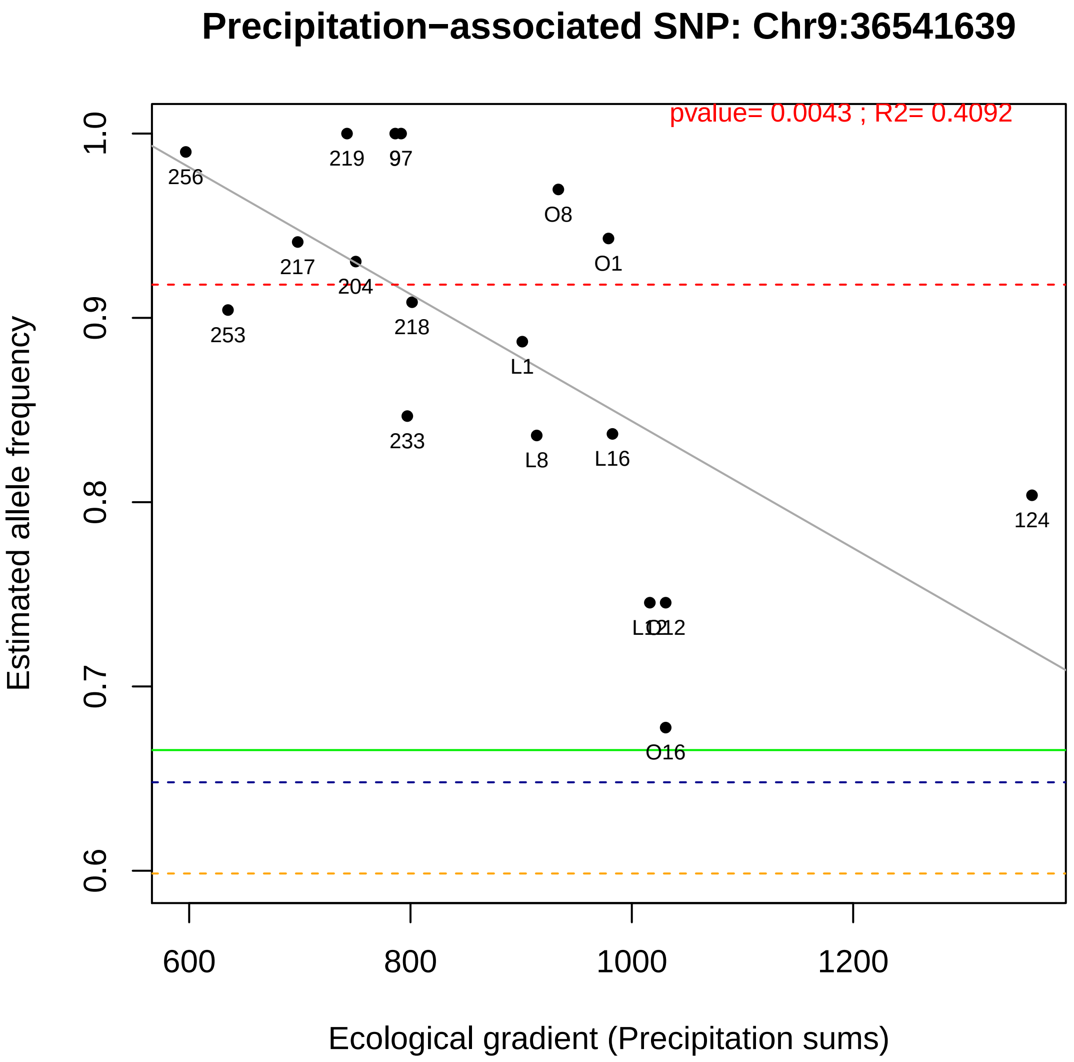

Precipitation−associated SNP: Chr9:36541639
pvalue= 0.0043 ; R2= 0.4092
256
219
97
O8
217
204
O1
253
218
L1
233
L8
L16
124
L10
O12
O16
1.0
0.9
0.8
0.7
0.6
600
800
1000
1200
Estimated allele frequency
Ecological gradient (Precipitation sums)

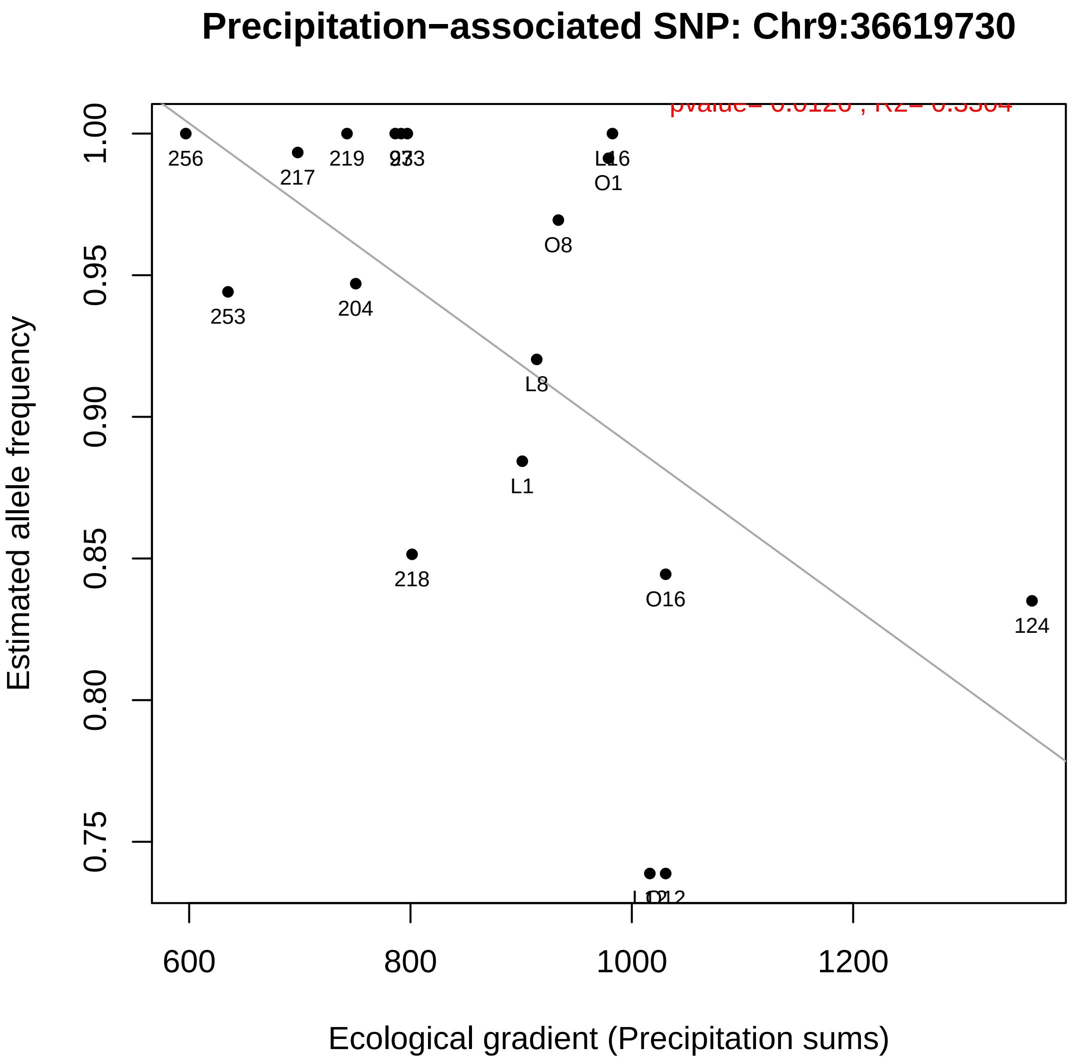
Precipitation−associated SNP: Chr9:36619730
Estimated allele frequency
Ecological gradient (Precipitation sums)
1.00
0.95
0.90
0.85
0.80
0.75
600
800
1000
1200
256
217
219
L16
O1
O8
253
204
L8
L1
218
O16
124

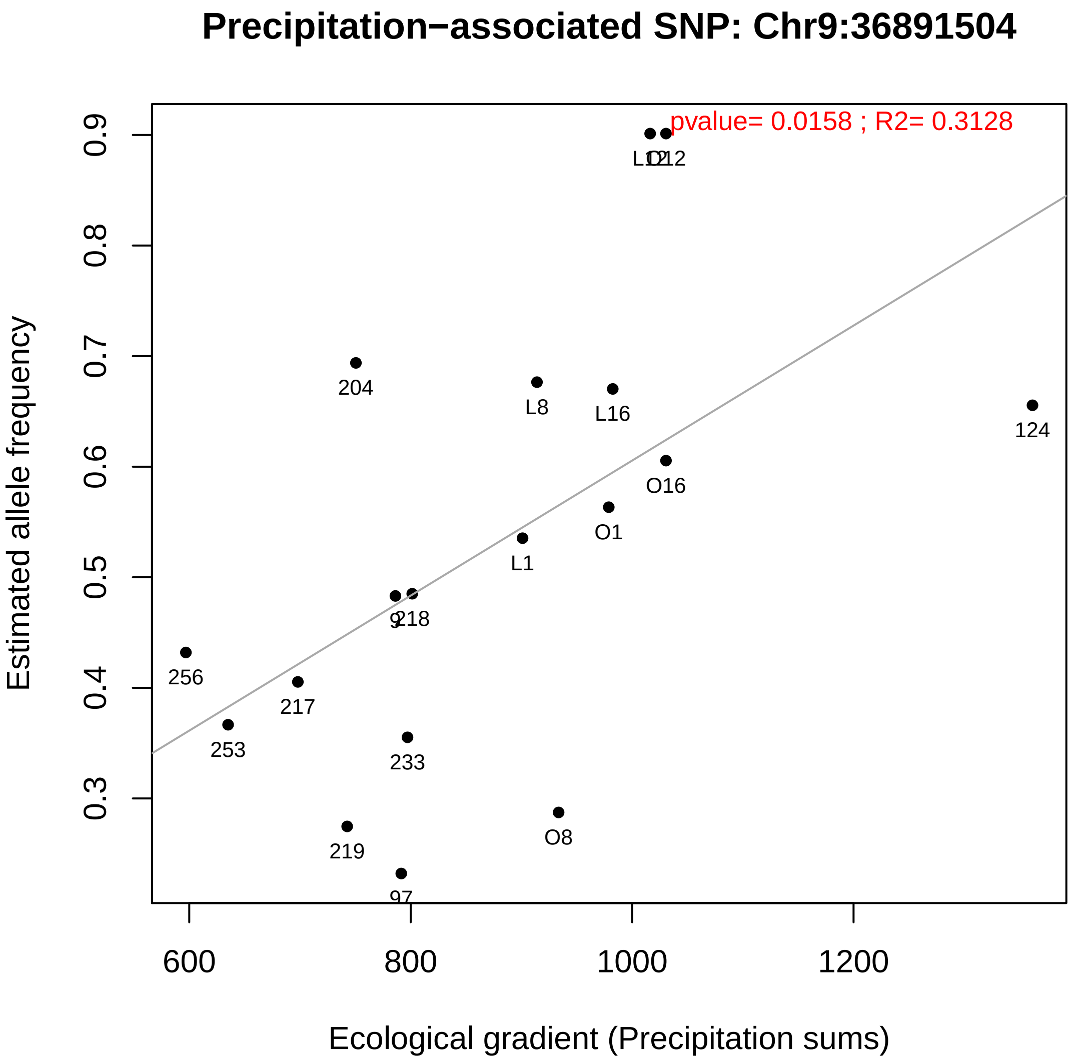
Precipitation−associated SNP: Chr9:36891504
pvalue= 0.0158 ; R2= 0.3128
Estimated allele frequency
Ecological gradient (Precipitation sums)
0.3
0.4
0.5
0.6
0.7
0.8
0.9
600
800
1000
1200
L10
O12
204
L8
L16
124
O16
O1
L1
9
218
256
217
253
233
O8
219
97

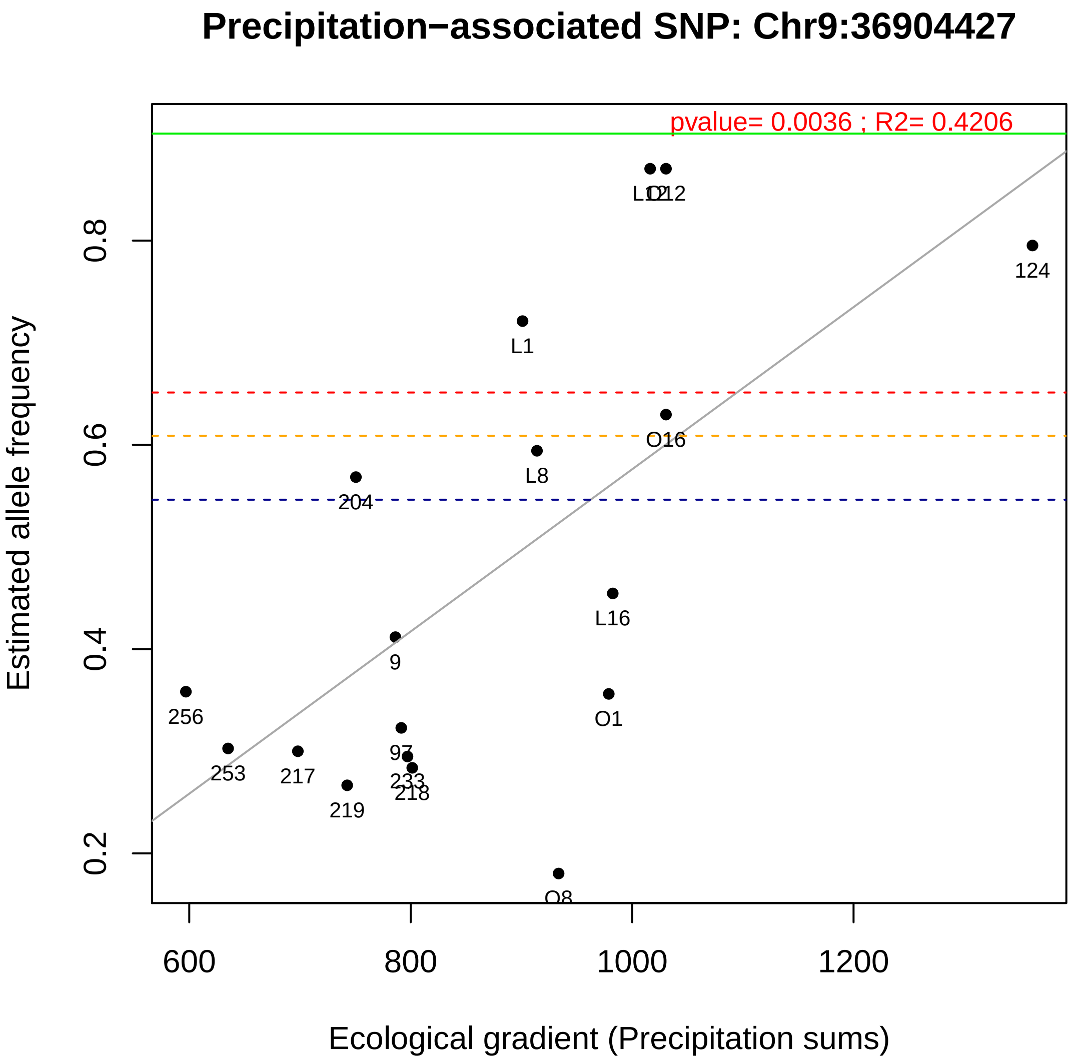

Precipitation−associated SNP: Chr9:36904427
pvalue= 0.0036 ; R2= 0.4206
Estimated allele frequency
0.2
0.4
0.6
0.8
600
800
1000
1200
Ecological gradient (Precipitation sums)
L1
L8
L16
O1
O8
O16
124
204
9
256
253
217
219
97
233
218

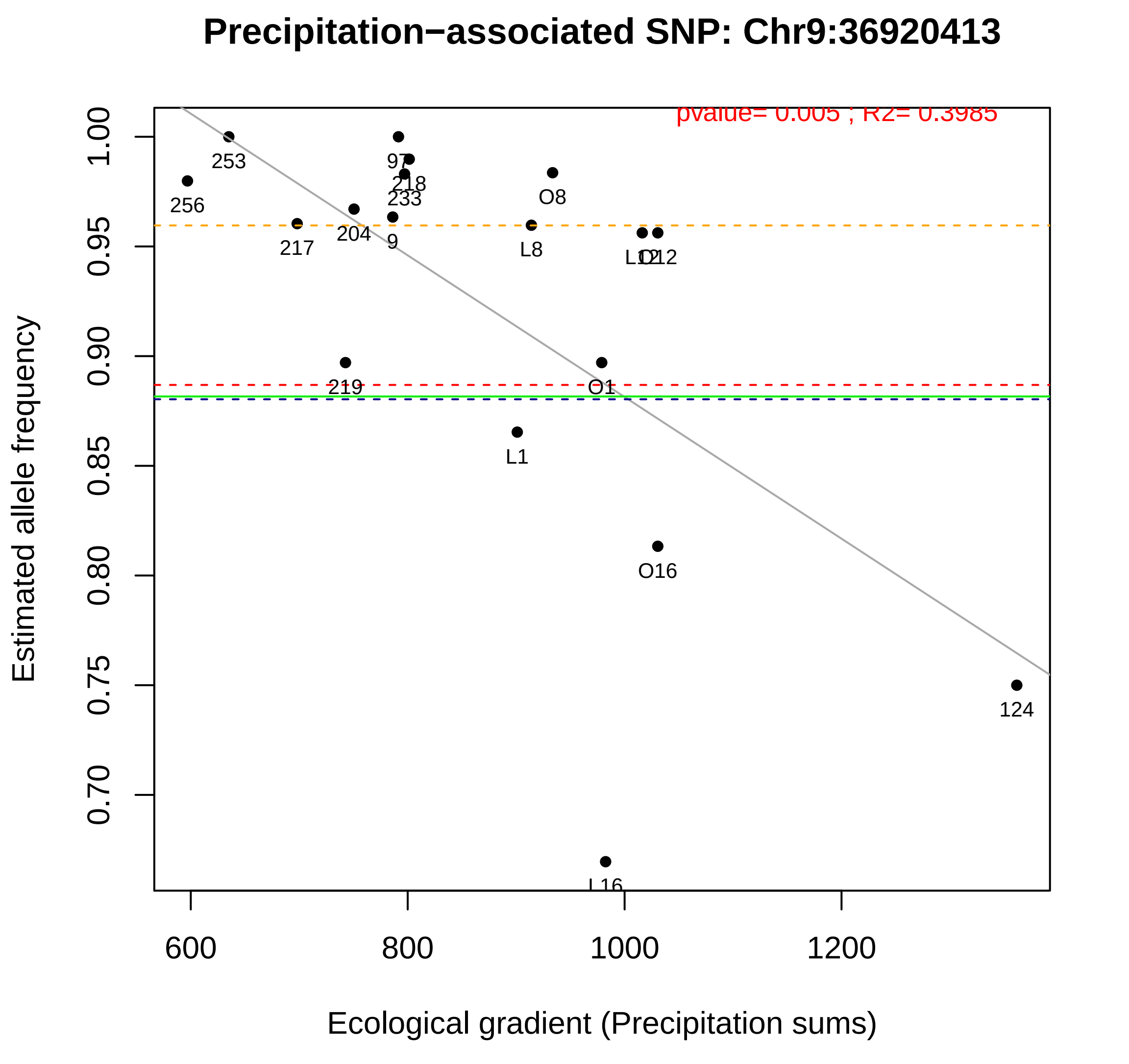

Precipitation−associated SNP: Chr9:36920413
pvalue= 0.005 ; R2= 0.3985
Estimated allele frequency
Ecological gradient (Precipitation sums)
1.00
0.95
0.90
0.85
0.80
0.75
0.70
600
800
1000
1200
253
256
217
204
219
97
218
233
9
O8
L8
L12
O12
O1
L1
O16
124
L16

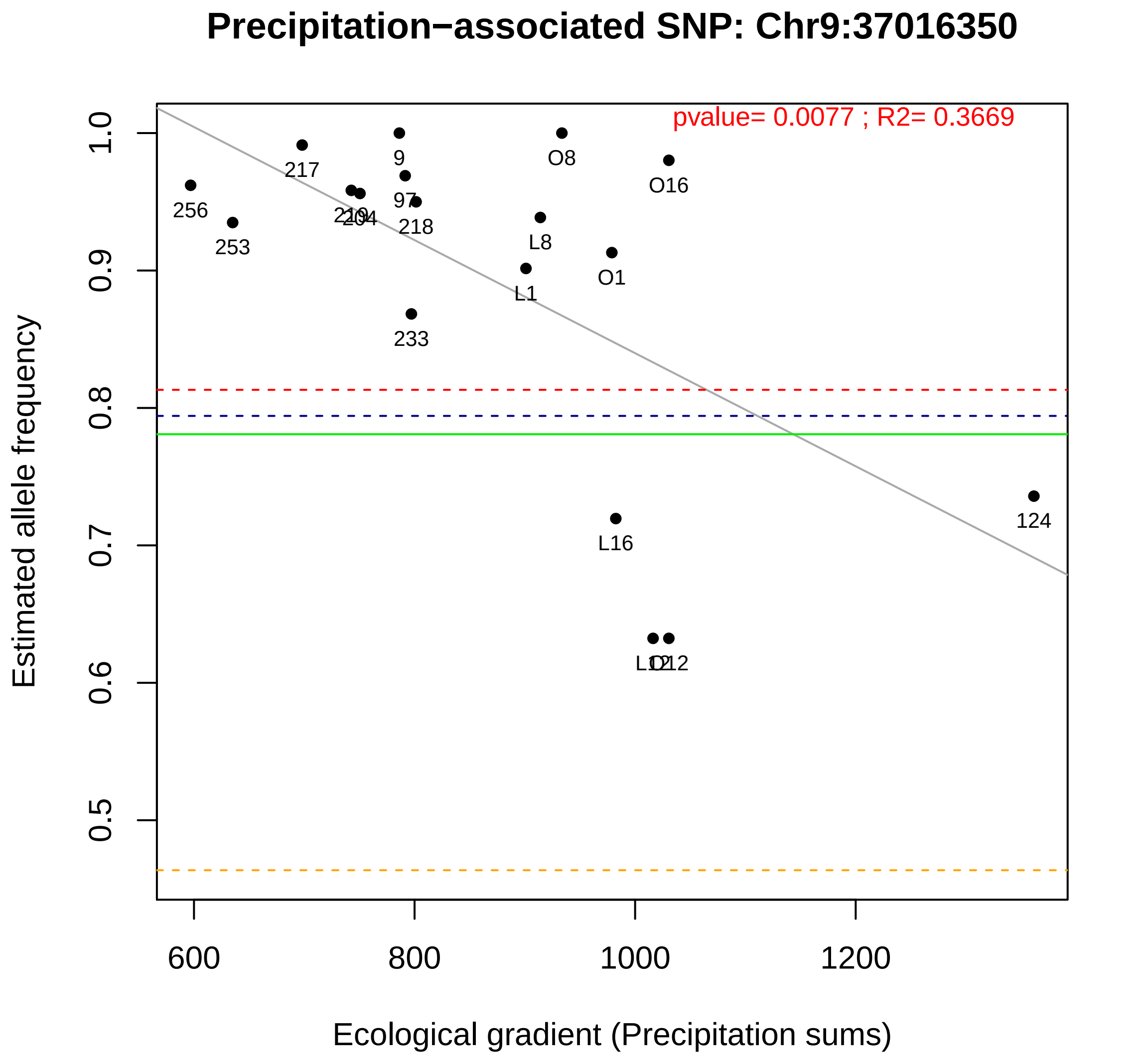

Precipitation−associated SNP: Chr9:37016350
pvalue= 0.0077 ; R2= 0.3669
Estimated allele frequency
Ecological gradient (Precipitation sums)
0.5
0.6
0.7
0.8
0.9
1.0
600
800
1000
1200
256
253
217
210
204
9
97
218
233
L1
L8
O8
O1
O16
L16
L12
O12
124

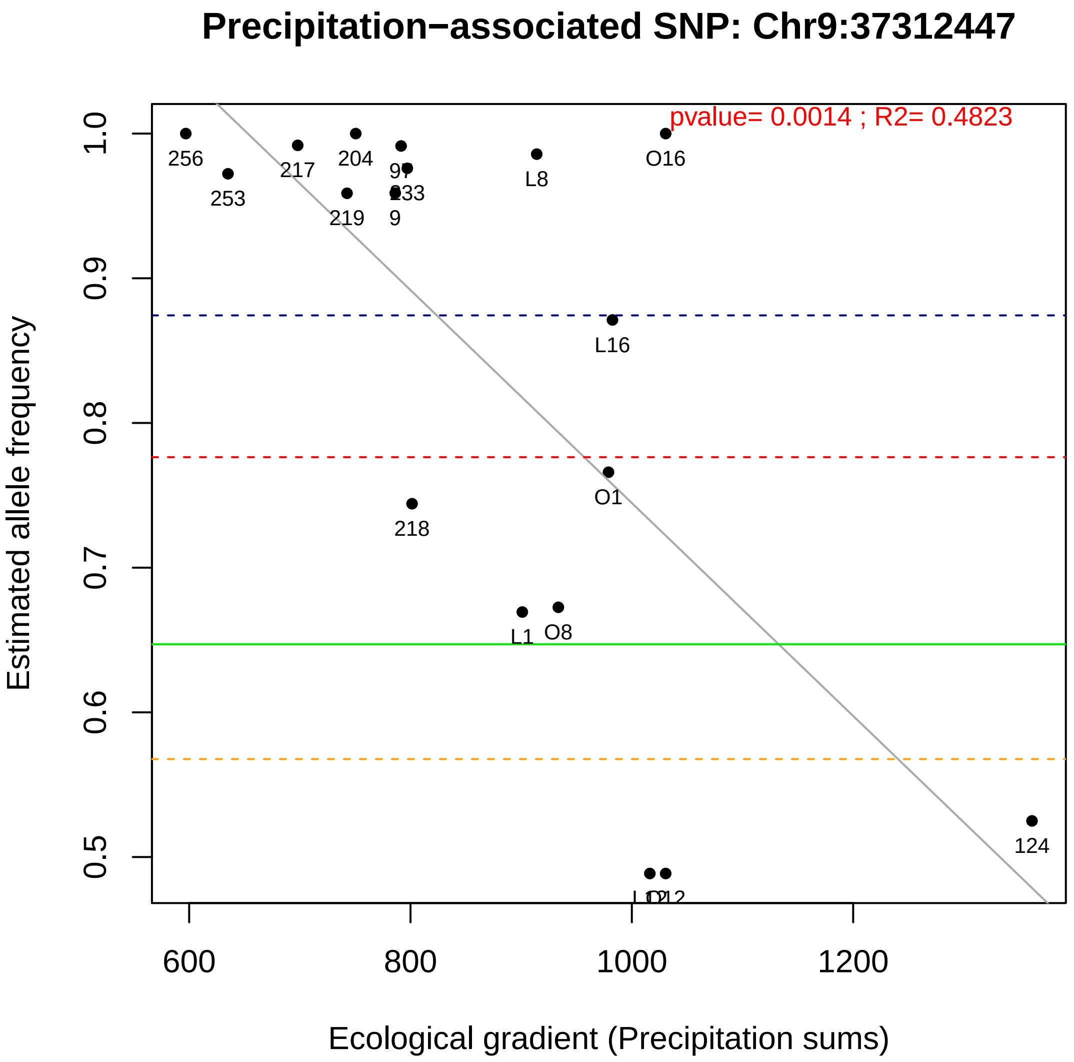
Precipitation−associated SNP: Chr9:37312447
pvalue= 0.0014 ; R2= 0.4823
Estimated allele frequency
Ecological gradient (Precipitation sums)
0.5
0.6
0.7
0.8
0.9
1.0
600
800
1000
1200
256
253
217
204
219
233
9
L8
O16
L16
O1
218
L1
O8
124

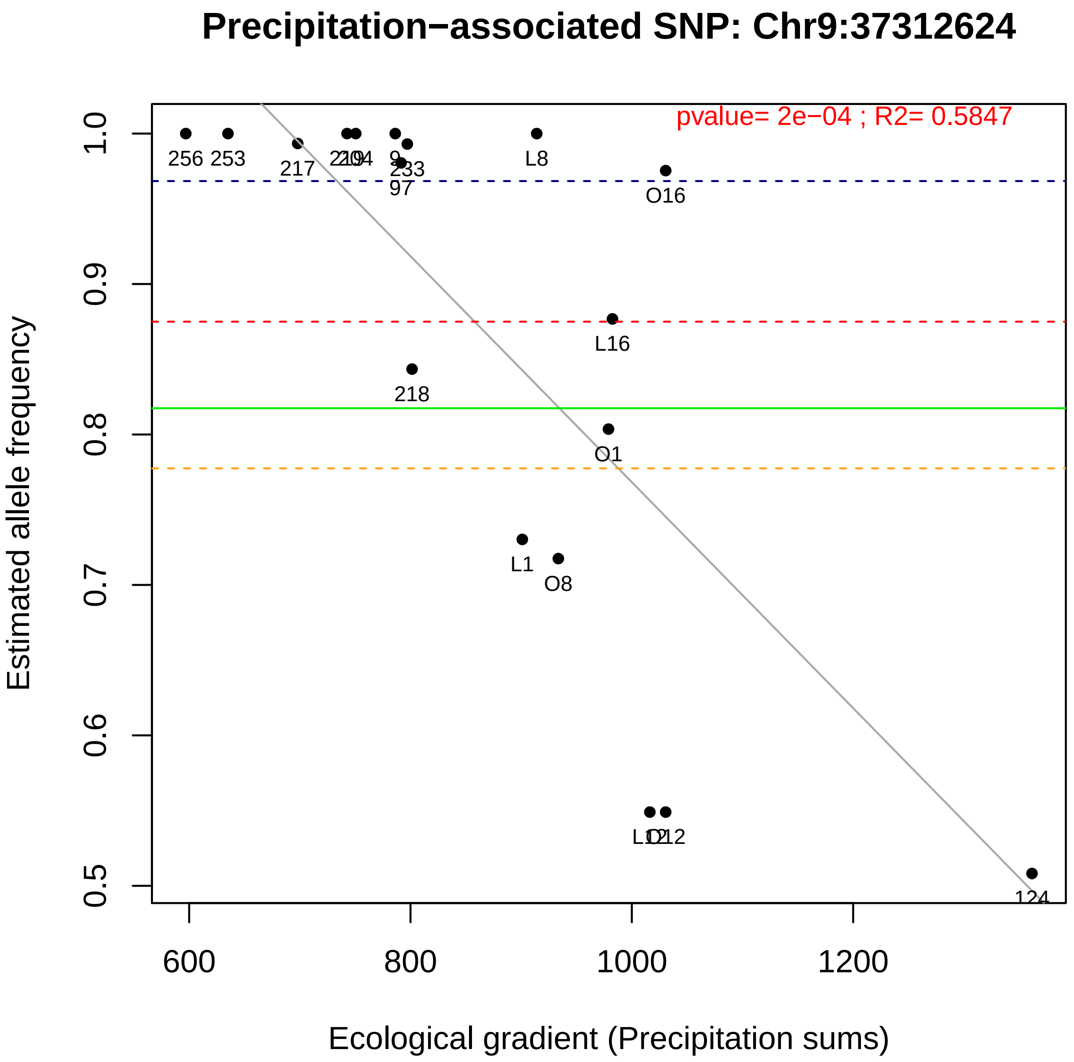
Precipitation−associated SNP: Chr9:37312624
pvalue= 2e−04 ; R2= 0.5847
Estimated allele frequency
Ecological gradient (Precipitation sums)
1.0
0.9
0.8
0.7
0.6
0.5
600
800
1000
1200
256
253
217
L8
O16
L16
218
O1
L1
O8
124

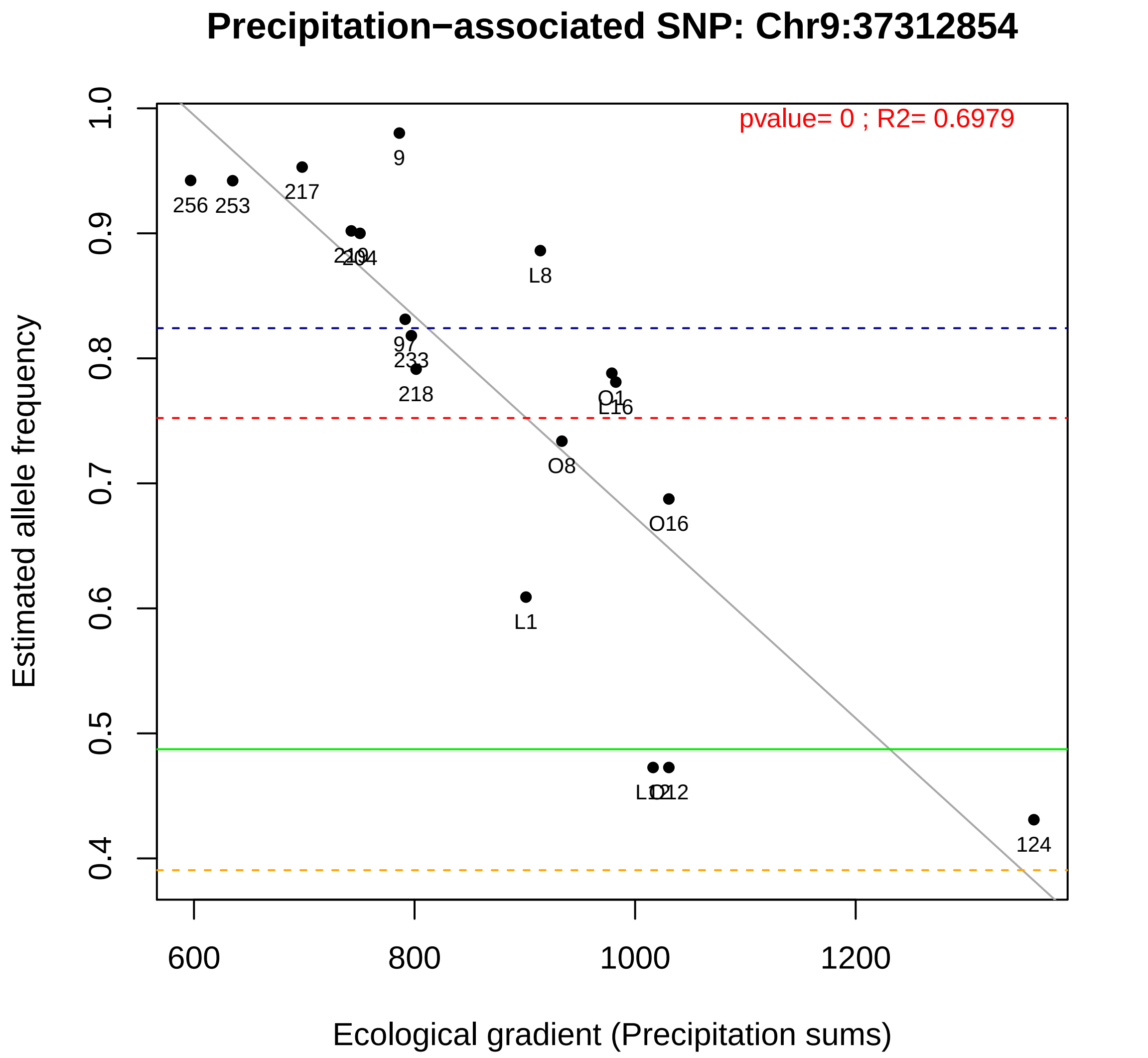

Precipitation−associated SNP: Chr9:37312854
pvalue= 0 ; R2= 0.6979
Estimated allele frequency
Ecological gradient (Precipitation sums)
1.0
0.9
0.8
0.7
0.6
0.5
0.4
600
800
1000
1200
9
256
253
217
L8
97
233
218
O8
O16
L1
124

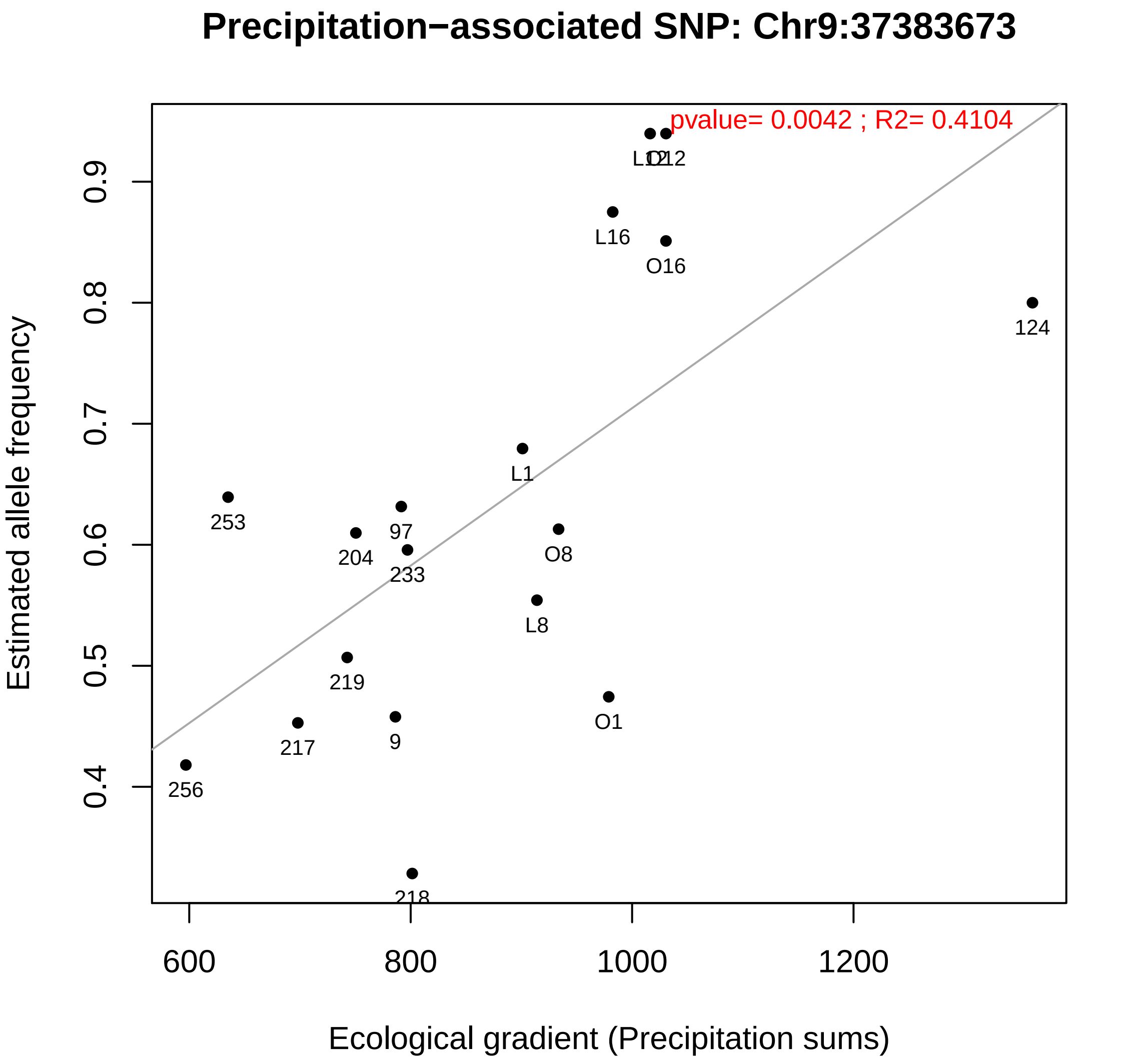
Precipitation−associated SNP: Chr9:37383673
pvalue= 0.0042 ; R2= 0.4104
Estimated allele frequency
Ecological gradient (Precipitation sums)
0.4
0.5
0.6
0.7
0.8
0.9
600
800
1000
1200
O12
L16
O16
124
L1
253
97
204
233
O8
L8
219
O1
217
9
256
218

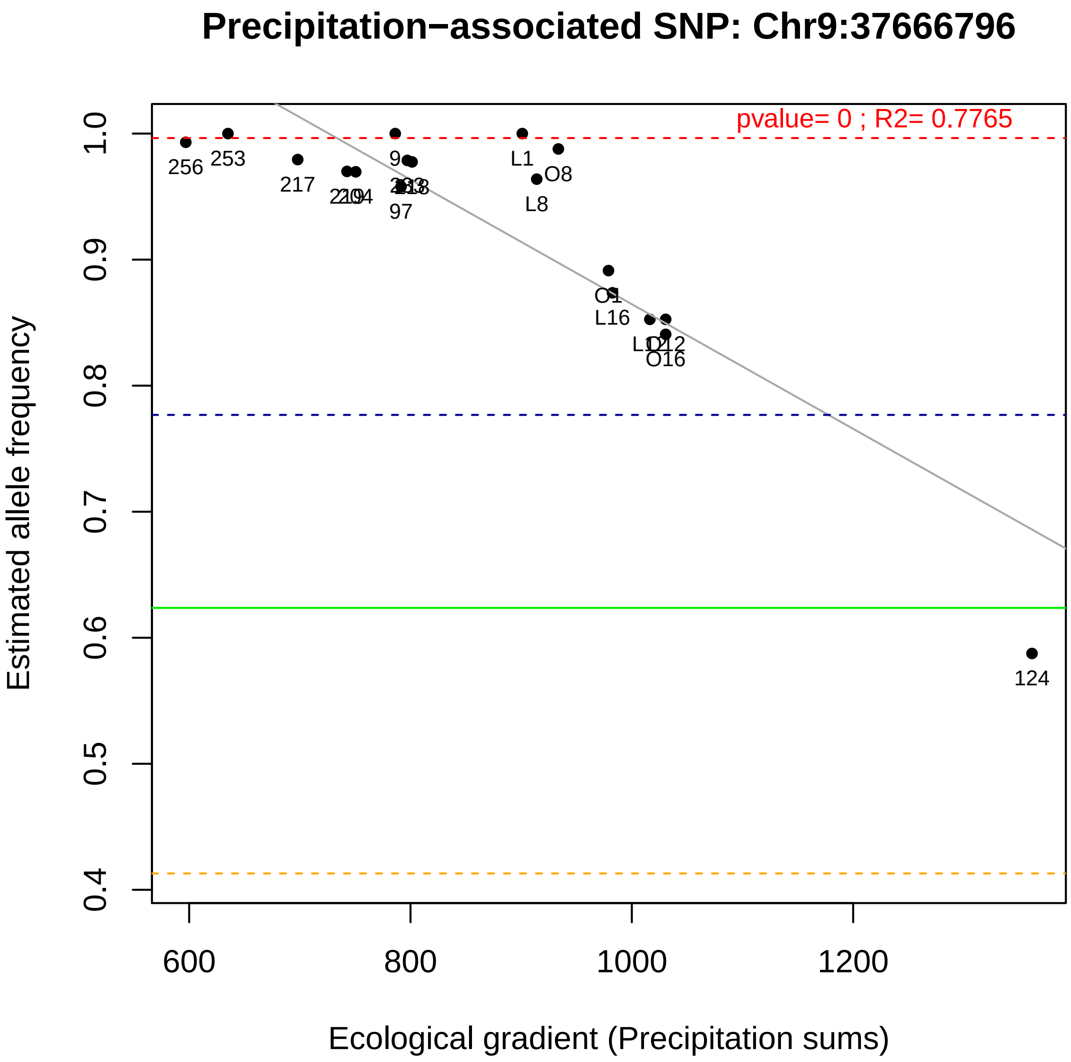
Precipitation−associated SNP: Chr9:37666796
pvalue= 0 ; R2= 0.7765
Estimated allele frequency
Ecological gradient (Precipitation sums)
1.0
0.9
0.8
0.7
0.6
0.5
0.4
600
800
1000
1200
256
253
217
L1
O8
L8
97
124
L16
O16

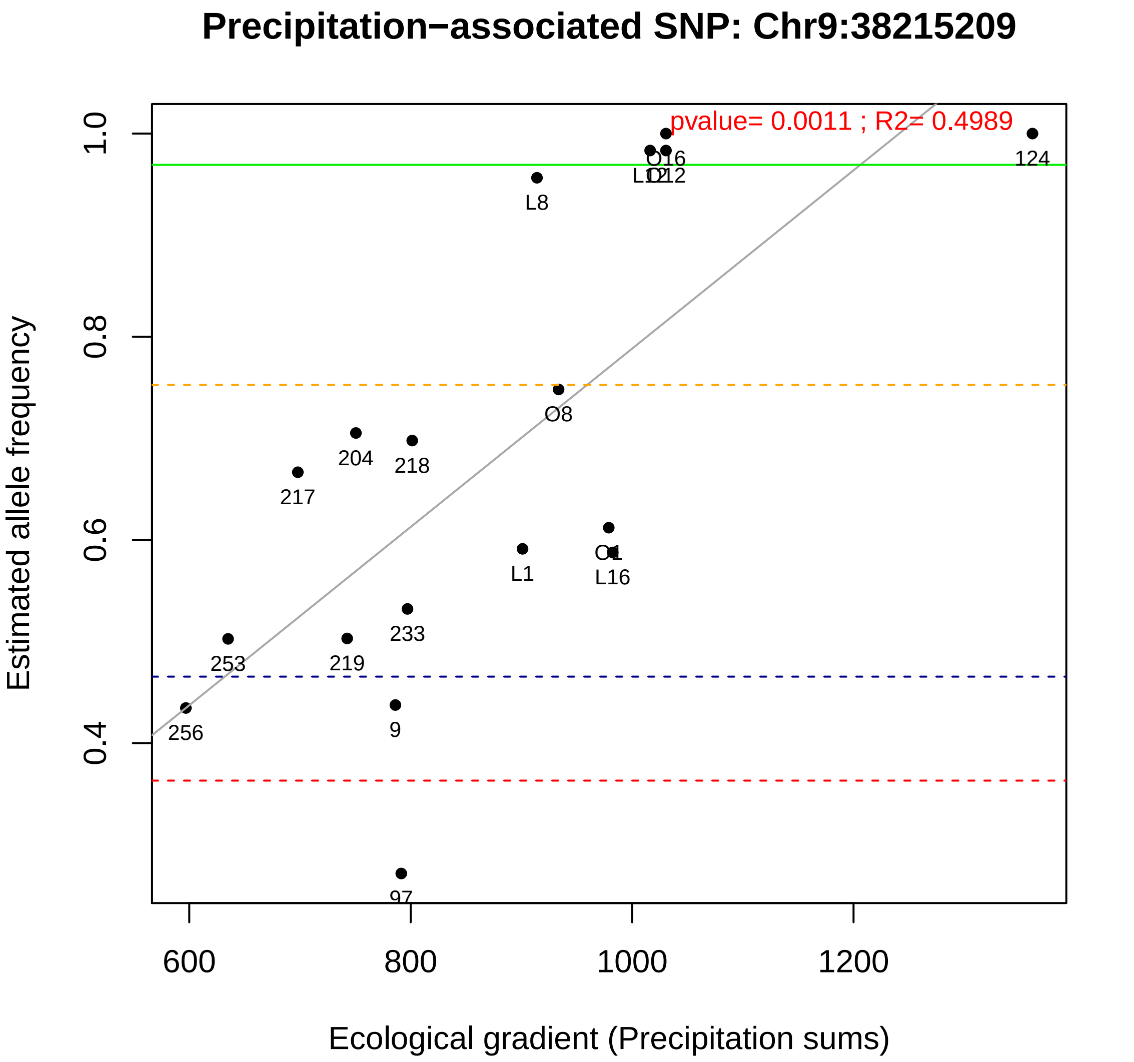
Precipitation−associated SNP: Chr9:38215209
pvalue= 0.0011 ; R2= 0.4989
Estimated allele frequency
0.4
0.6
0.8
1.0
600
800
1000
1200
Ecological gradient (Precipitation sums)
124
O16
L10
O12
L8
O8
204
218
217
O1
L1
L16
233
253
219
256
9
97

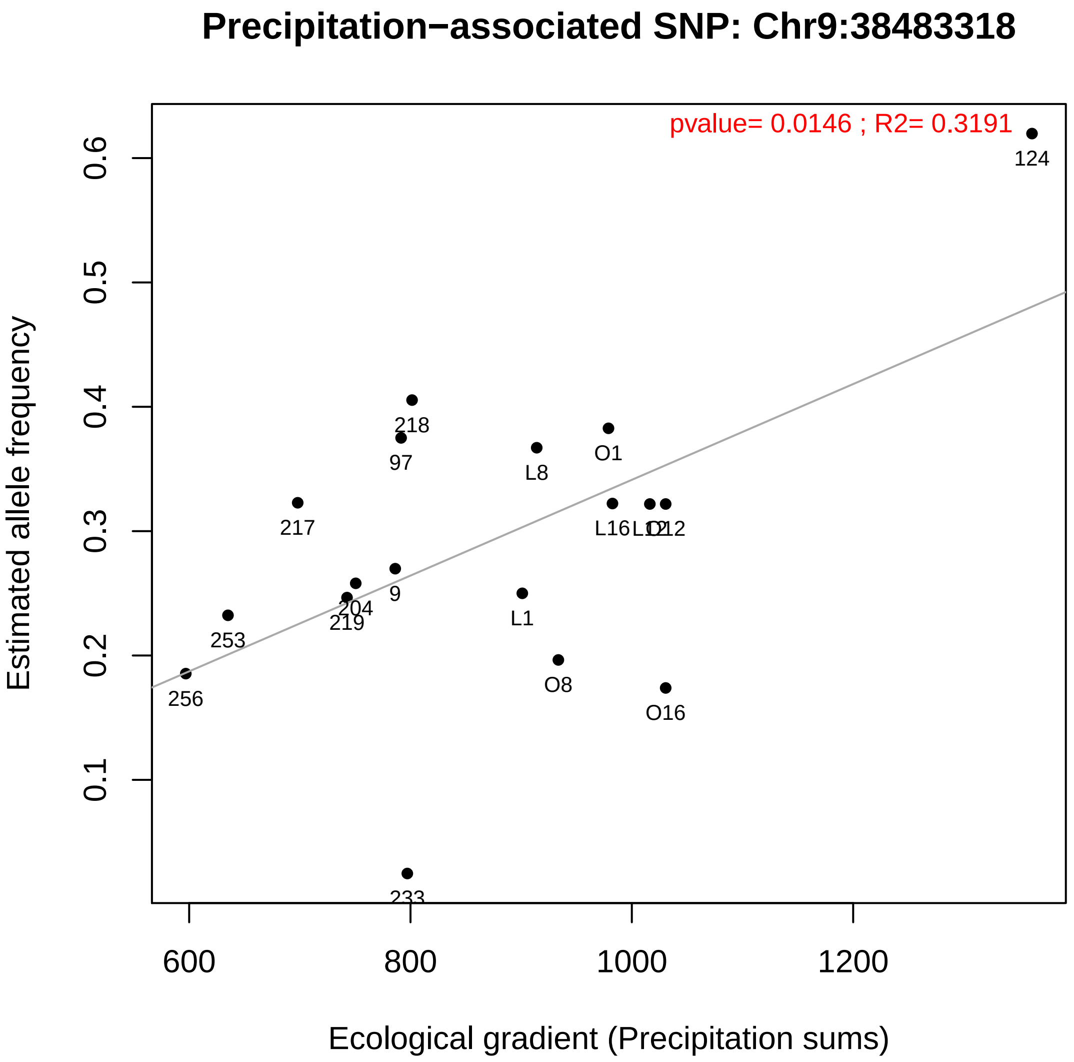
Precipitation−associated SNP: Chr9:38483318
pvalue= 0.0146 ; R2= 0.3191
124
218
97
L8
O1
217
L16
L12
O12
9
204
219
L1
253
256
O8
O16
233
0.1
0.2
0.3
0.4
0.5
0.6
600
800
1000
1200
Estimated allele frequency
Ecological gradient (Precipitation sums)

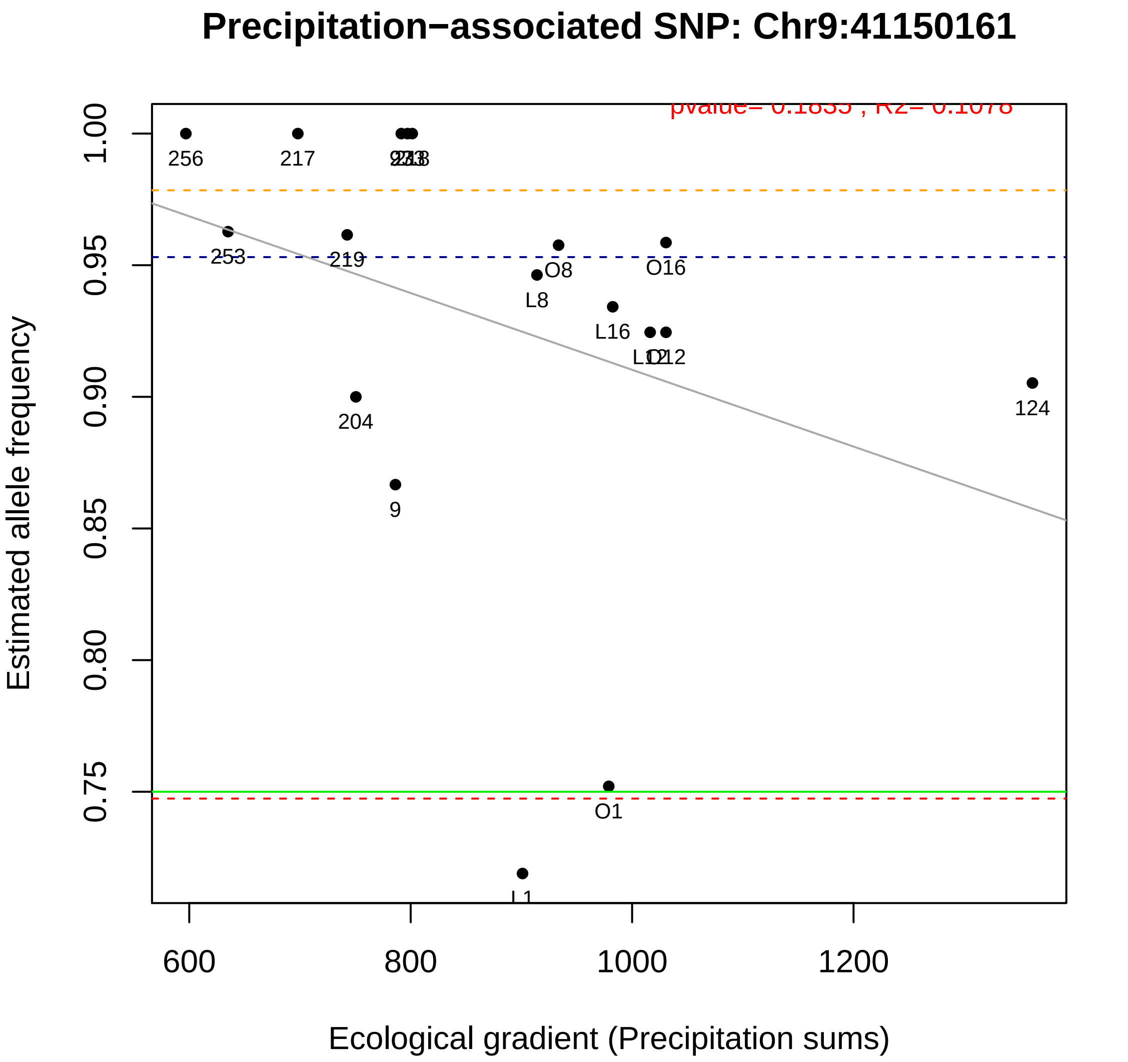
Precipitation−associated SNP: Chr9:41150161
pvalue= 0.1835 , R2= 0.1078
Estimated allele frequency
Ecological gradient (Precipitation sums)
0.75
0.80
0.85
0.90
0.95
1.00
600
800
1000
1200
256
217
253
219
O8
O16
L8
L16
O12
204
124
9
O1

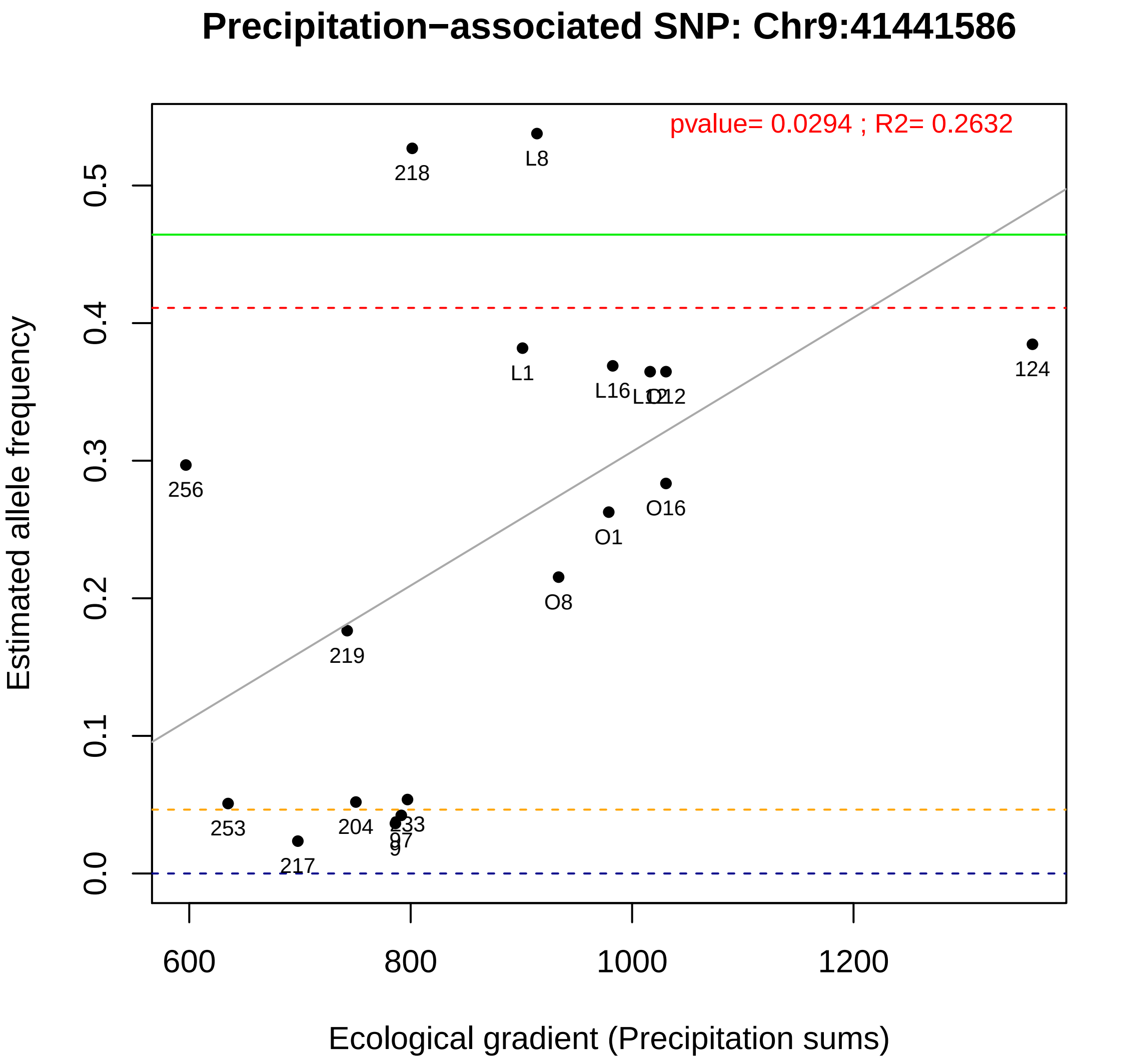

Precipitation−associated SNP: Chr9:41441586
pvalue= 0.0294 ; R2= 0.2632
Estimated allele frequency
0.0
0.1
0.2
0.3
0.4
0.5
Ecological gradient (Precipitation sums)
600
800
1000
1200
218
L8
L1
L16
O12
124
256
O16
O1
O8
219
253
204
233
97
9
217

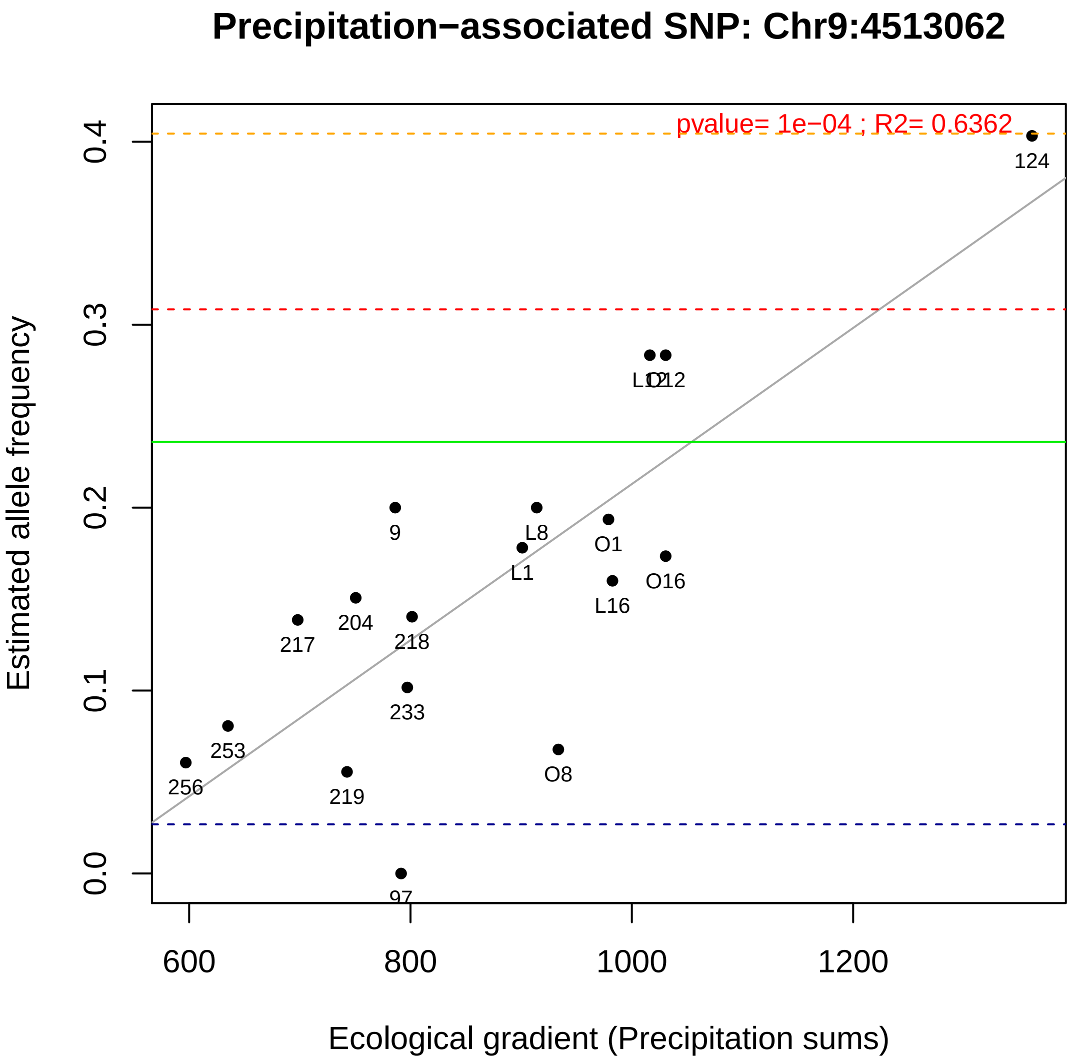
Precipitation−associated SNP: Chr9:4513062
pvalue= 1e−04 ; R2= 0.6362
124
L12
O12
9
L8
O1
L1
O16
L16
204
217
218
233
253
256
219
O8
97
0.0
0.1
0.2
0.3
0.4
600
800
1000
1200
Estimated allele frequency
Ecological gradient (Precipitation sums)

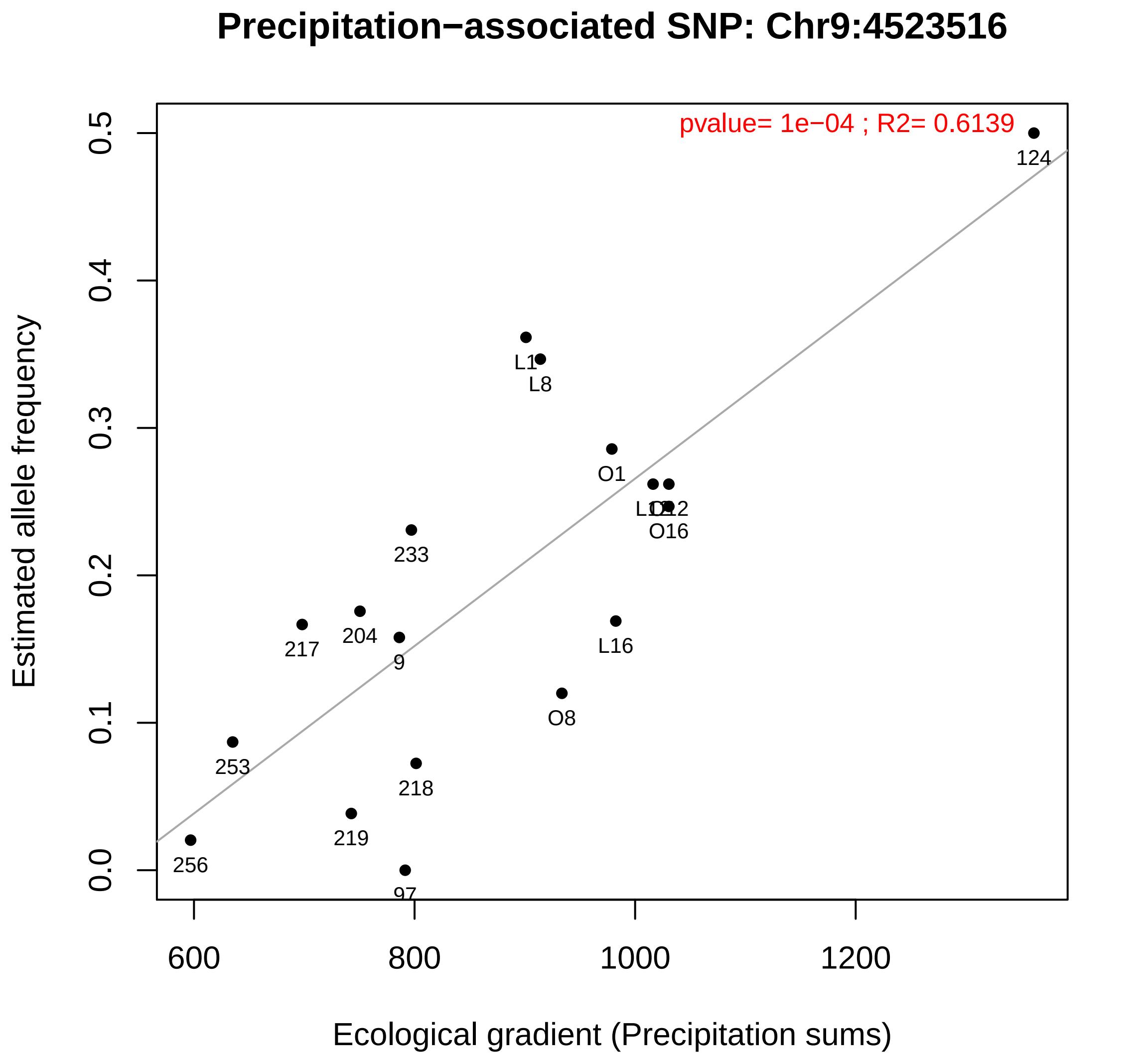

Precipitation−associated SNP: Chr9:4523516
pvalue= 1e−04 ; R2= 0.6139
Estimated allele frequency
Ecological gradient (Precipitation sums)
0.0
0.1
0.2
0.3
0.4
0.5
600
800
1000
1200
124
L1
L8
O1
O12
O16
233
204
217
9
L16
O8
253
218
219
256
97

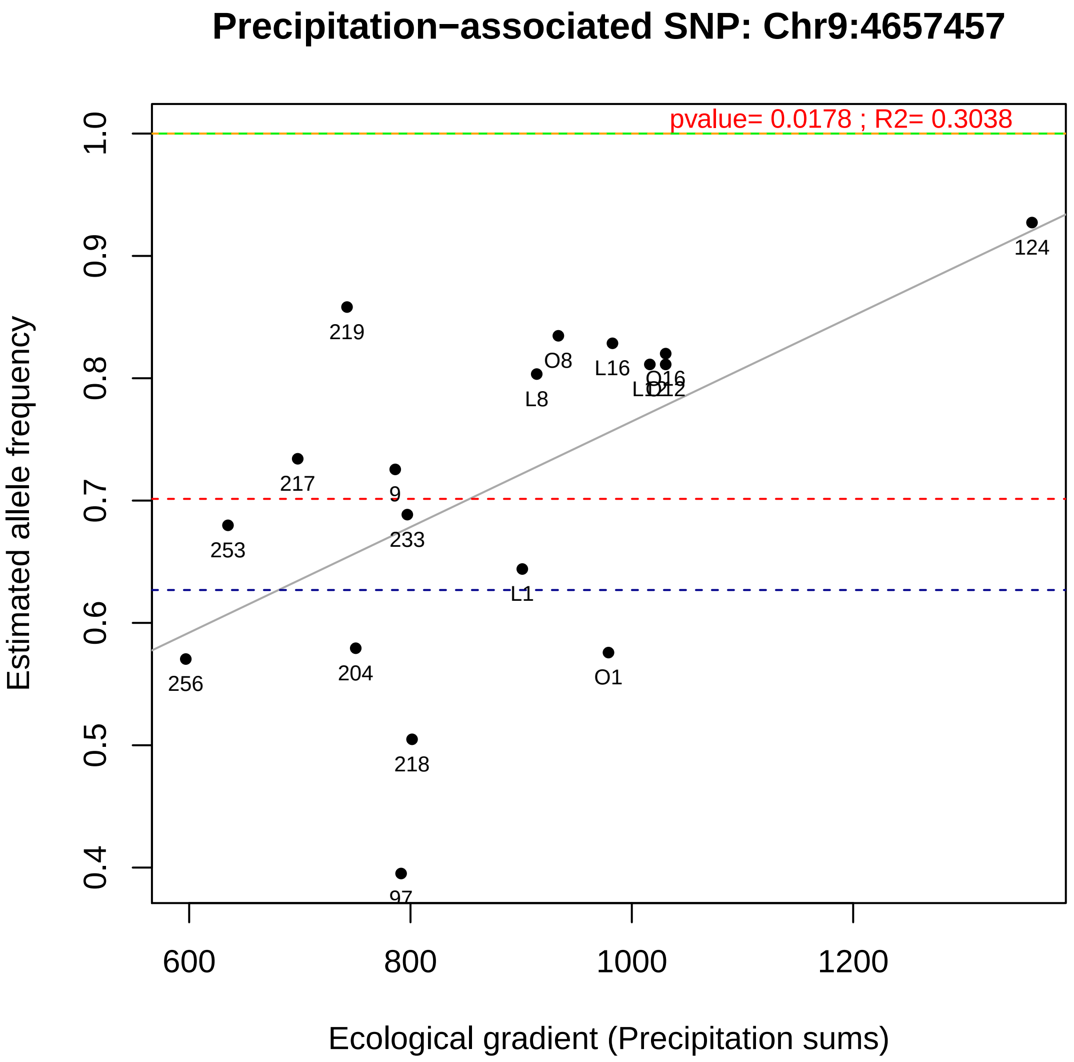
Precipitation−associated SNP: Chr9:4657457
pvalue= 0.0178 ; R2= 0.3038
Estimated allele frequency
Ecological gradient (Precipitation sums)
0.4
0.5
0.6
0.7
0.8
0.9
1.0
600
800
1000
1200
124
219
O8
L16
L8
217
9
253
233
L1
256
204
O1
218
97

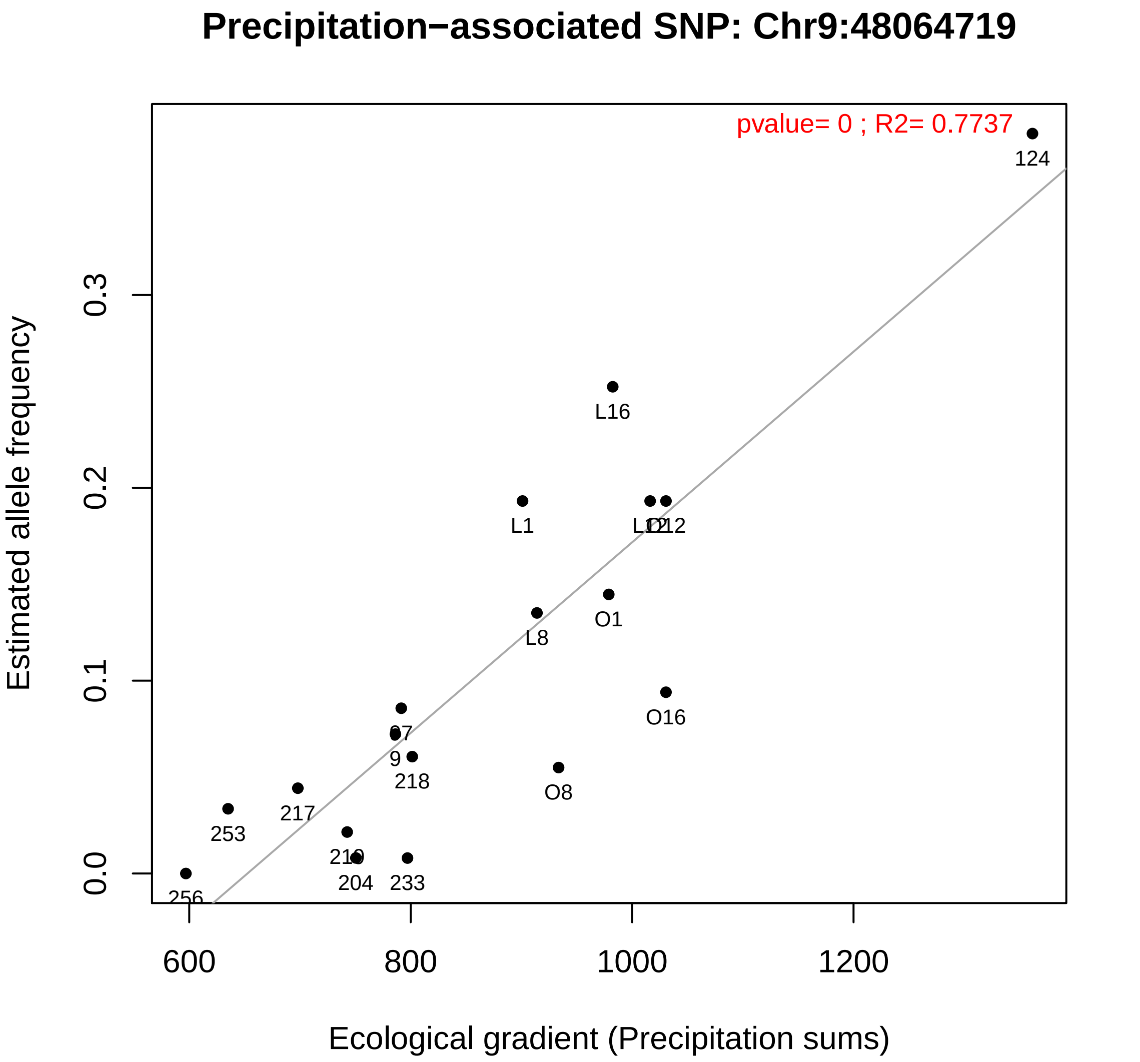
Precipitation−associated SNP: Chr9:48064719
pvalue= 0 ; R2= 0.7737
124
L16
L1
O12
O1
L8
O16
218
O8
217
253
204
233
256
0.0
0.1
0.2
0.3
600
800
1000
1200
Estimated allele frequency
Ecological gradient (Precipitation sums)

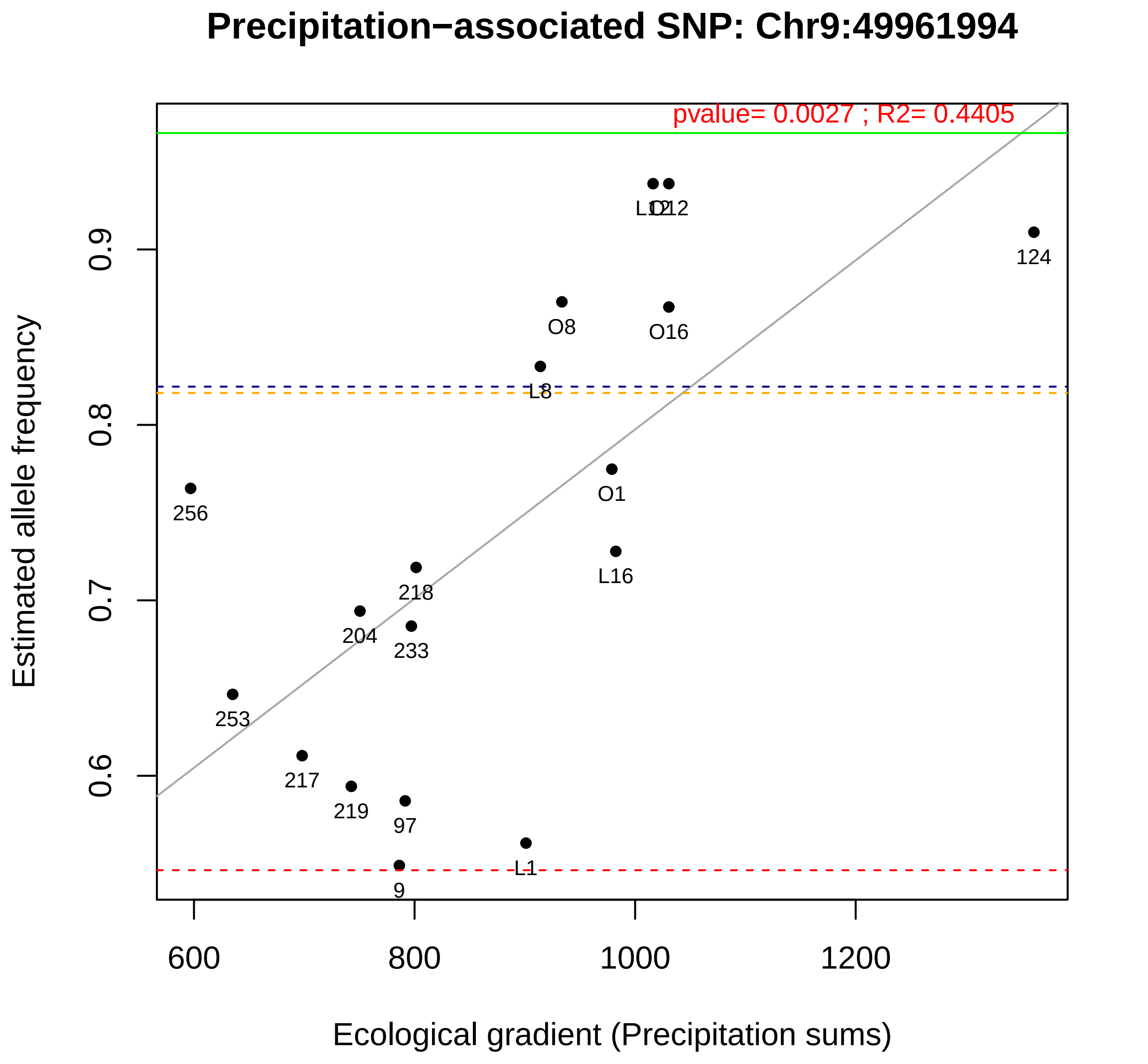
Precipitation−associated SNP: Chr9:49961994
pvalue= 0.0027 ; R2= 0.4405
Estimated allele frequency
0.6
0.7
0.8
0.9
Ecological gradient (Precipitation sums)
600
800
1000
1200
124
O8
O16
L8
256
O1
L16
218
204
233
253
217
219
97
L1
9

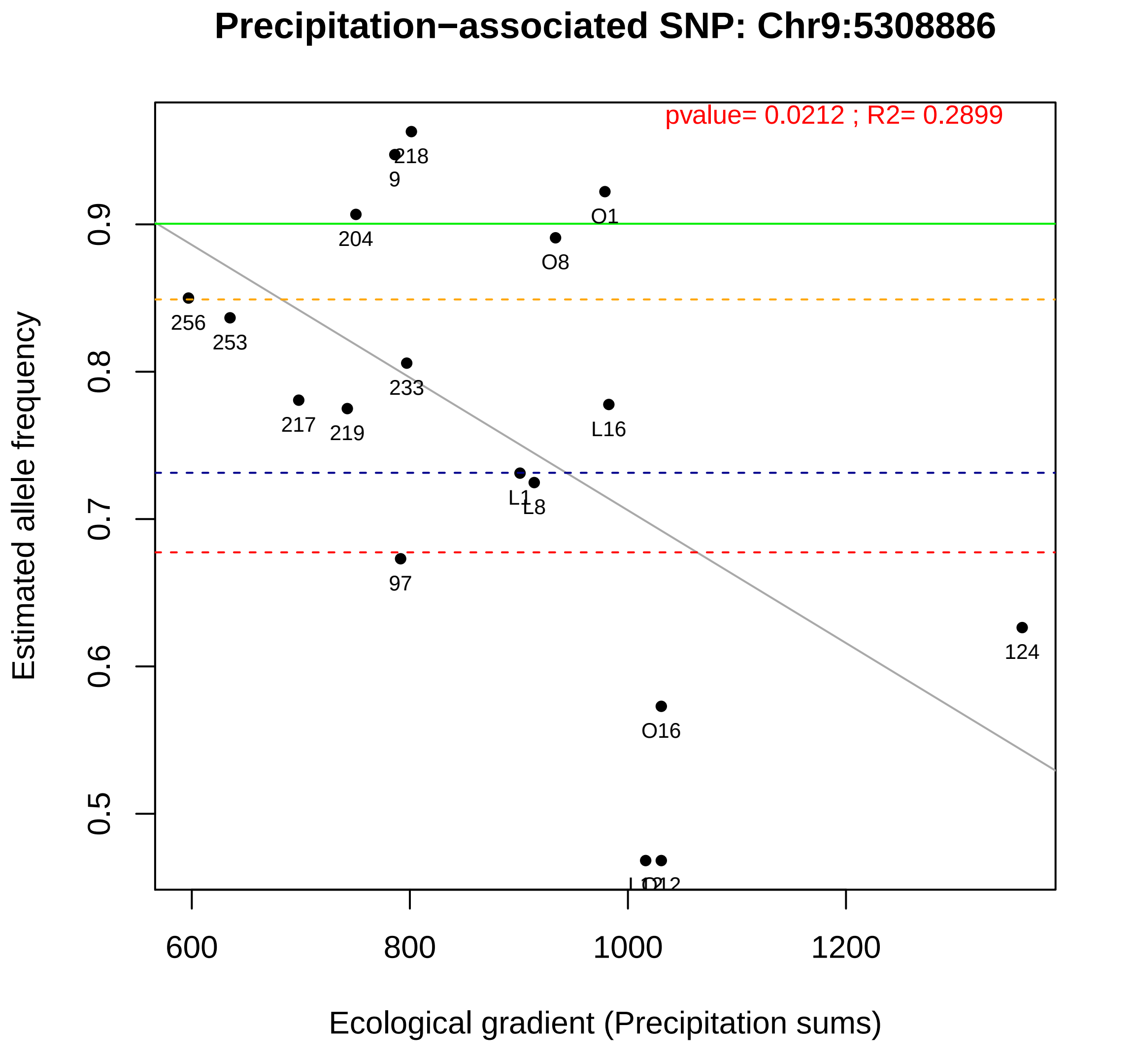

Precipitation−associated SNP: Chr9:5308886
pvalue= 0.0212 ; R2= 0.2899
Estimated allele frequency
Ecological gradient (Precipitation sums)
0.5
0.6
0.7
0.8
0.9
600
800
1000
1200
218
9
O1
204
O8
256
253
233
217
219
L16
L1
L8
97
124
O16
O12

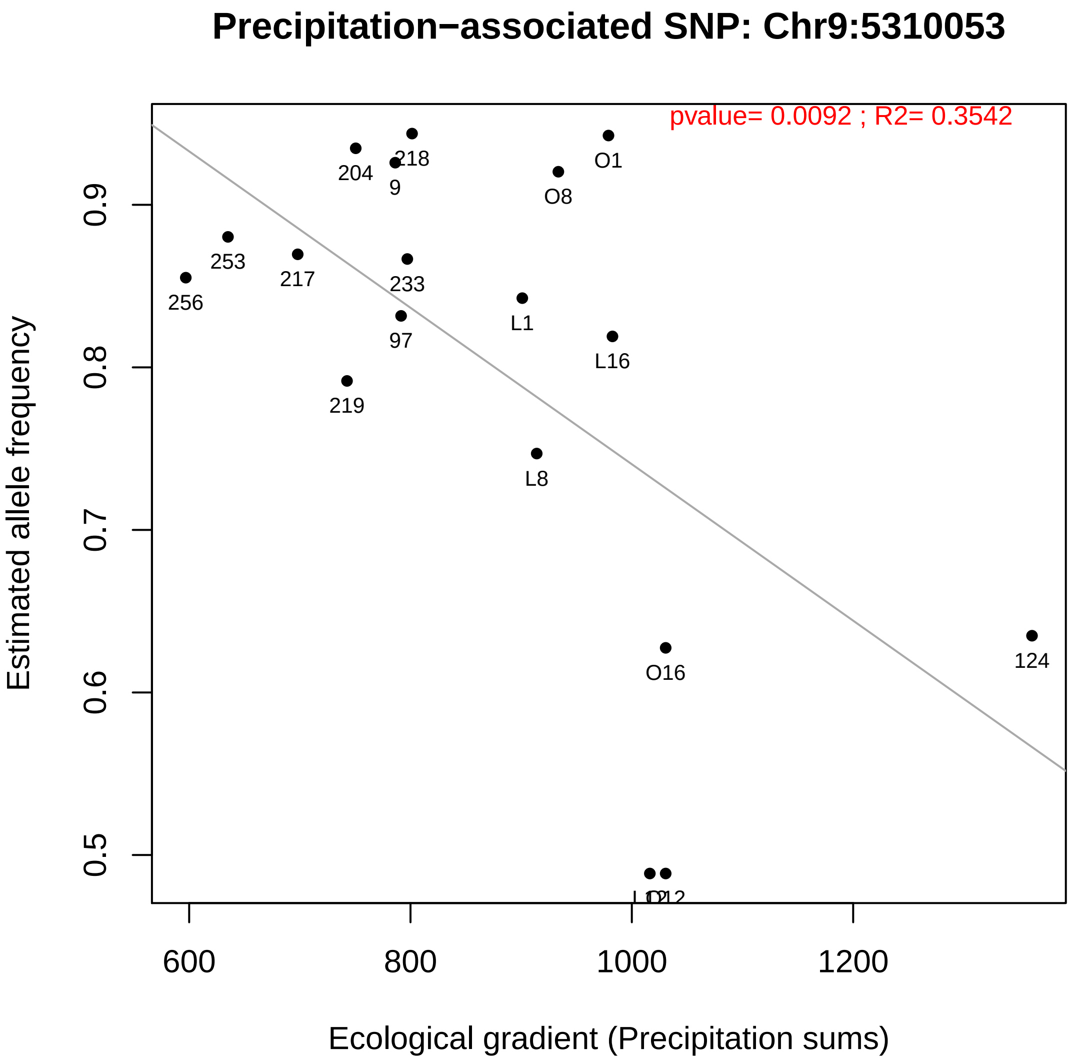
Precipitation−associated SNP: Chr9:5310053
pvalue= 0.0092 ; R2= 0.3542
Estimated allele frequency
0.5
0.6
0.7
0.8
0.9
600
800
1000
1200
Ecological gradient (Precipitation sums)
204
218
9
O1
O8
253
217
233
256
L1
97
L16
219
L8
O16
124

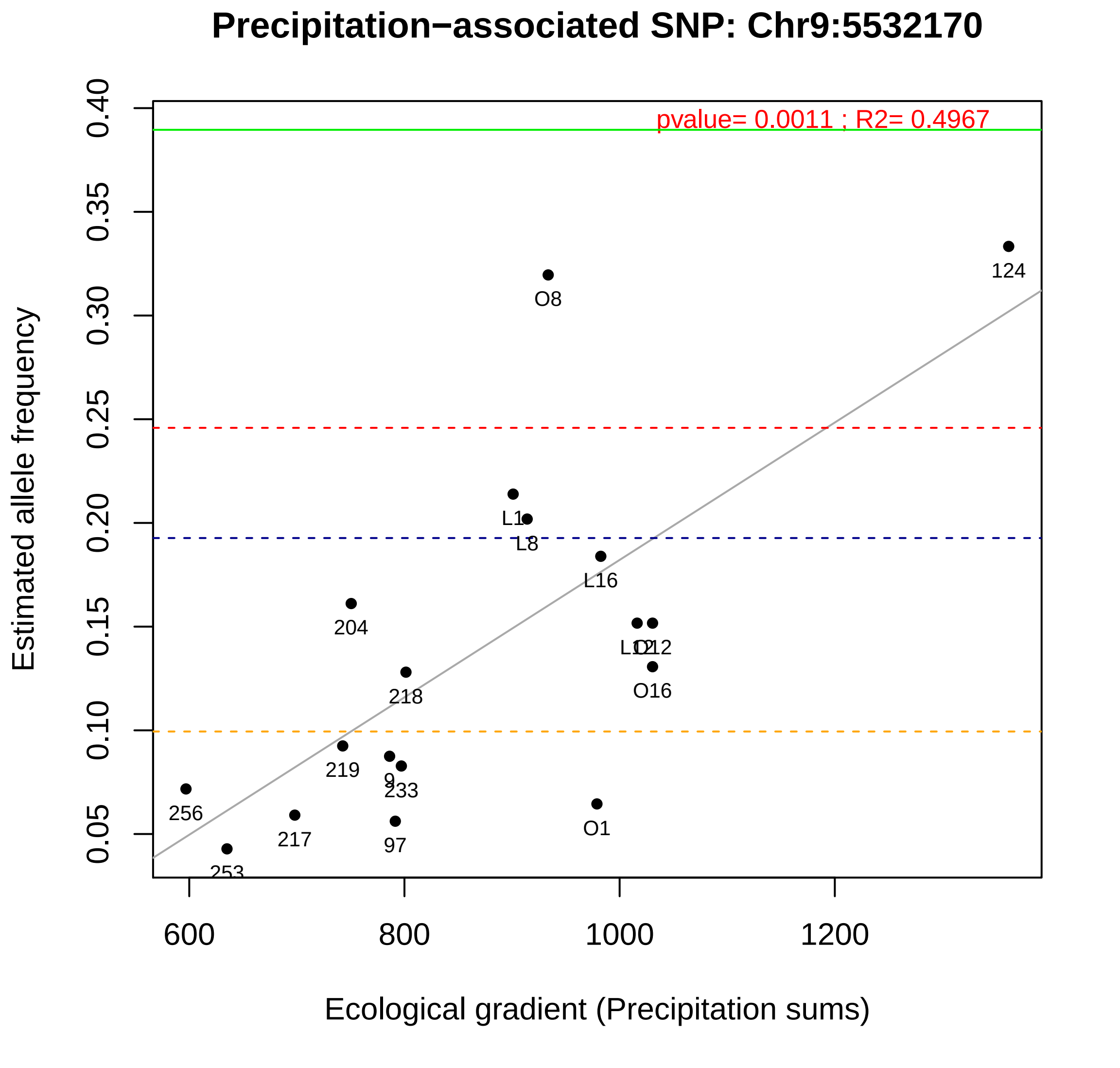

Precipitation−associated SNP: Chr9:5532170
pvalue= 0.0011 ; R2= 0.4967
Estimated allele frequency
0.05
0.10
0.15
0.20
0.25
0.30
0.35
0.40
600
800
1000
1200
Ecological gradient (Precipitation sums)
124
O8
L1
L8
L16
204
L12
O12
O16
218
219
9
233
256
217
97
O1
253

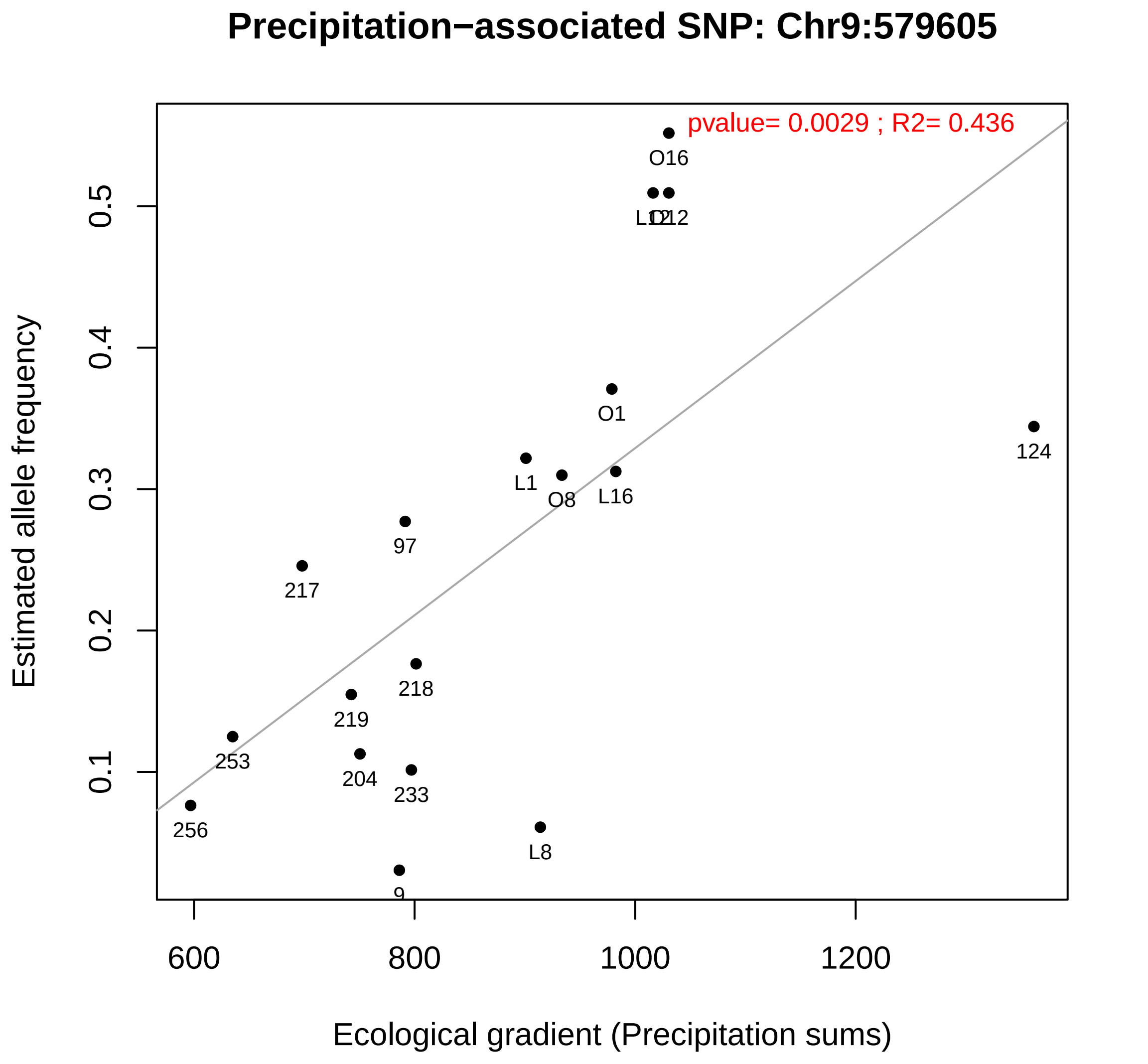

Precipitation−associated SNP: Chr9:579605
pvalue= 0.0029 ; R2= 0.436
Estimated allele frequency
0.1
0.2
0.3
0.4
0.5
600
800
1000
1200
Ecological gradient (Precipitation sums)
O16
L12
O12
O1
124
L1
O8
L16
97
217
218
219
253
204
233
256
L8
9

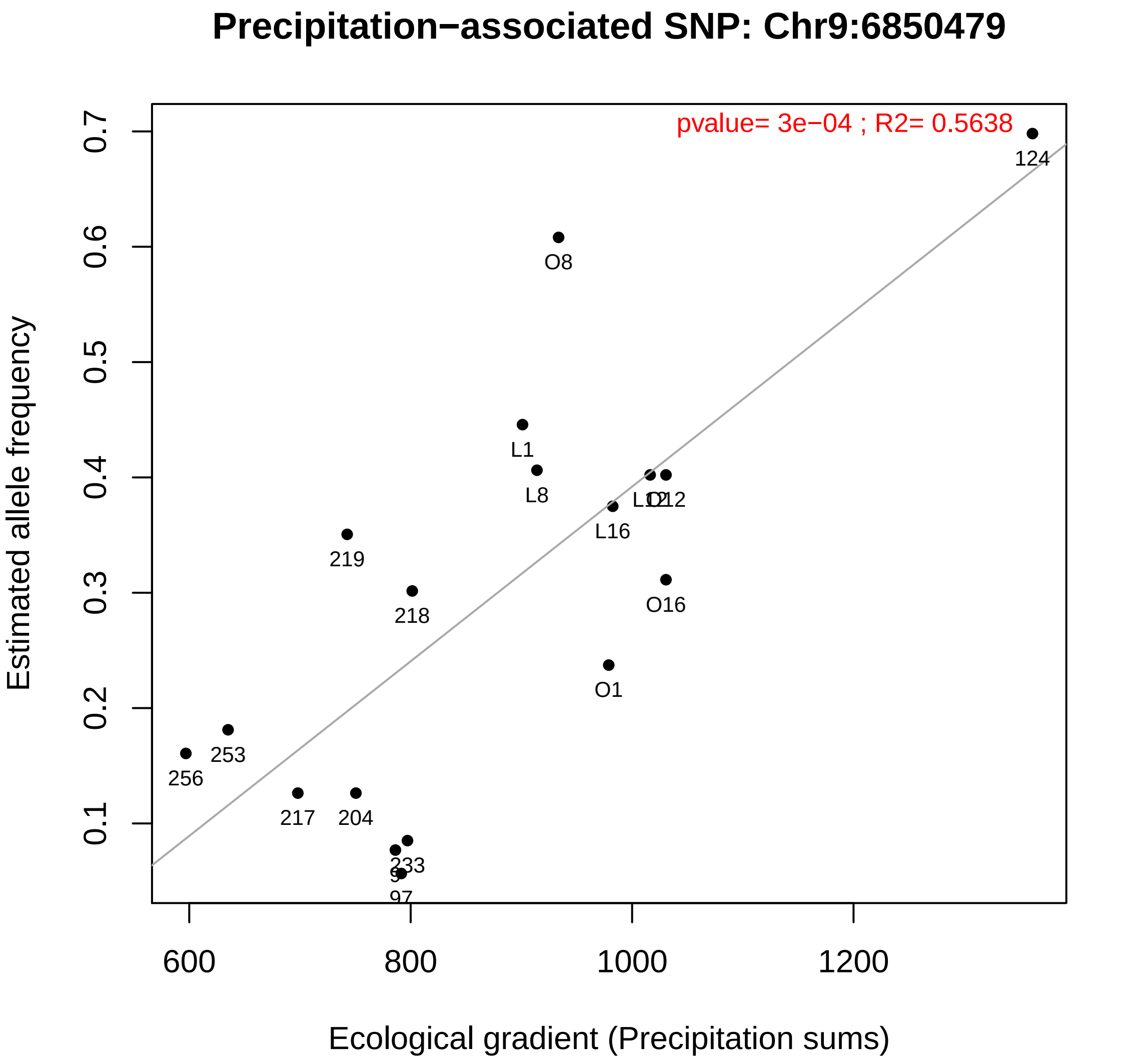
Precipitation−associated SNP: Chr9:6850479
pvalue= 3e−04 ; R2= 0.5638
Estimated allele frequency
Ecological gradient (Precipitation sums)
0.1
0.2
0.3
0.4
0.5
0.6
0.7
600
800
1000
1200
124
O8
L1
L8
L12
O12
L16
219
218
O16
O1
253
256
217
204
233
97

**Precipitation−associated SNP: Chr9:6954971**

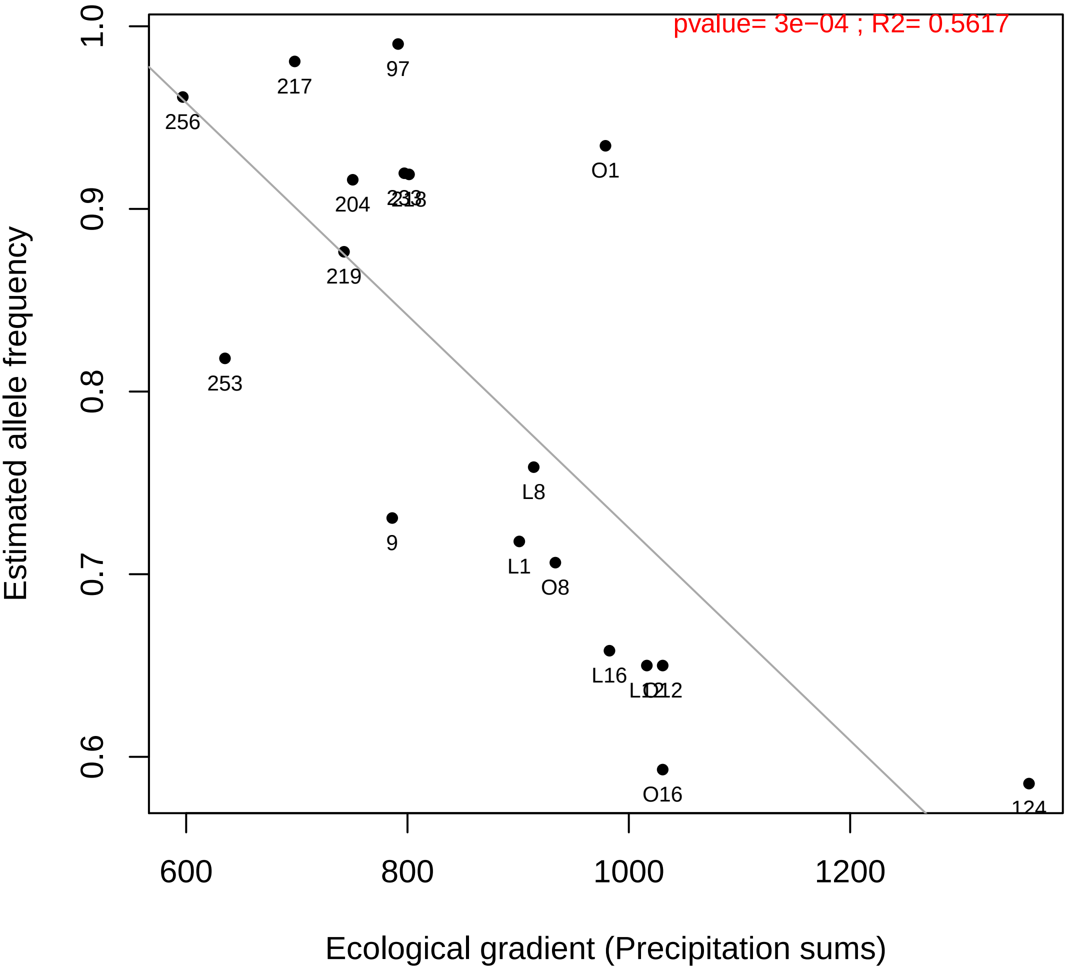

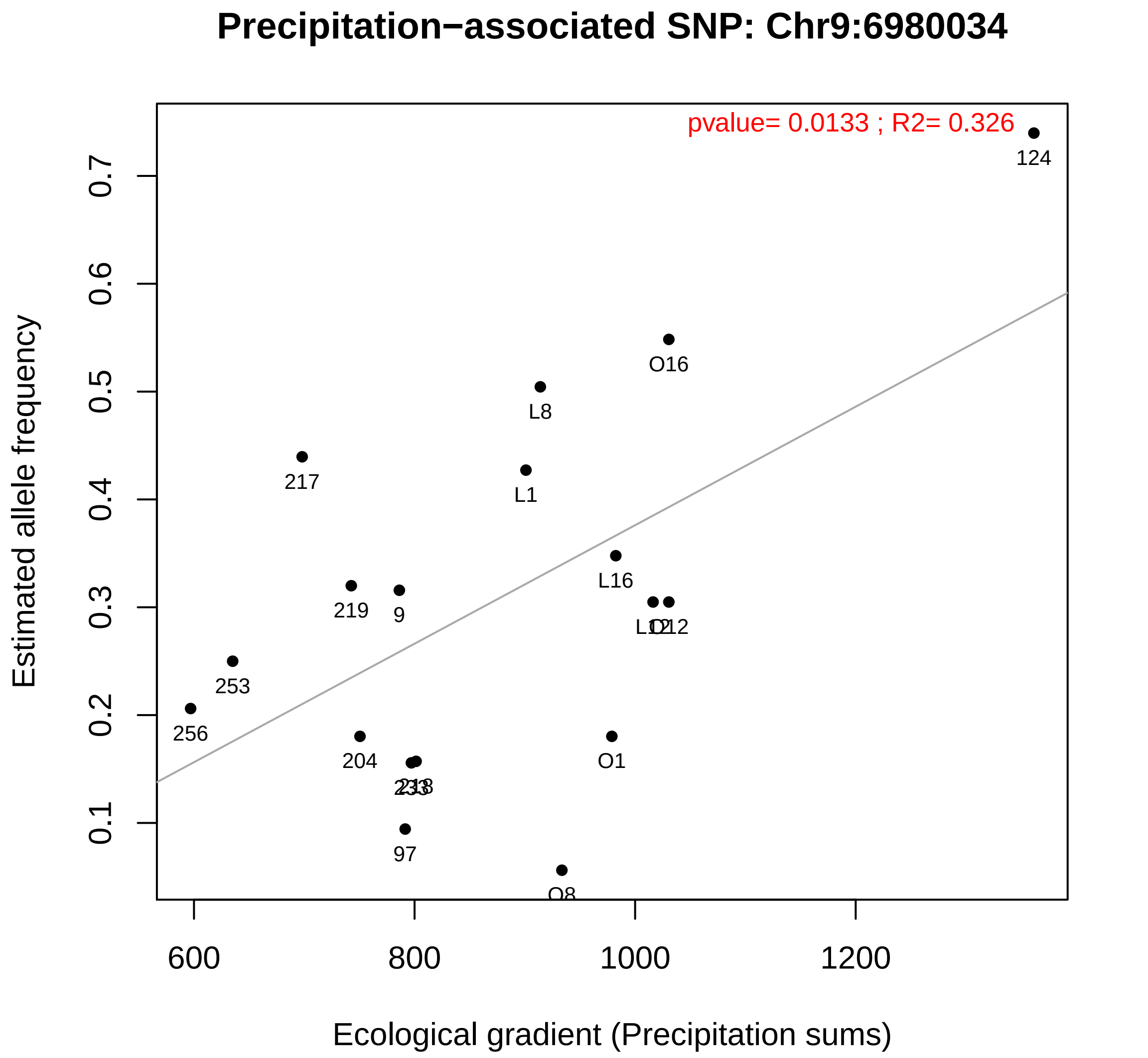

Precipitation−associated SNP: Chr9:6980034
pvalue= 0.0133 ; R2= 0.326
Estimated allele frequency
0.1
0.2
0.3
0.4
0.5
0.6
0.7
600
800
1000
1200
Ecological gradient (Precipitation sums)
124
O16
L8
217
L1
L16
219
9
253
256
204
O1
97

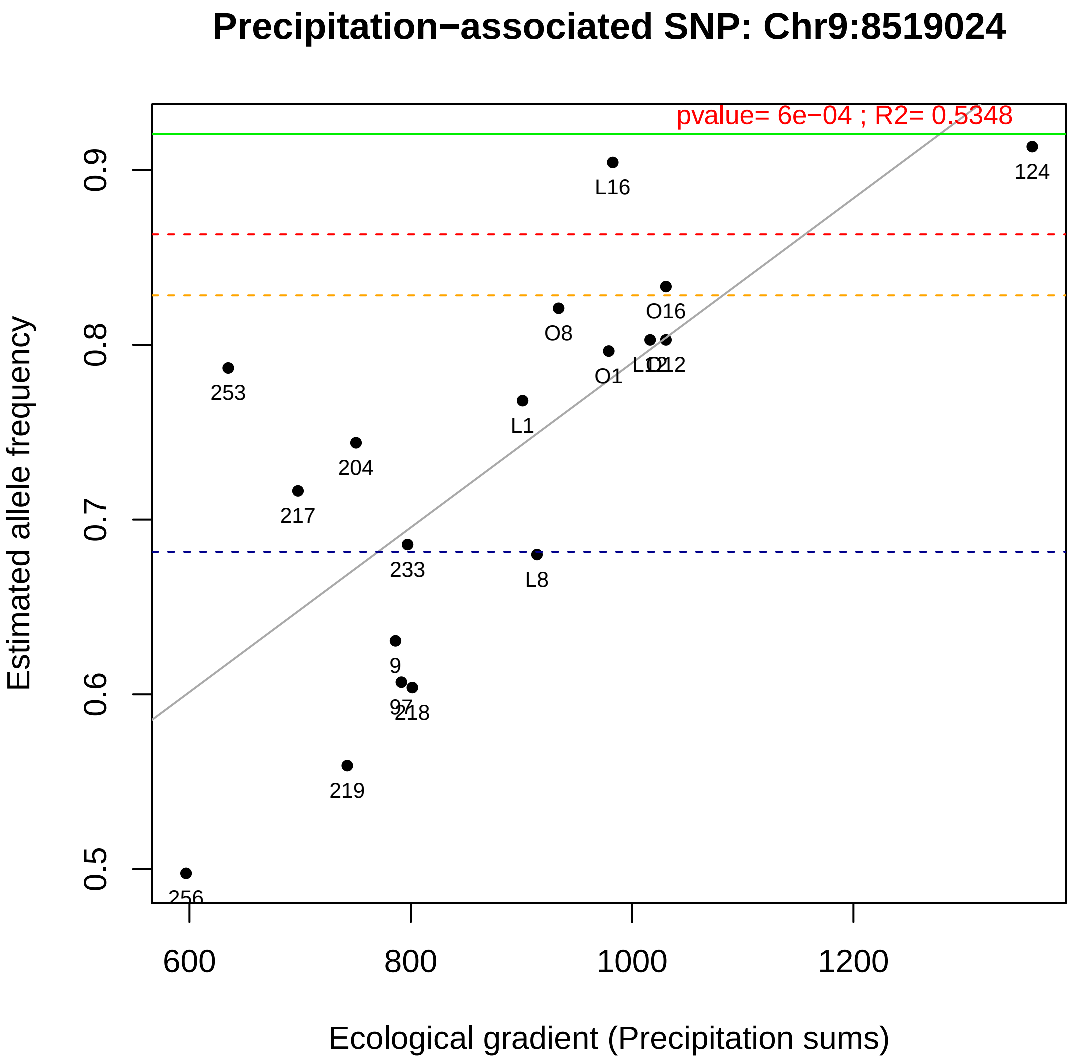

Precipitation−associated SNP: Chr9:8519024
pvalue= 6e−04 ; R2= 0.5348
Estimated allele frequency
Ecological gradient (Precipitation sums)
0.5
0.6
0.7
0.8
0.9
600
800
1000
1200
124
L16
O16
O8
L12
O12
O1
L1
253
204
217
233
L8
9
97
218
219
256

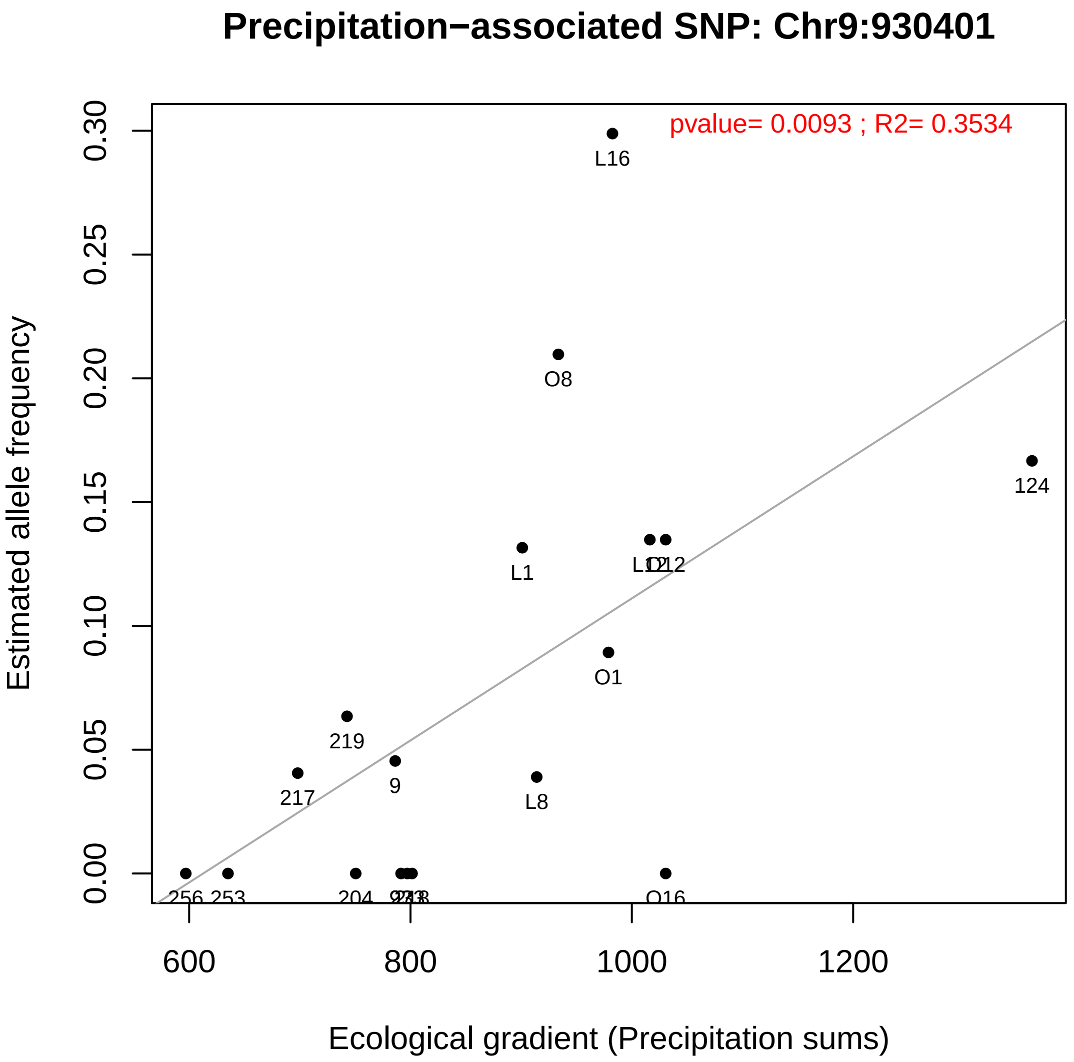

Precipitation−associated SNP: Chr9:930401
pvalue= 0.0093 ; R2= 0.3534
Estimated allele frequency
Ecological gradient (Precipitation sums)
0.00
0.05
0.10
0.15
0.20
0.25
0.30
600
800
1000
1200
L16
O8
124
L1
O1
219
217
9
L8
256
253
204
O16

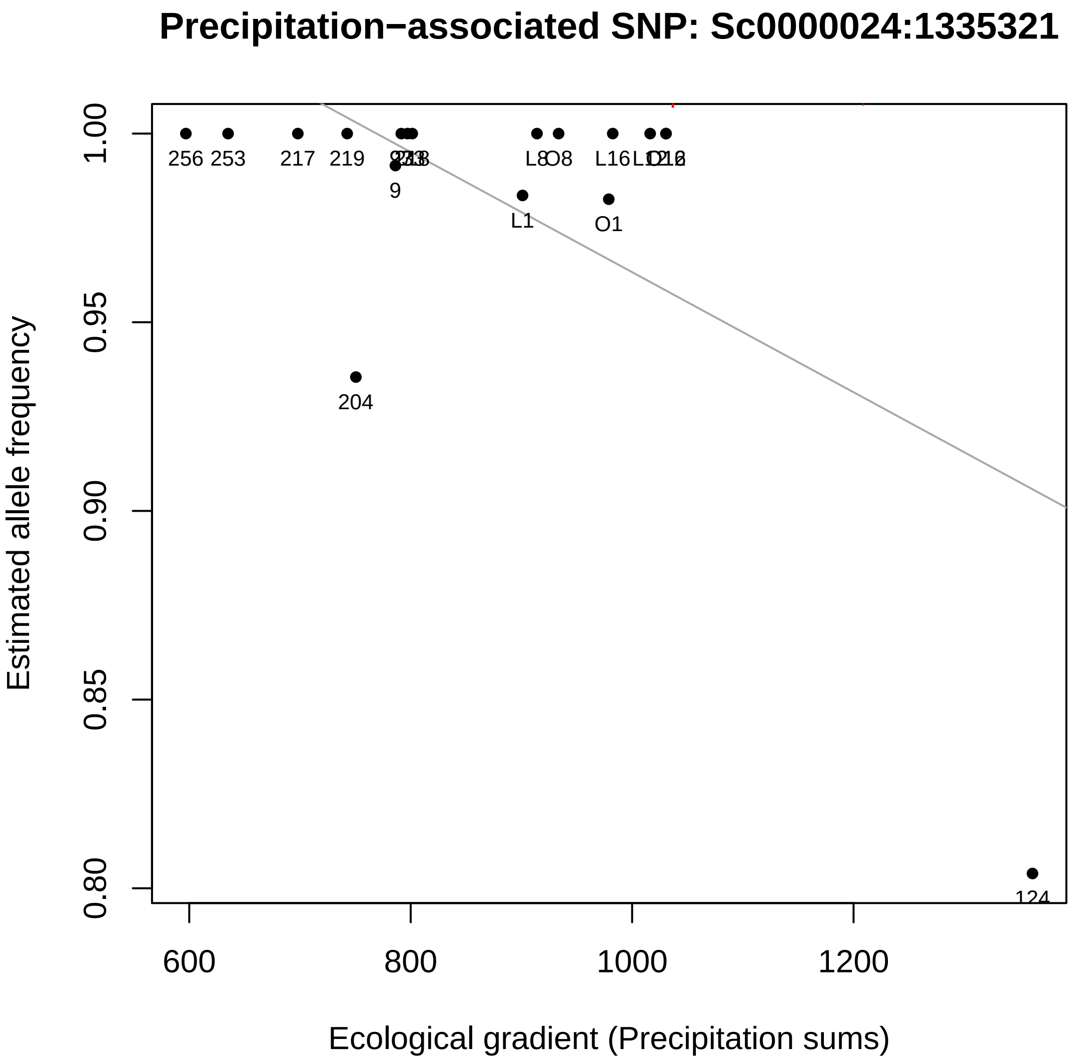

Precipitation−associated SNP: Sc0000024:1335321
Estimated allele frequency
Ecological gradient (Precipitation sums)
0.80
0.85
0.90
0.95
1.00
600
800
1000
1200
256
253
217
219
9
204
L1
L8
O8
L16
O1
124

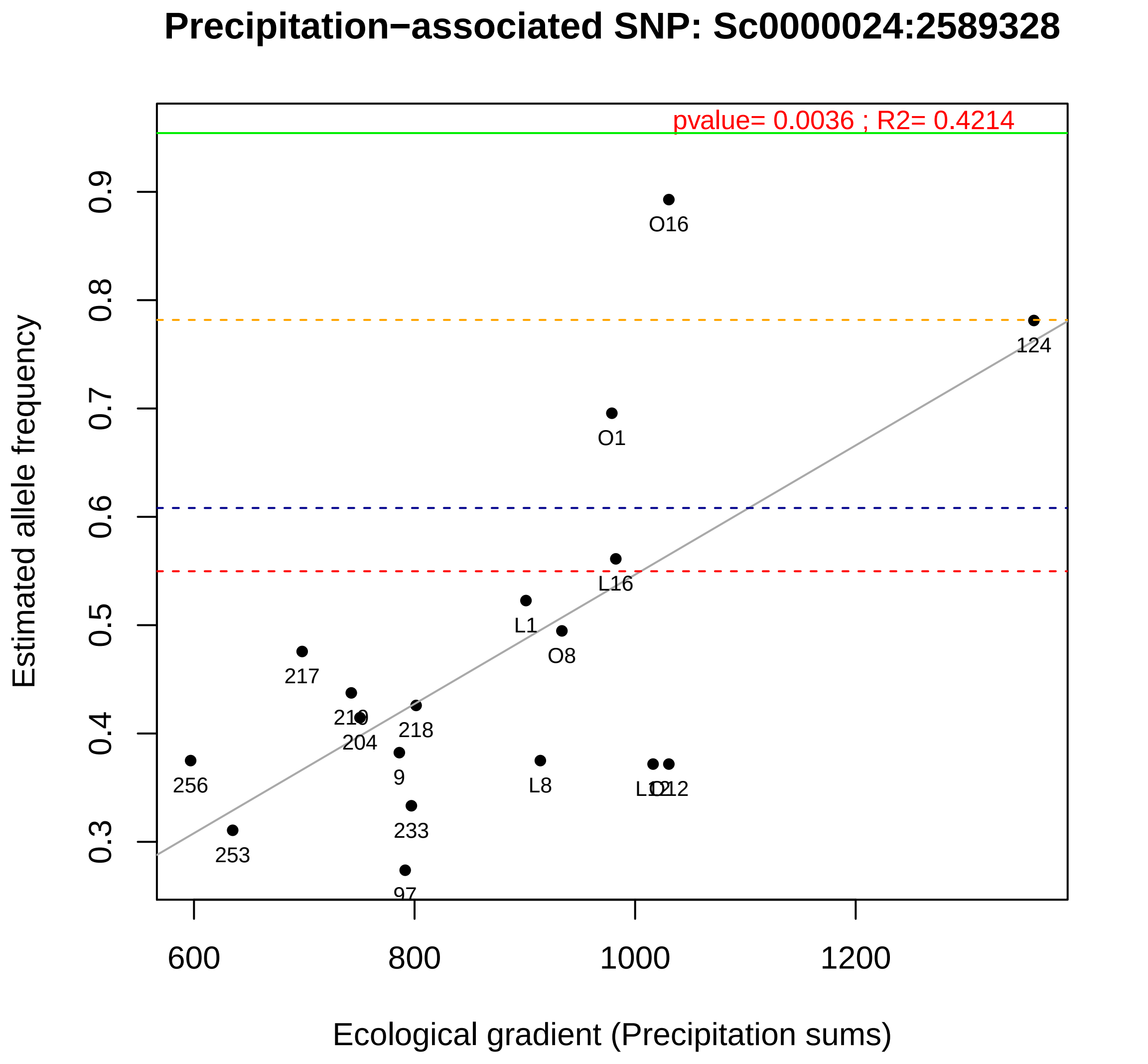

Precipitation−associated SNP: Sc0000024:2589328
pvalue= 0.0036 ; R2= 0.4214
Estimated allele frequency
0.3
0.4
0.5
0.6
0.7
0.8
0.9
600
800
1000
1200
Ecological gradient (Precipitation sums)
O16
124
O1
L16
L1
O8
217
210
204
218
256
9
L8
L12
O12
233
253
97

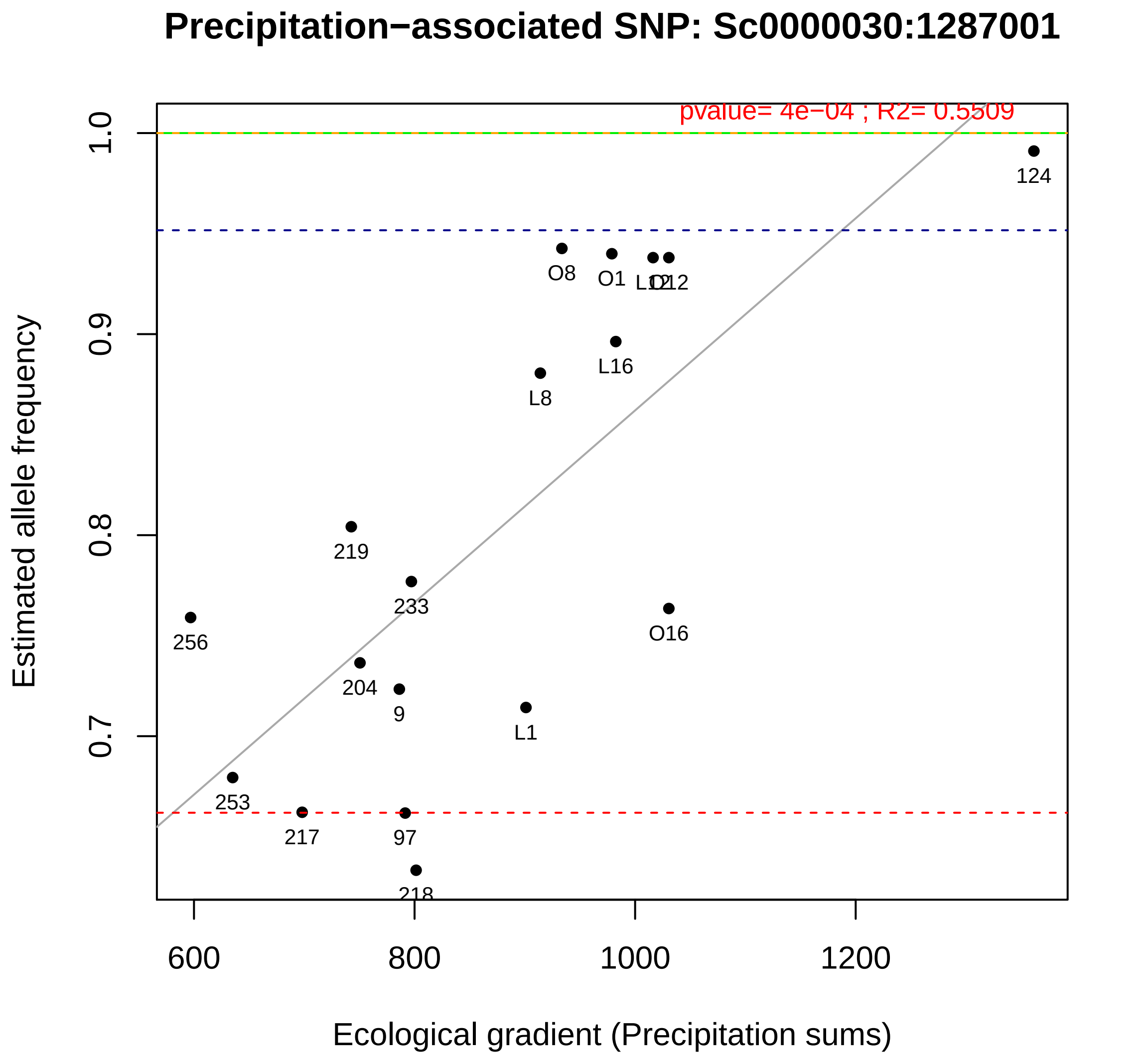
Precipitation−associated SNP: Sc0000030:1287001
pvalue= 4e−04 ; R2= 0.5509
Estimated allele frequency
Ecological gradient (Precipitation sums)
0.7
0.8
0.9
1.0
600
800
1000
1200
124
O8
O1
L12
O12
L16
L8
219
233
O16
256
204
9
L1
253
217
97
218

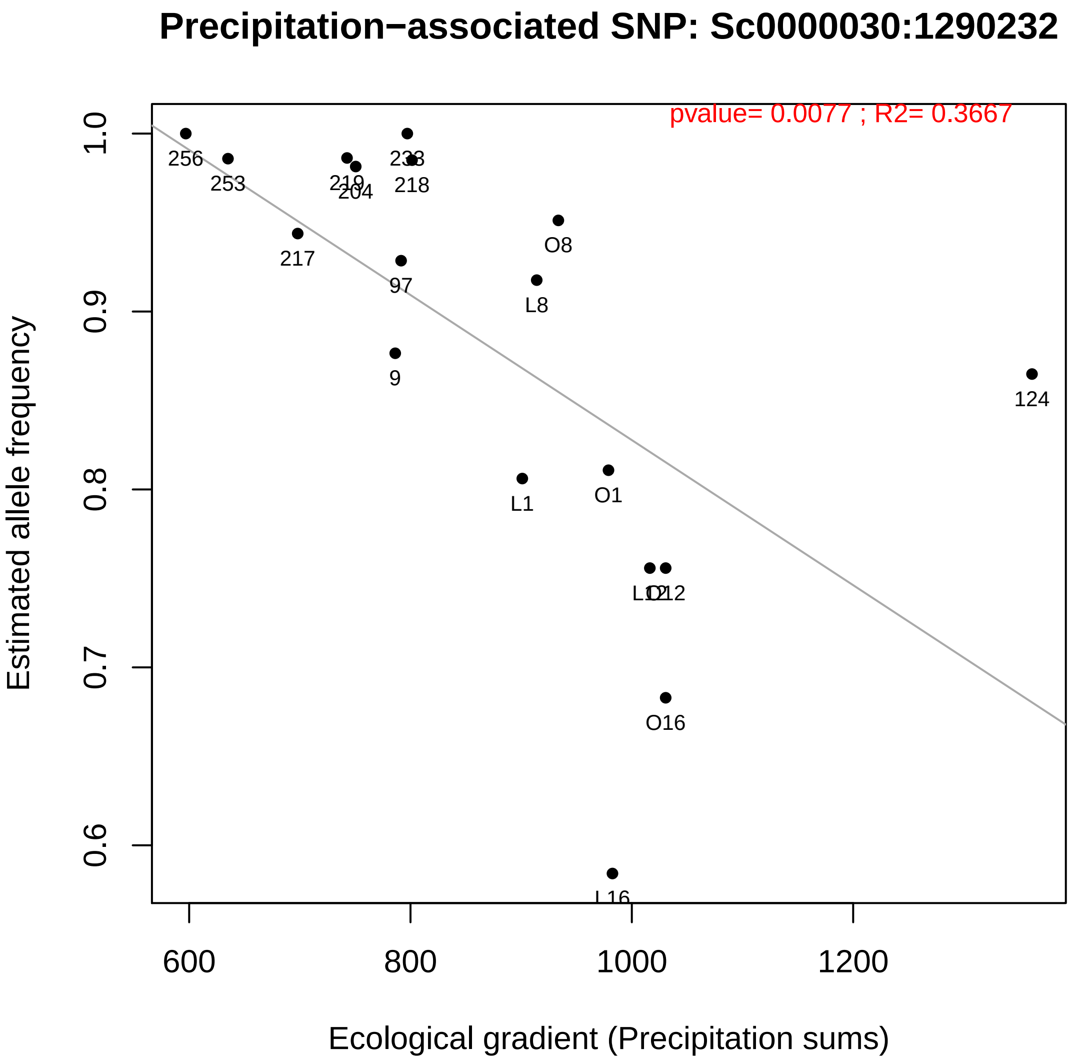

Precipitation−associated SNP: Sc0000030:1290232
pvalue= 0.0077 ; R2= 0.3667
Estimated allele frequency
1.0
0.9
0.8
0.7
0.6
600
800
1000
1200
Ecological gradient (Precipitation sums)
256
253
219
204
233
218
217
97
9
O8
L8
L1
O1
L12
O12
O16
L16
124

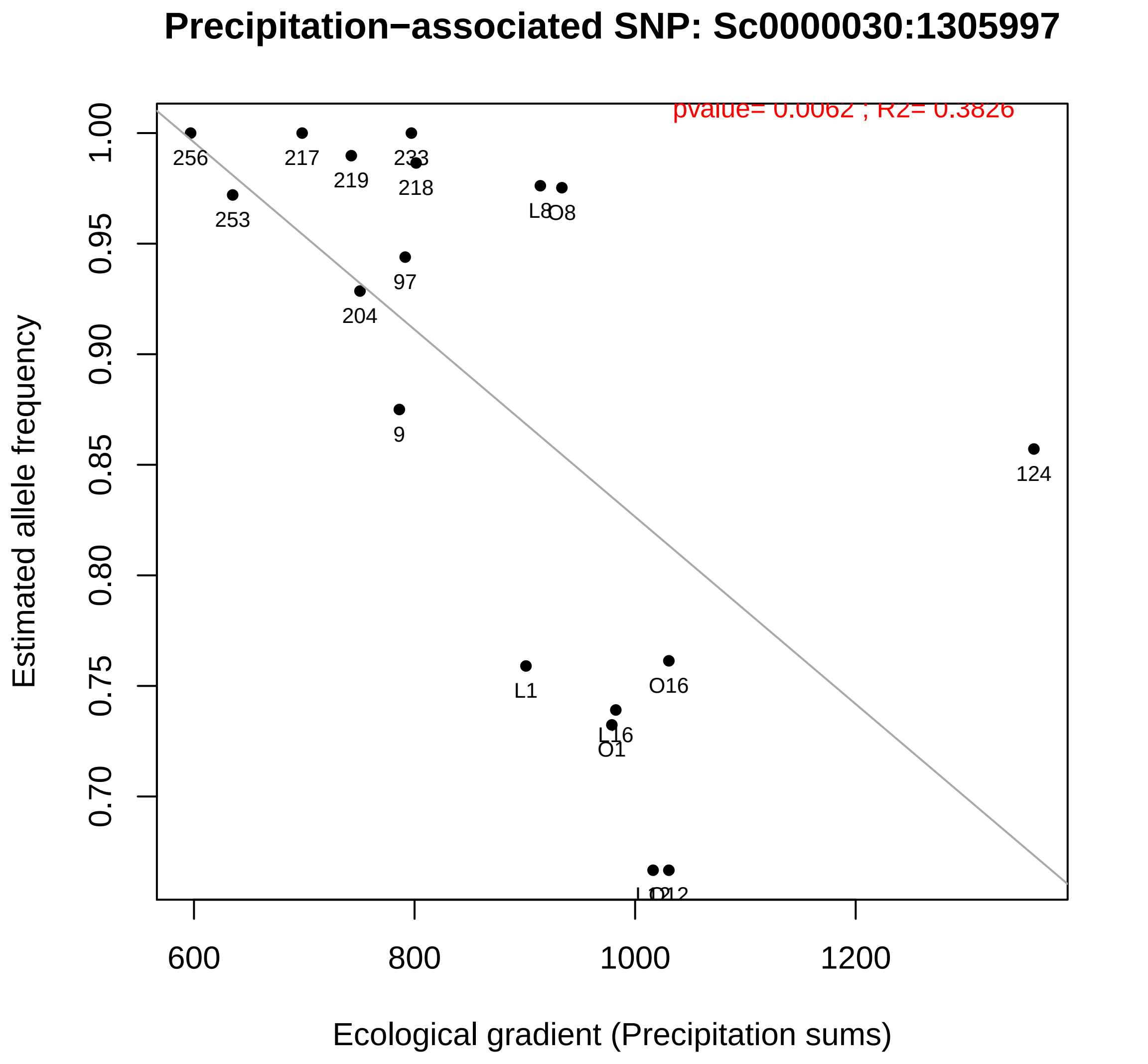

Precipitation−associated SNP: Sc0000030:1305997
pvalue= 0.0062 ; R2= 0.3826
Estimated allele frequency
Ecological gradient (Precipitation sums)
1.00
0.95
0.90
0.85
0.80
0.75
0.70
600
800
1000
1200
256
217
233
219
218
253
L8
O8
97
204
9
124
L1
O16
L16
O1

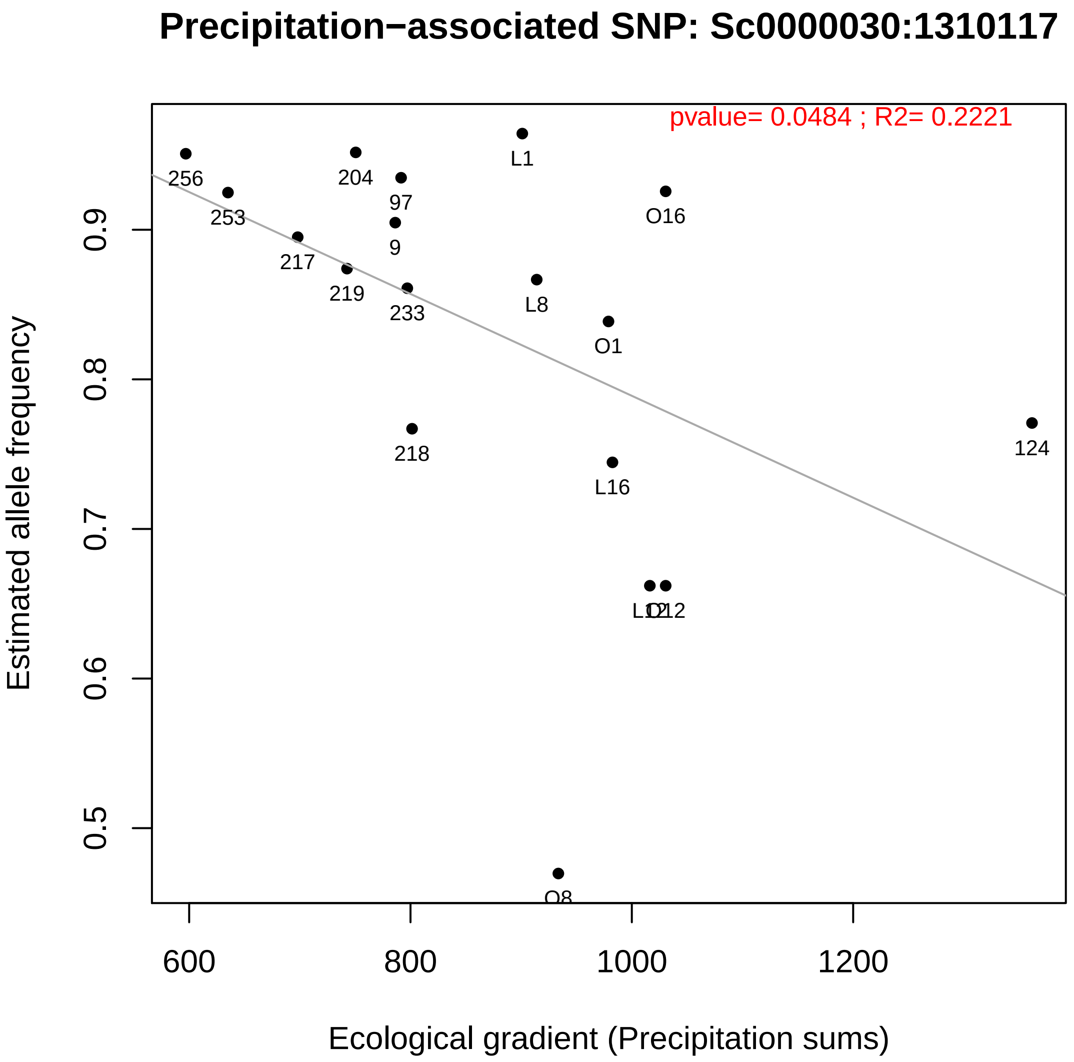

Precipitation−associated SNP: Sc0000030:1310117
pvalue= 0.0484 ; R2= 0.2221
256
253
217
219
204
97
9
233
218
L1
L8
O16
O1
L16
L12
O12
O8
124
0.5
0.6
0.7
0.8
0.9
600
800
1000
1200
Estimated allele frequency
Ecological gradient (Precipitation sums)

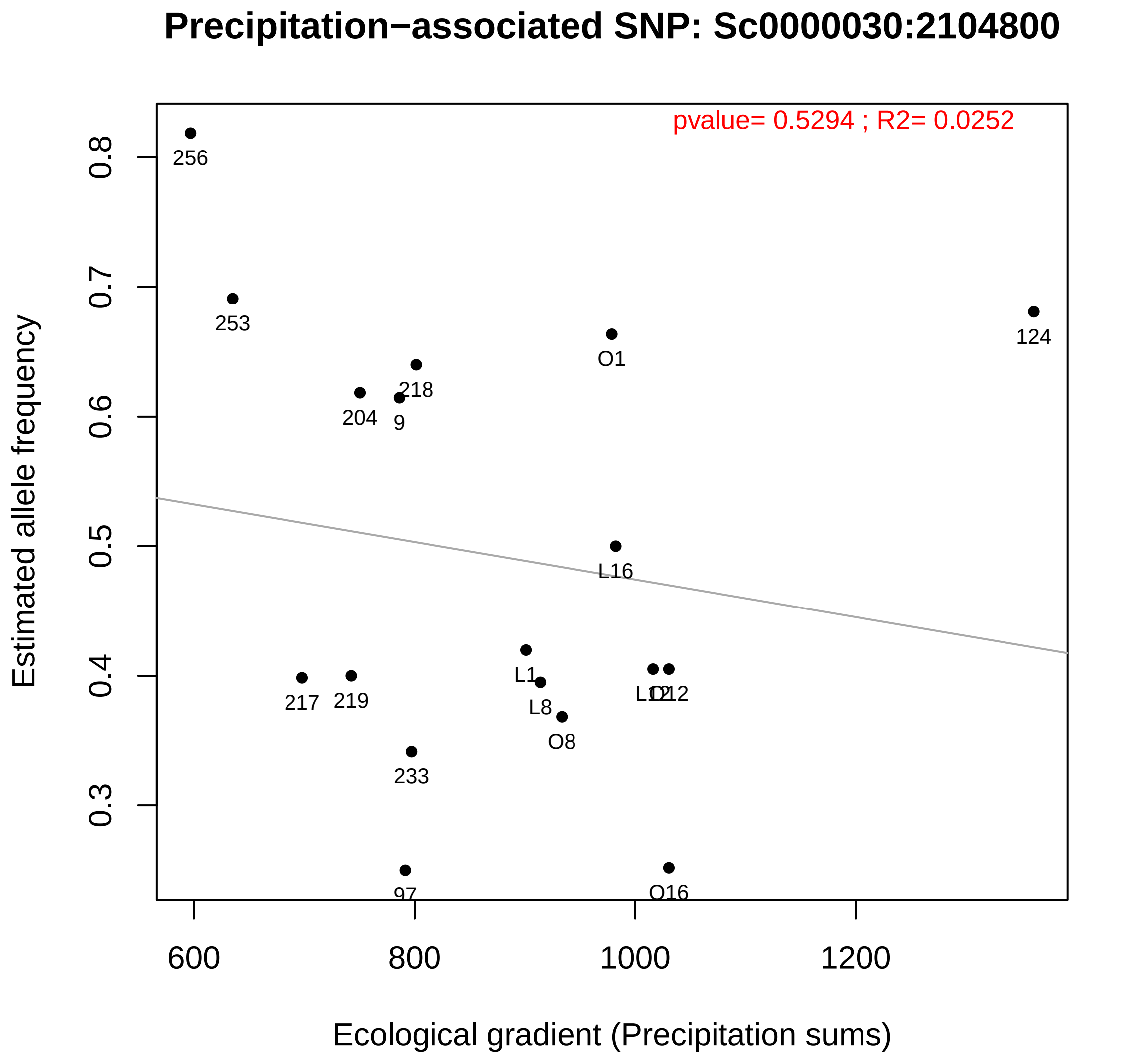

Precipitation−associated SNP: Sc0000030:2104800
pvalue= 0.5294 ; R2= 0.0252
Estimated allele frequency
0.3
0.4
0.5
0.6
0.7
0.8
600
800
1000
1200
Ecological gradient (Precipitation sums)
256
253
218
204
9
O1
124
L16
217
219
L1
L8
O8
233
97
O16

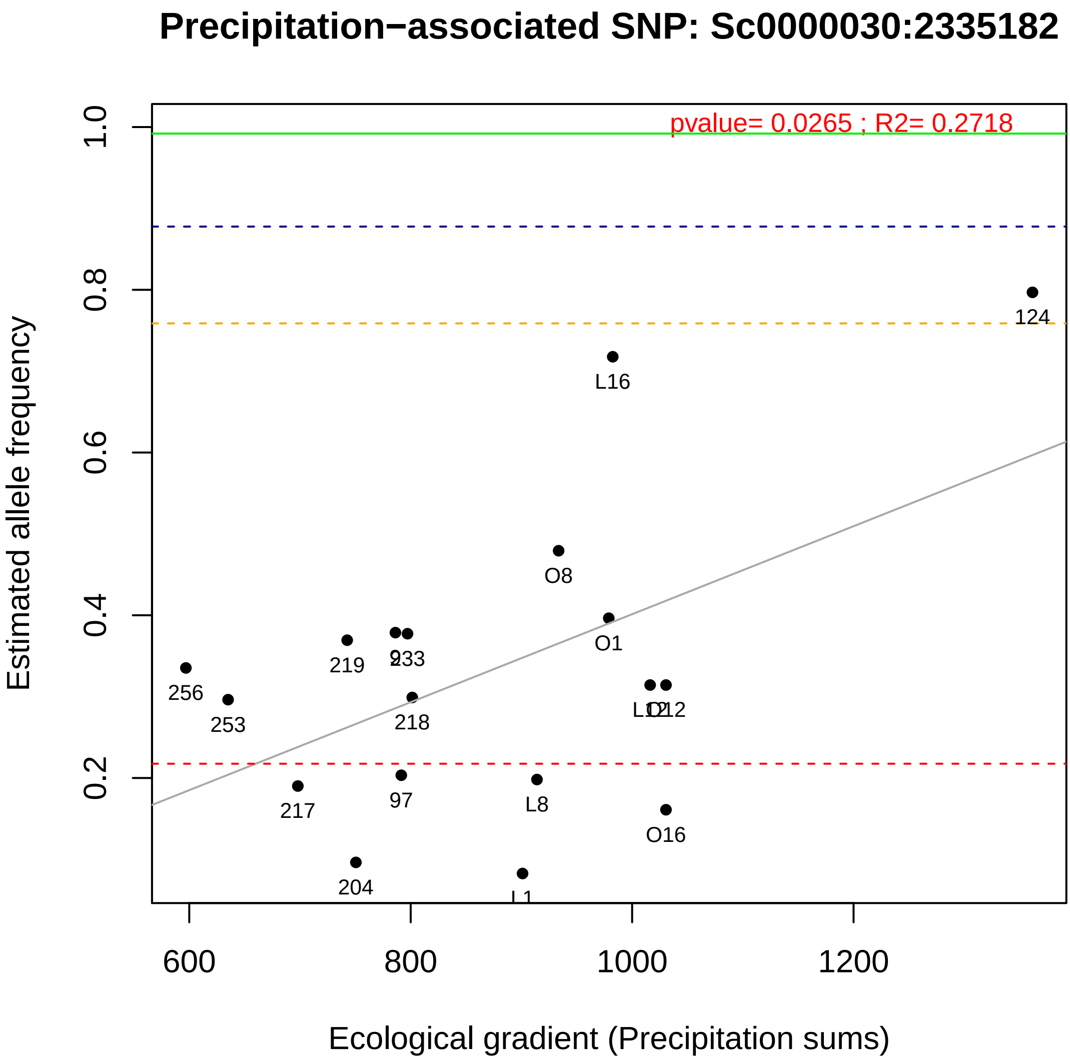

Precipitation−associated SNP: Sc0000030:2335182
pvalue= 0.0265 ; R2= 0.2718
Estimated allele frequency
Ecological gradient (Precipitation sums)
0.2
0.4
0.6
0.8
1.0
600
800
1000
1200
124
L16
O8
O1
219
233
256
253
218
217
97
L8
O16
204

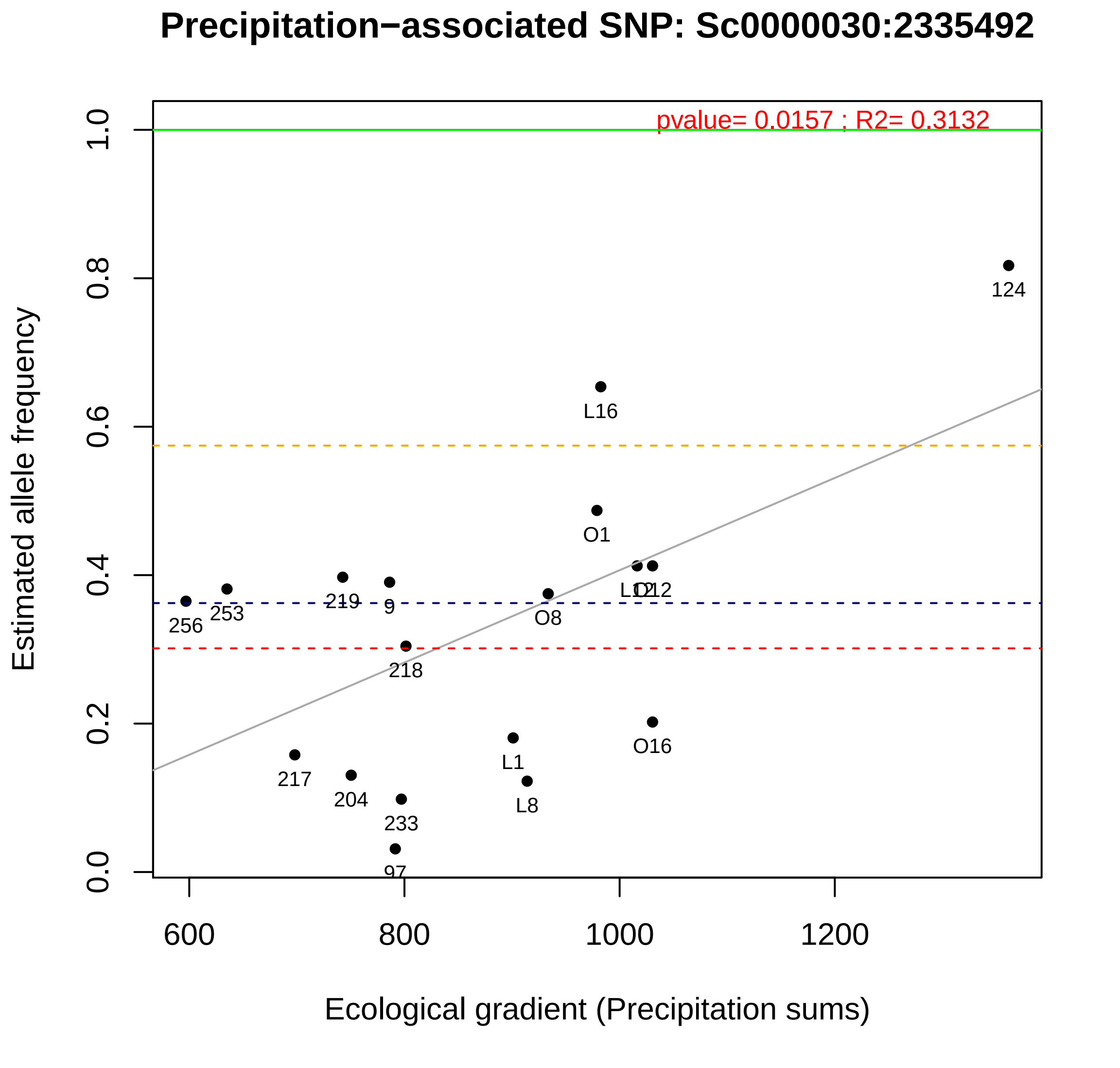

Precipitation−associated SNP: Sc0000030:2335492
pvalue= 0.0157 ; R2= 0.3132
Estimated allele frequency
0.0
0.2
0.4
0.6
0.8
1.0
600
800
1000
1200
Ecological gradient (Precipitation sums)
124
L16
O1
L10
O12
O8
219
9
253
256
218
O16
L1
L8
217
204
233
97

#### Precipitation−associated SNP: Sc0000030:2361805

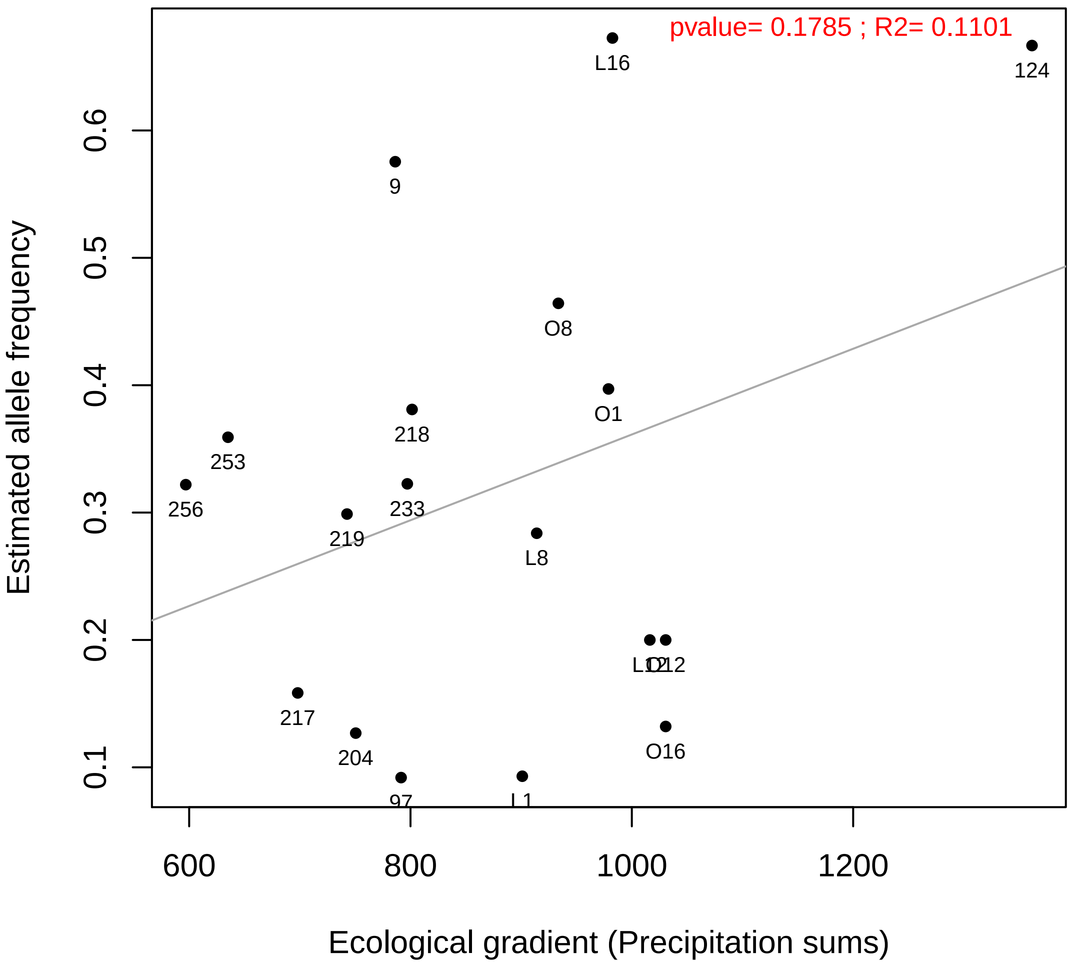

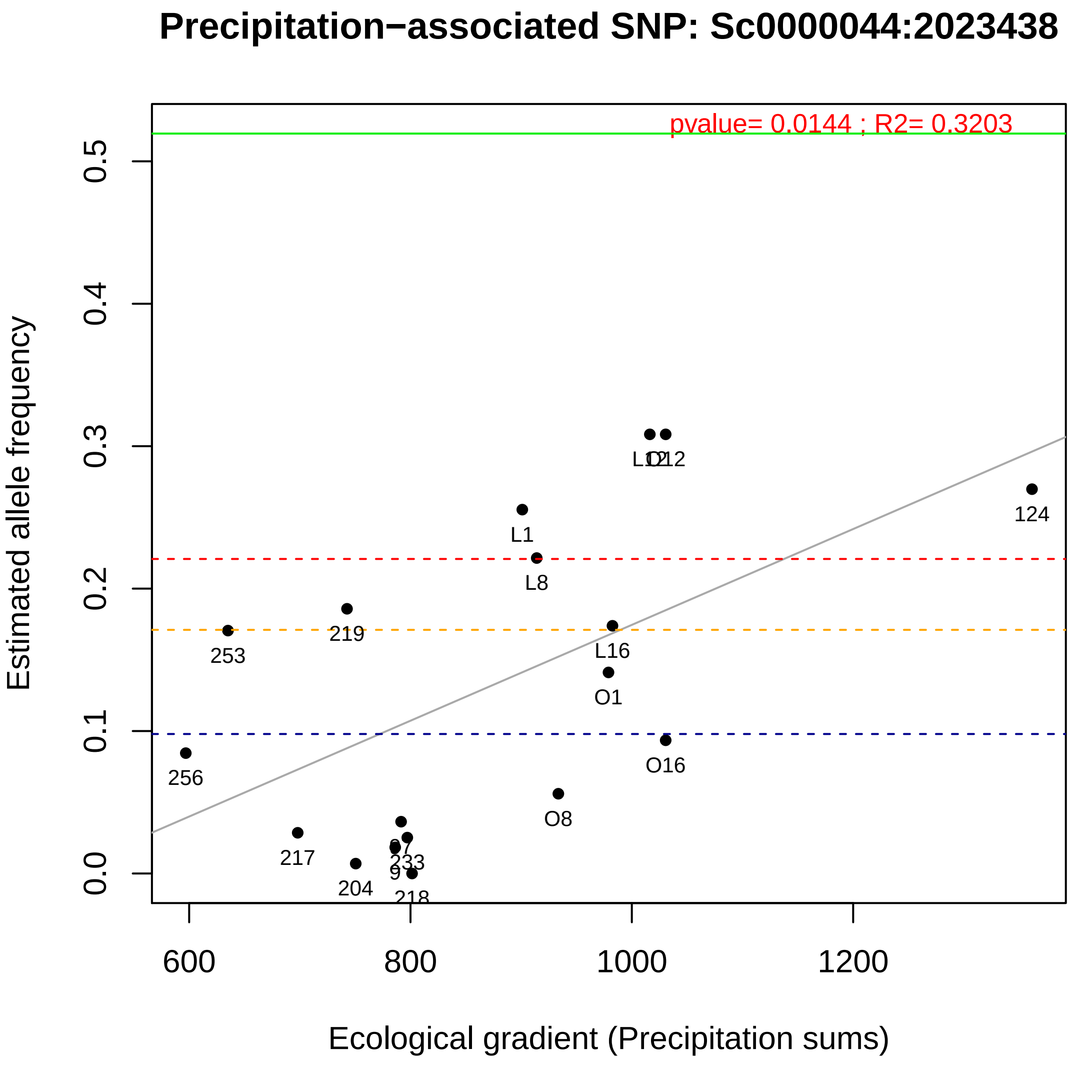
Precipitation−associated SNP: Sc0000044:2023438
pvalue= 0.0144 ; R2= 0.3203
Estimated allele frequency
0.0
0.1
0.2
0.3
0.4
0.5
Ecological gradient (Precipitation sums)
600
800
1000
1200
L10
O12
124
L1
L8
219
253
L16
O1
O16
256
O8
217
204
233
218

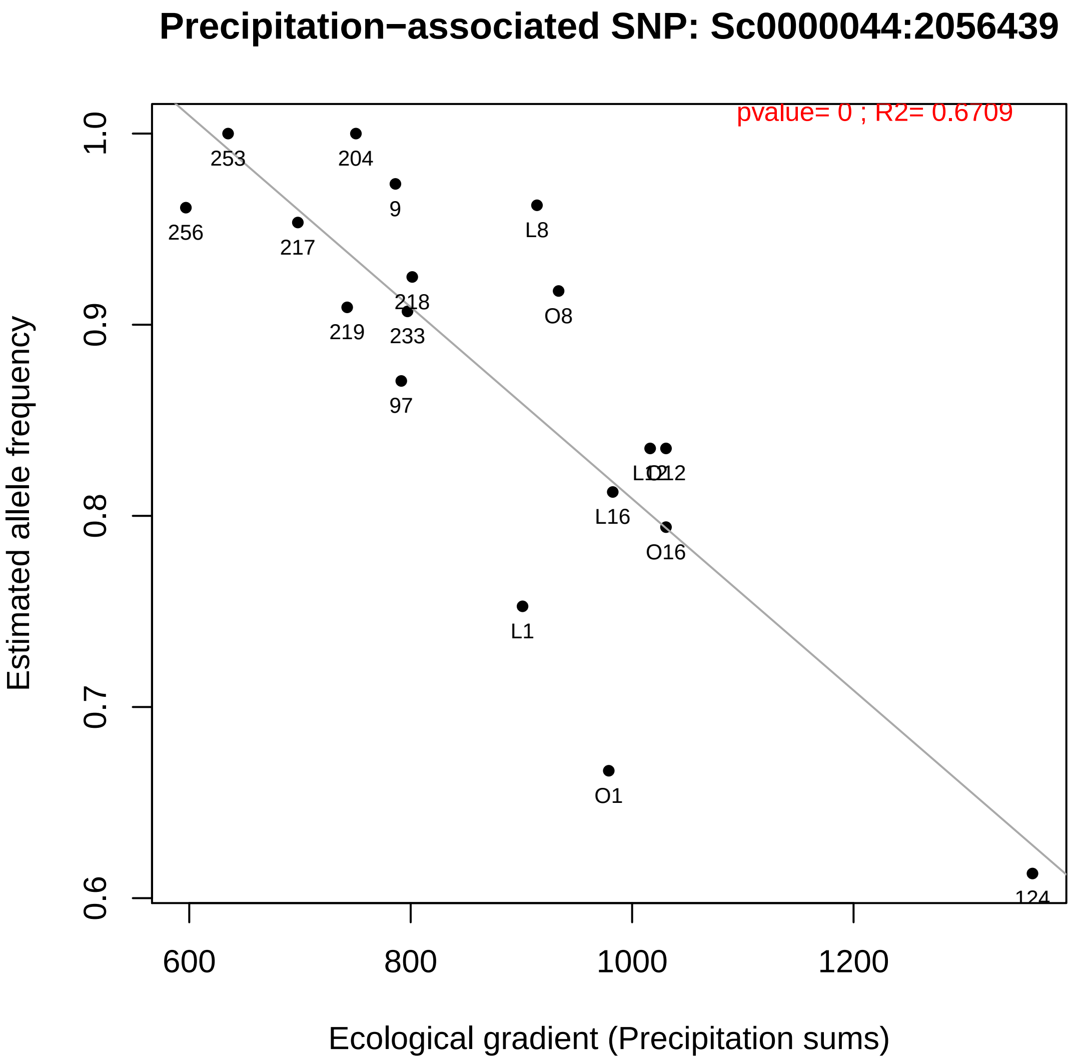
Precipitation−associated SNP: Sc0000044:2056439
pvalue= 0 ; R2= 0.6709
Estimated allele frequency
Ecological gradient (Precipitation sums)
0.6
0.7
0.8
0.9
1.0
600
800
1000
1200
253
204
9
256
217
L8
218
219
233
O8
97
L16
O16
L1
O1
124

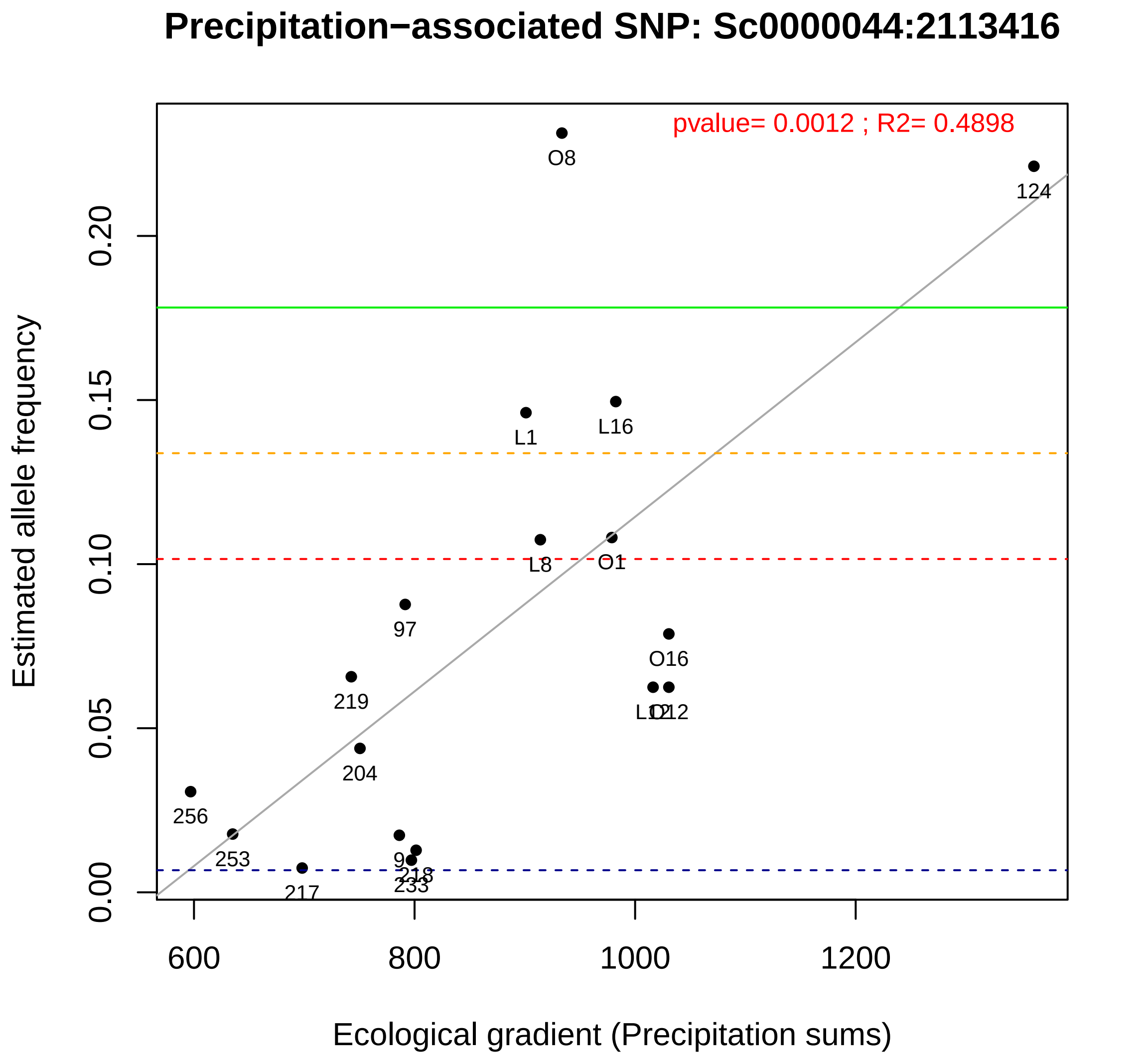

Precipitation−associated SNP: Sc0000044:2113416
pvalue= 0.0012 ; R2= 0.4898
O8
124
L1
L16
L8
O1
97
O16
219
L12
O12
204
256
253
217
218
233
0.00
0.05
0.10
0.15
0.20
600
800
1000
1200
Estimated allele frequency
Ecological gradient (Precipitation sums)

#### Precipitation−associated SNP: Sc0000044:2149038

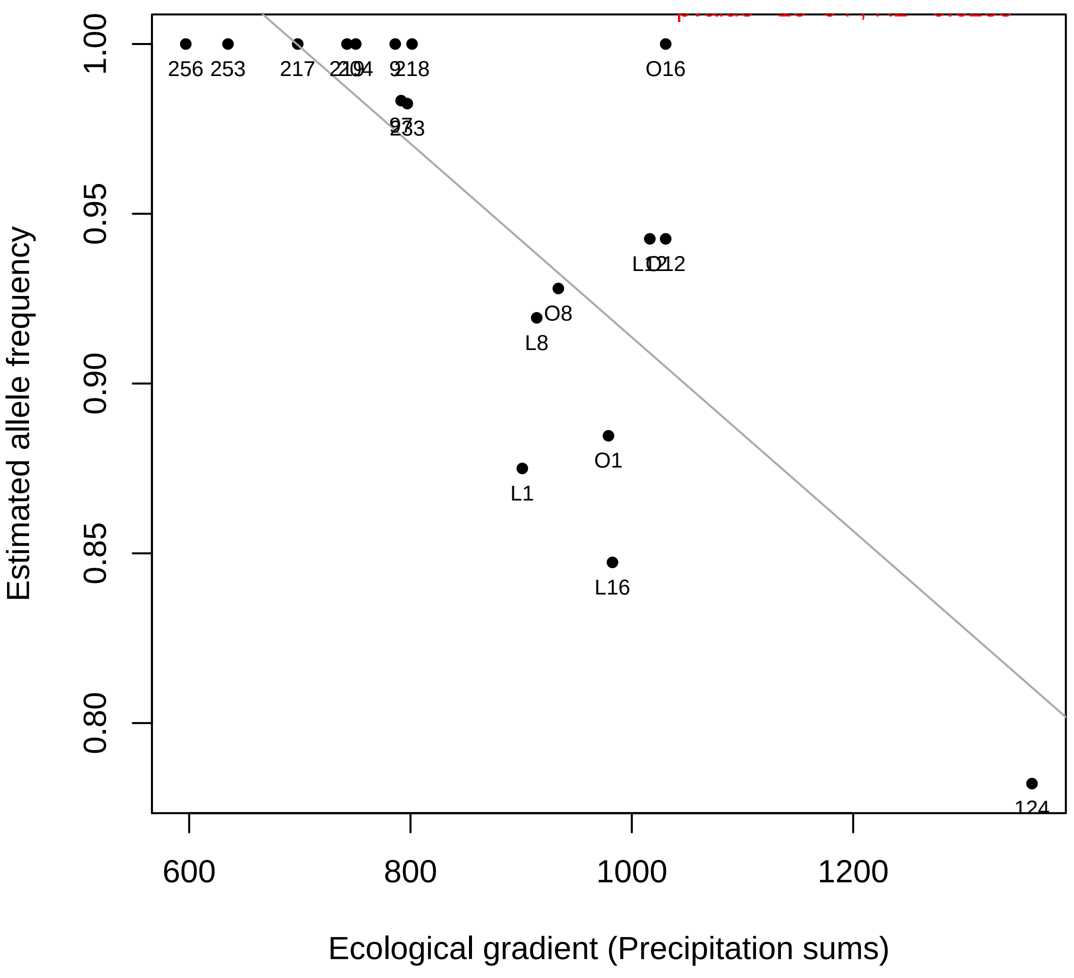

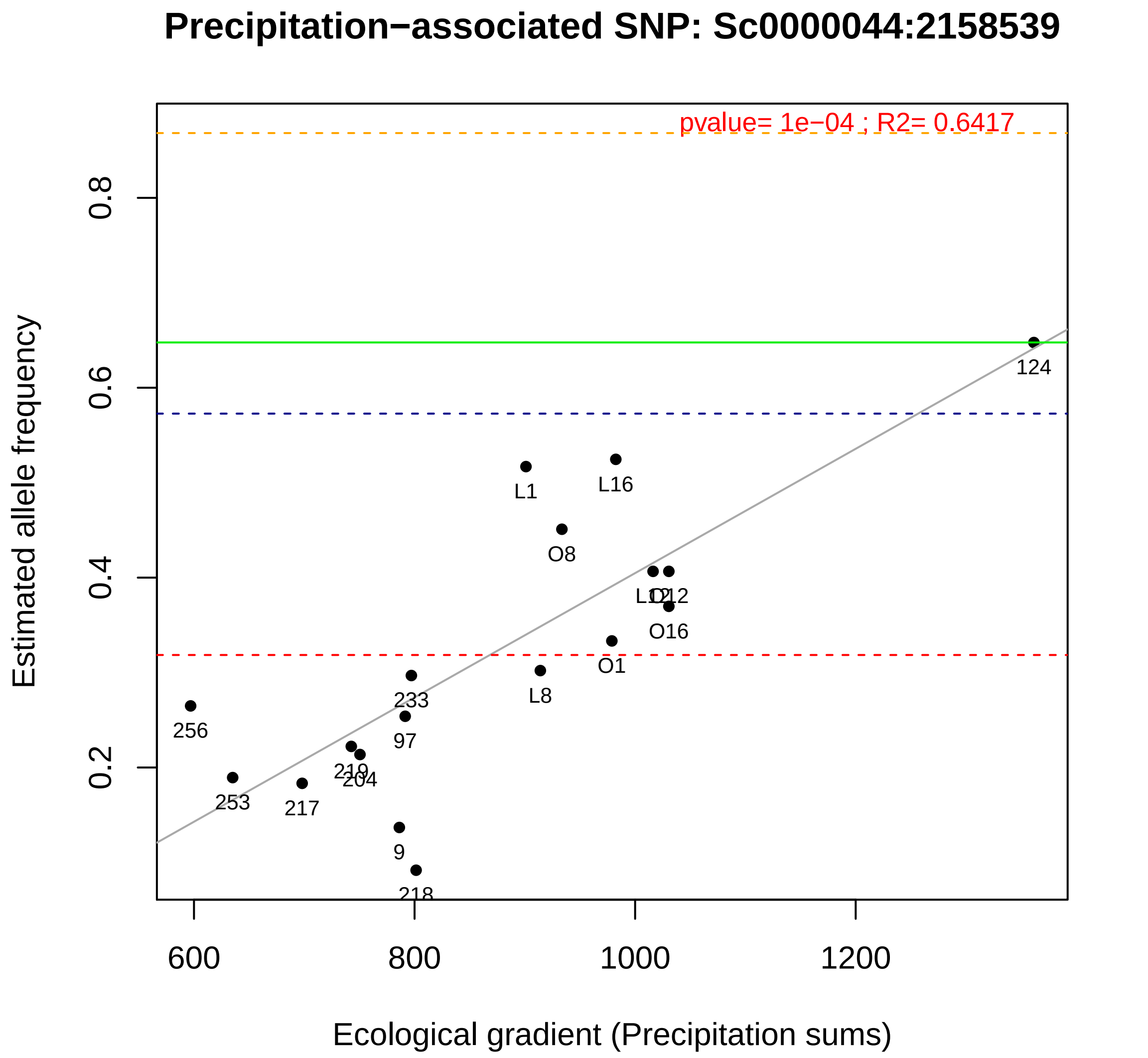

Precipitation−associated SNP: Sc0000044:2158539
pvalue= 1e−04 ; R2= 0.6417
Estimated allele frequency
0.2
0.4
0.6
0.8
Ecological gradient (Precipitation sums)
600
800
1000
1200
124
L1
L16
O8
O16
O1
L8
233
256
97
204
253
217
9
218

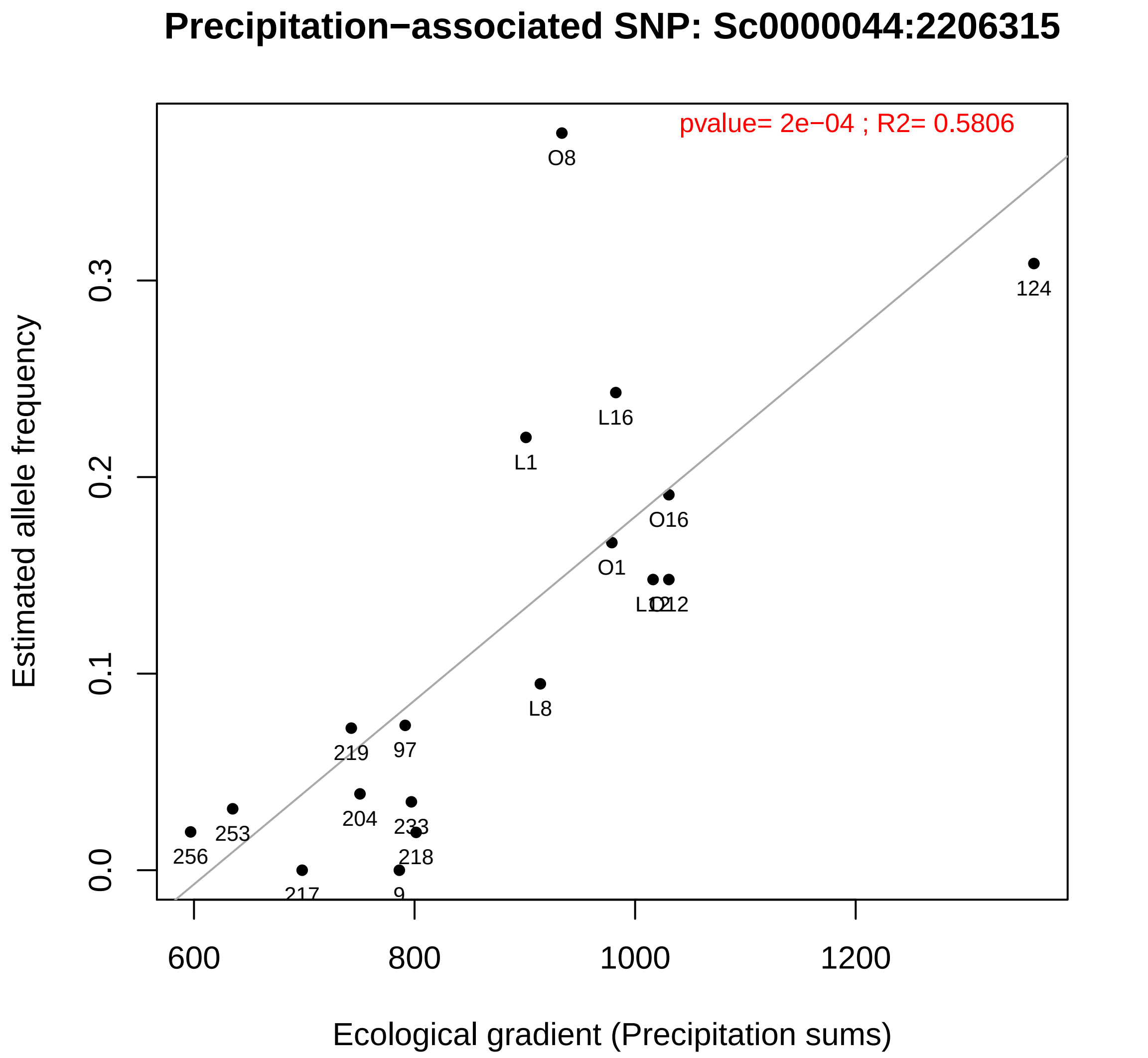

Precipitation−associated SNP: Sc0000044:2206315
pvalue= 2e−04 ; R2= 0.5806
Estimated allele frequency
0.0
0.1
0.2
0.3
600
800
1000
1200
Ecological gradient (Precipitation sums)
O8
124
L16
L1
O16
O1
L12
O12
L8
219
97
204
233
218
253
256
217
9

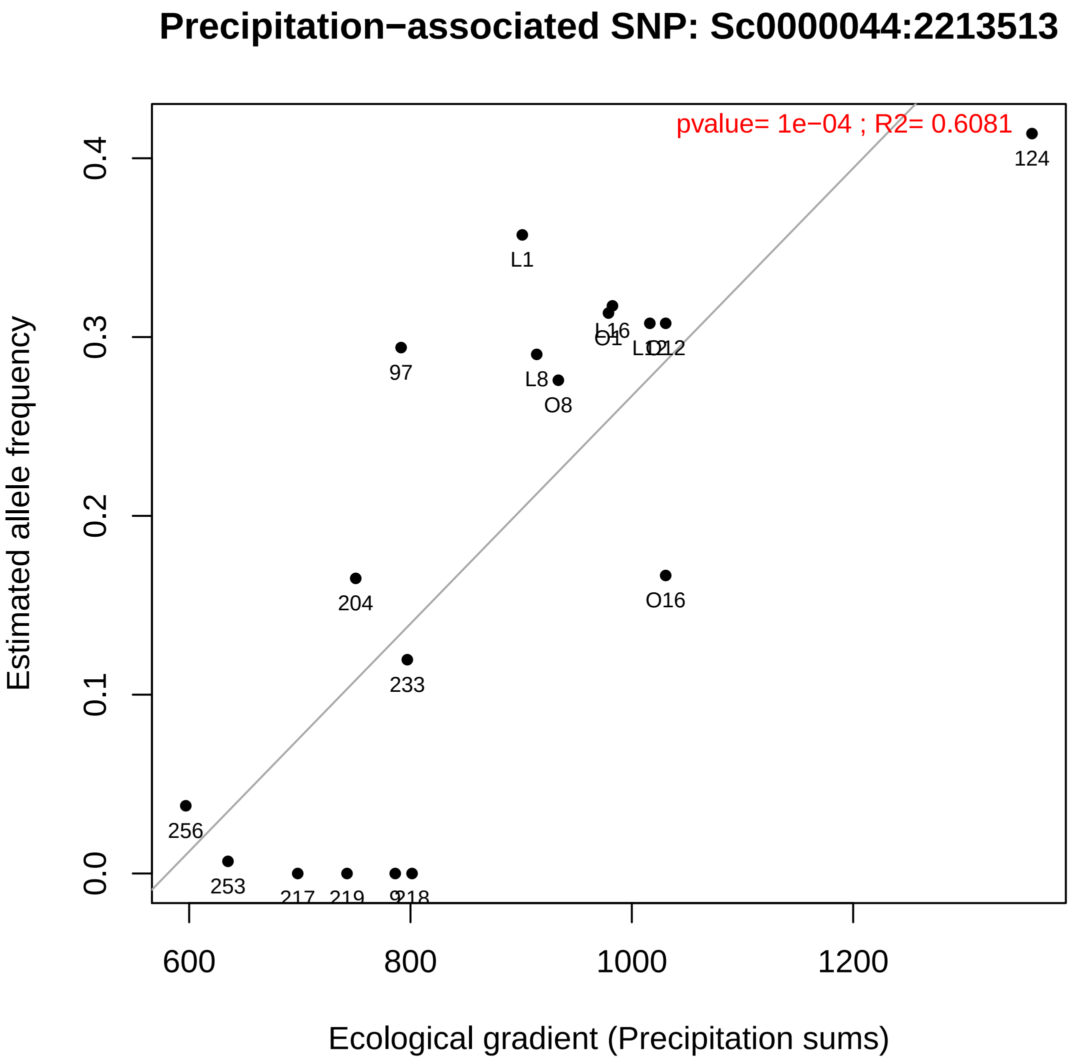

Precipitation−associated SNP: Sc0000044:2213513
pvalue= 1e−04 ; R2= 0.6081
Estimated allele frequency
0.0
0.1
0.2
0.3
0.4
600
800
1000
1200
Ecological gradient (Precipitation sums)
124
L1
L16
O1
L12
O12
97
L8
O8
204
O16
233
256
253
217
219
218

### Precipitation−associated SNP: Sc0000044:2224819

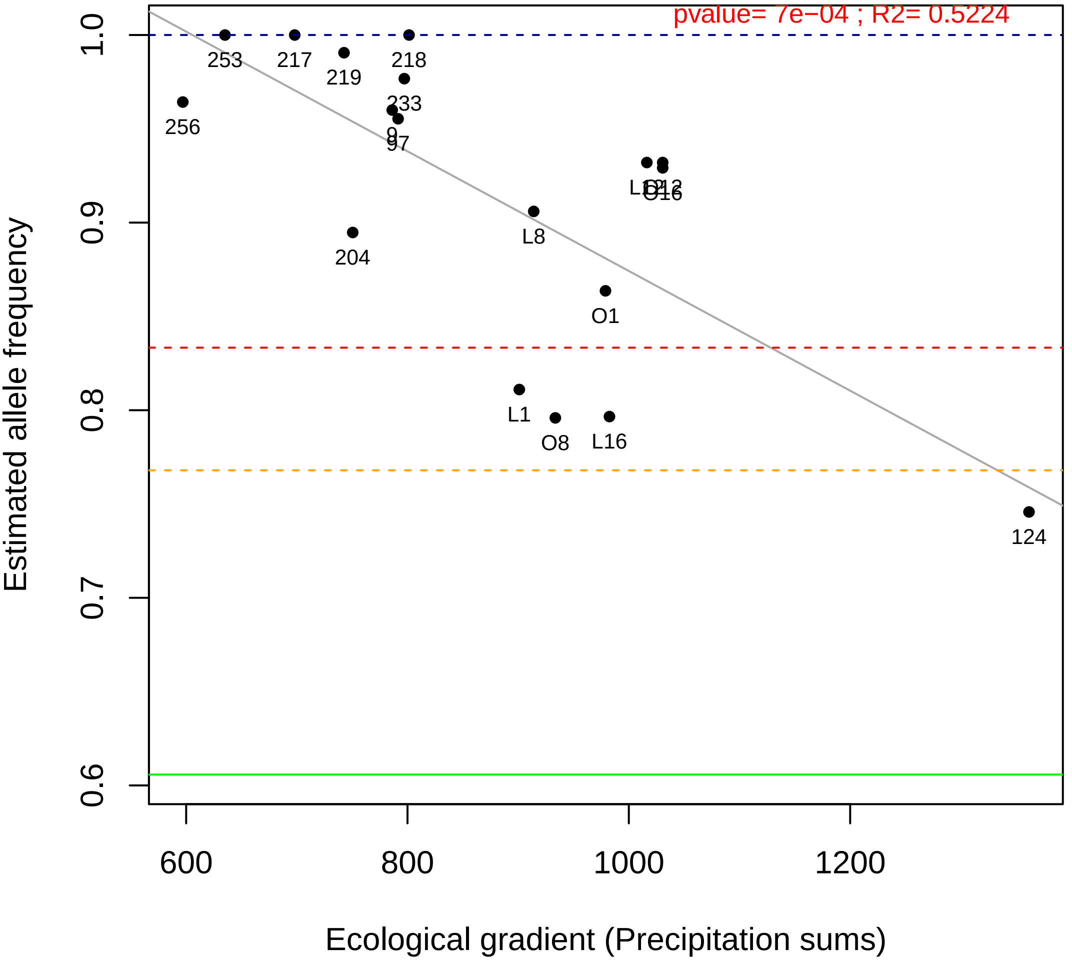

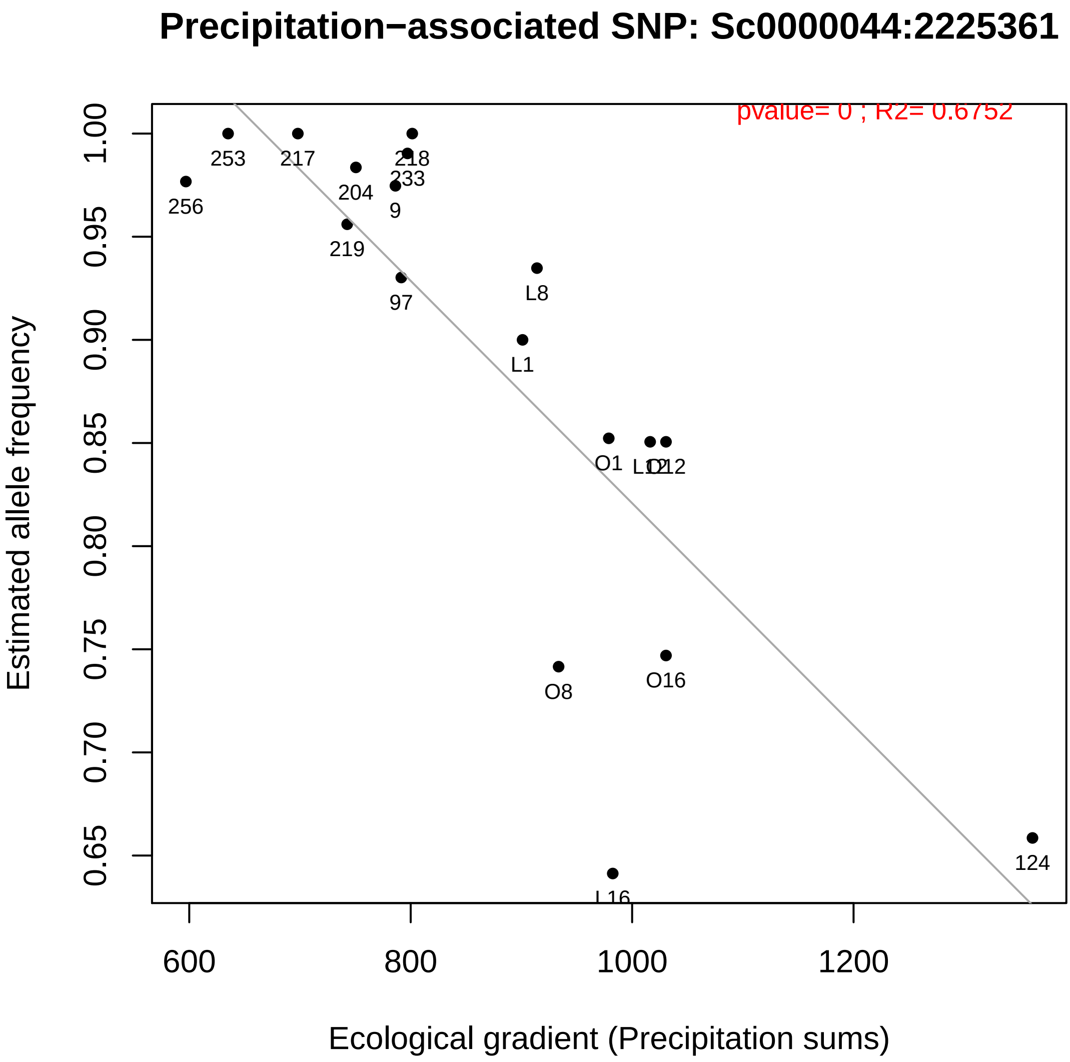
Precipitation−associated SNP: Sc0000044:2225361
pvalue= 0 ; R2= 0.6752
Estimated allele frequency
Ecological gradient (Precipitation sums)
0.65
0.70
0.75
0.80
0.85
0.90
0.95
1.00
600
800
1000
1200
253
217
218
233
204
9
256
219
97
L8
L1
O1
L12
O12
O8
O16
124
L16

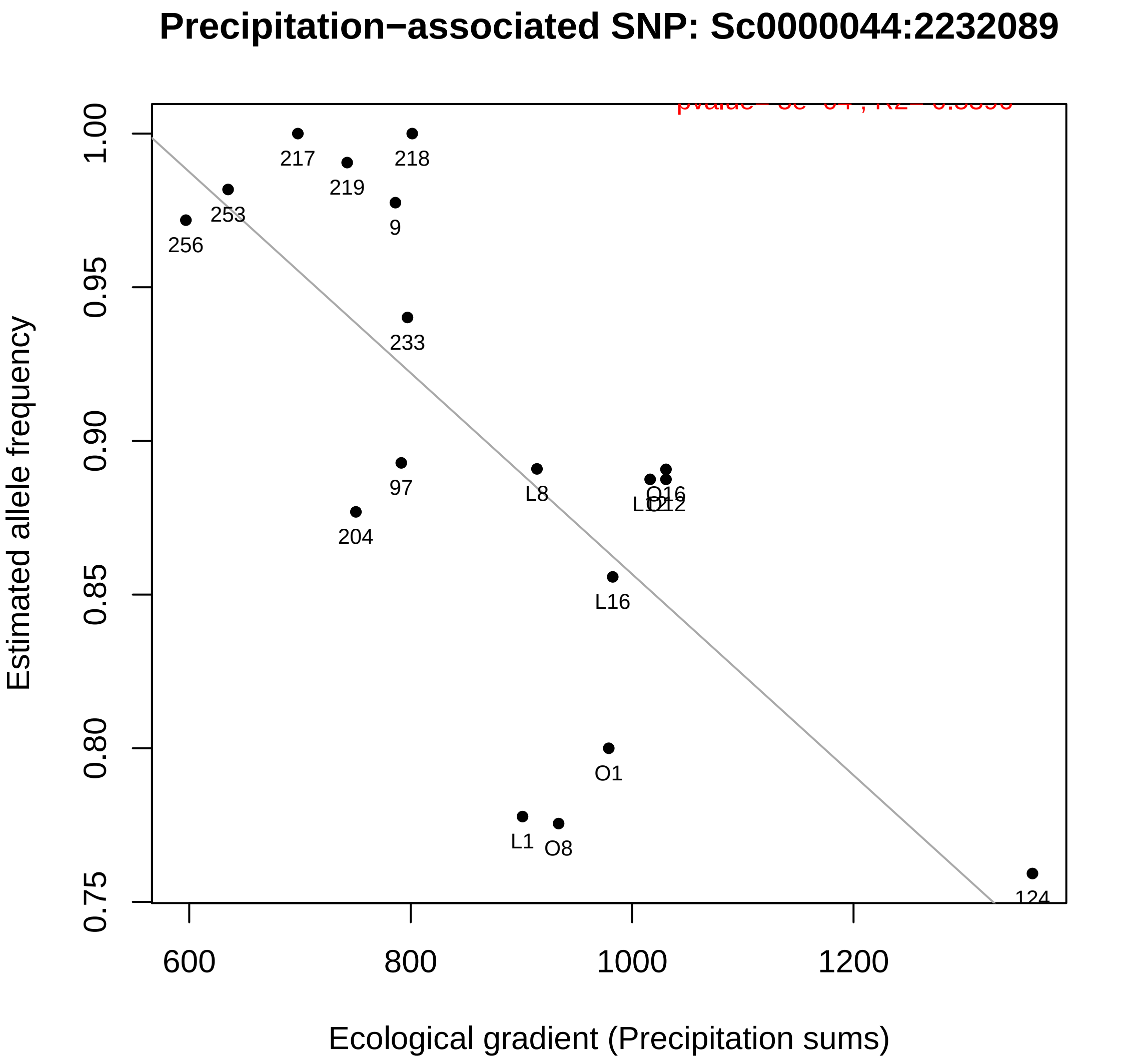

Precipitation−associated SNP: Sc0000044:2232089
Estimated allele frequency
Ecological gradient (Precipitation sums)
1.00
0.95
0.90
0.85
0.80
0.75
600
800
1000
1200
217
218
219
253
256
9
233
97
204
L8
O16
L12
L16
O1
L1
O8
124

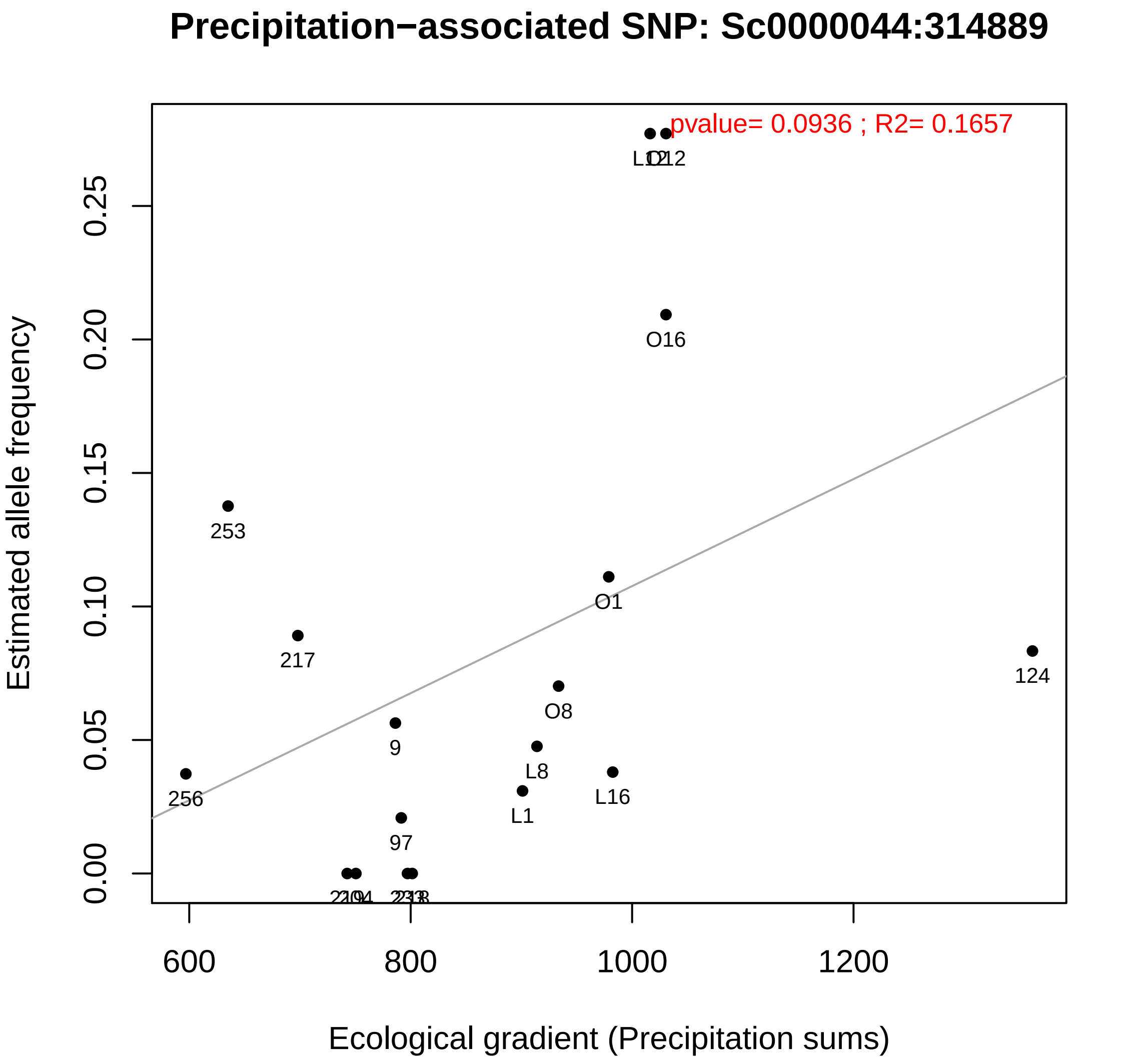

Precipitation−associated SNP: Sc0000044:314889
pvalue= 0.0936 ; R2= 0.1657
Estimated allele frequency
0.00
0.05
0.10
0.15
0.20
0.25
600
800
1000
1200
Ecological gradient (Precipitation sums)
O16
253
O1
217
124
O8
9
L8
L16
256
L1
97

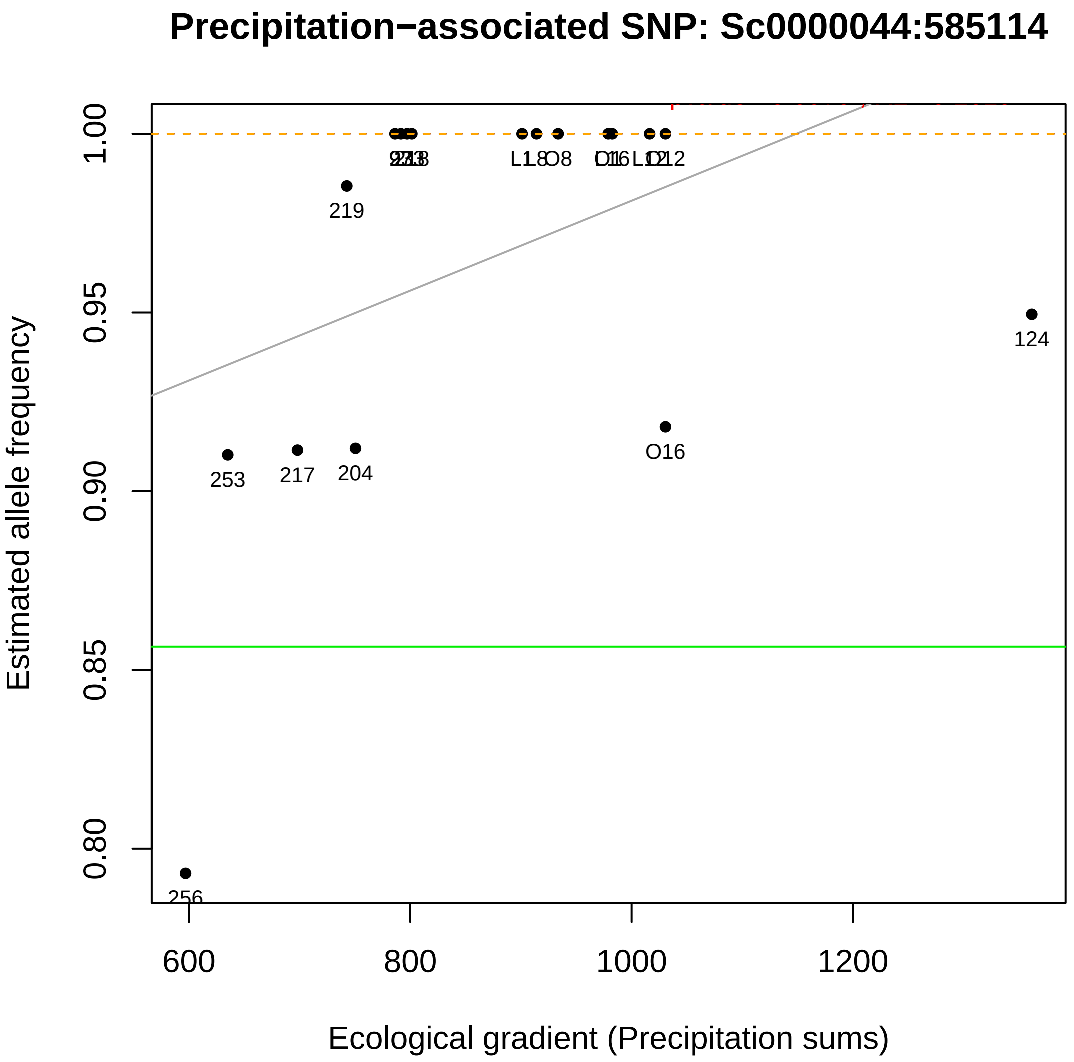
Precipitation−associated SNP: Sc0000044:585114
Estimated allele frequency
Ecological gradient (Precipitation sums)
1.00
0.95
0.90
0.85
0.80
600
800
1000
1200
219
253
217
204
O16
124
256
O8

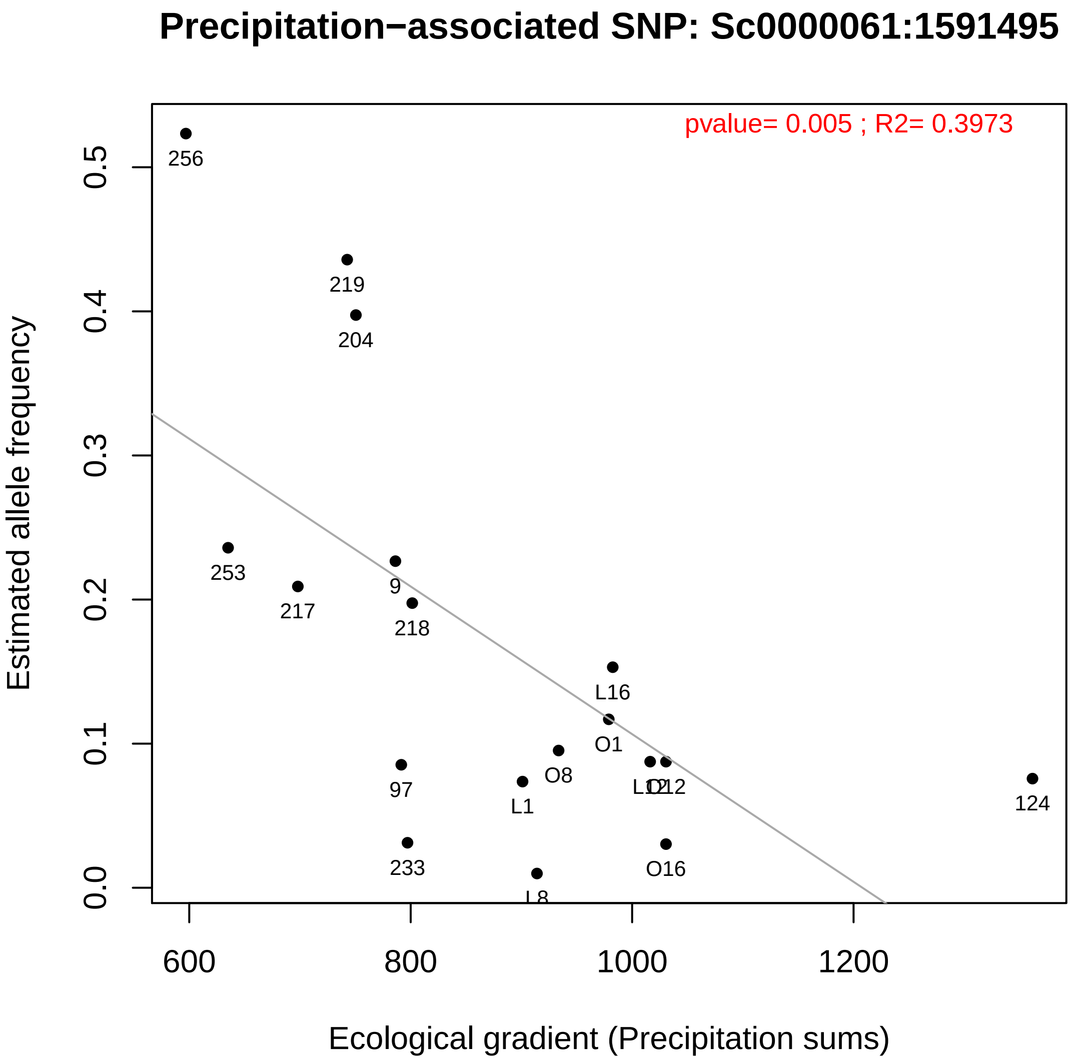

Precipitation−associated SNP: Sc0000061:1591495
pvalue= 0.005 ; R2= 0.3973
Estimated allele frequency
Ecological gradient (Precipitation sums)
0.0
0.1
0.2
0.3
0.4
0.5
600
800
1000
1200
256
219
204
253
217
9
218
L16
O1
O8
97
L1
L12
O12
124
233
O16
L8

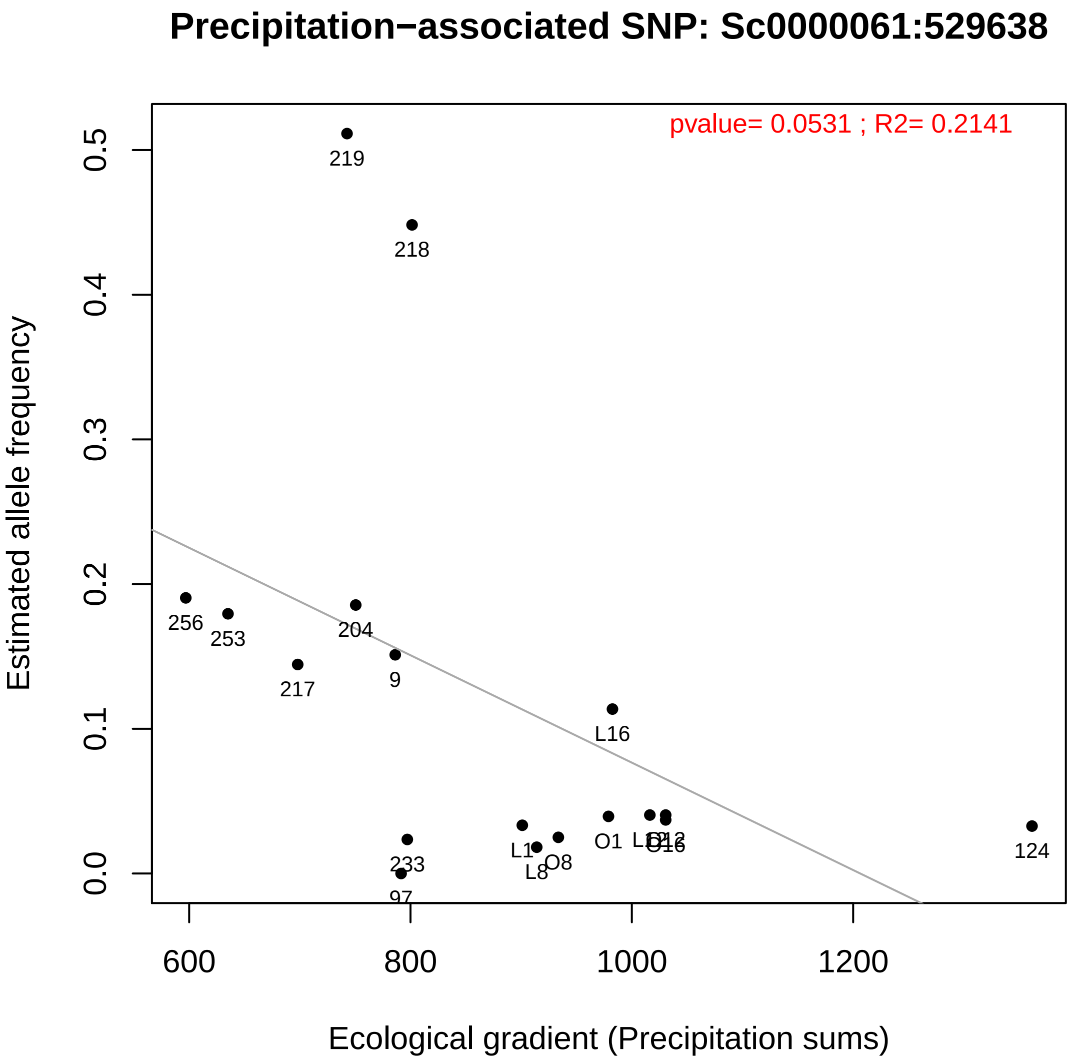
Precipitation−associated SNP: Sc0000061:529638
pvalue= 0.0531 ; R2= 0.2141
Estimated allele frequency
Ecological gradient (Precipitation sums)
0.0
0.1
0.2
0.3
0.4
0.5
600
800
1000
1200
219
218
256
253
204
217
9
L16
233
97
L1
L8
O8
O1
124

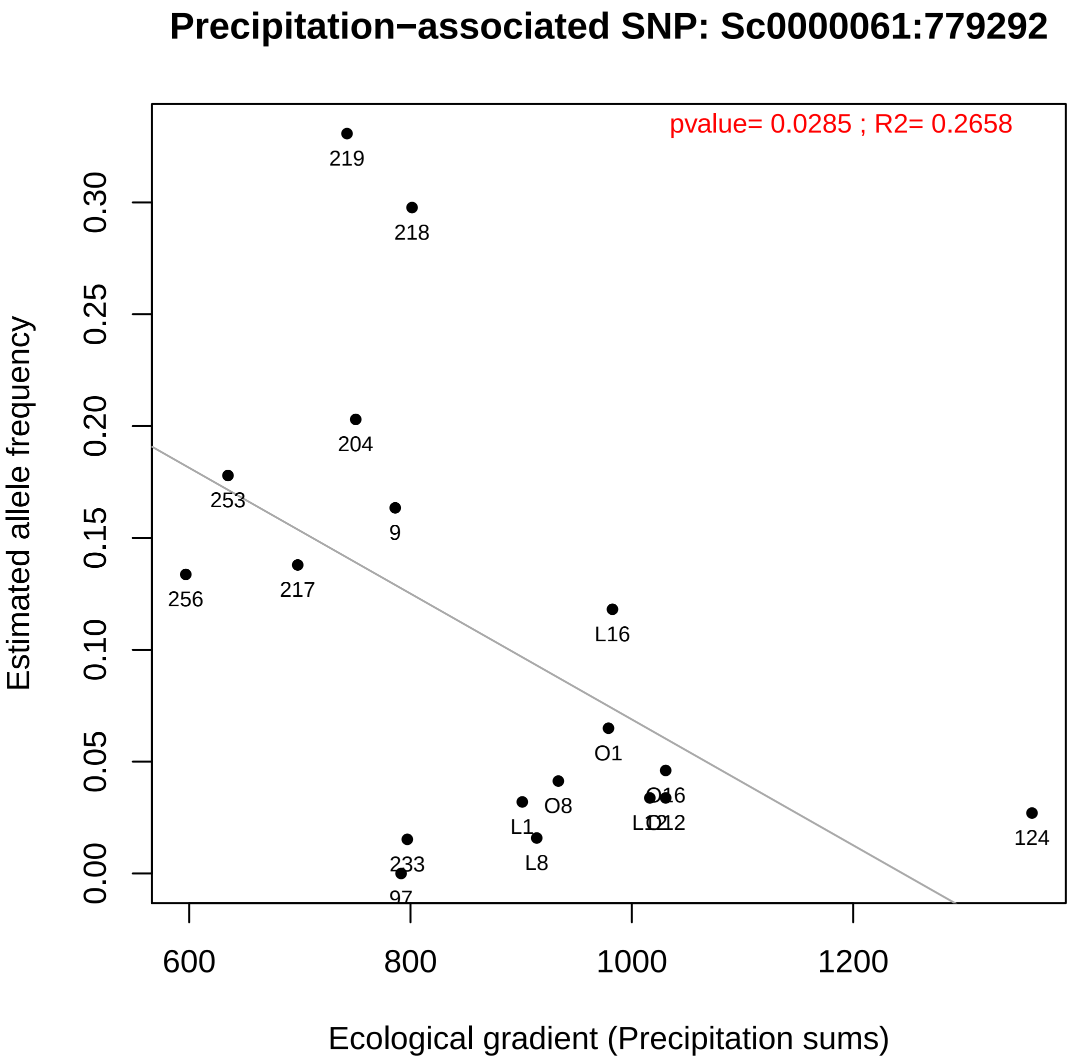

Precipitation−associated SNP: Sc0000061:779292
pvalue= 0.0285 ; R2= 0.2658
219
218
204
253
9
256
217
L16
O1
O8
O16
L1
L12
O12
124
L8
233
97
0.00
0.05
0.10
0.15
0.20
0.25
0.30
600
800
1000
1200
Estimated allele frequency
Ecological gradient (Precipitation sums)

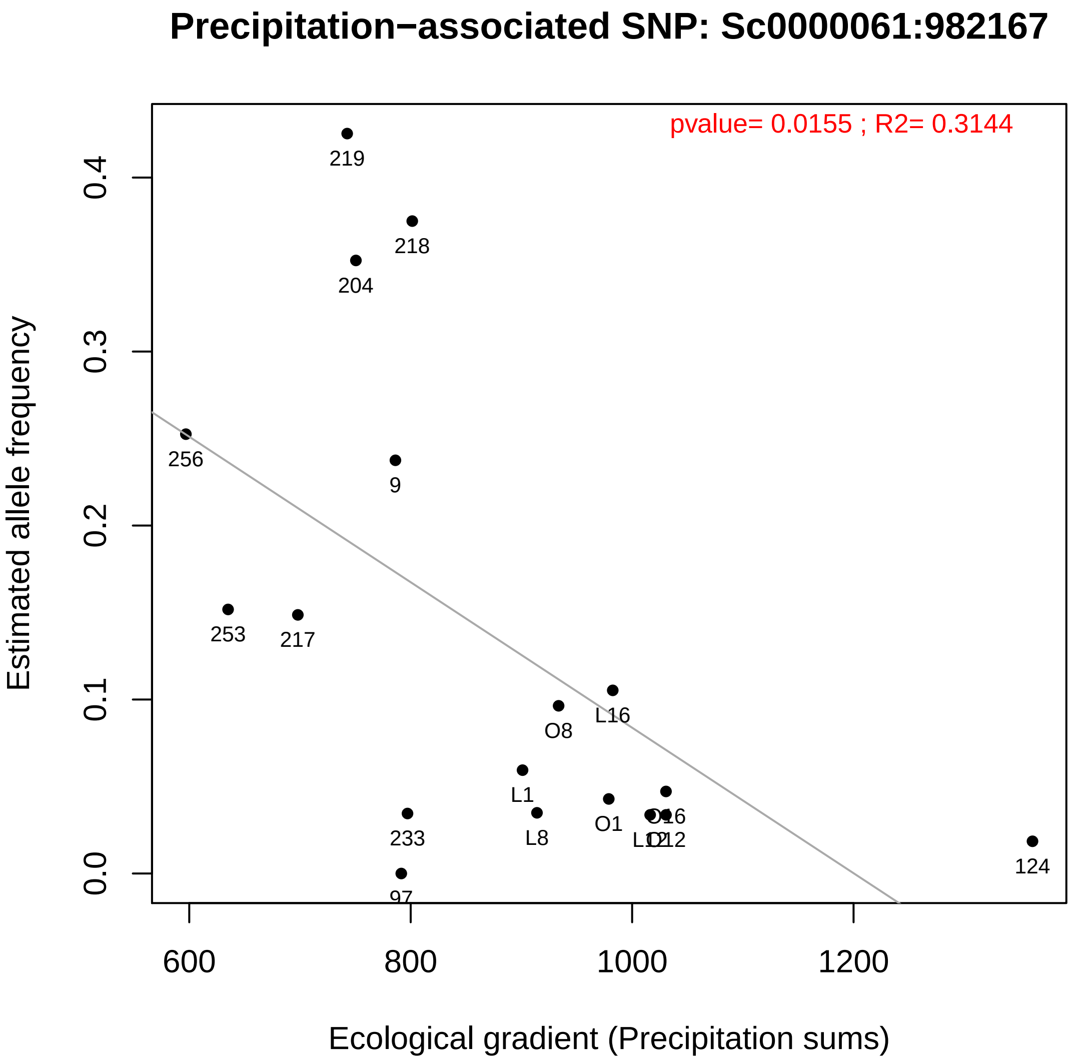
Precipitation−associated SNP: Sc0000061:982167
pvalue= 0.0155 ; R2= 0.3144
Estimated allele frequency
0.0
0.1
0.2
0.3
0.4
600
800
1000
1200
Ecological gradient (Precipitation sums)
219
218
204
256
9
253
217
L16
O8
L1
233
L8
O1
O16
L12
O12
124
97

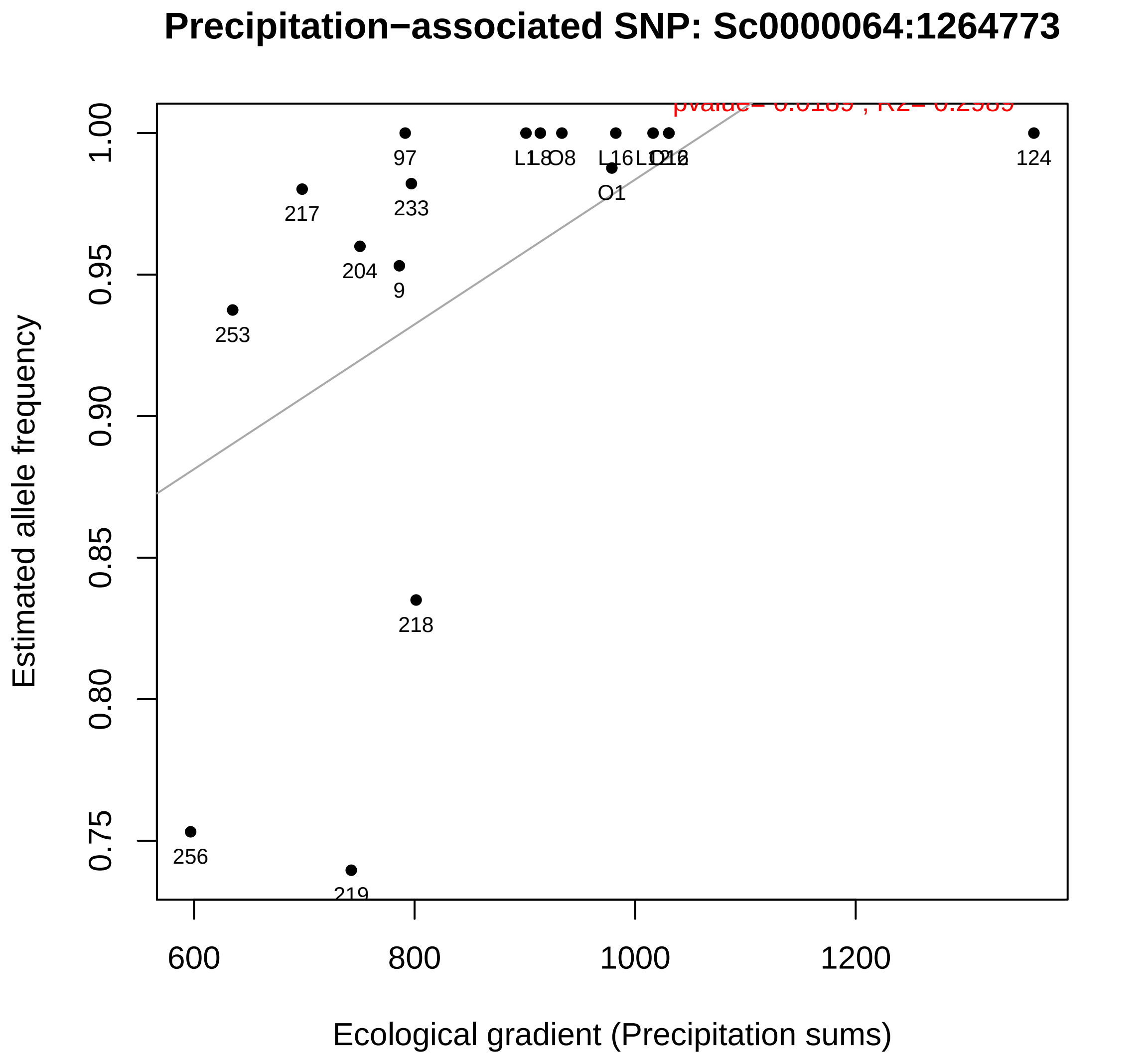
Precipitation−associated SNP: Sc0000064:1264773
pvalue= 0.0189 ; R2= 0.2989
Estimated allele frequency
Ecological gradient (Precipitation sums)
0.75
0.80
0.85
0.90
0.95
1.00
600
800
1000
1200
97
O8
L16
124
O1
217
233
204
9
253
218
256
219

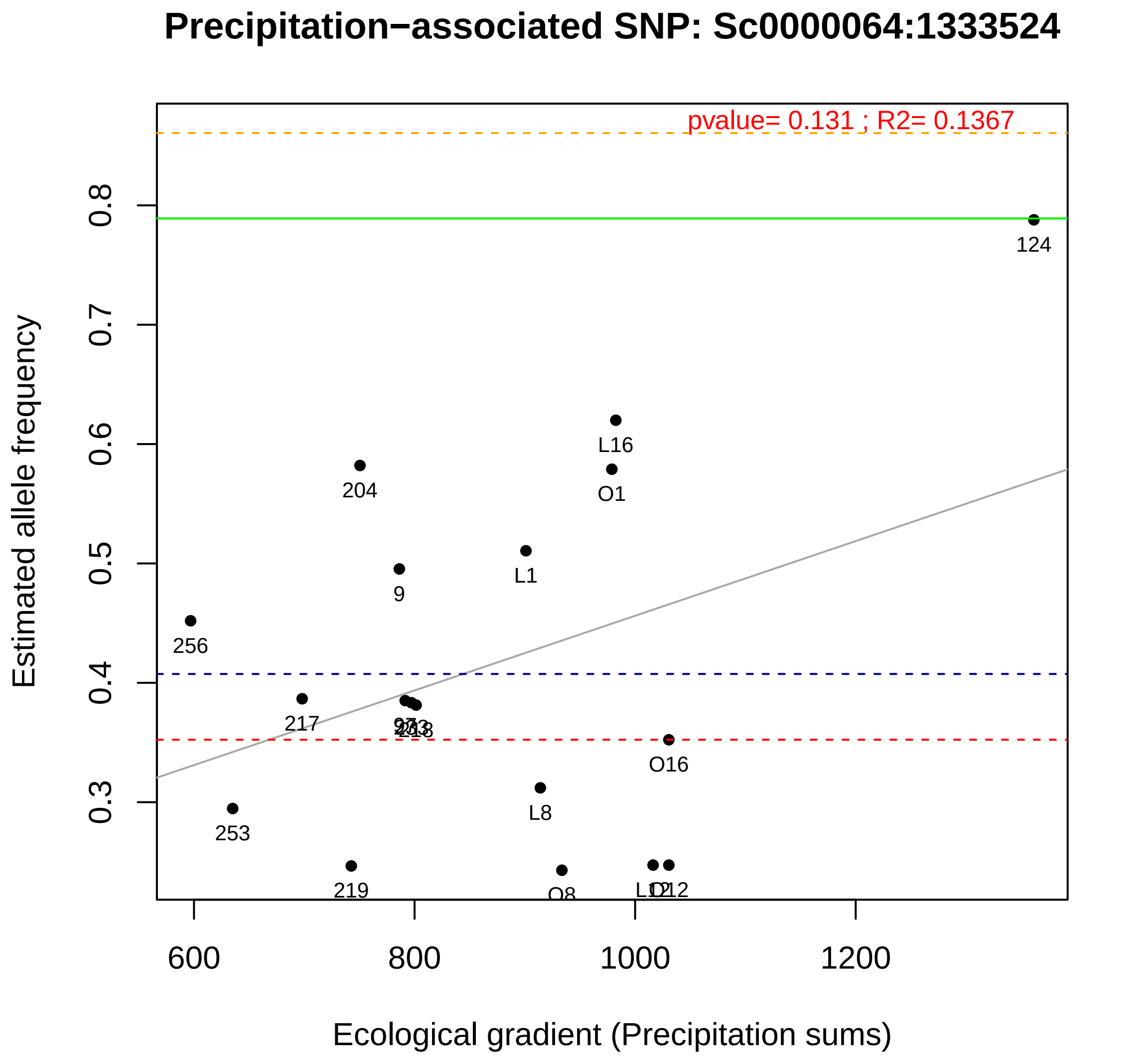

Precipitation−associated SNP: Sc0000064:1333524
pvalue= 0.131 ; R2= 0.1367
Estimated allele frequency
0.3
0.4
0.5
0.6
0.7
0.8
600
800
1000
1200
Ecological gradient (Precipitation sums)
124
L16
204
O1
L1
9
256
217
O16
L8
253
219

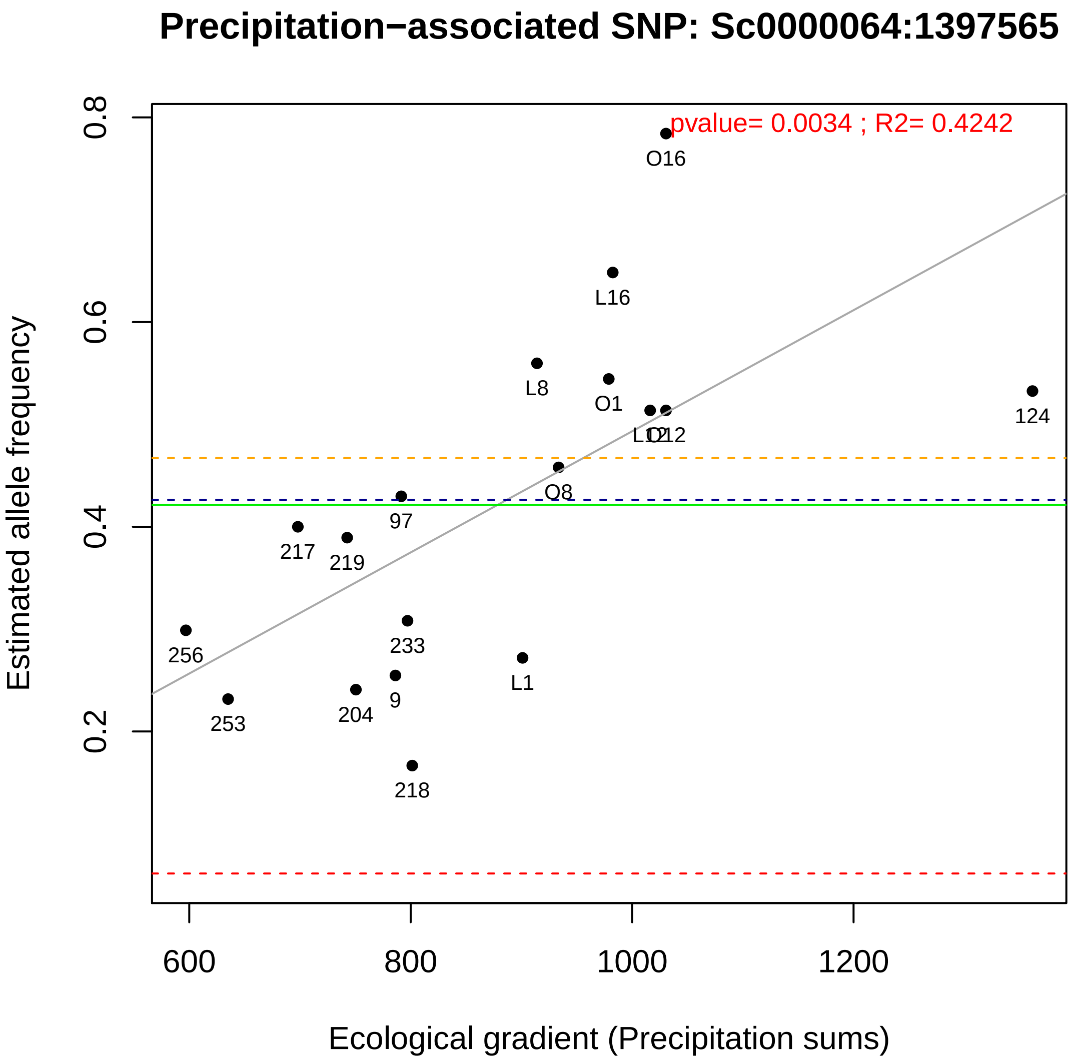
Precipitation−associated SNP: Sc0000064:1397565
pvalue= 0.0034 ; R2= 0.4242
O16
L16
L8
O1
L10
O12
124
Q8
97
217
219
256
233
L1
9
253
204
218
0.2
0.4
0.6
0.8
600
800
1000
1200
Estimated allele frequency
Ecological gradient (Precipitation sums)

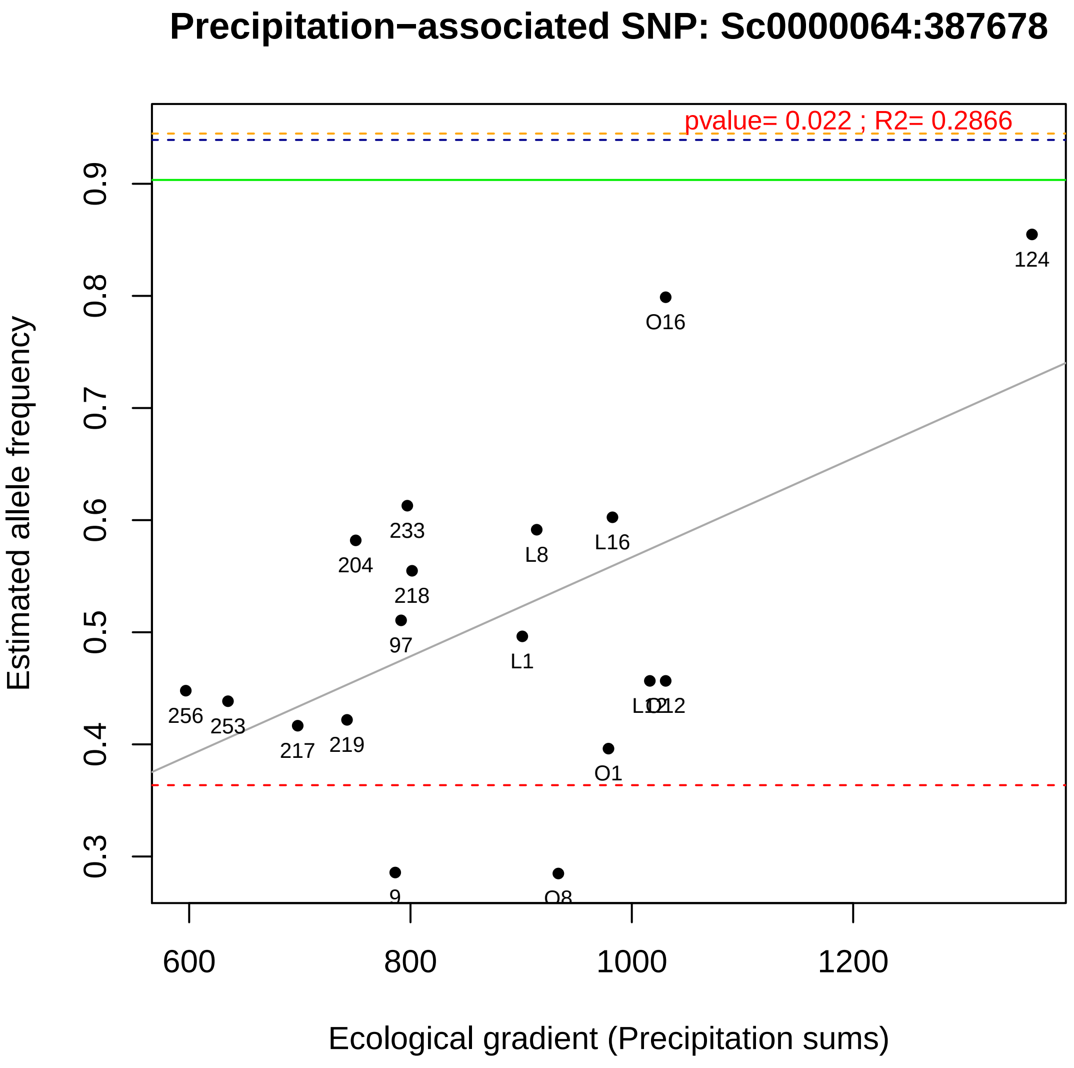

Precipitation−associated SNP: Sc0000064:387678
pvalue= 0.022 ; R2= 0.2866
Estimated allele frequency
0.3
0.4
0.5
0.6
0.7
0.8
0.9
600
800
1000
1200
Ecological gradient (Precipitation sums)
124
O16
233
204
218
L8
L16
97
L1
256
253
217
219
O1
9
O8

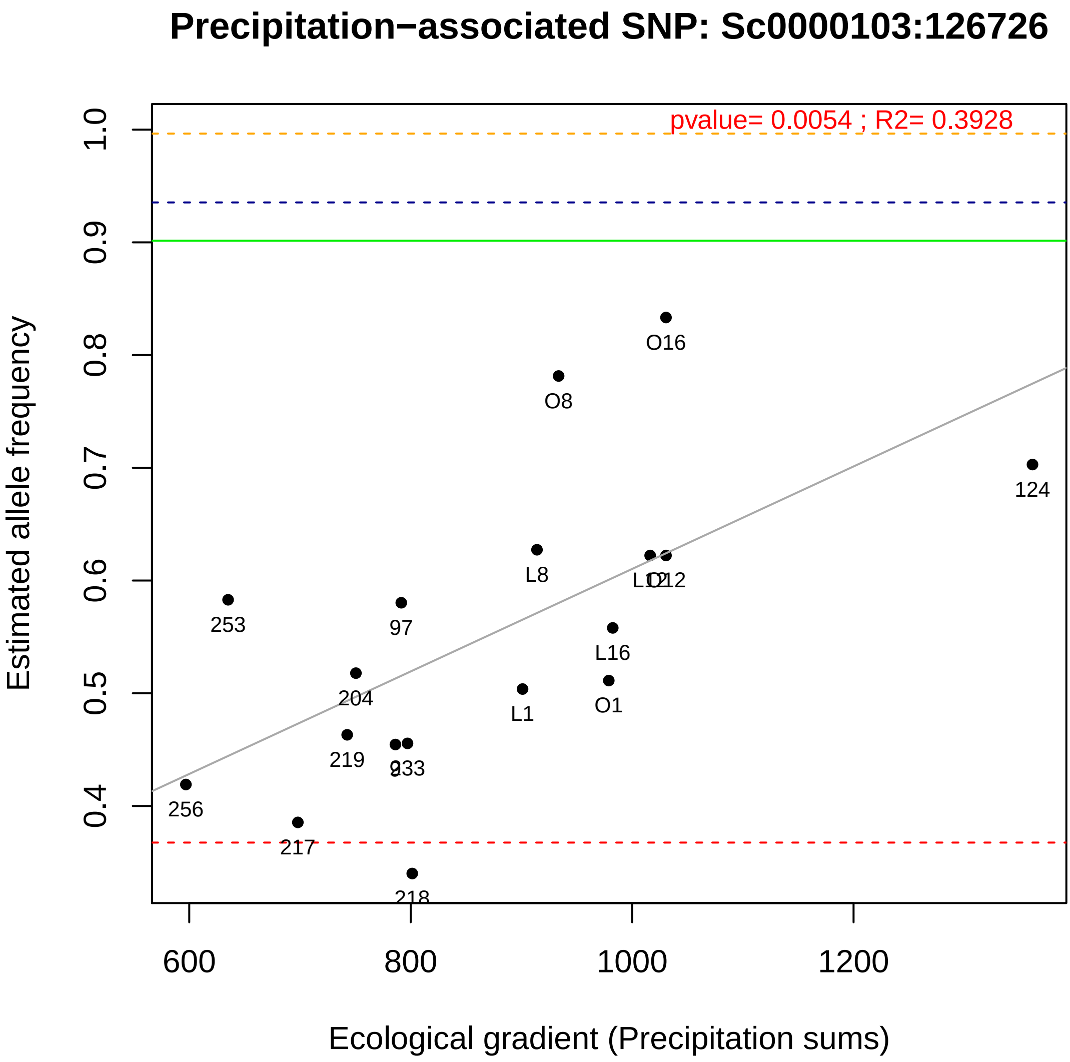
Precipitation−associated SNP: Sc0000103:126726
pvalue= 0.0054 ; R2= 0.3928
Estimated allele frequency
Ecological gradient (Precipitation sums)
1.0
0.9
0.8
0.7
0.6
0.5
0.4
600
800
1000
1200
O16
O8
124
L8
L12
O12
253
97
L16
204
L1
O1
219
2
233
256
217
218

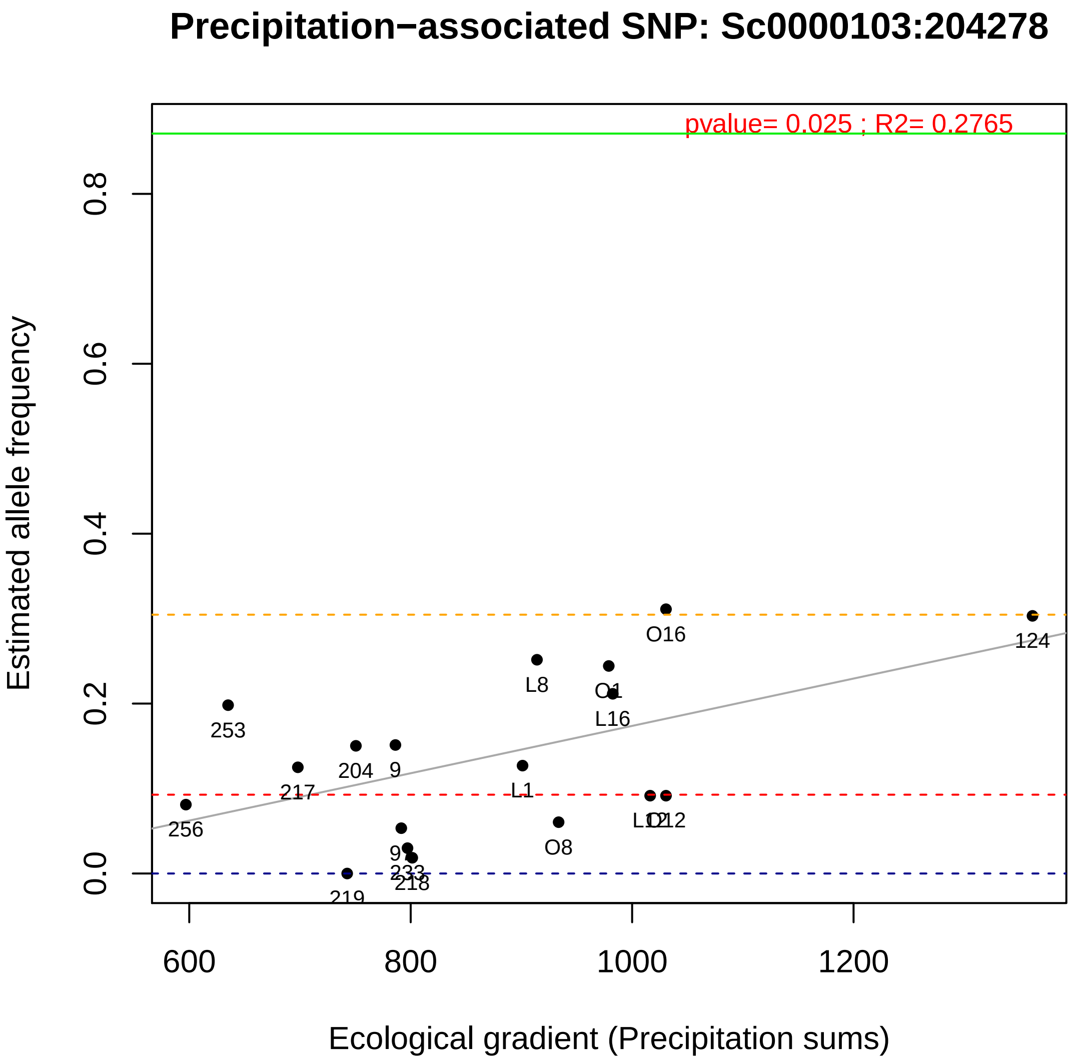

Precipitation−associated SNP: Sc0000103:204278
pvalue= 0.025 ; R2= 0.2765
Estimated allele frequency
0.0
0.2
0.4
0.6
0.8
600
800
1000
1200
Ecological gradient (Precipitation sums)
O16
124
L8
O1
L16
253
204
9
217
L1
256
O8
O12
97
233
218
219

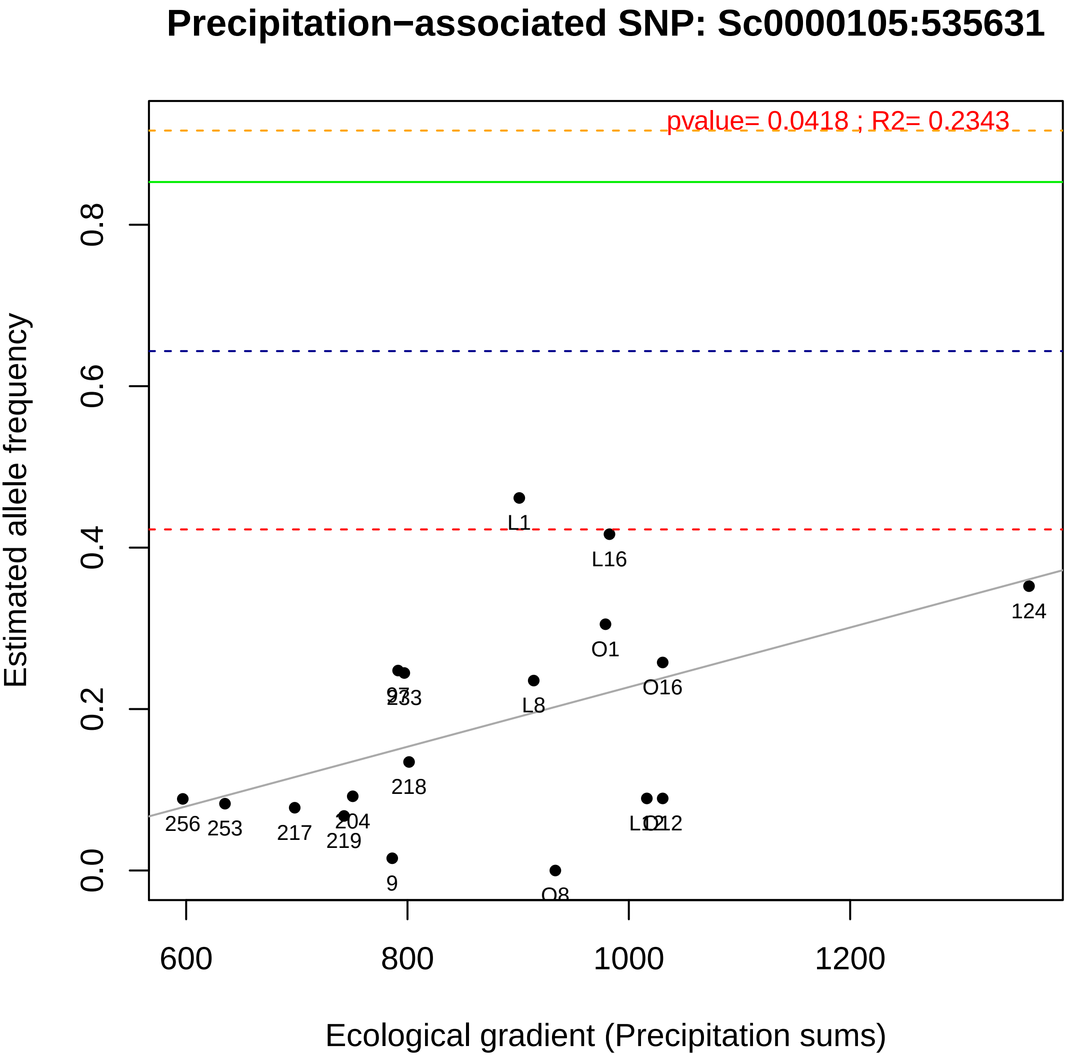

Precipitation−associated SNP: Sc0000105:535631
pvalue= 0.0418 ; R2= 0.2343
Estimated allele frequency
0.0
0.2
0.4
0.6
0.8
600
800
1000
1200
Ecological gradient (Precipitation sums)
L1
L16
124
O1
O16
L8
218
256
253
217
204
219
9

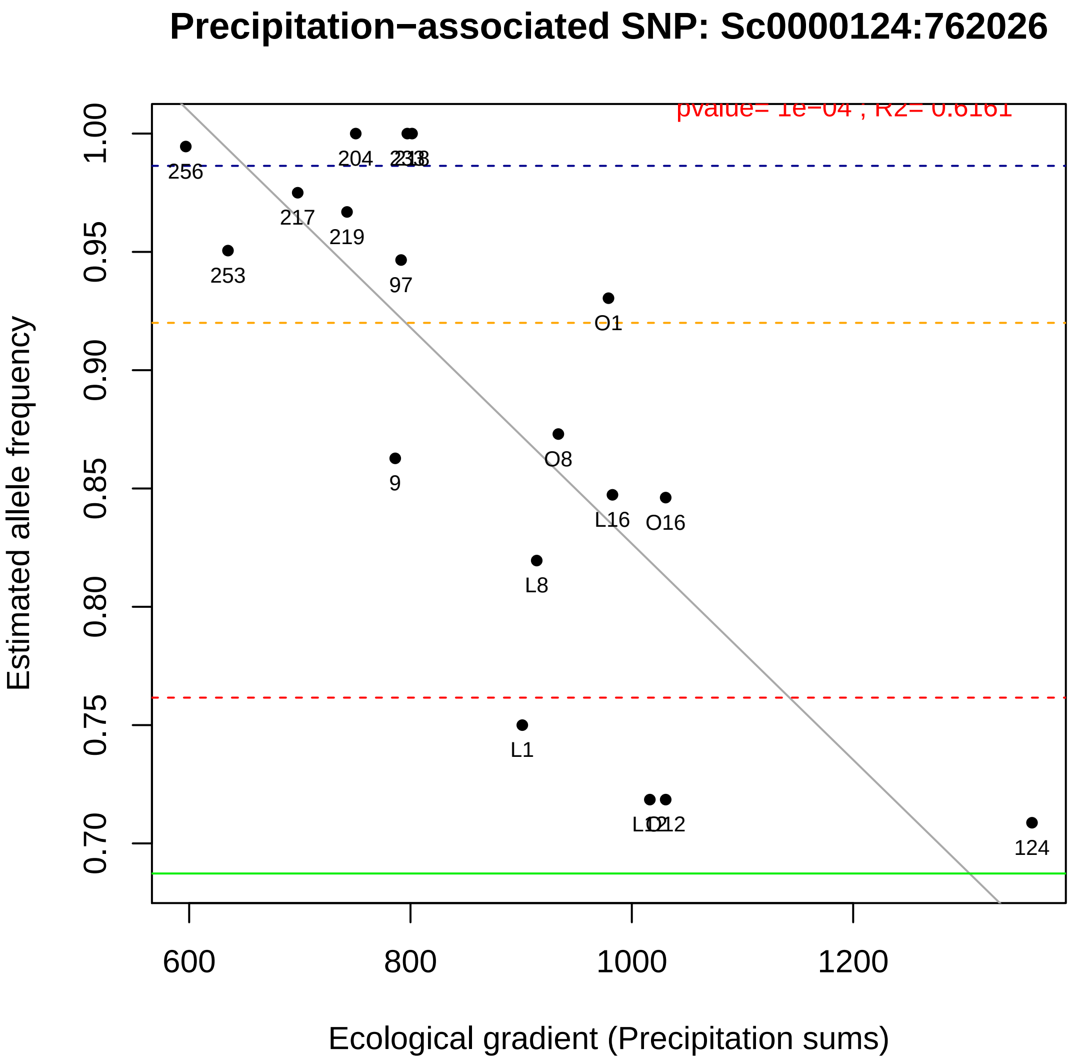

Precipitation−associated SNP: Sc0000124:762026
pvalue= 1e−04 ; R2= 0.6161
Estimated allele frequency
Ecological gradient (Precipitation sums)
0.70
0.75
0.80
0.85
0.90
0.95
1.00
600
800
1000
1200
256
204
217
219
253
97
9
O1
O8
L16
O16
L8
L1
124

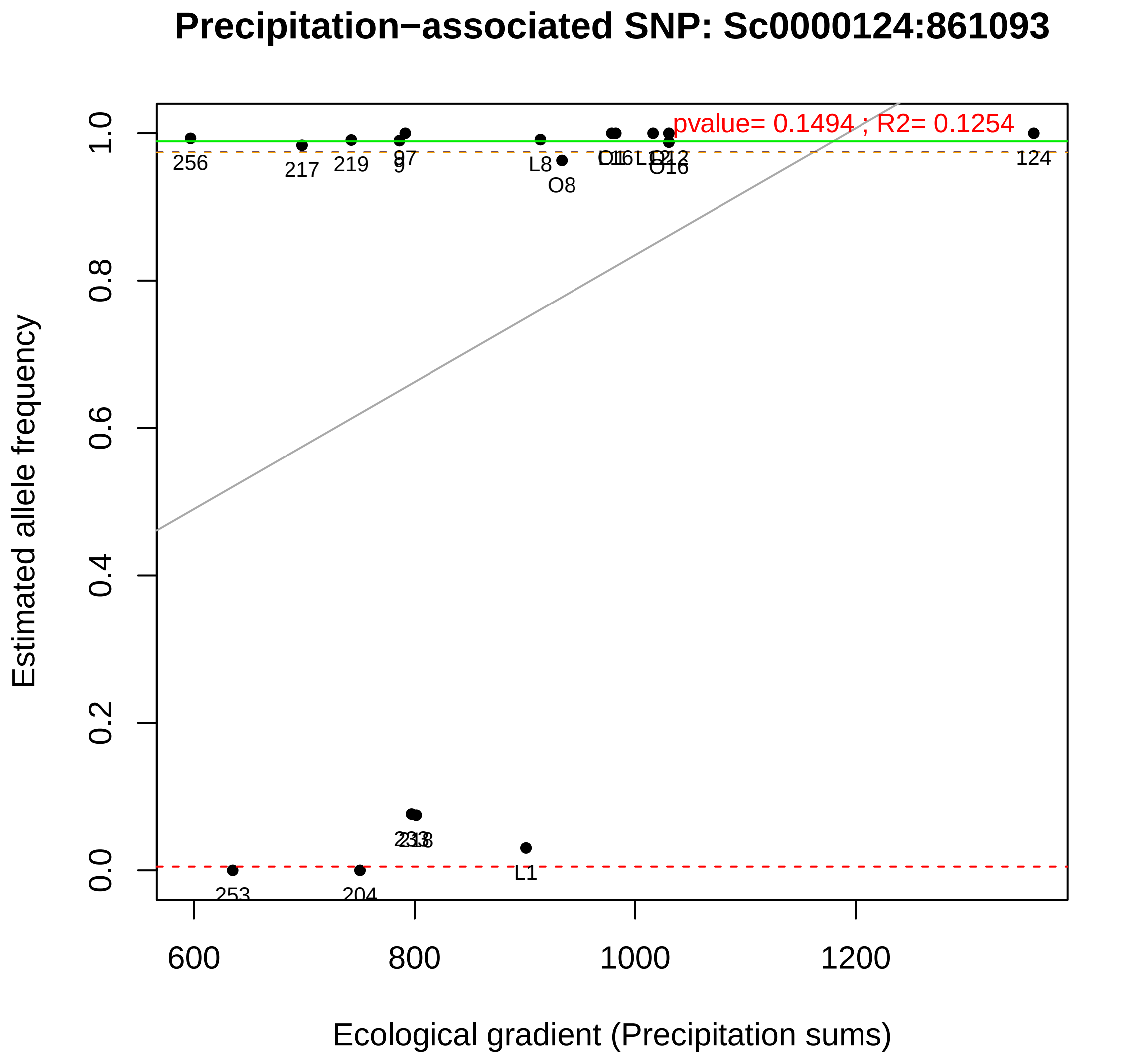

Precipitation−associated SNP: Sc0000124:861093
pvalue= 0.1494 ; R2= 0.1254
Estimated allele frequency
Ecological gradient (Precipitation sums)
0.0
0.2
0.4
0.6
0.8
1.0
600
800
1000
1200
256
217
219
97
9
L8
O8
124
O16
253
204
L1

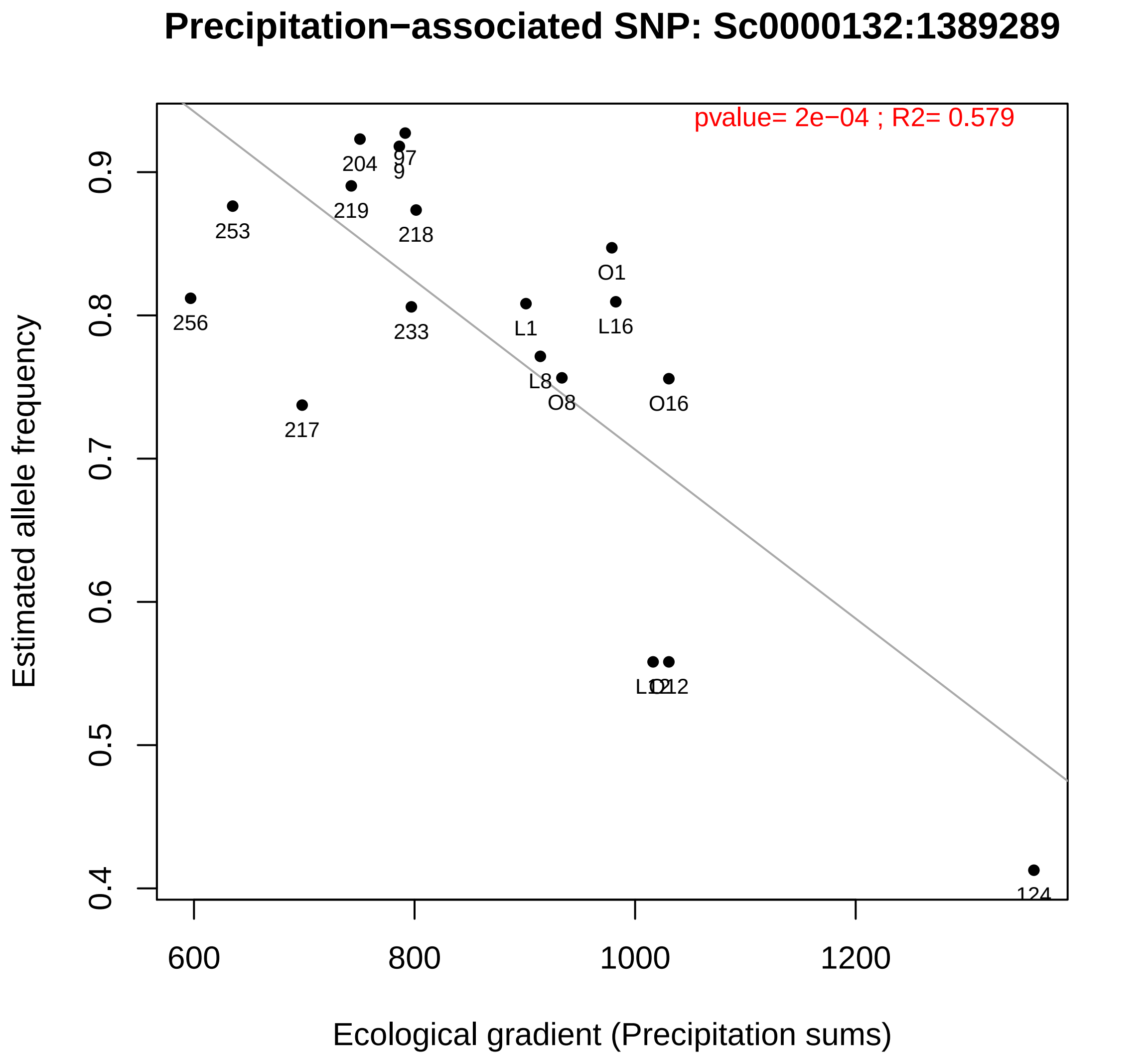
Precipitation−associated SNP: Sc0000132:1389289
pvalue= 2e−04 ; R2= 0.579
Estimated allele frequency
Ecological gradient (Precipitation sums)
0.4
0.5
0.6
0.7
0.8
0.9
600
800
1000
1200
204
97
9
219
253
218
O1
256
233
L1
L16
L8
O8
O16
217
O12
124

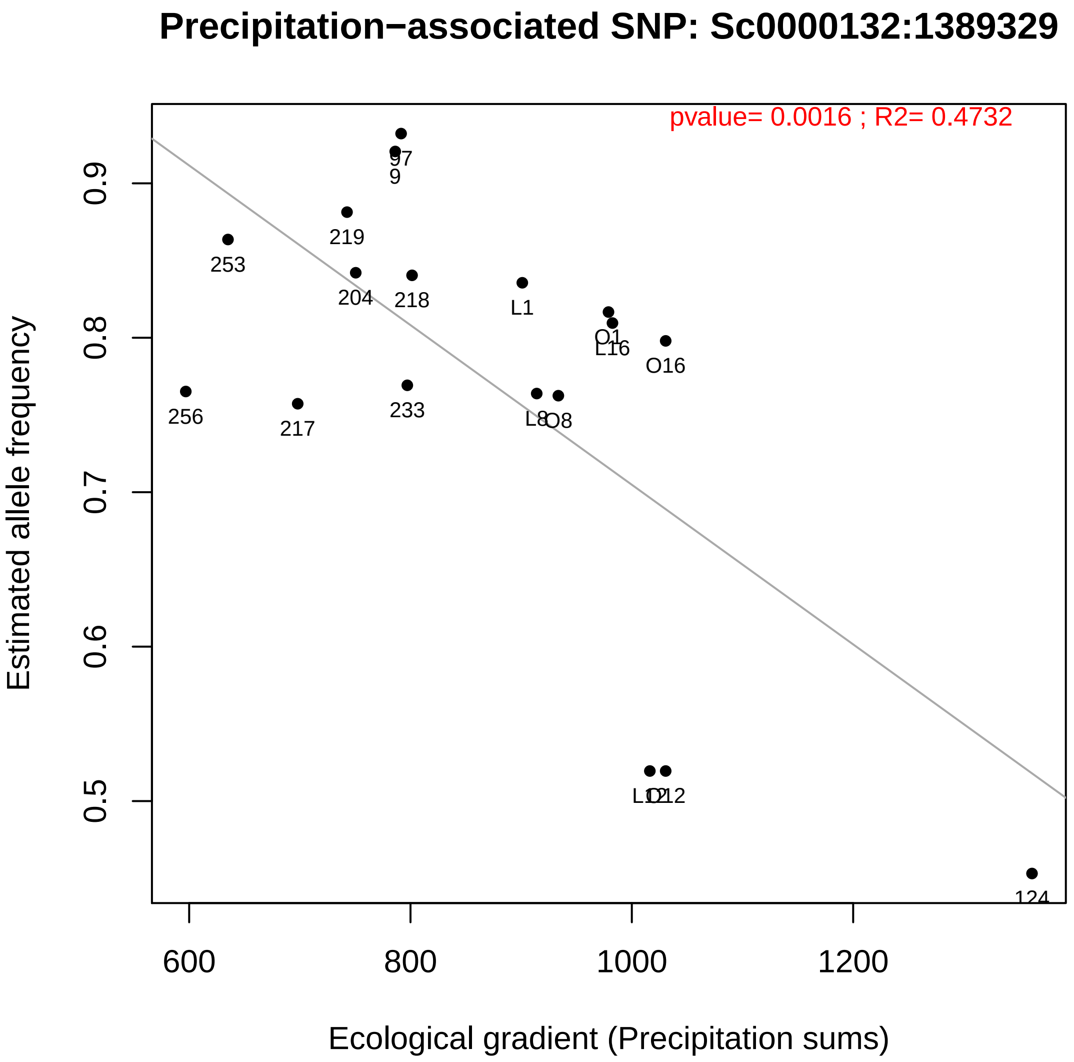
Precipitation−associated SNP: Sc0000132:1389329
pvalue= 0.0016 ; R2= 0.4732
Estimated allele frequency
0.5
0.6
0.7
0.8
0.9
600
800
1000
1200
Ecological gradient (Precipitation sums)
97
9
219
253
204
218
L1
O1
L16
O16
256
217
233
L8
O8
L12
O12
124

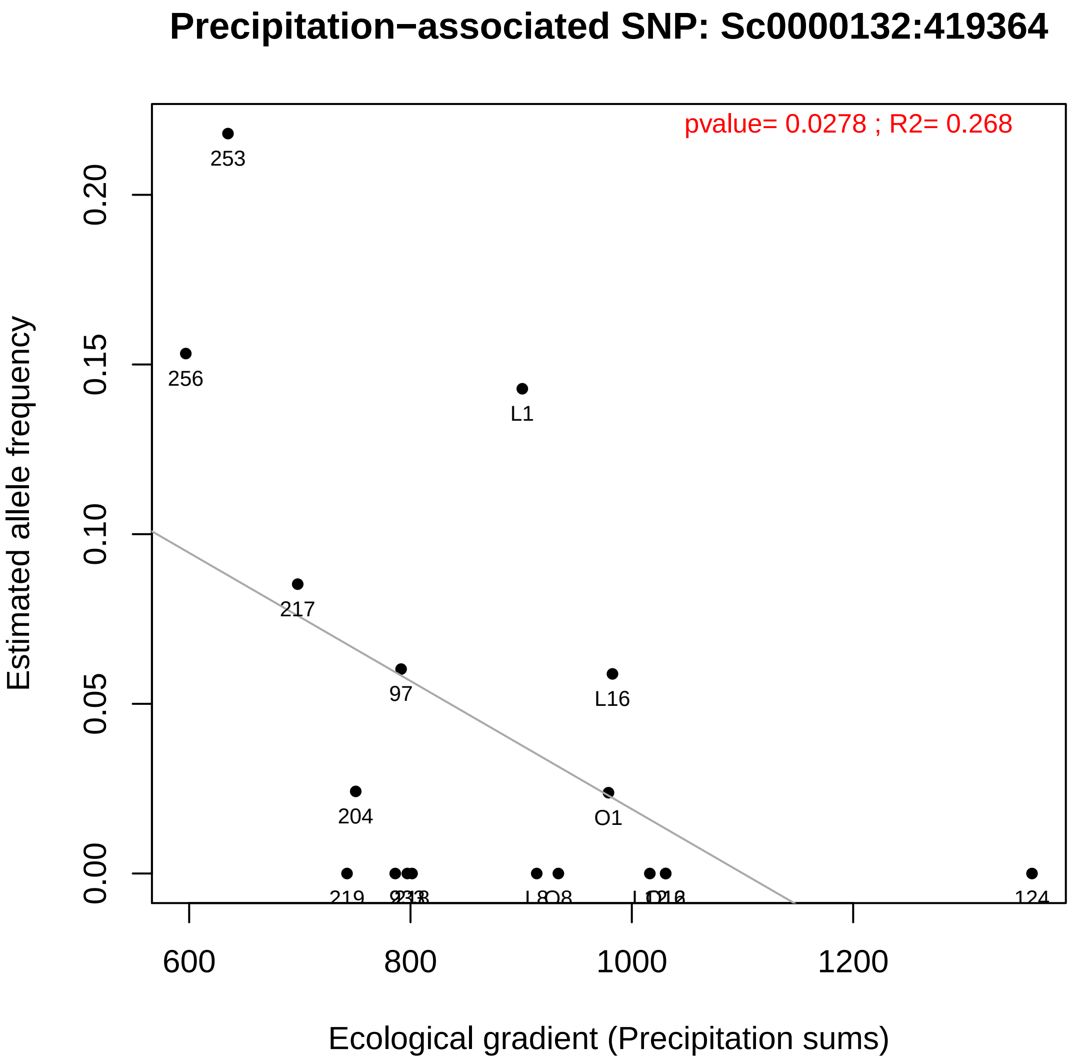

Precipitation−associated SNP: Sc0000132:419364
pvalue= 0.0278 ; R2= 0.268
Estimated allele frequency
Ecological gradient (Precipitation sums)
0.00
0.05
0.10
0.15
0.20
600
800
1000
1200
253
256
L1
217
97
L16
204
O1
219
124

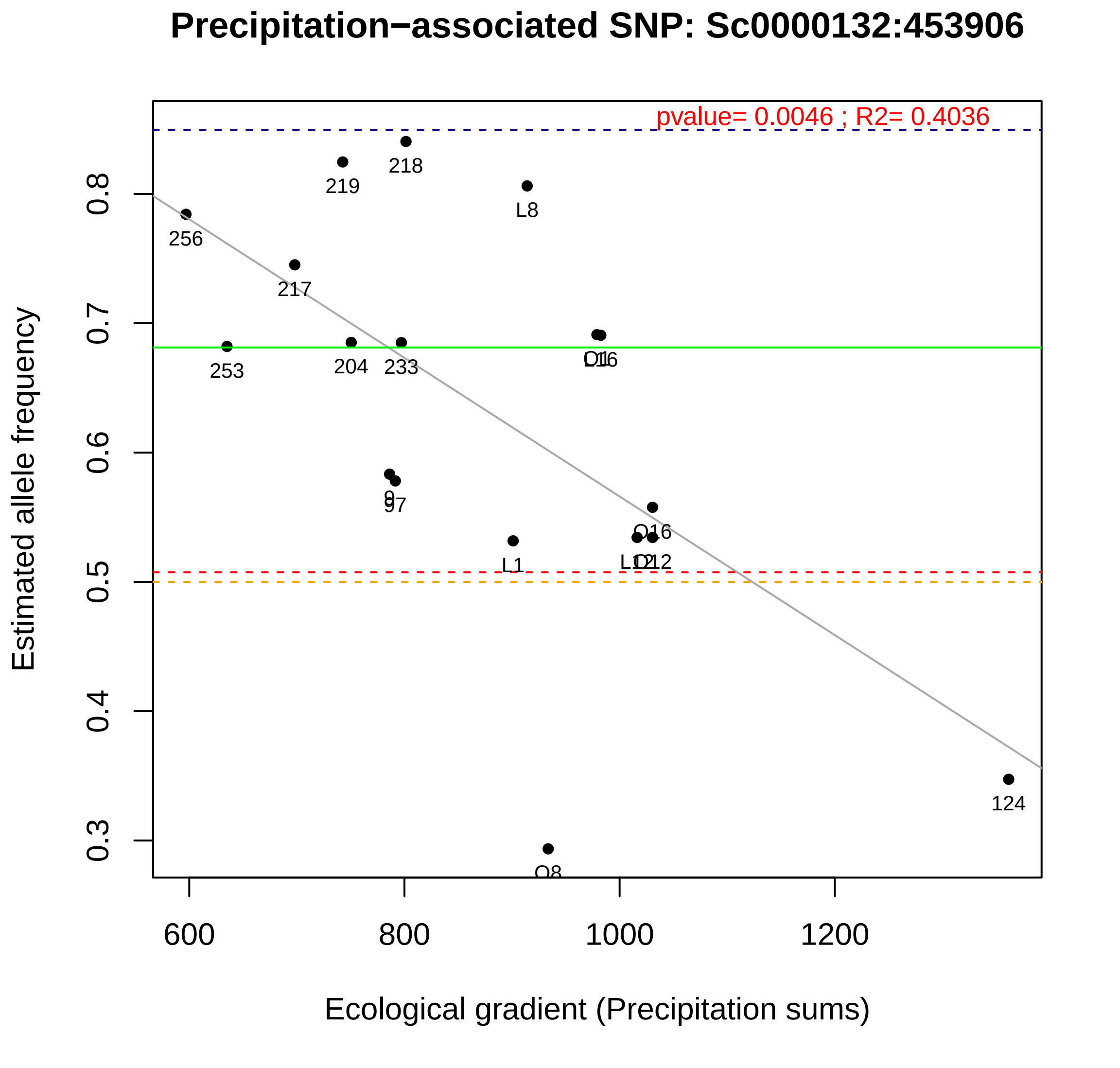
Precipitation−associated SNP: Sc0000132:453906
pvalue= 0.0046 ; R2= 0.4036
Estimated allele frequency
Ecological gradient (Precipitation sums)
0.3
0.4
0.5
0.6
0.7
0.8
600
800
1000
1200
218
219
L8
256
217
253
204
233
O16
97
L1
L12
124
O8

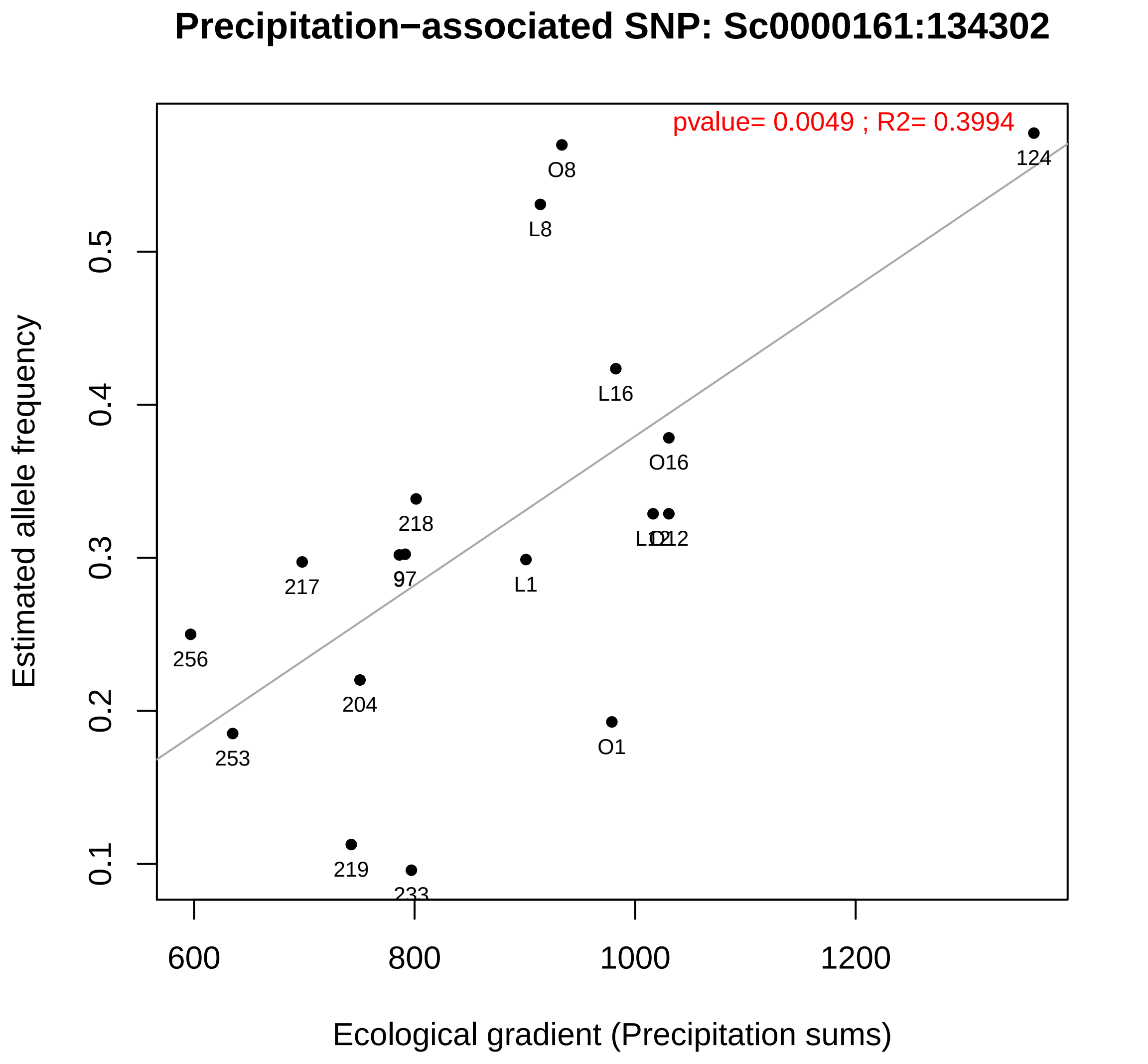
Precipitation−associated SNP: Sc0000161:134302
pvalue= 0.0049 ; R2= 0.3994
124
O8
L8
L16
O16
218
L12
O12
217
97
L1
256
204
253
O1
219
233
Estimated allele frequency
0.1
0.2
0.3
0.4
0.5
600
800
1000
1200
Ecological gradient (Precipitation sums)

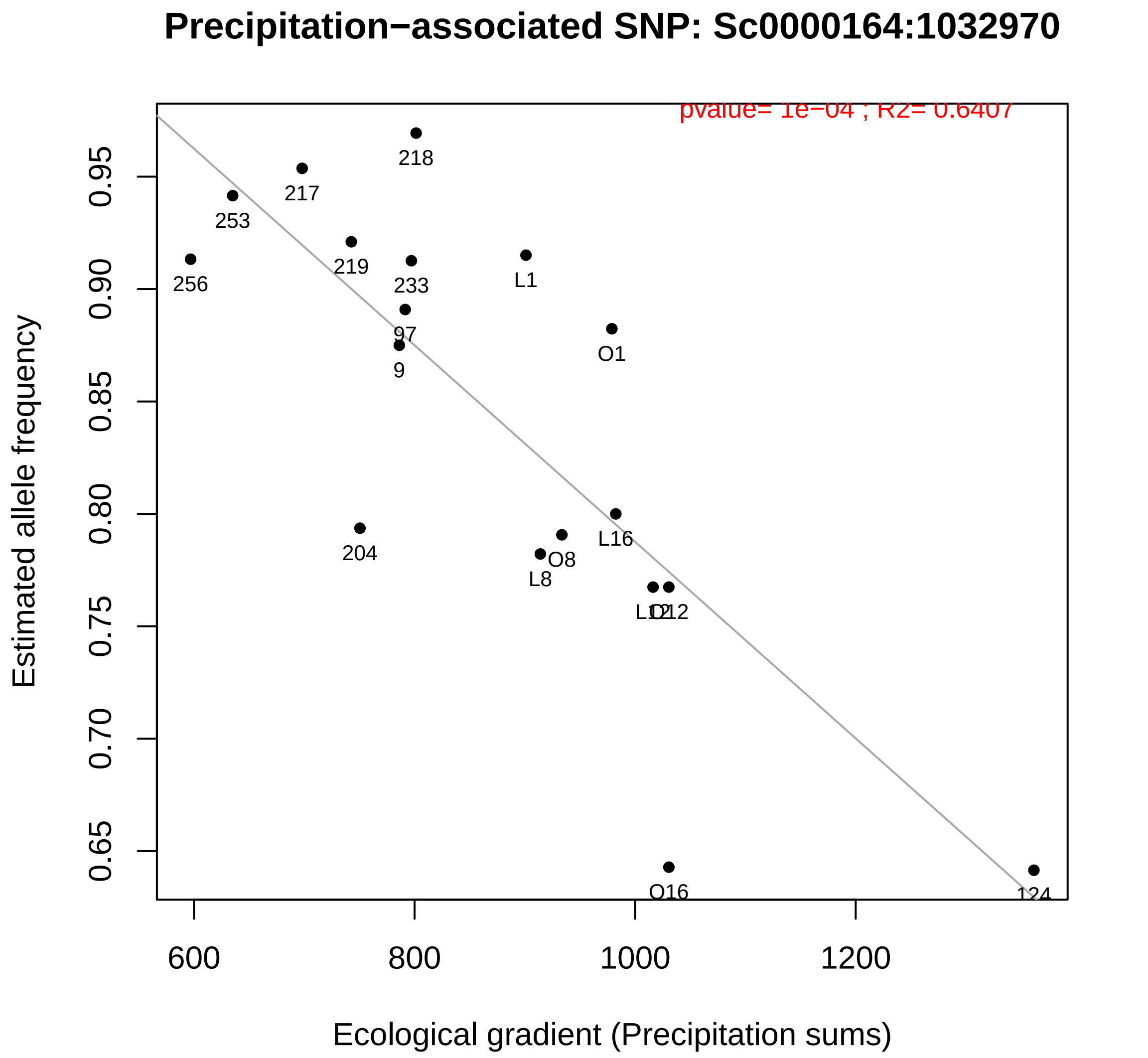

Precipitation−associated SNP: Sc0000164:1032970
pvalue= 1e−04 ; R2= 0.6407
Estimated allele frequency
0.65
0.70
0.75
0.80
0.85
0.90
0.95
600
800
1000
1200
Ecological gradient (Precipitation sums)
218
217
253
219
256
233
L1
97
9
O1
204
L16
O8
L8
O12
O16
124

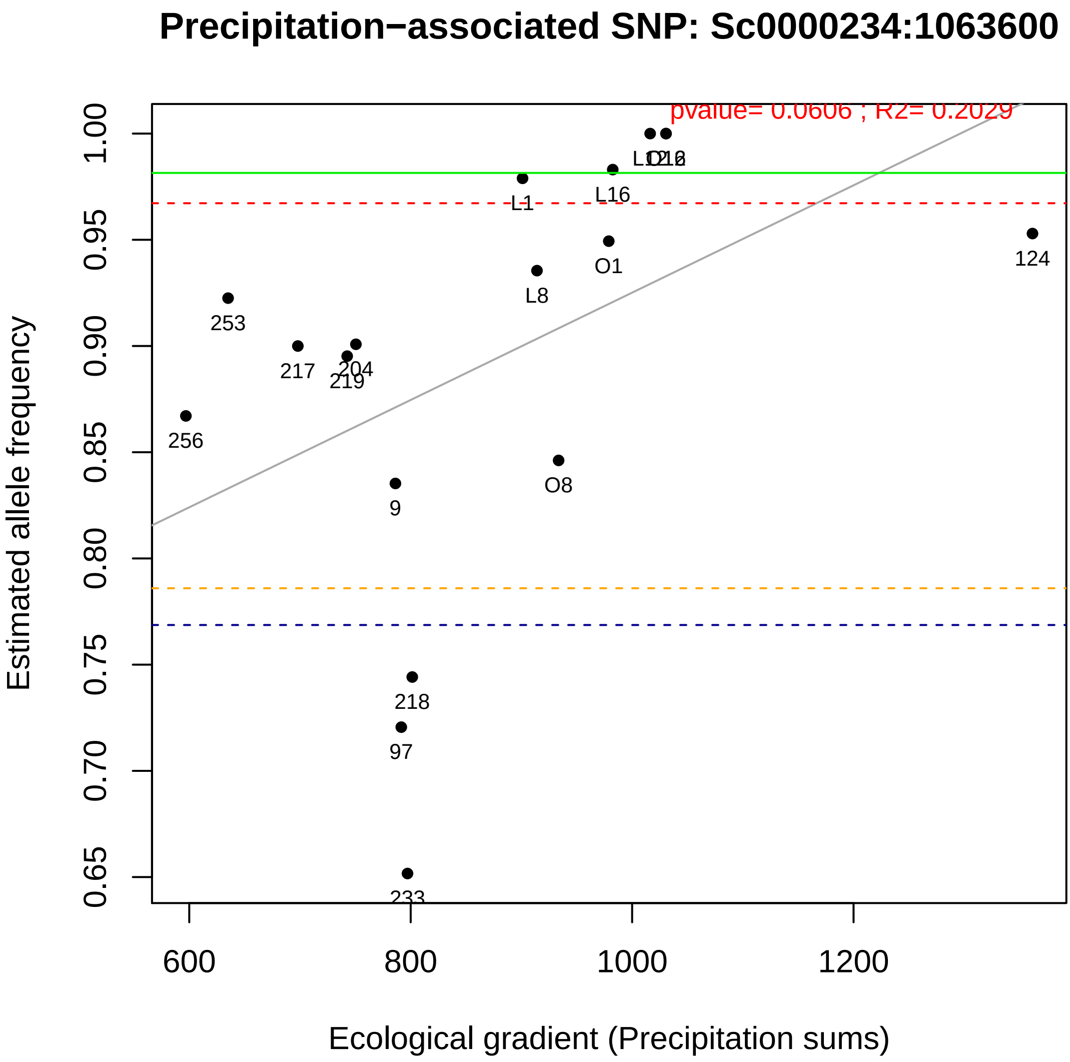

Precipitation−associated SNP: Sc0000234:1063600
pvalue= 0.0606 ; R2= 0.2029
Estimated allele frequency
Ecological gradient (Precipitation sums)
0.65
0.70
0.75
0.80
0.85
0.90
0.95
1.00
600
800
1000
1200
L1
L16
O1
L8
124
253
217
204
219
256
9
O8
218
97
233

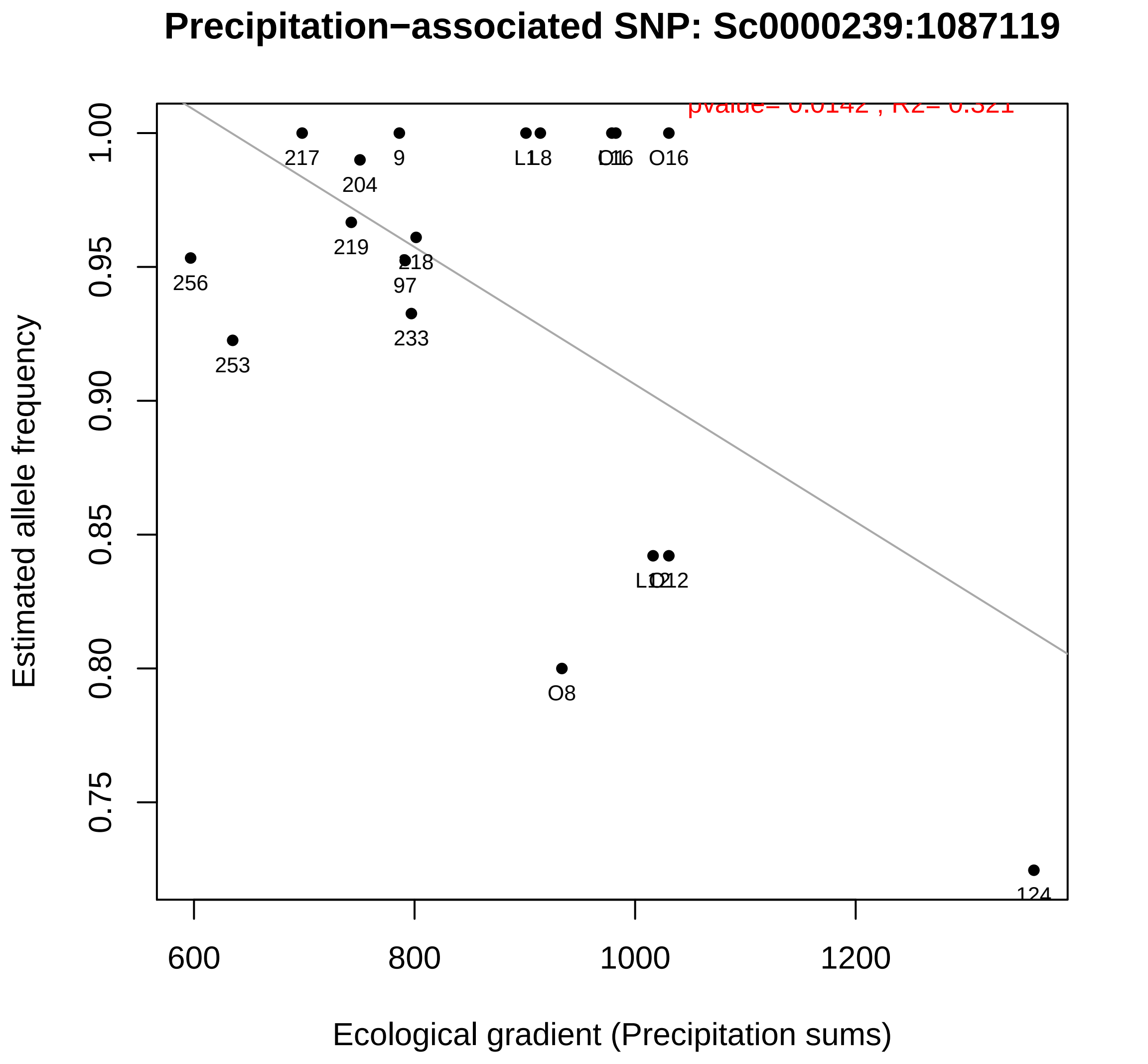

Precipitation−associated SNP: Sc0000239:1087119
pvalue= 0.0142 ; R2= 0.321
Estimated allele frequency
1.00
0.95
0.90
0.85
0.80
0.75
600
800
1000
1200
Ecological gradient (Precipitation sums)
217
204
9
219
218
97
256
233
253
O8
O16
124

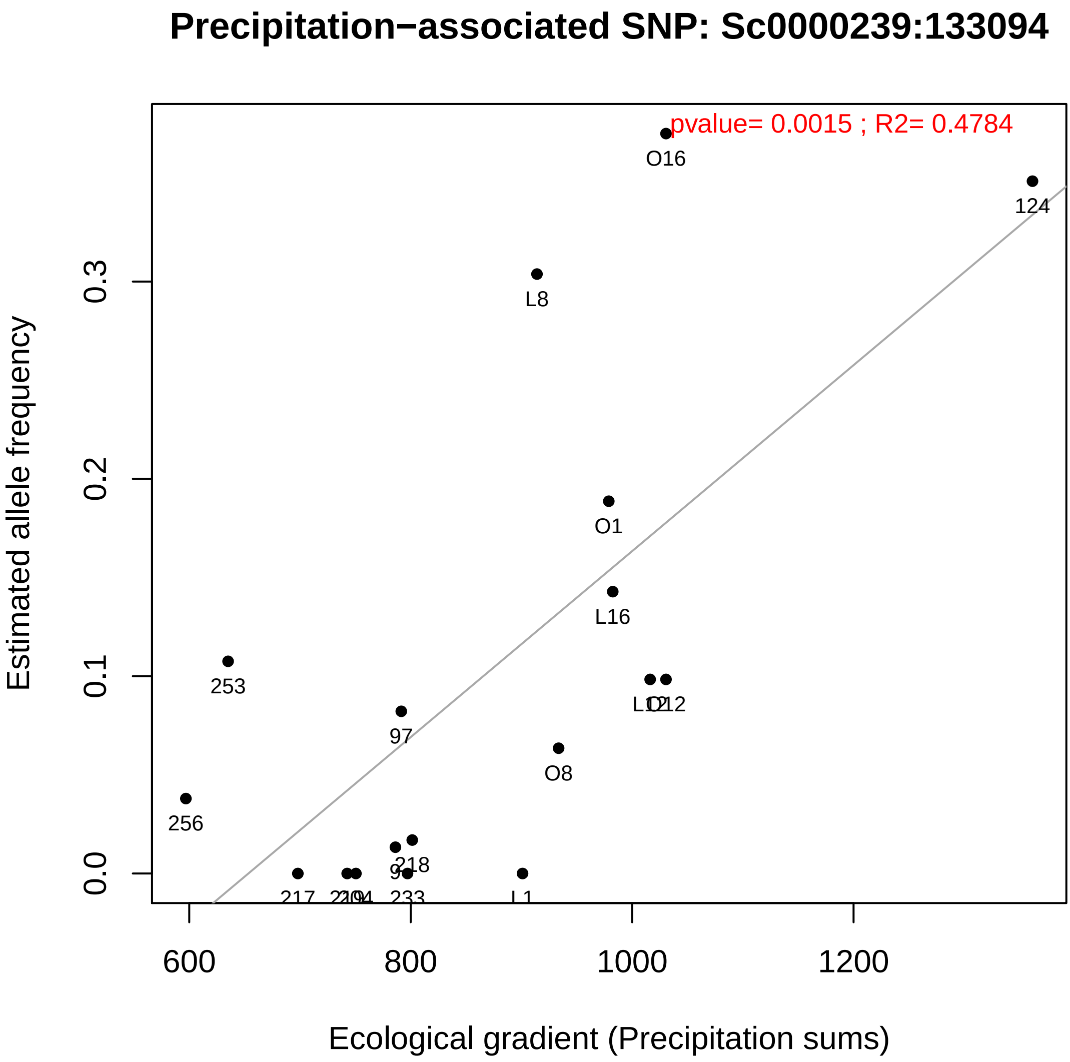

Precipitation−associated SNP: Sc0000239:133094
pvalue= 0.0015 ; R2= 0.4784
O16
124
L8
O1
L16
253
L12
O12
97
O8
256
218
217
233
L1
0.0
0.1
0.2
0.3
600
800
1000
1200
Estimated allele frequency
Ecological gradient (Precipitation sums)

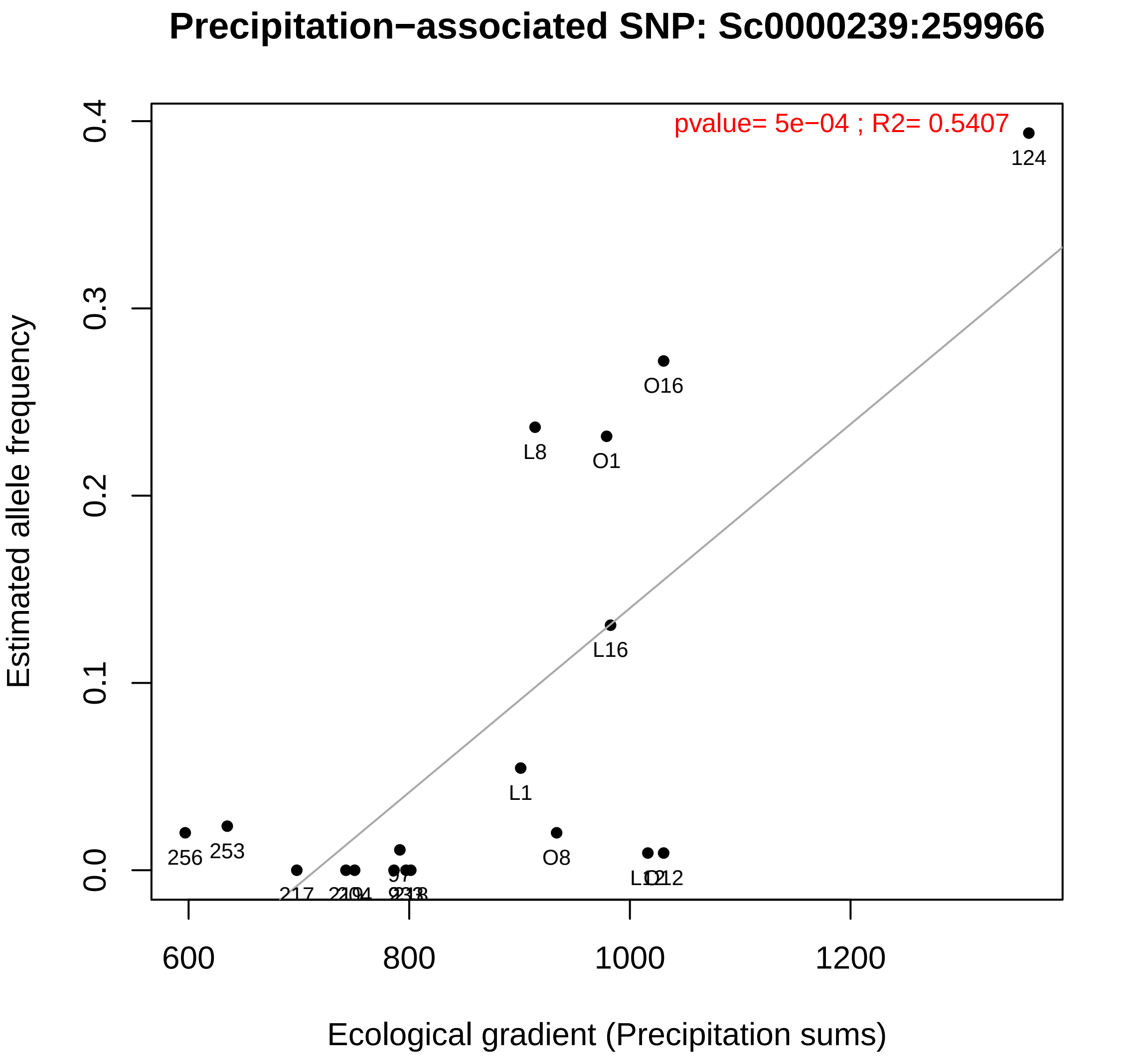

Precipitation−associated SNP: Sc0000239:259966
pvalue= 5e−04 ; R2= 0.5407
Estimated allele frequency
Ecological gradient (Precipitation sums)
0.0
0.1
0.2
0.3
0.4
600
800
1000
1200
124
O16
L8
O1
L16
L1
O8
256
253
217

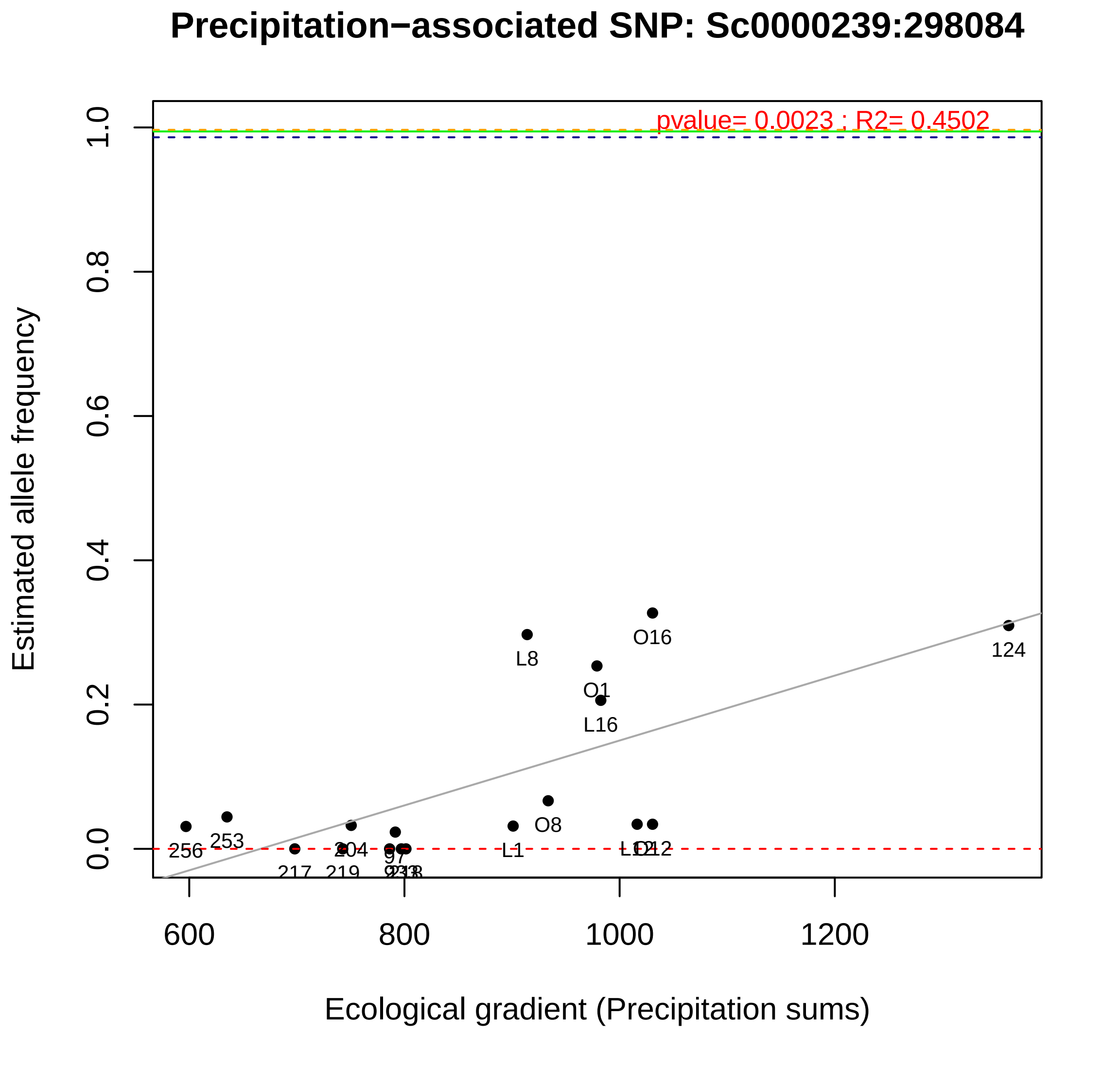
Precipitation−associated SNP: Sc0000239:298084
pvalue= 0.0023 ; R2= 0.4502
Estimated allele frequency
0.0
0.2
0.4
0.6
0.8
1.0
600
800
1000
1200
Ecological gradient (Precipitation sums)
256
253
217
219
204
97
L8
L1
O8
O1
L16
O16
124

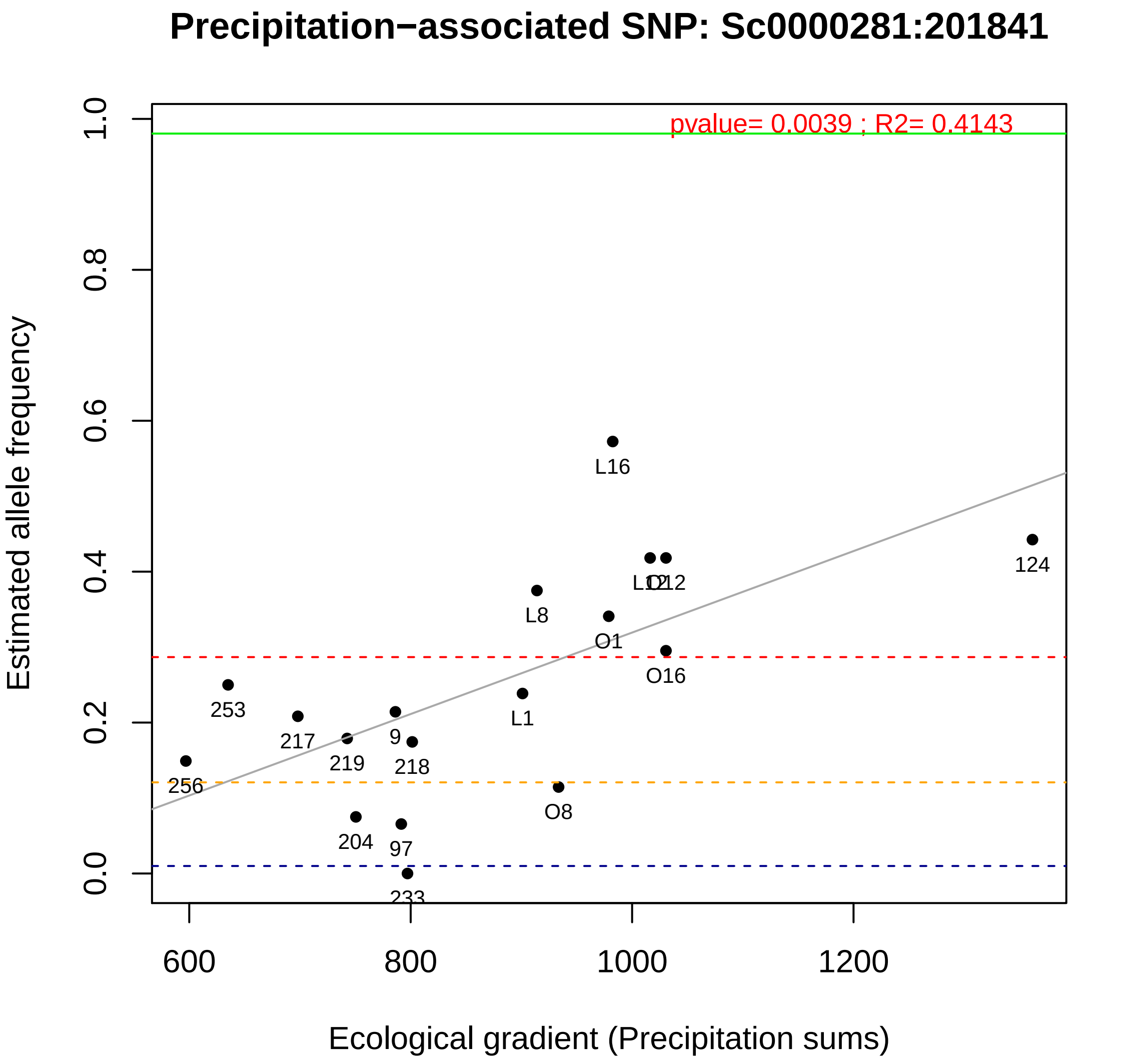
Precipitation−associated SNP: Sc0000281:201841
pvalue= 0.0039 ; R2= 0.4143
Estimated allele frequency
0.0
0.2
0.4
0.6
0.8
1.0
L16
124
L12
O12
L8
O1
O16
253
L1
217
9
219
218
256
O8
204
97
233
600
800
1000
1200
Ecological gradient (Precipitation sums)

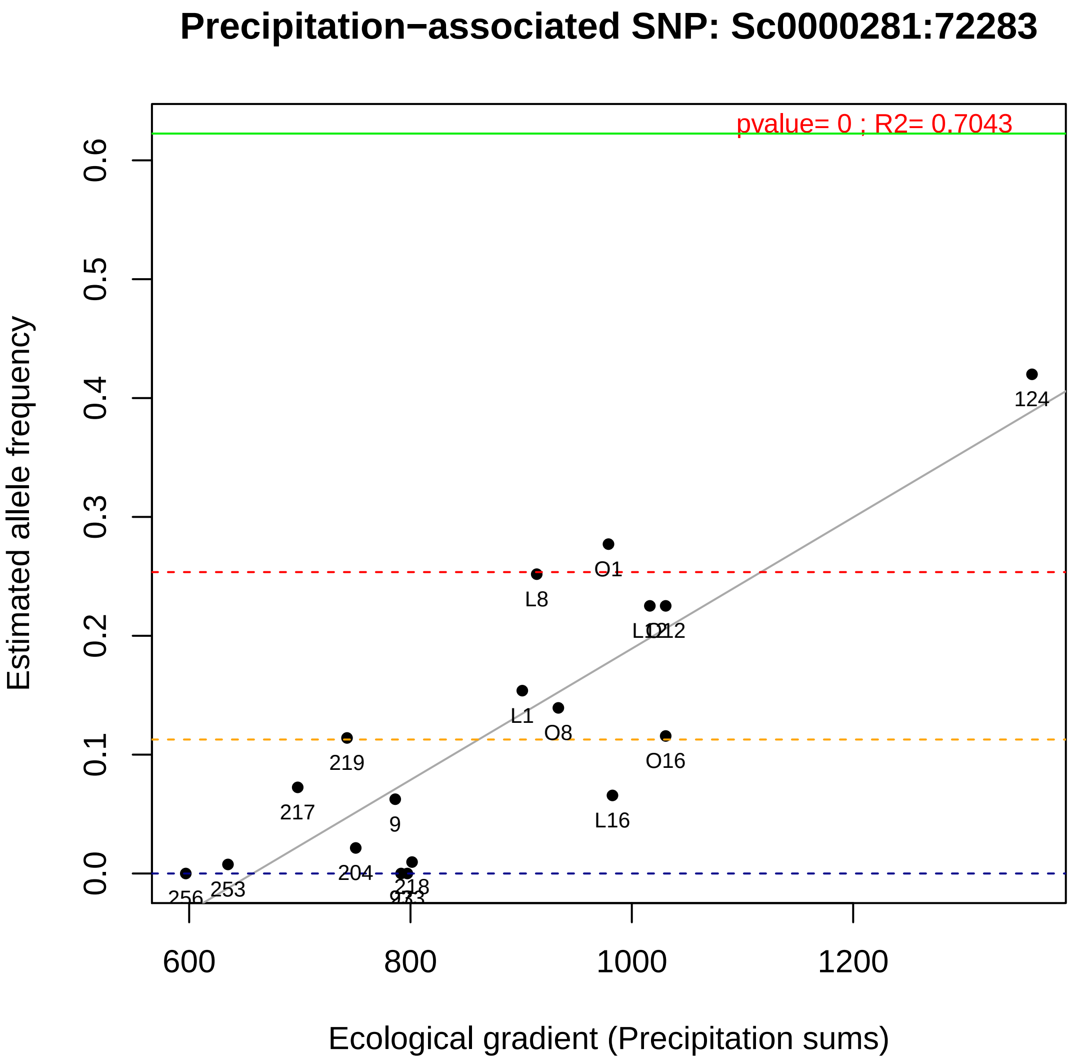

Precipitation−associated SNP: Sc0000281:72283
pvalue= 0 ; R2= 0.7043
Estimated allele frequency
0.0
0.1
0.2
0.3
0.4
0.5
0.6
600
800
1000
1200
Ecological gradient (Precipitation sums)
124
O1
L8
L1
O8
O16
219
217
9
L16
204
218
253
256

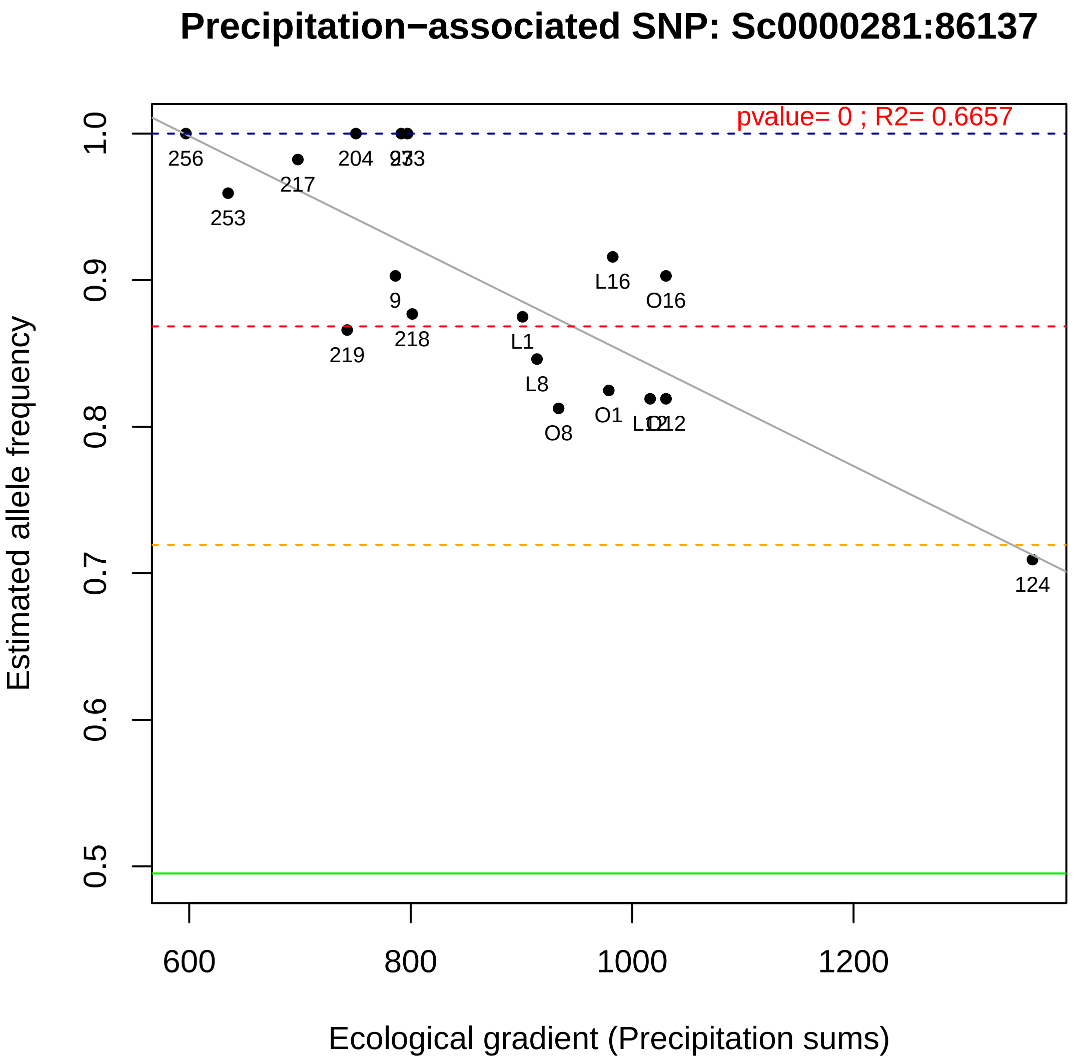
Precipitation−associated SNP: Sc0000281:86137
pvalue= 0 ; R2= 0.6657
Estimated allele frequency
1.0
0.9
0.8
0.7
0.6
0.5
600
800
1000
1200
Ecological gradient (Precipitation sums)
256
204
217
253
9
218
219
L1
L8
L16
O16
O1
O8
O12
124

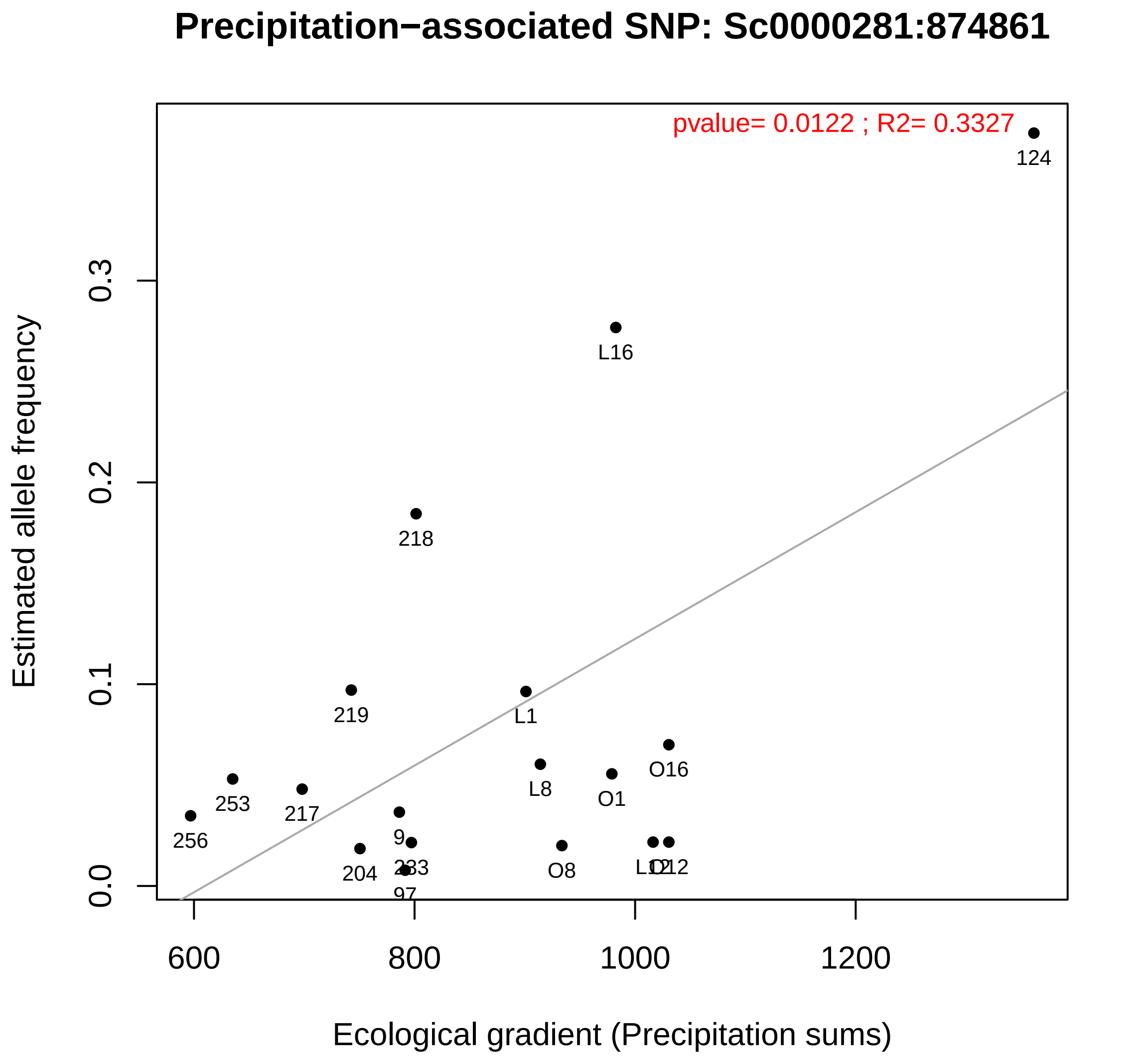

Precipitation−associated SNP: Sc0000281:874861
pvalue= 0.0122 ; R2= 0.3327
Estimated allele frequency
0.0
0.1
0.2
0.3
600
800
1000
1200
Ecological gradient (Precipitation sums)
124
L16
218
219
L1
O16
L8
O1
253
217
256
9
233
204
97
O8
L12
O12

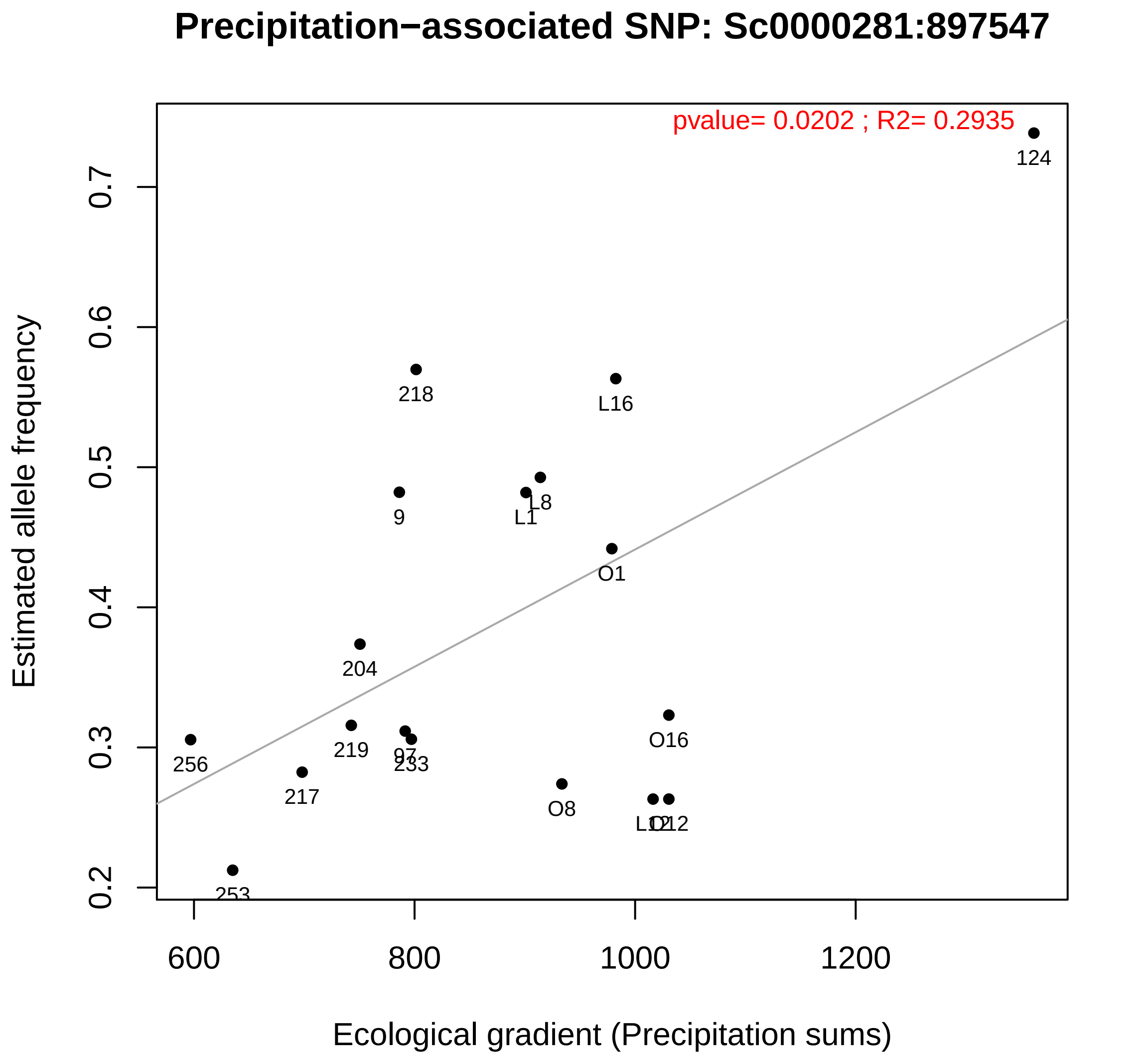

Precipitation−associated SNP: Sc0000281:897547
pvalue= 0.0202 ; R2= 0.2935
Estimated allele frequency
0.2
0.3
0.4
0.5
0.6
0.7
600
800
1000
1200
Ecological gradient (Precipitation sums)
124
218
L16
9
L1
L8
O1
204
256
219
97
233
217
O16
O8
L12
O12
253

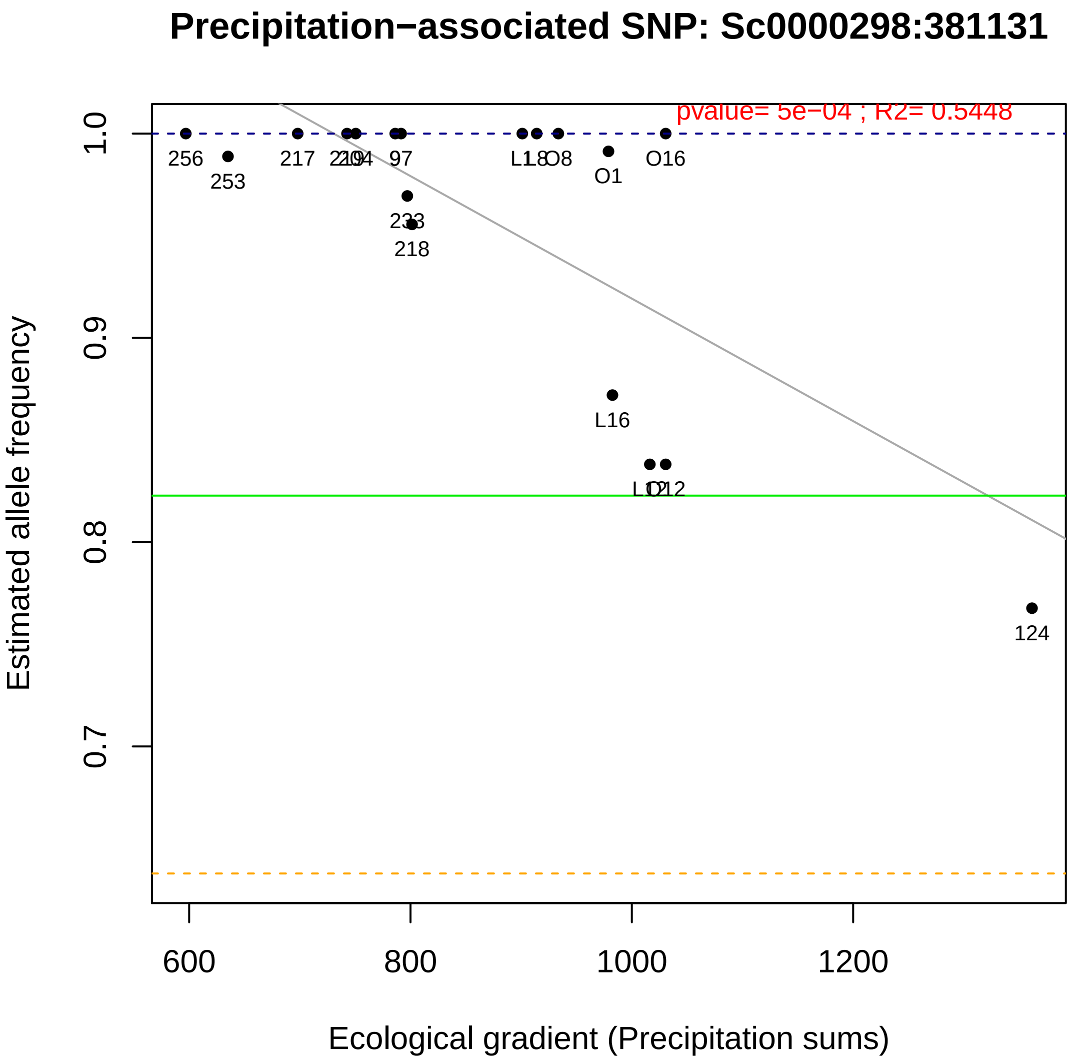

Precipitation−associated SNP: Sc0000298:381131
pvalue= 5e−04 ; R2= 0.5448
Estimated allele frequency
1.0
0.9
0.8
0.7
256
253
217
97
218
O8
O1
O16
L16
124
600
800
1000
1200
Ecological gradient (Precipitation sums)

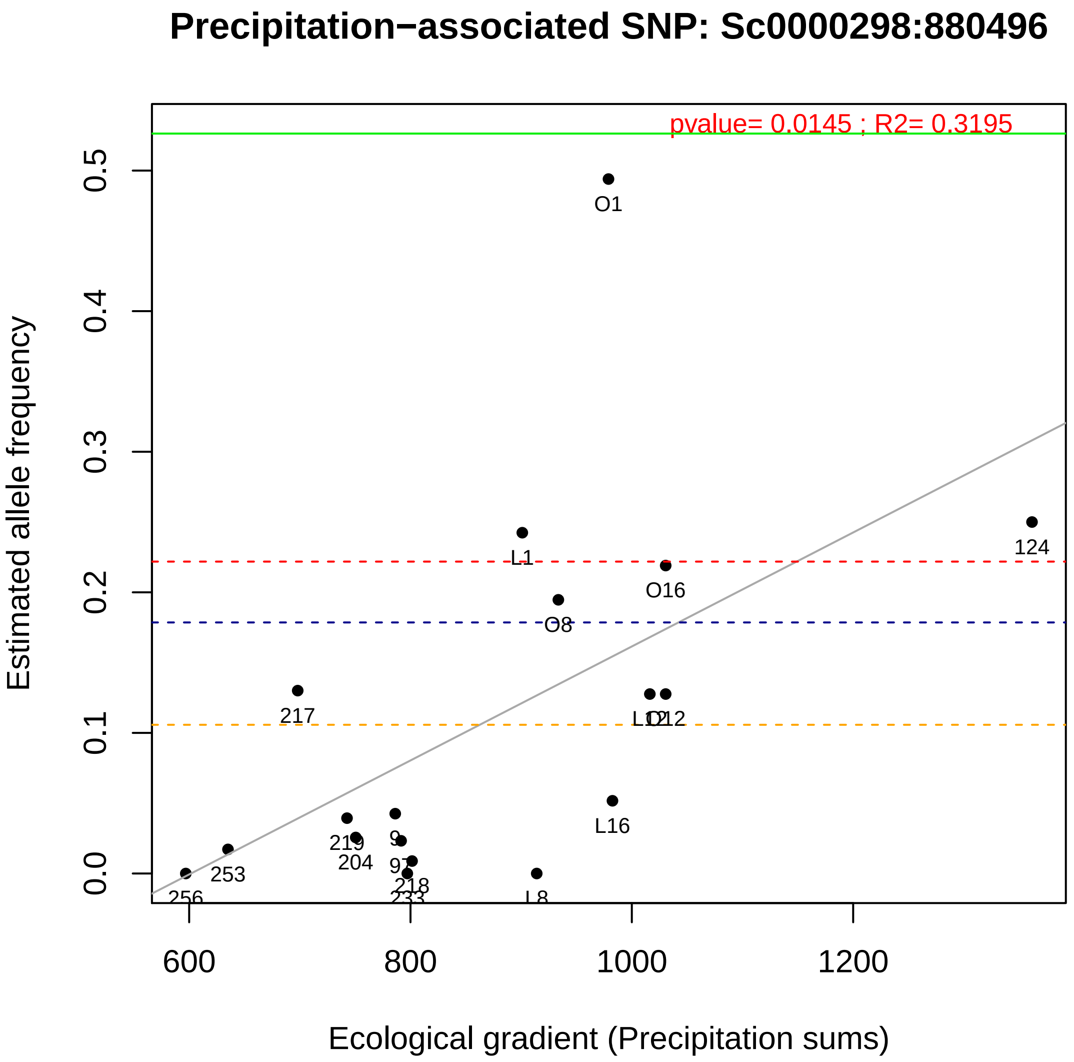

Precipitation−associated SNP: Sc0000298:880496
pvalue= 0.0145 ; R2= 0.3195
Estimated allele frequency
0.0
0.1
0.2
0.3
0.4
0.5
600
800
1000
1200
Ecological gradient (Precipitation sums)
O1
L1
124
O16
O8
217
O12
L16
219
204
253
218
L8

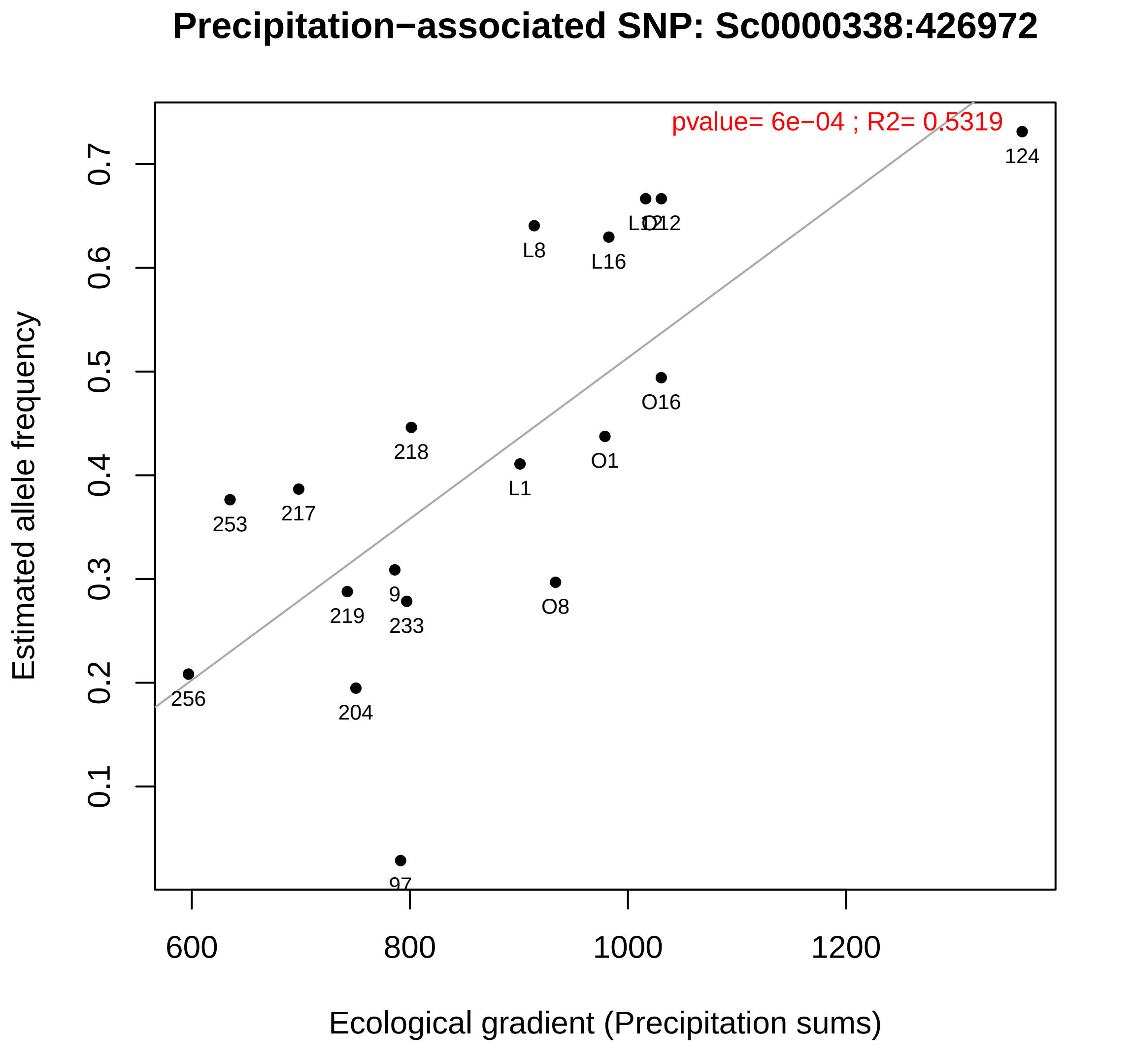

Precipitation−associated SNP: Sc0000338:426972
pvalue= 6e−04 ; R2= 0.5319
Estimated allele frequency
0.1
0.2
0.3
0.4
0.5
0.6
0.7
600
800
1000
1200
Ecological gradient (Precipitation sums)
124
L8
L16
L12
O12
O16
218
O1
L1
253
217
9
219
233
O8
256
204
97

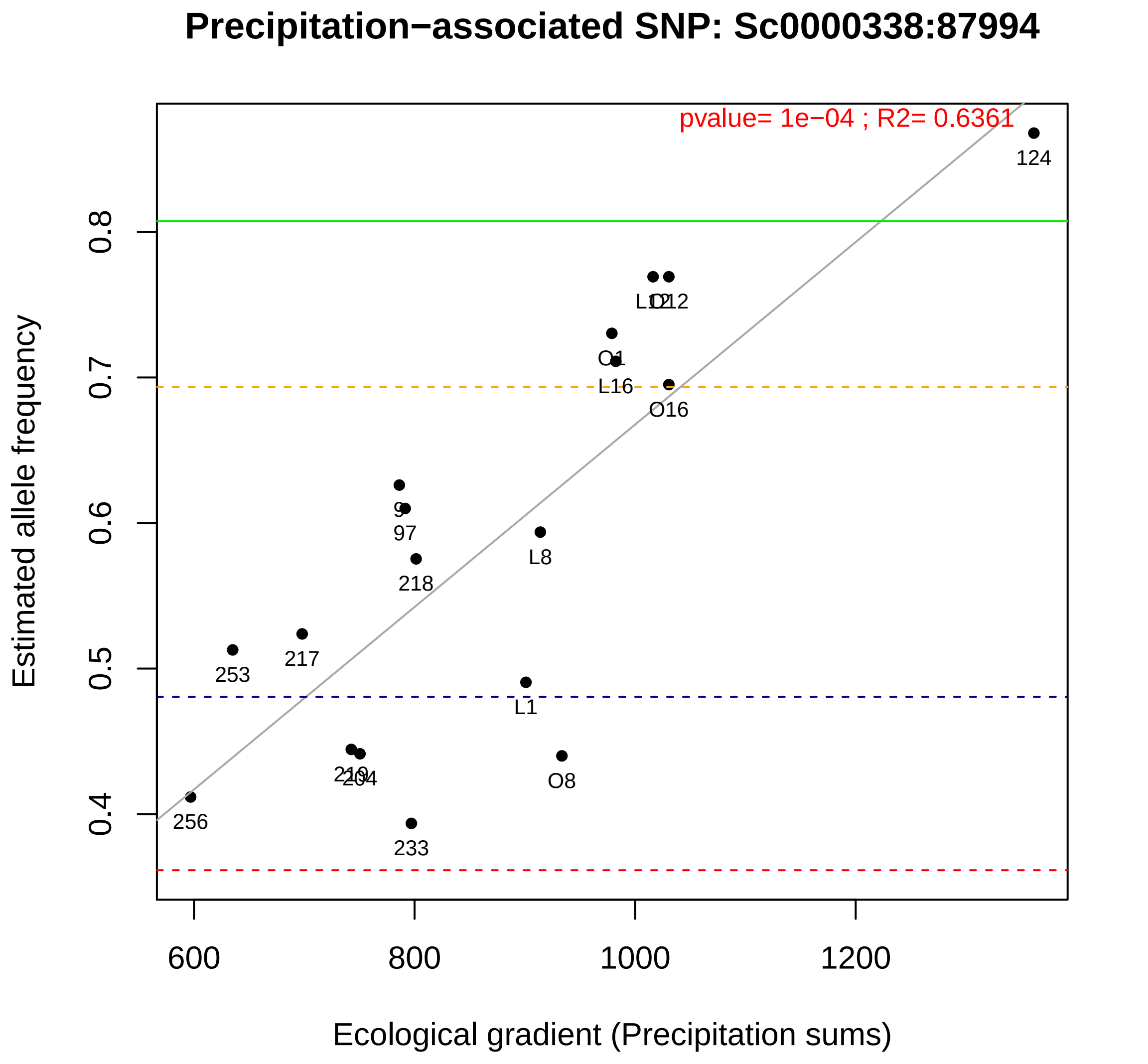
Precipitation−associated SNP: Sc0000338:87994
pvalue= 1e−04 ; R2= 0.6361
Estimated allele frequency
Ecological gradient (Precipitation sums)
0.4
0.5
0.6
0.7
0.8
600
800
1000
1200
124
O12
O1
L16
O16
97
L8
218
217
253
L1
O8
256
233

**Precipitation−associated SNP: Sc0000338:91769**

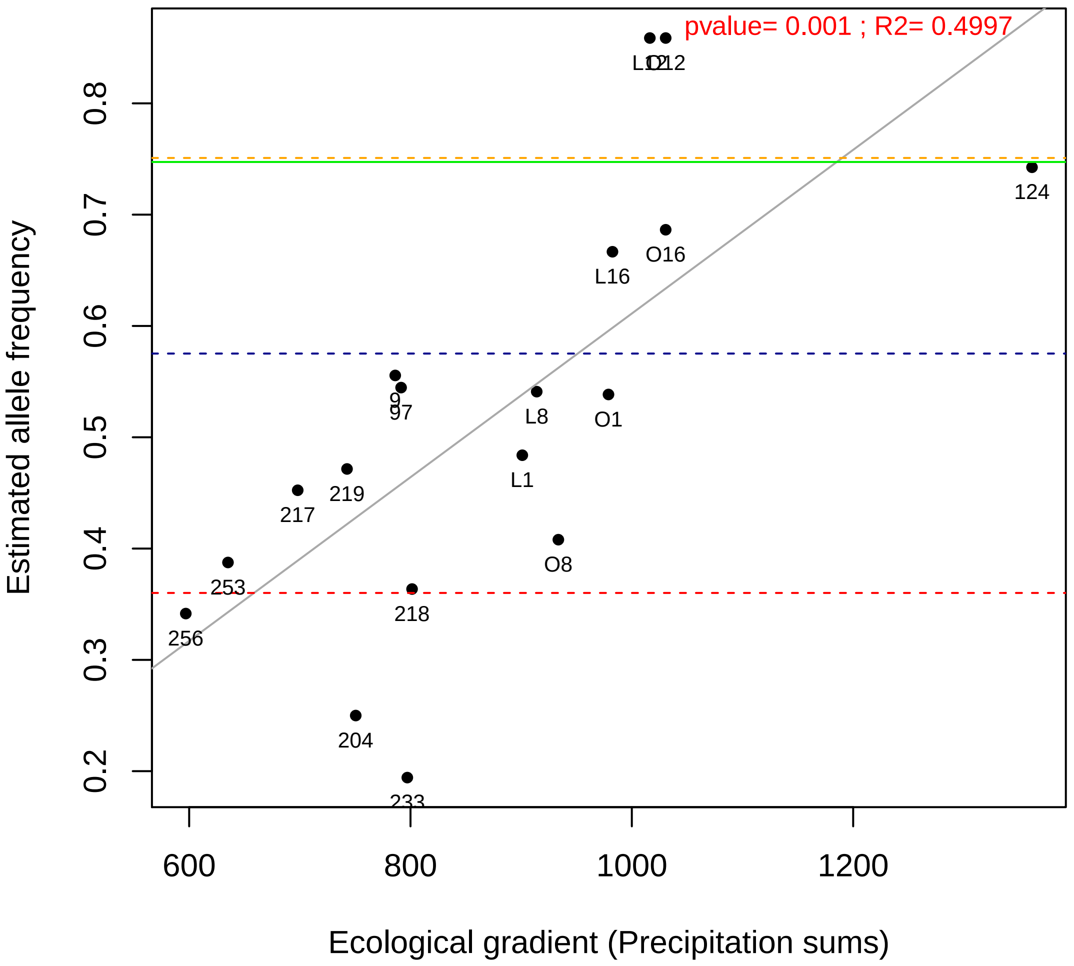

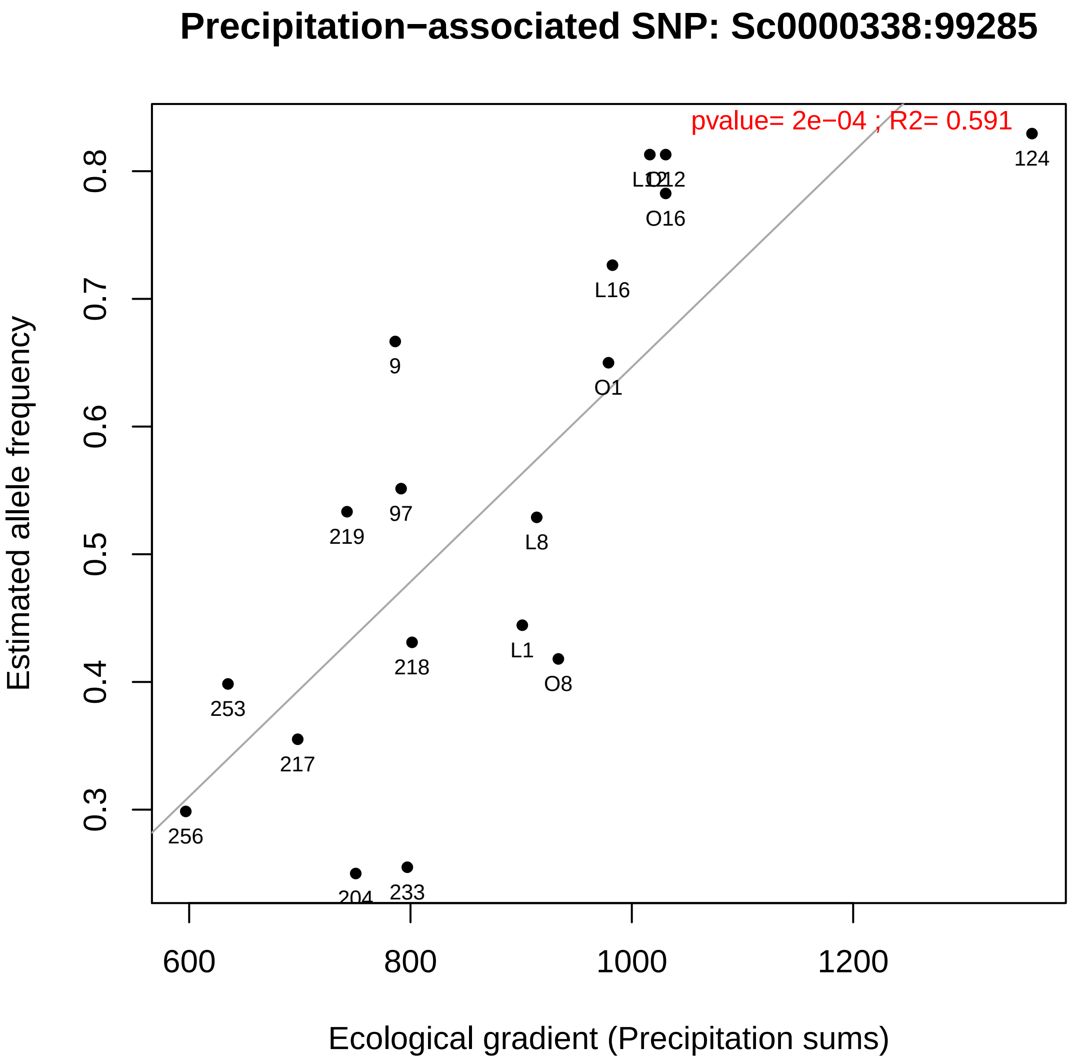

Precipitation−associated SNP: Sc0000338:99285
pvalue= 2e−04 ; R2= 0.591
Estimated allele frequency
0.3
0.4
0.5
0.6
0.7
0.8
Ecological gradient (Precipitation sums)
600
800
1000
1200
124
L12
O12
O16
L16
9
O1
97
219
L8
L1
218
O8
253
217
256
204
233

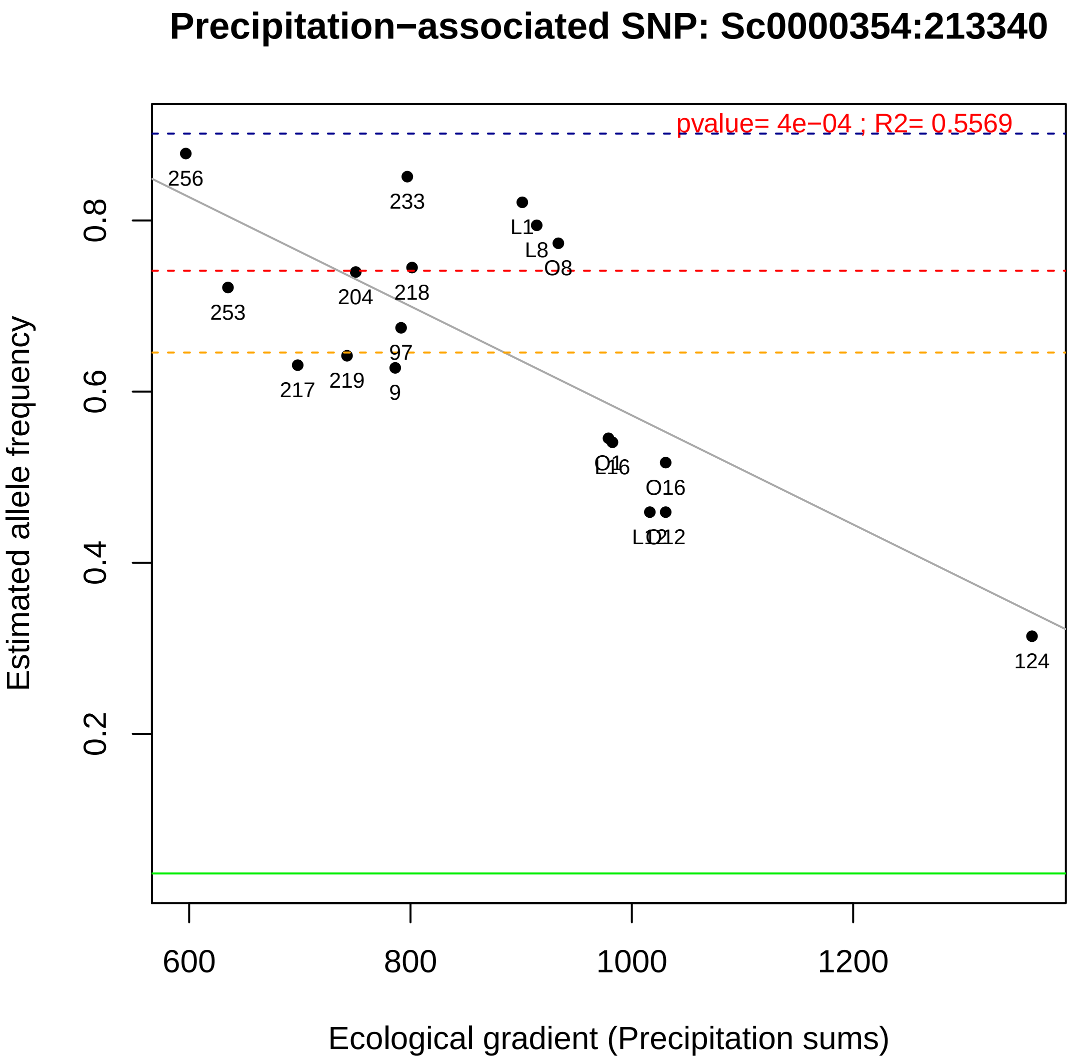
Precipitation−associated SNP: Sc0000354:213340
pvalue= 4e−04 ; R2= 0.5569
Estimated allele frequency
Ecological gradient (Precipitation sums)
0.2
0.4
0.6
0.8
600
800
1000
1200
256
233
L1
L8
O8
204
218
253
97
217
219
9
O1
L16
O16
L12
O12
124

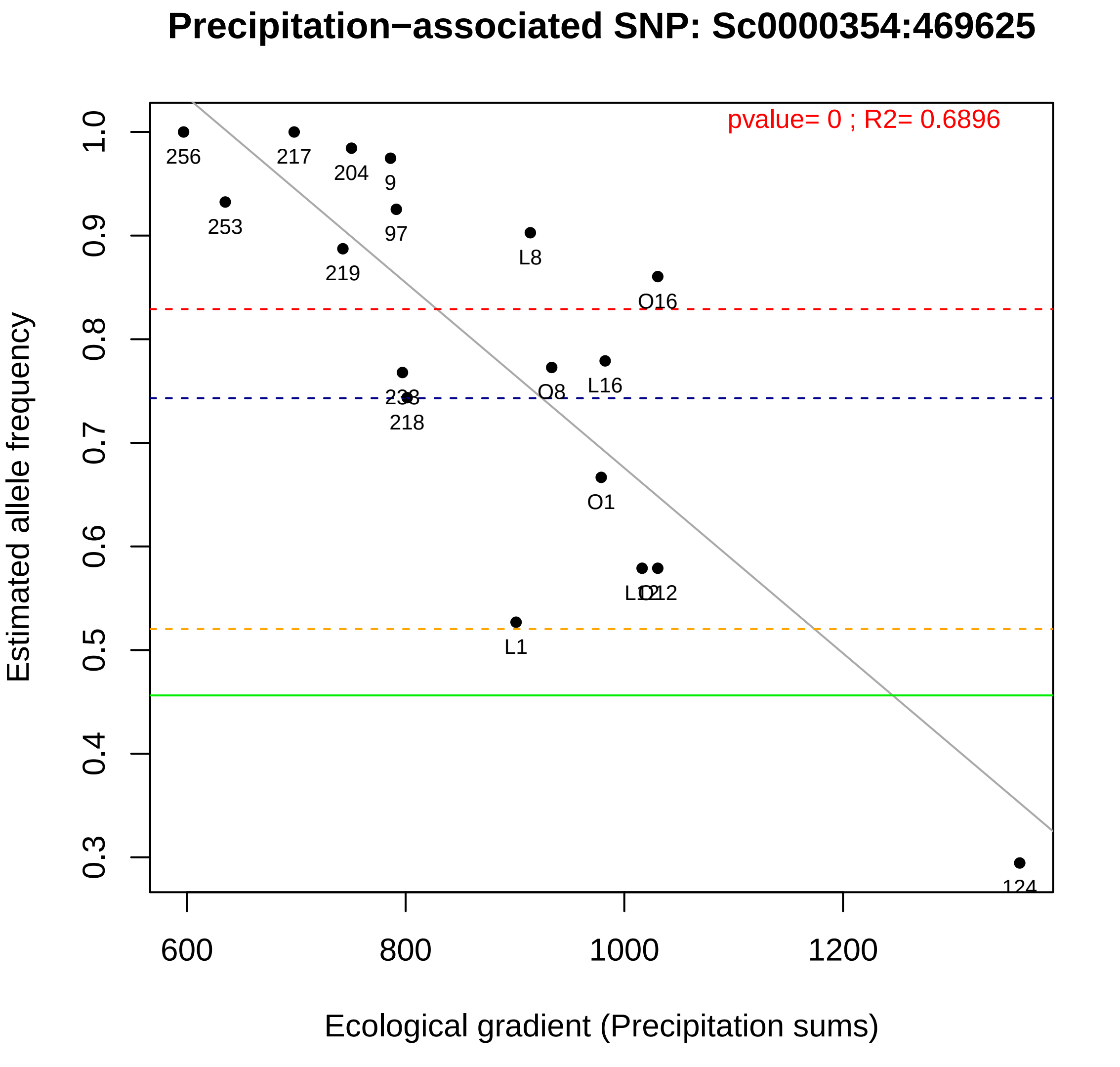
Precipitation−associated SNP: Sc0000354:469625
pvalue= 0 ; R2= 0.6896
Estimated allele frequency
0.3
0.4
0.5
0.6
0.7
0.8
0.9
1.0
600
800
1000
1200
Ecological gradient (Precipitation sums)
256
217
204
9
253
97
219
L8
O16
L16
O8
218
O1
L1
124

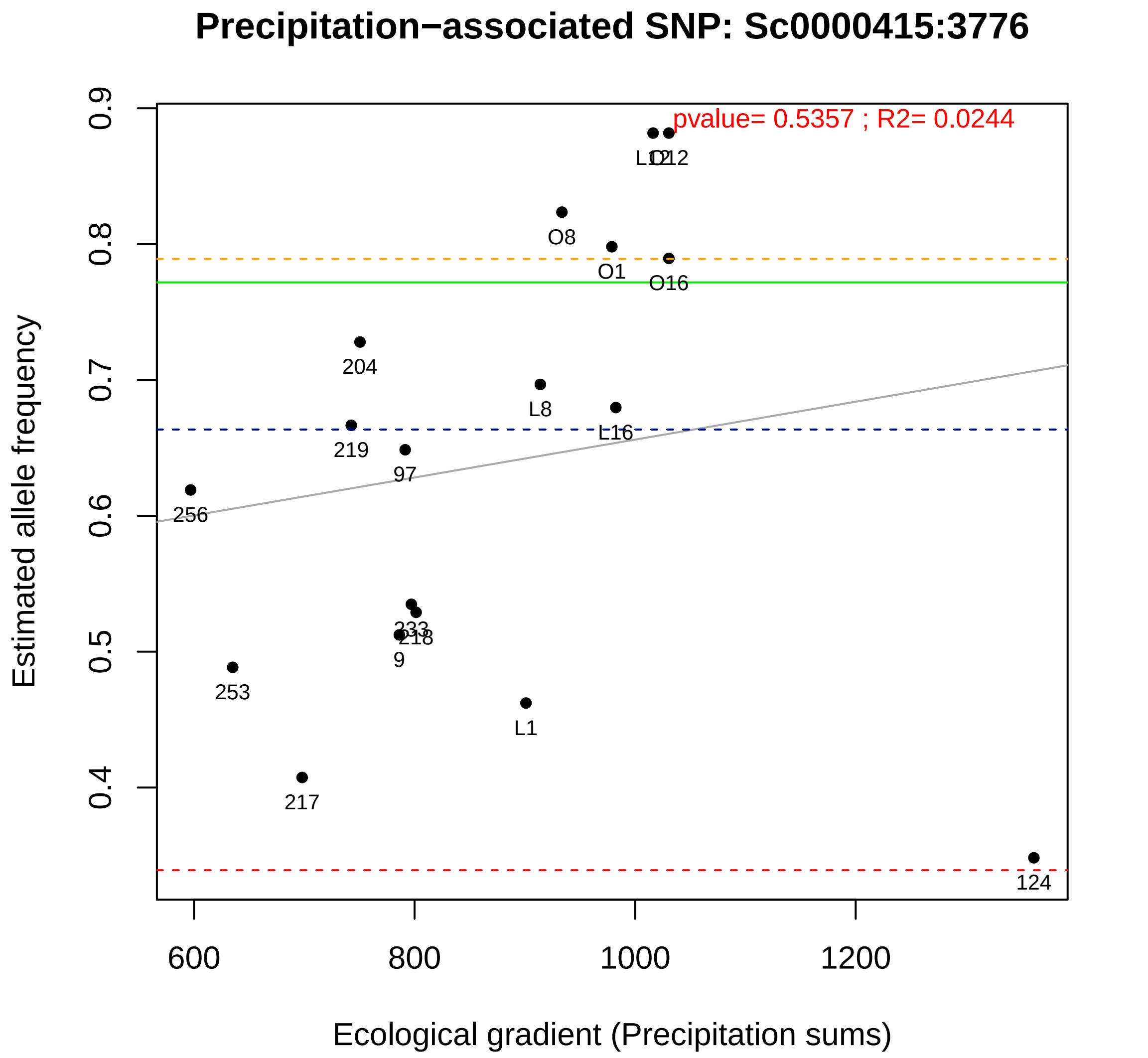

Precipitation−associated SNP: Sc0000415:3776
pvalue= 0.5357 ; R2= 0.0244
Estimated allele frequency
0.4
0.5
0.6
0.7
0.8
0.9
600
800
1000
1200
Ecological gradient (Precipitation sums)
L12
O12
O8
O1
O16
204
L8
L16
219
97
256
233
218
9
253
L1
217
124

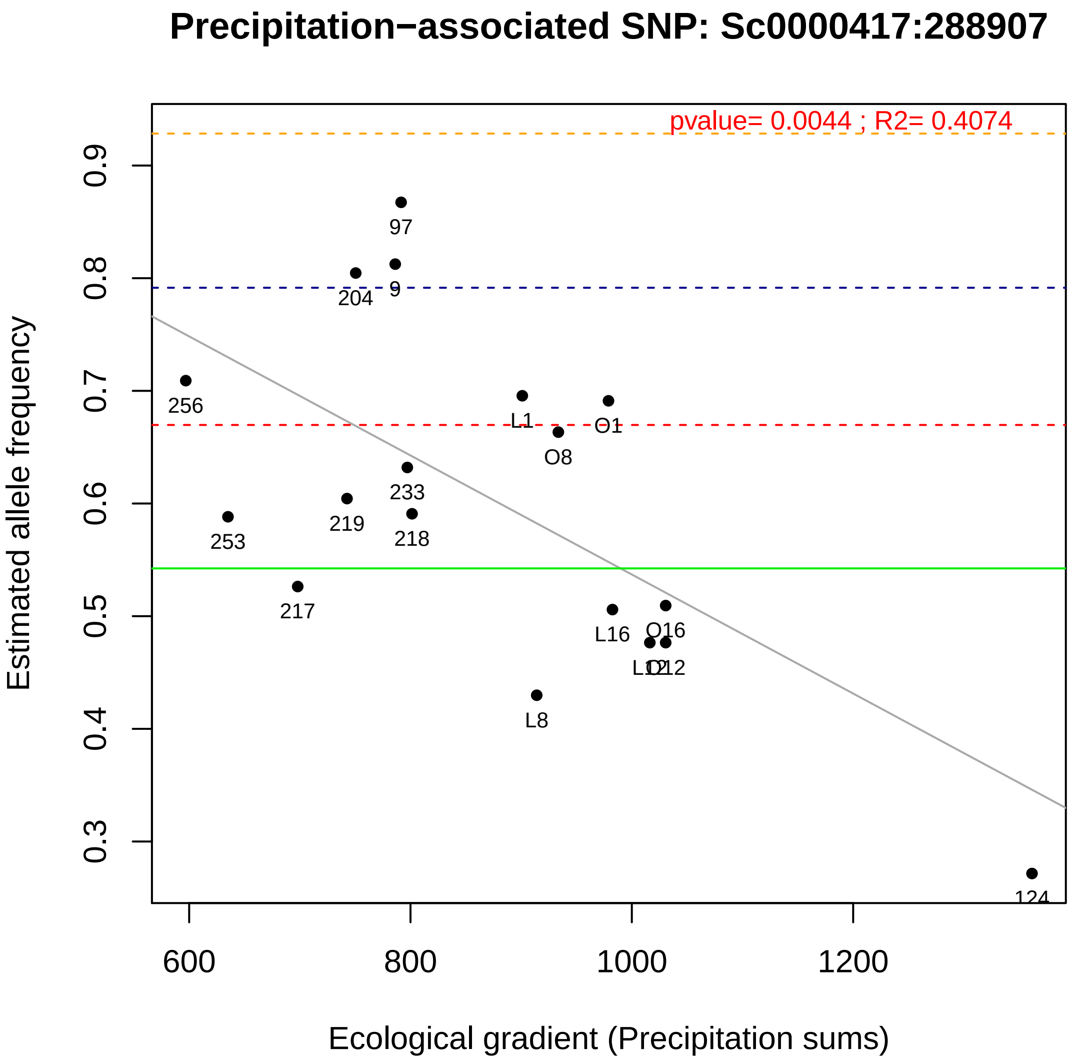

Precipitation−associated SNP: Sc0000417:288907
pvalue= 0.0044 ; R2= 0.4074
Estimated allele frequency
0.3
0.4
0.5
0.6
0.7
0.8
0.9
600
800
1000
1200
Ecological gradient (Precipitation sums)
97
204
9
256
L1
O1
O8
233
219
218
253
217
L16
O16
L8
124

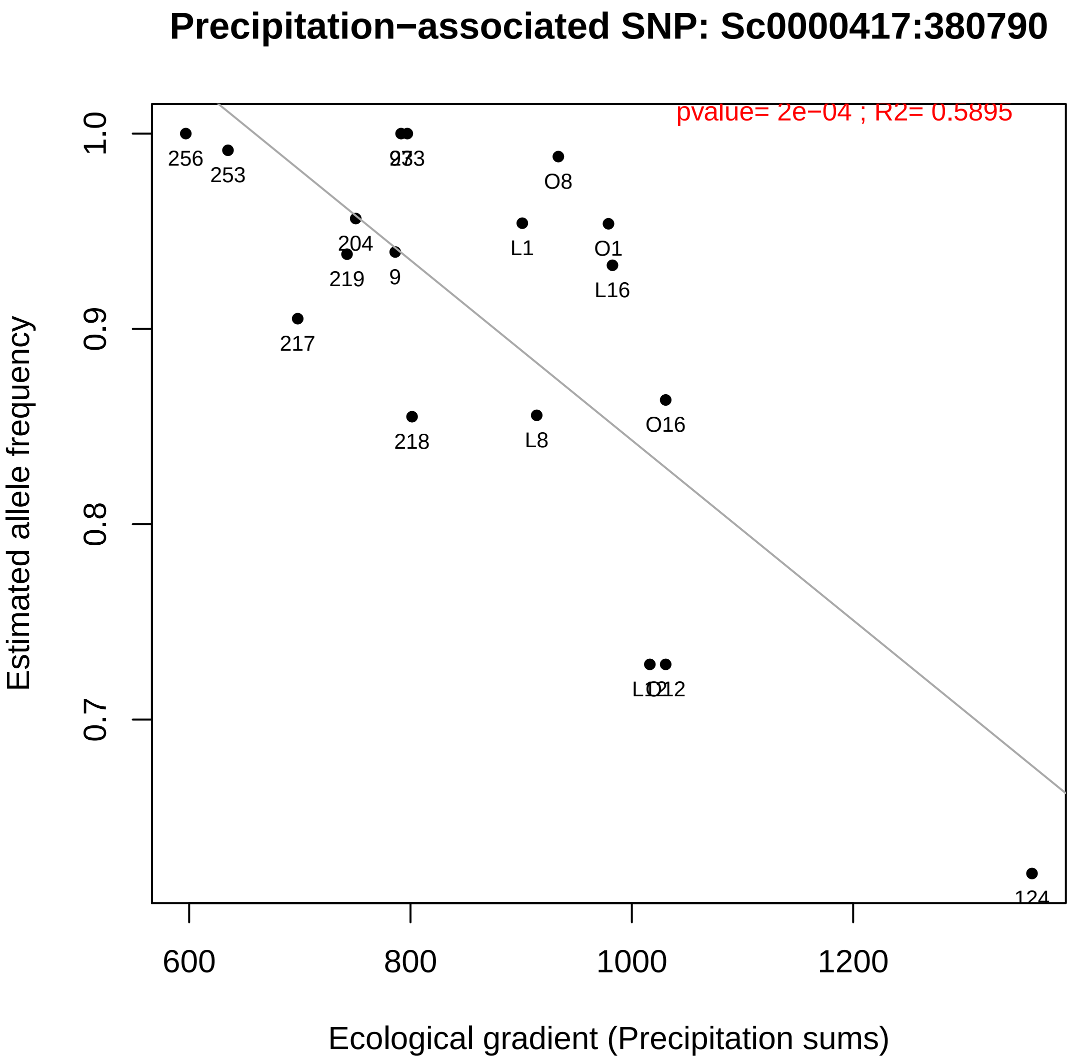
Precipitation−associated SNP: Sc0000417:380790
pvalue= 2e−04 ; R2= 0.5895
Estimated allele frequency
Ecological gradient (Precipitation sums)
0.7
0.8
0.9
1.0
600
800
1000
1200
256
253
204
219
9
217
218
L1
O8
O1
L16
L8
O16
124

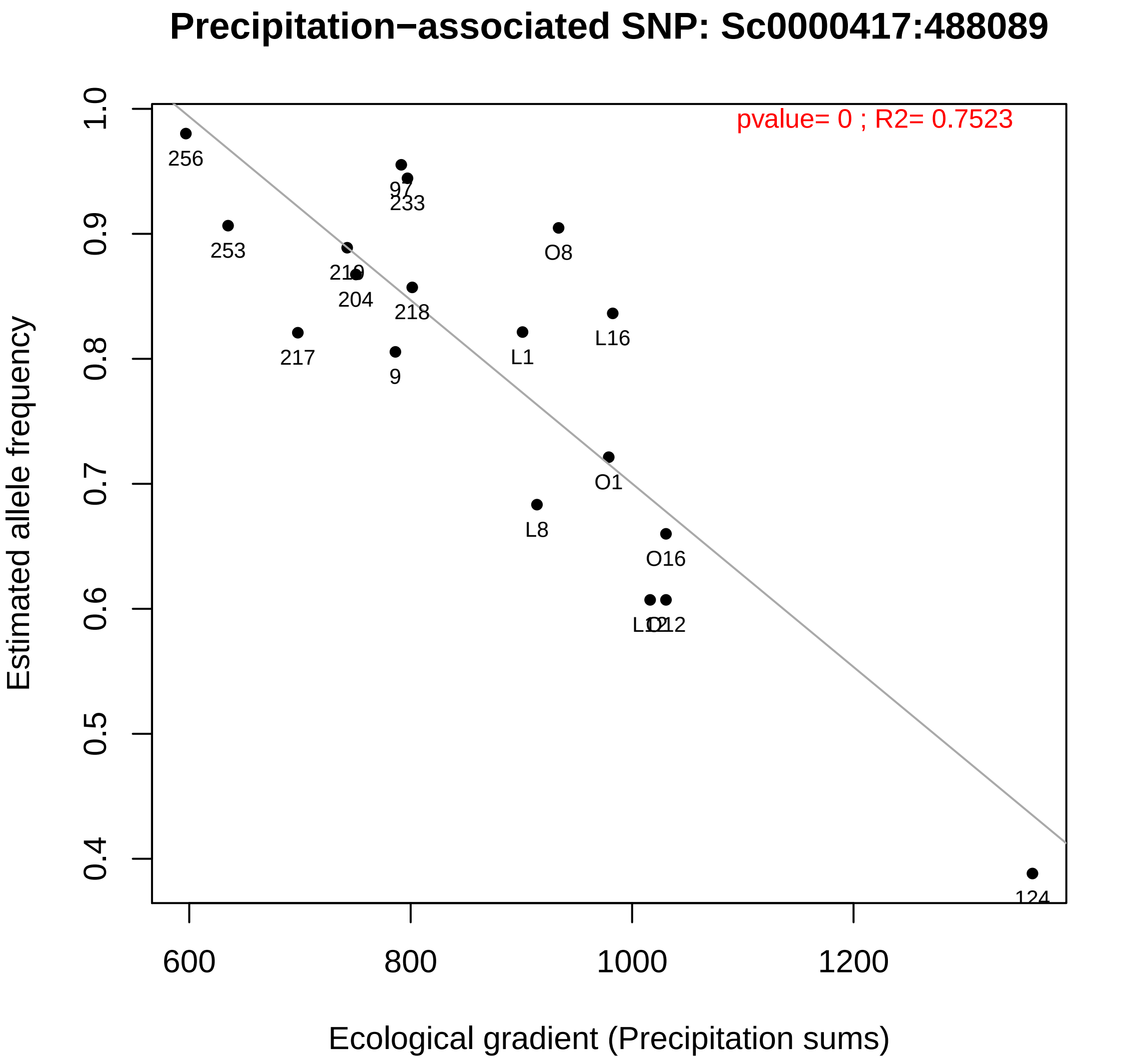
Precipitation−associated SNP: Sc0000417:488089
pvalue= 0 ; R2= 0.7523
Estimated allele frequency
Ecological gradient (Precipitation sums)
1.0
0.9
0.8
0.7
0.6
0.5
0.4
600
800
1000
1200
256
97
233
253
210
204
218
O8
217
9
L1
L16
O1
L8
O16
L12
O12
124

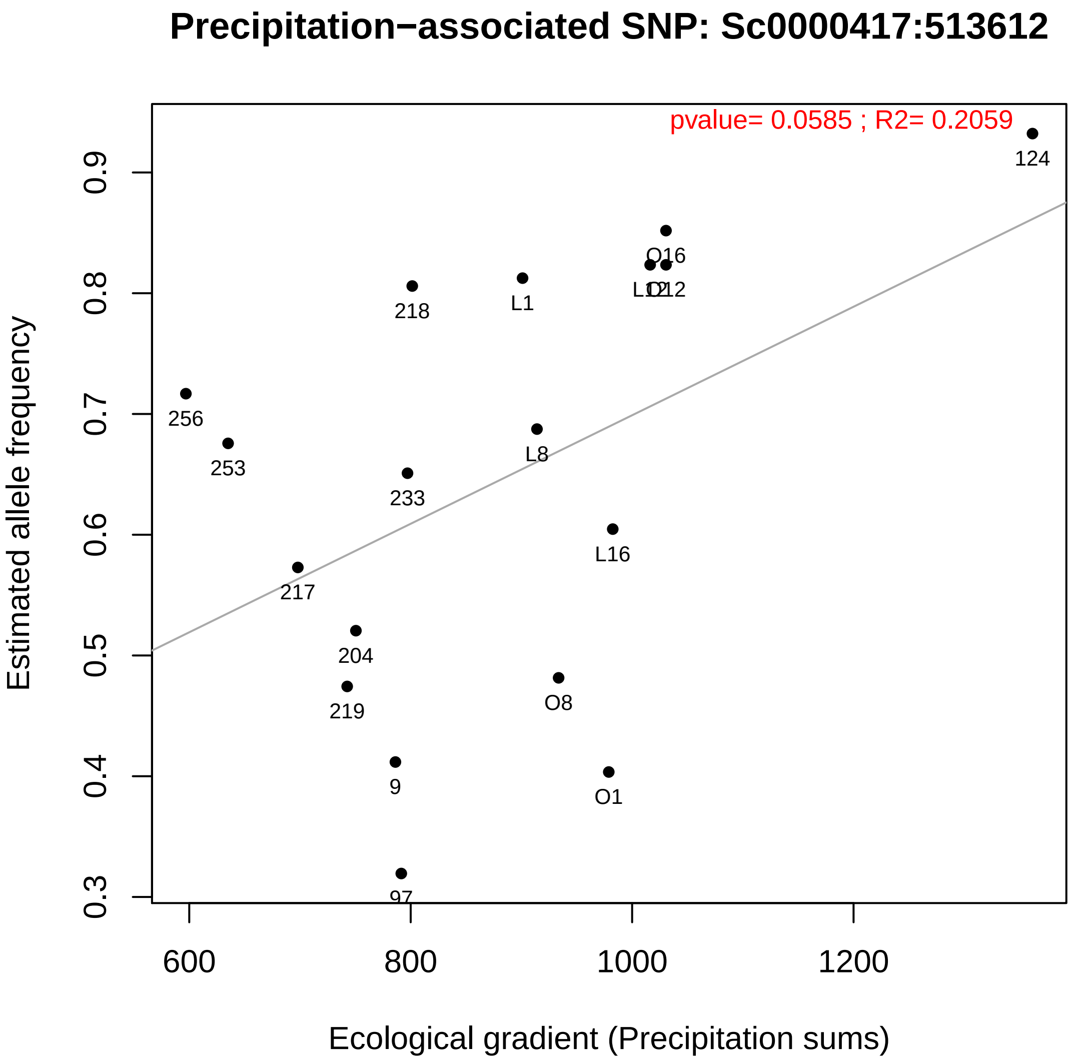

Precipitation−associated SNP: Sc0000417:513612
pvalue= 0.0585 ; R2= 0.2059
Estimated allele frequency
0.3
0.4
0.5
0.6
0.7
0.8
0.9
600
800
1000
1200
Ecological gradient (Precipitation sums)
124
O16
L12
O12
L1
218
256
253
L8
233
L16
217
204
O8
219
9
O1
97

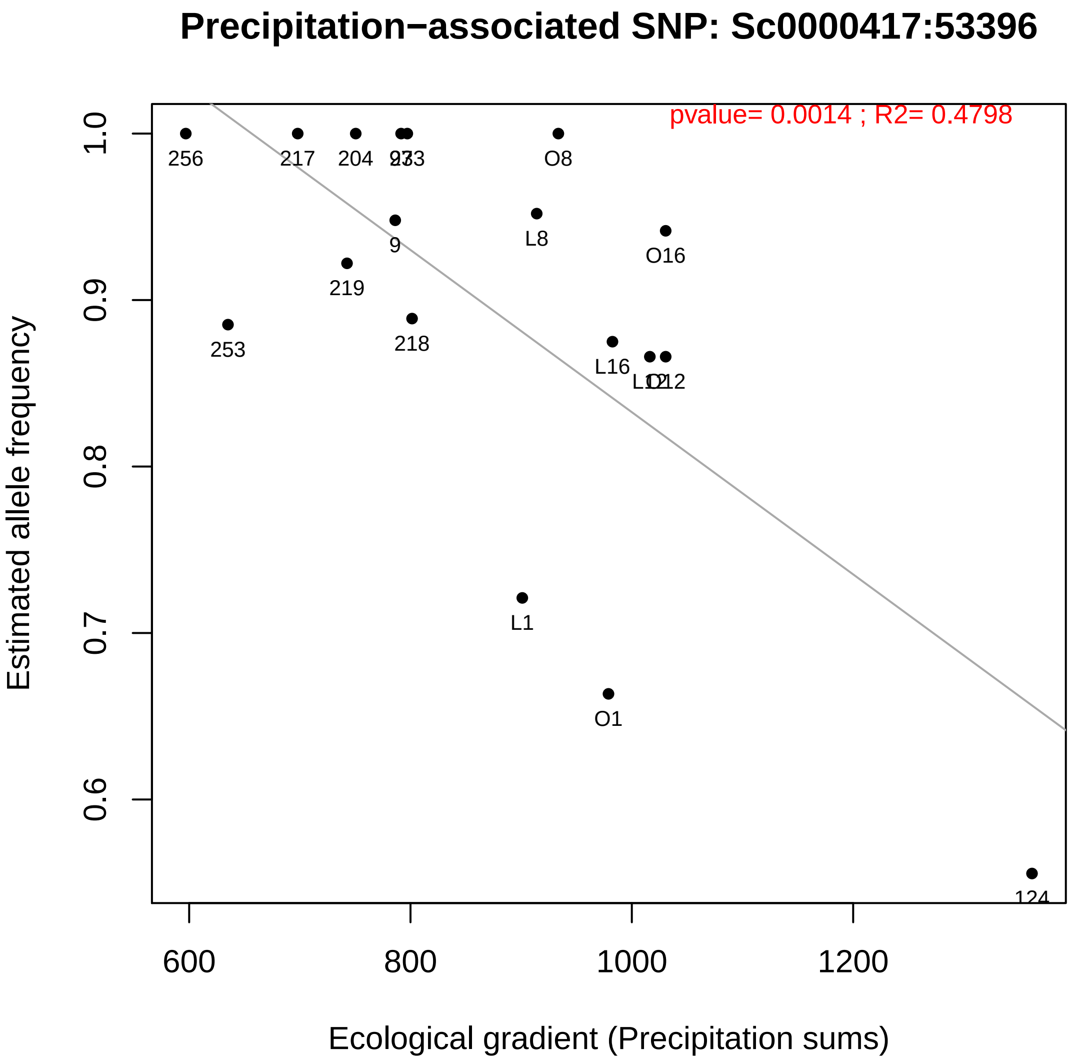

Precipitation−associated SNP: Sc0000417:53396
pvalue= 0.0014 ; R2= 0.4798
Estimated allele frequency
0.6
0.7
0.8
0.9
1.0
Ecological gradient (Precipitation sums)
600
800
1000
1200
256
217
204
O8
9
L8
O16
219
253
218
L16
O12
L1
O1
124

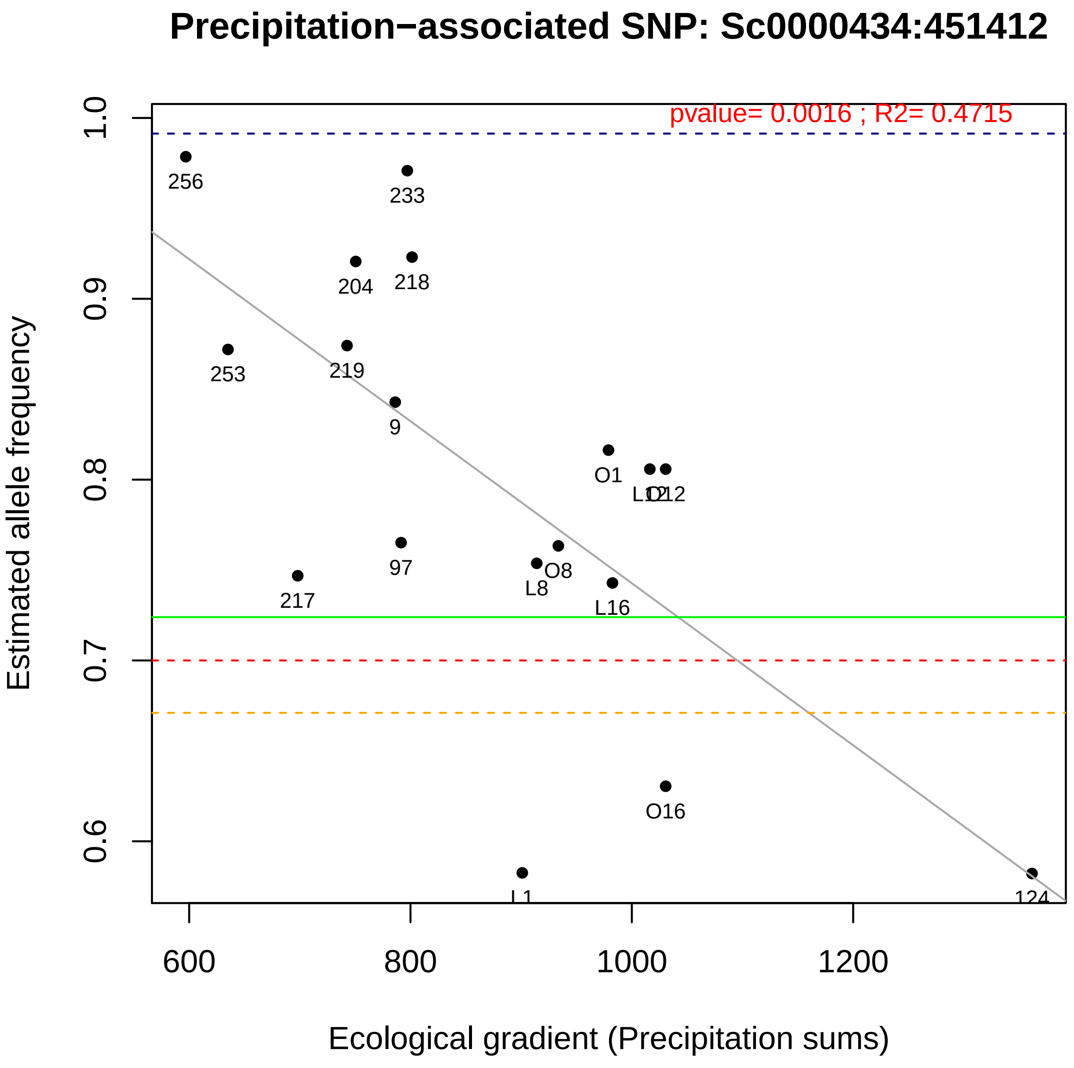
Precipitation−associated SNP: Sc0000434:451412
pvalue= 0.0016 ; R2= 0.4715
Estimated allele frequency
1.0
0.9
0.8
0.7
0.6
256
233
204
218
253
219
9
O1
L12
O12
97
O8
L8
217
L16
O16
L1
124
600
800
1000
1200
Ecological gradient (Precipitation sums)

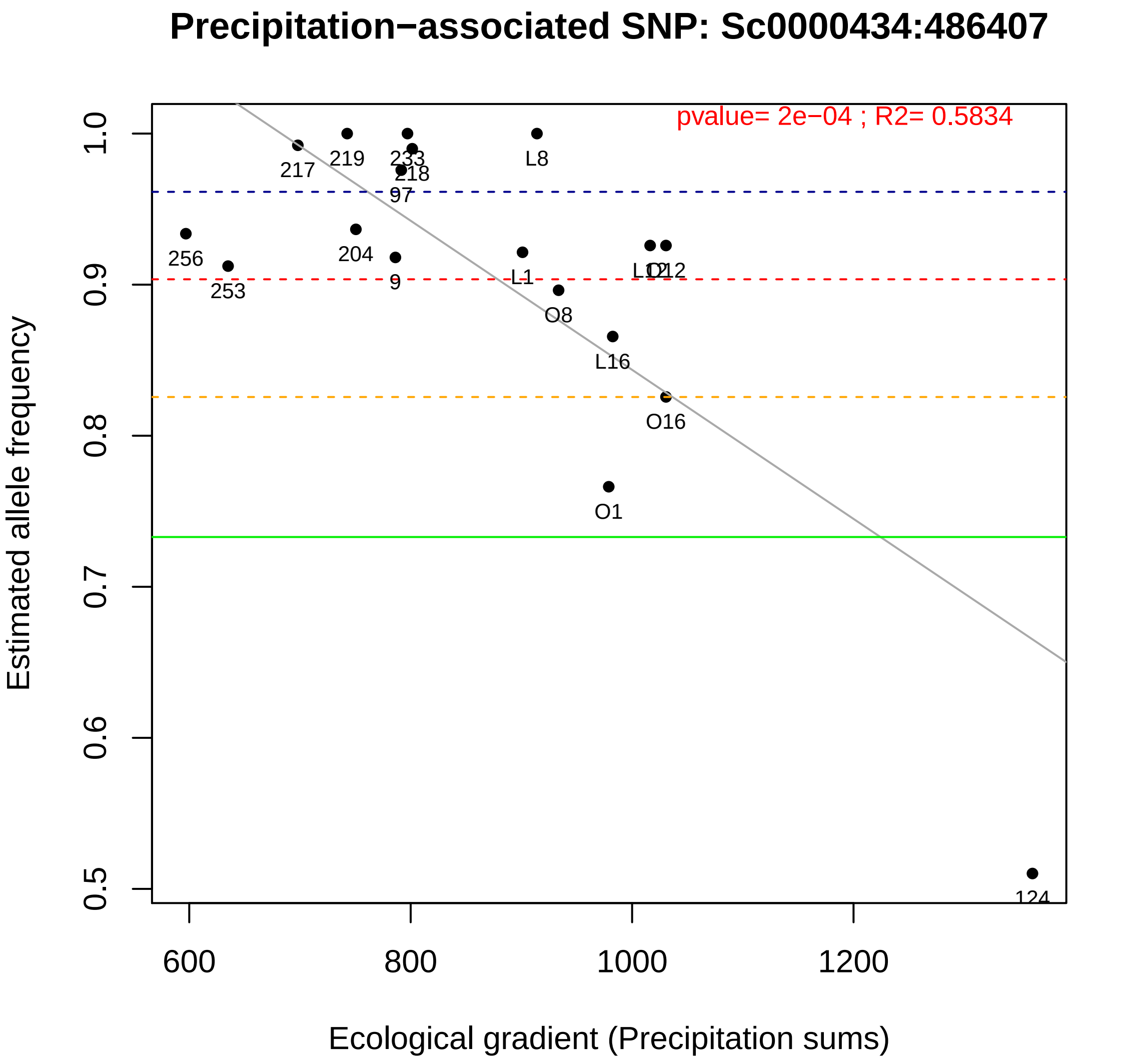

Precipitation−associated SNP: Sc0000434:486407
pvalue= 2e−04 ; R2= 0.5834
Estimated allele frequency
Ecological gradient (Precipitation sums)
0.5
0.6
0.7
0.8
0.9
1.0
600
800
1000
1200
217
219
233
218
97
L8
256
253
204
9
L1
L12
O12
O8
L16
O16
O1
124

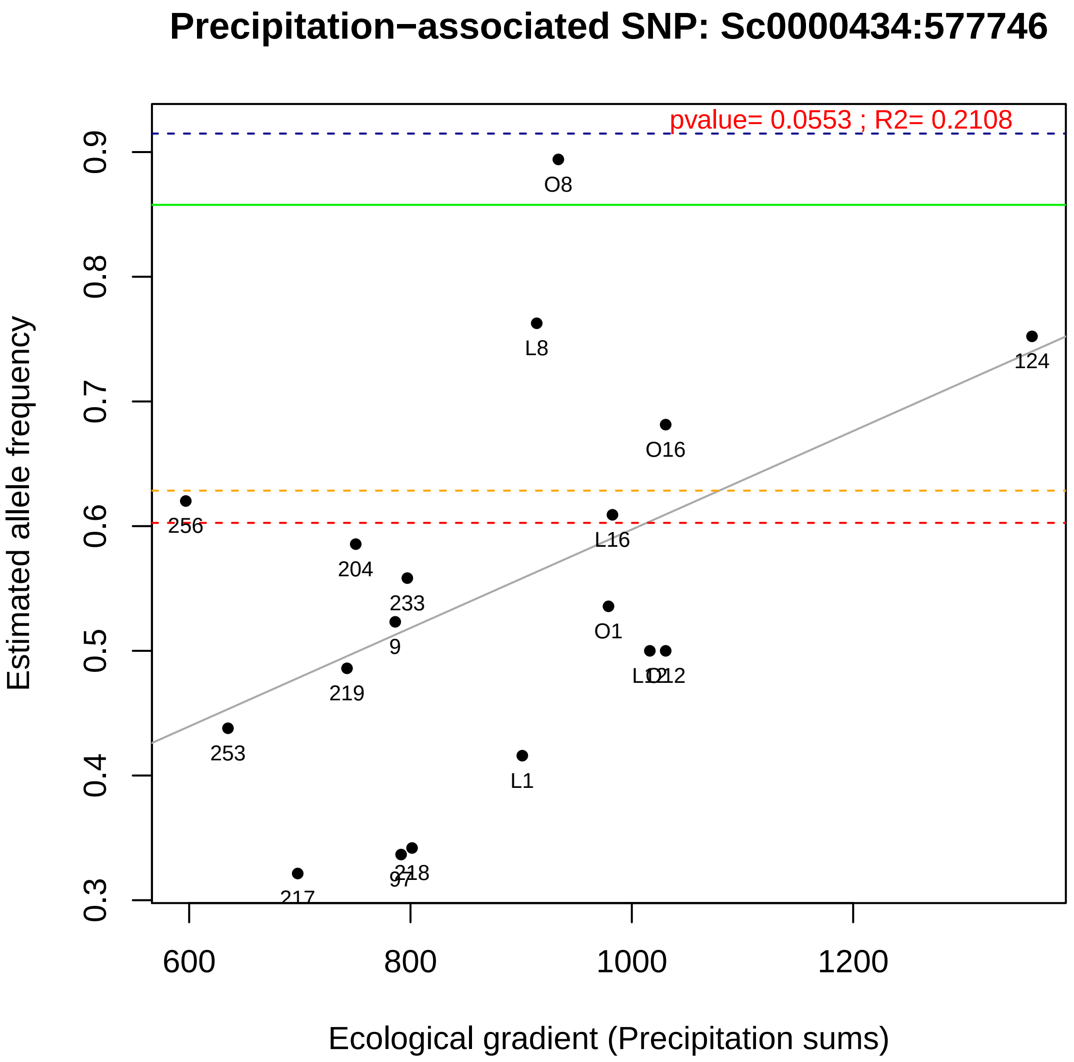

Precipitation−associated SNP: Sc0000434:577746
pvalue= 0.0553 ; R2= 0.2108
Estimated allele frequency
Ecological gradient (Precipitation sums)
0.3
0.4
0.5
0.6
0.7
0.8
0.9
600
800
1000
1200
O8
L8
124
O16
256
L16
204
233
9
O1
L12
O12
219
253
L1
217
97
218

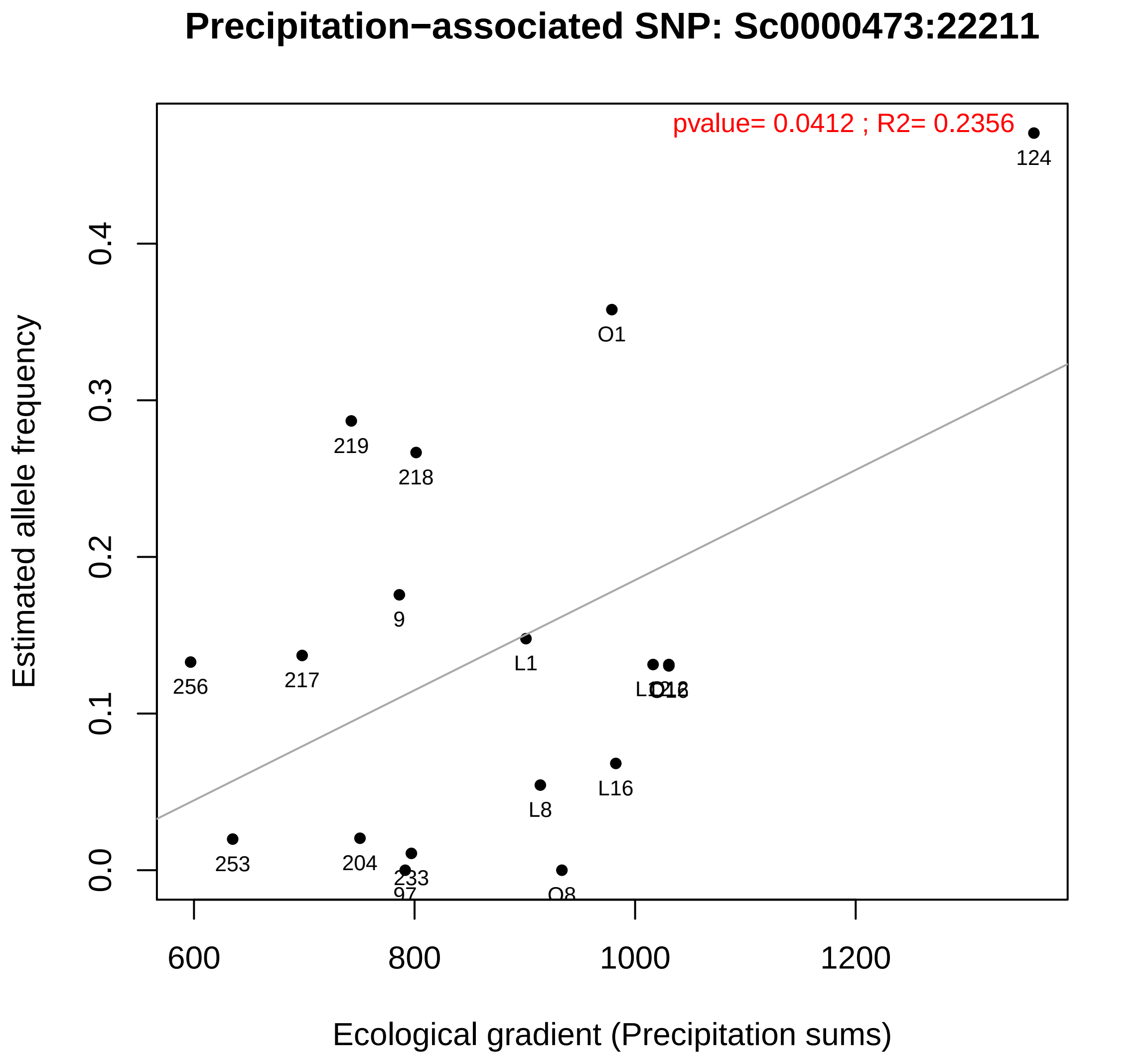

Precipitation−associated SNP: Sc0000473:22211
pvalue= 0.0412 ; R2= 0.2356
Estimated allele frequency
Ecological gradient (Precipitation sums)
0.0
0.1
0.2
0.3
0.4
600
800
1000
1200
124
O1
219
218
9
L1
256
217
L16
L8
253
204
233
97
O8

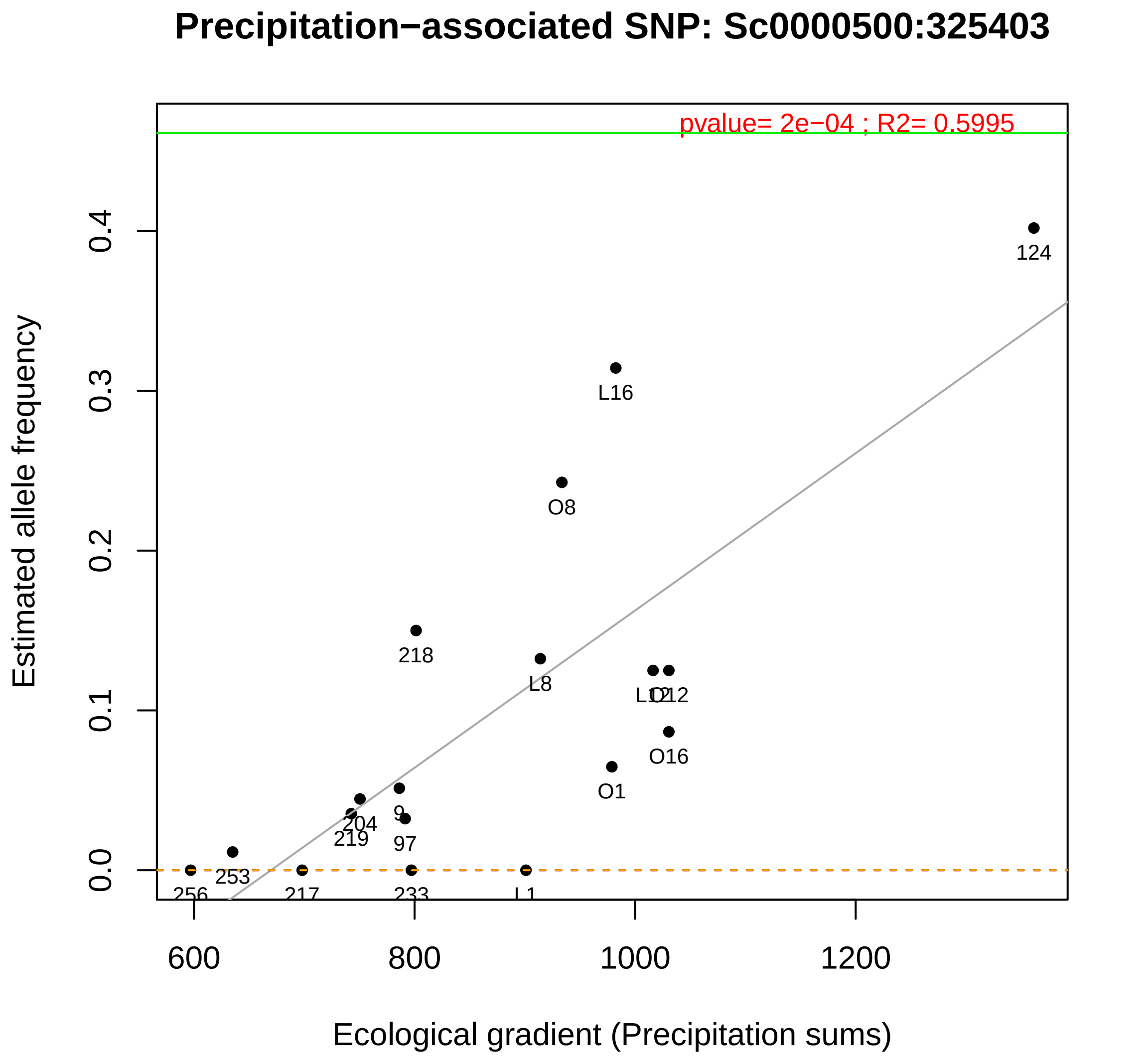
Precipitation−associated SNP: Sc0000500:325403
pvalue= 2e−04 ; R2= 0.5995
Estimated allele frequency
0.0
0.1
0.2
0.3
0.4
600
800
1000
1200
Ecological gradient (Precipitation sums)
124
L16
O8
218
L8
O12
O16
O1
204
219
9
97
253
256
217
233
L1

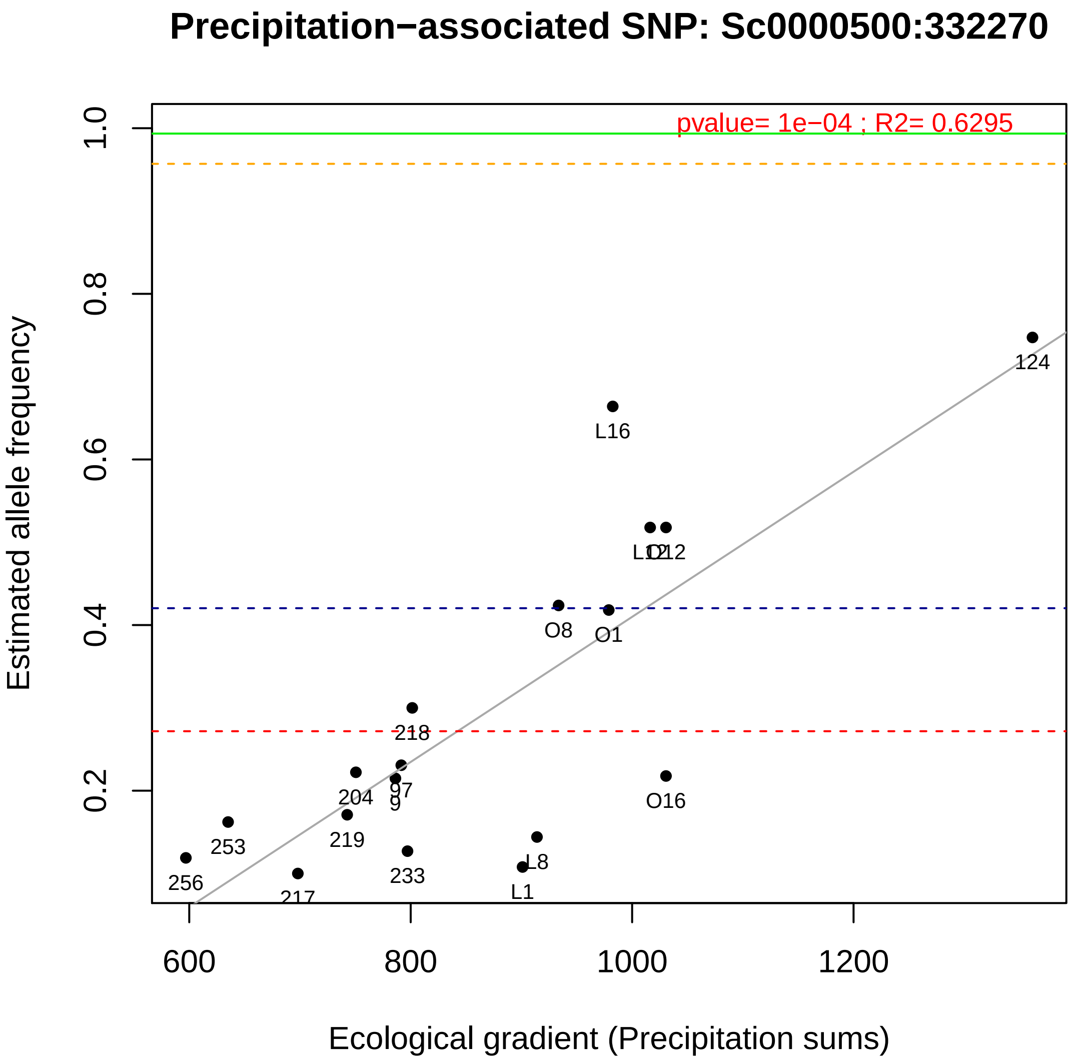

Precipitation−associated SNP: Sc0000500:332270
pvalue= 1e−04 ; R2= 0.6295
Estimated allele frequency
Ecological gradient (Precipitation sums)
0.2
0.4
0.6
0.8
1.0
600
800
1000
1200
124
L16
L12
O12
O8
O1
218
204
97
9
219
253
256
217
233
L8
L1
O16

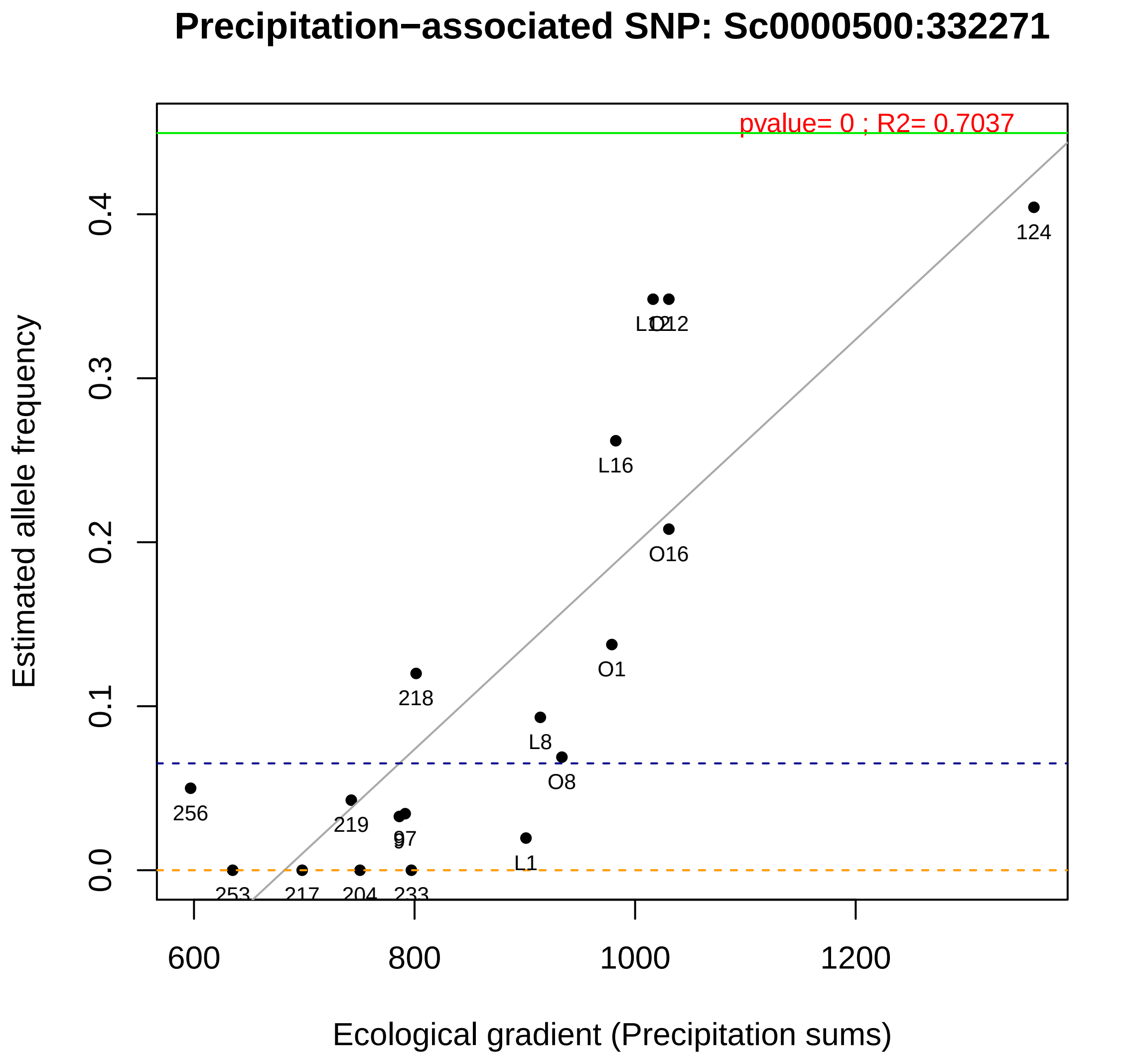

Precipitation−associated SNP: Sc0000500:332271
pvalue= 0 ; R2= 0.7037
124
L12
O12
L16
O16
O1
218
L8
O8
256
219
9
O7
L1
253
217
204
233
0.0
0.1
0.2
0.3
0.4
600
800
1000
1200
Estimated allele frequency
Ecological gradient (Precipitation sums)

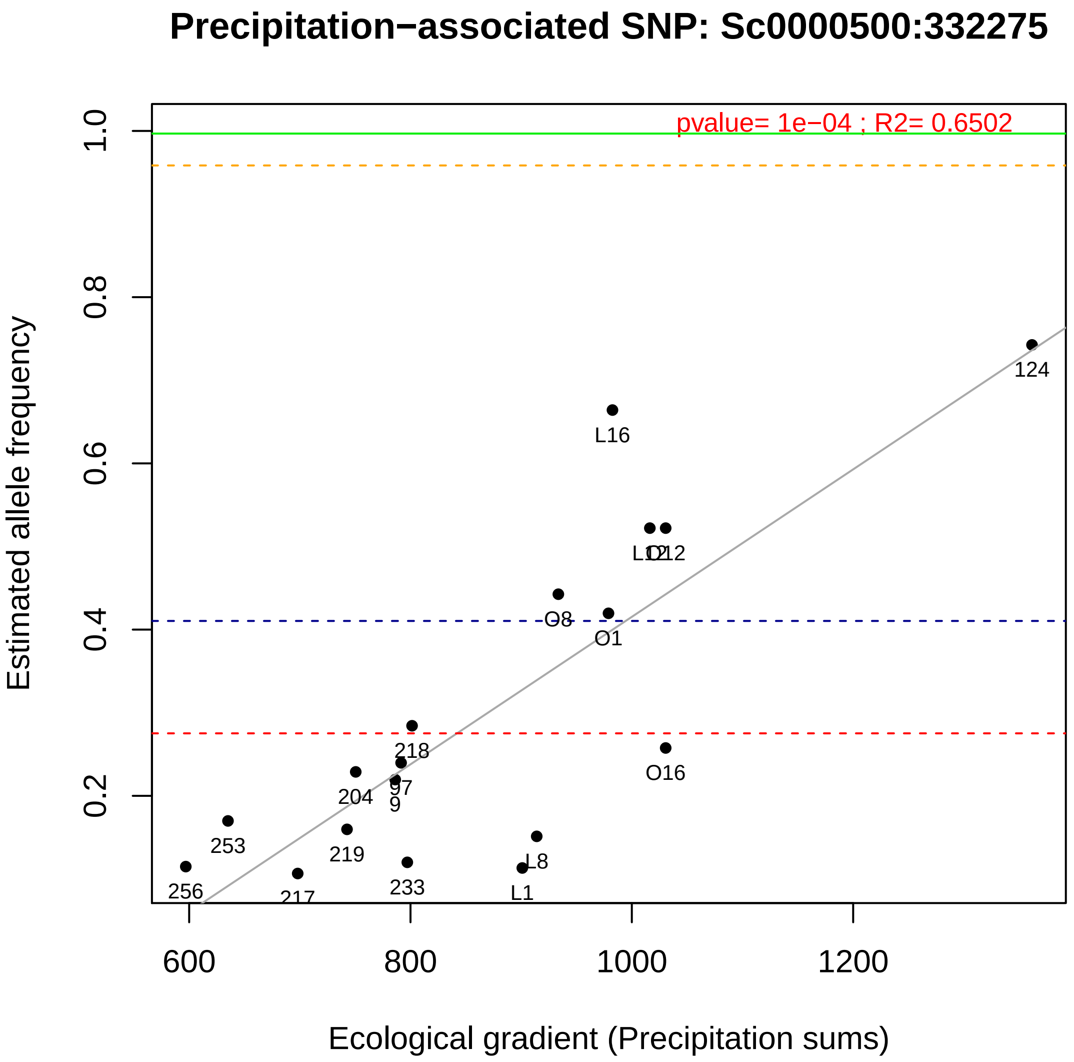

Precipitation−associated SNP: Sc0000500:332275
pvalue= 1e−04 ; R2= 0.6502
Estimated allele frequency
Ecological gradient (Precipitation sums)
0.2
0.4
0.6
0.8
1.0
600
800
1000
1200
124
L16
L12
O12
O8
O1
218
O16
204
97
9
253
219
256
217
233
L8
L1

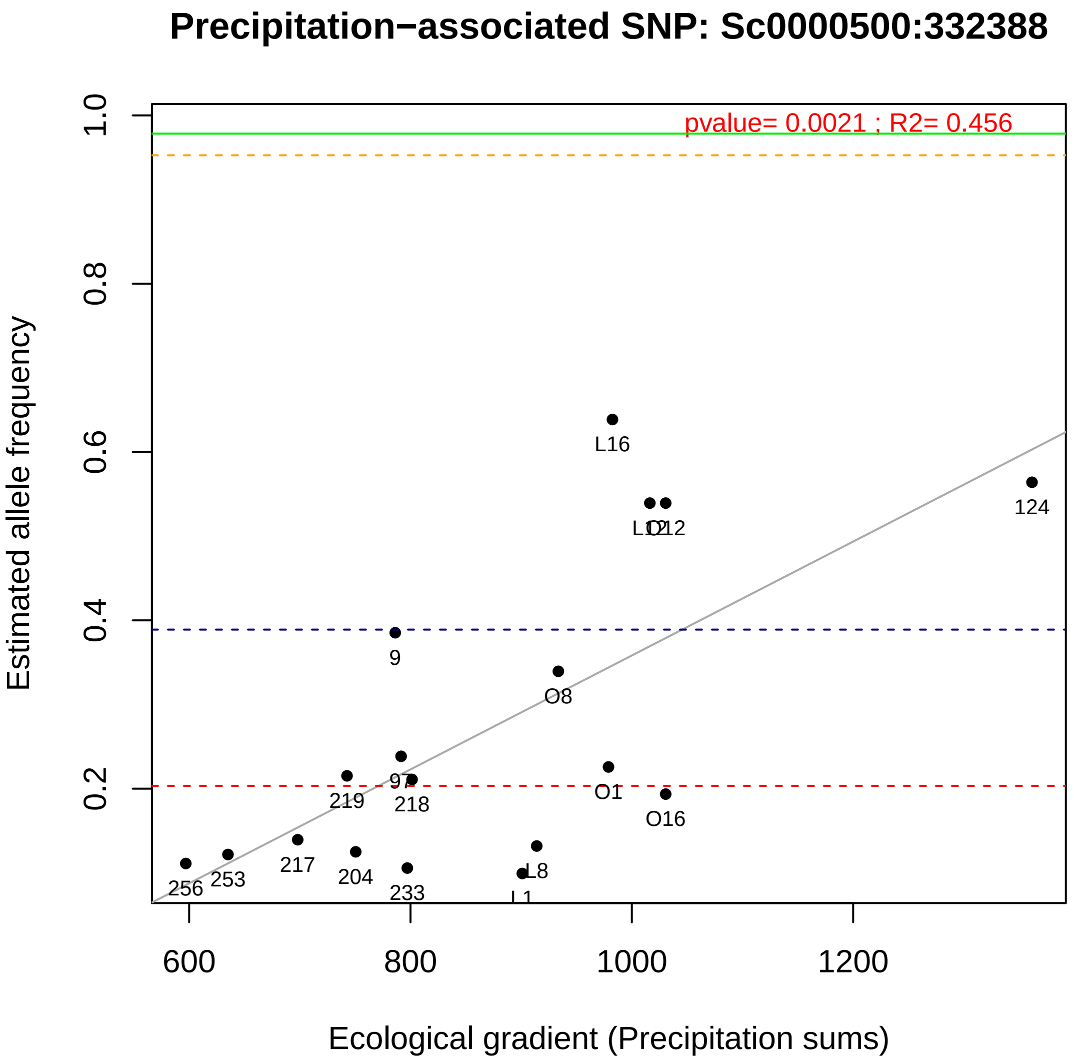

Precipitation−associated SNP: Sc0000500:332388
pvalue= 0.0021 ; R2= 0.456
Estimated allele frequency
Ecological gradient (Precipitation sums)
0.2
0.4
0.6
0.8
1.0
600
800
1000
1200
L16
124
O8
9
O1
O16
219
218
217
256
253
204
233
L8

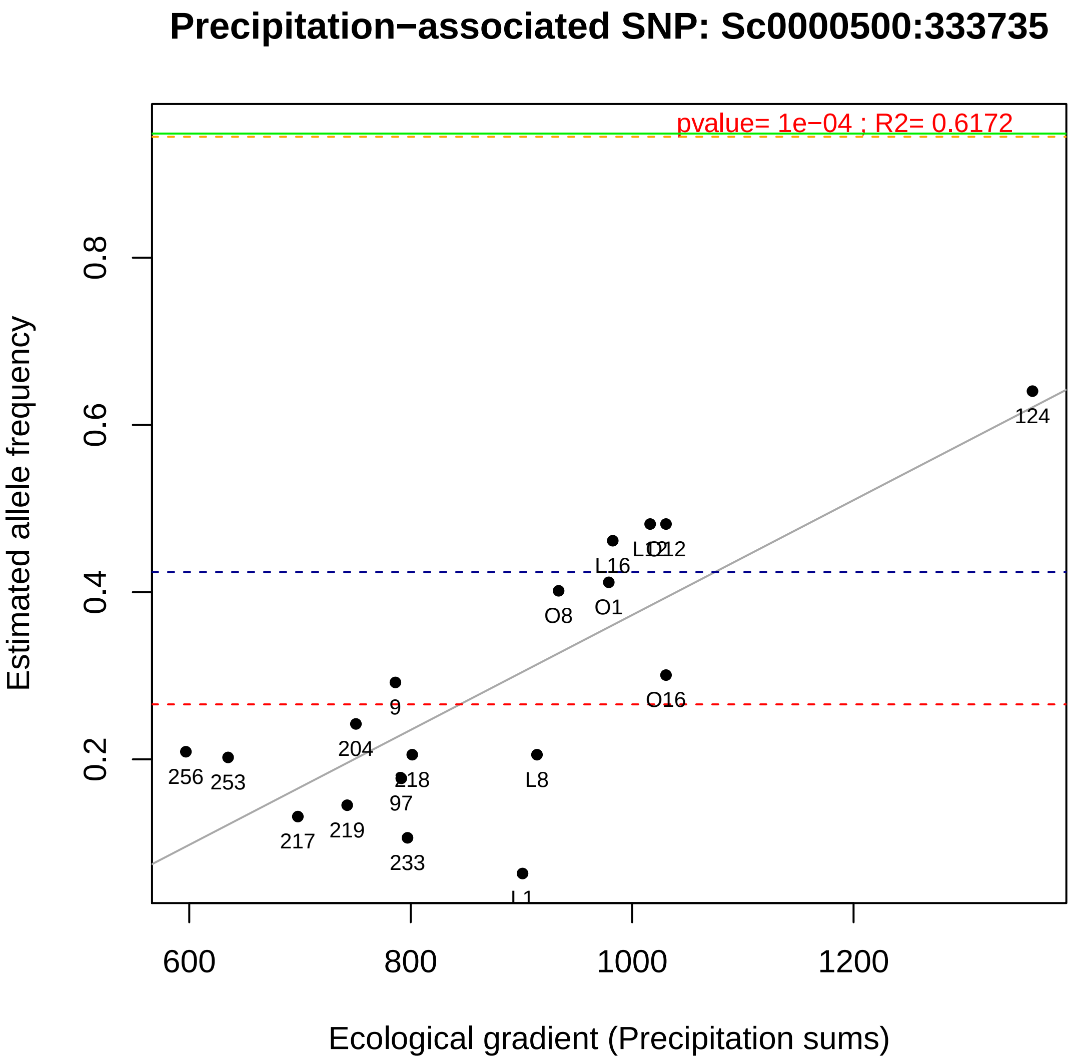

Precipitation−associated SNP: Sc0000500:333735
pvalue= 1e−04 ; R2= 0.6172
Estimated allele frequency
0.2
0.4
0.6
0.8
600
800
1000
1200
Ecological gradient (Precipitation sums)
124
L16
O8
O1
O16
9
204
256
253
218
97
L8
217
219
233

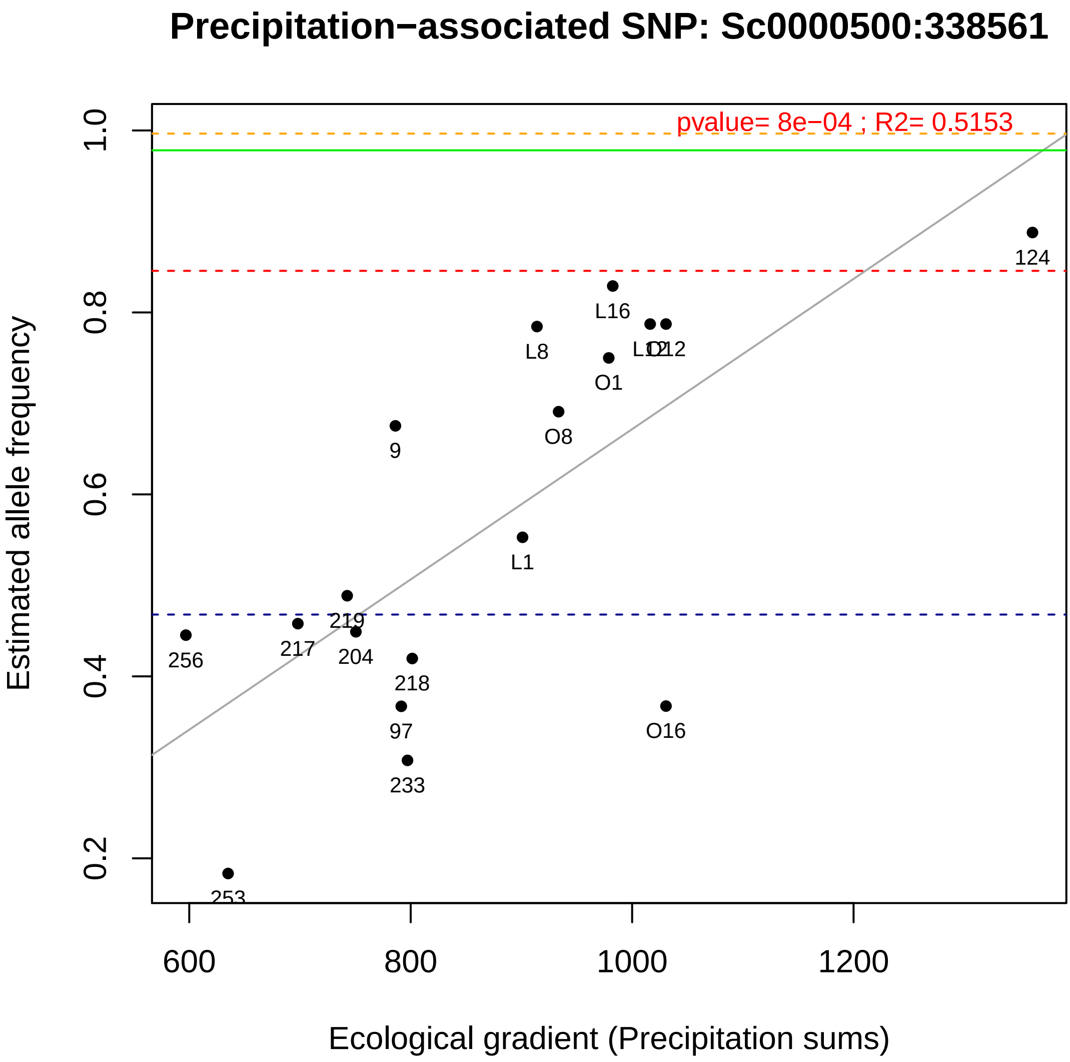
Precipitation−associated SNP: Sc0000500:338561
pvalue= 8e−04 ; R2= 0.5153
Estimated allele frequency
0.2
0.4
0.6
0.8
1.0
600
800
1000
1200
Ecological gradient (Precipitation sums)
124
L16
L8
L12
O12
O1
O8
9
L1
219
217
204
256
218
97
O16
233
253

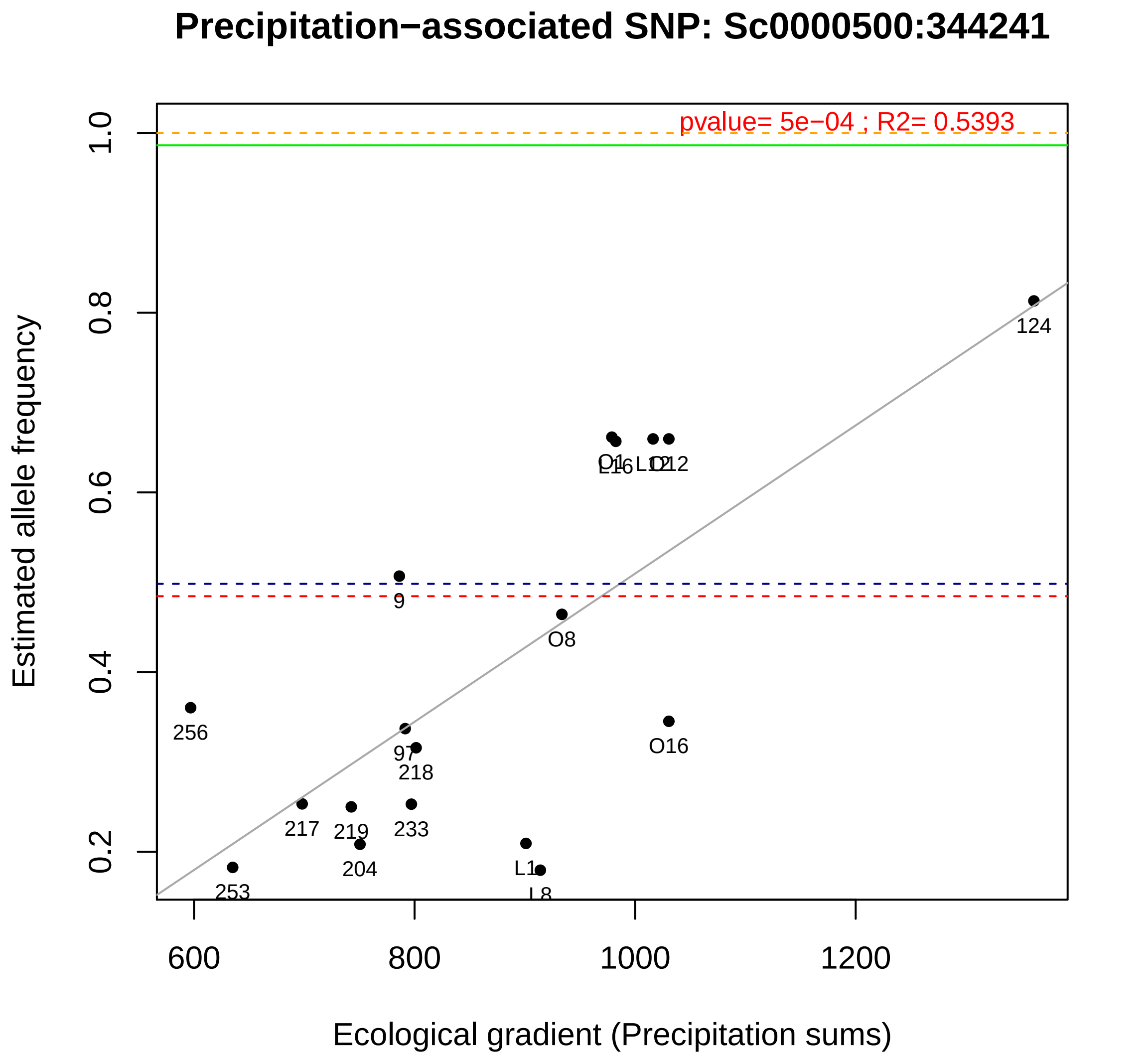

Precipitation−associated SNP: Sc0000500:344241
pvalue= 5e−04 ; R2= 0.5393
Estimated allele frequency
0.2
0.4
0.6
0.8
1.0
600
800
1000
1200
Ecological gradient (Precipitation sums)
124
O1
L16
L10
O12
9
O8
256
97
218
O16
217
219
233
204
253
L1
L8

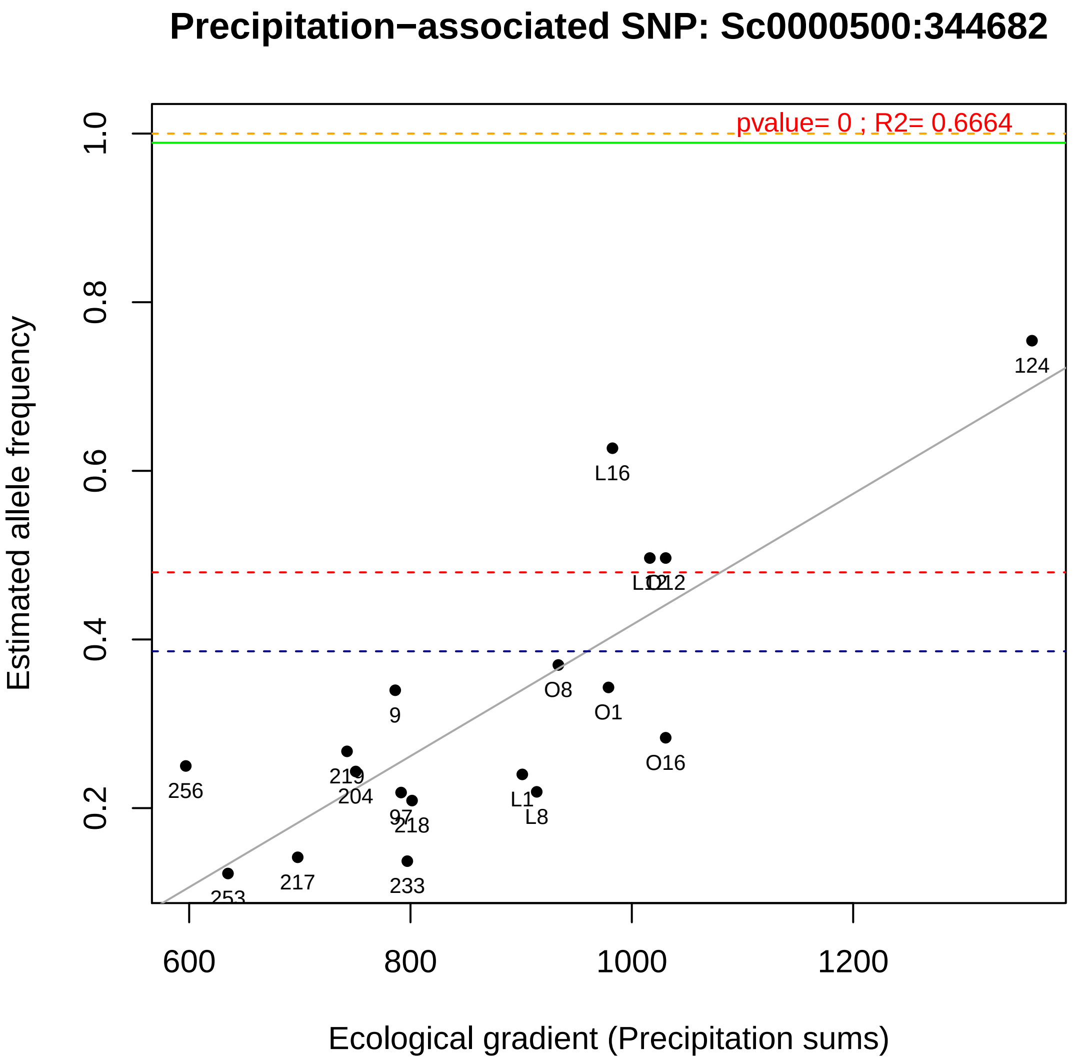
Precipitation−associated SNP: Sc0000500:344682
pvalue= 0 ; R2= 0.6664
Estimated allele frequency
1.0
0.8
0.6
0.4
0.2
124
L16
L10
O12
O8
O1
9
O16
219
204
256
97
218
L1
L8
217
233
253
600
800
1000
1200
Ecological gradient (Precipitation sums)

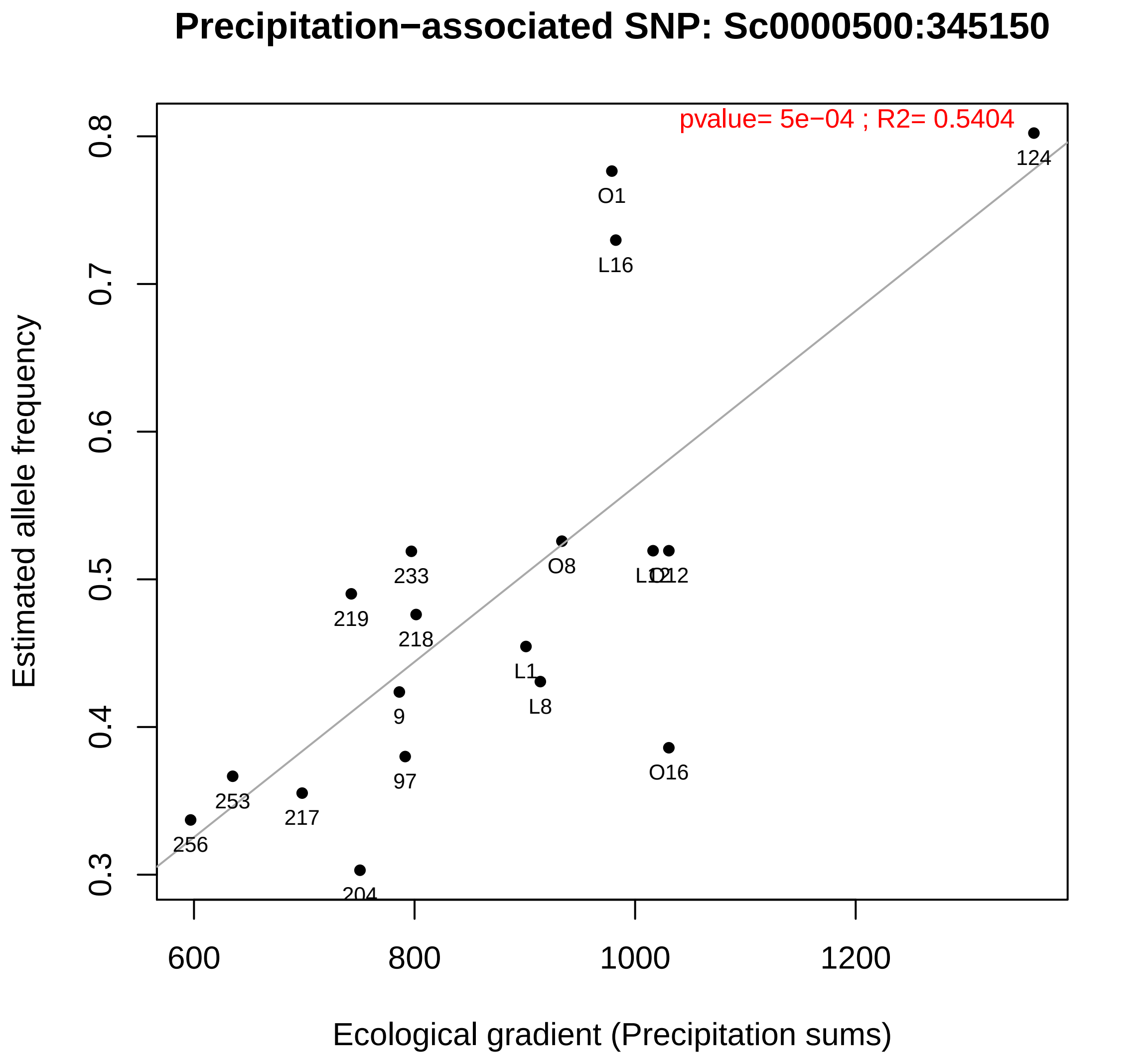

Precipitation−associated SNP: Sc0000500:345150
pvalue= 5e−04 ; R2= 0.5404
Estimated allele frequency
Ecological gradient (Precipitation sums)
0.3
0.4
0.5
0.6
0.7
0.8
600
800
1000
1200
124
O1
L16
O8
L12
O12
233
219
218
L1
L8
9
97
O16
253
217
256
204

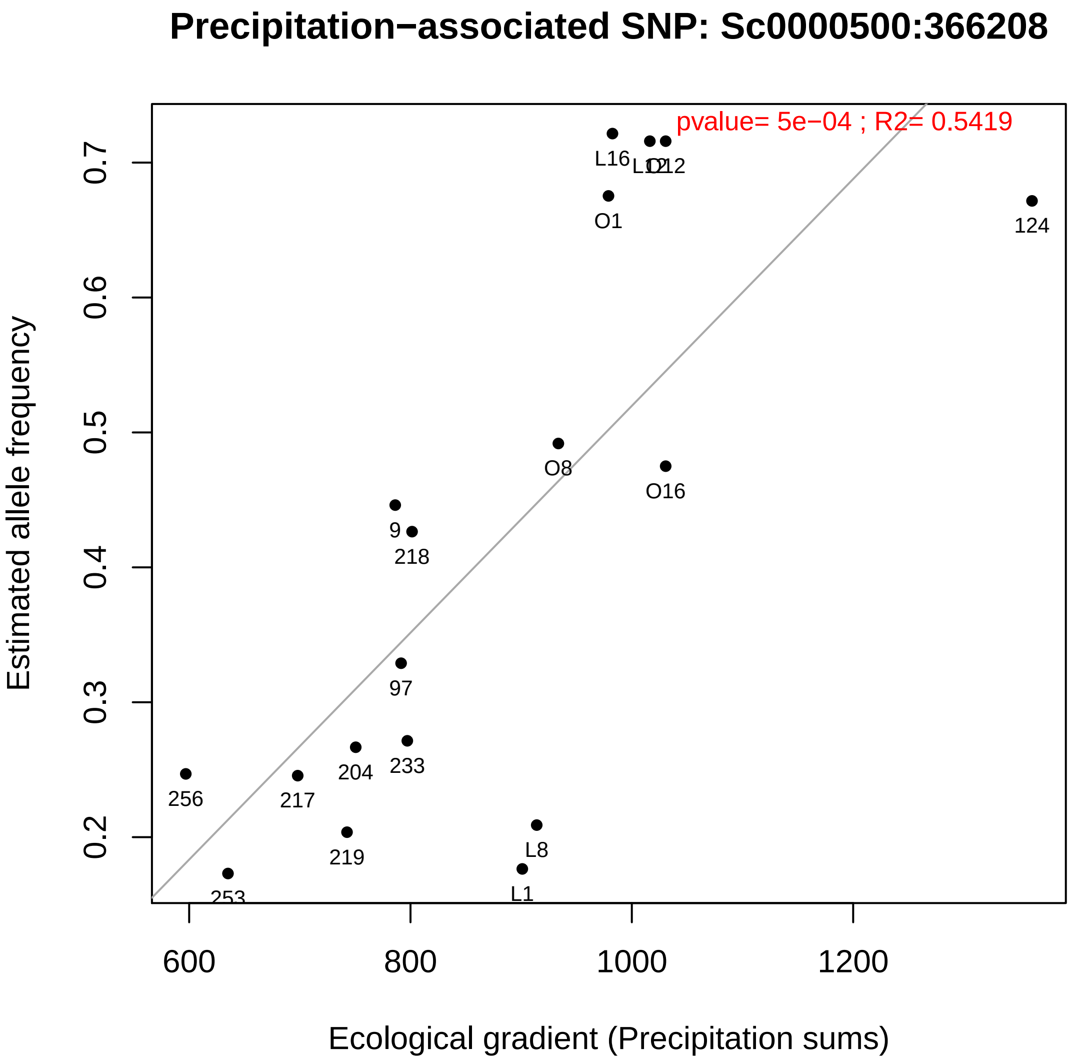
Precipitation−associated SNP: Sc0000500:366208
pvalue= 5e−04 ; R2= 0.5419
Estimated allele frequency
0.2
0.3
0.4
0.5
0.6
0.7
600
800
1000
1200
Ecological gradient (Precipitation sums)
L16
L12
O12
O1
124
O8
O16
9
218
97
204
233
256
217
219
L8
L1
253

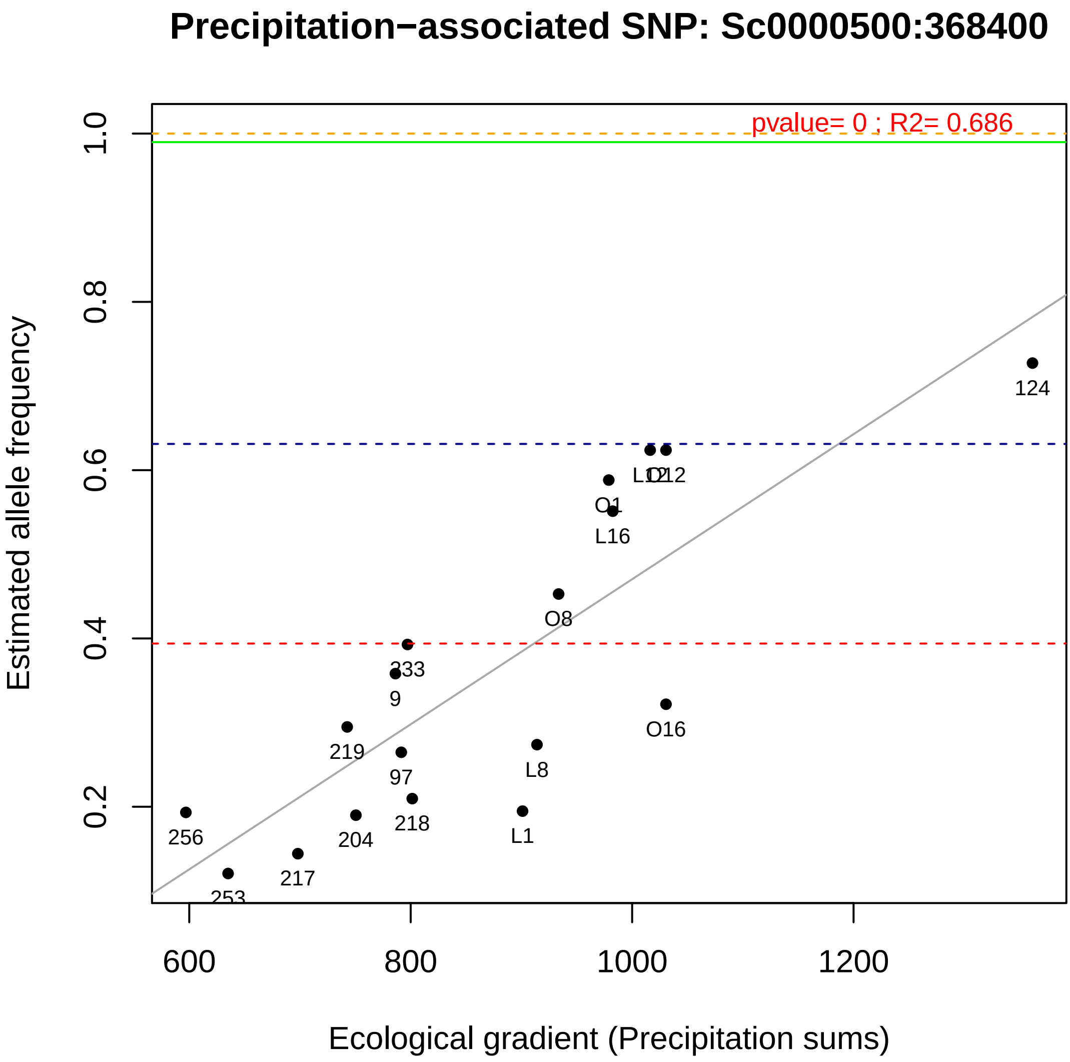

Precipitation−associated SNP: Sc0000500:368400
pvalue= 0 ; R2= 0.686
Estimated allele frequency
0.2
0.4
0.6
0.8
1.0
600
800
1000
1200
Ecological gradient (Precipitation sums)
124
L12
O12
O1
L16
O8
233
9
O16
219
L8
97
256
204
218
L1
217
253

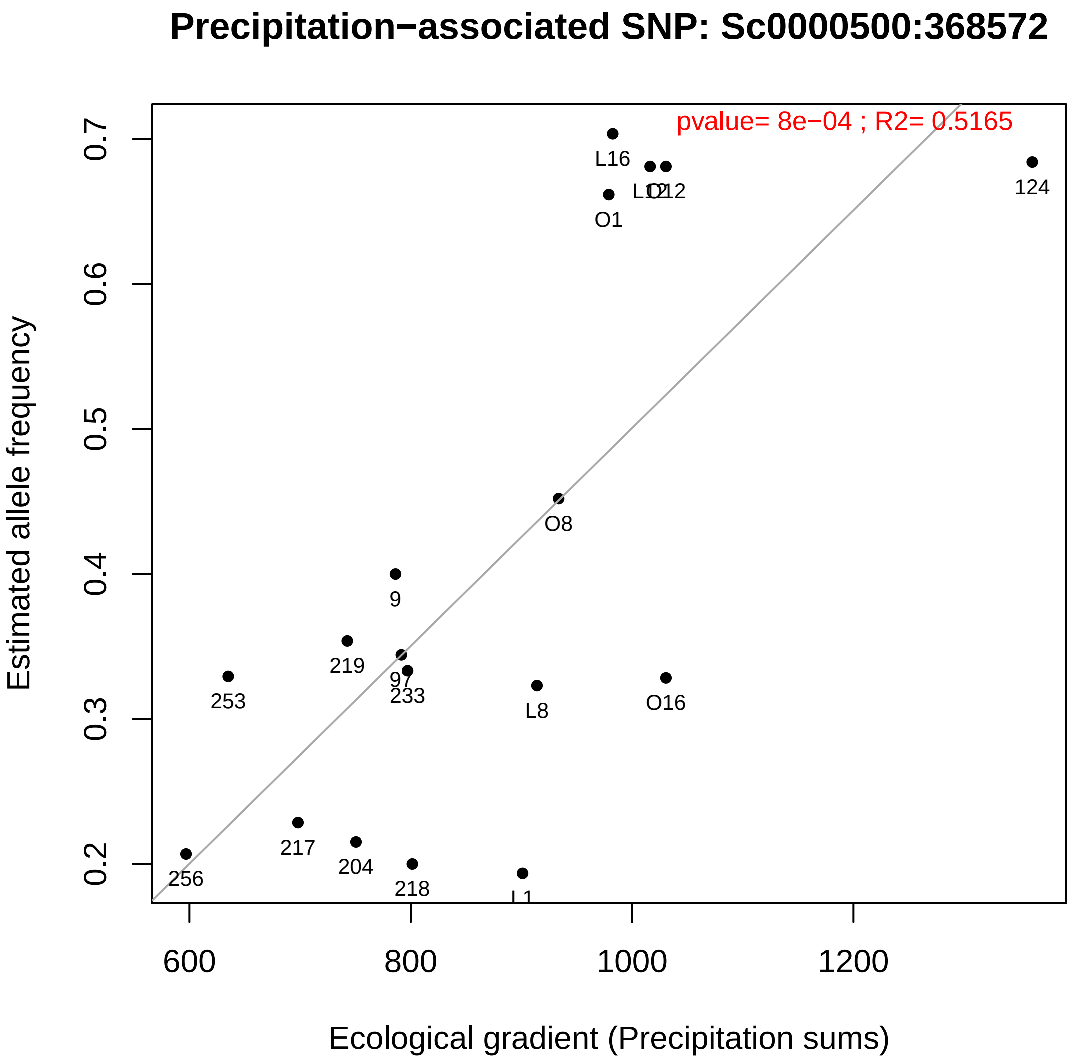

Precipitation−associated SNP: Sc0000500:368572
pvalue= 8e−04 ; R2= 0.5165
Estimated allele frequency
0.2
0.3
0.4
0.5
0.6
0.7
Ecological gradient (Precipitation sums)
600
800
1000
1200
L16
O1
O12
124
O8
9
219
97
233
253
L8
O16
217
204
218
256

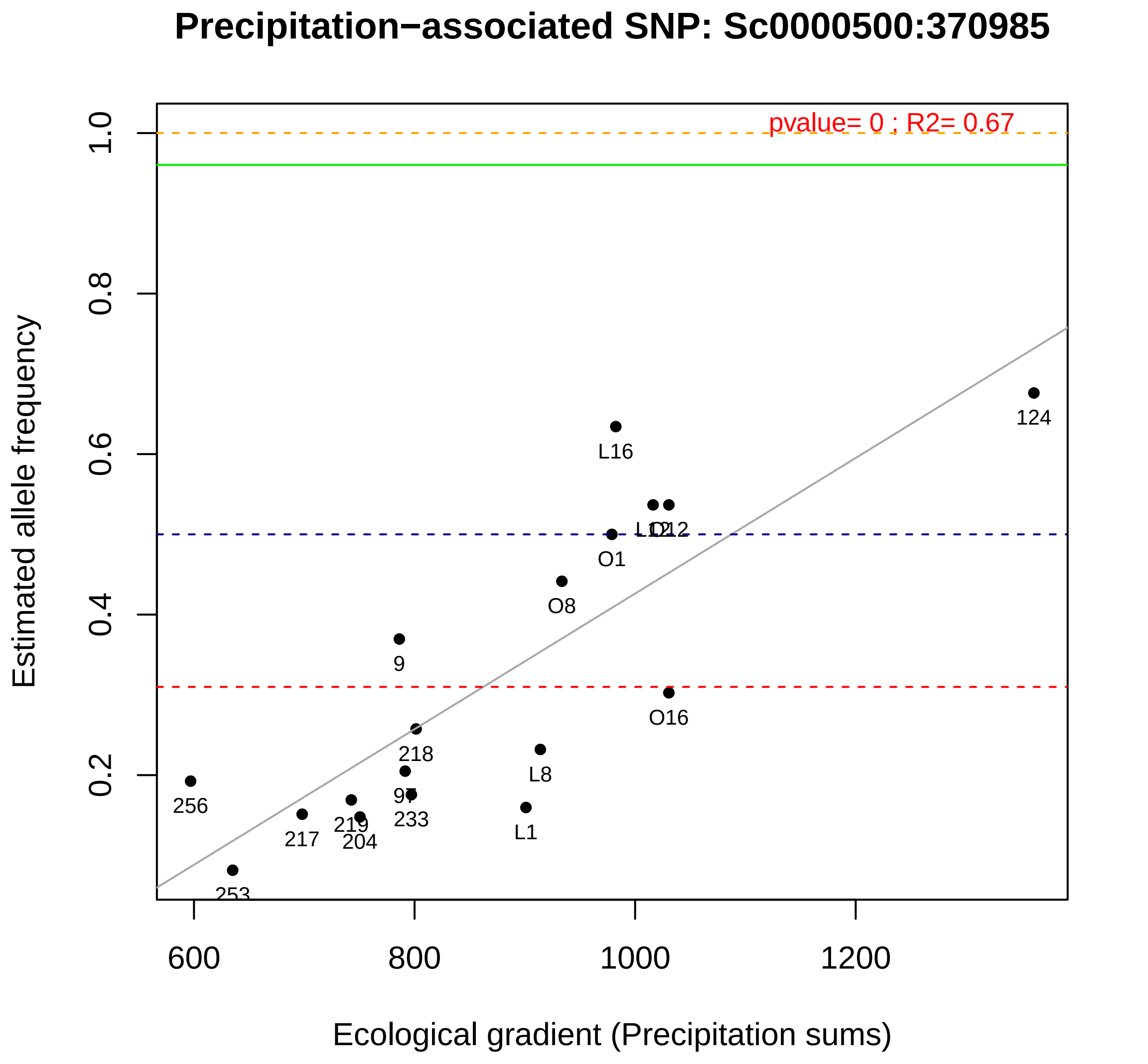

Precipitation−associated SNP: Sc0000500:370985
pvalue= 0 ; R2= 0.67
Estimated allele frequency
0.2
0.4
0.6
0.8
1.0
600
800
1000
1200
Ecological gradient (Precipitation sums)
124
L16
L12
O12
O1
O8
9
O16
218
L8
256
97
233
219
204
217
L1
253

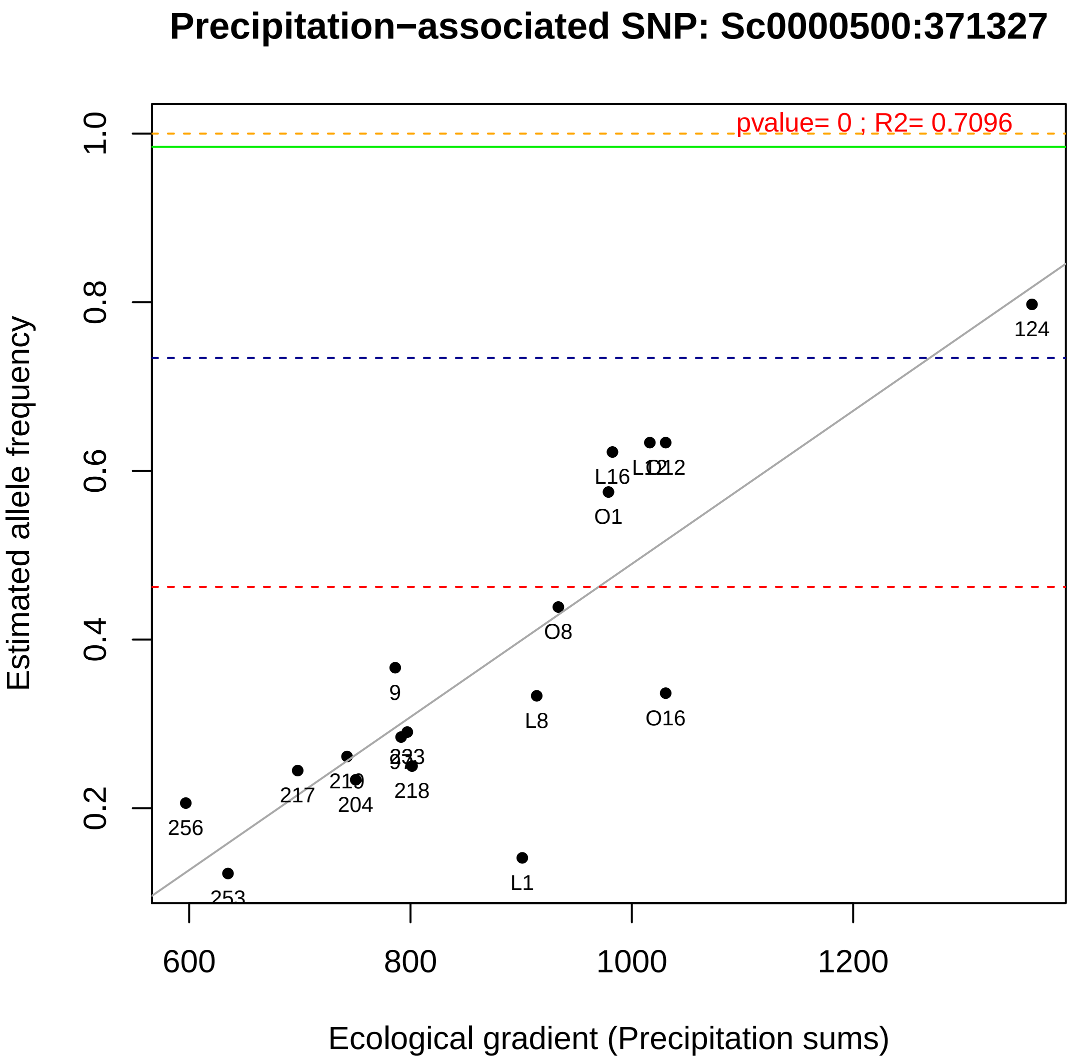
Precipitation−associated SNP: Sc0000500:371327
pvalue= 0 ; R2= 0.7096
Estimated allele frequency
Ecological gradient (Precipitation sums)
0.2
0.4
0.6
0.8
1.0
600
800
1000
1200
124
L16
O1
O8
9
L8
O16
217
204
218
256
253
L1

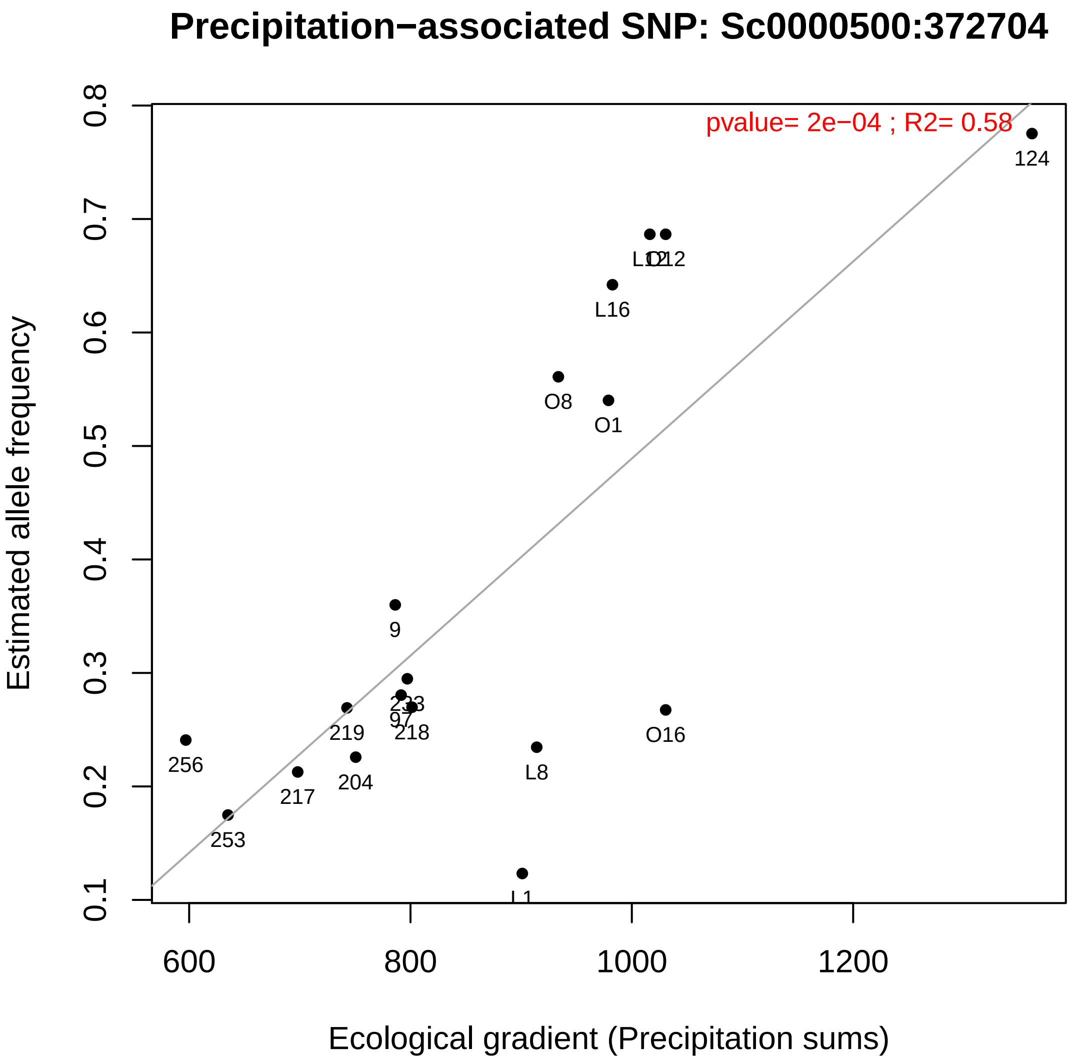

Precipitation−associated SNP: Sc0000500:372704
pvalue= 2e−04 ; R2= 0.58
Estimated allele frequency
Ecological gradient (Precipitation sums)
0.1
0.2
0.3
0.4
0.5
0.6
0.7
0.8
600
800
1000
1200
124
O12
L16
O8
O1
9
219
218
97
256
204
217
253
L8
O16

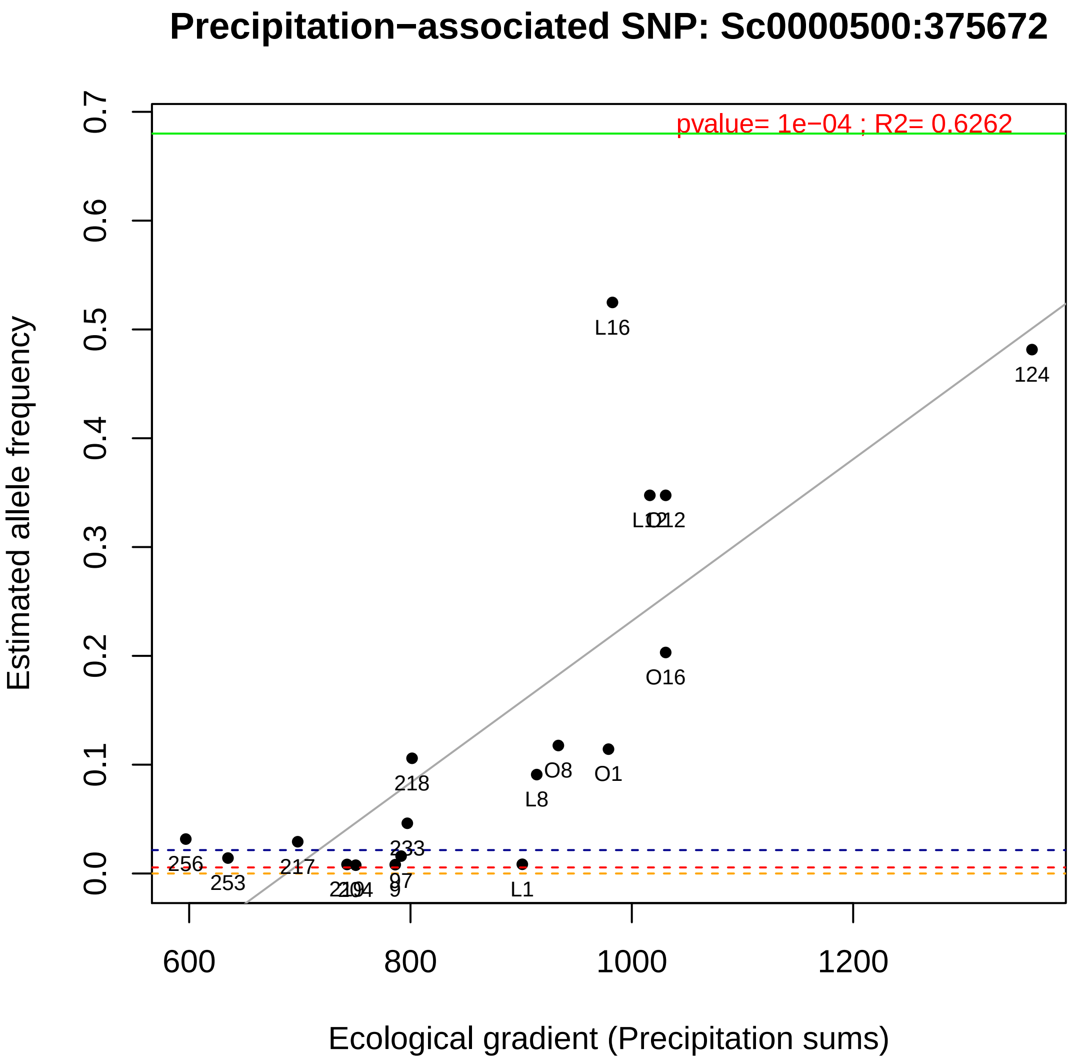
Precipitation−associated SNP: Sc0000500:375672
pvalue= 1e−04 ; R2= 0.6262
Estimated allele frequency
0.0
0.1
0.2
0.3
0.4
0.5
0.6
0.7
600
800
1000
1200
Ecological gradient (Precipitation sums)
L16
124
L12
O12
O16
O8
O1
218
L8
233
256
253
217
97
9
L1

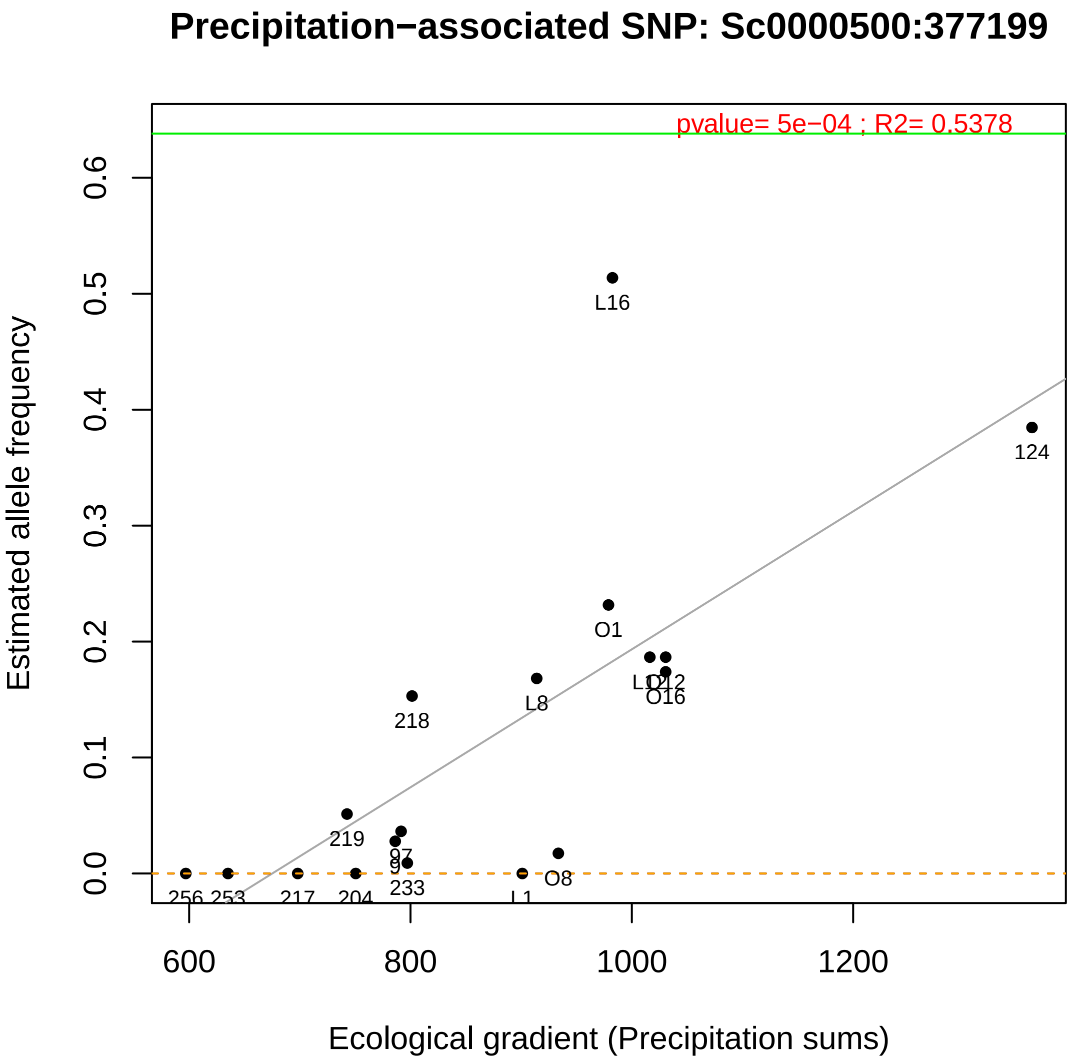
Precipitation−associated SNP: Sc0000500:377199
pvalue= 5e−04 ; R2= 0.5378
L16
124
O1
L12
O12
O16
L8
218
219
97
9
233
O8
256
253
217
204
L1
0.0
0.1
0.2
0.3
0.4
0.5
0.6
600
800
1000
1200
Estimated allele frequency
Ecological gradient (Precipitation sums)

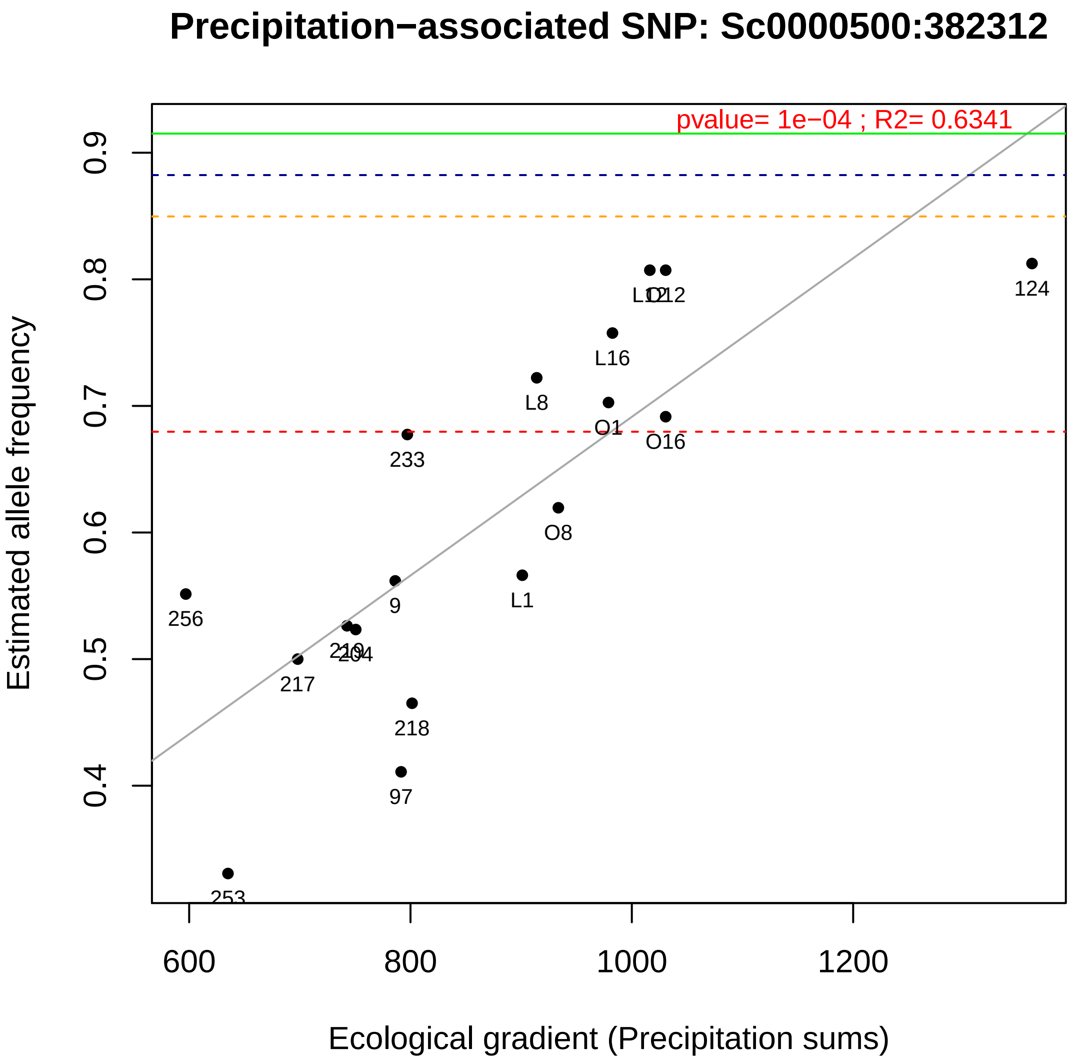
Precipitation−associated SNP: Sc0000500:382312
pvalue= 1e−04 ; R2= 0.6341
Estimated allele frequency
Ecological gradient (Precipitation sums)
0.4
0.5
0.6
0.7
0.8
0.9
600
800
1000
1200
124
L16
L8
O1
O16
233
O8
L1
9
256
217
218
97
253

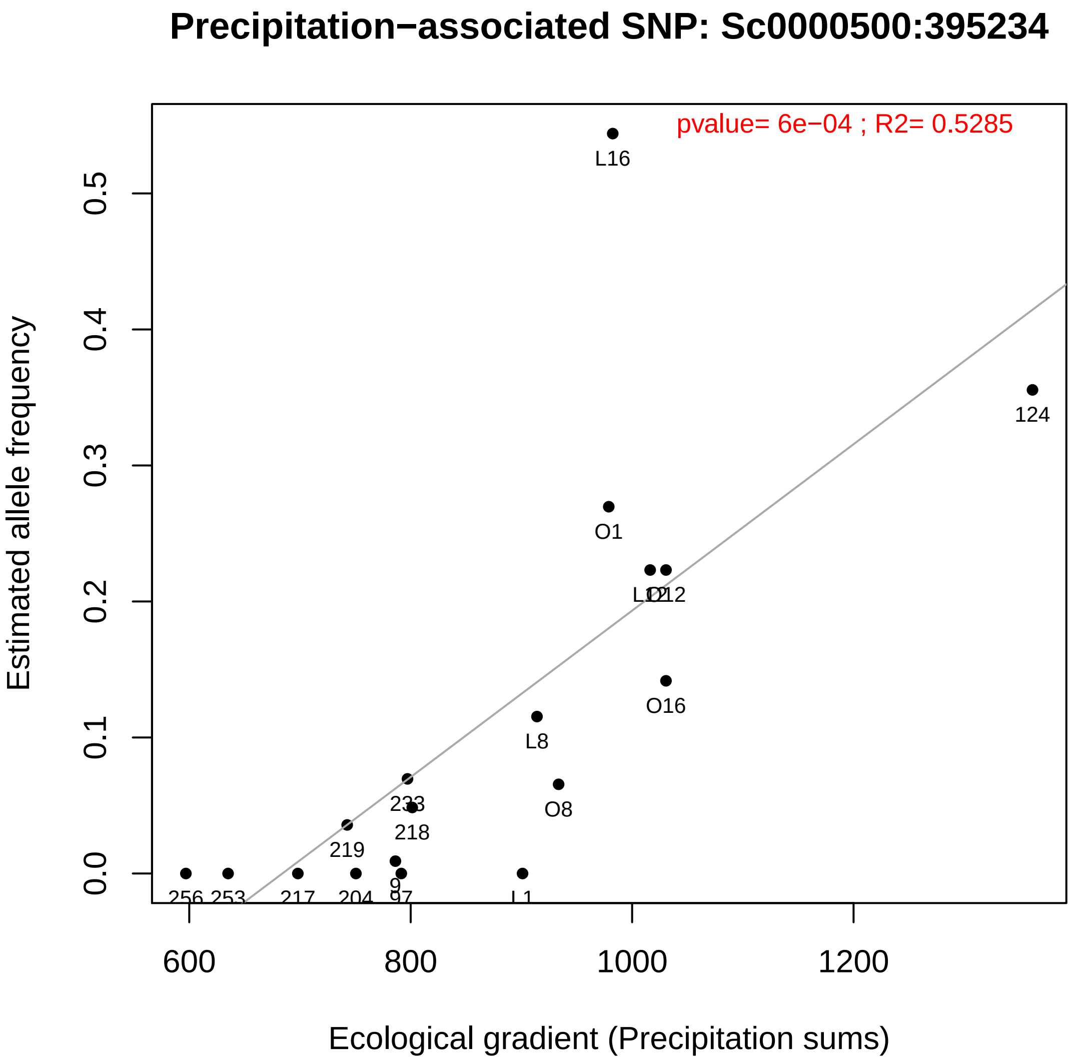

Precipitation−associated SNP: Sc0000500:395234
pvalue= 6e−04 ; R2= 0.5285
Estimated allele frequency
Ecological gradient (Precipitation sums)
0.0
0.1
0.2
0.3
0.4
0.5
600
800
1000
1200
L16
124
O1
O12
O16
L8
O8
233
218
219
9
256
253
217
204
97
L1

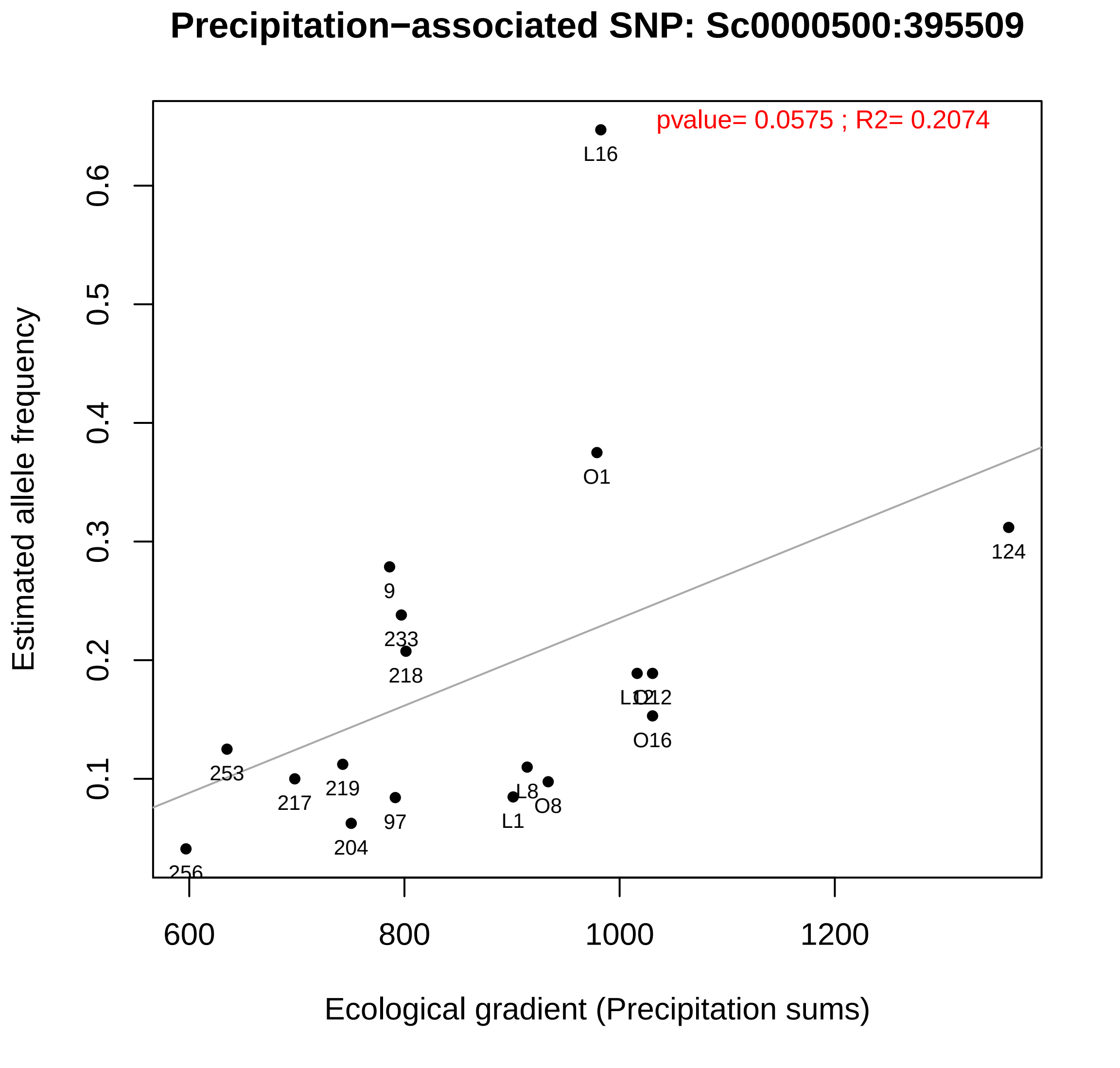

Precipitation−associated SNP: Sc0000500:395509
pvalue= 0.0575 ; R2= 0.2074
Estimated allele frequency
0.1
0.2
0.3
0.4
0.5
0.6
600
800
1000
1200
Ecological gradient (Precipitation sums)
L16
O1
124
9
233
218
L12
O12
O16
253
217
219
97
204
256
L8
O8
L1

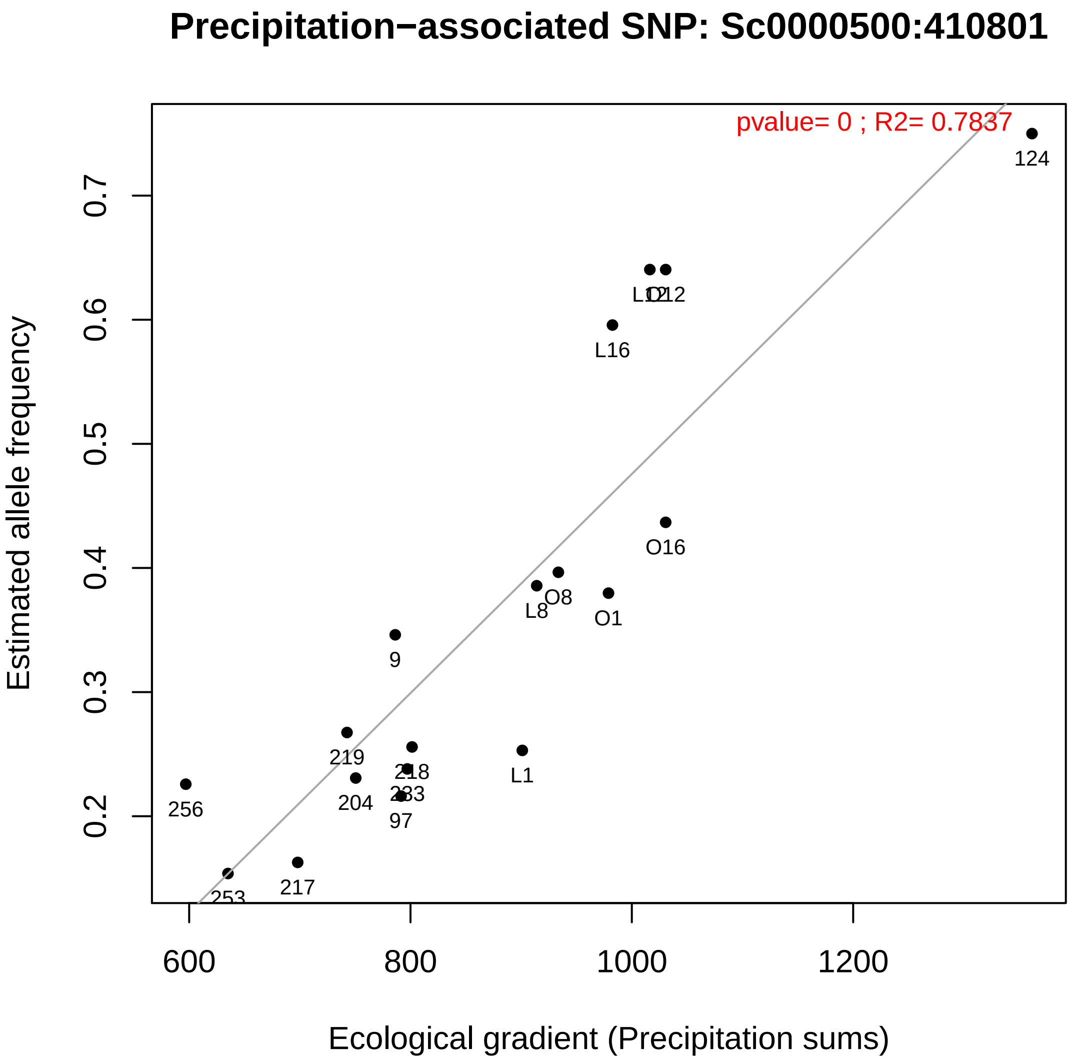

Precipitation−associated SNP: Sc0000500:410801
pvalue= 0 ; R2= 0.7837
Estimated allele frequency
Ecological gradient (Precipitation sums)
0.2
0.3
0.4
0.5
0.6
0.7
600
800
1000
1200
124
L12
O12
L16
O16
O8
L8
O1
9
219
218
233
204
97
L1
256
253
217

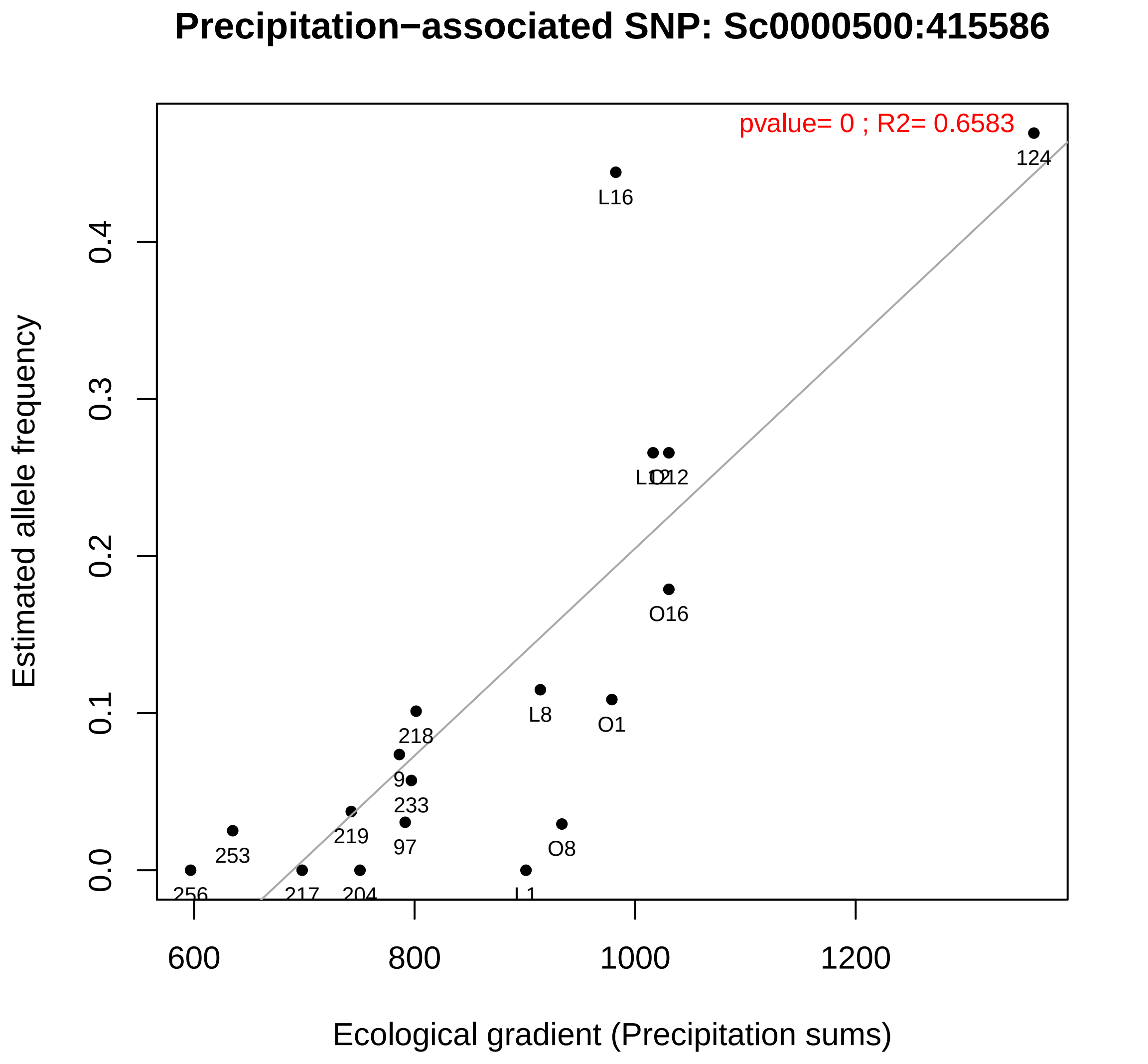
Precipitation−associated SNP: Sc0000500:415586
pvalue= 0 ; R2= 0.6583
124
L16
L10
O12
O16
L8
O1
218
9
233
219
97
O8
253
256
217
204
L1
0.0
0.1
0.2
0.3
0.4
600
800
1000
1200
Estimated allele frequency
Ecological gradient (Precipitation sums)

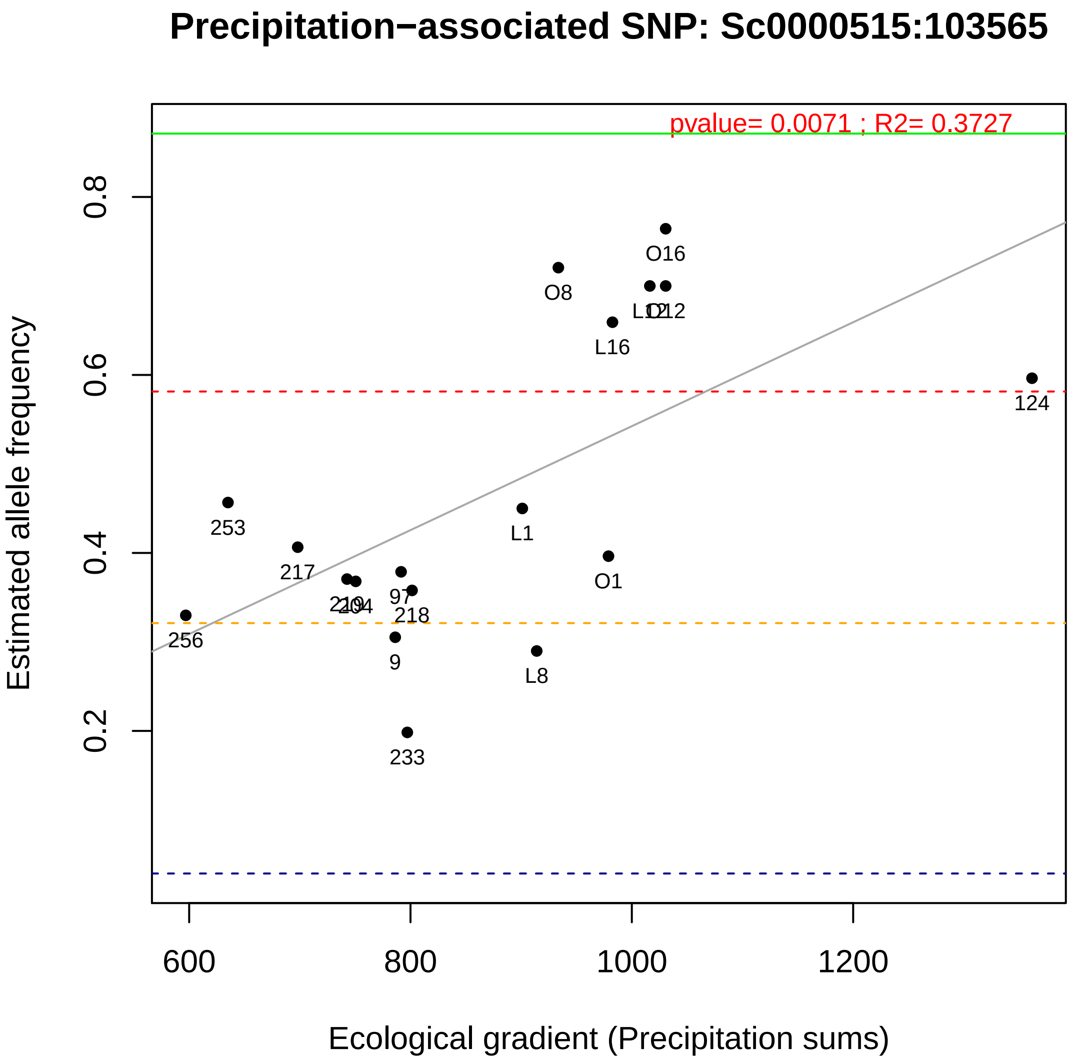

Precipitation−associated SNP: Sc0000515:103565
pvalue= 0.0071 ; R2= 0.3727
Estimated allele frequency
0.2
0.4
0.6
0.8
600
800
1000
1200
Ecological gradient (Precipitation sums)
O16
O8
L10
O12
L16
124
253
L1
217
O1
97
218
256
9
L8
233

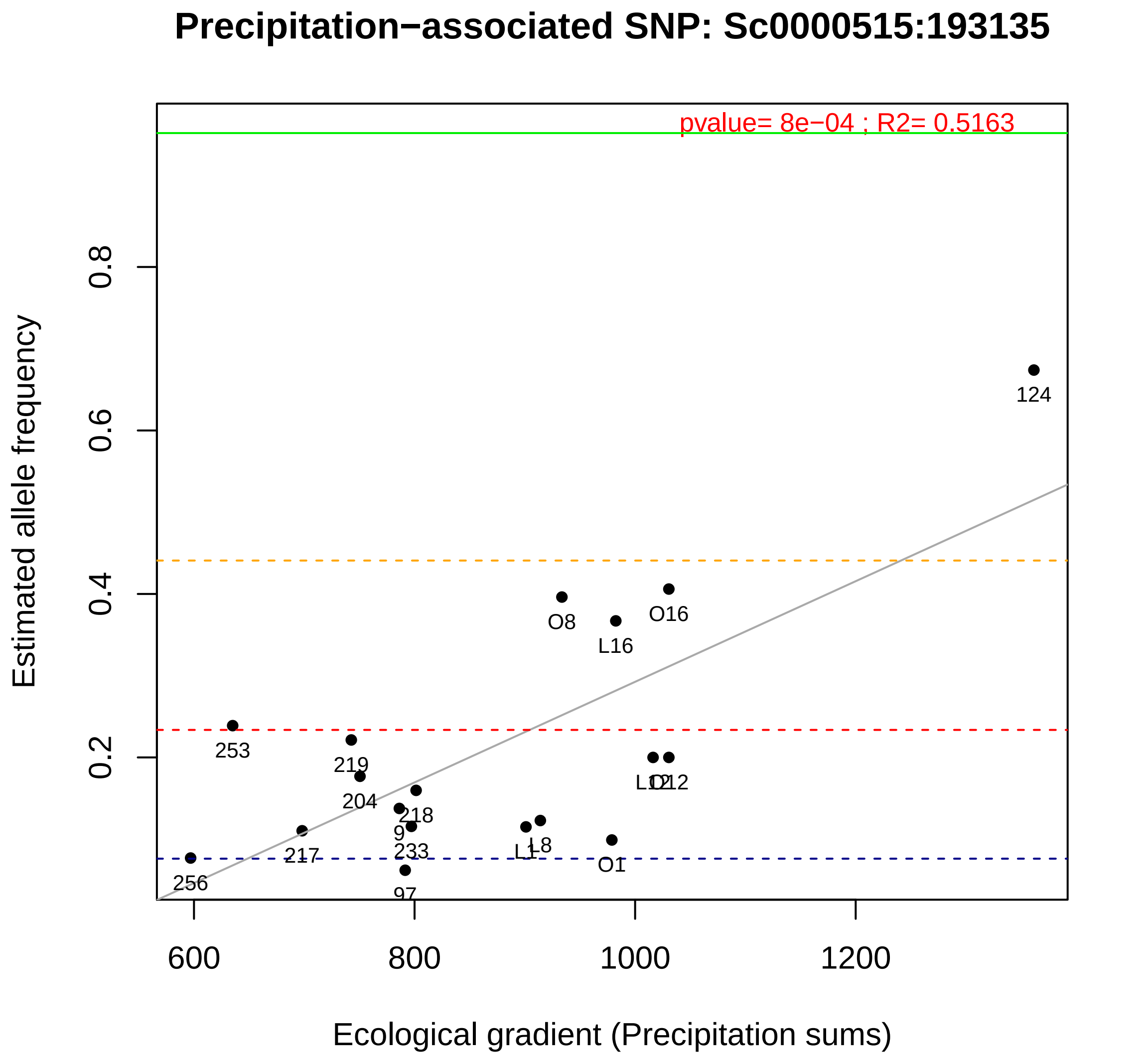

Precipitation−associated SNP: Sc0000515:193135
pvalue= 8e−04 ; R2= 0.5163
Estimated allele frequency
0.2
0.4
0.6
0.8
600
800
1000
1200
Ecological gradient (Precipitation sums)
124
O8
O16
L16
253
219
204
218
9
233
217
256
97
L1
L8
O1
L12
O12

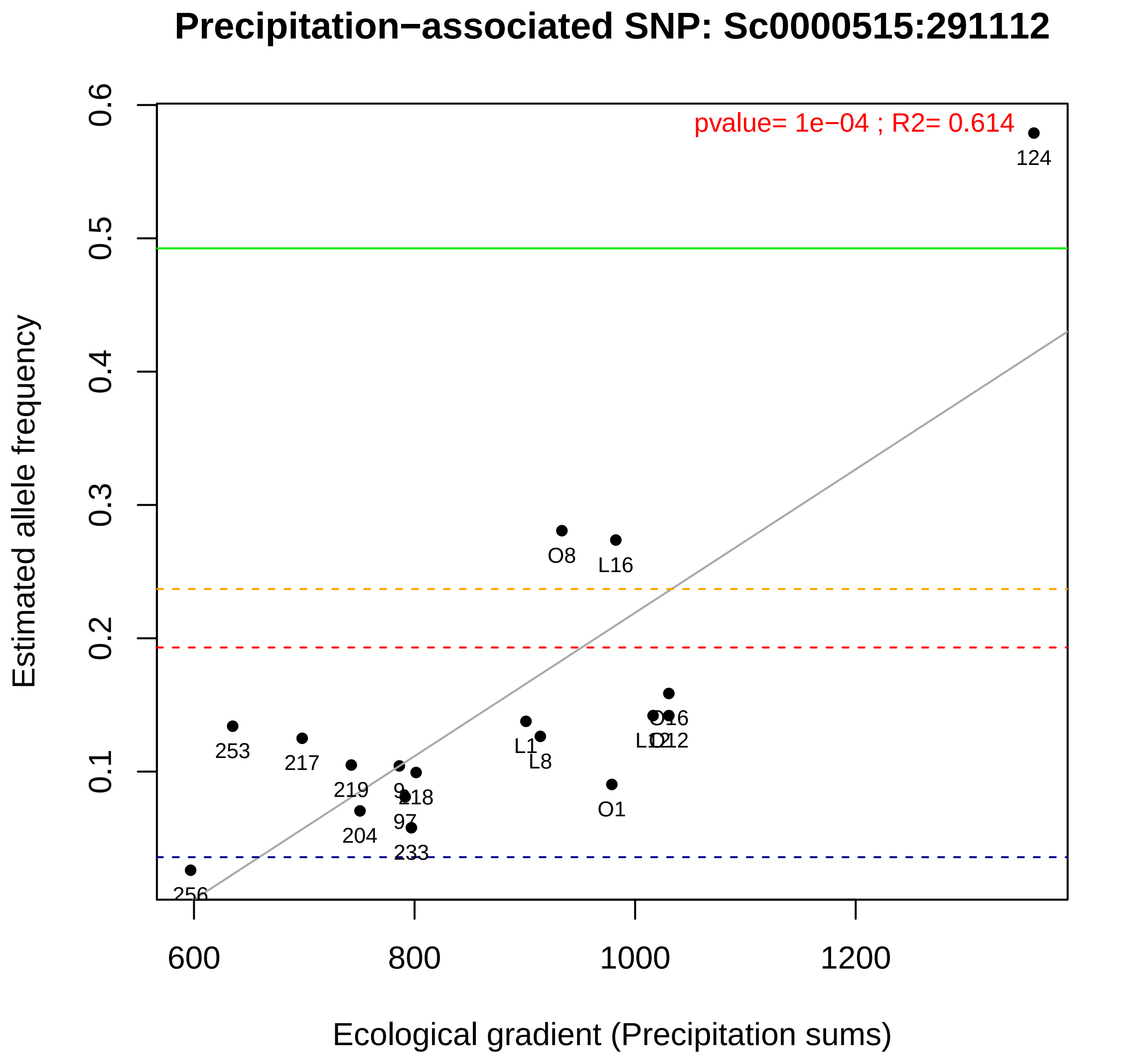
Precipitation−associated SNP: Sc0000515:291112
pvalue= 1e−04 ; R2= 0.614
Estimated allele frequency
0.1
0.2
0.3
0.4
0.5
0.6
Ecological gradient (Precipitation sums)
600
800
1000
1200
124
O8
L16
253
217
219
204
9
218
97
233
L1
L8
O1
O16
L10
O12
256

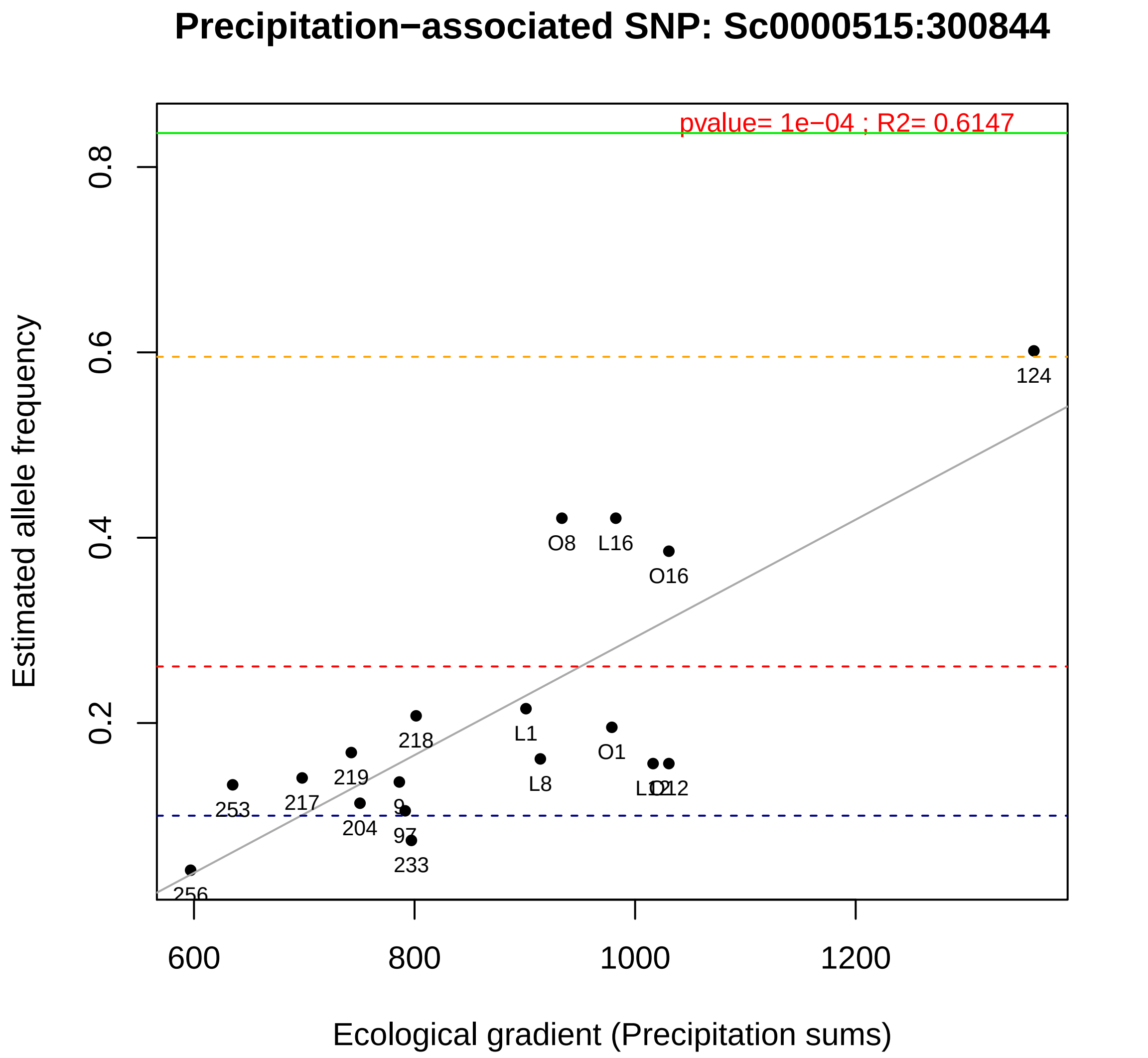

Precipitation−associated SNP: Sc0000515:300844
pvalue= 1e−04 ; R2= 0.6147
Estimated allele frequency
0.2
0.4
0.6
0.8
600
800
1000
1200
Ecological gradient (Precipitation sums)
124
O8
L16
O16
218
L1
O1
219
L8
253
217
204
9
97
233
256

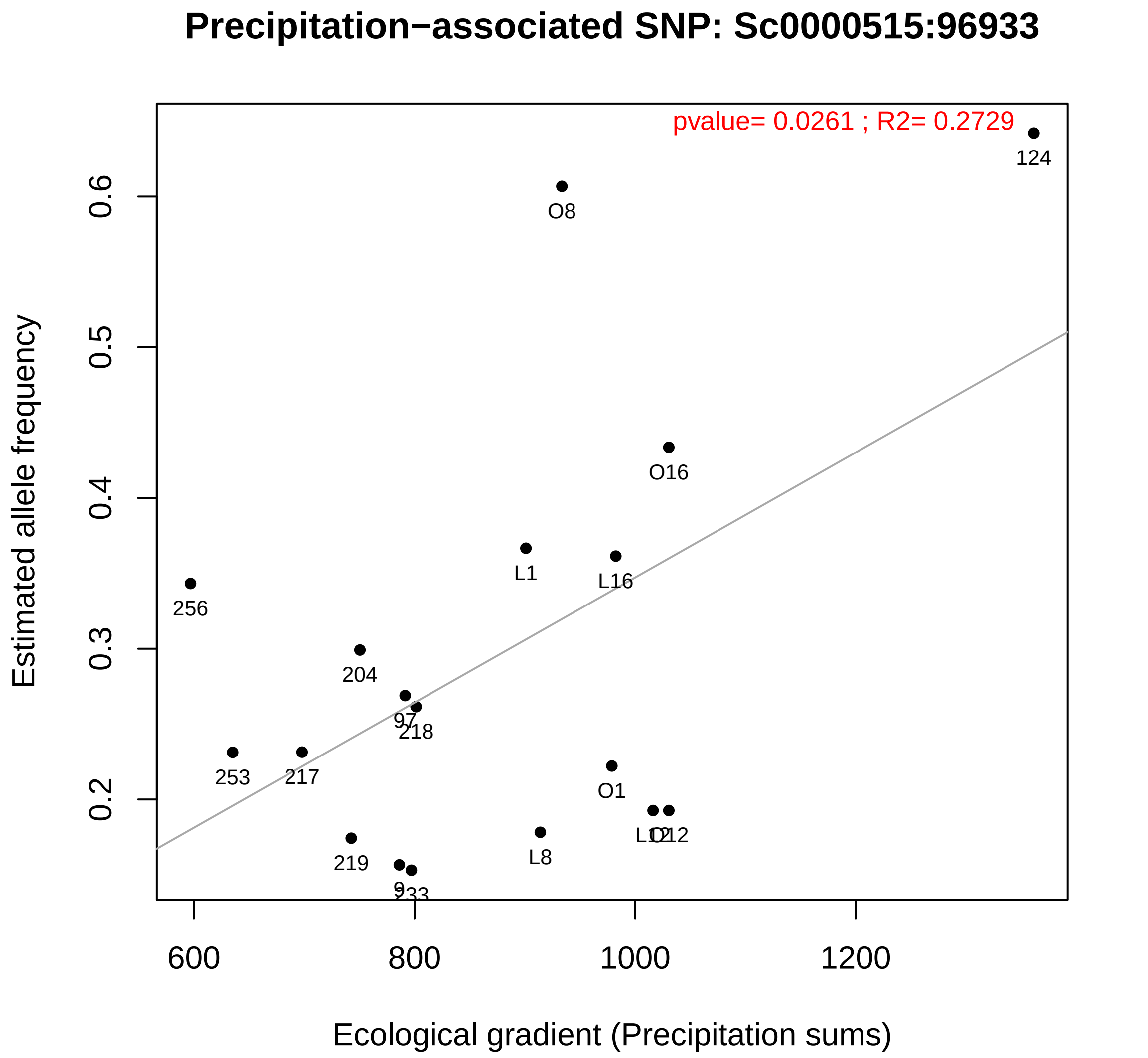
Precipitation−associated SNP: Sc0000515:96933
pvalue= 0.0261 ; R2= 0.2729
Estimated allele frequency
0.2
0.3
0.4
0.5
0.6
600
800
1000
1200
Ecological gradient (Precipitation sums)
124
O8
O16
L1
L16
256
204
97
218
253
217
O1
L8
219

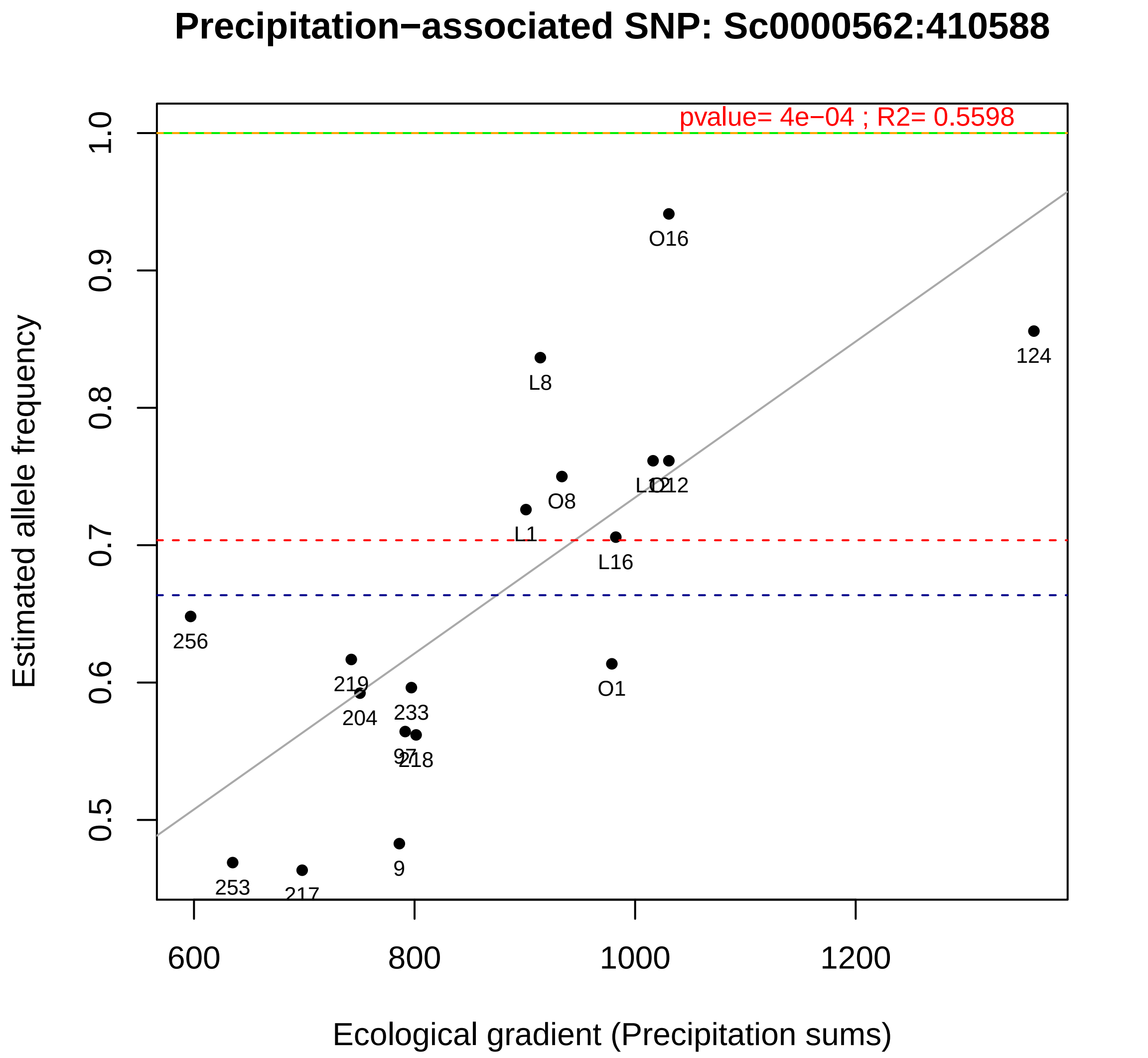

Precipitation−associated SNP: Sc0000562:410588
pvalue= 4e−04 ; R2= 0.5598
Estimated allele frequency
Ecological gradient (Precipitation sums)
0.5
0.6
0.7
0.8
0.9
1.0
600
800
1000
1200
O16
124
L8
O8
L1
L16
256
219
204
233
O1
9
253
217

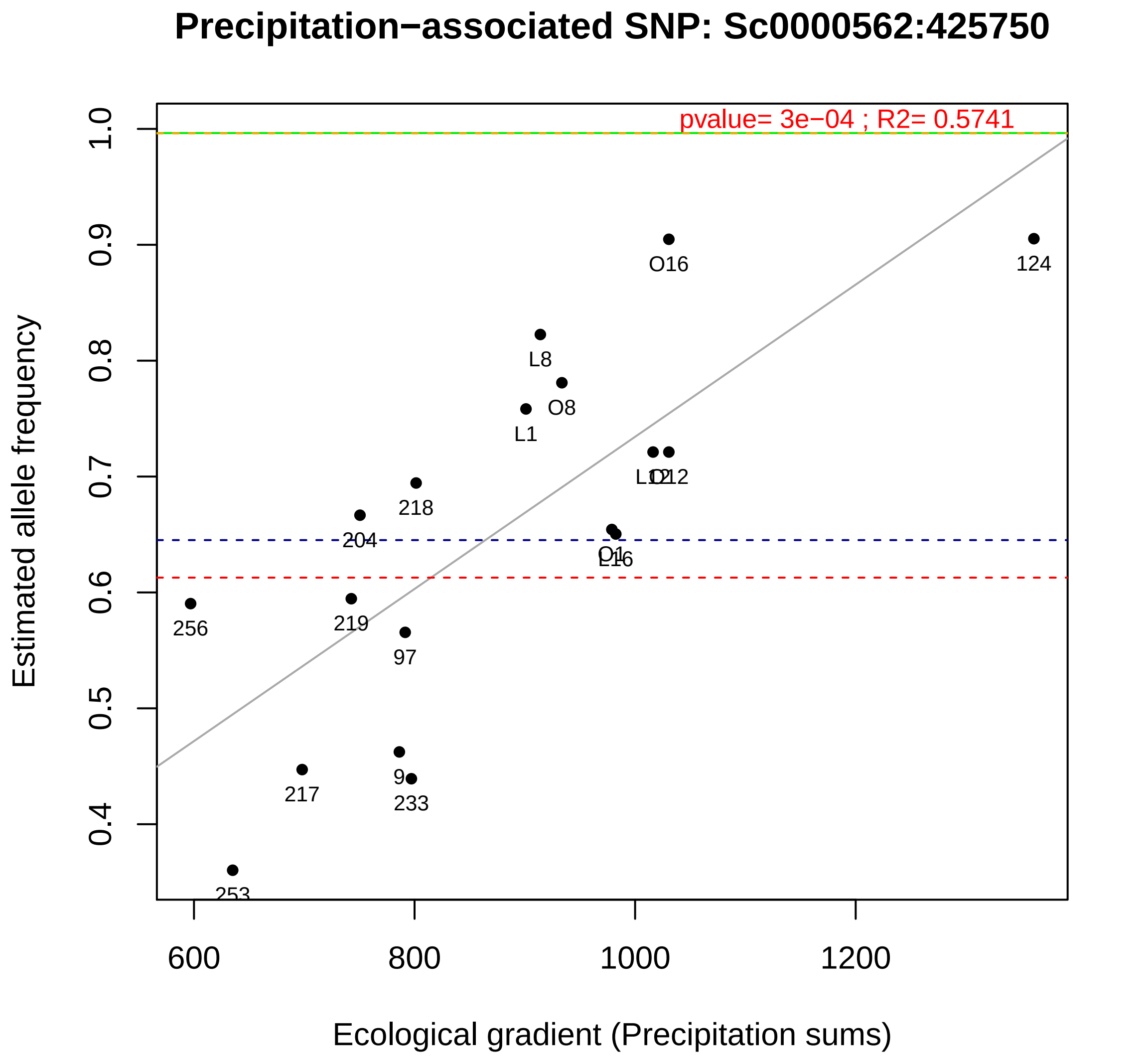

Precipitation−associated SNP: Sc0000562:425750
pvalue= 3e−04 ; R2= 0.5741
Estimated allele frequency
Ecological gradient (Precipitation sums)
0.4
0.5
0.6
0.7
0.8
0.9
1.0
600
800
1000
1200
O16
124
L8
O8
L1
L10
O12
218
204
O1
L16
256
219
97
217
9
233
253

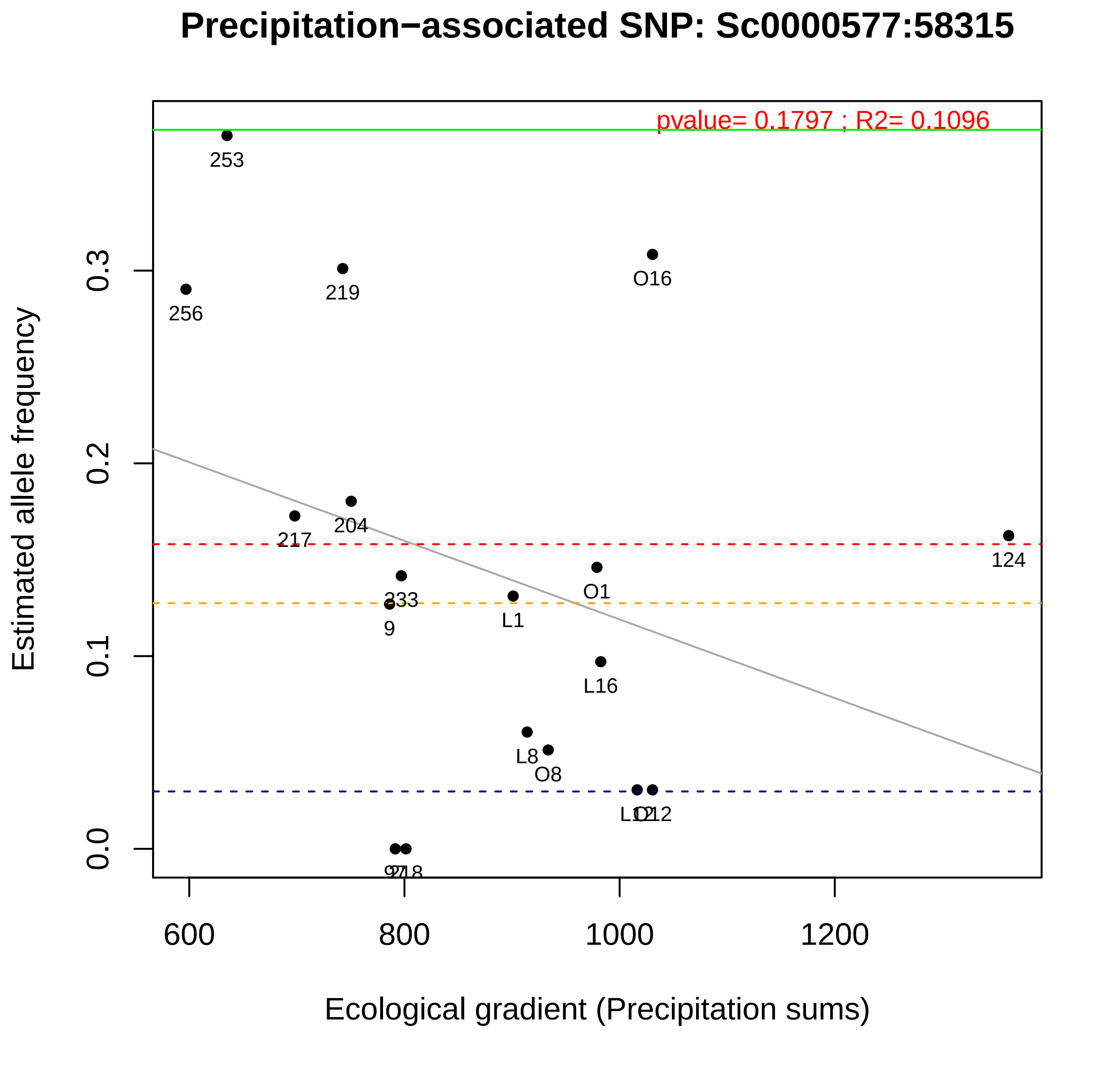

Precipitation−associated SNP: Sc0000577:58315
pvalue= 0.1797 ; R2= 0.1096
Estimated allele frequency
Ecological gradient (Precipitation sums)
0.0
0.1
0.2
0.3
600
800
1000
1200
253
256
219
O16
217
204
124
233
9
L1
O1
L16
L8
O8

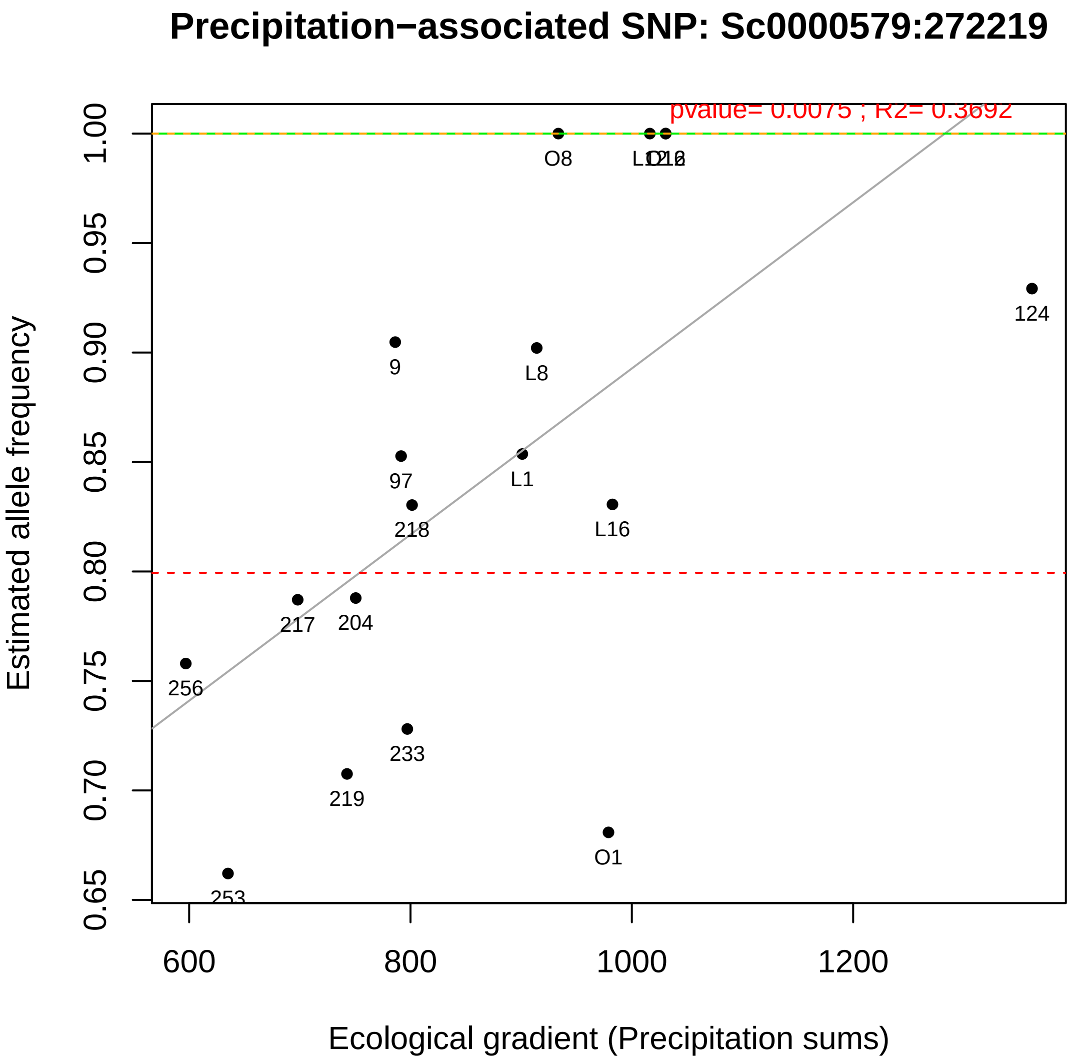
Precipitation−associated SNP: Sc0000579:272219
pvalue= 0.0075 ; R2= 0.3692
Estimated allele frequency
1.00
0.95
0.90
0.85
0.80
0.75
0.70
0.65
600
800
1000
1200
Ecological gradient (Precipitation sums)
O8
124
9
L8
97
L1
218
L16
217
204
256
233
219
O1
253

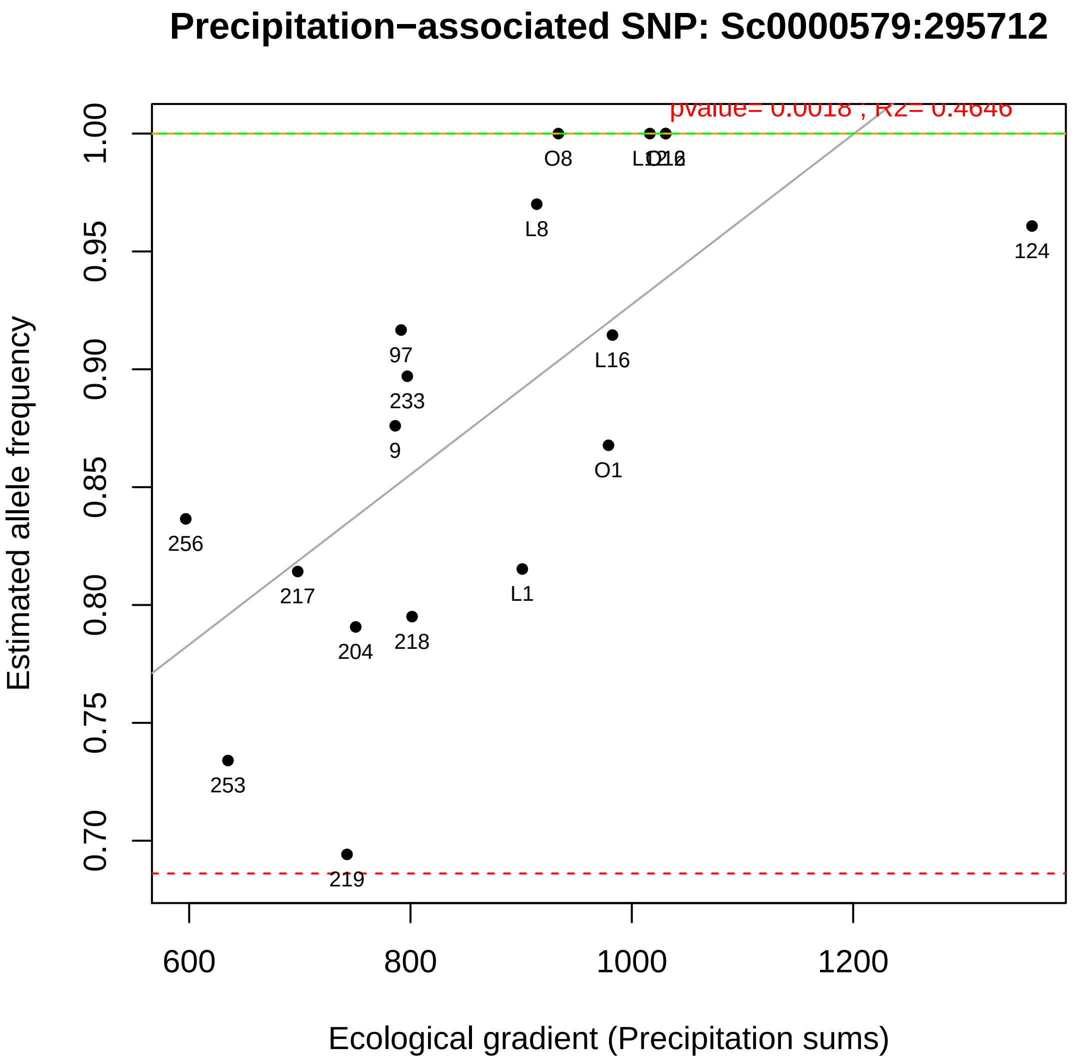

Precipitation−associated SNP: Sc0000579:295712
pvalue= 0.0018 ; R2= 0.4646
Estimated allele frequency
Ecological gradient (Precipitation sums)
1.00
0.95
0.90
0.85
0.80
0.75
0.70
600
800
1000
1200
O8
L8
124
97
233
9
L16
O1
256
217
204
218
L1
253
219

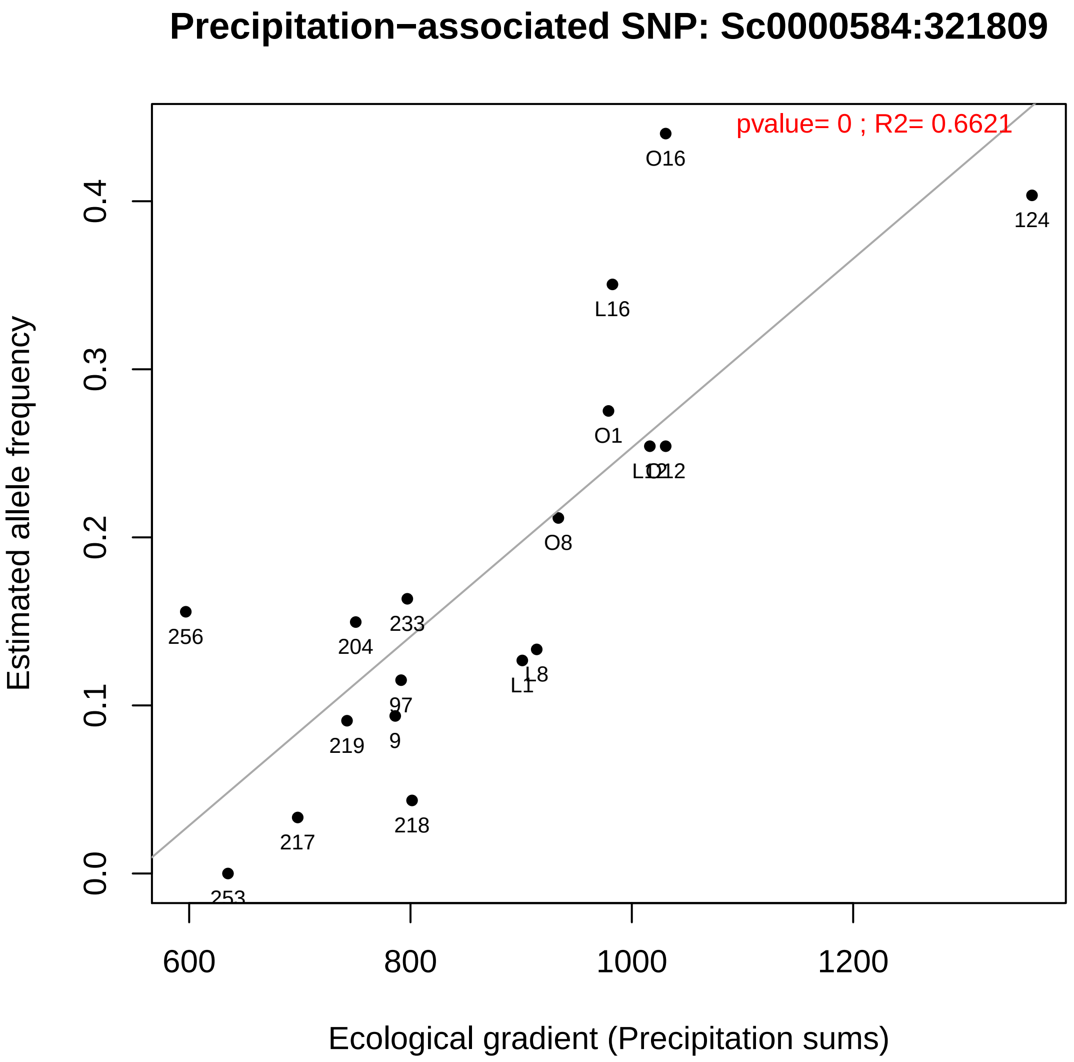
Precipitation−associated SNP: Sc0000584:321809
pvalue= 0 ; R2= 0.6621
O16
124
L16
O1
L12
O12
O8
233
256
204
L8
L1
97
219
9
218
217
253
0.0
0.1
0.2
0.3
0.4
Estimated allele frequency
600
800
1000
1200
Ecological gradient (Precipitation sums)

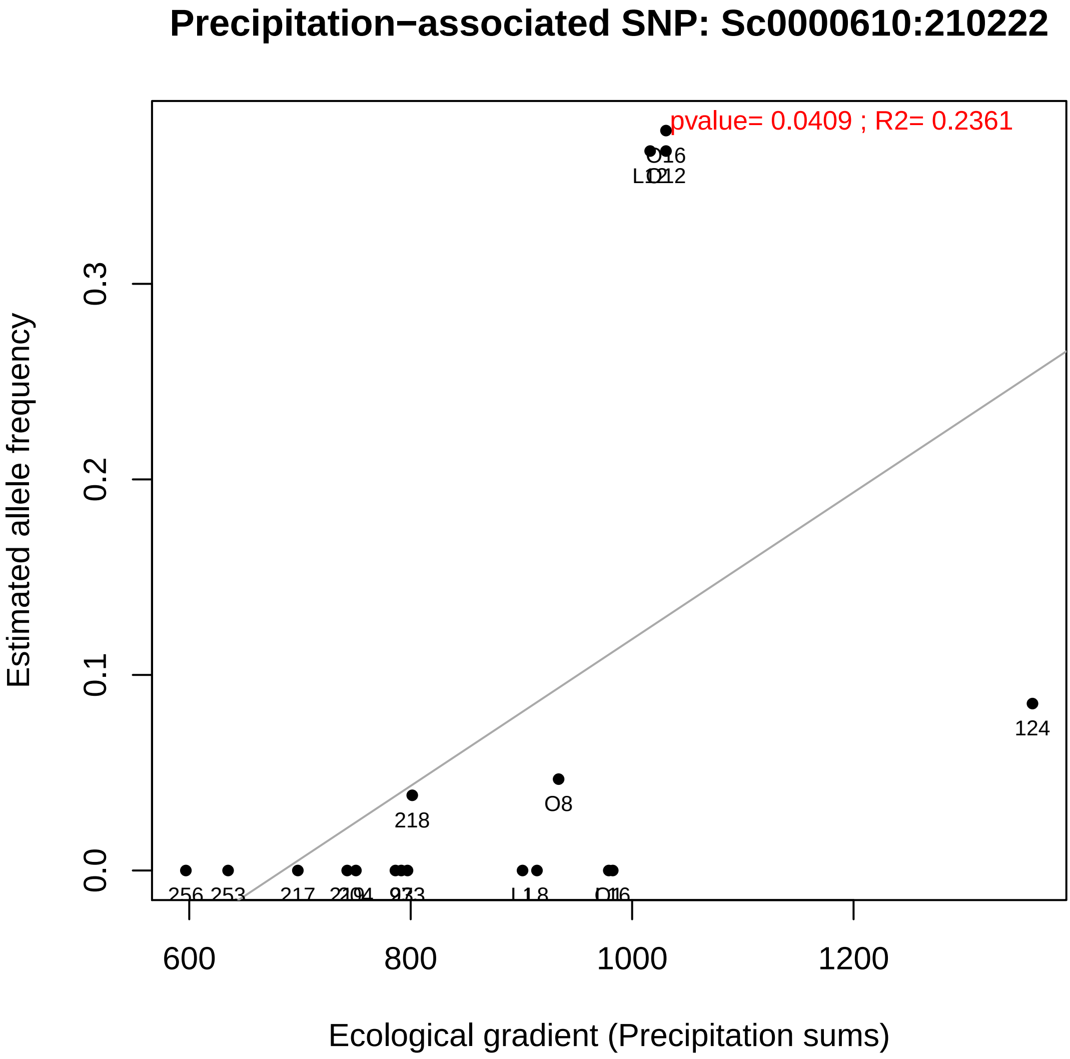
Precipitation−associated SNP: Sc0000610:210222
pvalue= 0.0409 ; R2= 0.2361
O16
124
218
O8
0.0
0.1
0.2
0.3
600
800
1000
1200
Estimated allele frequency
Ecological gradient (Precipitation sums)

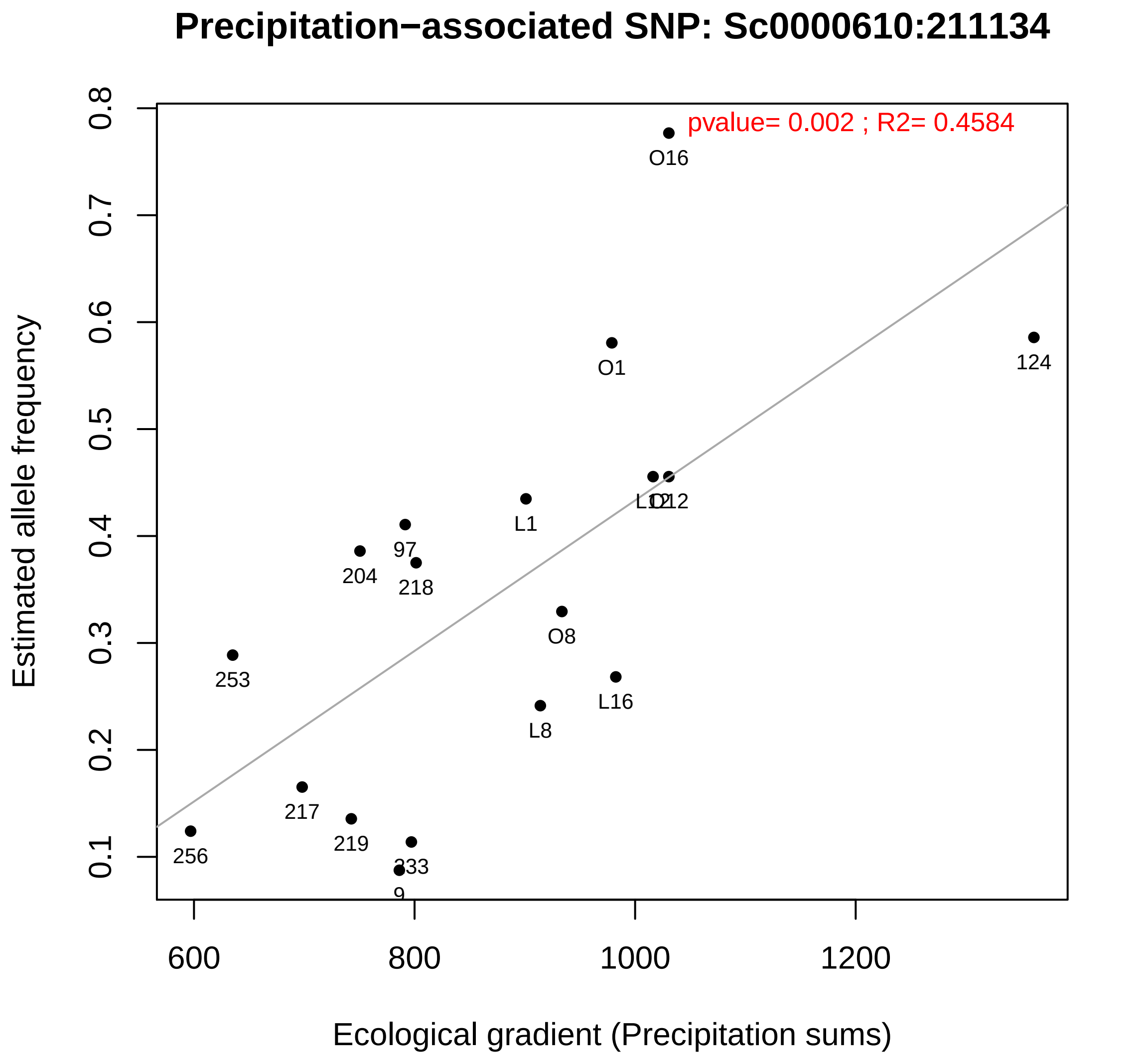
Precipitation−associated SNP: Sc0000610:211134
pvalue= 0.002 ; R2= 0.4584
O16
O1
124
L10
O12
L1
97
204
218
O8
253
L16
L8
217
219
256
233
9
Estimated allele frequency
0.1
0.2
0.3
0.4
0.5
0.6
0.7
0.8
600
800
1000
1200
Ecological gradient (Precipitation sums)

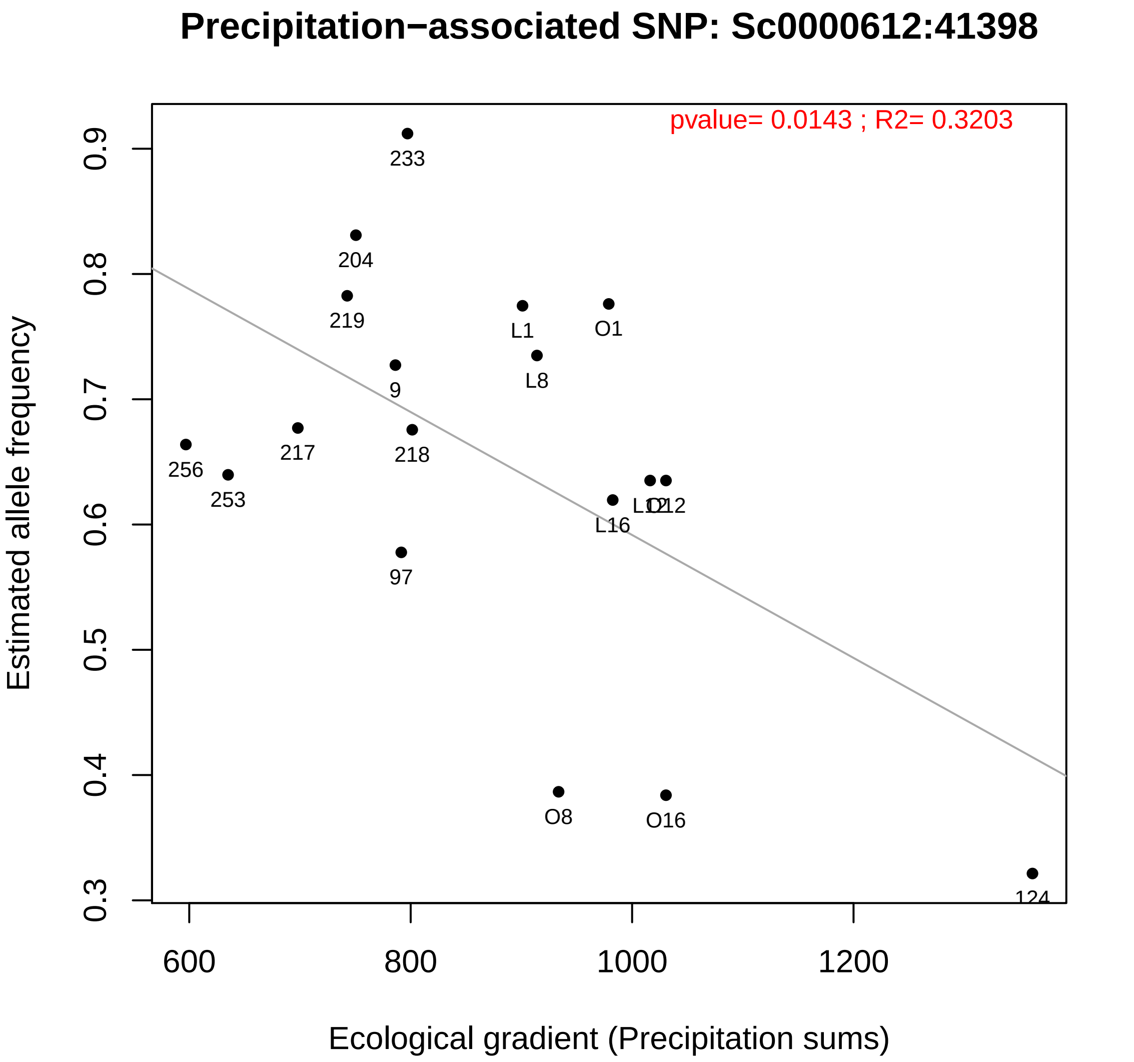

Precipitation−associated SNP: Sc0000612:41398
pvalue= 0.0143 ; R2= 0.3203
Estimated allele frequency
0.3
0.4
0.5
0.6
0.7
0.8
0.9
600
800
1000
1200
Ecological gradient (Precipitation sums)
233
204
219
L1
O1
L8
9
217
218
256
253
L16
L12
O12
97
O8
O16
124

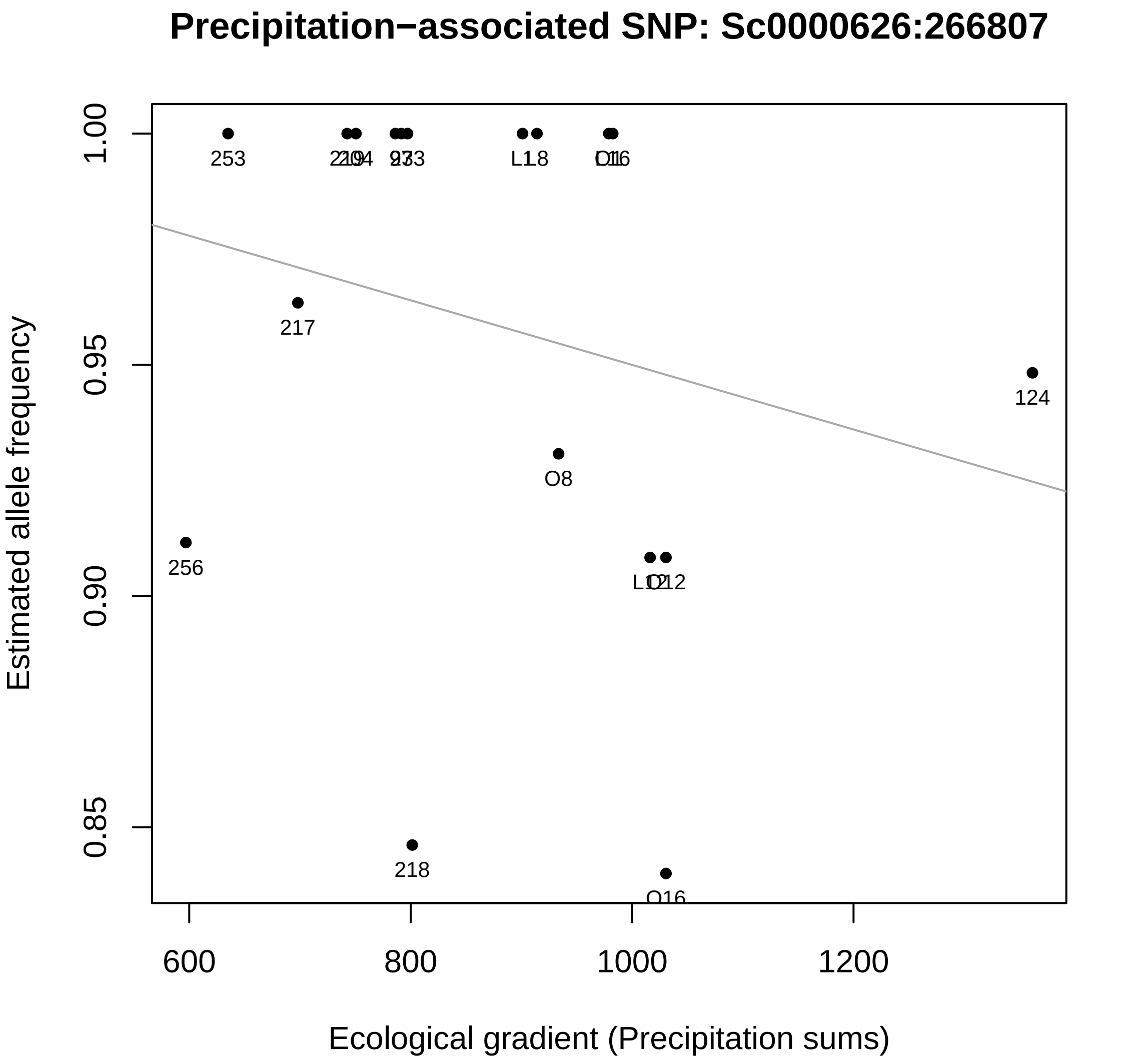
Precipitation−associated SNP: Sc0000626:266807
Estimated allele frequency
Ecological gradient (Precipitation sums)
1.00
0.95
0.90
0.85
600
800
1000
1200
253
217
256
218
O8
124

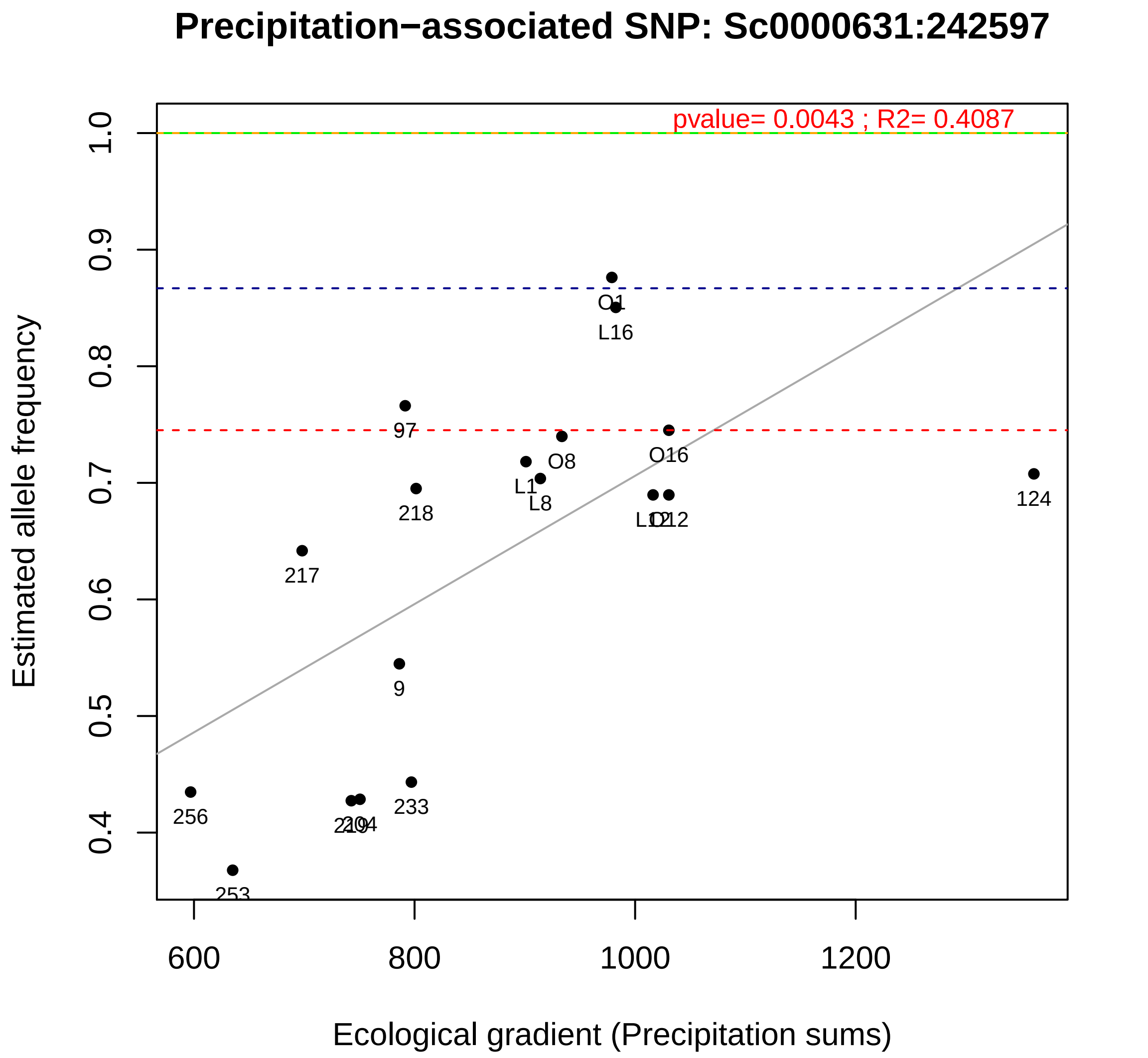

Precipitation−associated SNP: Sc0000631:242597
pvalue= 0.0043 ; R2= 0.4087
Estimated allele frequency
Ecological gradient (Precipitation sums)
0.4
0.5
0.6
0.7
0.8
0.9
1.0
600
800
1000
1200
O1
L16
97
O8
O16
L1
L8
218
124
L12
O12
217
9
256
233
209
204
253

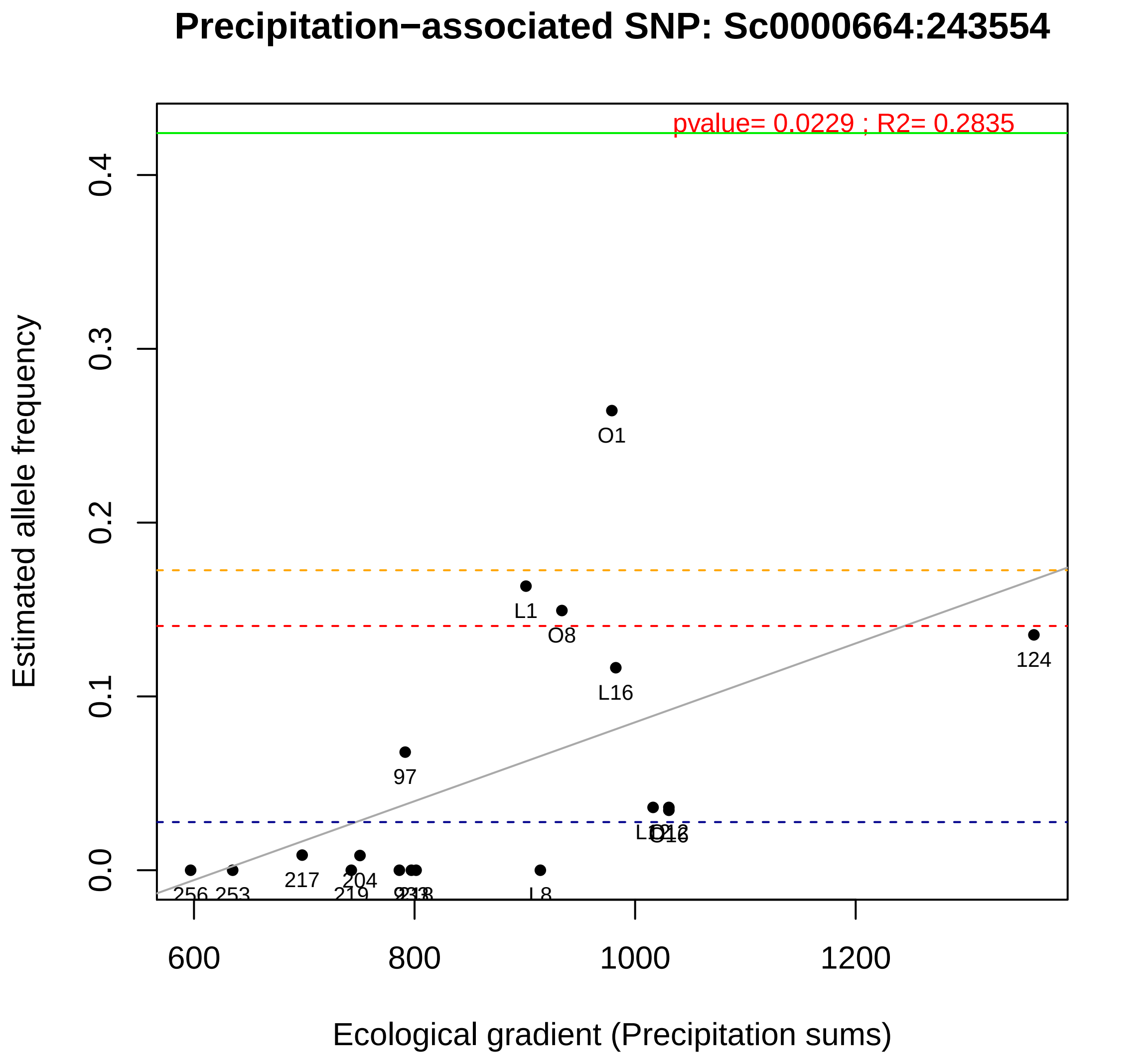

Precipitation−associated SNP: Sc0000664:243554
pvalue= 0.0229 ; R2= 0.2835
Estimated allele frequency
0.0
0.1
0.2
0.3
0.4
Ecological gradient (Precipitation sums)
600
800
1000
1200
O1
L1
O8
124
L16
97
217
204

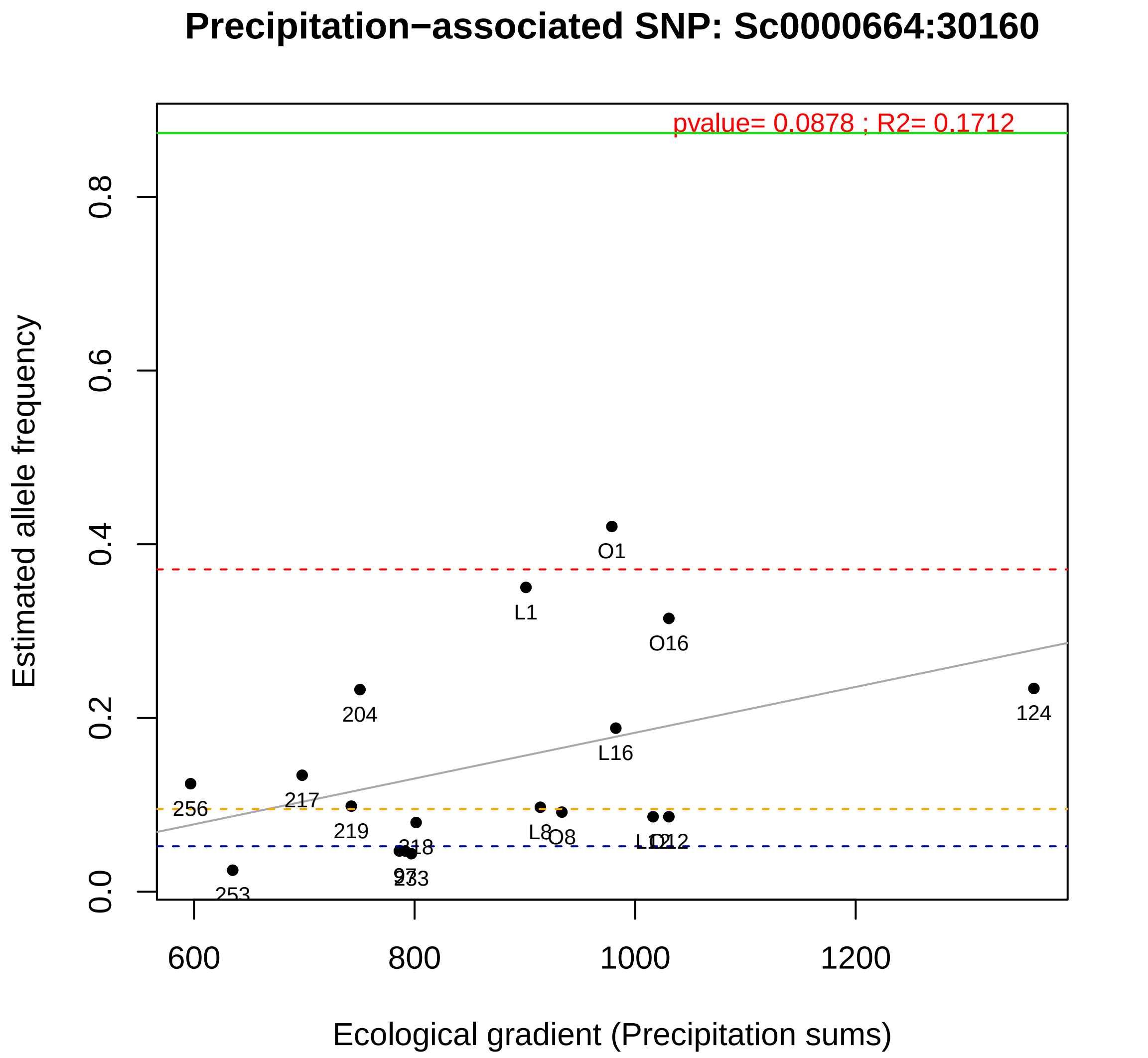

Precipitation−associated SNP: Sc0000664:30160
pvalue= 0.0878 ; R2= 0.1712
Estimated allele frequency
Ecological gradient (Precipitation sums)
0.0
0.2
0.4
0.6
0.8
600
800
1000
1200
O1
L1
O16
204
124
L16
256
217
219
L8
O8
218
253

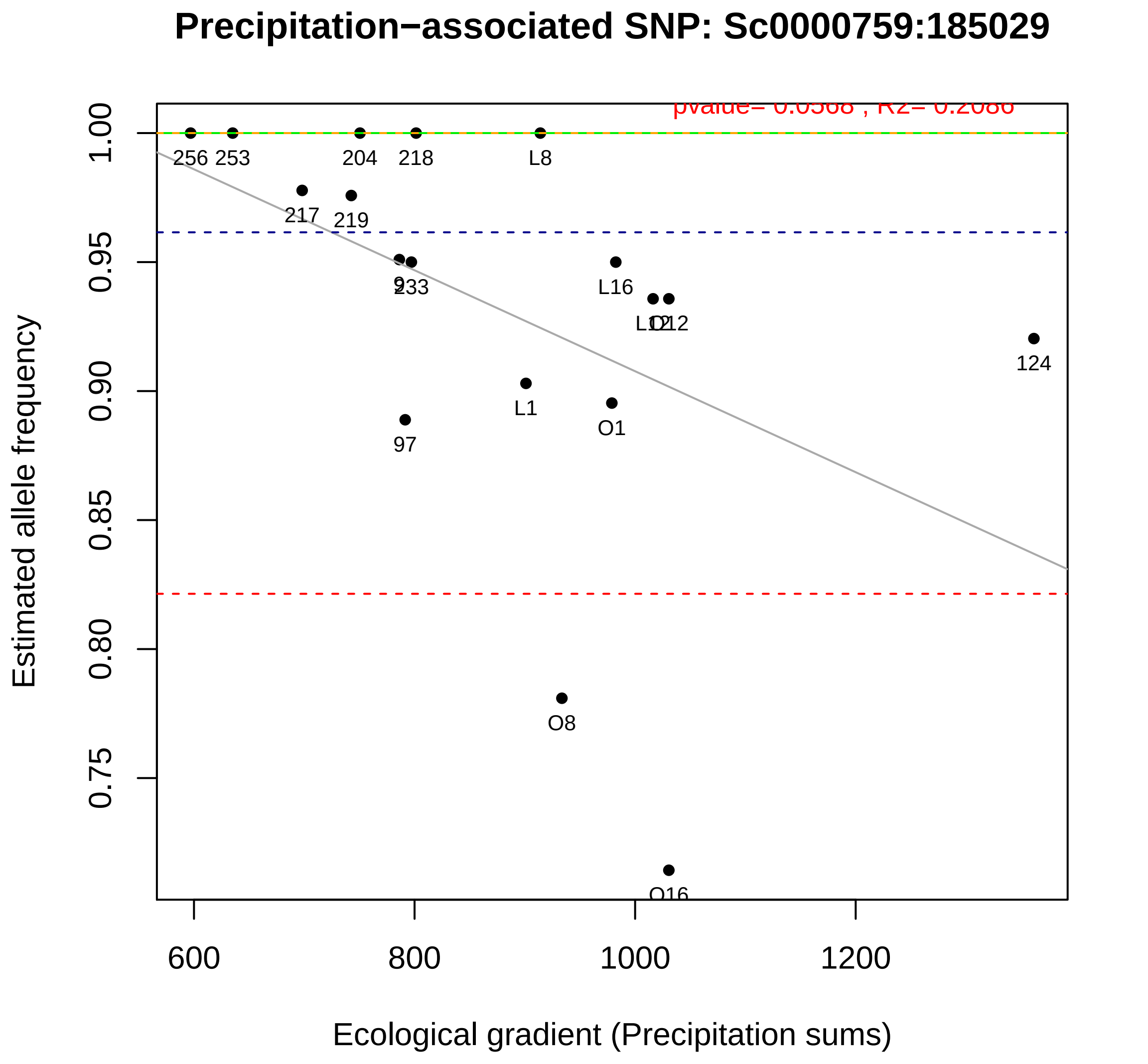

Precipitation−associated SNP: Sc0000759:185029
pvalue= 0.0568 , R2= 0.2086
Estimated allele frequency
1.00
0.95
0.90
0.85
0.80
0.75
256
253
204
218
L8
217
219
233
L16
L12
O12
124
L1
O1
97
O8
O16
600
800
1000
1200
Ecological gradient (Precipitation sums)

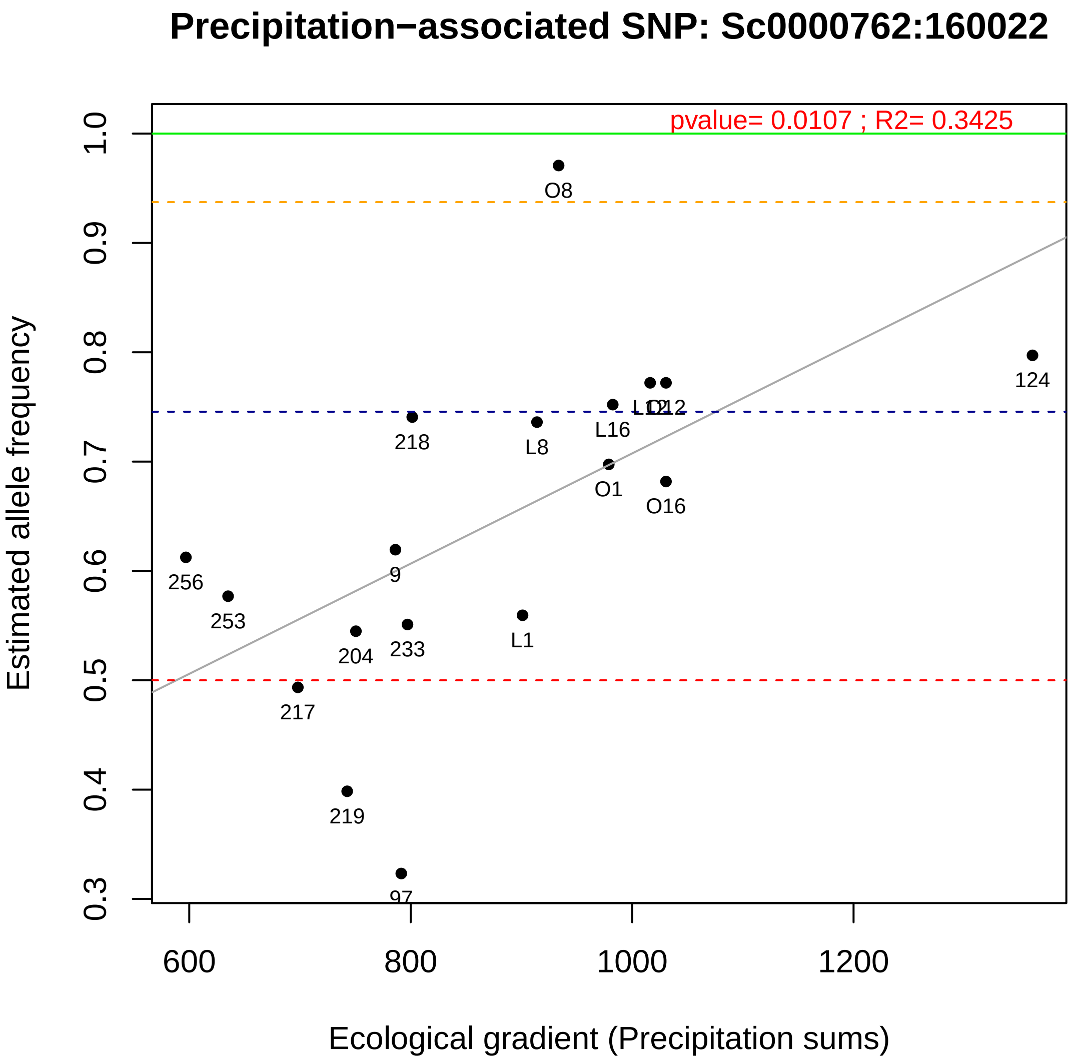

Precipitation−associated SNP: Sc0000762:160022
pvalue= 0.0107 ; R2= 0.3425
Estimated allele frequency
Ecological gradient (Precipitation sums)
0.3
0.4
0.5
0.6
0.7
0.8
0.9
1.0
600
800
1000
1200
O8
124
L12
O12
L16
218
L8
O1
O16
256
9
253
204
233
L1
217
219
97

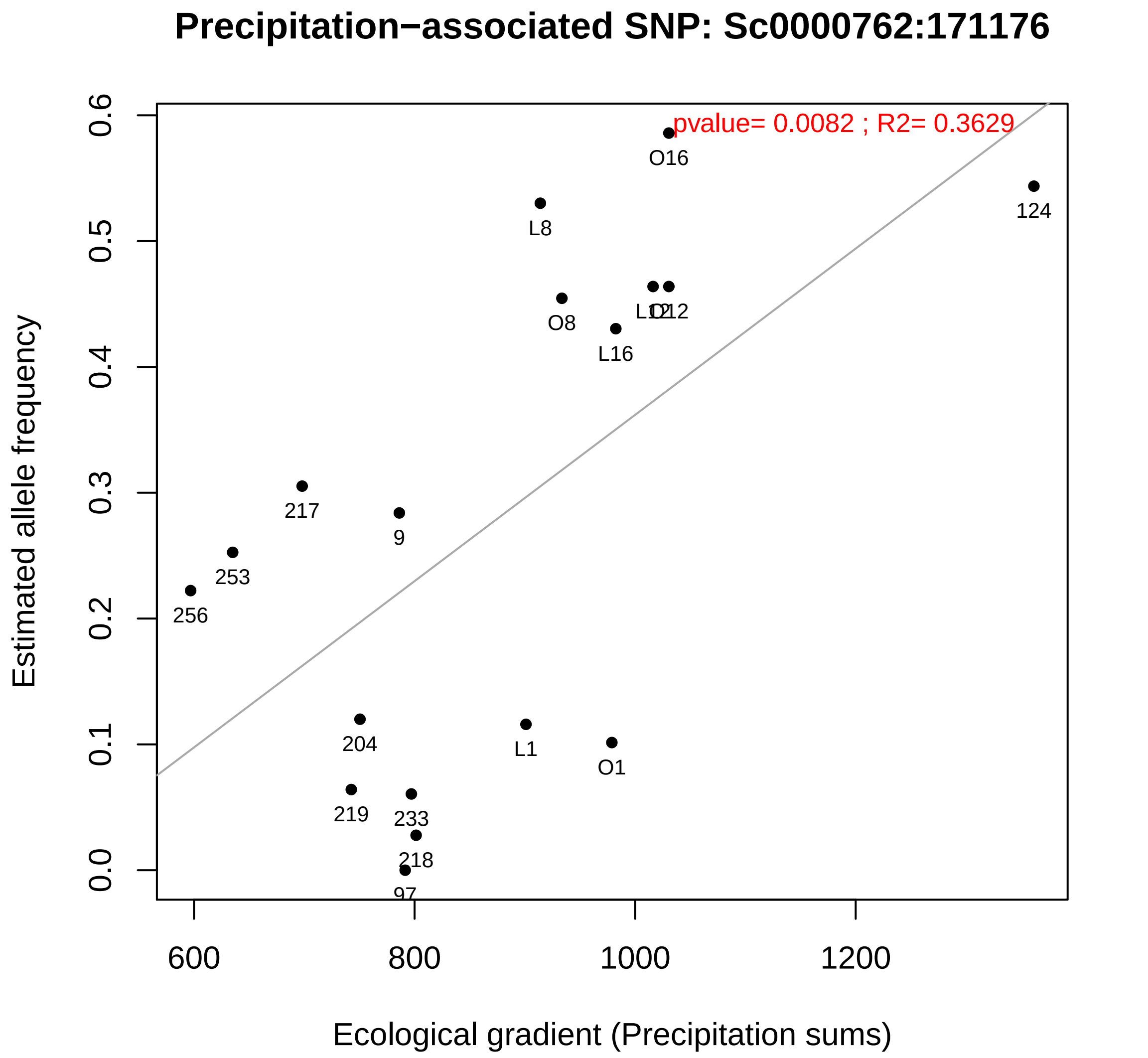

Precipitation−associated SNP: Sc0000762:171176
pvalue= 0.0082 ; R2= 0.3629
Estimated allele frequency
0.0
0.1
0.2
0.3
0.4
0.5
0.6
600
800
1000
1200
Ecological gradient (Precipitation sums)
O16
124
L8
O8
L12
O12
L16
217
9
253
256
204
L1
O1
219
233
218
97

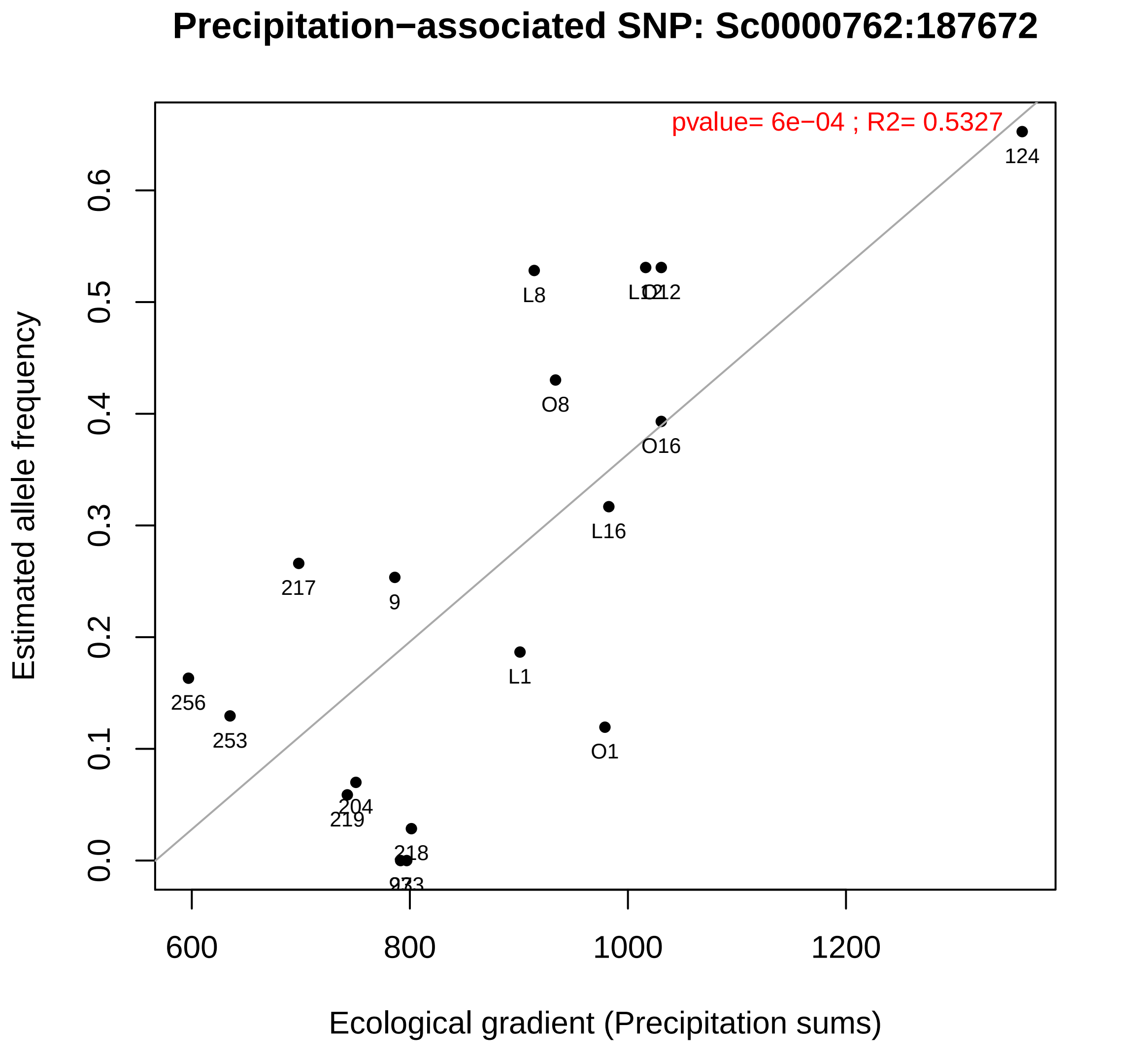
Precipitation−associated SNP: Sc0000762:187672
pvalue= 6e−04 ; R2= 0.5327
Estimated allele frequency
Ecological gradient (Precipitation sums)
0.0
0.1
0.2
0.3
0.4
0.5
0.6
600
800
1000
1200
124
L8
O12
O8
O16
L16
217
9
L1
256
253
O1
204
219
218

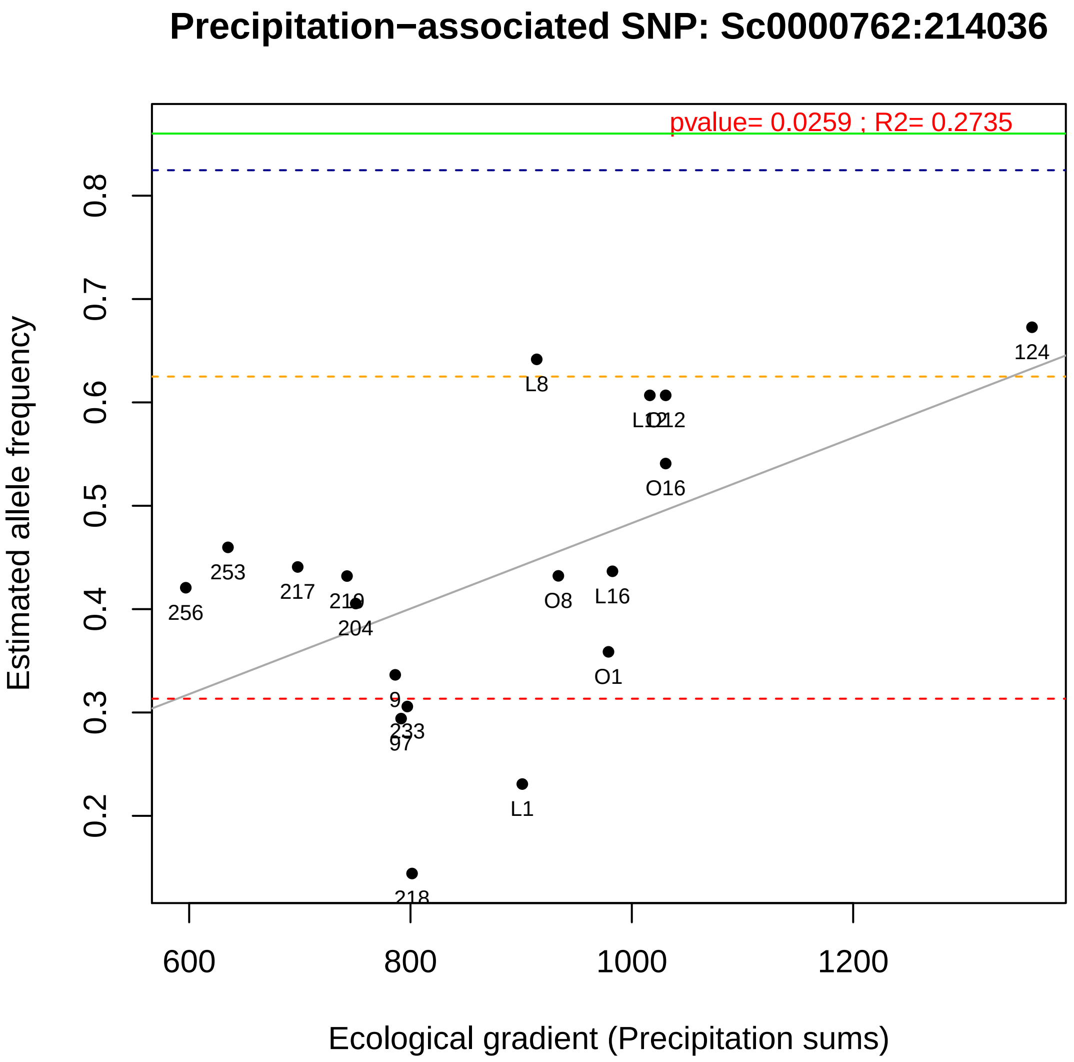

Precipitation−associated SNP: Sc0000762:214036
pvalue= 0.0259 ; R2= 0.2735
Estimated allele frequency
0.2
0.3
0.4
0.5
0.6
0.7
0.8
600
800
1000
1200
Ecological gradient (Precipitation sums)
124
L8
O16
253
217
256
204
O8
L16
O1
9
233
97
L1
218

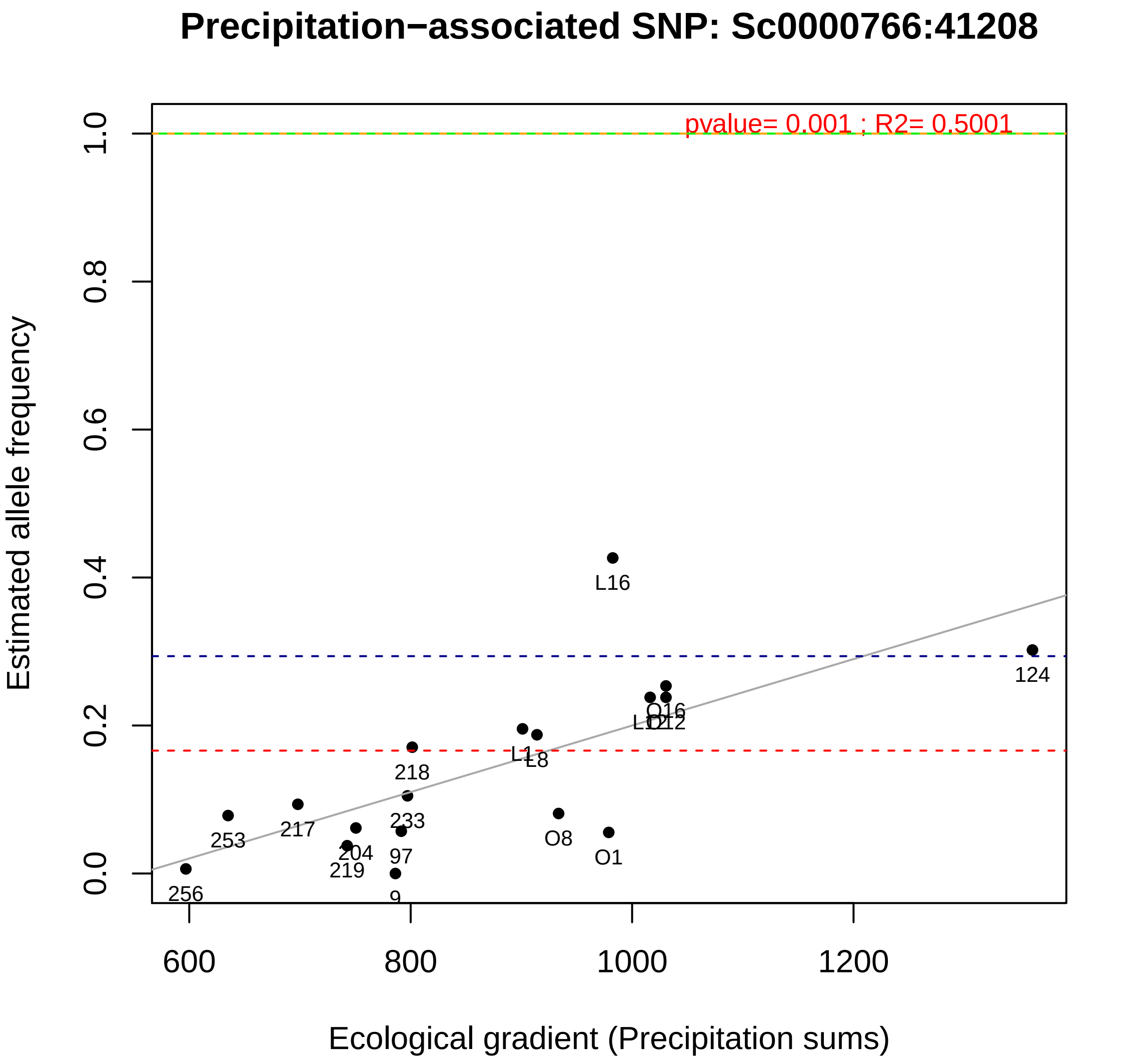
Precipitation−associated SNP: Sc0000766:41208
pvalue= 0.001 ; R2= 0.5001
Estimated allele frequency
0.0
0.2
0.4
0.6
0.8
1.0
600
800
1000
1200
Ecological gradient (Precipitation sums)
L16
124
O16
L12
O12
L1
L8
218
233
O8
O1
253
217
204
219
97
256
9

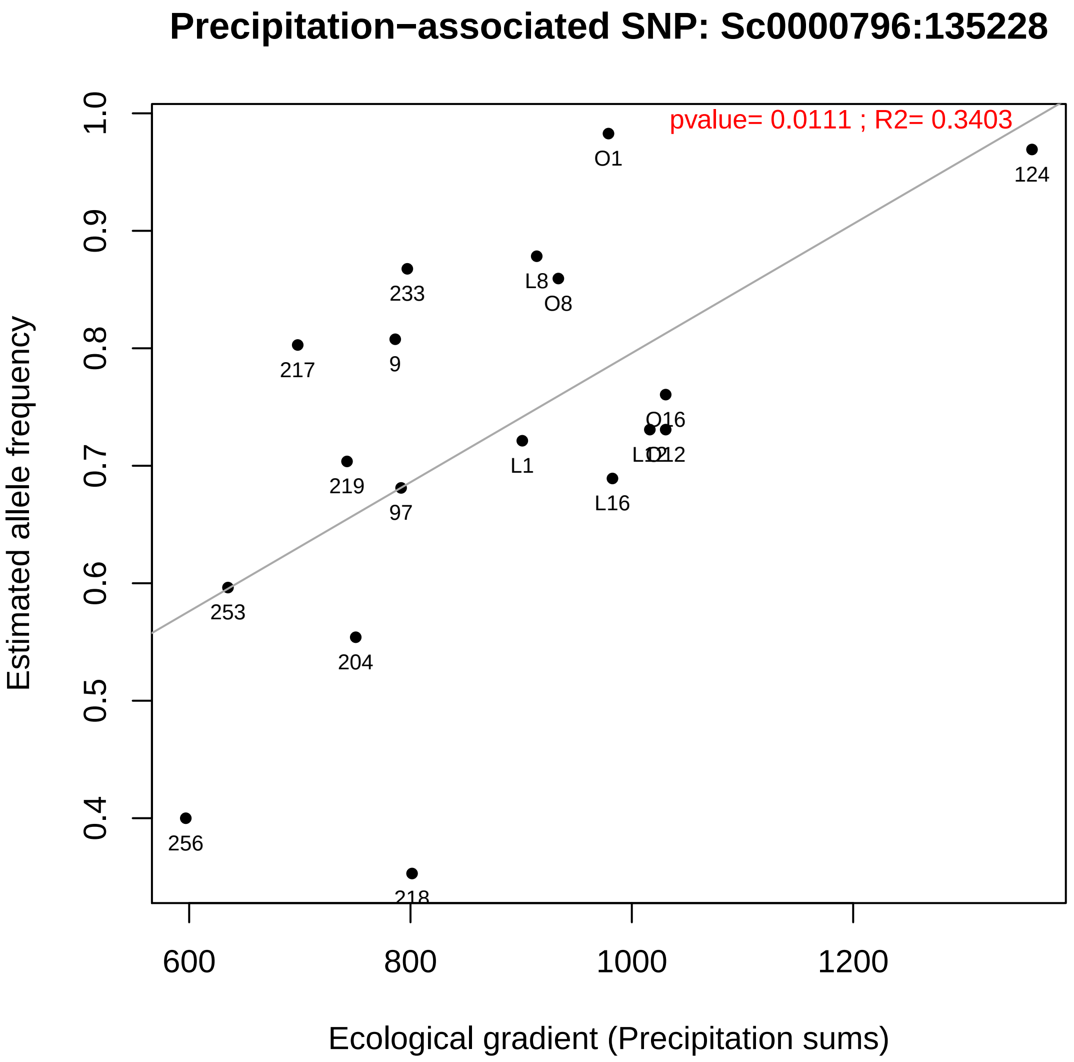

Precipitation−associated SNP: Sc0000796:135228
pvalue= 0.0111 ; R2= 0.3403
O1
124
233
L8
O8
217
9
O16
L12
O12
L1
219
97
L16
253
204
256
218
1.0
0.9
0.8
0.7
0.6
0.5
0.4
600
800
1000
1200
Estimated allele frequency
Ecological gradient (Precipitation sums)

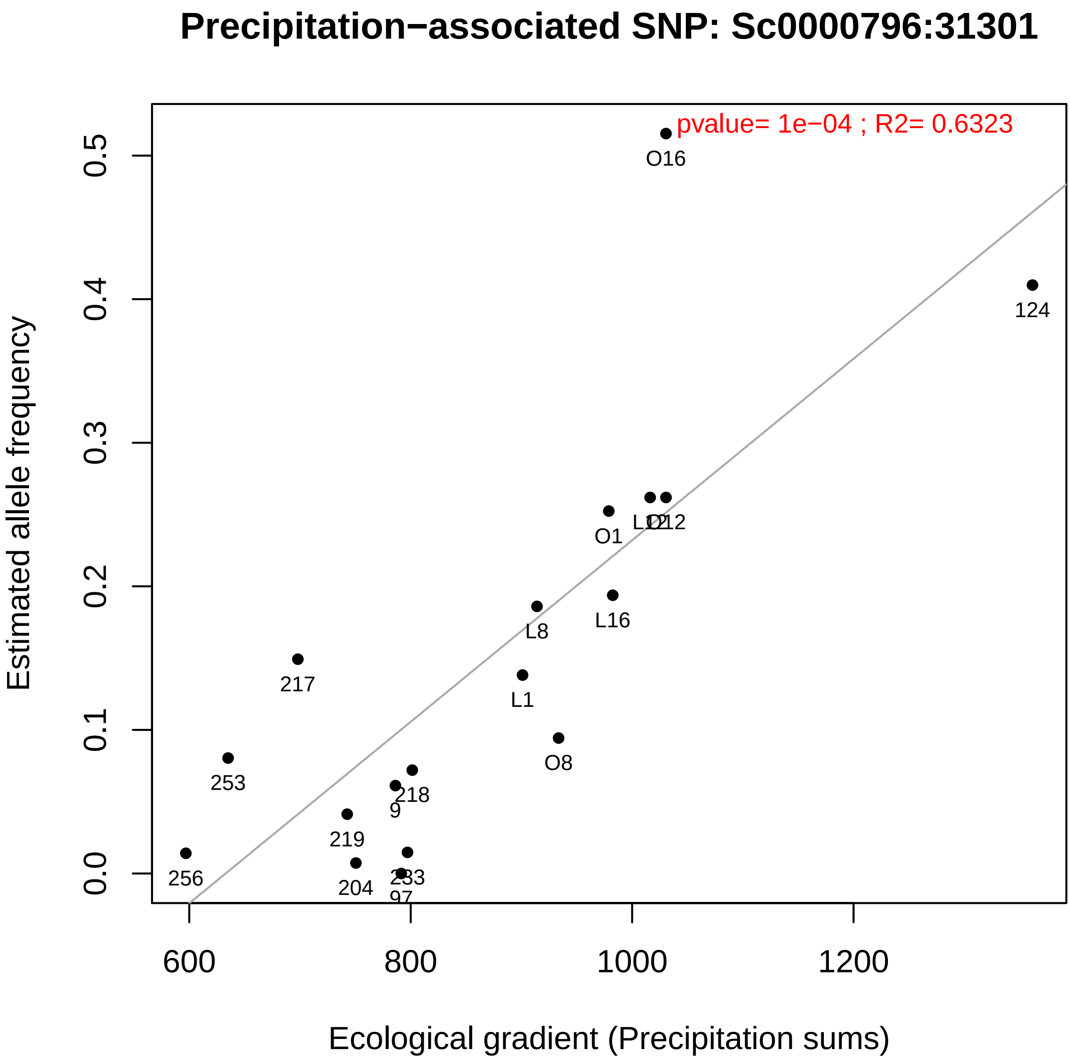

Precipitation−associated SNP: Sc0000796:31301
pvalue= 1e−04 ; R2= 0.6323
O16
124
O1
L12
O12
L8
L16
217
L1
O8
253
218
9
219
256
204
233
97
0.0
0.1
0.2
0.3
0.4
0.5
600
800
1000
1200
Estimated allele frequency
Ecological gradient (Precipitation sums)

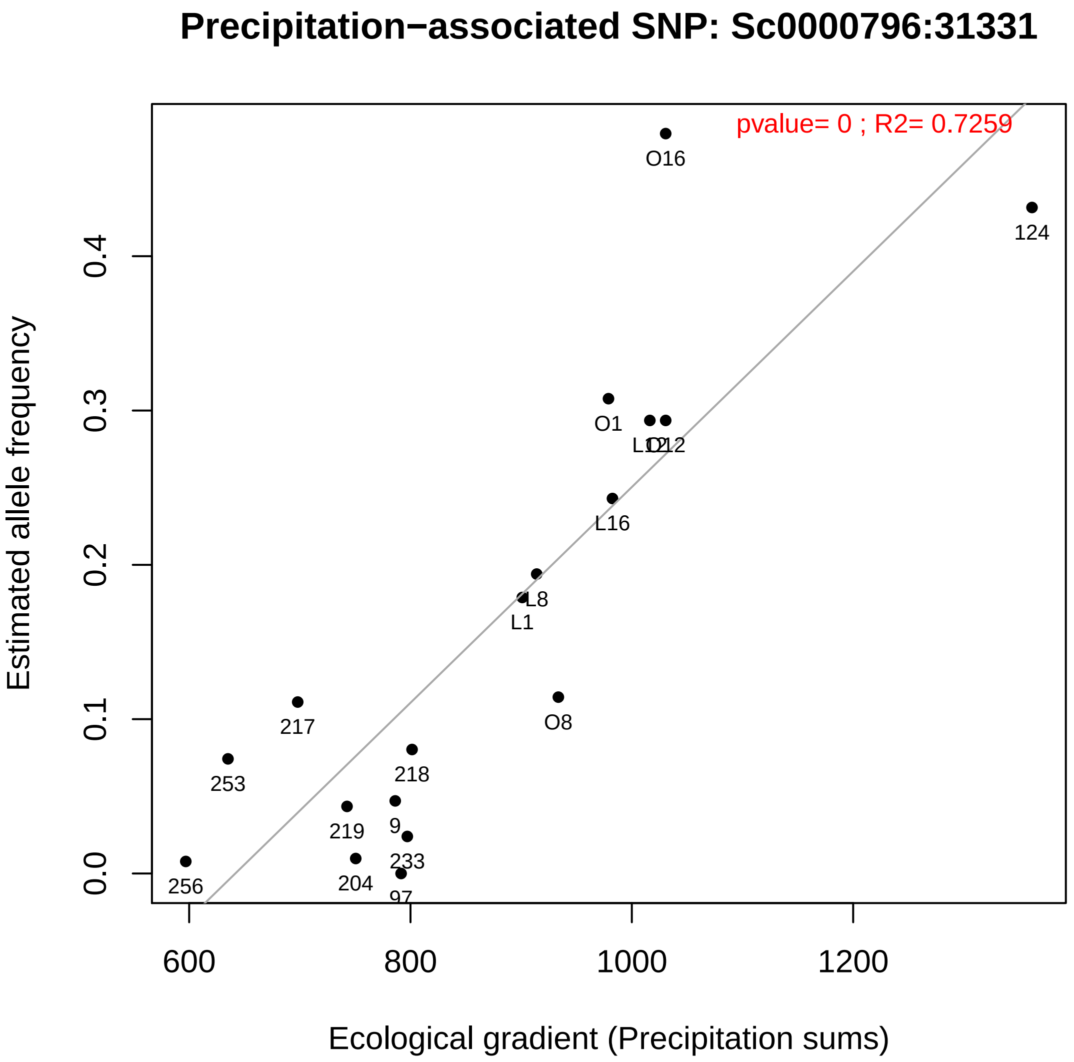
Precipitation−associated SNP: Sc0000796:31331
pvalue= 0 ; R2= 0.7259
O16
124
O1
L10
O12
L16
L8
L1
O8
217
253
218
219
9
233
256
204
97
0.0
0.1
0.2
0.3
0.4
600
800
1000
1200
Estimated allele frequency
Ecological gradient (Precipitation sums)

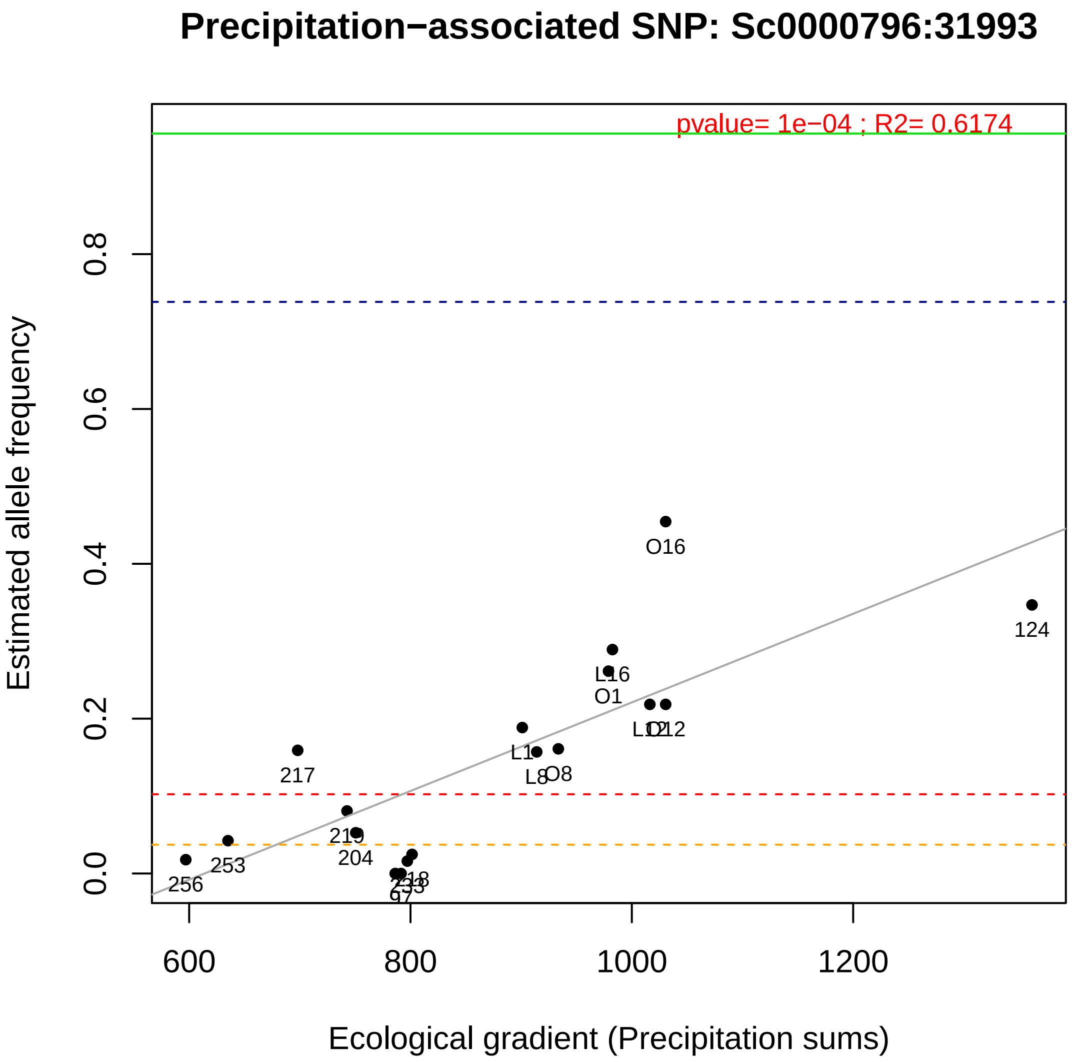
Precipitation−associated SNP: Sc0000796:31993
pvalue= 1e−04 ; R2= 0.6174
Estimated allele frequency
0.0
0.2
0.4
0.6
0.8
600
800
1000
1200
Ecological gradient (Precipitation sums)
O16
124
L16
O1
L12
O12
L1
L8
O8
217
219
204
253
256

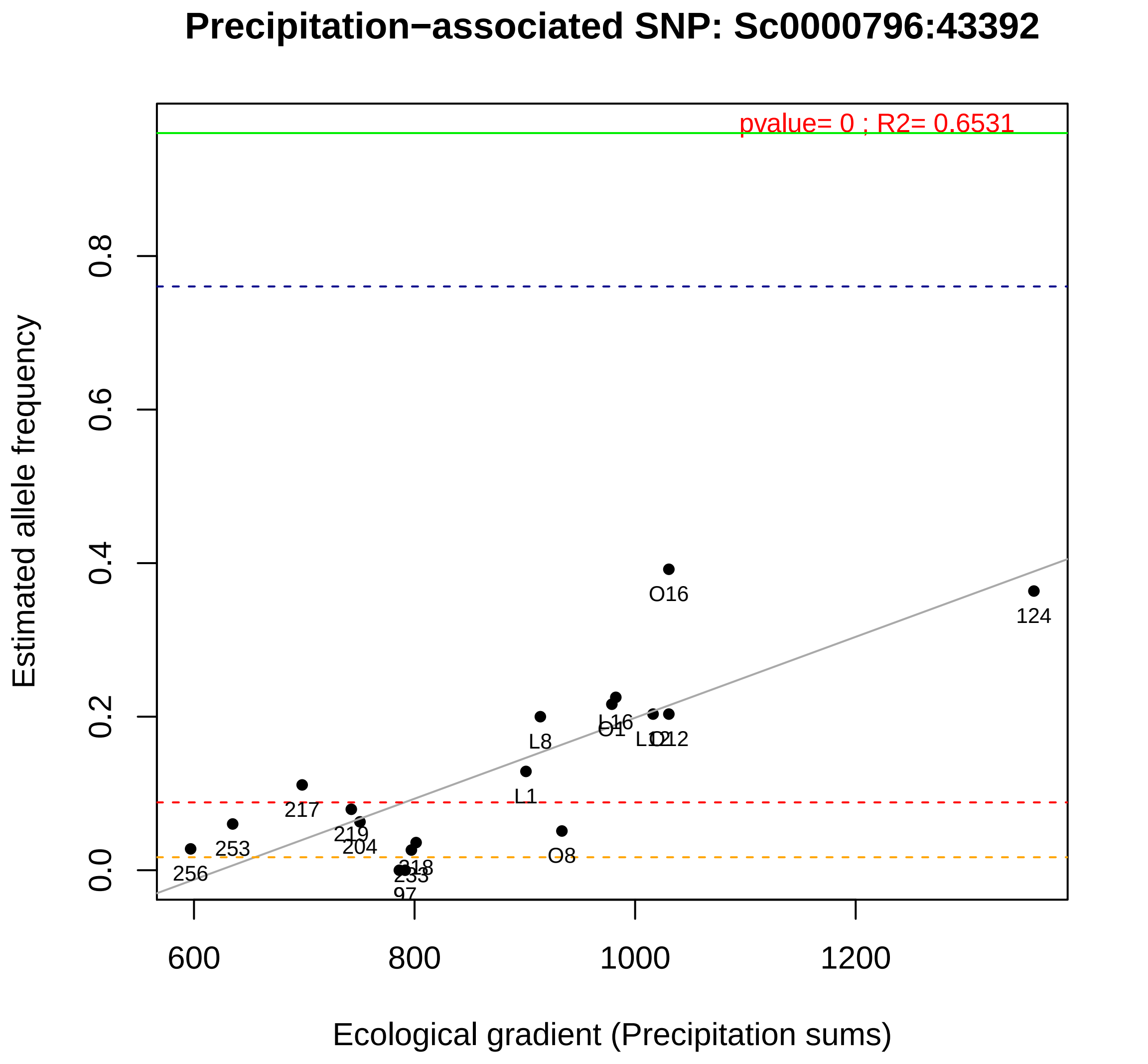

Precipitation−associated SNP: Sc0000796:43392
pvalue= 0 ; R2= 0.6531
Estimated allele frequency
0.0
0.2
0.4
0.6
0.8
600
800
1000
1200
Ecological gradient (Precipitation sums)
O16
124
L16
O1
L10
O12
L8
L1
217
219
204
253
256
O8
218
233
97

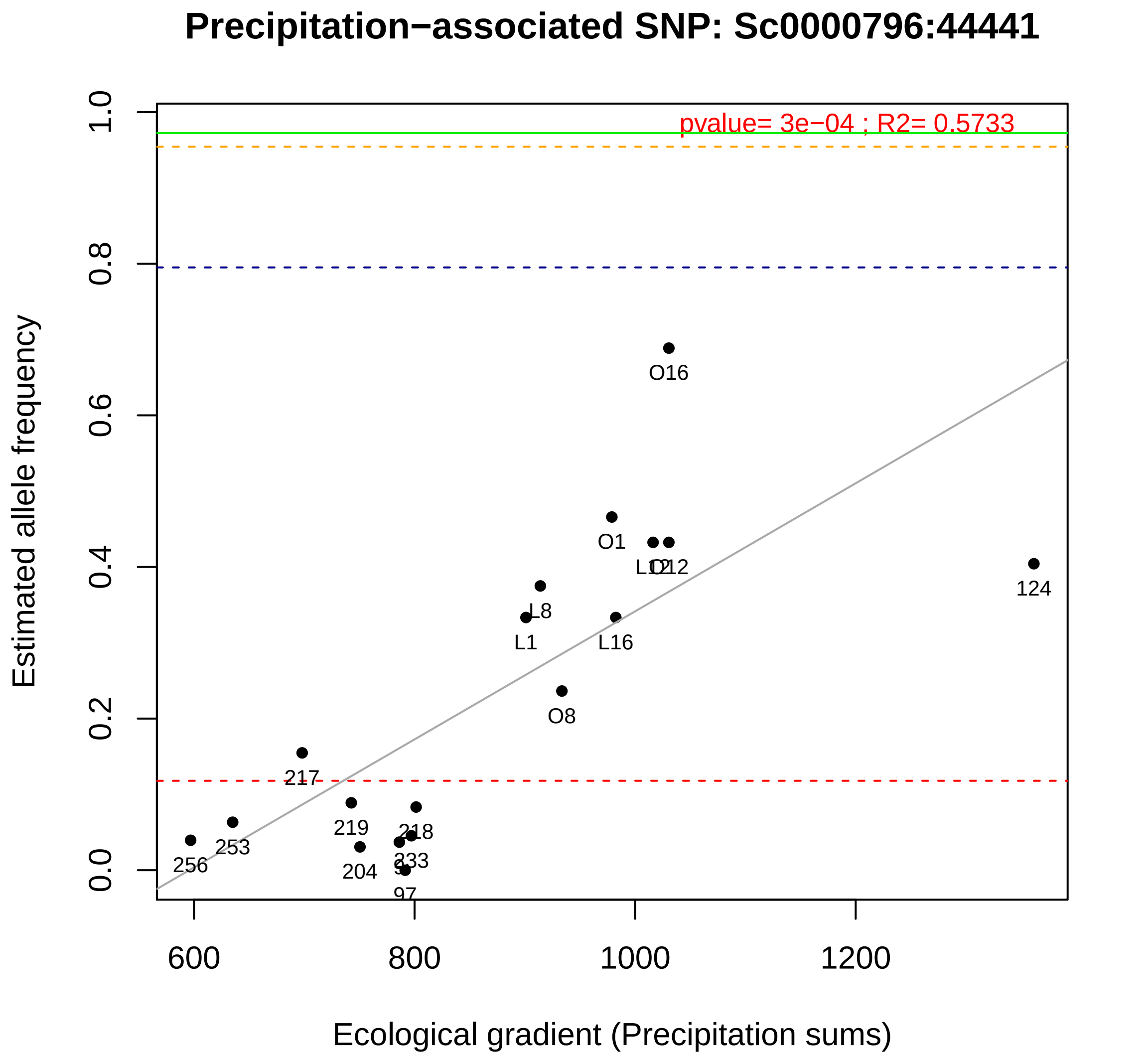
Precipitation−associated SNP: Sc0000796:44441
pvalue= 3e−04 ; R2= 0.5733
Estimated allele frequency
Ecological gradient (Precipitation sums)
0.0
0.2
0.4
0.6
0.8
1.0
600
800
1000
1200
O16
O1
L12
O12
124
L8
L1
L16
O8
217
219
218
253
256
204
233
97

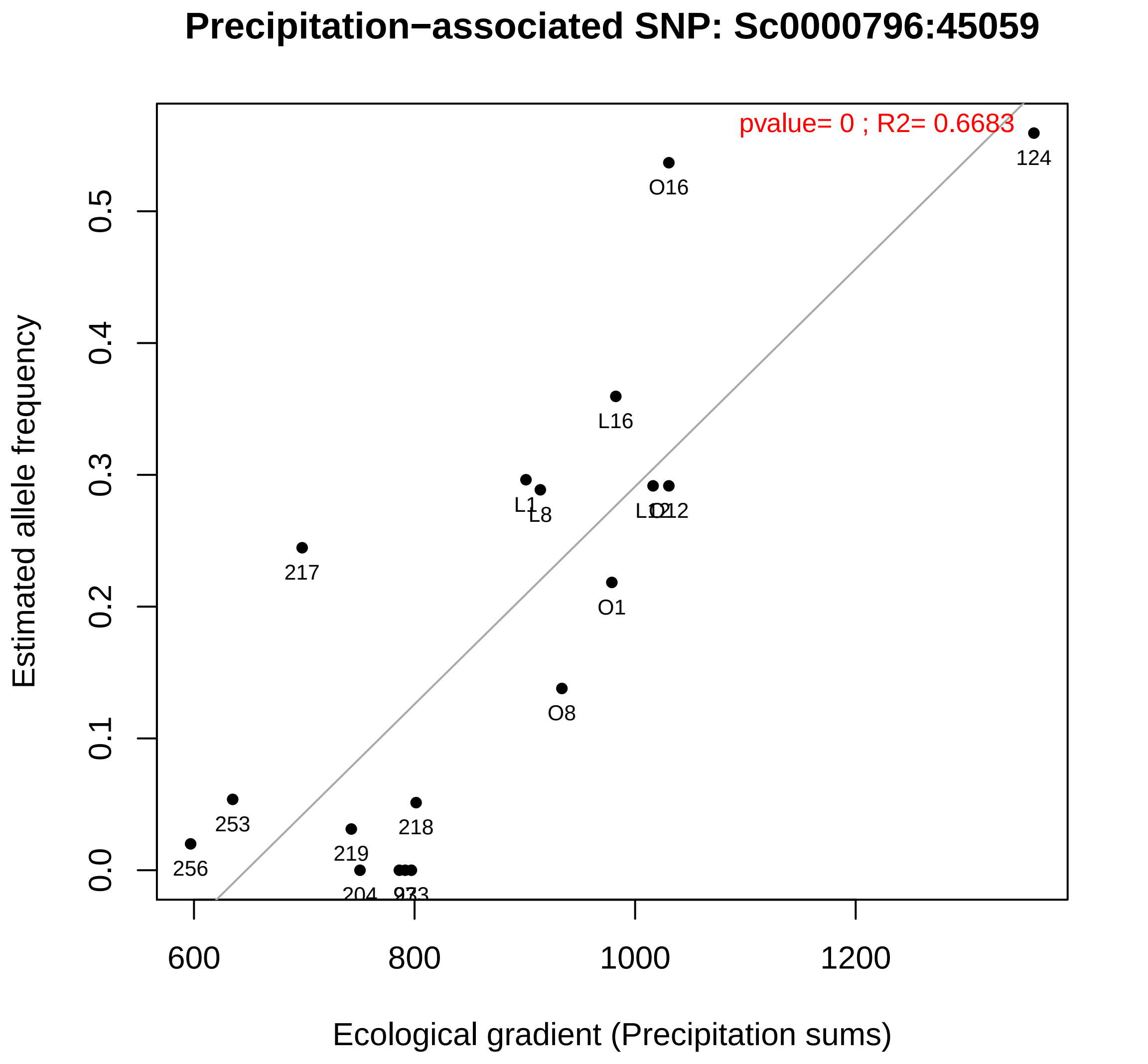

Precipitation−associated SNP: Sc0000796:45059
pvalue= 0 ; R2= 0.6683
Estimated allele frequency
Ecological gradient (Precipitation sums)
0.0
0.1
0.2
0.3
0.4
0.5
600
800
1000
1200
124
O16
L16
L1
L8
O12
O1
O8
217
253
256
219
218
204

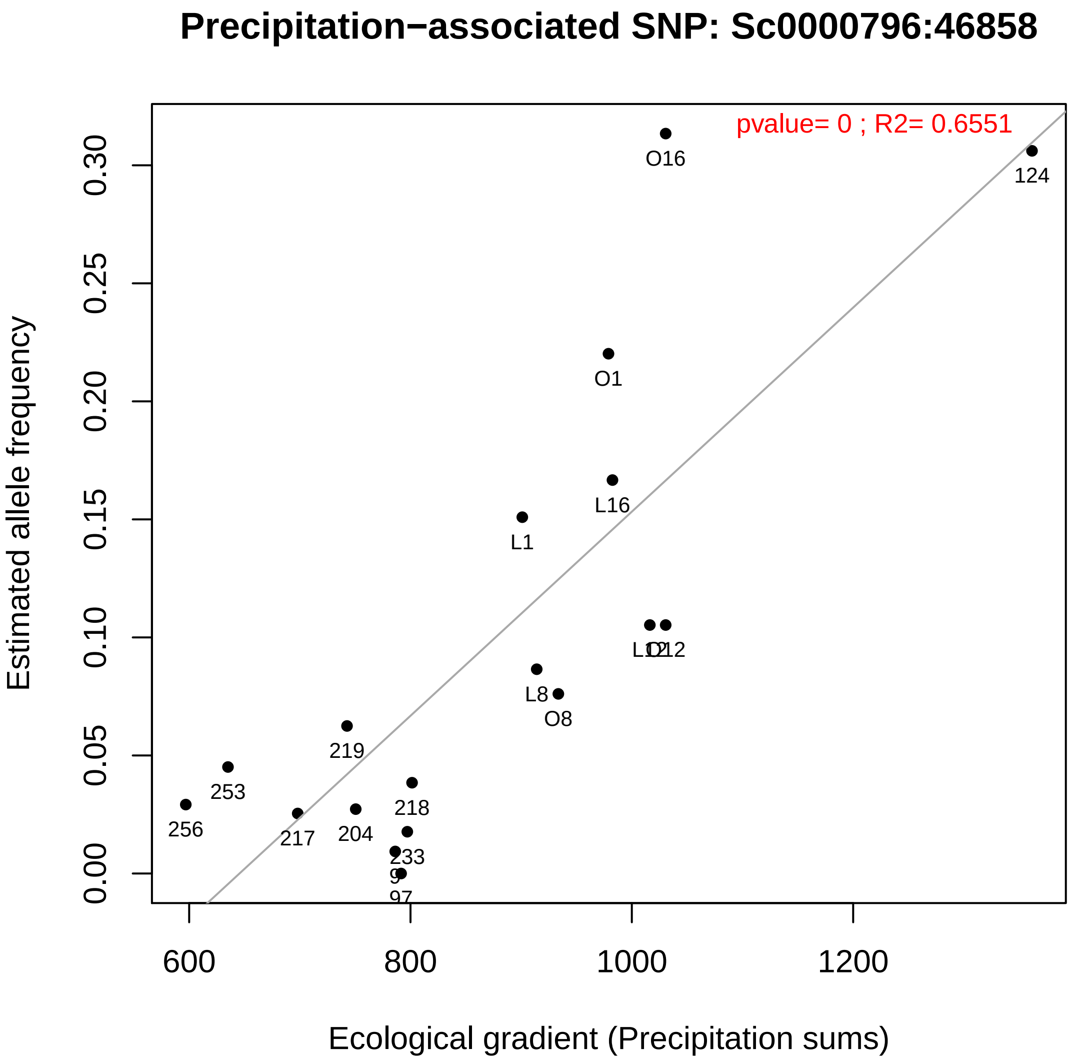

Precipitation−associated SNP: Sc0000796:46858
pvalue= 0 ; R2= 0.6551
Estimated allele frequency
0.00
0.05
0.10
0.15
0.20
0.25
0.30
600
800
1000
1200
Ecological gradient (Precipitation sums)
O16
124
O1
L16
L1
L8
O8
219
253
256
217
204
218
233
97

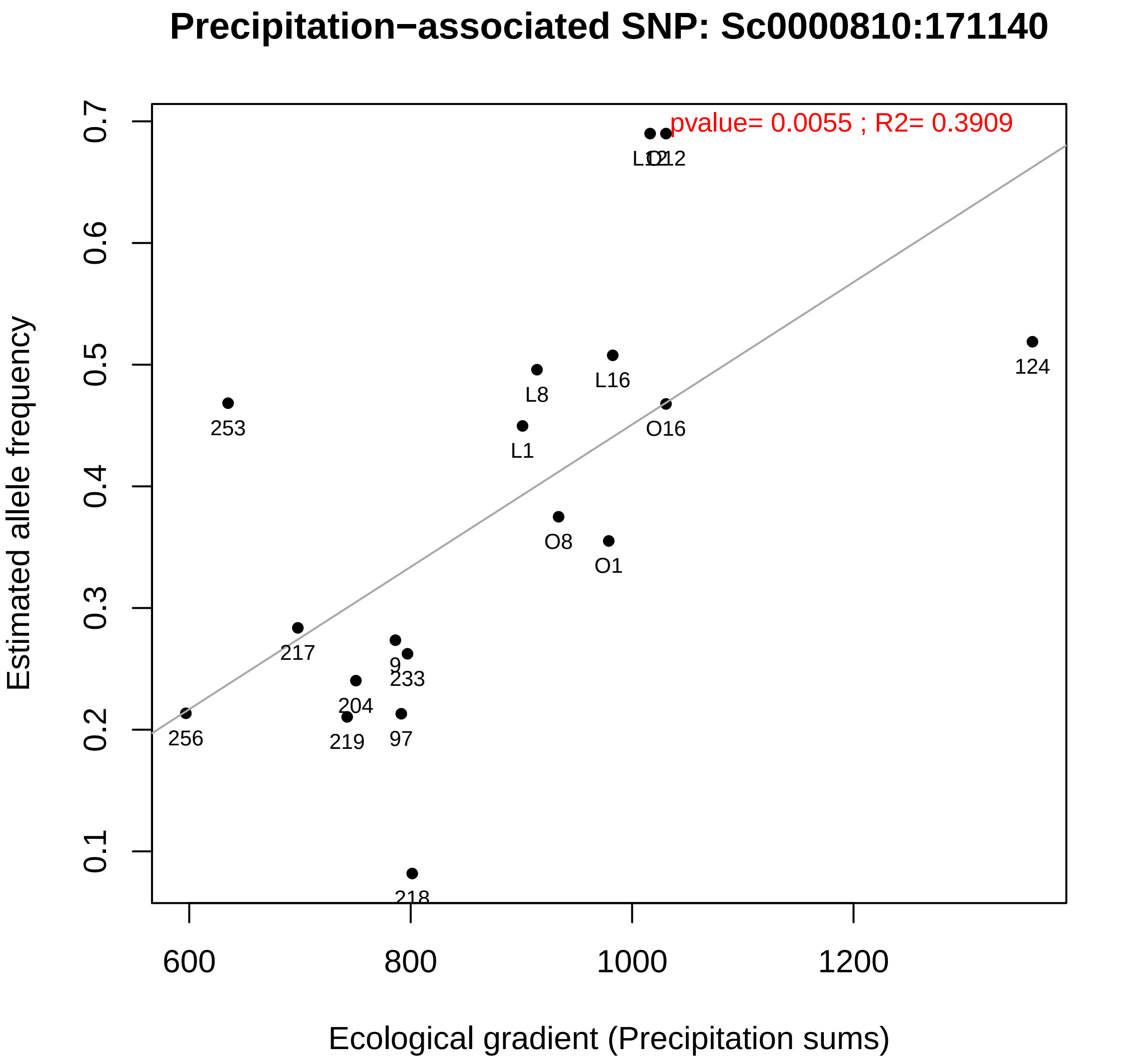
Precipitation−associated SNP: Sc0000810:171140
pvalue= 0.0055 ; R2= 0.3909
Estimated allele frequency
0.1
0.2
0.3
0.4
0.5
0.6
0.7
600
800
1000
1200
Ecological gradient (Precipitation sums)
L12
O12
L16
L8
124
253
O16
L1
O8
O1
217
9
233
204
256
219
97
218

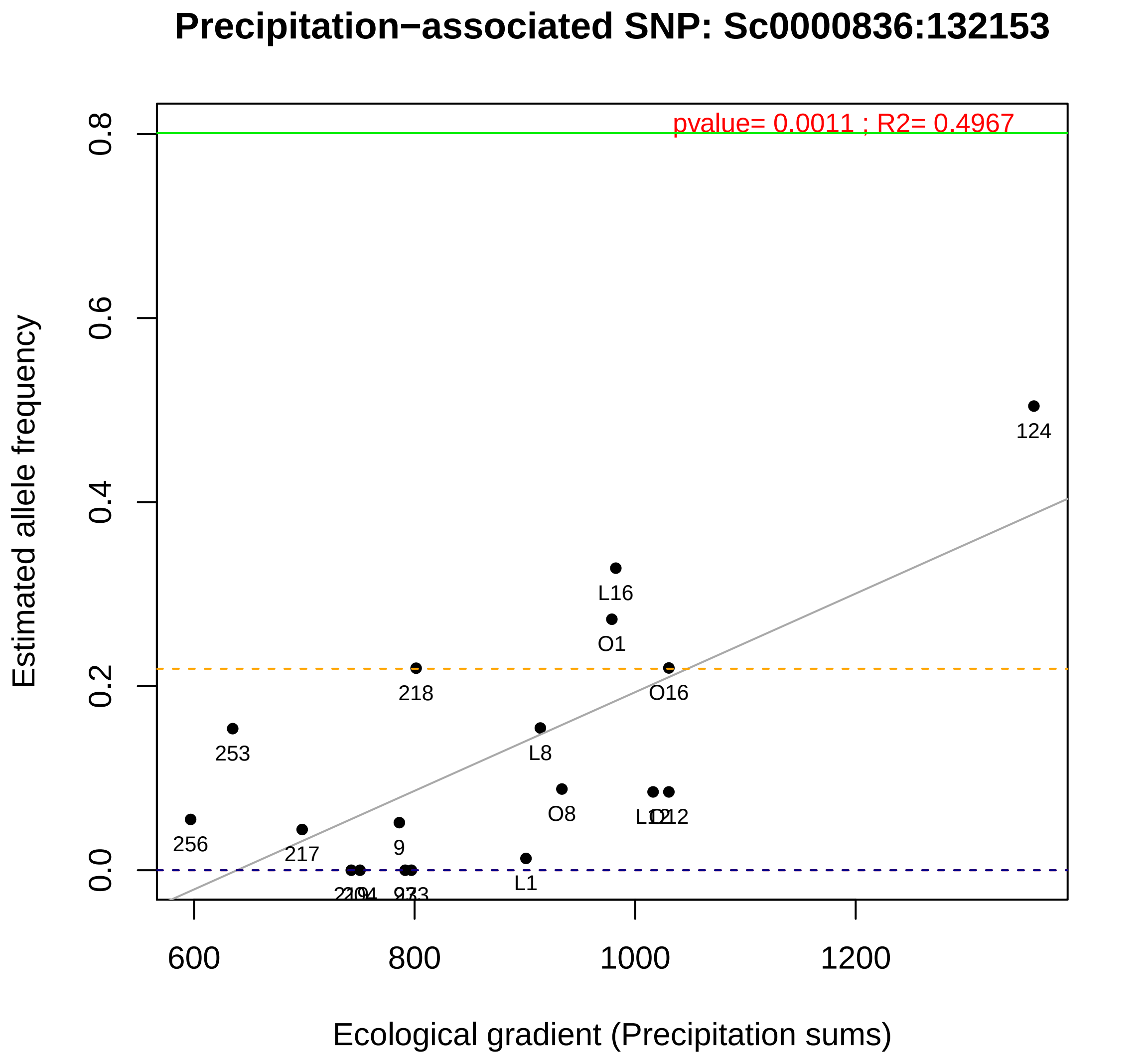
Precipitation−associated SNP: Sc0000836:132153
pvalue= 0.0011 ; R2= 0.4967
Estimated allele frequency
Ecological gradient (Precipitation sums)
0.0
0.2
0.4
0.6
0.8
600
800
1000
1200
124
L16
O1
O16
218
253
L8
O8
256
217
9
L1

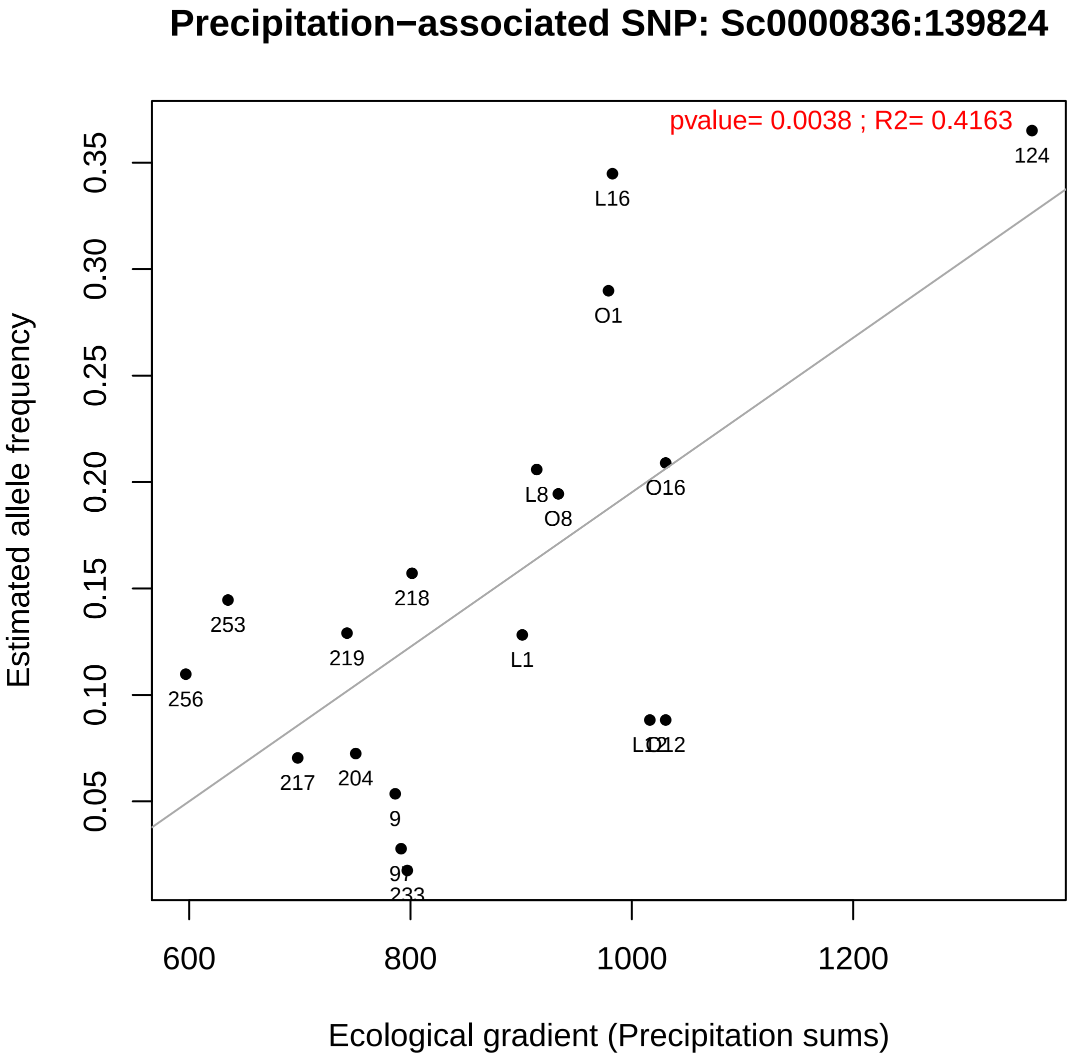
Precipitation−associated SNP: Sc0000836:139824
pvalue= 0.0038 ; R2= 0.4163
Estimated allele frequency
0.05
0.10
0.15
0.20
0.25
0.30
0.35
Ecological gradient (Precipitation sums)
600
800
1000
1200
124
L16
O1
L8
O8
O16
218
253
219
L1
256
L12
O12
217
204
9
97
233

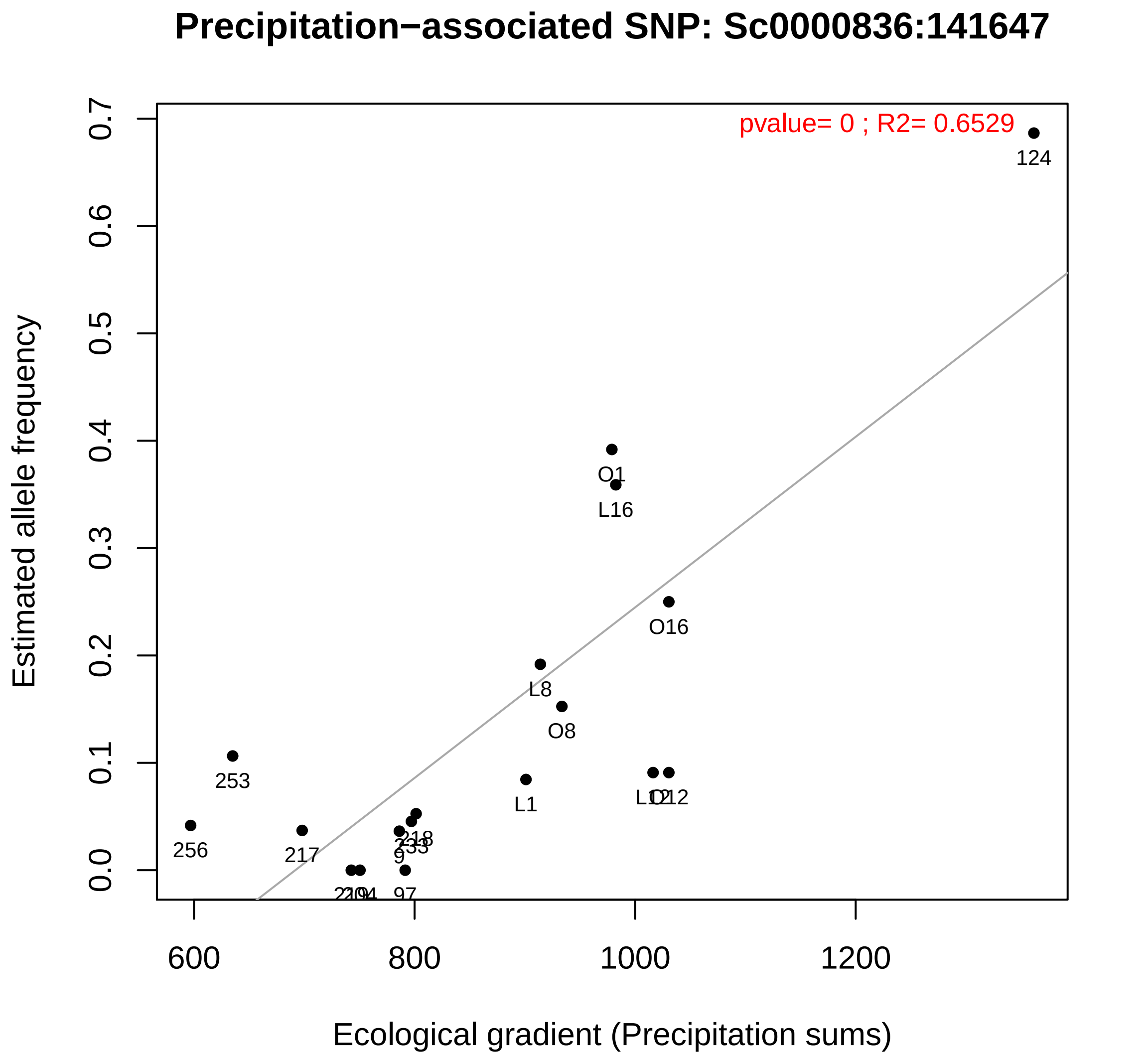

Precipitation−associated SNP: Sc0000836:141647
pvalue= 0 ; R2= 0.6529
Estimated allele frequency
0.0
0.1
0.2
0.3
0.4
0.5
0.6
0.7
Ecological gradient (Precipitation sums)
600
800
1000
1200
124
O1
L16
O16
L8
O8
253
L1
256
217
218
233
9
97

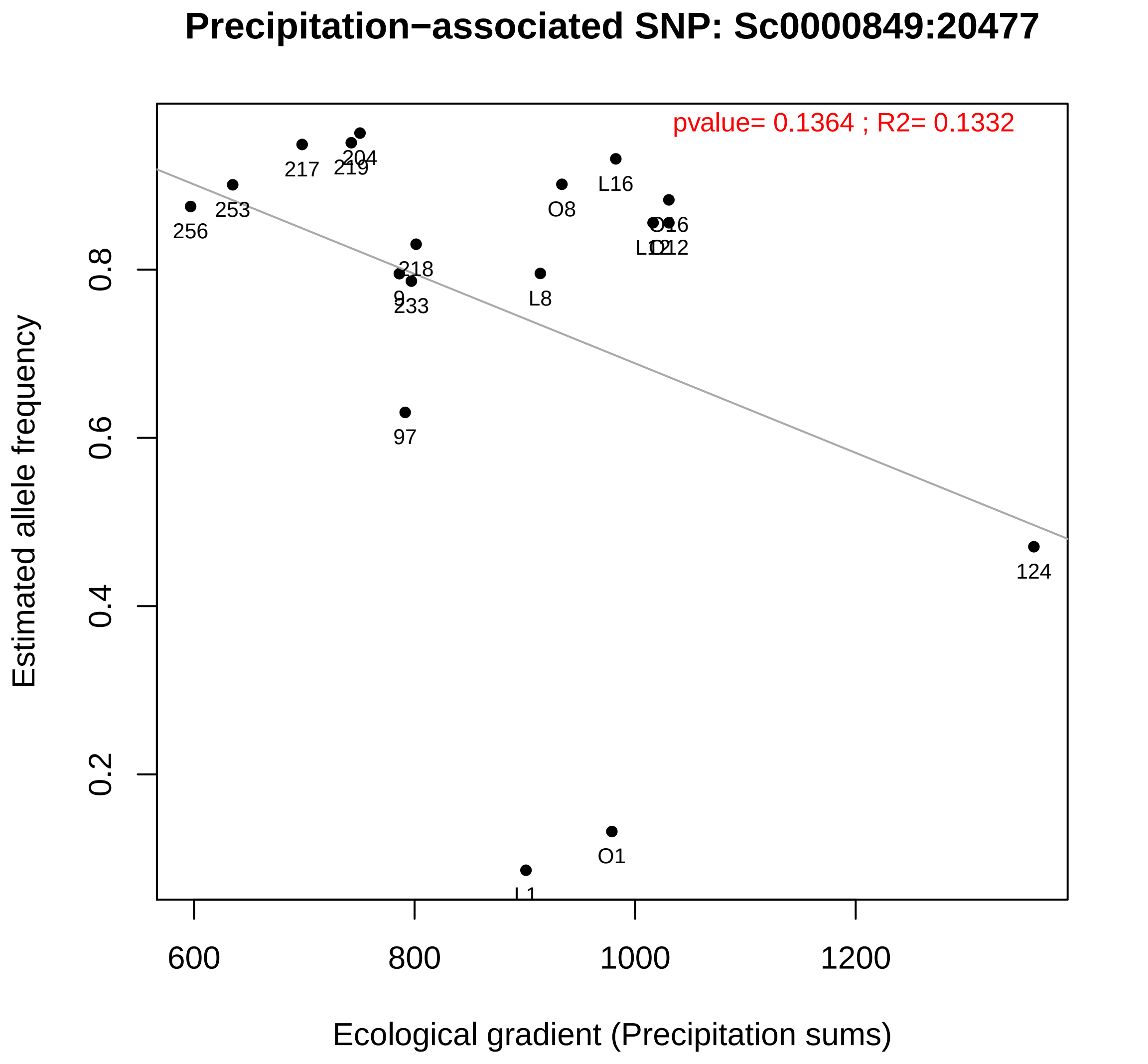

Precipitation−associated SNP: Sc0000849:20477
pvalue= 0.1364 ; R2= 0.1332
Estimated allele frequency
0.2
0.4
0.6
0.8
600
800
1000
1200
Ecological gradient (Precipitation sums)
217
219
204
253
256
218
9
233
97
L8
O8
L16
O16
L12
O12
124
O1
L1

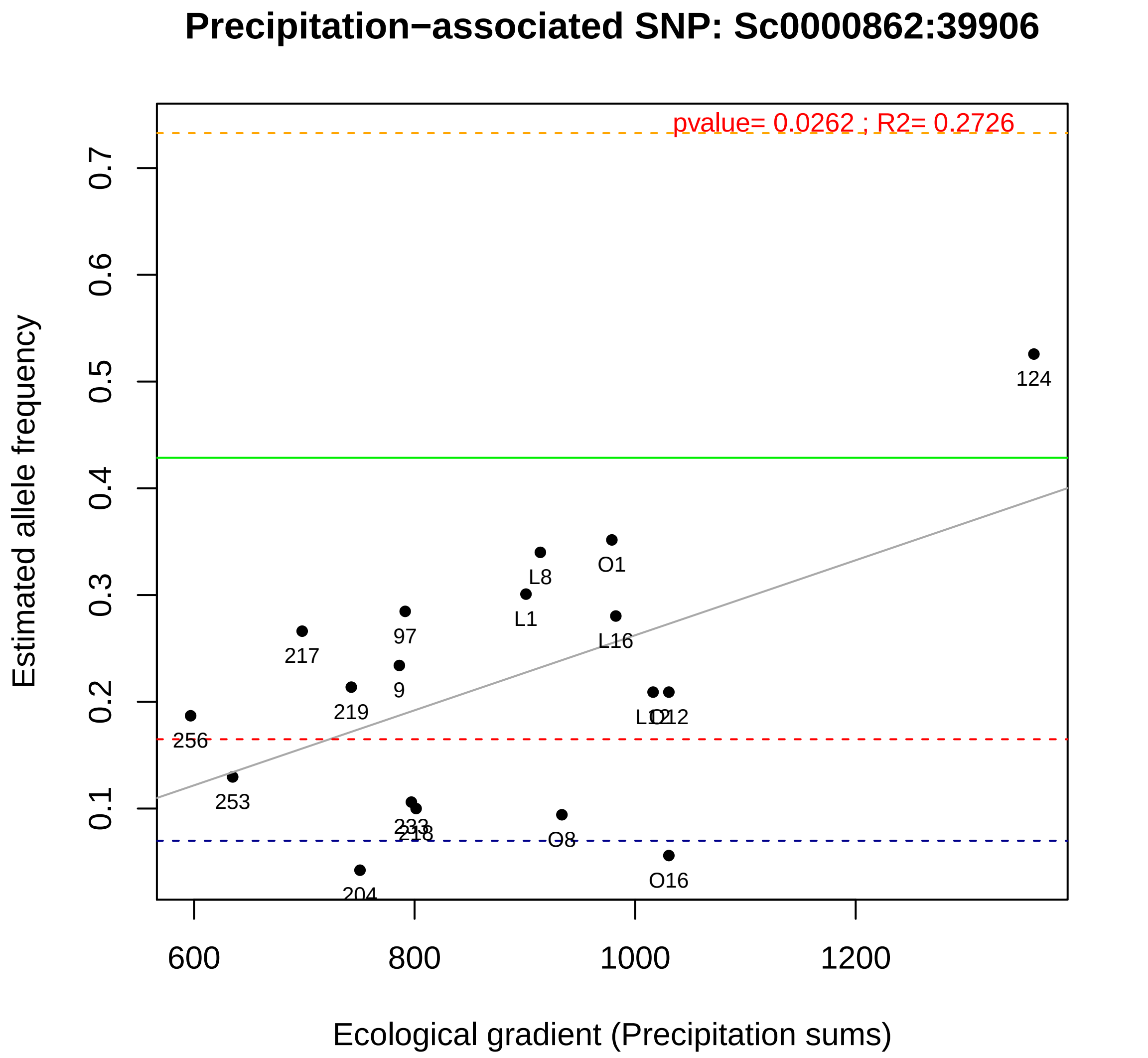
Precipitation−associated SNP: Sc0000862:39906
pvalue= 0.0262 ; R2= 0.2726
Estimated allele frequency
0.1
0.2
0.3
0.4
0.5
0.6
0.7
Ecological gradient (Precipitation sums)
600
800
1000
1200
124
O1
L8
L1
97
L16
217
9
219
L12
O12
256
253
233
218
O8
O16
204

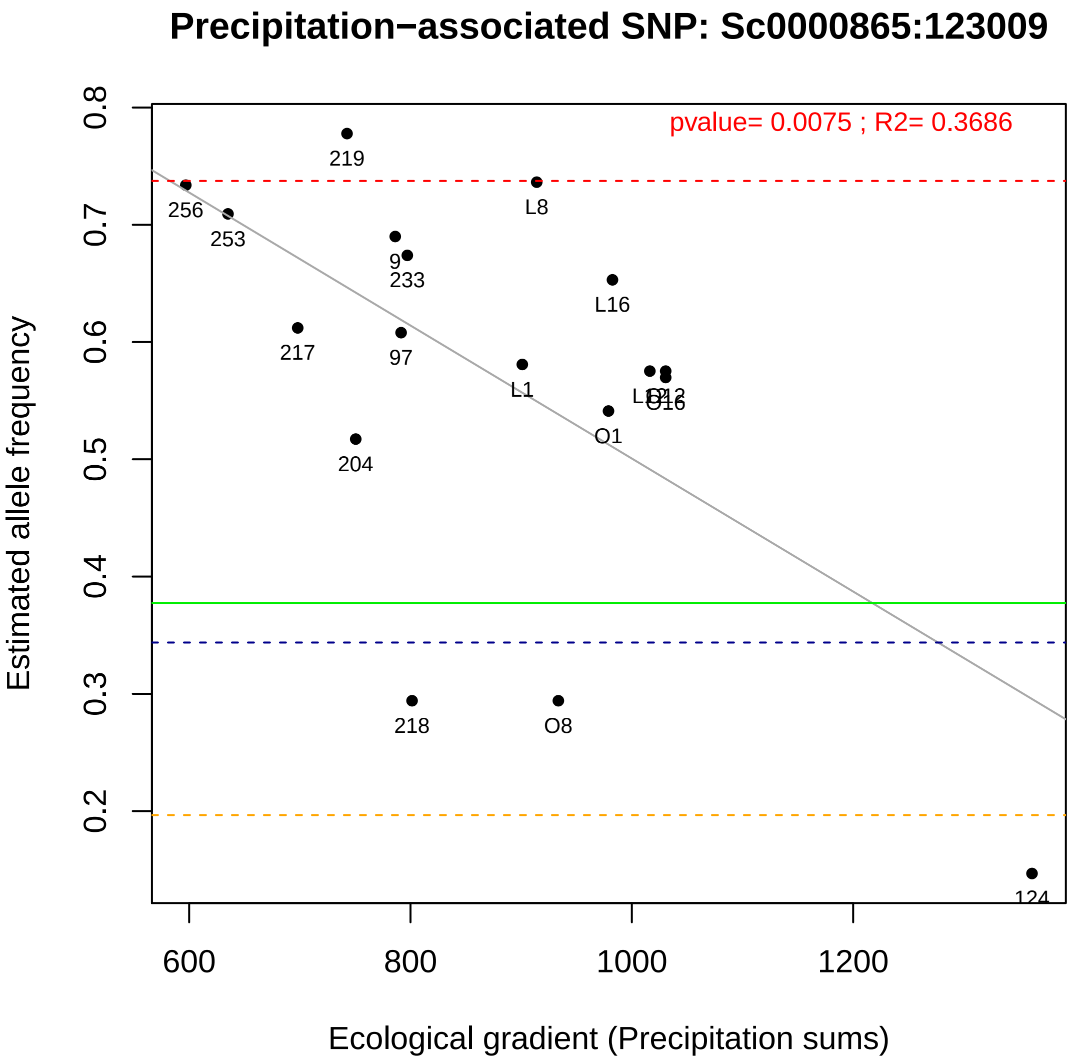
Precipitation−associated SNP: Sc0000865:123009
pvalue= 0.0075 ; R2= 0.3686
Estimated allele frequency
0.8
0.7
0.6
0.5
0.4
0.3
0.2
219
256
253
L8
9
233
L16
217
97
L1
L12
O12
O16
O1
204
218
O8
124
600
800
1000
1200
Ecological gradient (Precipitation sums)

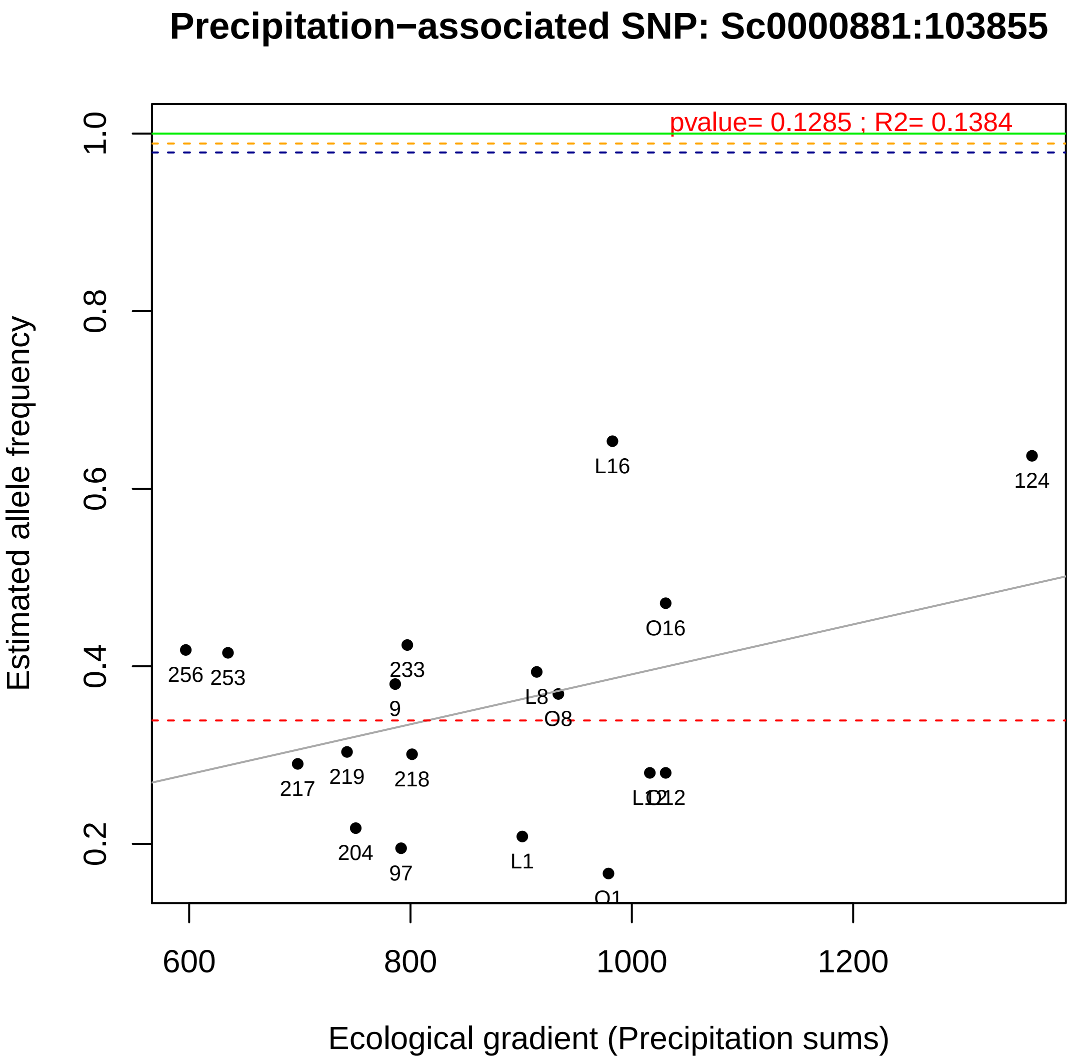

Precipitation−associated SNP: Sc0000881:103855
pvalue= 0.1285 ; R2= 0.1384
Estimated allele frequency
0.2
0.4
0.6
0.8
1.0
600
800
1000
1200
Ecological gradient (Precipitation sums)
L16
124
O16
256
253
233
9
L8
O8
217
219
218
L12
O12
204
97
L1

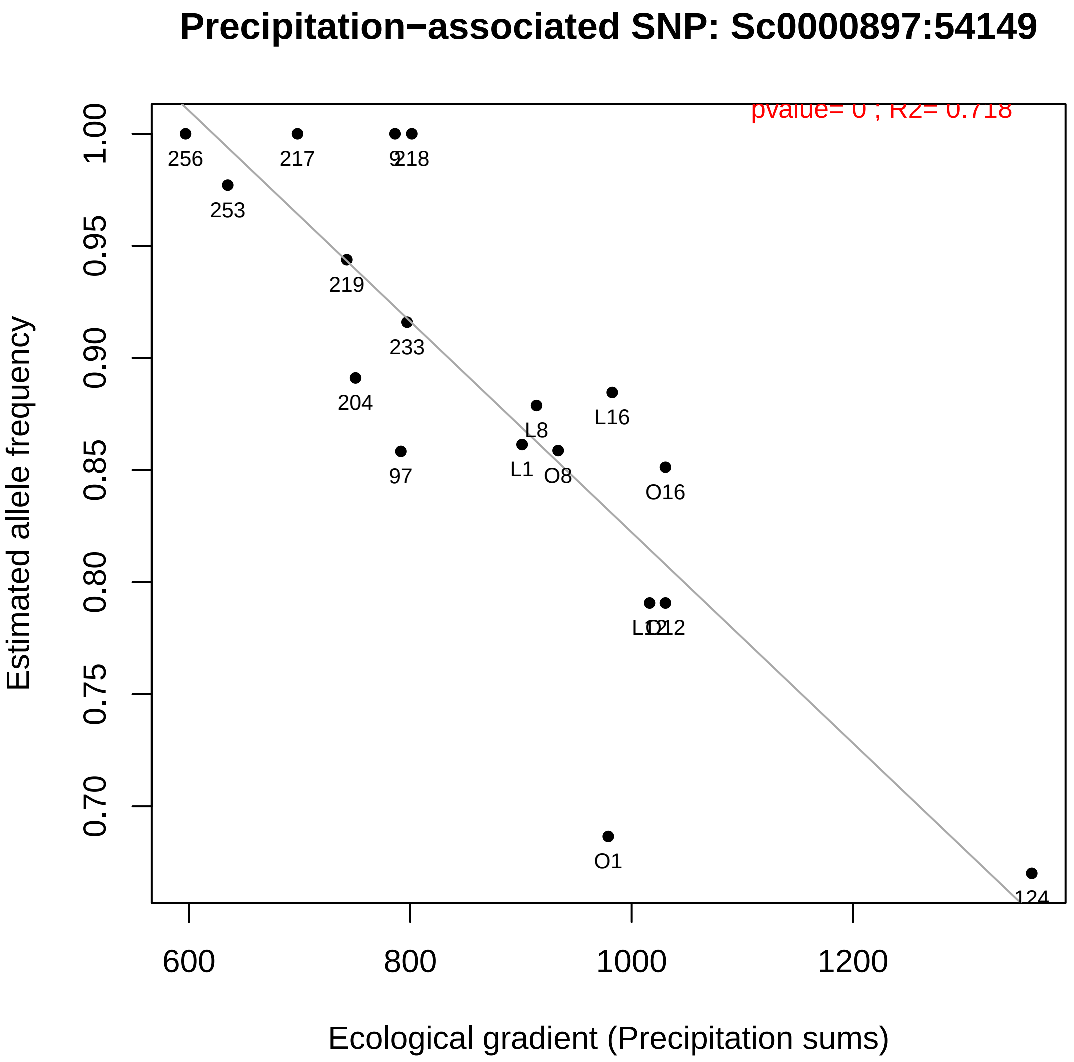

Precipitation−associated SNP: Sc0000897:54149
pvalue= 0 ; R2= 0.718
Estimated allele frequency
1.00
0.95
0.90
0.85
0.80
0.75
0.70
600
800
1000
1200
Ecological gradient (Precipitation sums)
256
217
218
253
219
233
204
97
L8
L1
O8
L16
O16
O12
O1
124

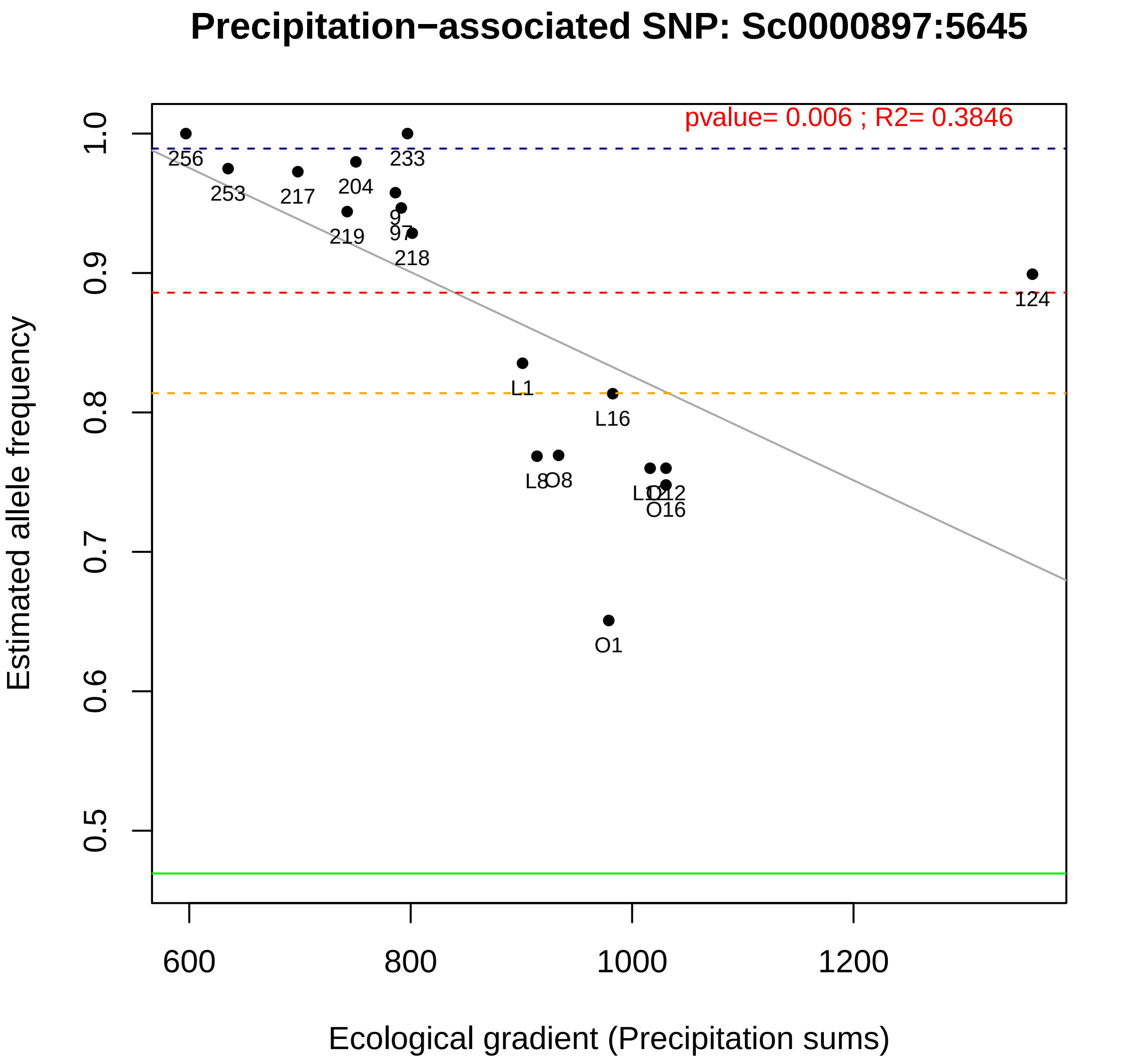

Precipitation−associated SNP: Sc0000897:5645
pvalue= 0.006 ; R2= 0.3846
Estimated allele frequency
Ecological gradient (Precipitation sums)
0.5
0.6
0.7
0.8
0.9
1.0
600
800
1000
1200
256
253
217
204
233
219
9
97
218
124
L1
L16
L8
O8
L12
O12
O16
O1

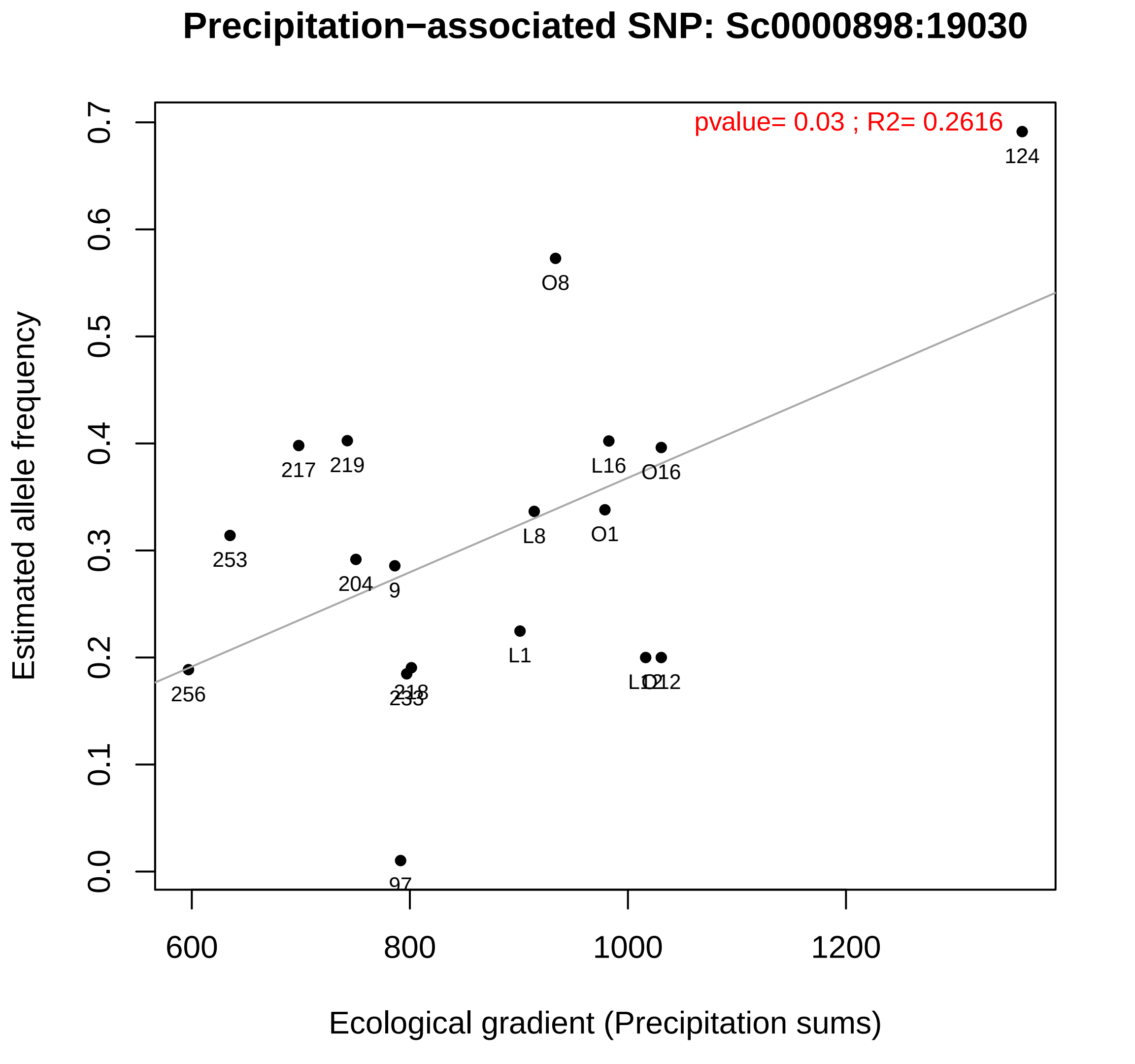

Precipitation−associated SNP: Sc0000898:19030
pvalue= 0.03 ; R2= 0.2616
Estimated allele frequency
0.0
0.1
0.2
0.3
0.4
0.5
0.6
0.7
600
800
1000
1200
Ecological gradient (Precipitation sums)
124
O8
217
219
L16
O16
L8
O1
253
204
9
L1
256
218
233
97

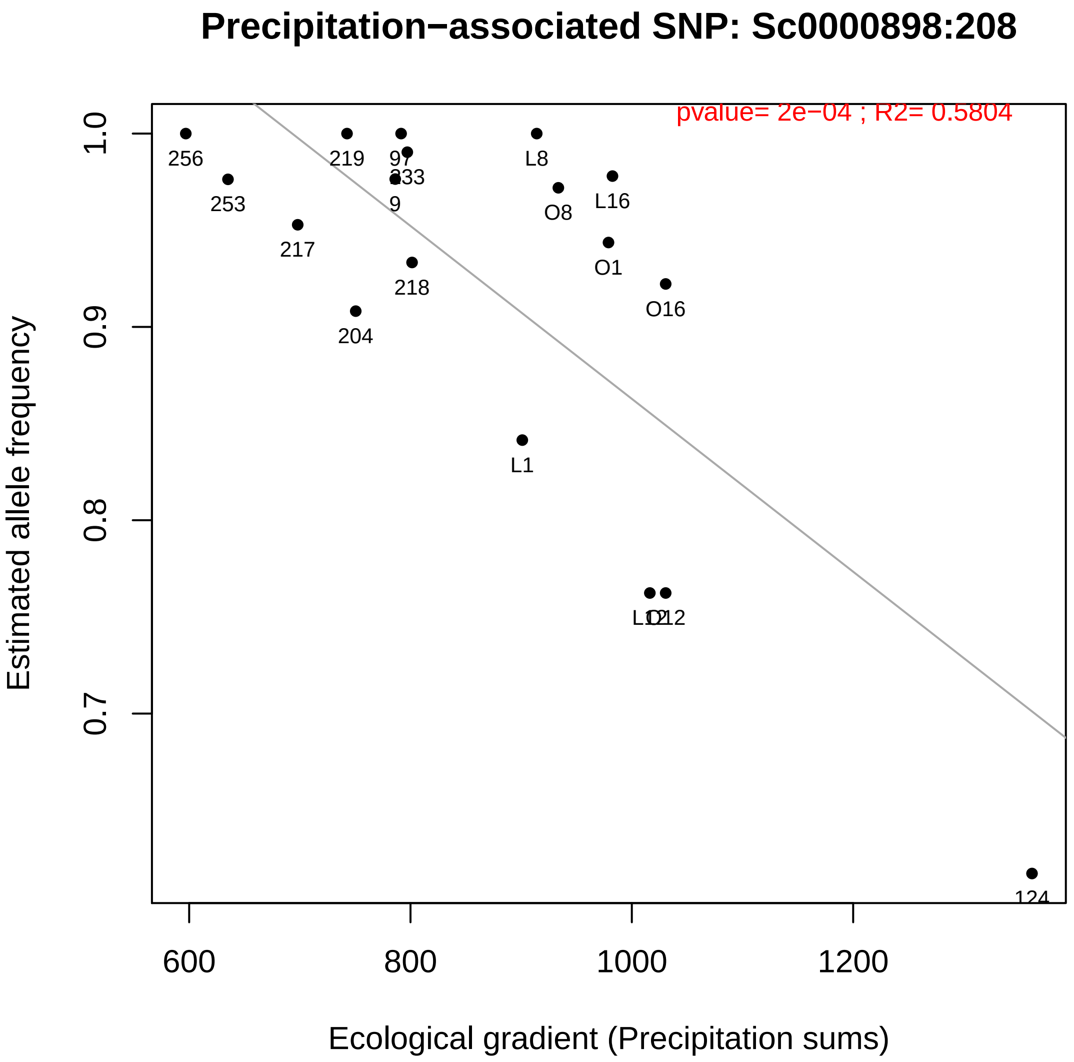
Precipitation−associated SNP: Sc0000898:208
pvalue= 2e−04 ; R2= 0.5804
Estimated allele frequency
1.0
0.9
0.8
0.7
600
800
1000
1200
Ecological gradient (Precipitation sums)
256
253
217
219
204
97
233
9
218
L8
O8
L16
O1
O16
L1
L12
O12
124

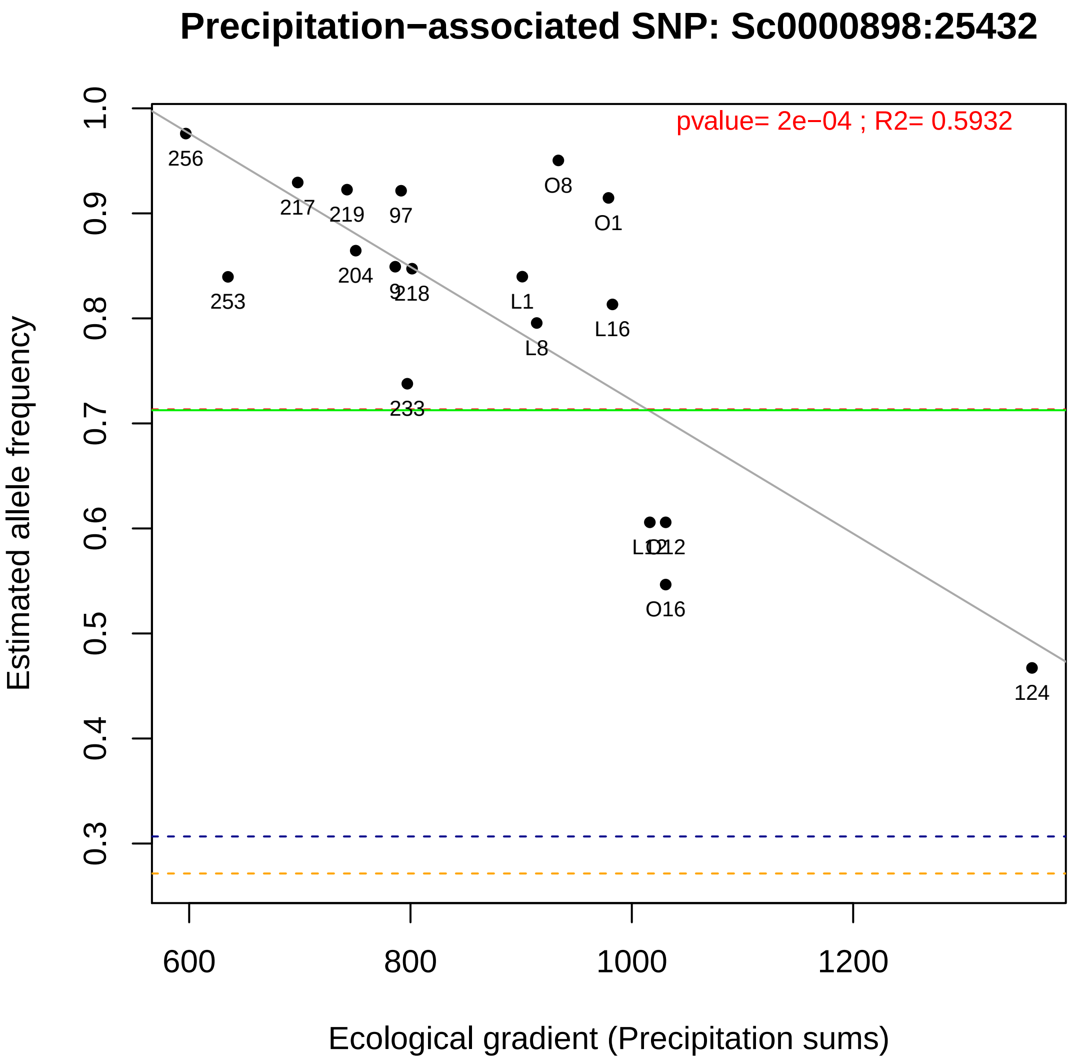

Precipitation−associated SNP: Sc0000898:25432
pvalue= 2e−04 ; R2= 0.5932
256
217
219
97
O8
O1
253
204
9
218
L1
L16
L8
233
L10
O12
O16
124
1.0
0.9
0.8
0.7
0.6
0.5
0.4
0.3
Estimated allele frequency
600
800
1000
1200
Ecological gradient (Precipitation sums)

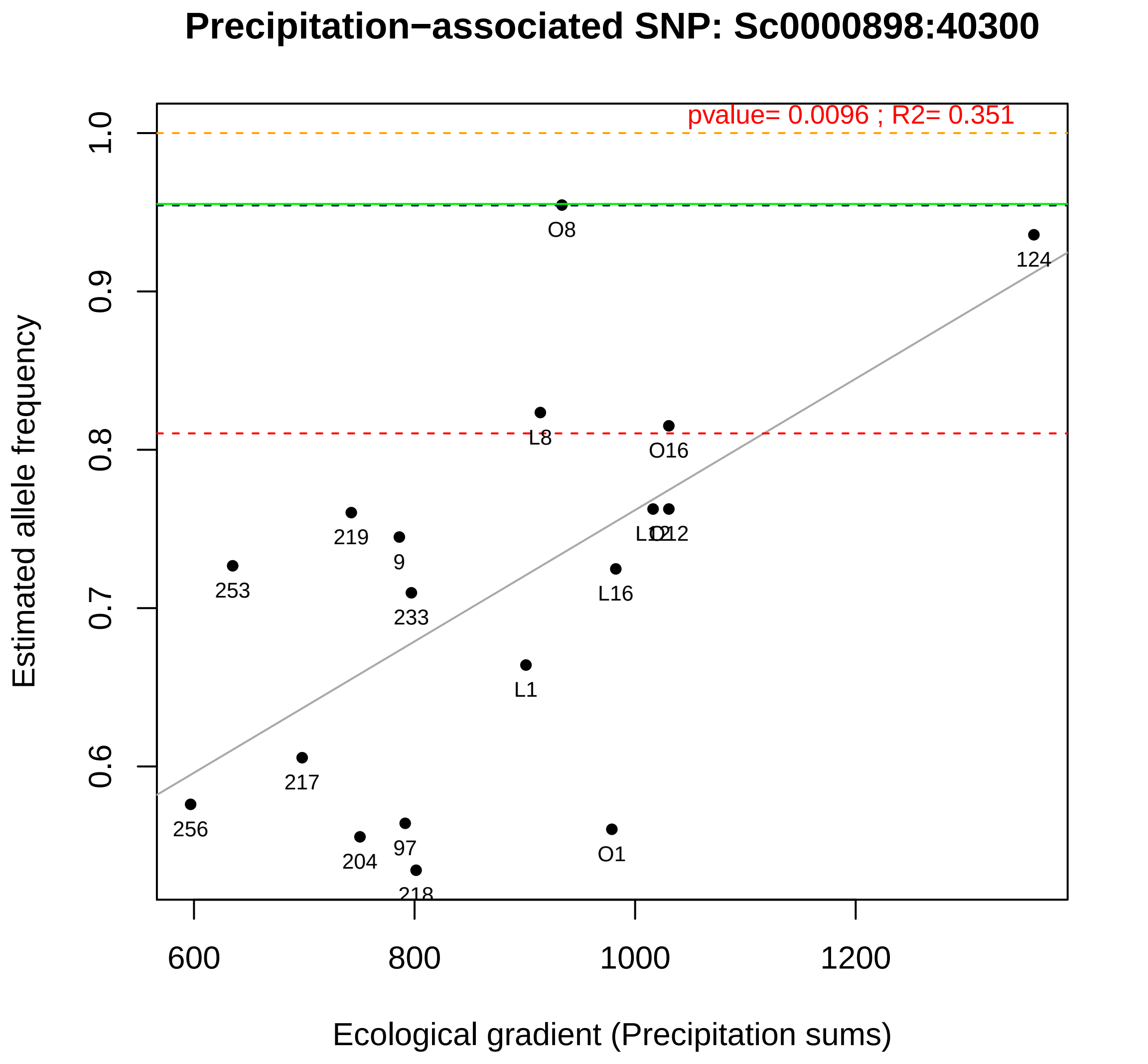

Precipitation−associated SNP: Sc0000898:40300
pvalue= 0.0096 ; R2= 0.351
Estimated allele frequency
0.6
0.7
0.8
0.9
1.0
600
800
1000
1200
Ecological gradient (Precipitation sums)
O8
124
L8
O16
219
9
L12
O12
253
233
L16
L1
217
256
204
97
218
O1

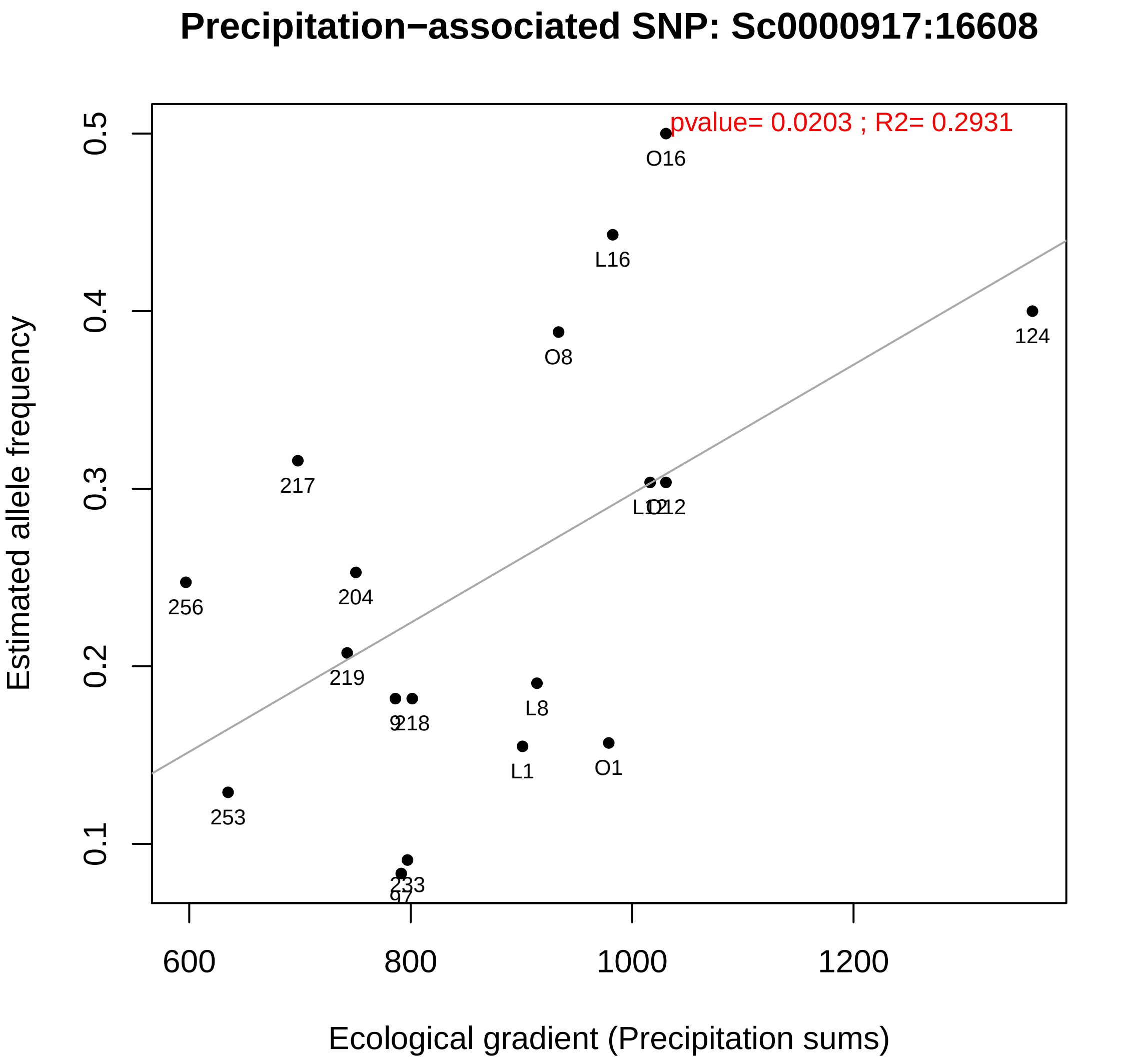

Precipitation−associated SNP: Sc0000917:16608
pvalue= 0.0203 ; R2= 0.2931
O16
L16
124
O8
217
L10
O12
256
204
219
9
218
L8
L1
O1
253
233
97
0.1
0.2
0.3
0.4
0.5
600
800
1000
1200
Estimated allele frequency
Ecological gradient (Precipitation sums)

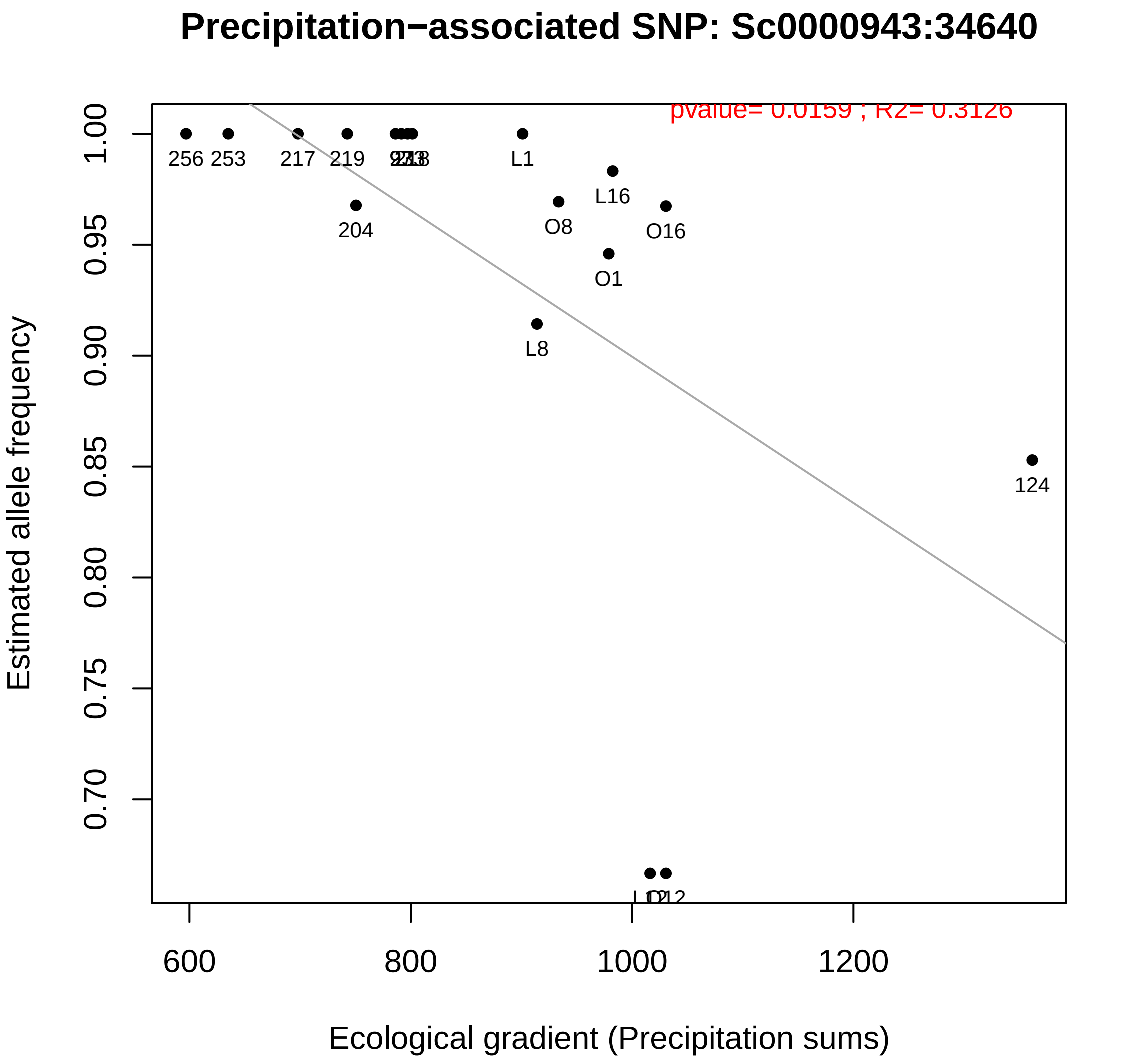
Precipitation−associated SNP: Sc0000943:34640
pvalue= 0.0159 ; R2= 0.3126
Estimated allele frequency
Ecological gradient (Precipitation sums)
1.00
0.95
0.90
0.85
0.80
0.75
0.70
600
800
1000
1200
256
253
217
219
204
L1
L16
O8
O16
O1
L8
124

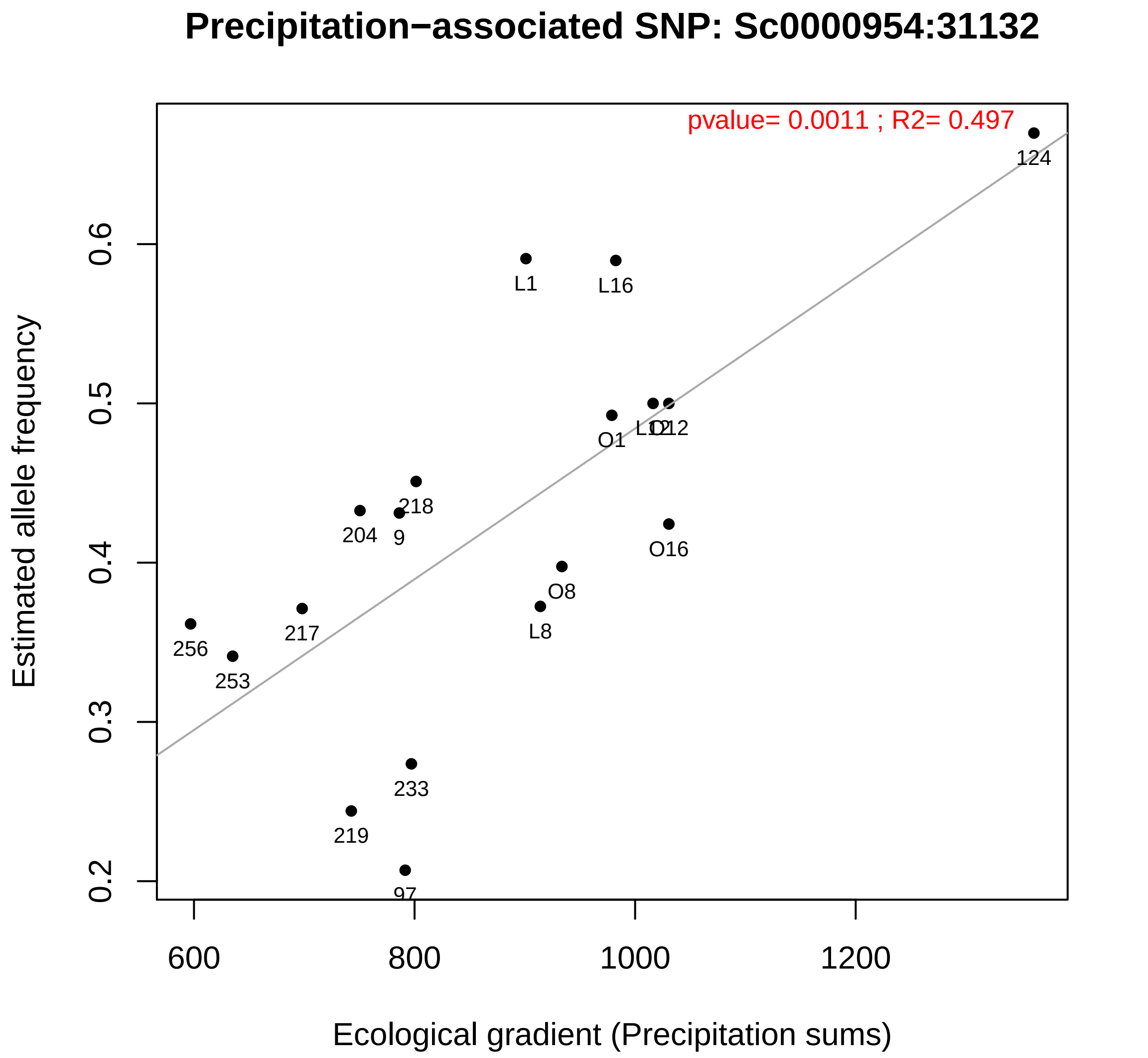
Precipitation−associated SNP: Sc0000954:31132
pvalue= 0.0011 ; R2= 0.497
124
L1
L16
L12
O12
O1
218
204
9
O16
O8
L8
217
256
253
233
219
97
0.2
0.3
0.4
0.5
0.6
600
800
1000
1200
Estimated allele frequency
Ecological gradient (Precipitation sums)

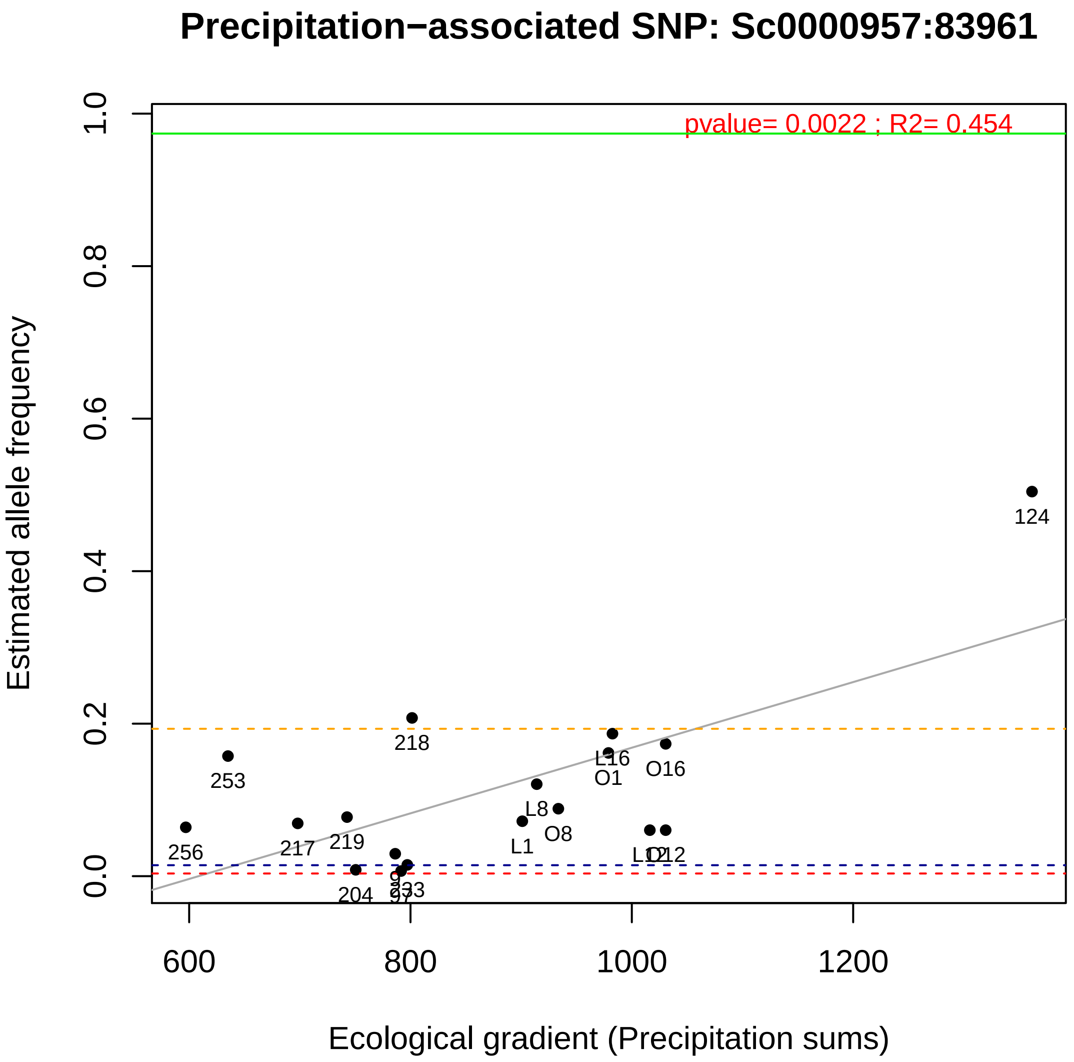

Precipitation−associated SNP: Sc0000957:83961
pvalue= 0.0022 ; R2= 0.454
Estimated allele frequency
Ecological gradient (Precipitation sums)
0.0
0.2
0.4
0.6
0.8
1.0
600
800
1000
1200
124
218
253
256
217
219
204
L16
O1
O16
L8
O8
L1

**Precipitation−associated SNP: Sc0001007:67512**

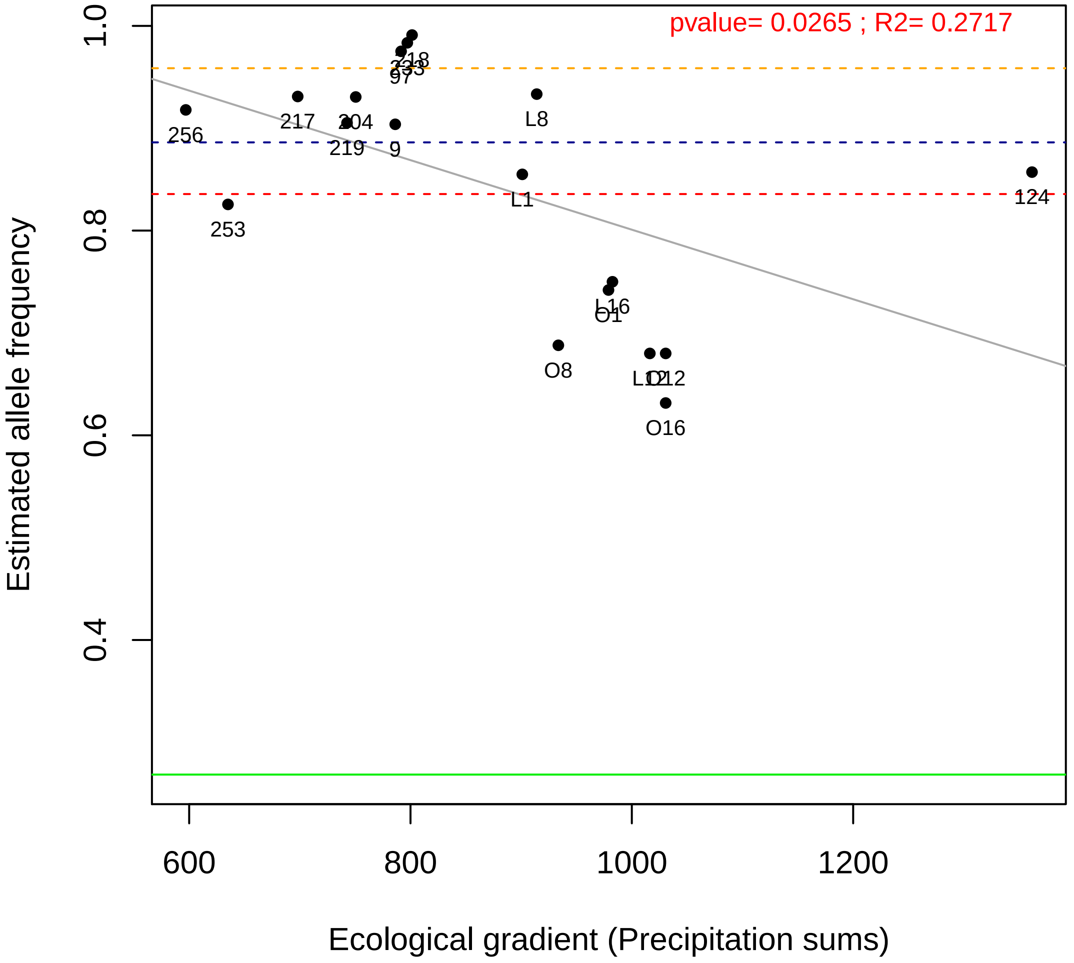

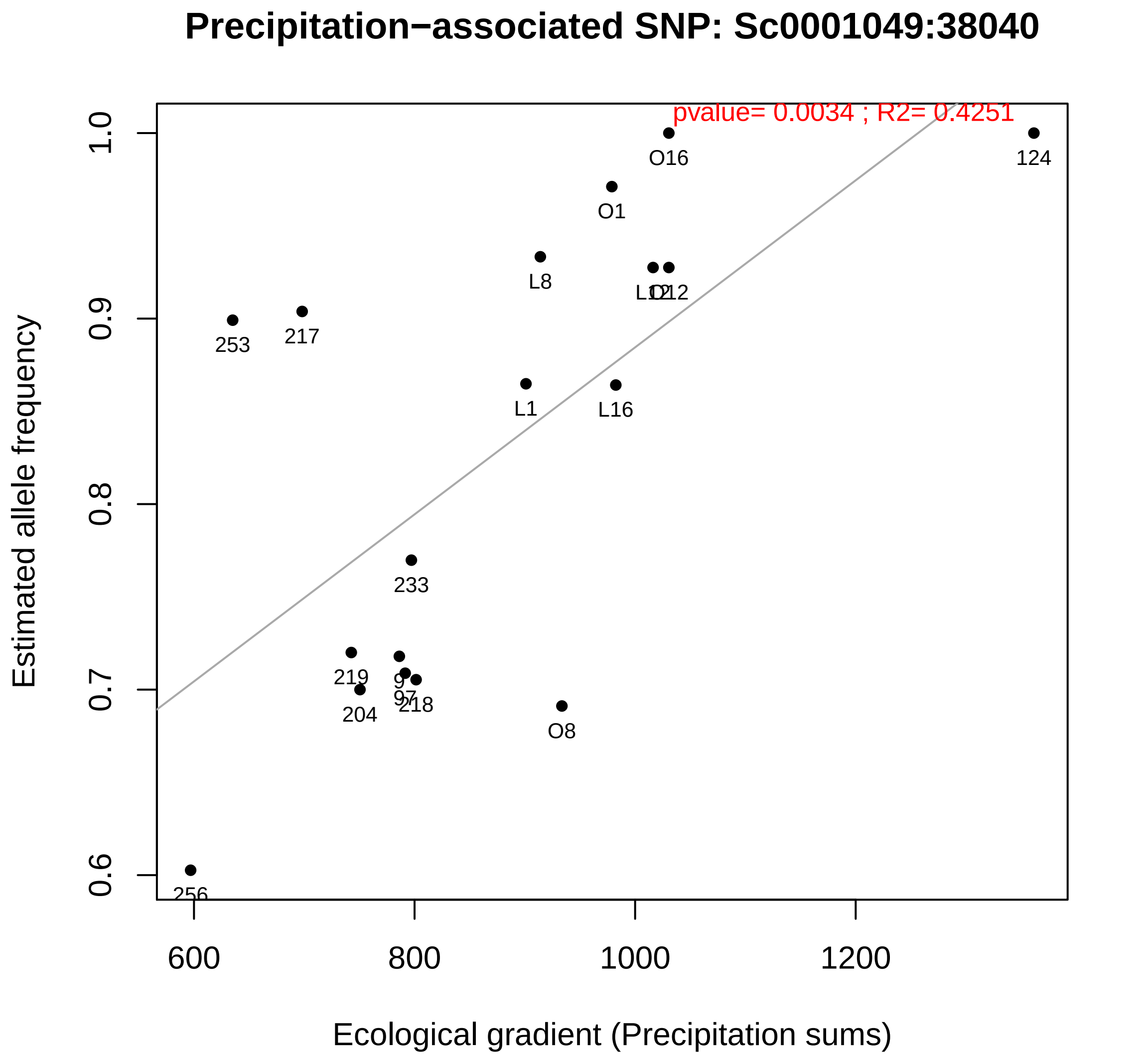

Precipitation−associated SNP: Sc0001049:38040
pvalue= 0.0034 ; R2= 0.4251
Estimated allele frequency
Ecological gradient (Precipitation sums)
1.0
0.9
0.8
0.7
0.6
600
800
1000
1200
O16
124
O1
L8
253
217
L1
L16
233
219
204
218
O8

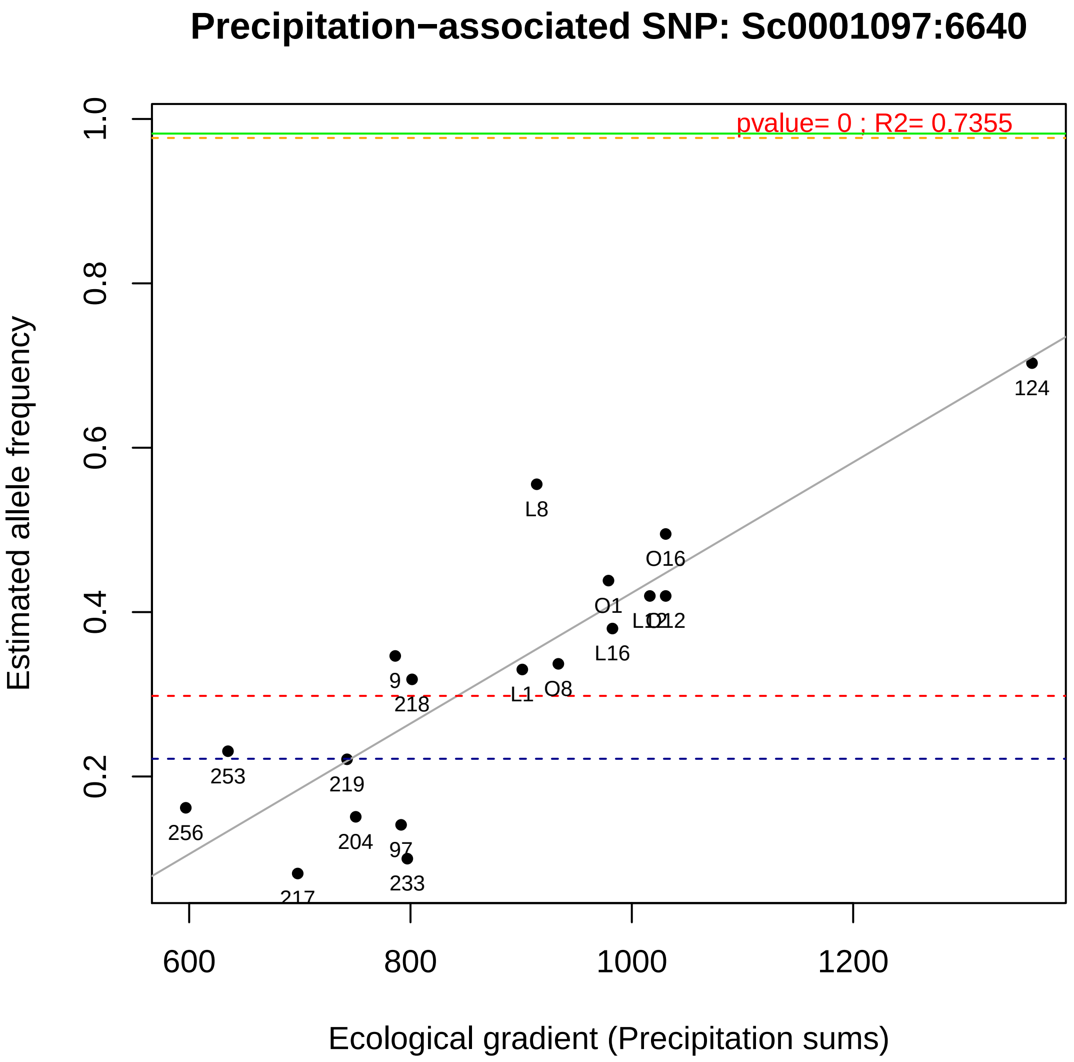
Precipitation−associated SNP: Sc0001097:6640
pvalue= 0 ; R2= 0.7355
Estimated allele frequency
0.2
0.4
0.6
0.8
1.0
Ecological gradient (Precipitation sums)
600
800
1000
1200
124
L8
O16
O1
L16
O8
L1
9
218
253
219
256
204
97
233
217

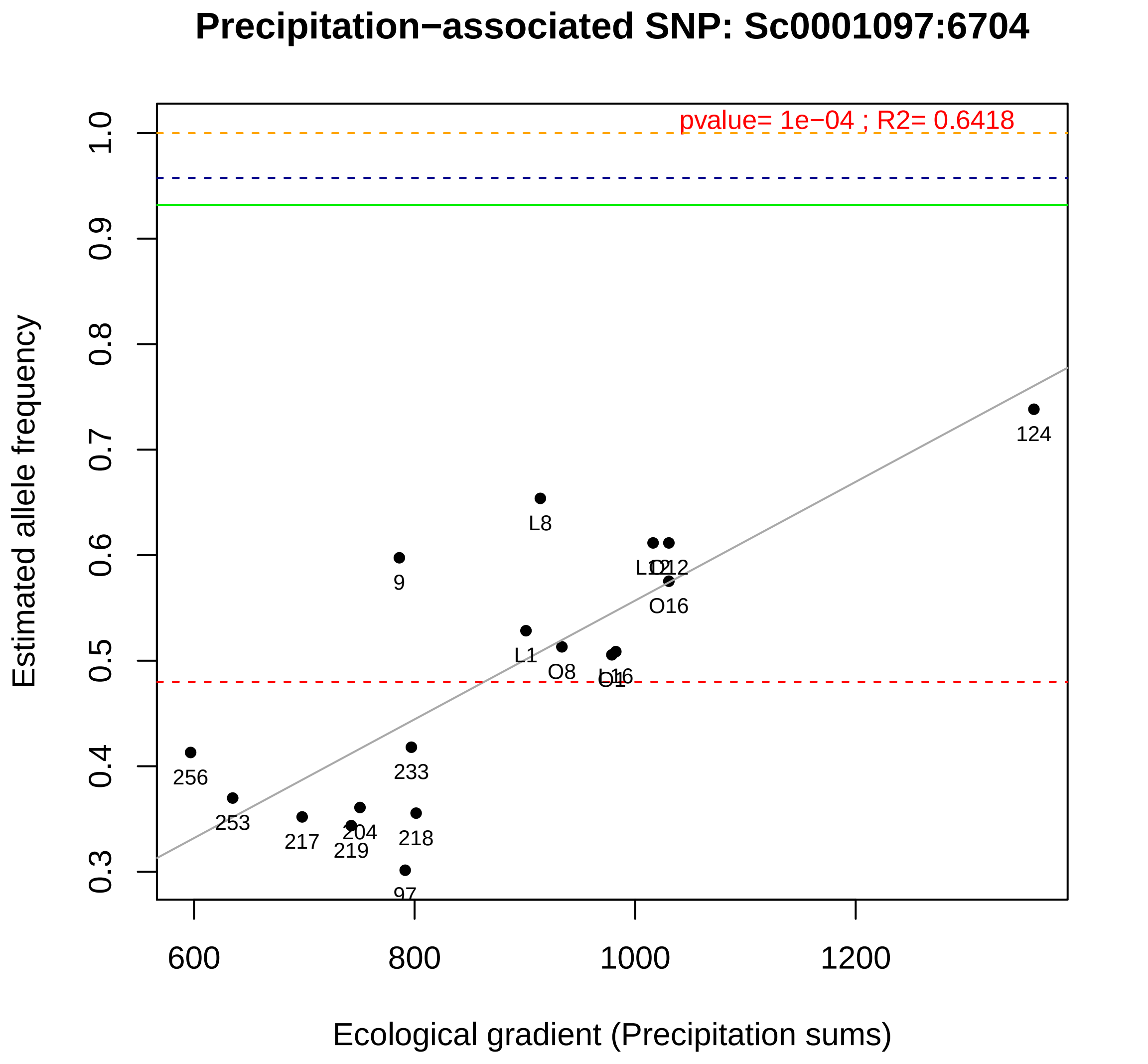

Precipitation−associated SNP: Sc0001097:6704
pvalue= 1e−04 ; R2= 0.6418
Estimated allele frequency
Ecological gradient (Precipitation sums)
1.0
0.9
0.8
0.7
0.6
0.5
0.4
0.3
600
800
1000
1200
124
L8
9
L12
O12
O16
L1
O8
L16
O1
256
233
253
217
204
219
218
97

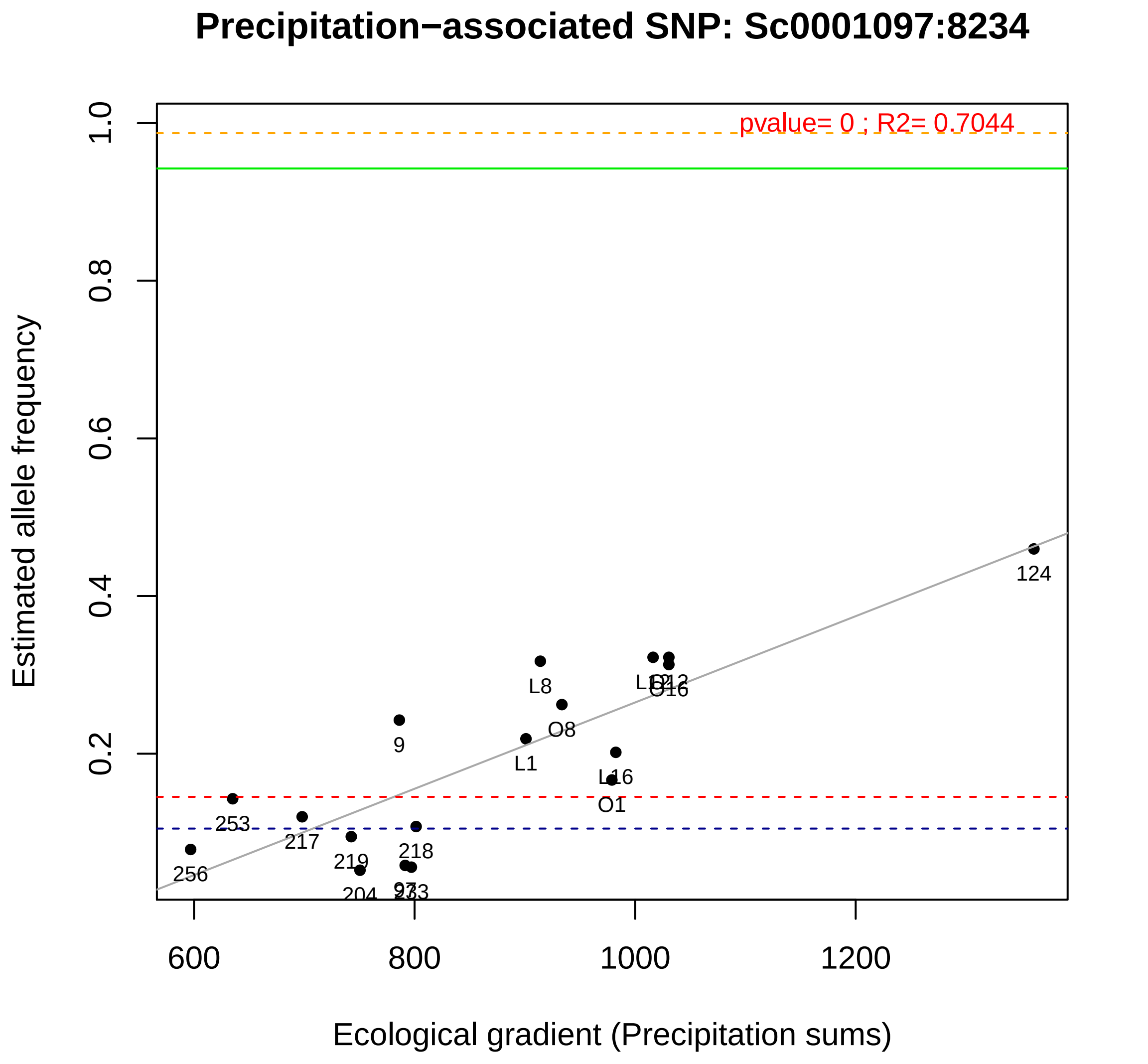
Precipitation−associated SNP: Sc0001097:8234
pvalue= 0 ; R2= 0.7044
Estimated allele frequency
Ecological gradient (Precipitation sums)
0.2
0.4
0.6
0.8
1.0
600
800
1000
1200
124
L8
O8
9
L1
L16
O1
253
217
219
218
256
204

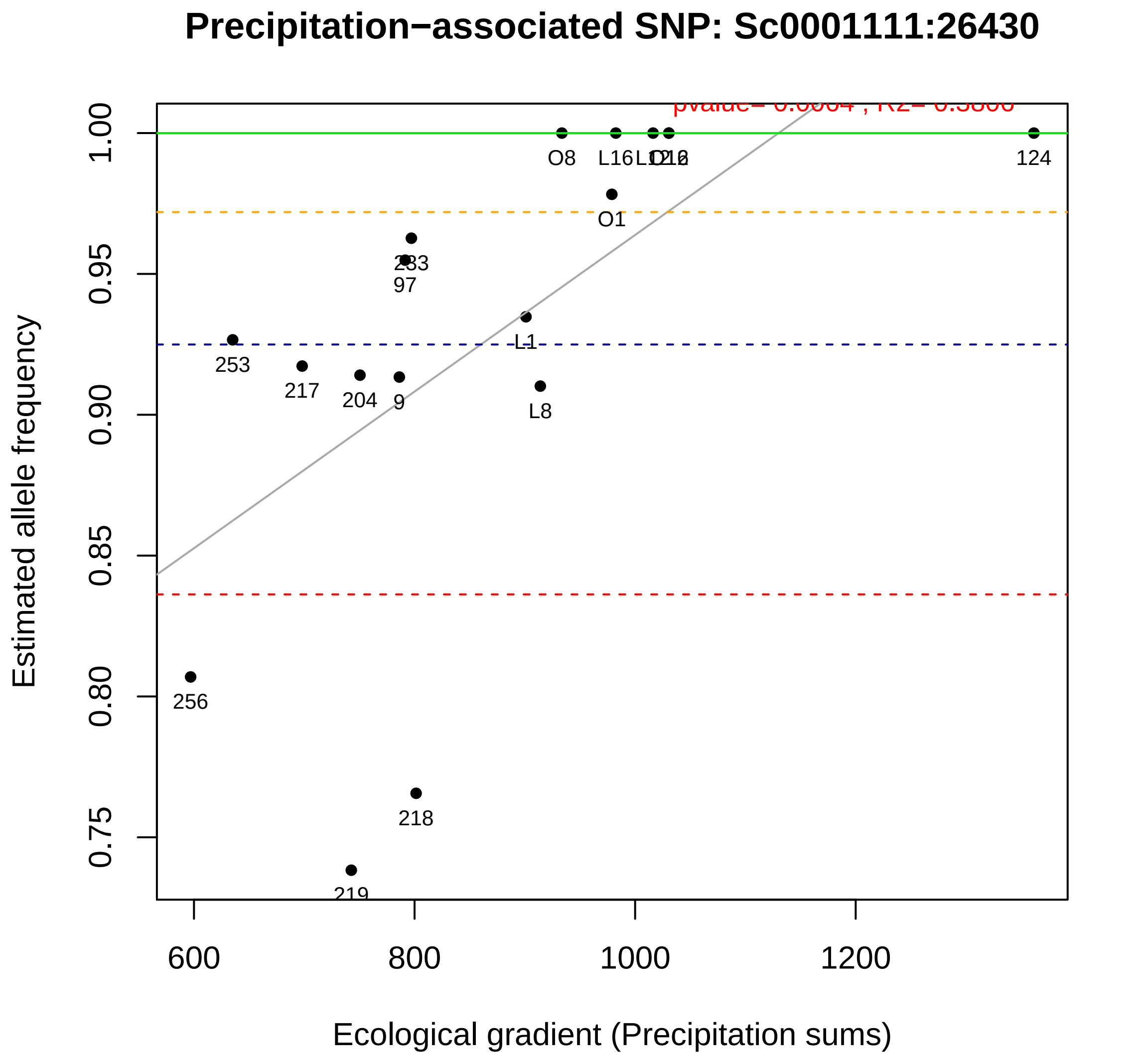
Precipitation−associated SNP: Sc0001111:26430
pvalue= 0.0004 ; R2= 0.3806
Estimated allele frequency
Ecological gradient (Precipitation sums)
1.00
0.95
0.90
0.85
0.80
0.75
600
800
1000
1200
O8
L16
L12
O2
124
O1
233
97
L1
253
217
204
9
L8
256
218
219

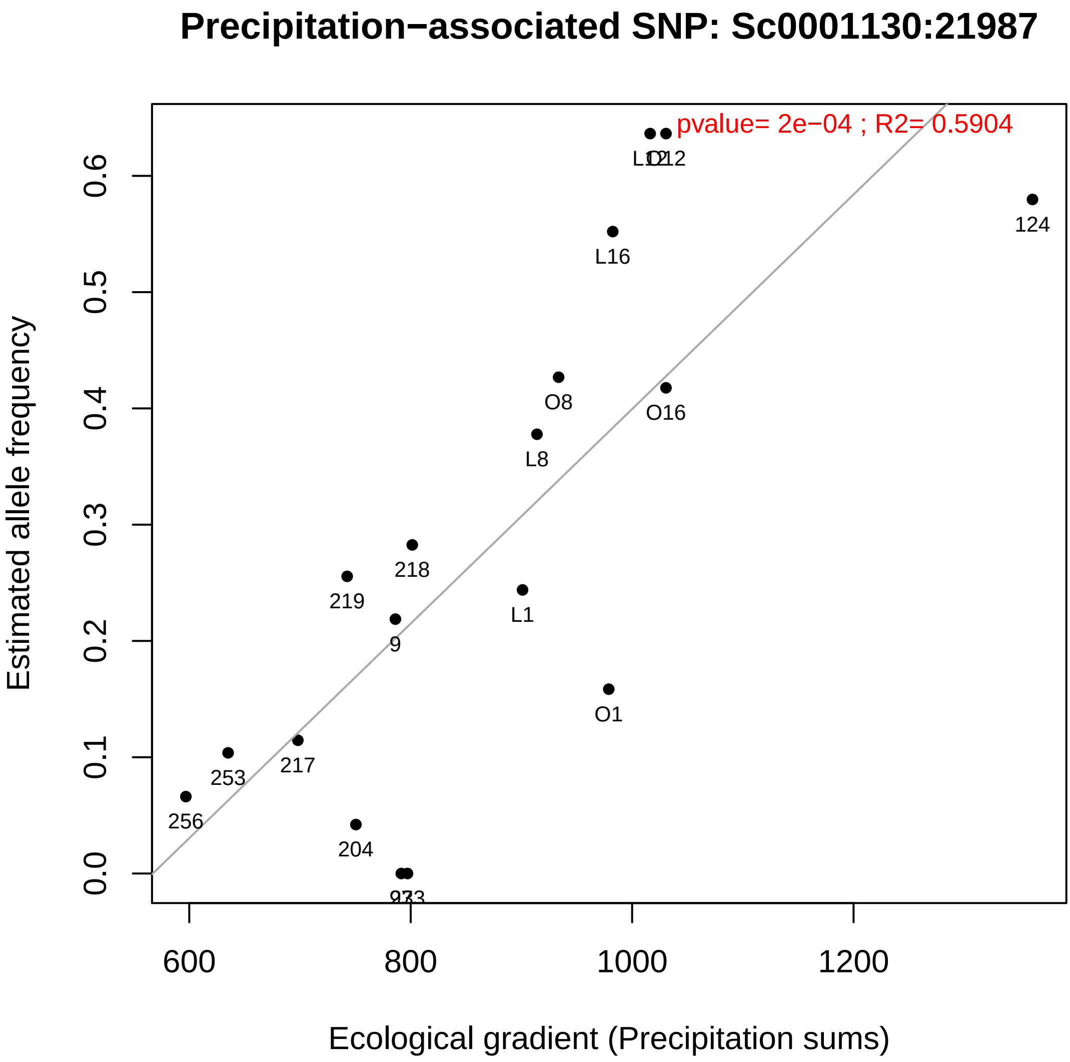

Precipitation−associated SNP: Sc0001130:21987
pvalue= 2e−04 ; R2= 0.5904
Estimated allele frequency
Ecological gradient (Precipitation sums)
0.0
0.1
0.2
0.3
0.4
0.5
0.6
600
800
1000
1200
124
L16
O8
O16
L8
218
219
9
L1
O1
217
253
256
204

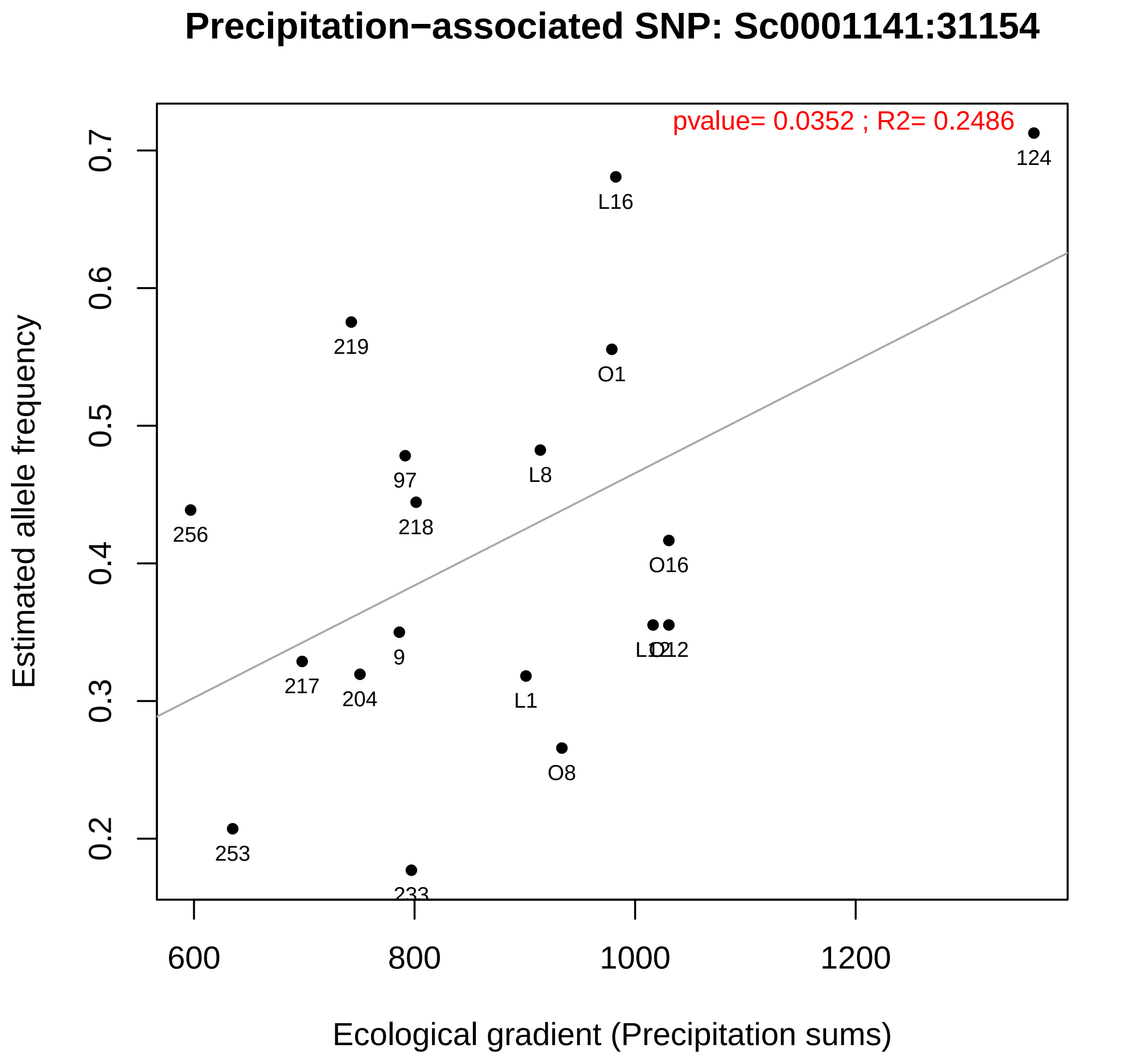

Precipitation−associated SNP: Sc0001141:31154
pvalue= 0.0352 ; R2= 0.2486
Estimated allele frequency
0.2
0.3
0.4
0.5
0.6
0.7
600
800
1000
1200
Ecological gradient (Precipitation sums)
124
L16
219
O1
97
L8
218
256
O16
L12
O12
9
217
204
L1
O8
253
233

**Precipitation−associated SNP: Sc0001189:11447**

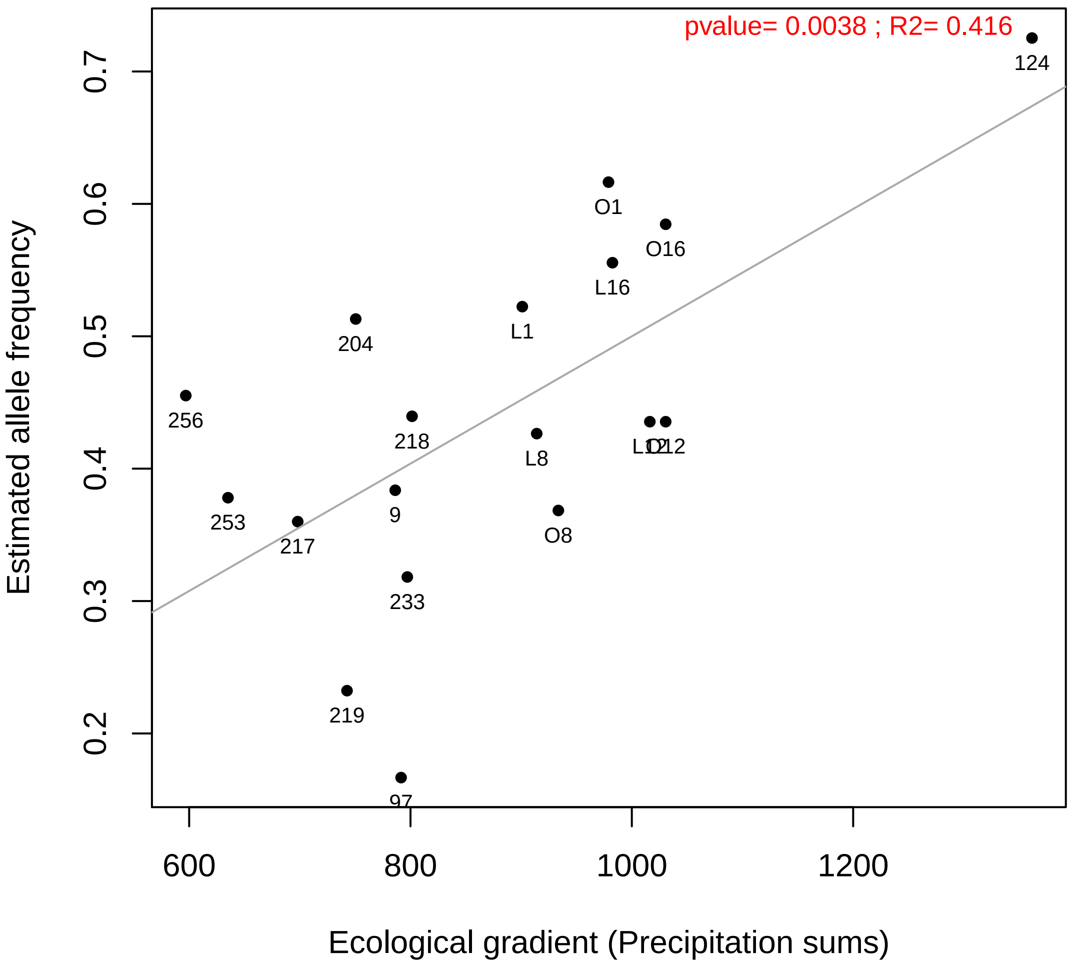

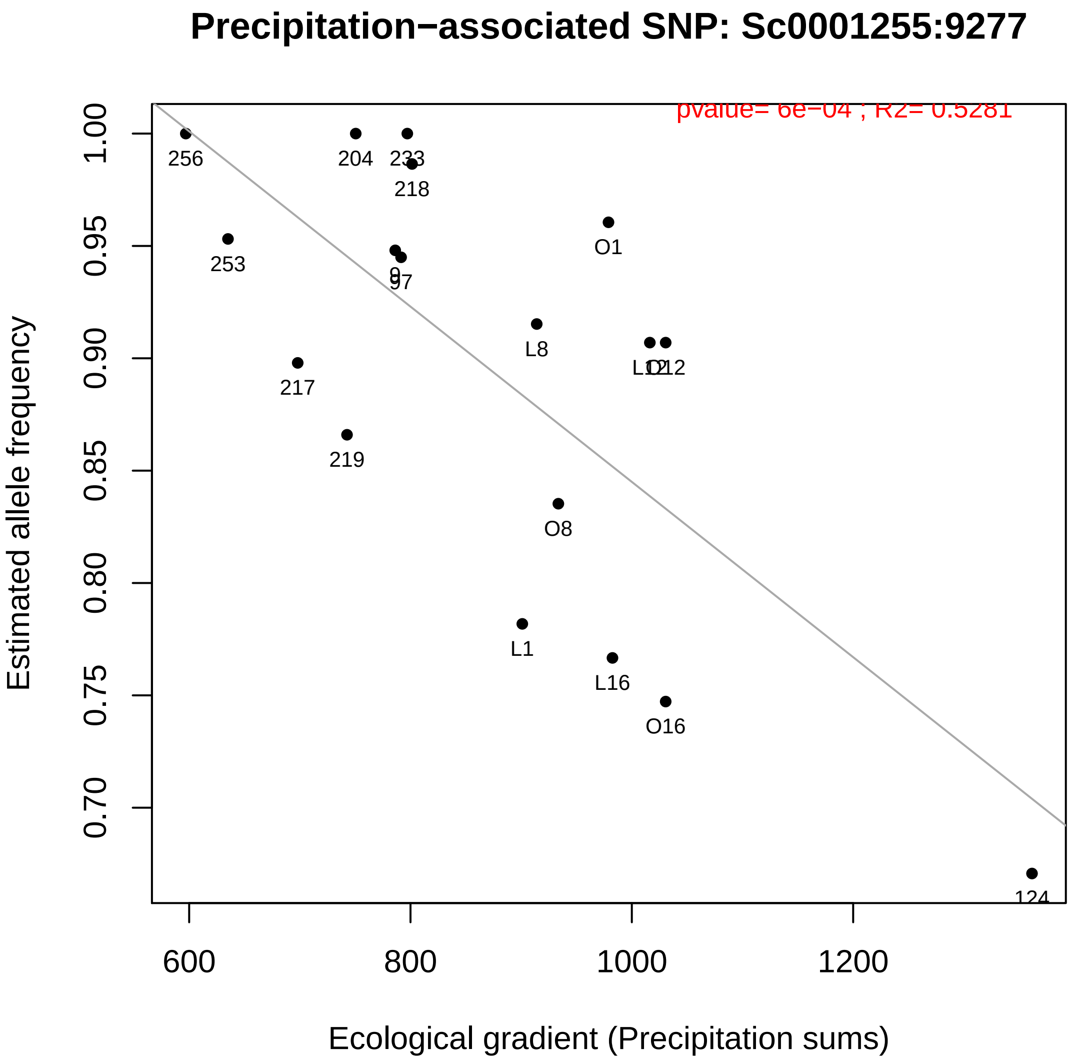
Precipitation−associated SNP: Sc0001255:9277
pvalue= 6e−04 ; R2= 0.5281
Estimated allele frequency
Ecological gradient (Precipitation sums)
1.00
0.95
0.90
0.85
0.80
0.75
0.70
600
800
1000
1200
256
204
233
218
253
O1
9
97
L8
217
219
O8
L1
L16
O16
124

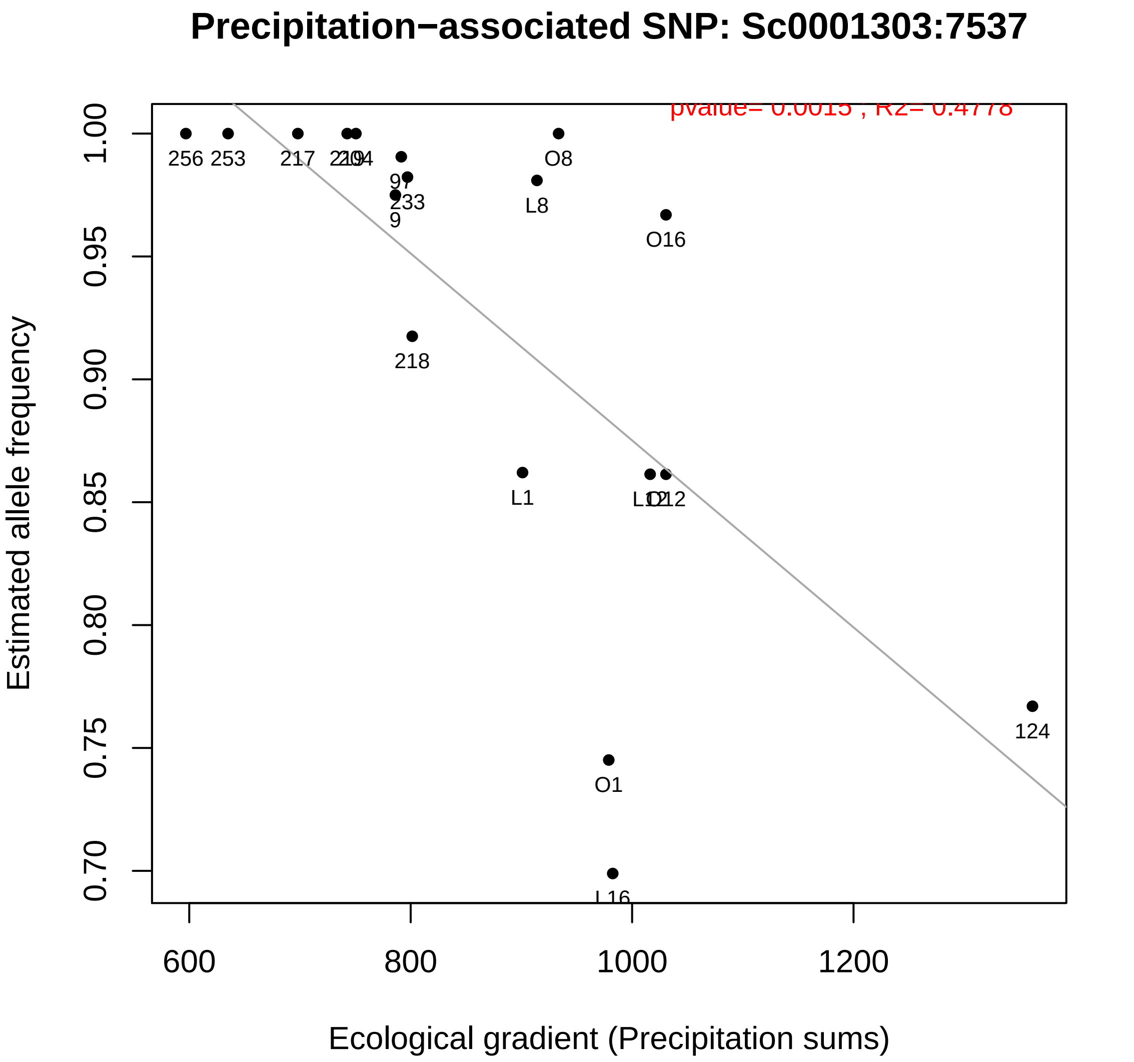

Precipitation−associated SNP: Sc0001303:7537
pvalue= 0.0015 , R2= 0.4778
Estimated allele frequency
1.00
0.95
0.90
0.85
0.80
0.75
0.70
600
800
1000
1200
Ecological gradient (Precipitation sums)
256
253
217
O8
97
233
9
L8
O16
218
L1
O12
124
O1

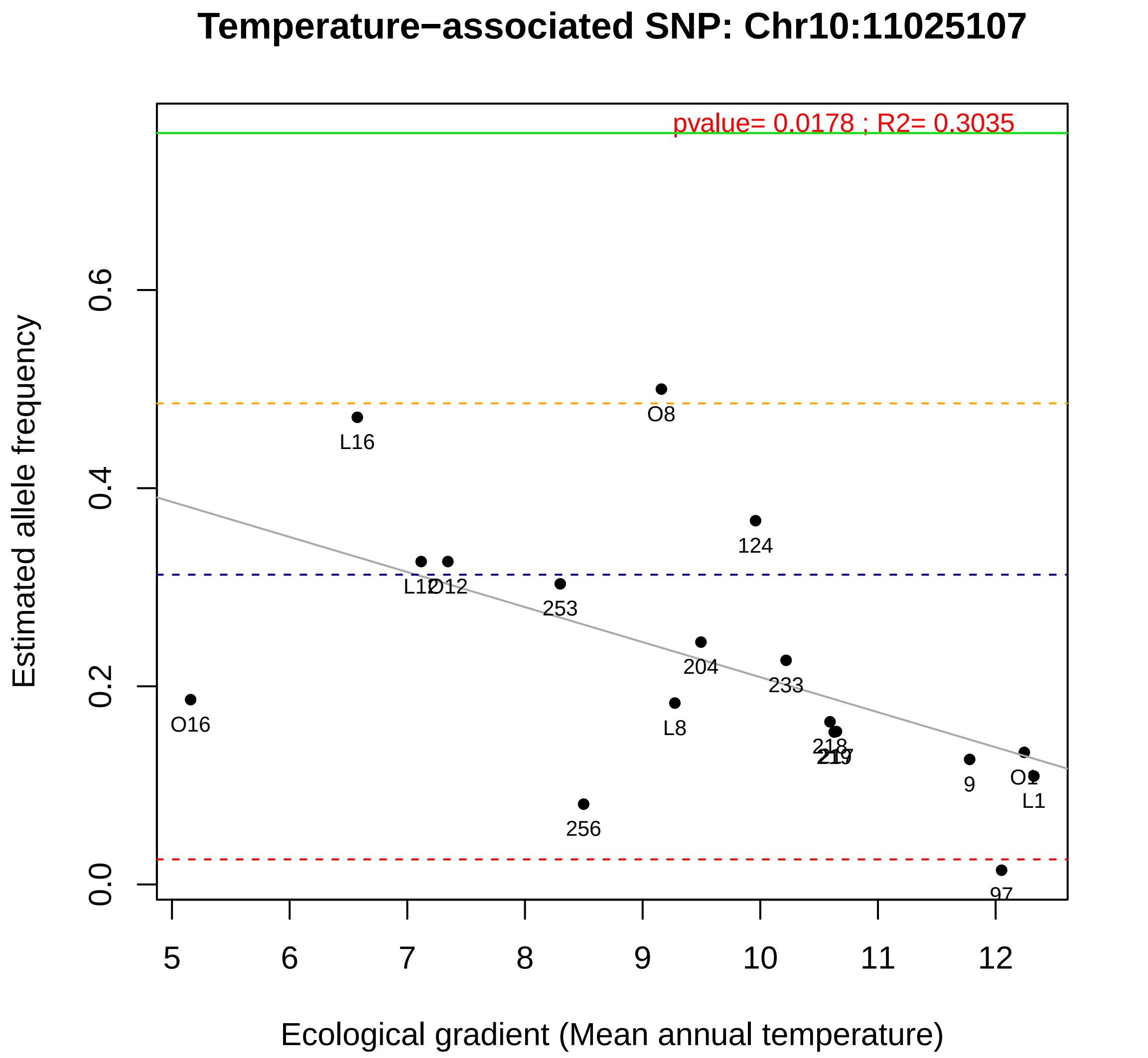

Temperature−associated SNP: Chr10:11025107
pvalue= 0.0178 ; R2= 0.3035
Estimated allele frequency
0.0
0.2
0.4
0.6
5
6
7
8
9
10
11
12
Ecological gradient (Mean annual temperature)
O16
L16
L12
O12
253
256
O8
L8
204
124
233
218
9
O1
L1
97

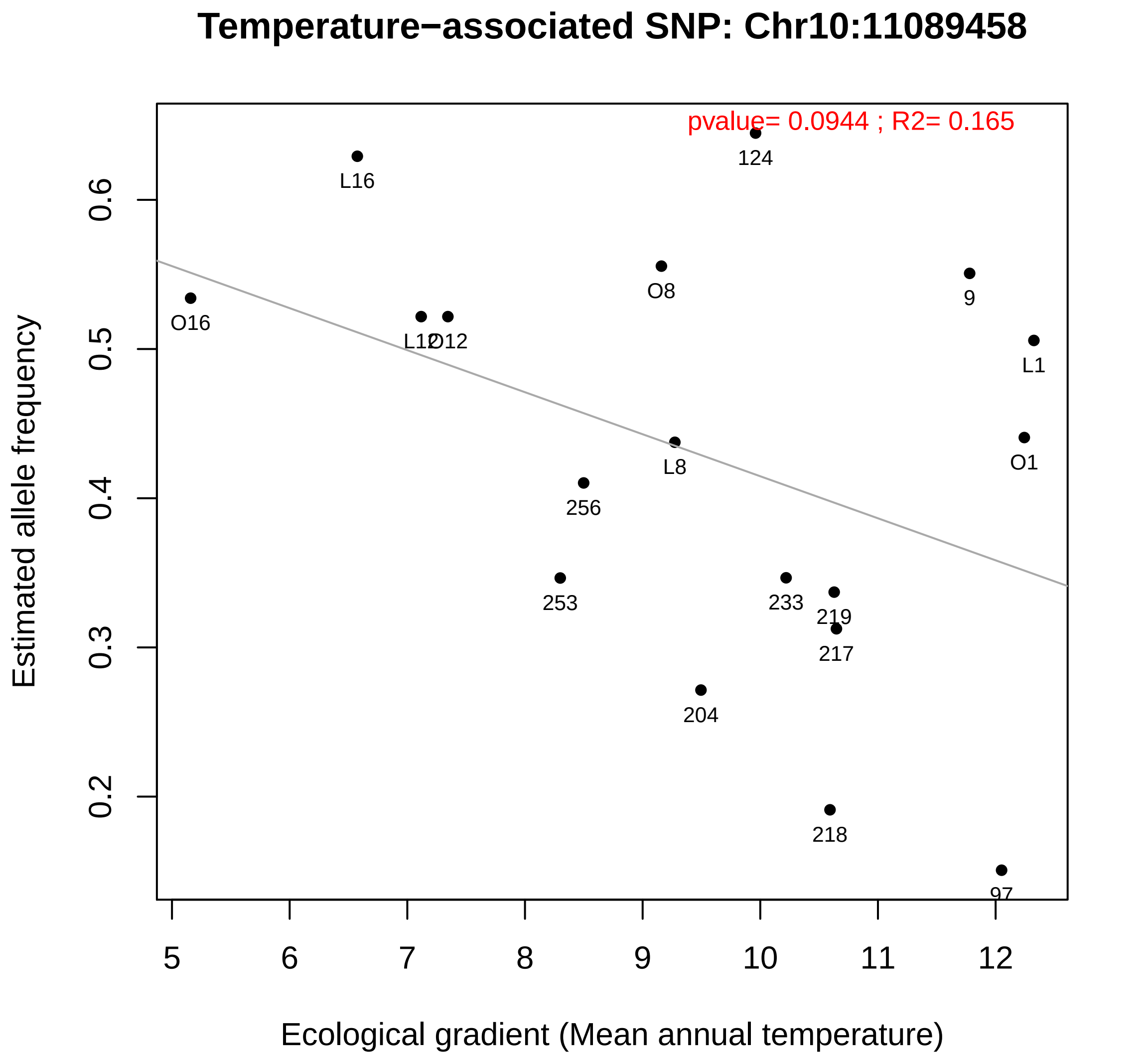
Temperature−associated SNP: Chr10:11089458
pvalue= 0.0944 ; R2= 0.165
L16
124
O8
9
O16
L12
O12
L1
L8
O1
256
253
233
219
217
204
218
97
0.2
0.3
0.4
0.5
0.6
Estimated allele frequency
5
6
7
8
9
10
11
12
Ecological gradient (Mean annual temperature)

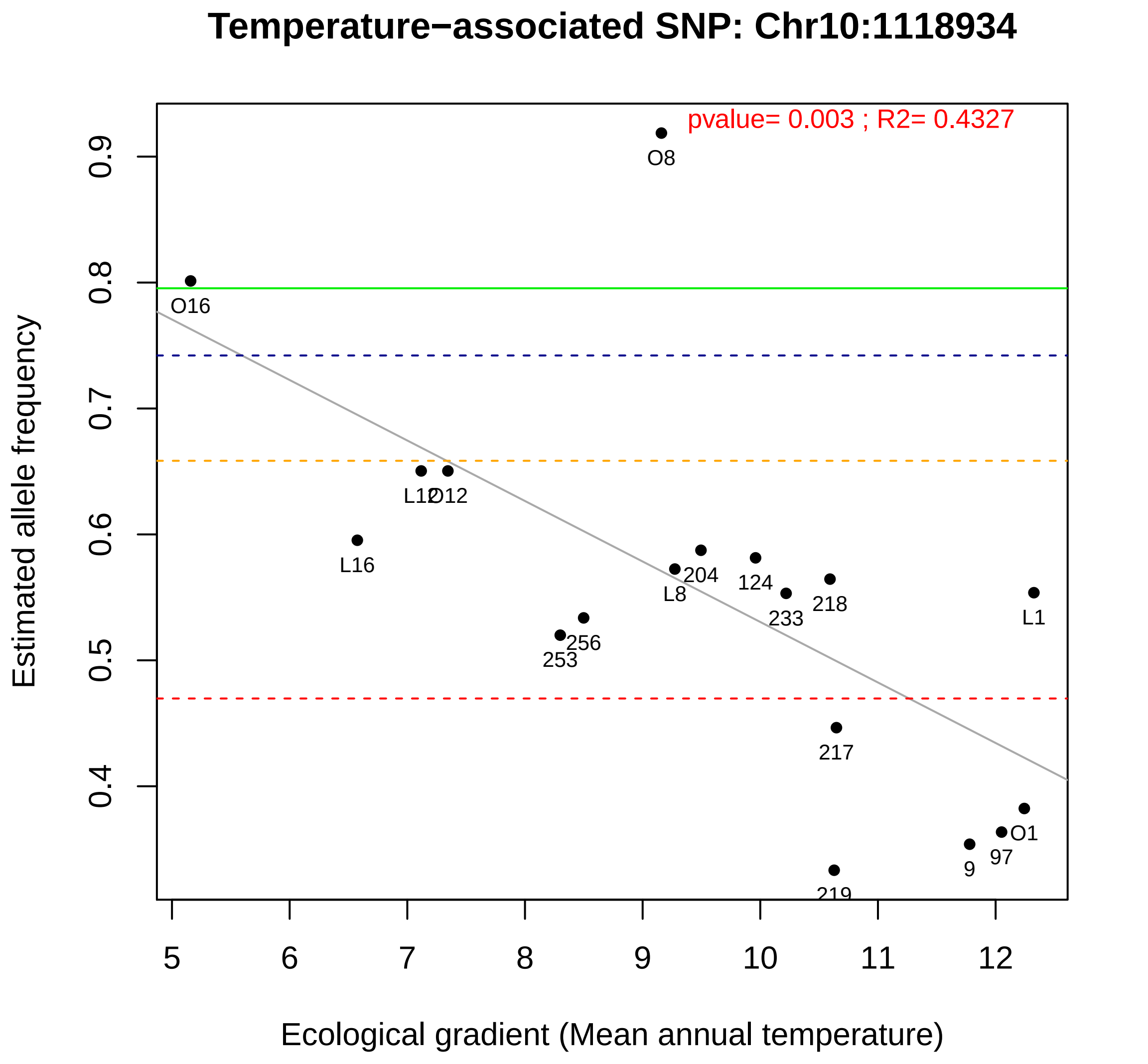

Temperature−associated SNP: Chr10:1118934
pvalue= 0.003 ; R2= 0.4327
Estimated allele frequency
0.4
0.5
0.6
0.7
0.8
0.9
5
6
7
8
9
10
11
12
Ecological gradient (Mean annual temperature)
O8
O16
L12
O12
L16
L8
204
124
233
218
L1
256
253
217
O1
97
9
219

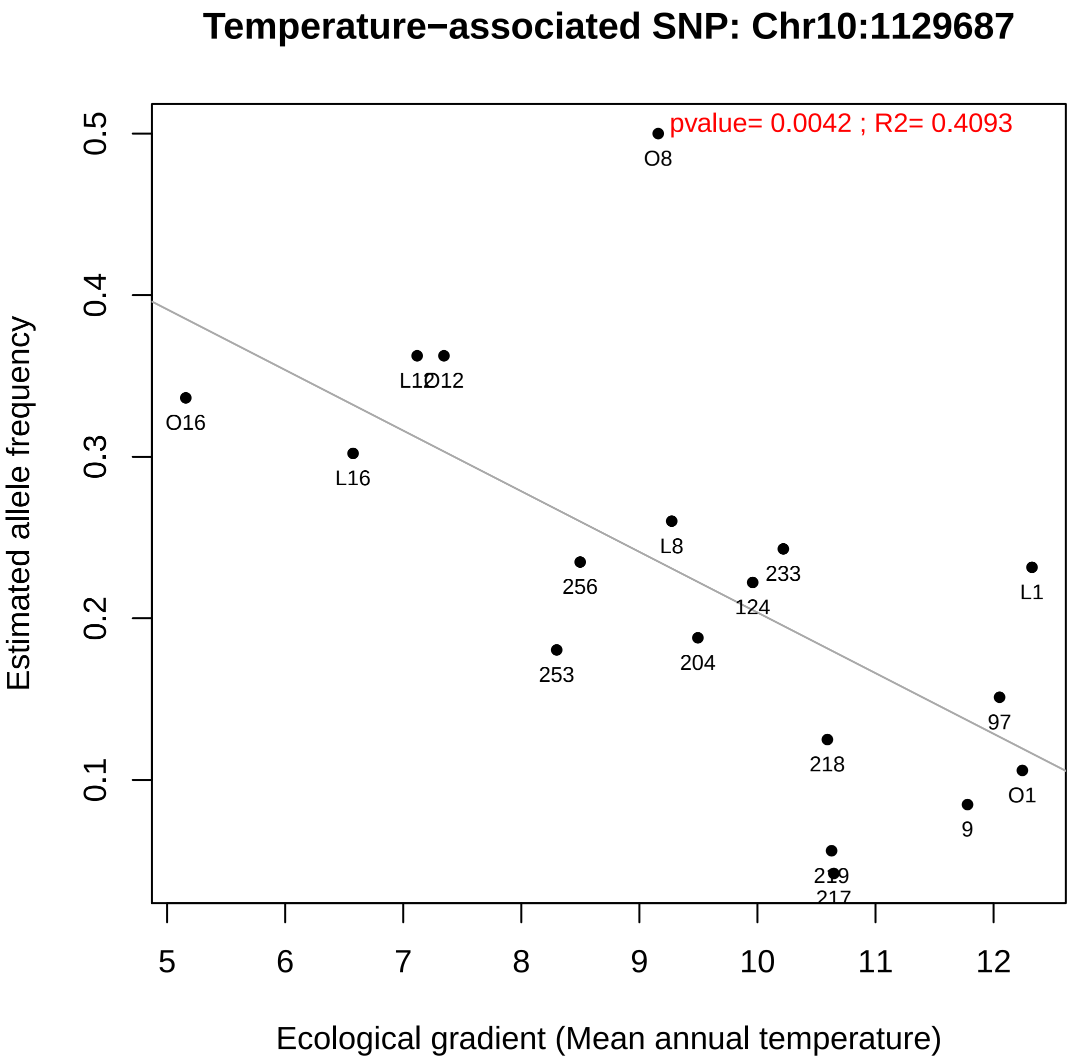

Temperature−associated SNP: Chr10:1129687
pvalue= 0.0042 ; R2= 0.4093
Estimated allele frequency
0.1
0.2
0.3
0.4
0.5
Ecological gradient (Mean annual temperature)
5
6
7
8
9
10
11
12
O8
L12
O12
O16
L16
L8
256
233
124
L1
253
204
97
218
O1
9
219
217

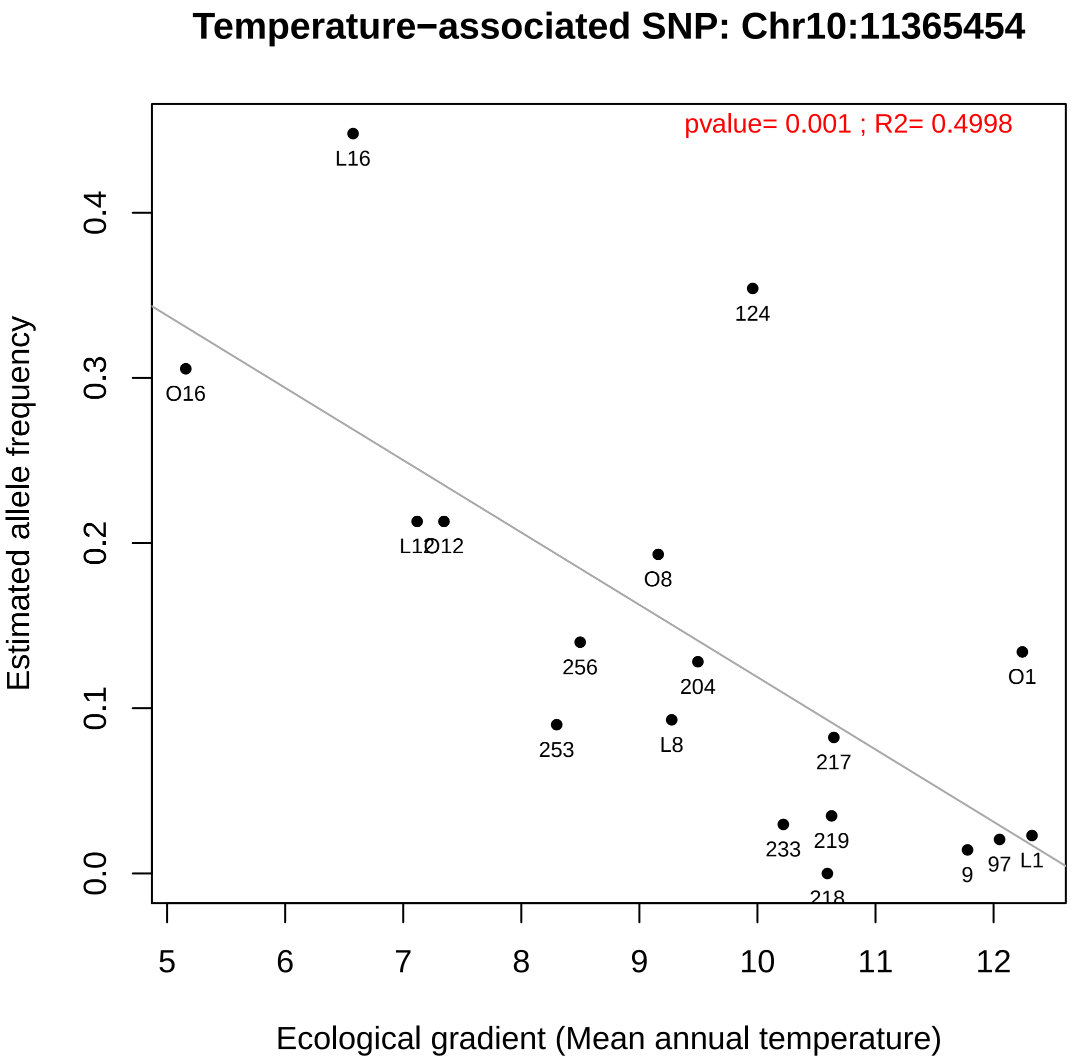
Temperature−associated SNP: Chr10:11365454
pvalue= 0.001 ; R2= 0.4998
Estimated allele frequency
0.0
0.1
0.2
0.3
0.4
5
6
7
8
9
10
11
12
Ecological gradient (Mean annual temperature)
L16
124
O16
L12
O12
O8
256
204
O1
253
L8
217
233
219
218
9
97
L1

**Temperature−associated SNP: Chr10:13849277**

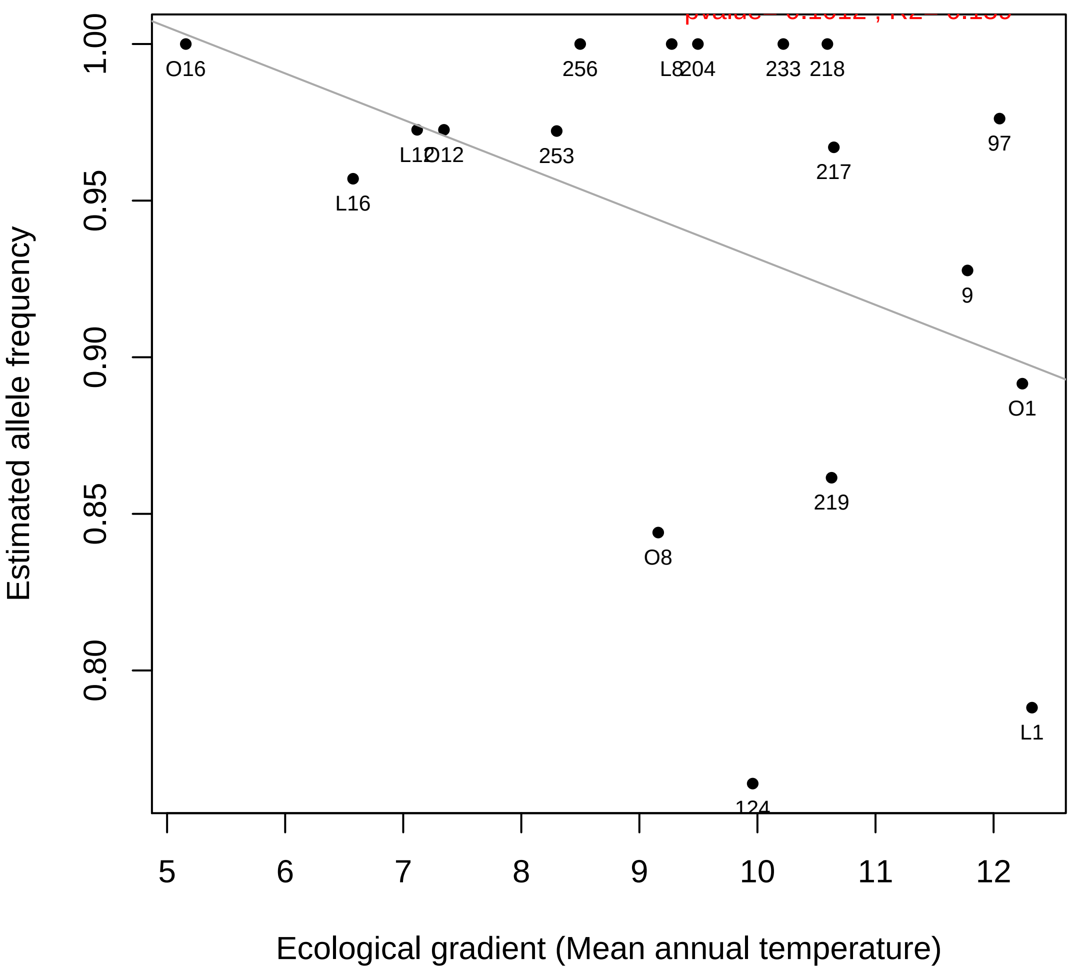

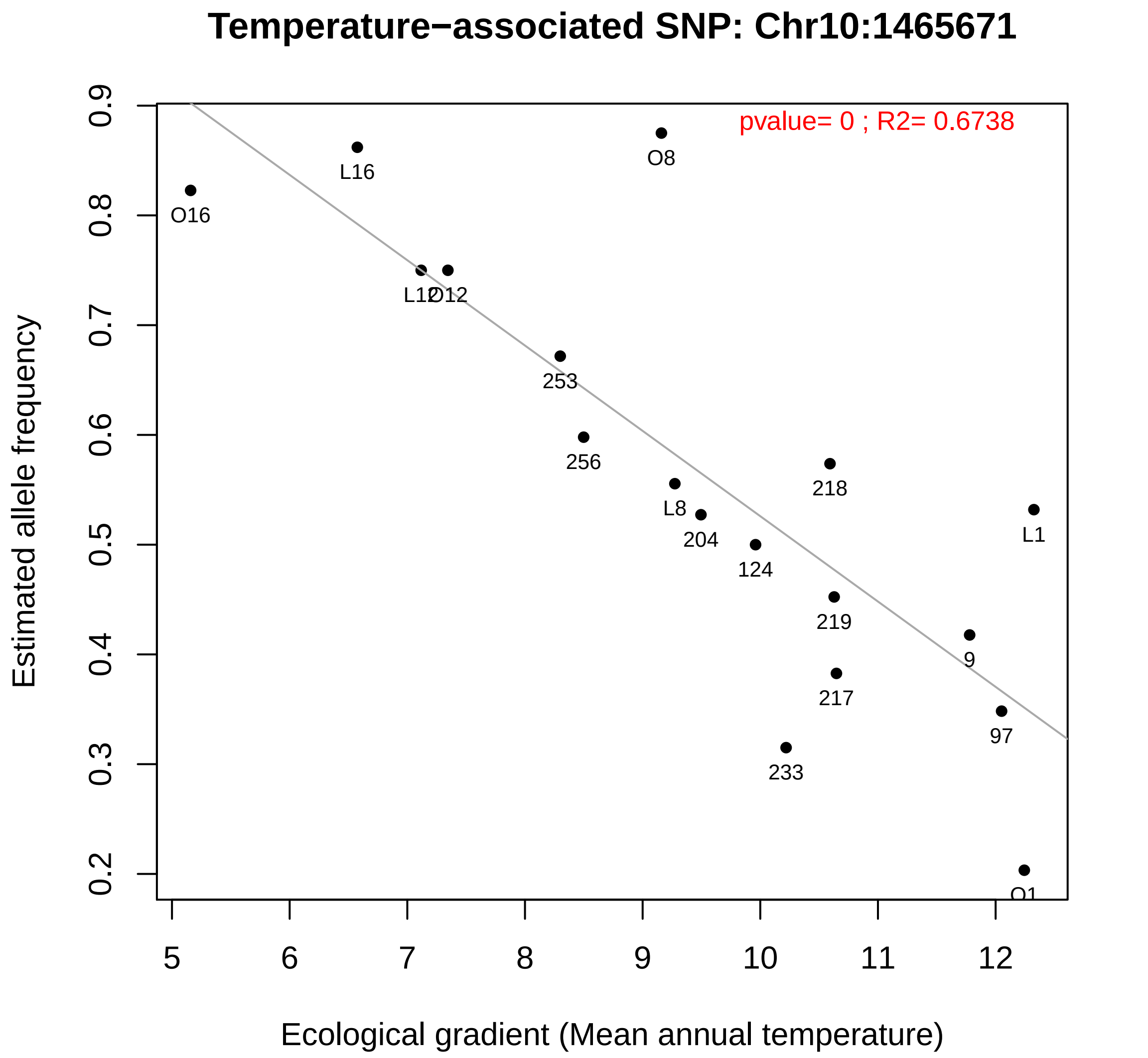

Temperature−associated SNP: Chr10:1465671
pvalue= 0 ; R2= 0.6738
Estimated allele frequency
0.2
0.3
0.4
0.5
0.6
0.7
0.8
0.9
5
6
7
8
9
10
11
12
Ecological gradient (Mean annual temperature)
O16
L16
O8
L12
O12
253
256
218
L8
204
L1
124
219
9
217
97
233
O1

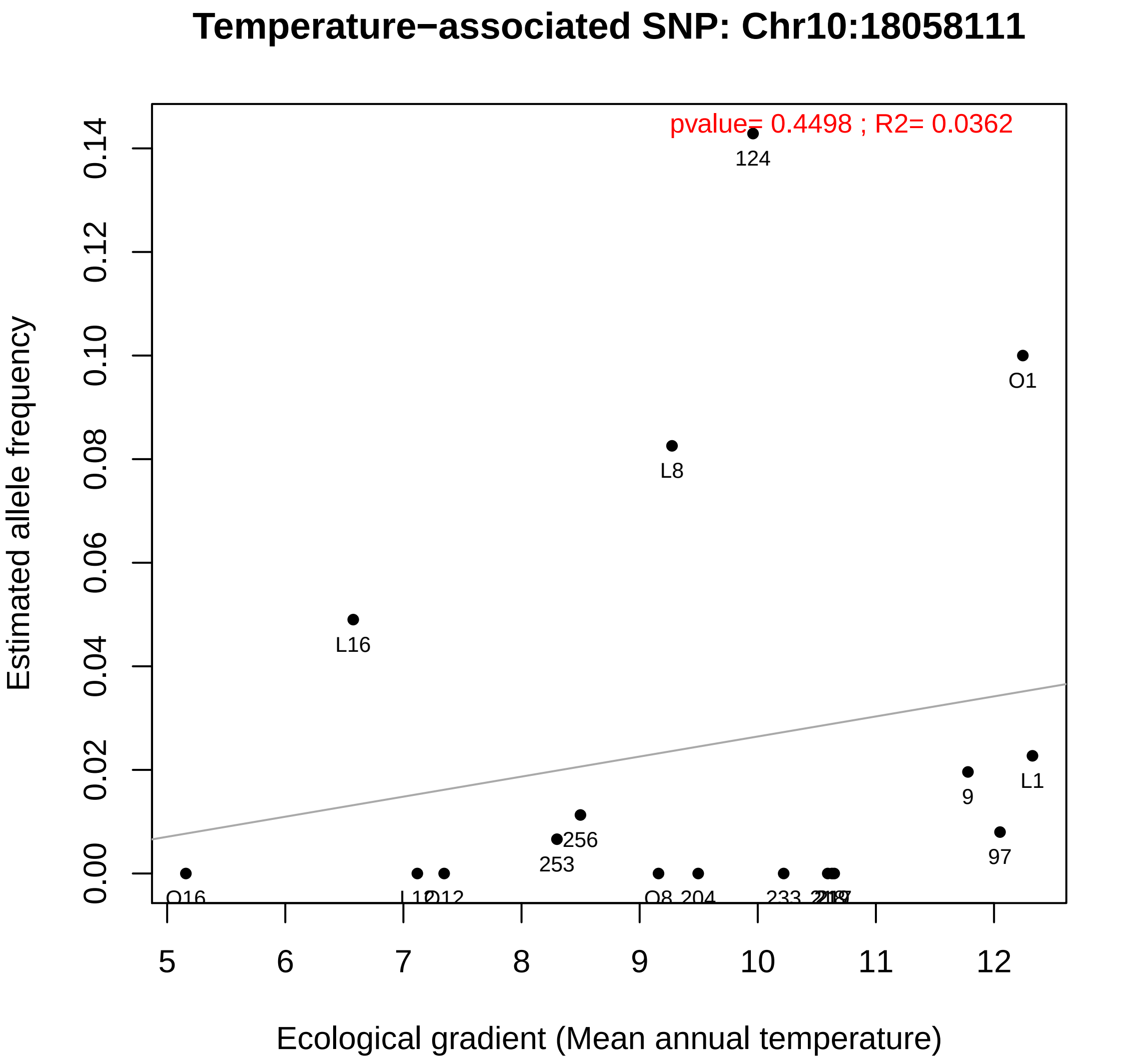
Temperature−associated SNP: Chr10:18058111
pvalue= 0.4498 ; R2= 0.0362
Estimated allele frequency
Ecological gradient (Mean annual temperature)
0.00
0.02
0.04
0.06
0.08
0.10
0.12
0.14
5
6
7
8
9
10
11
12
124
O1
L8
L16
L1
9
256
253
97
O16
O12
O8
204
233

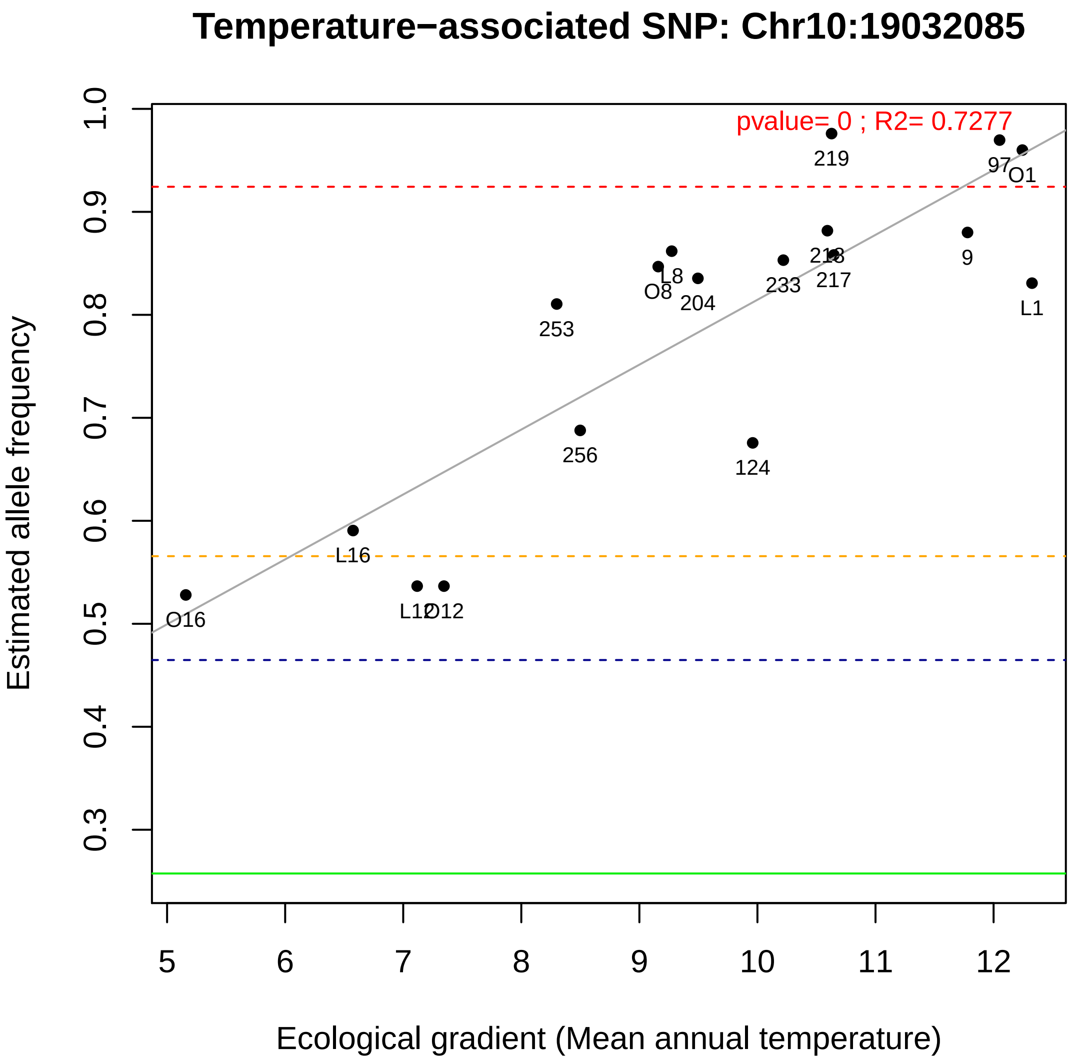
Temperature−associated SNP: Chr10:19032085
pvalue= 0 ; R2= 0.7277
Estimated allele frequency
Ecological gradient (Mean annual temperature)
1.0
0.9
0.8
0.7
0.6
0.5
0.4
0.3
5
6
7
8
9
10
11
12
219
97
O1
218
217
9
L8
O8
204
233
L1
253
256
124
L16
O16
L12
O12

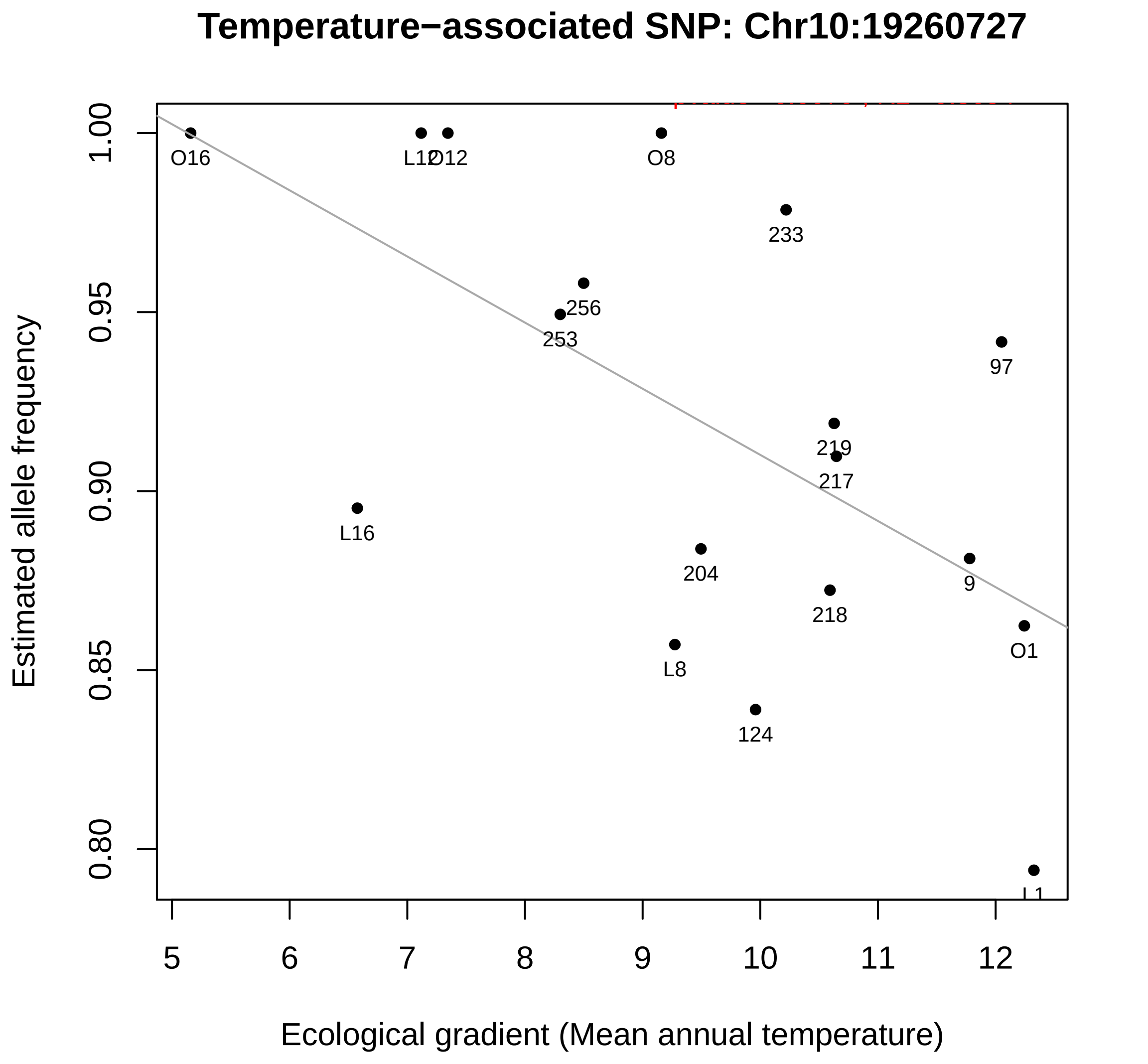
Temperature−associated SNP: Chr10:19260727
Estimated allele frequency
Ecological gradient (Mean annual temperature)
1.00
0.95
0.90
0.85
0.80
5
6
7
8
9
10
11
12
O16
L12
O12
O8
233
256
253
97
219
217
L16
204
9
218
O1
L8
124

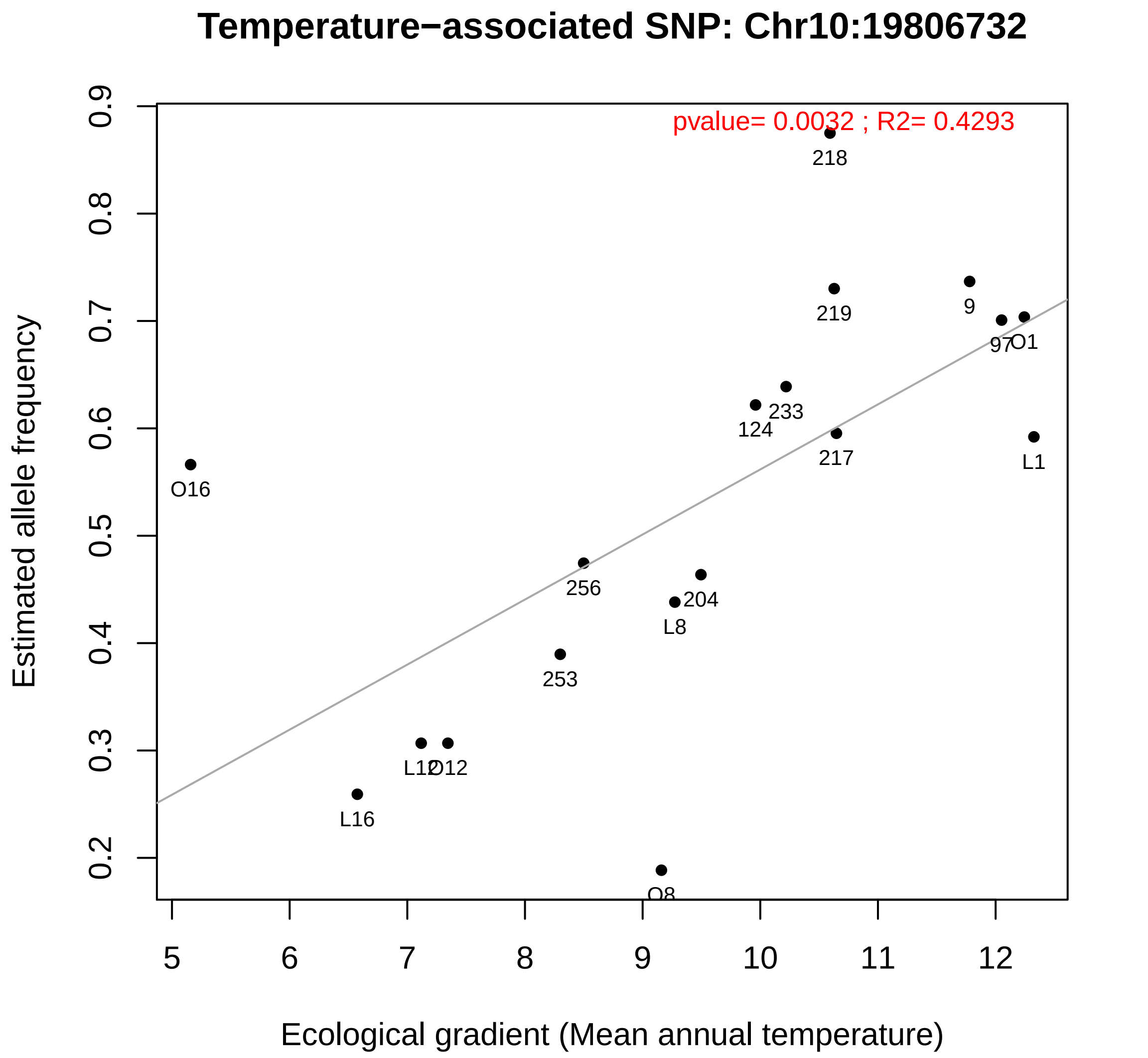

Temperature−associated SNP: Chr10:19806732
pvalue= 0.0032 ; R2= 0.4293
Estimated allele frequency
Ecological gradient (Mean annual temperature)
0.2
0.3
0.4
0.5
0.6
0.7
0.8
0.9
5
6
7
8
9
10
11
12
218
219
9
97
O1
233
124
217
L1
O16
256
204
L8
253
L12
O12
L16
O8

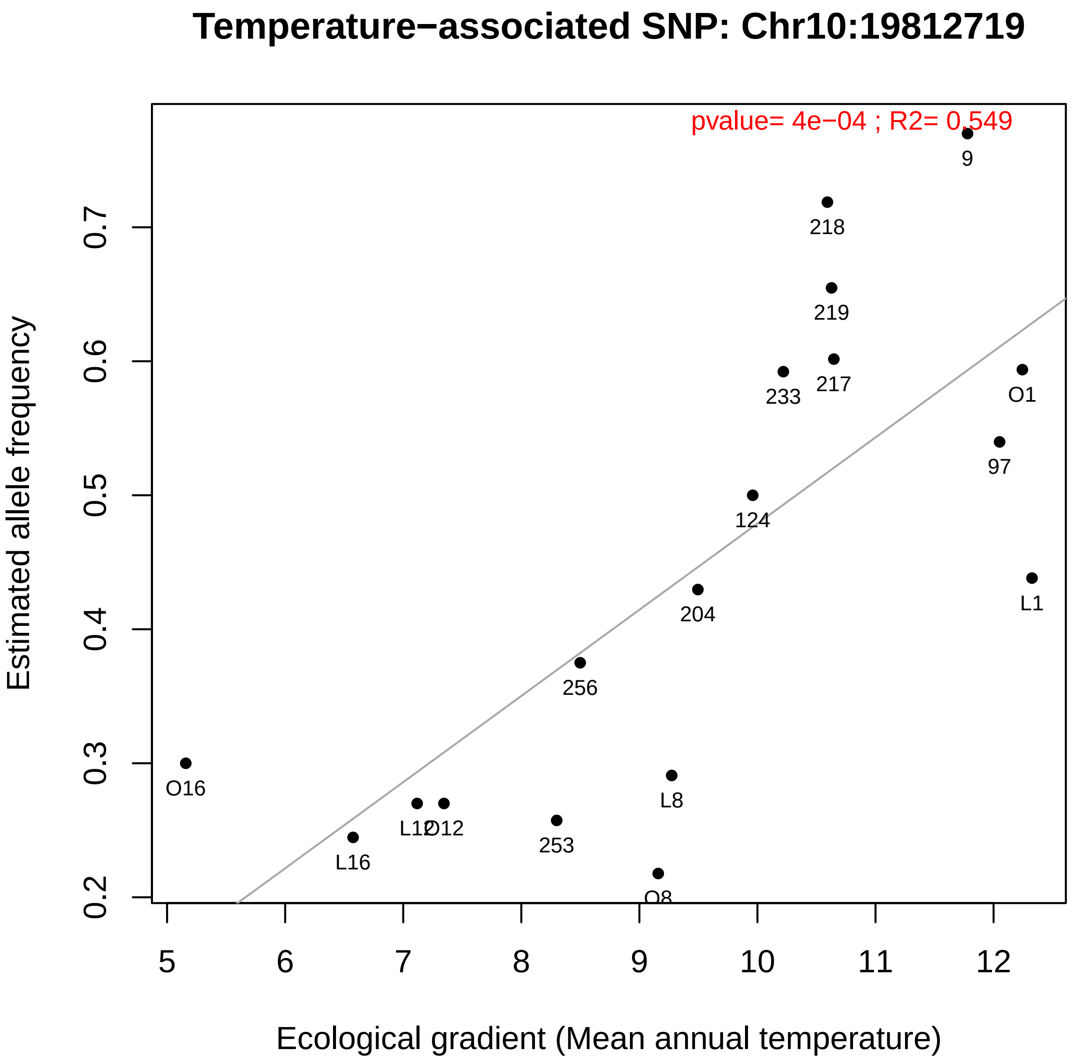

Temperature−associated SNP: Chr10:19812719
pvalue= 4e−04 ; R2= 0.549
Estimated allele frequency
Ecological gradient (Mean annual temperature)
0.2
0.3
0.4
0.5
0.6
0.7
5
6
7
8
9
10
11
12
9
218
219
217
233
O1
97
124
L1
204
256
O16
L8
L12
O12
253
L16
O8

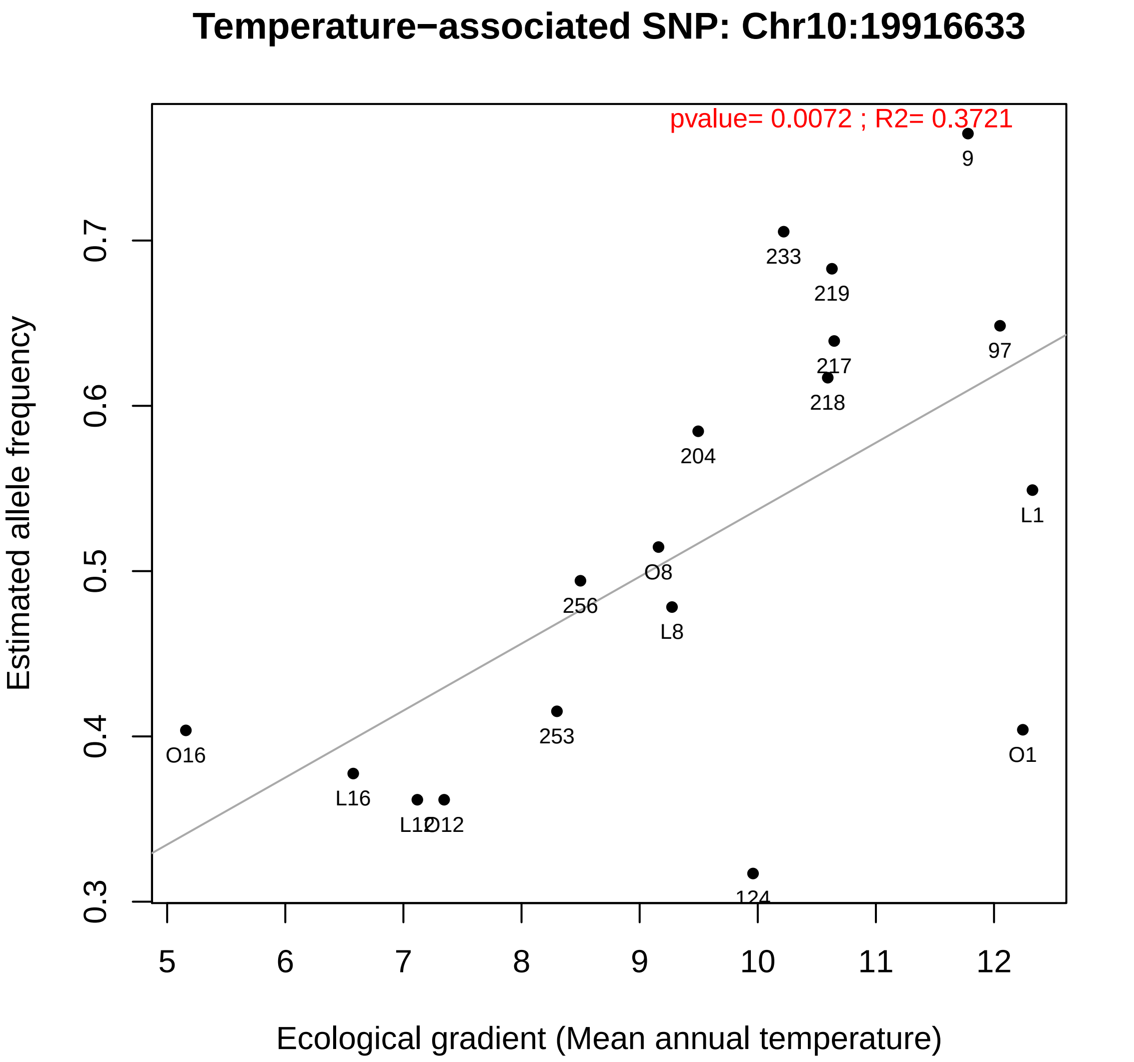
Temperature−associated SNP: Chr10:19916633
pvalue= 0.0072 ; R2= 0.3721
Estimated allele frequency
0.3
0.4
0.5
0.6
0.7
5
6
7
8
9
10
11
12
Ecological gradient (Mean annual temperature)
9
233
219
97
217
218
204
L1
O8
256
L8
253
O16
O1
L16
L12
O12
124

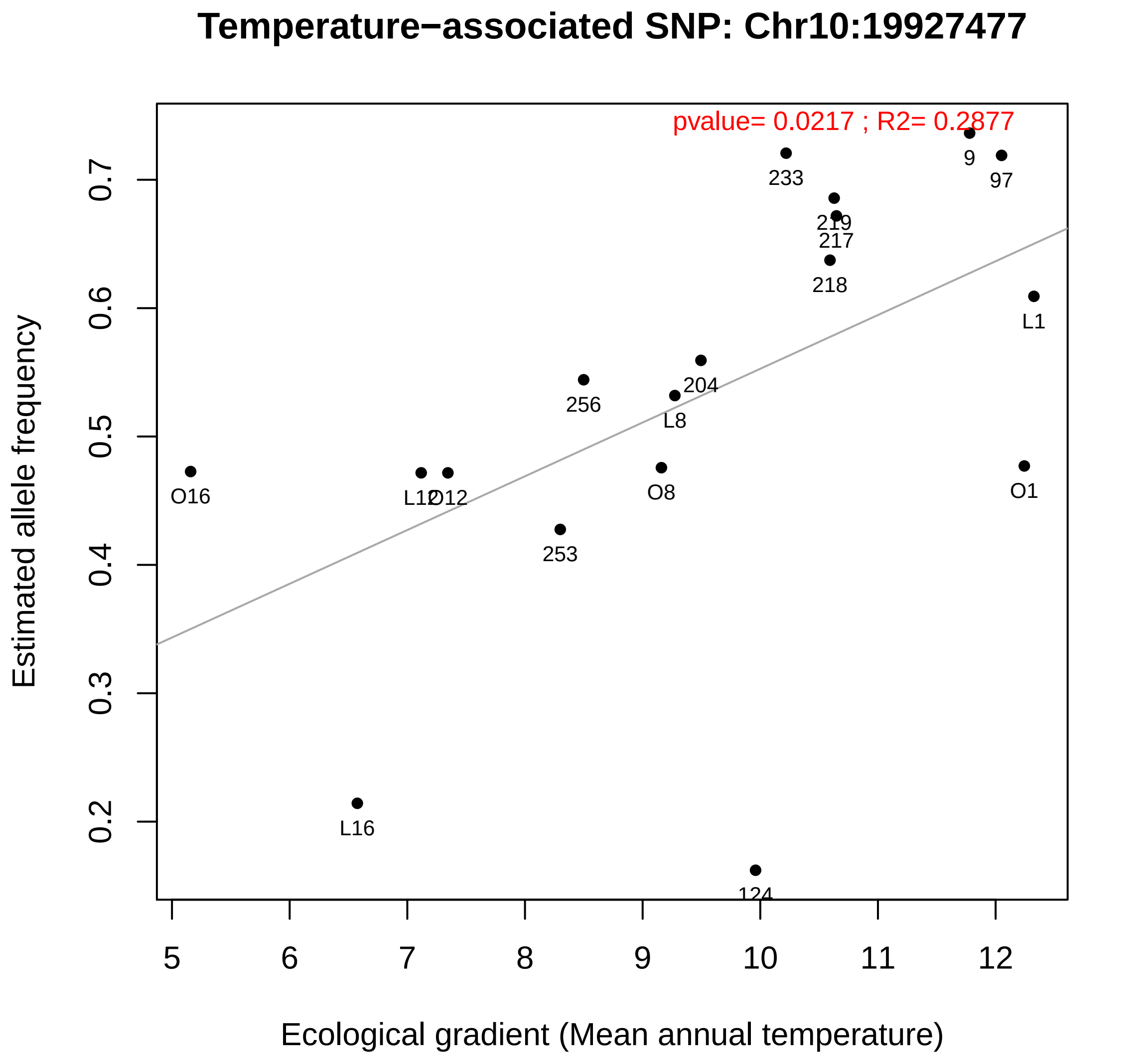

Temperature−associated SNP: Chr10:19927477
pvalue= 0.0217 ; R2= 0.2877
Estimated allele frequency
0.2
0.3
0.4
0.5
0.6
0.7
5
6
7
8
9
10
11
12
Ecological gradient (Mean annual temperature)
O16
L16
L12
O12
253
256
O8
L8
204
124
233
219
217
218
9
97
L1
O1

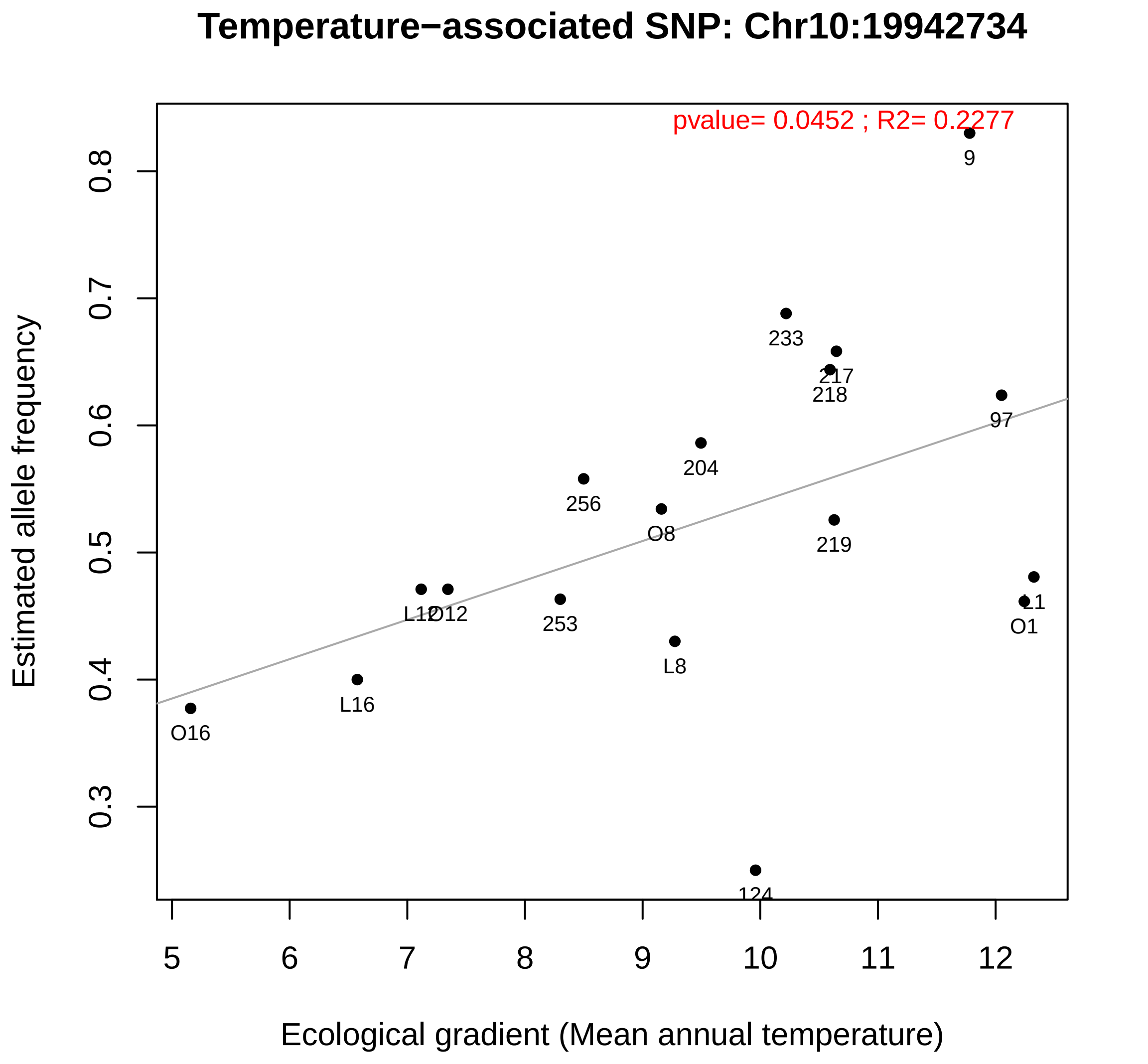
Temperature−associated SNP: Chr10:19942734
pvalue= 0.0452 ; R2= 0.2277
Estimated allele frequency
Ecological gradient (Mean annual temperature)
0.3
0.4
0.5
0.6
0.7
0.8
5
6
7
8
9
10
11
12
9
233
217
218
97
204
256
O8
219
L12
O12
253
L8
L1
O1
L16
O16
124

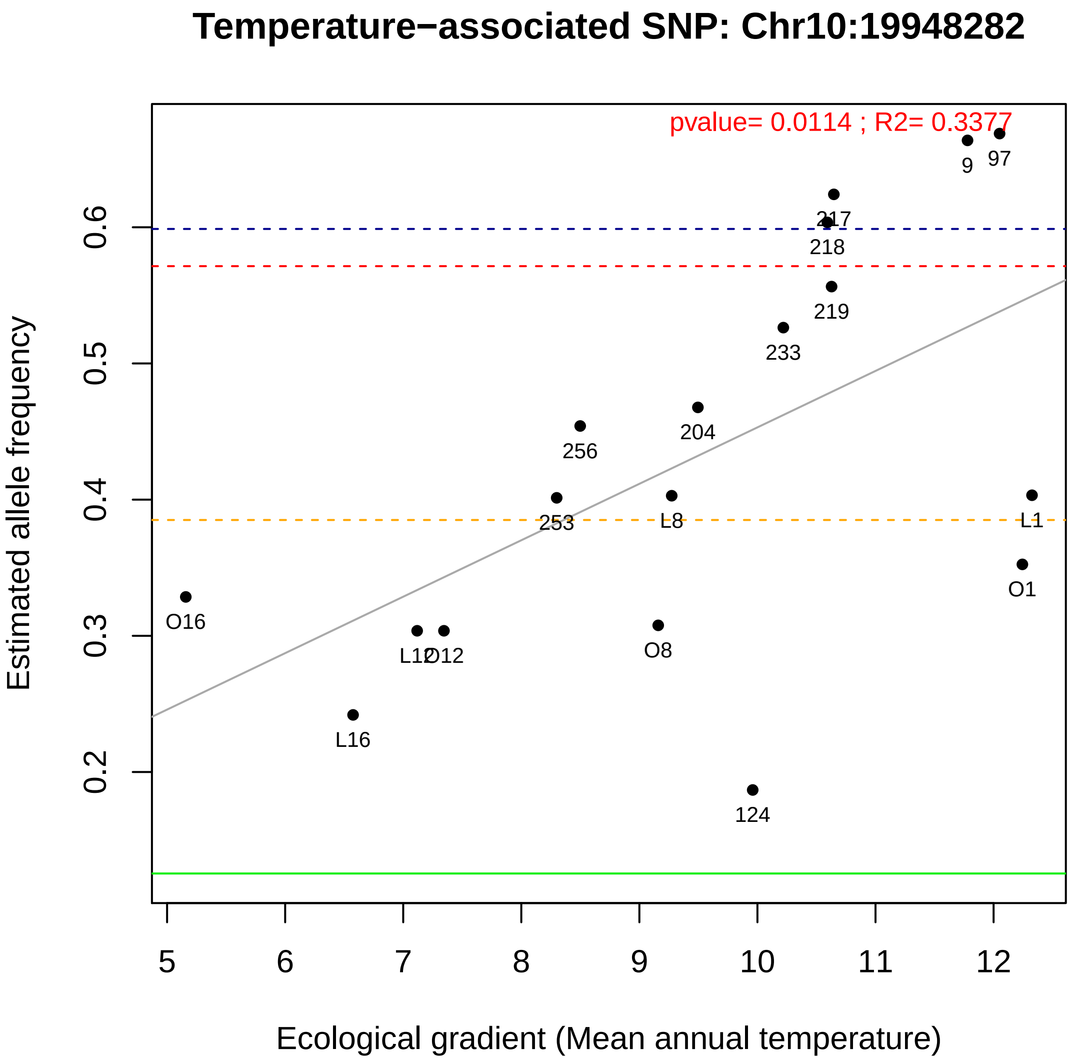

Temperature−associated SNP: Chr10:19948282
pvalue= 0.0114 ; R2= 0.3377
Estimated allele frequency
0.2
0.3
0.4
0.5
0.6
Ecological gradient (Mean annual temperature)
5
6
7
8
9
10
11
12
9
97
217
218
219
233
204
256
253
L8
L1
O1
O16
L12
O12
O8
L16
124

### Temperature−associated SNP: Chr10:19955355

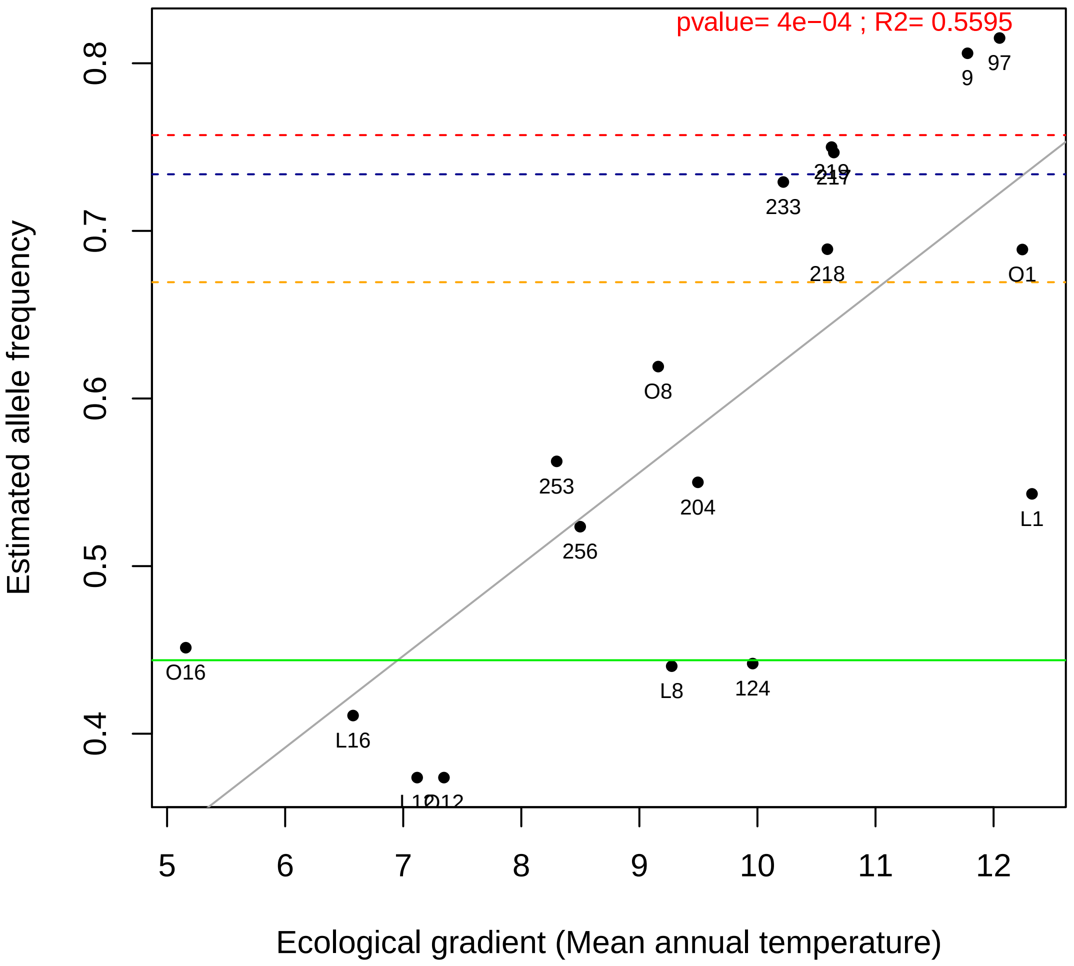

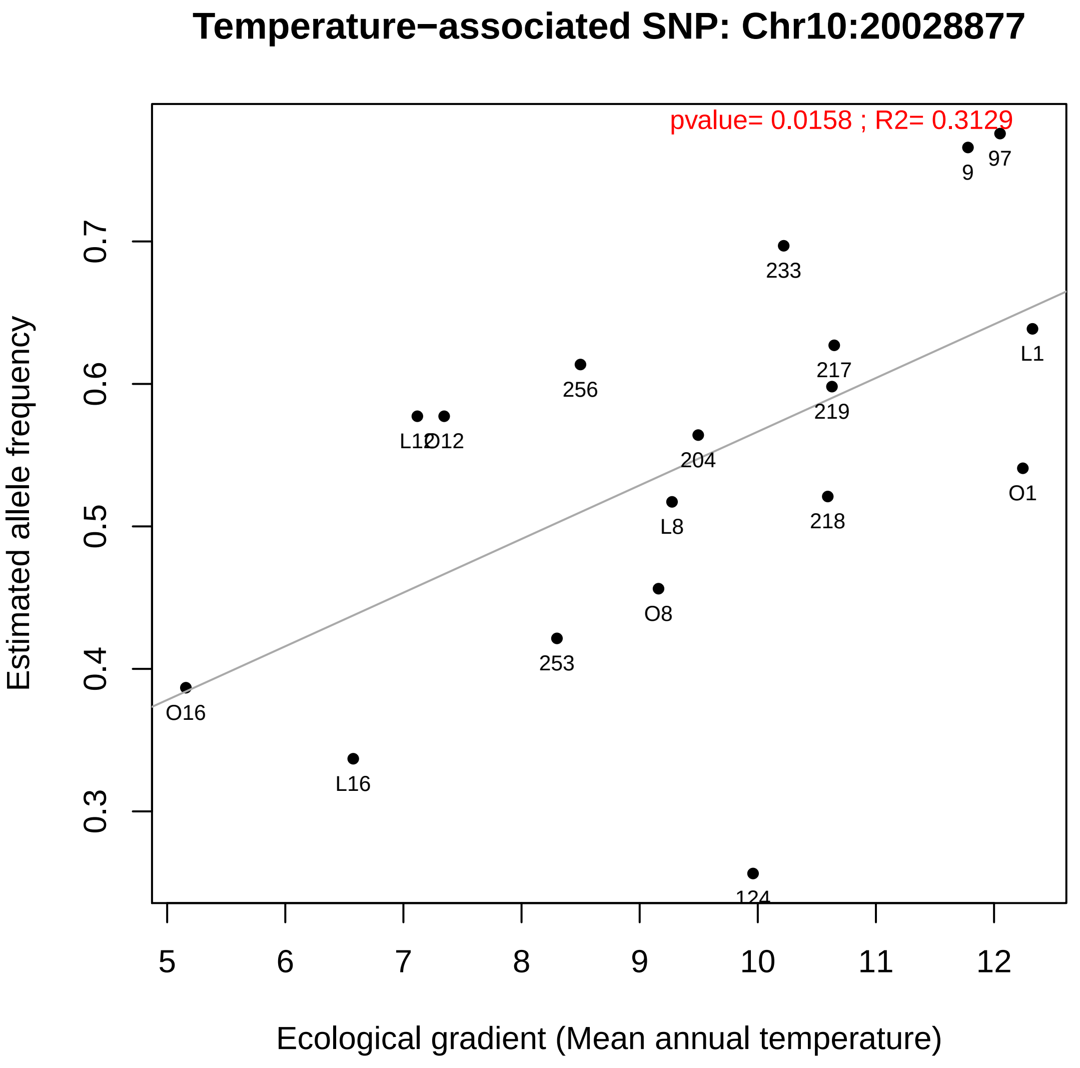

Temperature−associated SNP: Chr10:20028877
pvalue= 0.0158 ; R2= 0.3129
Estimated allele frequency
0.3
0.4
0.5
0.6
0.7
5
6
7
8
9
10
11
12
Ecological gradient (Mean annual temperature)
9
97
233
L1
217
219
256
L12
O12
204
O1
L8
218
O8
253
O16
L16
124

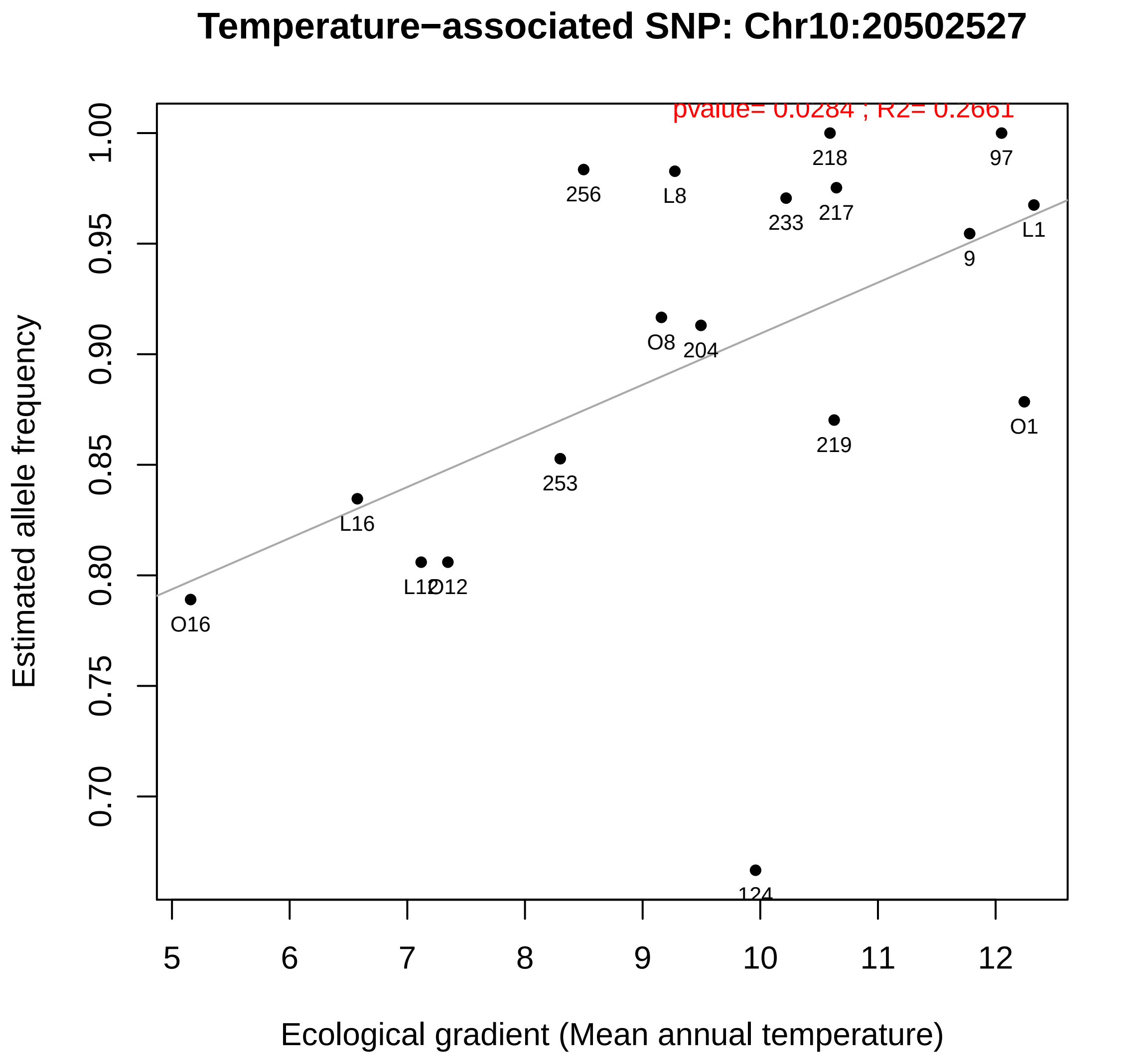

Temperature−associated SNP: Chr10:20502527
pvalue= 0.0284 ; R2= 0.2661
Estimated allele frequency
1.00
0.95
0.90
0.85
0.80
0.75
0.70
5
6
7
8
9
10
11
12
Ecological gradient (Mean annual temperature)
218
97
256
L8
233
217
L1
9
O8
204
O1
219
253
L16
L12
O12
O16
124

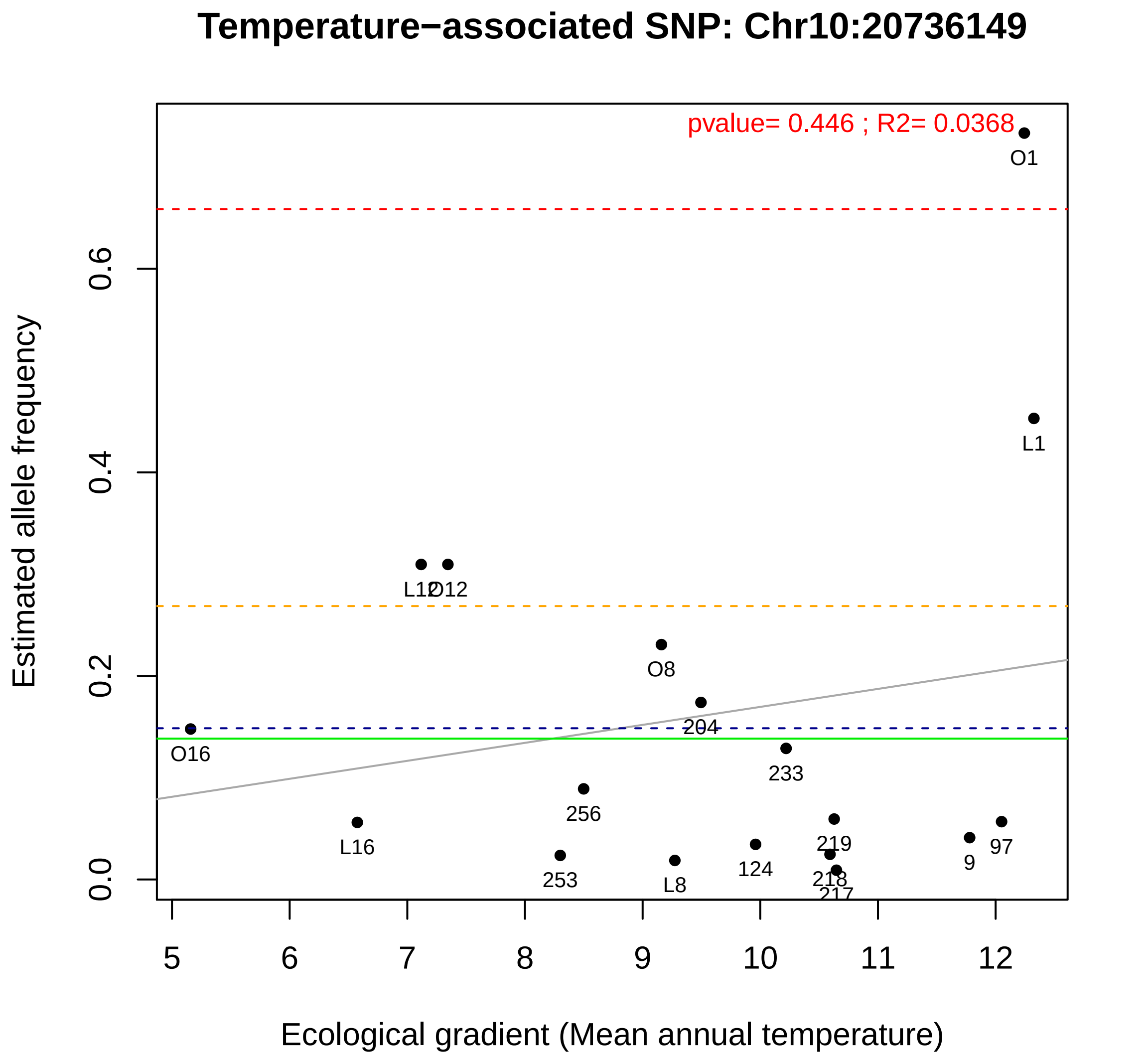
Temperature−associated SNP: Chr10:20736149
pvalue= 0.446 ; R2= 0.0368
Estimated allele frequency
0.0
0.2
0.4
0.6
5
6
7
8
9
10
11
12
Ecological gradient (Mean annual temperature)
O1
L1
L12
O12
O8
204
O16
233
256
L16
219
97
9
124
253
L8
218
217

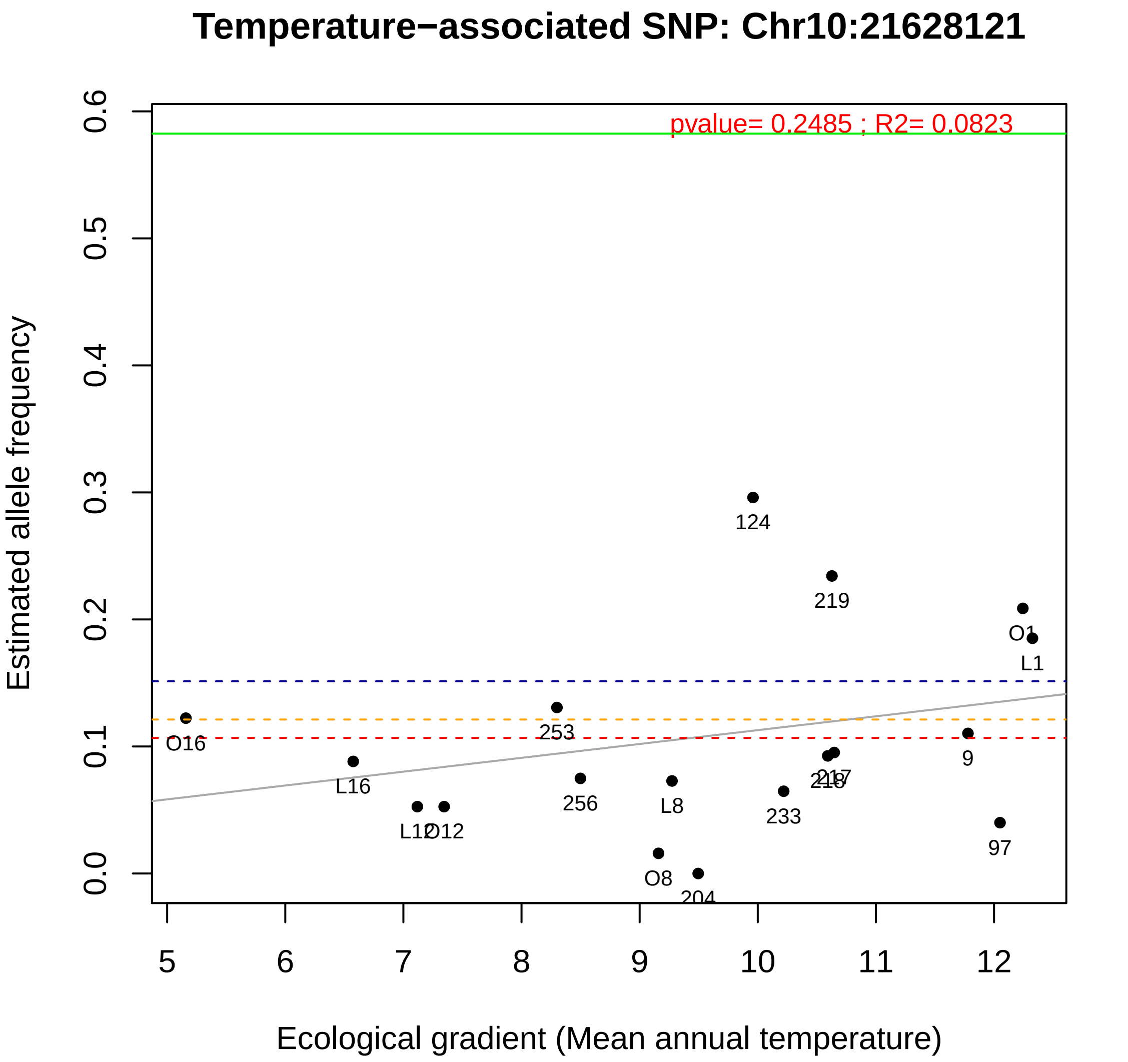
Temperature−associated SNP: Chr10:21628121
pvalue= 0.2485 ; R2= 0.0823
Estimated allele frequency
0.0
0.1
0.2
0.3
0.4
0.5
0.6
124
219
O1
L1
O16
253
9
L16
256
L8
233
L12
O12
97
O8
204
5
6
7
8
9
10
11
12
Ecological gradient (Mean annual temperature)

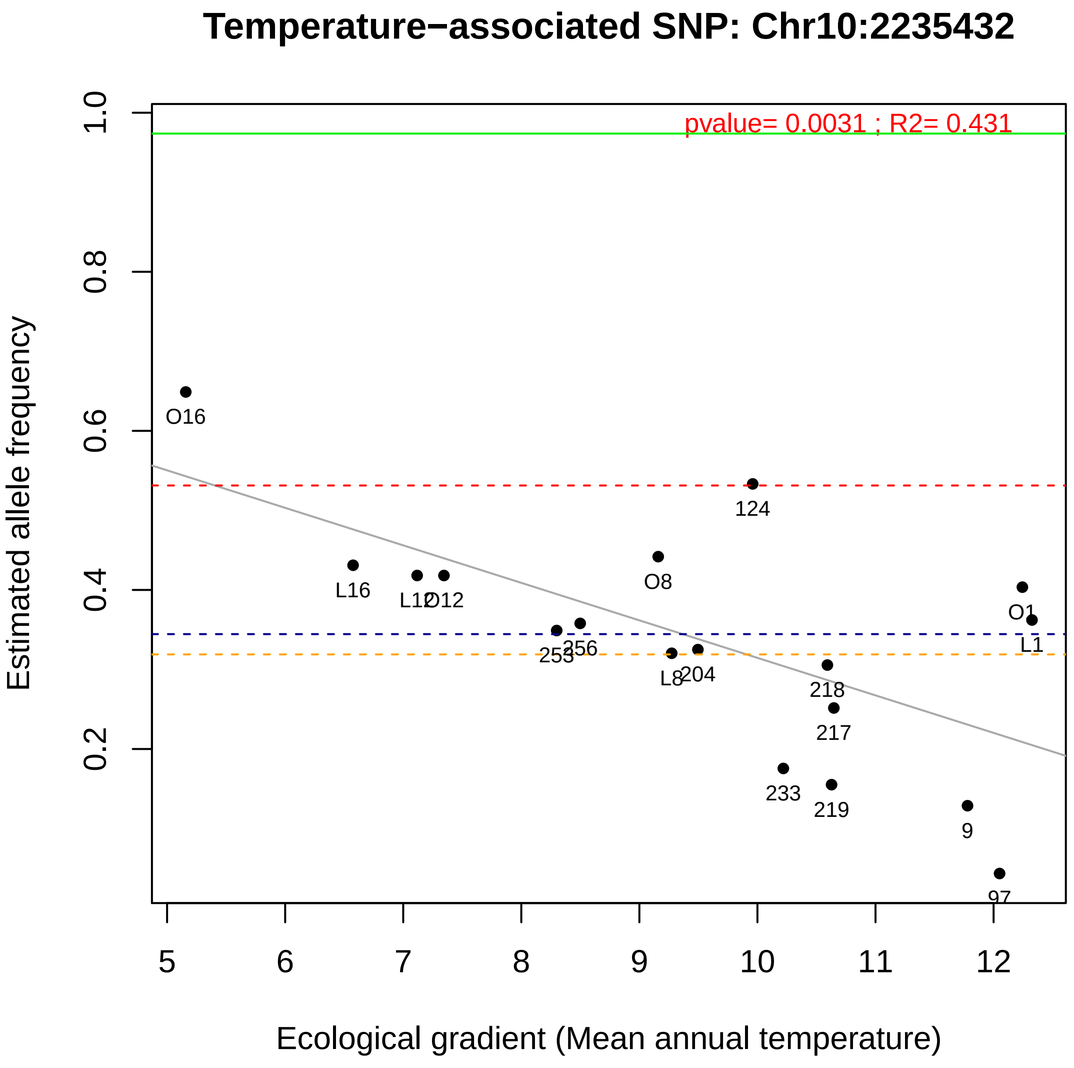

Temperature−associated SNP: Chr10:2235432
pvalue= 0.0031 ; R2= 0.431
Estimated allele frequency
1.0
0.8
0.6
0.4
0.2
O16
124
L16
L12
O12
O8
O1
L1
253
256
L8
204
218
217
233
219
9
97
5
6
7
8
9
10
11
12
Ecological gradient (Mean annual temperature)

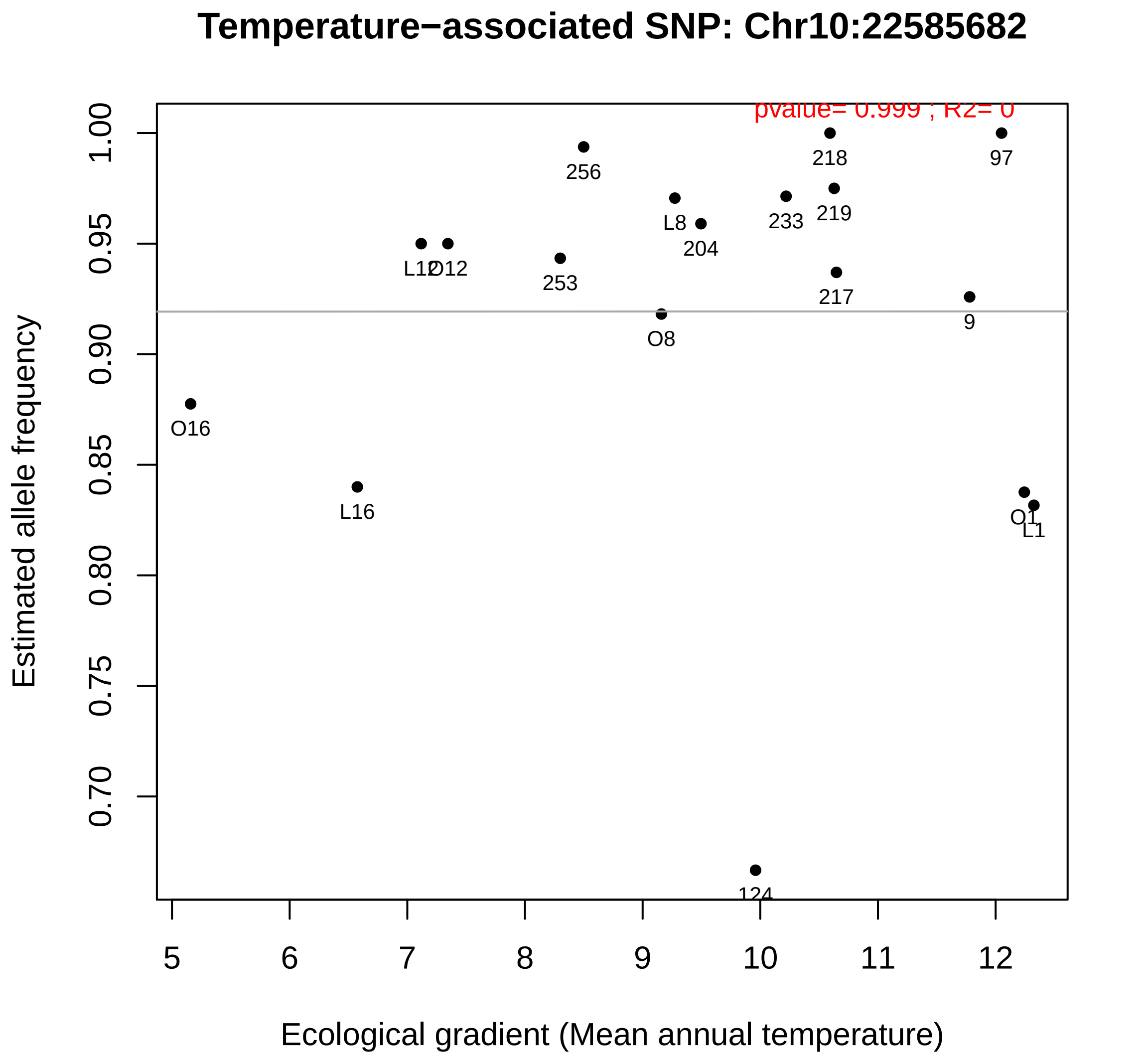
Temperature−associated SNP: Chr10:22585682
pvalue= 0.999 ; R2= 0
Estimated allele frequency
1.00
0.95
0.90
0.85
0.80
0.75
0.70
256
218
97
L8
204
233
219
L12
O12
253
217
O8
9
O16
L16
O1
L1
124
5
6
7
8
9
10
11
12
Ecological gradient (Mean annual temperature)

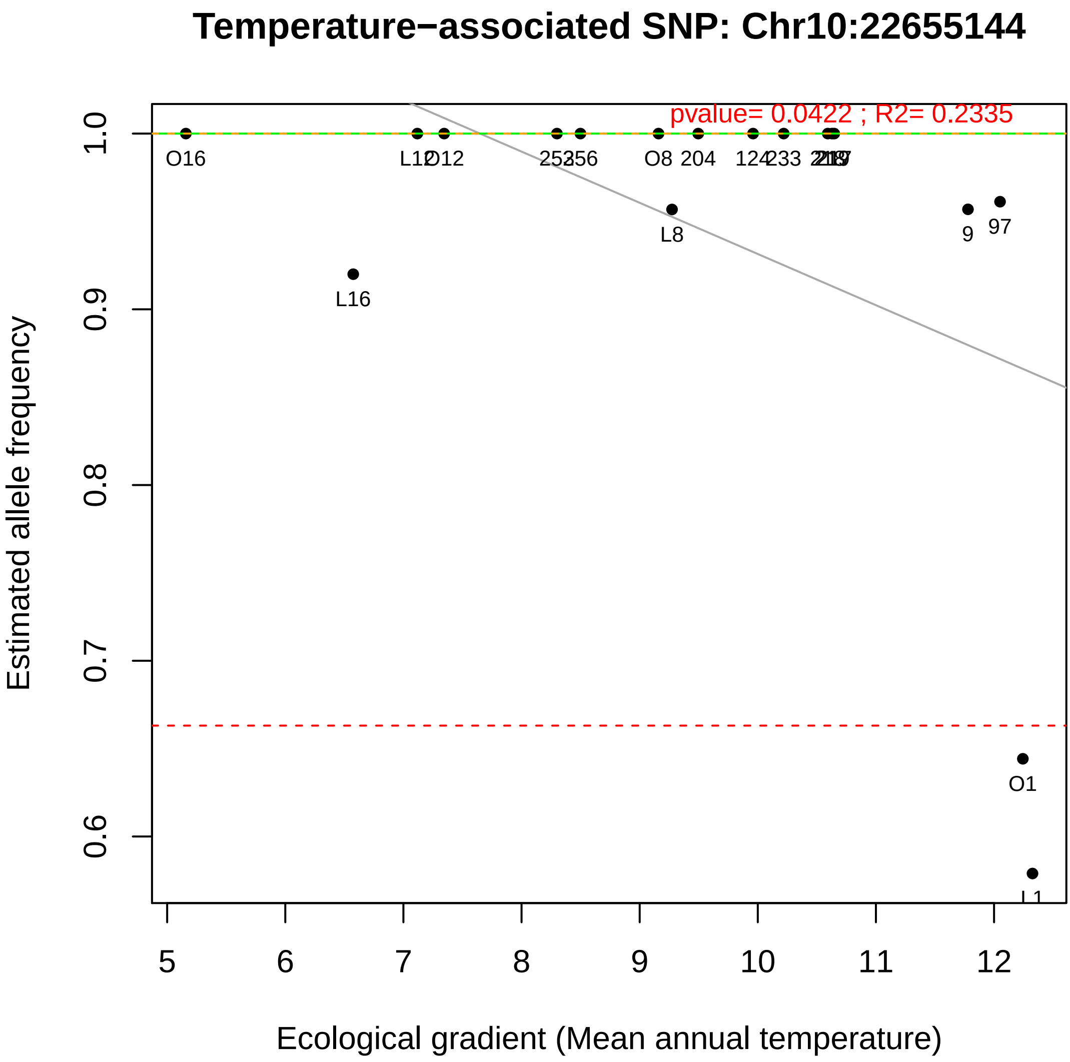

Temperature−associated SNP: Chr10:22655144
pvalue= 0.0422 ; R2= 0.2335
Estimated allele frequency
Ecological gradient (Mean annual temperature)
1.0
0.9
0.8
0.7
0.6
5
6
7
8
9
10
11
12
O16
L16
O12
L8
204
124
233
9
97
O1

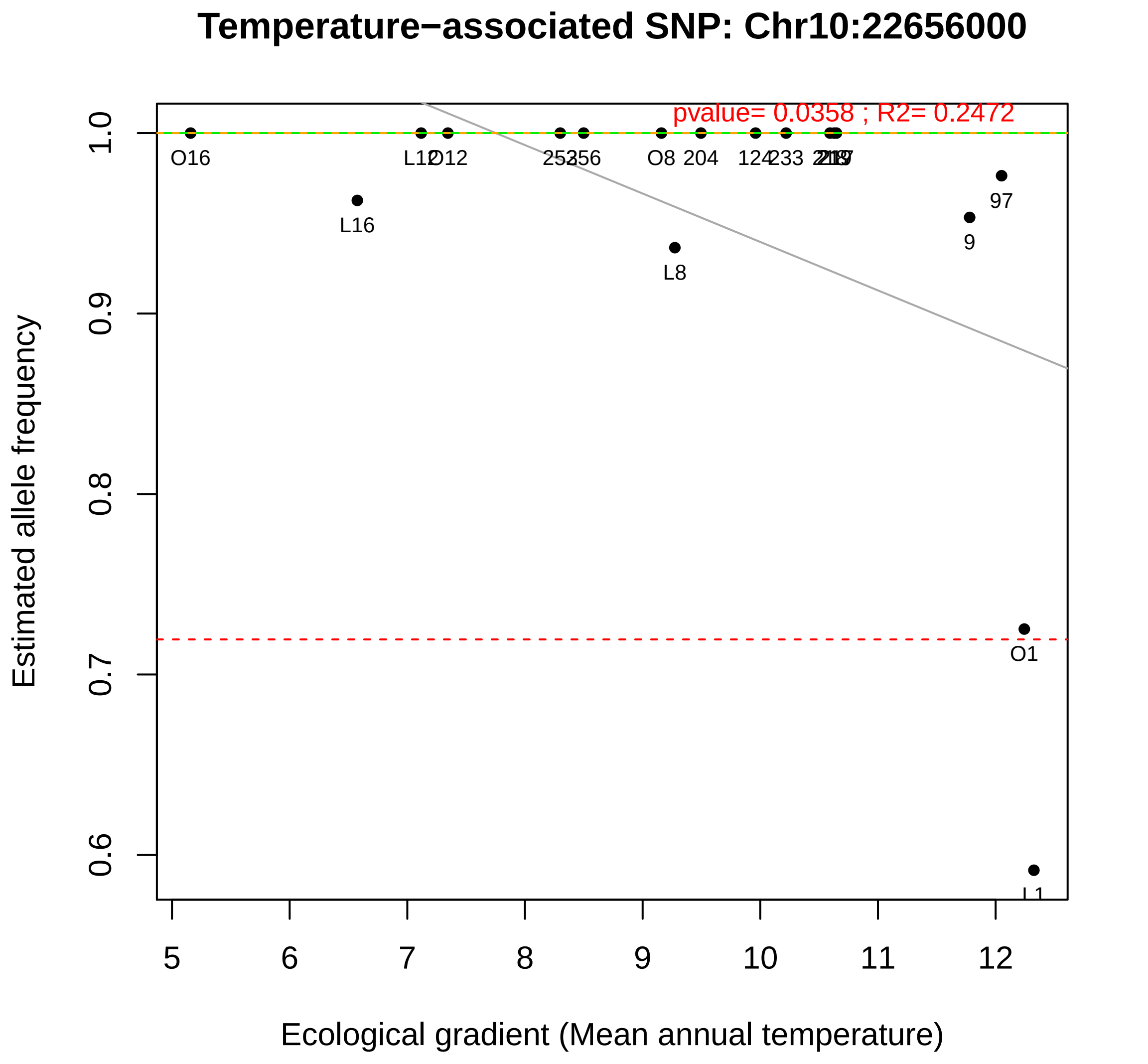

Temperature−associated SNP: Chr10:22656000
pvalue= 0.0358 ; R2= 0.2472
Estimated allele frequency
Ecological gradient (Mean annual temperature)
1.0
0.9
0.8
0.7
0.6
5
6
7
8
9
10
11
12
O16
L16
L12
O12
253
256
O8
204
L8
124
233
97
9
O1

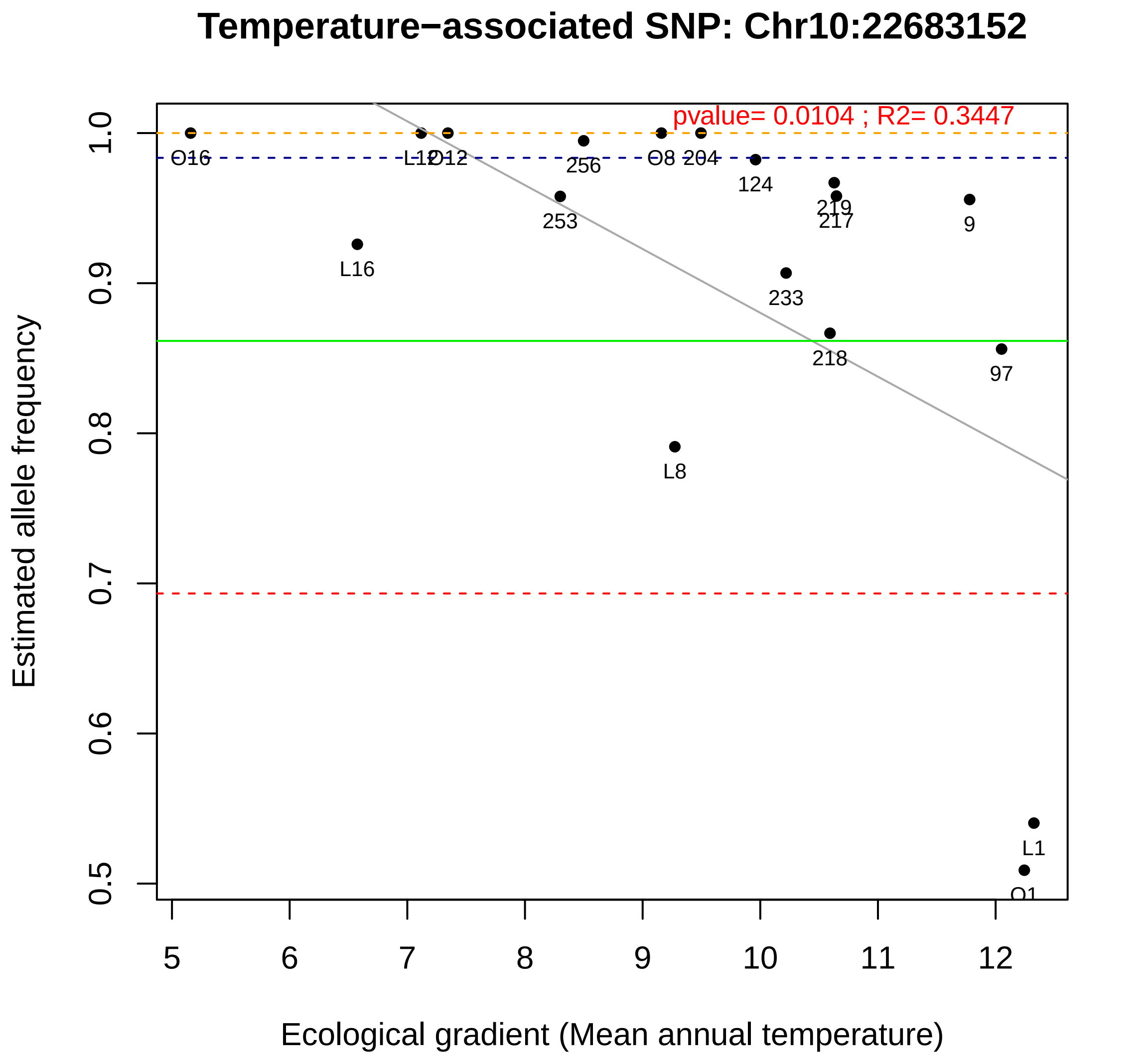
Temperature−associated SNP: Chr10:22683152
pvalue= 0.0104 ; R2= 0.3447
Estimated allele frequency
Ecological gradient (Mean annual temperature)
1.0
0.9
0.8
0.7
0.6
0.5
5
6
7
8
9
10
11
12
O16
L12
O12
256
O8
204
124
253
219
217
9
L16
233
218
97
L8
L1
O1

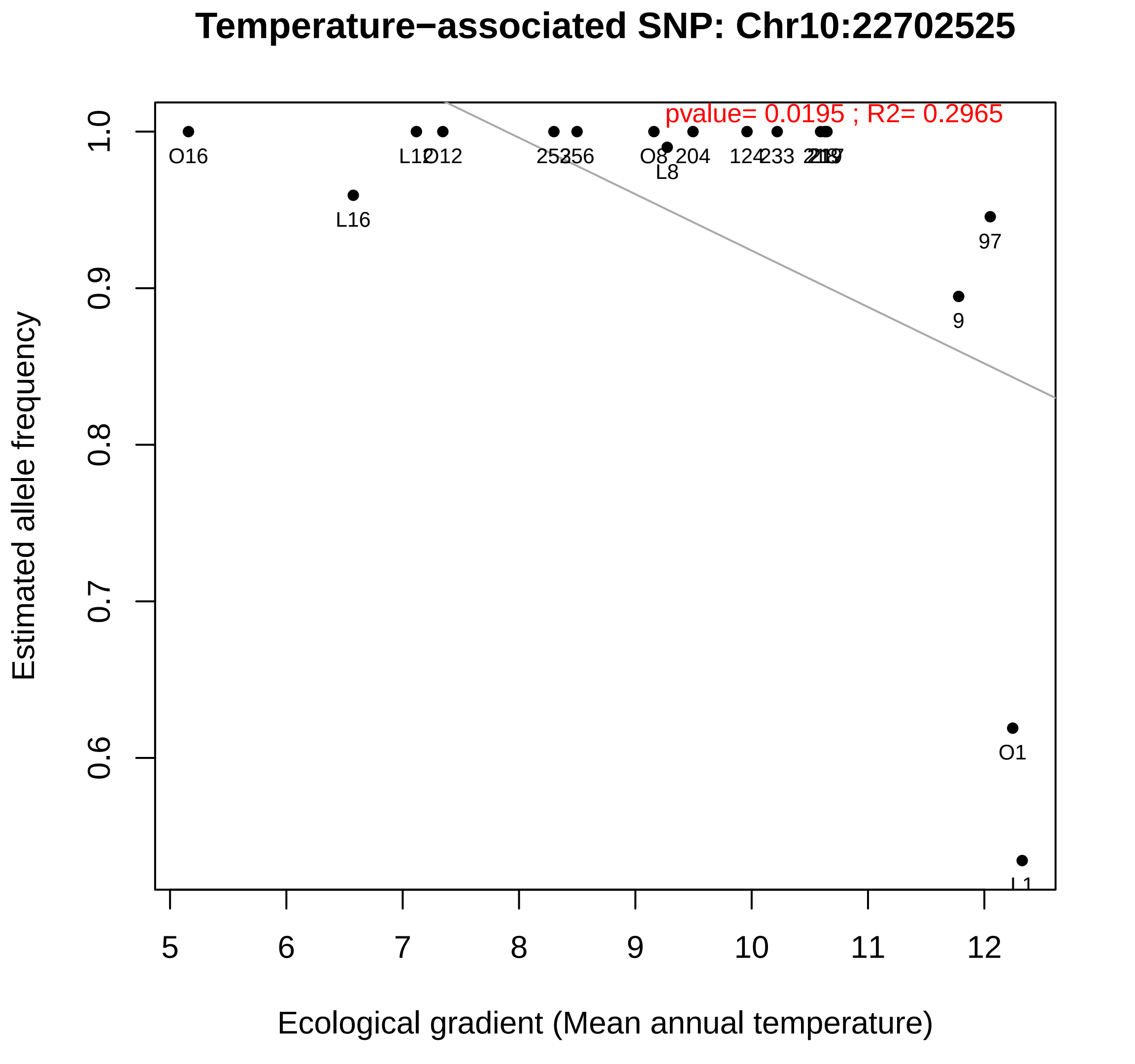

Temperature−associated SNP: Chr10:22702525
pvalue= 0.0195 ; R2= 0.2965
Estimated allele frequency
Ecological gradient (Mean annual temperature)
0.6
0.7
0.8
0.9
1.0
5
6
7
8
9
10
11
12
O16
L16
L12
O12
204
124
233
97
9
O1
L8
O8

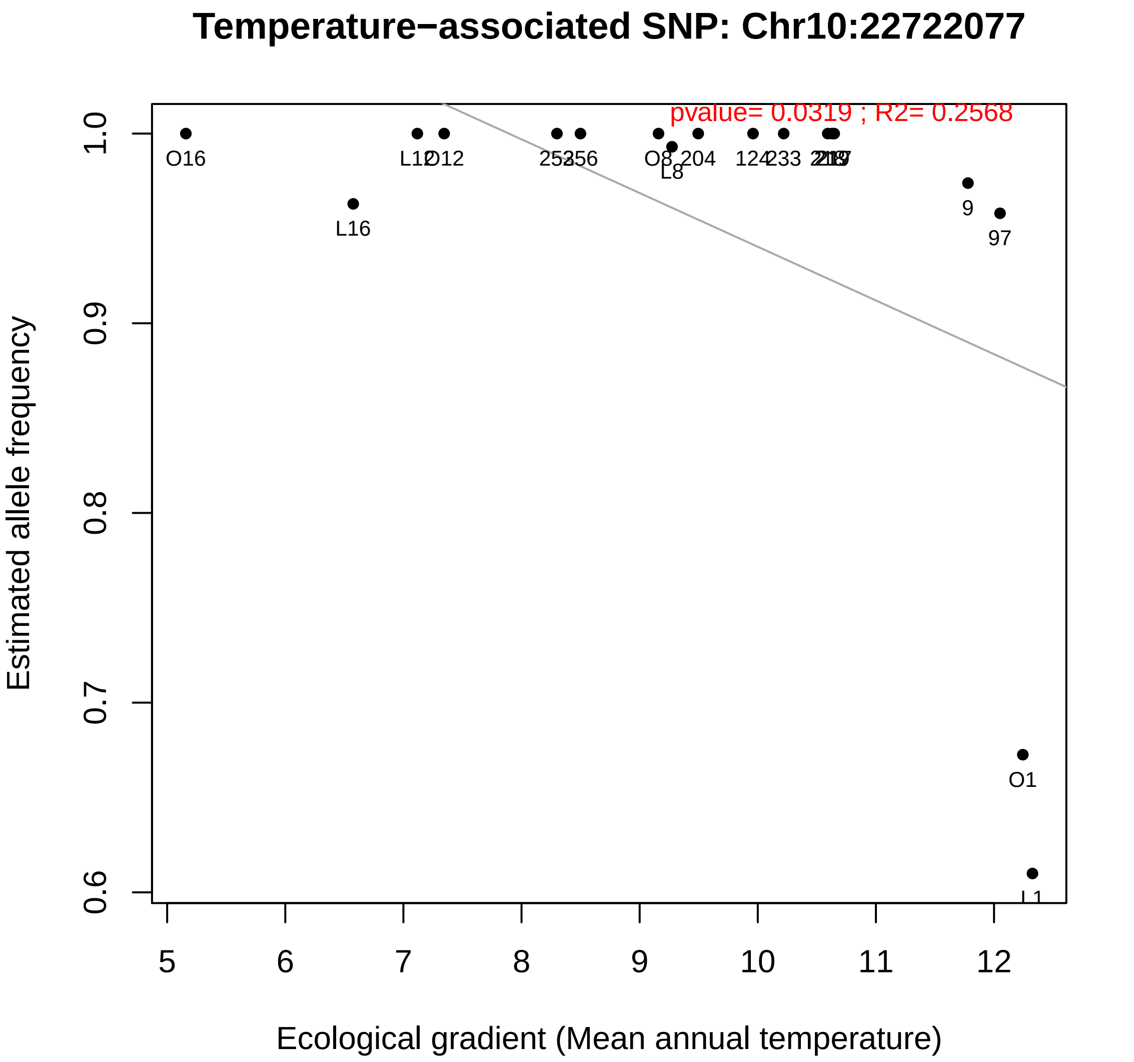

Temperature−associated SNP: Chr10:22722077
pvalue= 0.0319 ; R2= 0.2568
Estimated allele frequency
Ecological gradient (Mean annual temperature)
1.0
0.9
0.8
0.7
0.6
5
6
7
8
9
10
11
12
O16
L16
L12
O12
O8
L8
204
124
233
9
97
O1

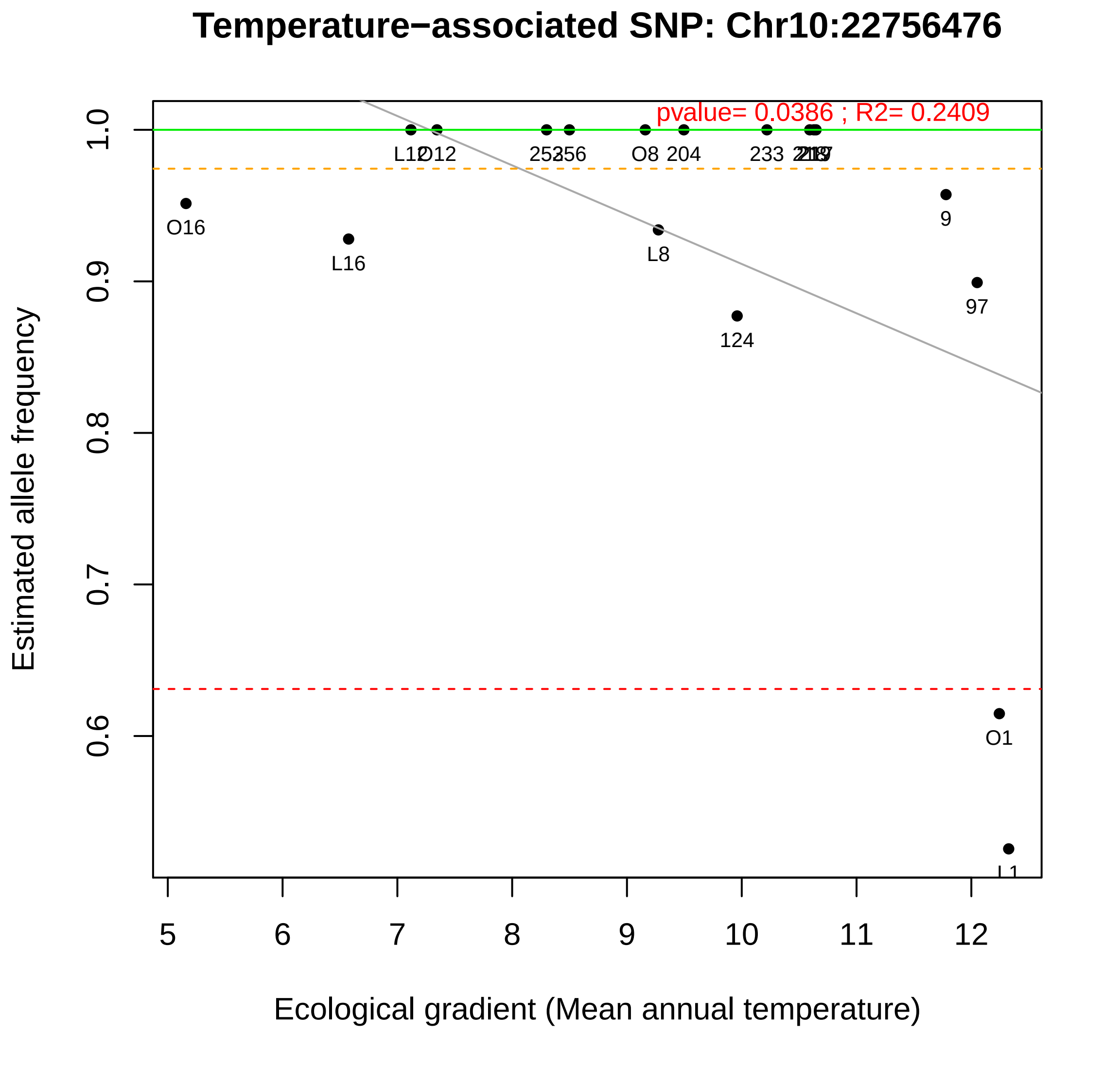

Temperature−associated SNP: Chr10:22756476
pvalue= 0.0386 ; R2= 0.2409
Estimated allele frequency
Ecological gradient (Mean annual temperature)
1.0
0.9
0.8
0.7
0.6
5
6
7
8
9
10
11
12
O16
L16
L12
O12
256
O8
204
233
L8
124
9
97
O1

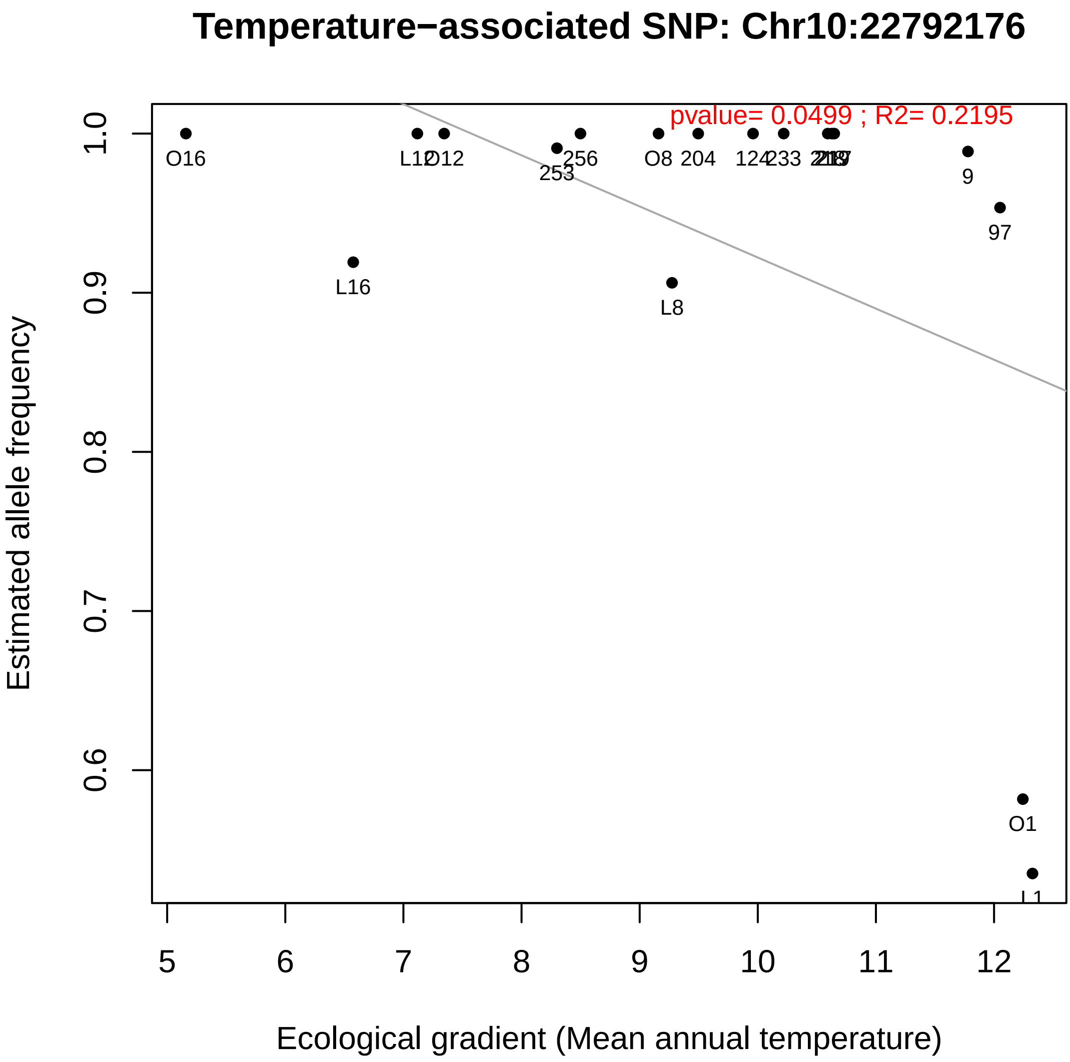
Temperature−associated SNP: Chr10:22792176
pvalue= 0.0499 ; R2= 0.2195
Estimated allele frequency
Ecological gradient (Mean annual temperature)
1.0
0.9
0.8
0.7
0.6
5
6
7
8
9
10
11
12
O16
L12
O12
253
256
O8
204
124
233
9
97
L16
L8
O1

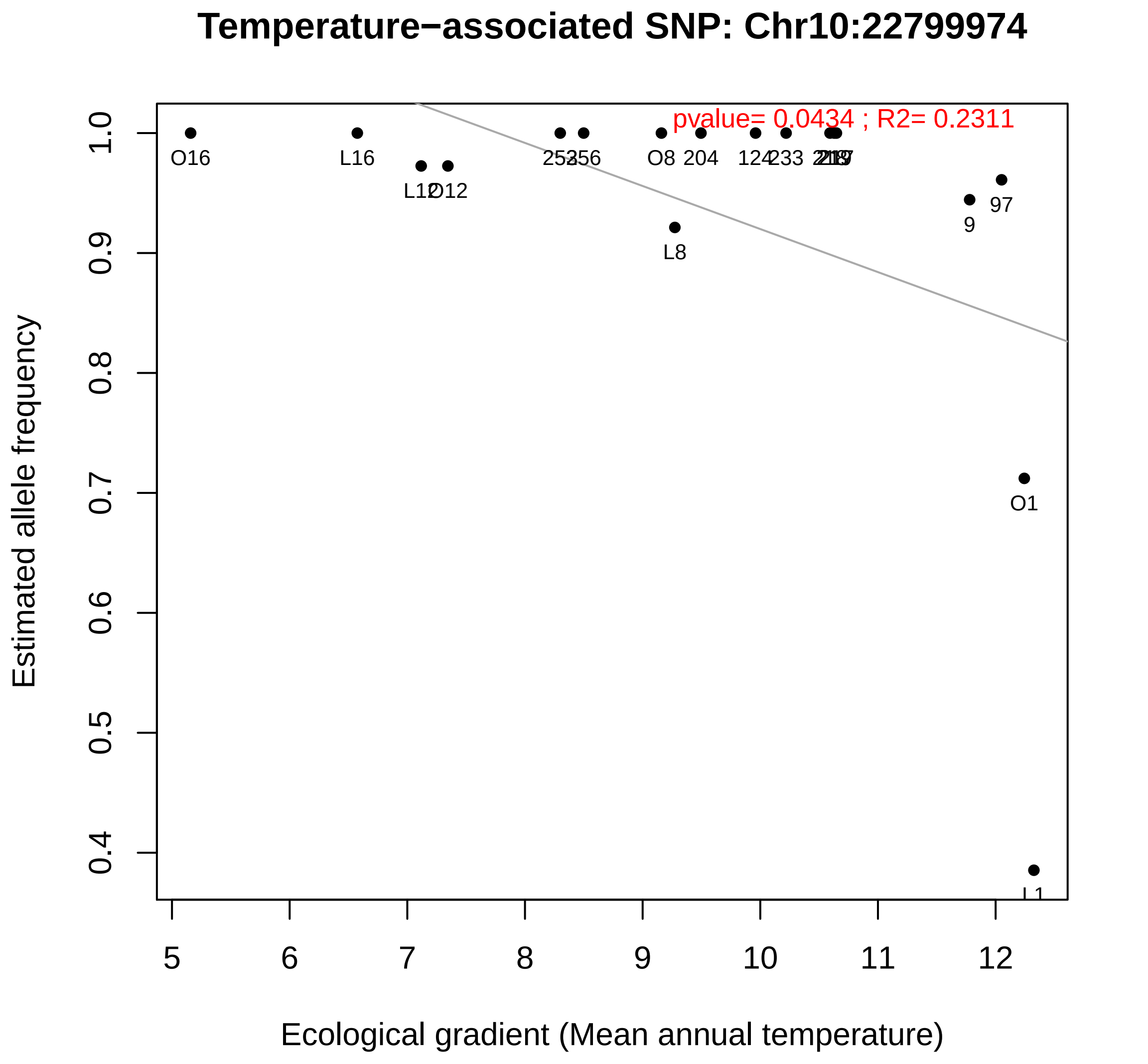

Temperature−associated SNP: Chr10:22799974
pvalue= 0.0434 ; R2= 0.2311
Estimated allele frequency
Ecological gradient (Mean annual temperature)
1.0
0.9
0.8
0.7
0.6
0.5
0.4
5
6
7
8
9
10
11
12
O16
L16
L12
O12
O8
204
L8
97
9
O1

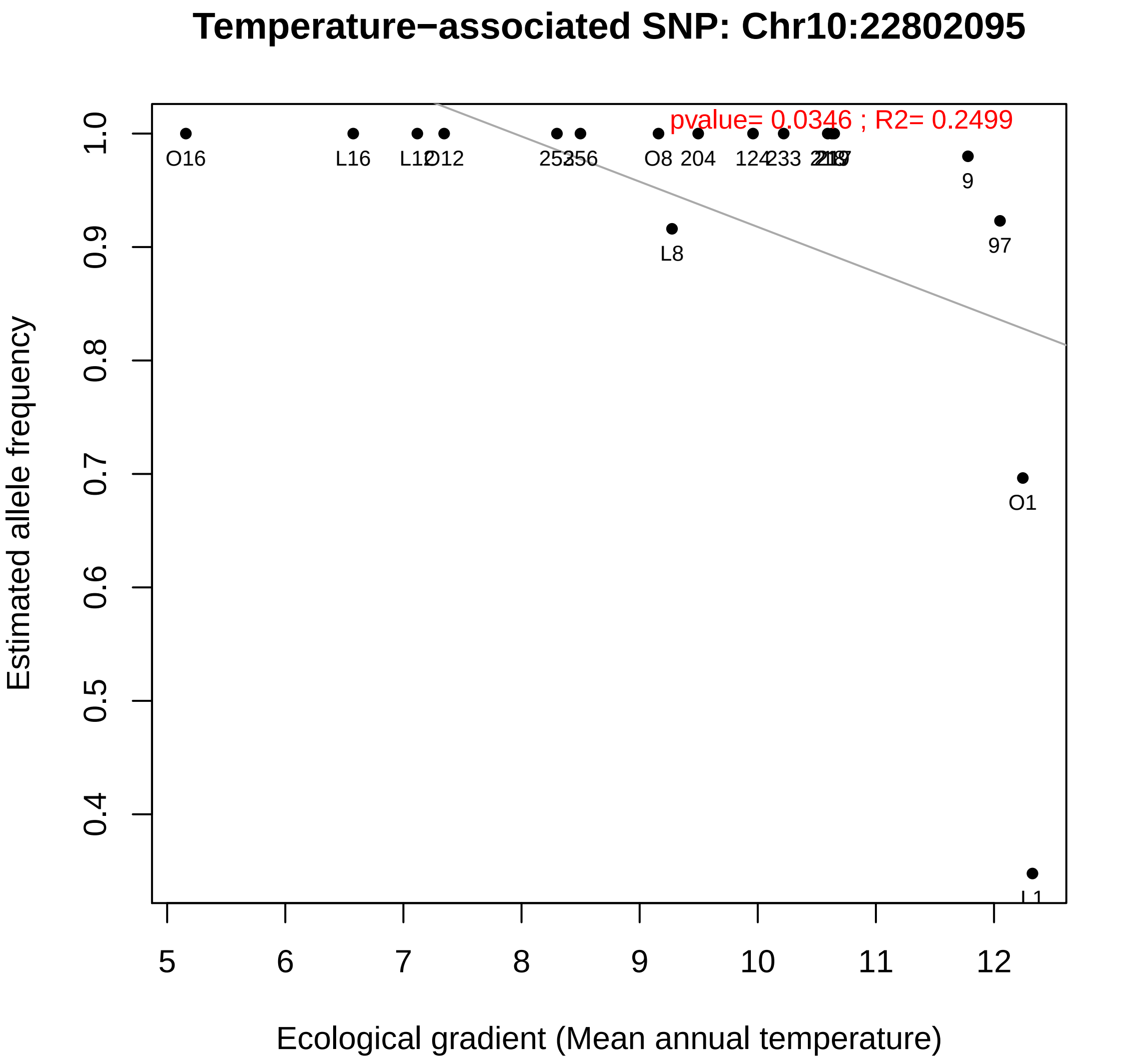

Temperature−associated SNP: Chr10:22802095
pvalue= 0.0346 ; R2= 0.2499
Estimated allele frequency
Ecological gradient (Mean annual temperature)
0.4
0.5
0.6
0.7
0.8
0.9
1.0
5
6
7
8
9
10
11
12
O16
L16
L12
O12
O8
204
L8
9
97
O1

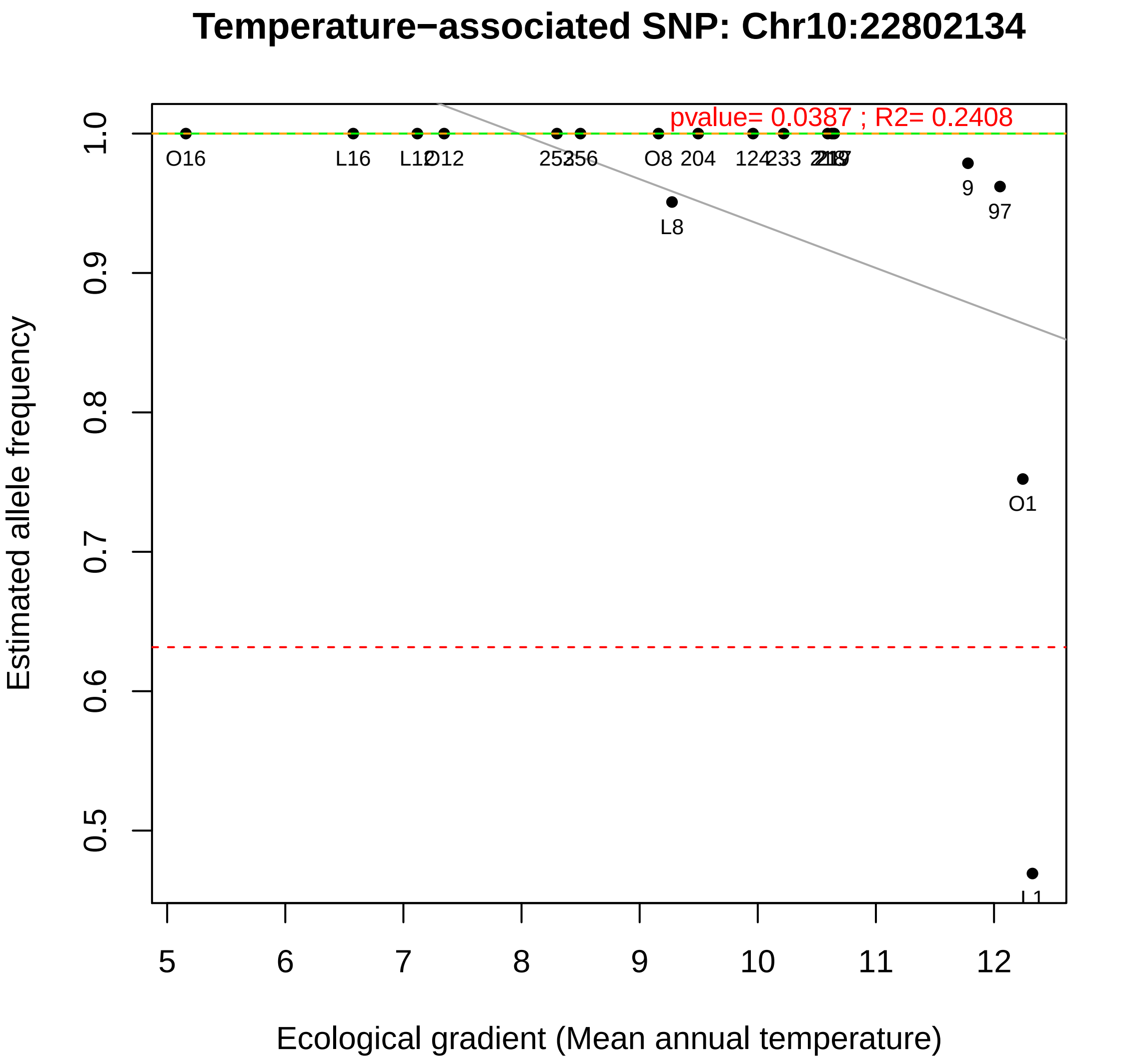

Temperature−associated SNP: Chr10:22802134
pvalue= 0.0387 ; R2= 0.2408
O16
L16
O12
204
124
233
9
97
L8
O1
Estimated allele frequency
1.0
0.9
0.8
0.7
0.6
0.5
5
6
7
8
9
10
11
12
Ecological gradient (Mean annual temperature)

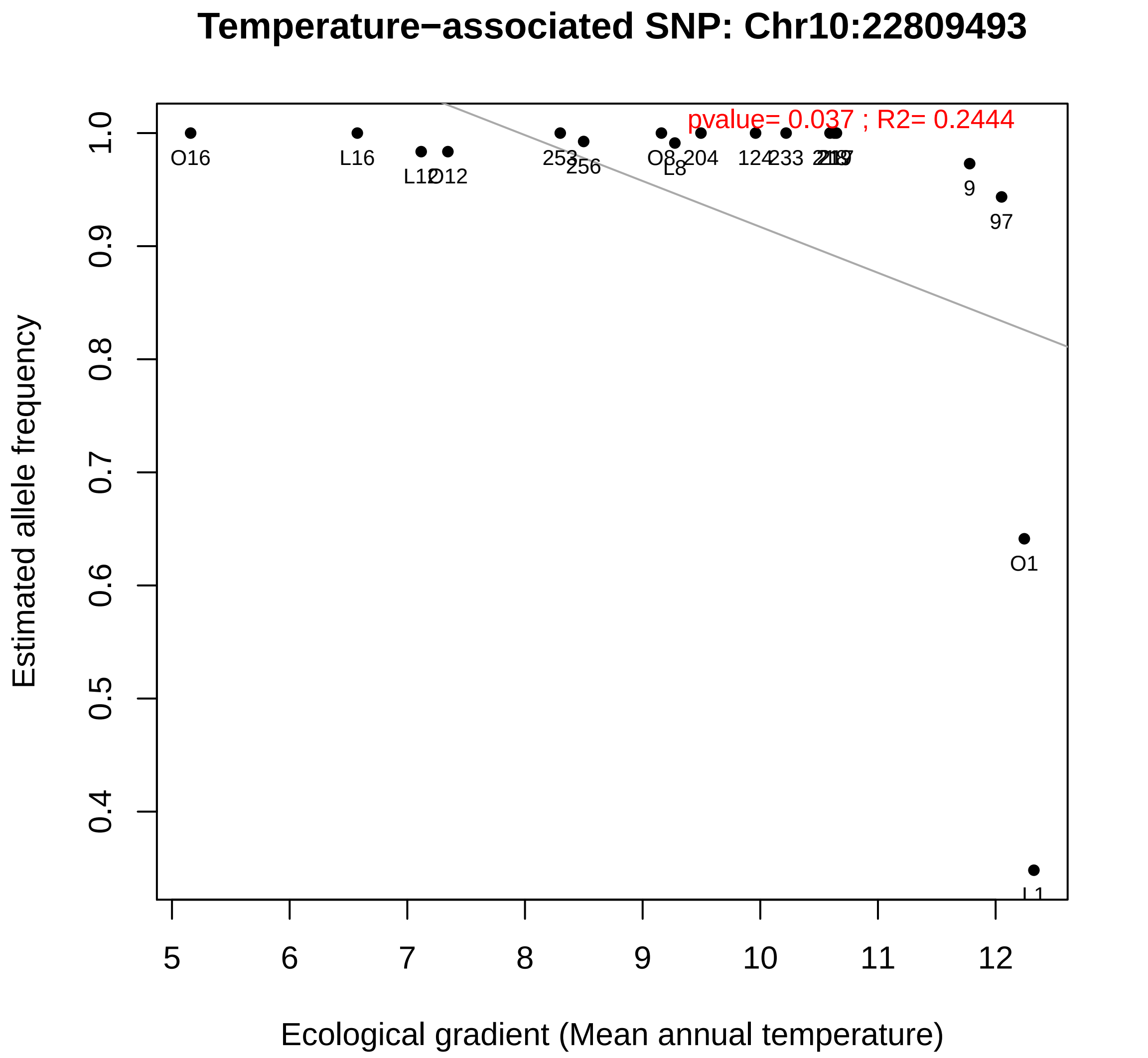

Temperature−associated SNP: Chr10:22809493
pvalue= 0.037 ; R2= 0.2444
O16
L16
L12
O12
253
256
O8
L8
204
124
233
9
97
O1
Estimated allele frequency
1.0
0.9
0.8
0.7
0.6
0.5
0.4
5
6
7
8
9
10
11
12
Ecological gradient (Mean annual temperature)

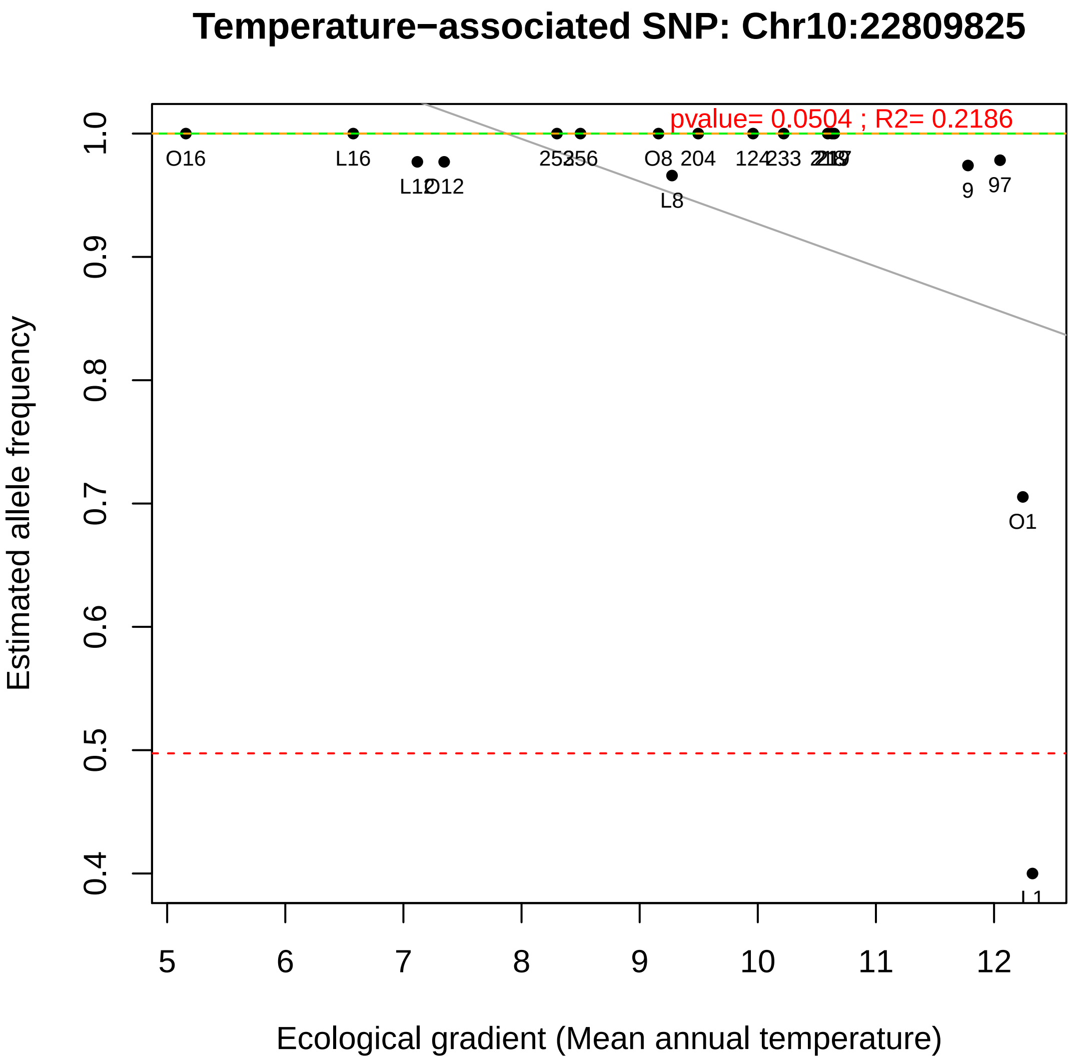

Temperature−associated SNP: Chr10:22809825
pvalue= 0.0504 ; R2= 0.2186
Estimated allele frequency
Ecological gradient (Mean annual temperature)
0.4
0.5
0.6
0.7
0.8
0.9
1.0
5
6
7
8
9
10
11
12
O16
L16
L12
O12
O8
204
124
233
L8
9
97
O1

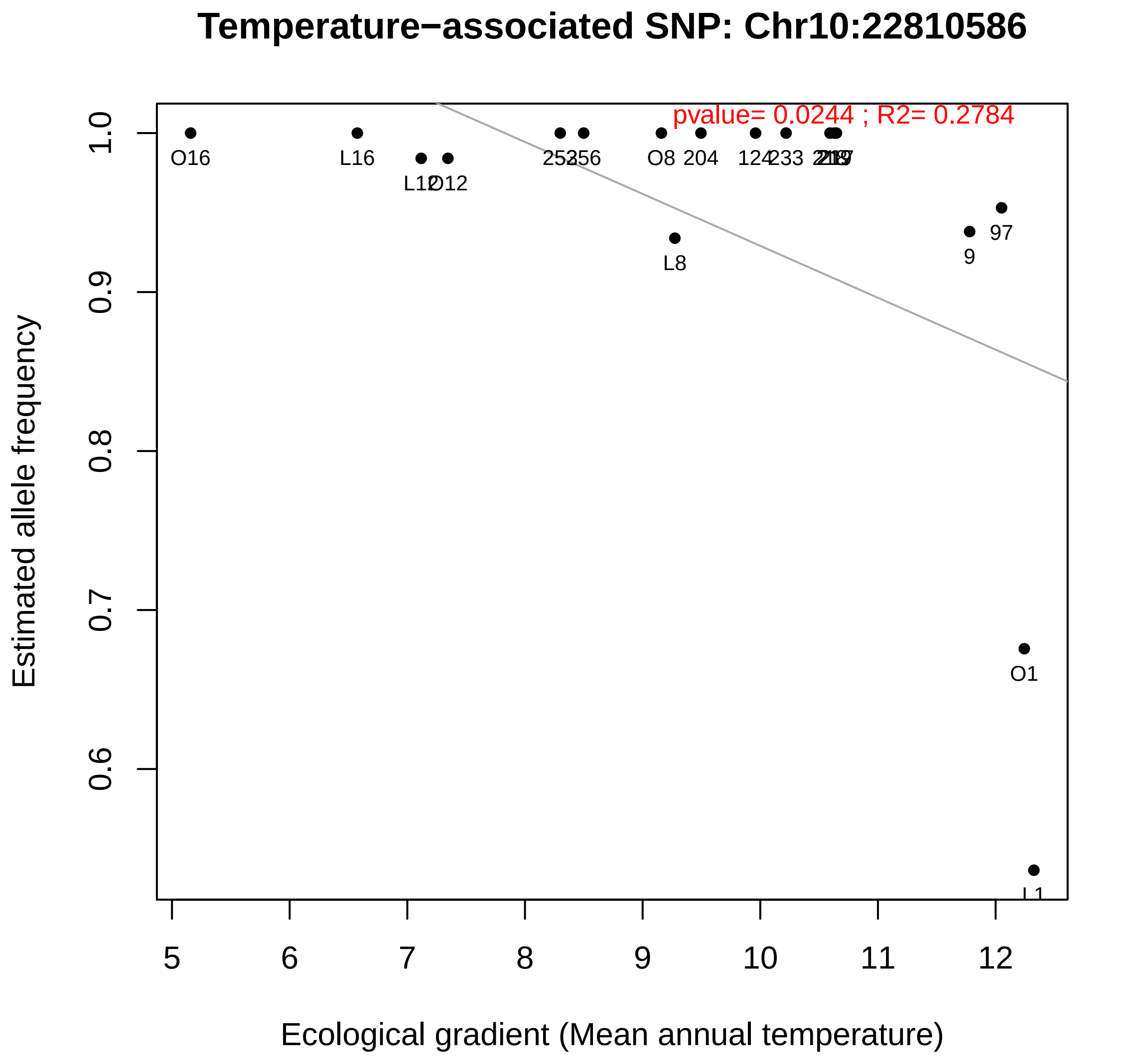
Temperature−associated SNP: Chr10:22810586
pvalue= 0.0244 ; R2= 0.2784
Estimated allele frequency
Ecological gradient (Mean annual temperature)
1.0
0.9
0.8
0.7
0.6
5
6
7
8
9
10
11
12
O16
L16
L12
O12
253
256
O8
204
124
233
L8
9
97
O1

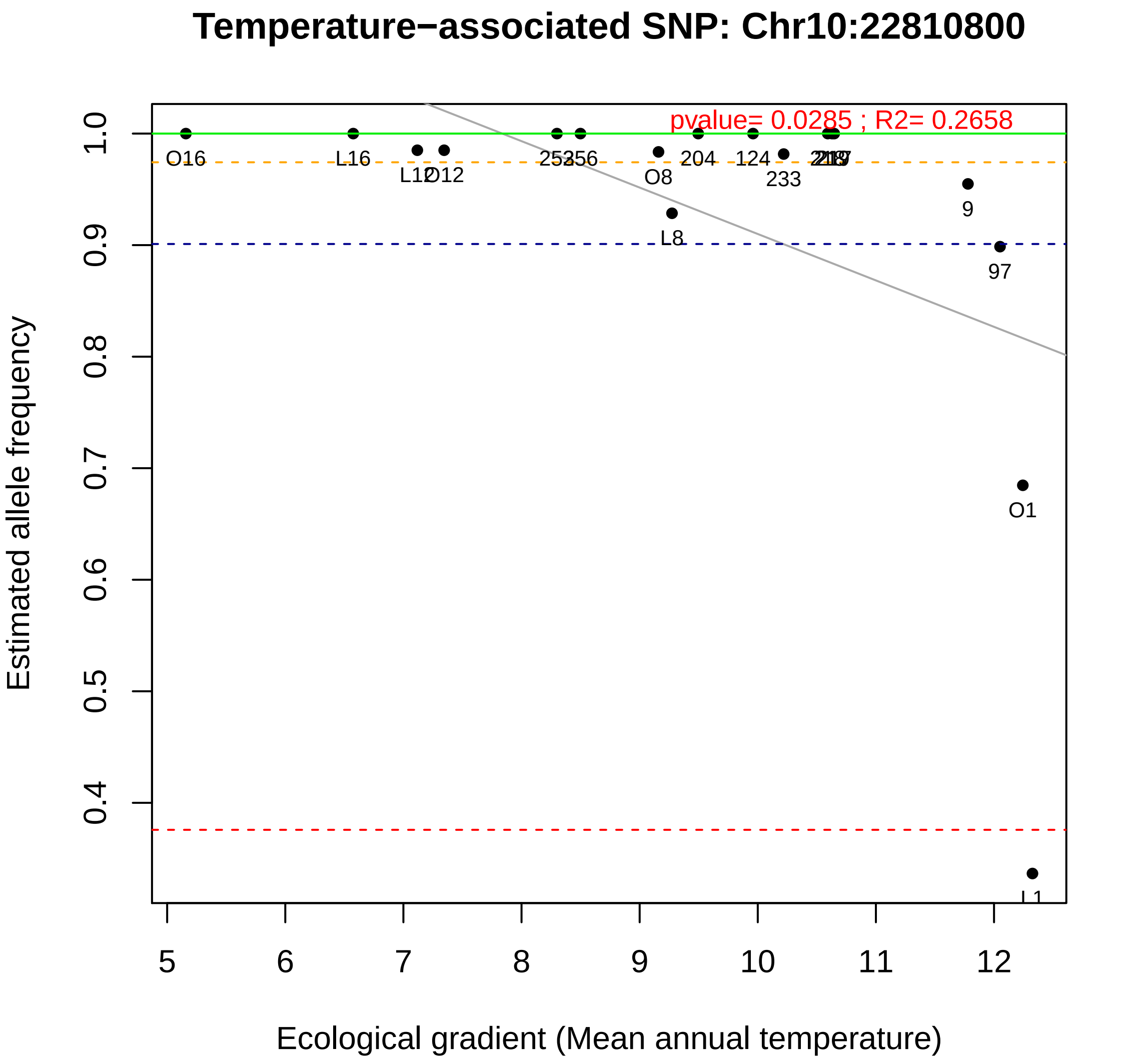

Temperature−associated SNP: Chr10:22810800
pvalue= 0.0285 ; R2= 0.2658
Estimated allele frequency
Ecological gradient (Mean annual temperature)
1.0
0.9
0.8
0.7
0.6
0.5
0.4
5
6
7
8
9
10
11
12
O16
L16
L12
O12
253
256
O8
L8
204
124
233
9
97
O1

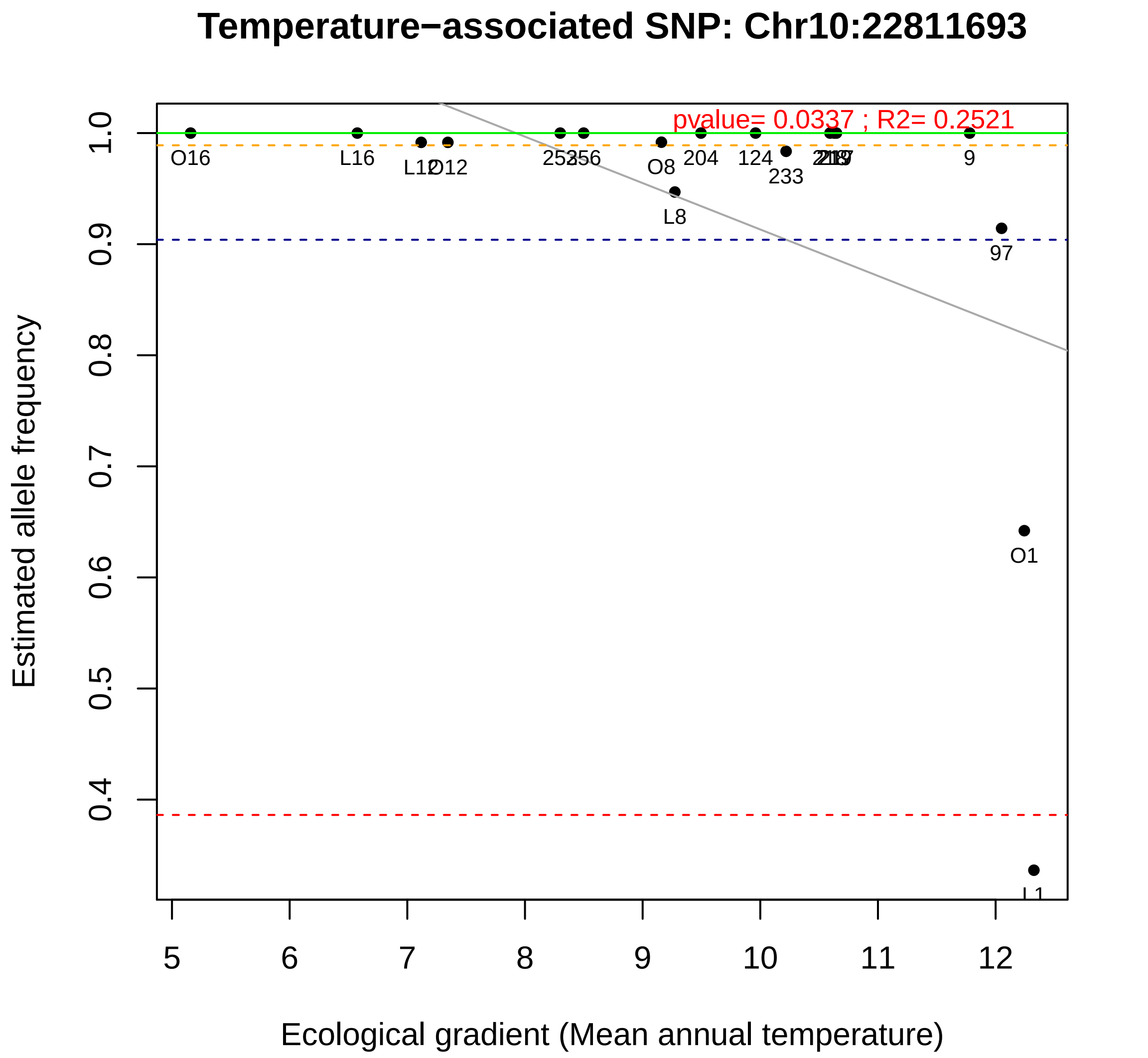

Temperature−associated SNP: Chr10:22811693
pvalue= 0.0337 ; R2= 0.2521
Estimated allele frequency
Ecological gradient (Mean annual temperature)
1.0
0.9
0.8
0.7
0.6
0.5
0.4
5
6
7
8
9
10
11
12
O16
L16
L12
O12
253
256
O8
204
124
233
L8
9
97
O1

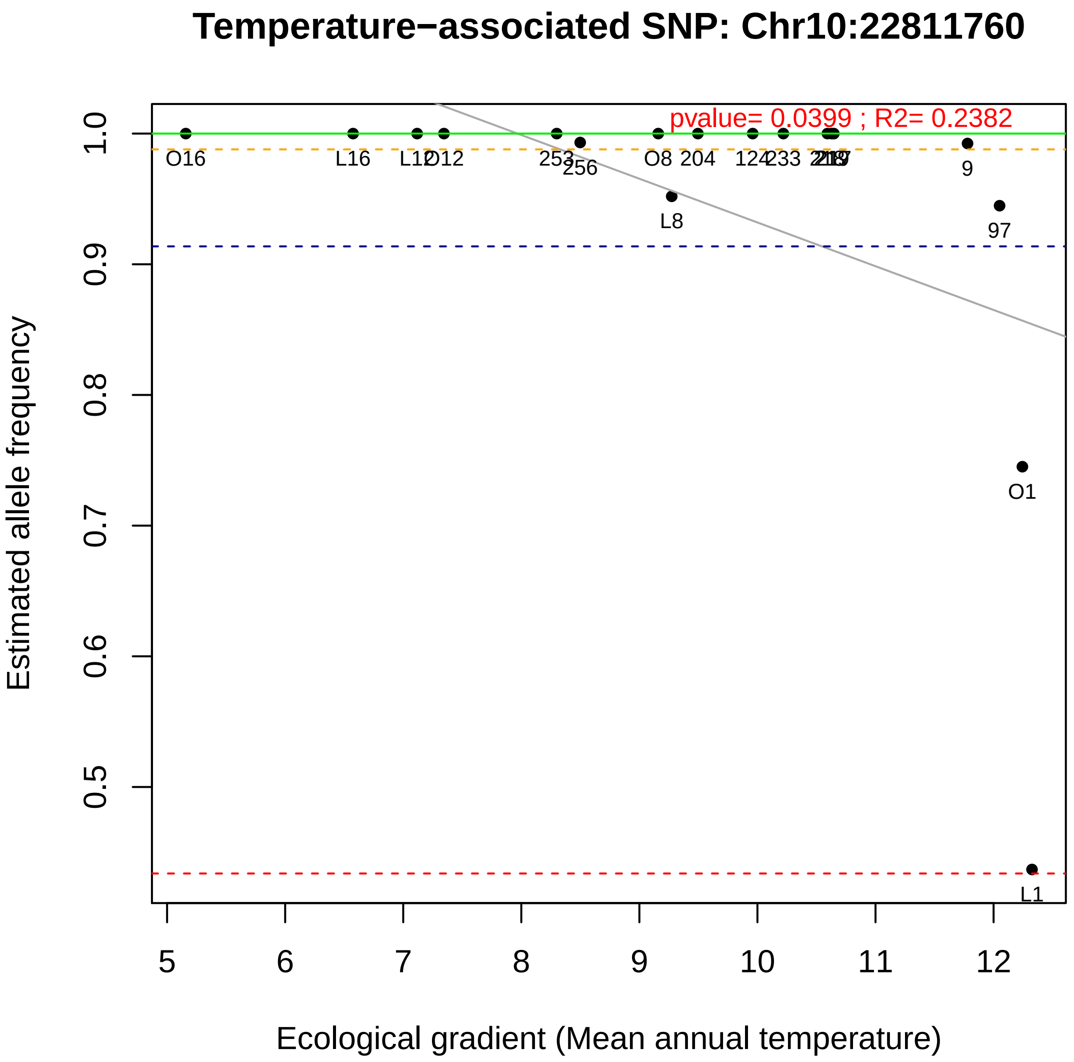

Temperature−associated SNP: Chr10:22811760
pvalue= 0.0399 ; R2= 0.2382
O16
L16
L12
O12
253
256
O8
204
L8
124
233
9
97
O1
L1
1.0
0.9
0.8
0.7
0.6
0.5
5
6
7
8
9
10
11
12
Estimated allele frequency
Ecological gradient (Mean annual temperature)

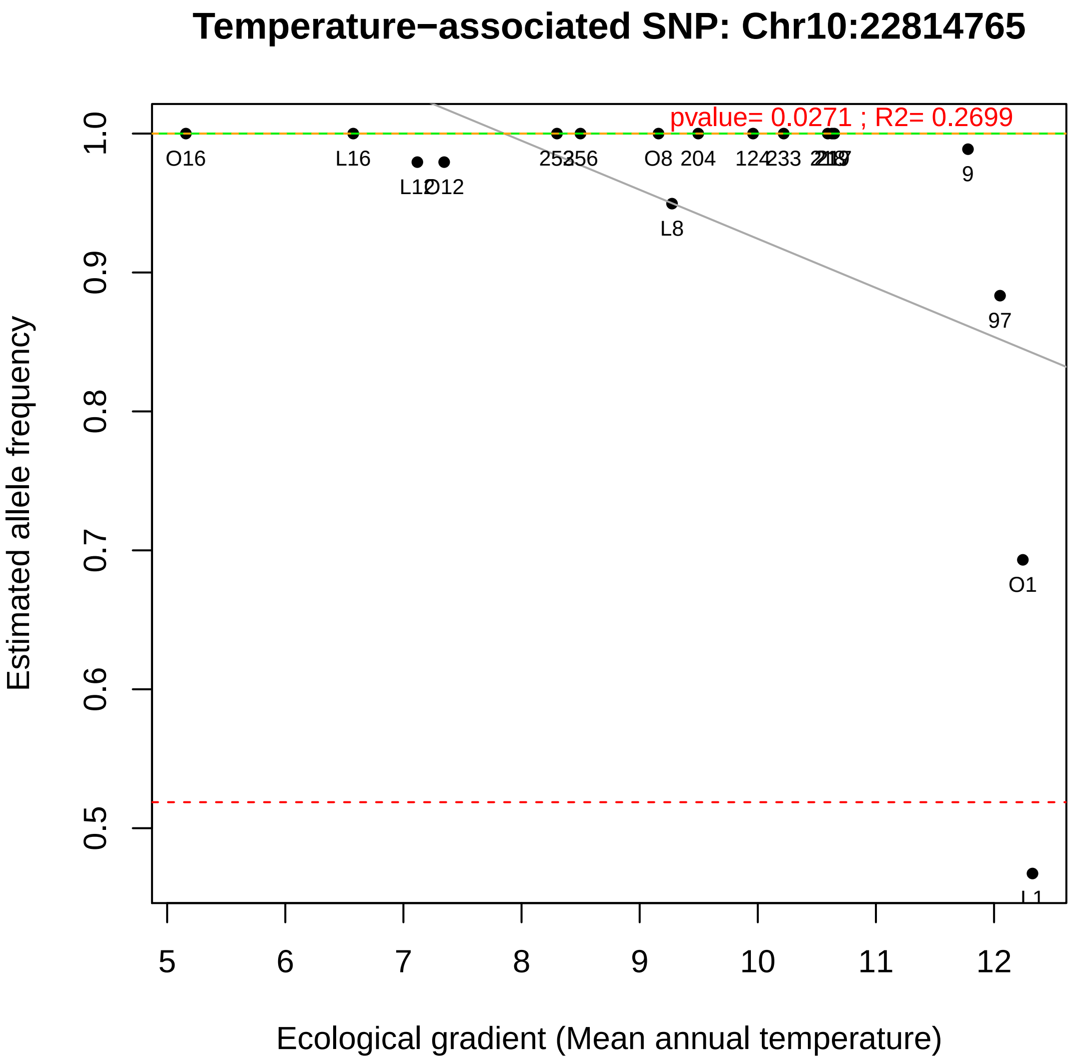

Temperature−associated SNP: Chr10:22814765
pvalue= 0.0271 ; R2= 0.2699
O16
L16
L12
O12
O8
204
L8
9
97
O1
Estimated allele frequency
Ecological gradient (Mean annual temperature)

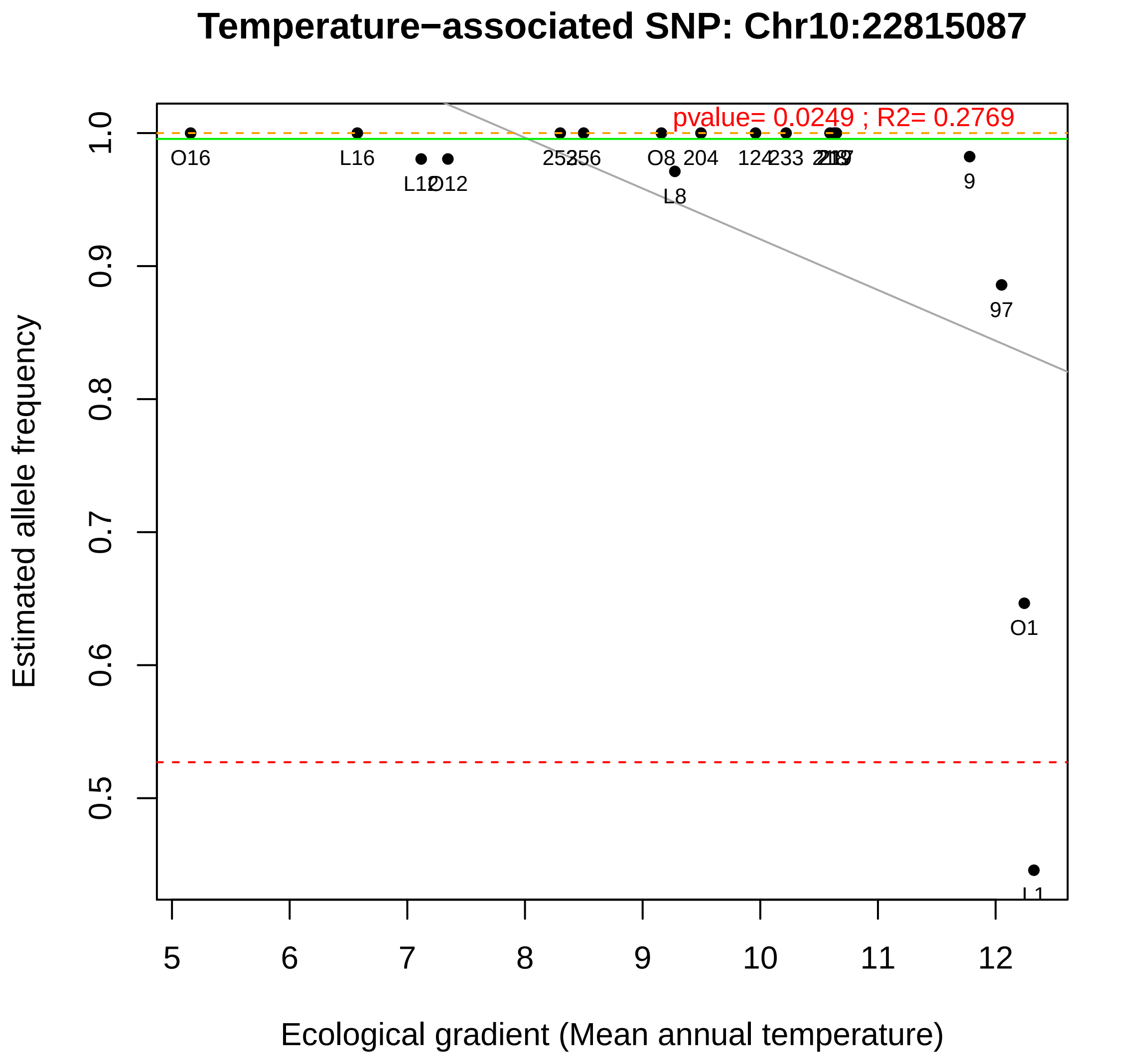

Temperature−associated SNP: Chr10:22815087
pvalue= 0.0249 ; R2= 0.2769
Estimated allele frequency
Ecological gradient (Mean annual temperature)
1.0
0.9
0.8
0.7
0.6
0.5
5
6
7
8
9
10
11
12
O16
L16
L12
O12
O8
204
L8
9
97
O1

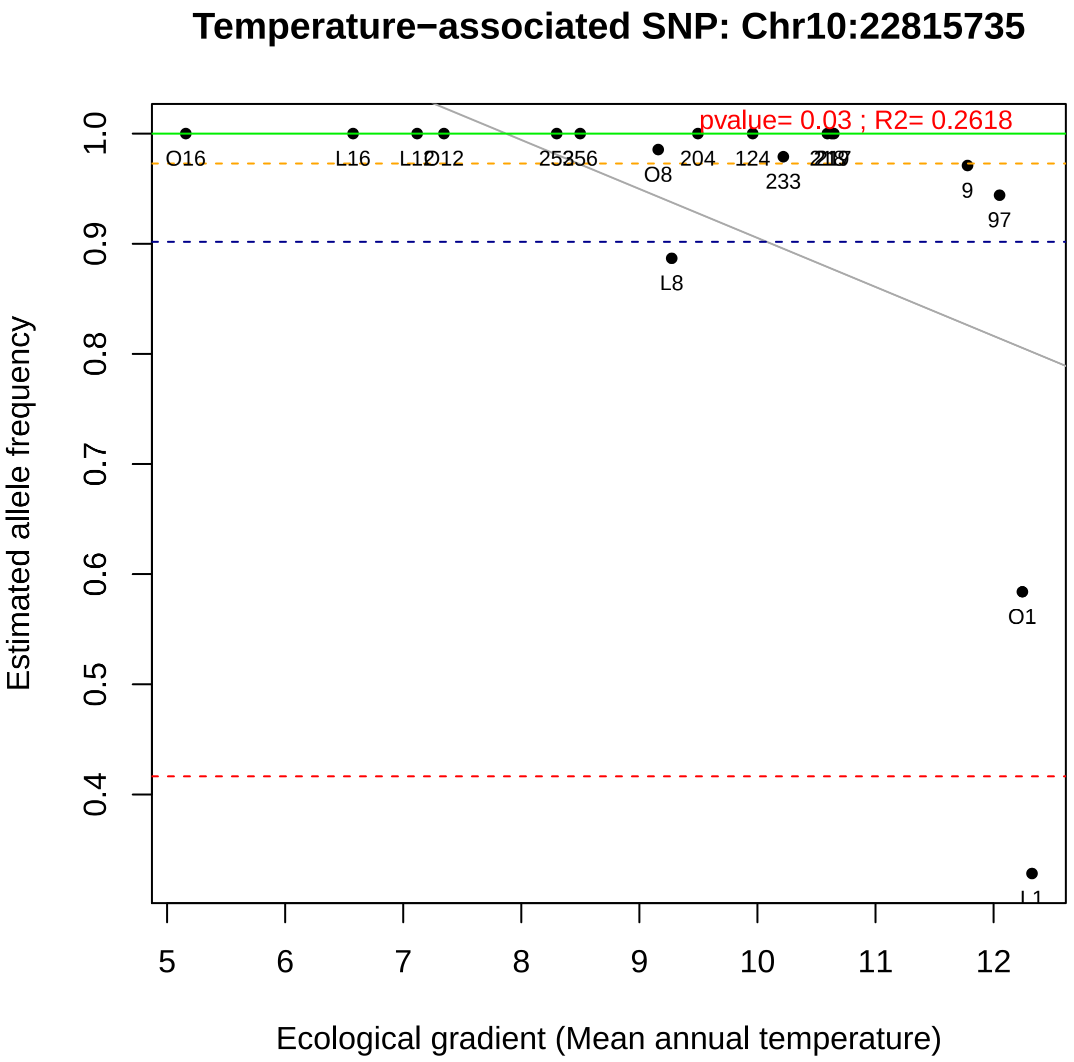

Temperature−associated SNP: Chr10:22815735
pvalue= 0.03 ; R2= 0.2618
Estimated allele frequency
Ecological gradient (Mean annual temperature)
1.0
0.9
0.8
0.7
0.6
0.5
0.4
5
6
7
8
9
10
11
12
O16
L16
O12
204
124
O8
233
9
97
L8
O1

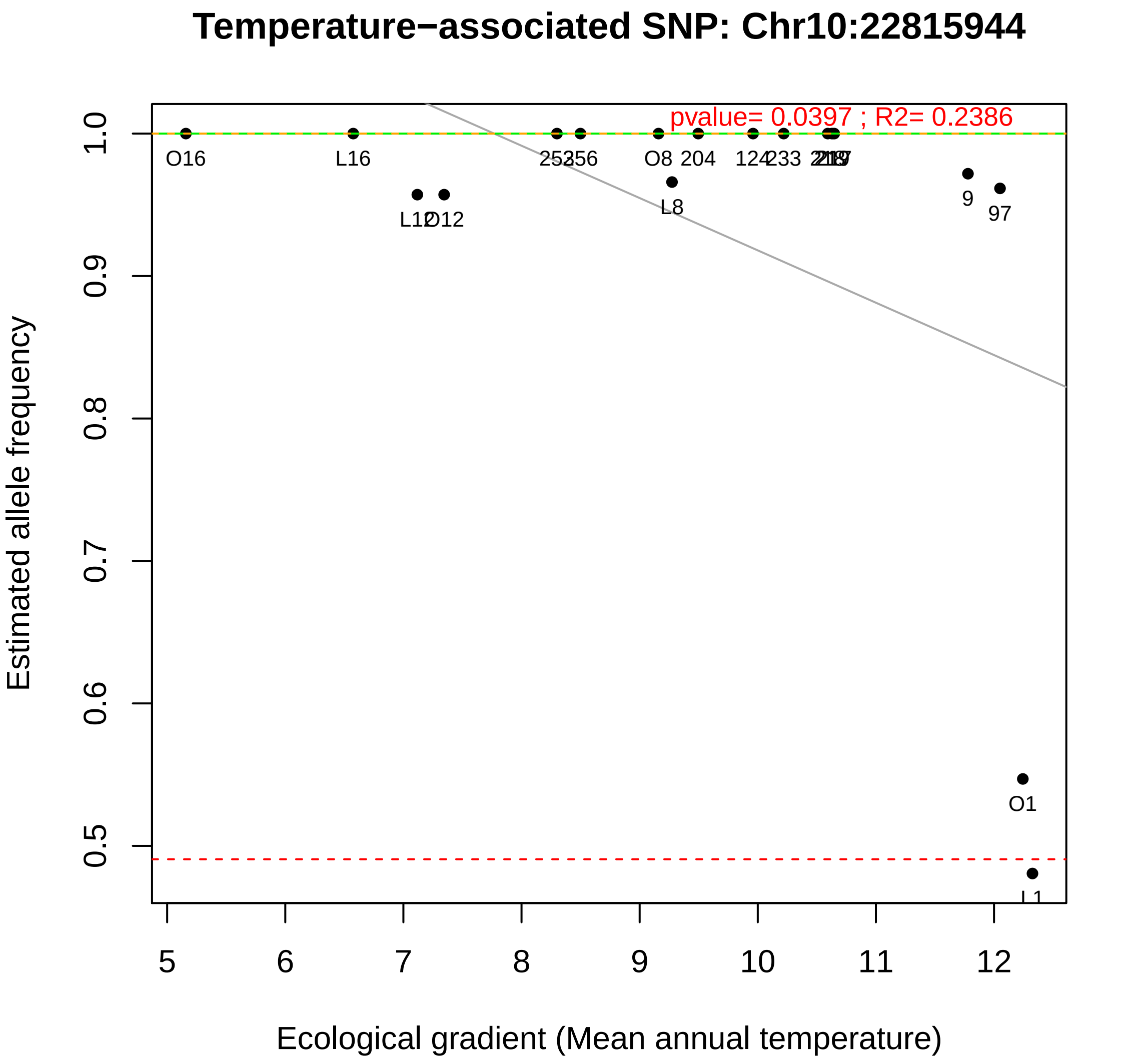

Temperature−associated SNP: Chr10:22815944
pvalue= 0.0397 ; R2= 0.2386
Estimated allele frequency
Ecological gradient (Mean annual temperature)
0.5
0.6
0.7
0.8
0.9
1.0
5
6
7
8
9
10
11
12
O16
L16
L12
O12
253
256
O8
204
L8
124
233
9
97
O1

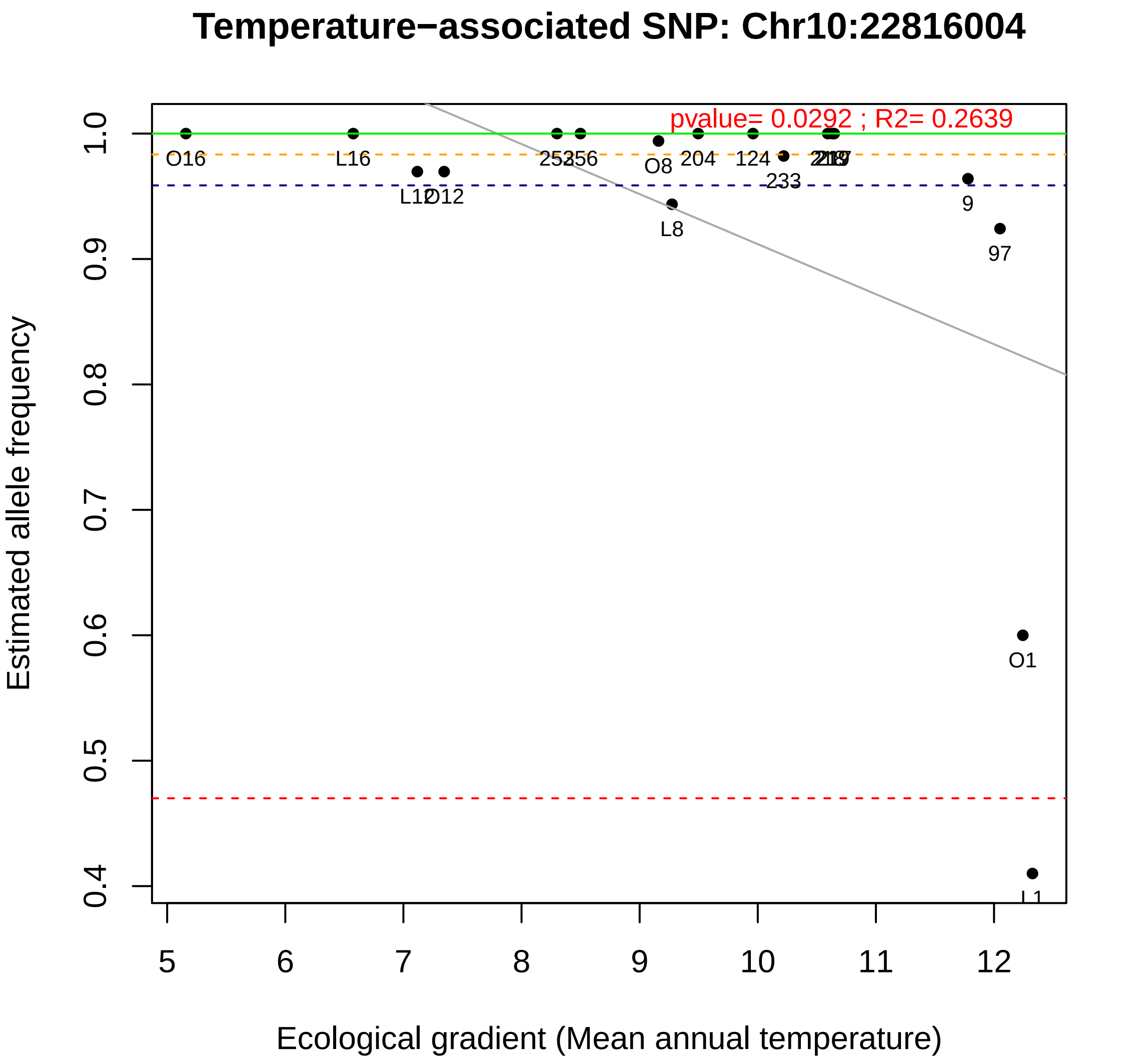

Temperature−associated SNP: Chr10:22816004
pvalue= 0.0292 ; R2= 0.2639
Estimated allele frequency
Ecological gradient (Mean annual temperature)
O16
L16
L12
O12
O8
204
124
233
9
L8
97
O1

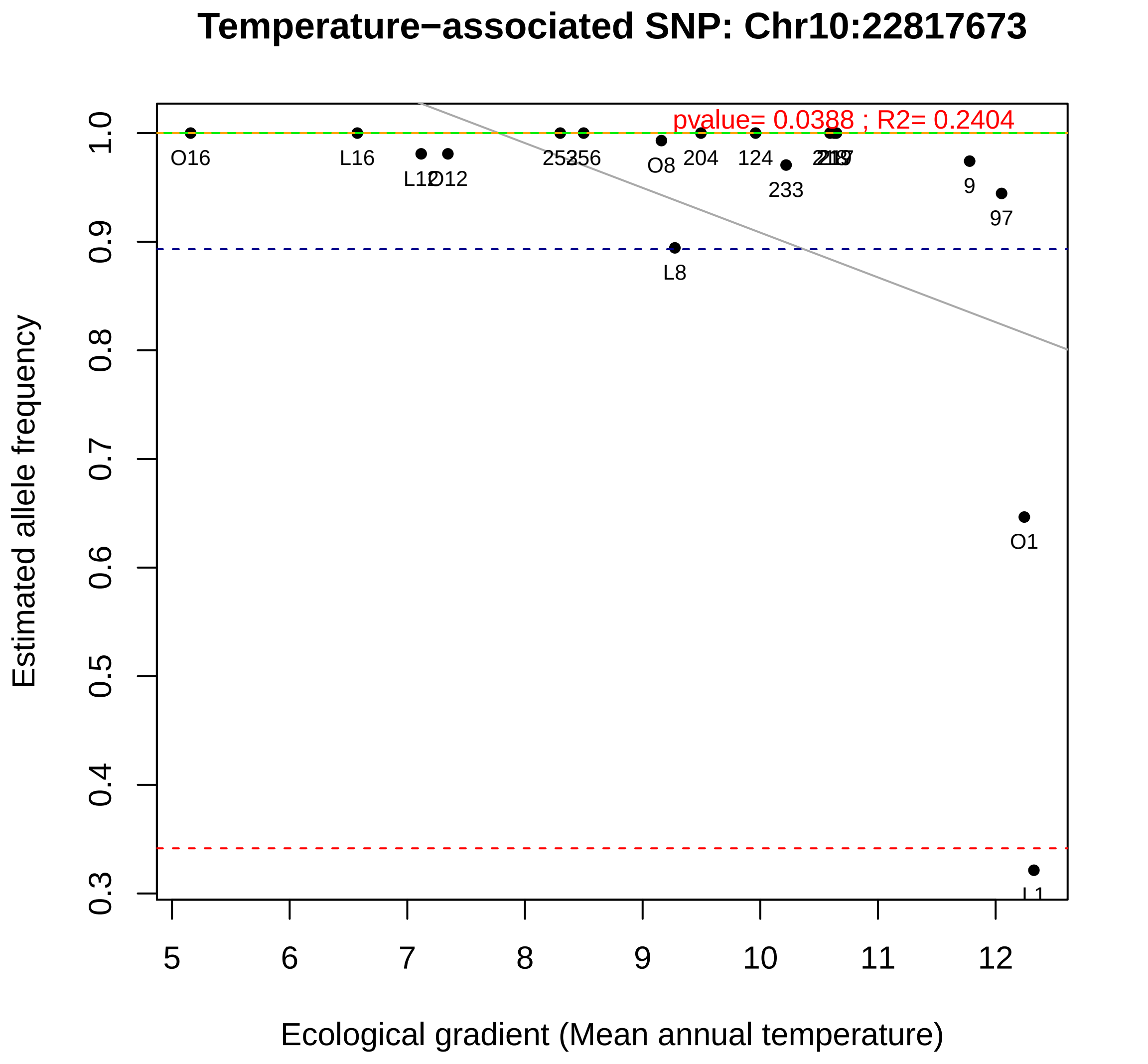

Temperature−associated SNP: Chr10:22817673
pvalue= 0.0388 ; R2= 0.2404
Estimated allele frequency
Ecological gradient (Mean annual temperature)
1.0
0.9
0.8
0.7
0.6
0.5
0.4
0.3
5
6
7
8
9
10
11
12
O16
L16
L12
O12
O8
204
124
233
9
97
L8
O1

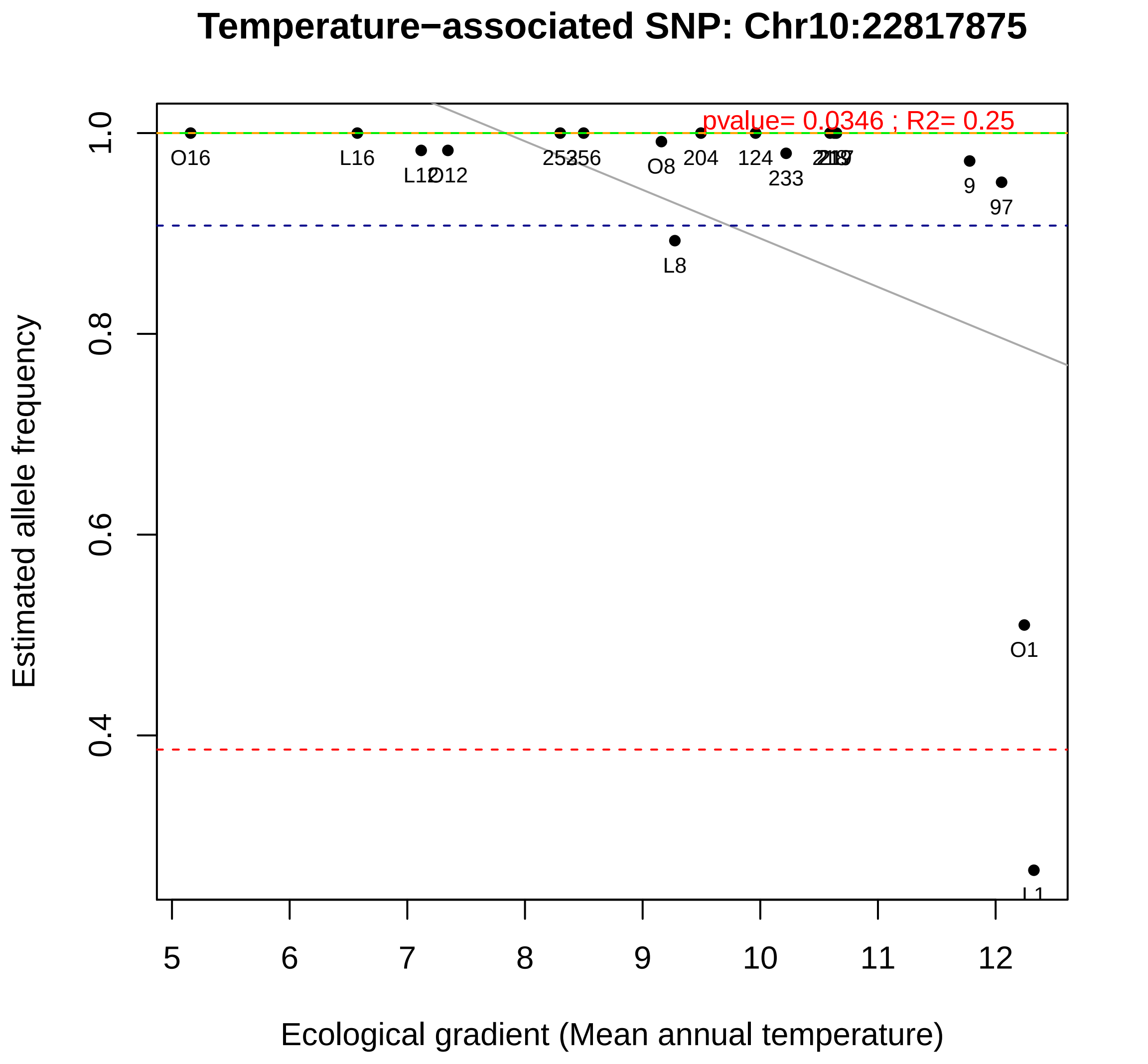

Temperature−associated SNP: Chr10:22817875
pvalue= 0.0346 ; R2= 0.25
Estimated allele frequency
Ecological gradient (Mean annual temperature)
1.0
0.8
0.6
0.4
5
6
7
8
9
10
11
12
O16
L16
L12
O12
O8
204
124
233
9
97
L8
O1

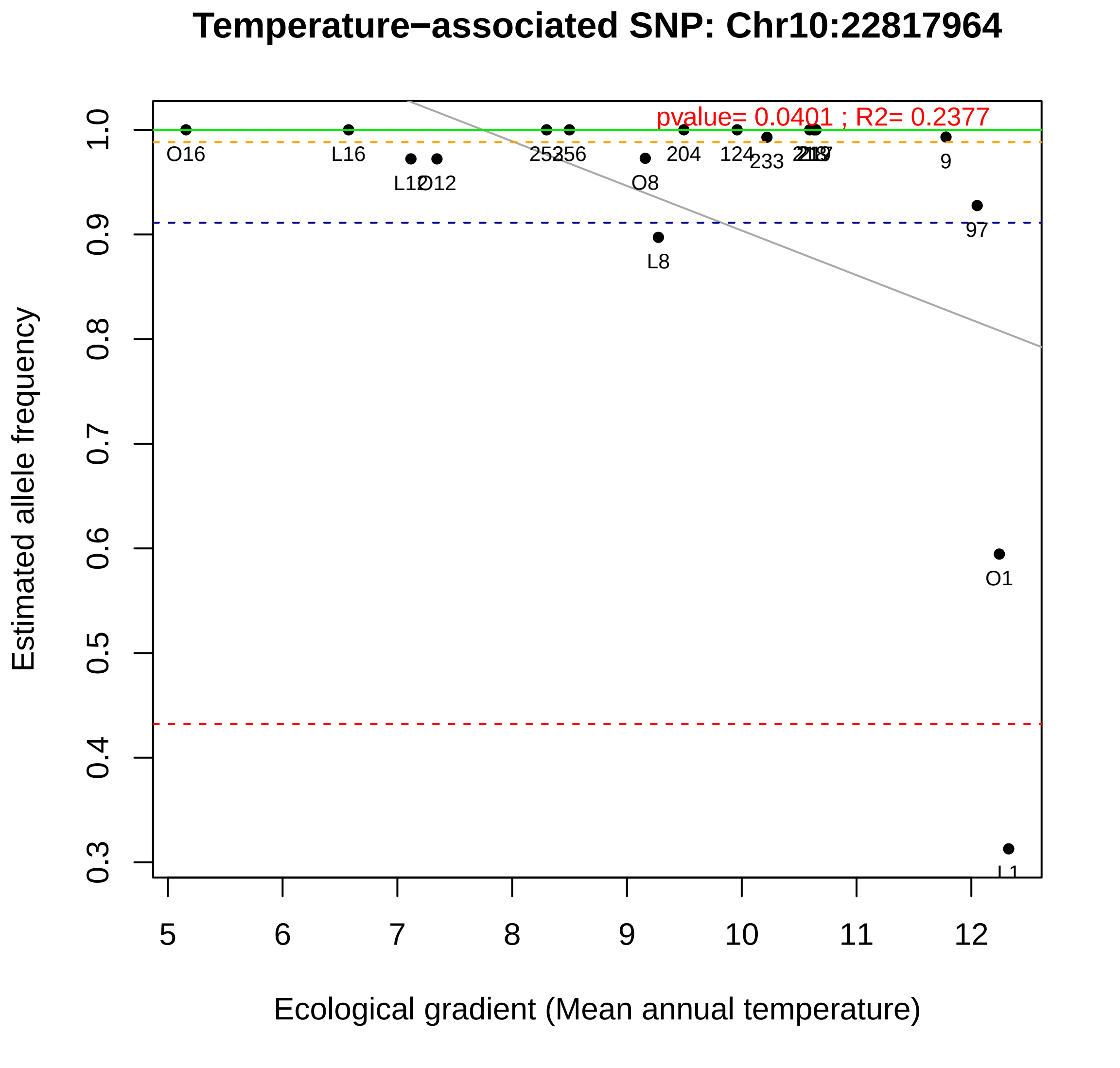
Temperature−associated SNP: Chr10:22817964
pvalue= 0.0401 ; R2= 0.2377
Estimated allele frequency
1.0
0.9
0.8
0.7
0.6
0.5
0.4
0.3
5
6
7
8
9
10
11
12
Ecological gradient (Mean annual temperature)
O16
L16
L12
O12
O8
204
124
233
9
97
L8
O1

**Temperature−associated SNP: Chr10:22818078**

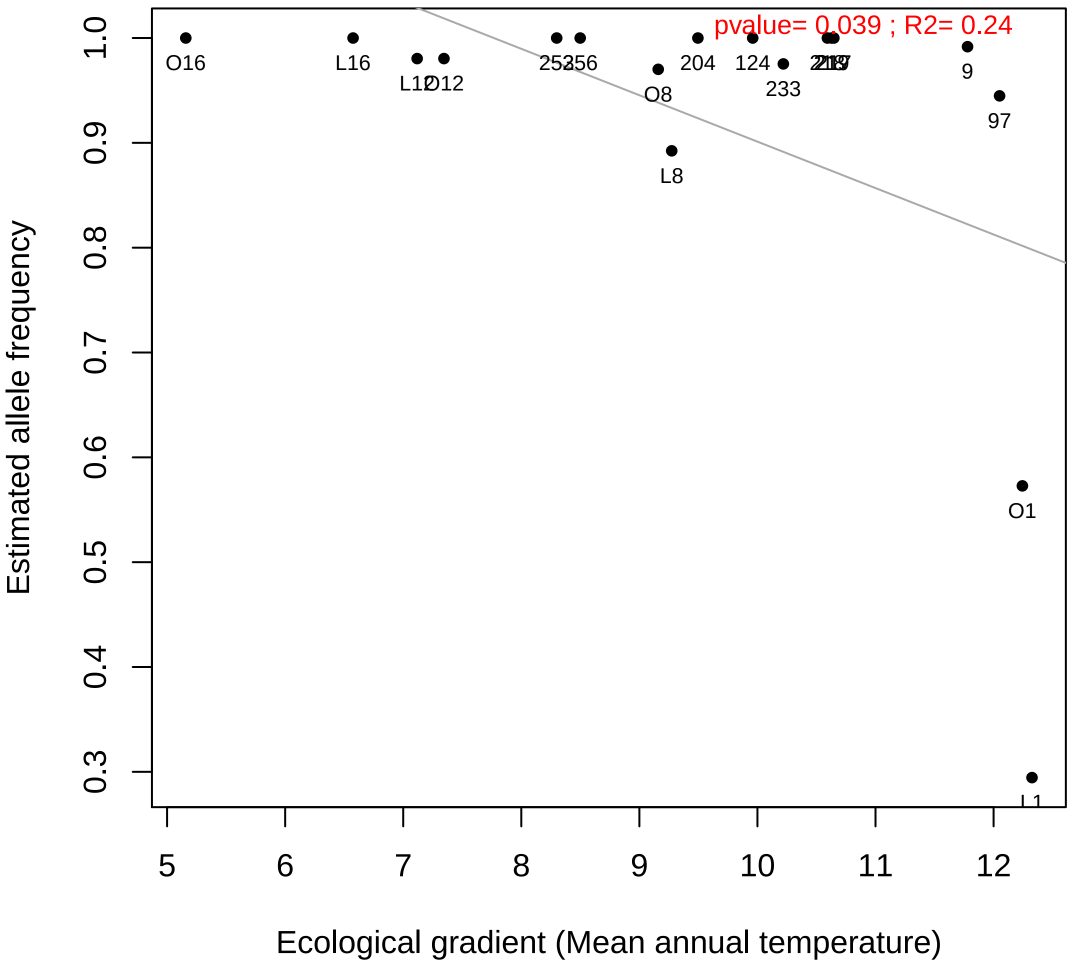

**Temperature−associated SNP: Chr10:22818334**

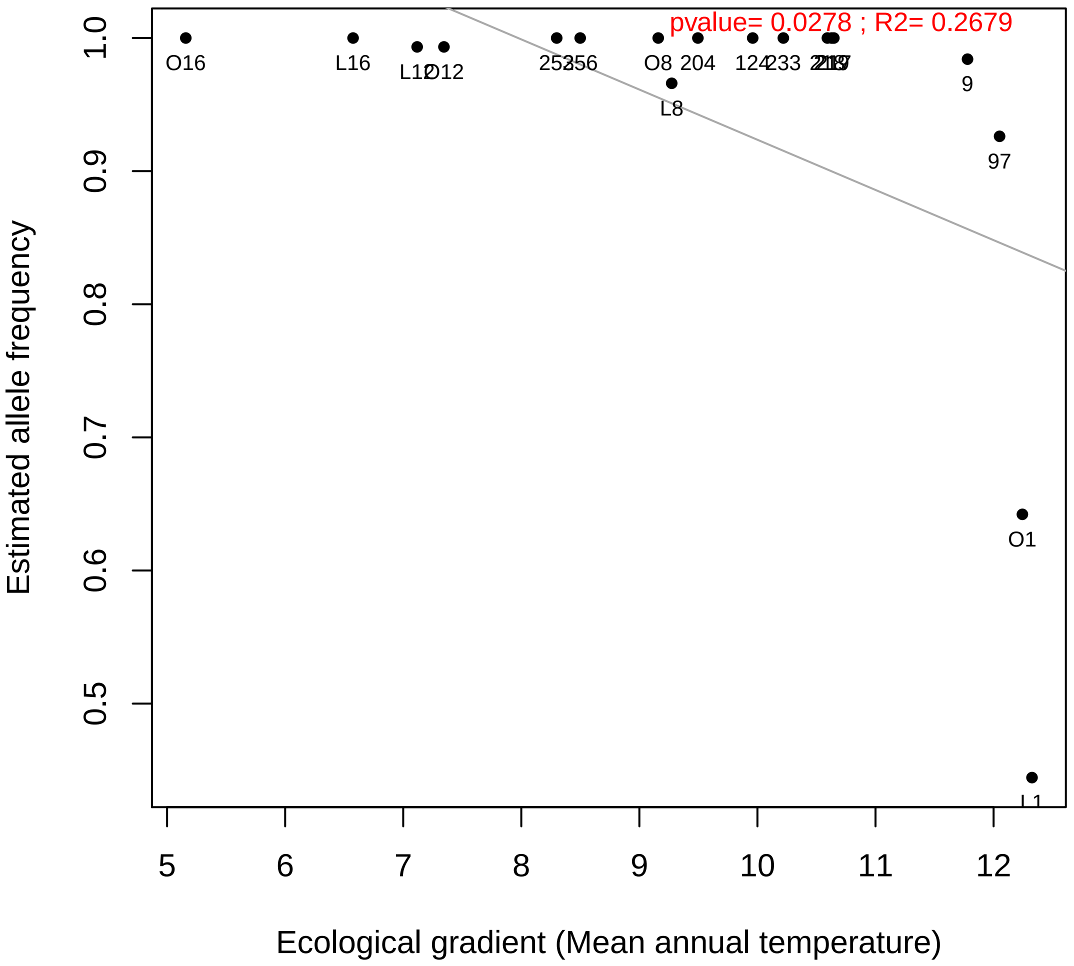

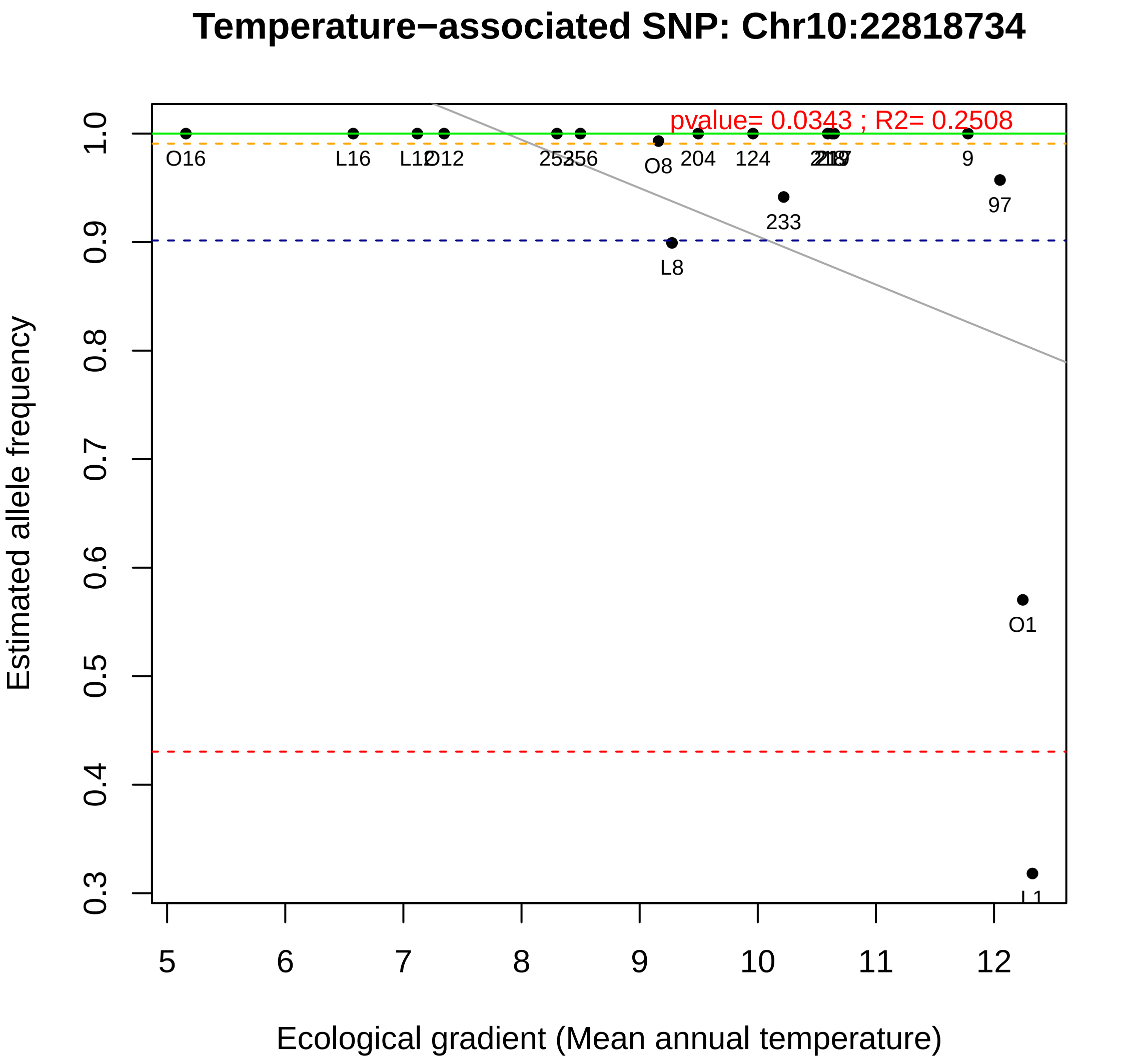

Temperature−associated SNP: Chr10:22818734
pvalue= 0.0343 ; R2= 0.2508
O16
L16
L12
O12
253
256
O8
204
124
9
97
233
L8
O1
1.0
0.9
0.8
0.7
0.6
0.5
0.4
0.3
5
6
7
8
9
10
11
12
Estimated allele frequency
Ecological gradient (Mean annual temperature)

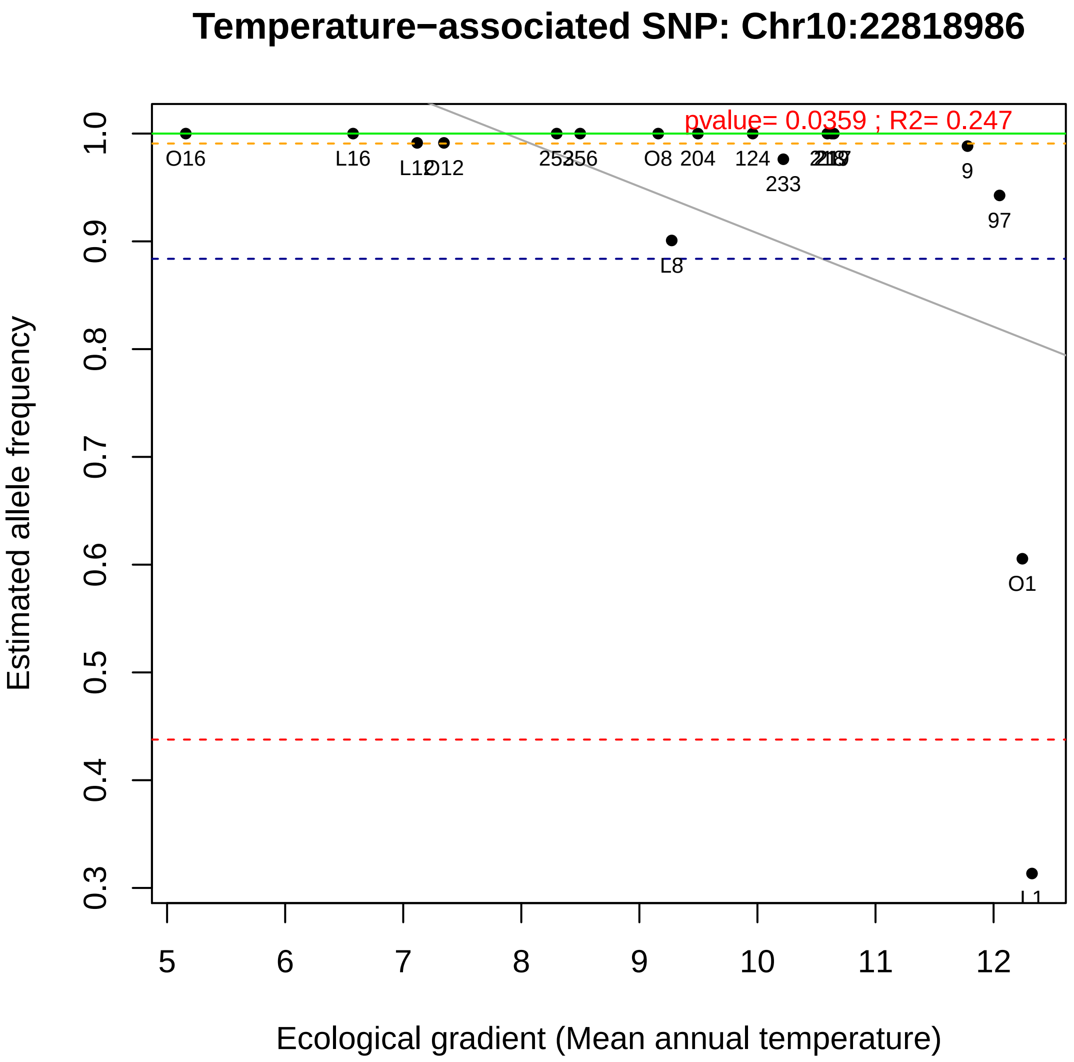

Temperature−associated SNP: Chr10:22818986
pvalue= 0.0359 ; R2= 0.247
Estimated allele frequency
Ecological gradient (Mean annual temperature)
1.0
0.9
0.8
0.7
0.6
0.5
0.4
0.3
5
6
7
8
9
10
11
12
O16
L16
L12
O12
O8
204
124
233
9
97
L8
O1

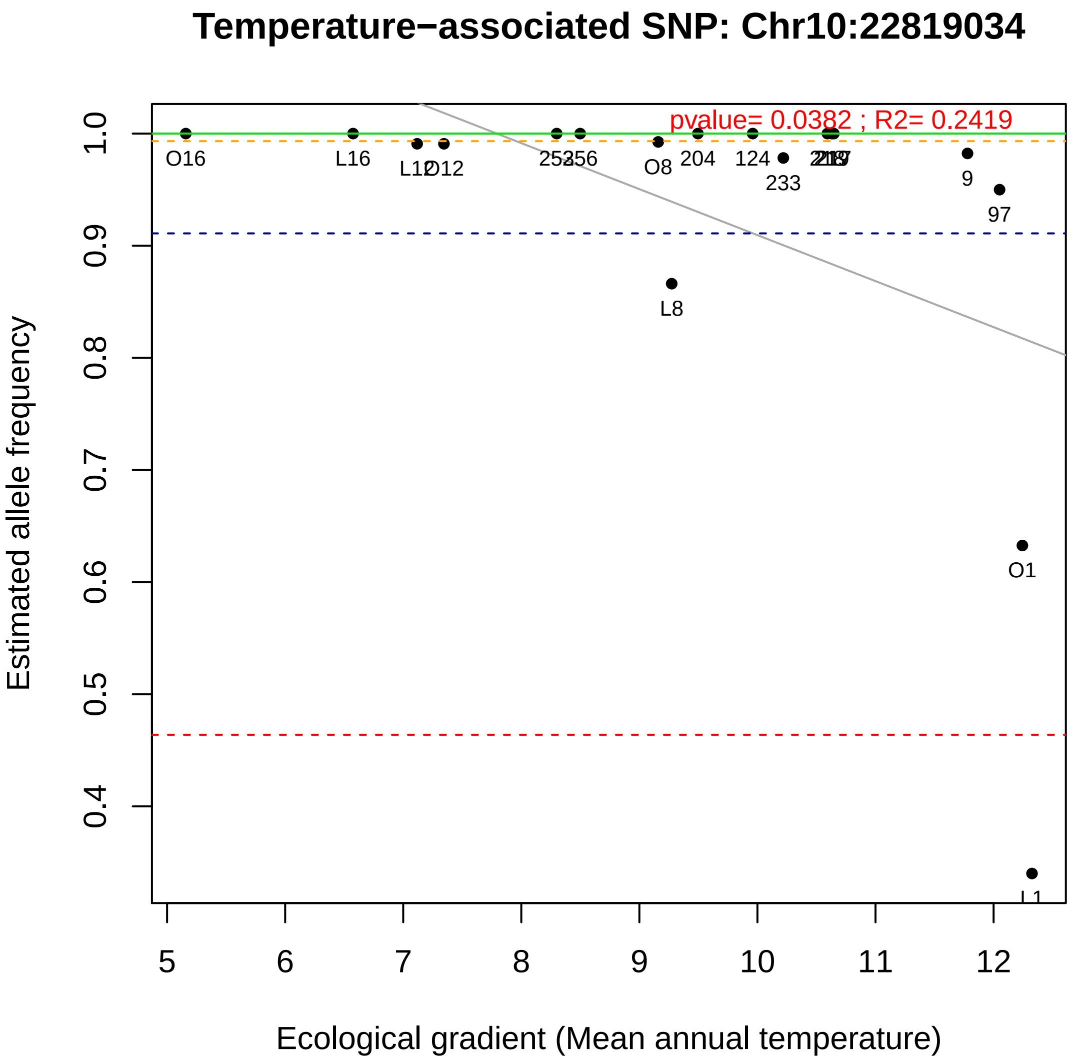
Temperature−associated SNP: Chr10:22819034
pvalue= 0.0382 ; R2= 0.2419
Estimated allele frequency
Ecological gradient (Mean annual temperature)
1.0
0.9
0.8
0.7
0.6
0.5
0.4
5
6
7
8
9
10
11
12
O16
L16
L12
O12
O8
204
124
233
9
97
L8
O1

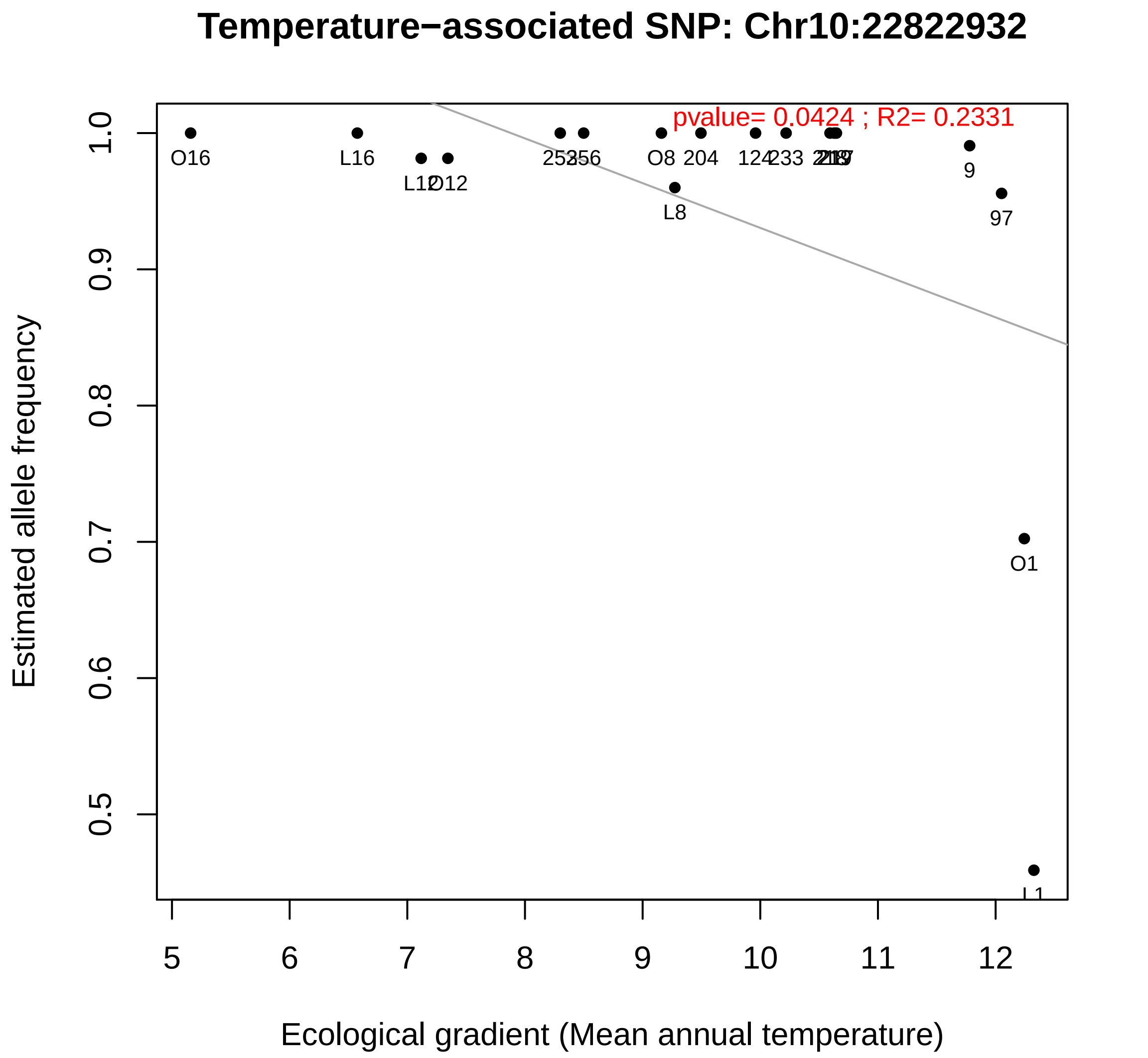
Temperature−associated SNP: Chr10:22822932
pvalue= 0.0424 ; R2= 0.2331
Estimated allele frequency
Ecological gradient (Mean annual temperature)
1.0
0.9
0.8
0.7
0.6
0.5
5
6
7
8
9
10
11
12
O16
L16
L12
O12
O8
204
L8
9
97
O1

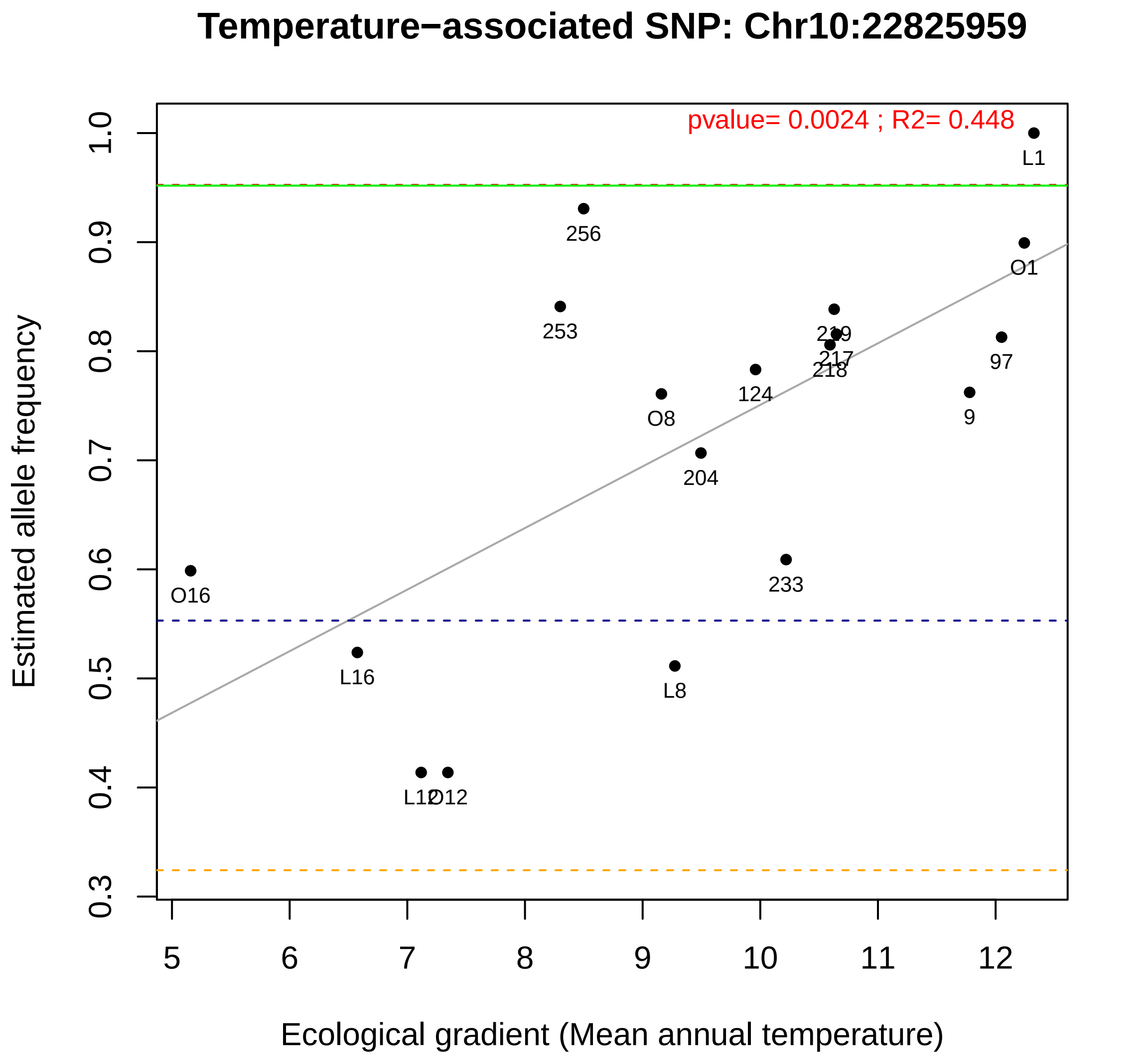
Temperature−associated SNP: Chr10:22825959
pvalue= 0.0024 ; R2= 0.448
Estimated allele frequency
0.3
0.4
0.5
0.6
0.7
0.8
0.9
1.0
5
6
7
8
9
10
11
12
Ecological gradient (Mean annual temperature)
L1
256
O1
253
219
217
218
97
124
O8
9
204
233
O16
L16
L8
L12
O12

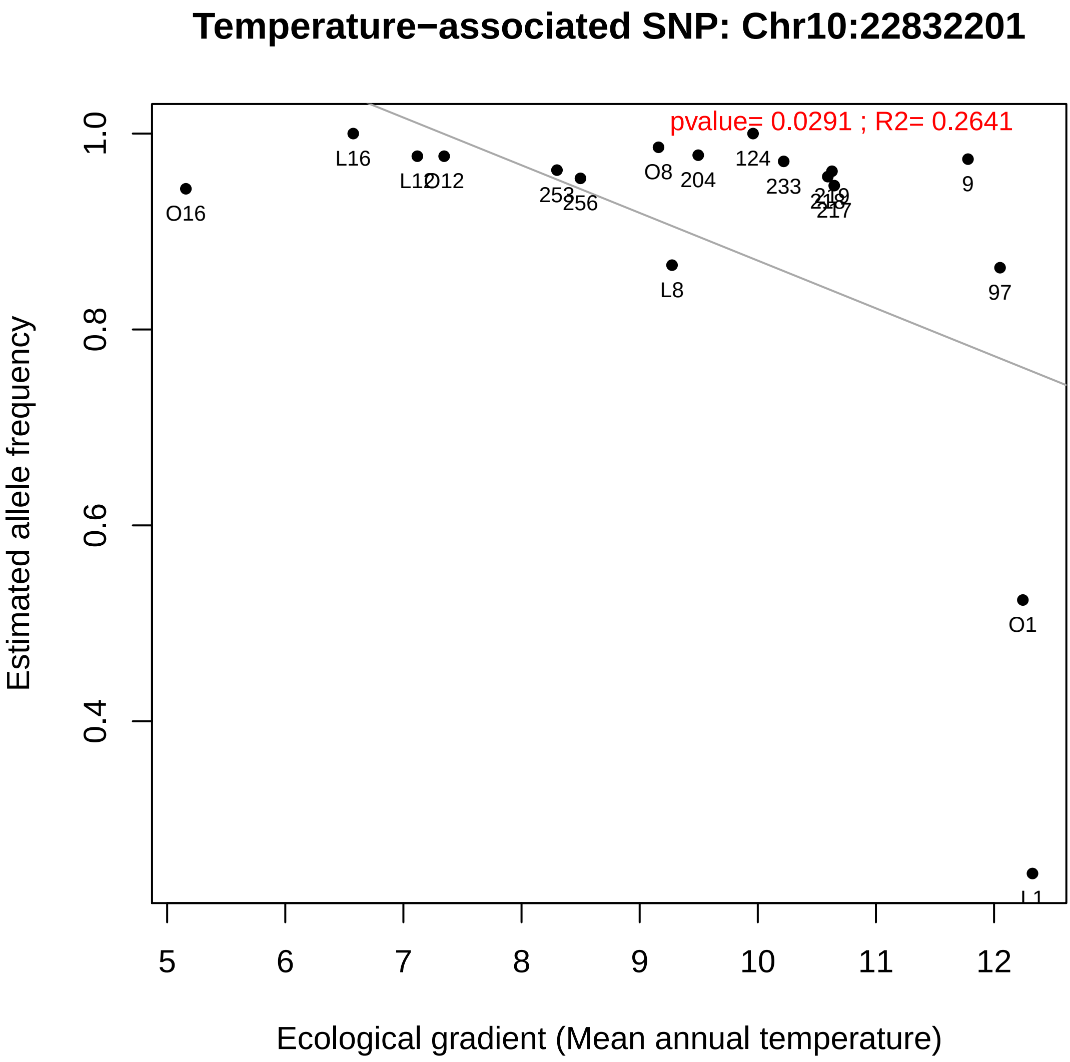

Temperature−associated SNP: Chr10:22832201
pvalue= 0.0291 ; R2= 0.2641
Estimated allele frequency
Ecological gradient (Mean annual temperature)
1.0
0.8
0.6
0.4
5
6
7
8
9
10
11
12
O16
L16
L12
O12
253
256
O8
L8
204
124
233
219
218
217
9
97
O1

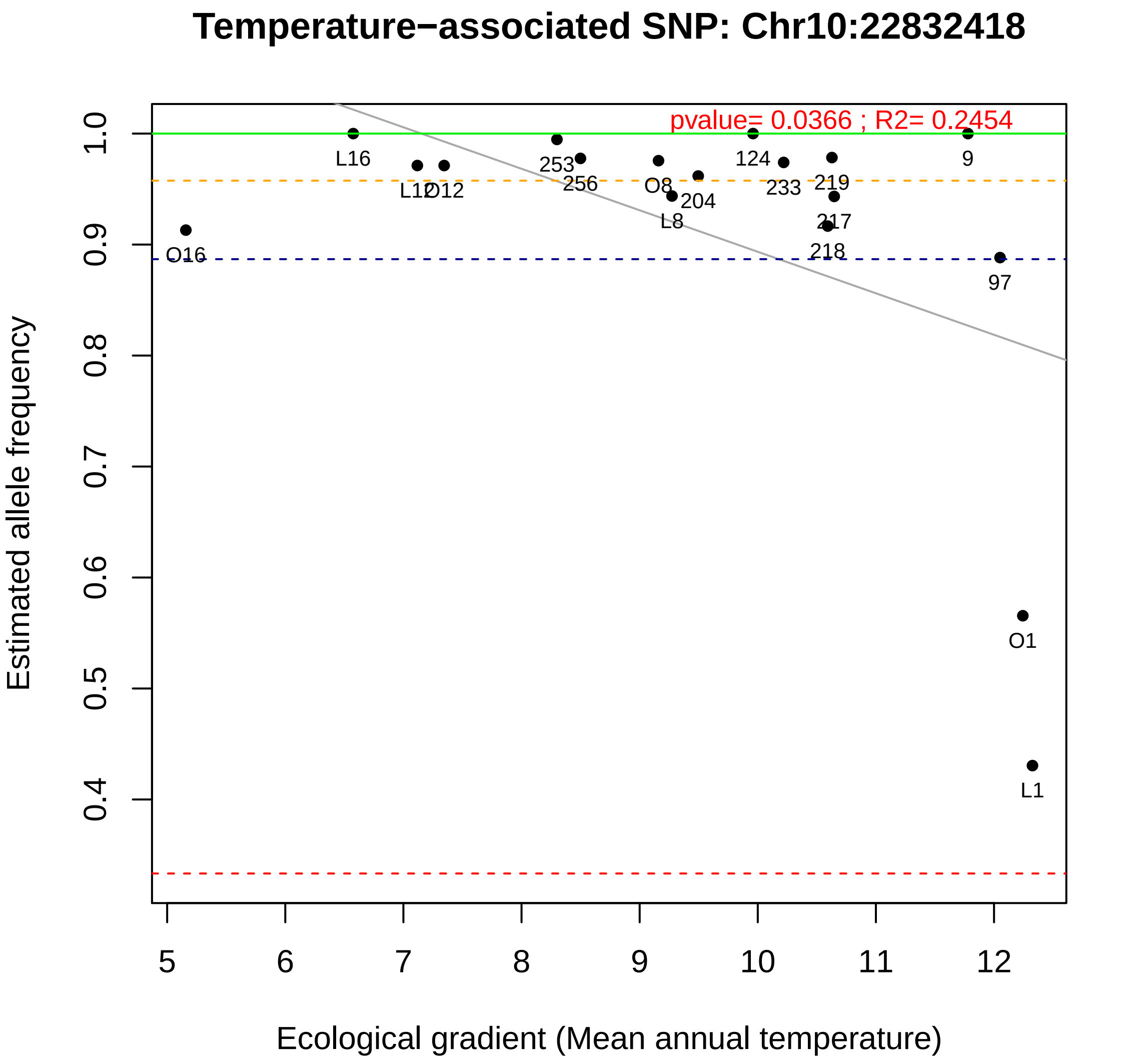
Temperature−associated SNP: Chr10:22832418
pvalue= 0.0366 ; R2= 0.2454
Estimated allele frequency
Ecological gradient (Mean annual temperature)
0.4
0.5
0.6
0.7
0.8
0.9
1.0
5
6
7
8
9
10
11
12
O16
L16
L12
O12
253
256
O8
L8
204
124
233
219
217
218
9
97
O1
L1

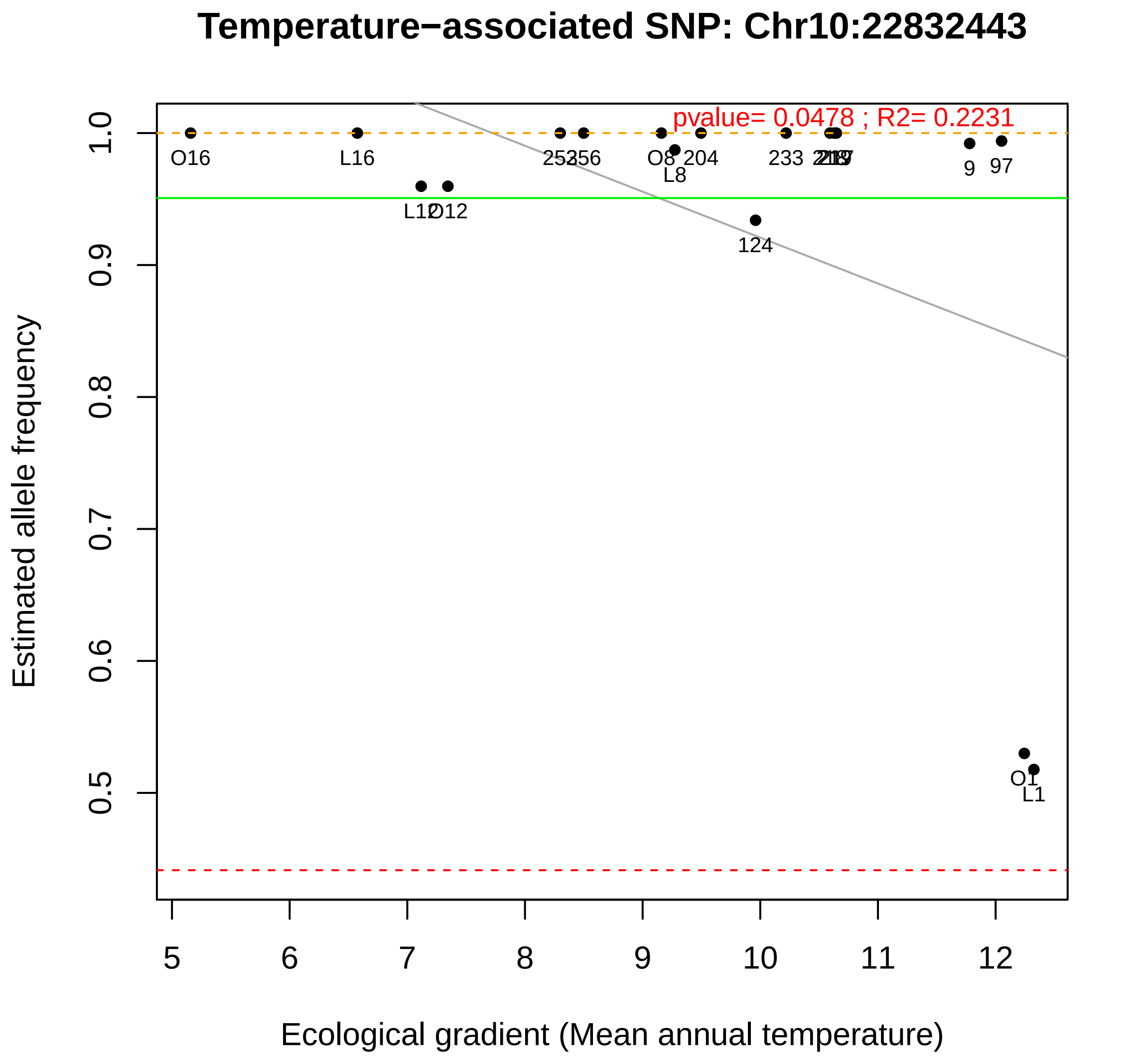
Temperature−associated SNP: Chr10:22832443
pvalue= 0.0478 ; R2= 0.2231
Estimated allele frequency
Ecological gradient (Mean annual temperature)
1.0
0.9
0.8
0.7
0.6
0.5
5
6
7
8
9
10
11
12
O16
L16
L12
O12
O8
L8
204
124
233
9
97
O1
L1

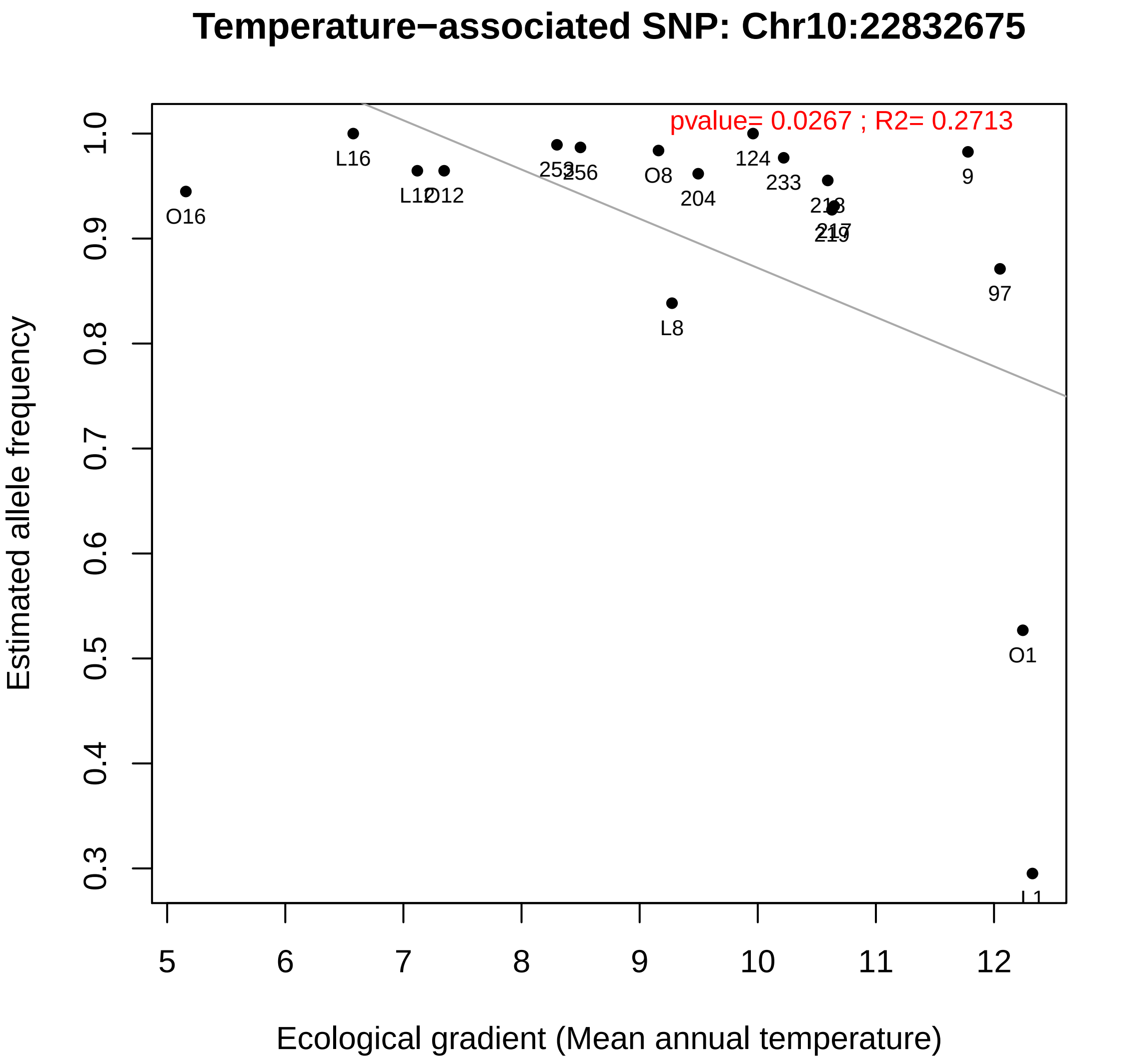

Temperature−associated SNP: Chr10:22832675
pvalue= 0.0267 ; R2= 0.2713
Estimated allele frequency
Ecological gradient (Mean annual temperature)
0.3
0.4
0.5
0.6
0.7
0.8
0.9
1.0
5
6
7
8
9
10
11
12
O16
L16
L12
O12
256
O8
204
124
233
9
97
L8
O1

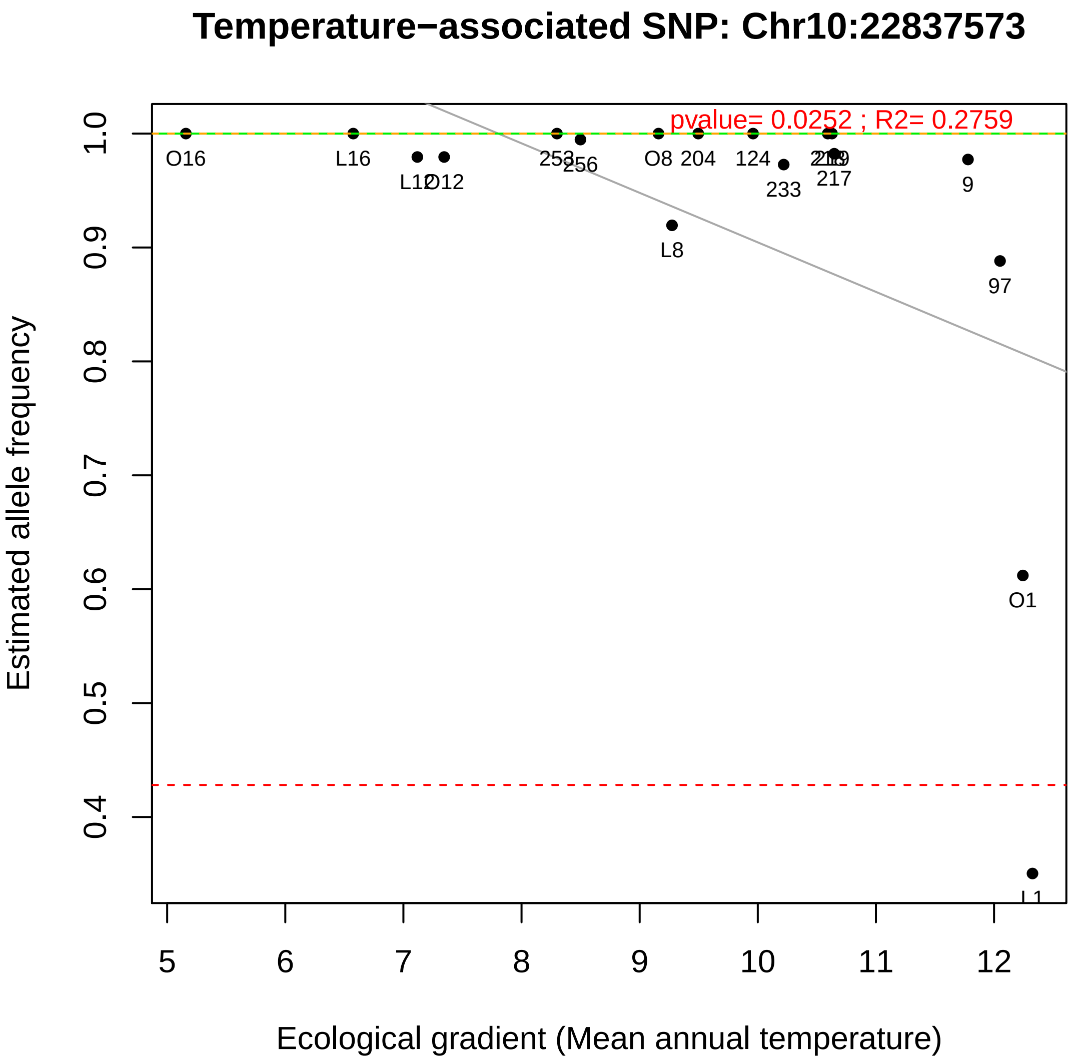

Temperature−associated SNP: Chr10:22837573
pvalue= 0.0252 ; R2= 0.2759
Estimated allele frequency
Ecological gradient (Mean annual temperature)
1.0
0.9
0.8
0.7
0.6
0.5
0.4
5
6
7
8
9
10
11
12
O16
L16
L12
O12
253
256
O8
204
124
L8
233
217
9
97
O1

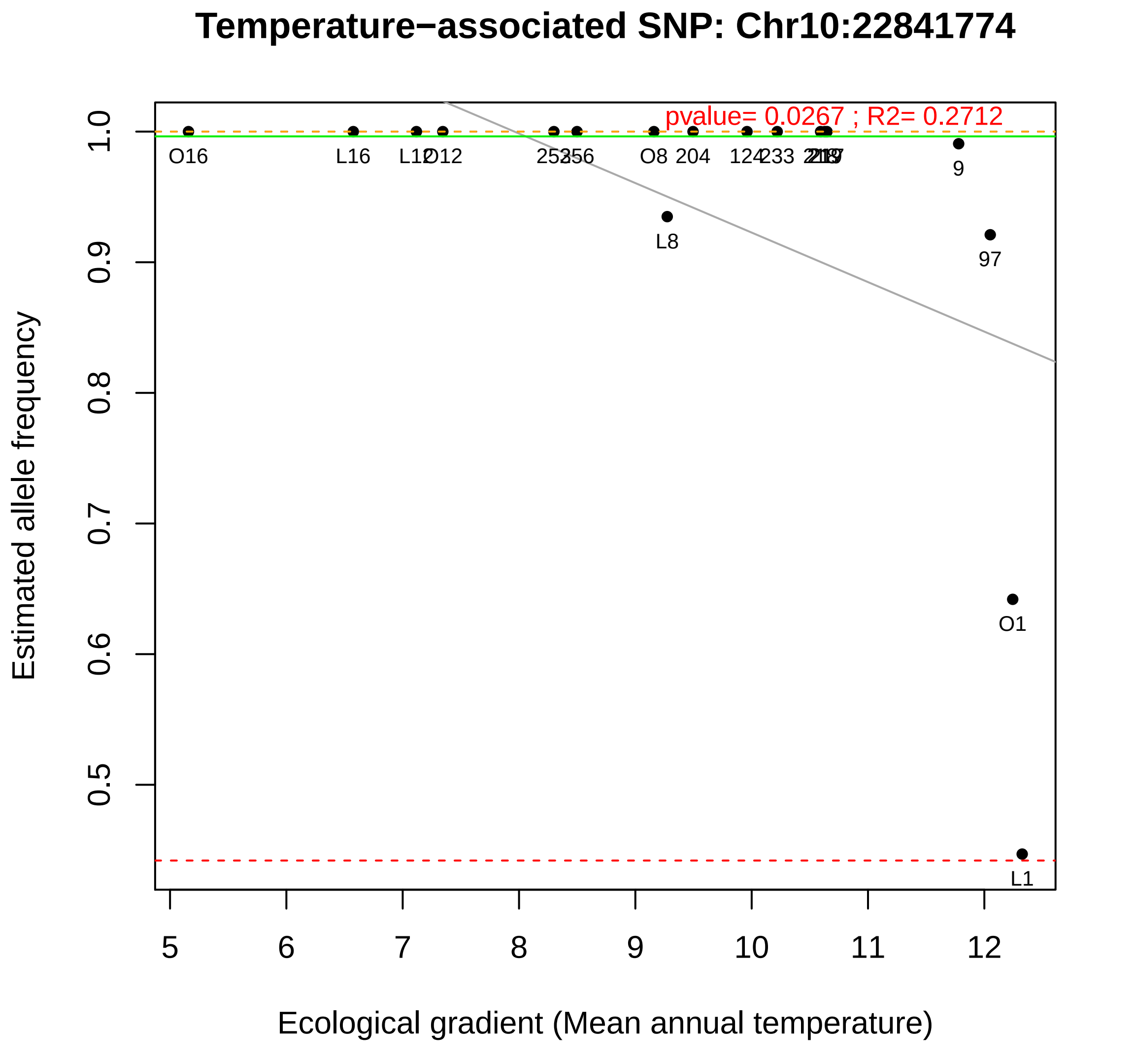

Temperature−associated SNP: Chr10:22841774
pvalue= 0.0267 ; R2= 0.2712
O16
L16
L12
O12
253
256
O8
204
124
233
218
197
9
L8
97
O1
L1
1.0
0.9
0.8
0.7
0.6
0.5
5
6
7
8
9
10
11
12
Estimated allele frequency
Ecological gradient (Mean annual temperature)

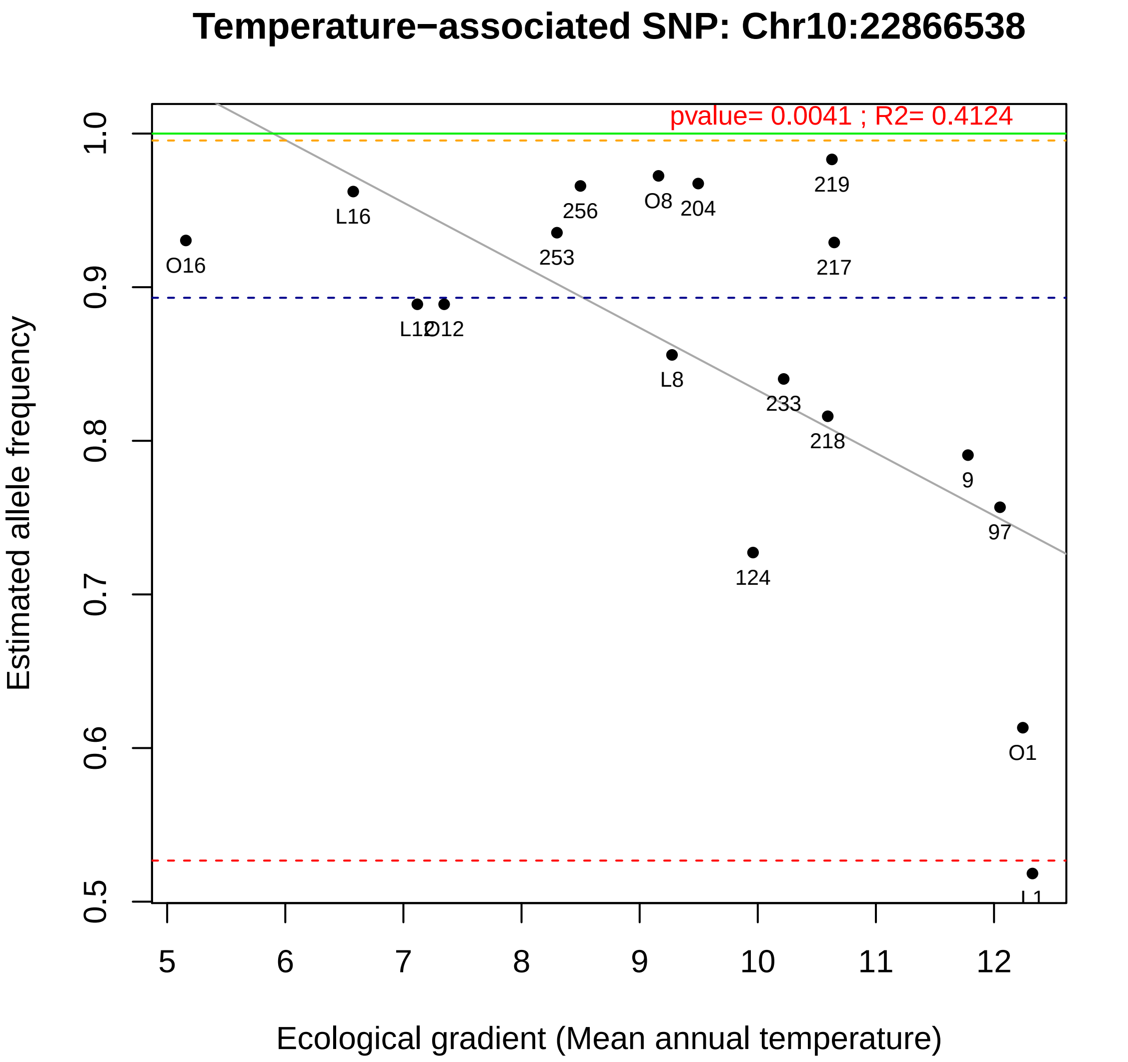
Temperature−associated SNP: Chr10:22866538
pvalue= 0.0041 ; R2= 0.4124
Estimated allele frequency
Ecological gradient (Mean annual temperature)
1.0
0.9
0.8
0.7
0.6
0.5
5
6
7
8
9
10
11
12
O16
L16
L12
O12
253
256
O8
204
219
217
L8
233
218
124
9
97
O1

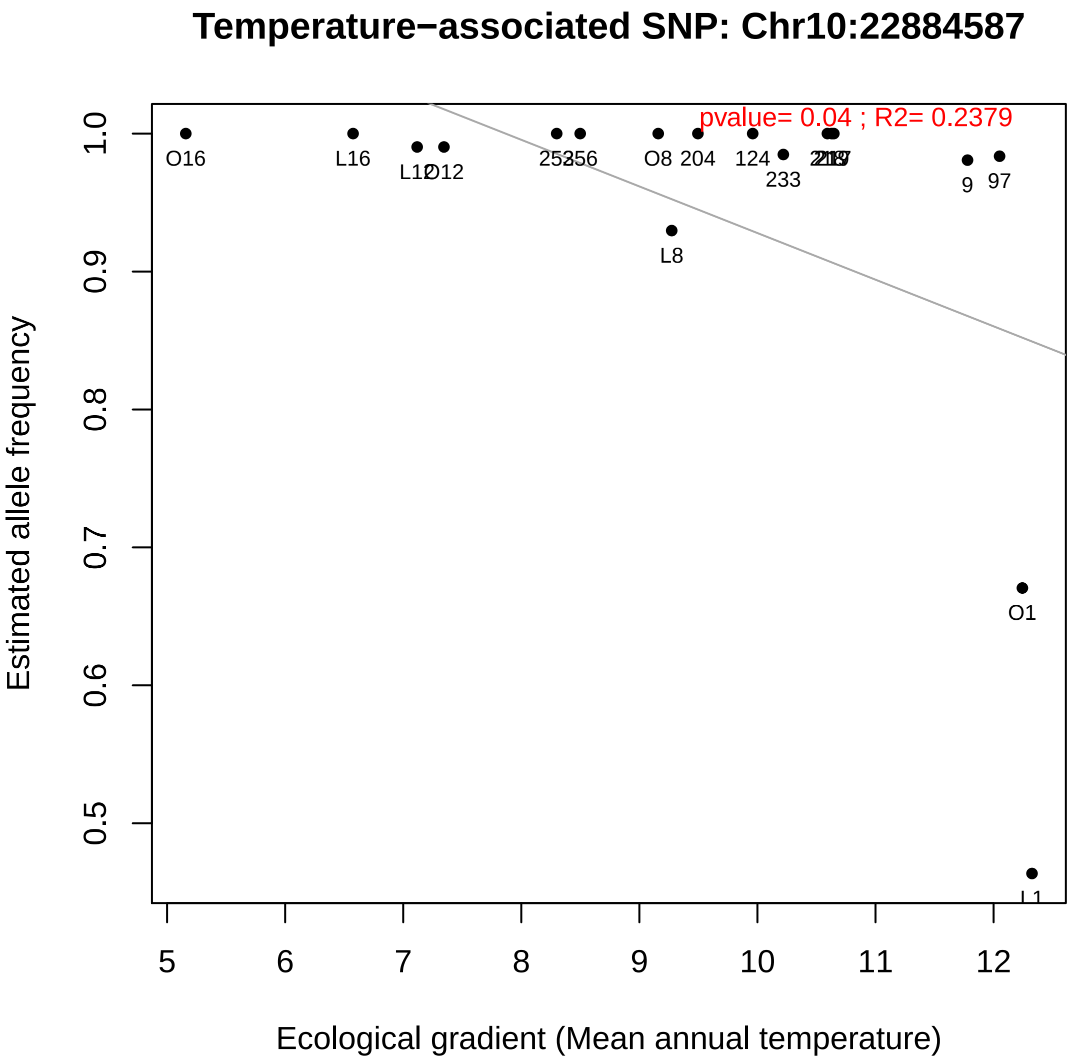

Temperature−associated SNP: Chr10:22884587
pvalue= 0.04 ; R2= 0.2379
Estimated allele frequency
Ecological gradient (Mean annual temperature)
1.0
0.9
0.8
0.7
0.6
0.5
5
6
7
8
9
10
11
12
O16
L16
L12
O12
253
256
O8
204
124
233
218
217
9
97
L8
O1

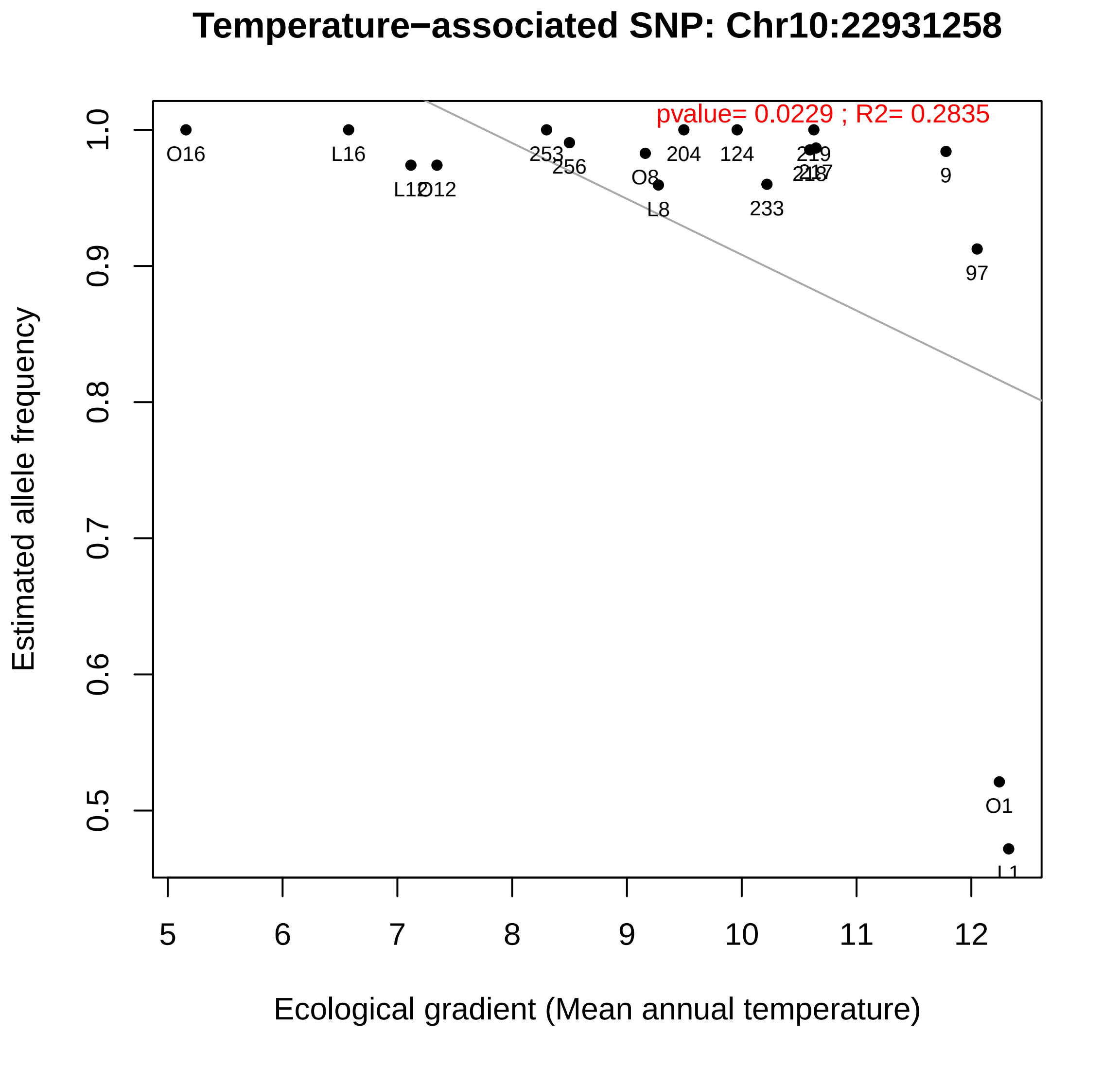

Temperature−associated SNP: Chr10:22931258
pvalue= 0.0229 ; R2= 0.2835
Estimated allele frequency
Ecological gradient (Mean annual temperature)
O16
L16
L12
O12
253
256
O8
L8
204
124
233
219
217
218
9
97
O1
L1
0.5
0.6
0.7
0.8
0.9
1.0
5
6
7
8
9
10
11
12

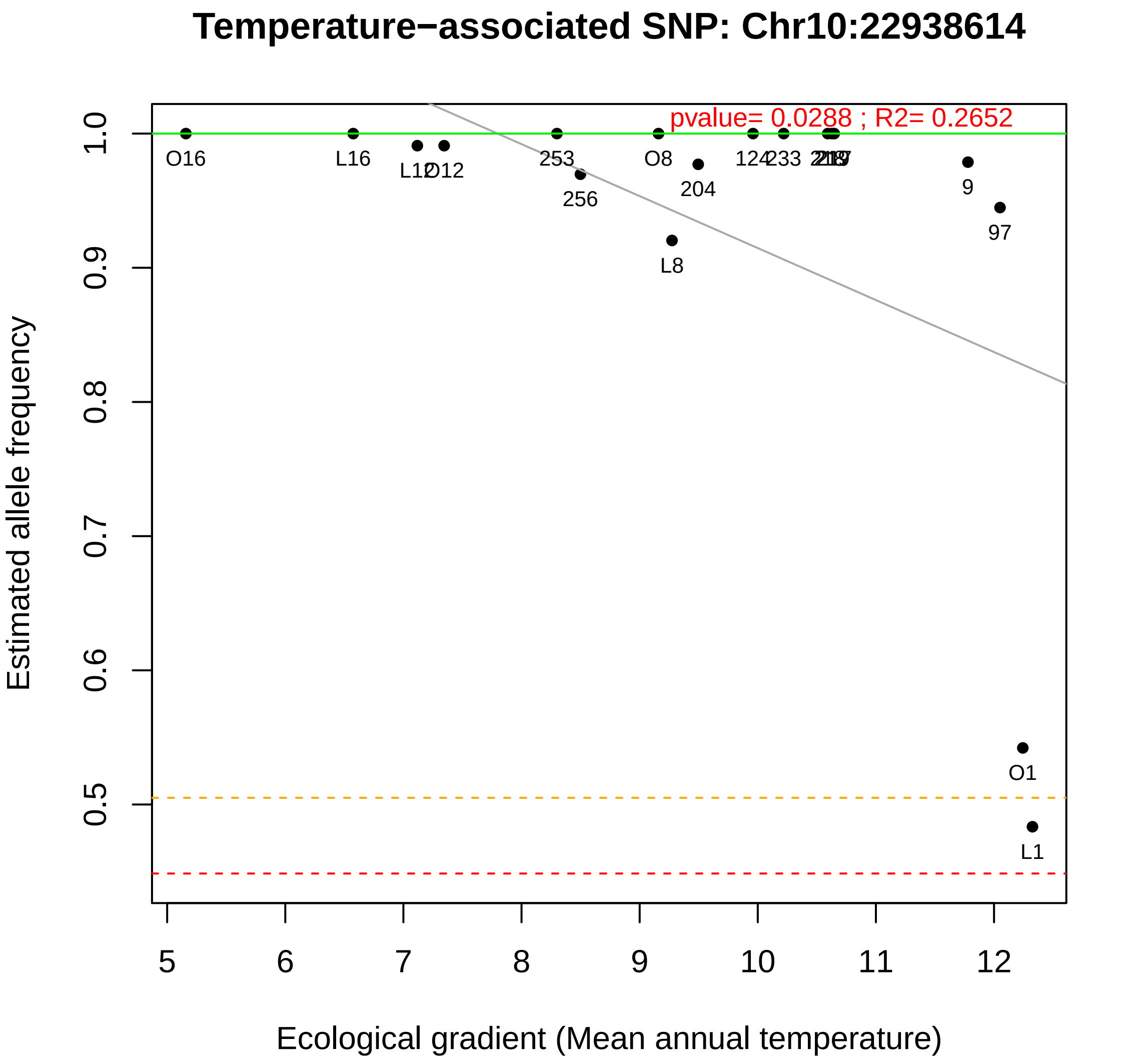
Temperature−associated SNP: Chr10:22938614
pvalue= 0.0288 ; R2= 0.2652
Estimated allele frequency
Ecological gradient (Mean annual temperature)
0.5
0.6
0.7
0.8
0.9
1.0
5
6
7
8
9
10
11
12
O16
L16
L12
O12
253
256
O8
L8
204
124
233
9
97
O1
L1

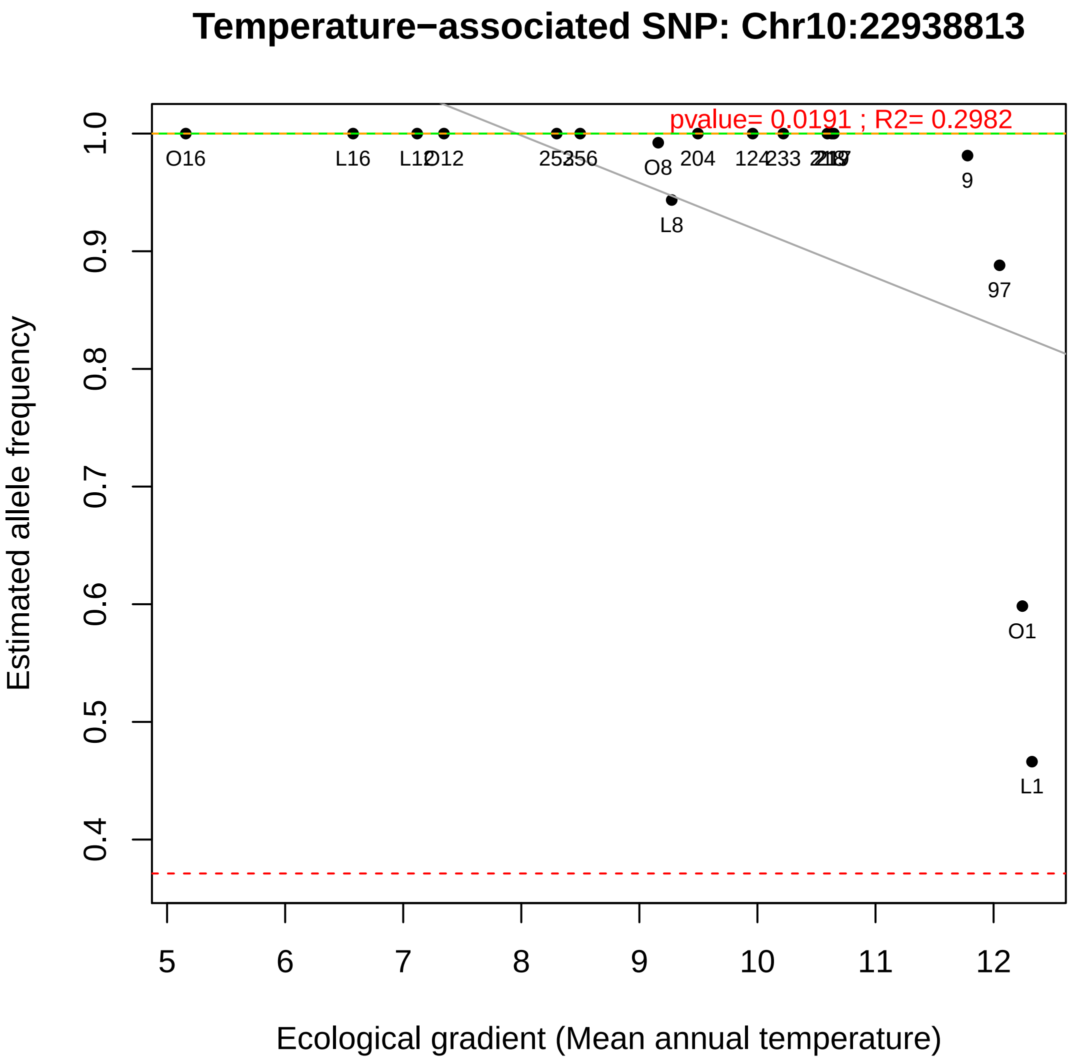
Temperature−associated SNP: Chr10:22938813
pvalue= 0.0191 ; R2= 0.2982
O16
L16
L12
O12
253
256
O8
L8
204
124
233
9
97
O1
L1
1.0
0.9
0.8
0.7
0.6
0.5
0.4
5
6
7
8
9
10
11
12
Estimated allele frequency
Ecological gradient (Mean annual temperature)

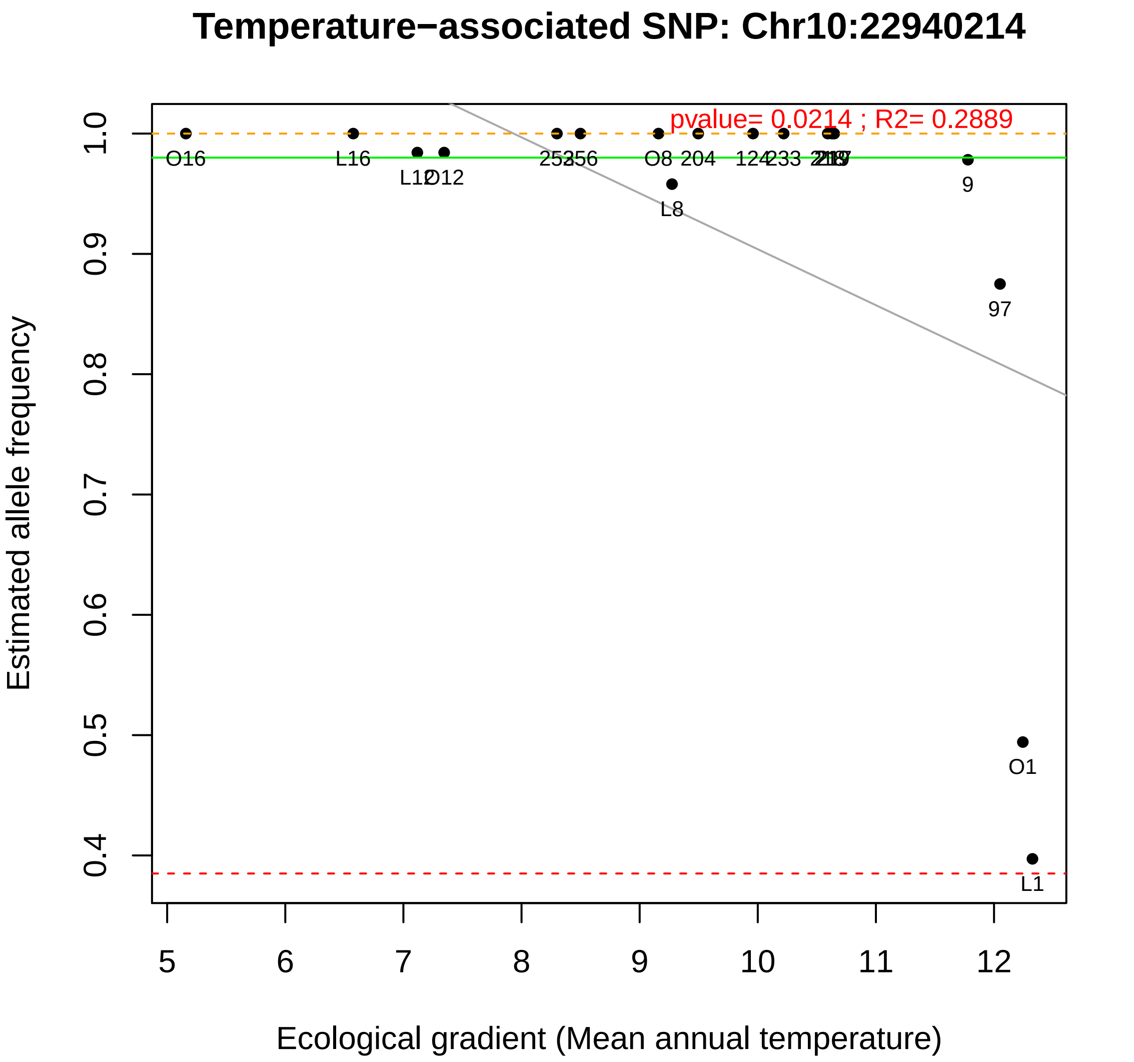
Temperature−associated SNP: Chr10:22940214
pvalue= 0.0214 ; R2= 0.2889
Estimated allele frequency
Ecological gradient (Mean annual temperature)
O16
L16
L12
O12
O8
204
L8
9
97
O1
L1
1.0
0.9
0.8
0.7
0.6
0.5
0.4
5
6
7
8
9
10
11
12

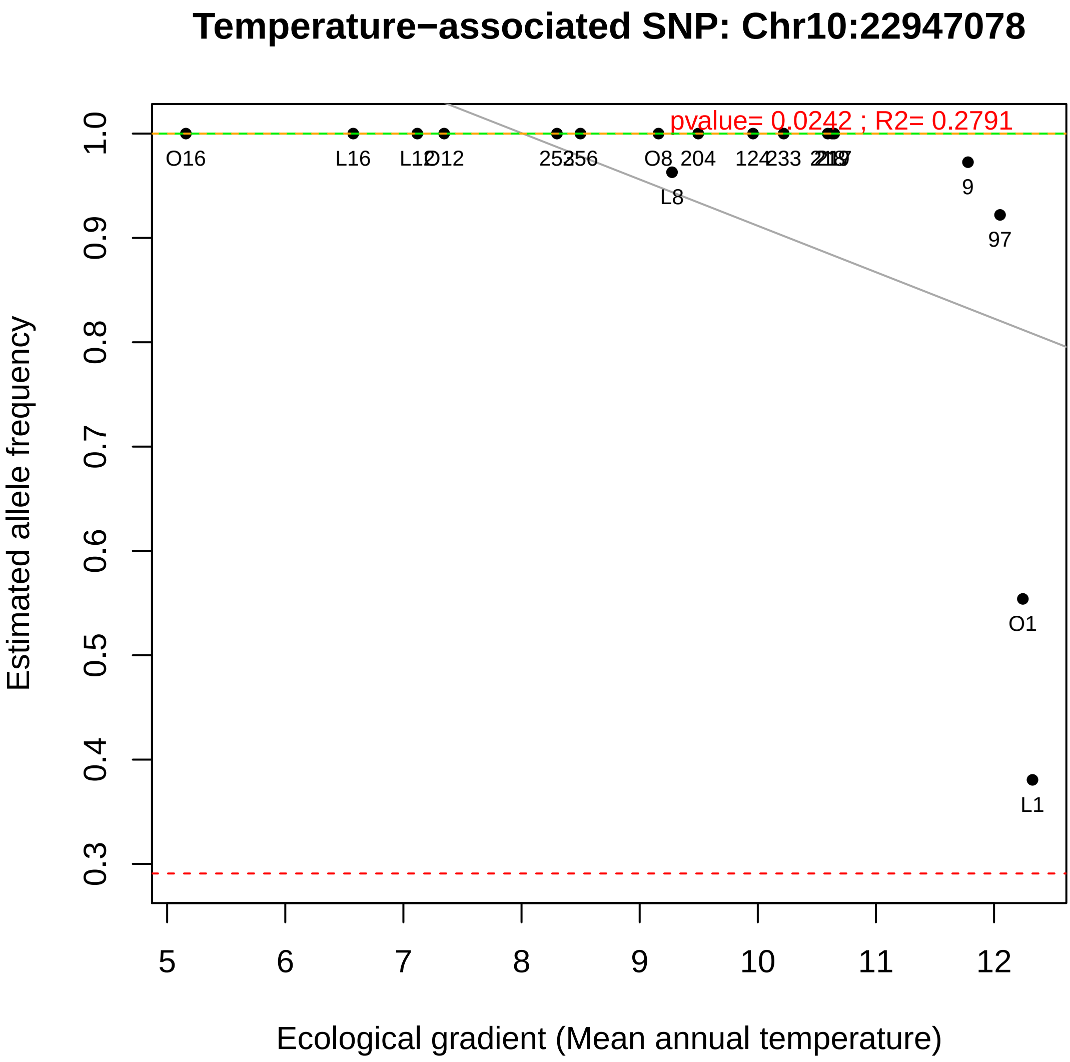
Temperature−associated SNP: Chr10:22947078
pvalue= 0.0242 ; R2= 0.2791
Estimated allele frequency
Ecological gradient (Mean annual temperature)
1.0
0.9
0.8
0.7
0.6
0.5
0.4
0.3
5
6
7
8
9
10
11
12
O16
L16
L12
O12
253
256
O8
204
124
233
L8
9
97
O1
L1

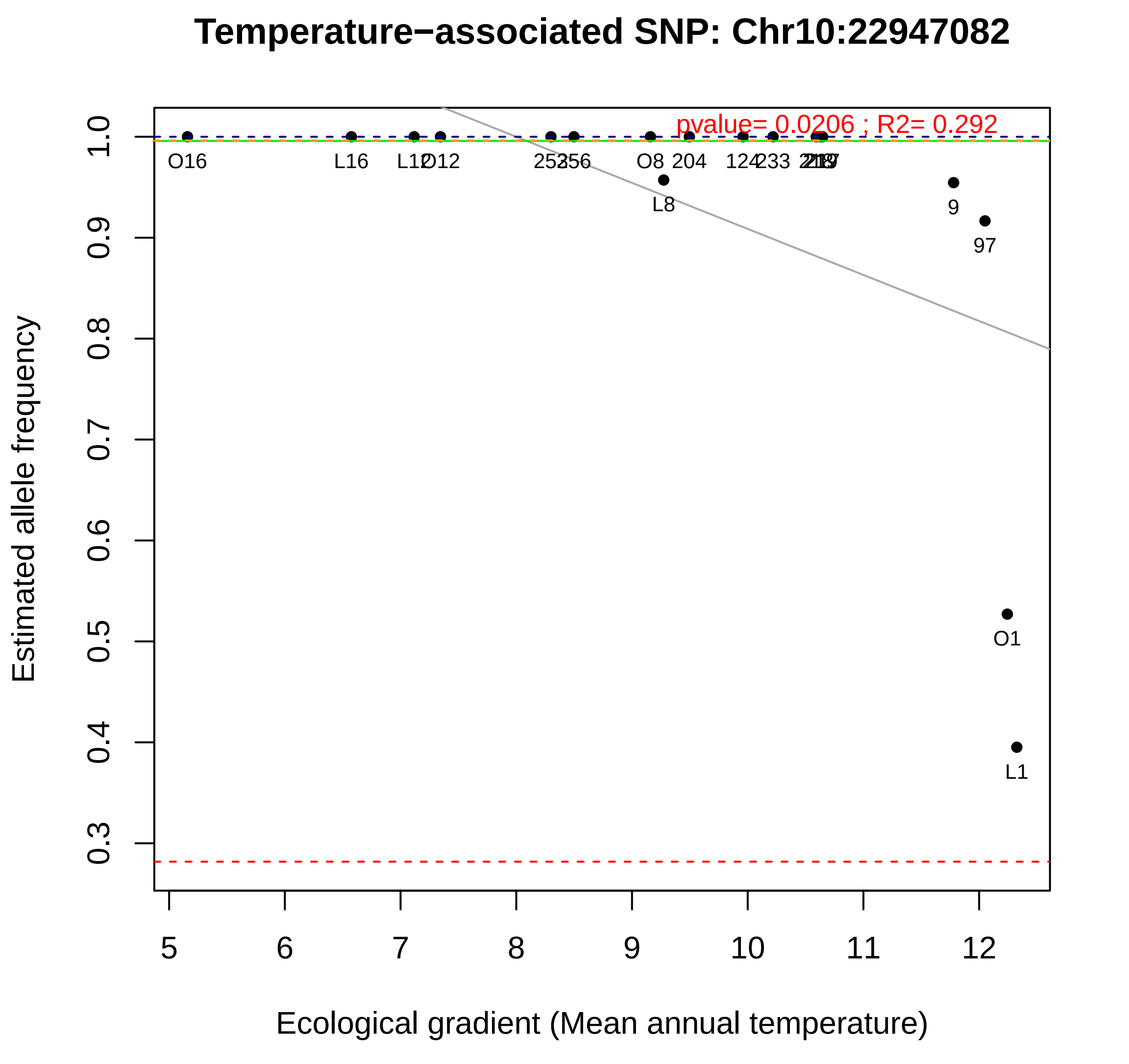
Temperature−associated SNP: Chr10:22947082
pvalue= 0.0206 ; R2= 0.292
Estimated allele frequency
Ecological gradient (Mean annual temperature)
O16
L16
L12
O12
253
256
O8
204
124
233
L8
9
97
O1
L1
1.0
0.9
0.8
0.7
0.6
0.5
0.4
0.3
5
6
7
8
9
10
11
12

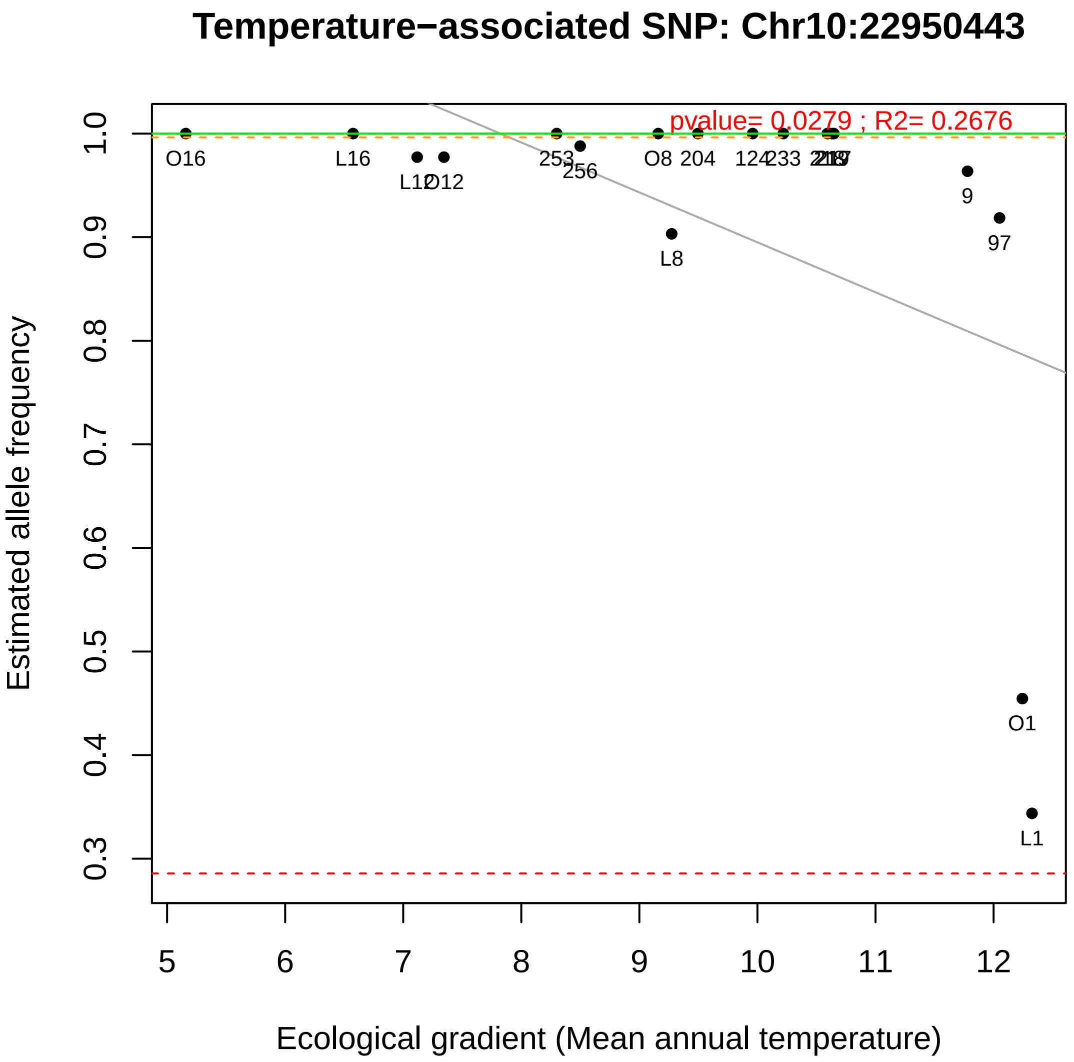
Temperature−associated SNP: Chr10:22950443
pvalue= 0.0279 ; R2= 0.2676
O16
L16
L12
O12
253
256
O8
204
124
233
L8
9
97
O1
L1
Estimated allele frequency
1.0
0.9
0.8
0.7
0.6
0.5
0.4
0.3
5
6
7
8
9
10
11
12
Ecological gradient (Mean annual temperature)

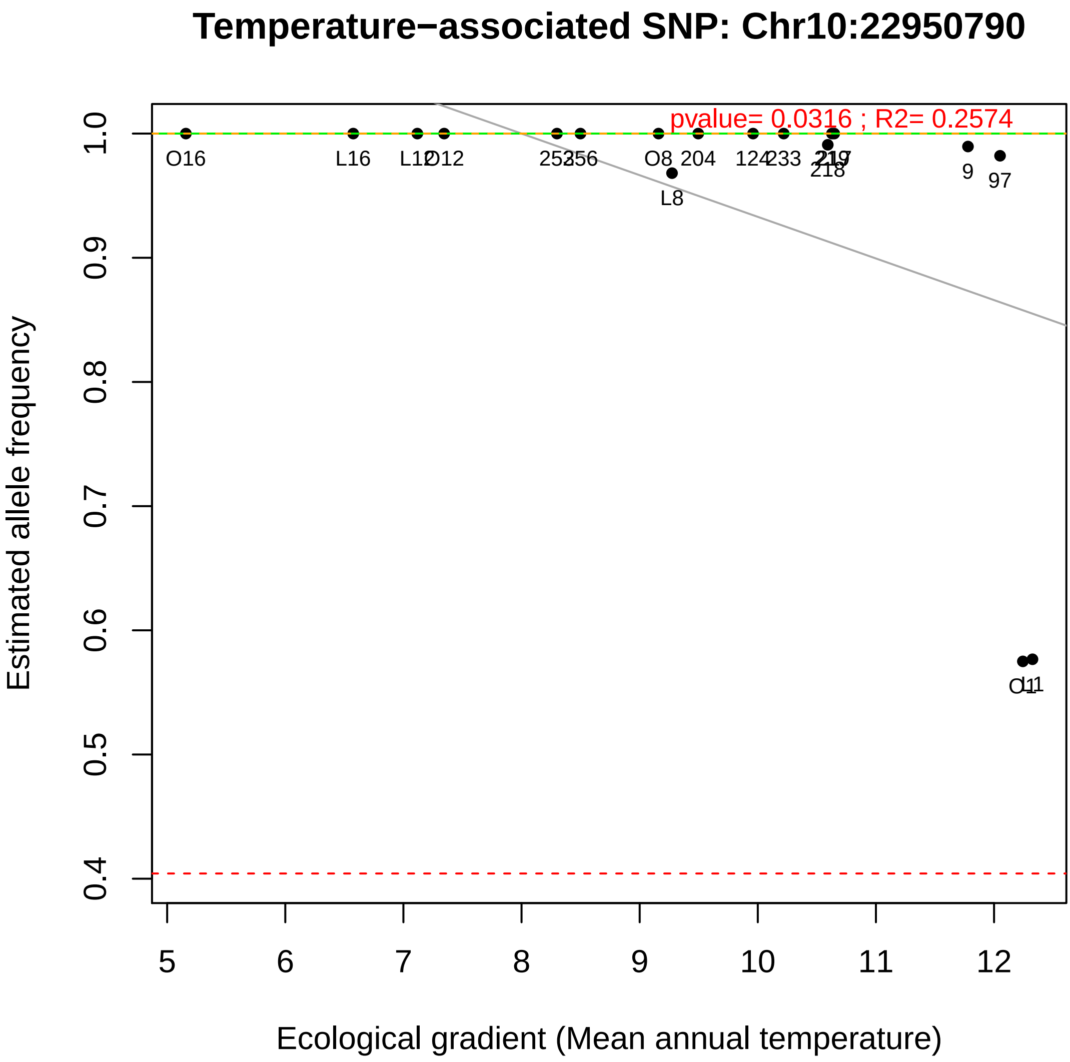

Temperature−associated SNP: Chr10:22950790
pvalue= 0.0316 ; R2= 0.2574
Estimated allele frequency
Ecological gradient (Mean annual temperature)
1.0
0.9
0.8
0.7
0.6
0.5
0.4
5
6
7
8
9
10
11
12
O16
L16
L12
O12
O8
204
L8
233
218
9
97

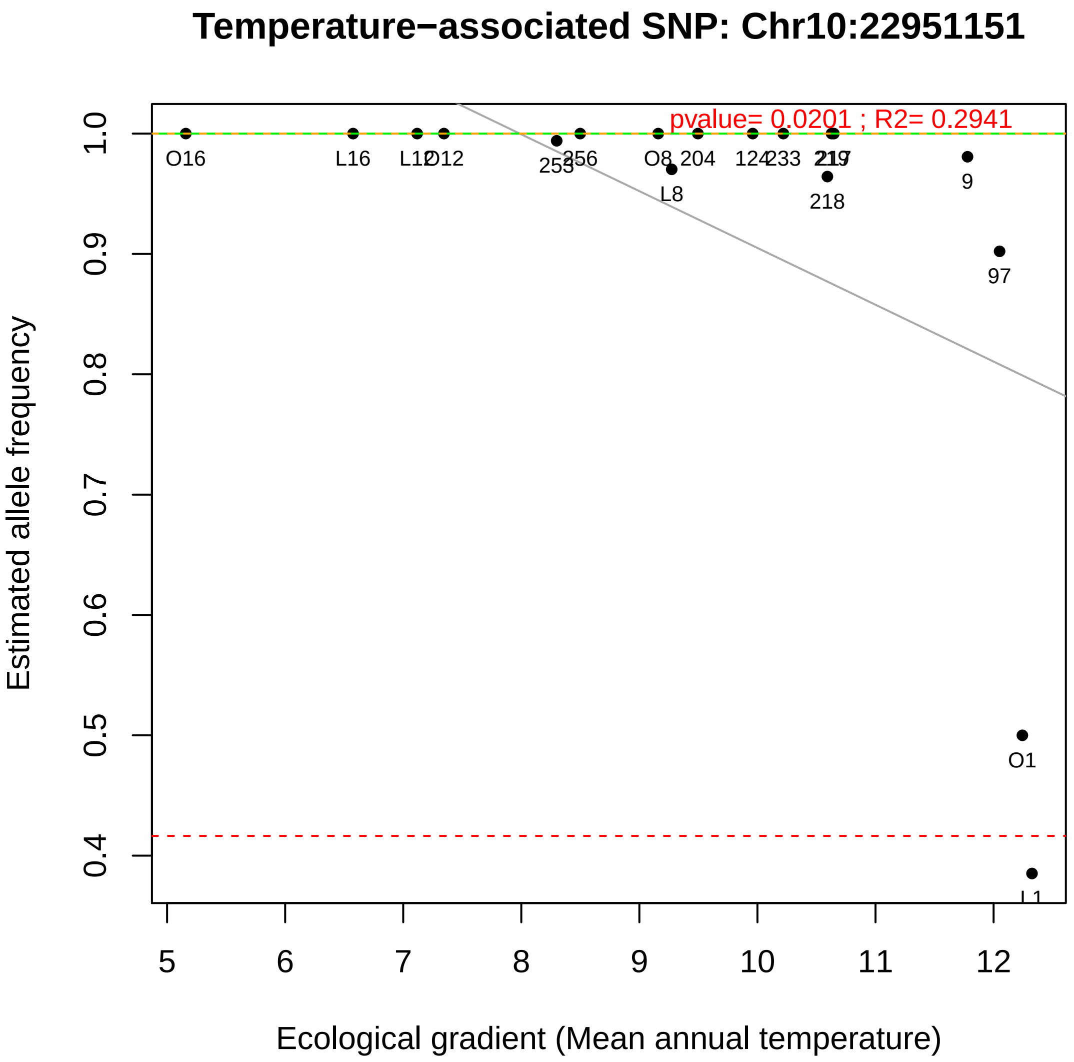
Temperature−associated SNP: Chr10:22951151
pvalue= 0.0201 ; R2= 0.2941
Estimated allele frequency
Ecological gradient (Mean annual temperature)
1.0
0.9
0.8
0.7
0.6
0.5
0.4
5
6
7
8
9
10
11
12
O16
L16
L12
O12
253
256
O8
204
L8
124
233
217
219
218
9
97
O1
L1

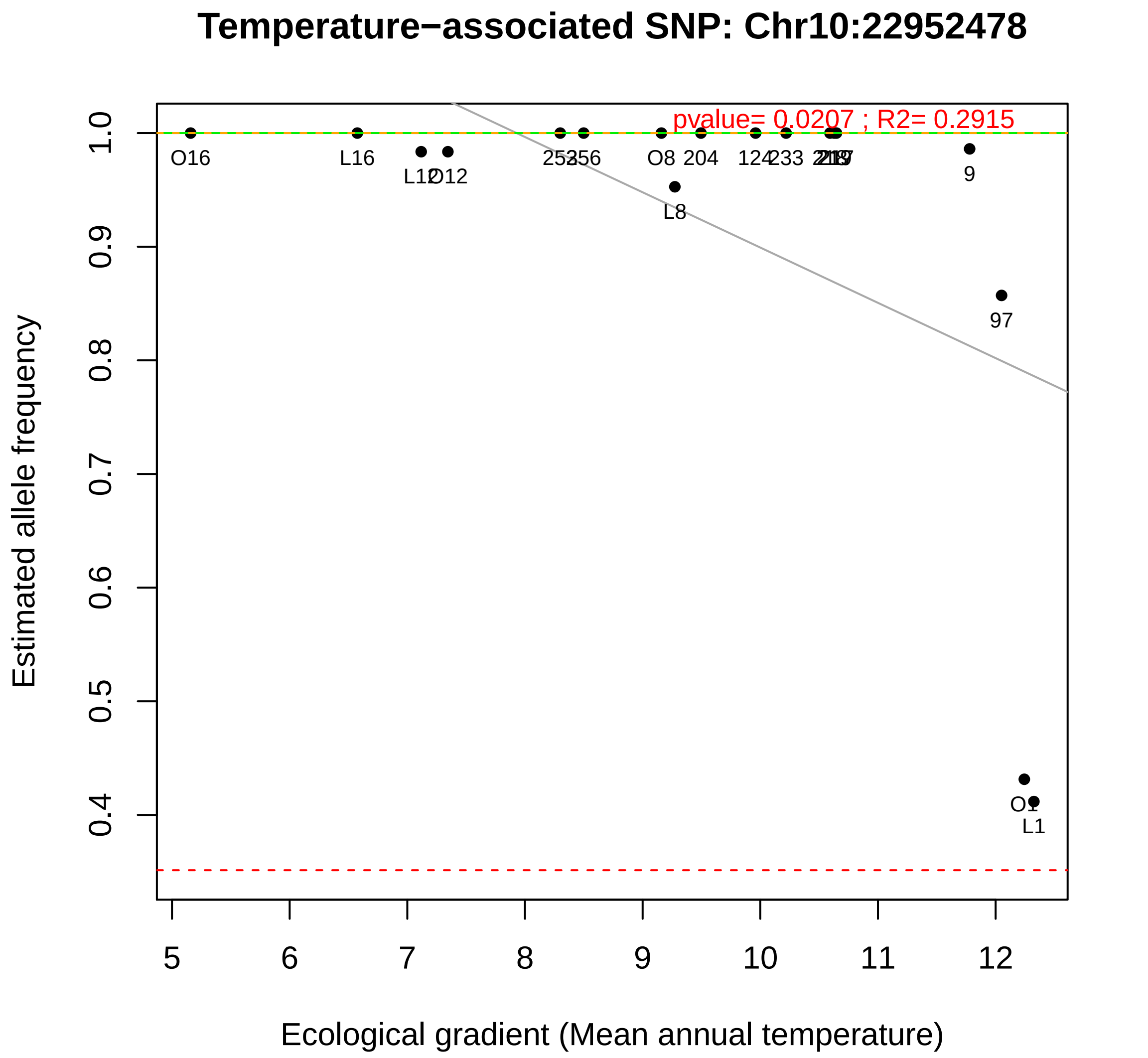

Temperature−associated SNP: Chr10:22952478
pvalue= 0.0207 ; R2= 0.2915
O16
L16
L12
O12
O8
204
L8
9
97
O1
L1
1.0
0.9
0.8
0.7
0.6
0.5
0.4
5
6
7
8
9
10
11
12
Estimated allele frequency
Ecological gradient (Mean annual temperature)

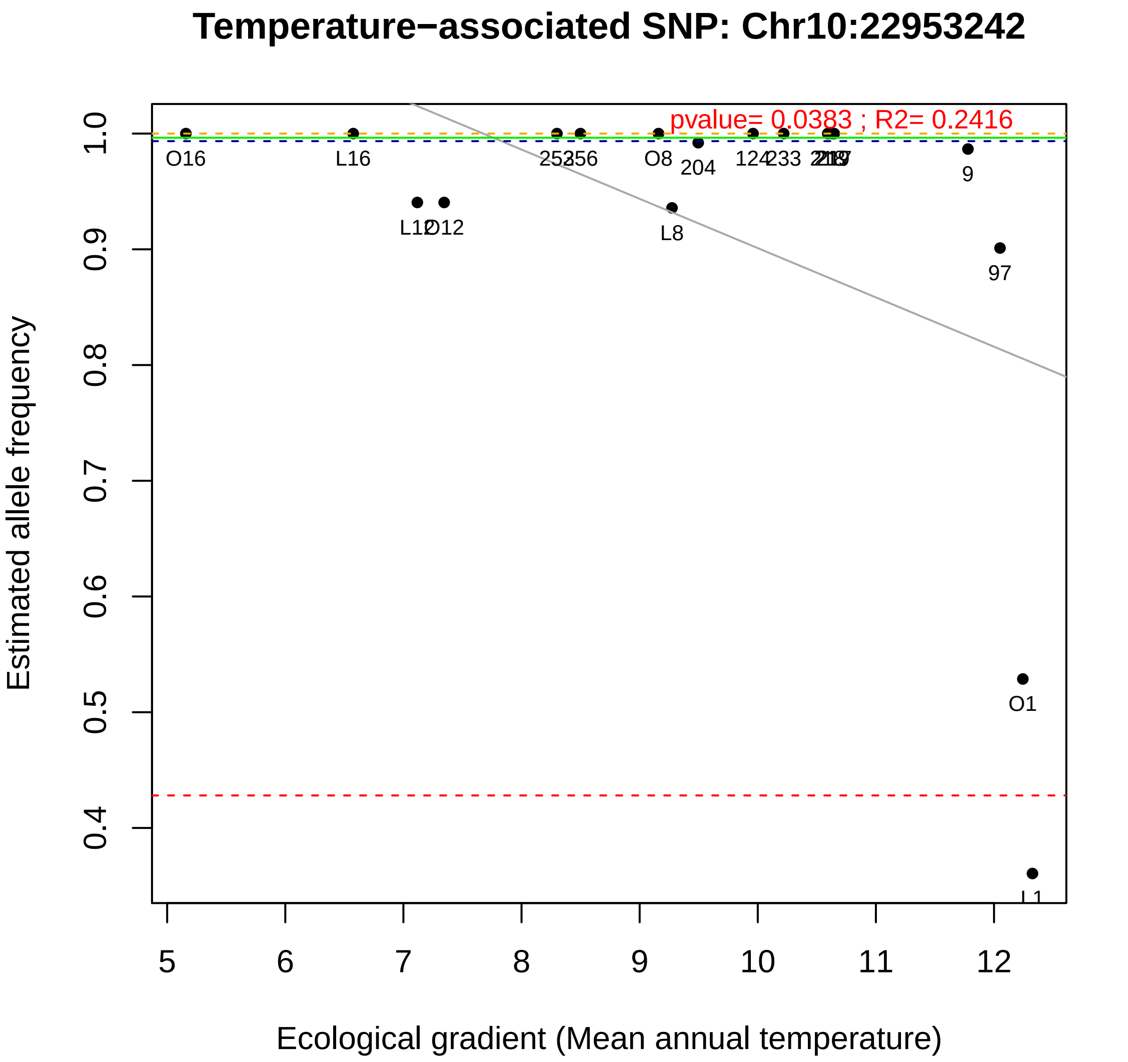

Temperature−associated SNP: Chr10:22953242
pvalue= 0.0383 ; R2= 0.2416
Estimated allele frequency
Ecological gradient (Mean annual temperature)
1.0
0.9
0.8
0.7
0.6
0.5
0.4
5
6
7
8
9
10
11
12
O16
L16
L12
O12
256
O8
204
L8
124
233
9
97
O1

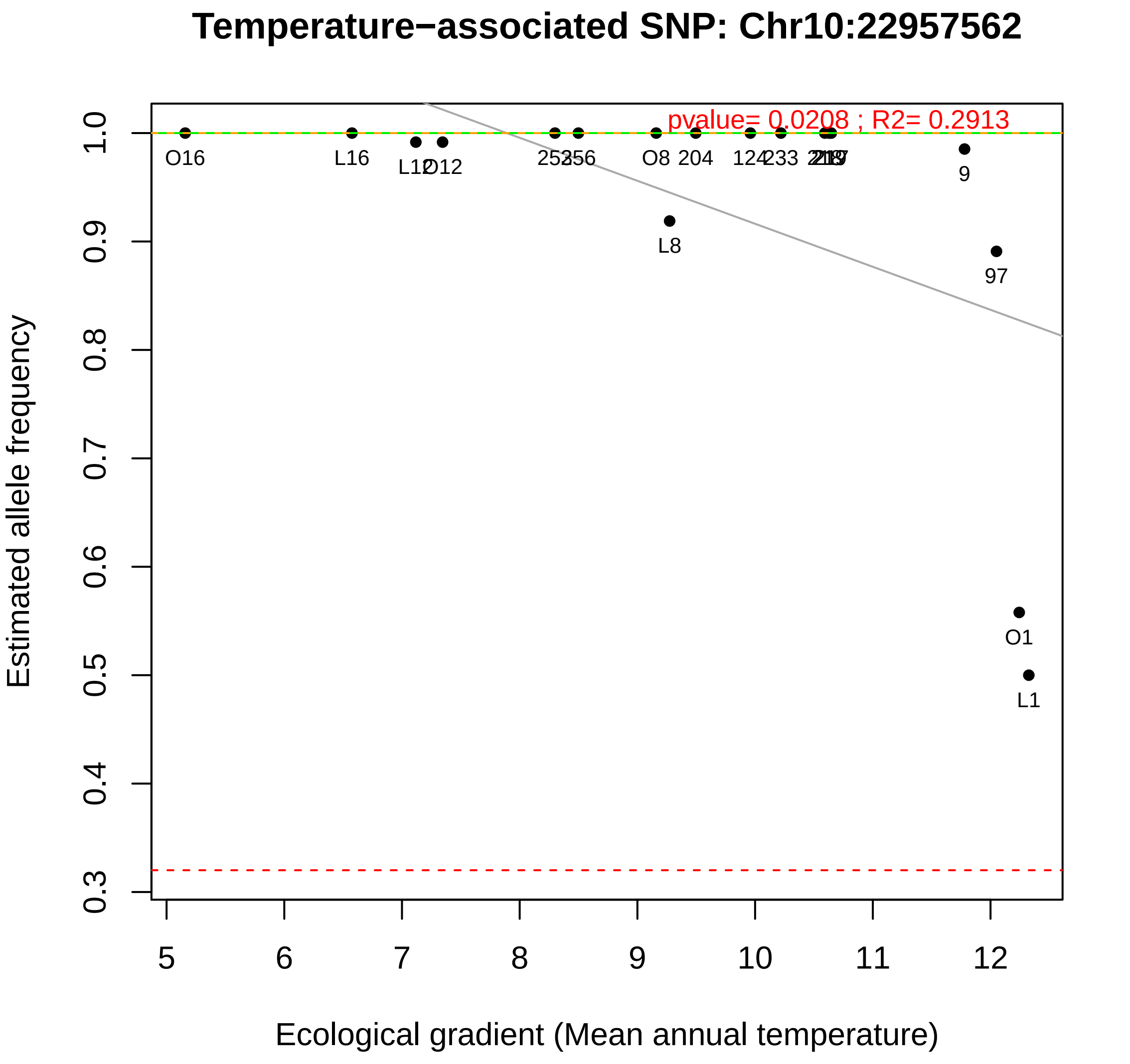

Temperature−associated SNP: Chr10:22957562
pvalue= 0.0208 ; R2= 0.2913
Estimated allele frequency
Ecological gradient (Mean annual temperature)
1.0
0.9
0.8
0.7
0.6
0.5
0.4
0.3
5
6
7
8
9
10
11
12
O16
L16
O12
O8
204
233
9
L8
97
O1
L1

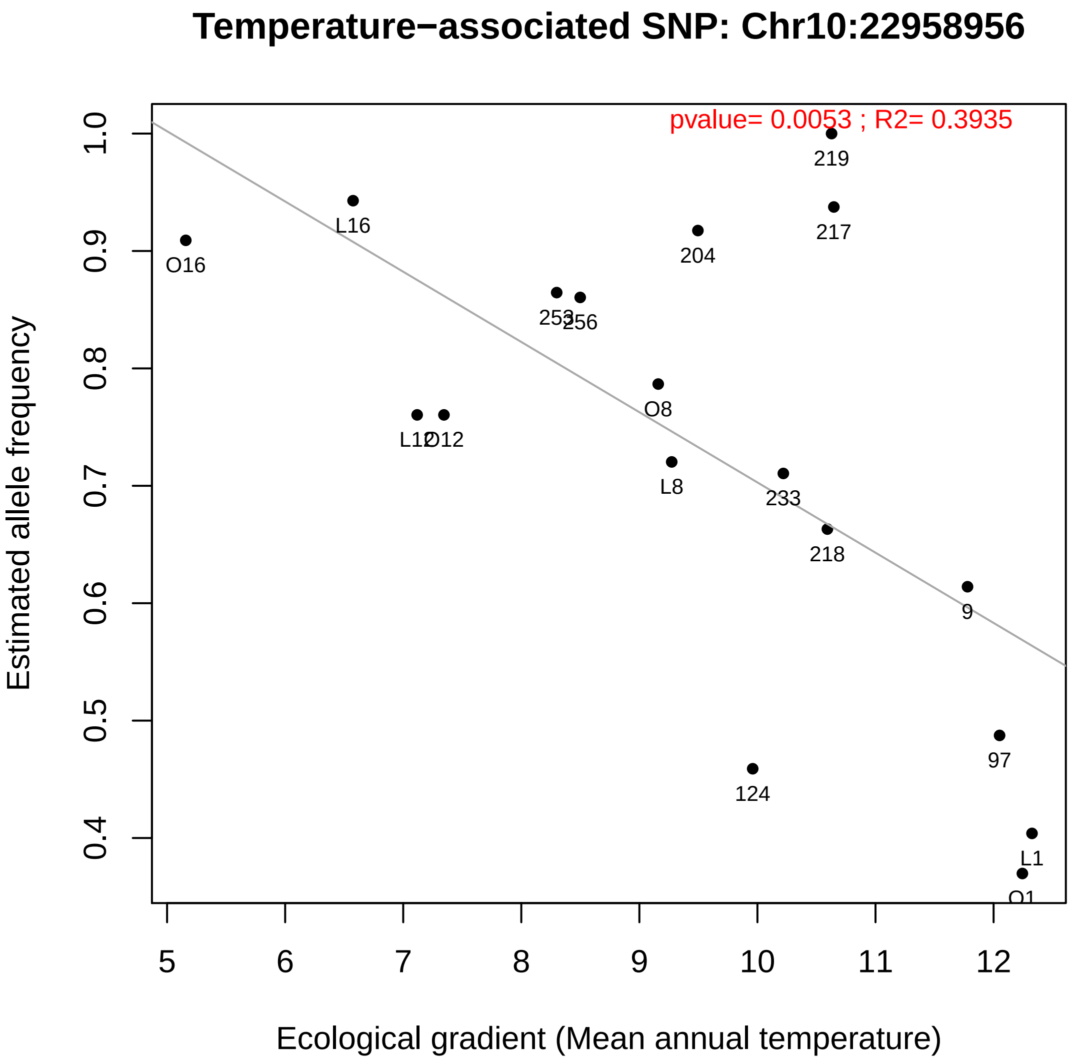
Temperature−associated SNP: Chr10:22958956
pvalue= 0.0053 ; R2= 0.3935
Estimated allele frequency
Ecological gradient (Mean annual temperature)
1.0
0.9
0.8
0.7
0.6
0.5
0.4
5
6
7
8
9
10
11
12
219
L16
217
O16
204
253
256
O8
L12
O12
L8
233
218
9
97
124
L1
O1

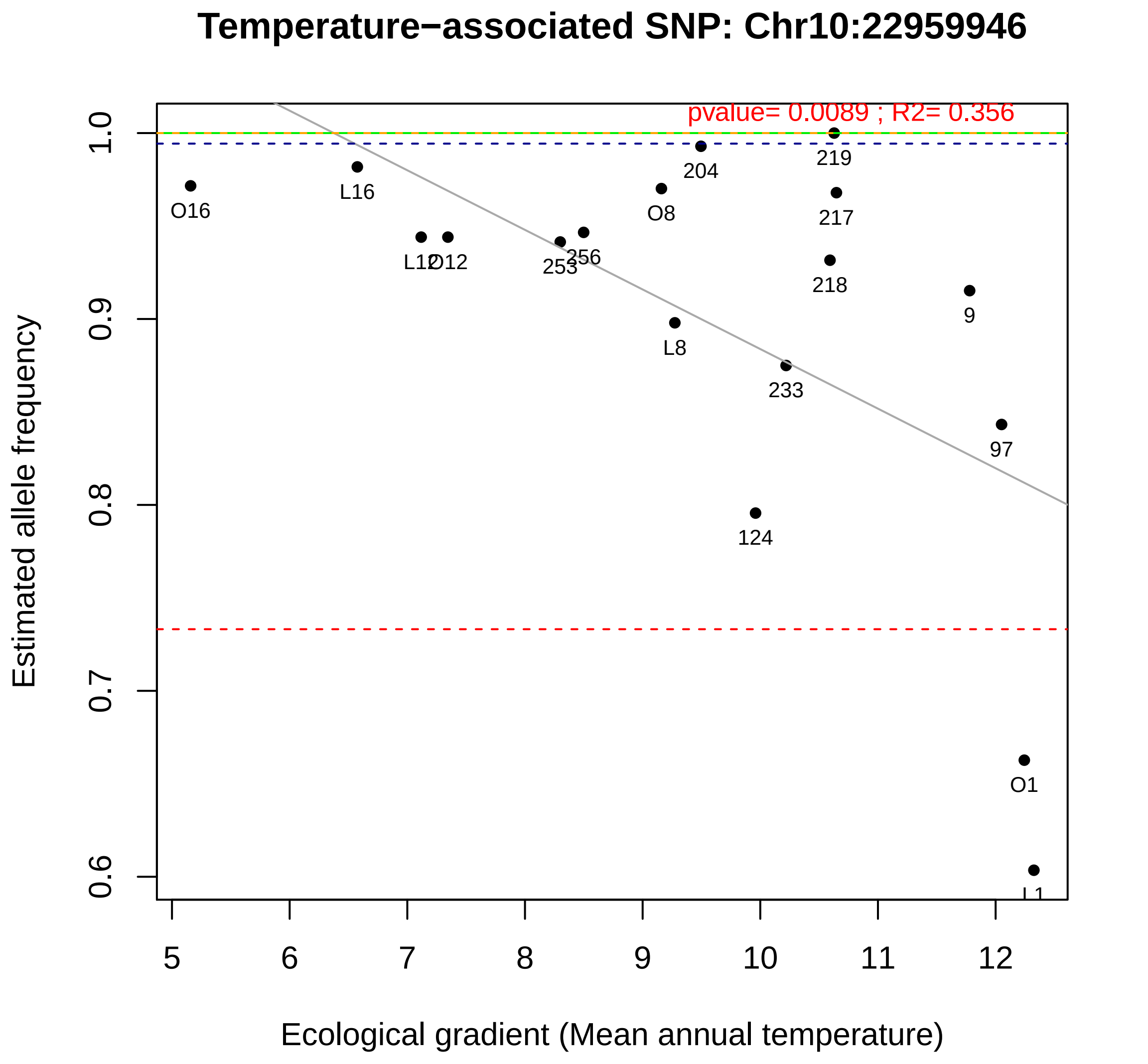
Temperature−associated SNP: Chr10:22959946
pvalue= 0.0089 ; R2= 0.356
Estimated allele frequency
Ecological gradient (Mean annual temperature)
1.0
0.9
0.8
0.7
0.6
5
6
7
8
9
10
11
12
O16
L16
L12
O12
253
256
O8
204
219
217
218
L8
233
124
9
97
O1
L1

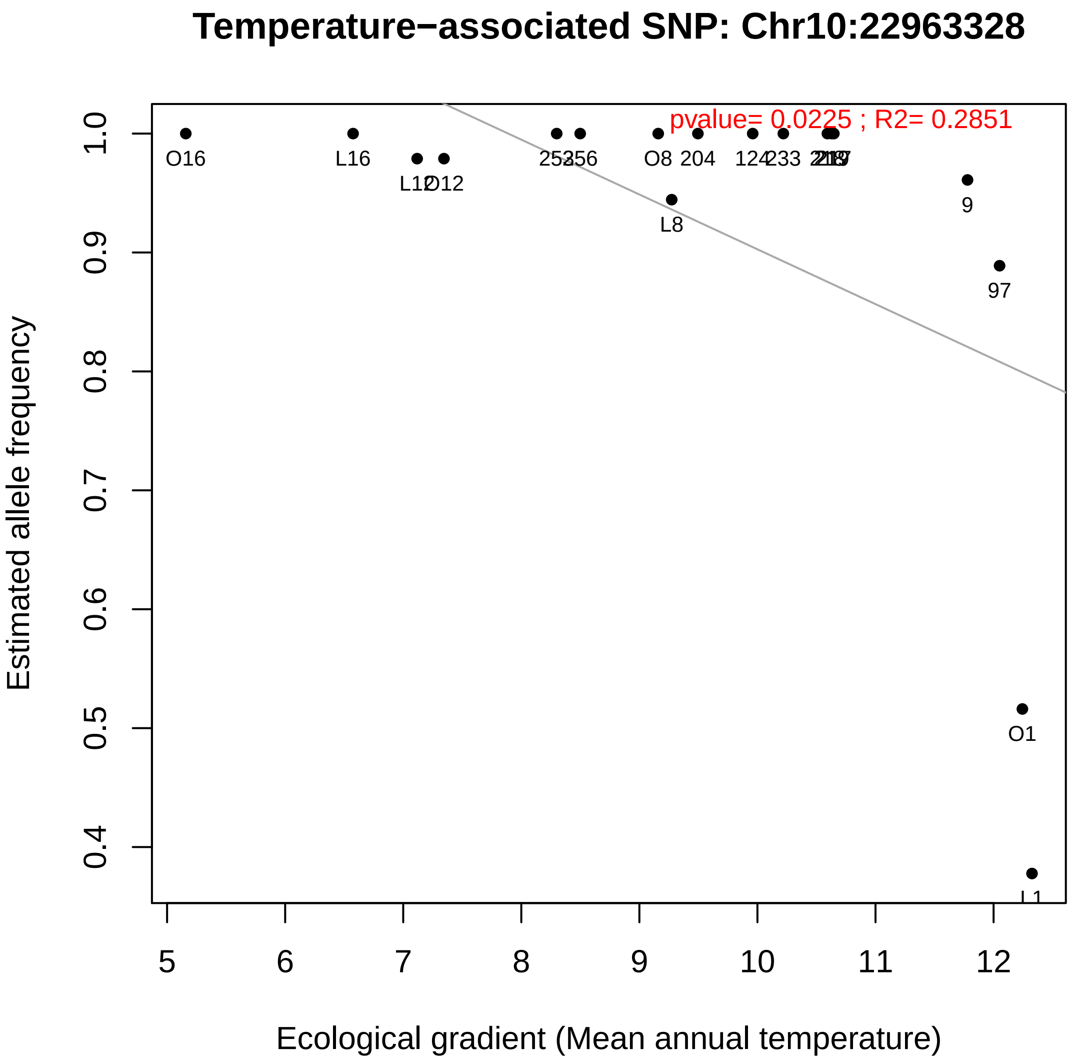
Temperature−associated SNP: Chr10:22963328
pvalue= 0.0225 ; R2= 0.2851
Estimated allele frequency
Ecological gradient (Mean annual temperature)
1.0
0.9
0.8
0.7
0.6
0.5
0.4
5
6
7
8
9
10
11
12
O16
L16
L12
O12
O8
204
L8
9
97
O1

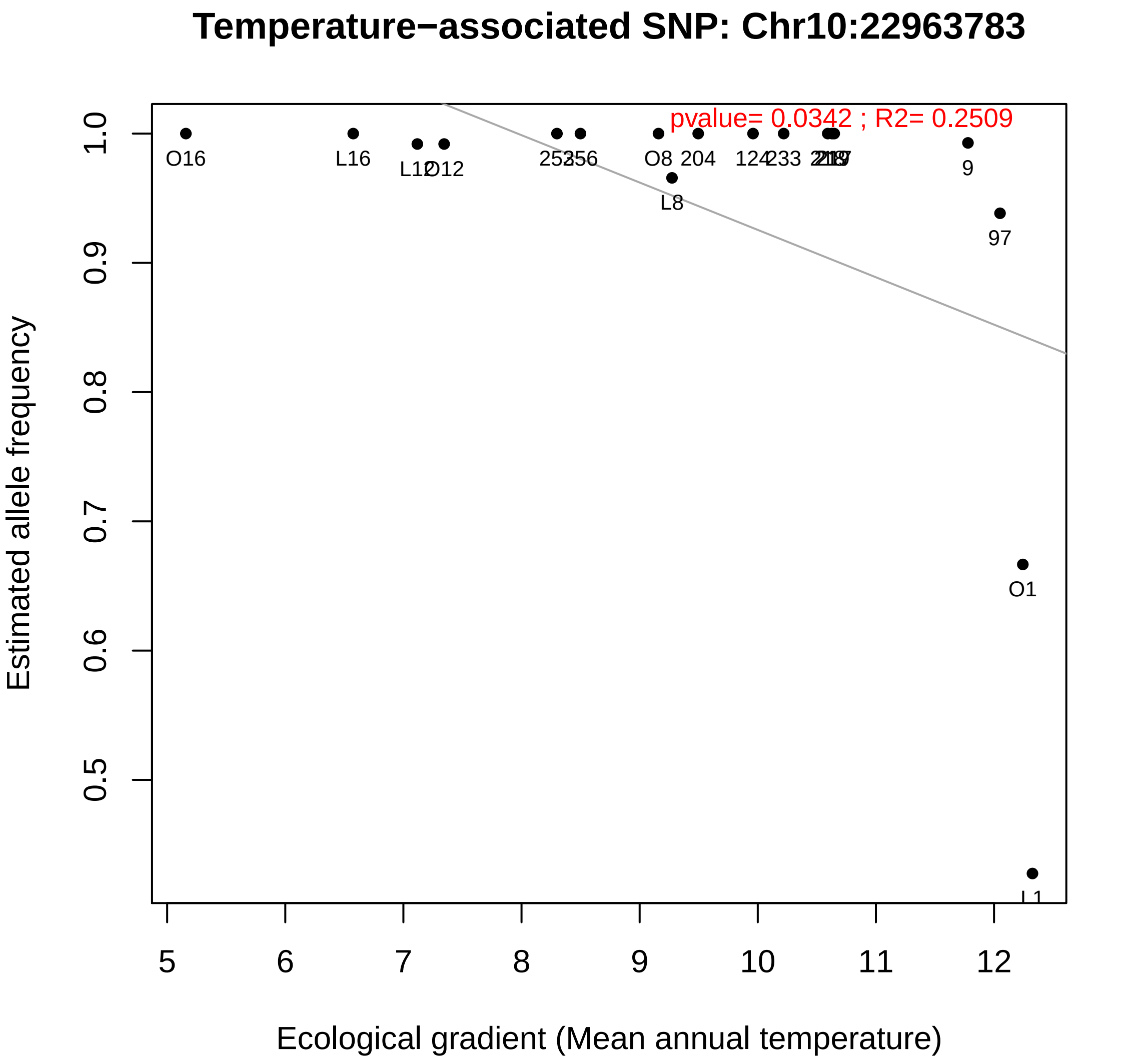

Temperature−associated SNP: Chr10:22963783
pvalue= 0.0342 ; R2= 0.2509
Estimated allele frequency
Ecological gradient (Mean annual temperature)
1.0
0.9
0.8
0.7
0.6
0.5
5
6
7
8
9
10
11
12
O16
L16
L12
O12
O8
204
L8
9
97
O1

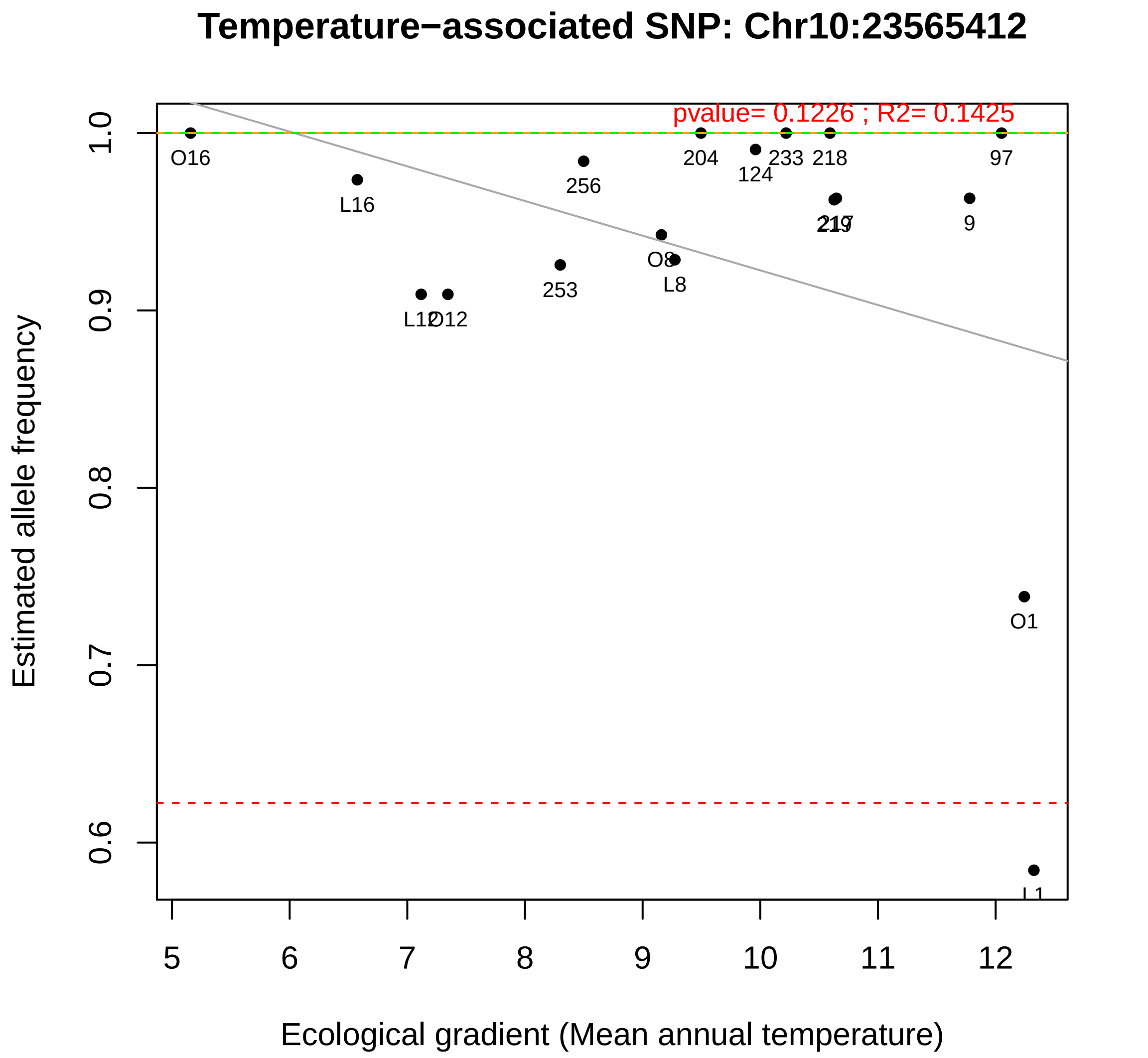

Temperature−associated SNP: Chr10:23565412
pvalue= 0.1226 ; R2= 0.1425
Estimated allele frequency
Ecological gradient (Mean annual temperature)
1.0
0.9
0.8
0.7
0.6
5
6
7
8
9
10
11
12
O16
L16
L12
O12
253
256
O8
L8
204
124
233
218
97
9
O1

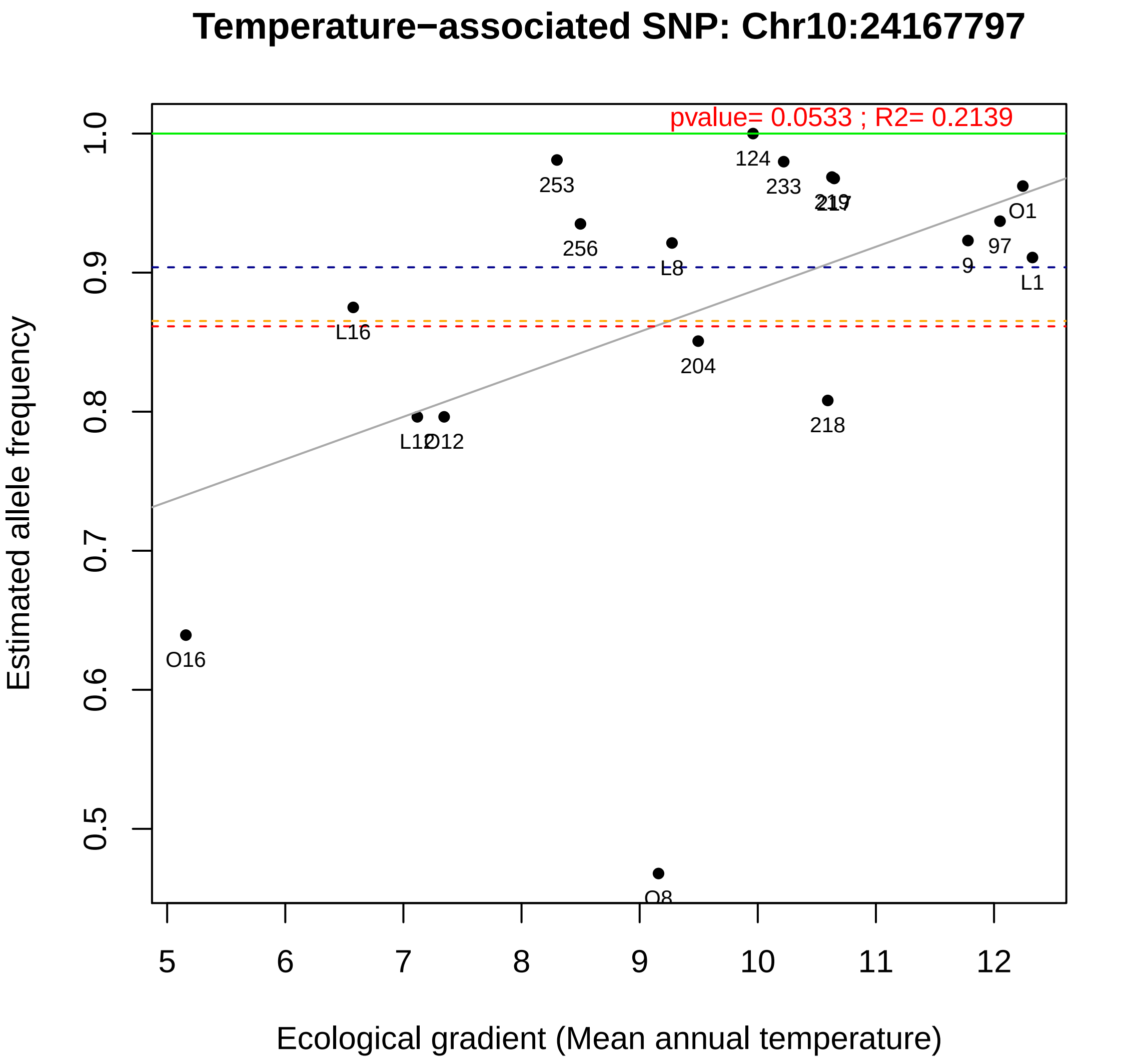

Temperature−associated SNP: Chr10:24167797
pvalue= 0.0533 ; R2= 0.2139
Estimated allele frequency
Ecological gradient (Mean annual temperature)
1.0
0.9
0.8
0.7
0.6
0.5
5
6
7
8
9
10
11
12
124
253
233
O1
256
L8
97
9
L1
L16
204
218
L12
O12
O16
O8

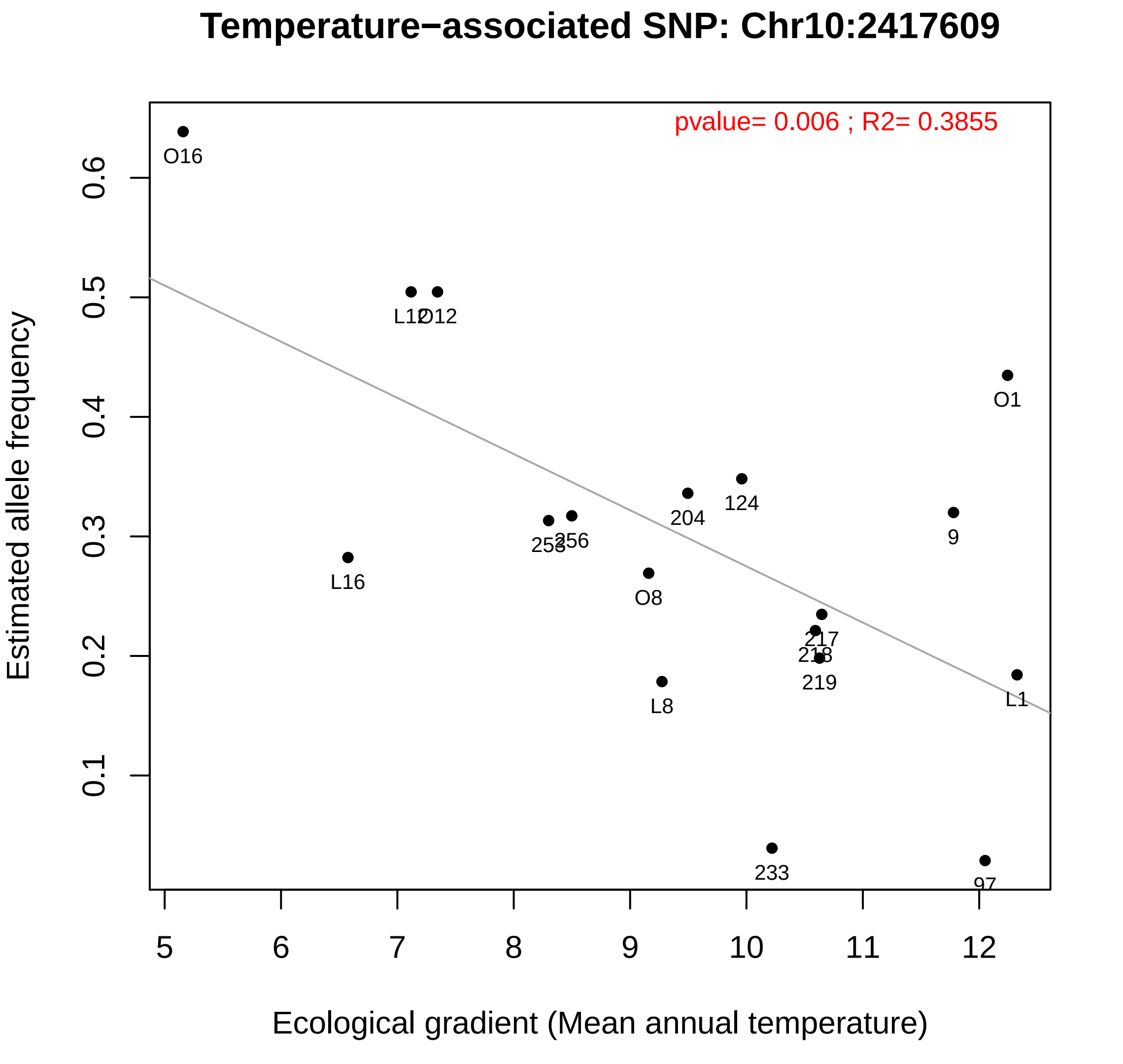

Temperature−associated SNP: Chr10:2417609
pvalue= 0.006 ; R2= 0.3855
Estimated allele frequency
Ecological gradient (Mean annual temperature)
0.1
0.2
0.3
0.4
0.5
0.6
5
6
7
8
9
10
11
12
O16
L12
O12
O1
124
204
253
256
9
L16
O8
217
218
219
L8
L1
233
97

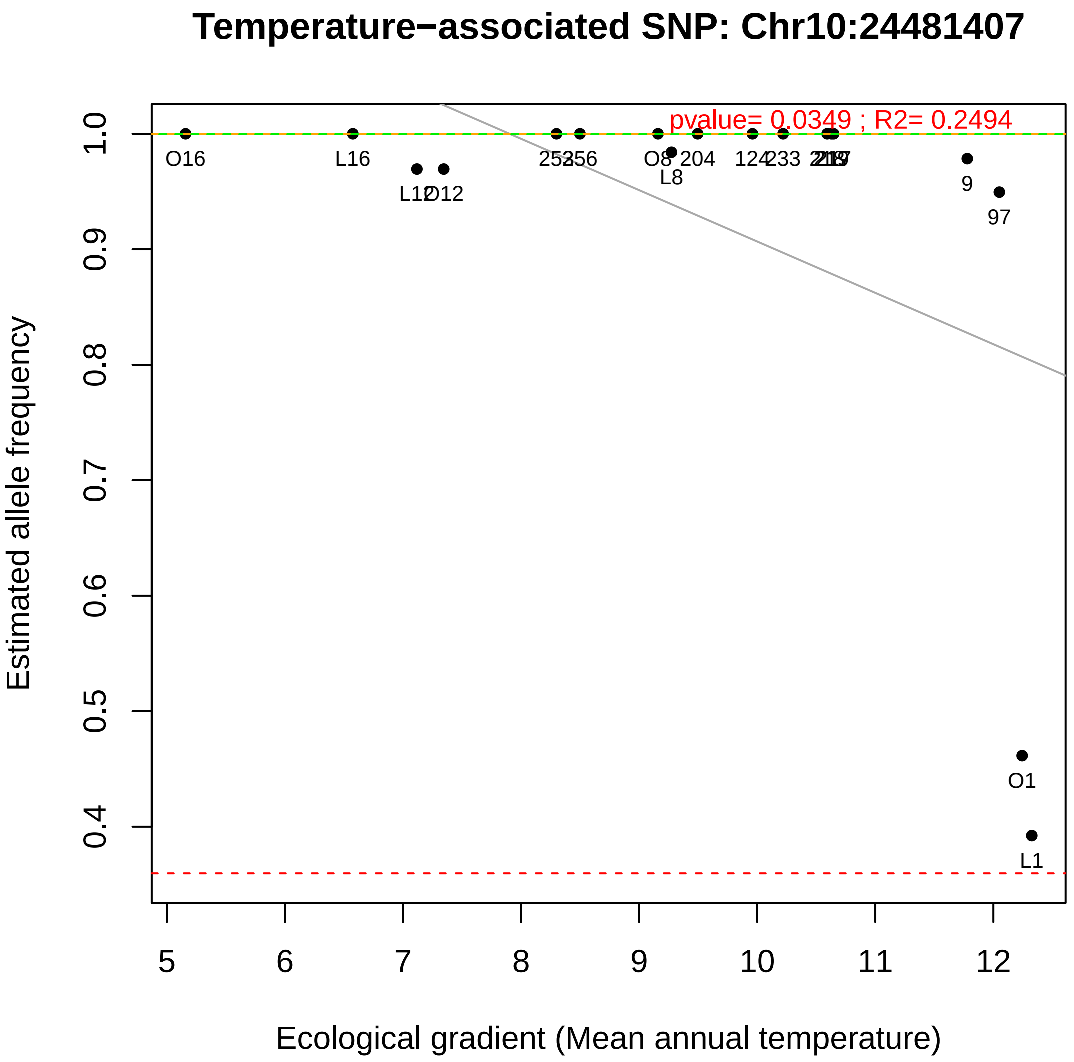

Temperature−associated SNP: Chr10:24481407
pvalue= 0.0349 ; R2= 0.2494
O16
L16
L12
O12
253
256
O8
L8
204
124
233
9
97
O1
L1
1.0
0.9
0.8
0.7
0.6
0.5
0.4
Estimated allele frequency
5
6
7
8
9
10
11
12
Ecological gradient (Mean annual temperature)

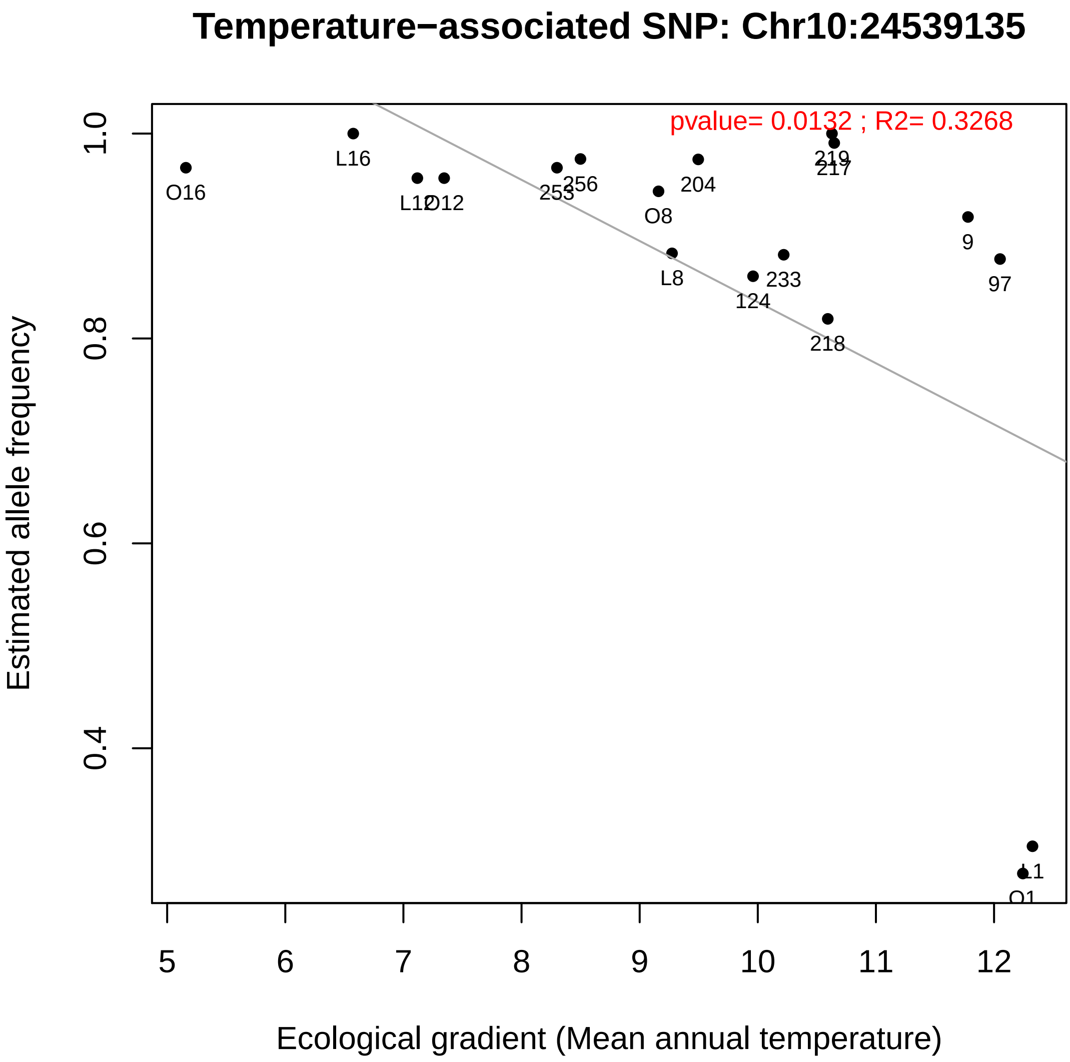
Temperature−associated SNP: Chr10:24539135
pvalue= 0.0132 ; R2= 0.3268
Estimated allele frequency
Ecological gradient (Mean annual temperature)
1.0
0.8
0.6
0.4
5
6
7
8
9
10
11
12
O16
L16
L12
O12
253
256
O8
204
219
217
L8
124
233
218
9
97
L1
O1

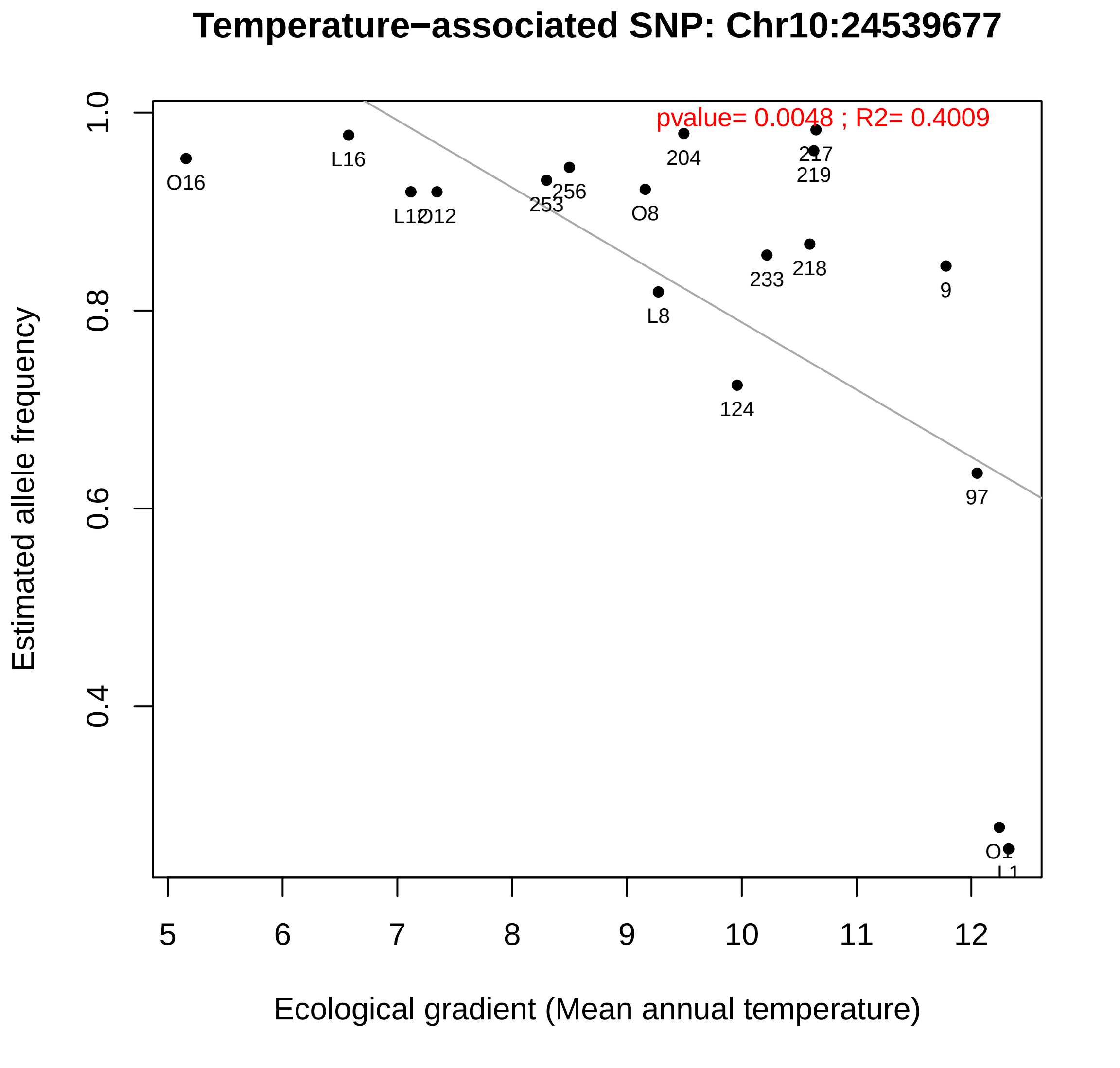

Temperature−associated SNP: Chr10:24539677
pvalue= 0.0048 ; R2= 0.4009
O16
L16
L12
O12
253
256
O8
204
217
219
L8
233
218
9
124
97
O1
L1
Estimated allele frequency
0.4
0.6
0.8
1.0
5
6
7
8
9
10
11
12
Ecological gradient (Mean annual temperature)

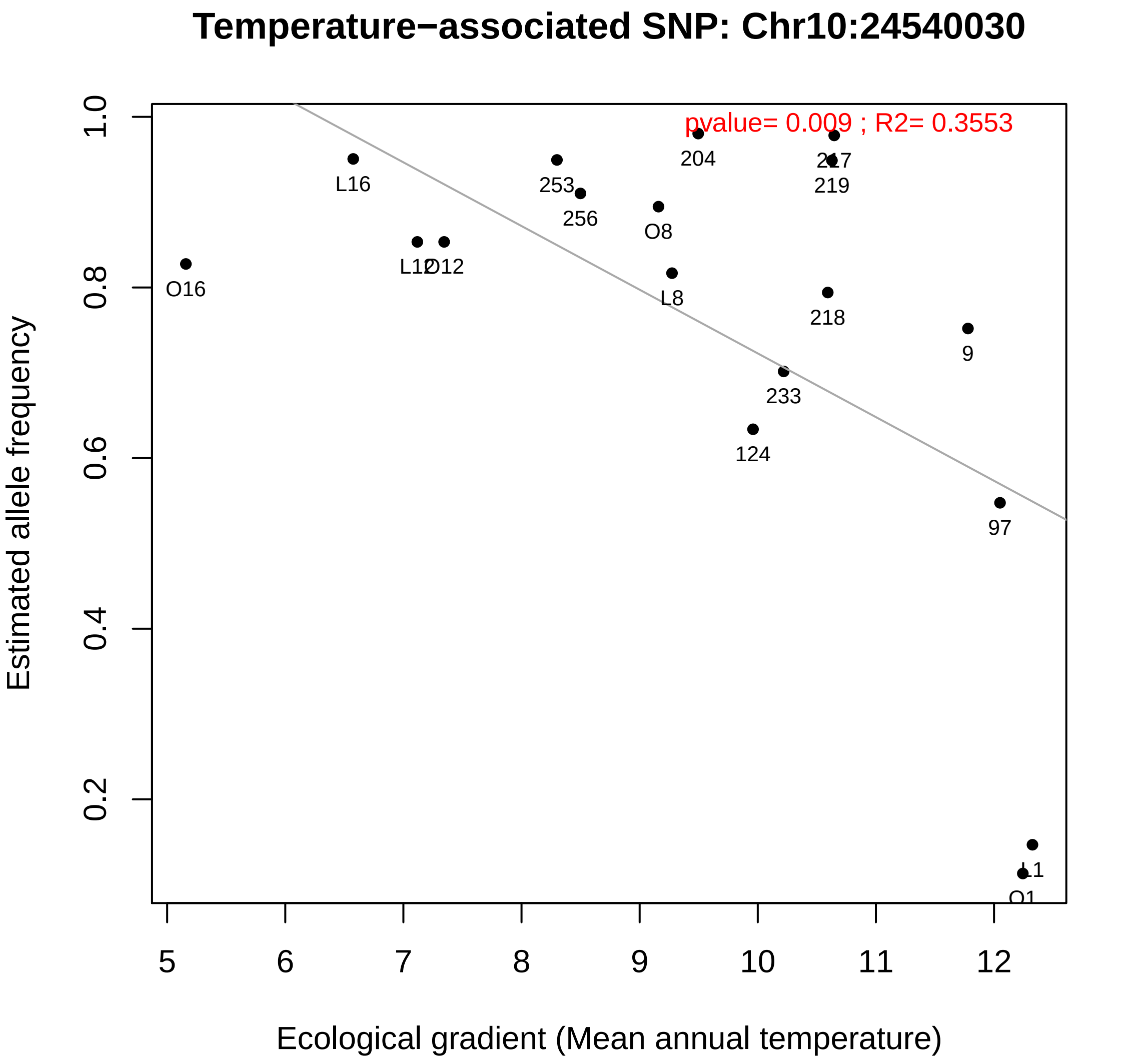

Temperature−associated SNP: Chr10:24540030
pvalue= 0.009 ; R2= 0.3553
Estimated allele frequency
Ecological gradient (Mean annual temperature)
0.2
0.4
0.6
0.8
1.0
5
6
7
8
9
10
11
12
O16
L16
L12
O12
253
256
204
O8
L8
217
219
218
9
233
124
97
L1
O1

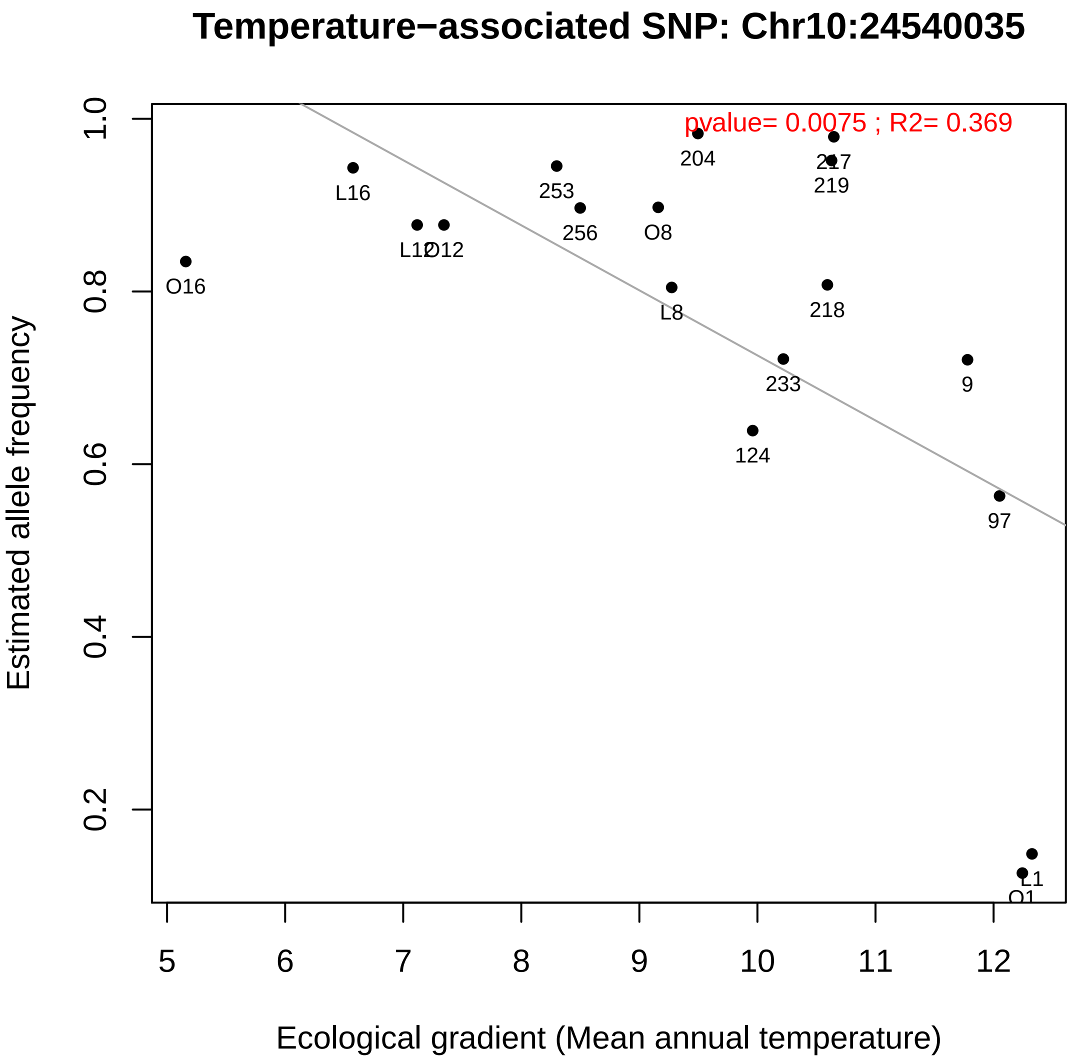

Temperature−associated SNP: Chr10:24540035
pvalue= 0.0075 ; R2= 0.369
Estimated allele frequency
Ecological gradient (Mean annual temperature)
0.2
0.4
0.6
0.8
1.0
5
6
7
8
9
10
11
12
O16
L16
L12
O12
253
256
O8
204
217
219
L8
218
233
124
9
97
L1
O1

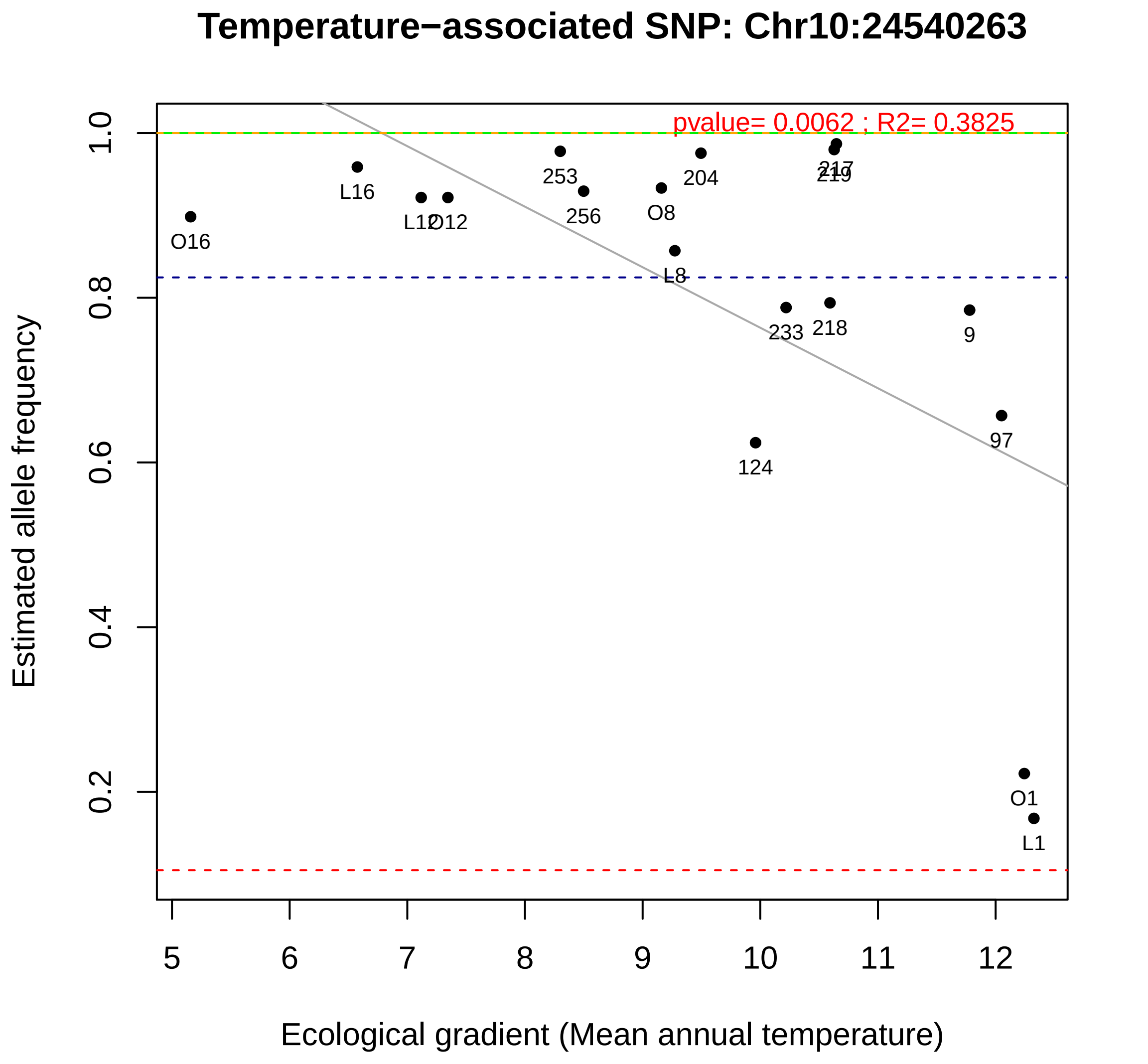

Temperature−associated SNP: Chr10:24540263
pvalue= 0.0062 ; R2= 0.3825
Estimated allele frequency
Ecological gradient (Mean annual temperature)
0.2
0.4
0.6
0.8
1.0
5
6
7
8
9
10
11
12
O16
L16
L12
O12
253
256
O8
204
L8
217
219
233
218
124
9
97
O1
L1

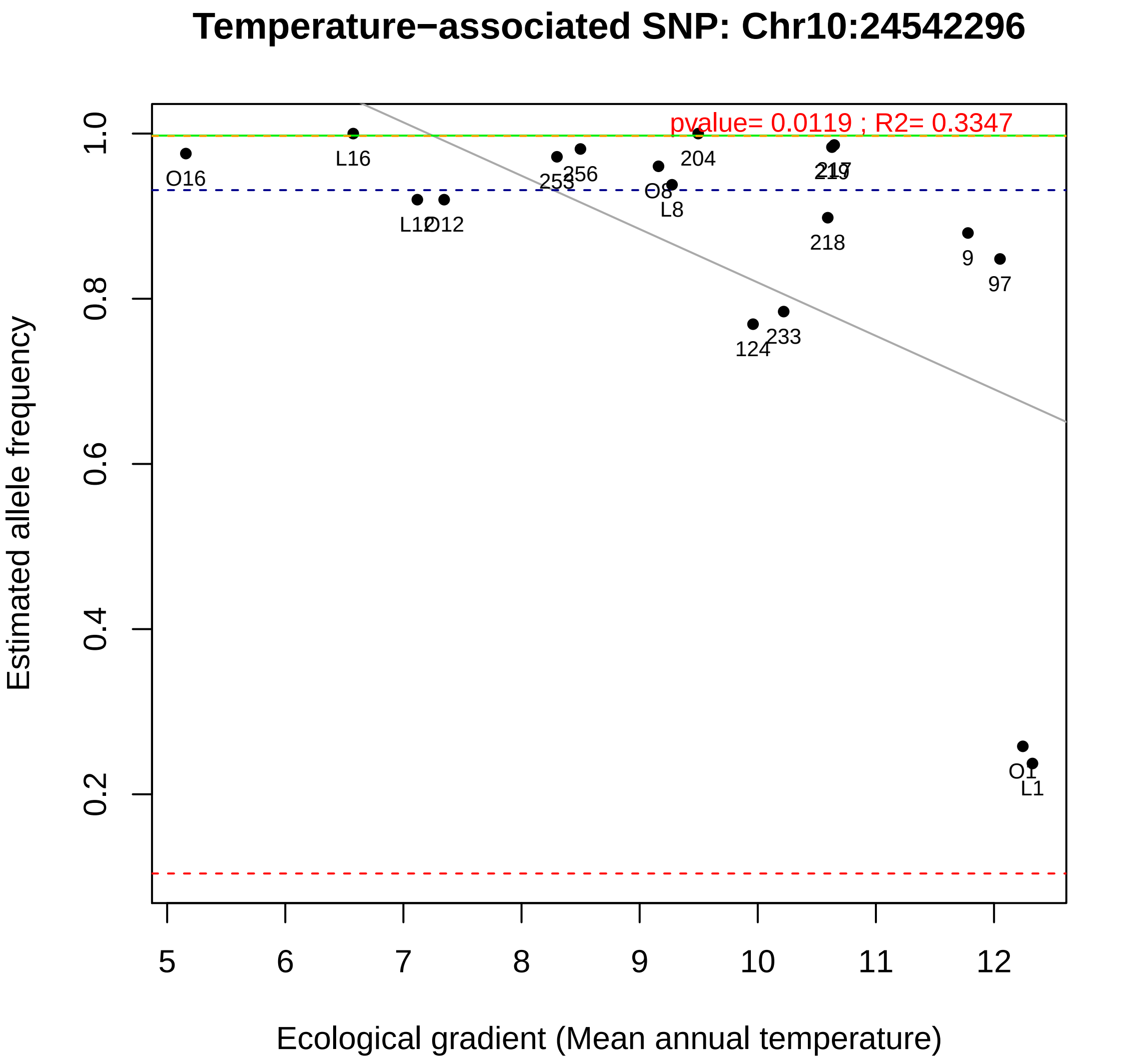

Temperature−associated SNP: Chr10:24542296
pvalue= 0.0119 ; R2= 0.3347
Estimated allele frequency
Ecological gradient (Mean annual temperature)
0.2
0.4
0.6
0.8
1.0
5
6
7
8
9
10
11
12
O16
L16
L12
O12
253
256
O8
L8
204
124
233
218
9
97
O1
L1

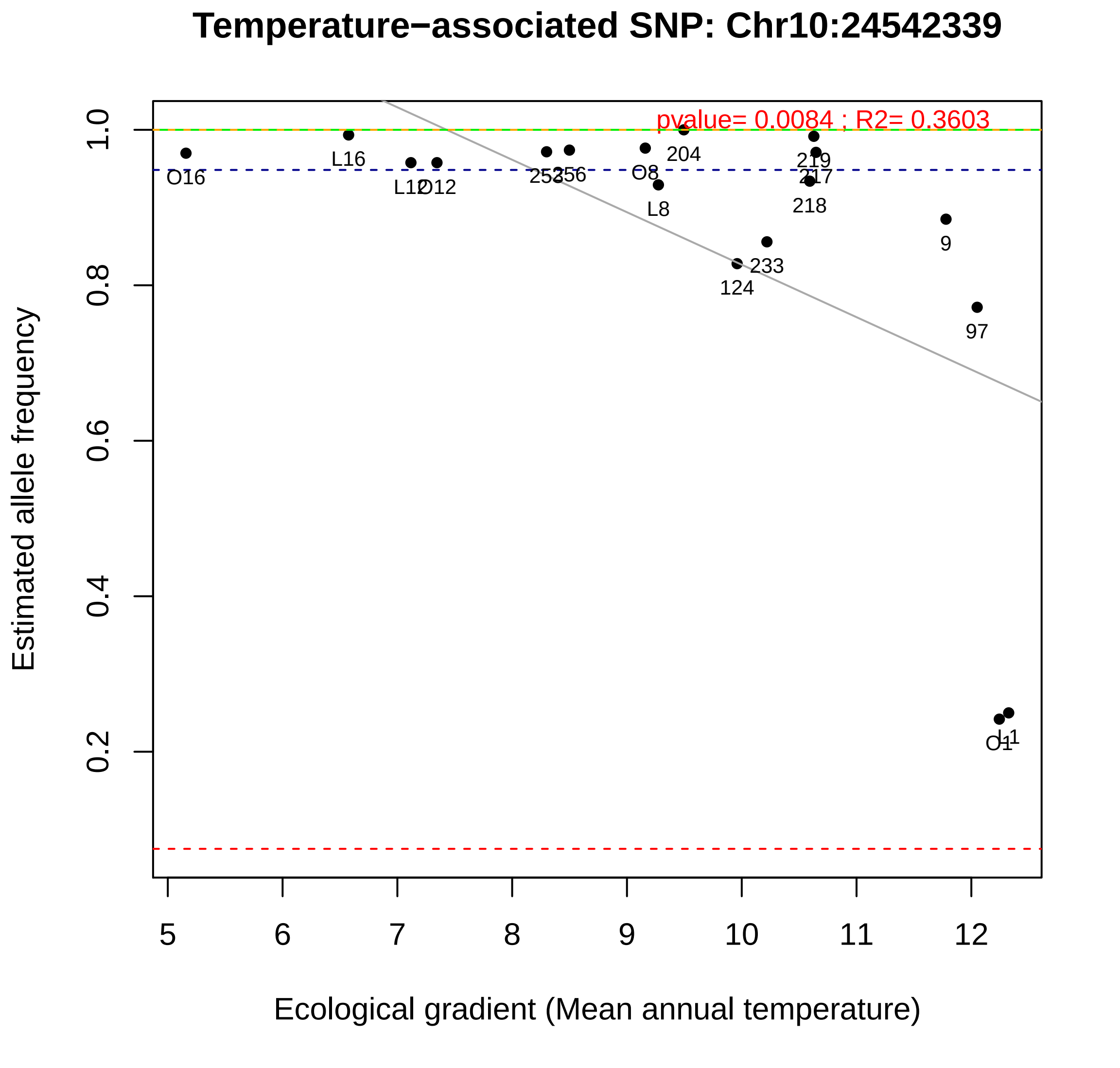

Temperature−associated SNP: Chr10:24542339
pvalue= 0.0084 ; R2= 0.3603
Estimated allele frequency
0.2
0.4
0.6
0.8
1.0
5
6
7
8
9
10
11
12
Ecological gradient (Mean annual temperature)
O16
L16
L12
O12
253
256
O8
L8
204
219
217
218
124
233
9
97
O1
L1

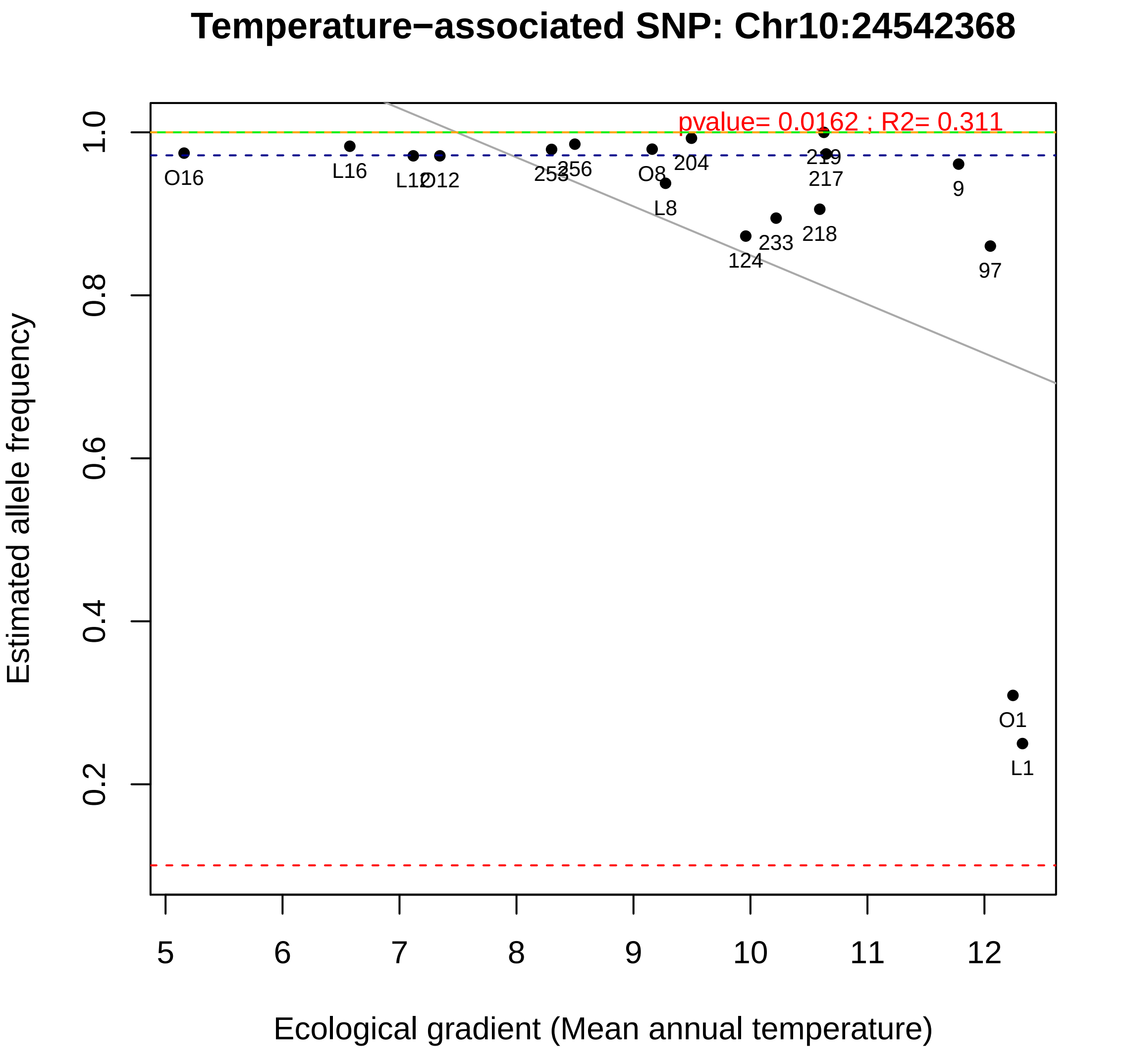

Temperature−associated SNP: Chr10:24542368
pvalue= 0.0162 ; R2= 0.311
Estimated allele frequency
Ecological gradient (Mean annual temperature)
0.2
0.4
0.6
0.8
1.0
5
6
7
8
9
10
11
12
O16
L16
L12
O12
253
256
O8
L8
204
219
217
218
233
124
9
97
O1
L1

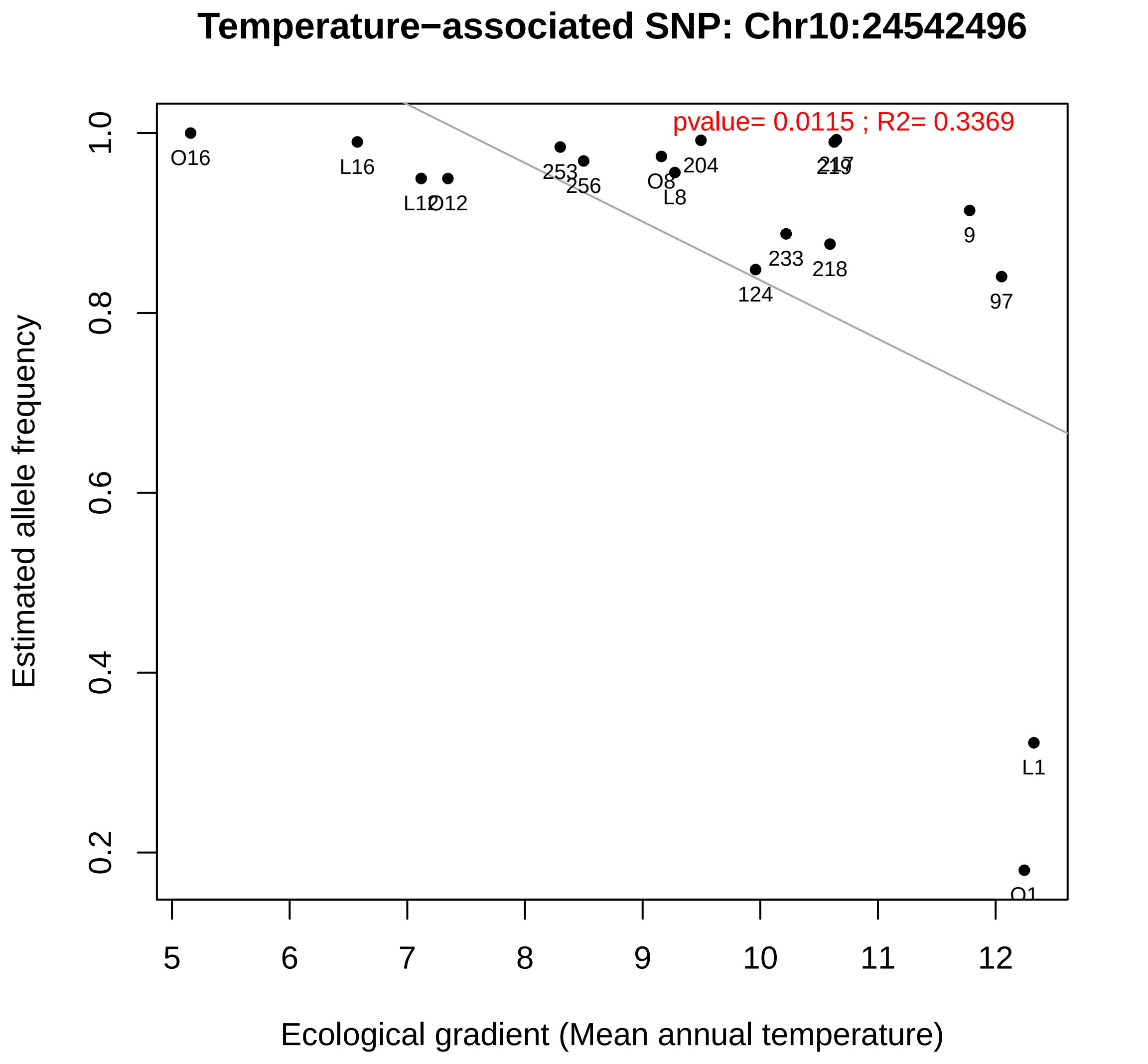

Temperature−associated SNP: Chr10:24542496
pvalue= 0.0115 ; R2= 0.3369
Estimated allele frequency
Ecological gradient (Mean annual temperature)
1.0
0.8
0.6
0.4
0.2
5
6
7
8
9
10
11
12
O16
L16
L12
O12
253
256
O8
L8
204
217
219
233
218
124
9
97
L1
O1

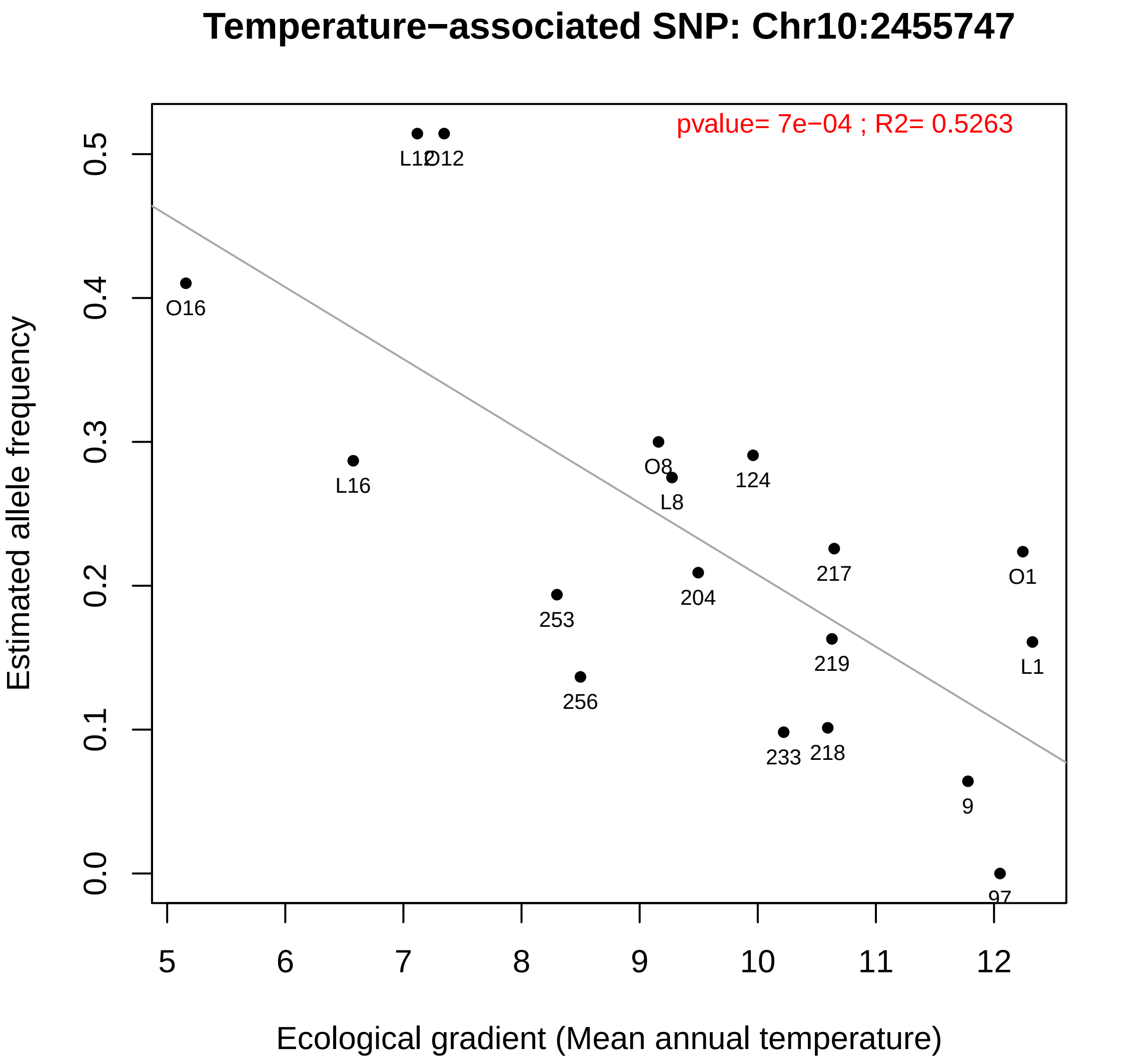
Temperature−associated SNP: Chr10:2455747
pvalue= 7e−04 ; R2= 0.5263
L12
O12
O16
L16
O8
L8
124
217
O1
204
253
219
L1
256
233
218
9
97
0.0
0.1
0.2
0.3
0.4
0.5
5
6
7
8
9
10
11
12
Estimated allele frequency
Ecological gradient (Mean annual temperature)

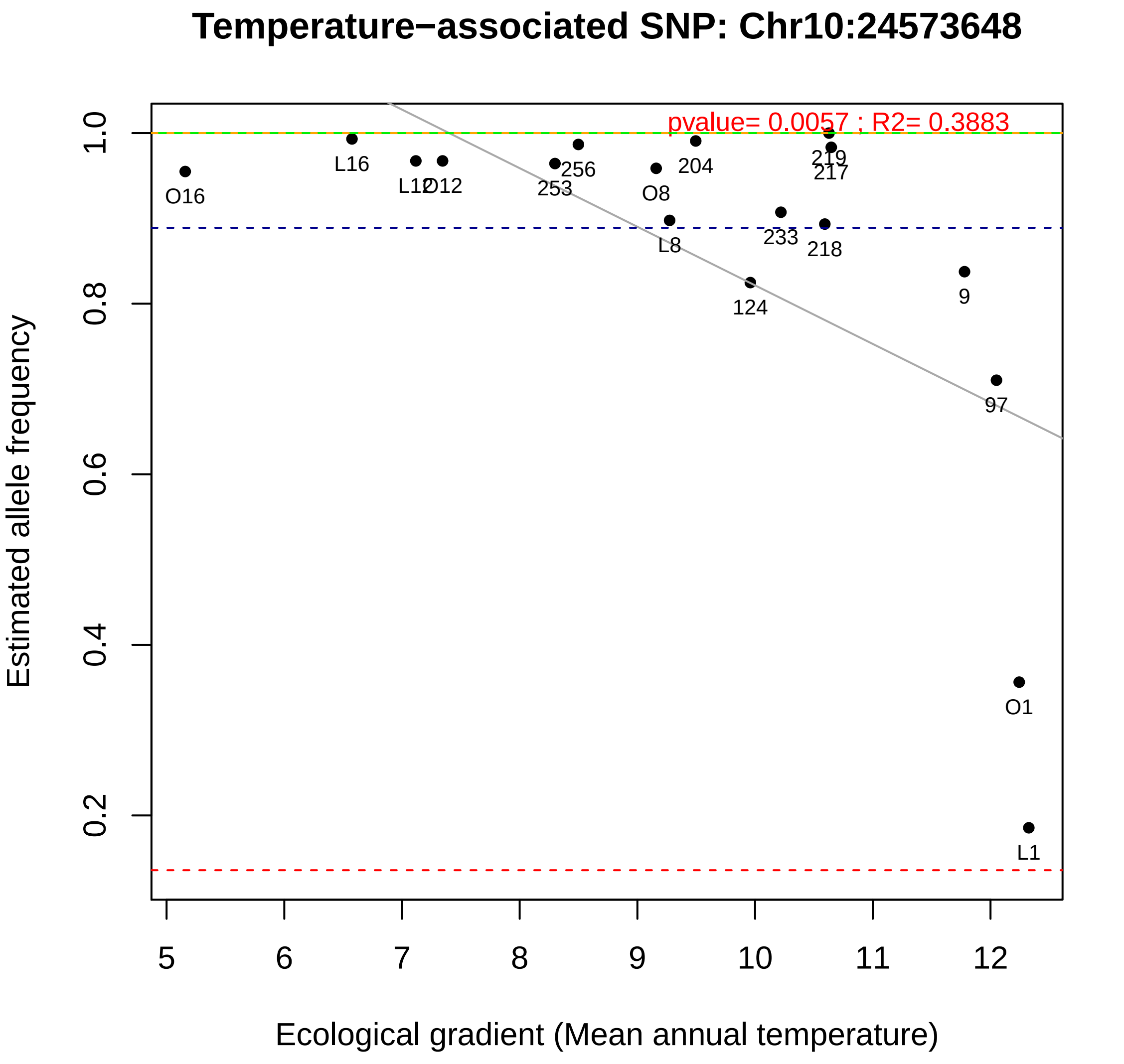
Temperature−associated SNP: Chr10:24573648
pvalue= 0.0057 ; R2= 0.3883
Estimated allele frequency
0.2
0.4
0.6
0.8
1.0
5
6
7
8
9
10
11
12
Ecological gradient (Mean annual temperature)
O16
L16
L12
O12
253
256
O8
204
219
217
L8
233
218
124
9
97
O1
L1

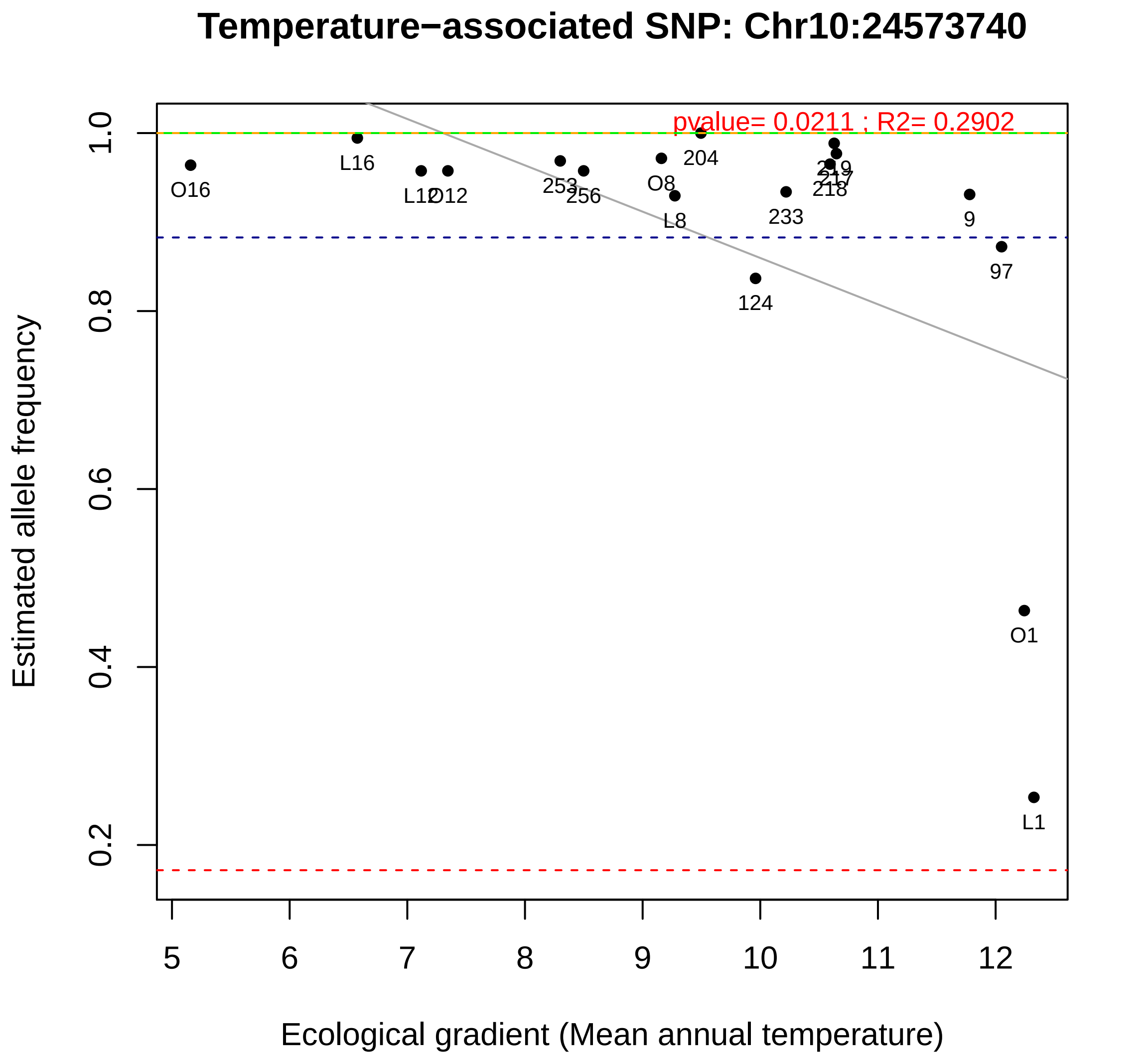

Temperature−associated SNP: Chr10:24573740
pvalue= 0.0211 ; R2= 0.2902
Estimated allele frequency
Ecological gradient (Mean annual temperature)
1.0
0.8
0.6
0.4
0.2
5
6
7
8
9
10
11
12
O16
L16
L12
O12
253
256
O8
L8
204
124
233
219
217
218
9
97
O1
L1

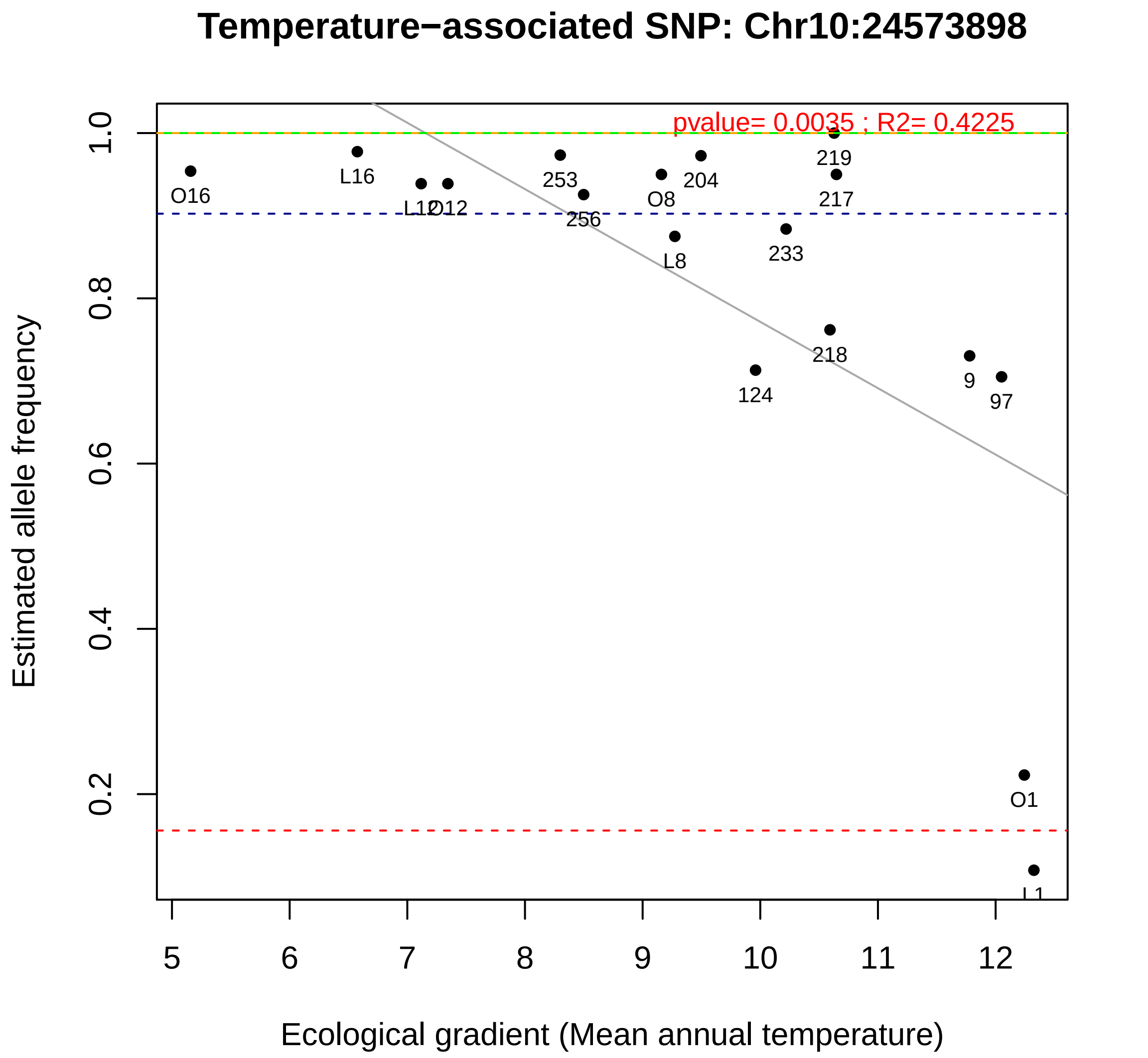

Temperature−associated SNP: Chr10:24573898
pvalue= 0.0035 ; R2= 0.4225
Estimated allele frequency
1.0
0.8
0.6
0.4
0.2
O16
L16
L12
O12
253
256
O8
204
L8
233
219
217
218
124
9
97
O1
L1
5
6
7
8
9
10
11
12
Ecological gradient (Mean annual temperature)

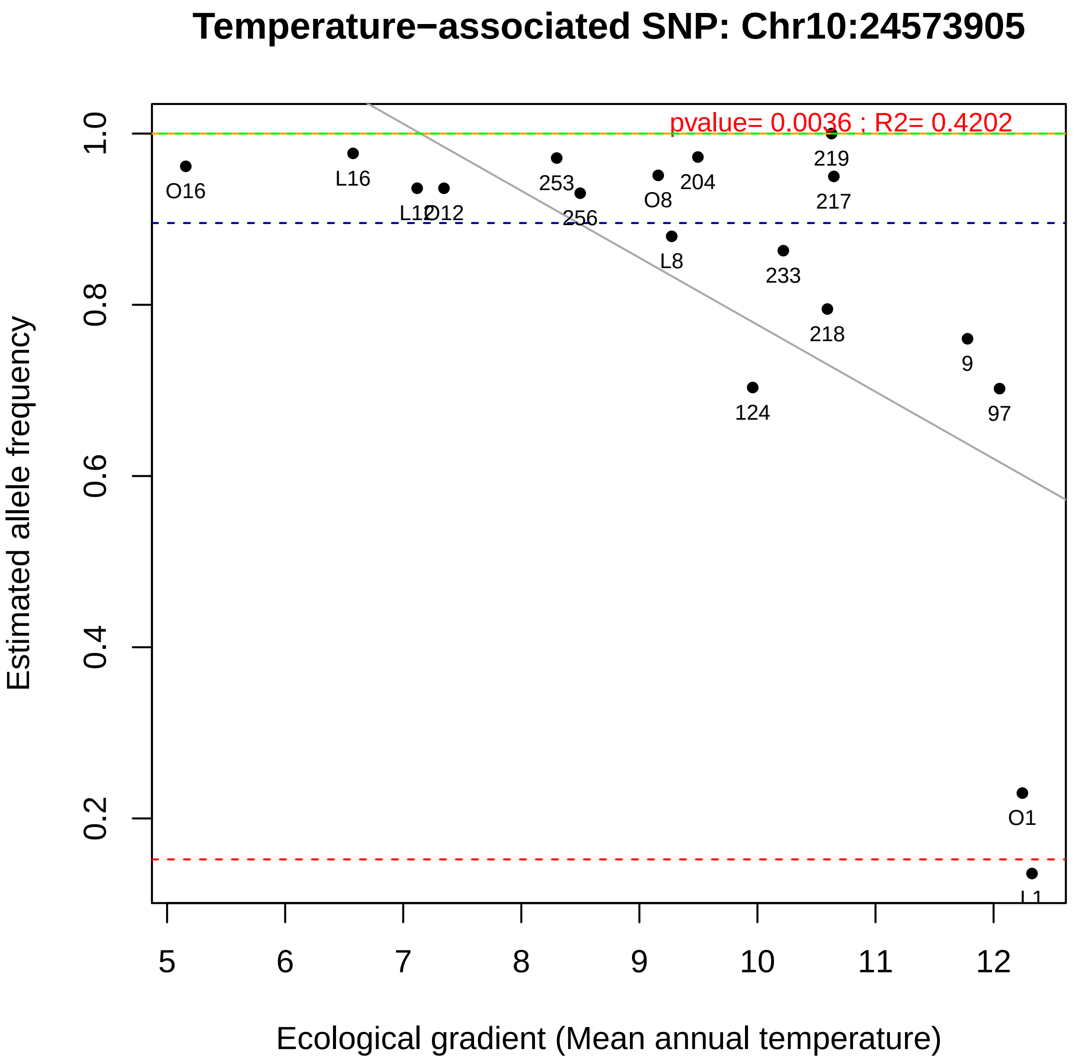

Temperature−associated SNP: Chr10:24573905
pvalue= 0.0036 ; R2= 0.4202
Estimated allele frequency
Ecological gradient (Mean annual temperature)
1.0
0.8
0.6
0.4
0.2
5
6
7
8
9
10
11
12
O16
L16
L12
O12
253
256
O8
204
L8
219
217
233
218
124
9
97
O1

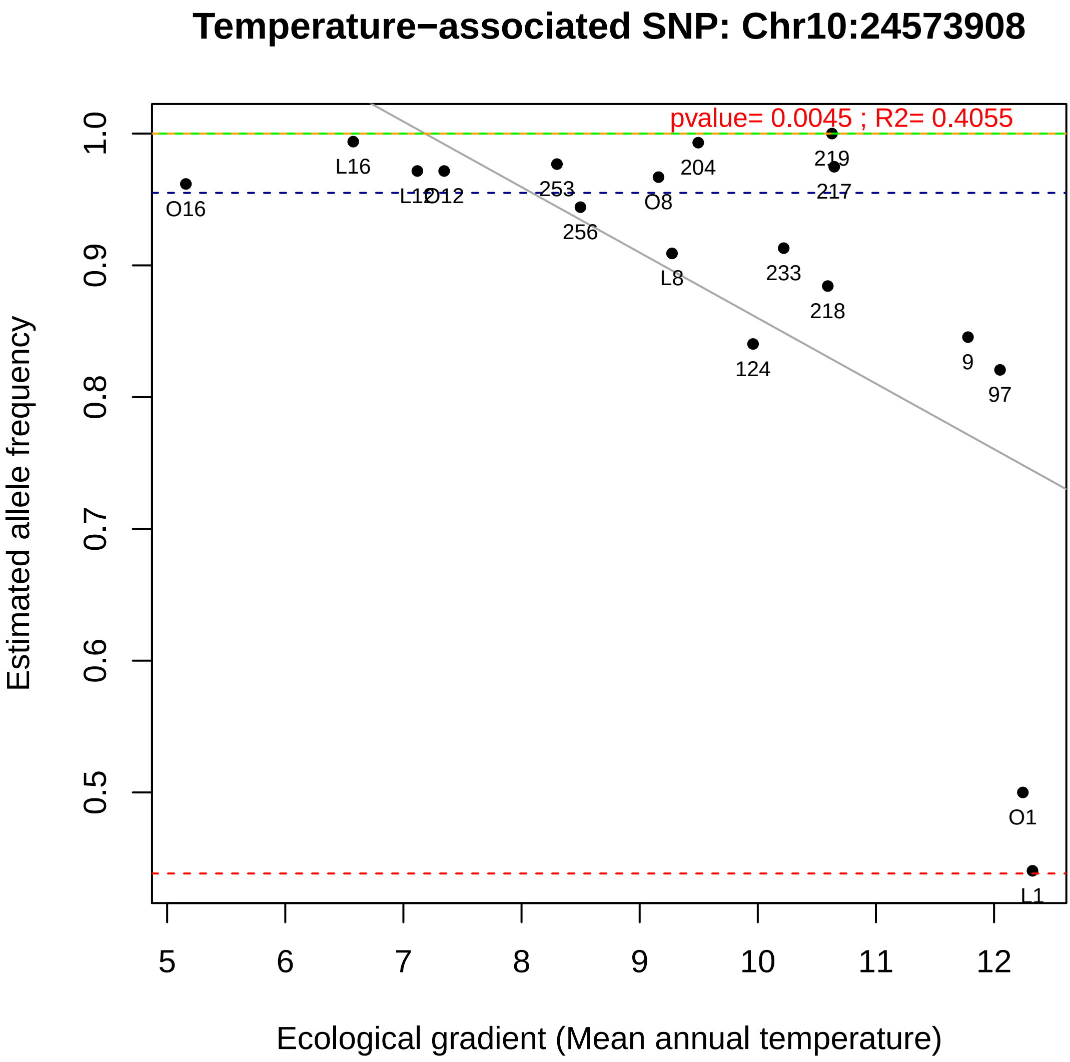

Temperature−associated SNP: Chr10:24573908
pvalue= 0.0045 ; R2= 0.4055
Estimated allele frequency
Ecological gradient (Mean annual temperature)
1.0
0.9
0.8
0.7
0.6
0.5
5
6
7
8
9
10
11
12
O16
L16
L12
O12
253
256
O8
204
L8
233
219
217
218
124
9
97
O1
L1

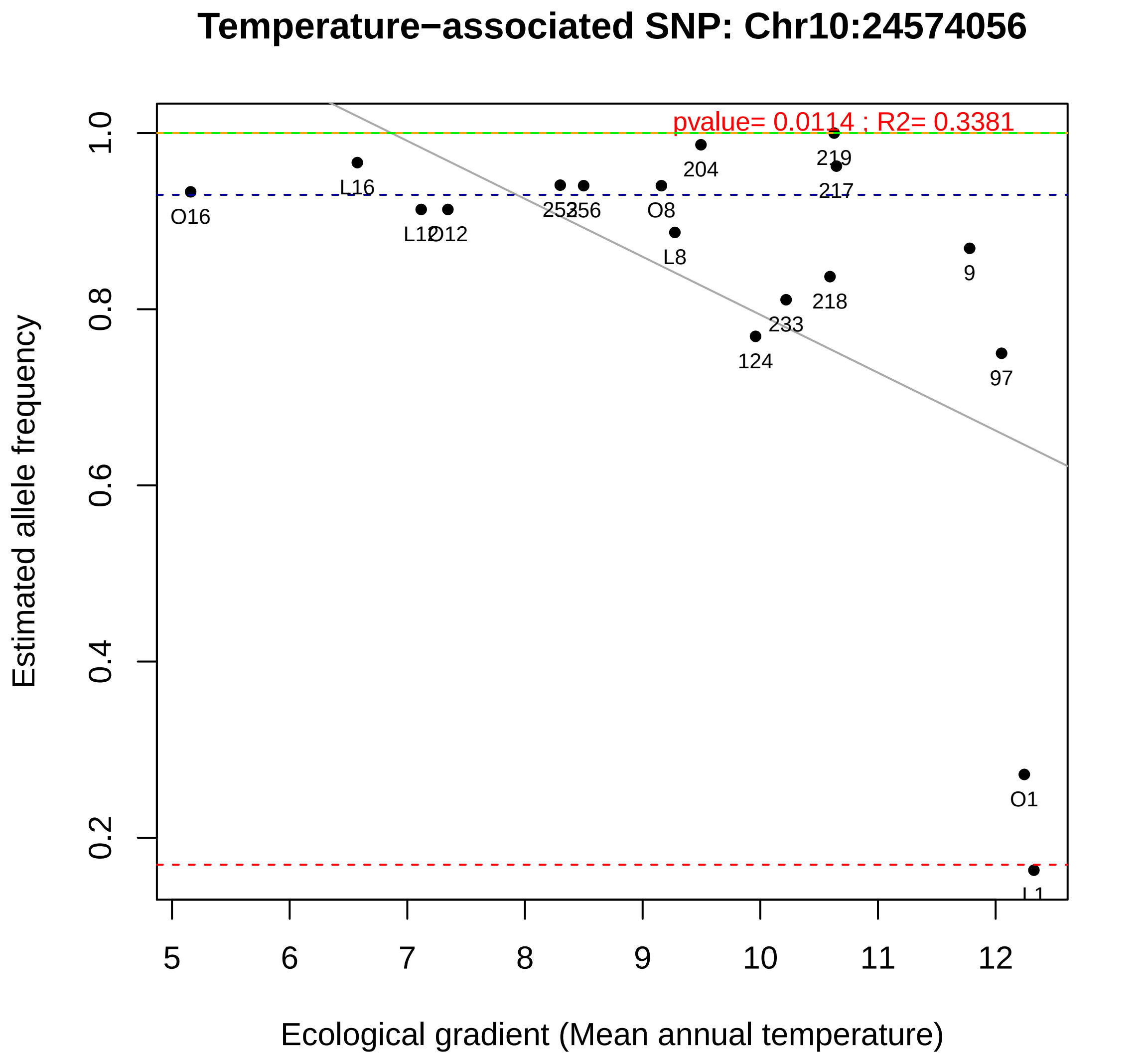

Temperature−associated SNP: Chr10:24574056
pvalue= 0.0114 ; R2= 0.3381
Estimated allele frequency
Ecological gradient (Mean annual temperature)
0.2
0.4
0.6
0.8
1.0
5
6
7
8
9
10
11
12
O16
L16
L12
O12
252
256
O8
L8
204
219
217
218
233
124
9
97
O1

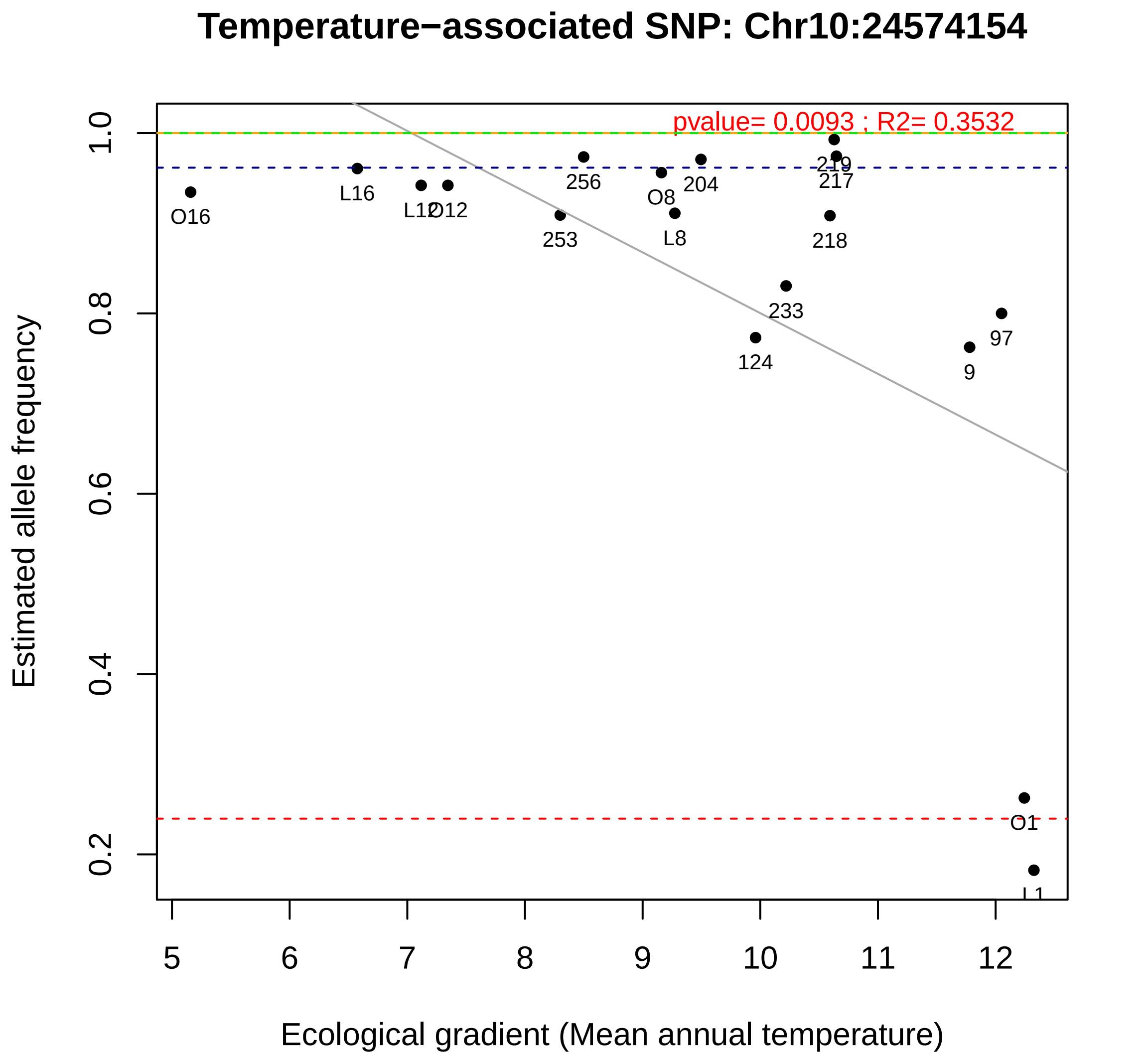
Temperature−associated SNP: Chr10:24574154
pvalue= 0.0093 ; R2= 0.3532
Estimated allele frequency
0.2
0.4
0.6
0.8
1.0
Ecological gradient (Mean annual temperature)
5
6
7
8
9
10
11
12
O16
L16
L12
O12
253
256
O8
L8
204
219
217
218
233
124
9
97
O1

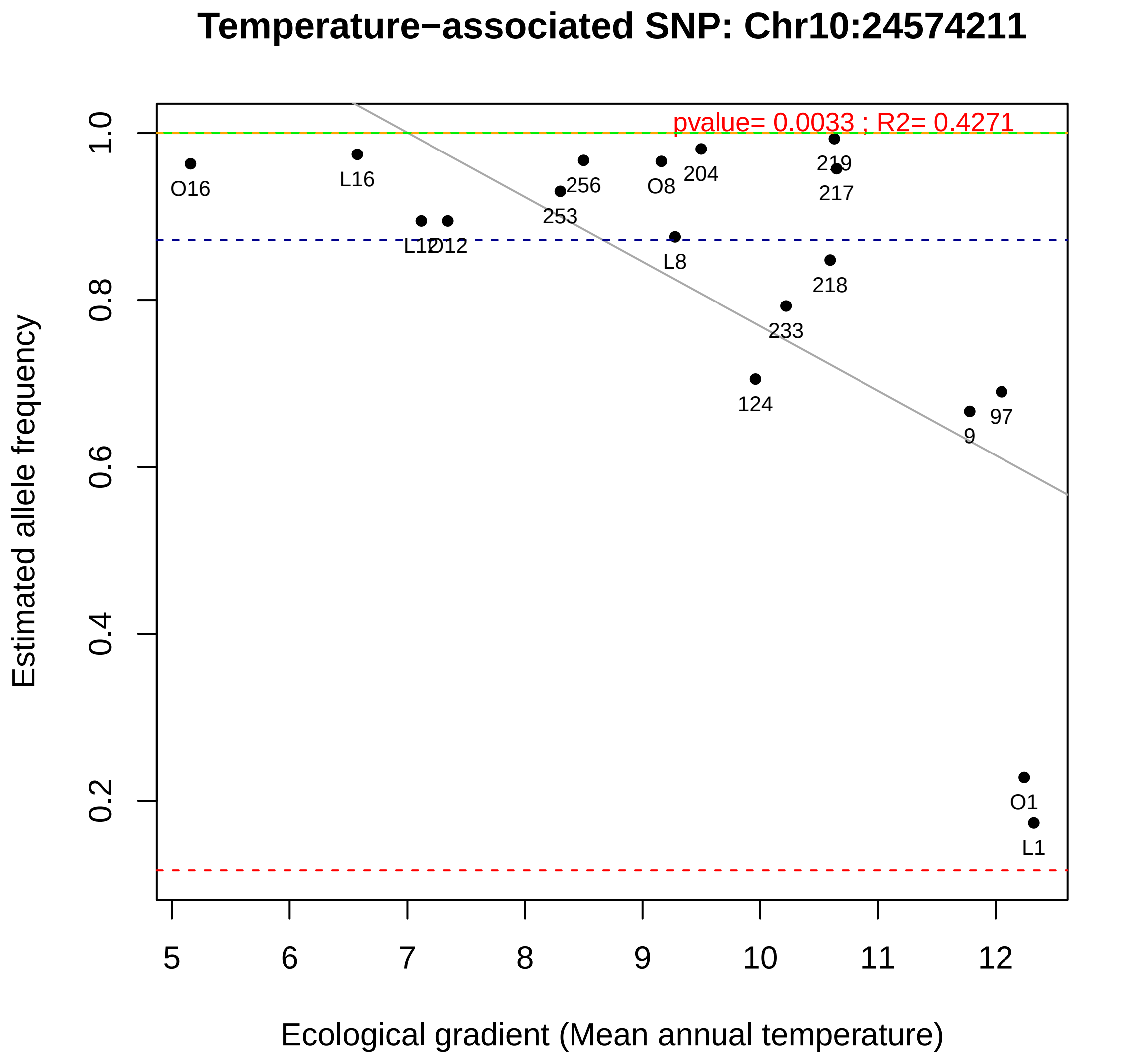

Temperature−associated SNP: Chr10:24574211
pvalue= 0.0033 ; R2= 0.4271
Estimated allele frequency
Ecological gradient (Mean annual temperature)
0.2
0.4
0.6
0.8
1.0
5
6
7
8
9
10
11
12
O16
L16
L12
O12
253
256
O8
204
L8
219
217
218
233
124
9
97
O1
L1

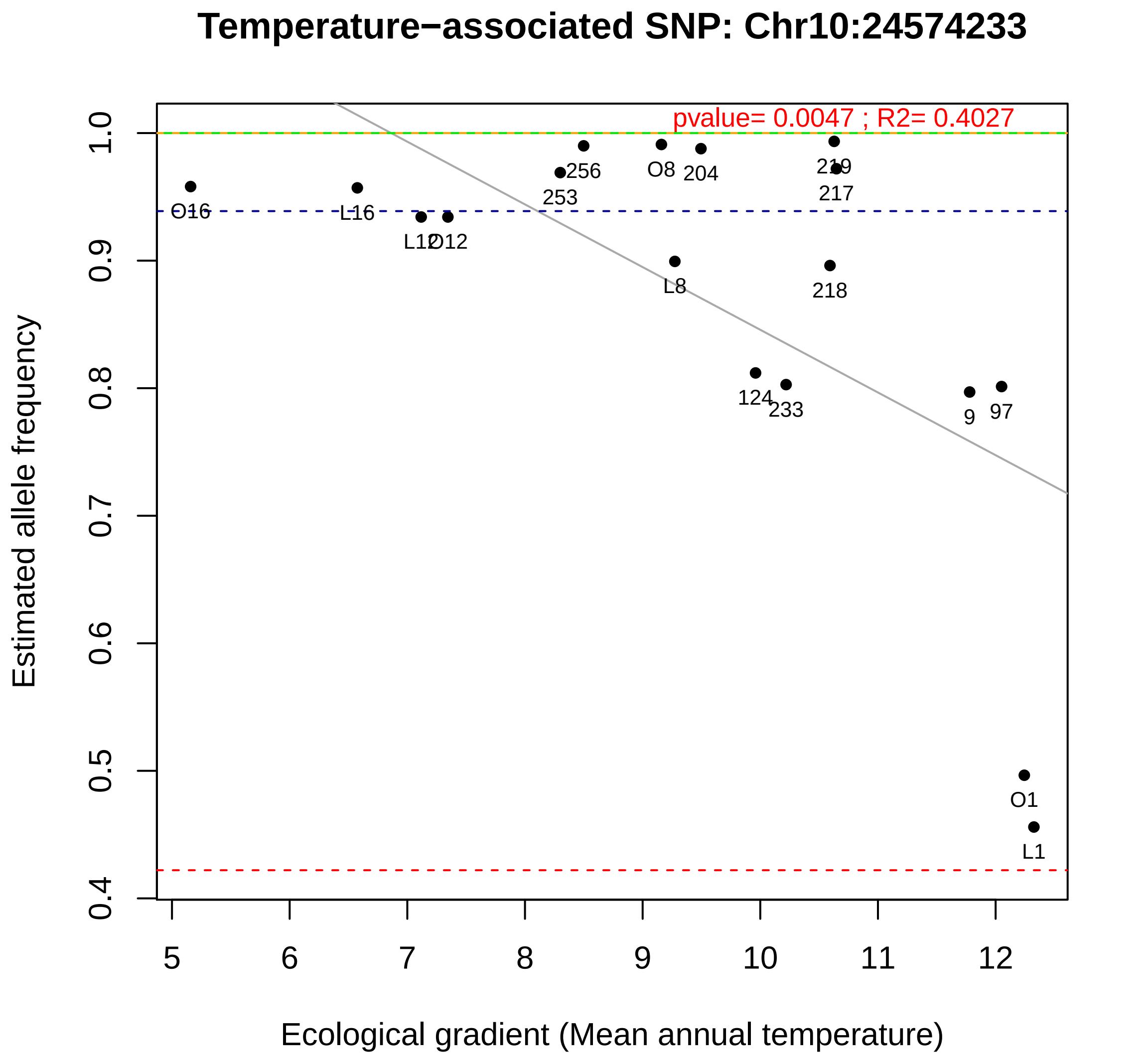
Temperature−associated SNP: Chr10:24574233
pvalue= 0.0047 ; R2= 0.4027
Estimated allele frequency
Ecological gradient (Mean annual temperature)
0.4
0.5
0.6
0.7
0.8
0.9
1.0
5
6
7
8
9
10
11
12
O16
L16
L12
O12
253
256
O8
204
219
217
L8
218
124
233
9
97
O1
L1

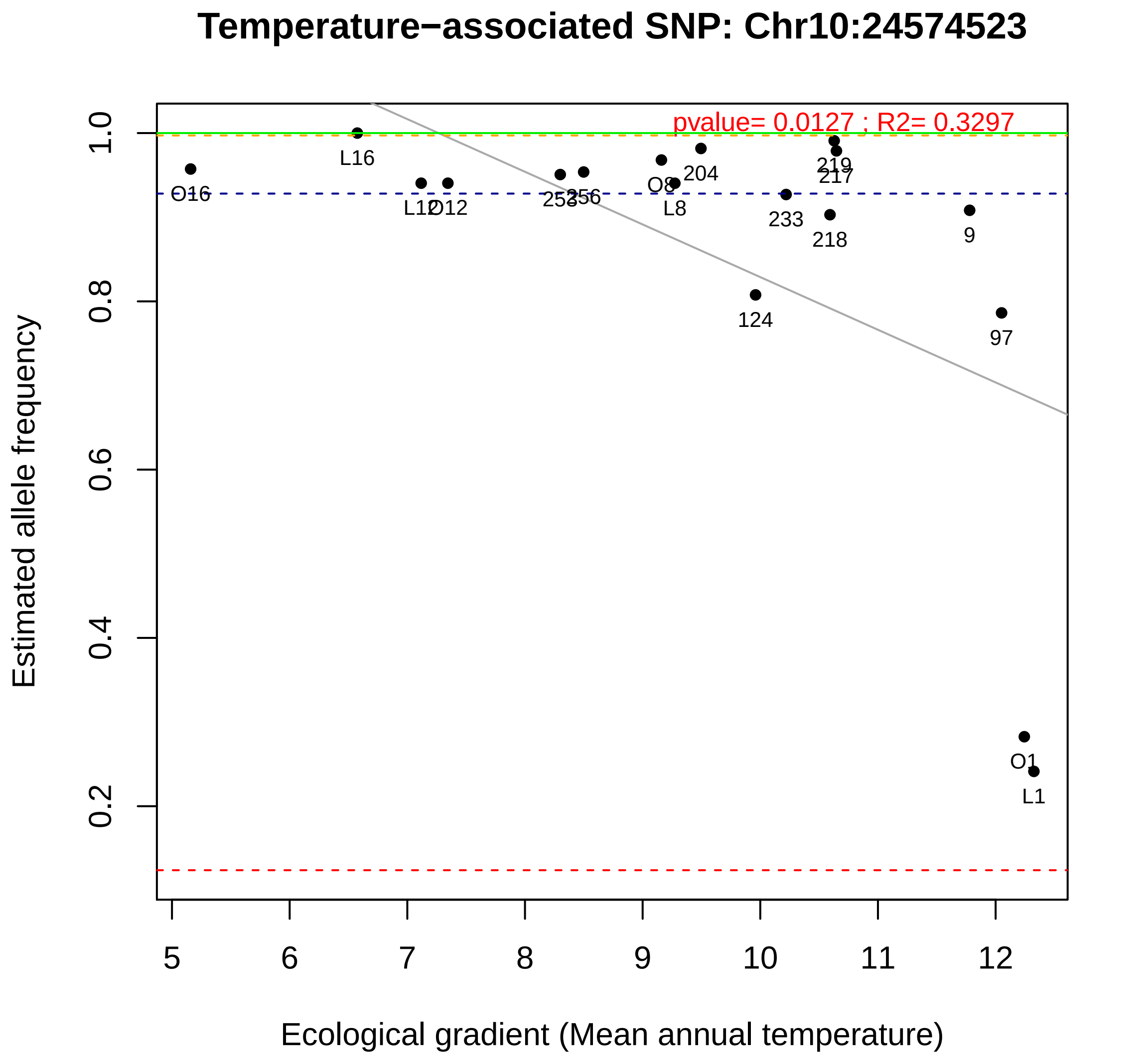
Temperature−associated SNP: Chr10:24574523
pvalue= 0.0127 ; R2= 0.3297
Estimated allele frequency
Ecological gradient (Mean annual temperature)
1.0
0.8
0.6
0.4
0.2
5
6
7
8
9
10
11
12
L16
O16
L12
O12
256
O8
L8
204
219
217
233
218
9
124
97
O1
L1

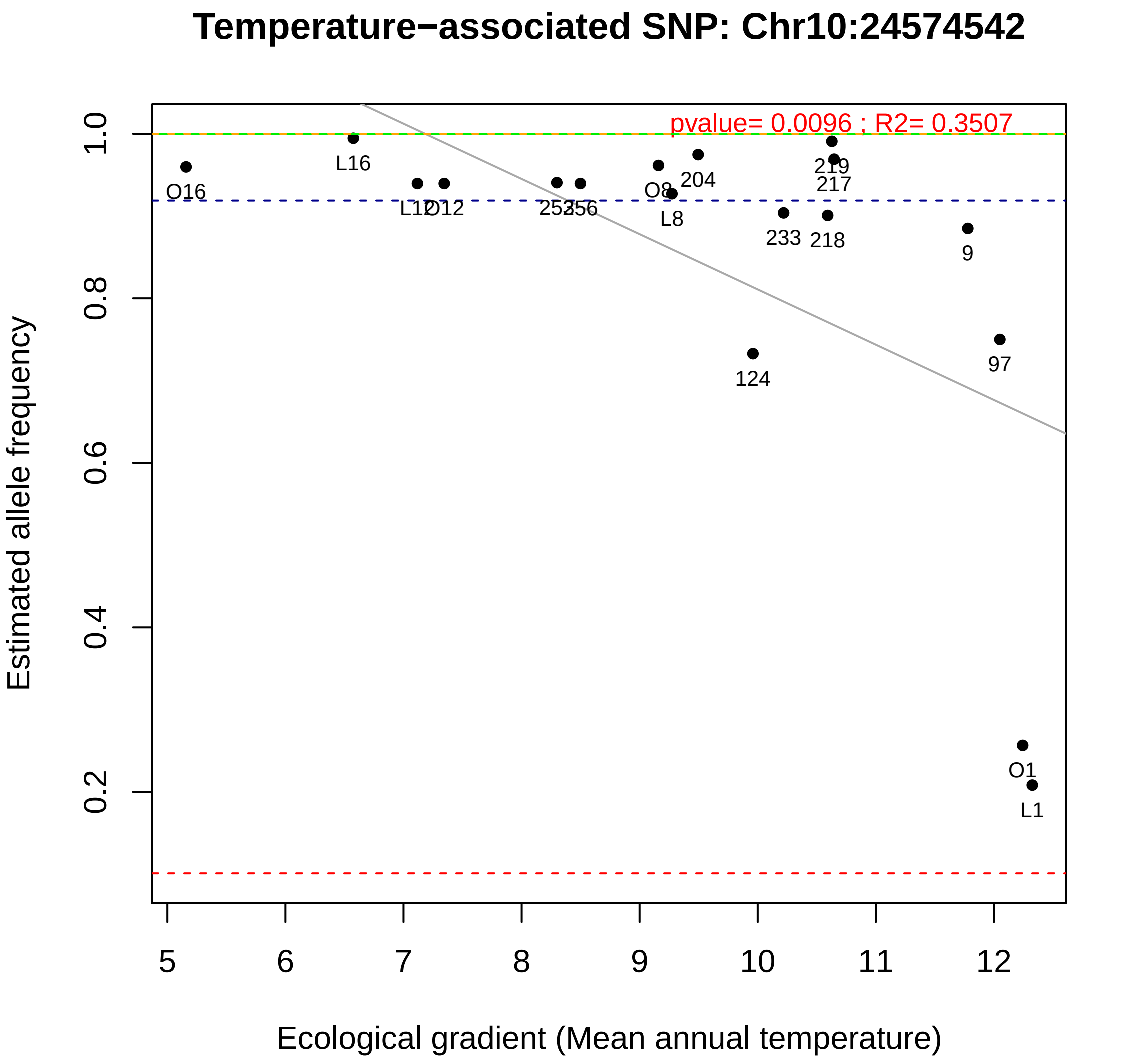

Temperature−associated SNP: Chr10:24574542
pvalue= 0.0096 ; R2= 0.3507
Estimated allele frequency
Ecological gradient (Mean annual temperature)
0.2
0.4
0.6
0.8
1.0
5
6
7
8
9
10
11
12
O16
L16
L12
O12
253
256
O8
L8
204
124
233
219
217
218
9
97
O1
L1

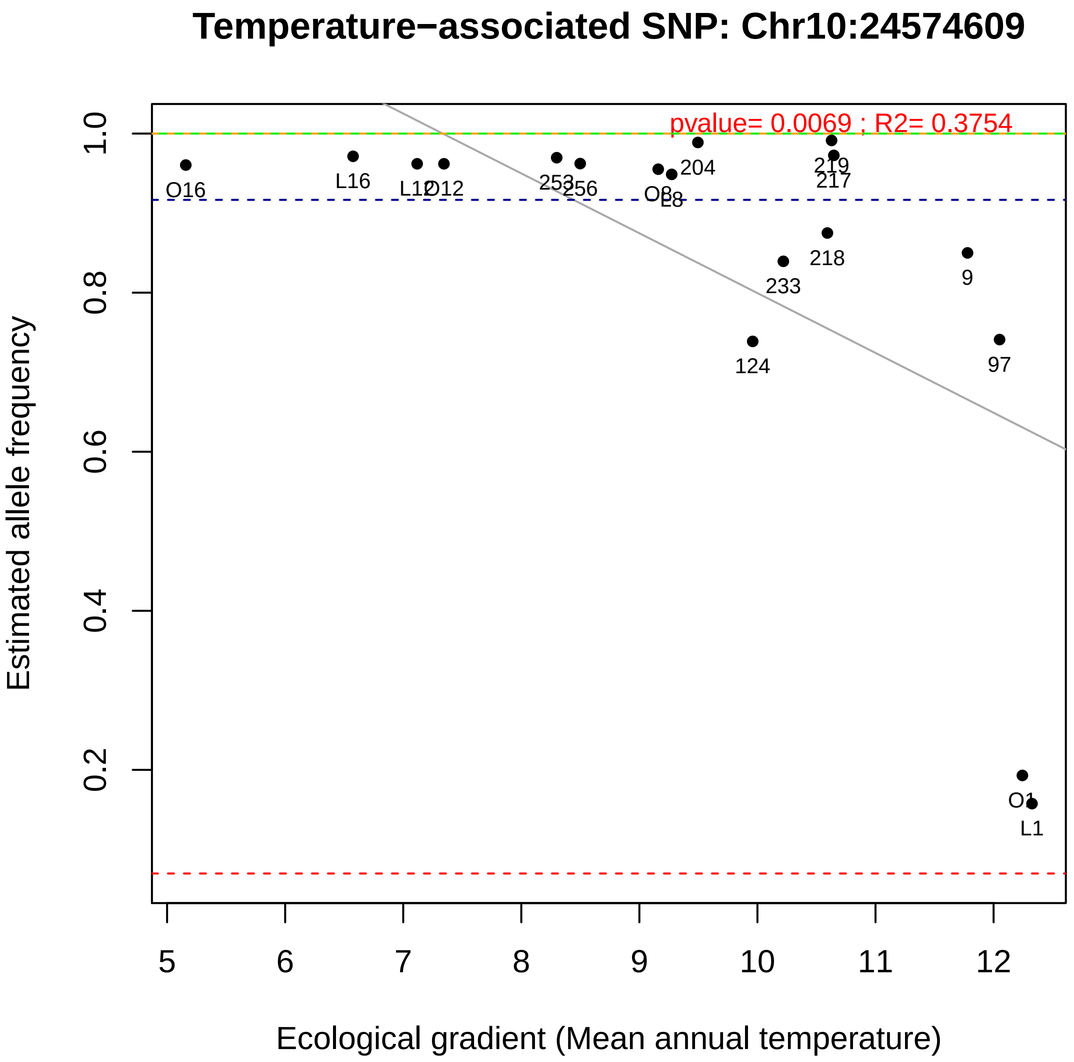

Temperature−associated SNP: Chr10:24574609
pvalue= 0.0069 ; R2= 0.3754
Estimated allele frequency
Ecological gradient (Mean annual temperature)
0.2
0.4
0.6
0.8
1.0
5
6
7
8
9
10
11
12
O16
L16
L12
O12
253
256
O8
L8
204
219
217
218
233
124
9
97
O1
L1

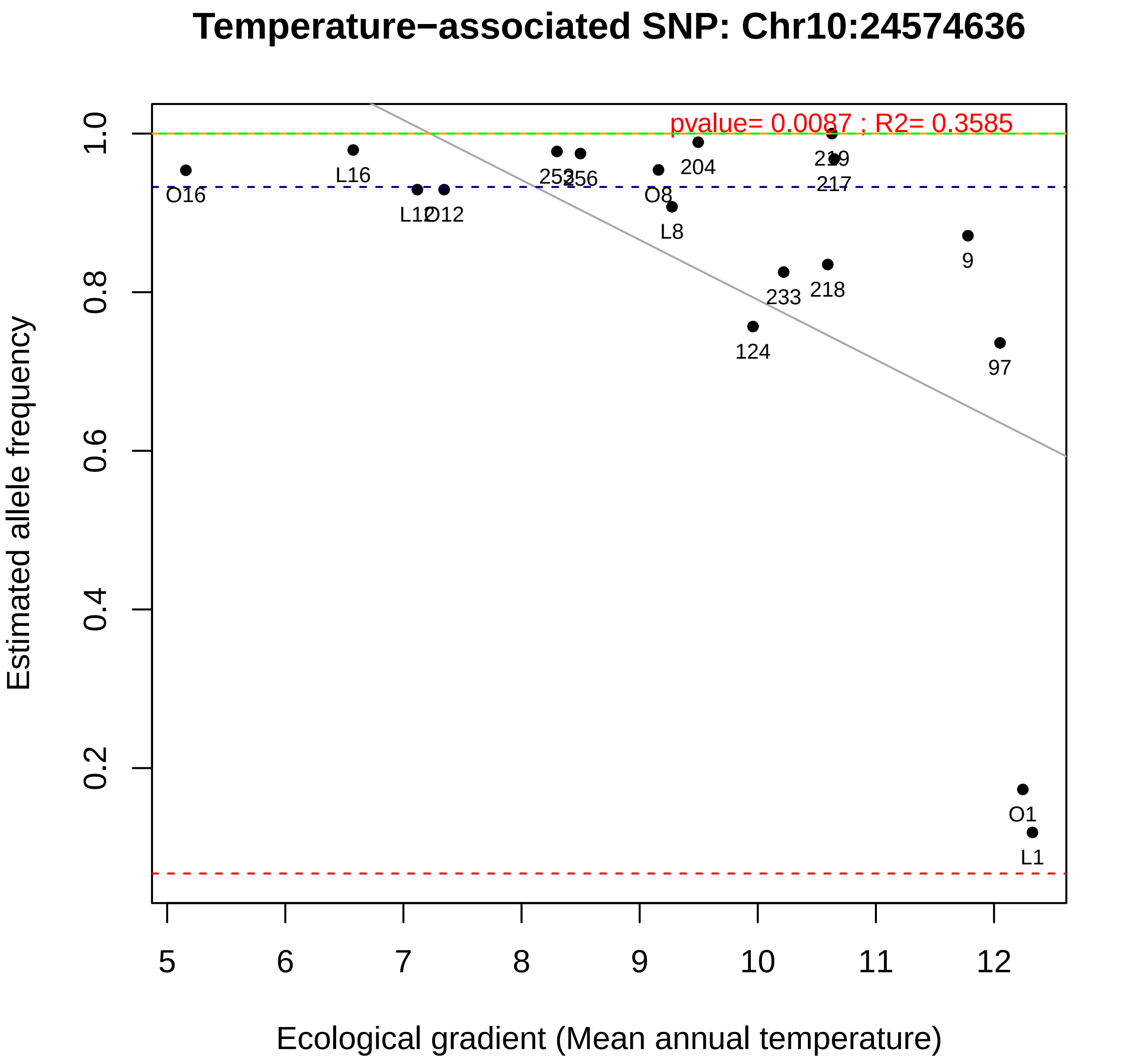

Temperature−associated SNP: Chr10:24574636
pvalue= 0.0087 ; R2= 0.3585
Estimated allele frequency
Ecological gradient (Mean annual temperature)
0.2
0.4
0.6
0.8
1.0
5
6
7
8
9
10
11
12
O16
L16
L12
O12
252
256
O8
L8
204
219
217
124
233
218
9
97
O1
L1

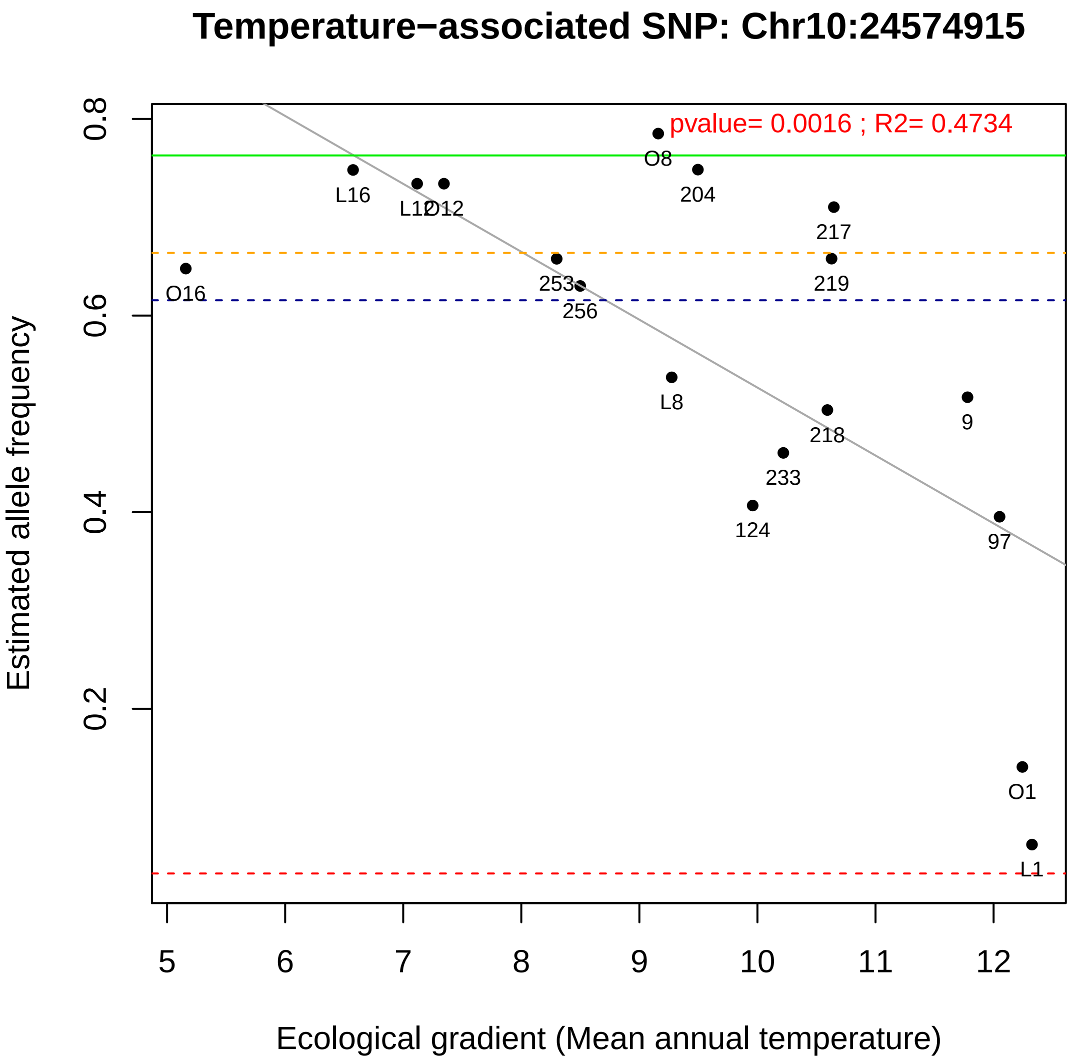
Temperature−associated SNP: Chr10:24574915
pvalue= 0.0016 ; R2= 0.4734
Estimated allele frequency
0.2
0.4
0.6
0.8
Ecological gradient (Mean annual temperature)
5
6
7
8
9
10
11
12
O16
L16
L12
O12
253
256
O8
204
L8
124
233
218
217
219
9
97
O1
L1

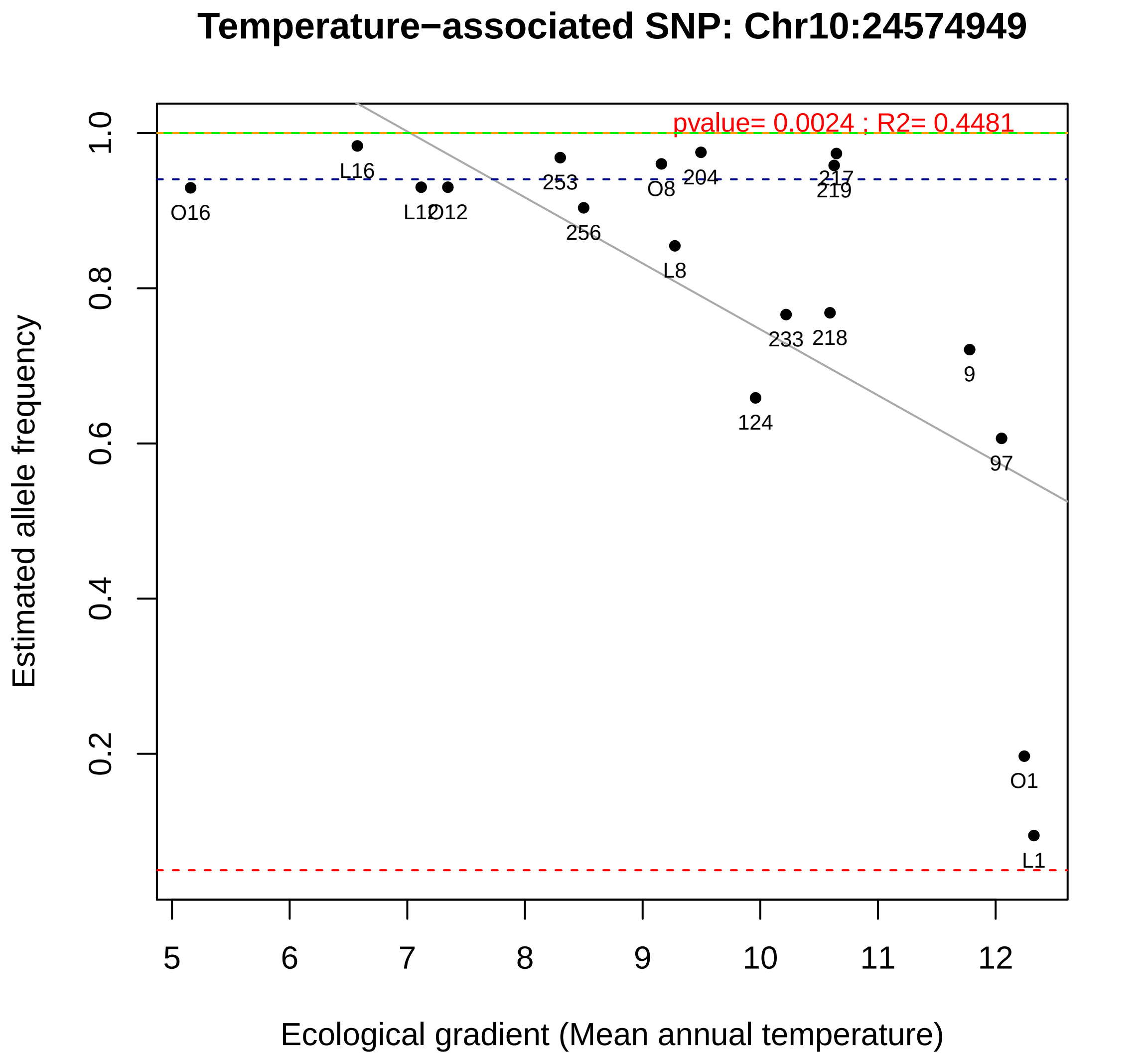

Temperature−associated SNP: Chr10:24574949
pvalue= 0.0024 ; R2= 0.4481
Estimated allele frequency
0.2
0.4
0.6
0.8
1.0
5
6
7
8
9
10
11
12
Ecological gradient (Mean annual temperature)
O16
L16
L12
O12
253
256
O8
204
L8
217
219
233
218
124
9
97
O1
L1

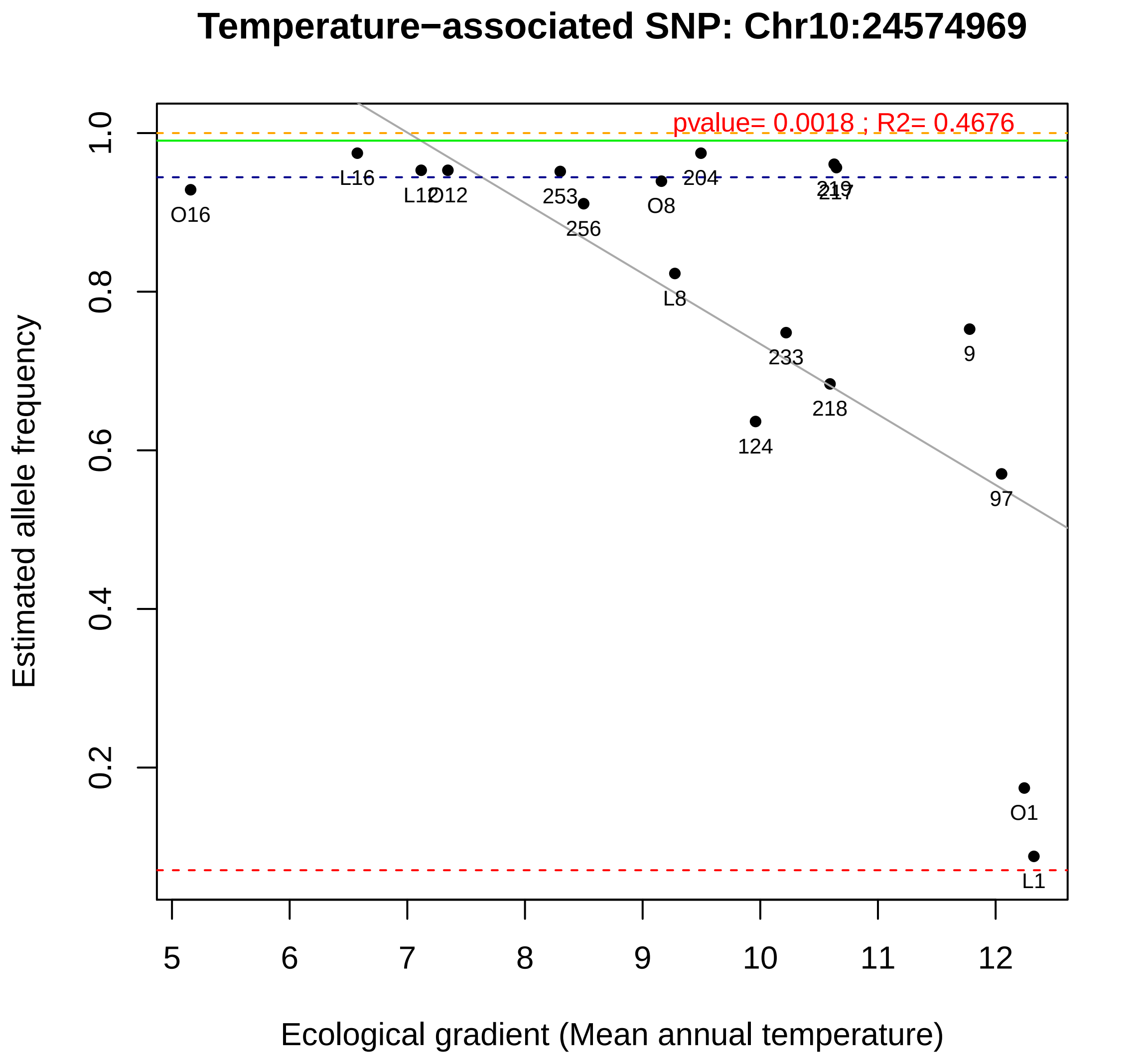

Temperature−associated SNP: Chr10:24574969
pvalue= 0.0018 ; R2= 0.4676
Estimated allele frequency
Ecological gradient (Mean annual temperature)
0.2
0.4
0.6
0.8
1.0
5
6
7
8
9
10
11
12
O16
L16
L12
O12
253
256
O8
204
219
217
L8
233
9
218
124
97
O1
L1

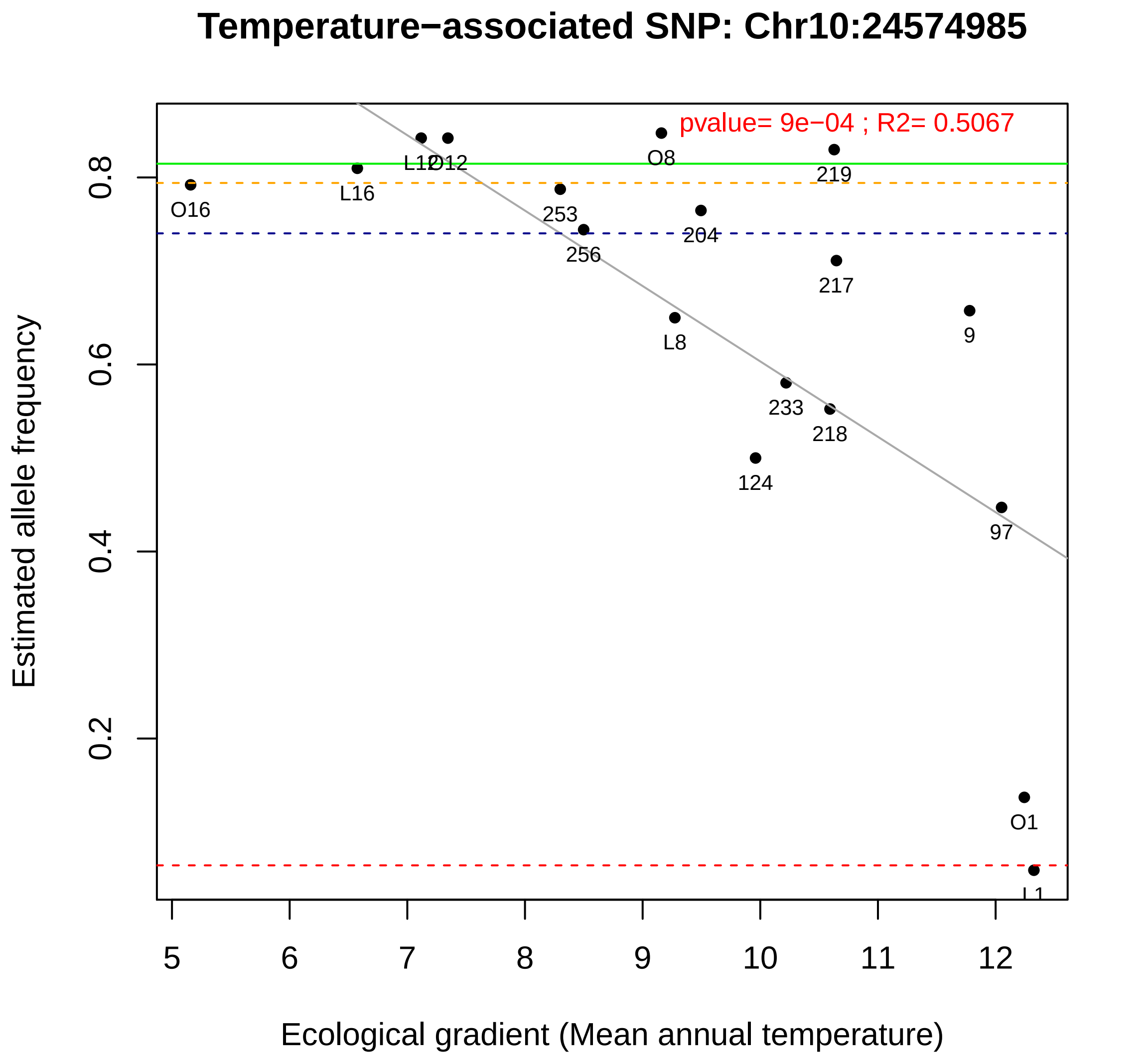
Temperature−associated SNP: Chr10:24574985
pvalue= 9e−04 ; R2= 0.5067
Estimated allele frequency
Ecological gradient (Mean annual temperature)
0.2
0.4
0.6
0.8
5
6
7
8
9
10
11
12
O16
L16
L12
O12
253
256
O8
204
219
217
L8
9
233
218
124
97
O1
L1

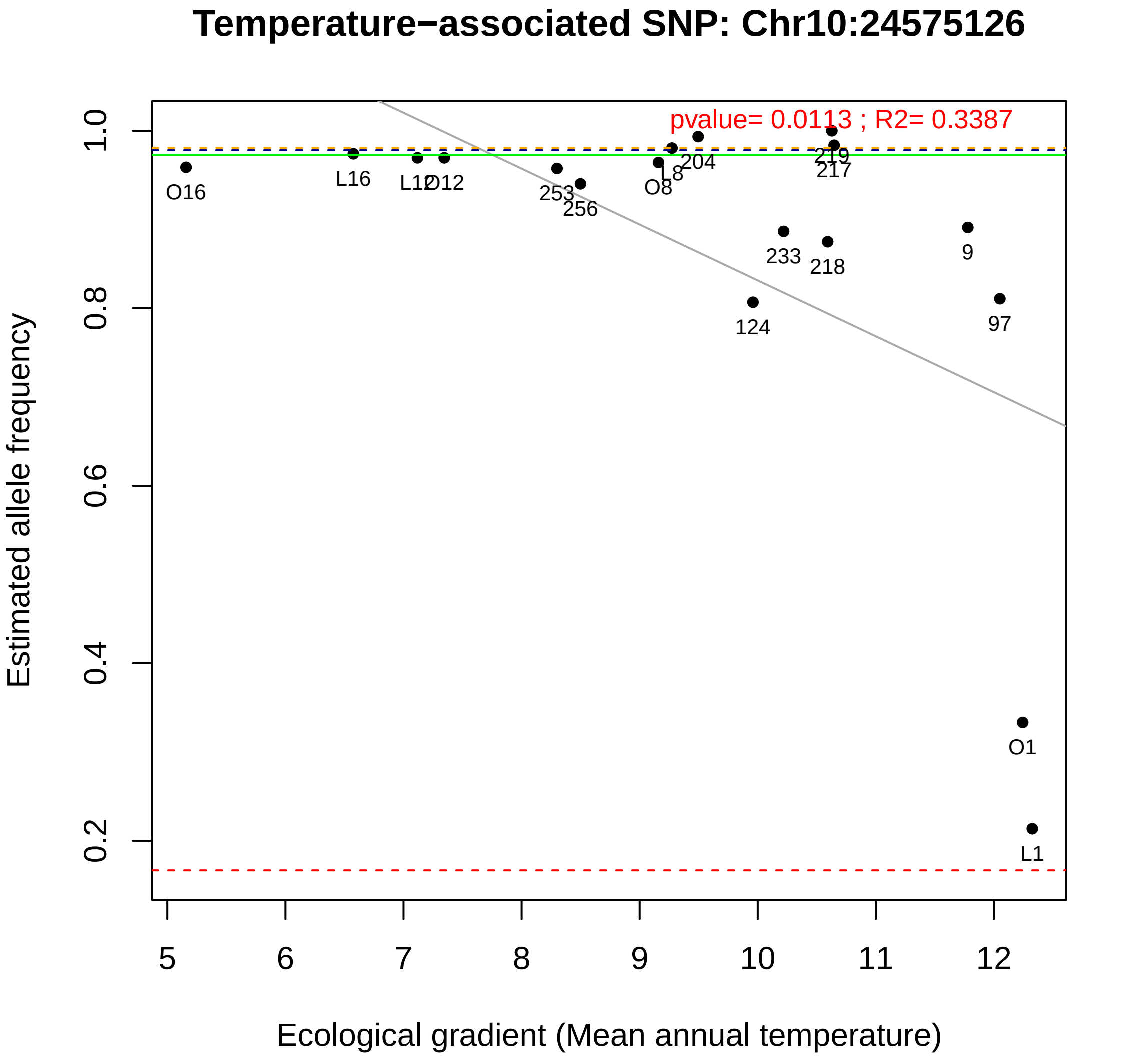
Temperature−associated SNP: Chr10:24575126
pvalue= 0.0113 ; R2= 0.3387
Estimated allele frequency
0.2
0.4
0.6
0.8
1.0
5
6
7
8
9
10
11
12
Ecological gradient (Mean annual temperature)
O16
L16
253
256
O8
204
233
218
9
124
97
O1
L1
217

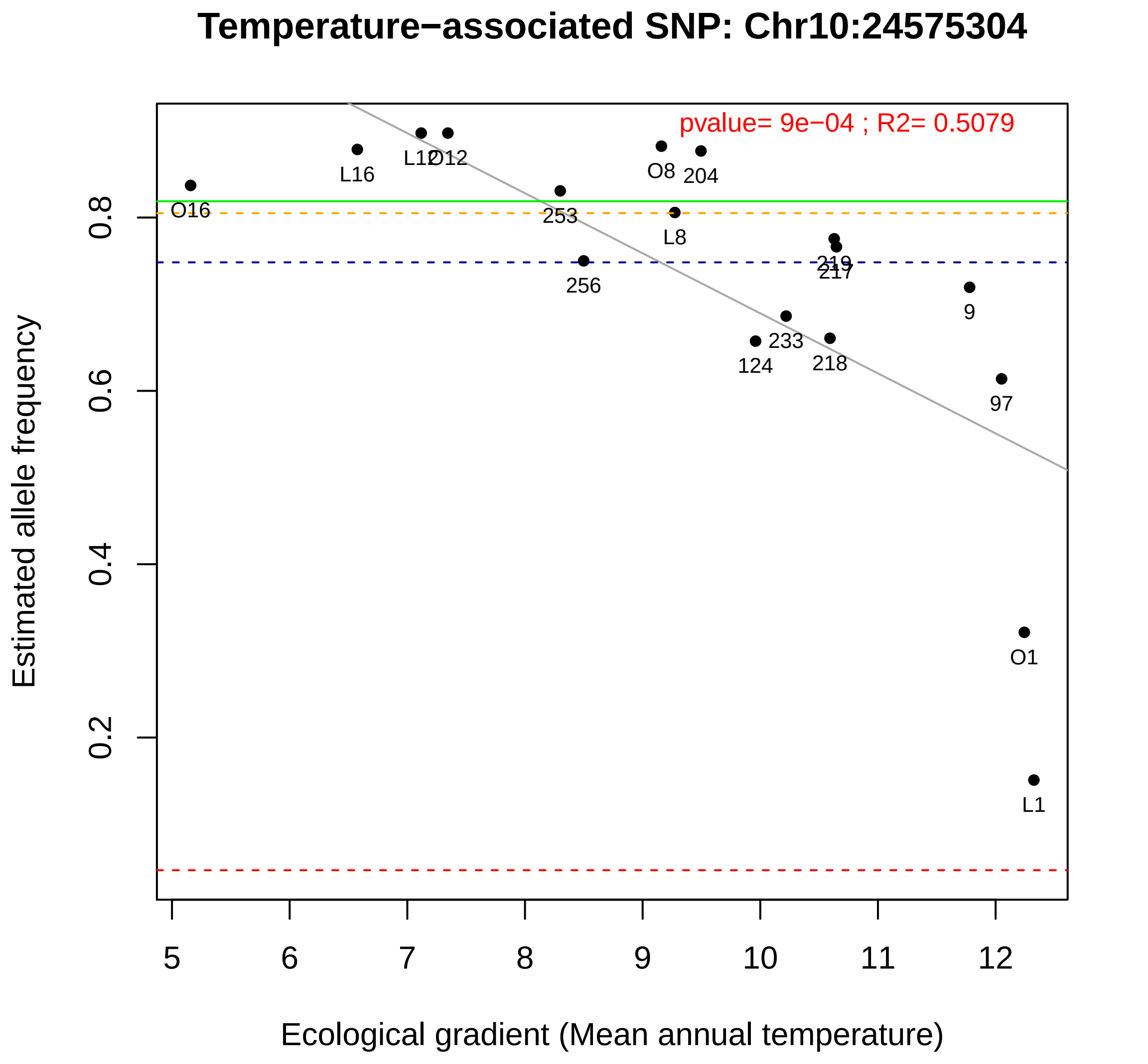

Temperature−associated SNP: Chr10:24575304
pvalue= 9e−04 ; R2= 0.5079
O16
L16
L12
O12
253
256
O8
204
L8
219
217
233
124
218
9
97
O1
L1
0.2
0.4
0.6
0.8
5
6
7
8
9
10
11
12
Estimated allele frequency
Ecological gradient (Mean annual temperature)

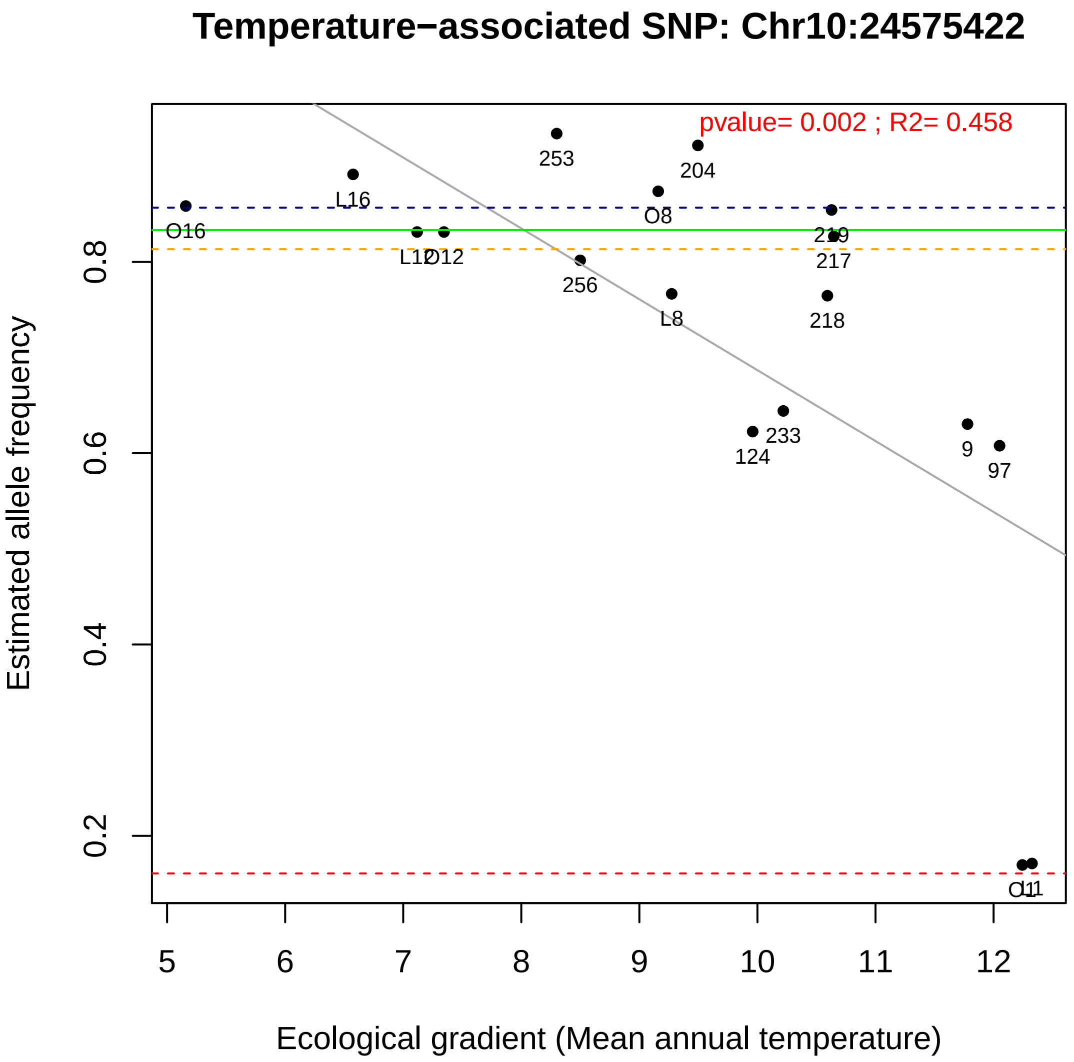
Temperature−associated SNP: Chr10:24575422
pvalue= 0.002 ; R2= 0.458
Estimated allele frequency
0.2
0.4
0.6
0.8
5
6
7
8
9
10
11
12
Ecological gradient (Mean annual temperature)
O16
L16
253
204
O8
256
L8
217
218
233
124
9
97

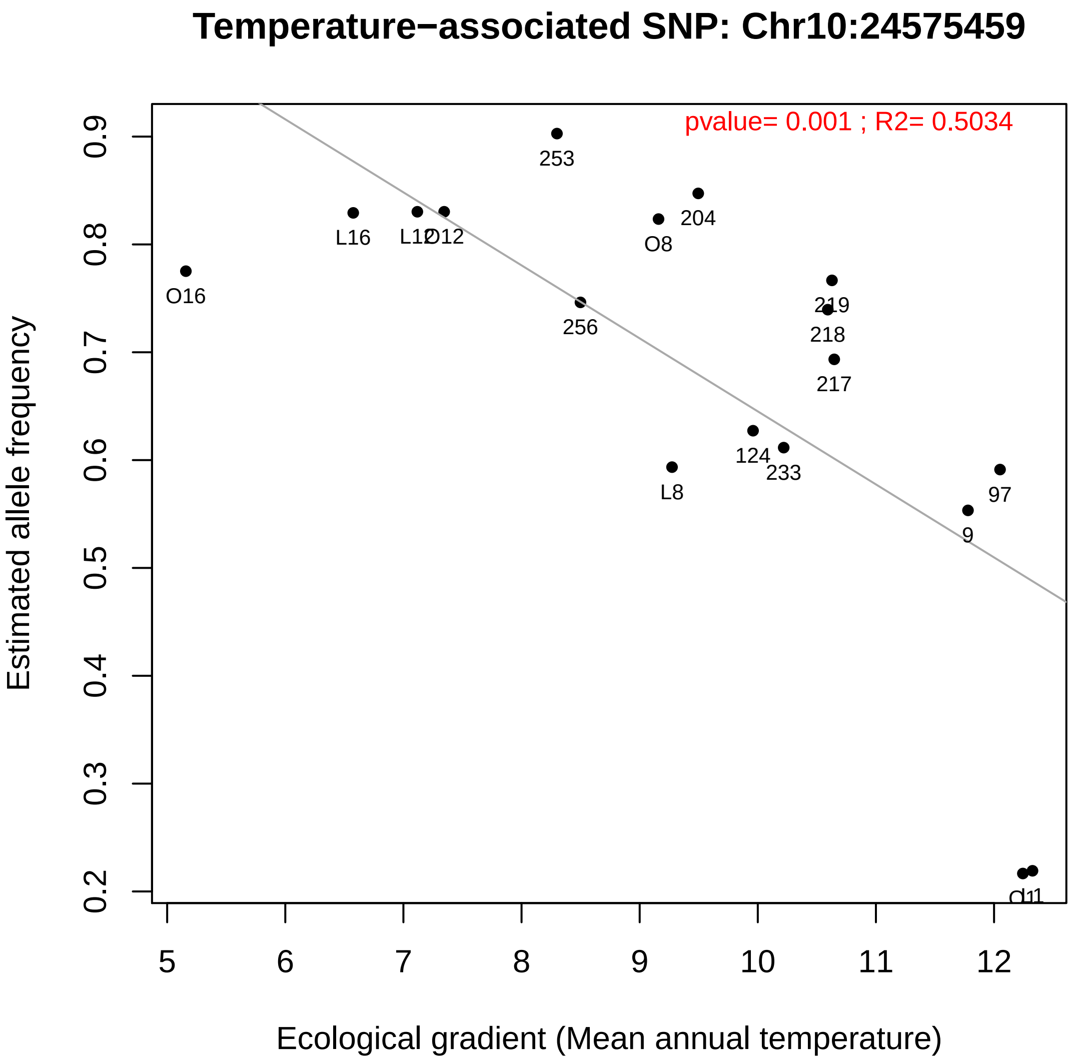

Temperature−associated SNP: Chr10:24575459
pvalue= 0.001 ; R2= 0.5034
Estimated allele frequency
Ecological gradient (Mean annual temperature)
0.2
0.3
0.4
0.5
0.6
0.7
0.8
0.9
5
6
7
8
9
10
11
12
O16
L16
L12
O12
253
256
O8
204
L8
124
233
219
218
217
9
97

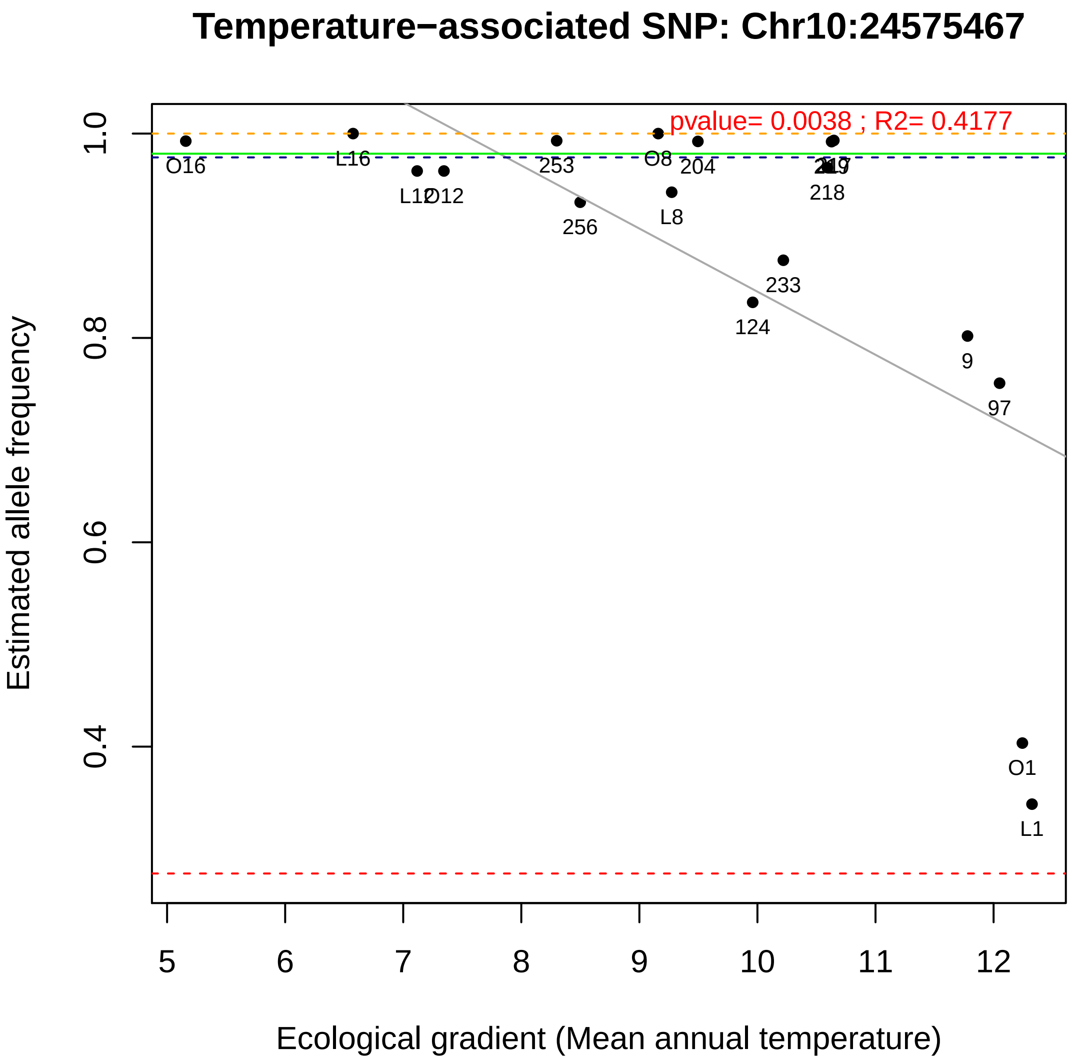
Temperature−associated SNP: Chr10:24575467
pvalue= 0.0038 ; R2= 0.4177
Estimated allele frequency
0.4
0.6
0.8
1.0
Ecological gradient (Mean annual temperature)
5
6
7
8
9
10
11
12
O16
L16
L12
O12
253
256
O8
204
L8
218
233
124
9
97
O1
L1

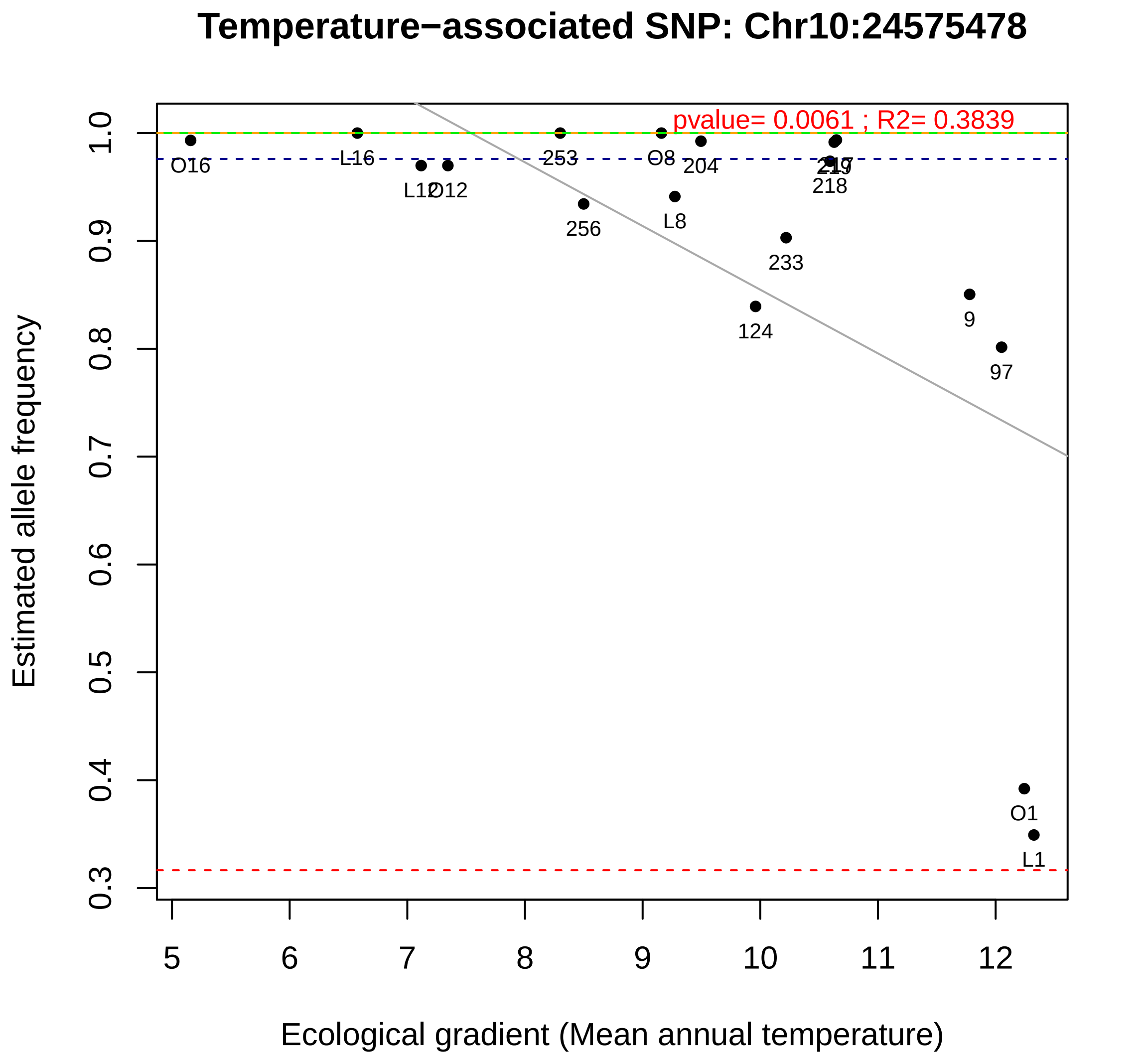

Temperature−associated SNP: Chr10:24575478
pvalue= 0.0061 ; R2= 0.3839
Estimated allele frequency
Ecological gradient (Mean annual temperature)
0.3
0.4
0.5
0.6
0.7
0.8
0.9
1.0
5
6
7
8
9
10
11
12
O16
L16
L12
O12
253
256
O8
204
L8
233
124
217
219
218
9
97
O1
L1

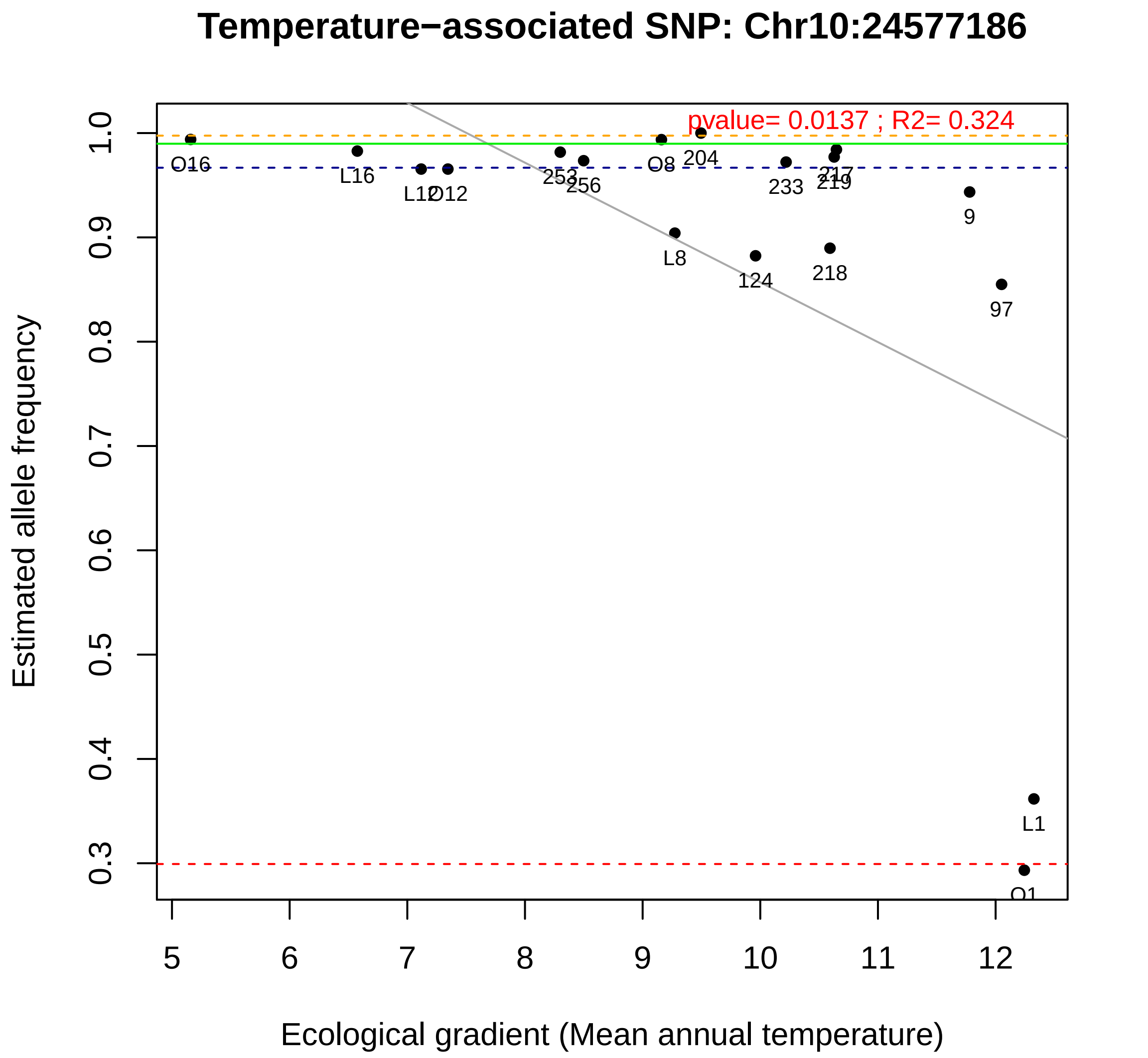
Temperature−associated SNP: Chr10:24577186
pvalue= 0.0137 ; R2= 0.324
Estimated allele frequency
Ecological gradient (Mean annual temperature)
1.0
0.9
0.8
0.7
0.6
0.5
0.4
0.3
5
6
7
8
9
10
11
12
O16
L16
L12
O12
253
256
O8
204
L8
124
233
217
219
218
9
97
L1
O1

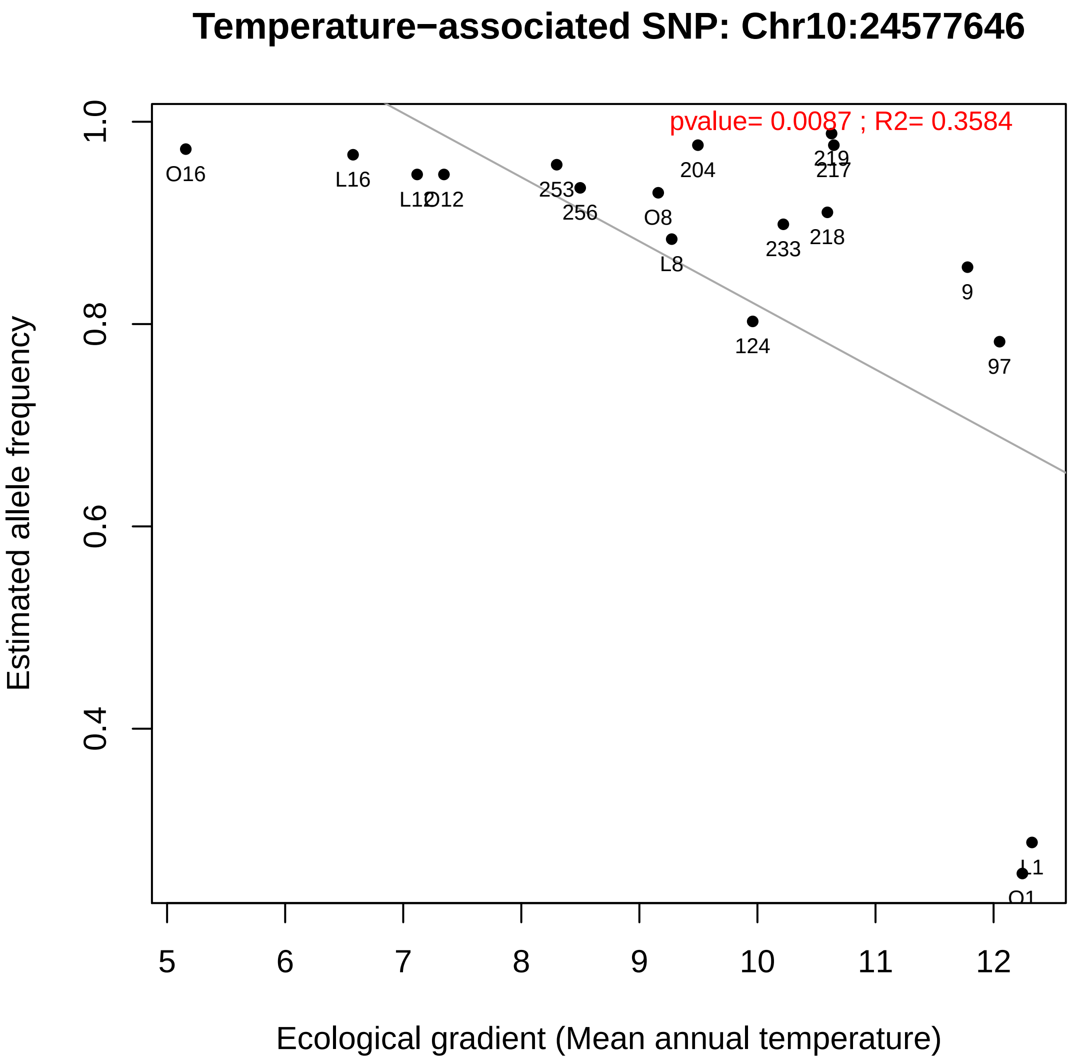

Temperature−associated SNP: Chr10:24577646
pvalue= 0.0087 ; R2= 0.3584
Estimated allele frequency
Ecological gradient (Mean annual temperature)
0.4
0.6
0.8
1.0
5
6
7
8
9
10
11
12
O16
L16
L12
O12
253
256
O8
L8
204
124
233
219
217
218
9
97
L1
O1

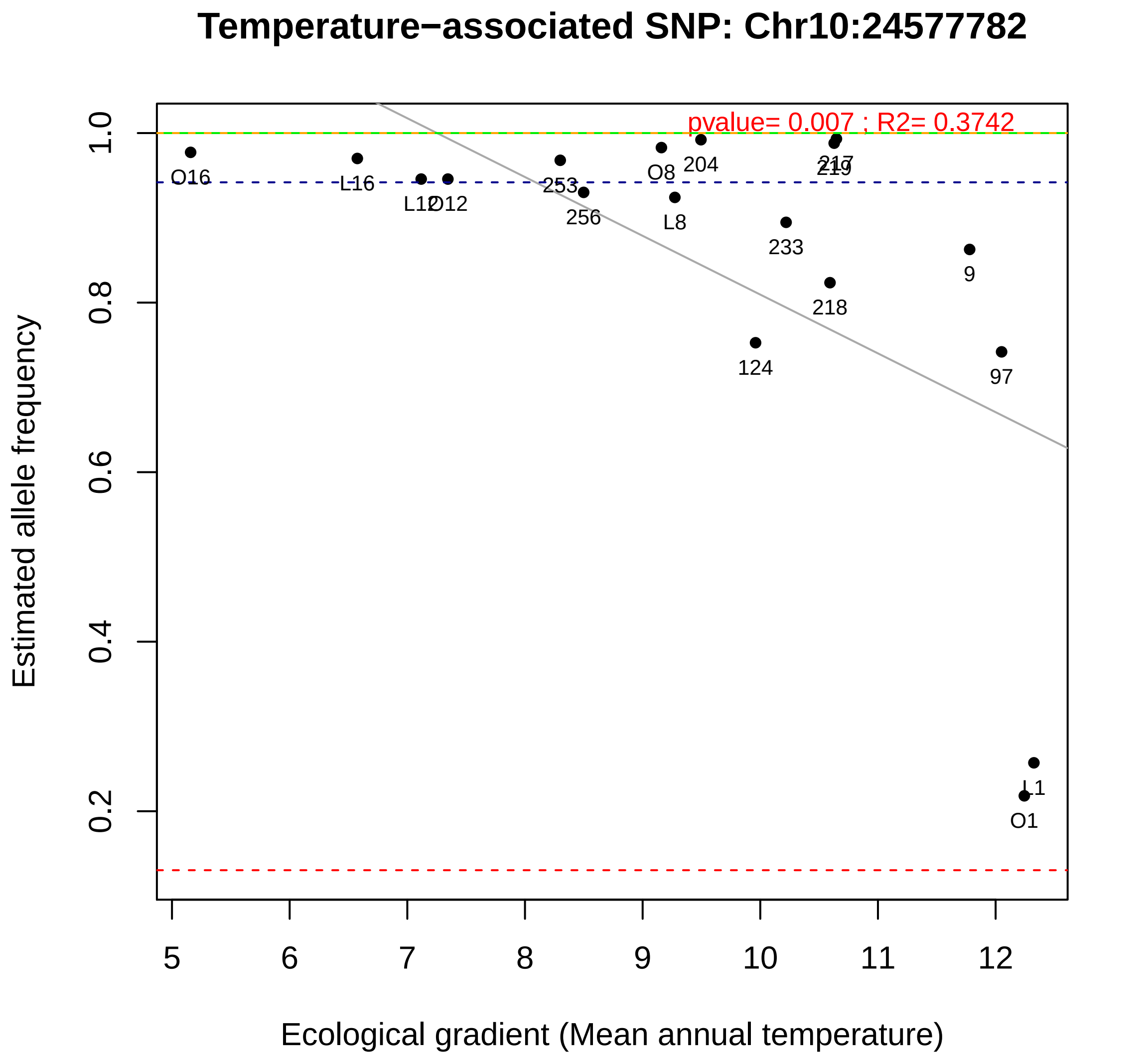

Temperature−associated SNP: Chr10:24577782
pvalue= 0.007 ; R2= 0.3742
Estimated allele frequency
Ecological gradient (Mean annual temperature)
0.2
0.4
0.6
0.8
1.0
5
6
7
8
9
10
11
12
O16
L16
L12
O12
253
256
O8
204
L8
233
217
219
124
218
9
97
L1
O1

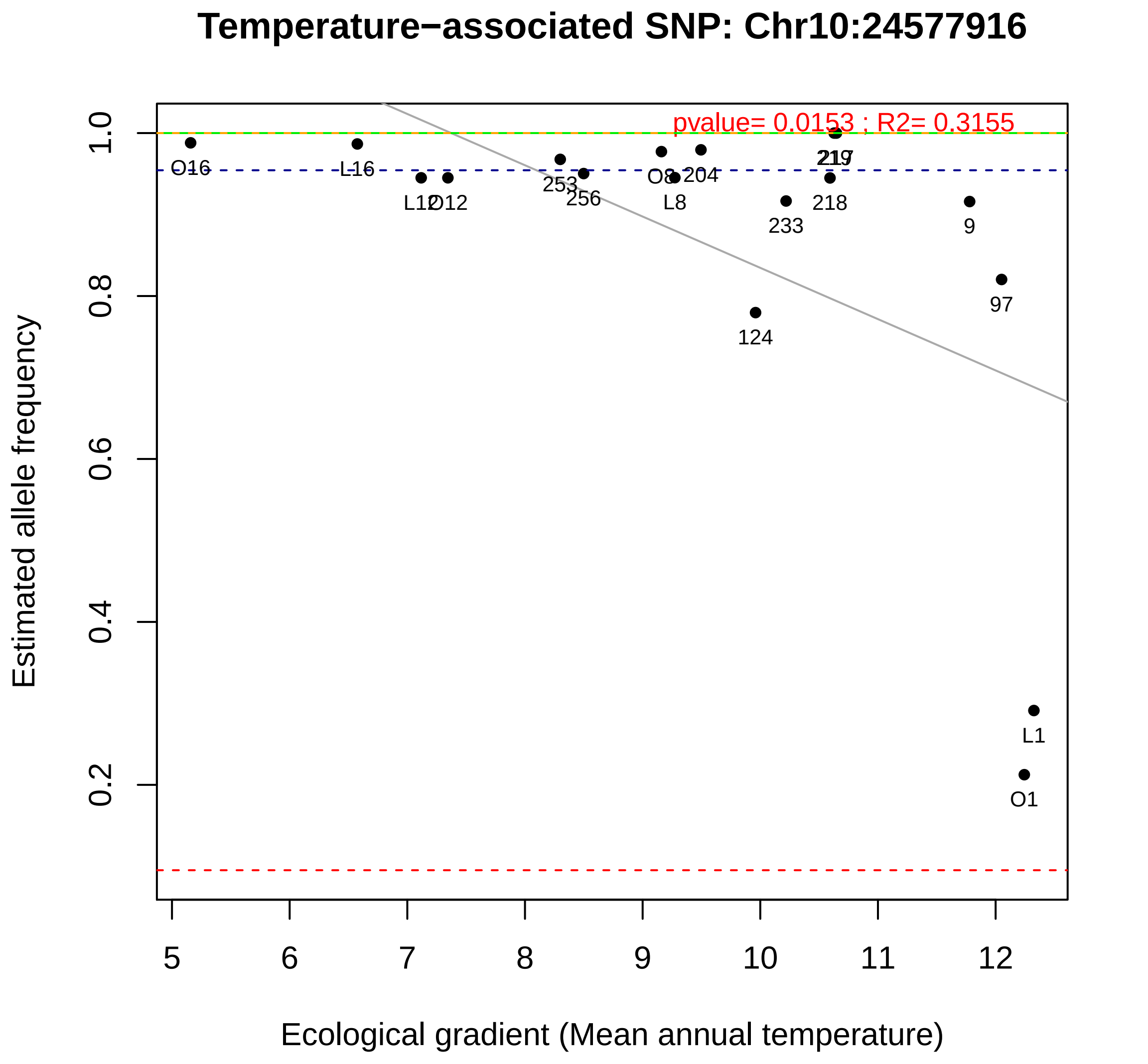

Temperature−associated SNP: Chr10:24577916
pvalue= 0.0153 ; R2= 0.3155
Estimated allele frequency
Ecological gradient (Mean annual temperature)
1.0
0.8
0.6
0.4
0.2
5
6
7
8
9
10
11
12
O16
L16
L12
O12
253
256
O8
L8
204
124
233
217
219
218
9
97
L1
O1

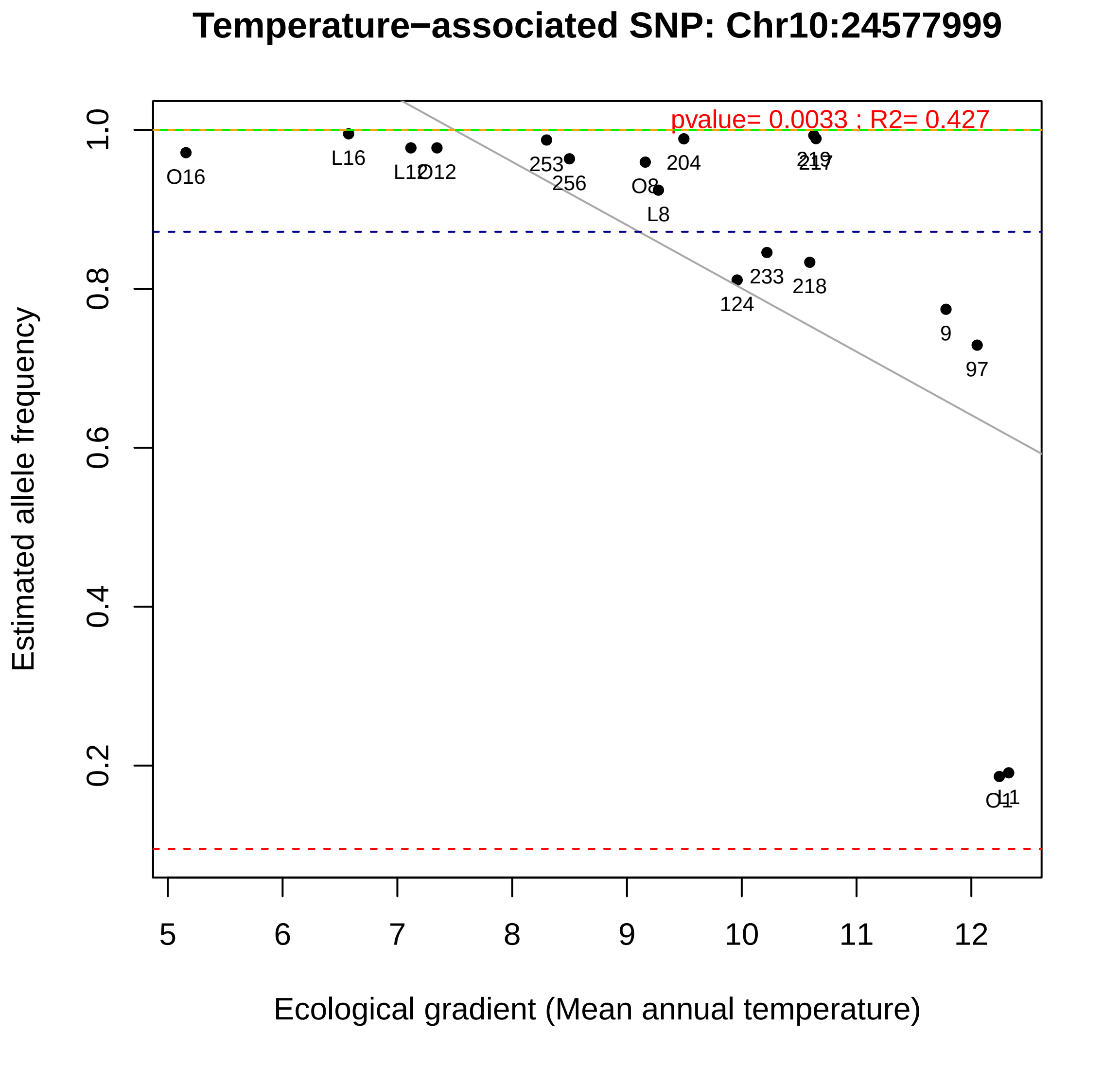
Temperature−associated SNP: Chr10:24577999
pvalue= 0.0033 ; R2= 0.427
Estimated allele frequency
0.2
0.4
0.6
0.8
1.0
5
6
7
8
9
10
11
12
Ecological gradient (Mean annual temperature)
O16
L16
L12
O12
253
256
O8
L8
204
219
217
233
124
218
9
97
O1
L1

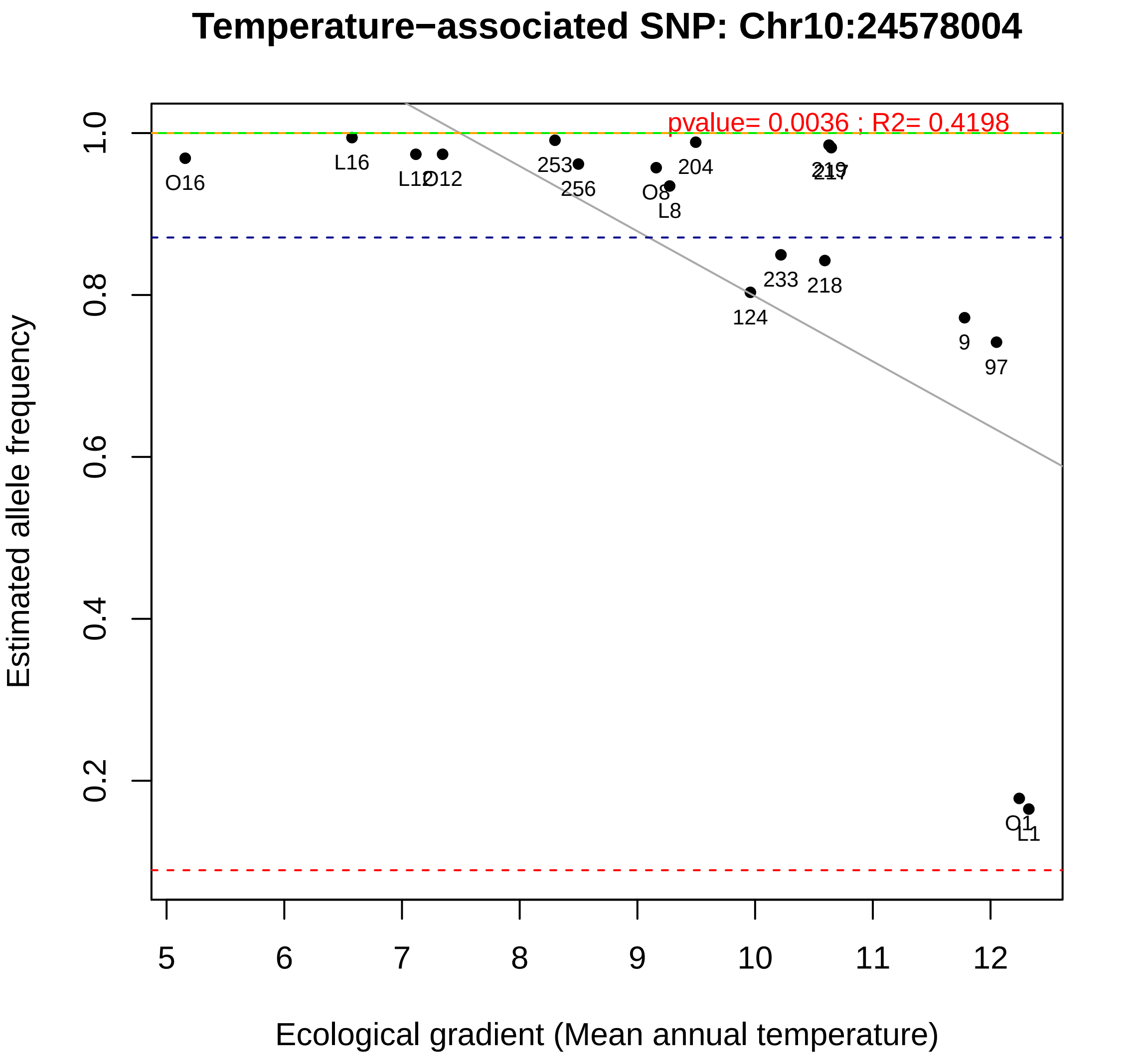

Temperature−associated SNP: Chr10:24578004
pvalue= 0.0036 ; R2= 0.4198
Estimated allele frequency
1.0
0.8
0.6
0.4
0.2
O16
L16
L12
O12
253
256
O8
L8
204
219
217
233
218
124
9
97
O1
L1
5
6
7
8
9
10
11
12
Ecological gradient (Mean annual temperature)

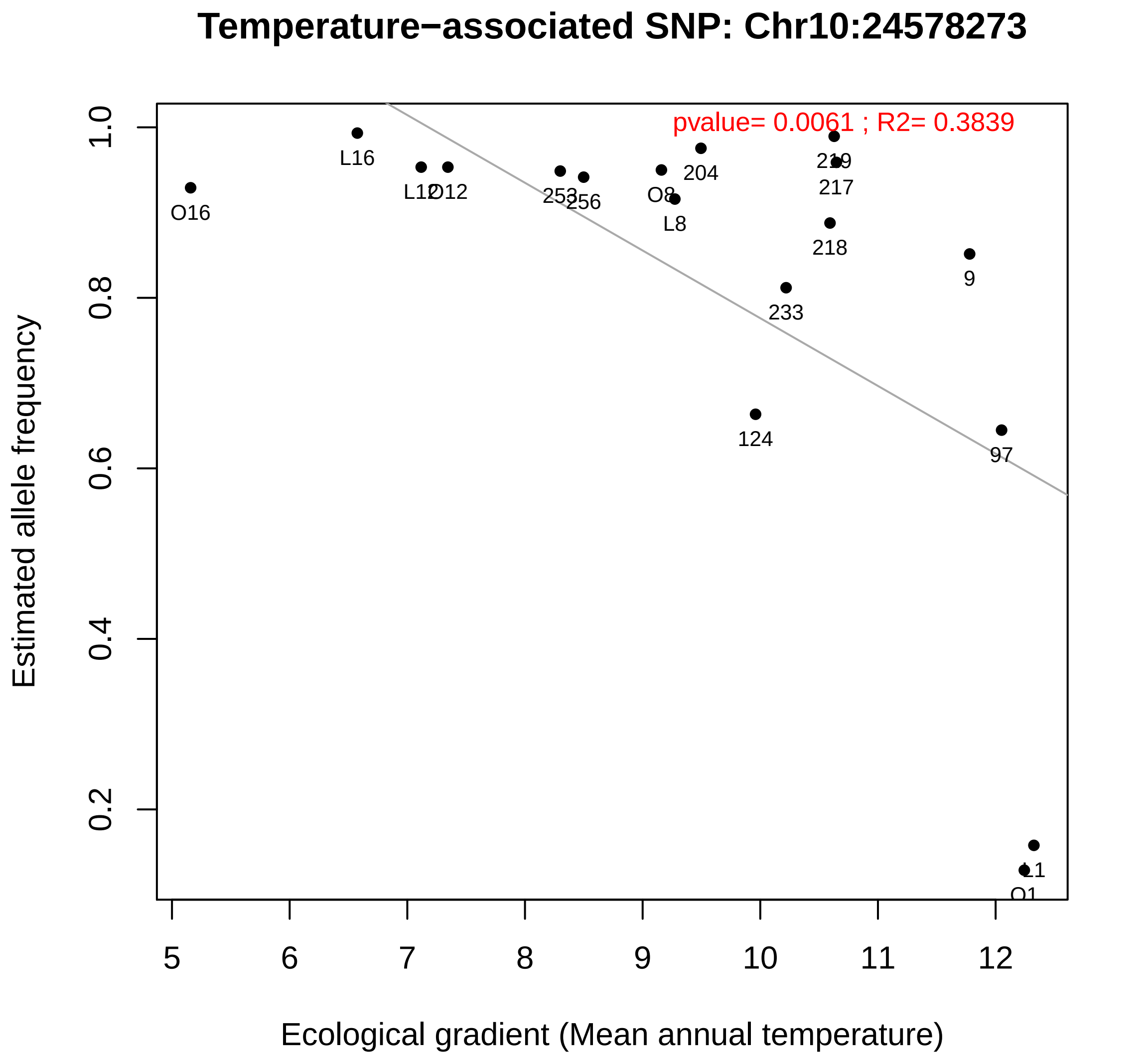

Temperature−associated SNP: Chr10:24578273
pvalue= 0.0061 ; R2= 0.3839
Estimated allele frequency
Ecological gradient (Mean annual temperature)
0.2
0.4
0.6
0.8
1.0
5
6
7
8
9
10
11
12
O16
L16
L12
O12
253
256
O8
L8
204
219
217
218
233
124
9
97
L1
O1

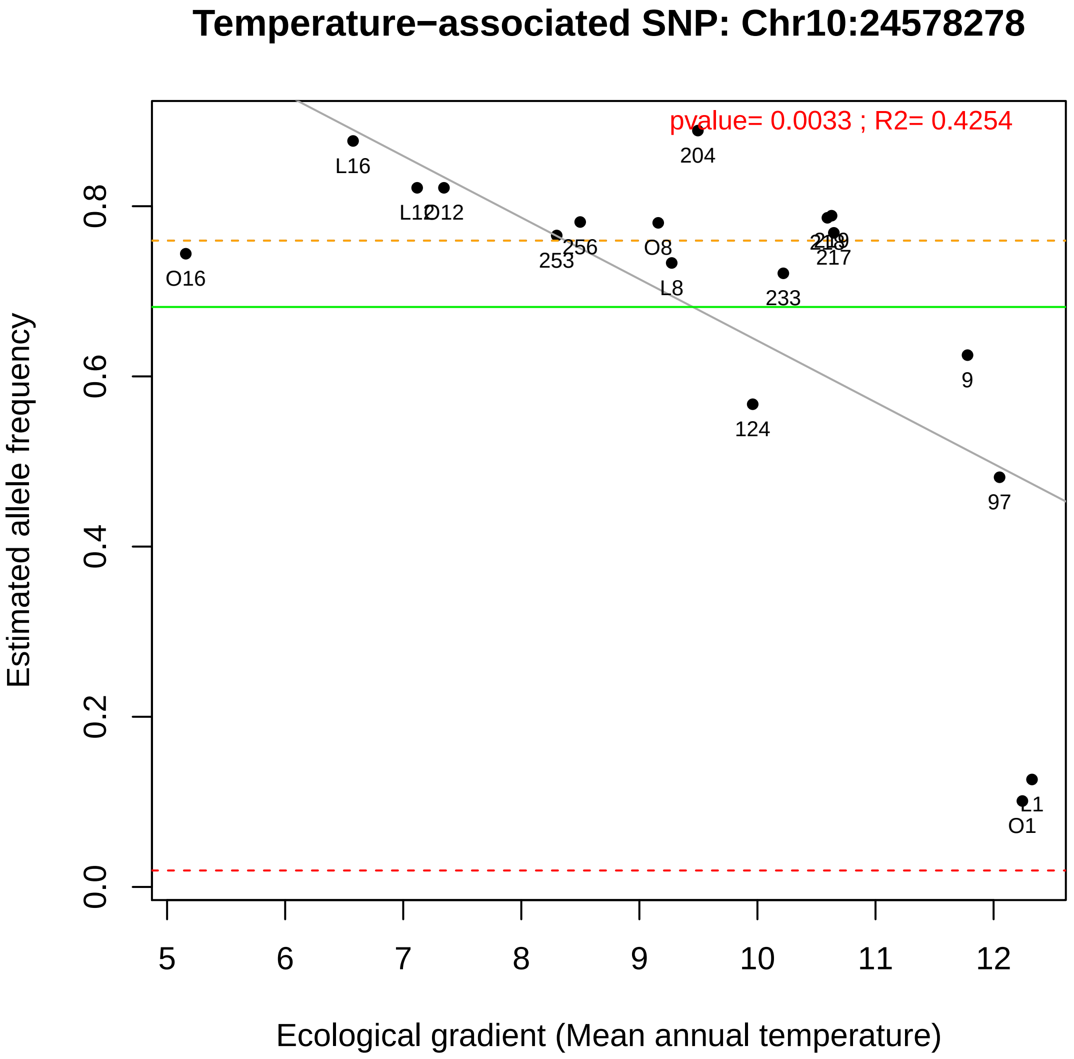

Temperature−associated SNP: Chr10:24578278
pvalue= 0.0033 ; R2= 0.4254
Estimated allele frequency
Ecological gradient (Mean annual temperature)
0.0
0.2
0.4
0.6
0.8
5
6
7
8
9
10
11
12
L16
204
O12
256
253
O8
O16
L8
217
233
9
124
97
L1
O1

**Temperature−associated SNP: Chr10:24578299**

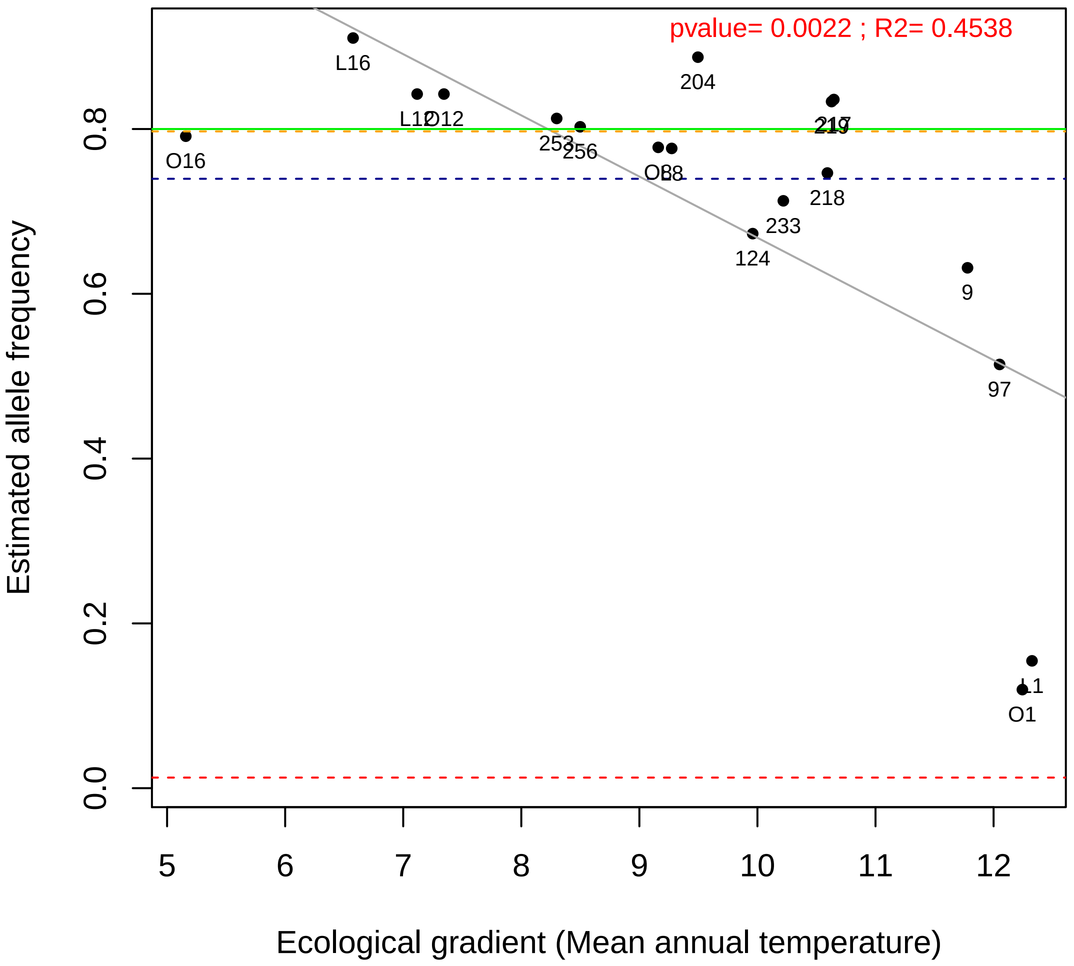

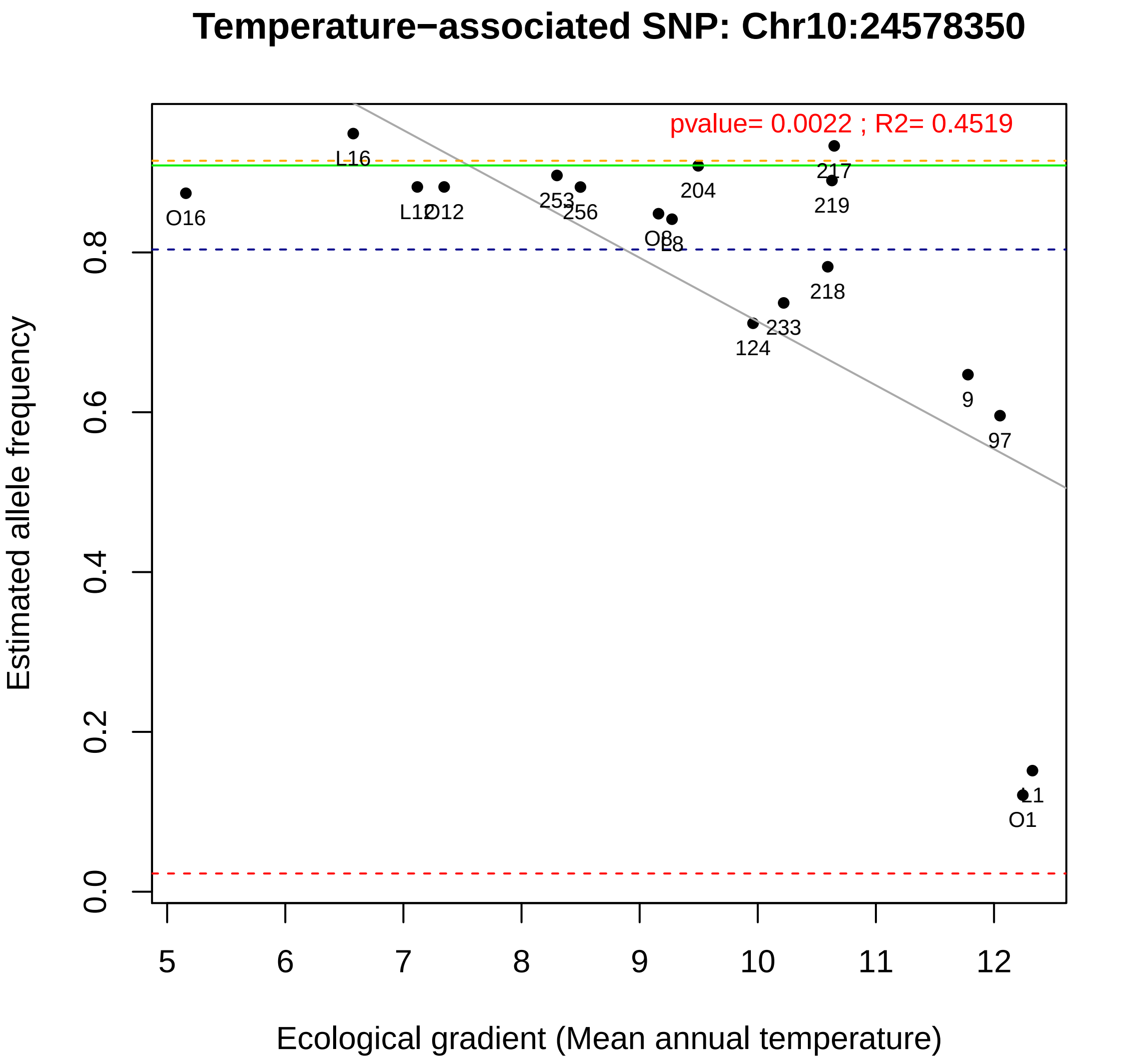
Temperature−associated SNP: Chr10:24578350
pvalue= 0.0022 ; R2= 0.4519
Estimated allele frequency
0.0
0.2
0.4
0.6
0.8
5
6
7
8
9
10
11
12
Ecological gradient (Mean annual temperature)
O16
L16
L12
O12
253
256
O8
L8
204
124
233
217
219
218
9
97
L1
O1

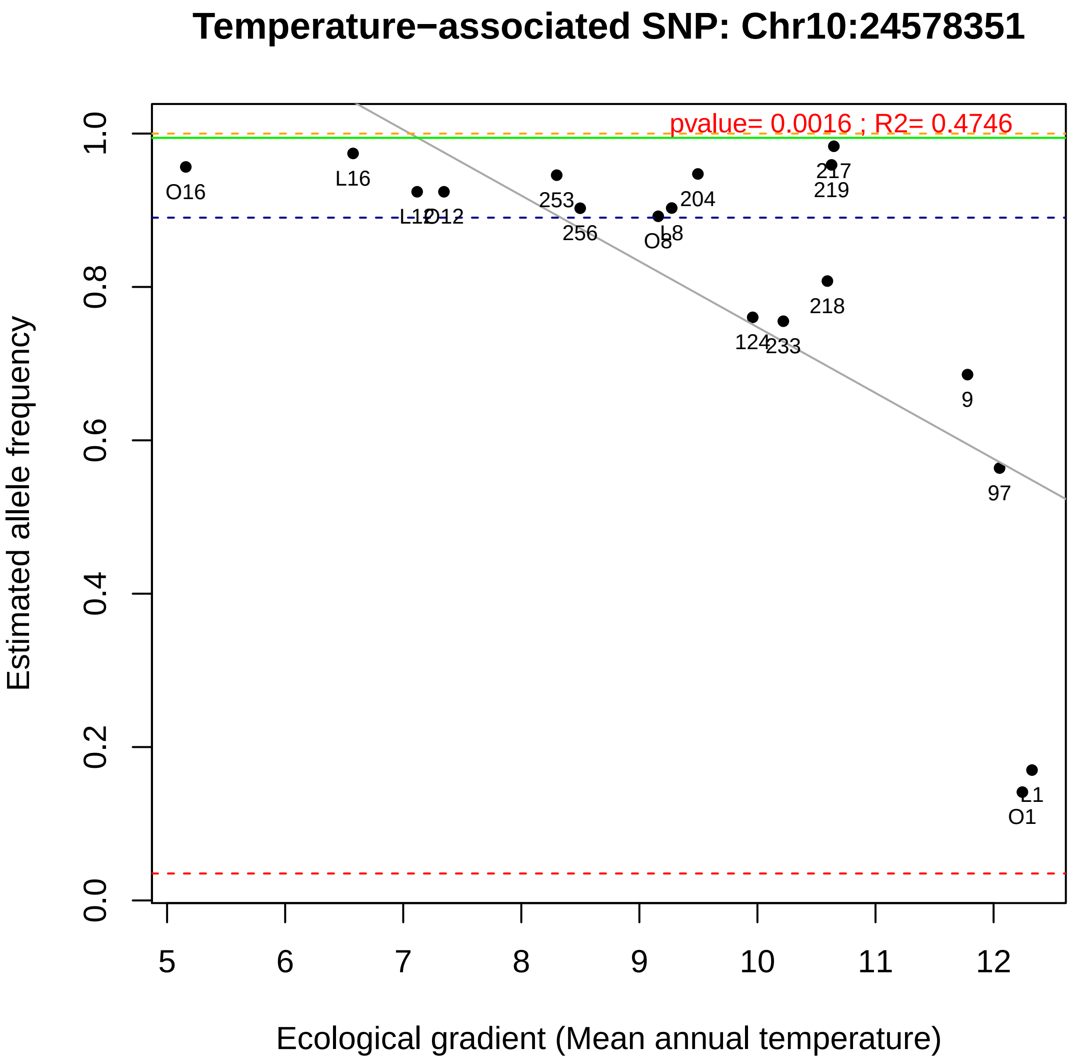
Temperature−associated SNP: Chr10:24578351
pvalue= 0.0016 ; R2= 0.4746
Estimated allele frequency
0.0
0.2
0.4
0.6
0.8
1.0
5
6
7
8
9
10
11
12
Ecological gradient (Mean annual temperature)
O16
L16
L12
O12
253
256
O8
L8
204
217
219
218
124
233
9
97
L1
O1

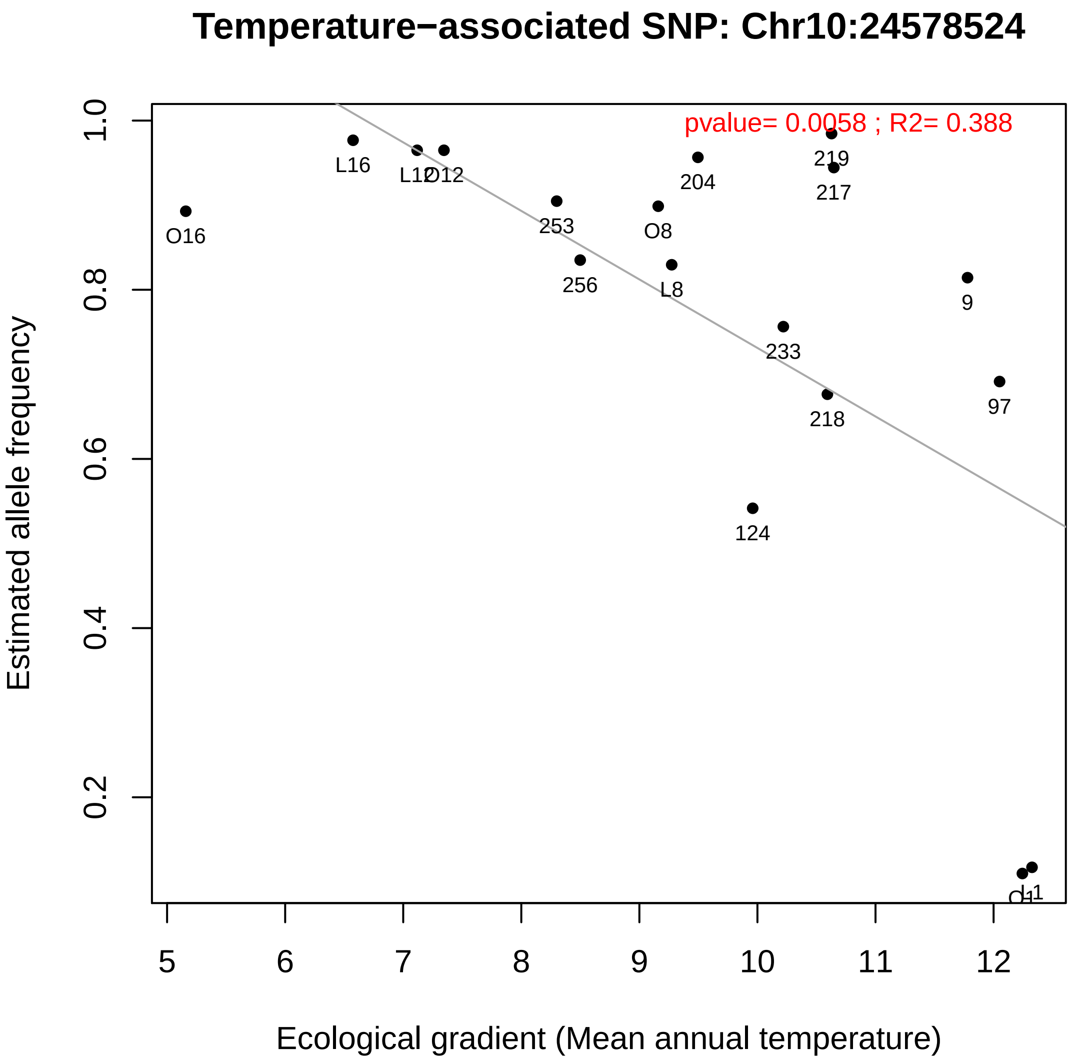

Temperature−associated SNP: Chr10:24578524
pvalue= 0.0058 ; R2= 0.388
Estimated allele frequency
Ecological gradient (Mean annual temperature)
0.2
0.4
0.6
0.8
1.0
5
6
7
8
9
10
11
12
O16
L16
L12
O12
253
256
O8
L8
204
219
217
233
218
124
9
97

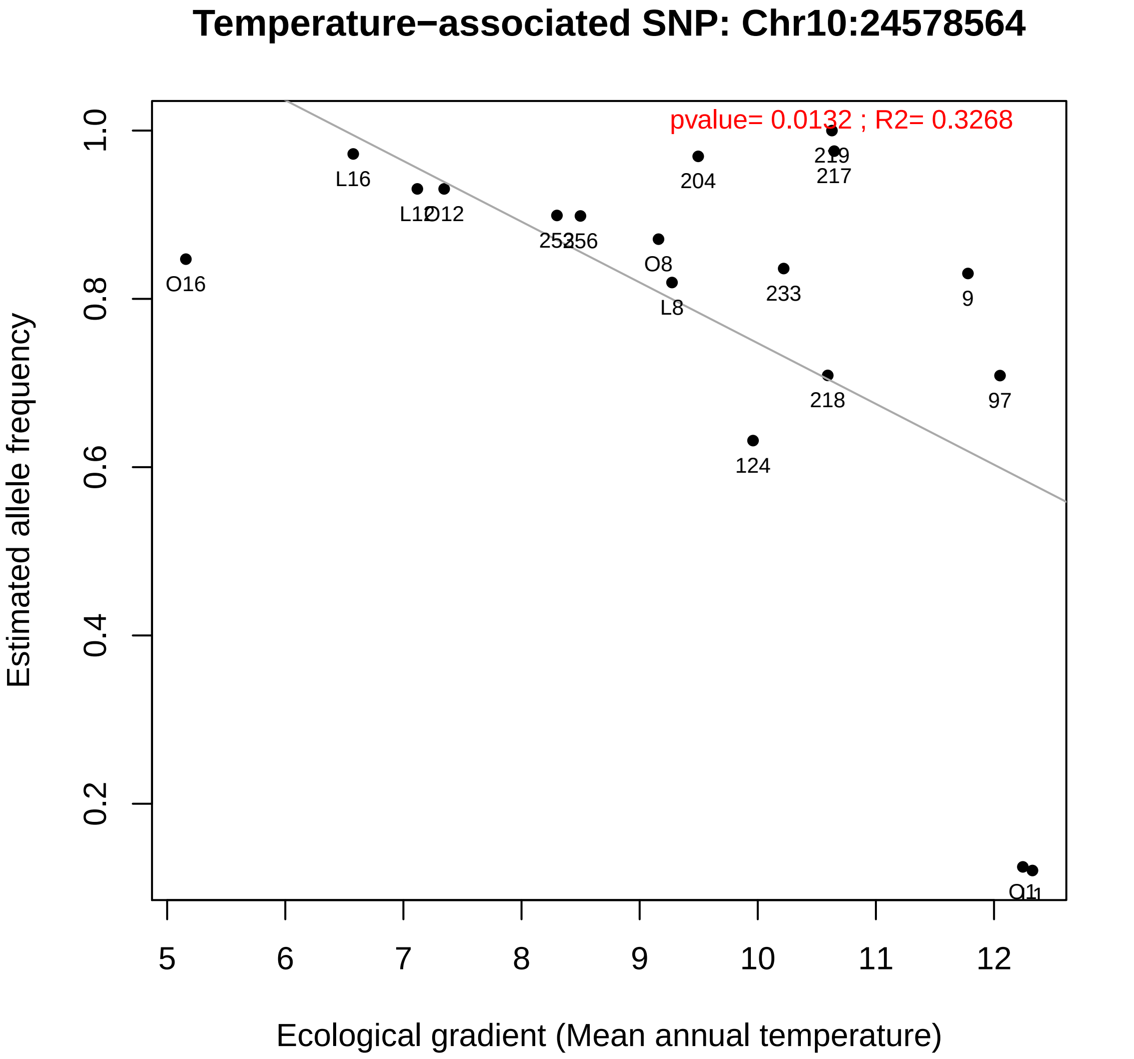
Temperature−associated SNP: Chr10:24578564
pvalue= 0.0132 ; R2= 0.3268
Estimated allele frequency
Ecological gradient (Mean annual temperature)
0.2
0.4
0.6
0.8
1.0
5
6
7
8
9
10
11
12
O16
L16
O12
256
O8
L8
204
233
217
9
218
97
124

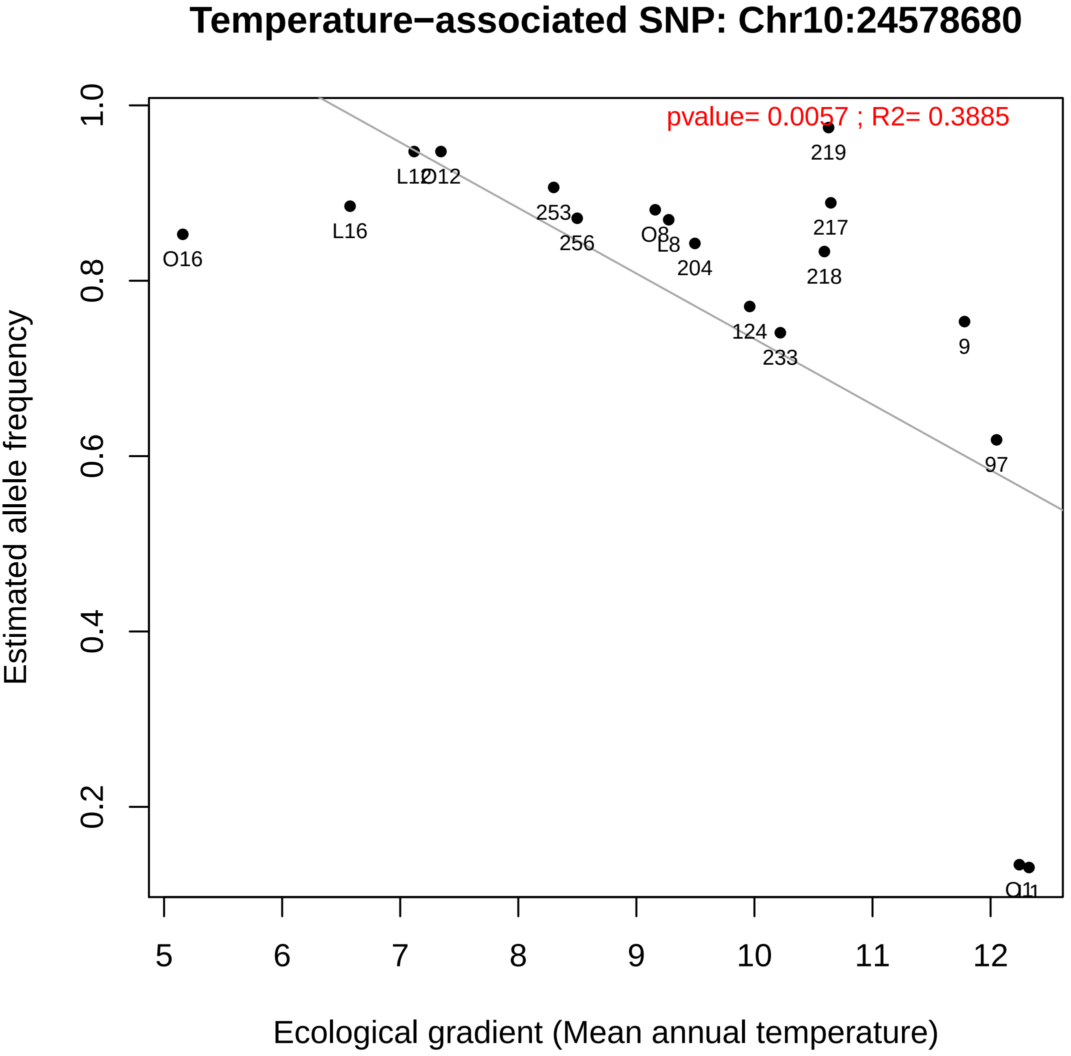

Temperature−associated SNP: Chr10:24578680
pvalue= 0.0057 ; R2= 0.3885
Estimated allele frequency
Ecological gradient (Mean annual temperature)
0.2
0.4
0.6
0.8
1.0
5
6
7
8
9
10
11
12
O16
L16
L12
O12
253
256
O8
L8
204
124
233
219
217
218
9
97

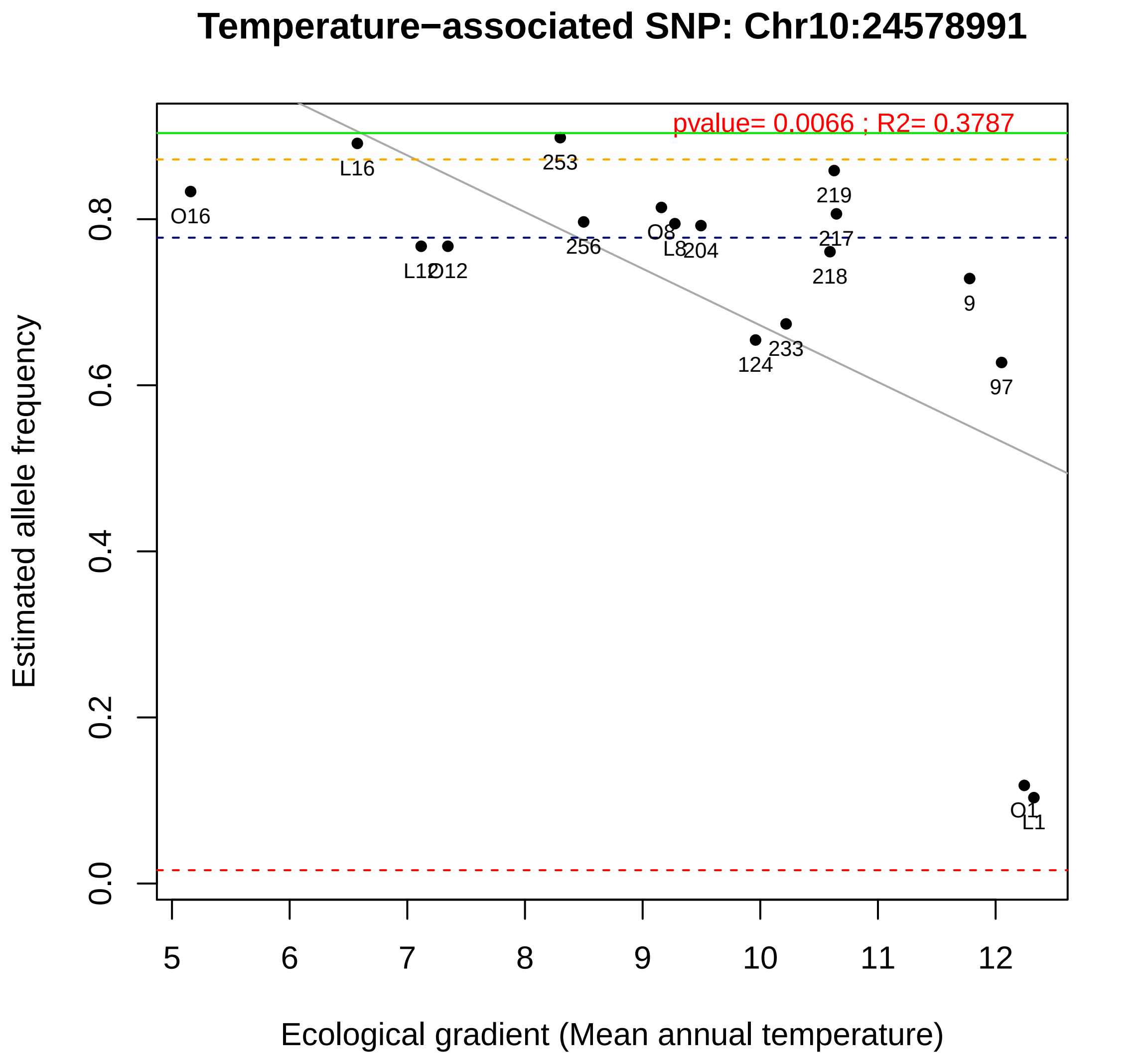
Temperature−associated SNP: Chr10:24578991
pvalue= 0.0066 ; R2= 0.3787
Estimated allele frequency
0.0
0.2
0.4
0.6
0.8
5
6
7
8
9
10
11
12
Ecological gradient (Mean annual temperature)
O16
L16
253
256
L12
O12
O8
L8
204
219
217
218
124
233
9
97
O1
L1

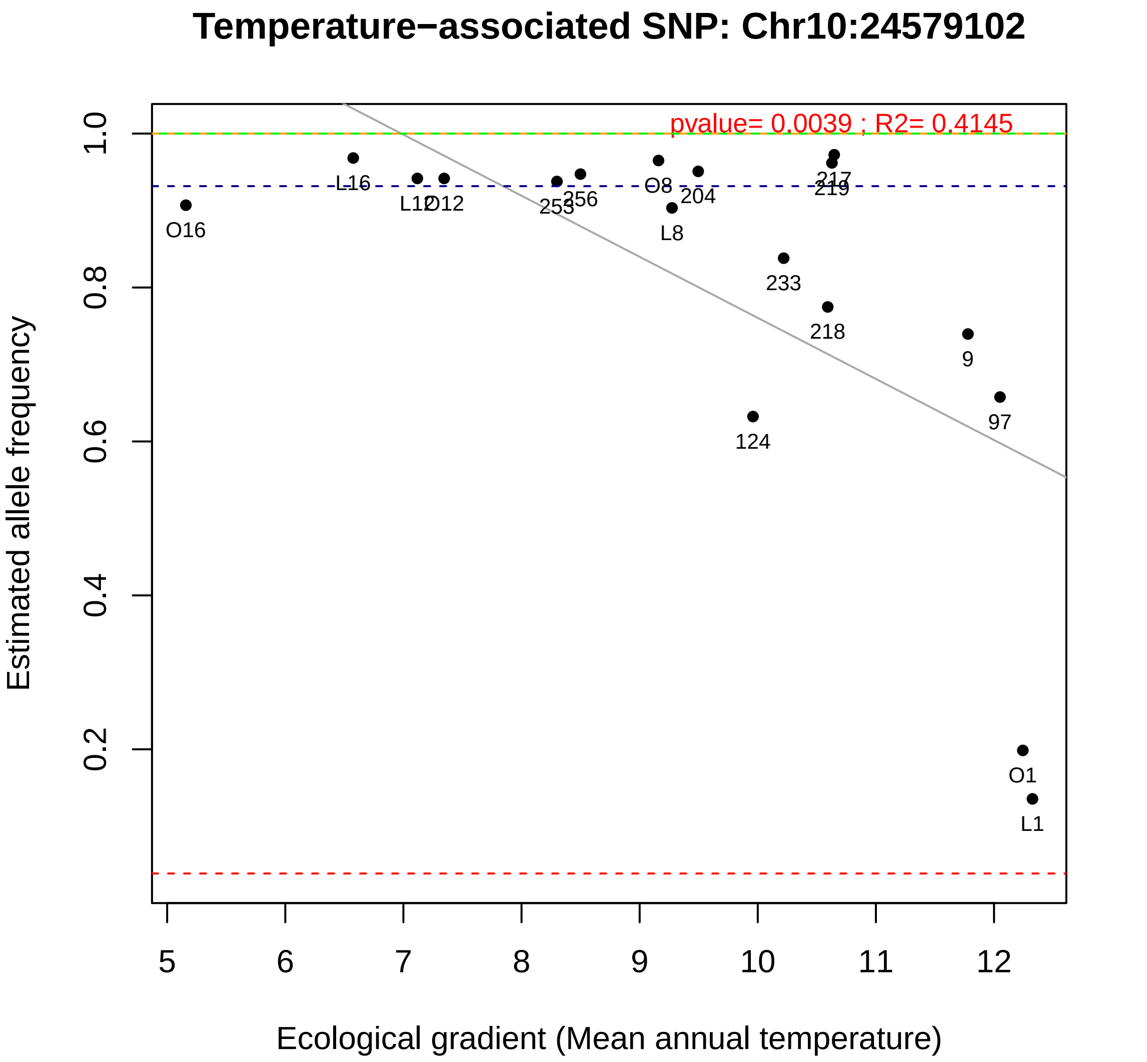
Temperature−associated SNP: Chr10:24579102
pvalue= 0.0039 ; R2= 0.4145
Estimated allele frequency
Ecological gradient (Mean annual temperature)
1.0
0.8
0.6
0.4
0.2
5
6
7
8
9
10
11
12
O16
L16
L12
O12
253
256
O8
L8
204
217
219
233
218
124
9
97
O1
L1

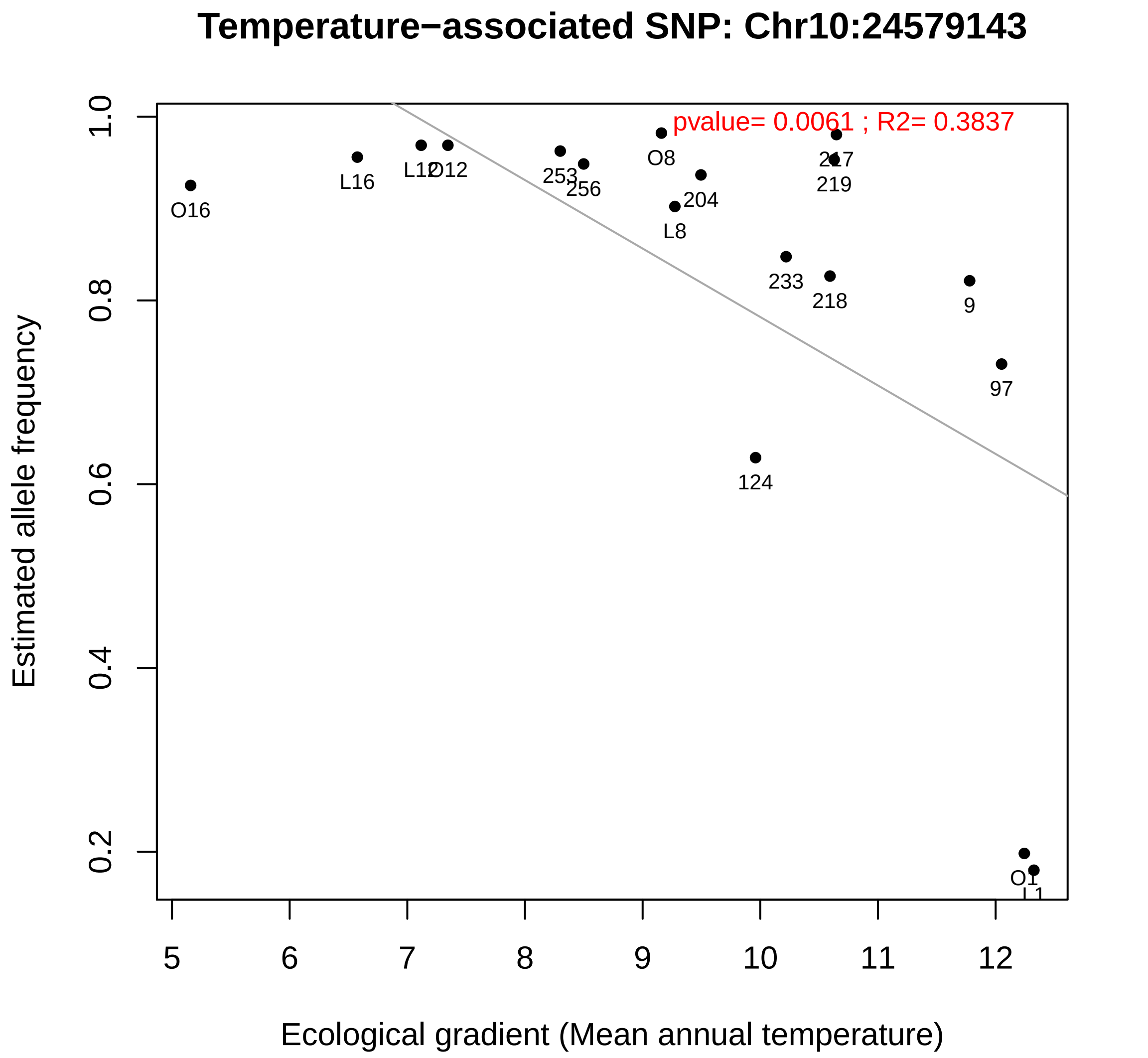

Temperature−associated SNP: Chr10:24579143
pvalue= 0.0061 ; R2= 0.3837
Estimated allele frequency
Ecological gradient (Mean annual temperature)
0.2
0.4
0.6
0.8
1.0
5
6
7
8
9
10
11
12
O16
L16
L12
O12
253
256
O8
204
L8
217
219
233
218
124
9
97
O1

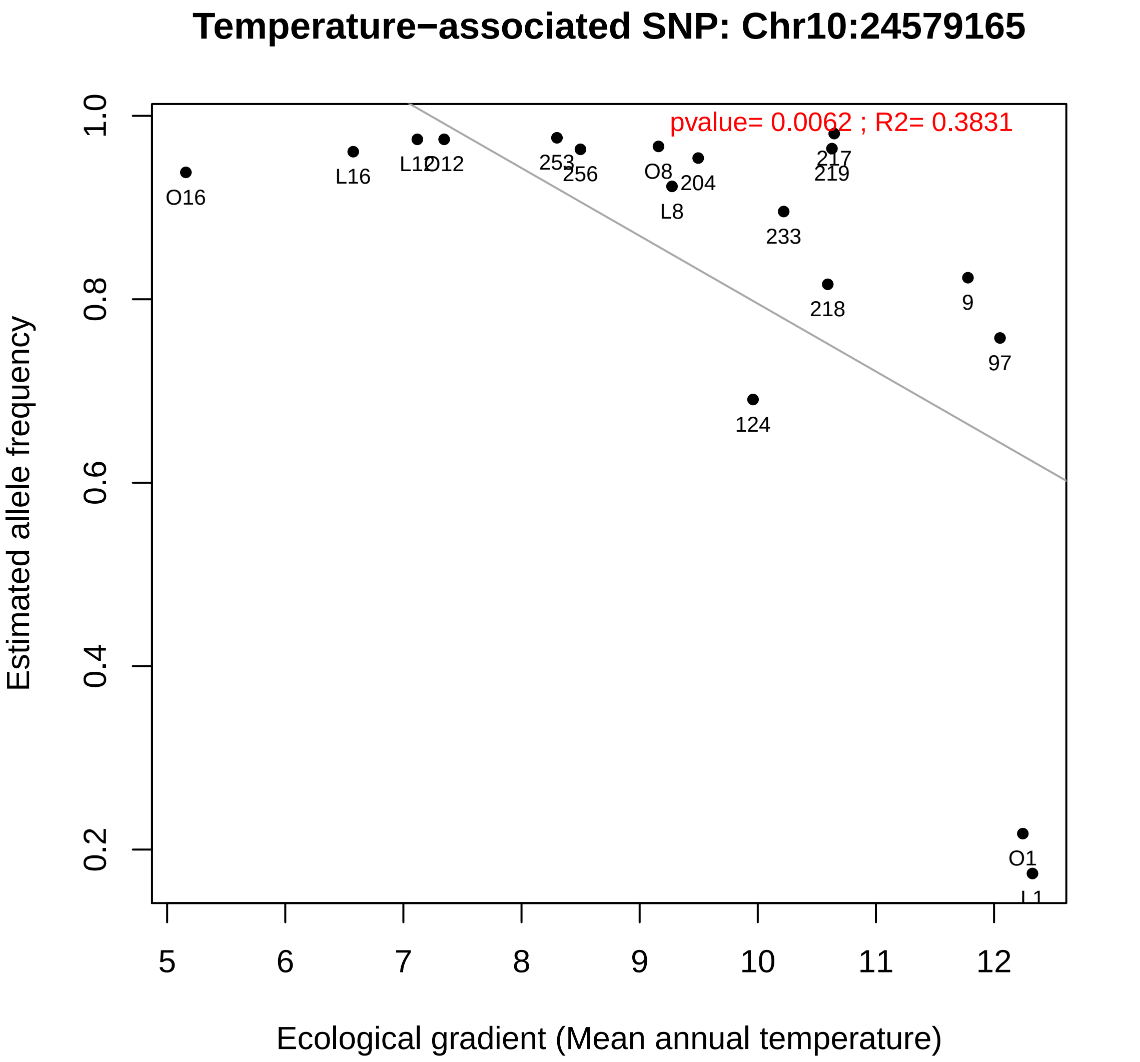

Temperature−associated SNP: Chr10:24579165
pvalue= 0.0062 ; R2= 0.3831
Estimated allele frequency
Ecological gradient (Mean annual temperature)
0.2
0.4
0.6
0.8
1.0
5
6
7
8
9
10
11
12
O16
L16
L12
O12
253
256
O8
204
L8
217
219
233
218
9
97
124
O1

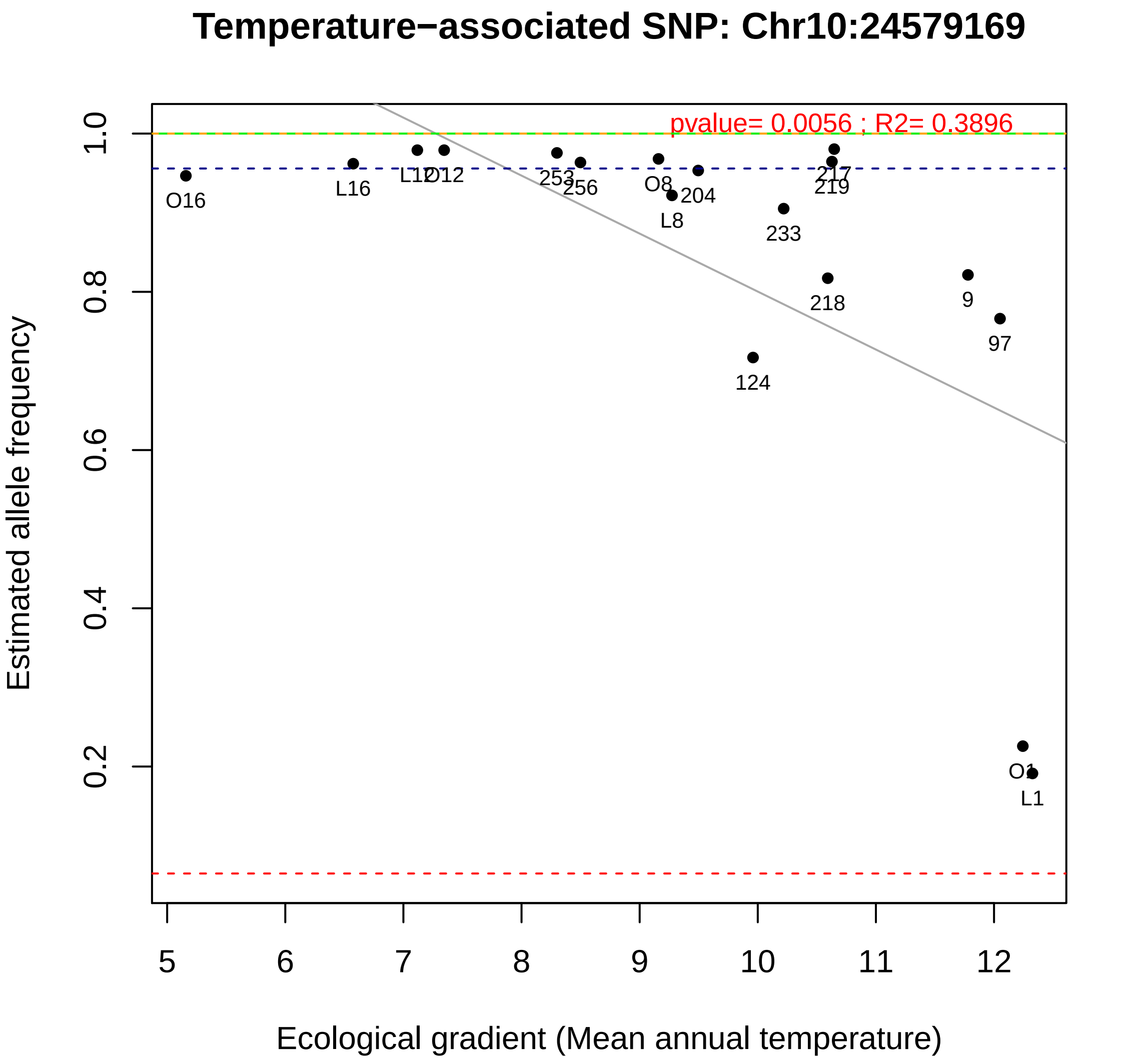
Temperature−associated SNP: Chr10:24579169
pvalue= 0.0056 ; R2= 0.3896
Estimated allele frequency
Ecological gradient (Mean annual temperature)
0.2
0.4
0.6
0.8
1.0
5
6
7
8
9
10
11
12
O16
L16
L12
O12
253
256
O8
L8
204
233
217
219
218
124
9
97
O1
L1

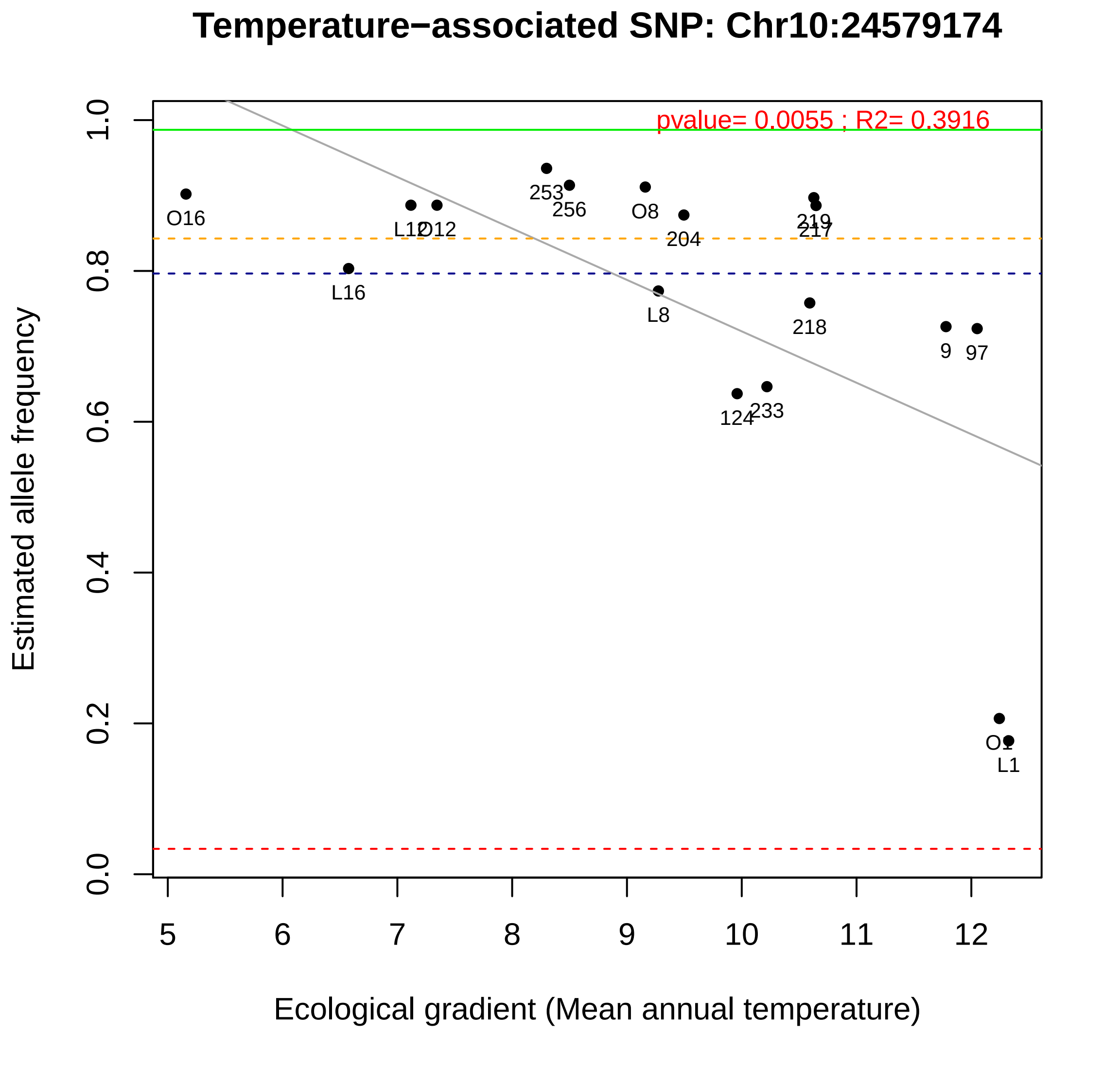
Temperature−associated SNP: Chr10:24579174
pvalue= 0.0055 ; R2= 0.3916
Estimated allele frequency
0.0
0.2
0.4
0.6
0.8
1.0
Ecological gradient (Mean annual temperature)
5
6
7
8
9
10
11
12
O16
L16
L12
O12
253
256
O8
204
L8
219
217
218
124
233
9
97
O1
L1

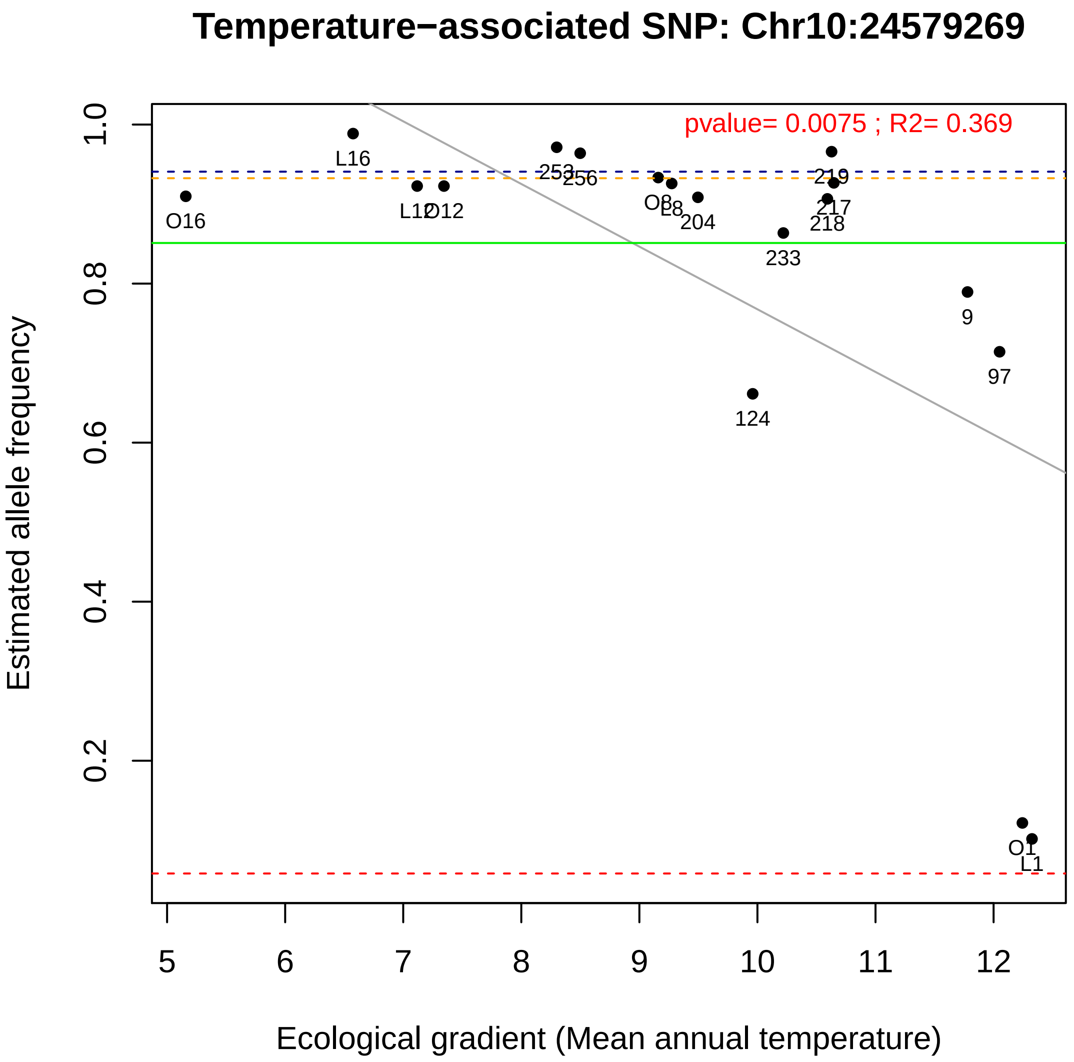
Temperature−associated SNP: Chr10:24579269
pvalue= 0.0075 ; R2= 0.369
Estimated allele frequency
Ecological gradient (Mean annual temperature)
1.0
0.8
0.6
0.4
0.2
5
6
7
8
9
10
11
12
O16
L16
L12
O12
253
256
O8
L8
204
124
233
219
217
218
9
97
O1
L1

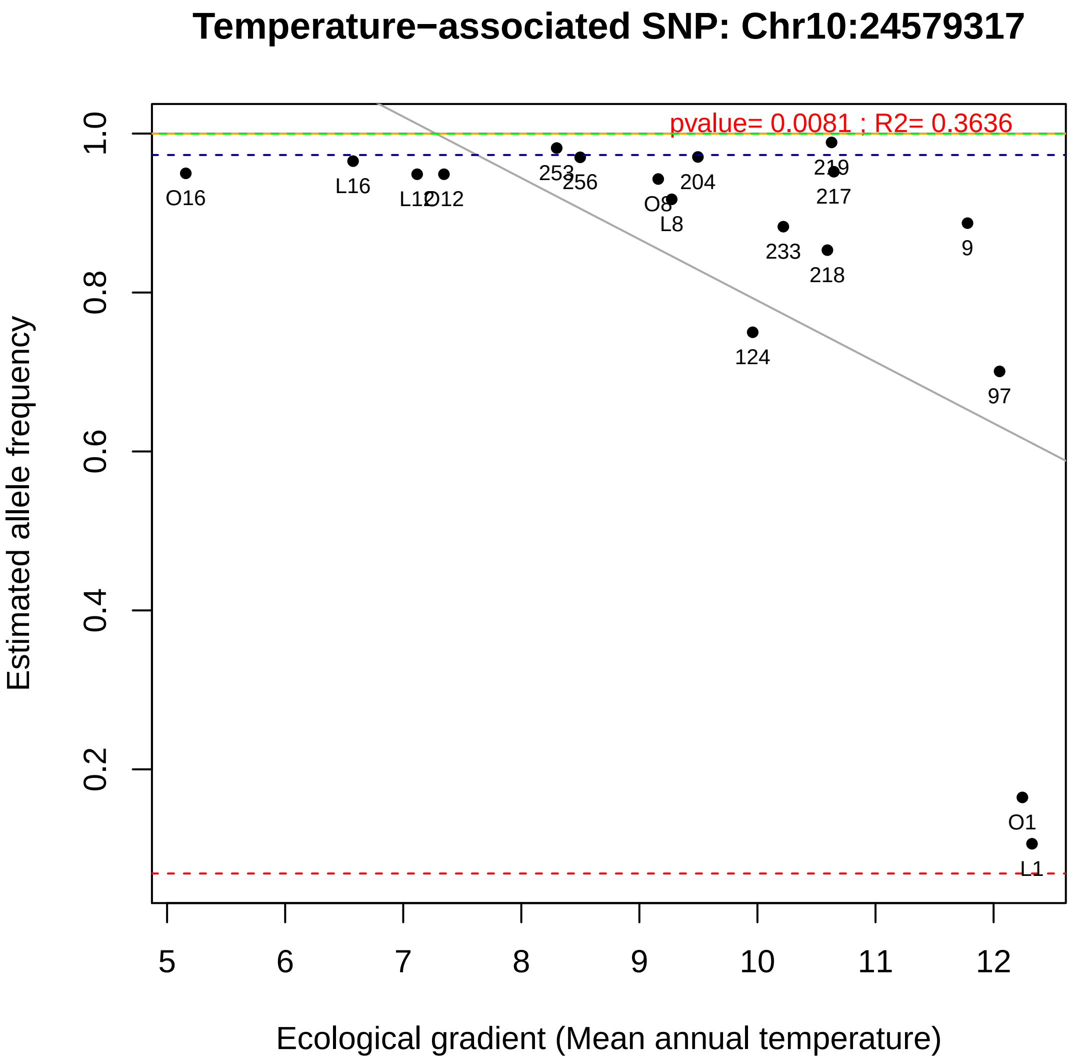

Temperature−associated SNP: Chr10:24579317
pvalue= 0.0081 ; R2= 0.3636
Estimated allele frequency
Ecological gradient (Mean annual temperature)
1.0
0.8
0.6
0.4
0.2
5
6
7
8
9
10
11
12
O16
L16
L12
O12
253
256
O8
L8
204
219
217
233
218
124
9
97
O1
L1

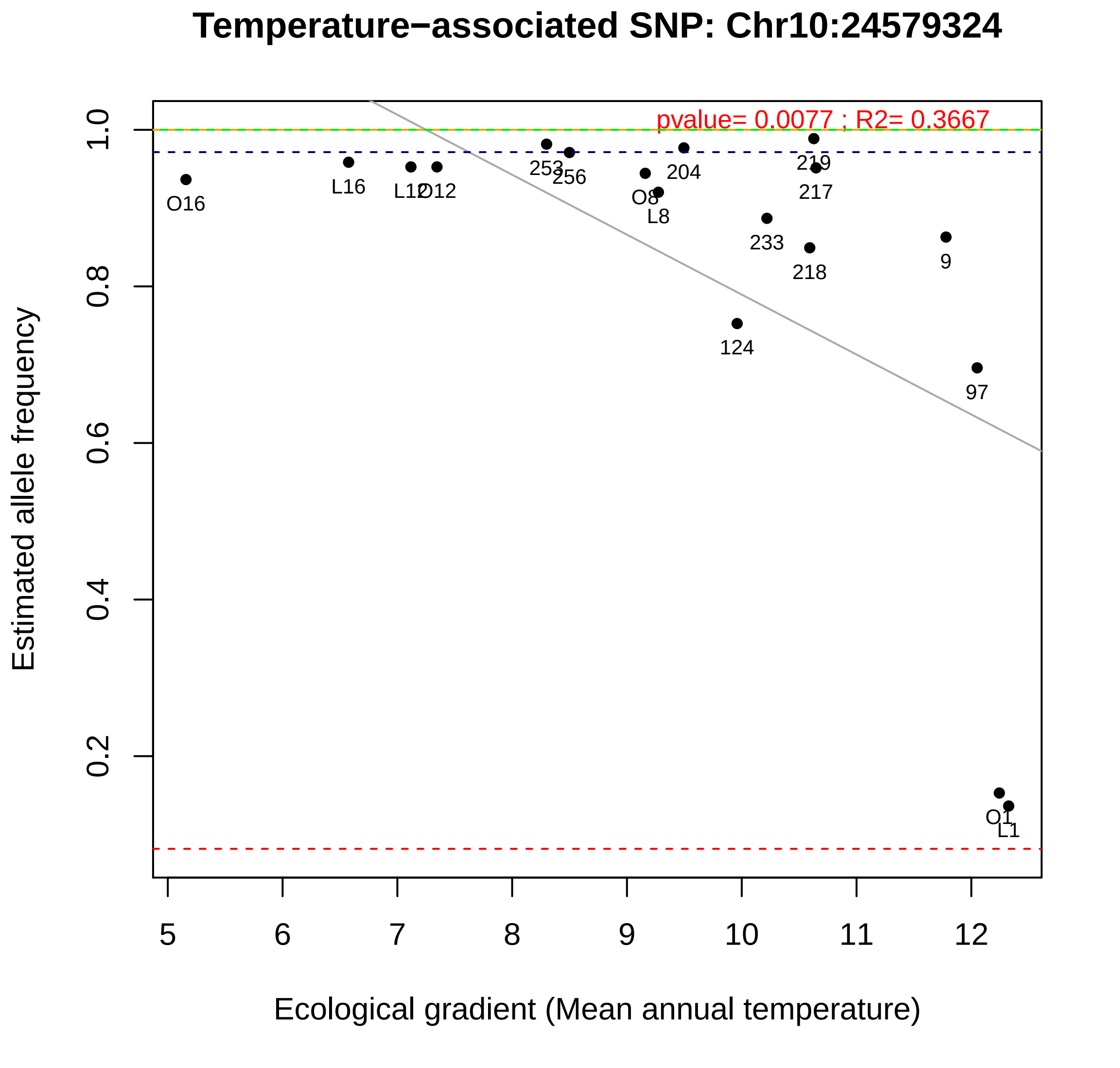
Temperature−associated SNP: Chr10:24579324
pvalue= 0.0077 ; R2= 0.3667
Estimated allele frequency
0.2
0.4
0.6
0.8
1.0
Ecological gradient (Mean annual temperature)
5
6
7
8
9
10
11
12
O16
L16
L12
O12
253
256
O8
L8
204
219
217
233
218
124
9
97
O1
L1

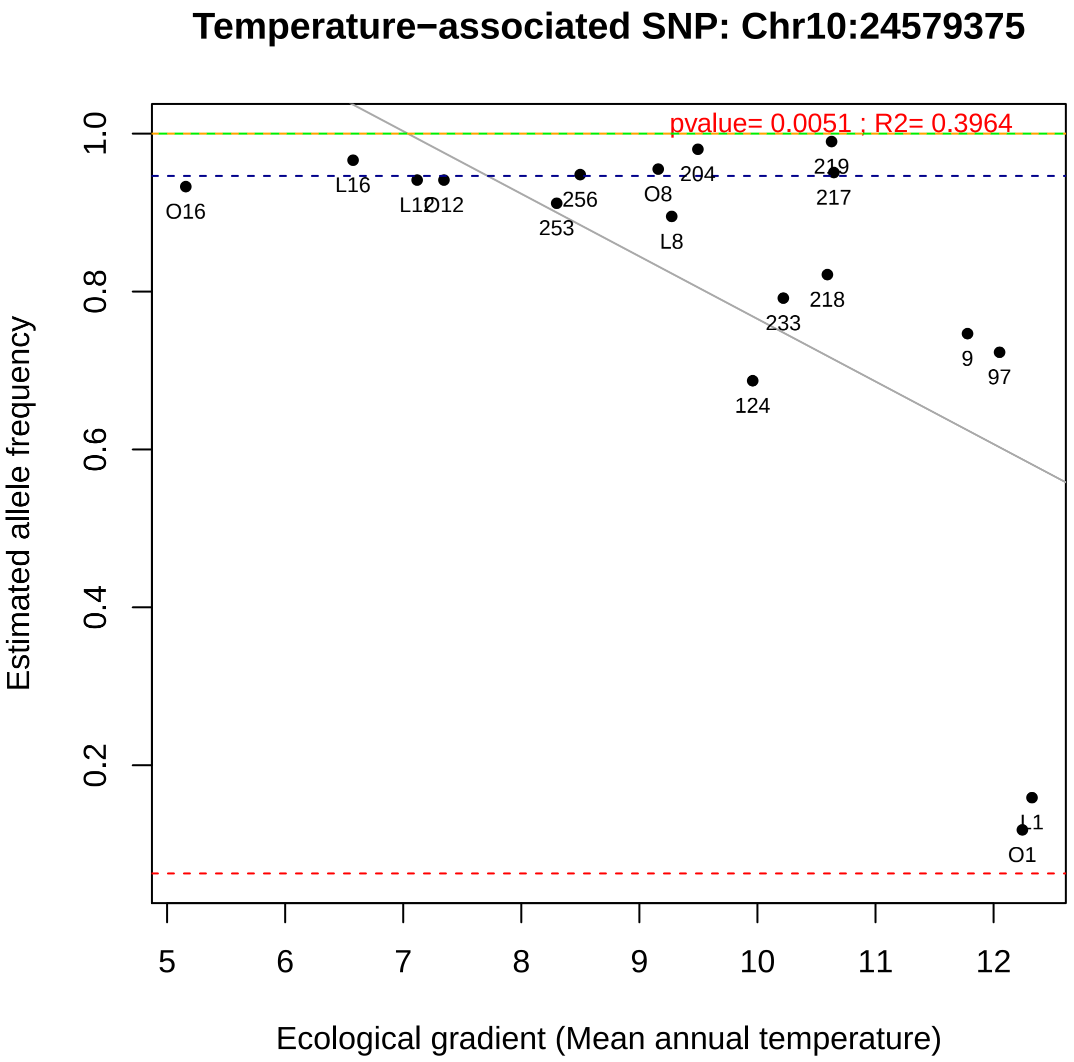
Temperature−associated SNP: Chr10:24579375
pvalue= 0.0051 ; R2= 0.3964
Estimated allele frequency
0.2
0.4
0.6
0.8
1.0
5
6
7
8
9
10
11
12
Ecological gradient (Mean annual temperature)
O16
L16
L12
O12
253
256
O8
L8
204
219
217
218
233
124
9
97
L1
O1

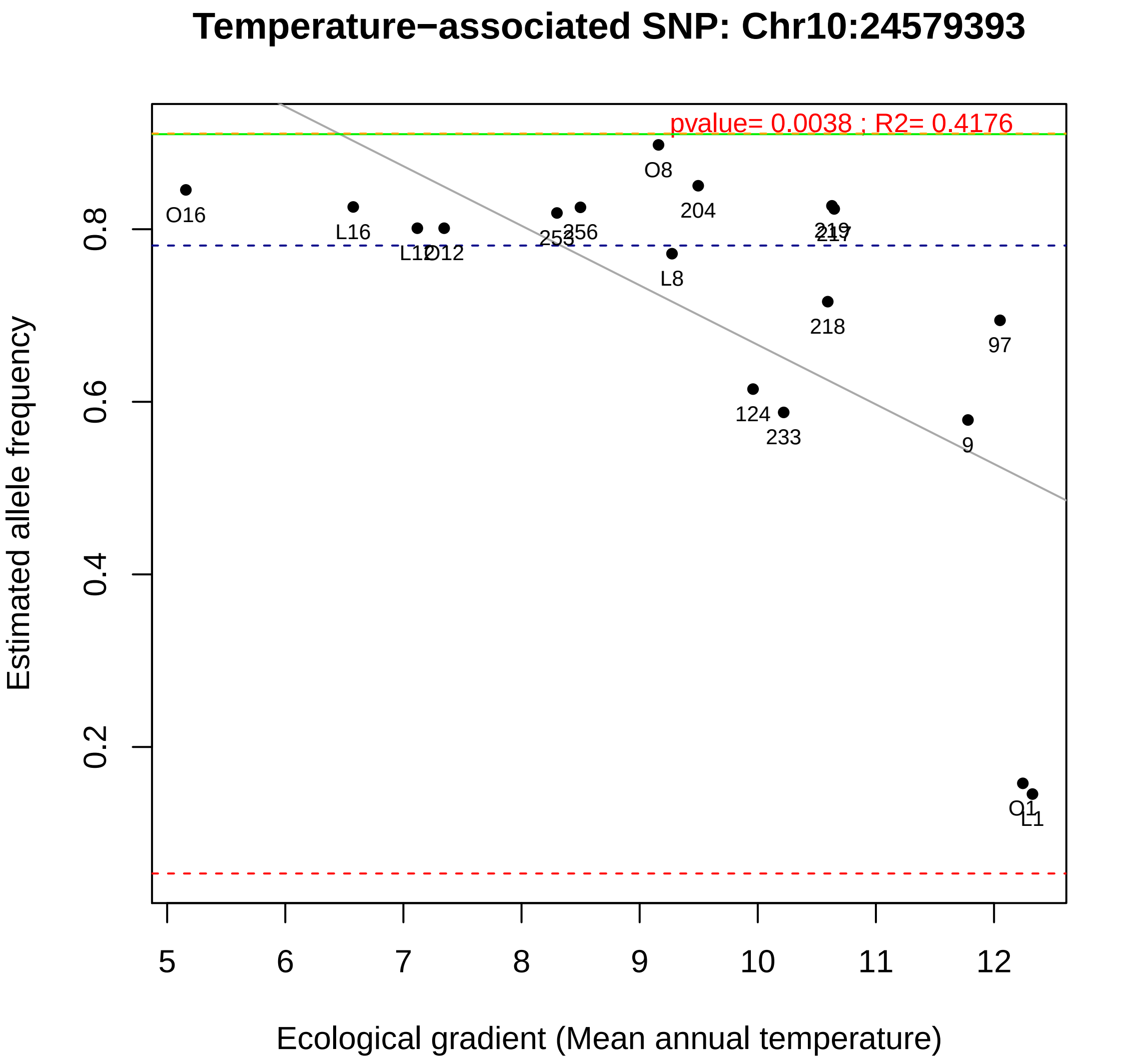
Temperature−associated SNP: Chr10:24579393
pvalue= 0.0038 ; R2= 0.4176
Estimated allele frequency
0.2
0.4
0.6
0.8
5
6
7
8
9
10
11
12
Ecological gradient (Mean annual temperature)
O16
L16
L12
O12
253
256
O8
204
L8
219
217
218
97
124
233
9
O1
L1

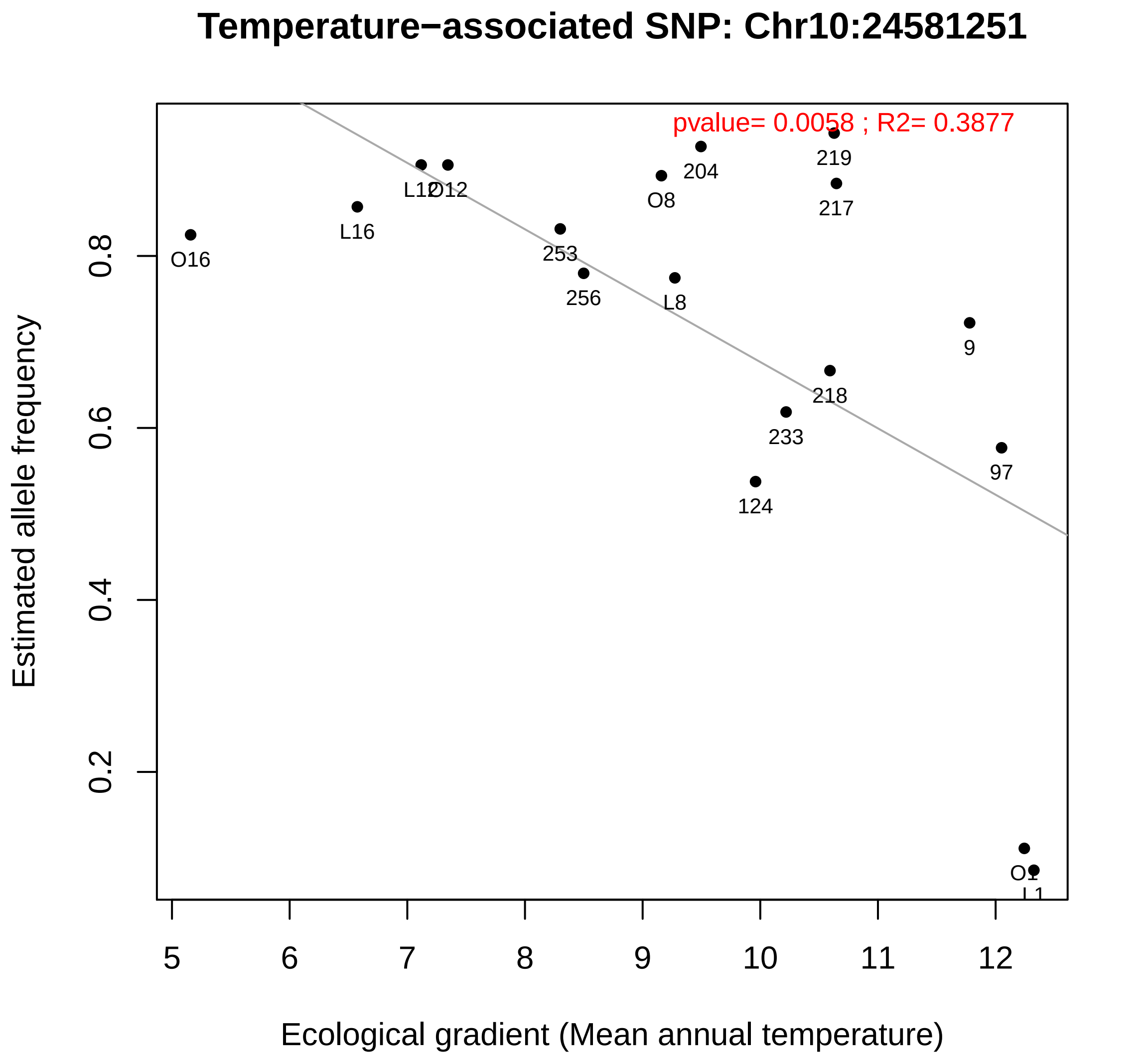

Temperature−associated SNP: Chr10:24581251
pvalue= 0.0058 ; R2= 0.3877
O16
L16
L12
O12
253
256
O8
204
L8
219
217
9
218
233
124
97
0.2
0.4
0.6
0.8
5
6
7
8
9
10
11
12
Estimated allele frequency
Ecological gradient (Mean annual temperature)

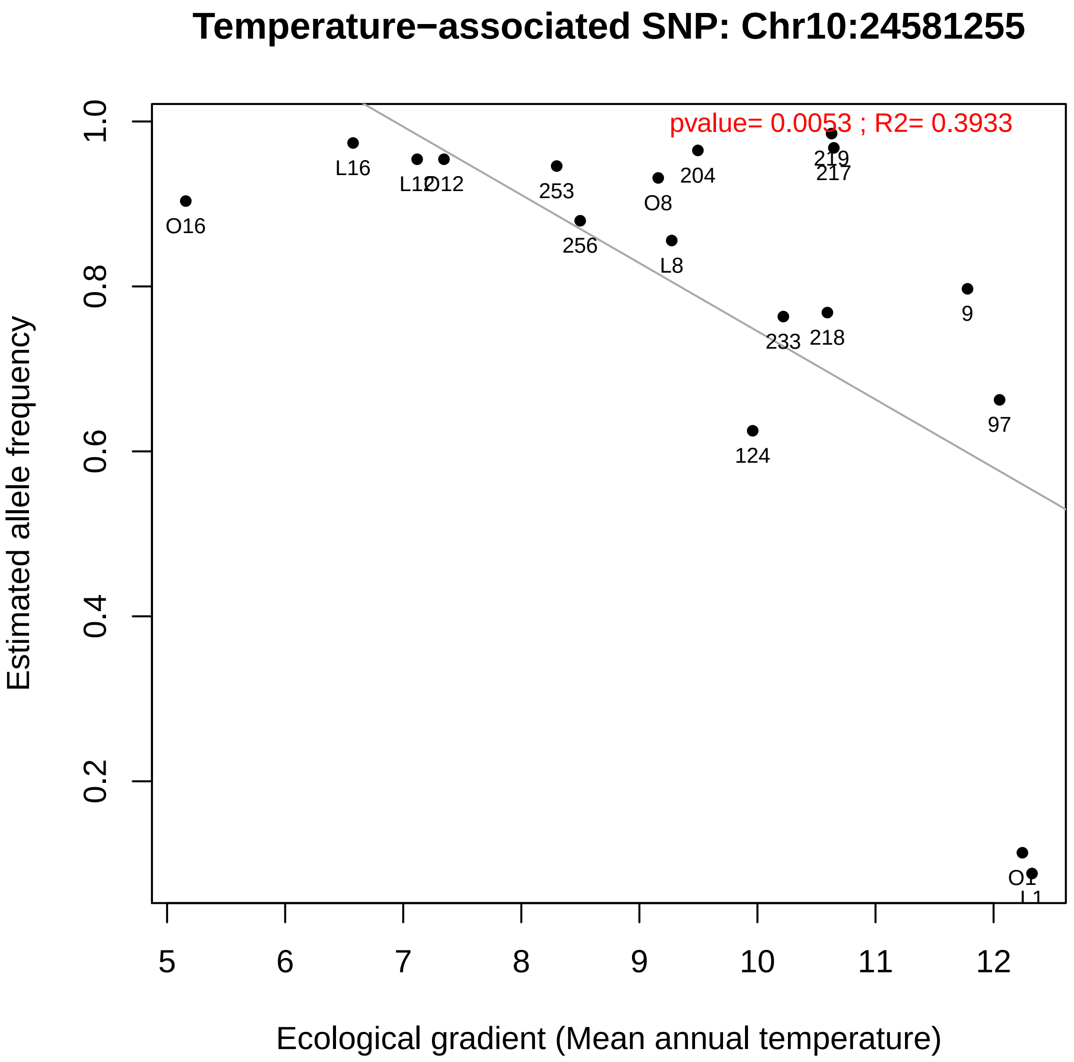
Temperature−associated SNP: Chr10:24581255
pvalue= 0.0053 ; R2= 0.3933
Estimated allele frequency
1.0
0.8
0.6
0.4
0.2
5
6
7
8
9
10
11
12
Ecological gradient (Mean annual temperature)
O16
L16
L12
O12
253
256
O8
L8
204
219
217
233
218
124
9
97

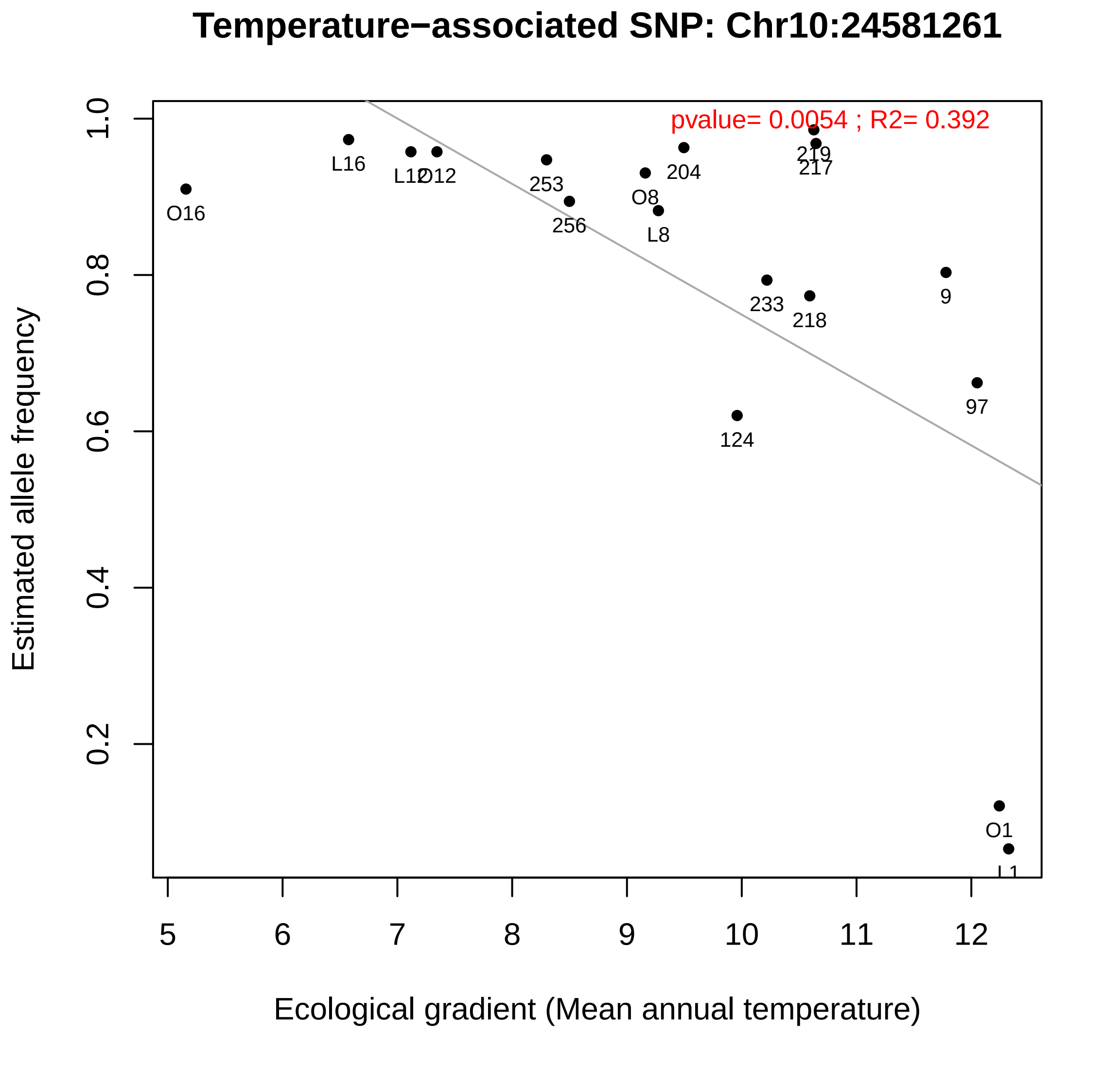

Temperature−associated SNP: Chr10:24581261
pvalue= 0.0054 ; R2= 0.392
Estimated allele frequency
Ecological gradient (Mean annual temperature)
0.2
0.4
0.6
0.8
1.0
5
6
7
8
9
10
11
12
O16
L16
L12
O12
253
256
O8
L8
204
219
217
233
218
124
9
97
O1

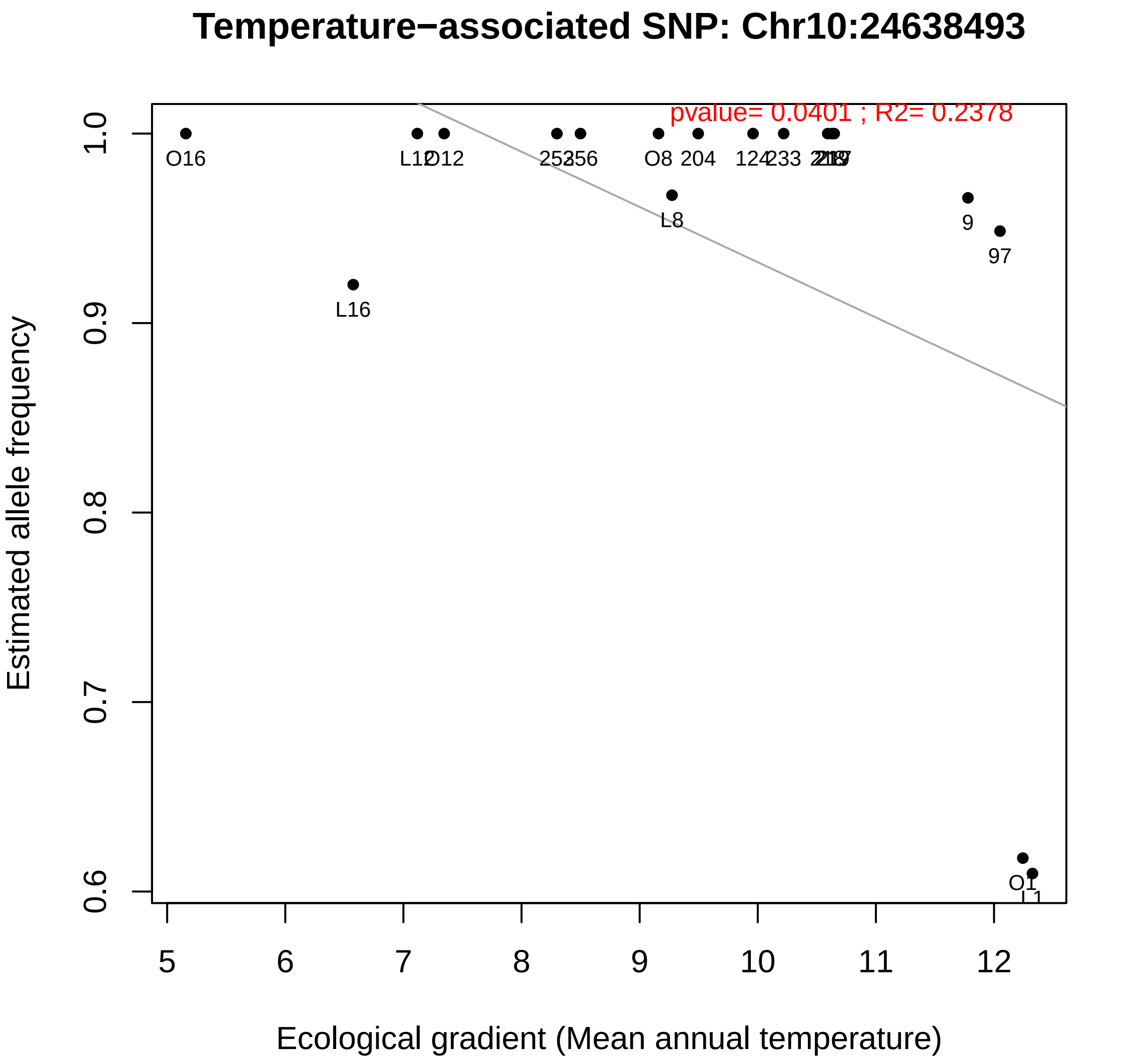

Temperature−associated SNP: Chr10:24638493
pvalue= 0.0401 ; R2= 0.2378
Estimated allele frequency
1.0
0.9
0.8
0.7
0.6
O16
L12
O12
253
256
O8
204
124
233
L8
9
97
L16
O1
L1
5
6
7
8
9
10
11
12
Ecological gradient (Mean annual temperature)

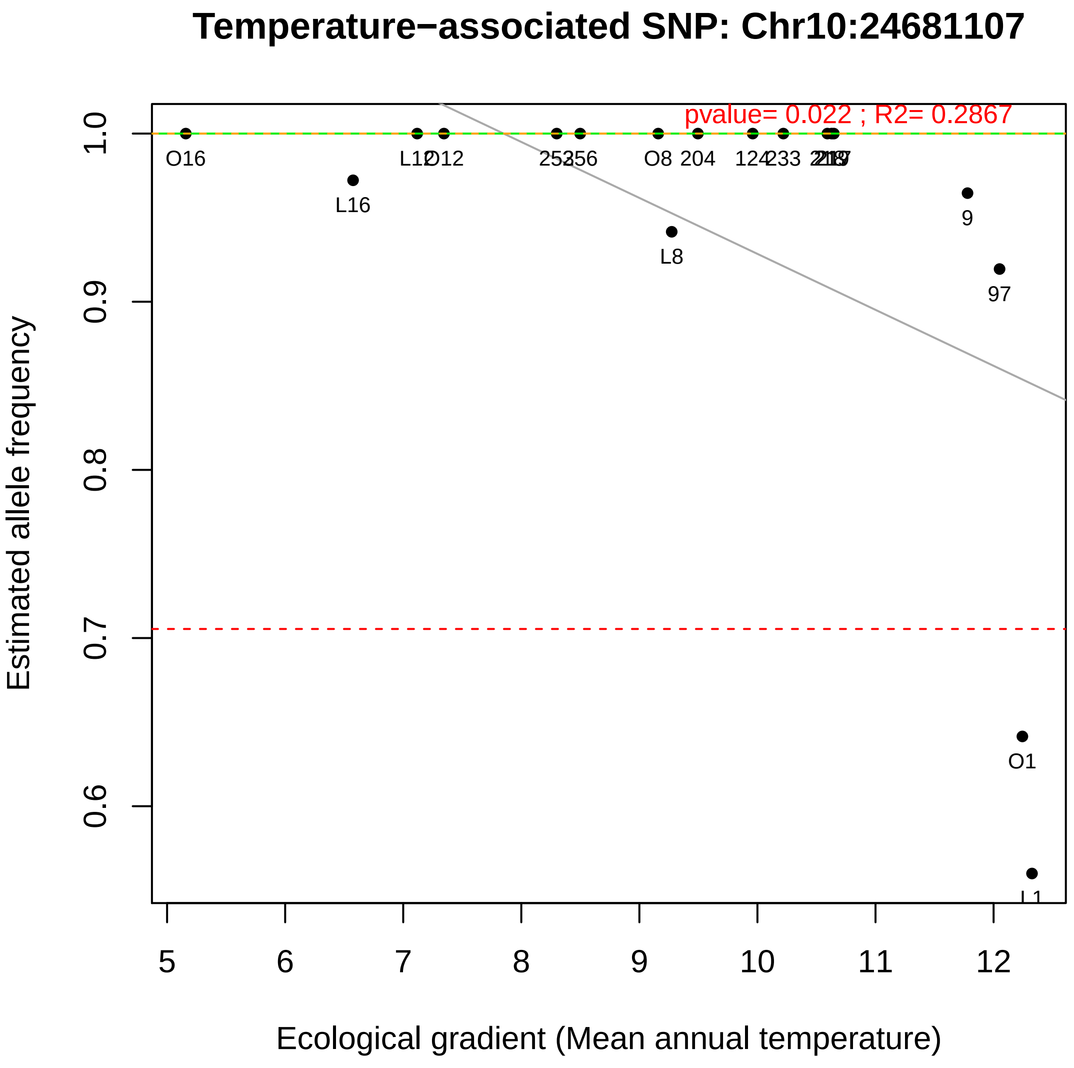

Temperature−associated SNP: Chr10:24681107
pvalue= 0.022 ; R2= 0.2867
Estimated allele frequency
Ecological gradient (Mean annual temperature)
1.0
0.9
0.8
0.7
0.6
5
6
7
8
9
10
11
12
O16
L16
L12
O12
253
256
O8
204
L8
124
233
9
97
O1

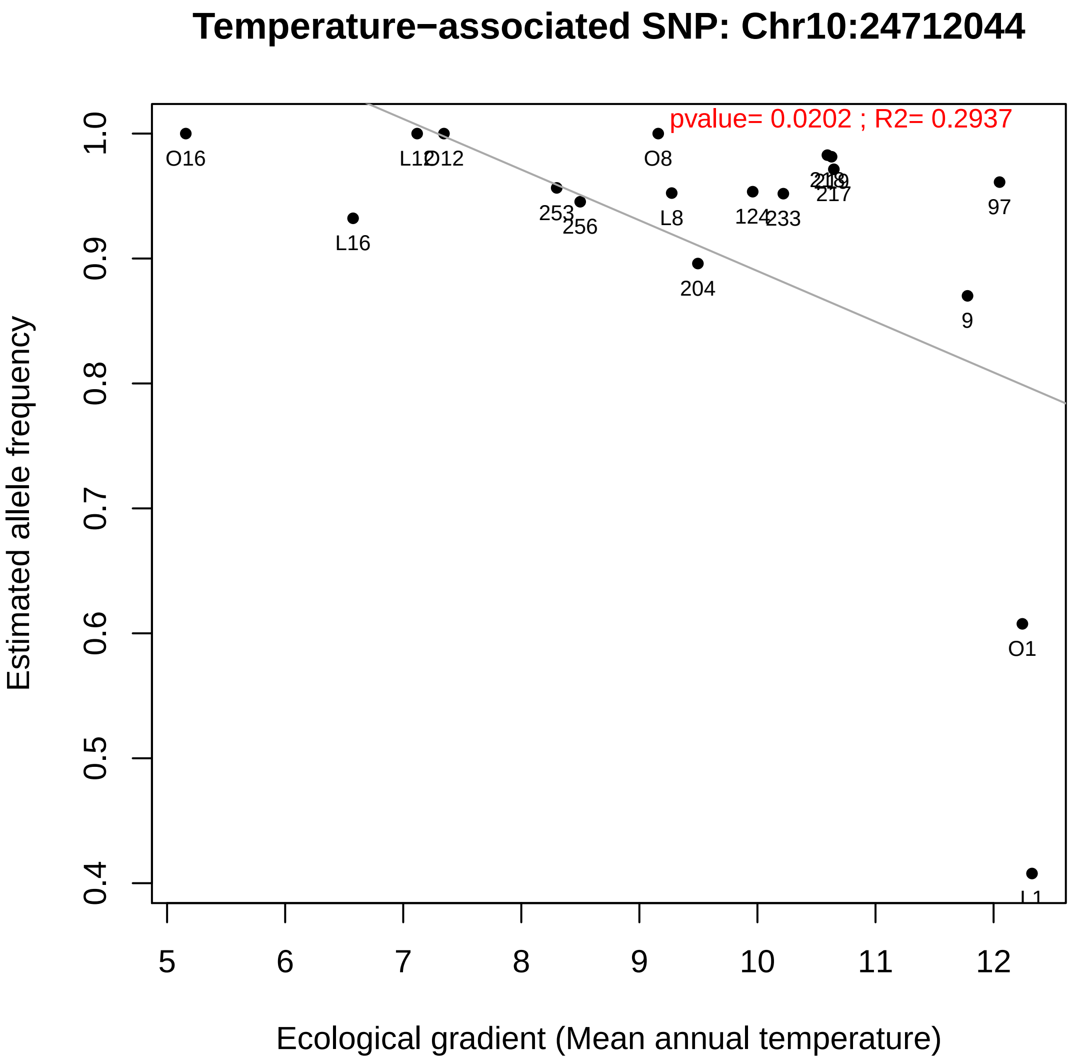

Temperature−associated SNP: Chr10:24712044
pvalue= 0.0202 ; R2= 0.2937
O16
L16
L12
O12
253
256
O8
L8
204
124
233
219
218
217
97
9
O1
L1
1.0
0.9
0.8
0.7
0.6
0.5
0.4
5
6
7
8
9
10
11
12
Estimated allele frequency
Ecological gradient (Mean annual temperature)

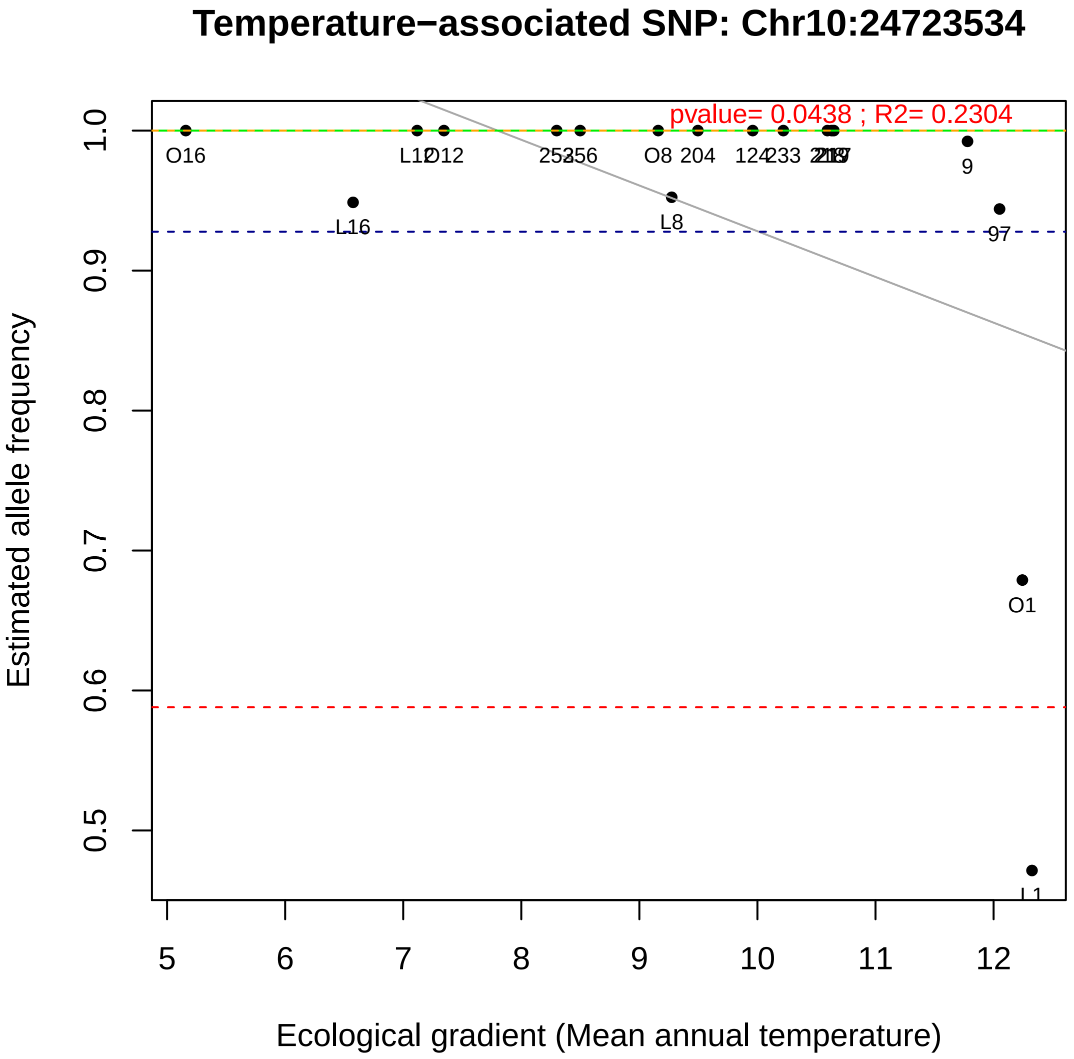
Temperature−associated SNP: Chr10:24723534
pvalue= 0.0438 ; R2= 0.2304
Estimated allele frequency
Ecological gradient (Mean annual temperature)
0.5
0.6
0.7
0.8
0.9
1.0
5
6
7
8
9
10
11
12
O16
L16
O8
204
L8
9
97
O1

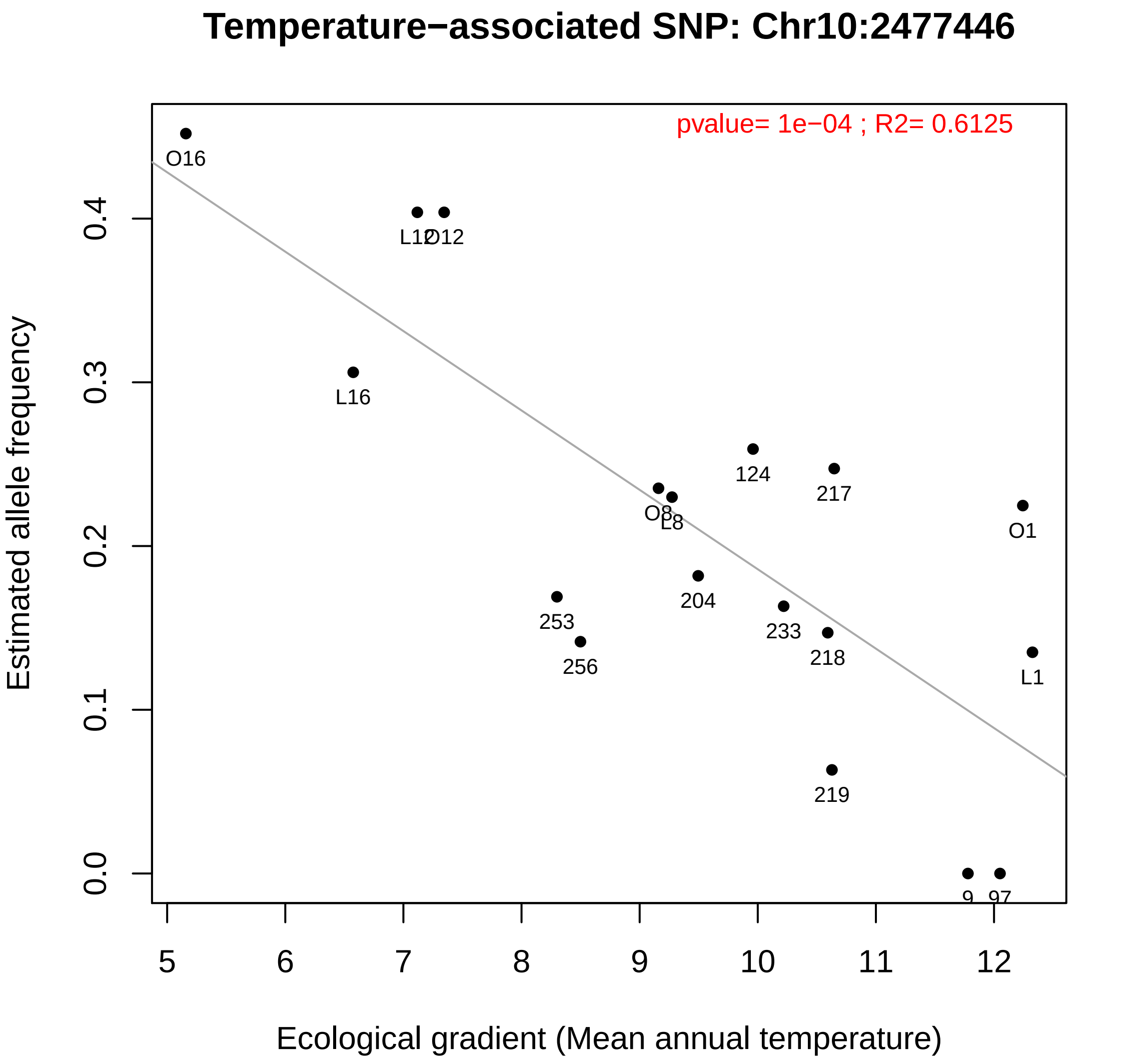
Temperature−associated SNP: Chr10:2477446
pvalue= 1e−04 ; R2= 0.6125
Estimated allele frequency
0.0
0.1
0.2
0.3
0.4
5
6
7
8
9
10
11
12
Ecological gradient (Mean annual temperature)
O16
L12
O12
L16
124
217
O8
L8
O1
204
253
233
218
256
L1
219
9
97

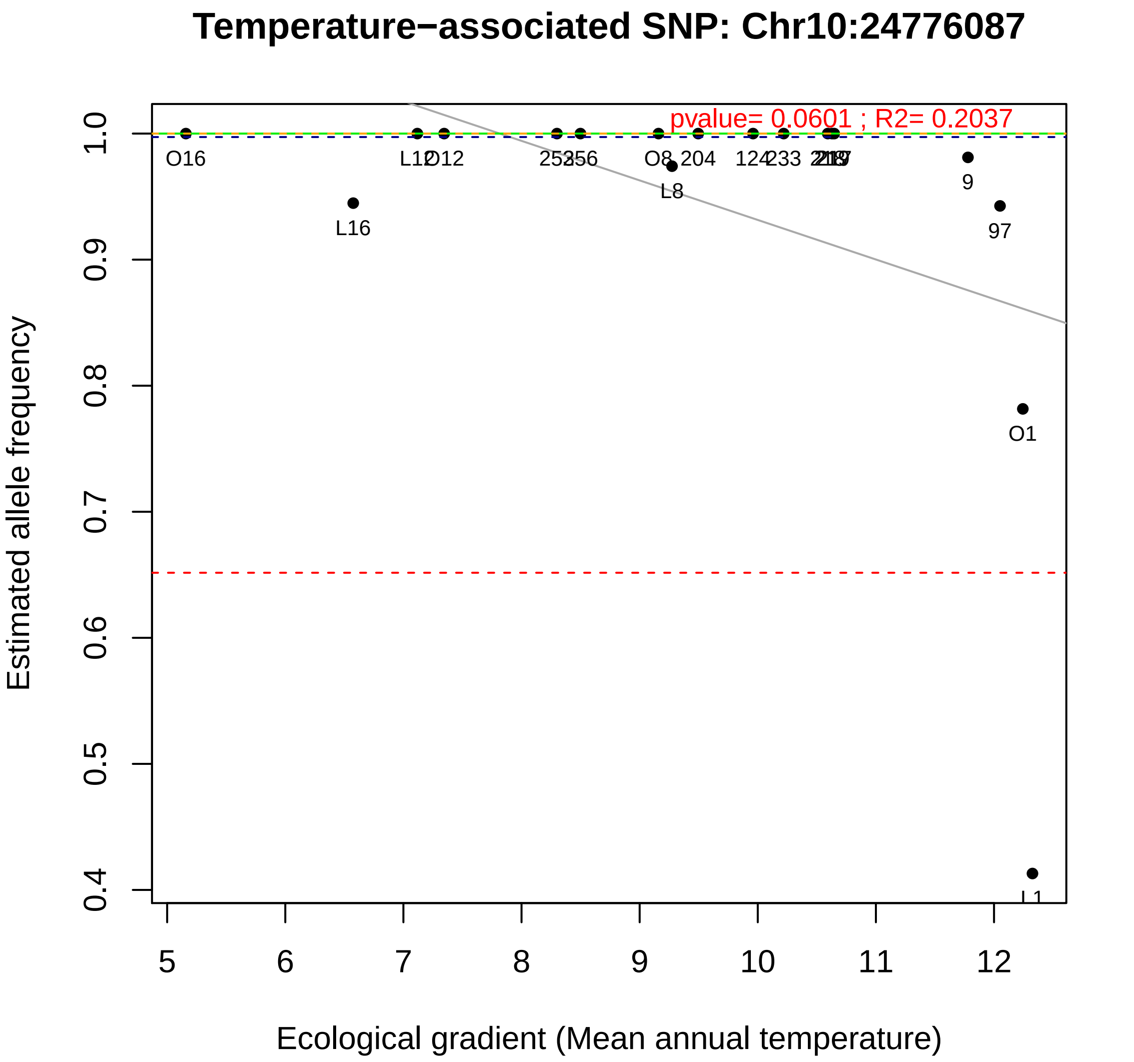

Temperature−associated SNP: Chr10:24776087
pvalue= 0.0601 ; R2= 0.2037
Estimated allele frequency
Ecological gradient (Mean annual temperature)
O16
L16
L12
O12
9
97
O1
L8
204
124
233
1.0
0.9
0.8
0.7
0.6
0.5
0.4
5
6
7
8
9
10
11
12

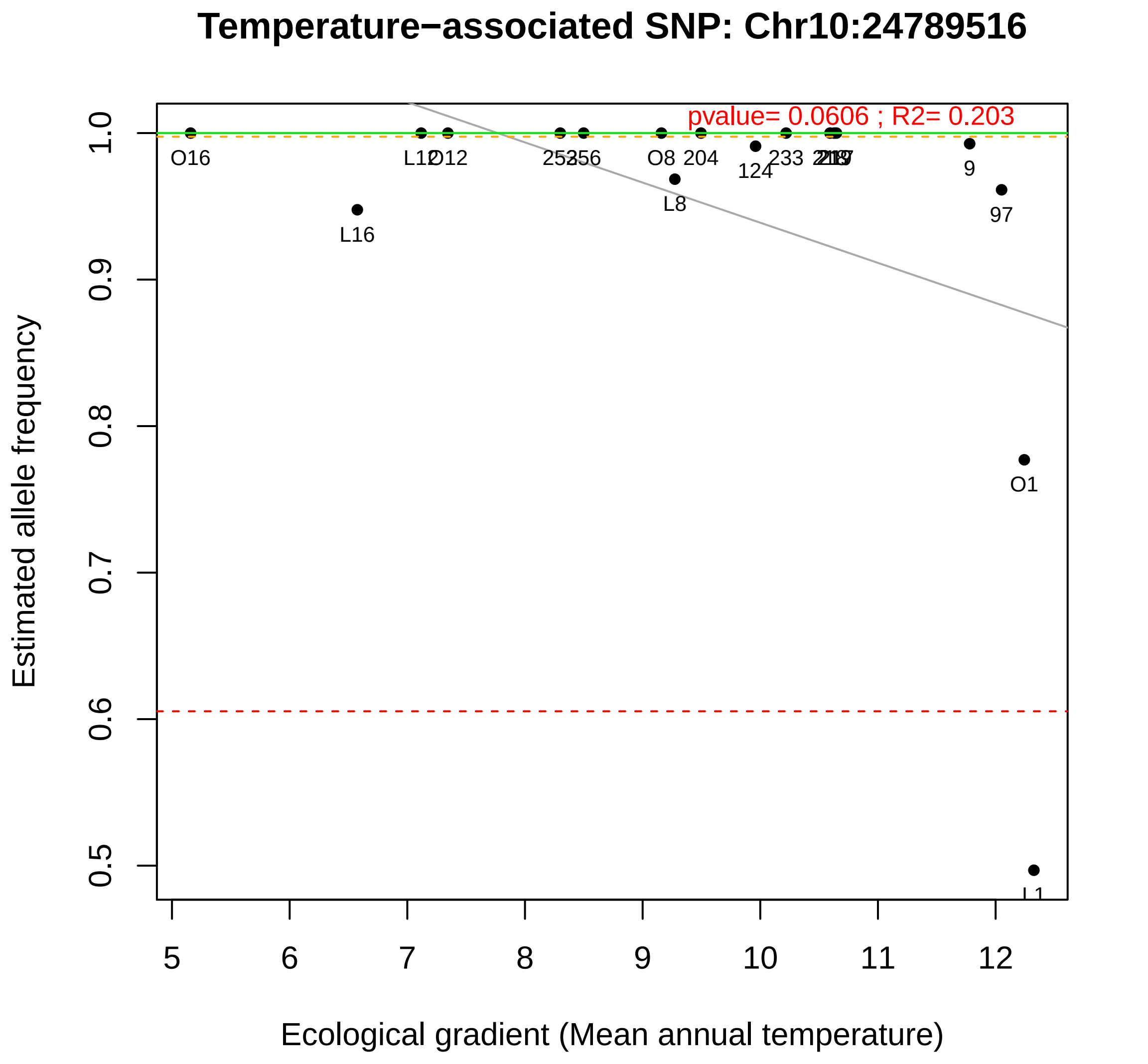
Temperature−associated SNP: Chr10:24789516
pvalue= 0.0606 ; R2= 0.203
Estimated allele frequency
Ecological gradient (Mean annual temperature)
1.0
0.9
0.8
0.7
0.6
0.5
5
6
7
8
9
10
11
12
O16
L16
L12
O12
O8
204
233
124
L8
9
97
O1

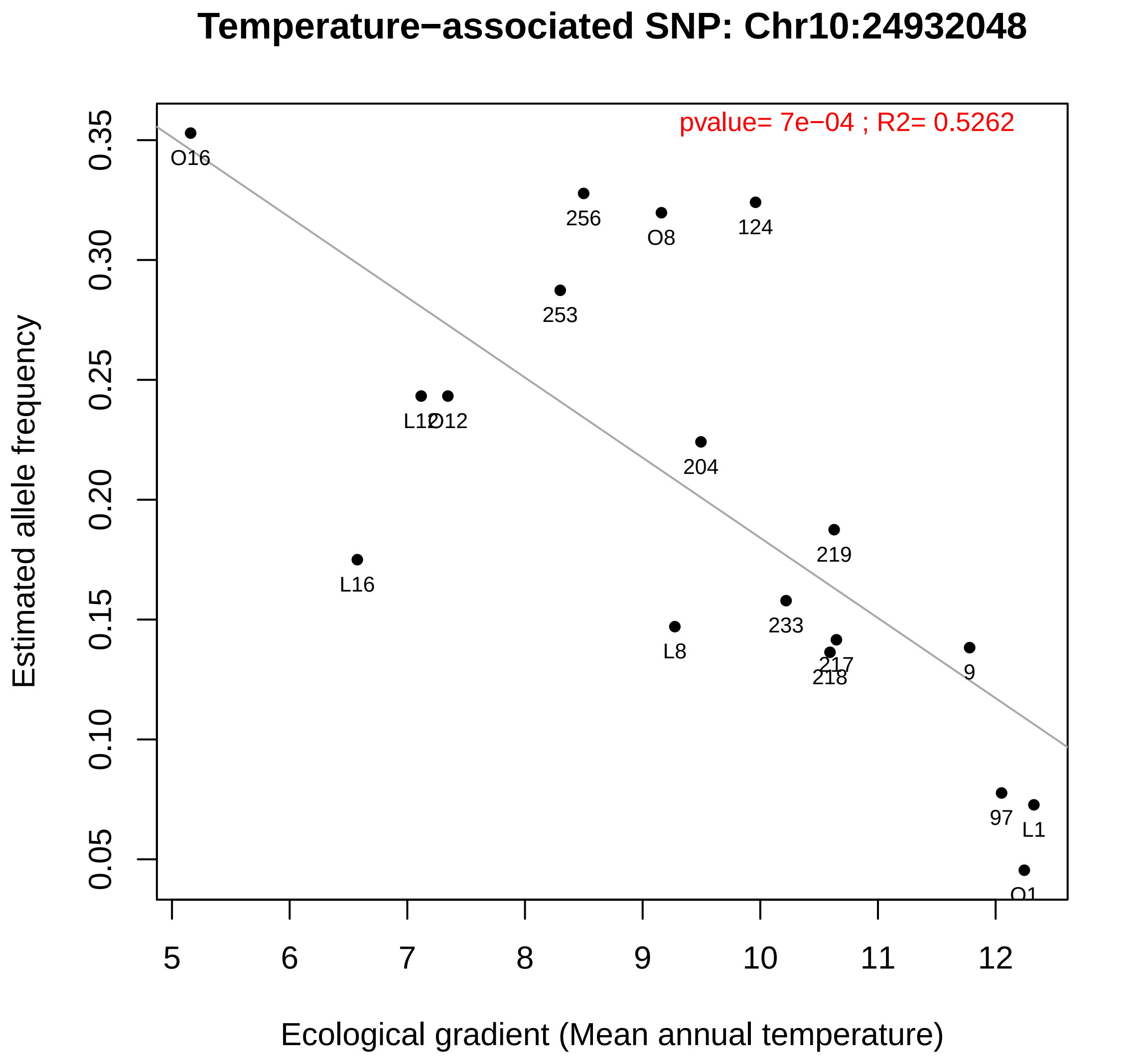
Temperature−associated SNP: Chr10:24932048
pvalue= 7e−04 ; R2= 0.5262
O16
256
O8
124
253
L12
O12
204
219
L16
233
L8
217
218
9
97
L1
O1
0.05
0.10
0.15
0.20
0.25
0.30
0.35
Estimated allele frequency
5
6
7
8
9
10
11
12
Ecological gradient (Mean annual temperature)

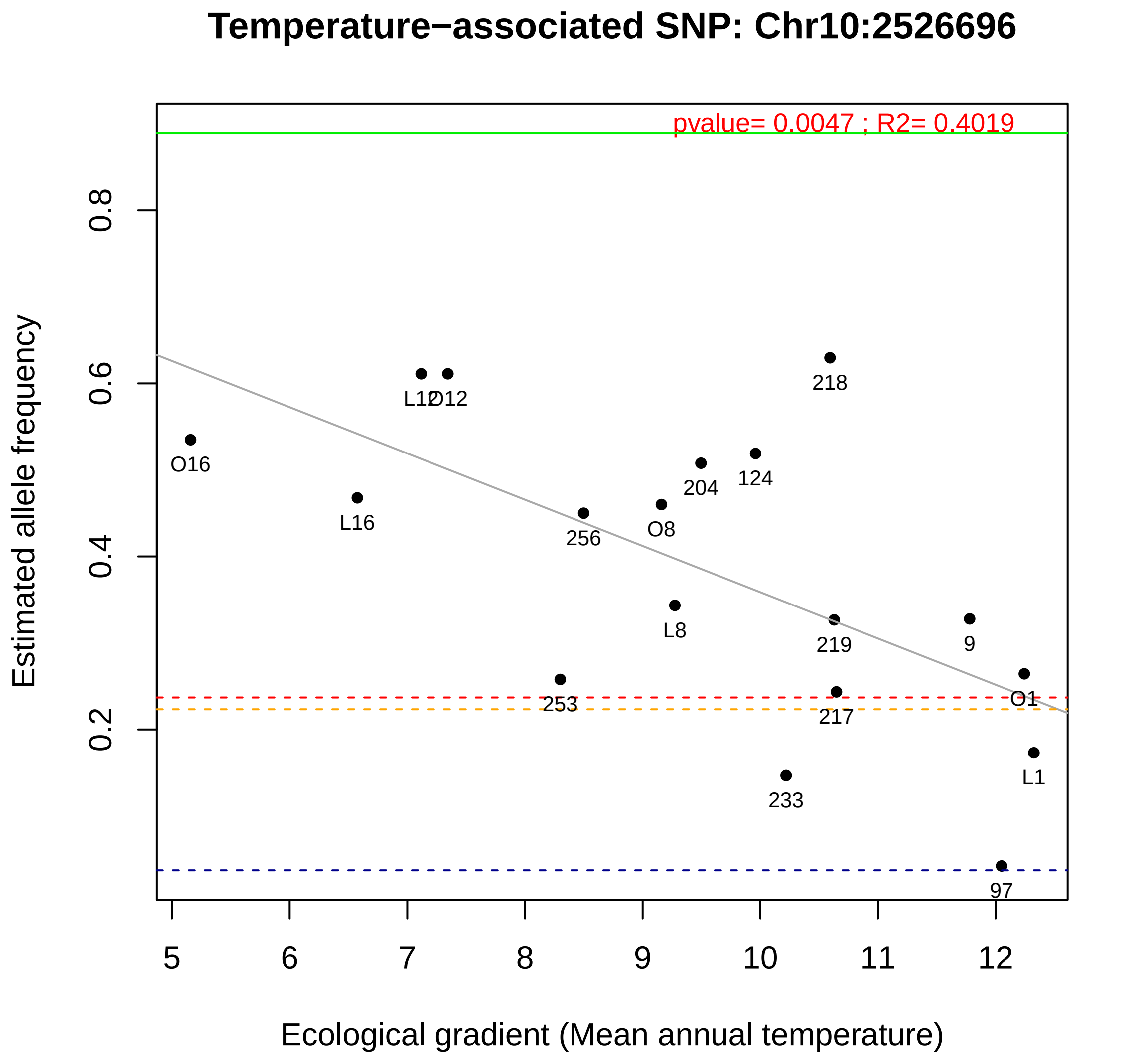

Temperature−associated SNP: Chr10:2526696
pvalue= 0.0047 ; R2= 0.4019
Estimated allele frequency
0.2
0.4
0.6
0.8
O16
L16
L12
O12
253
256
L8
O8
204
124
233
218
219
217
9
O1
L1
97
5
6
7
8
9
10
11
12
Ecological gradient (Mean annual temperature)

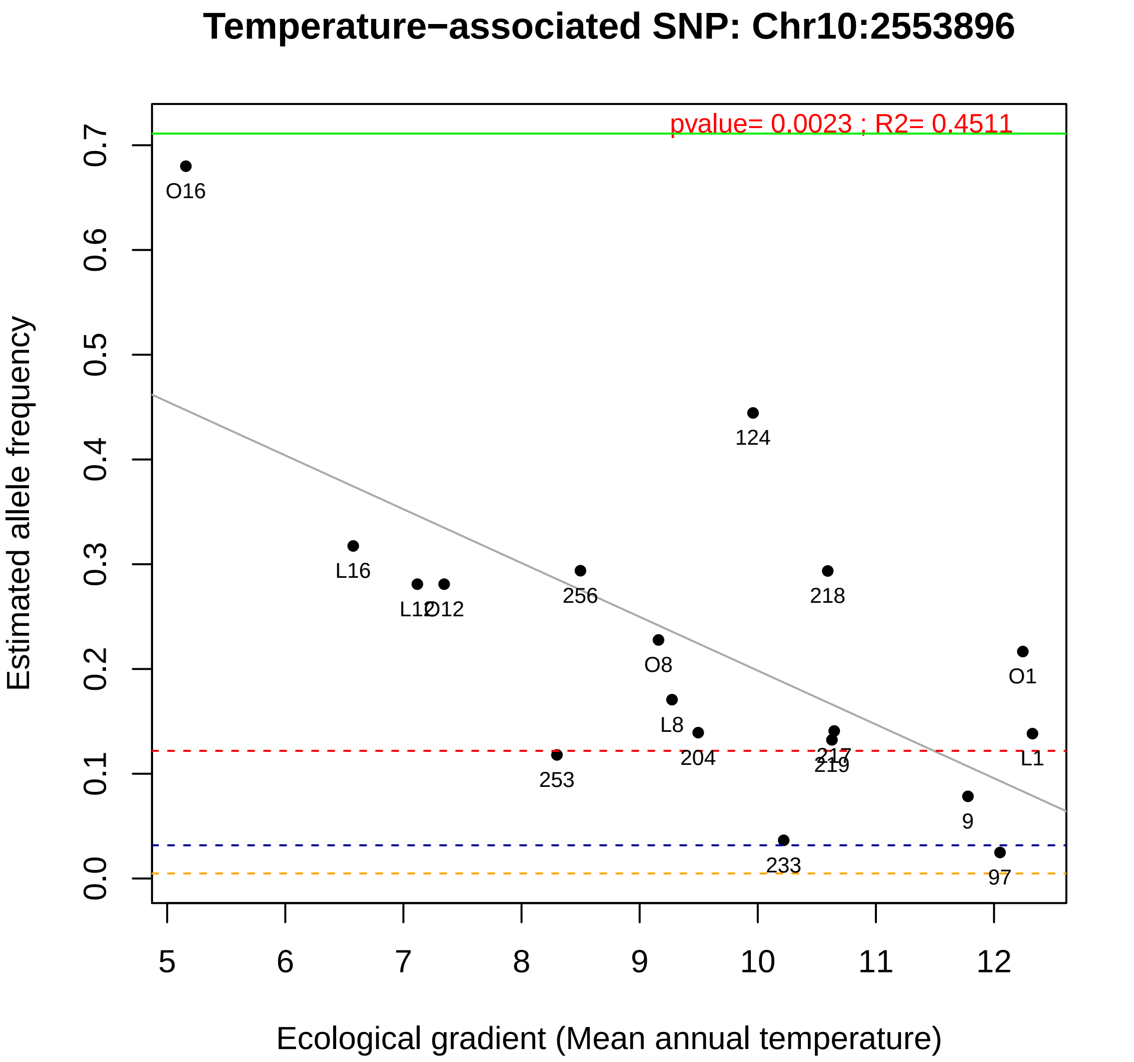

Temperature−associated SNP: Chr10:2553896
pvalue= 0.0023 ; R2= 0.4511
Estimated allele frequency
0.0
0.1
0.2
0.3
0.4
0.5
0.6
0.7
Ecological gradient (Mean annual temperature)
5
6
7
8
9
10
11
12
O16
L16
L12
O12
256
124
218
O8
O1
L8
204
217
219
L1
253
9
233
97

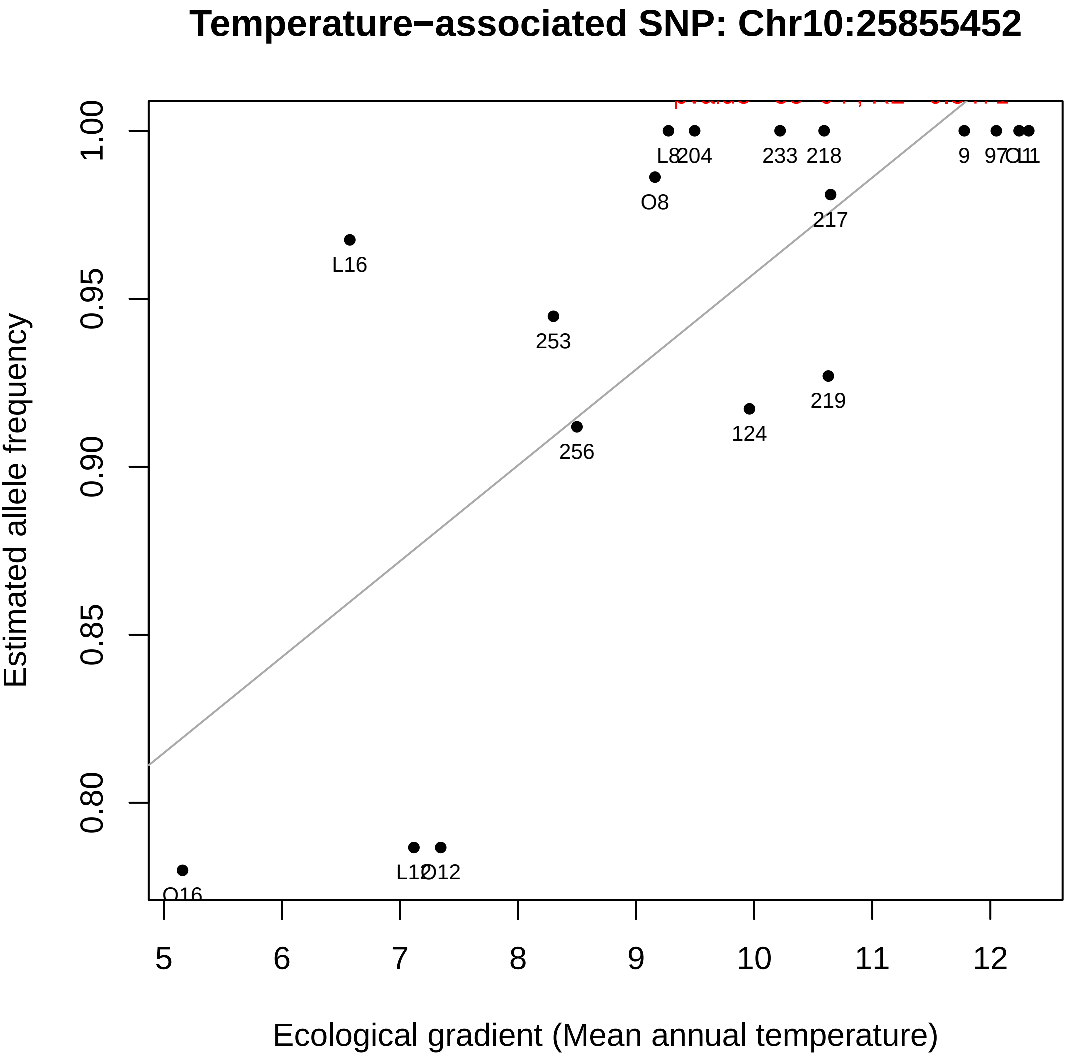

Temperature−associated SNP: Chr10:25855452
Estimated allele frequency
1.00
0.95
0.90
0.85
0.80
5
6
7
8
9
10
11
12
Ecological gradient (Mean annual temperature)
L8
204
233
218
9
97
O8
217
L16
253
256
124
219
L12
O12
O16

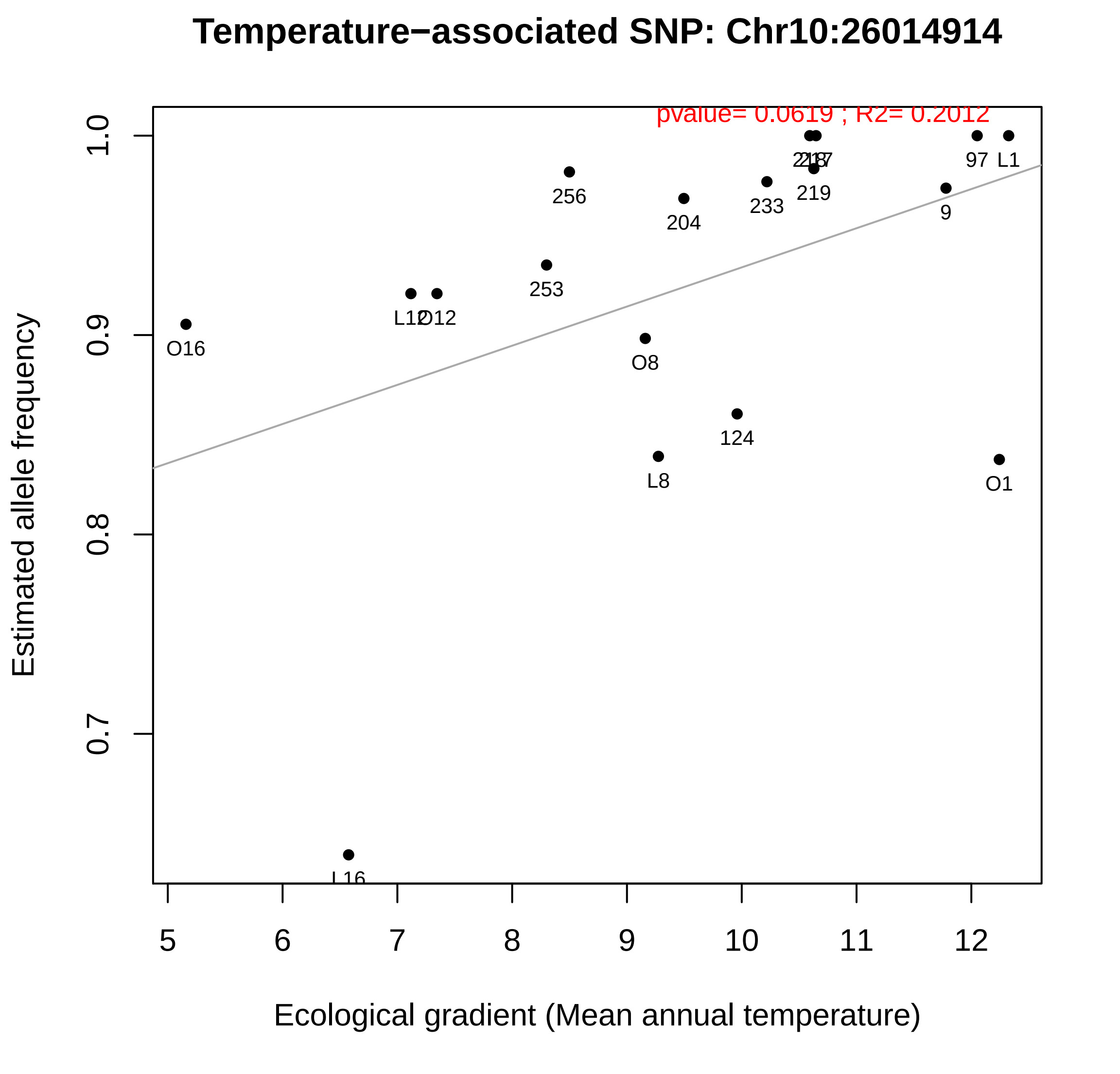

Temperature−associated SNP: Chr10:26014914
pvalue= 0.0619 ; R2= 0.2012
Estimated allele frequency
Ecological gradient (Mean annual temperature)
1.0
0.9
0.8
0.7
5
6
7
8
9
10
11
12
O16
L12
O12
253
256
204
233
219
9
97
L1
O8
124
L8
O1
L16

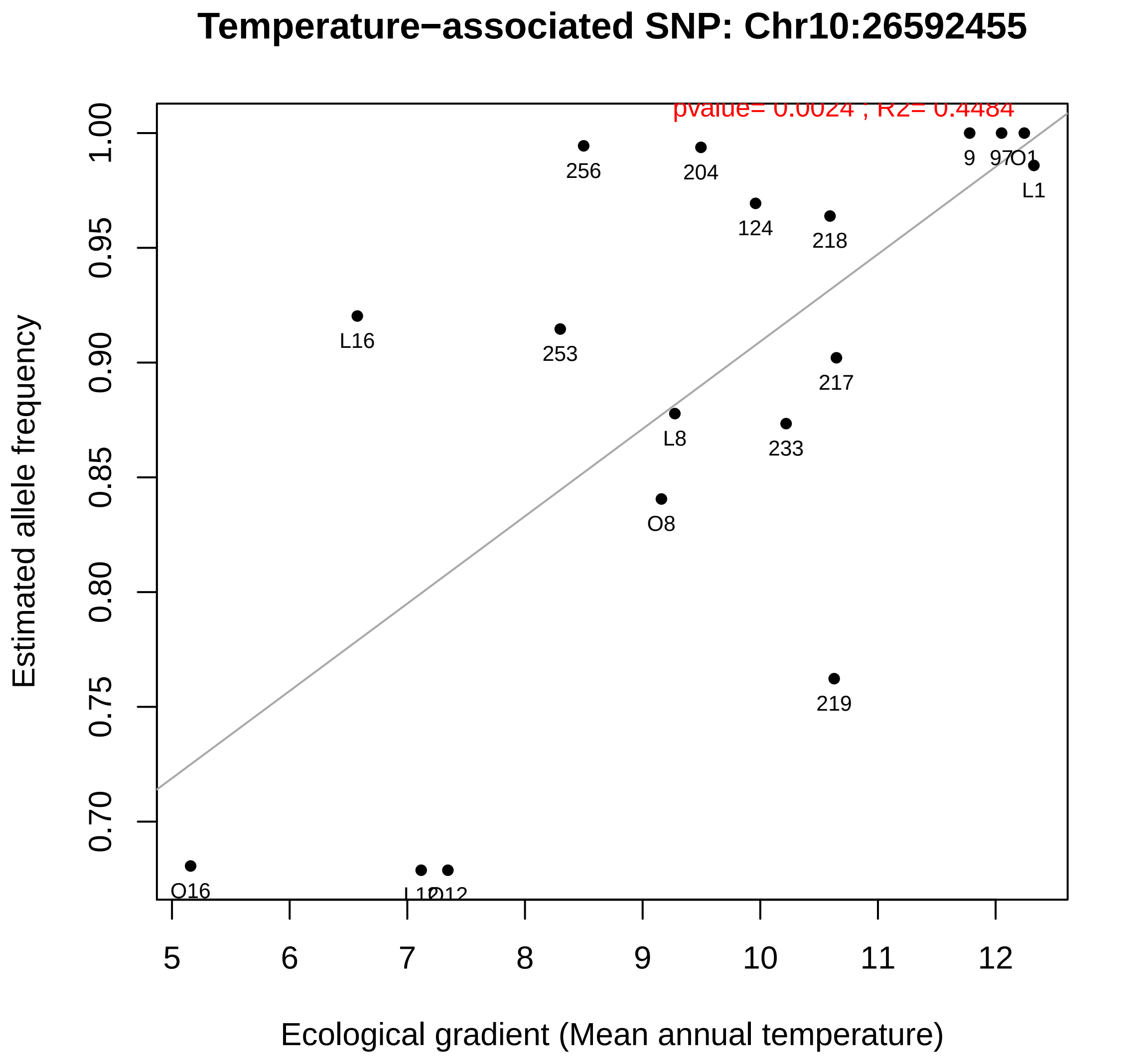
Temperature−associated SNP: Chr10:26592455
pvalue= 0.0024 ; R2= 0.4484
Estimated allele frequency
1.00
0.95
0.90
0.85
0.80
0.75
0.70
256
204
9
97
O1
L1
124
218
L16
253
217
L8
233
O8
219
O16
O12
5
6
7
8
9
10
11
12
Ecological gradient (Mean annual temperature)

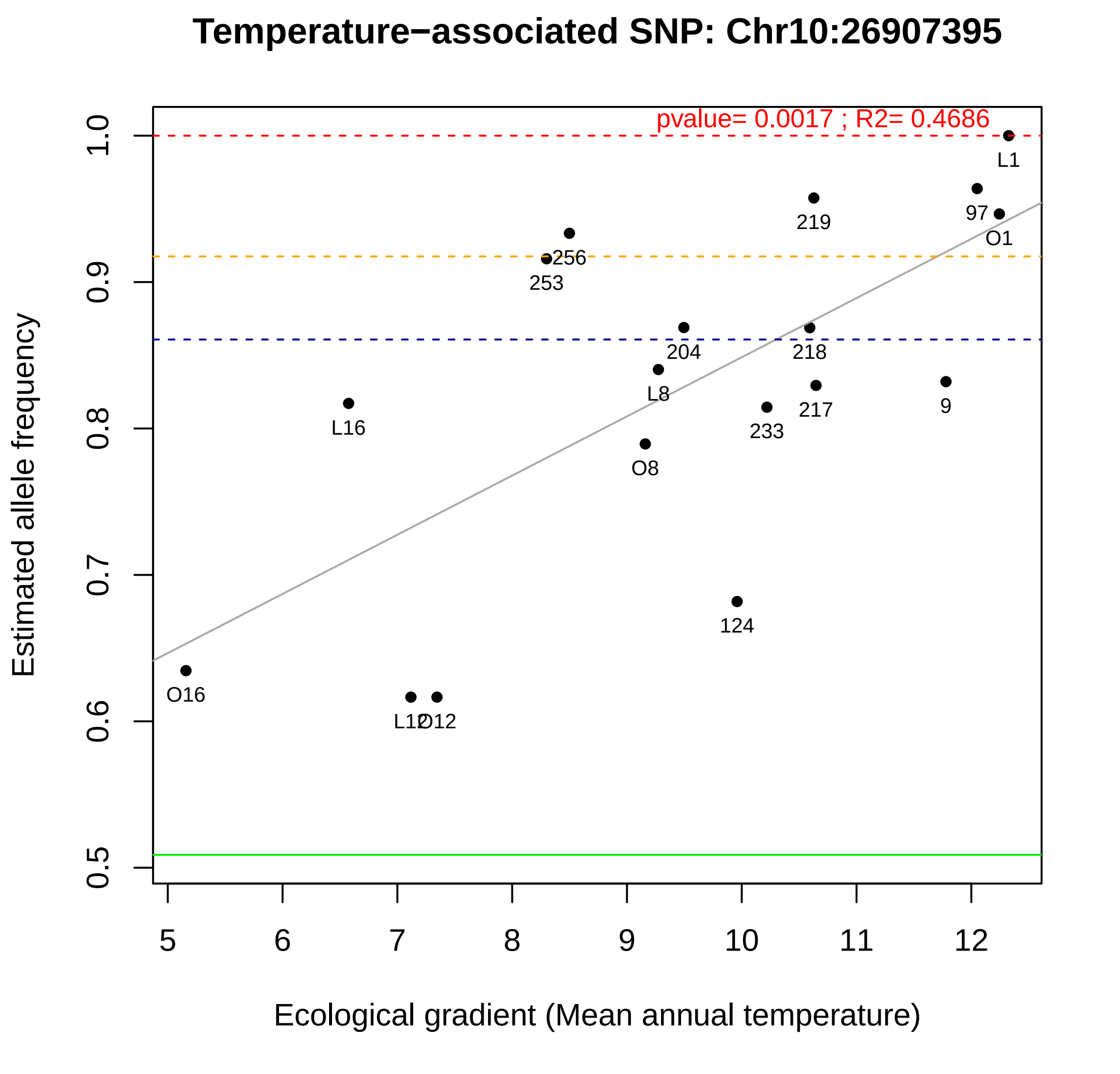

Temperature−associated SNP: Chr10:26907395
pvalue= 0.0017 ; R2= 0.4686
Estimated allele frequency
Ecological gradient (Mean annual temperature)
1.0
0.9
0.8
0.7
0.6
0.5
5
6
7
8
9
10
11
12
L1
97
O1
219
256
253
204
218
L8
217
9
L16
233
O8
124
O16
L12
O12

### Temperature−associated SNP: Chr10:27779422

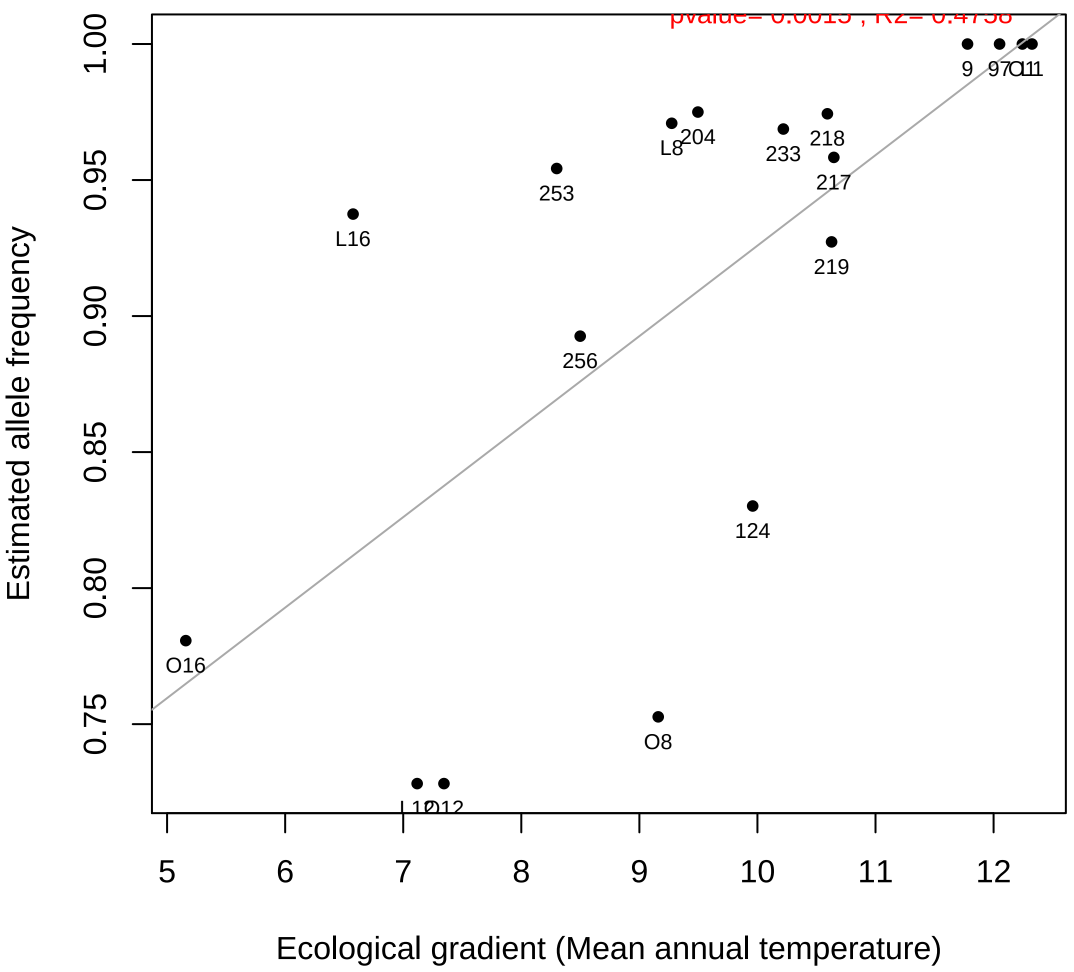

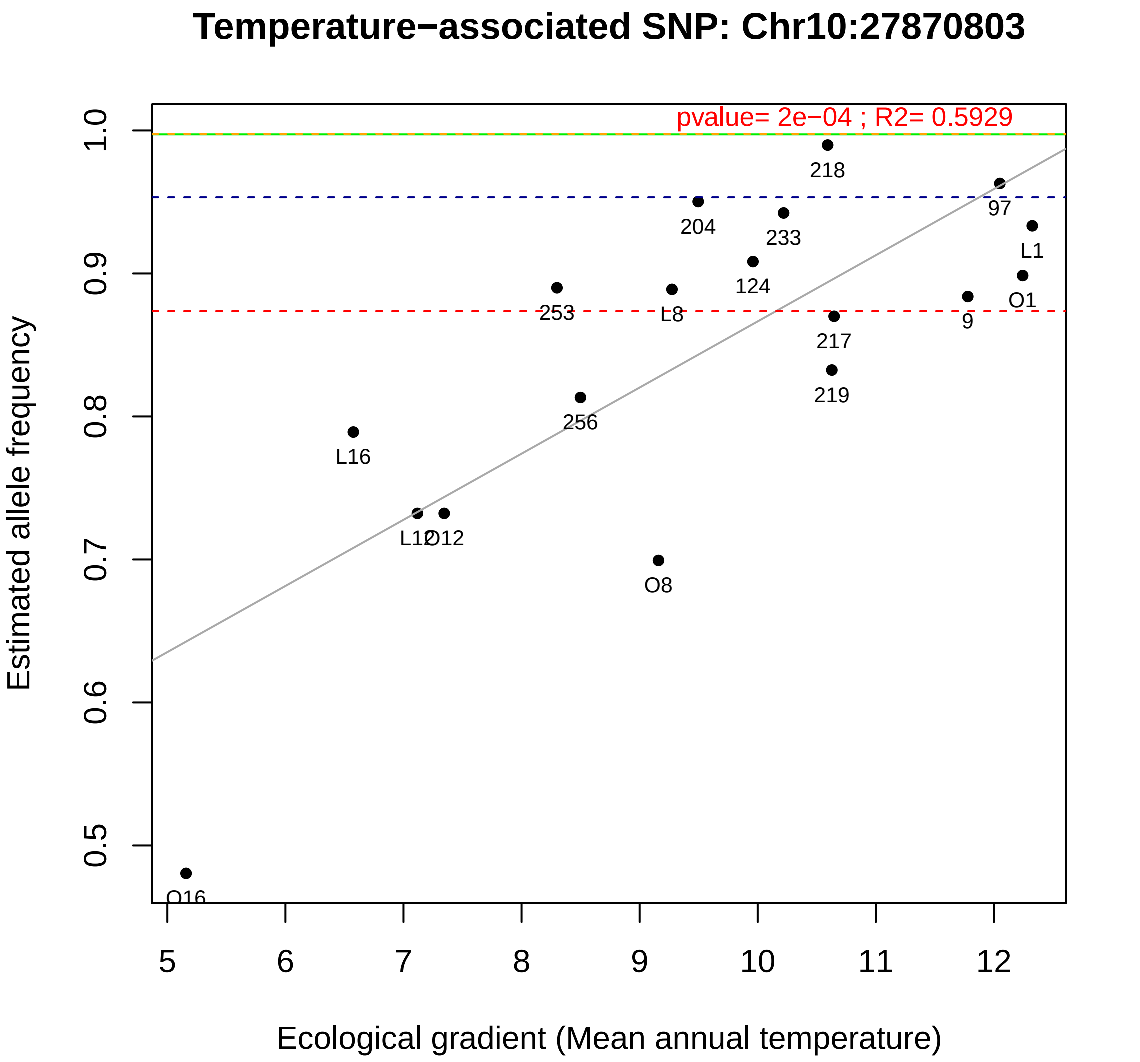
Temperature−associated SNP: Chr10:27870803
pvalue= 2e−04 ; R2= 0.5929
Estimated allele frequency
Ecological gradient (Mean annual temperature)
0.5
0.6
0.7
0.8
0.9
1.0
5
6
7
8
9
10
11
12
218
97
204
233
L1
124
O1
253
L8
9
217
219
256
L16
L12
O12
O8
O16

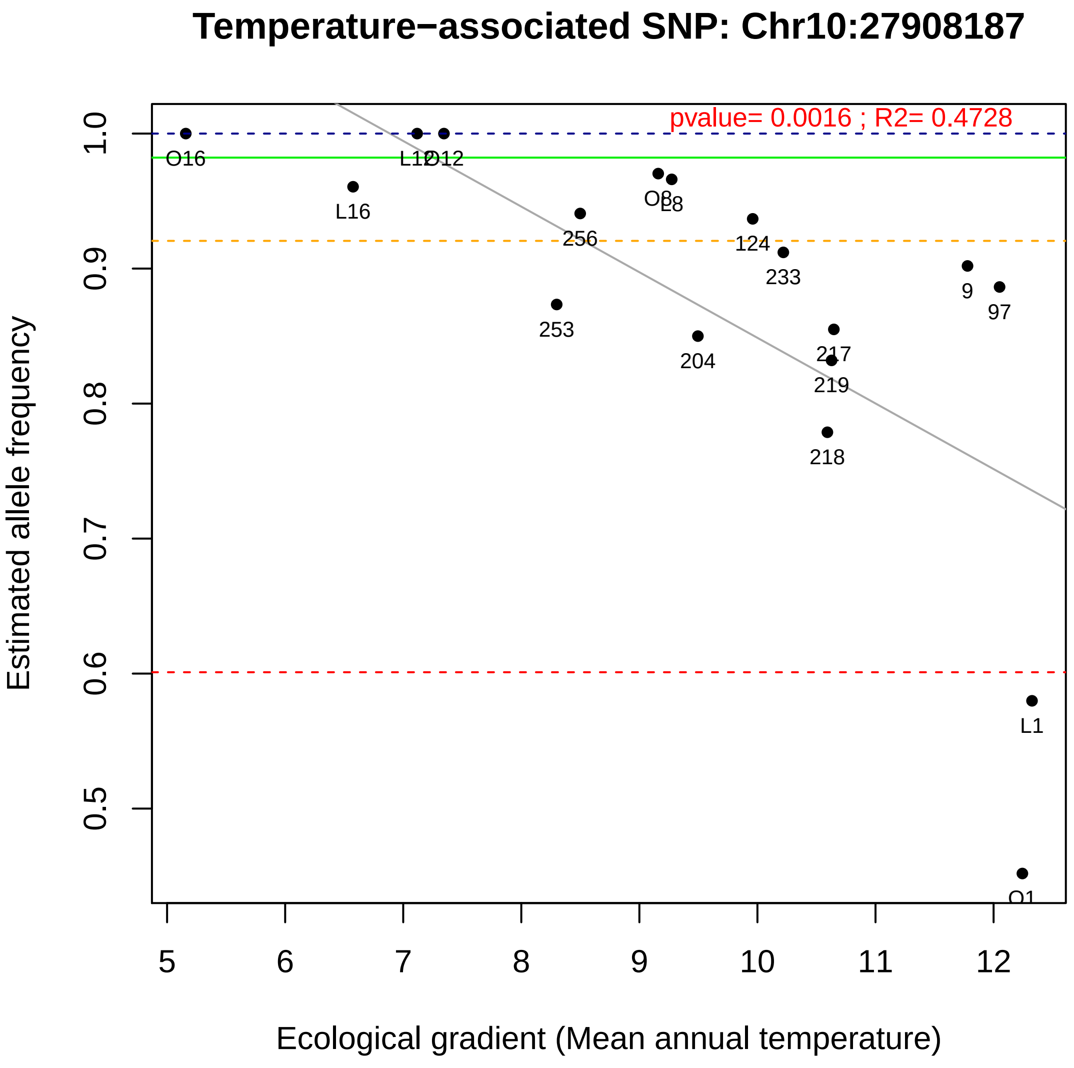

Temperature−associated SNP: Chr10:27908187
pvalue= 0.0016 ; R2= 0.4728
O16
L12
O12
L16
O8
L8
256
124
233
253
204
217
219
218
9
97
L1
O1
1.0
0.9
0.8
0.7
0.6
0.5
5
6
7
8
9
10
11
12
Estimated allele frequency
Ecological gradient (Mean annual temperature)

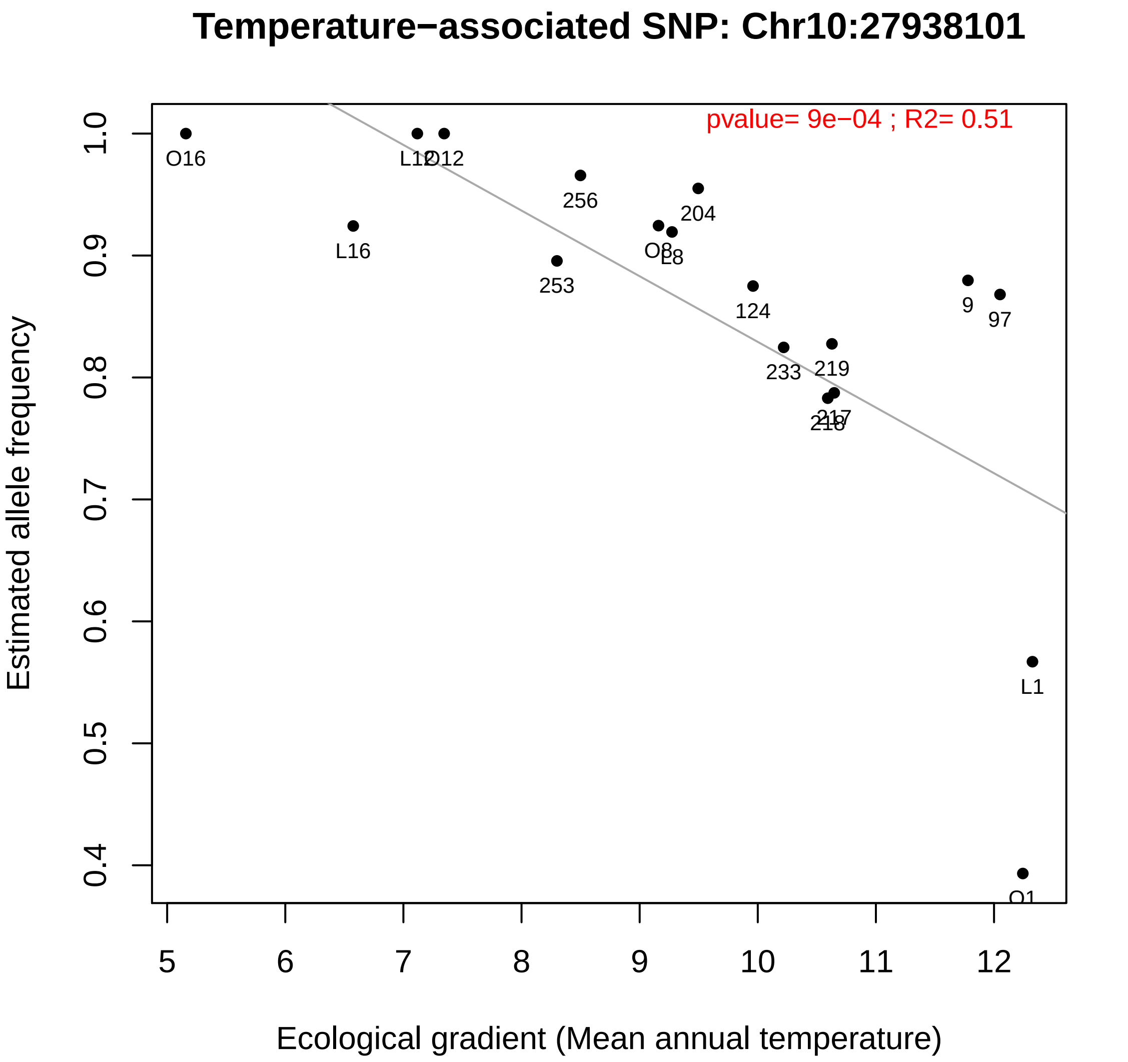

Temperature−associated SNP: Chr10:27938101
pvalue= 9e−04 ; R2= 0.51
Estimated allele frequency
Ecological gradient (Mean annual temperature)
1.0
0.9
0.8
0.7
0.6
0.5
0.4
5
6
7
8
9
10
11
12
O16
L12
O12
L16
256
204
253
O8
L8
124
9
97
233
219
217
218
L1
O1

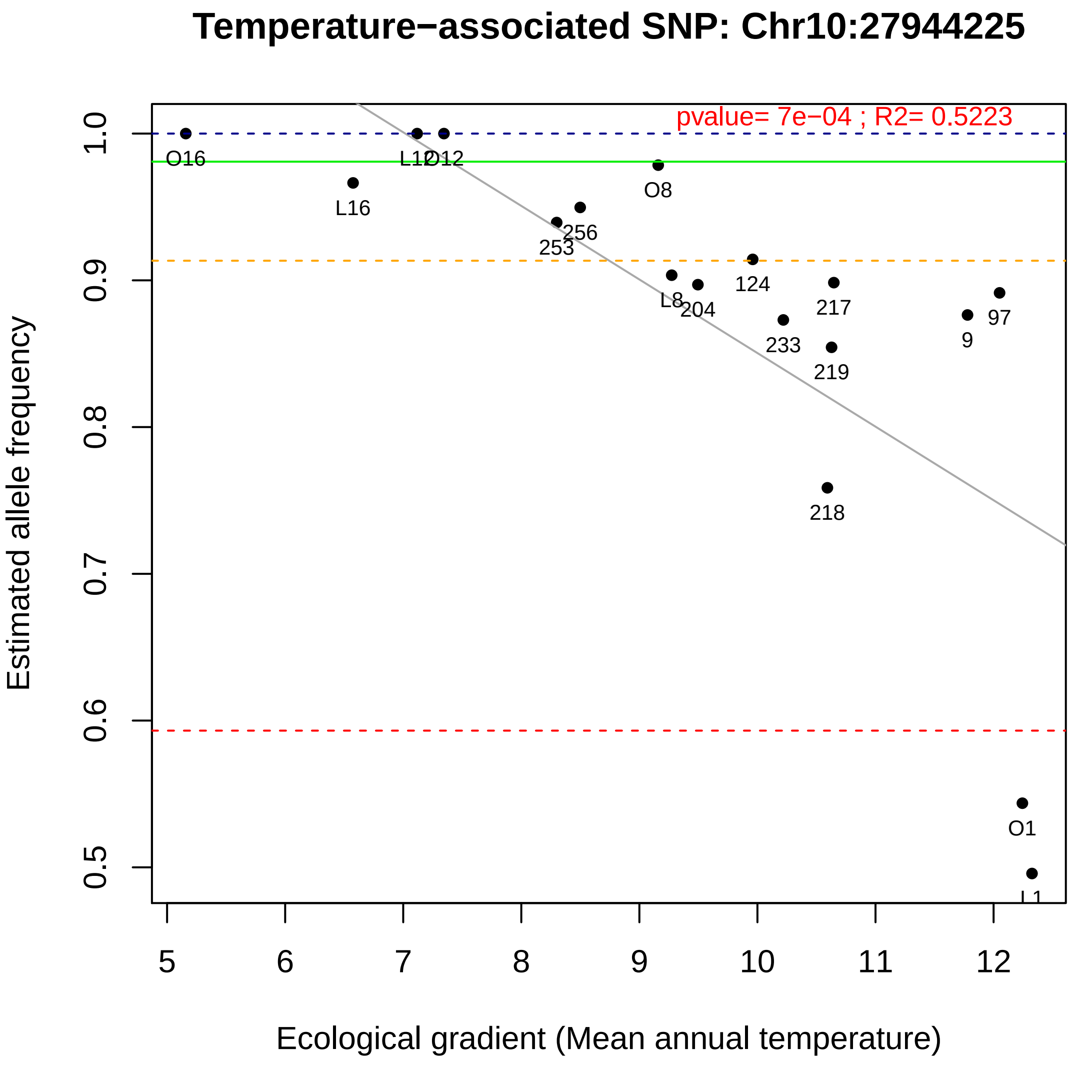

Temperature−associated SNP: Chr10:27944225
pvalue= 7e−04 ; R2= 0.5223
Estimated allele frequency
Ecological gradient (Mean annual temperature)
1.0
0.9
0.8
0.7
0.6
0.5
5
6
7
8
9
10
11
12
O16
L12
O12
L16
O8
256
253
124
L8
204
217
233
9
97
219
218
O1

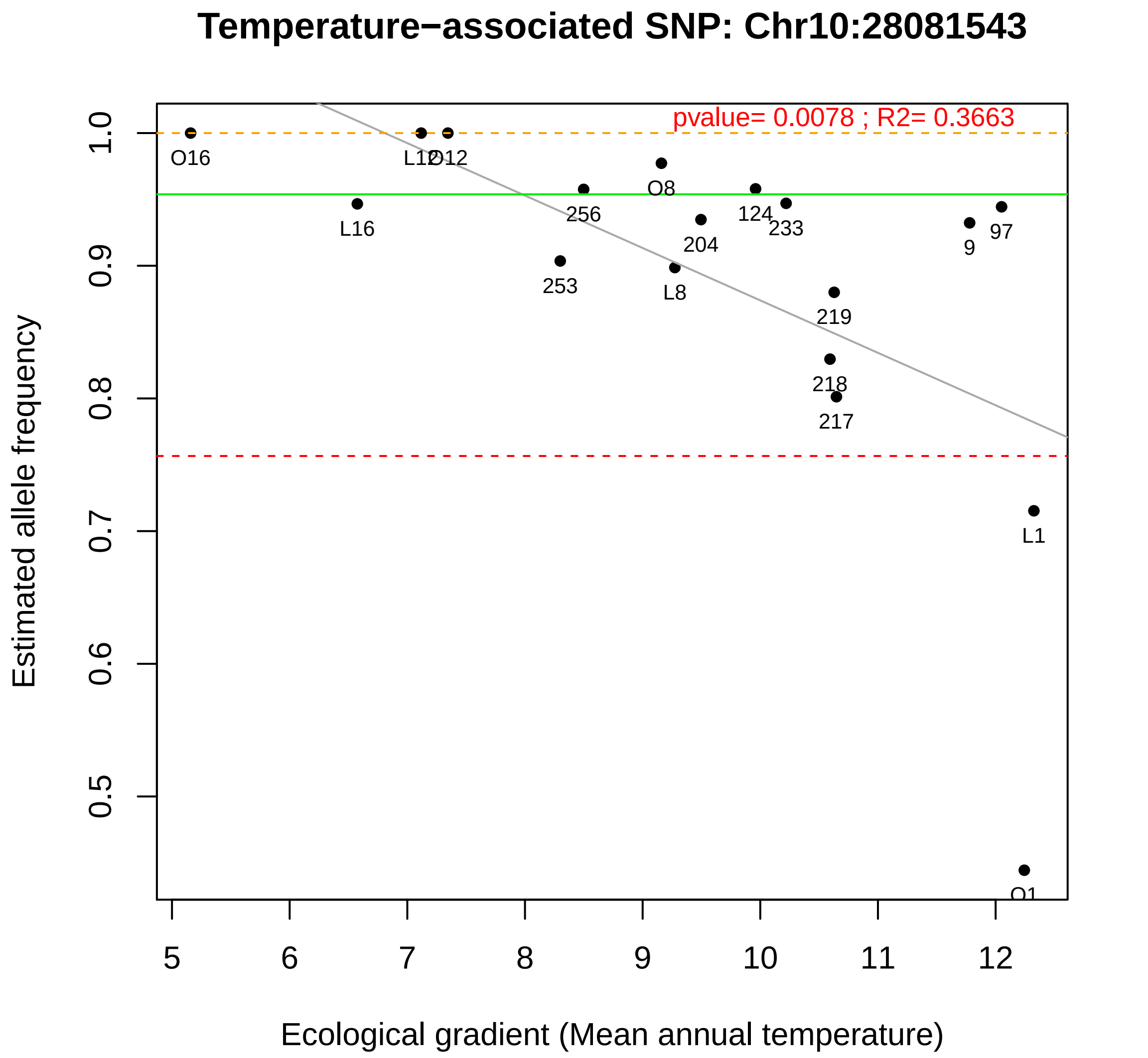

Temperature−associated SNP: Chr10:28081543
pvalue= 0.0078 ; R2= 0.3663
Estimated allele frequency
Ecological gradient (Mean annual temperature)
1.0
0.9
0.8
0.7
0.6
0.5
5
6
7
8
9
10
11
12
O16
L12
O12
L16
256
O8
124
233
204
253
L8
219
218
217
9
97
L1

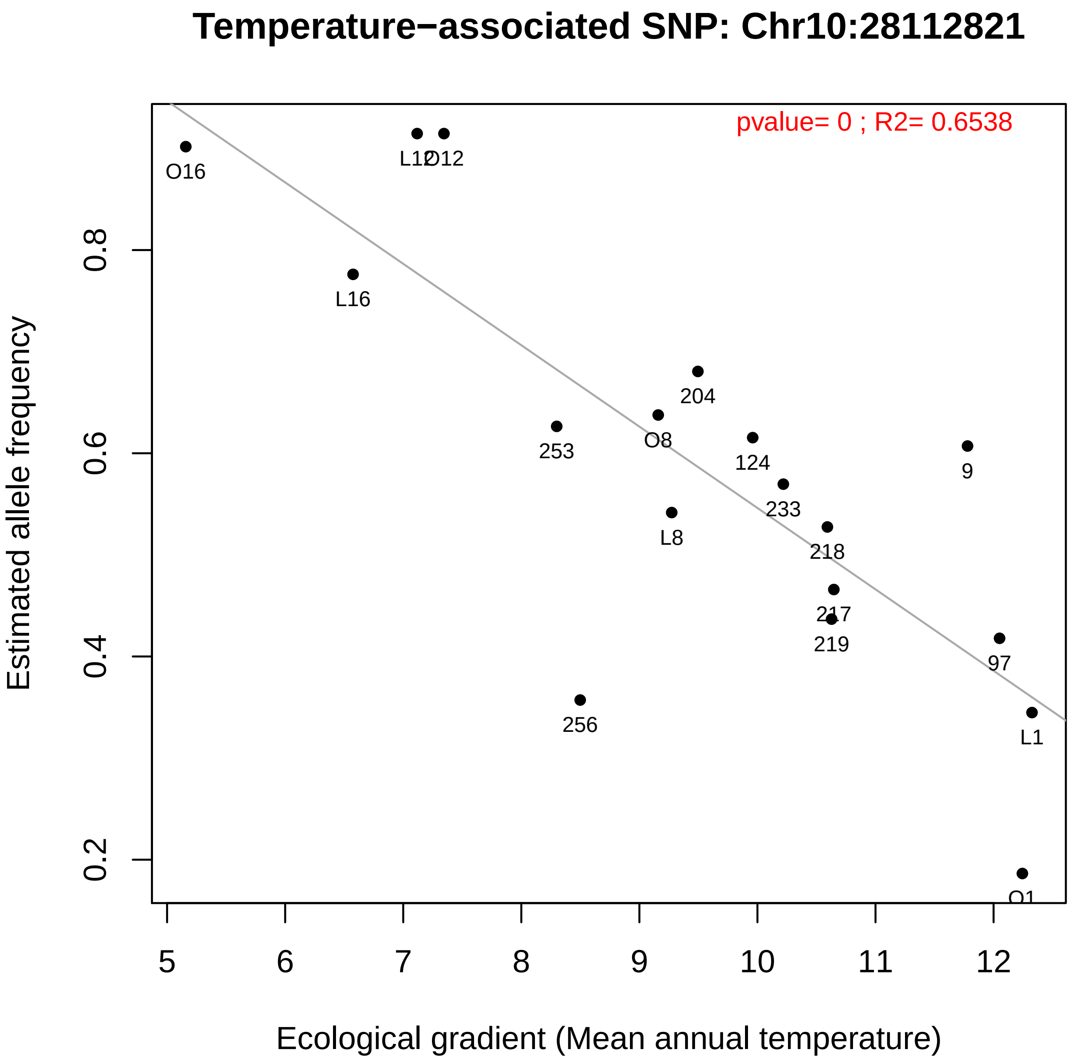
Temperature−associated SNP: Chr10:28112821
pvalue= 0 ; R2= 0.6538
O16
L12
O12
L16
204
253
O8
124
9
233
L8
218
217
219
97
256
L1
O1
0.2
0.4
0.6
0.8
5
6
7
8
9
10
11
12
Estimated allele frequency
Ecological gradient (Mean annual temperature)

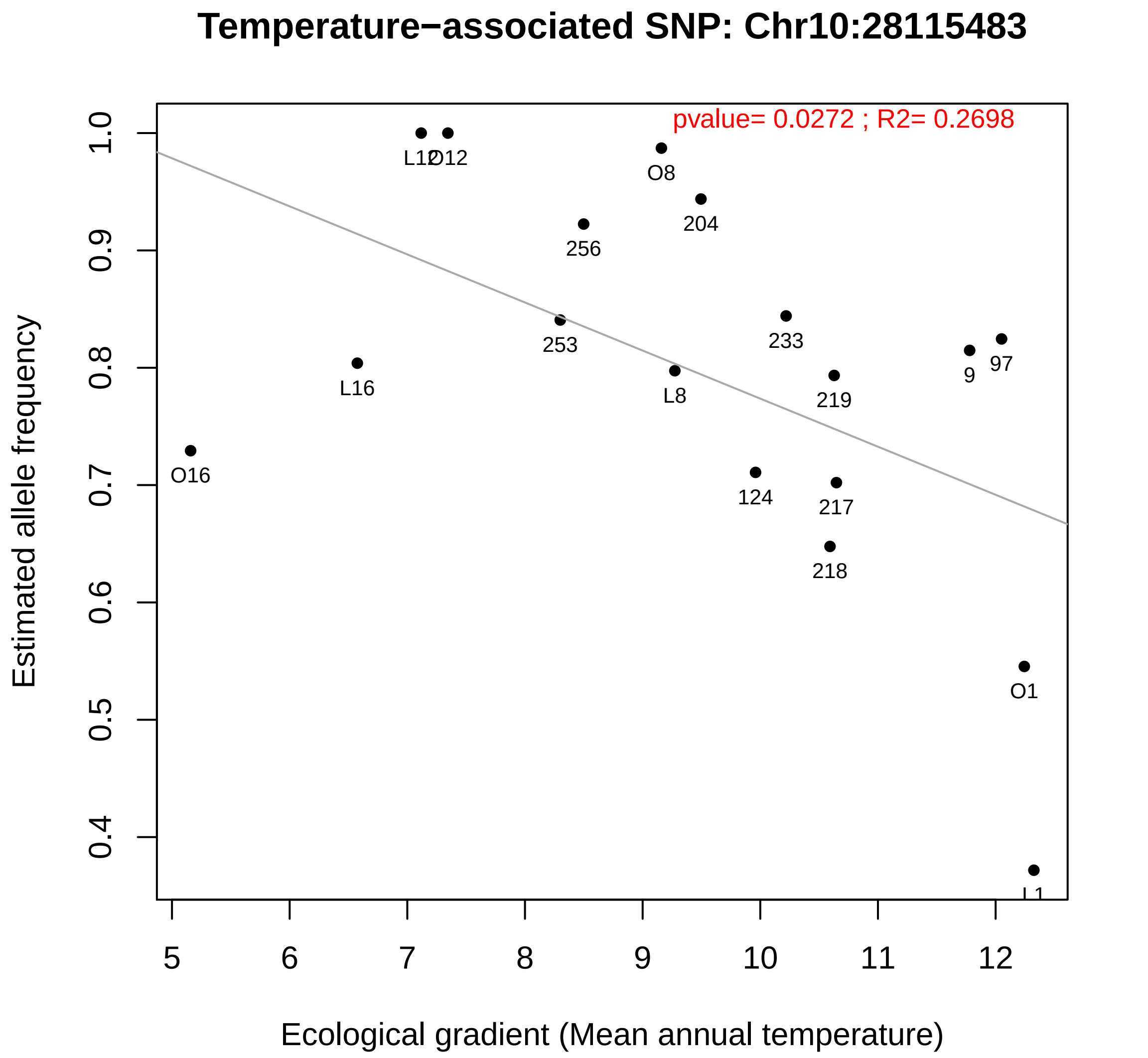
Temperature−associated SNP: Chr10:28115483
pvalue= 0.0272 ; R2= 0.2698
Estimated allele frequency
Ecological gradient (Mean annual temperature)
1.0
0.9
0.8
0.7
0.6
0.5
0.4
5
6
7
8
9
10
11
12
L12
O12
O8
204
256
253
233
L16
L8
219
9
97
O16
124
217
218
O1

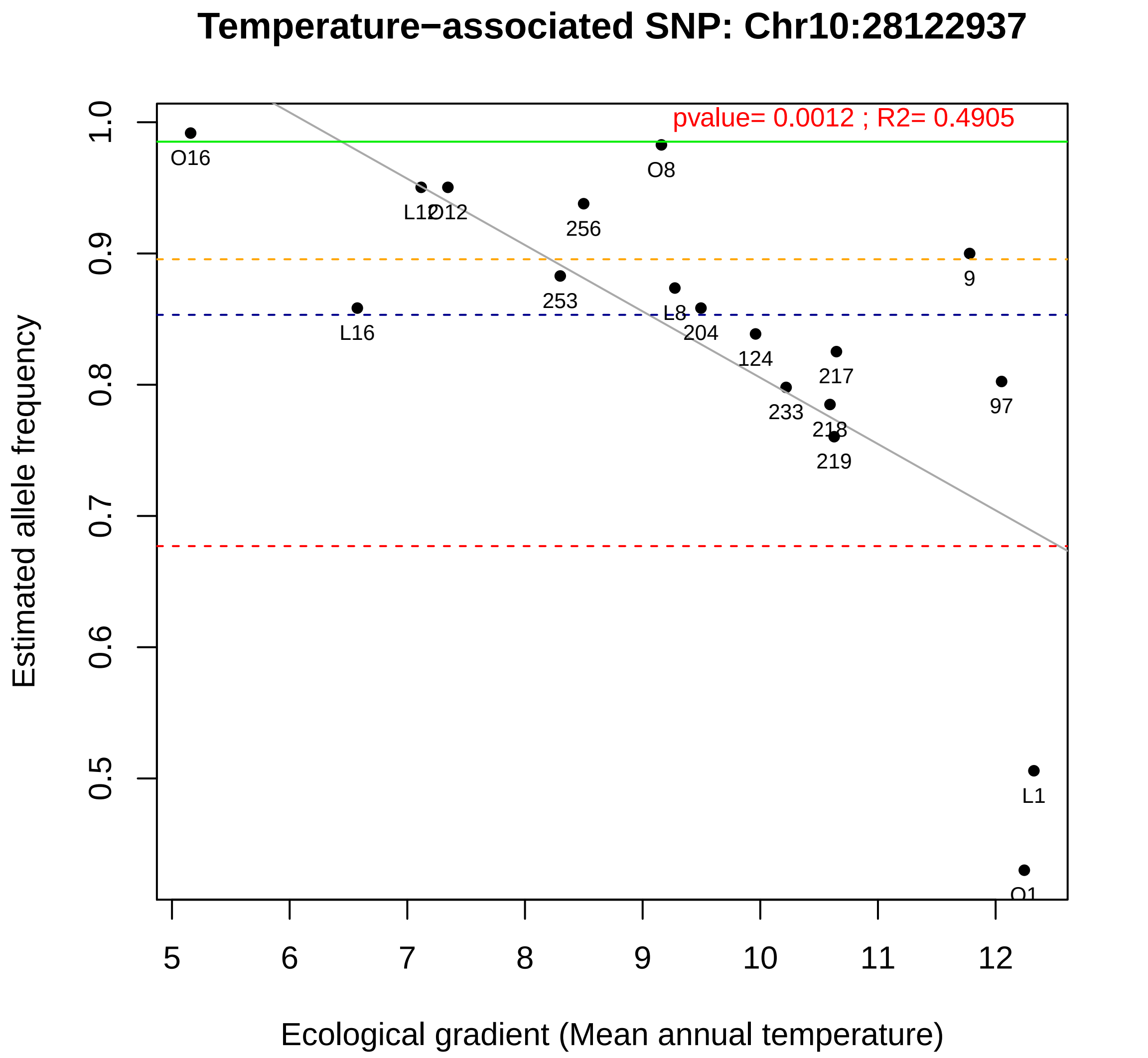

Temperature−associated SNP: Chr10:28122937
pvalue= 0.0012 ; R2= 0.4905
O16
O8
L12
O12
256
9
253
L8
204
L16
124
217
233
97
218
219
L1
O1
1.0
0.9
0.8
0.7
0.6
0.5
Estimated allele frequency
5
6
7
8
9
10
11
12
Ecological gradient (Mean annual temperature)

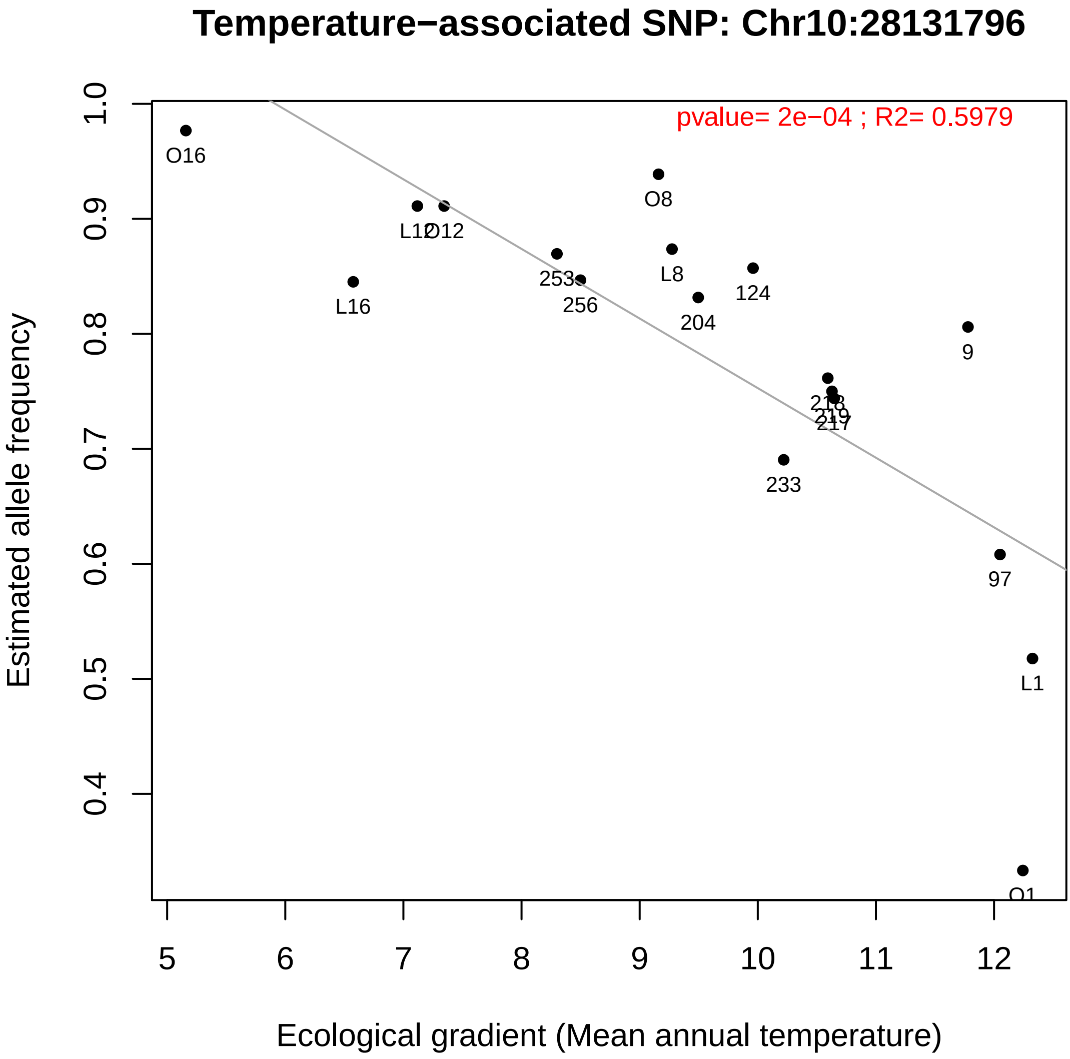
Temperature−associated SNP: Chr10:28131796
pvalue= 2e−04 ; R2= 0.5979
Estimated allele frequency
Ecological gradient (Mean annual temperature)
1.0
0.9
0.8
0.7
0.6
0.5
0.4
5
6
7
8
9
10
11
12
O16
L12
O12
L16
253
256
O8
L8
204
124
9
218
219
217
233
97
L1

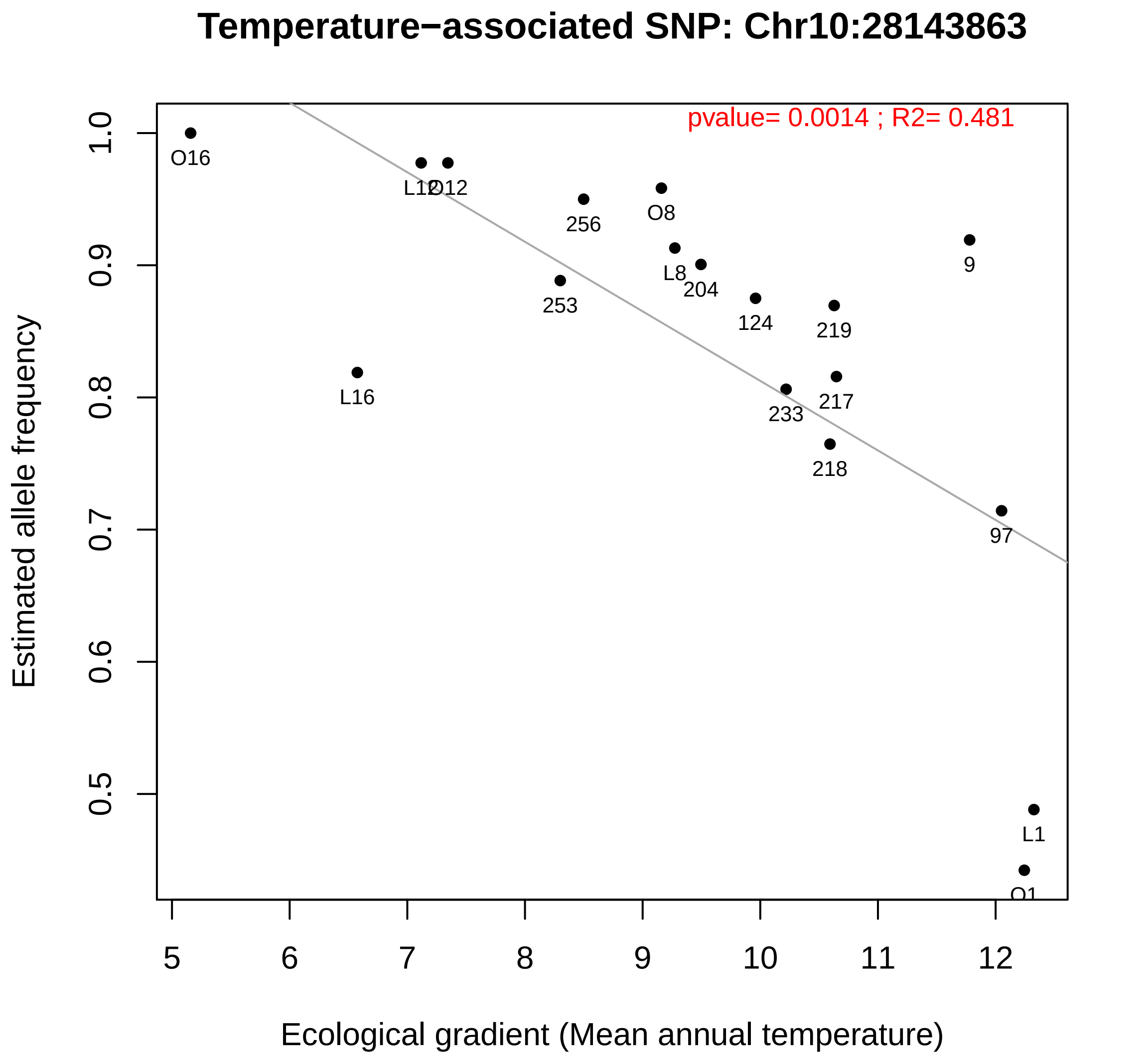

Temperature−associated SNP: Chr10:28143863
pvalue= 0.0014 ; R2= 0.481
Estimated allele frequency
Ecological gradient (Mean annual temperature)
1.0
0.9
0.8
0.7
0.6
0.5
5
6
7
8
9
10
11
12
O16
L12
O12
256
O8
L8
204
253
124
219
9
L16
233
217
218
97
L1
O1

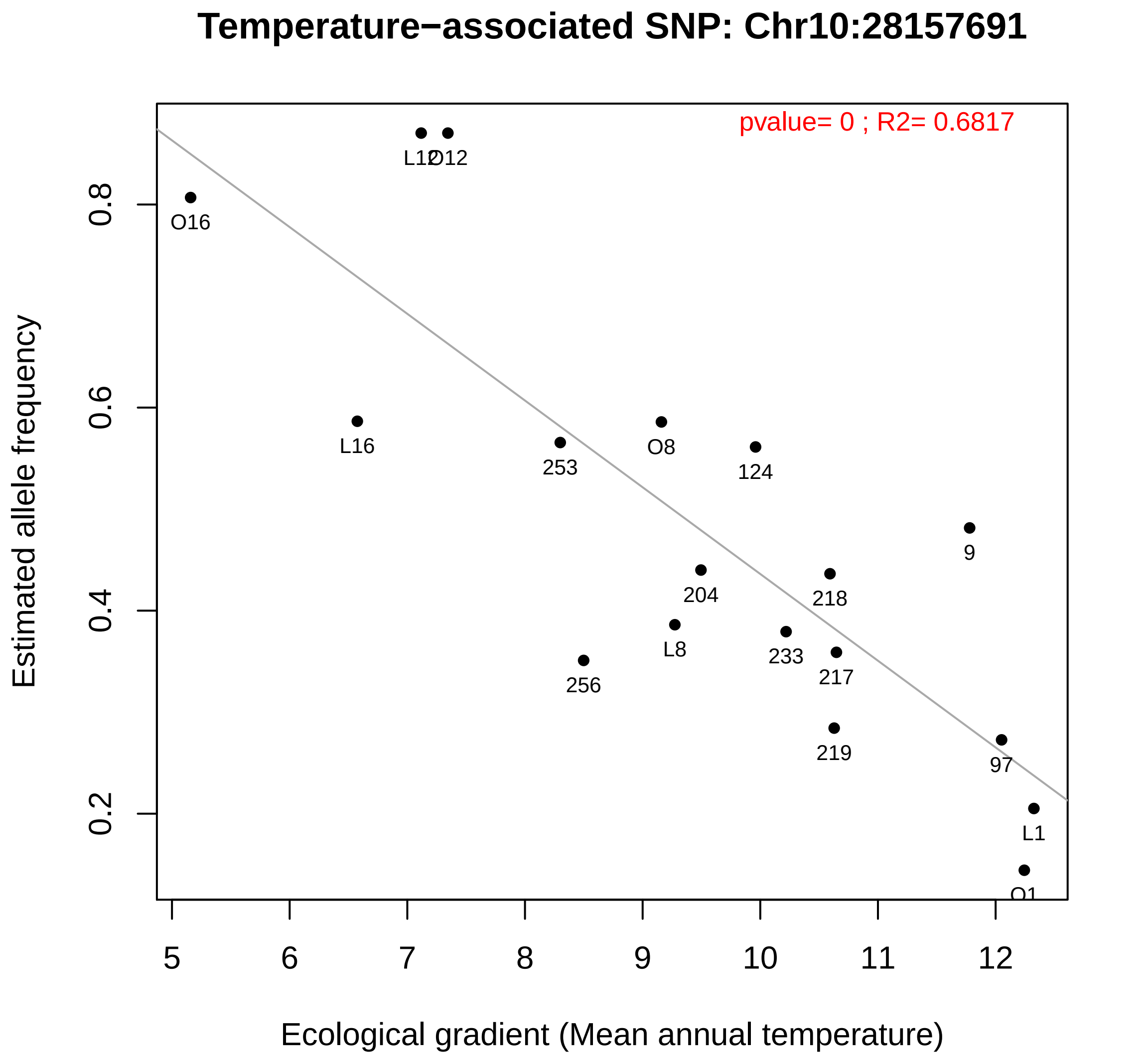
Temperature−associated SNP: Chr10:28157691
pvalue= 0 ; R2= 0.6817
Estimated allele frequency
0.2
0.4
0.6
0.8
5
6
7
8
9
10
11
12
Ecological gradient (Mean annual temperature)
O16
L12
O12
L16
253
O8
124
9
204
218
L8
233
217
256
219
97
L1
O1

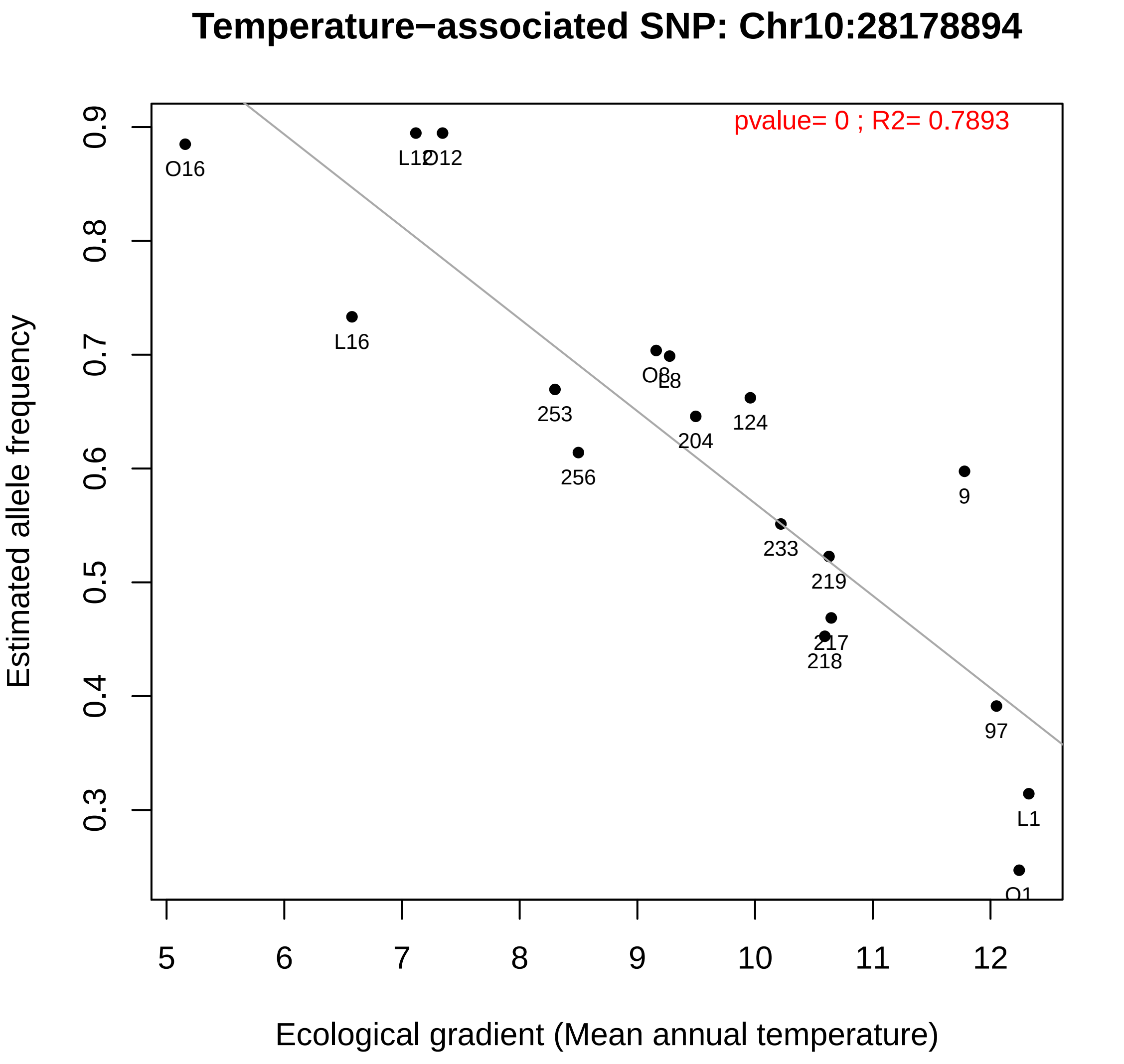

Temperature−associated SNP: Chr10:28178894
pvalue= 0 ; R2= 0.7893
Estimated allele frequency
Ecological gradient (Mean annual temperature)
O16
L12
O12
L16
O8
L8
253
256
204
124
9
233
219
217
218
97
L1
O1
0.3
0.4
0.5
0.6
0.7
0.8
0.9
5
6
7
8
9
10
11
12

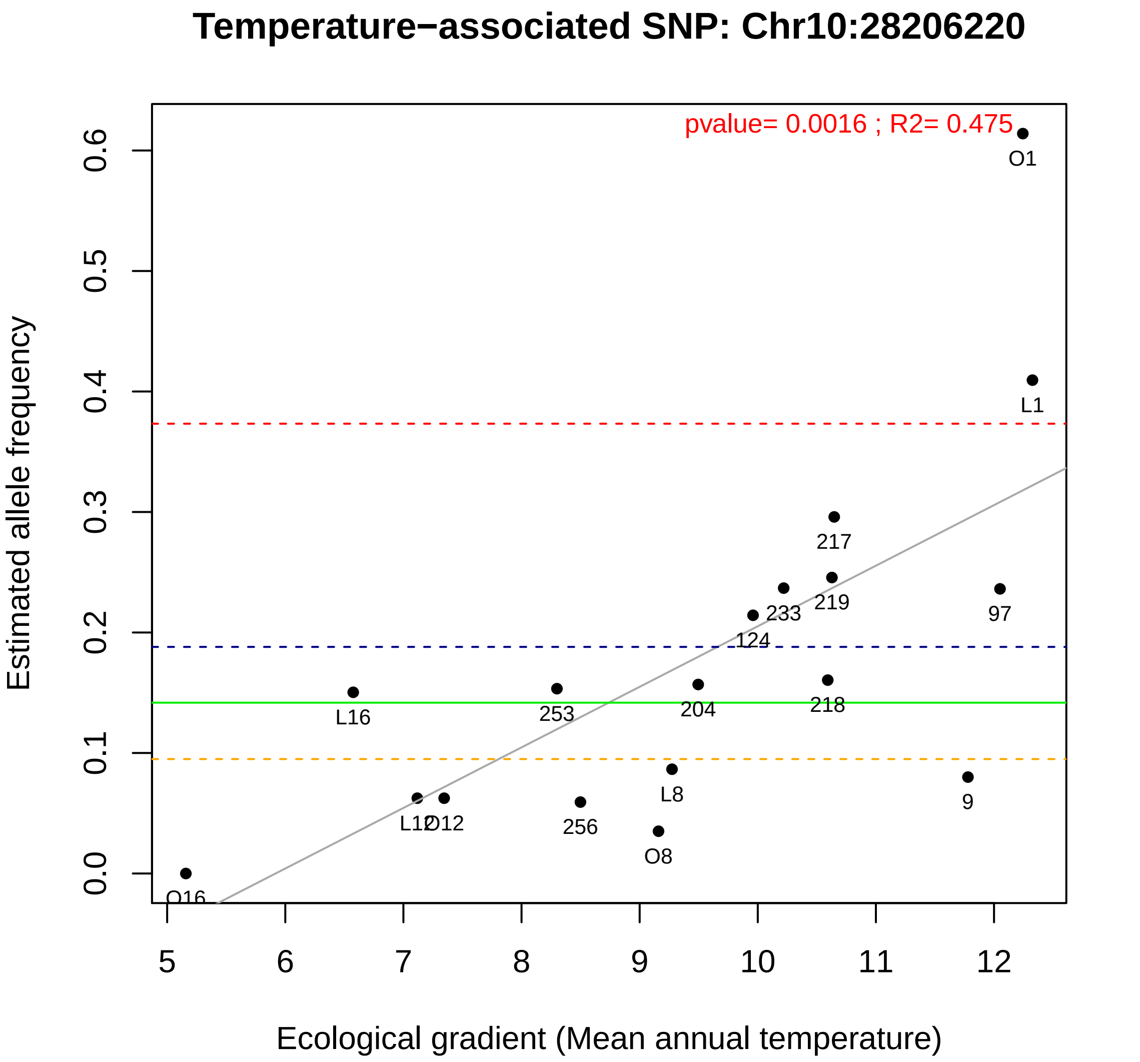

Temperature−associated SNP: Chr10:28206220
pvalue= 0.0016 ; R2= 0.475
Estimated allele frequency
Ecological gradient (Mean annual temperature)
0.0
0.1
0.2
0.3
0.4
0.5
0.6
5
6
7
8
9
10
11
12
O1
L1
217
219
233
124
97
204
218
253
L16
L8
9
L12
O12
256
O8
O16

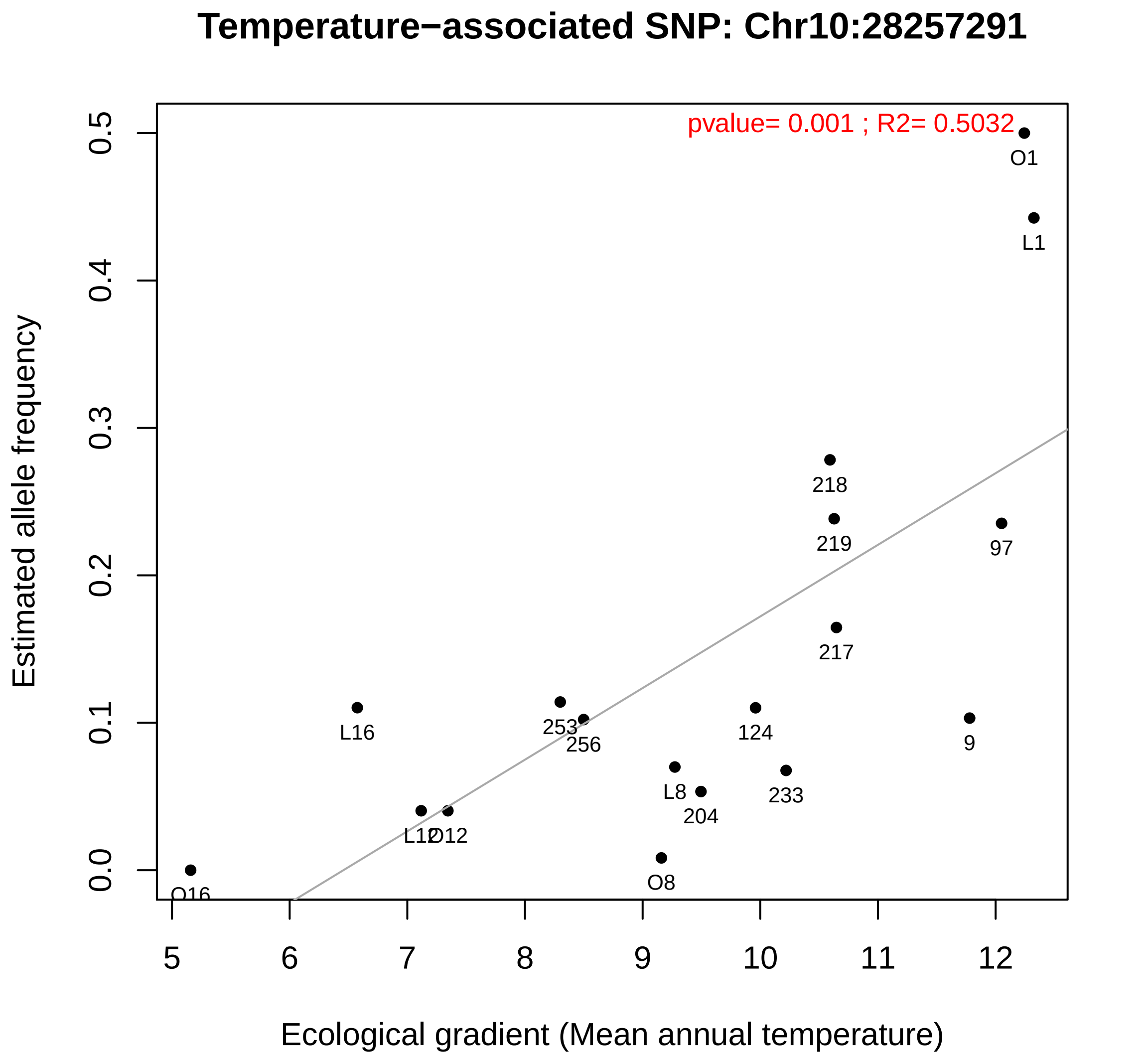
Temperature−associated SNP: Chr10:28257291
pvalue= 0.001 ; R2= 0.5032
Estimated allele frequency
0.0
0.1
0.2
0.3
0.4
0.5
5
6
7
8
9
10
11
12
Ecological gradient (Mean annual temperature)
O1
L1
218
219
97
217
L16
253
256
124
9
L8
204
233
L12
O12
O8
O16

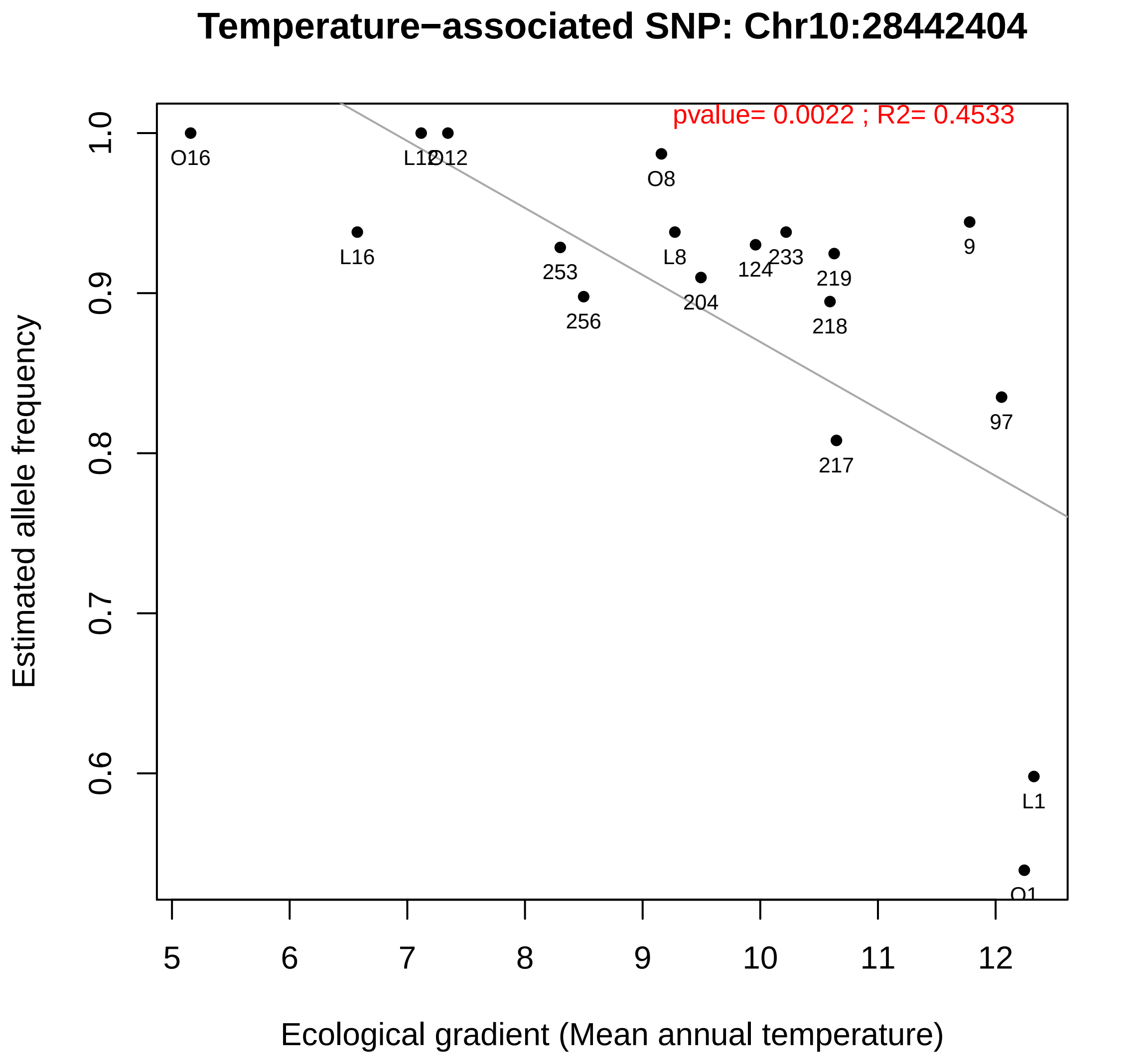
Temperature−associated SNP: Chr10:28442404
pvalue= 0.0022 ; R2= 0.4533
Estimated allele frequency
Ecological gradient (Mean annual temperature)
1.0
0.9
0.8
0.7
0.6
5
6
7
8
9
10
11
12
O16
L12
O12
O8
L16
253
L8
124
233
219
9
204
256
218
97
217
L1
O1

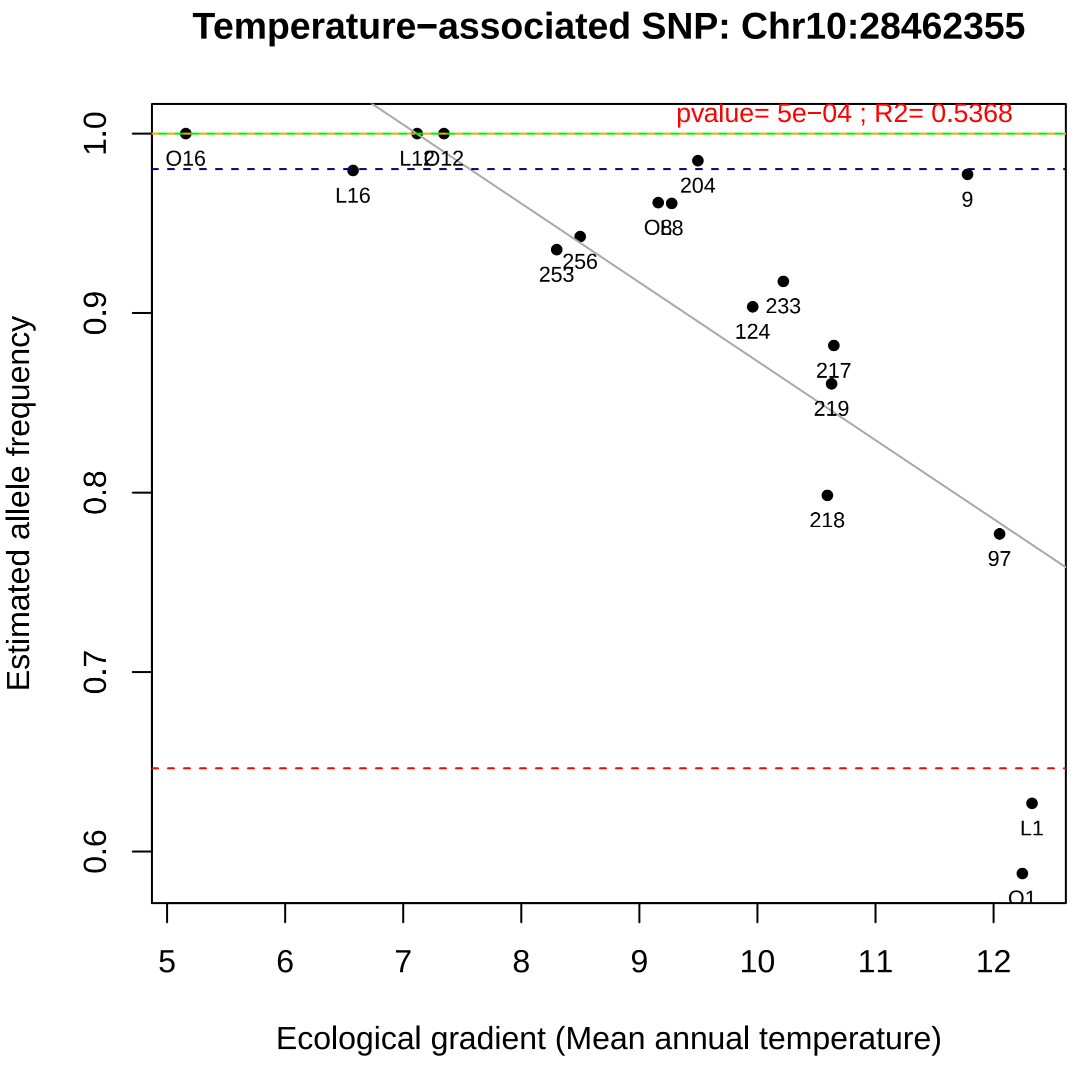
Temperature−associated SNP: Chr10:28462355
pvalue= 5e−04 ; R2= 0.5368
Estimated allele frequency
Ecological gradient (Mean annual temperature)
0.6
0.7
0.8
0.9
1.0
5
6
7
8
9
10
11
12
O16
L16
L12
O12
204
9
O8
L8
256
253
233
124
217
219
218
97
L1
O1

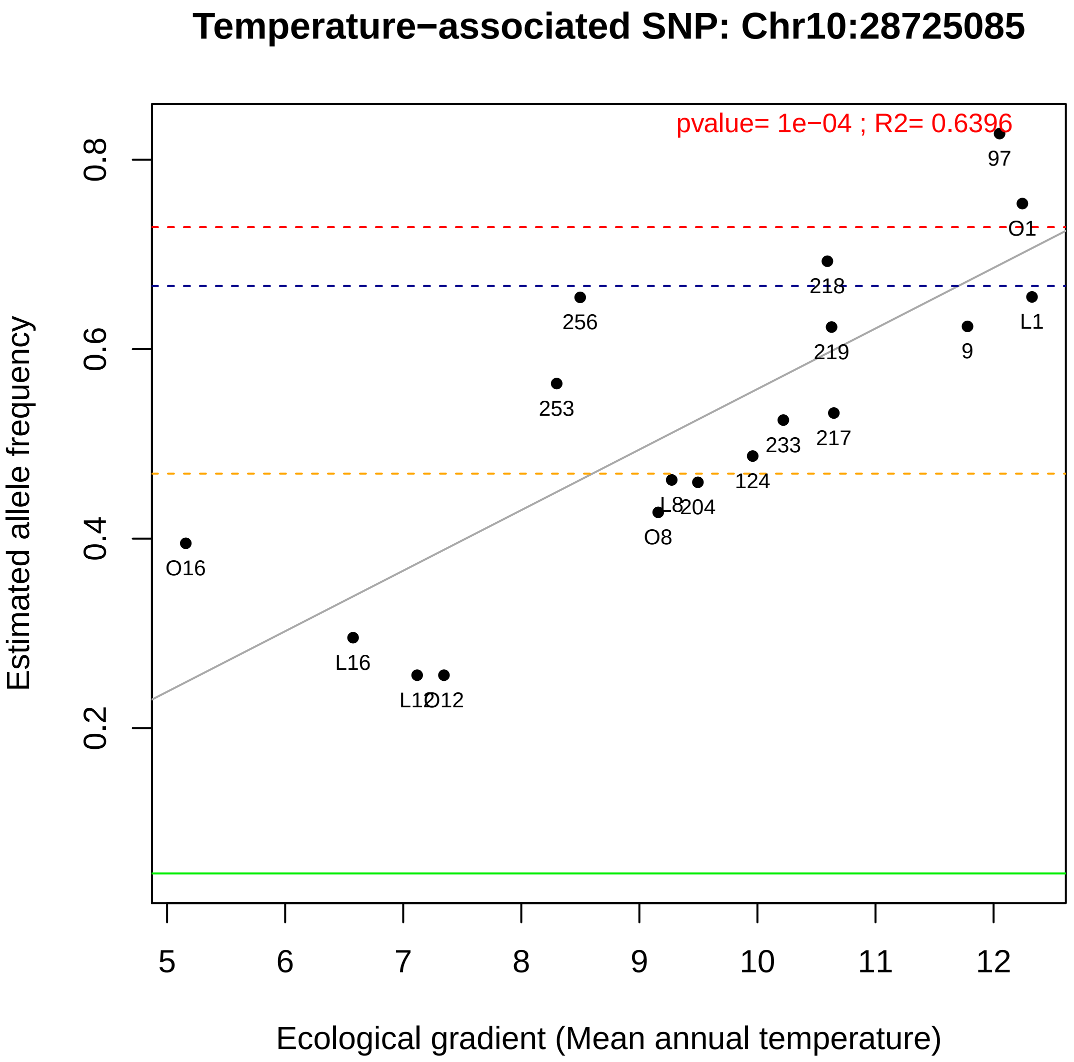

Temperature−associated SNP: Chr10:28725085
pvalue= 1e−04 ; R2= 0.6396
97
O1
218
256
L1
219
9
253
233
217
124
L8
204
O8
O16
L16
L12
O12
0.2
0.4
0.6
0.8
Estimated allele frequency
5
6
7
8
9
10
11
12
Ecological gradient (Mean annual temperature)

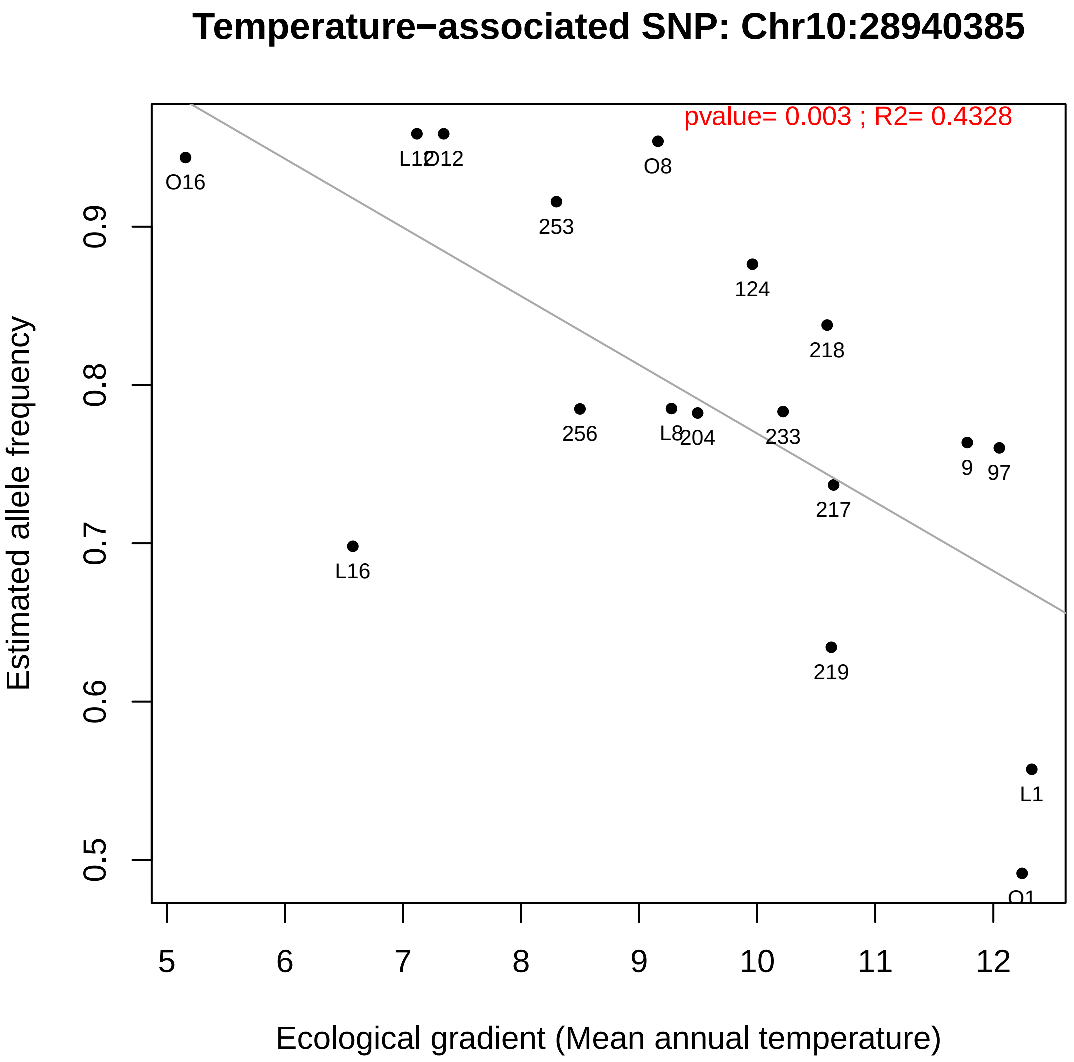

Temperature−associated SNP: Chr10:28940385
pvalue= 0.003 ; R2= 0.4328
O16
L12
O12
O8
253
124
218
256
L8
204
233
9
97
217
L16
219
L1
O1
0.5
0.6
0.7
0.8
0.9
Estimated allele frequency
5
6
7
8
9
10
11
12
Ecological gradient (Mean annual temperature)

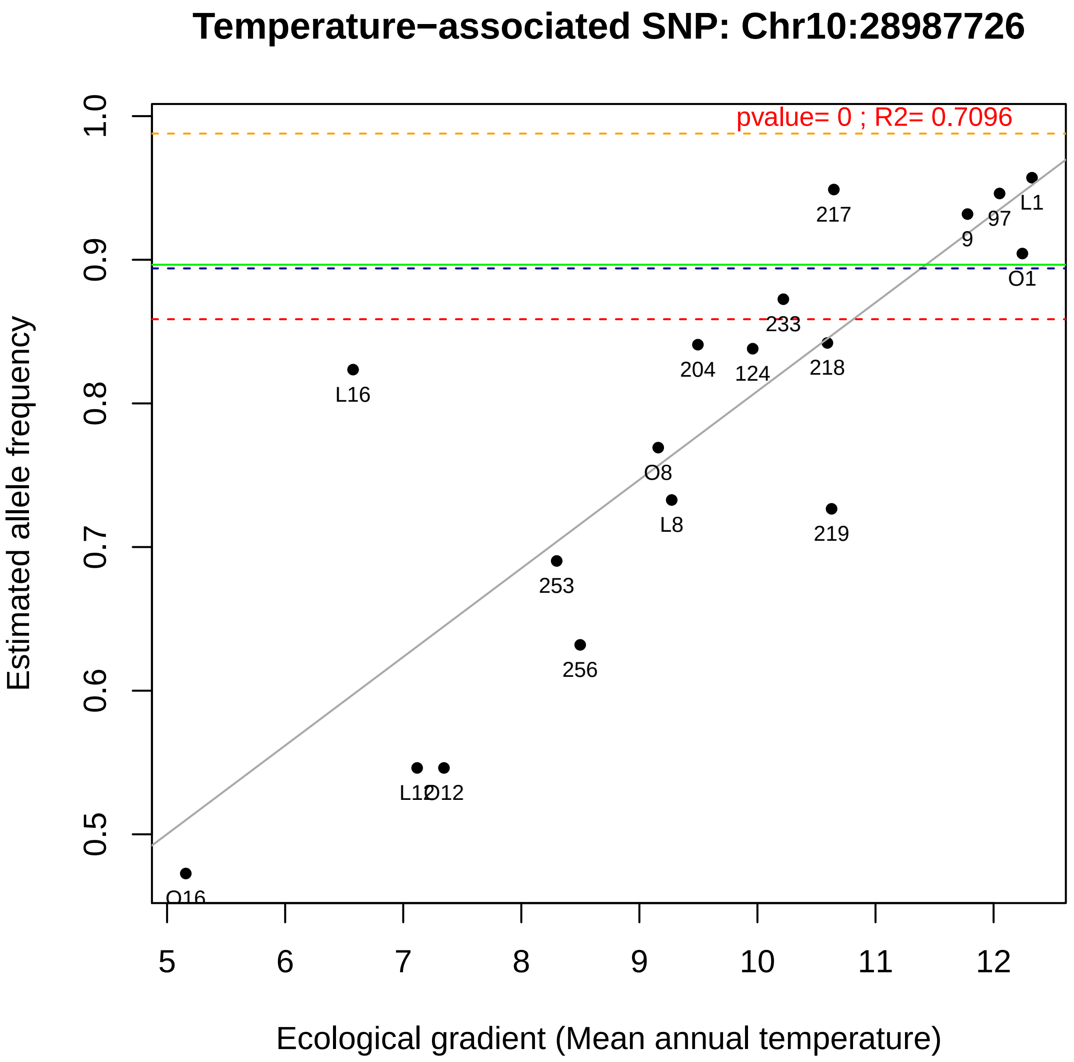

Temperature−associated SNP: Chr10:28987726
pvalue= 0 ; R2= 0.7096
Estimated allele frequency
1.0
0.9
0.8
0.7
0.6
0.5
217
97
L1
9
O1
233
204
124
218
L16
O8
L8
219
253
256
L12
O12
O16
5
6
7
8
9
10
11
12
Ecological gradient (Mean annual temperature)

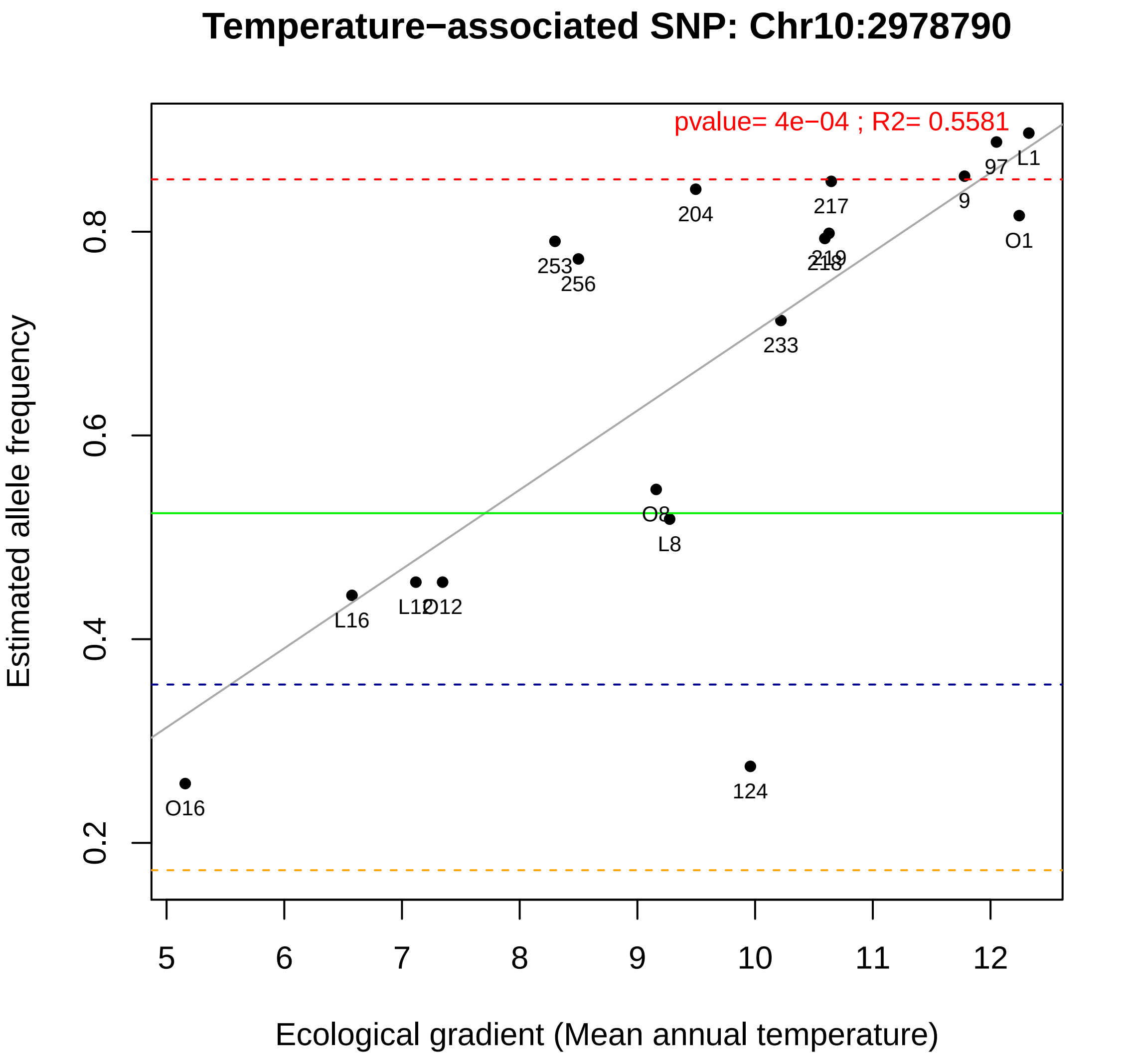

Temperature−associated SNP: Chr10:2978790
pvalue= 4e−04 ; R2= 0.5581
Estimated allele frequency
0.2
0.4
0.6
0.8
Ecological gradient (Mean annual temperature)
5
6
7
8
9
10
11
12
L1
97
9
O1
217
204
219
218
253
256
233
O8
L8
L16
L12
O12
O16
124

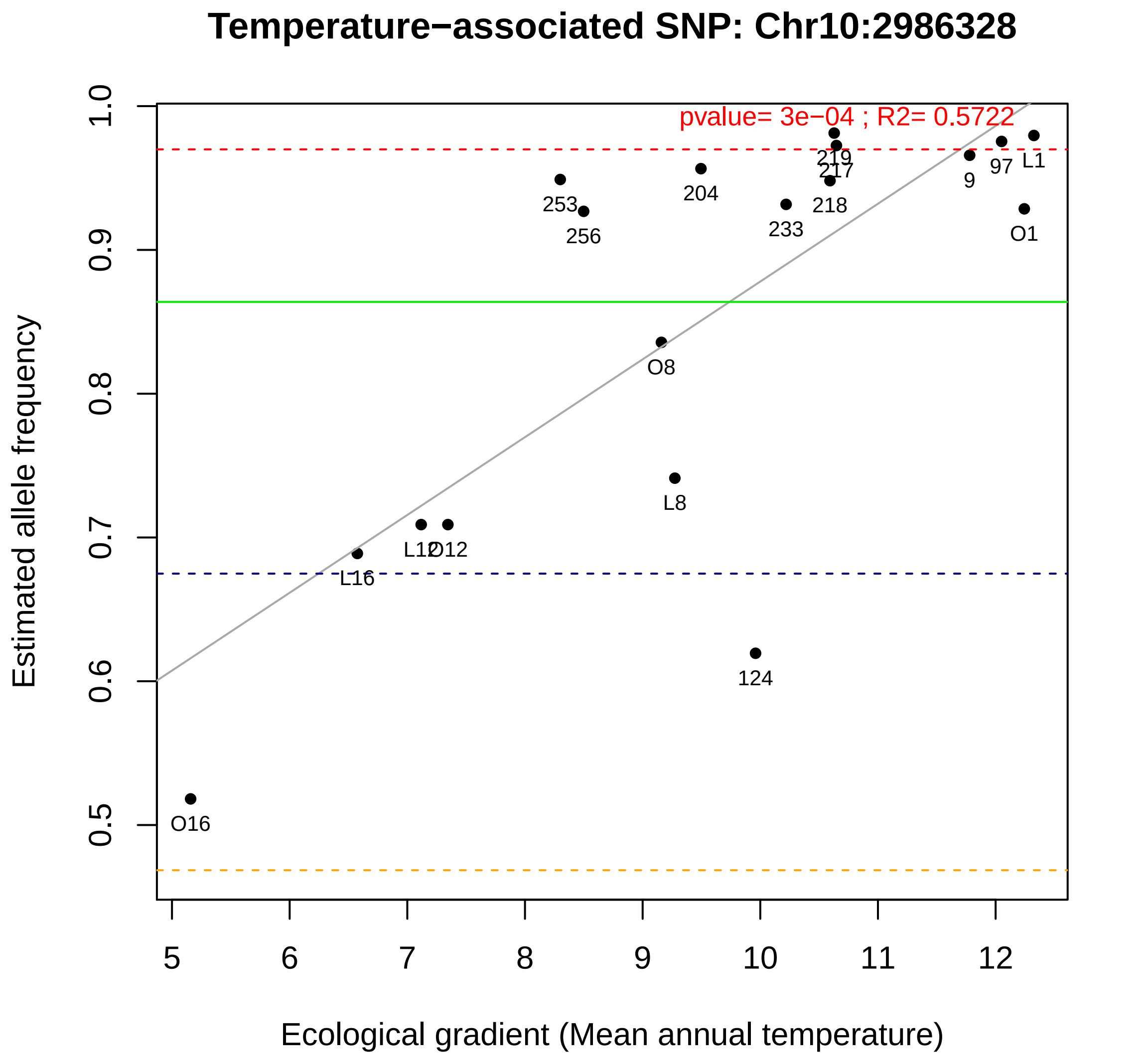
Temperature−associated SNP: Chr10:2986328
pvalue= 3e−04 ; R2= 0.5722
Estimated allele frequency
0.5
0.6
0.7
0.8
0.9
1.0
5
6
7
8
9
10
11
12
Ecological gradient (Mean annual temperature)
O16
L16
L12
O12
253
256
O8
L8
204
124
233
219
217
218
9
97
L1
O1

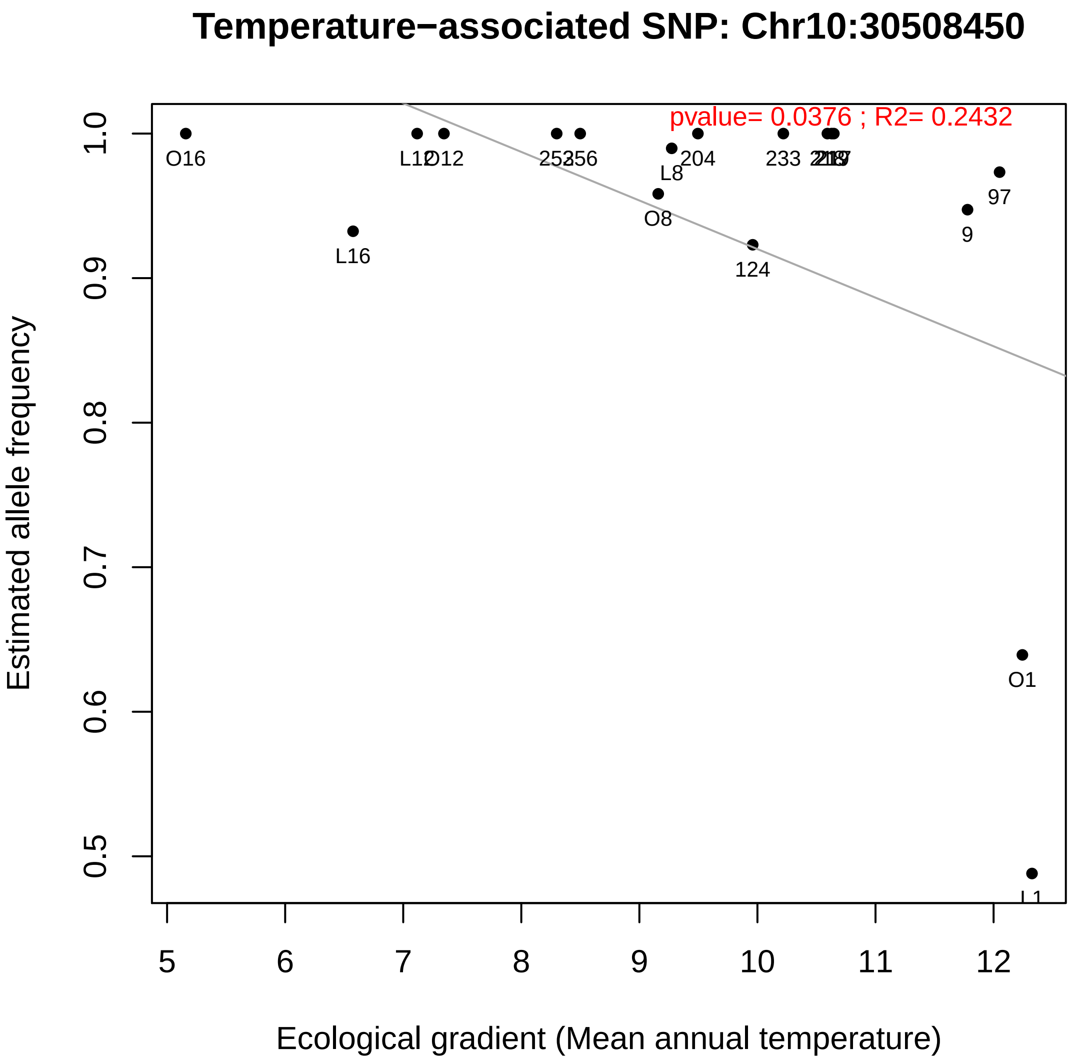

Temperature−associated SNP: Chr10:30508450
pvalue= 0.0376 ; R2= 0.2432
Estimated allele frequency
Ecological gradient (Mean annual temperature)
1.0
0.9
0.8
0.7
0.6
0.5
5
6
7
8
9
10
11
12
O16
L16
L12
O12
253
256
L8
O8
204
124
233
97
9
O1

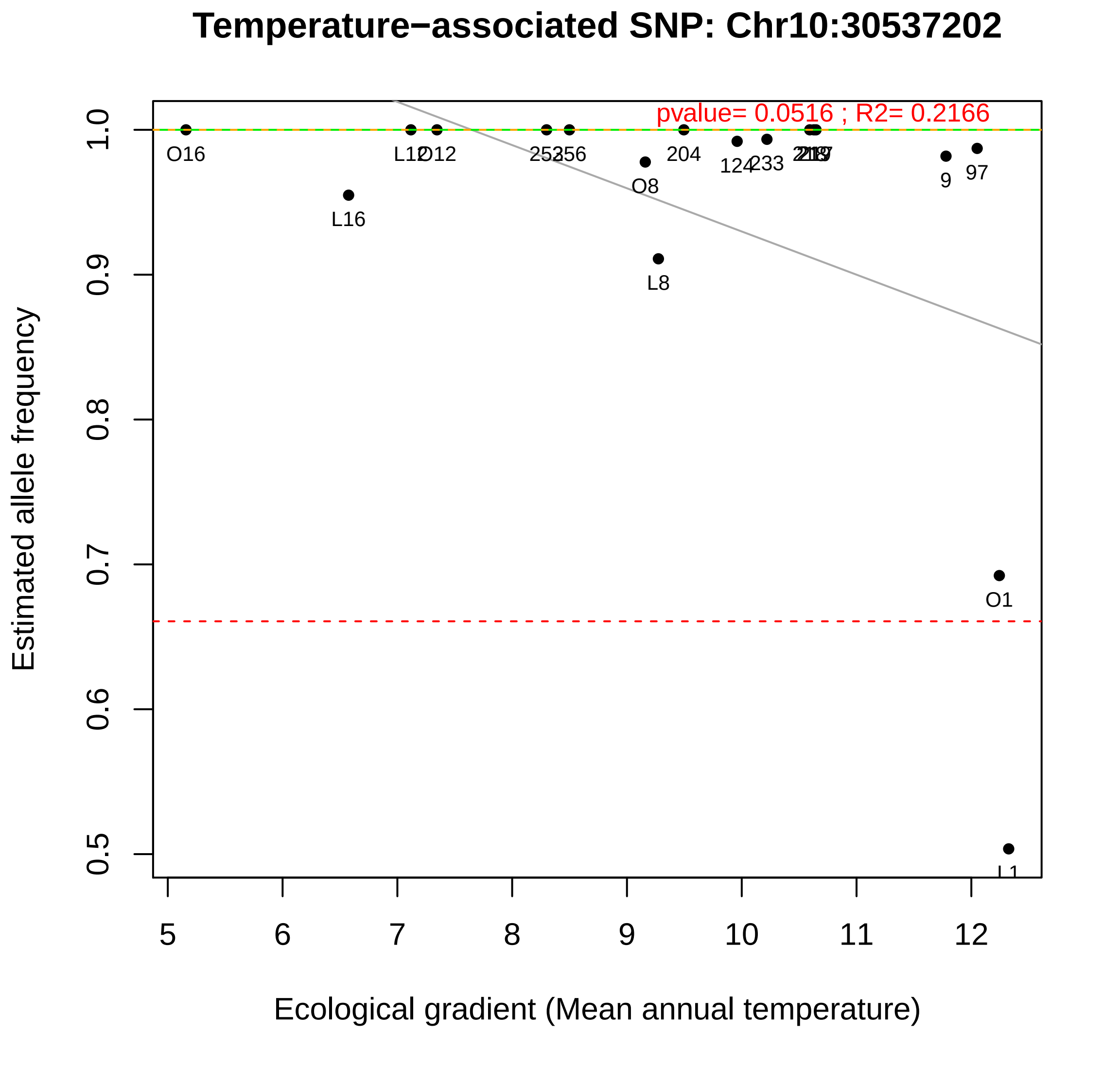

Temperature−associated SNP: Chr10:30537202
pvalue= 0.0516 ; R2= 0.2166
Estimated allele frequency
Ecological gradient (Mean annual temperature)
0.5
0.6
0.7
0.8
0.9
1.0
5
6
7
8
9
10
11
12
O16
L16
L12
O12
204
O8
L8
124
233
97
9
O1
L1

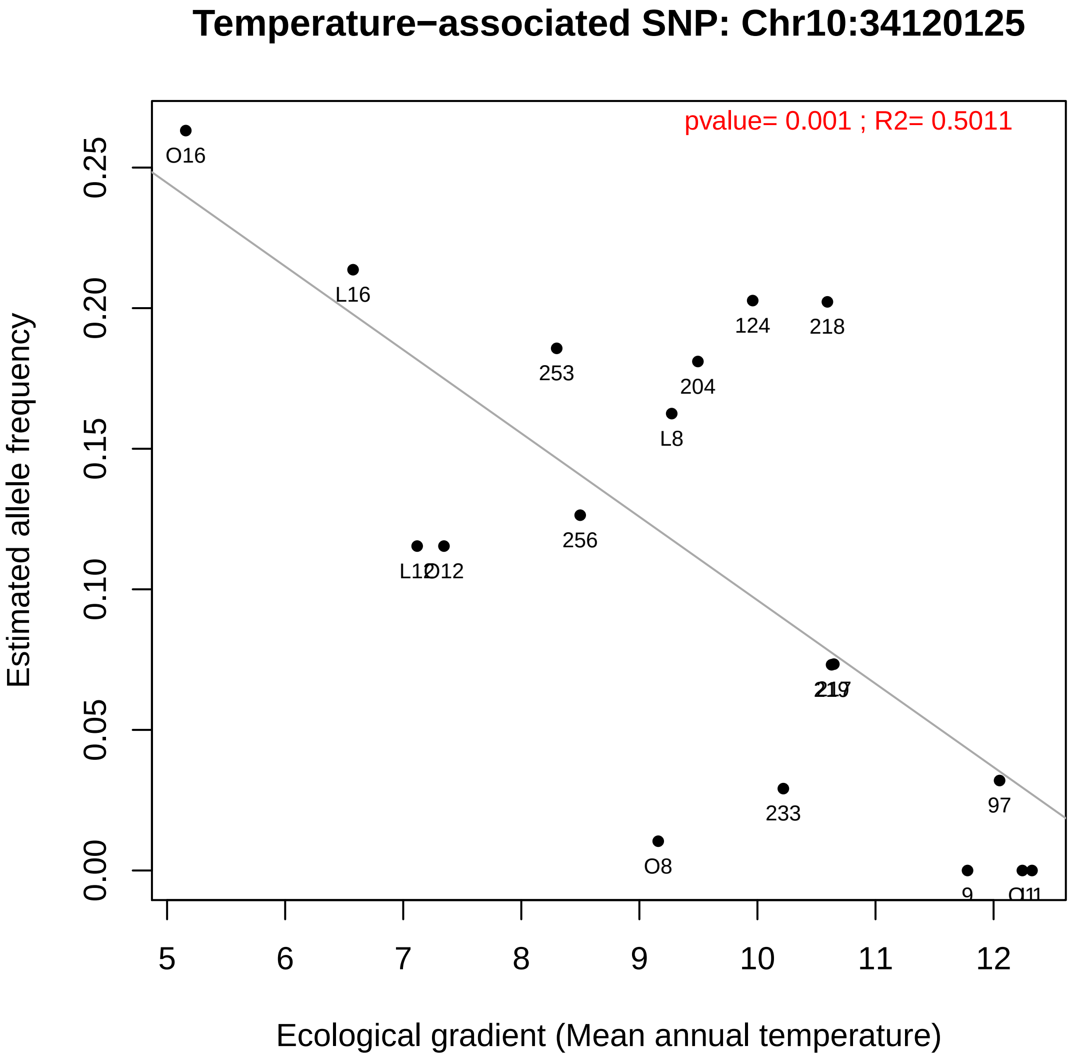
Temperature−associated SNP: Chr10:34120125
pvalue= 0.001 ; R2= 0.5011
Estimated allele frequency
0.00
0.05
0.10
0.15
0.20
0.25
5
6
7
8
9
10
11
12
Ecological gradient (Mean annual temperature)
O16
L16
253
204
124
218
L8
256
O12
233
O8
97
9

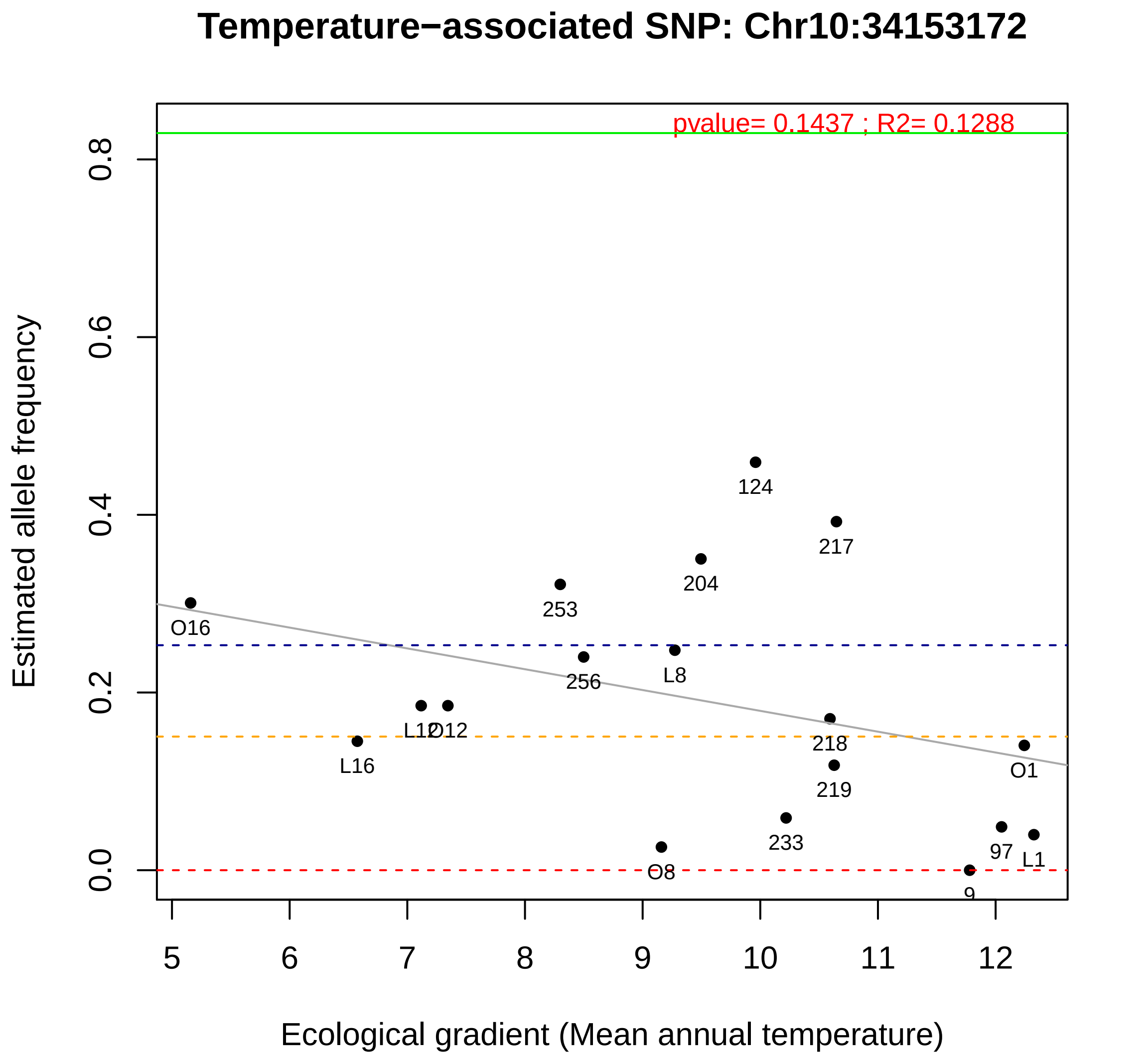

Temperature−associated SNP: Chr10:34153172
pvalue= 0.1437 ; R2= 0.1288
Estimated allele frequency
0.0
0.2
0.4
0.6
0.8
5
6
7
8
9
10
11
12
Ecological gradient (Mean annual temperature)
O16
L16
L12
O12
253
256
O8
L8
204
124
233
217
218
219
9
97
L1
O1

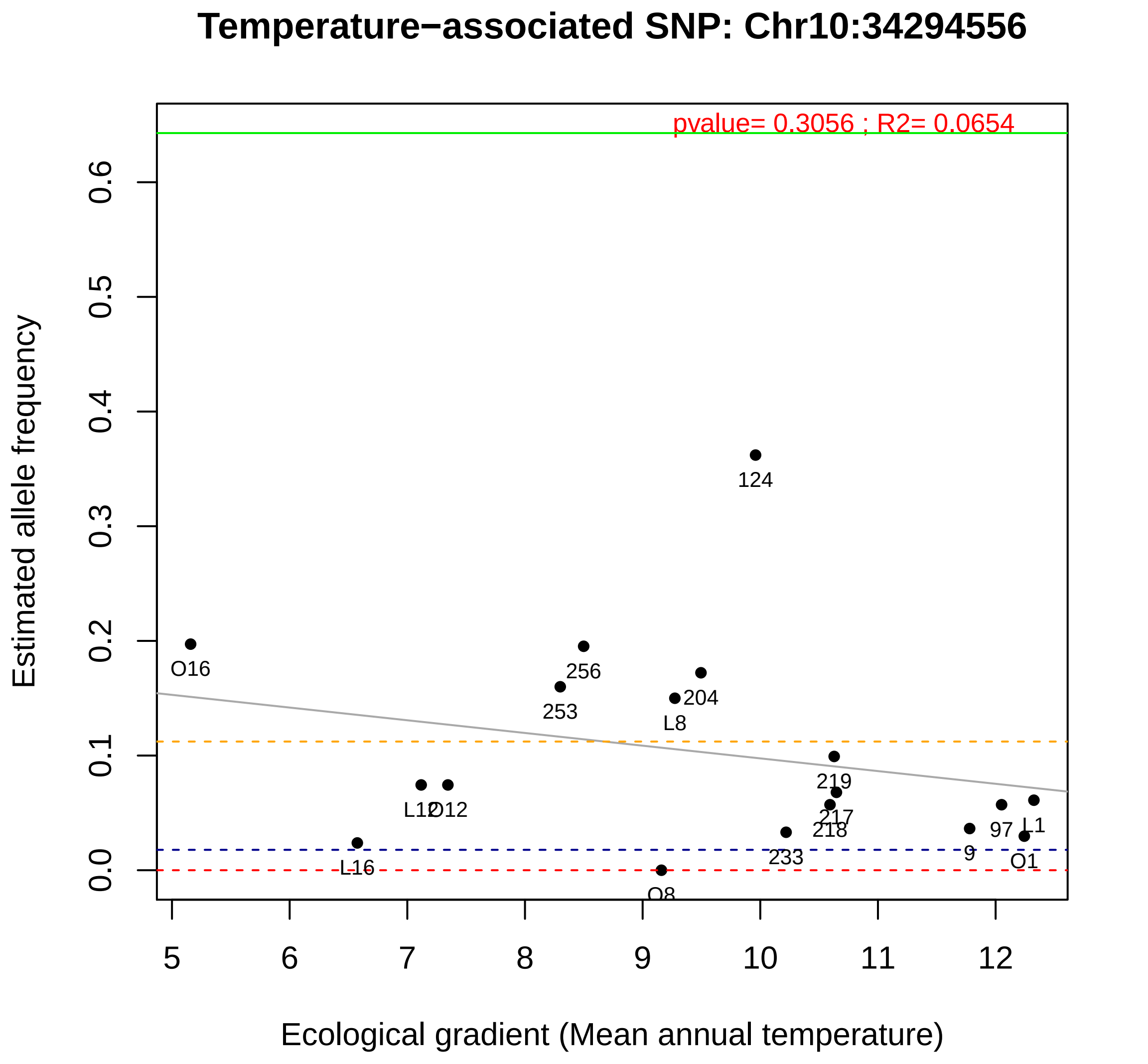

Temperature−associated SNP: Chr10:34294556
pvalue= 0.3056 ; R2= 0.0654
Estimated allele frequency
Ecological gradient (Mean annual temperature)
0.0
0.1
0.2
0.3
0.4
0.5
0.6
5
6
7
8
9
10
11
12
O16
L16
L12
O12
253
256
O8
L8
204
124
233
219
217
218
9
97
L1
O1

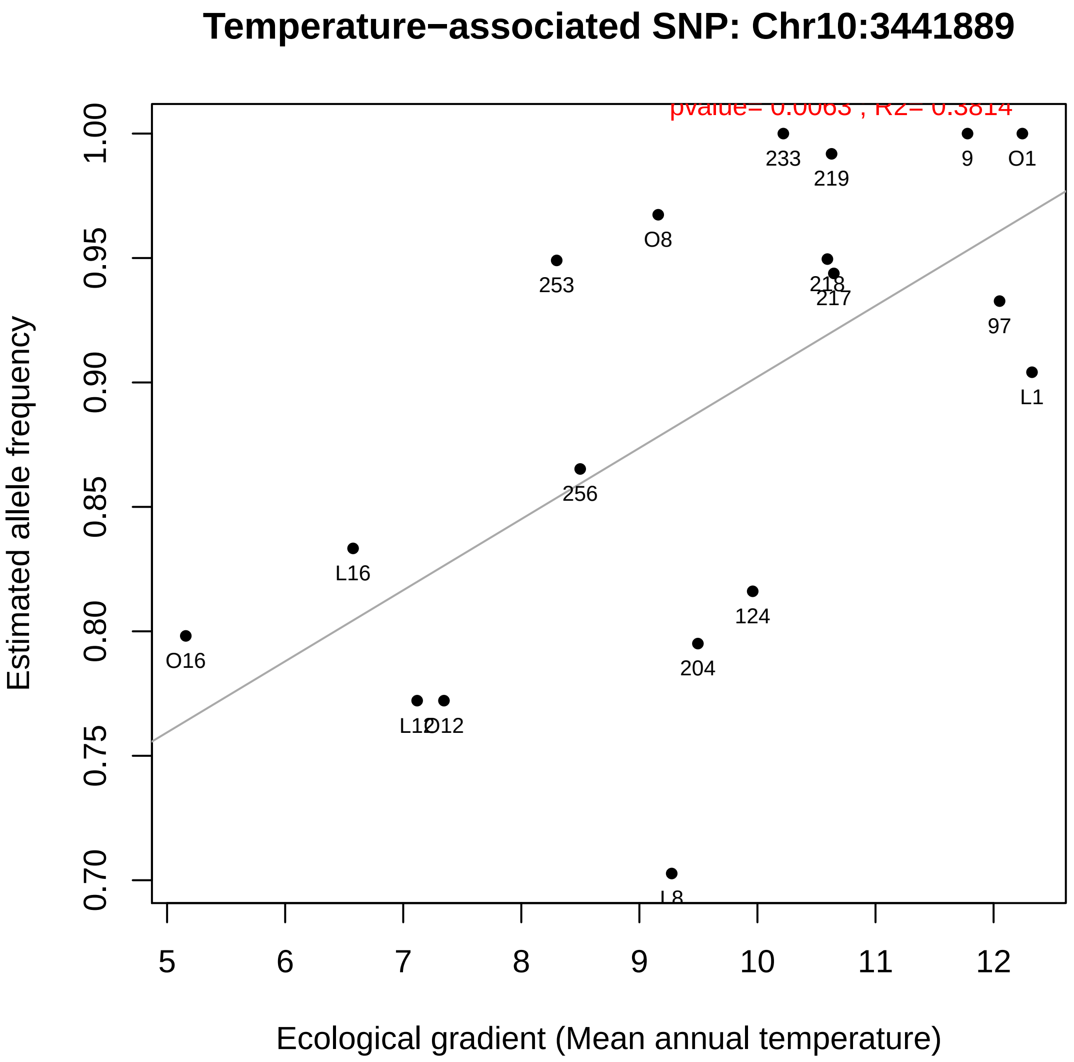

Temperature−associated SNP: Chr10:3441889
pvalue= 0.0063 , R2= 0.3814
Estimated allele frequency
1.00
0.95
0.90
0.85
0.80
0.75
0.70
5
6
7
8
9
10
11
12
Ecological gradient (Mean annual temperature)
233
219
9
O1
O8
253
218
217
97
L1
256
L16
124
O16
204
L12
O12

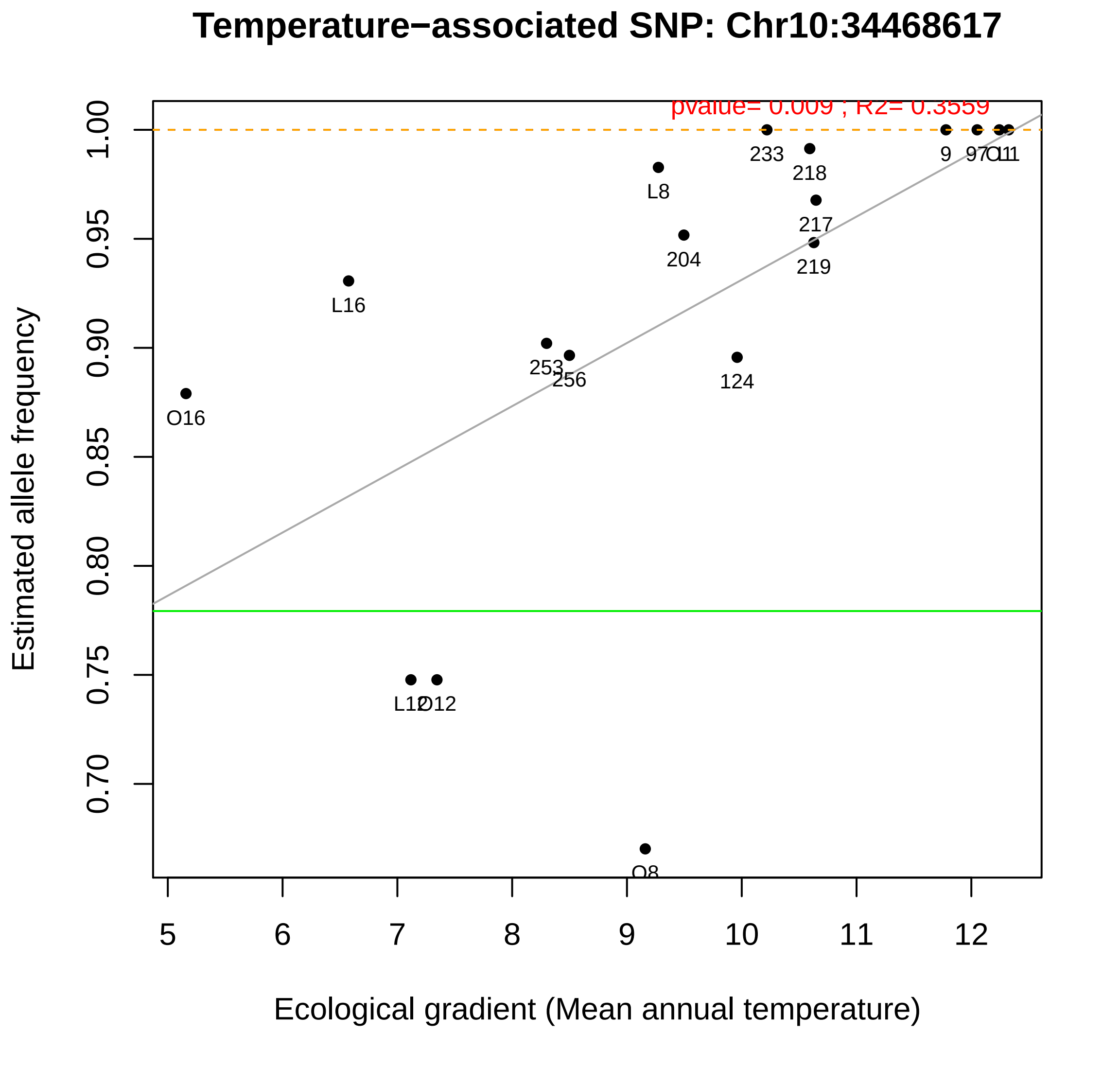

Temperature−associated SNP: Chr10:34468617
pvalue= 0.009 ; R2= 0.3559
Estimated allele frequency
Ecological gradient (Mean annual temperature)
1.00
0.95
0.90
0.85
0.80
0.75
0.70
5
6
7
8
9
10
11
12
233
218
L8
217
204
219
L16
253
256
124
O16
L12
O12
O8
9
97
O11

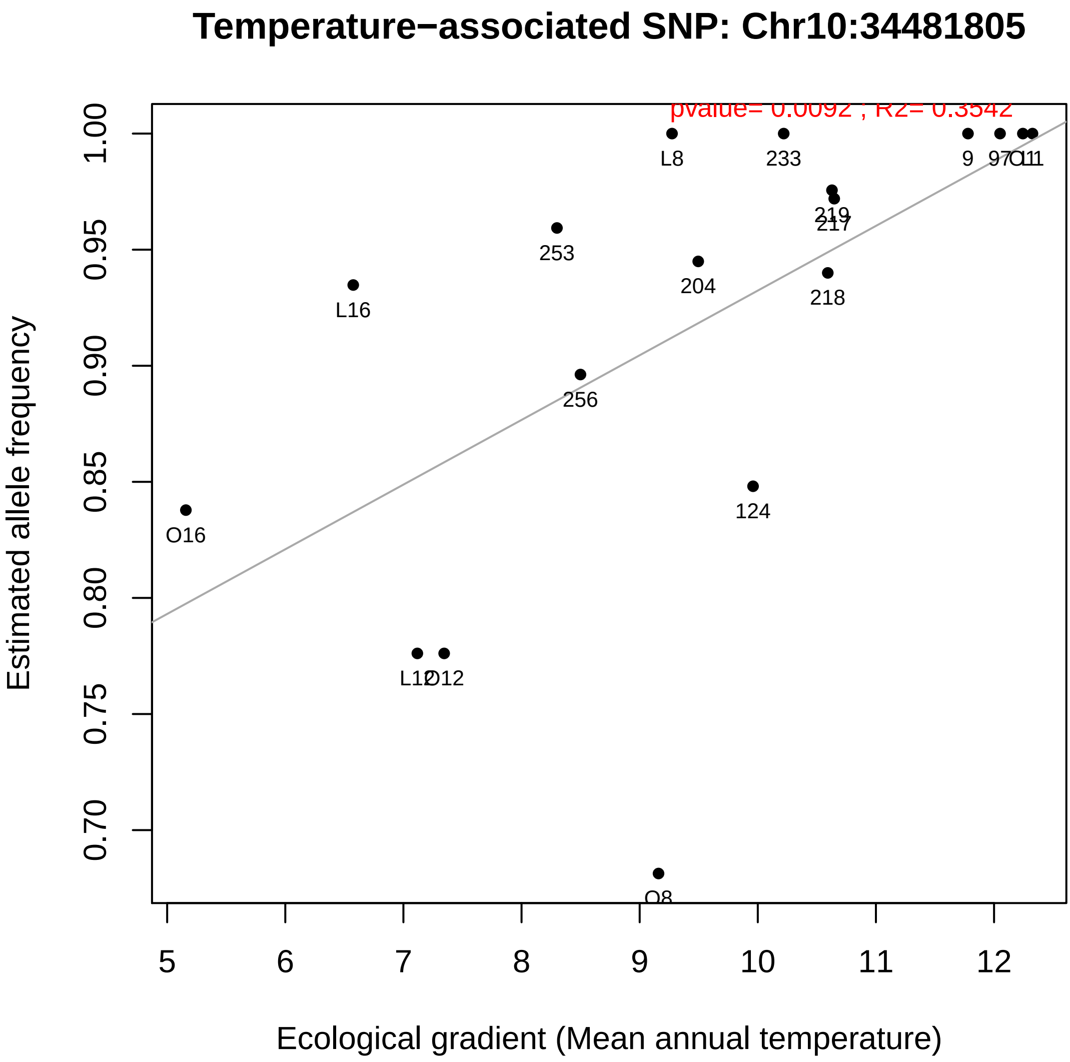
Temperature−associated SNP: Chr10:34481805
pvalue= 0.0092 ; R2= 0.3542
Estimated allele frequency
Ecological gradient (Mean annual temperature)
1.00
0.95
0.90
0.85
0.80
0.75
0.70
5
6
7
8
9
10
11
12
L8
233
9
97
O11
219
217
253
204
218
L16
256
124
O16
L12
O12
O8

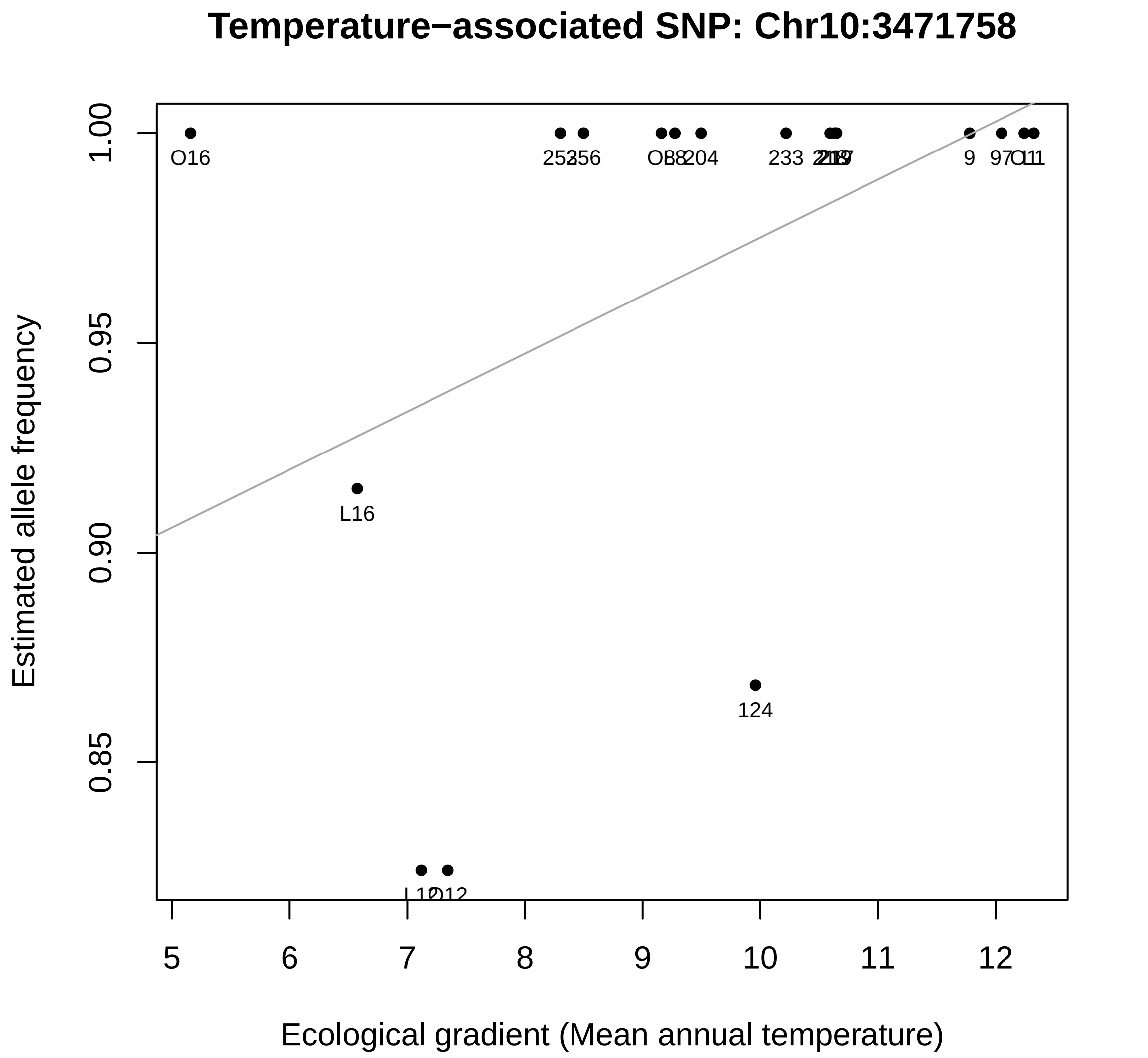
Temperature−associated SNP: Chr10:3471758
Estimated allele frequency
Ecological gradient (Mean annual temperature)
0.85
0.90
0.95
1.00
5
6
7
8
9
10
11
12
O16
L16
124
233
204
9

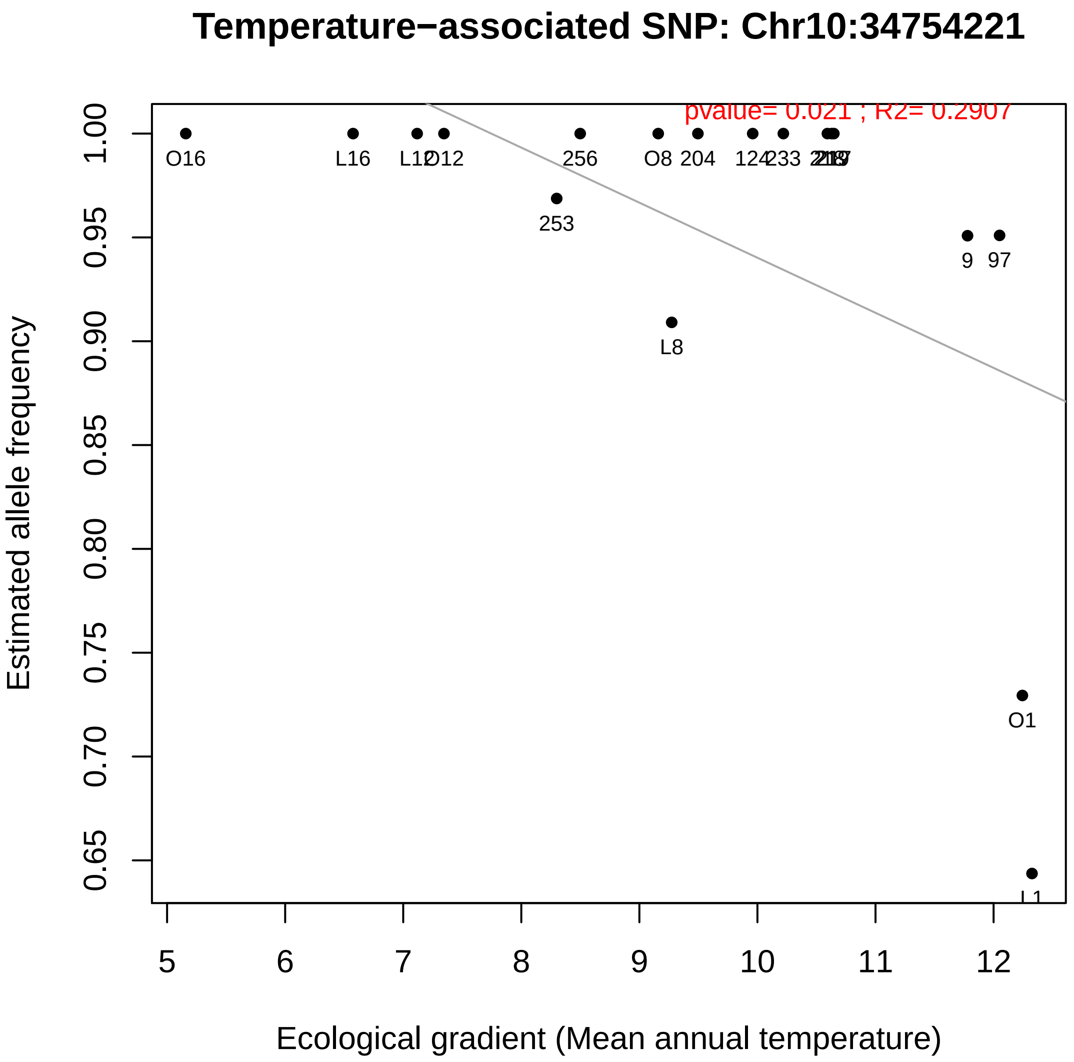

Temperature−associated SNP: Chr10:34754221
pvalue= 0.021 ; R2= 0.2907
O16
L16
L12
O12
256
O8
204
124
233
253
9
97
L8
O1
Estimated allele frequency
Ecological gradient (Mean annual temperature)
1.00
0.95
0.90
0.85
0.80
0.75
0.70
0.65
5
6
7
8
9
10
11
12

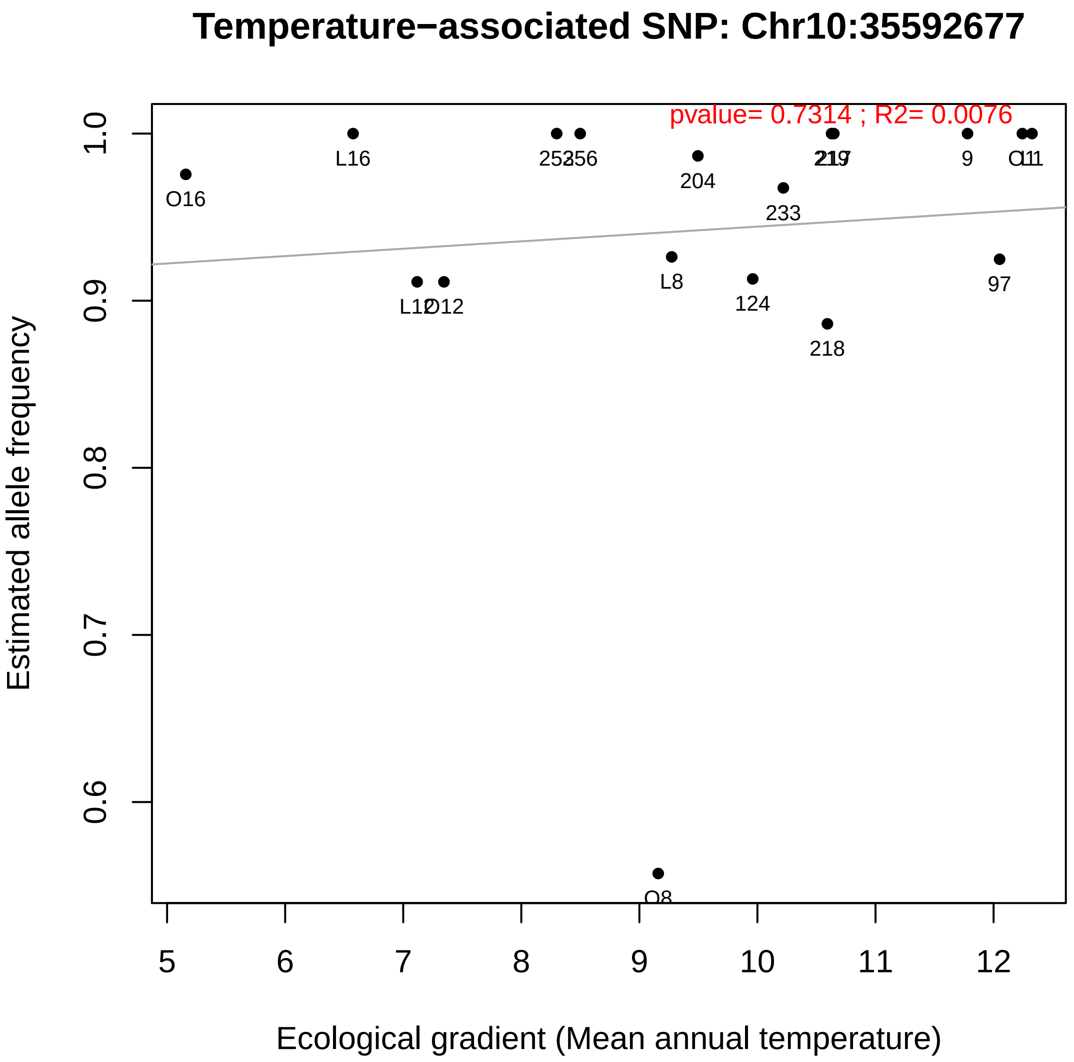
Temperature−associated SNP: Chr10:35592677
pvalue= 0.7314 ; R2= 0.0076
Estimated allele frequency
Ecological gradient (Mean annual temperature)
1.0
0.9
0.8
0.7
0.6
5
6
7
8
9
10
11
12
O16
L16
204
233
L8
124
218
9
97
O8

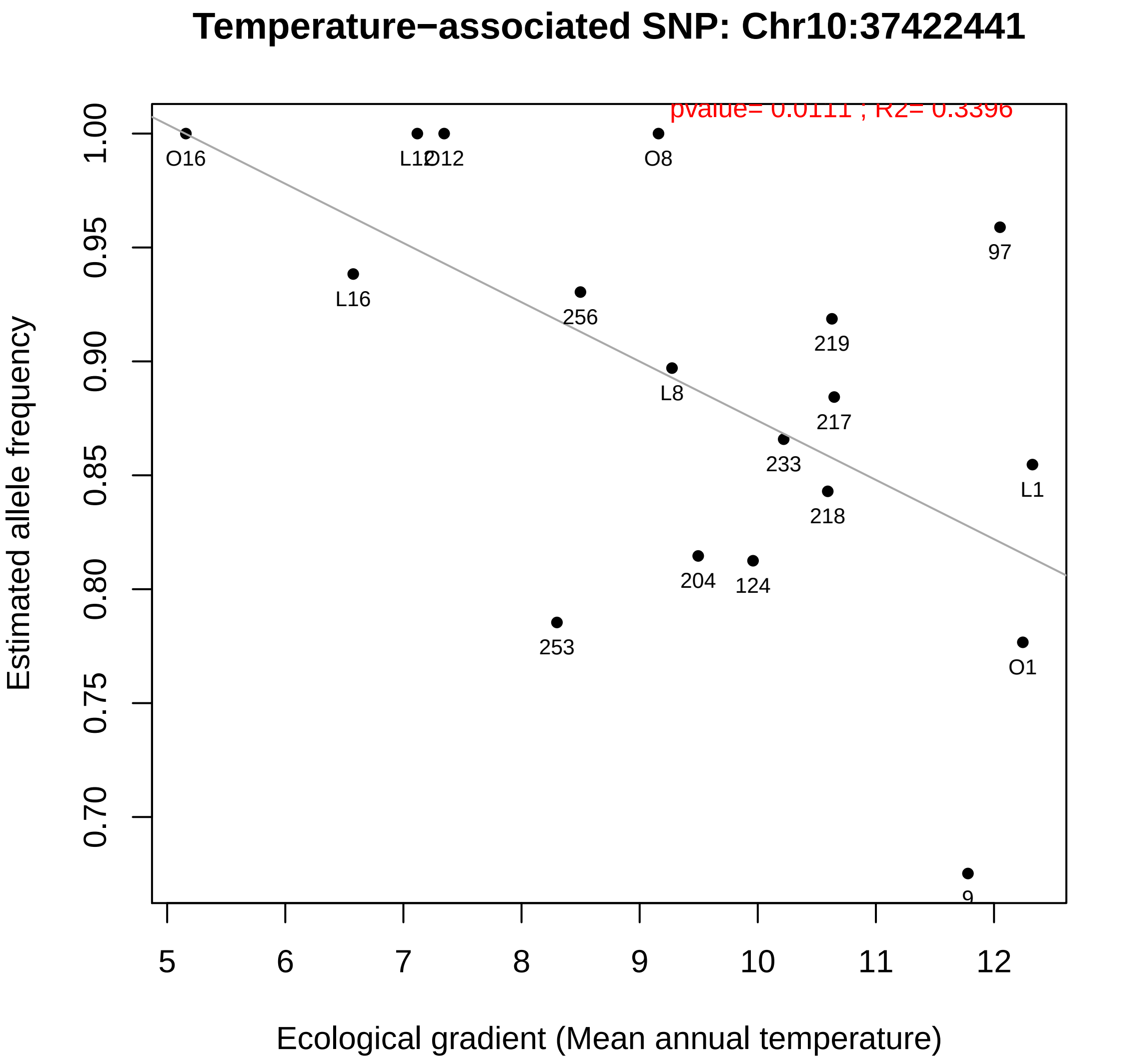

Temperature−associated SNP: Chr10:37422441
pvalue= 0.0111 ; R2= 0.3396
Estimated allele frequency
Ecological gradient (Mean annual temperature)
1.00
0.95
0.90
0.85
0.80
0.75
0.70
5
6
7
8
9
10
11
12
O16
L12
O12
O8
97
L16
256
219
L8
217
233
L1
218
204
124
253
O1
9

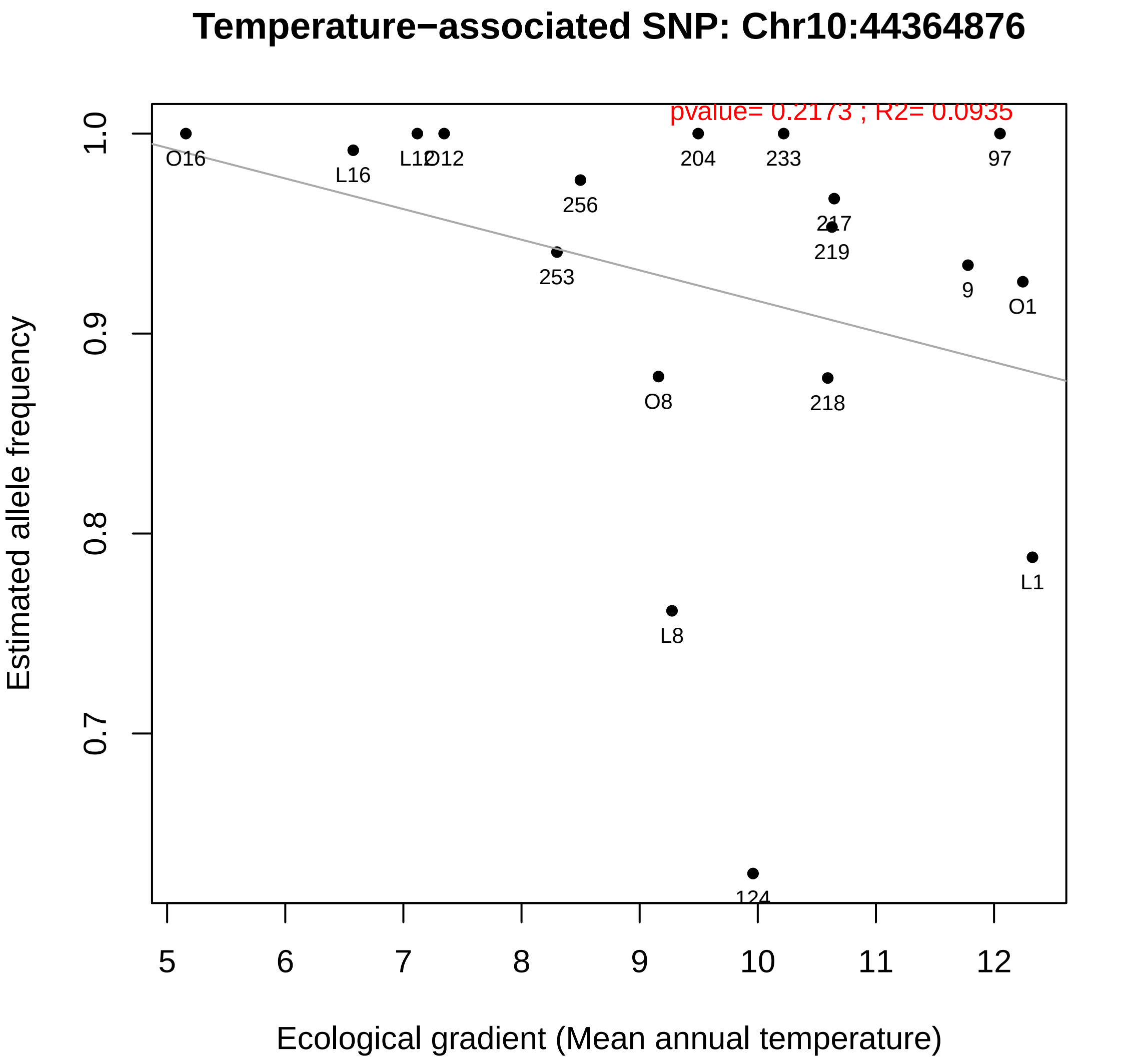

Temperature−associated SNP: Chr10:44364876
pvalue= 0.2173 ; R2= 0.0935
Estimated allele frequency
Ecological gradient (Mean annual temperature)
1.0
0.9
0.8
0.7
5
6
7
8
9
10
11
12
O16
L16
L12
O12
256
253
204
233
217
219
97
9
O1
O8
218
L8
L1
124

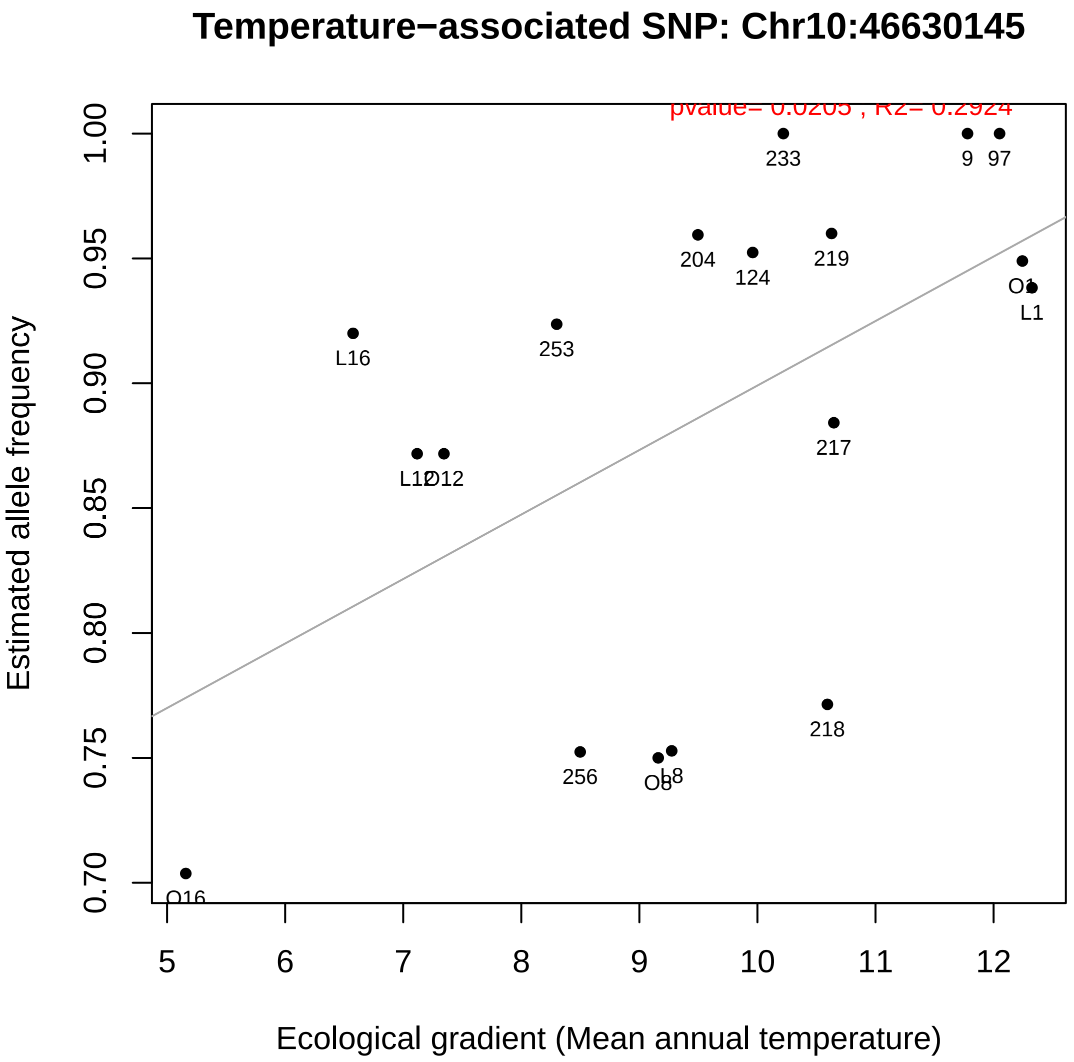
Temperature−associated SNP: Chr10:46630145
pvalue= 0.0205 , R2= 0.2924
Estimated allele frequency
Ecological gradient (Mean annual temperature)
1.00
0.95
0.90
0.85
0.80
0.75
0.70
5
6
7
8
9
10
11
12
233
9
97
204
124
219
O1
L1
L16
253
L12
O12
217
218
256
O8
L8
O16

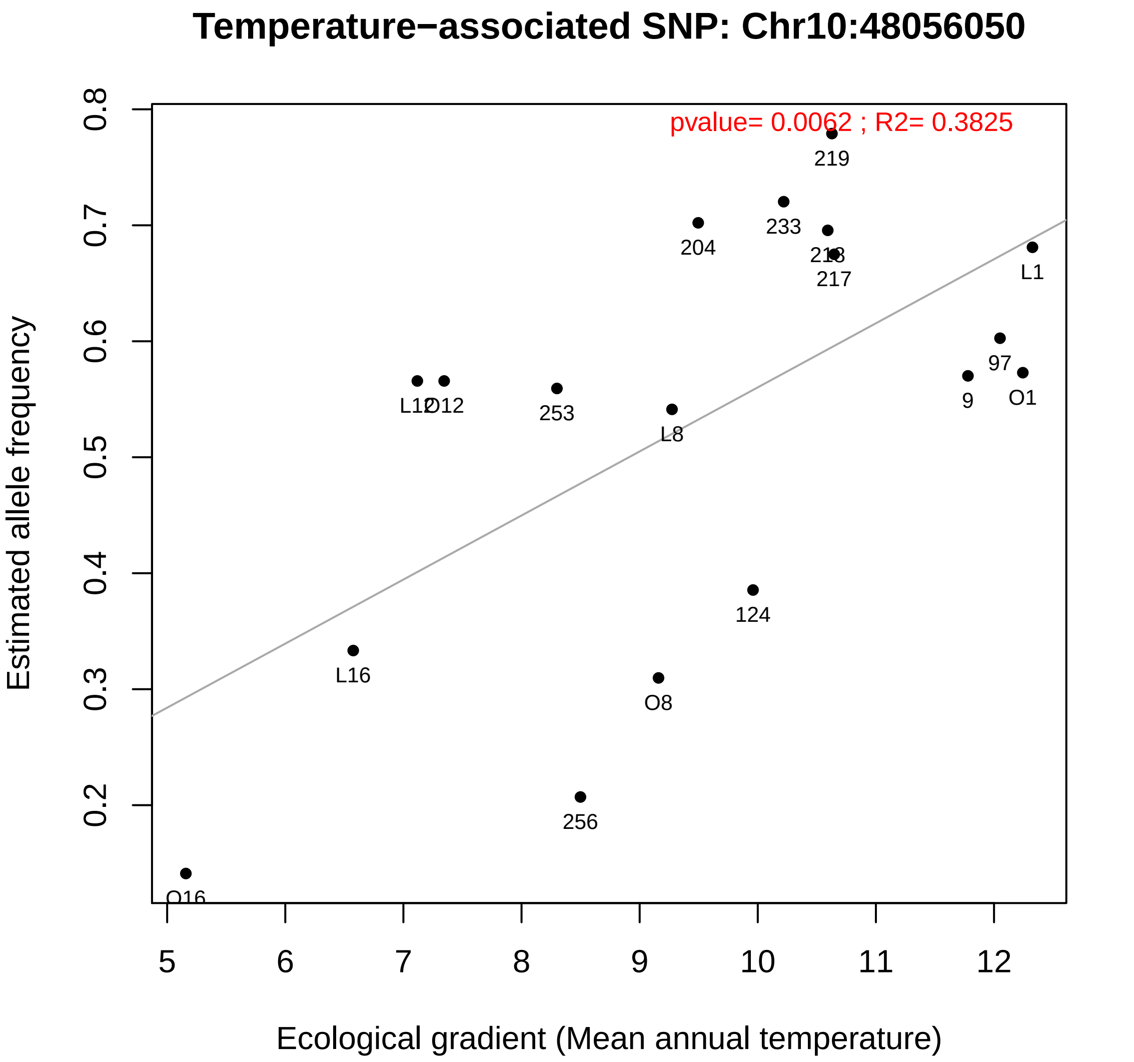

Temperature−associated SNP: Chr10:48056050
pvalue= 0.0062 ; R2= 0.3825
Estimated allele frequency
Ecological gradient (Mean annual temperature)
0.2
0.3
0.4
0.5
0.6
0.7
0.8
5
6
7
8
9
10
11
12
219
233
204
218
217
L1
97
9
O1
L12
O12
253
L8
124
L16
O8
256
O16

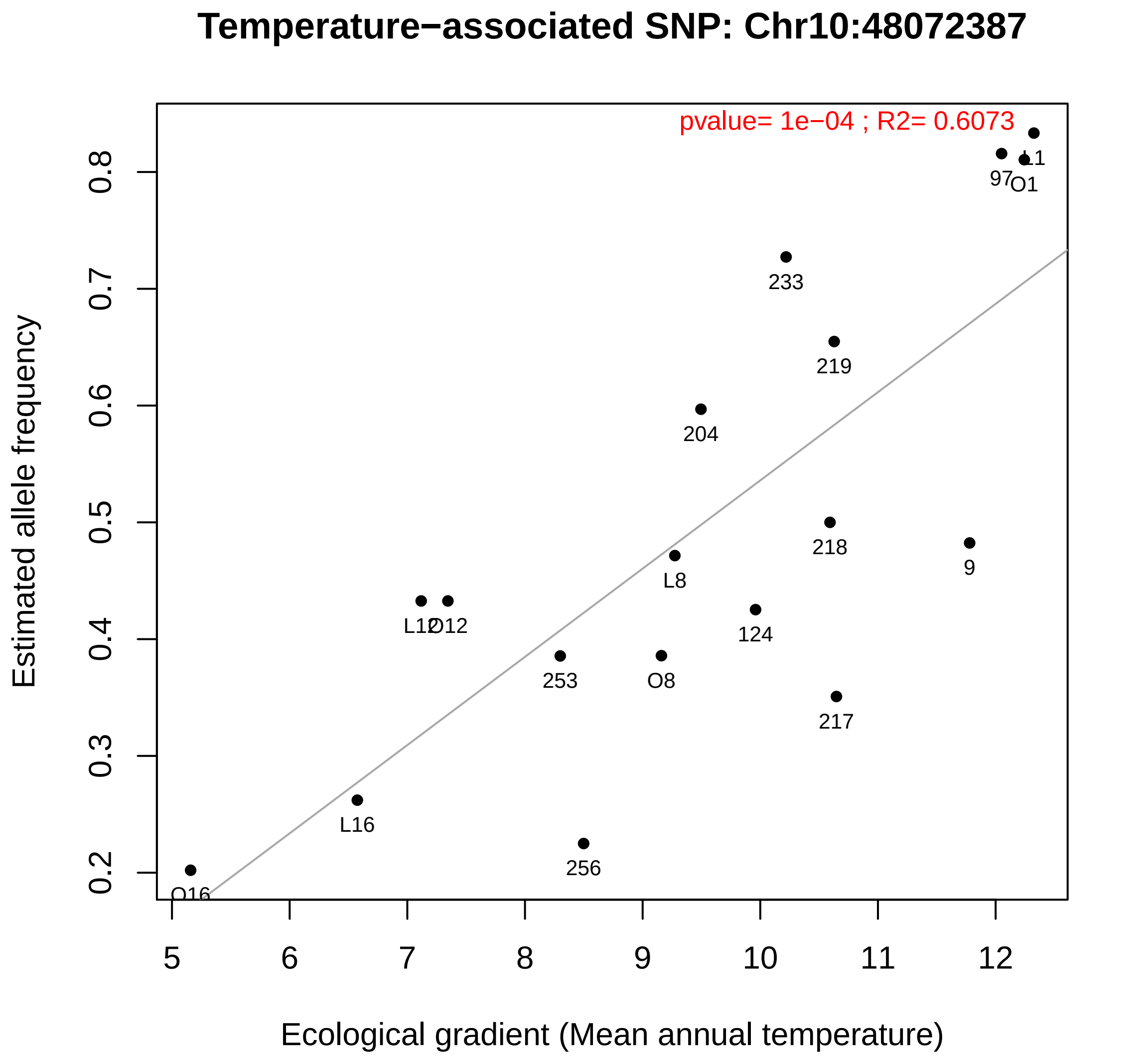

Temperature−associated SNP: Chr10:48072387
pvalue= 1e−04 ; R2= 0.6073
Estimated allele frequency
0.2
0.3
0.4
0.5
0.6
0.7
0.8
5
6
7
8
9
10
11
12
Ecological gradient (Mean annual temperature)
L1
97
O1
233
219
204
218
9
L8
L12
O12
124
253
O8
217
L16
256
O16

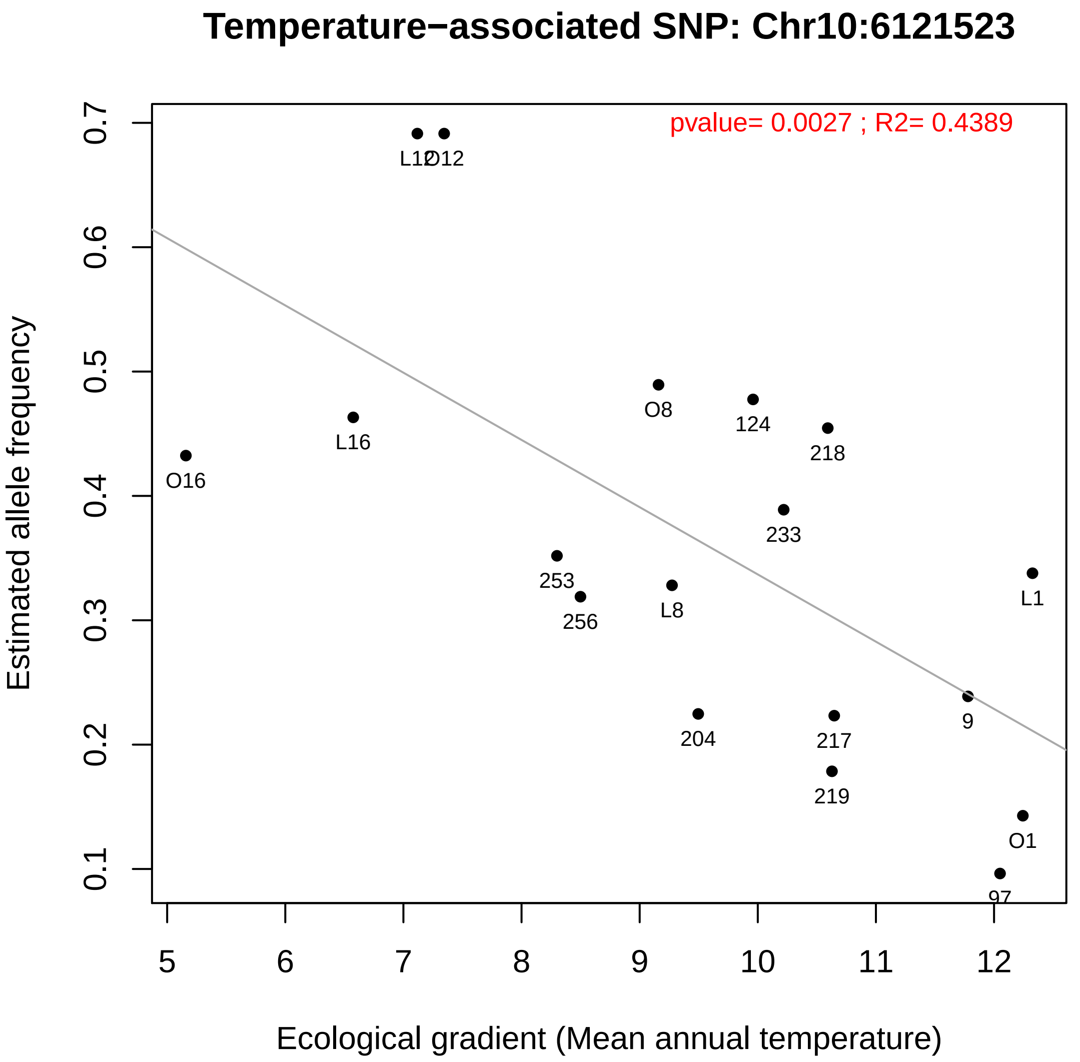

Temperature−associated SNP: Chr10:6121523
pvalue= 0.0027 ; R2= 0.4389
Estimated allele frequency
0.1
0.2
0.3
0.4
0.5
0.6
0.7
Ecological gradient (Mean annual temperature)
5
6
7
8
9
10
11
12
L12
O12
O16
L16
253
256
O8
L8
124
204
218
233
217
219
9
L1
O1
97

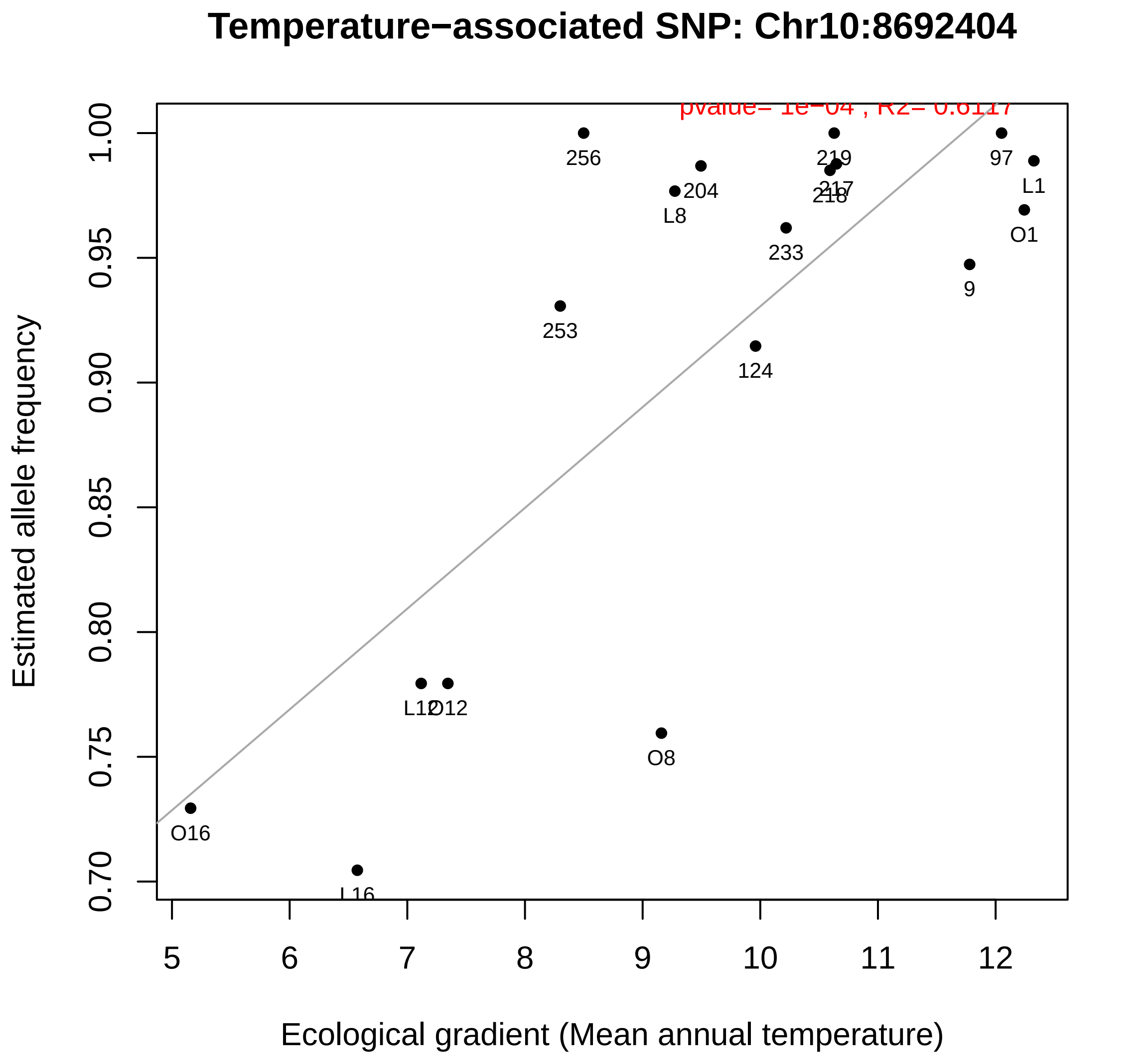
Temperature−associated SNP: Chr10:8692404
pvalue= 1e−04 , R2= 0.6117
Estimated allele frequency
Ecological gradient (Mean annual temperature)
0.70
0.75
0.80
0.85
0.90
0.95
1.00
5
6
7
8
9
10
11
12
256
219
97
204
L1
L8
217
218
O1
233
9
253
124
L12
O12
O8
O16
L16

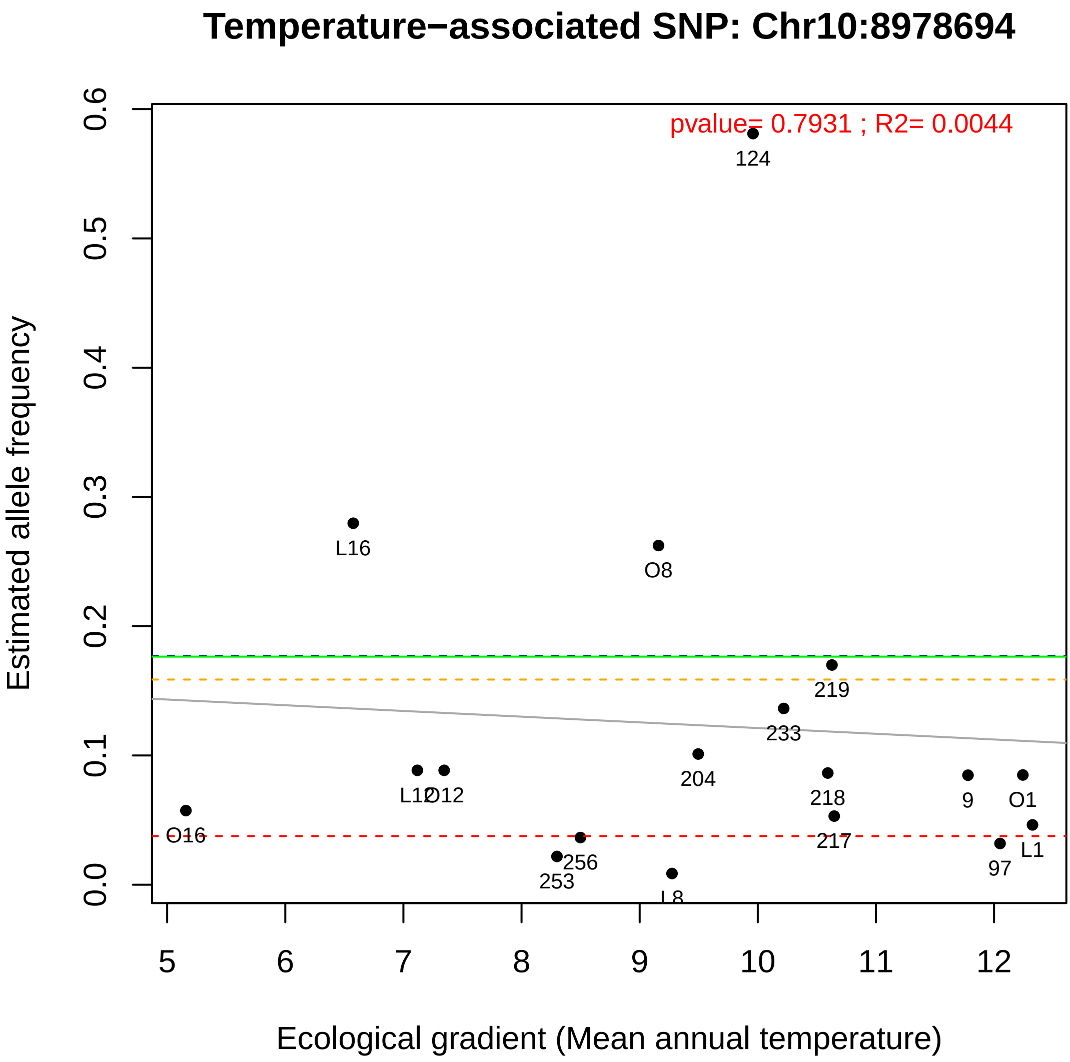

Temperature−associated SNP: Chr10:8978694
pvalue= 0.7931 ; R2= 0.0044
Estimated allele frequency
Ecological gradient (Mean annual temperature)
0.0
0.1
0.2
0.3
0.4
0.5
0.6
5
6
7
8
9
10
11
12
124
L16
O8
219
233
204
L12
O12
218
9
O1
O16
217
L1
256
253
97
L8

### Temperature−associated SNP: Chr10:8978809

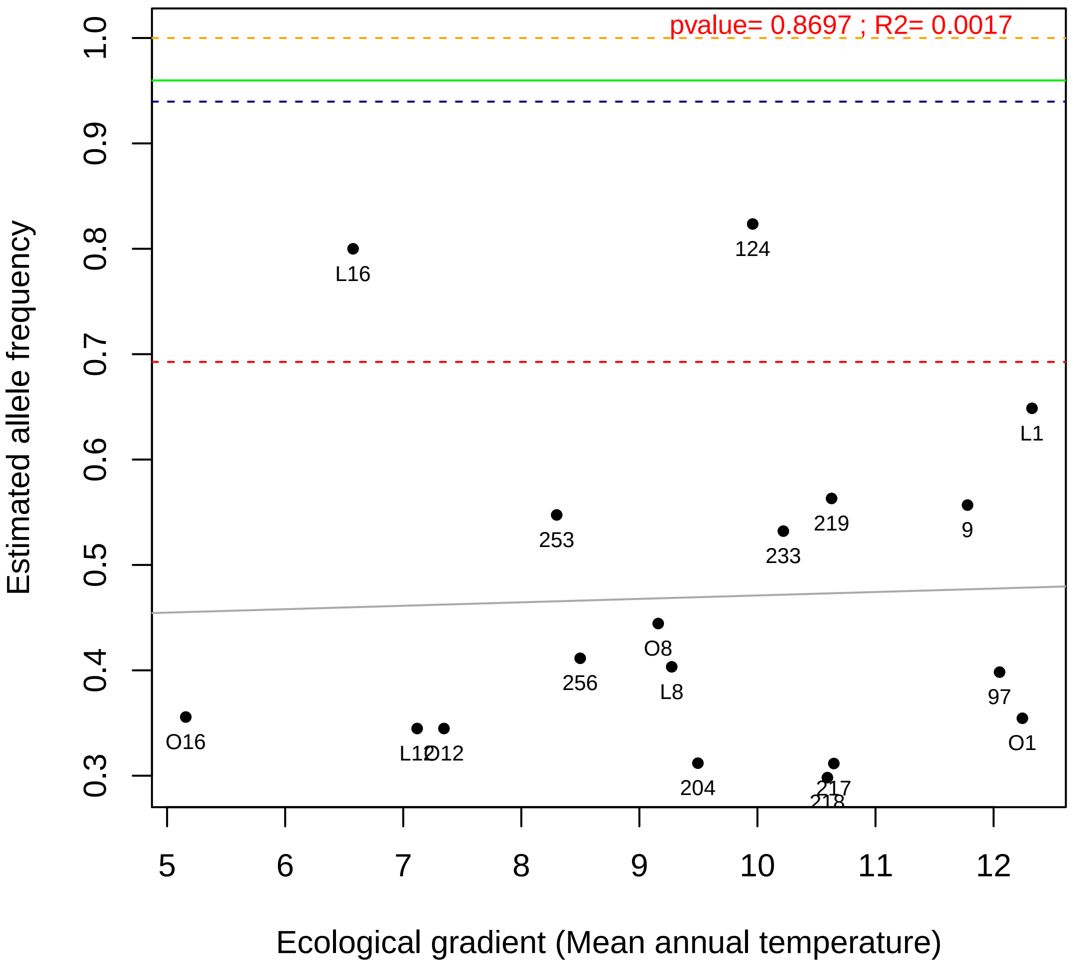

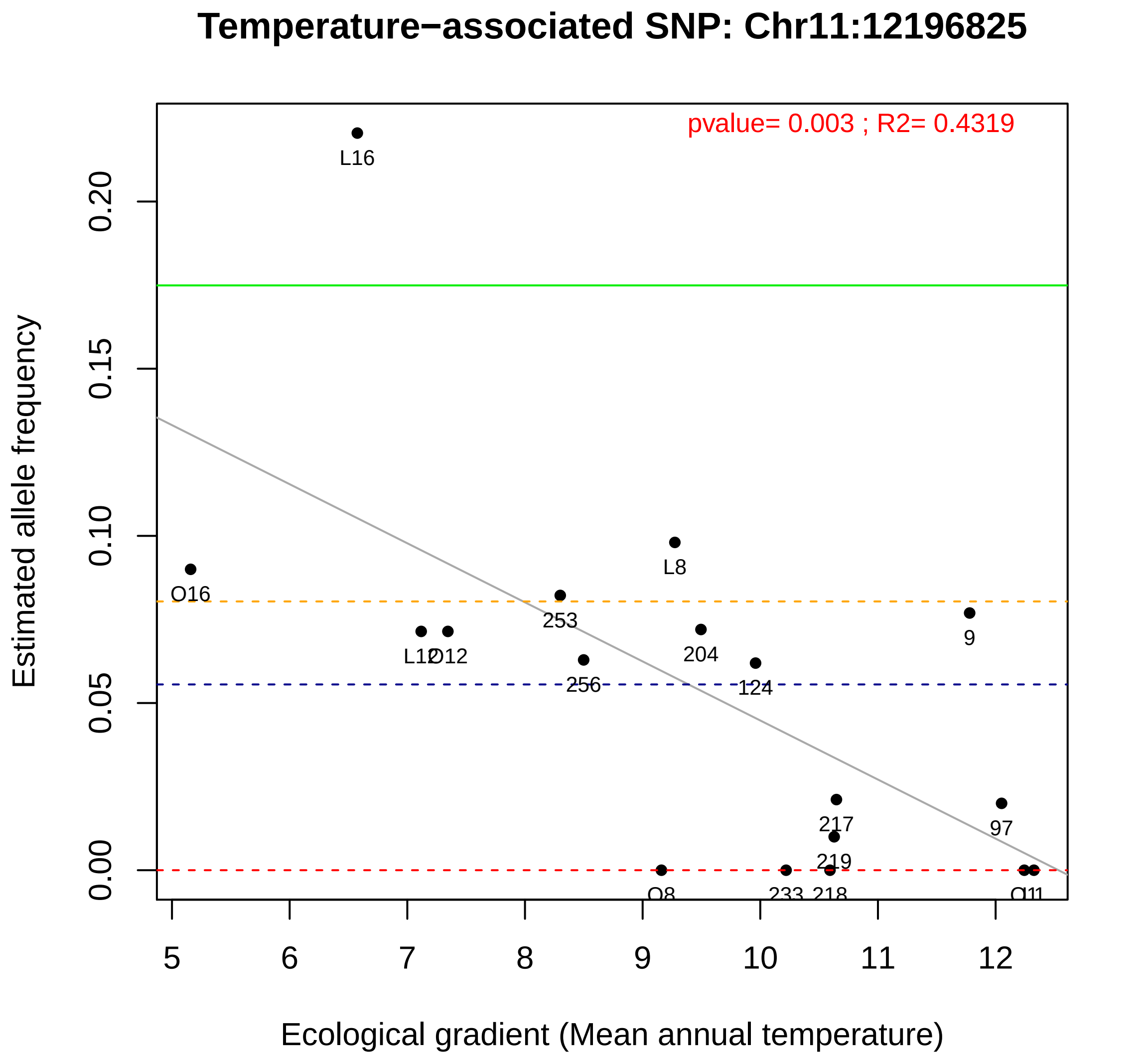
Temperature−associated SNP: Chr11:12196825
pvalue= 0.003 ; R2= 0.4319
Estimated allele frequency
0.00
0.05
0.10
0.15
0.20
5
6
7
8
9
10
11
12
Ecological gradient (Mean annual temperature)
L16
O16
L12
O12
253
256
L8
204
124
9
217
219
97
O8
233
218

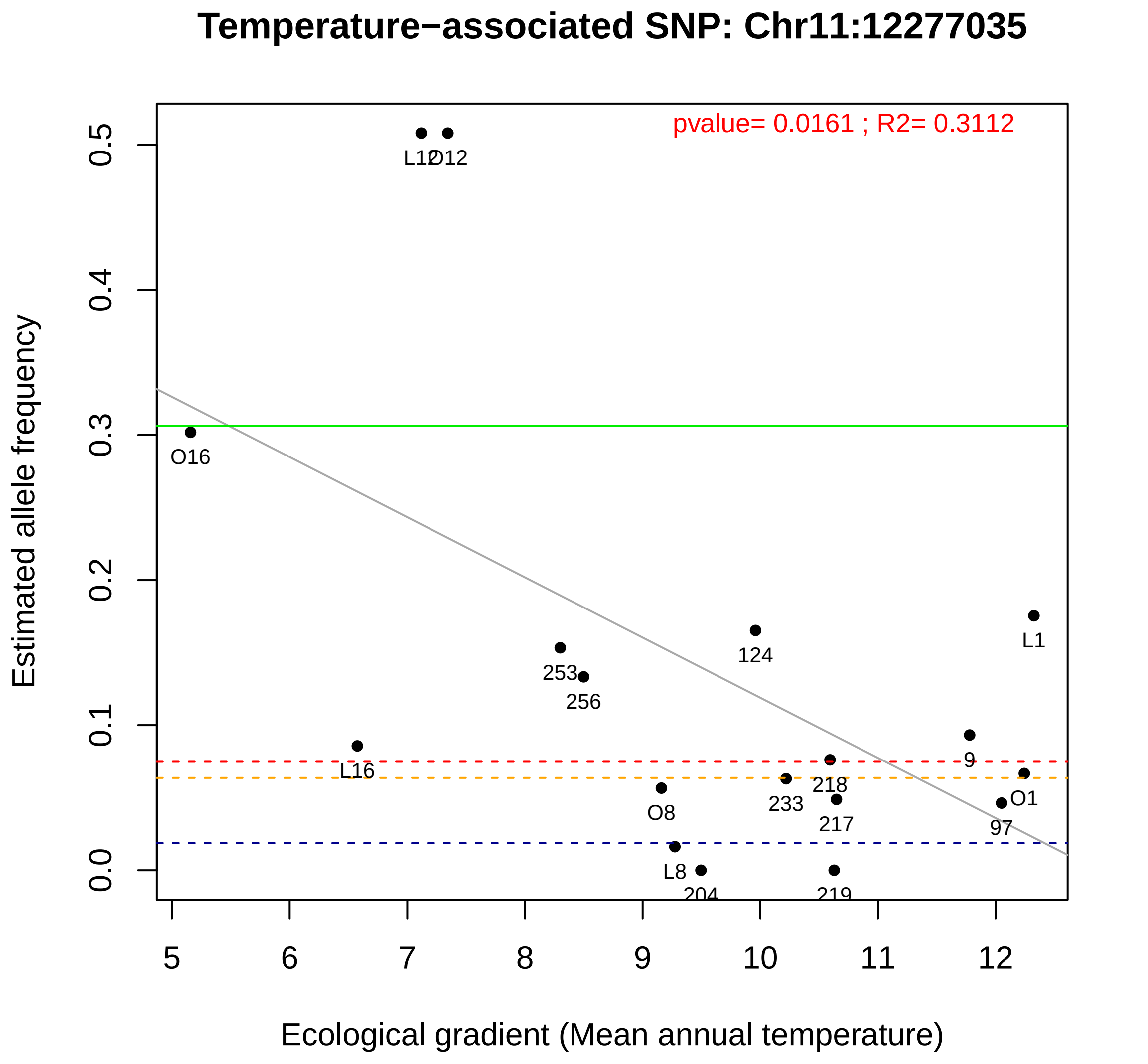

Temperature−associated SNP: Chr11:12277035
pvalue= 0.0161 ; R2= 0.3112
Estimated allele frequency
Ecological gradient (Mean annual temperature)
0.0
0.1
0.2
0.3
0.4
0.5
5
6
7
8
9
10
11
12
L12
O12
O16
253
256
124
L1
L16
9
218
233
O1
O8
217
97
L8
204
219

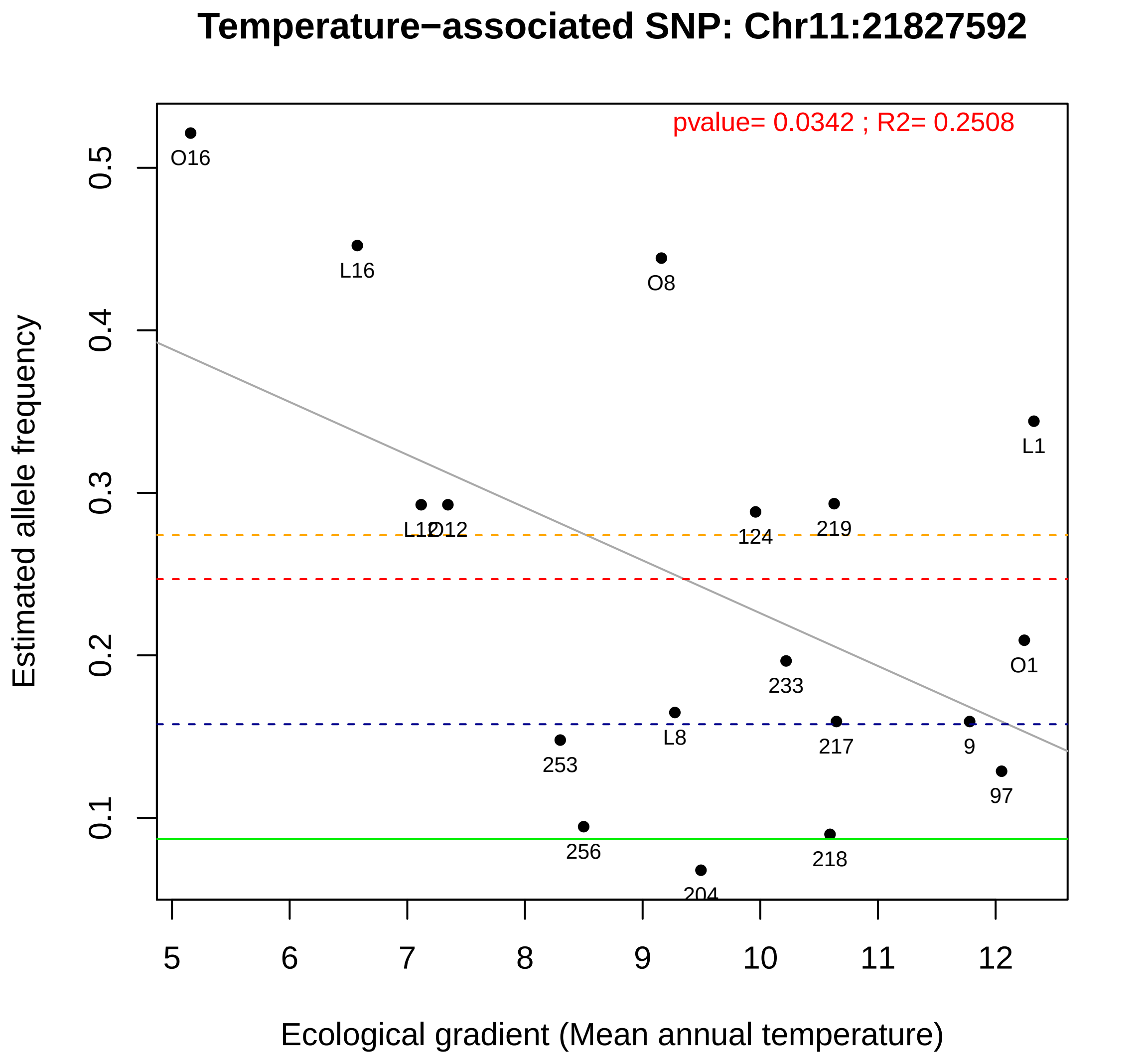
Temperature−associated SNP: Chr11:21827592
pvalue= 0.0342 ; R2= 0.2508
Estimated allele frequency
0.1
0.2
0.3
0.4
0.5
5
6
7
8
9
10
11
12
Ecological gradient (Mean annual temperature)
O16
L16
O8
L1
L12
O12
124
219
O1
233
L8
217
9
253
97
256
218
204

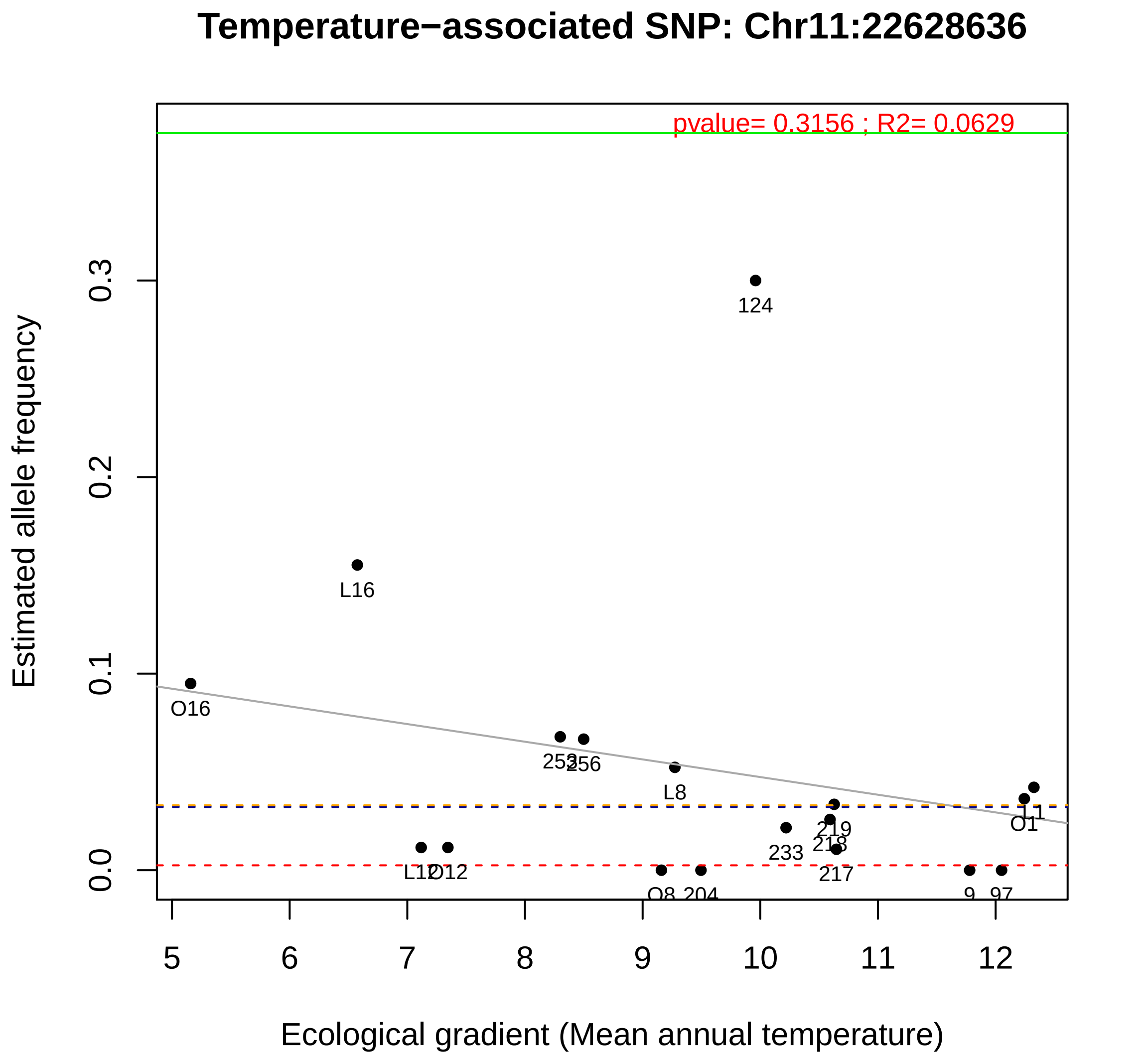

Temperature−associated SNP: Chr11:22628636
pvalue= 0.3156 ; R2= 0.0629
Estimated allele frequency
0.0
0.1
0.2
0.3
Ecological gradient (Mean annual temperature)
5
6
7
8
9
10
11
12
124
L16
O16
256
L8
219
218
233
217
O1
L1
O8
204
9
97

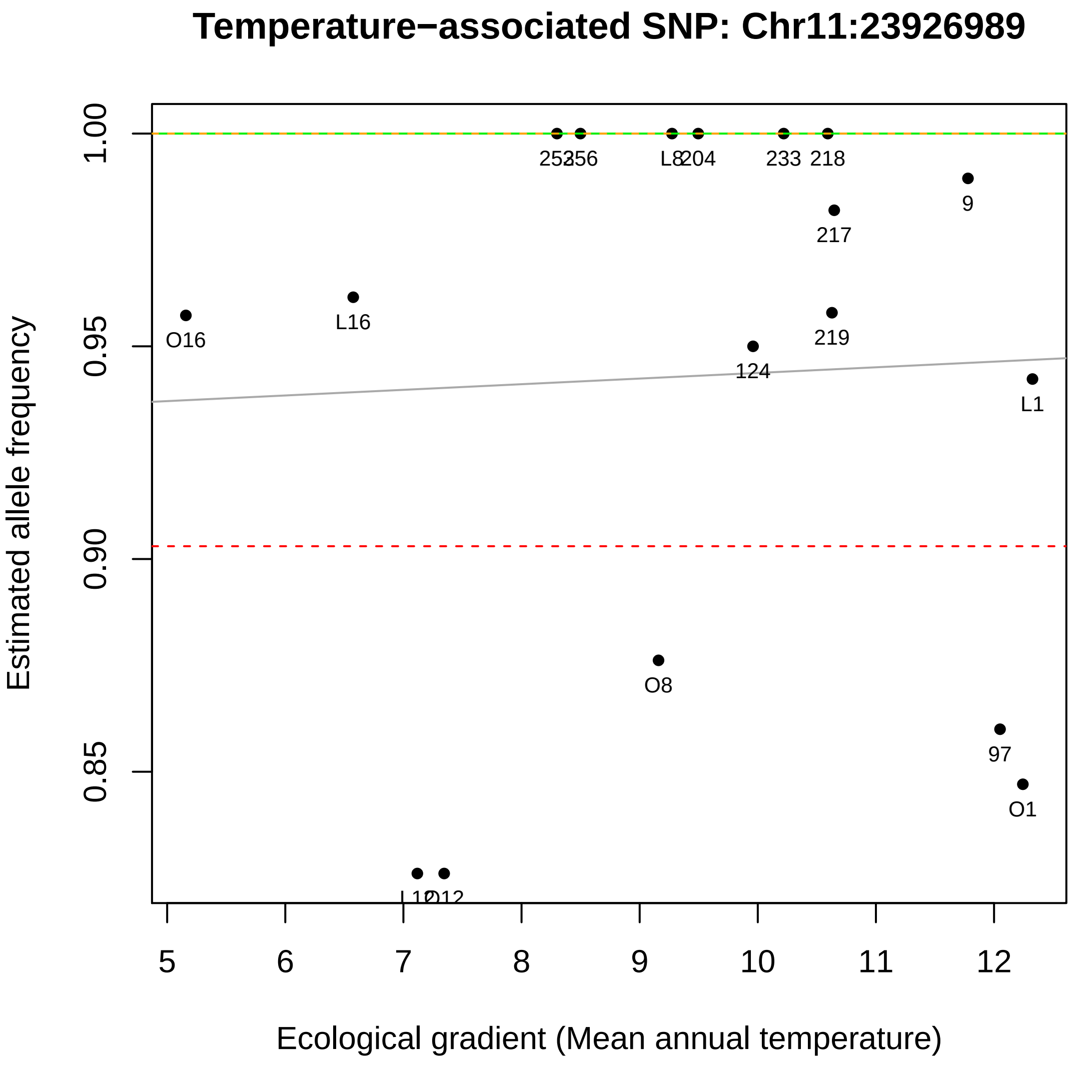
Temperature−associated SNP: Chr11:23926989
Estimated allele frequency
Ecological gradient (Mean annual temperature)
1.00
0.95
0.90
0.85
5
6
7
8
9
10
11
12
O16
L16
L12
O12
253
256
L8
204
O8
124
233
218
217
219
9
97
O1
L1

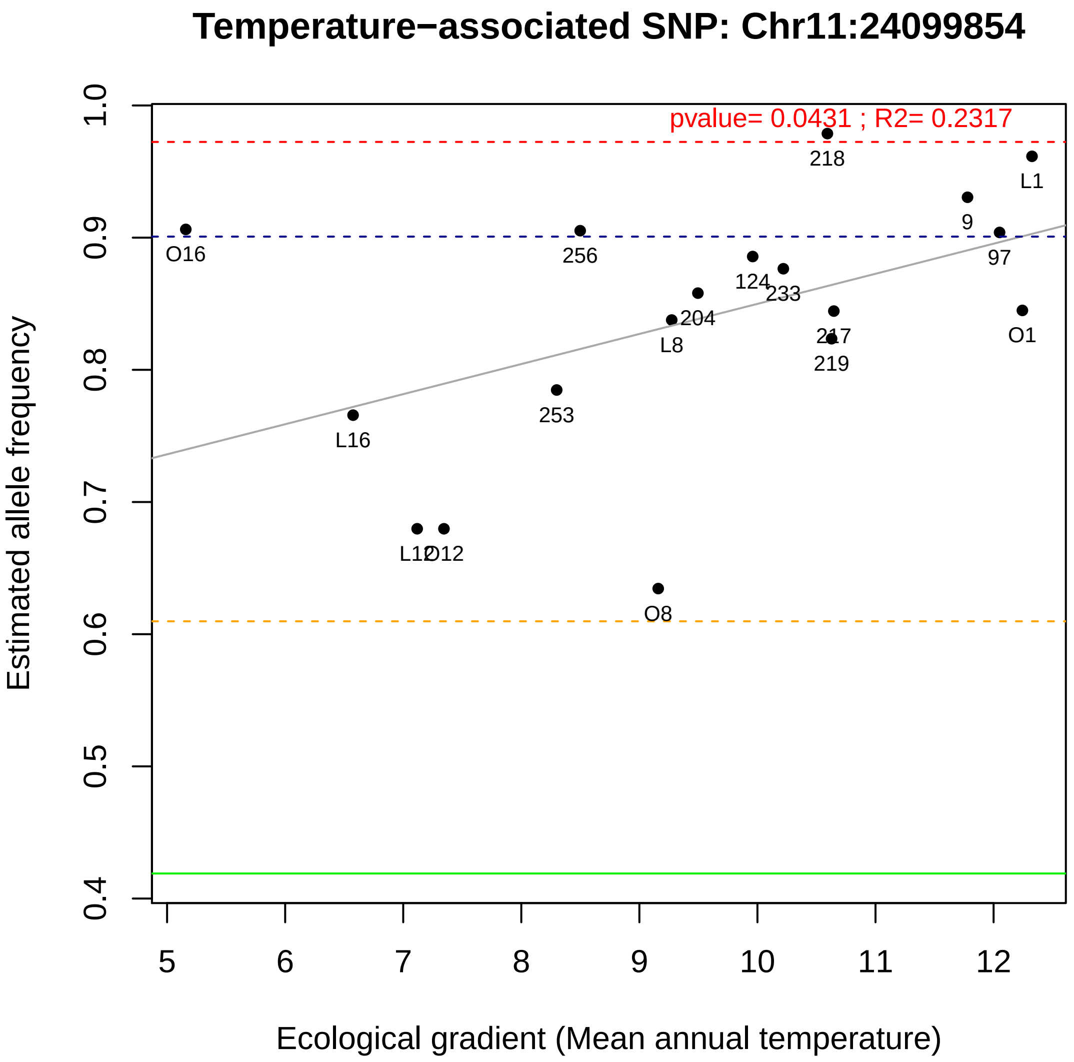
Temperature−associated SNP: Chr11:24099854
pvalue= 0.0431 ; R2= 0.2317
Estimated allele frequency
Ecological gradient (Mean annual temperature)
1.0
0.9
0.8
0.7
0.6
0.5
0.4
5
6
7
8
9
10
11
12
218
L1
9
97
O16
256
124
233
204
L8
217
219
O1
253
L16
L12
O12
O8

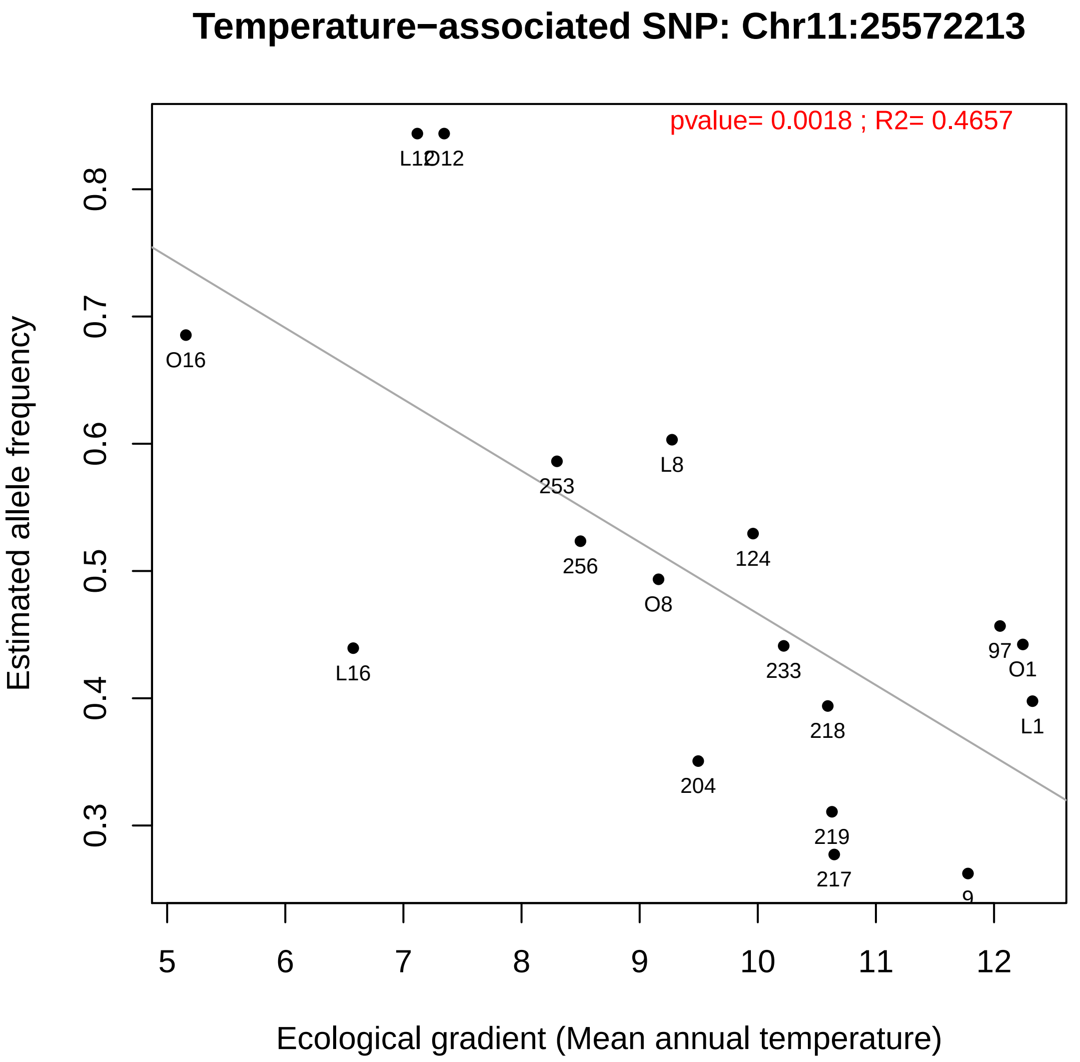
Temperature−associated SNP: Chr11:25572213
pvalue= 0.0018 ; R2= 0.4657
Estimated allele frequency
0.3
0.4
0.5
0.6
0.7
0.8
5
6
7
8
9
10
11
12
Ecological gradient (Mean annual temperature)
L12
O12
O16
253
L8
256
124
O8
L16
233
97
O1
218
L1
204
219
217
9

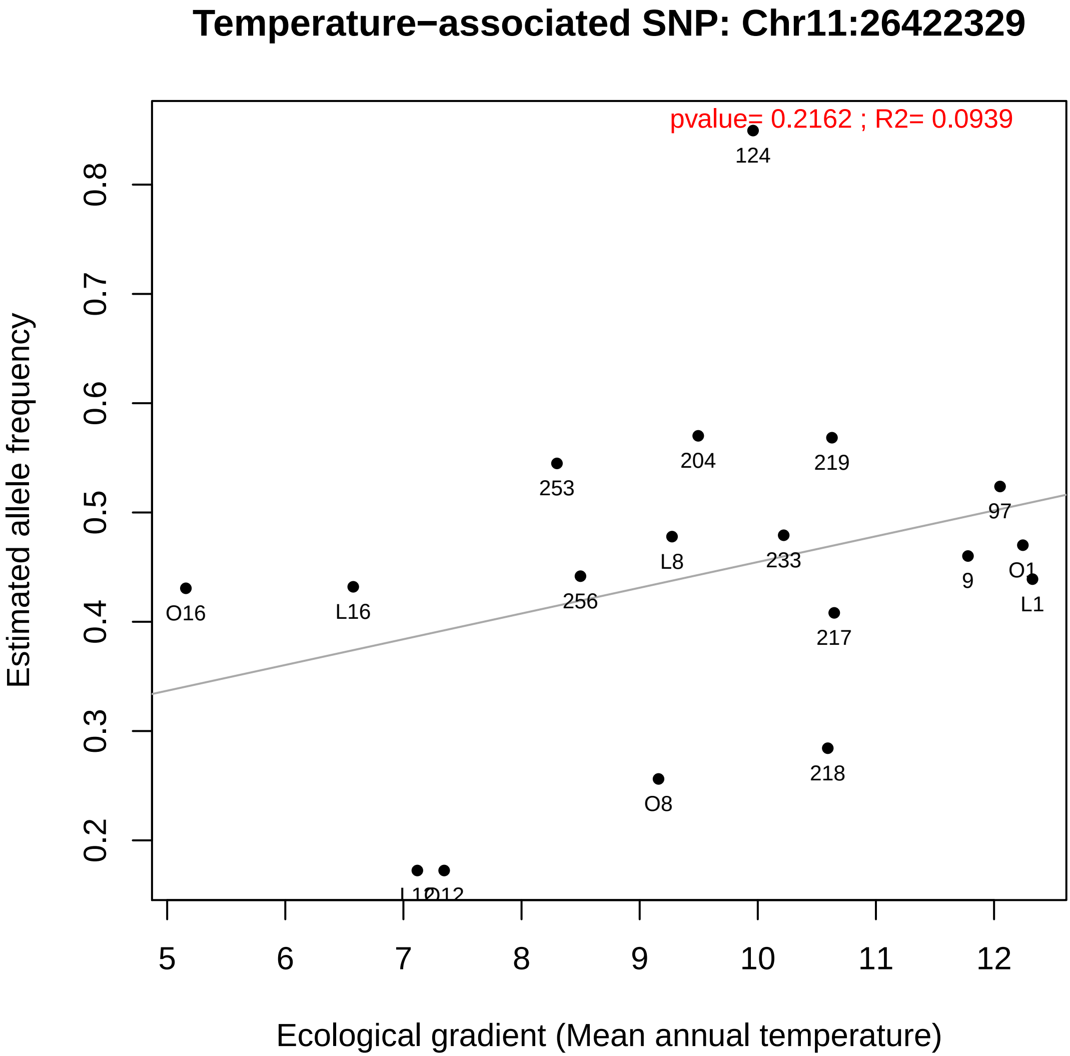
Temperature−associated SNP: Chr11:26422329
pvalue= 0.2162 ; R2= 0.0939
Estimated allele frequency
0.2
0.3
0.4
0.5
0.6
0.7
0.8
Ecological gradient (Mean annual temperature)
5
6
7
8
9
10
11
12
124
204
219
253
97
L8
233
256
9
O1
L1
O16
L16
217
218
O8
O12

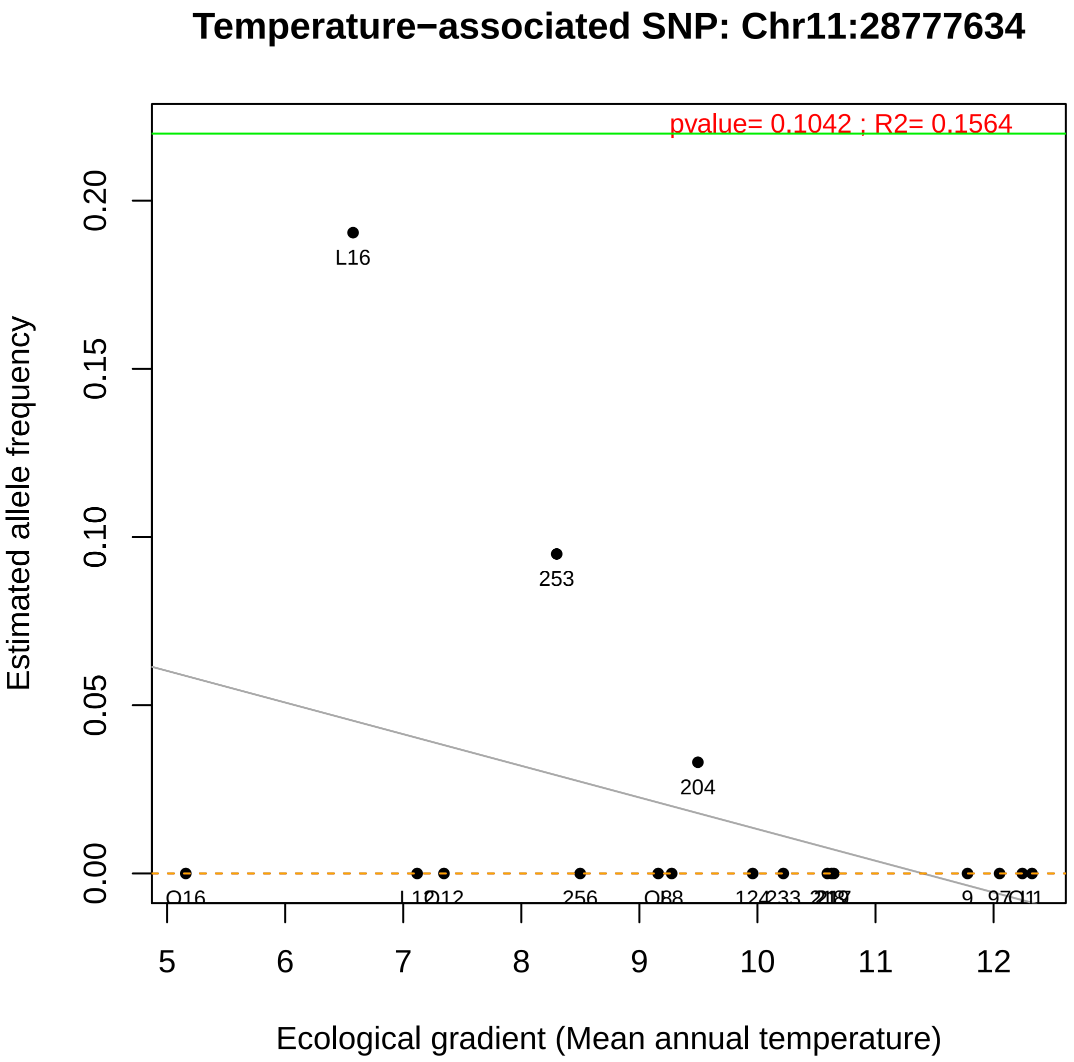

Temperature−associated SNP: Chr11:28777634
pvalue= 0.1042 ; R2= 0.1564
Estimated allele frequency
0.00
0.05
0.10
0.15
0.20
L16
253
204
O16
256
9
5
6
7
8
9
10
11
12
Ecological gradient (Mean annual temperature)

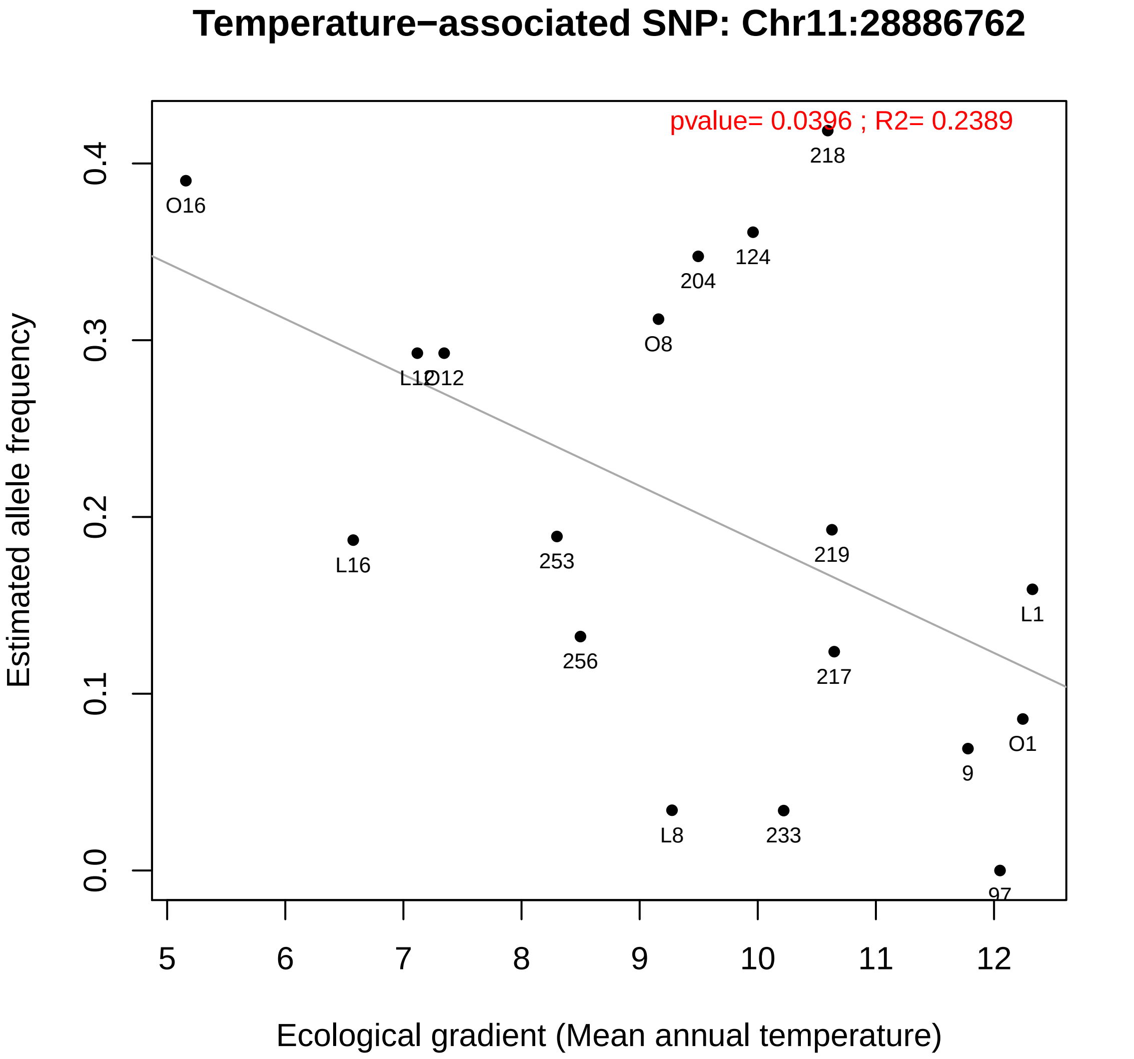
Temperature−associated SNP: Chr11:28886762
pvalue= 0.0396 ; R2= 0.2389
Estimated allele frequency
0.0
0.1
0.2
0.3
0.4
Ecological gradient (Mean annual temperature)
5
6
7
8
9
10
11
12
O16
218
124
204
O8
L12
O12
L16
253
219
L1
256
217
O1
9
L8
233
97

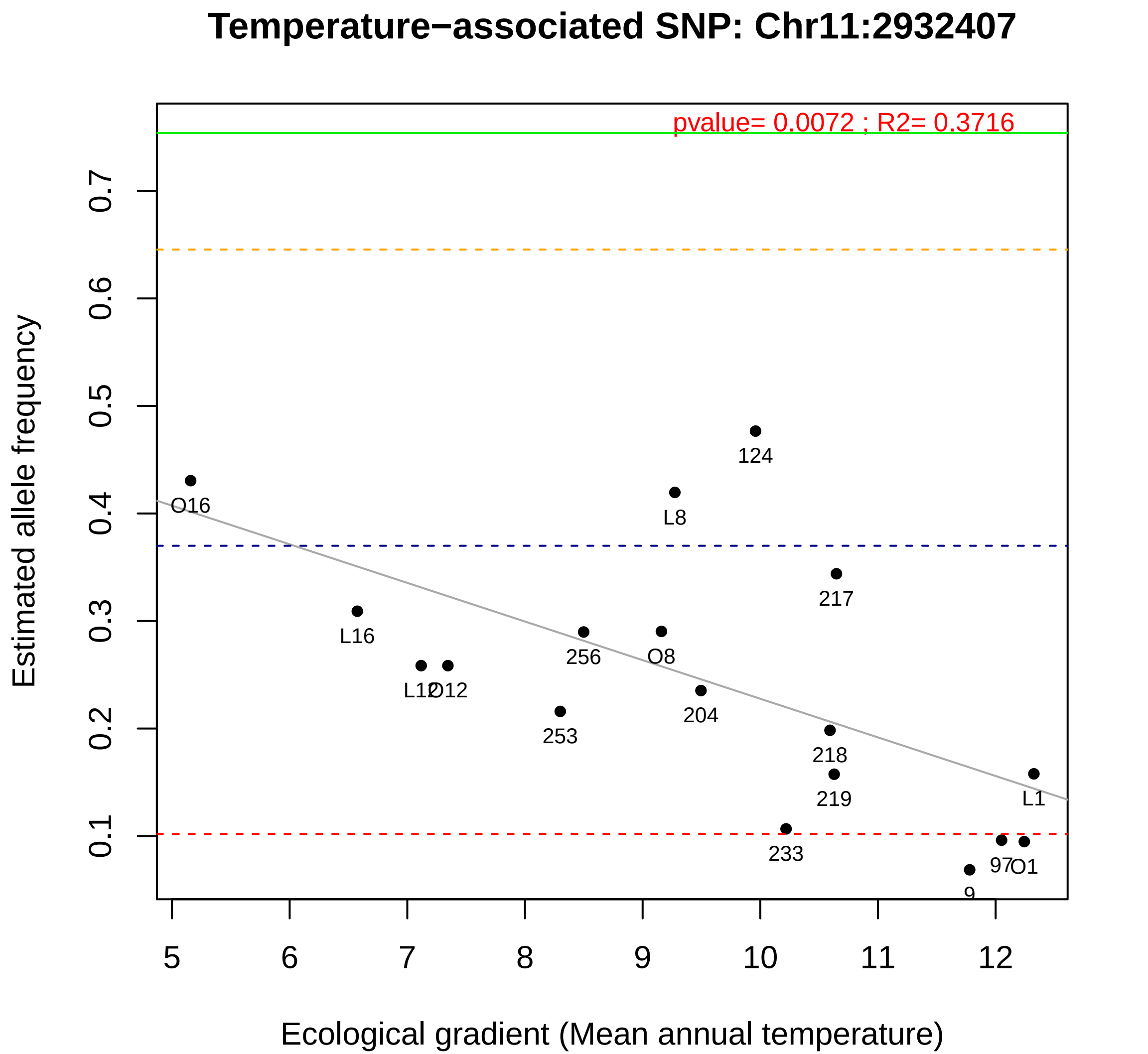

Temperature−associated SNP: Chr11:2932407
pvalue= 0.0072 ; R2= 0.3716
Estimated allele frequency
Ecological gradient (Mean annual temperature)
0.1
0.2
0.3
0.4
0.5
0.6
0.7
5
6
7
8
9
10
11
12
O16
L16
L12
O12
253
256
O8
L8
204
124
233
218
219
217
9
97
O1
L1

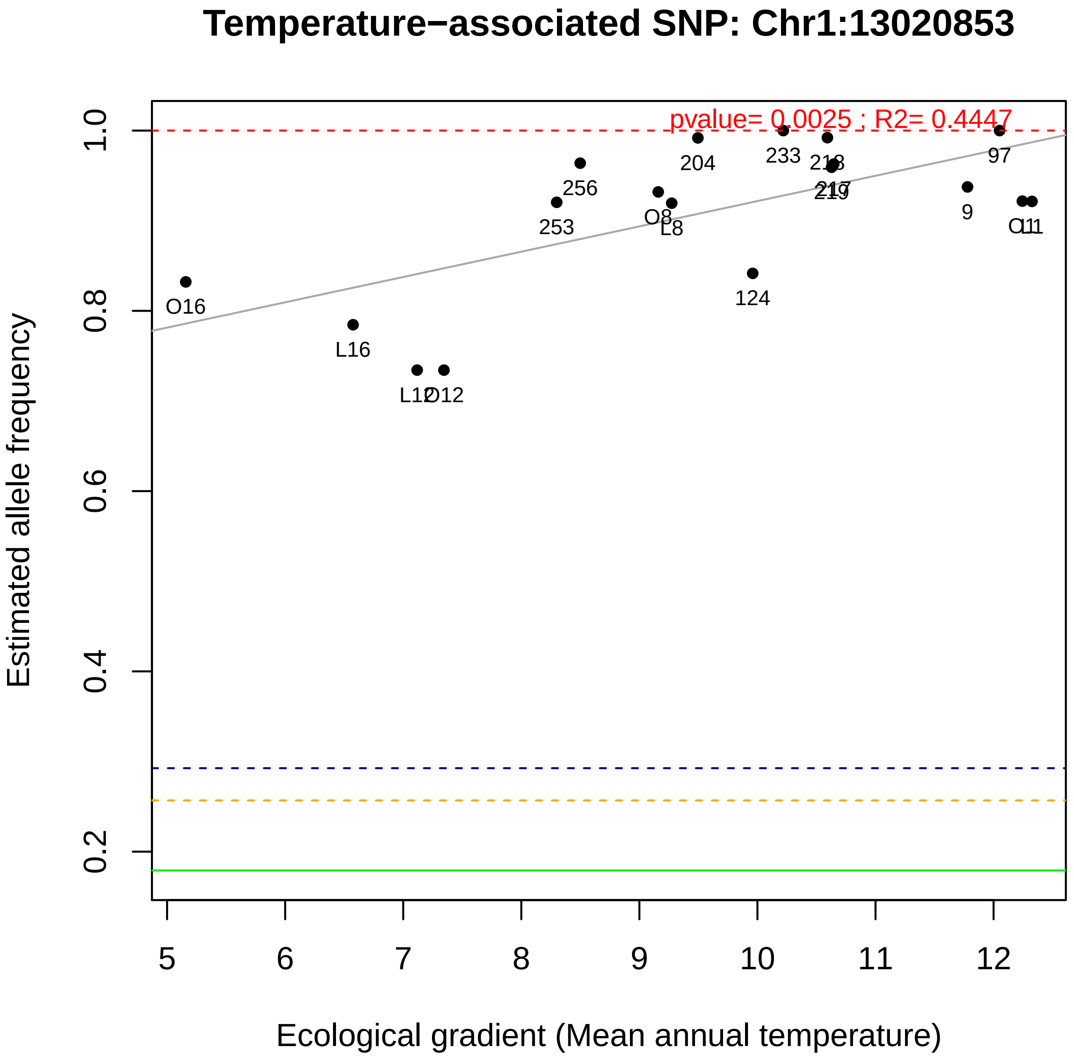

Temperature−associated SNP: Chr1:13020853
pvalue= 0.0025 ; R2= 0.4447
Estimated allele frequency
0.2
0.4
0.6
0.8
1.0
Ecological gradient (Mean annual temperature)
5
6
7
8
9
10
11
12
O16
L16
L12
O12
253
256
O8
L8
204
124
233
9
97

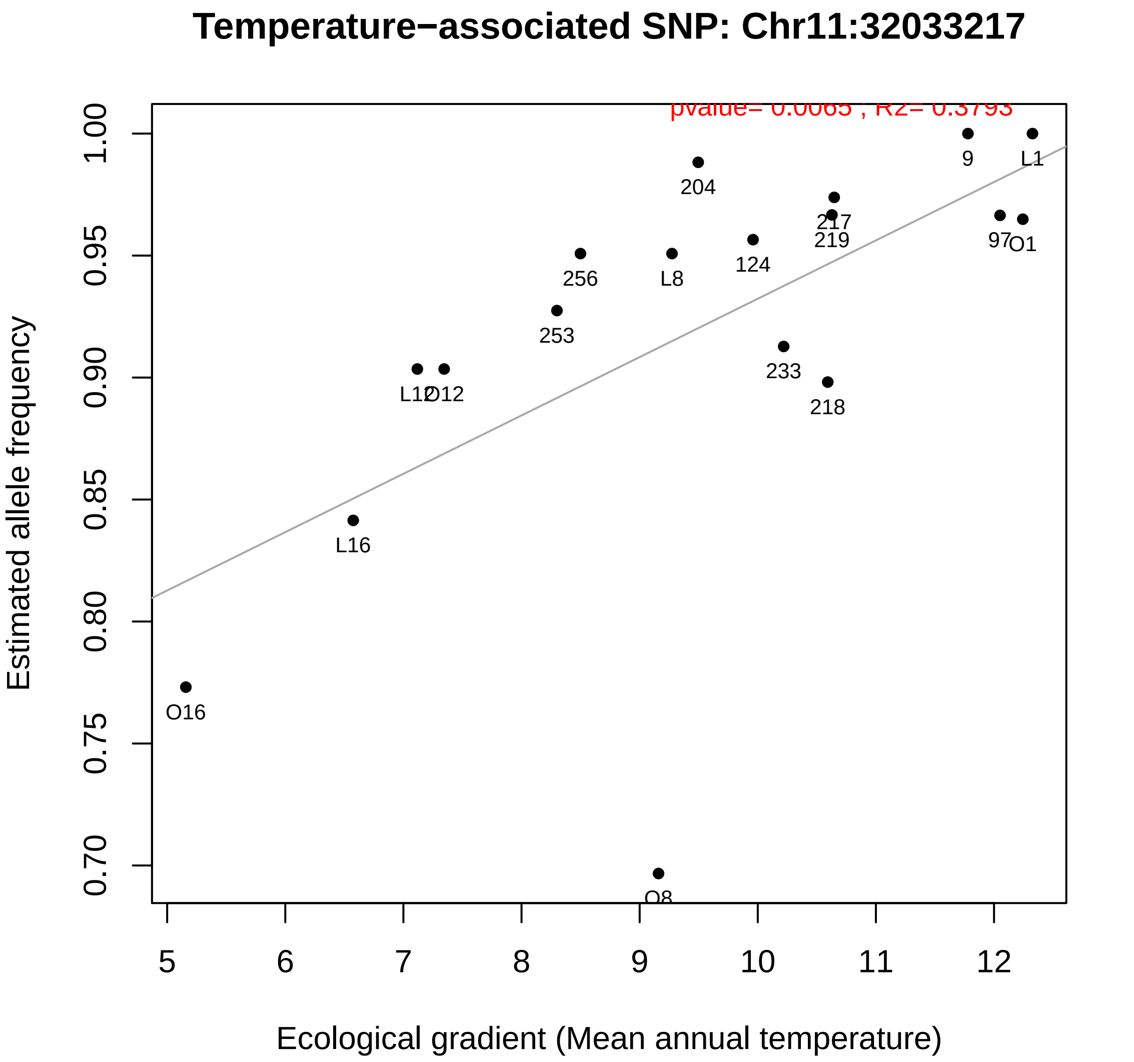

Temperature−associated SNP: Chr11:32033217
pvalue= 0.0065 ; R2= 0.3793
Estimated allele frequency
Ecological gradient (Mean annual temperature)
1.00
0.95
0.90
0.85
0.80
0.75
0.70
5
6
7
8
9
10
11
12
9
L1
204
217
219
97
O1
256
L8
124
253
233
218
L12
O12
L16
O16
O8

**Temperature−associated SNP: Chr11:34740463**

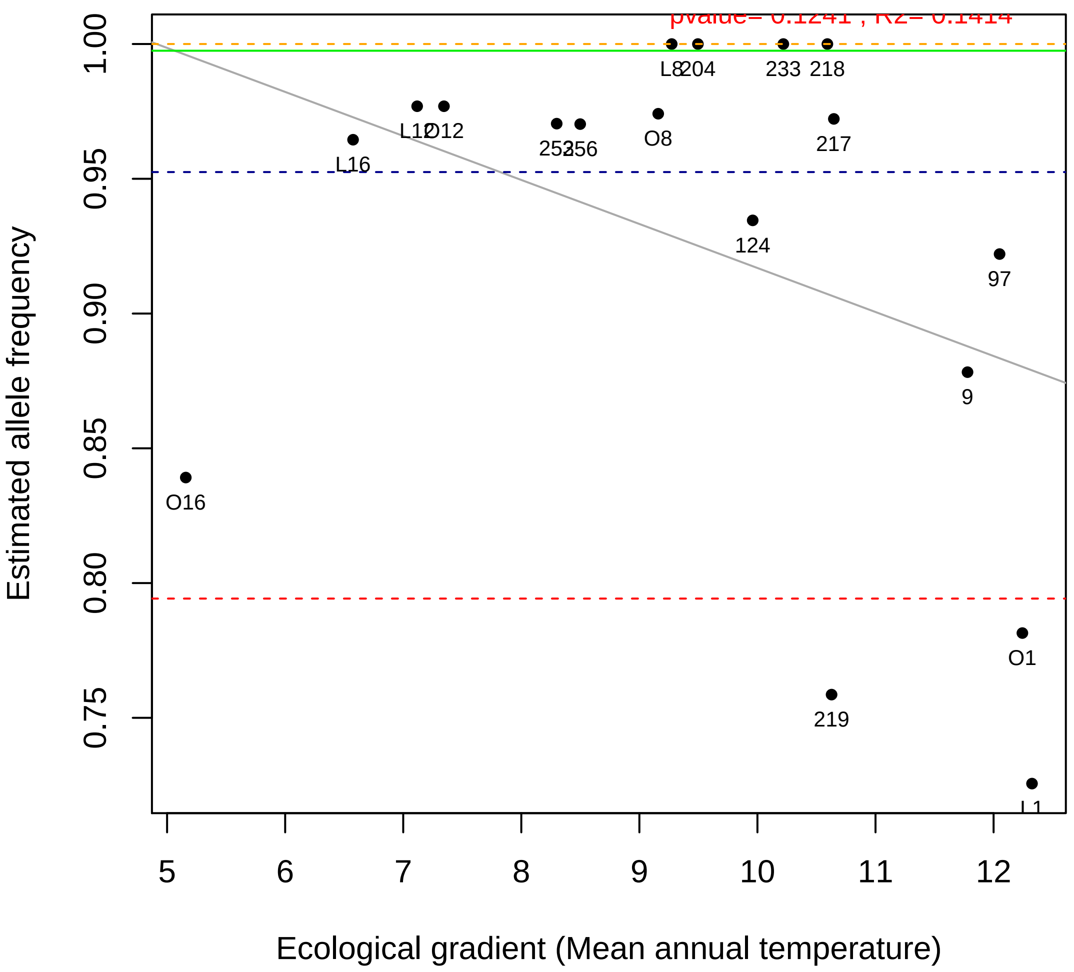

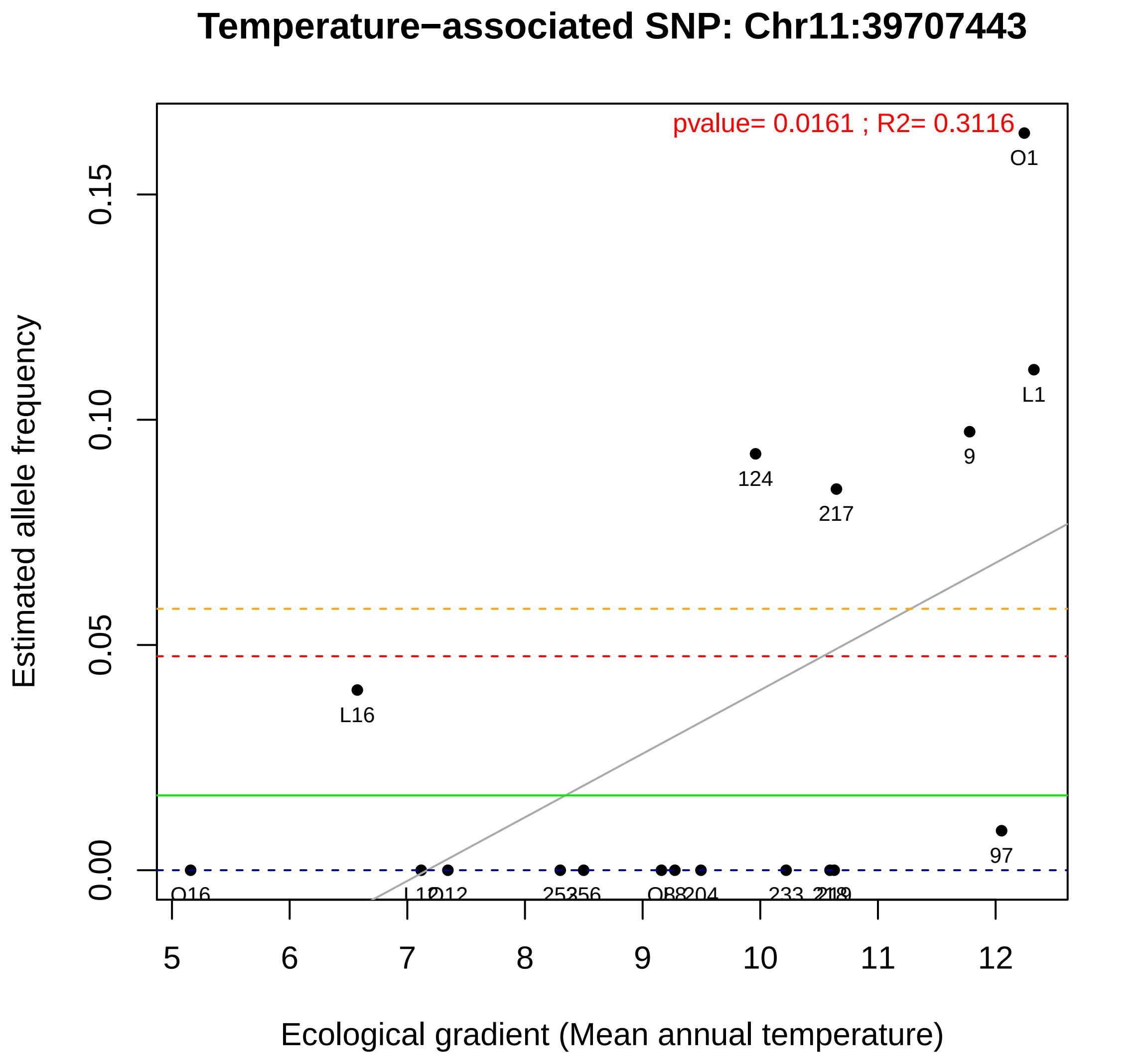

Temperature−associated SNP: Chr11:39707443
pvalue= 0.0161 ; R2= 0.3116
Estimated allele frequency
0.00
0.05
0.10
0.15
Ecological gradient (Mean annual temperature)
5
6
7
8
9
10
11
12
O1
L1
9
124
217
L16
97
O16
233
204

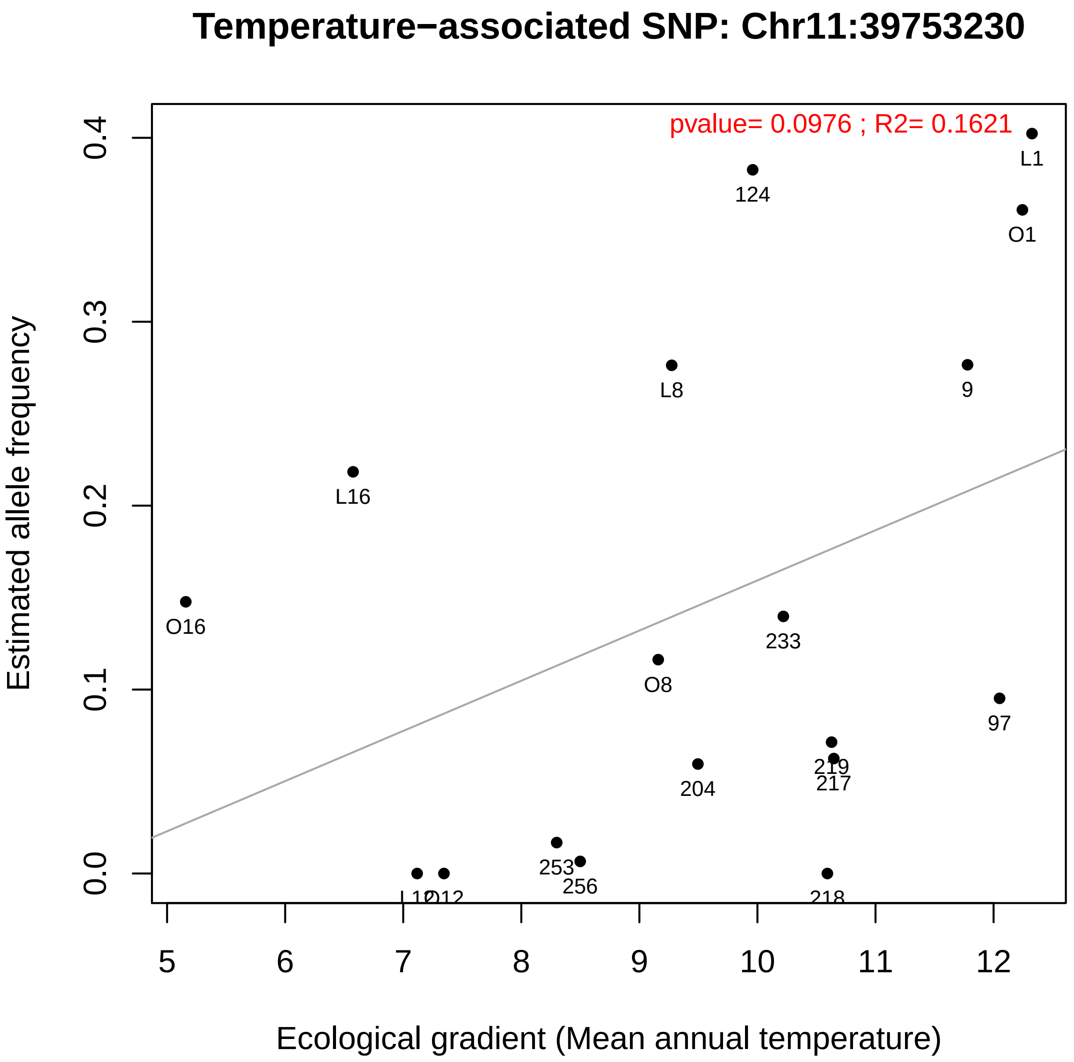
Temperature−associated SNP: Chr11:39753230
pvalue= 0.0976 ; R2= 0.1621
Estimated allele frequency
0.0
0.1
0.2
0.3
0.4
Ecological gradient (Mean annual temperature)
5
6
7
8
9
10
11
12
L1
124
O1
L8
9
L16
O16
233
O8
97
219
217
204
253
256
218

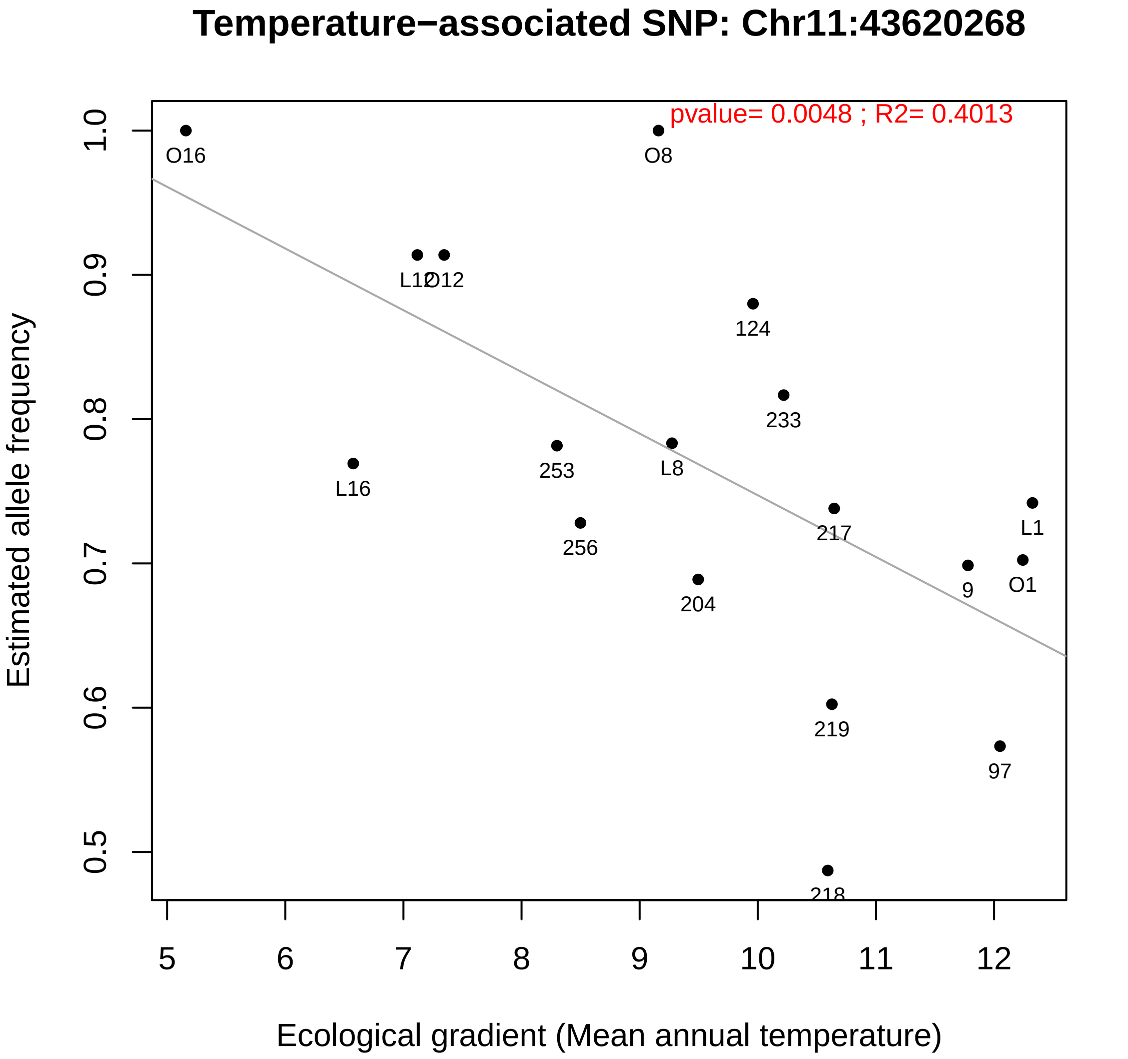
Temperature−associated SNP: Chr11:43620268
pvalue= 0.0048 ; R2= 0.4013
Estimated allele frequency
Ecological gradient (Mean annual temperature)
1.0
0.9
0.8
0.7
0.6
0.5
5
6
7
8
9
10
11
12
O16
O8
L12
O12
124
233
L16
253
L8
256
217
L1
204
9
O1
219
97
218

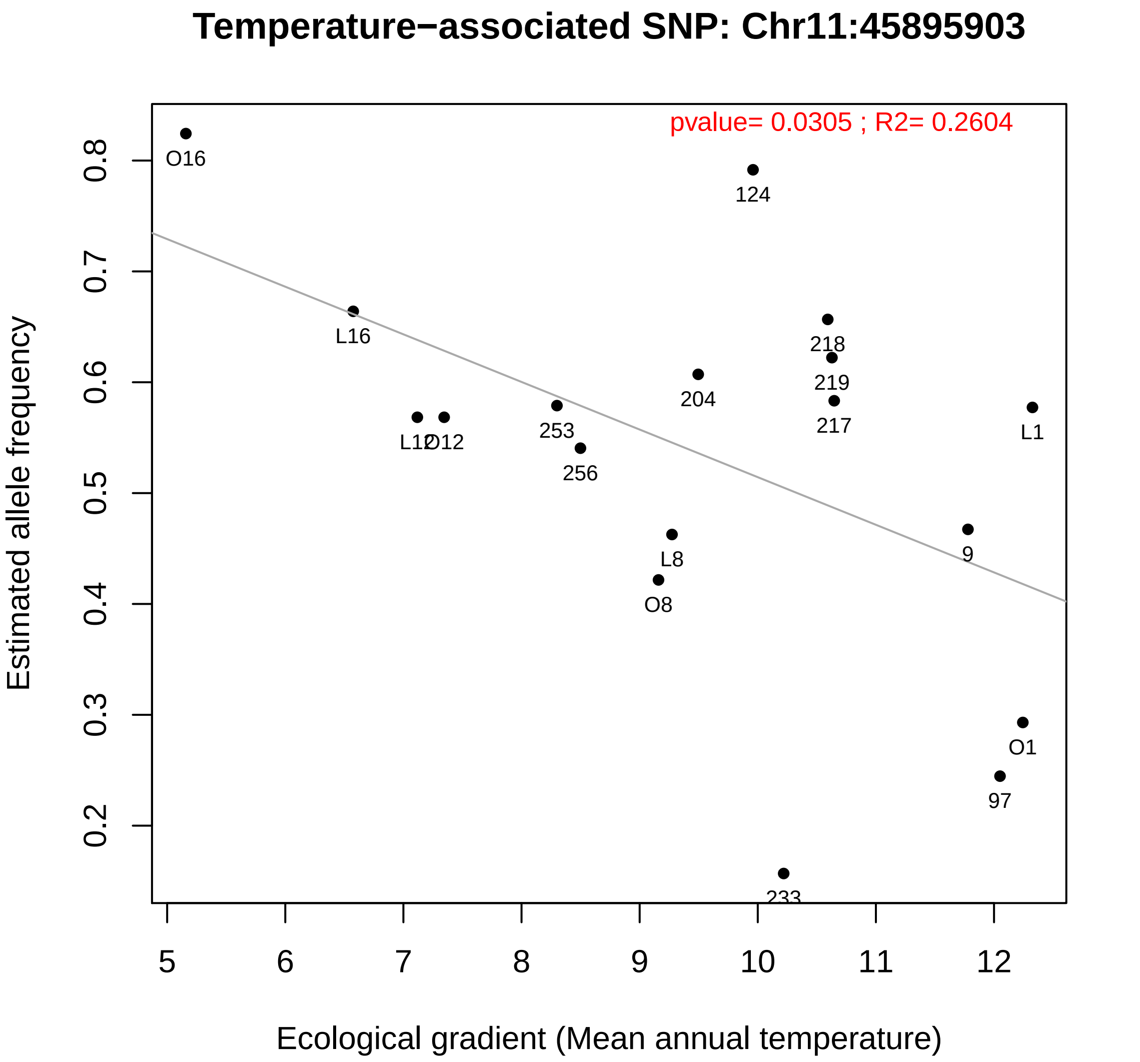
Temperature−associated SNP: Chr11:45895903
pvalue= 0.0305 ; R2= 0.2604
O16
124
L16
218
219
204
217
L1
L12
O12
253
256
L8
9
O8
O1
97
233
0.2
0.3
0.4
0.5
0.6
0.7
0.8
5
6
7
8
9
10
11
12
Estimated allele frequency
Ecological gradient (Mean annual temperature)

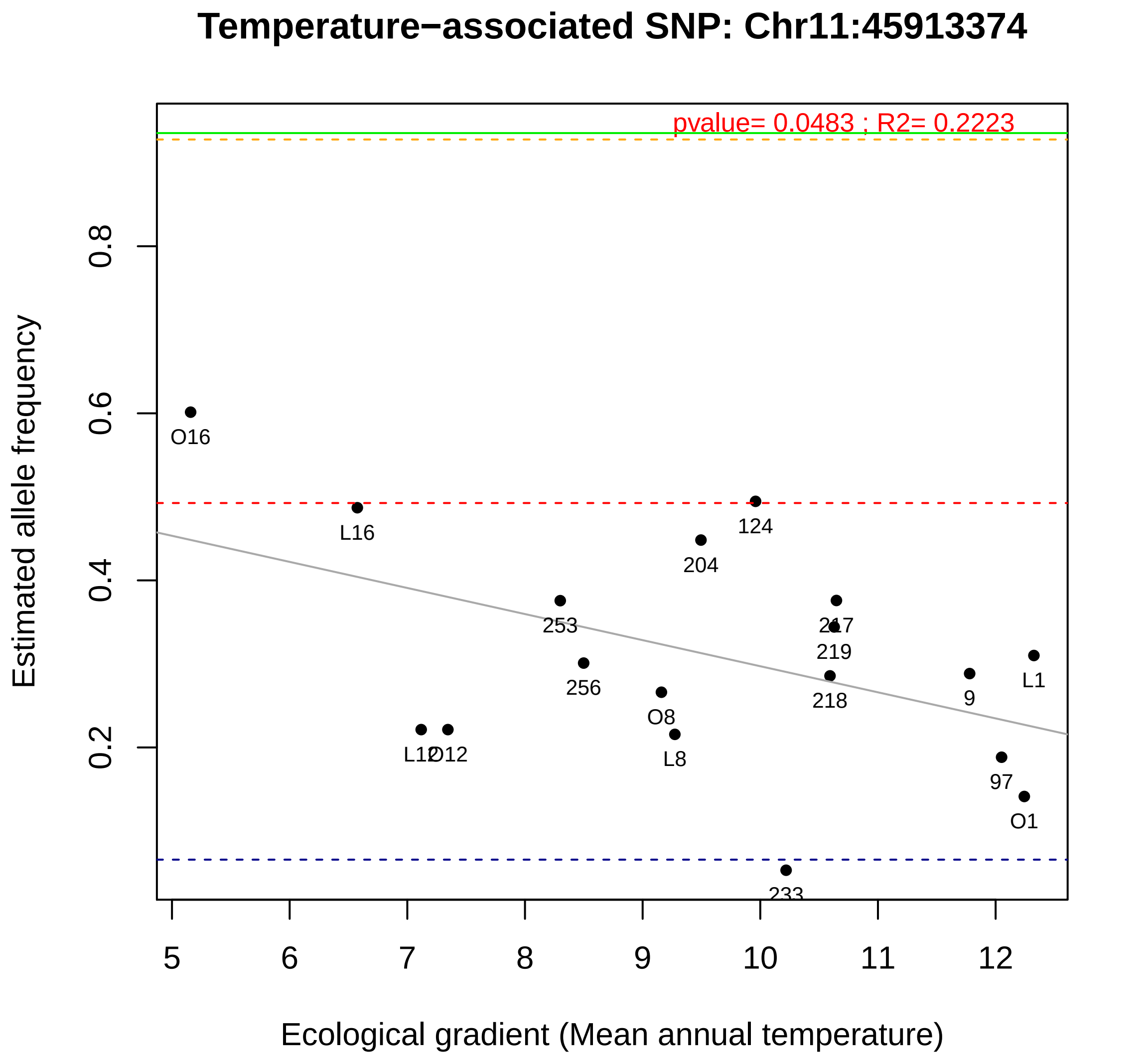
Temperature−associated SNP: Chr11:45913374
pvalue= 0.0483 ; R2= 0.2223
Estimated allele frequency
0.2
0.4
0.6
0.8
Ecological gradient (Mean annual temperature)
5
6
7
8
9
10
11
12
O16
L16
124
204
253
217
219
256
218
9
L1
O8
L12
O12
L8
97
O1
233

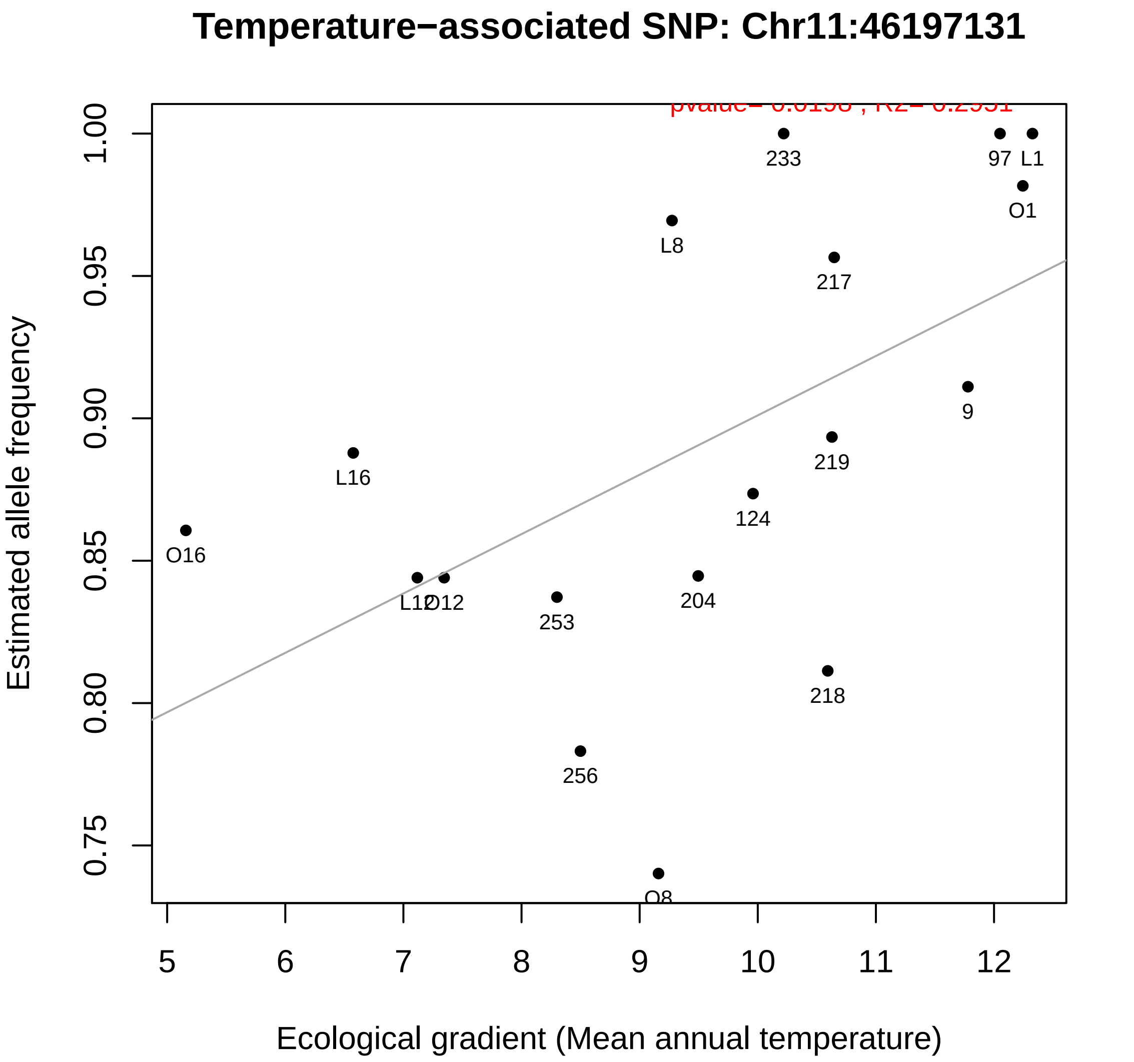
Temperature−associated SNP: Chr11:46197131
Estimated allele frequency
Ecological gradient (Mean annual temperature)
1.00
0.95
0.90
0.85
0.80
0.75
5
6
7
8
9
10
11
12
233
97
L1
O1
L8
217
9
219
L16
124
O16
L12
O12
253
204
218
256
O8

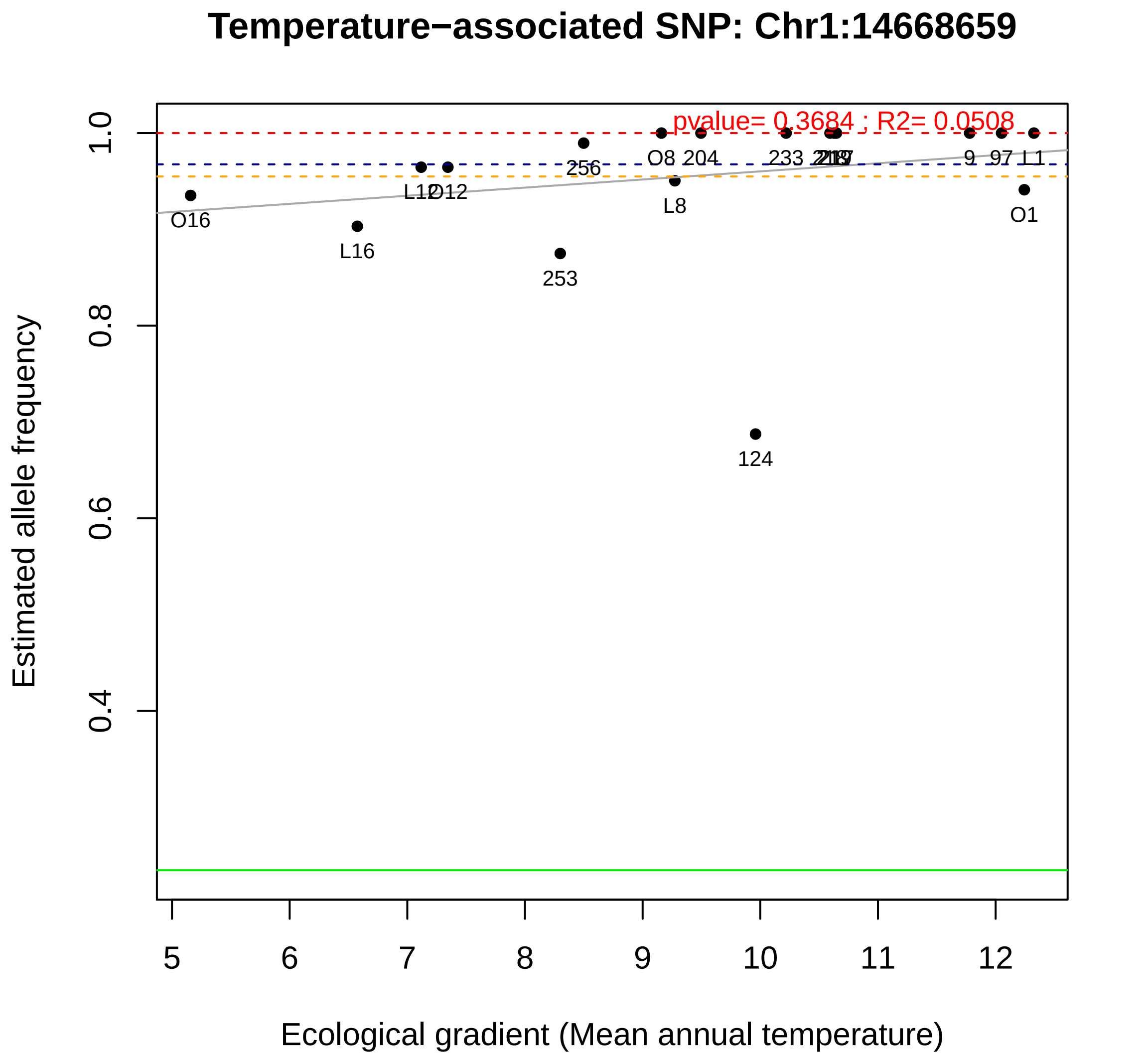
Temperature−associated SNP: Chr1:14668659
pvalue= 0.3684 ; R2= 0.0508
Estimated allele frequency
1.0
0.8
0.6
0.4
5
6
7
8
9
10
11
12
Ecological gradient (Mean annual temperature)
O16
L16
L12
O12
253
256
O8
204
L8
124
233
9
97
L1
O1

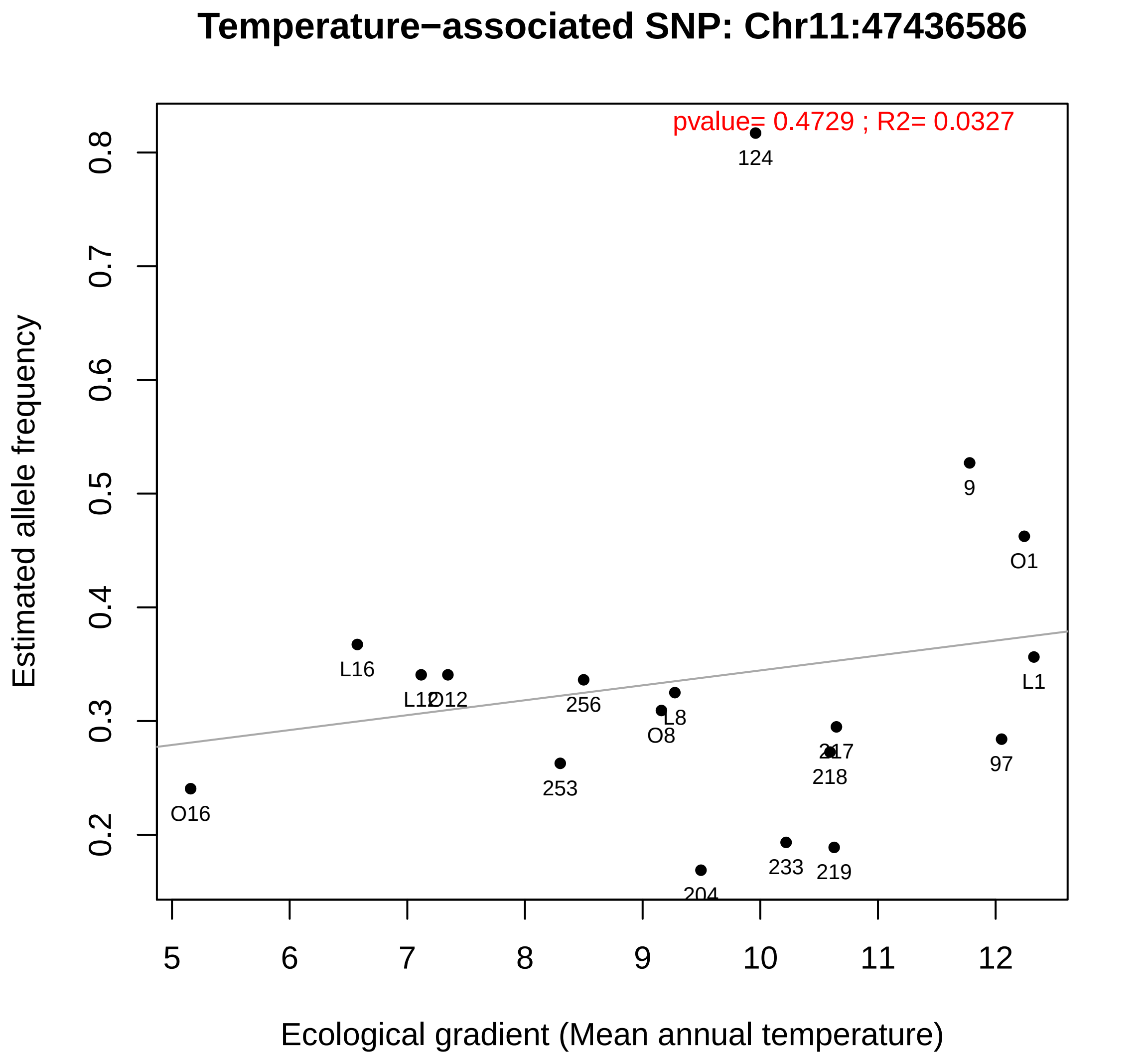

Temperature−associated SNP: Chr11:47436586
pvalue= 0.4729 ; R2= 0.0327
Estimated allele frequency
0.2
0.3
0.4
0.5
0.6
0.7
0.8
5
6
7
8
9
10
11
12
Ecological gradient (Mean annual temperature)
124
9
O1
L1
97
L16
L12
O12
256
253
O16
L8
O8
217
218
233
219
204

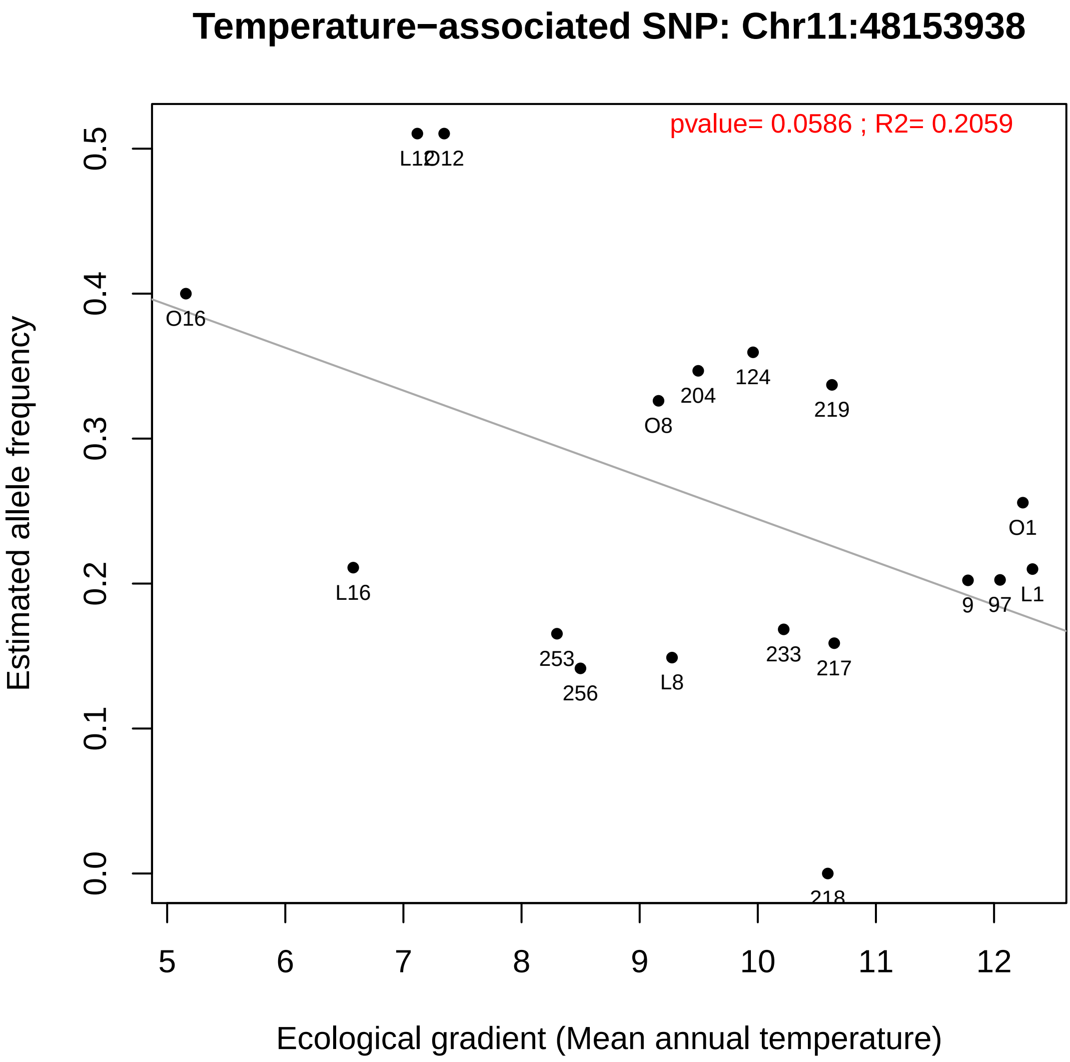
Temperature−associated SNP: Chr11:48153938
pvalue= 0.0586 ; R2= 0.2059
Estimated allele frequency
0.0
0.1
0.2
0.3
0.4
0.5
5
6
7
8
9
10
11
12
Ecological gradient (Mean annual temperature)
O16
L12
O12
L16
253
256
O8
204
124
219
L8
233
217
218
O1
9
97
L1

#### Temperature−associated SNP: Chr11:48283827

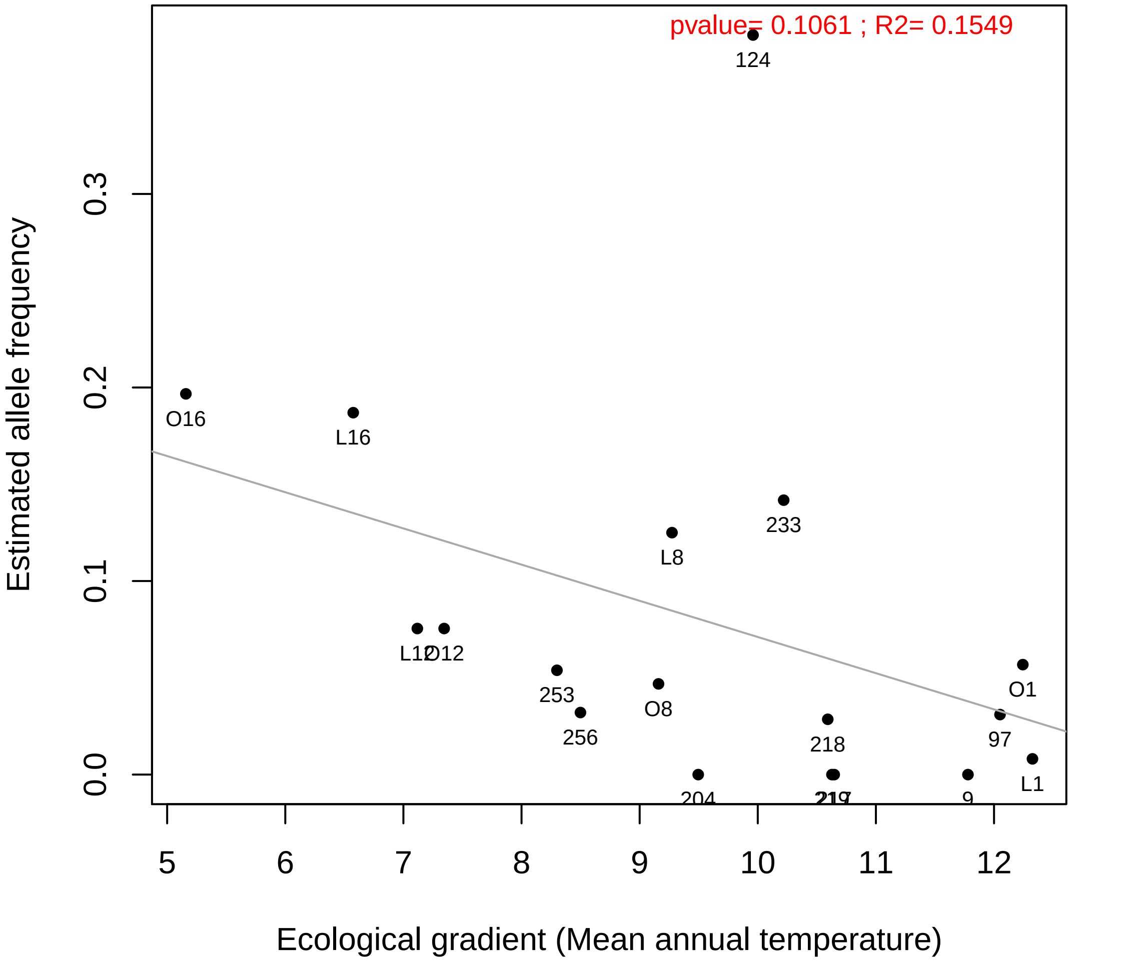

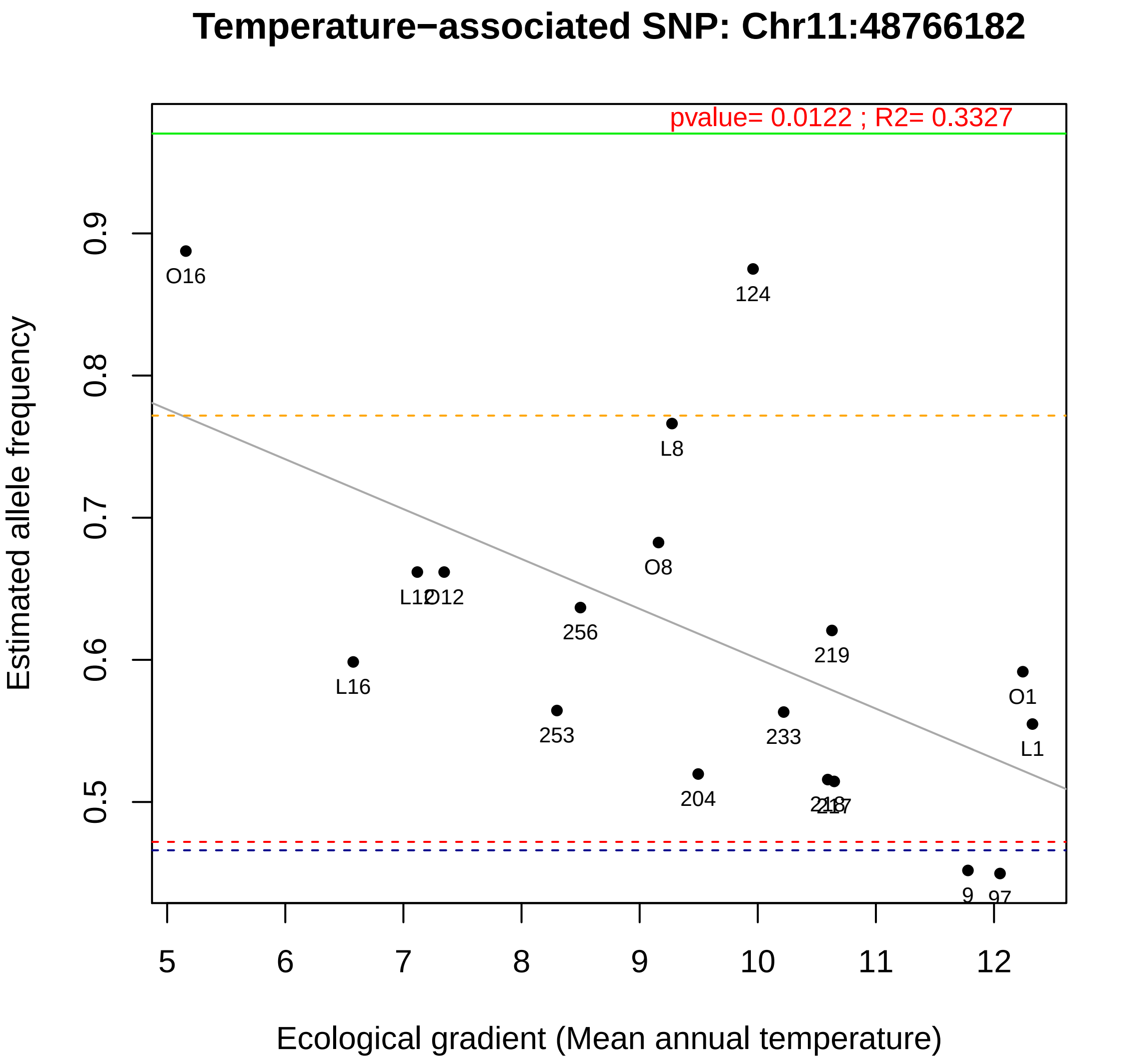

Temperature−associated SNP: Chr11:48766182
pvalue= 0.0122 ; R2= 0.3327
Estimated allele frequency
0.5
0.6
0.7
0.8
0.9
Ecological gradient (Mean annual temperature)
5
6
7
8
9
10
11
12
O16
124
L8
O8
O12
256
219
L16
O1
253
233
L1
204
9
97

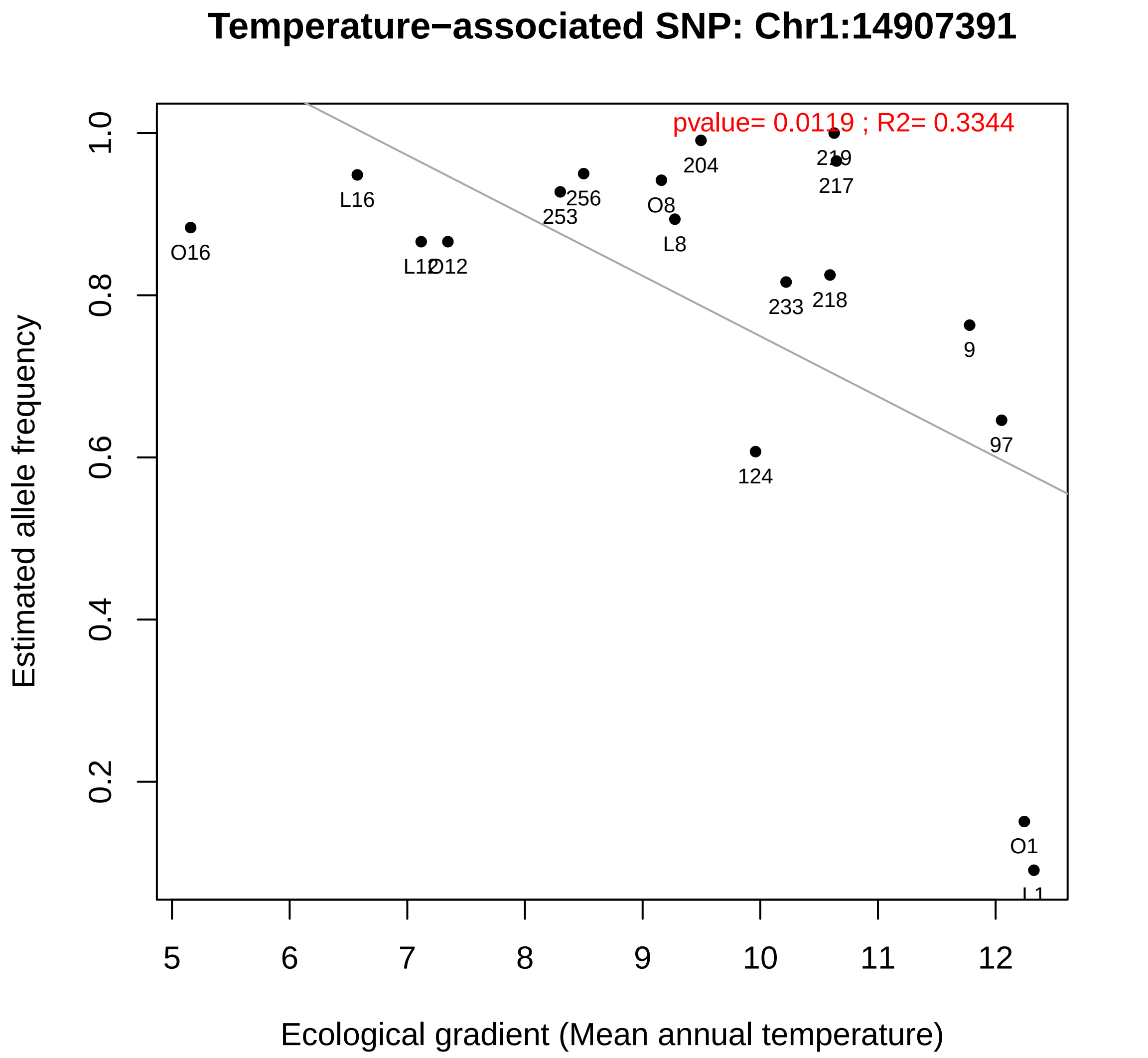
Temperature−associated SNP: Chr1:14907391
pvalue= 0.0119 ; R2= 0.3344
O16
L16
L12
O12
253
256
O8
L8
204
219
217
233
218
124
9
97
O1
L1
Estimated allele frequency
0.2
0.4
0.6
0.8
1.0
5
6
7
8
9
10
11
12
Ecological gradient (Mean annual temperature)

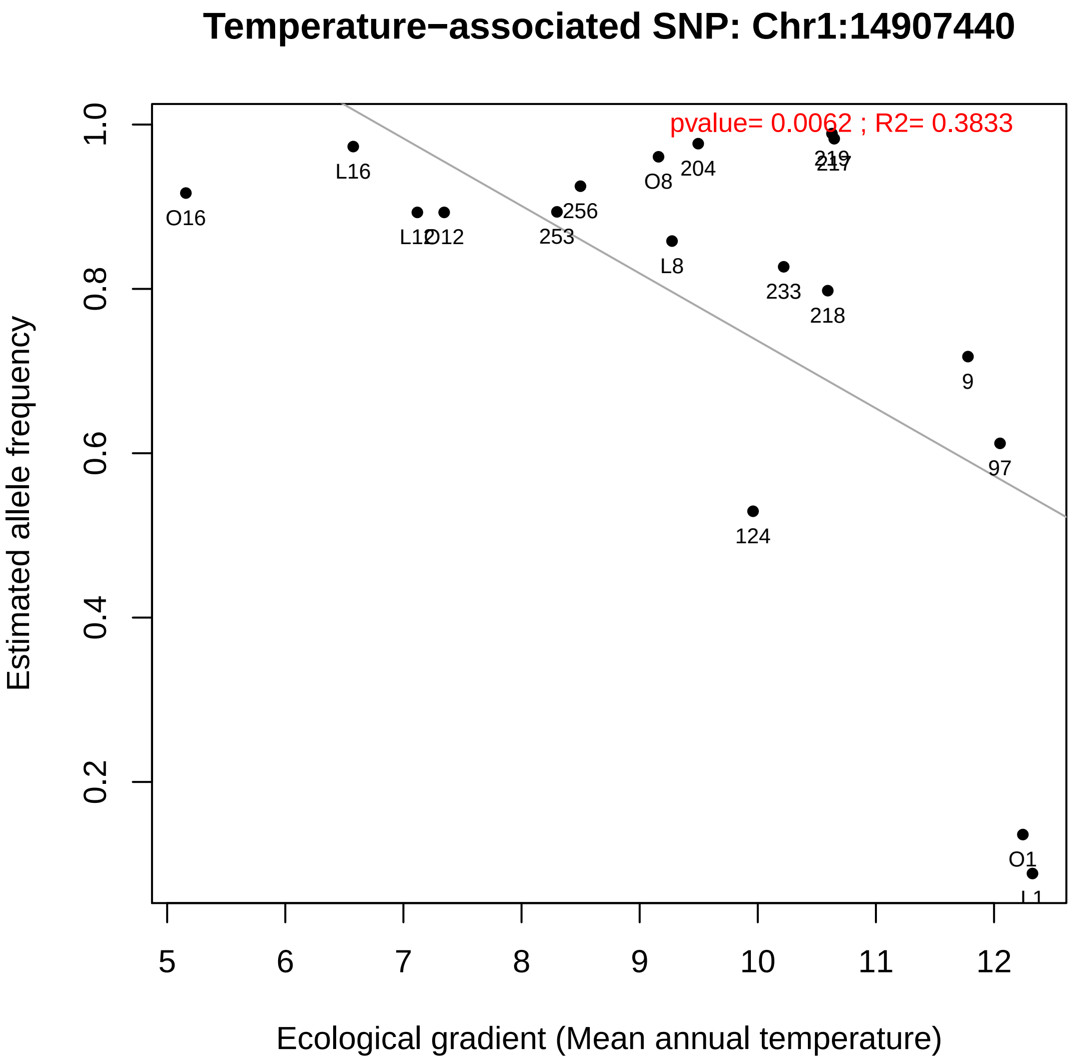

Temperature−associated SNP: Chr1:14907440
pvalue= 0.0062 ; R2= 0.3833
O16
L16
L12
O12
253
256
O8
204
L8
219
217
233
218
124
9
97
O1
L1
1.0
0.8
0.6
0.4
0.2
5
6
7
8
9
10
11
12
Estimated allele frequency
Ecological gradient (Mean annual temperature)

**Temperature−associated SNP: Chr1:14907522**

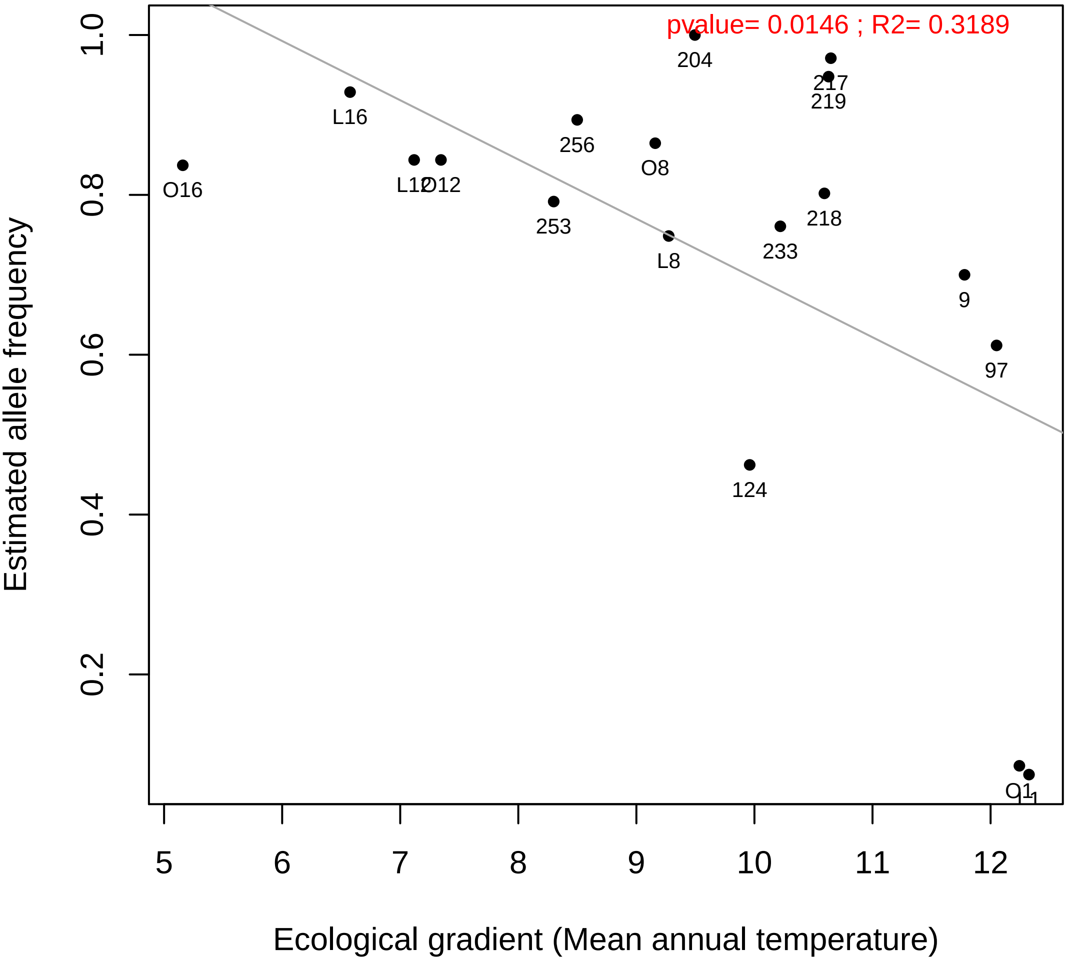

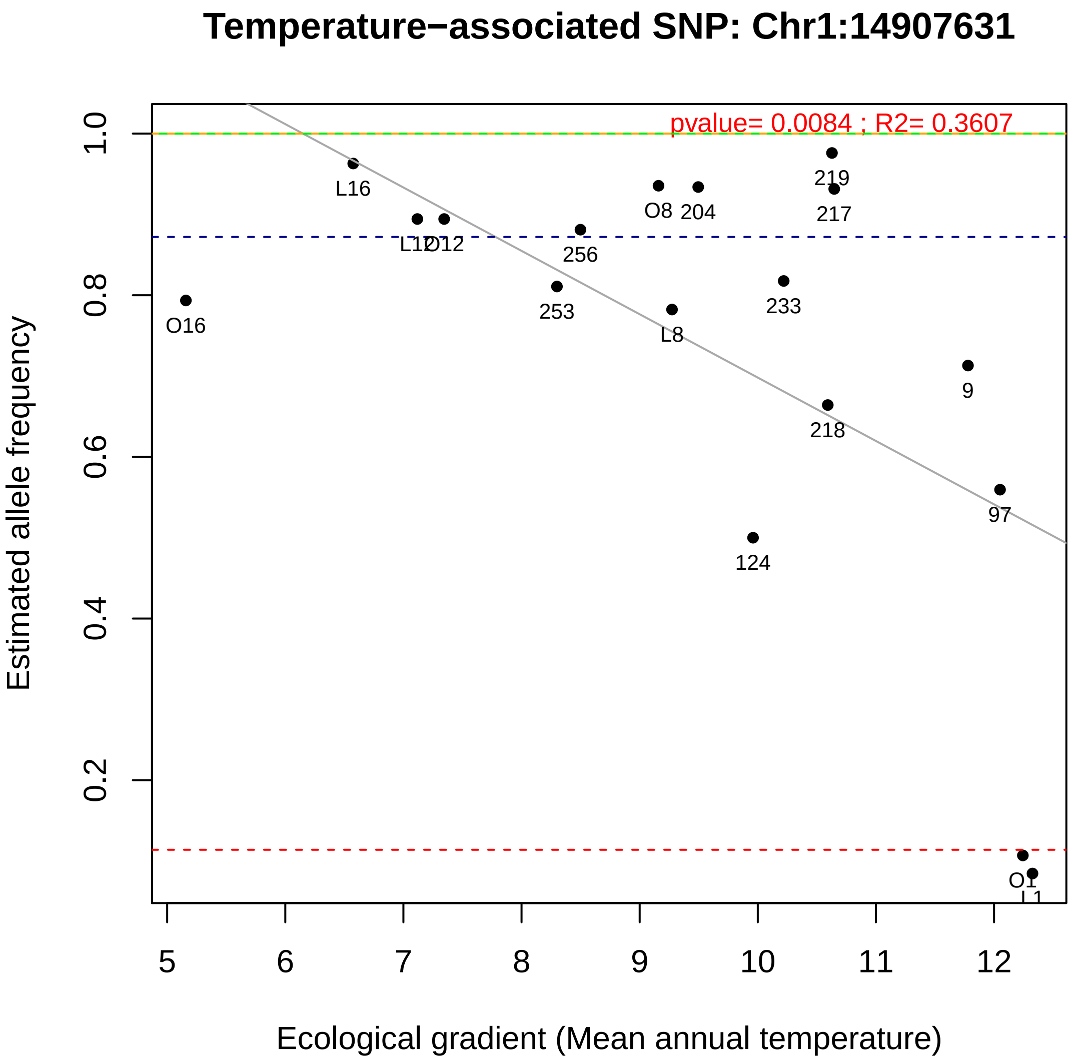
Temperature−associated SNP: Chr1:14907631
pvalue= 0.0084 ; R2= 0.3607
Estimated allele frequency
0.2
0.4
0.6
0.8
1.0
5
6
7
8
9
10
11
12
Ecological gradient (Mean annual temperature)
L16
L12
O12
256
253
O16
O8
204
L8
219
217
233
218
9
97
124
O1
L1

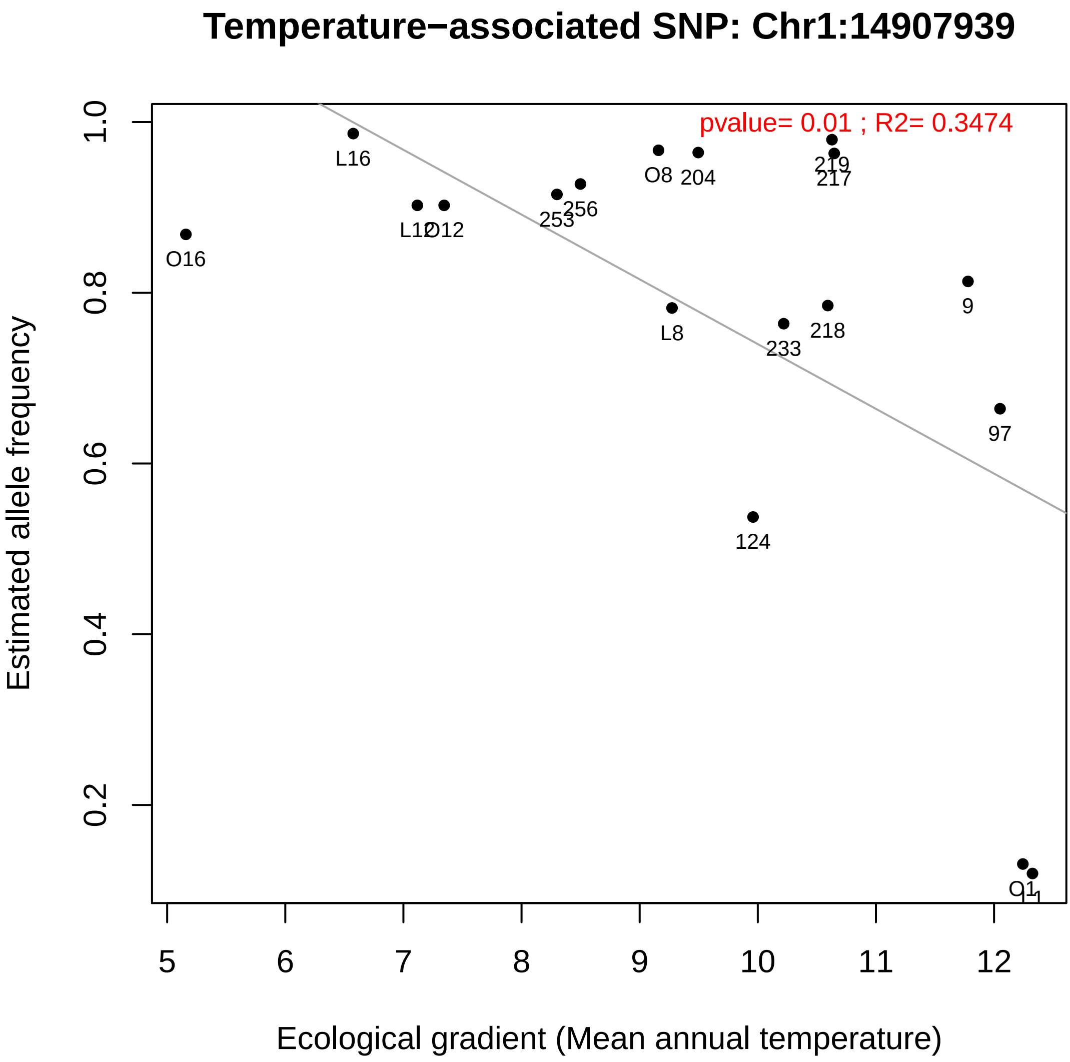
Temperature−associated SNP: Chr1:14907939
pvalue= 0.01 ; R2= 0.3474
Estimated allele frequency
Ecological gradient (Mean annual temperature)
0.2
0.4
0.6
0.8
1.0
5
6
7
8
9
10
11
12
L16
O16
L12
O12
253
256
O8
204
219
217
L8
233
218
9
97
124
O1

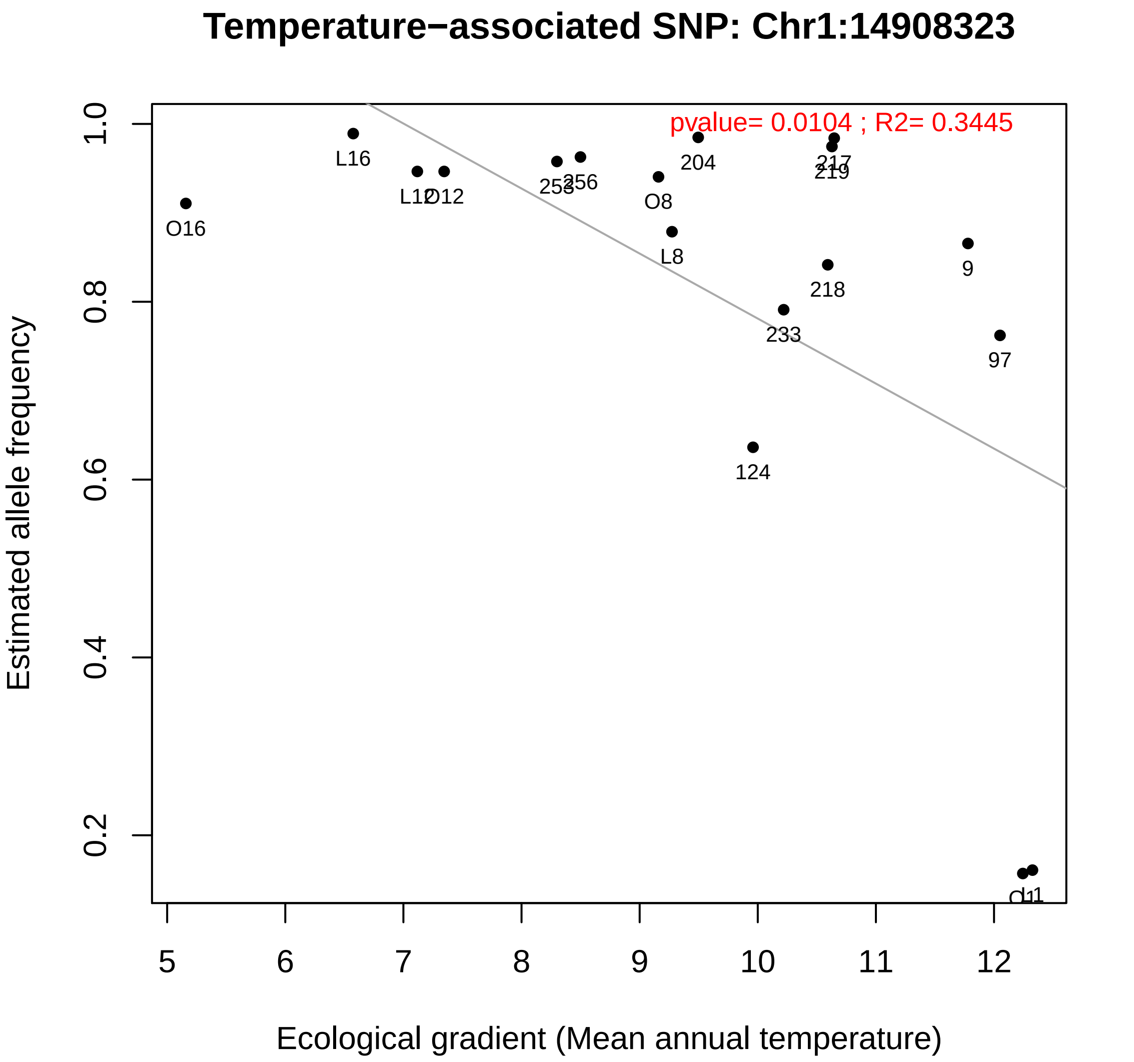
Temperature−associated SNP: Chr1:14908323
pvalue= 0.0104 ; R2= 0.3445
Estimated allele frequency
Ecological gradient (Mean annual temperature)
0.2
0.4
0.6
0.8
1.0
5
6
7
8
9
10
11
12
O16
L16
L12
O12
253
256
O8
204
L8
217
219
9
218
233
97
124

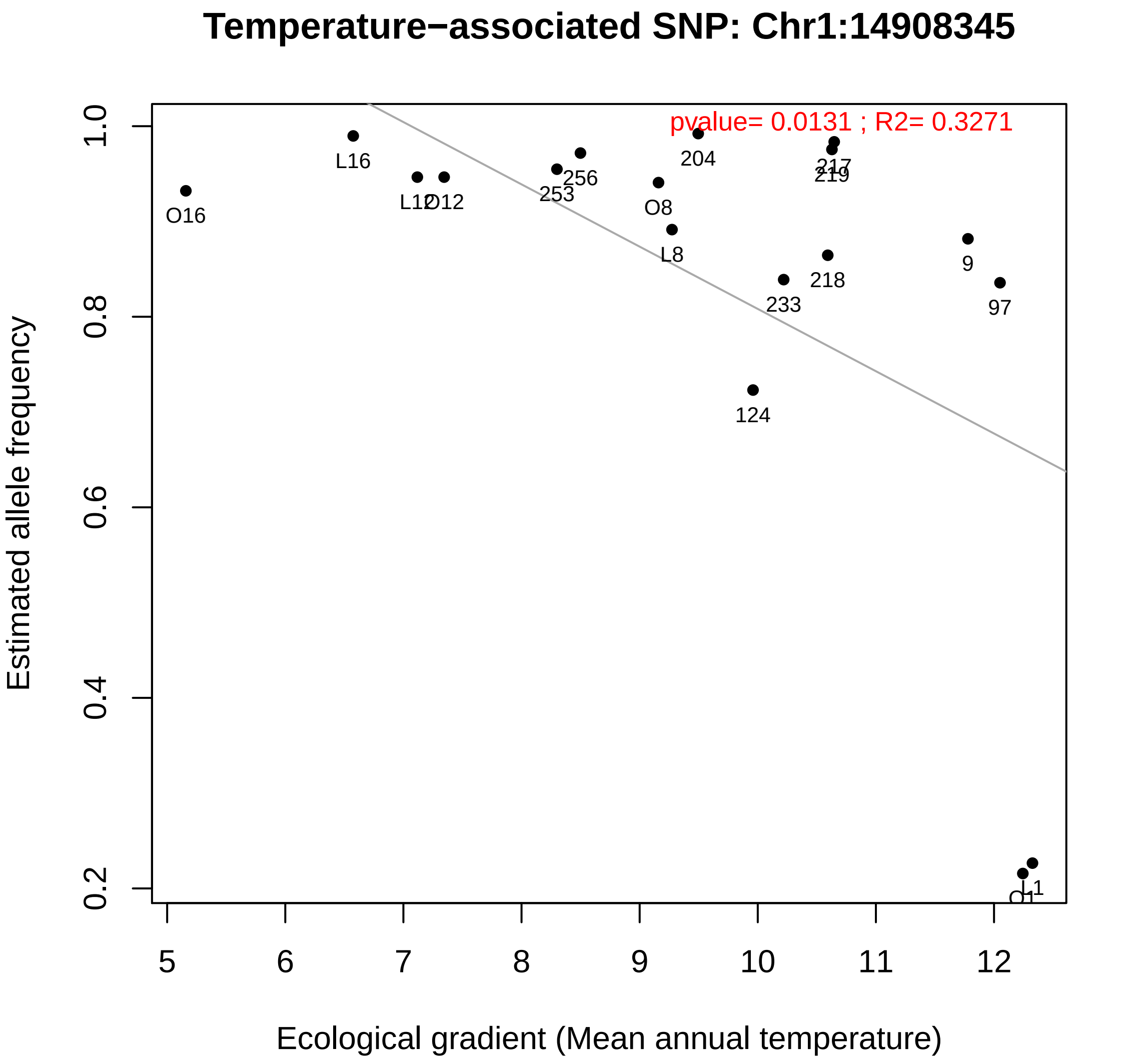

Temperature−associated SNP: Chr1:14908345
pvalue= 0.0131 ; R2= 0.3271
Estimated allele frequency
0.2
0.4
0.6
0.8
1.0
5
6
7
8
9
10
11
12
Ecological gradient (Mean annual temperature)
O16
L16
L12
O12
253
256
O8
L8
204
217
219
233
218
124
9
97
L1
O1

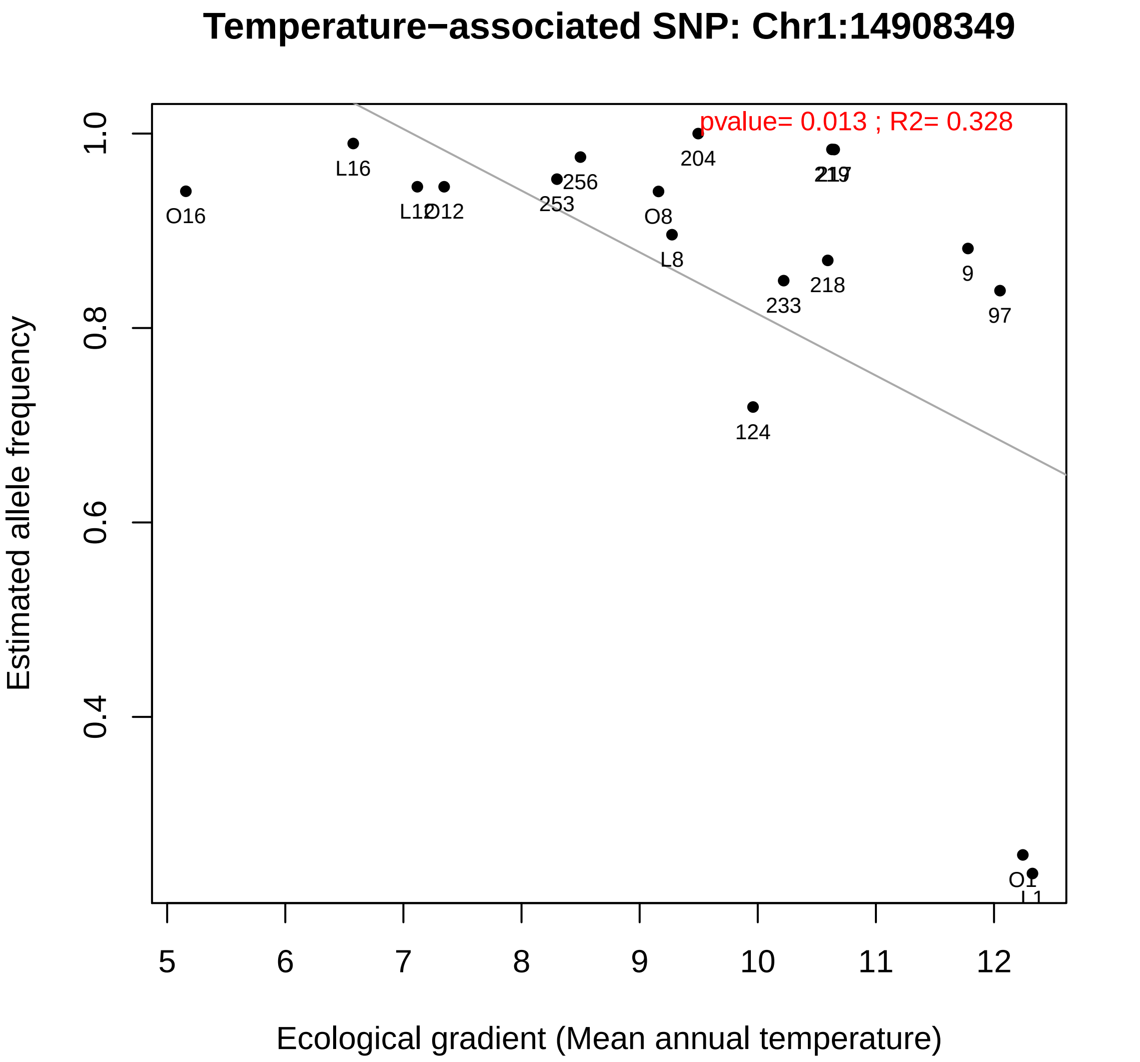

Temperature−associated SNP: Chr1:14908349
pvalue= 0.013 ; R2= 0.328
Estimated allele frequency
1.0
0.8
0.6
0.4
5
6
7
8
9
10
11
12
Ecological gradient (Mean annual temperature)
O16
L16
L12
O12
253
256
O8
204
L8
233
124
218
9
97
O1
L1

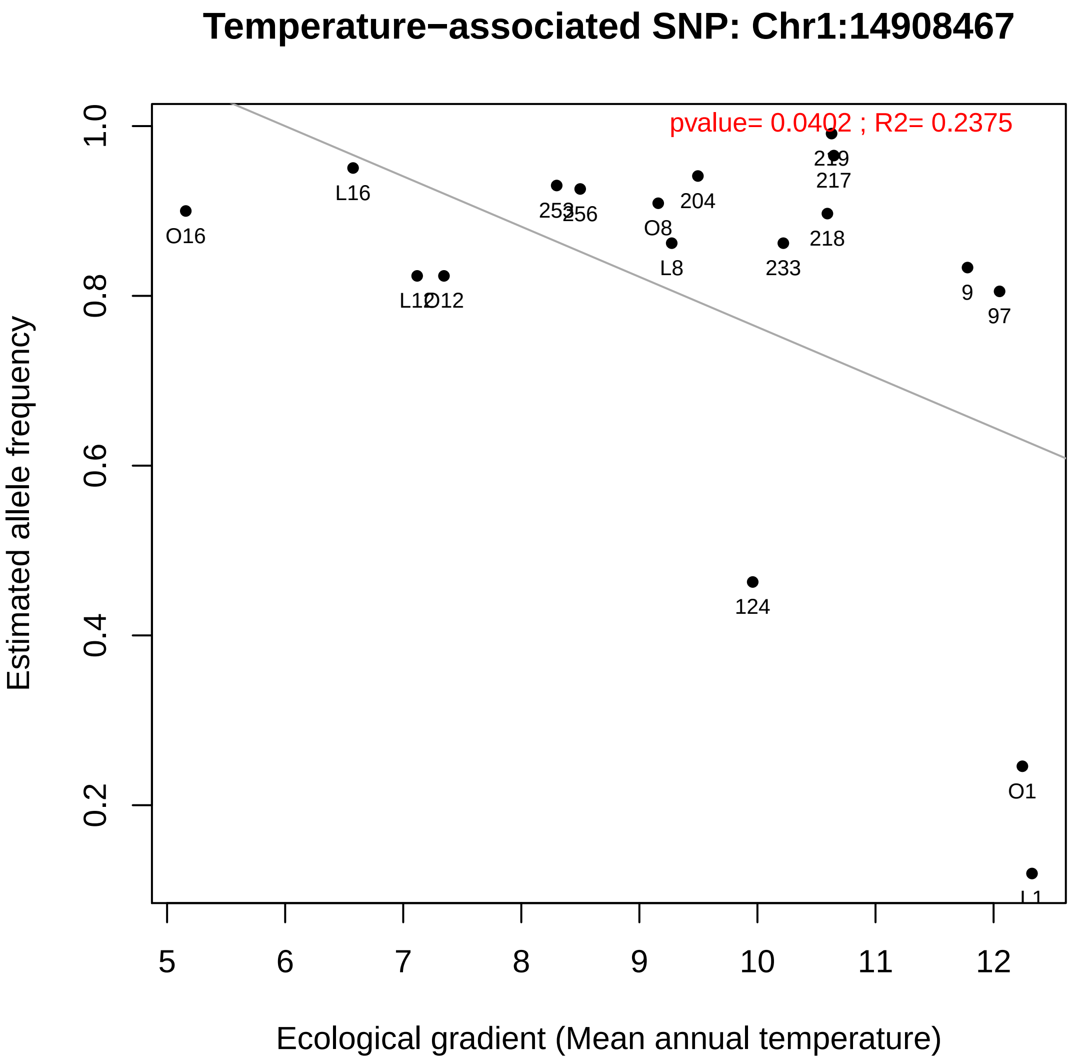

Temperature−associated SNP: Chr1:14908467
pvalue= 0.0402 ; R2= 0.2375
Estimated allele frequency
1.0
0.8
0.6
0.4
0.2
5
6
7
8
9
10
11
12
Ecological gradient (Mean annual temperature)
O16
L16
256
O8
204
217
218
L8
233
9
97
124
O1

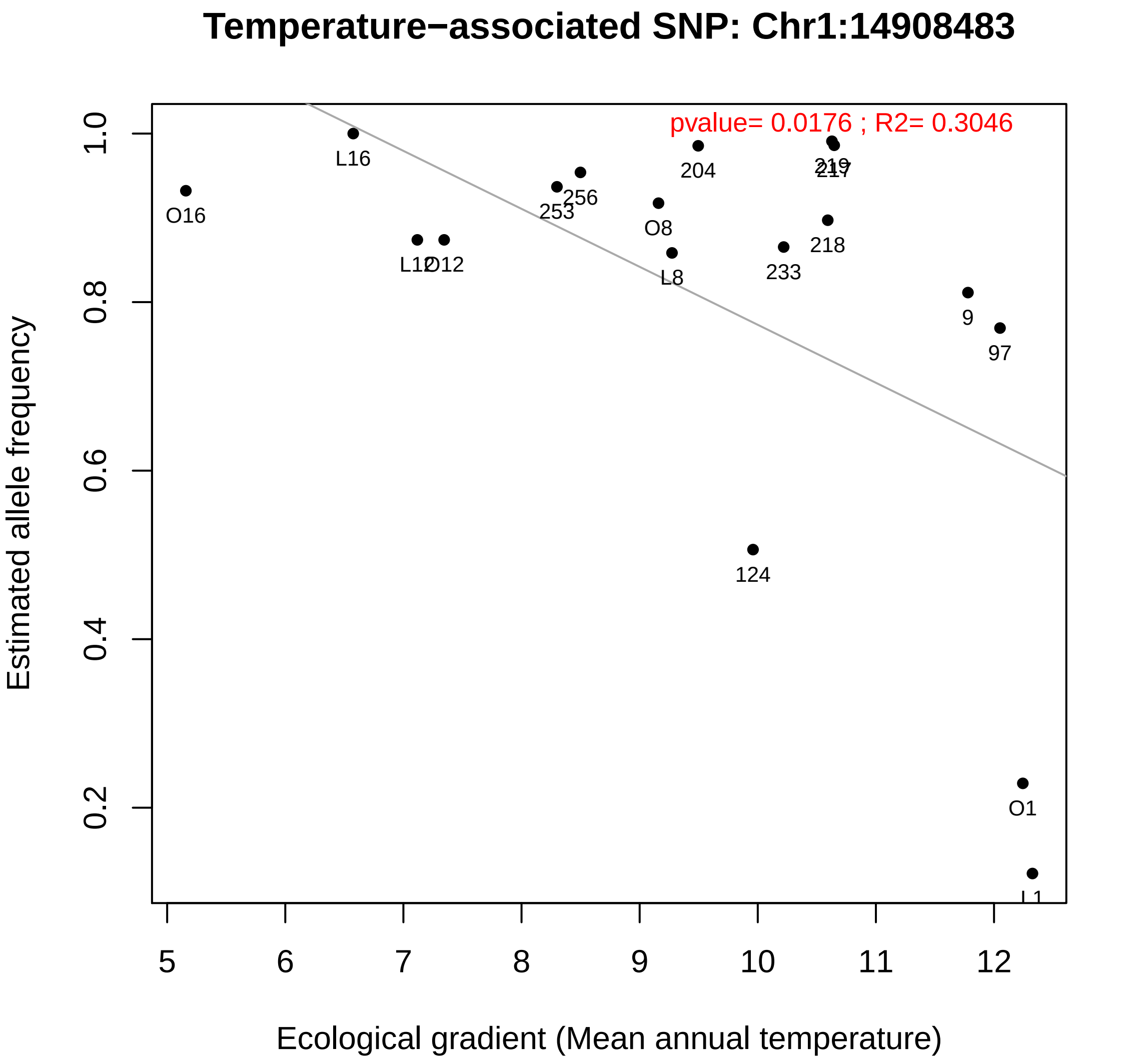
Temperature−associated SNP: Chr1:14908483
pvalue= 0.0176 ; R2= 0.3046
Estimated allele frequency
Ecological gradient (Mean annual temperature)
0.2
0.4
0.6
0.8
1.0
5
6
7
8
9
10
11
12
O16
L16
L12
O12
253
256
O8
L8
204
124
233
218
219
217
9
97
O1
L1

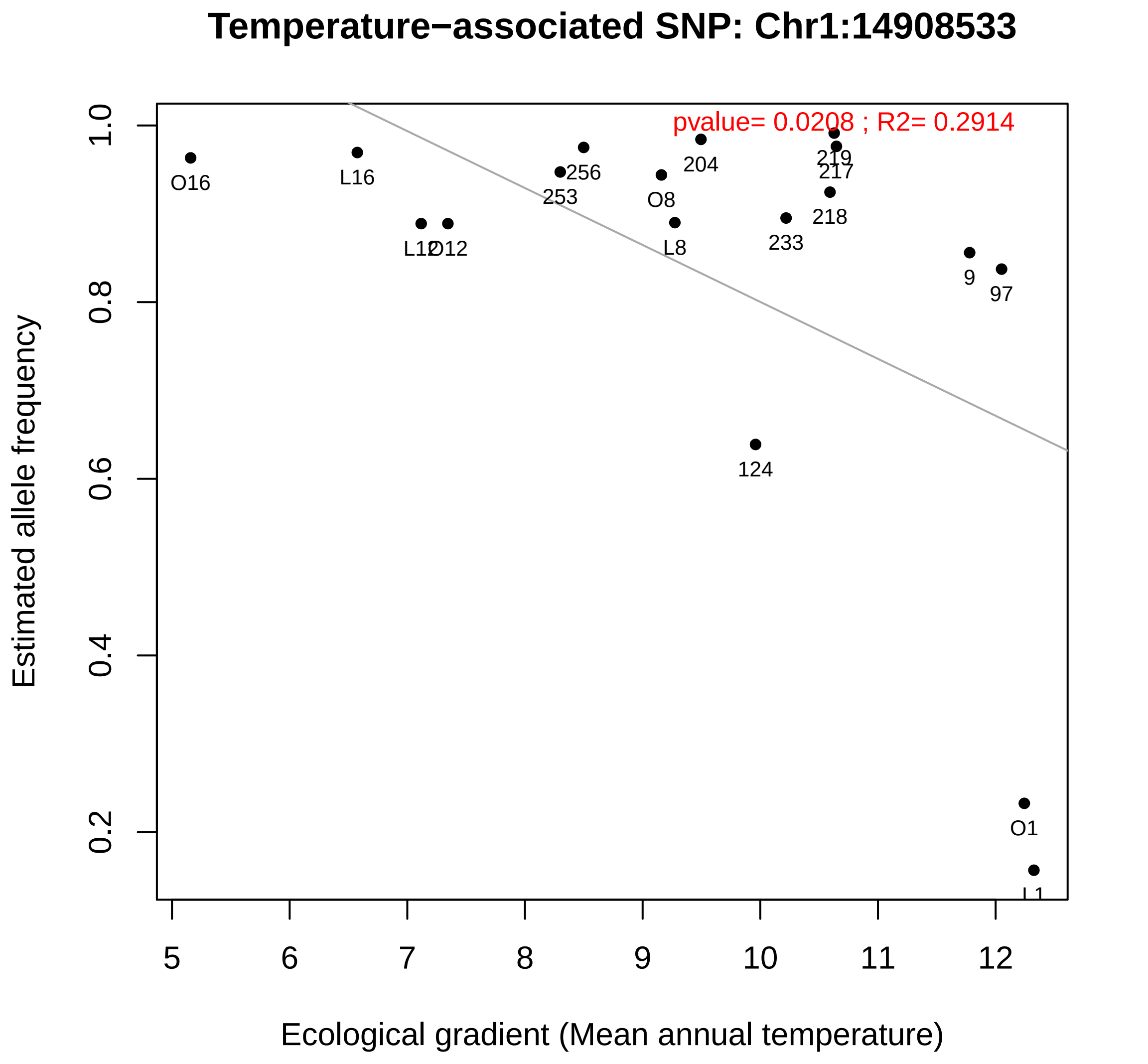
Temperature−associated SNP: Chr1:14908533
pvalue= 0.0208 ; R2= 0.2914
O16
L16
L12
O12
253
256
O8
L8
204
124
233
219
217
218
9
97
O1
L1
1.0
0.8
0.6
0.4
0.2
5
6
7
8
9
10
11
12
Estimated allele frequency
Ecological gradient (Mean annual temperature)

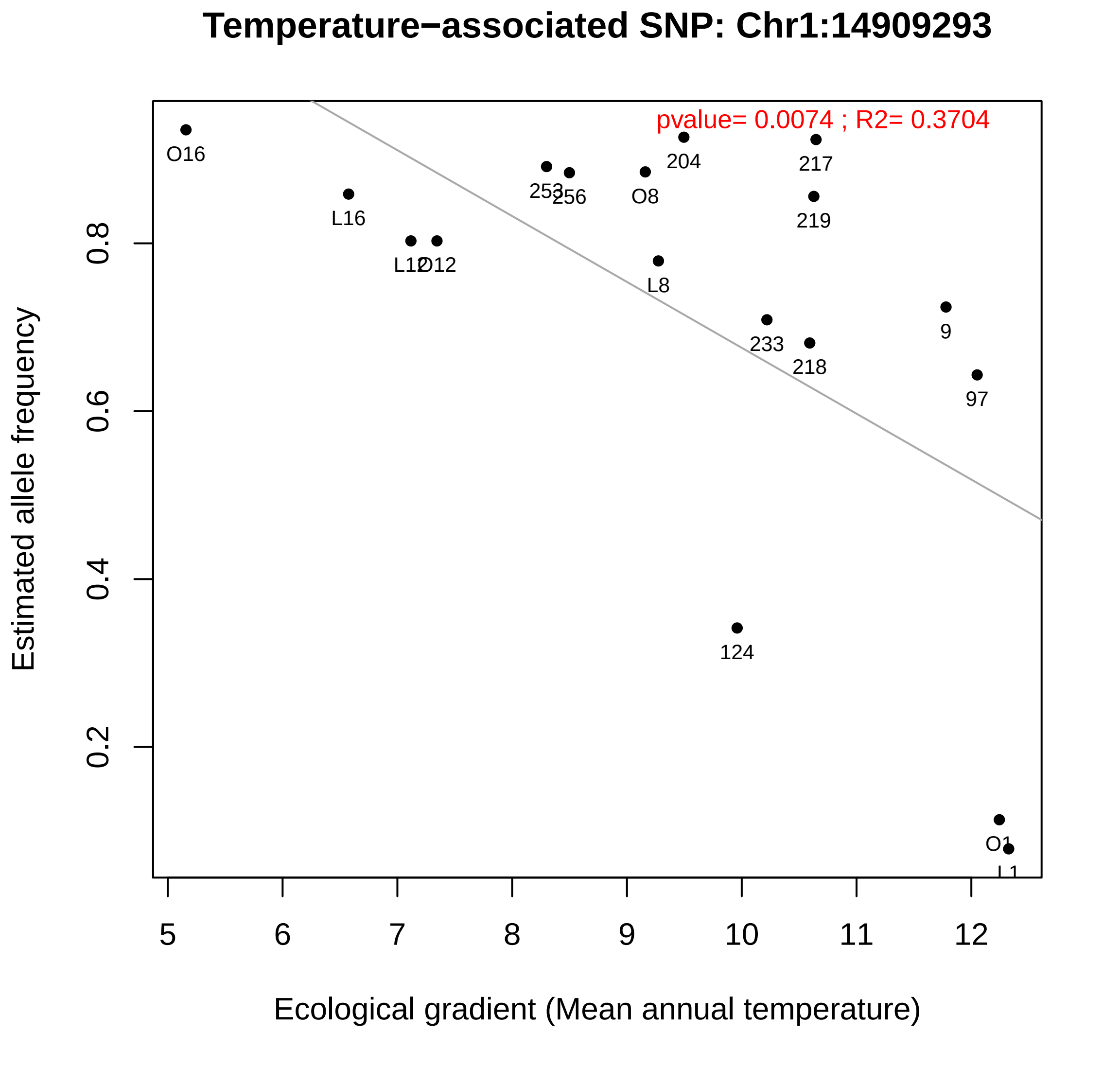

Temperature−associated SNP: Chr1:14909293
pvalue= 0.0074 ; R2= 0.3704
Estimated allele frequency
0.2
0.4
0.6
0.8
Ecological gradient (Mean annual temperature)
5
6
7
8
9
10
11
12
O16
L16
L12
O12
253
256
O8
204
217
219
L8
233
218
9
97
124
O1
L1

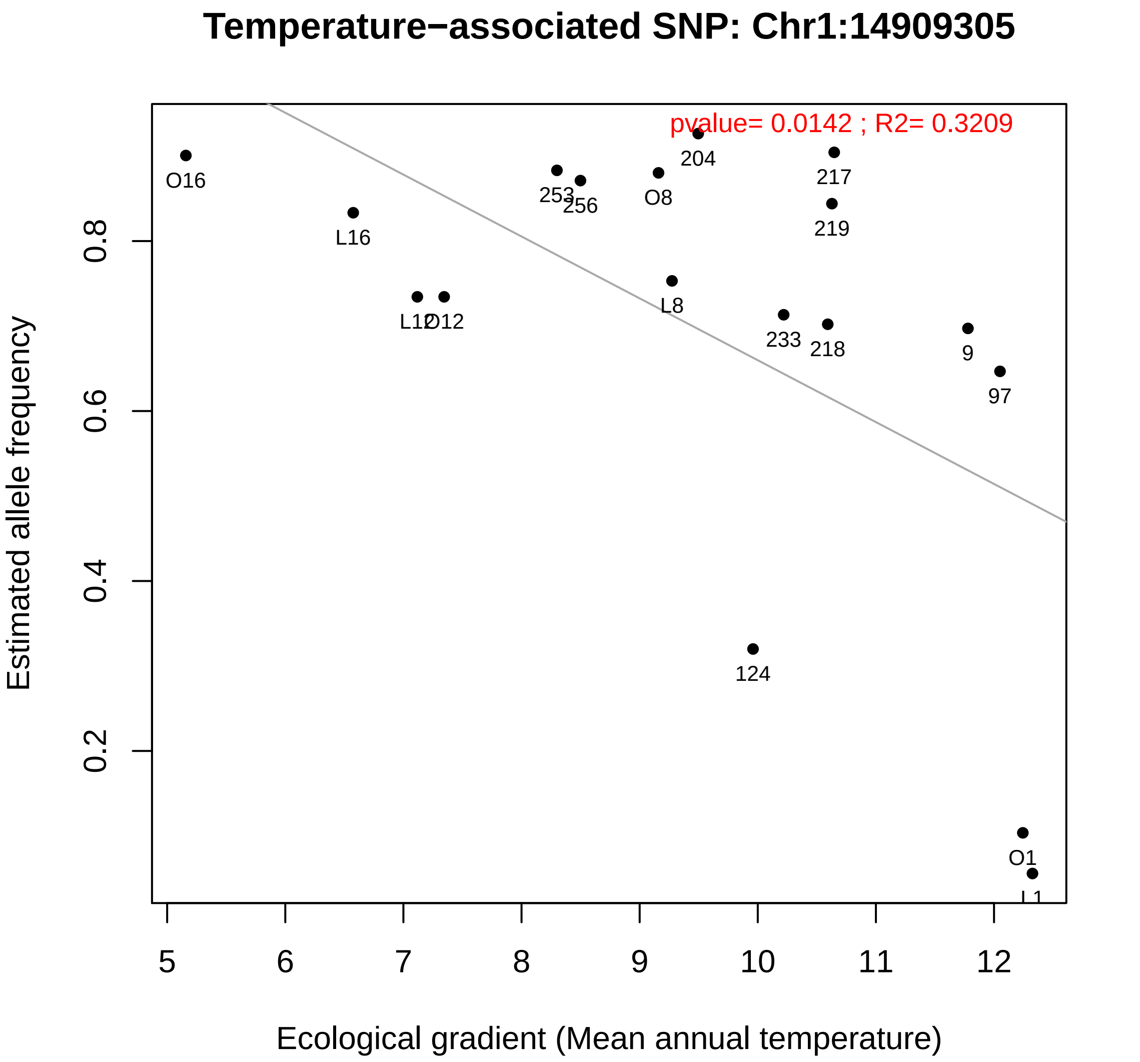

Temperature−associated SNP: Chr1:14909305
pvalue= 0.0142 ; R2= 0.3209
Estimated allele frequency
0.2
0.4
0.6
0.8
5
6
7
8
9
10
11
12
Ecological gradient (Mean annual temperature)
O16
L16
L12
O12
253
256
O8
204
L8
217
219
233
218
124
9
97
O1

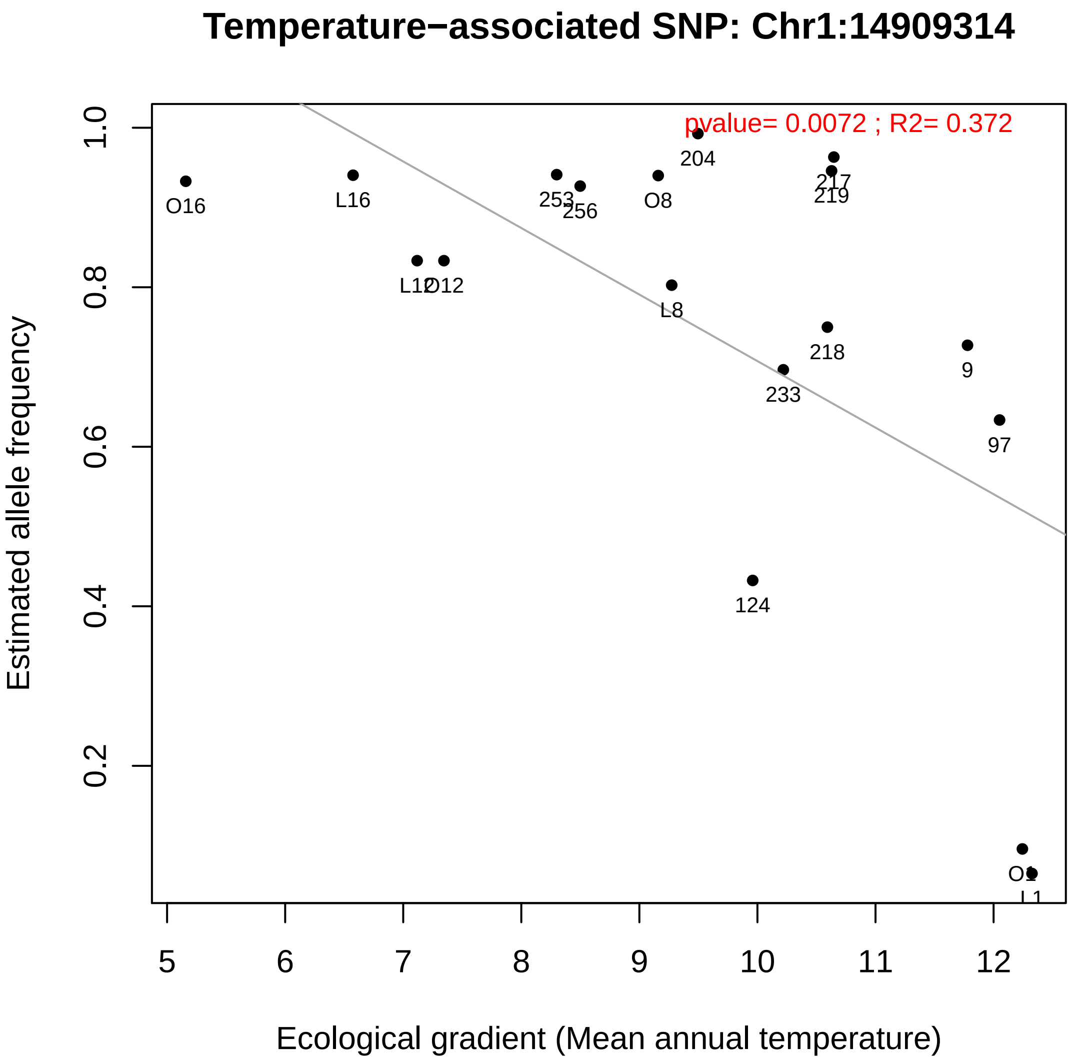

Temperature−associated SNP: Chr1:14909314
pvalue= 0.0072 ; R2= 0.372
Estimated allele frequency
Ecological gradient (Mean annual temperature)
0.2
0.4
0.6
0.8
1.0
5
6
7
8
9
10
11
12
O16
L16
L12
O12
253
256
O8
204
217
219
L8
218
233
9
97
124
O1
L1

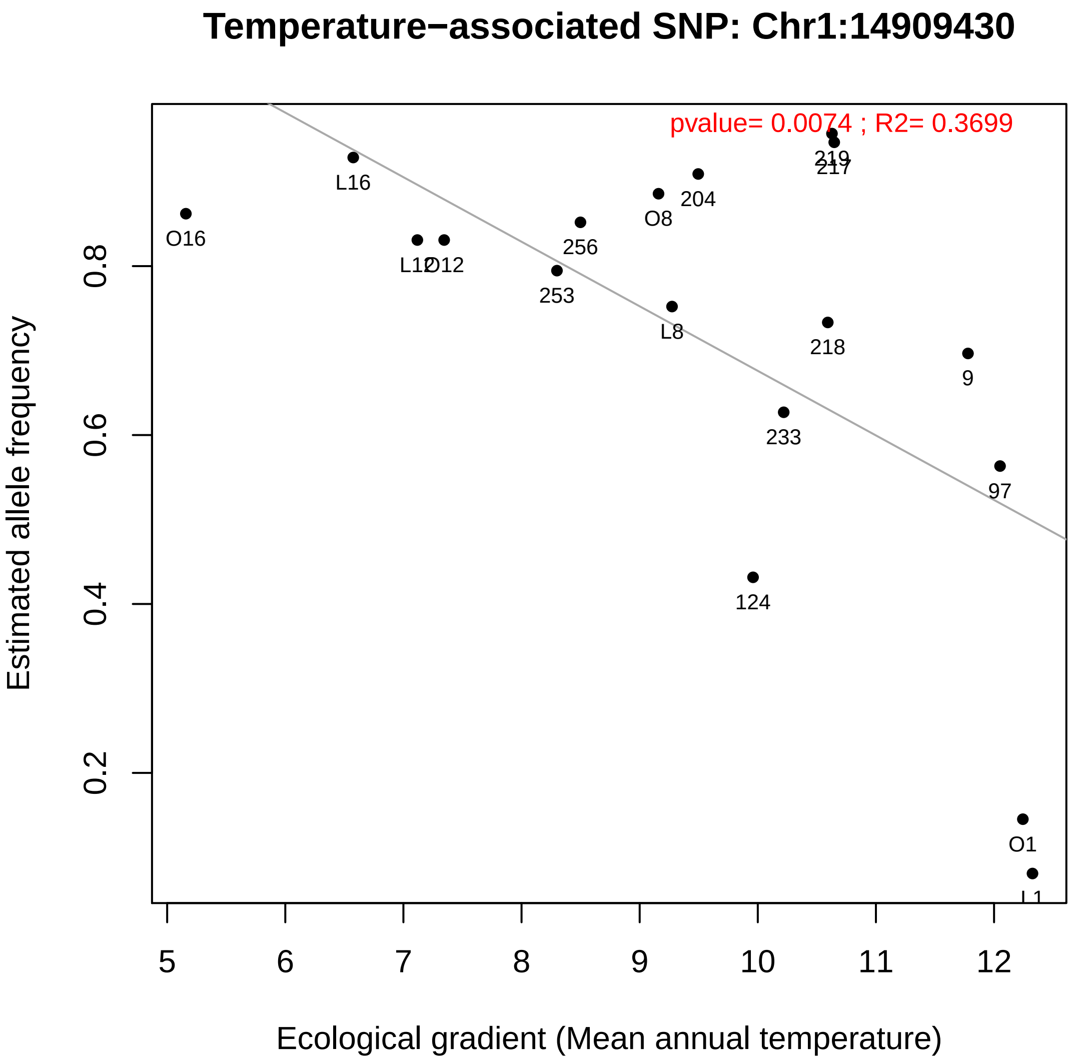

Temperature−associated SNP: Chr1:14909430
pvalue= 0.0074 ; R2= 0.3699
Estimated allele frequency
0.2
0.4
0.6
0.8
5
6
7
8
9
10
11
12
Ecological gradient (Mean annual temperature)
O16
L16
L12
O12
253
256
O8
204
L8
219
217
218
233
124
9
97
O1

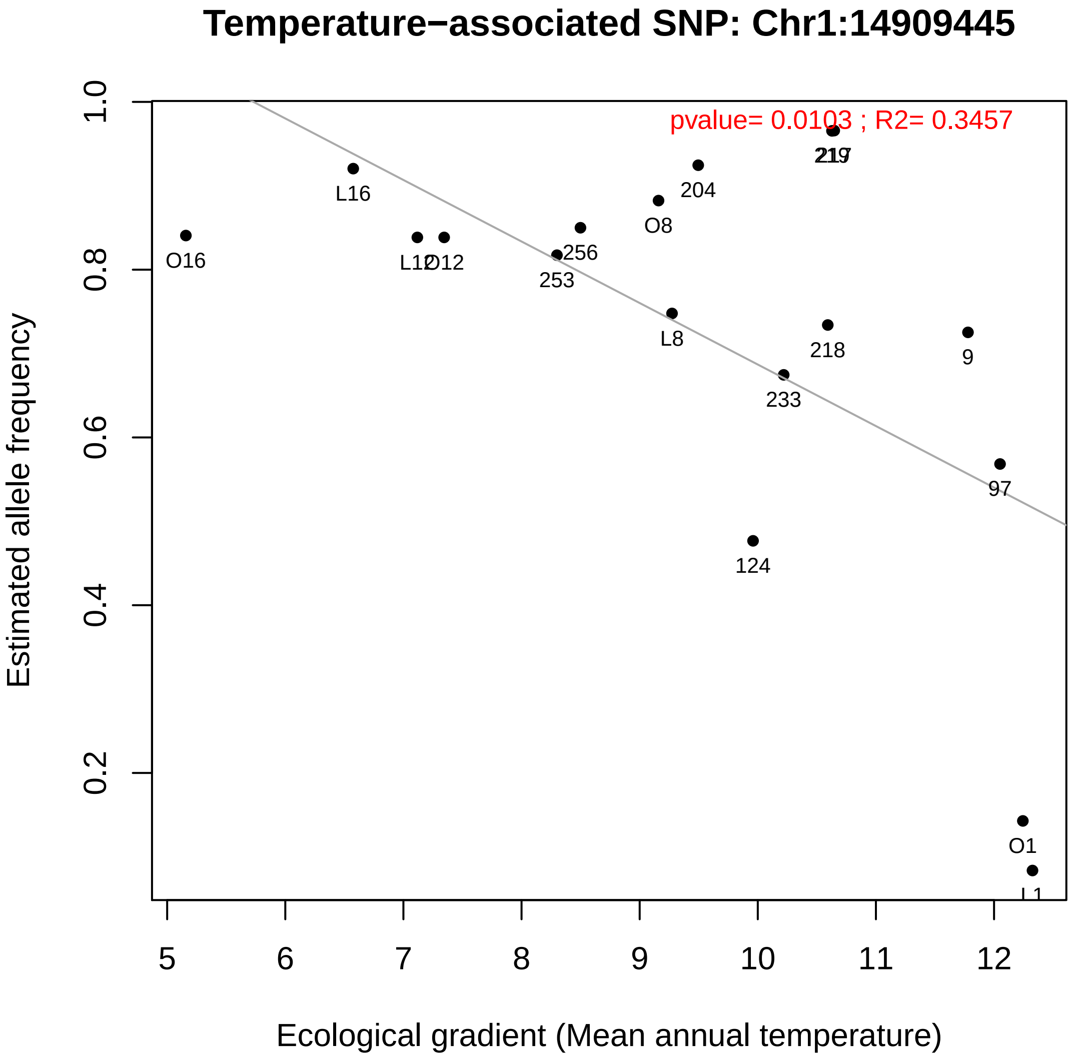
Temperature−associated SNP: Chr1:14909445
pvalue= 0.0103 ; R2= 0.3457
Estimated allele frequency
0.2
0.4
0.6
0.8
1.0
Ecological gradient (Mean annual temperature)
5
6
7
8
9
10
11
12
O16
L16
L12
O12
253
256
O8
204
L8
233
218
9
97
124
O1

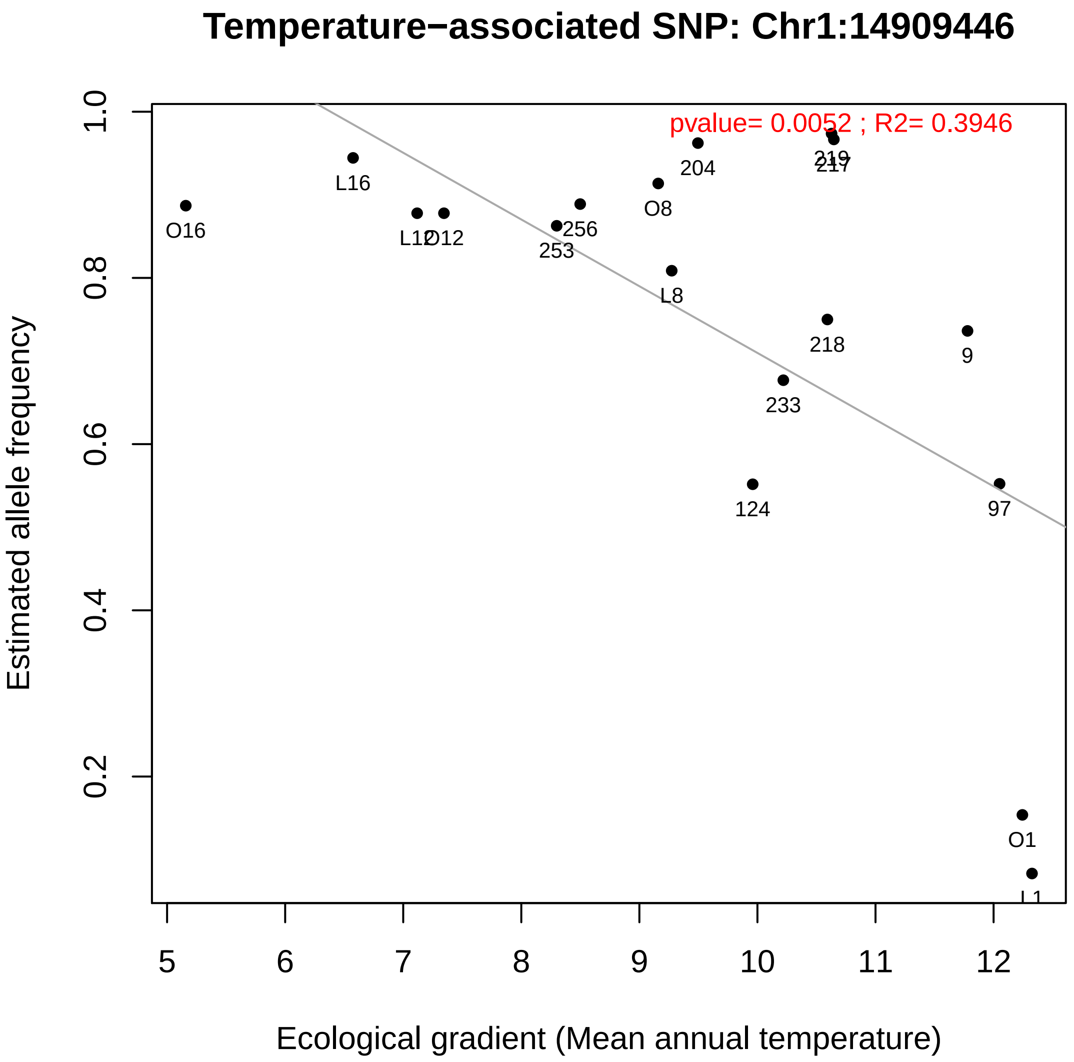
Temperature−associated SNP: Chr1:14909446
pvalue= 0.0052 ; R2= 0.3946
Estimated allele frequency
Ecological gradient (Mean annual temperature)
0.2
0.4
0.6
0.8
1.0
5
6
7
8
9
10
11
12
O16
L16
L12
O12
253
256
O8
204
L8
219
217
218
233
124
9
97
O1

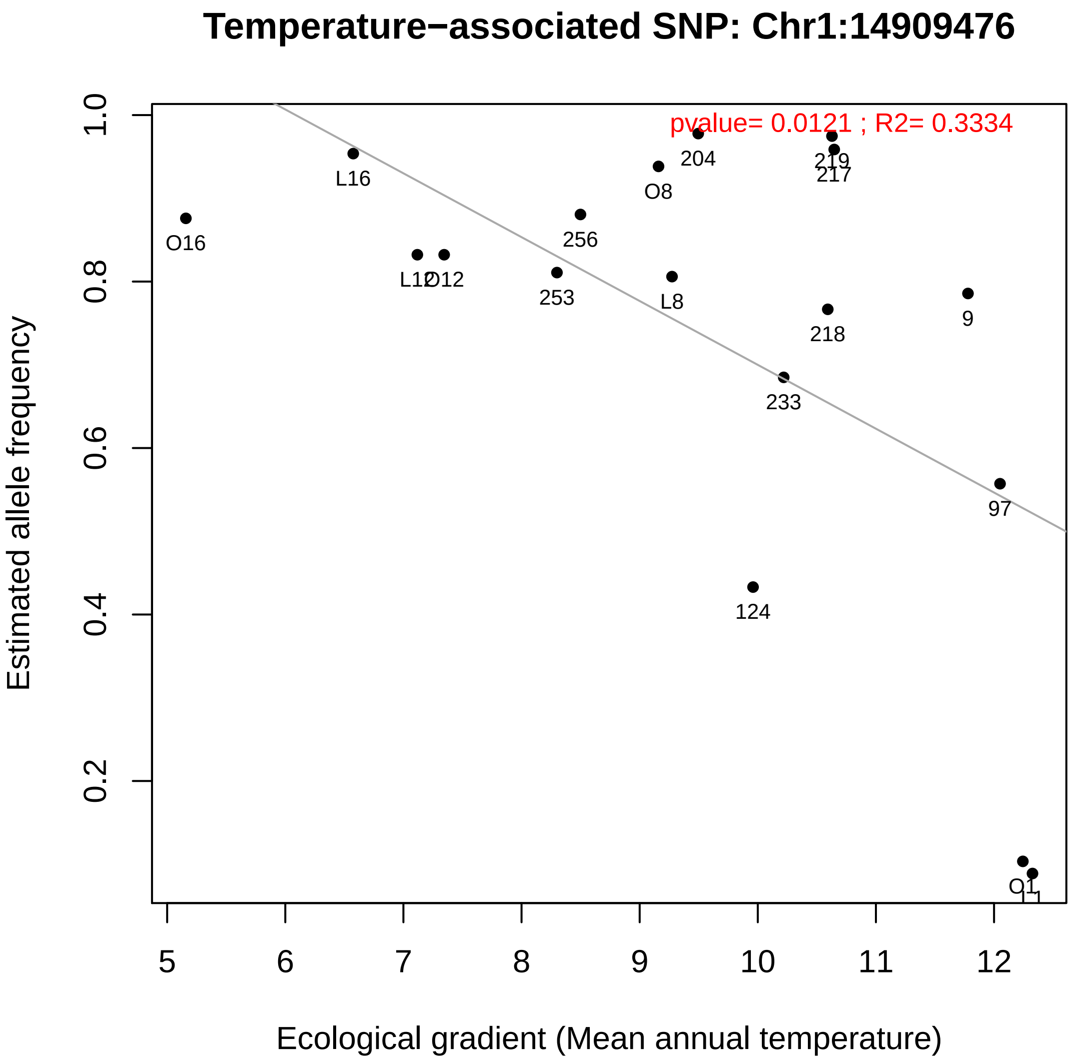
Temperature−associated SNP: Chr1:14909476
pvalue= 0.0121 ; R2= 0.3334
Estimated allele frequency
Ecological gradient (Mean annual temperature)
1.0
0.8
0.6
0.4
0.2
5
6
7
8
9
10
11
12
O16
L16
L12
O12
253
256
O8
204
L8
219
217
218
233
124
9
97
O1
L1

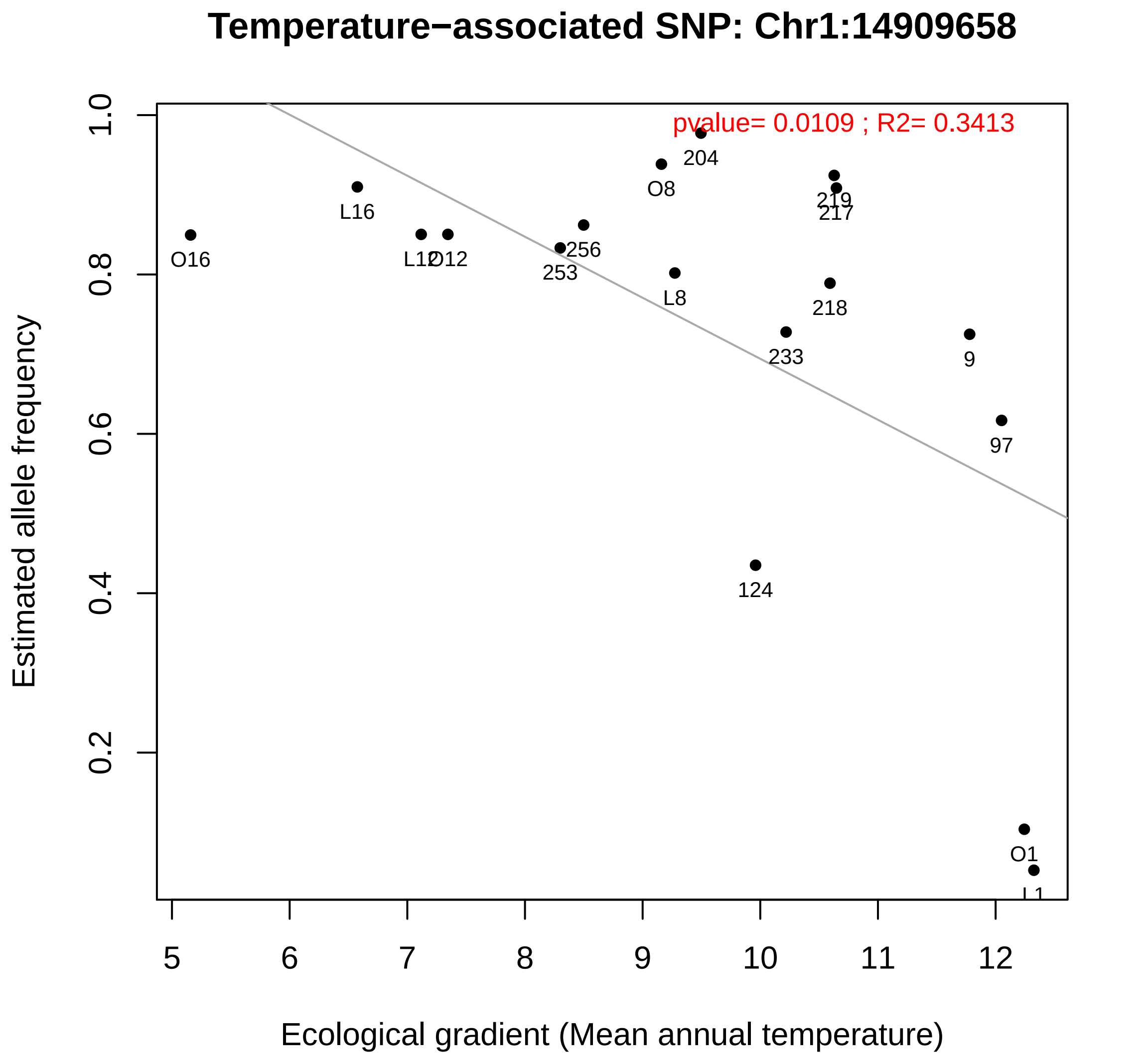

Temperature−associated SNP: Chr1:14909658
pvalue= 0.0109 ; R2= 0.3413
O16
L16
L12
O12
253
256
O8
204
L8
219
217
218
233
124
9
97
O1
L1
1.0
0.8
0.6
0.4
0.2
5
6
7
8
9
10
11
12
Estimated allele frequency
Ecological gradient (Mean annual temperature)

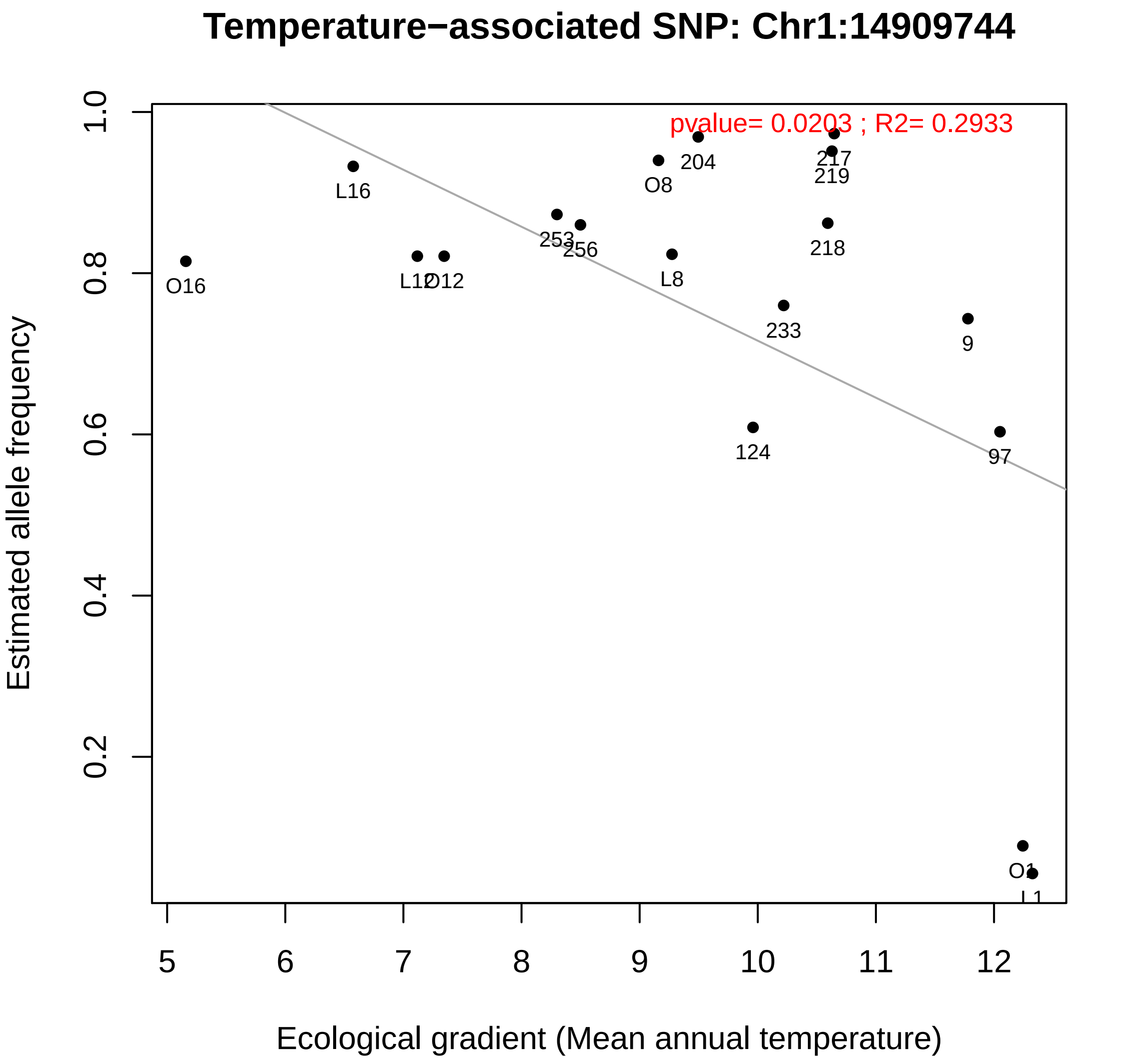

Temperature−associated SNP: Chr1:14909744
pvalue= 0.0203 ; R2= 0.2933
Estimated allele frequency
Ecological gradient (Mean annual temperature)
0.2
0.4
0.6
0.8
1.0
5
6
7
8
9
10
11
12
O16
L16
L12
O12
253
256
O8
204
L8
217
219
218
233
124
9
97
O1
L1

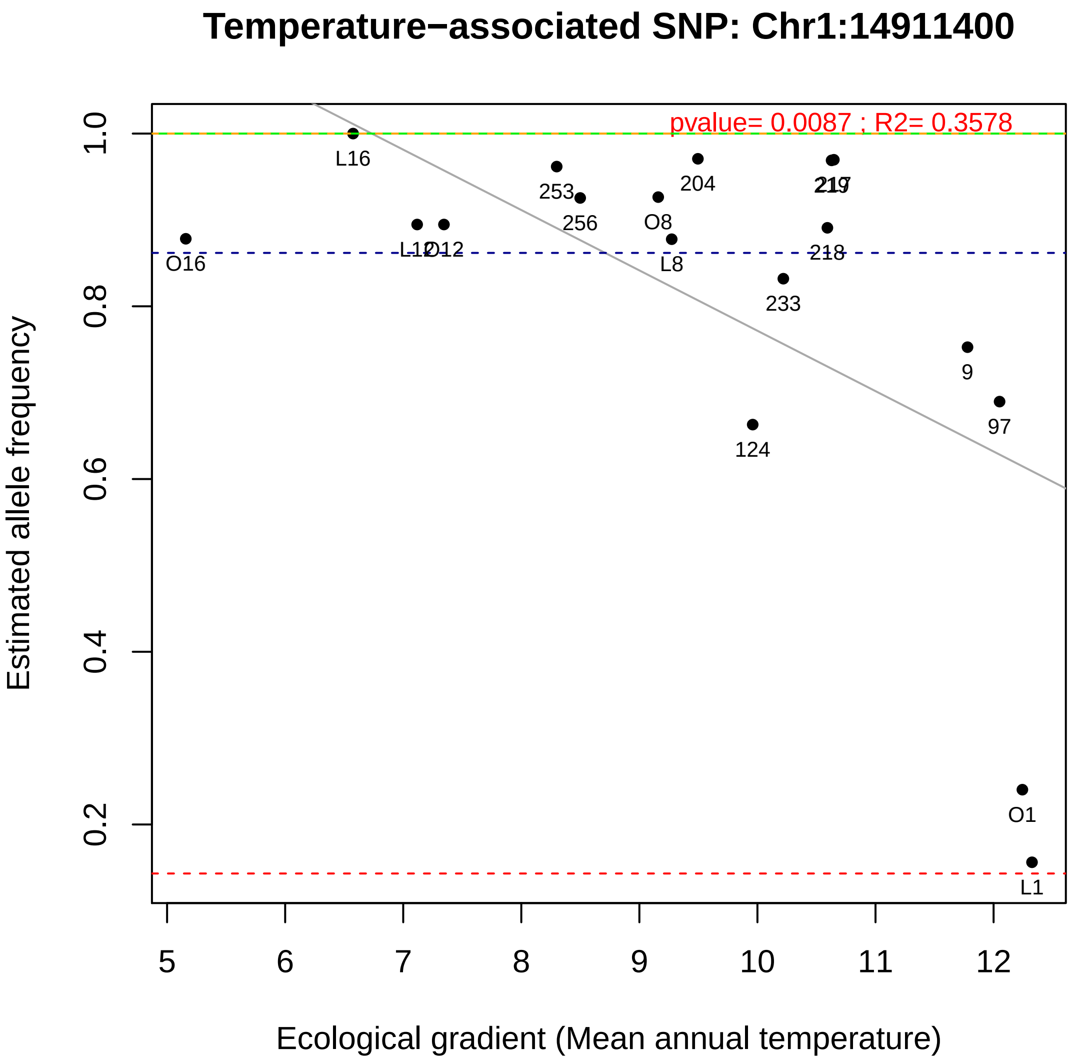

Temperature−associated SNP: Chr1:14911400
pvalue= 0.0087 ; R2= 0.3578
Estimated allele frequency
Ecological gradient (Mean annual temperature)
1.0
0.8
0.6
0.4
0.2
5
6
7
8
9
10
11
12
L16
253
256
204
217
219
O8
O16
L12
O12
L8
218
233
9
97
124
O1
L1

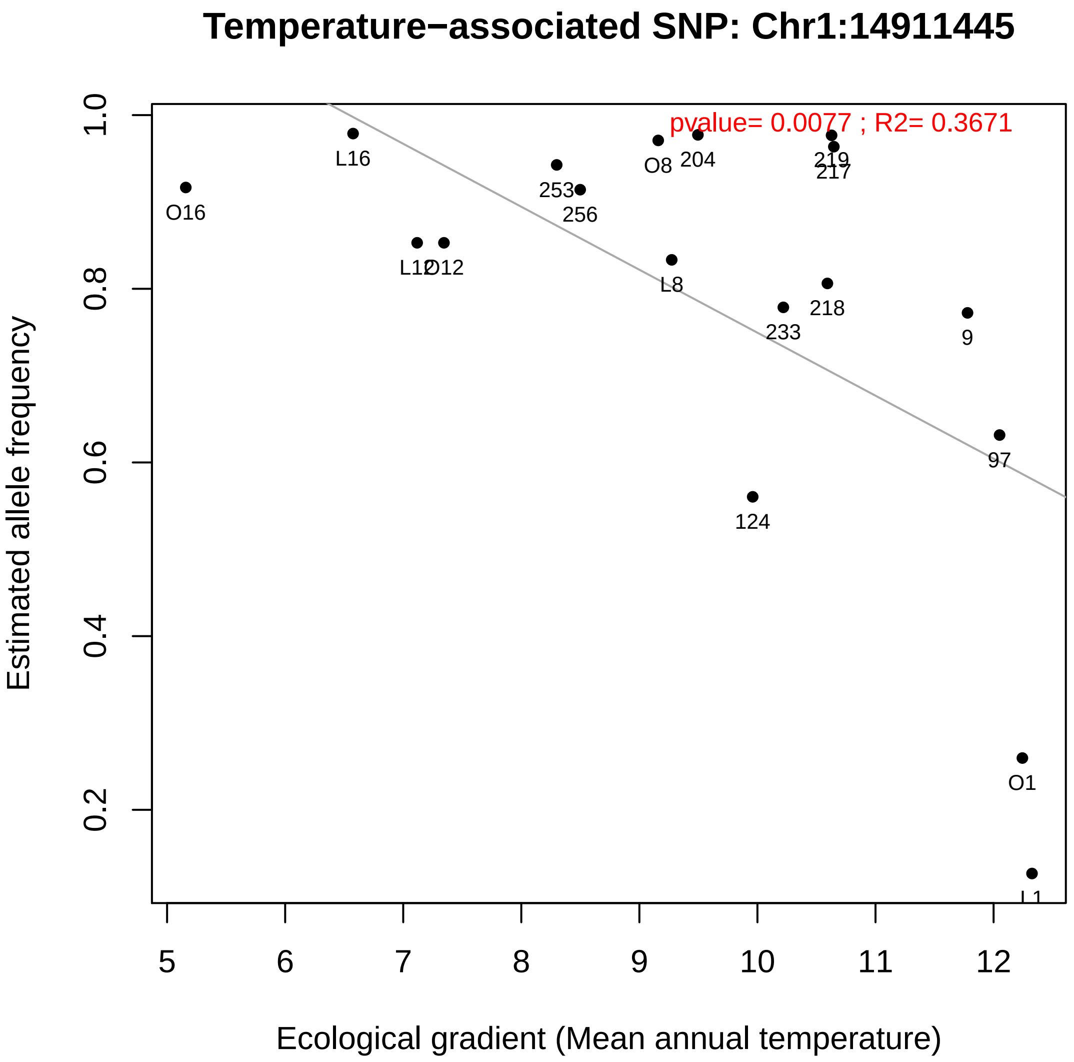

Temperature−associated SNP: Chr1:14911445
pvalue= 0.0077 ; R2= 0.3671
Estimated allele frequency
Ecological gradient (Mean annual temperature)
0.2
0.4
0.6
0.8
1.0
5
6
7
8
9
10
11
12
O16
L16
L12
O12
253
256
O8
204
L8
219
217
218
233
9
97
124
O1

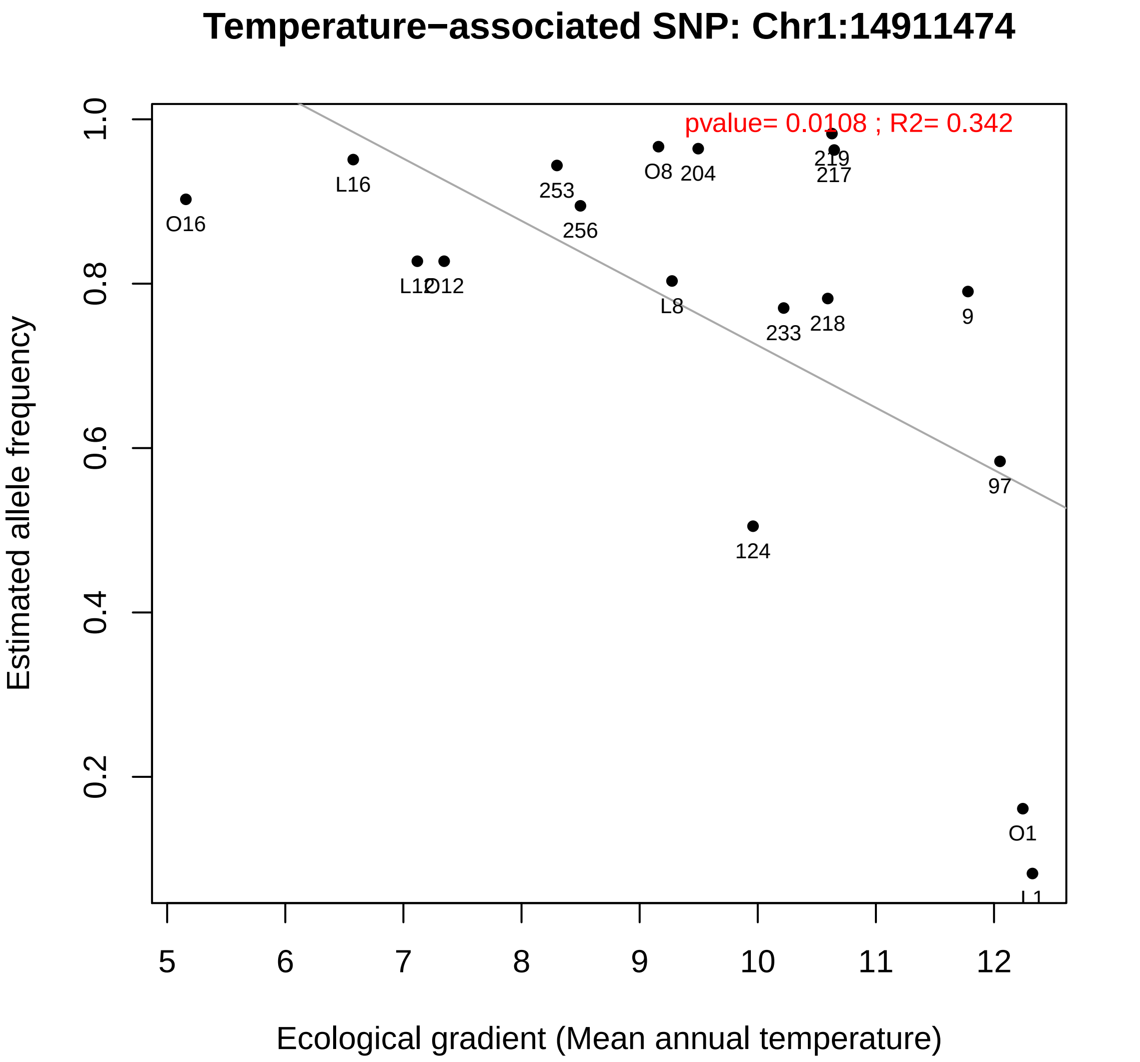
Temperature−associated SNP: Chr1:14911474
pvalue= 0.0108 ; R2= 0.342
Estimated allele frequency
Ecological gradient (Mean annual temperature)
1.0
0.8
0.6
0.4
0.2
5
6
7
8
9
10
11
12
O16
L16
L12
O12
253
256
O8
204
219
217
L8
233
218
9
97
124
O1

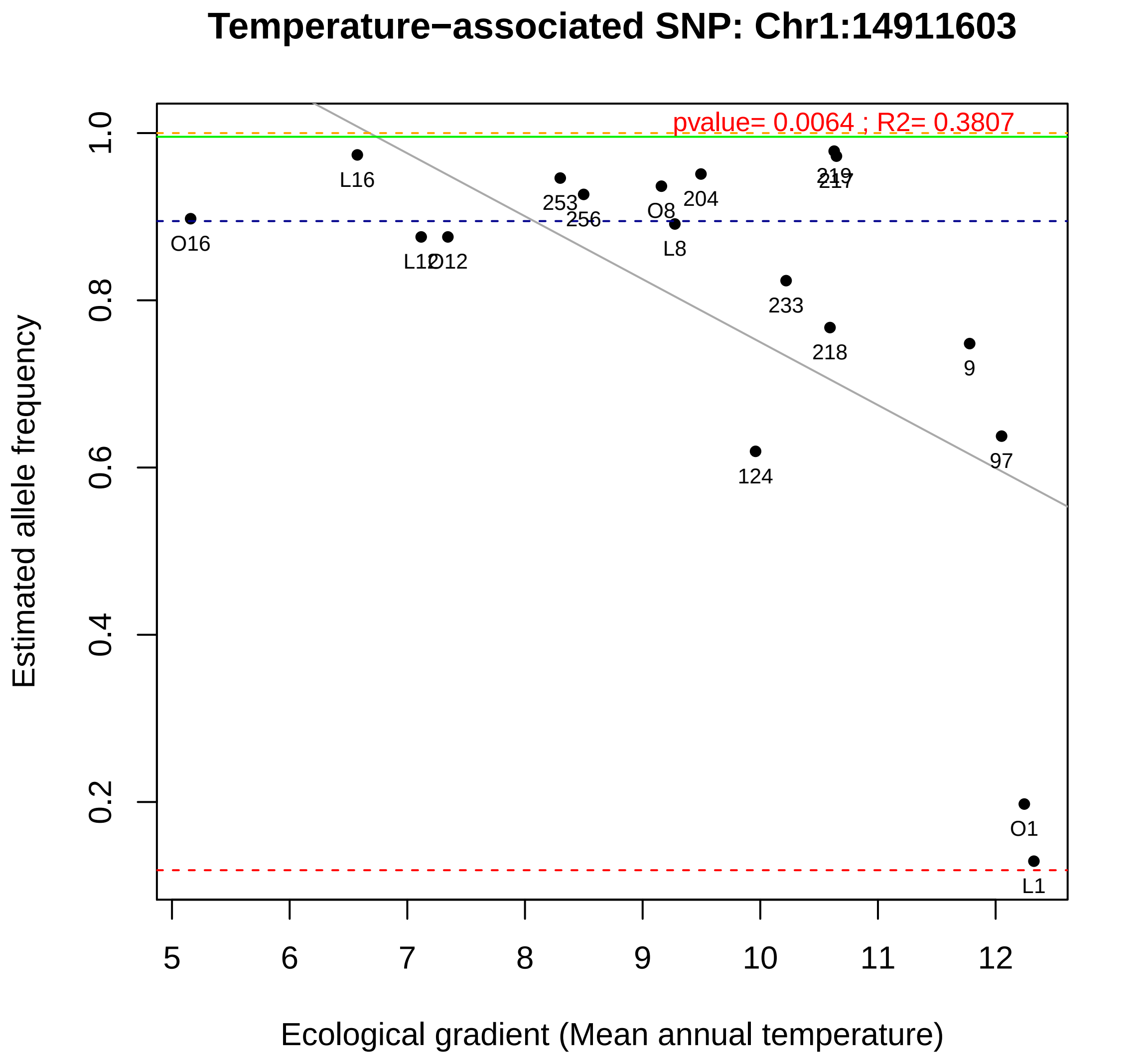

Temperature−associated SNP: Chr1:14911603
pvalue= 0.0064 ; R2= 0.3807
Estimated allele frequency
1.0
0.8
0.6
0.4
0.2
5
6
7
8
9
10
11
12
Ecological gradient (Mean annual temperature)
L16
O16
L12
O12
253
256
O8
204
L8
219
217
233
218
9
124
97
O1
L1

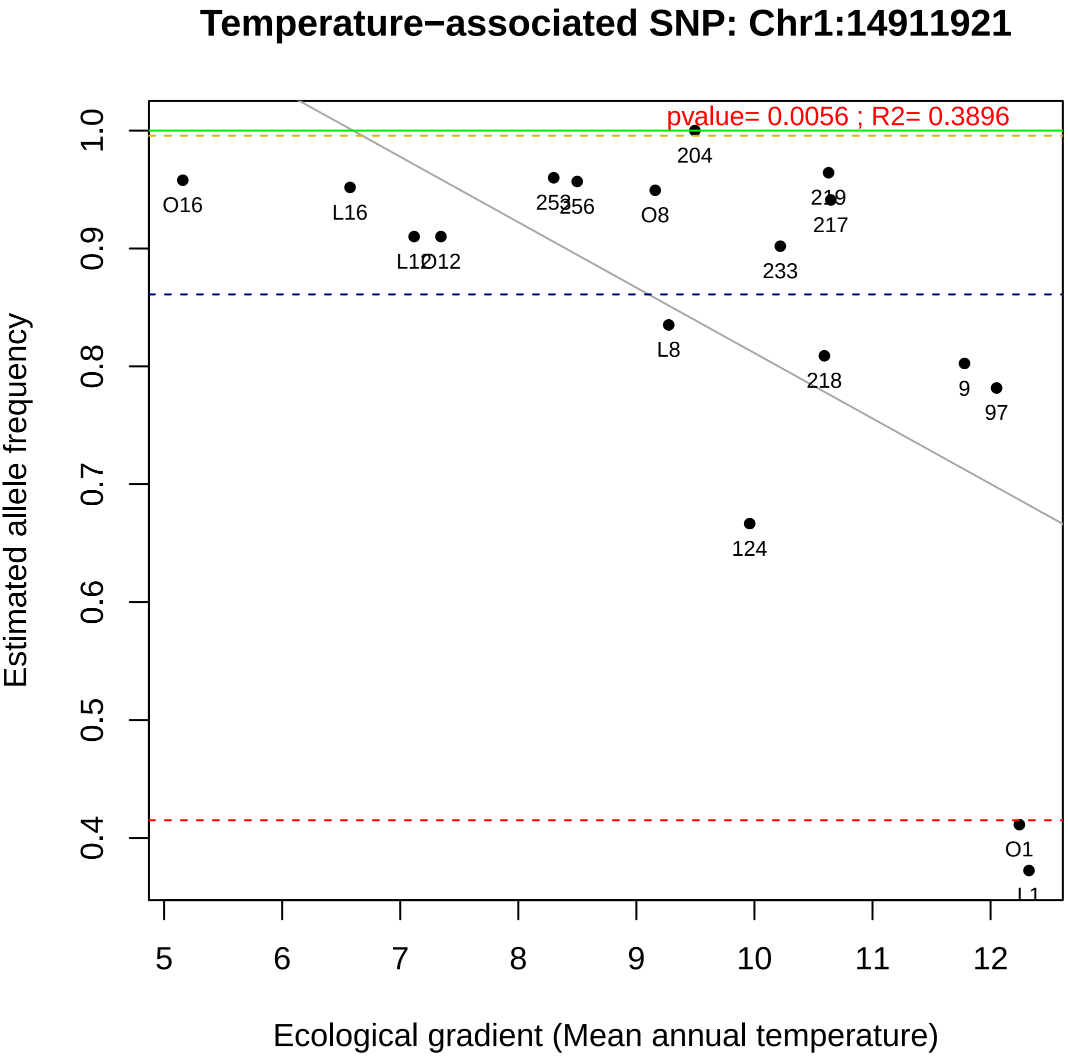

Temperature−associated SNP: Chr1:14911921
pvalue= 0.0056 ; R2= 0.3896
Estimated allele frequency
Ecological gradient (Mean annual temperature)
0.4
0.5
0.6
0.7
0.8
0.9
1.0
5
6
7
8
9
10
11
12
O16
L16
L12
O12
253
256
O8
204
219
217
233
L8
218
124
9
97
O1

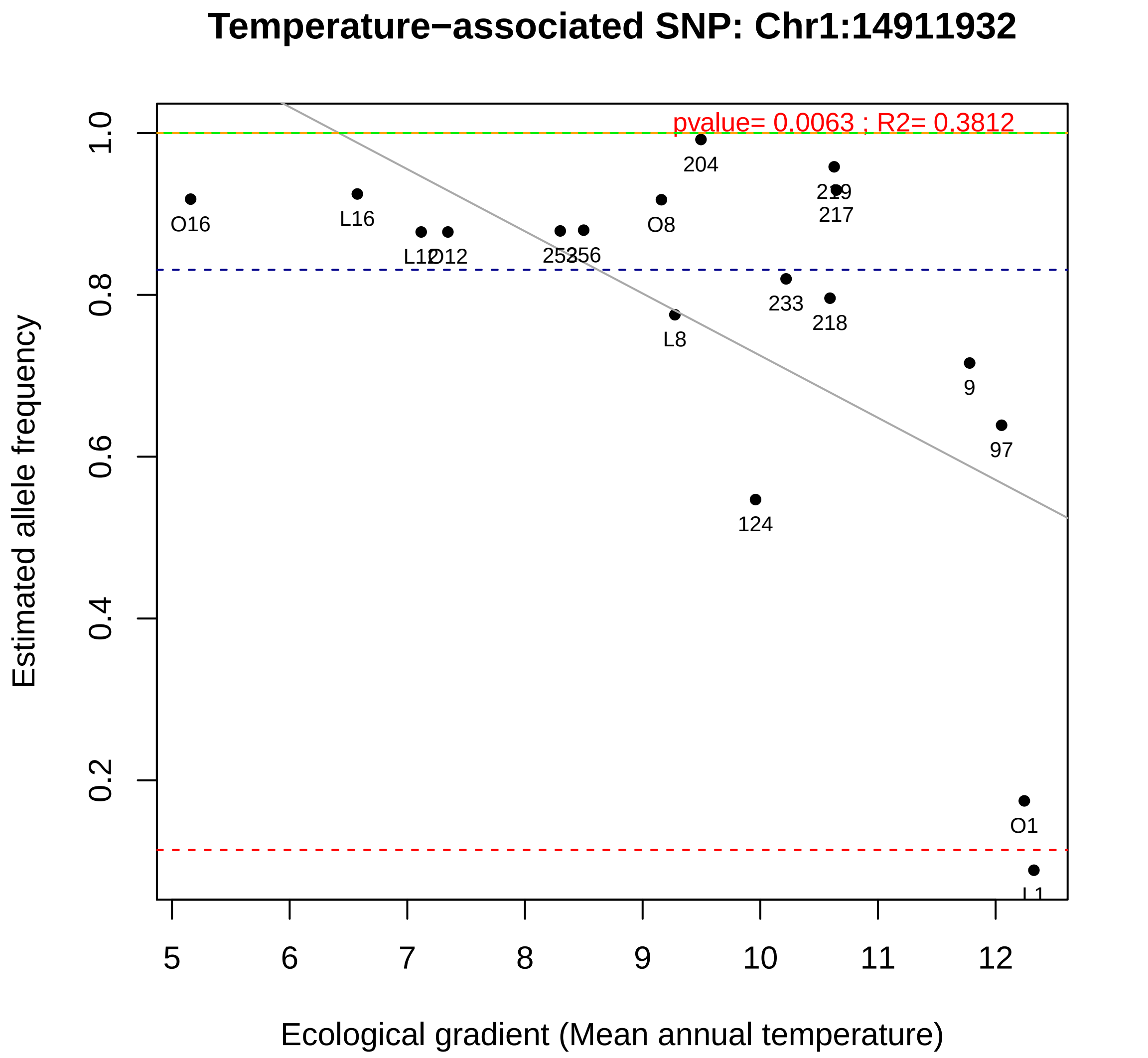

Temperature−associated SNP: Chr1:14911932
pvalue= 0.0063 ; R2= 0.3812
Estimated allele frequency
Ecological gradient (Mean annual temperature)
0.2
0.4
0.6
0.8
1.0
5
6
7
8
9
10
11
12
O16
L16
O8
204
217
233
218
L8
9
97
124
O1

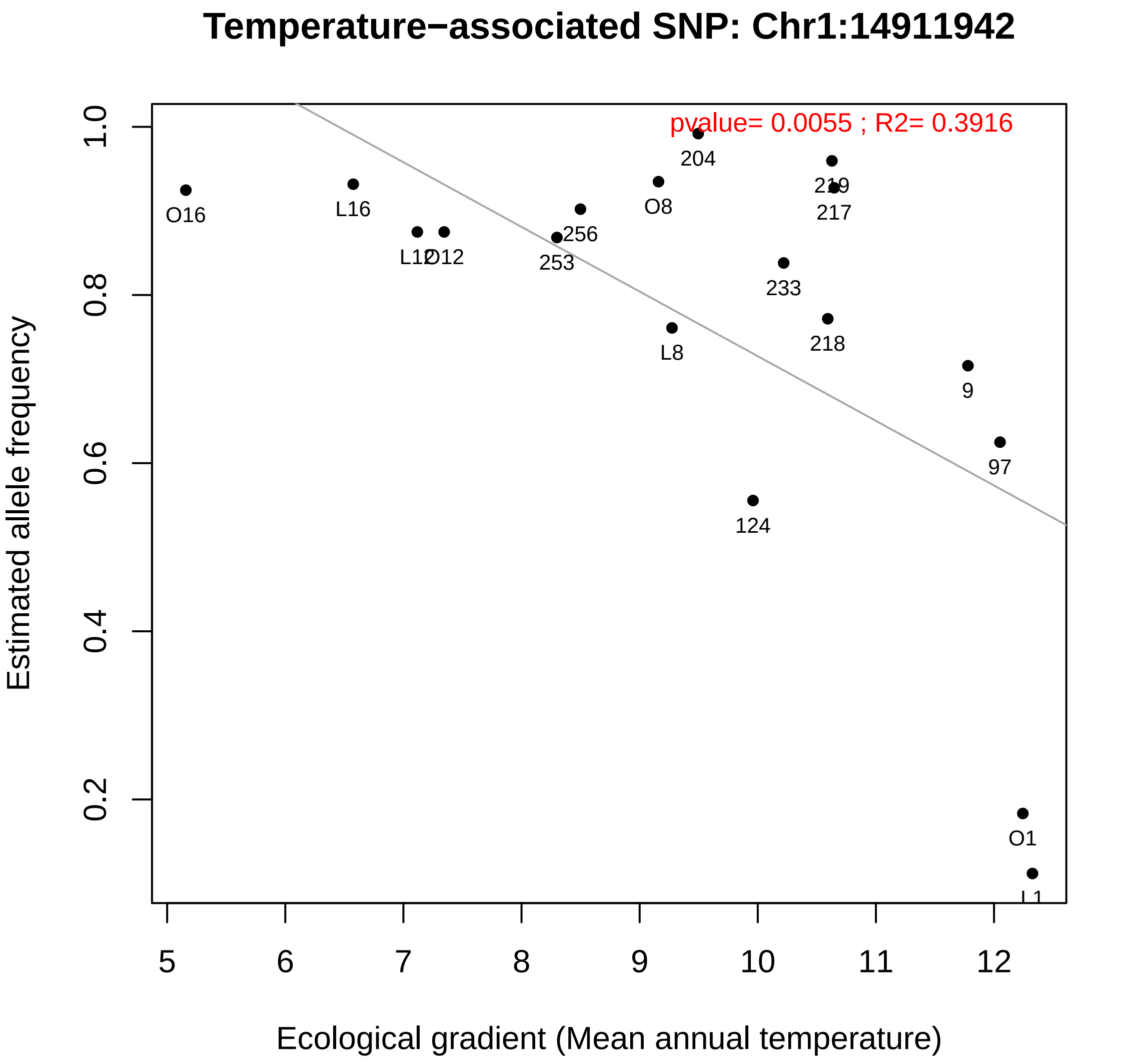
Temperature−associated SNP: Chr1:14911942
pvalue= 0.0055 ; R2= 0.3916
Estimated allele frequency
Ecological gradient (Mean annual temperature)
0.2
0.4
0.6
0.8
1.0
5
6
7
8
9
10
11
12
O16
L16
L12
O12
253
256
O8
204
L8
219
217
233
218
124
9
97
O1

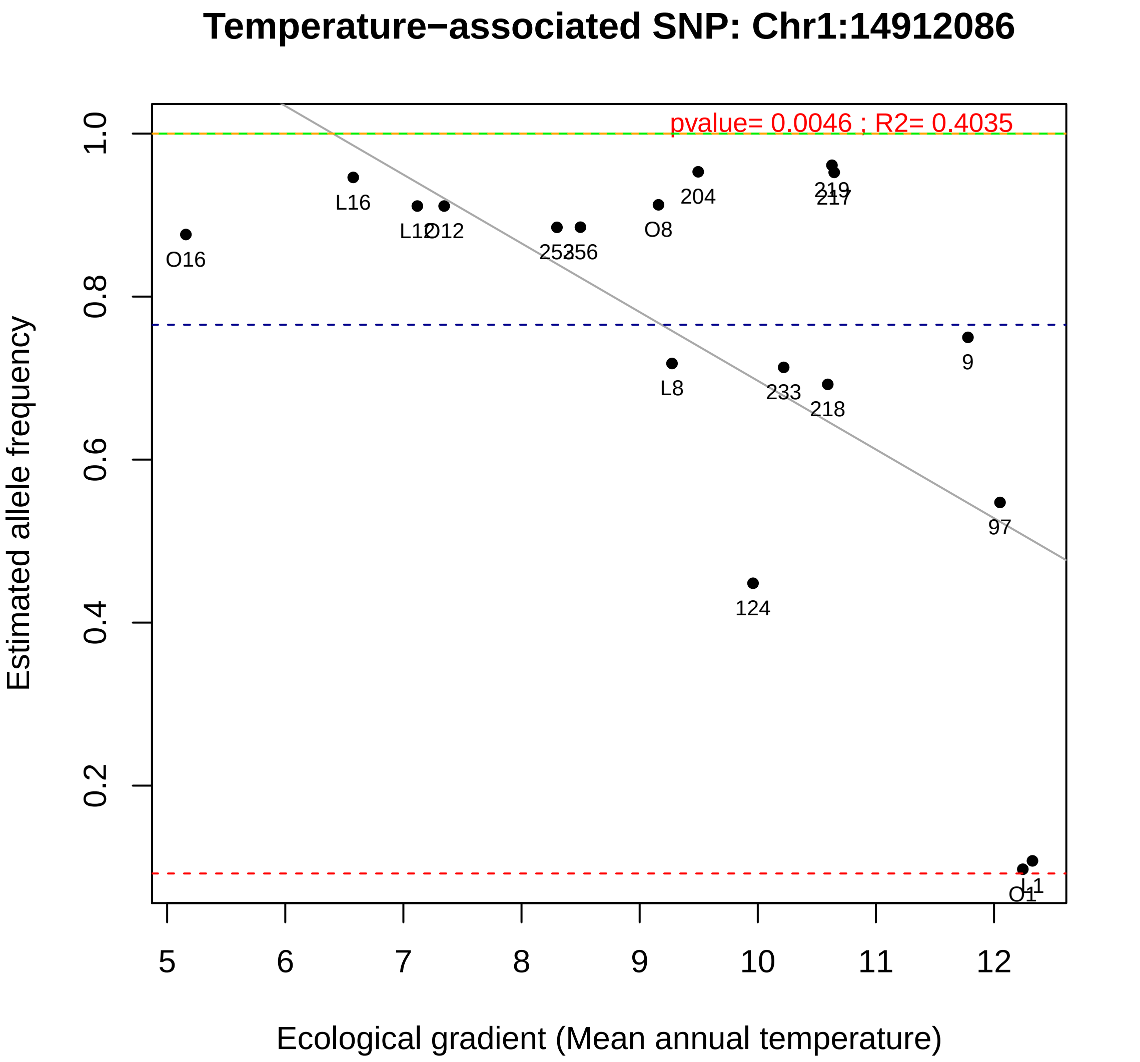

Temperature−associated SNP: Chr1:14912086
pvalue= 0.0046 ; R2= 0.4035
Estimated allele frequency
Ecological gradient (Mean annual temperature)
0.2
0.4
0.6
0.8
1.0
5
6
7
8
9
10
11
12
O16
L16
L12
O12
252
256
O8
204
219
217
L8
233
218
9
97
124
O1
L1

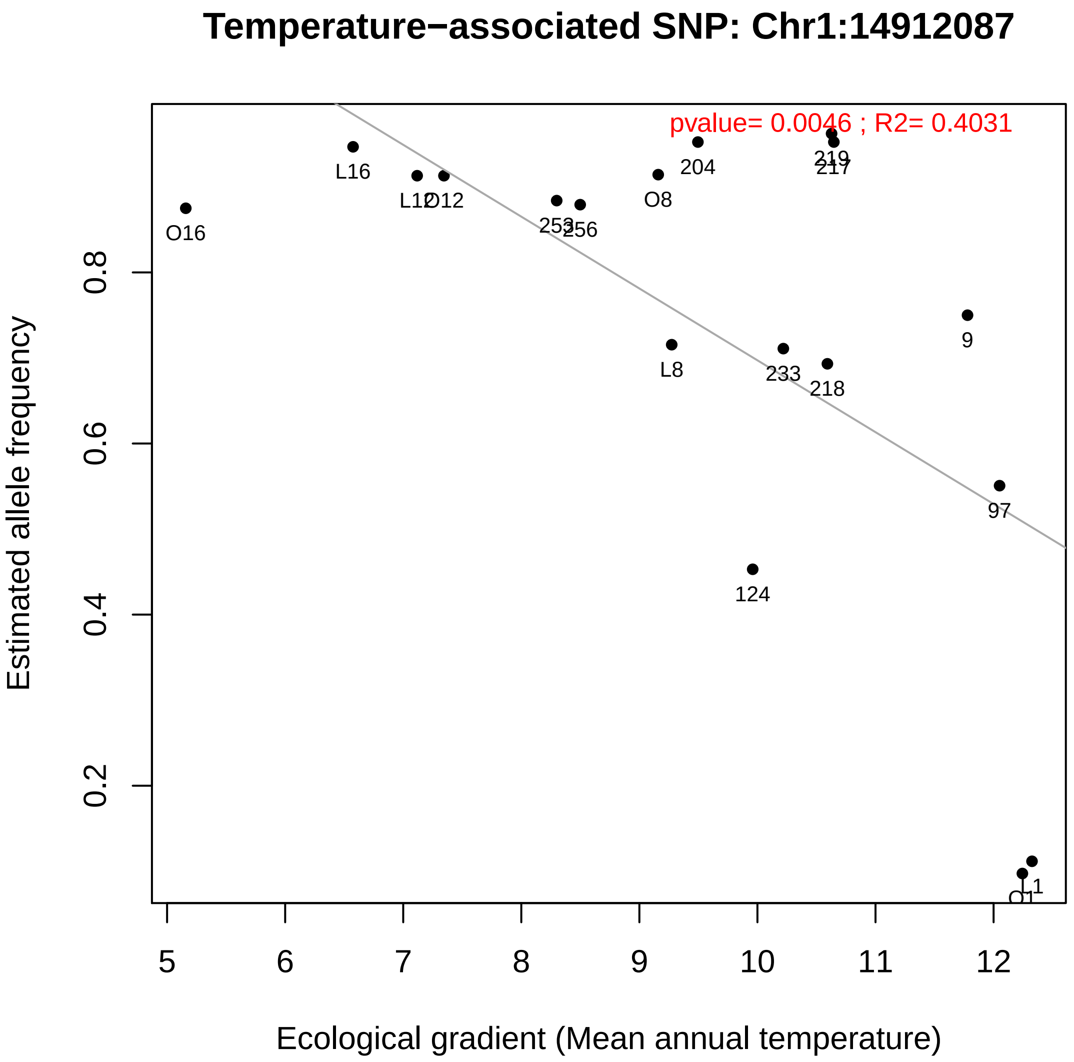
Temperature−associated SNP: Chr1:14912087
pvalue= 0.0046 ; R2= 0.4031
Estimated allele frequency
0.2
0.4
0.6
0.8
5
6
7
8
9
10
11
12
Ecological gradient (Mean annual temperature)
O16
L16
L12
O12
253
256
O8
204
219
217
L8
233
218
9
97
124
L1
O1

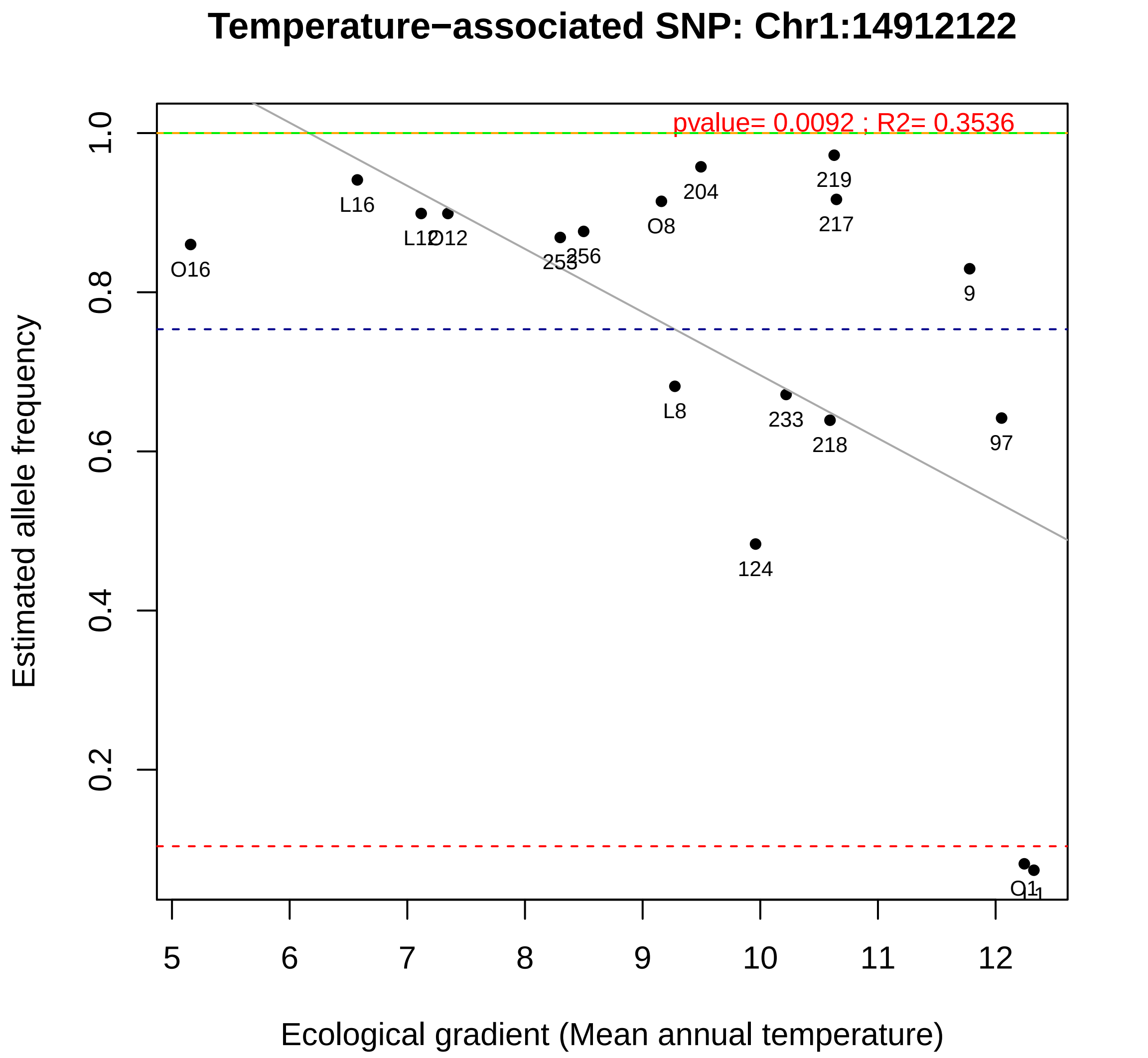

Temperature−associated SNP: Chr1:14912122
pvalue= 0.0092 ; R2= 0.3536
Estimated allele frequency
0.2
0.4
0.6
0.8
1.0
5
6
7
8
9
10
11
12
Ecological gradient (Mean annual temperature)
O16
L16
L12
O12
253
256
O8
204
219
217
9
L8
233
218
97
124

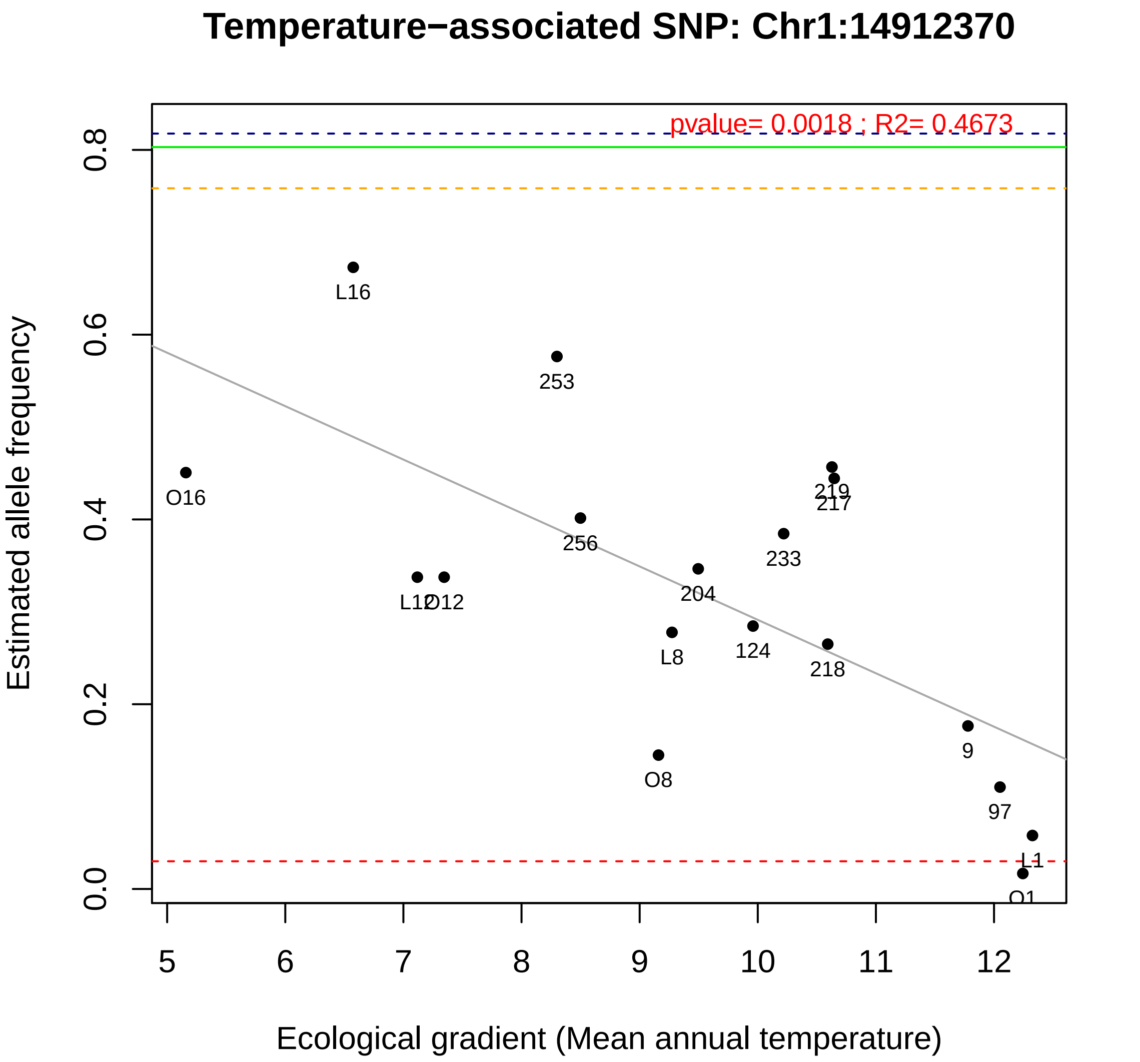
Temperature−associated SNP: Chr1:14912370
pvalue= 0.0018 ; R2= 0.4673
Estimated allele frequency
0.0
0.2
0.4
0.6
0.8
5
6
7
8
9
10
11
12
Ecological gradient (Mean annual temperature)
L16
253
O16
256
219
217
233
204
L12
O12
L8
124
218
9
O8
97
L1
O1

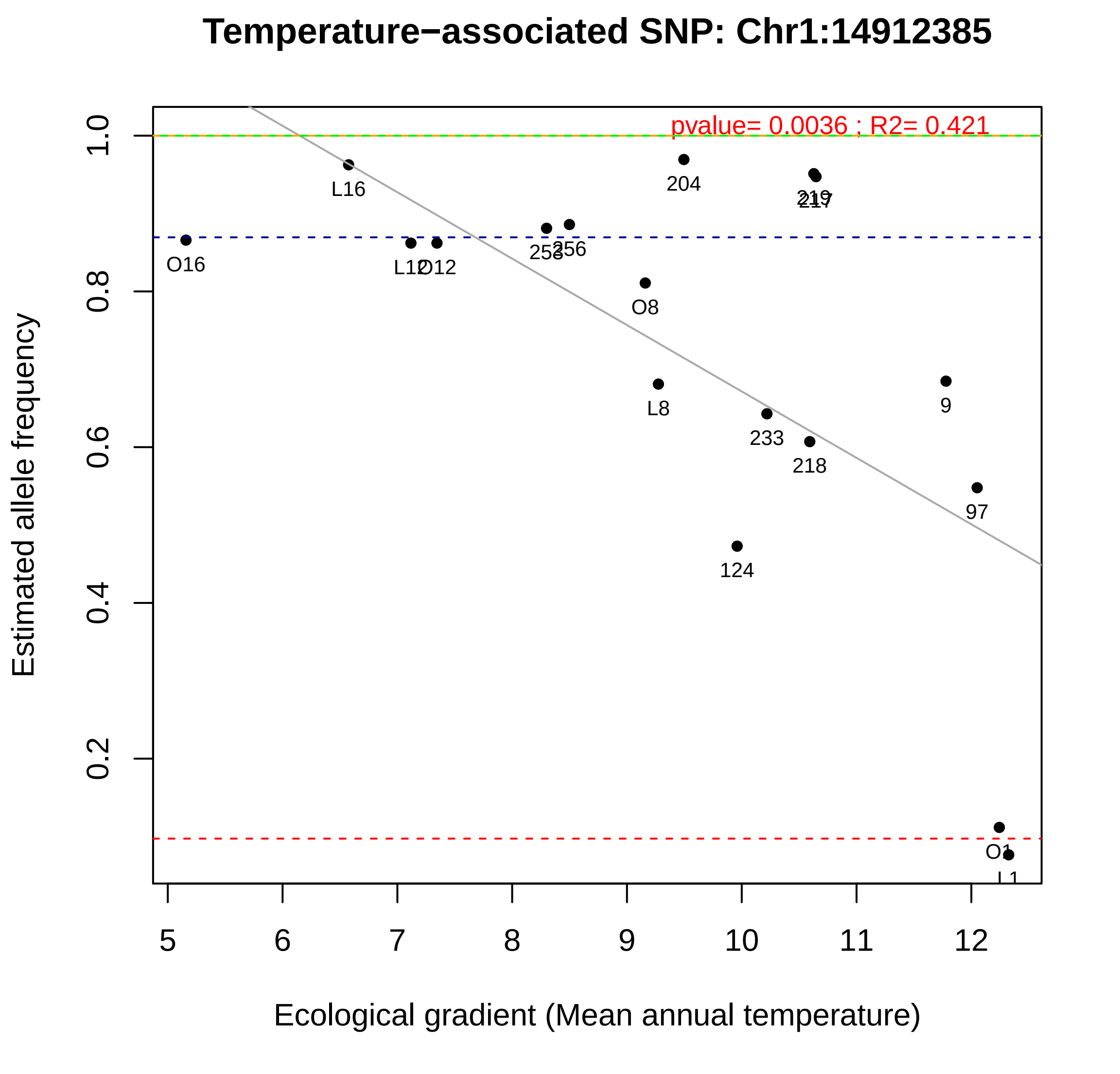

Temperature−associated SNP: Chr1:14912385
pvalue= 0.0036 ; R2= 0.421
Estimated allele frequency
Ecological gradient (Mean annual temperature)
0.2
0.4
0.6
0.8
1.0
5
6
7
8
9
10
11
12
O16
L16
L12
O12
253
256
204
O8
L8
219
217
233
218
124
9
97

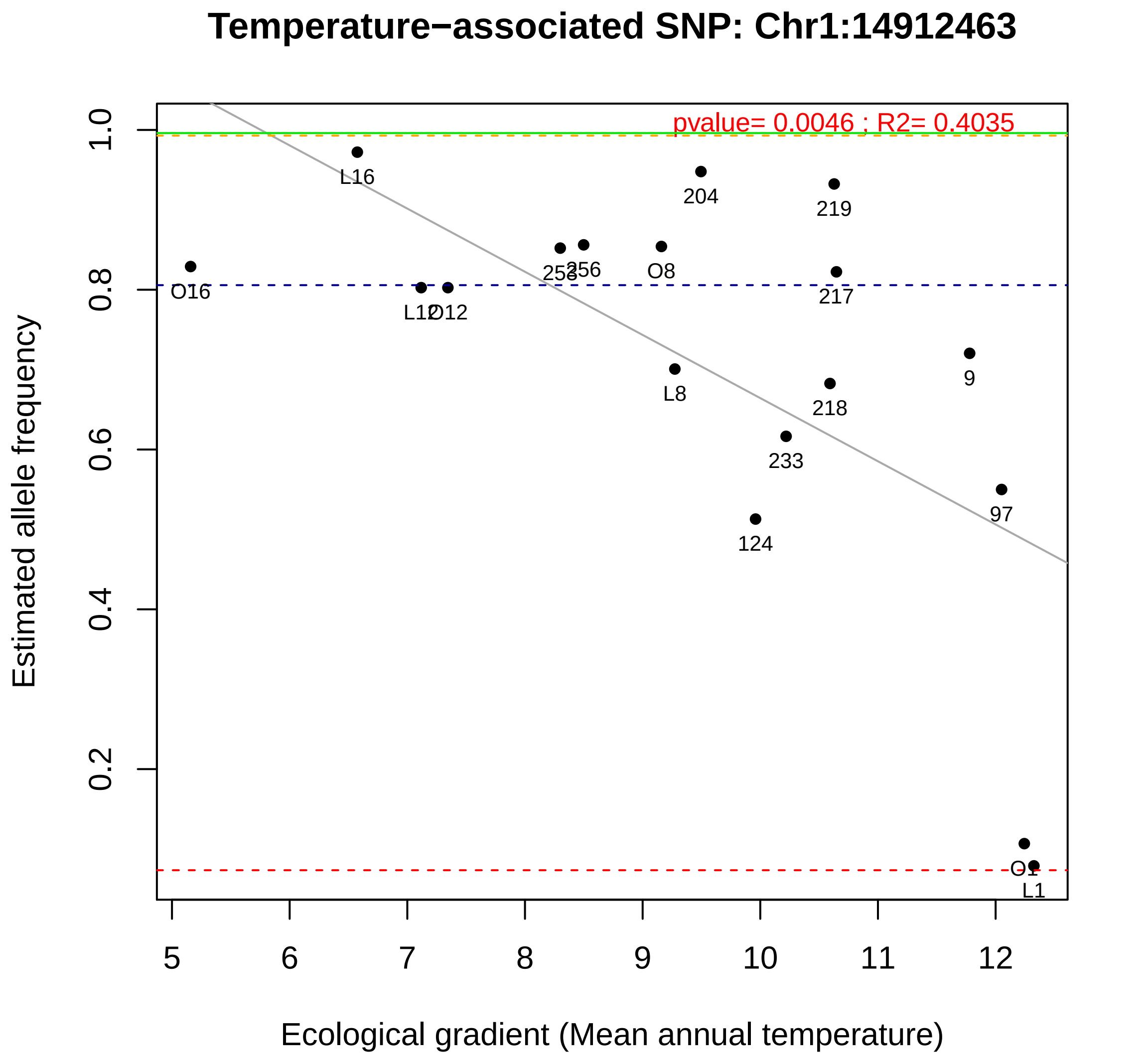

Temperature−associated SNP: Chr1:14912463
pvalue= 0.0046 ; R2= 0.4035
Estimated allele frequency
Ecological gradient (Mean annual temperature)
1.0
0.8
0.6
0.4
0.2
5
6
7
8
9
10
11
12
L16
204
219
O16
256
O8
217
L12
O12
9
L8
218
233
97
124
L1

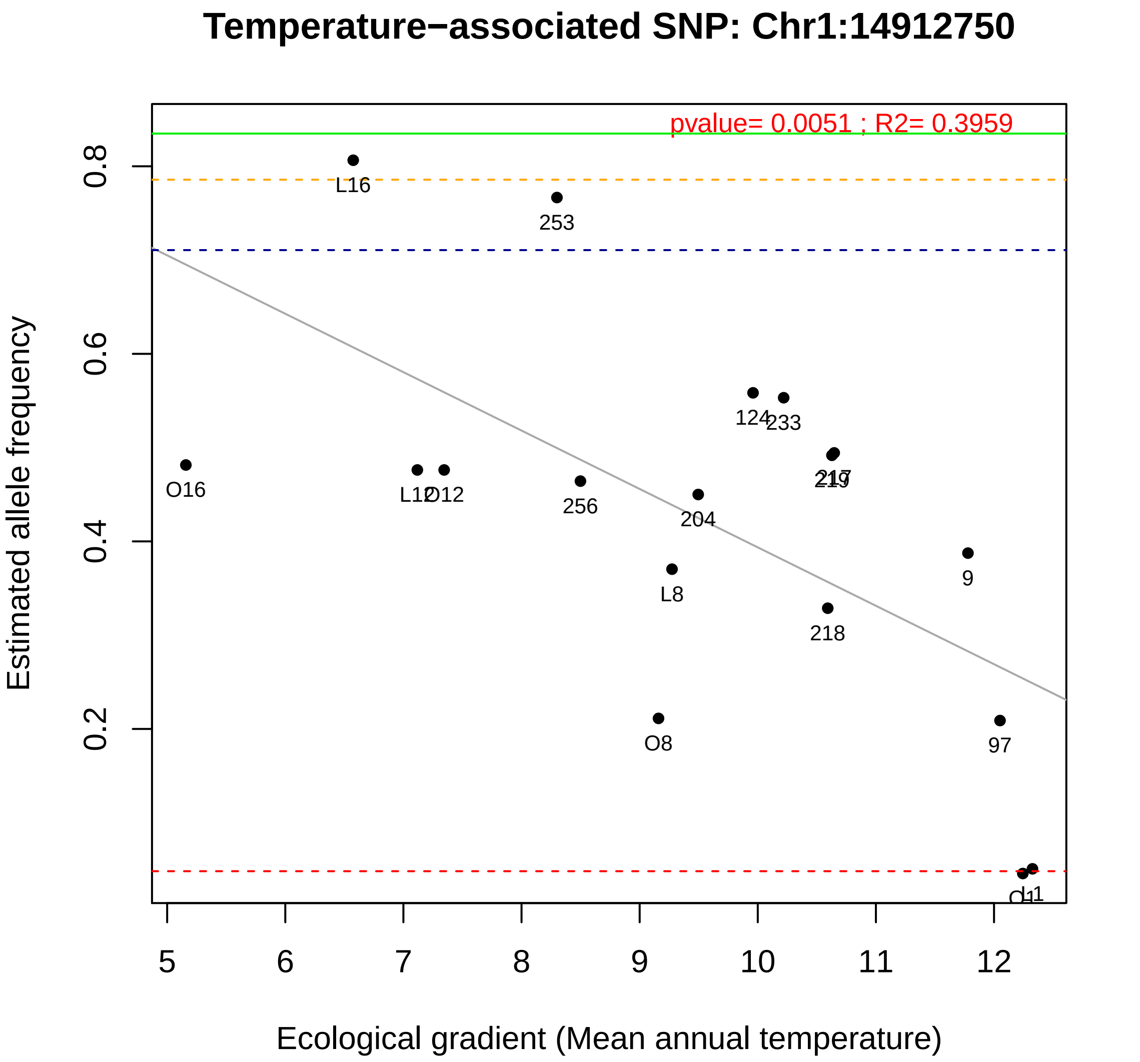

Temperature−associated SNP: Chr1:14912750
pvalue= 0.0051 ; R2= 0.3959
Estimated allele frequency
0.2
0.4
0.6
0.8
5
6
7
8
9
10
11
12
Ecological gradient (Mean annual temperature)
L16
253
124
233
217
219
O16
L12
O12
256
204
L8
9
218
O8
97

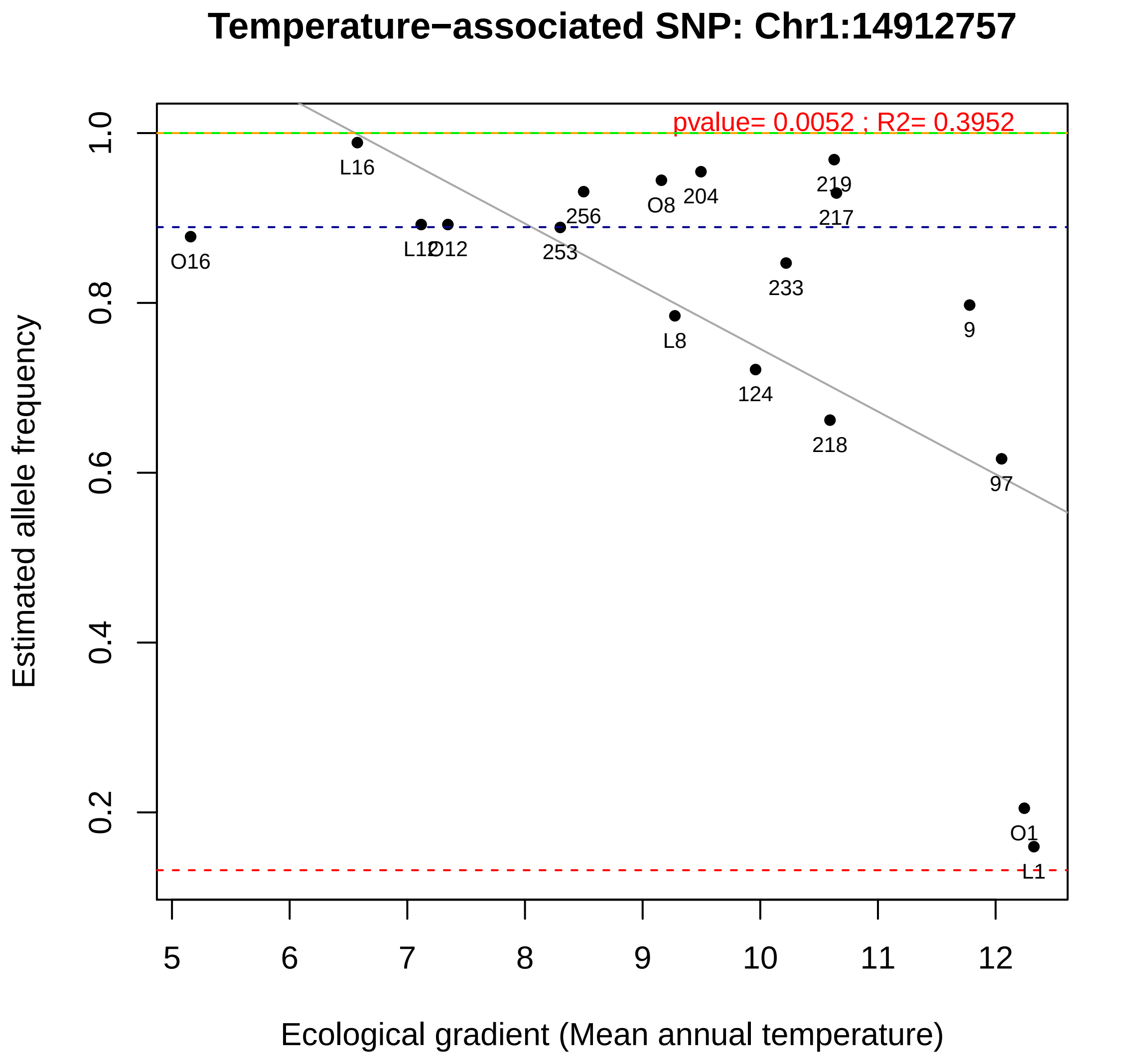
Temperature−associated SNP: Chr1:14912757
pvalue= 0.0052 ; R2= 0.3952
Estimated allele frequency
Ecological gradient (Mean annual temperature)
0.2
0.4
0.6
0.8
1.0
5
6
7
8
9
10
11
12
O16
L16
L12
O12
253
256
O8
204
219
217
233
L8
124
218
9
97
O1
L1

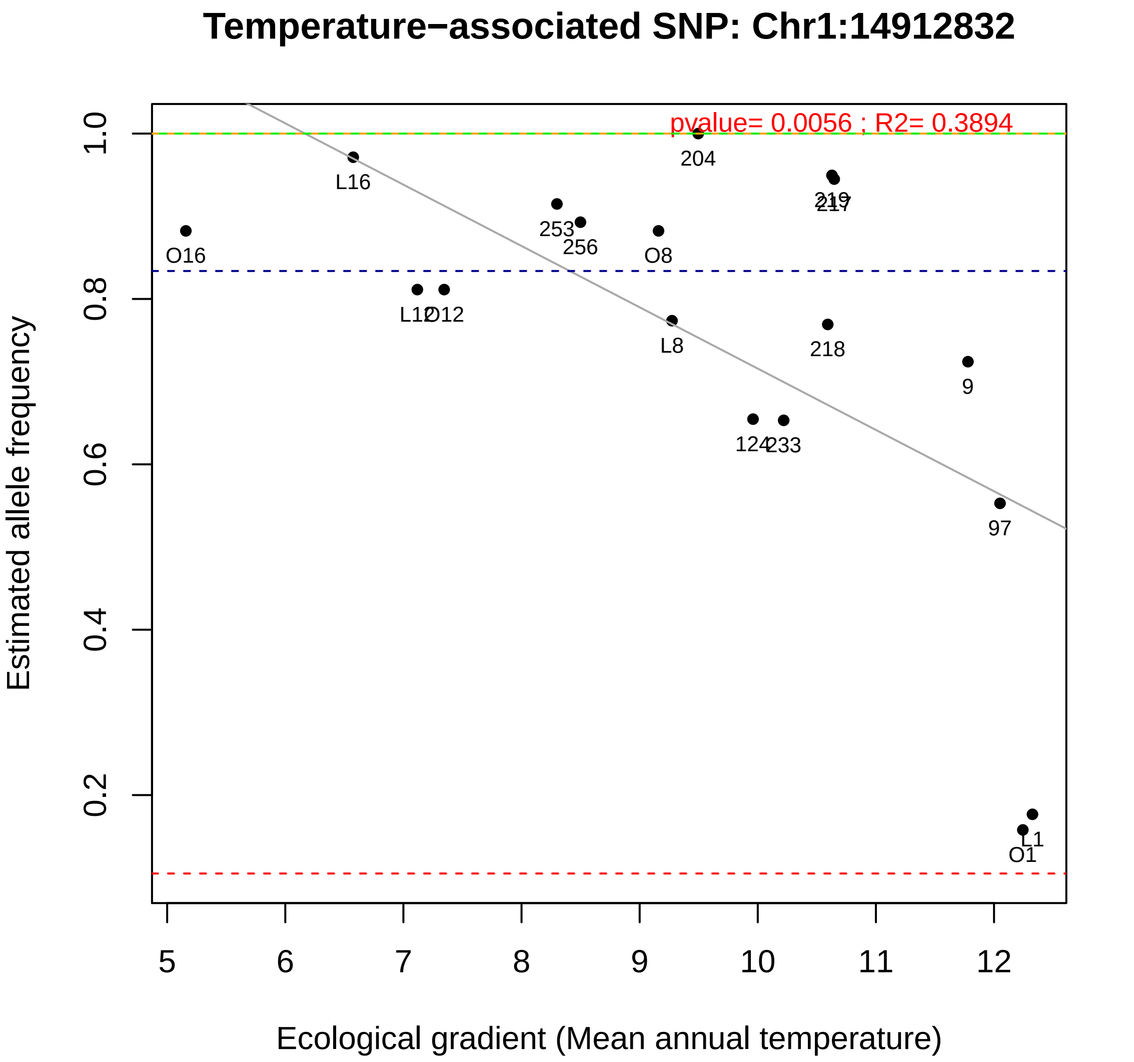

Temperature−associated SNP: Chr1:14912832
pvalue= 0.0056 ; R2= 0.3894
Estimated allele frequency
Ecological gradient (Mean annual temperature)
0.2
0.4
0.6
0.8
1.0
5
6
7
8
9
10
11
12
L16
204
219
217
O16
253
256
O8
L12
O12
L8
218
9
124
233
97
L1
O1

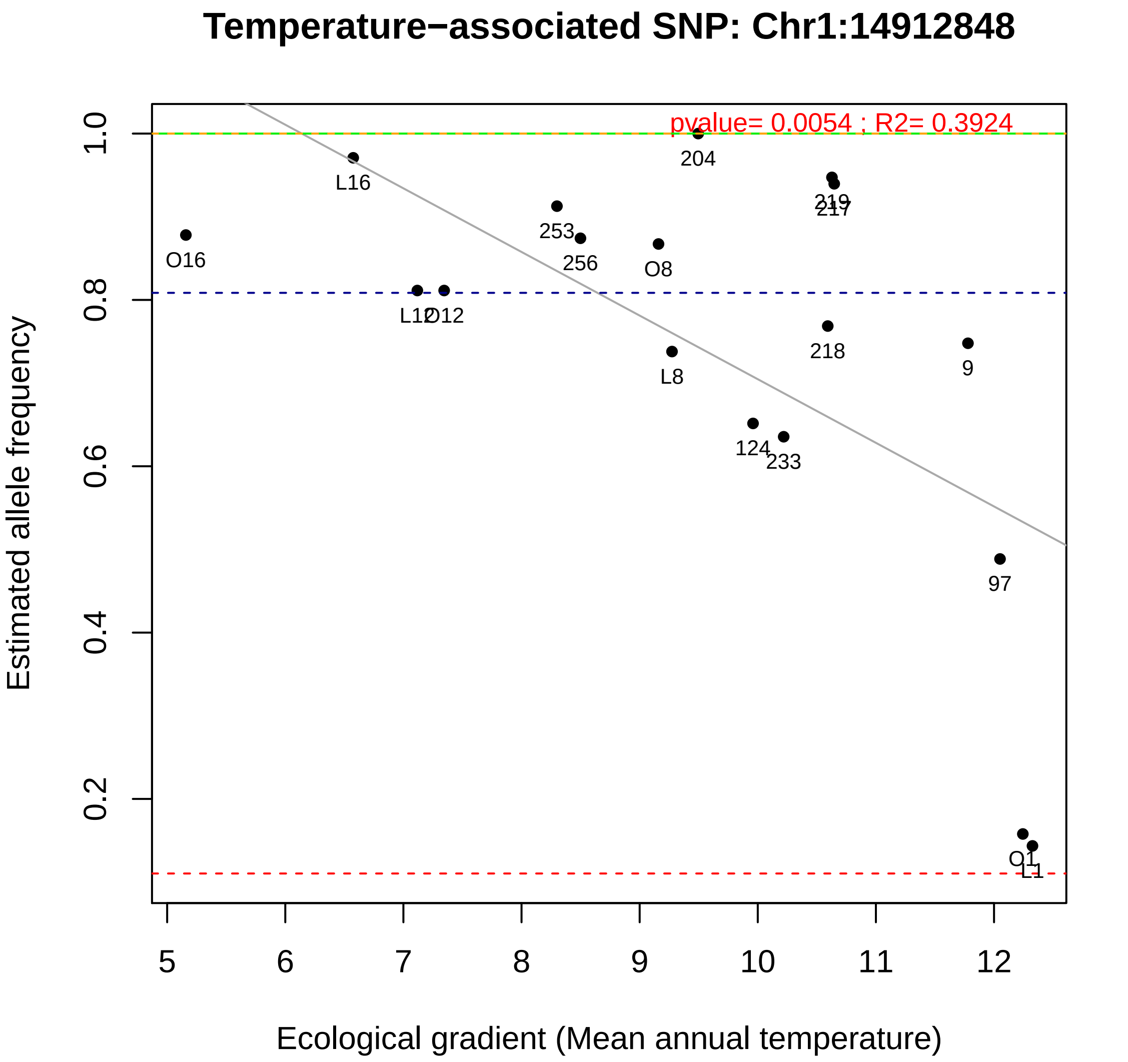

Temperature−associated SNP: Chr1:14912848
pvalue= 0.0054 ; R2= 0.3924
Estimated allele frequency
0.2
0.4
0.6
0.8
1.0
5
6
7
8
9
10
11
12
Ecological gradient (Mean annual temperature)
O16
L16
L12
O12
253
256
O8
204
L8
124
233
219
217
218
9
97
O1
L1

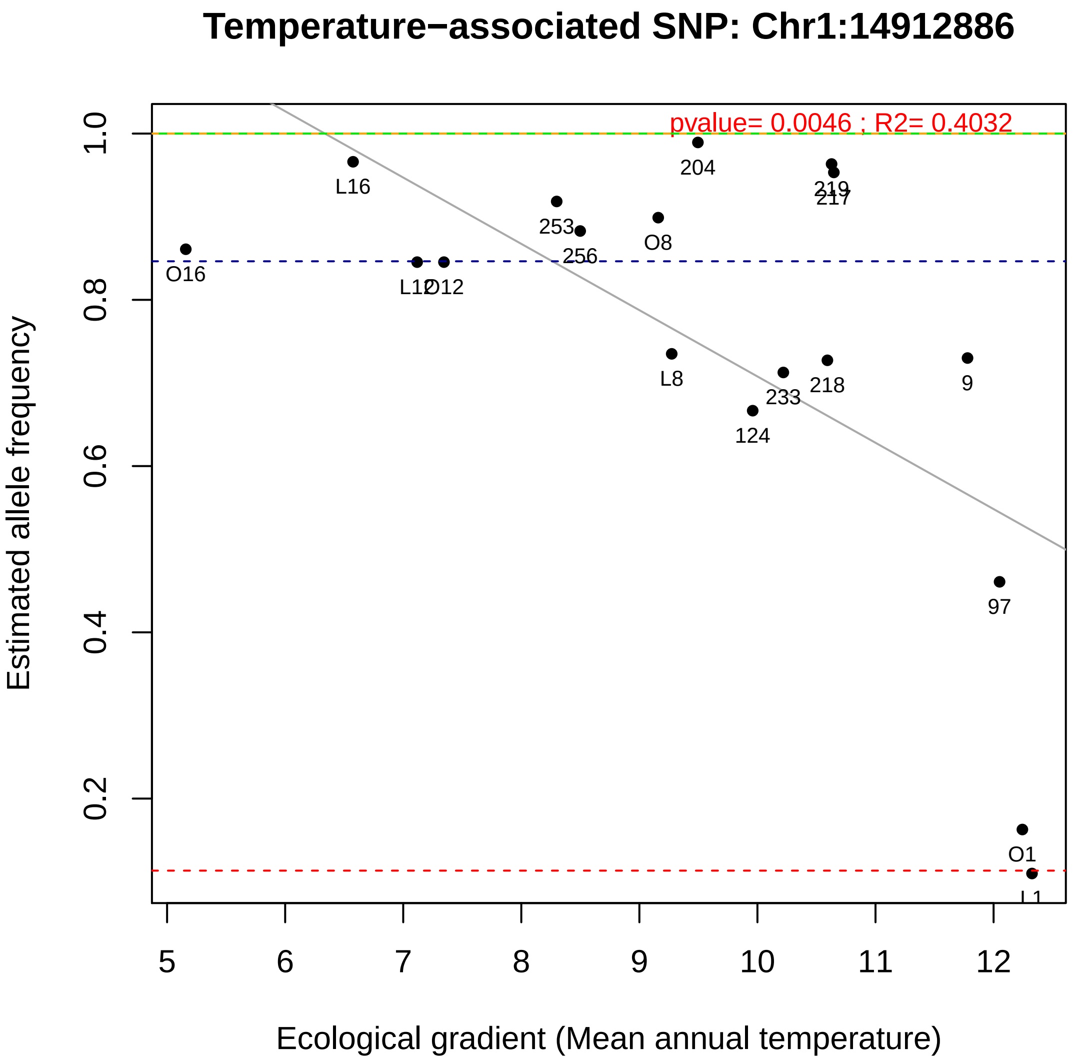

Temperature−associated SNP: Chr1:14912886
pvalue= 0.0046 ; R2= 0.4032
Estimated allele frequency
1.0
0.8
0.6
0.4
0.2
O16
L16
L12
O12
253
256
O8
204
L8
124
233
218
219
217
9
97
O1
L1
5
6
7
8
9
10
11
12
Ecological gradient (Mean annual temperature)

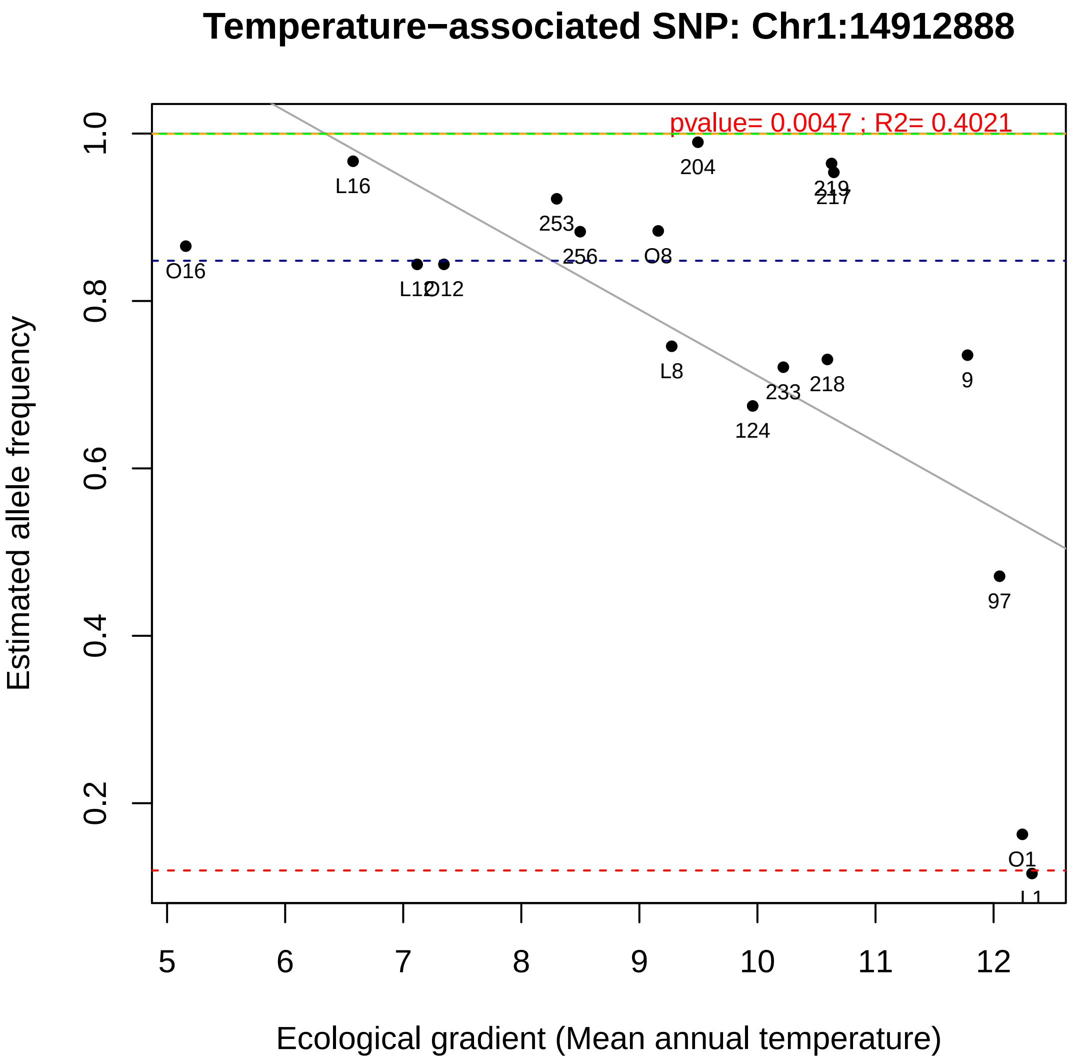

Temperature−associated SNP: Chr1:14912888
pvalue= 0.0047 ; R2= 0.4021
Estimated allele frequency
Ecological gradient (Mean annual temperature)
1.0
0.8
0.6
0.4
0.2
5
6
7
8
9
10
11
12
O16
L16
L12
O12
253
256
O8
204
219
217
L8
233
218
124
9
97
O1

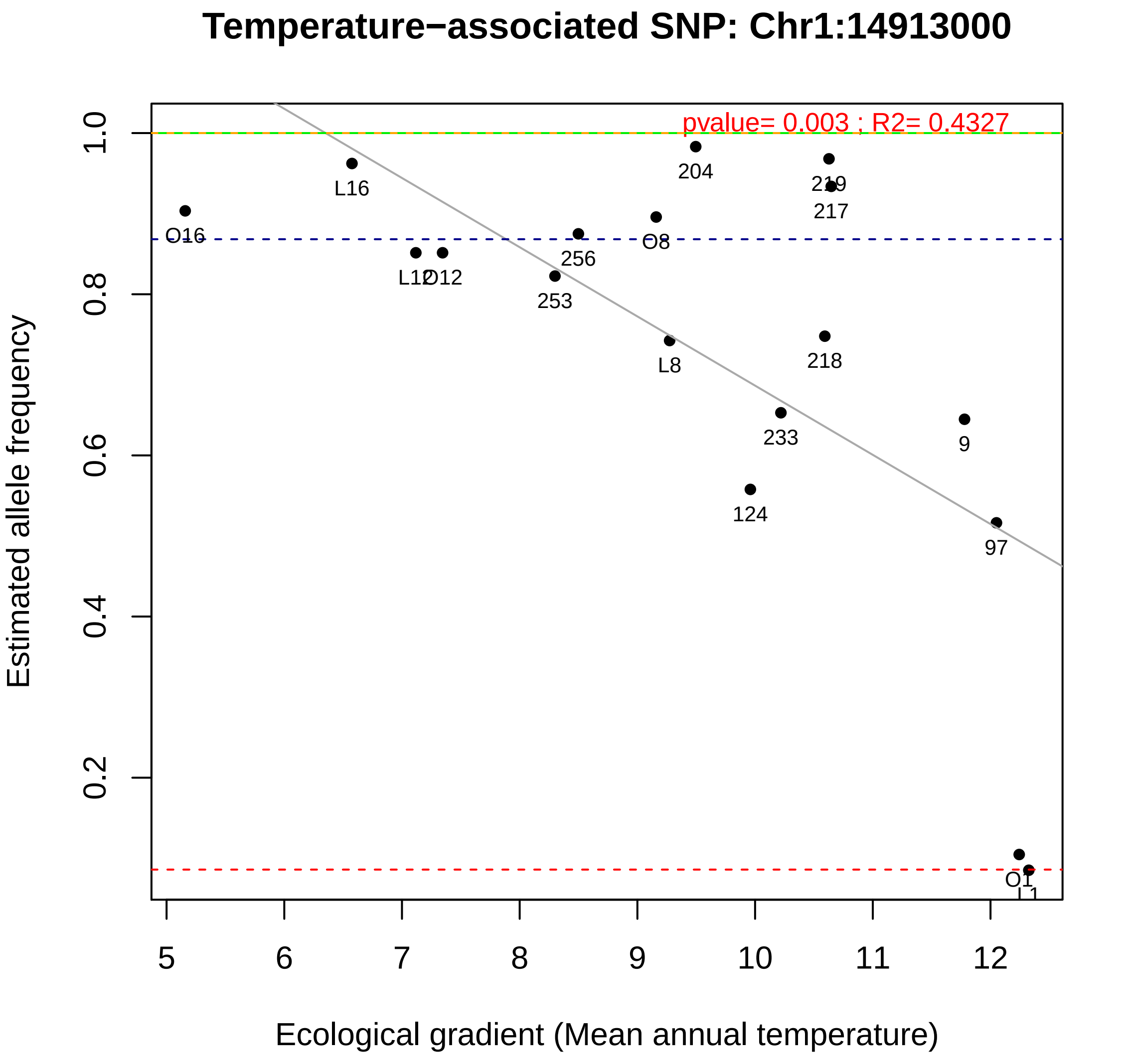

Temperature−associated SNP: Chr1:14913000
pvalue= 0.003 ; R2= 0.4327
Estimated allele frequency
Ecological gradient (Mean annual temperature)
0.2
0.4
0.6
0.8
1.0
5
6
7
8
9
10
11
12
O16
L16
L12
O12
253
256
O8
204
219
217
L8
218
233
124
9
97
O1
L1

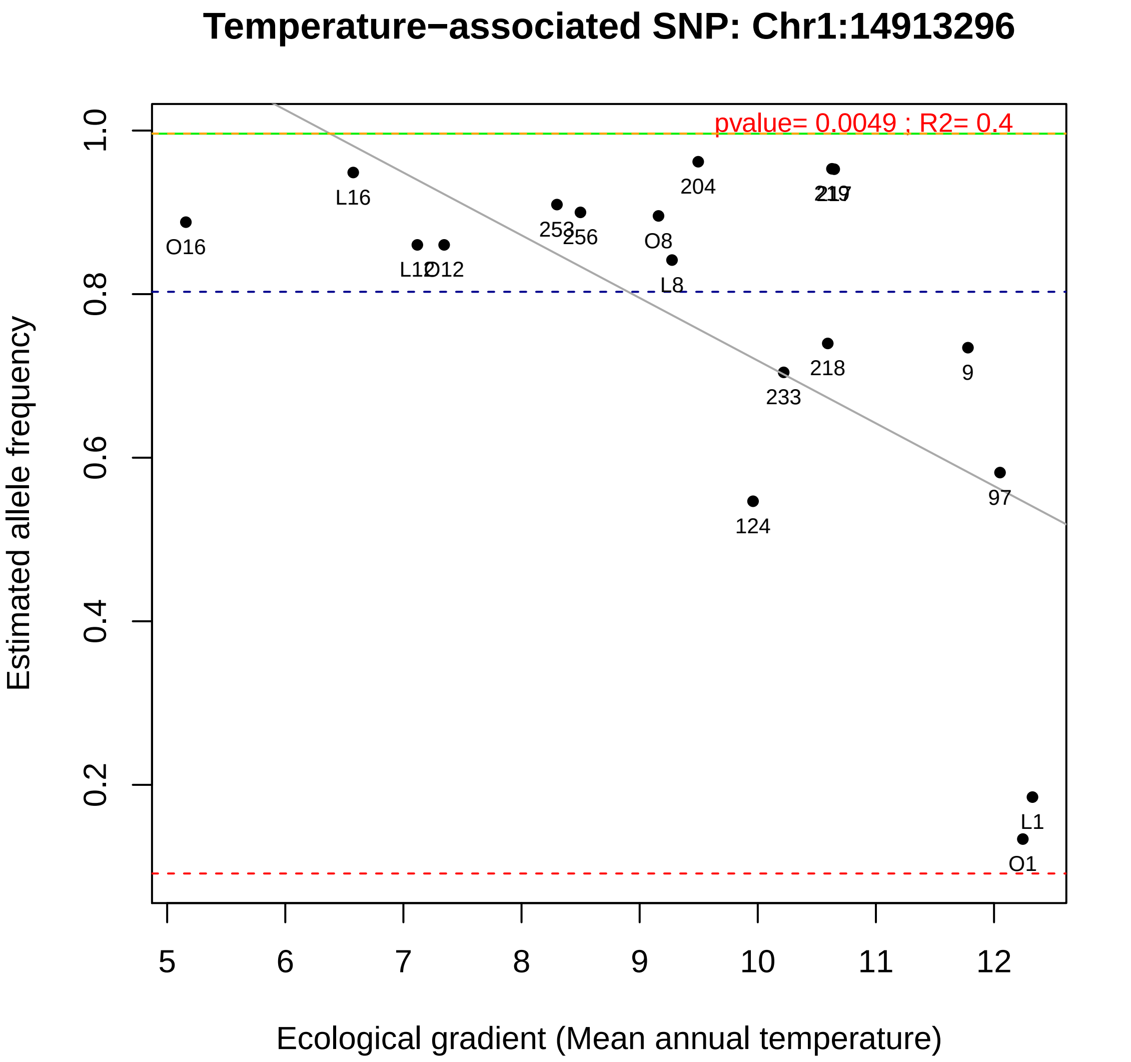
Temperature−associated SNP: Chr1:14913296
pvalue= 0.0049 ; R2= 0.4
Estimated allele frequency
Ecological gradient (Mean annual temperature)
1.0
0.8
0.6
0.4
0.2
5
6
7
8
9
10
11
12
O16
L16
L12
O12
253
256
O8
L8
204
217
219
233
218
124
9
97
L1
O1

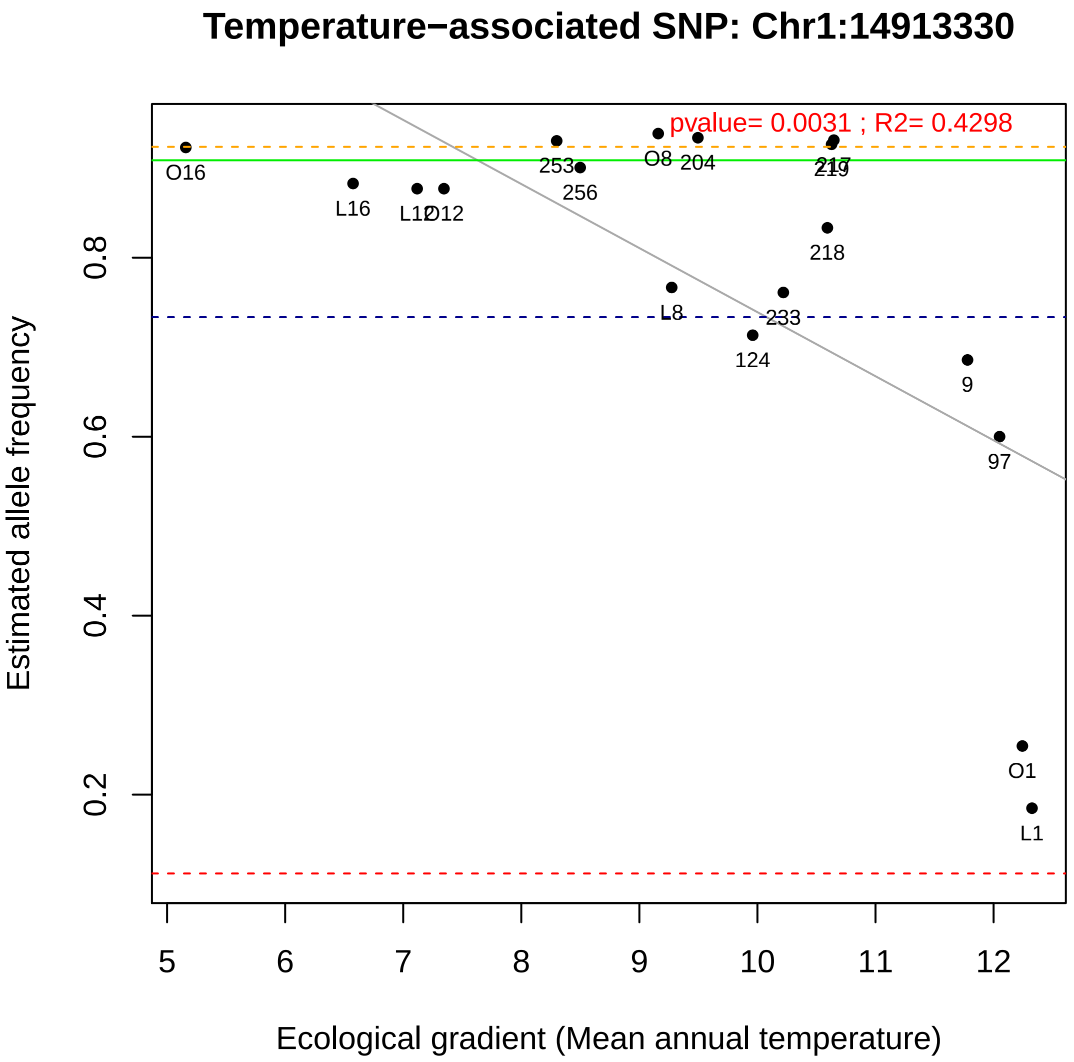
Temperature−associated SNP: Chr1:14913330
pvalue= 0.0031 ; R2= 0.4298
Estimated allele frequency
0.2
0.4
0.6
0.8
Ecological gradient (Mean annual temperature)
5
6
7
8
9
10
11
12
O16
L16
L12
O12
253
256
O8
204
217
219
218
L8
233
124
9
97
O1
L1

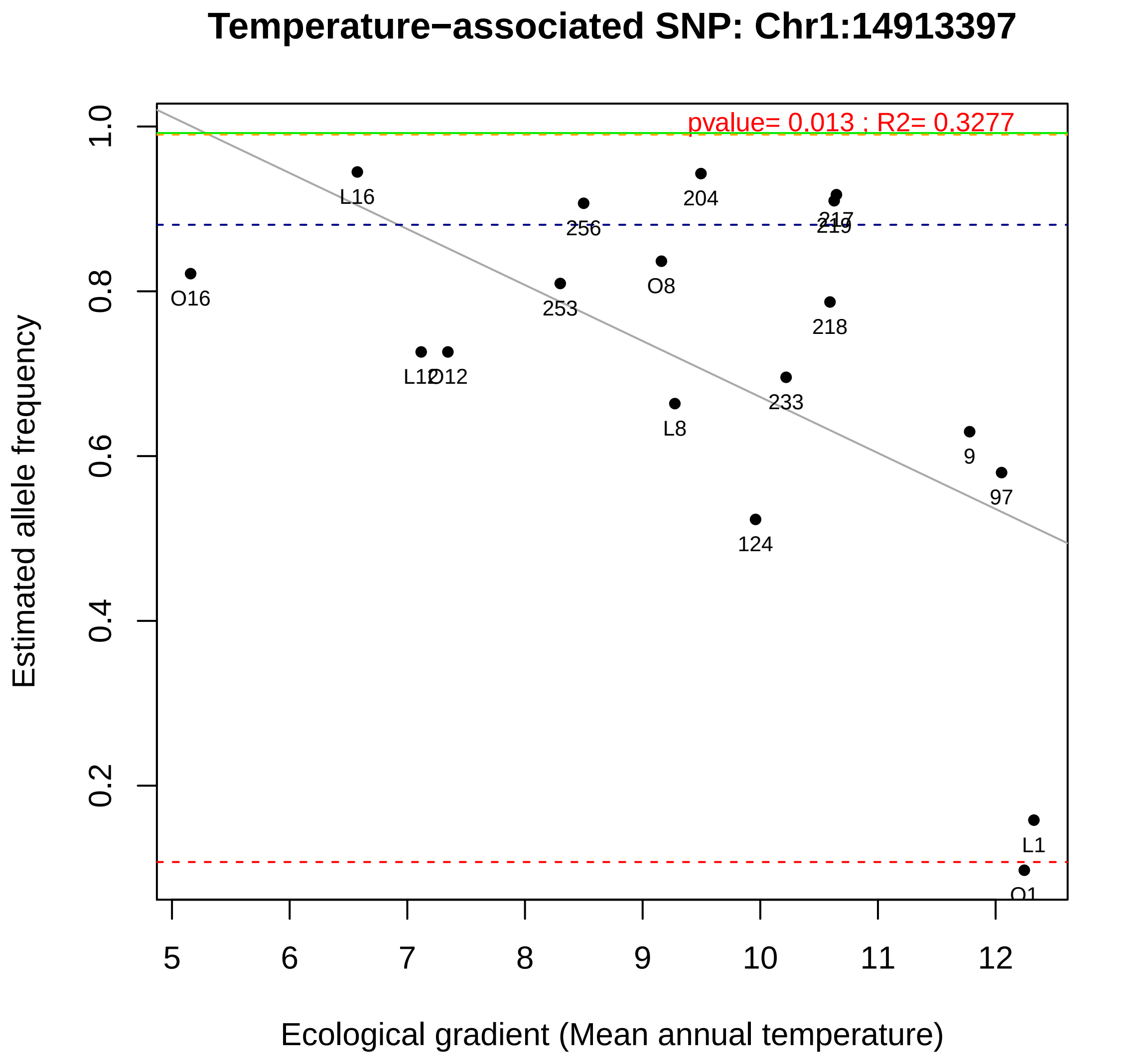
Temperature−associated SNP: Chr1:14913397
pvalue= 0.013 ; R2= 0.3277
Estimated allele frequency
Ecological gradient (Mean annual temperature)
1.0
0.8
0.6
0.4
0.2
5
6
7
8
9
10
11
12
L16
O16
256
253
204
O8
217
219
218
L12
O12
L8
233
124
9
97
L1
O1

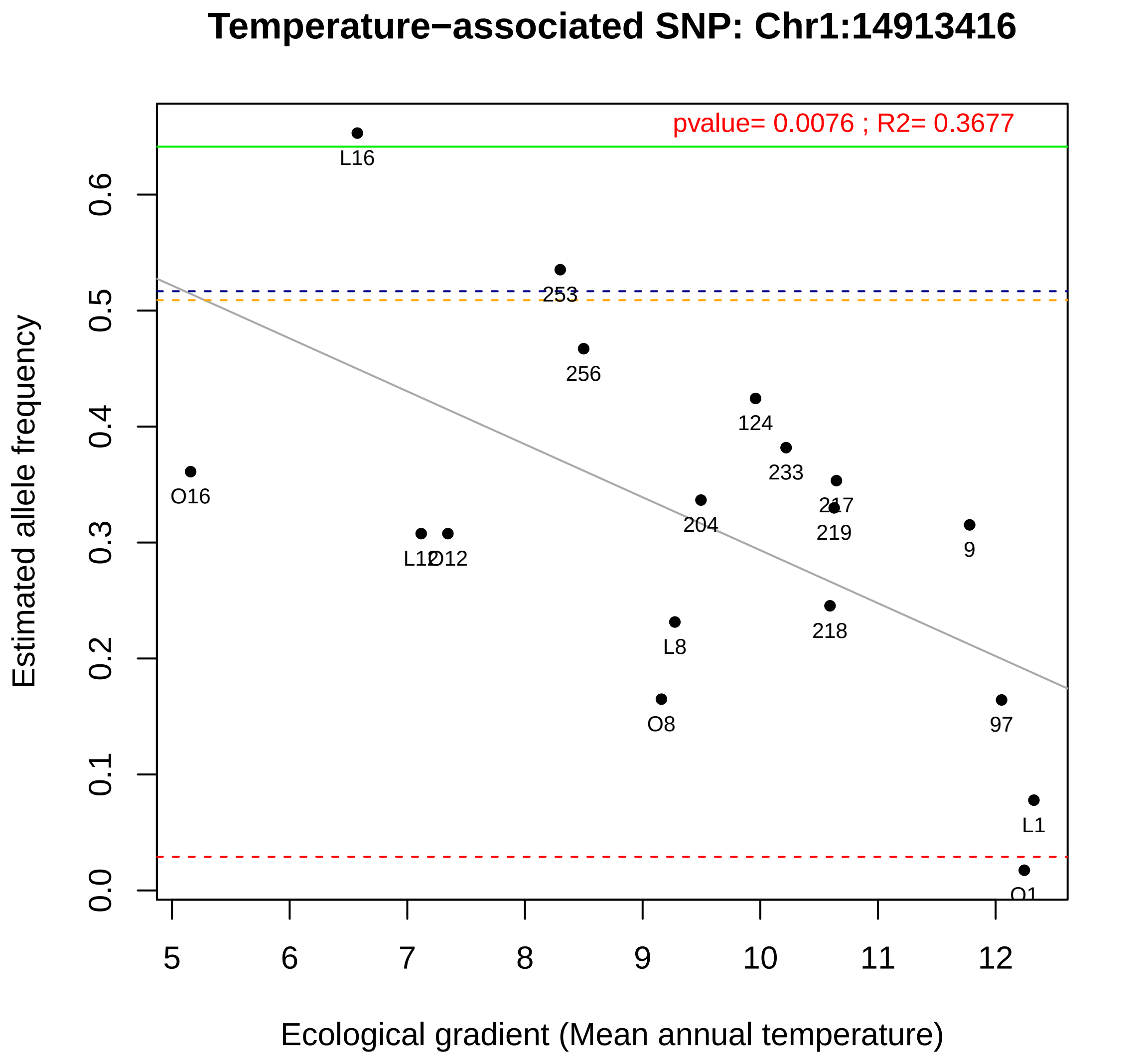
Temperature−associated SNP: Chr1:14913416
pvalue= 0.0076 ; R2= 0.3677
Estimated allele frequency
Ecological gradient (Mean annual temperature)
0.0
0.1
0.2
0.3
0.4
0.5
0.6
5
6
7
8
9
10
11
12
L16
253
256
124
233
217
219
204
O16
L12
O12
9
L8
218
O8
97
L1
O1

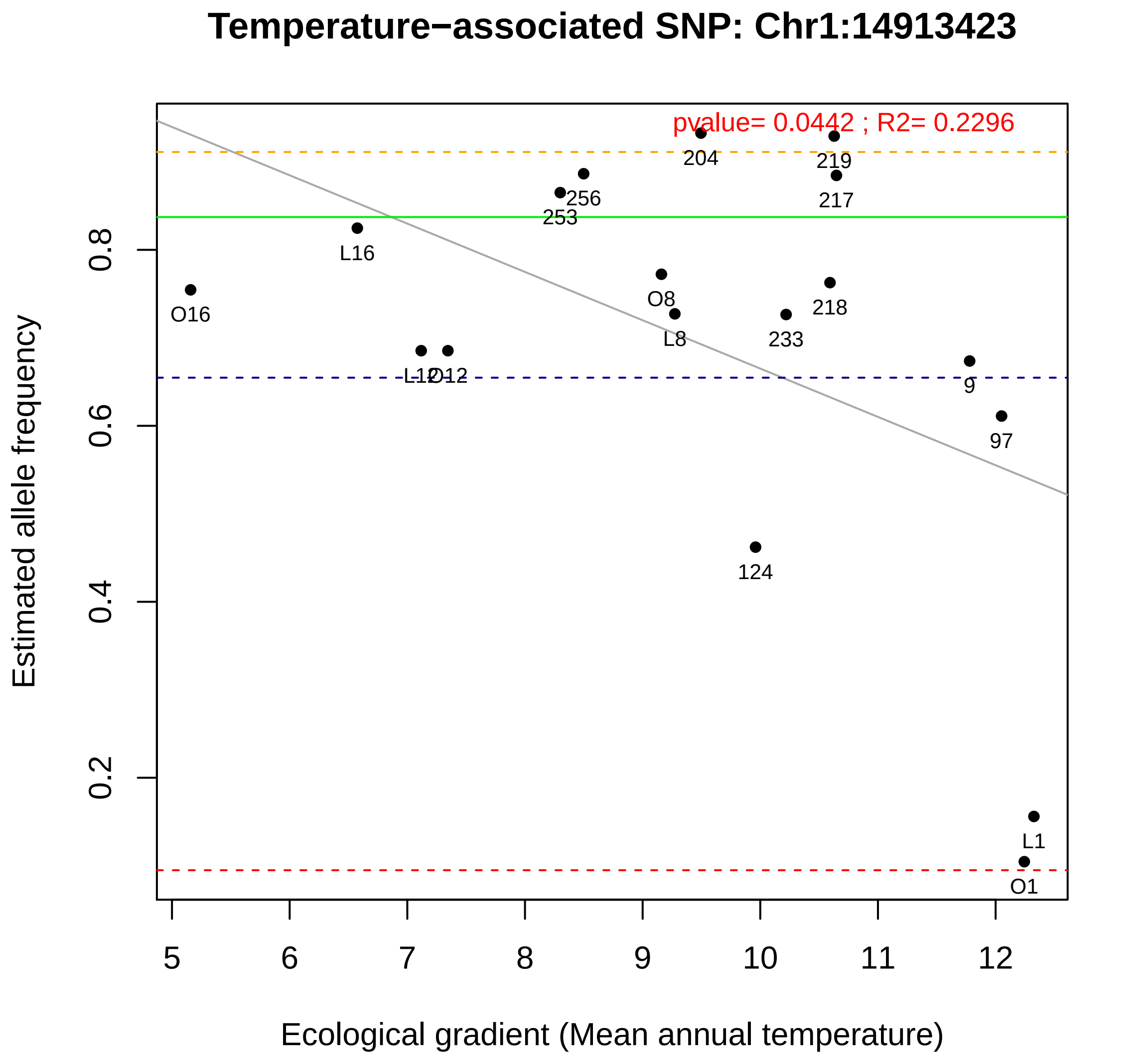

Temperature−associated SNP: Chr1:14913423
pvalue= 0.0442 ; R2= 0.2296
Estimated allele frequency
0.2
0.4
0.6
0.8
5
6
7
8
9
10
11
12
Ecological gradient (Mean annual temperature)
O16
L16
L12
O12
253
256
204
O8
L8
124
233
218
219
217
9
97
L1
O1

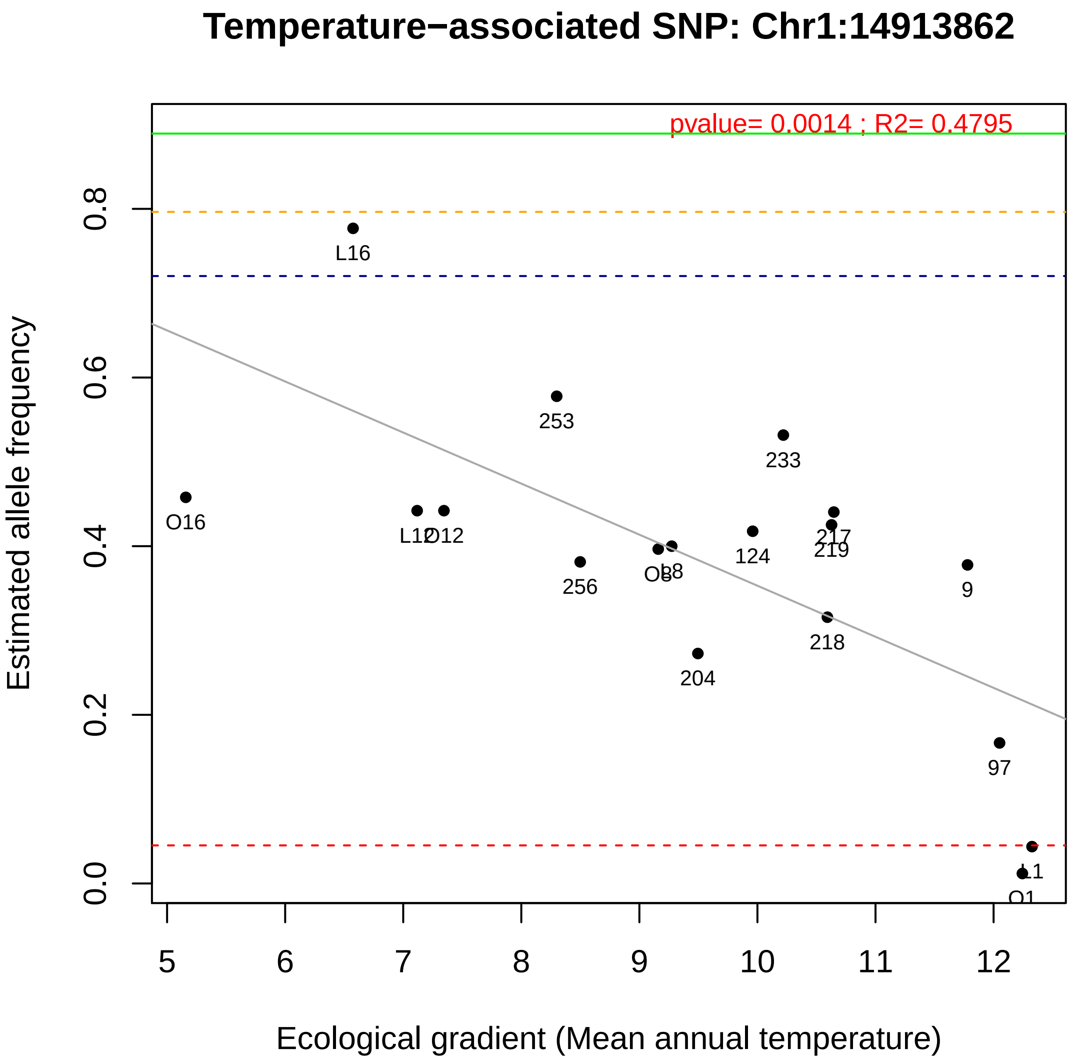
Temperature−associated SNP: Chr1:14913862
pvalue= 0.0014 ; R2= 0.4795
Estimated allele frequency
0.0
0.2
0.4
0.6
0.8
L16
253
233
O16
L12
O12
217
219
124
O8
L8
256
9
218
204
97
L1
O1
5
6
7
8
9
10
11
12
Ecological gradient (Mean annual temperature)

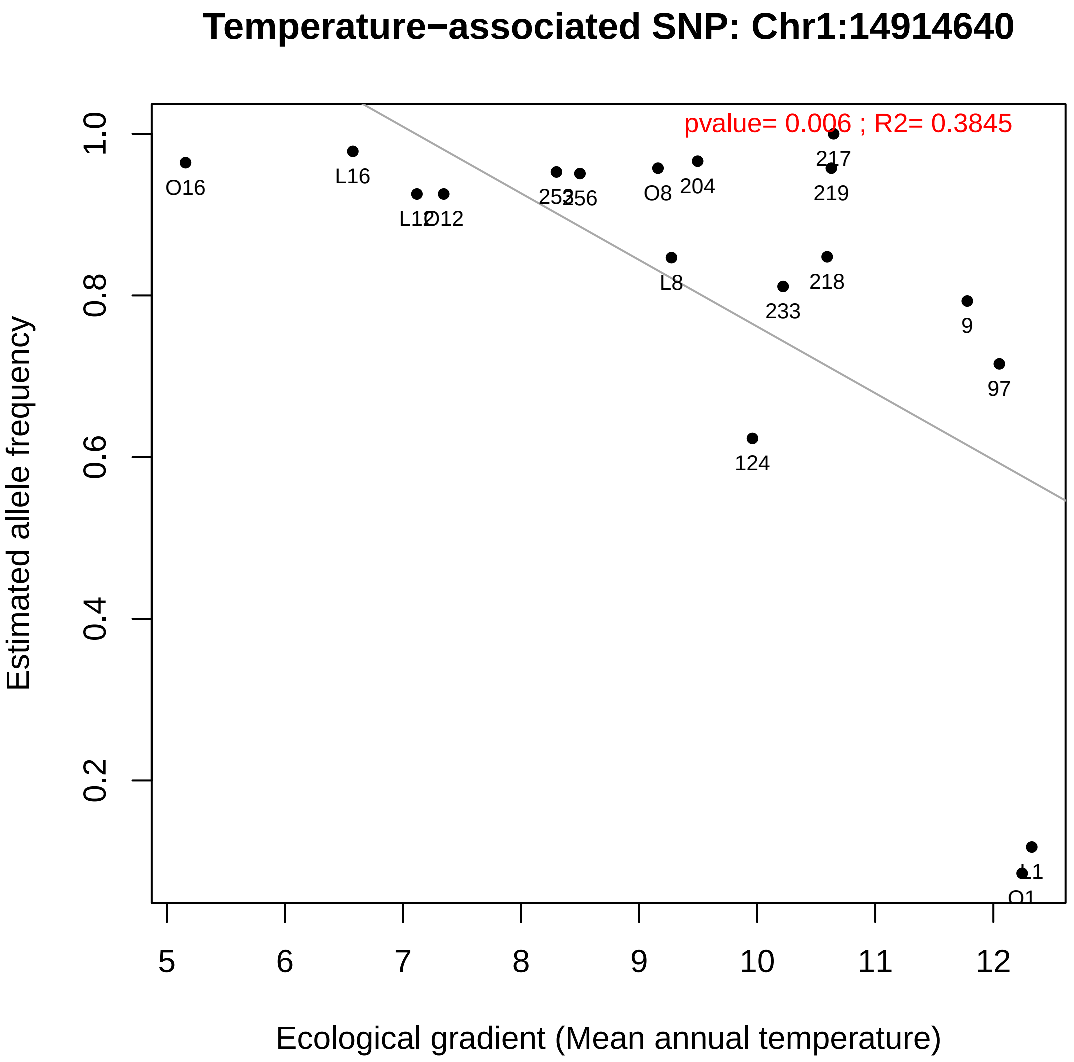

Temperature−associated SNP: Chr1:14914640
pvalue= 0.006 ; R2= 0.3845
Estimated allele frequency
Ecological gradient (Mean annual temperature)
0.2
0.4
0.6
0.8
1.0
5
6
7
8
9
10
11
12
O16
L16
L12
O12
252
256
O8
204
217
219
L8
218
233
9
97
124
L1
O1

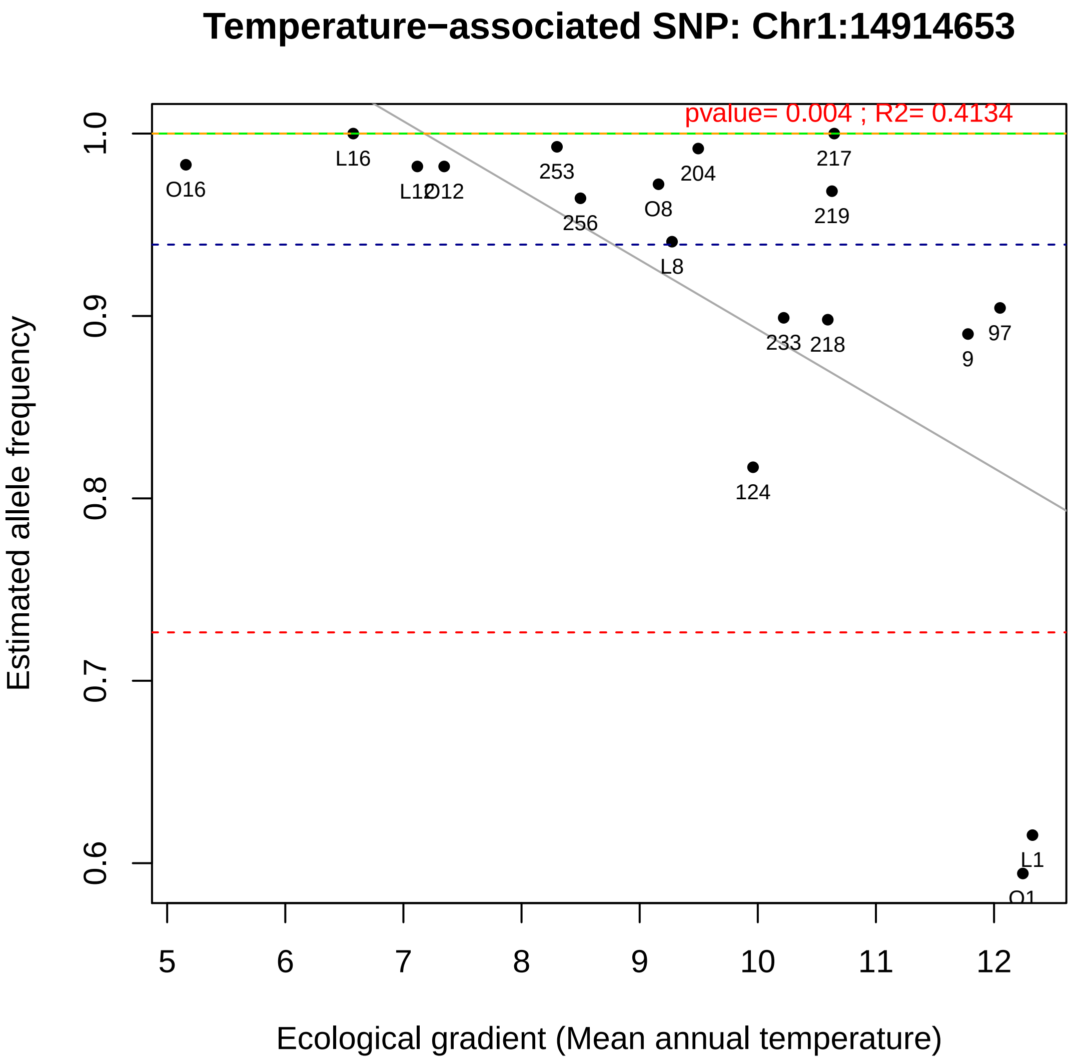

Temperature−associated SNP: Chr1:14914653
pvalue= 0.004 ; R2= 0.4134
Estimated allele frequency
1.0
0.9
0.8
0.7
0.6
Ecological gradient (Mean annual temperature)
5
6
7
8
9
10
11
12
O16
L16
L12
O12
253
256
O8
204
L8
217
219
124
233
218
9
97
L1
O1

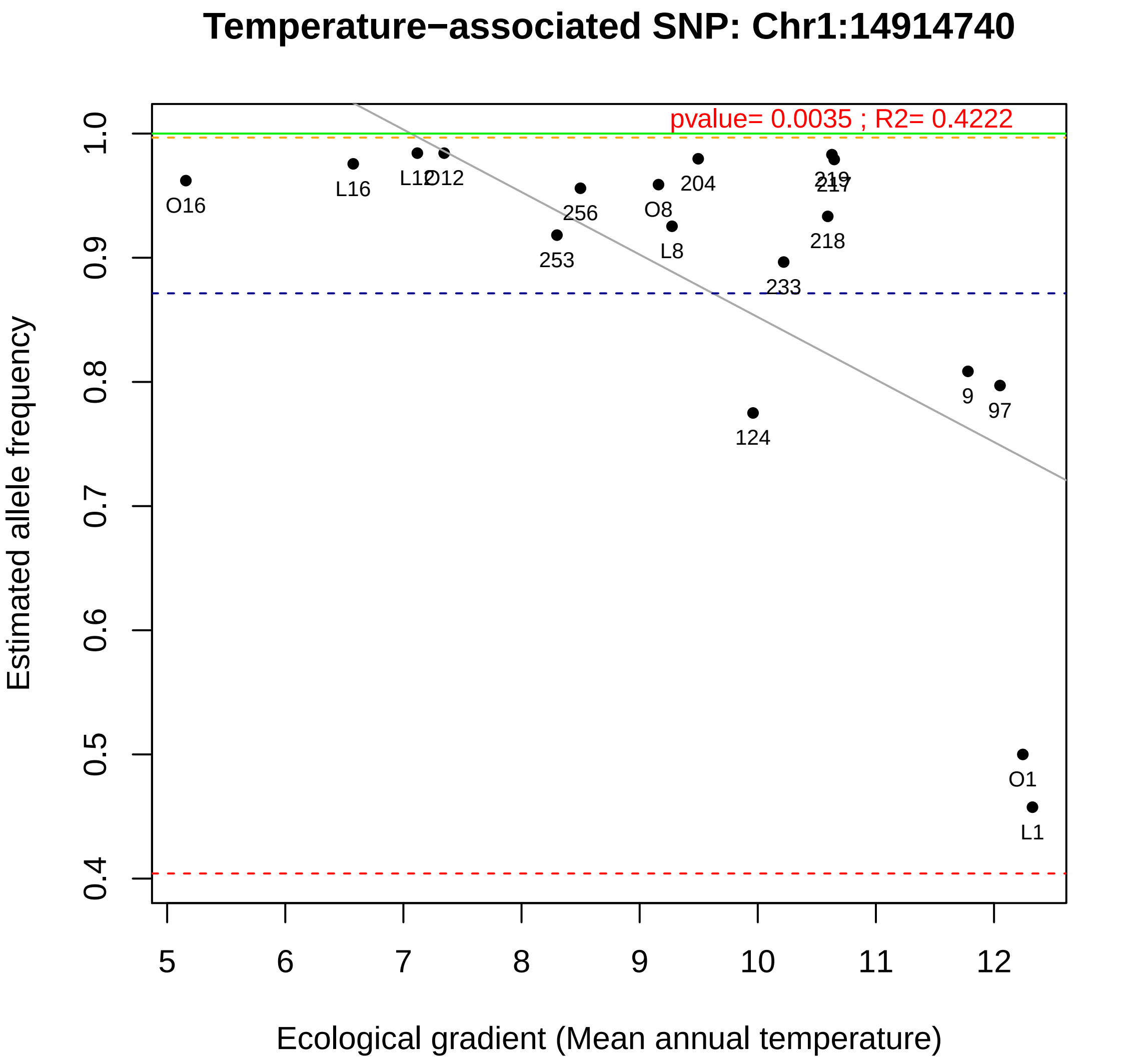

Temperature−associated SNP: Chr1:14914740
pvalue= 0.0035 ; R2= 0.4222
Estimated allele frequency
1.0
0.9
0.8
0.7
0.6
0.5
0.4
5
6
7
8
9
10
11
12
Ecological gradient (Mean annual temperature)
O16
L16
L12
O12
256
253
O8
204
L8
219
217
218
233
124
9
97
O1
L1

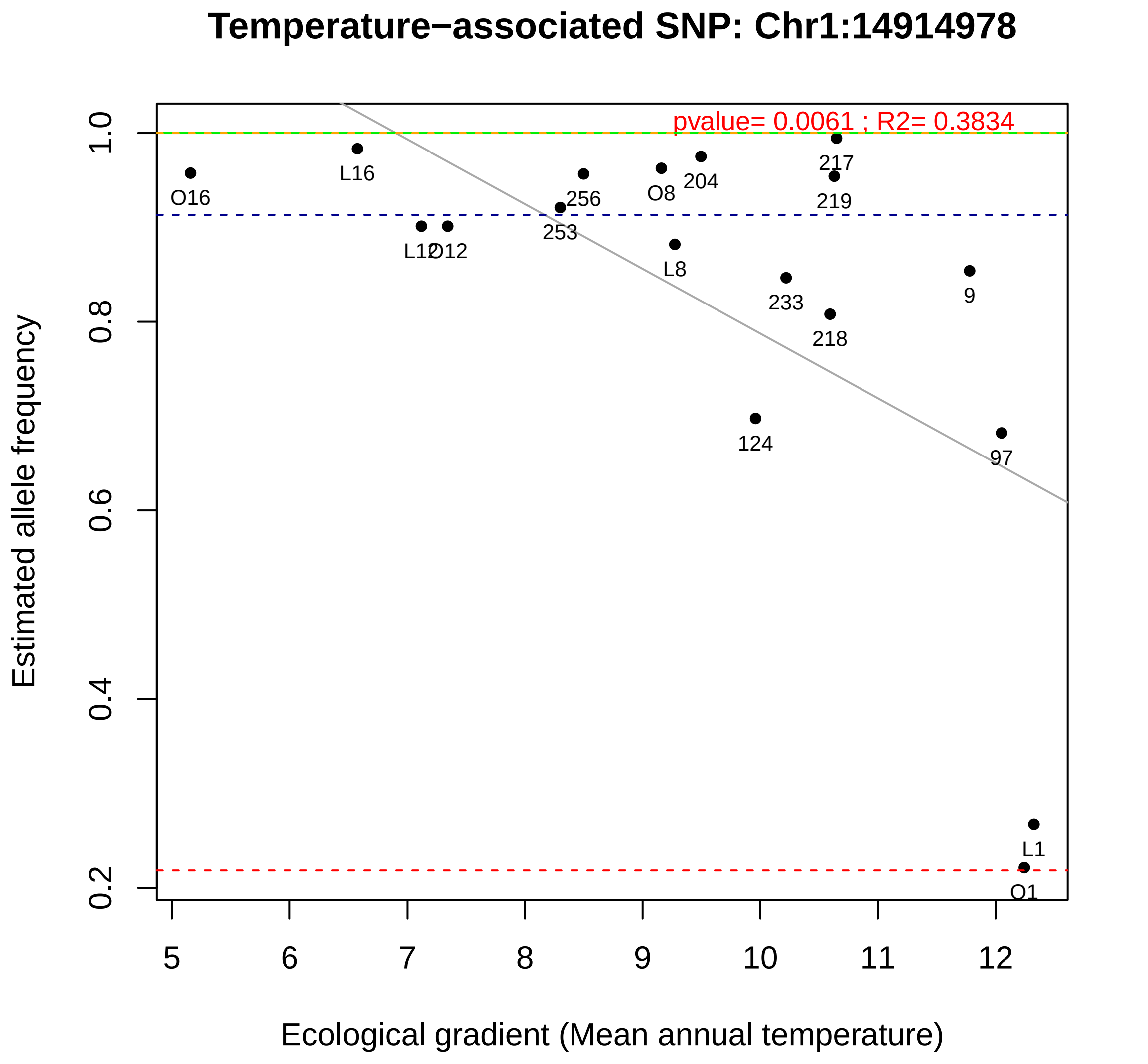
Temperature−associated SNP: Chr1:14914978
pvalue= 0.0061 ; R2= 0.3834
Estimated allele frequency
0.2
0.4
0.6
0.8
1.0
5
6
7
8
9
10
11
12
Ecological gradient (Mean annual temperature)
O16
L16
L12
O12
253
256
O8
204
L8
124
233
217
219
218
9
97
L1
O1

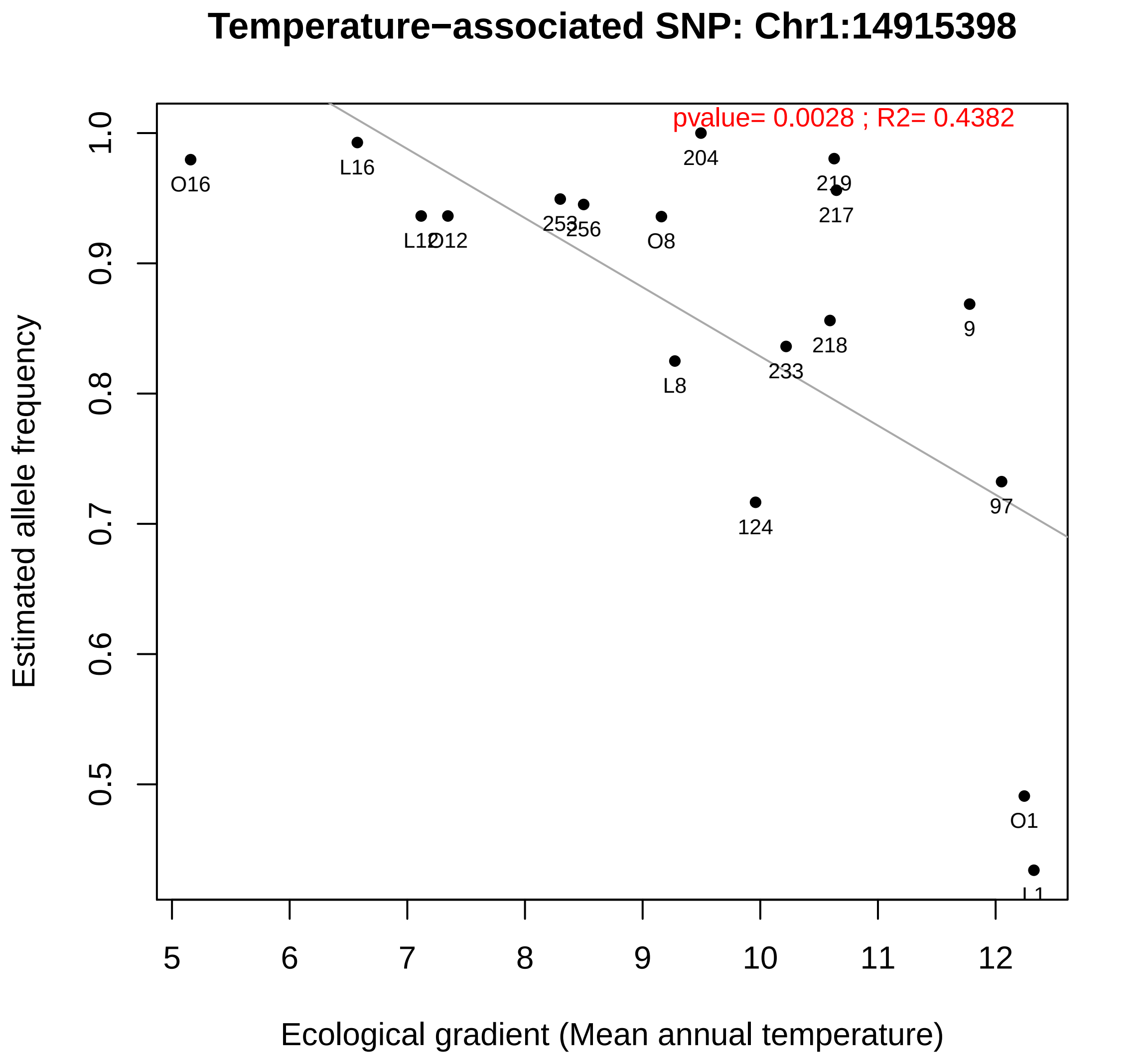

Temperature−associated SNP: Chr1:14915398
pvalue= 0.0028 ; R2= 0.4382
Estimated allele frequency
Ecological gradient (Mean annual temperature)
1.0
0.9
0.8
0.7
0.6
0.5
5
6
7
8
9
10
11
12
O16
L16
L12
O12
253
256
O8
204
219
217
L8
233
218
9
124
97
O1

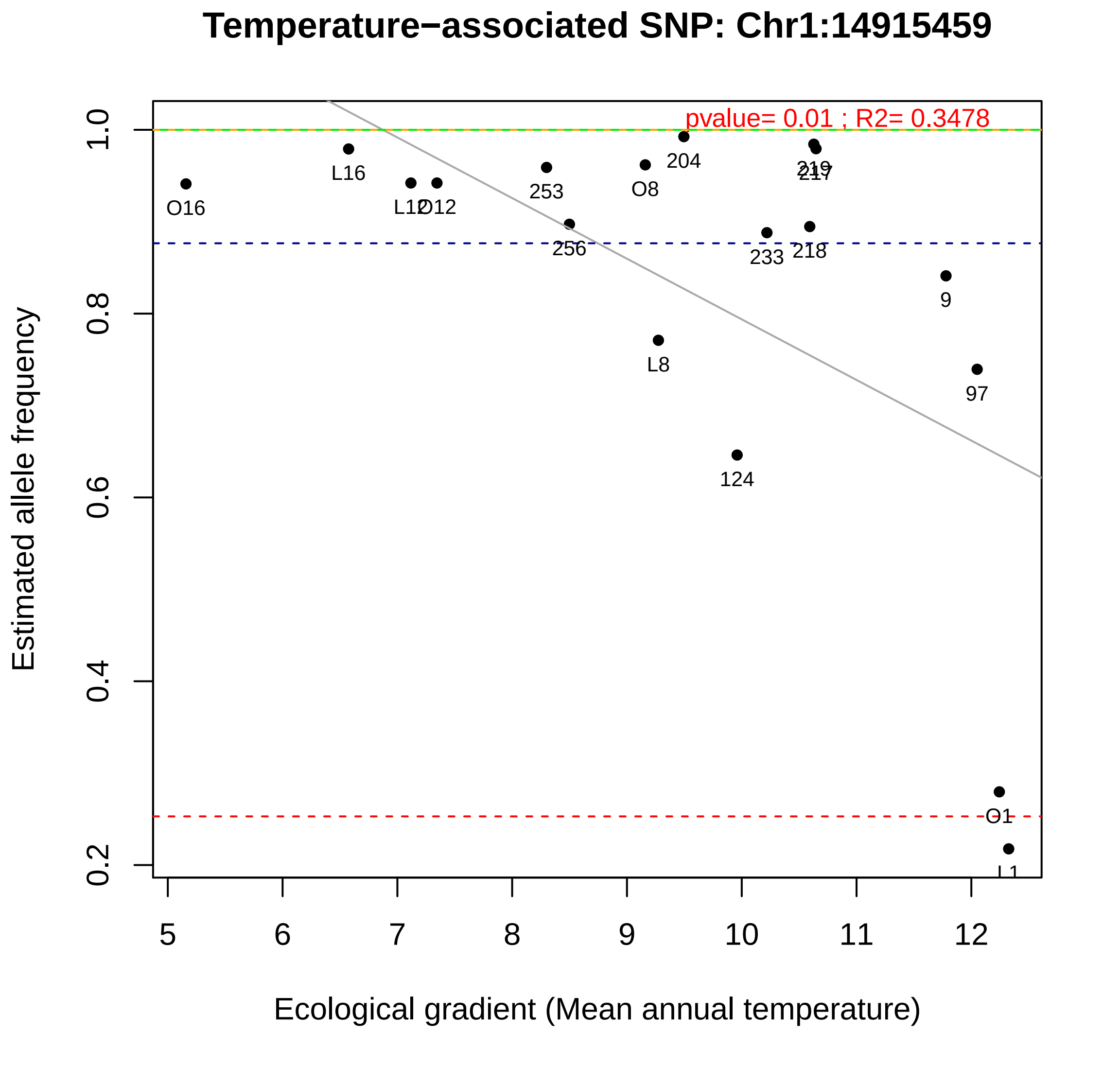
Temperature−associated SNP: Chr1:14915459
pvalue= 0.01 ; R2= 0.3478
Estimated allele frequency
0.2
0.4
0.6
0.8
1.0
5
6
7
8
9
10
11
12
Ecological gradient (Mean annual temperature)
O16
L16
L12
O12
253
256
O8
204
L8
124
233
218
219
217
9
97
O1
L1

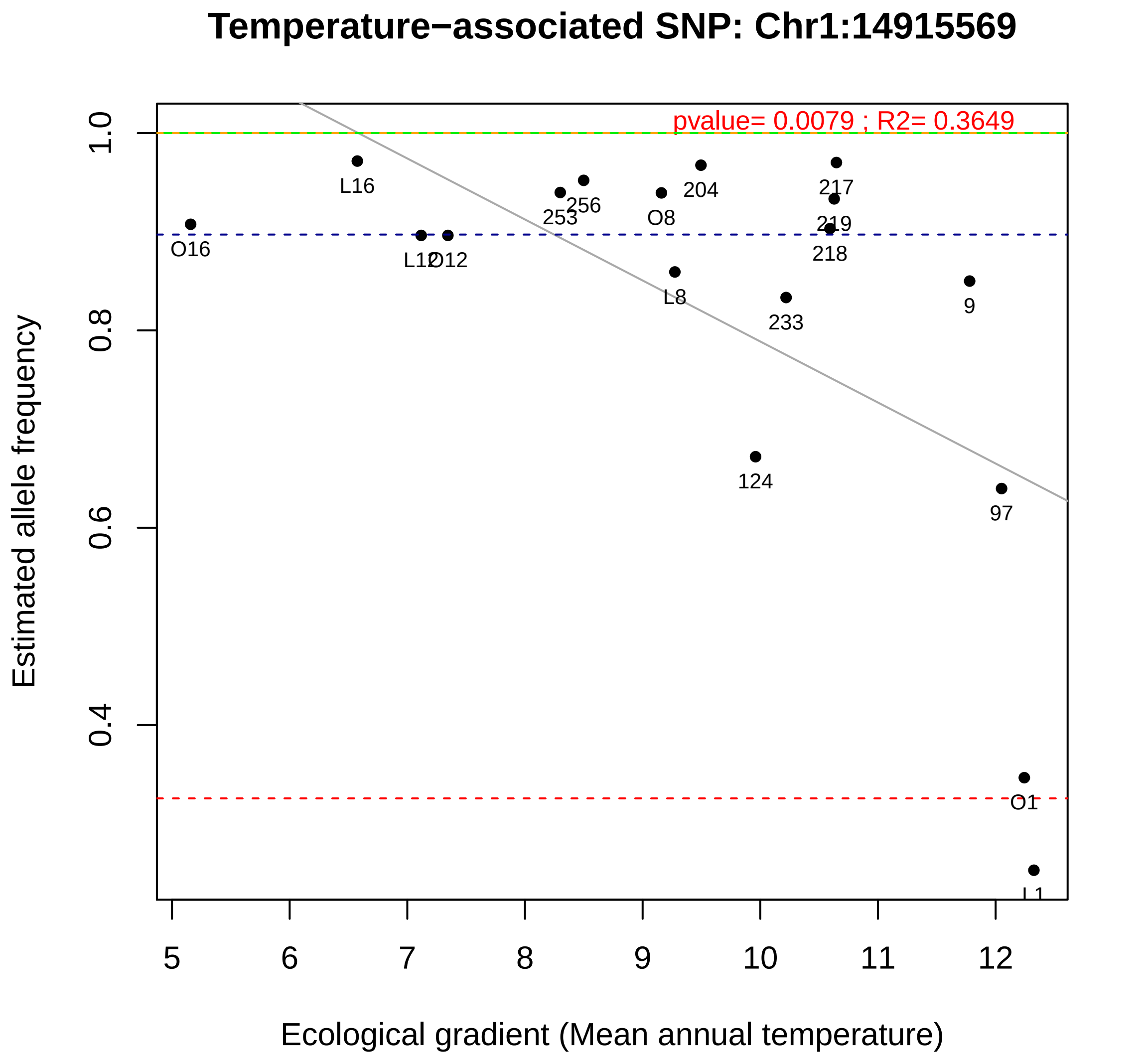
Temperature−associated SNP: Chr1:14915569
pvalue= 0.0079 ; R2= 0.3649
Estimated allele frequency
Ecological gradient (Mean annual temperature)
1.0
0.8
0.6
0.4
5
6
7
8
9
10
11
12
O16
L16
L12
O12
253
256
O8
204
L8
217
219
218
233
124
9
97
O1

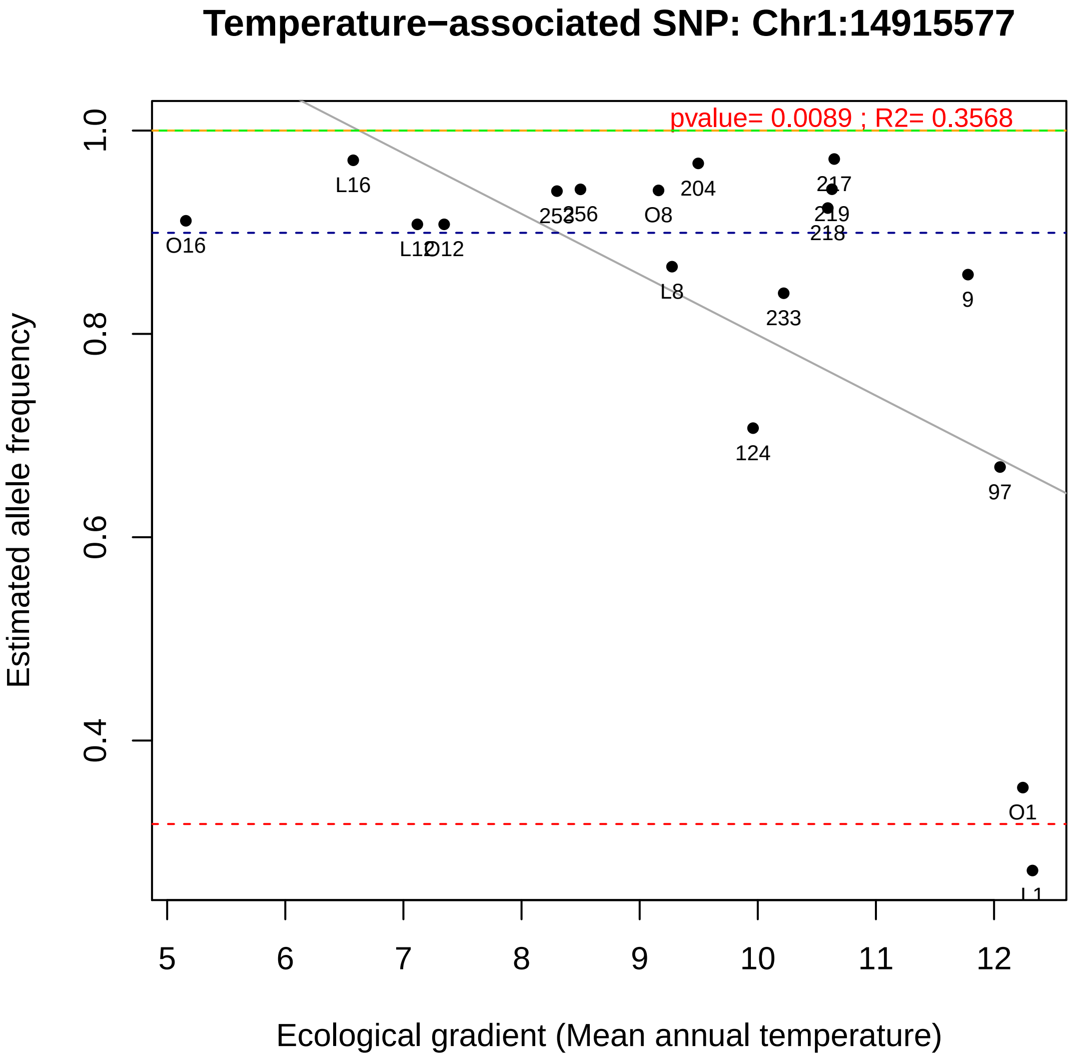
Temperature−associated SNP: Chr1:14915577
pvalue= 0.0089 ; R2= 0.3568
Estimated allele frequency
Ecological gradient (Mean annual temperature)
1.0
0.8
0.6
0.4
5
6
7
8
9
10
11
12
O16
L16
L12
O12
253
256
O8
204
L8
124
233
217
219
218
9
97
O1
L1

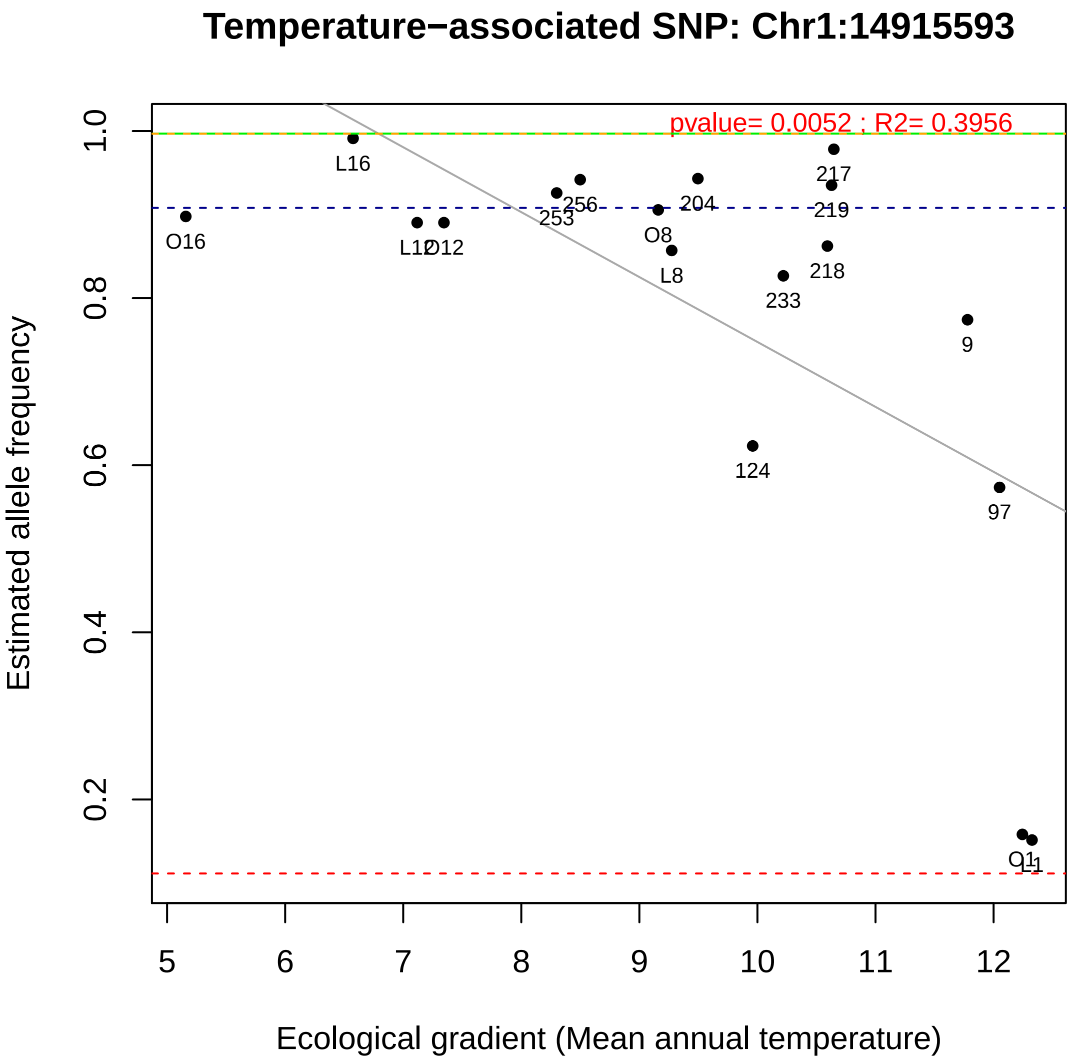

Temperature−associated SNP: Chr1:14915593
pvalue= 0.0052 ; R2= 0.3956
Estimated allele frequency
Ecological gradient (Mean annual temperature)
1.0
0.8
0.6
0.4
0.2
5
6
7
8
9
10
11
12
L16
O16
L12
O12
253
256
O8
L8
204
217
219
218
233
9
124
97
O1
L1

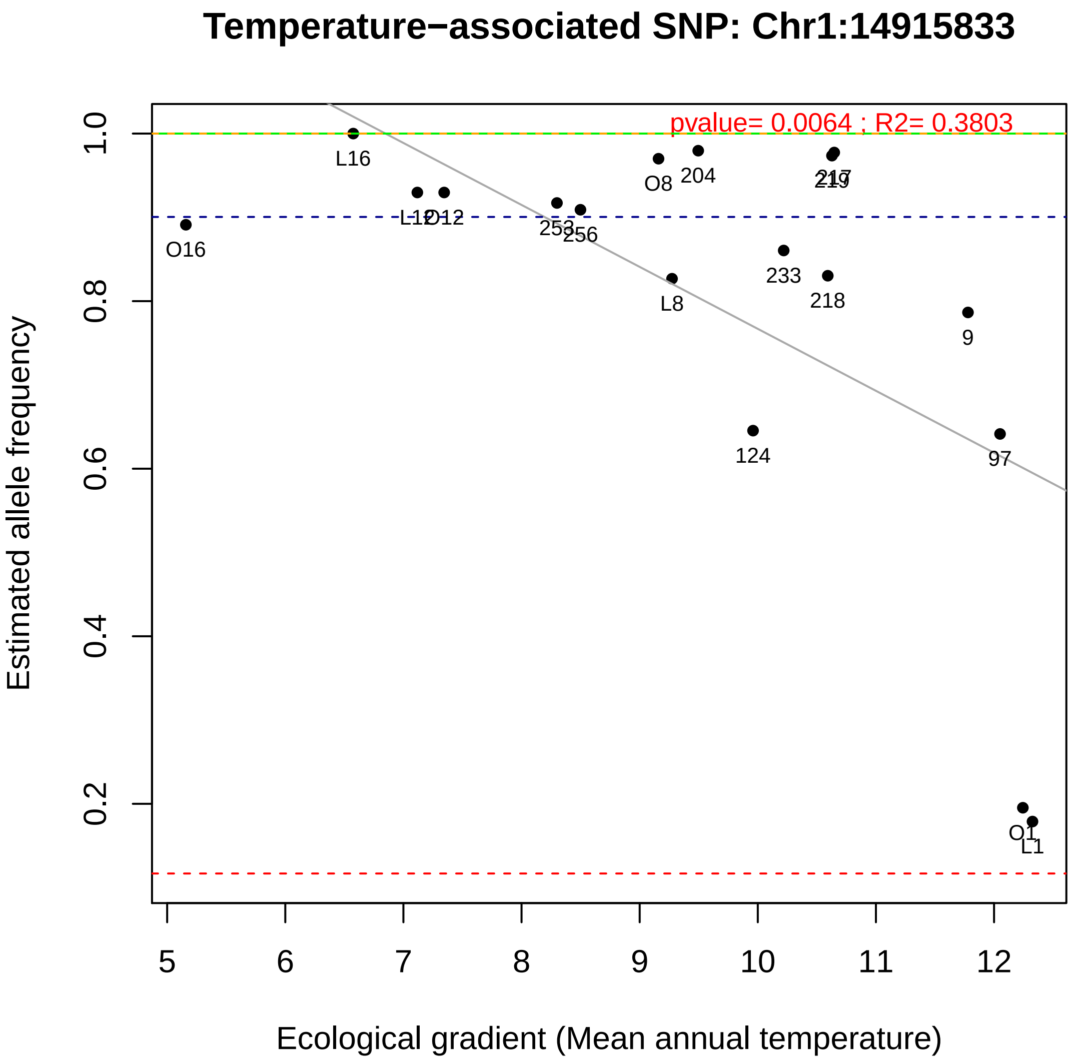

Temperature−associated SNP: Chr1:14915833
pvalue= 0.0064 ; R2= 0.3803
Estimated allele frequency
Ecological gradient (Mean annual temperature)
1.0
0.8
0.6
0.4
0.2
5
6
7
8
9
10
11
12
L16
O16
L12
O12
253
256
O8
204
L8
217
219
233
218
124
9
97
O1
L1

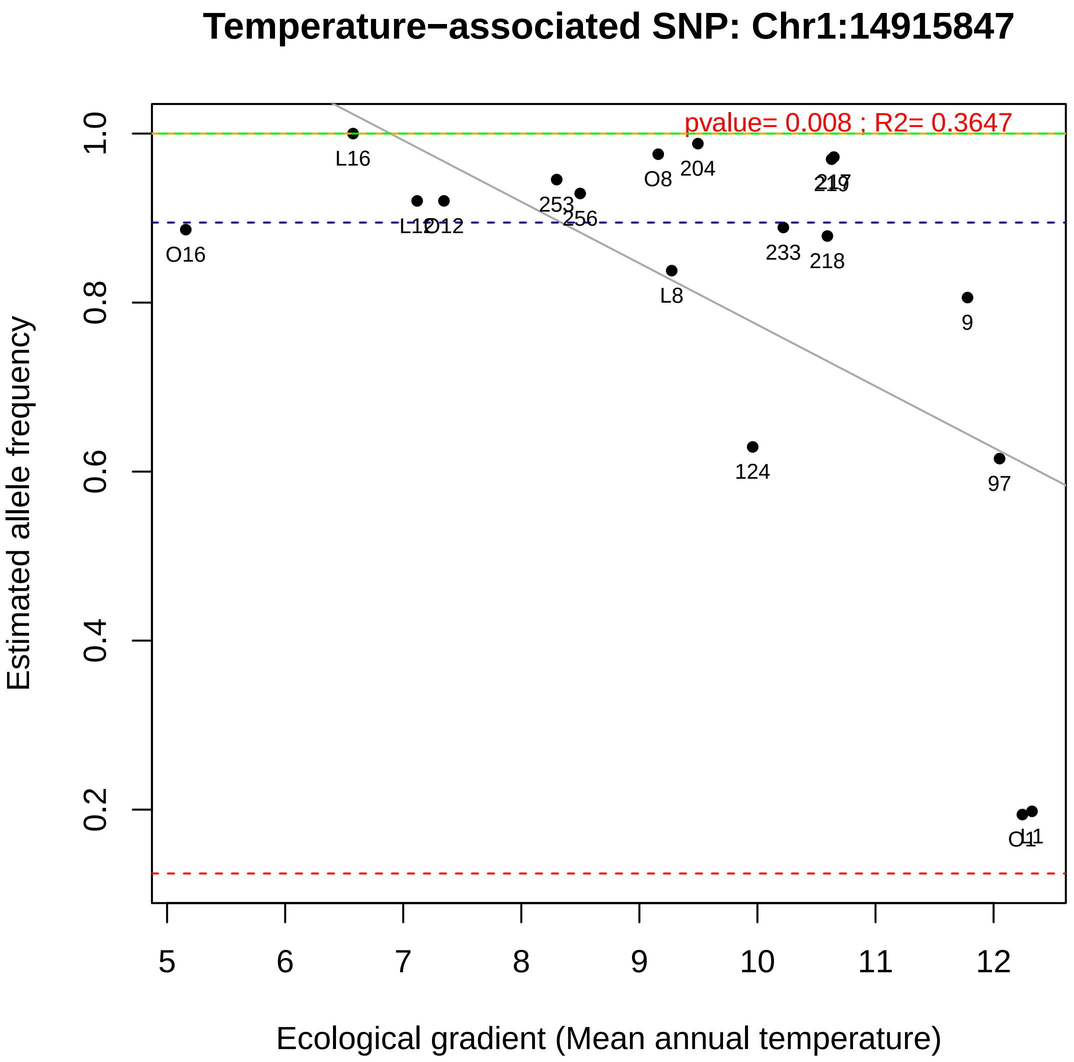
Temperature−associated SNP: Chr1:14915847
pvalue= 0.008 ; R2= 0.3647
Estimated allele frequency
Ecological gradient (Mean annual temperature)
1.0
0.8
0.6
0.4
0.2
5
6
7
8
9
10
11
12
L16
O16
253
256
O8
204
L8
233
218
124
9
97

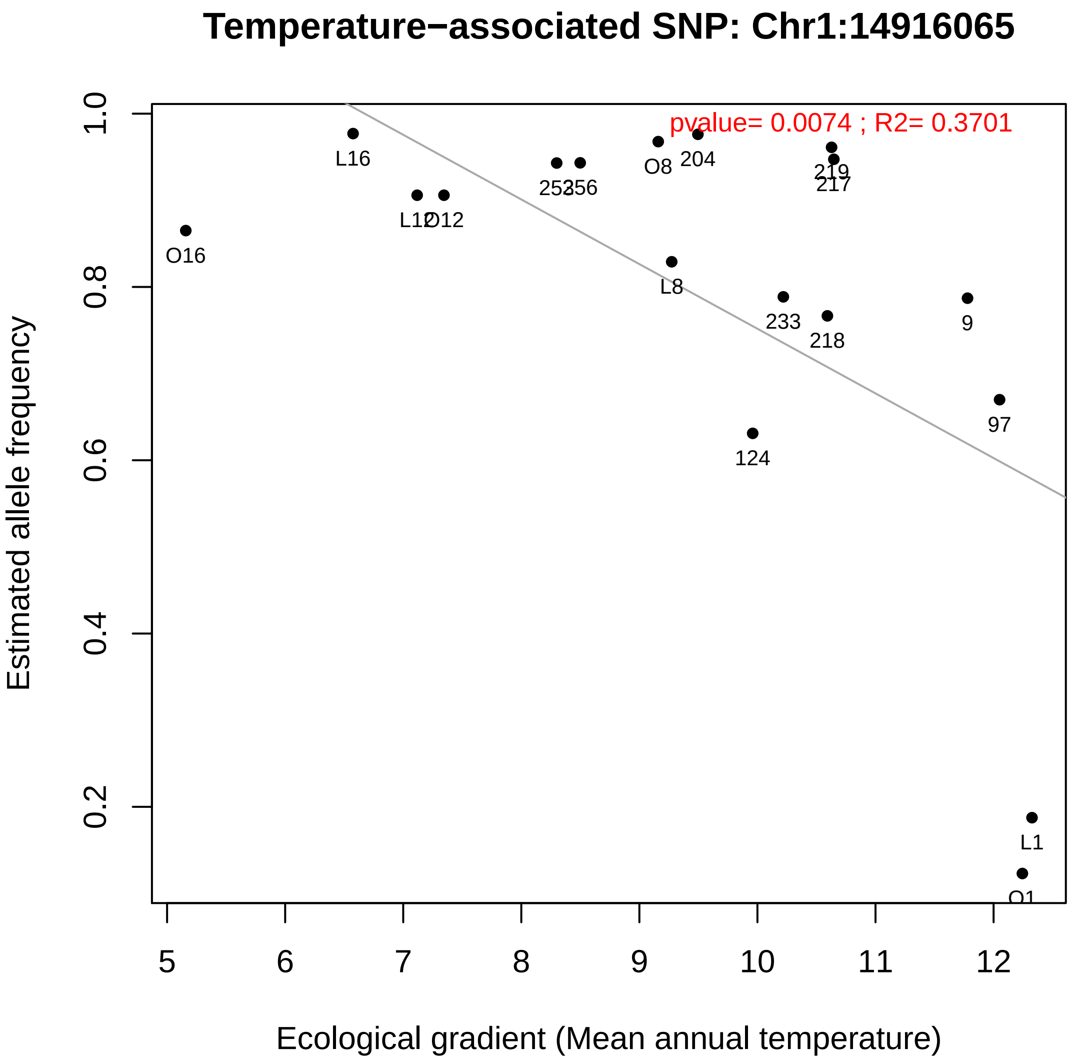
Temperature−associated SNP: Chr1:14916065
pvalue= 0.0074 ; R2= 0.3701
Estimated allele frequency
Ecological gradient (Mean annual temperature)
0.2
0.4
0.6
0.8
1.0
5
6
7
8
9
10
11
12
O16
L16
L12
O12
O8
204
219
217
233
218
9
97
124
L8
L1

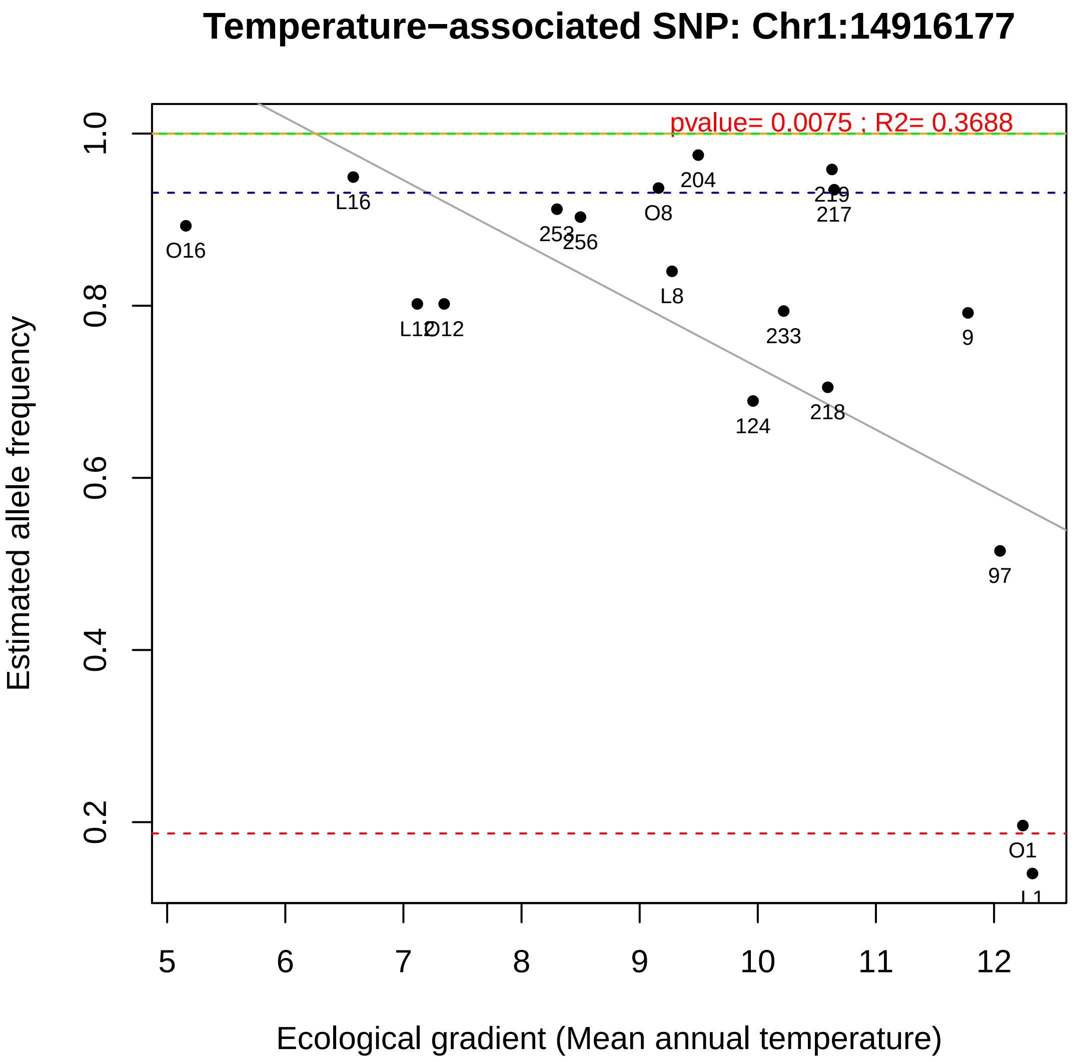

Temperature−associated SNP: Chr1:14916177
pvalue= 0.0075 ; R2= 0.3688
Estimated allele frequency
Ecological gradient (Mean annual temperature)
0.2
0.4
0.6
0.8
1.0
5
6
7
8
9
10
11
12
O16
L16
L12
O12
253
256
O8
204
L8
219
217
233
124
218
9
97
O1

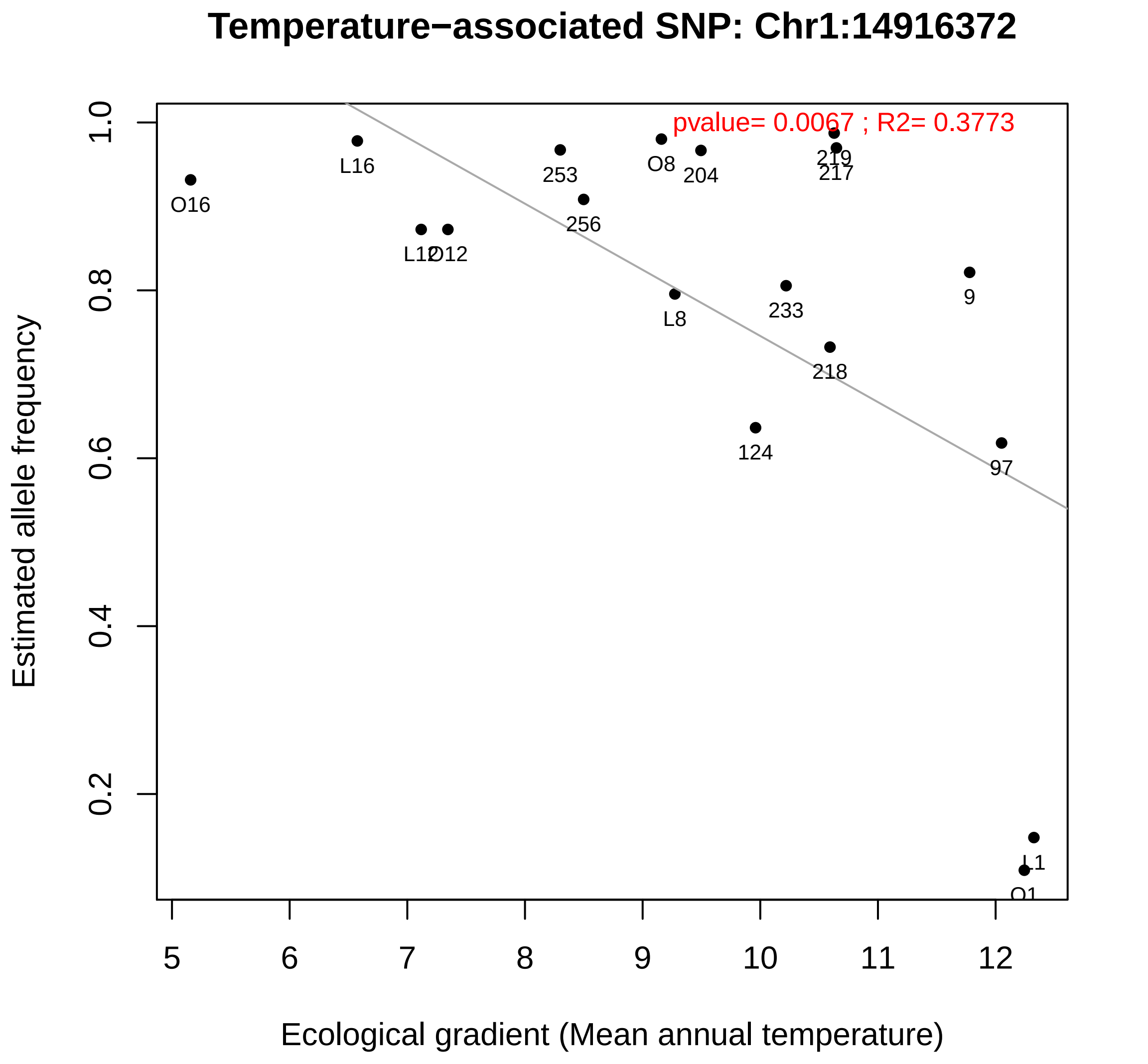
Temperature−associated SNP: Chr1:14916372
pvalue= 0.0067 ; R2= 0.3773
Estimated allele frequency
0.2
0.4
0.6
0.8
1.0
Ecological gradient (Mean annual temperature)
5
6
7
8
9
10
11
12
O16
L16
L12
O12
253
256
O8
204
L8
233
124
218
219
217
9
97
L1
O1

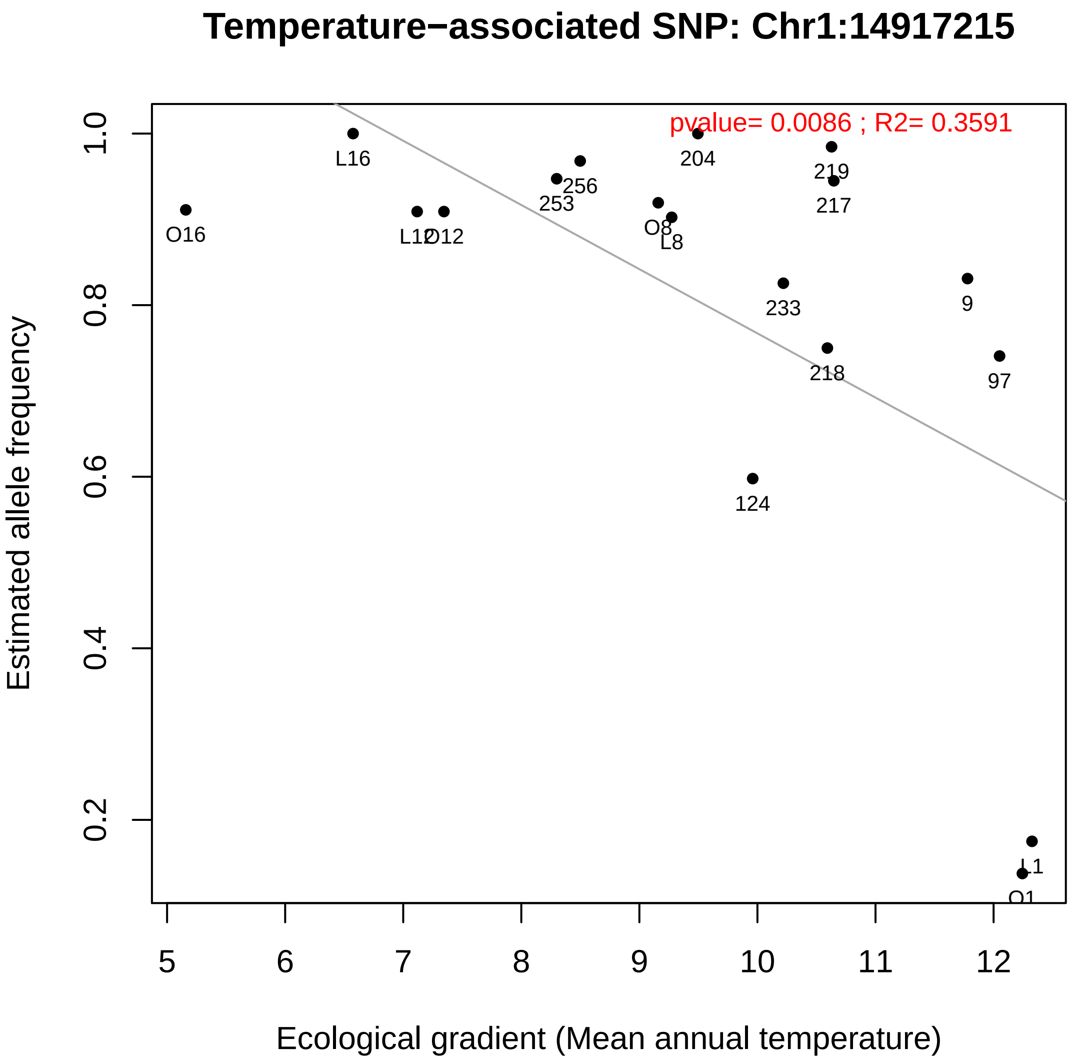
Temperature−associated SNP: Chr1:14917215
pvalue= 0.0086 ; R2= 0.3591
Estimated allele frequency
Ecological gradient (Mean annual temperature)
0.2
0.4
0.6
0.8
1.0
5
6
7
8
9
10
11
12
O16
L16
L12
O12
253
256
O8
L8
204
219
217
233
218
124
9
97
L1
O1

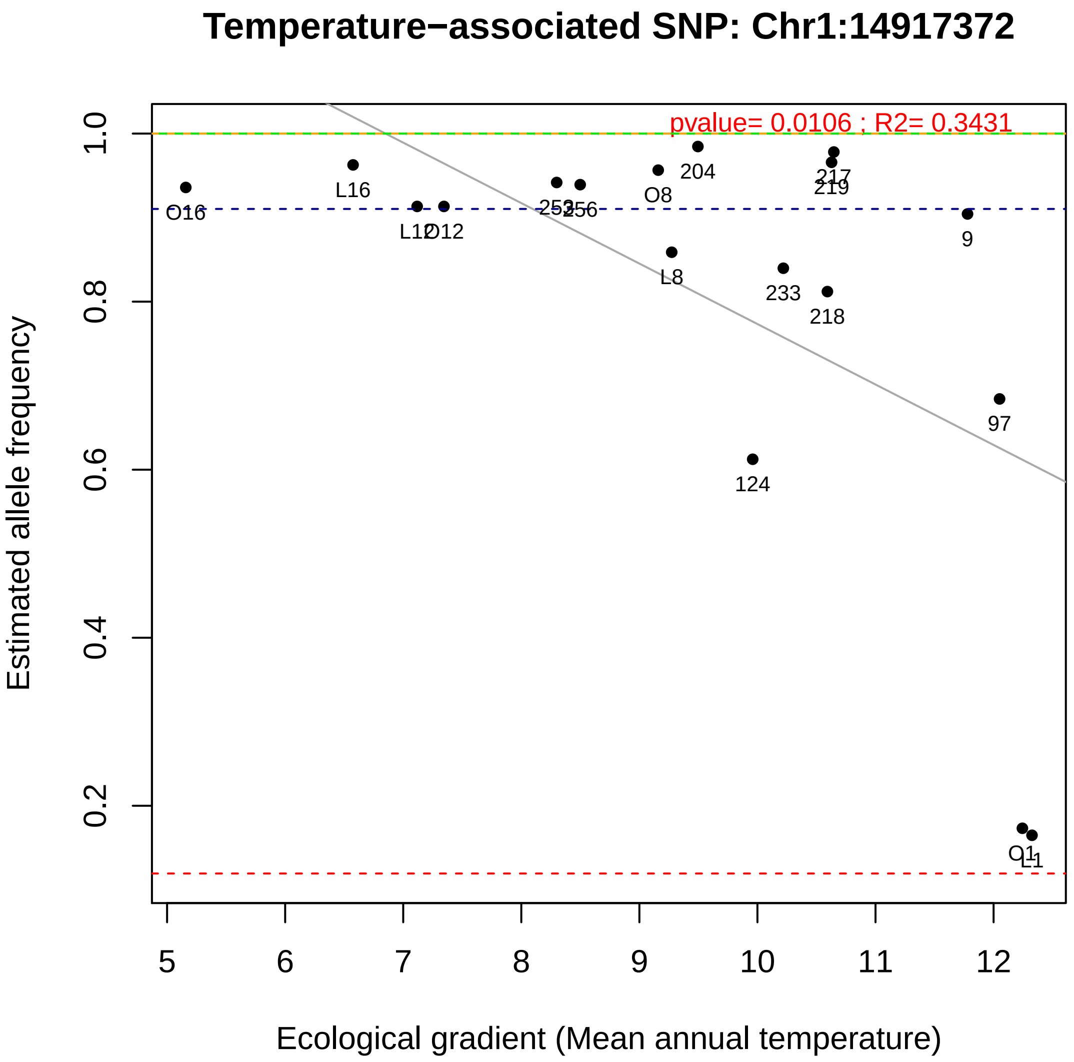

Temperature−associated SNP: Chr1:14917372
pvalue= 0.0106 ; R2= 0.3431
Estimated allele frequency
1.0
0.8
0.6
0.4
0.2
5
6
7
8
9
10
11
12
Ecological gradient (Mean annual temperature)
O16
L16
L12
O12
252
256
O8
204
L8
217
219
233
218
124
9
97
O1
L1

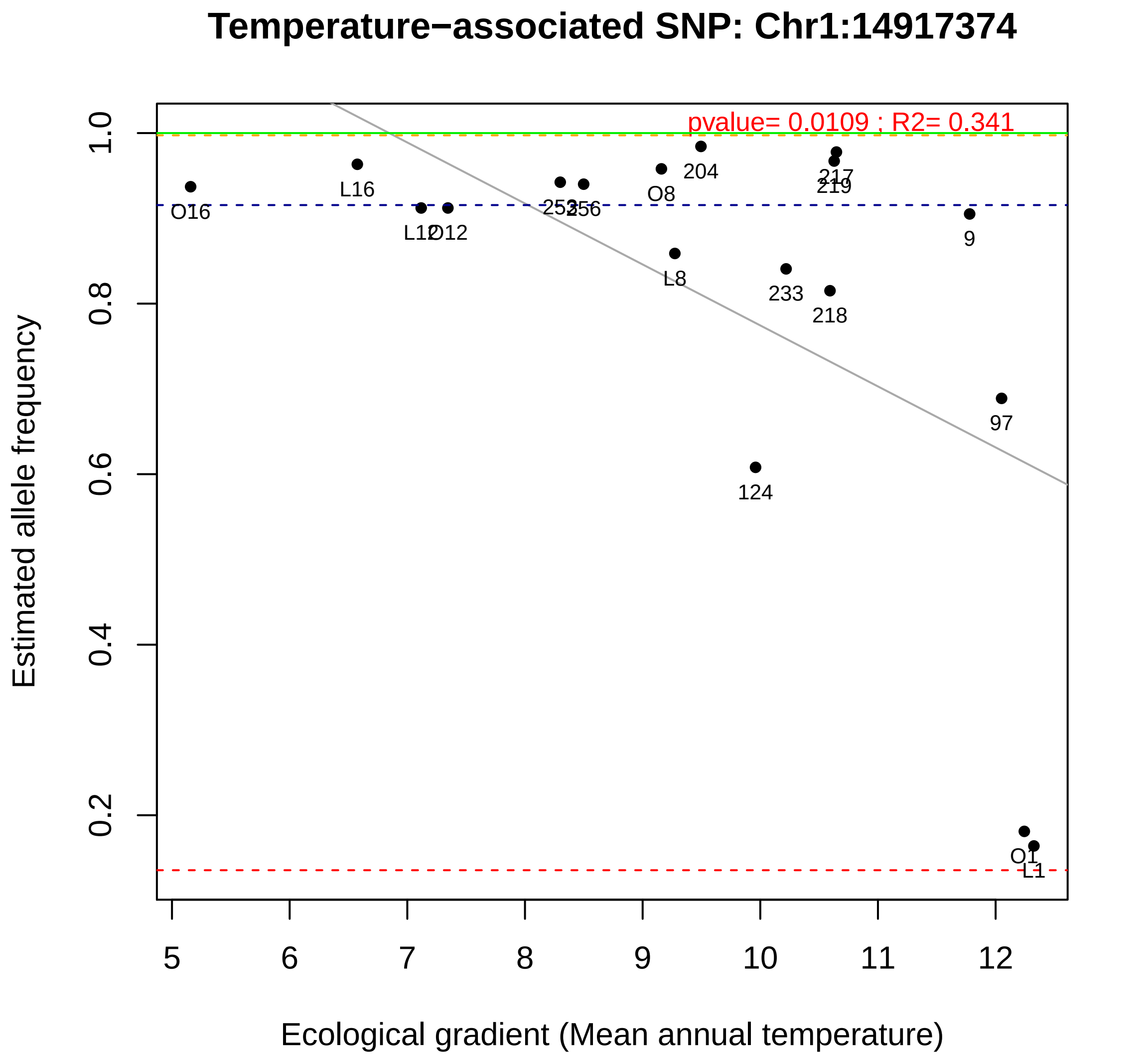

Temperature−associated SNP: Chr1:14917374
pvalue= 0.0109 ; R2= 0.341
Estimated allele frequency
Ecological gradient (Mean annual temperature)
0.2
0.4
0.6
0.8
1.0
5
6
7
8
9
10
11
12
O16
L16
L12
O12
O8
204
217
219
9
L8
233
218
97
124
O1
L1

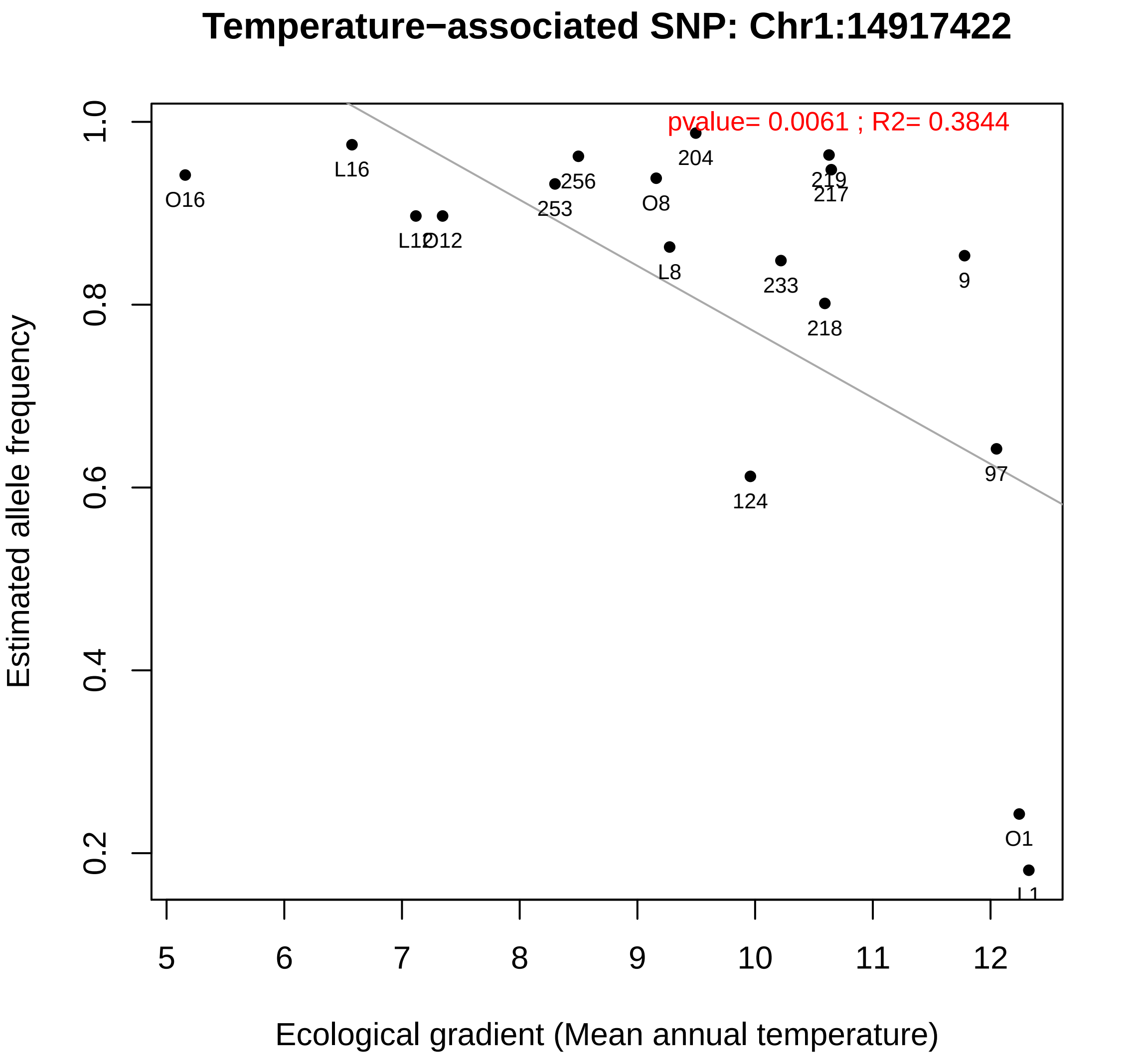

Temperature−associated SNP: Chr1:14917422
pvalue= 0.0061 ; R2= 0.3844
Estimated allele frequency
Ecological gradient (Mean annual temperature)
0.2
0.4
0.6
0.8
1.0
5
6
7
8
9
10
11
12
O16
L16
L12
O12
253
256
O8
204
L8
233
219
217
218
9
124
97
O1

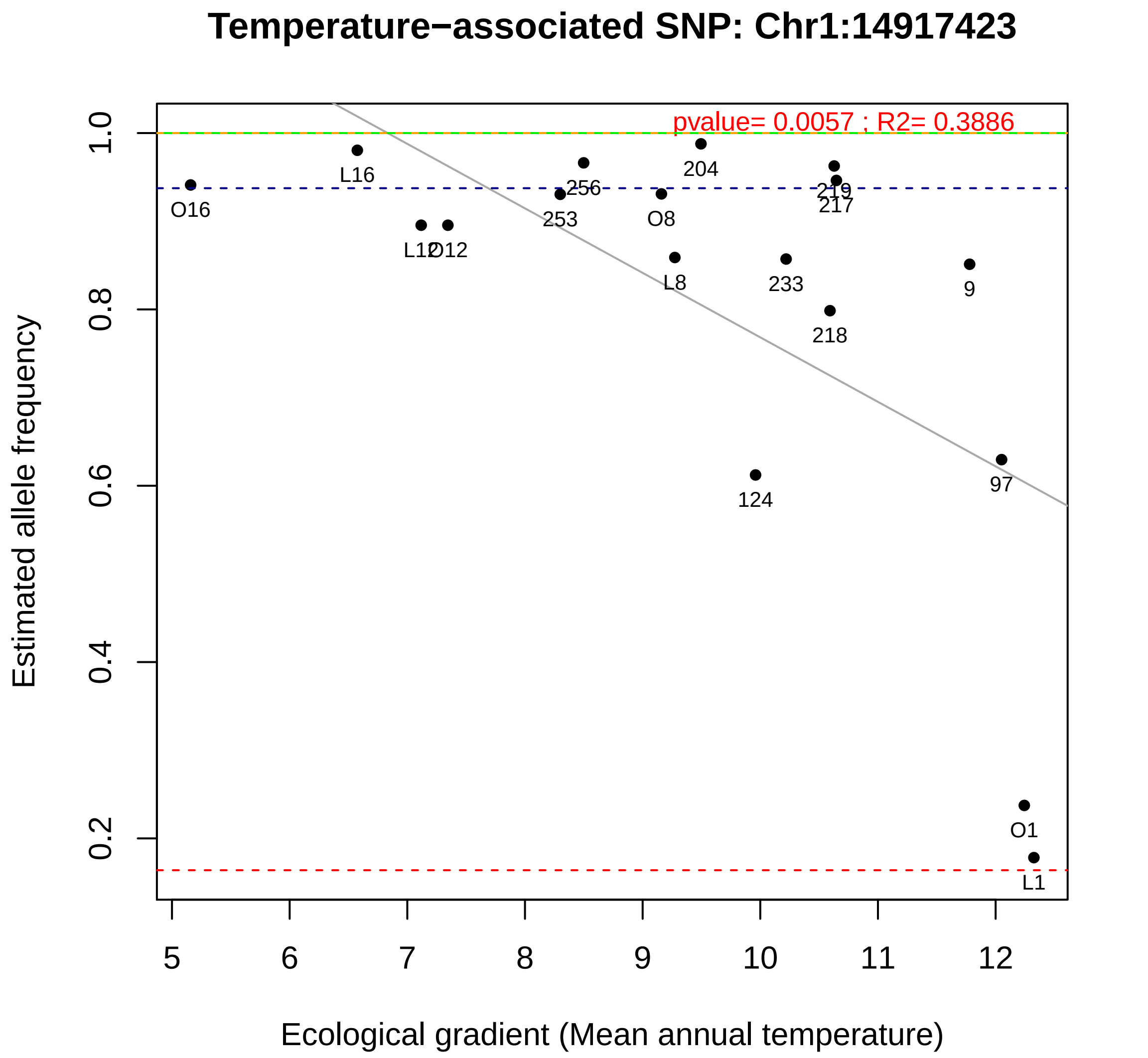
Temperature−associated SNP: Chr1:14917423
pvalue= 0.0057 ; R2= 0.3886
Estimated allele frequency
Ecological gradient (Mean annual temperature)
0.2
0.4
0.6
0.8
1.0
5
6
7
8
9
10
11
12
O16
L16
L12
O12
253
256
O8
204
L8
233
219
217
218
9
124
97
O1
L1

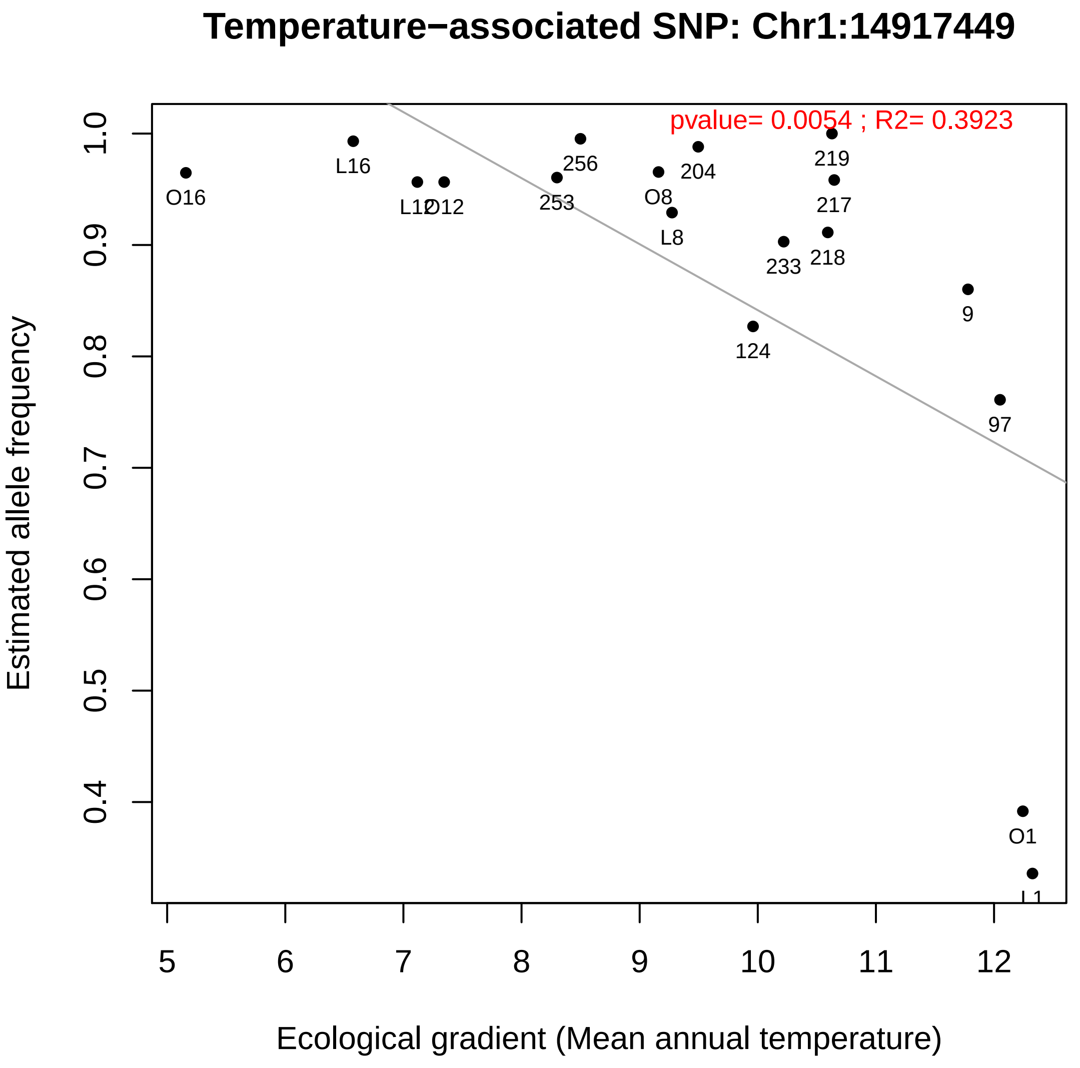
Temperature−associated SNP: Chr1:14917449
pvalue= 0.0054 ; R2= 0.3923
Estimated allele frequency
Ecological gradient (Mean annual temperature)
1.0
0.9
0.8
0.7
0.6
0.5
0.4
5
6
7
8
9
10
11
12
O16
L16
L12
O12
253
256
O8
L8
204
219
217
218
233
124
9
97
O1

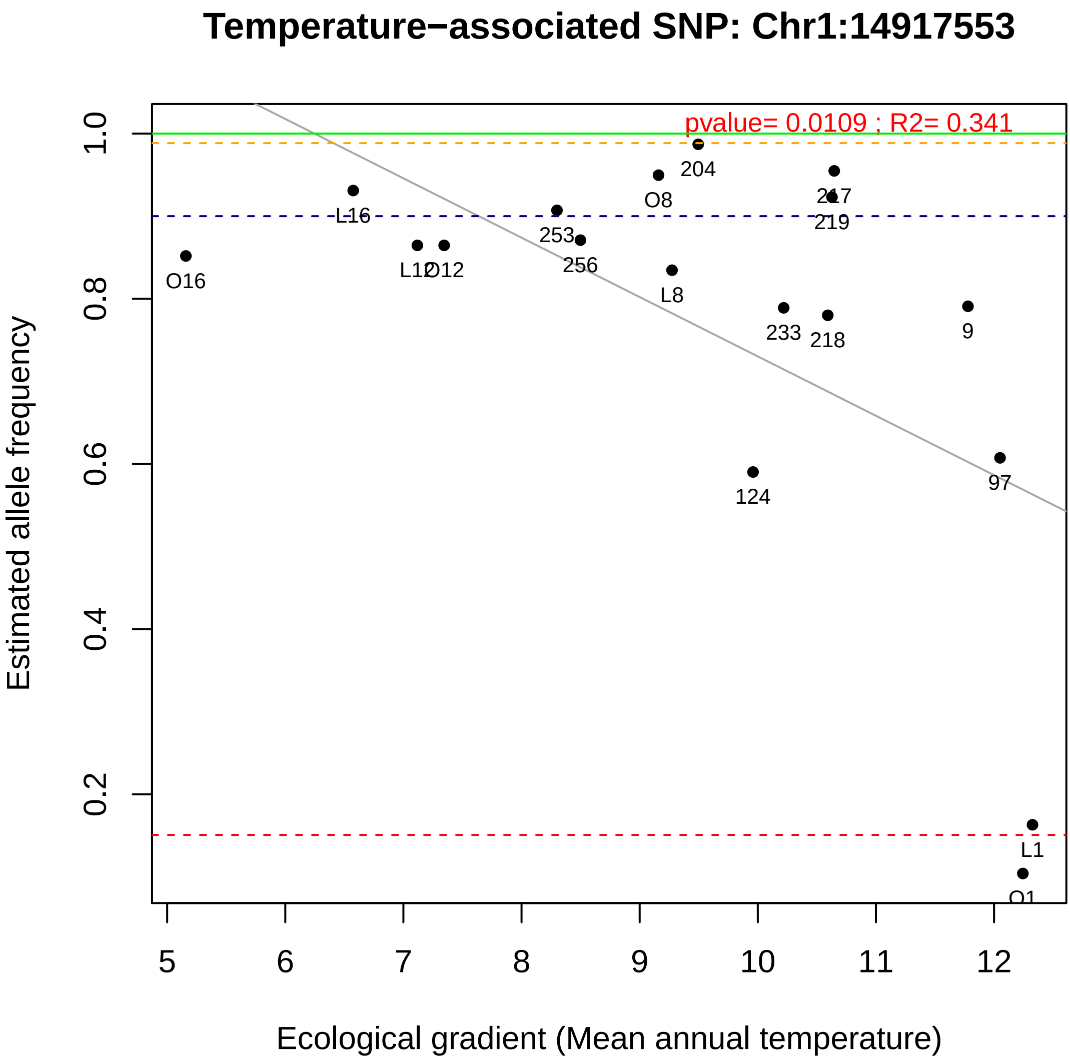

Temperature−associated SNP: Chr1:14917553
pvalue= 0.0109 ; R2= 0.341
Estimated allele frequency
Ecological gradient (Mean annual temperature)
1.0
0.8
0.6
0.4
0.2
5
6
7
8
9
10
11
12
O16
L16
L12
O12
253
256
O8
204
L8
217
219
233
218
124
9
97
L1
O1

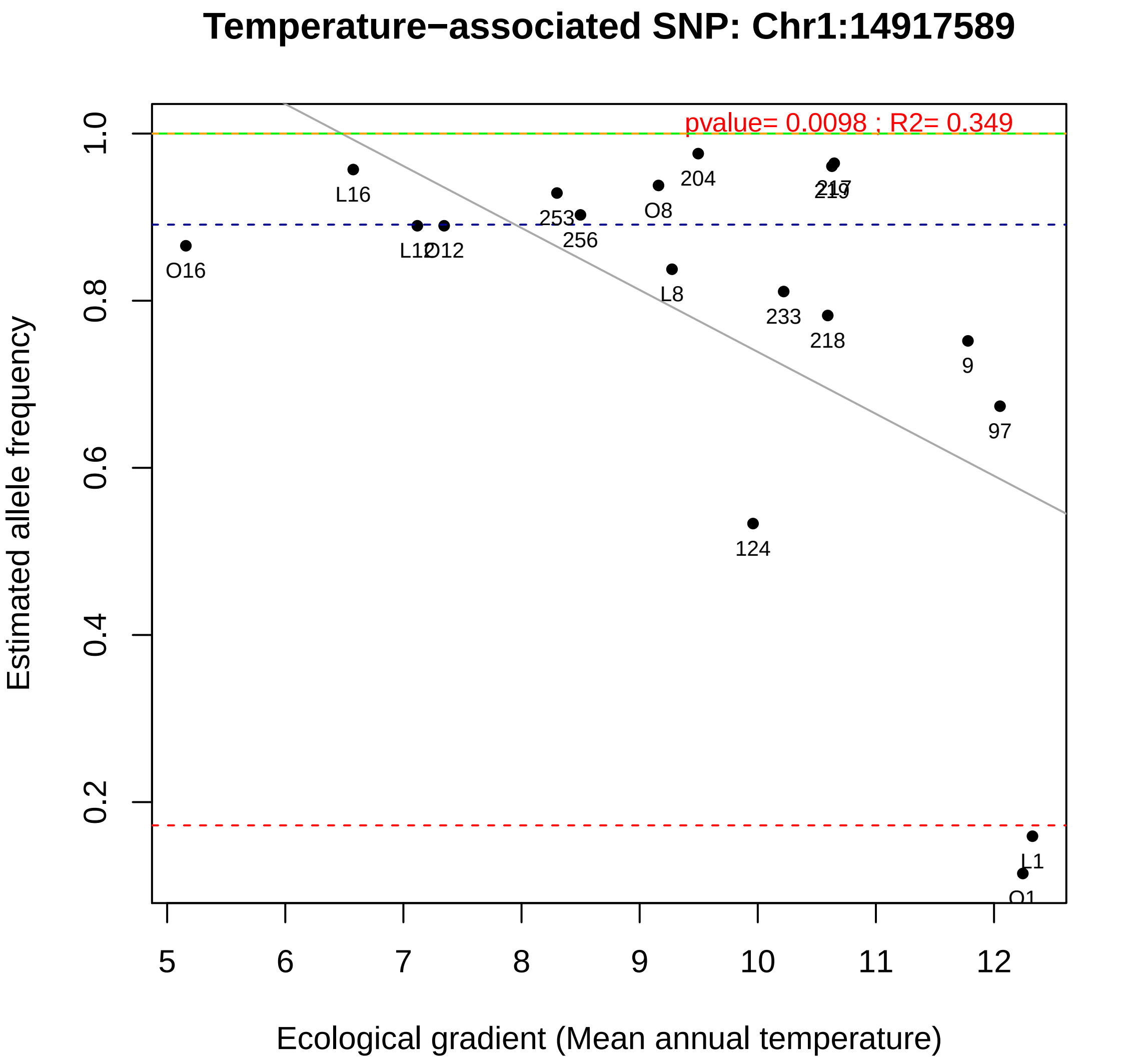

Temperature−associated SNP: Chr1:14917589
pvalue= 0.0098 ; R2= 0.349
Estimated allele frequency
Ecological gradient (Mean annual temperature)
1.0
0.8
0.6
0.4
0.2
5
6
7
8
9
10
11
12
O16
L16
L12
O12
253
256
O8
204
L8
124
233
217
219
218
9
97
L1
O1

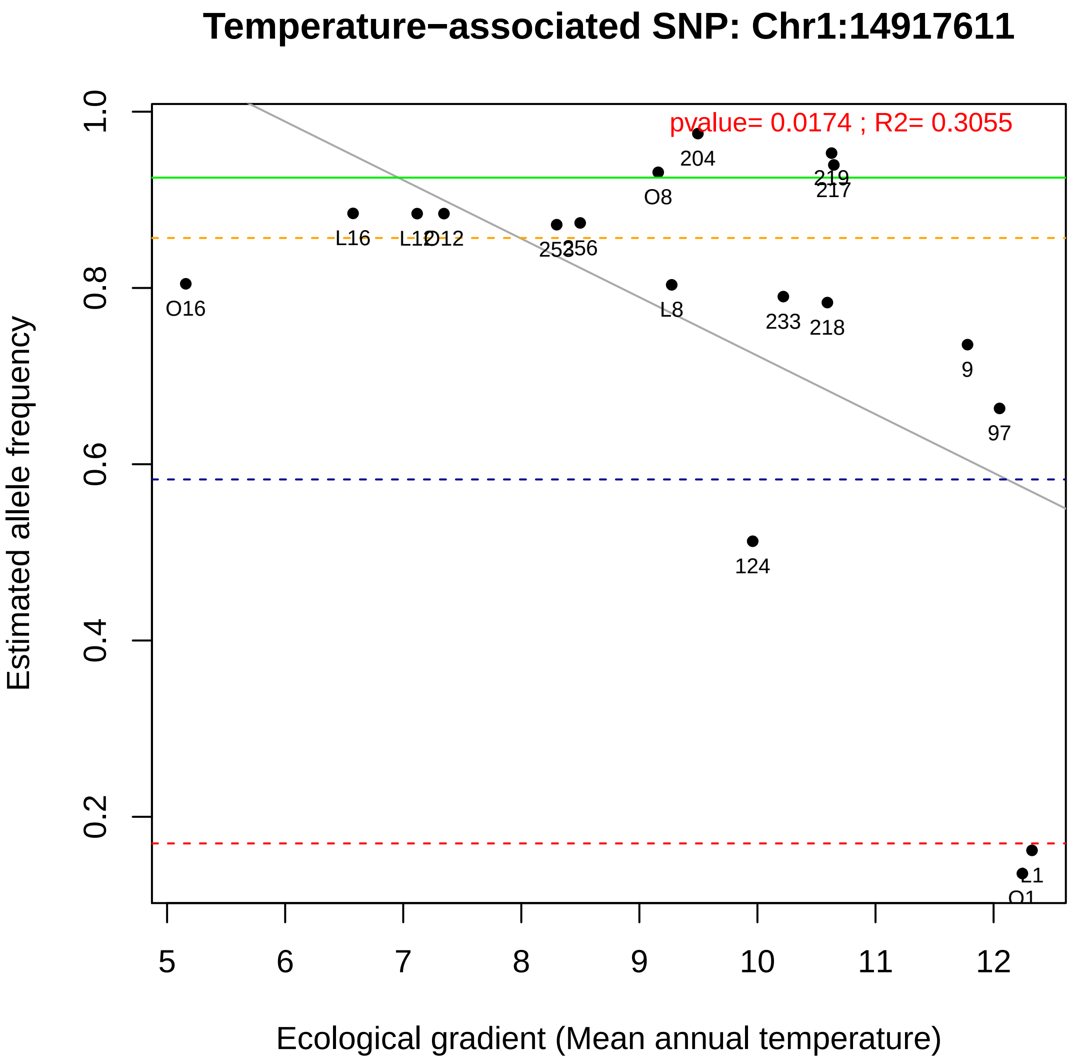

Temperature−associated SNP: Chr1:14917611
pvalue= 0.0174 ; R2= 0.3055
Estimated allele frequency
Ecological gradient (Mean annual temperature)
0.2
0.4
0.6
0.8
1.0
5
6
7
8
9
10
11
12
O16
L16
L12
O12
O8
204
219
217
L8
233
218
124
9
97
L1
O1

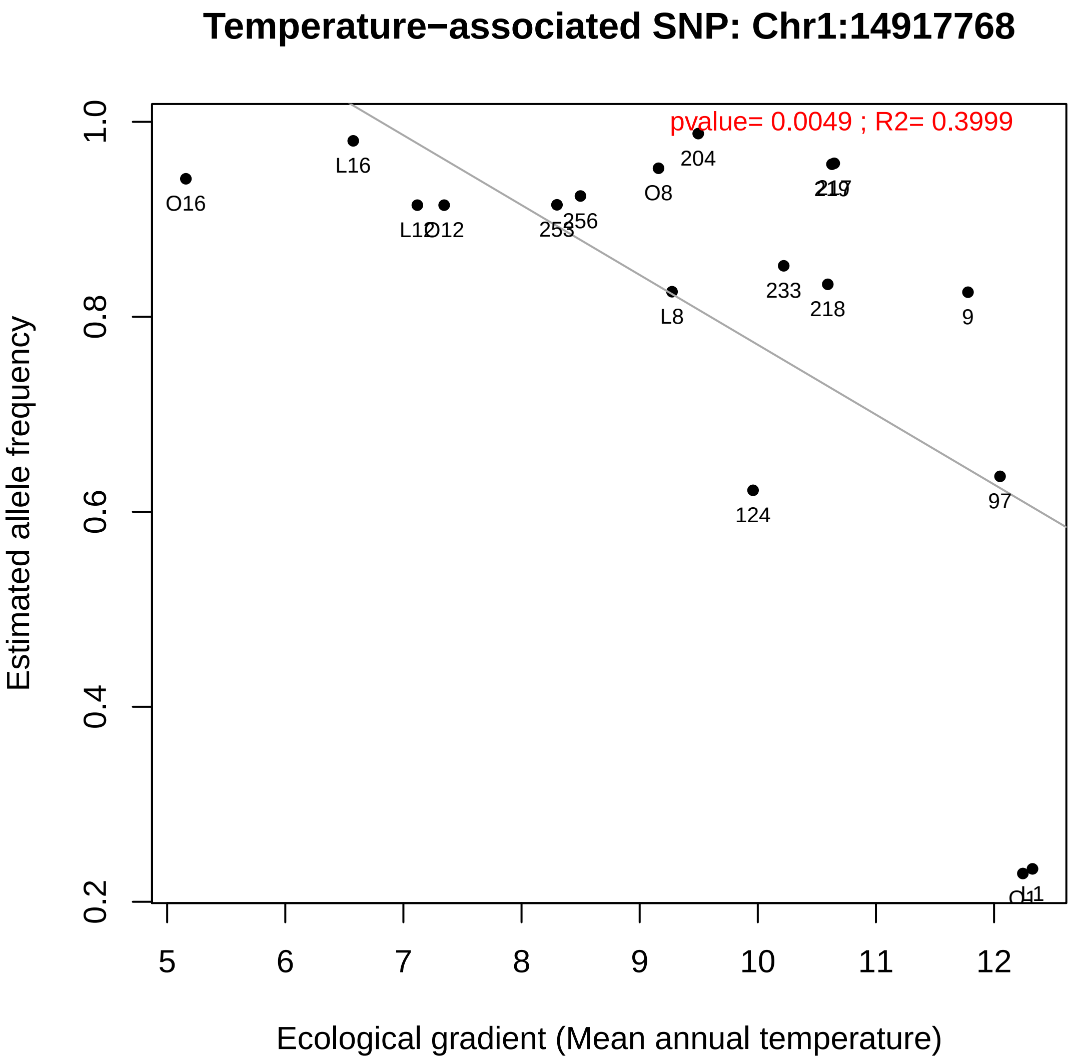
Temperature−associated SNP: Chr1:14917768
pvalue= 0.0049 ; R2= 0.3999
Estimated allele frequency
Ecological gradient (Mean annual temperature)
1.0
0.8
0.6
0.4
0.2
5
6
7
8
9
10
11
12
O16
L16
L12
O12
253
256
O8
204
233
218
L8
9
124
97

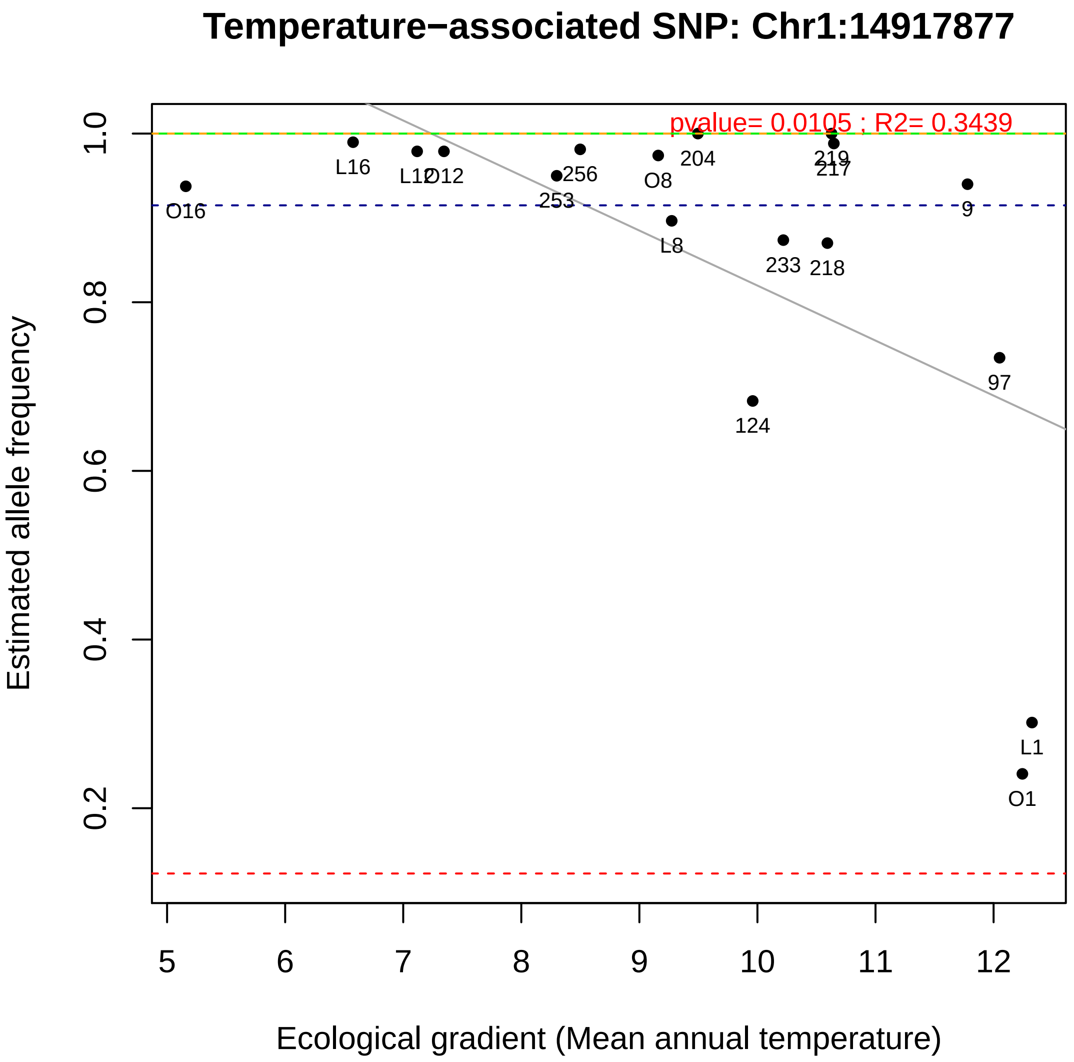

Temperature−associated SNP: Chr1:14917877
pvalue= 0.0105 ; R2= 0.3439
Estimated allele frequency
0.2
0.4
0.6
0.8
1.0
5
6
7
8
9
10
11
12
Ecological gradient (Mean annual temperature)
O16
L16
L12
O12
253
256
O8
204
L8
124
233
219
217
218
9
97
L1
O1

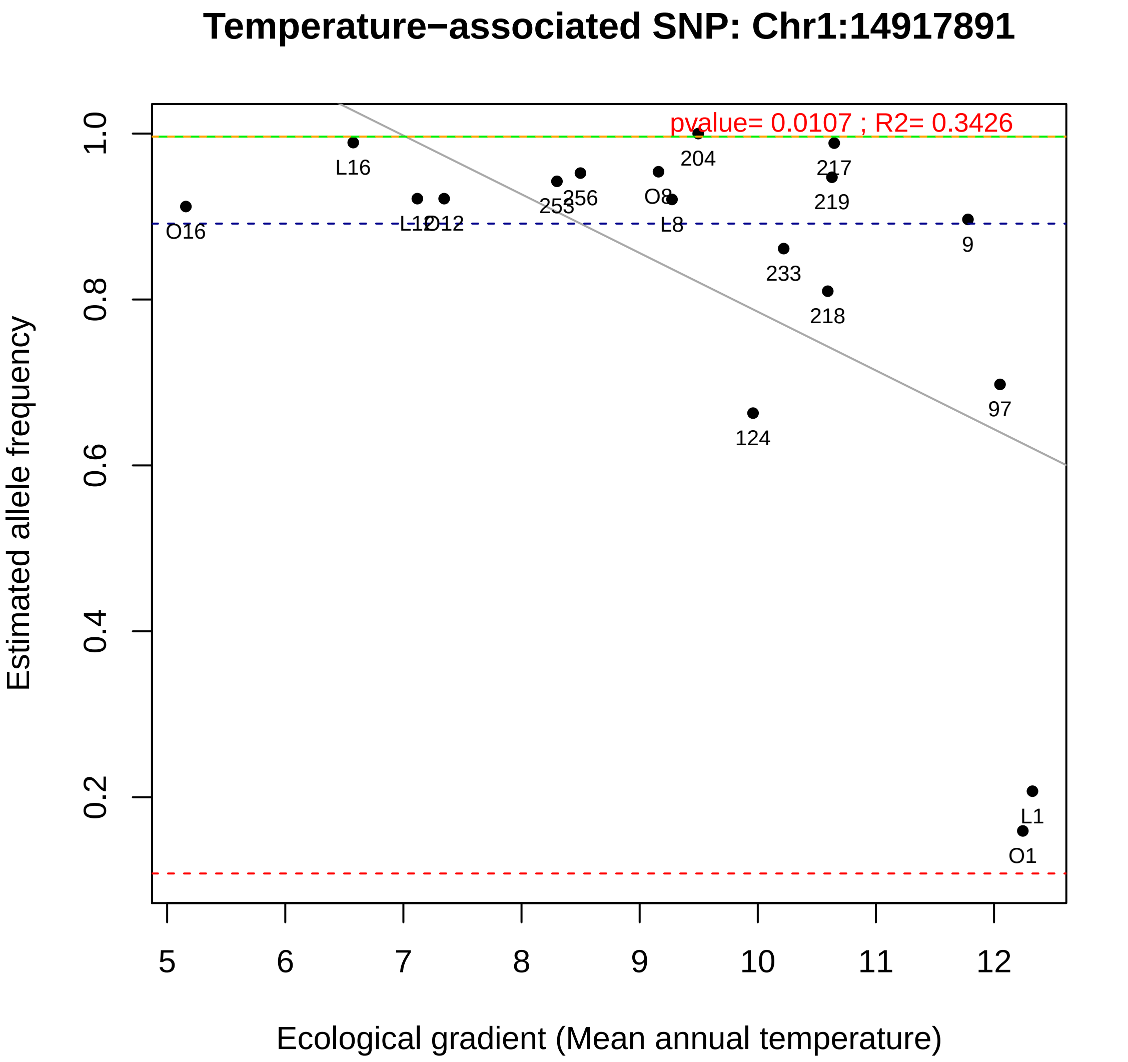
Temperature−associated SNP: Chr1:14917891
pvalue= 0.0107 ; R2= 0.3426
Estimated allele frequency
Ecological gradient (Mean annual temperature)
0.2
0.4
0.6
0.8
1.0
5
6
7
8
9
10
11
12
O16
L16
L12
O12
253
256
O8
L8
204
217
219
233
218
124
9
97
L1
O1

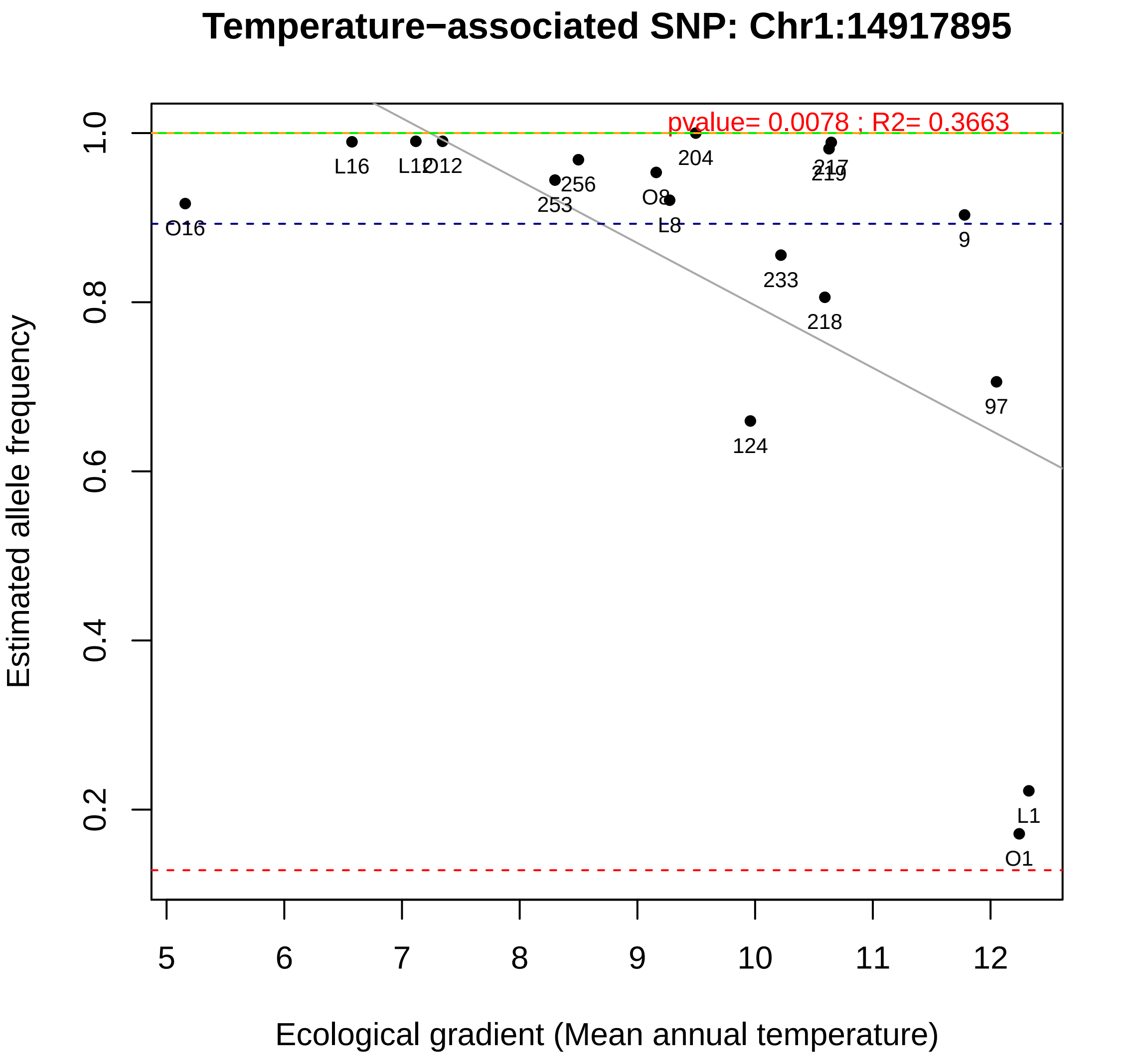

Temperature−associated SNP: Chr1:14917895
pvalue= 0.0078 ; R2= 0.3663
Estimated allele frequency
Ecological gradient (Mean annual temperature)
0.2
0.4
0.6
0.8
1.0
5
6
7
8
9
10
11
12
O16
L16
L12
O12
253
256
O8
L8
204
233
218
124
217
219
9
97
L1
O1

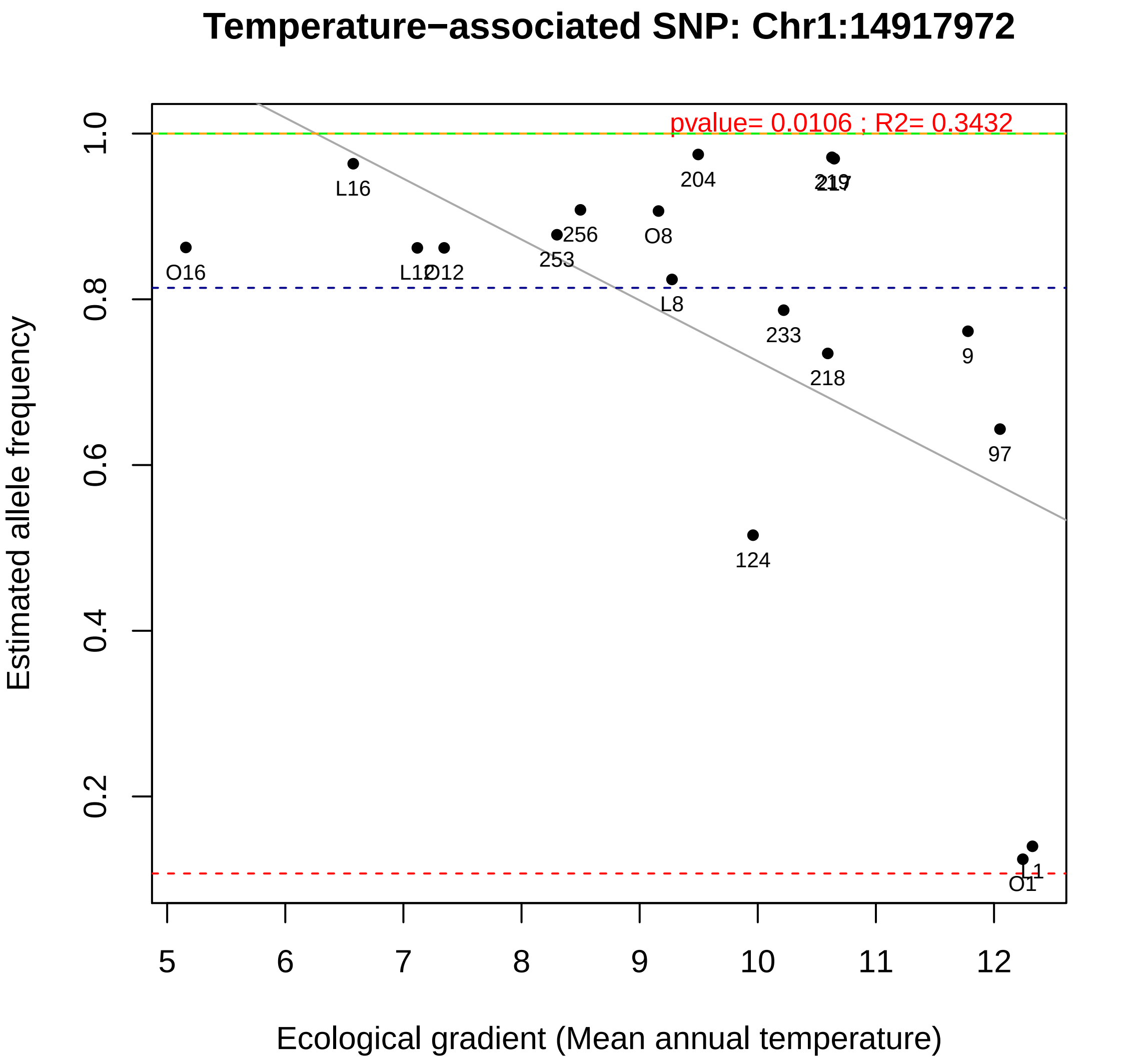

Temperature−associated SNP: Chr1:14917972
pvalue= 0.0106 ; R2= 0.3432
Estimated allele frequency
0.2
0.4
0.6
0.8
1.0
5
6
7
8
9
10
11
12
Ecological gradient (Mean annual temperature)
O16
L16
L12
O12
253
256
O8
204
L8
233
218
9
97
124
O1
L1

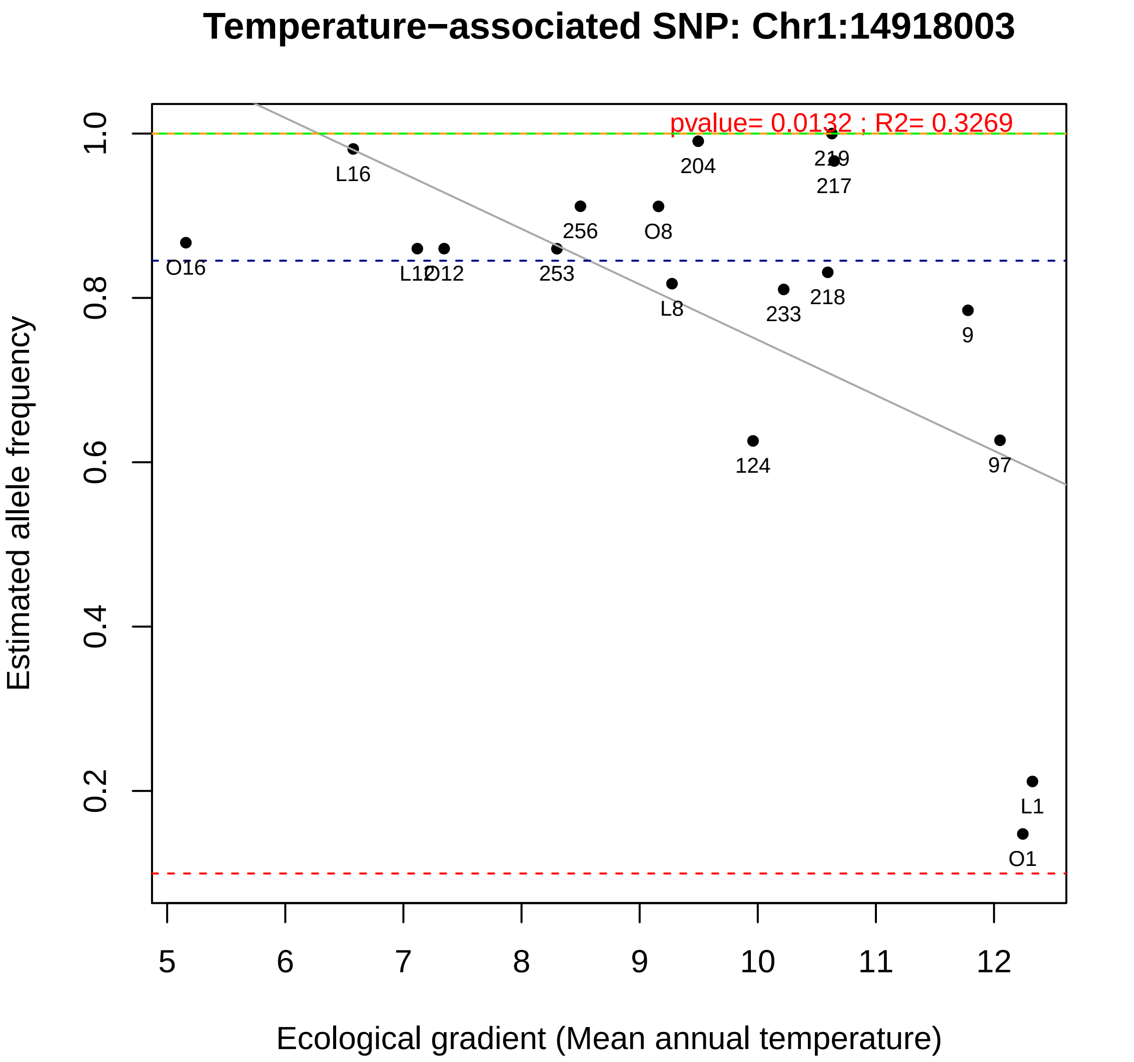
Temperature−associated SNP: Chr1:14918003
pvalue= 0.0132 ; R2= 0.3269
Estimated allele frequency
Ecological gradient (Mean annual temperature)
0.2
0.4
0.6
0.8
1.0
5
6
7
8
9
10
11
12
O16
L16
L12
O12
253
256
O8
L8
204
124
233
218
219
217
9
97
L1
O1

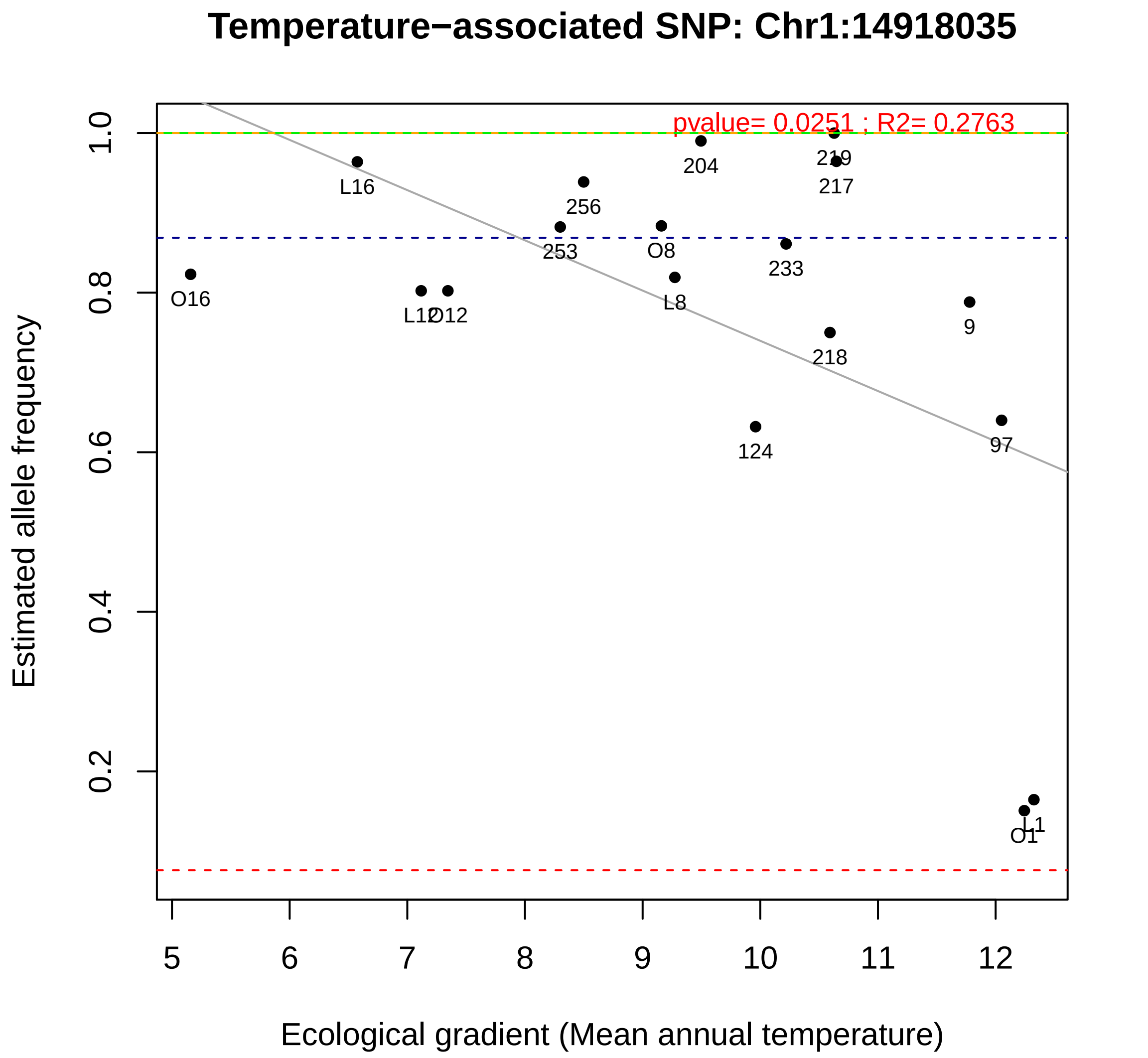

Temperature−associated SNP: Chr1:14918035
pvalue= 0.0251 ; R2= 0.2763
Estimated allele frequency
Ecological gradient (Mean annual temperature)
1.0
0.8
0.6
0.4
0.2
5
6
7
8
9
10
11
12
O16
L16
L12
O12
253
256
O8
L8
204
124
233
219
217
218
9
97
O1
L1

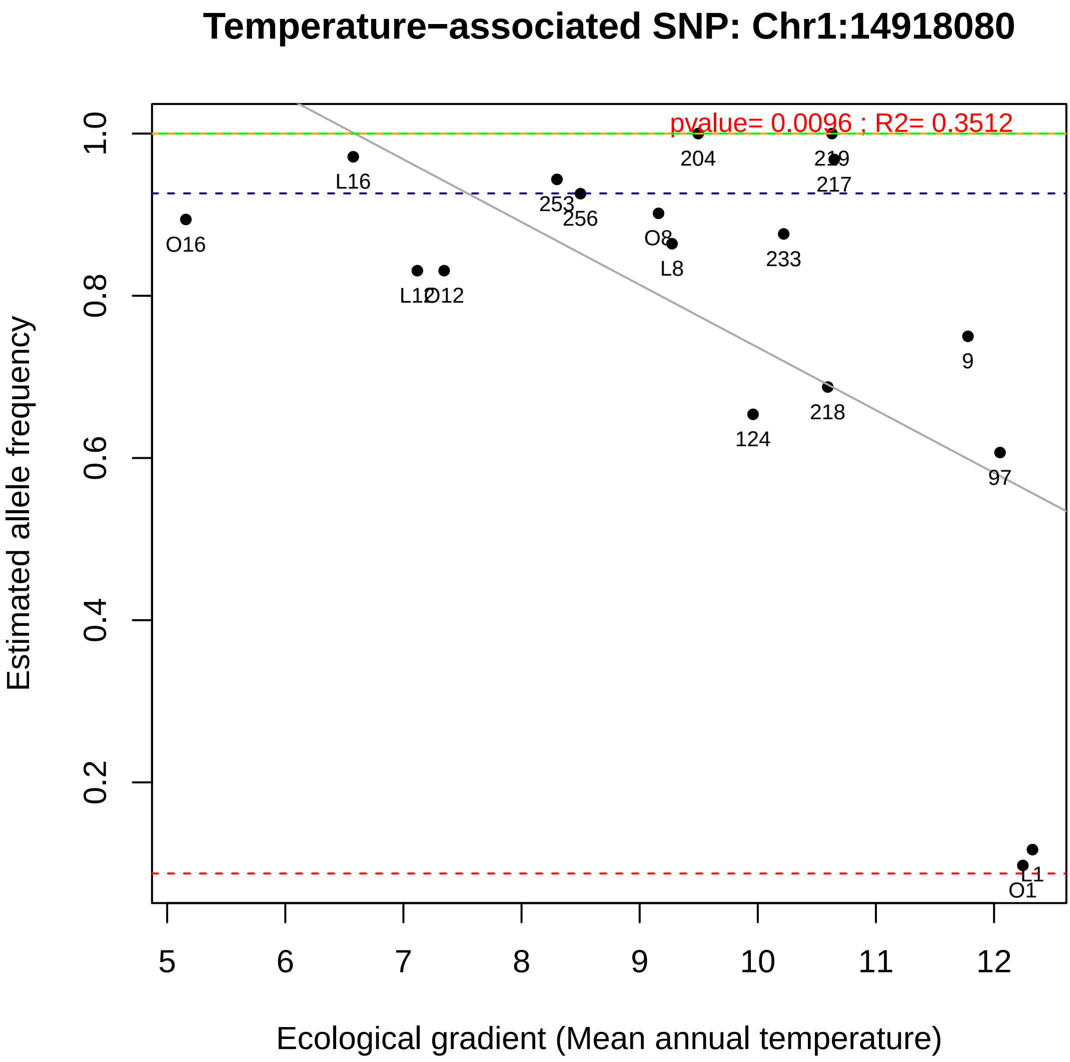

Temperature−associated SNP: Chr1:14918080
pvalue= 0.0096 ; R2= 0.3512
Estimated allele frequency
0.2
0.4
0.6
0.8
1.0
5
6
7
8
9
10
11
12
Ecological gradient (Mean annual temperature)
O16
L16
L12
O12
253
256
O8
L8
204
233
219
217
124
218
9
97
L1
O1

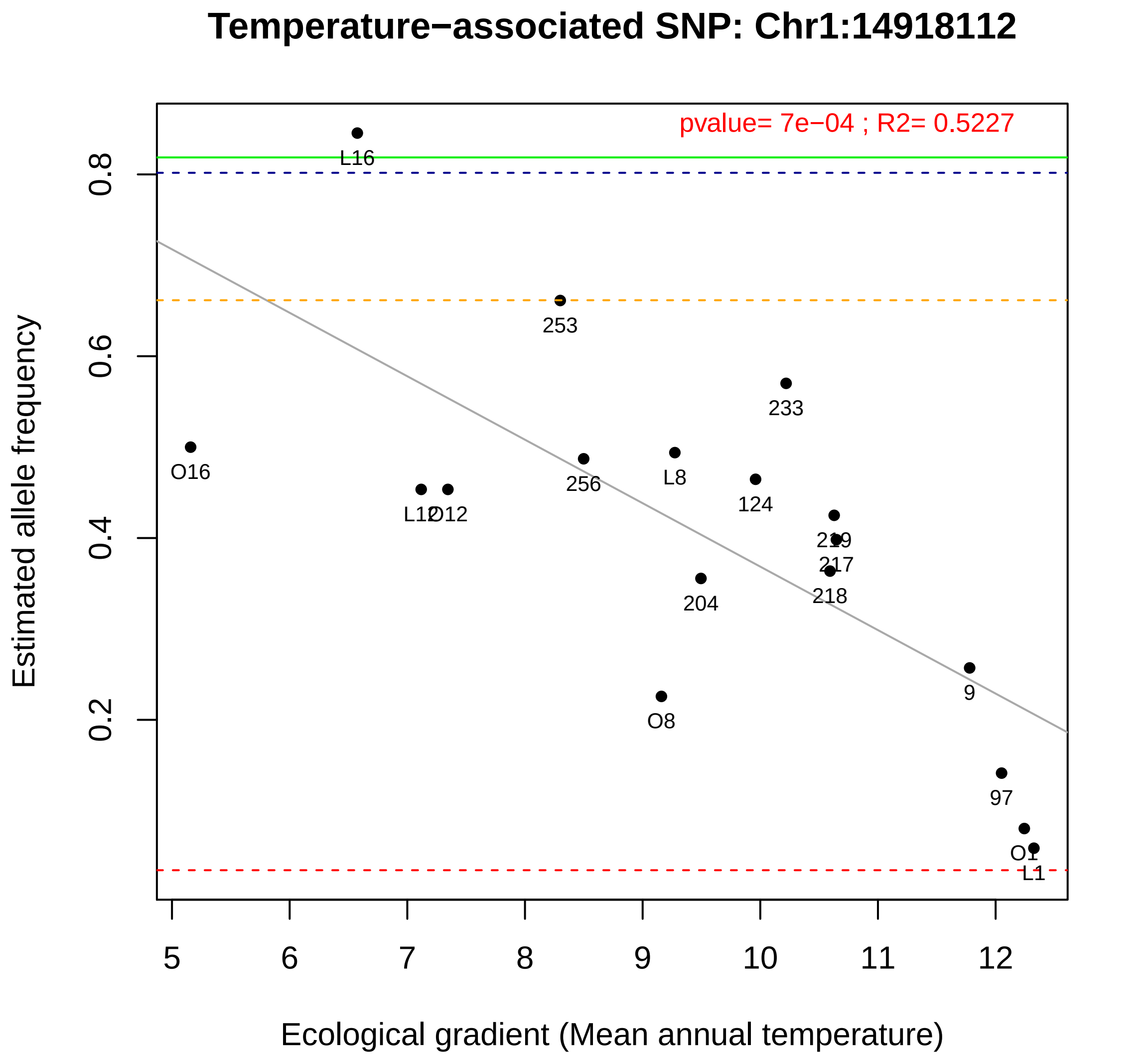
Temperature−associated SNP: Chr1:14918112
pvalue= 7e−04 ; R2= 0.5227
Estimated allele frequency
0.2
0.4
0.6
0.8
L16
253
233
O16
L12
O12
256
L8
124
219
217
218
204
9
O8
97
O1
L1
5
6
7
8
9
10
11
12
Ecological gradient (Mean annual temperature)

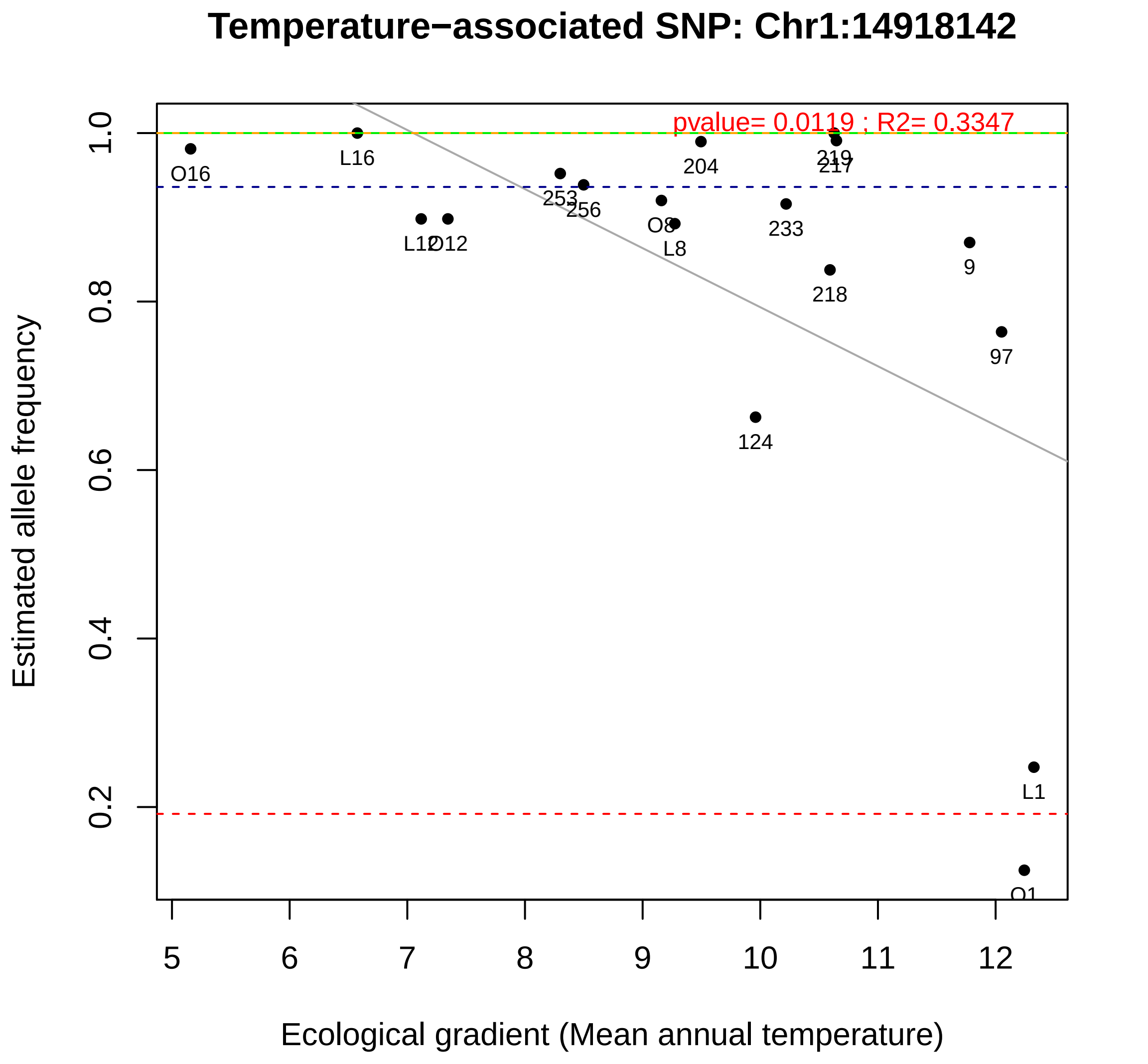
Temperature−associated SNP: Chr1:14918142
pvalue= 0.0119 ; R2= 0.3347
Estimated allele frequency
Ecological gradient (Mean annual temperature)
1.0
0.8
0.6
0.4
0.2
5
6
7
8
9
10
11
12
O16
L16
L12
O12
253
256
O8
L8
204
233
219
217
218
124
9
97
L1
O1

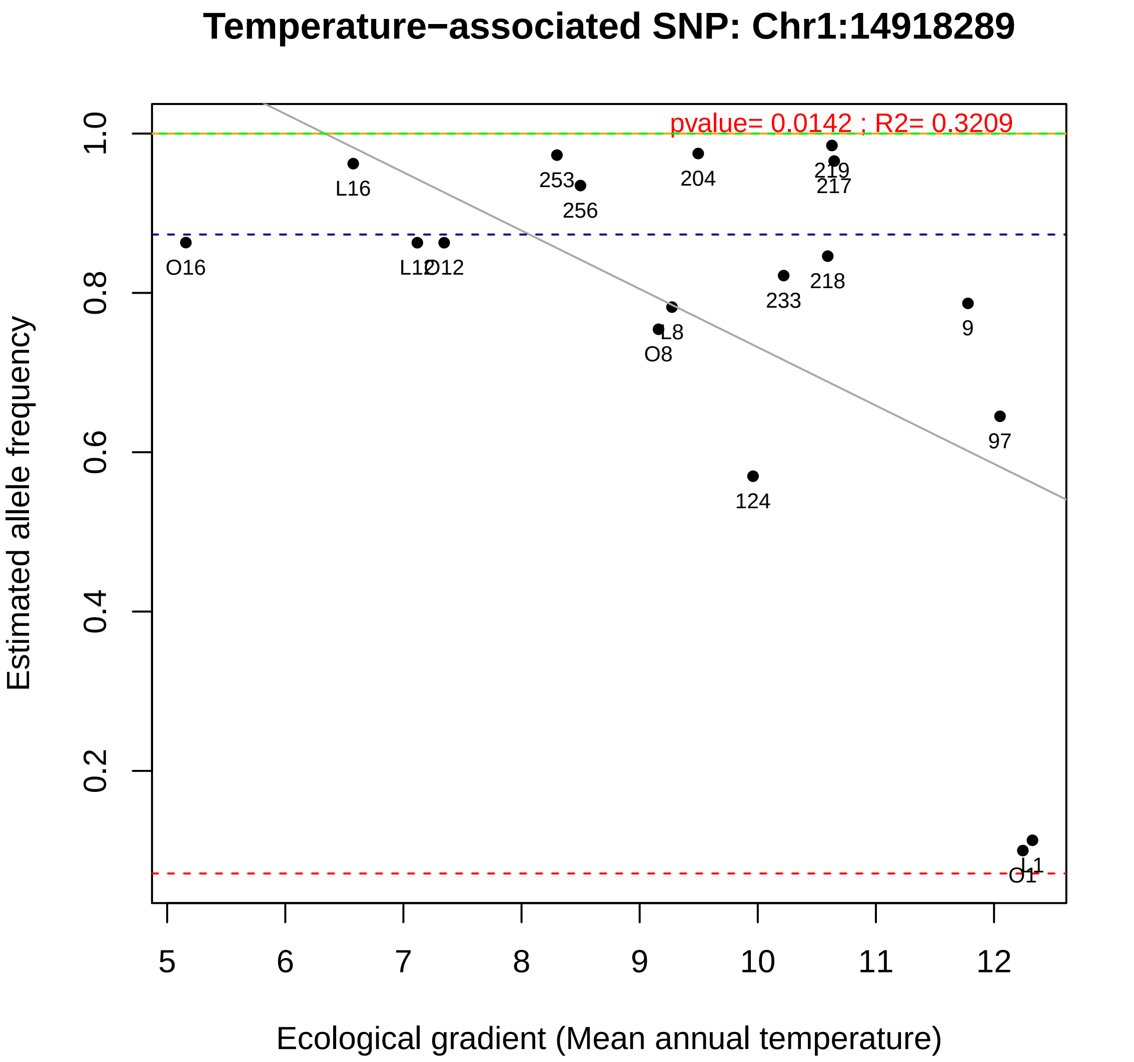

Temperature−associated SNP: Chr1:14918289
pvalue= 0.0142 ; R2= 0.3209
Estimated allele frequency
Ecological gradient (Mean annual temperature)
1.0
0.8
0.6
0.4
0.2
5
6
7
8
9
10
11
12
O16
L16
L12
O12
253
256
204
219
217
218
233
L8
O8
9
97
124
L1
O1

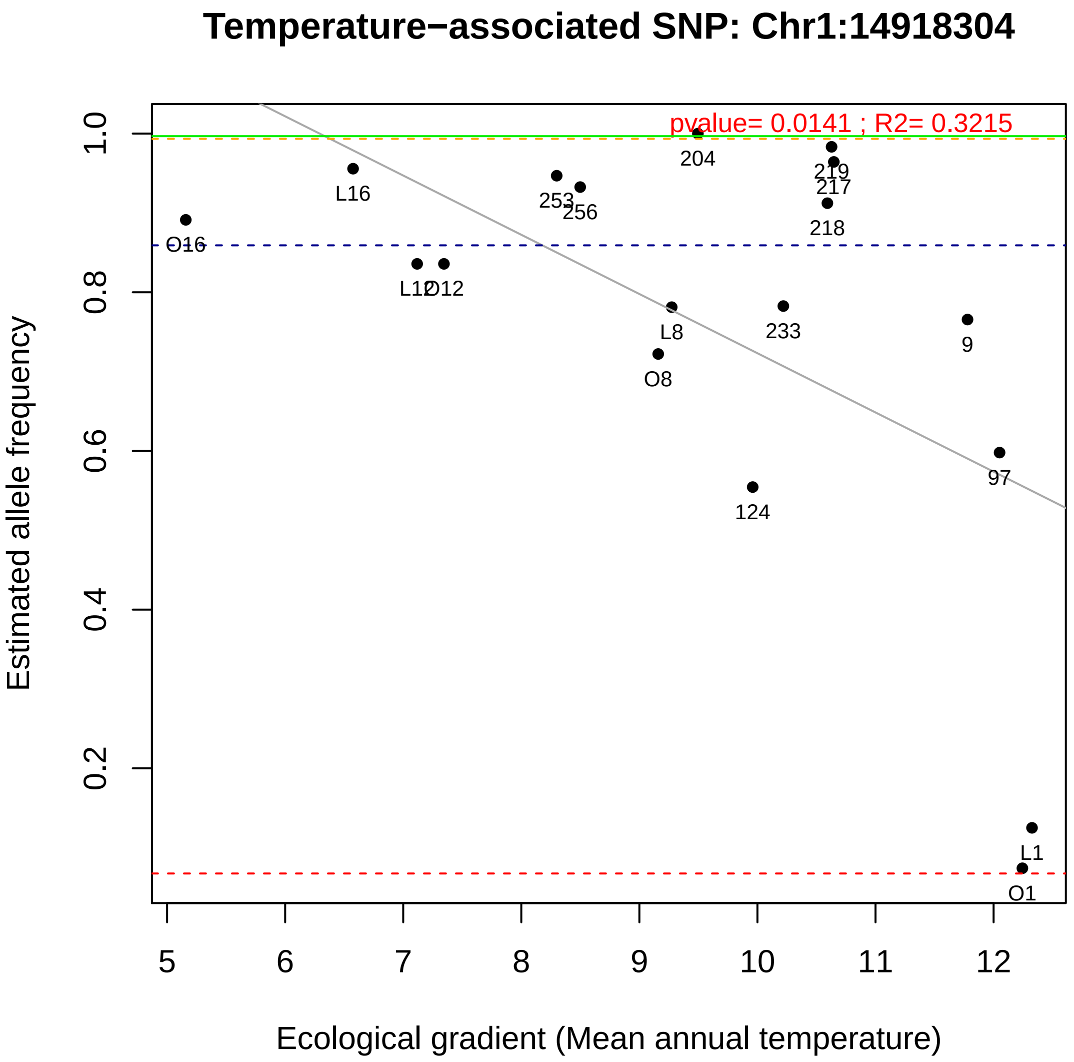

Temperature−associated SNP: Chr1:14918304
pvalue= 0.0141 ; R2= 0.3215
Estimated allele frequency
Ecological gradient (Mean annual temperature)
0.2
0.4
0.6
0.8
1.0
5
6
7
8
9
10
11
12
O16
L16
L12
O12
253
256
204
219
217
218
L8
O8
233
9
124
97
L1
O1

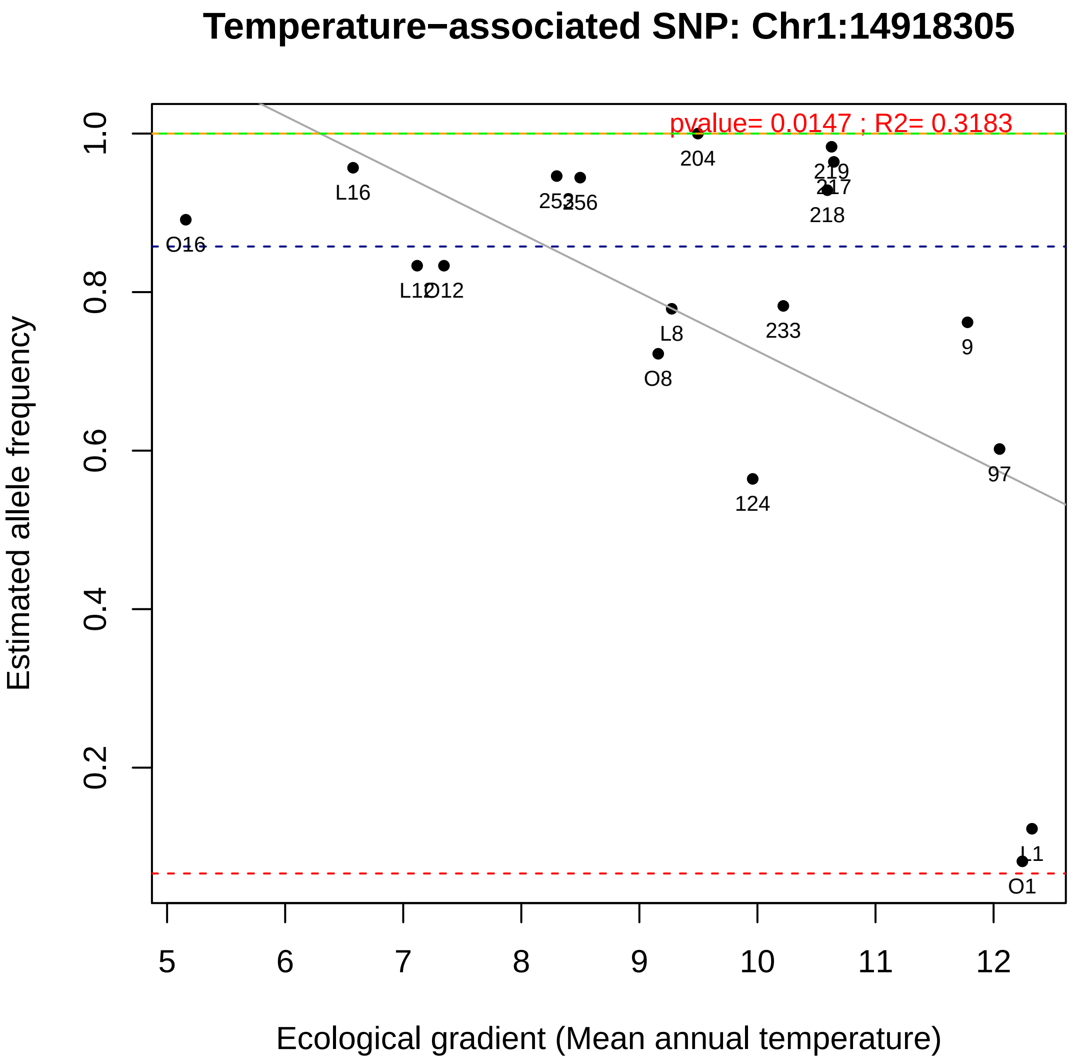
Temperature−associated SNP: Chr1:14918305
pvalue= 0.0147 ; R2= 0.3183
Estimated allele frequency
Ecological gradient (Mean annual temperature)
1.0
0.8
0.6
0.4
0.2
5
6
7
8
9
10
11
12
O16
L16
L12
O12
252
256
204
219
217
218
L8
O8
233
124
9
97
L1
O1

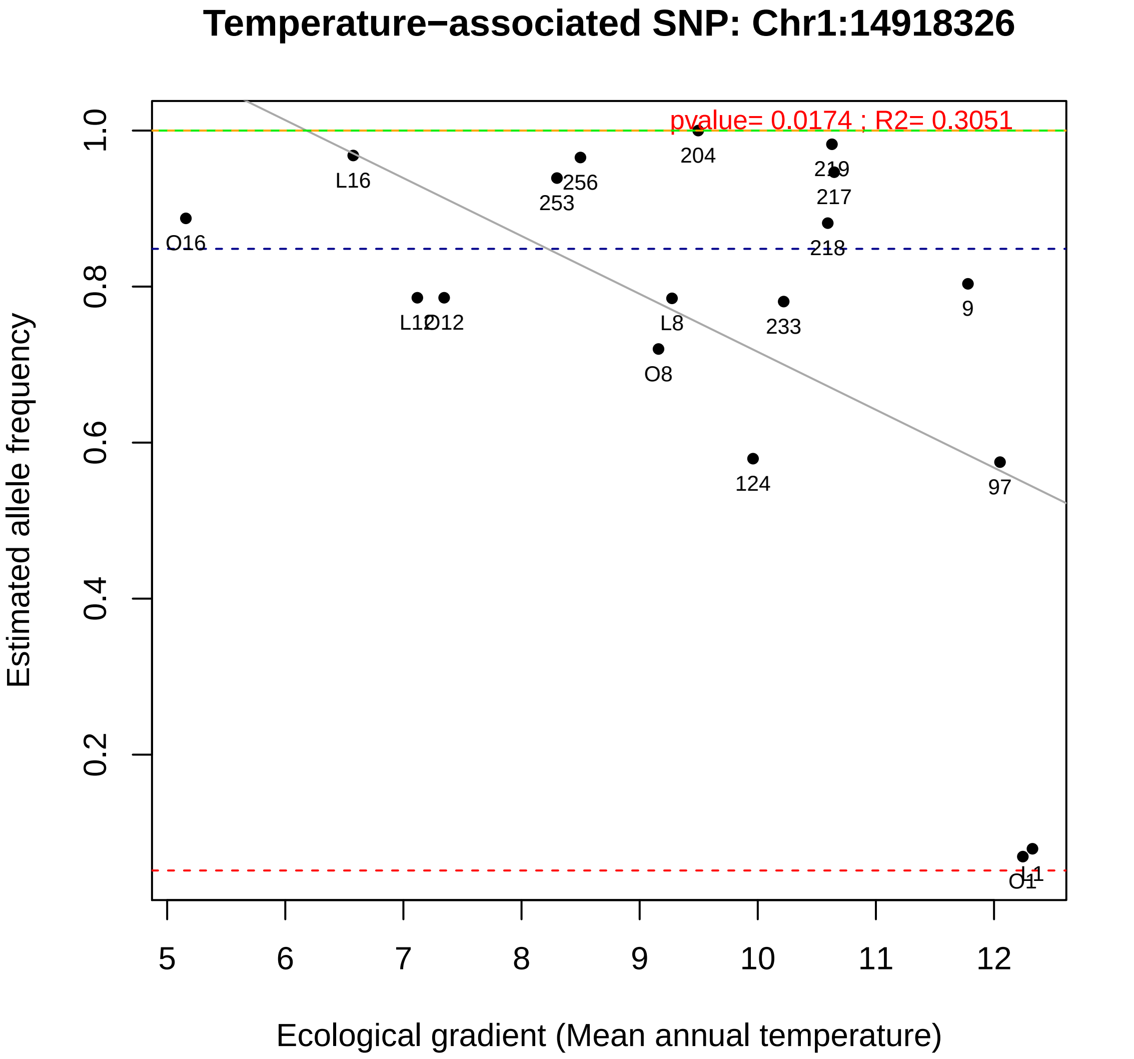
Temperature−associated SNP: Chr1:14918326
pvalue= 0.0174 ; R2= 0.3051
Estimated allele frequency
Ecological gradient (Mean annual temperature)
0.2
0.4
0.6
0.8
1.0
5
6
7
8
9
10
11
12
O16
L16
L12
O12
253
256
204
O8
L8
124
233
219
217
218
9
97
O1
L1

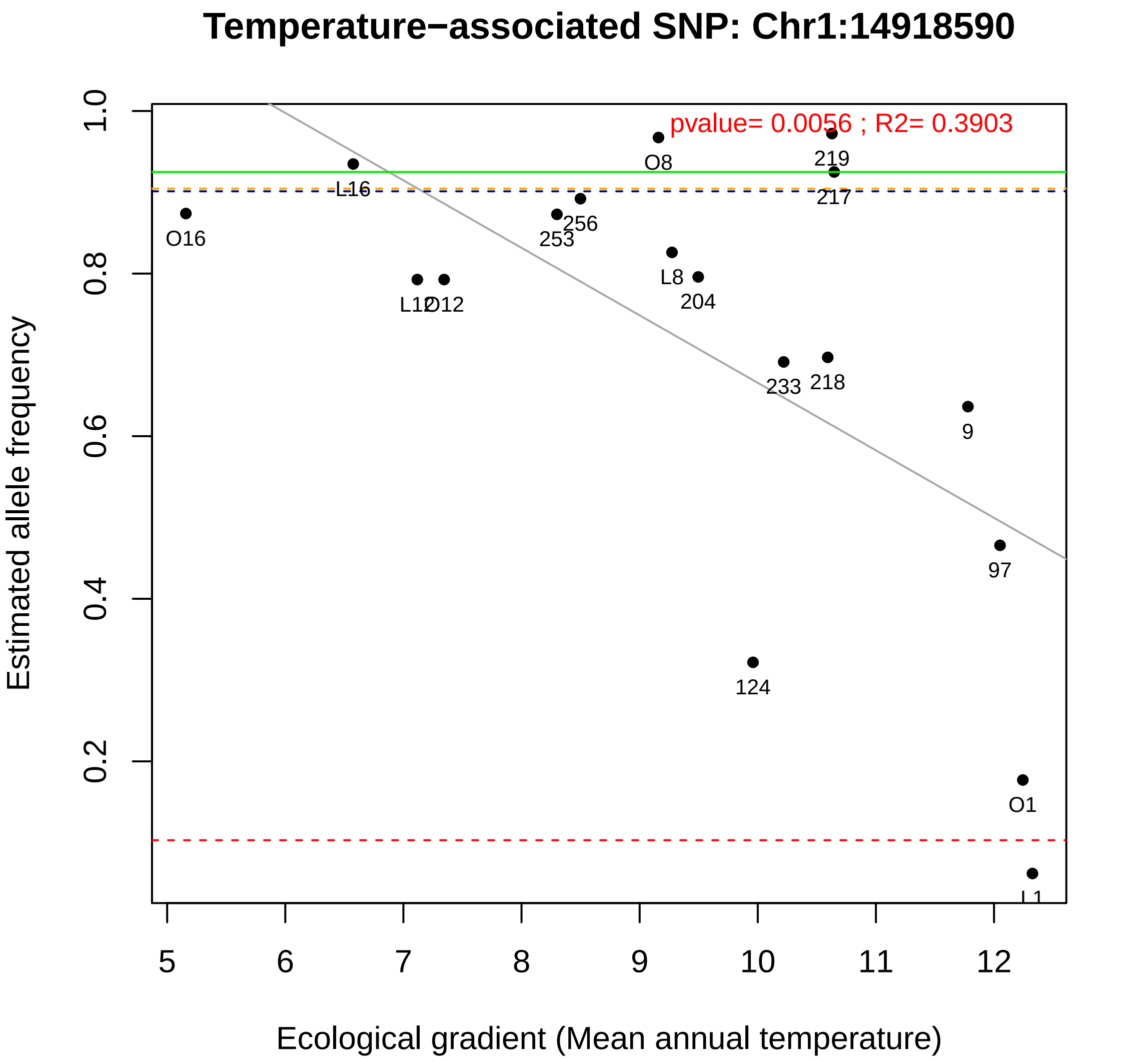
Temperature−associated SNP: Chr1:14918590
pvalue= 0.0056 ; R2= 0.3903
Estimated allele frequency
Ecological gradient (Mean annual temperature)
1.0
0.8
0.6
0.4
0.2
5
6
7
8
9
10
11
12
O16
L16
L12
O12
253
256
O8
L8
204
219
217
233
218
124
9
97
O1
L1

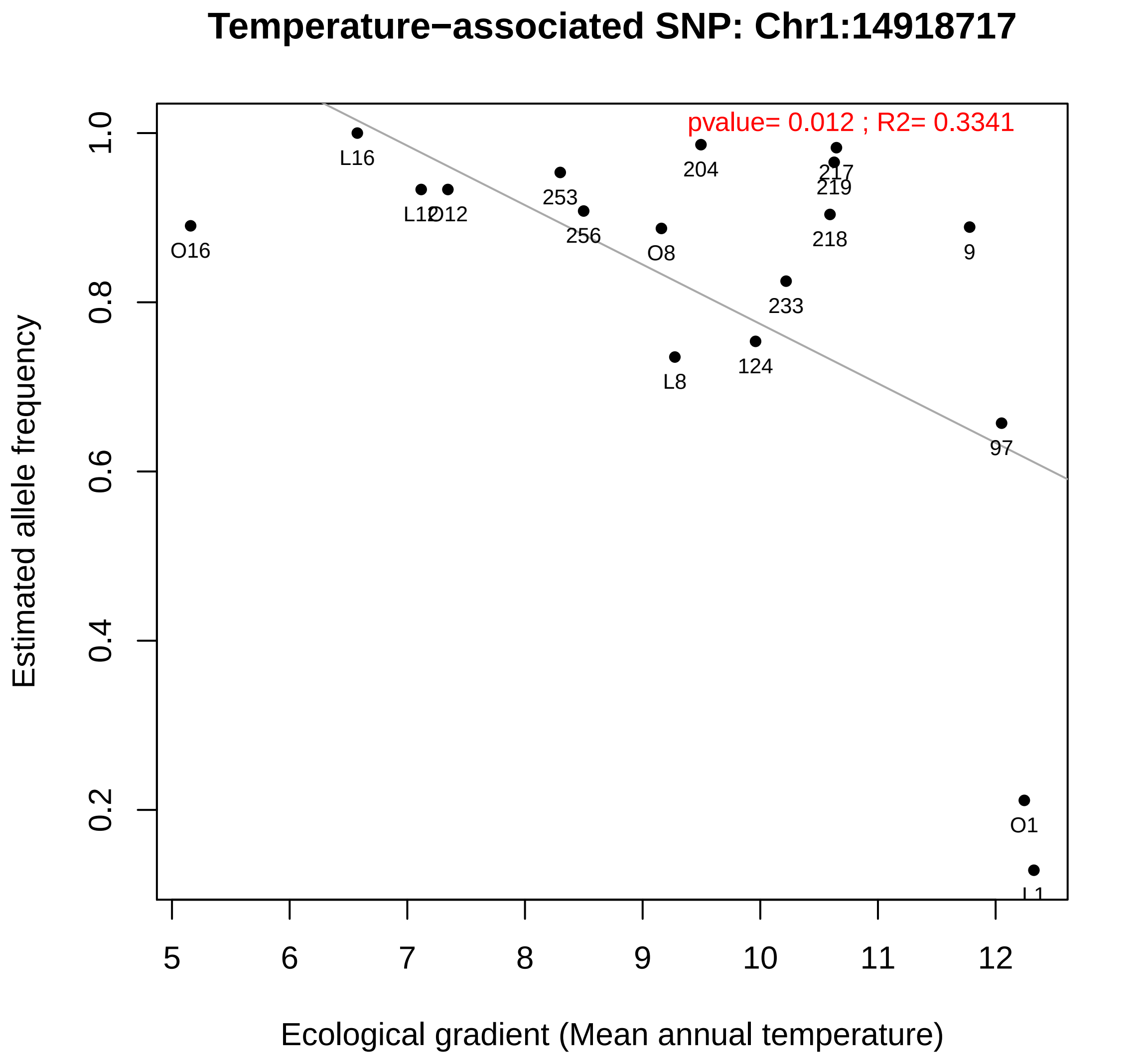

Temperature−associated SNP: Chr1:14918717
pvalue= 0.012 ; R2= 0.3341
Estimated allele frequency
Ecological gradient (Mean annual temperature)
0.2
0.4
0.6
0.8
1.0
5
6
7
8
9
10
11
12
L16
O16
L12
O12
253
256
O8
L8
204
124
233
217
219
218
9
97
O1
L1

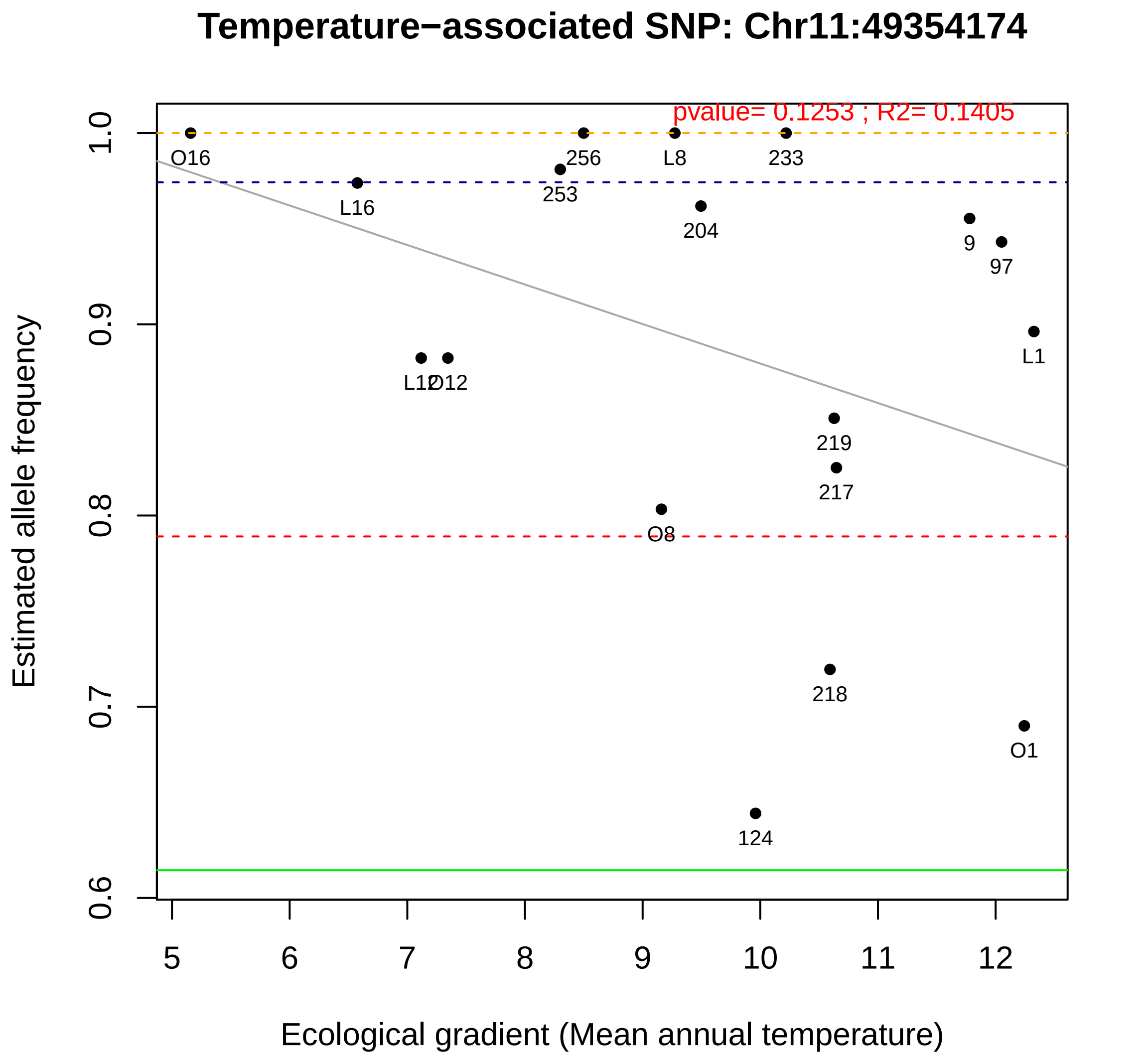

Temperature−associated SNP: Chr11:49354174
pvalue= 0.1253 ; R2= 0.1405
Estimated allele frequency
Ecological gradient (Mean annual temperature)
0.6
0.7
0.8
0.9
1.0
5
6
7
8
9
10
11
12
O16
L16
256
253
L8
233
204
9
97
L1
L12
O12
219
217
O8
218
O1
124

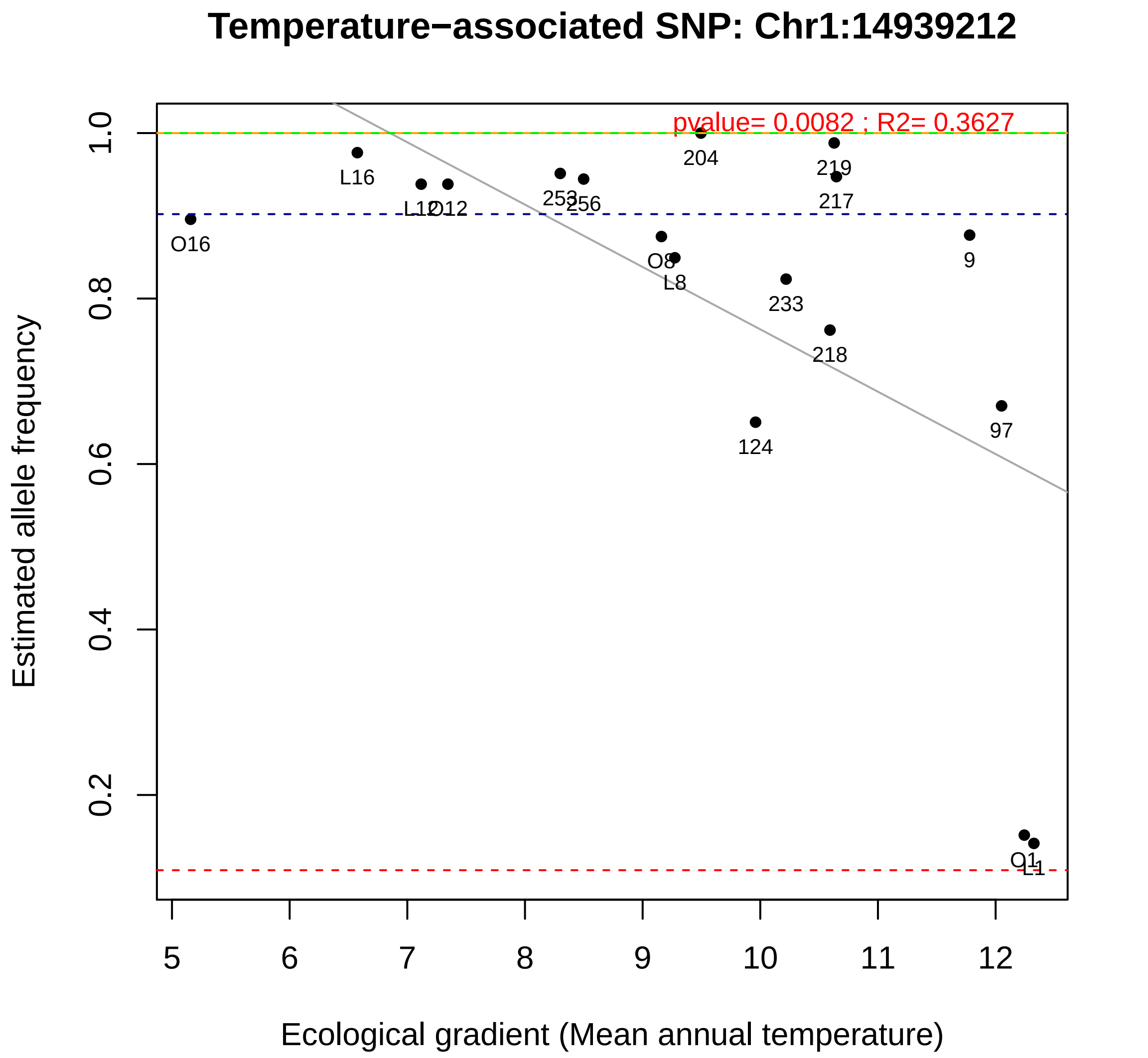

Temperature−associated SNP: Chr1:14939212
pvalue= 0.0082 ; R2= 0.3627
Estimated allele frequency
1.0
0.8
0.6
0.4
0.2
O16
L16
L12
O12
253
256
204
O8
L8
233
124
218
219
217
9
97
O1
L1
5
6
7
8
9
10
11
12
Ecological gradient (Mean annual temperature)

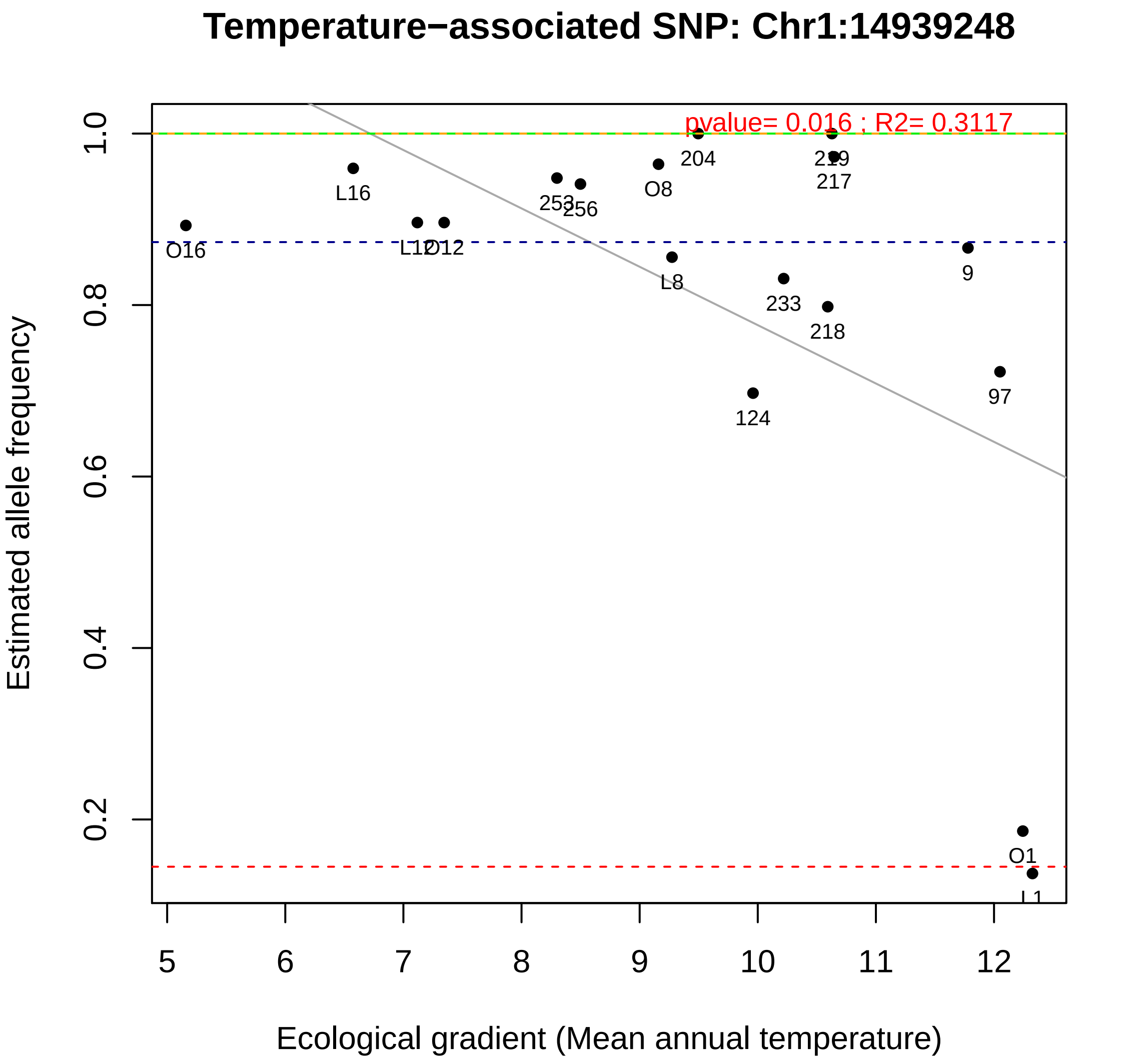

Temperature−associated SNP: Chr1:14939248
pvalue= 0.016 ; R2= 0.3117
Estimated allele frequency
Ecological gradient (Mean annual temperature)
1.0
0.8
0.6
0.4
0.2
5
6
7
8
9
10
11
12
O16
L16
L12
O12
253
256
O8
204
219
217
L8
233
218
9
124
97
O1
L1

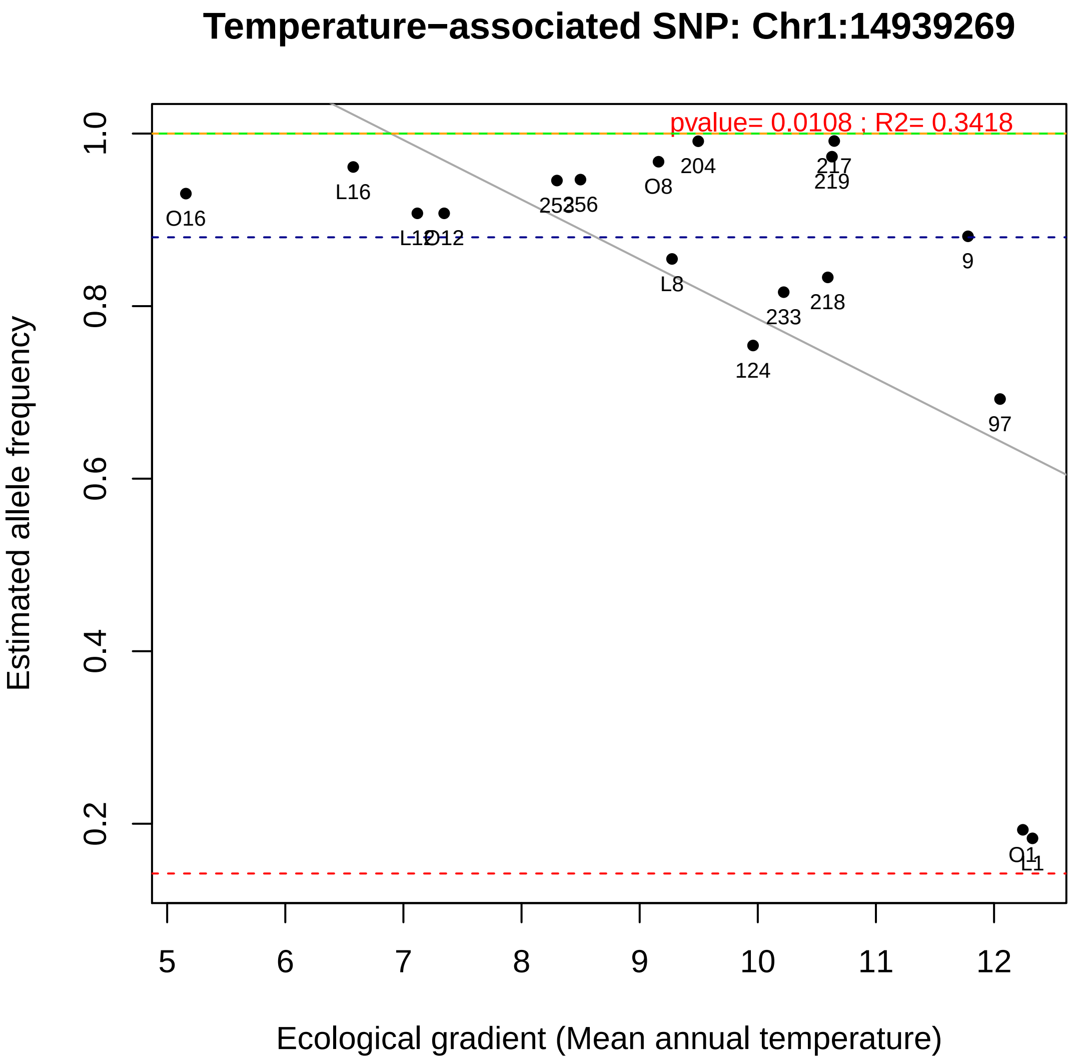

Temperature−associated SNP: Chr1:14939269
pvalue= 0.0108 ; R2= 0.3418
Estimated allele frequency
Ecological gradient (Mean annual temperature)
1.0
0.8
0.6
0.4
0.2
5
6
7
8
9
10
11
12
O16
L16
L12
O12
253
256
O8
204
217
219
L8
233
218
124
9
97
O1
L1

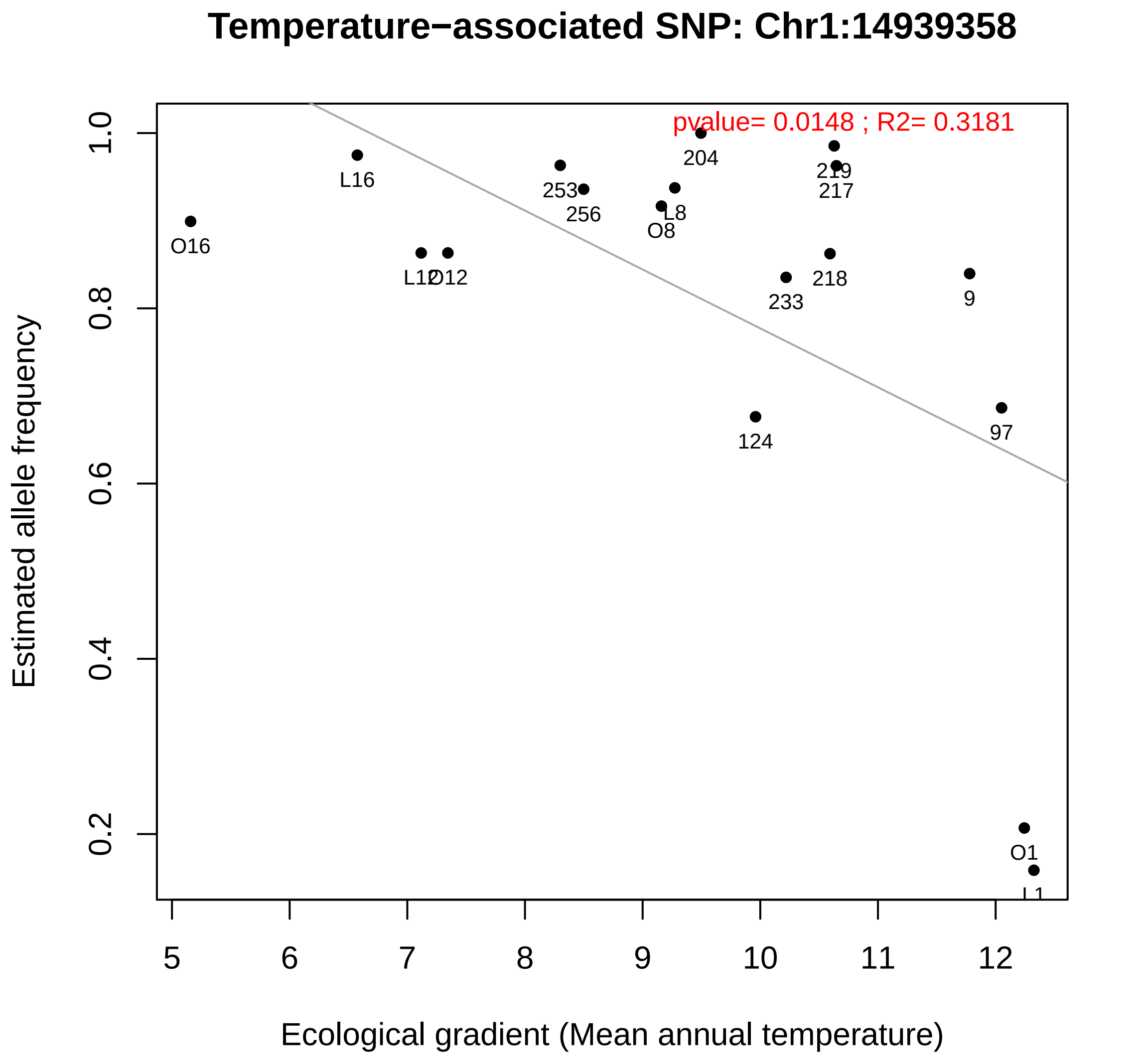
Temperature−associated SNP: Chr1:14939358
pvalue= 0.0148 ; R2= 0.3181
Estimated allele frequency
Ecological gradient (Mean annual temperature)
0.2
0.4
0.6
0.8
1.0
5
6
7
8
9
10
11
12
O16
L16
L12
O12
253
256
O8
L8
204
219
217
233
218
124
9
97
O1

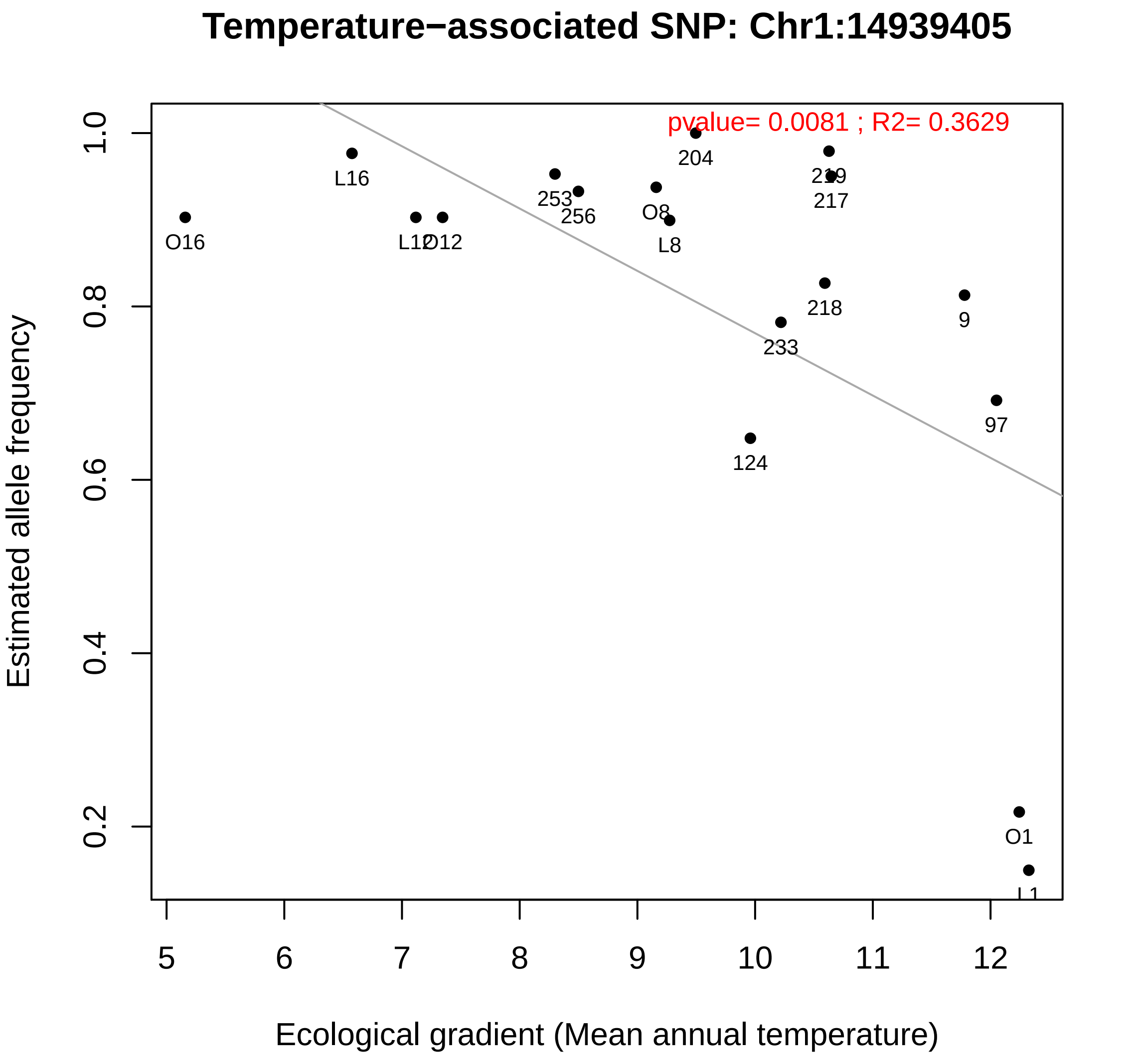

Temperature−associated SNP: Chr1:14939405
pvalue= 0.0081 ; R2= 0.3629
Estimated allele frequency
Ecological gradient (Mean annual temperature)
1.0
0.8
0.6
0.4
0.2
5
6
7
8
9
10
11
12
L16
O16
L12
O12
253
256
O8
L8
204
219
217
218
233
124
9
97
O1

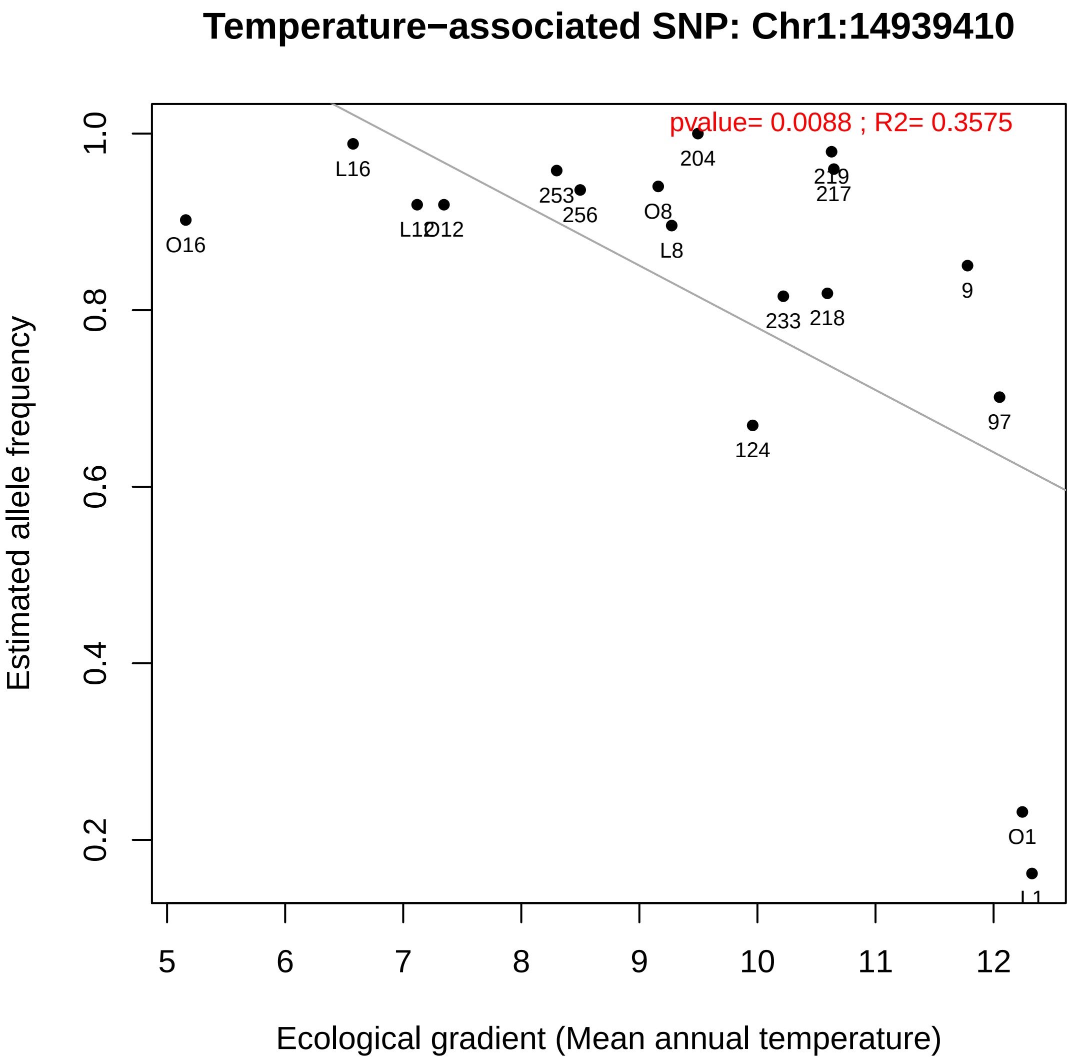
Temperature−associated SNP: Chr1:14939410
pvalue= 0.0088 ; R2= 0.3575
Estimated allele frequency
0.2
0.4
0.6
0.8
1.0
Ecological gradient (Mean annual temperature)
5
6
7
8
9
10
11
12
O16
L16
L12
O12
253
256
O8
L8
204
219
217
233
218
124
9
97
O1

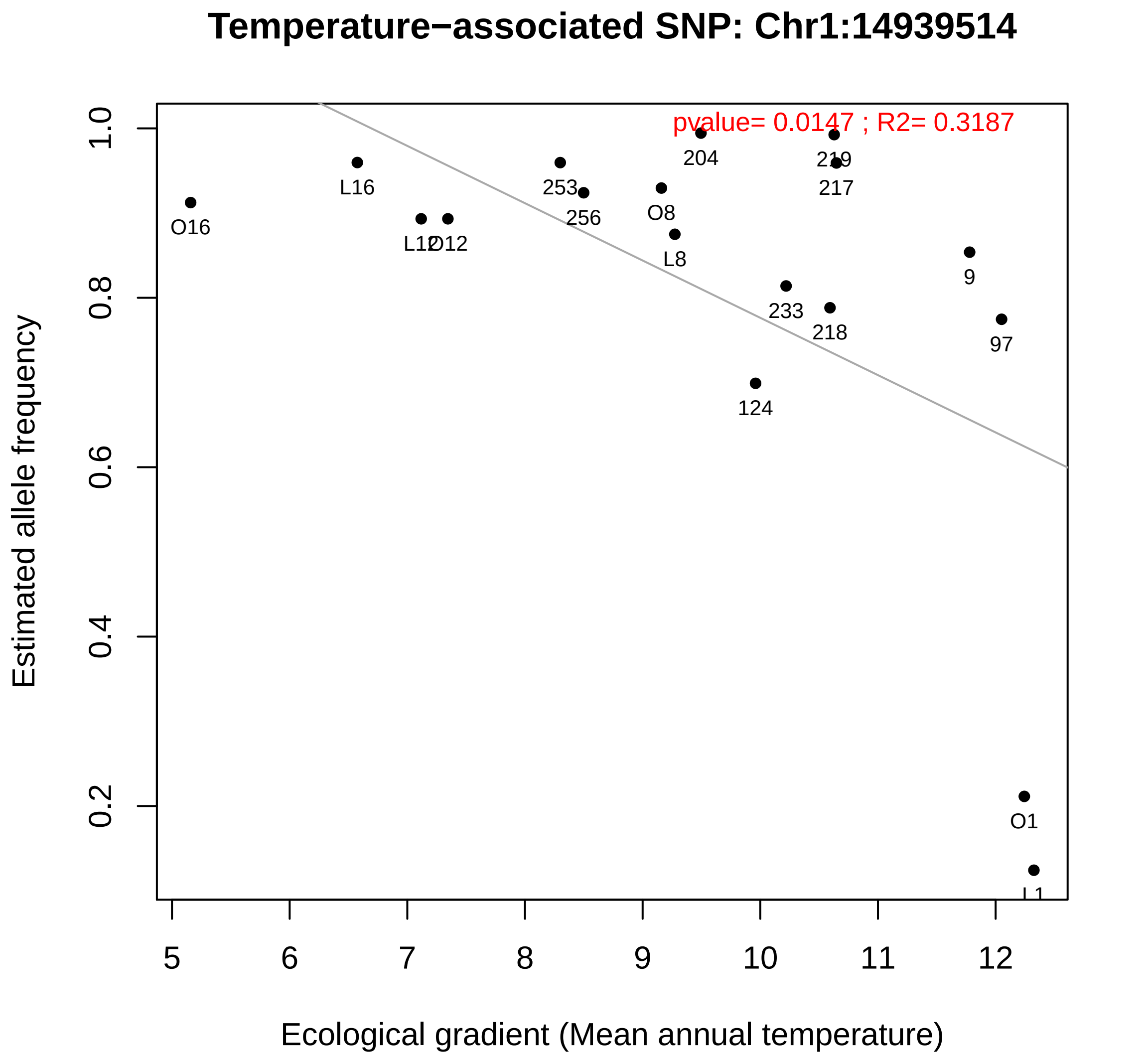
Temperature−associated SNP: Chr1:14939514
pvalue= 0.0147 ; R2= 0.3187
Estimated allele frequency
Ecological gradient (Mean annual temperature)
1.0
0.8
0.6
0.4
0.2
5
6
7
8
9
10
11
12
O16
L16
L12
O12
253
256
O8
L8
204
219
217
233
218
124
9
97
O1
L1

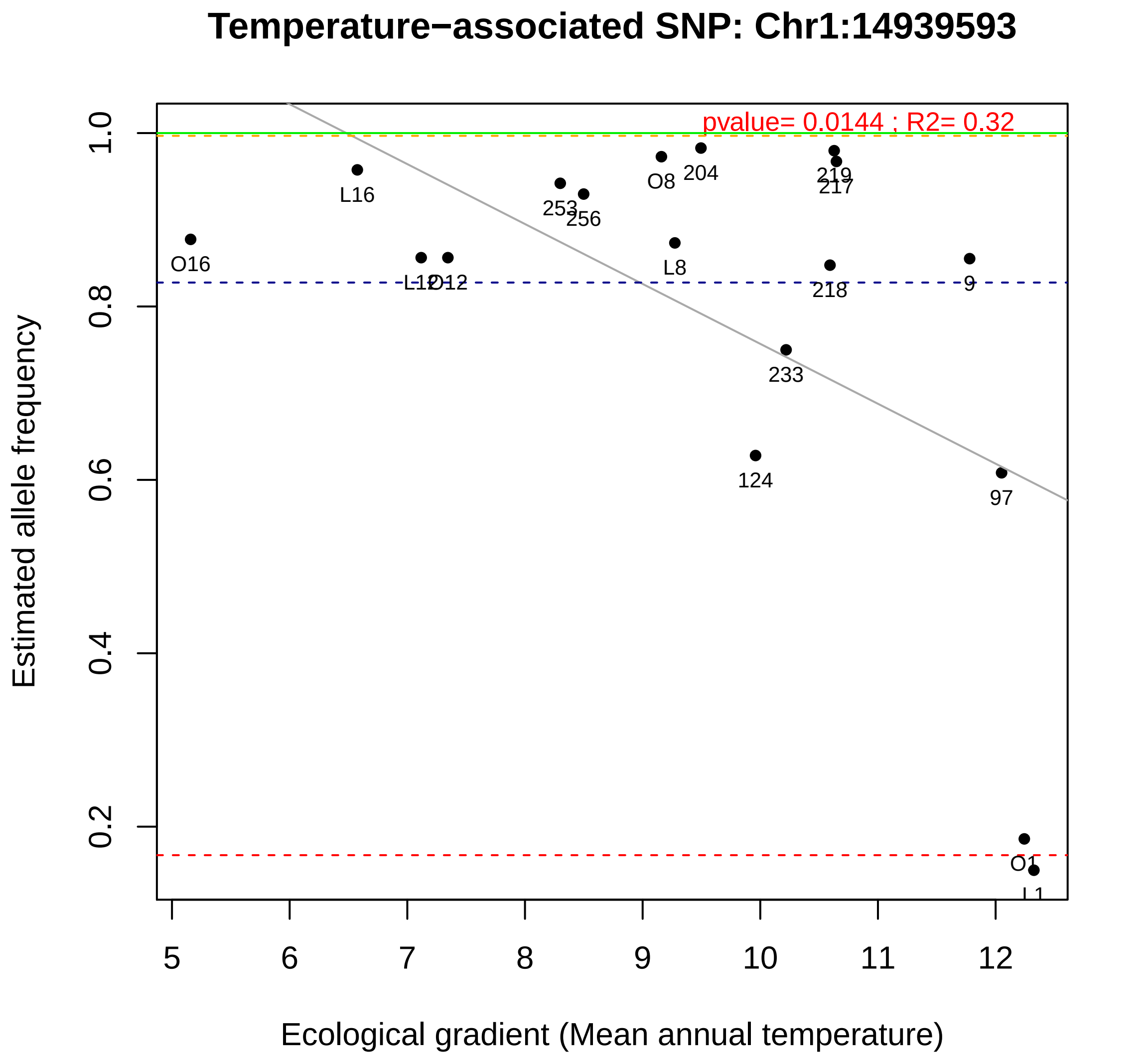
Temperature−associated SNP: Chr1:14939593
pvalue= 0.0144 ; R2= 0.32
Estimated allele frequency
0.2
0.4
0.6
0.8
1.0
5
6
7
8
9
10
11
12
Ecological gradient (Mean annual temperature)
O16
L16
L12
O12
253
256
O8
204
L8
219
217
218
9
233
124
97
O1
L1

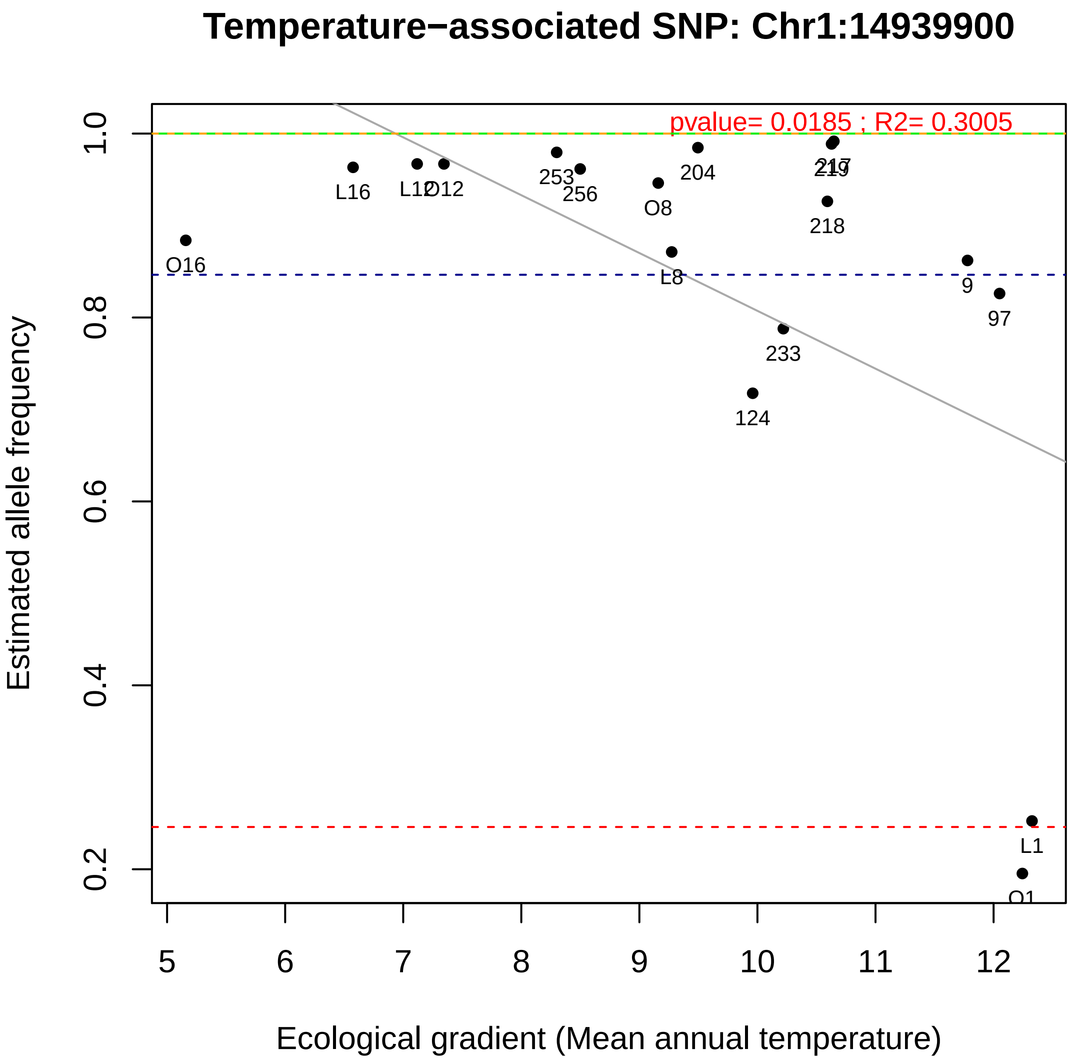

Temperature−associated SNP: Chr1:14939900
pvalue= 0.0185 ; R2= 0.3005
Estimated allele frequency
Ecological gradient (Mean annual temperature)
1.0
0.8
0.6
0.4
0.2
5
6
7
8
9
10
11
12
O16
L16
L12
O12
253
256
O8
204
L8
217
219
218
233
124
9
97
L1
O1

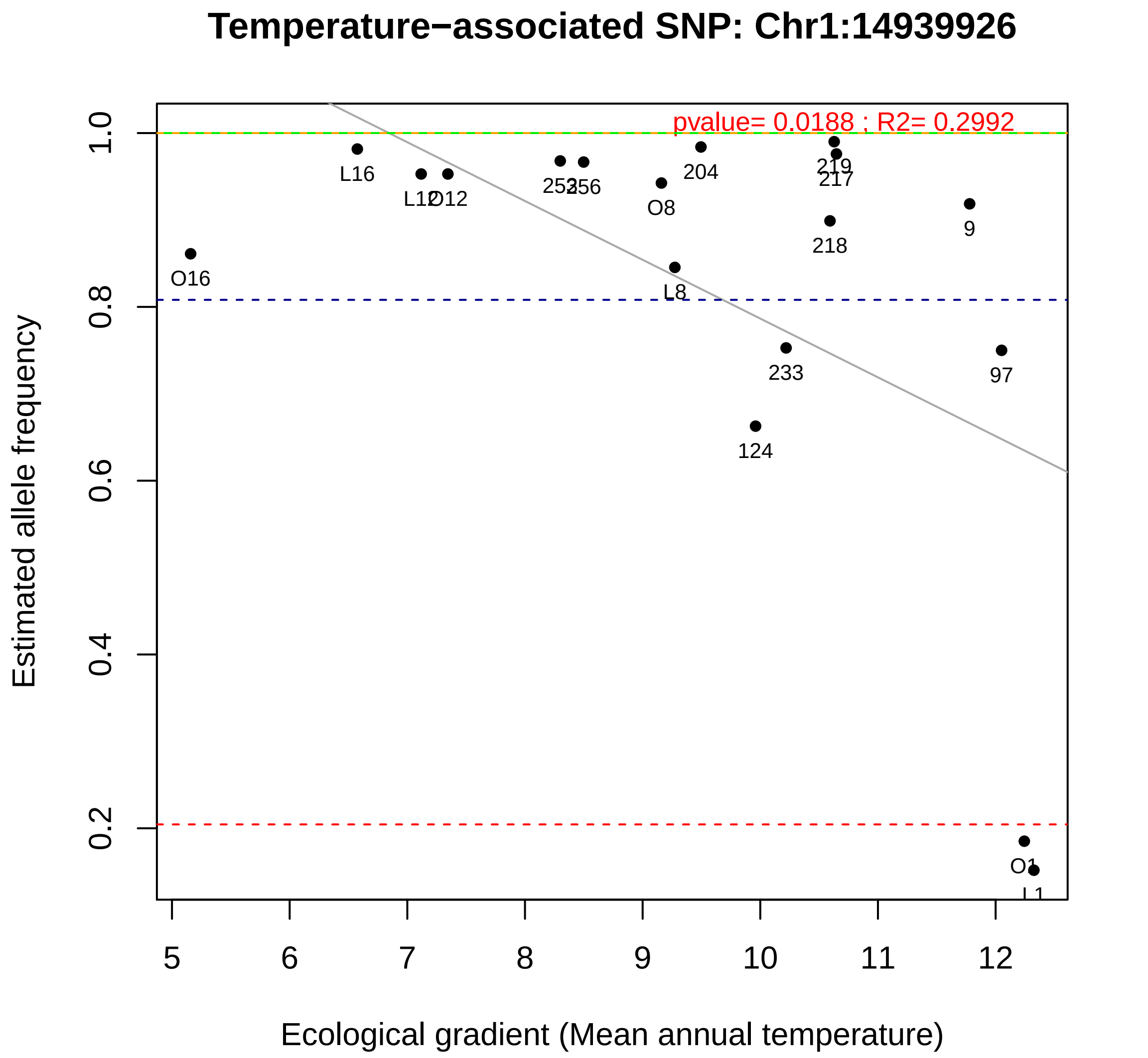

Temperature−associated SNP: Chr1:14939926
pvalue= 0.0188 ; R2= 0.2992
Estimated allele frequency
Ecological gradient (Mean annual temperature)
0.2
0.4
0.6
0.8
1.0
5
6
7
8
9
10
11
12
O16
L16
L12
O12
252
256
O8
L8
204
124
233
219
217
218
9
97
O1
L1

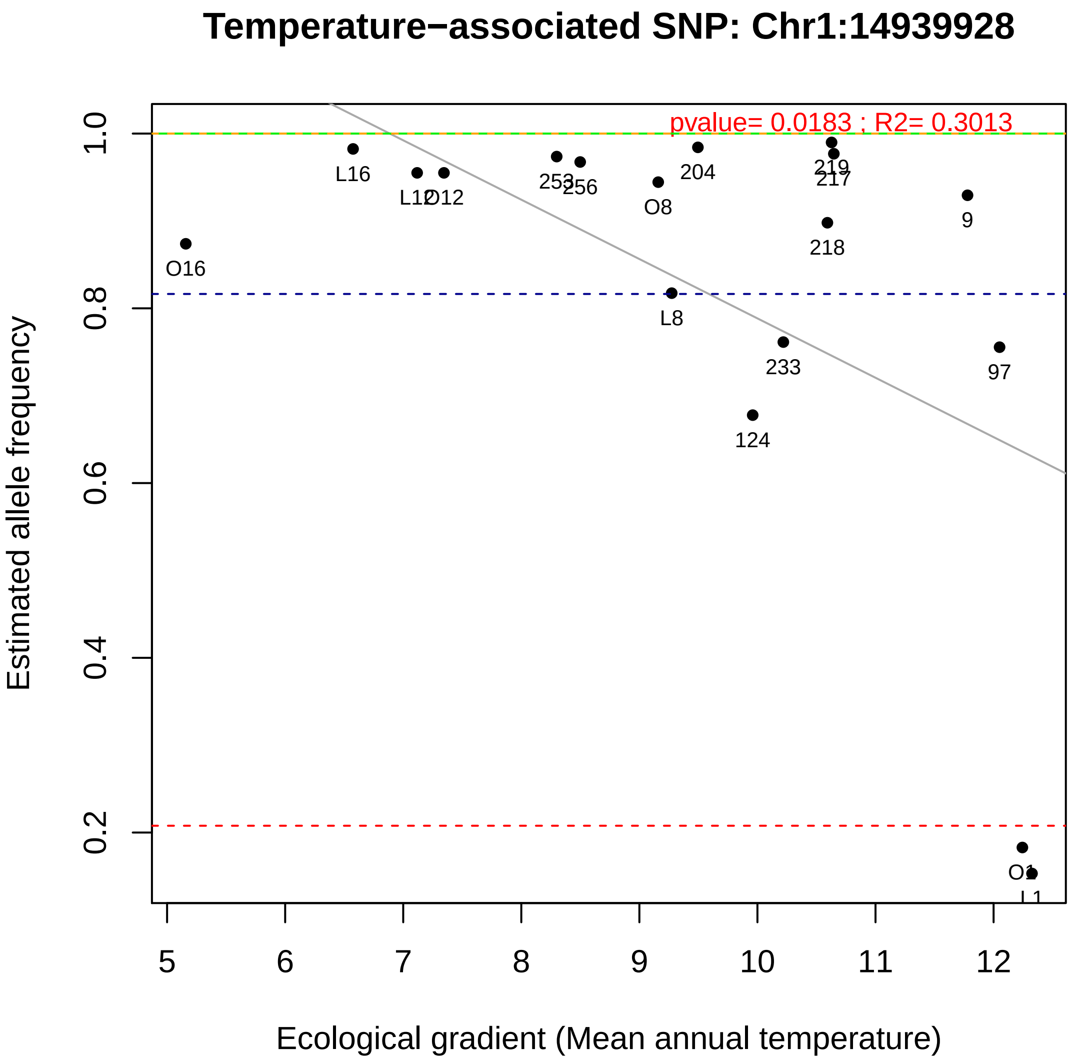
Temperature−associated SNP: Chr1:14939928
pvalue= 0.0183 ; R2= 0.3013
Estimated allele frequency
1.0
0.8
0.6
0.4
0.2
5
6
7
8
9
10
11
12
Ecological gradient (Mean annual temperature)
O16
L16
L12
O12
253
256
O8
204
219
217
218
9
L8
233
97
124
O1
L1

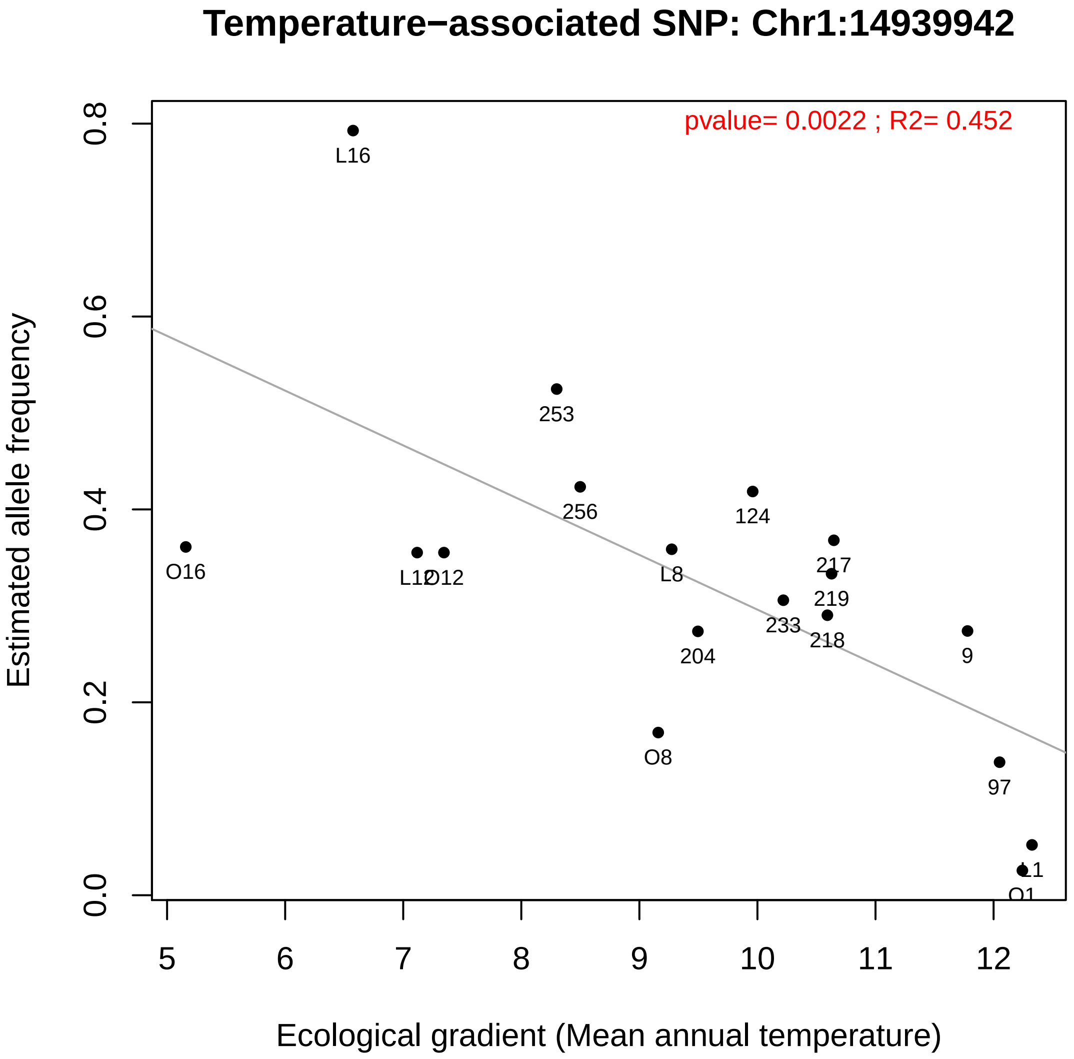

Temperature−associated SNP: Chr1:14939942
pvalue= 0.0022 ; R2= 0.452
Estimated allele frequency
0.0
0.2
0.4
0.6
0.8
5
6
7
8
9
10
11
12
Ecological gradient (Mean annual temperature)
L16
253
256
124
O16
L12
O12
L8
217
219
233
218
204
9
O8
97
L1
O1

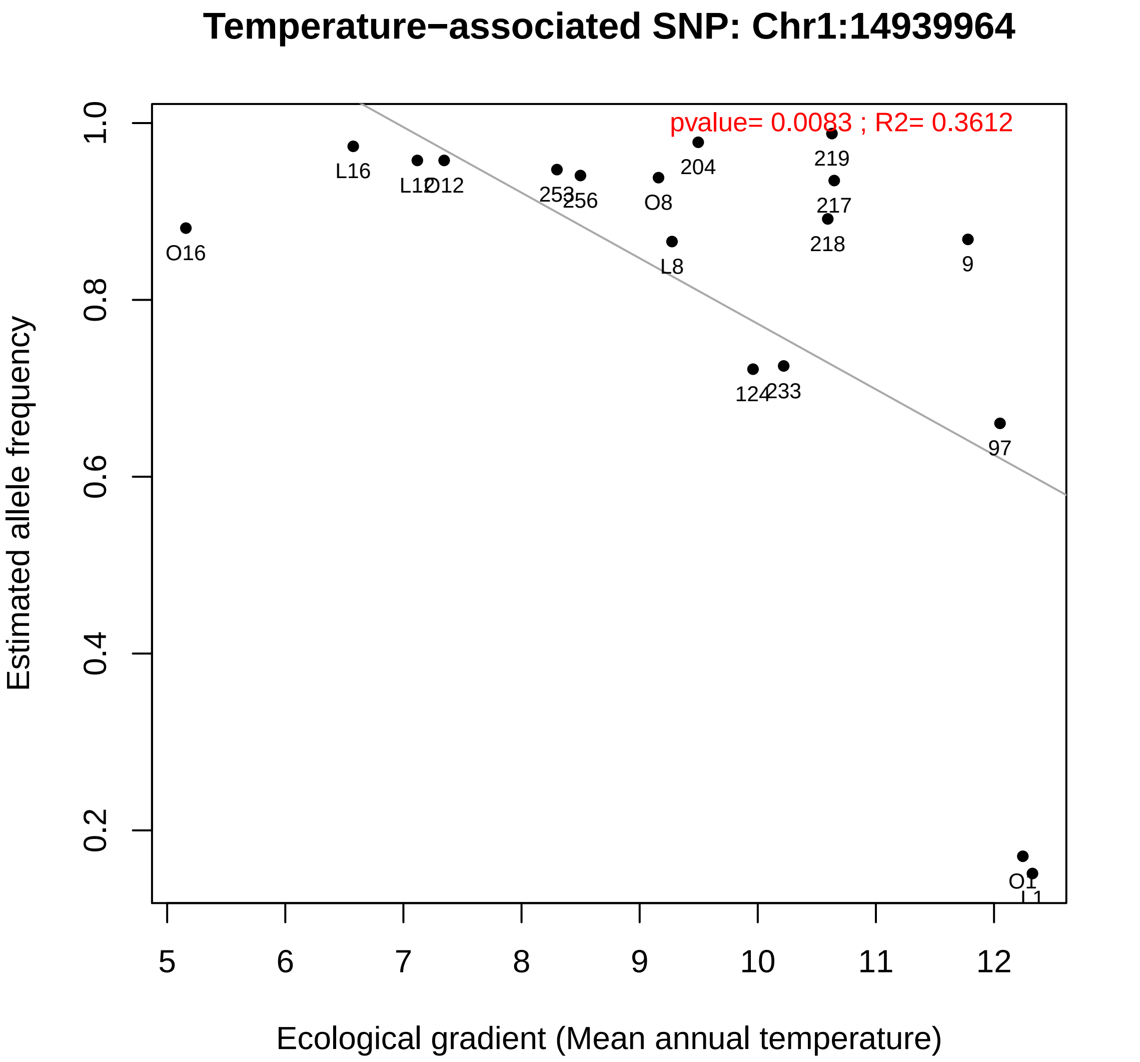
Temperature−associated SNP: Chr1:14939964
pvalue= 0.0083 ; R2= 0.3612
Estimated allele frequency
Ecological gradient (Mean annual temperature)
1.0
0.8
0.6
0.4
0.2
5
6
7
8
9
10
11
12
O16
L16
L12
O12
253
256
O8
204
L8
219
217
218
9
124
233
97
O1

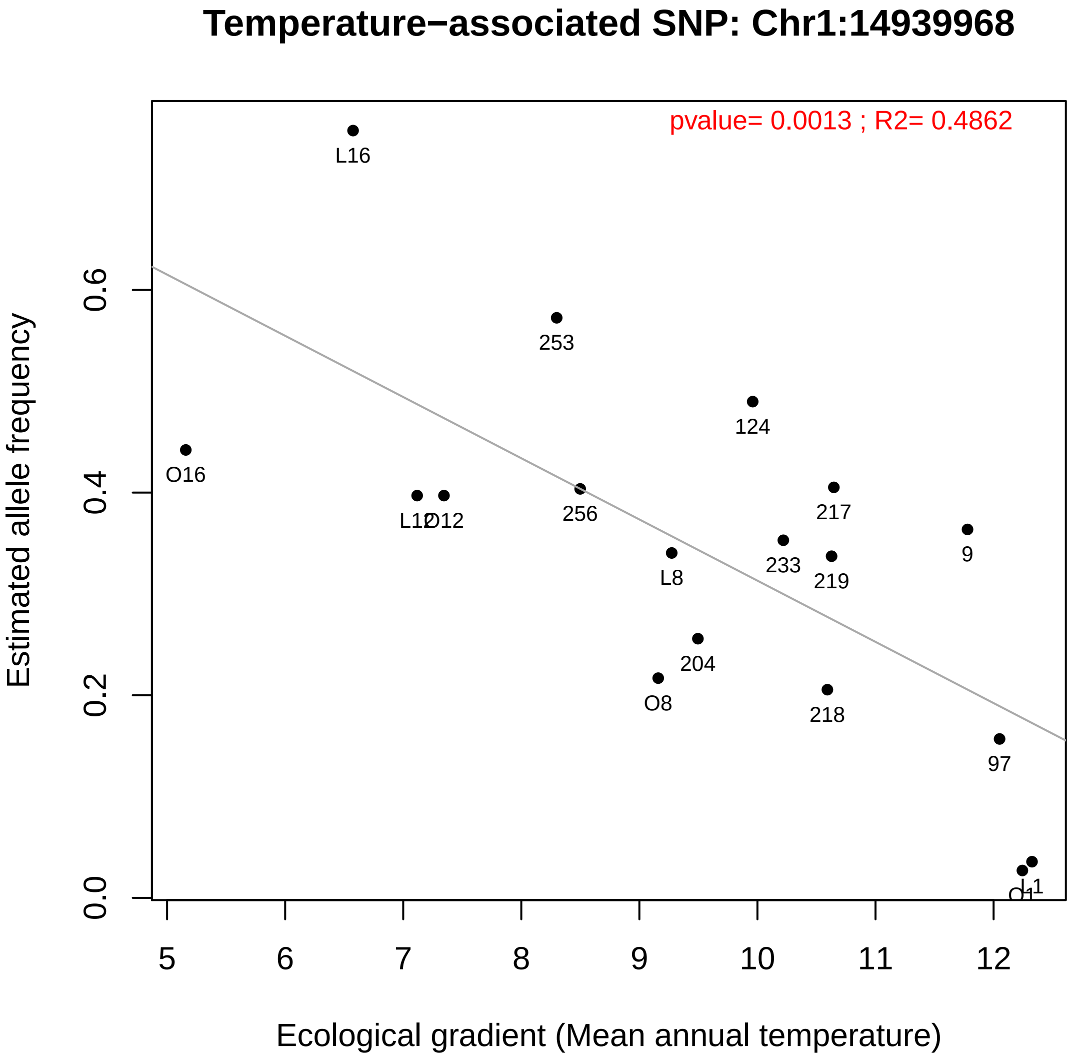
Temperature−associated SNP: Chr1:14939968
pvalue= 0.0013 ; R2= 0.4862
L16
253
124
O16
L12
O12
256
217
233
9
L8
219
204
O8
218
97
L1
O1
0.0
0.2
0.4
0.6
Estimated allele frequency
5
6
7
8
9
10
11
12
Ecological gradient (Mean annual temperature)

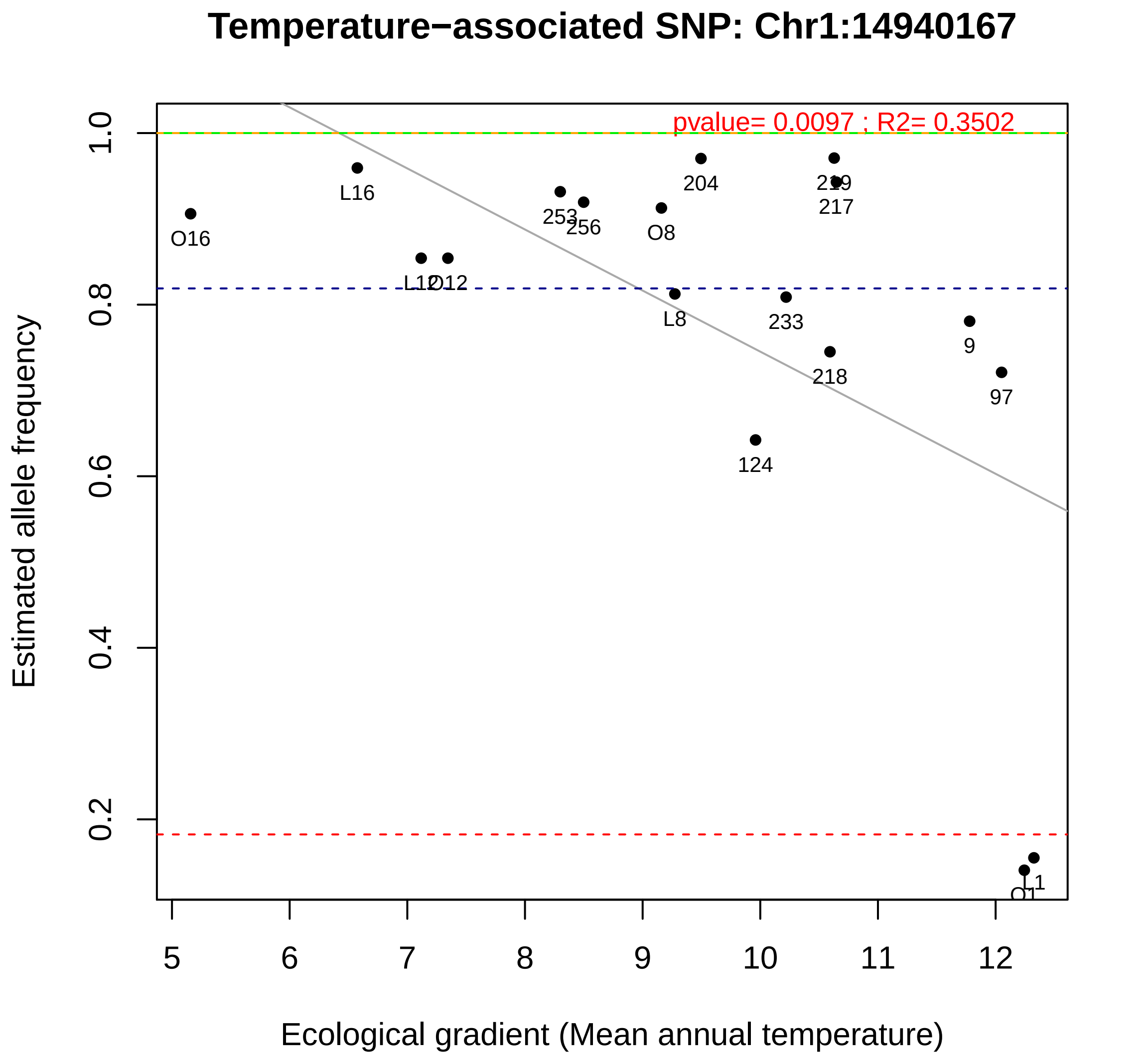
Temperature−associated SNP: Chr1:14940167
pvalue= 0.0097 ; R2= 0.3502
Estimated allele frequency
Ecological gradient (Mean annual temperature)
0.2
0.4
0.6
0.8
1.0
5
6
7
8
9
10
11
12
O16
L16
L12
O12
253
256
O8
204
219
217
L8
233
218
124
9
97
L1
O1

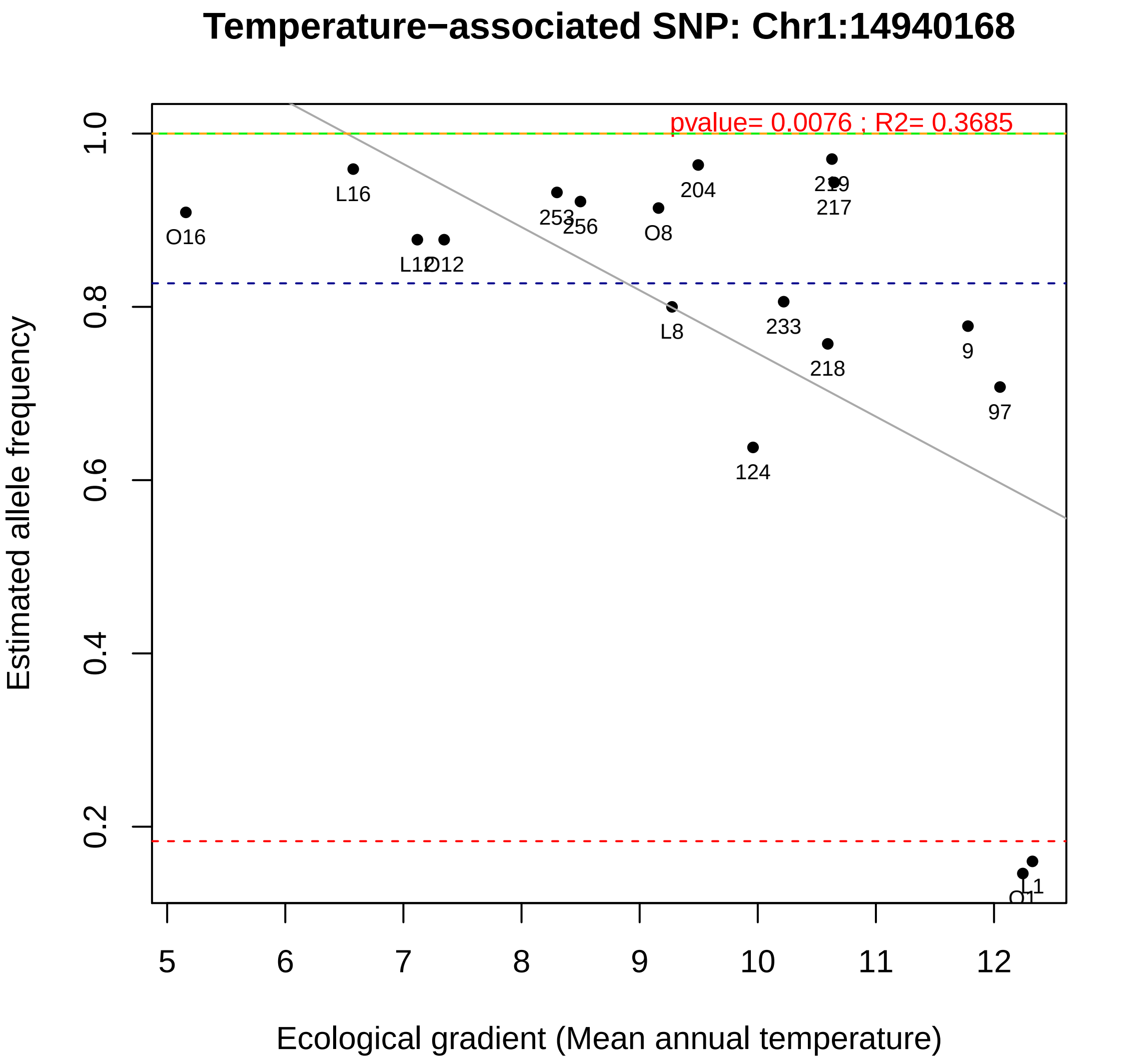
Temperature−associated SNP: Chr1:14940168
pvalue= 0.0076 ; R2= 0.3685
Estimated allele frequency
Ecological gradient (Mean annual temperature)
1.0
0.8
0.6
0.4
0.2
5
6
7
8
9
10
11
12
O16
L16
L12
O12
253
256
O8
204
219
217
L8
233
218
124
9
97
L1
O1

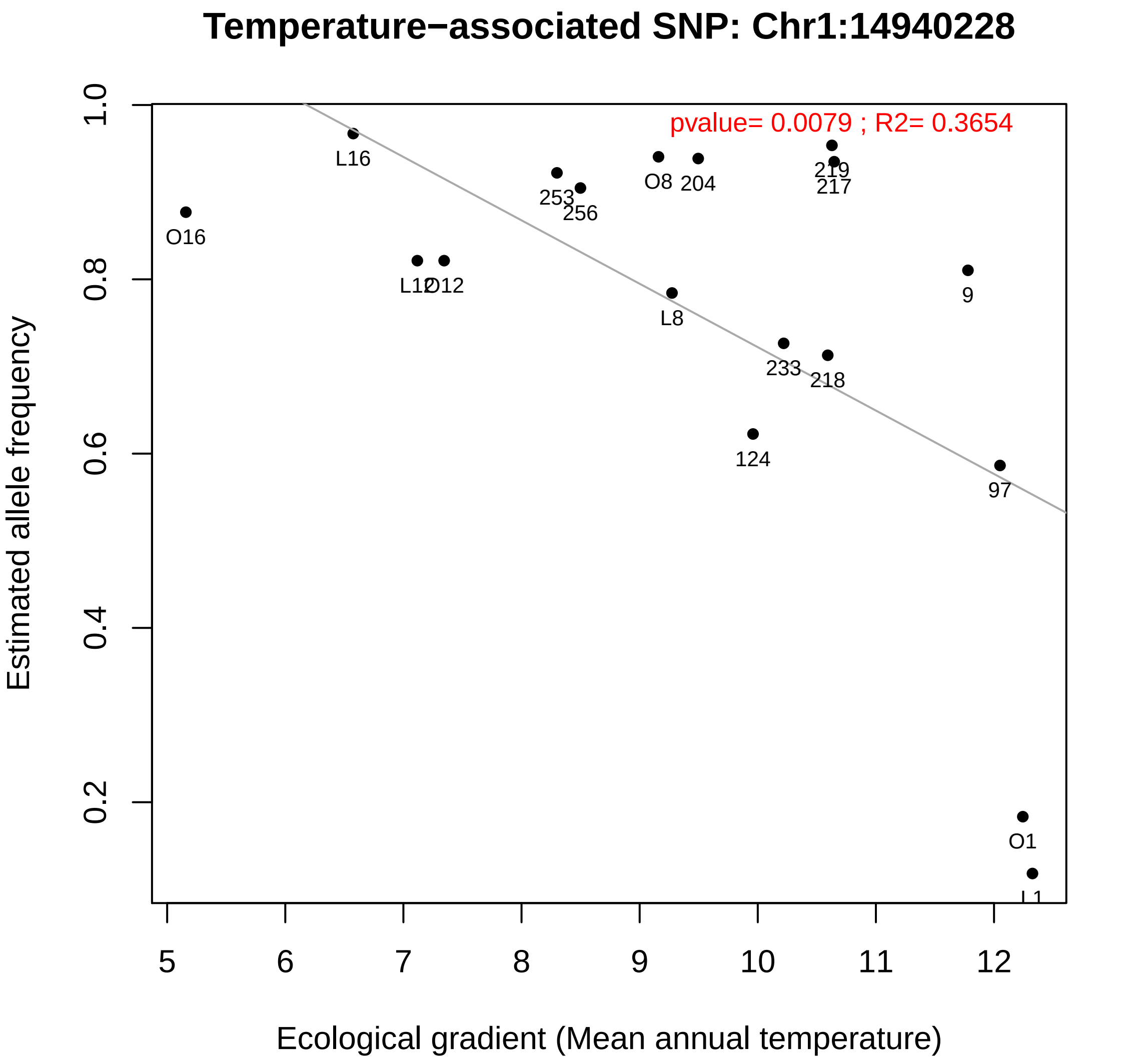
Temperature−associated SNP: Chr1:14940228
pvalue= 0.0079 ; R2= 0.3654
Estimated allele frequency
1.0
0.8
0.6
0.4
0.2
5
6
7
8
9
10
11
12
Ecological gradient (Mean annual temperature)
L16
O16
253
256
O8
204
219
217
L12
O12
L8
9
233
218
124
97
O1

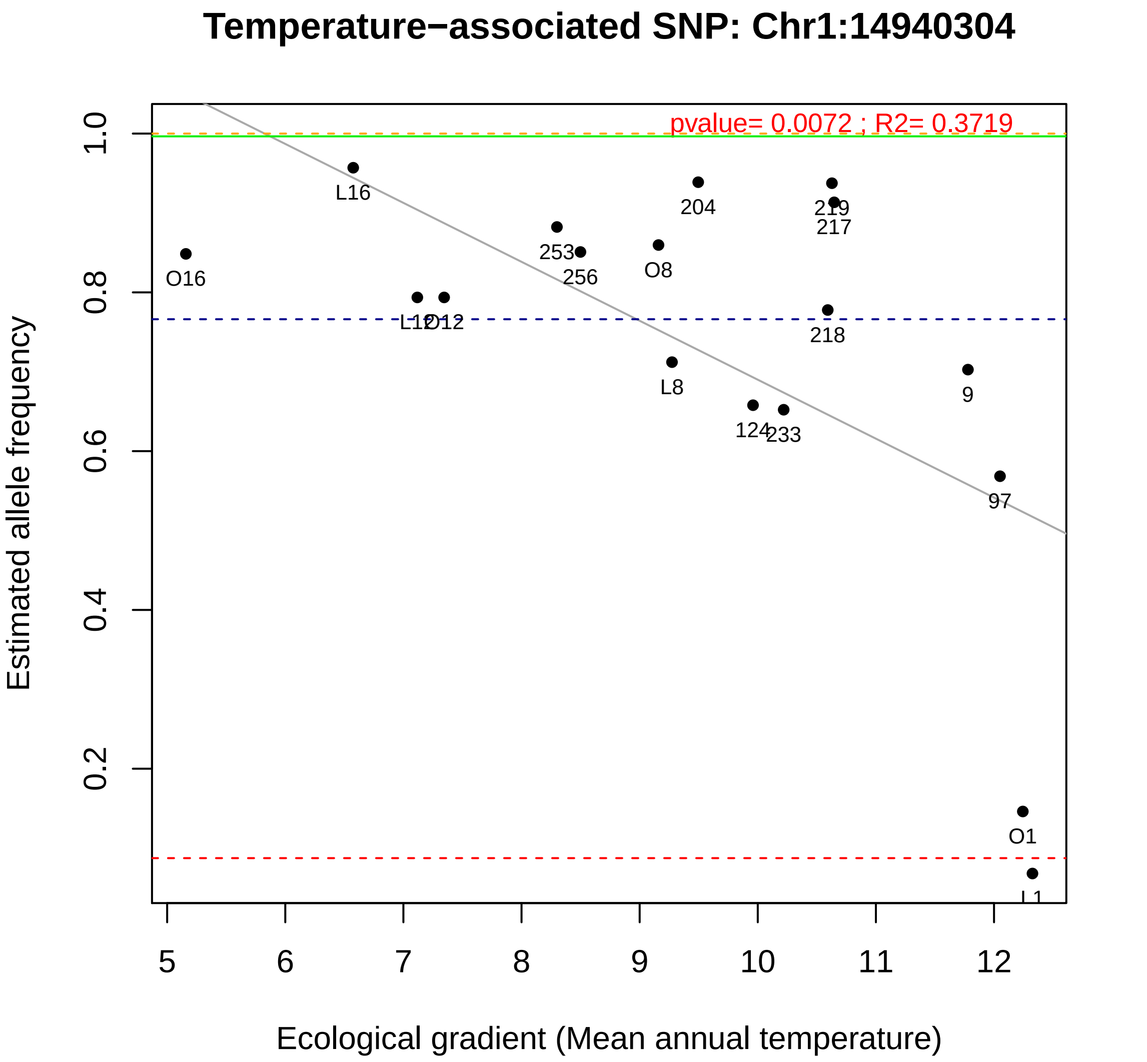

Temperature−associated SNP: Chr1:14940304
pvalue= 0.0072 ; R2= 0.3719
Estimated allele frequency
Ecological gradient (Mean annual temperature)
0.2
0.4
0.6
0.8
1.0
5
6
7
8
9
10
11
12
O16
L16
L12
O12
253
256
O8
204
L8
219
217
218
124
233
9
97
O1

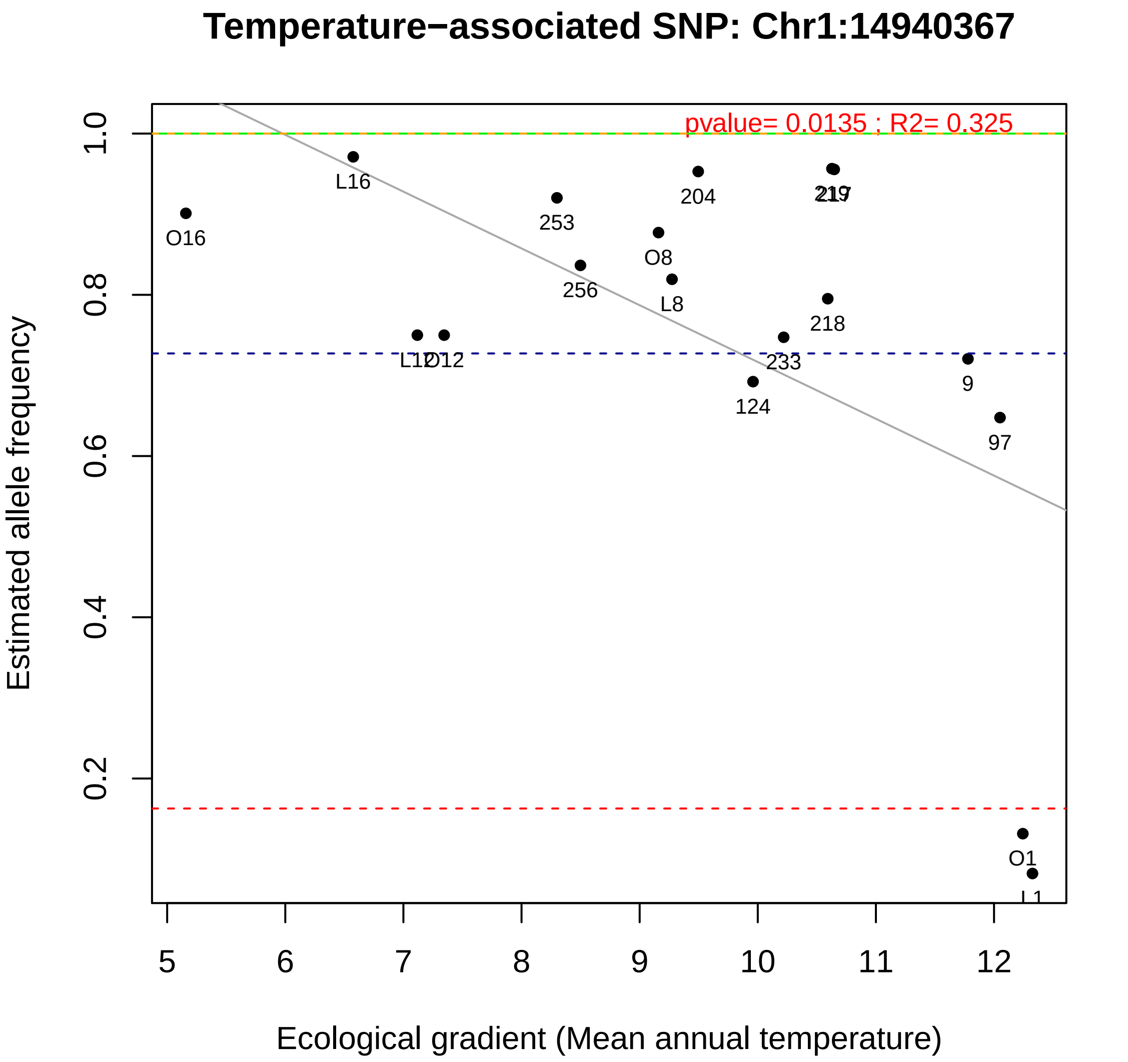

Temperature−associated SNP: Chr1:14940367
pvalue= 0.0135 ; R2= 0.325
Estimated allele frequency
Ecological gradient (Mean annual temperature)
1.0
0.8
0.6
0.4
0.2
5
6
7
8
9
10
11
12
L16
O16
253
256
204
O8
L8
218
233
124
9
97
O1

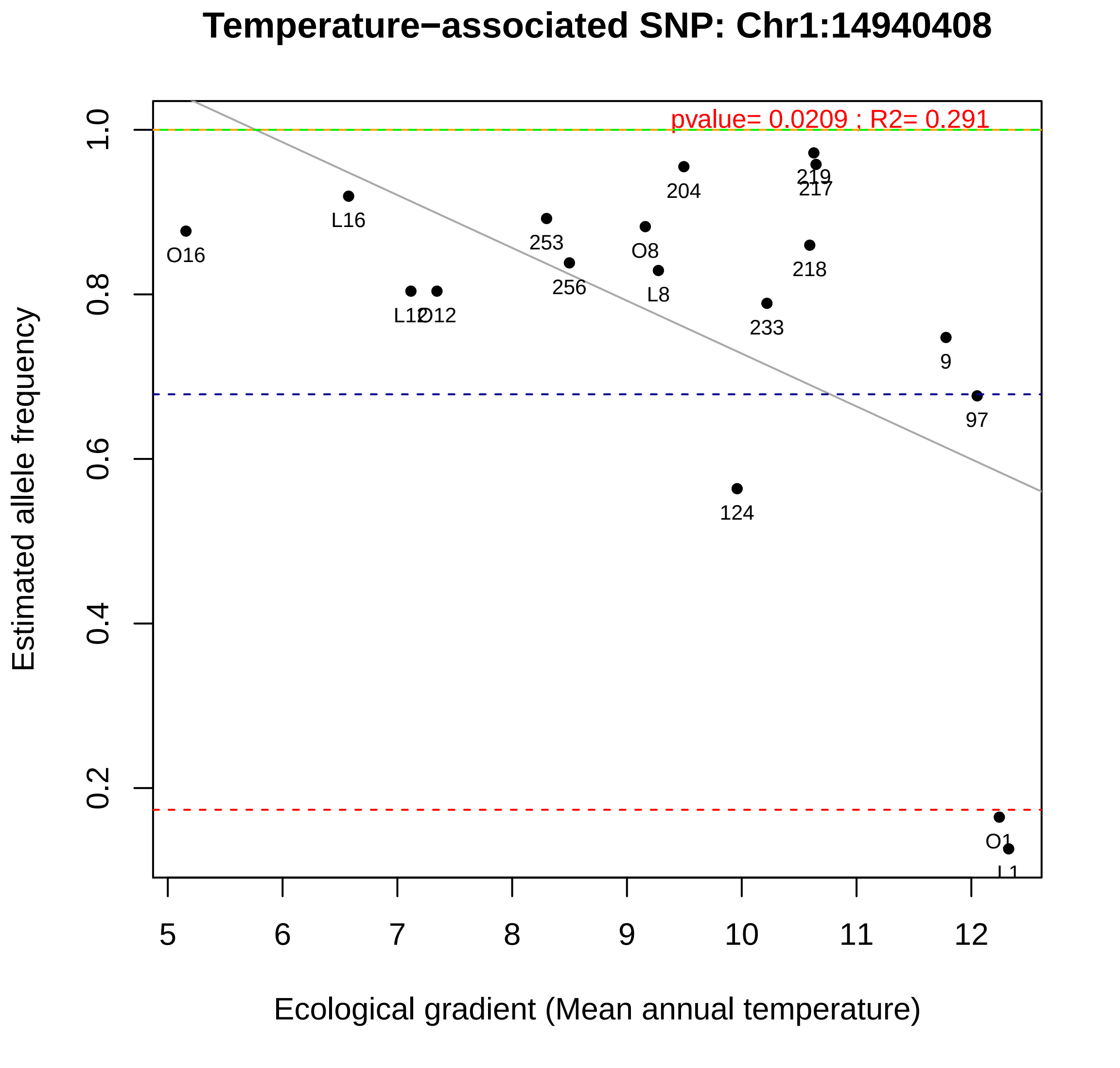
Temperature−associated SNP: Chr1:14940408
pvalue= 0.0209 ; R2= 0.291
Estimated allele frequency
0.2
0.4
0.6
0.8
1.0
5
6
7
8
9
10
11
12
Ecological gradient (Mean annual temperature)
O16
L16
L12
O12
253
256
O8
L8
204
219
217
218
233
124
9
97
O1

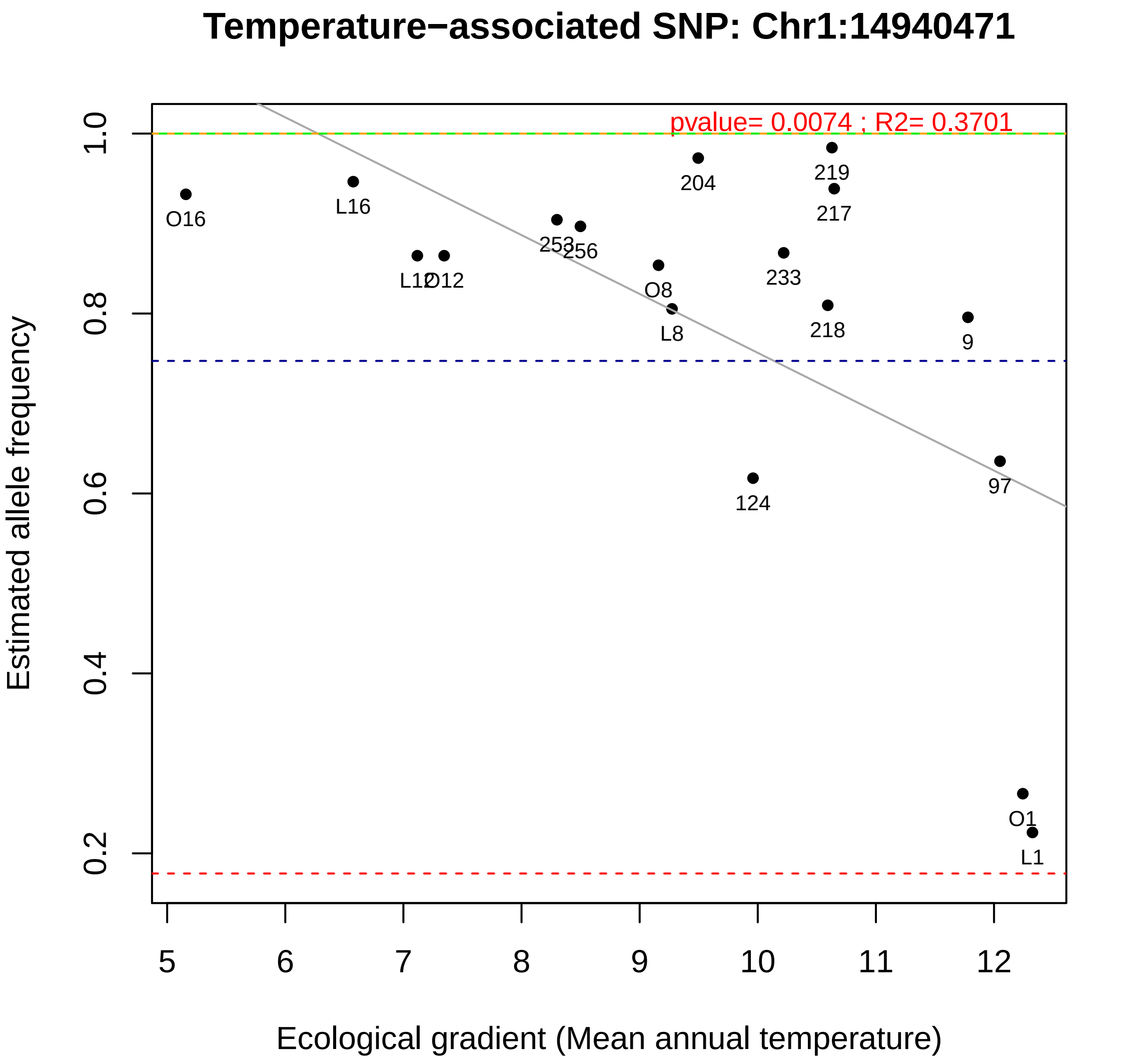

Temperature−associated SNP: Chr1:14940471
pvalue= 0.0074 ; R2= 0.3701
Estimated allele frequency
Ecological gradient (Mean annual temperature)
0.2
0.4
0.6
0.8
1.0
5
6
7
8
9
10
11
12
O16
L16
L12
O12
253
256
O8
L8
204
124
233
219
217
218
9
97
O1
L1

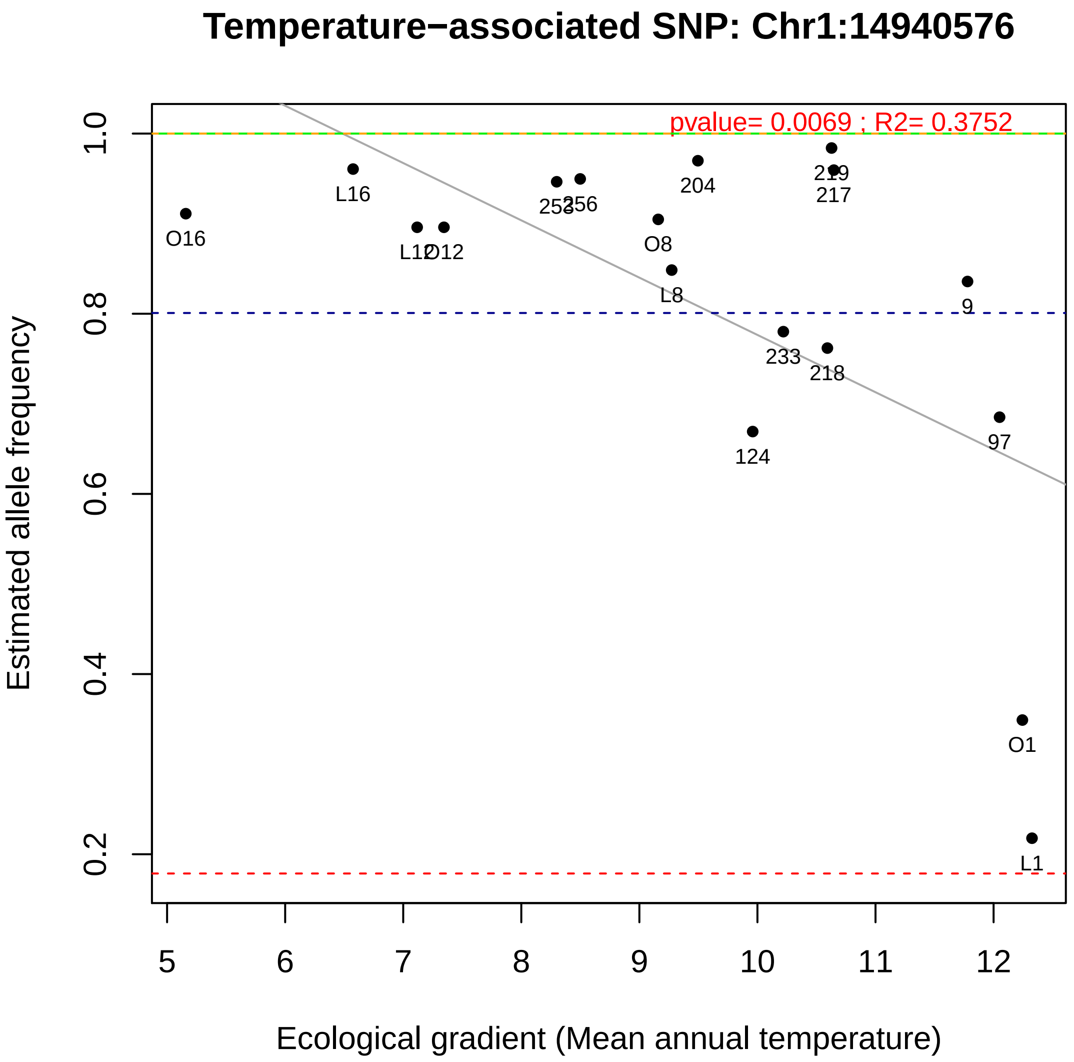

Temperature−associated SNP: Chr1:14940576
pvalue= 0.0069 ; R2= 0.3752
Estimated allele frequency
Ecological gradient (Mean annual temperature)
0.2
0.4
0.6
0.8
1.0
5
6
7
8
9
10
11
12
O16
L16
L12
O12
256
204
219
217
O8
L8
9
233
218
124
97
O1
L1

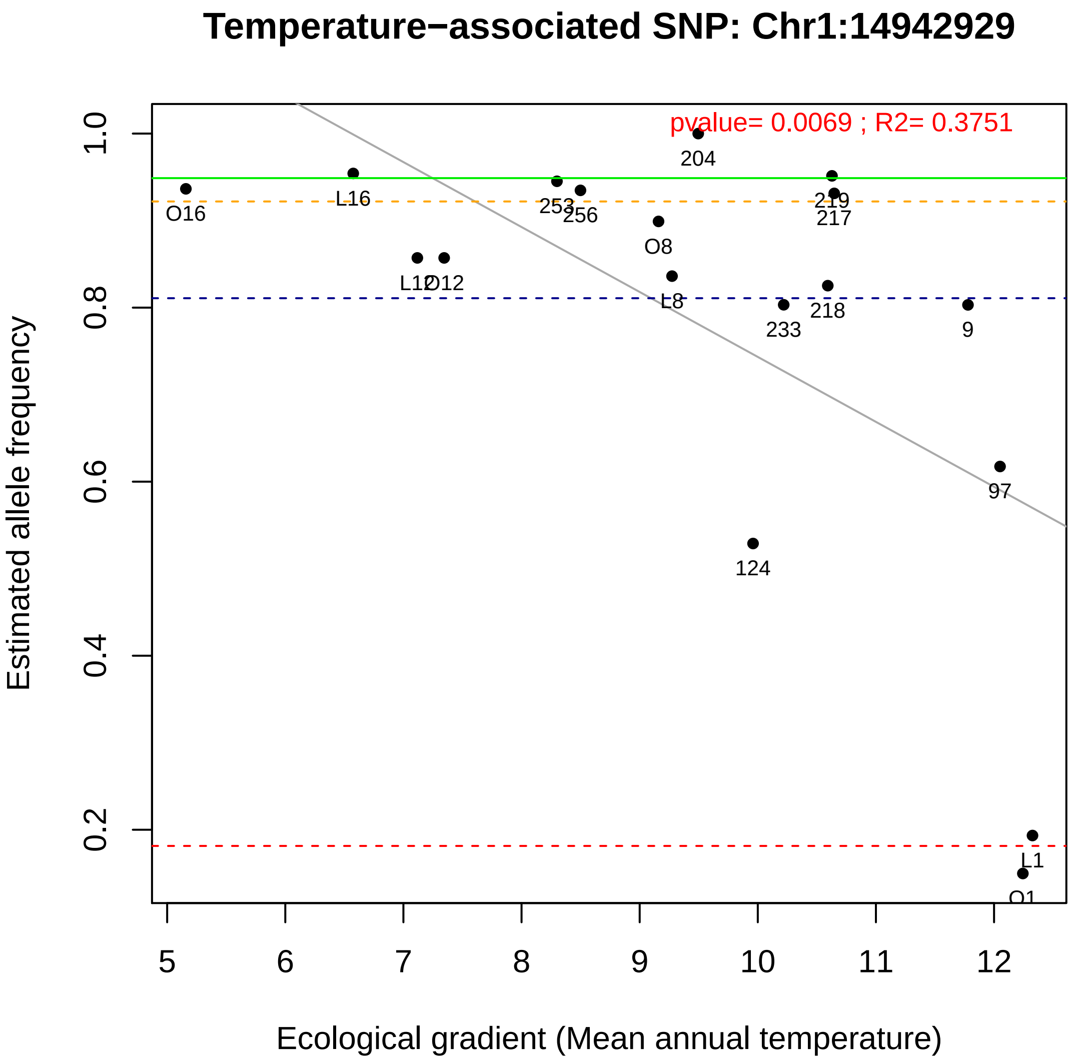
Temperature−associated SNP: Chr1:14942929
pvalue= 0.0069 ; R2= 0.3751
Estimated allele frequency
1.0
0.8
0.6
0.4
0.2
5
6
7
8
9
10
11
12
Ecological gradient (Mean annual temperature)
O16
L16
L12
O12
253
256
O8
L8
204
219
217
233
218
124
9
97
L1
O1

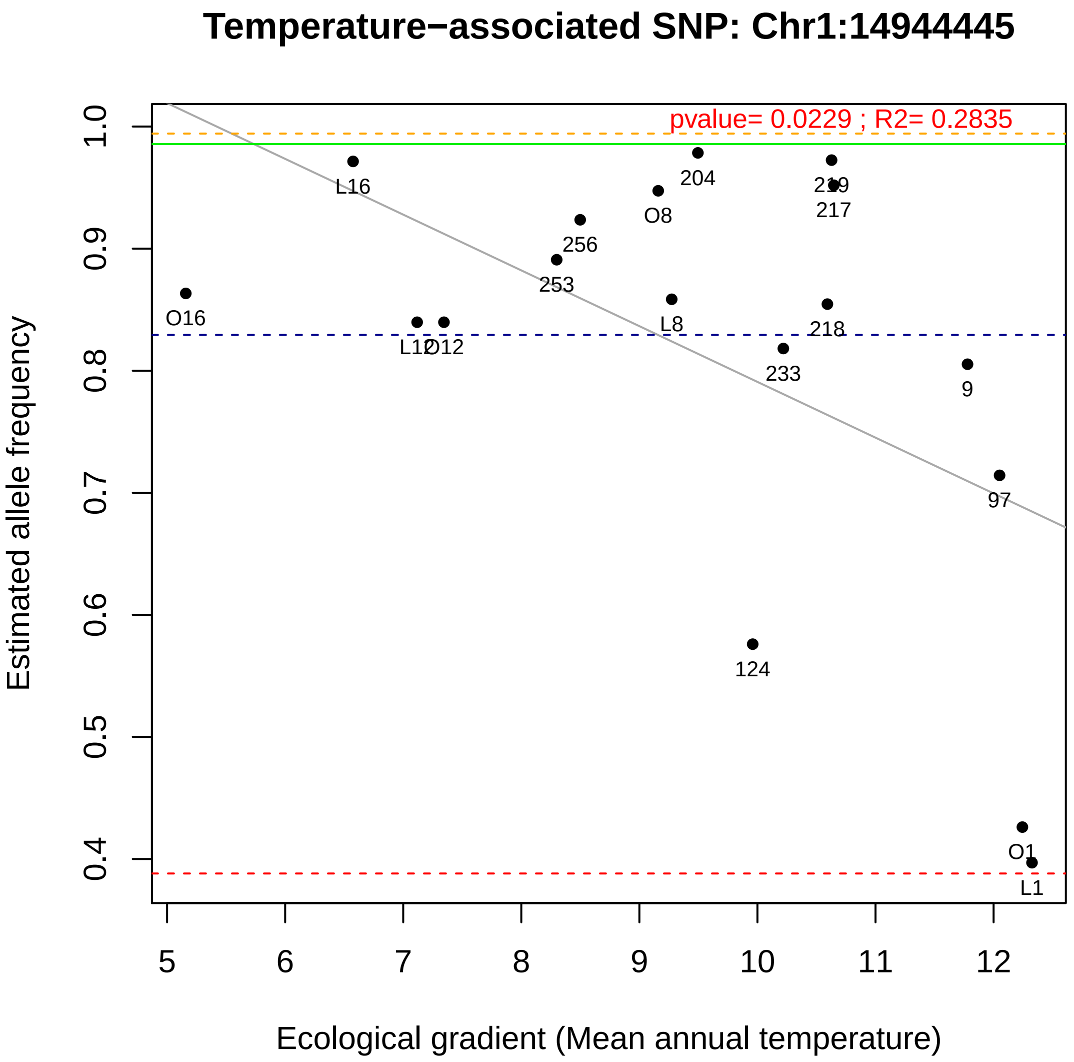

Temperature−associated SNP: Chr1:14944445
pvalue= 0.0229 ; R2= 0.2835
Estimated allele frequency
Ecological gradient (Mean annual temperature)
0.4
0.5
0.6
0.7
0.8
0.9
1.0
5
6
7
8
9
10
11
12
O16
L16
L12
O12
253
256
O8
L8
204
124
233
219
217
218
9
97
O1
L1

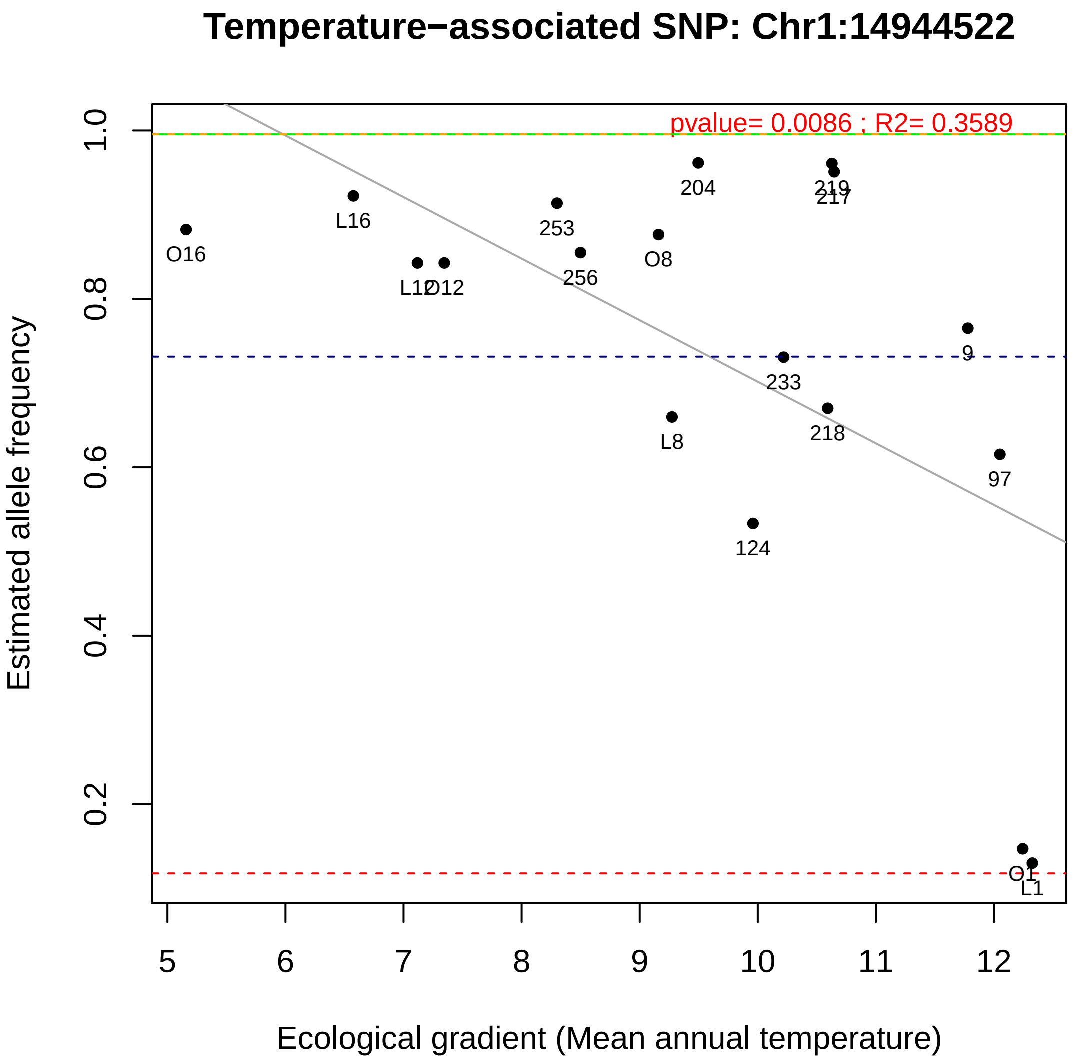

Temperature−associated SNP: Chr1:14944522
pvalue= 0.0086 ; R2= 0.3589
Estimated allele frequency
Ecological gradient (Mean annual temperature)
0.2
0.4
0.6
0.8
1.0
5
6
7
8
9
10
11
12
O16
L16
L12
O12
253
256
O8
204
219
217
L8
124
233
218
9
97
O1
L1

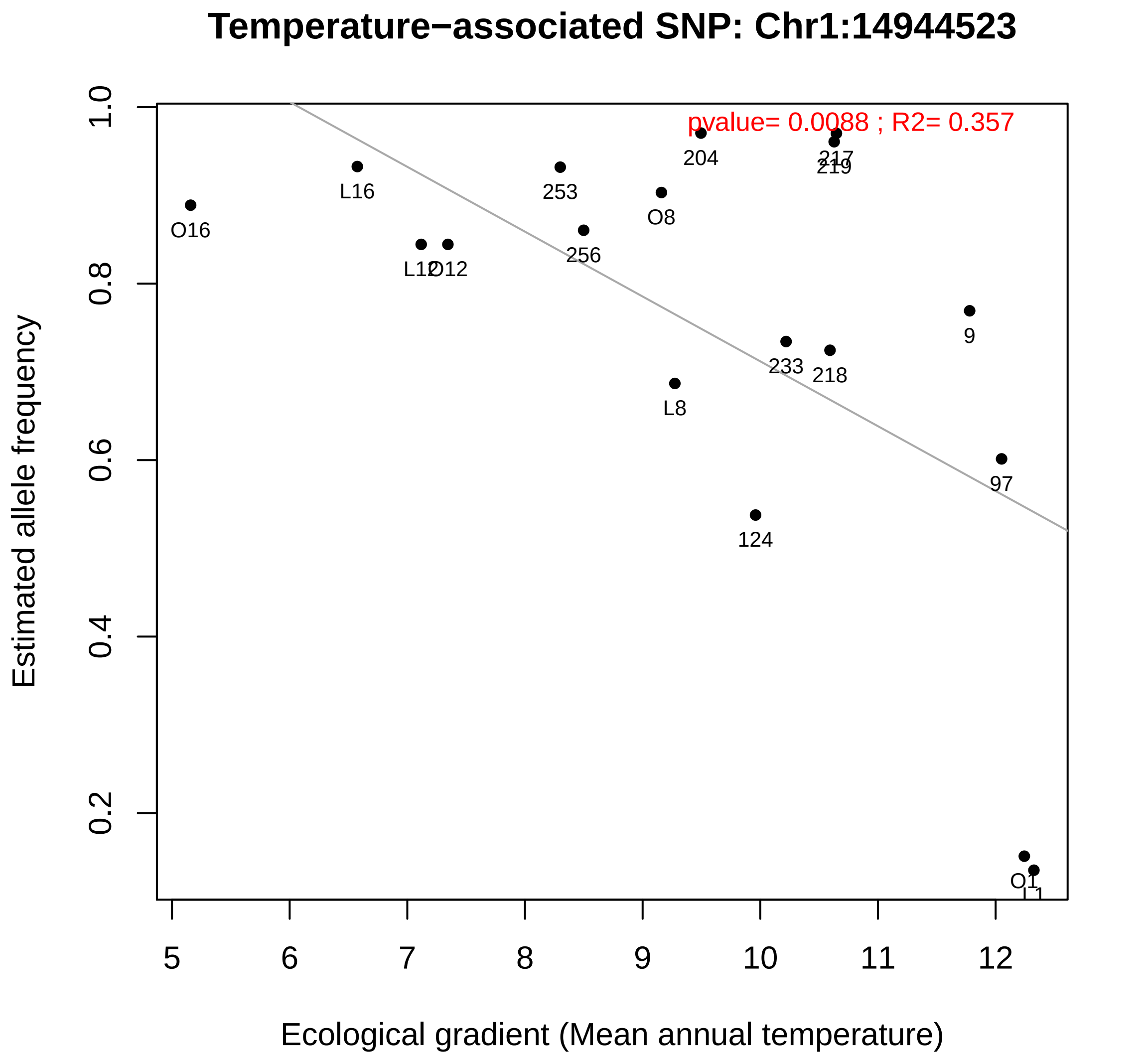
Temperature−associated SNP: Chr1:14944523
pvalue= 0.0088 ; R2= 0.357
Estimated allele frequency
Ecological gradient (Mean annual temperature)
1.0
0.8
0.6
0.4
0.2
5
6
7
8
9
10
11
12
O16
L16
L12
O12
253
256
O8
204
217
219
L8
233
218
124
9
97
O1
L1

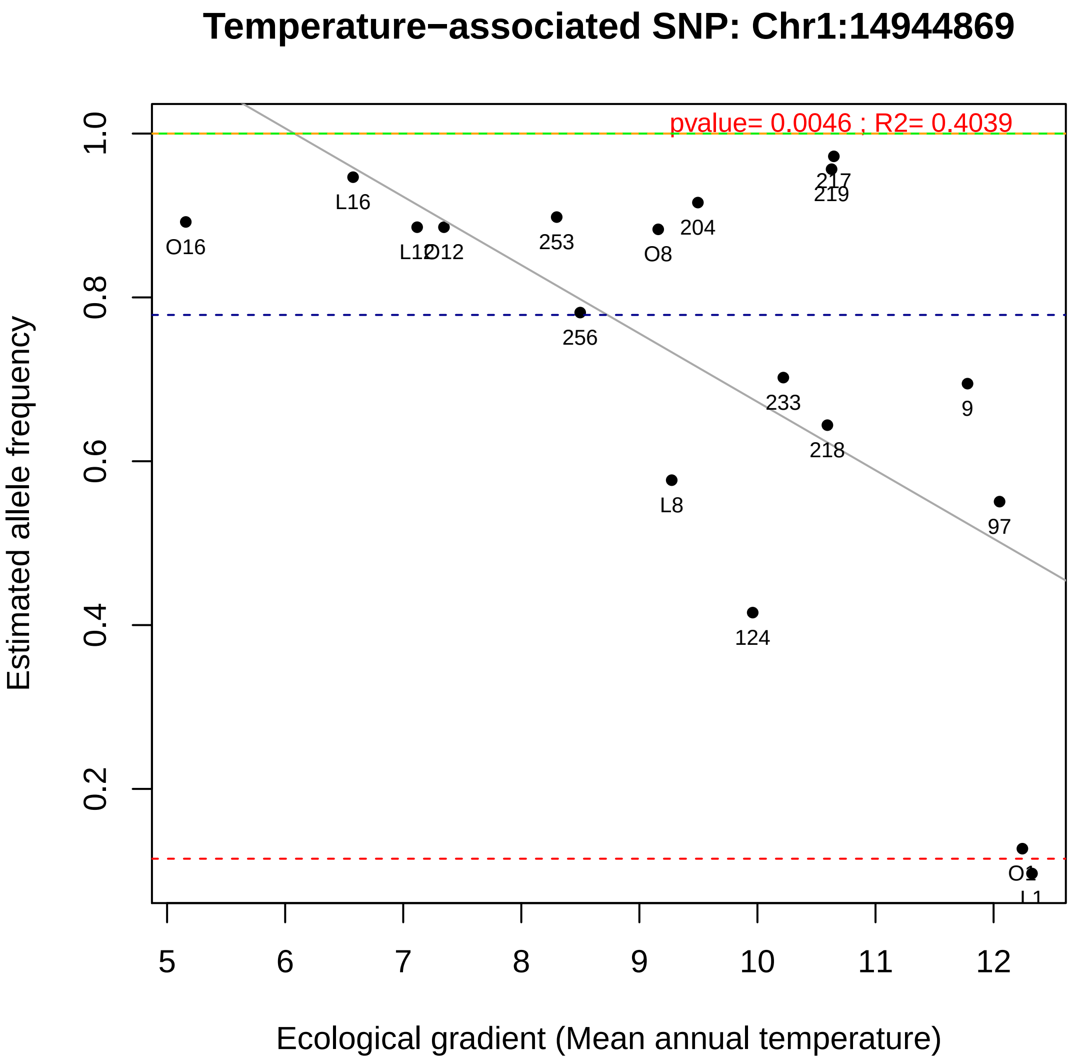

Temperature−associated SNP: Chr1:14944869
pvalue= 0.0046 ; R2= 0.4039
Estimated allele frequency
0.2
0.4
0.6
0.8
1.0
5
6
7
8
9
10
11
12
Ecological gradient (Mean annual temperature)
O16
L16
L12
O12
253
256
O8
204
217
219
L8
124
233
218
9
97
O1
L1

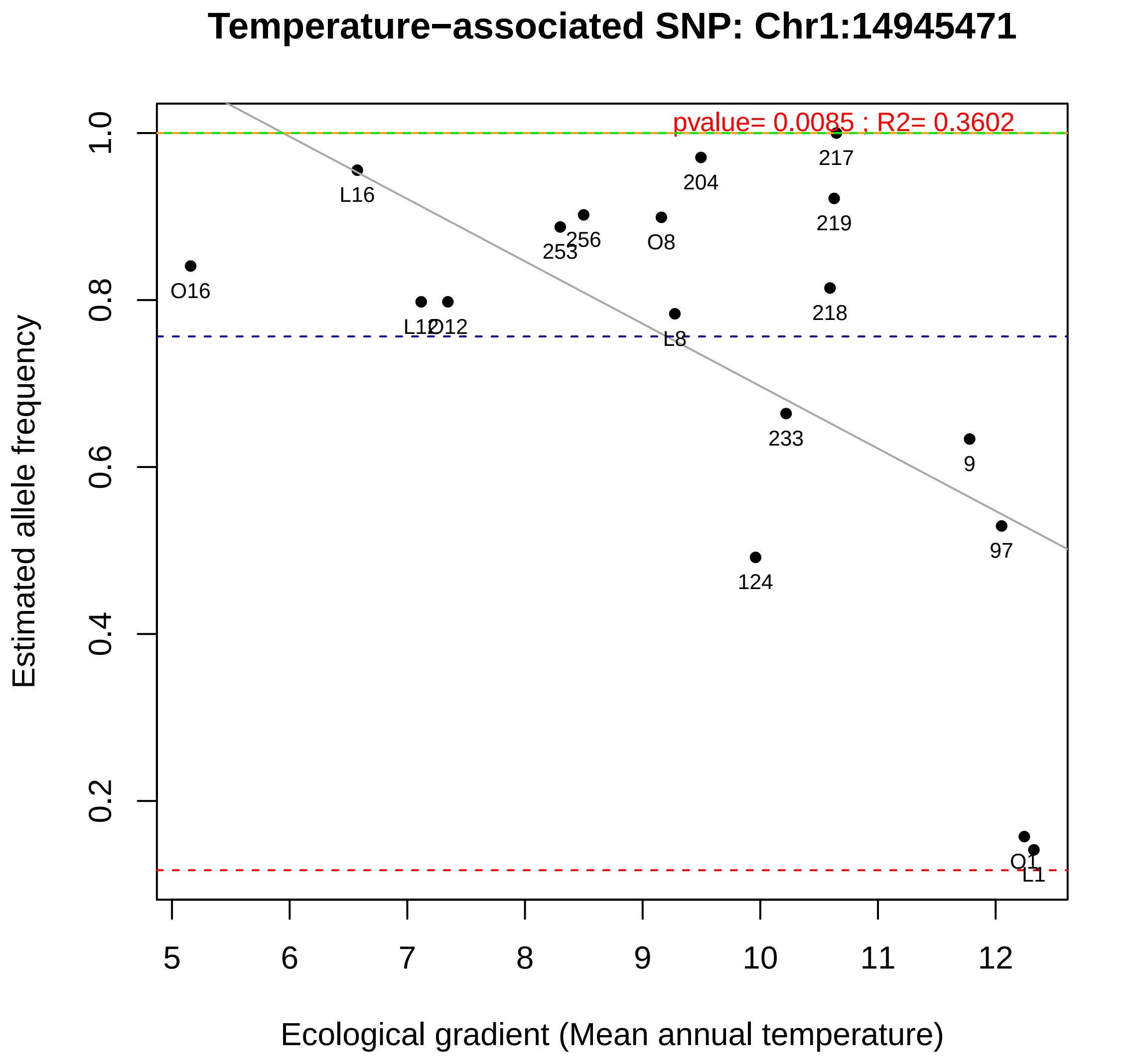

Temperature−associated SNP: Chr1:14945471
pvalue= 0.0085 ; R2= 0.3602
Estimated allele frequency
Ecological gradient (Mean annual temperature)
0.2
0.4
0.6
0.8
1.0
5
6
7
8
9
10
11
12
O16
L16
L12
O12
253
256
O8
L8
204
233
124
217
219
218
9
97
O1
L1

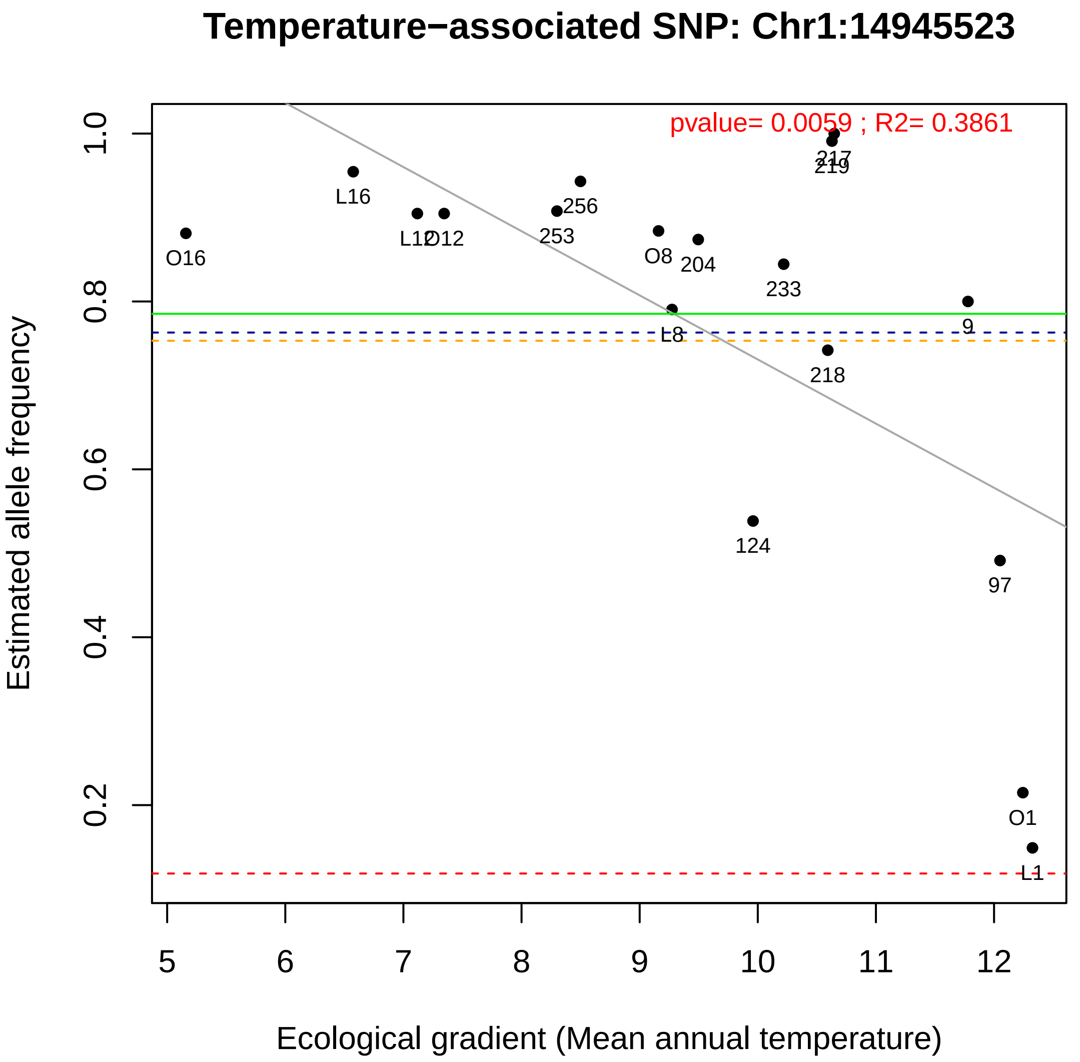

Temperature−associated SNP: Chr1:14945523
pvalue= 0.0059 ; R2= 0.3861
Estimated allele frequency
Ecological gradient (Mean annual temperature)
1.0
0.8
0.6
0.4
0.2
5
6
7
8
9
10
11
12
O16
L16
L12
O12
253
256
O8
204
L8
233
217
219
218
124
9
97
O1
L1

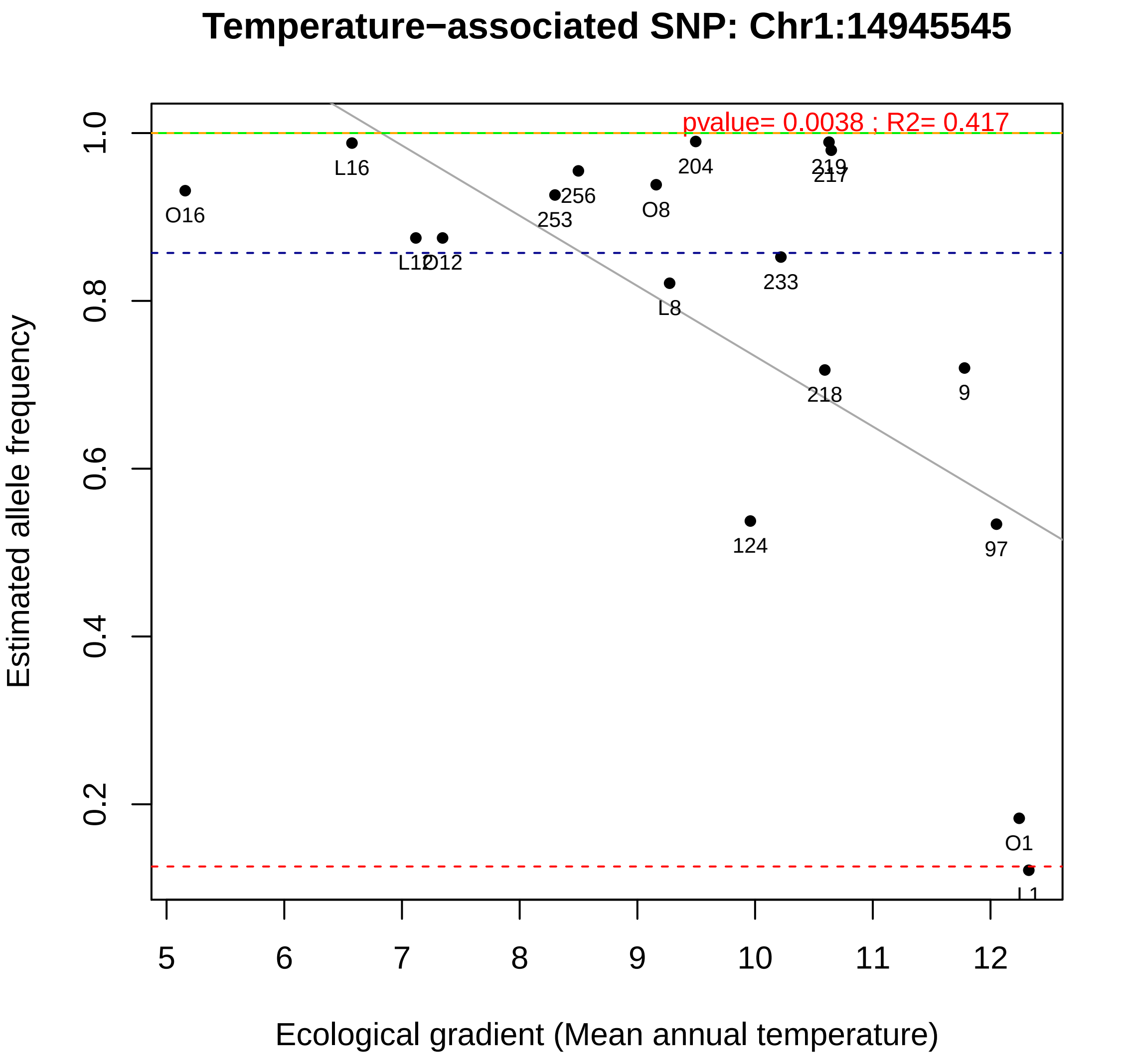

Temperature−associated SNP: Chr1:14945545
pvalue= 0.0038 ; R2= 0.417
Estimated allele frequency
Ecological gradient (Mean annual temperature)
1.0
0.8
0.6
0.4
0.2
5
6
7
8
9
10
11
12
O16
L16
L12
O12
253
256
O8
204
219
217
L8
233
218
9
124
97
O1

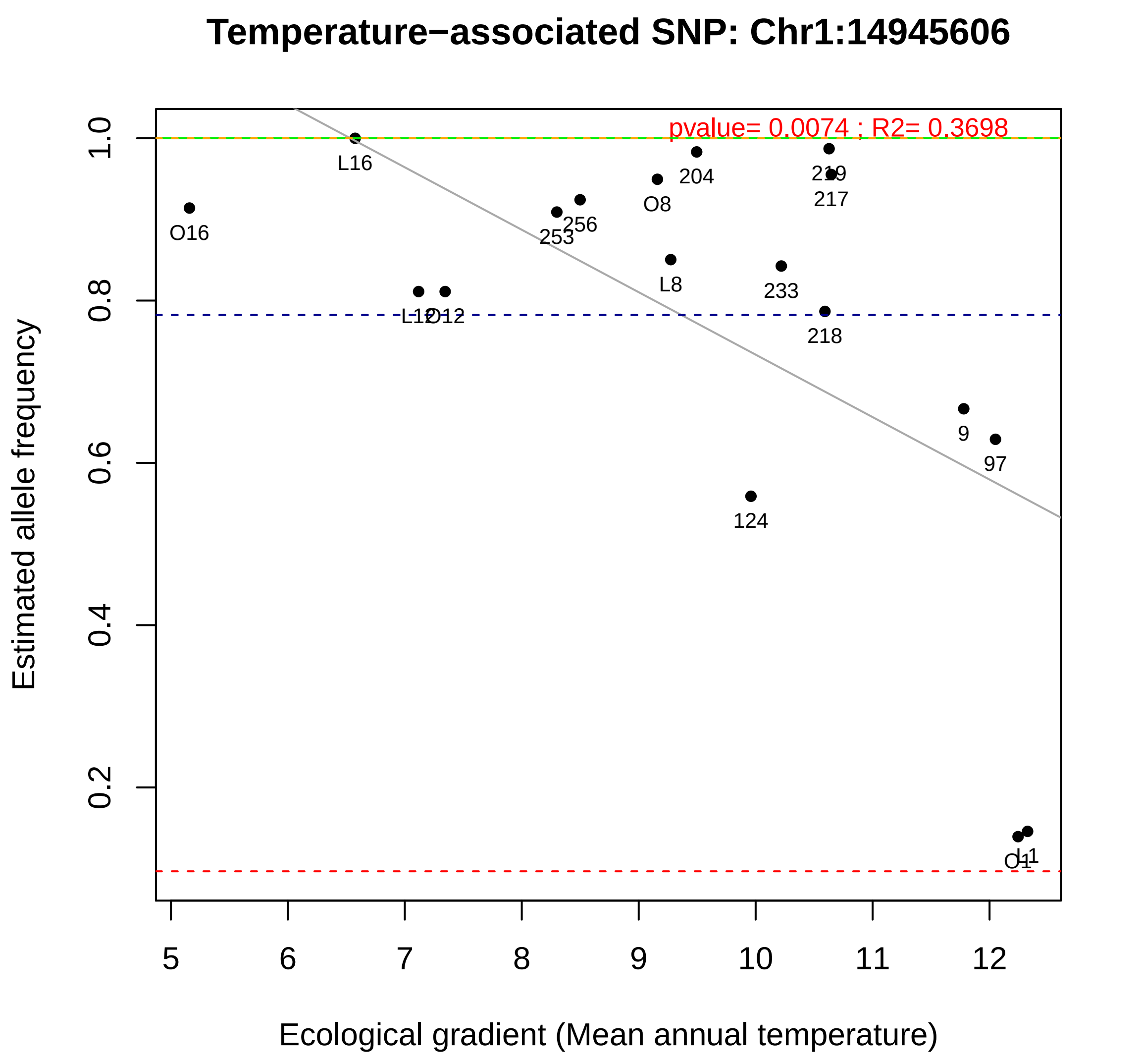
Temperature−associated SNP: Chr1:14945606
pvalue= 0.0074 ; R2= 0.3698
Estimated allele frequency
Ecological gradient (Mean annual temperature)
0.2
0.4
0.6
0.8
1.0
5
6
7
8
9
10
11
12
L16
O16
L12
O12
253
256
O8
204
L8
233
219
217
218
124
9
97
O1
L1

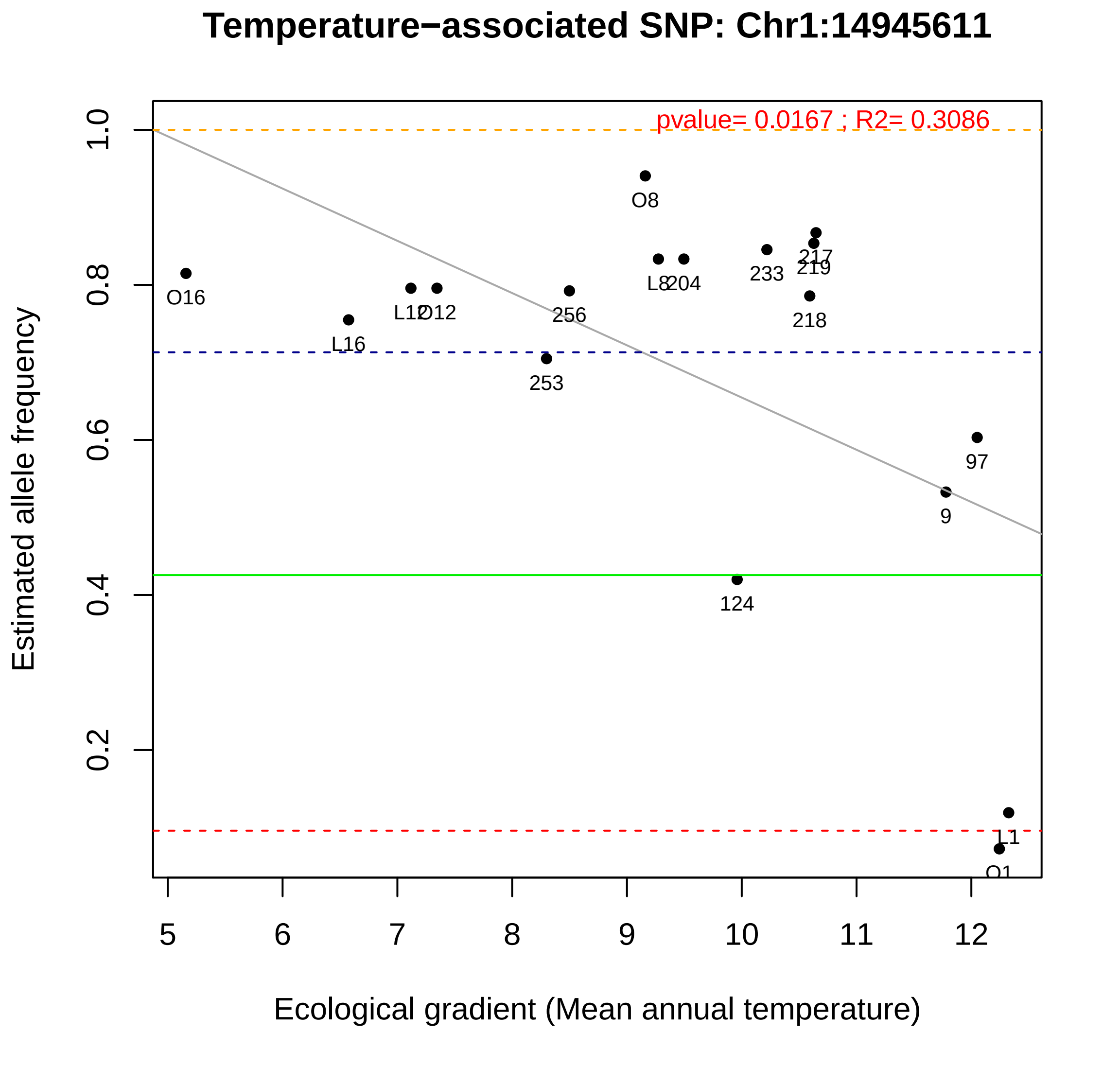
Temperature−associated SNP: Chr1:14945611
pvalue= 0.0167 ; R2= 0.3086
Estimated allele frequency
Ecological gradient (Mean annual temperature)
1.0
0.8
0.6
0.4
0.2
5
6
7
8
9
10
11
12
O16
L16
L12
O12
253
256
O8
L8
204
233
217
219
218
124
97
9
L1
O1

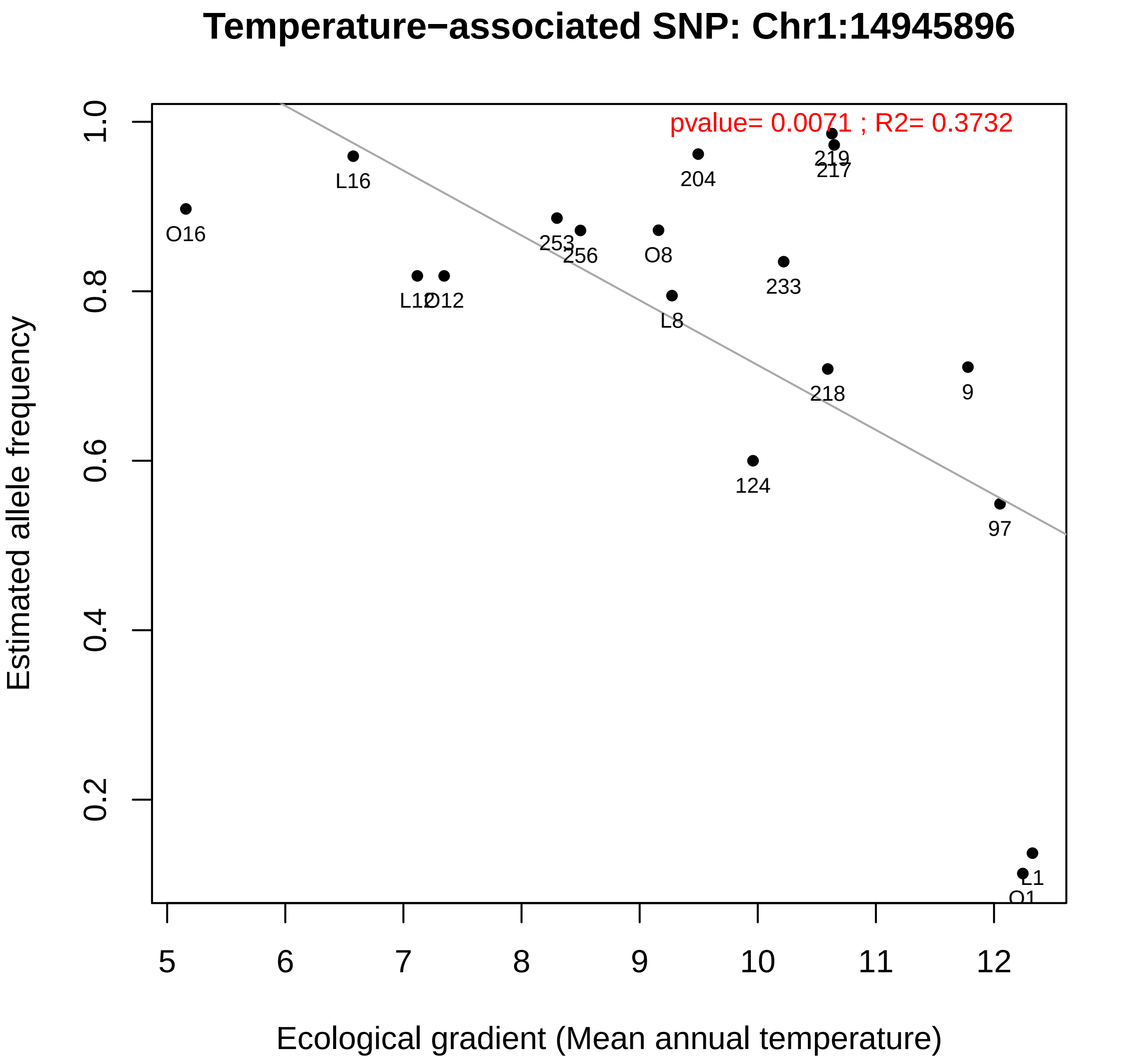
Temperature−associated SNP: Chr1:14945896
pvalue= 0.0071 ; R2= 0.3732
Estimated allele frequency
0.2
0.4
0.6
0.8
1.0
5
6
7
8
9
10
11
12
Ecological gradient (Mean annual temperature)
O16
L16
L12
O12
253
256
O8
L8
204
219
217
233
218
124
9
97
L1
O1

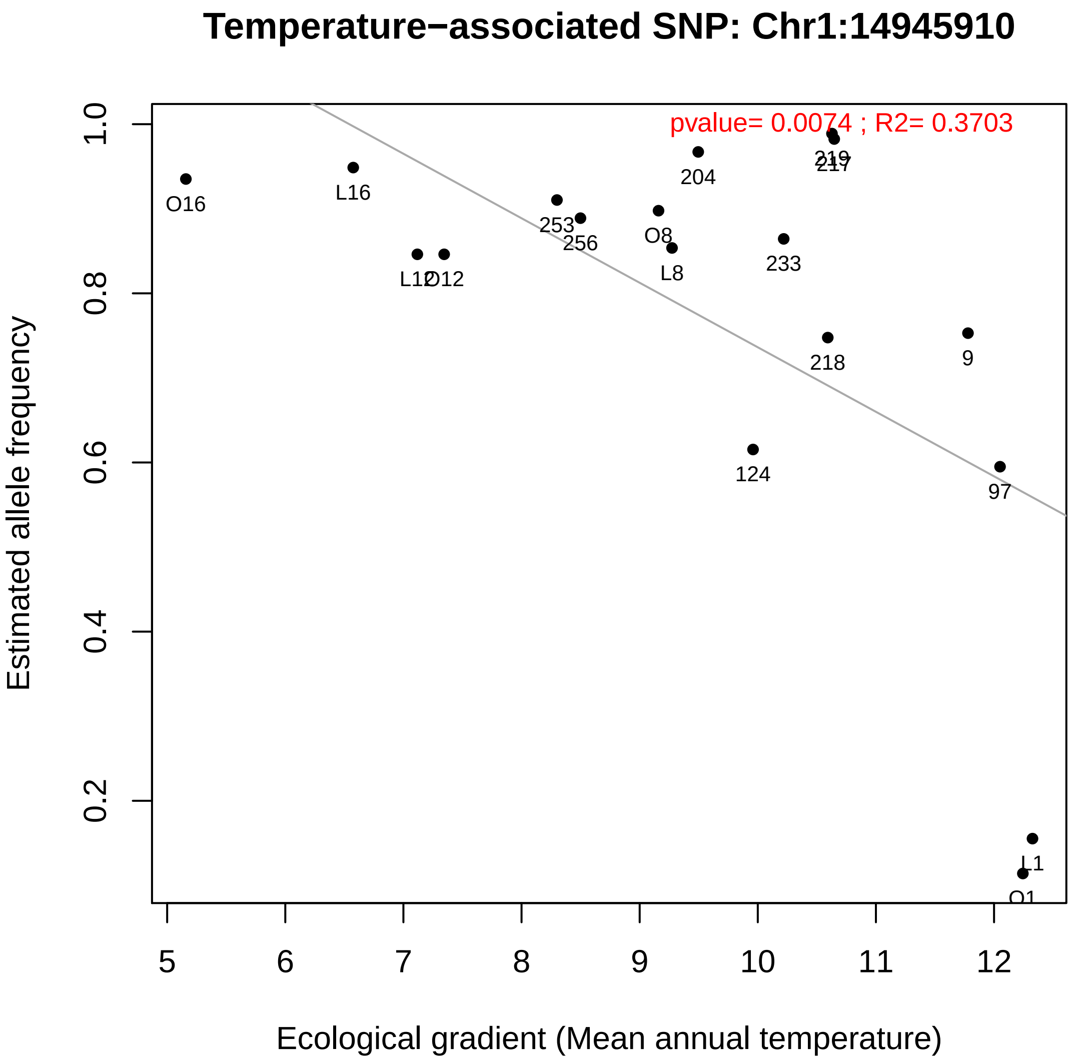

Temperature−associated SNP: Chr1:14945910
pvalue= 0.0074 ; R2= 0.3703
O16
L16
L12
O12
253
256
O8
L8
204
233
219
217
218
9
124
97
L1
O1
1.0
0.8
0.6
0.4
0.2
Estimated allele frequency
5
6
7
8
9
10
11
12
Ecological gradient (Mean annual temperature)

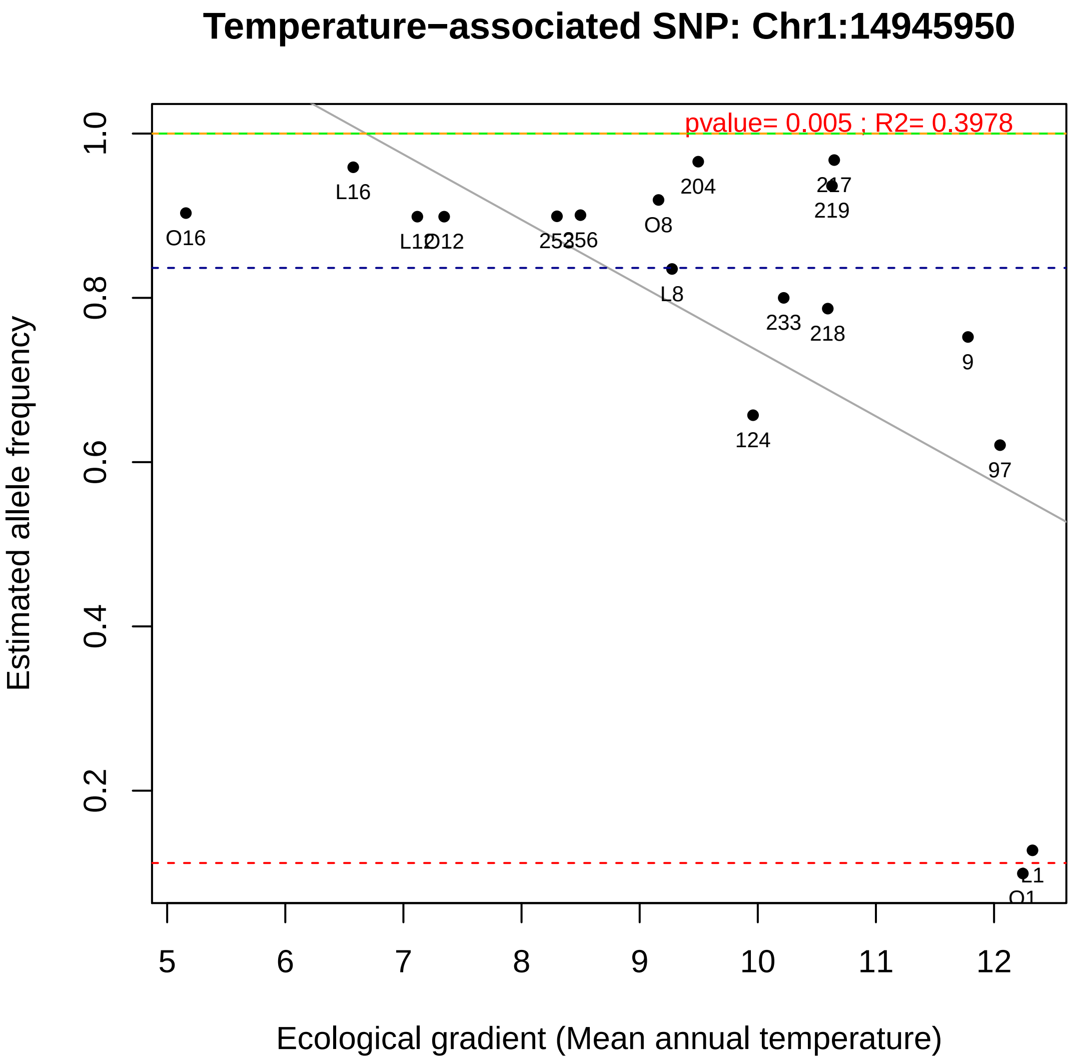

Temperature−associated SNP: Chr1:14945950
pvalue= 0.005 ; R2= 0.3978
Estimated allele frequency
Ecological gradient (Mean annual temperature)
1.0
0.8
0.6
0.4
0.2
5
6
7
8
9
10
11
12
O16
L16
L12
O12
256
204
O8
217
219
L8
233
218
9
124
97
L1

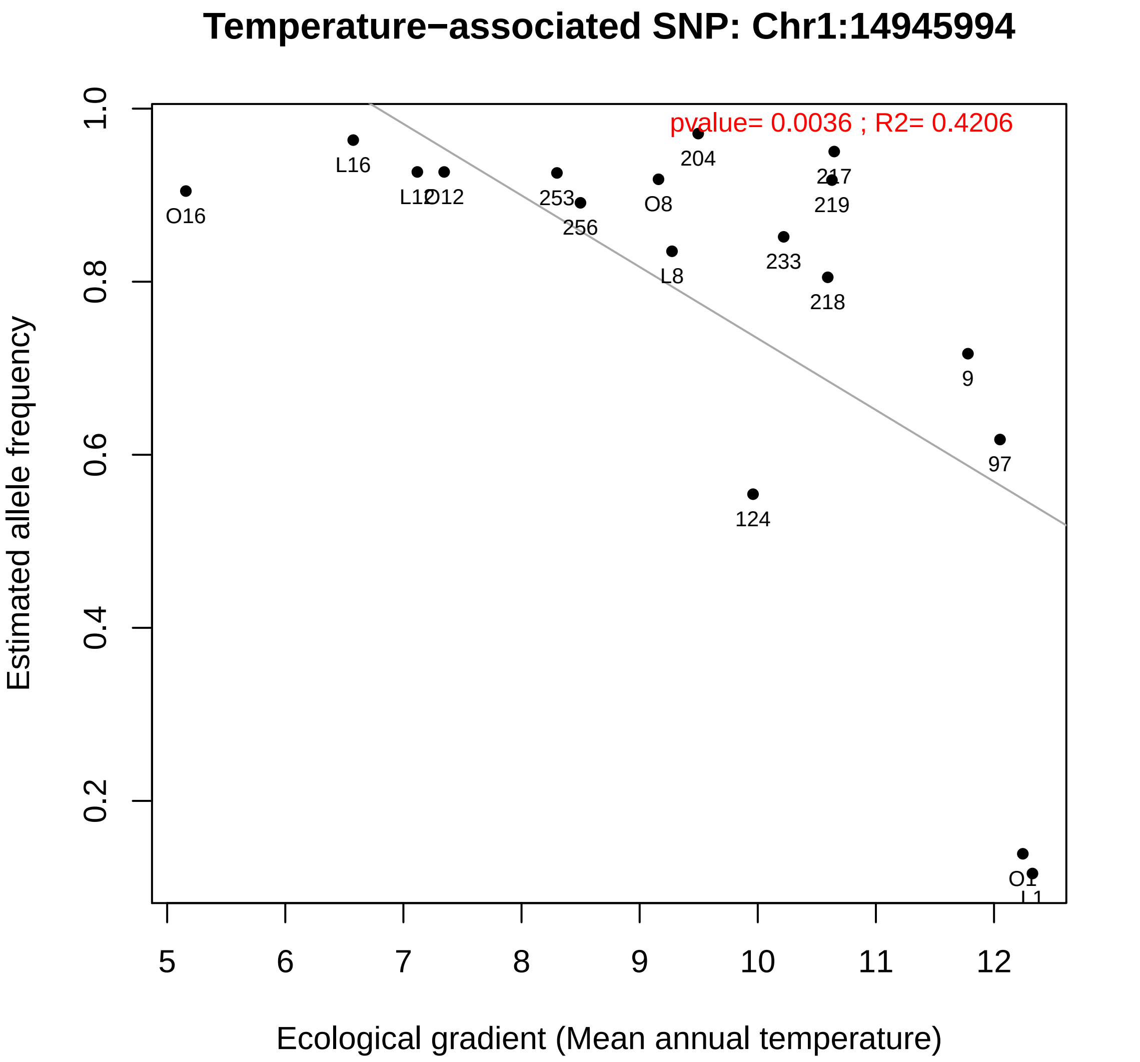

Temperature−associated SNP: Chr1:14945994
pvalue= 0.0036 ; R2= 0.4206
Estimated allele frequency
Ecological gradient (Mean annual temperature)
1.0
0.8
0.6
0.4
0.2
5
6
7
8
9
10
11
12
O16
L16
L12
O12
253
256
O8
204
L8
233
217
219
218
124
9
97
O1

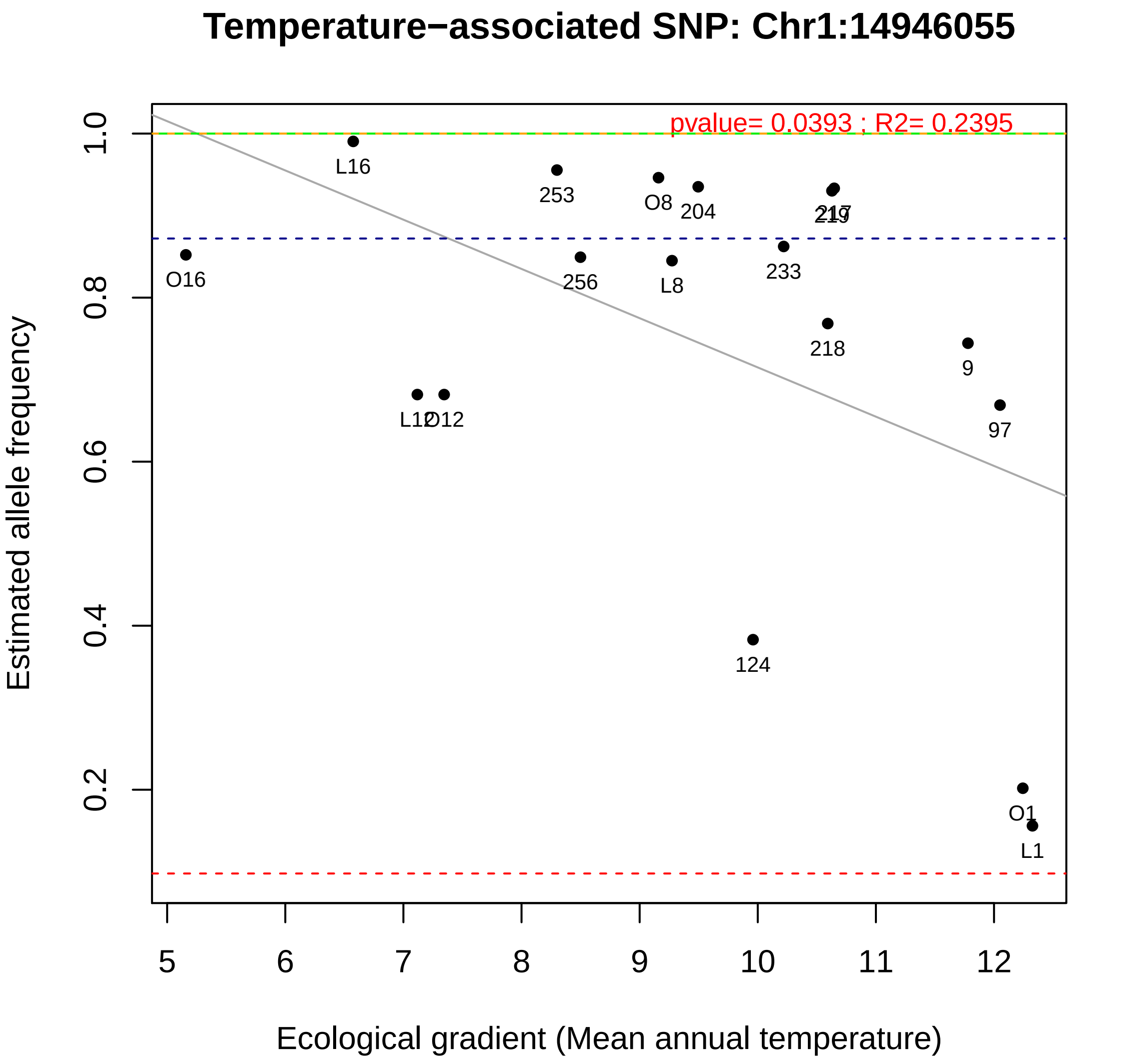

Temperature−associated SNP: Chr1:14946055
pvalue= 0.0393 ; R2= 0.2395
Estimated allele frequency
Ecological gradient (Mean annual temperature)
1.0
0.8
0.6
0.4
0.2
5
6
7
8
9
10
11
12
L16
253
O8
204
217
219
O16
256
L8
233
218
9
L12
O12
97
124
O1
L1

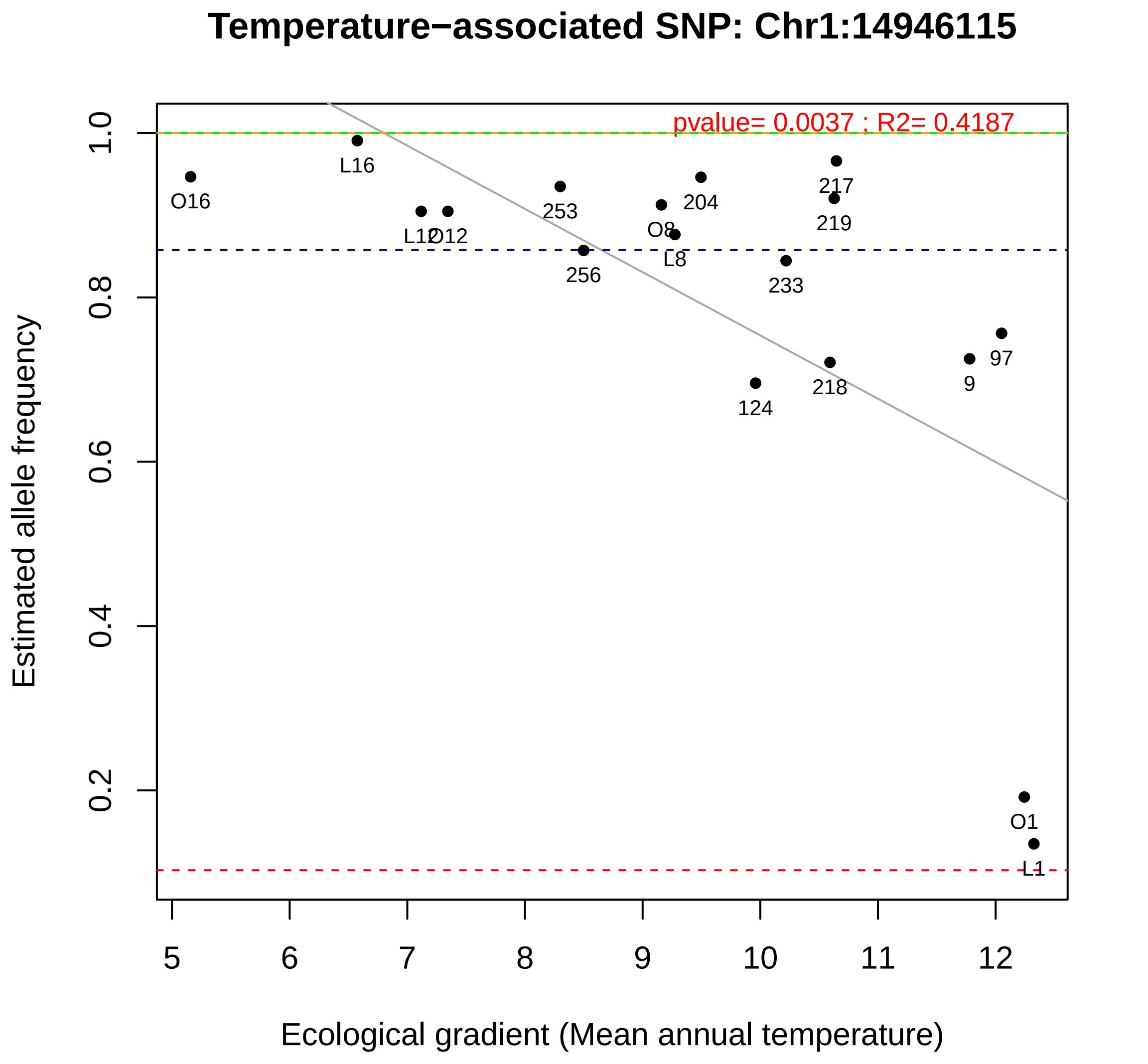

Temperature−associated SNP: Chr1:14946115
pvalue= 0.0037 ; R2= 0.4187
Estimated allele frequency
1.0
0.8
0.6
0.4
0.2
O16
L16
L12
O12
253
256
O8
L8
204
233
217
219
124
218
9
97
O1
L1
5
6
7
8
9
10
11
12
Ecological gradient (Mean annual temperature)

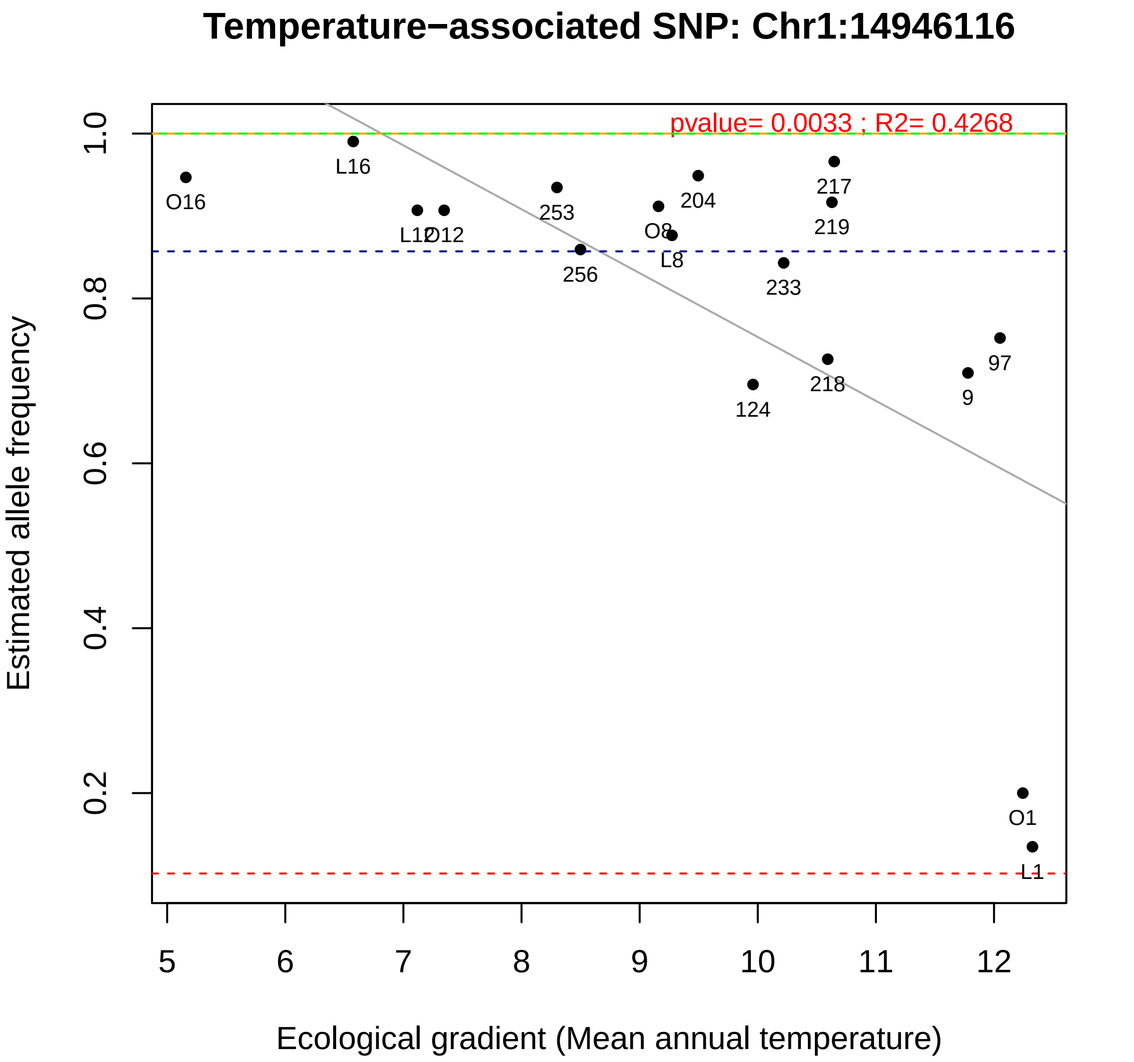

Temperature−associated SNP: Chr1:14946116
pvalue= 0.0033 ; R2= 0.4268
Estimated allele frequency
0.2
0.4
0.6
0.8
1.0
5
6
7
8
9
10
11
12
Ecological gradient (Mean annual temperature)
O16
L16
L12
O12
253
256
O8
L8
204
217
219
233
124
218
9
97
O1
L1

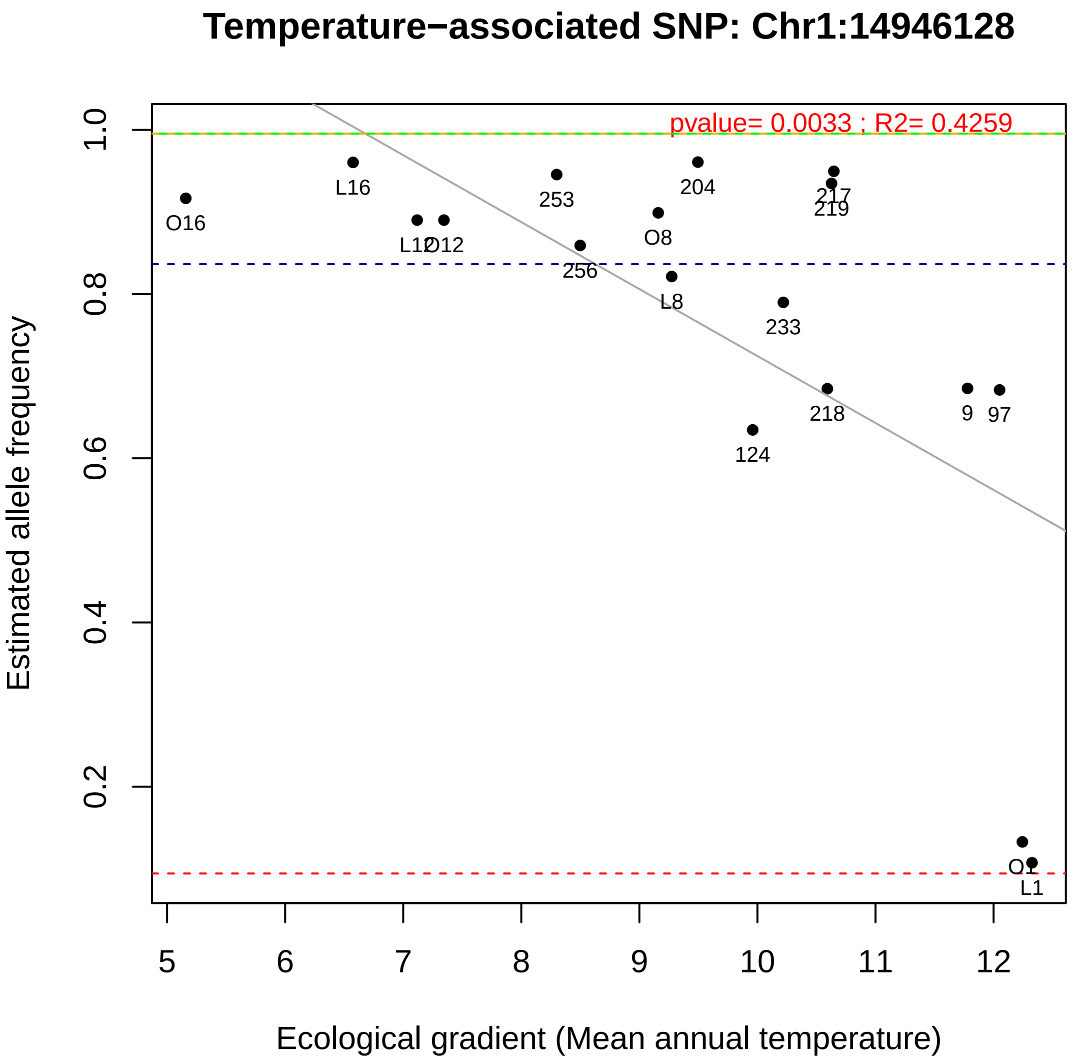

Temperature−associated SNP: Chr1:14946128
pvalue= 0.0033 ; R2= 0.4259
Estimated allele frequency
0.2
0.4
0.6
0.8
1.0
O16
L16
L12
O12
253
256
O8
L8
204
233
124
217
219
218
9
97
O1
L1
5
6
7
8
9
10
11
12
Ecological gradient (Mean annual temperature)

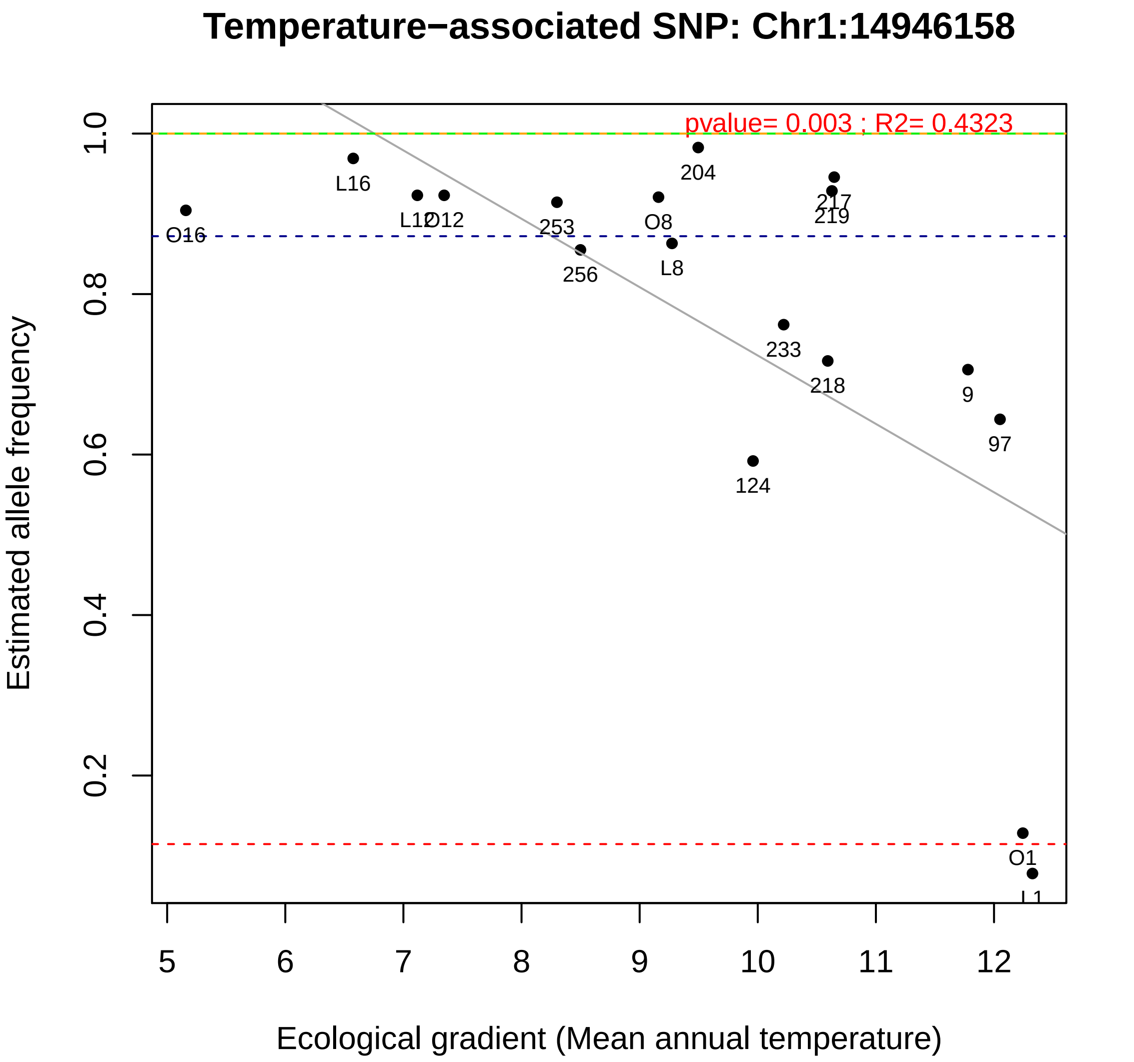
Temperature−associated SNP: Chr1:14946158
pvalue= 0.003 ; R2= 0.4323
Estimated allele frequency
0.2
0.4
0.6
0.8
1.0
Ecological gradient (Mean annual temperature)
5
6
7
8
9
10
11
12
O16
L16
L12
O12
253
256
O8
L8
204
217
219
233
218
124
9
97
O1

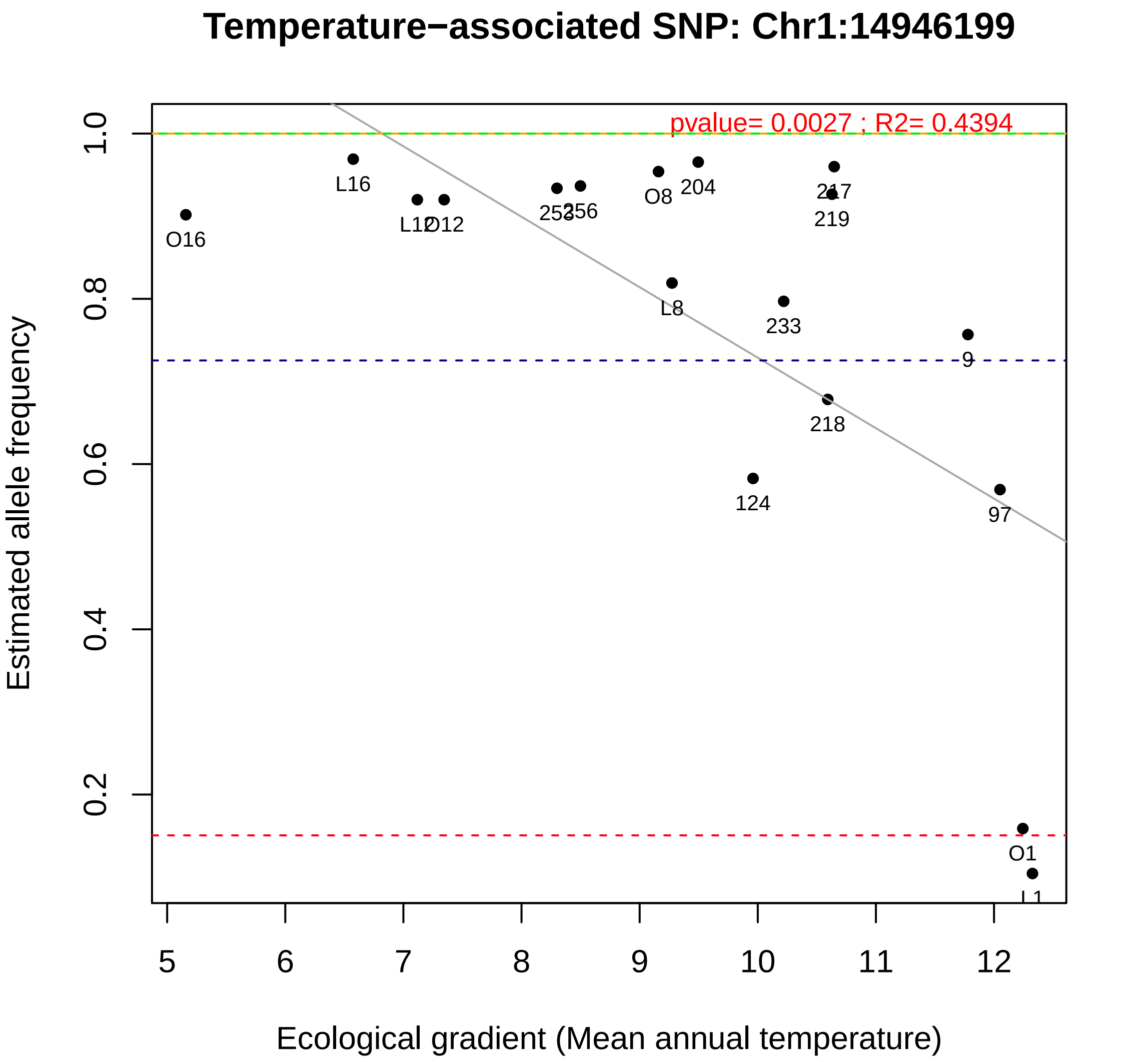
Temperature−associated SNP: Chr1:14946199
pvalue= 0.0027 ; R2= 0.4394
Estimated allele frequency
Ecological gradient (Mean annual temperature)
0.2
0.4
0.6
0.8
1.0
5
6
7
8
9
10
11
12
O16
L16
L12
O12
252
256
O8
204
217
219
L8
233
9
218
124
97
O1

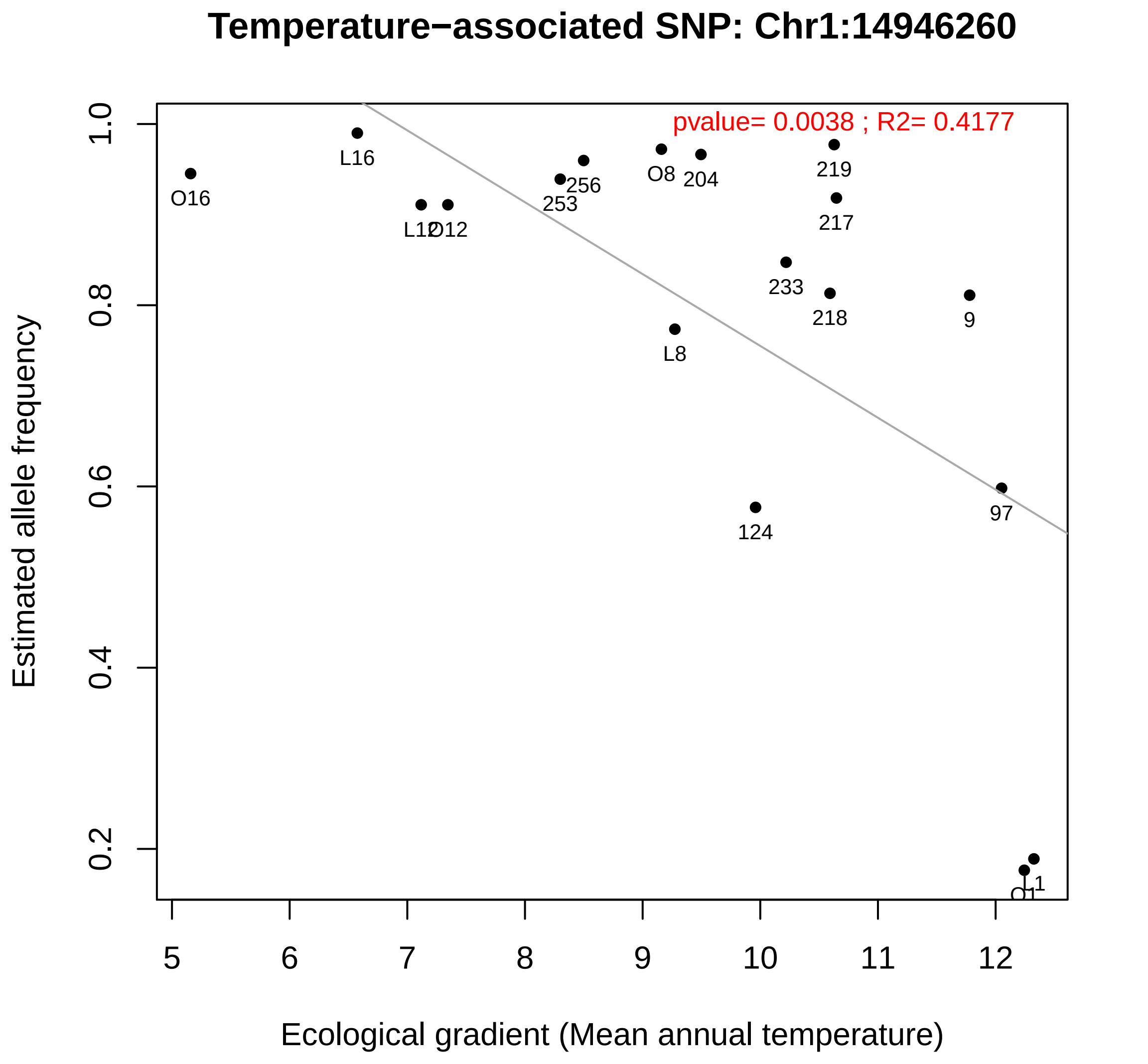

Temperature−associated SNP: Chr1:14946260
pvalue= 0.0038 ; R2= 0.4177
Estimated allele frequency
Ecological gradient (Mean annual temperature)
0.2
0.4
0.6
0.8
1.0
5
6
7
8
9
10
11
12
O16
L16
L12
O12
253
256
O8
204
219
217
233
218
9
L8
124
97
L1
O1

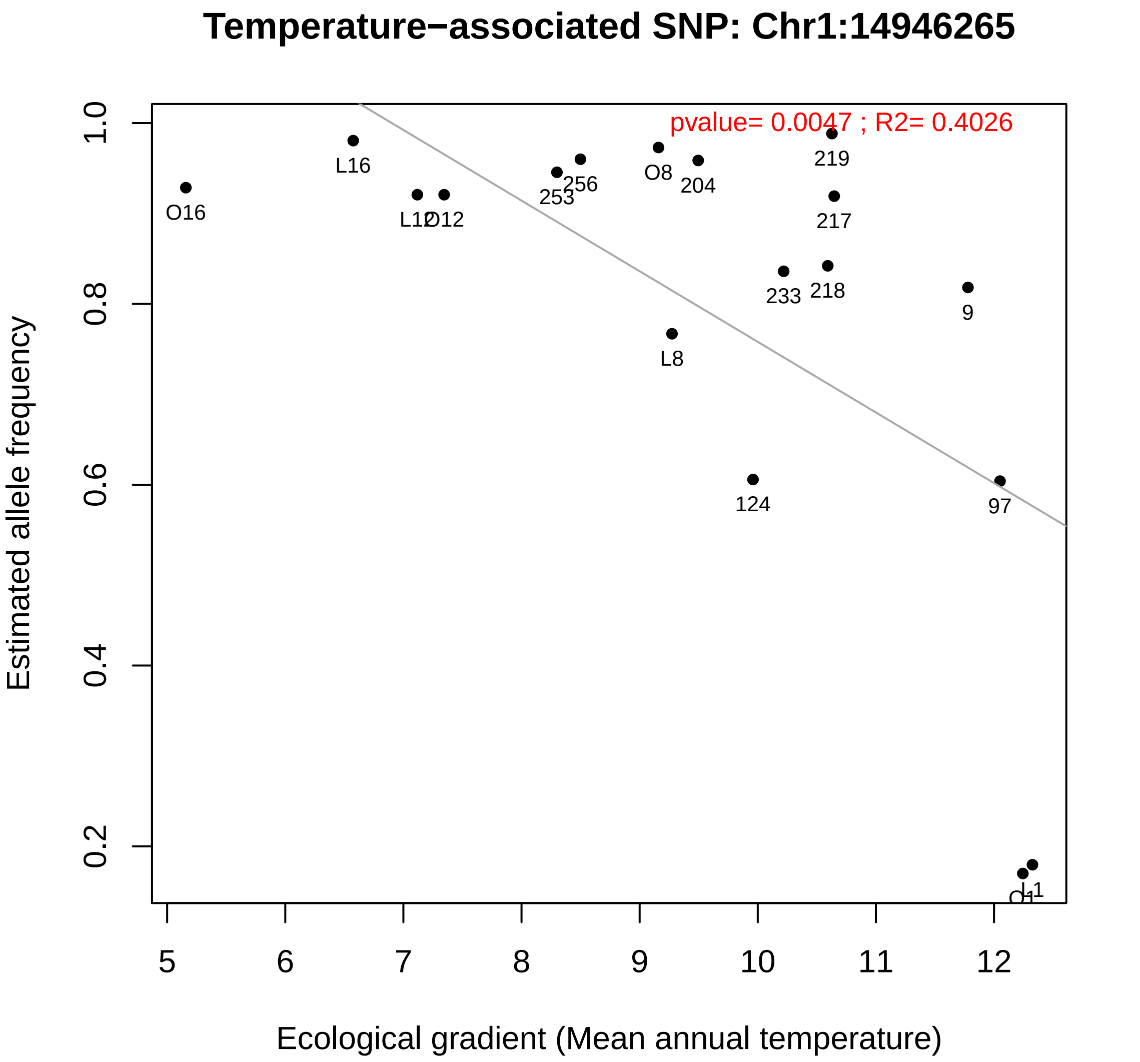
Temperature−associated SNP: Chr1:14946265
pvalue= 0.0047 ; R2= 0.4026
Estimated allele frequency
Ecological gradient (Mean annual temperature)
0.2
0.4
0.6
0.8
1.0
5
6
7
8
9
10
11
12
O16
L16
L12
O12
253
256
O8
204
219
217
233
218
9
L8
124
97
L1

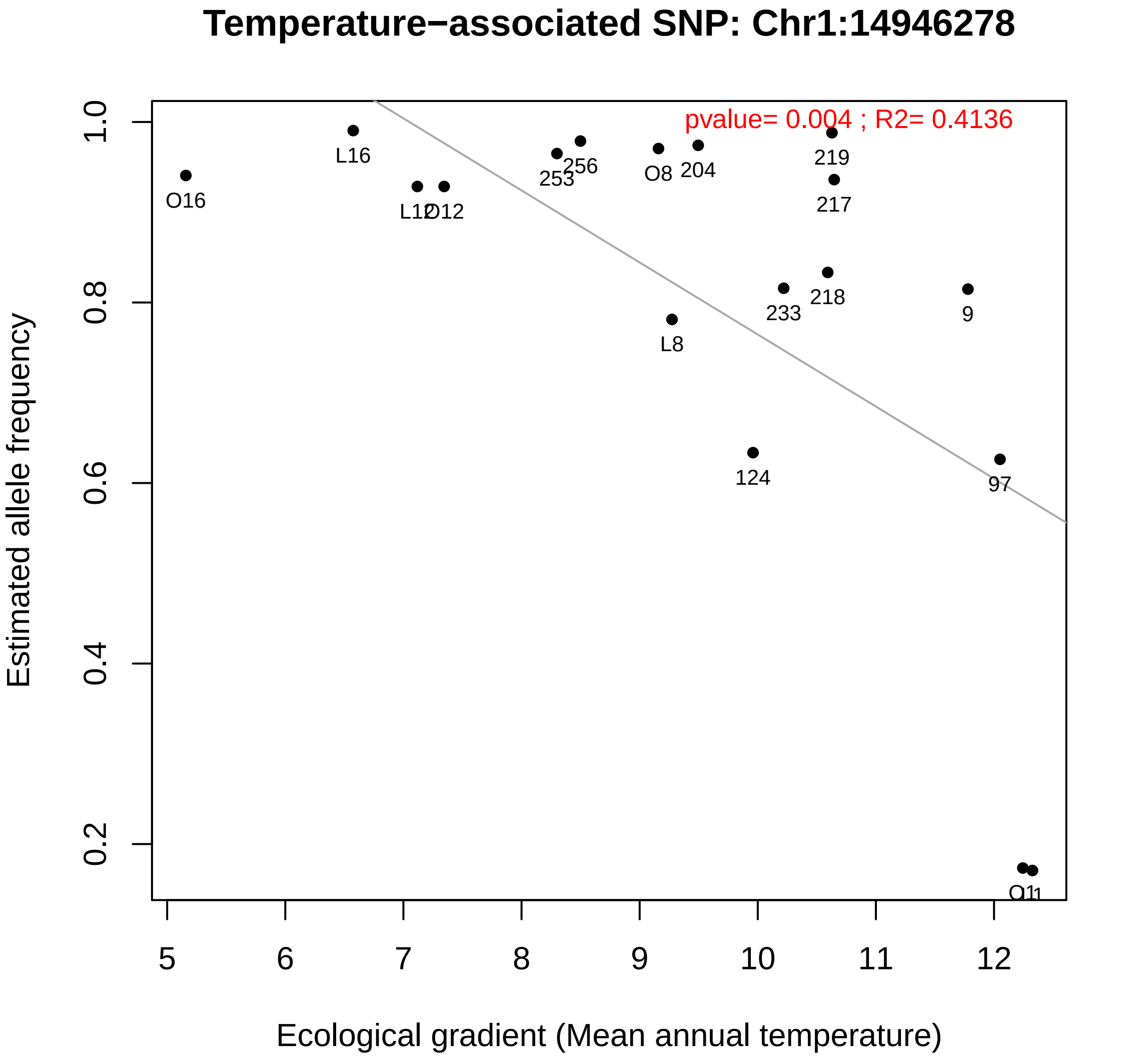
Temperature−associated SNP: Chr1:14946278
pvalue= 0.004 ; R2= 0.4136
Estimated allele frequency
0.2
0.4
0.6
0.8
1.0
5
6
7
8
9
10
11
12
Ecological gradient (Mean annual temperature)
O16
L16
O12
253
256
O8
204
219
217
218
233
9
L8
124
97

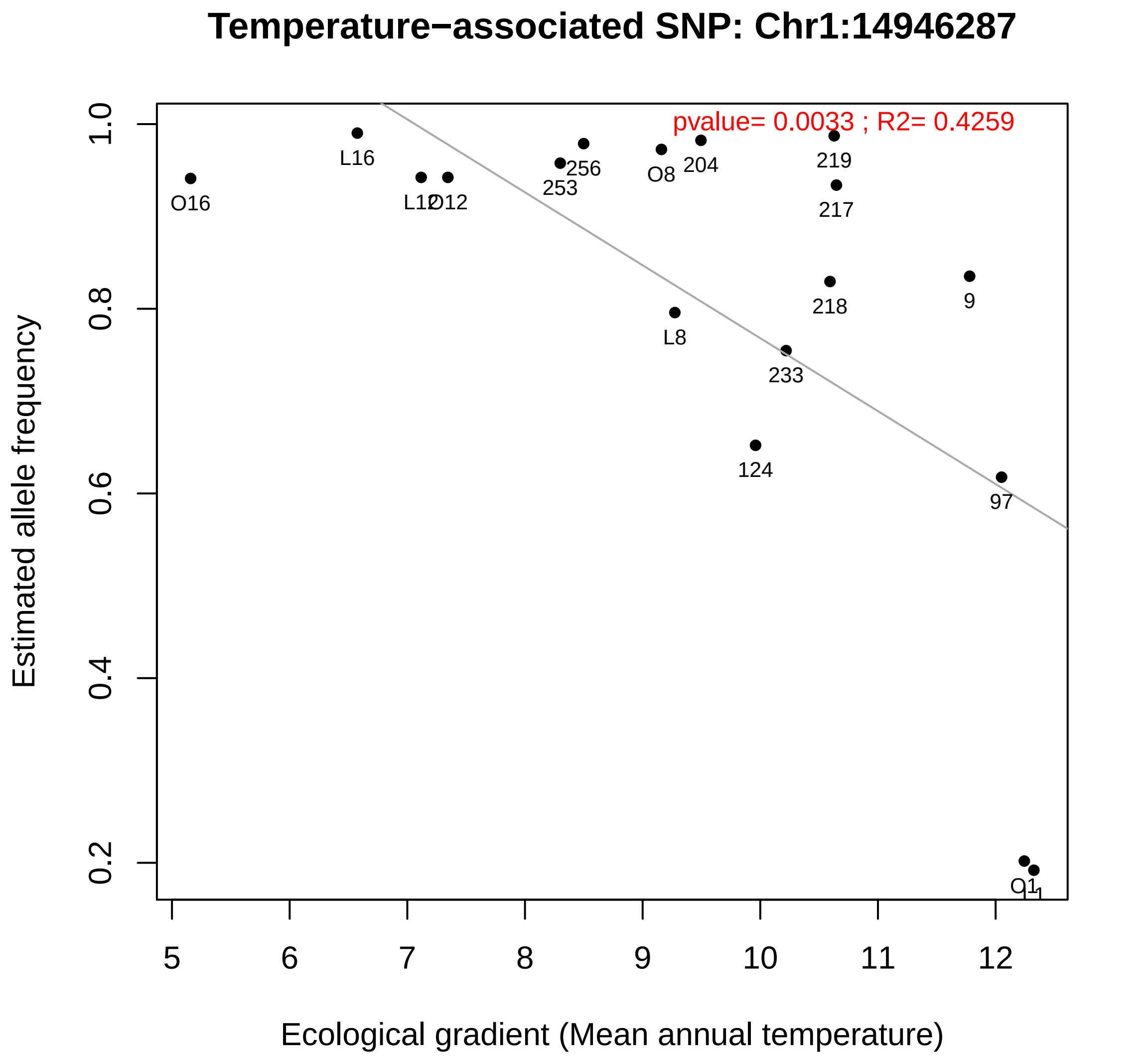

Temperature−associated SNP: Chr1:14946287
pvalue= 0.0033 ; R2= 0.4259
Estimated allele frequency
Ecological gradient (Mean annual temperature)
0.2
0.4
0.6
0.8
1.0
5
6
7
8
9
10
11
12
O16
L16
L12
O12
253
256
O8
204
219
217
218
9
L8
233
124
97
O1

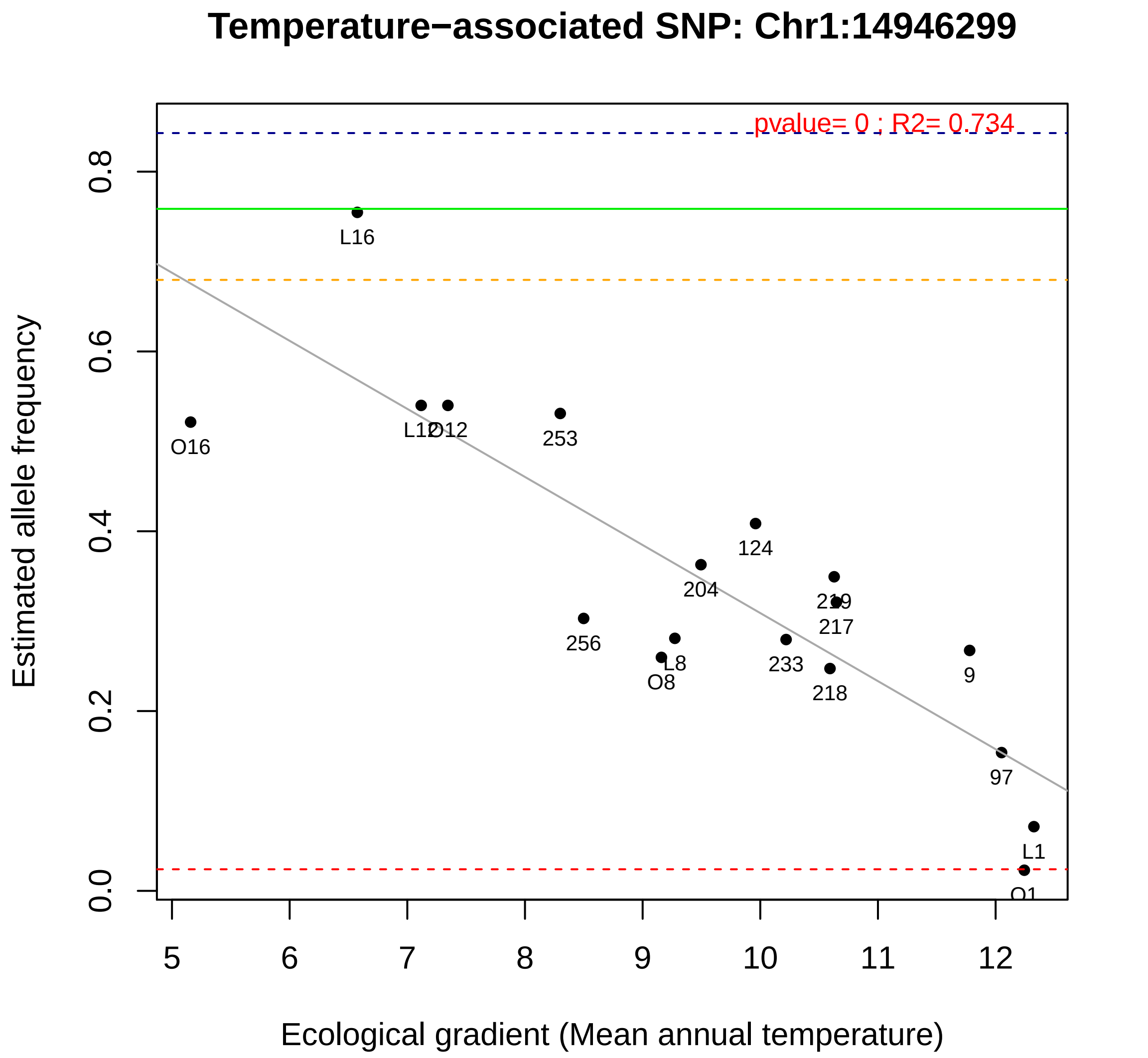

Temperature−associated SNP: Chr1:14946299
pvalue= 0 ; R2= 0.734
Estimated allele frequency
0.0
0.2
0.4
0.6
0.8
5
6
7
8
9
10
11
12
Ecological gradient (Mean annual temperature)
L16
O16
L12
O12
253
256
124
204
219
217
233
L8
O8
218
9
97
L1
O1

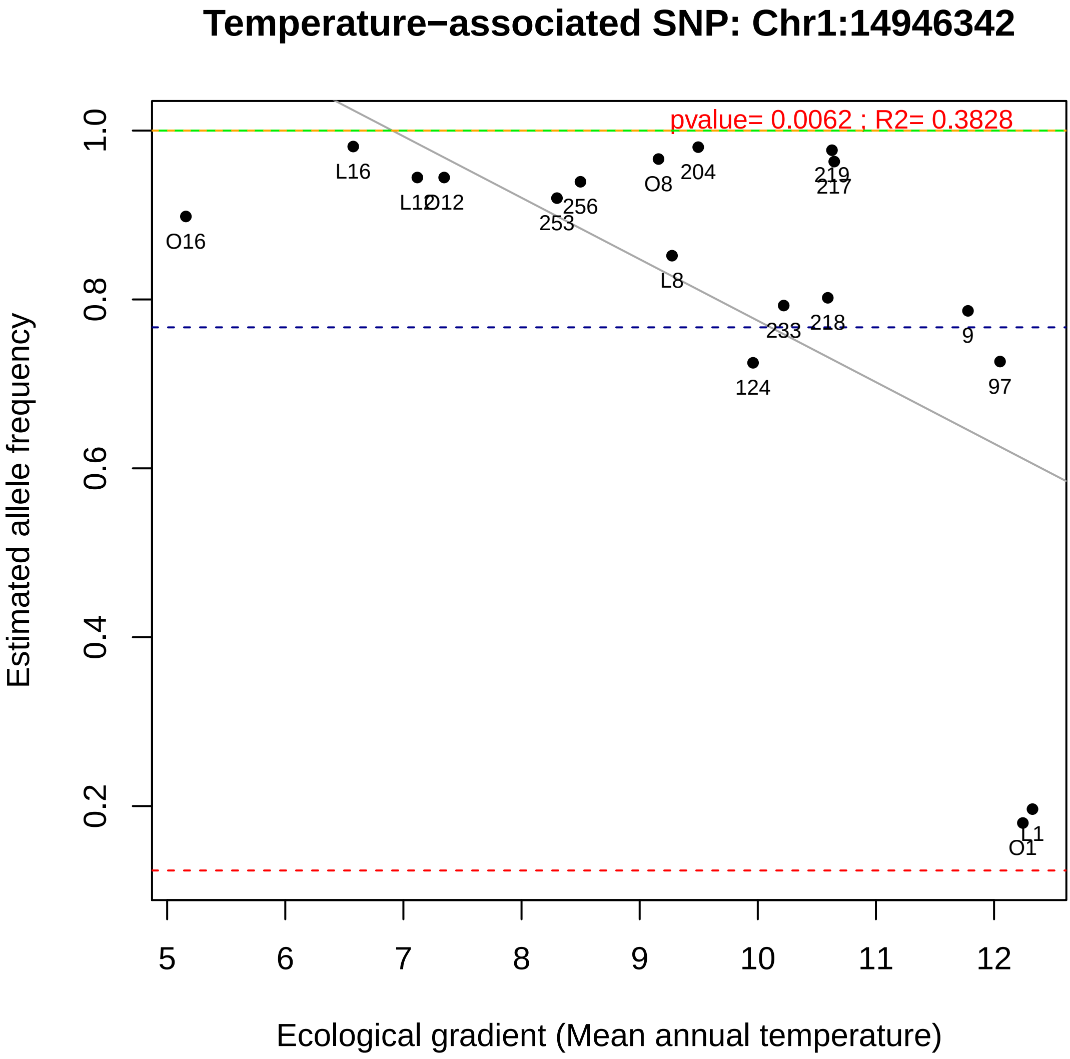

Temperature−associated SNP: Chr1:14946342
pvalue= 0.0062 ; R2= 0.3828
Estimated allele frequency
Ecological gradient (Mean annual temperature)
0.2
0.4
0.6
0.8
1.0
5
6
7
8
9
10
11
12
O16
L16
L12
O12
253
256
O8
204
L8
219
217
233
218
124
9
97
L1
O1

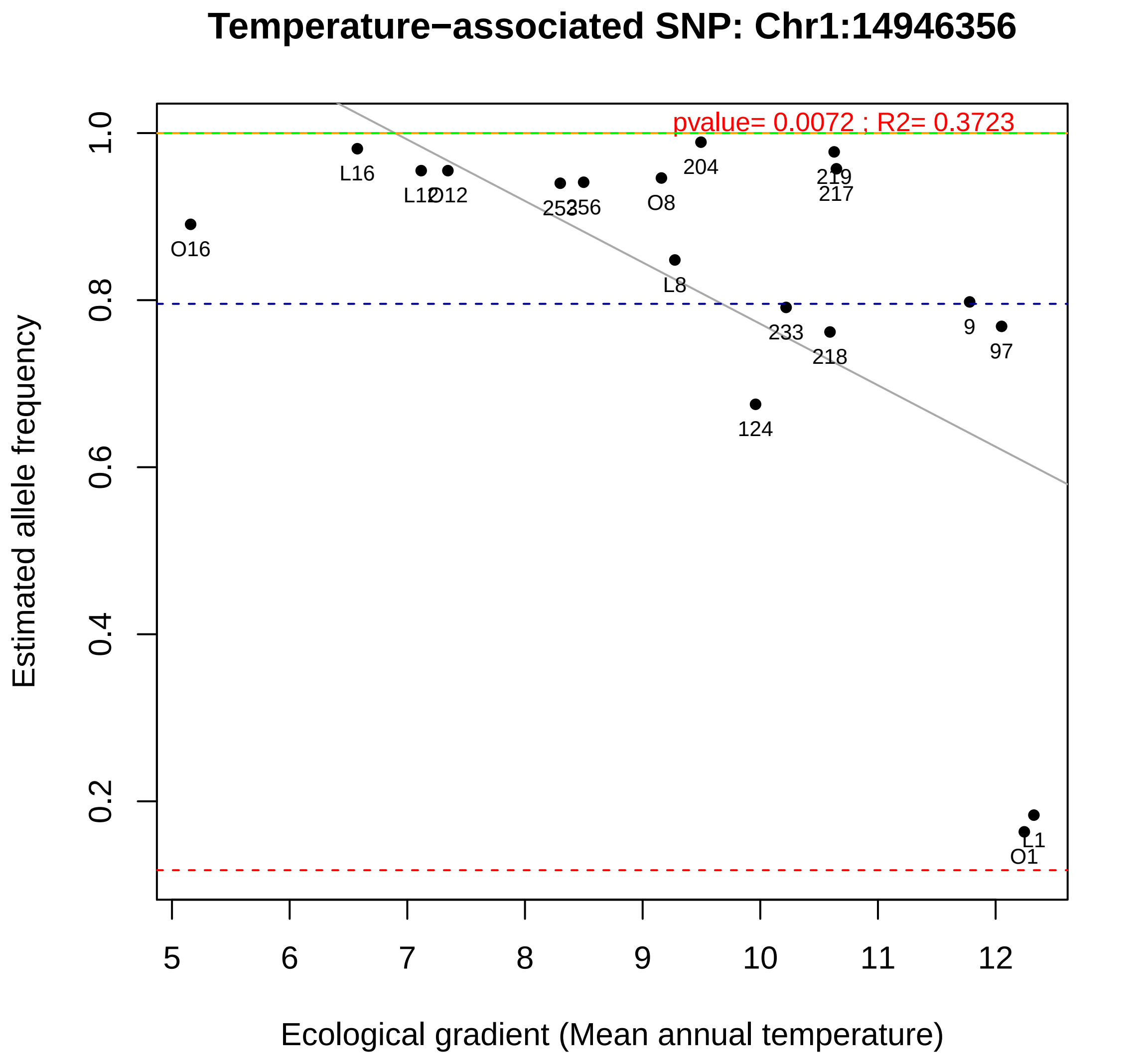

Temperature−associated SNP: Chr1:14946356
pvalue= 0.0072 ; R2= 0.3723
Estimated allele frequency
0.2
0.4
0.6
0.8
1.0
5
6
7
8
9
10
11
12
Ecological gradient (Mean annual temperature)
O16
L16
L12
O12
253
256
O8
204
L8
219
217
233
218
124
9
97
L1
O1

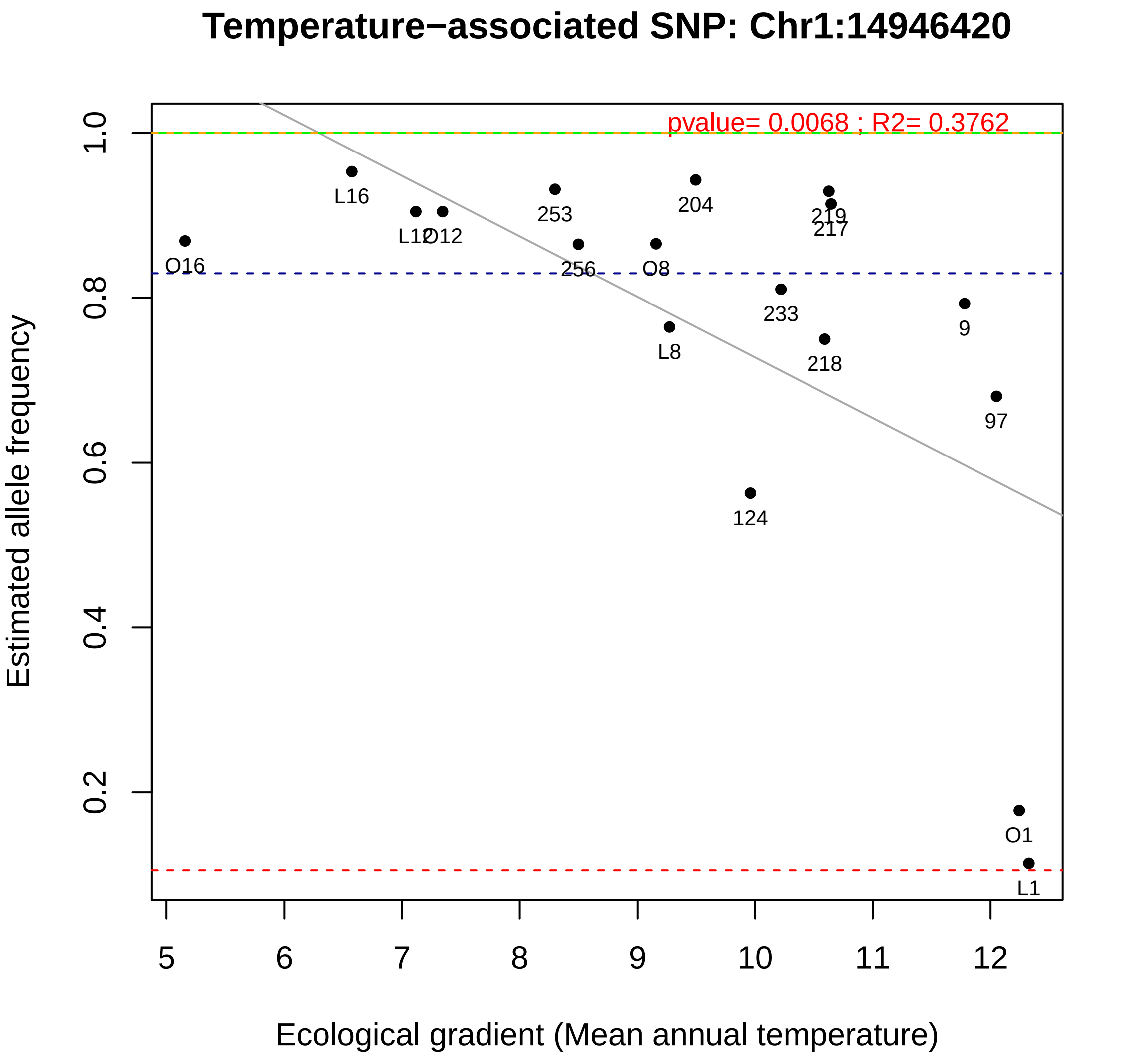

Temperature−associated SNP: Chr1:14946420
pvalue= 0.0068 ; R2= 0.3762
Estimated allele frequency
0.2
0.4
0.6
0.8
1.0
5
6
7
8
9
10
11
12
Ecological gradient (Mean annual temperature)
O16
L16
L12
O12
253
256
O8
204
219
217
233
L8
218
9
124
97
O1
L1

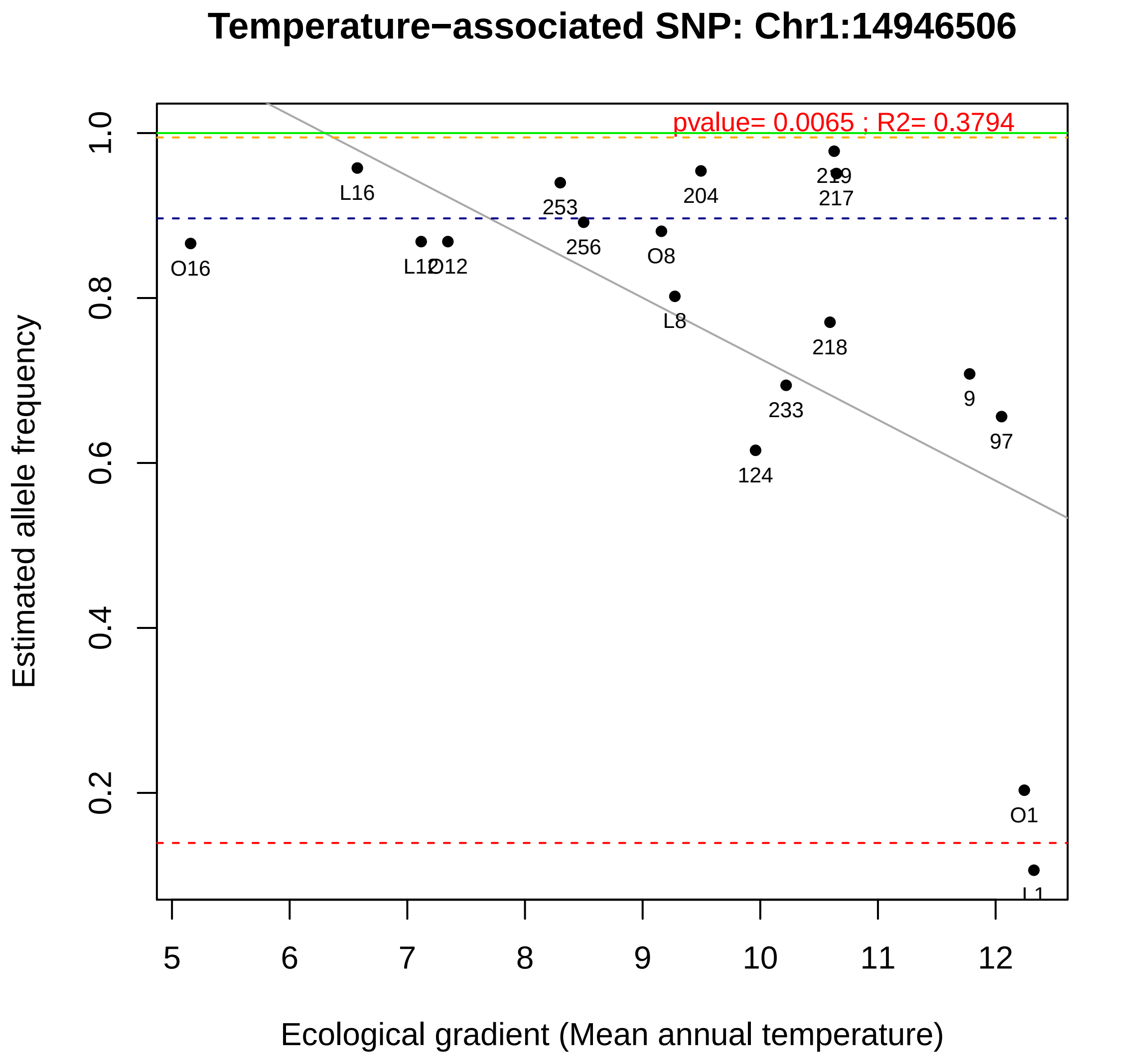
Temperature−associated SNP: Chr1:14946506
pvalue= 0.0065 ; R2= 0.3794
Estimated allele frequency
0.2
0.4
0.6
0.8
1.0
5
6
7
8
9
10
11
12
Ecological gradient (Mean annual temperature)
O16
L16
L12
O12
253
256
204
O8
L8
219
217
218
233
124
9
97
O1

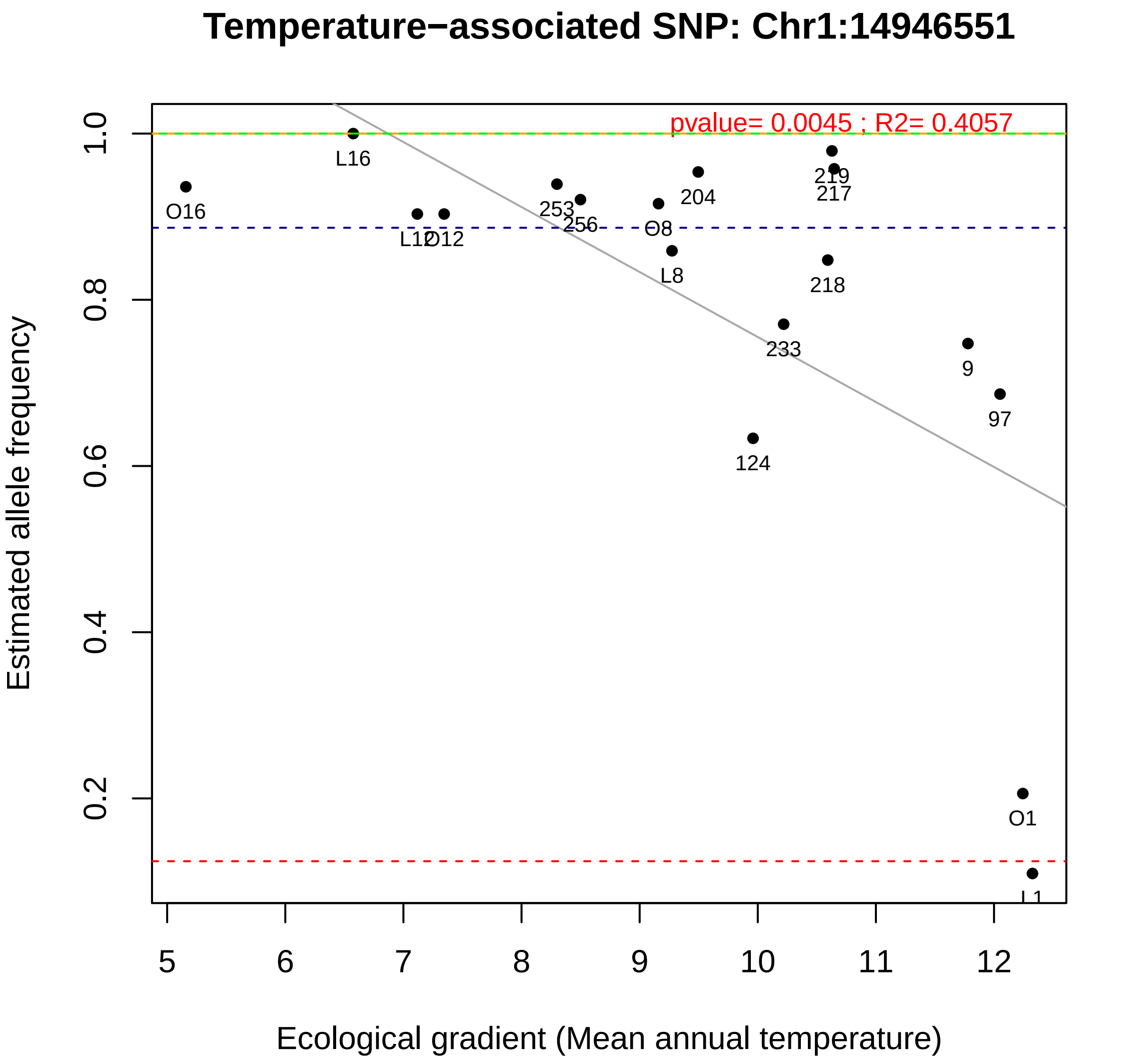
Temperature−associated SNP: Chr1:14946551
pvalue= 0.0045 ; R2= 0.4057
Estimated allele frequency
Ecological gradient (Mean annual temperature)
0.2
0.4
0.6
0.8
1.0
5
6
7
8
9
10
11
12
O16
L16
L12
O12
253
256
O8
204
L8
219
217
218
233
124
9
97
O1

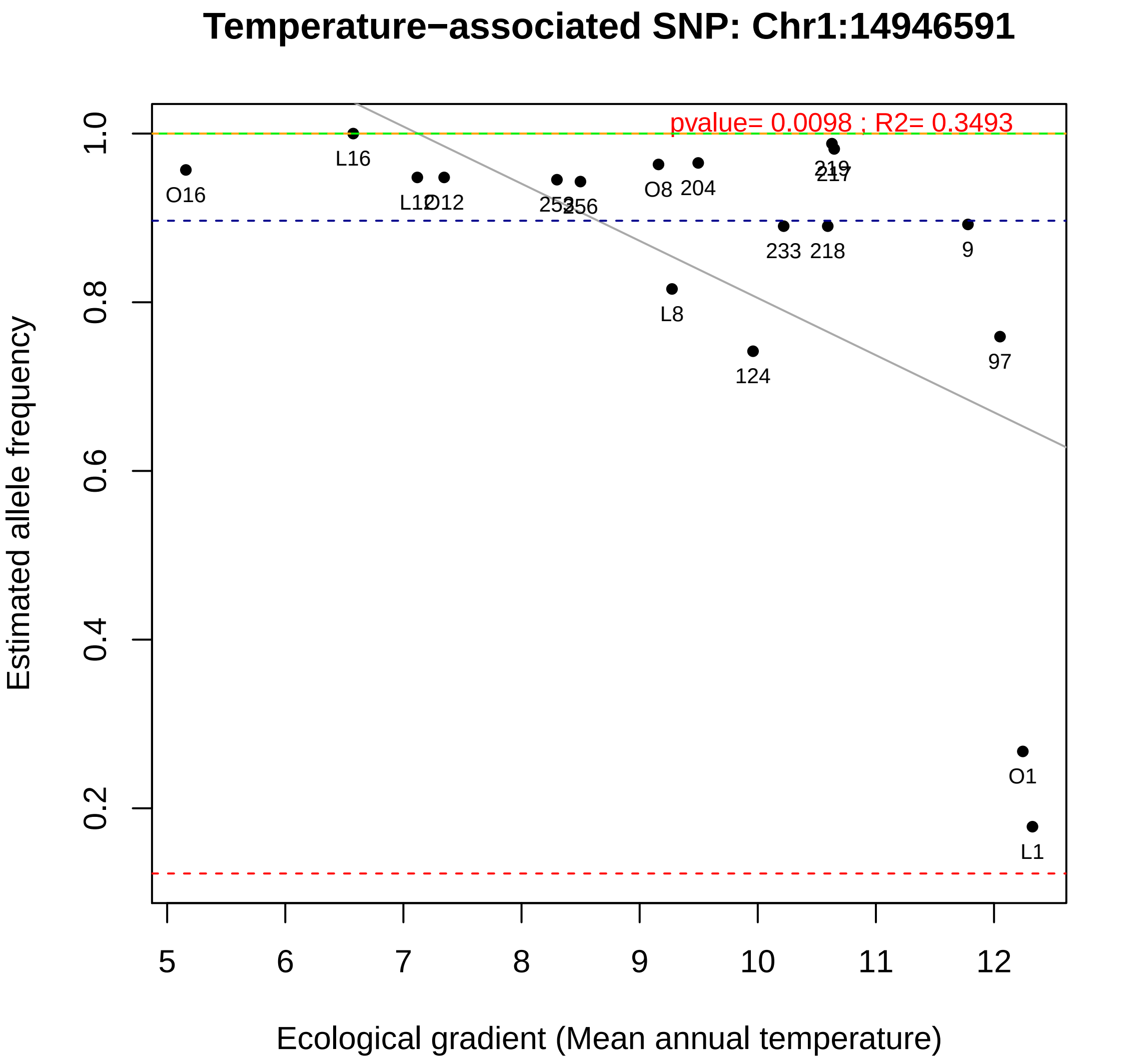
Temperature−associated SNP: Chr1:14946591
pvalue= 0.0098 ; R2= 0.3493
Estimated allele frequency
1.0
0.8
0.6
0.4
0.2
5
6
7
8
9
10
11
12
Ecological gradient (Mean annual temperature)
O16
L16
L12
O12
O8
204
219
217
233
218
9
L8
124
97
O1
L1
256

**Temperature−associated SNP: Chr1:14946622**

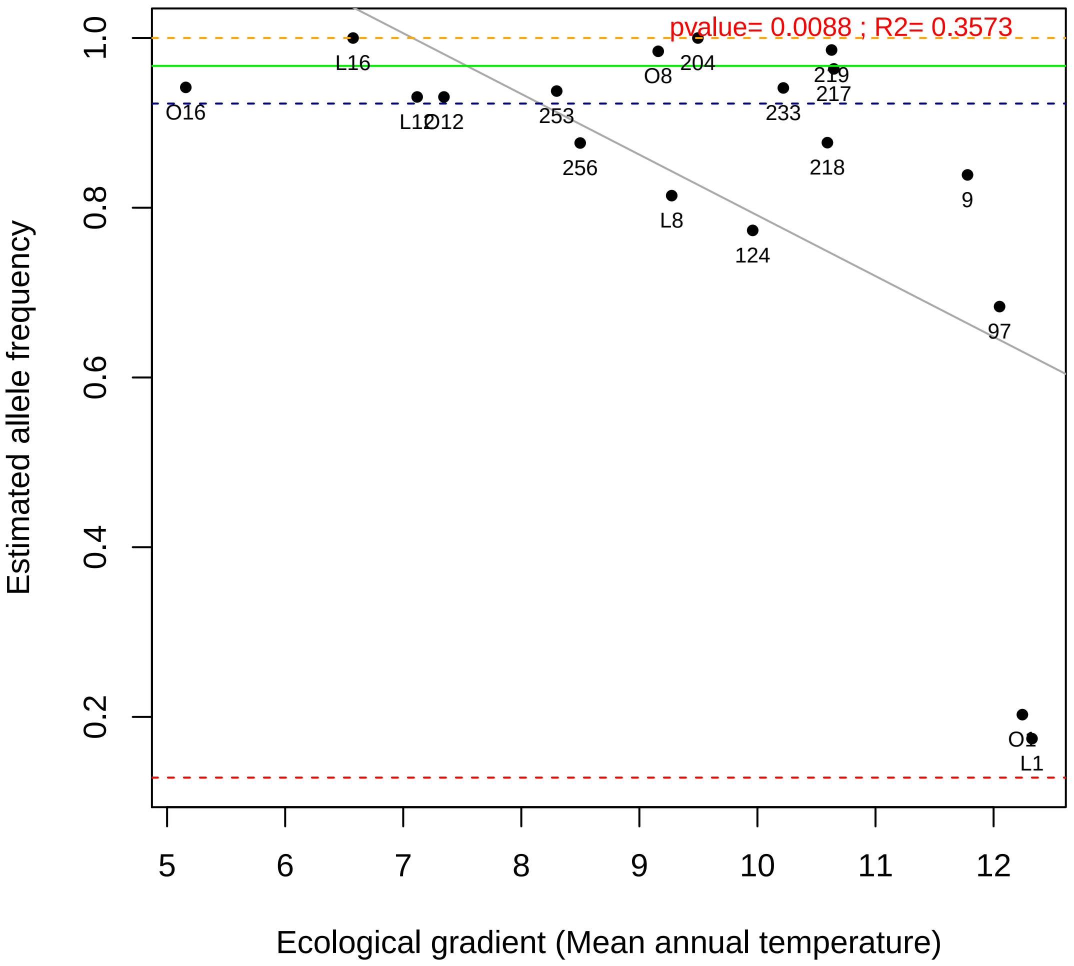

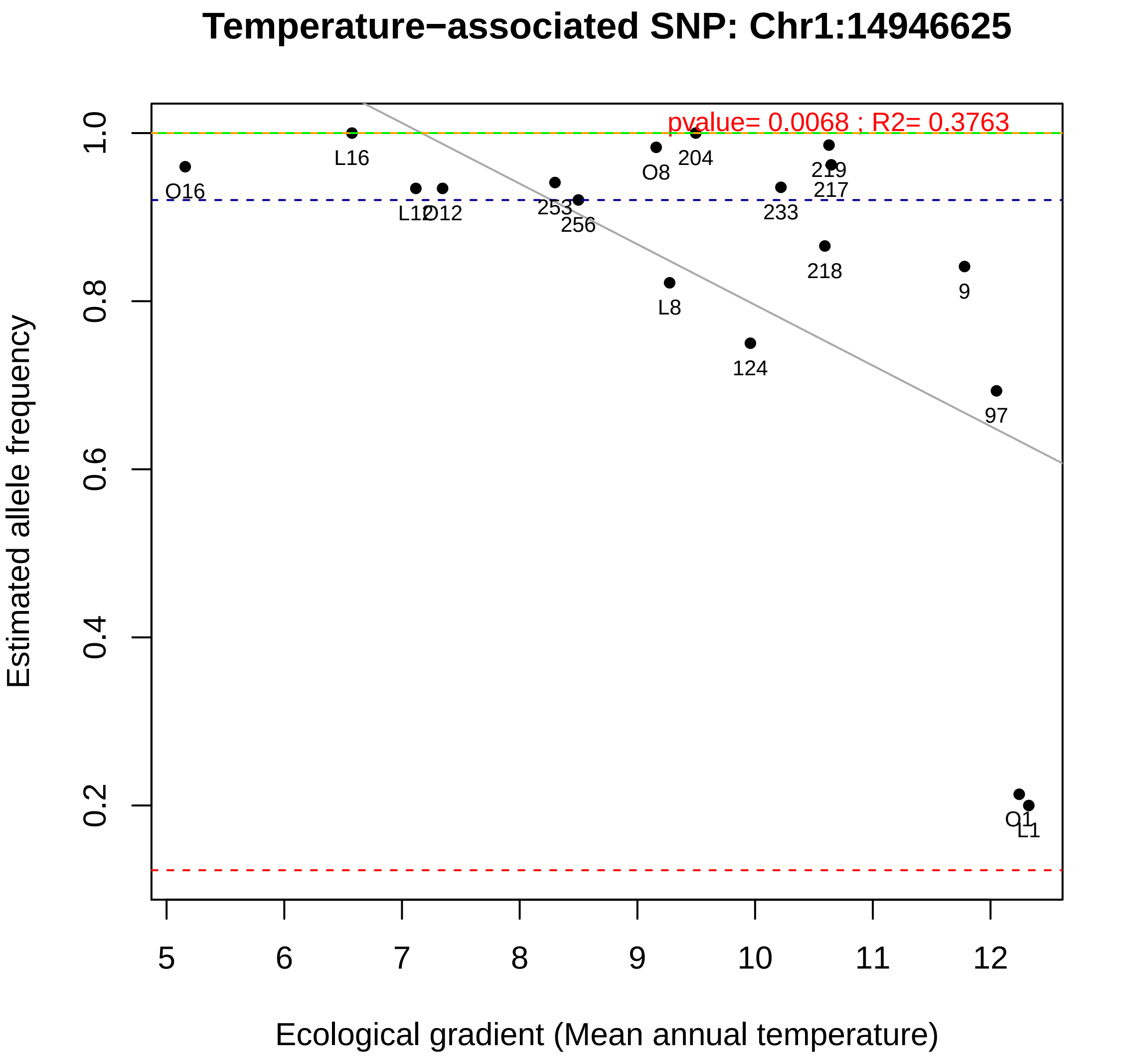

Temperature−associated SNP: Chr1:14946625
pvalue= 0.0068 ; R2= 0.3763
Estimated allele frequency
0.2
0.4
0.6
0.8
1.0
5
6
7
8
9
10
11
12
Ecological gradient (Mean annual temperature)
O16
L16
L12
O12
253
256
O8
204
L8
124
233
219
217
218
9
97
O1
L1

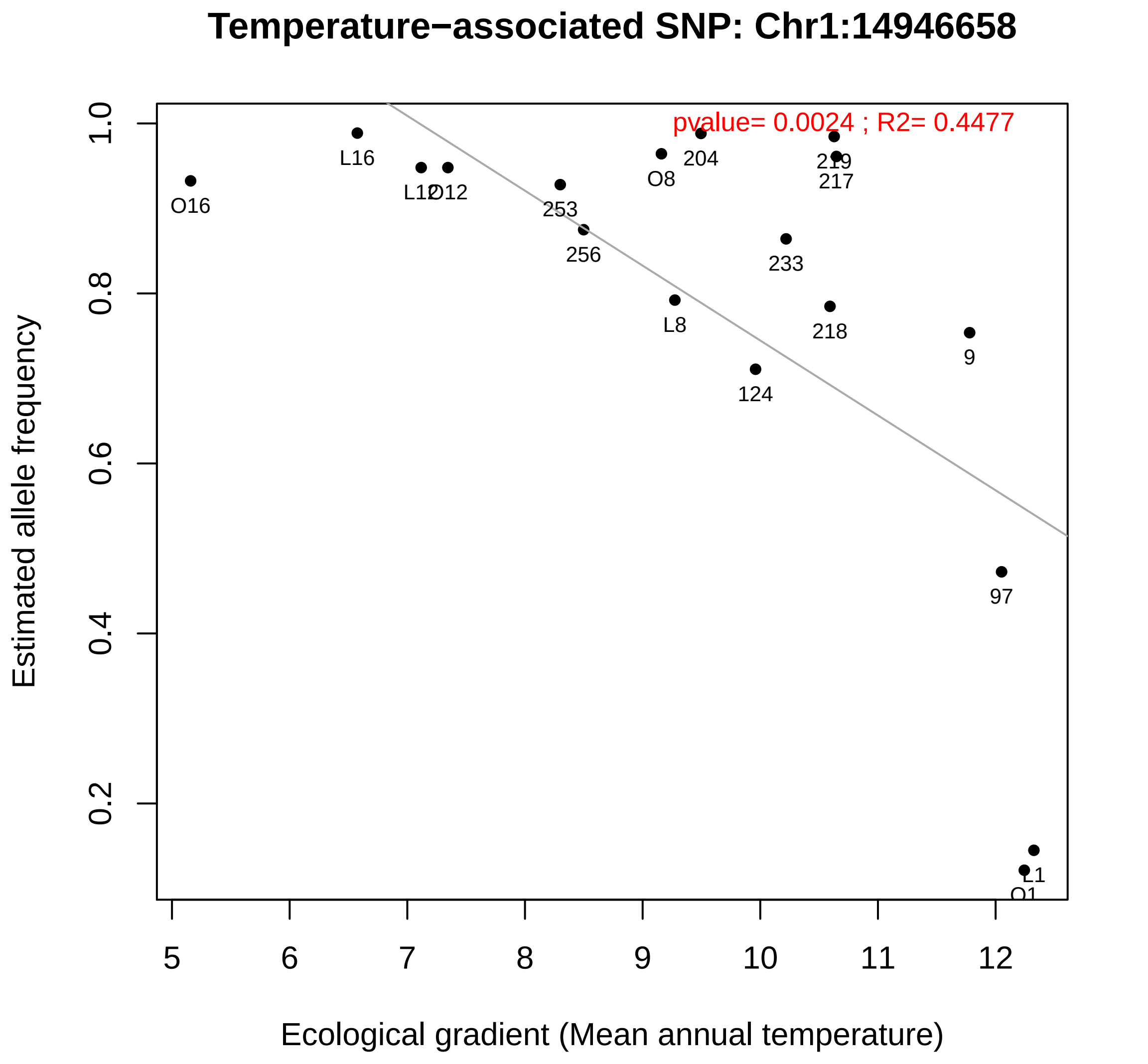

Temperature−associated SNP: Chr1:14946658
pvalue= 0.0024 ; R2= 0.4477
Estimated allele frequency
0.2
0.4
0.6
0.8
1.0
5
6
7
8
9
10
11
12
Ecological gradient (Mean annual temperature)
O16
L16
L12
O12
253
256
O8
204
L8
124
233
219
217
218
9
97
L1
O1

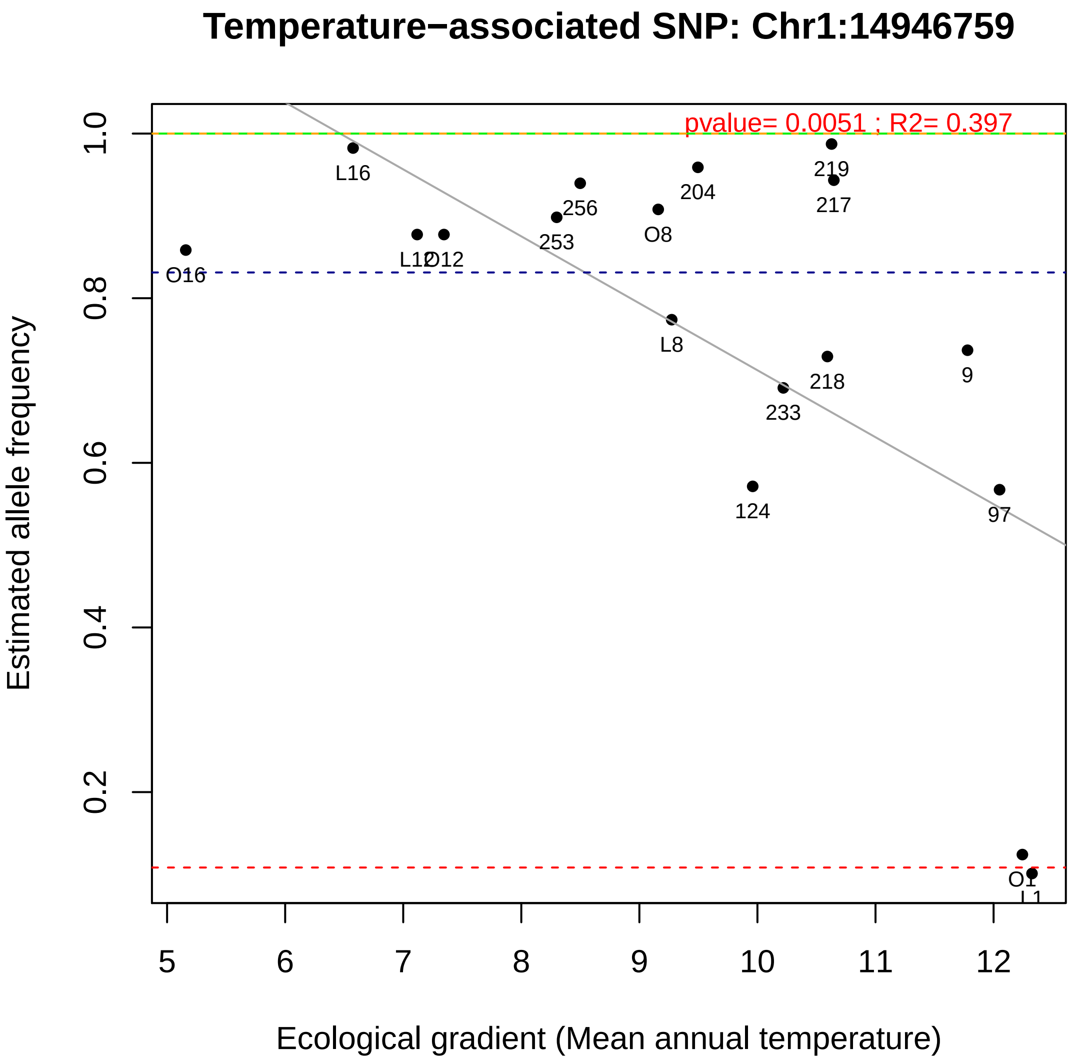
Temperature−associated SNP: Chr1:14946759
pvalue= 0.0051 ; R2= 0.397
Estimated allele frequency
Ecological gradient (Mean annual temperature)
0.2
0.4
0.6
0.8
1.0
5
6
7
8
9
10
11
12
L16
O16
L12
O12
253
256
O8
204
219
217
L8
218
9
233
124
97
O1

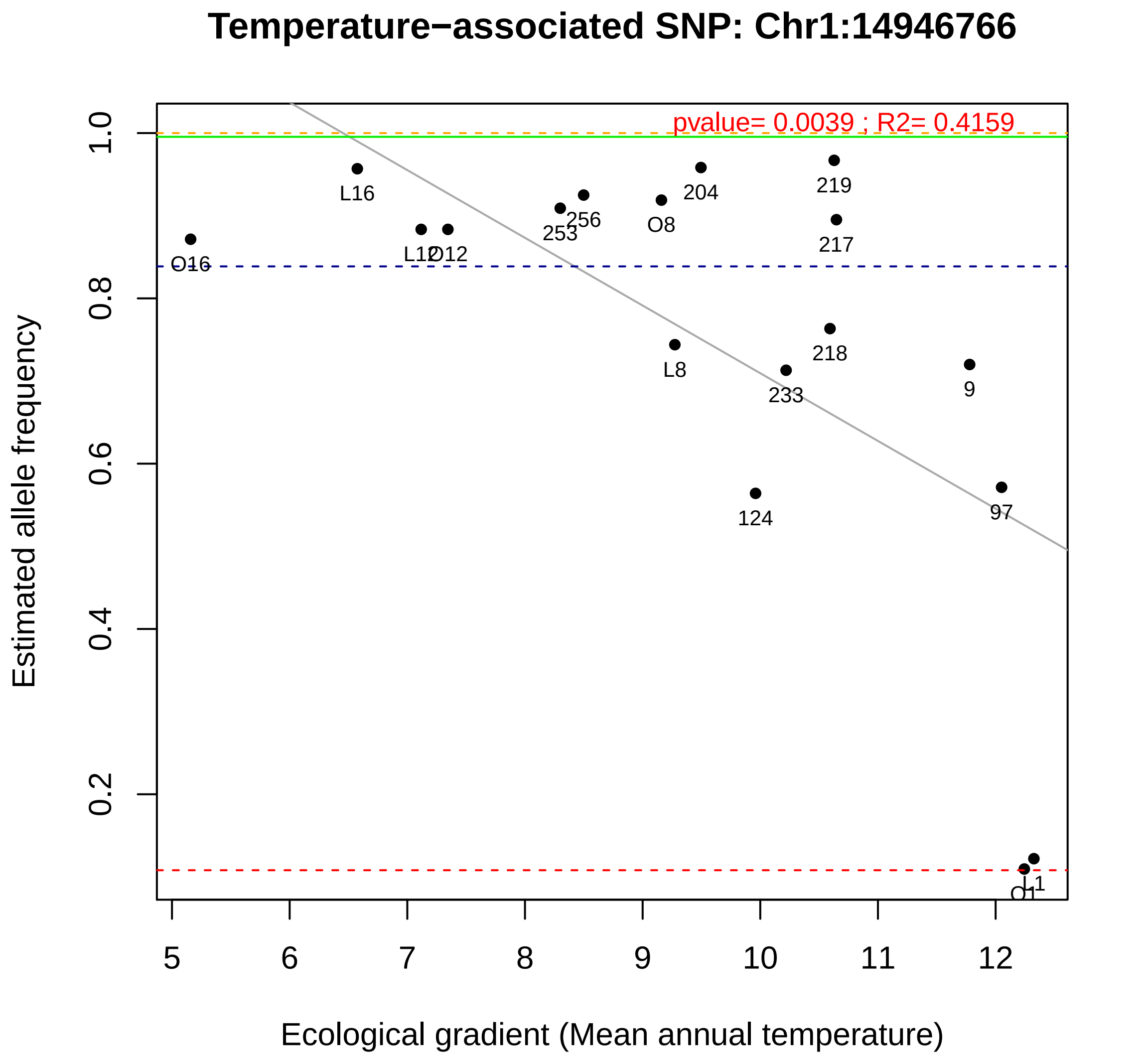

Temperature−associated SNP: Chr1:14946766
pvalue= 0.0039 ; R2= 0.4159
Estimated allele frequency
0.2
0.4
0.6
0.8
1.0
5
6
7
8
9
10
11
12
Ecological gradient (Mean annual temperature)
O16
L16
L12
O12
253
256
O8
204
219
217
L8
233
218
124
9
97
L1
O1

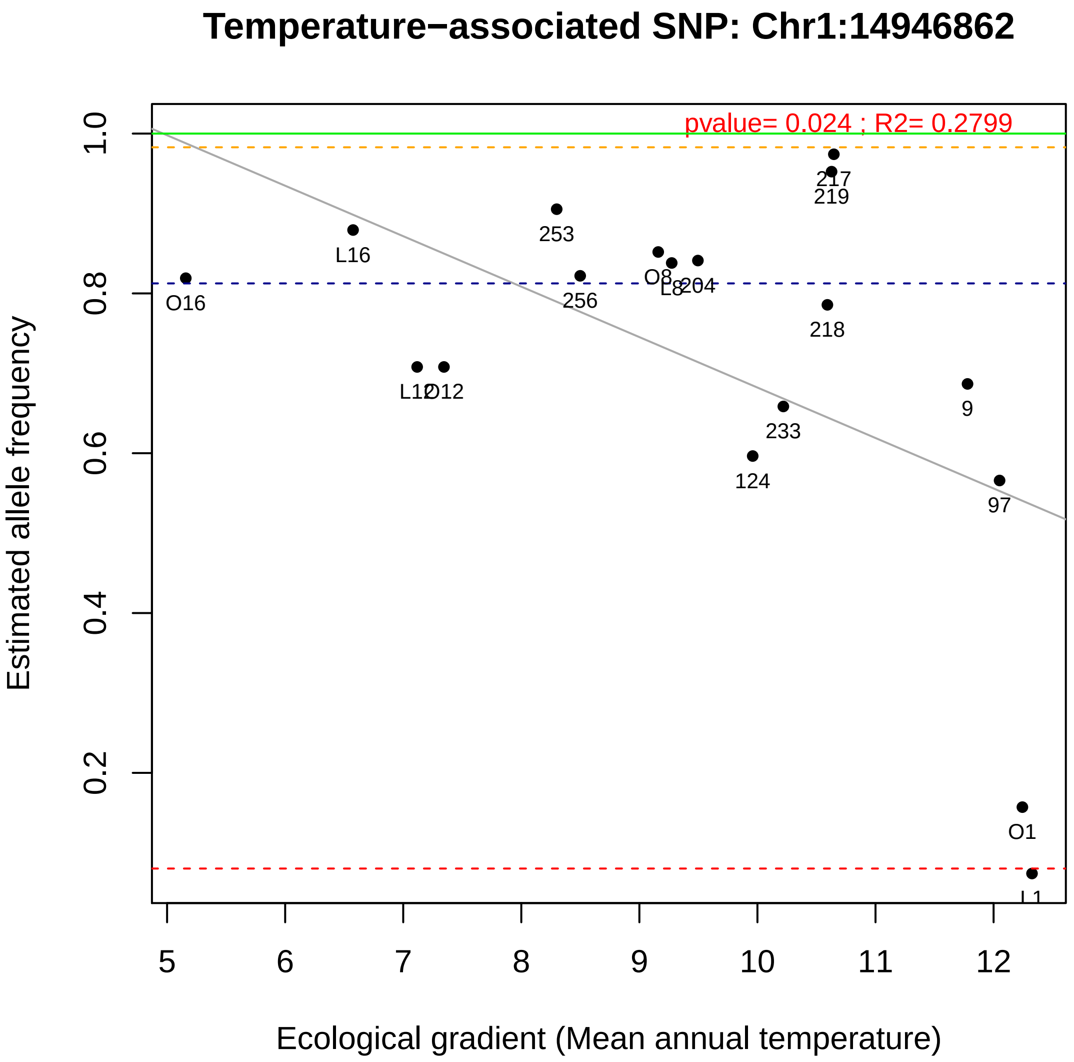

Temperature−associated SNP: Chr1:14946862
pvalue= 0.024 ; R2= 0.2799
Estimated allele frequency
Ecological gradient (Mean annual temperature)
0.2
0.4
0.6
0.8
1.0
5
6
7
8
9
10
11
12
O16
L16
L12
O12
253
256
O8
L8
204
217
219
218
233
124
9
97
O1
L1

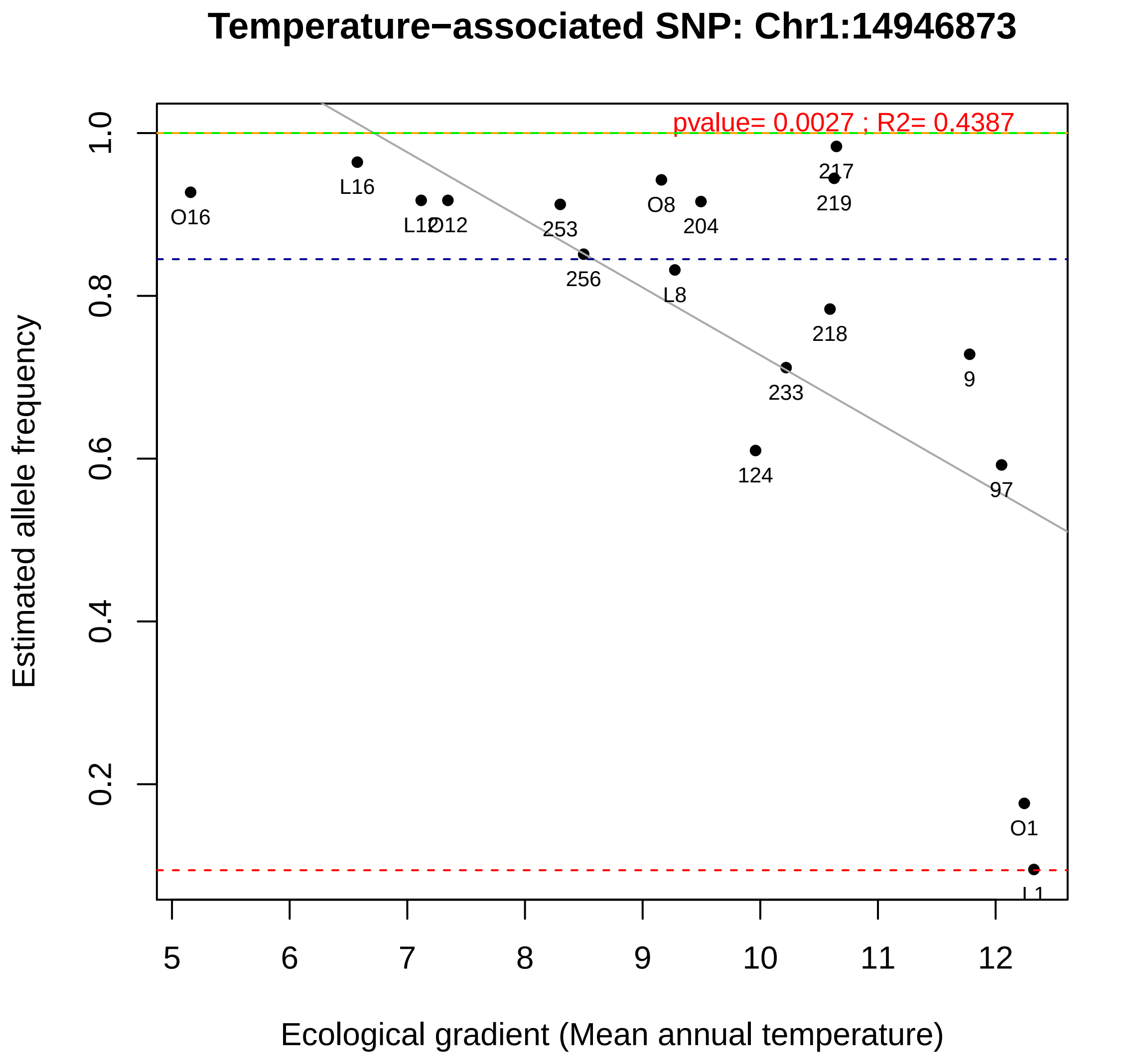

Temperature−associated SNP: Chr1:14946873
pvalue= 0.0027 ; R2= 0.4387
Estimated allele frequency
Ecological gradient (Mean annual temperature)
0.2
0.4
0.6
0.8
1.0
5
6
7
8
9
10
11
12
O16
L16
L12
O12
253
256
O8
204
L8
217
219
218
233
124
9
97
O1

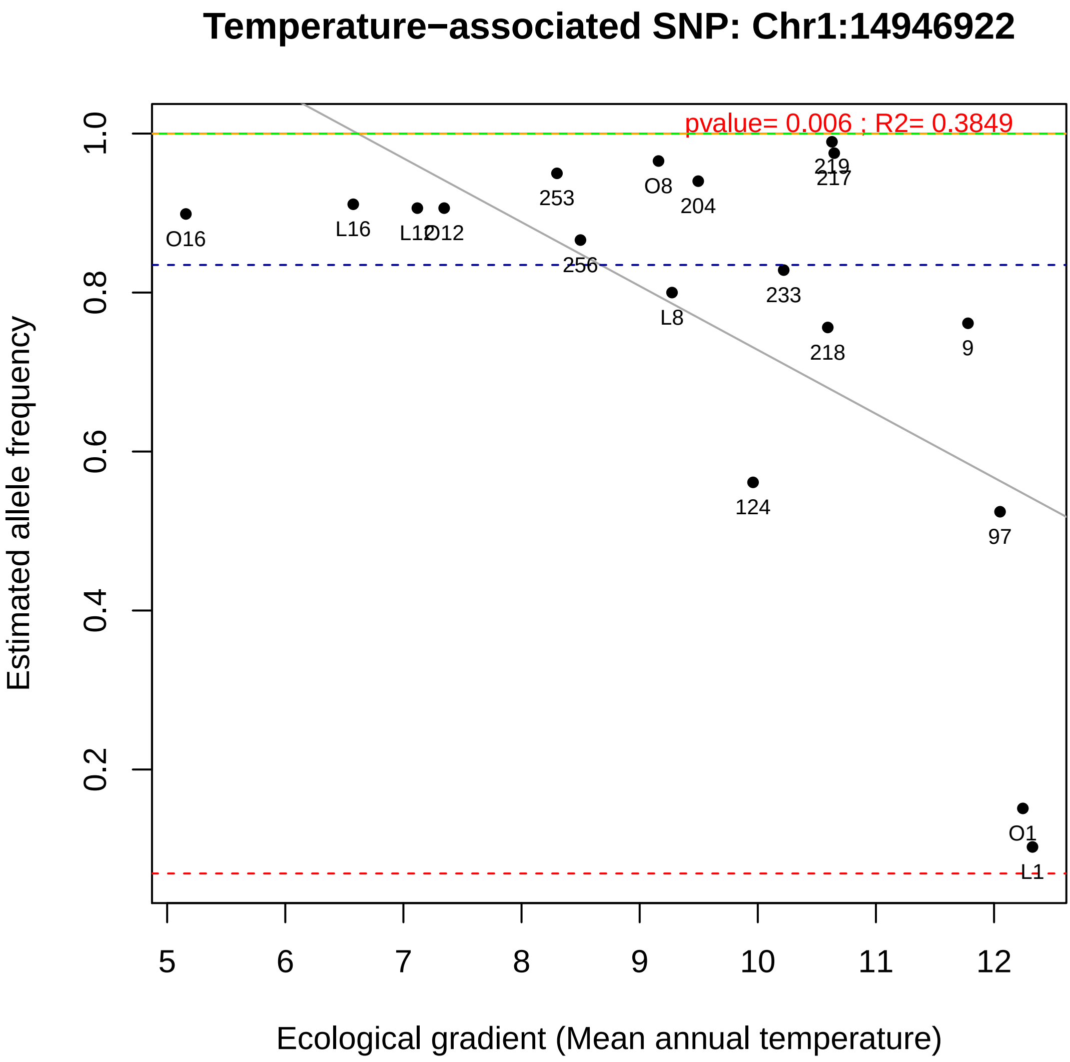

Temperature−associated SNP: Chr1:14946922
pvalue= 0.006 ; R2= 0.3849
Estimated allele frequency
0.2
0.4
0.6
0.8
1.0
5
6
7
8
9
10
11
12
Ecological gradient (Mean annual temperature)
O16
L16
L12
O12
253
256
O8
204
L8
219
217
233
218
9
124
97
O1
L1

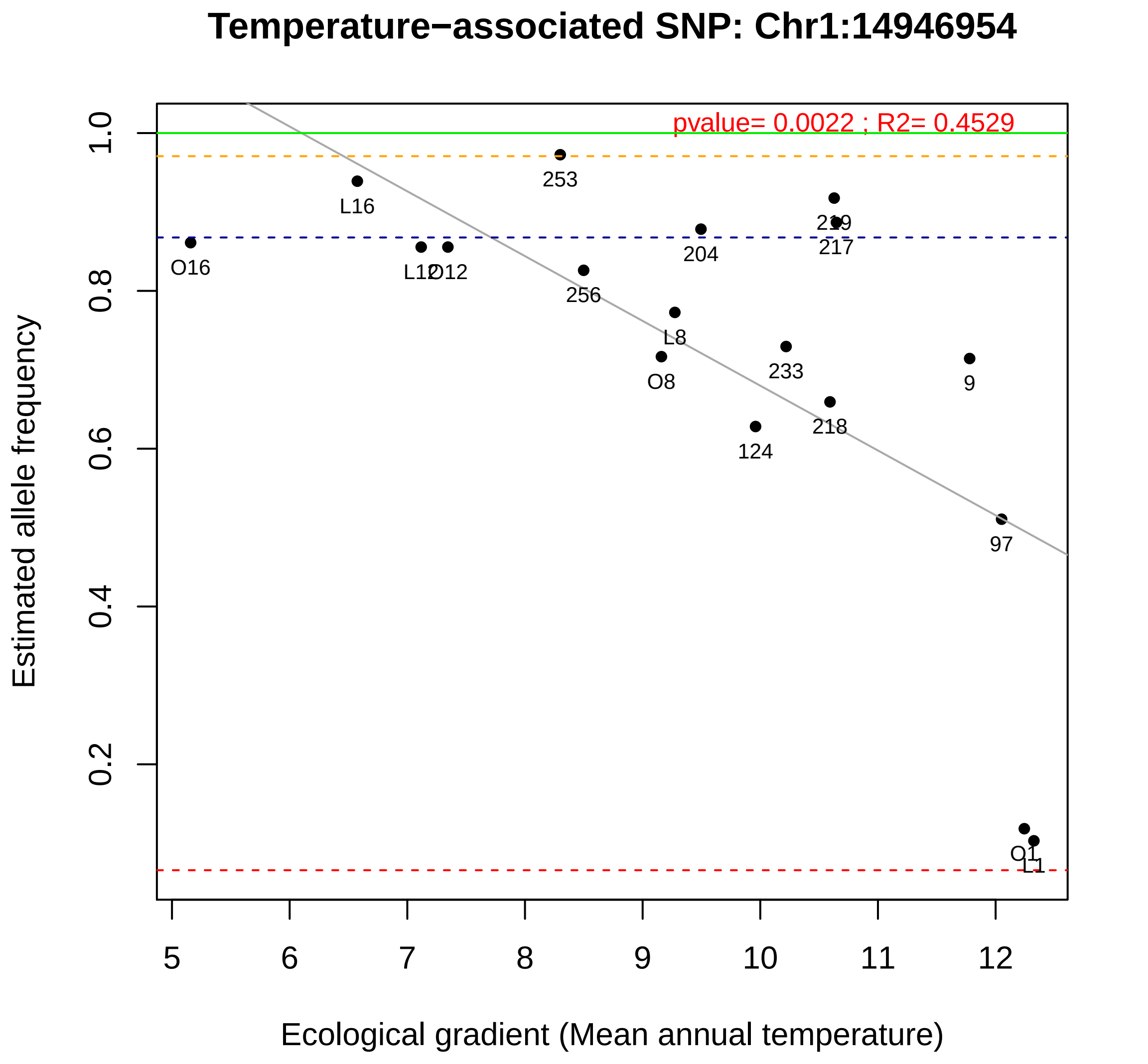
Temperature−associated SNP: Chr1:14946954
pvalue= 0.0022 ; R2= 0.4529
Estimated allele frequency
0.2
0.4
0.6
0.8
1.0
5
6
7
8
9
10
11
12
Ecological gradient (Mean annual temperature)
O16
L16
L12
O12
253
256
204
219
217
L8
O8
233
124
218
9
97
O1
L1

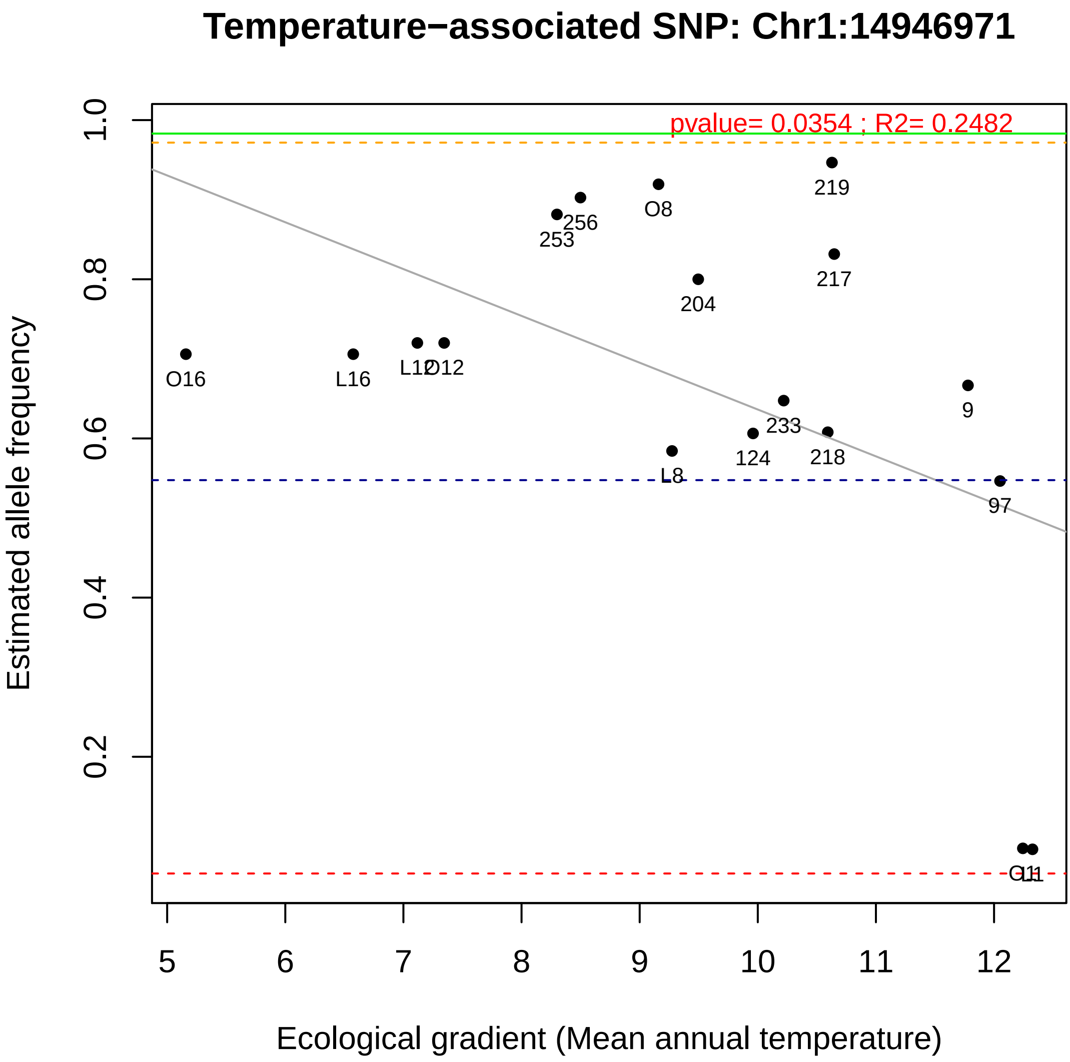

Temperature−associated SNP: Chr1:14946971
pvalue= 0.0354 ; R2= 0.2482
Estimated allele frequency
Ecological gradient (Mean annual temperature)
0.2
0.4
0.6
0.8
1.0
5
6
7
8
9
10
11
12
O16
L16
L12
O12
253
256
O8
219
204
217
233
124
218
L8
9
97
O11
L11

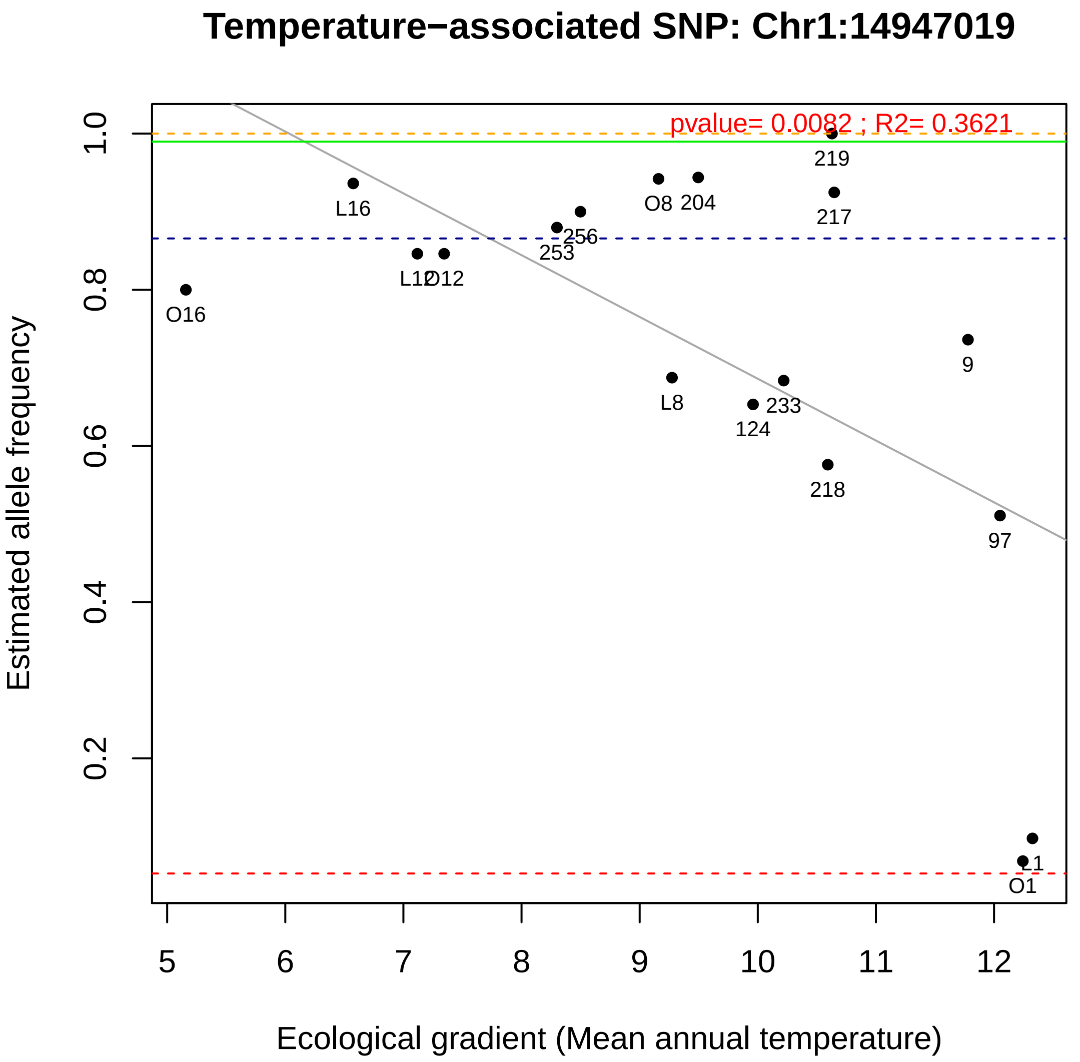

Temperature−associated SNP: Chr1:14947019
pvalue= 0.0082 ; R2= 0.3621
Estimated allele frequency
Ecological gradient (Mean annual temperature)
0.2
0.4
0.6
0.8
1.0
5
6
7
8
9
10
11
12
O16
L16
L12
O12
253
256
O8
204
219
217
L8
124
233
218
9
97
L1
O1

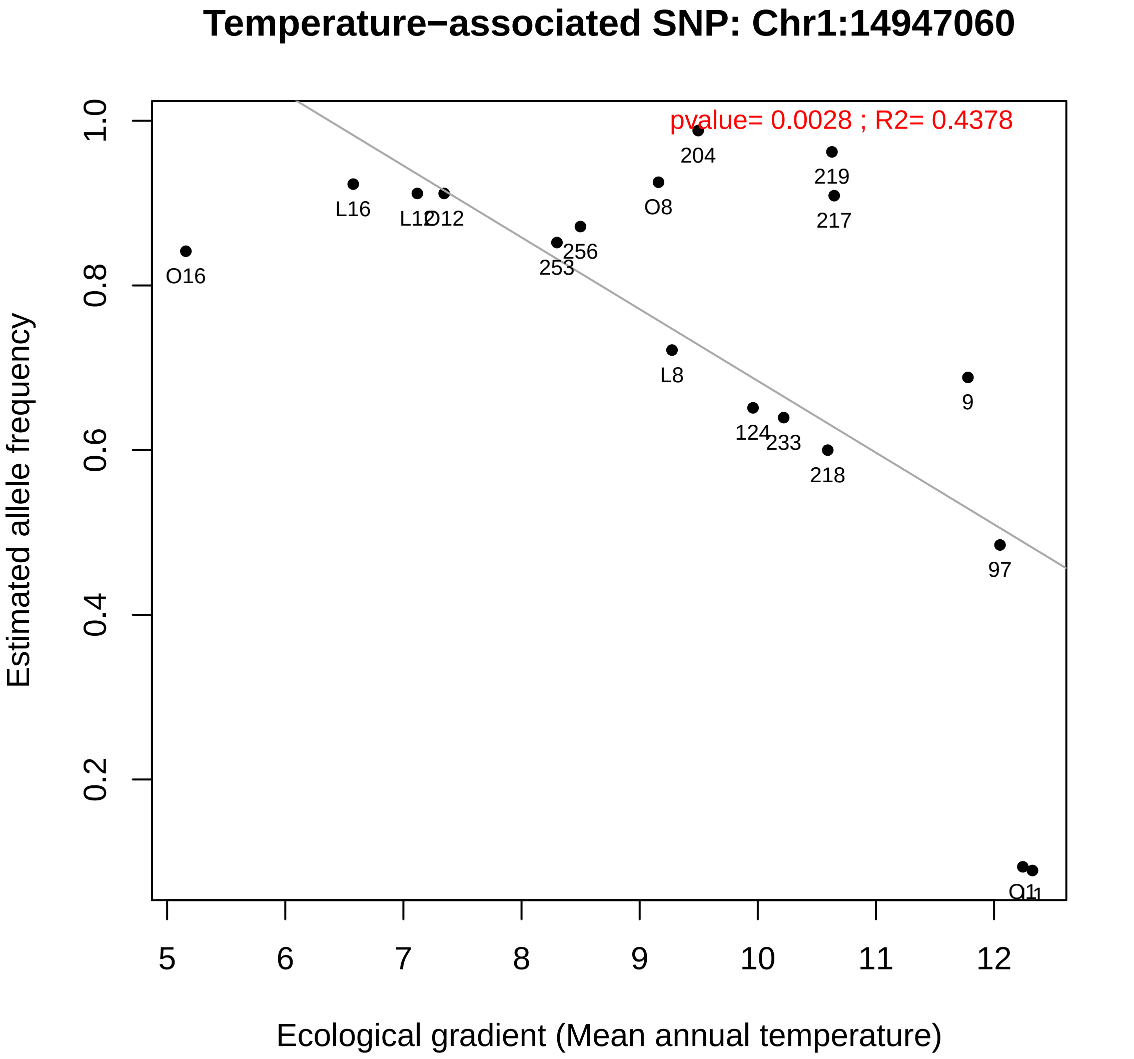
Temperature−associated SNP: Chr1:14947060
pvalue= 0.0028 ; R2= 0.4378
Estimated allele frequency
0.2
0.4
0.6
0.8
1.0
Ecological gradient (Mean annual temperature)
5
6
7
8
9
10
11
12
O16
L16
L12
O12
253
256
O8
204
L8
124
233
219
217
218
9
97

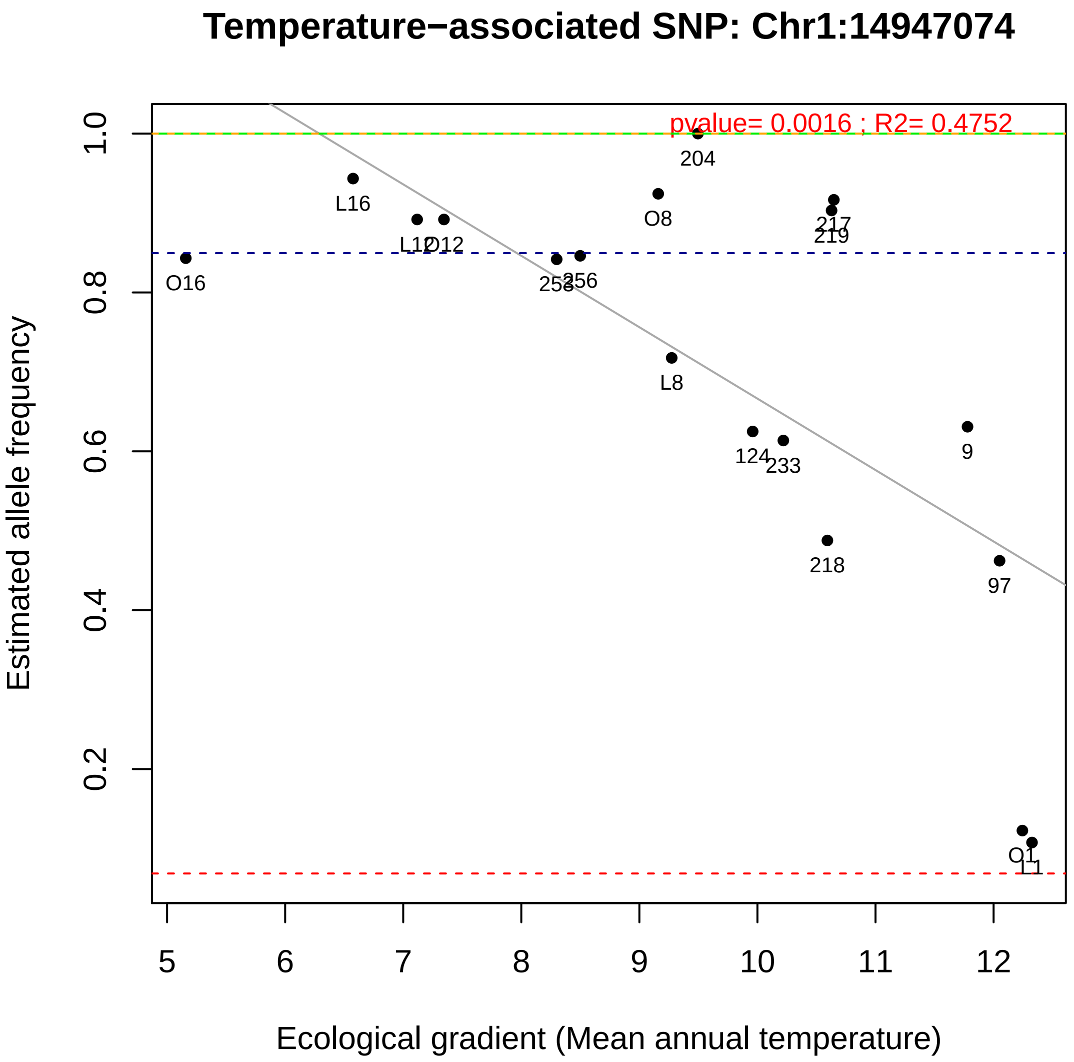
Temperature−associated SNP: Chr1:14947074
pvalue= 0.0016 ; R2= 0.4752
Estimated allele frequency
Ecological gradient (Mean annual temperature)
0.2
0.4
0.6
0.8
1.0
5
6
7
8
9
10
11
12
O16
L16
L12
O12
253
256
O8
204
L8
124
233
217
219
218
9
97
O1
L1

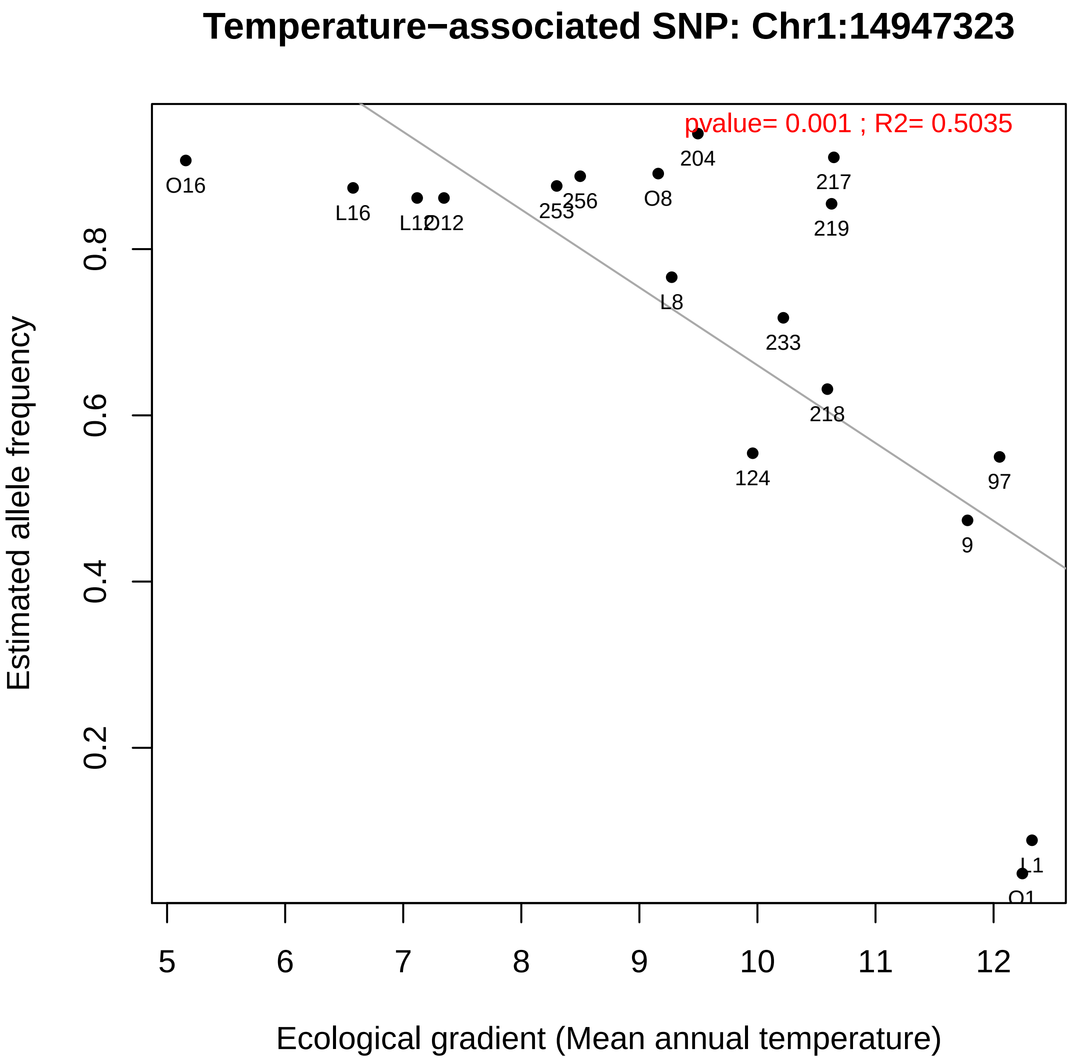
Temperature−associated SNP: Chr1:14947323
pvalue= 0.001 ; R2= 0.5035
Estimated allele frequency
0.2
0.4
0.6
0.8
Ecological gradient (Mean annual temperature)
5
6
7
8
9
10
11
12
O16
L16
L12
O12
253
256
O8
204
217
219
L8
233
218
124
97
9
L1
O1

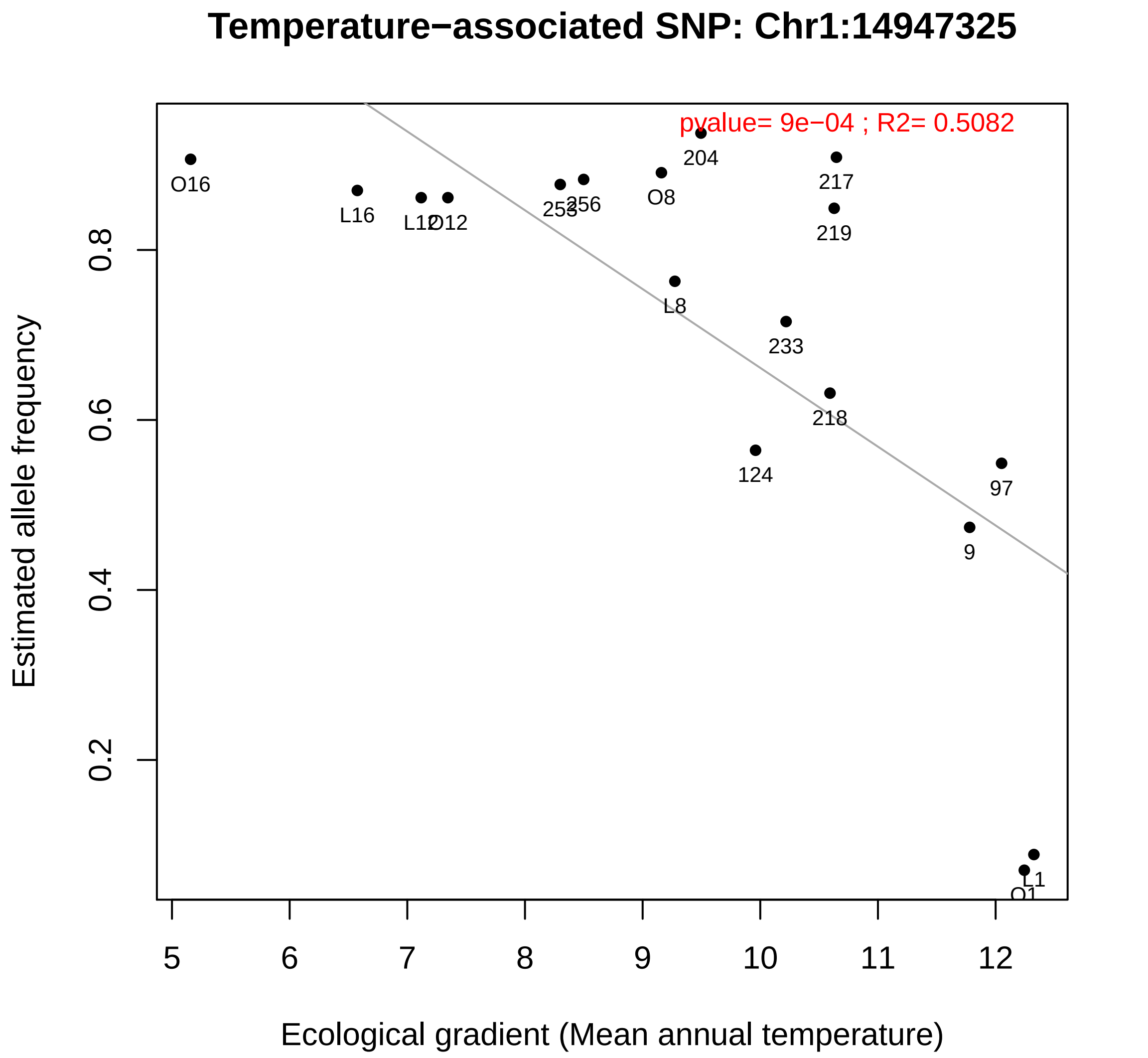

Temperature−associated SNP: Chr1:14947325
pvalue= 9e−04 ; R2= 0.5082
Estimated allele frequency
0.2
0.4
0.6
0.8
Ecological gradient (Mean annual temperature)
5
6
7
8
9
10
11
12
O16
L16
L12
O12
253
256
O8
204
217
219
L8
233
218
124
97
9
L1
O1

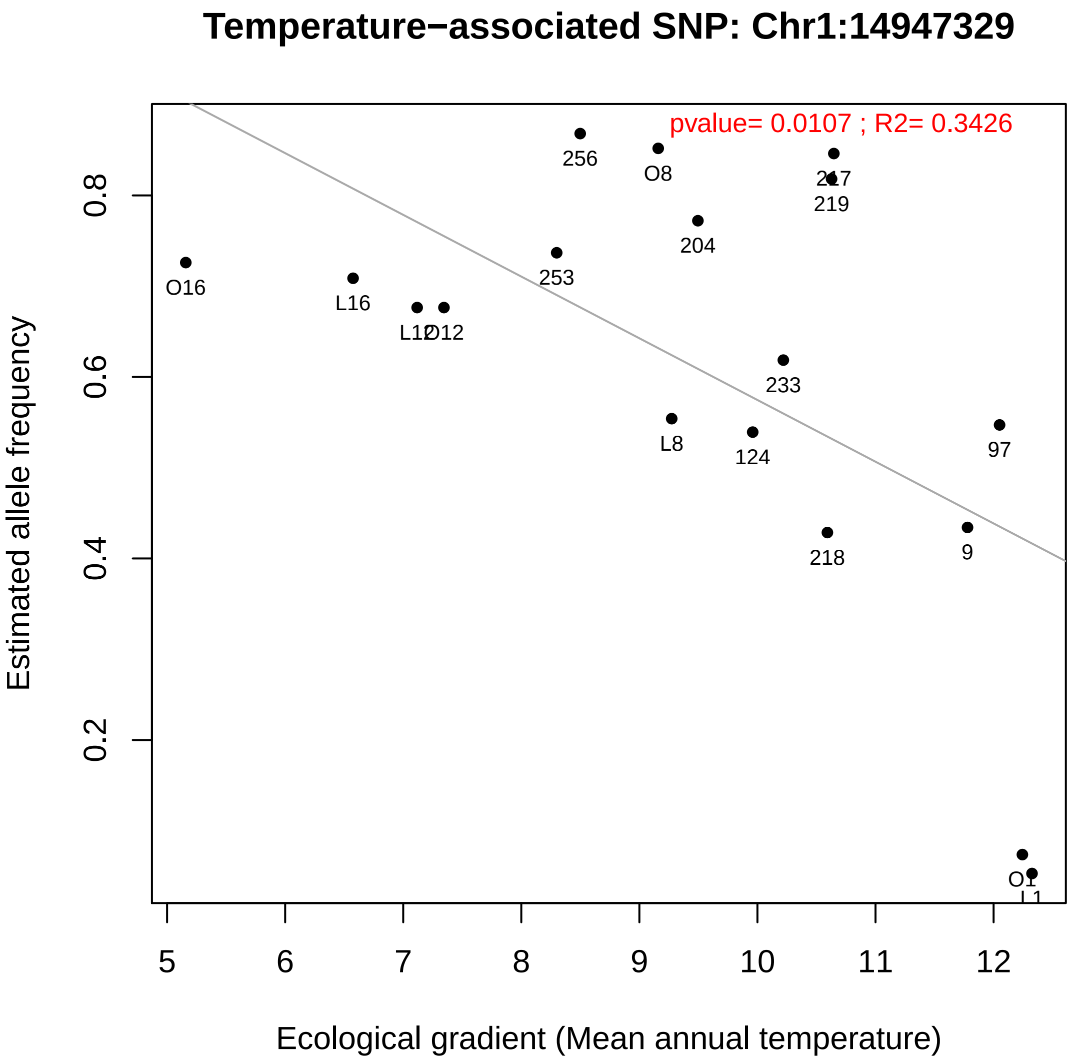

Temperature−associated SNP: Chr1:14947329
pvalue= 0.0107 ; R2= 0.3426
Estimated allele frequency
0.2
0.4
0.6
0.8
5
6
7
8
9
10
11
12
Ecological gradient (Mean annual temperature)
O16
L16
L12
O12
253
256
O8
217
219
204
233
L8
124
97
218
9
O1

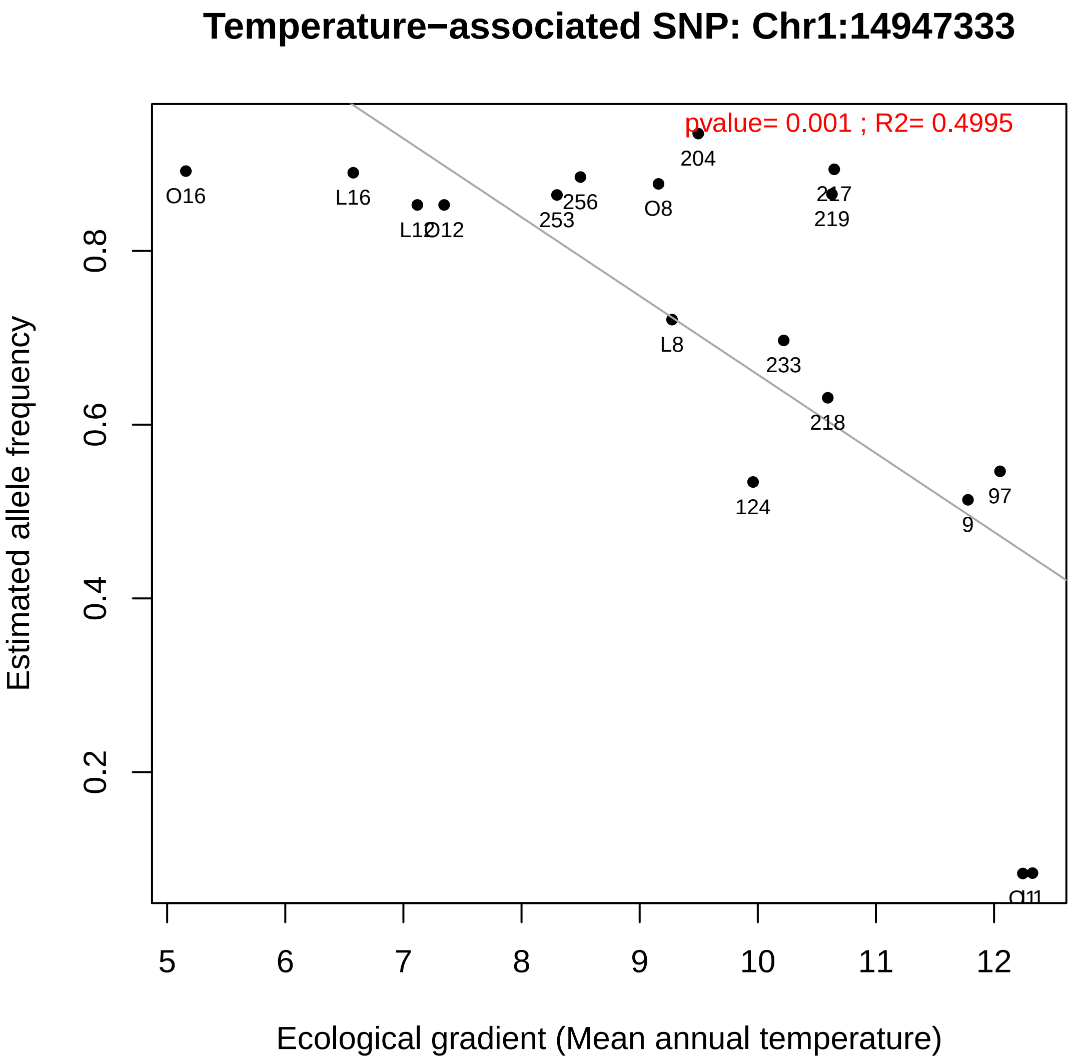
Temperature−associated SNP: Chr1:14947333
pvalue= 0.001 ; R2= 0.4995
O16
L16
L12
O12
253
256
O8
204
217
219
L8
233
218
124
9
97
0.2
0.4
0.6
0.8
5
6
7
8
9
10
11
12
Estimated allele frequency
Ecological gradient (Mean annual temperature)

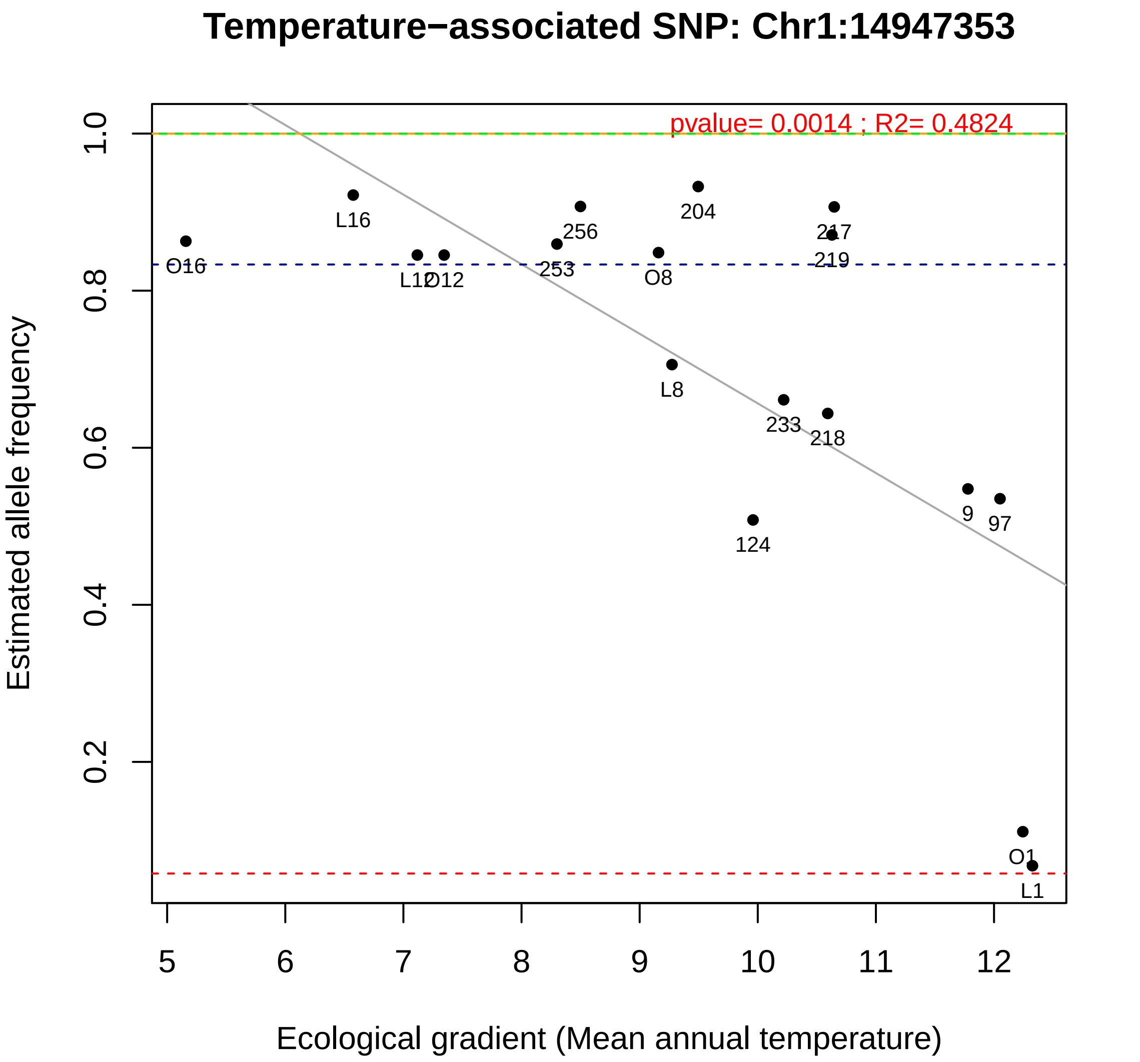
Temperature−associated SNP: Chr1:14947353
pvalue= 0.0014 ; R2= 0.4824
Estimated allele frequency
Ecological gradient (Mean annual temperature)
0.2
0.4
0.6
0.8
1.0
5
6
7
8
9
10
11
12
O16
L16
L12
O12
253
256
O8
204
217
219
L8
233
218
124
9
97
O1
L1

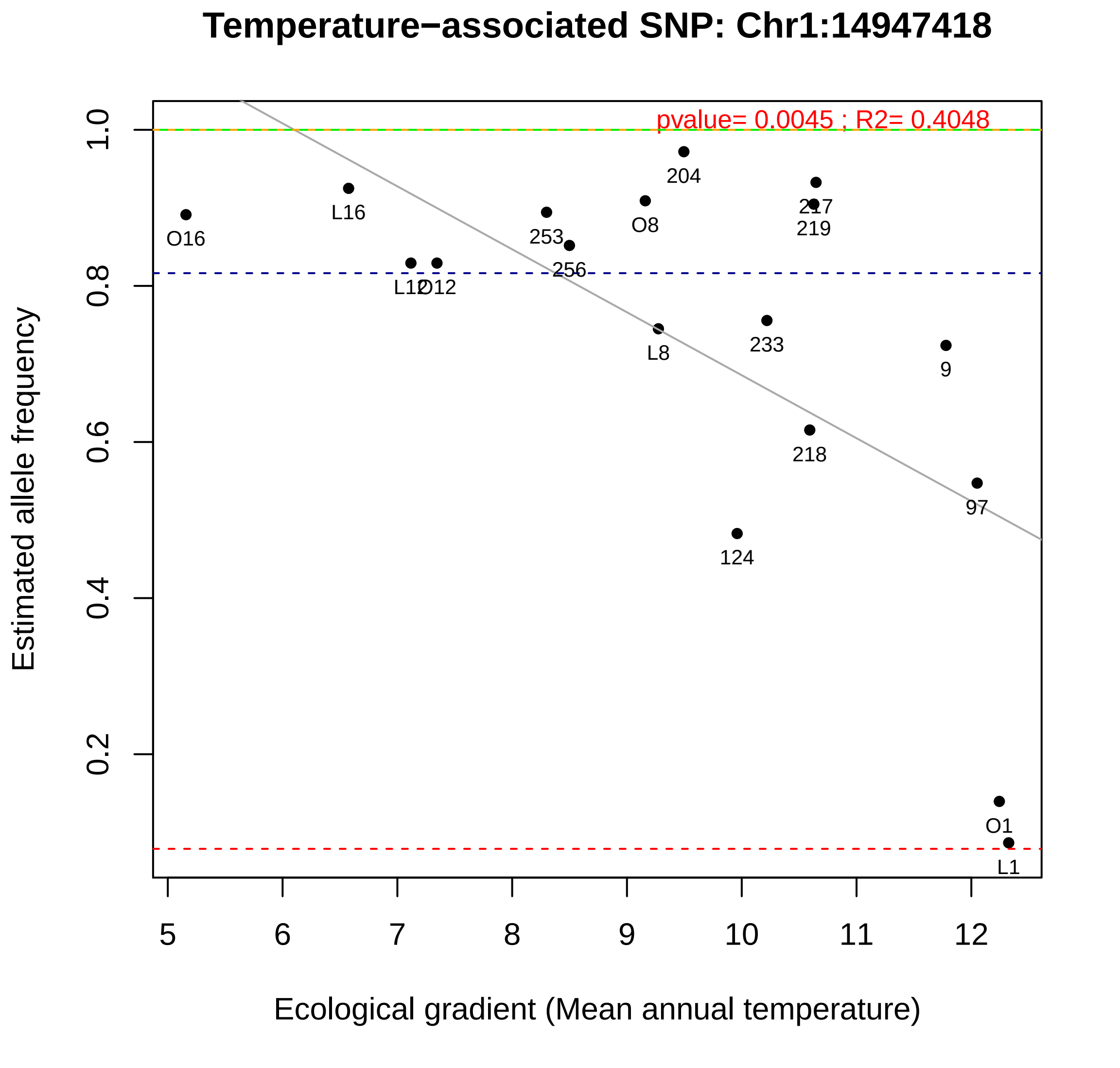

Temperature−associated SNP: Chr1:14947418
pvalue= 0.0045 ; R2= 0.4048
Estimated allele frequency
0.2
0.4
0.6
0.8
1.0
5
6
7
8
9
10
11
12
Ecological gradient (Mean annual temperature)
O16
L16
L12
O12
253
256
O8
204
L8
217
219
233
124
218
9
97
O1
L1

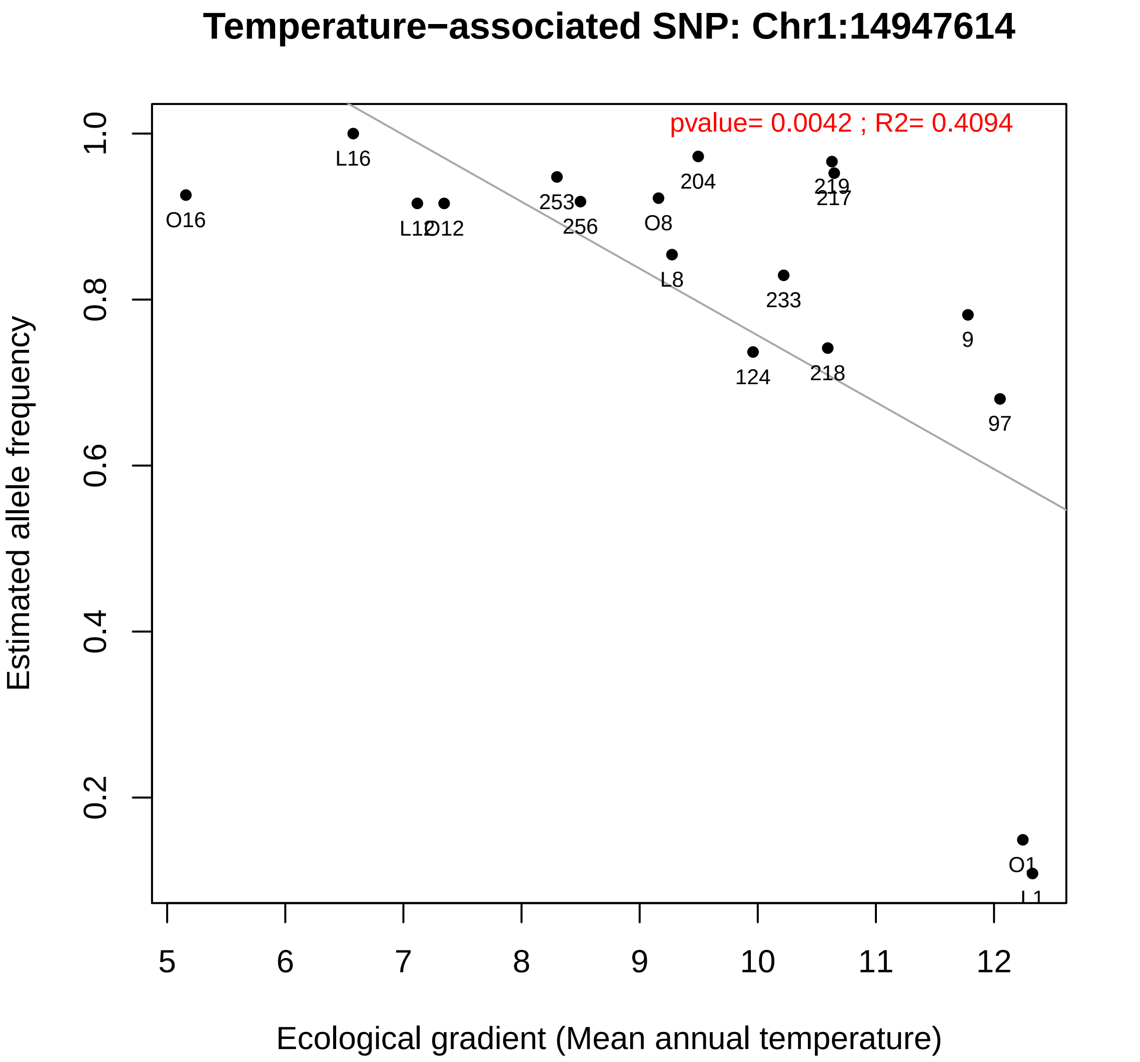

Temperature−associated SNP: Chr1:14947614
pvalue= 0.0042 ; R2= 0.4094
Estimated allele frequency
Ecological gradient (Mean annual temperature)
0.2
0.4
0.6
0.8
1.0
5
6
7
8
9
10
11
12
O16
L16
L12
O12
253
256
O8
204
L8
219
217
233
124
218
9
97
O1

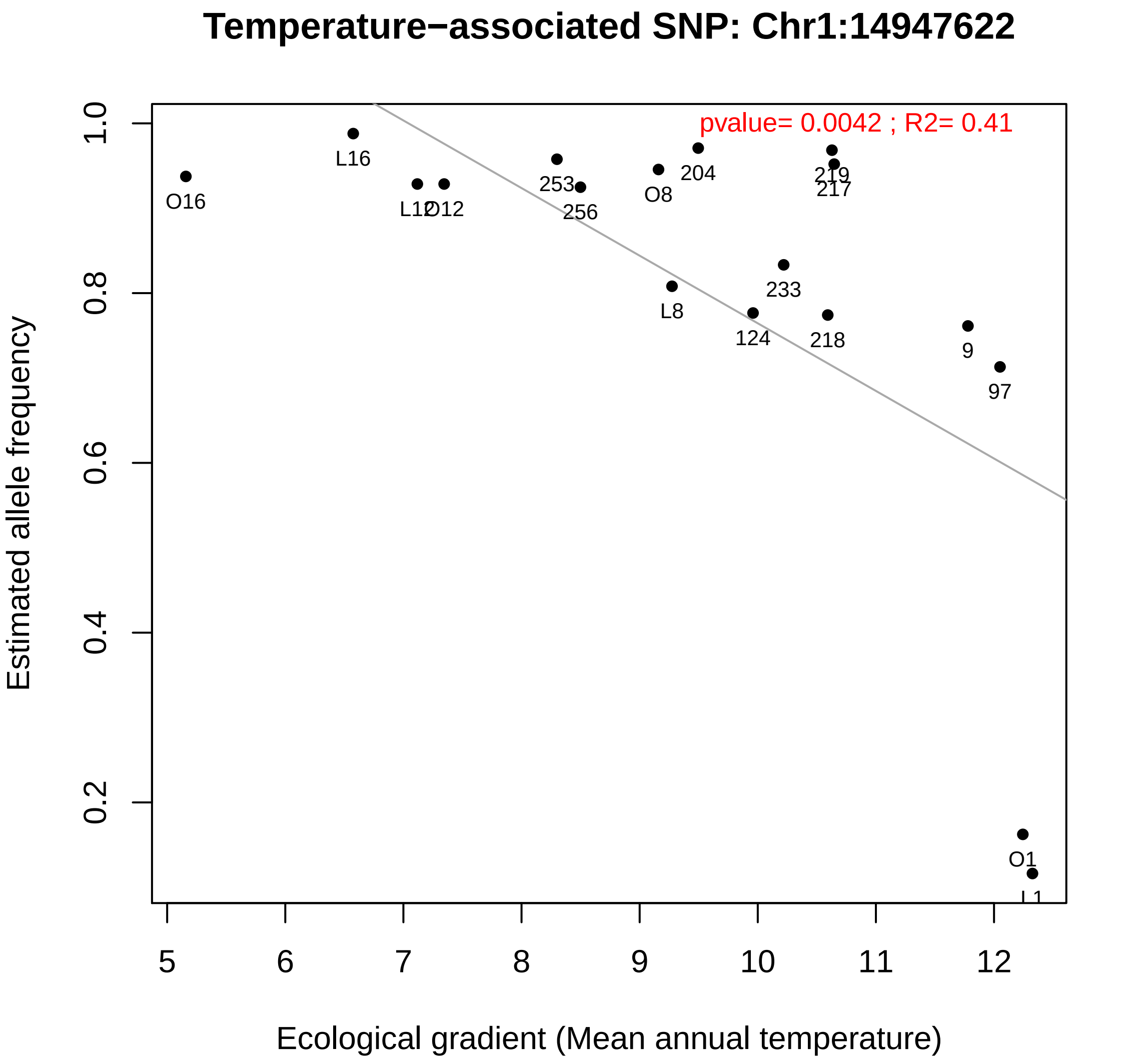
Temperature−associated SNP: Chr1:14947622
pvalue= 0.0042 ; R2= 0.41
Estimated allele frequency
Ecological gradient (Mean annual temperature)
0.2
0.4
0.6
0.8
1.0
5
6
7
8
9
10
11
12
O16
L16
L12
O12
253
256
O8
204
219
217
L8
233
124
218
9
97
O1

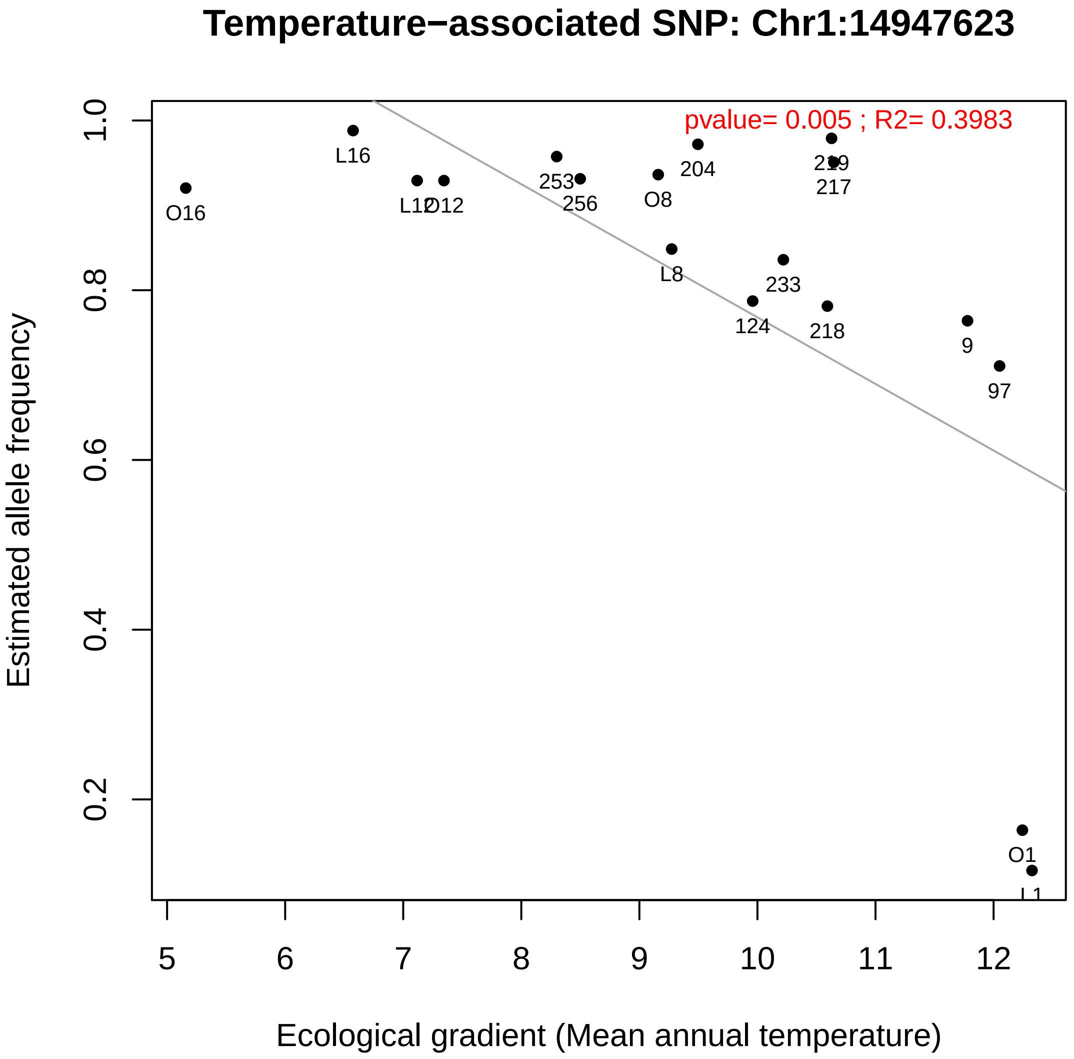

Temperature−associated SNP: Chr1:14947623
pvalue= 0.005 ; R2= 0.3983
Estimated allele frequency
0.2
0.4
0.6
0.8
1.0
Ecological gradient (Mean annual temperature)
5
6
7
8
9
10
11
12
O16
L16
L12
O12
253
256
O8
204
219
217
L8
233
124
218
9
97
O1

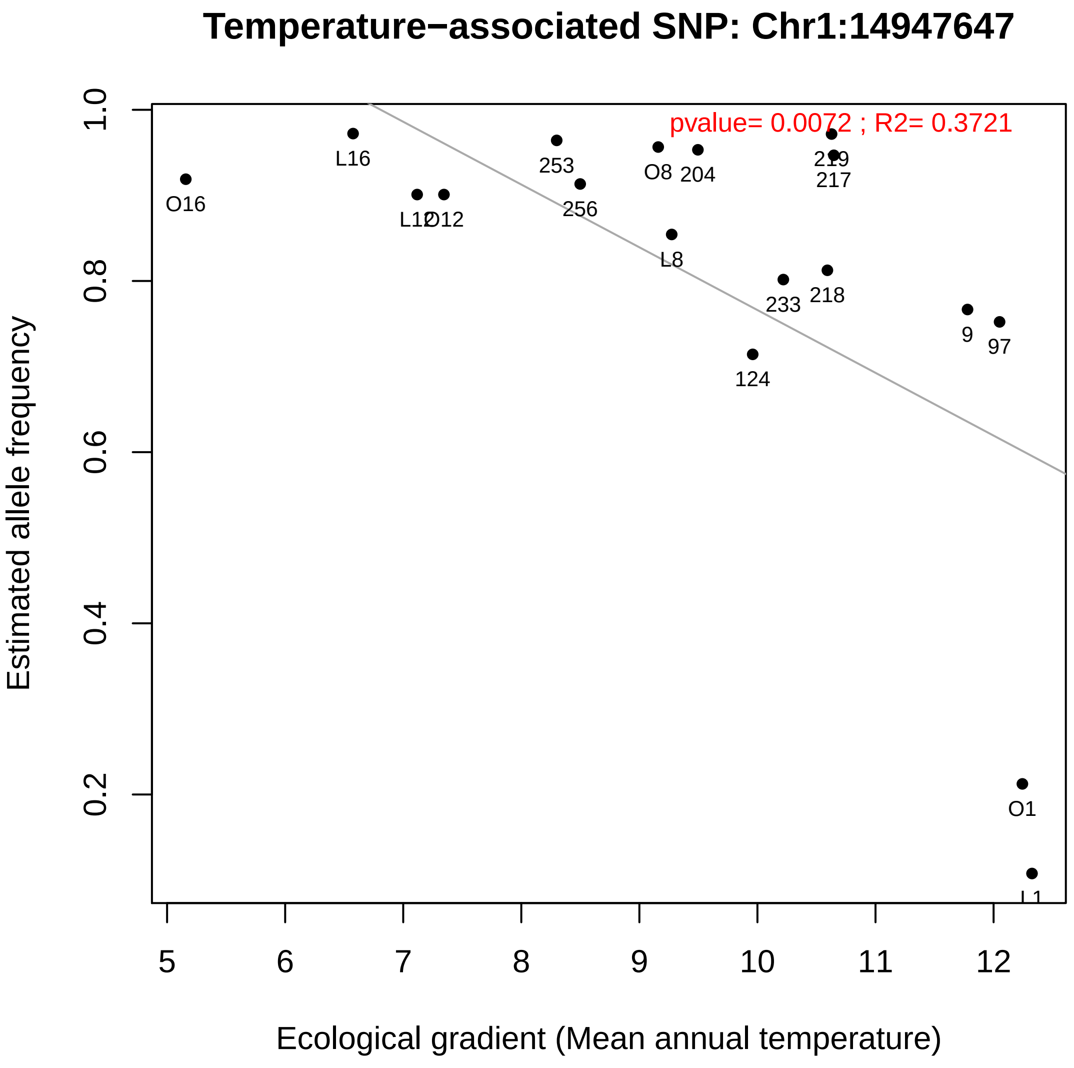

Temperature−associated SNP: Chr1:14947647
pvalue= 0.0072 ; R2= 0.3721
Estimated allele frequency
Ecological gradient (Mean annual temperature)
0.2
0.4
0.6
0.8
1.0
5
6
7
8
9
10
11
12
O16
L16
L12
O12
253
256
O8
204
L8
219
217
233
218
124
9
97
O1

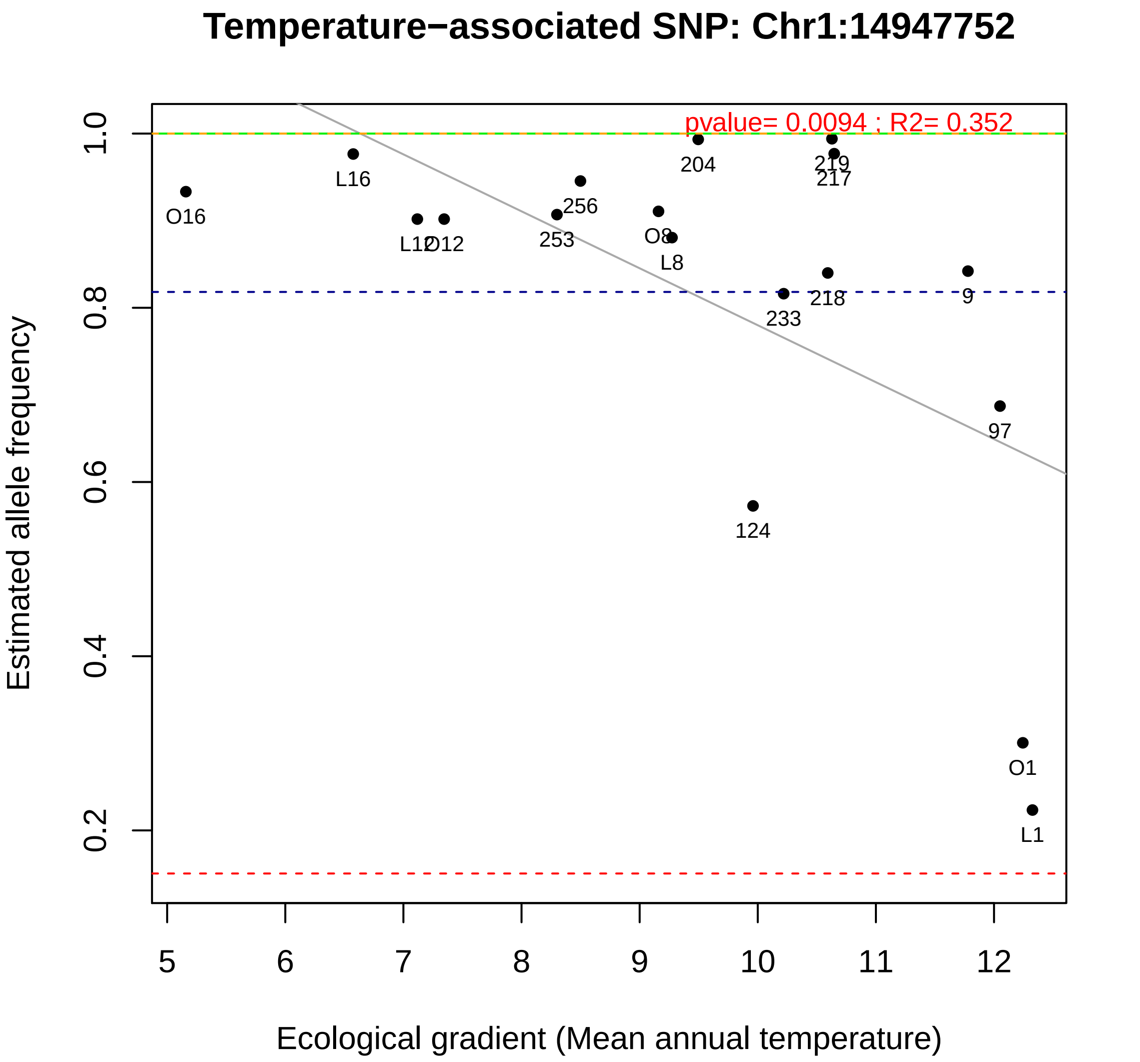
Temperature−associated SNP: Chr1:14947752
pvalue= 0.0094 ; R2= 0.352
Estimated allele frequency
0.2
0.4
0.6
0.8
1.0
5
6
7
8
9
10
11
12
Ecological gradient (Mean annual temperature)
O16
L16
L12
O12
253
256
O8
L8
204
219
217
233
218
124
9
97
O1
L1

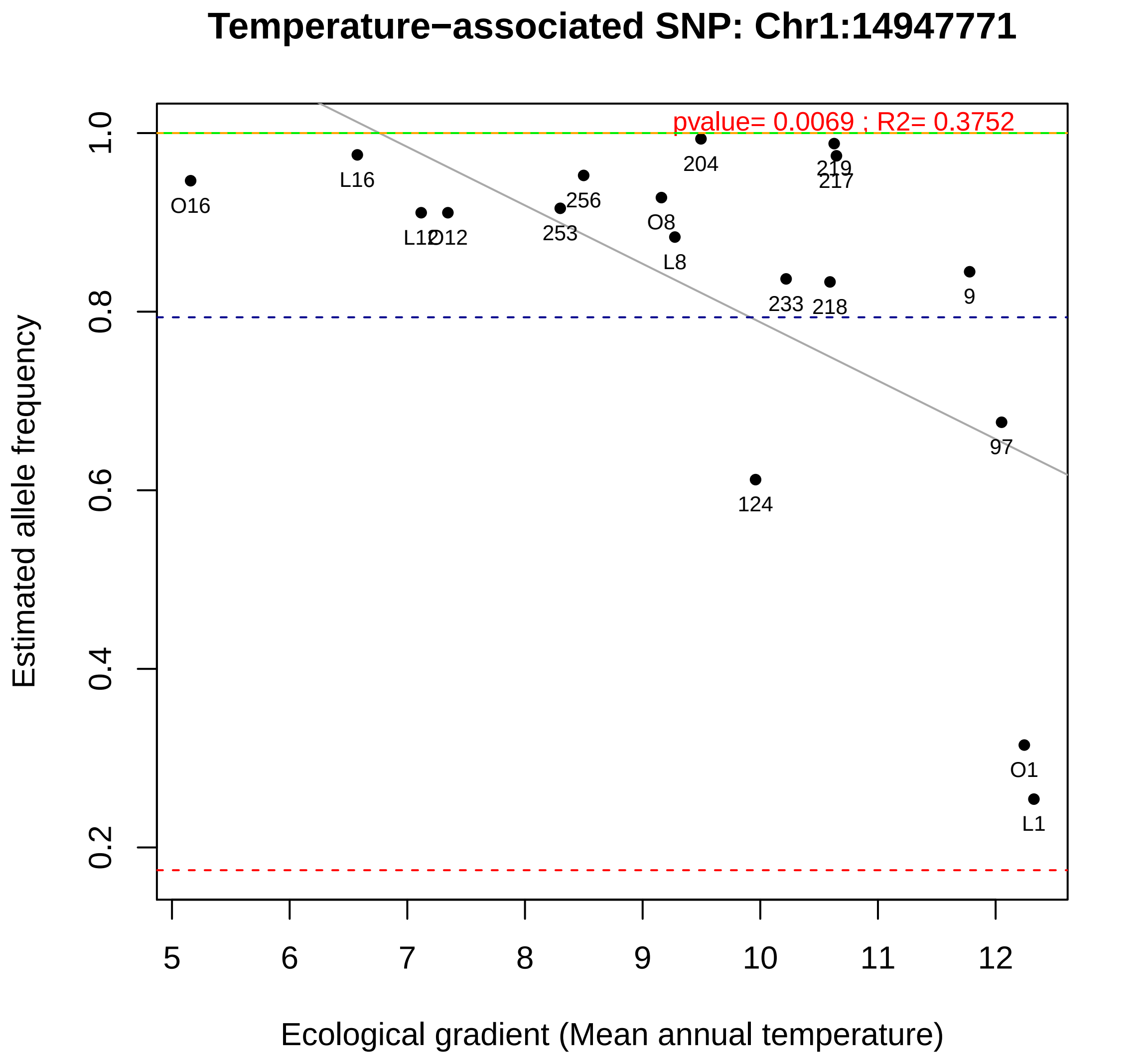

Temperature−associated SNP: Chr1:14947771
pvalue= 0.0069 ; R2= 0.3752
Estimated allele frequency
0.2
0.4
0.6
0.8
1.0
Ecological gradient (Mean annual temperature)
5
6
7
8
9
10
11
12
O16
L16
L12
O12
253
256
O8
L8
204
219
217
233
218
124
9
97
O1
L1

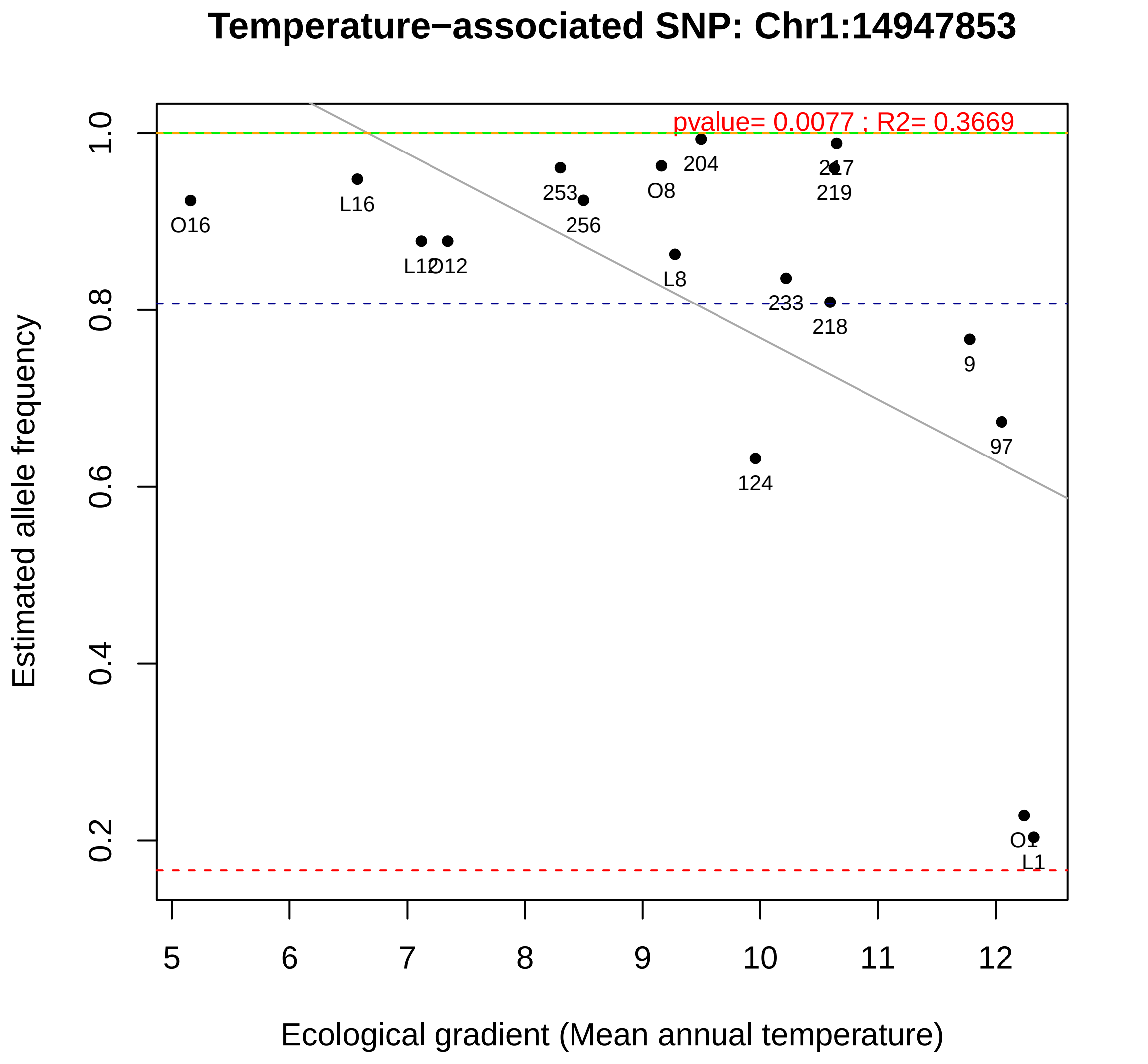

Temperature−associated SNP: Chr1:14947853
pvalue= 0.0077 ; R2= 0.3669
Estimated allele frequency
Ecological gradient (Mean annual temperature)
1.0
0.8
0.6
0.4
0.2
5
6
7
8
9
10
11
12
O16
L16
L12
O12
253
256
O8
204
L8
217
219
233
218
124
9
97
O1
L1

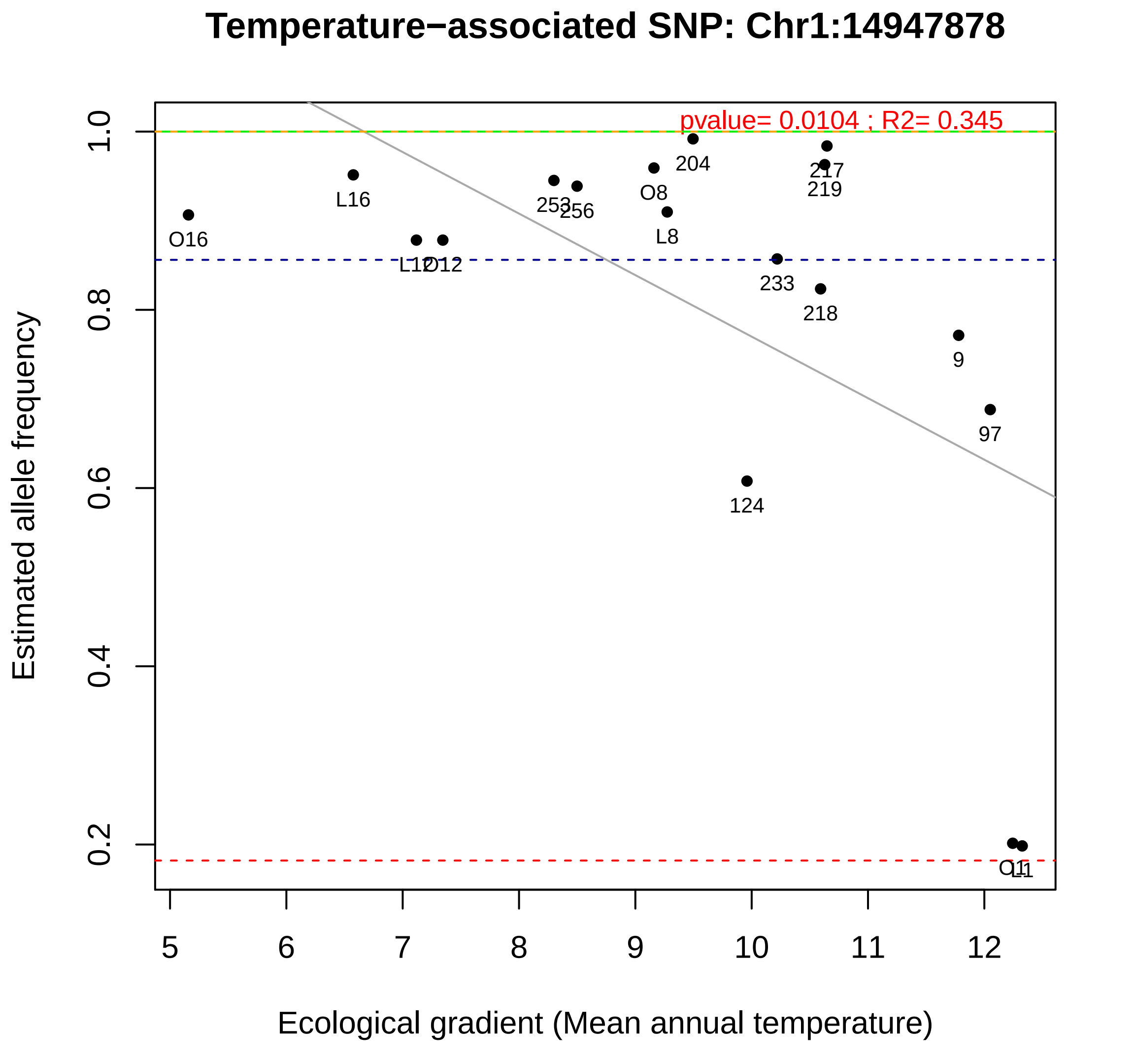

Temperature−associated SNP: Chr1:14947878
pvalue= 0.0104 ; R2= 0.345
Estimated allele frequency
0.2
0.4
0.6
0.8
1.0
5
6
7
8
9
10
11
12
Ecological gradient (Mean annual temperature)
O16
L16
L12
O12
253
256
O8
L8
204
217
219
233
218
124
9
97

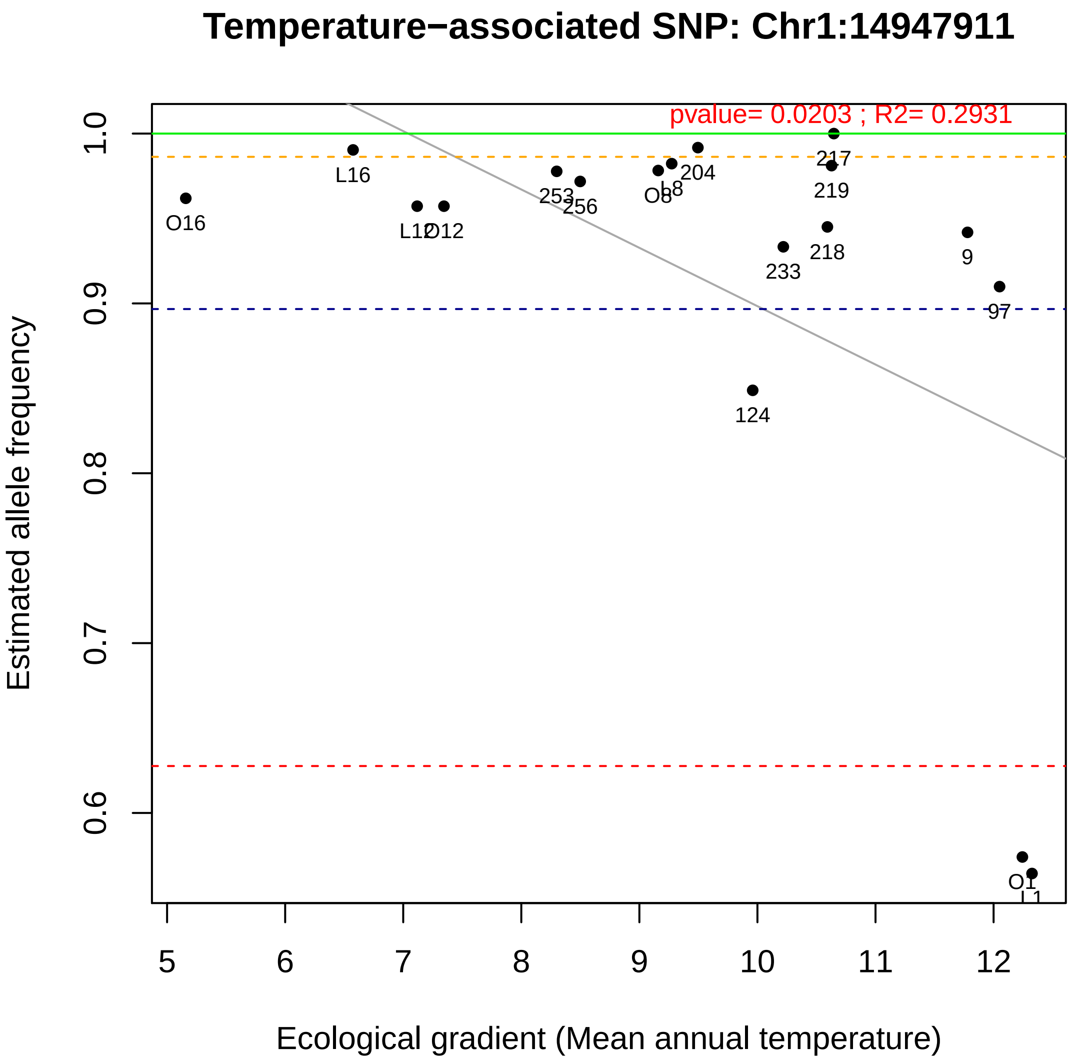

Temperature−associated SNP: Chr1:14947911
pvalue= 0.0203 ; R2= 0.2931
Estimated allele frequency
0.6
0.7
0.8
0.9
1.0
5
6
7
8
9
10
11
12
Ecological gradient (Mean annual temperature)
O16
L16
L12
O12
253
256
O8
L8
204
217
219
218
233
124
9
97
O1
L1

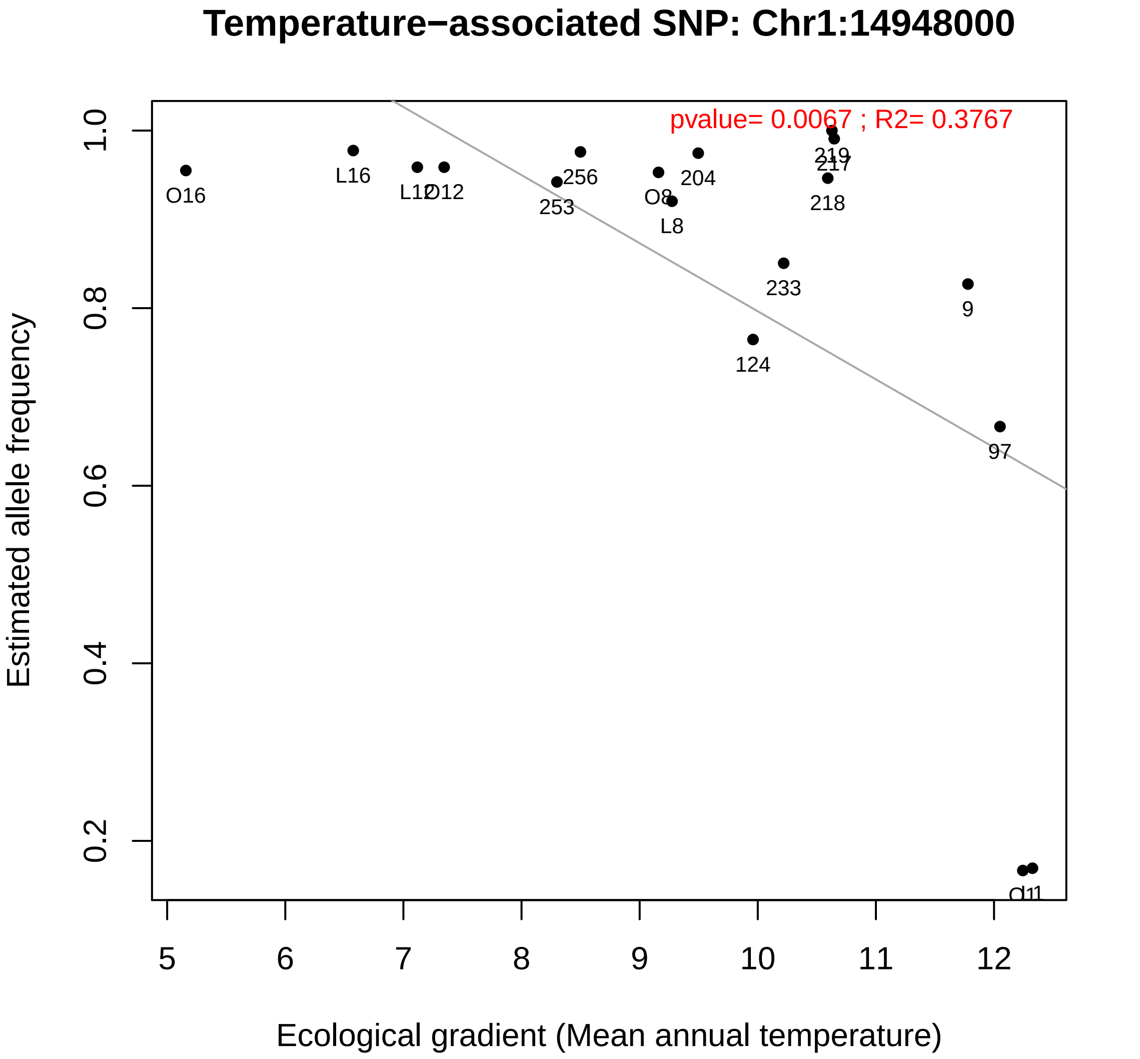

Temperature−associated SNP: Chr1:14948000
pvalue= 0.0067 ; R2= 0.3767
Estimated allele frequency
Ecological gradient (Mean annual temperature)
0.2
0.4
0.6
0.8
1.0
5
6
7
8
9
10
11
12
O16
L16
L12
O12
253
256
O8
L8
204
219
217
218
233
124
9
97

**Temperature−associated SNP: Chr1:14948032**

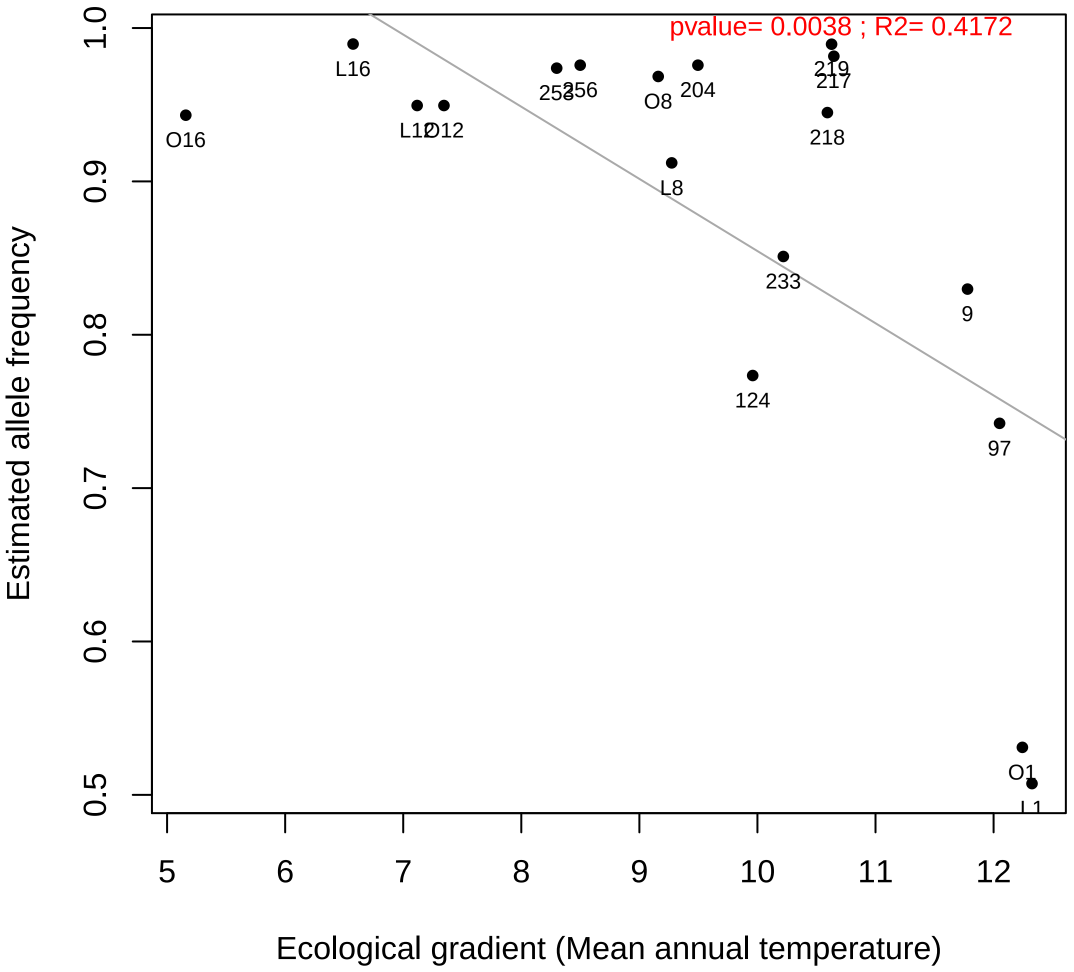

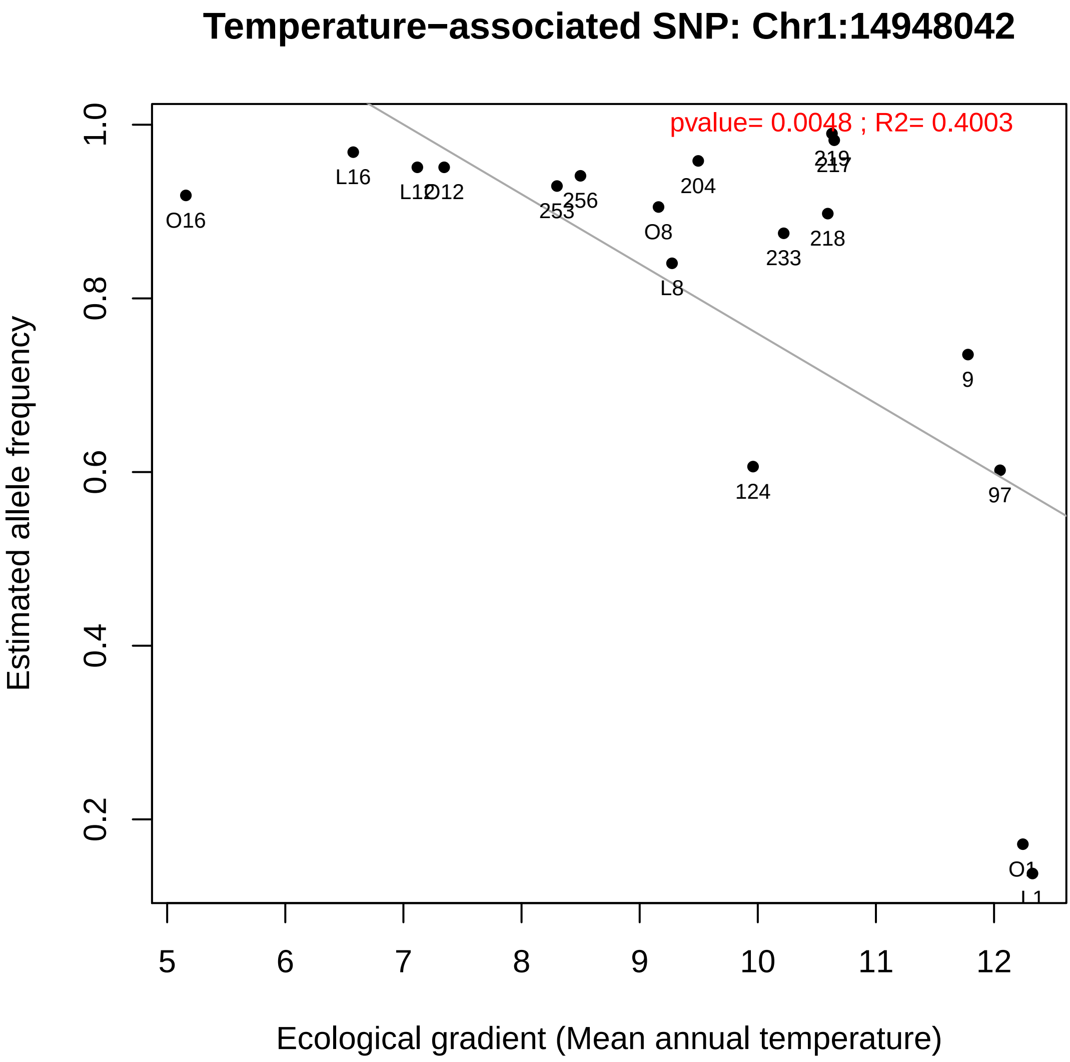

Temperature−associated SNP: Chr1:14948042
pvalue= 0.0048 ; R2= 0.4003
O16
L16
L12
O12
253
256
O8
L8
204
233
219
217
218
9
124
97
O1
L1
1.0
0.8
0.6
0.4
0.2
5
6
7
8
9
10
11
12
Estimated allele frequency
Ecological gradient (Mean annual temperature)

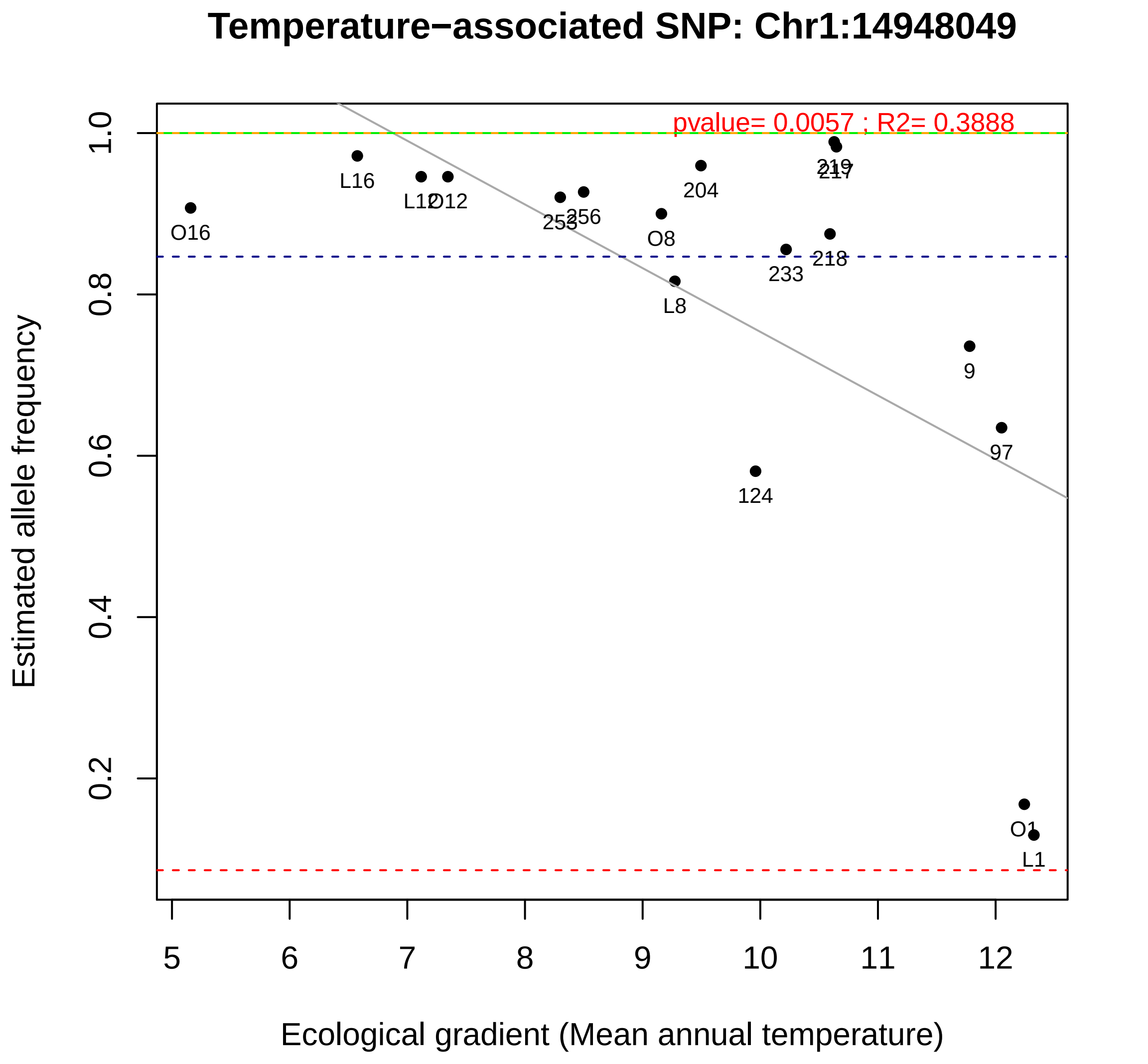
Temperature−associated SNP: Chr1:14948049
pvalue= 0.0057 ; R2= 0.3888
Estimated allele frequency
Ecological gradient (Mean annual temperature)
0.2
0.4
0.6
0.8
1.0
5
6
7
8
9
10
11
12
O16
L16
L12
O12
253
256
O8
204
L8
233
218
219
217
124
9
97
O1
L1

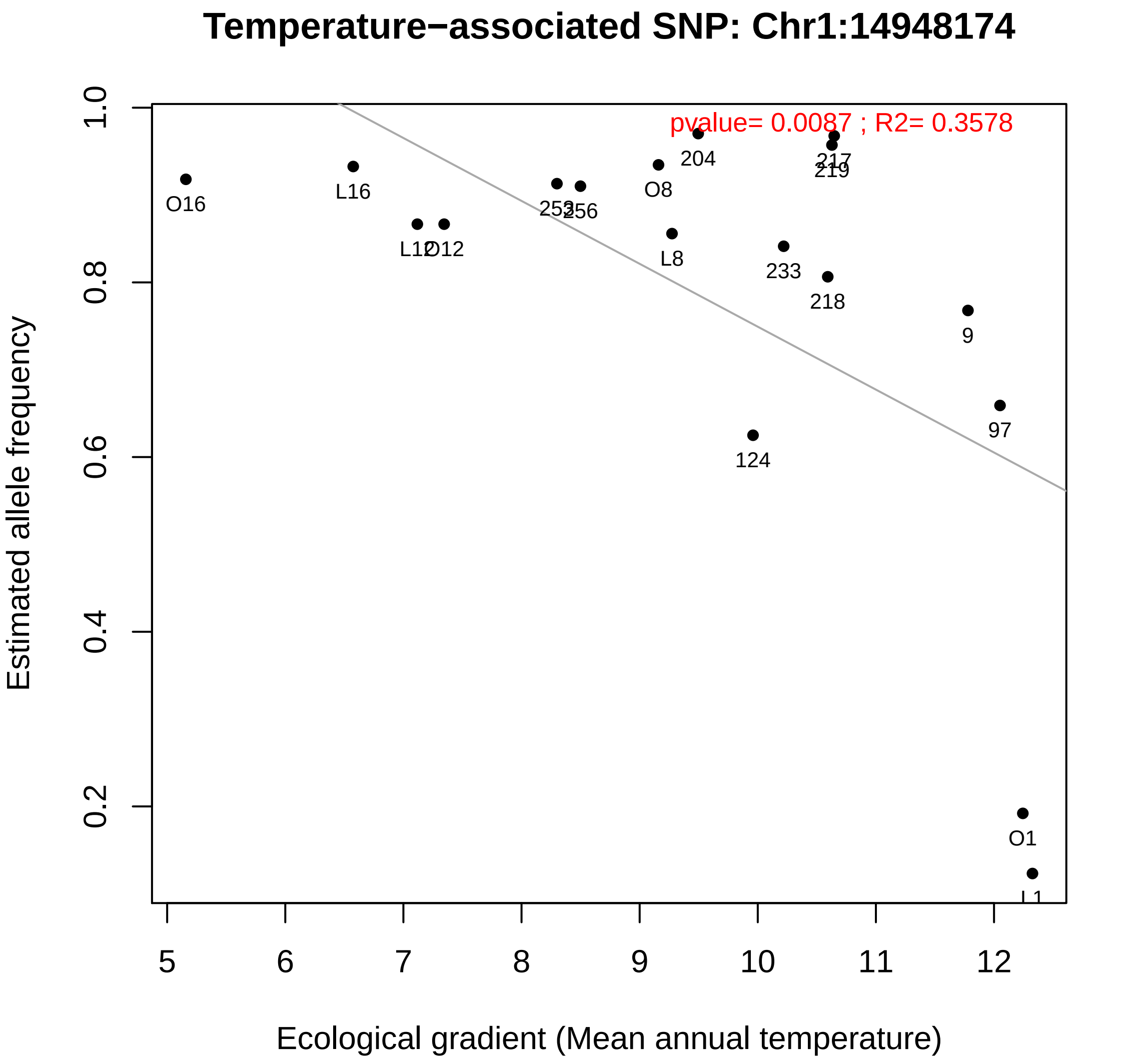

Temperature−associated SNP: Chr1:14948174
pvalue= 0.0087 ; R2= 0.3578
Estimated allele frequency
0.2
0.4
0.6
0.8
1.0
5
6
7
8
9
10
11
12
Ecological gradient (Mean annual temperature)
O16
L16
L12
O12
252
256
O8
204
L8
233
217
219
218
9
97
124
O1

**Temperature−associated SNP: Chr1:14948291**

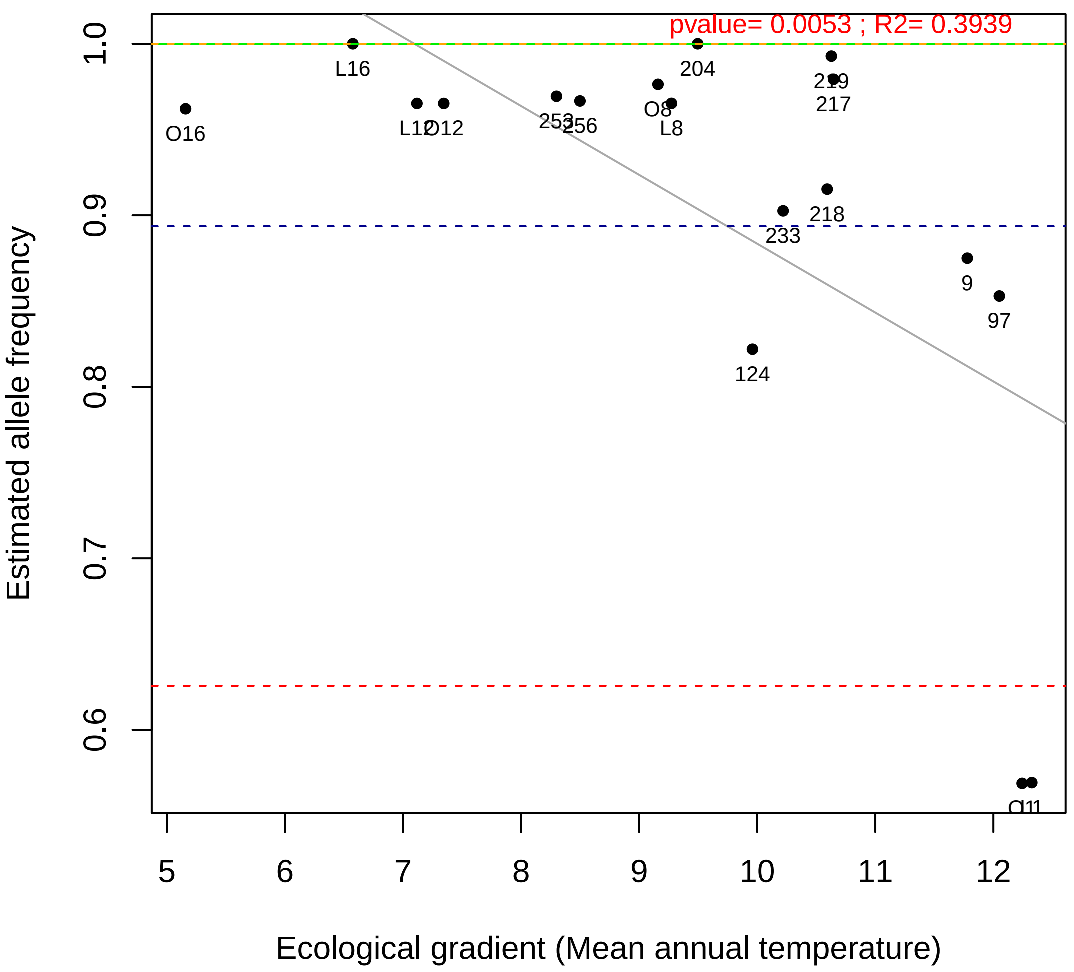

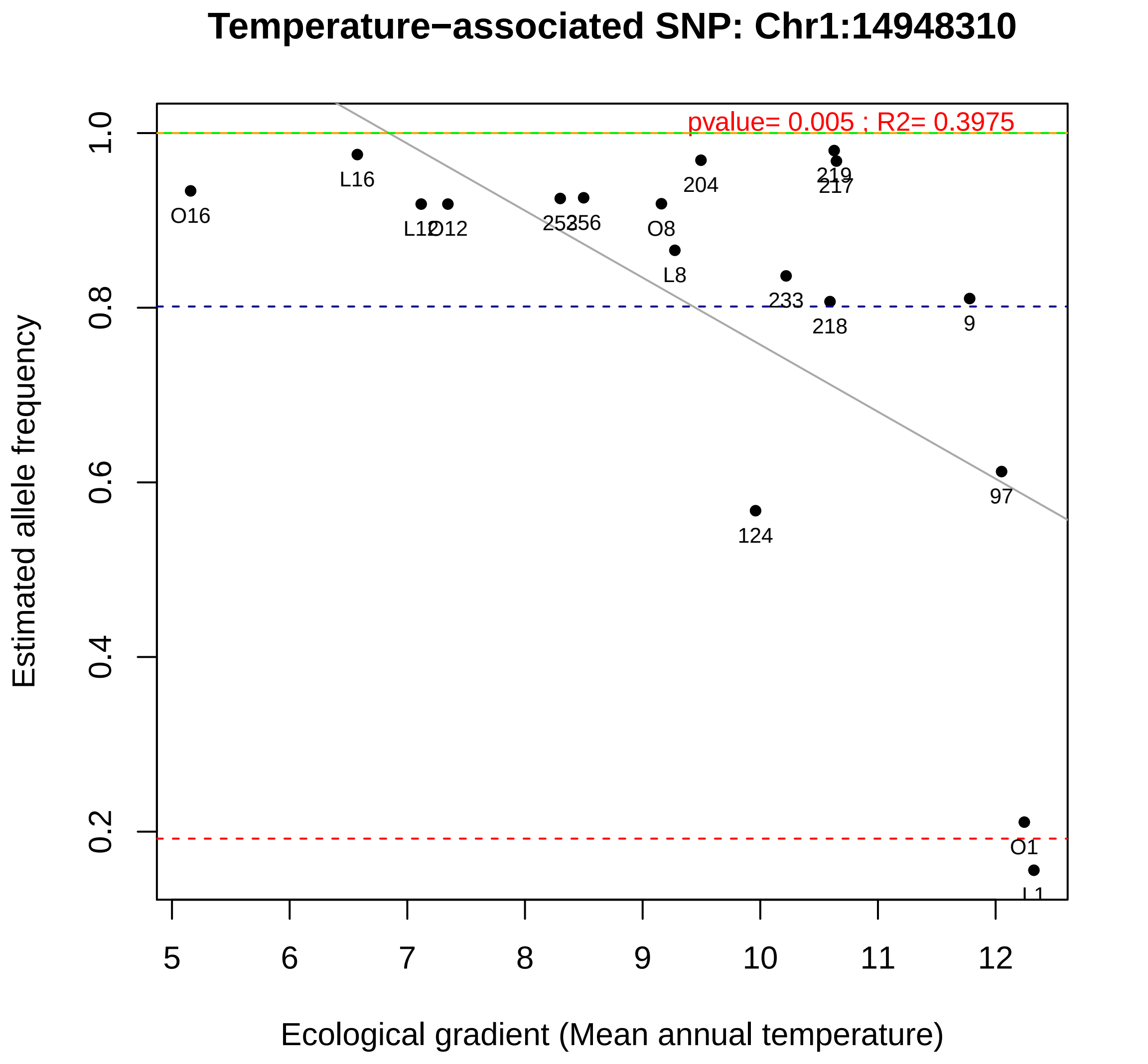
Temperature−associated SNP: Chr1:14948310
pvalue= 0.005 ; R2= 0.3975
Estimated allele frequency
Ecological gradient (Mean annual temperature)
1.0
0.8
0.6
0.4
0.2
5
6
7
8
9
10
11
12
O16
L16
L12
O12
253
256
O8
204
219
217
L8
233
218
9
124
97
O1
L1

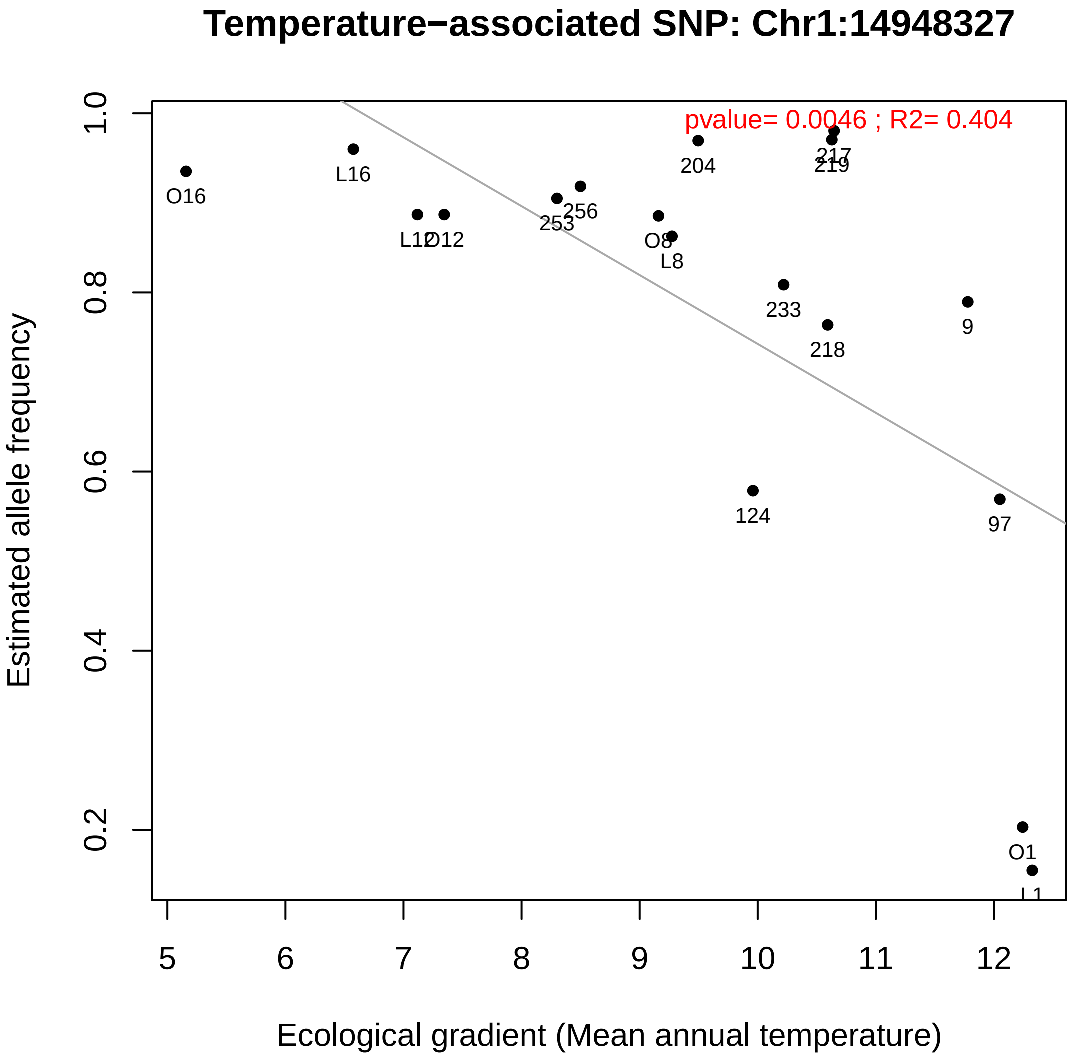
Temperature−associated SNP: Chr1:14948327
pvalue= 0.0046 ; R2= 0.404
Estimated allele frequency
Ecological gradient (Mean annual temperature)
0.2
0.4
0.6
0.8
1.0
5
6
7
8
9
10
11
12
O16
L16
L12
O12
253
256
O8
L8
204
217
219
233
218
9
124
97
O1

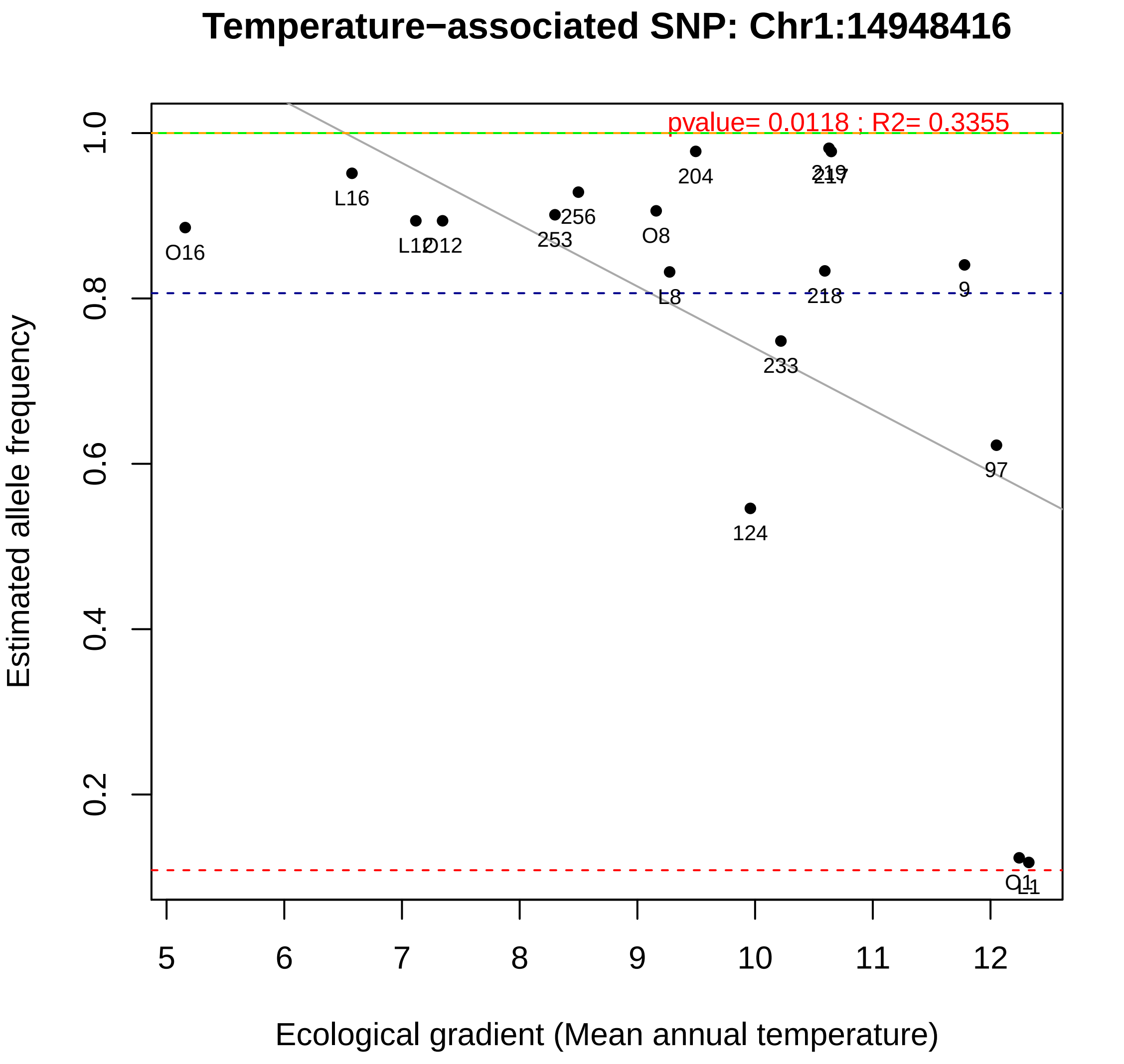

Temperature−associated SNP: Chr1:14948416
pvalue= 0.0118 ; R2= 0.3355
O16
L16
L12
O12
253
256
O8
204
L8
219
217
218
233
9
124
97
O1
L1
1.0
0.8
0.6
0.4
0.2
5
6
7
8
9
10
11
12
Estimated allele frequency
Ecological gradient (Mean annual temperature)

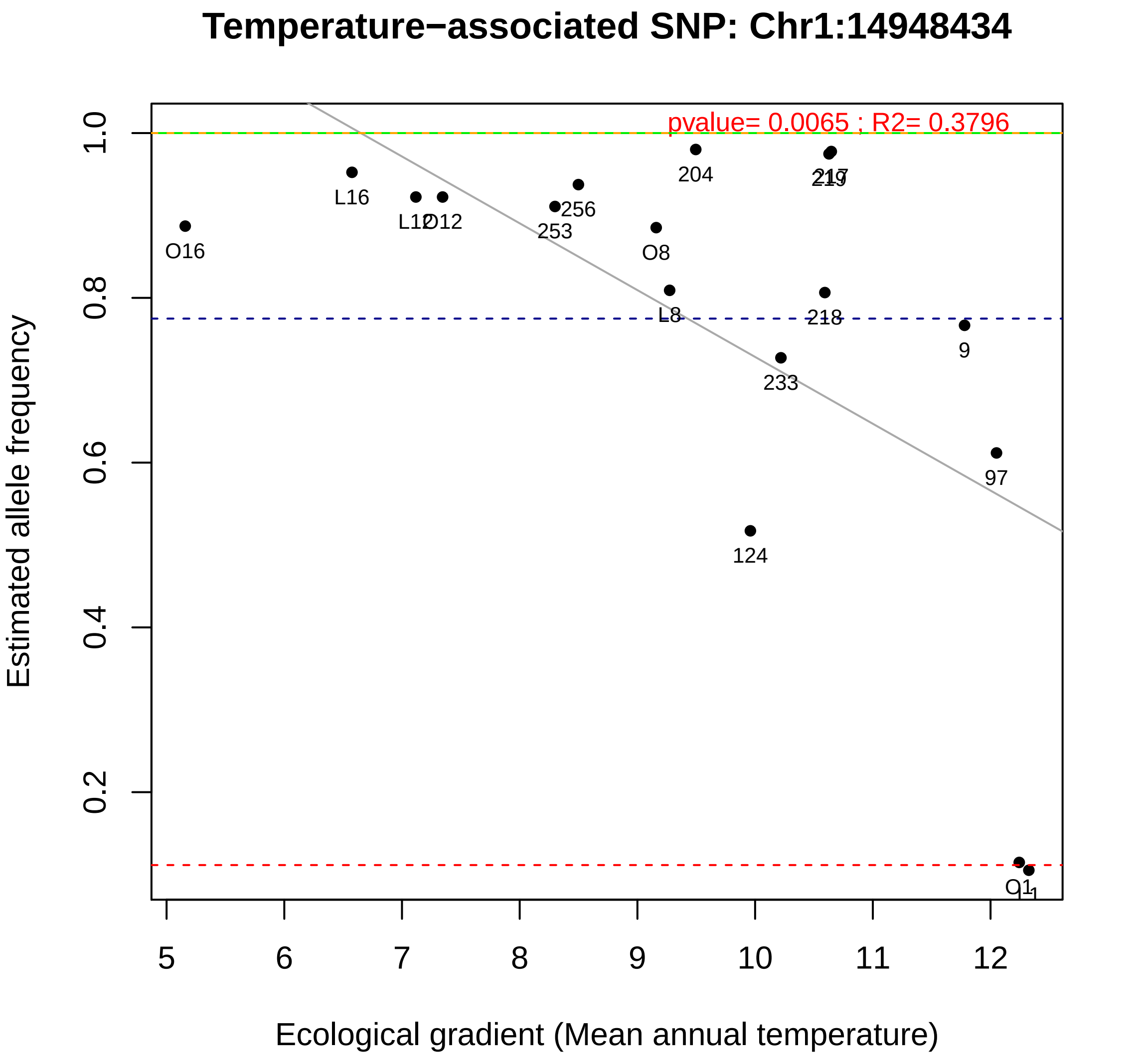
Temperature−associated SNP: Chr1:14948434
pvalue= 0.0065 ; R2= 0.3796
Estimated allele frequency
Ecological gradient (Mean annual temperature)
1.0
0.8
0.6
0.4
0.2
5
6
7
8
9
10
11
12
O16
L16
L12
O12
253
256
O8
L8
204
217
219
218
233
124
9
97
O1
L1

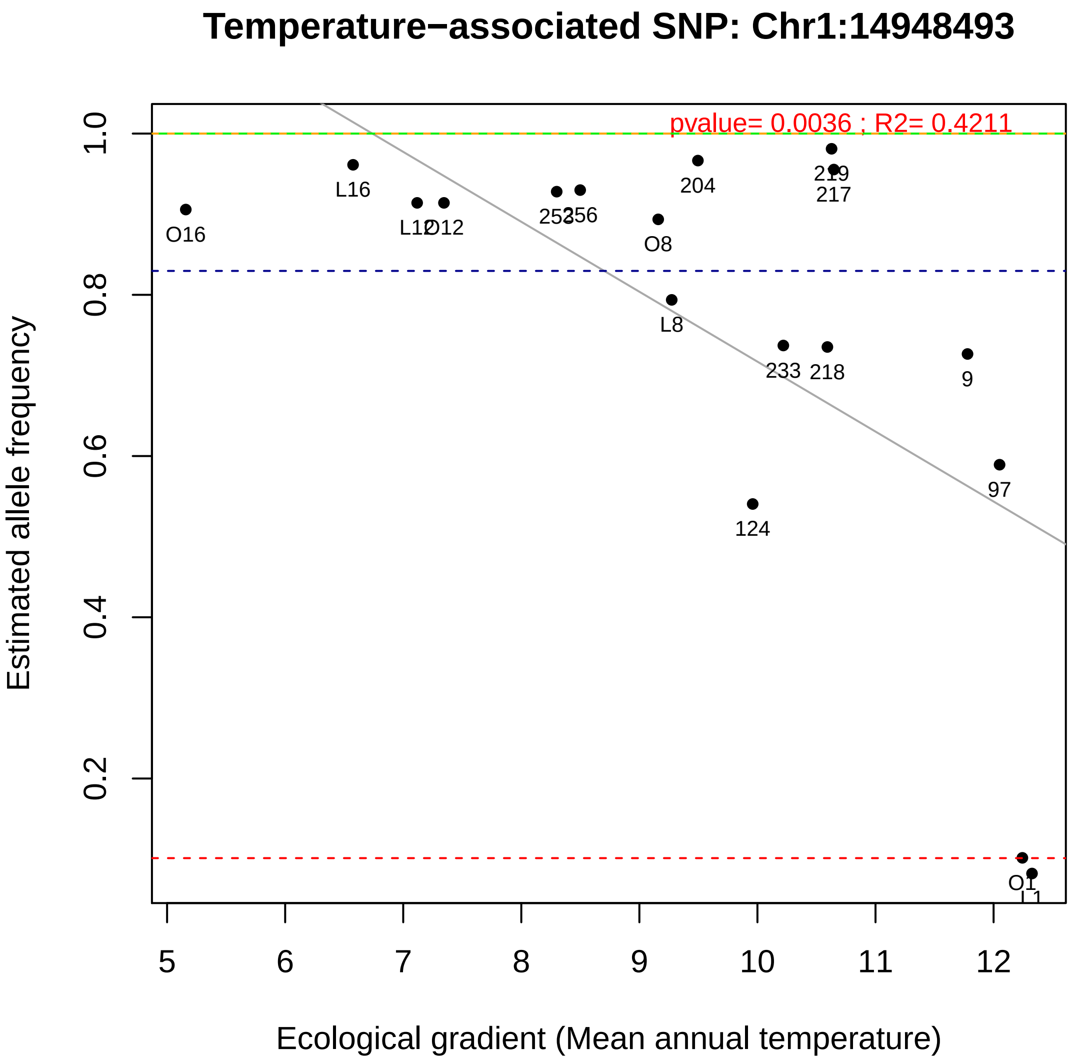

Temperature−associated SNP: Chr1:14948493
pvalue= 0.0036 ; R2= 0.4211
Estimated allele frequency
Ecological gradient (Mean annual temperature)
0.2
0.4
0.6
0.8
1.0
5
6
7
8
9
10
11
12
O16
L16
L12
O12
256
O8
204
219
217
L8
233
218
9
97
124
O1

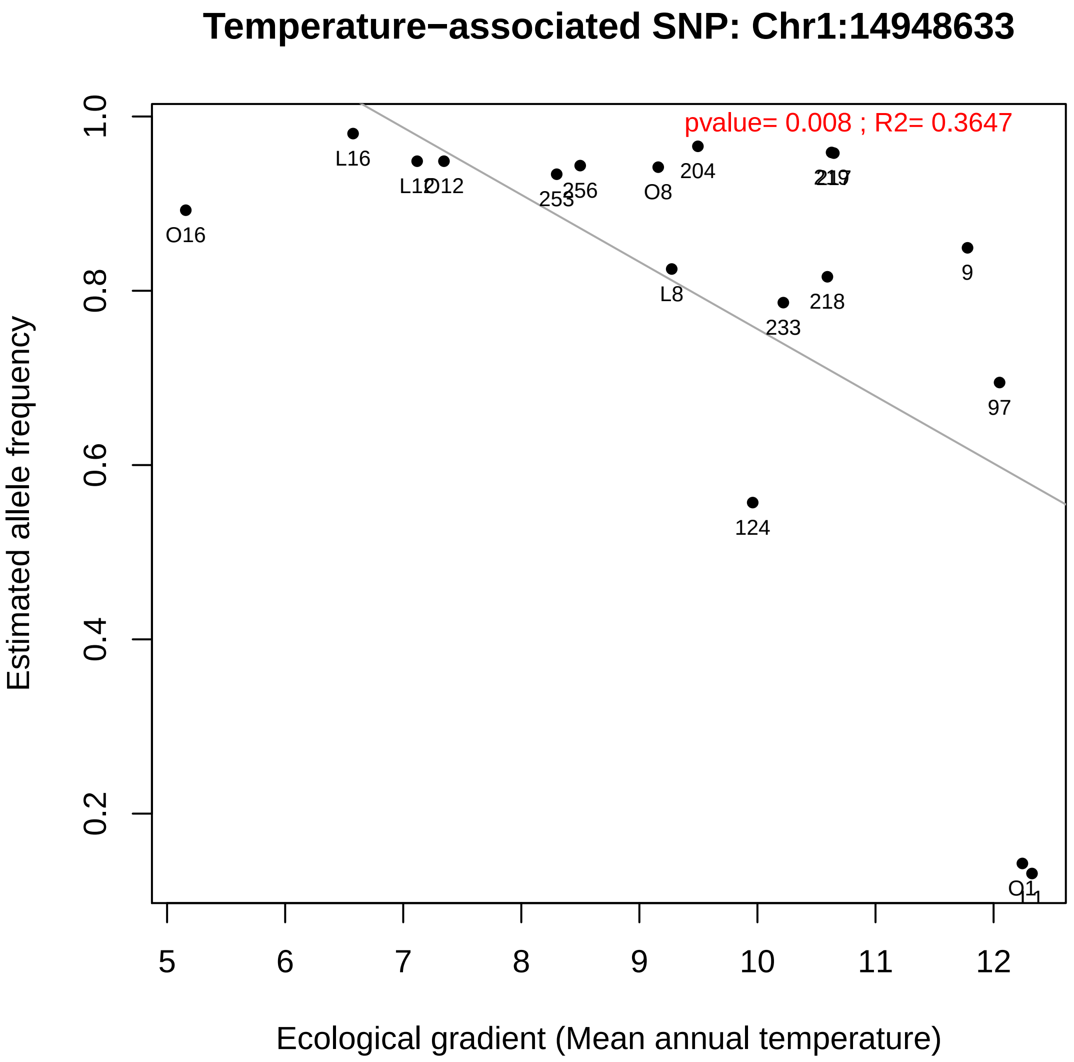

Temperature−associated SNP: Chr1:14948633
pvalue= 0.008 ; R2= 0.3647
Estimated allele frequency
Ecological gradient (Mean annual temperature)
0.2
0.4
0.6
0.8
1.0
5
6
7
8
9
10
11
12
O16
L16
L12
O12
253
256
O8
204
219
217
L8
233
218
9
97
124

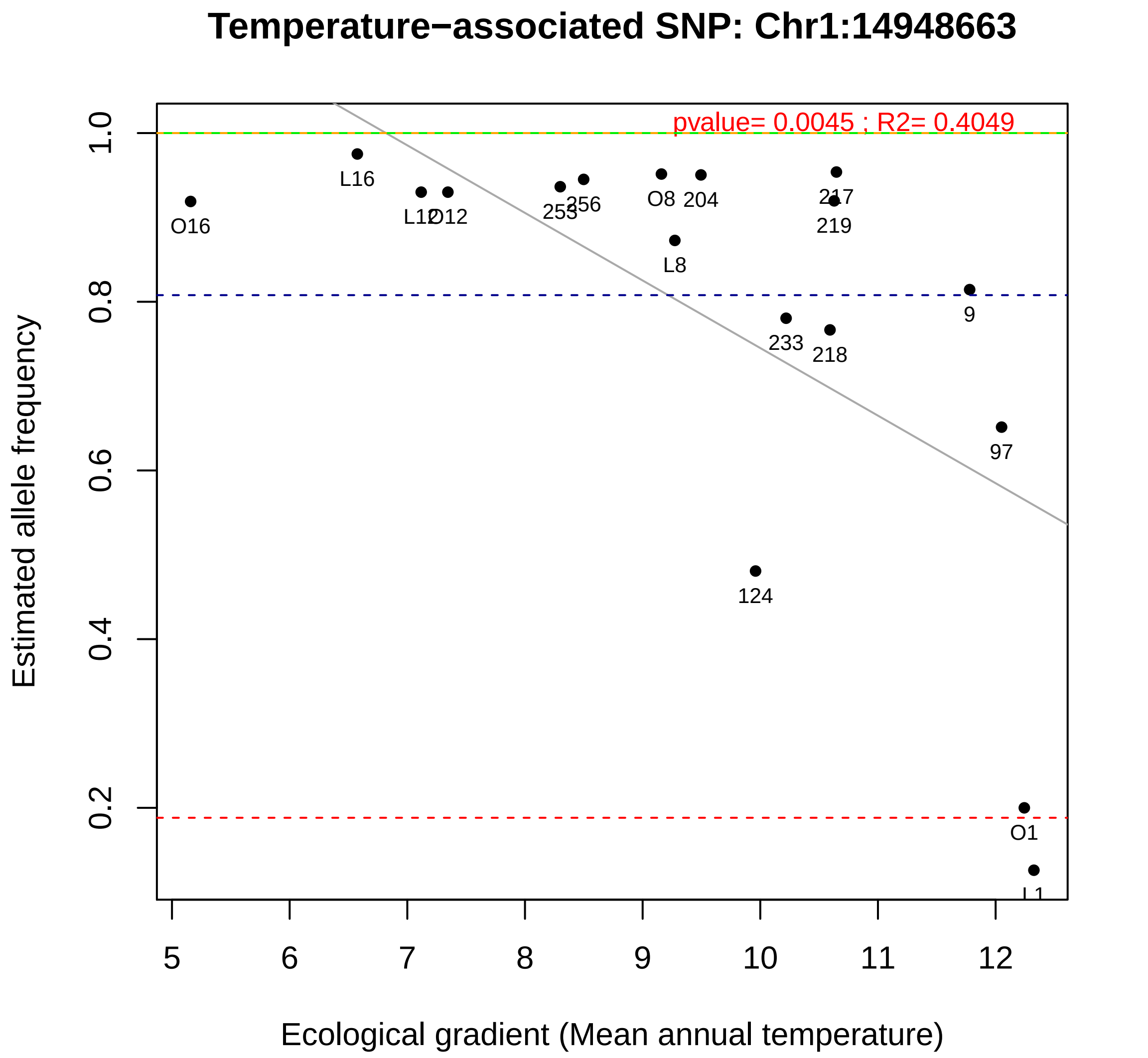

Temperature−associated SNP: Chr1:14948663
pvalue= 0.0045 ; R2= 0.4049
Estimated allele frequency
Ecological gradient (Mean annual temperature)
1.0
0.8
0.6
0.4
0.2
5
6
7
8
9
10
11
12
O16
L16
L12
O12
253
256
O8
204
217
219
L8
9
233
218
97
124
O1

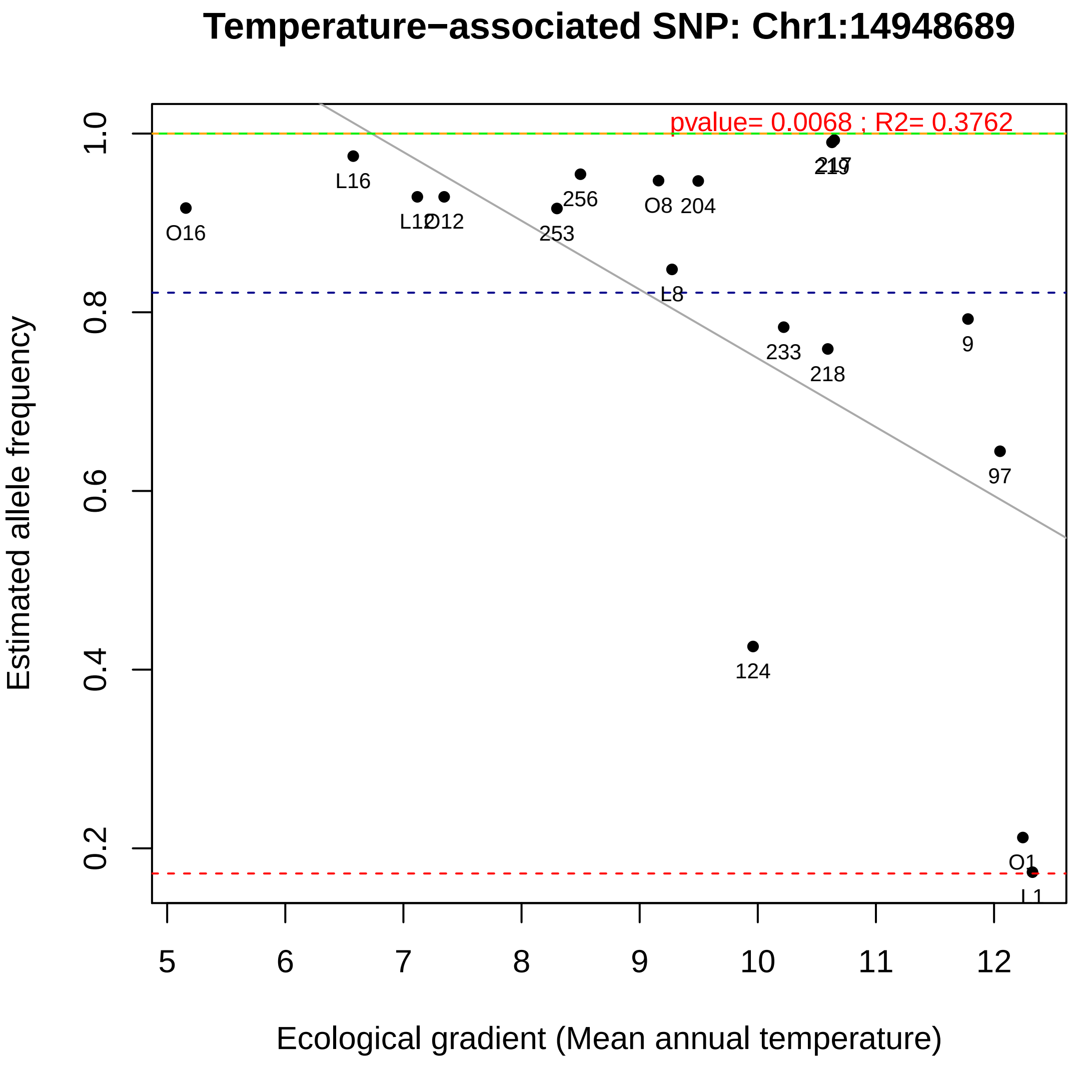
Temperature−associated SNP: Chr1:14948689
pvalue= 0.0068 ; R2= 0.3762
Estimated allele frequency
0.2
0.4
0.6
0.8
1.0
5
6
7
8
9
10
11
12
Ecological gradient (Mean annual temperature)
O16
L16
O12
256
253
O8
204
L8
233
218
9
97
124
O1

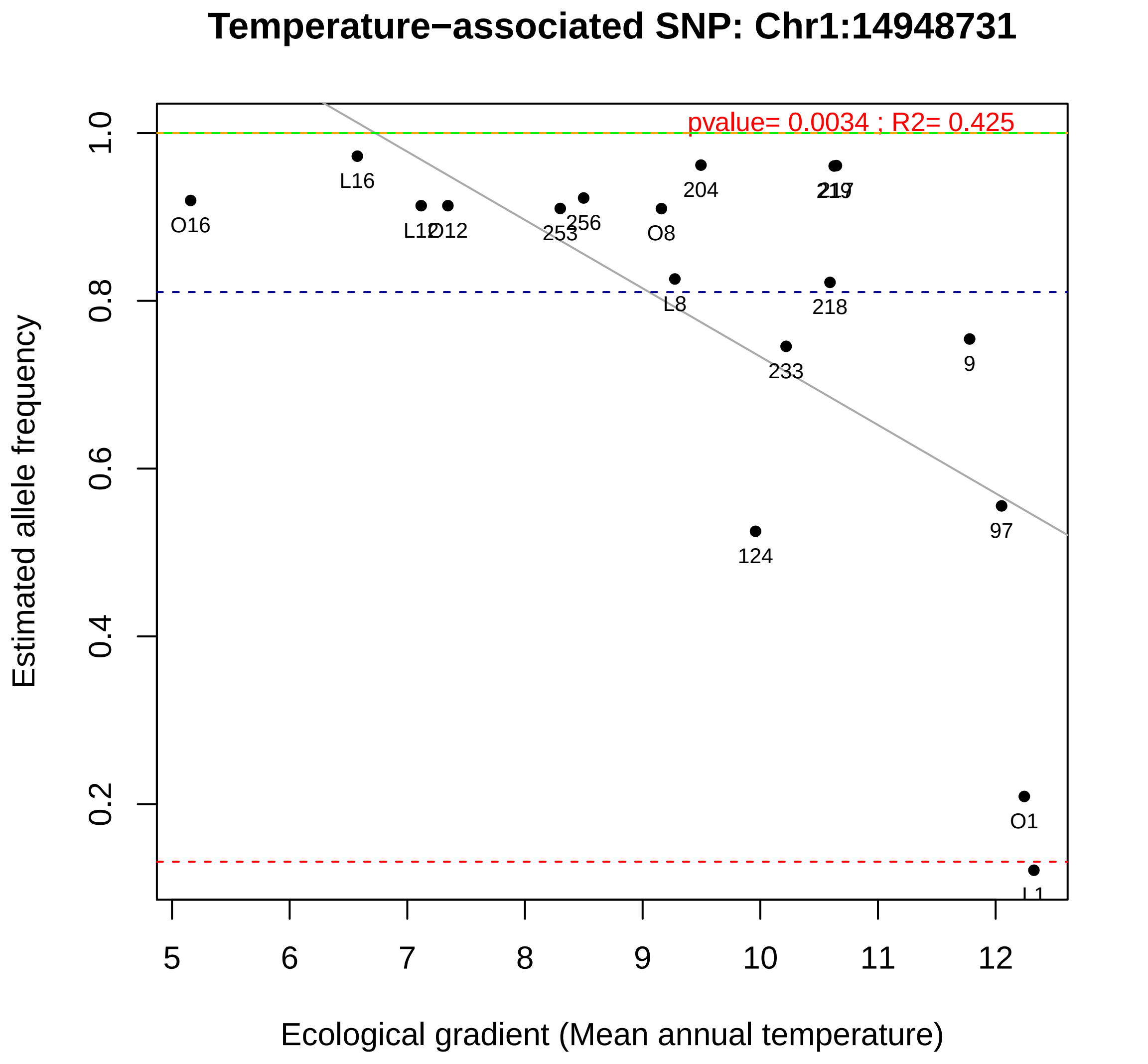
Temperature−associated SNP: Chr1:14948731
pvalue= 0.0034 ; R2= 0.425
Estimated allele frequency
0.2
0.4
0.6
0.8
1.0
Ecological gradient (Mean annual temperature)
5
6
7
8
9
10
11
12
O16
L16
L12
O12
253
256
O8
204
L8
217
219
218
233
9
124
97
O1
L1

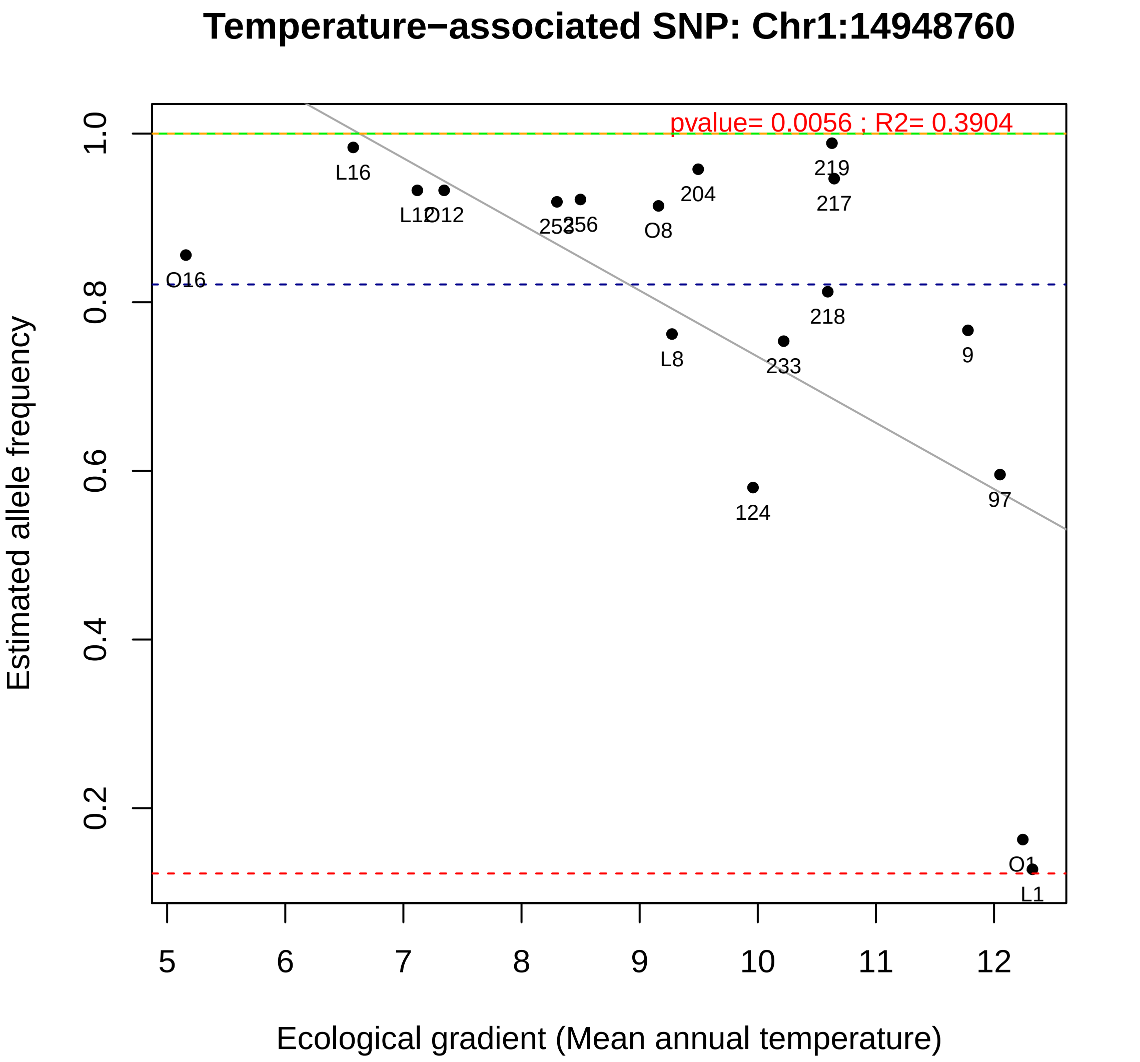
Temperature−associated SNP: Chr1:14948760
pvalue= 0.0056 ; R2= 0.3904
Estimated allele frequency
Ecological gradient (Mean annual temperature)
1.0
0.8
0.6
0.4
0.2
5
6
7
8
9
10
11
12
O16
L16
L12
O12
253
256
O8
204
219
217
218
L8
233
9
124
97
O1
L1

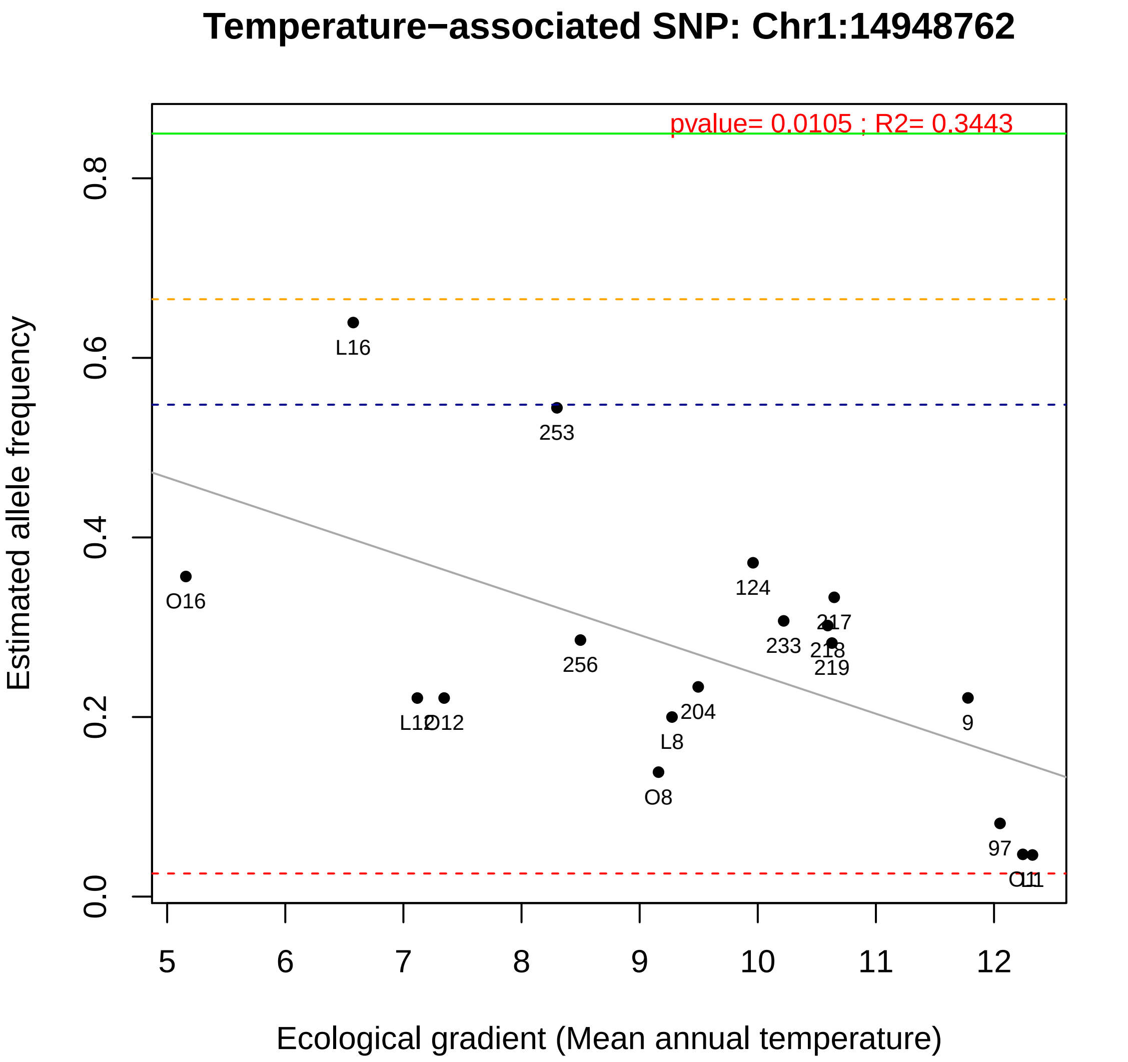

Temperature−associated SNP: Chr1:14948762
pvalue= 0.0105 ; R2= 0.3443
Estimated allele frequency
0.0
0.2
0.4
0.6
0.8
5
6
7
8
9
10
11
12
Ecological gradient (Mean annual temperature)
L16
253
O16
256
L12
O12
124
233
217
218
219
204
L8
O8
9
97
O11
L11

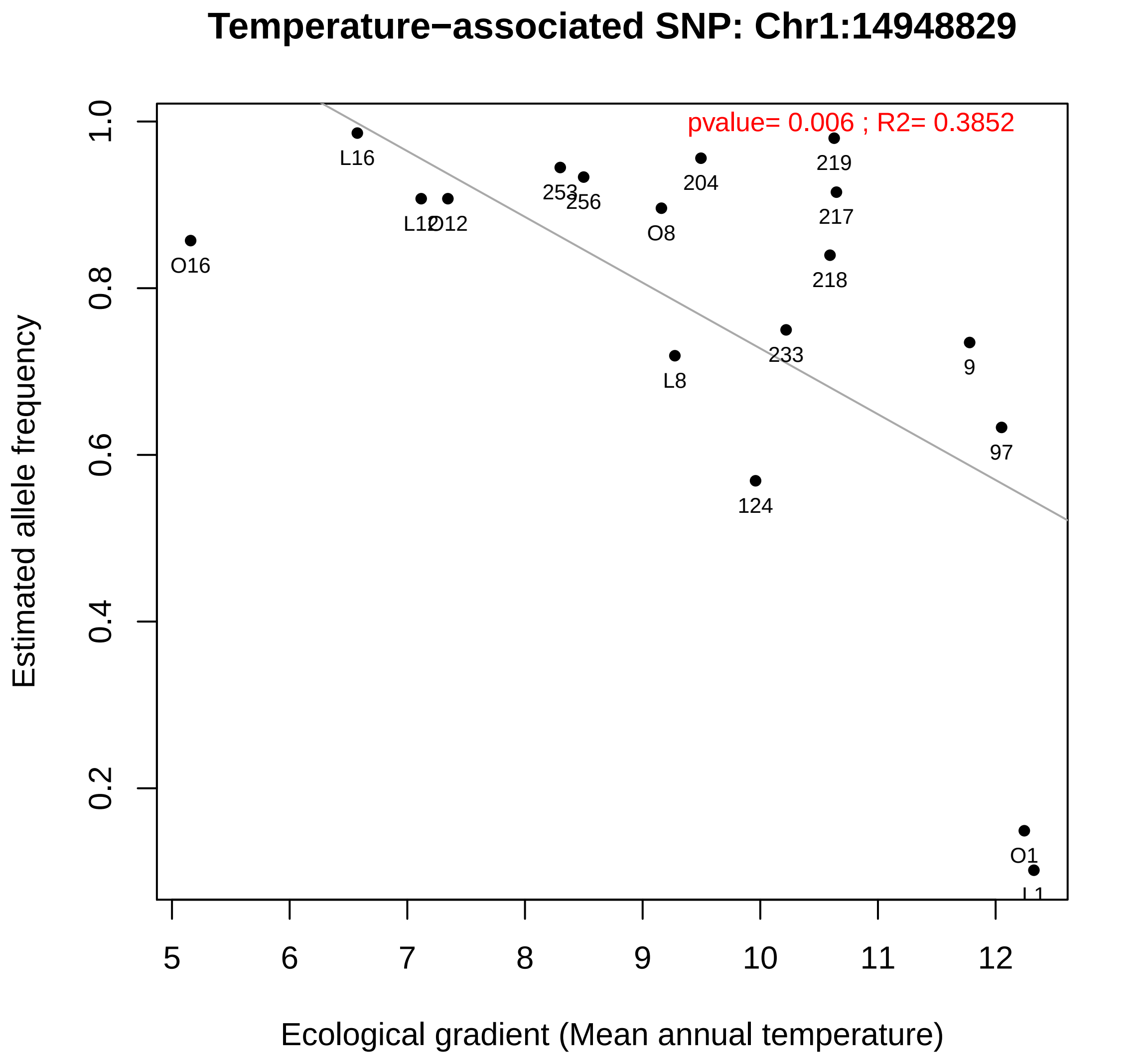

Temperature−associated SNP: Chr1:14948829
pvalue= 0.006 ; R2= 0.3852
Estimated allele frequency
0.2
0.4
0.6
0.8
1.0
Ecological gradient (Mean annual temperature)
5
6
7
8
9
10
11
12
O16
L16
L12
O12
253
256
O8
L8
204
124
233
219
217
218
9
97
O1

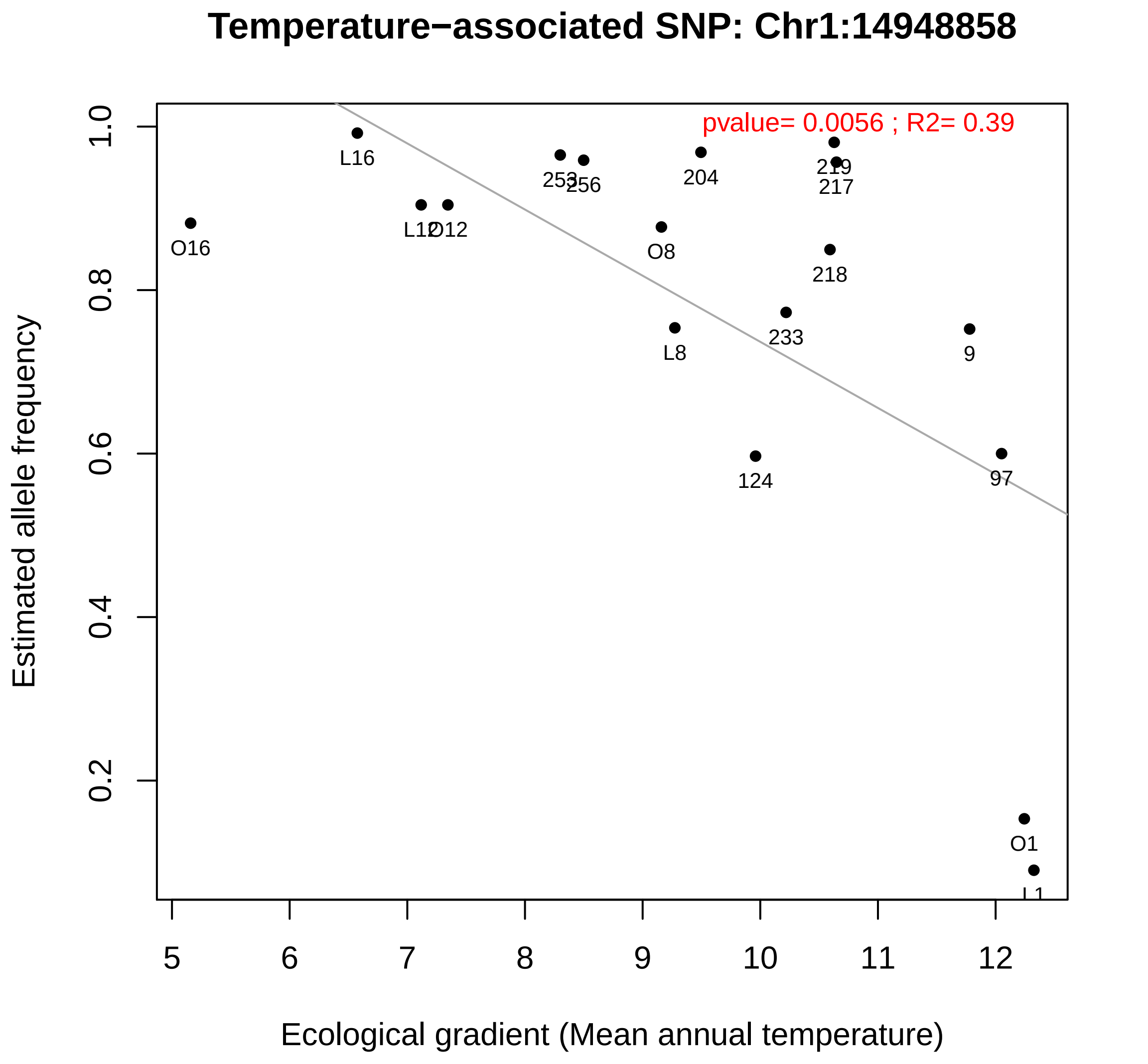
Temperature−associated SNP: Chr1:14948858
pvalue= 0.0056 ; R2= 0.39
Estimated allele frequency
Ecological gradient (Mean annual temperature)
1.0
0.8
0.6
0.4
0.2
5
6
7
8
9
10
11
12
L16
O16
L12
O12
253
256
204
O8
219
217
218
233
L8
9
124
97
O1
L1

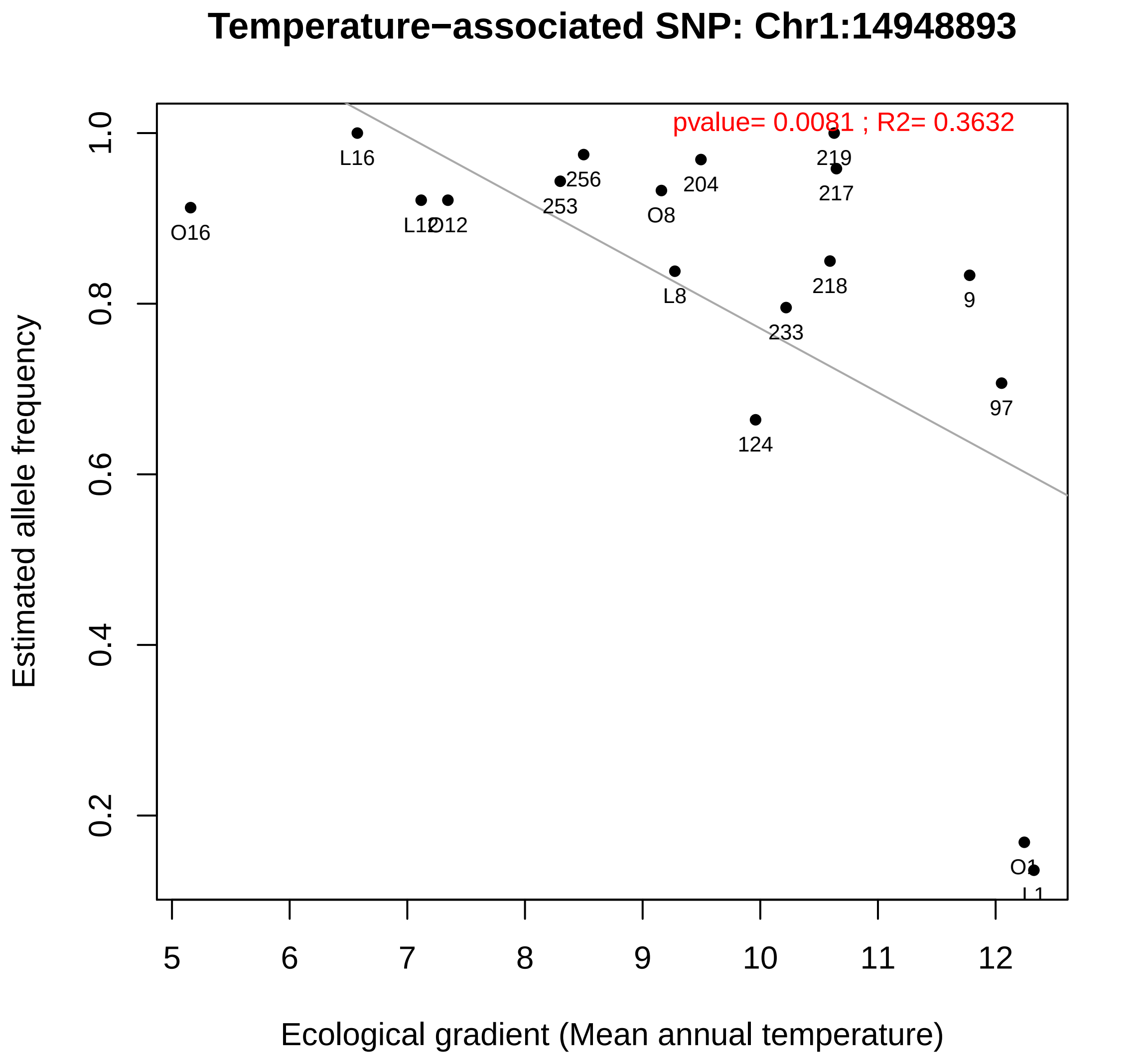

Temperature−associated SNP: Chr1:14948893
pvalue= 0.0081 ; R2= 0.3632
L16
O16
L12
O12
256
253
204
O8
219
217
L8
218
9
233
97
124
Estimated allele frequency
1.0
0.8
0.6
0.4
0.2
5
6
7
8
9
10
11
12
Ecological gradient (Mean annual temperature)

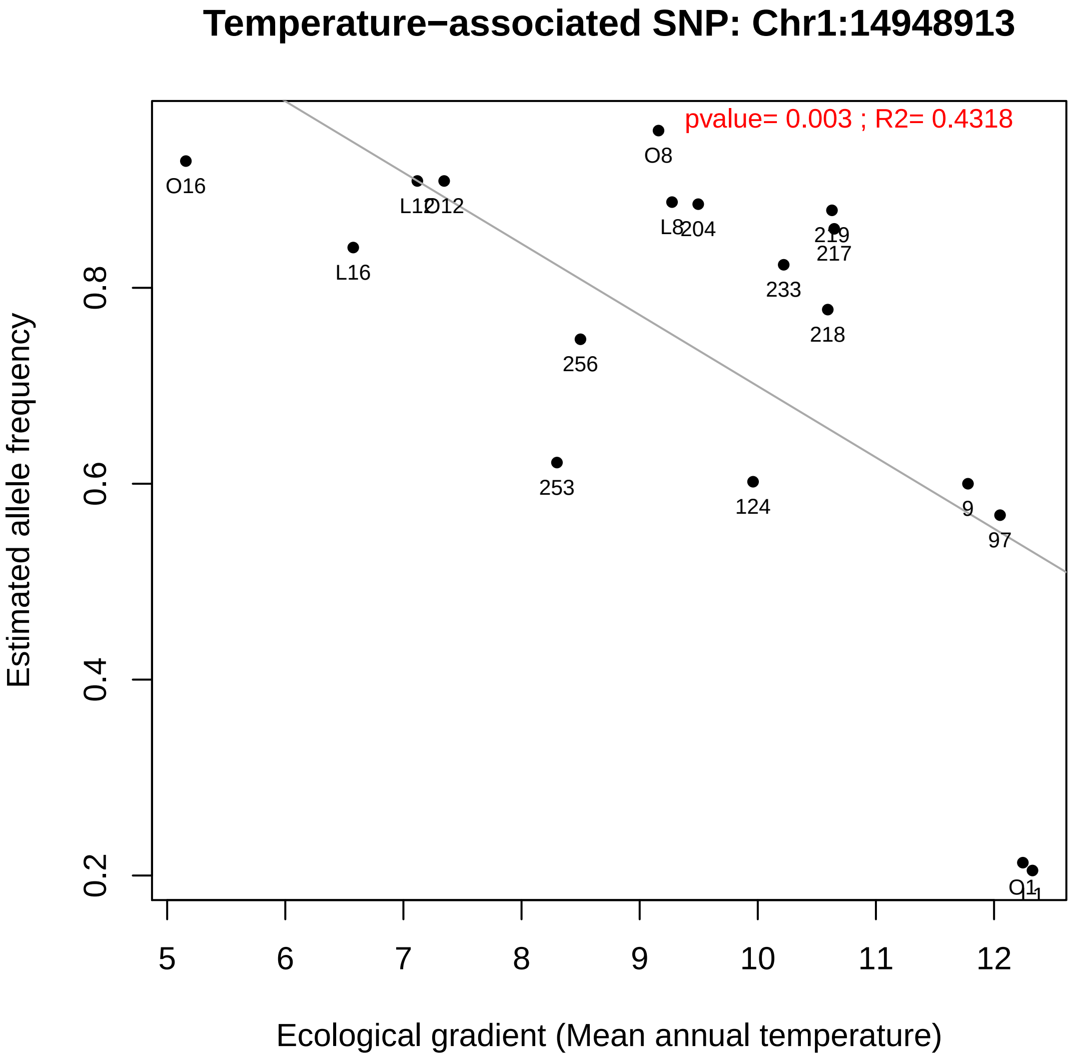
Temperature−associated SNP: Chr1:14948913
pvalue= 0.003 ; R2= 0.4318
Estimated allele frequency
0.2
0.4
0.6
0.8
5
6
7
8
9
10
11
12
Ecological gradient (Mean annual temperature)
O16
L16
L12
O12
253
256
O8
L8
204
124
233
219
217
218
9
97

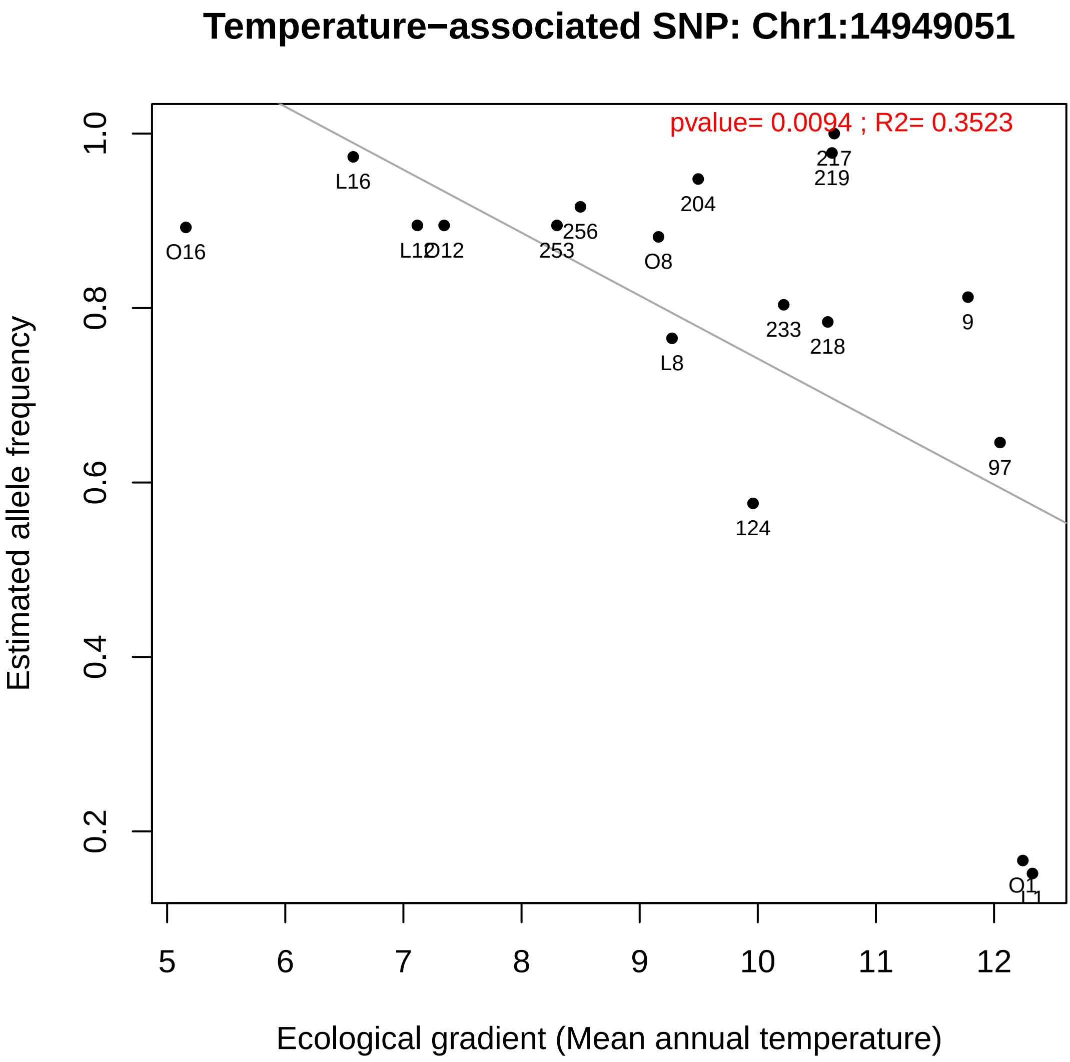

Temperature−associated SNP: Chr1:14949051
pvalue= 0.0094 ; R2= 0.3523
Estimated allele frequency
Ecological gradient (Mean annual temperature)
1.0
0.8
0.6
0.4
0.2
5
6
7
8
9
10
11
12
O16
L16
L12
O12
253
256
O8
L8
204
217
219
233
218
124
9
97
O1

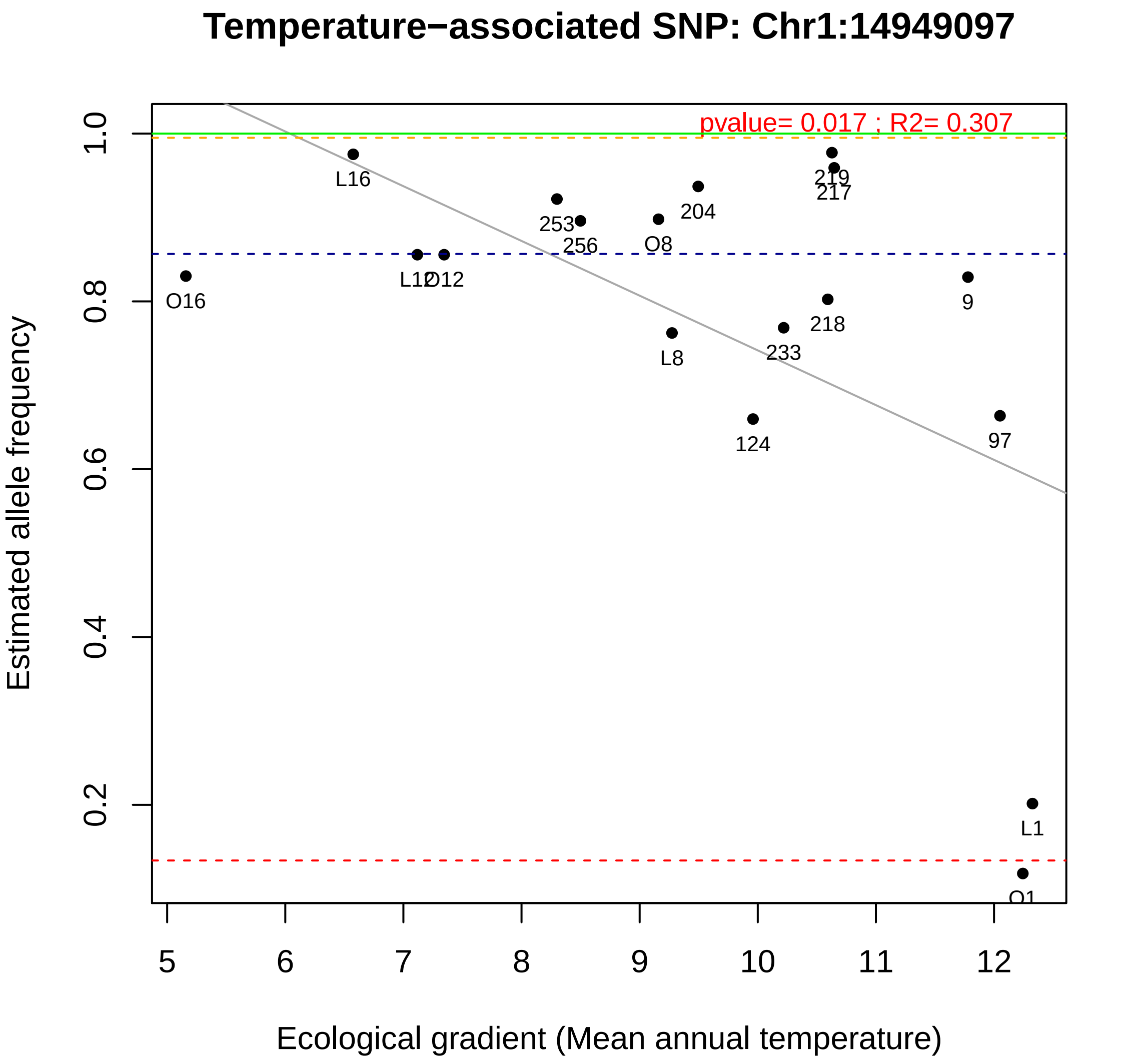

Temperature−associated SNP: Chr1:14949097
pvalue= 0.017 ; R2= 0.307
Estimated allele frequency
Ecological gradient (Mean annual temperature)
1.0
0.8
0.6
0.4
0.2
5
6
7
8
9
10
11
12
L16
O16
L12
O12
253
256
O8
204
219
217
L8
233
218
124
9
97
L1
O1

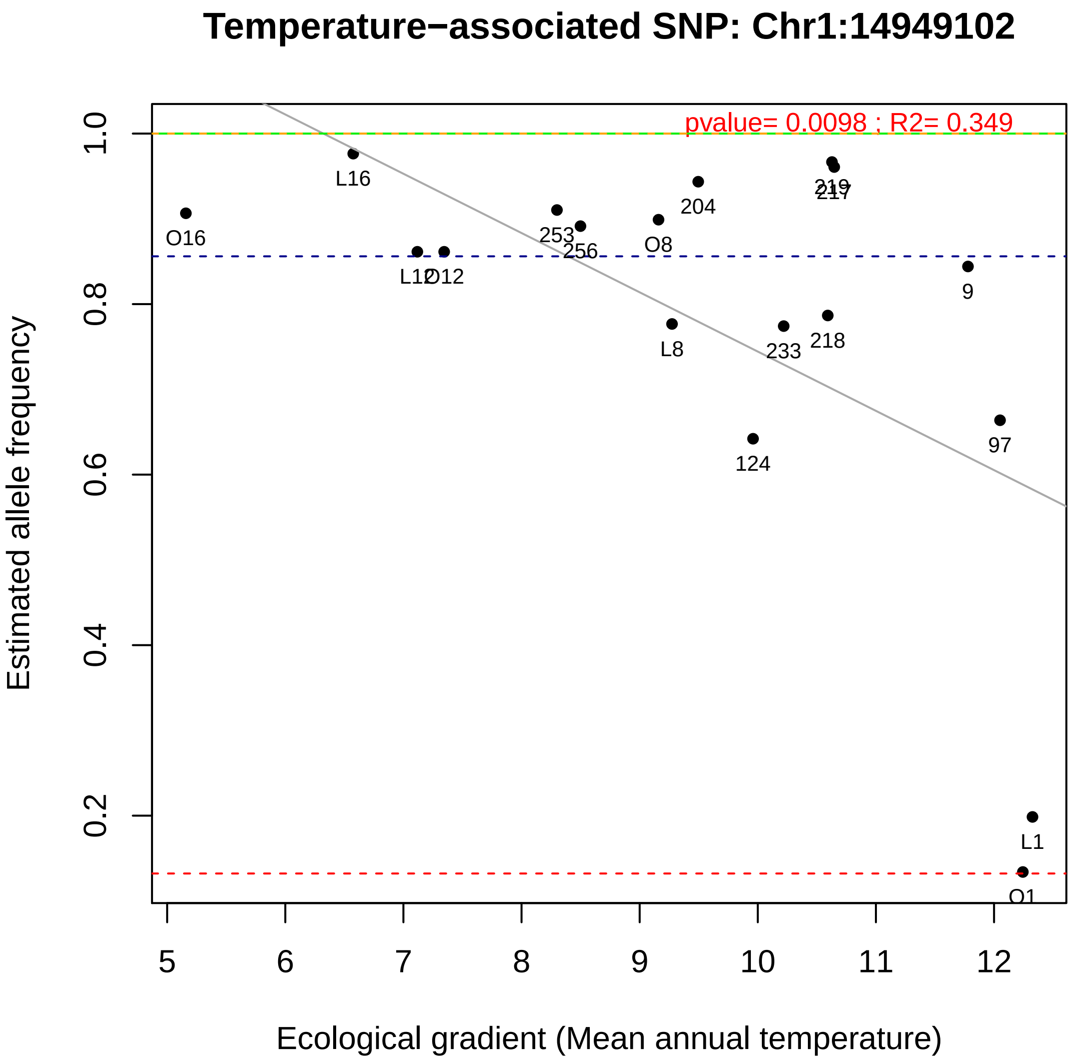

Temperature−associated SNP: Chr1:14949102
pvalue= 0.0098 ; R2= 0.349
Estimated allele frequency
Ecological gradient (Mean annual temperature)
1.0
0.8
0.6
0.4
0.2
5
6
7
8
9
10
11
12
O16
L16
L12
O12
253
256
O8
204
L8
124
233
218
219
217
9
97
L1
O1

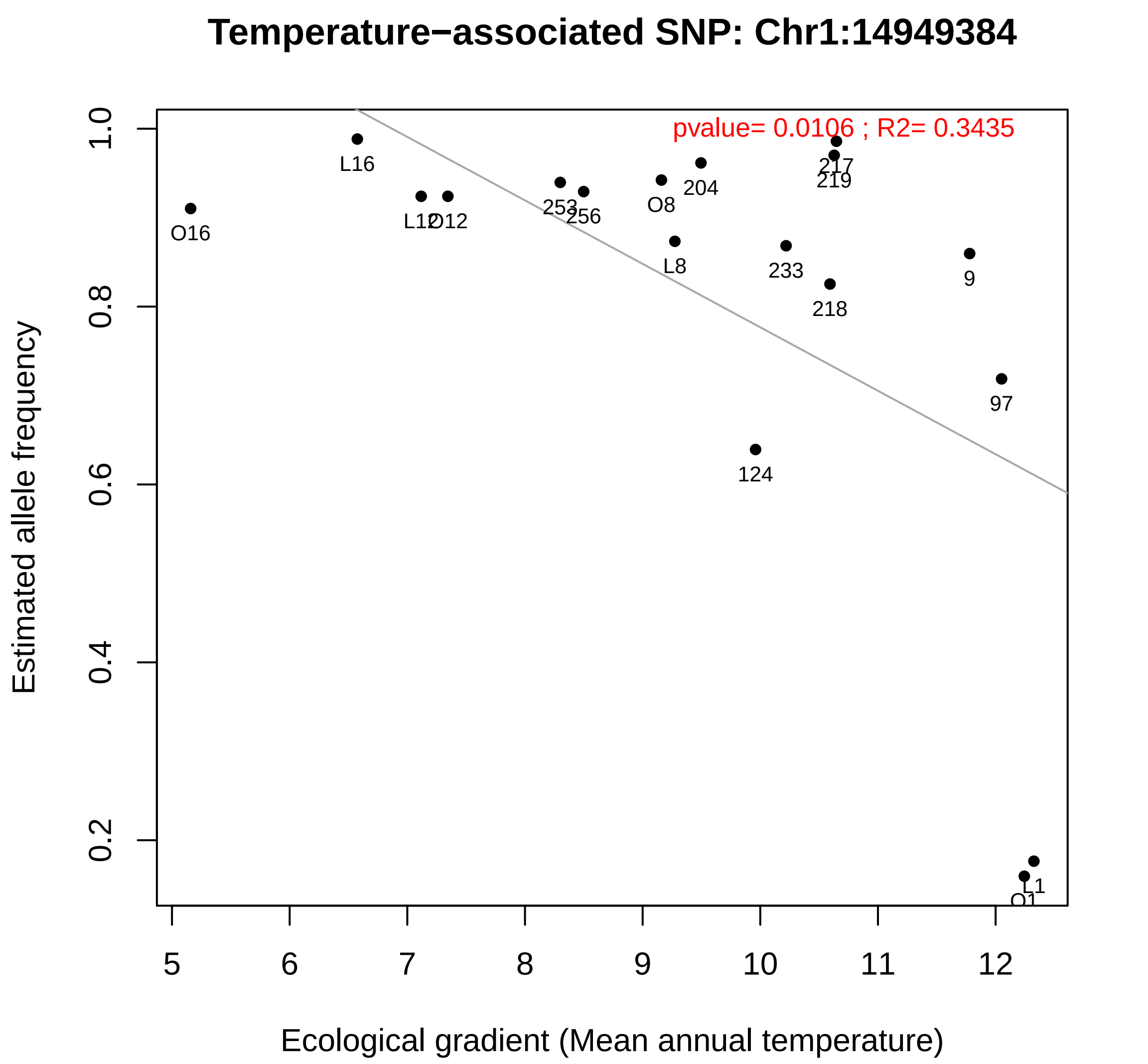
Temperature−associated SNP: Chr1:14949384
pvalue= 0.0106 ; R2= 0.3435
Estimated allele frequency
Ecological gradient (Mean annual temperature)
1.0
0.8
0.6
0.4
0.2
5
6
7
8
9
10
11
12
O16
L16
L12
O12
253
256
O8
204
217
219
L8
233
218
9
97
124
L1
O1

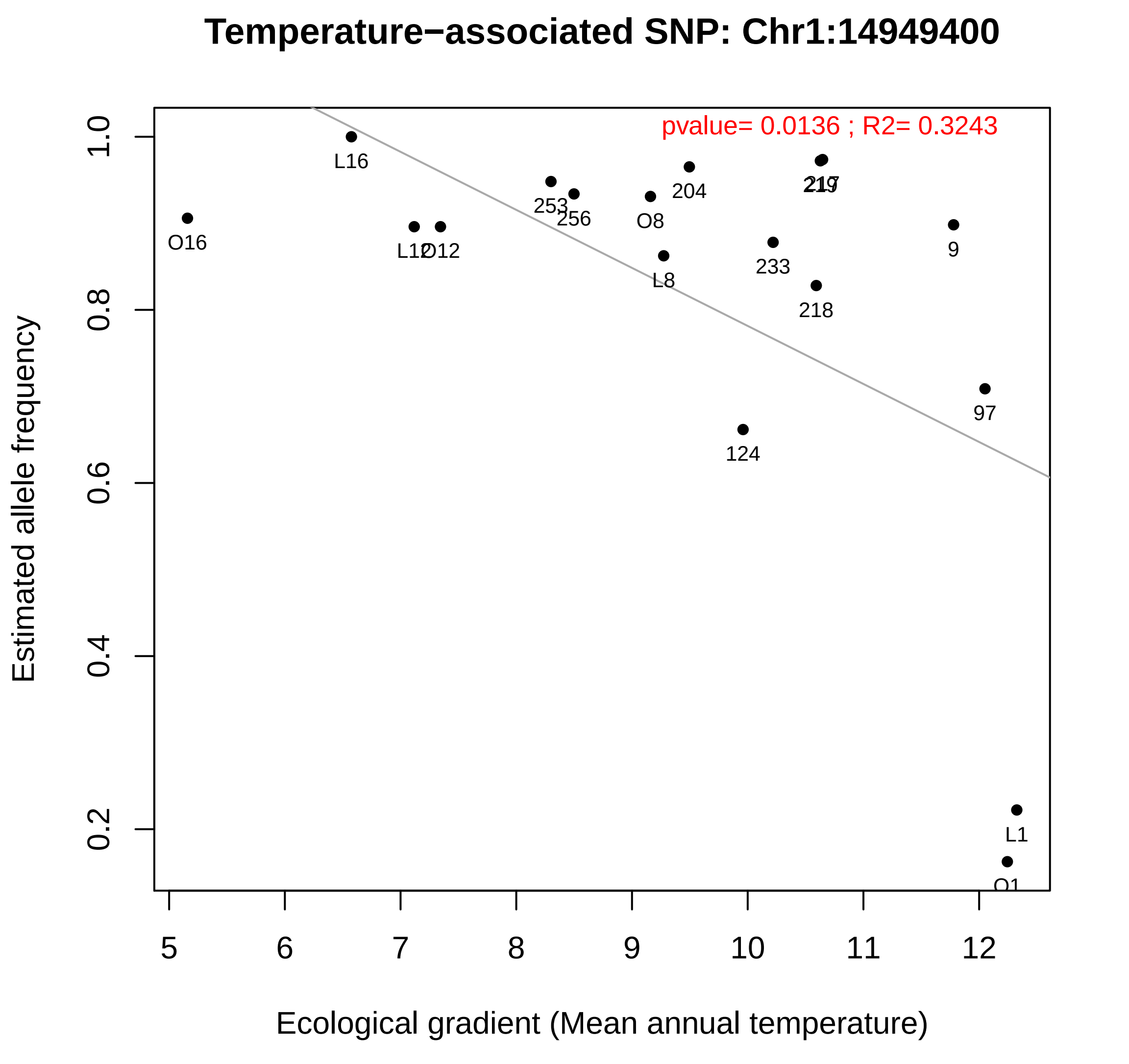
Temperature−associated SNP: Chr1:14949400
pvalue= 0.0136 ; R2= 0.3243
Estimated allele frequency
Ecological gradient (Mean annual temperature)
0.2
0.4
0.6
0.8
1.0
5
6
7
8
9
10
11
12
O16
L16
L12
O12
253
256
O8
204
L8
233
218
9
97
124
L1
O1

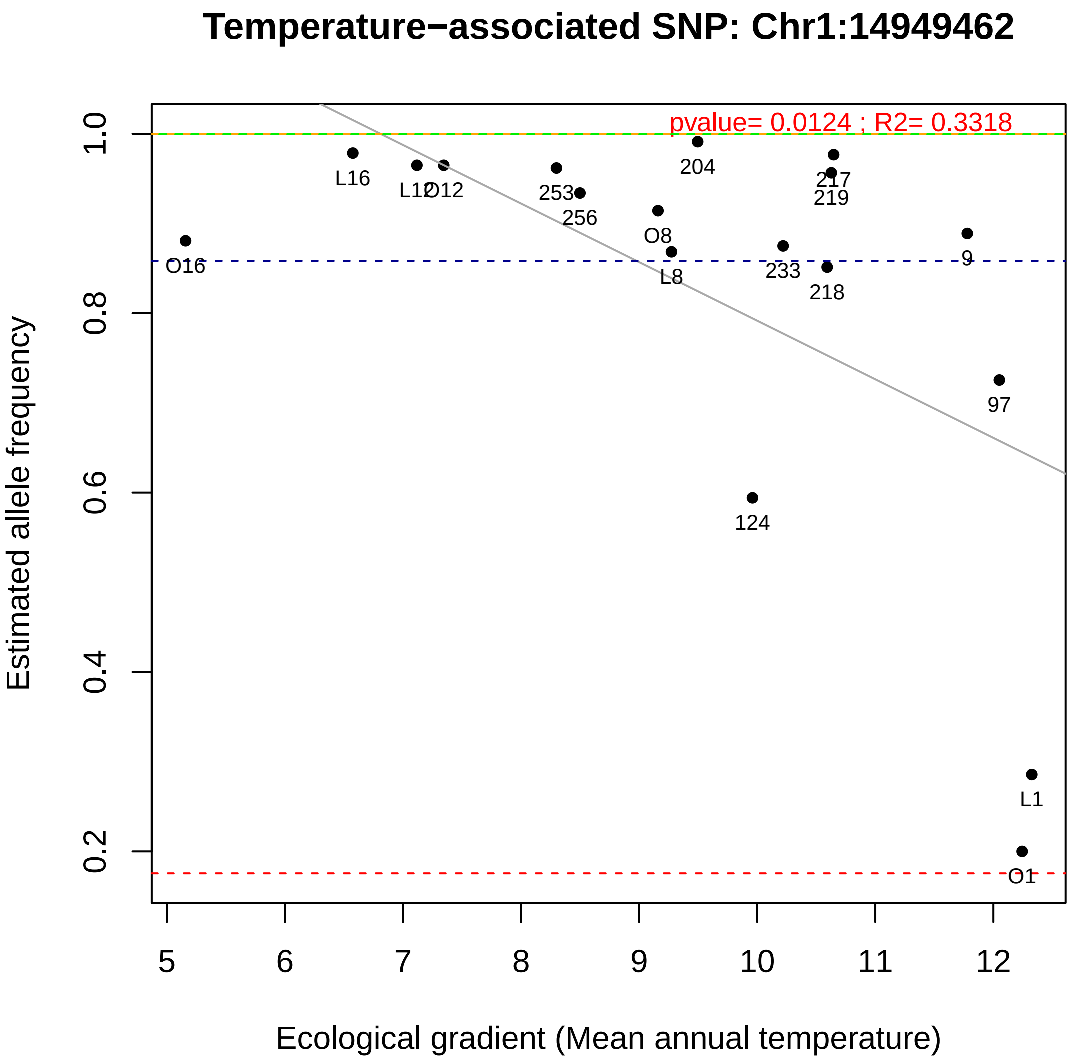

Temperature−associated SNP: Chr1:14949462
pvalue= 0.0124 ; R2= 0.3318
Estimated allele frequency
0.2
0.4
0.6
0.8
1.0
5
6
7
8
9
10
11
12
Ecological gradient (Mean annual temperature)
O16
L16
L12
O12
253
256
204
O8
L8
124
233
217
219
218
9
97
L1
O1

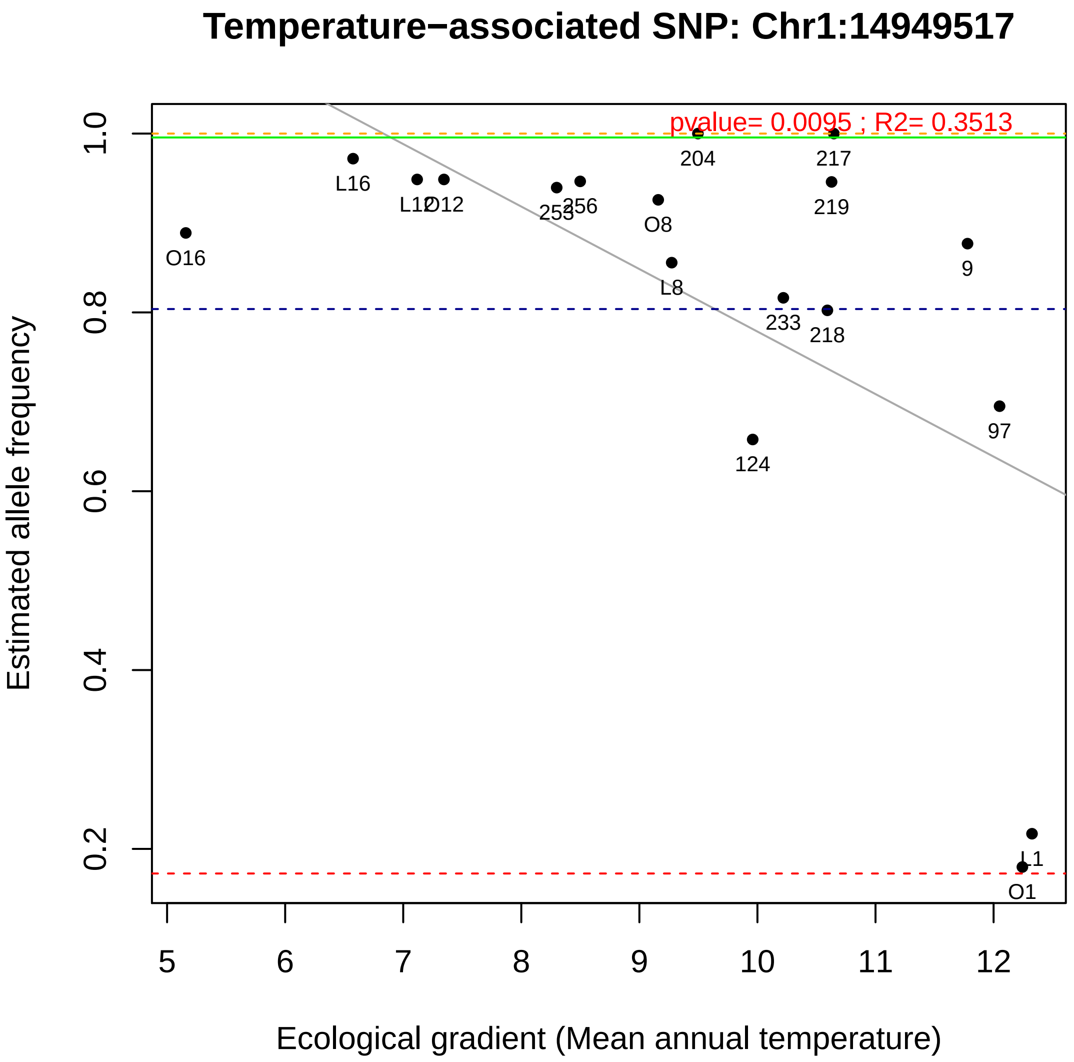

Temperature−associated SNP: Chr1:14949517
pvalue= 0.0095 ; R2= 0.3513
Estimated allele frequency
Ecological gradient (Mean annual temperature)
1.0
0.8
0.6
0.4
0.2
5
6
7
8
9
10
11
12
O16
L16
L12
O12
253
256
O8
204
L8
124
233
217
219
218
9
97
L1
O1

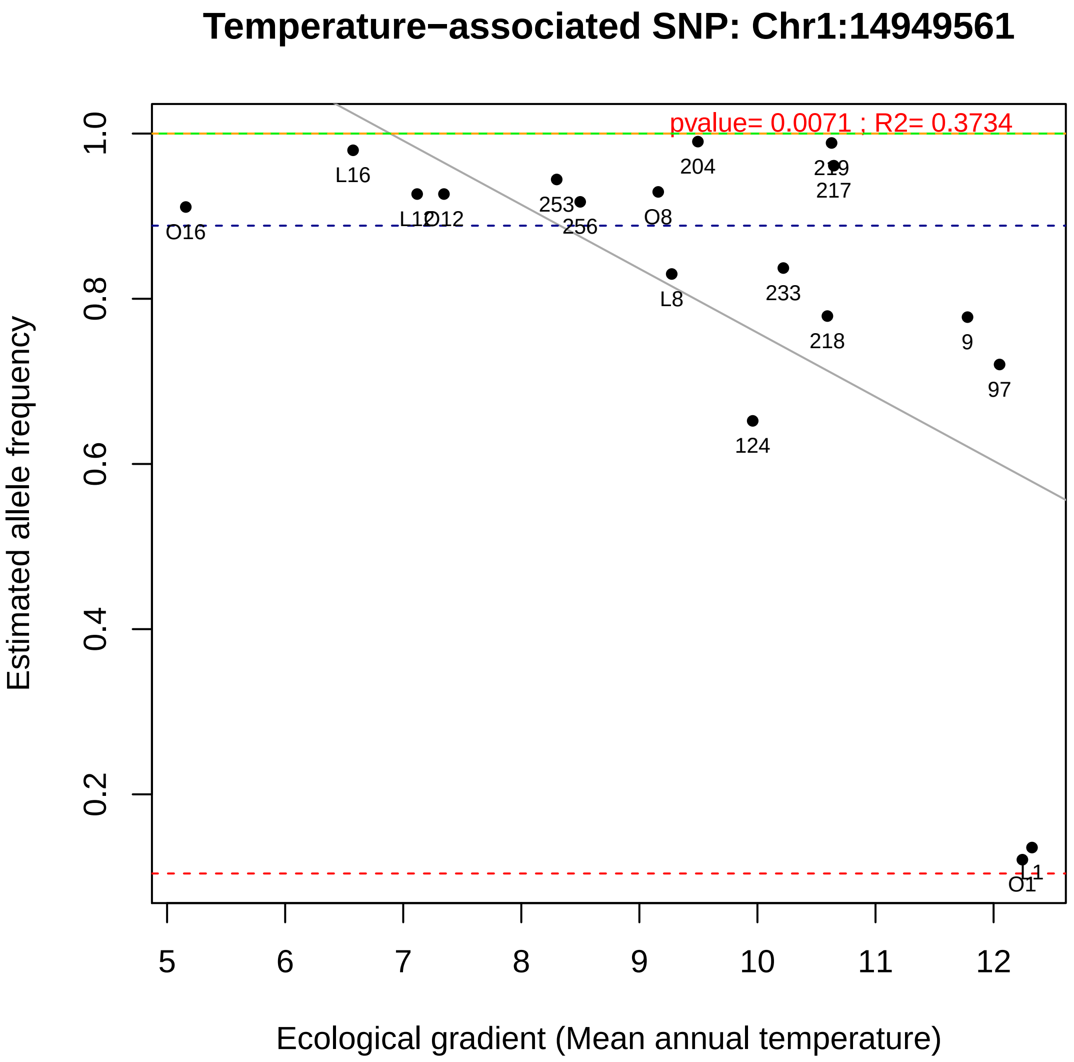

Temperature−associated SNP: Chr1:14949561
pvalue= 0.0071 ; R2= 0.3734
Estimated allele frequency
Ecological gradient (Mean annual temperature)
0.2
0.4
0.6
0.8
1.0
5
6
7
8
9
10
11
12
O16
L16
L12
O12
253
256
O8
L8
204
124
233
219
217
218
9
97
L1
O1

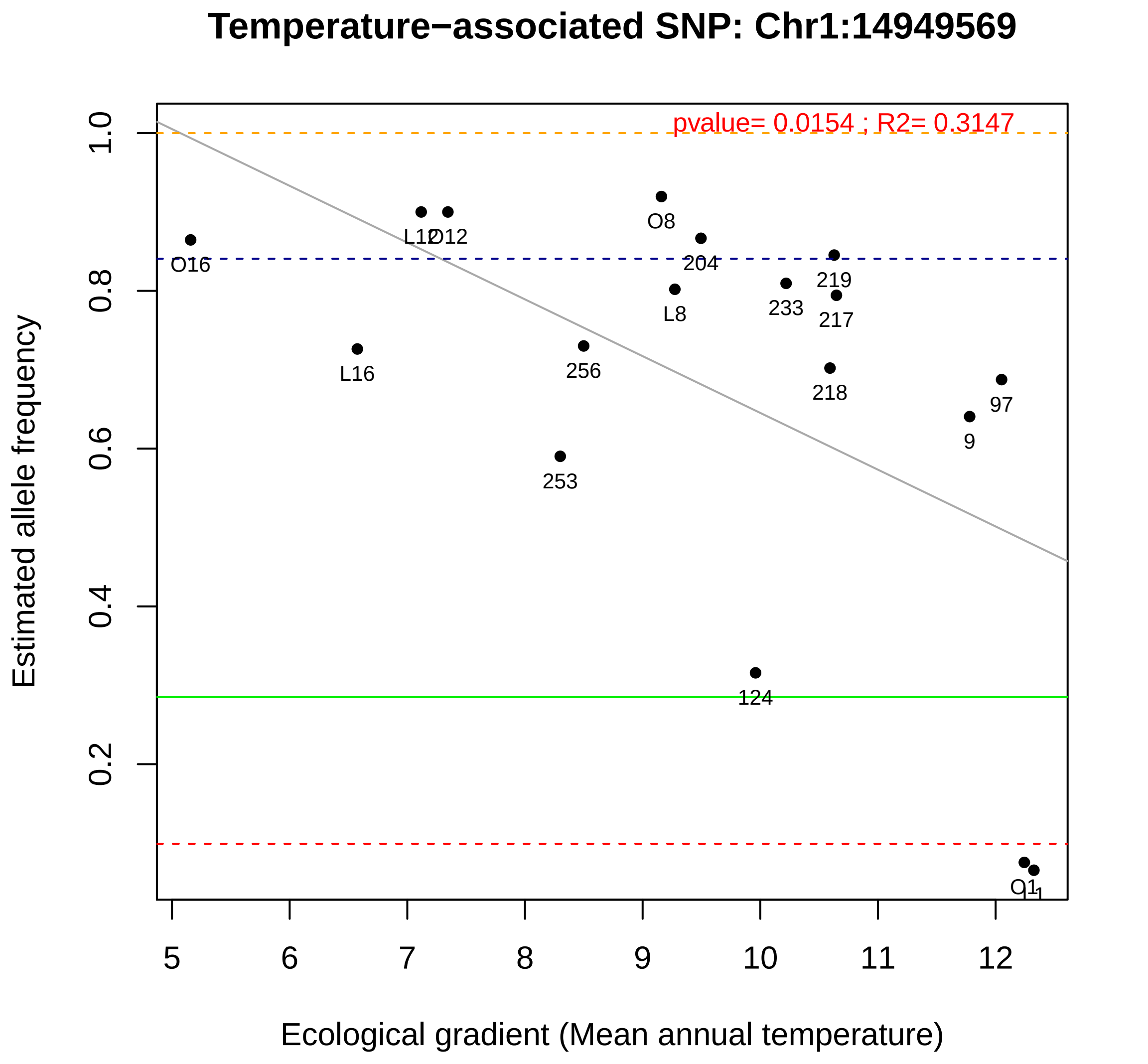

Temperature−associated SNP: Chr1:14949569
pvalue= 0.0154 ; R2= 0.3147
Estimated allele frequency
Ecological gradient (Mean annual temperature)
0.2
0.4
0.6
0.8
1.0
5
6
7
8
9
10
11
12
O16
L16
L12
O12
256
253
O8
204
L8
233
219
217
218
124
9
97

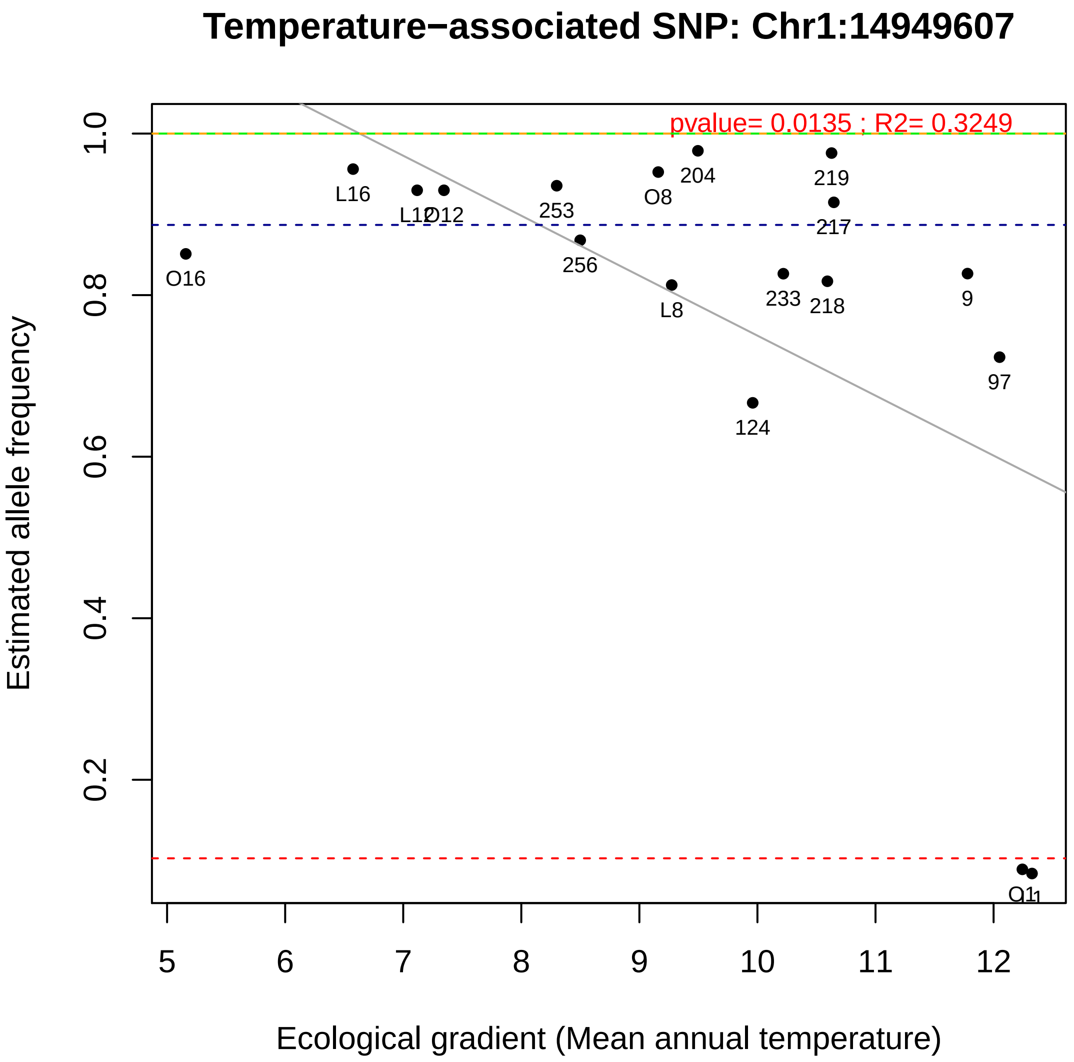
Temperature−associated SNP: Chr1:14949607
pvalue= 0.0135 ; R2= 0.3249
Estimated allele frequency
Ecological gradient (Mean annual temperature)
0.2
0.4
0.6
0.8
1.0
5
6
7
8
9
10
11
12
O16
L16
L12
O12
253
256
O8
204
L8
124
233
219
217
218
9
97

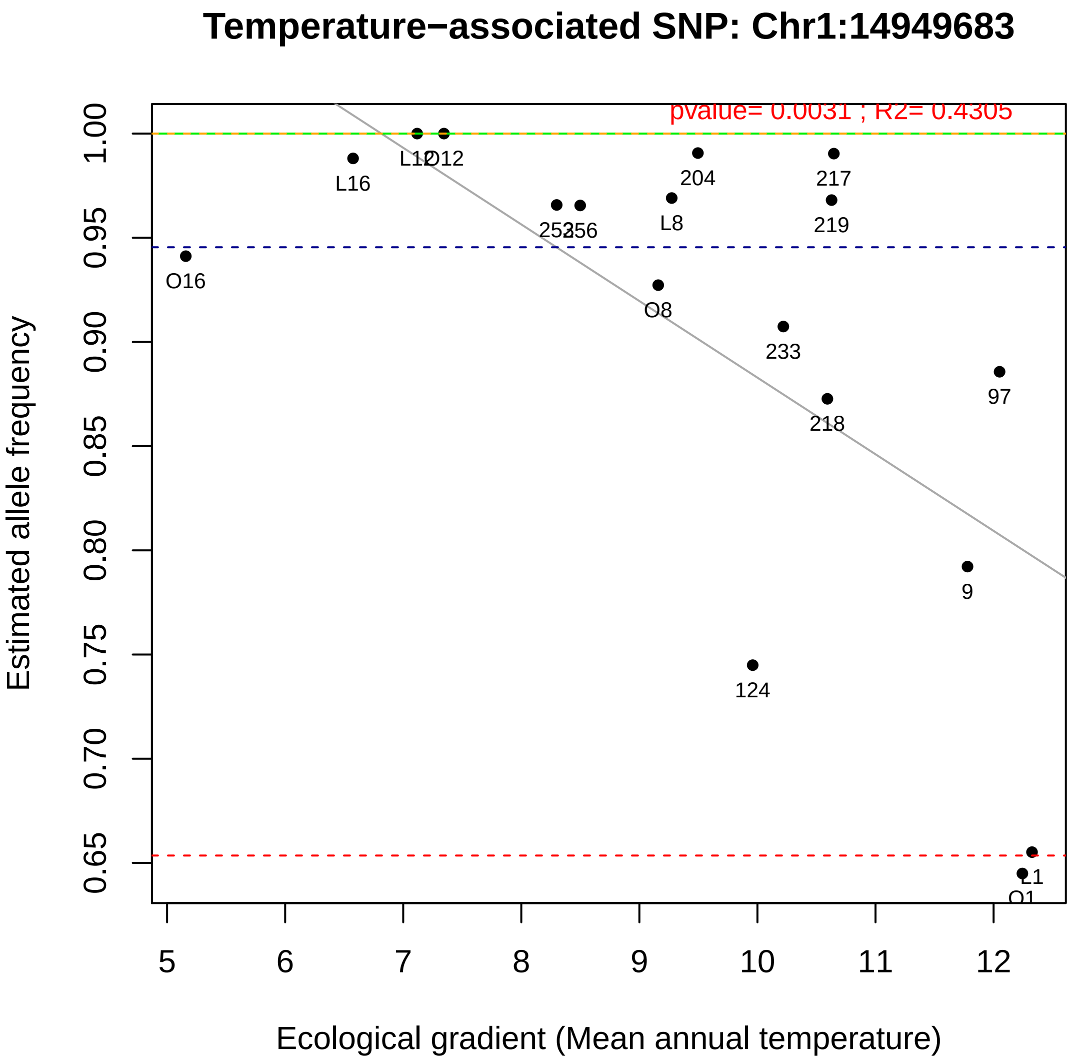

Temperature−associated SNP: Chr1:14949683
pvalue= 0.0031 ; R2= 0.4305
Estimated allele frequency
Ecological gradient (Mean annual temperature)
1.00
0.95
0.90
0.85
0.80
0.75
0.70
0.65
5
6
7
8
9
10
11
12
O16
L16
L12
O12
253
256
L8
204
217
219
O8
233
218
97
9
124
L1
O1

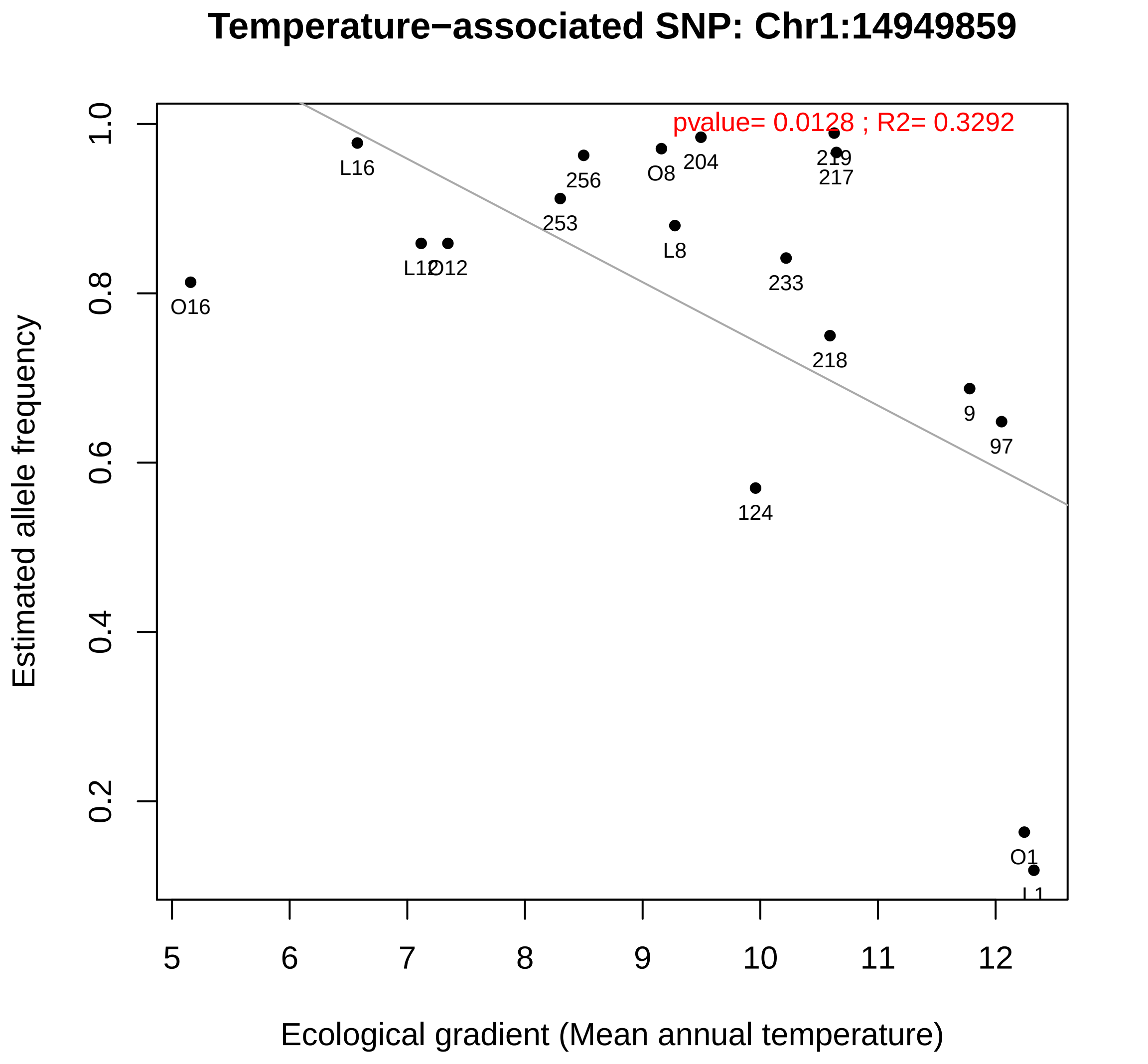
Temperature−associated SNP: Chr1:14949859
pvalue= 0.0128 ; R2= 0.3292
L16
256
O8
204
219
217
253
L8
L12
O12
233
O16
218
9
97
124
O1
L1
1.0
0.8
0.6
0.4
0.2
Estimated allele frequency
5
6
7
8
9
10
11
12
Ecological gradient (Mean annual temperature)

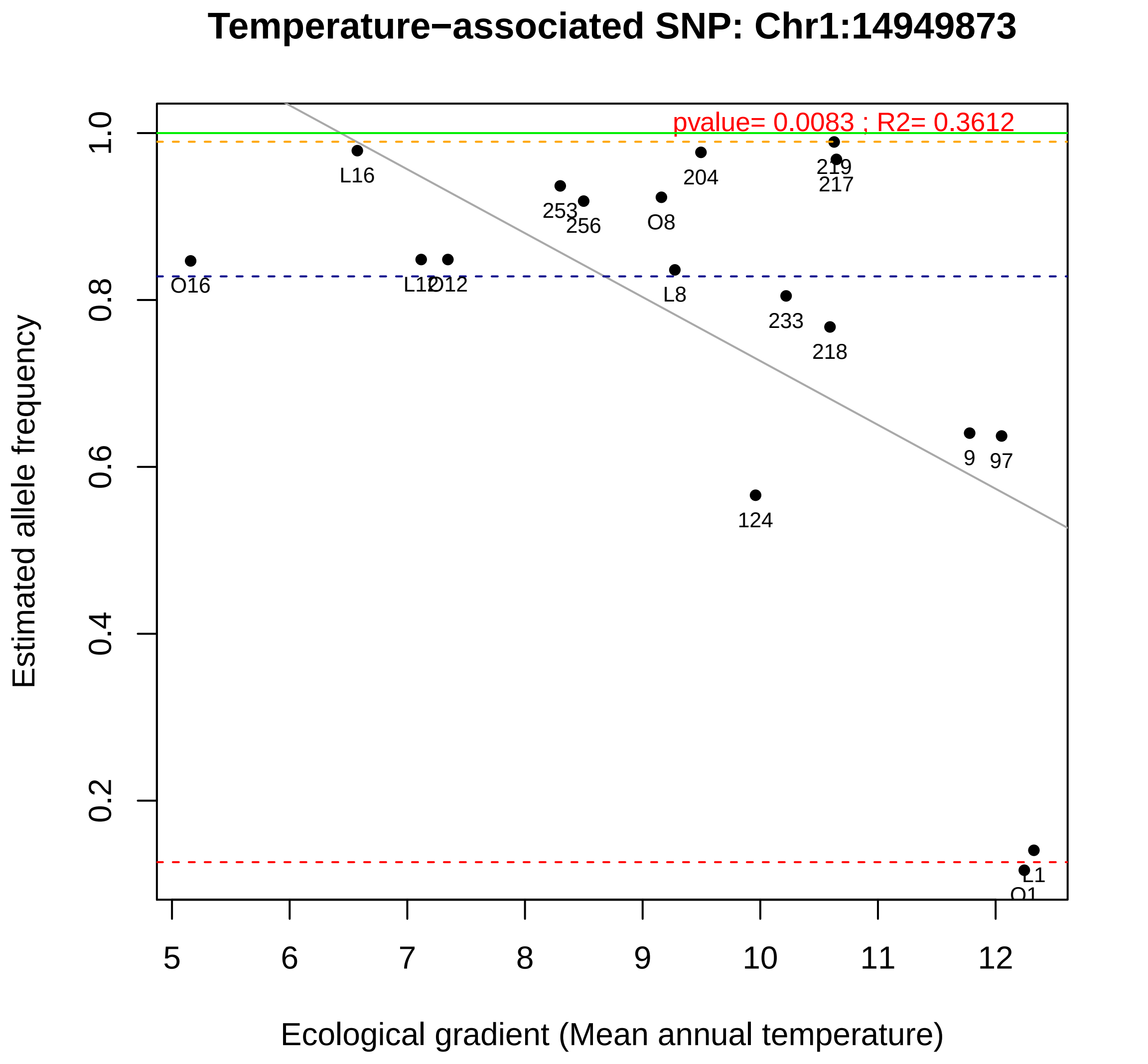

Temperature−associated SNP: Chr1:14949873
pvalue= 0.0083 ; R2= 0.3612
Estimated allele frequency
Ecological gradient (Mean annual temperature)
0.2
0.4
0.6
0.8
1.0
5
6
7
8
9
10
11
12
O16
L16
L12
O12
253
256
O8
L8
204
233
218
219
217
124
9
97
L1
O1

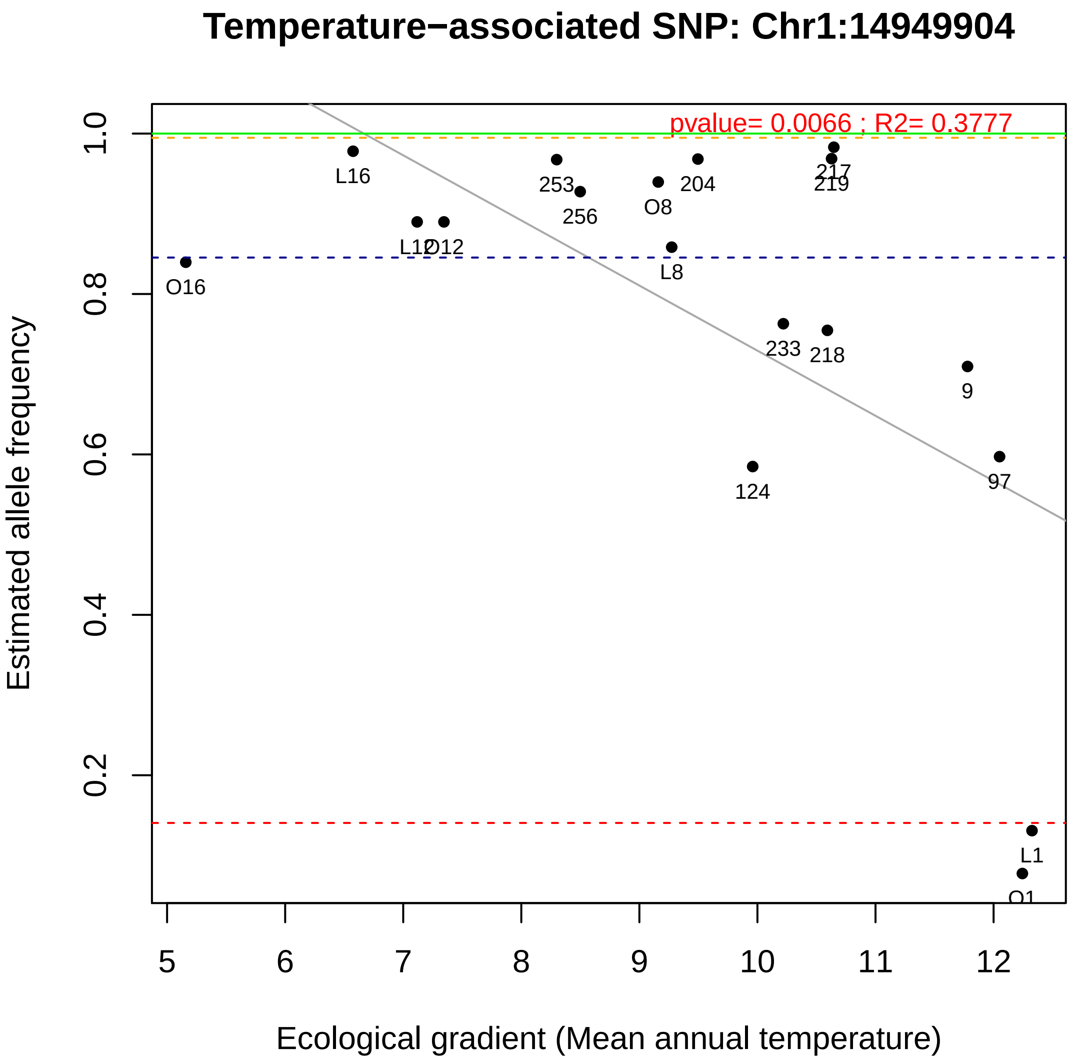
Temperature−associated SNP: Chr1:14949904
pvalue= 0.0066 ; R2= 0.3777
Estimated allele frequency
Ecological gradient (Mean annual temperature)
1.0
0.8
0.6
0.4
0.2
5
6
7
8
9
10
11
12
O16
L16
L12
O12
253
256
O8
204
L8
217
219
233
218
124
9
97
L1
O1

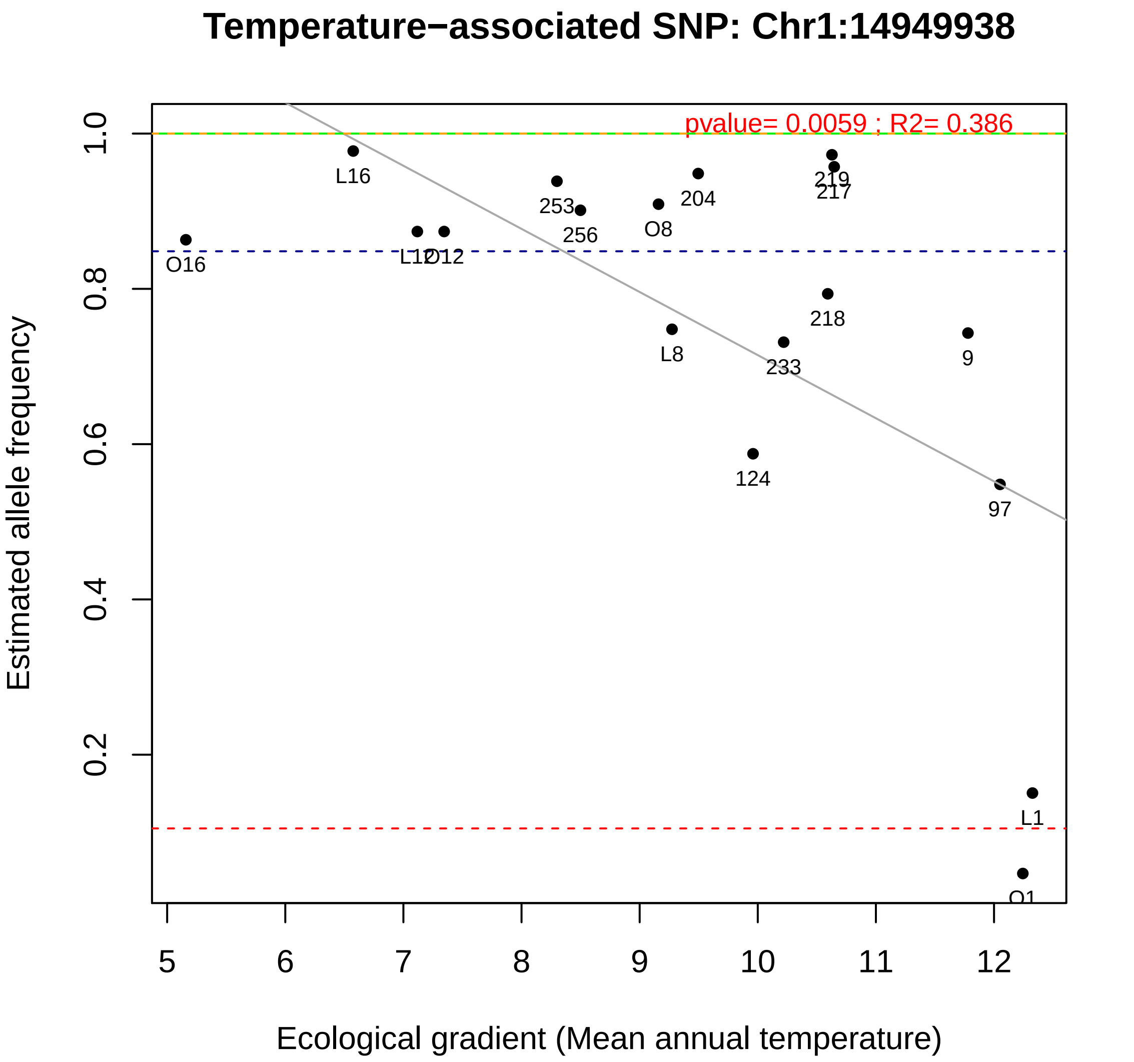

Temperature−associated SNP: Chr1:14949938
pvalue= 0.0059 ; R2= 0.386
Estimated allele frequency
Ecological gradient (Mean annual temperature)
0.2
0.4
0.6
0.8
1.0
5
6
7
8
9
10
11
12
O16
L16
L12
O12
253
256
O8
204
219
217
L8
233
218
124
9
97
L1
O1

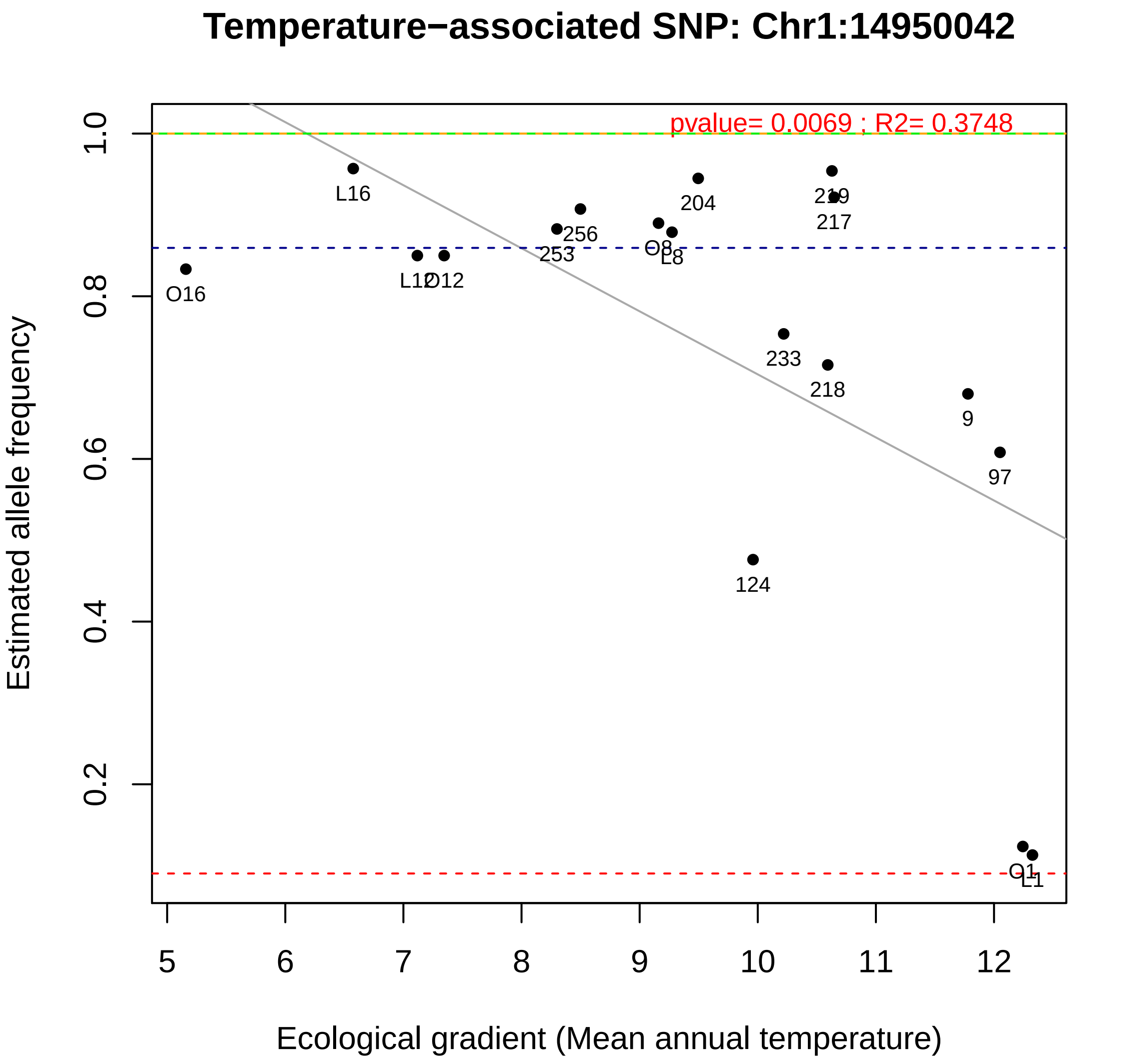
Temperature−associated SNP: Chr1:14950042
pvalue= 0.0069 ; R2= 0.3748
Estimated allele frequency
0.2
0.4
0.6
0.8
1.0
5
6
7
8
9
10
11
12
Ecological gradient (Mean annual temperature)
L16
O16
L12
O12
253
256
O8
L8
204
219
217
233
218
9
97
124
O1
L1

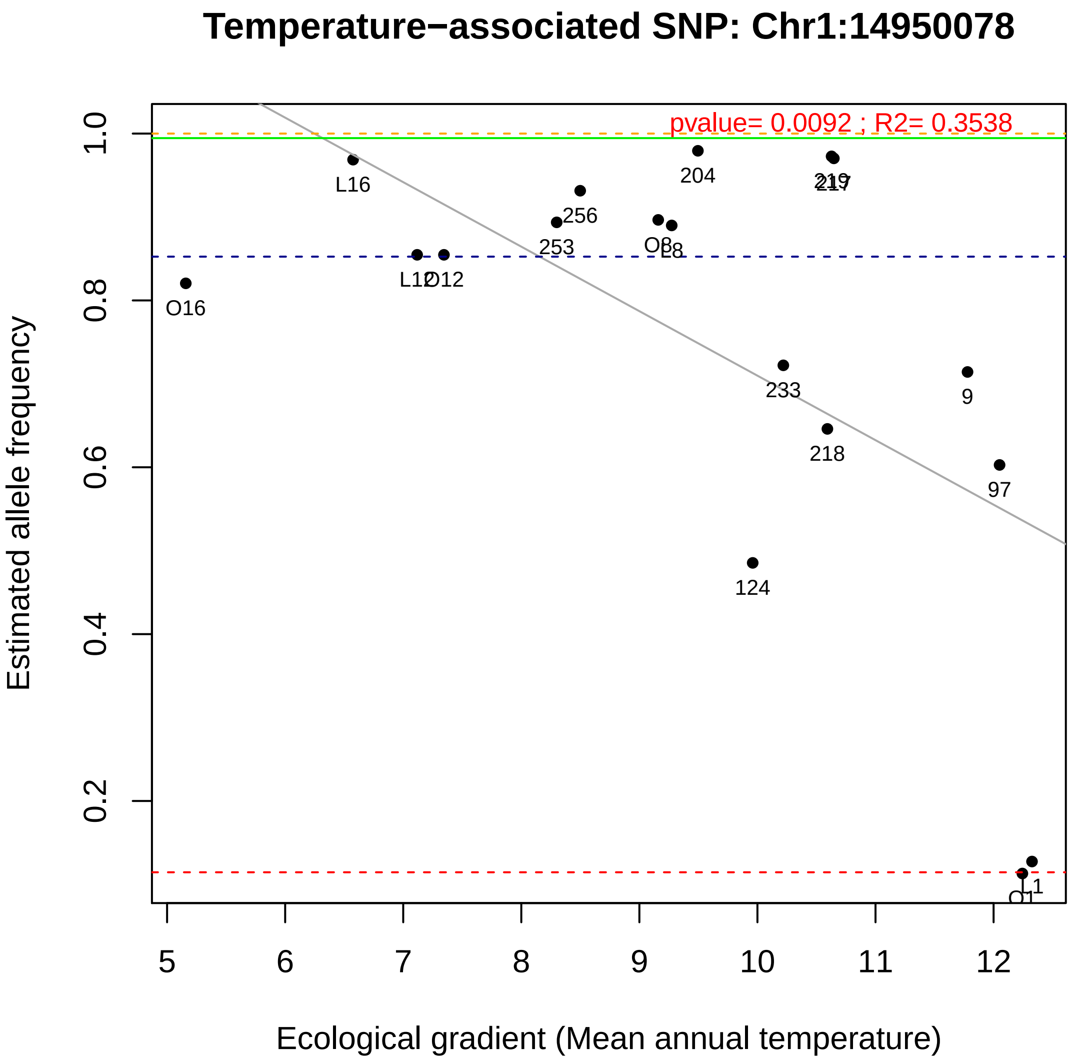
Temperature−associated SNP: Chr1:14950078
pvalue= 0.0092 ; R2= 0.3538
Estimated allele frequency
Ecological gradient (Mean annual temperature)
0.2
0.4
0.6
0.8
1.0
5
6
7
8
9
10
11
12
O16
L16
L12
O12
253
256
O8
L8
204
219
217
233
218
124
9
97
L1
O1

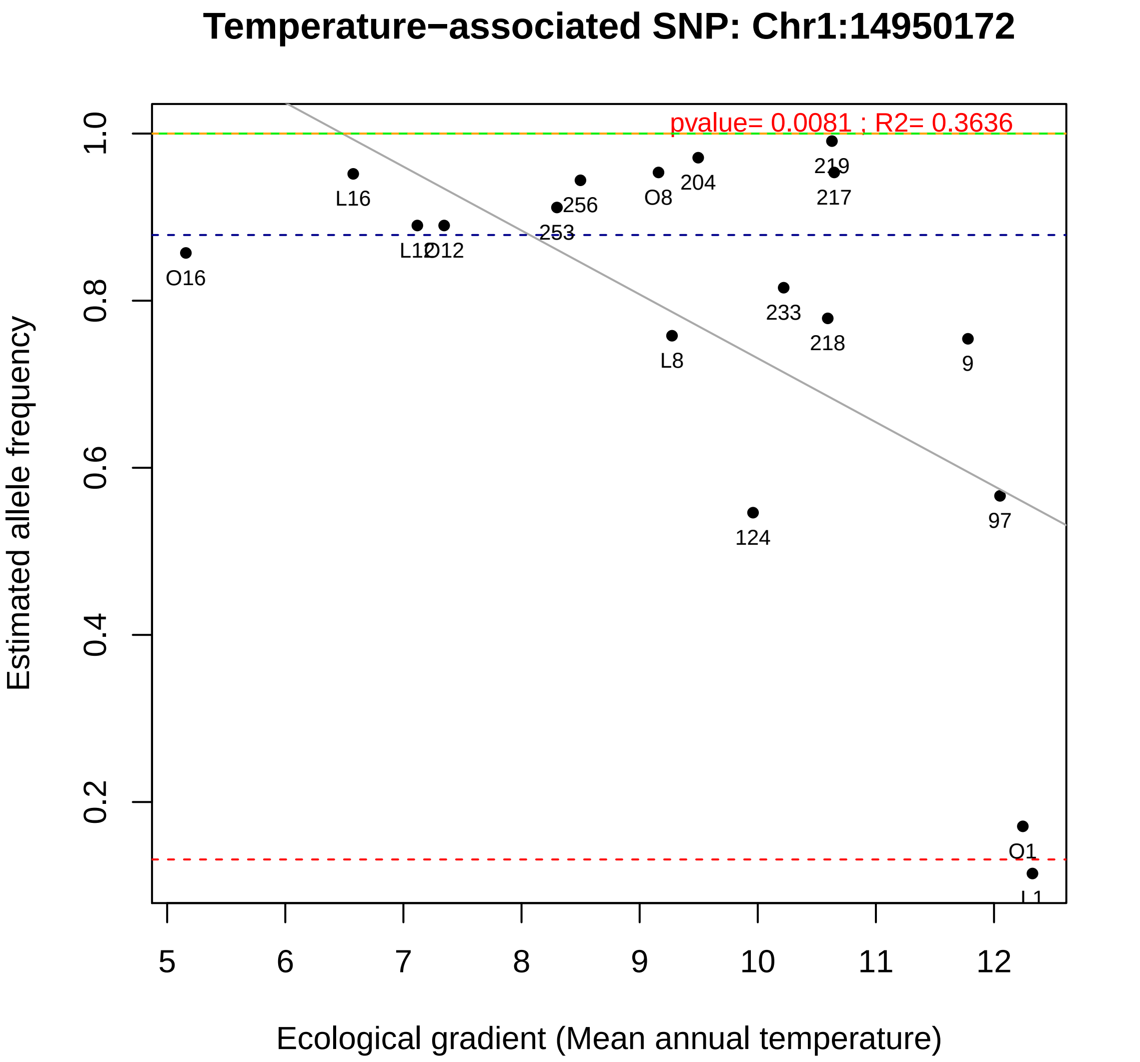

Temperature−associated SNP: Chr1:14950172
pvalue= 0.0081 ; R2= 0.3636
Estimated allele frequency
Ecological gradient (Mean annual temperature)
0.2
0.4
0.6
0.8
1.0
5
6
7
8
9
10
11
12
O16
L16
L12
O12
253
256
O8
204
219
217
L8
233
218
124
9
97
O1

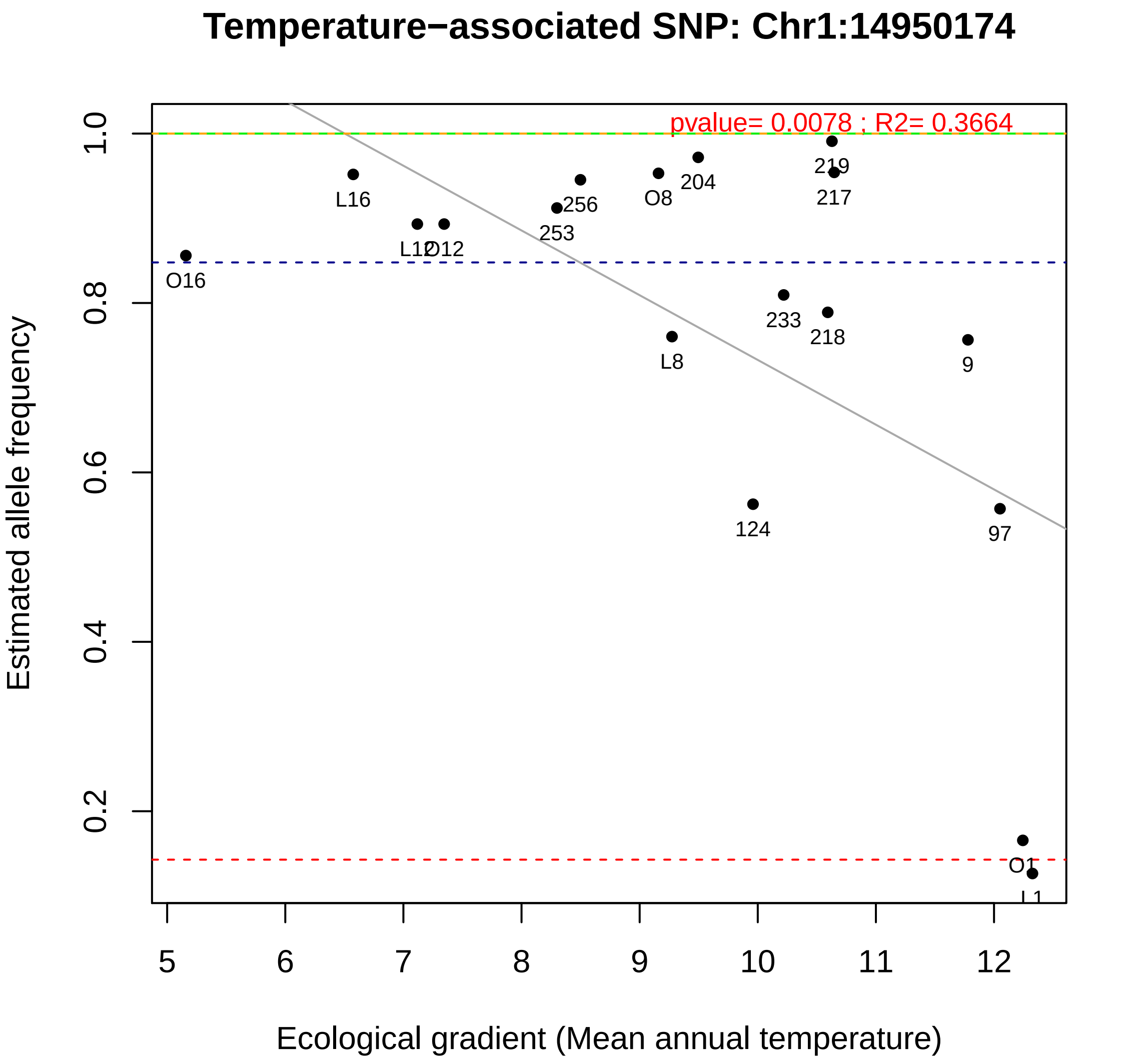

Temperature−associated SNP: Chr1:14950174
pvalue= 0.0078 ; R2= 0.3664
Estimated allele frequency
Ecological gradient (Mean annual temperature)
1.0
0.8
0.6
0.4
0.2
5
6
7
8
9
10
11
12
O16
L16
L12
O12
253
256
O8
204
219
217
233
218
L8
9
124
97
O1

**Temperature−associated SNP: Chr1:14950233**

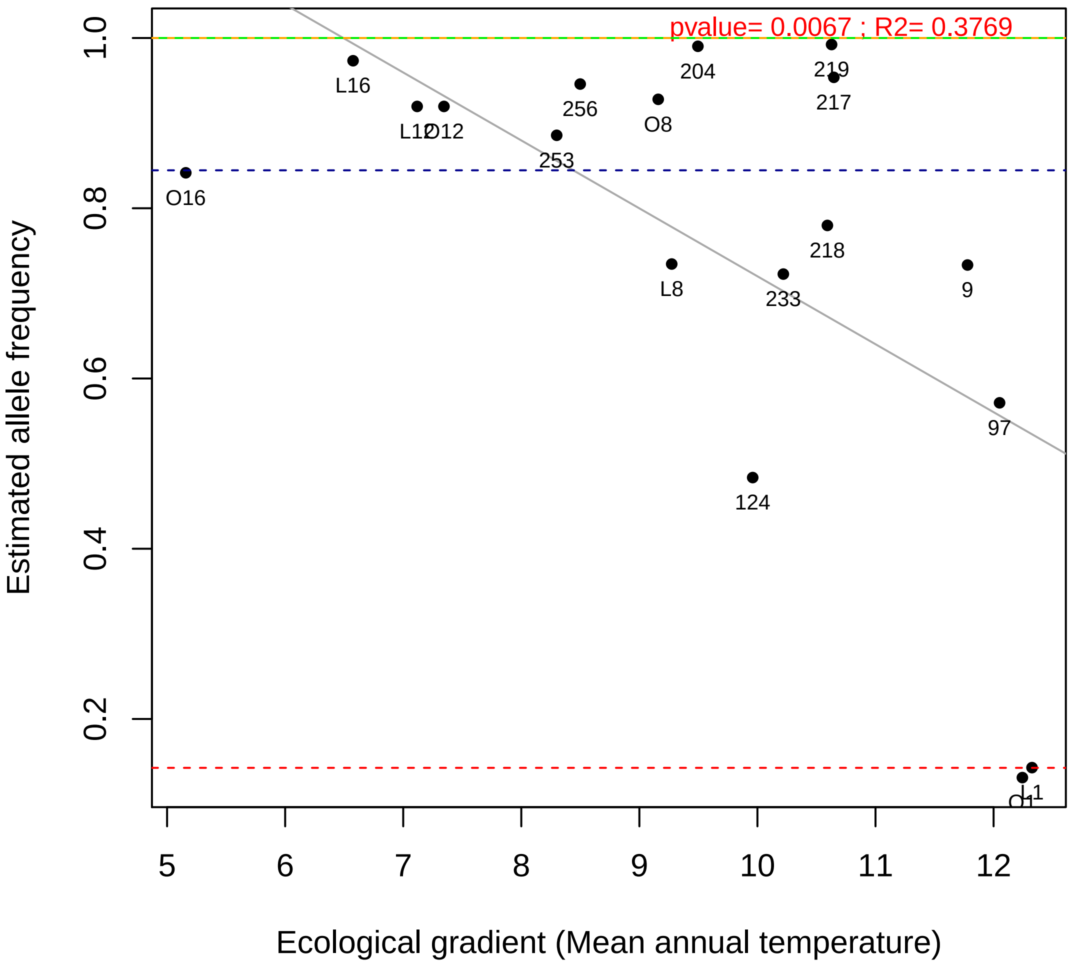

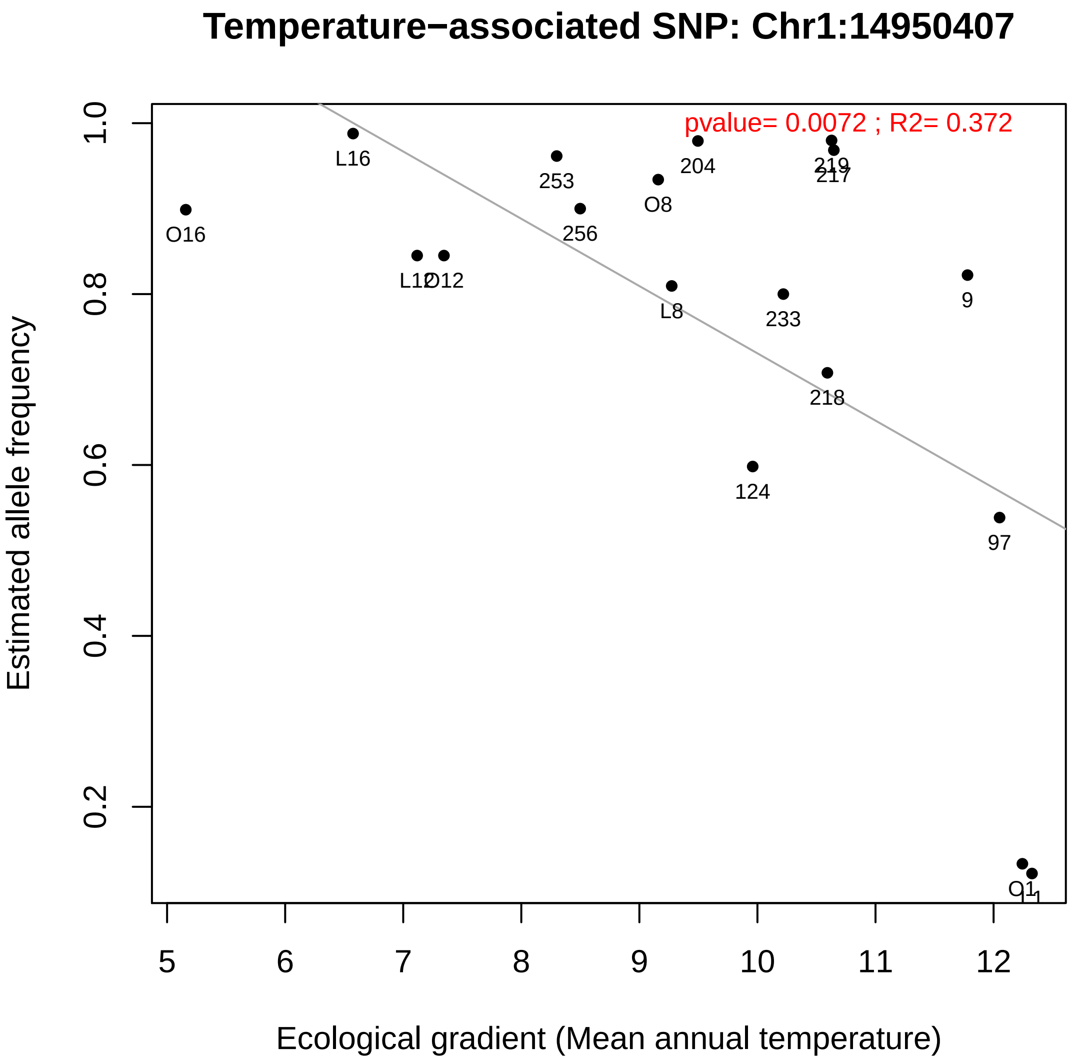
Temperature−associated SNP: Chr1:14950407
pvalue= 0.0072 ; R2= 0.372
Estimated allele frequency
0.2
0.4
0.6
0.8
1.0
Ecological gradient (Mean annual temperature)
5
6
7
8
9
10
11
12
O16
L16
L12
O12
253
256
O8
204
L8
233
219
217
218
124
9
97
O1
L1

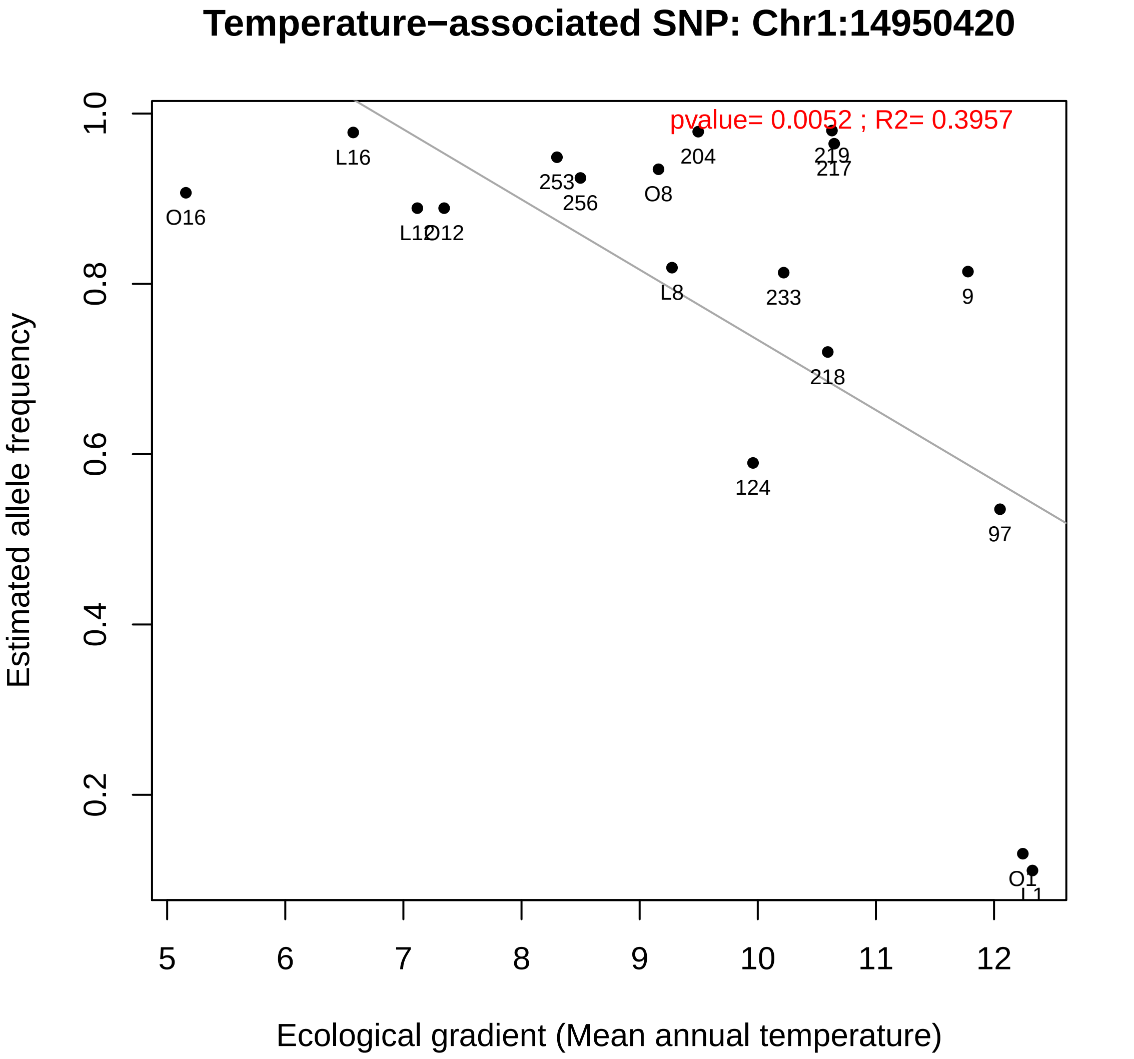
Temperature−associated SNP: Chr1:14950420
pvalue= 0.0052 ; R2= 0.3957
Estimated allele frequency
0.2
0.4
0.6
0.8
1.0
5
6
7
8
9
10
11
12
Ecological gradient (Mean annual temperature)
O16
L16
L12
O12
253
256
O8
204
219
217
L8
233
9
218
124
97
O1

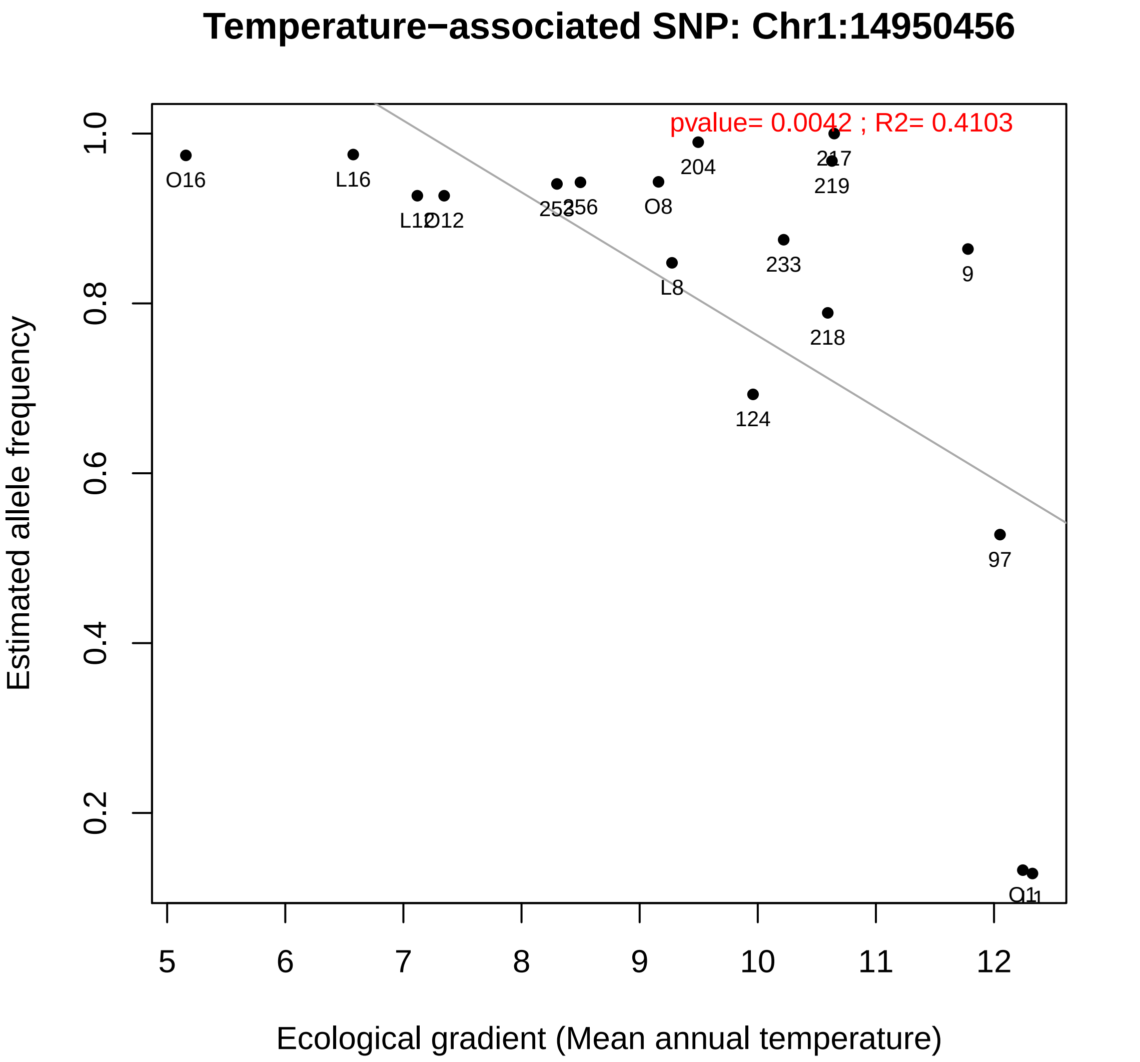
Temperature−associated SNP: Chr1:14950456
pvalue= 0.0042 ; R2= 0.4103
Estimated allele frequency
Ecological gradient (Mean annual temperature)
0.2
0.4
0.6
0.8
1.0
5
6
7
8
9
10
11
12
O16
L16
L12
O12
256
O8
204
217
219
L8
233
9
218
124
97

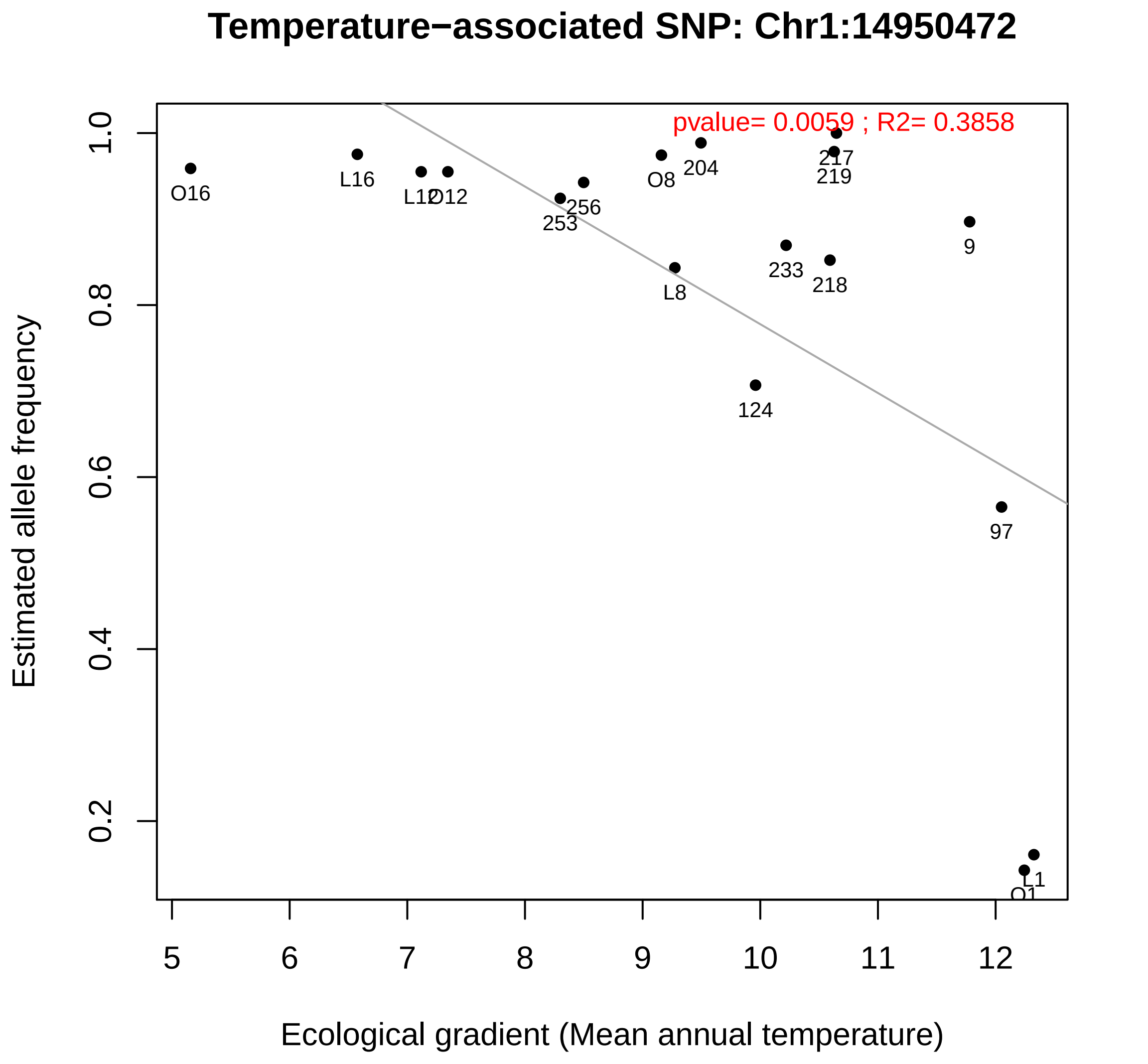
Temperature−associated SNP: Chr1:14950472
pvalue= 0.0059 ; R2= 0.3858
Estimated allele frequency
Ecological gradient (Mean annual temperature)
0.2
0.4
0.6
0.8
1.0
5
6
7
8
9
10
11
12
O16
L16
L12
O12
253
256
O8
204
217
219
L8
233
218
9
124
97
L1
O1

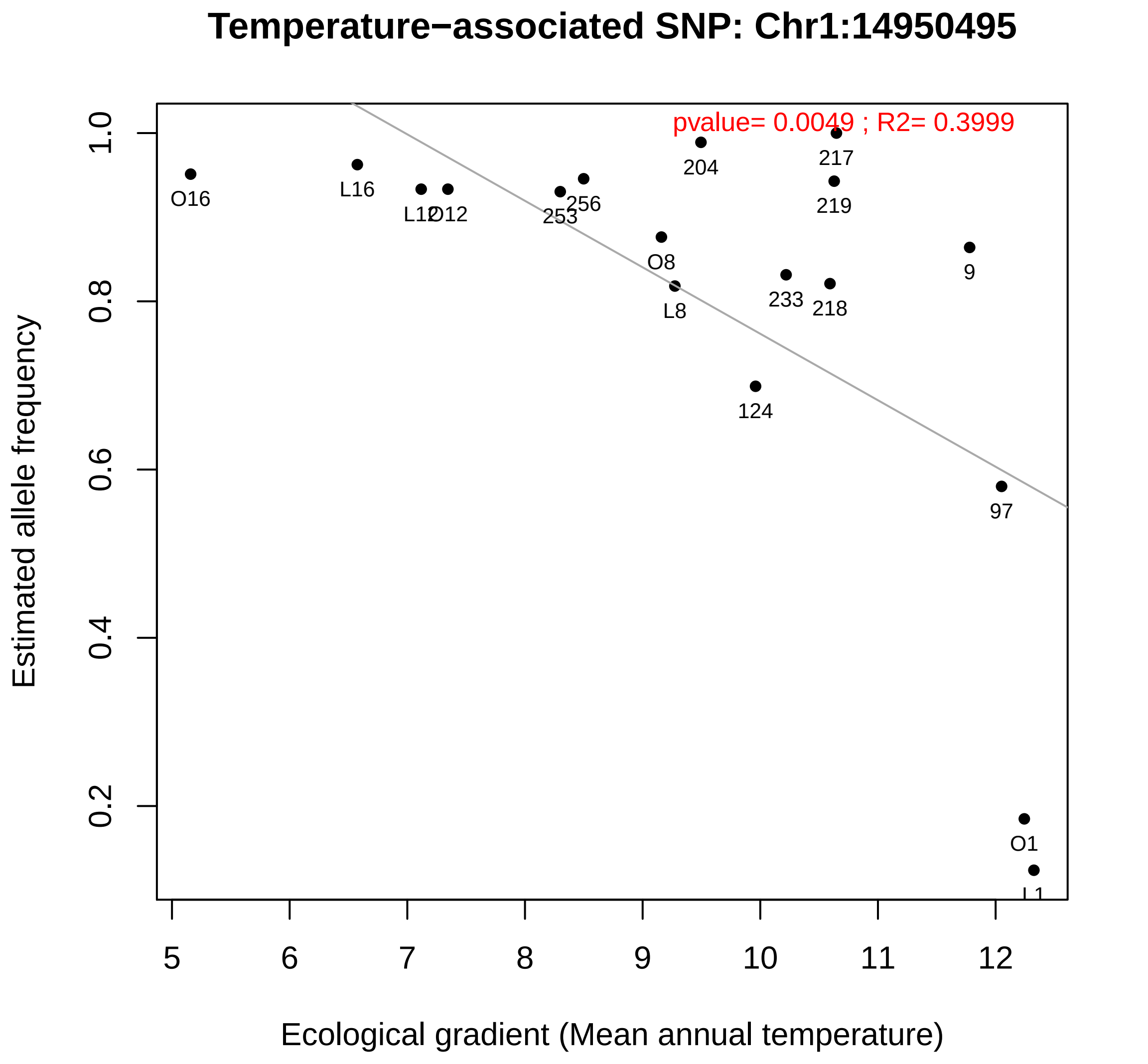

Temperature−associated SNP: Chr1:14950495
pvalue= 0.0049 ; R2= 0.3999
Estimated allele frequency
Ecological gradient (Mean annual temperature)
0.2
0.4
0.6
0.8
1.0
5
6
7
8
9
10
11
12
O16
L16
L12
O12
253
256
204
217
219
O8
L8
233
218
9
124
97
O1

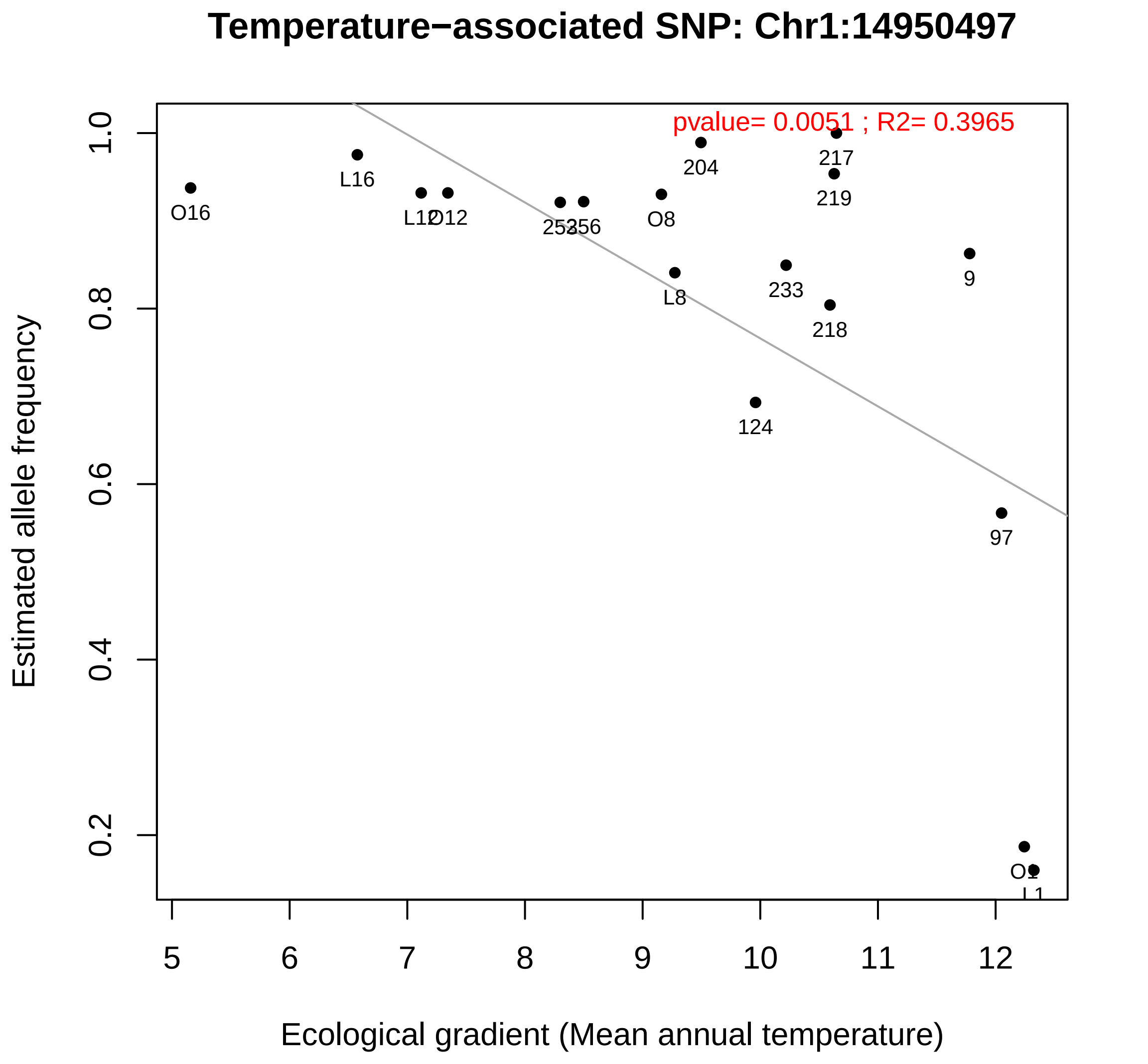
Temperature−associated SNP: Chr1:14950497
pvalue= 0.0051 ; R2= 0.3965
Estimated allele frequency
0.2
0.4
0.6
0.8
1.0
5
6
7
8
9
10
11
12
Ecological gradient (Mean annual temperature)
O16
L16
L12
O12
256
O8
204
217
219
L8
233
218
9
124
97

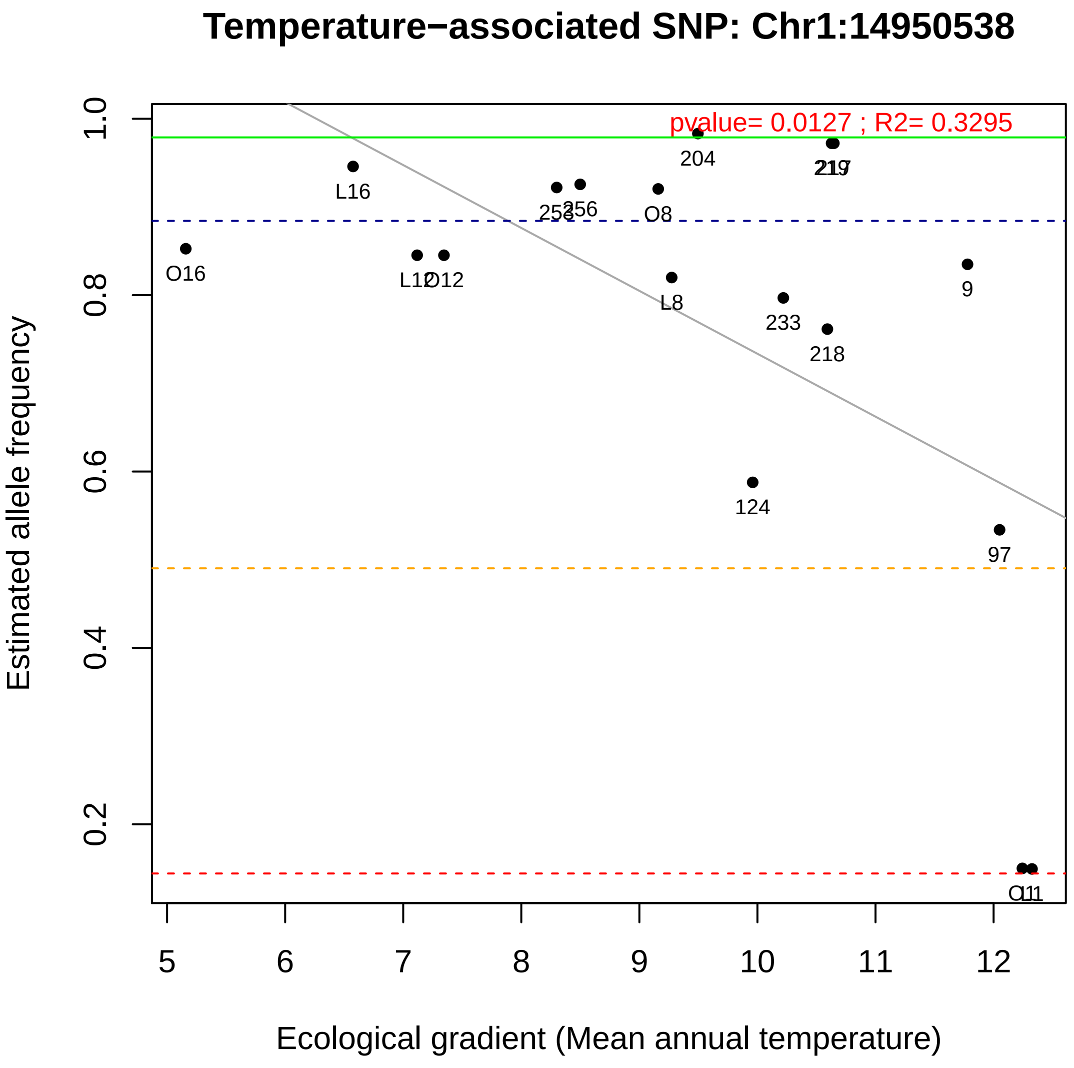

Temperature−associated SNP: Chr1:14950538
pvalue= 0.0127 ; R2= 0.3295
Estimated allele frequency
Ecological gradient (Mean annual temperature)
0.2
0.4
0.6
0.8
1.0
5
6
7
8
9
10
11
12
O16
L16
L12
O12
253
256
O8
204
L8
233
217
219
218
124
9
97

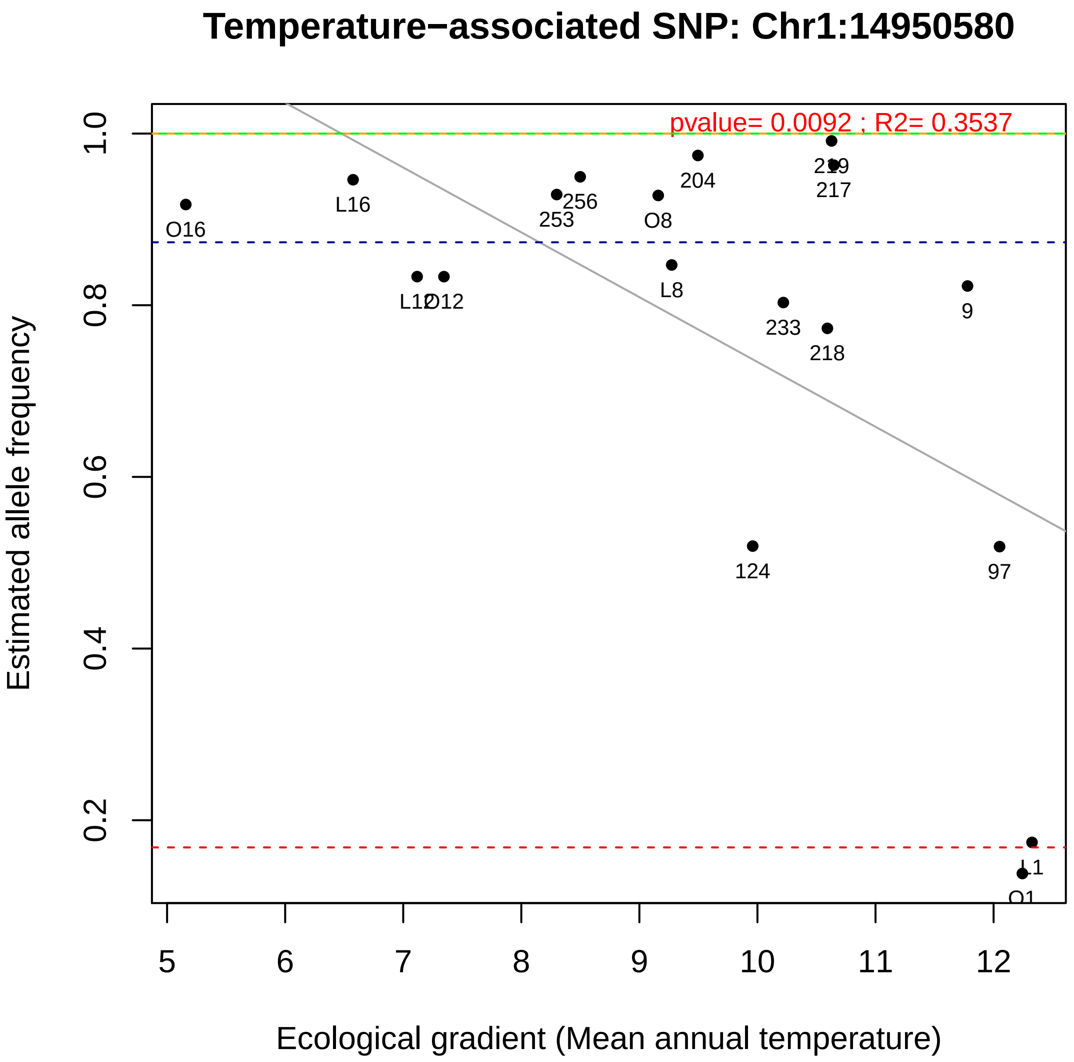
Temperature−associated SNP: Chr1:14950580
pvalue= 0.0092 ; R2= 0.3537
Estimated allele frequency
Ecological gradient (Mean annual temperature)
0.2
0.4
0.6
0.8
1.0
5
6
7
8
9
10
11
12
O16
L16
L12
O12
253
256
O8
L8
204
219
217
233
218
124
9
97
L1
O1

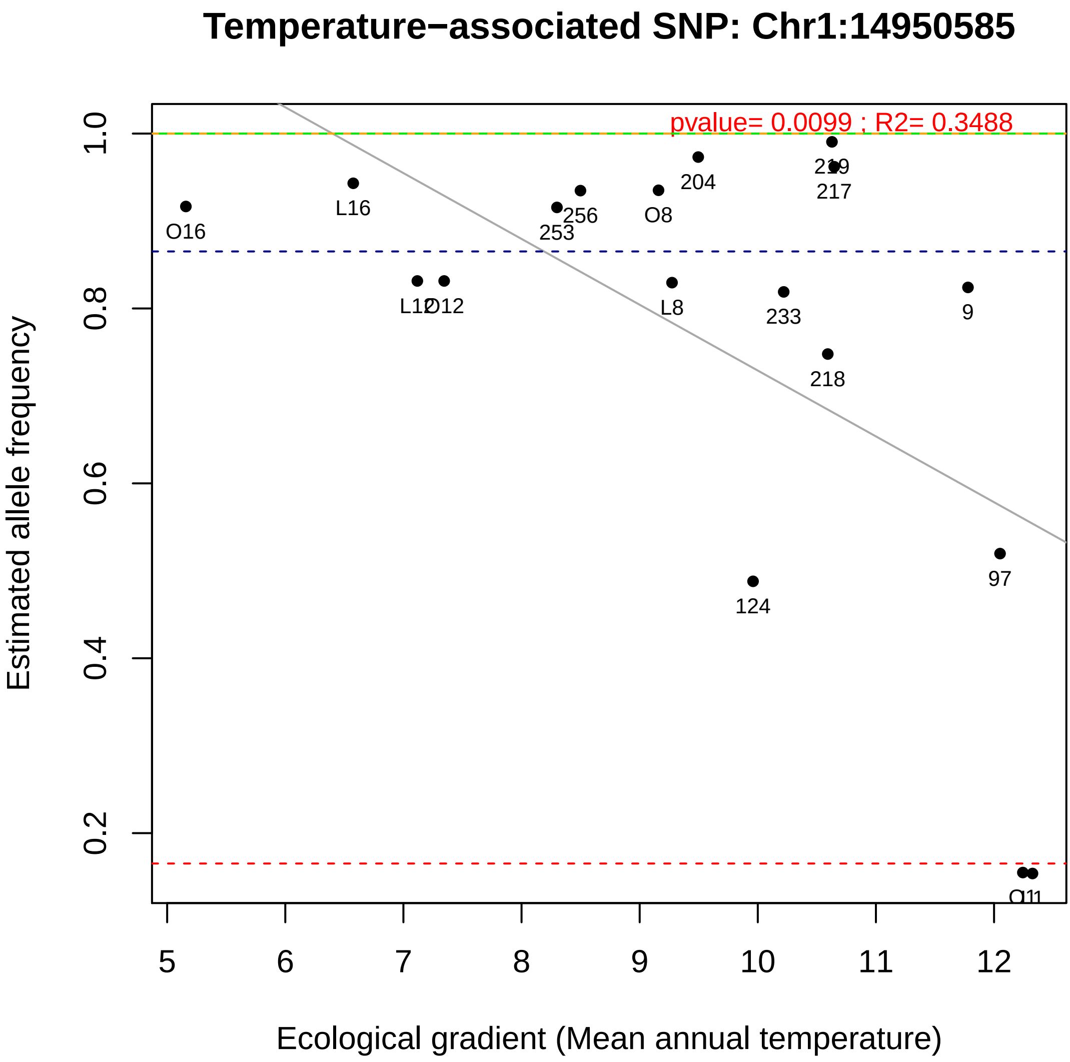

Temperature−associated SNP: Chr1:14950585
pvalue= 0.0099 ; R2= 0.3488
Estimated allele frequency
Ecological gradient (Mean annual temperature)
0.2
0.4
0.6
0.8
1.0
5
6
7
8
9
10
11
12
O16
L16
L12
O12
253
256
O8
204
219
217
L8
233
218
9
124
97

### Temperature−associated SNP: Chr1:14950668

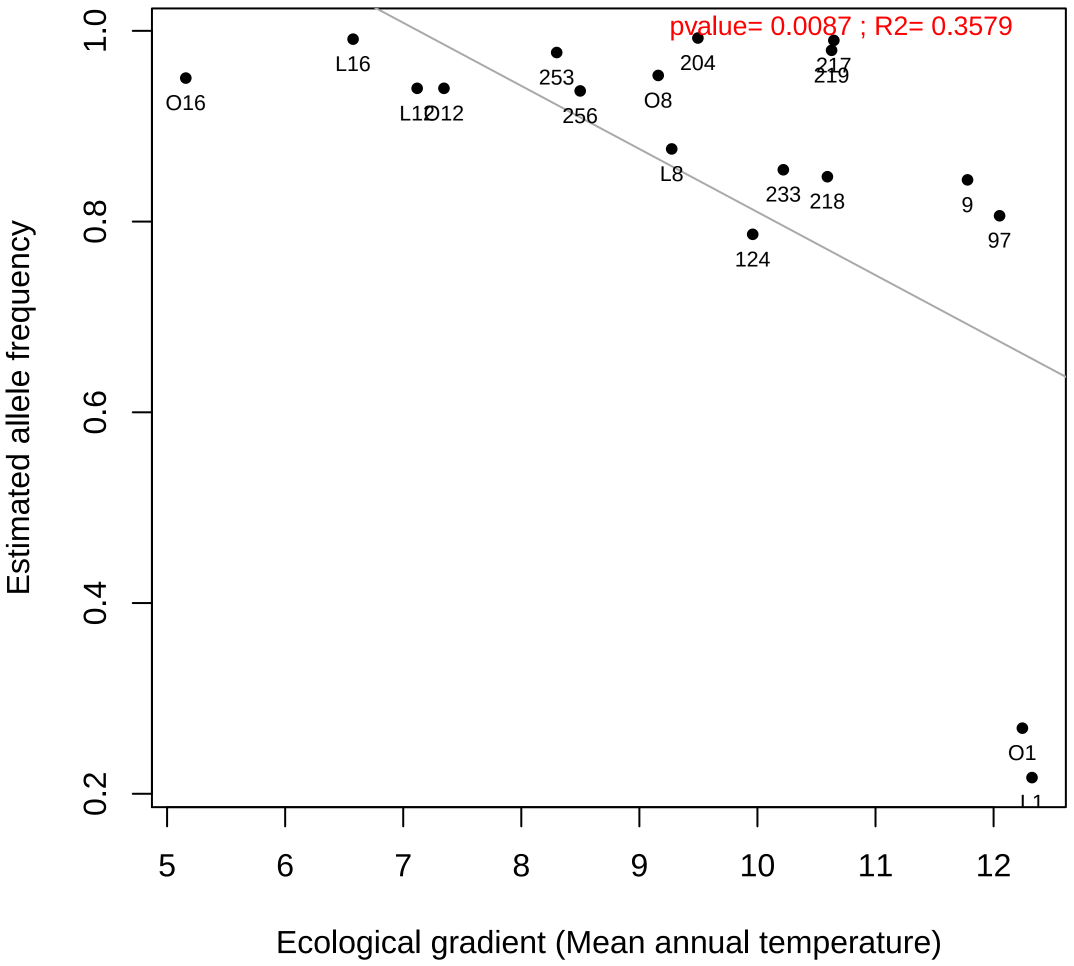

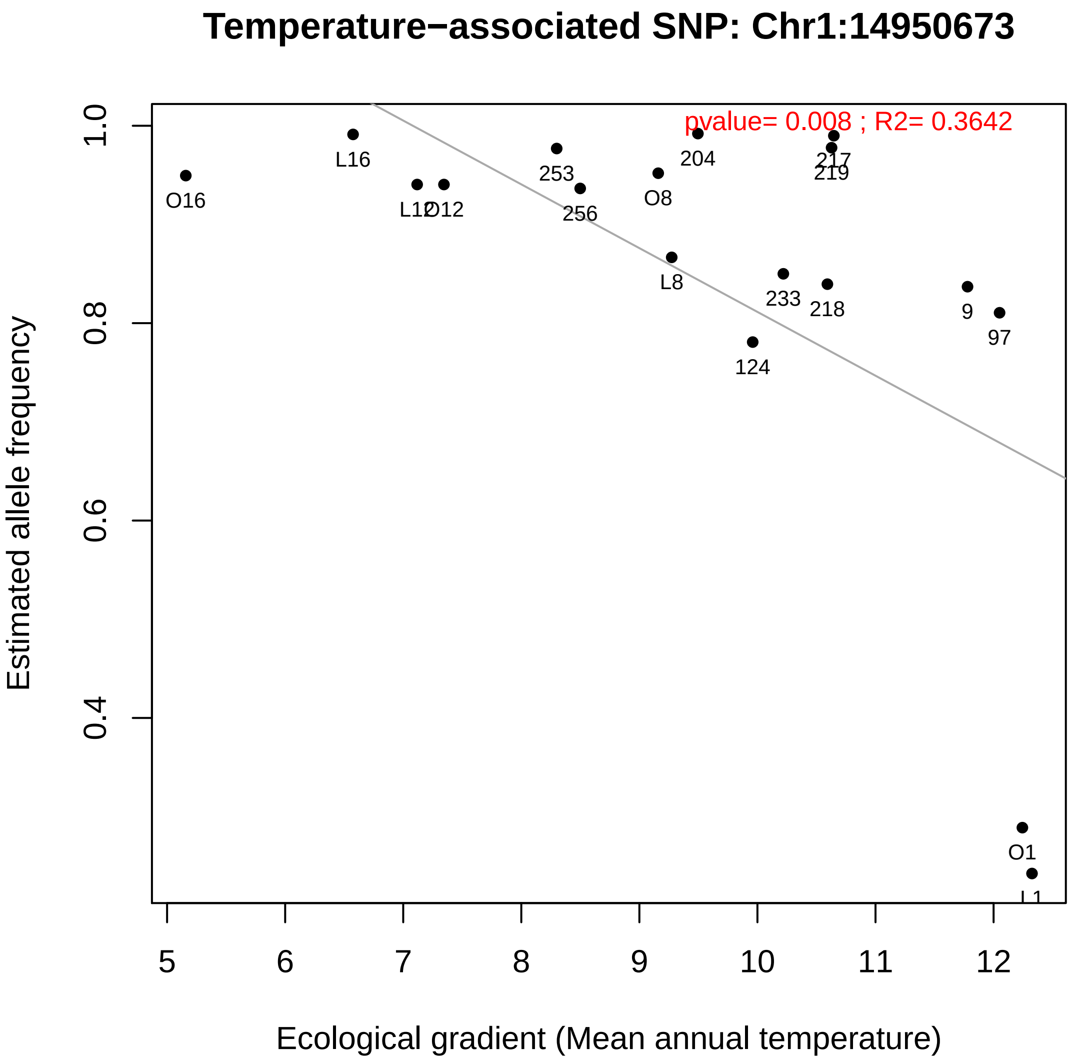

Temperature−associated SNP: Chr1:14950673
pvalue= 0.008 ; R2= 0.3642
Estimated allele frequency
0.4
0.6
0.8
1.0
5
6
7
8
9
10
11
12
Ecological gradient (Mean annual temperature)
O16
L16
L12
O12
253
256
O8
204
217
219
L8
233
218
124
9
97
O1

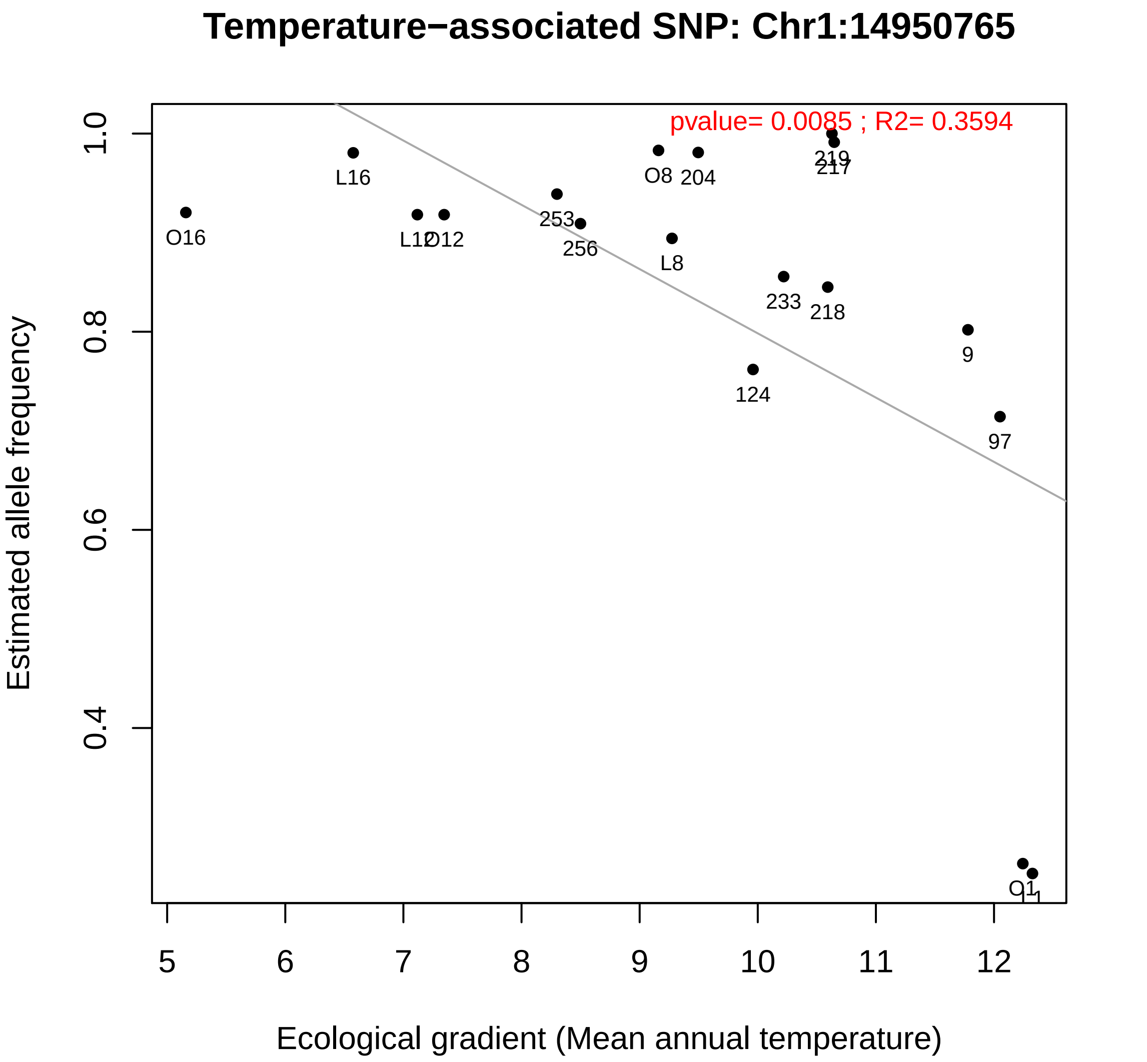

Temperature−associated SNP: Chr1:14950765
pvalue= 0.0085 ; R2= 0.3594
O16
L16
L12
O12
253
256
O8
204
L8
219
217
233
218
124
9
97
O1
L1
1.0
0.8
0.6
0.4
5
6
7
8
9
10
11
12
Estimated allele frequency
Ecological gradient (Mean annual temperature)

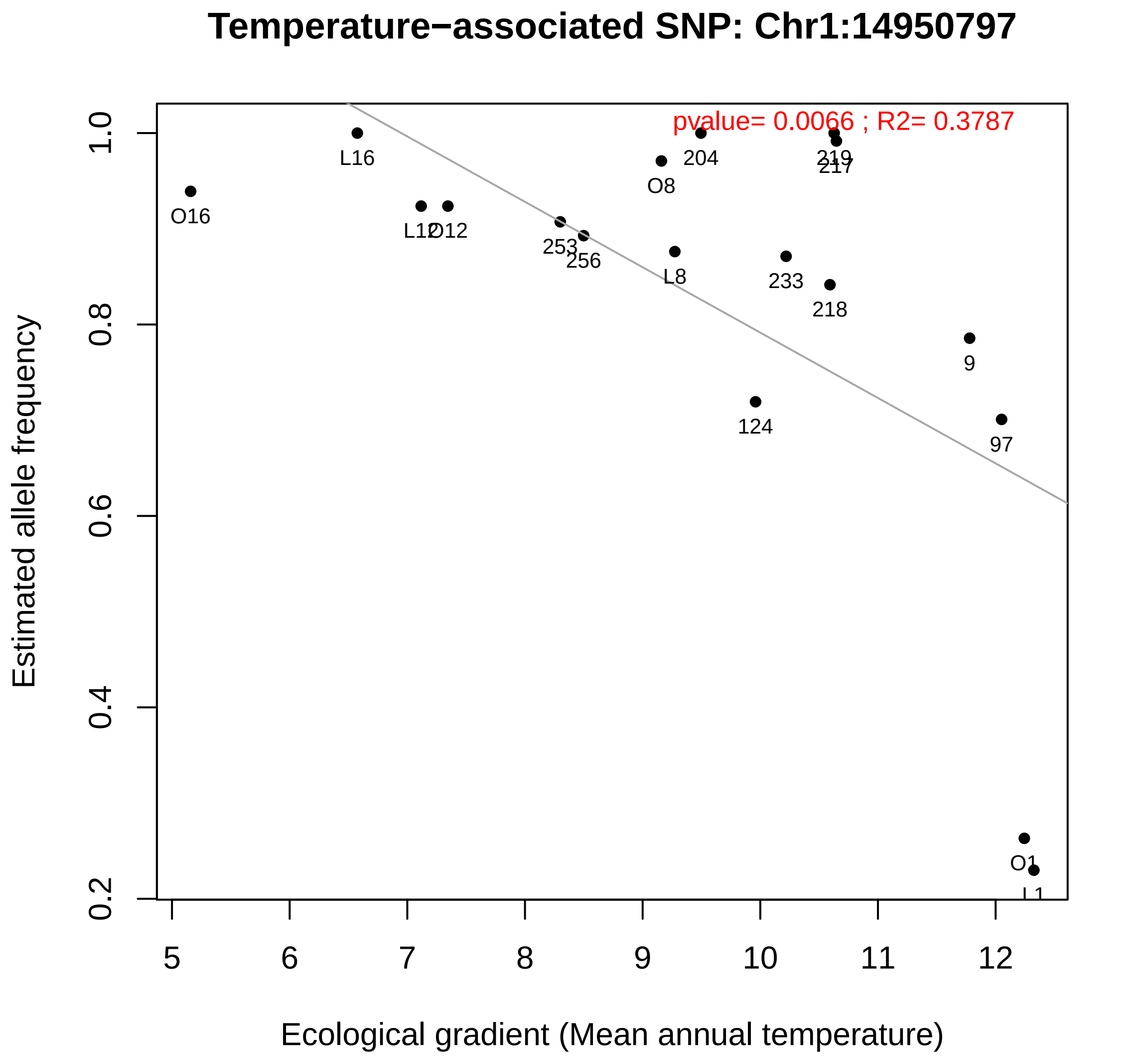

Temperature−associated SNP: Chr1:14950797
pvalue= 0.0066 ; R2= 0.3787
Estimated allele frequency
0.2
0.4
0.6
0.8
1.0
5
6
7
8
9
10
11
12
Ecological gradient (Mean annual temperature)
L16
O16
L12
O12
253
256
O8
204
L8
233
219
217
218
124
9
97
O1
L1

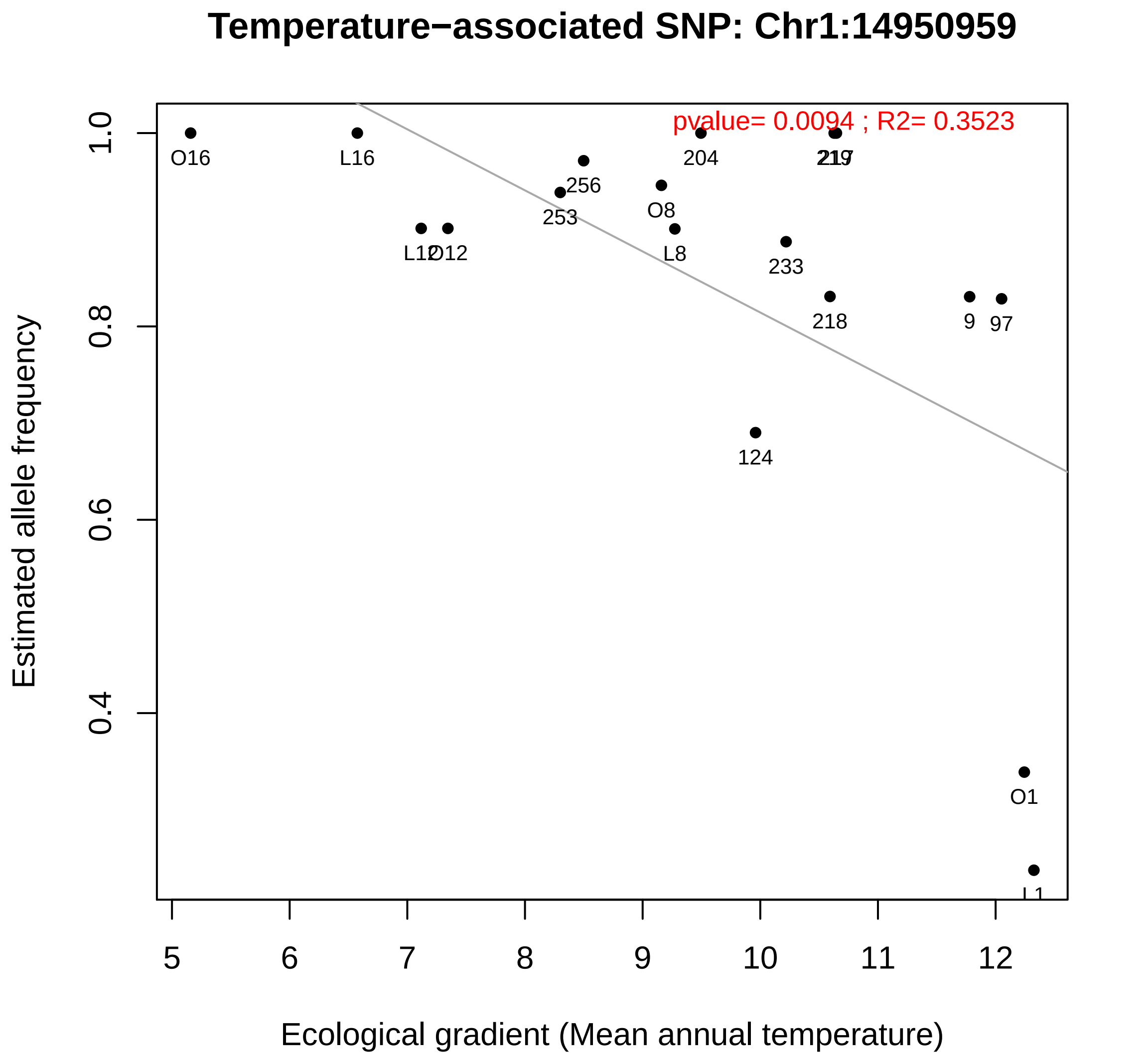

Temperature−associated SNP: Chr1:14950959
pvalue= 0.0094 ; R2= 0.3523
O16
L16
L12
O12
253
256
O8
L8
204
233
218
124
9
97
O1
Estimated allele frequency
1.0
0.8
0.6
0.4
5
6
7
8
9
10
11
12
Ecological gradient (Mean annual temperature)

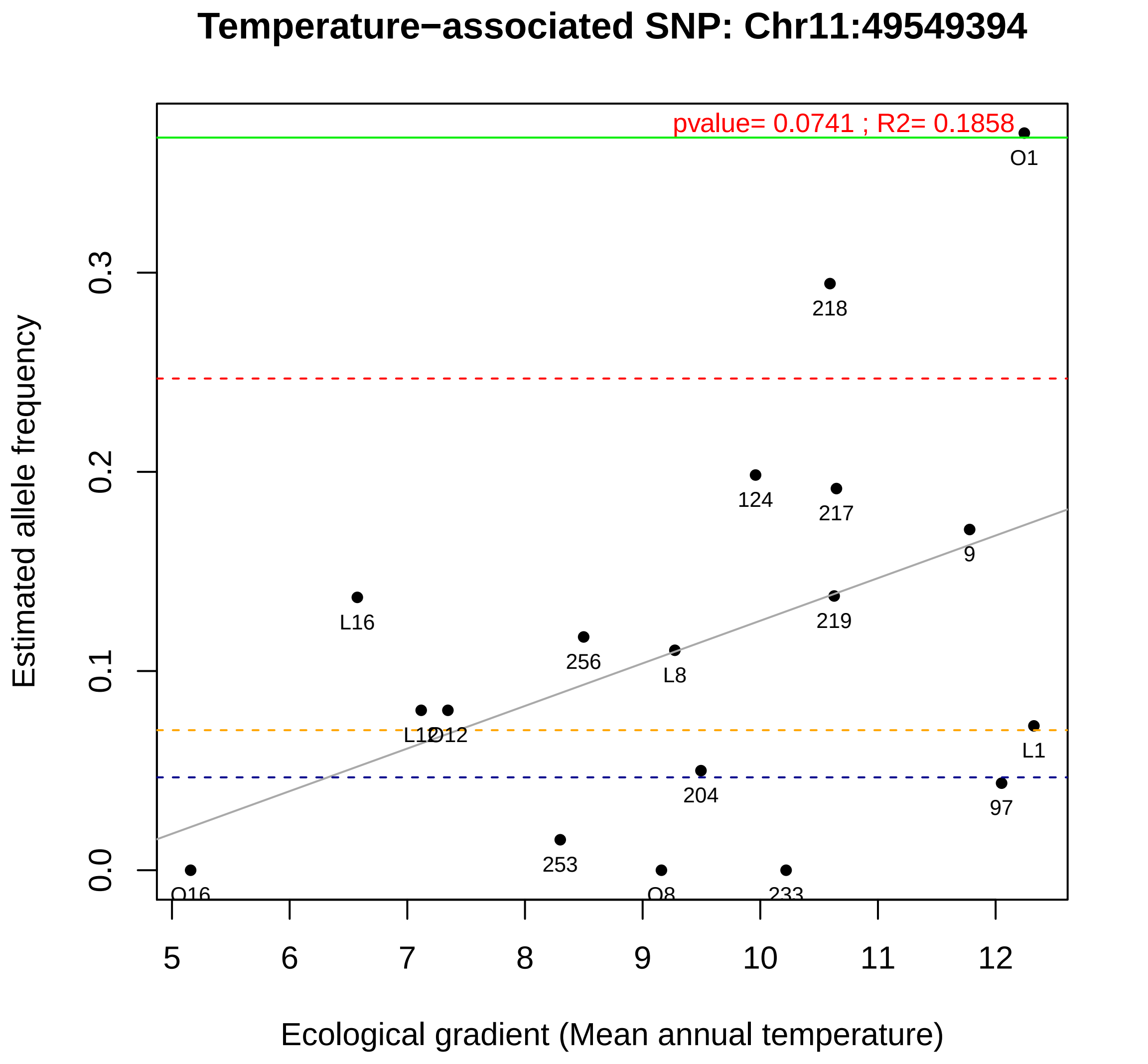

Temperature−associated SNP: Chr11:49549394
pvalue= 0.0741 ; R2= 0.1858
Estimated allele frequency
0.0
0.1
0.2
0.3
5
6
7
8
9
10
11
12
Ecological gradient (Mean annual temperature)
O1
218
124
217
9
L16
219
256
L8
L12
O12
L1
204
97
253
O16
O8
233

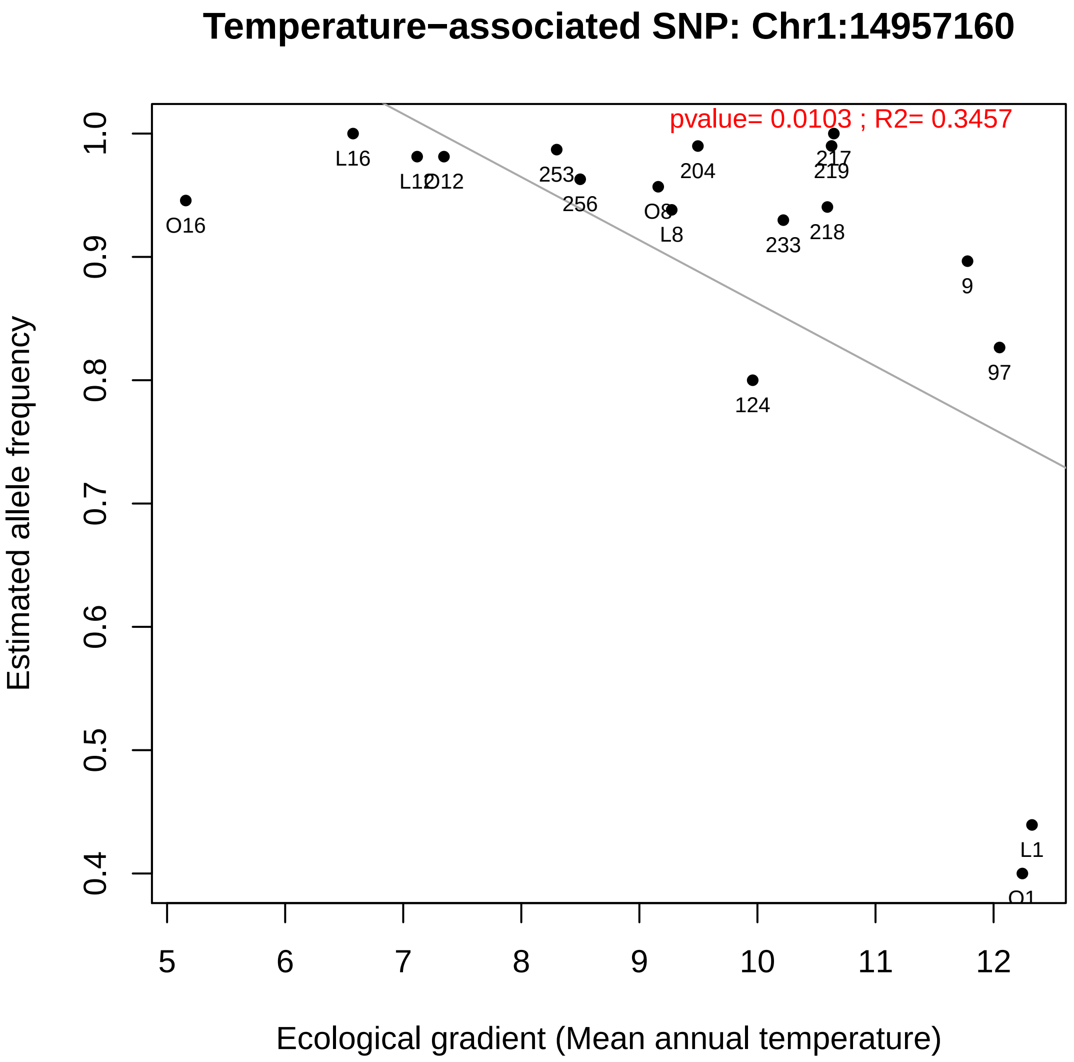

Temperature−associated SNP: Chr1:14957160
pvalue= 0.0103 ; R2= 0.3457
Estimated allele frequency
1.0
0.9
0.8
0.7
0.6
0.5
0.4
5
6
7
8
9
10
11
12
Ecological gradient (Mean annual temperature)
O16
L16
L12
O12
253
256
O8
L8
204
233
124
217
219
218
9
97
L1
O1

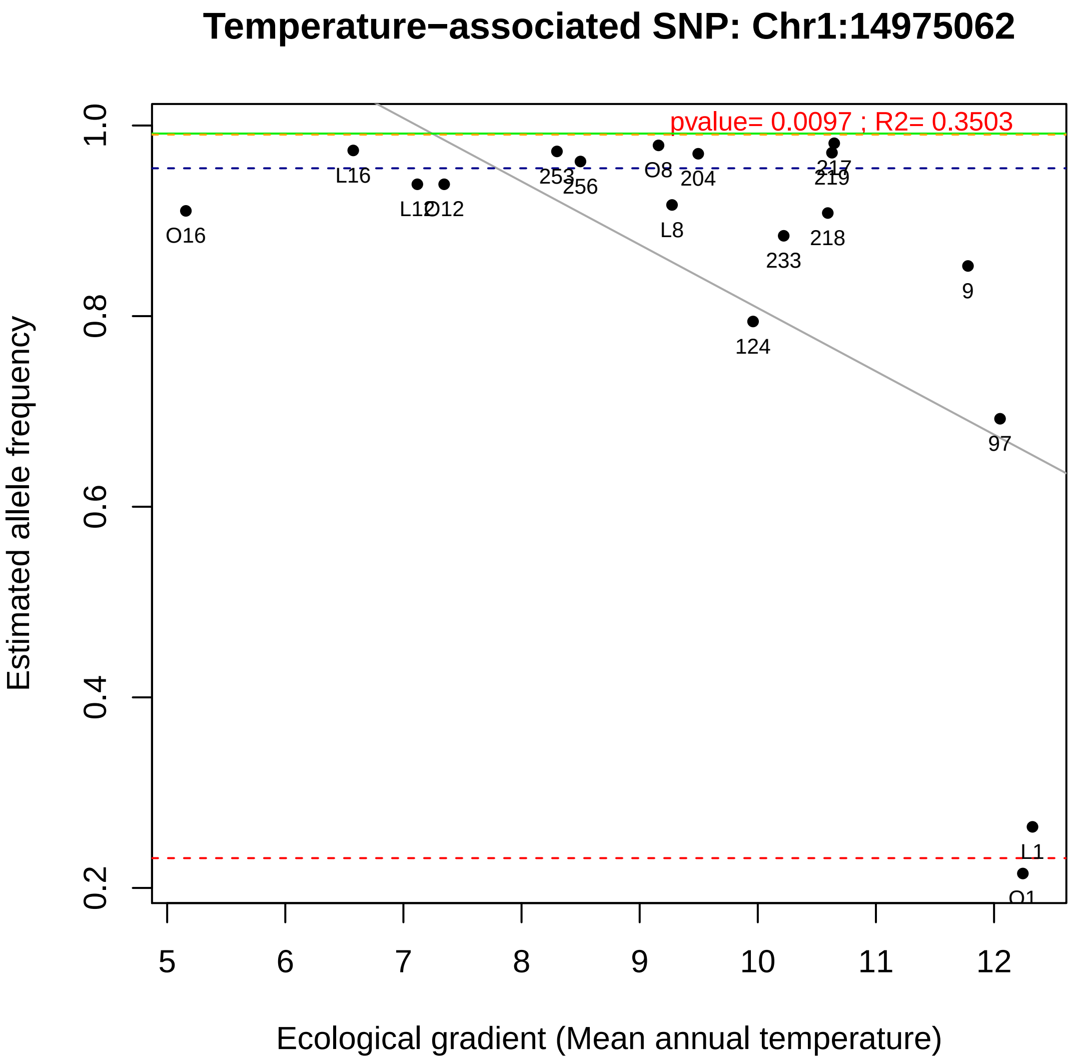

Temperature−associated SNP: Chr1:14975062
pvalue= 0.0097 ; R2= 0.3503
Estimated allele frequency
0.2
0.4
0.6
0.8
1.0
Ecological gradient (Mean annual temperature)
5
6
7
8
9
10
11
12
O16
L16
L12
O12
253
256
O8
204
L8
124
233
217
219
218
9
97
L1
O1

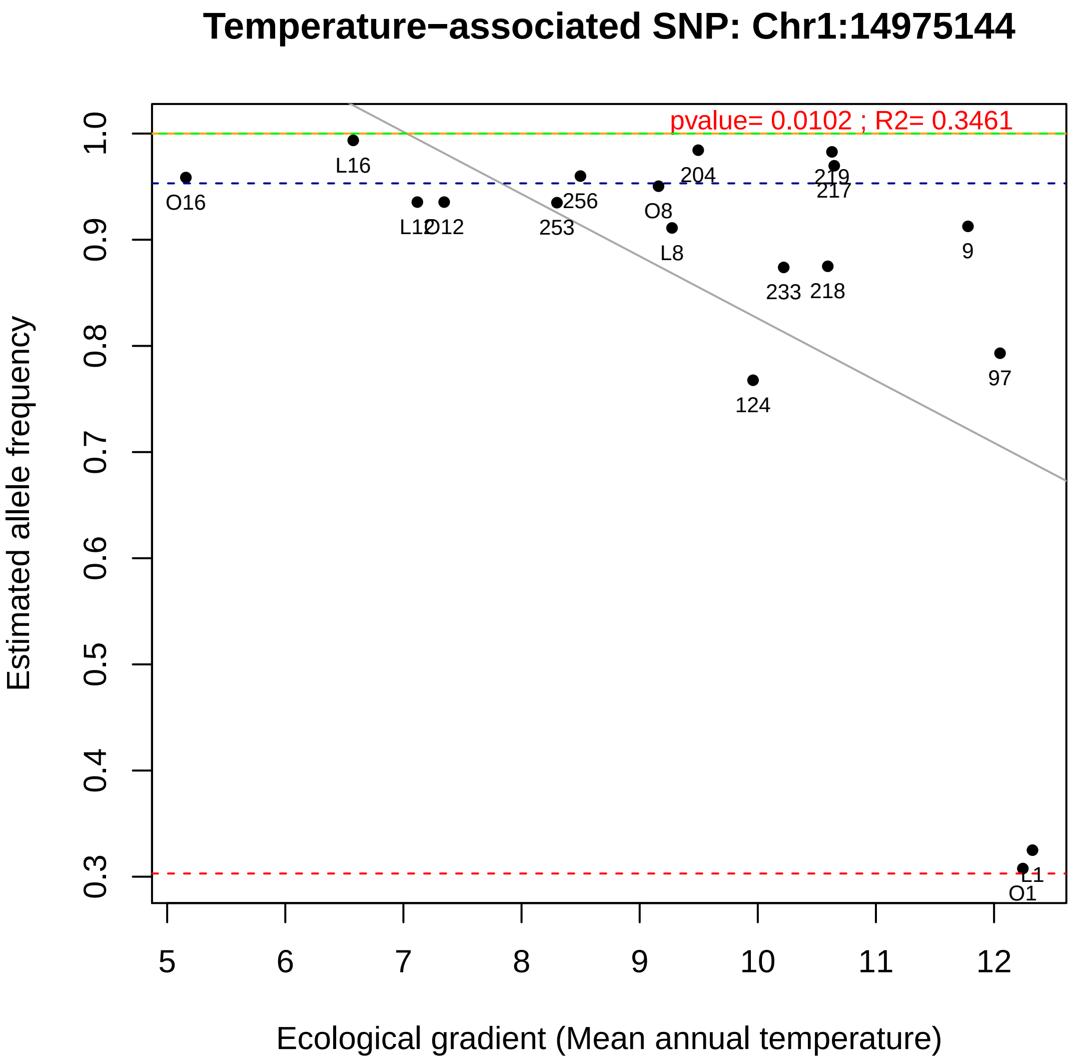

Temperature−associated SNP: Chr1:14975144
pvalue= 0.0102 ; R2= 0.3461
Estimated allele frequency
Ecological gradient (Mean annual temperature)
1.0
0.9
0.8
0.7
0.6
0.5
0.4
0.3
5
6
7
8
9
10
11
12
O16
L16
L12
O12
253
256
O8
L8
204
124
233
218
219
217
9
97
L1
O1

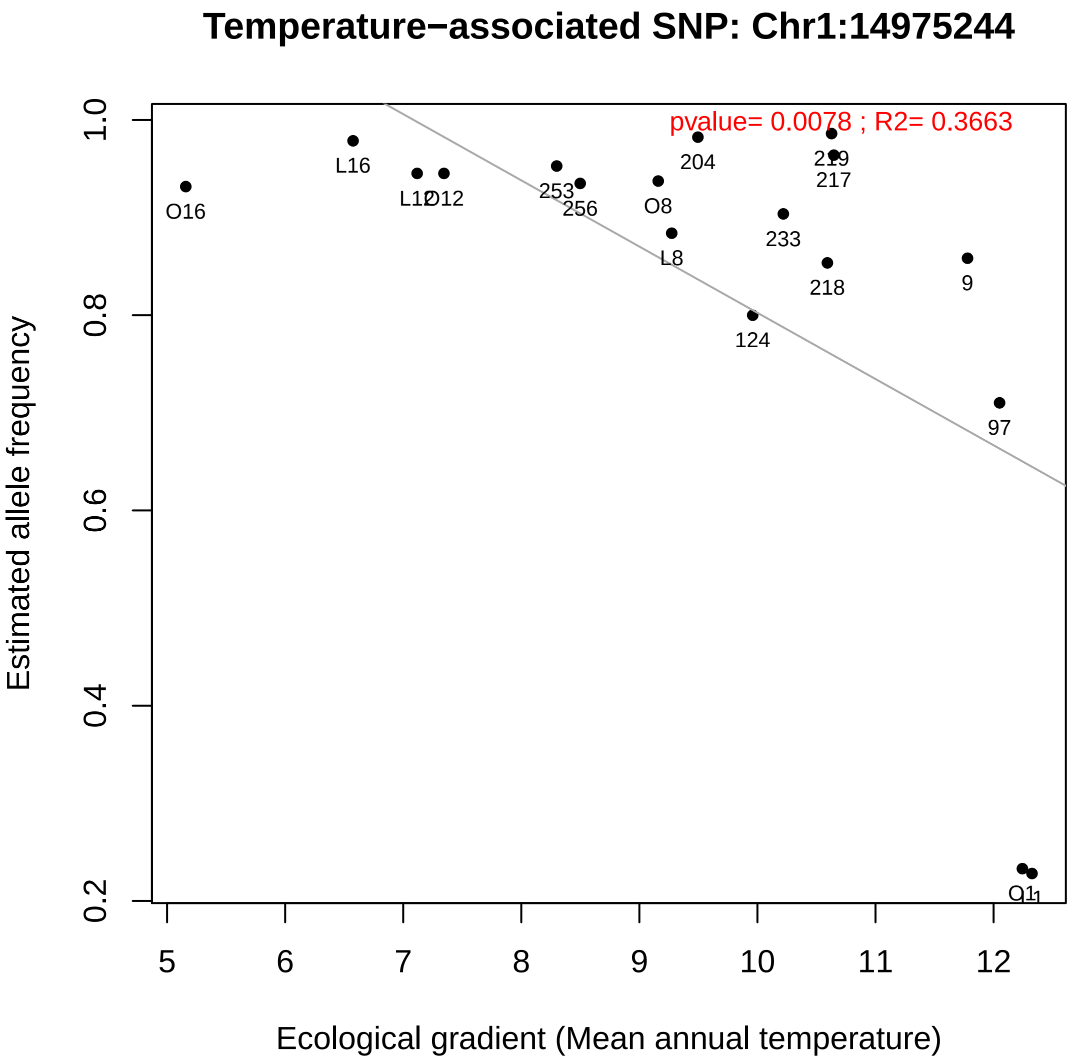
Temperature−associated SNP: Chr1:14975244
pvalue= 0.0078 ; R2= 0.3663
Estimated allele frequency
Ecological gradient (Mean annual temperature)
0.2
0.4
0.6
0.8
1.0
5
6
7
8
9
10
11
12
O16
L16
L12
O12
253
256
O8
204
L8
124
233
219
217
218
9
97

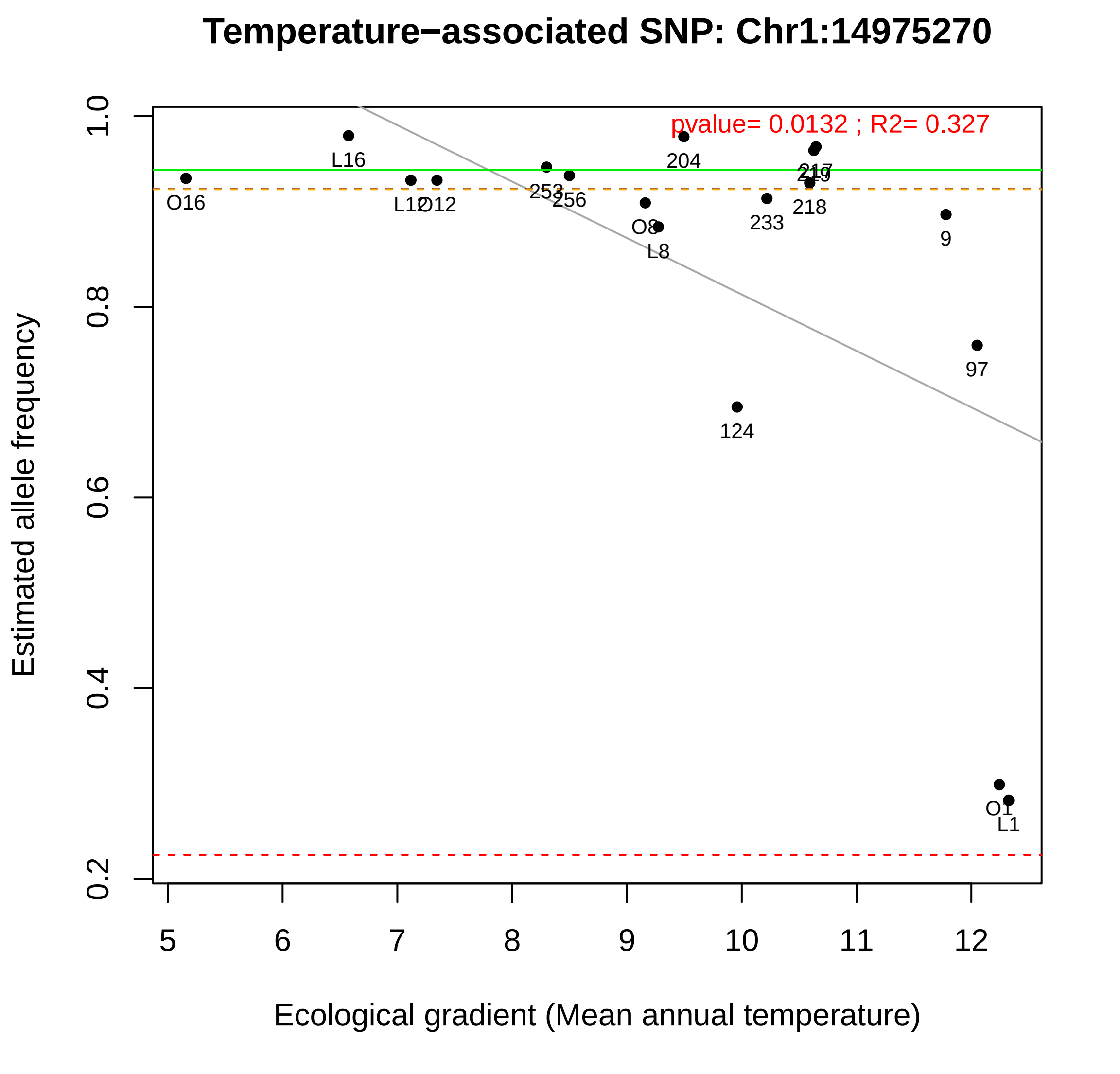

Temperature−associated SNP: Chr1:14975270
pvalue= 0.0132 ; R2= 0.327
Estimated allele frequency
Ecological gradient (Mean annual temperature)
0.2
0.4
0.6
0.8
1.0
5
6
7
8
9
10
11
12
O16
L16
L12
O12
253
256
O8
L8
204
124
233
217
219
218
9
97
O1
L1

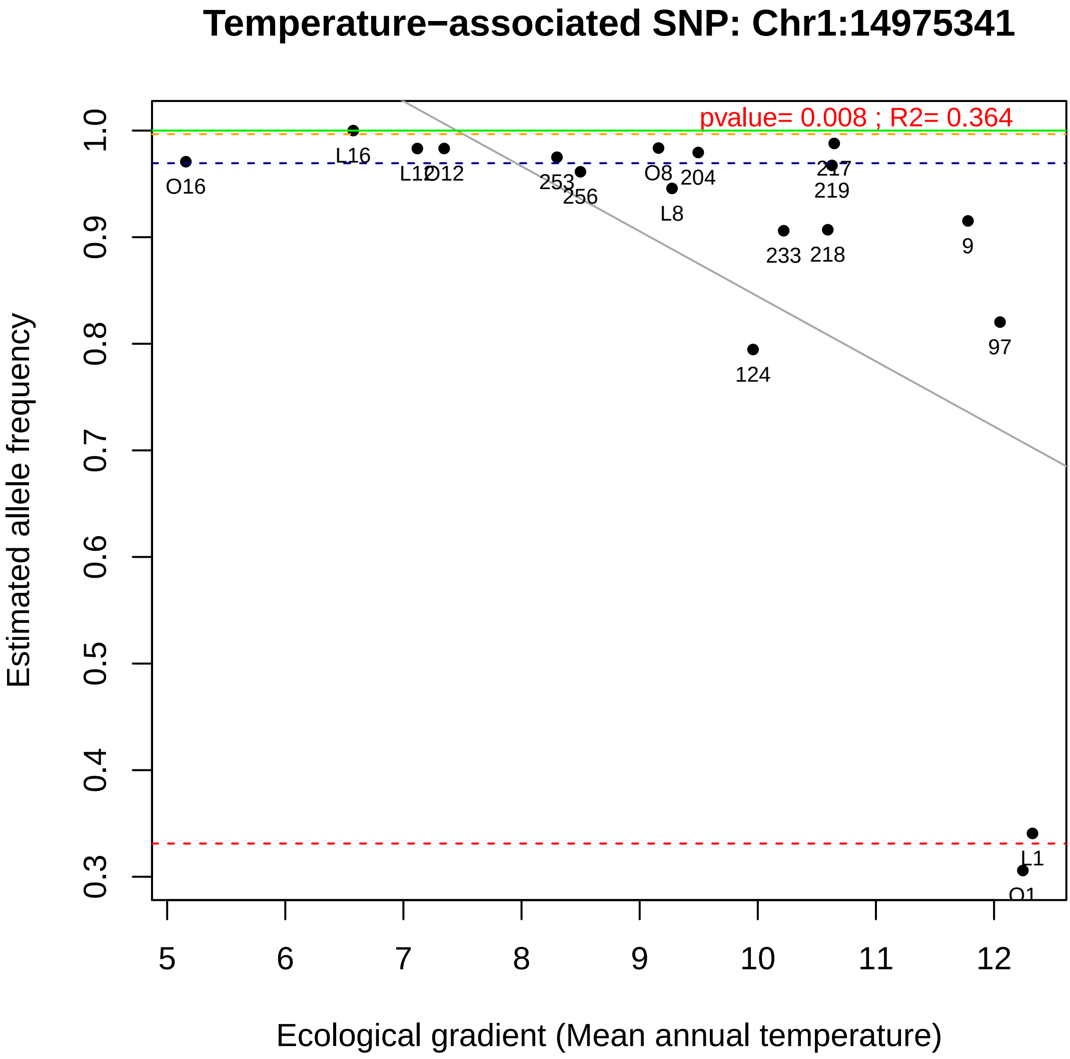

Temperature−associated SNP: Chr1:14975341
pvalue= 0.008 ; R2= 0.364
Estimated allele frequency
Ecological gradient (Mean annual temperature)
1.0
0.9
0.8
0.7
0.6
0.5
0.4
0.3
5
6
7
8
9
10
11
12
O16
L16
L12
O12
253
256
O8
204
L8
217
219
233
218
124
9
97
L1
O1

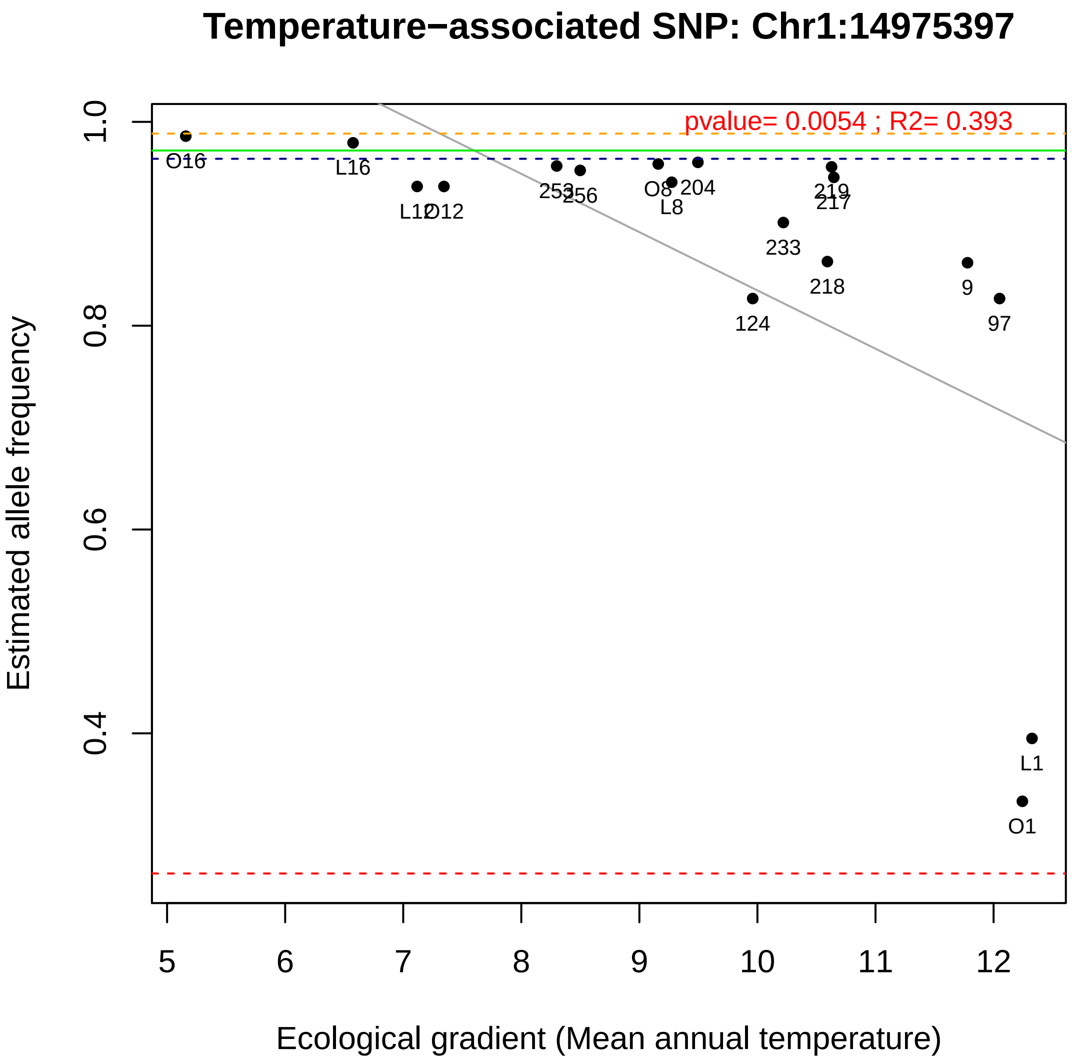

Temperature−associated SNP: Chr1:14975397
pvalue= 0.0054 ; R2= 0.393
Estimated allele frequency
1.0
0.8
0.6
0.4
Ecological gradient (Mean annual temperature)
5
6
7
8
9
10
11
12
O16
L16
L12
O12
253
256
O8
L8
204
219
217
233
218
124
9
97
L1
O1

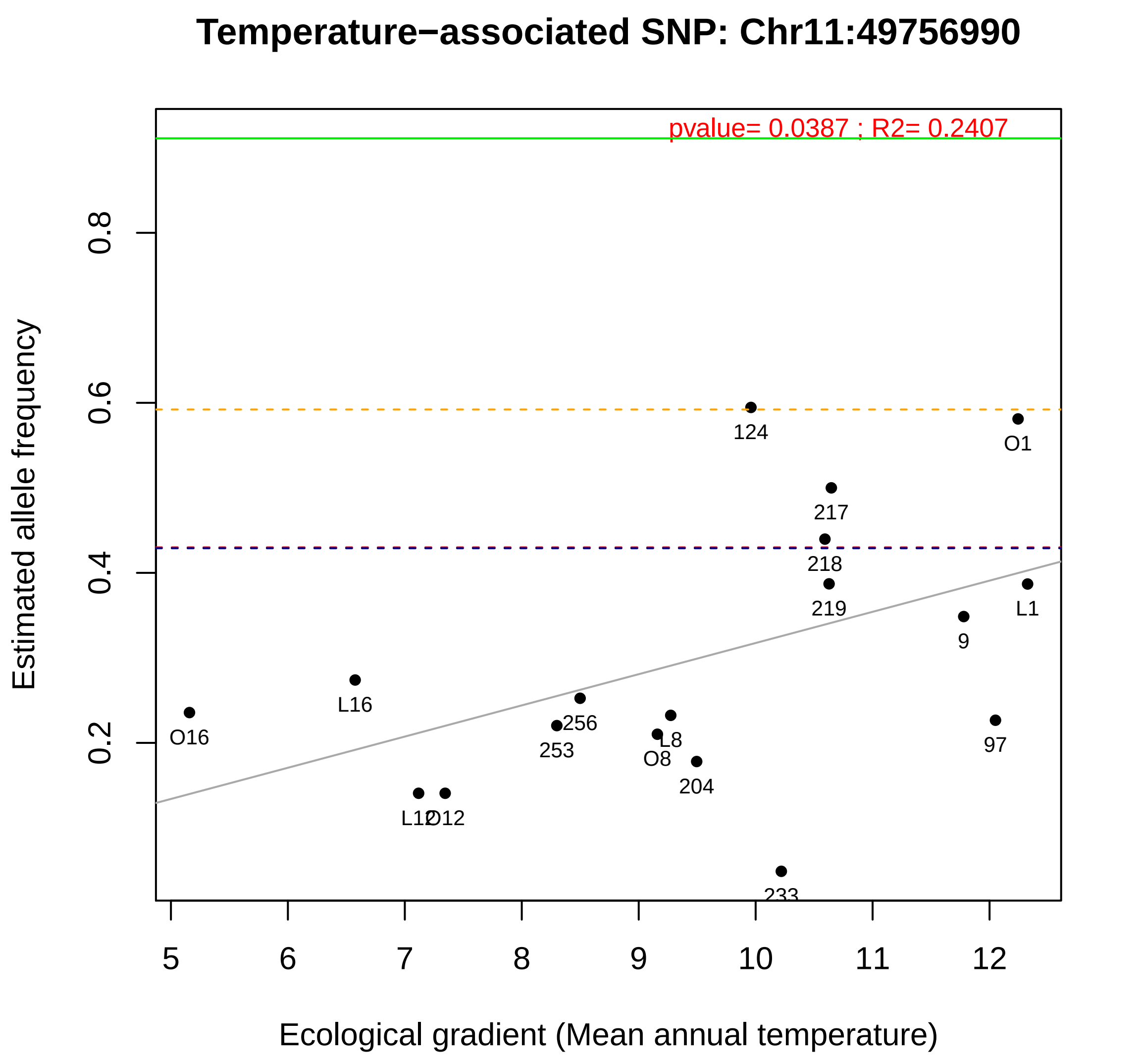
Temperature−associated SNP: Chr11:49756990
pvalue= 0.0387 ; R2= 0.2407
Estimated allele frequency
0.2
0.4
0.6
0.8
124
O1
217
218
219
L1
9
L16
O16
256
253
L8
O8
204
97
L12
O12
233
5
6
7
8
9
10
11
12
Ecological gradient (Mean annual temperature)

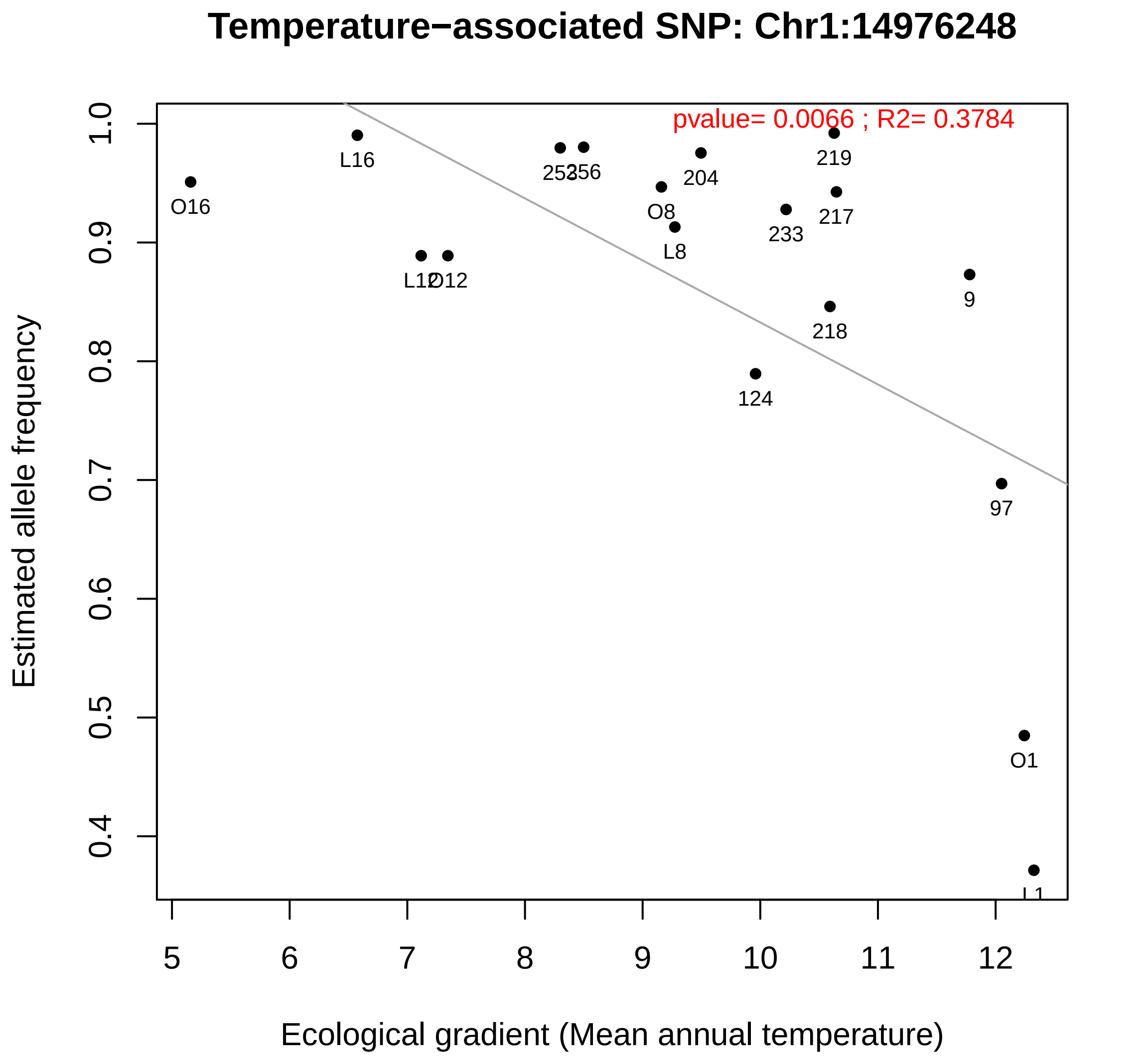

Temperature−associated SNP: Chr1:14976248
pvalue= 0.0066 ; R2= 0.3784
Estimated allele frequency
Ecological gradient (Mean annual temperature)
1.0
0.9
0.8
0.7
0.6
0.5
0.4
5
6
7
8
9
10
11
12
O16
L16
L12
O12
256
204
O8
L8
233
219
217
9
218
124
97
O1

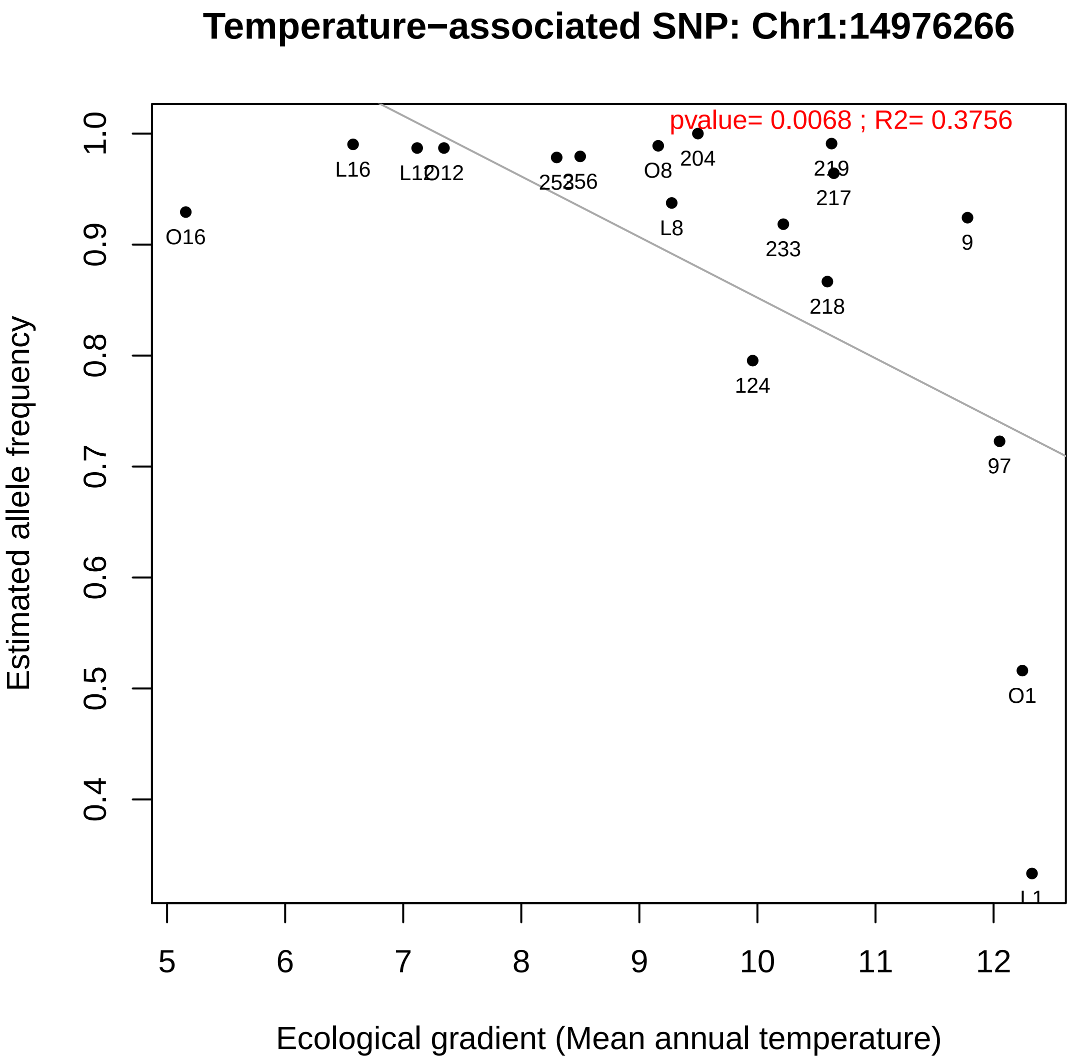

Temperature−associated SNP: Chr1:14976266
pvalue= 0.0068 ; R2= 0.3756
Estimated allele frequency
Ecological gradient (Mean annual temperature)
O16
L16
L12
O12
253
256
O8
204
L8
219
217
233
218
124
9
97
O1

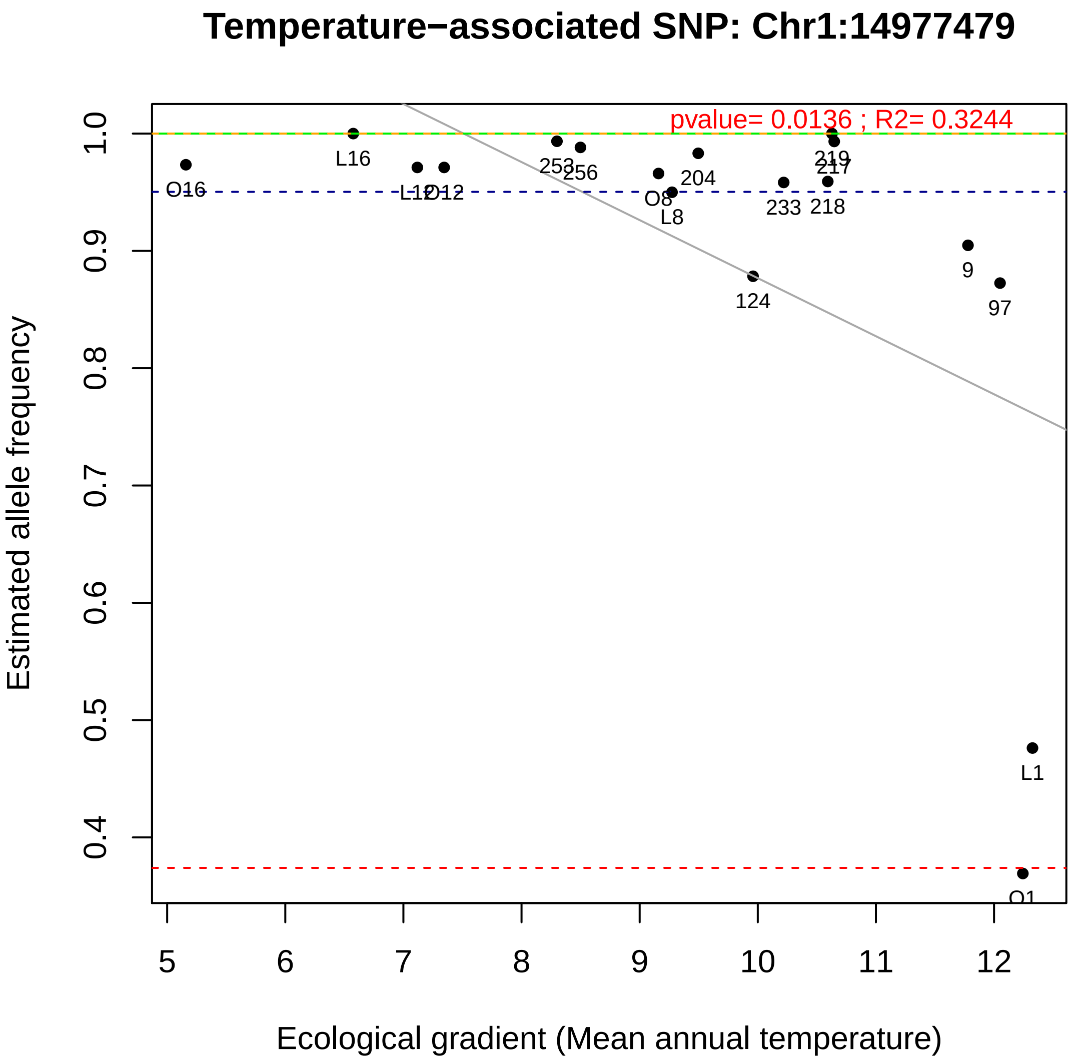
Temperature−associated SNP: Chr1:14977479
pvalue= 0.0136 ; R2= 0.3244
Estimated allele frequency
Ecological gradient (Mean annual temperature)
1.0
0.9
0.8
0.7
0.6
0.5
0.4
5
6
7
8
9
10
11
12
O16
L16
L12
O12
253
256
O8
L8
204
124
233
219
217
218
9
97
L1
O1

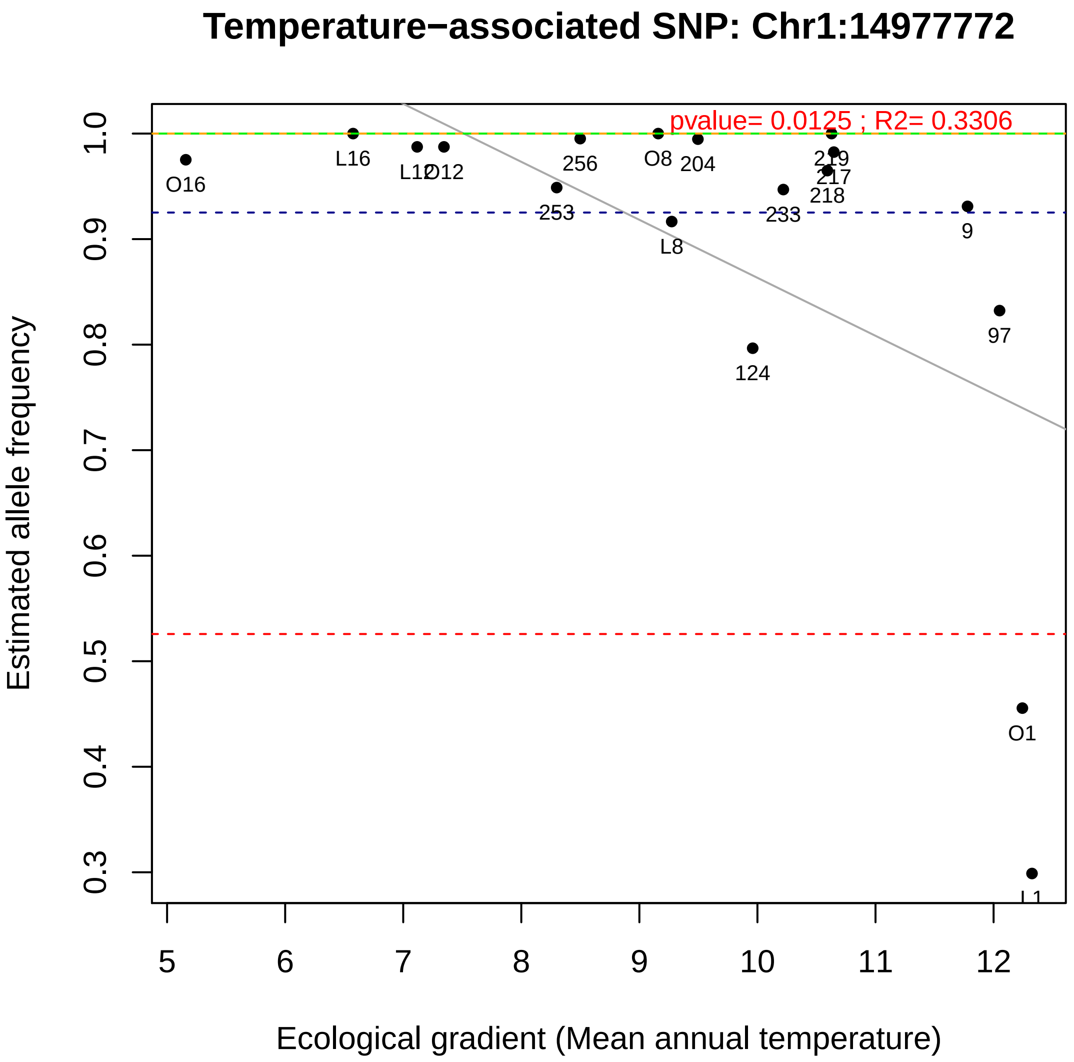
Temperature−associated SNP: Chr1:14977772
pvalue= 0.0125 ; R2= 0.3306
Estimated allele frequency
Ecological gradient (Mean annual temperature)
0.3
0.4
0.5
0.6
0.7
0.8
0.9
1.0
5
6
7
8
9
10
11
12
O16
L16
L12
O12
253
256
O8
L8
204
124
233
219
217
218
9
97
O1

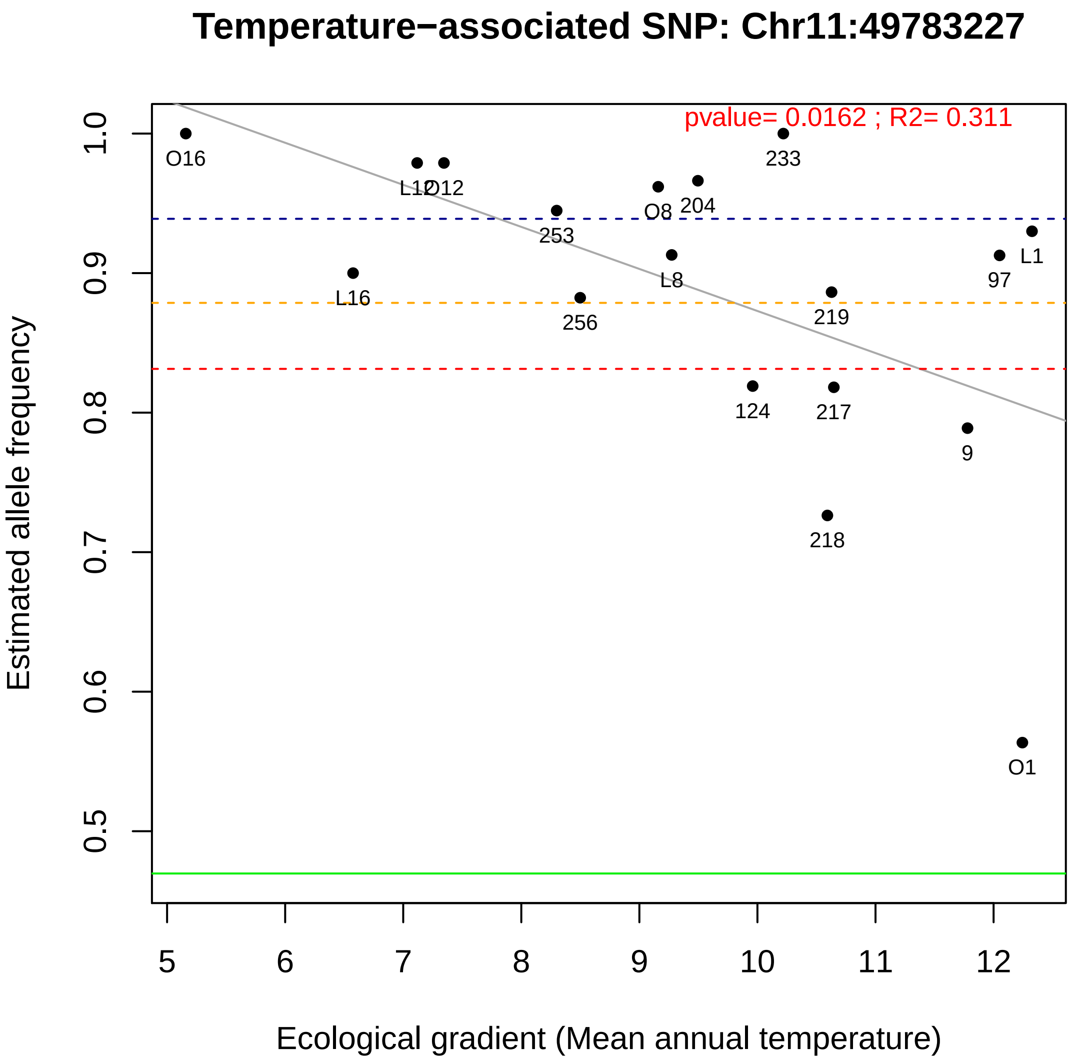

Temperature−associated SNP: Chr11:49783227
pvalue= 0.0162 ; R2= 0.311
O16
L12
O12
233
204
O8
253
L16
256
L8
219
L1
97
124
217
9
218
O1
1.0
0.9
0.8
0.7
0.6
0.5
5
6
7
8
9
10
11
12
Estimated allele frequency
Ecological gradient (Mean annual temperature)

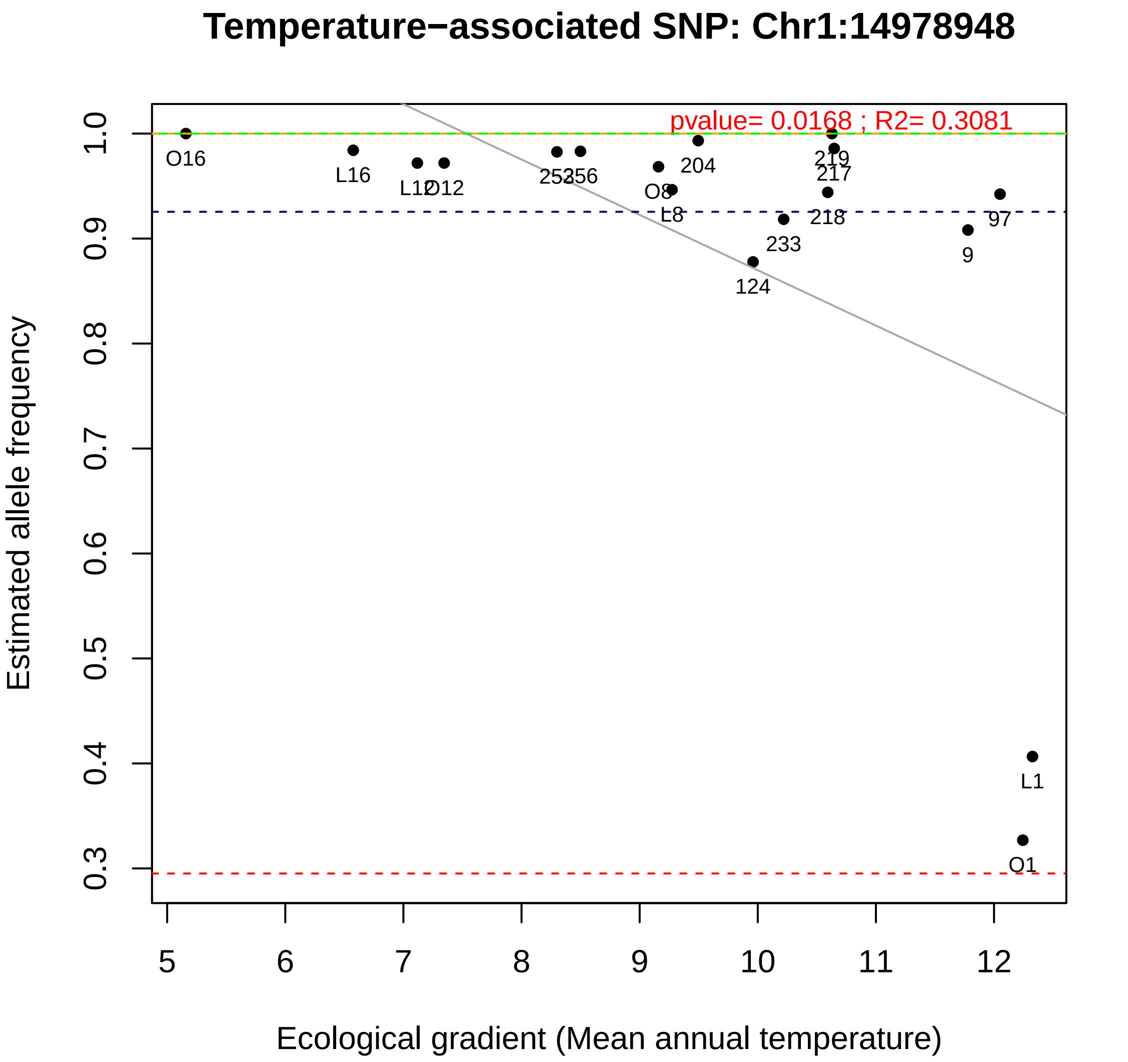
Temperature−associated SNP: Chr1:14978948
pvalue= 0.0168 ; R2= 0.3081
Estimated allele frequency
Ecological gradient (Mean annual temperature)
O16
L16
L12
O12
253
256
O8
L8
204
219
217
218
233
124
9
97
L1
O1
1.0
0.9
0.8
0.7
0.6
0.5
0.4
0.3
5
6
7
8
9
10
11
12

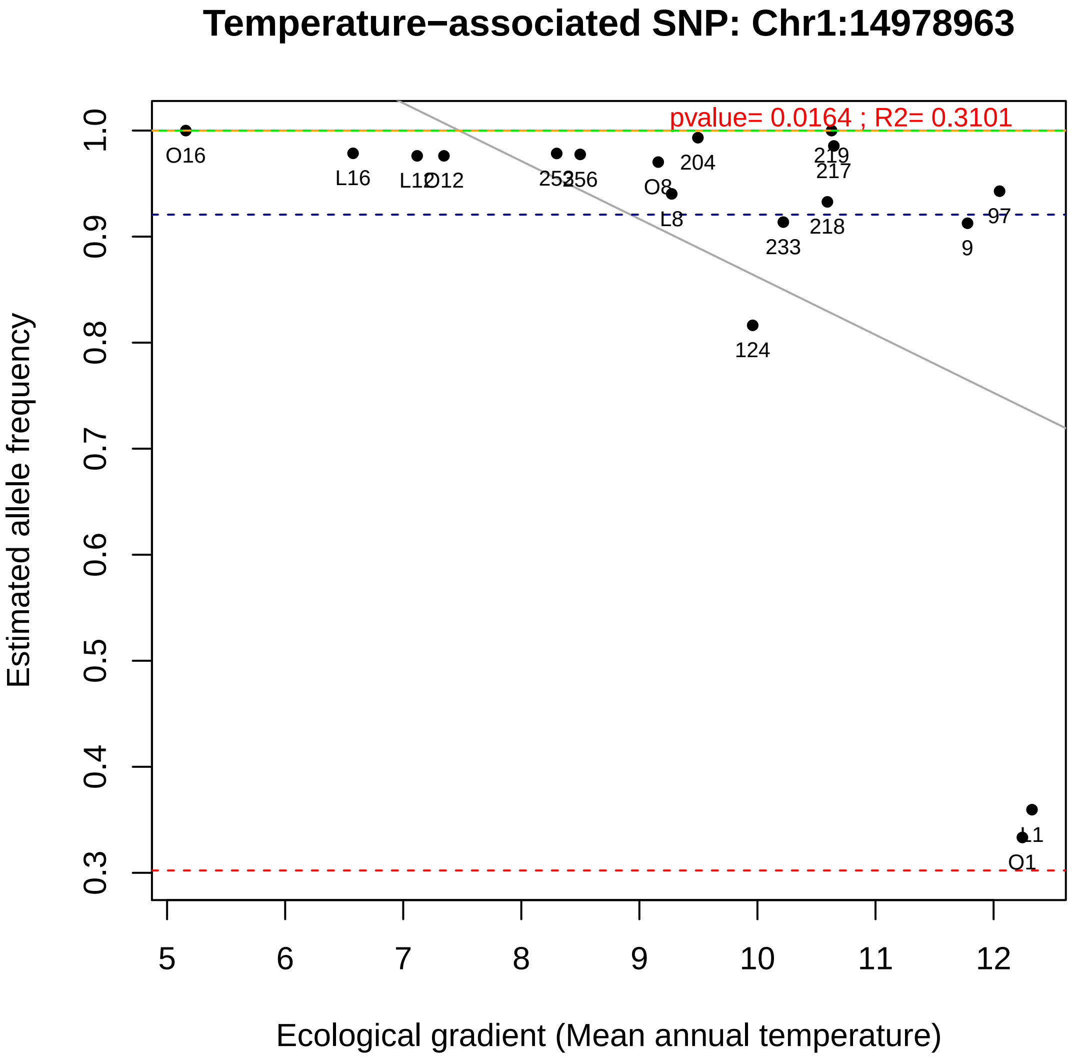

Temperature−associated SNP: Chr1:14978963
pvalue= 0.0164 ; R2= 0.3101
Estimated allele frequency
Ecological gradient (Mean annual temperature)
1.0
0.9
0.8
0.7
0.6
0.5
0.4
0.3
5
6
7
8
9
10
11
12
O16
L16
L12
O12
256
O8
204
L8
219
217
218
233
124
9
97
L1
O1

**Temperature−associated SNP: Chr1:14981282**

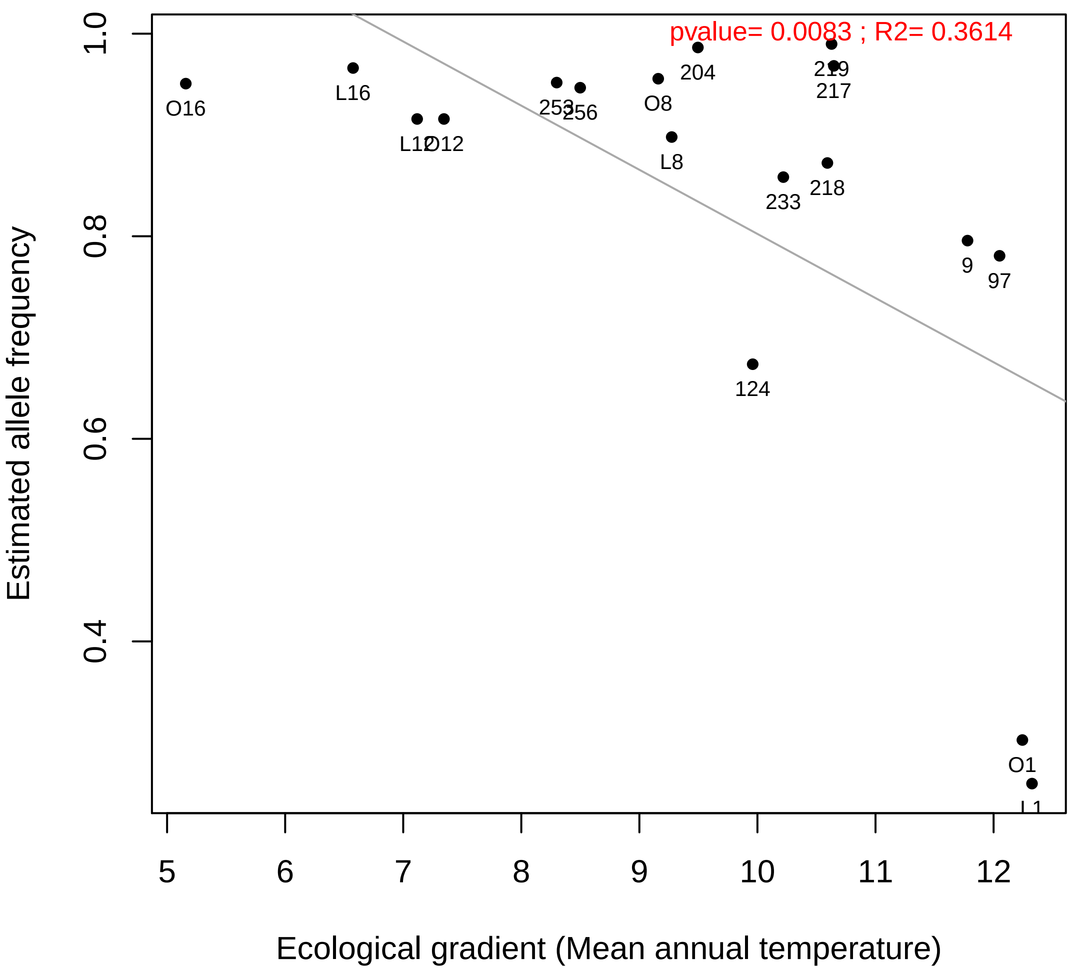

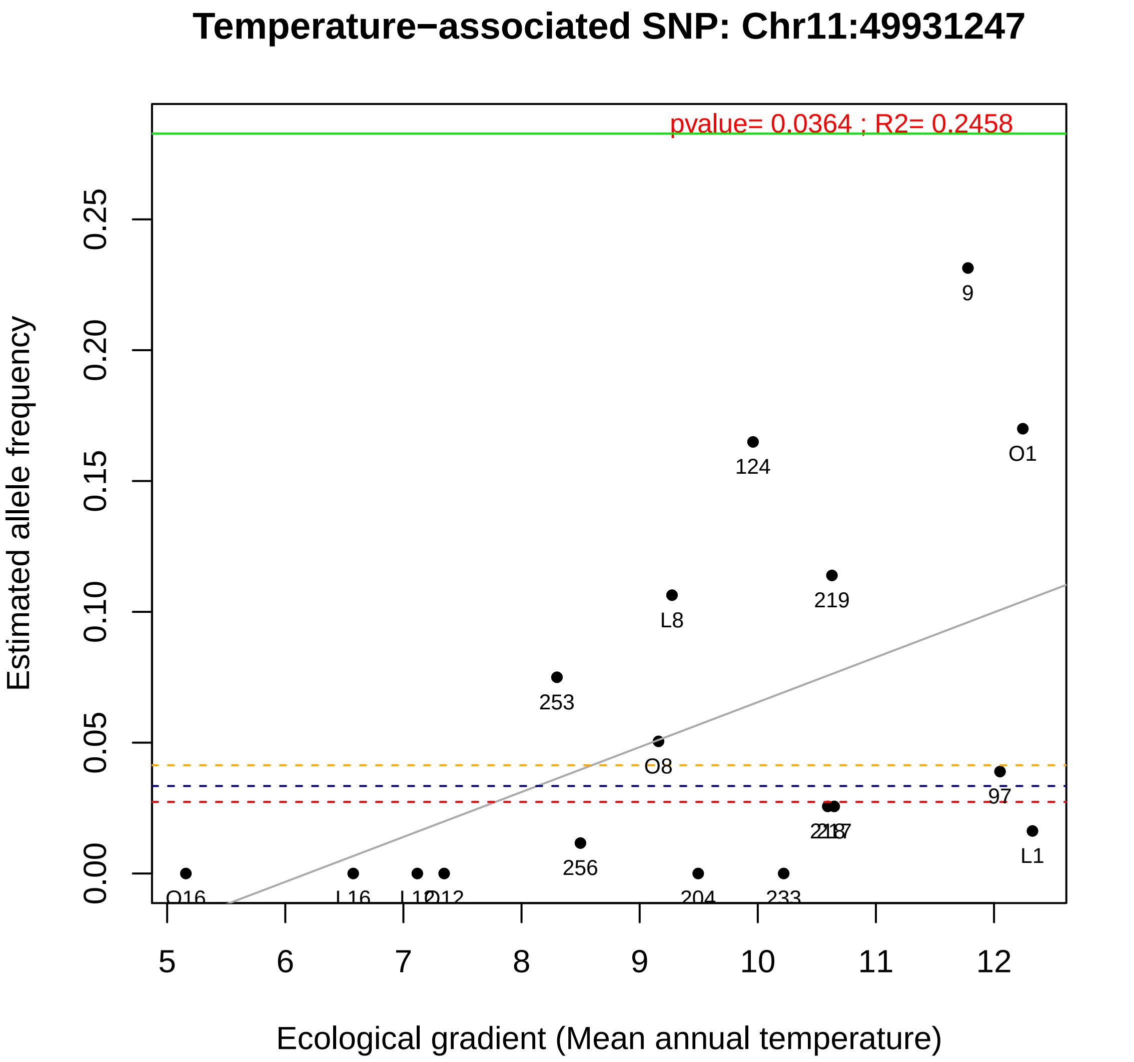
Temperature−associated SNP: Chr11:49931247
pvalue= 0.0364 ; R2= 0.2458
Estimated allele frequency
0.00
0.05
0.10
0.15
0.20
0.25
Ecological gradient (Mean annual temperature)
5
6
7
8
9
10
11
12
9
O1
124
L8
219
253
O8
97
L1
256
O16
L16
O12
204
233

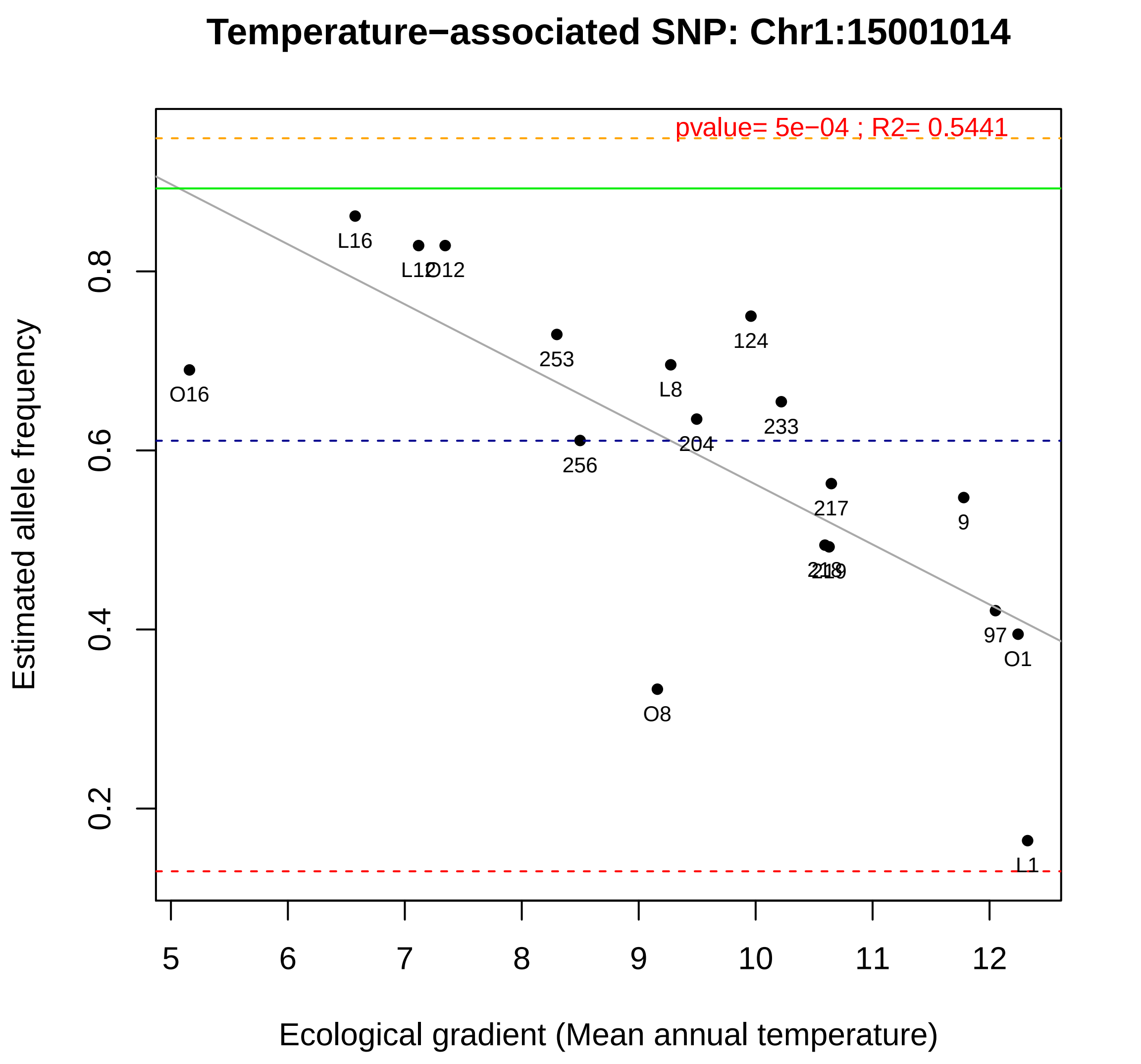
Temperature−associated SNP: Chr1:15001014
pvalue= 5e−04 ; R2= 0.5441
Estimated allele frequency
0.2
0.4
0.6
0.8
5
6
7
8
9
10
11
12
Ecological gradient (Mean annual temperature)
L16
L12
O12
253
O16
L8
124
233
204
256
217
9
97
O1
O8
L1

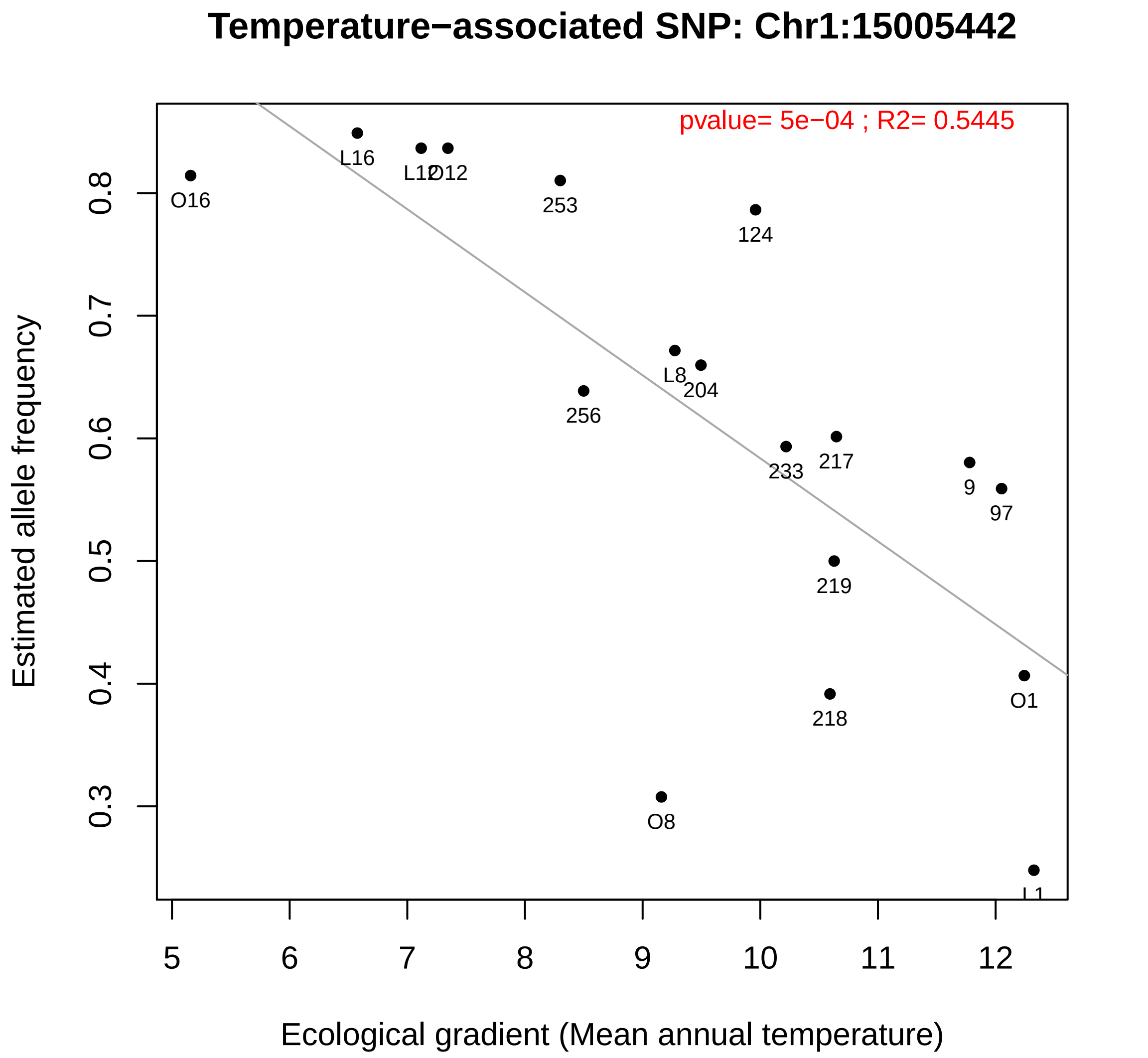

Temperature−associated SNP: Chr1:15005442
pvalue= 5e−04 ; R2= 0.5445
Estimated allele frequency
0.3
0.4
0.5
0.6
0.7
0.8
5
6
7
8
9
10
11
12
Ecological gradient (Mean annual temperature)
O16
L16
L12
O12
253
124
L8
204
256
233
217
9
97
219
O1
218
O8

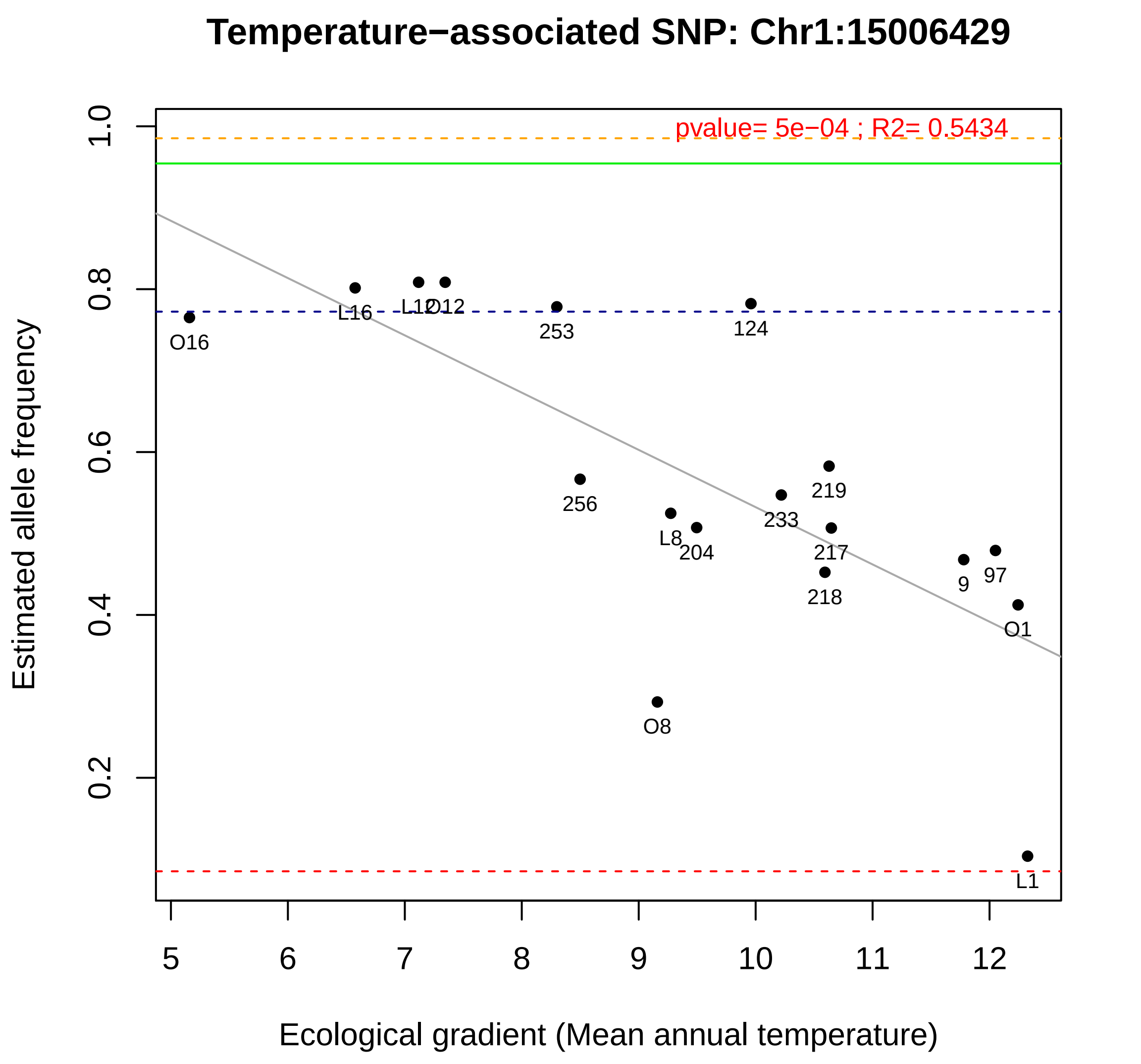

Temperature−associated SNP: Chr1:15006429
pvalue= 5e−04 ; R2= 0.5434
Estimated allele frequency
Ecological gradient (Mean annual temperature)
1.0
0.8
0.6
0.4
0.2
5
6
7
8
9
10
11
12
O16
L16
L12
O12
253
124
256
L8
204
233
219
217
218
9
97
O1
O8
L1

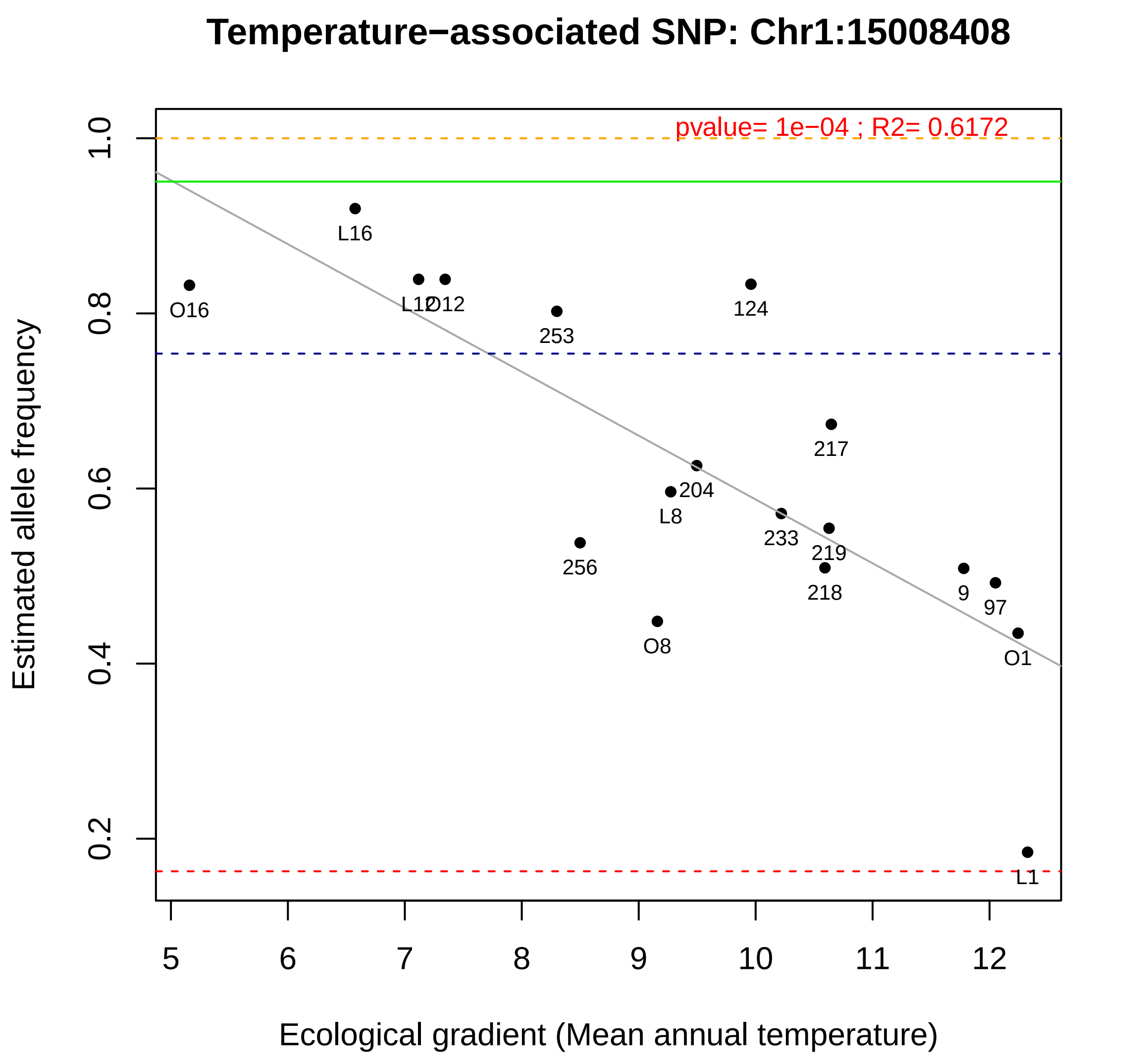

Temperature−associated SNP: Chr1:15008408
pvalue= 1e−04 ; R2= 0.6172
Estimated allele frequency
Ecological gradient (Mean annual temperature)
0.2
0.4
0.6
0.8
1.0
5
6
7
8
9
10
11
12
L16
O16
L12
O12
253
124
217
204
L8
233
219
256
218
9
97
O8
O1
L1

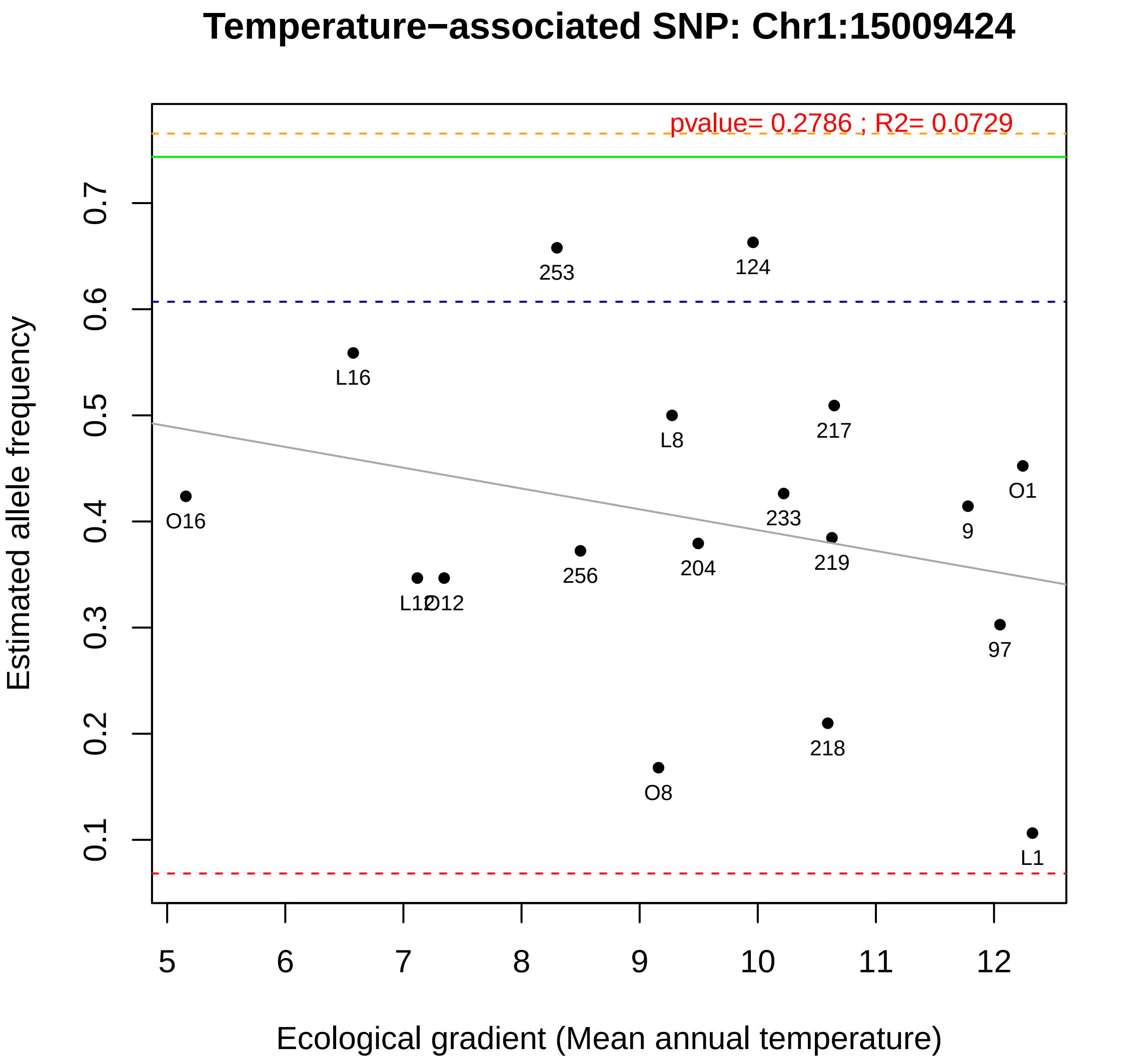

Temperature−associated SNP: Chr1:15009424
pvalue= 0.2786 ; R2= 0.0729
Estimated allele frequency
0.1
0.2
0.3
0.4
0.5
0.6
0.7
5
6
7
8
9
10
11
12
Ecological gradient (Mean annual temperature)
253
124
L16
L8
217
O1
O16
233
9
204
219
256
L12
O12
97
218
O8
L1

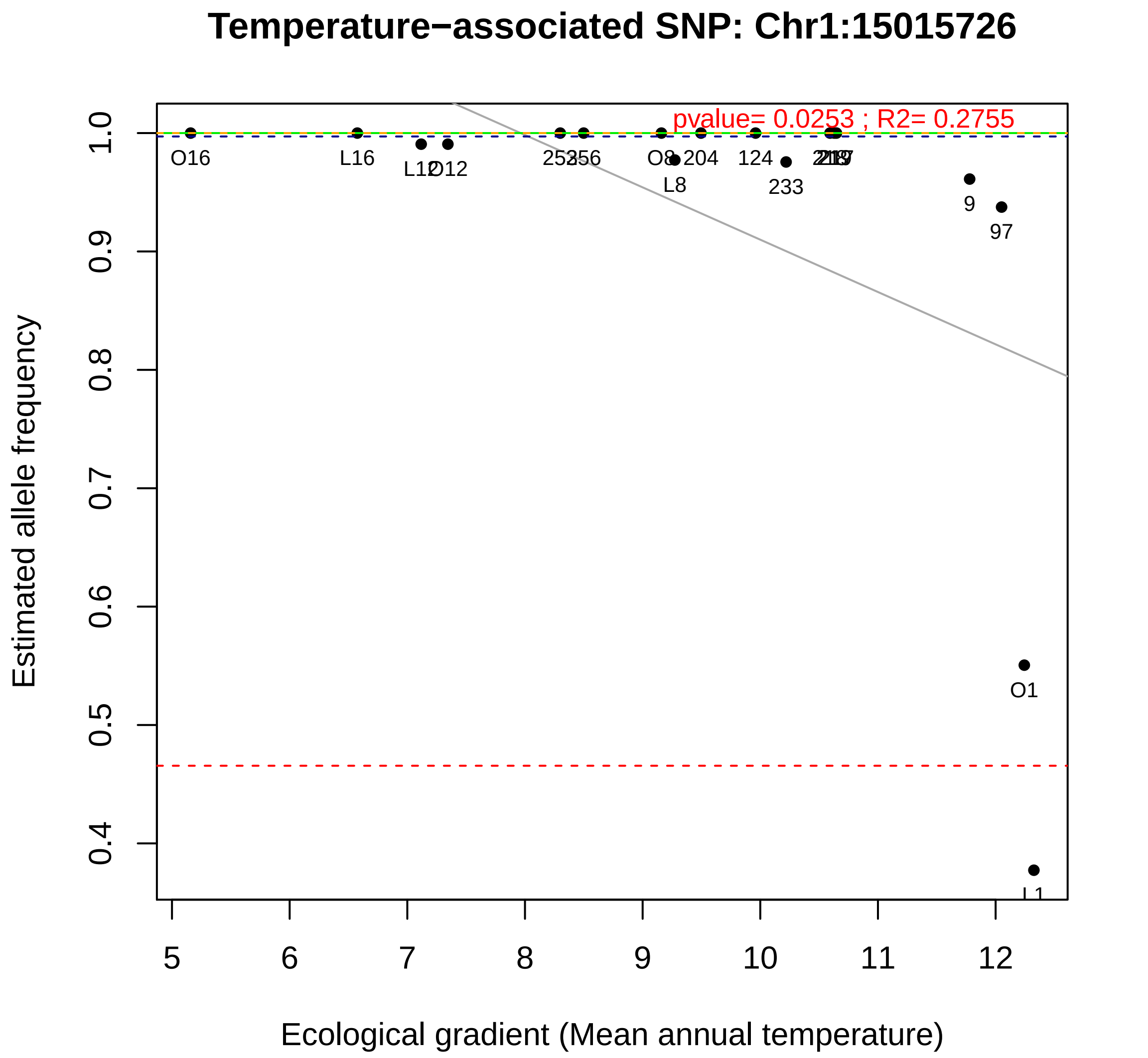

Temperature−associated SNP: Chr1:15015726
pvalue= 0.0253 ; R2= 0.2755
O16
L16
L12
O12
253
256
O8
L8
204
124
233
218
217
9
97
O1
L1
1.0
0.9
0.8
0.7
0.6
0.5
0.4
5
6
7
8
9
10
11
12
Estimated allele frequency
Ecological gradient (Mean annual temperature)

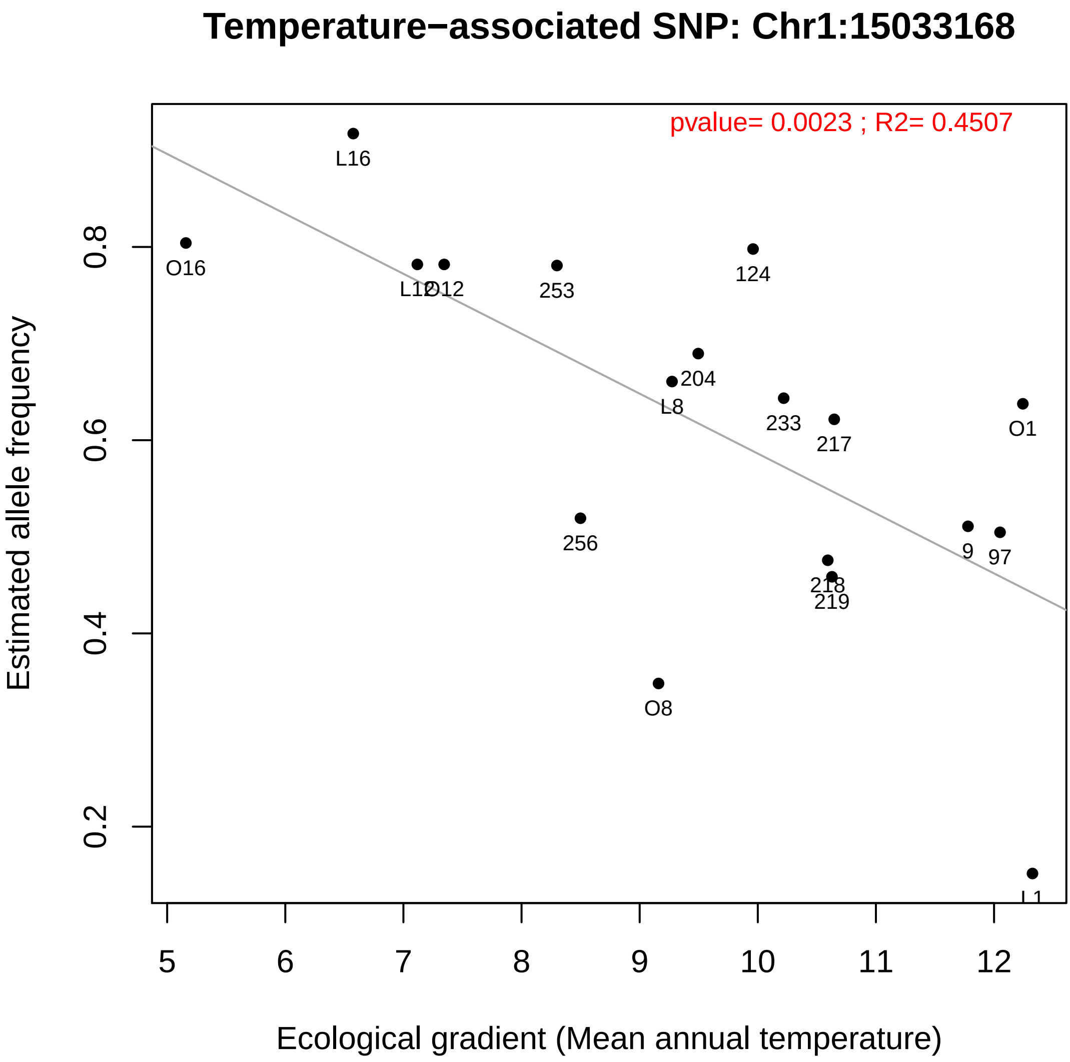

Temperature−associated SNP: Chr1:15033168
pvalue= 0.0023 ; R2= 0.4507
L16
O16
L12
O12
253
124
204
L8
233
217
O1
256
9
97
218
219
O8
L1
0.2
0.4
0.6
0.8
5
6
7
8
9
10
11
12
Estimated allele frequency
Ecological gradient (Mean annual temperature)

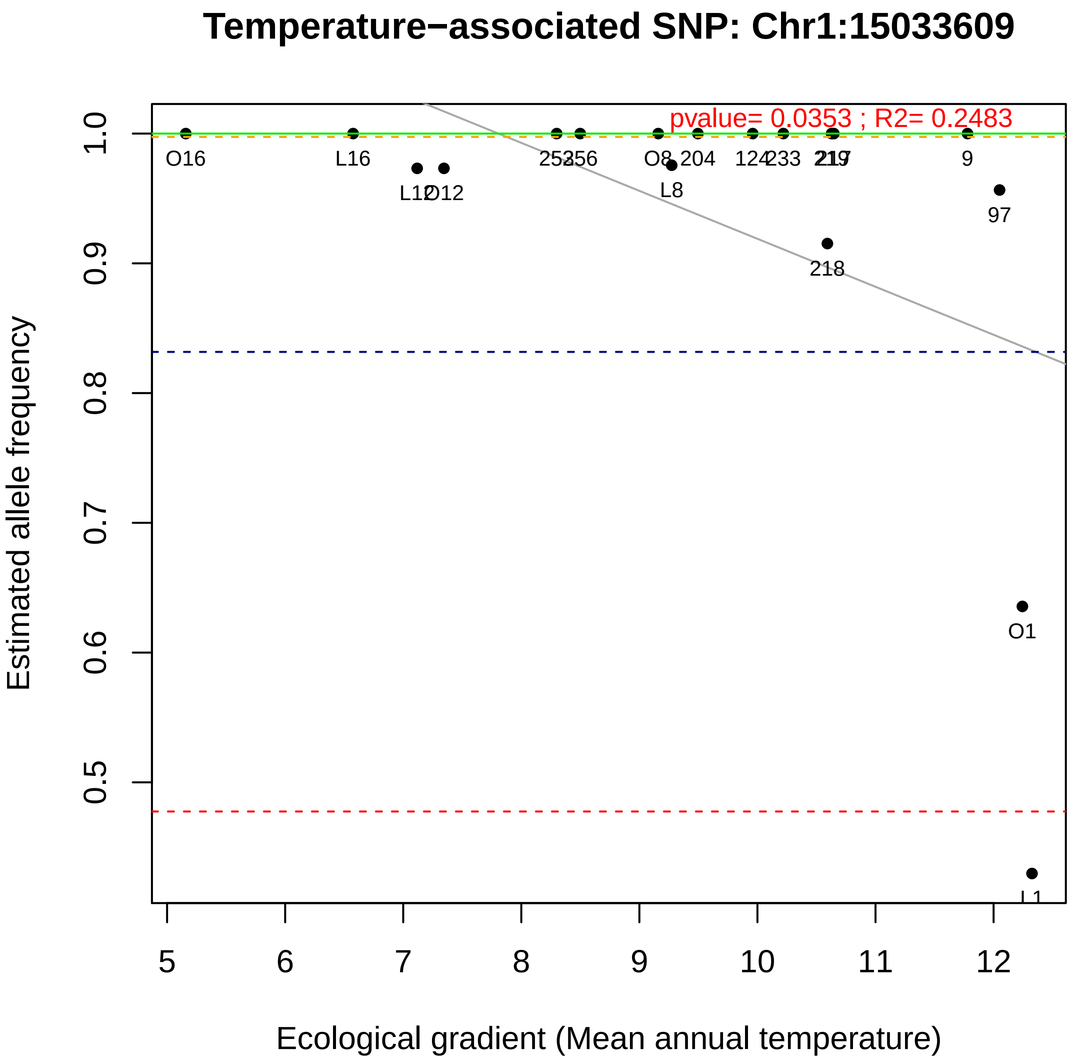

Temperature−associated SNP: Chr1:15033609
pvalue= 0.0353 ; R2= 0.2483
Estimated allele frequency
Ecological gradient (Mean annual temperature)
1.0
0.9
0.8
0.7
0.6
0.5
5
6
7
8
9
10
11
12
O16
L16
L12
O12
253
256
O8
204
L8
124
233
217
219
9
97
218
O1
L1

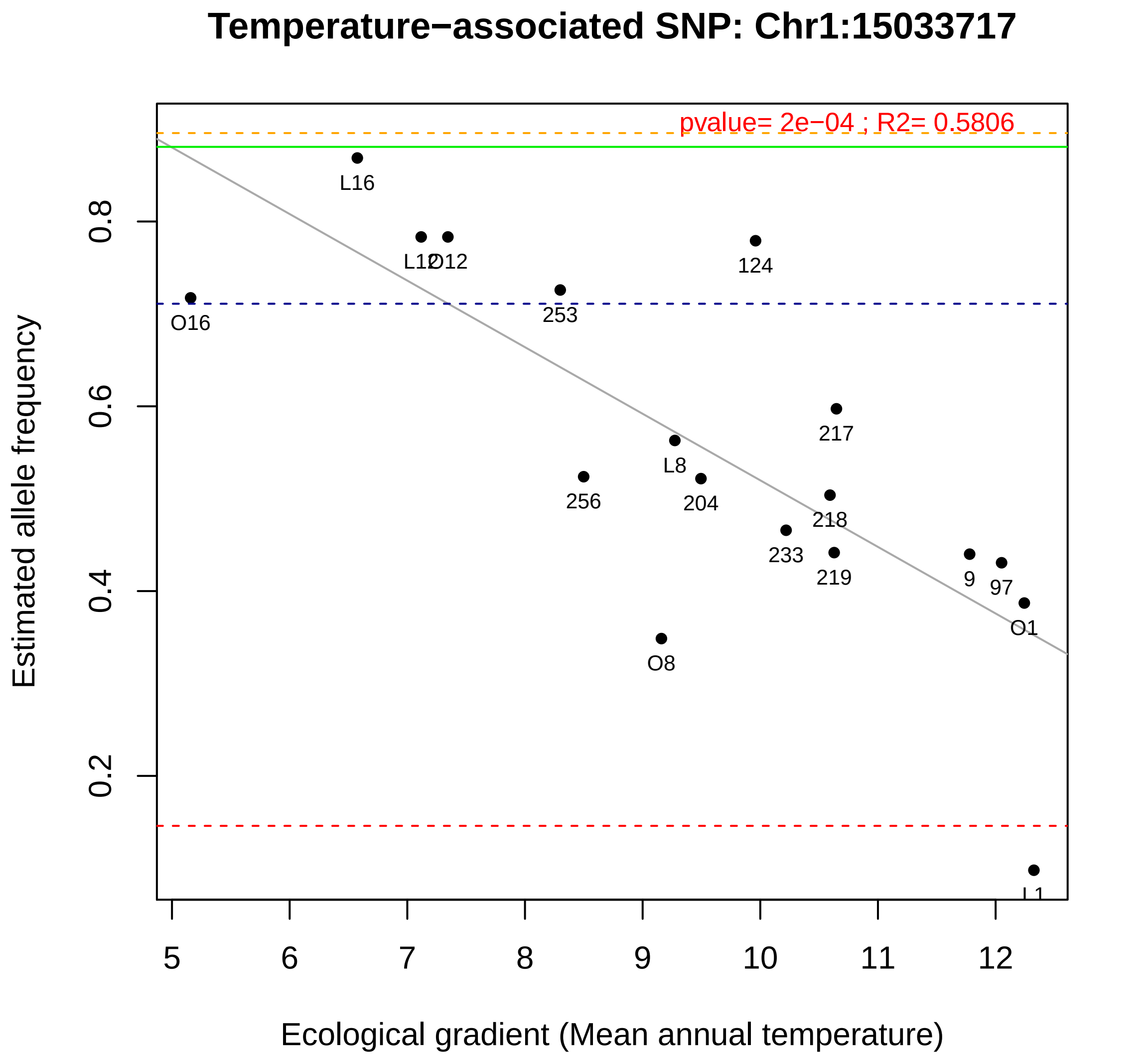
Temperature−associated SNP: Chr1:15033717
pvalue= 2e−04 ; R2= 0.5806
Estimated allele frequency
0.2
0.4
0.6
0.8
Ecological gradient (Mean annual temperature)
5
6
7
8
9
10
11
12
L16
L12
O12
124
O16
253
217
L8
256
204
218
233
219
9
97
O1
O8

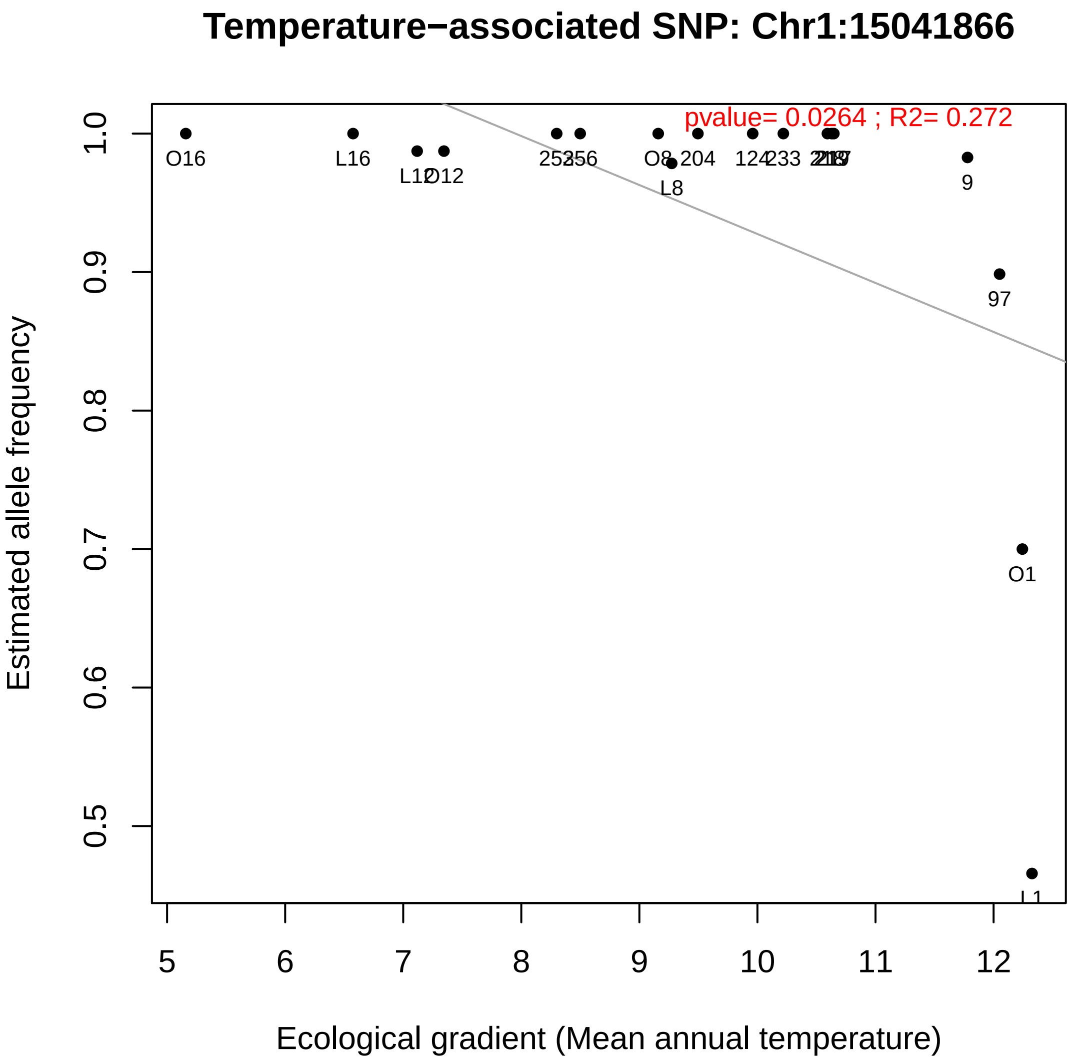

Temperature−associated SNP: Chr1:15041866
pvalue= 0.0264 ; R2= 0.272
O16
L16
L12
O12
253
256
O8
L8
204
124
233
9
97
O1
L1
1.0
0.9
0.8
0.7
0.6
0.5
Estimated allele frequency
5
6
7
8
9
10
11
12
Ecological gradient (Mean annual temperature)

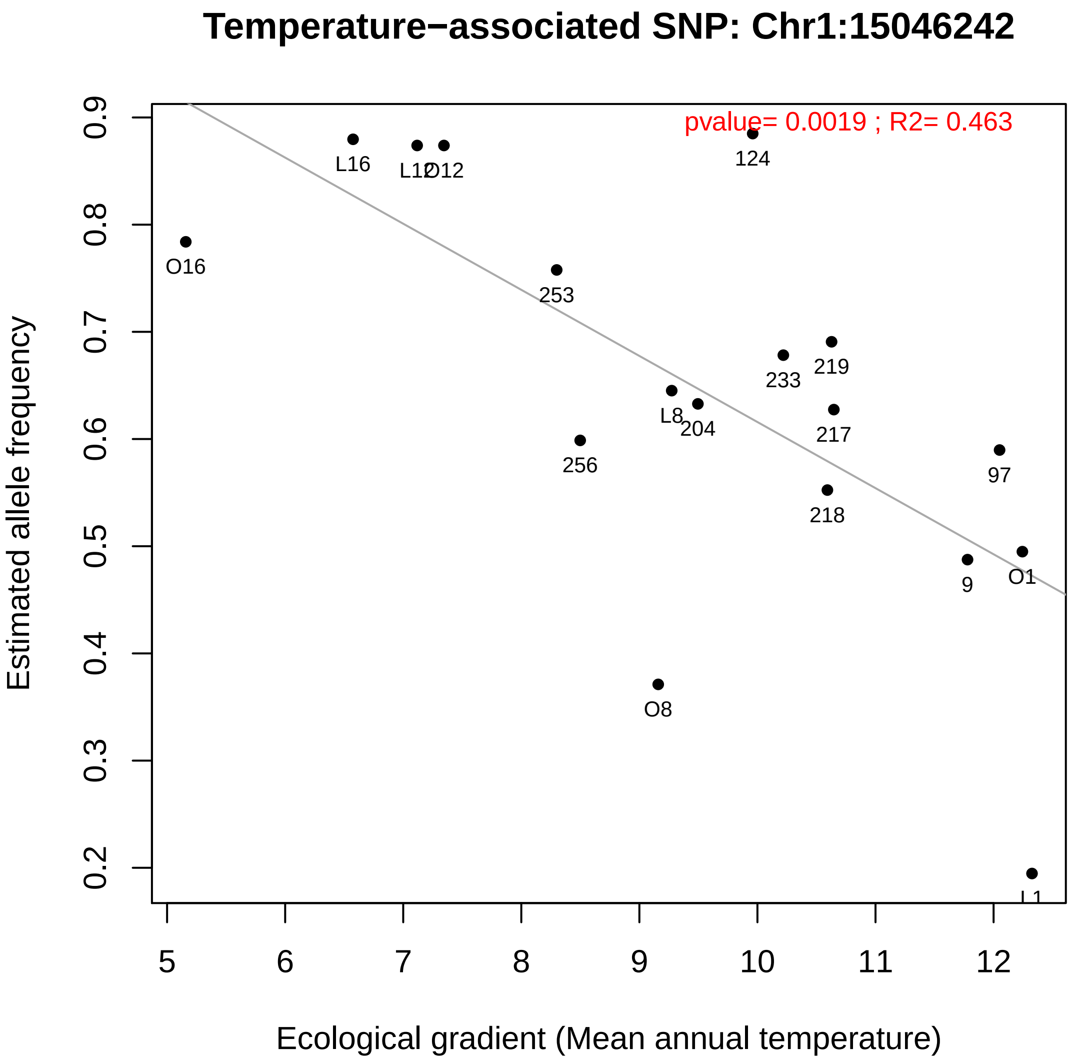
Temperature−associated SNP: Chr1:15046242
pvalue= 0.0019 ; R2= 0.463
Estimated allele frequency
0.2
0.3
0.4
0.5
0.6
0.7
0.8
0.9
5
6
7
8
9
10
11
12
Ecological gradient (Mean annual temperature)
O16
L16
L12
O12
124
253
256
L8
204
233
219
217
218
O8
97
9
O1

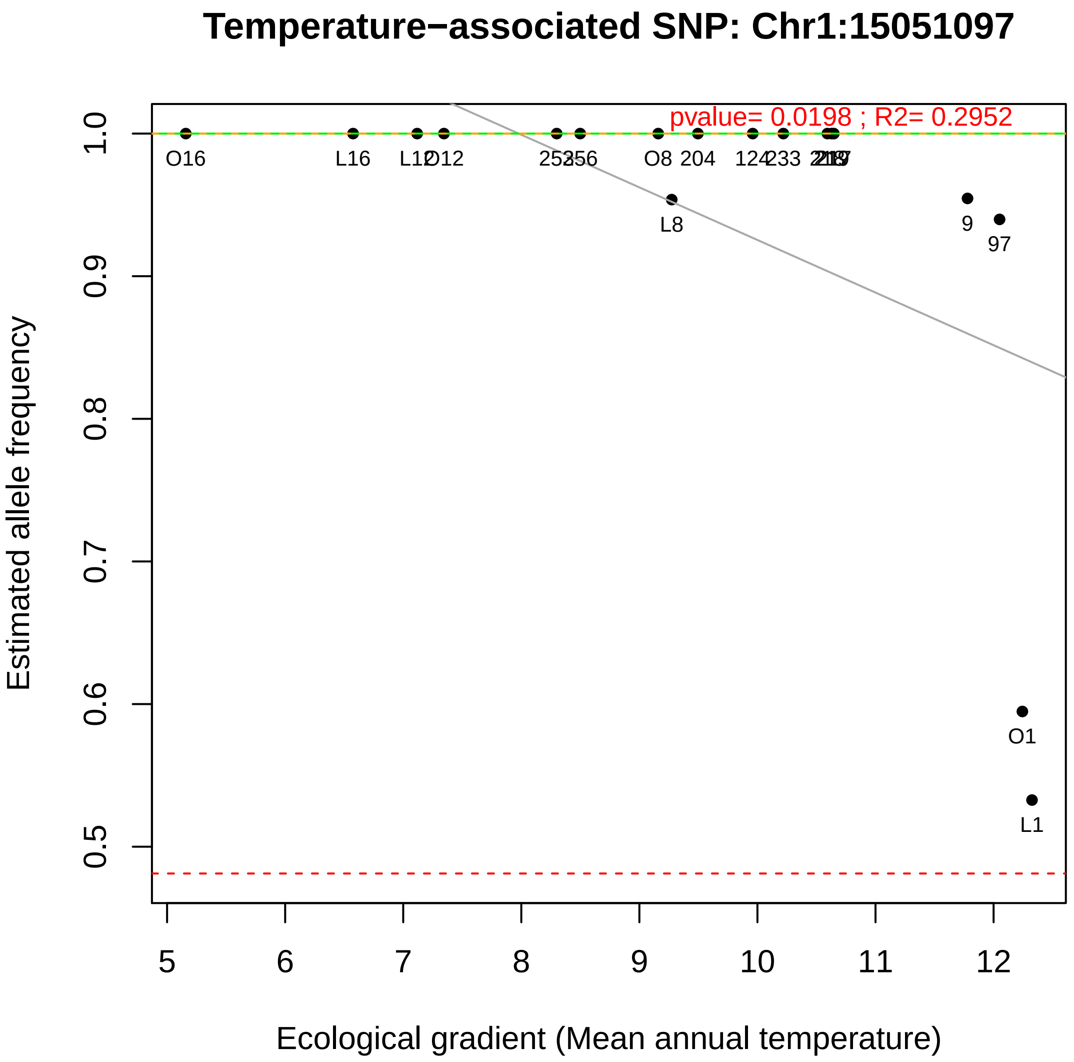
Temperature−associated SNP: Chr1:15051097
pvalue= 0.0198 ; R2= 0.2952
Estimated allele frequency
1.0
0.9
0.8
0.7
0.6
0.5
O16
L16
O12
256
O8
204
233
L8
9
97
O1
L1
5
6
7
8
9
10
11
12
Ecological gradient (Mean annual temperature)

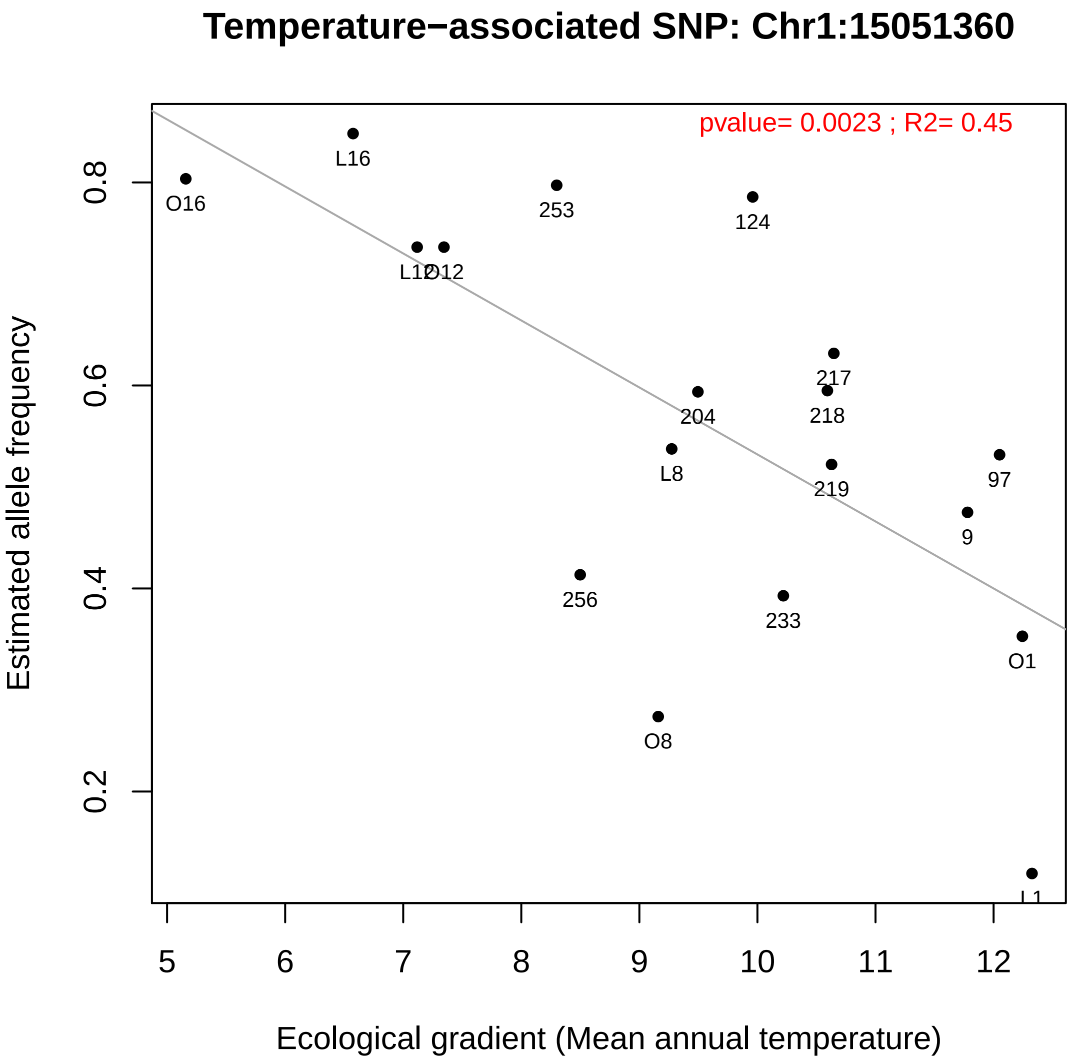

Temperature−associated SNP: Chr1:15051360
pvalue= 0.0023 ; R2= 0.45
Estimated allele frequency
Ecological gradient (Mean annual temperature)
0.2
0.4
0.6
0.8
5
6
7
8
9
10
11
12
O16
L16
253
124
L12
O12
217
204
218
L8
219
97
9
256
233
O1
O8

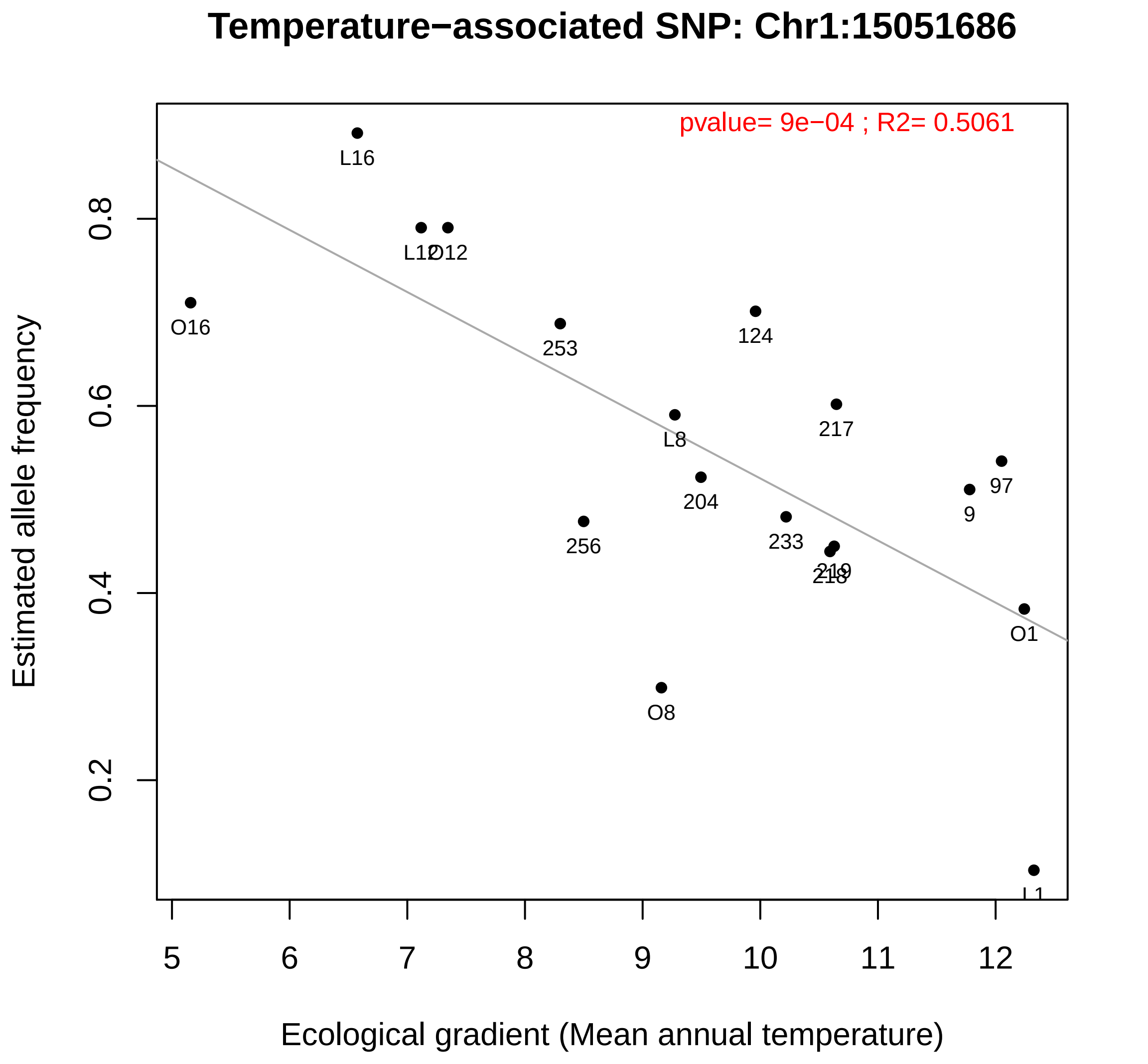

Temperature−associated SNP: Chr1:15051686
pvalue= 9e−04 ; R2= 0.5061
L16
L12
O12
O16
253
124
L8
217
204
97
9
256
233
219
218
O1
O8
L1
0.2
0.4
0.6
0.8
5
6
7
8
9
10
11
12
Estimated allele frequency
Ecological gradient (Mean annual temperature)

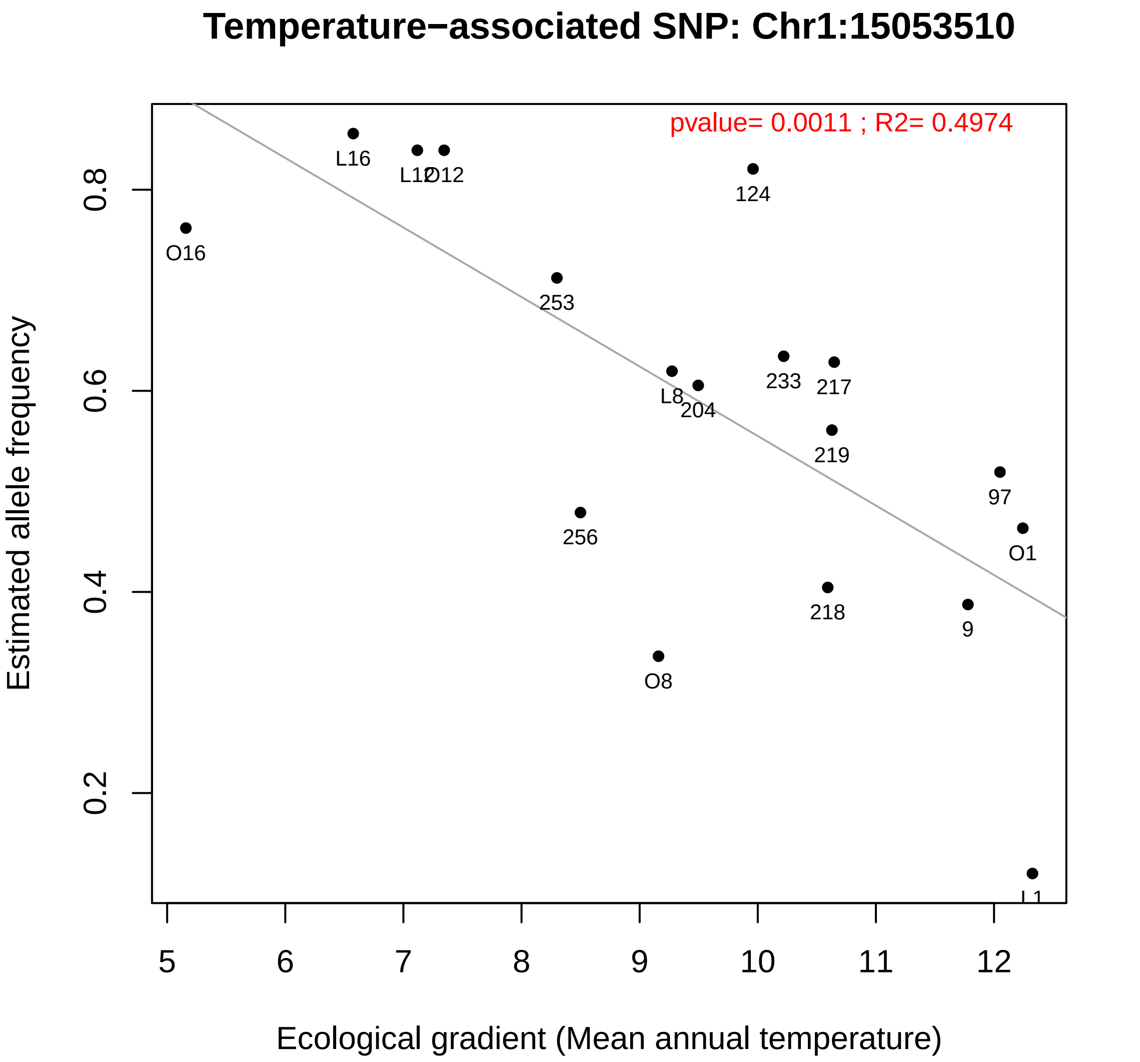

Temperature−associated SNP: Chr1:15053510
pvalue= 0.0011 ; R2= 0.4974
Estimated allele frequency
0.2
0.4
0.6
0.8
5
6
7
8
9
10
11
12
Ecological gradient (Mean annual temperature)
O16
L16
L12
O12
124
253
233
217
L8
204
219
97
O1
256
218
9
O8

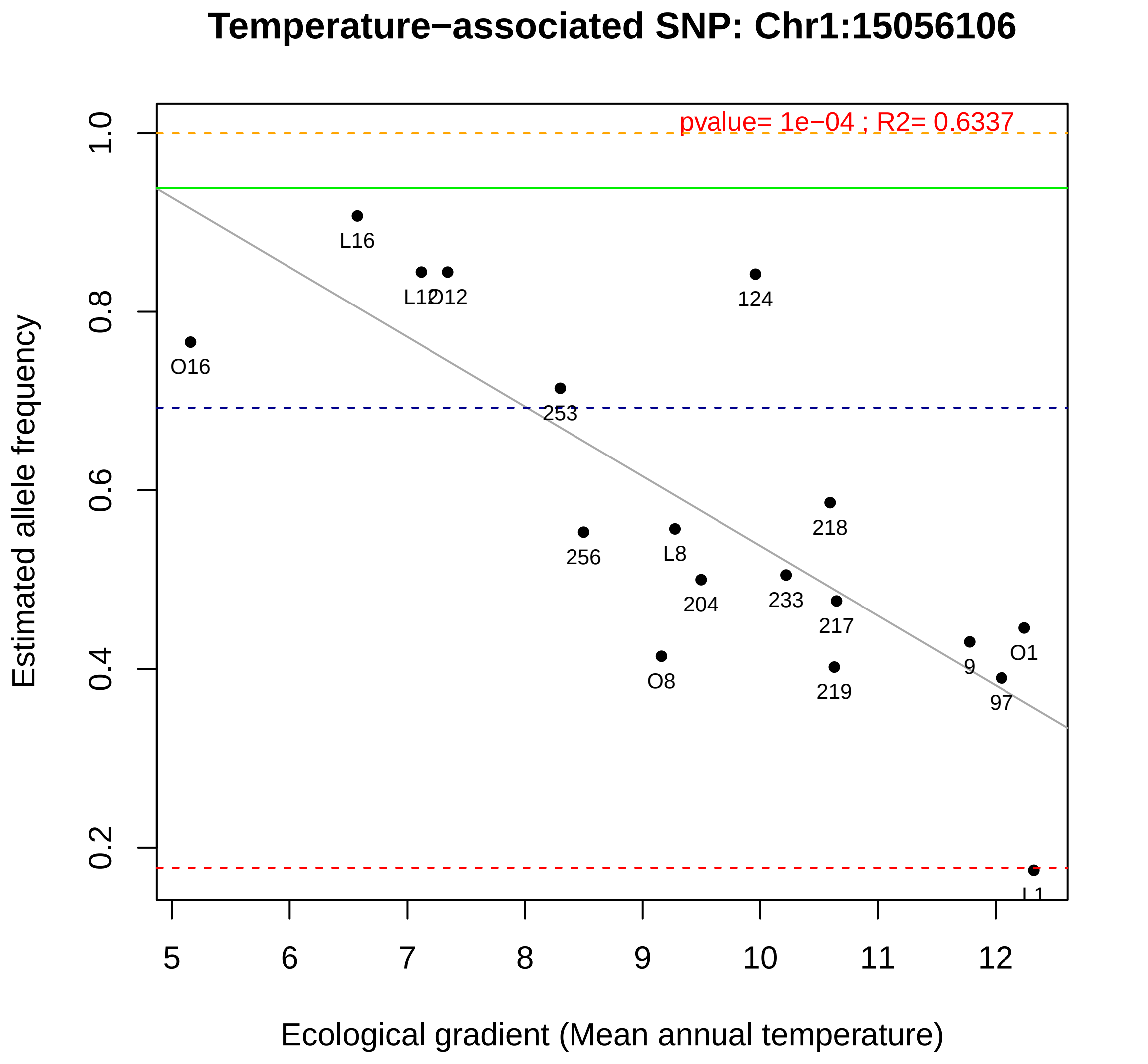

Temperature−associated SNP: Chr1:15056106
pvalue= 1e−04 ; R2= 0.6337
Estimated allele frequency
Ecological gradient (Mean annual temperature)
0.2
0.4
0.6
0.8
1.0
5
6
7
8
9
10
11
12
L16
L12
O12
124
O16
253
218
256
L8
204
233
217
O1
9
O8
219
97

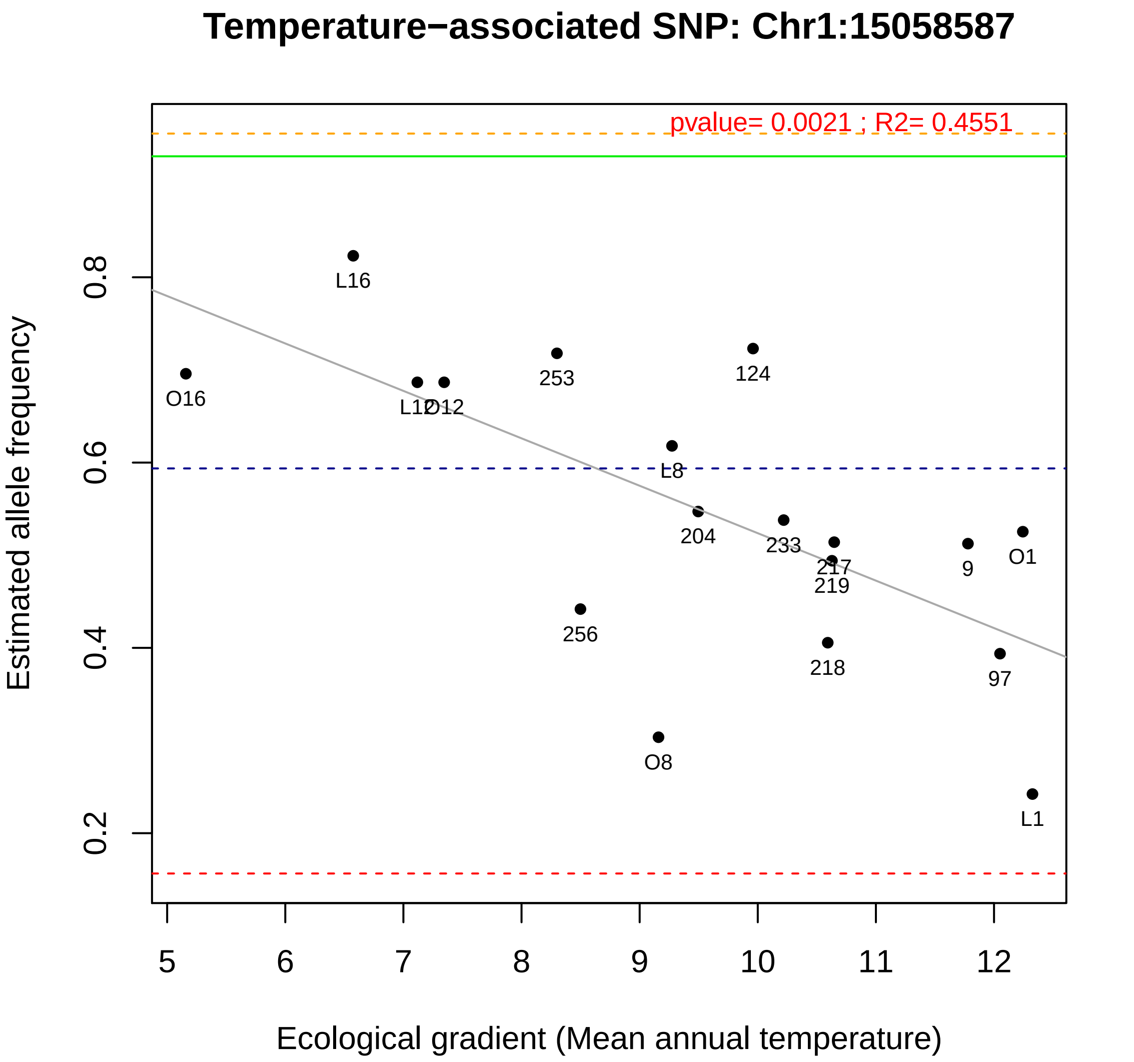
Temperature−associated SNP: Chr1:15058587
pvalue= 0.0021 ; R2= 0.4551
Estimated allele frequency
0.2
0.4
0.6
0.8
5
6
7
8
9
10
11
12
Ecological gradient (Mean annual temperature)
L16
O16
L12
O12
253
124
L8
204
233
217
219
9
O1
256
218
97
O8
L1

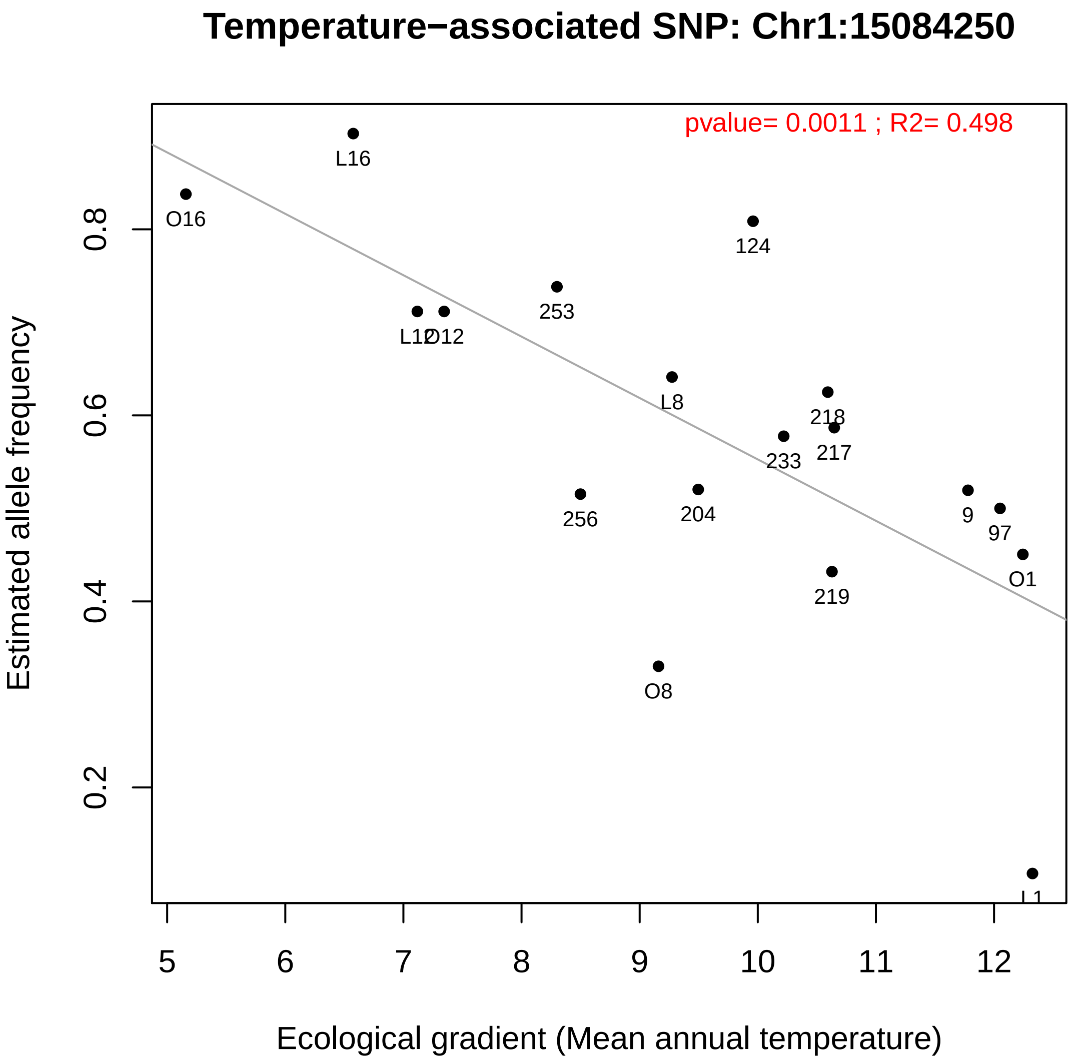

Temperature−associated SNP: Chr1:15084250
pvalue= 0.0011 ; R2= 0.498
Estimated allele frequency
0.2
0.4
0.6
0.8
5
6
7
8
9
10
11
12
Ecological gradient (Mean annual temperature)
L16
O16
124
253
L12
O12
L8
218
217
233
256
204
9
97
O1
219
O8

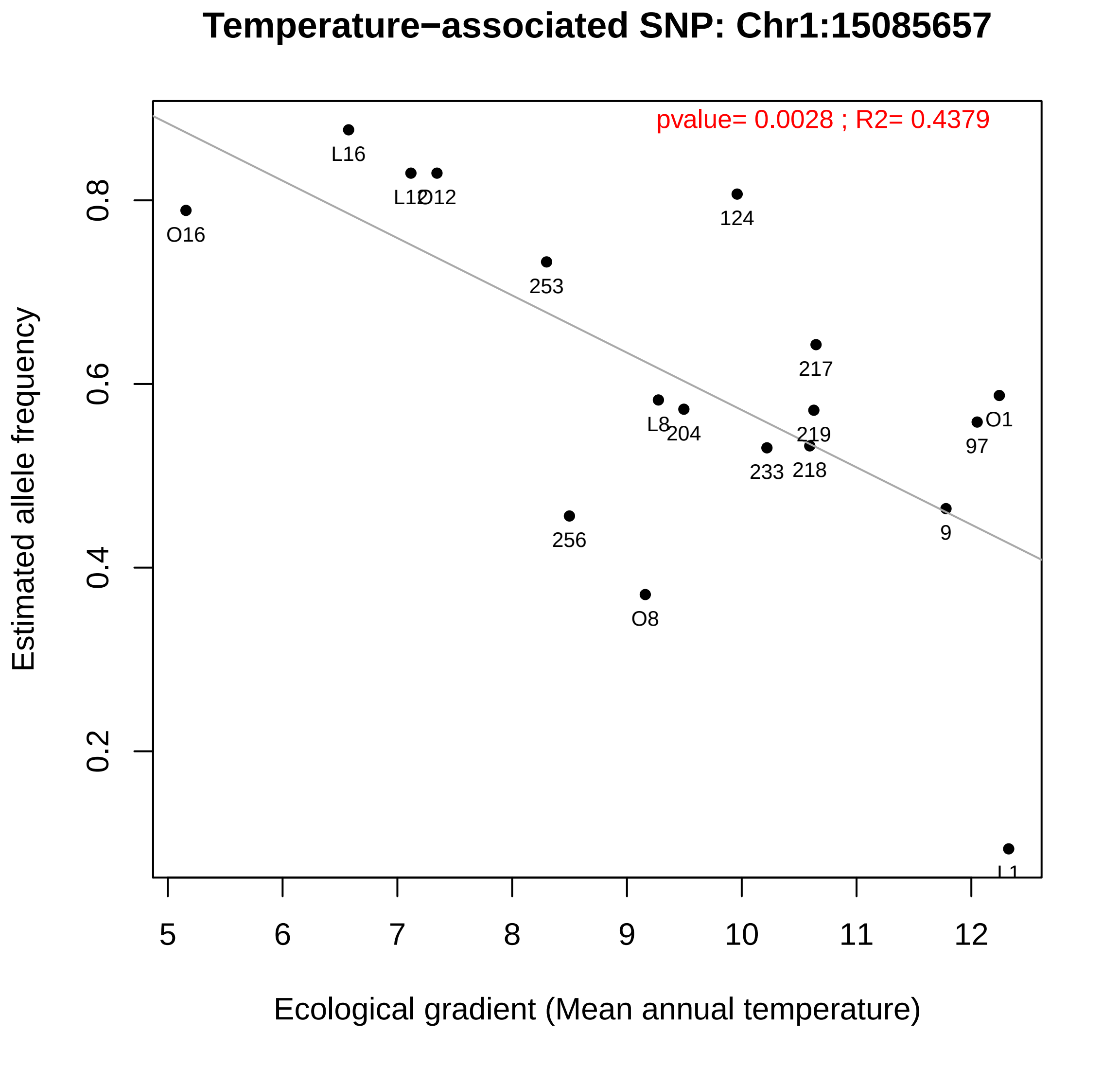

Temperature−associated SNP: Chr1:15085657
pvalue= 0.0028 ; R2= 0.4379
L16
L12
O12
O16
124
253
217
L8
204
O1
219
97
233
218
256
9
O8
L1
0.8
0.6
0.4
0.2
5
6
7
8
9
10
11
12
Estimated allele frequency
Ecological gradient (Mean annual temperature)

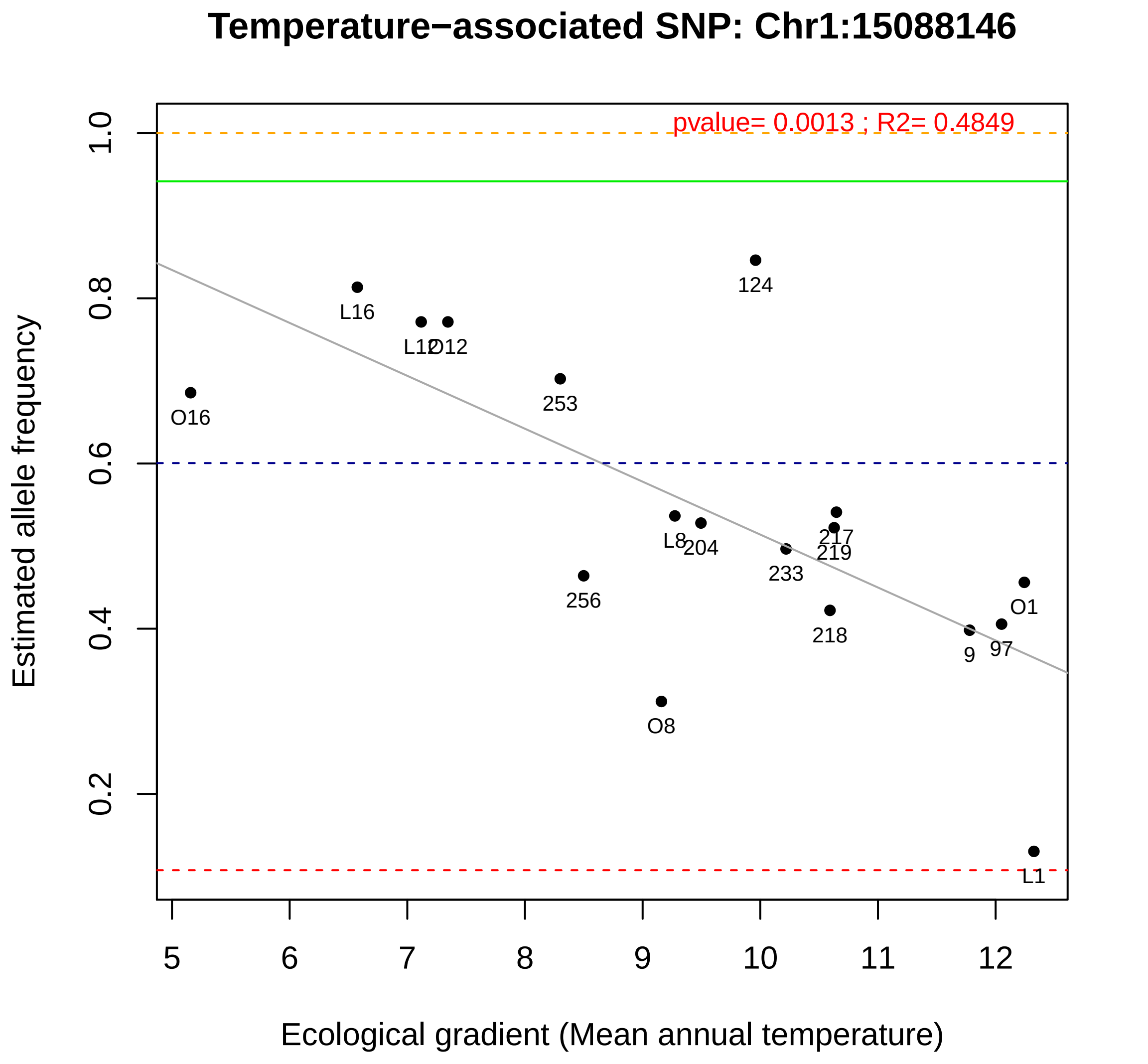
Temperature−associated SNP: Chr1:15088146
pvalue= 0.0013 ; R2= 0.4849
Estimated allele frequency
Ecological gradient (Mean annual temperature)
1.0
0.8
0.6
0.4
0.2
5
6
7
8
9
10
11
12
O16
L16
L12
O12
253
124
256
L8
204
O8
233
217
219
218
O1
9
97
L1

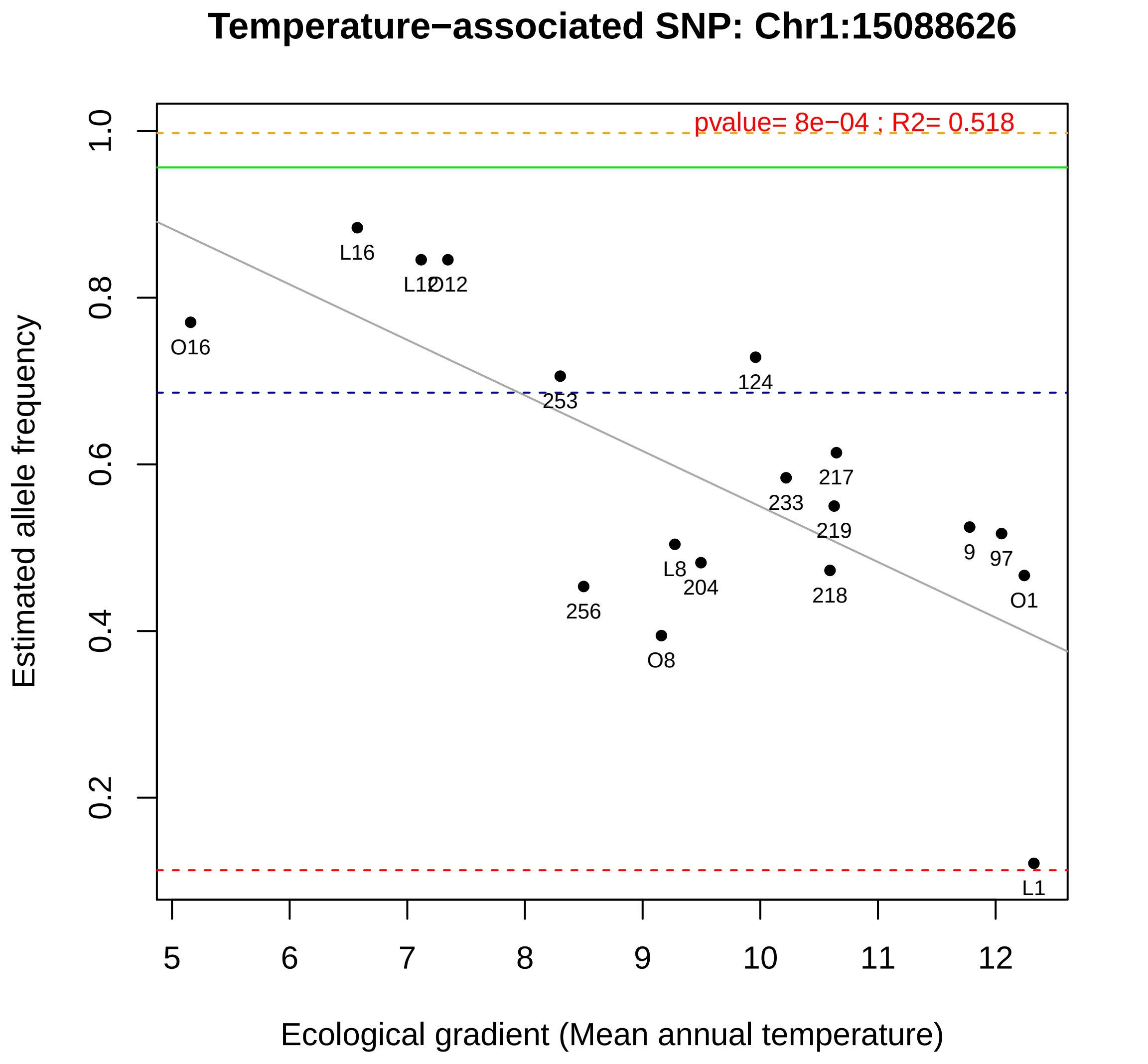

Temperature−associated SNP: Chr1:15088626
pvalue= 8e−04 ; R2= 0.518
Estimated allele frequency
0.2
0.4
0.6
0.8
1.0
5
6
7
8
9
10
11
12
Ecological gradient (Mean annual temperature)
O16
L16
L12
O12
253
124
256
L8
204
O8
233
217
219
218
9
97
O1
L1

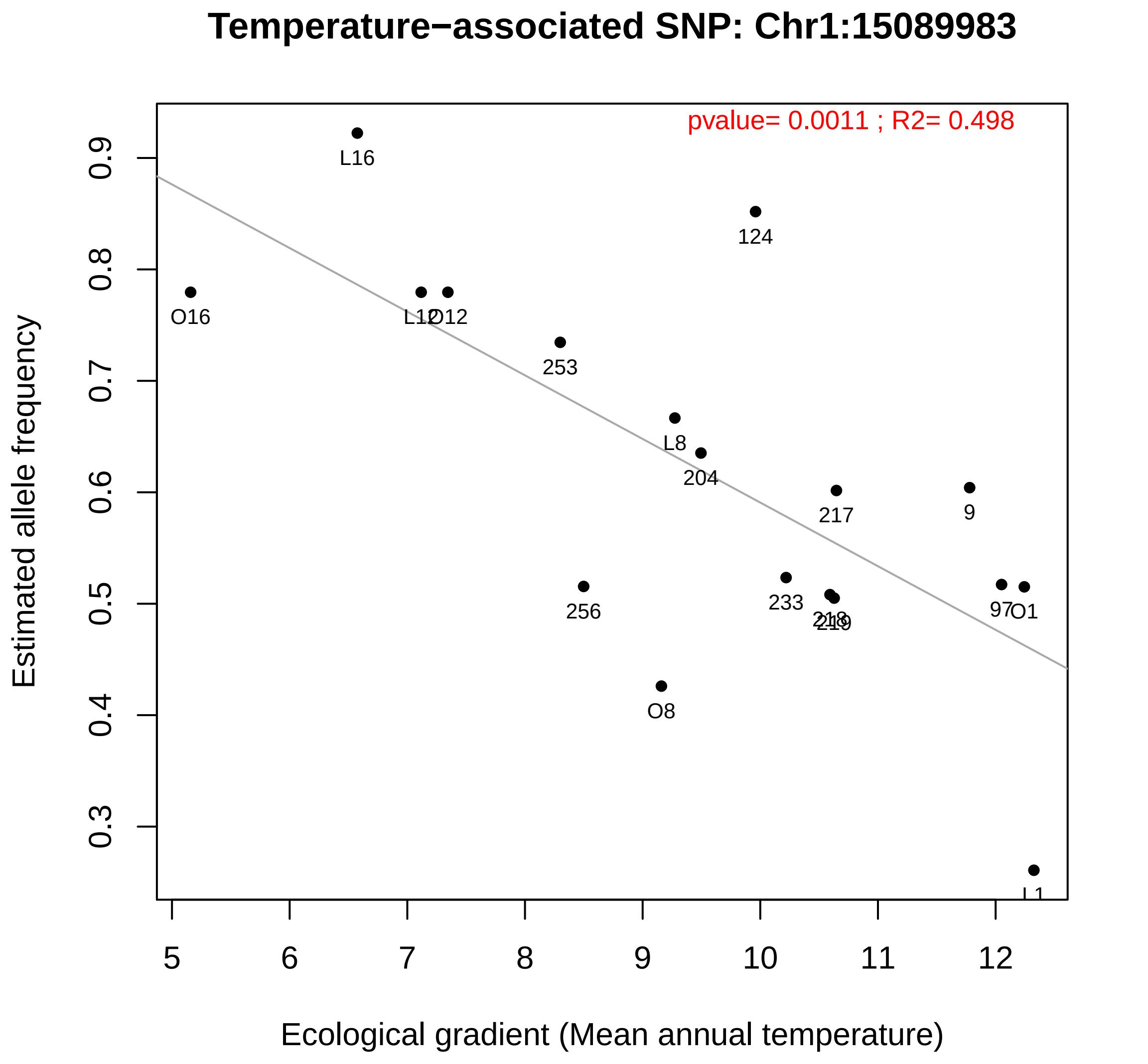
Temperature−associated SNP: Chr1:15089983
pvalue= 0.0011 ; R2= 0.498
Estimated allele frequency
Ecological gradient (Mean annual temperature)
0.3
0.4
0.5
0.6
0.7
0.8
0.9
5
6
7
8
9
10
11
12
L16
124
O16
L12
O12
253
L8
204
217
9
256
233
97
O1
O8

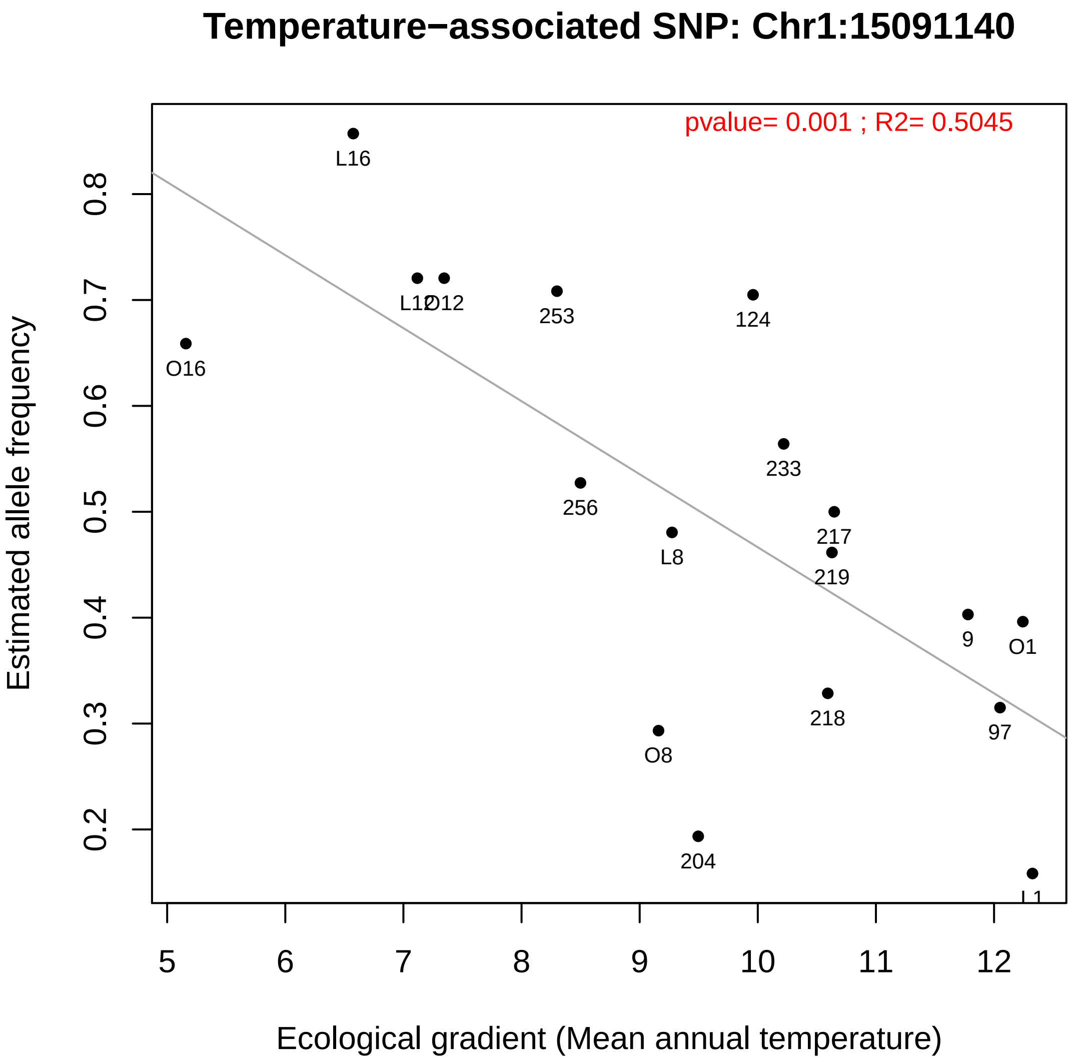

Temperature−associated SNP: Chr1:15091140
pvalue= 0.001 ; R2= 0.5045
Estimated allele frequency
0.2
0.3
0.4
0.5
0.6
0.7
0.8
Ecological gradient (Mean annual temperature)
5
6
7
8
9
10
11
12
L16
O16
L12
O12
253
124
256
233
217
219
L8
9
O1
218
97
O8
204

### Temperature−associated SNP: Chr1:15098057

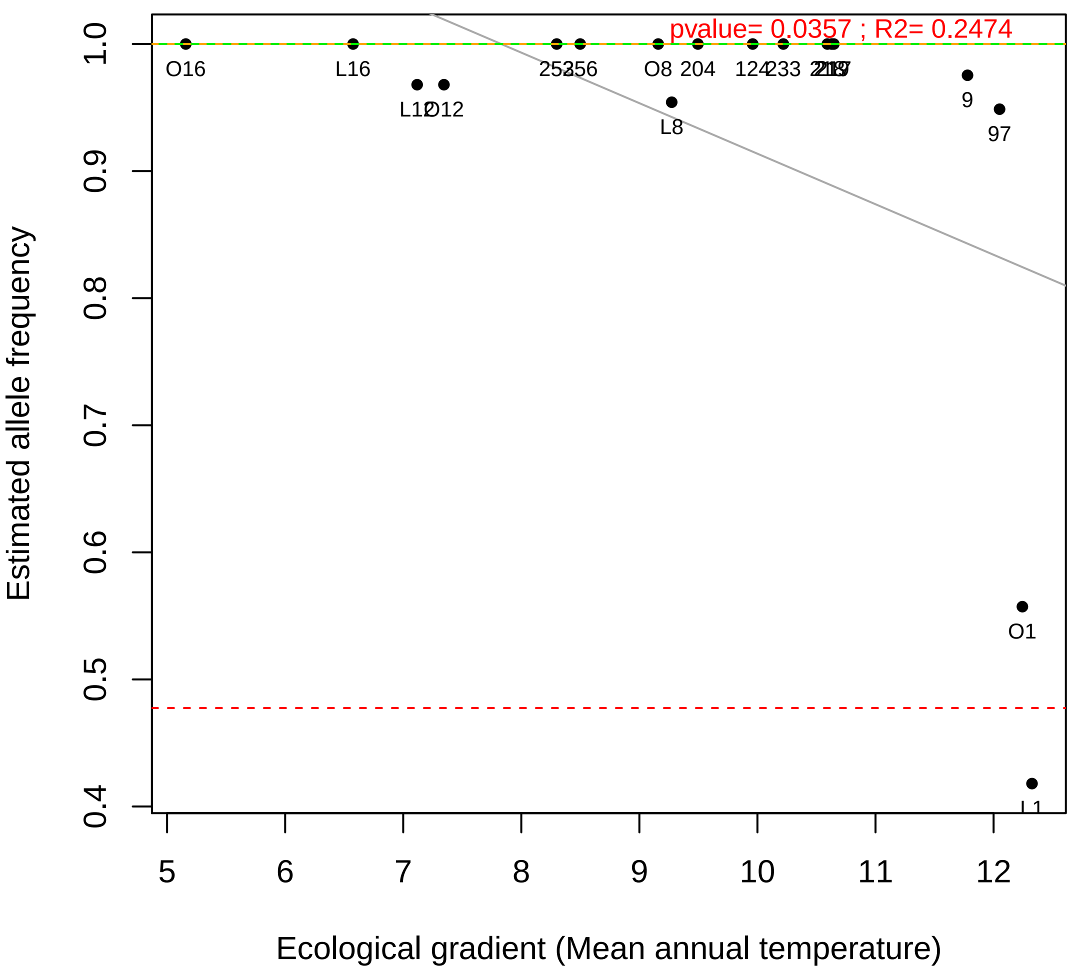

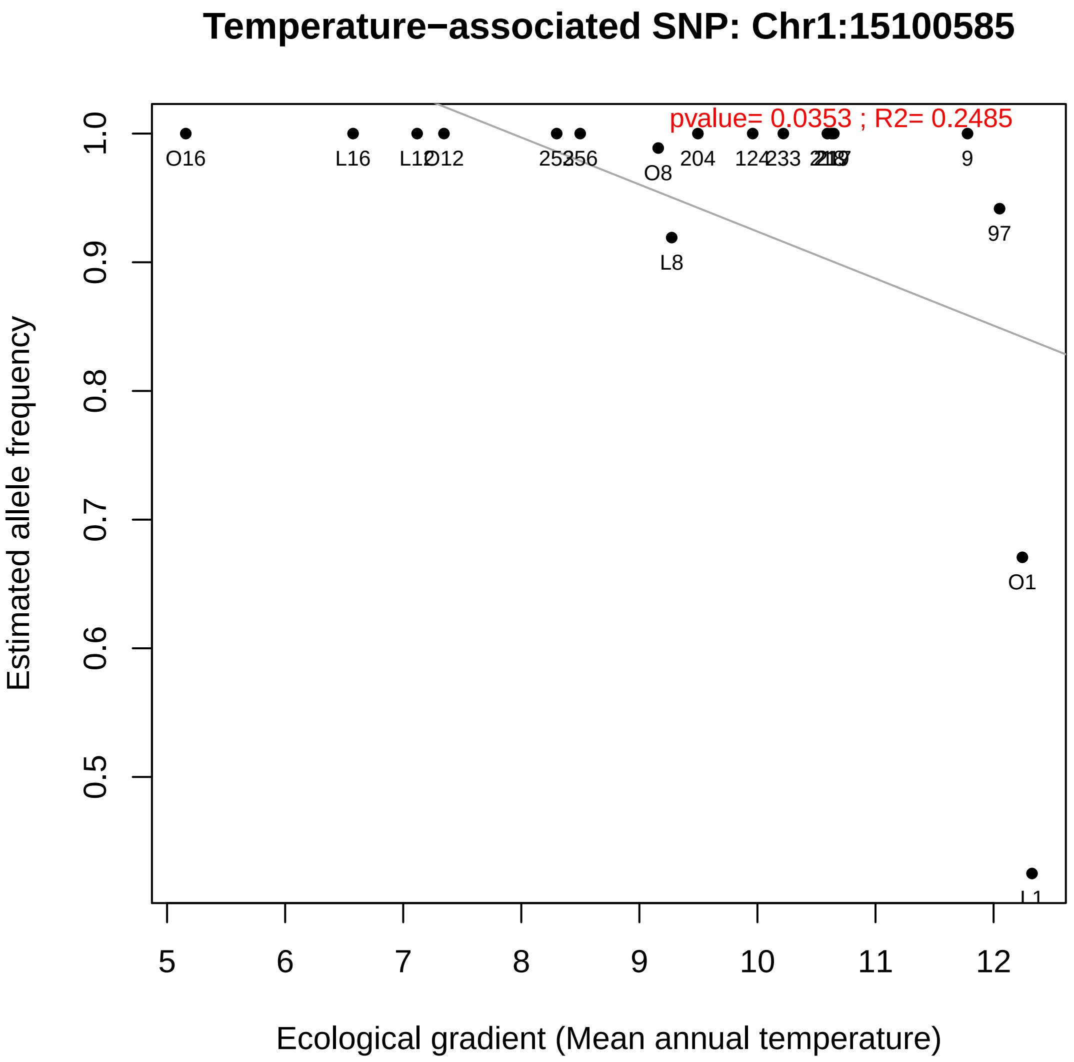
Temperature−associated SNP: Chr1:15100585
pvalue= 0.0353 ; R2= 0.2485
Estimated allele frequency
Ecological gradient (Mean annual temperature)
O16
L16
L12
O12
204
124
233
9
O8
L8
97
O1
0.5
0.6
0.7
0.8
0.9
1.0
5
6
7
8
9
10
11
12

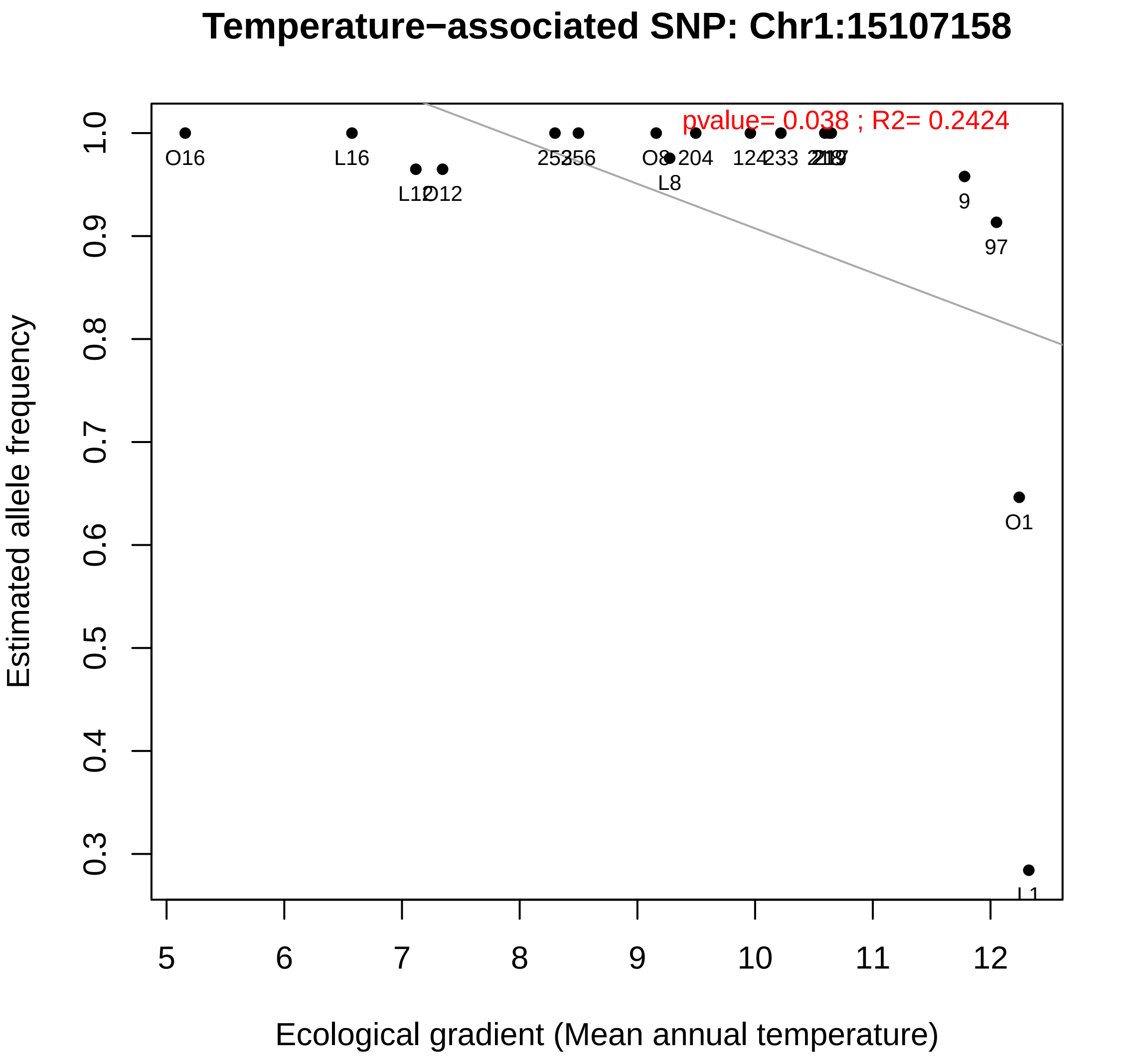

Temperature−associated SNP: Chr1:15107158
pvalue= 0.038 ; R2= 0.2424
Estimated allele frequency
Ecological gradient (Mean annual temperature)
1.0
0.9
0.8
0.7
0.6
0.5
0.4
0.3
5
6
7
8
9
10
11
12
O16
L16
L12
O12
253
256
O8
L8
204
124
233
9
97
O1
L1

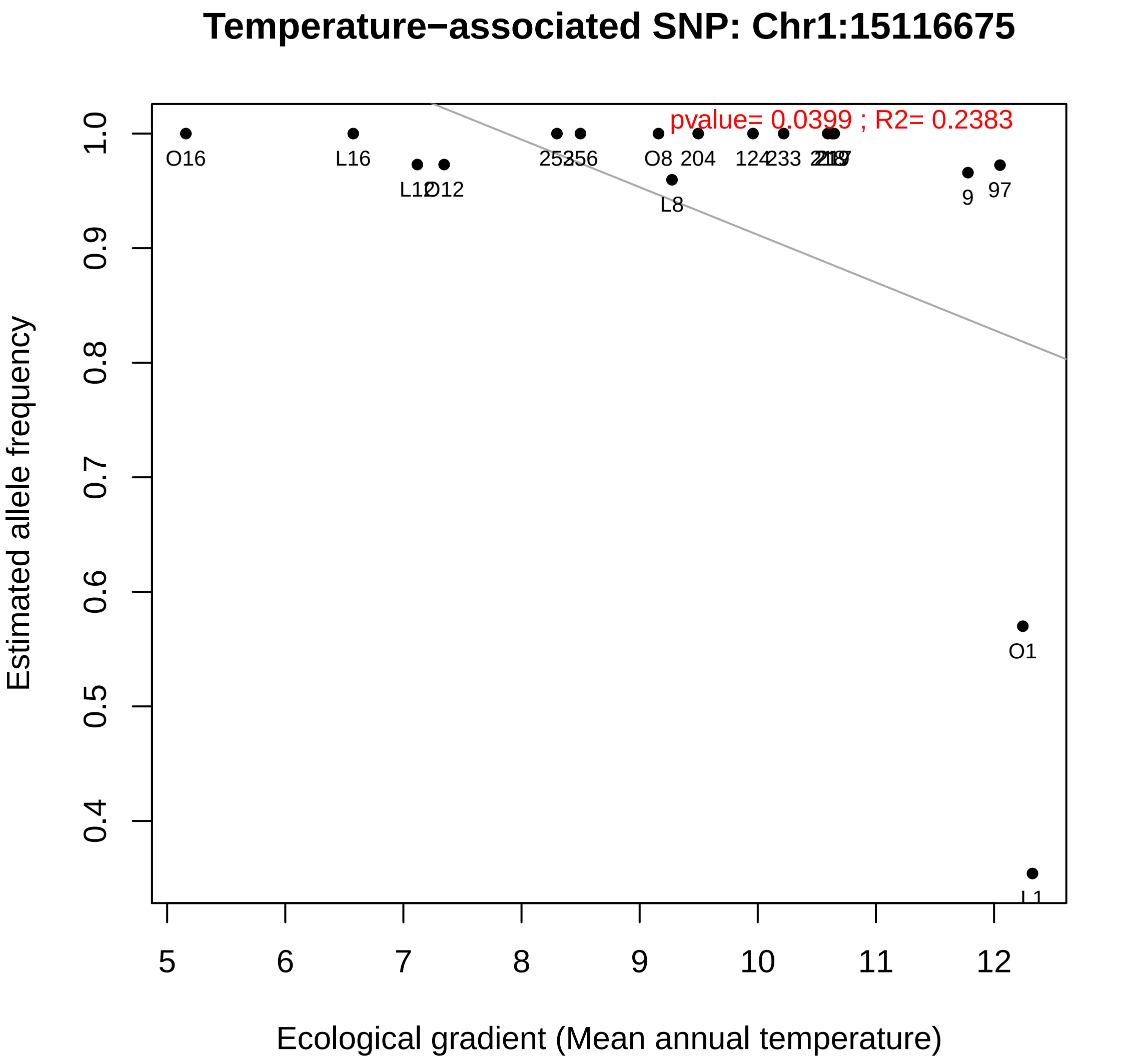
Temperature−associated SNP: Chr1:15116675
pvalue= 0.0399 ; R2= 0.2383
Estimated allele frequency
Ecological gradient (Mean annual temperature)
1.0
0.9
0.8
0.7
0.6
0.5
0.4
5
6
7
8
9
10
11
12
O16
L16
O12
O8
204
L8
9
97
O1

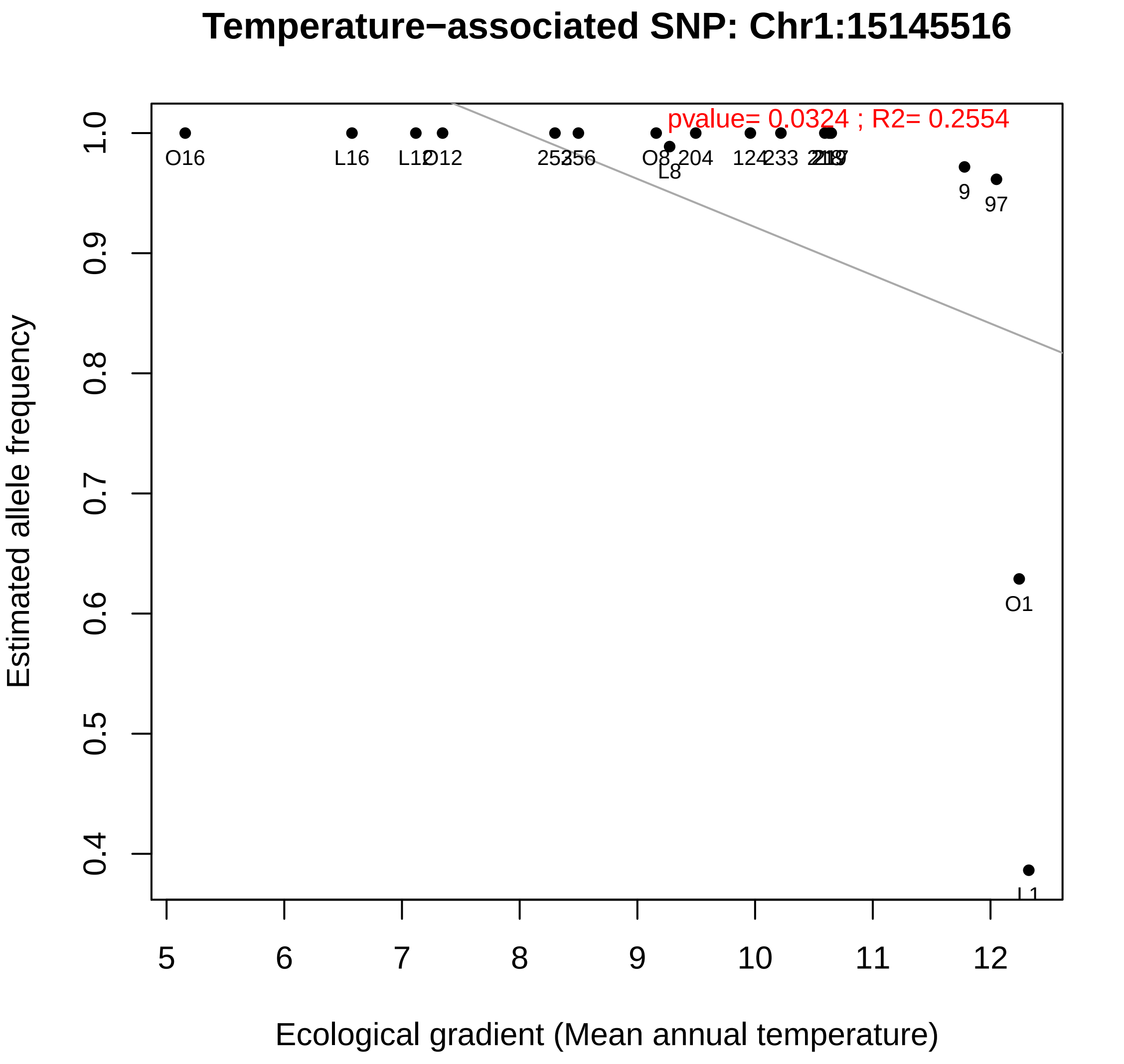

Temperature−associated SNP: Chr1:15145516
pvalue= 0.0324 ; R2= 0.2554
O16
L16
L12
O12
253
256
O8
L8
204
124
233
9
97
O1
L1
Estimated allele frequency
1.0
0.9
0.8
0.7
0.6
0.5
0.4
5
6
7
8
9
10
11
12
Ecological gradient (Mean annual temperature)

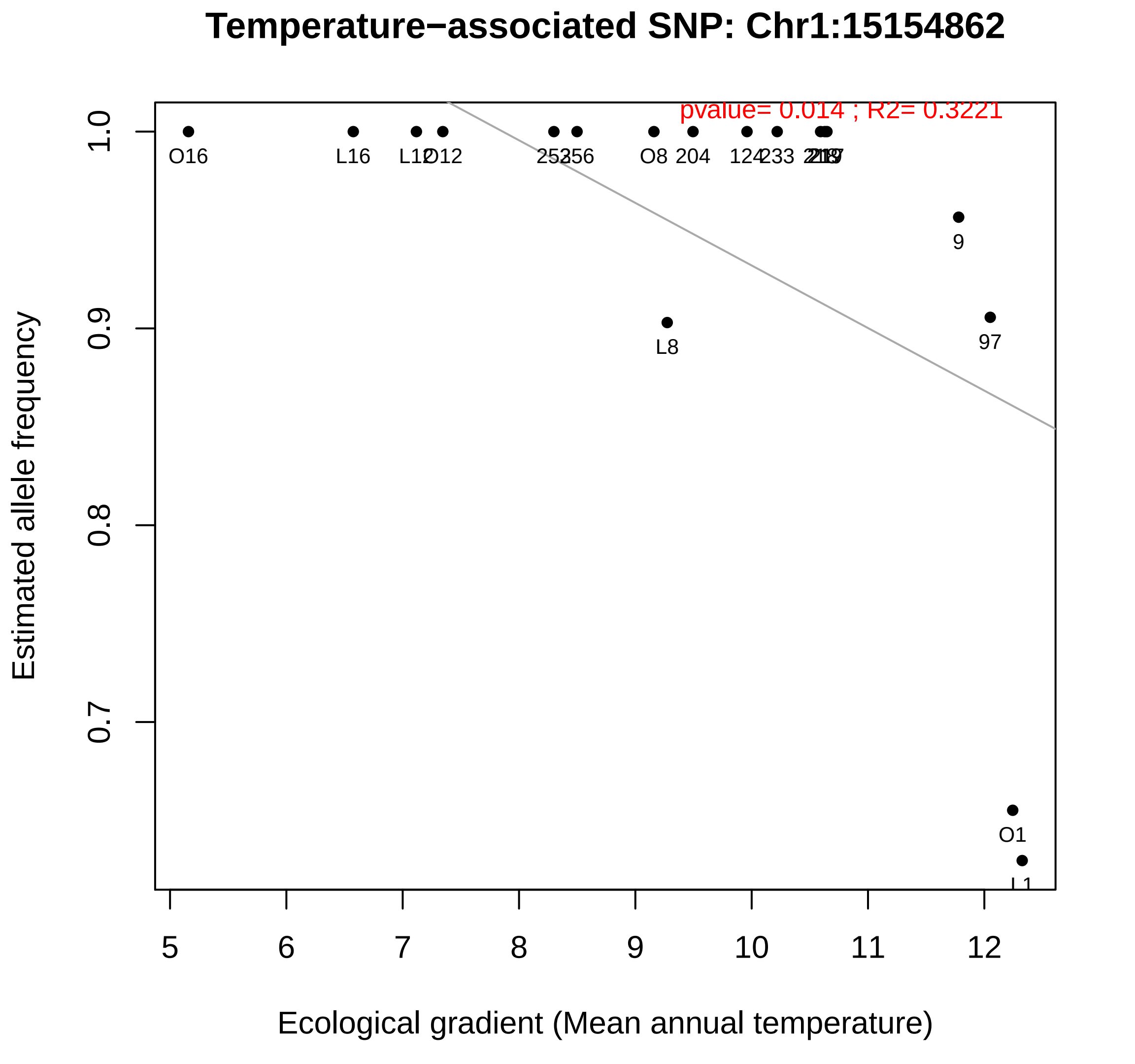
Temperature−associated SNP: Chr1:15154862
pvalue= 0.014 ; R2= 0.3221
Estimated allele frequency
Ecological gradient (Mean annual temperature)
1.0
0.9
0.8
0.7
5
6
7
8
9
10
11
12
O16
L16
O12
O8
204
124
233
9
L8
97
O1

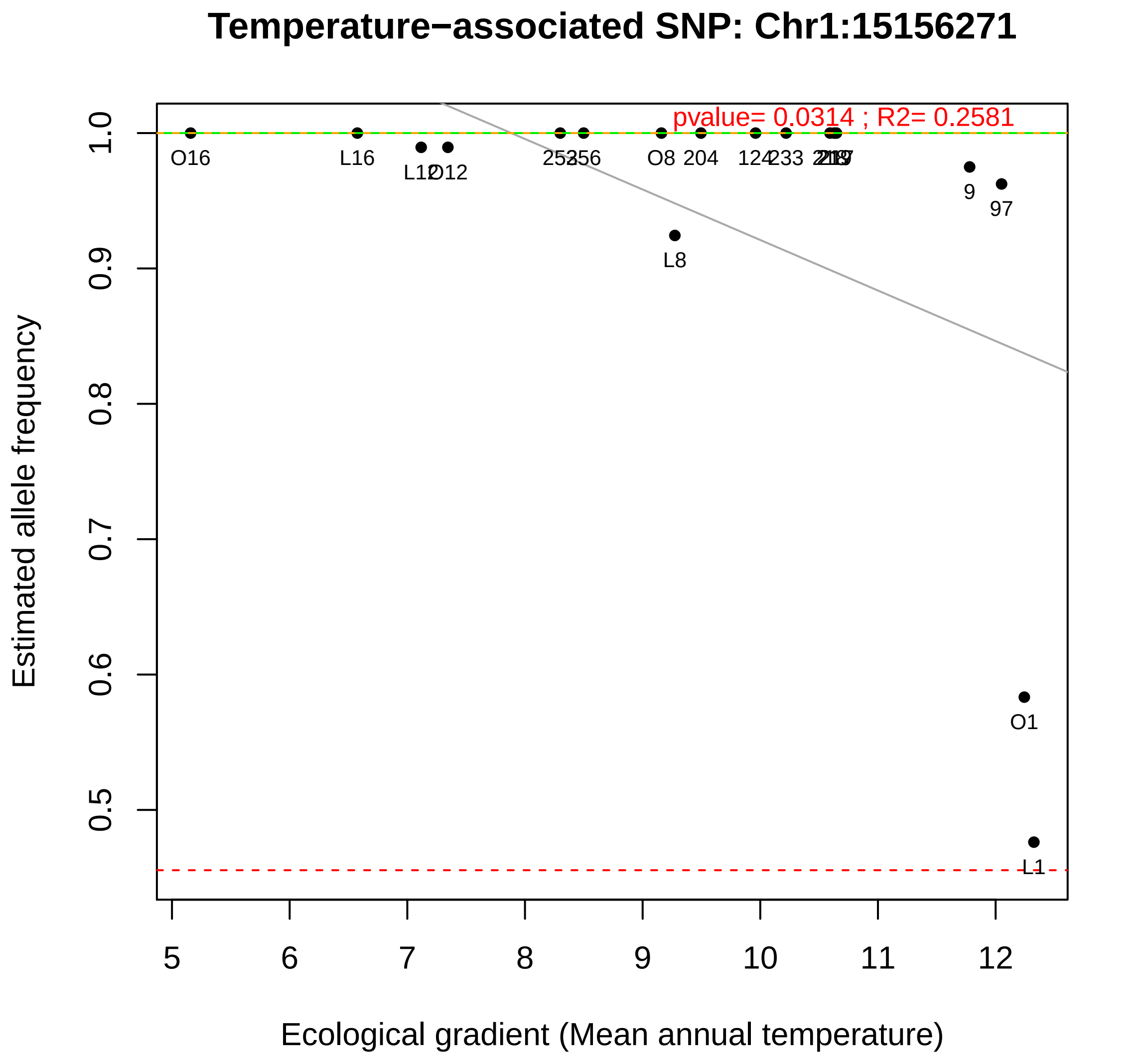

Temperature−associated SNP: Chr1:15156271
pvalue= 0.0314 ; R2= 0.2581
Estimated allele frequency
Ecological gradient (Mean annual temperature)
O16
L16
L12
O12
O8
204
124
233
9
97
L8
O1
L1
0.5
0.6
0.7
0.8
0.9
1.0
5
6
7
8
9
10
11
12

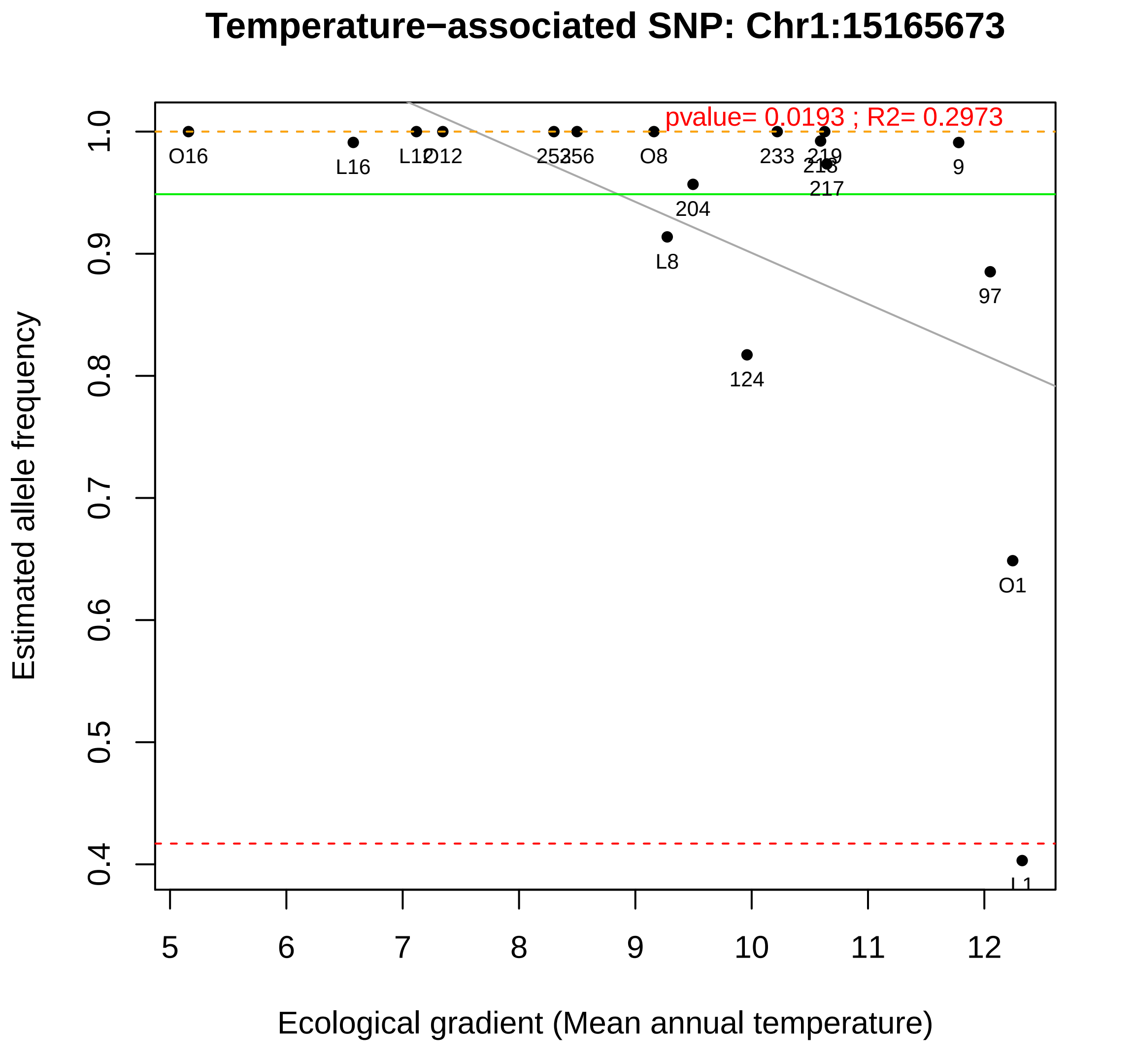

Temperature−associated SNP: Chr1:15165673
pvalue= 0.0193 ; R2= 0.2973
Estimated allele frequency
Ecological gradient (Mean annual temperature)
1.0
0.9
0.8
0.7
0.6
0.5
0.4
5
6
7
8
9
10
11
12
O16
L16
L12
O12
O8
233
219
217
204
L8
9
97
124
O1

**Temperature−associated SNP: Chr1:15173835**

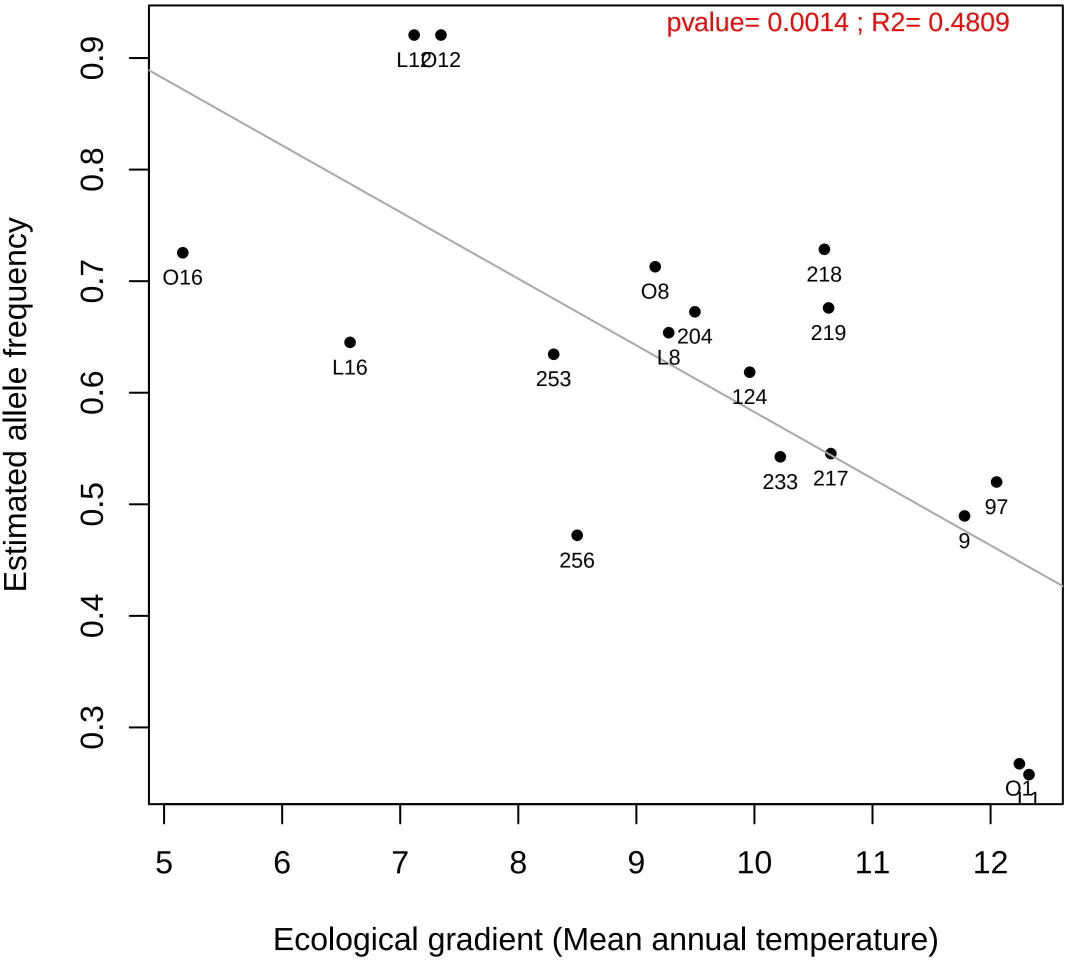

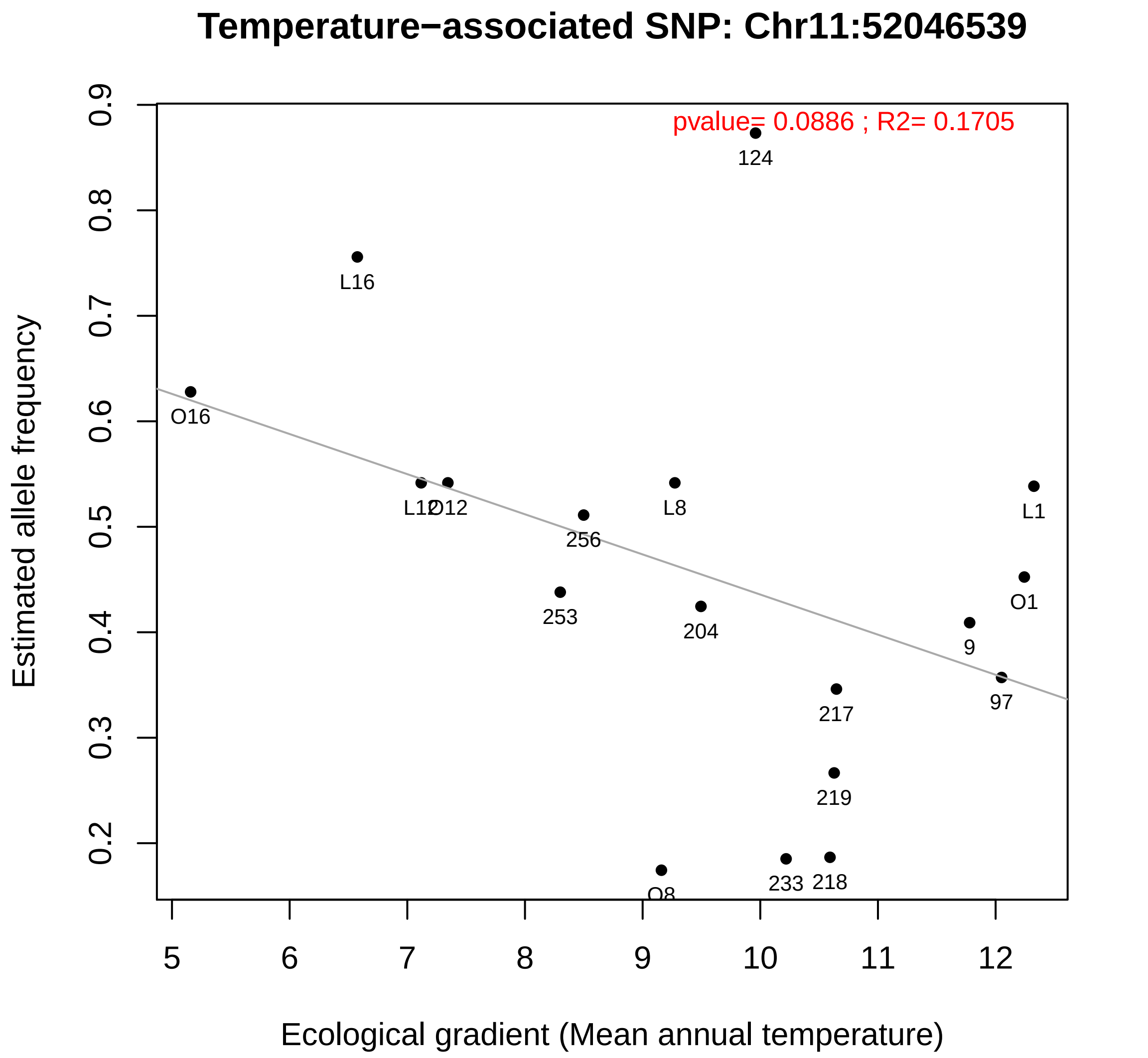
Temperature−associated SNP: Chr11:52046539
pvalue= 0.0886 ; R2= 0.1705
Estimated allele frequency
0.2
0.3
0.4
0.5
0.6
0.7
0.8
0.9
5
6
7
8
9
10
11
12
Ecological gradient (Mean annual temperature)
124
L16
O16
L12
O12
256
L8
L1
253
204
O1
9
97
217
219
233
218
O8

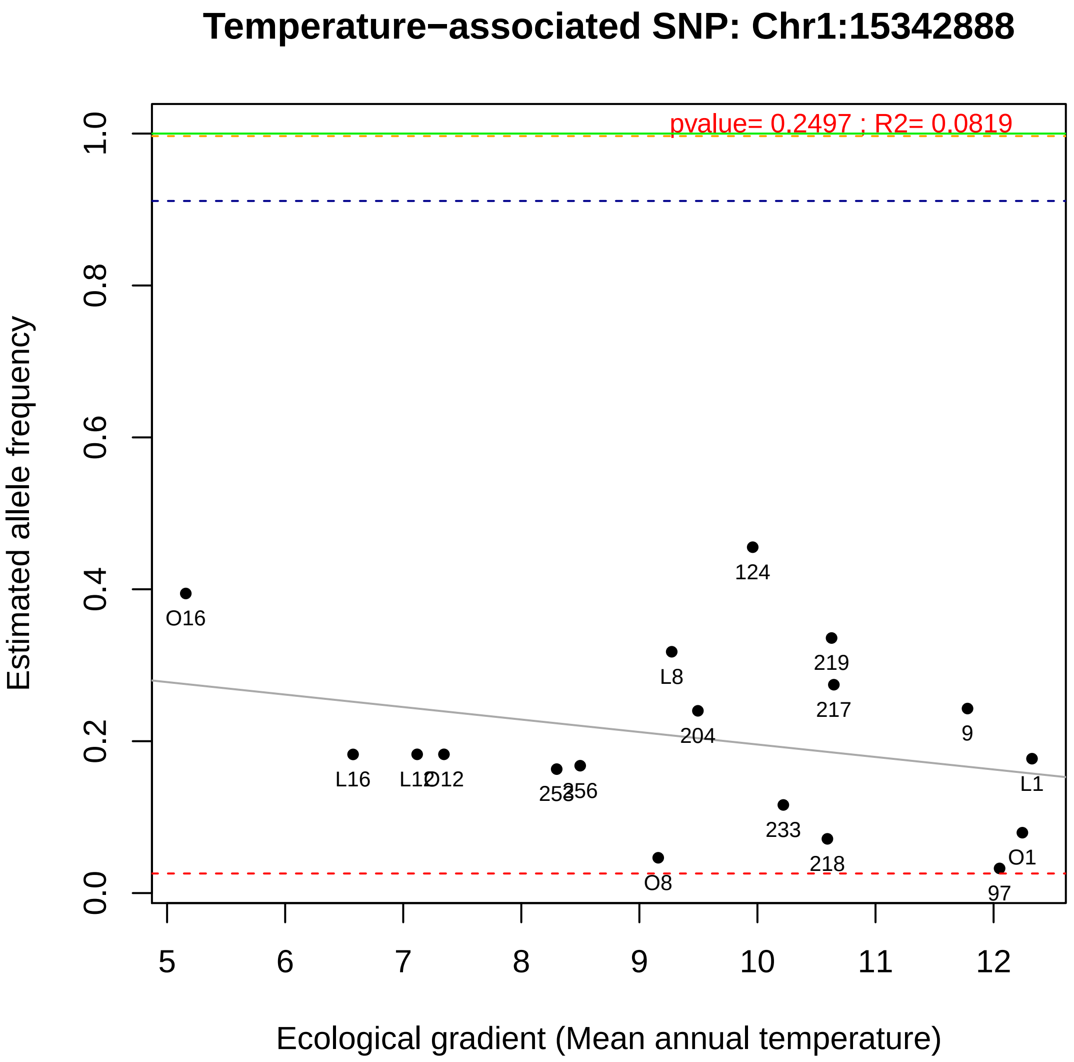

Temperature−associated SNP: Chr1:15342888
pvalue= 0.2497 ; R2= 0.0819
Estimated allele frequency
0.0
0.2
0.4
0.6
0.8
1.0
O16
L16
L12
O12
253
256
O8
L8
204
124
233
219
217
218
9
L1
O1
97
5
6
7
8
9
10
11
12
Ecological gradient (Mean annual temperature)

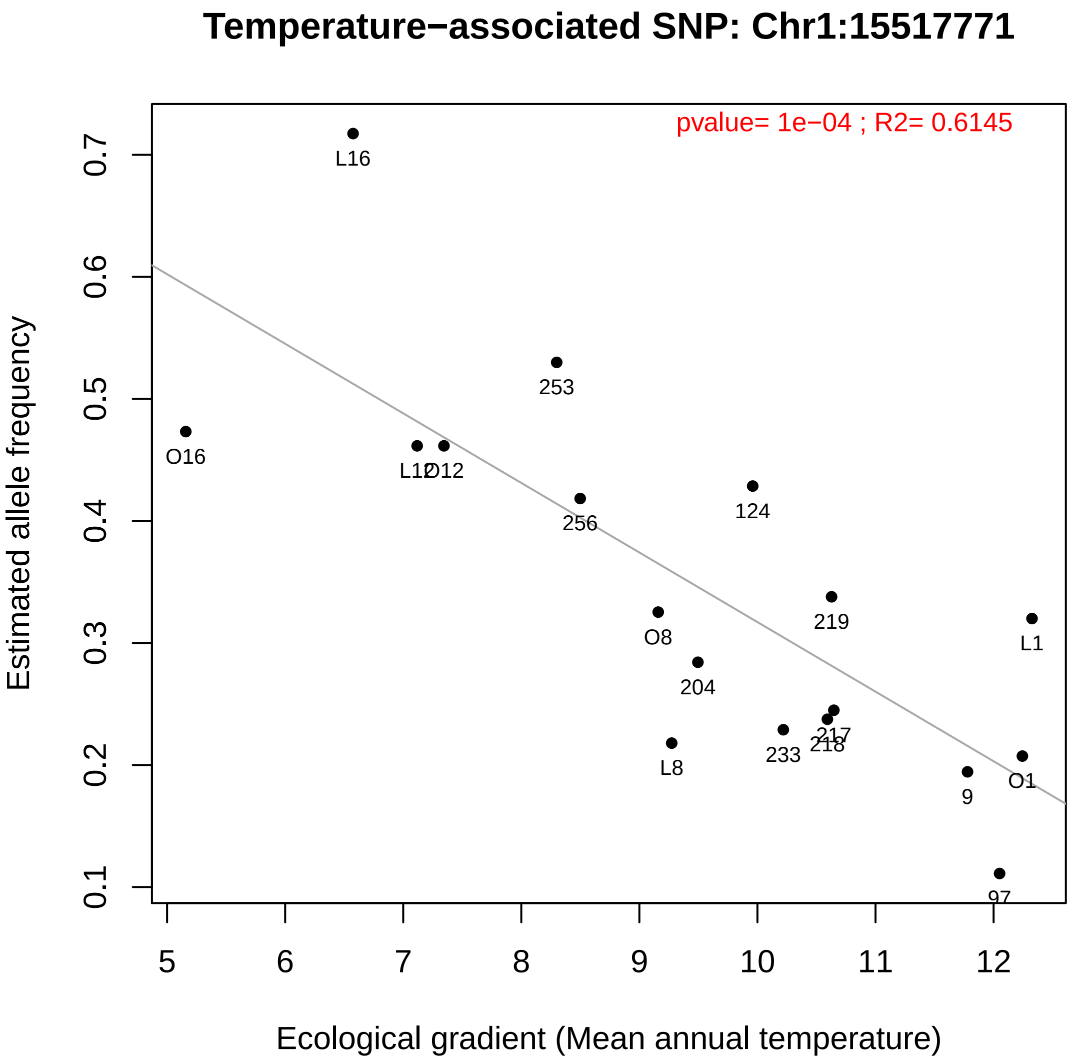
Temperature−associated SNP: Chr1:15517771
pvalue= 1e−04 ; R2= 0.6145
Estimated allele frequency
Ecological gradient (Mean annual temperature)
0.1
0.2
0.3
0.4
0.5
0.6
0.7
5
6
7
8
9
10
11
12
L16
253
O16
L12
O12
256
124
O8
219
L1
204
217
218
233
L8
O1
9
97

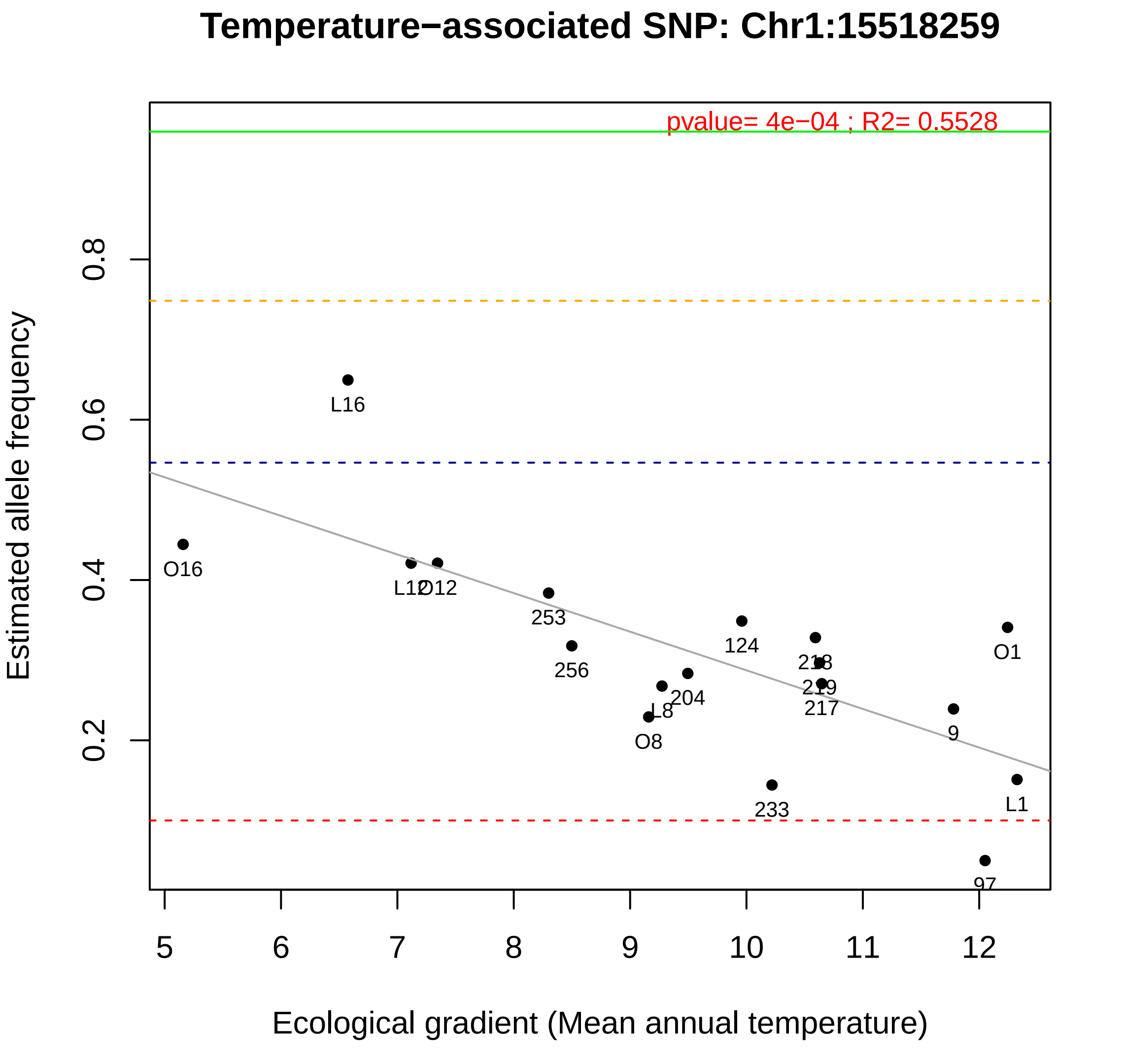
Temperature−associated SNP: Chr1:15518259
pvalue= 4e−04 ; R2= 0.5528
Estimated allele frequency
0.2
0.4
0.6
0.8
5
6
7
8
9
10
11
12
Ecological gradient (Mean annual temperature)
L16
O16
L12
O12
253
256
124
218
219
217
204
L8
O8
233
9
O1
L1
97

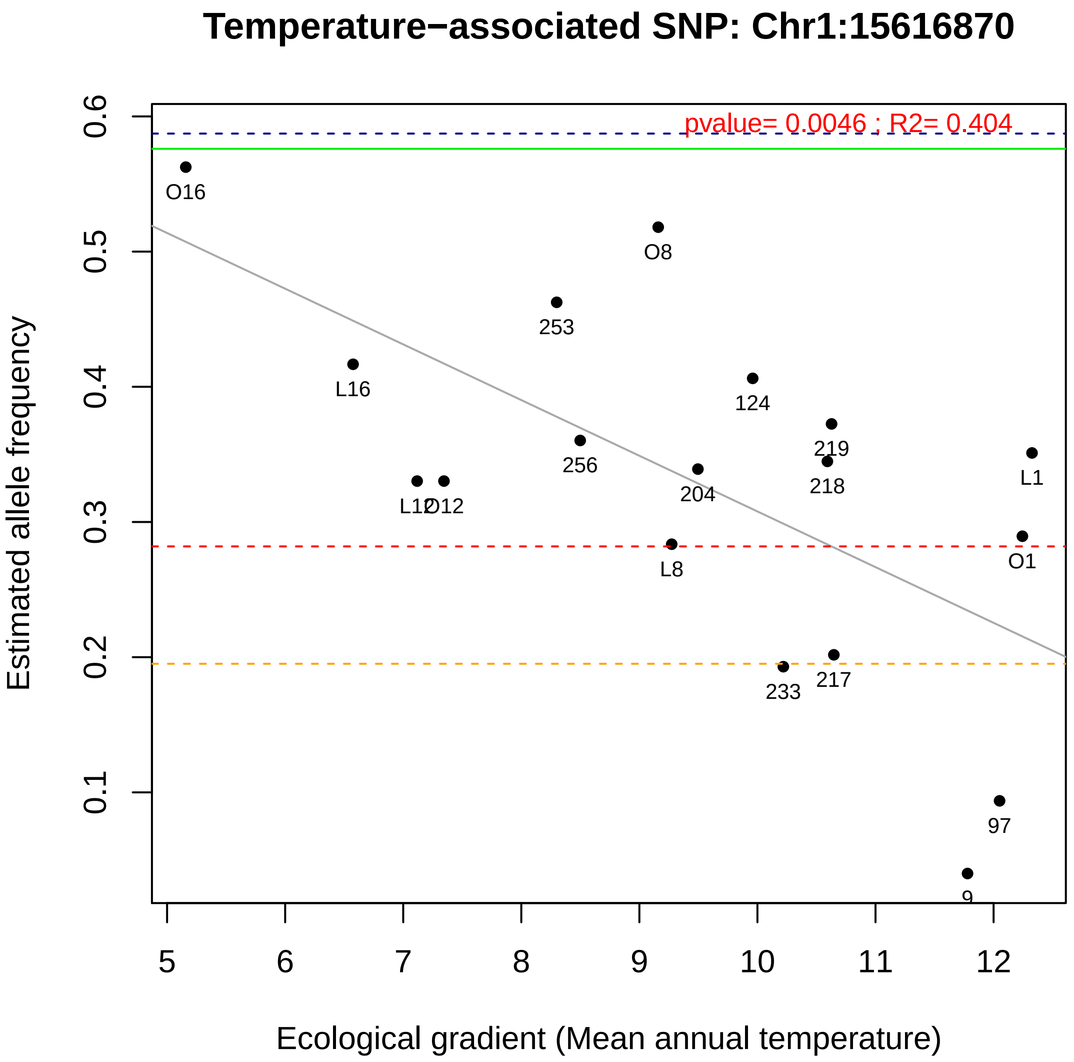
Temperature−associated SNP: Chr1:15616870
pvalue= 0.0046 ; R2= 0.404
Estimated allele frequency
0.1
0.2
0.3
0.4
0.5
0.6
O16
O8
253
L16
124
219
256
218
L1
L12
O12
204
L8
O1
233
217
97
9
5
6
7
8
9
10
11
12
Ecological gradient (Mean annual temperature)

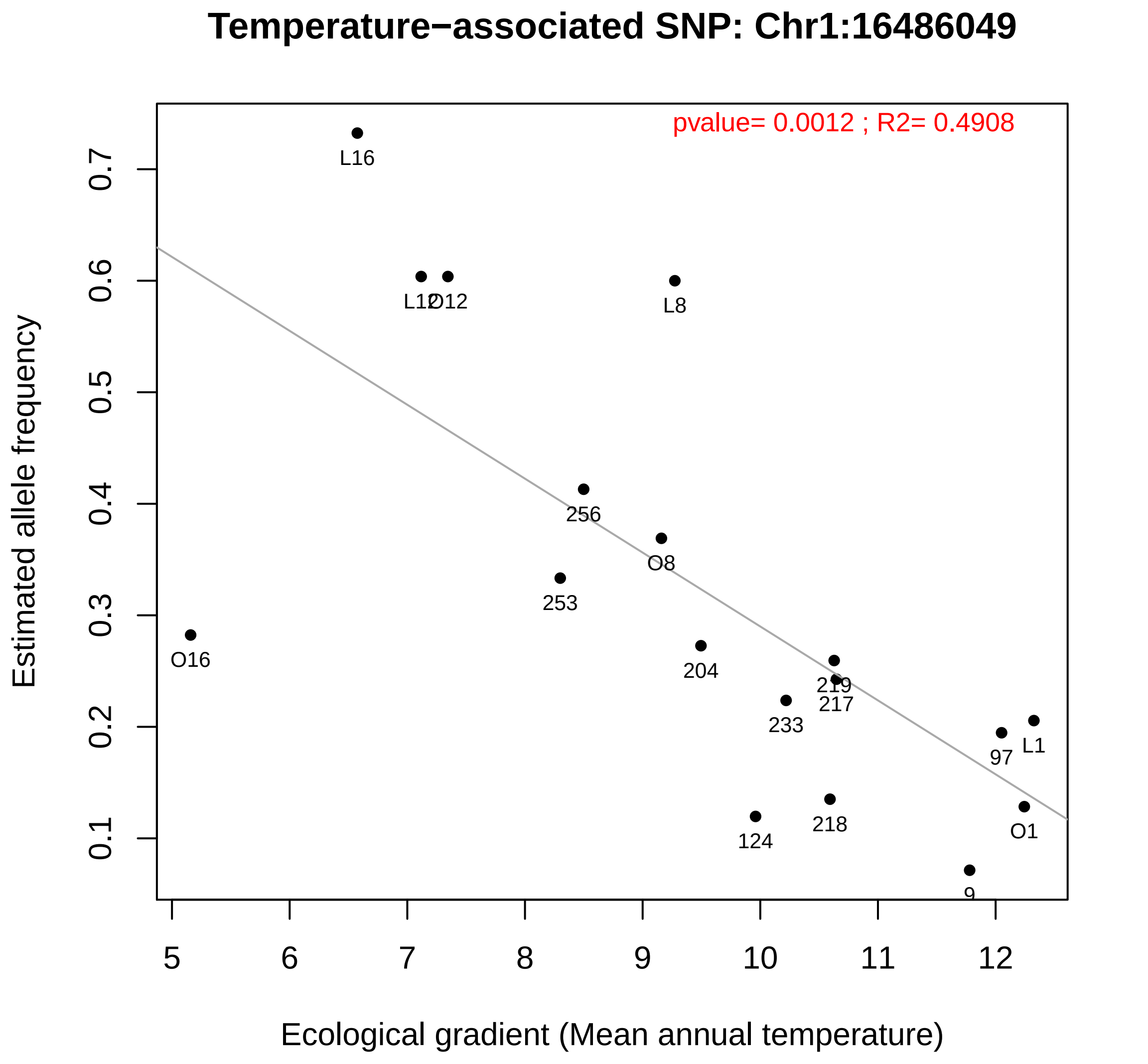

Temperature−associated SNP: Chr1:16486049
pvalue= 0.0012 ; R2= 0.4908
Estimated allele frequency
0.1
0.2
0.3
0.4
0.5
0.6
0.7
5
6
7
8
9
10
11
12
Ecological gradient (Mean annual temperature)
L16
L12
O12
L8
256
O8
253
O16
204
219
217
233
97
L1
218
124
O1
9

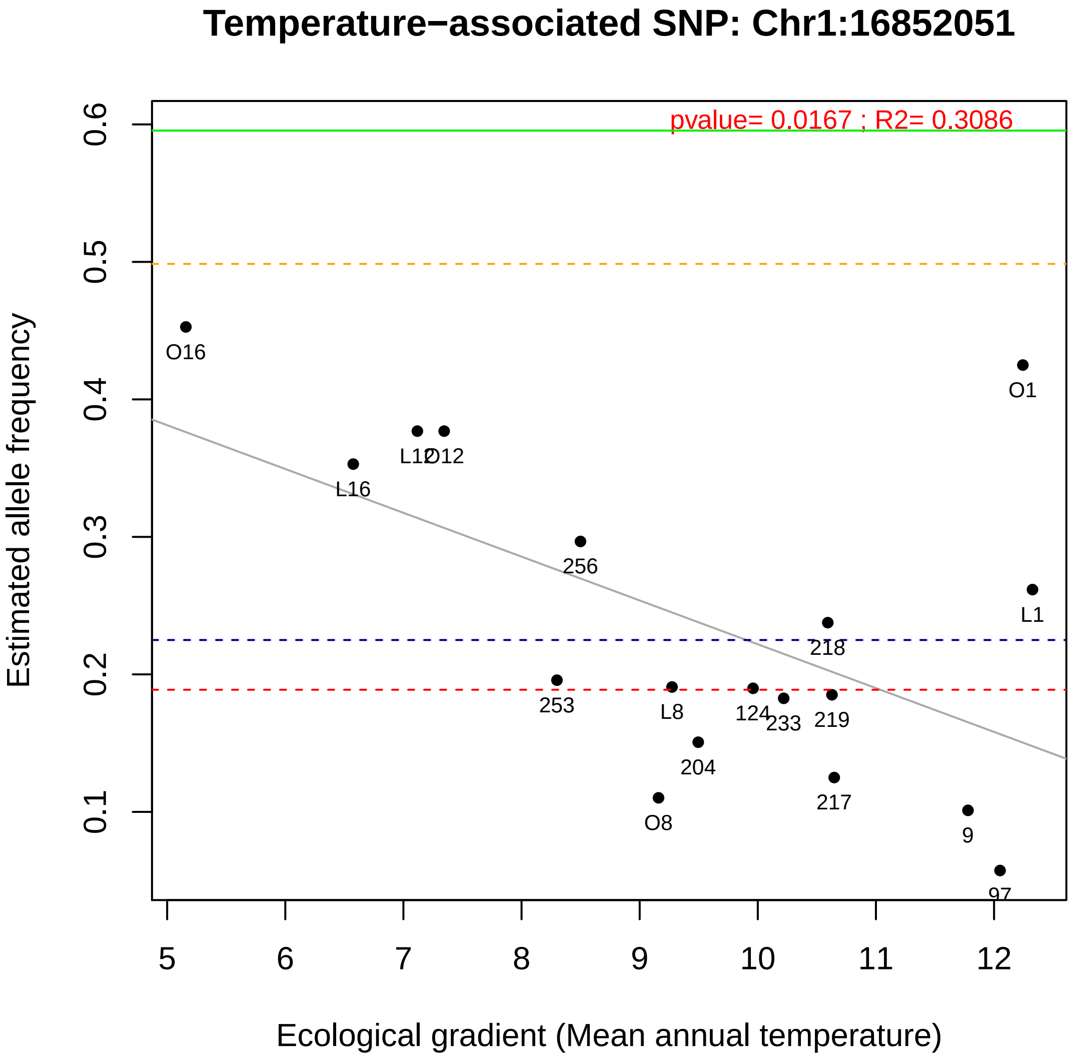

Temperature−associated SNP: Chr1:16852051
pvalue= 0.0167 ; R2= 0.3086
Estimated allele frequency
0.1
0.2
0.3
0.4
0.5
0.6
5
6
7
8
9
10
11
12
Ecological gradient (Mean annual temperature)
O16
O1
L12
O12
L16
256
L1
218
253
L8
124
233
219
204
217
O8
9
97

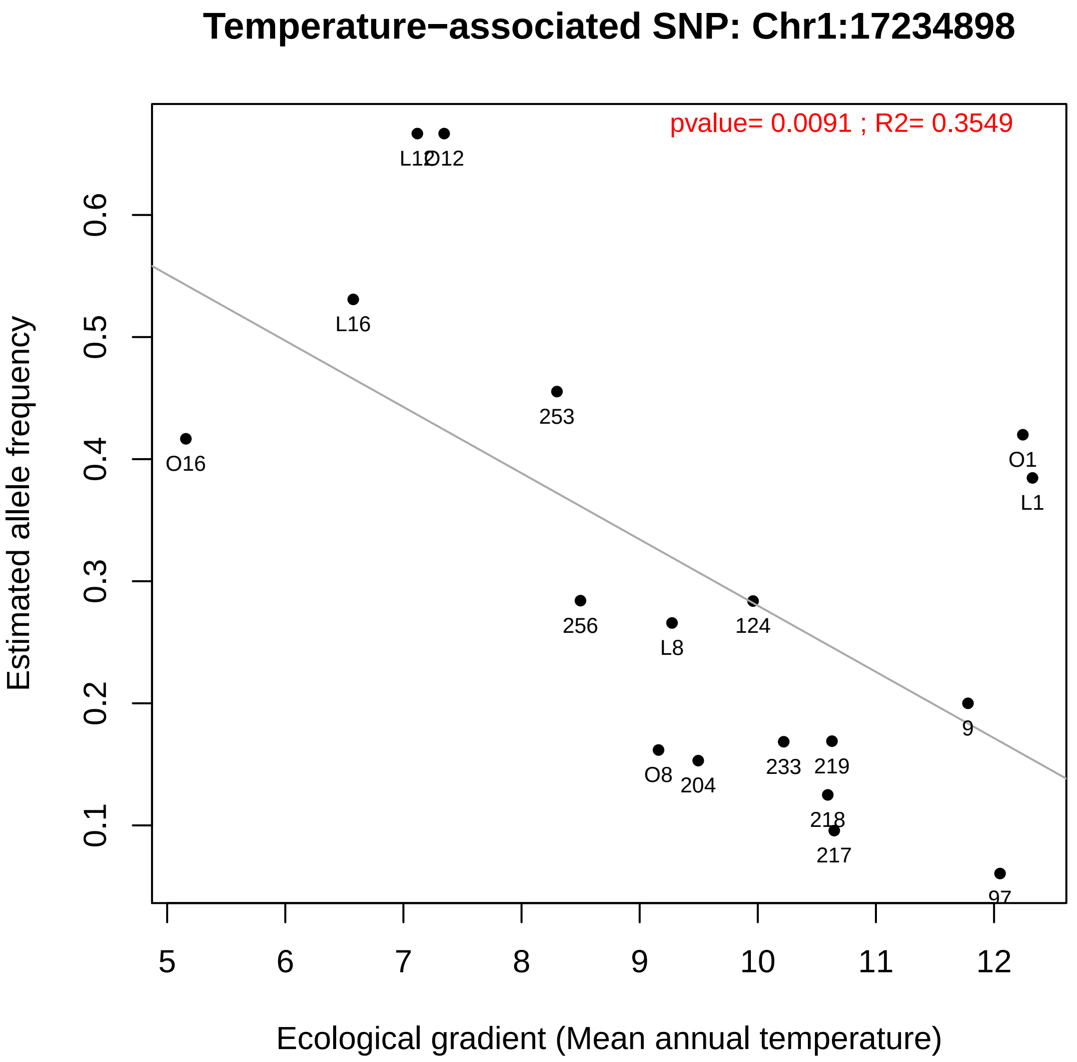

Temperature−associated SNP: Chr1:17234898
pvalue= 0.0091 ; R2= 0.3549
Estimated allele frequency
Ecological gradient (Mean annual temperature)
0.1
0.2
0.3
0.4
0.5
0.6
5
6
7
8
9
10
11
12
L12
O12
L16
253
O16
O1
L1
256
L8
124
9
O8
204
233
219
218
217
97

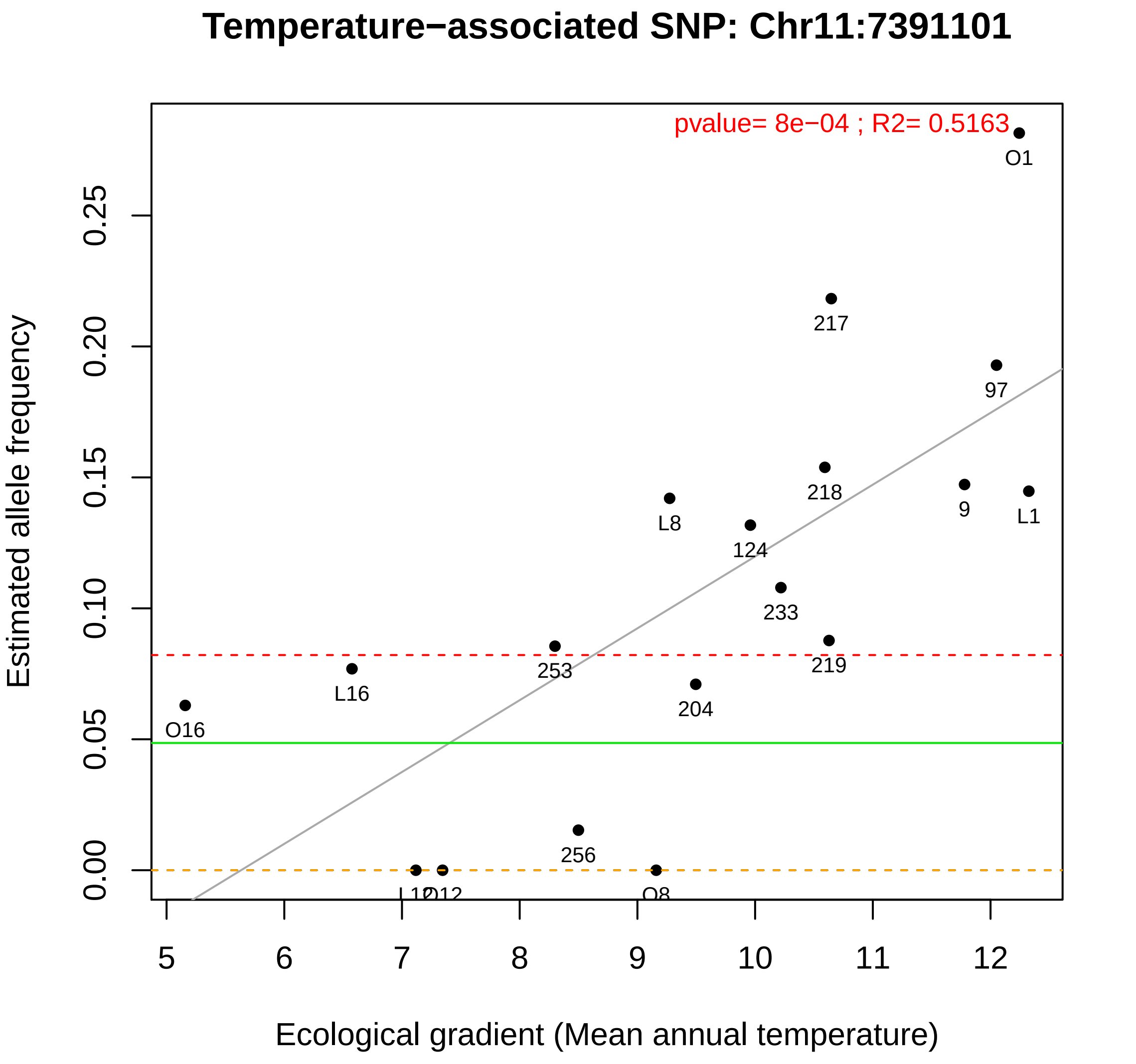
Temperature−associated SNP: Chr11:7391101
pvalue= 8e−04 ; R2= 0.5163
Estimated allele frequency
0.00
0.05
0.10
0.15
0.20
0.25
Ecological gradient (Mean annual temperature)
5
6
7
8
9
10
11
12
O1
217
97
218
9
L1
L8
124
233
219
253
L16
204
O16
256
O12
O8

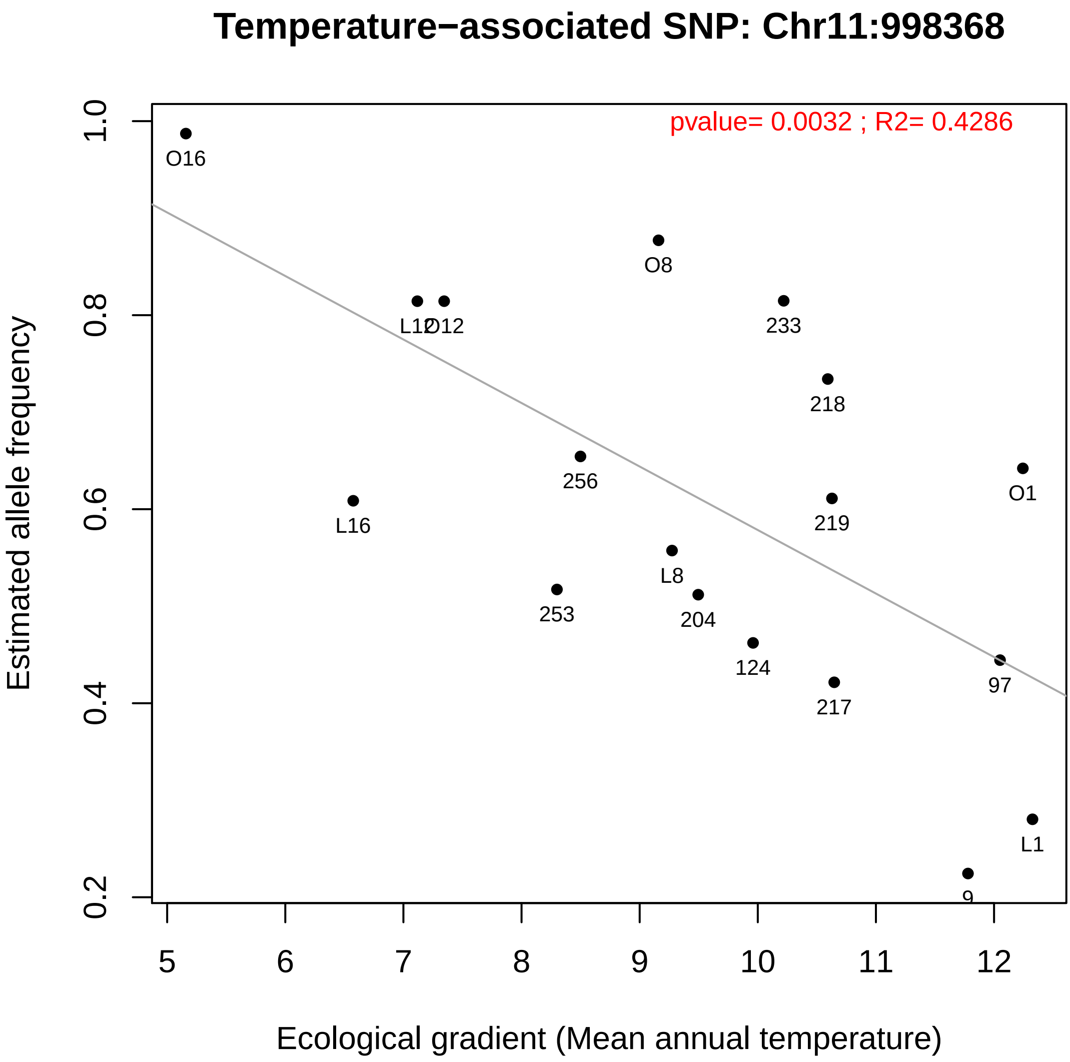

Temperature−associated SNP: Chr11:998368
pvalue= 0.0032 ; R2= 0.4286
Estimated allele frequency
Ecological gradient (Mean annual temperature)
0.2
0.4
0.6
0.8
1.0
5
6
7
8
9
10
11
12
O16
O8
L12
O12
233
218
256
O1
L16
219
L8
253
204
124
217
97
L1
9

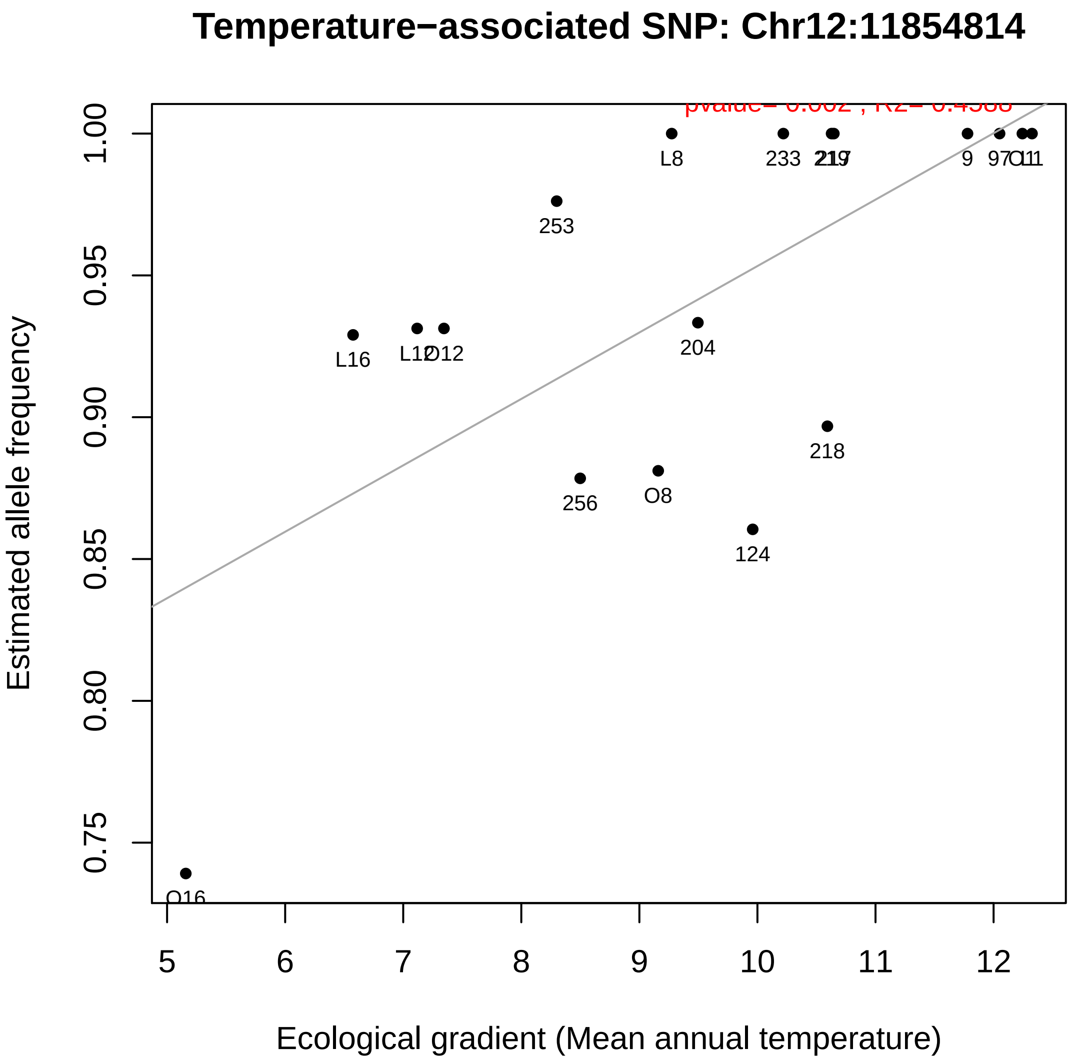

Temperature−associated SNP: Chr12:11854814
Estimated allele frequency
Ecological gradient (Mean annual temperature)
0.75
0.80
0.85
0.90
0.95
1.00
5
6
7
8
9
10
11
12
L8
233
9
253
L16
204
218
256
O8
124
O16

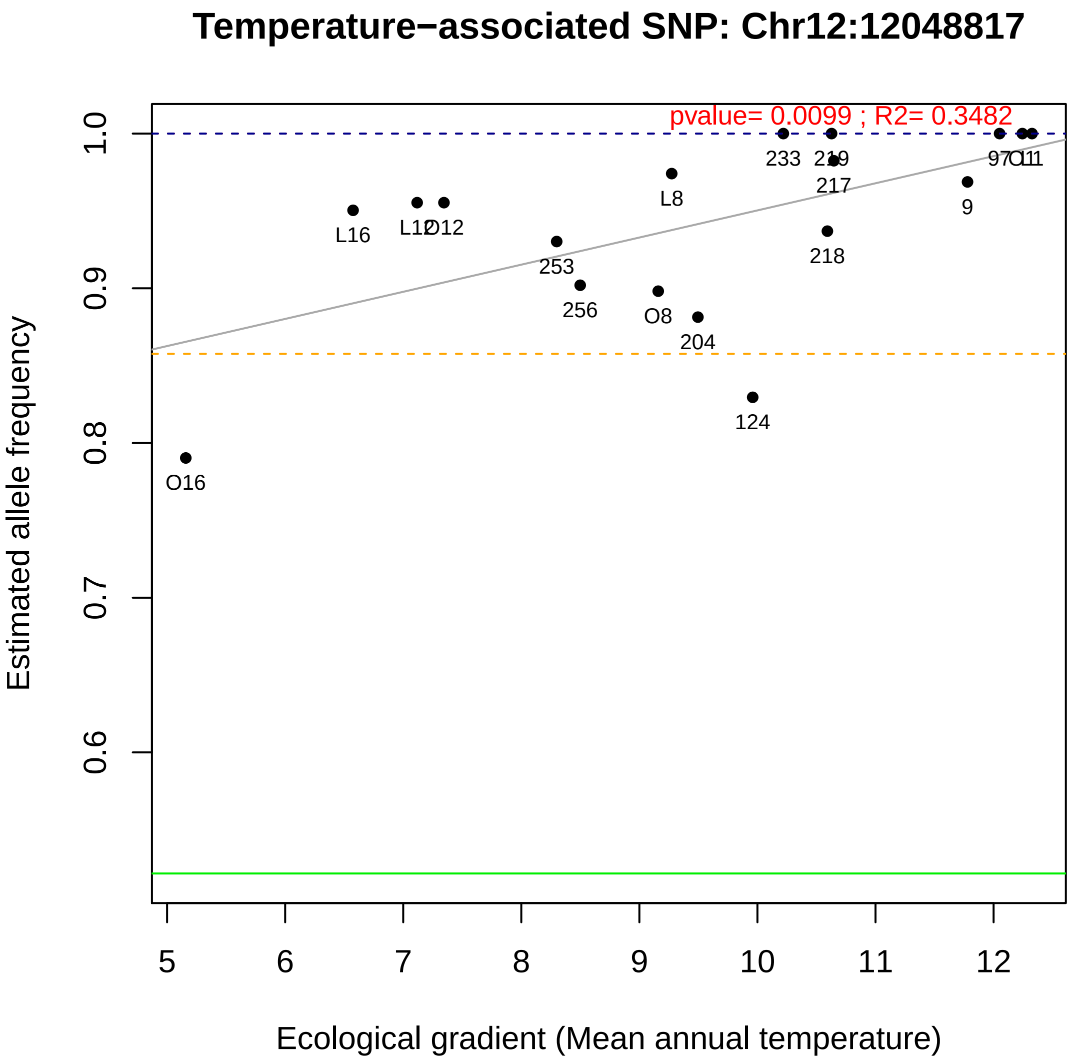

Temperature−associated SNP: Chr12:12048817
pvalue= 0.0099 ; R2= 0.3482
Estimated allele frequency
Ecological gradient (Mean annual temperature)
1.0
0.9
0.8
0.7
0.6
5
6
7
8
9
10
11
12
O16
L16
L12
O12
253
256
O8
204
124
L8
233
219
217
218
9
97
O11

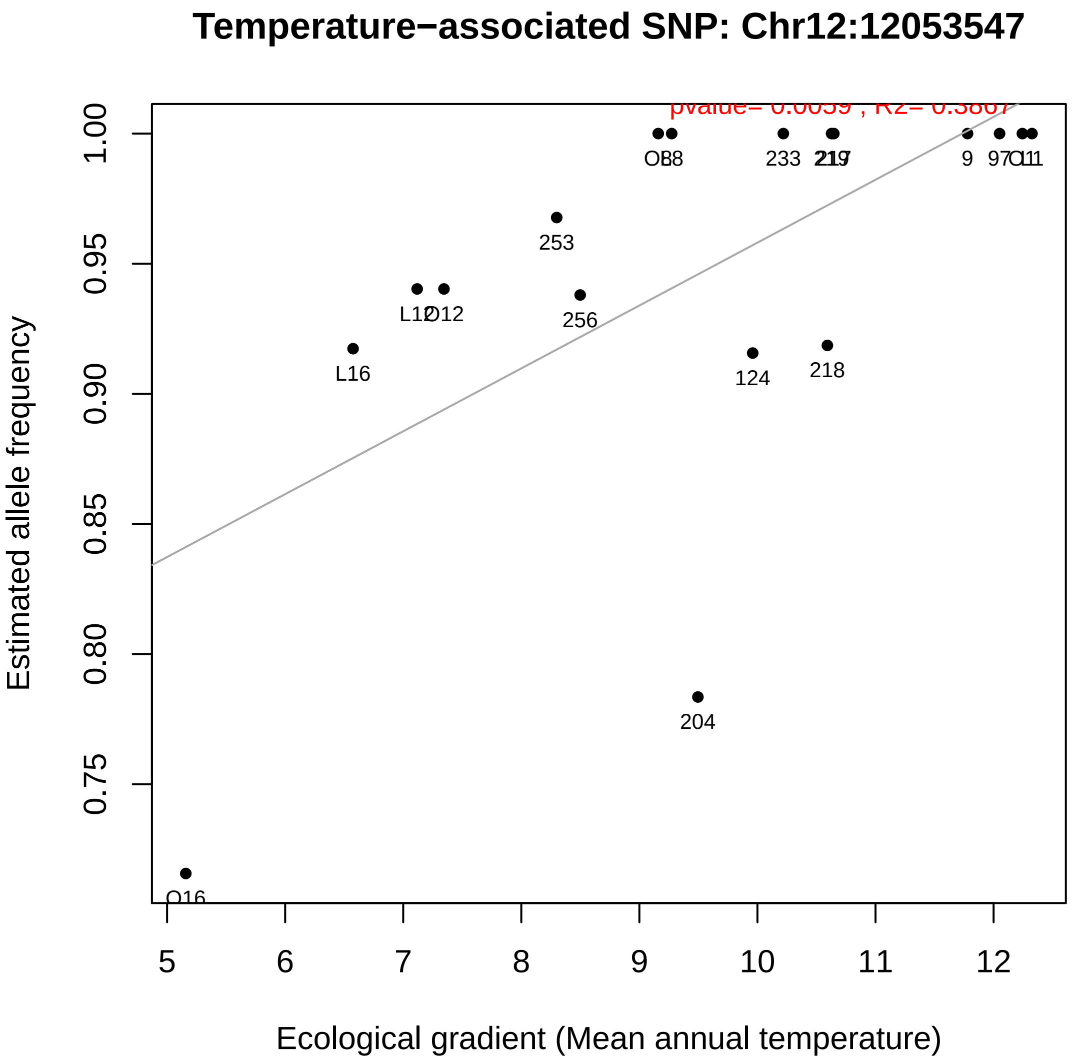
Temperature−associated SNP: Chr12:12053547
pvalue= 0.0059 , R2= 0.3867
Estimated allele frequency
Ecological gradient (Mean annual temperature)
1.00
0.95
0.90
0.85
0.80
0.75
5
6
7
8
9
10
11
12
253
L12
O12
256
L16
124
218
204
O16
233
9

Temperature−associated SNP: Chr12:12065198

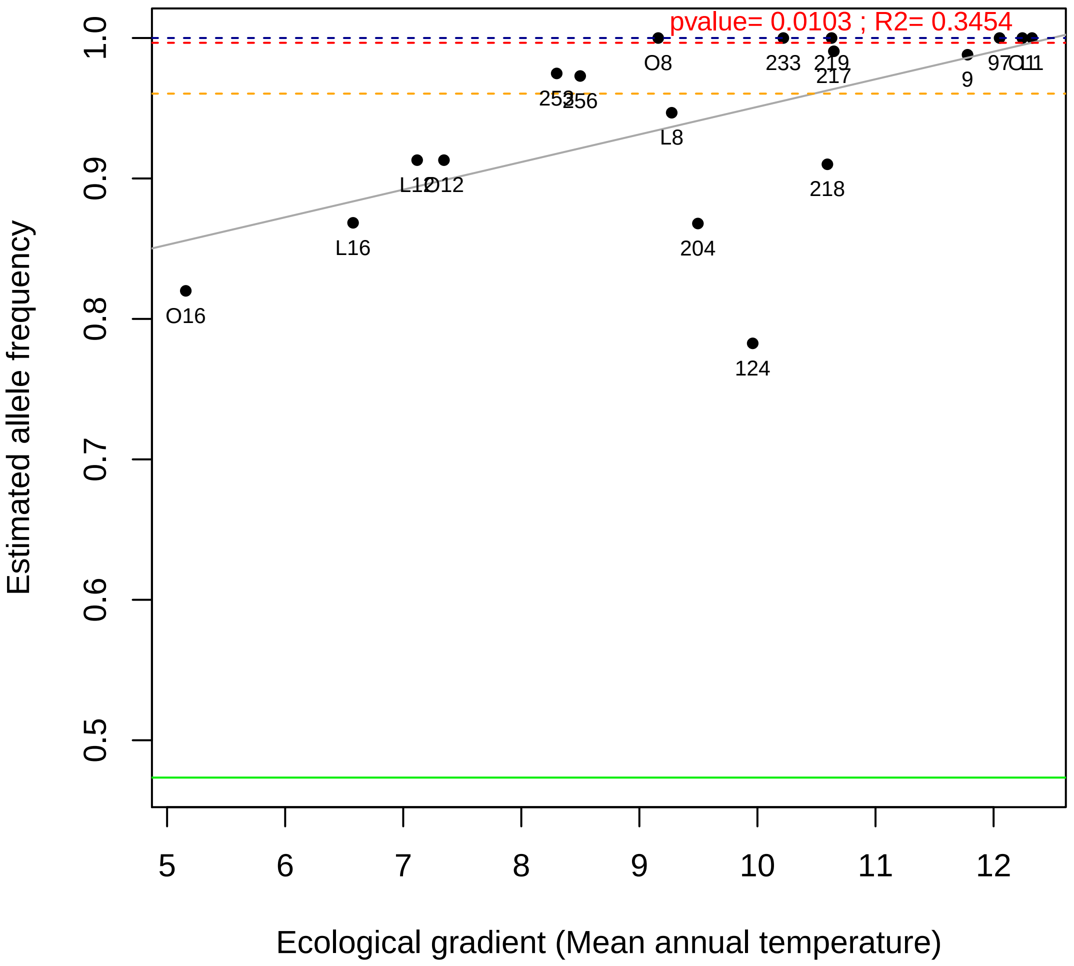

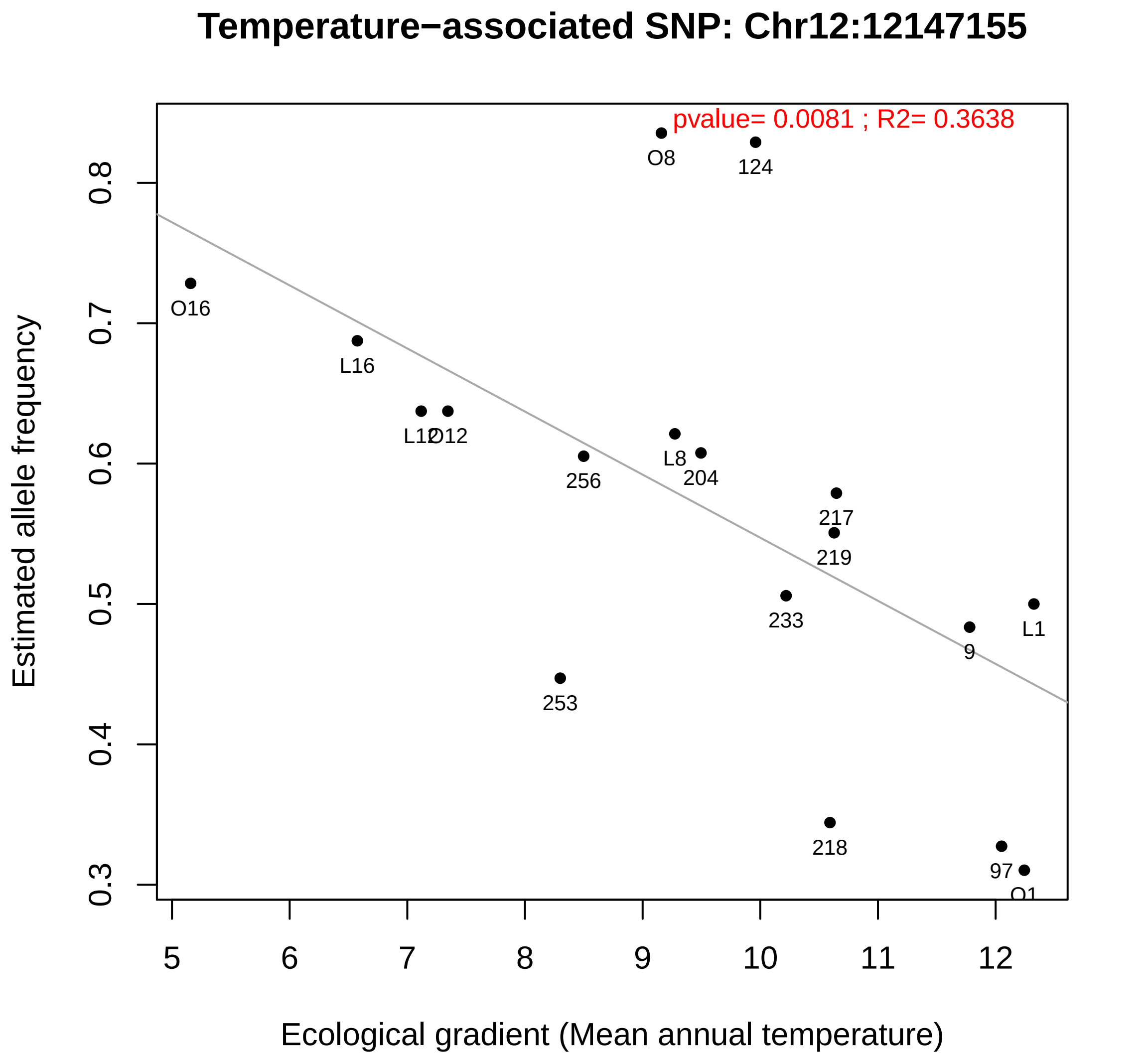

Temperature−associated SNP: Chr12:12147155
pvalue= 0.0081 ; R2= 0.3638
O8
124
O16
L16
L12
O12
256
L8
204
217
219
233
9
L1
253
218
97
O1
0.3
0.4
0.5
0.6
0.7
0.8
5
6
7
8
9
10
11
12
Estimated allele frequency
Ecological gradient (Mean annual temperature)

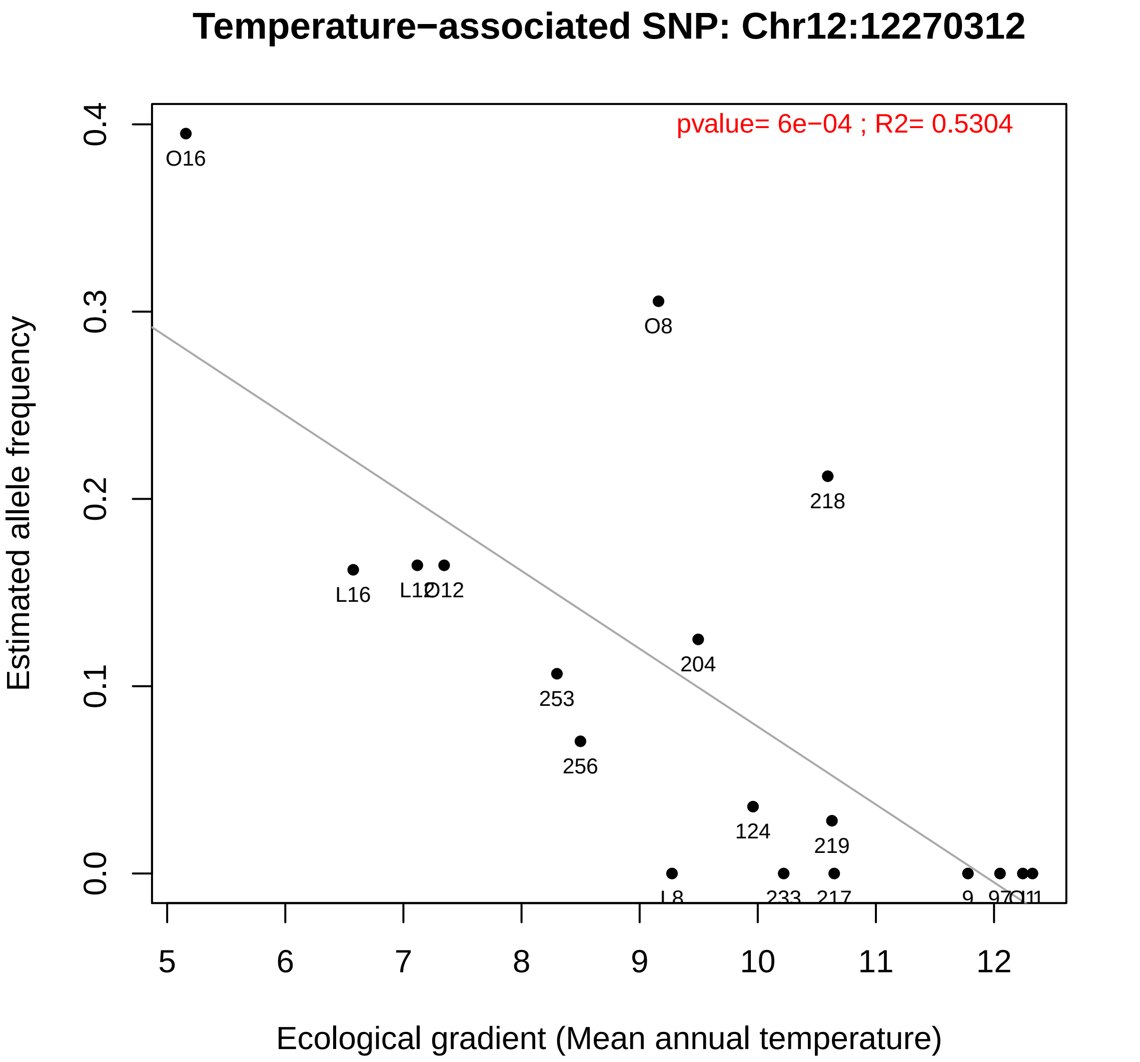
Temperature−associated SNP: Chr12:12270312
pvalue= 6e−04 ; R2= 0.5304
Estimated allele frequency
Ecological gradient (Mean annual temperature)
0.0
0.1
0.2
0.3
0.4
5
6
7
8
9
10
11
12
O16
O8
218
L16
L12
O12
204
253
256
124
219
L8
233
217
9
97
O11

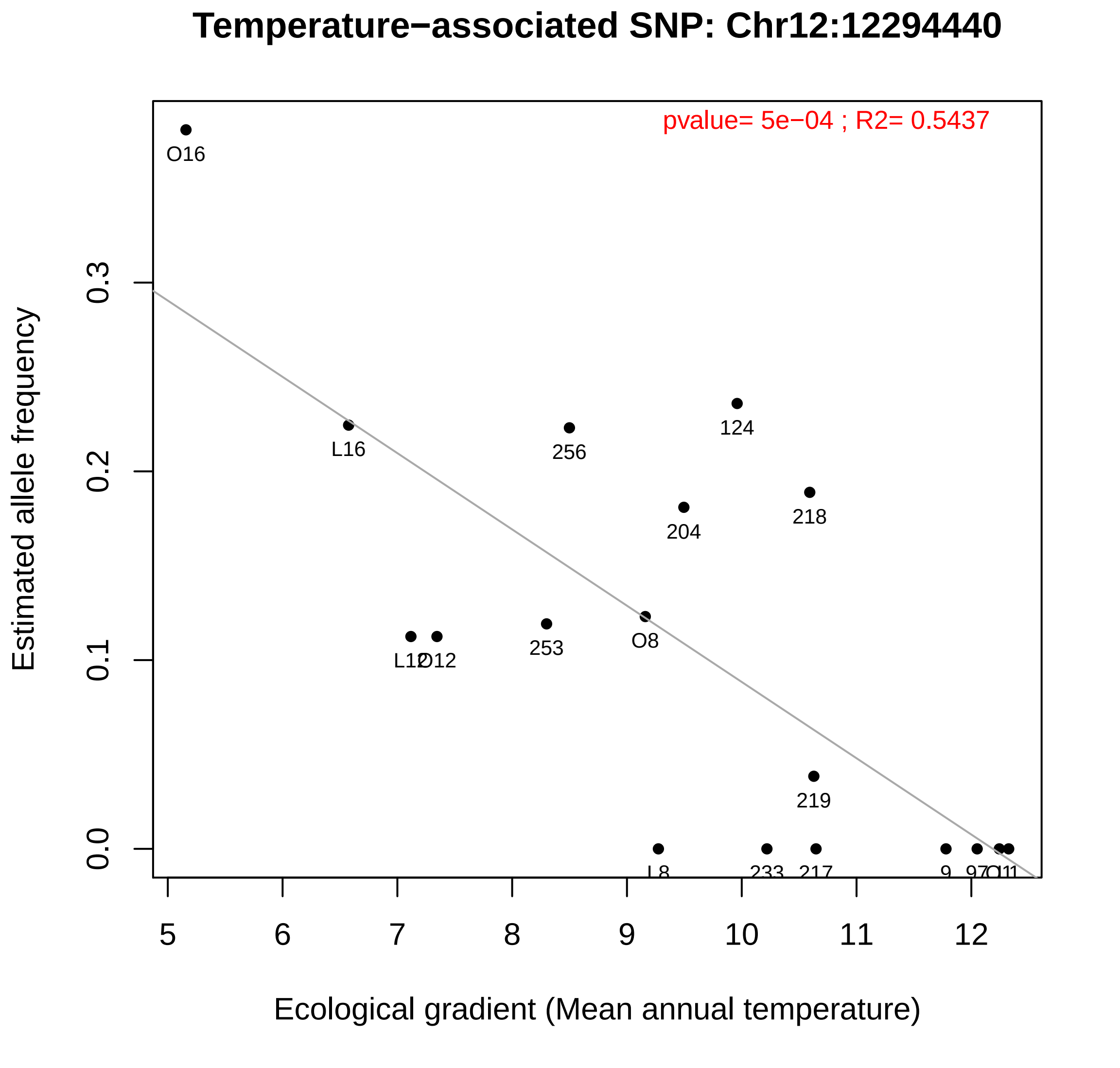

Temperature−associated SNP: Chr12:12294440
pvalue= 5e−04 ; R2= 0.5437
Estimated allele frequency
Ecological gradient (Mean annual temperature)
0.0
0.1
0.2
0.3
5
6
7
8
9
10
11
12
O16
L16
256
124
204
218
L12
O12
253
O8
219
L8
233
217
9
97

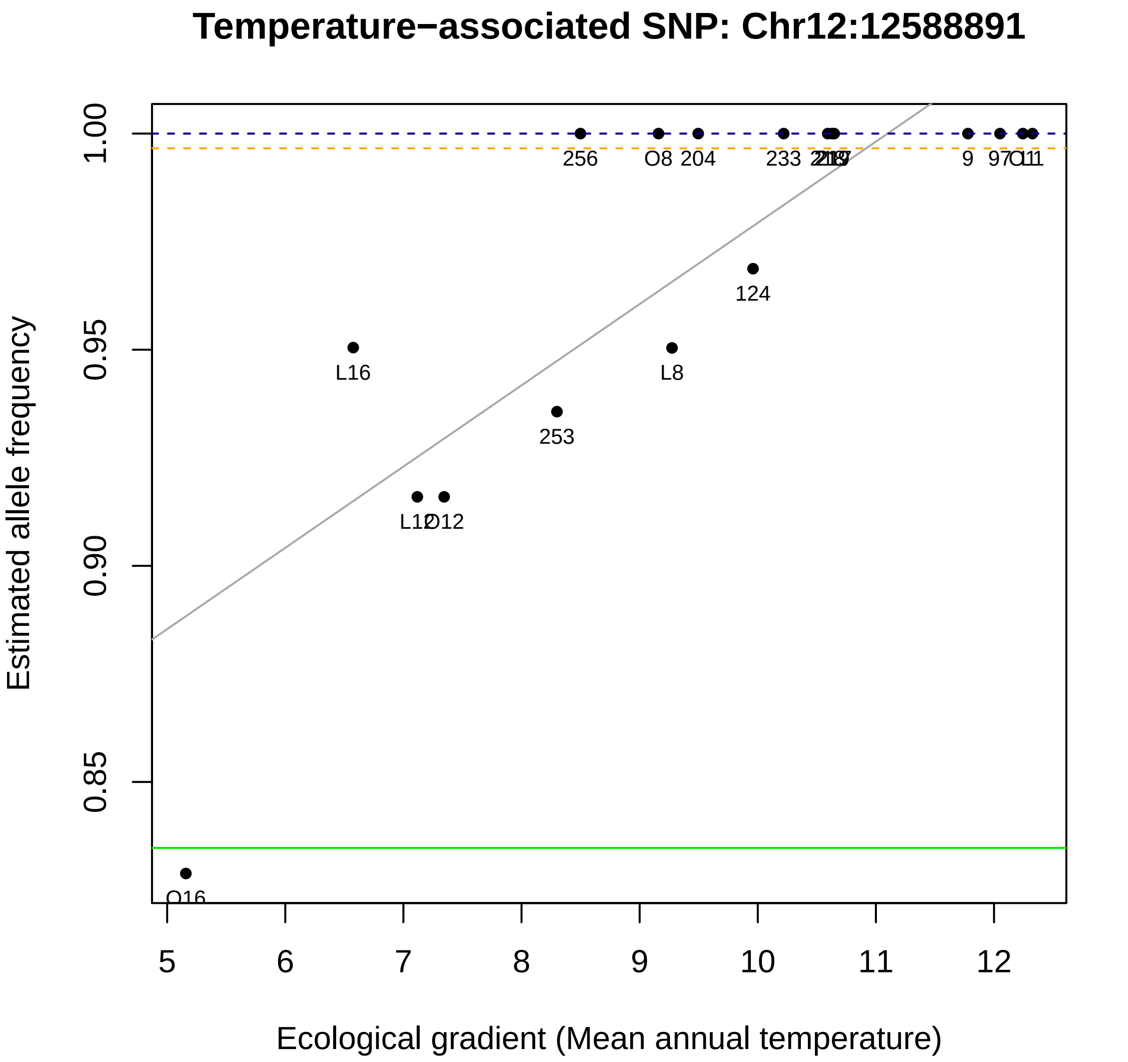
Temperature−associated SNP: Chr12:12588891
Estimated allele frequency
Ecological gradient (Mean annual temperature)
0.85
0.90
0.95
1.00
5
6
7
8
9
10
11
12
256
O8
204
233
9
124
L16
L8
253
L12
O12
O16

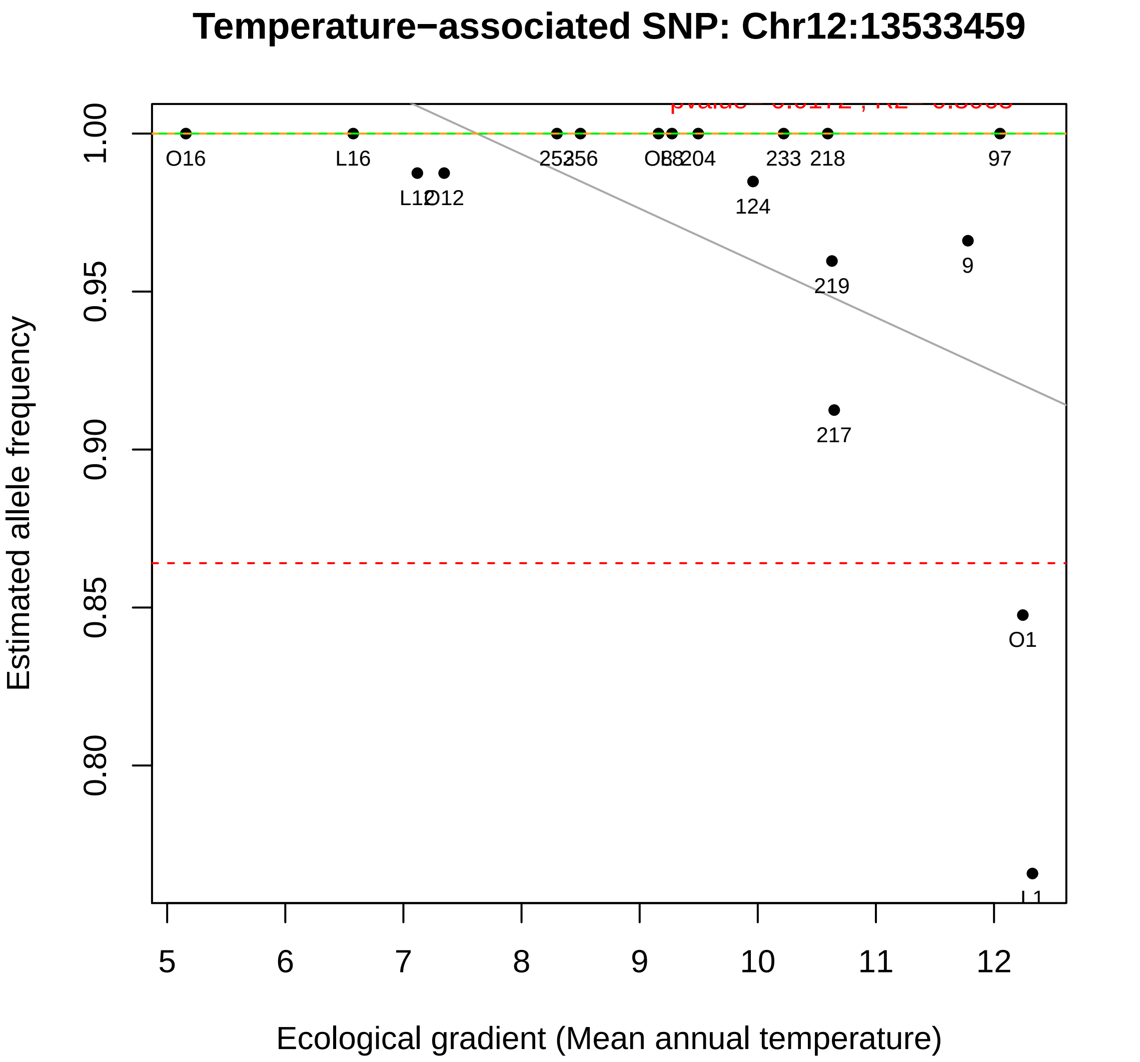

Temperature−associated SNP: Chr12:13533459
Estimated allele frequency
Ecological gradient (Mean annual temperature)
1.00
0.95
0.90
0.85
0.80
5
6
7
8
9
10
11
12
O16
L16
97
233
218
124
219
9
217
O1

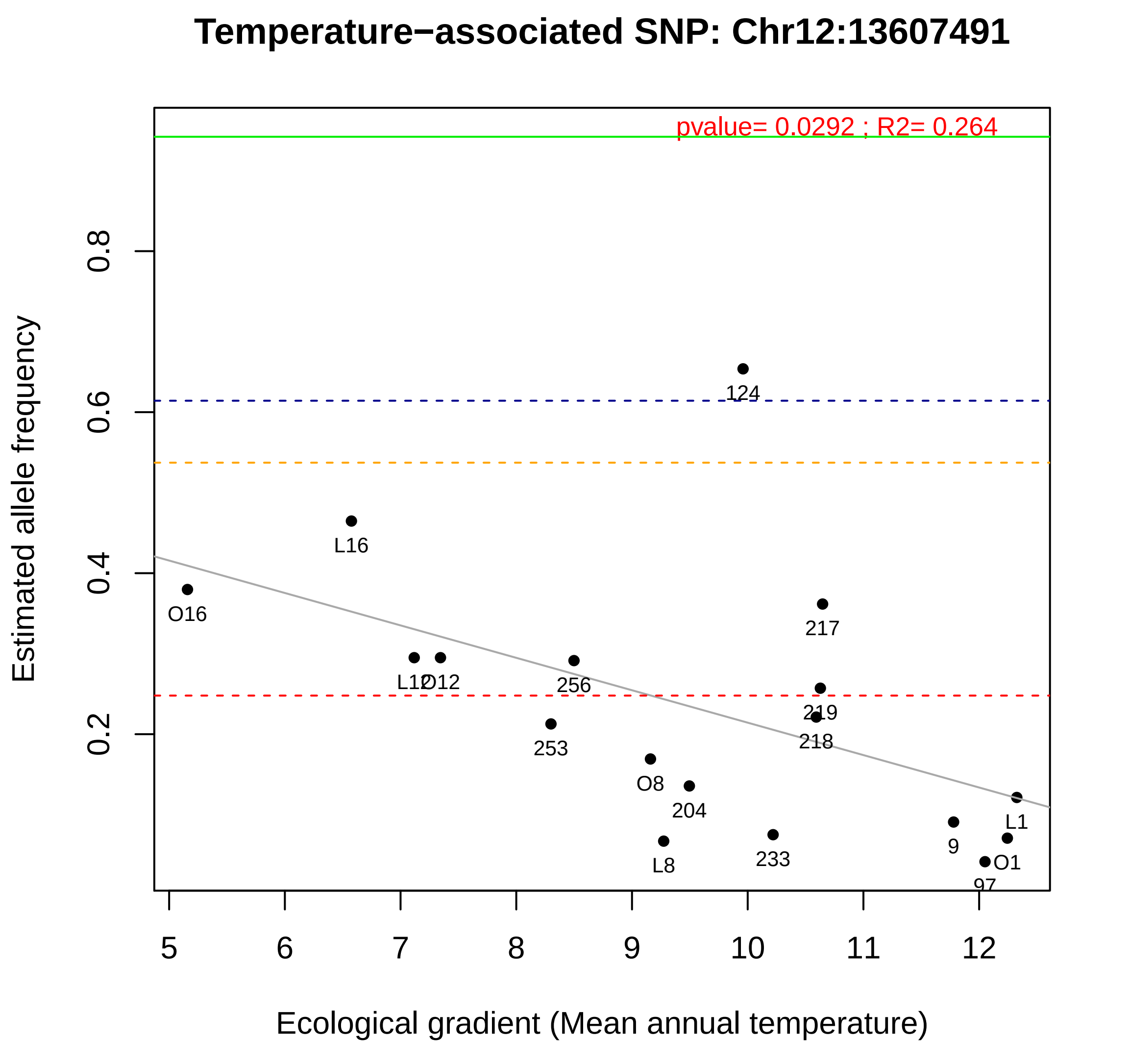

Temperature−associated SNP: Chr12:13607491
pvalue= 0.0292 ; R2= 0.264
Estimated allele frequency
0.2
0.4
0.6
0.8
Ecological gradient (Mean annual temperature)
5
6
7
8
9
10
11
12
124
L16
O16
217
L12
O12
256
219
218
253
O8
204
L1
L8
233
9
O1
97

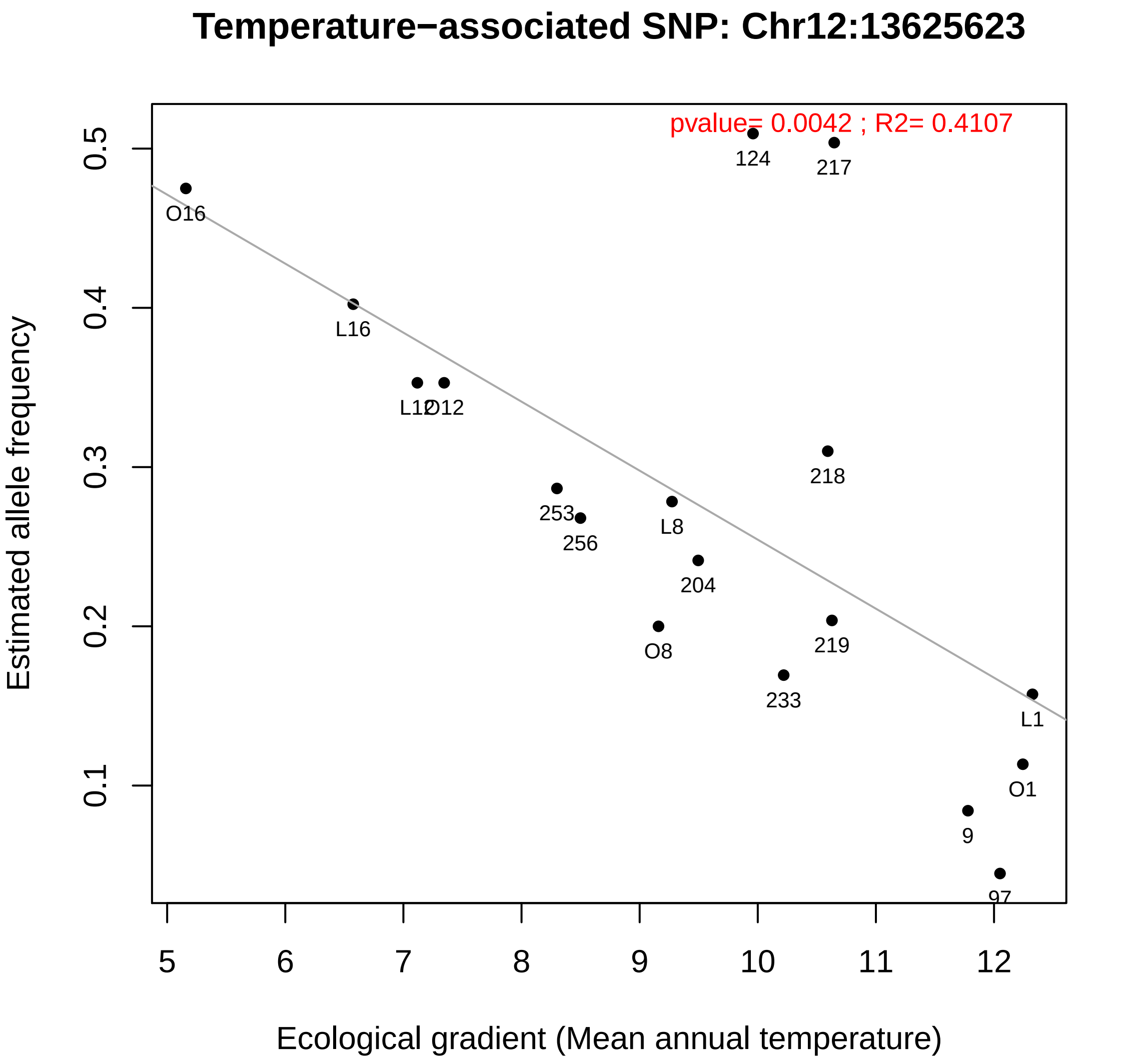

Temperature−associated SNP: Chr12:13625623
pvalue= 0.0042 ; R2= 0.4107
Estimated allele frequency
Ecological gradient (Mean annual temperature)
0.1
0.2
0.3
0.4
0.5
5
6
7
8
9
10
11
12
O16
L16
L12
O12
253
256
L8
204
O8
124
217
218
219
233
L1
O1
9
97

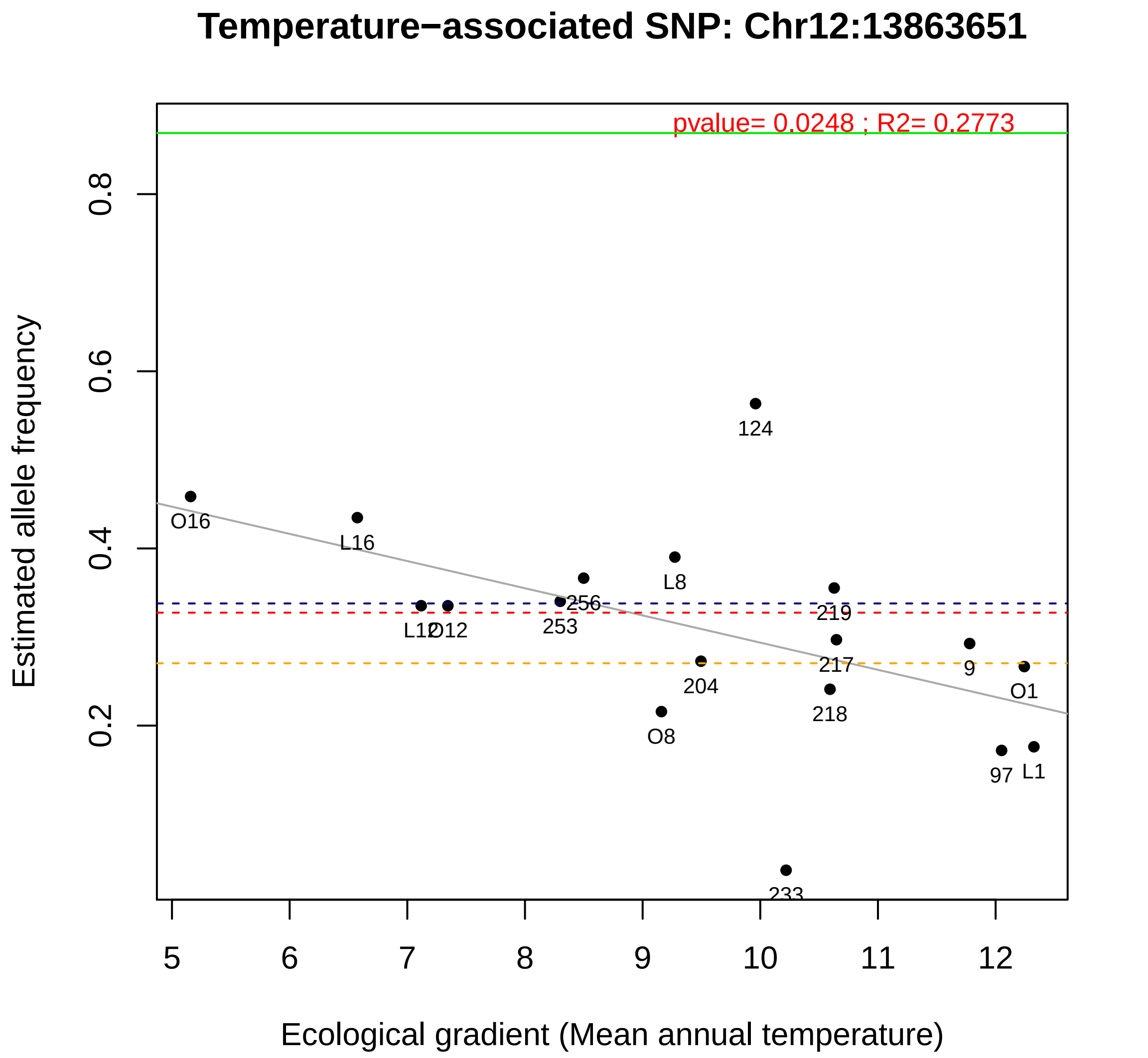

Temperature−associated SNP: Chr12:13863651
pvalue= 0.0248 ; R2= 0.2773
Estimated allele frequency
0.2
0.4
0.6
0.8
5
6
7
8
9
10
11
12
Ecological gradient (Mean annual temperature)
O16
L16
L12
O12
253
256
L8
124
204
O8
233
219
217
218
9
O1
97
L1

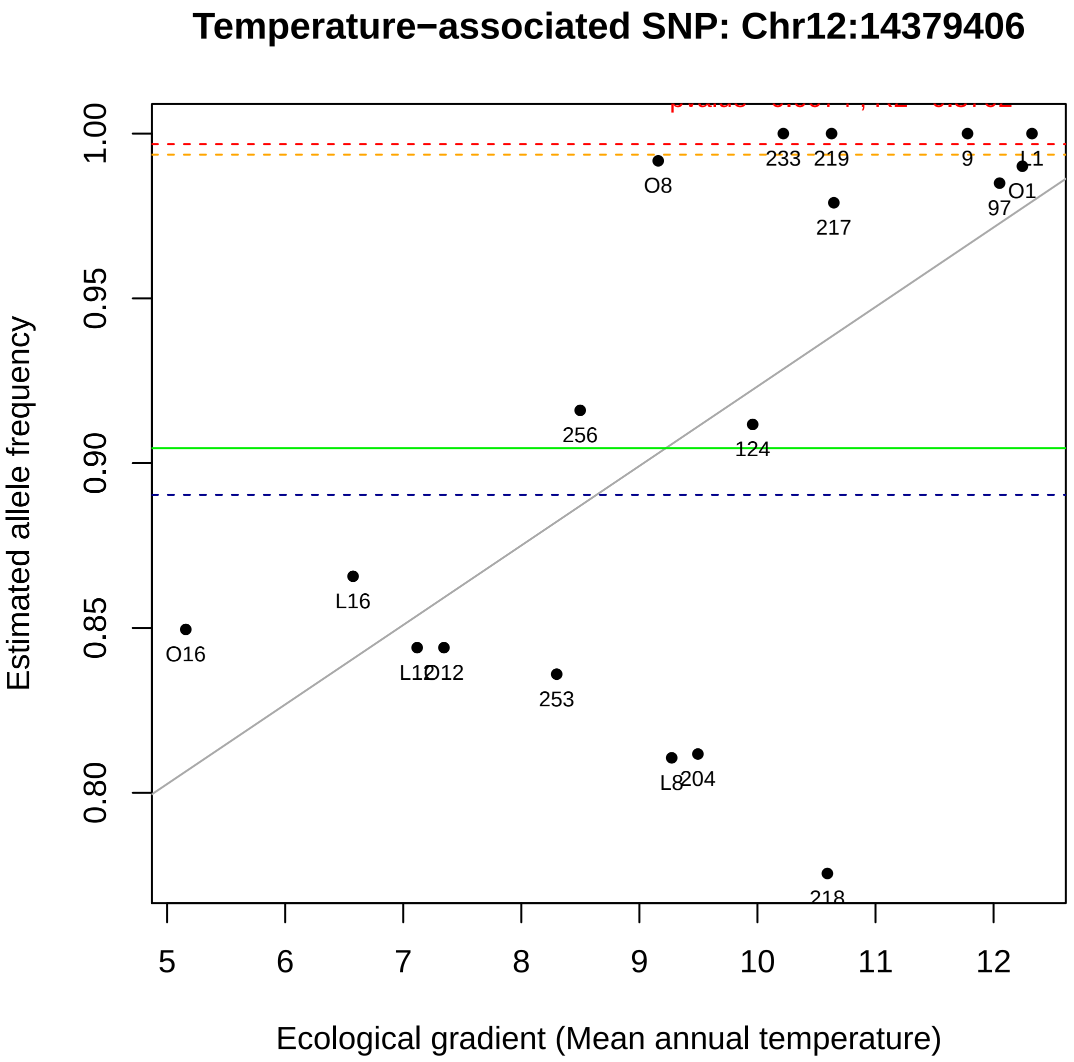
Temperature−associated SNP: Chr12:14379406
Estimated allele frequency
Ecological gradient (Mean annual temperature)
0.80
0.85
0.90
0.95
1.00
5
6
7
8
9
10
11
12
233
219
9
L1
O8
O1
97
217
256
124
L16
O16
L12
O12
253
L8
204
218

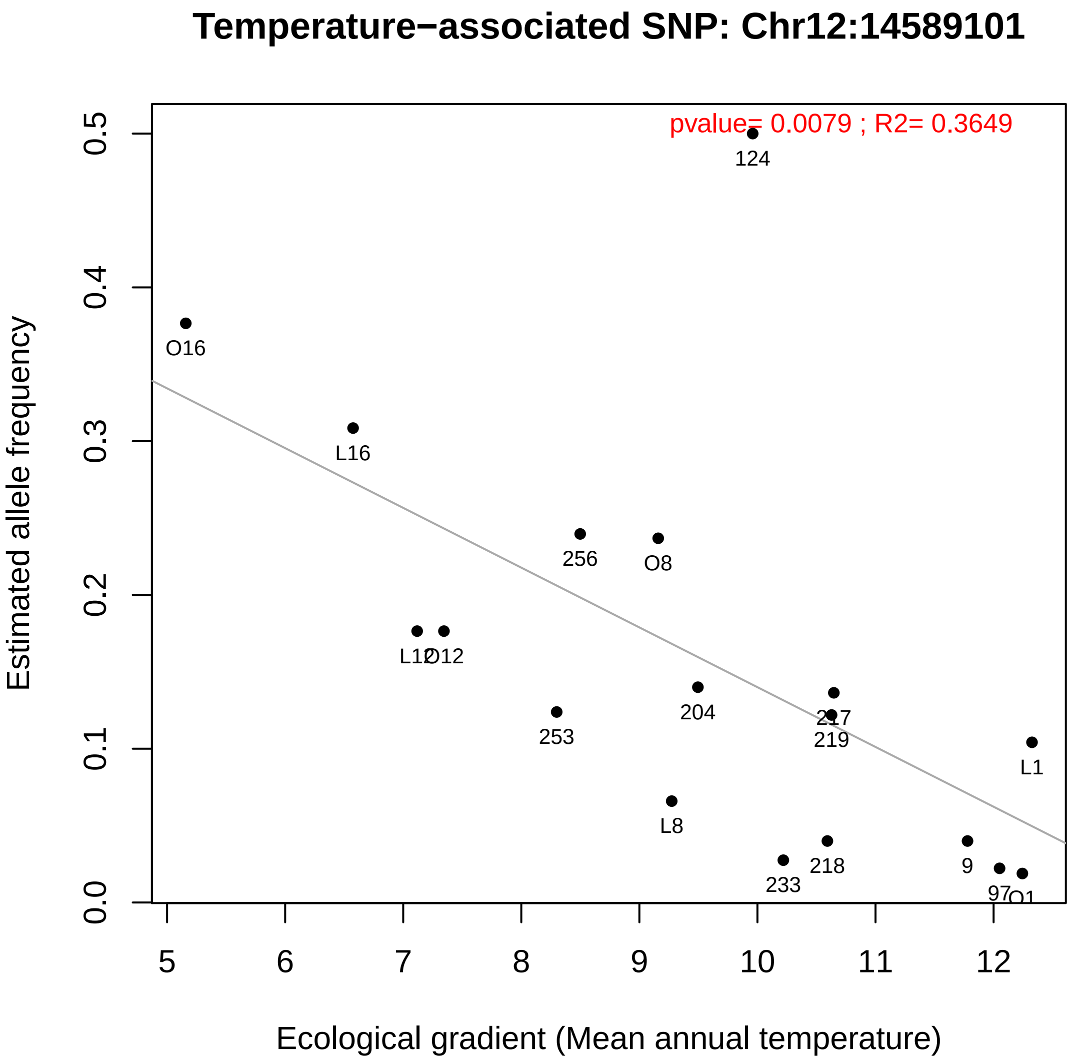

Temperature−associated SNP: Chr12:14589101
pvalue= 0.0079 ; R2= 0.3649
Estimated allele frequency
Ecological gradient (Mean annual temperature)
0.0
0.1
0.2
0.3
0.4
0.5
5
6
7
8
9
10
11
12
124
O16
L16
256
O8
L12
O12
253
204
217
219
L1
L8
233
218
9
97
O1

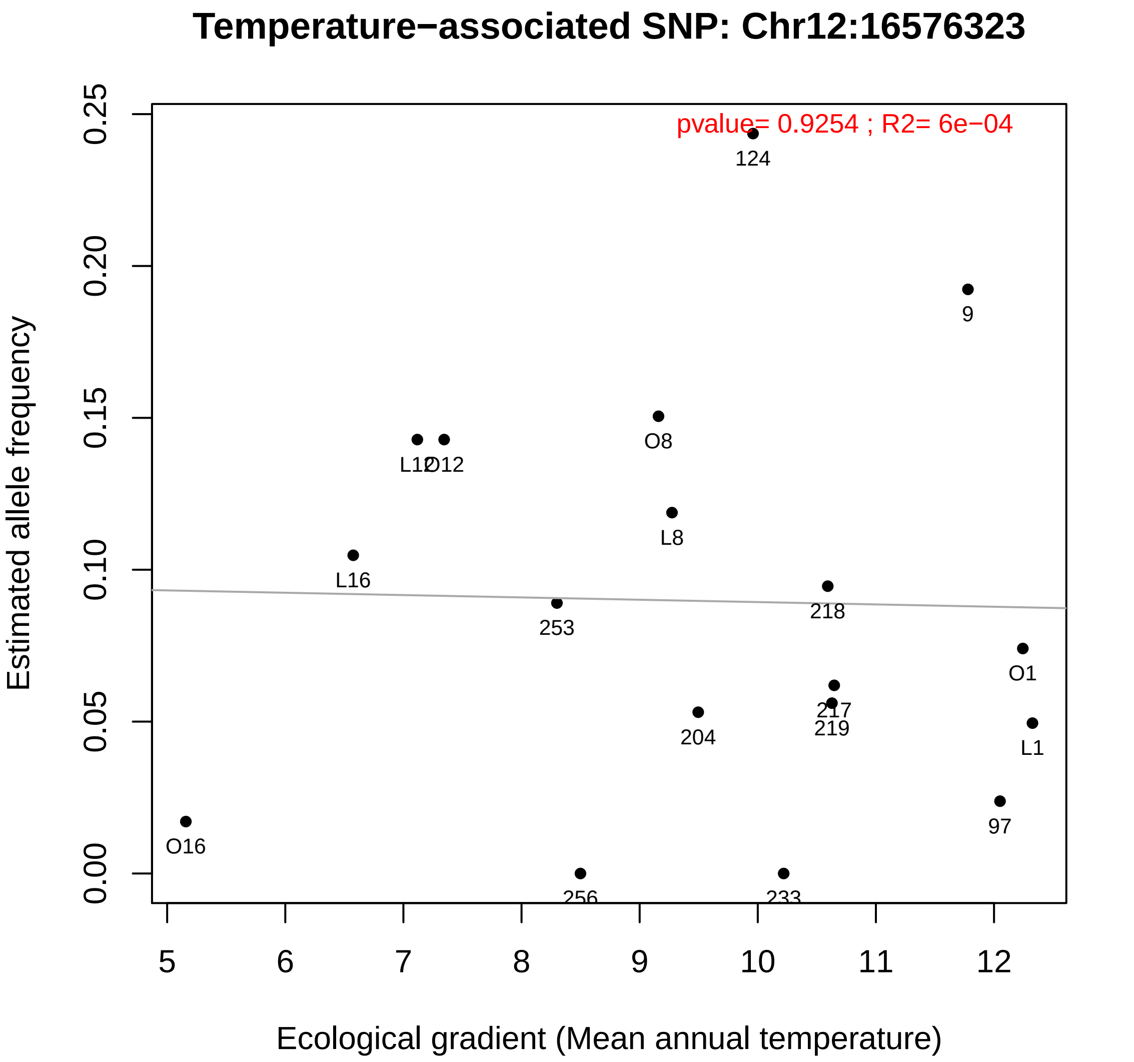
Temperature−associated SNP: Chr12:16576323
pvalue= 0.9254 ; R2= 6e−04
Estimated allele frequency
0.00
0.05
0.10
0.15
0.20
0.25
5
6
7
8
9
10
11
12
Ecological gradient (Mean annual temperature)
124
9
O8
L12
O12
L8
L16
218
253
O1
217
219
204
L1
97
O16
256
233

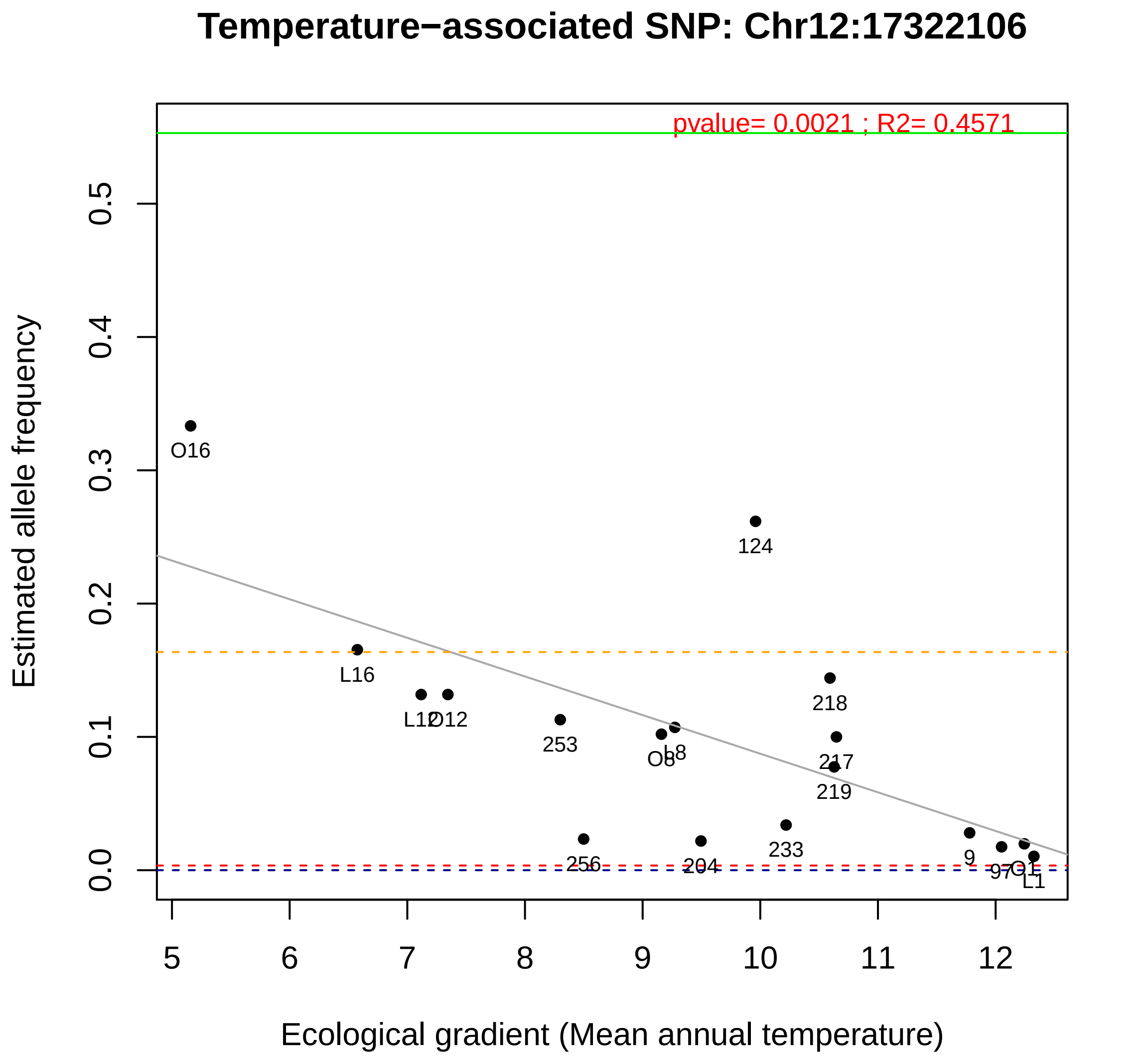
Temperature−associated SNP: Chr12:17322106
pvalue= 0.0021 ; R2= 0.4571
Estimated allele frequency
Ecological gradient (Mean annual temperature)
0.0
0.1
0.2
0.3
0.4
0.5
5
6
7
8
9
10
11
12
O16
L16
L12
O12
253
256
O8
L8
204
124
233
218
217
219
9
97
O1
L1

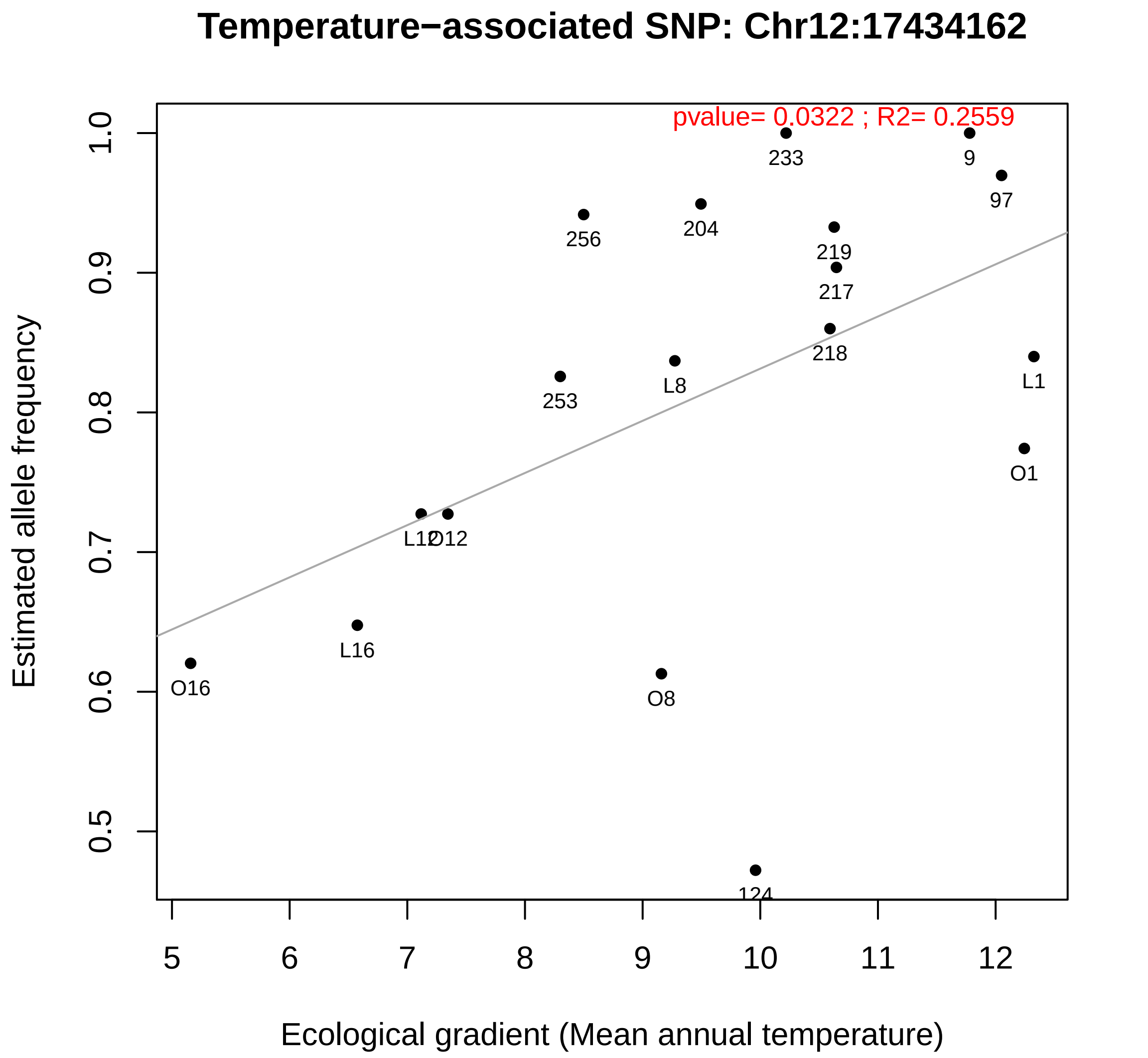
Temperature−associated SNP: Chr12:17434162
pvalue= 0.0322 ; R2= 0.2559
Estimated allele frequency
Ecological gradient (Mean annual temperature)
0.5
0.6
0.7
0.8
0.9
1.0
5
6
7
8
9
10
11
12
233
9
97
256
204
219
217
218
L1
253
L8
O1
L12
O12
L16
O16
O8
124

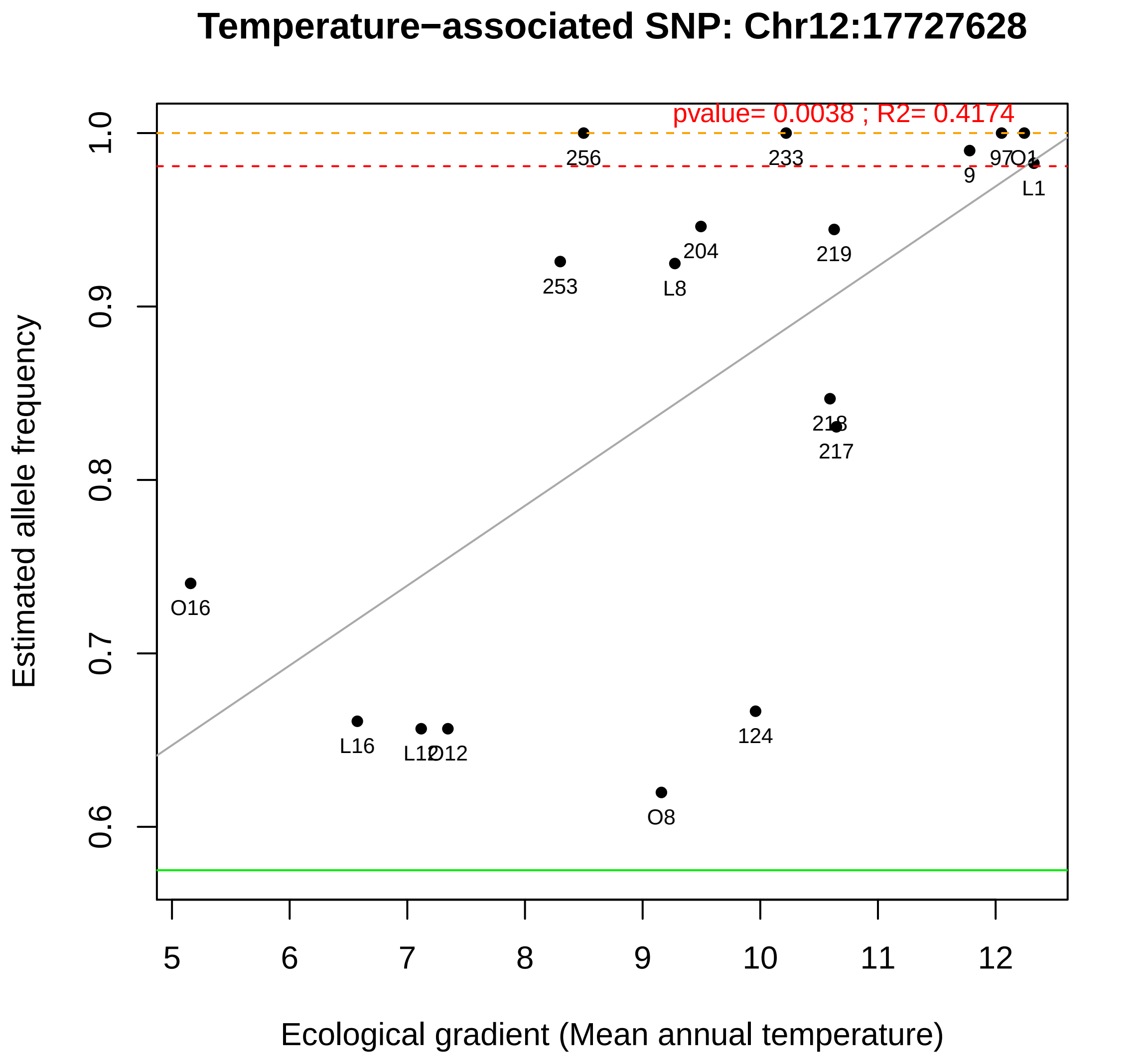
Temperature−associated SNP: Chr12:17727628
pvalue= 0.0038 ; R2= 0.4174
Estimated allele frequency
0.6
0.7
0.8
0.9
1.0
5
6
7
8
9
10
11
12
Ecological gradient (Mean annual temperature)
256
233
97
O1
9
L1
204
219
253
L8
218
217
O16
L16
L12
O12
124
O8

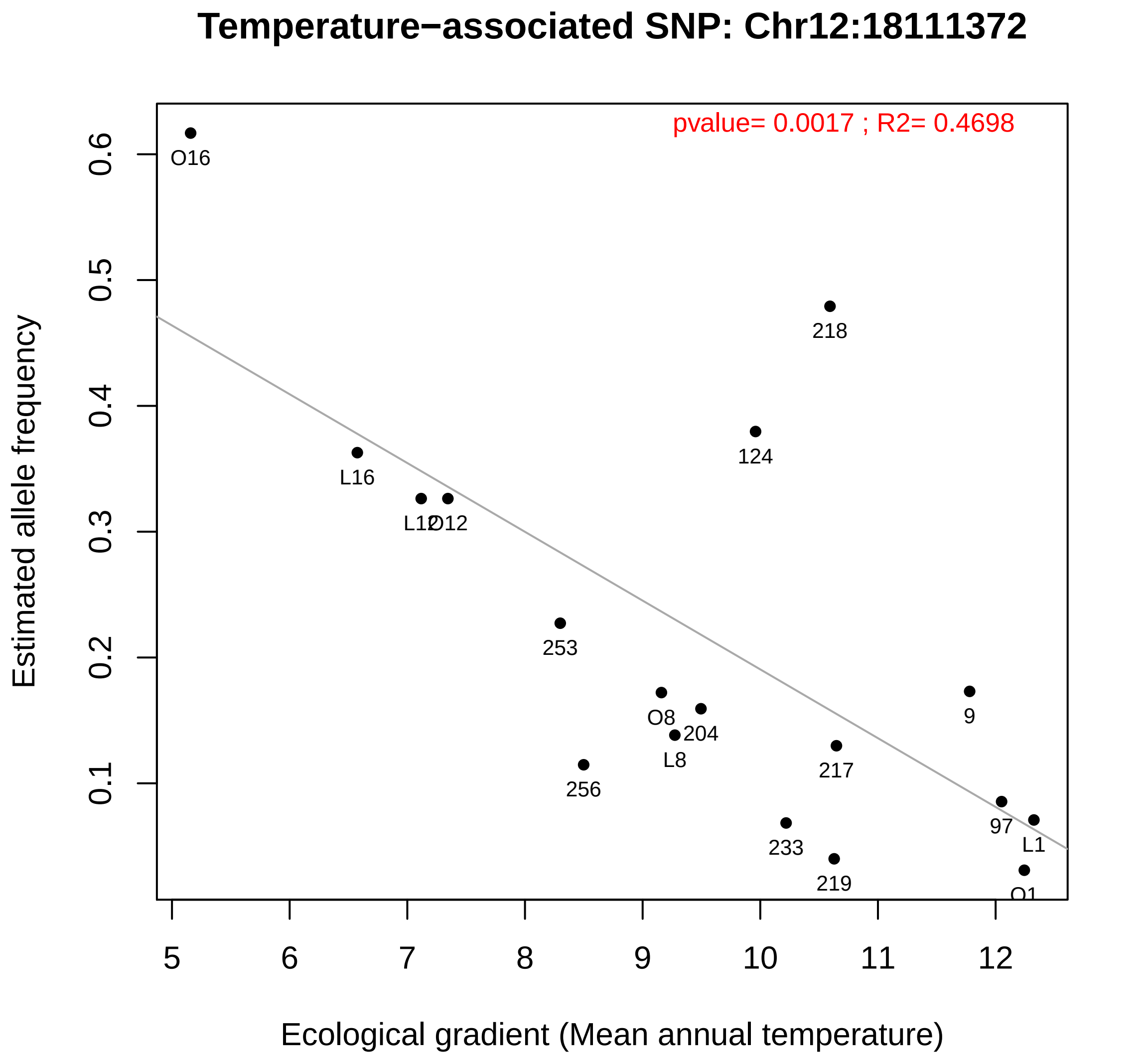

Temperature−associated SNP: Chr12:18111372
pvalue= 0.0017 ; R2= 0.4698
Estimated allele frequency
0.1
0.2
0.3
0.4
0.5
0.6
Ecological gradient (Mean annual temperature)
5
6
7
8
9
10
11
12
O16
L16
L12
O12
253
256
O8
204
L8
124
218
233
219
217
9
97
L1
O1

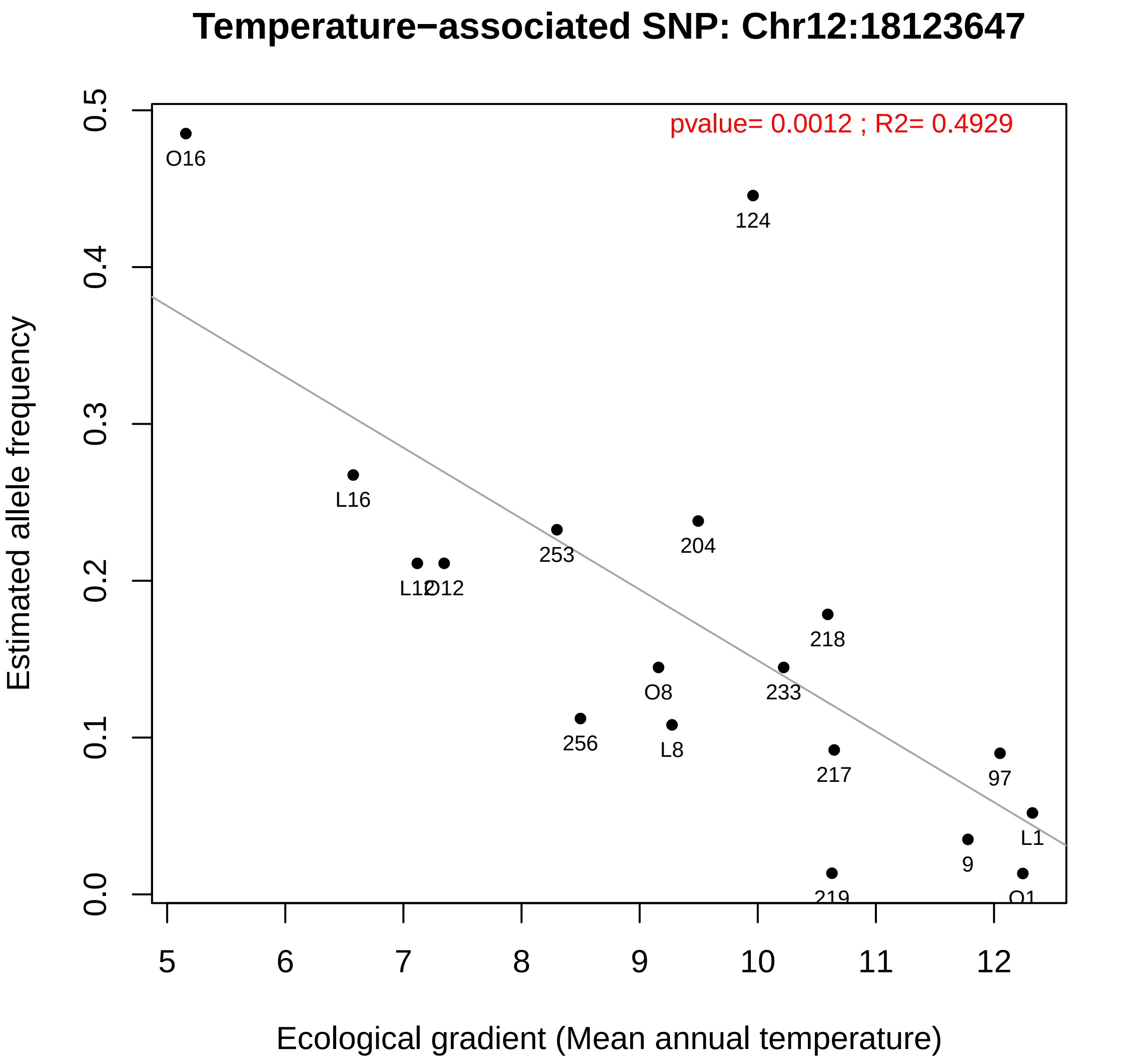
Temperature−associated SNP: Chr12:18123647
pvalue= 0.0012 ; R2= 0.4929
Estimated allele frequency
Ecological gradient (Mean annual temperature)
0.0
0.1
0.2
0.3
0.4
0.5
5
6
7
8
9
10
11
12
O16
124
L16
253
204
L12
O12
218
O8
233
256
L8
217
97
L1
9
219
O1

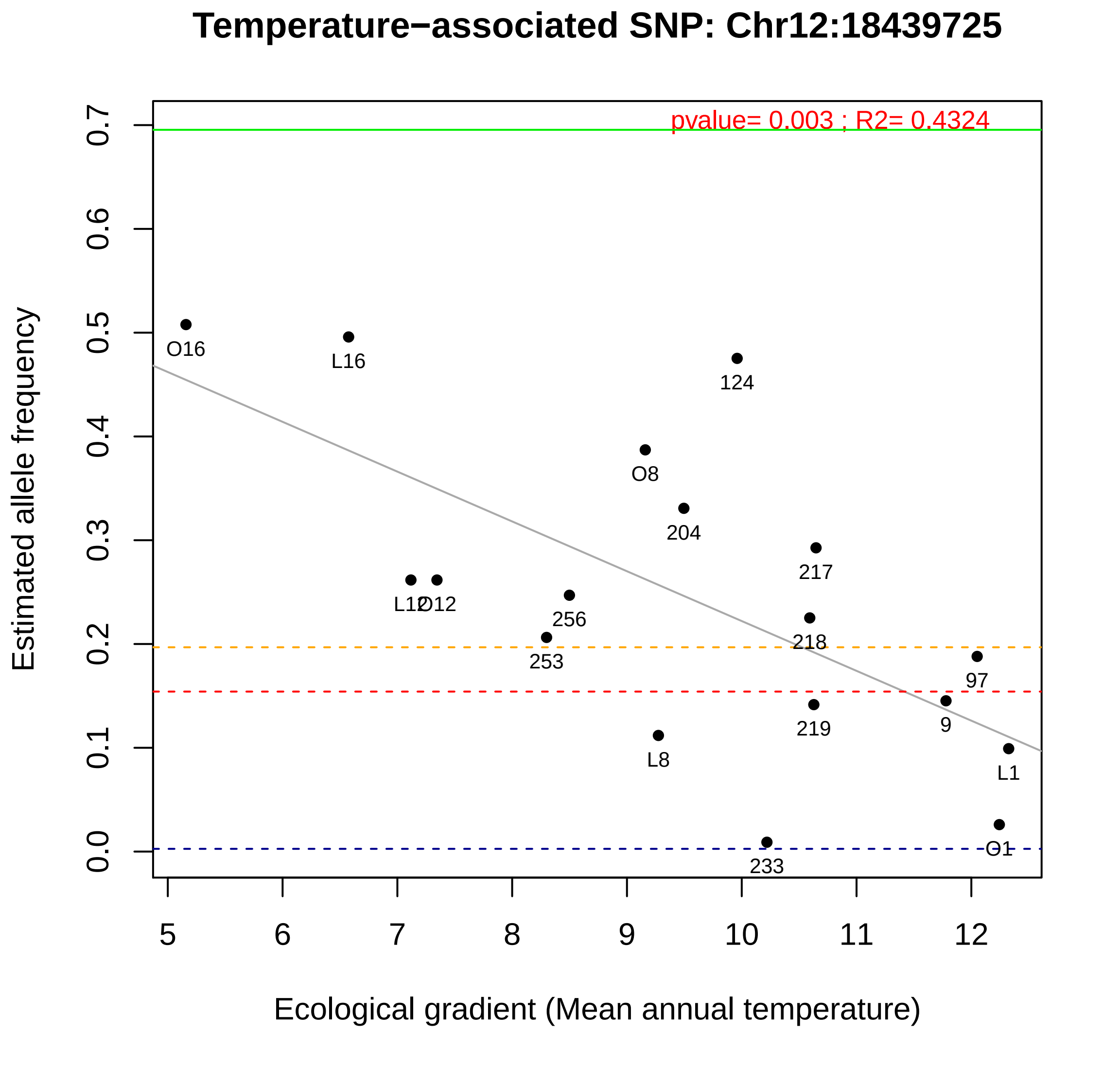
Temperature−associated SNP: Chr12:18439725
pvalue= 0.003 ; R2= 0.4324
Estimated allele frequency
0.0
0.1
0.2
0.3
0.4
0.5
0.6
0.7
5
6
7
8
9
10
11
12
Ecological gradient (Mean annual temperature)
O16
L16
124
O8
204
217
L12
O12
256
218
253
97
9
219
L8
L1
O1
233

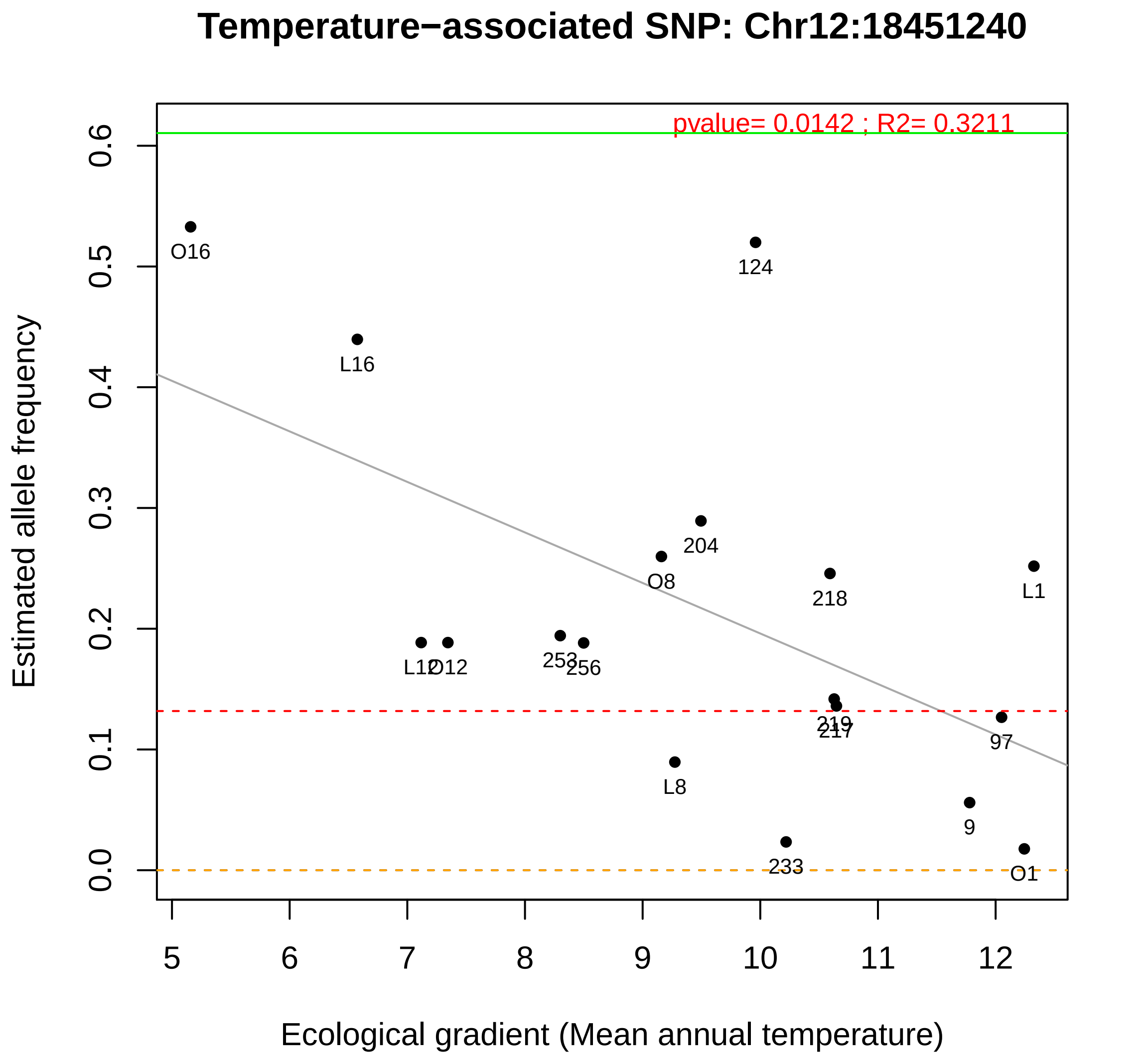
Temperature−associated SNP: Chr12:18451240
pvalue= 0.0142 ; R2= 0.3211
Estimated allele frequency
0.0
0.1
0.2
0.3
0.4
0.5
0.6
5
6
7
8
9
10
11
12
Ecological gradient (Mean annual temperature)
O16
L16
124
204
O8
218
L1
L12
O12
253
256
219
217
97
L8
9
233
O1

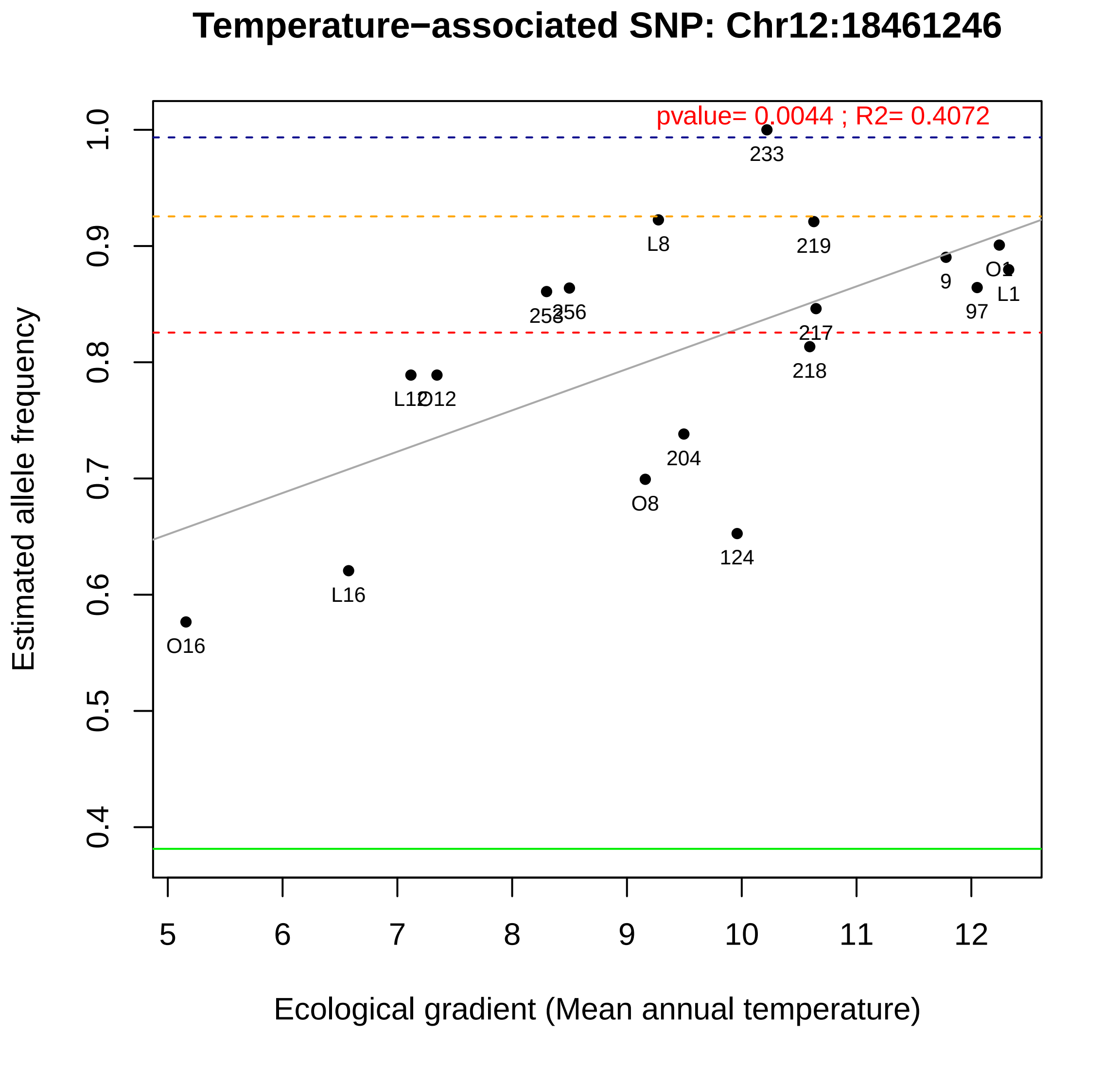

Temperature−associated SNP: Chr12:18461246
pvalue= 0.0044 ; R2= 0.4072
Estimated allele frequency
Ecological gradient (Mean annual temperature)
1.0
0.9
0.8
0.7
0.6
0.5
0.4
5
6
7
8
9
10
11
12
233
L8
219
9
97
L1
217
218
256
L12
O12
204
O8
124
L16
O16

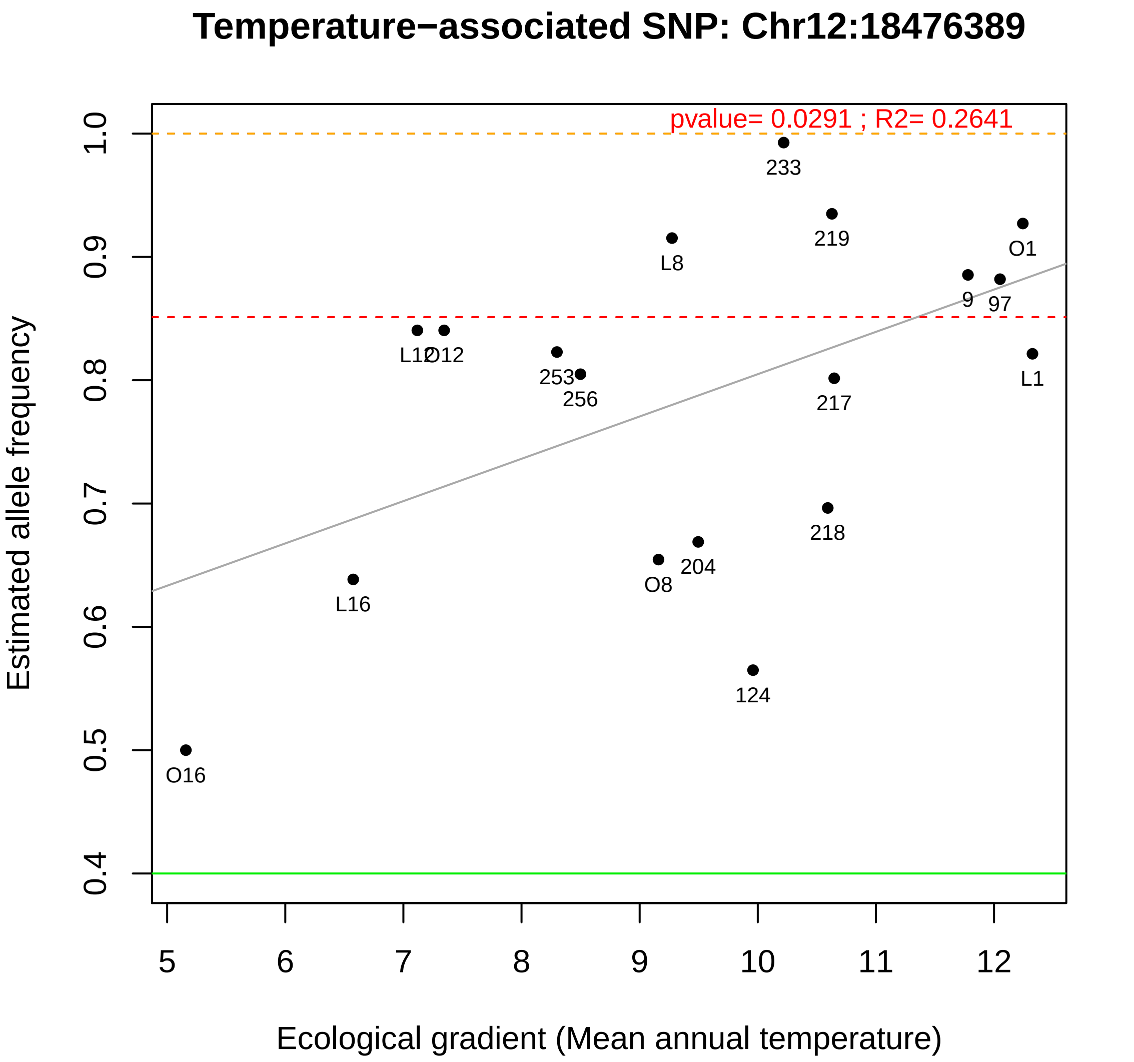

Temperature−associated SNP: Chr12:18476389
pvalue= 0.0291 ; R2= 0.2641
Estimated allele frequency
Ecological gradient (Mean annual temperature)
0.4
0.5
0.6
0.7
0.8
0.9
1.0
5
6
7
8
9
10
11
12
233
219
O1
L8
9
97
L12
O12
253
256
L1
217
218
204
O8
L16
124
O16

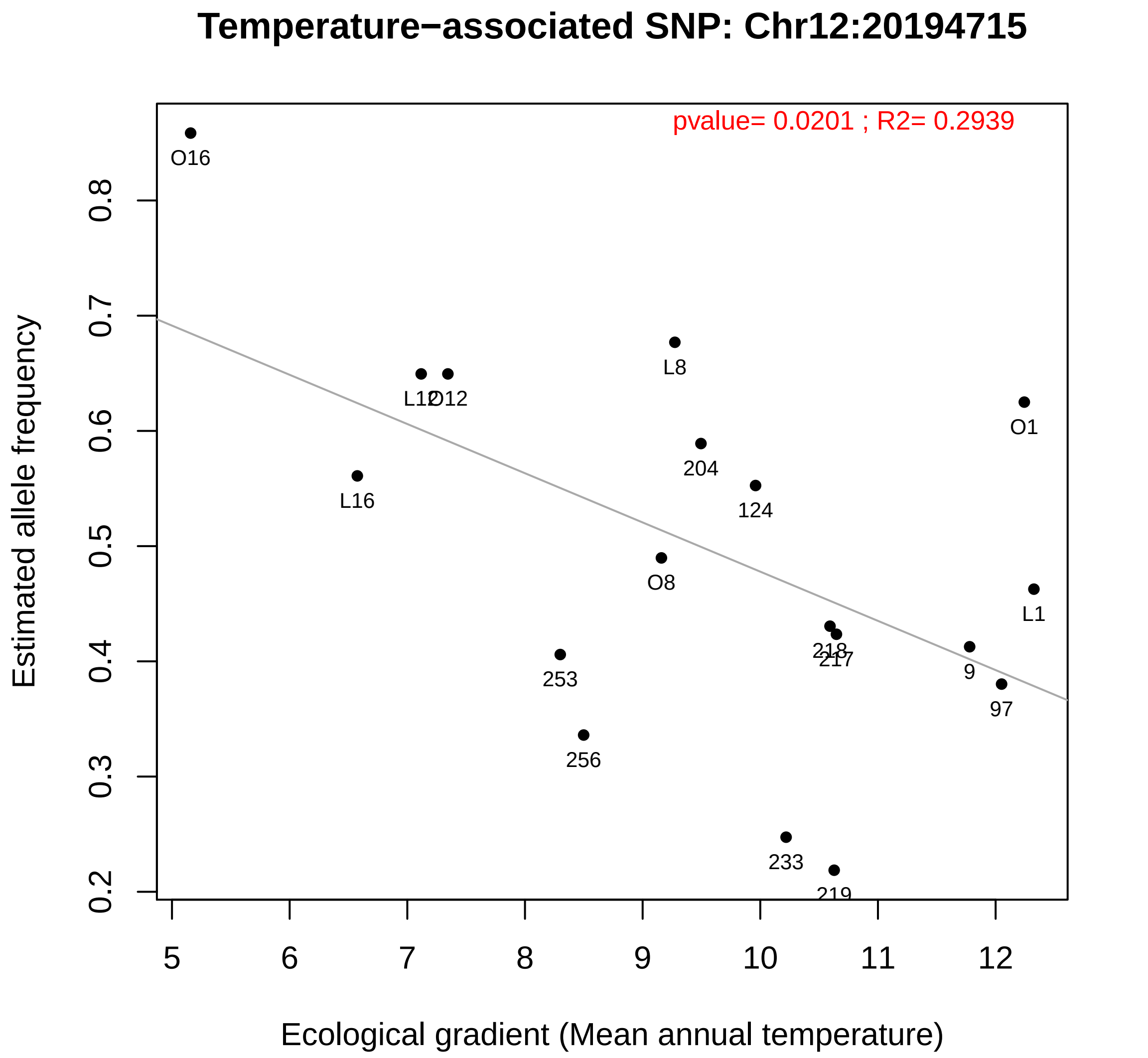
Temperature−associated SNP: Chr12:20194715
pvalue= 0.0201 ; R2= 0.2939
Estimated allele frequency
0.2
0.3
0.4
0.5
0.6
0.7
0.8
5
6
7
8
9
10
11
12
Ecological gradient (Mean annual temperature)
O16
L12
O12
L8
O1
204
L16
124
O8
L1
218
217
9
253
97
256
233
219

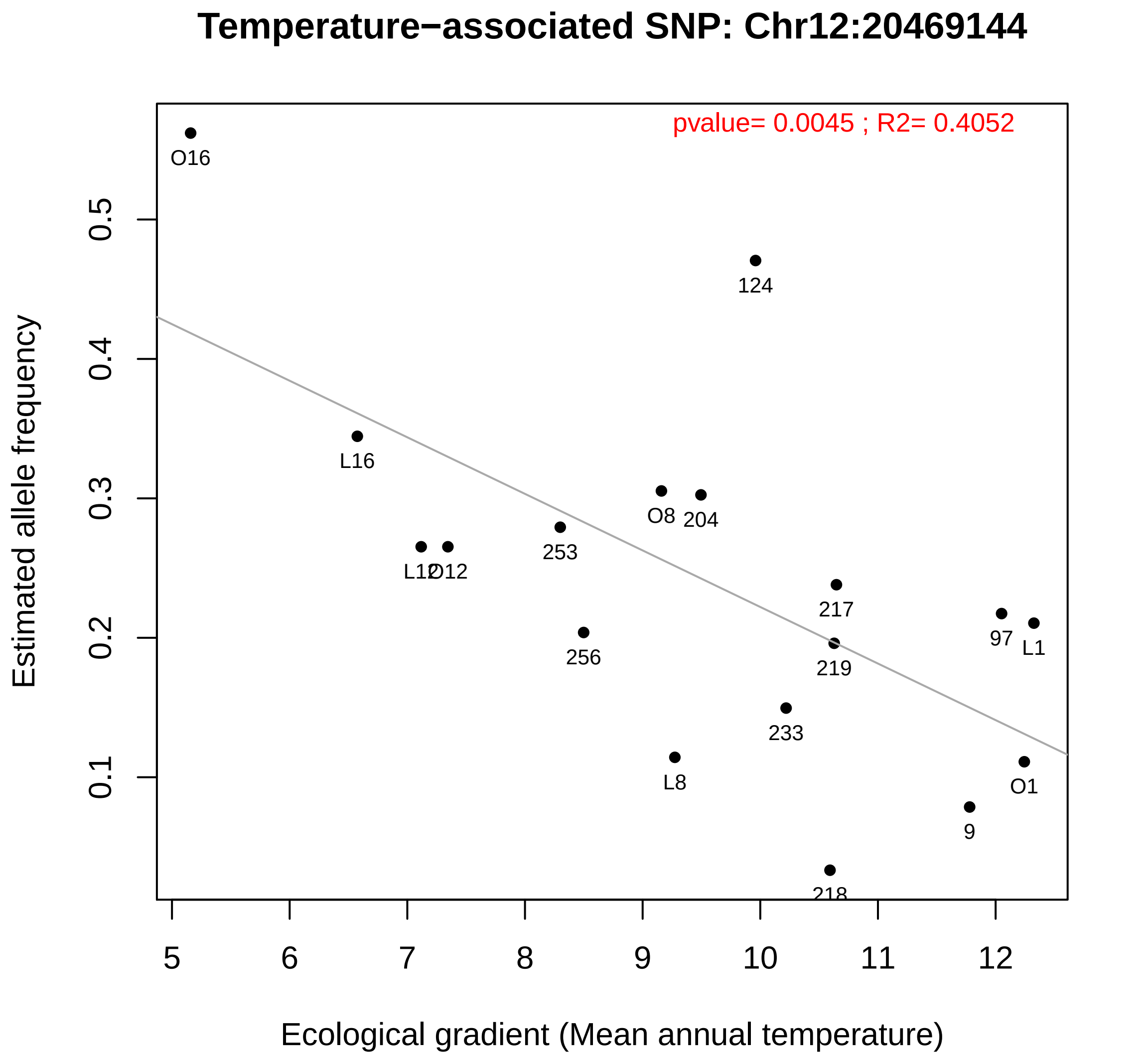
Temperature−associated SNP: Chr12:20469144
pvalue= 0.0045 ; R2= 0.4052
Estimated allele frequency
Ecological gradient (Mean annual temperature)
0.1
0.2
0.3
0.4
0.5
5
6
7
8
9
10
11
12
O16
124
L16
O8
204
253
L12
O12
217
97
L1
219
256
233
L8
O1
9
218

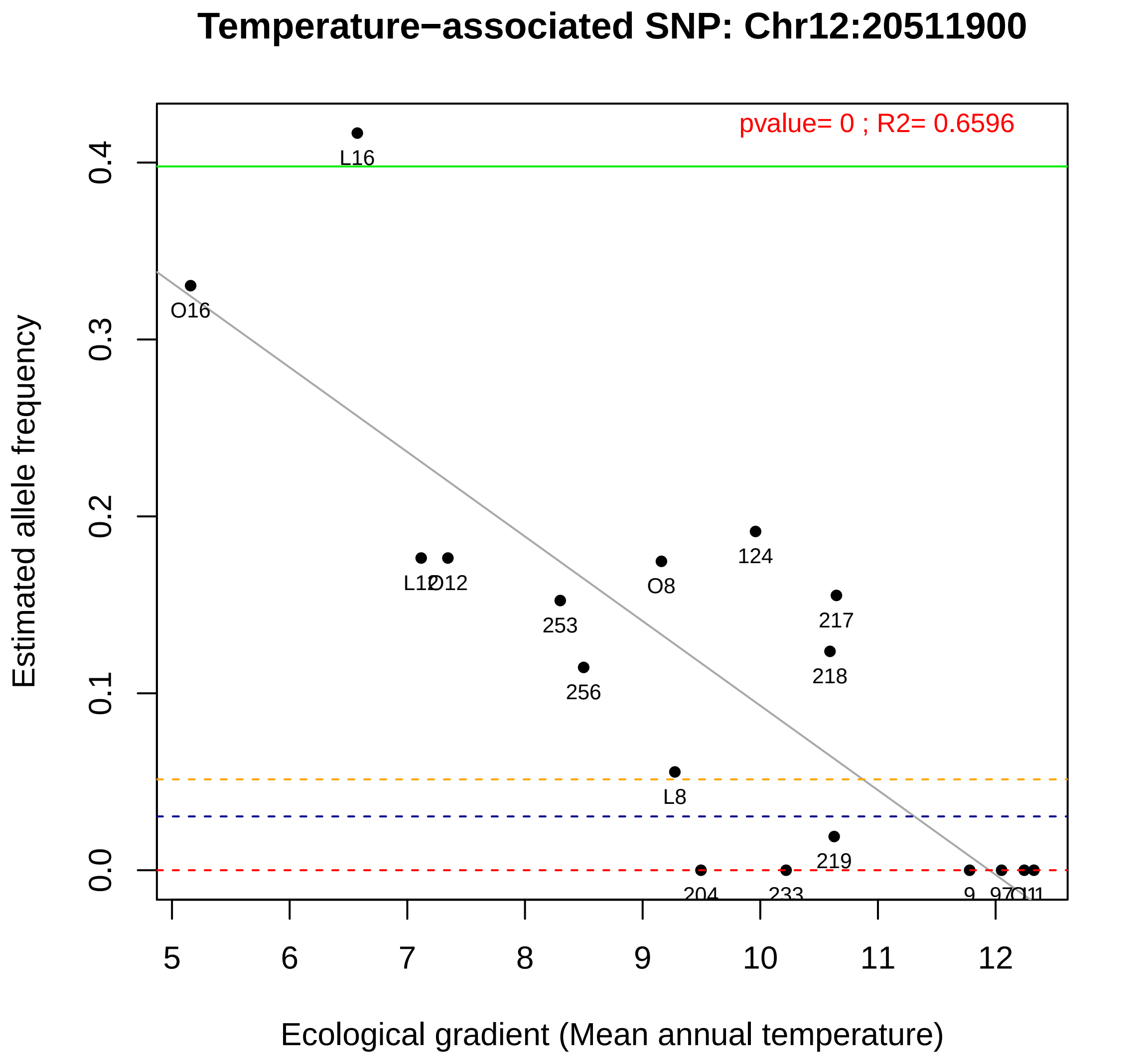

Temperature−associated SNP: Chr12:20511900
pvalue= 0 ; R2= 0.6596
Estimated allele frequency
0.0
0.1
0.2
0.3
0.4
5
6
7
8
9
10
11
12
Ecological gradient (Mean annual temperature)
L16
O16
L12
O12
253
256
O8
124
217
218
L8
219
204
233

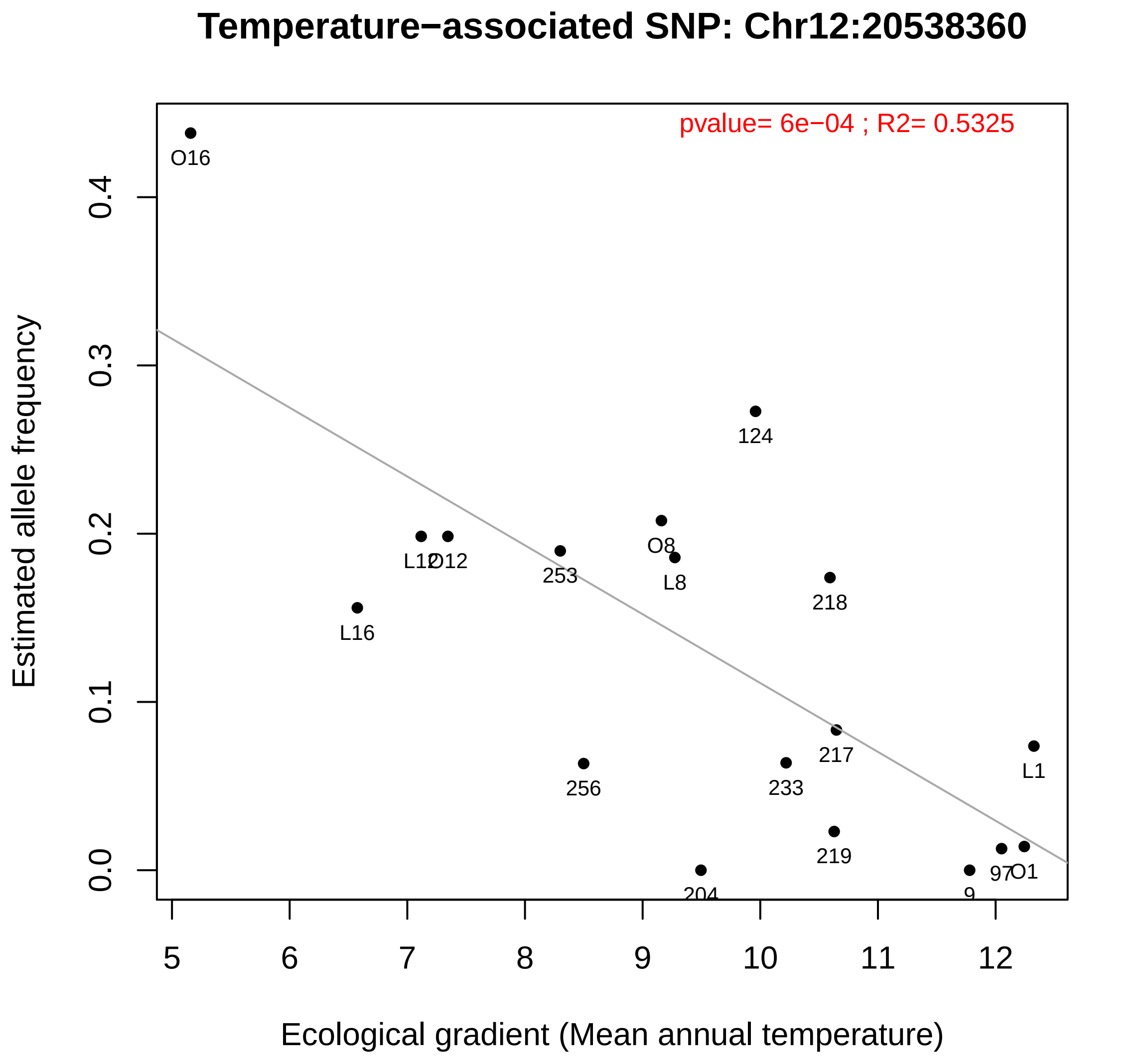
Temperature−associated SNP: Chr12:20538360
pvalue= 6e−04 ; R2= 0.5325
Estimated allele frequency
0.0
0.1
0.2
0.3
0.4
Ecological gradient (Mean annual temperature)
5
6
7
8
9
10
11
12
O16
124
O8
L12
O12
253
L8
218
L16
217
L1
256
233
219
204
9
97
O1

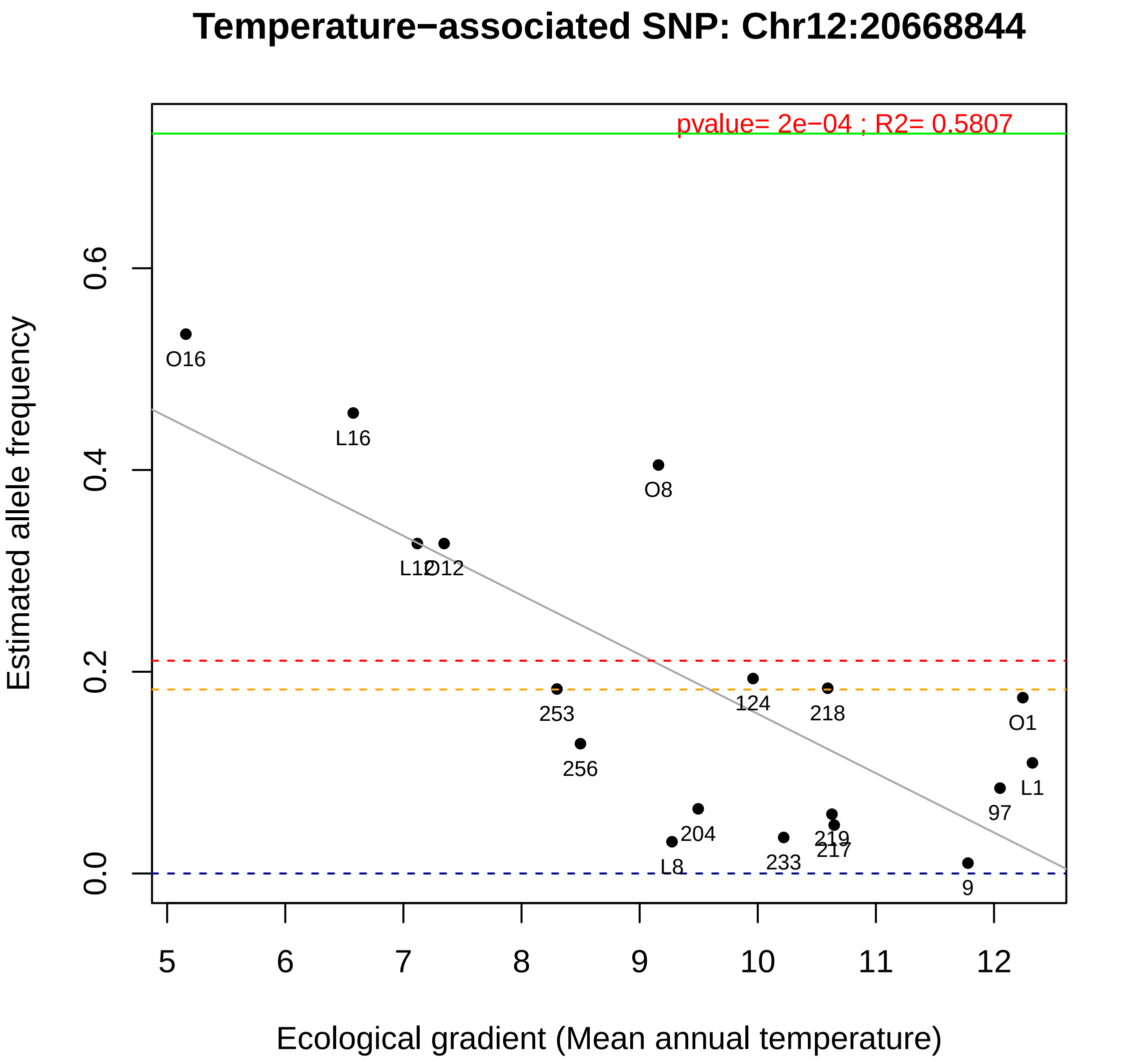
Temperature−associated SNP: Chr12:20668844
pvalue= 2e−04 ; R2= 0.5807
Estimated allele frequency
0.0
0.2
0.4
0.6
O16
L16
O8
L12
O12
253
124
218
O1
256
L1
97
204
219
217
L8
233
9
5
6
7
8
9
10
11
12
Ecological gradient (Mean annual temperature)

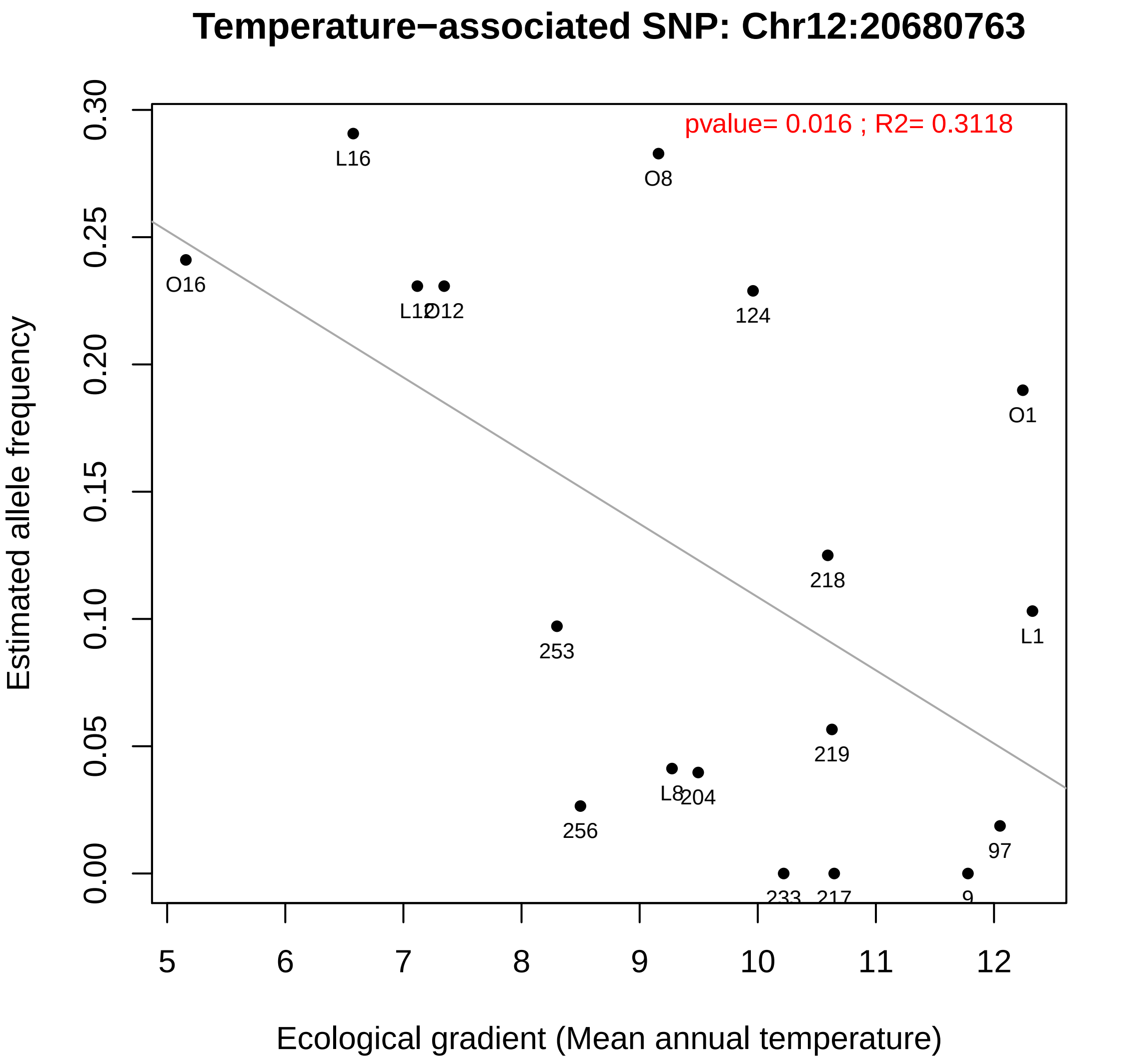

Temperature−associated SNP: Chr12:20680763
pvalue= 0.016 ; R2= 0.3118
L16
O8
O16
L12
O12
124
O1
218
L1
253
219
L8
204
256
97
233
217
9
Estimated allele frequency
0.00
0.05
0.10
0.15
0.20
0.25
0.30
5
6
7
8
9
10
11
12
Ecological gradient (Mean annual temperature)

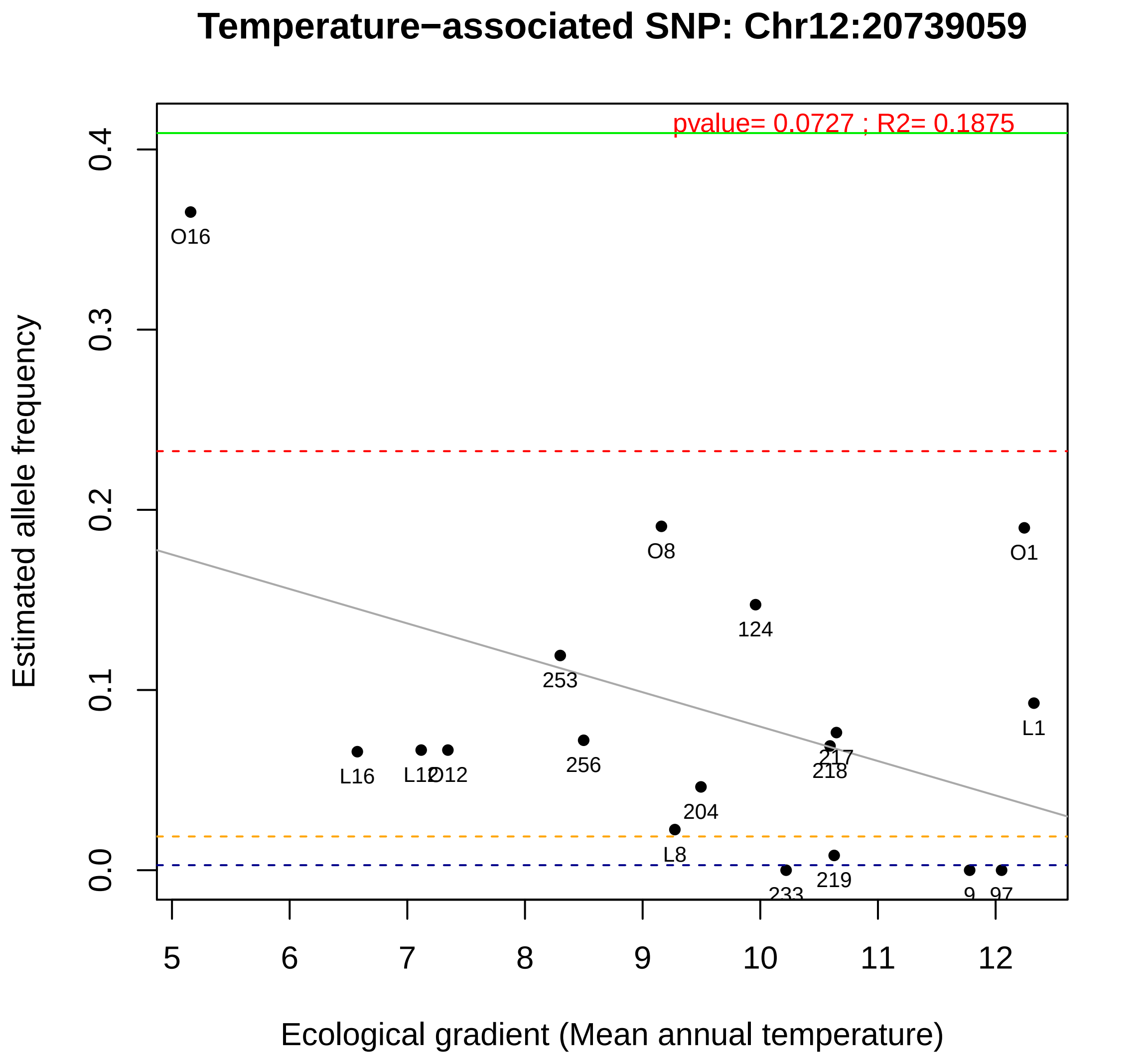

Temperature−associated SNP: Chr12:20739059
pvalue= 0.0727 ; R2= 0.1875
Estimated allele frequency
0.0
0.1
0.2
0.3
0.4
O16
O8
O1
124
253
L1
217
218
L16
L12
O12
256
204
L8
219
233
9
97
5
6
7
8
9
10
11
12
Ecological gradient (Mean annual temperature)

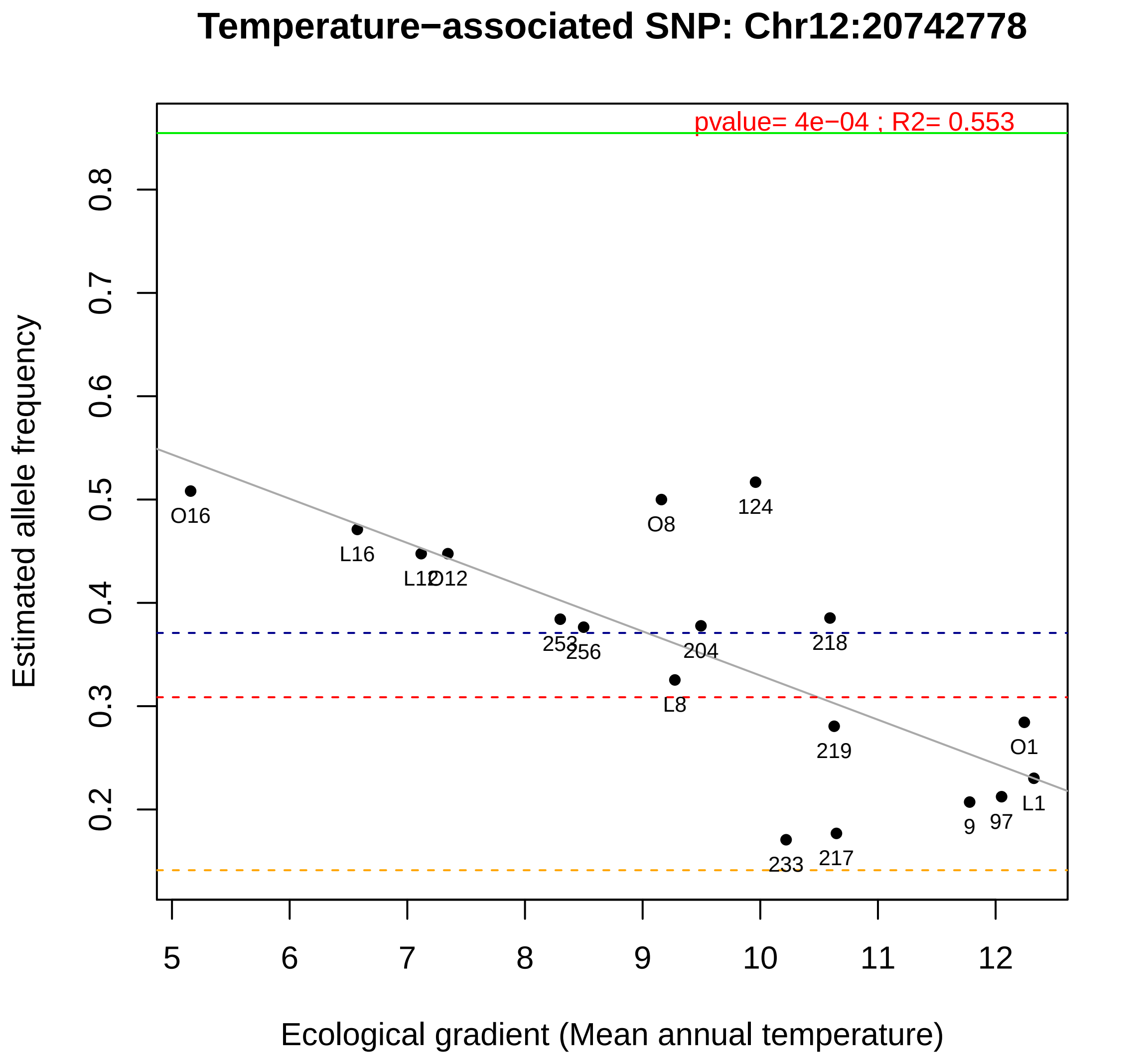
Temperature−associated SNP: Chr12:20742778
pvalue= 4e−04 ; R2= 0.553
Estimated allele frequency
0.2
0.3
0.4
0.5
0.6
0.7
0.8
O16
L16
L12
O12
253
256
O8
124
204
L8
218
219
O1
L1
9
97
233
217
5
6
7
8
9
10
11
12
Ecological gradient (Mean annual temperature)

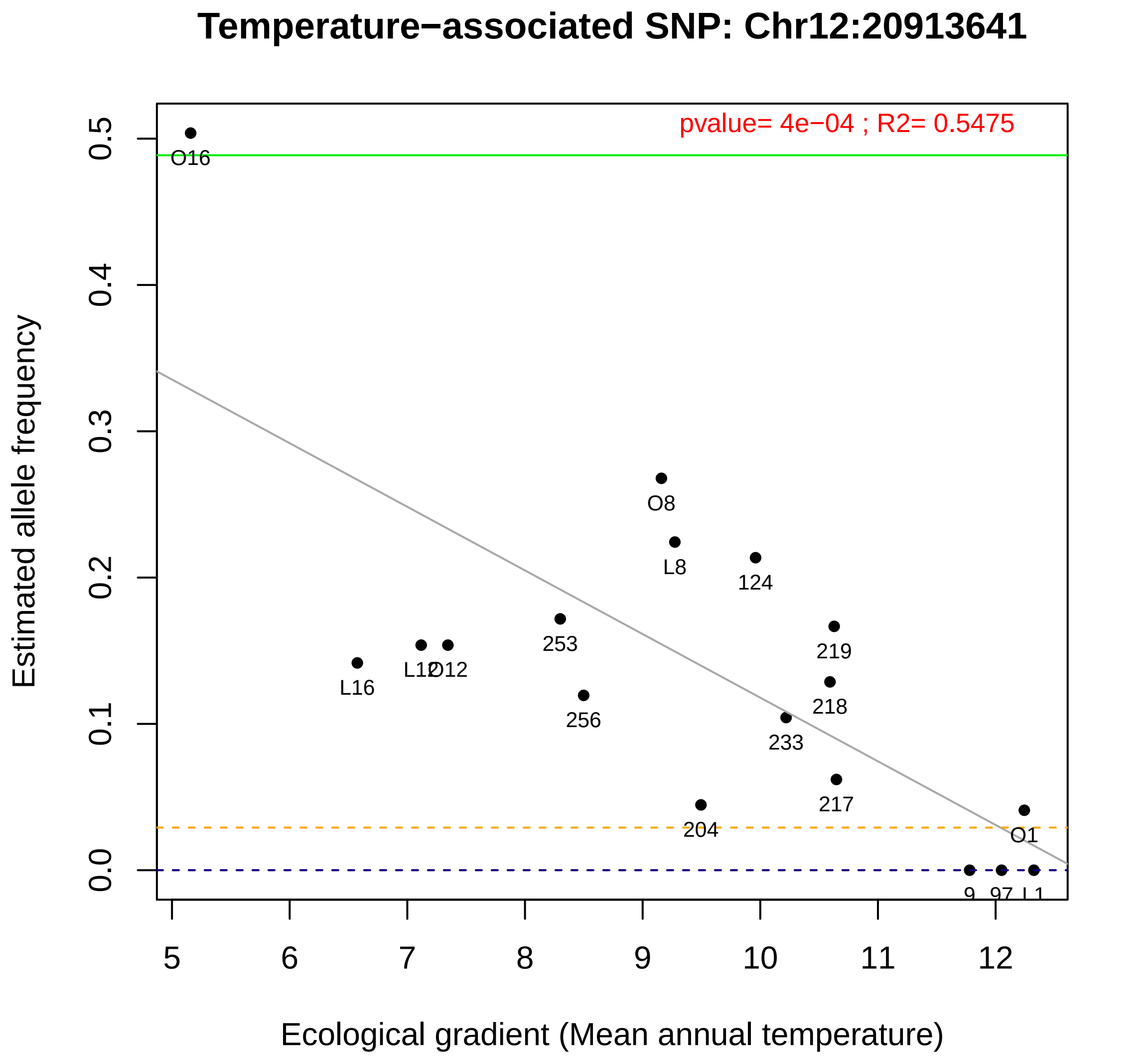

Temperature−associated SNP: Chr12:20913641
pvalue= 4e−04 ; R2= 0.5475
Estimated allele frequency
0.0
0.1
0.2
0.3
0.4
0.5
Ecological gradient (Mean annual temperature)
5
6
7
8
9
10
11
12
O16
O8
L8
124
253
219
L16
L12
O12
256
218
233
217
204
O1

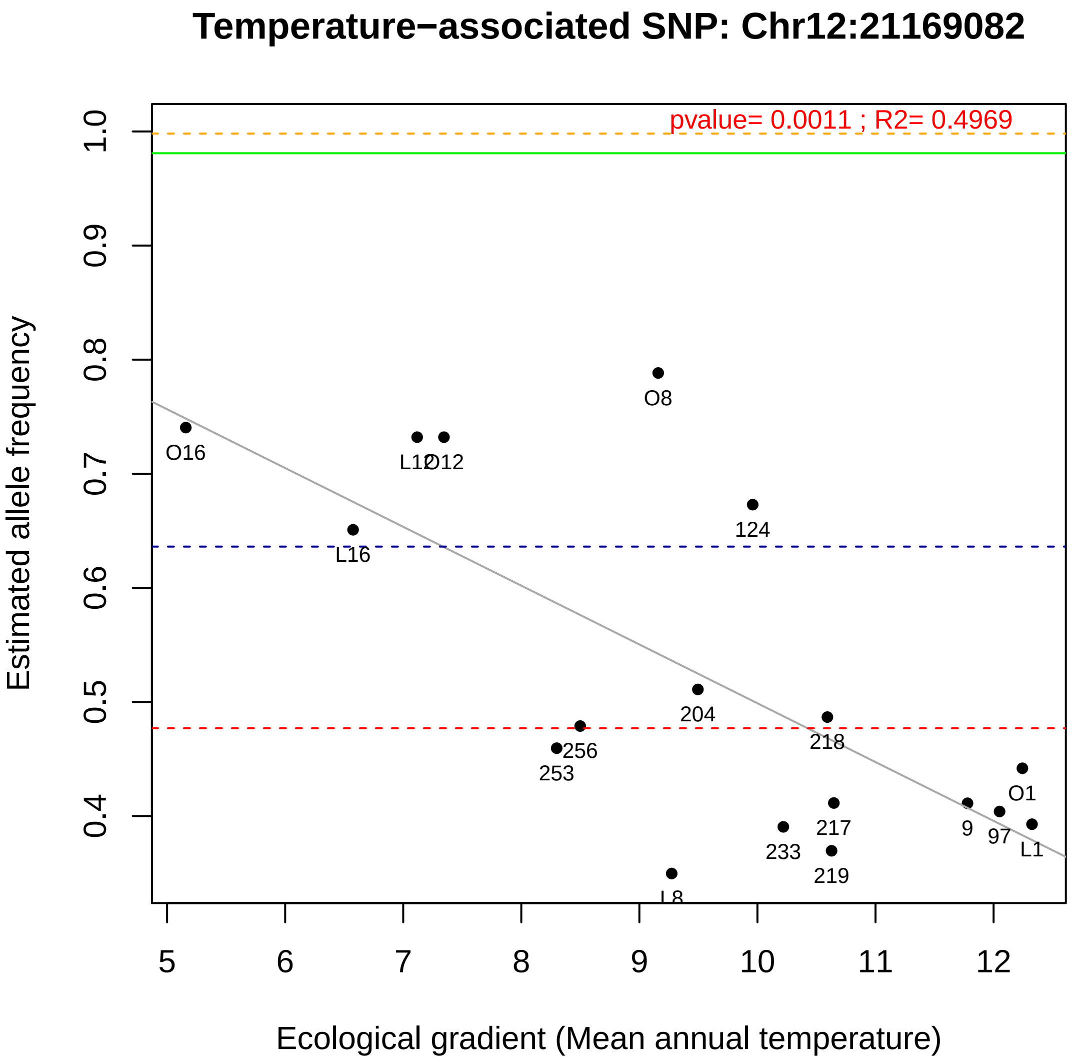

Temperature−associated SNP: Chr12:21169082
pvalue= 0.0011 ; R2= 0.4969
Estimated allele frequency
Ecological gradient (Mean annual temperature)
0.4
0.5
0.6
0.7
0.8
0.9
1.0
5
6
7
8
9
10
11
12
O16
L12
O12
O8
L16
124
204
218
256
253
O1
217
9
97
233
L1
219
L8

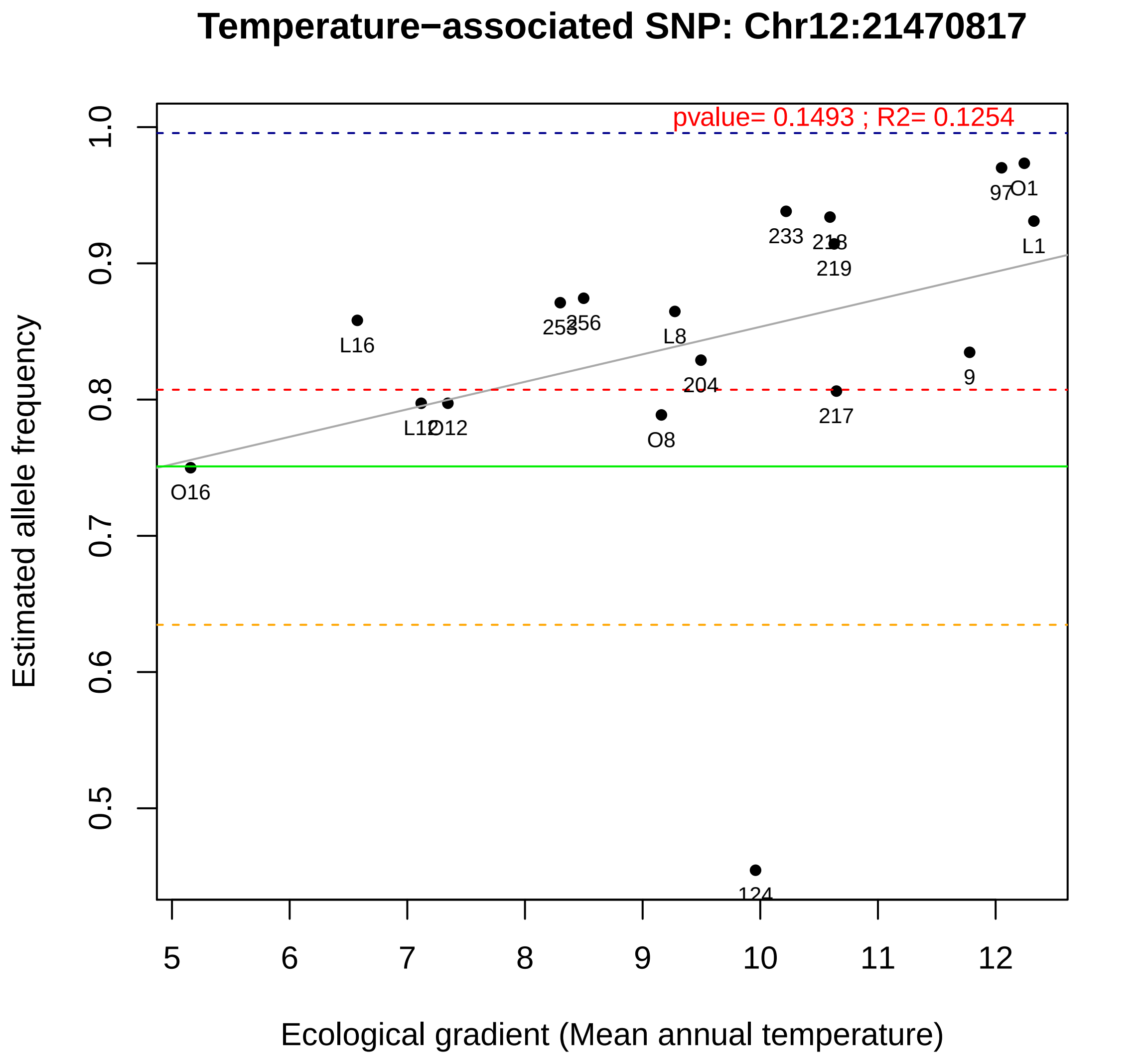
Temperature−associated SNP: Chr12:21470817
pvalue= 0.1493 ; R2= 0.1254
Estimated allele frequency
Ecological gradient (Mean annual temperature)
1.0
0.9
0.8
0.7
0.6
0.5
5
6
7
8
9
10
11
12
97
O1
233
218
219
L1
253
256
L16
L8
204
9
217
L12
O12
O8
O16
124

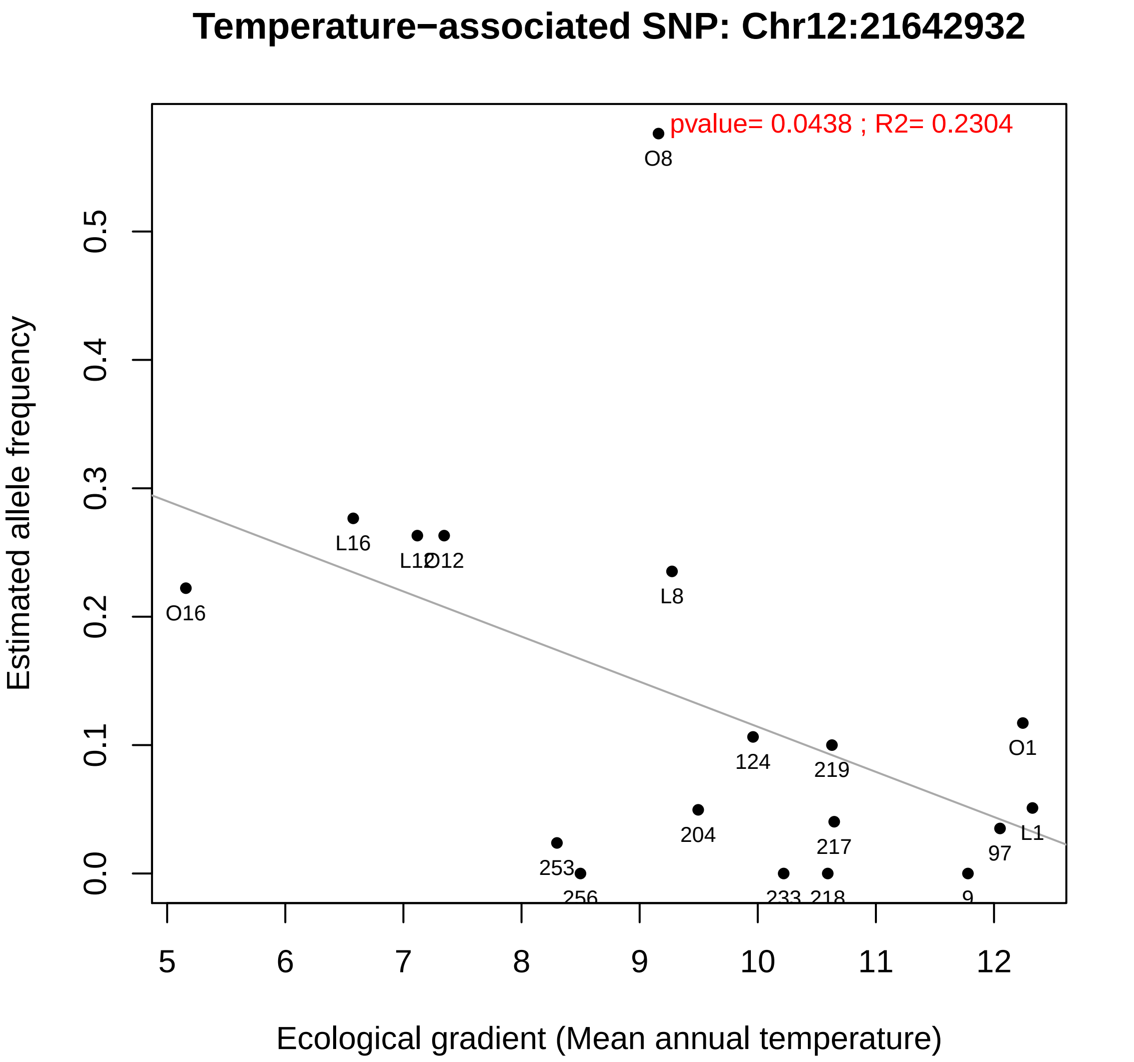

Temperature−associated SNP: Chr12:21642932
pvalue= 0.0438 ; R2= 0.2304
O8
L16
L12
O12
L8
O16
124
219
O1
204
217
L1
97
253
256
233
218
9
Estimated allele frequency
0.0
0.1
0.2
0.3
0.4
0.5
5
6
7
8
9
10
11
12
Ecological gradient (Mean annual temperature)

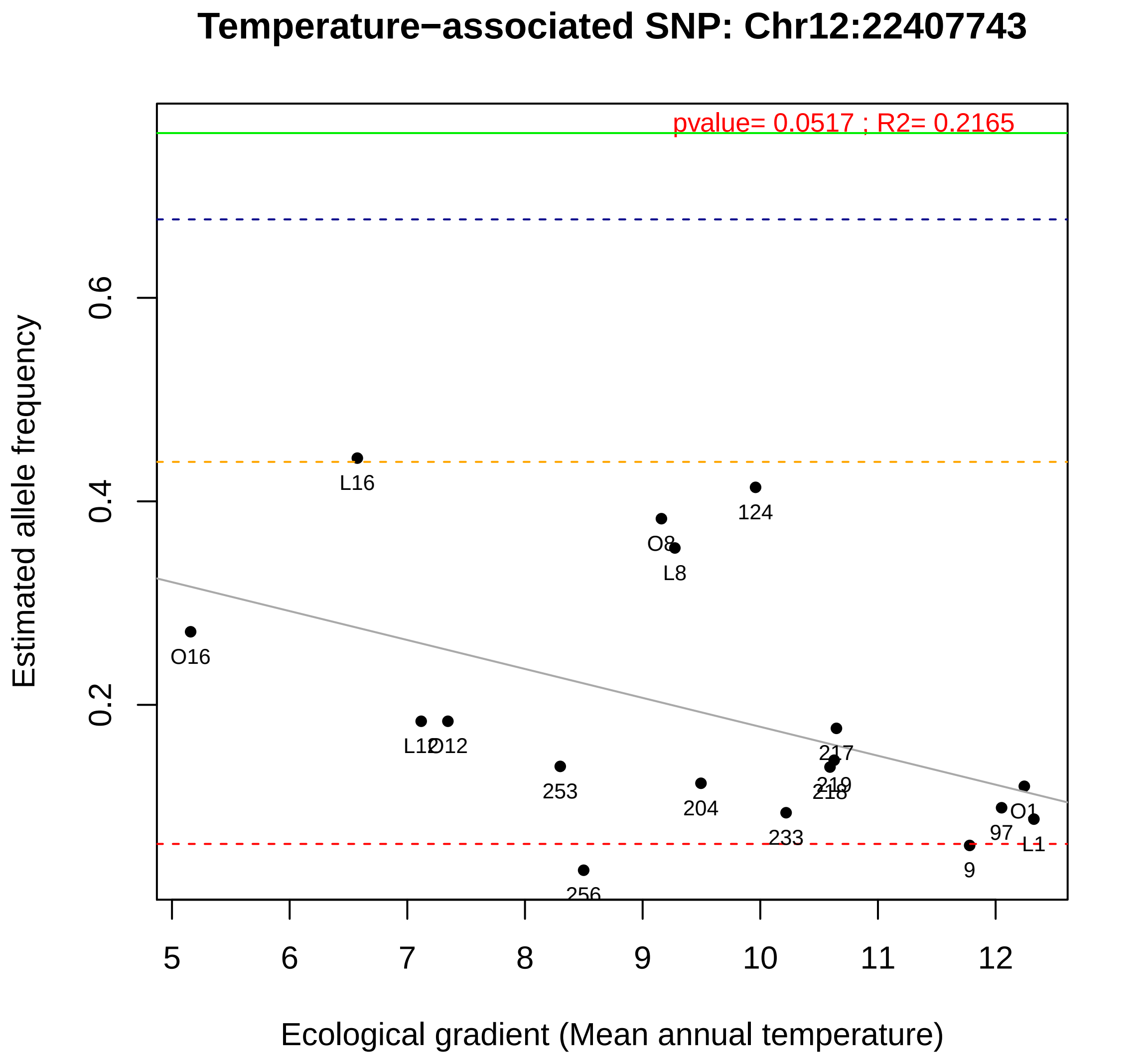
Temperature−associated SNP: Chr12:22407743
pvalue= 0.0517 ; R2= 0.2165
Estimated allele frequency
0.2
0.4
0.6
Ecological gradient (Mean annual temperature)
5
6
7
8
9
10
11
12
L16
124
O8
L8
O16
L12
O12
253
256
204
233
217
219
218
9
97
O1
L1

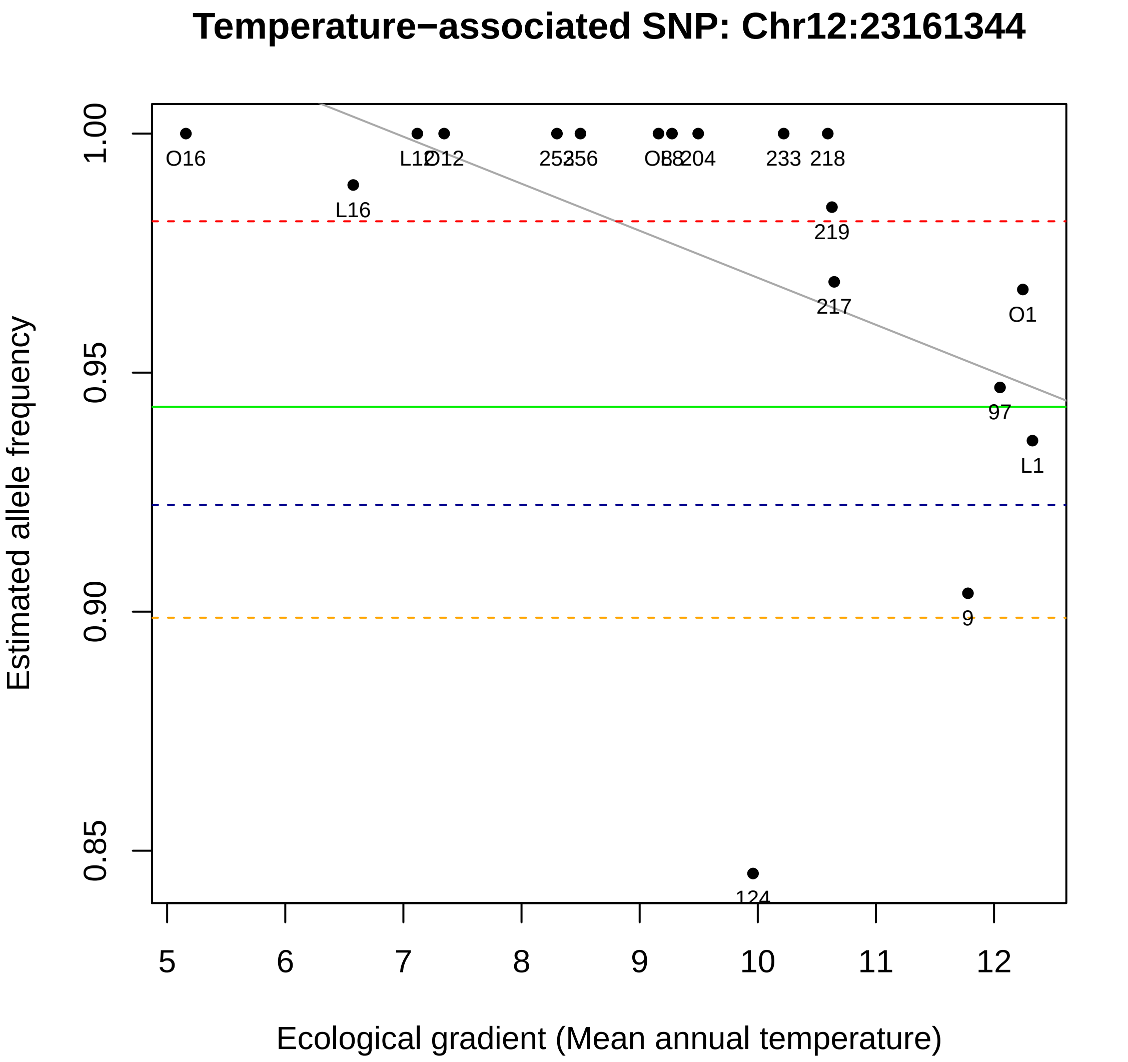

Temperature−associated SNP: Chr12:23161344
Estimated allele frequency
Ecological gradient (Mean annual temperature)
1.00
0.95
0.90
0.85
5
6
7
8
9
10
11
12
O16
L16
O12
233
218
219
217
O1
97
L1
9
124

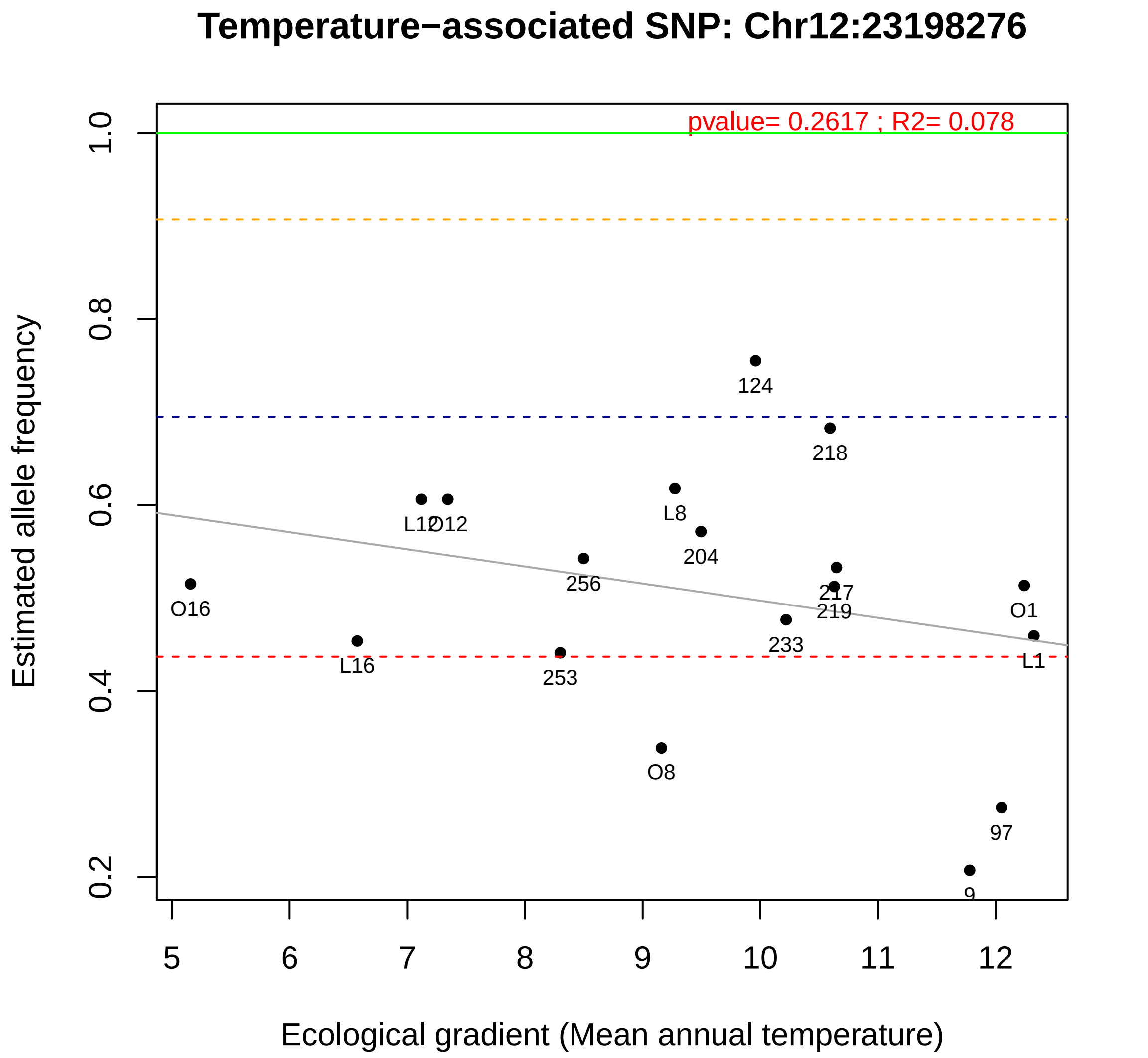

Temperature−associated SNP: Chr12:23198276
pvalue= 0.2617 ; R2= 0.078
Estimated allele frequency
1.0
0.8
0.6
0.4
0.2
124
218
L12
O12
L8
204
256
217
219
O1
O16
L16
253
233
L1
O8
97
9
5
6
7
8
9
10
11
12
Ecological gradient (Mean annual temperature)

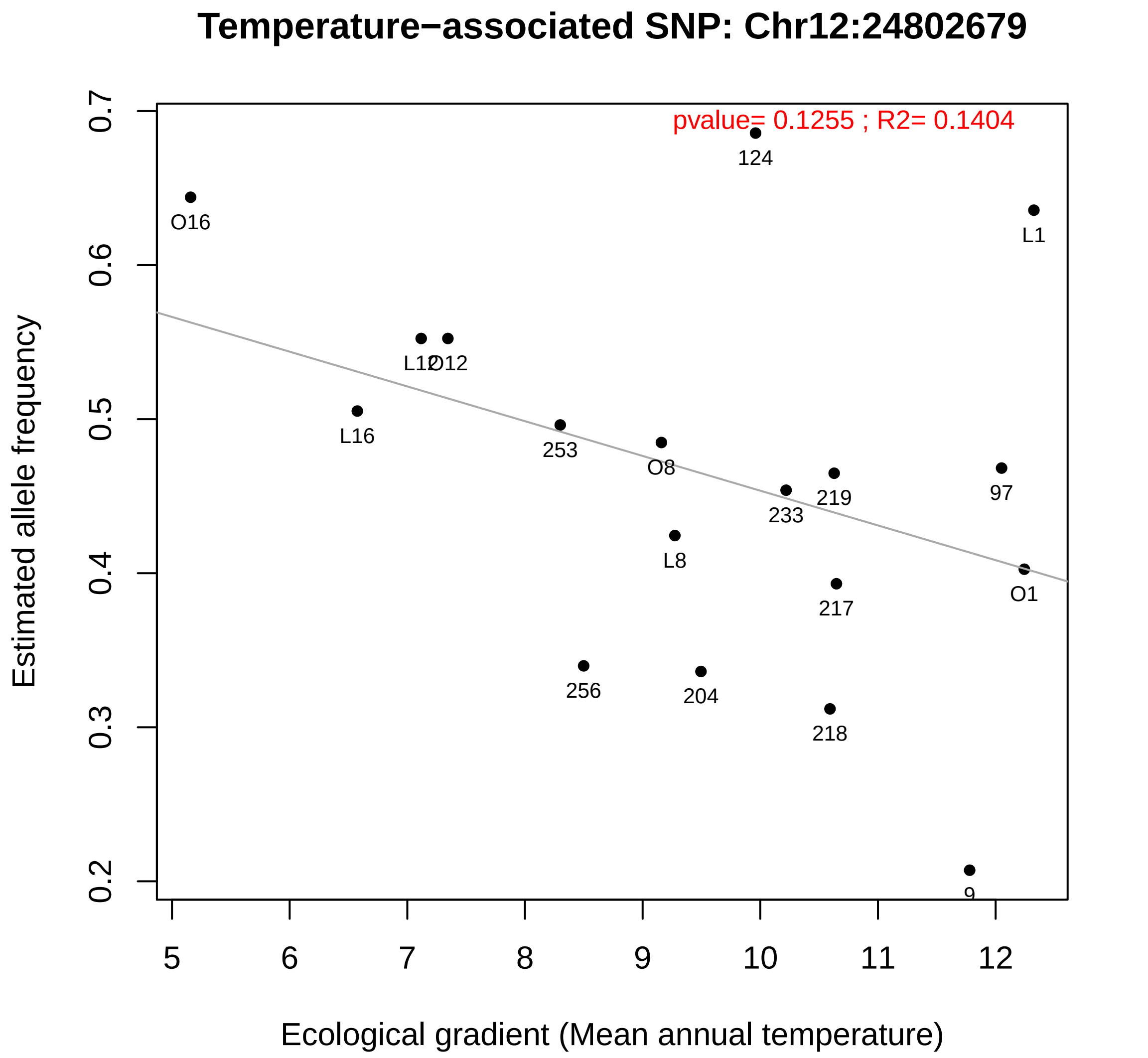
Temperature−associated SNP: Chr12:24802679
pvalue= 0.1255 ; R2= 0.1404
Estimated allele frequency
Ecological gradient (Mean annual temperature)
0.2
0.3
0.4
0.5
0.6
0.7
5
6
7
8
9
10
11
12
124
O16
L1
L12
O12
L16
253
O8
219
97
233
L8
O1
217
256
204
218
9

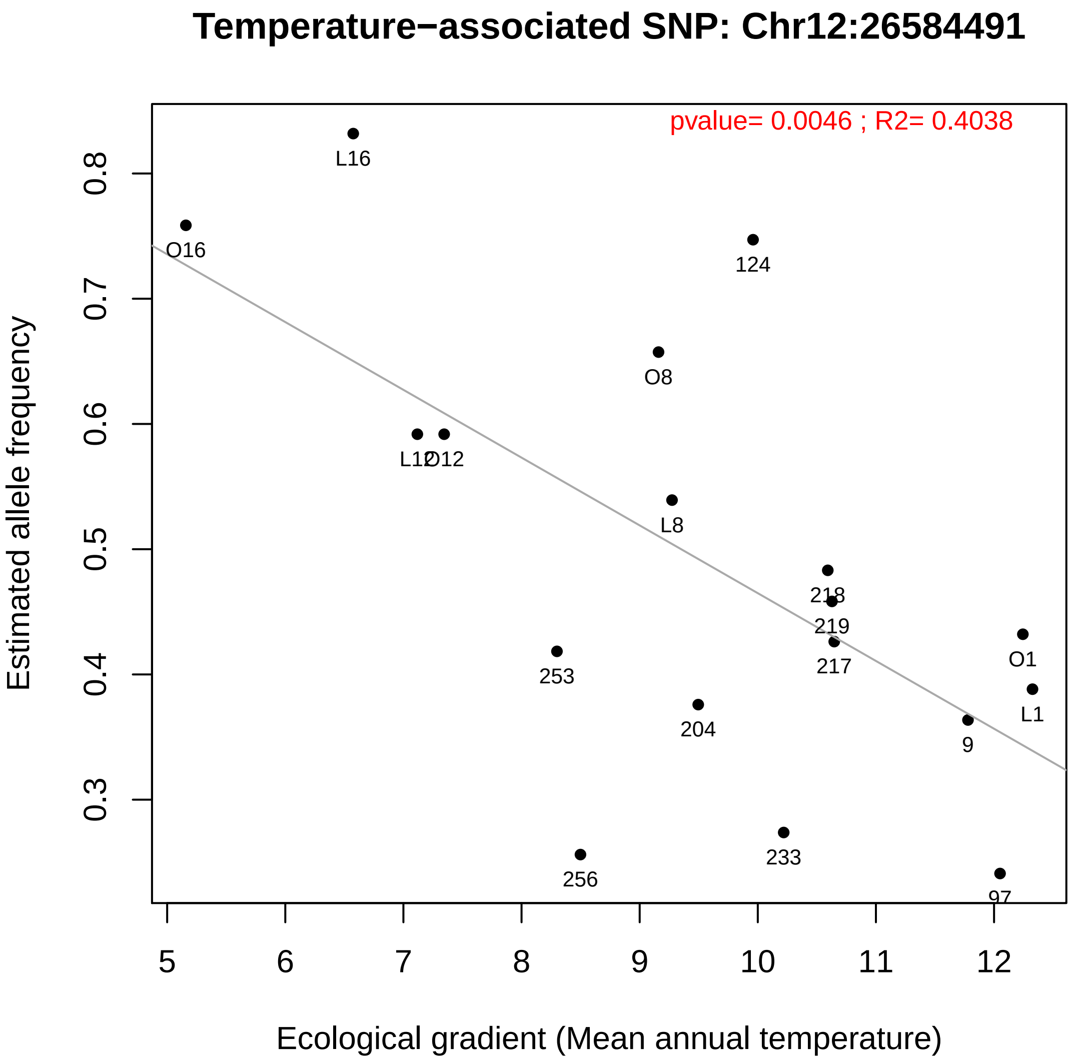

Temperature−associated SNP: Chr12:26584491
pvalue= 0.0046 ; R2= 0.4038
Estimated allele frequency
0.3
0.4
0.5
0.6
0.7
0.8
5
6
7
8
9
10
11
12
Ecological gradient (Mean annual temperature)
L16
O16
124
O8
L12
O12
L8
218
219
217
O1
L1
253
204
9
256
233
97

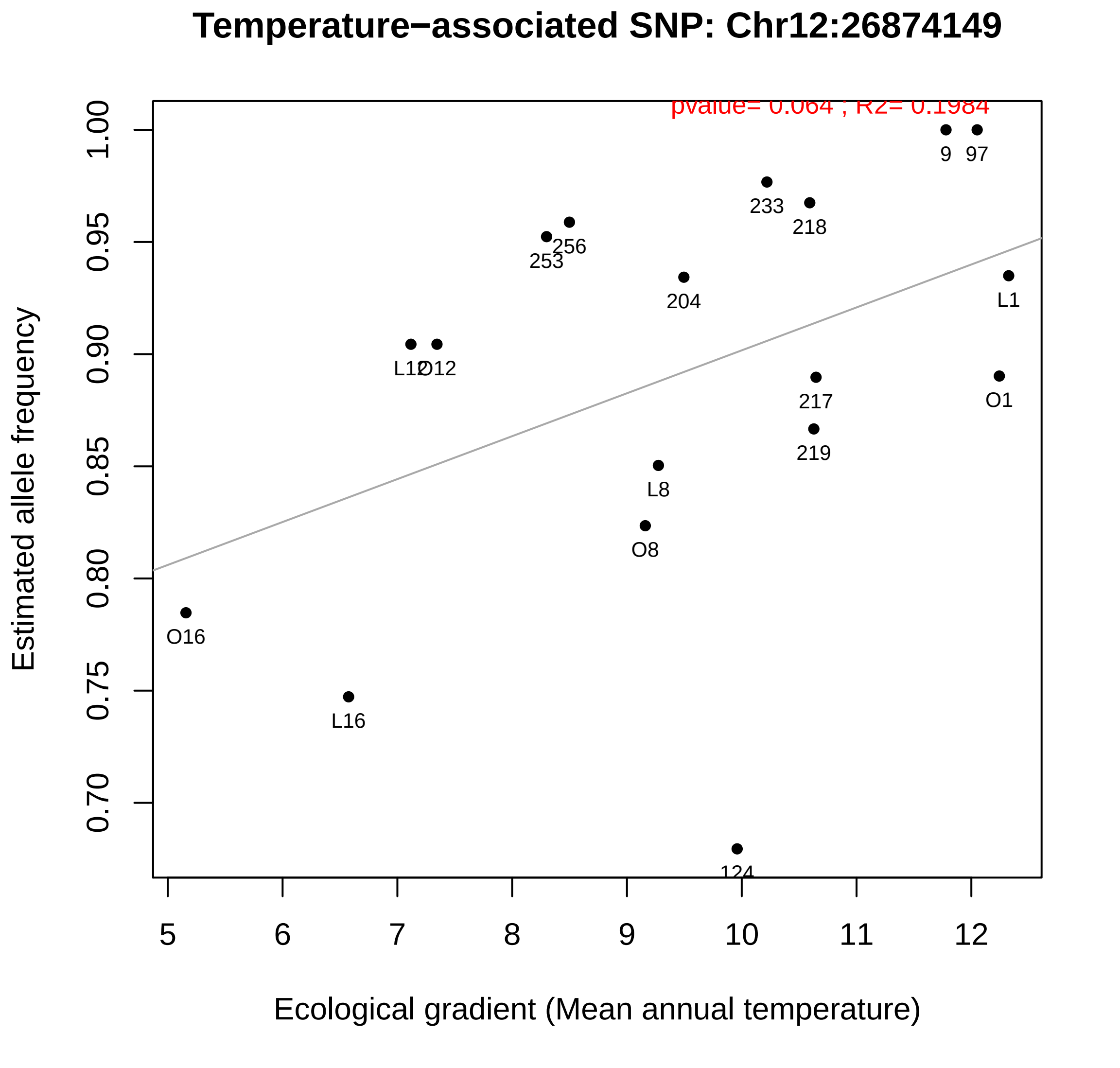
Temperature−associated SNP: Chr12:26874149
pvalue= 0.064 ; R2= 0.1984
Estimated allele frequency
Ecological gradient (Mean annual temperature)
1.00
0.95
0.90
0.85
0.80
0.75
0.70
5
6
7
8
9
10
11
12
9
97
233
218
256
253
204
L1
L12
O12
217
O1
219
L8
O8
O16
L16
124

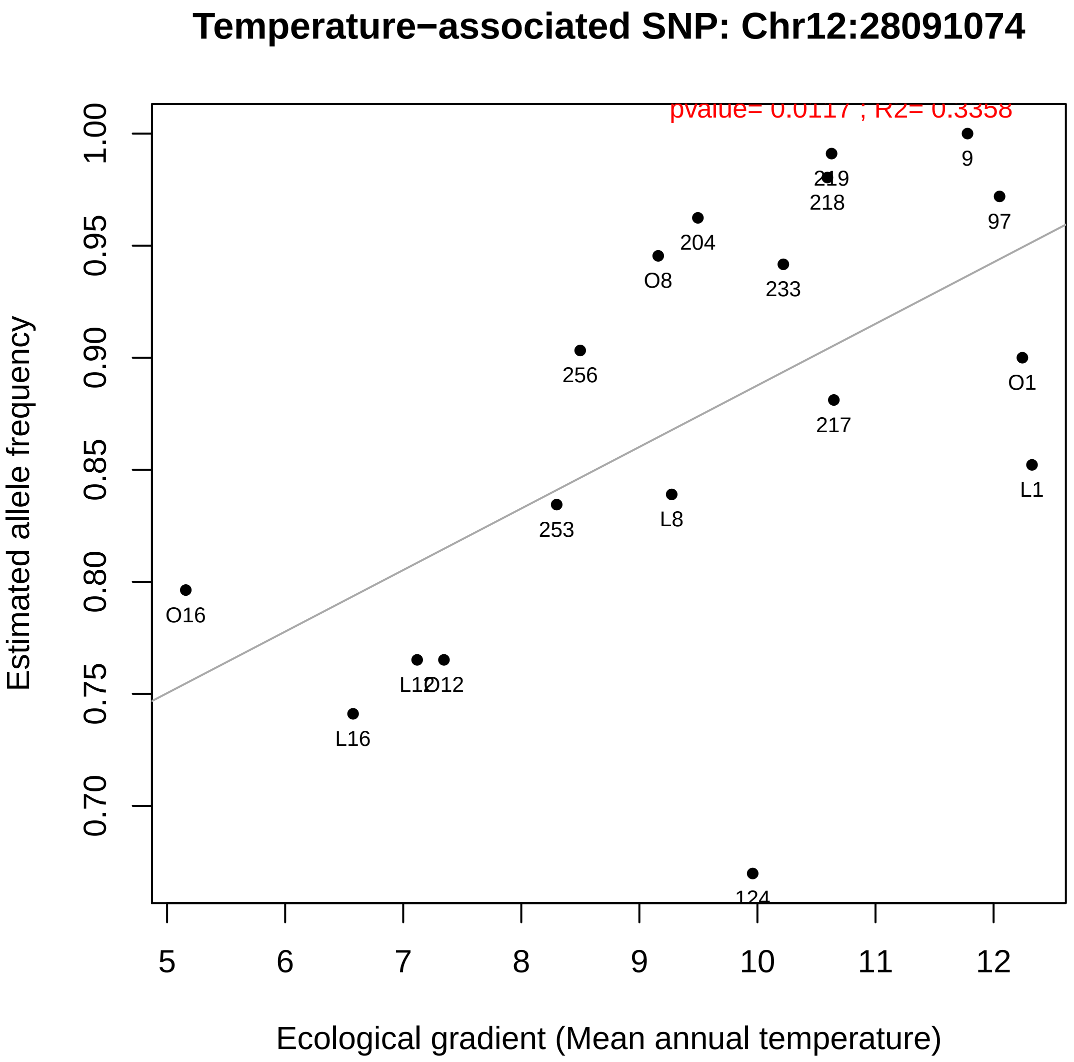

Temperature−associated SNP: Chr12:28091074
pvalue= 0.0117 ; R2= 0.3358
Estimated allele frequency
1.00
0.95
0.90
0.85
0.80
0.75
0.70
5
6
7
8
9
10
11
12
Ecological gradient (Mean annual temperature)
9
219
218
97
204
O8
233
256
O1
217
L1
253
L8
O16
L12
O12
L16
124

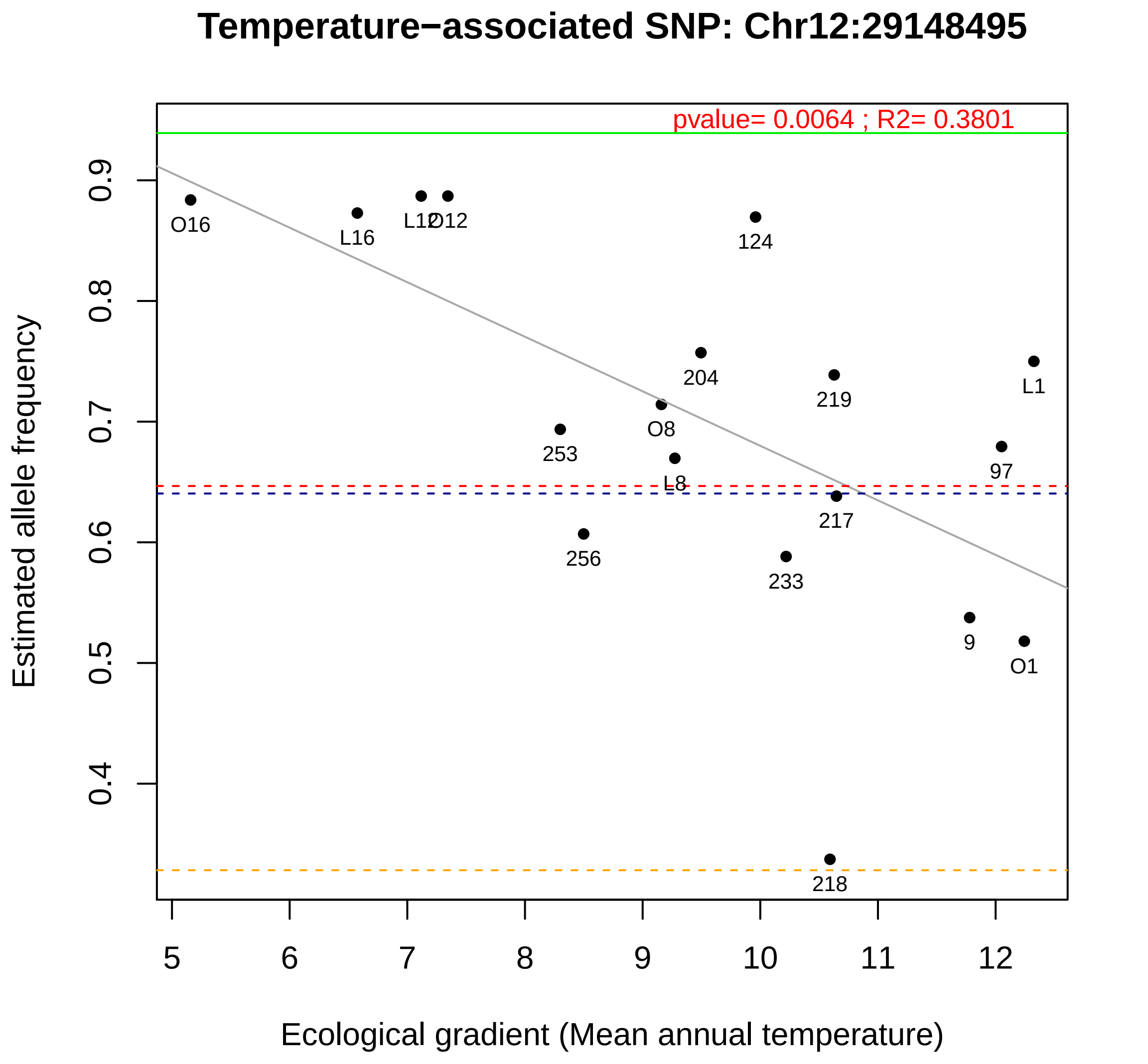
Temperature−associated SNP: Chr12:29148495
pvalue= 0.0064 ; R2= 0.3801
Estimated allele frequency
Ecological gradient (Mean annual temperature)
0.4
0.5
0.6
0.7
0.8
0.9
5
6
7
8
9
10
11
12
O16
L16
L12
O12
124
204
219
L1
O8
253
L8
97
217
256
233
9
O1
218

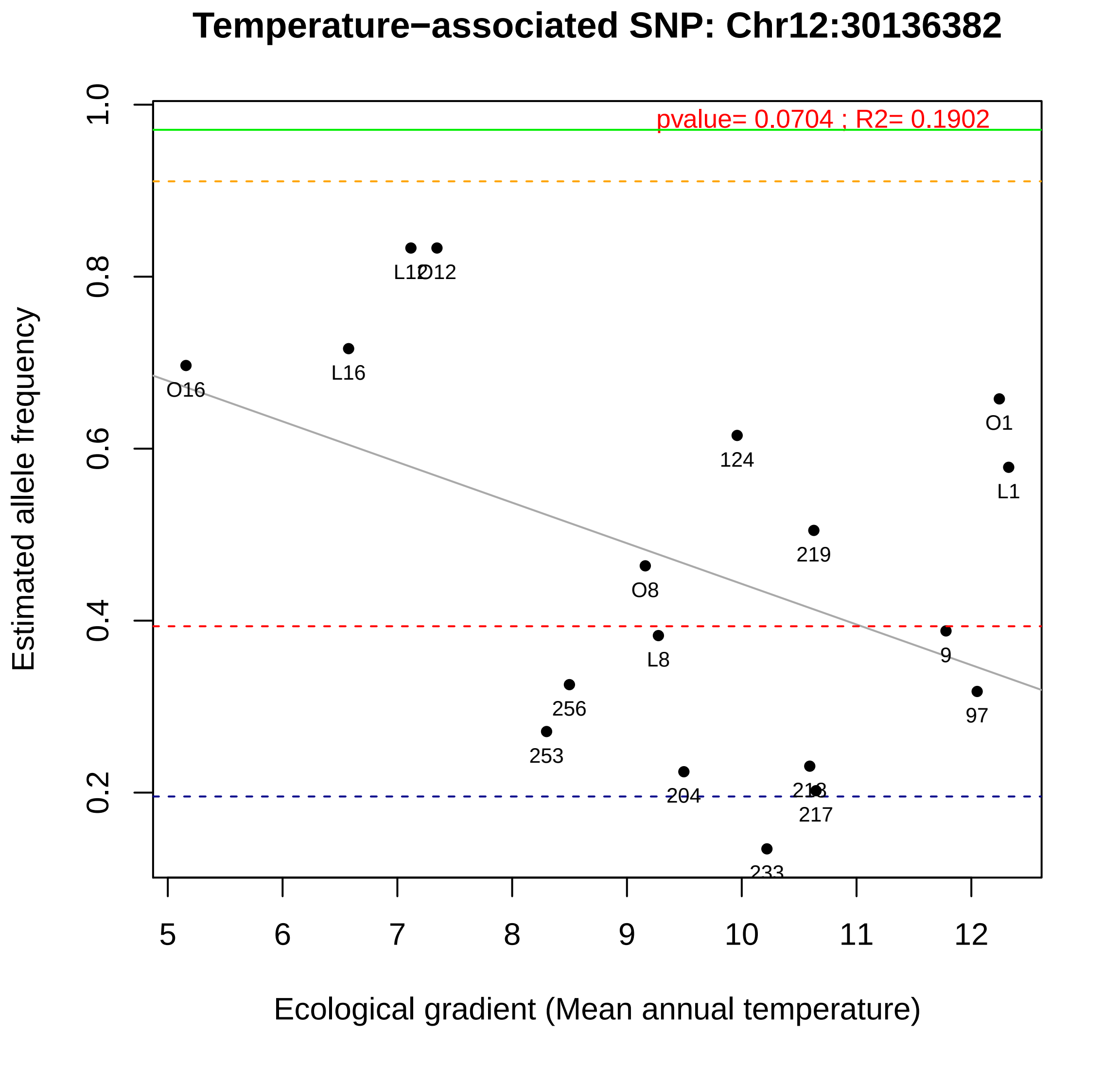

Temperature−associated SNP: Chr12:30136382
pvalue= 0.0704 ; R2= 0.1902
Estimated allele frequency
Ecological gradient (Mean annual temperature)
1.0
0.8
0.6
0.4
0.2
5
6
7
8
9
10
11
12
O16
L16
L12
O12
O8
L8
124
219
O1
L1
9
97
256
253
204
218
217
233

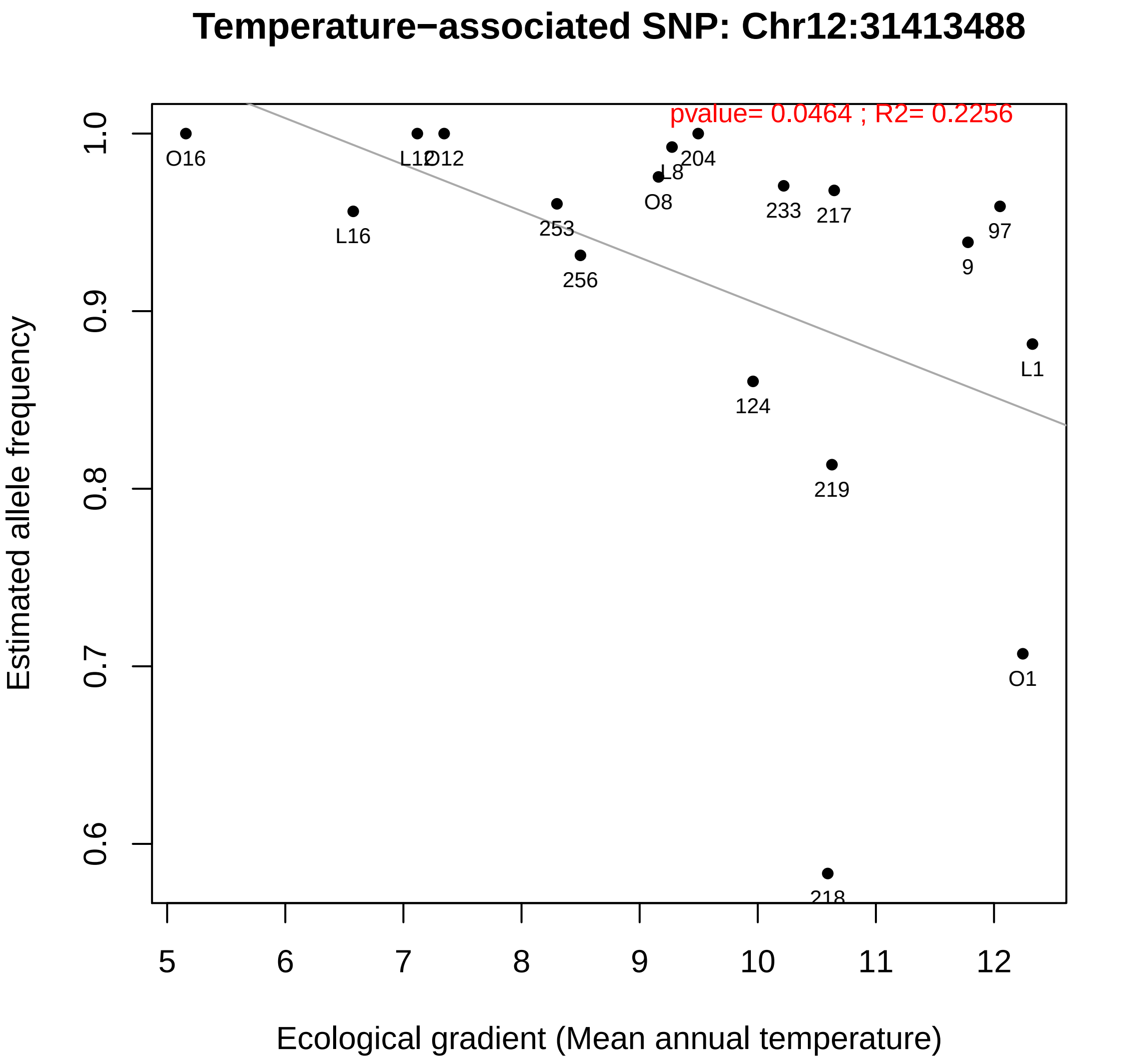

Temperature−associated SNP: Chr12:31413488
pvalue= 0.0464 ; R2= 0.2256
O16
L12
O12
204
L8
O8
L16
253
256
233
217
97
9
L1
124
219
O1
218
1.0
0.9
0.8
0.7
0.6
5
6
7
8
9
10
11
12
Estimated allele frequency
Ecological gradient (Mean annual temperature)

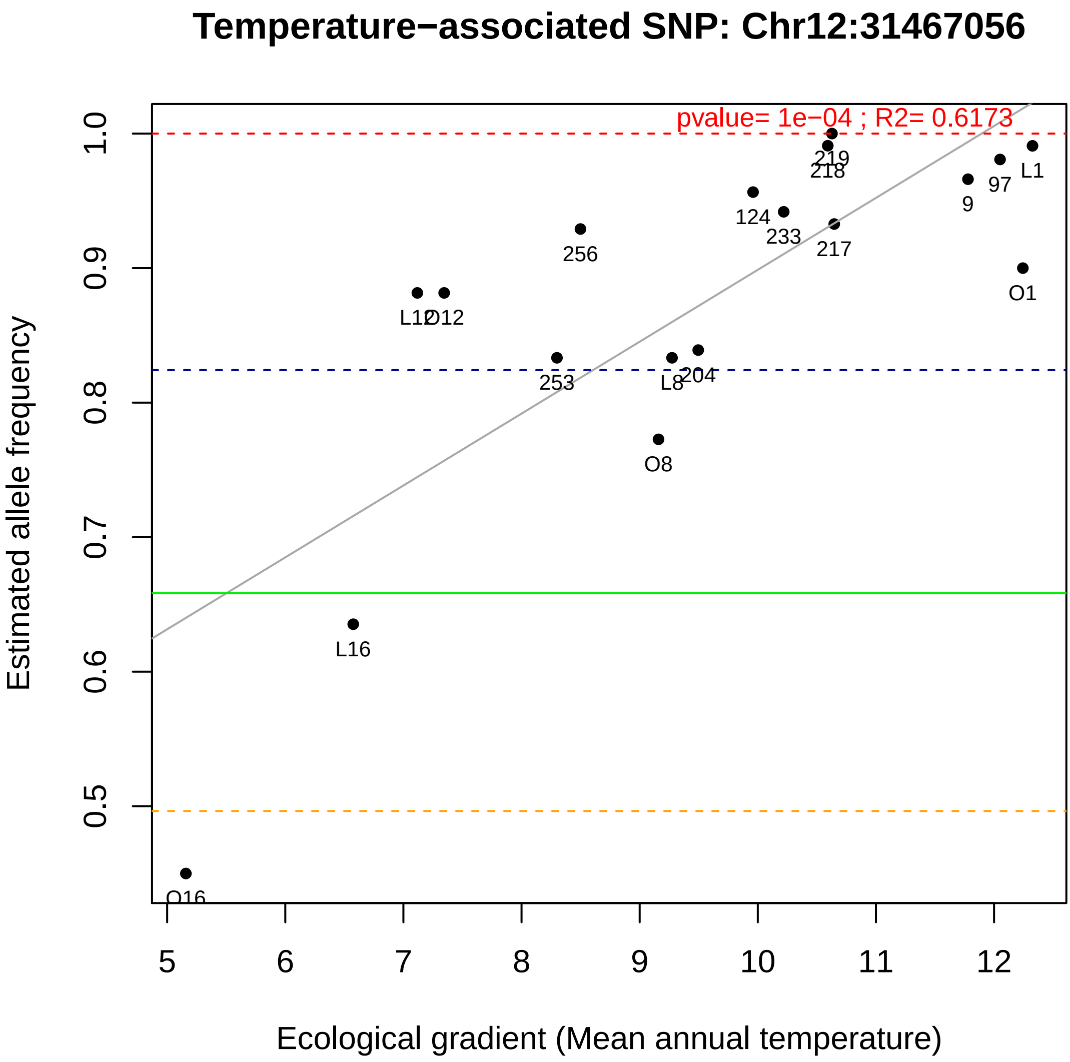

Temperature−associated SNP: Chr12:31467056
pvalue= 1e−04 ; R2= 0.6173
Estimated allele frequency
Ecological gradient (Mean annual temperature)
0.5
0.6
0.7
0.8
0.9
1.0
5
6
7
8
9
10
11
12
219
218
L1
97
9
124
233
217
256
O1
L12
O12
253
L8
204
O8
L16
O16

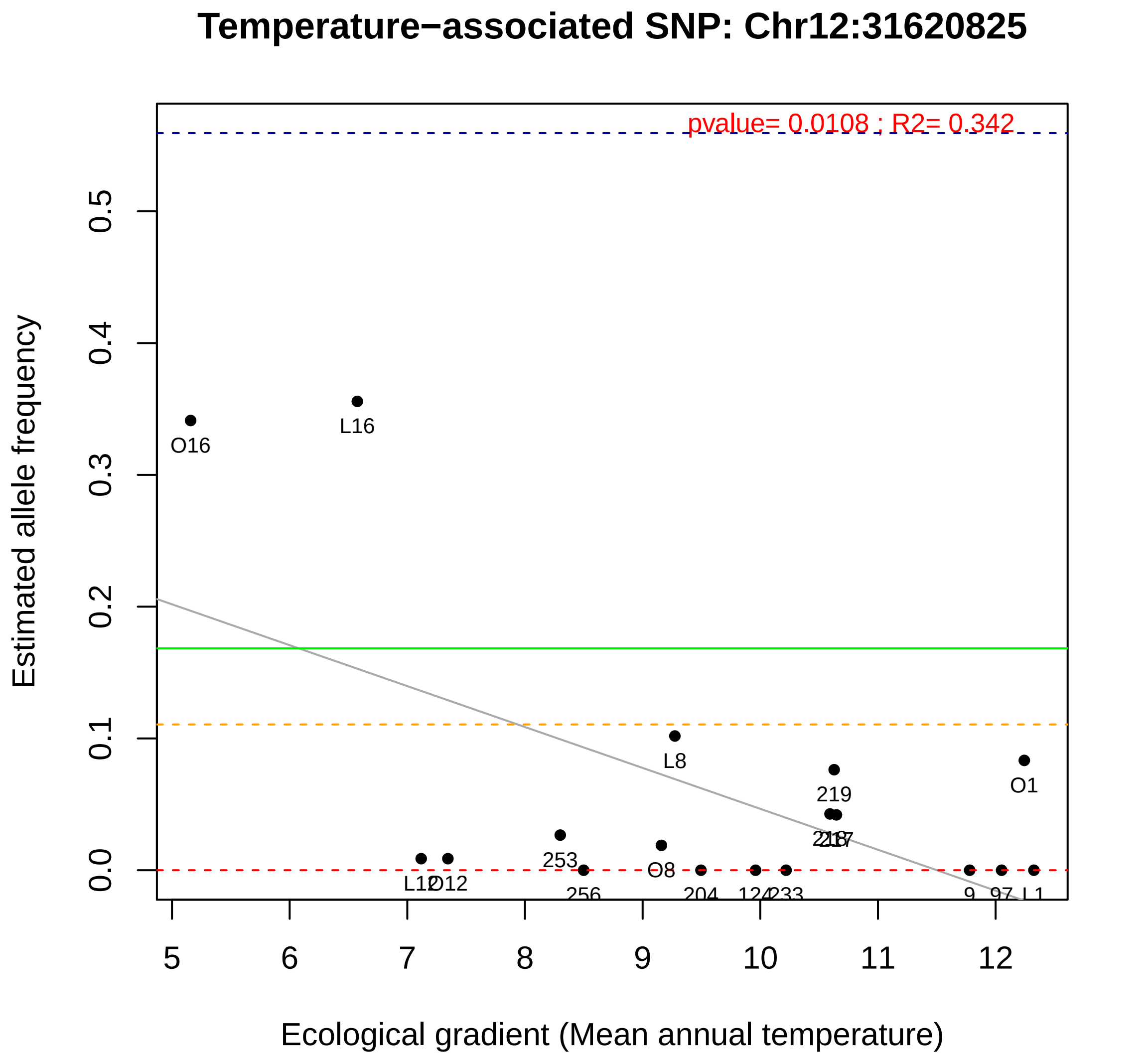

Temperature−associated SNP: Chr12:31620825
pvalue= 0.0108 ; R2= 0.342
Estimated allele frequency
0.0
0.1
0.2
0.3
0.4
0.5
Ecological gradient (Mean annual temperature)
5
6
7
8
9
10
11
12
O16
L16
L8
219
O1
253
O8
L12
O12
256
204
233
9
97
L1

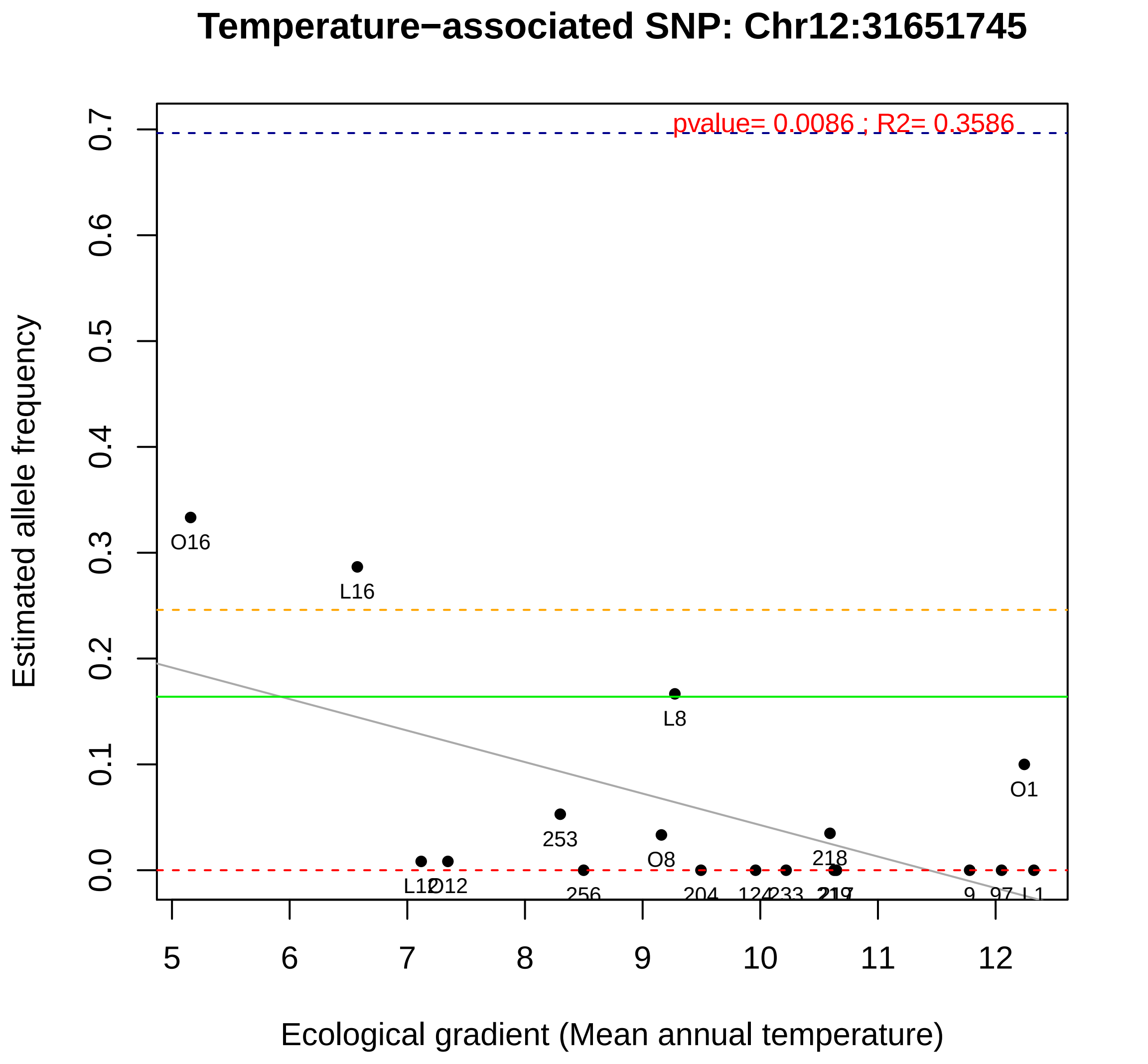

Temperature−associated SNP: Chr12:31651745
pvalue= 0.0086 ; R2= 0.3586
Estimated allele frequency
Ecological gradient (Mean annual temperature)
0.0
0.1
0.2
0.3
0.4
0.5
0.6
0.7
5
6
7
8
9
10
11
12
O16
L16
L8
O1
253
O8
218
L12
O12
256
204
9
97
L1

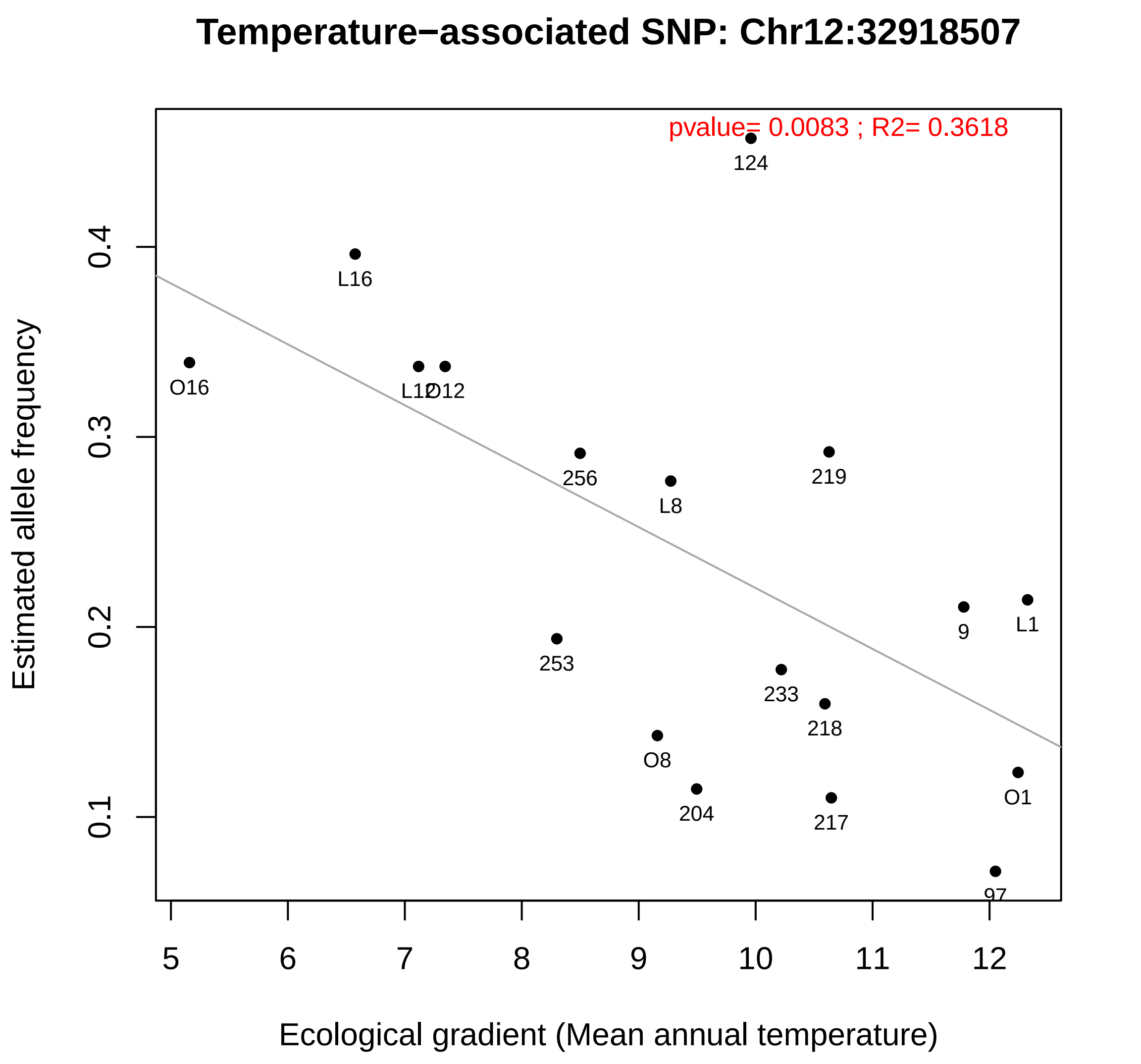

Temperature−associated SNP: Chr12:32918507
pvalue= 0.0083 ; R2= 0.3618
Estimated allele frequency
Ecological gradient (Mean annual temperature)
0.1
0.2
0.3
0.4
5
6
7
8
9
10
11
12
124
L16
O16
L12
O12
256
L8
219
9
L1
253
233
218
O8
O1
204
217
97

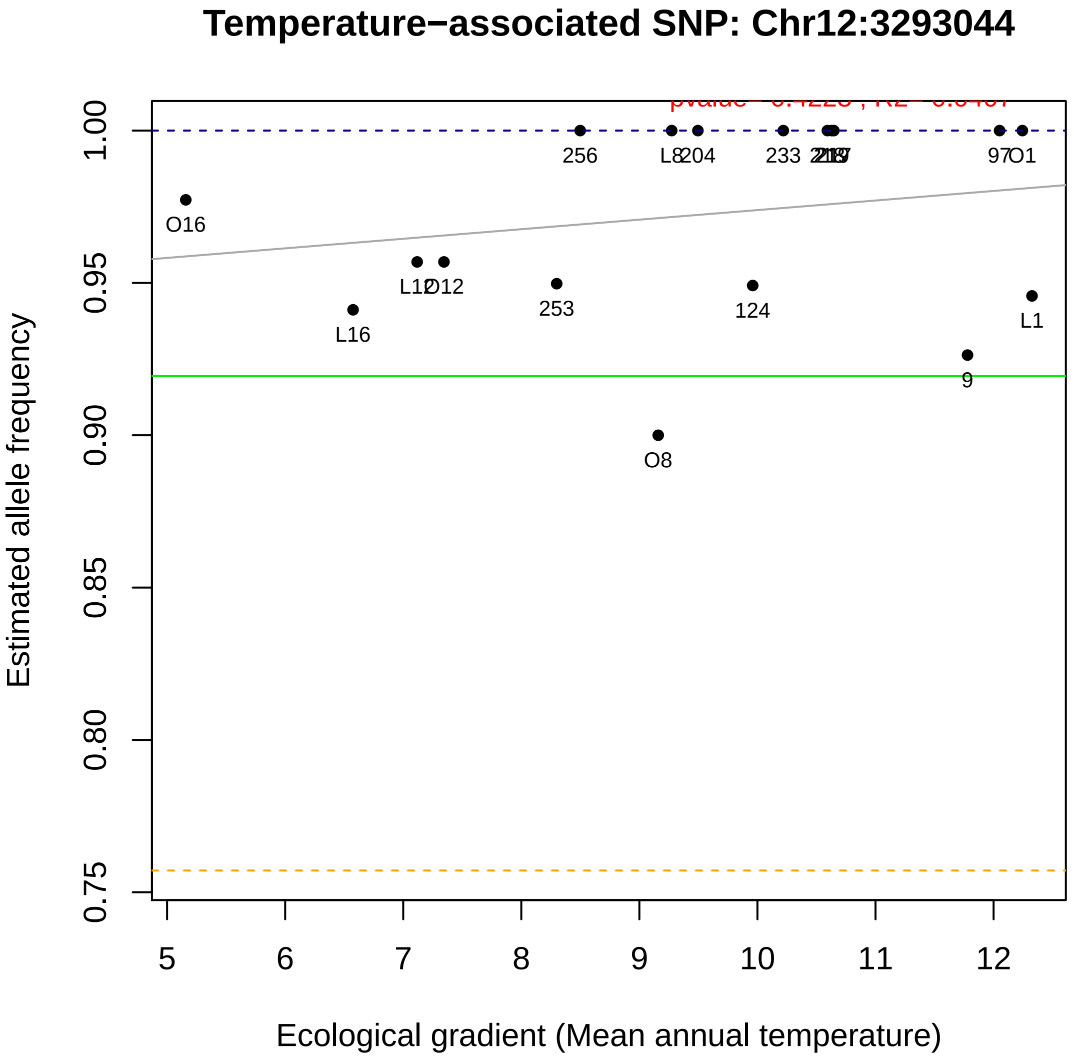
Temperature−associated SNP: Chr12:3293044
Estimated allele frequency
Ecological gradient (Mean annual temperature)
0.75
0.80
0.85
0.90
0.95
1.00
5
6
7
8
9
10
11
12
O16
L16
L12
O12
253
256
O8
L8
204
124
233
9
97
O1
L1

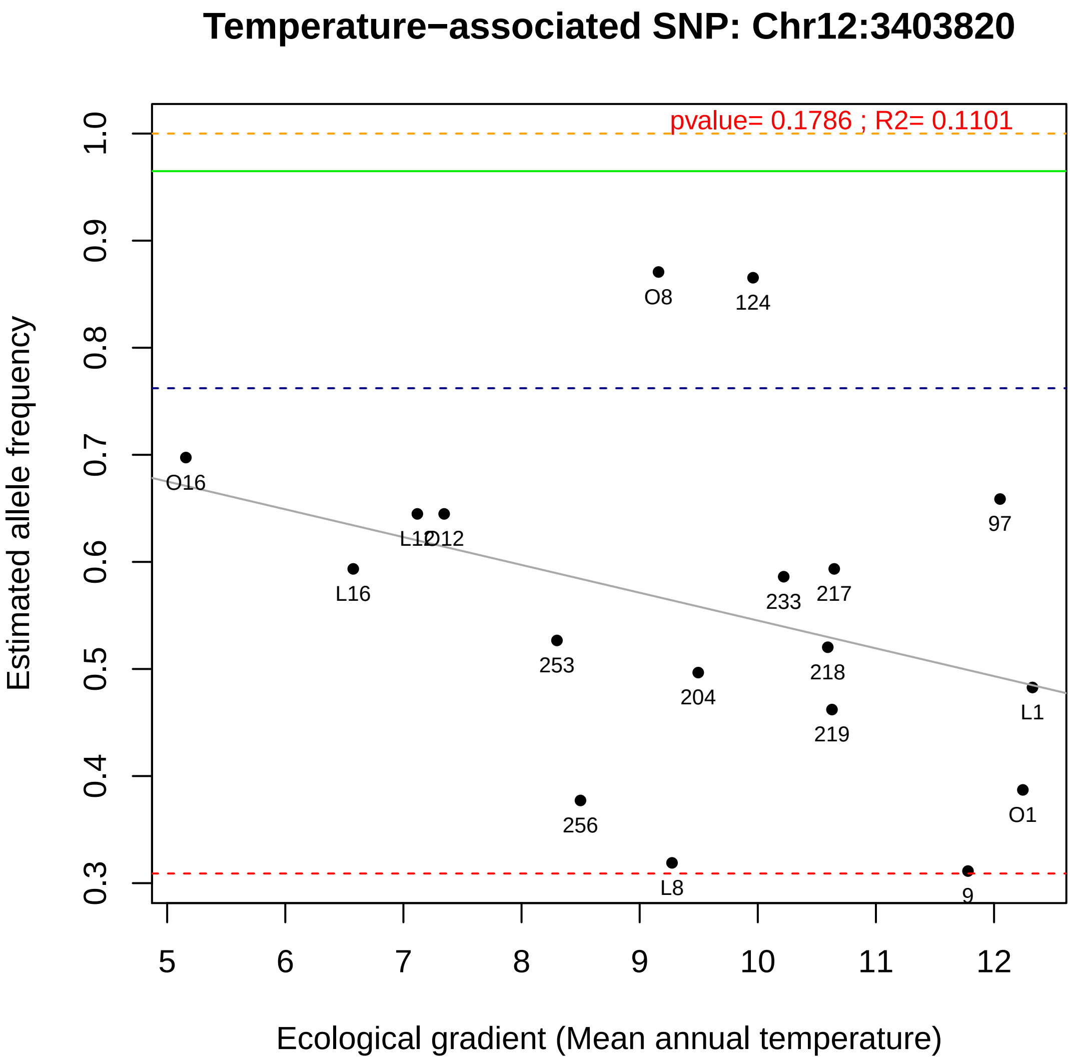
Temperature−associated SNP: Chr12:3403820
pvalue= 0.1786 ; R2= 0.1101
Estimated allele frequency
Ecological gradient (Mean annual temperature)
1.0
0.9
0.8
0.7
0.6
0.5
0.4
0.3
5
6
7
8
9
10
11
12
O8
124
O16
L12
O12
97
L16
233
217
253
218
204
L1
219
256
O1
L8
9

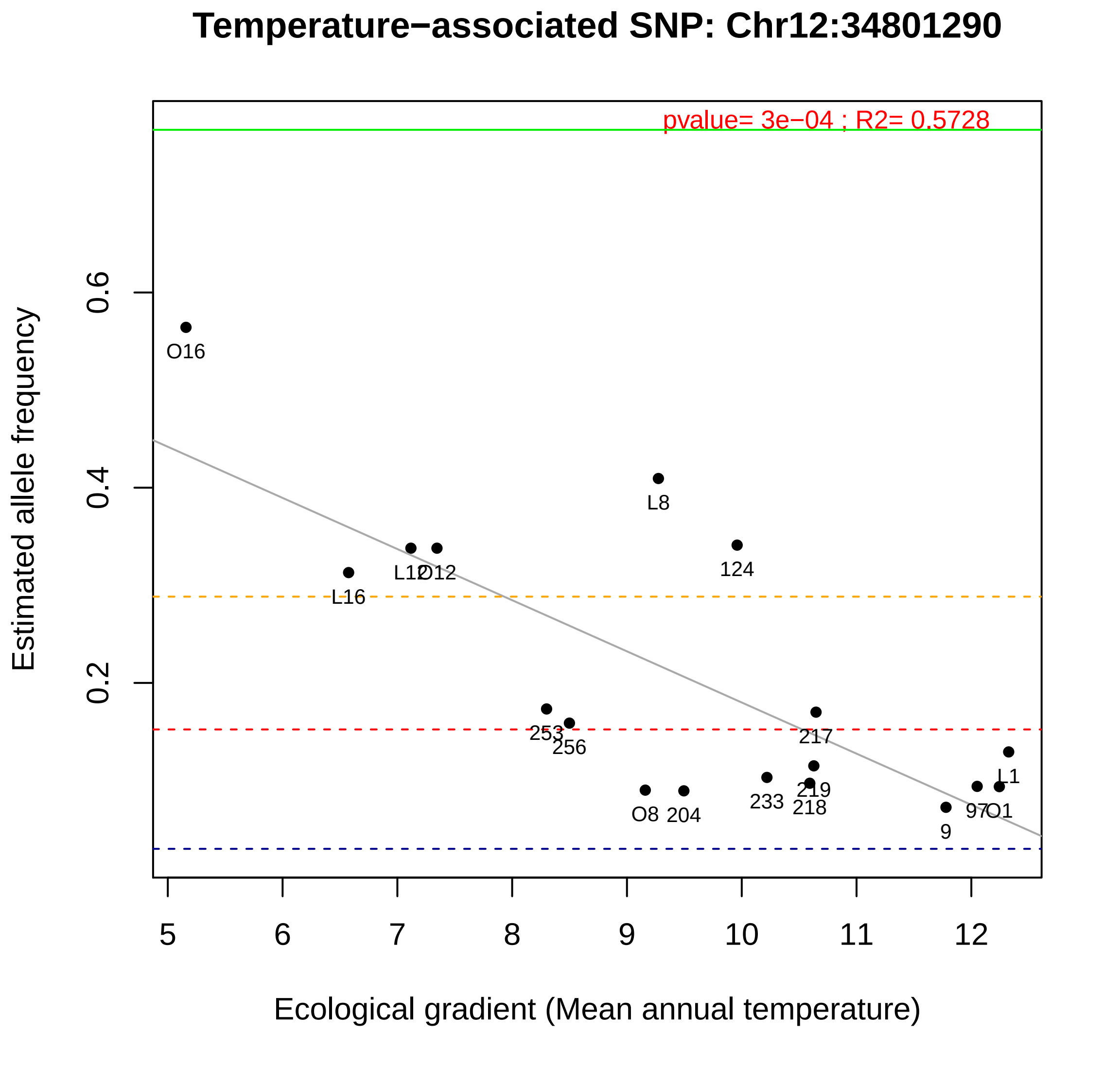
Temperature−associated SNP: Chr12:34801290
pvalue= 3e−04 ; R2= 0.5728
Estimated allele frequency
0.2
0.4
0.6
O16
L8
L12
O12
124
L16
253
256
217
L1
219
233
218
O8
204
97
O1
9
5
6
7
8
9
10
11
12
Ecological gradient (Mean annual temperature)

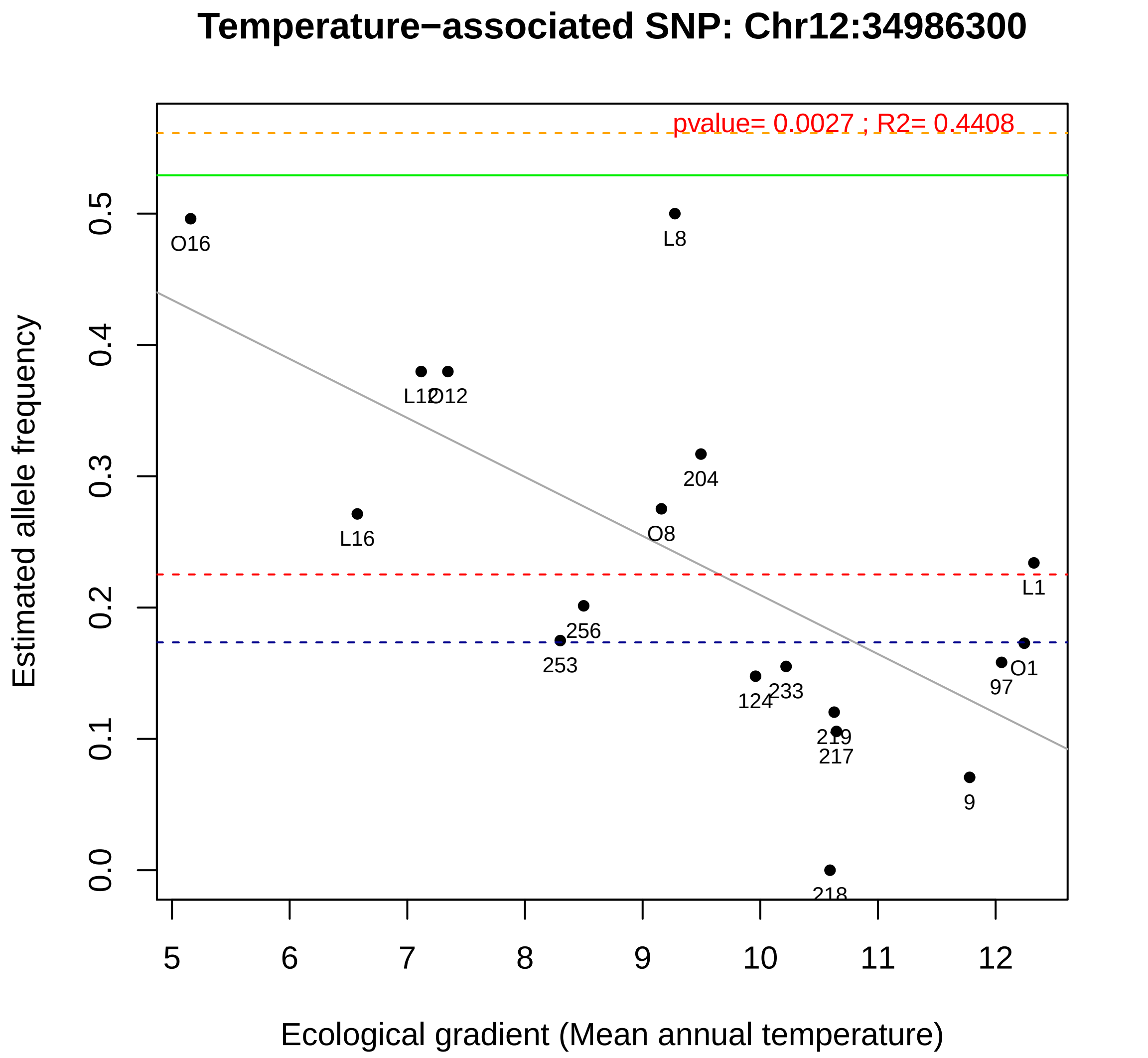

Temperature−associated SNP: Chr12:34986300
pvalue= 0.0027 ; R2= 0.4408
Estimated allele frequency
0.0
0.1
0.2
0.3
0.4
0.5
5
6
7
8
9
10
11
12
Ecological gradient (Mean annual temperature)
O16
L8
L12
O12
204
L16
O8
L1
256
253
O1
97
124
233
219
217
9
218

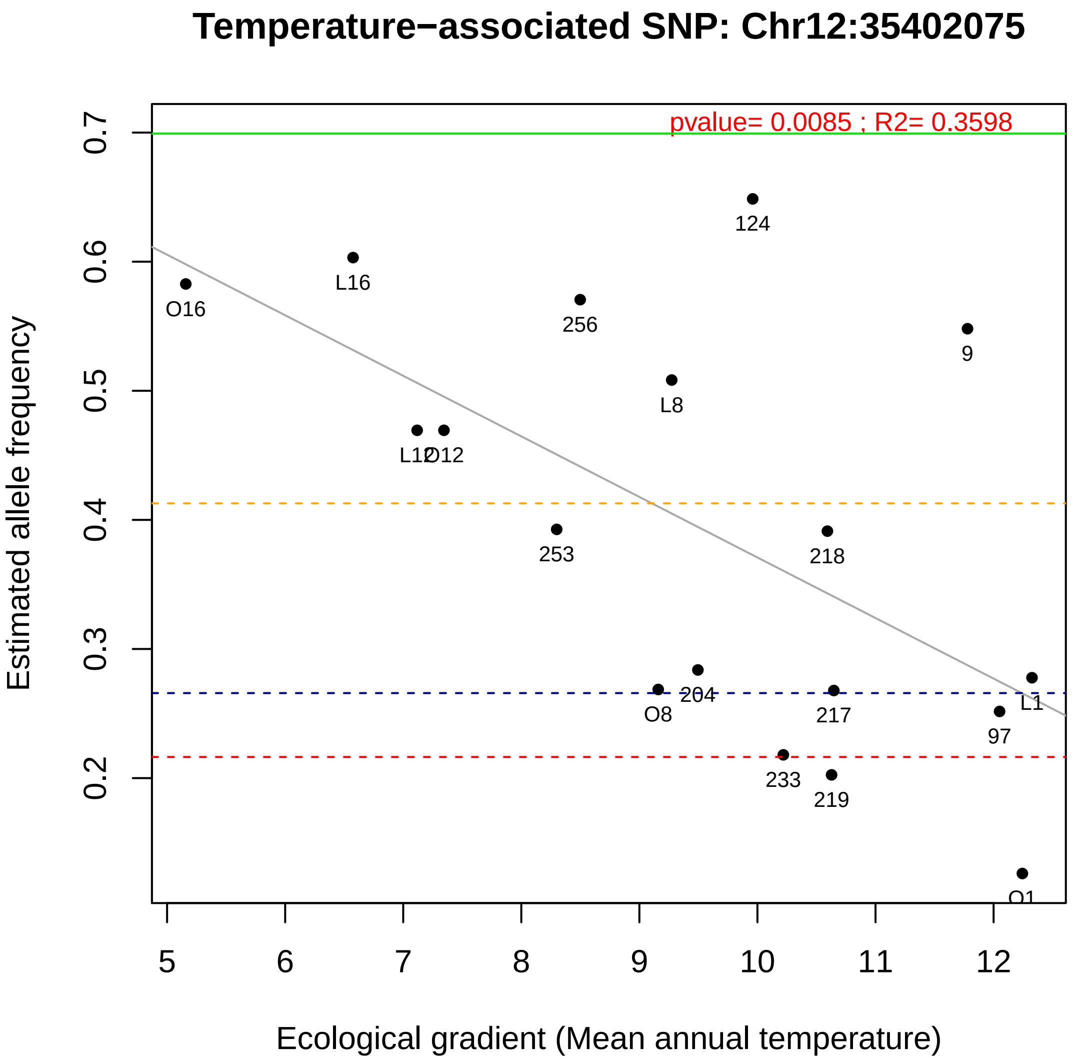

Temperature−associated SNP: Chr12:35402075
pvalue= 0.0085 ; R2= 0.3598
Estimated allele frequency
Ecological gradient (Mean annual temperature)
0.2
0.3
0.4
0.5
0.6
0.7
5
6
7
8
9
10
11
12
O16
L16
256
124
9
L8
L12
O12
253
218
O8
204
217
L1
97
233
219
O1

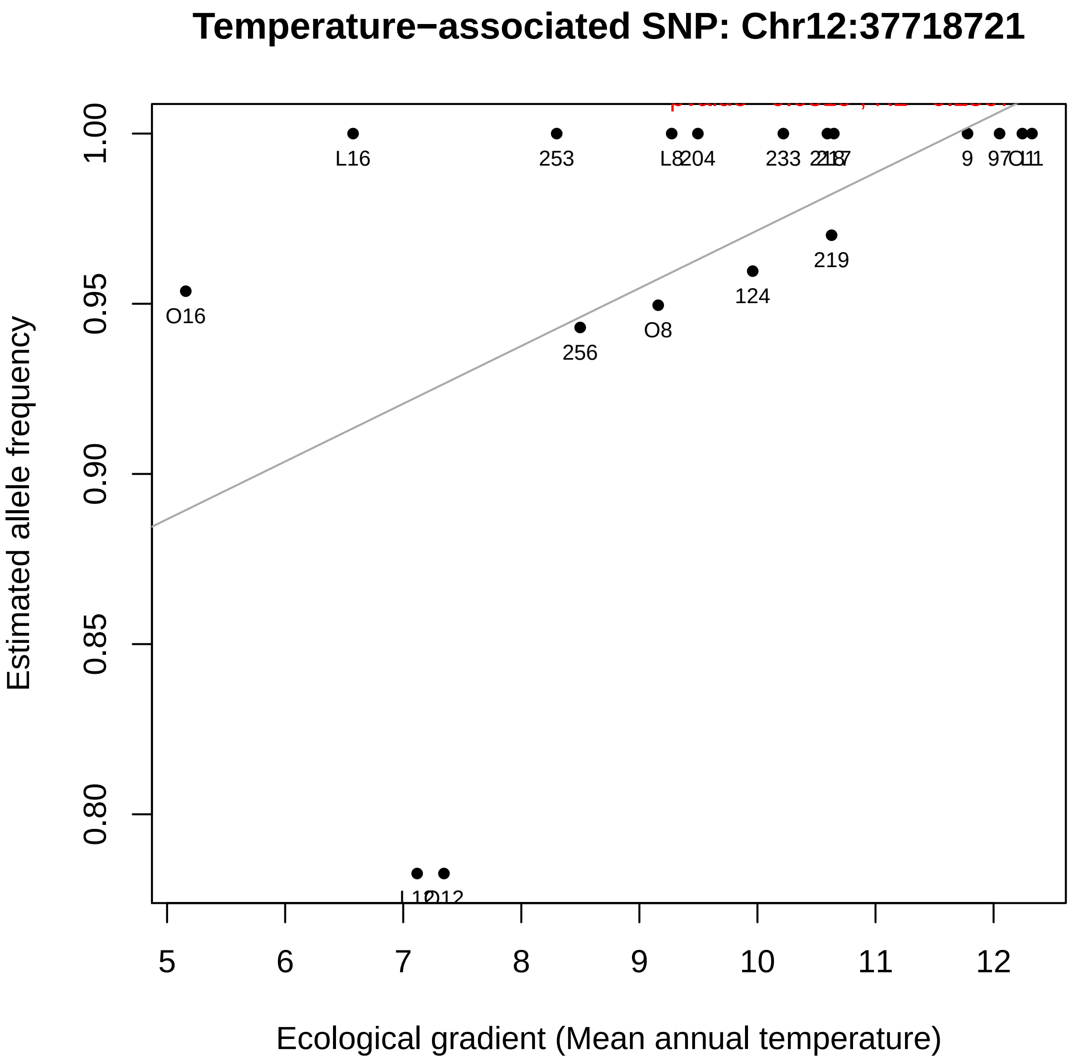
Temperature−associated SNP: Chr12:37718721
Estimated allele frequency
Ecological gradient (Mean annual temperature)
0.80
0.85
0.90
0.95
1.00
5
6
7
8
9
10
11
12
L16
253
L8
204
233
9
O16
256
O8
124
219
O12

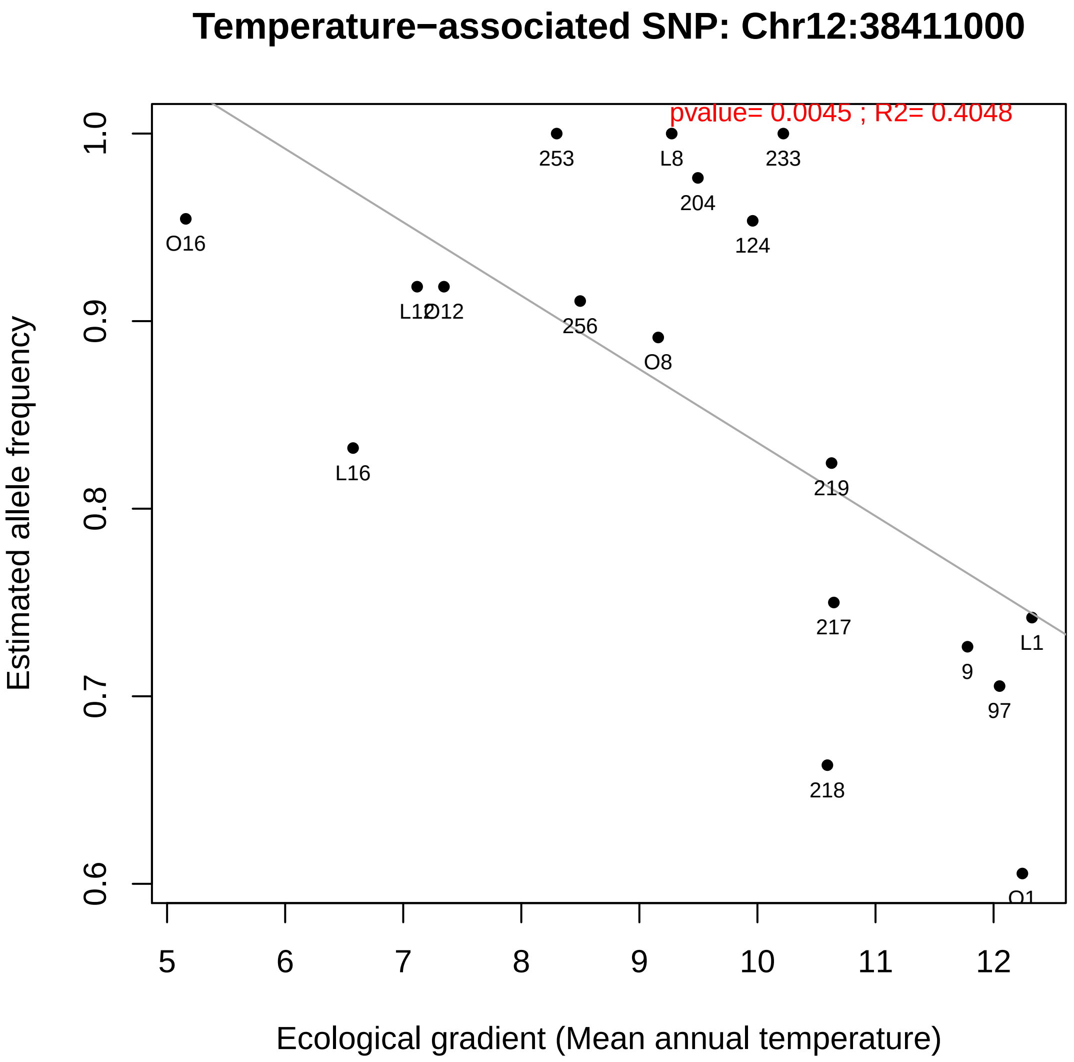

Temperature−associated SNP: Chr12:38411000
pvalue= 0.0045 ; R2= 0.4048
Estimated allele frequency
Ecological gradient (Mean annual temperature)
1.0
0.9
0.8
0.7
0.6
5
6
7
8
9
10
11
12
O16
L16
L12
O12
253
256
L8
204
O8
233
124
219
217
218
9
97
L1
O1

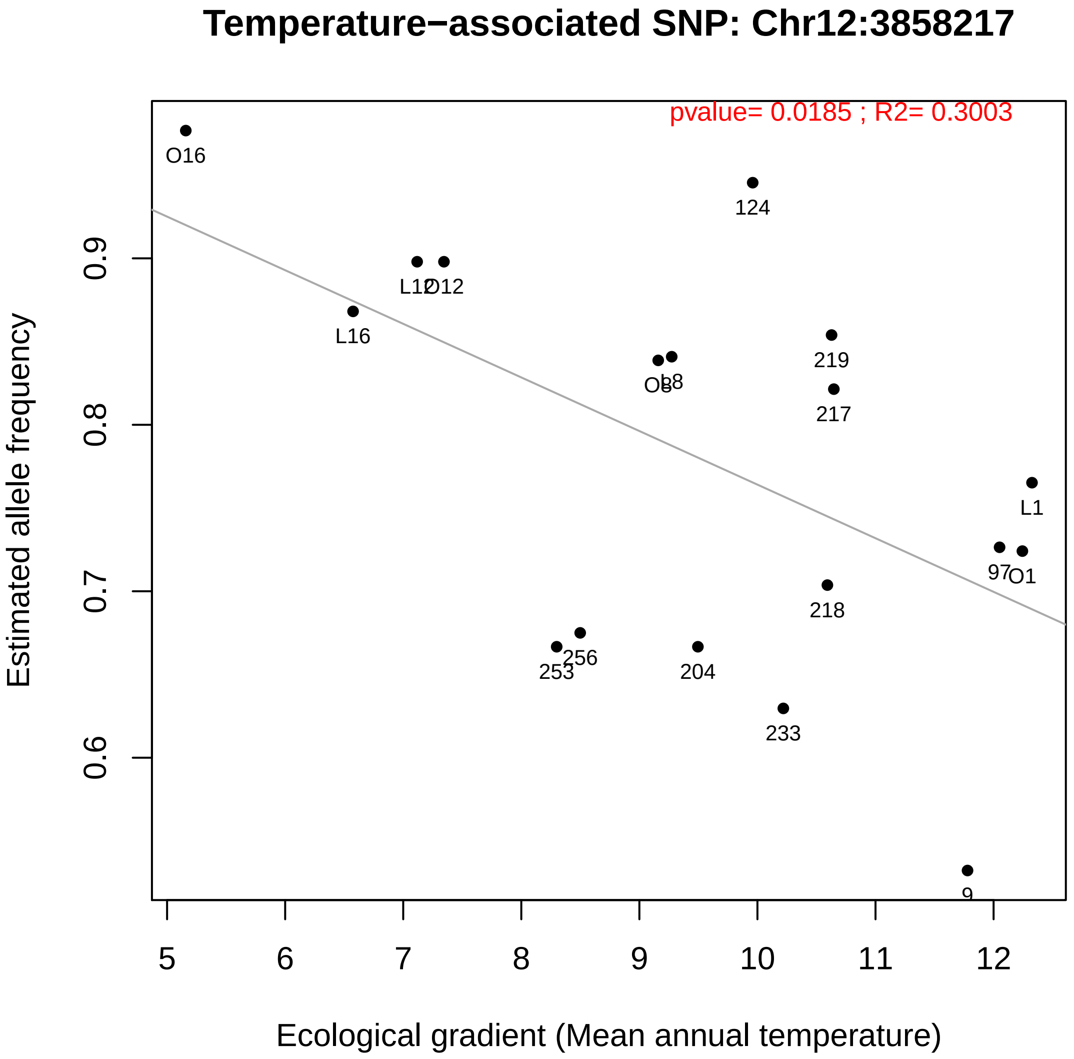

Temperature−associated SNP: Chr12:3858217
pvalue= 0.0185 ; R2= 0.3003
O16
124
L12
O12
L16
219
O8
L8
217
L1
97
O1
218
256
253
204
233
9
0.6
0.7
0.8
0.9
5
6
7
8
9
10
11
12
Estimated allele frequency
Ecological gradient (Mean annual temperature)

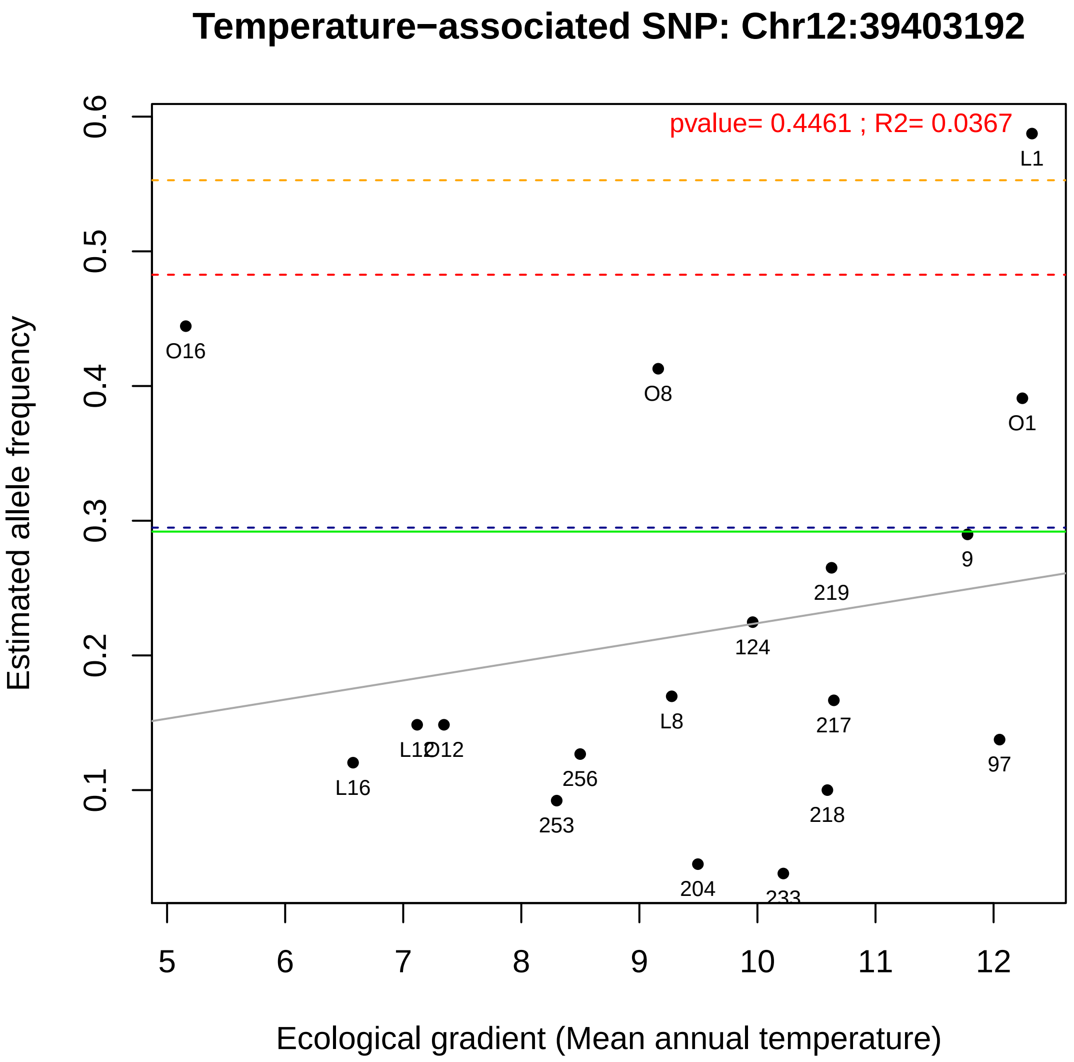

Temperature−associated SNP: Chr12:39403192
pvalue= 0.4461 ; R2= 0.0367
Estimated allele frequency
0.1
0.2
0.3
0.4
0.5
0.6
Ecological gradient (Mean annual temperature)
5
6
7
8
9
10
11
12
L1
O16
O8
O1
9
219
124
L8
217
97
L12
O12
L16
256
253
218
204
233

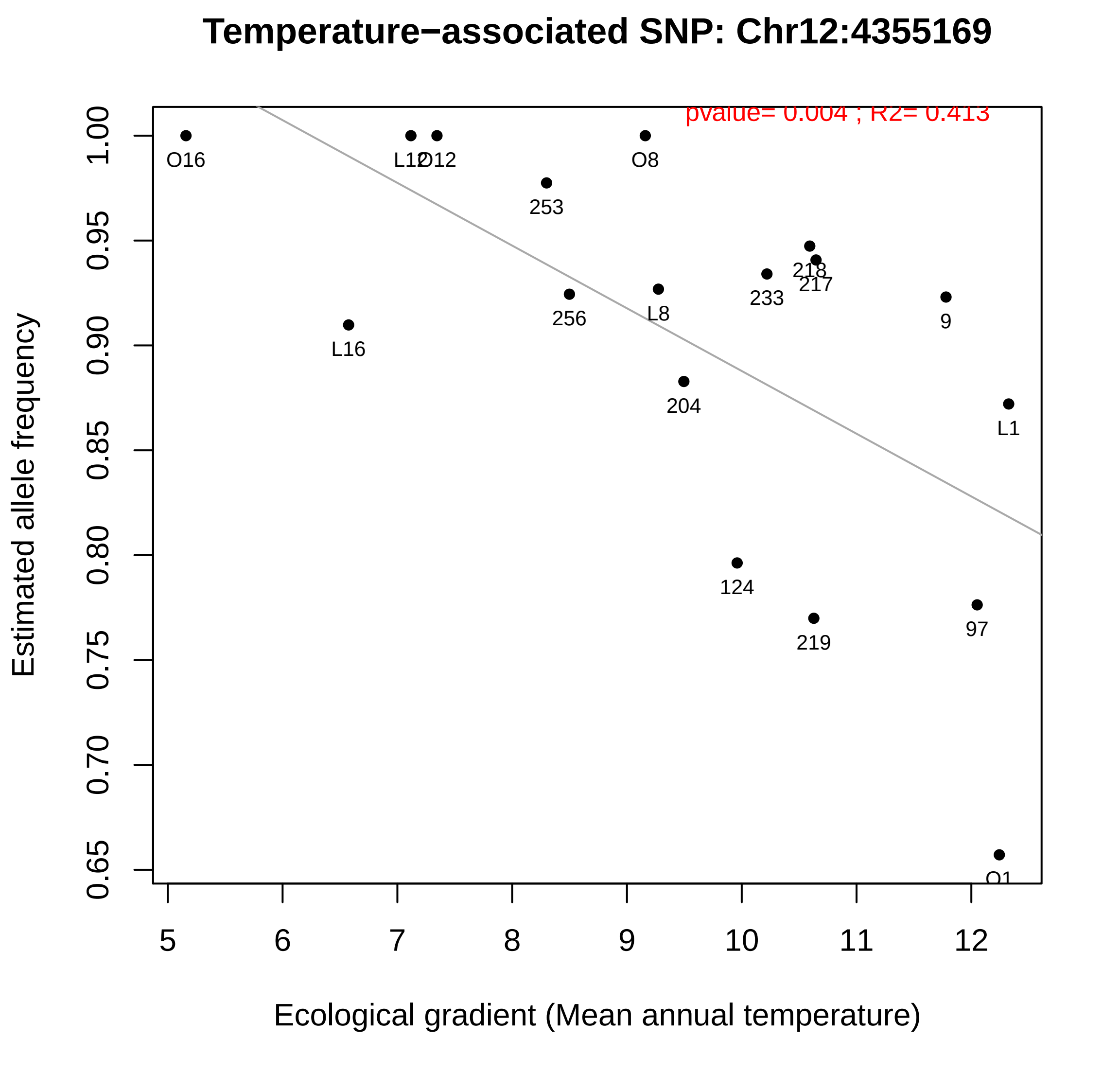
Temperature−associated SNP: Chr12:4355169
pvalue= 0.004 ; R2= 0.413
Estimated allele frequency
1.00
0.95
0.90
0.85
0.80
0.75
0.70
0.65
5
6
7
8
9
10
11
12
Ecological gradient (Mean annual temperature)
O16
L12
O12
O8
253
218
217
233
256
L8
9
L16
204
L1
124
97
219

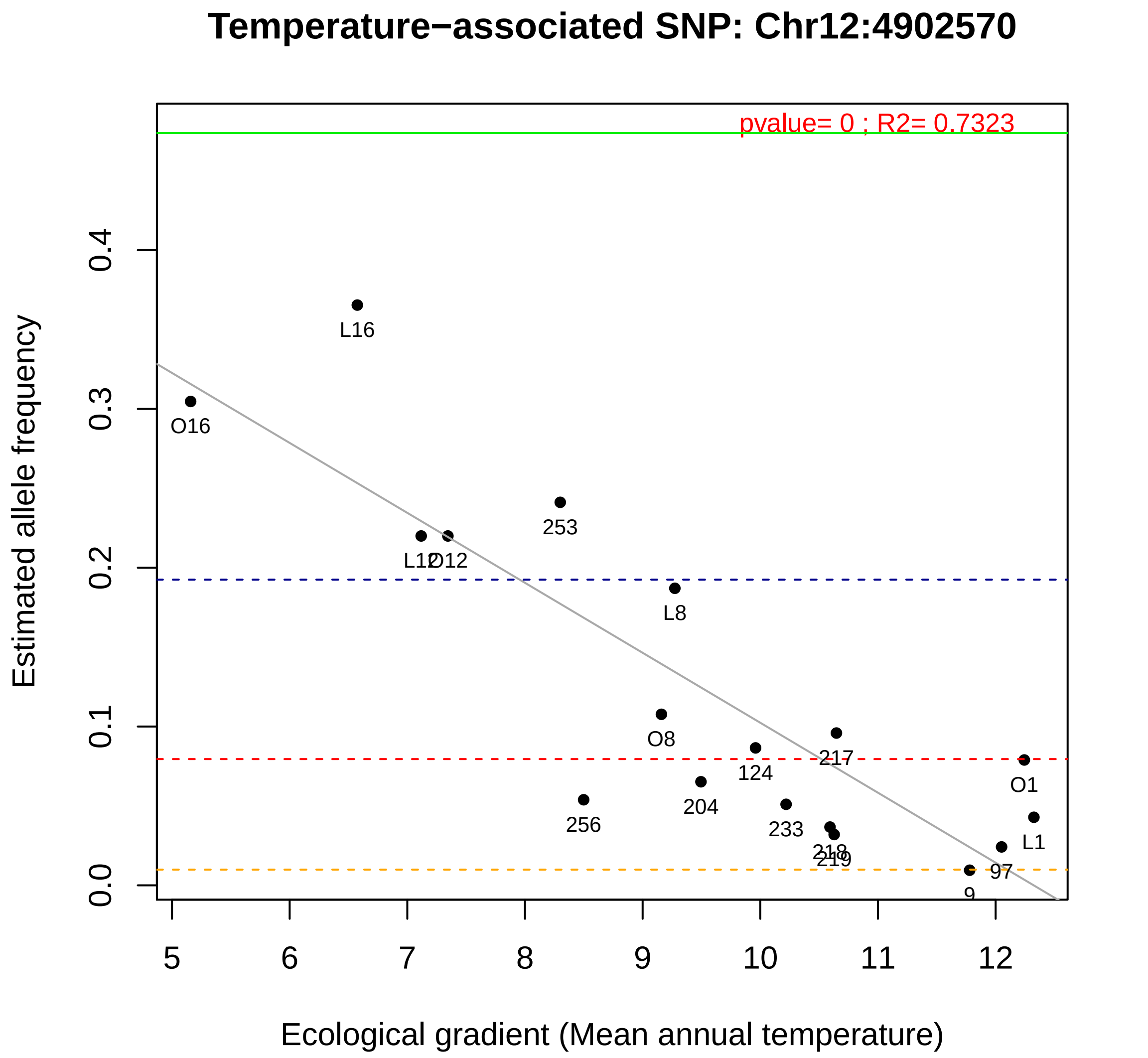

Temperature−associated SNP: Chr12:4902570
pvalue= 0 ; R2= 0.7323
Estimated allele frequency
0.0
0.1
0.2
0.3
0.4
5
6
7
8
9
10
11
12
Ecological gradient (Mean annual temperature)
L16
O16
253
L12
O12
L8
O8
217
124
O1
204
256
233
L1
218
219
97
9

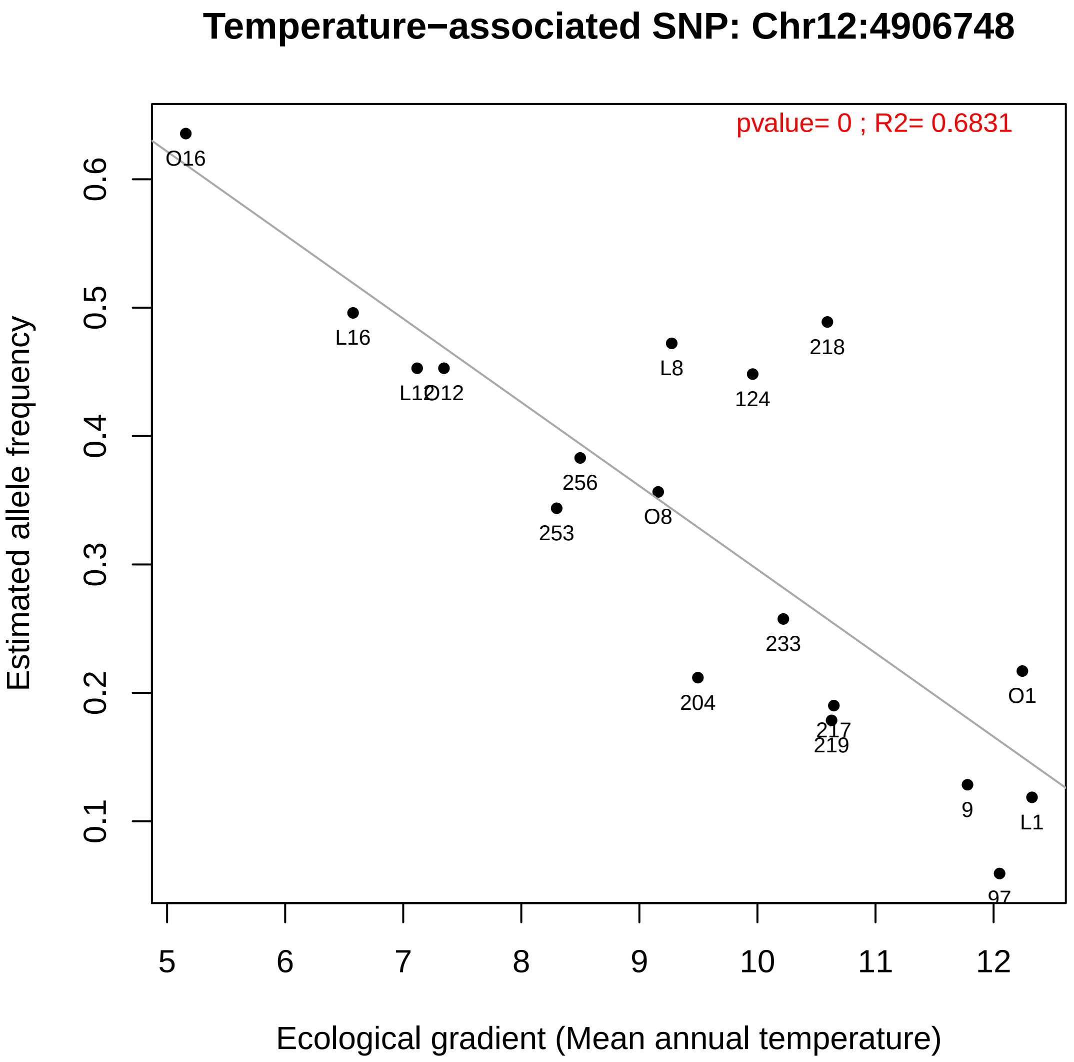
Temperature−associated SNP: Chr12:4906748
pvalue= 0 ; R2= 0.6831
Estimated allele frequency
Ecological gradient (Mean annual temperature)
0.1
0.2
0.3
0.4
0.5
0.6
5
6
7
8
9
10
11
12
O16
L16
L12
O12
256
253
O8
L8
124
218
233
204
217
219
O1
9
L1
97

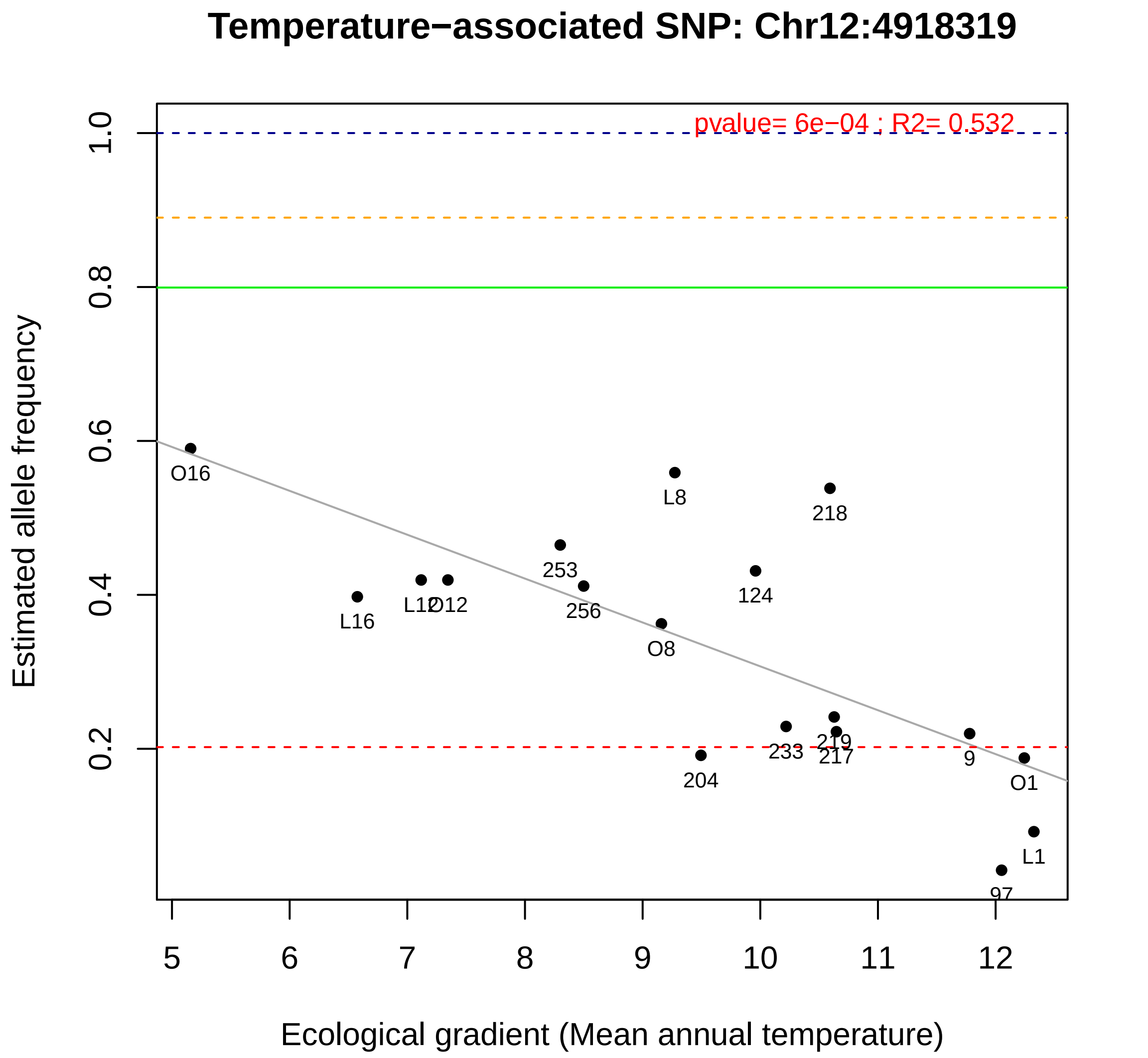

Temperature−associated SNP: Chr12:4918319
pvalue= 6e−04 ; R2= 0.532
Estimated allele frequency
1.0
0.8
0.6
0.4
0.2
O16
L8
218
253
124
L16
L12
O12
256
O8
233
219
217
9
204
O1
L1
97
5
6
7
8
9
10
11
12
Ecological gradient (Mean annual temperature)

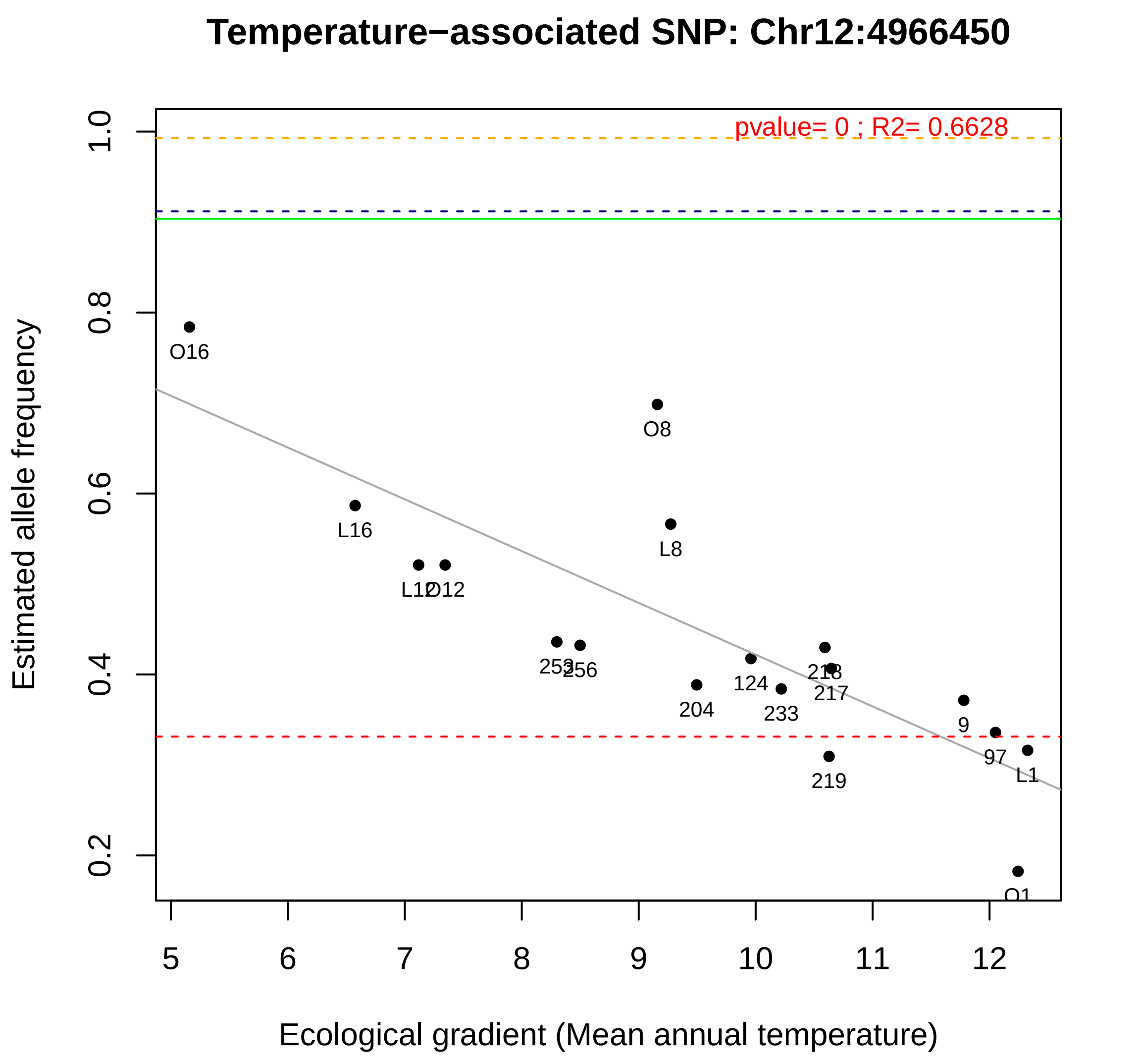

Temperature−associated SNP: Chr12:4966450
pvalue= 0 ; R2= 0.6628
Estimated allele frequency
0.2
0.4
0.6
0.8
1.0
5
6
7
8
9
10
11
12
Ecological gradient (Mean annual temperature)
O16
O8
L16
L8
L12
O12
253
256
204
124
233
218
217
219
9
97
L1
O1

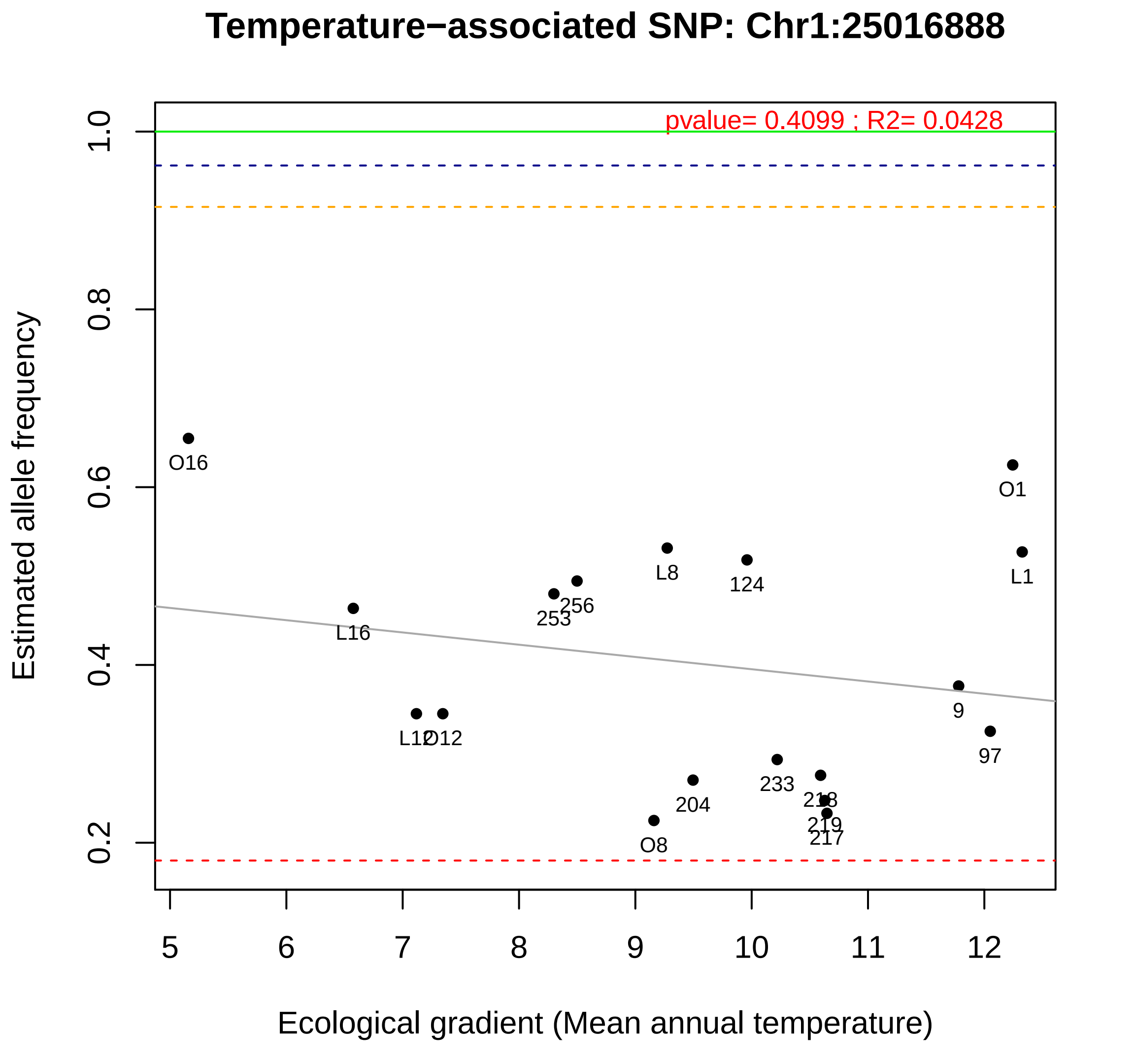

Temperature−associated SNP: Chr1:25016888
pvalue= 0.4099 ; R2= 0.0428
Estimated allele frequency
0.2
0.4
0.6
0.8
1.0
Ecological gradient (Mean annual temperature)
5
6
7
8
9
10
11
12
O16
O1
L1
L8
124
256
253
L16
9
97
L12
O12
233
218
204
219
217
O8

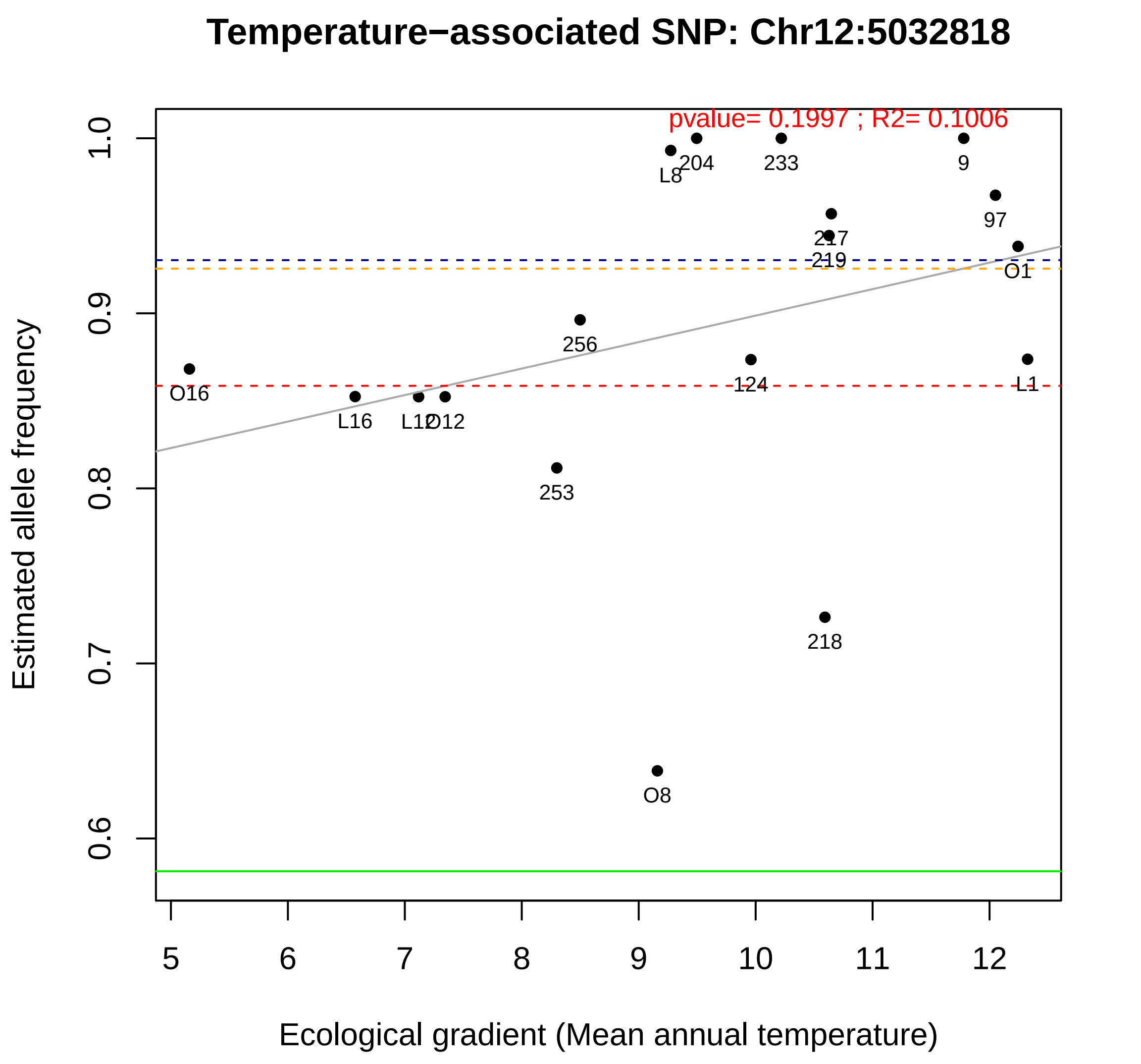
Temperature−associated SNP: Chr12:5032818
pvalue= 0.1997 ; R2= 0.1006
Estimated allele frequency
Ecological gradient (Mean annual temperature)
0.6
0.7
0.8
0.9
1.0
5
6
7
8
9
10
11
12
L8
204
233
9
97
217
219
O1
256
124
L1
O16
L16
L12
O12
253
218
O8

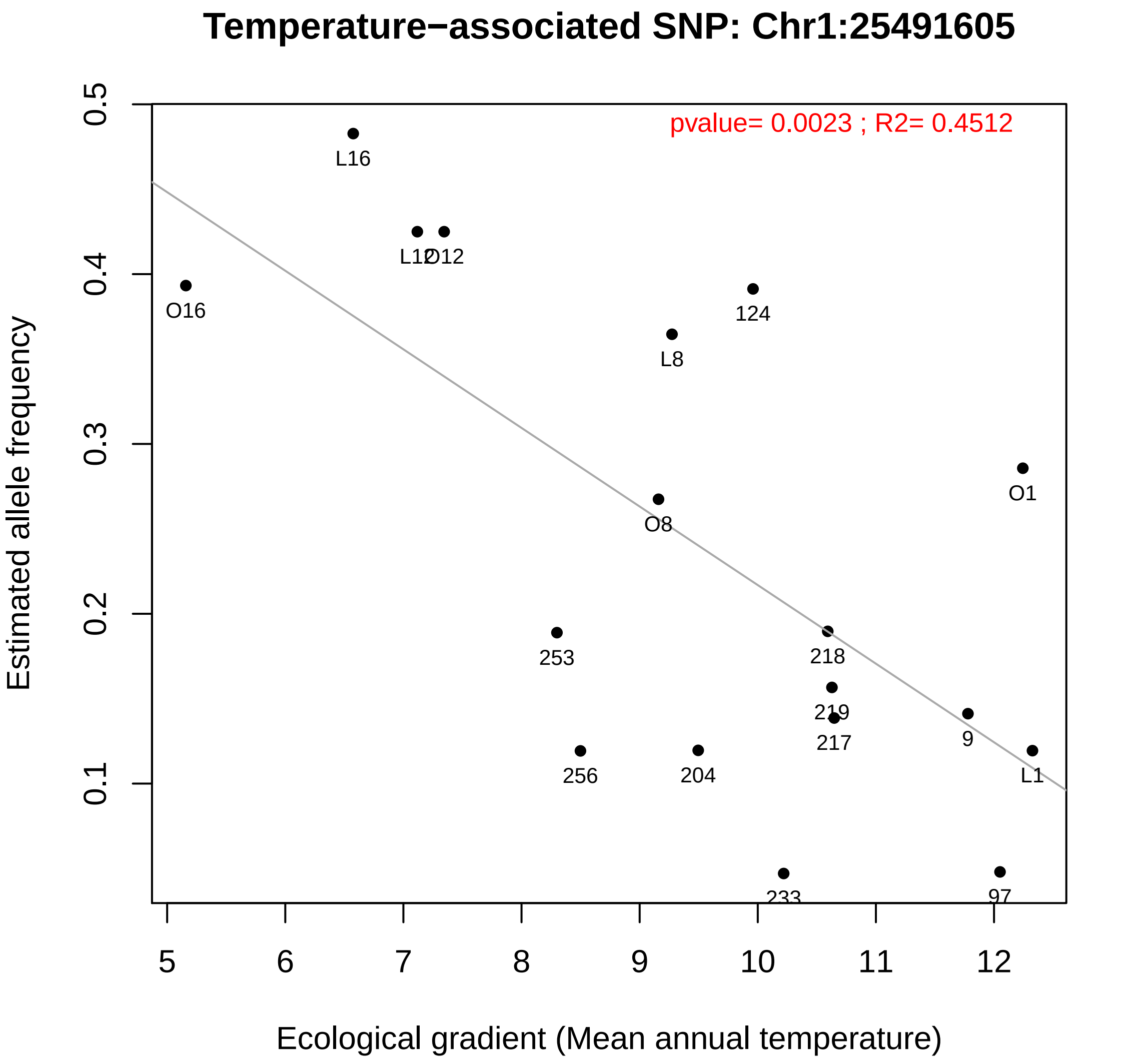
Temperature−associated SNP: Chr1:25491605
pvalue= 0.0023 ; R2= 0.4512
Estimated allele frequency
0.1
0.2
0.3
0.4
0.5
Ecological gradient (Mean annual temperature)
5
6
7
8
9
10
11
12
L16
O12
O16
124
L8
O1
O8
253
218
219
217
9
256
204
L1
97

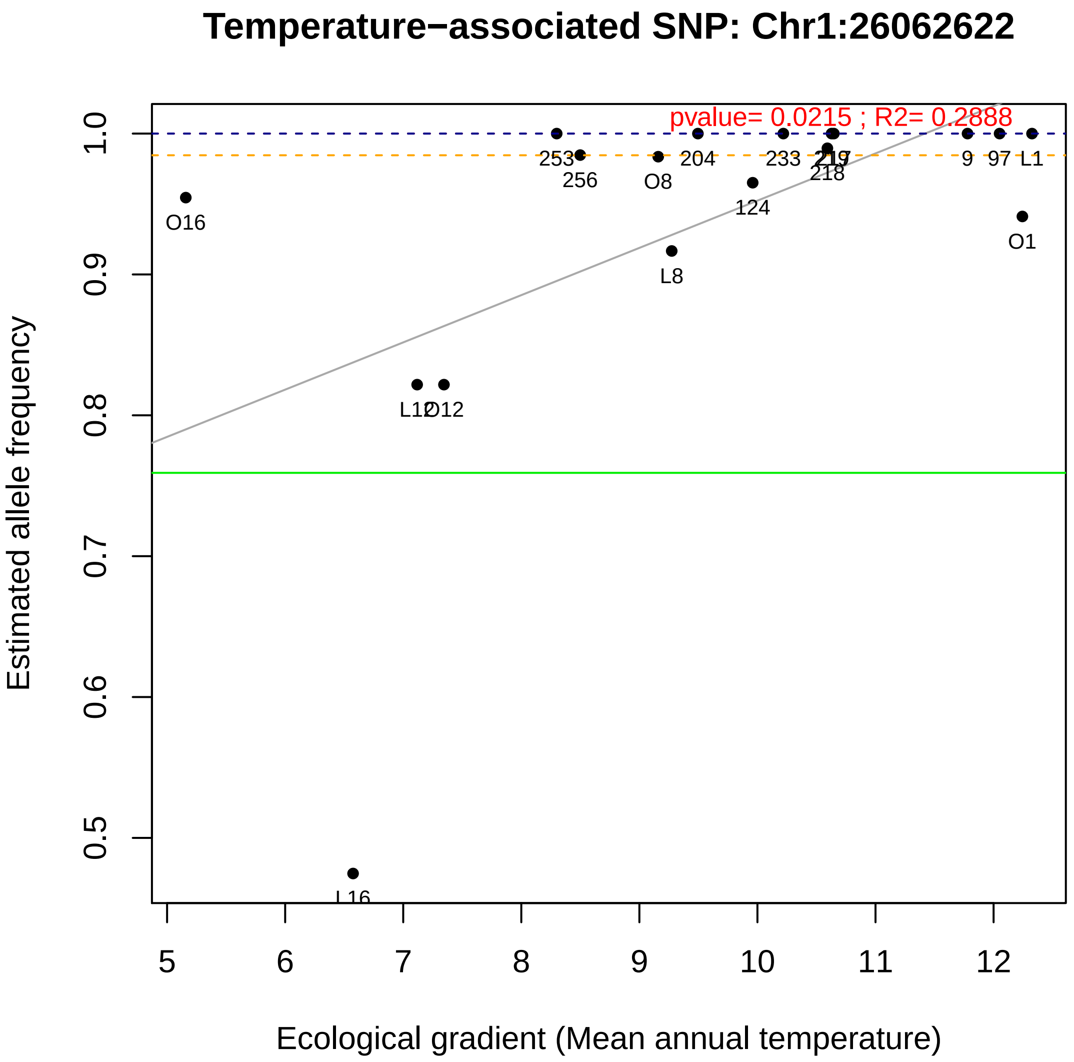

Temperature−associated SNP: Chr1:26062622
pvalue= 0.0215 ; R2= 0.2888
Estimated allele frequency
Ecological gradient (Mean annual temperature)
1.0
0.9
0.8
0.7
0.6
0.5
5
6
7
8
9
10
11
12
O16
253
256
O8
204
L8
124
233
217
219
218
9
97
L1
O1
L12
O12
L16

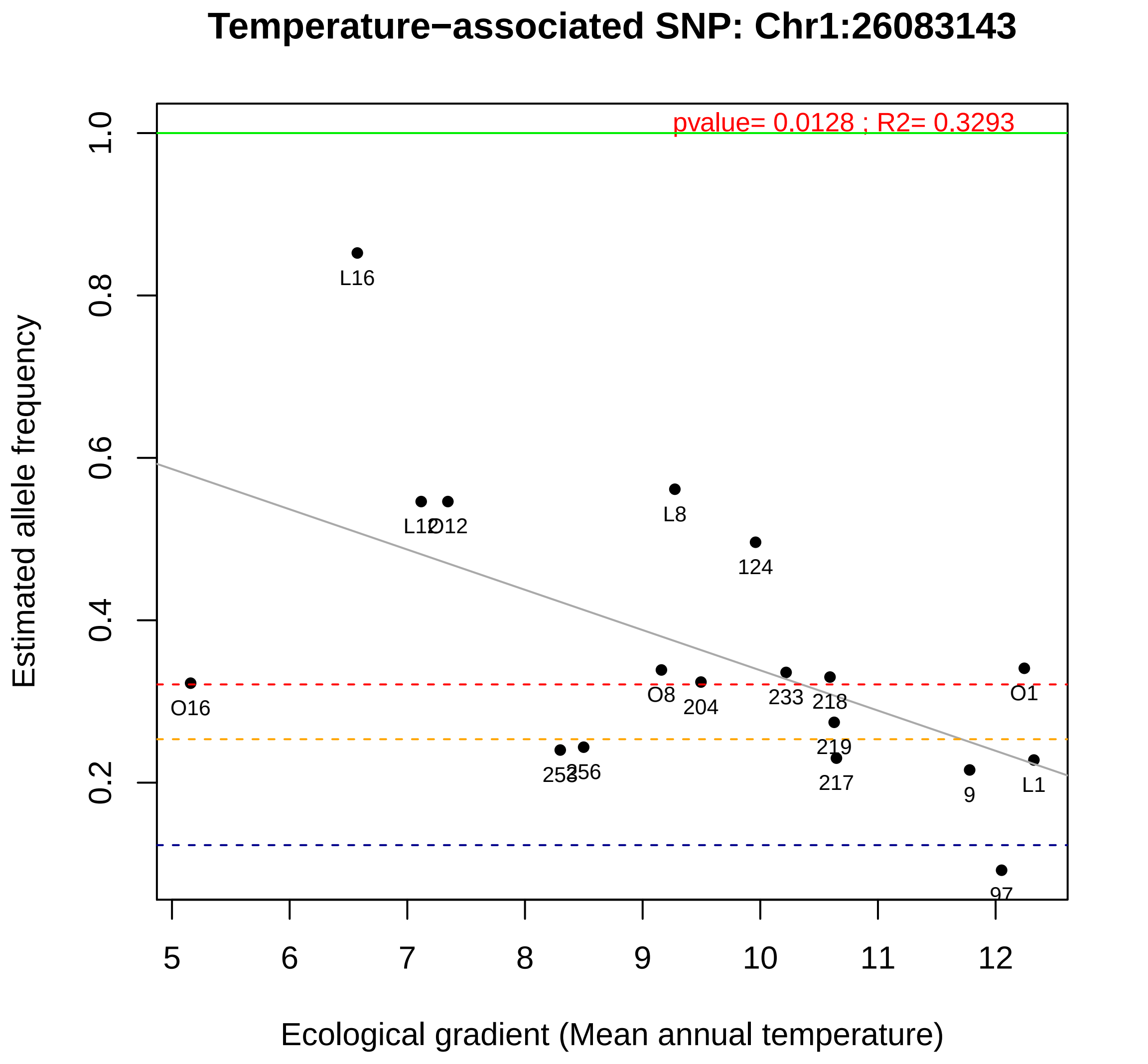

Temperature−associated SNP: Chr1:26083143
pvalue= 0.0128 ; R2= 0.3293
Estimated allele frequency
Ecological gradient (Mean annual temperature)
1.0
0.8
0.6
0.4
0.2
5
6
7
8
9
10
11
12
L16
L12
O12
L8
124
O16
O8
204
233
218
O1
219
217
256
9
L1
97

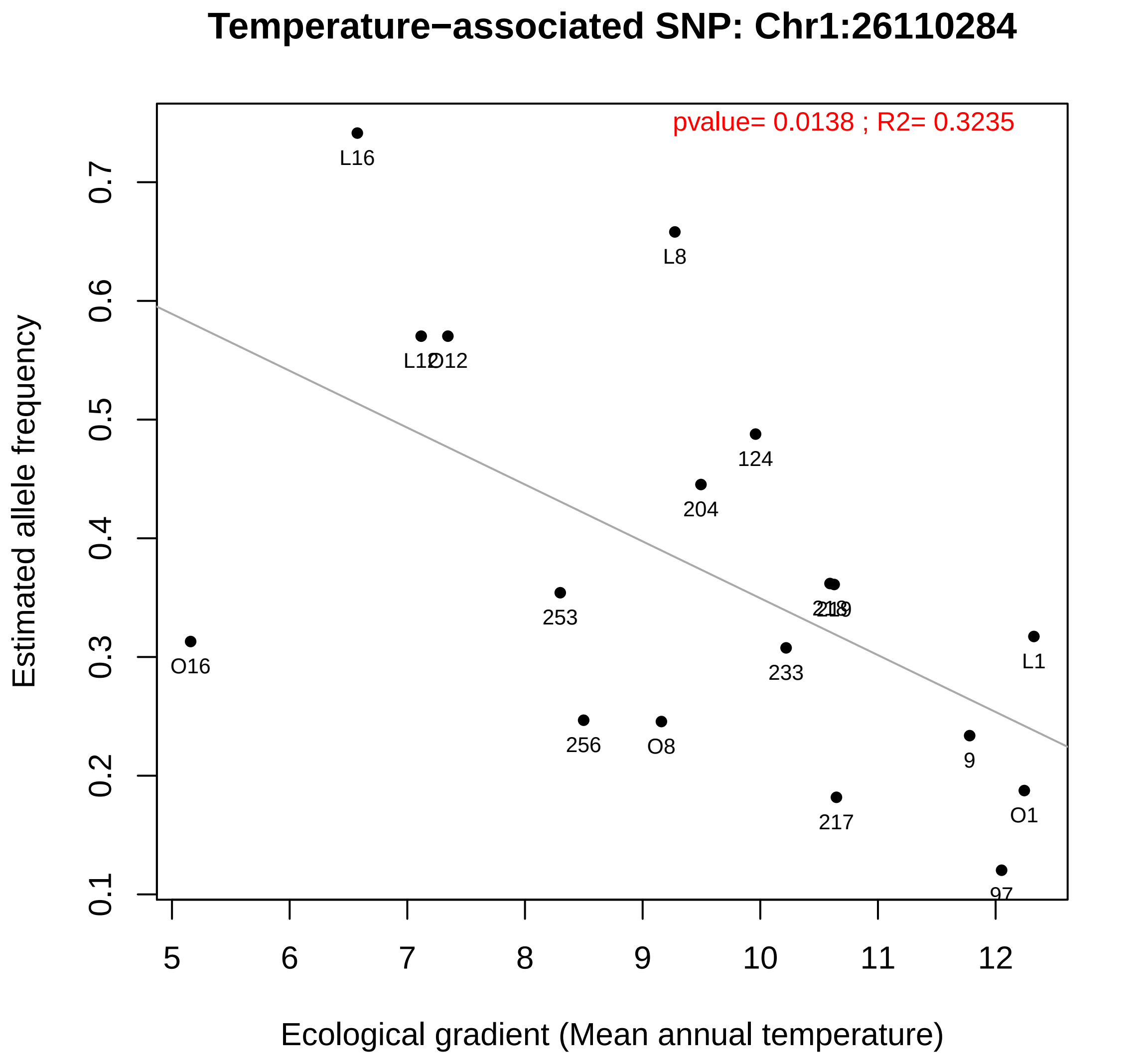

Temperature−associated SNP: Chr1:26110284
pvalue= 0.0138 ; R2= 0.3235
L16
L8
L12
O12
124
204
253
O16
233
L1
256
O8
9
217
O1
97
0.1
0.2
0.3
0.4
0.5
0.6
0.7
5
6
7
8
9
10
11
12
Estimated allele frequency
Ecological gradient (Mean annual temperature)

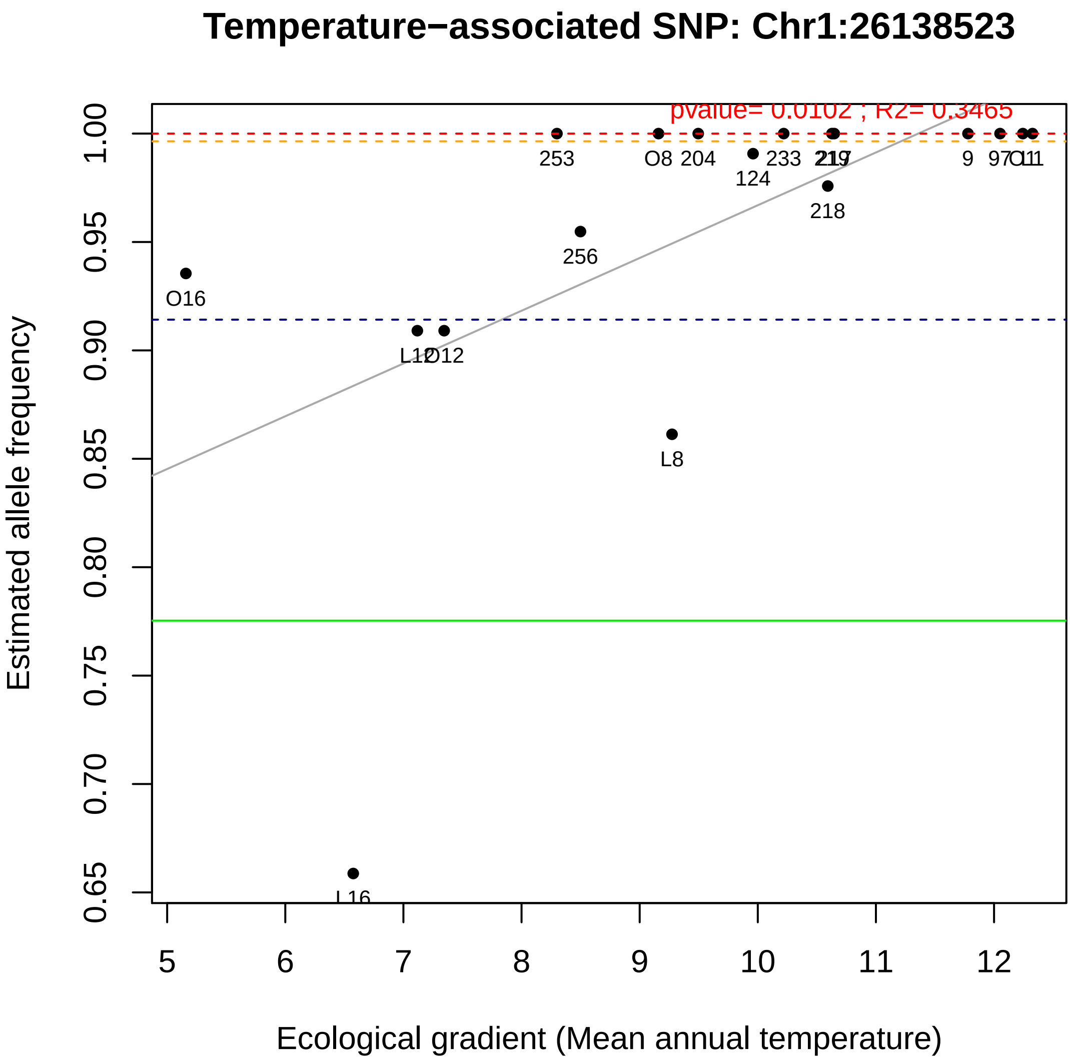
Temperature−associated SNP: Chr1:26138523
pvalue= 0.0102 ; R2= 0.3465
Estimated allele frequency
Ecological gradient (Mean annual temperature)
0.65
0.70
0.75
0.80
0.85
0.90
0.95
1.00
5
6
7
8
9
10
11
12
253
O8
204
233
124
218
256
O16
L12
O12
L8
L16
9

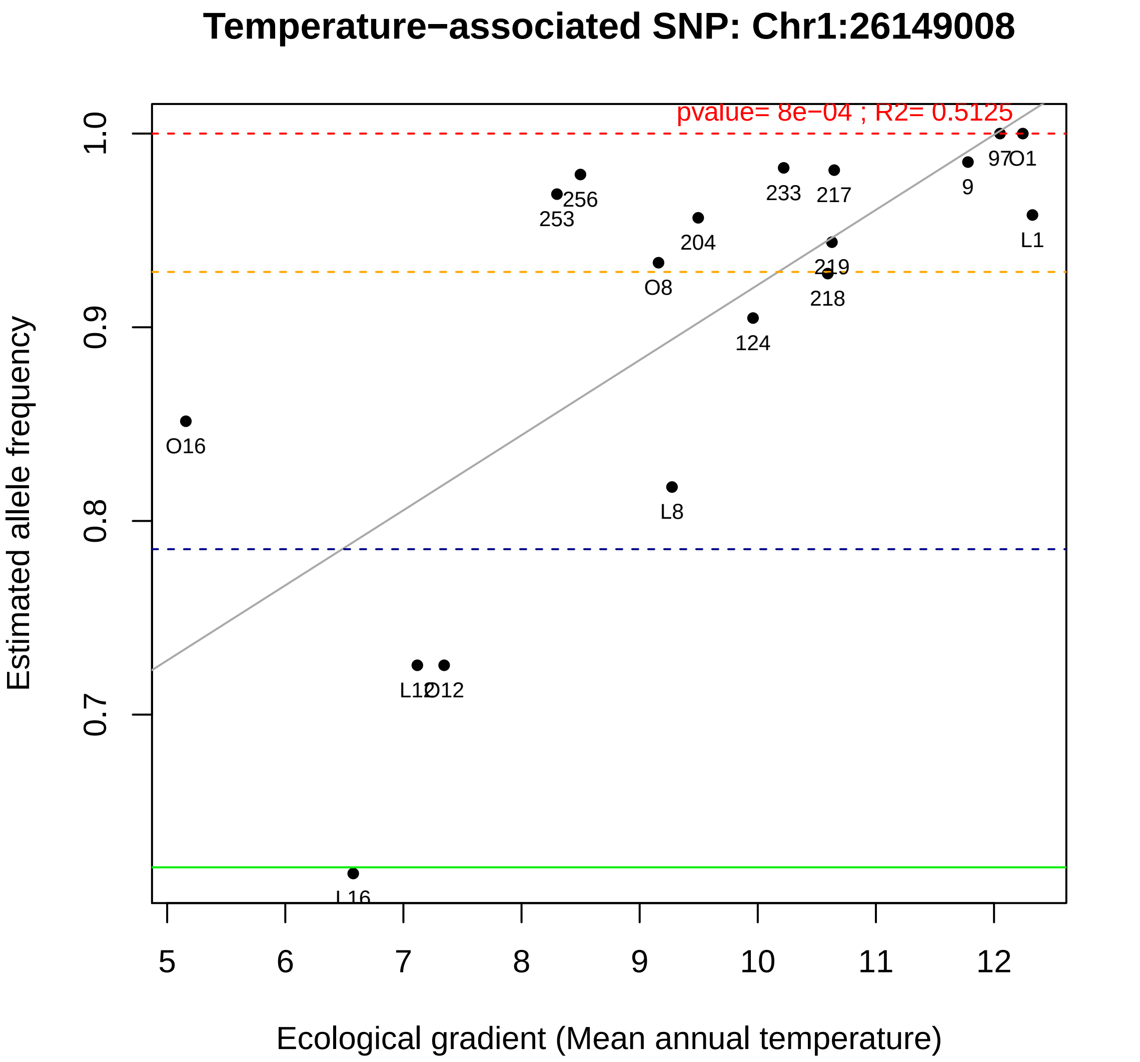
Temperature−associated SNP: Chr1:26149008
pvalue= 8e−04 ; R2= 0.5125
Estimated allele frequency
Ecological gradient (Mean annual temperature)
1.0
0.9
0.8
0.7
5
6
7
8
9
10
11
12
O16
L16
L12
O12
253
256
O8
L8
204
124
233
217
219
218
9
97
O1
L1

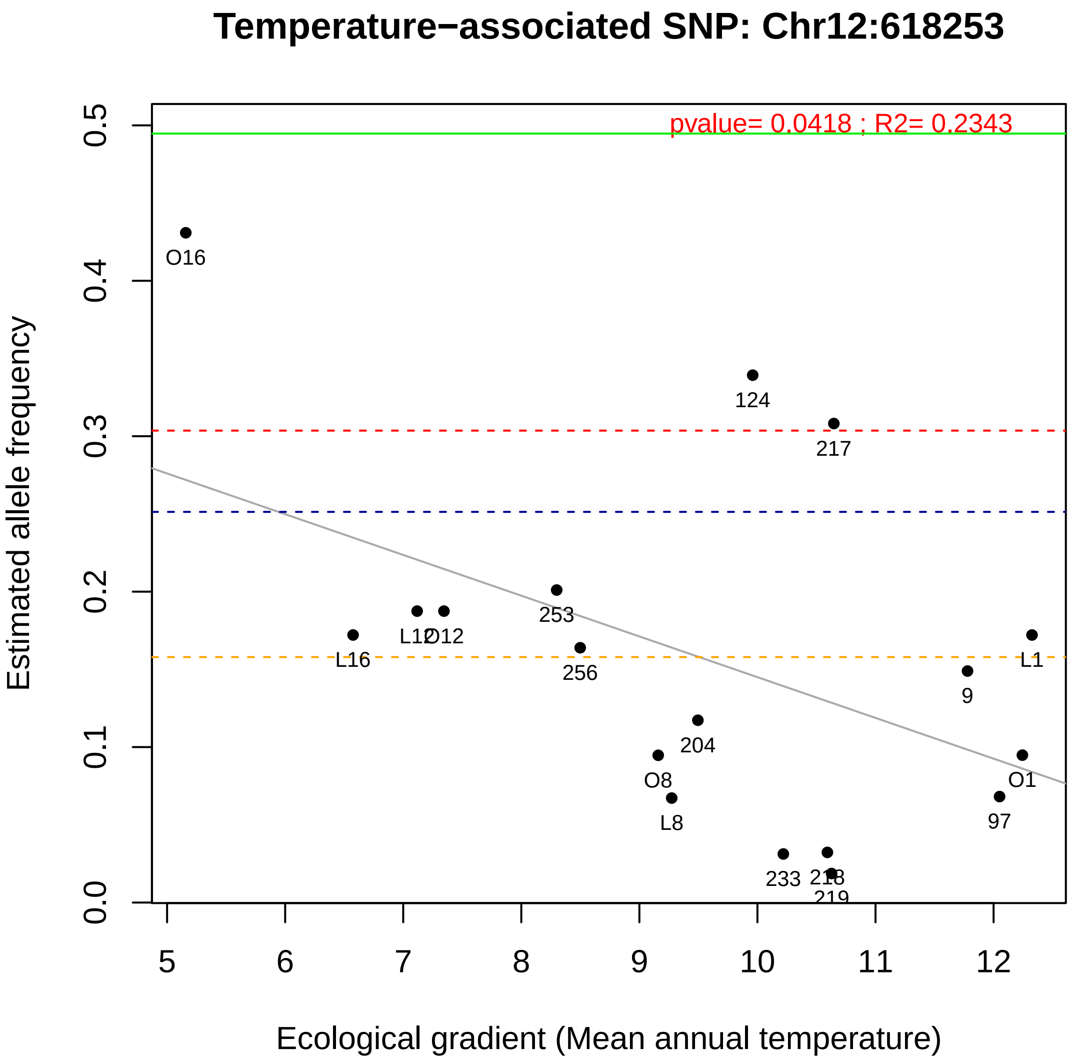

Temperature−associated SNP: Chr12:618253
pvalue= 0.0418 ; R2= 0.2343
Estimated allele frequency
Ecological gradient (Mean annual temperature)
0.0
0.1
0.2
0.3
0.4
0.5
5
6
7
8
9
10
11
12
O16
124
217
253
L12
O12
L16
256
L1
9
204
O8
O1
L8
97
233
218
219

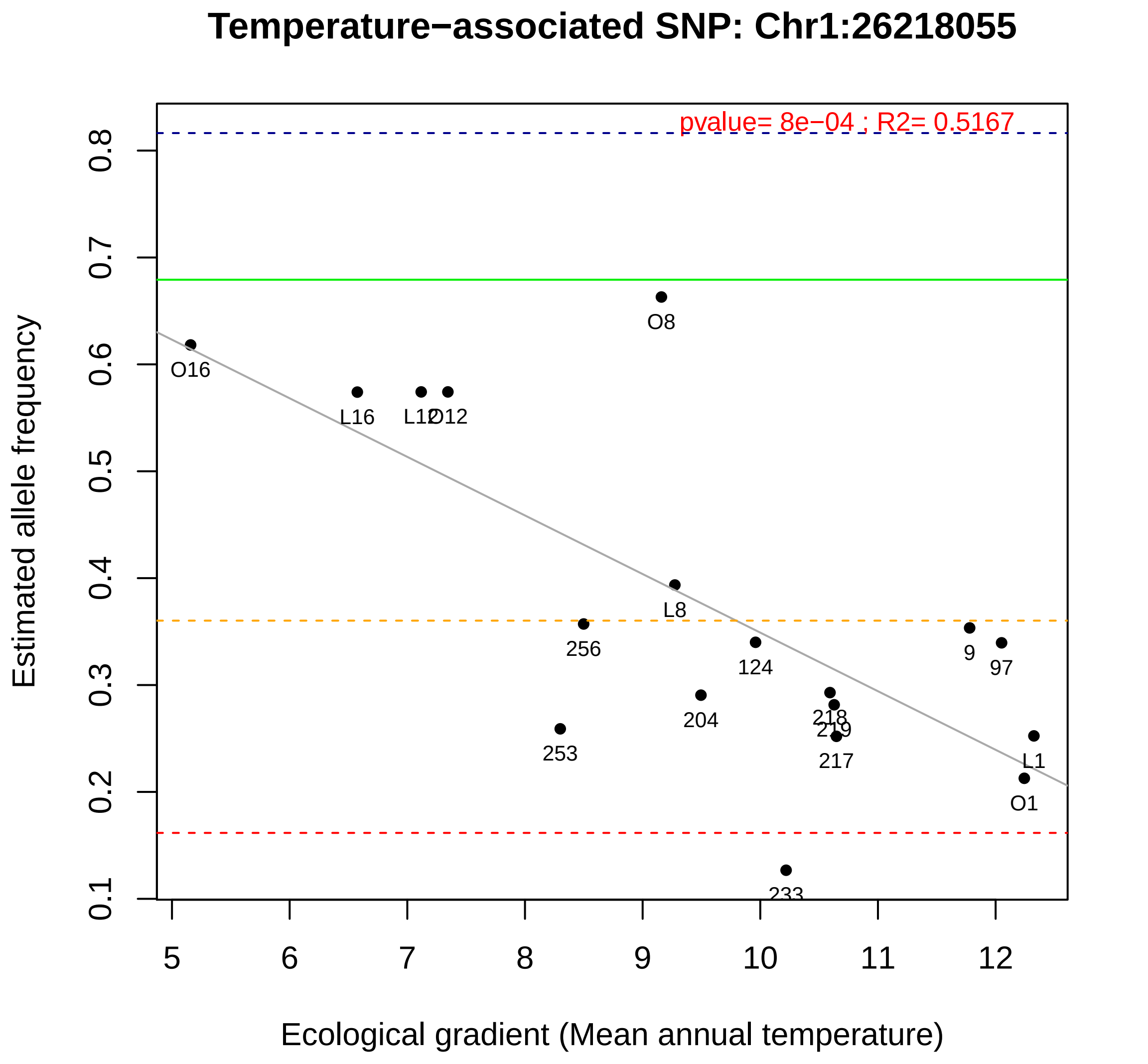
Temperature−associated SNP: Chr1:26218055
pvalue= 8e−04 ; R2= 0.5167
Estimated allele frequency
0.1
0.2
0.3
0.4
0.5
0.6
0.7
0.8
5
6
7
8
9
10
11
12
Ecological gradient (Mean annual temperature)
O16
L16
L12
O12
O8
256
253
L8
204
124
218
219
217
233
9
97
L1
O1

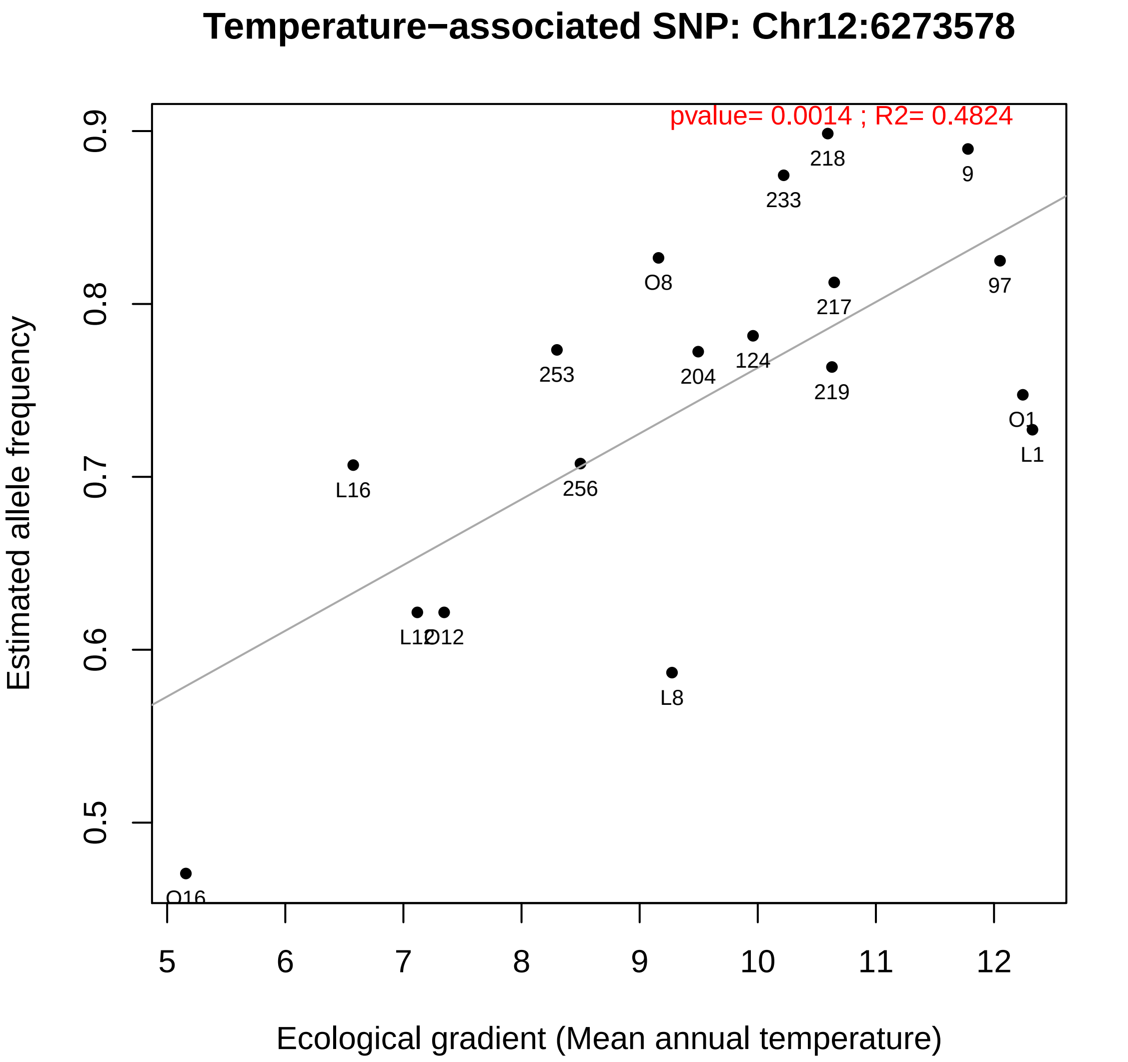

Temperature−associated SNP: Chr12:6273578
pvalue= 0.0014 ; R2= 0.4824
Estimated allele frequency
Ecological gradient (Mean annual temperature)
0.5
0.6
0.7
0.8
0.9
5
6
7
8
9
10
11
12
218
9
233
O8
97
217
253
204
124
219
O1
L1
L16
256
L12
O12
L8
O16

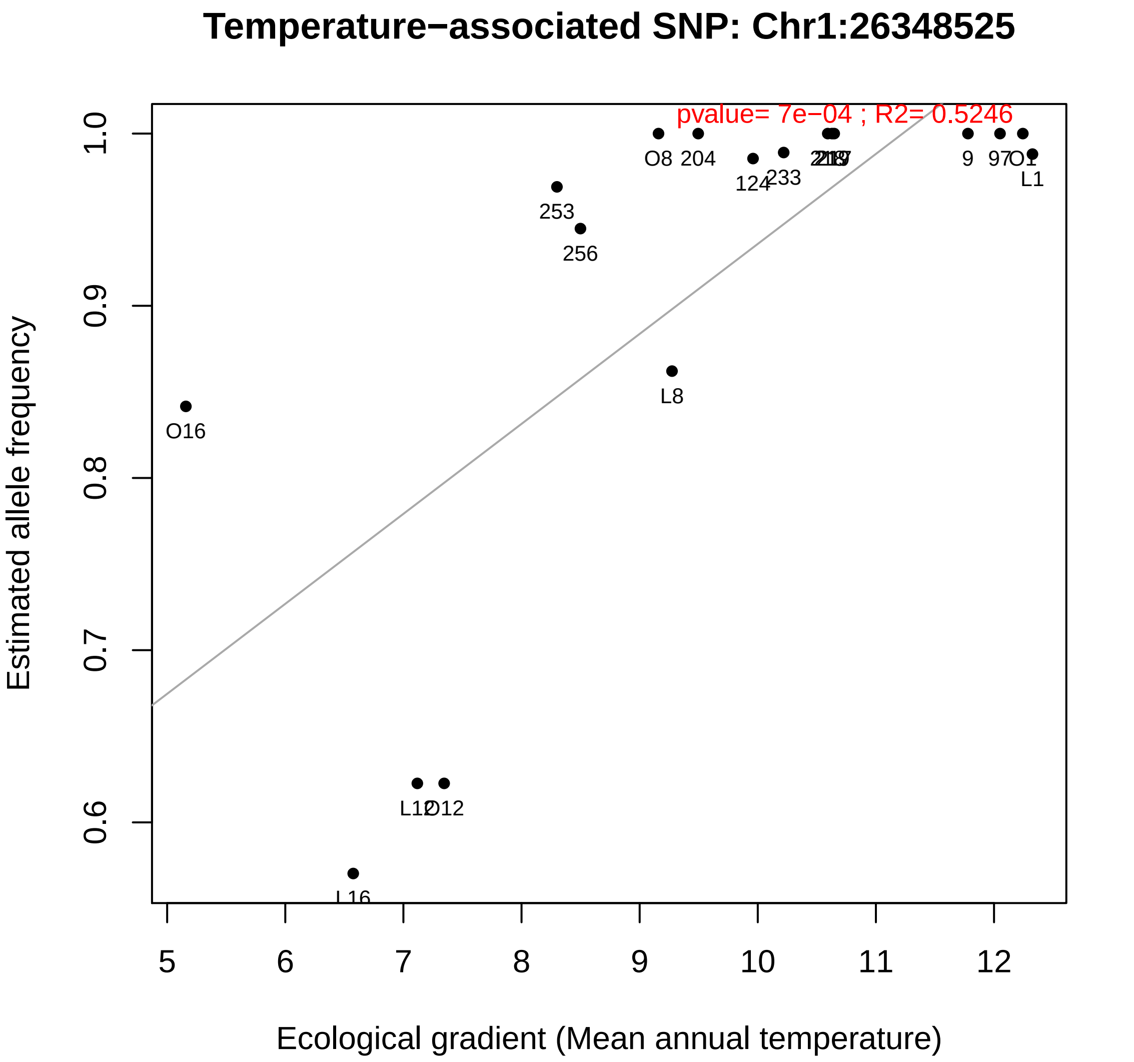

Temperature−associated SNP: Chr1:26348525
pvalue= 7e−04 ; R2= 0.5246
Estimated allele frequency
Ecological gradient (Mean annual temperature)
0.6
0.7
0.8
0.9
1.0
5
6
7
8
9
10
11
12
O16
L16
L12
O12
253
256
L8
O8
204
124
233
9
97
O1
L1

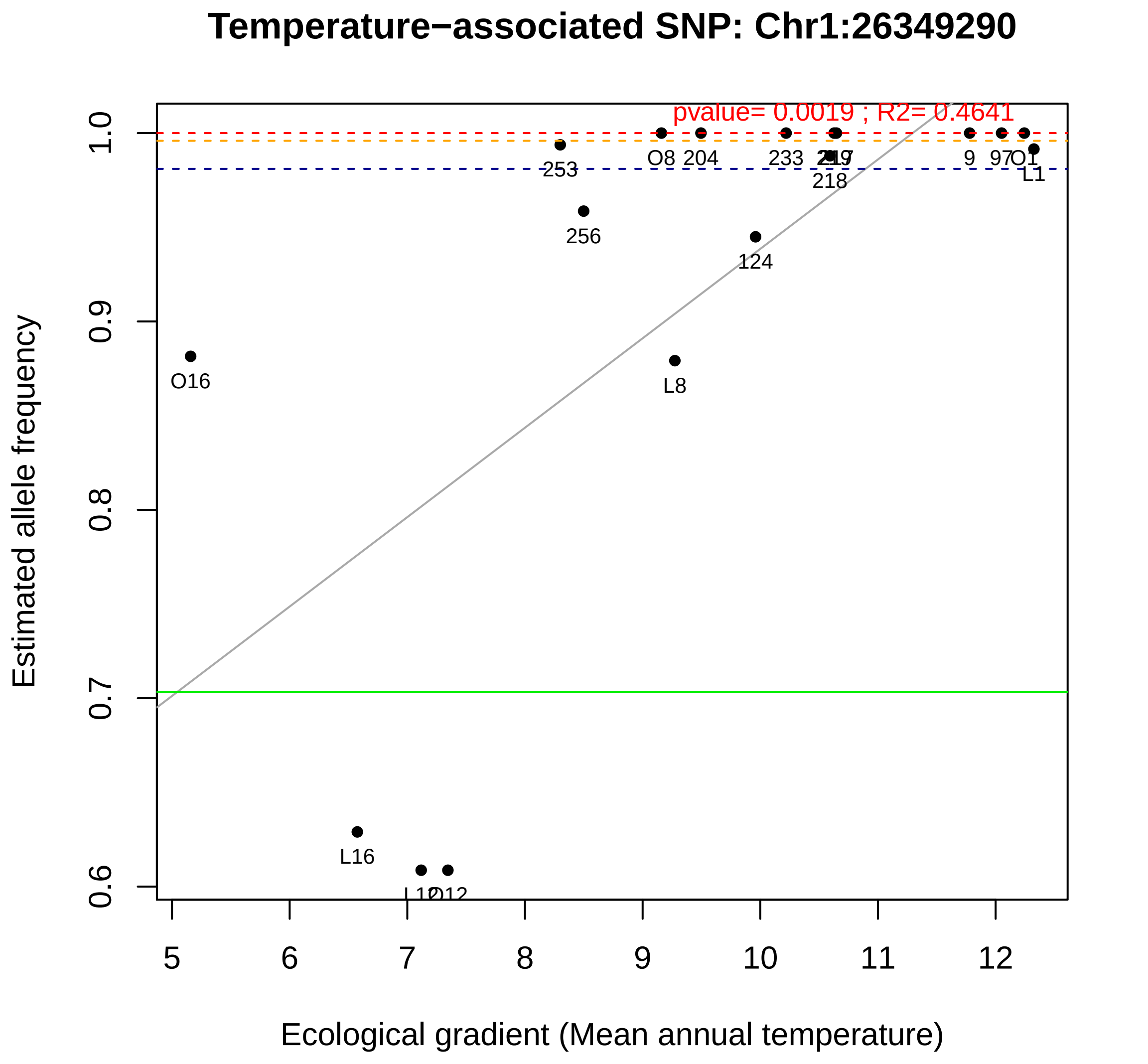

Temperature−associated SNP: Chr1:26349290
pvalue= 0.0019 ; R2= 0.4641
Estimated allele frequency
Ecological gradient (Mean annual temperature)
0.6
0.7
0.8
0.9
1.0
5
6
7
8
9
10
11
12
O16
L16
253
256
O8
204
L8
124
233
218
9
97
O1
L1

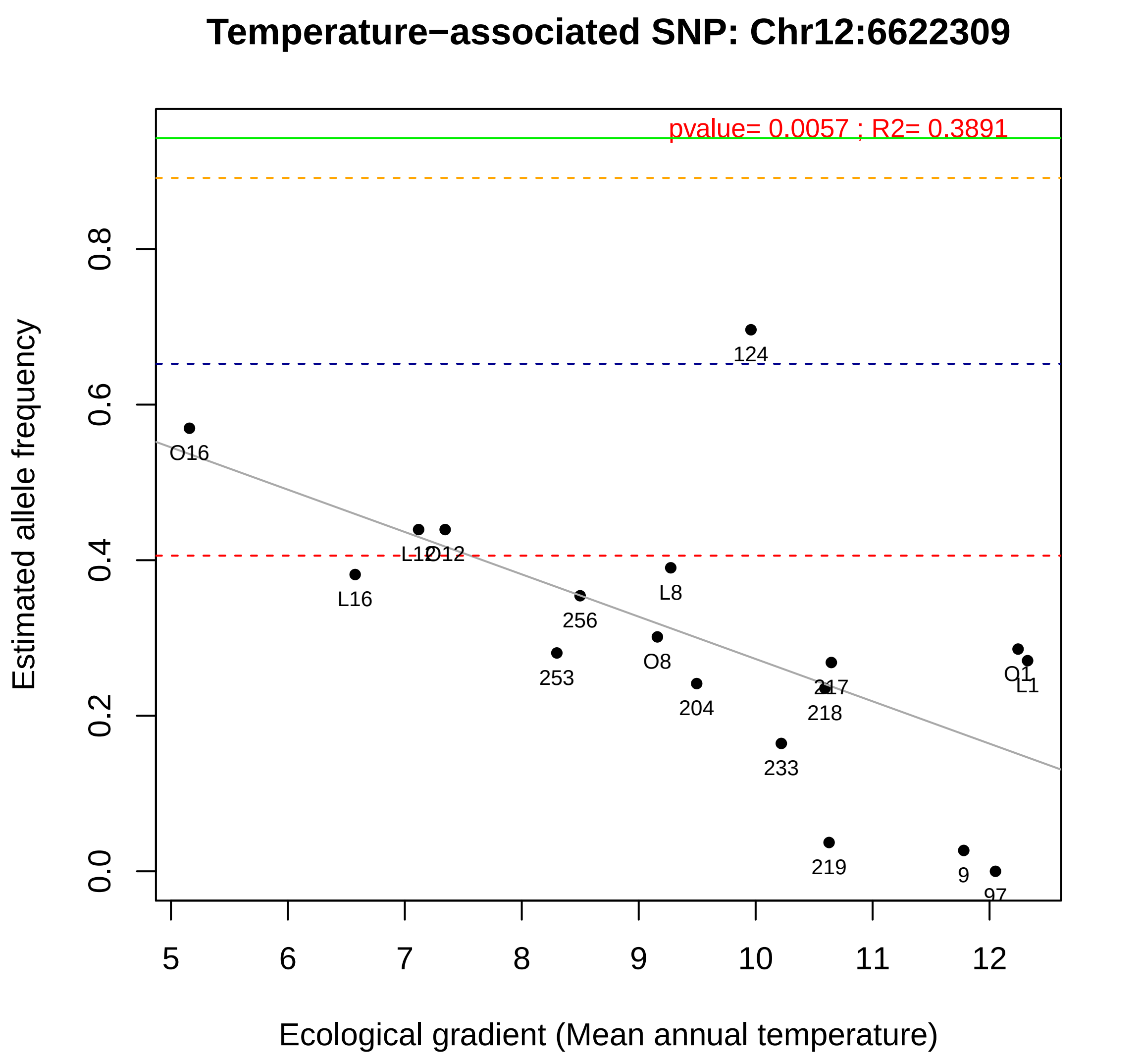
Temperature−associated SNP: Chr12:6622309
pvalue= 0.0057 ; R2= 0.3891
Estimated allele frequency
0.0
0.2
0.4
0.6
0.8
124
O16
L12
O12
L16
L8
256
O8
253
204
217
218
O1
L1
233
219
9
97
5
6
7
8
9
10
11
12
Ecological gradient (Mean annual temperature)

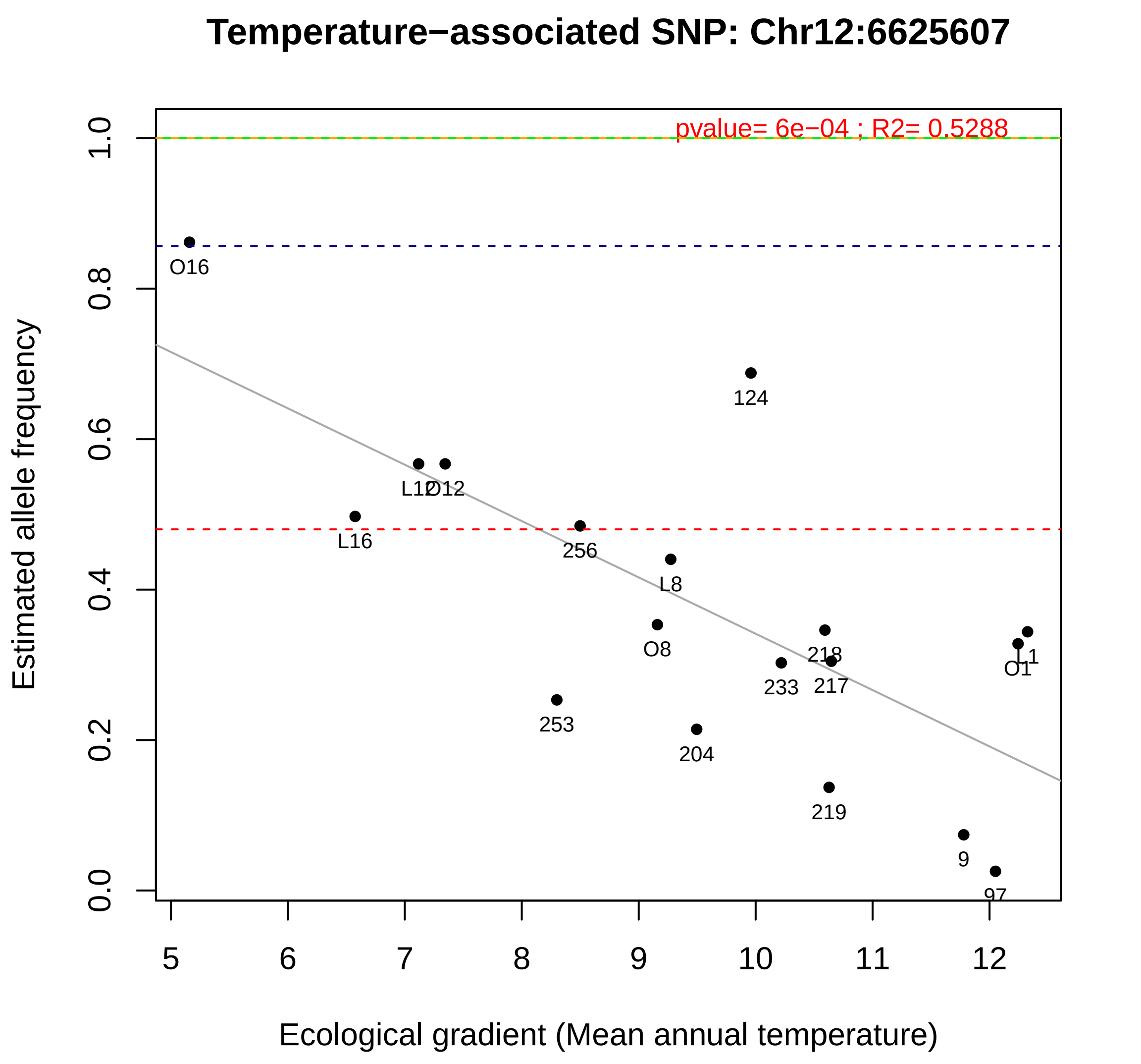

Temperature−associated SNP: Chr12:6625607
pvalue= 6e−04 ; R2= 0.5288
Estimated allele frequency
Ecological gradient (Mean annual temperature)
0.0
0.2
0.4
0.6
0.8
1.0
5
6
7
8
9
10
11
12
O16
124
L12
O12
L16
256
L8
O8
218
217
233
L1
O1
253
204
219
9
97

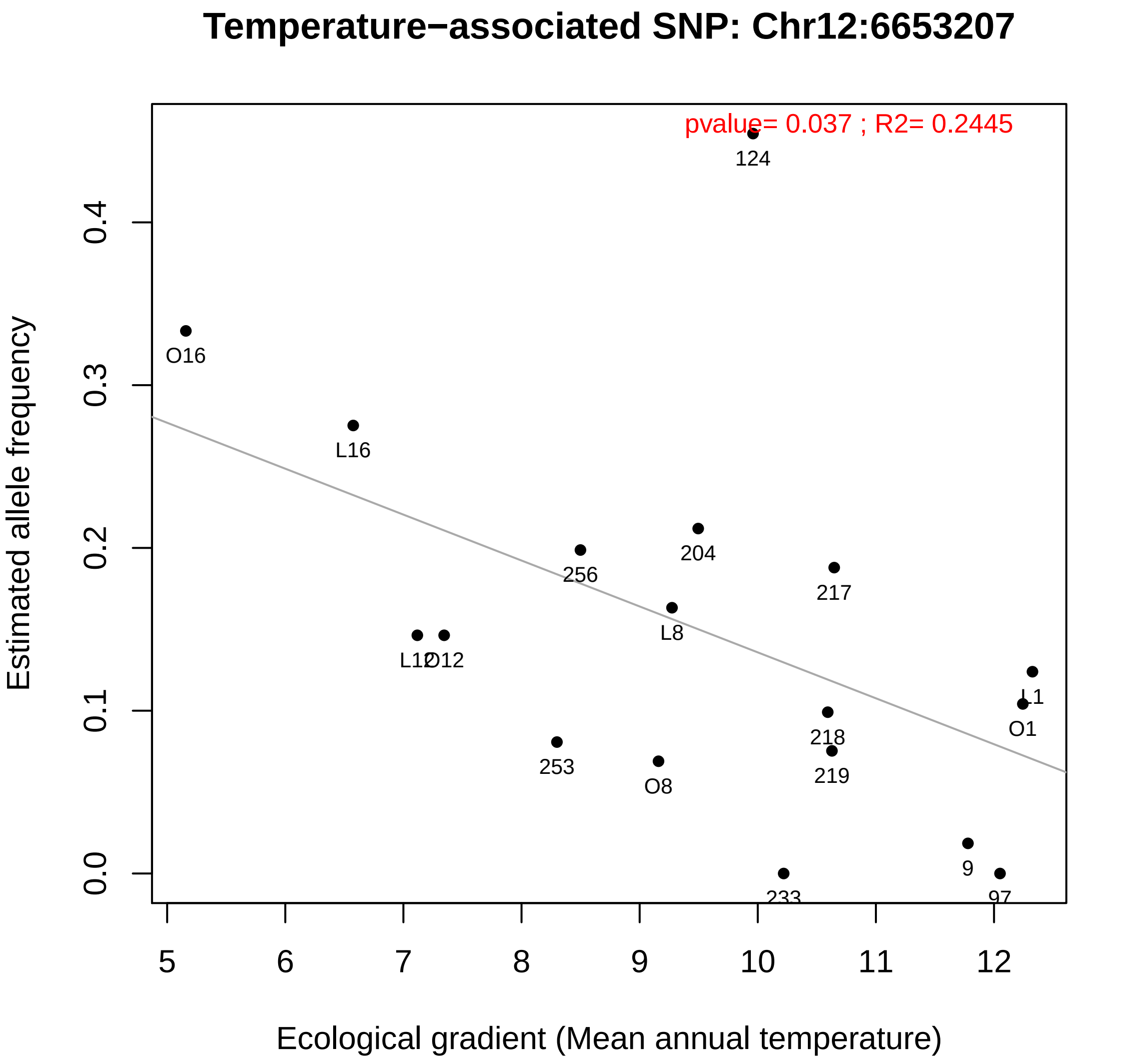

Temperature−associated SNP: Chr12:6653207
pvalue= 0.037 ; R2= 0.2445
Estimated allele frequency
Ecological gradient (Mean annual temperature)
0.0
0.1
0.2
0.3
0.4
5
6
7
8
9
10
11
12
O16
L16
L12
O12
256
204
217
L8
253
O8
218
219
L1
O1
124
9
233
97

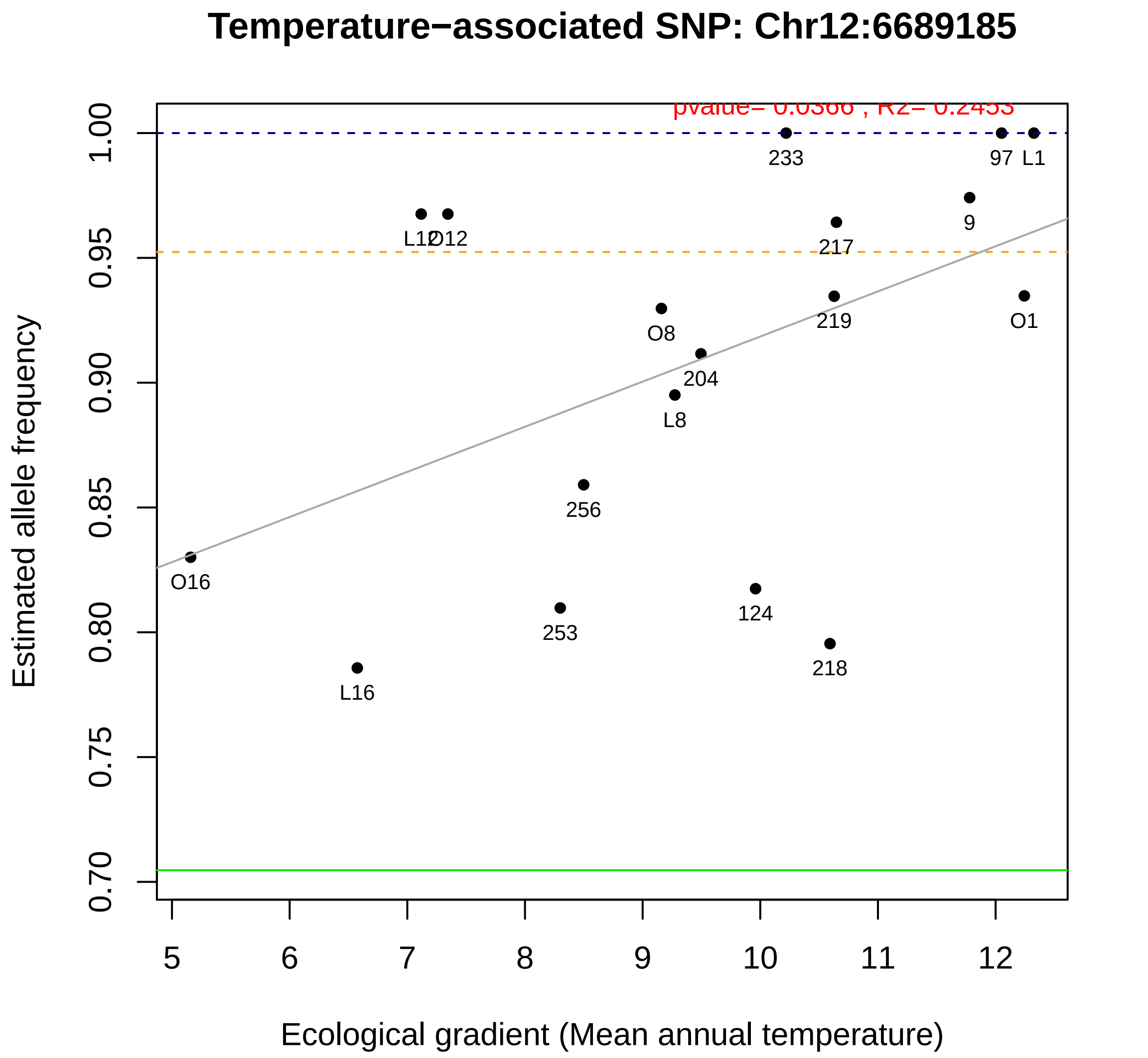

Temperature−associated SNP: Chr12:6689185
pvalue= 0.0366 , R2= 0.2453
Estimated allele frequency
Ecological gradient (Mean annual temperature)
1.00
0.95
0.90
0.85
0.80
0.75
0.70
5
6
7
8
9
10
11
12
233
97
L1
9
L12
O12
217
219
O1
O8
204
L8
256
O16
253
124
218
L16

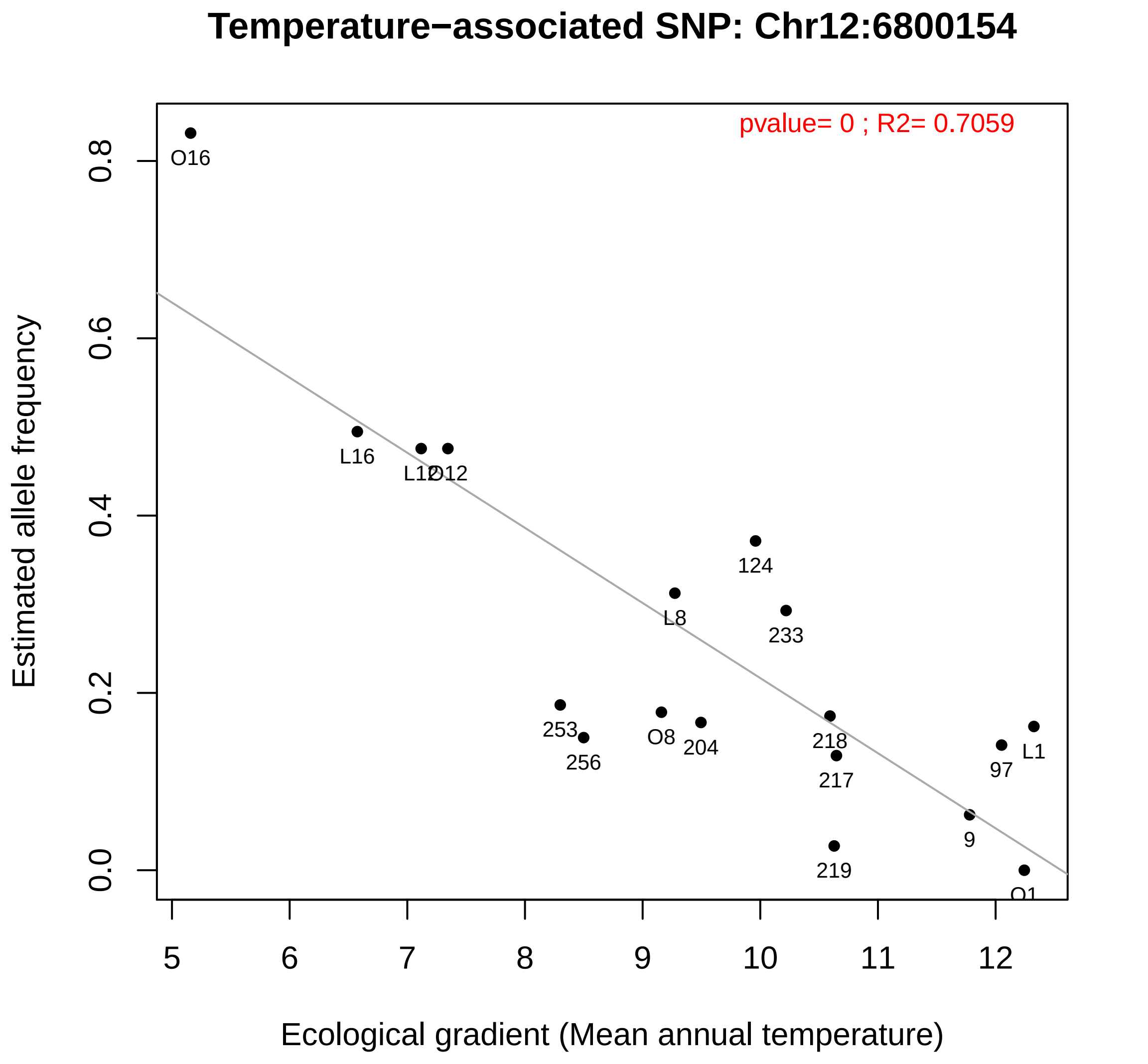
Temperature−associated SNP: Chr12:6800154
pvalue= 0 ; R2= 0.7059
Estimated allele frequency
0.0
0.2
0.4
0.6
0.8
Ecological gradient (Mean annual temperature)
5
6
7
8
9
10
11
12
O16
L16
L12
O12
124
L8
233
253
256
O8
204
218
217
219
9
97
L1
O1

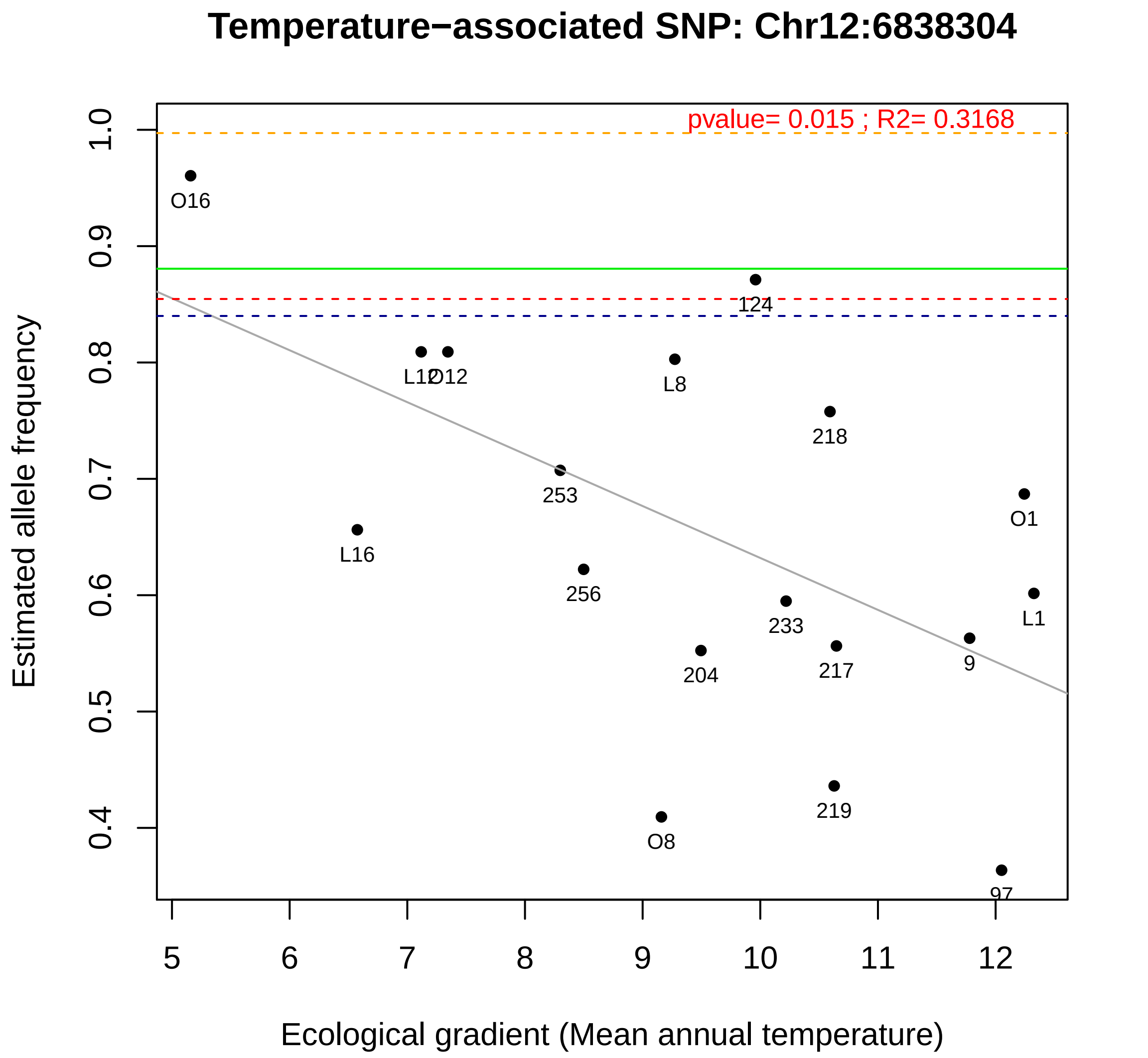

Temperature−associated SNP: Chr12:6838304
pvalue= 0.015 ; R2= 0.3168
Estimated allele frequency
0.4
0.5
0.6
0.7
0.8
0.9
1.0
5
6
7
8
9
10
11
12
Ecological gradient (Mean annual temperature)
O16
124
L12
O12
L8
218
253
O1
L16
256
L1
233
204
217
9
219
O8
97

### Temperature−associated SNP: Chr12:9665422

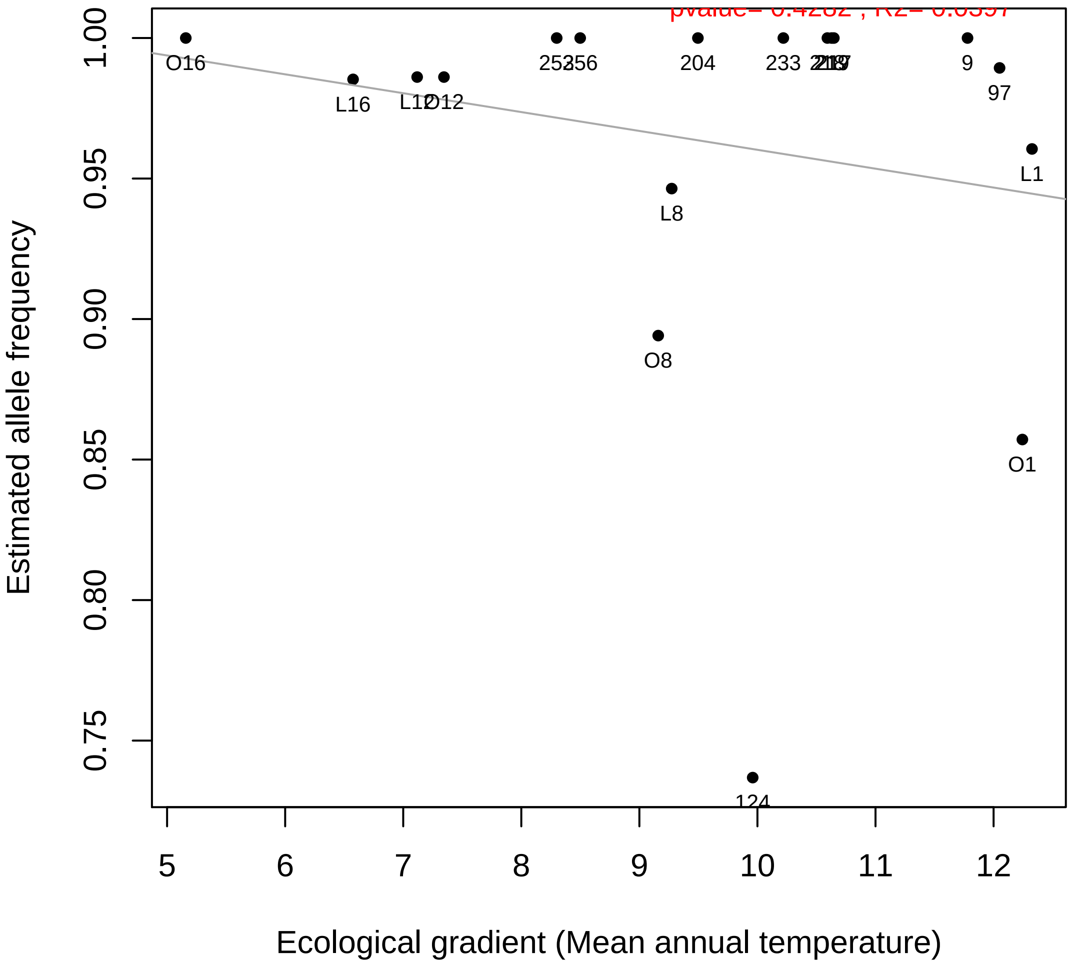

#### Temperature−associated SNP: Chr12:9913656

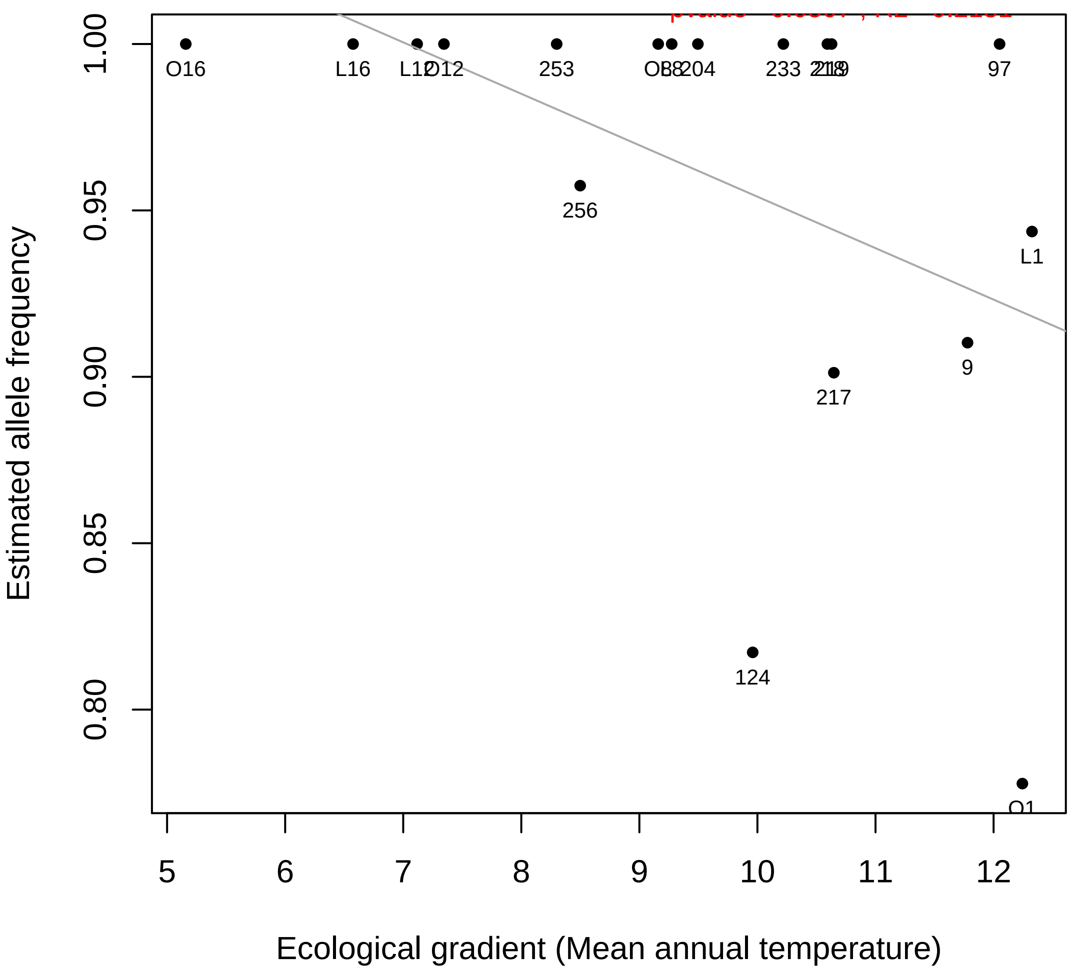

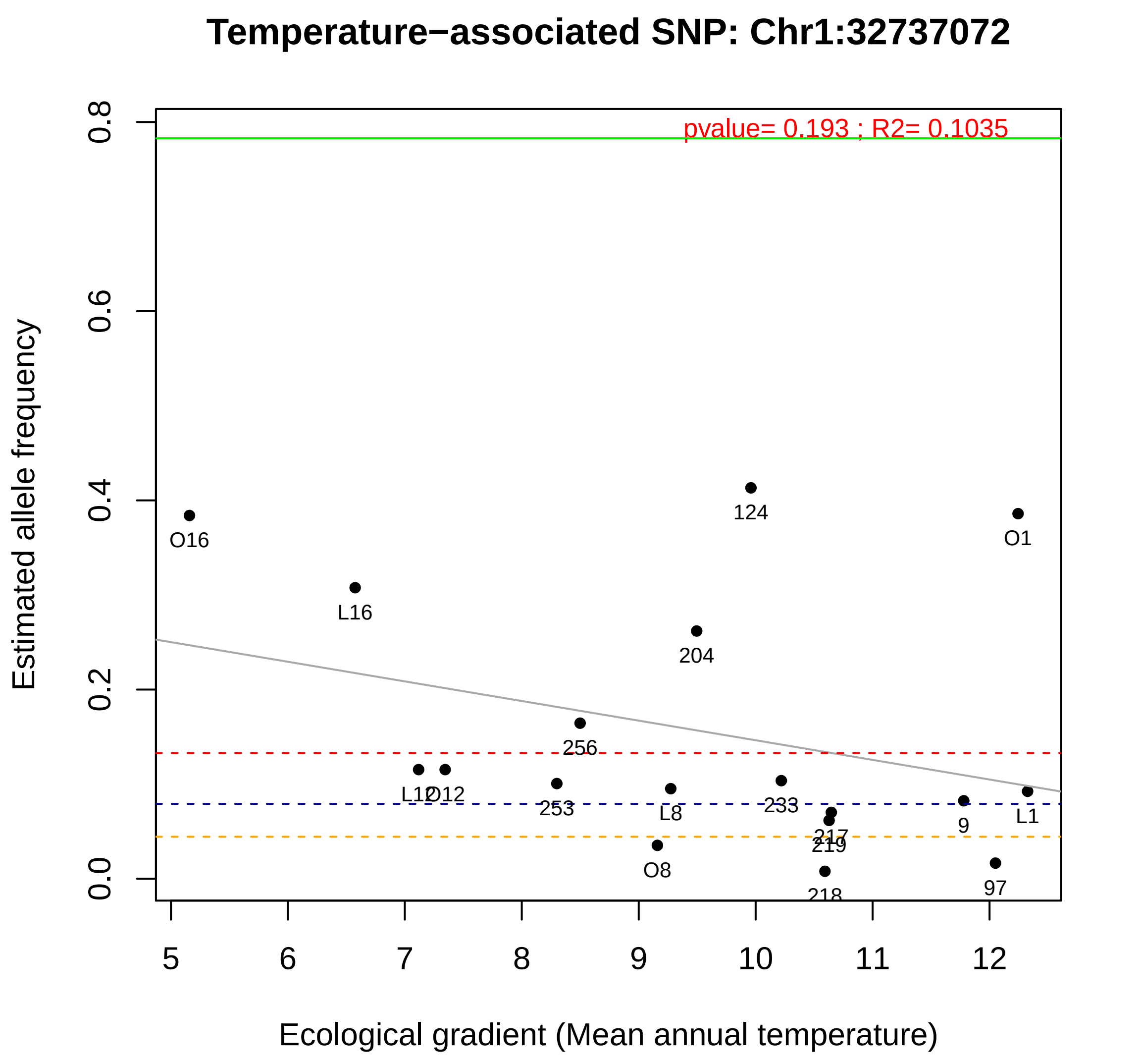

Temperature−associated SNP: Chr1:32737072
pvalue= 0.193 ; R2= 0.1035
Estimated allele frequency
0.0
0.2
0.4
0.6
0.8
5
6
7
8
9
10
11
12
Ecological gradient (Mean annual temperature)
O16
L16
L12
O12
253
256
O8
L8
204
124
233
217
219
218
9
97
O1
L1

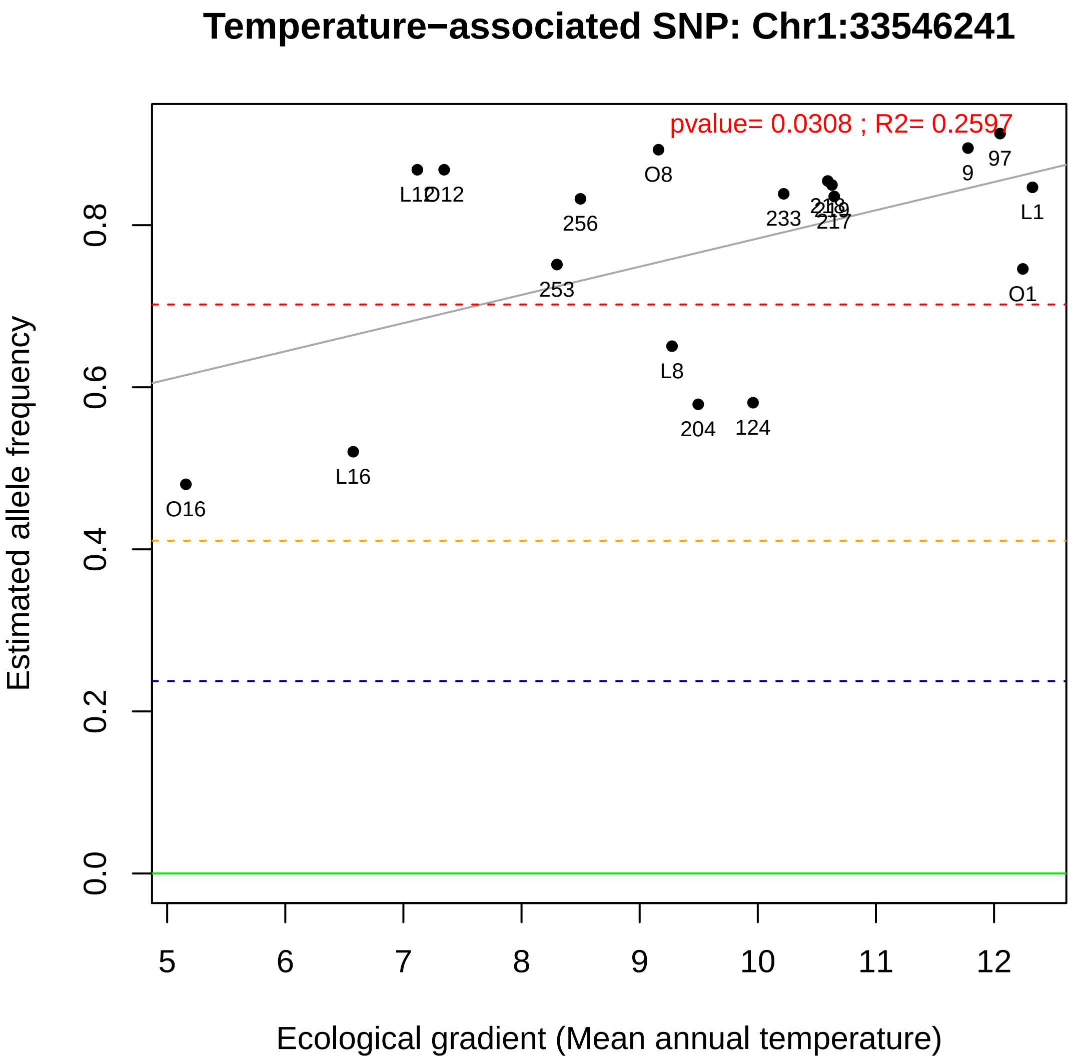

Temperature−associated SNP: Chr1:33546241
pvalue= 0.0308 ; R2= 0.2597
Estimated allele frequency
0.0
0.2
0.4
0.6
0.8
5
6
7
8
9
10
11
12
Ecological gradient (Mean annual temperature)
O16
L16
L12
O12
253
256
O8
L8
204
124
233
218
219
217
9
97
L1
O1

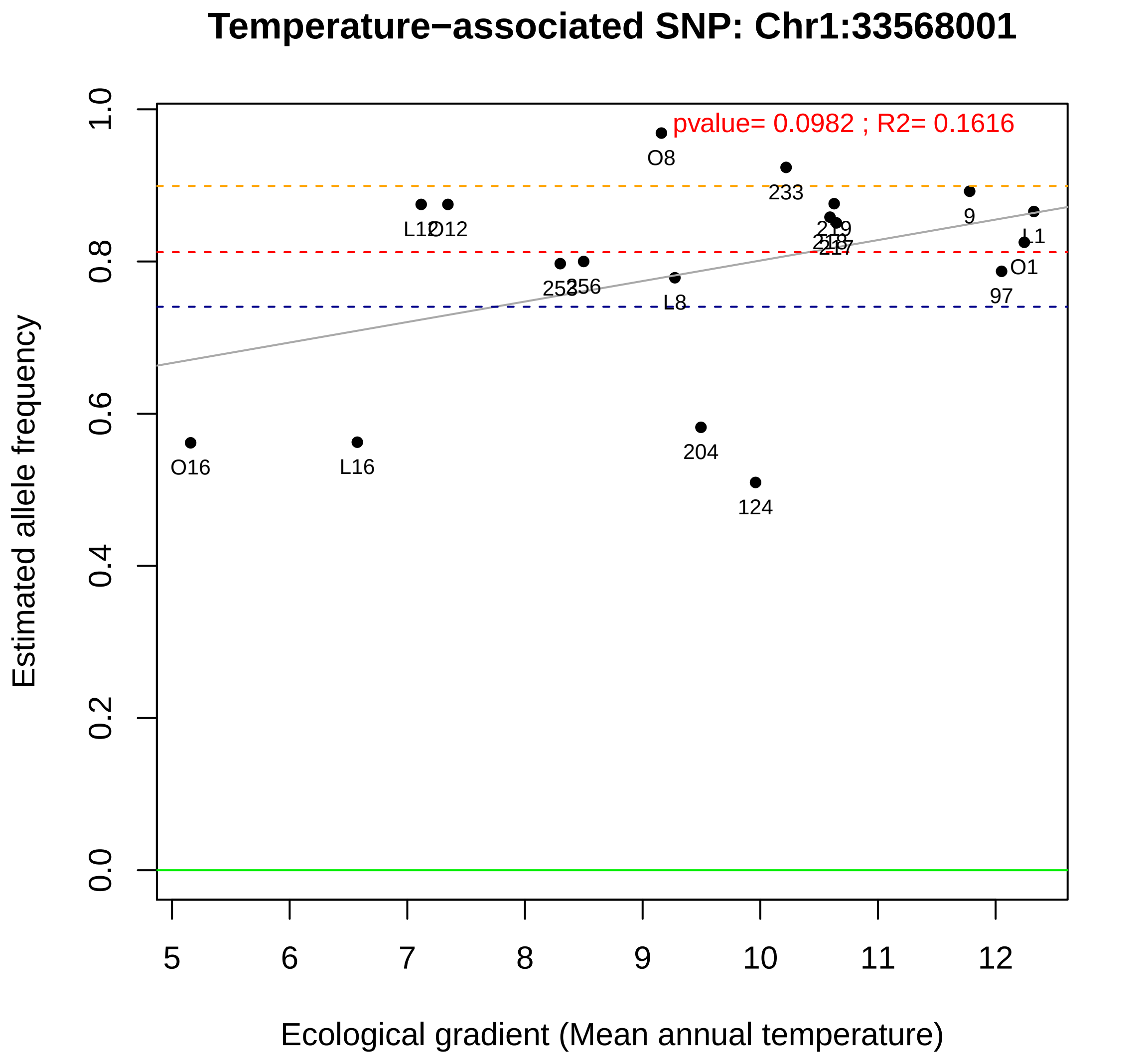

Temperature−associated SNP: Chr1:33568001
pvalue= 0.0982 ; R2= 0.1616
Estimated allele frequency
Ecological gradient (Mean annual temperature)
0.0
0.2
0.4
0.6
0.8
1.0
5
6
7
8
9
10
11
12
O8
233
L12
O12
9
253
256
L8
L1
O1
97
O16
L16
204
124

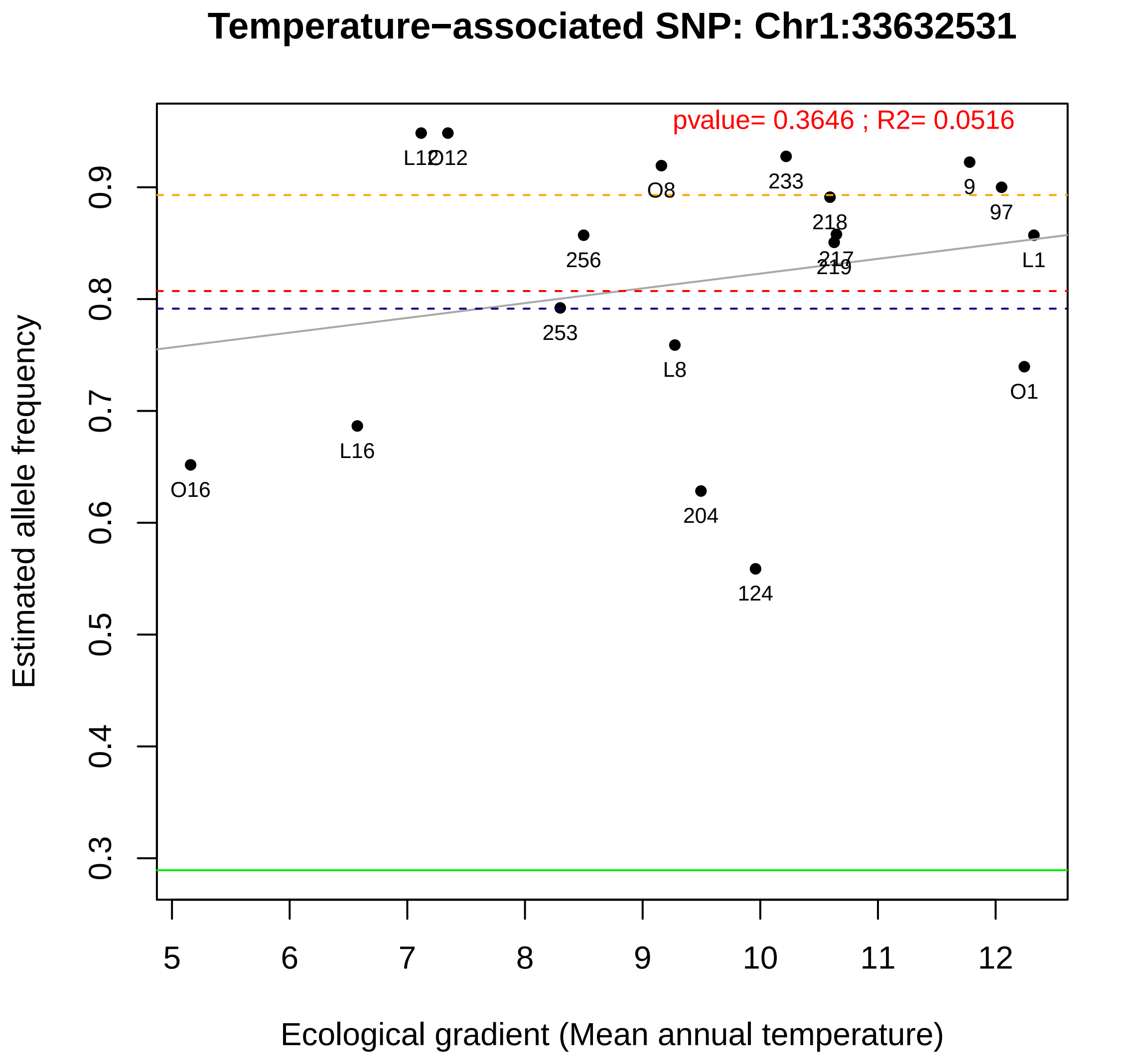

Temperature−associated SNP: Chr1:33632531
pvalue= 0.3646 ; R2= 0.0516
Estimated allele frequency
Ecological gradient (Mean annual temperature)
0.3
0.4
0.5
0.6
0.7
0.8
0.9
5
6
7
8
9
10
11
12
L12
O12
O8
233
9
97
218
256
217
219
L1
253
L8
O1
L16
O16
204
124

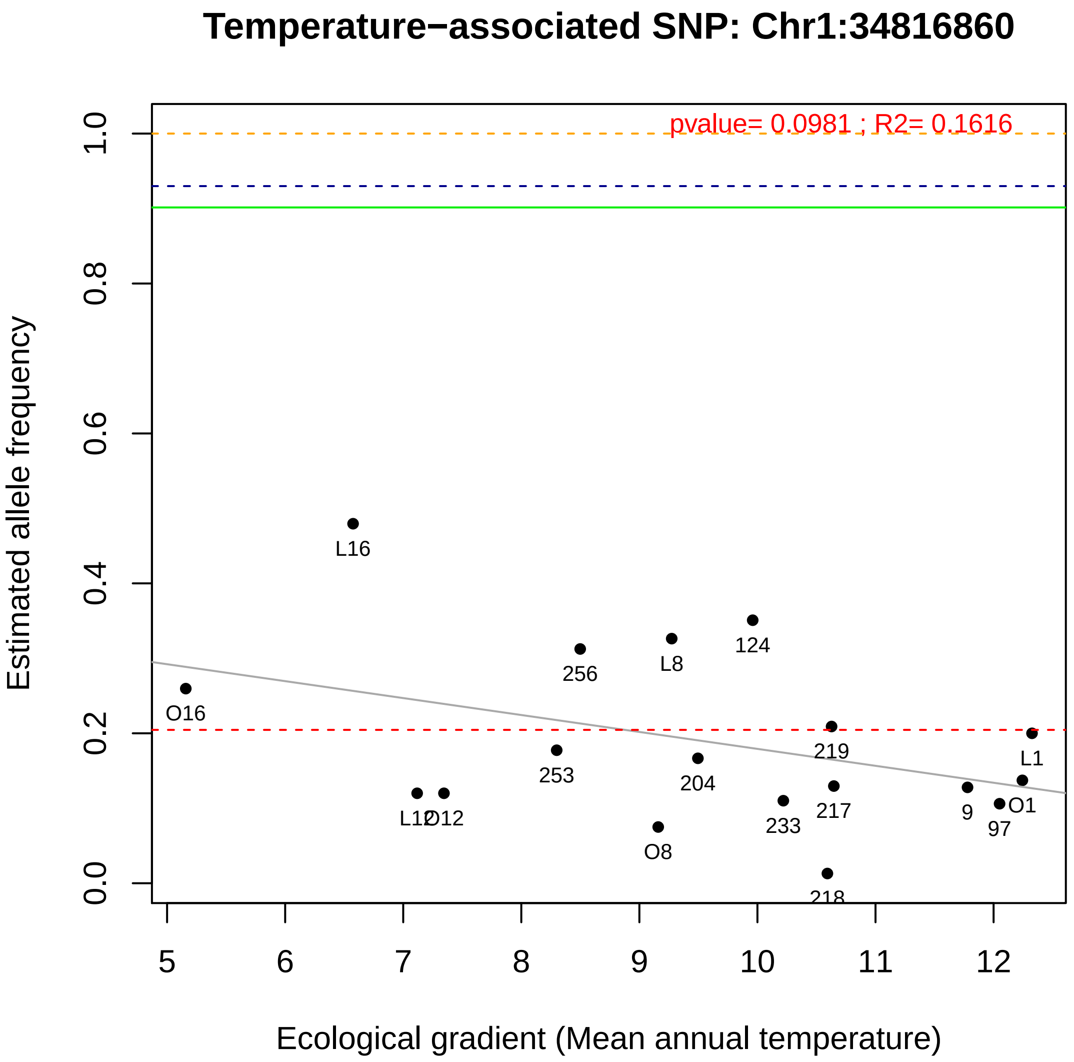

Temperature−associated SNP: Chr1:34816860
pvalue= 0.0981 ; R2= 0.1616
Estimated allele frequency
Ecological gradient (Mean annual temperature)
0.0
0.2
0.4
0.6
0.8
1.0
5
6
7
8
9
10
11
12
L16
256
L8
124
O16
219
L1
253
204
217
L12
O12
233
9
O1
97
O8
218

**Temperature−associated SNP: Chr1:35348319**

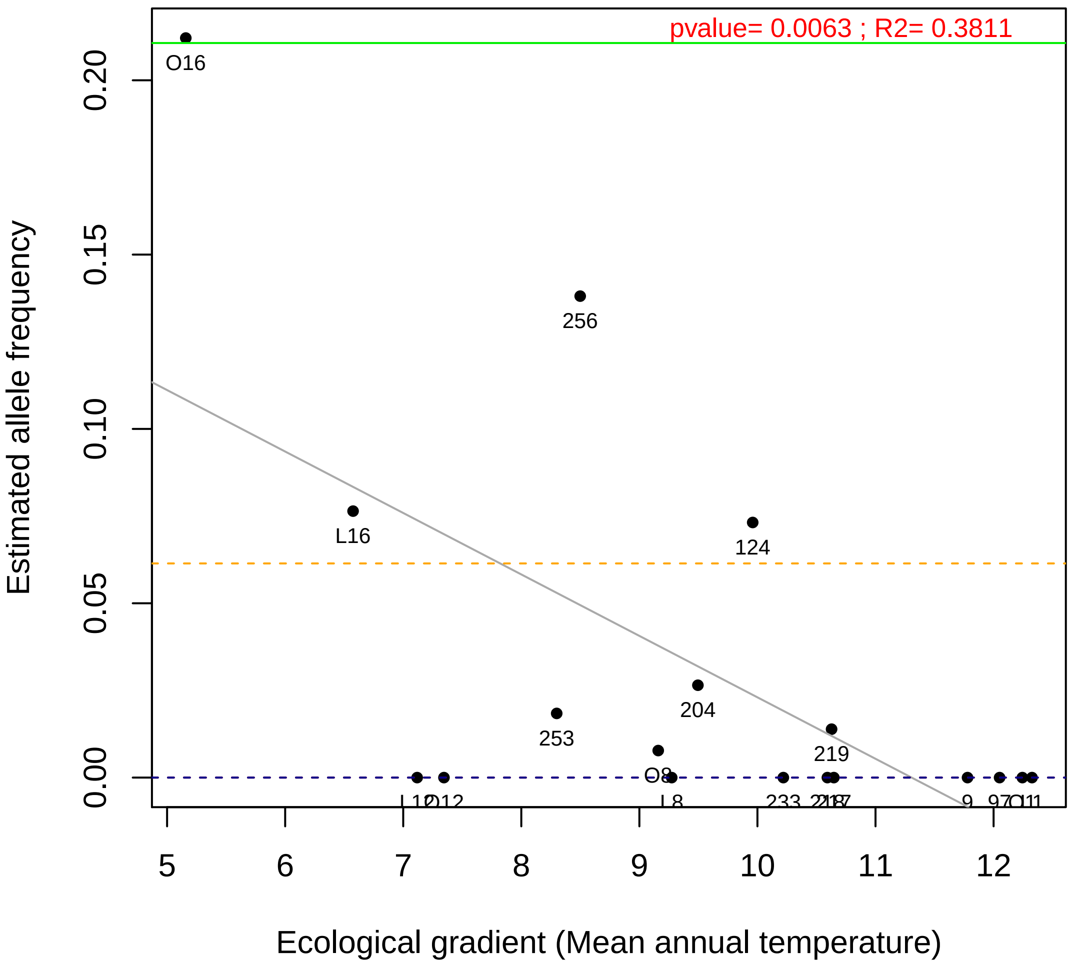

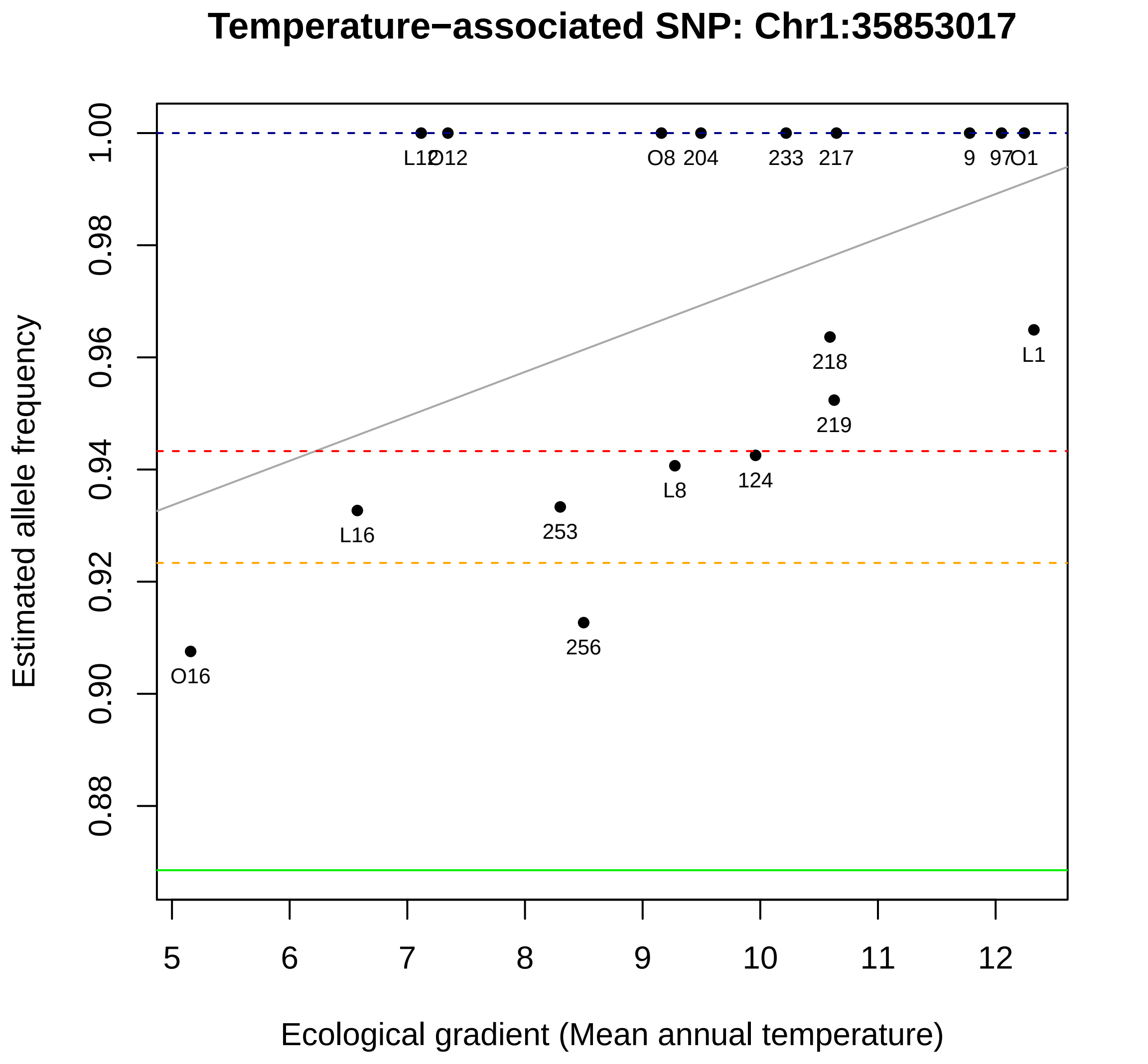
Temperature−associated SNP: Chr1:35853017
Estimated allele frequency
Ecological gradient (Mean annual temperature)
1.00
0.98
0.96
0.94
0.92
0.90
0.88
5
6
7
8
9
10
11
12
L12
O12
O8
204
233
217
9
97
O1
218
L1
219
124
L8
L16
253
256
O16

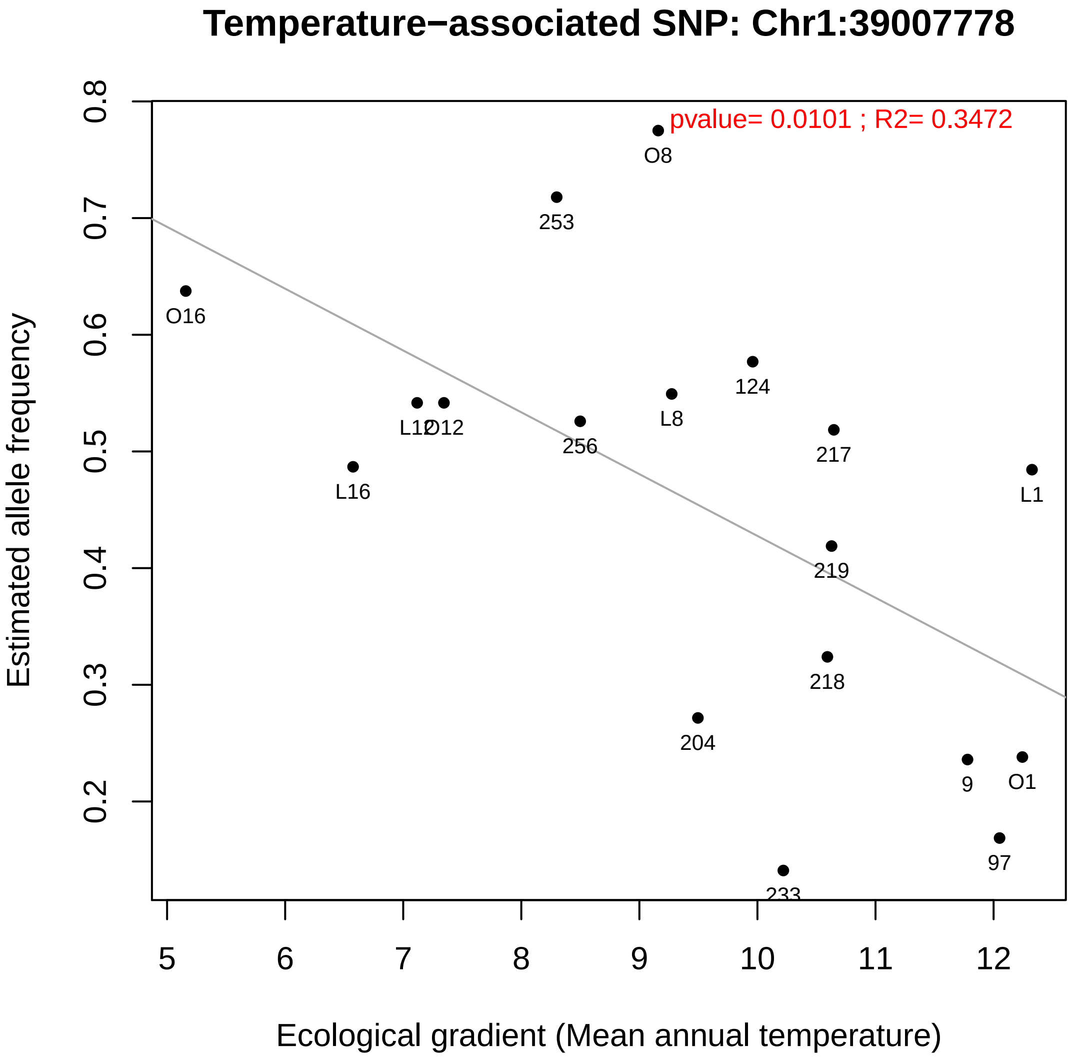
Temperature−associated SNP: Chr1:39007778
pvalue= 0.0101 ; R2= 0.3472
O8
253
O16
124
L8
L12
O12
256
217
L16
L1
219
218
204
9
O1
97
233
0.8
0.7
0.6
0.5
0.4
0.3
0.2
Estimated allele frequency
5
6
7
8
9
10
11
12
Ecological gradient (Mean annual temperature)

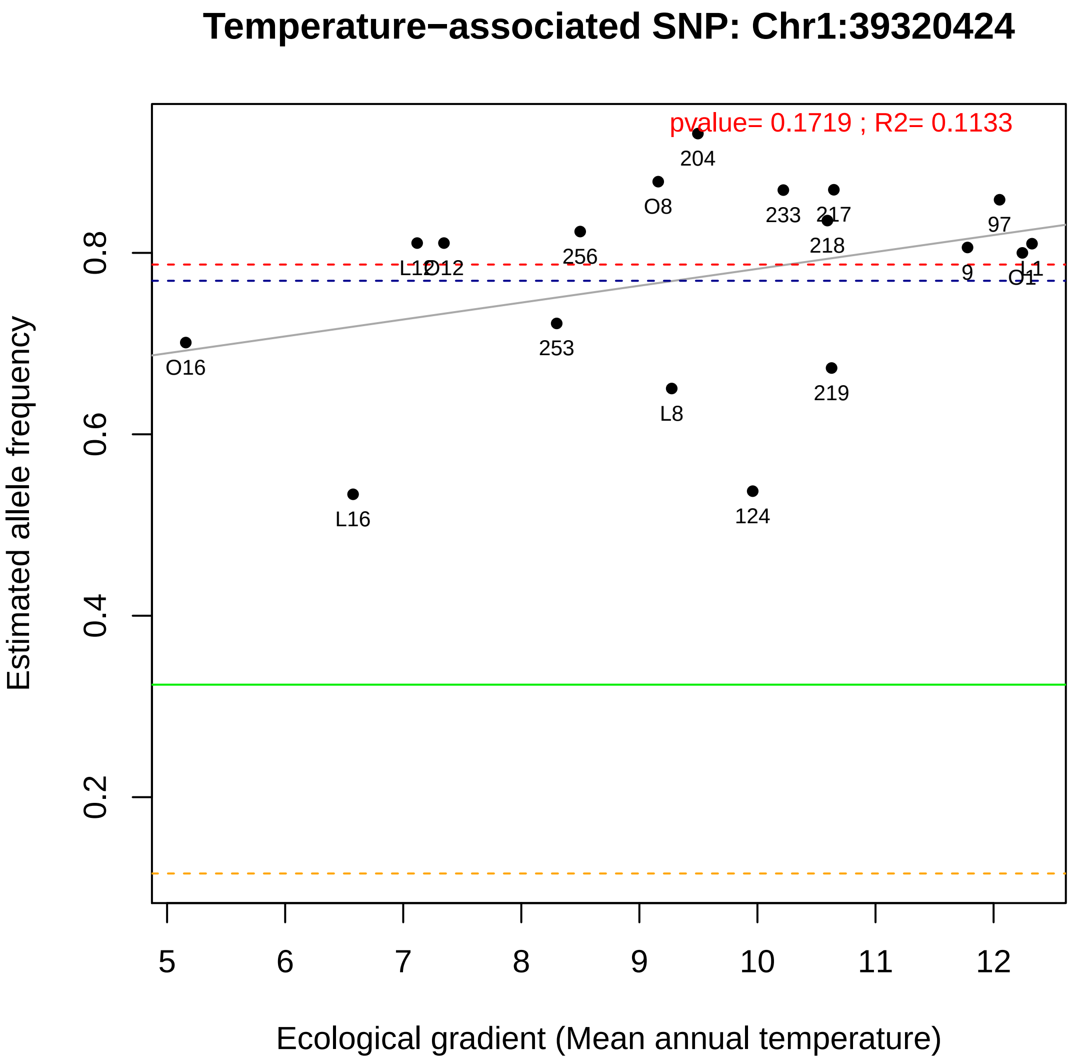
Temperature−associated SNP: Chr1:39320424
pvalue= 0.1719 ; R2= 0.1133
Estimated allele frequency
Ecological gradient (Mean annual temperature)
0.2
0.4
0.6
0.8
5
6
7
8
9
10
11
12
O16
L16
L12
O12
253
256
O8
L8
204
124
233
217
218
219
9
97
O1
L1

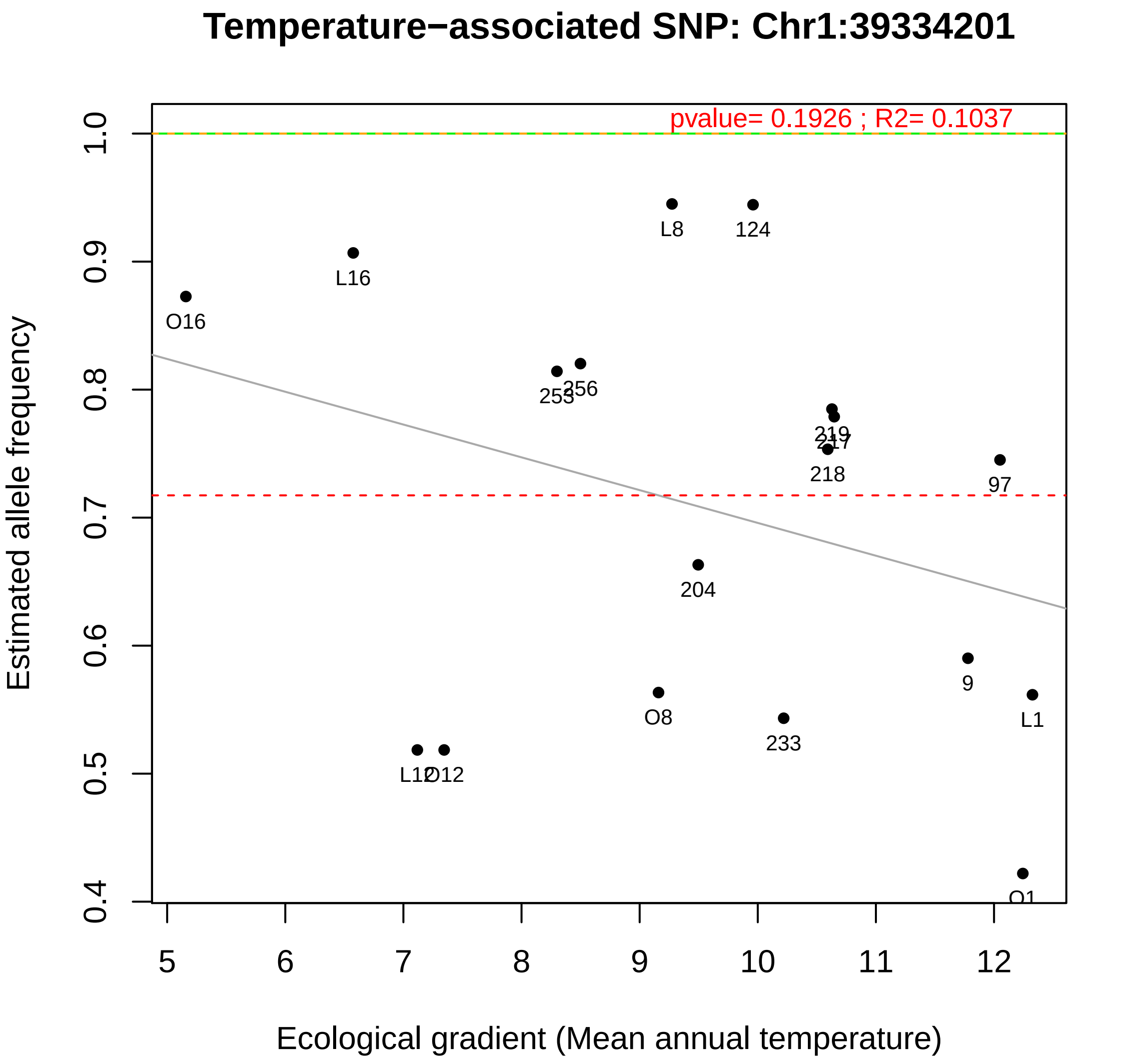

Temperature−associated SNP: Chr1:39334201
pvalue= 0.1926 ; R2= 0.1037
Estimated allele frequency
Ecological gradient (Mean annual temperature)
1.0
0.9
0.8
0.7
0.6
0.5
0.4
5
6
7
8
9
10
11
12
O16
L16
L8
124
253
256
219
217
218
97
204
9
O8
233
L1
L12
O12
O1

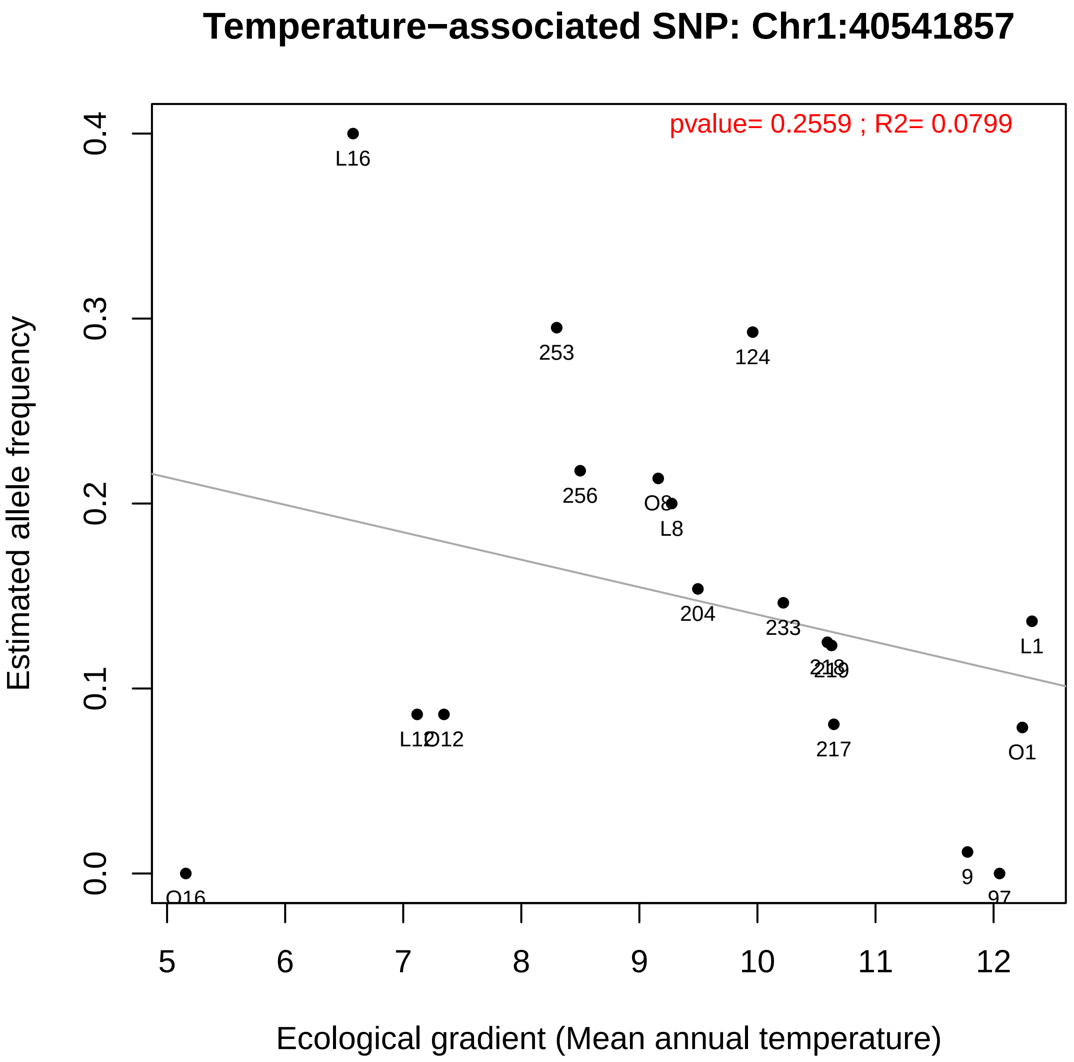

Temperature−associated SNP: Chr1:40541857
pvalue= 0.2559 ; R2= 0.0799
Estimated allele frequency
Ecological gradient (Mean annual temperature)
0.0
0.1
0.2
0.3
0.4
5
6
7
8
9
10
11
12
L16
253
124
256
O8
L8
204
233
L1
218
219
L12
O12
217
O1
9
97
O16

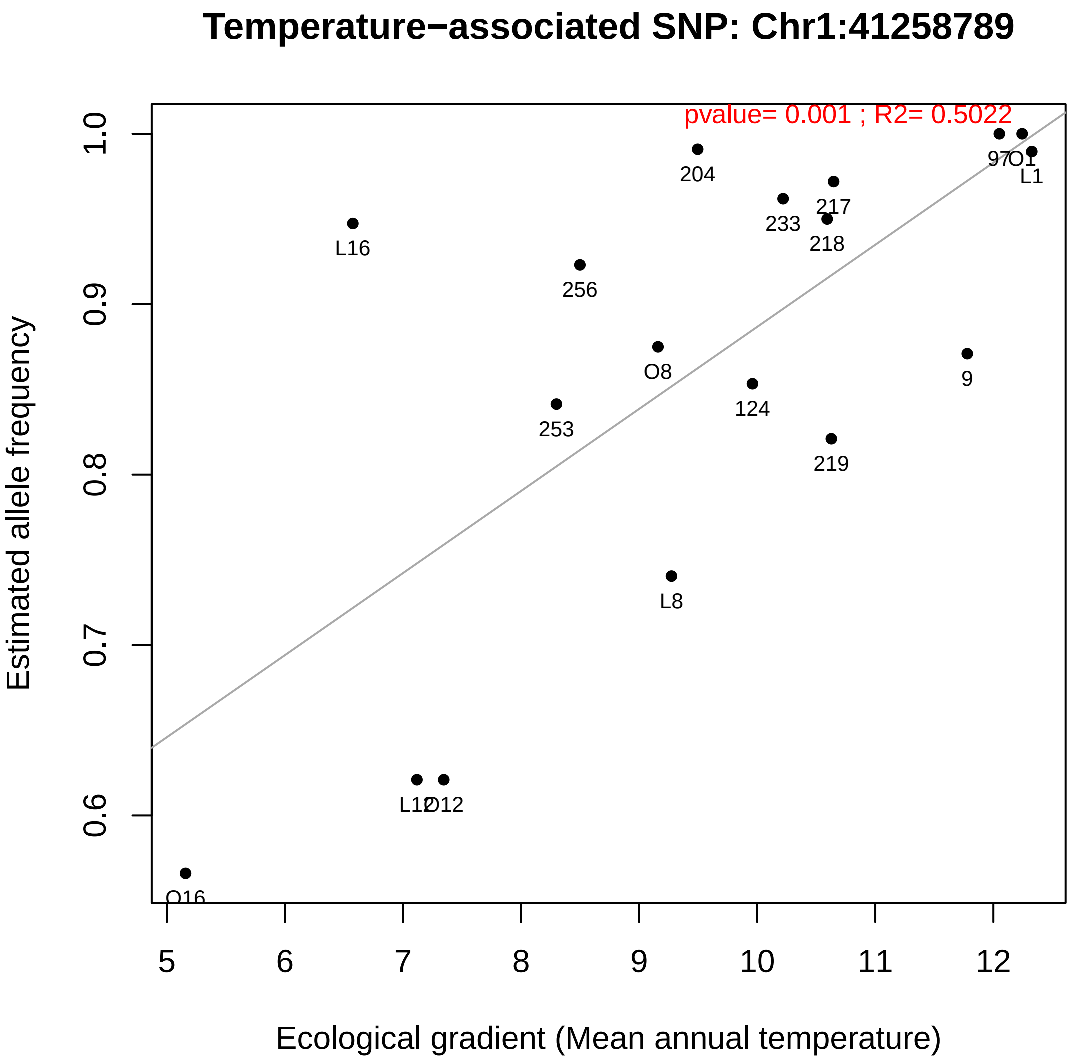
Temperature−associated SNP: Chr1:41258789
pvalue= 0.001 ; R2= 0.5022
Estimated allele frequency
Ecological gradient (Mean annual temperature)
0.6
0.7
0.8
0.9
1.0
5
6
7
8
9
10
11
12
204
97
O1
L1
233
217
218
L16
256
O8
9
124
253
219
L8
O12
O16

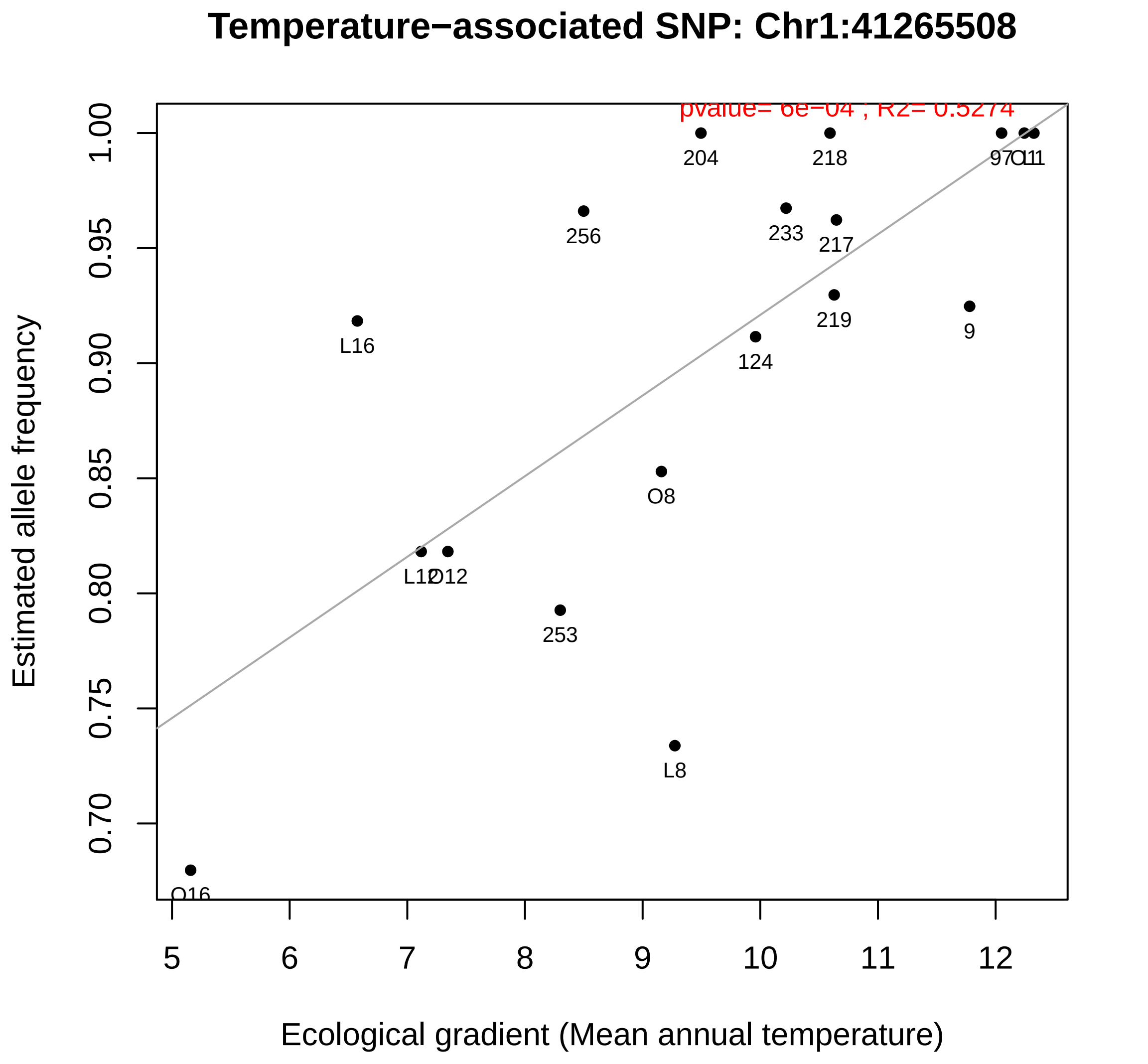
Temperature−associated SNP: Chr1:41265508
pvalue= 6e−04 ; R2= 0.5274
Estimated allele frequency
Ecological gradient (Mean annual temperature)
1.00
0.95
0.90
0.85
0.80
0.75
0.70
5
6
7
8
9
10
11
12
204
218
97
O1
1
256
233
217
219
9
L16
124
O8
L12
O12
253
L8
O16

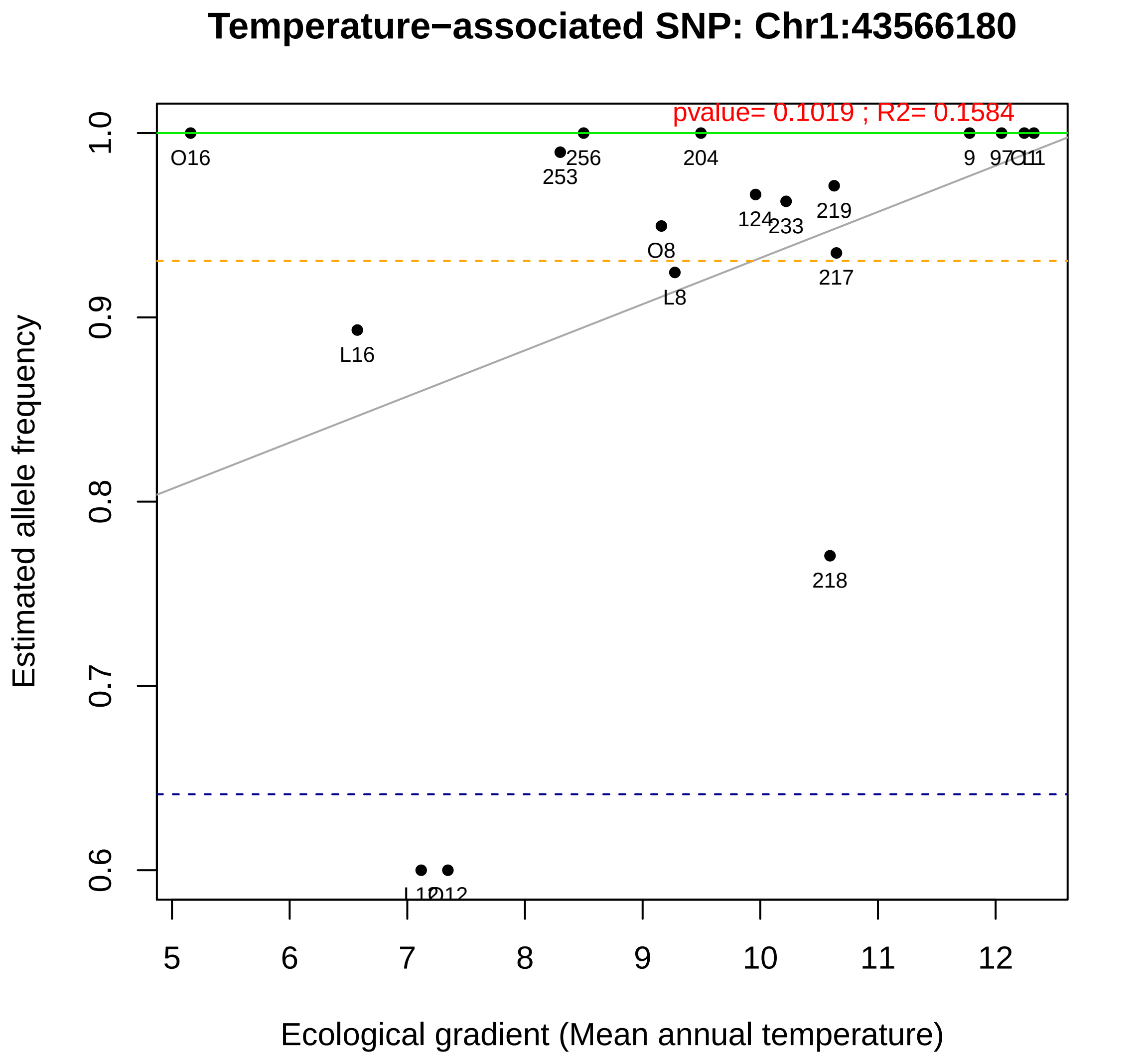

Temperature−associated SNP: Chr1:43566180
pvalue= 0.1019 ; R2= 0.1584
Estimated allele frequency
Ecological gradient (Mean annual temperature)
1.0
0.9
0.8
0.7
0.6
5
6
7
8
9
10
11
12
O16
256
253
204
9
124
233
219
O8
217
L8
L16
218

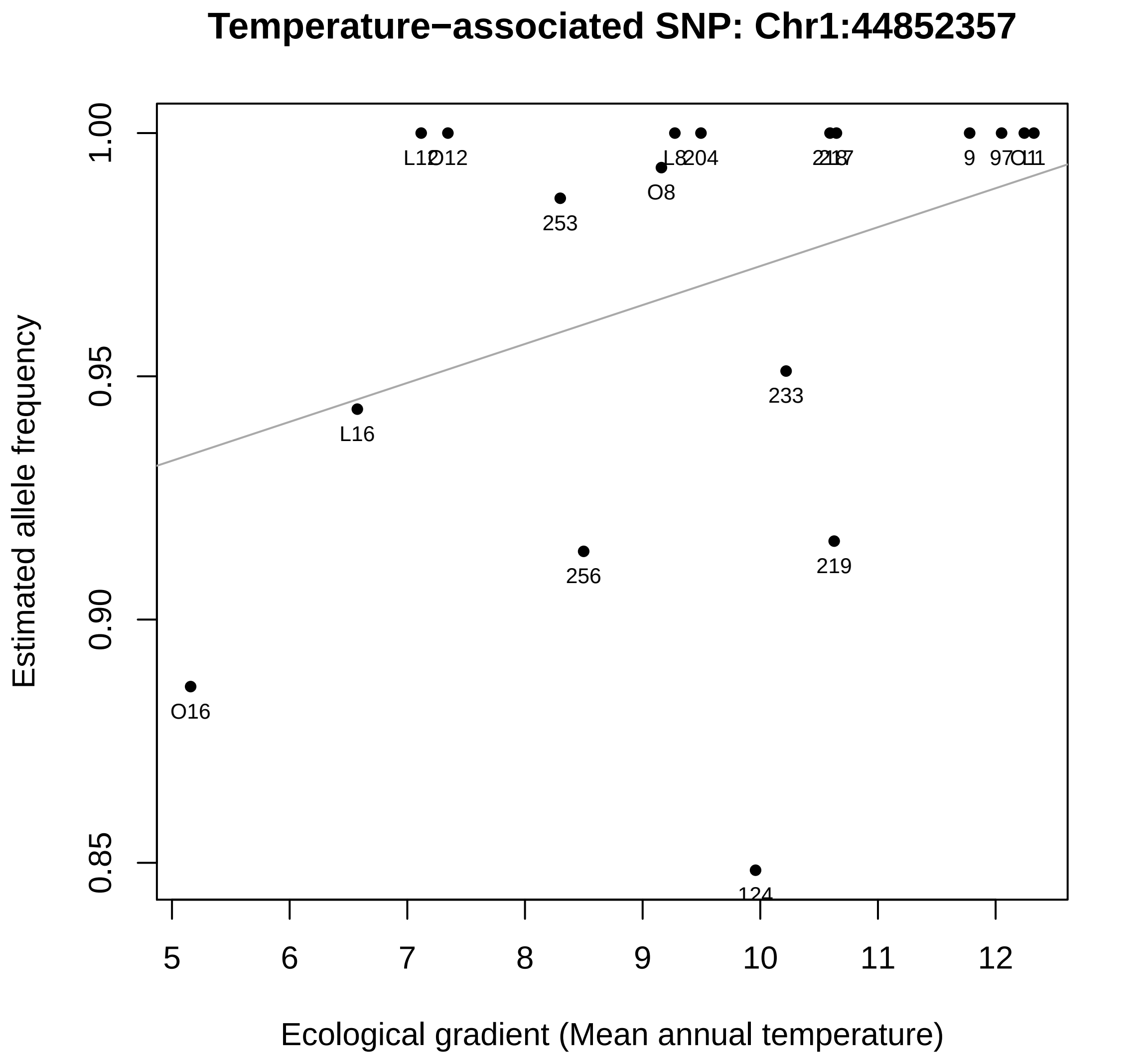

Temperature−associated SNP: Chr1:44852357
Estimated allele frequency
Ecological gradient (Mean annual temperature)
0.85
0.90
0.95
1.00
5
6
7
8
9
10
11
12
L12
O12
L8
204
218
217
9
97
O11
O8
253
L16
233
256
219
O16
124

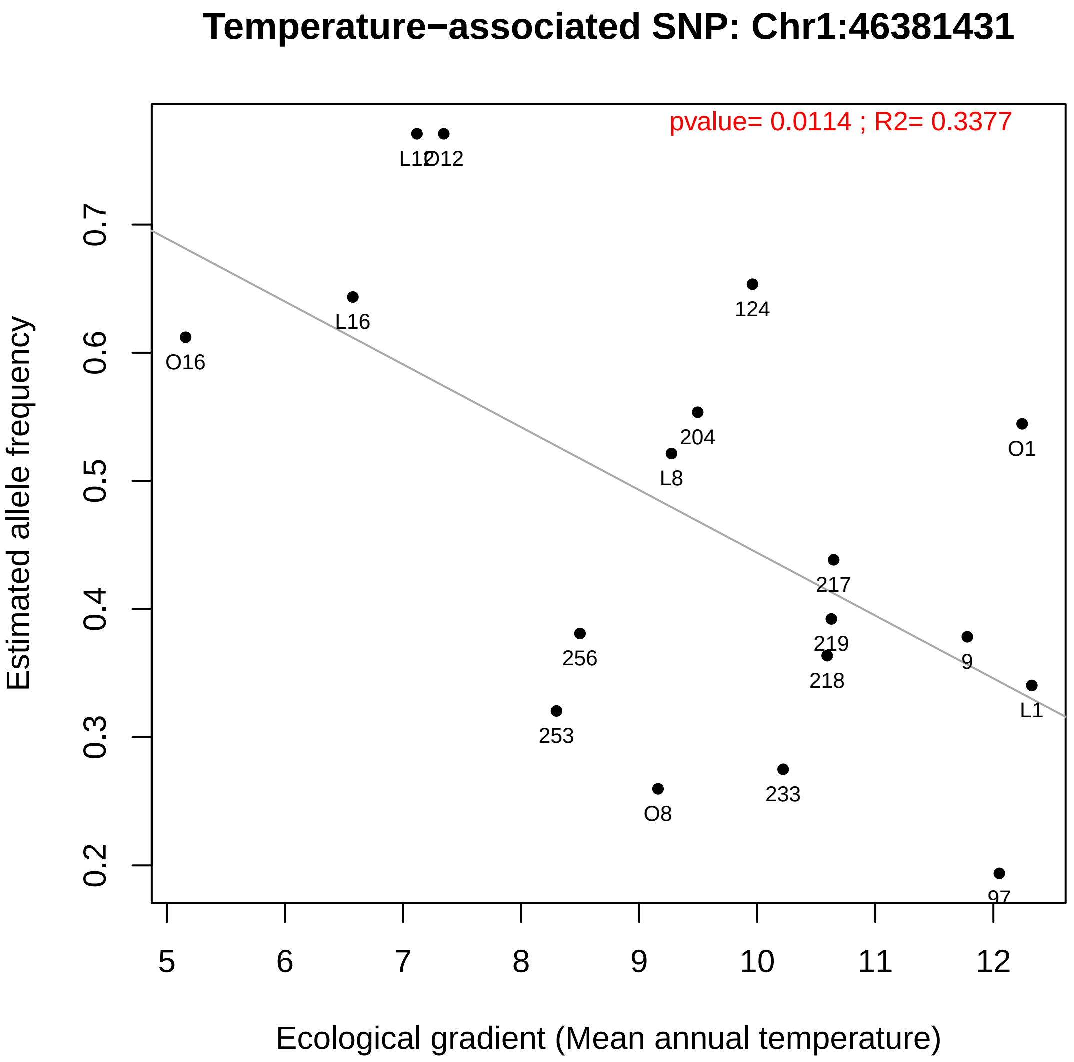
Temperature−associated SNP: Chr1:46381431
pvalue= 0.0114 ; R2= 0.3377
Estimated allele frequency
Ecological gradient (Mean annual temperature)
0.2
0.3
0.4
0.5
0.6
0.7
5
6
7
8
9
10
11
12
L12
O12
L16
O16
124
204
L8
O1
217
219
218
9
L1
256
253
O8
233
97

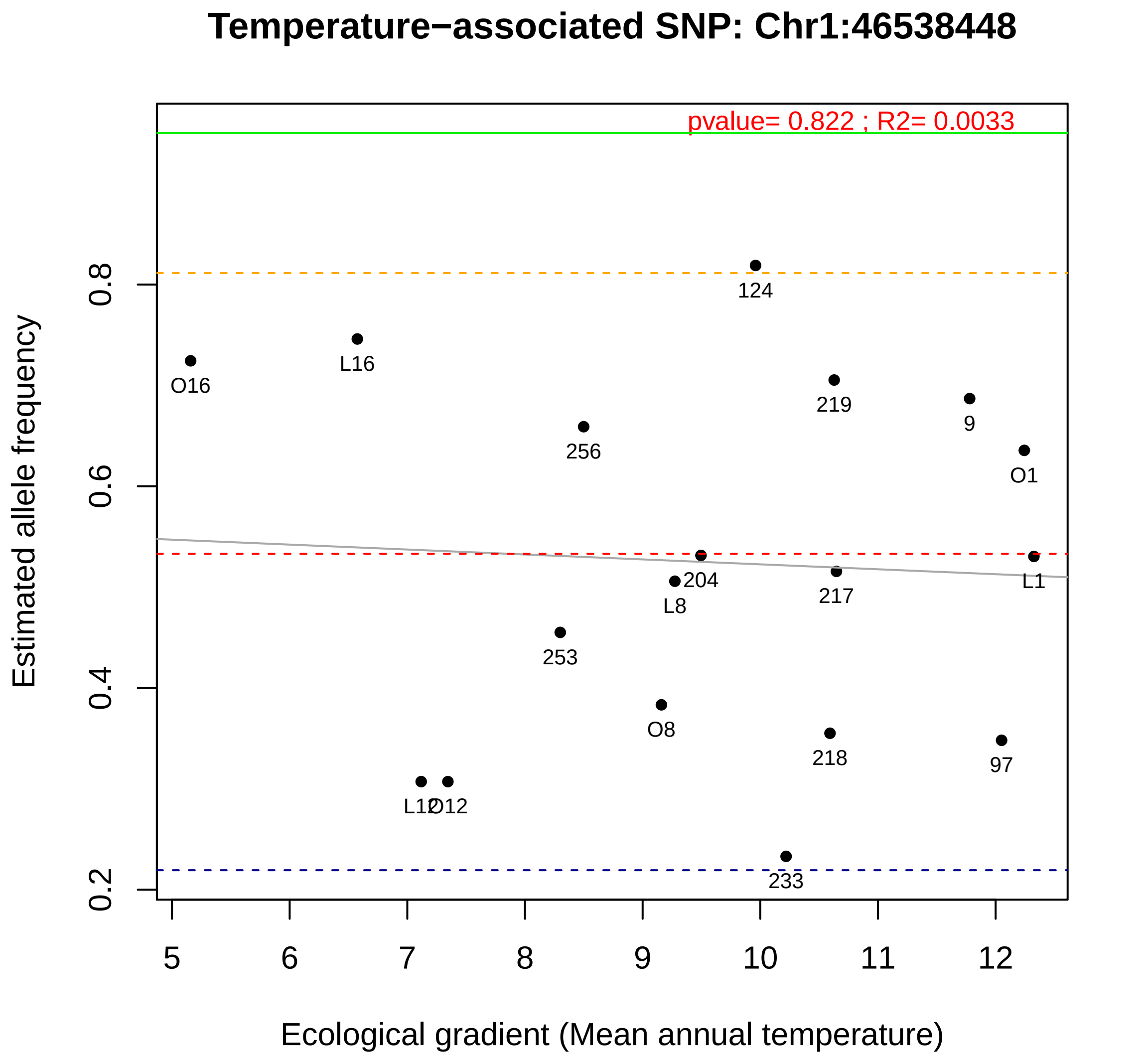
Temperature−associated SNP: Chr1:46538448
pvalue= 0.822 ; R2= 0.0033
Estimated allele frequency
0.2
0.4
0.6
0.8
Ecological gradient (Mean annual temperature)
5
6
7
8
9
10
11
12
O16
L16
124
219
9
256
O1
204
L8
217
L1
253
O8
218
97
L12
O12
233

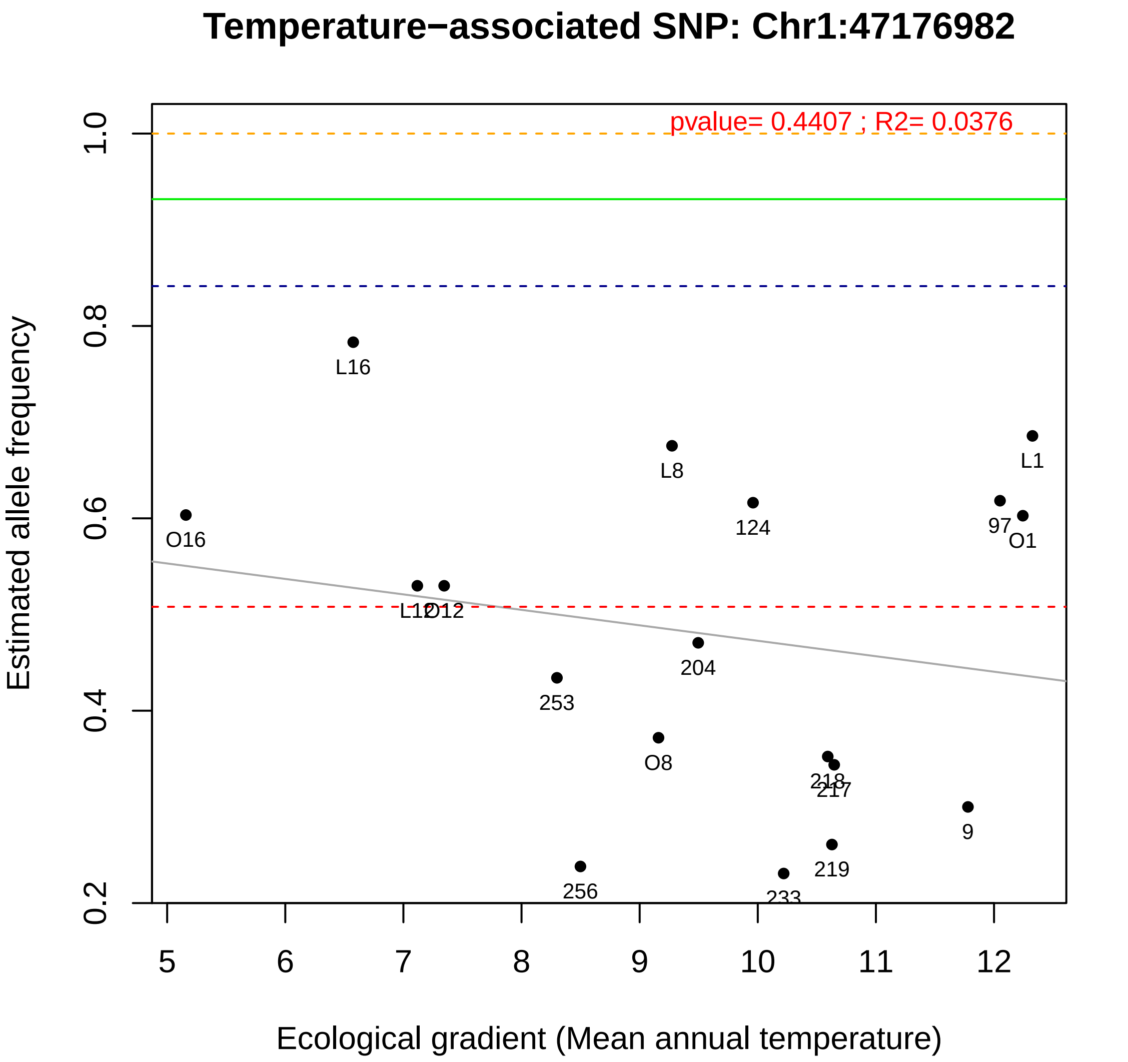

Temperature−associated SNP: Chr1:47176982
pvalue= 0.4407 ; R2= 0.0376
Estimated allele frequency
0.2
0.4
0.6
0.8
1.0
5
6
7
8
9
10
11
12
Ecological gradient (Mean annual temperature)
L16
L8
L1
O16
124
97
O1
L12
O12
204
253
O8
218
217
9
219
256
233

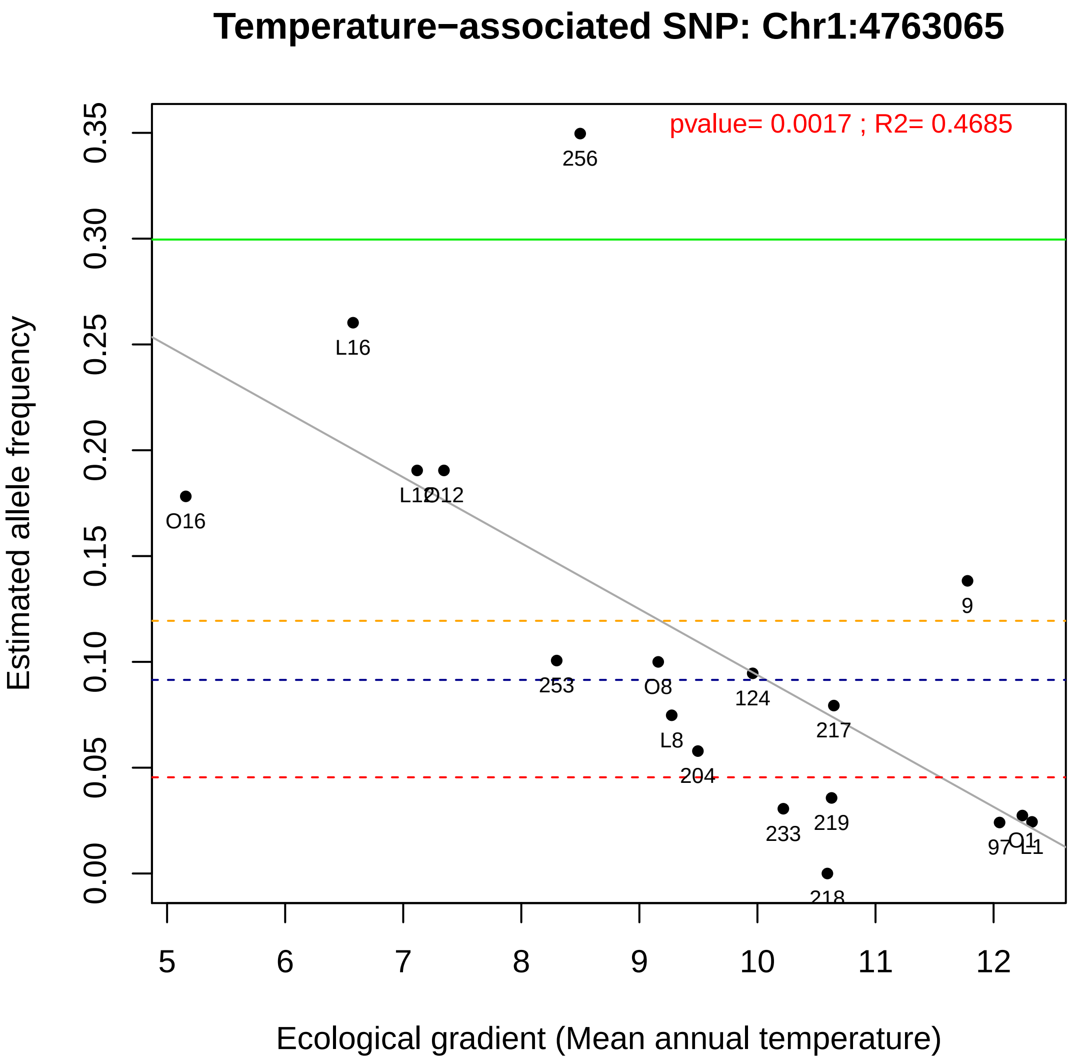

Temperature−associated SNP: Chr1:4763065
pvalue= 0.0017 ; R2= 0.4685
Estimated allele frequency
Ecological gradient (Mean annual temperature)
0.00
0.05
0.10
0.15
0.20
0.25
0.30
0.35
5
6
7
8
9
10
11
12
256
L16
L12
O12
O16
9
253
O8
124
217
L8
204
219
233
218
97
O1
L1

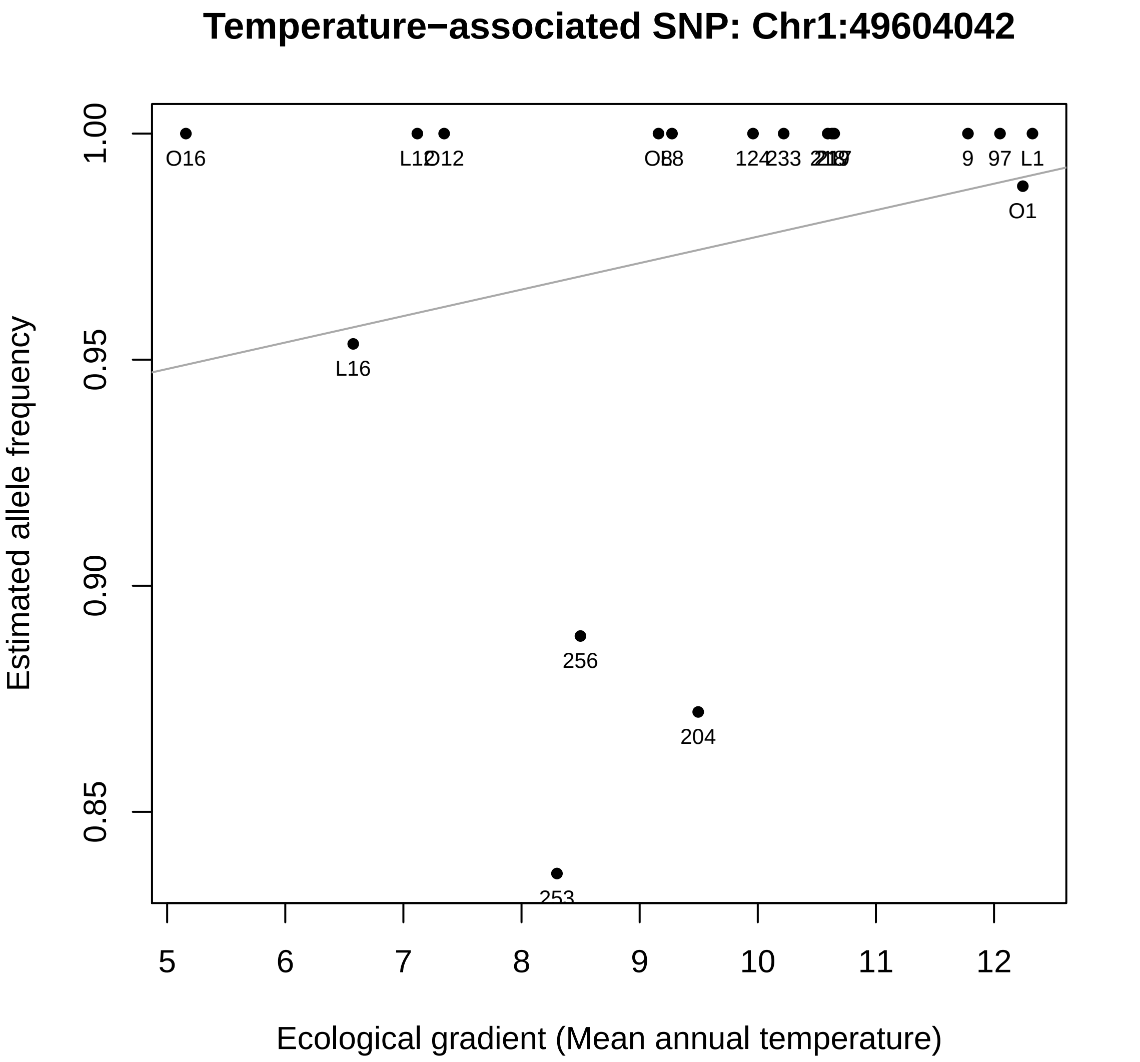
Temperature−associated SNP: Chr1:49604042
Estimated allele frequency
Ecological gradient (Mean annual temperature)
1.00
0.95
0.90
0.85
5
6
7
8
9
10
11
12
O16
L12
O12
O8
L8
124
233
9
97
L1
O1
L16
256
204
253

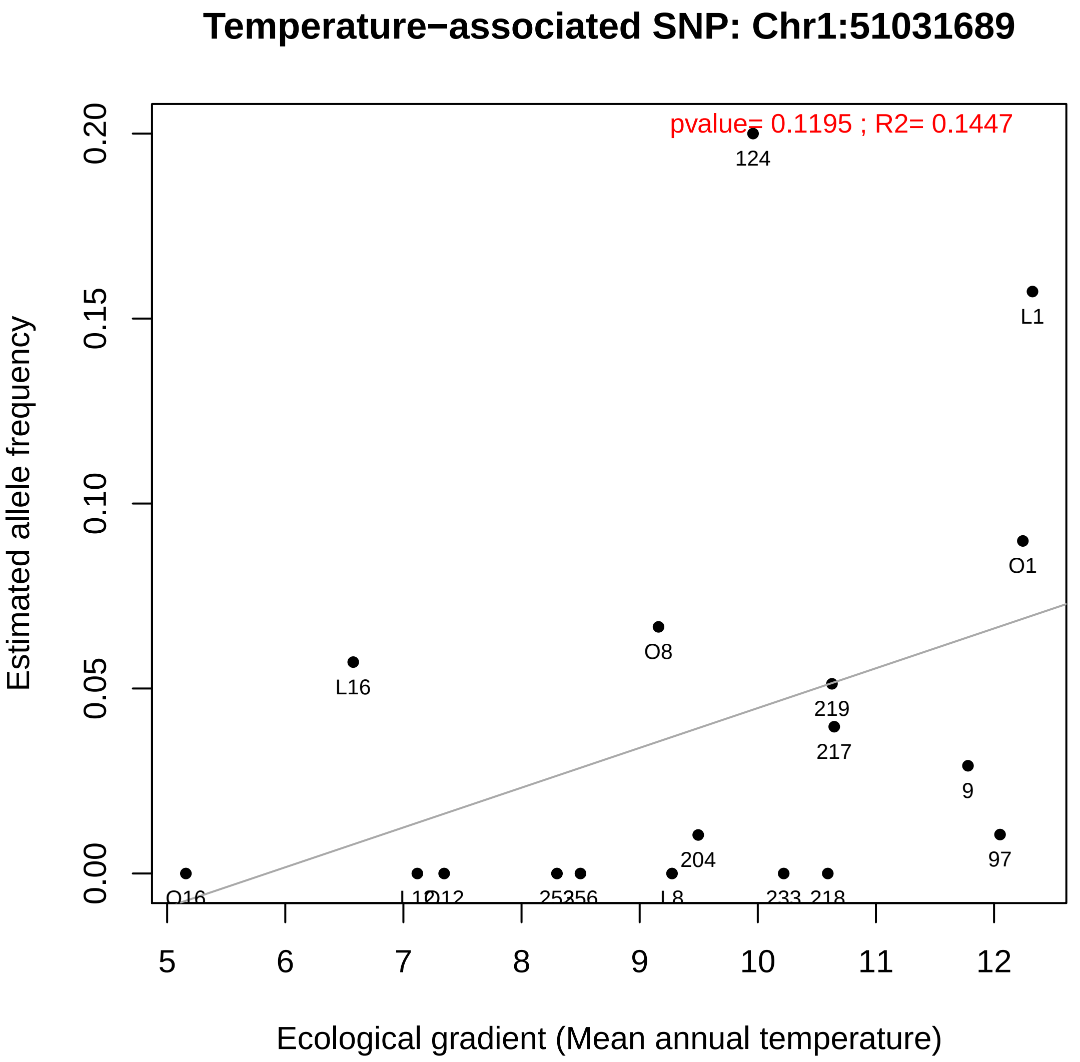

Temperature−associated SNP: Chr1:51031689
pvalue= 0.1195 ; R2= 0.1447
Estimated allele frequency
0.00
0.05
0.10
0.15
0.20
5
6
7
8
9
10
11
12
Ecological gradient (Mean annual temperature)
124
L1
O1
O8
L16
219
217
9
204
97
233
218

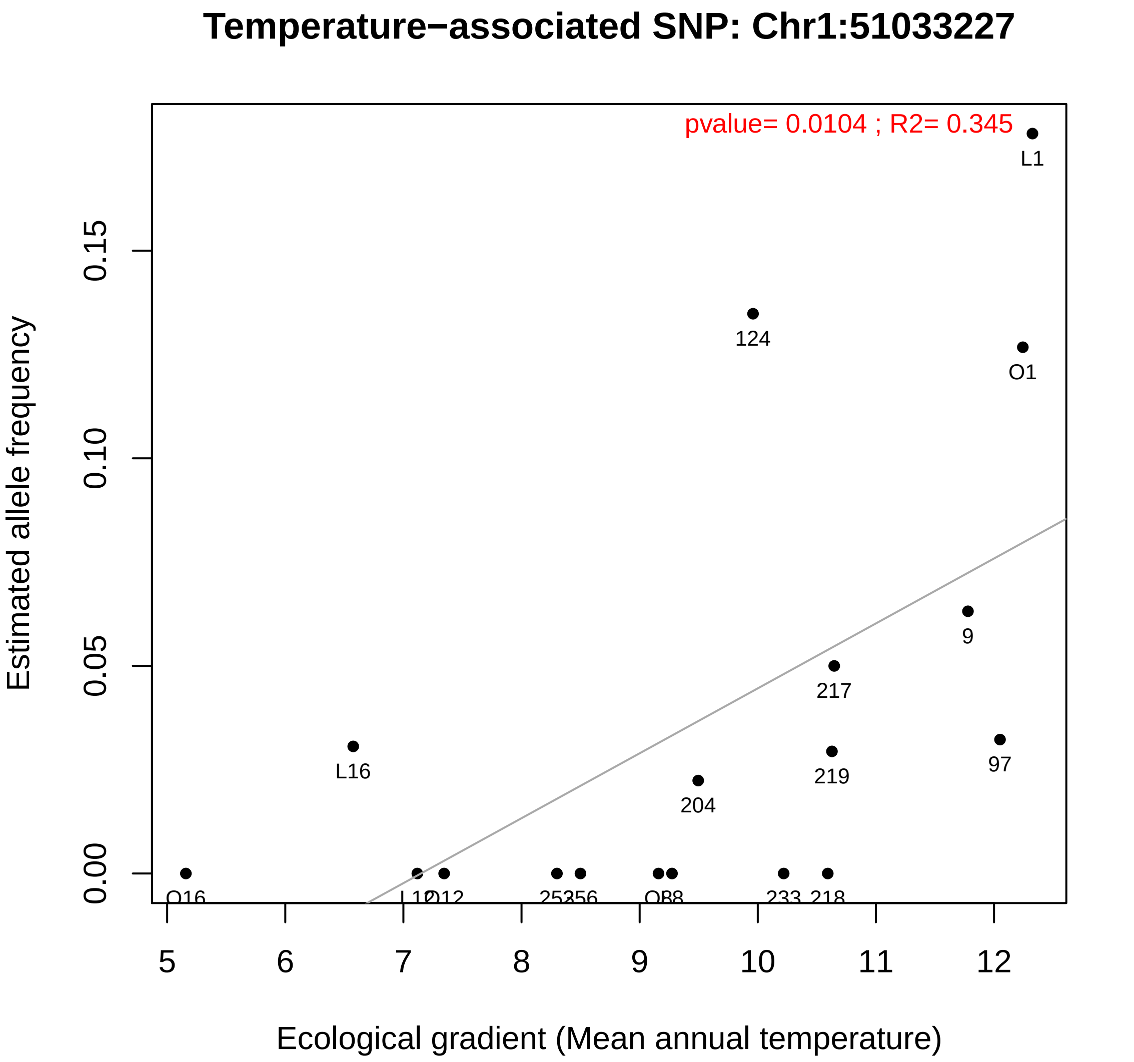
Temperature−associated SNP: Chr1:51033227
pvalue= 0.0104 ; R2= 0.345
Estimated allele frequency
0.00
0.05
0.10
0.15
Ecological gradient (Mean annual temperature)
5
6
7
8
9
10
11
12
L1
124
O1
9
217
97
219
L16
204

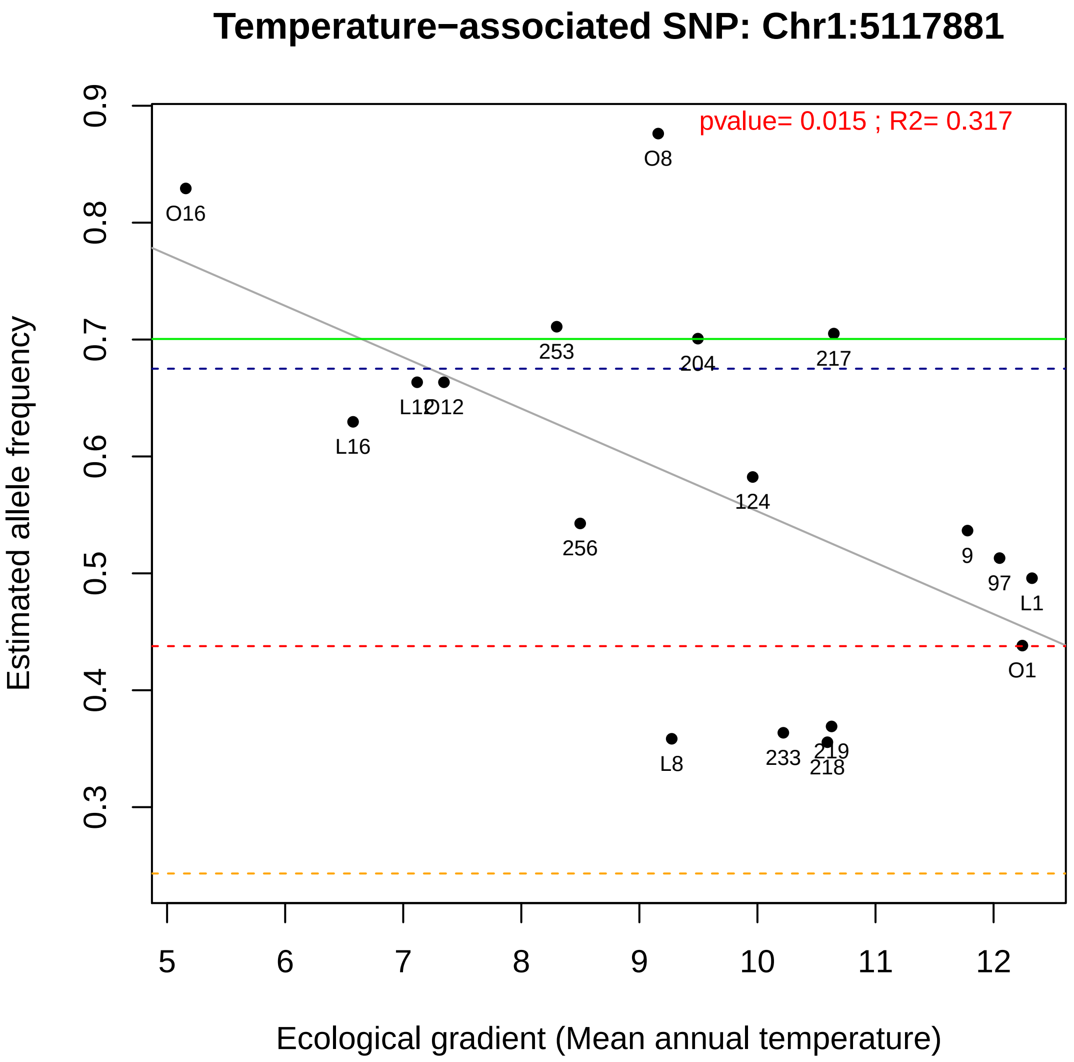

Temperature−associated SNP: Chr1:5117881
pvalue= 0.015 ; R2= 0.317
Estimated allele frequency
Ecological gradient (Mean annual temperature)
0.3
0.4
0.5
0.6
0.7
0.8
0.9
5
6
7
8
9
10
11
12
O16
O8
253
204
217
L12
O12
L16
124
256
9
97
L1
O1
L8
233
219
218

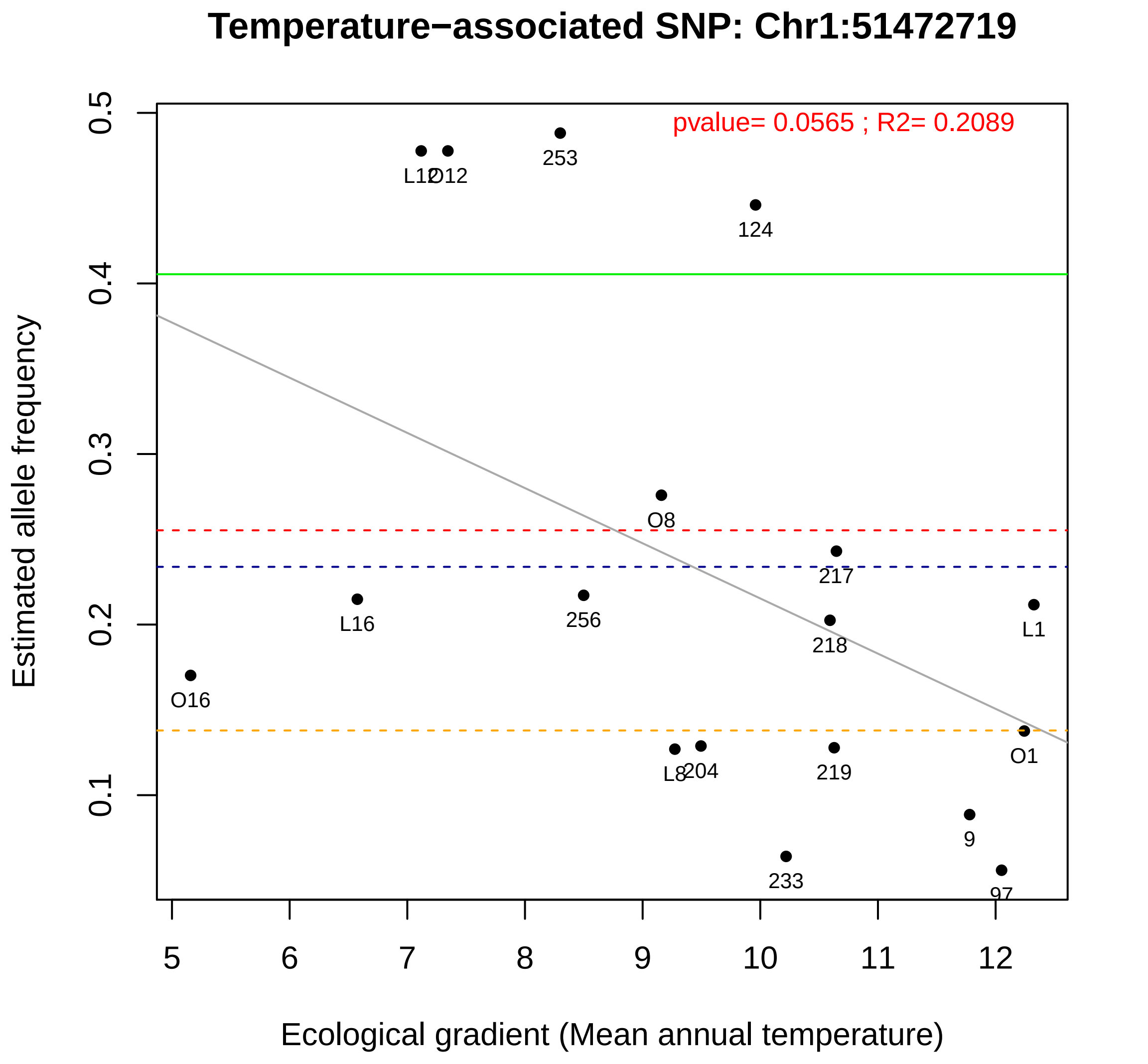
Temperature−associated SNP: Chr1:51472719
pvalue= 0.0565 ; R2= 0.2089
Estimated allele frequency
Ecological gradient (Mean annual temperature)
0.1
0.2
0.3
0.4
0.5
5
6
7
8
9
10
11
12
L12
O12
253
124
O8
217
L16
256
218
L1
O16
O1
L8
204
219
9
233
97

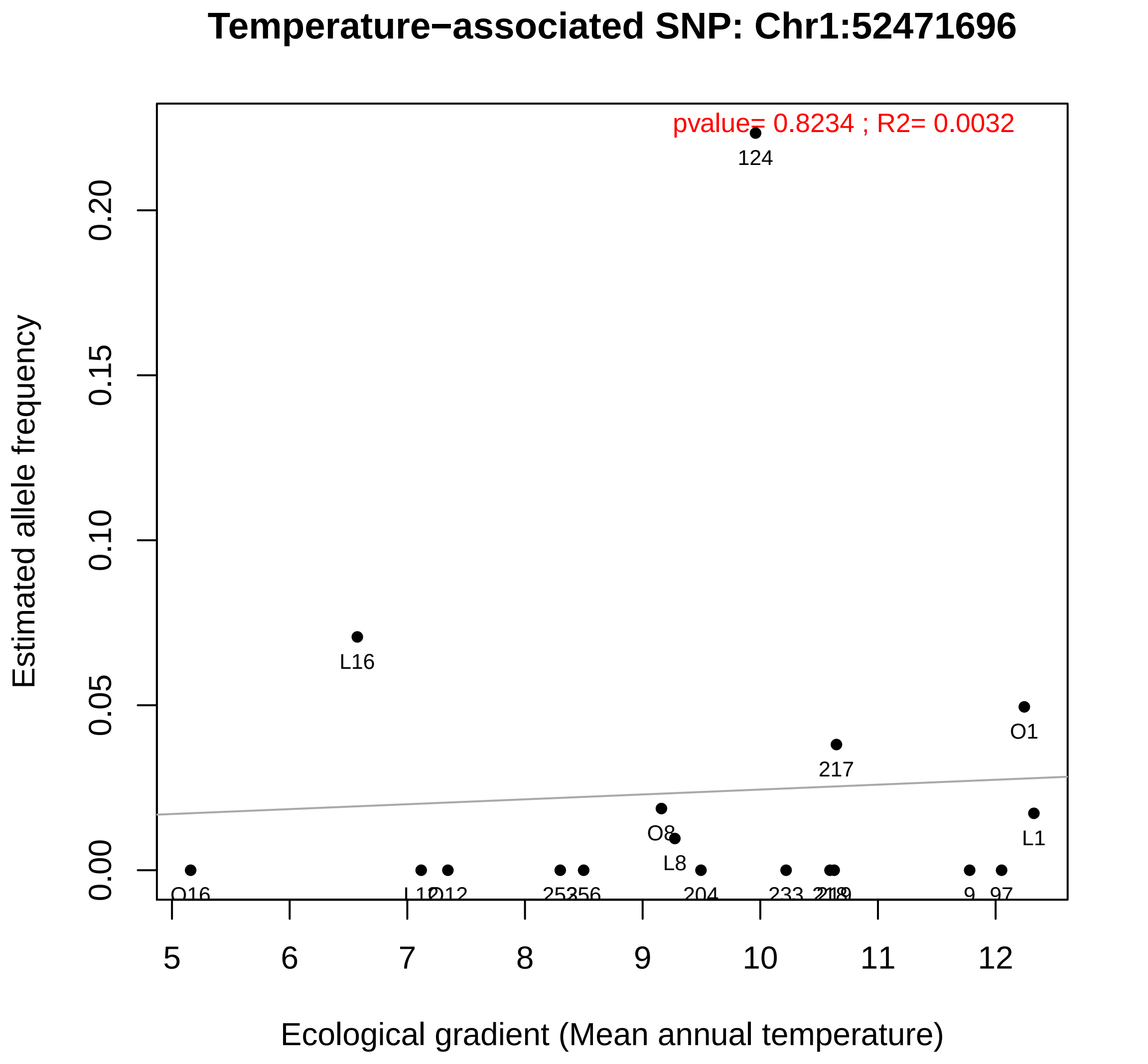
Temperature−associated SNP: Chr1:52471696
pvalue= 0.8234 ; R2= 0.0032
Estimated allele frequency
0.00
0.05
0.10
0.15
0.20
124
L16
O1
217
L1
O8
L8
O16
204
233
9
97
5
6
7
8
9
10
11
12
Ecological gradient (Mean annual temperature)

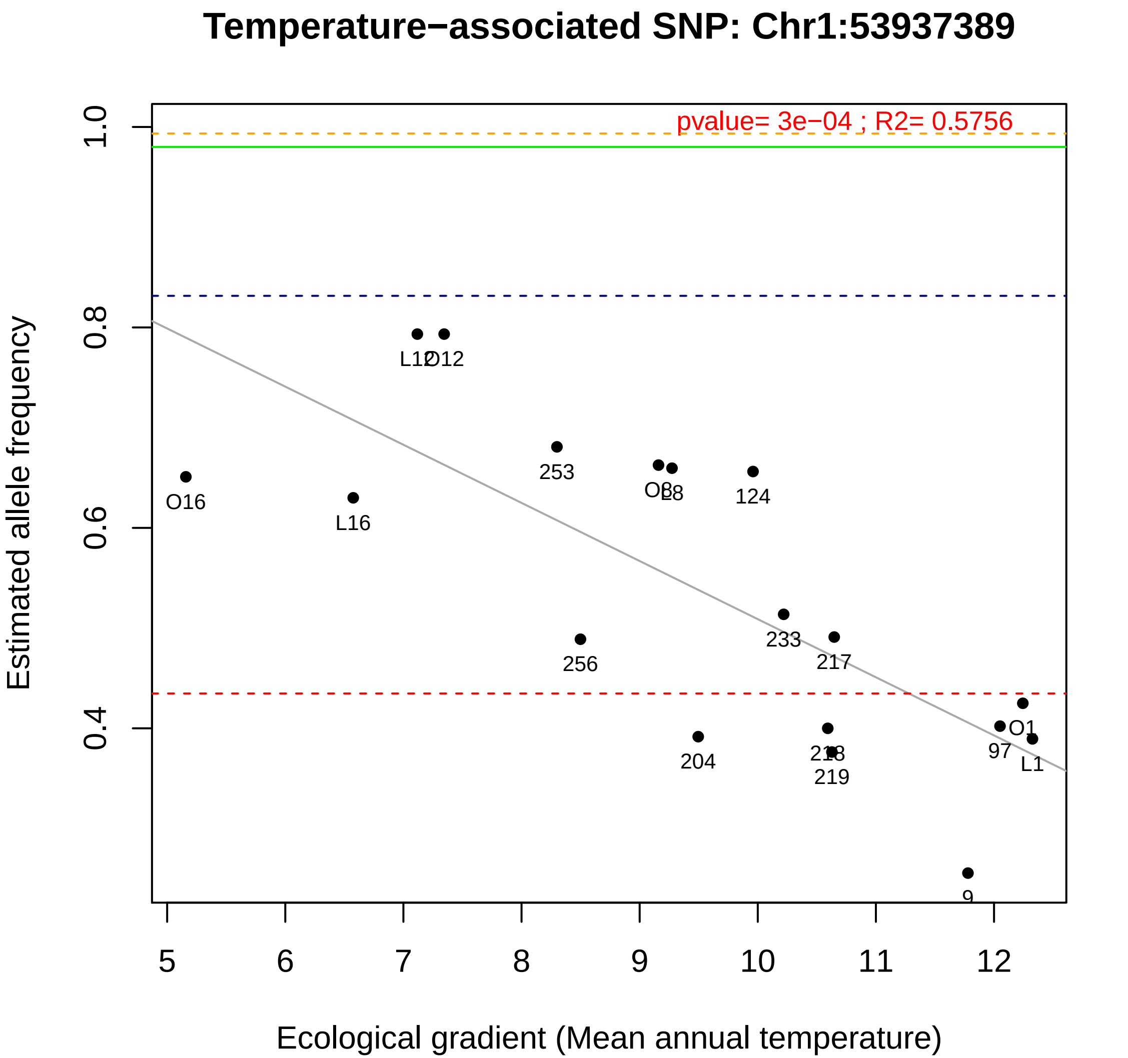

Temperature−associated SNP: Chr1:53937389
pvalue= 3e−04 ; R2= 0.5756
Estimated allele frequency
Ecological gradient (Mean annual temperature)
0.4
0.6
0.8
1.0
5
6
7
8
9
10
11
12
O16
L16
L12
O12
253
256
O8
L8
124
204
233
217
218
219
9
97
O1
L1

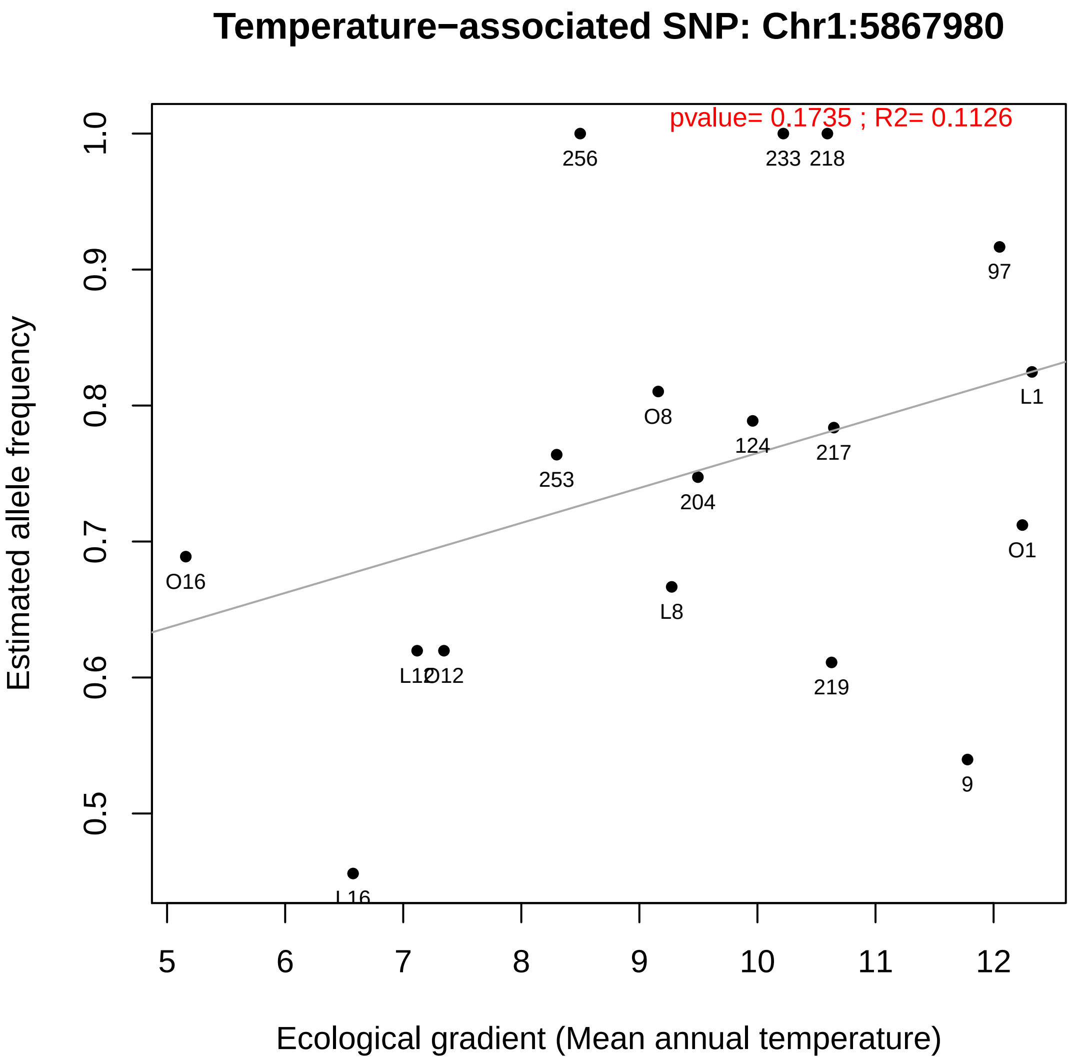

Temperature−associated SNP: Chr1:5867980
pvalue= 0.1735 ; R2= 0.1126
Estimated allele frequency
Ecological gradient (Mean annual temperature)
1.0
0.9
0.8
0.7
0.6
0.5
5
6
7
8
9
10
11
12
256
233
218
97
L1
O8
124
217
253
204
O1
O16
L8
L12
O12
219
9
L16

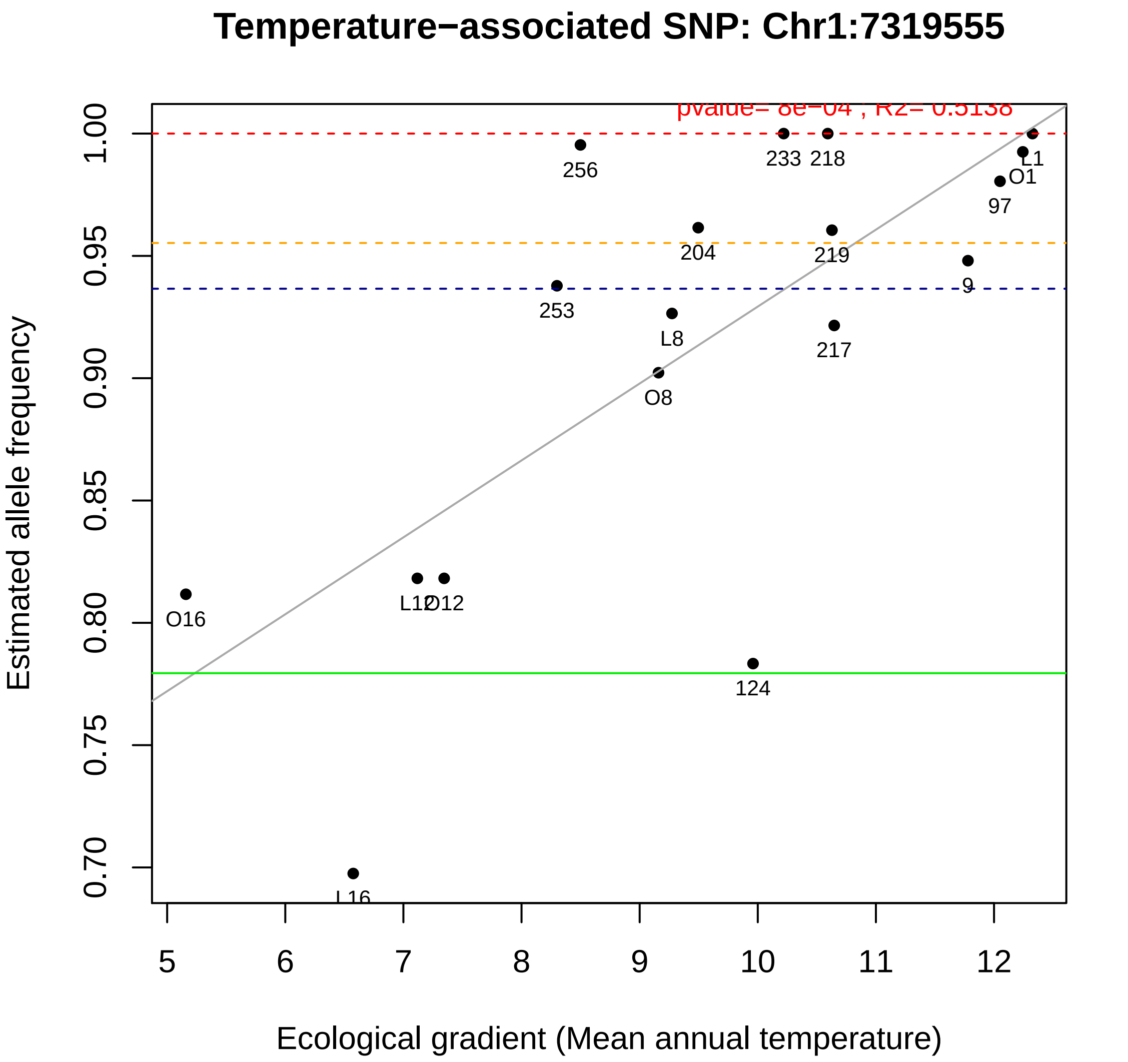
Temperature−associated SNP: Chr1:7319555
pvalue= 8e−04 , R2= 0.5138
Estimated allele frequency
1.00
0.95
0.90
0.85
0.80
0.75
0.70
256
233
218
L1
O1
97
204
219
9
253
L8
217
O8
L12
O12
O16
124
L16
5
6
7
8
9
10
11
12
Ecological gradient (Mean annual temperature)

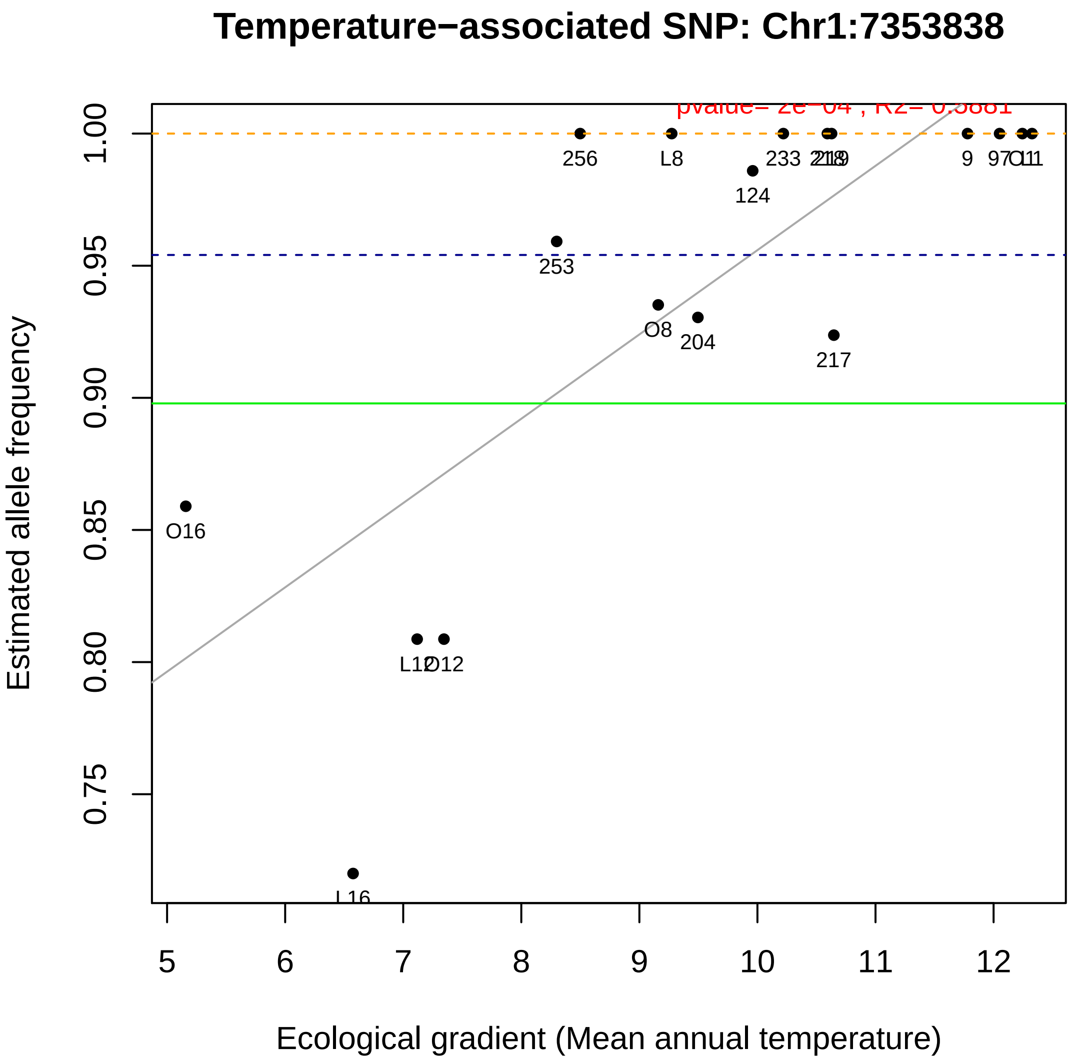

Temperature−associated SNP: Chr1:7353838
pvalue= 2e−04 , R2= 0.5881
Estimated allele frequency
Ecological gradient (Mean annual temperature)
1.00
0.95
0.90
0.85
0.80
0.75
5
6
7
8
9
10
11
12
256
L8
233
124
253
O8
204
217
9
O16
L12
O12
L16

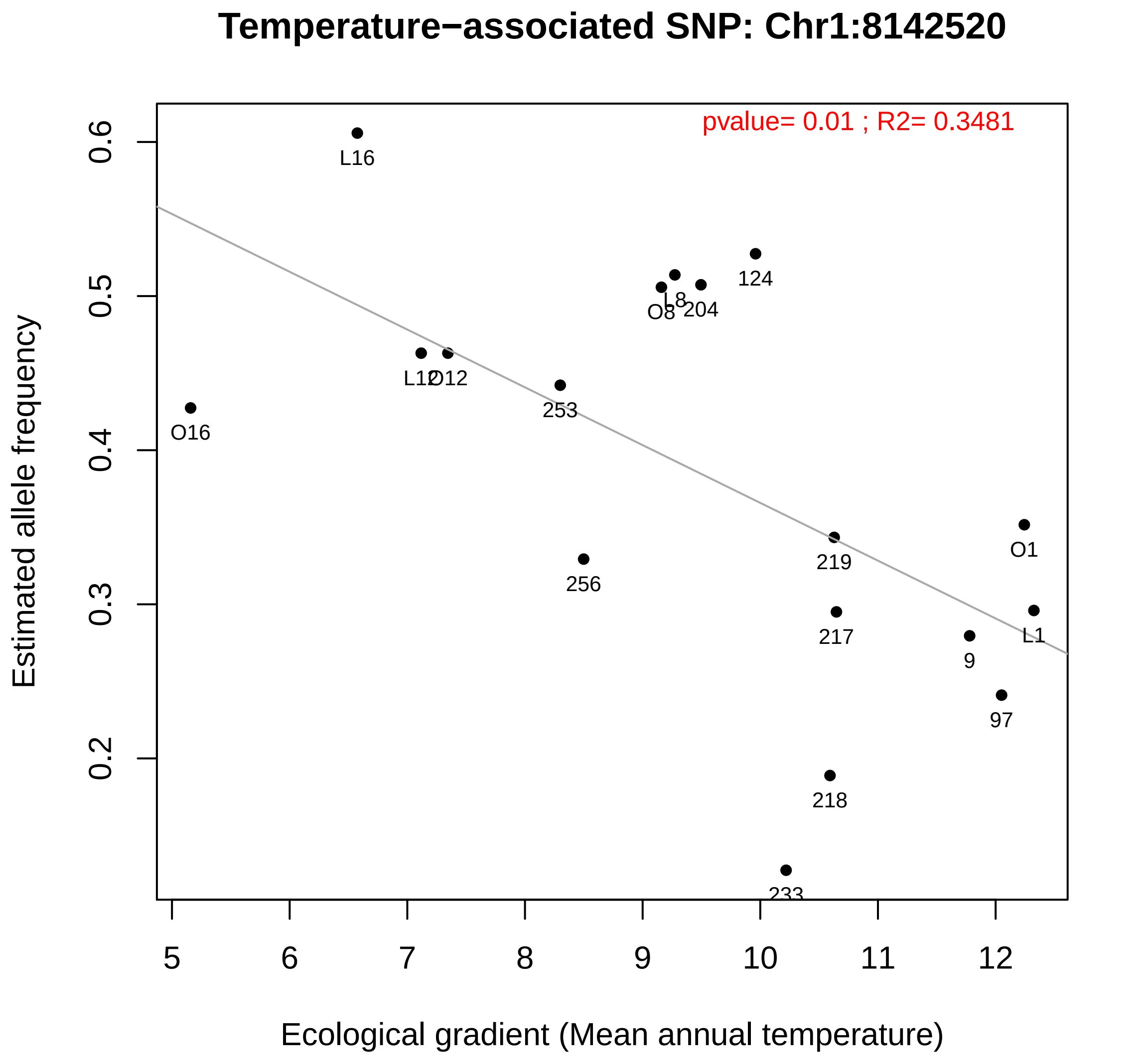
Temperature−associated SNP: Chr1:8142520
pvalue= 0.01 ; R2= 0.3481
Estimated allele frequency
Ecological gradient (Mean annual temperature)
0.2
0.3
0.4
0.5
0.6
5
6
7
8
9
10
11
12
L16
124
O8
L8
204
L12
O12
253
O16
219
O1
256
217
L1
9
97
218
233

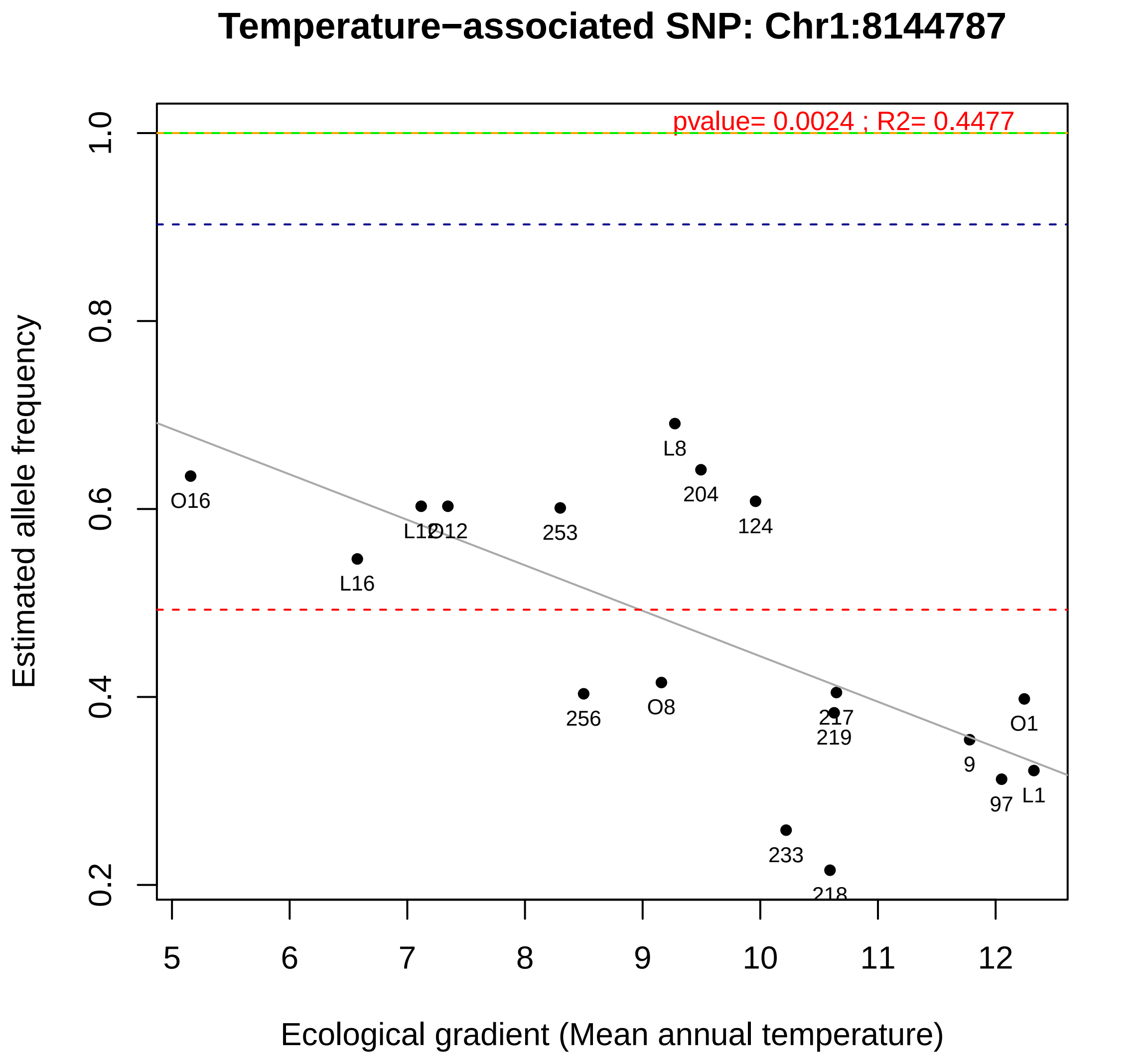
Temperature−associated SNP: Chr1:8144787
pvalue= 0.0024 ; R2= 0.4477
Estimated allele frequency
Ecological gradient (Mean annual temperature)
0.2
0.4
0.6
0.8
1.0
5
6
7
8
9
10
11
12
O16
L16
L12
O12
253
256
L8
204
O8
124
233
217
219
218
9
O1
97
L1

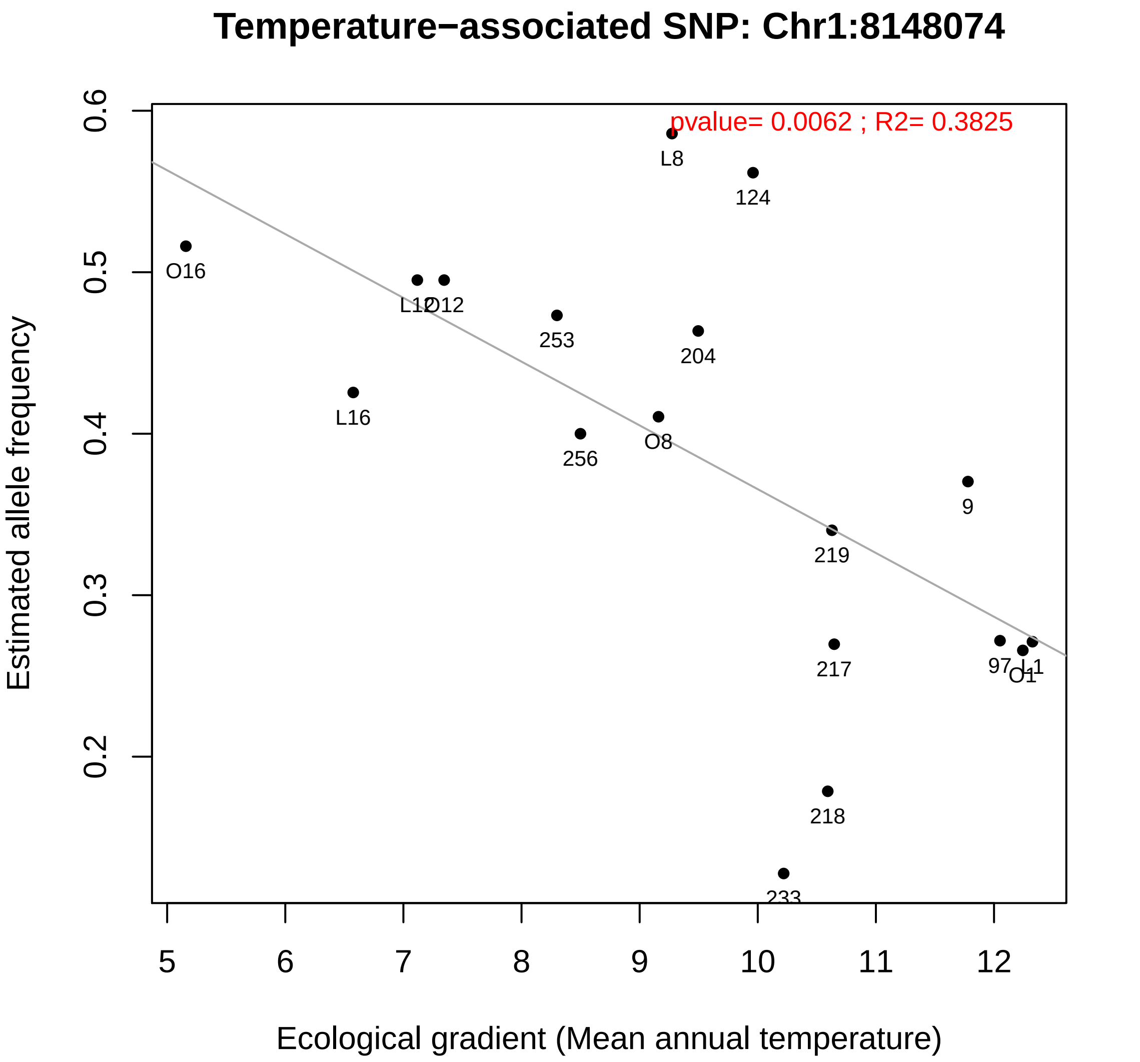

Temperature−associated SNP: Chr1:8148074
pvalue= 0.0062 ; R2= 0.3825
Estimated allele frequency
Ecological gradient (Mean annual temperature)
0.2
0.3
0.4
0.5
0.6
5
6
7
8
9
10
11
12
L8
124
O16
L12
O12
253
204
L16
256
O8
9
219
217
97
L1
O1
218
233

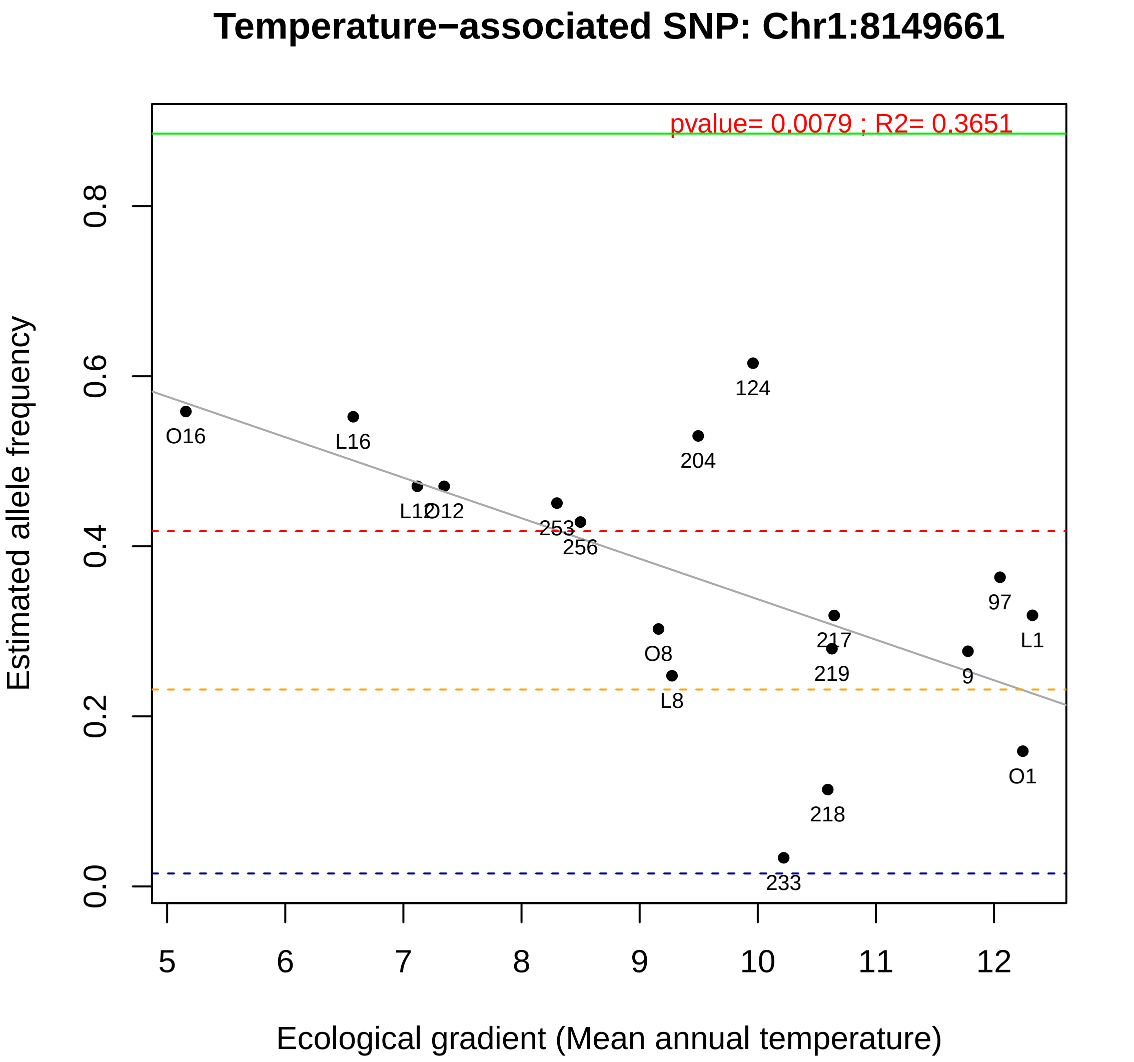

Temperature−associated SNP: Chr1:8149661
pvalue= 0.0079 ; R2= 0.3651
Estimated allele frequency
0.0
0.2
0.4
0.6
0.8
Ecological gradient (Mean annual temperature)
5
6
7
8
9
10
11
12
O16
L16
L12
O12
253
256
204
124
O8
L8
217
219
218
233
9
97
L1
O1

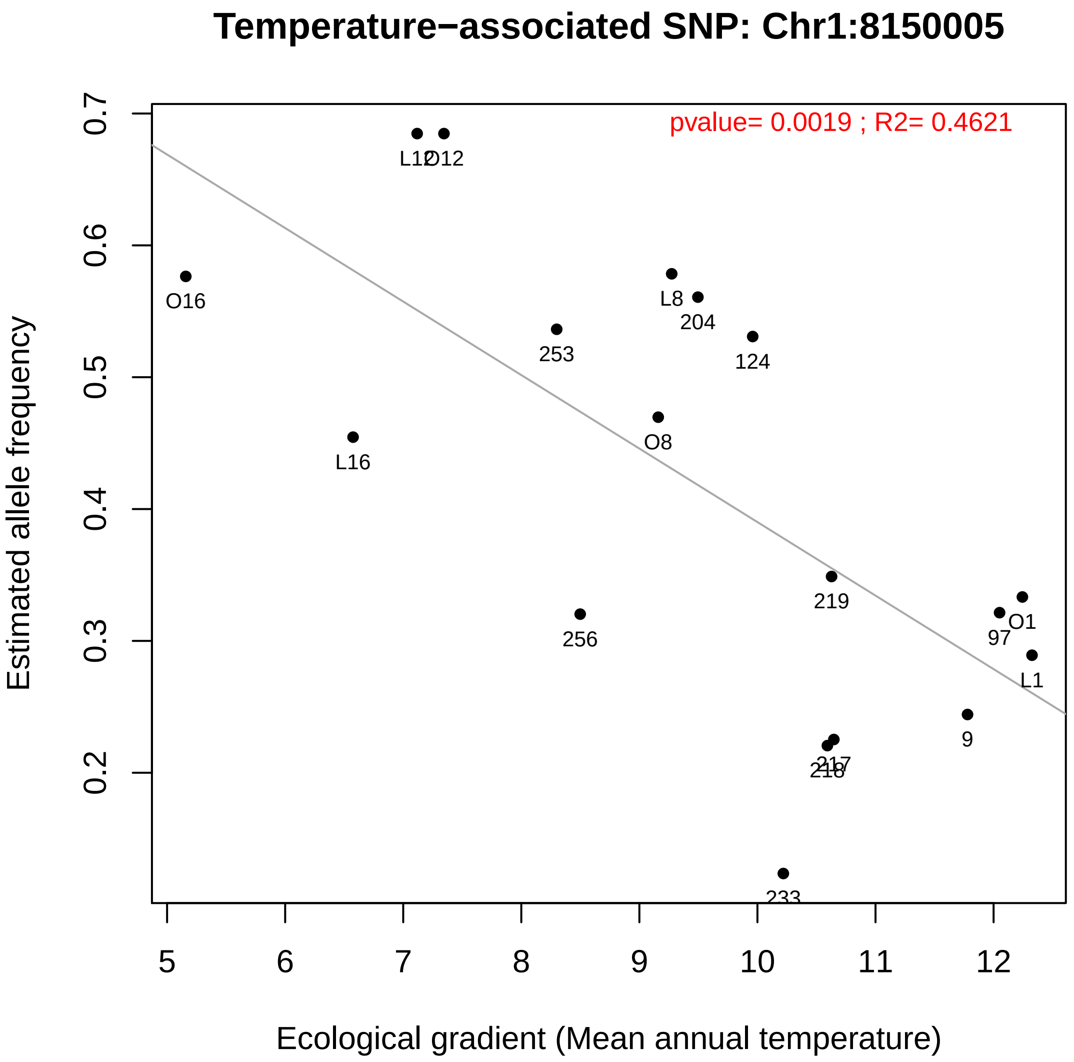

Temperature−associated SNP: Chr1:8150005
pvalue= 0.0019 ; R2= 0.4621
Estimated allele frequency
0.7
0.6
0.5
0.4
0.3
0.2
5
6
7
8
9
10
11
12
Ecological gradient (Mean annual temperature)
L12
O12
O16
L8
204
253
124
O8
L16
219
256
O1
97
L1
9
217
218
233

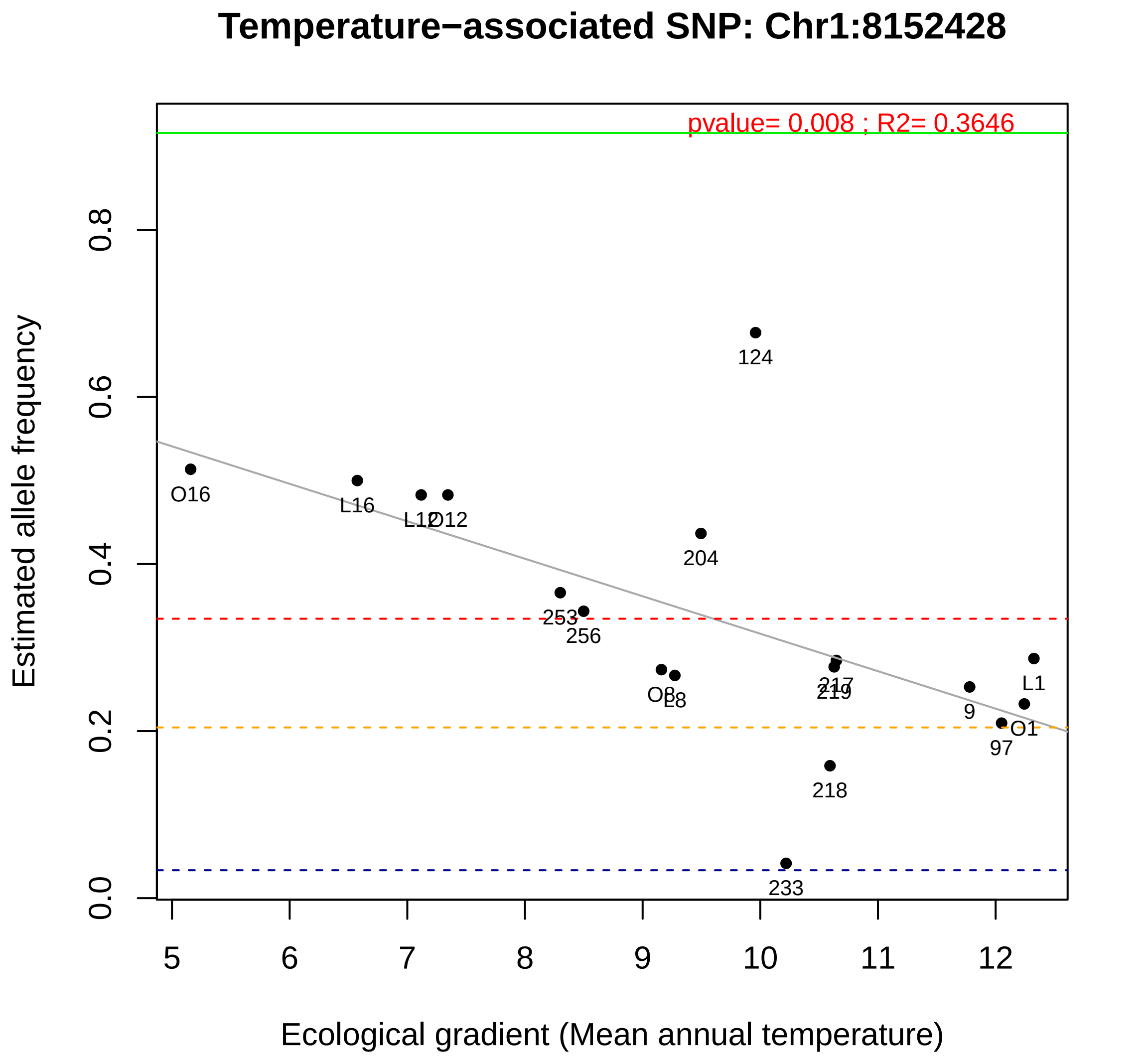
Temperature−associated SNP: Chr1:8152428
pvalue= 0.008 ; R2= 0.3646
Estimated allele frequency
0.0
0.2
0.4
0.6
0.8
5
6
7
8
9
10
11
12
Ecological gradient (Mean annual temperature)
O16
L16
L12
O12
124
204
253
256
O8
L8
217
219
218
233
9
L1
O1
97

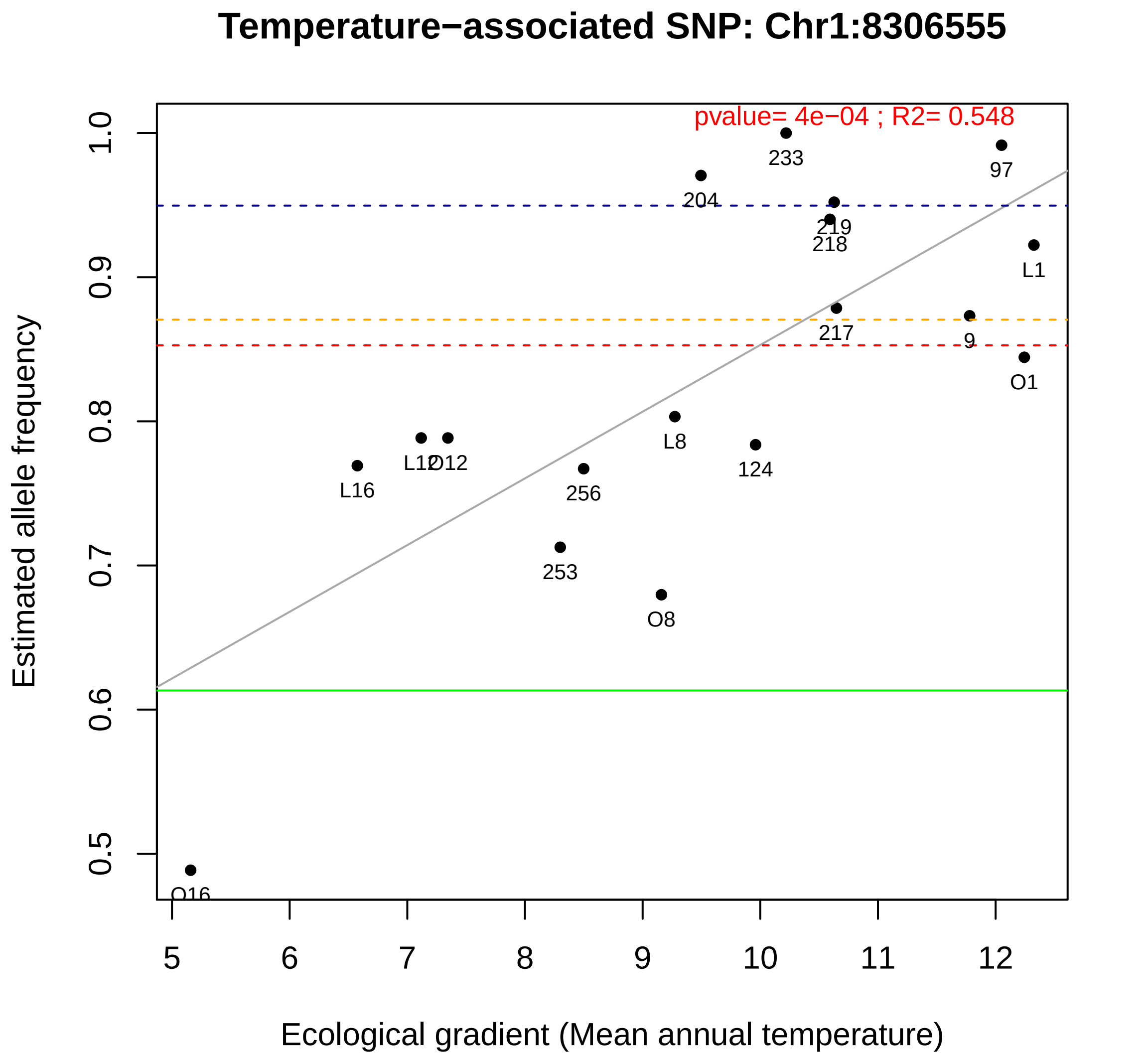
Temperature−associated SNP: Chr1:8306555
pvalue= 4e−04 ; R2= 0.548
Estimated allele frequency
Ecological gradient (Mean annual temperature)
0.5
0.6
0.7
0.8
0.9
1.0
5
6
7
8
9
10
11
12
233
97
204
219
218
L1
217
9
O1
L8
L12
O12
124
L16
256
253
O8
O16

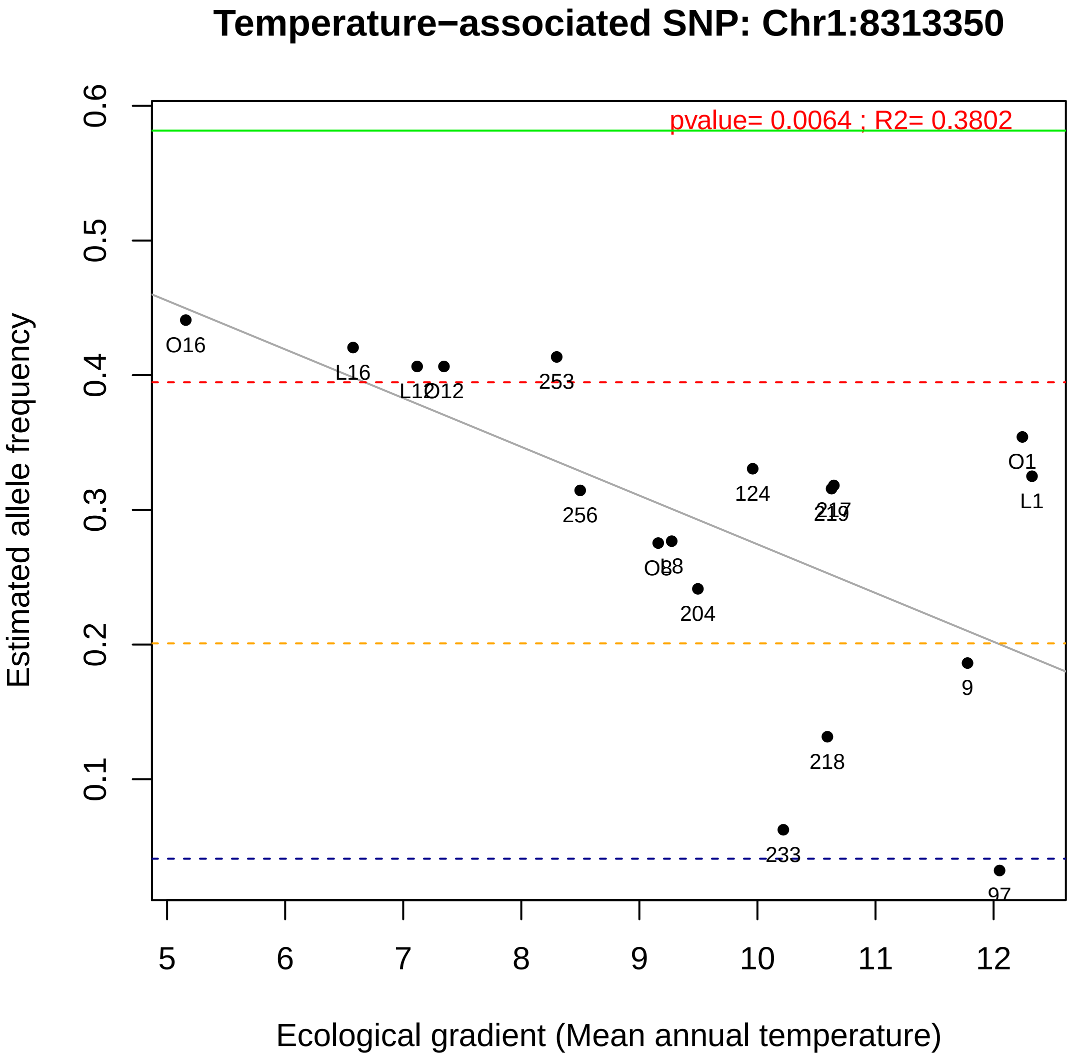
Temperature−associated SNP: Chr1:8313350
pvalue= 0.0064 ; R2= 0.3802
Estimated allele frequency
Ecological gradient (Mean annual temperature)
0.1
0.2
0.3
0.4
0.5
0.6
5
6
7
8
9
10
11
12
O16
L16
253
256
124
O1
L1
204
9
218
233
97

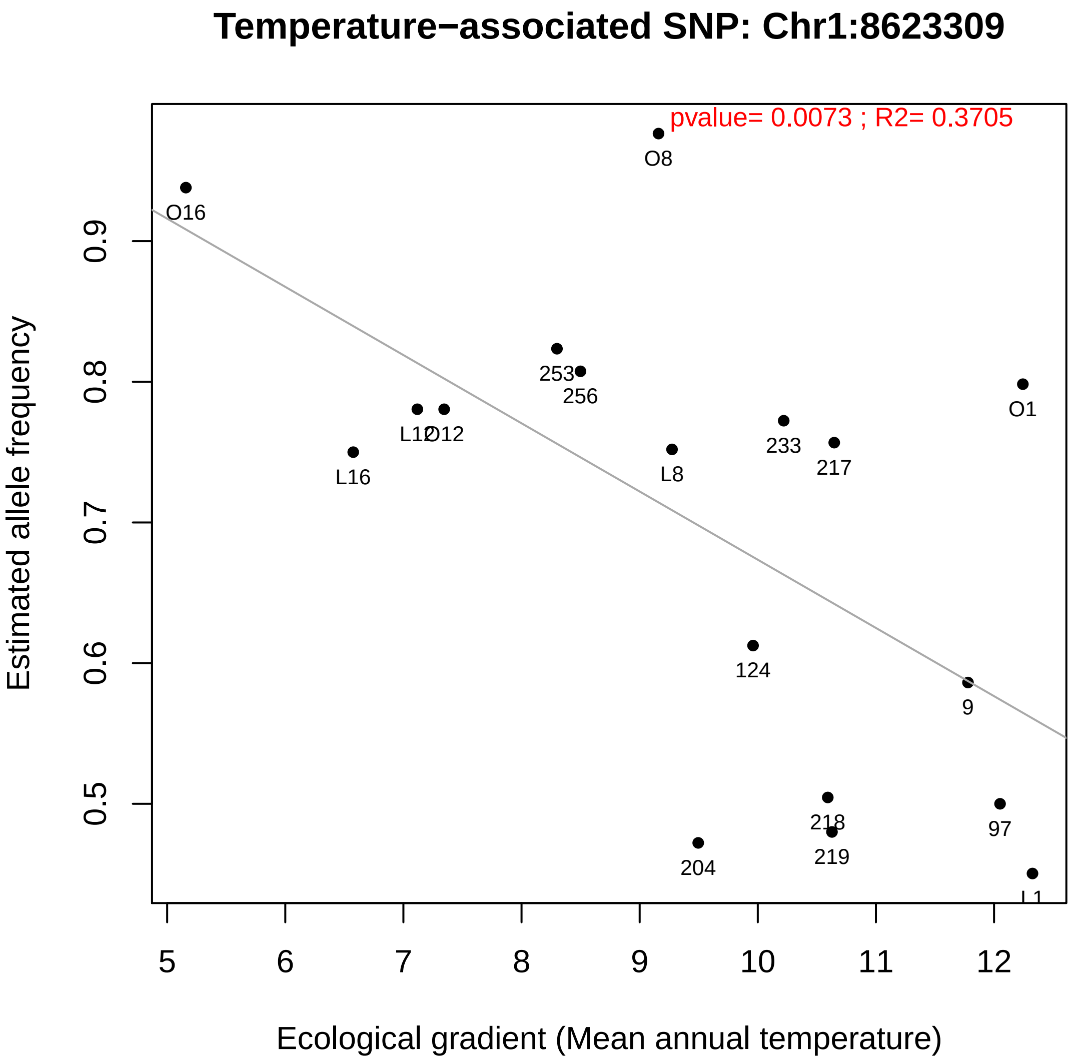
Temperature−associated SNP: Chr1:8623309
pvalue= 0.0073 ; R2= 0.3705
Estimated allele frequency
Ecological gradient (Mean annual temperature)
0.5
0.6
0.7
0.8
0.9
5
6
7
8
9
10
11
12
O16
O8
253
256
L12
O12
L16
L8
233
217
O1
124
9
218
219
204
97

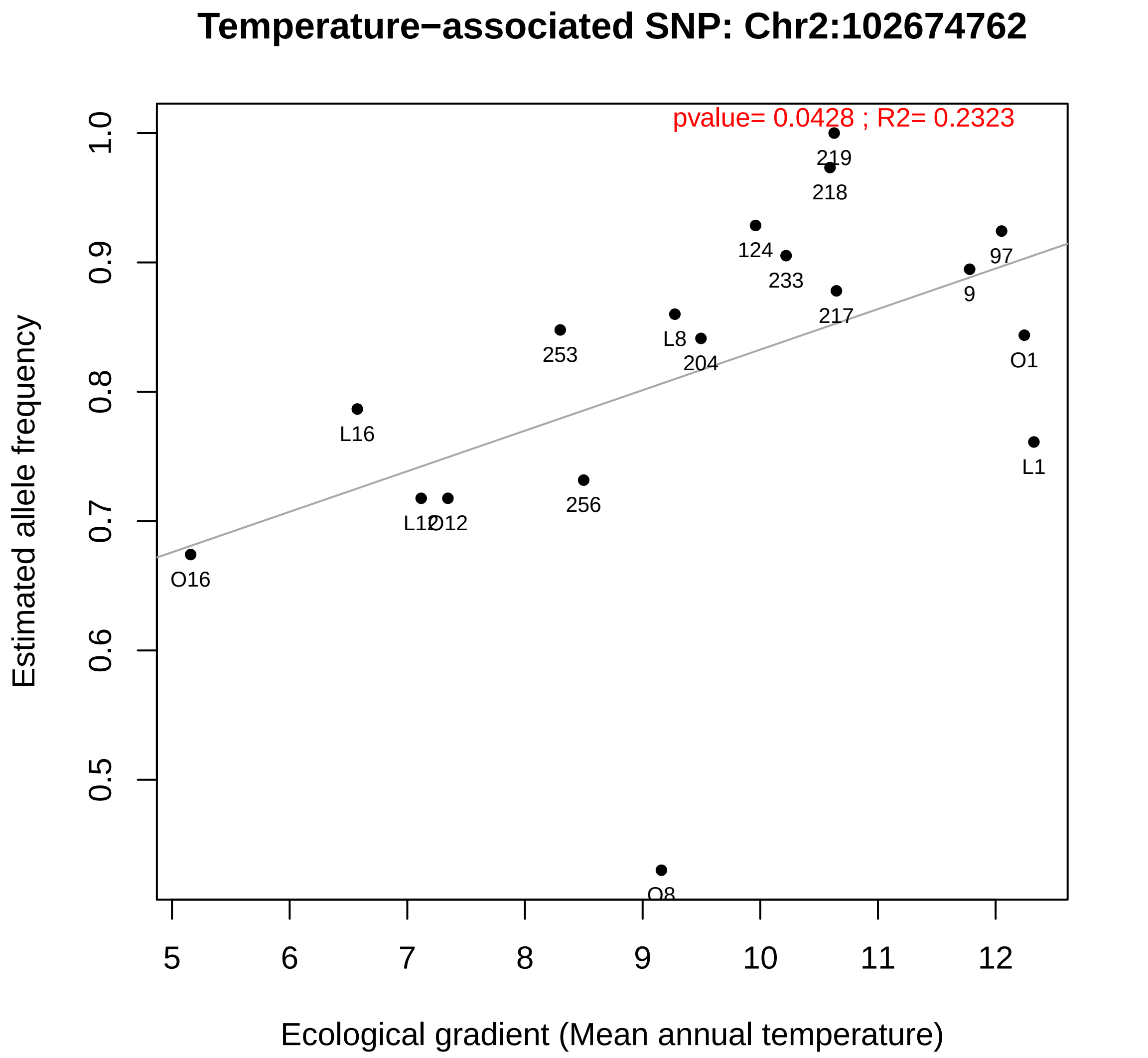
Temperature−associated SNP: Chr2:102674762
pvalue= 0.0428 ; R2= 0.2323
Estimated allele frequency
0.5
0.6
0.7
0.8
0.9
1.0
5
6
7
8
9
10
11
12
Ecological gradient (Mean annual temperature)
219
218
124
233
97
9
217
L8
204
253
O1
L16
L1
256
L12
O12
O16
O8

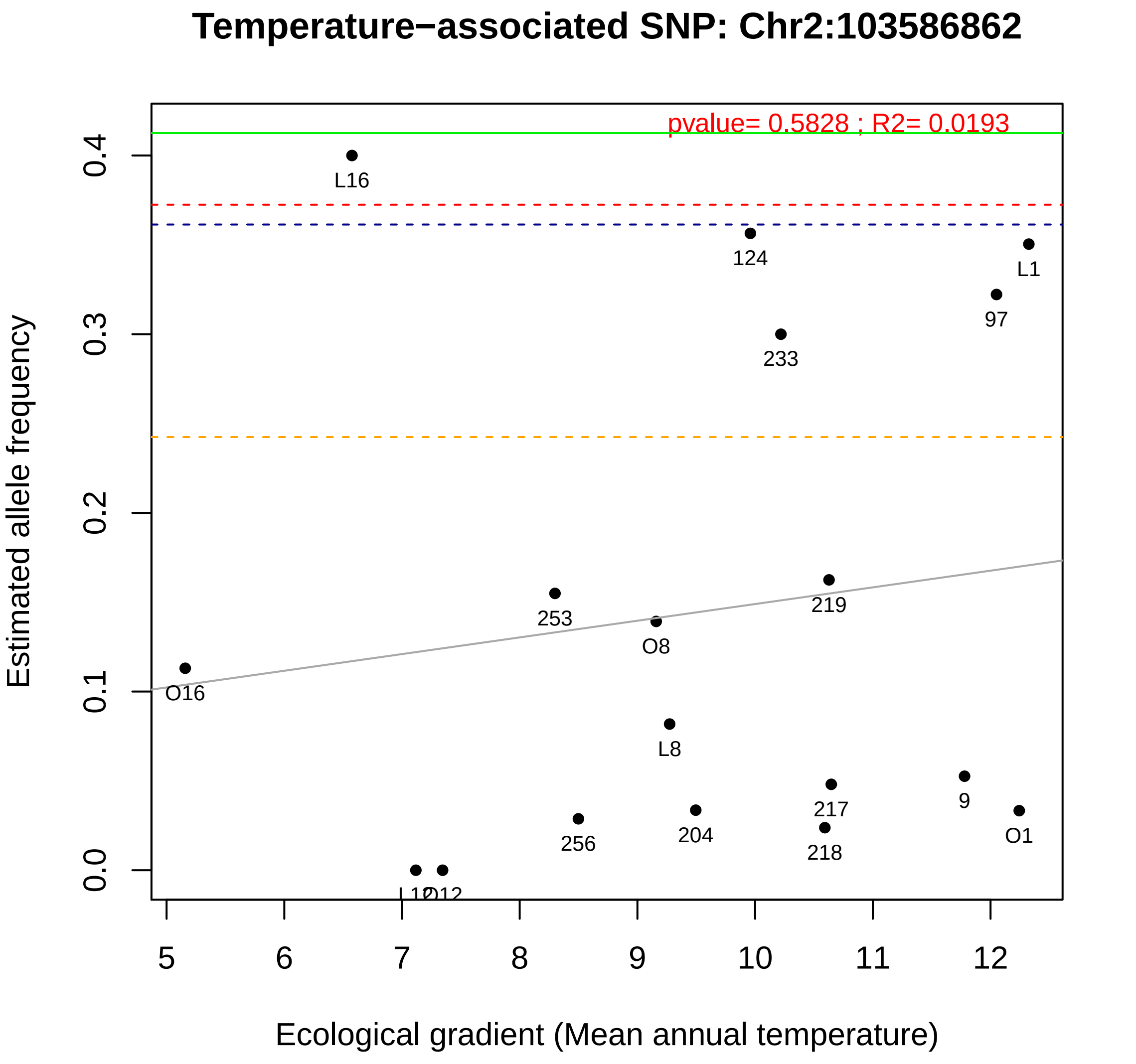
Temperature−associated SNP: Chr2:103586862
pvalue= 0.5828 ; R2= 0.0193
Estimated allele frequency
0.0
0.1
0.2
0.3
0.4
Ecological gradient (Mean annual temperature)
5
6
7
8
9
10
11
12
L16
124
L1
97
233
219
253
O8
O16
L8
9
217
O1
204
256
218
O12

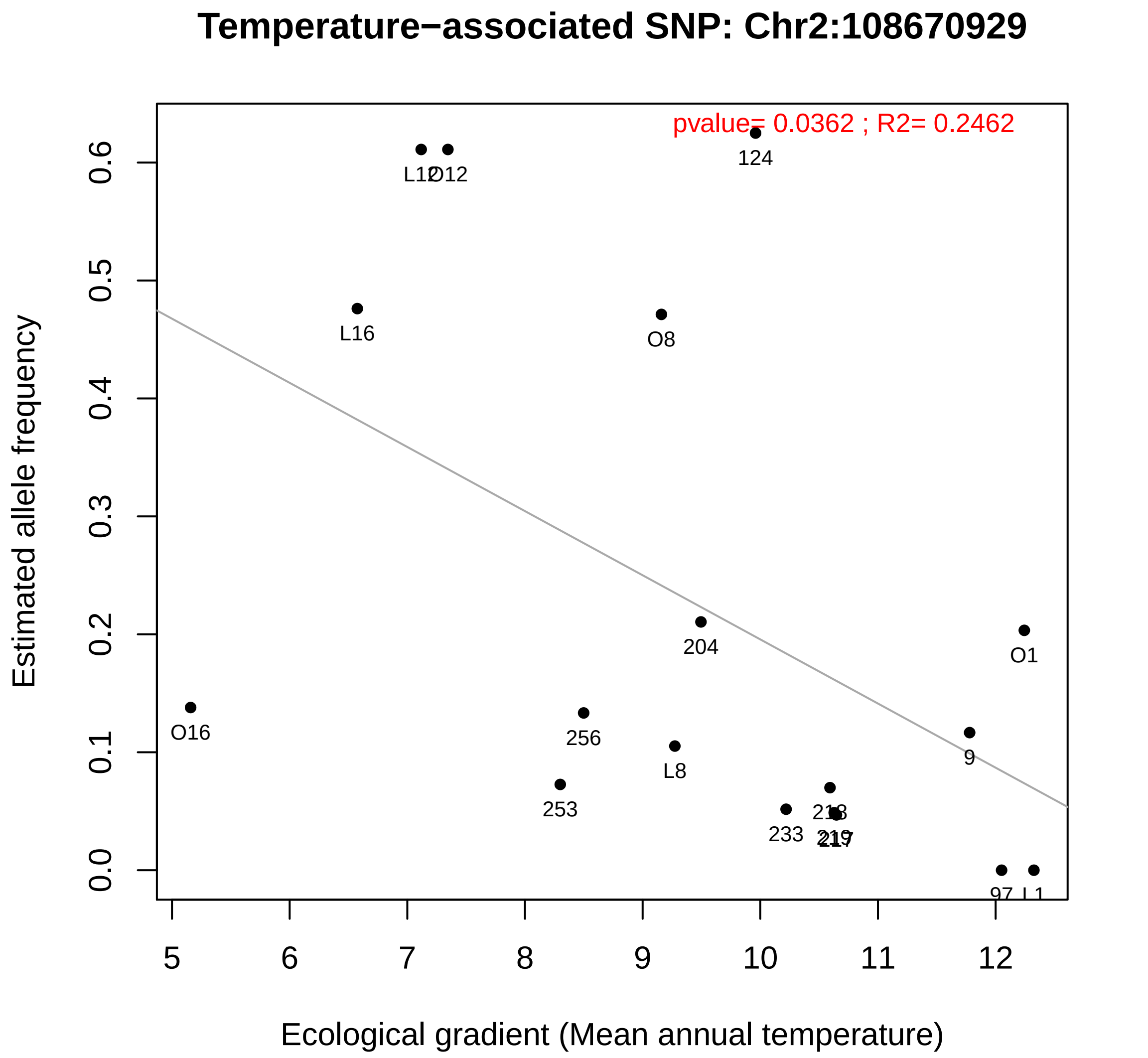
Temperature−associated SNP: Chr2:108670929
pvalue= 0.0362 ; R2= 0.2462
Estimated allele frequency
Ecological gradient (Mean annual temperature)
0.0
0.1
0.2
0.3
0.4
0.5
0.6
5
6
7
8
9
10
11
12
124
L12
O12
L16
O8
204
O1
O16
256
L8
9
253
233
218
217
97
L1

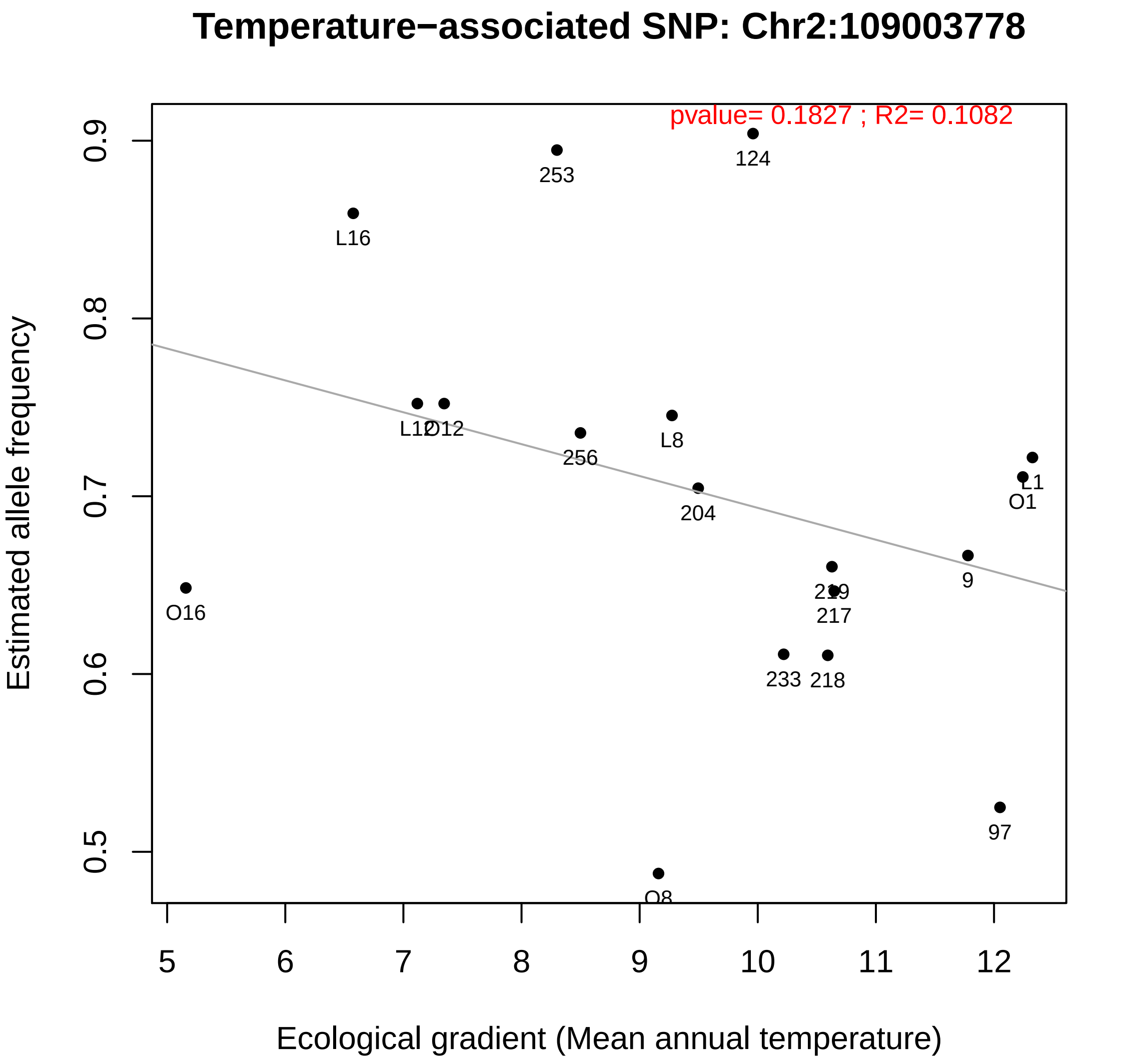
Temperature−associated SNP: Chr2:109003778
pvalue= 0.1827 ; R2= 0.1082
Estimated allele frequency
0.5
0.6
0.7
0.8
0.9
5
6
7
8
9
10
11
12
Ecological gradient (Mean annual temperature)
253
124
L16
L12
O12
256
L8
204
L1
O1
9
219
217
233
218
O16
97
O8

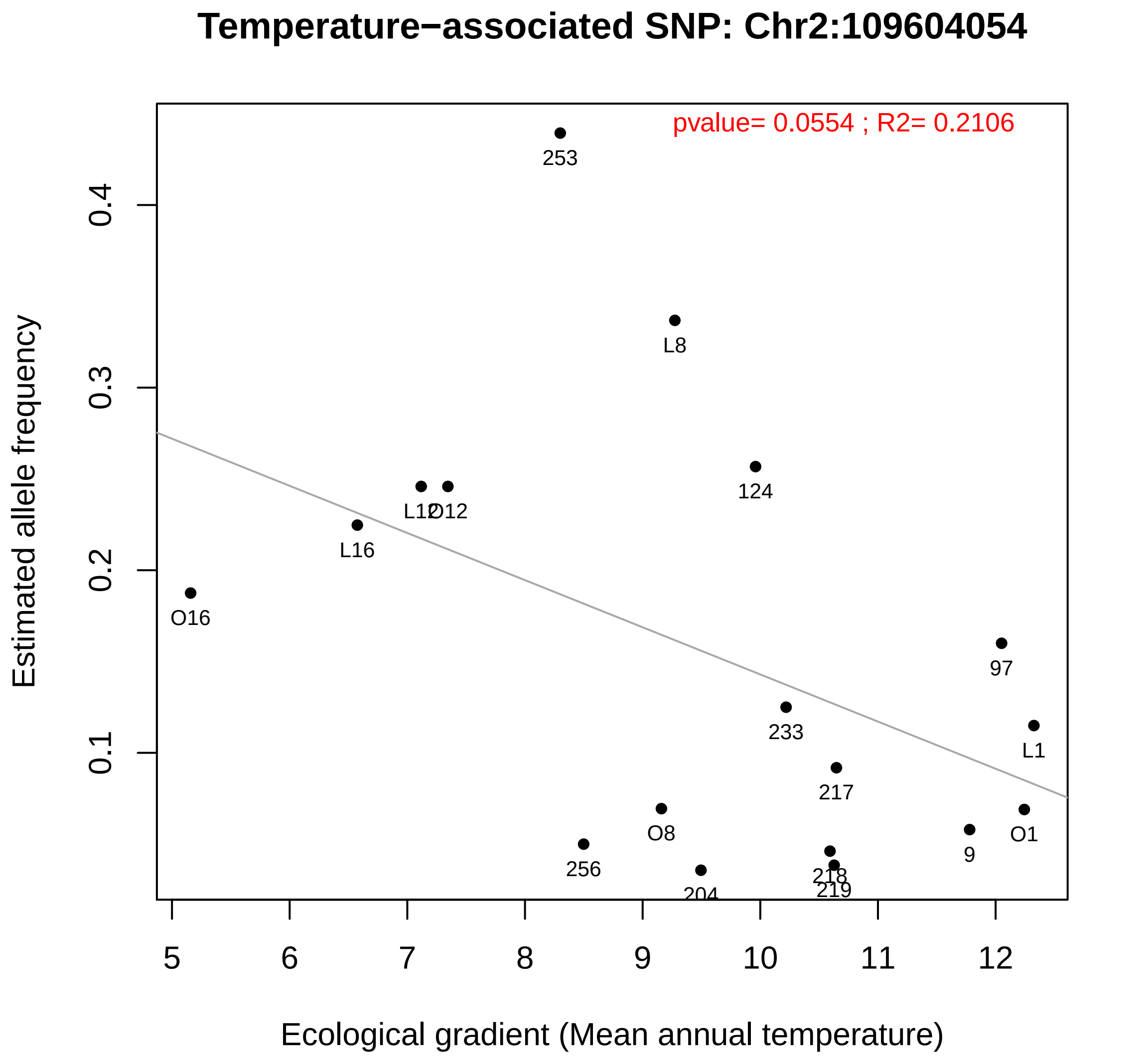

Temperature−associated SNP: Chr2:109604054
pvalue= 0.0554 ; R2= 0.2106
Estimated allele frequency
Ecological gradient (Mean annual temperature)
0.1
0.2
0.3
0.4
5
6
7
8
9
10
11
12
253
L8
124
L12
O12
L16
O16
97
233
L1
217
O8
O1
9
256
204
218
219

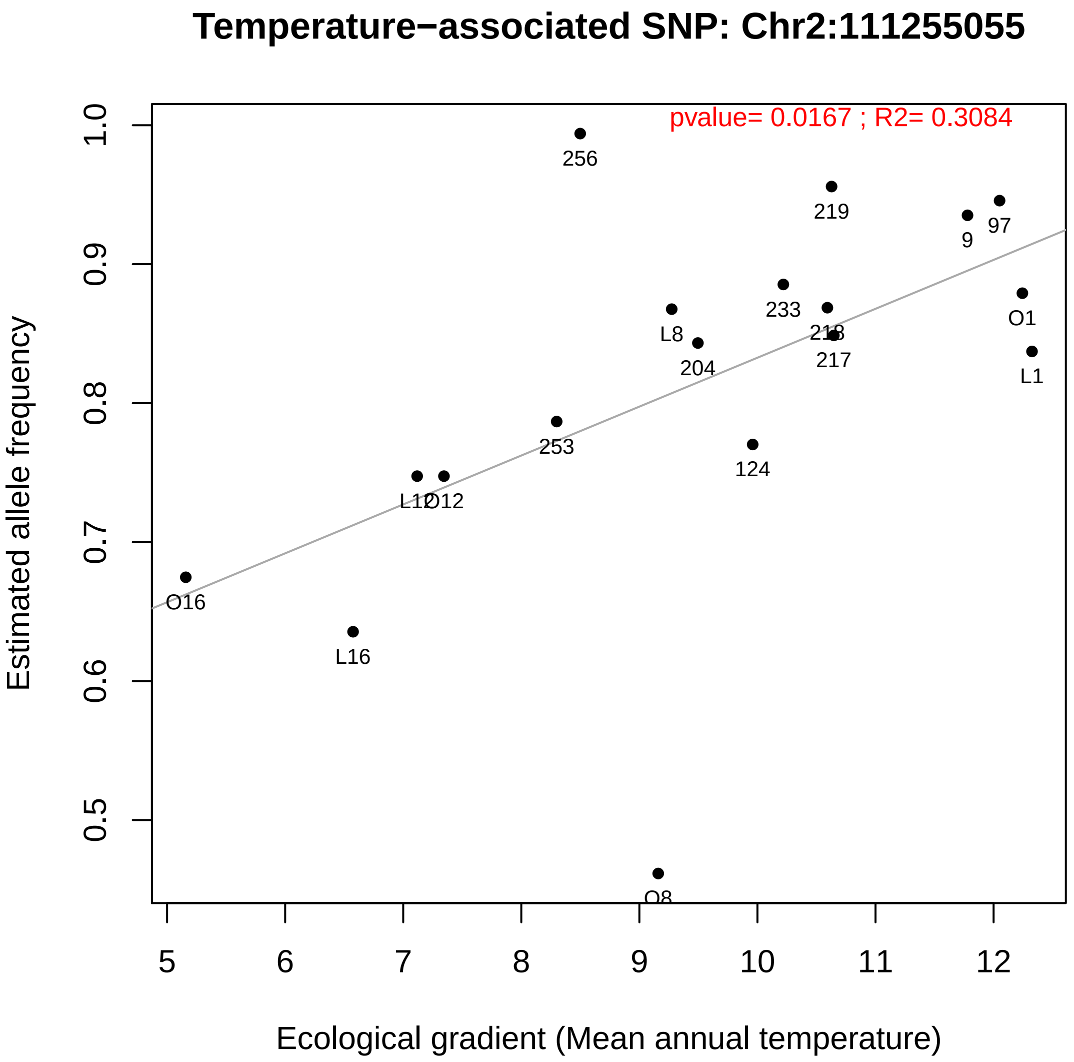
Temperature−associated SNP: Chr2:111255055
pvalue= 0.0167 ; R2= 0.3084
Estimated allele frequency
Ecological gradient (Mean annual temperature)
256
219
97
9
233
O1
L8
218
217
204
L1
253
124
L12
O12
O16
L16
O8
0.5
0.6
0.7
0.8
0.9
1.0
5
6
7
8
9
10
11
12

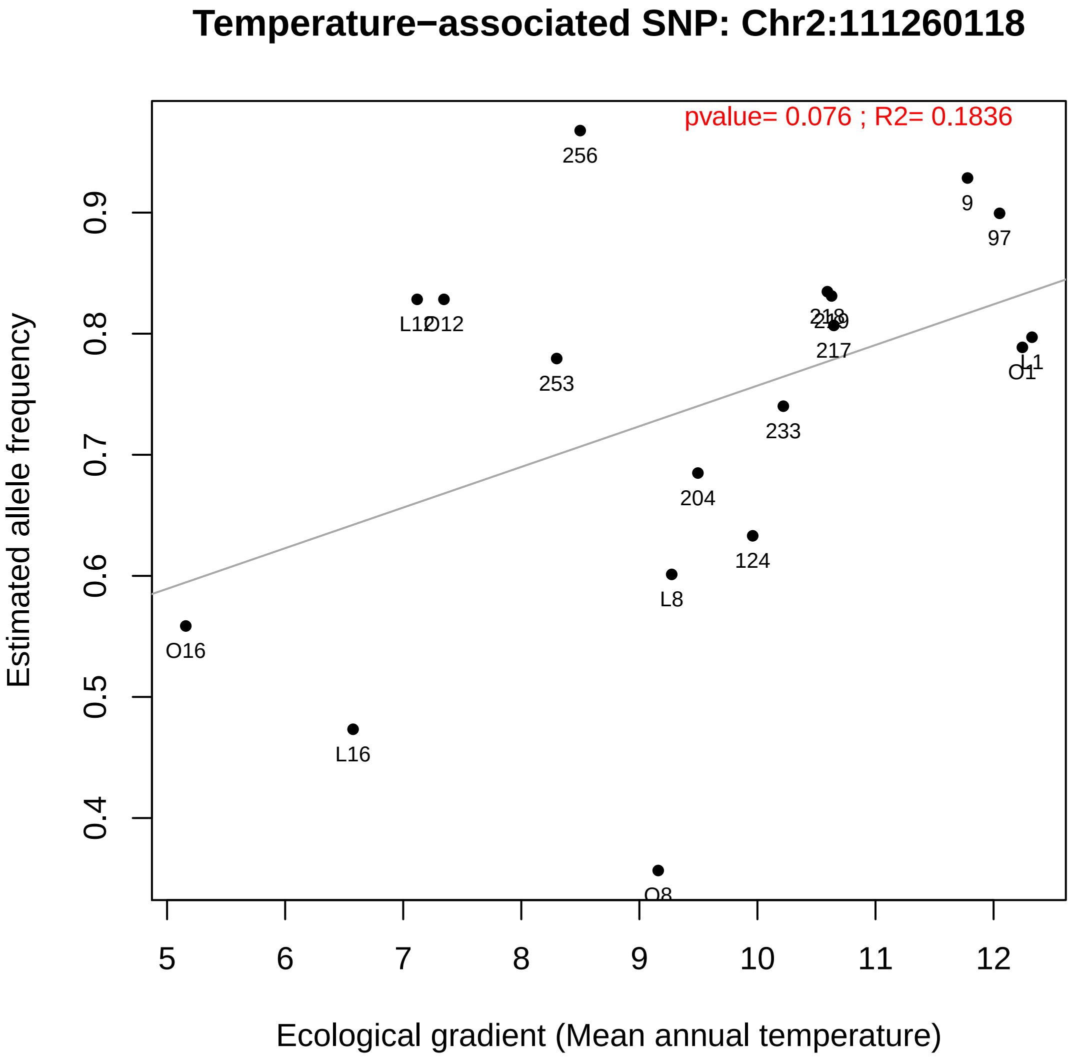
Temperature−associated SNP: Chr2:111260118
pvalue= 0.076 ; R2= 0.1836
Estimated allele frequency
0.4
0.5
0.6
0.7
0.8
0.9
5
6
7
8
9
10
11
12
Ecological gradient (Mean annual temperature)
256
9
97
L12
O12
253
218
217
O1
233
204
124
L8
O16
L16
O8

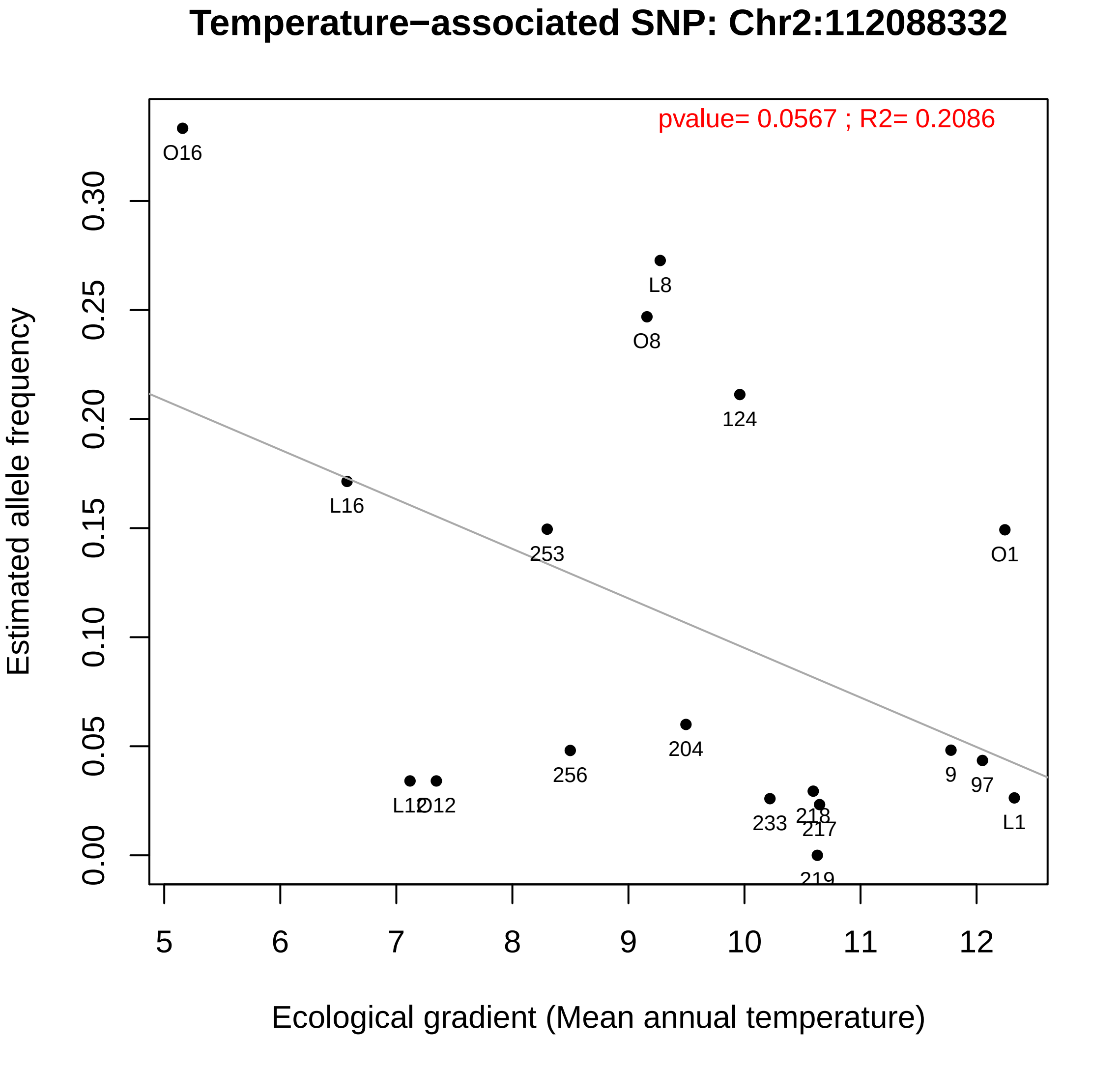

Temperature−associated SNP: Chr2:112088332
pvalue= 0.0567 ; R2= 0.2086
Estimated allele frequency
0.00
0.05
0.10
0.15
0.20
0.25
0.30
Ecological gradient (Mean annual temperature)
5
6
7
8
9
10
11
12
O16
L8
O8
124
L16
253
O1
204
256
9
97
L12
O12
233
218
217
L1
219

**Temperature−associated SNP: Chr2:112829511**

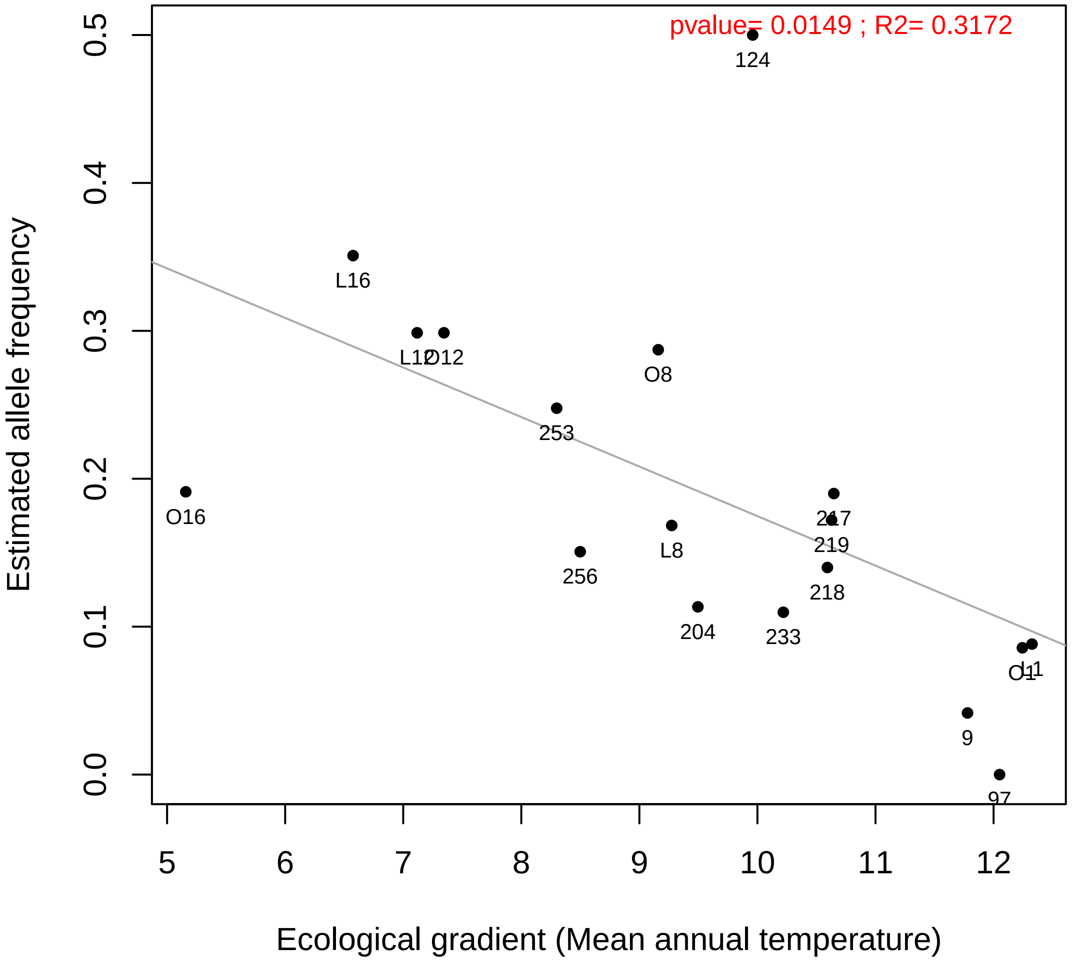

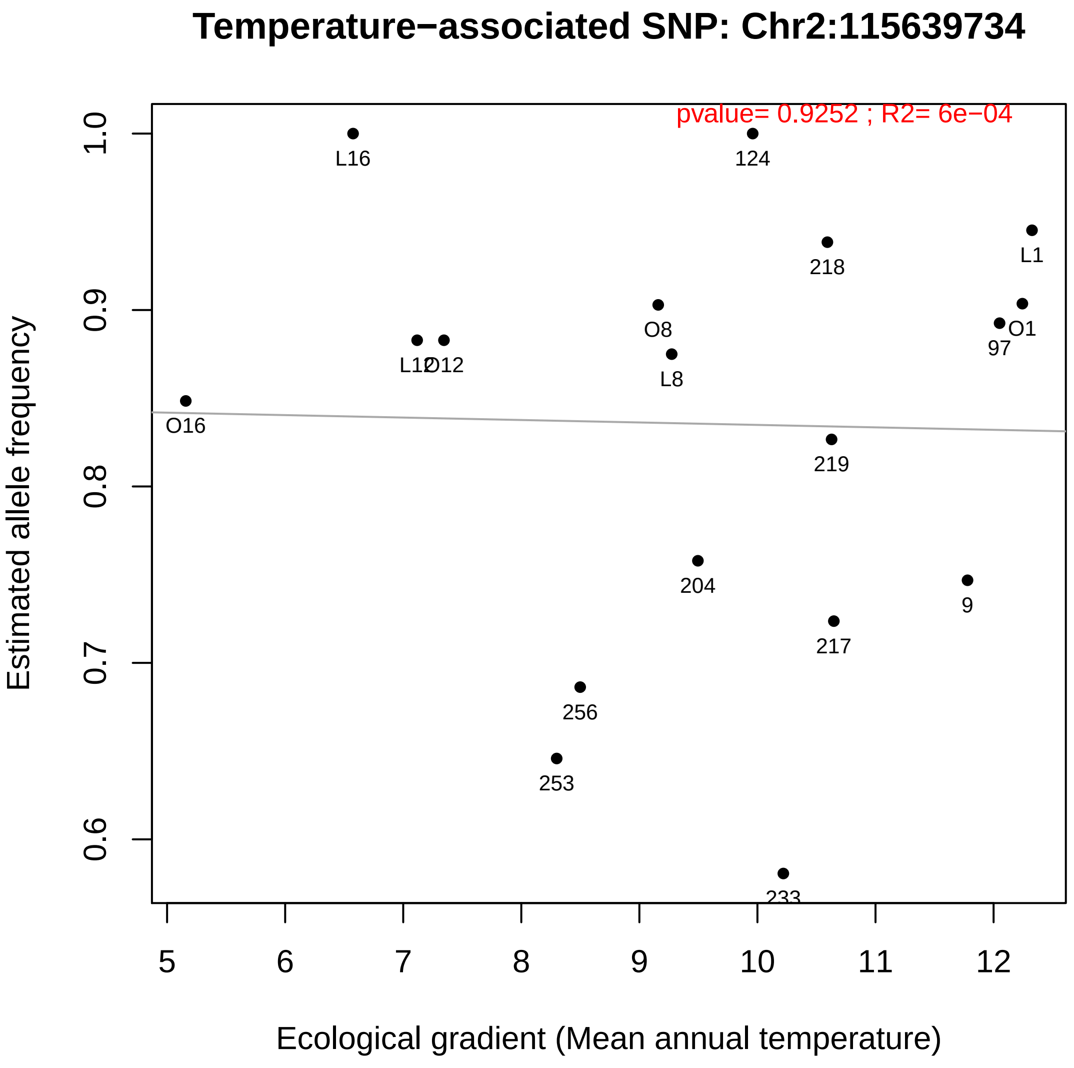
Temperature−associated SNP: Chr2:115639734
pvalue= 0.9252 ; R2= 6e−04
Estimated allele frequency
0.6
0.7
0.8
0.9
1.0
5
6
7
8
9
10
11
12
Ecological gradient (Mean annual temperature)
L16
124
218
L1
O8
O1
97
L12
O12
L8
O16
219
204
9
217
256
253
233

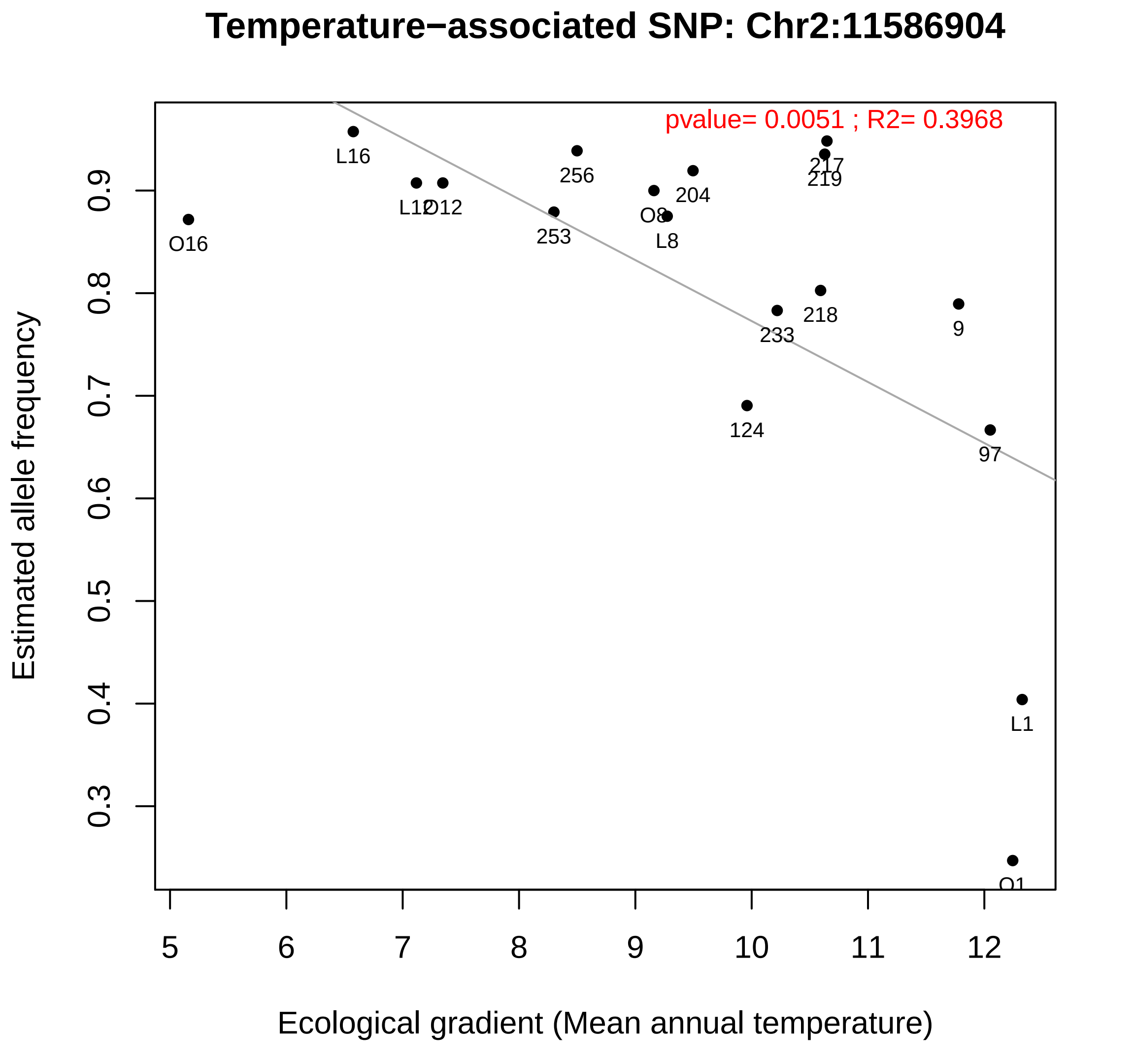
Temperature−associated SNP: Chr2:11586904
pvalue= 0.0051 ; R2= 0.3968
Estimated allele frequency
Ecological gradient (Mean annual temperature)
0.3
0.4
0.5
0.6
0.7
0.8
0.9
5
6
7
8
9
10
11
12
L16
256
217
219
204
L12
O12
O16
253
O8
L8
218
9
233
124
97
L1
O1

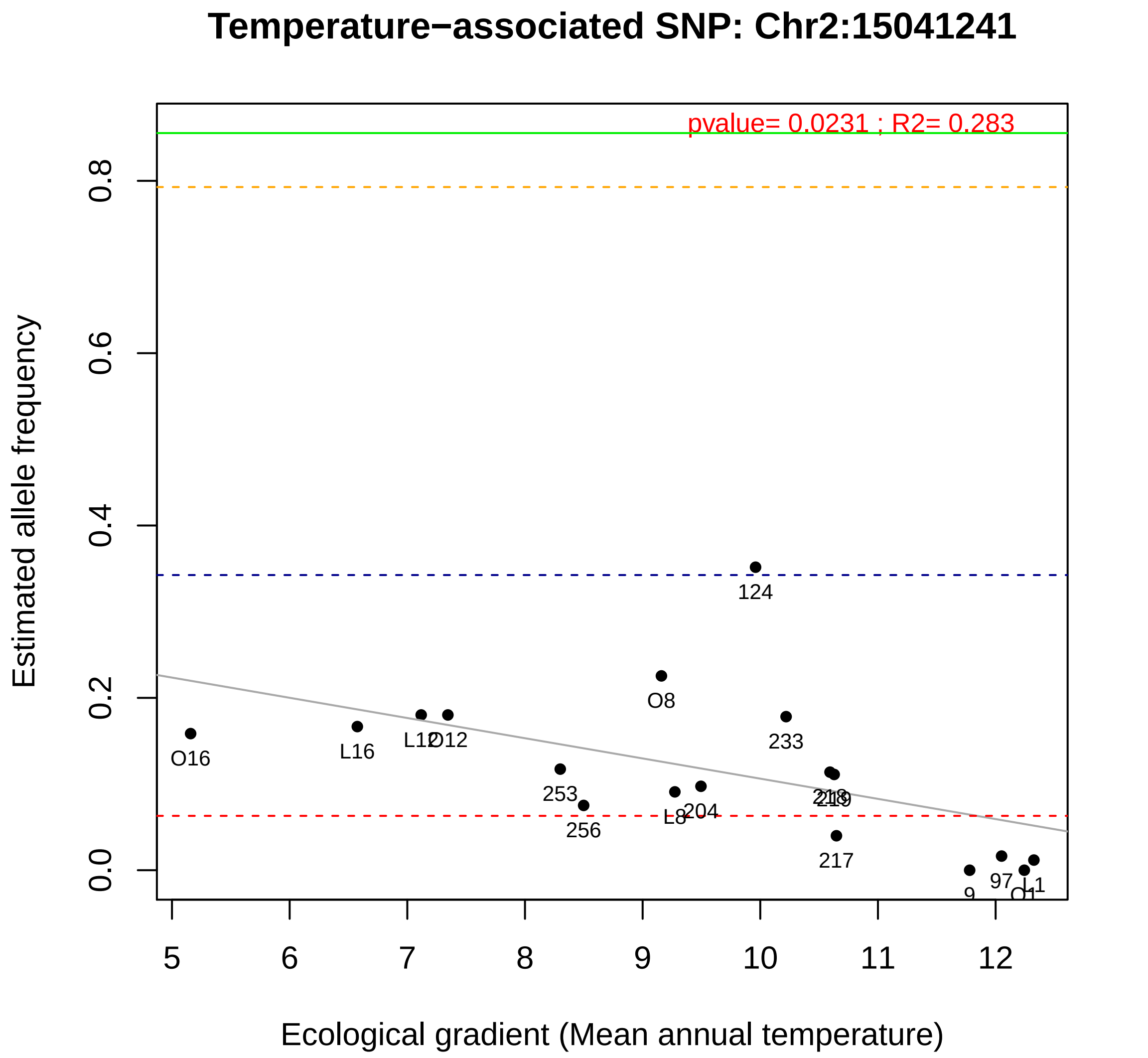

Temperature−associated SNP: Chr2:15041241
pvalue= 0.0231 ; R2= 0.283
Estimated allele frequency
0.0
0.2
0.4
0.6
0.8
5
6
7
8
9
10
11
12
Ecological gradient (Mean annual temperature)
O16
L16
L12
O12
253
256
O8
L8
204
124
233
218
219
217
9
97
O1
L1

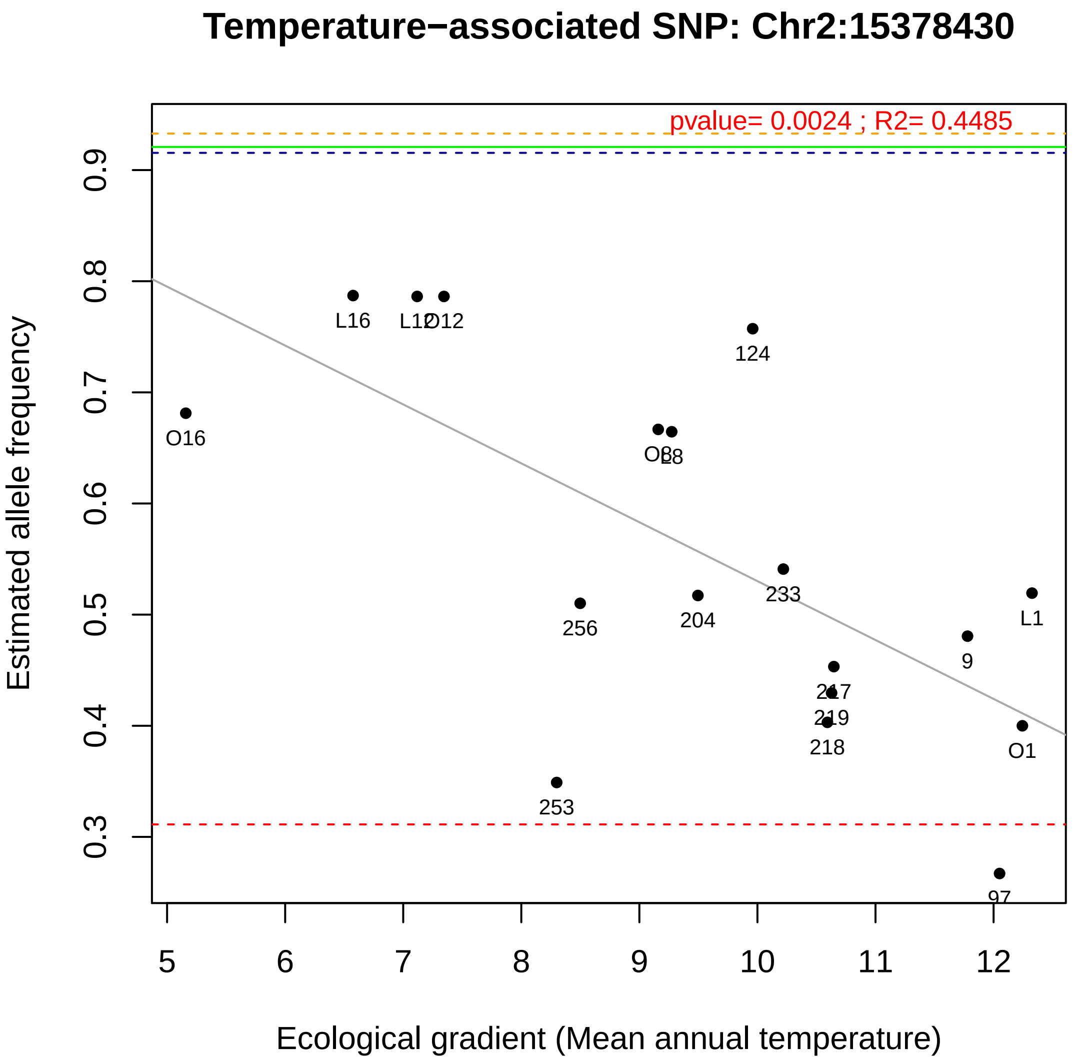
Temperature−associated SNP: Chr2:15378430
pvalue= 0.0024 ; R2= 0.4485
Estimated allele frequency
0.3
0.4
0.5
0.6
0.7
0.8
0.9
5
6
7
8
9
10
11
12
Ecological gradient (Mean annual temperature)
L16
L12
O12
124
O16
O8
L8
233
256
204
L1
9
217
219
218
O1
253
97

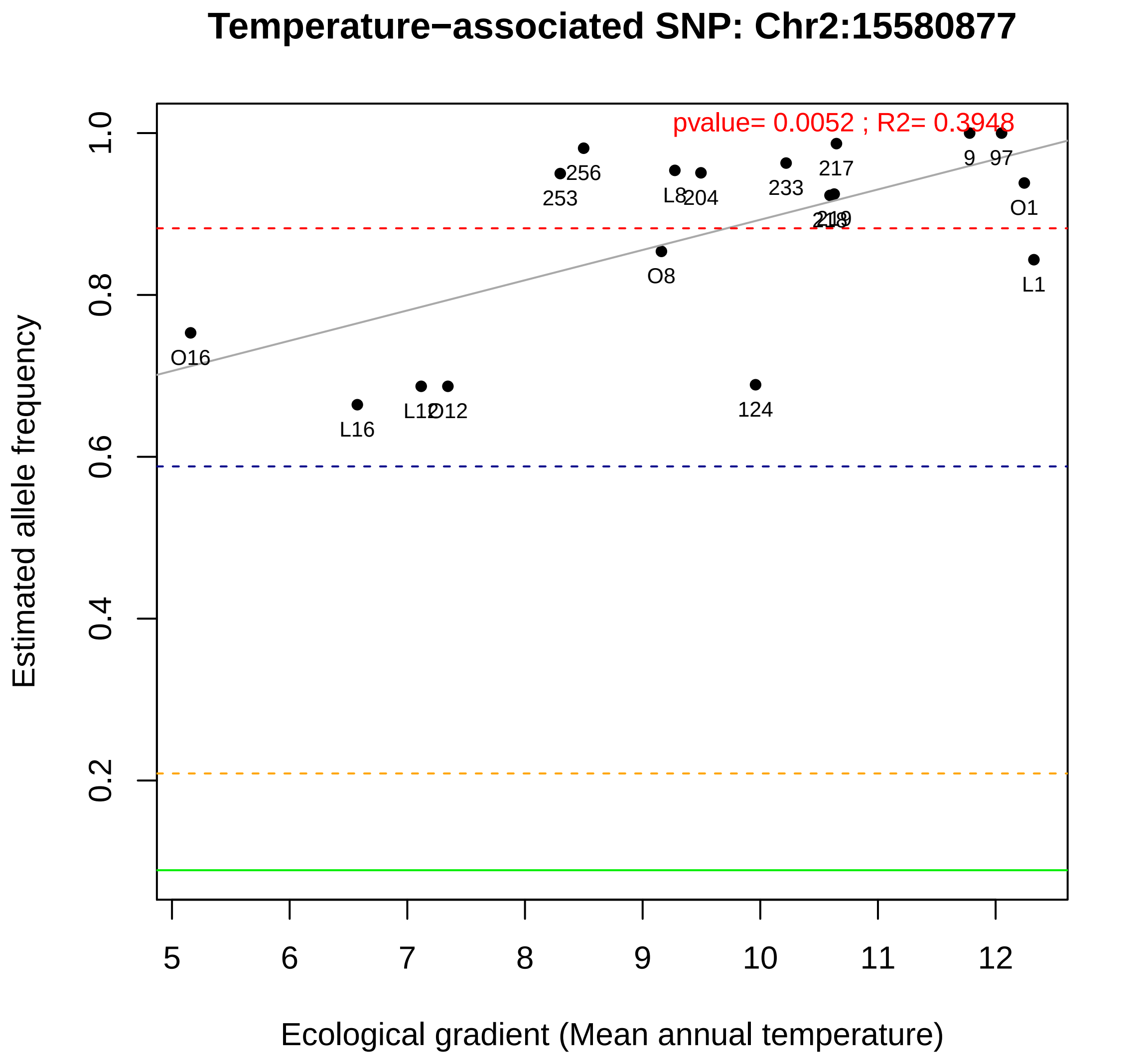

Temperature−associated SNP: Chr2:15580877
pvalue= 0.0052 ; R2= 0.3948
256
253
L8
204
233
217
9
97
O1
219
O8
L1
O16
L16
L12
O12
124
Estimated allele frequency
0.2
0.4
0.6
0.8
1.0
5
6
7
8
9
10
11
12
Ecological gradient (Mean annual temperature)

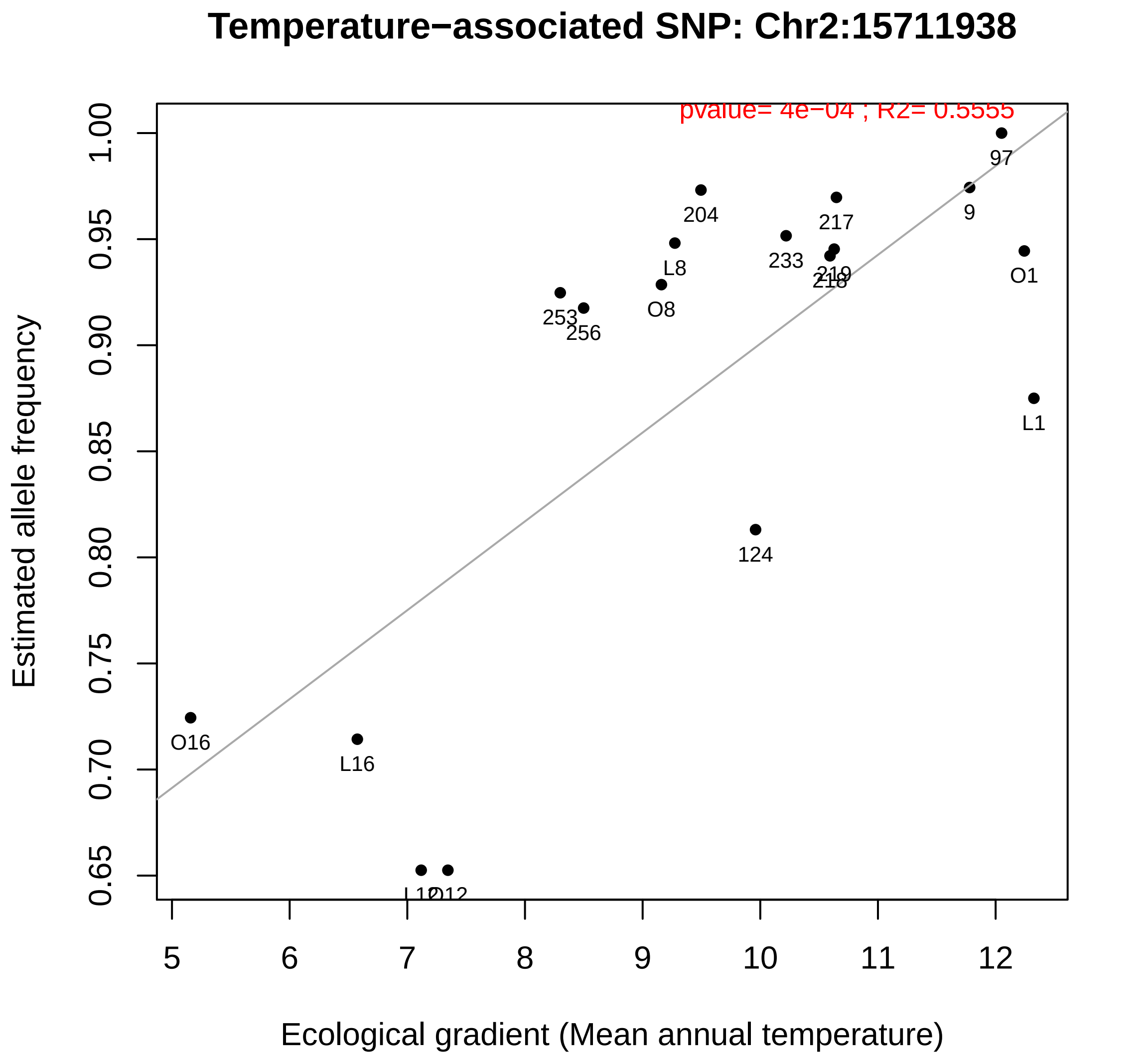

Temperature−associated SNP: Chr2:15711938
pvalue= 4e−04 ; R2= 0.5555
Estimated allele frequency
Ecological gradient (Mean annual temperature)
0.65
0.70
0.75
0.80
0.85
0.90
0.95
1.00
5
6
7
8
9
10
11
12
97
9
204
217
233
219
218
O1
L8
O8
253
256
L1
124
O16
L16
O12

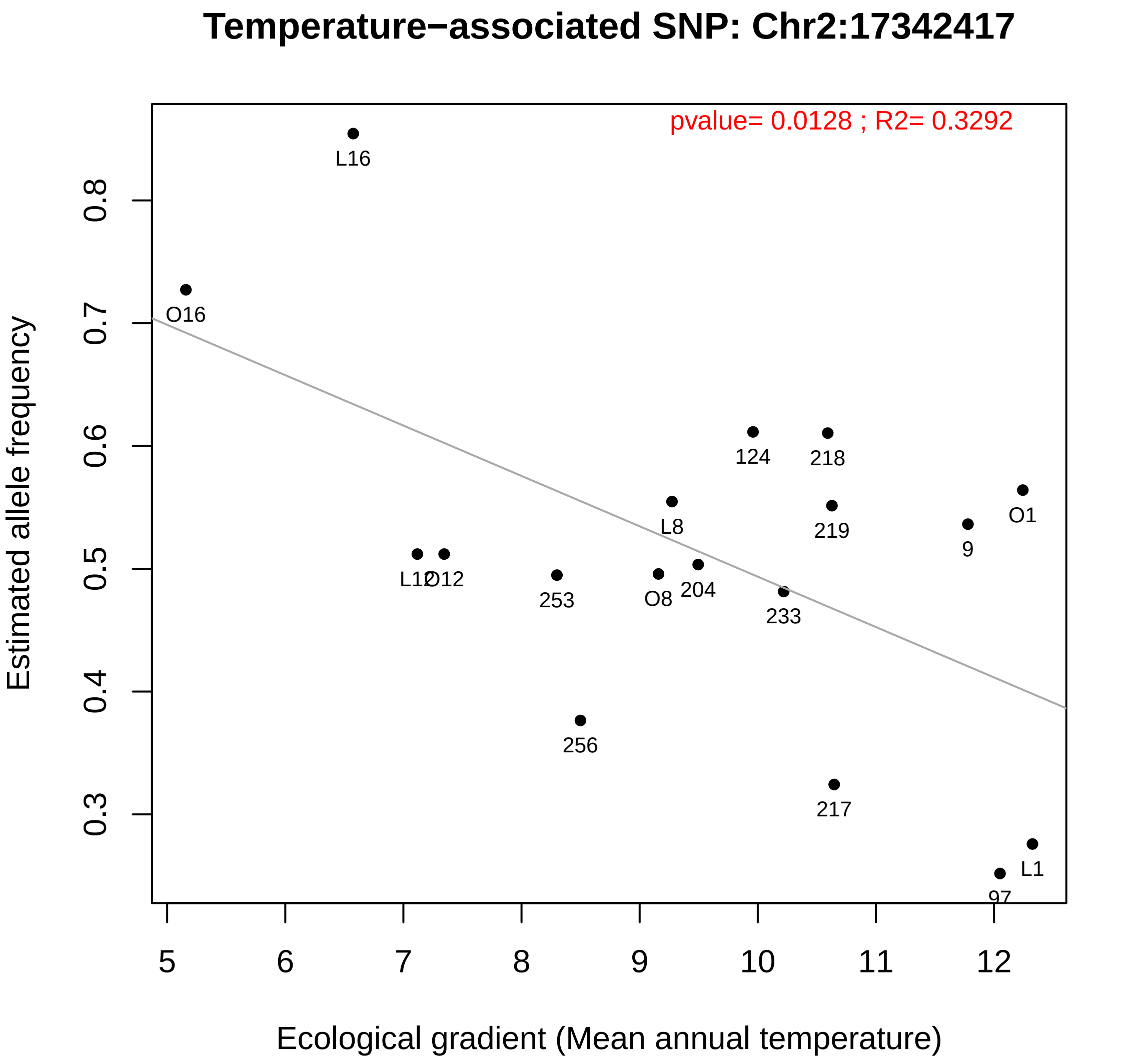
Temperature−associated SNP: Chr2:17342417
pvalue= 0.0128 ; R2= 0.3292
Estimated allele frequency
Ecological gradient (Mean annual temperature)
0.3
0.4
0.5
0.6
0.7
0.8
5
6
7
8
9
10
11
12
L16
O16
124
218
L8
219
O1
9
L12
O12
253
O8
204
233
256
217
L1
97

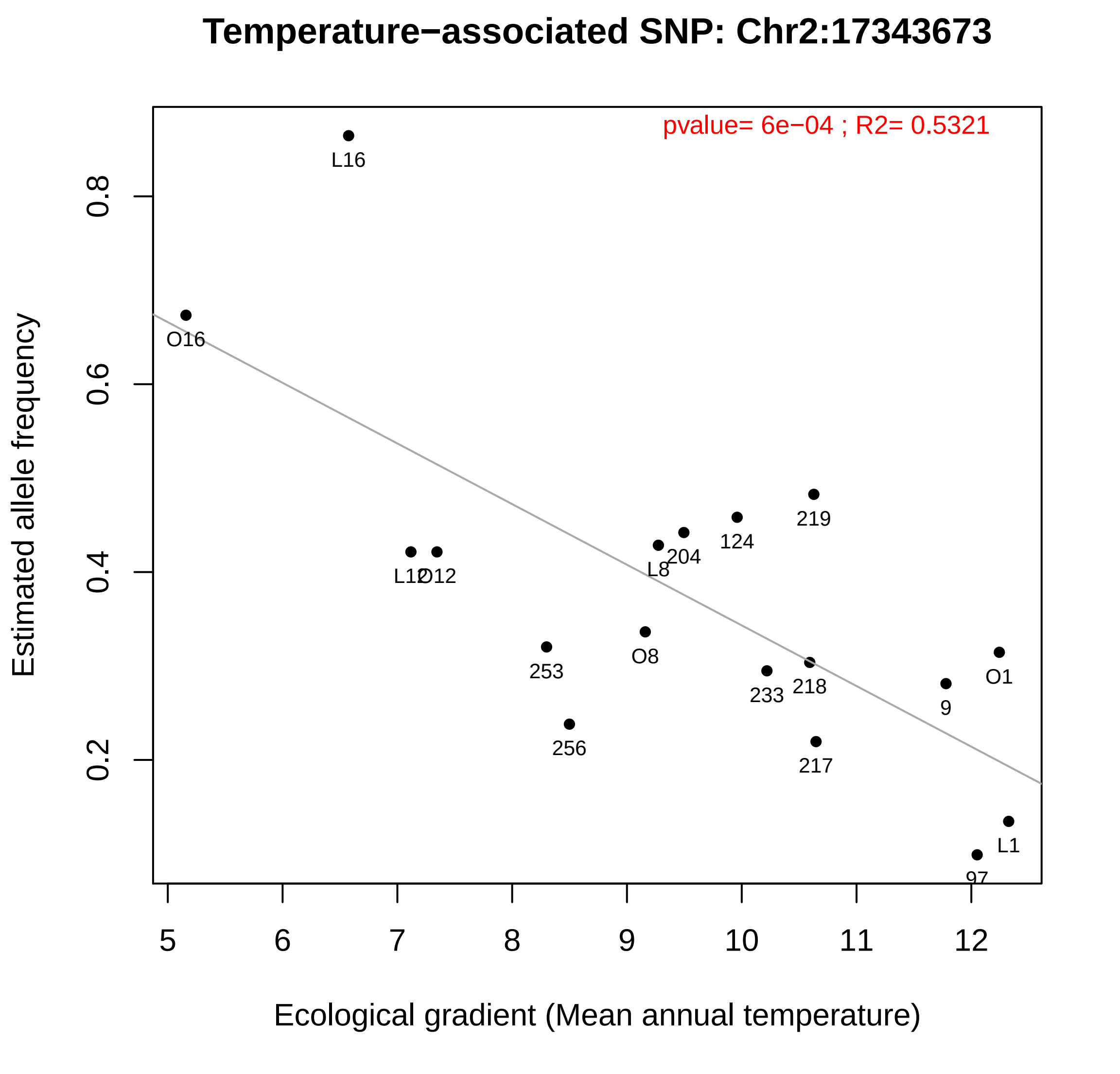
Temperature−associated SNP: Chr2:17343673
pvalue= 6e−04 ; R2= 0.5321
Estimated allele frequency
0.2
0.4
0.6
0.8
Ecological gradient (Mean annual temperature)
5
6
7
8
9
10
11
12
L16
O16
L12
O12
253
256
O8
L8
204
124
219
233
218
217
9
O1
L1
97

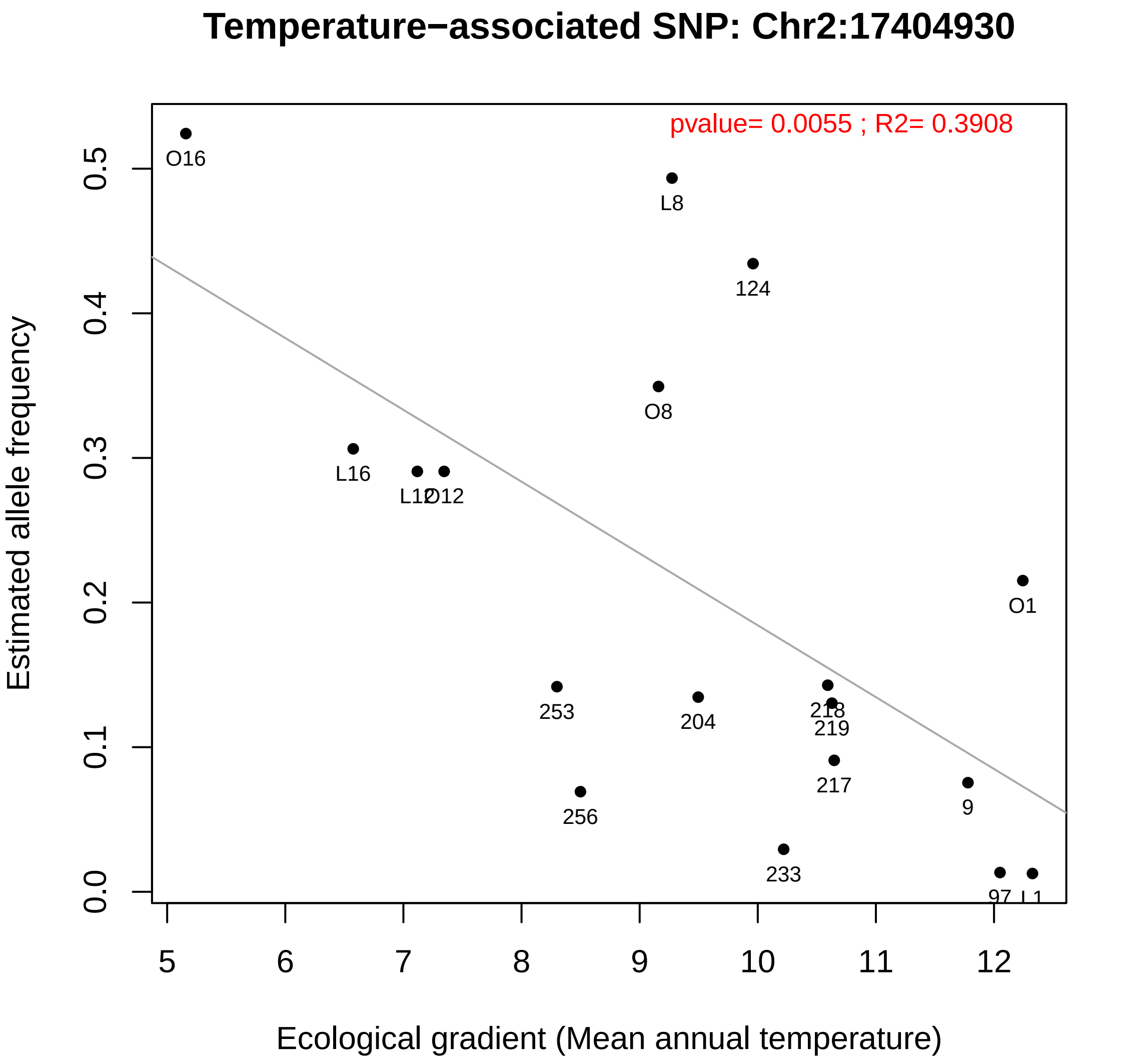
Temperature−associated SNP: Chr2:17404930
pvalue= 0.0055 ; R2= 0.3908
Estimated allele frequency
Ecological gradient (Mean annual temperature)
0.0
0.1
0.2
0.3
0.4
0.5
5
6
7
8
9
10
11
12
O16
L8
124
O8
L16
L12
O12
O1
253
204
218
219
217
256
9
233
97
L1

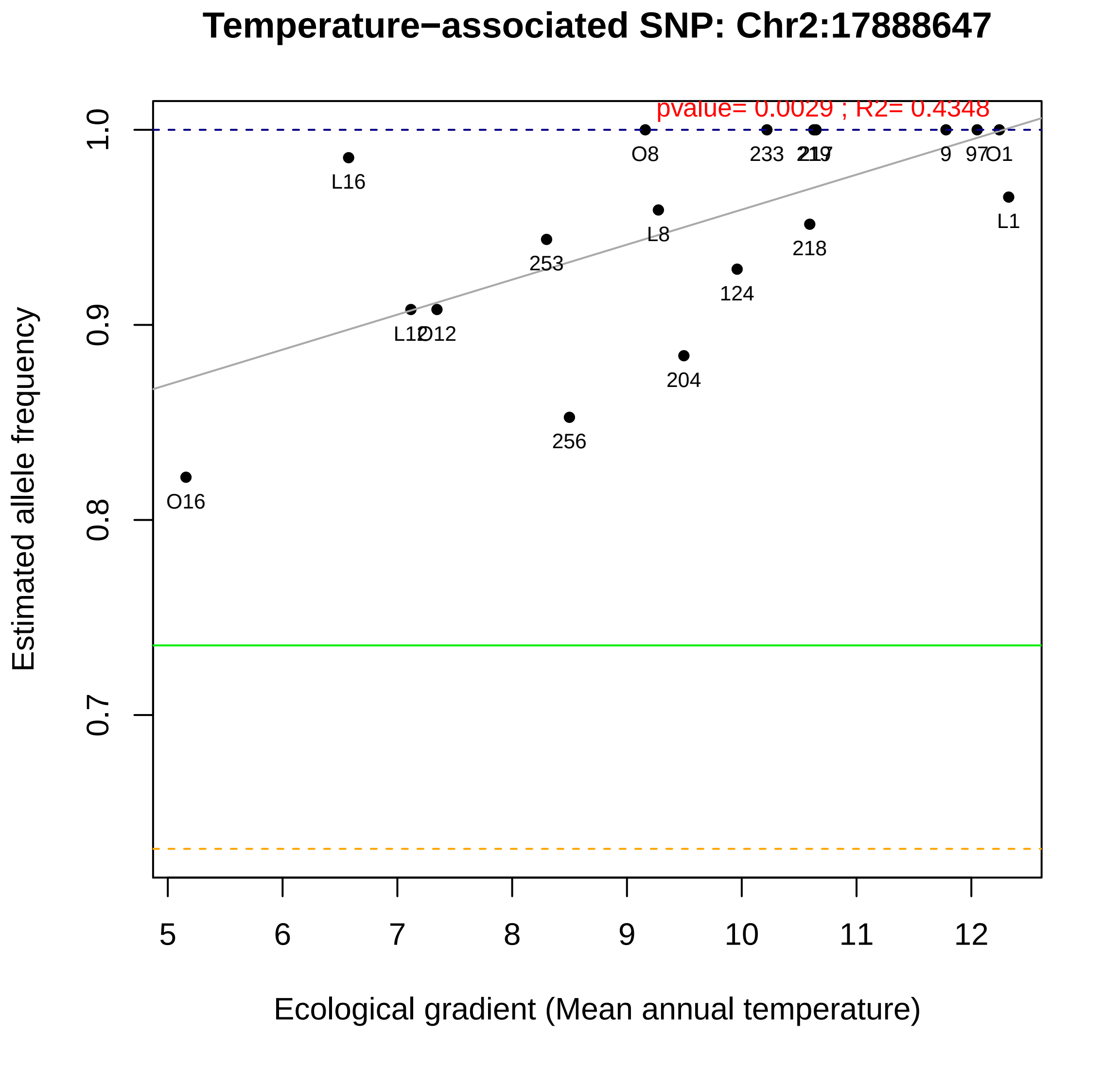

Temperature−associated SNP: Chr2:17888647
pvalue= 0.0029 ; R2= 0.4348
Estimated allele frequency
Ecological gradient (Mean annual temperature)
1.0
0.9
0.8
0.7
5
6
7
8
9
10
11
12
L16
O8
233
9
O1
L1
L8
218
253
124
O12
204
256
O16

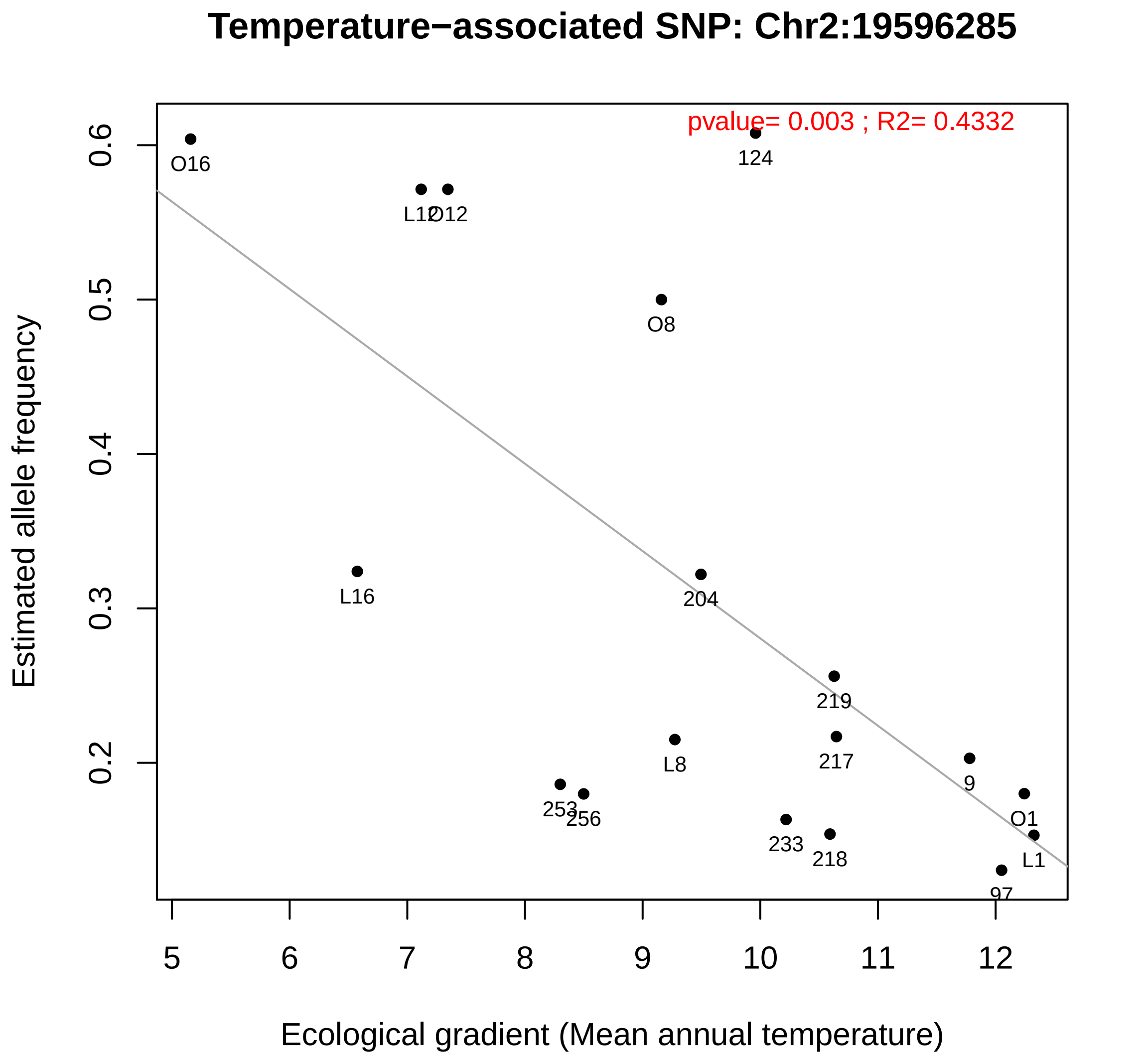
Temperature−associated SNP: Chr2:19596285
pvalue= 0.003 ; R2= 0.4332
Estimated allele frequency
Ecological gradient (Mean annual temperature)
0.2
0.3
0.4
0.5
0.6
5
6
7
8
9
10
11
12
O16
L12
O12
124
O8
L16
204
219
L8
217
9
253
256
O1
233
218
L1
97

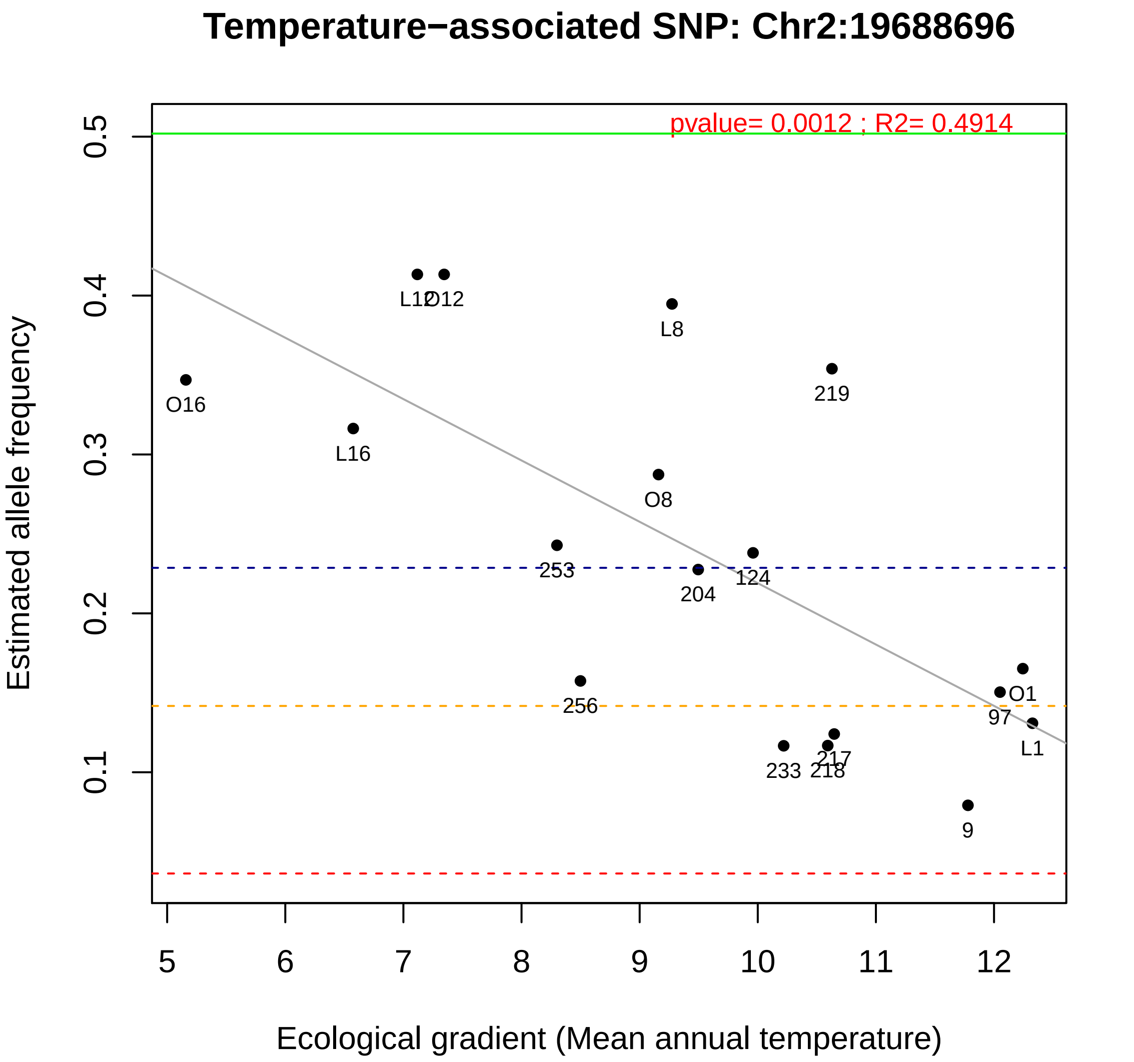

Temperature−associated SNP: Chr2:19688696
pvalue= 0.0012 ; R2= 0.4914
Estimated allele frequency
0.1
0.2
0.3
0.4
0.5
5
6
7
8
9
10
11
12
Ecological gradient (Mean annual temperature)
O16
L16
L12
O12
L8
219
O8
253
204
124
256
233
218
217
9
97
O1
L1

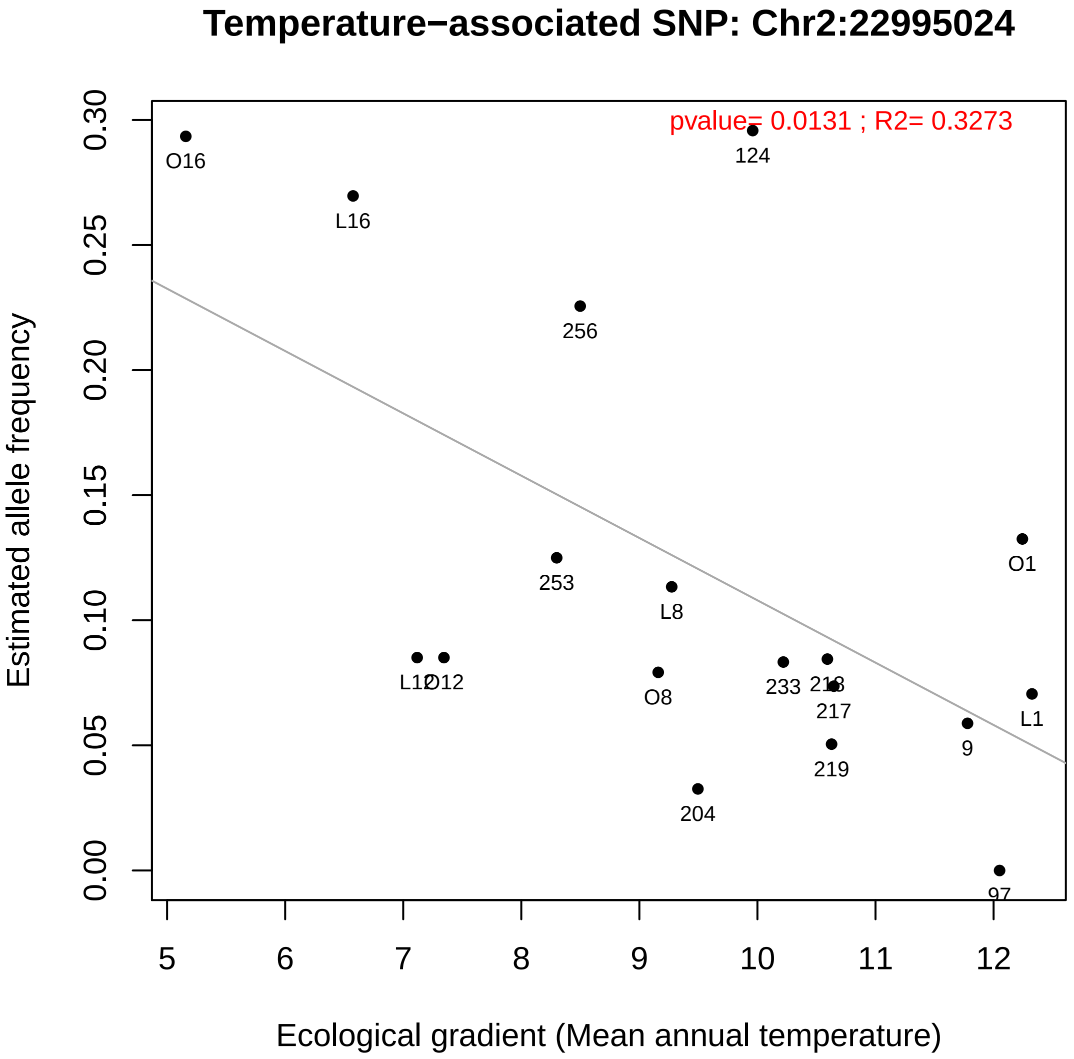

Temperature−associated SNP: Chr2:22995024
pvalue= 0.0131 ; R2= 0.3273
Estimated allele frequency
0.00
0.05
0.10
0.15
0.20
0.25
0.30
Ecological gradient (Mean annual temperature)
5
6
7
8
9
10
11
12
O16
L16
124
256
253
L8
O1
L12
O12
O8
233
218
217
L1
9
219
204
97

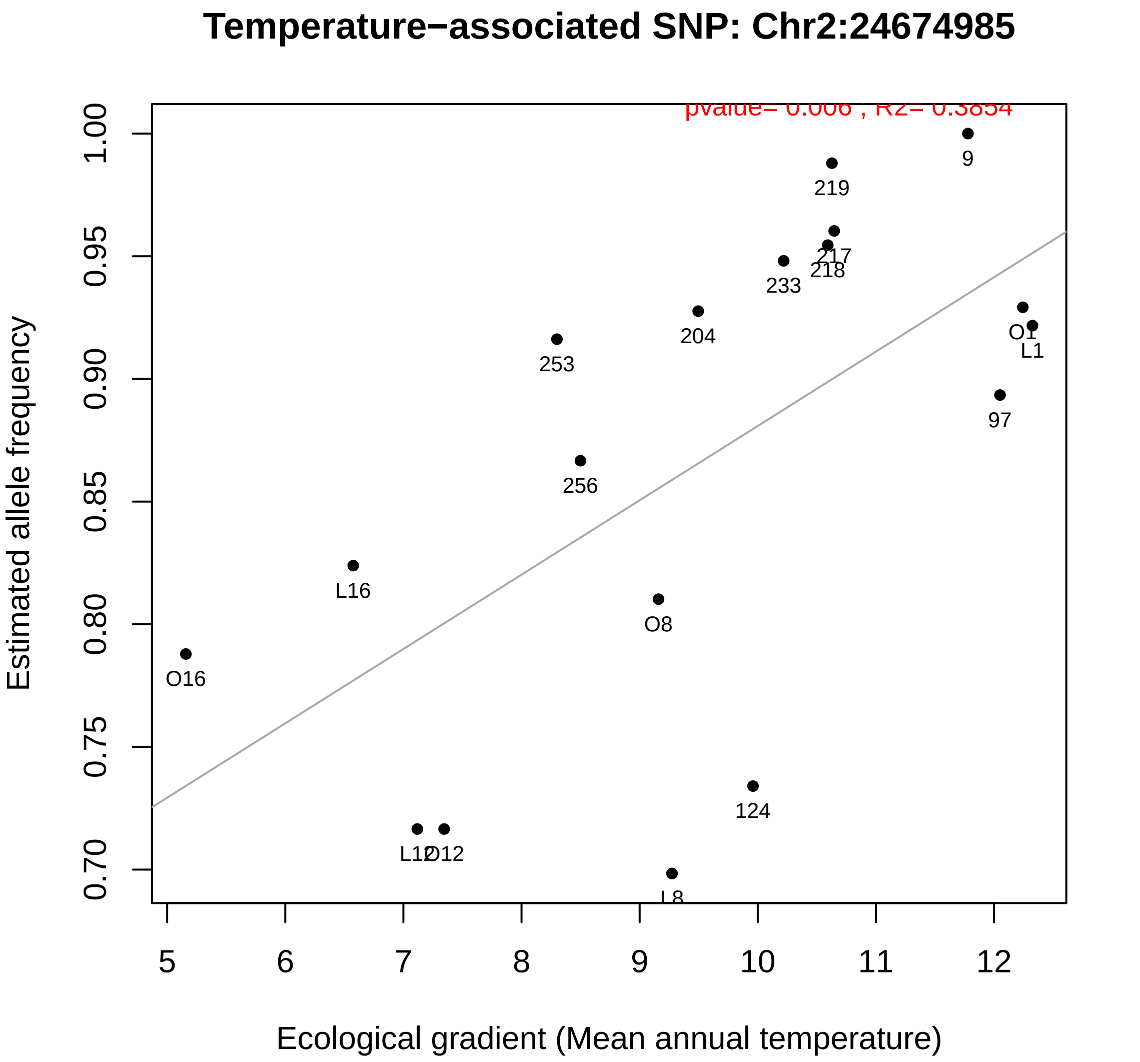
Temperature−associated SNP: Chr2:24674985
pvalue= 0.006 , R2= 0.3854
Estimated allele frequency
1.00
0.95
0.90
0.85
0.80
0.75
0.70
5
6
7
8
9
10
11
12
Ecological gradient (Mean annual temperature)
9
219
217
218
233
204
O1
L1
253
97
256
L16
O8
O16
124
O12

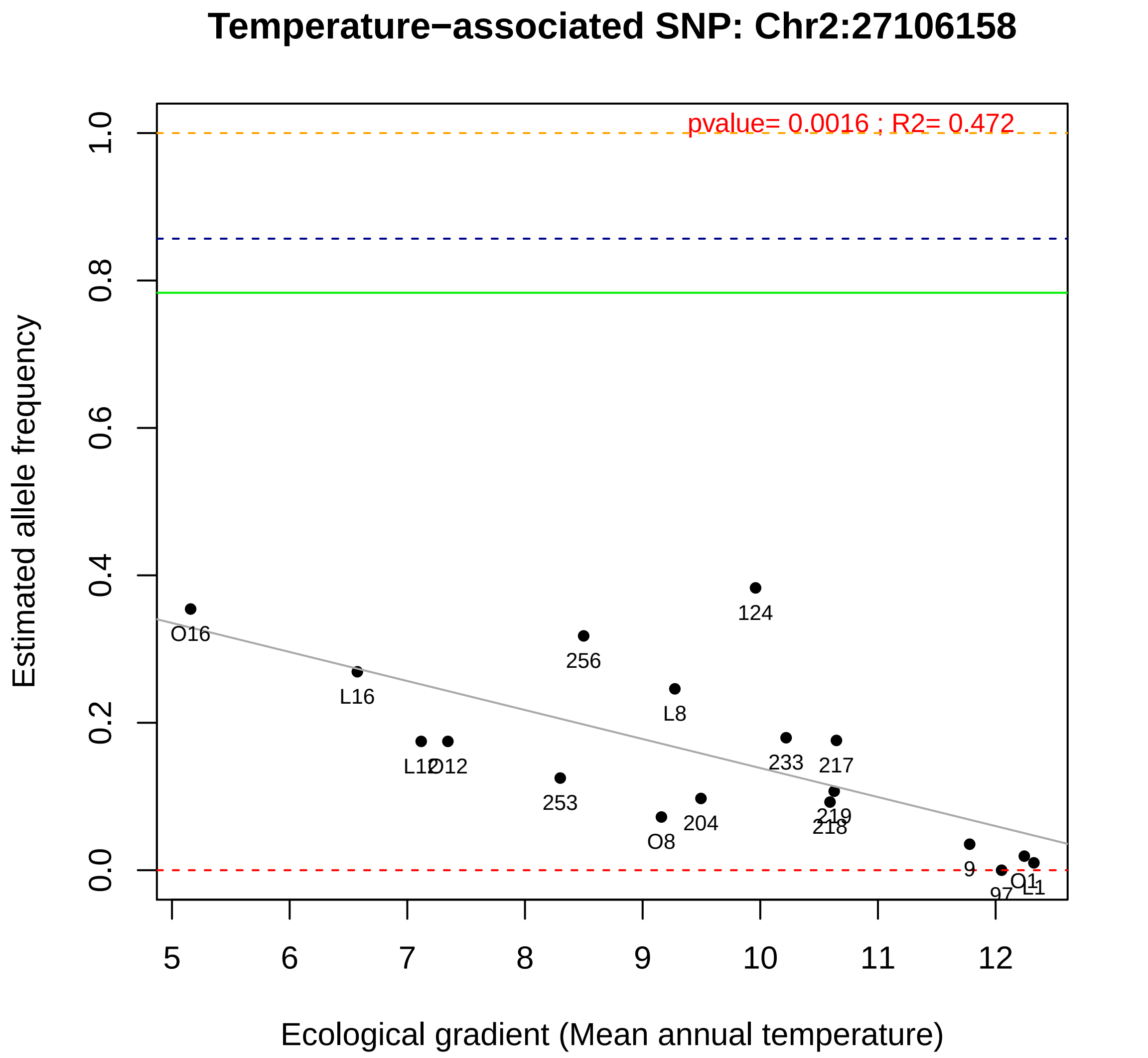
Temperature−associated SNP: Chr2:27106158
pvalue= 0.0016 ; R2= 0.472
Estimated allele frequency
Ecological gradient (Mean annual temperature)
0.0
0.2
0.4
0.6
0.8
1.0
5
6
7
8
9
10
11
12
O16
L16
L12
O12
253
256
O8
L8
204
124
233
217
219
218
9
97
O1
L1

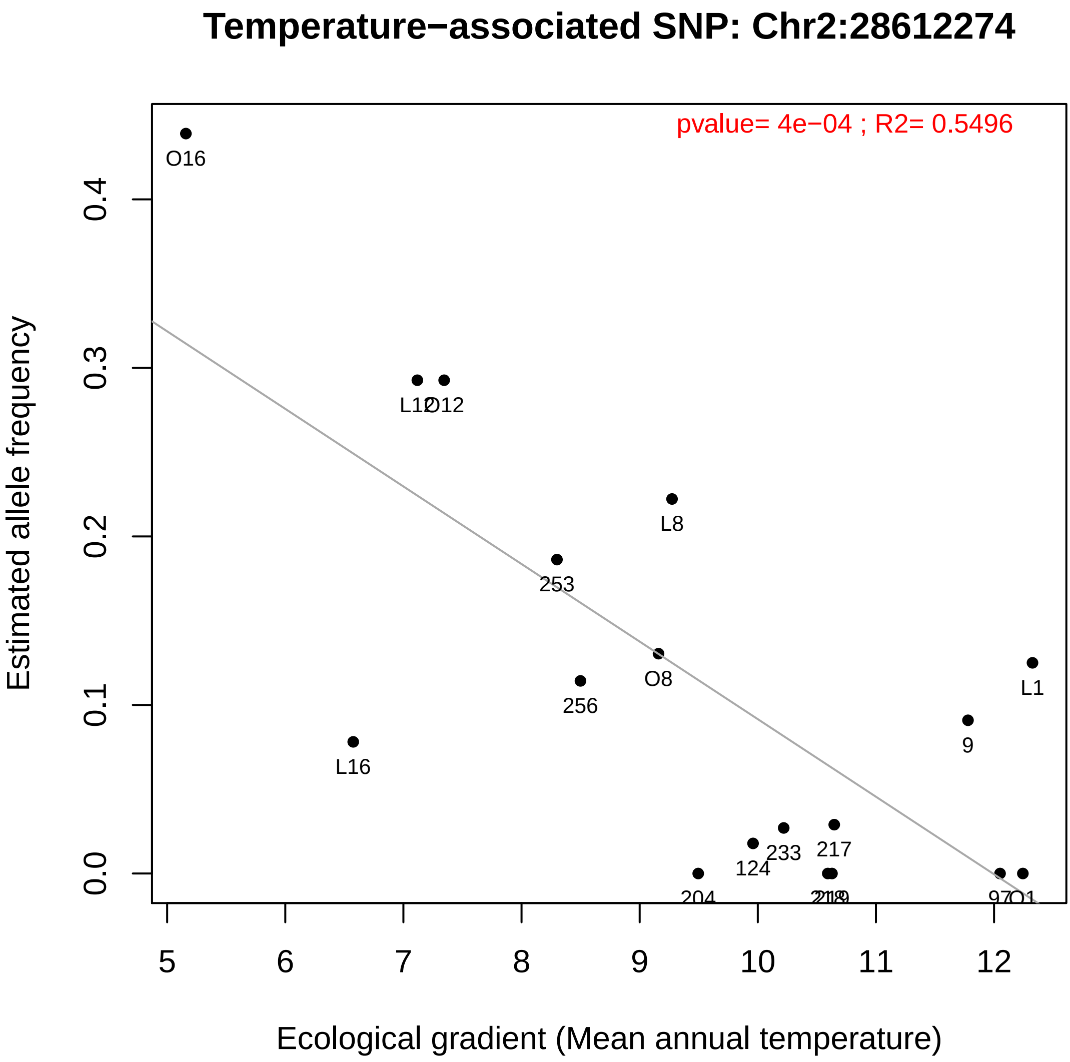
Temperature−associated SNP: Chr2:28612274
pvalue= 4e−04 ; R2= 0.5496
Estimated allele frequency
Ecological gradient (Mean annual temperature)
0.0
0.1
0.2
0.3
0.4
5
6
7
8
9
10
11
12
O16
L12
O12
L8
253
O8
256
L1
9
L16
233
217
124
204

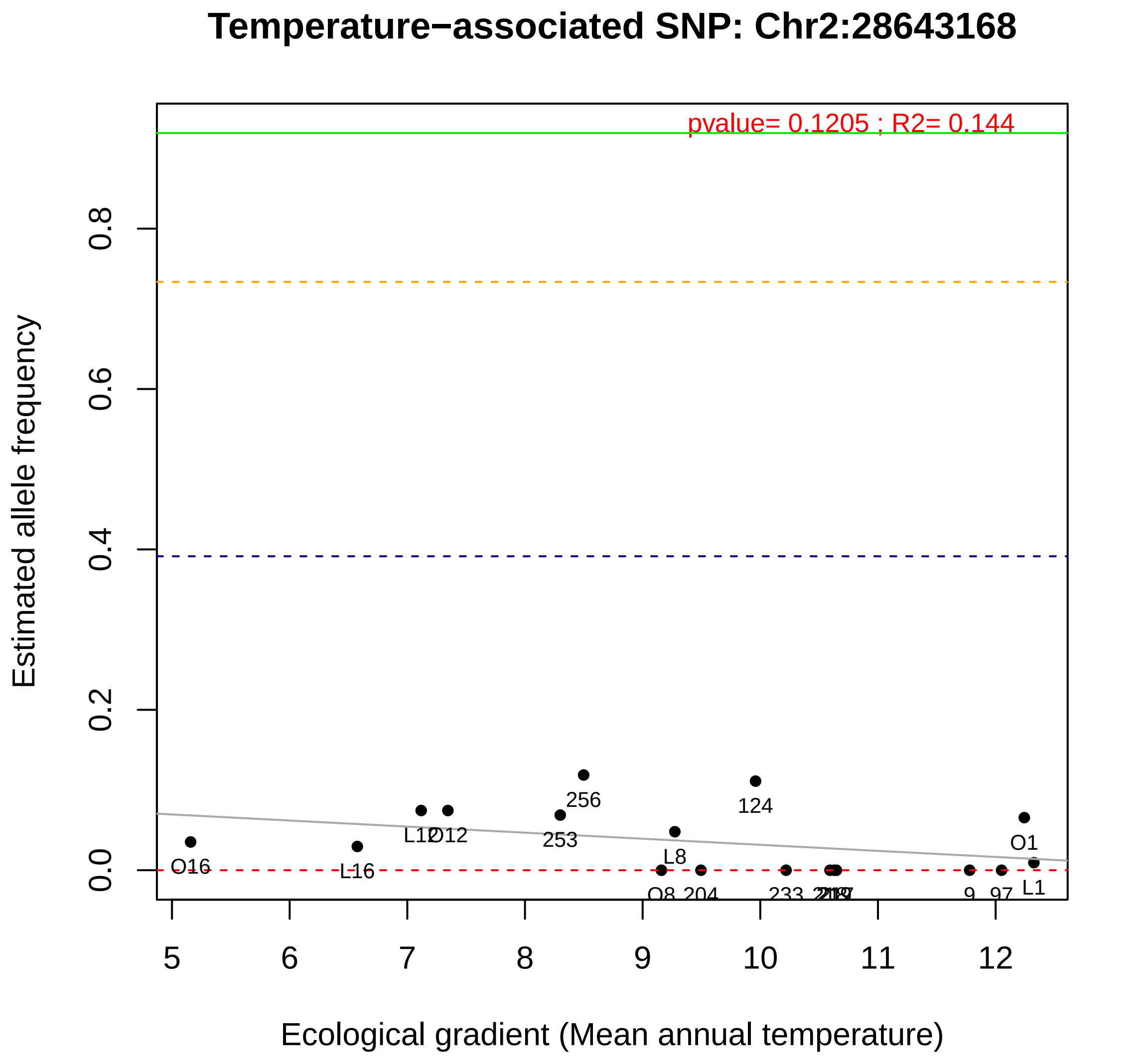

Temperature−associated SNP: Chr2:28643168
pvalue= 0.1205 ; R2= 0.144
Estimated allele frequency
0.0
0.2
0.4
0.6
0.8
5
6
7
8
9
10
11
12
Ecological gradient (Mean annual temperature)
O16
L16
L12
O12
253
256
L8
O8
204
124
233
O1
L1
9
97

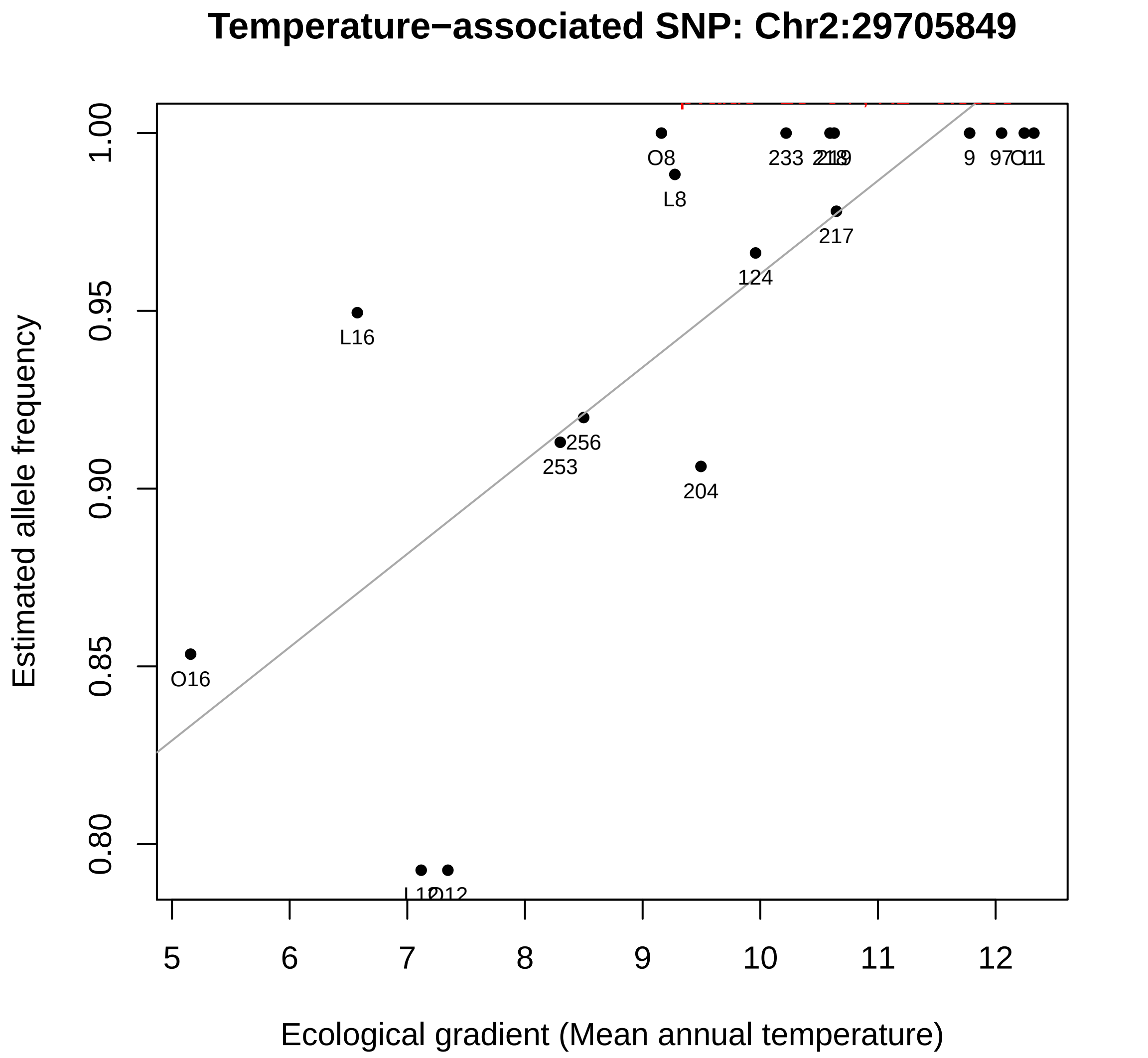
Temperature−associated SNP: Chr2:29705849
Estimated allele frequency
Ecological gradient (Mean annual temperature)
1.00
0.95
0.90
0.85
0.80
5
6
7
8
9
10
11
12
O8
233
9
L8
217
124
L16
256
253
204
O16
O12

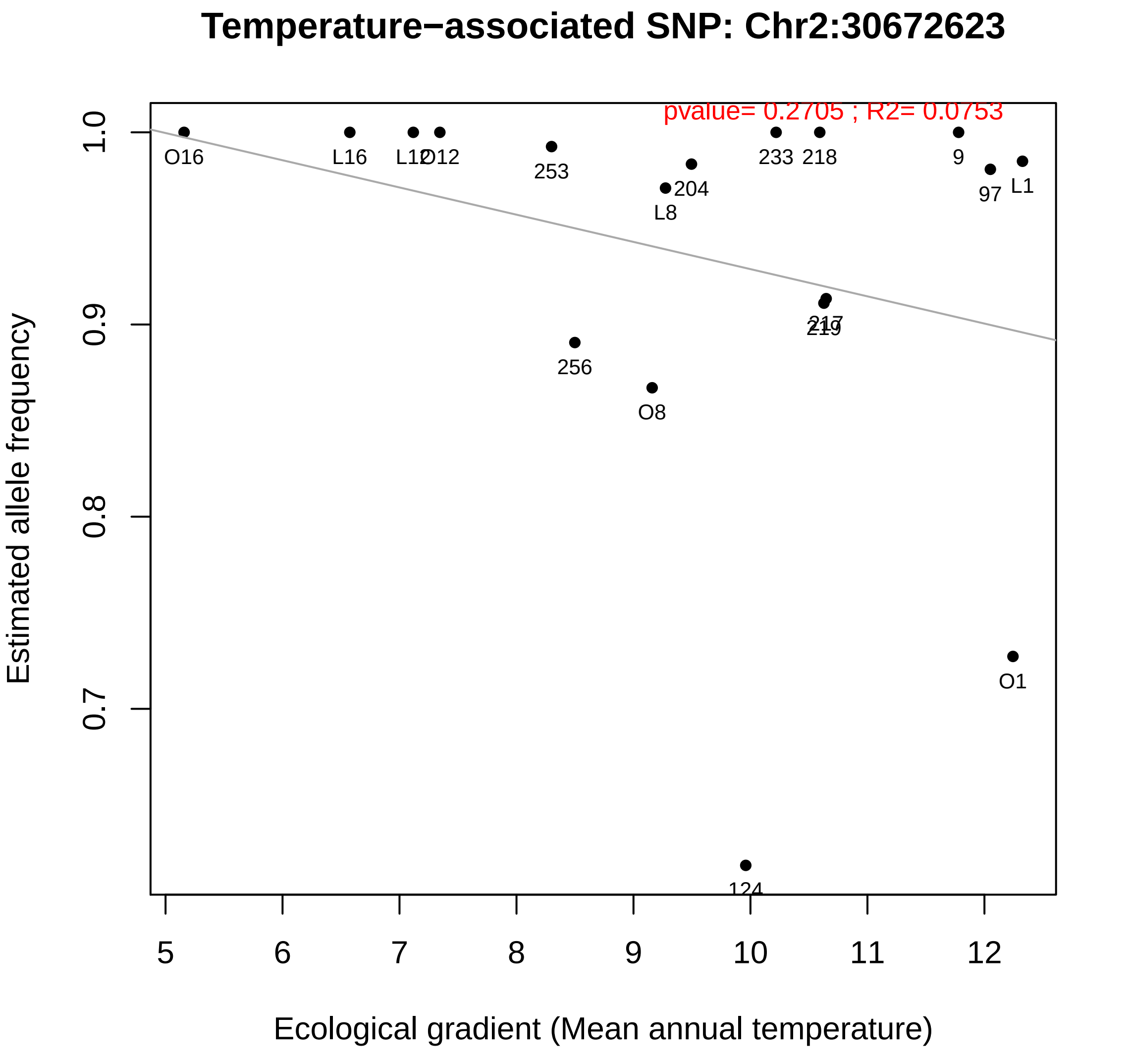

Temperature−associated SNP: Chr2:30672623
pvalue= 0.2705 ; R2= 0.0753
Estimated allele frequency
Ecological gradient (Mean annual temperature)
1.0
0.9
0.8
0.7
5
6
7
8
9
10
11
12
O16
L16
L12
O12
253
204
L8
233
218
9
97
L1
217
219
256
O8
O1
124

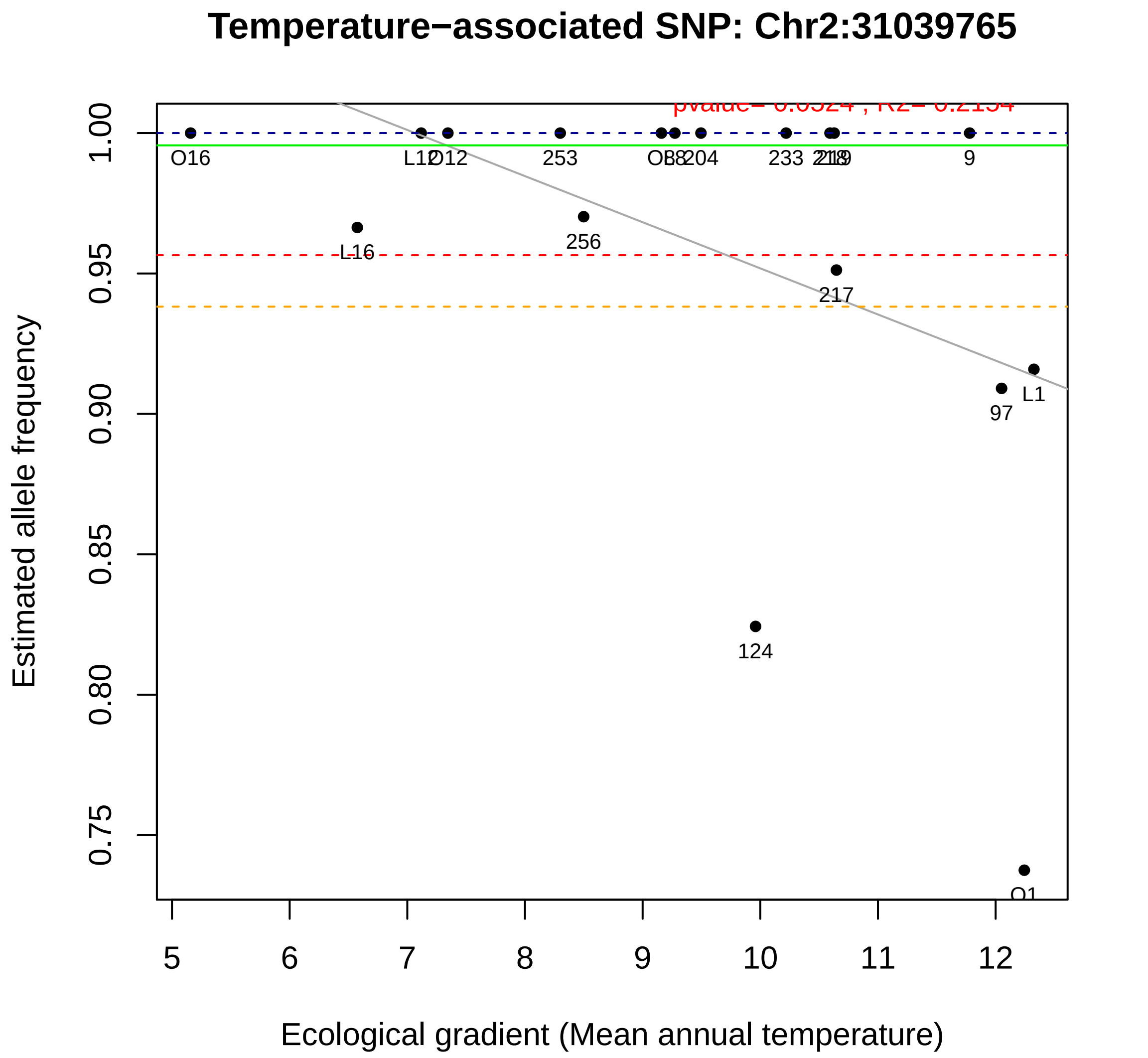

Temperature−associated SNP: Chr2:31039765
pvalue= 0.0324 ; R2= 0.2134
Estimated allele frequency
Ecological gradient (Mean annual temperature)
1.00
0.95
0.90
0.85
0.80
0.75
5
6
7
8
9
10
11
12
O16
L12
O12
253
204
233
9
L16
256
217
L1
97
124
O1

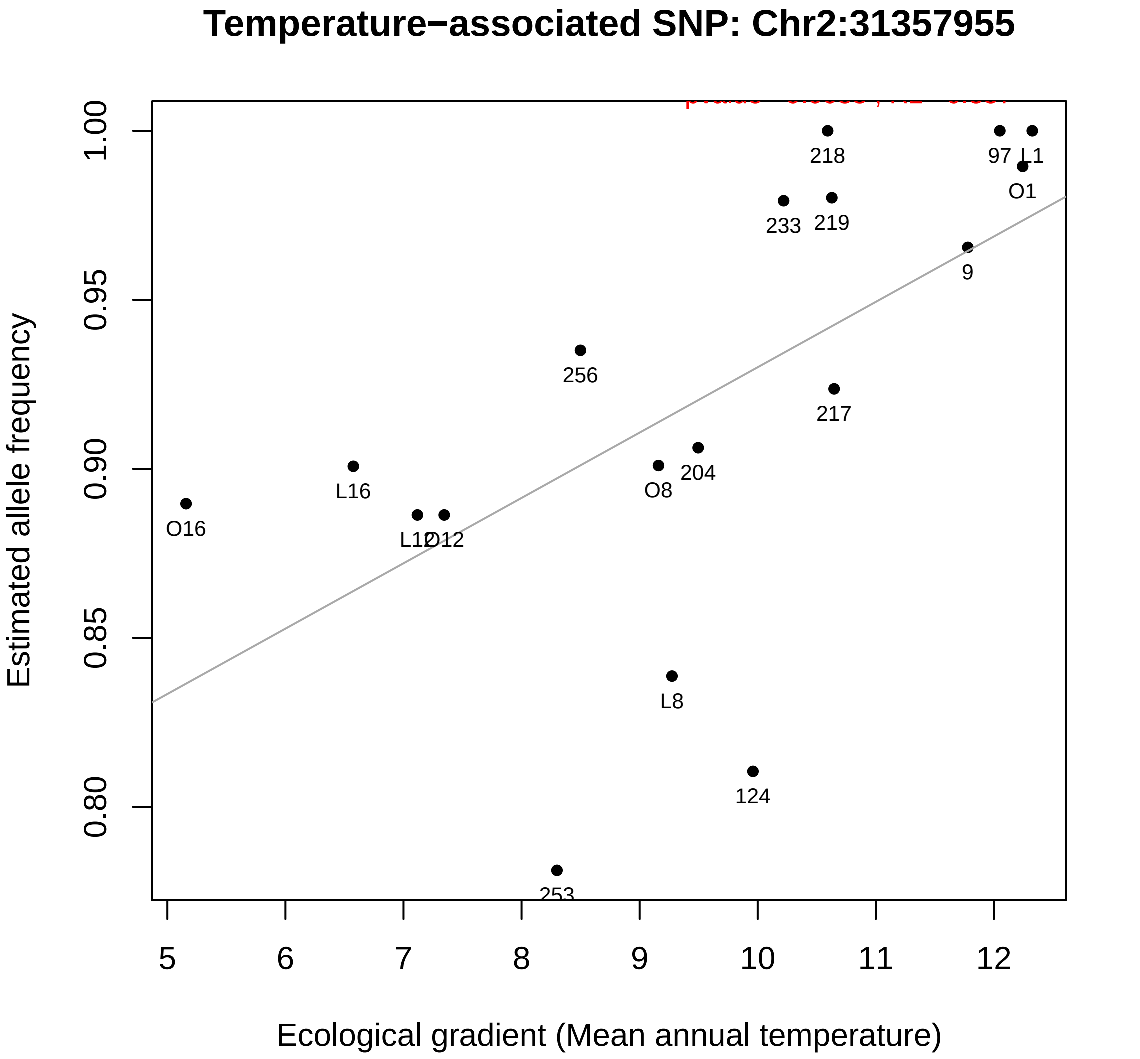

Temperature−associated SNP: Chr2:31357955
Estimated allele frequency
1.00
0.95
0.90
0.85
0.80
5
6
7
8
9
10
11
12
Ecological gradient (Mean annual temperature)
218
97
L1
O1
233
219
9
256
217
204
O8
L16
O16
O12
L8
124

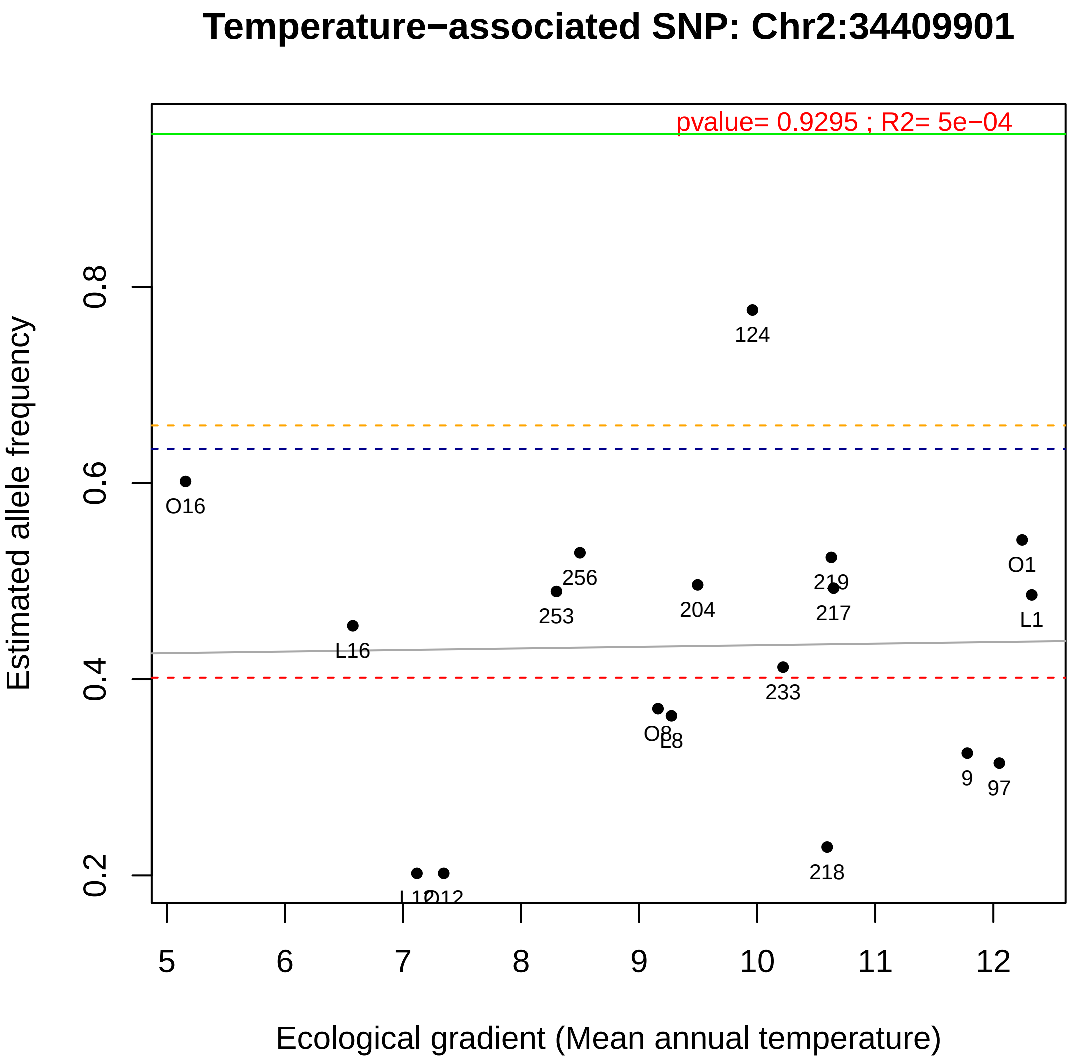

Temperature−associated SNP: Chr2:34409901
pvalue= 0.9295 ; R2= 5e−04
Estimated allele frequency
0.2
0.4
0.6
0.8
5
6
7
8
9
10
11
12
Ecological gradient (Mean annual temperature)
124
O16
256
253
204
219
217
O1
L1
L16
233
O8
L8
9
97
218

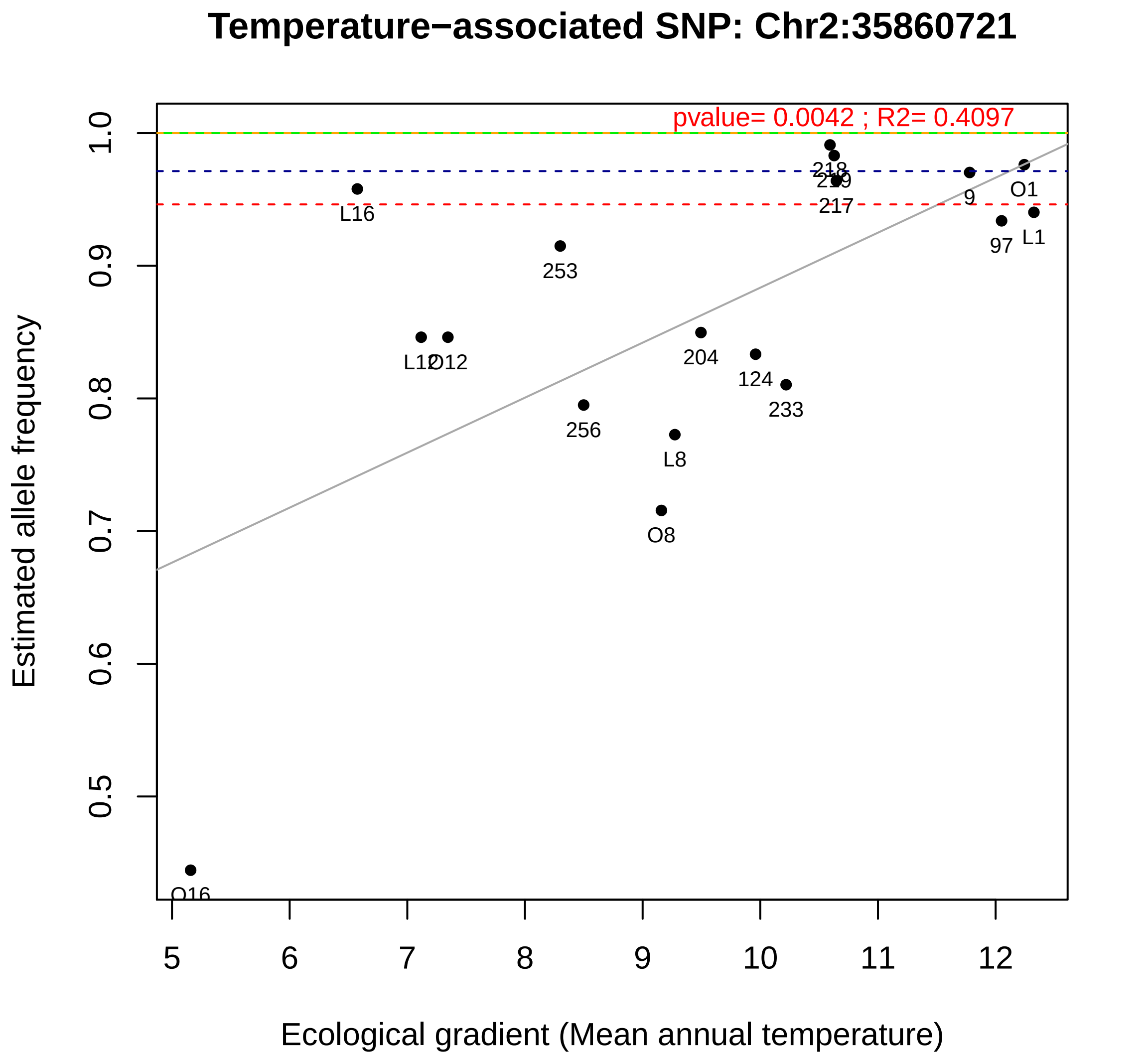

Temperature−associated SNP: Chr2:35860721
pvalue= 0.0042 ; R2= 0.4097
Estimated allele frequency
Ecological gradient (Mean annual temperature)
1.0
0.9
0.8
0.7
0.6
0.5
5
6
7
8
9
10
11
12
L16
218
219
217
9
O1
97
L1
253
L12
O12
204
124
233
256
L8
O8
O16

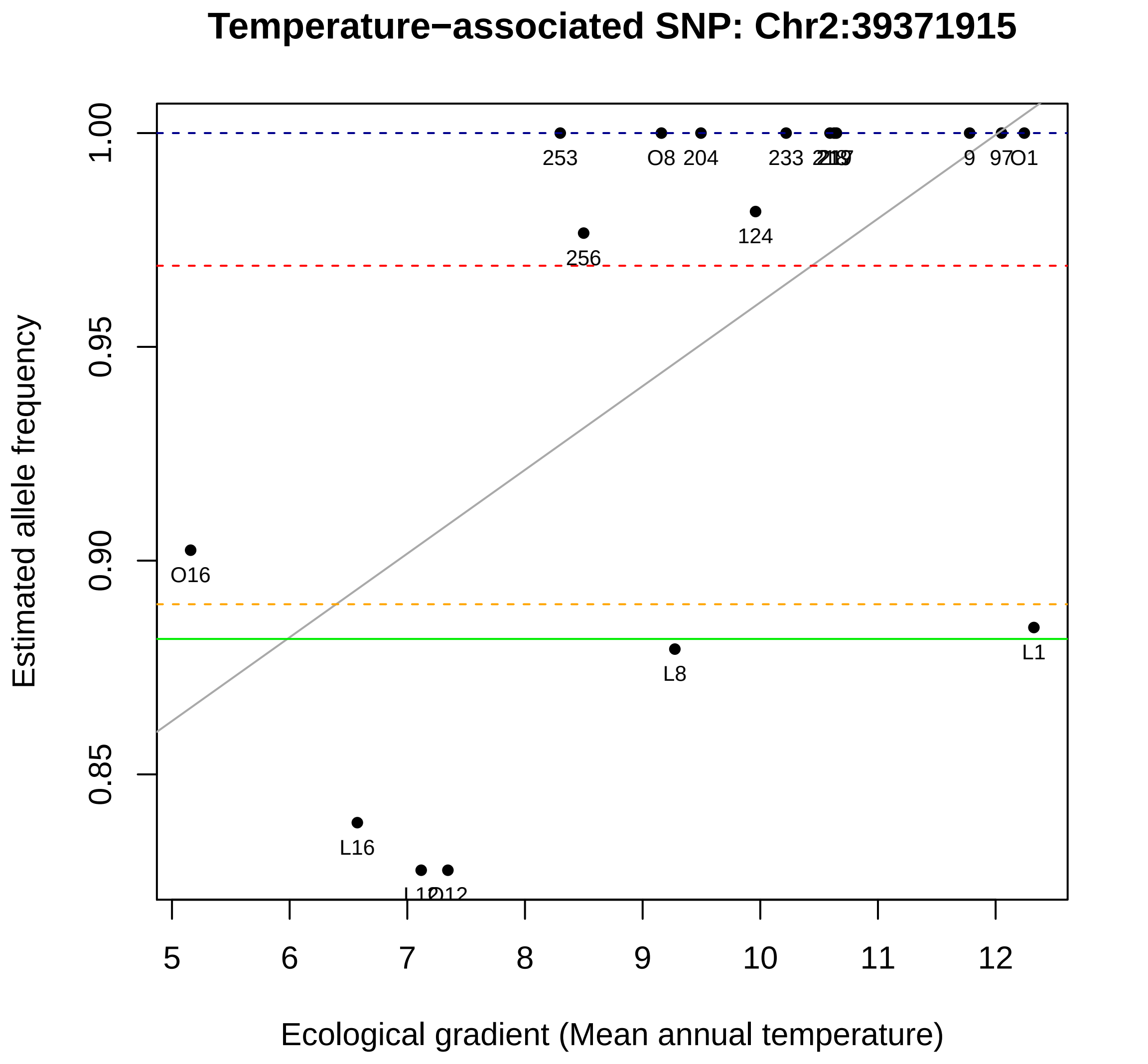

Temperature−associated SNP: Chr2:39371915
Estimated allele frequency
Ecological gradient (Mean annual temperature)
1.00
0.95
0.90
0.85
5
6
7
8
9
10
11
12
253
O8
204
233
9
97
O1
256
124
O16
L8
L1
L16

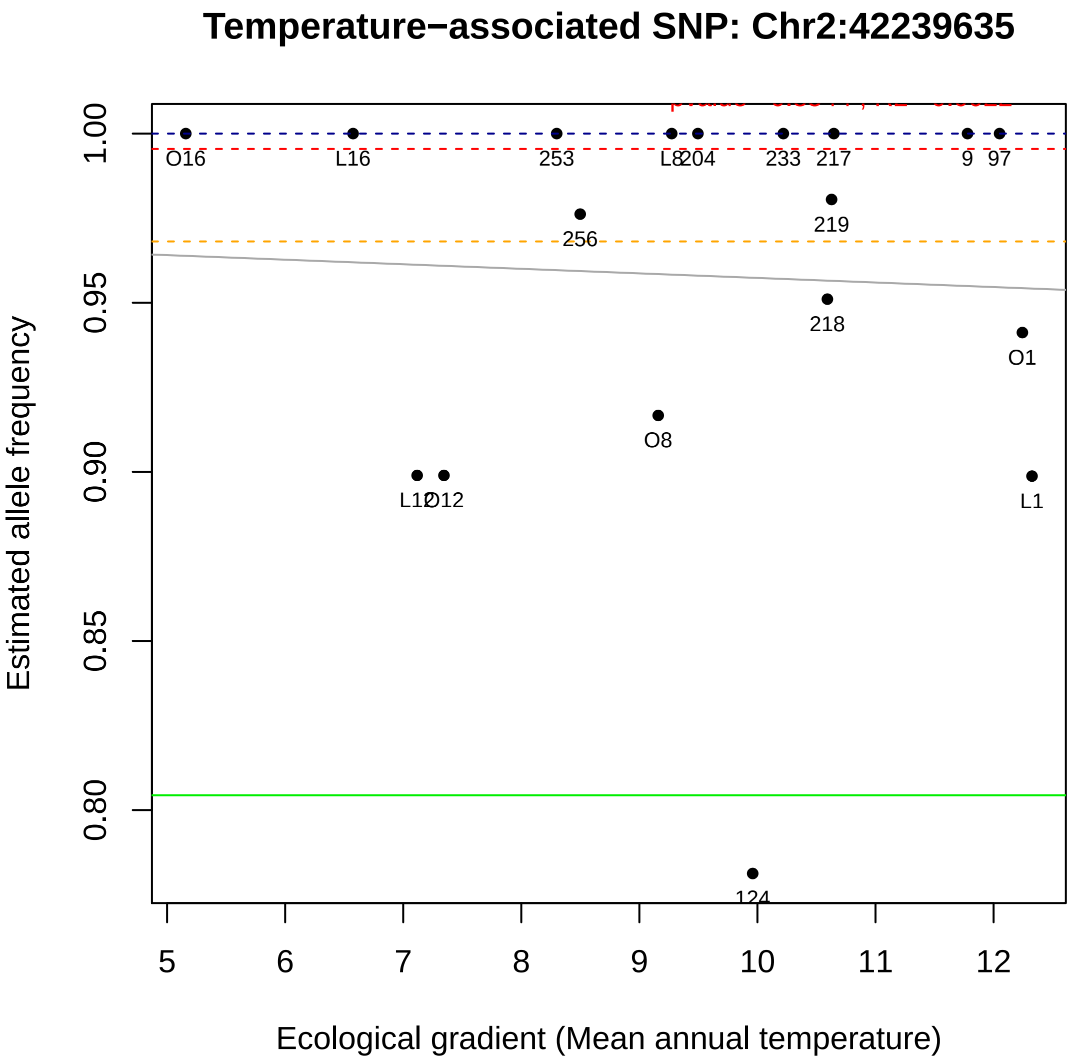

Temperature−associated SNP: Chr2:42239635
Estimated allele frequency
Ecological gradient (Mean annual temperature)
0.80
0.85
0.90
0.95
1.00
5
6
7
8
9
10
11
12
O16
L16
253
L8
204
233
217
9
97
256
219
218
O1
O8
L12
O12
L1
124

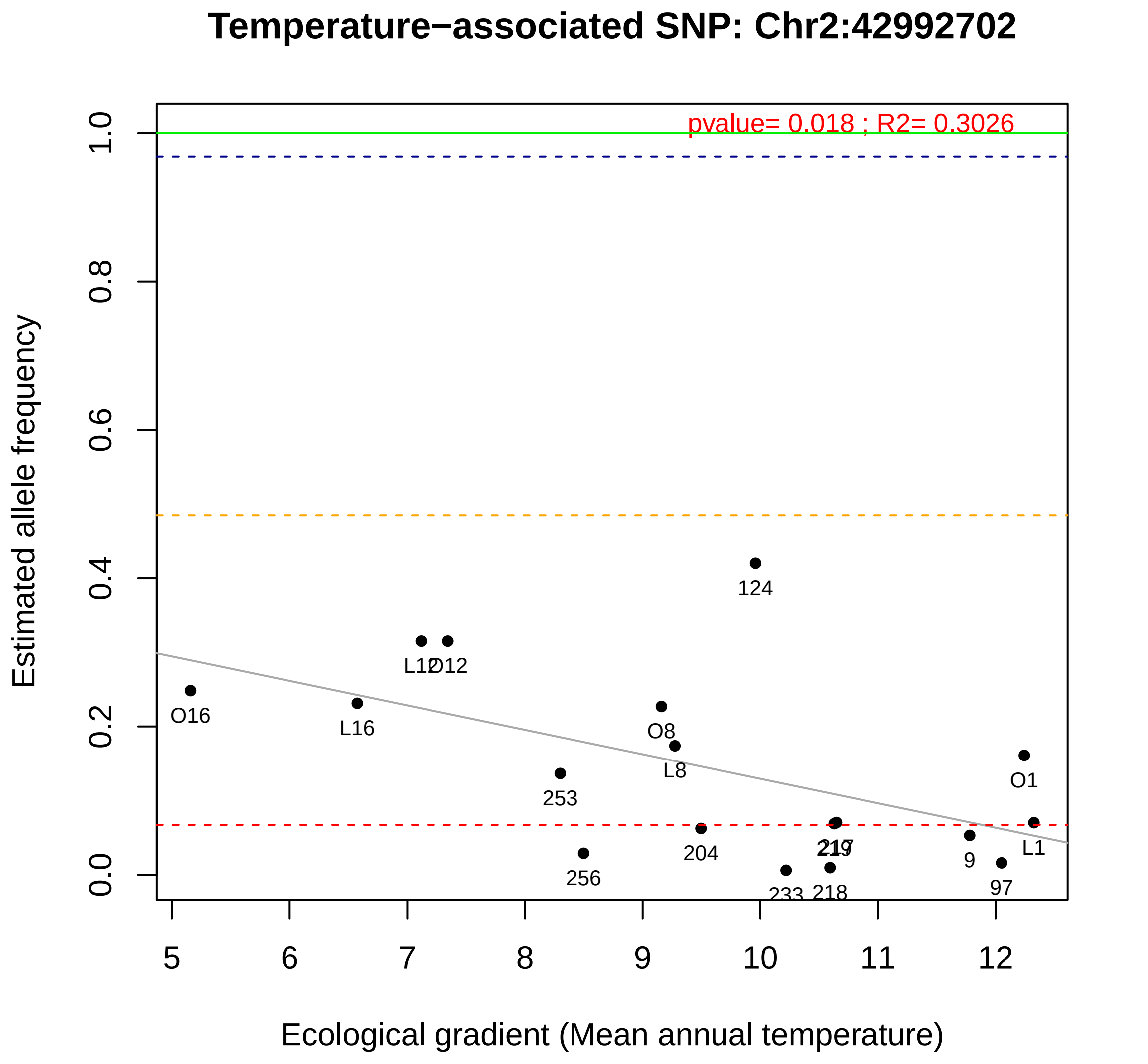
Temperature−associated SNP: Chr2:42992702
pvalue= 0.018 ; R2= 0.3026
Estimated allele frequency
Ecological gradient (Mean annual temperature)
0.0
0.2
0.4
0.6
0.8
1.0
5
6
7
8
9
10
11
12
O16
L16
L12
O12
253
256
O8
L8
204
124
233
218
9
97
O1
L1

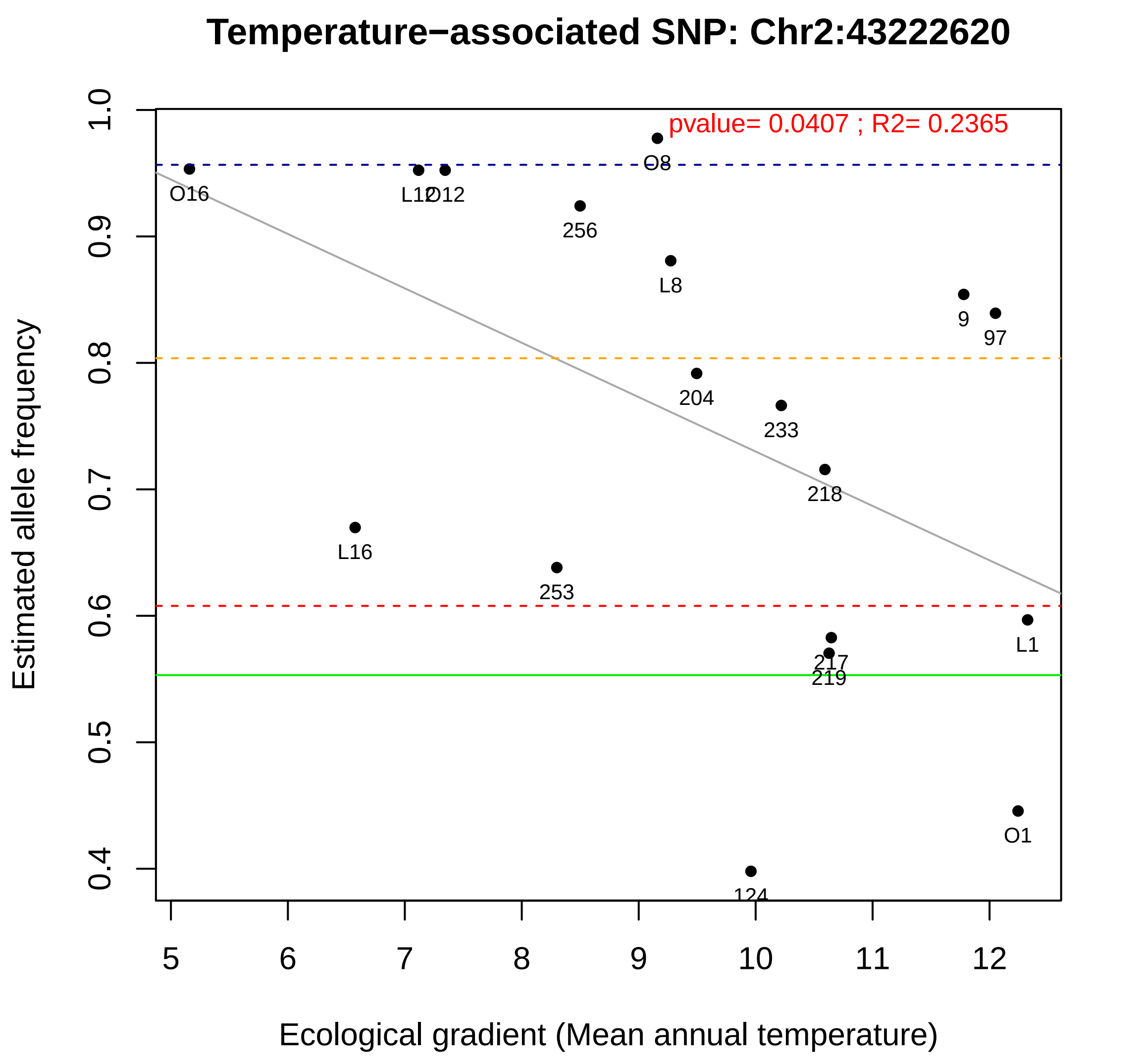
Temperature−associated SNP: Chr2:43222620
pvalue= 0.0407 ; R2= 0.2365
Estimated allele frequency
Ecological gradient (Mean annual temperature)
1.0
0.9
0.8
0.7
0.6
0.5
0.4
5
6
7
8
9
10
11
12
O16
L12
O12
O8
256
L8
9
97
204
233
218
L16
253
L1
217
219
O1
124

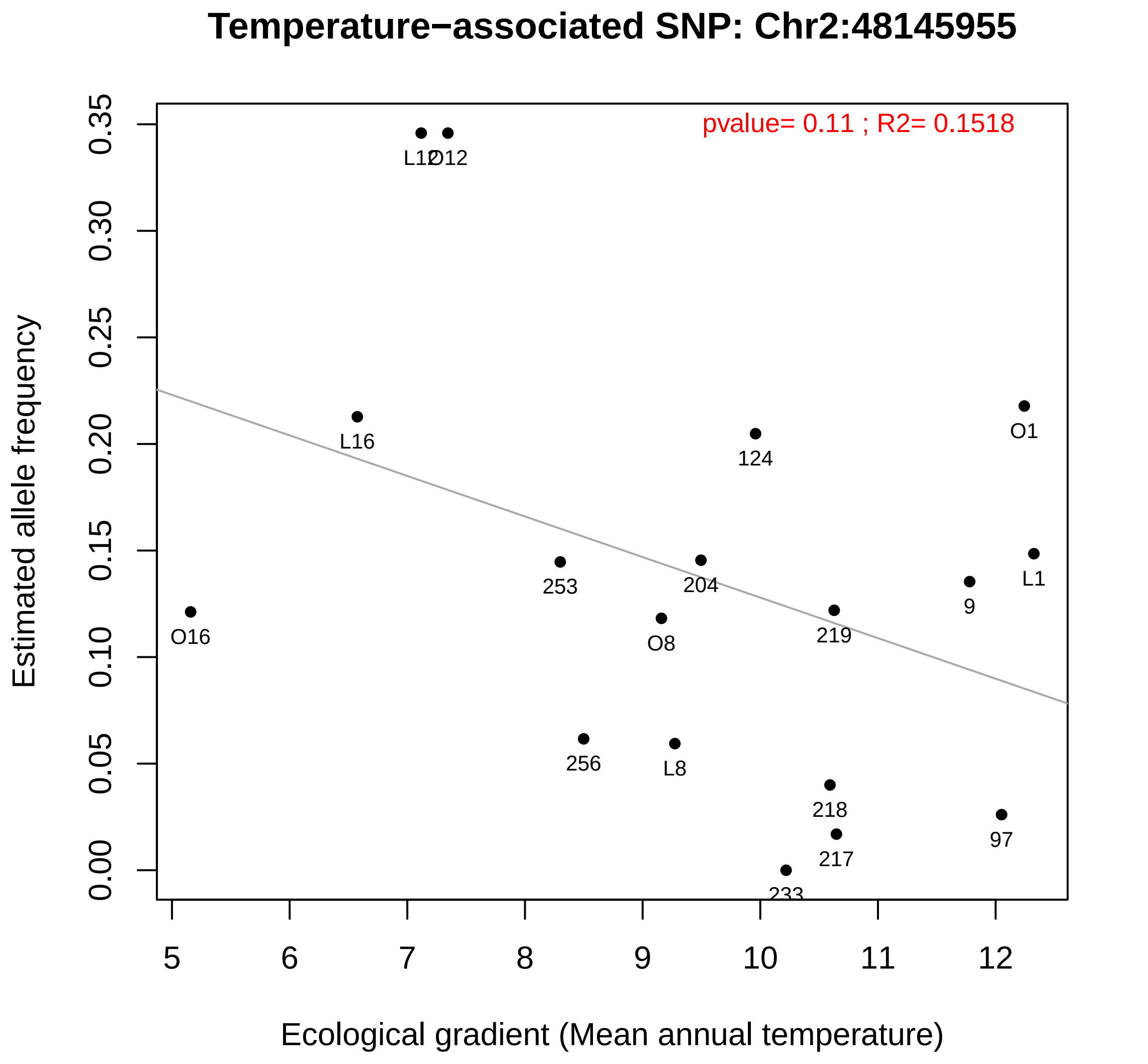
Temperature−associated SNP: Chr2:48145955
pvalue= 0.11 ; R2= 0.1518
Estimated allele frequency
0.00
0.05
0.10
0.15
0.20
0.25
0.30
0.35
Ecological gradient (Mean annual temperature)
5
6
7
8
9
10
11
12
L12
O12
L16
O1
124
253
204
L1
9
O16
O8
219
256
L8
218
97
217
233

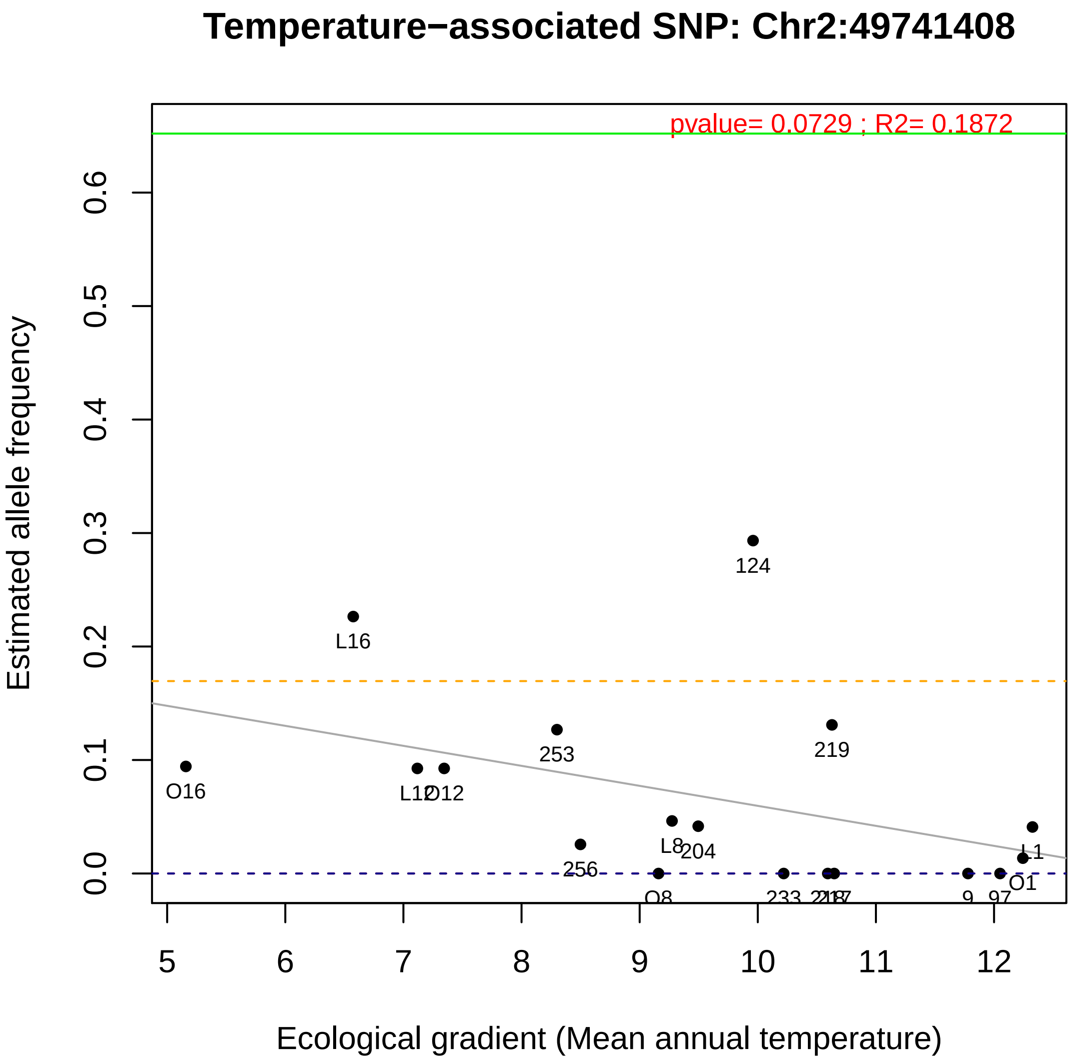

Temperature−associated SNP: Chr2:49741408
pvalue= 0.0729 ; R2= 0.1872
Estimated allele frequency
0.0
0.1
0.2
0.3
0.4
0.5
0.6
124
L16
253
219
O16
L12
O12
L8
204
256
L1
O8
233
O1
9
97
5
6
7
8
9
10
11
12
Ecological gradient (Mean annual temperature)

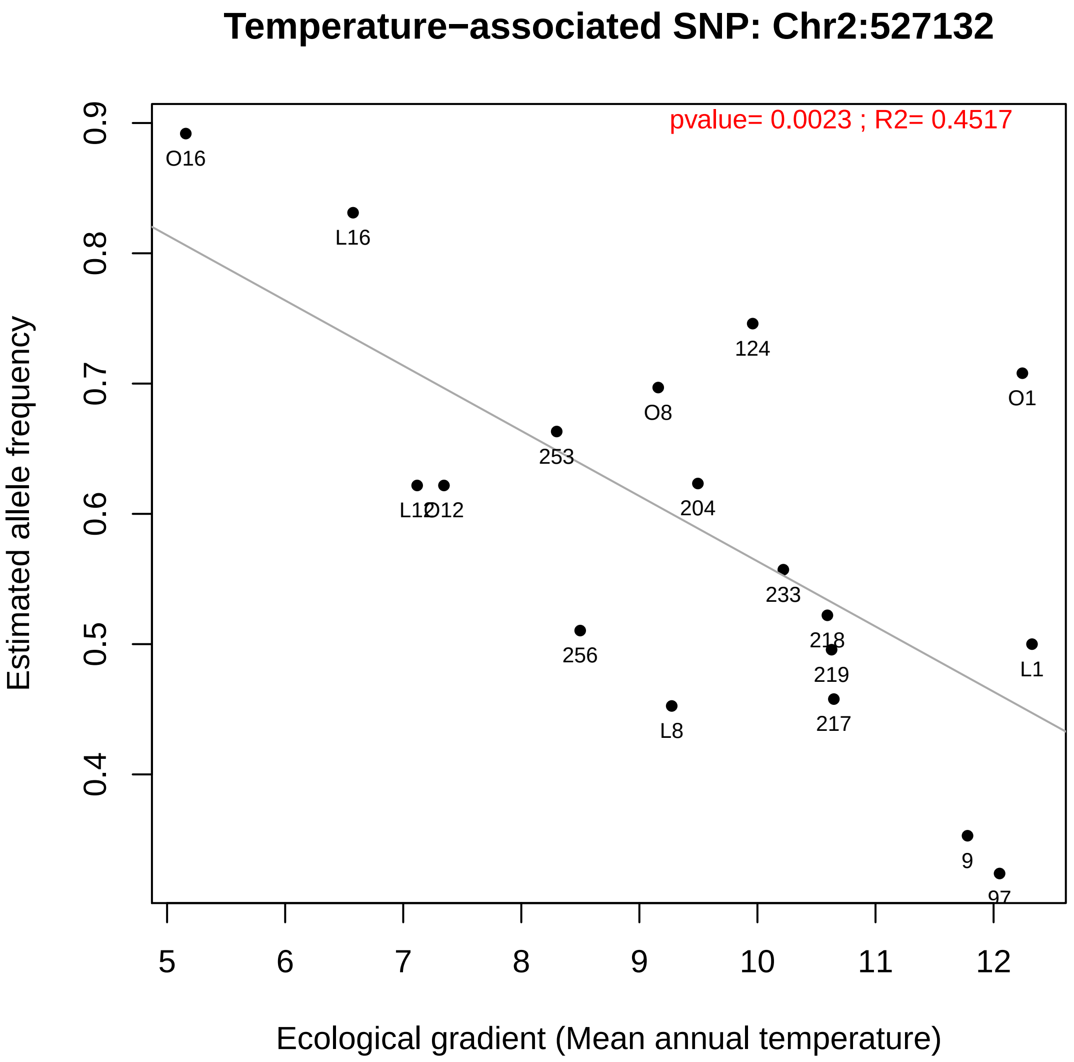
Temperature−associated SNP: Chr2:527132
pvalue= 0.0023 ; R2= 0.4517
O16
L16
124
O1
O8
253
L12
O12
204
233
218
256
219
L1
L8
217
9
97
0.9
0.8
0.7
0.6
0.5
0.4
5
6
7
8
9
10
11
12
Estimated allele frequency
Ecological gradient (Mean annual temperature)

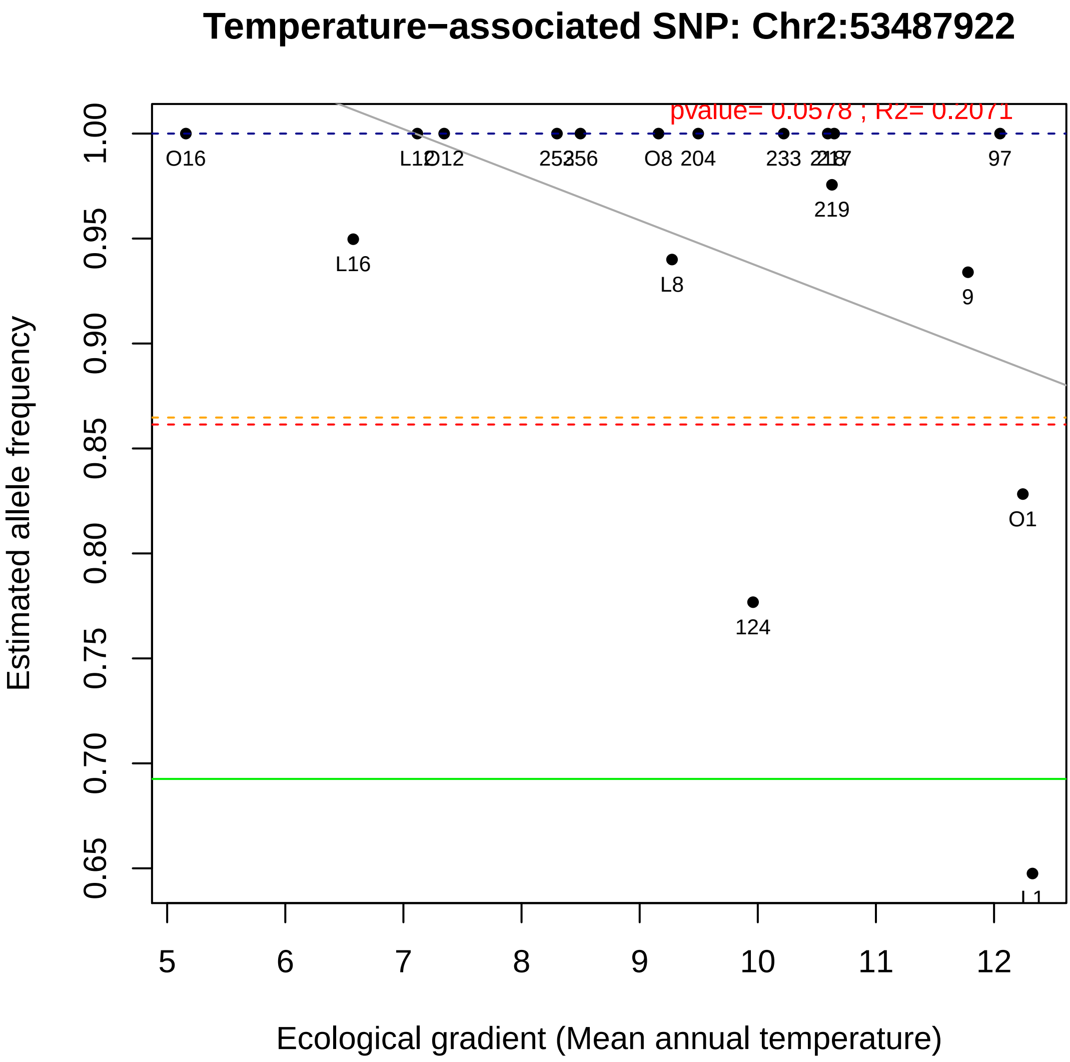
Temperature−associated SNP: Chr2:53487922
pvalue= 0.0578 ; R2= 0.2071
Estimated allele frequency
1.00
0.95
0.90
0.85
0.80
0.75
0.70
0.65
O16
L12
O12
253
256
O8
204
233
218
217
97
219
L16
L8
9
O1
124
L1
5
6
7
8
9
10
11
12
Ecological gradient (Mean annual temperature)

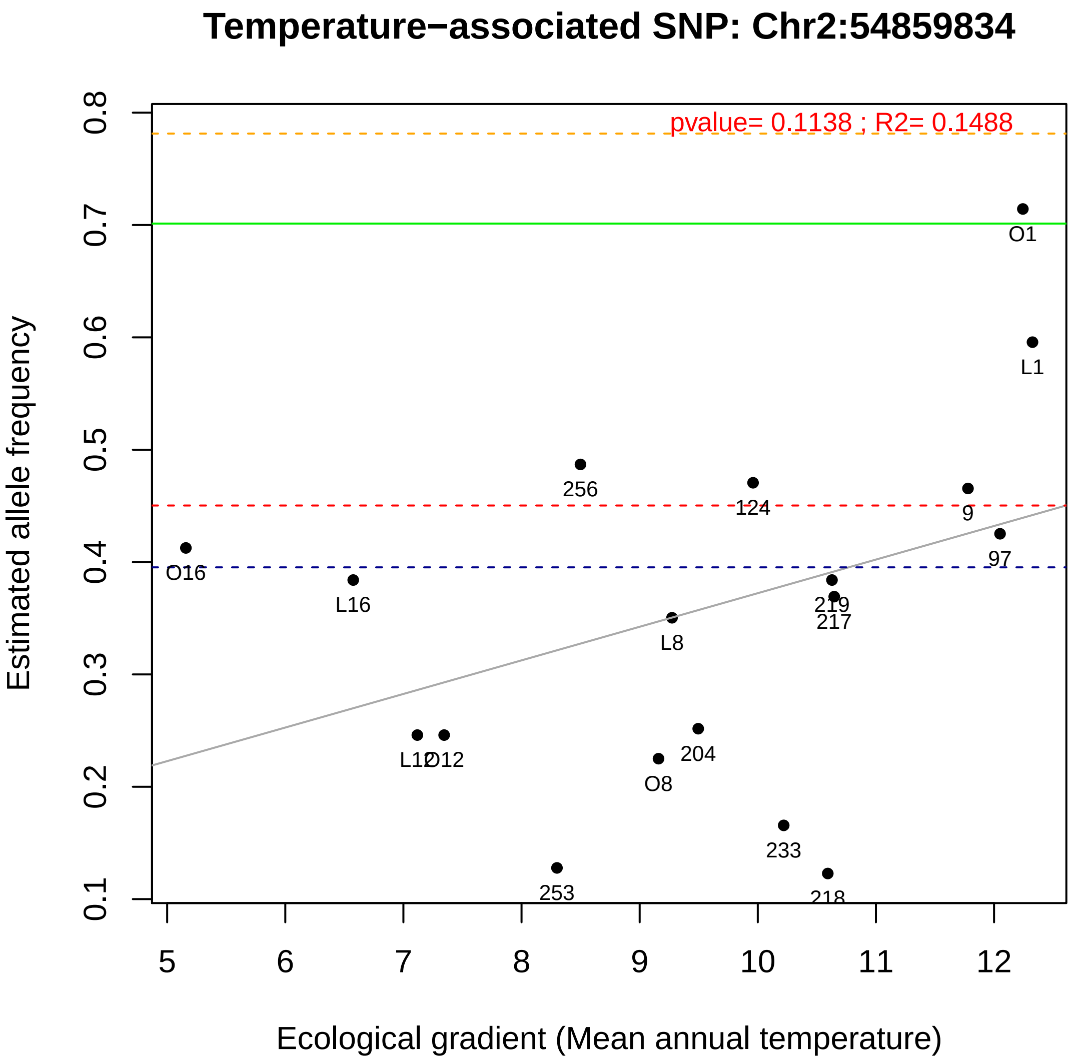
Temperature−associated SNP: Chr2:54859834
pvalue= 0.1138 ; R2= 0.1488
Estimated allele frequency
0.1
0.2
0.3
0.4
0.5
0.6
0.7
0.8
5
6
7
8
9
10
11
12
Ecological gradient (Mean annual temperature)
O1
L1
256
124
9
97
O16
L16
219
217
L8
L12
O12
204
O8
233
253
218

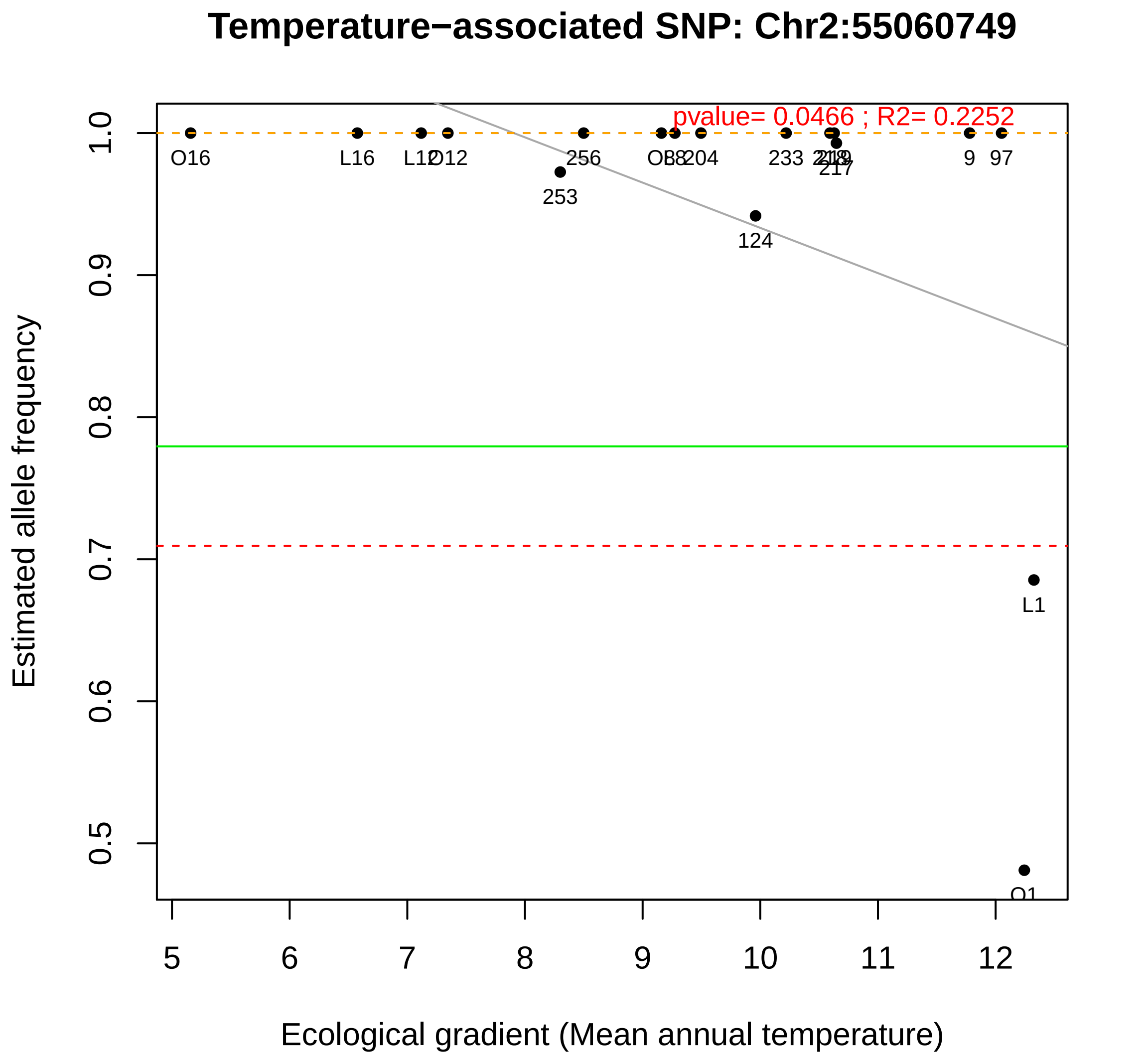

Temperature−associated SNP: Chr2:55060749
pvalue= 0.0466 ; R2= 0.2252
O16
L16
L12
O12
256
253
O8
L8
204
124
233
218
219
217
9
97
L1
O1
Estimated allele frequency
1.0
0.9
0.8
0.7
0.6
0.5
5
6
7
8
9
10
11
12
Ecological gradient (Mean annual temperature)

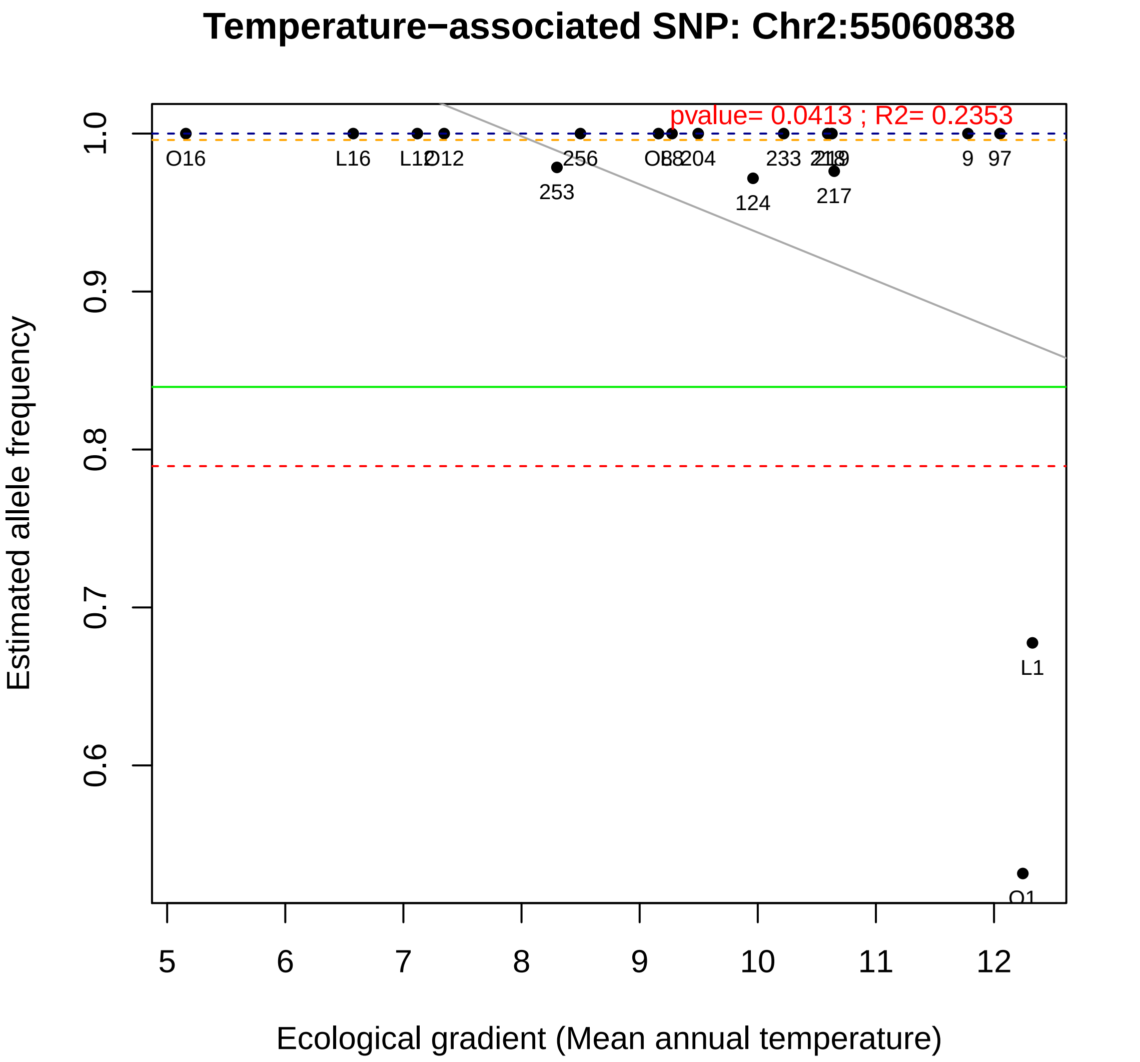

Temperature−associated SNP: Chr2:55060838
pvalue= 0.0413 ; R2= 0.2353
Estimated allele frequency
Ecological gradient (Mean annual temperature)
1.0
0.9
0.8
0.7
0.6
5
6
7
8
9
10
11
12
O16
L16
L12
O12
256
253
204
233
124
217
9
97
L1
O1

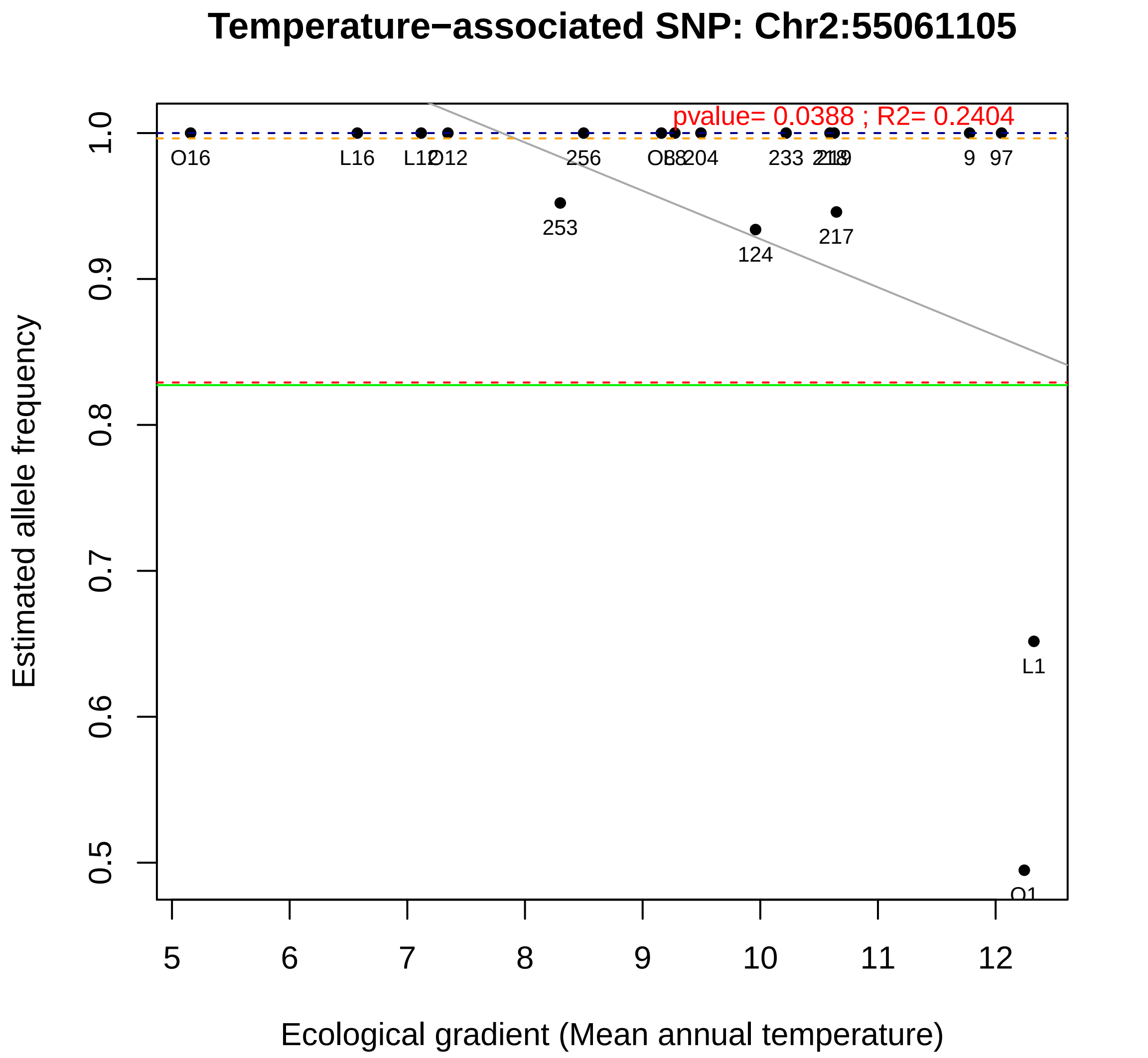

Temperature−associated SNP: Chr2:55061105
pvalue= 0.0388 ; R2= 0.2404
Estimated allele frequency
Ecological gradient (Mean annual temperature)
0.5
0.6
0.7
0.8
0.9
1.0
5
6
7
8
9
10
11
12
O16
L16
O12
256
204
233
9
97
253
124
217
L1

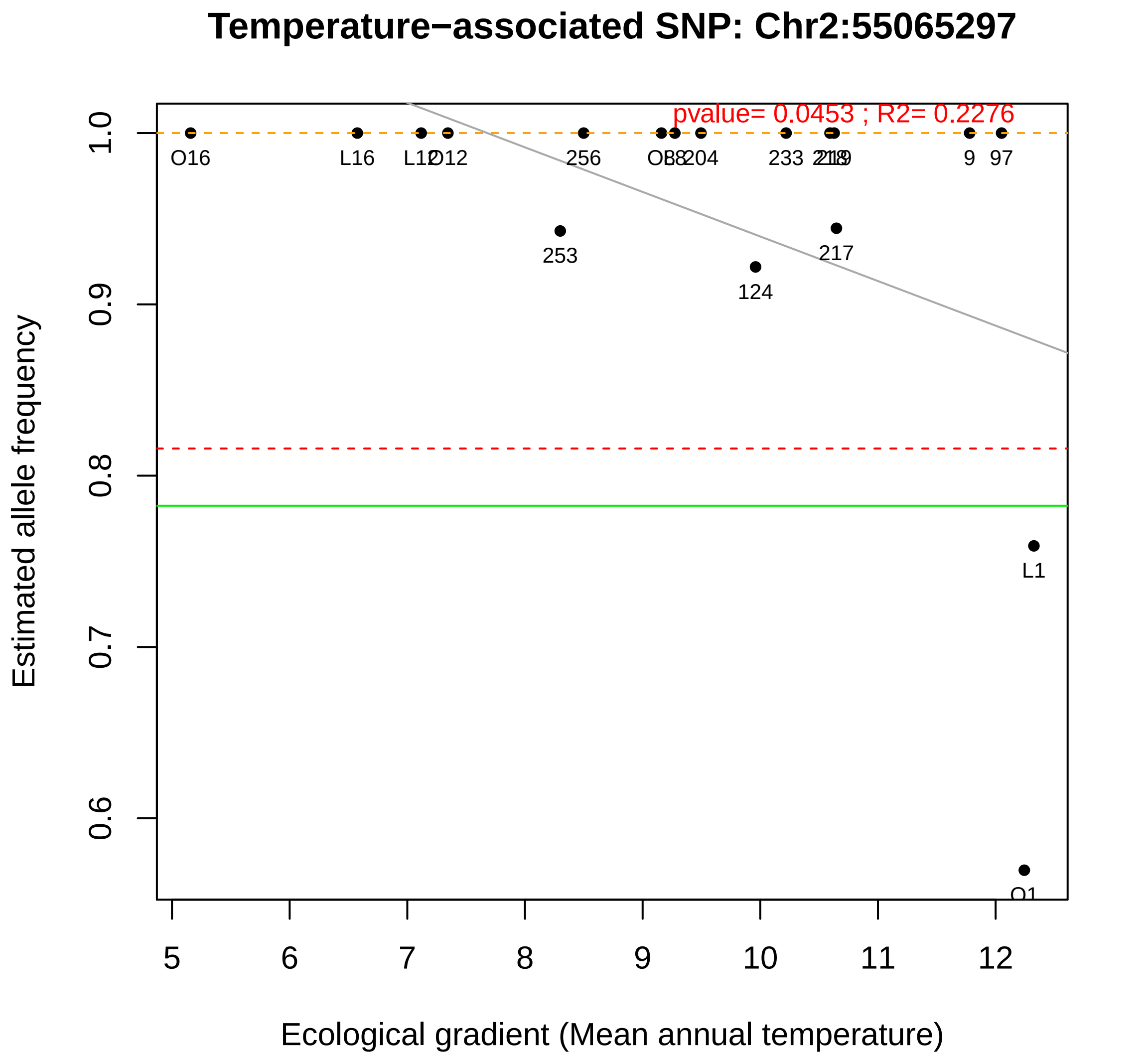
Temperature−associated SNP: Chr2:55065297
pvalue= 0.0453 ; R2= 0.2276
O16
L16
L12
O12
256
253
204
233
124
217
9
97
L1
O1
Estimated allele frequency
0.6
0.7
0.8
0.9
1.0
5
6
7
8
9
10
11
12
Ecological gradient (Mean annual temperature)

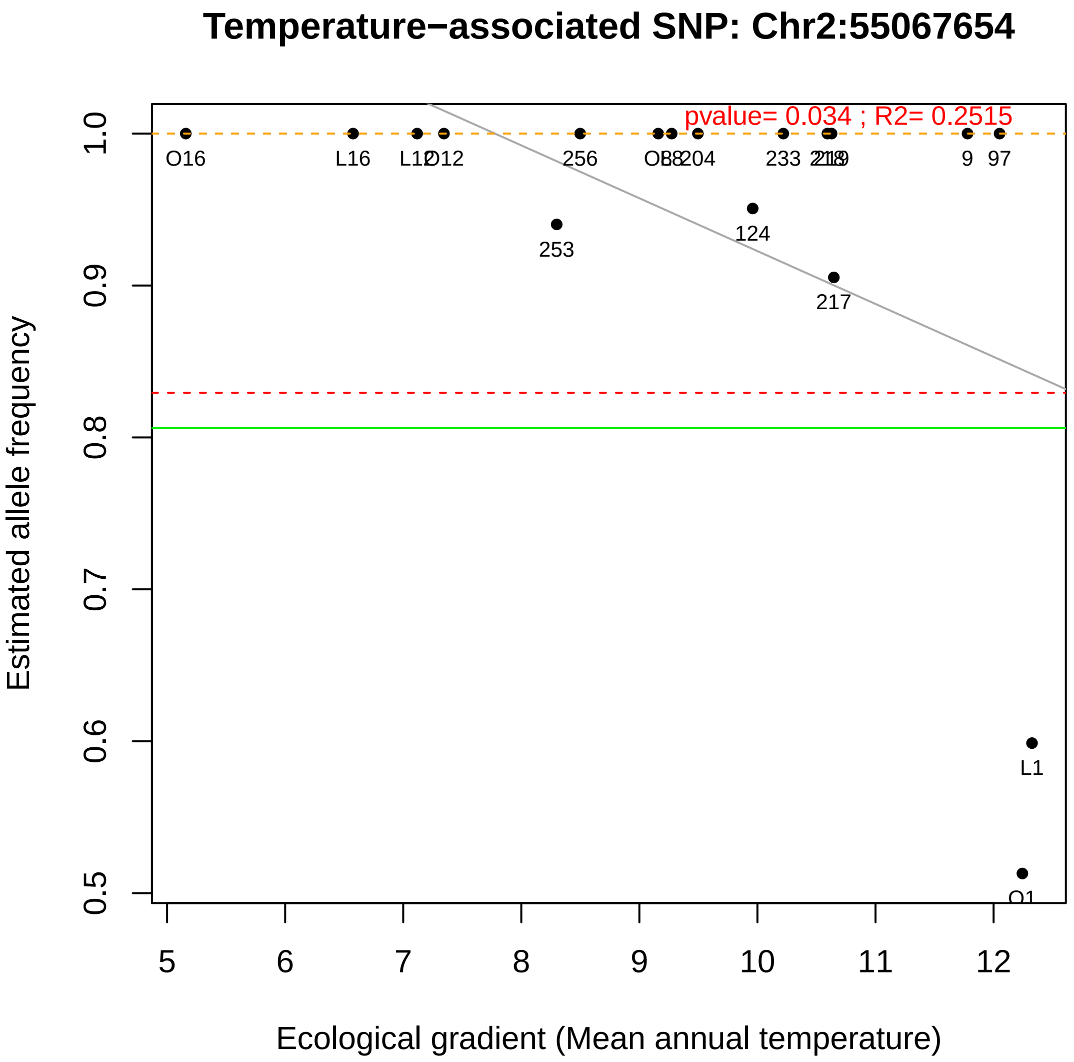
Temperature−associated SNP: Chr2:55067654
pvalue= 0.034 ; R2= 0.2515
Estimated allele frequency
Ecological gradient (Mean annual temperature)
0.5
0.6
0.7
0.8
0.9
1.0
5
6
7
8
9
10
11
12
O16
L16
256
204
233
9
97
253
124
217
L1

**Temperature−associated SNP: Chr2:55176046**

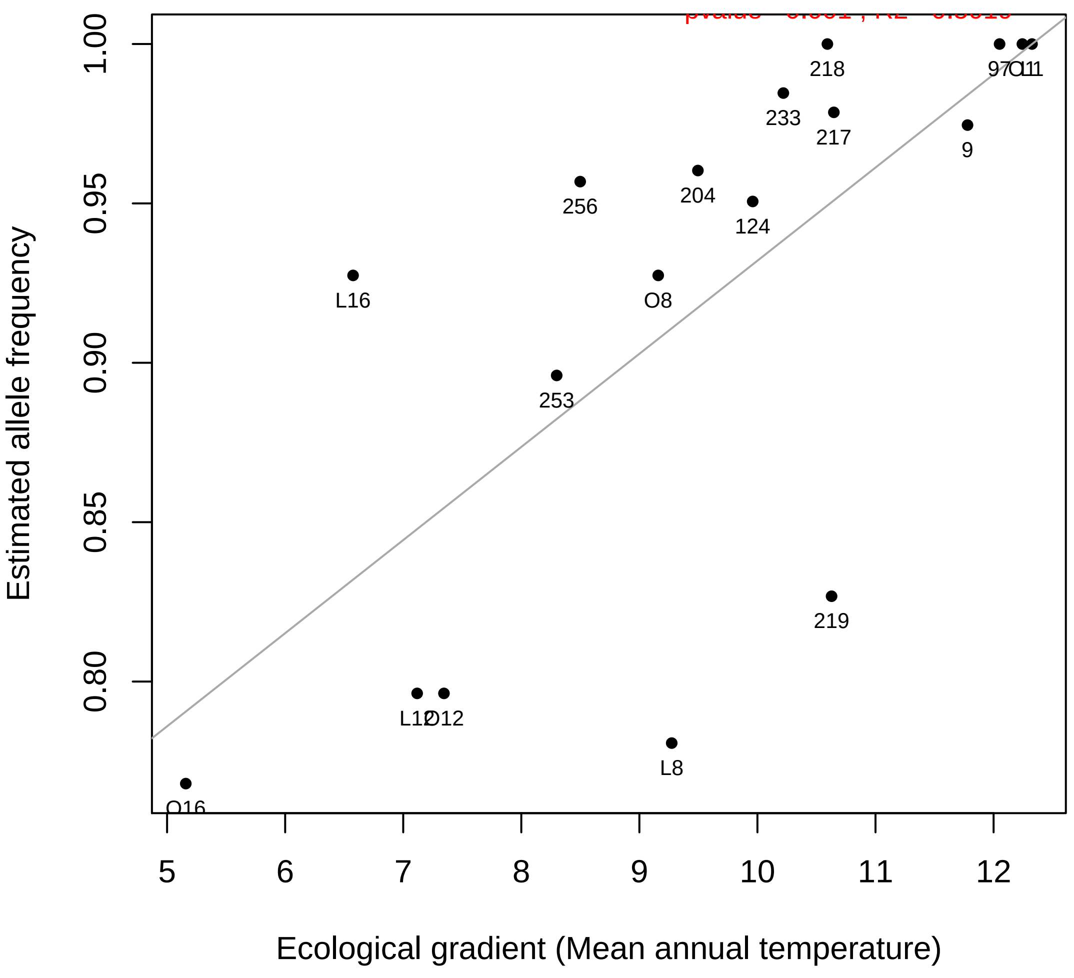

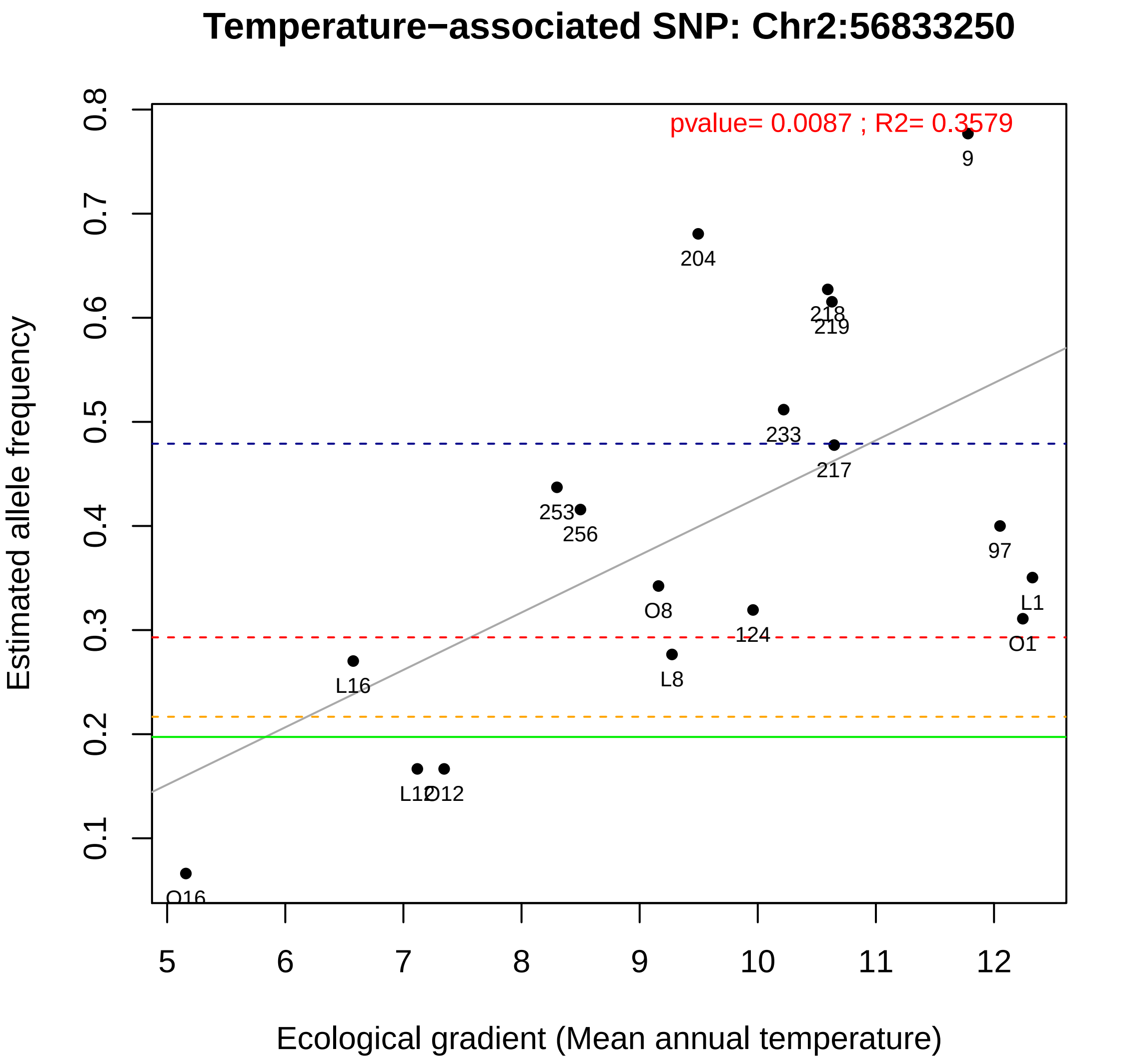
Temperature−associated SNP: Chr2:56833250
pvalue= 0.0087 ; R2= 0.3579
Estimated allele frequency
0.1
0.2
0.3
0.4
0.5
0.6
0.7
0.8
Ecological gradient (Mean annual temperature)
5
6
7
8
9
10
11
12
9
204
218
219
233
217
253
256
97
L1
O8
124
O1
L16
L8
L12
O12
O16

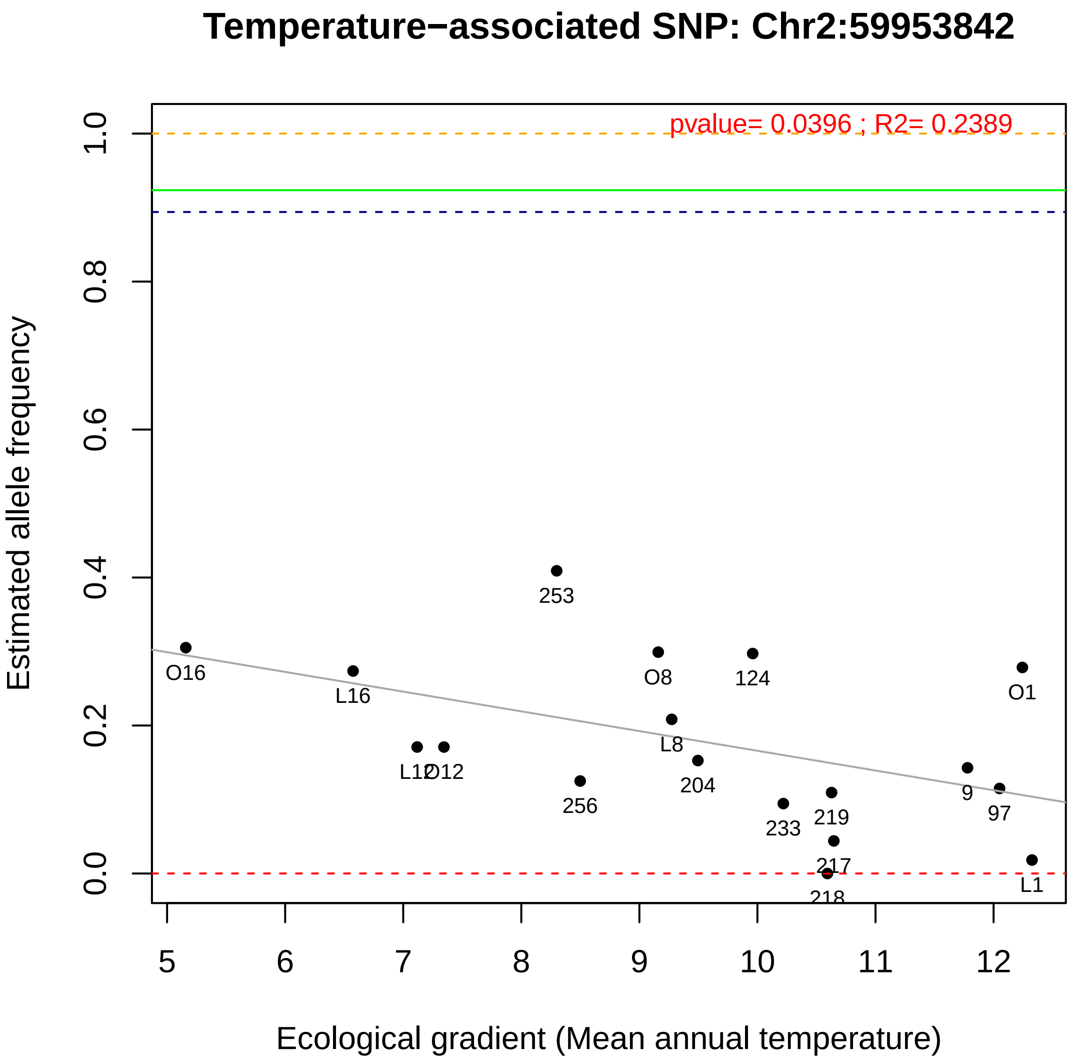

Temperature−associated SNP: Chr2:59953842
pvalue= 0.0396 ; R2= 0.2389
Estimated allele frequency
0.0
0.2
0.4
0.6
0.8
1.0
O16
L16
L12
O12
253
256
O8
L8
204
124
233
219
217
218
9
97
O1
L1
5
6
7
8
9
10
11
12
Ecological gradient (Mean annual temperature)

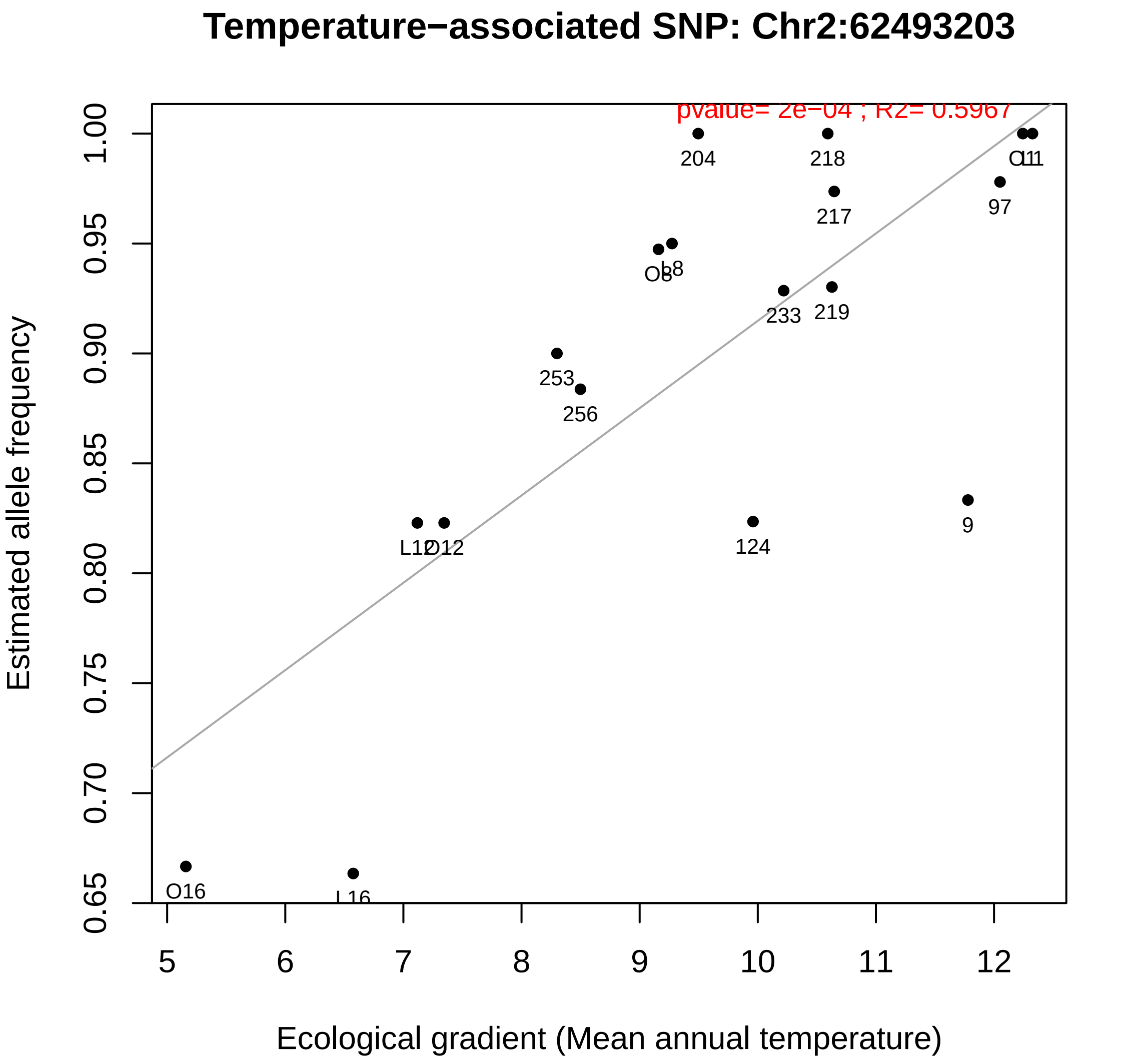

Temperature−associated SNP: Chr2:62493203
pvalue= 2e−04 ; R2= 0.5967
Estimated allele frequency
Ecological gradient (Mean annual temperature)
204
218
217
97
O8
L8
233
219
253
256
L12
O12
124
9
O16
L16
0.65
0.70
0.75
0.80
0.85
0.90
0.95
1.00
5
6
7
8
9
10
11
12

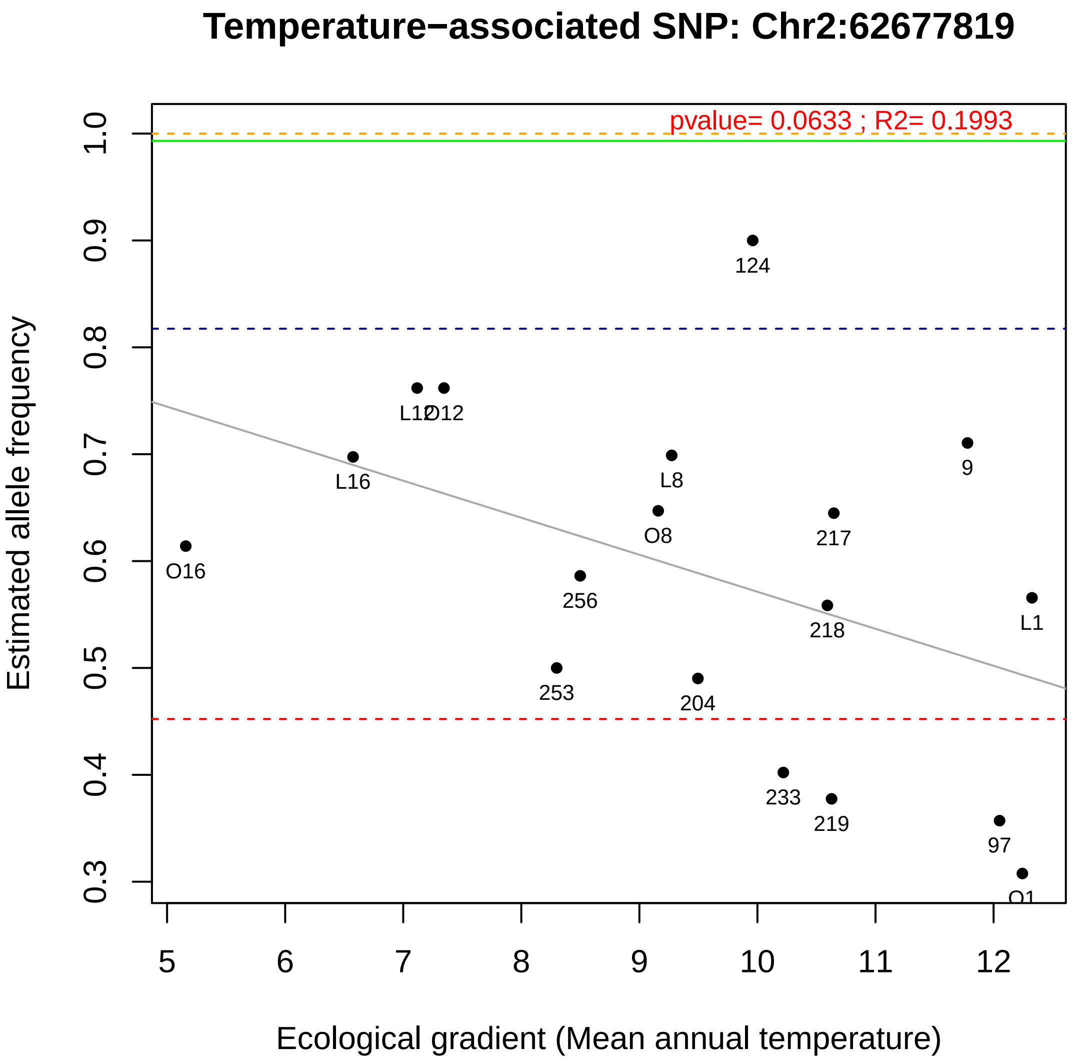
Temperature−associated SNP: Chr2:62677819
pvalue= 0.0633 ; R2= 0.1993
Estimated allele frequency
Ecological gradient (Mean annual temperature)
0.3
0.4
0.5
0.6
0.7
0.8
0.9
1.0
5
6
7
8
9
10
11
12
124
L12
O12
L16
L8
9
O8
217
O16
256
218
L1
253
204
233
219
97
O1

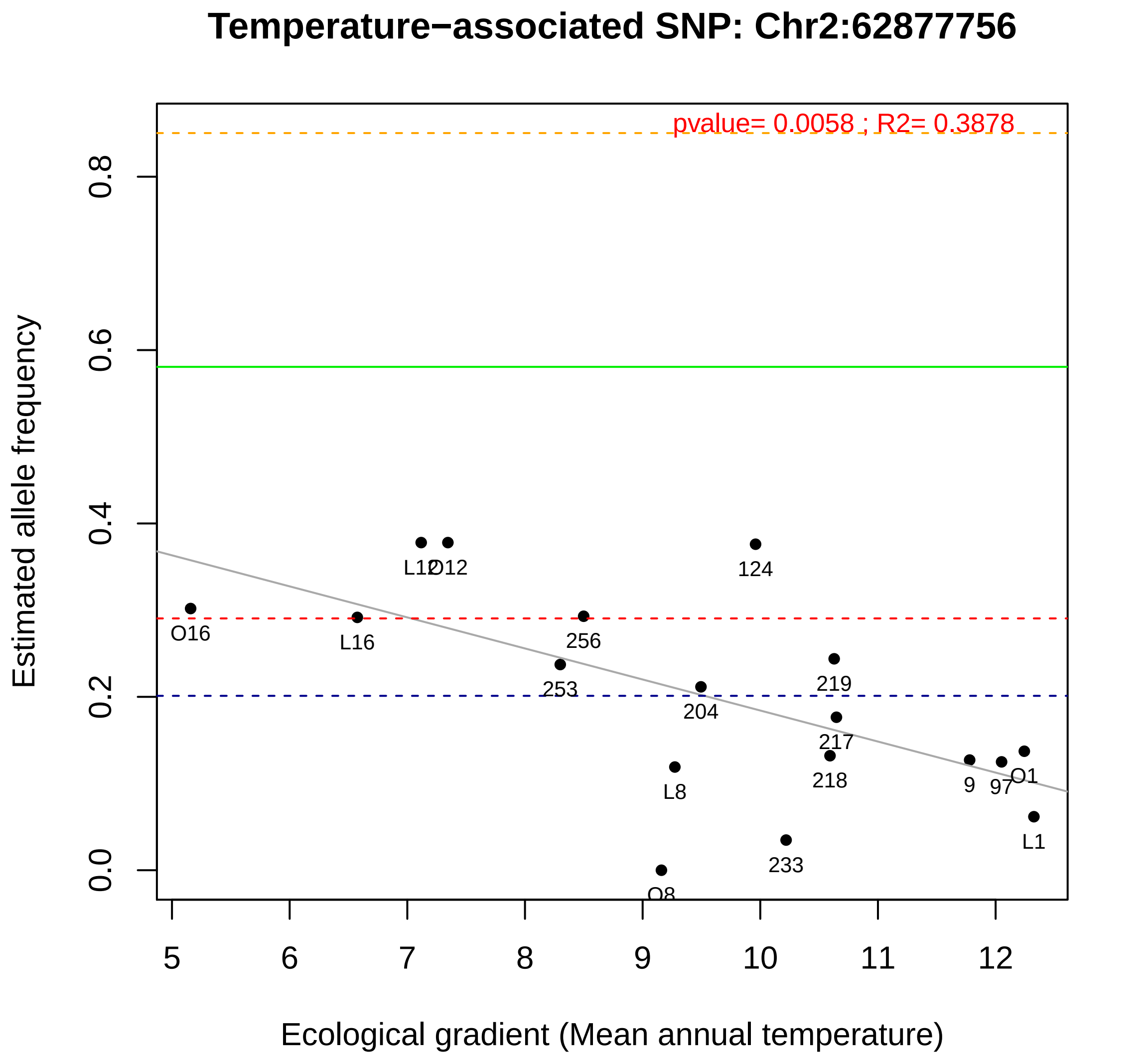

Temperature−associated SNP: Chr2:62877756
pvalue= 0.0058 ; R2= 0.3878
Estimated allele frequency
0.0
0.2
0.4
0.6
0.8
5
6
7
8
9
10
11
12
Ecological gradient (Mean annual temperature)
O16
L16
L12
O12
256
253
124
204
219
217
218
L8
O8
233
9
97
O1
L1

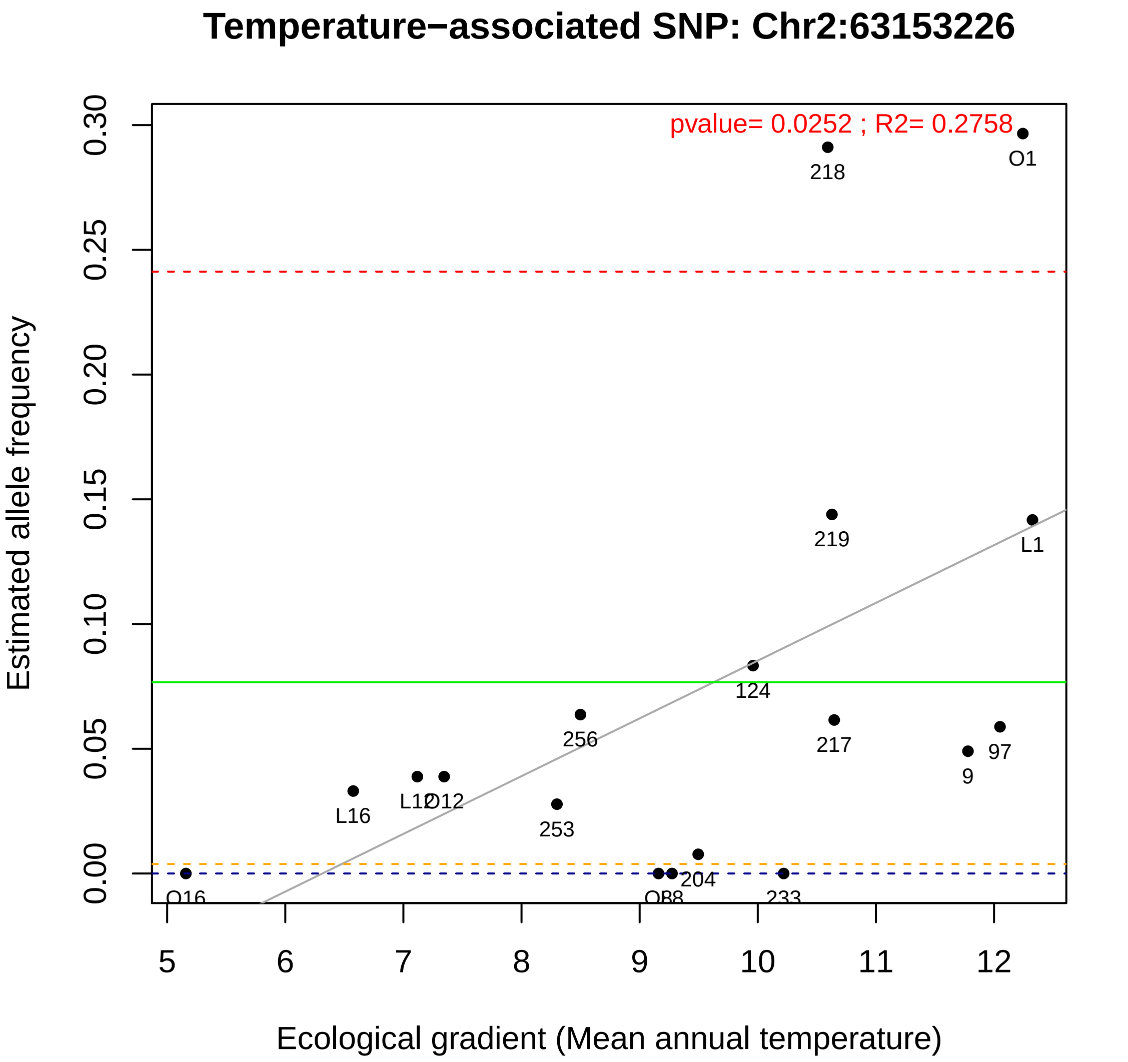

Temperature−associated SNP: Chr2:63153226
pvalue= 0.0252 ; R2= 0.2758
Estimated allele frequency
0.00
0.05
0.10
0.15
0.20
0.25
0.30
Ecological gradient (Mean annual temperature)
5
6
7
8
9
10
11
12
218
O1
219
L1
124
256
217
97
9
L16
L12
O12
253
204
O16
O8
233

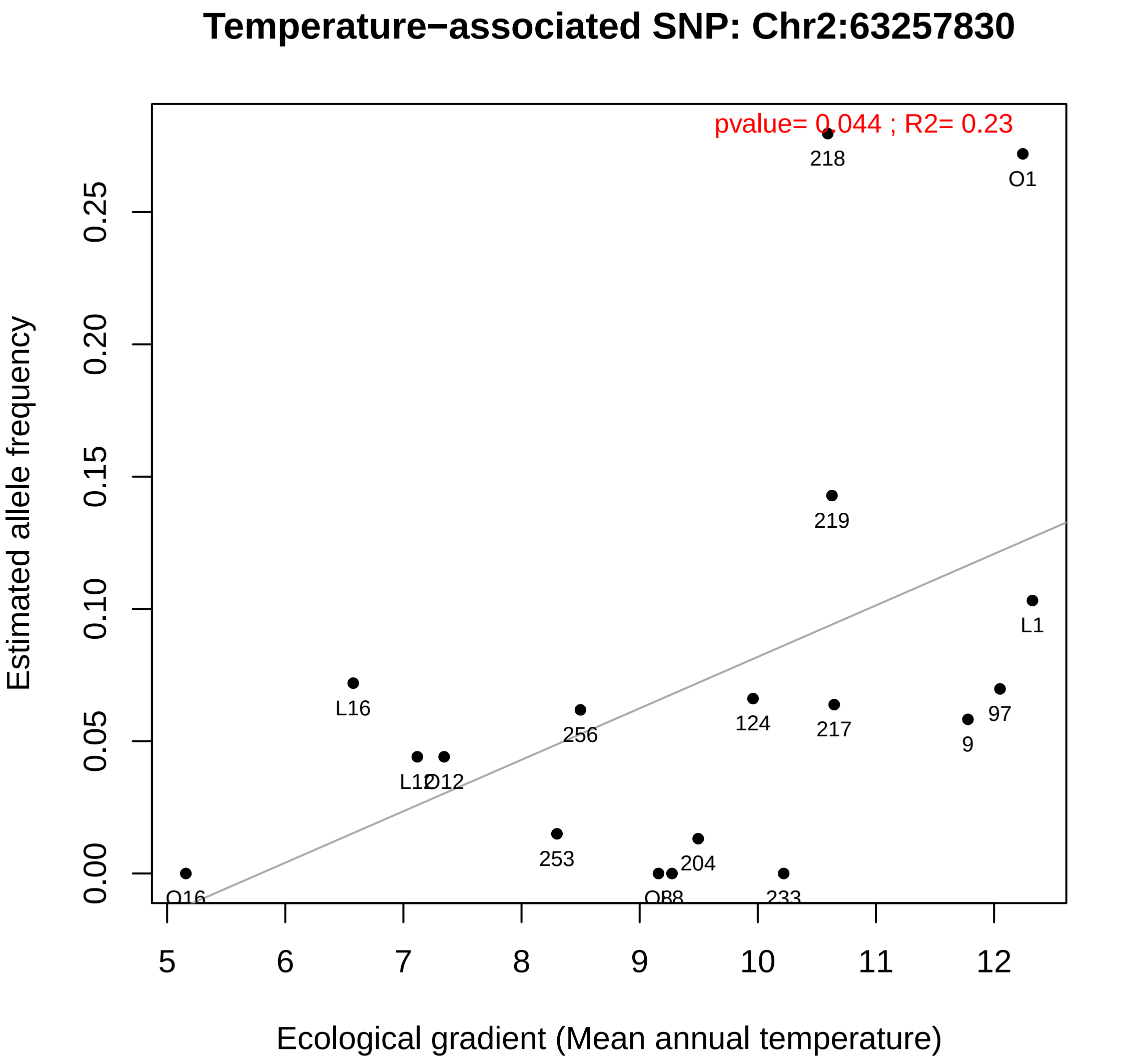

Temperature−associated SNP: Chr2:63257830
pvalue= 0.044 ; R2= 0.23
218
O1
219
L1
L16
256
124
217
97
9
L12
O12
253
204
O16
O8
8
233
0.00
0.05
0.10
0.15
0.20
0.25
5
6
7
8
9
10
11
12
Estimated allele frequency
Ecological gradient (Mean annual temperature)

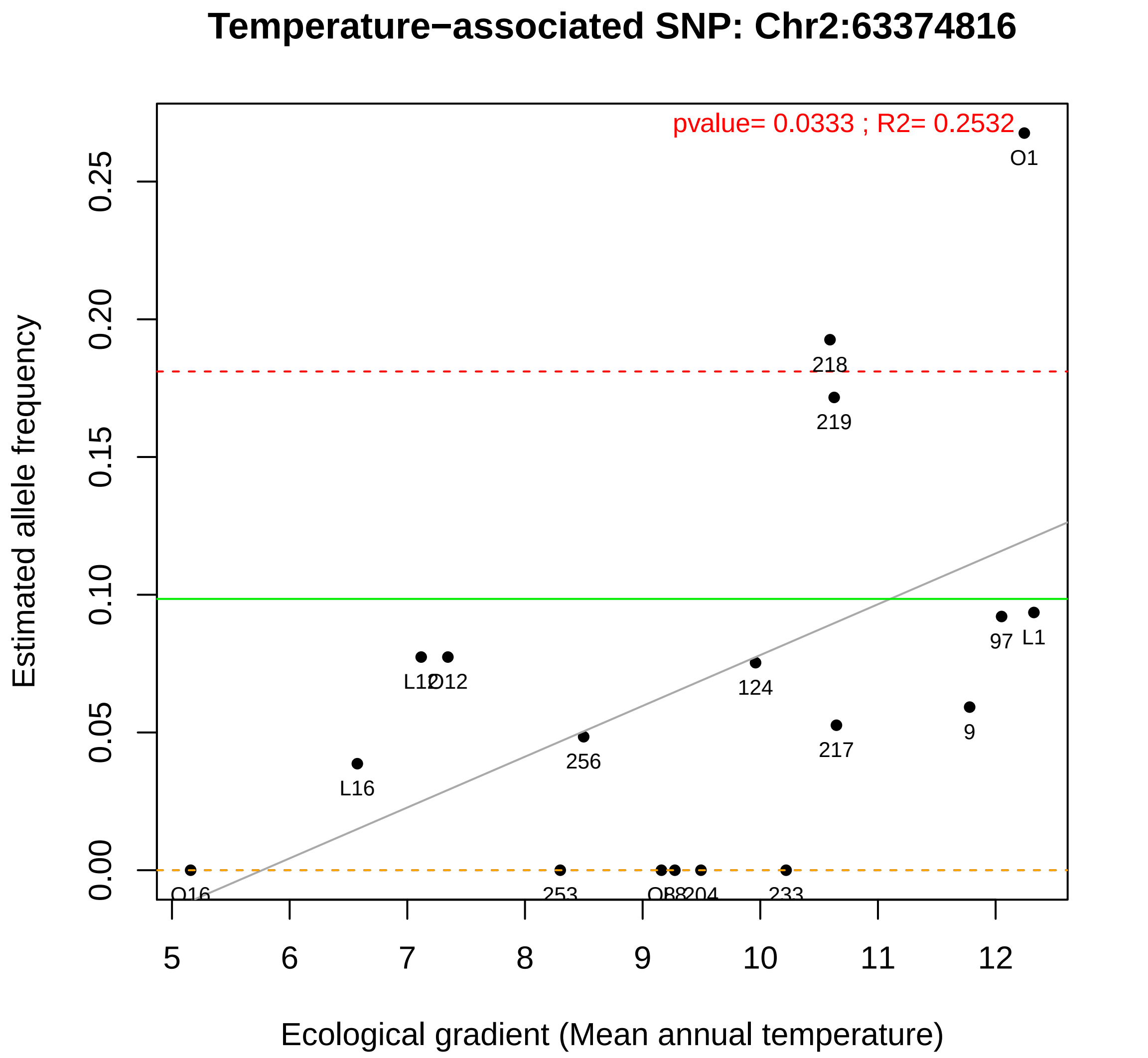
Temperature−associated SNP: Chr2:63374816
pvalue= 0.0333 ; R2= 0.2532
Estimated allele frequency
Ecological gradient (Mean annual temperature)
0.00
0.05
0.10
0.15
0.20
0.25
5
6
7
8
9
10
11
12
O1
218
219
97
L1
L12
O12
124
9
217
256
L16
O16
253
204
233

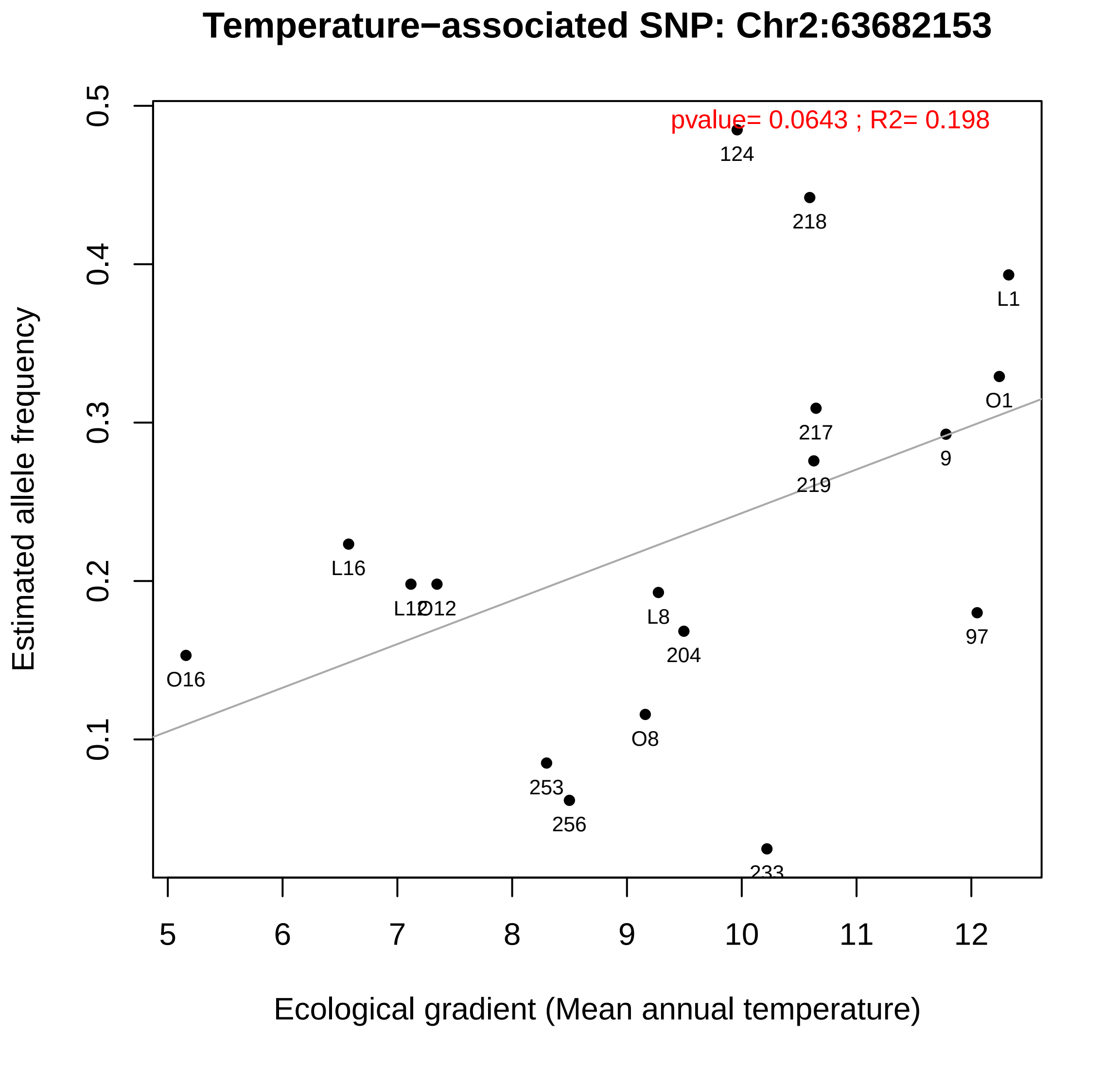

Temperature−associated SNP: Chr2:63682153
pvalue= 0.0643 ; R2= 0.198
Estimated allele frequency
0.1
0.2
0.3
0.4
0.5
5
6
7
8
9
10
11
12
Ecological gradient (Mean annual temperature)
124
218
L1
O1
217
9
219
L16
L12
O12
L8
204
97
O16
O8
253
256
233

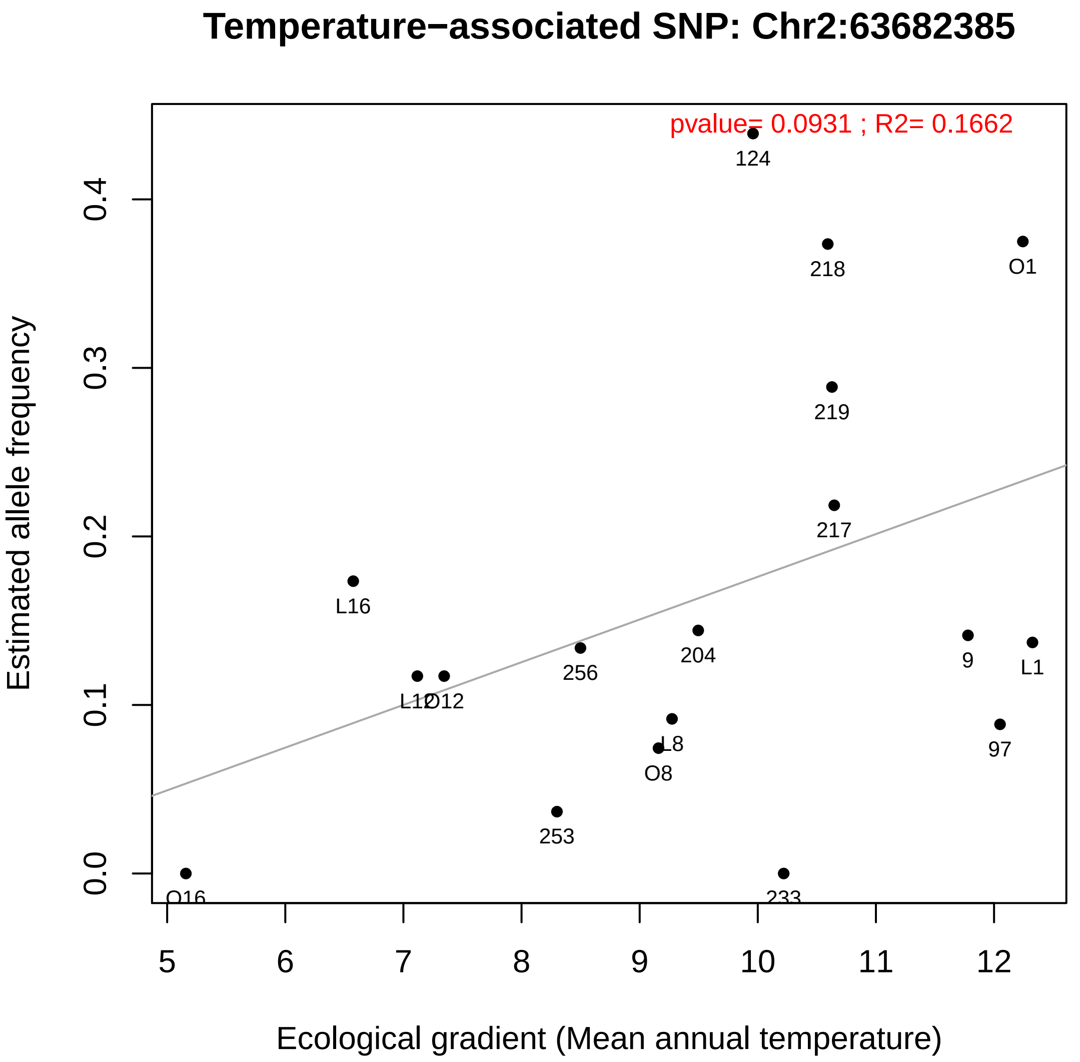
Temperature−associated SNP: Chr2:63682385
pvalue= 0.0931 ; R2= 0.1662
Estimated allele frequency
Ecological gradient (Mean annual temperature)
0.0
0.1
0.2
0.3
0.4
5
6
7
8
9
10
11
12
124
218
O1
219
217
L16
204
9
L1
256
L12
O12
L8
O8
97
253
O16
233

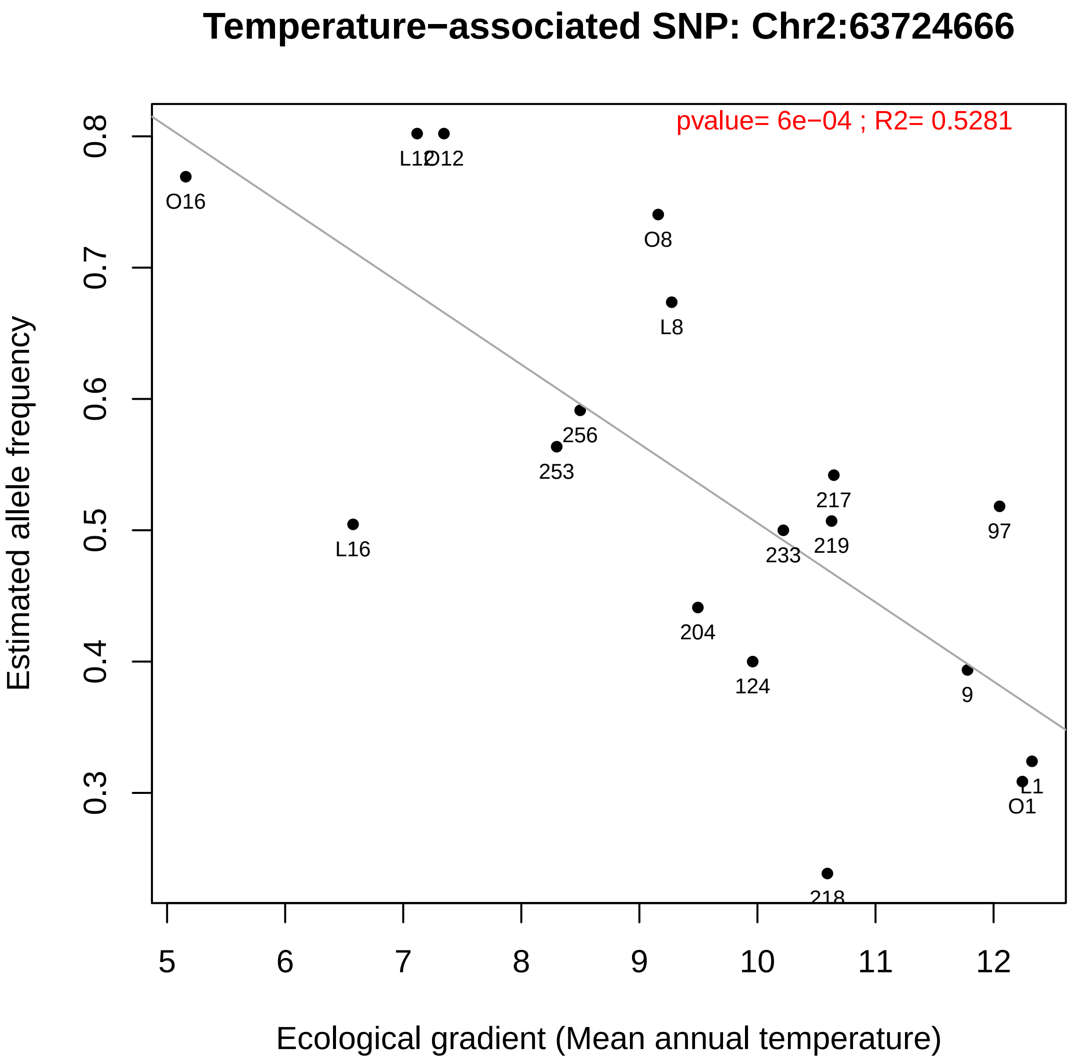

Temperature−associated SNP: Chr2:63724666
pvalue= 6e−04 ; R2= 0.5281
Estimated allele frequency
Ecological gradient (Mean annual temperature)
0.3
0.4
0.5
0.6
0.7
0.8
5
6
7
8
9
10
11
12
O16
L12
O12
O8
L8
256
253
L16
217
219
233
97
204
124
9
L1
O1
218

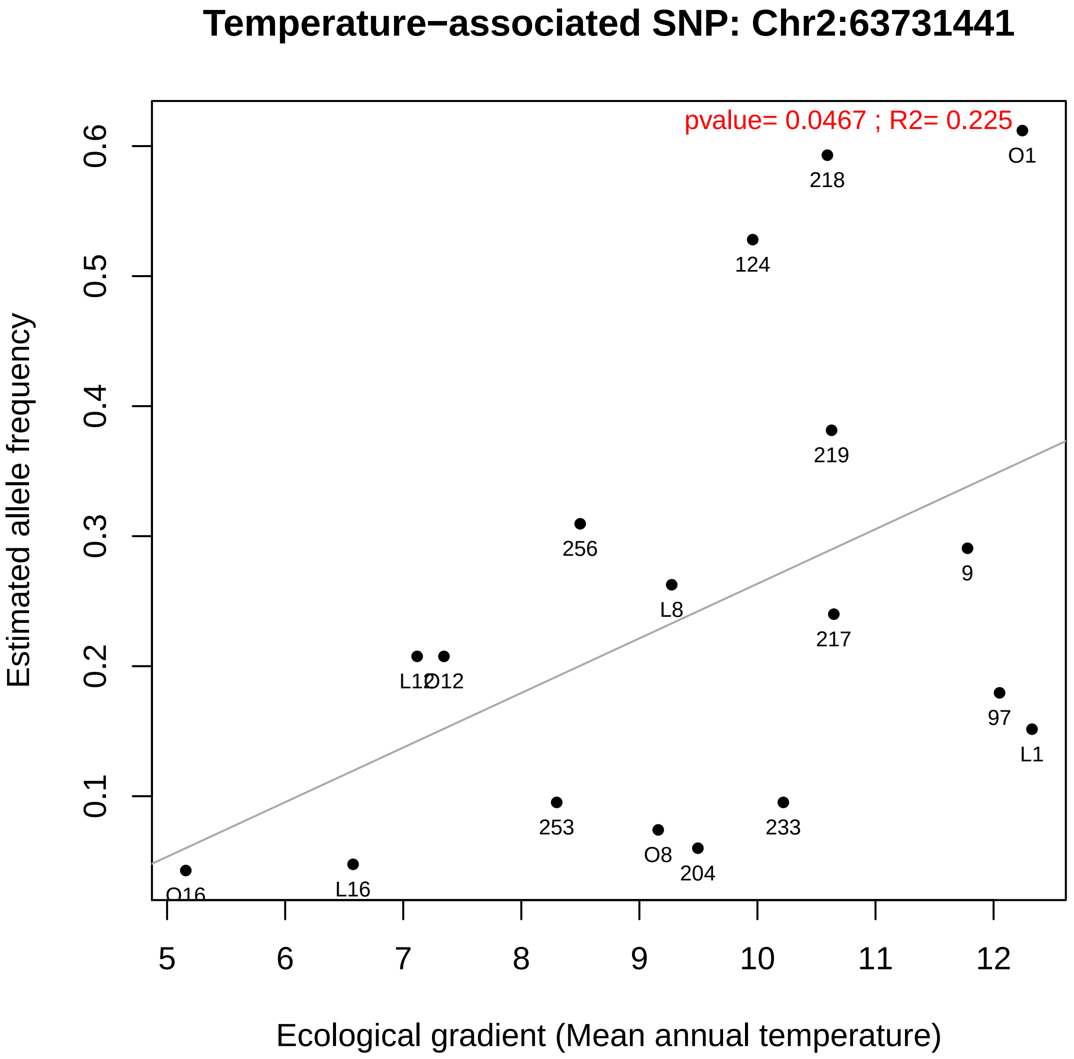

Temperature−associated SNP: Chr2:63731441
pvalue= 0.0467 ; R2= 0.225
Estimated allele frequency
0.1
0.2
0.3
0.4
0.5
0.6
Ecological gradient (Mean annual temperature)
5
6
7
8
9
10
11
12
O1
218
124
219
256
9
L8
217
L12
O12
97
L1
253
233
O8
204
O16
L16

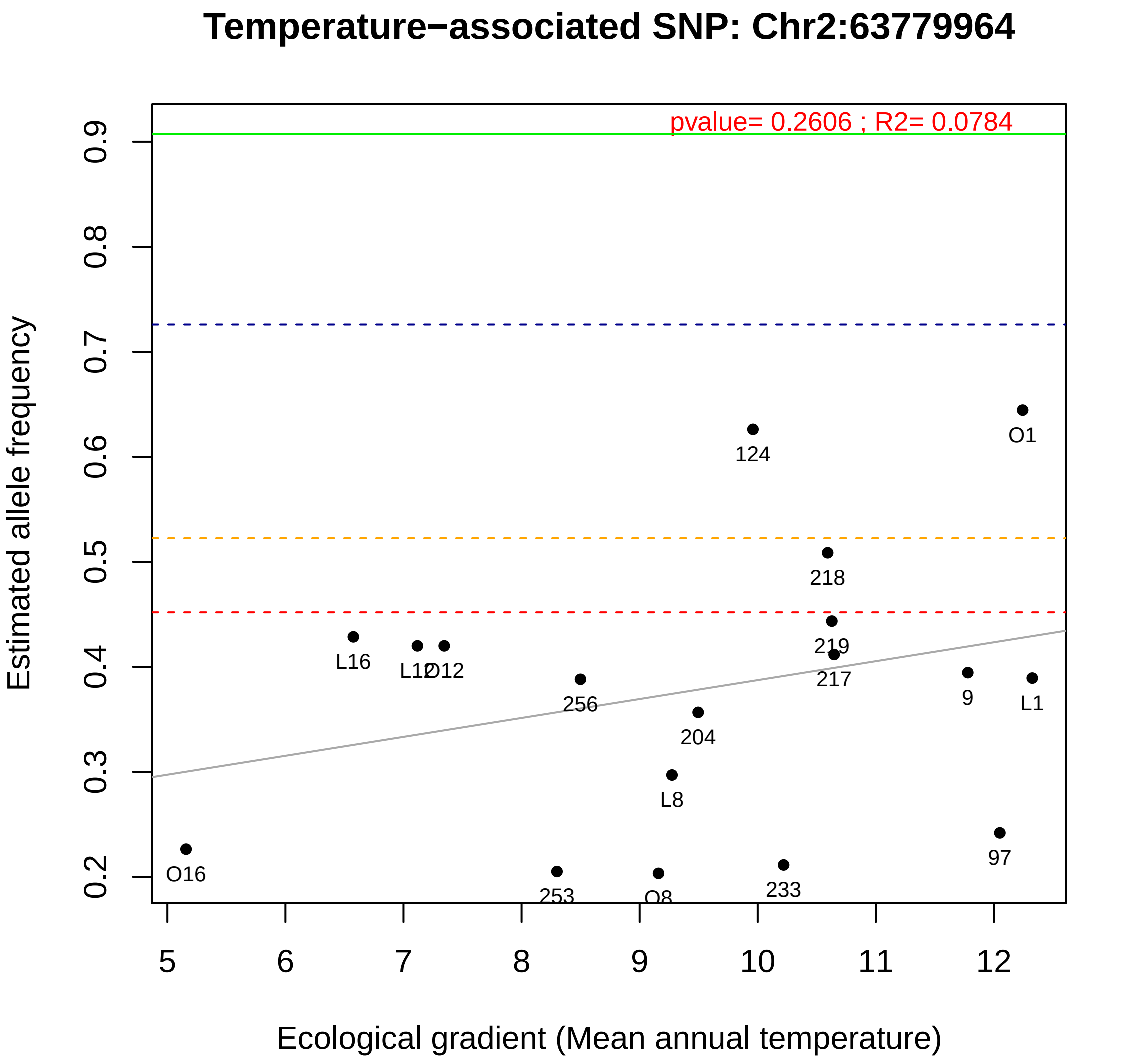

Temperature−associated SNP: Chr2:63779964
pvalue= 0.2606 ; R2= 0.0784
Estimated allele frequency
0.2
0.3
0.4
0.5
0.6
0.7
0.8
0.9
5
6
7
8
9
10
11
12
Ecological gradient (Mean annual temperature)
O16
L16
L12
O12
253
256
O8
L8
204
124
233
218
219
217
9
97
O1
L1

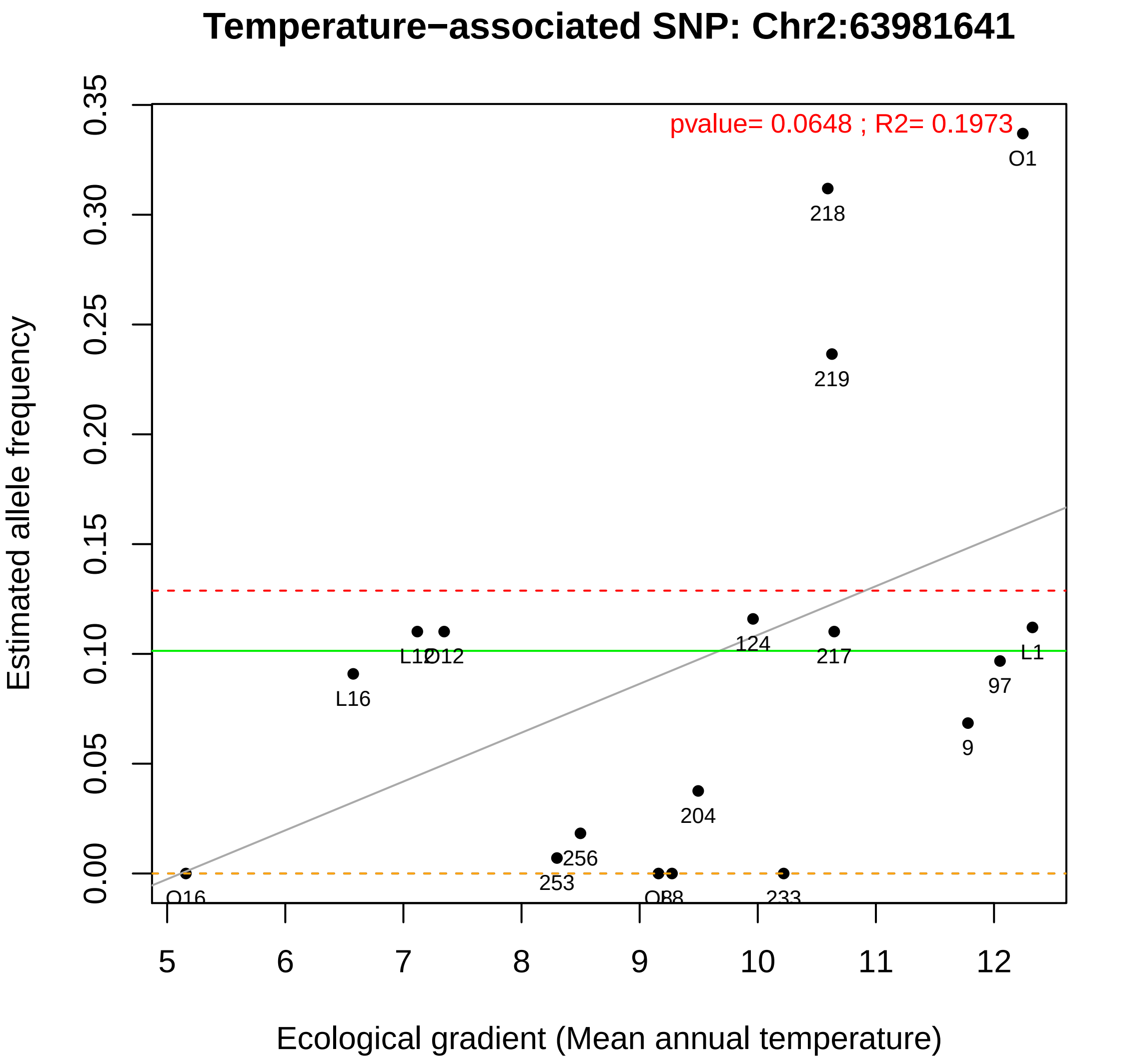

Temperature−associated SNP: Chr2:63981641
pvalue= 0.0648 ; R2= 0.1973
Estimated allele frequency
Ecological gradient (Mean annual temperature)
0.00
0.05
0.10
0.15
0.20
0.25
0.30
0.35
5
6
7
8
9
10
11
12
O1
218
219
124
217
L1
97
9
L16
L12
O12
204
256
253
233
O16

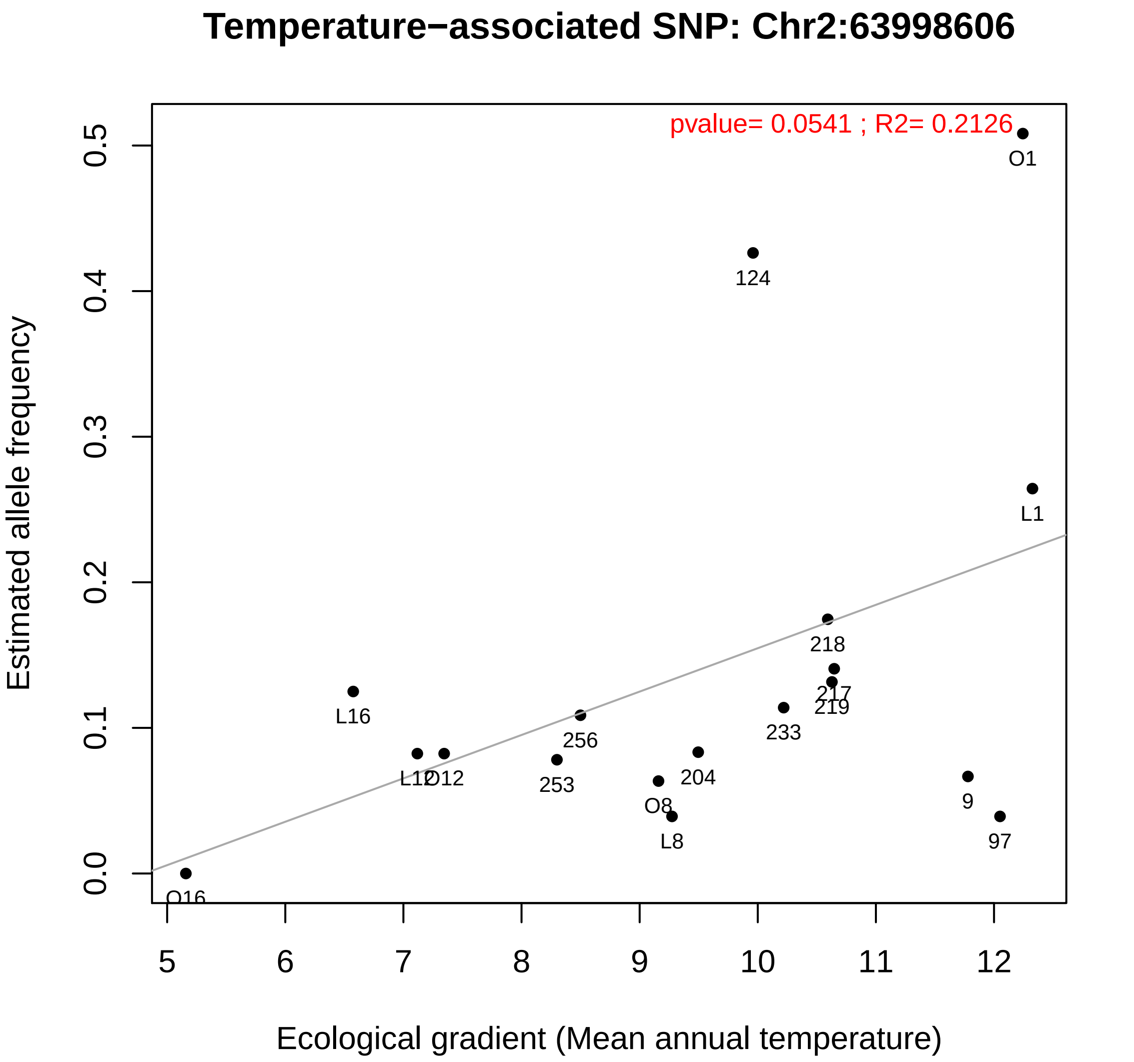

Temperature−associated SNP: Chr2:63998606
pvalue= 0.0541 ; R2= 0.2126
Estimated allele frequency
0.0
0.1
0.2
0.3
0.4
0.5
O1
124
L1
218
217
219
233
L16
256
L12
O12
253
204
O8
L8
9
97
O16
5
6
7
8
9
10
11
12
Ecological gradient (Mean annual temperature)

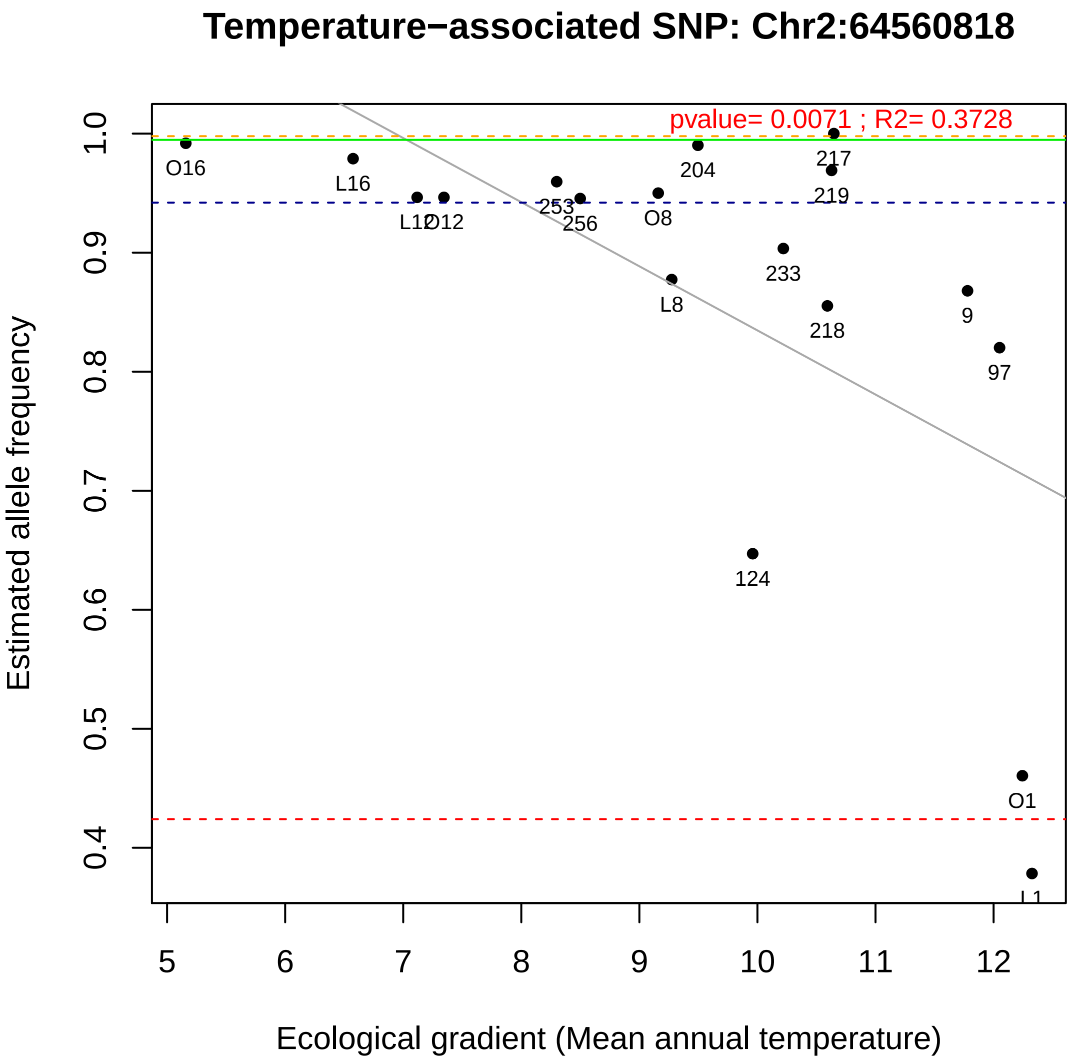
Temperature−associated SNP: Chr2:64560818
pvalue= 0.0071 ; R2= 0.3728
O16
L16
L12
O12
253
256
O8
204
217
219
233
L8
218
9
97
124
O1
L1
1.0
0.9
0.8
0.7
0.6
0.5
0.4
5
6
7
8
9
10
11
12
Estimated allele frequency
Ecological gradient (Mean annual temperature)

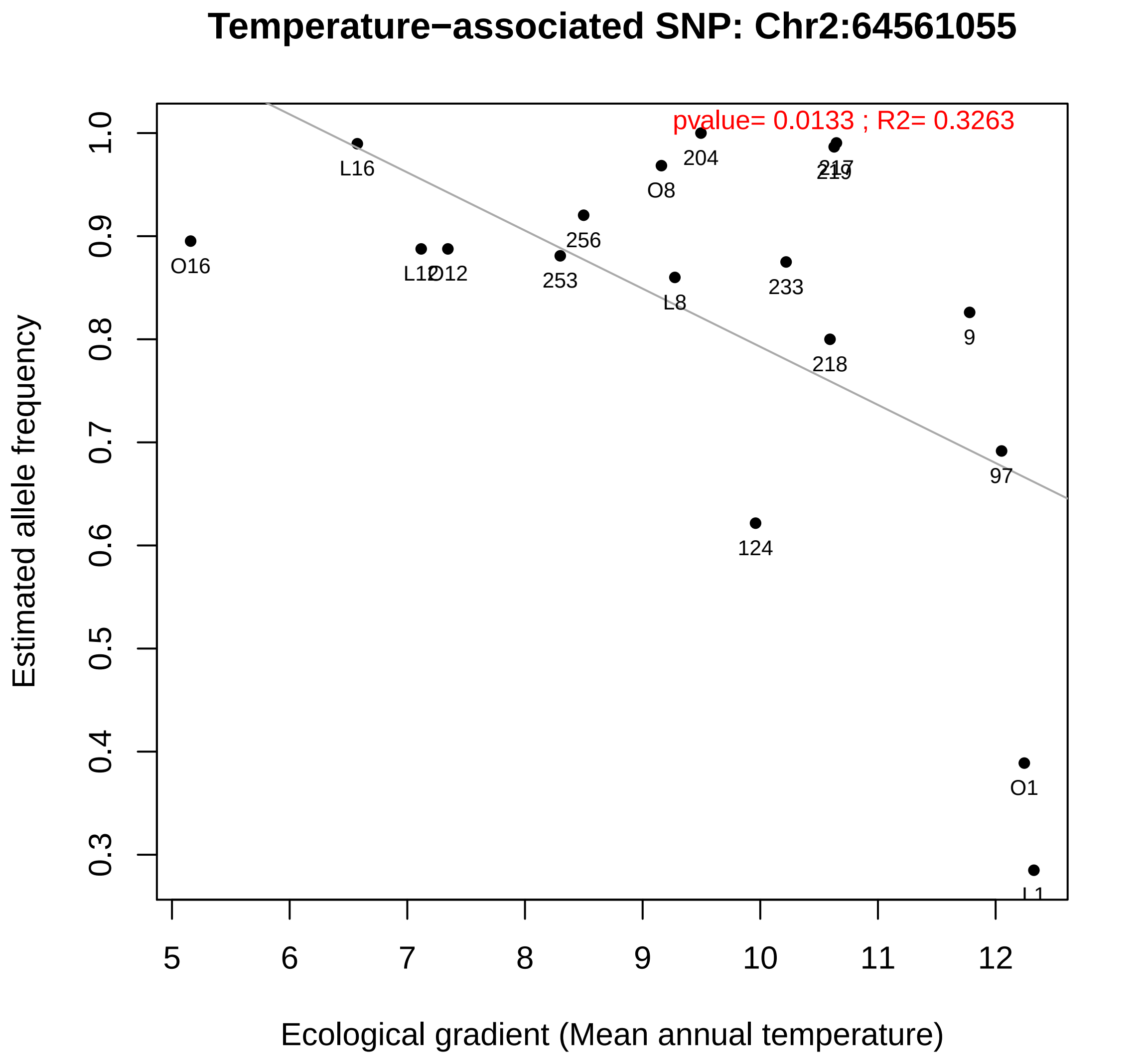

Temperature−associated SNP: Chr2:64561055
pvalue= 0.0133 ; R2= 0.3263
Estimated allele frequency
Ecological gradient (Mean annual temperature)
0.3
0.4
0.5
0.6
0.7
0.8
0.9
1.0
5
6
7
8
9
10
11
12
L16
204
O8
217
219
256
O16
L12
O12
253
L8
233
9
218
97
124
O1
L1

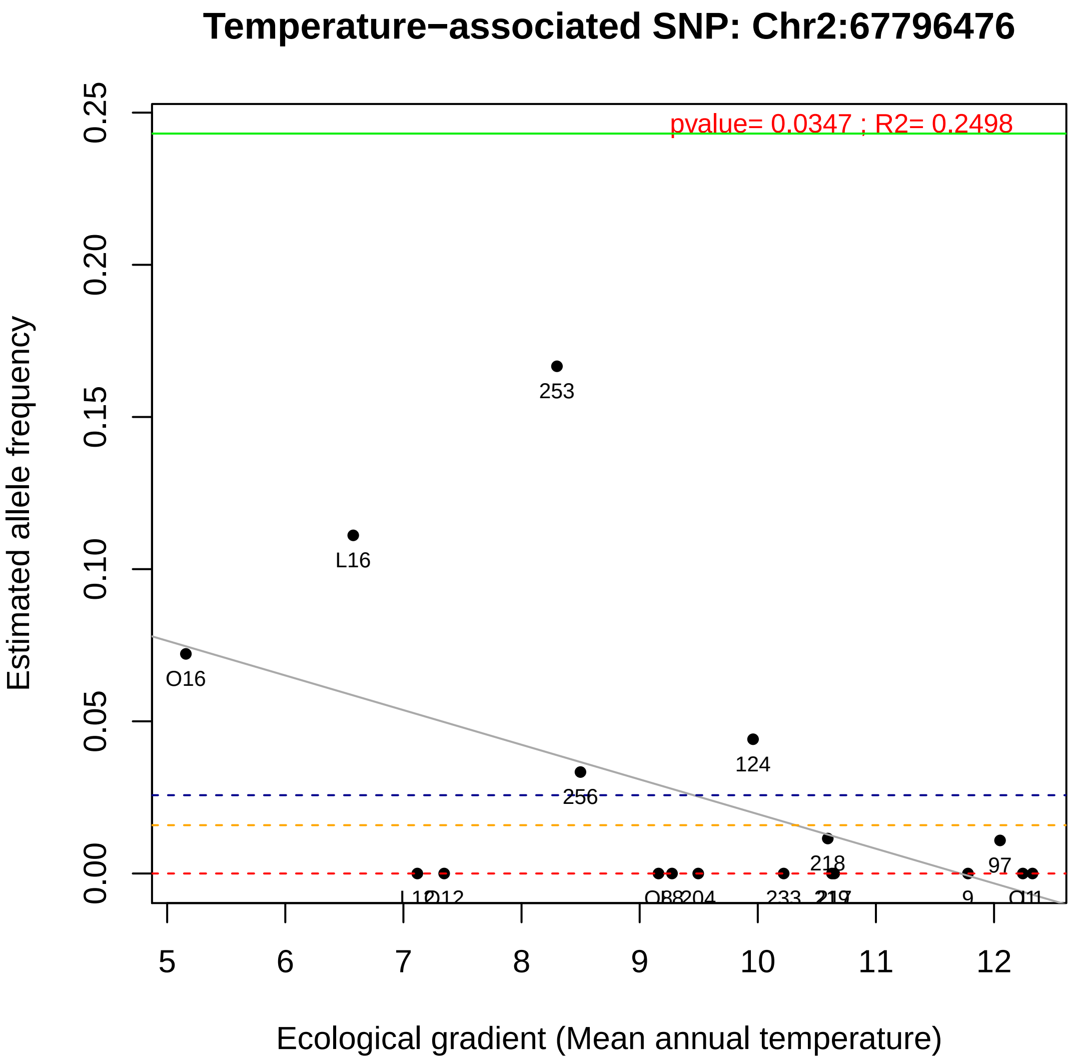
Temperature−associated SNP: Chr2:67796476
pvalue= 0.0347 ; R2= 0.2498
Estimated allele frequency
0.00
0.05
0.10
0.15
0.20
0.25
5
6
7
8
9
10
11
12
Ecological gradient (Mean annual temperature)
253
L16
O16
256
124
218
97
O12
204
233
9

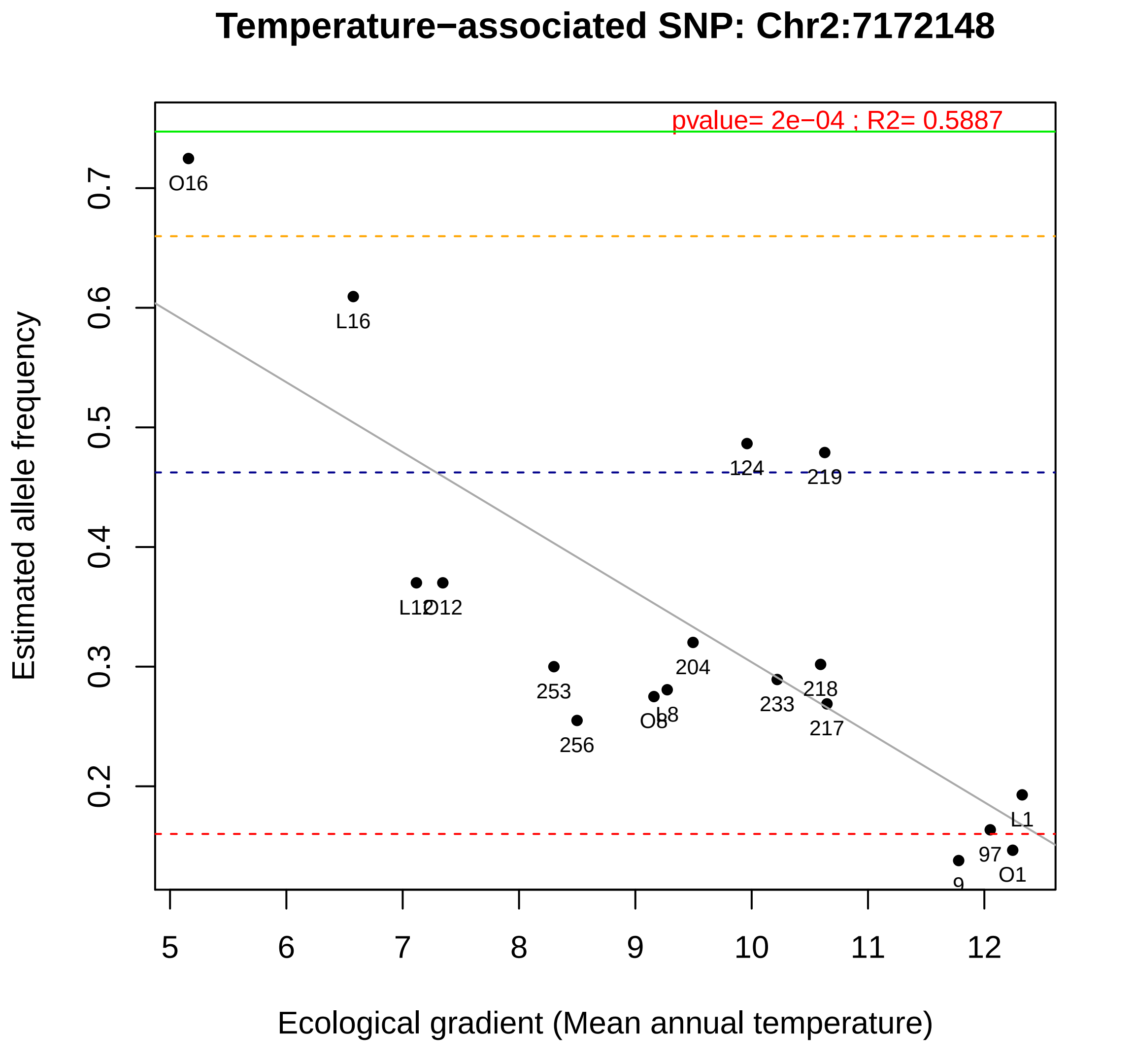
Temperature−associated SNP: Chr2:7172148
pvalue= 2e−04 ; R2= 0.5887
Estimated allele frequency
0.2
0.3
0.4
0.5
0.6
0.7
Ecological gradient (Mean annual temperature)
5
6
7
8
9
10
11
12
O16
L16
124
219
L12
O12
204
253
218
233
L8
O8
217
256
L1
97
O1
9

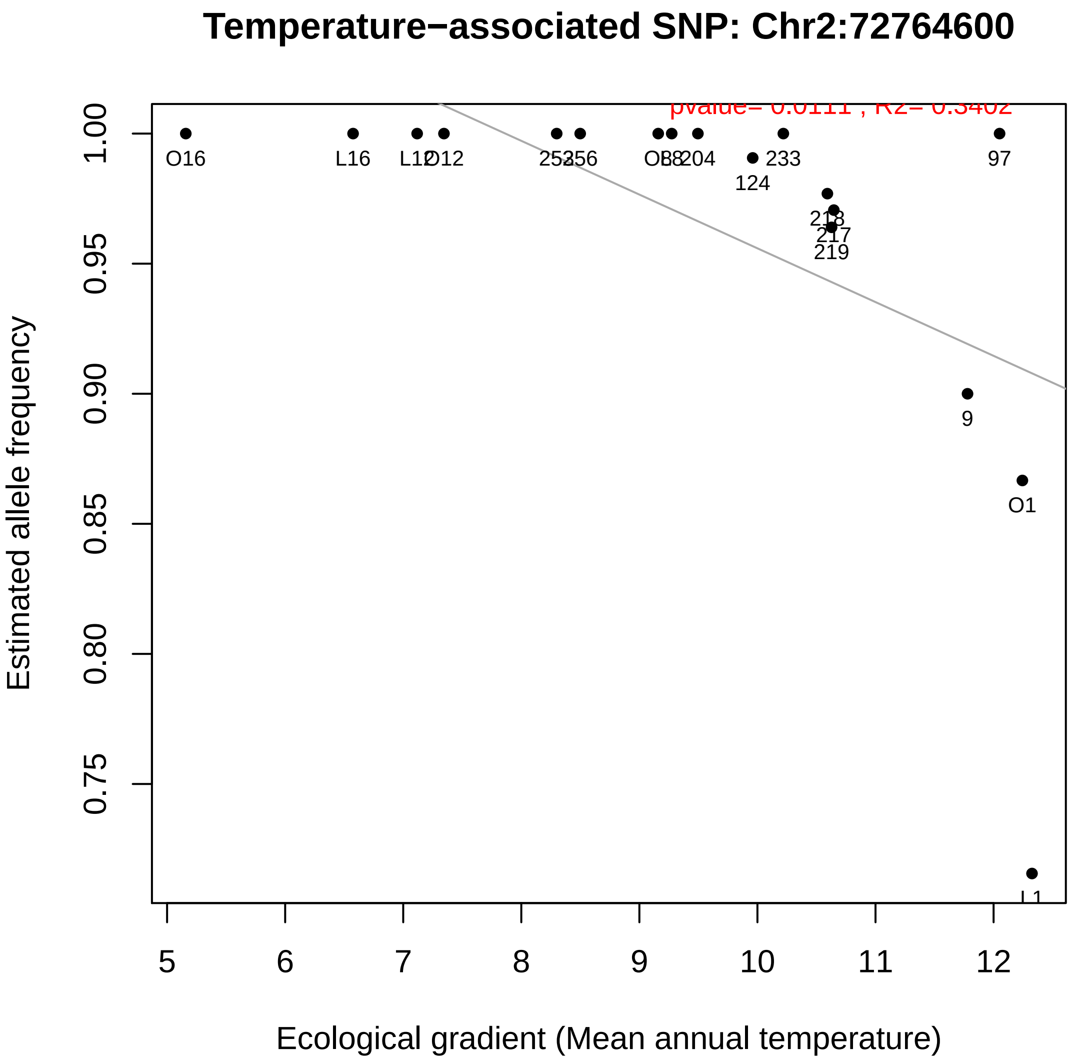
Temperature−associated SNP: Chr2:72764600
pvalue= 0.0111 , R2= 0.3402
Estimated allele frequency
Ecological gradient (Mean annual temperature)
O16
L16
L12
O12
253
256
O8
L8
204
124
233
218
217
219
97
9
O1
L1
1.00
0.95
0.90
0.85
0.80
0.75
5
6
7
8
9
10
11
12

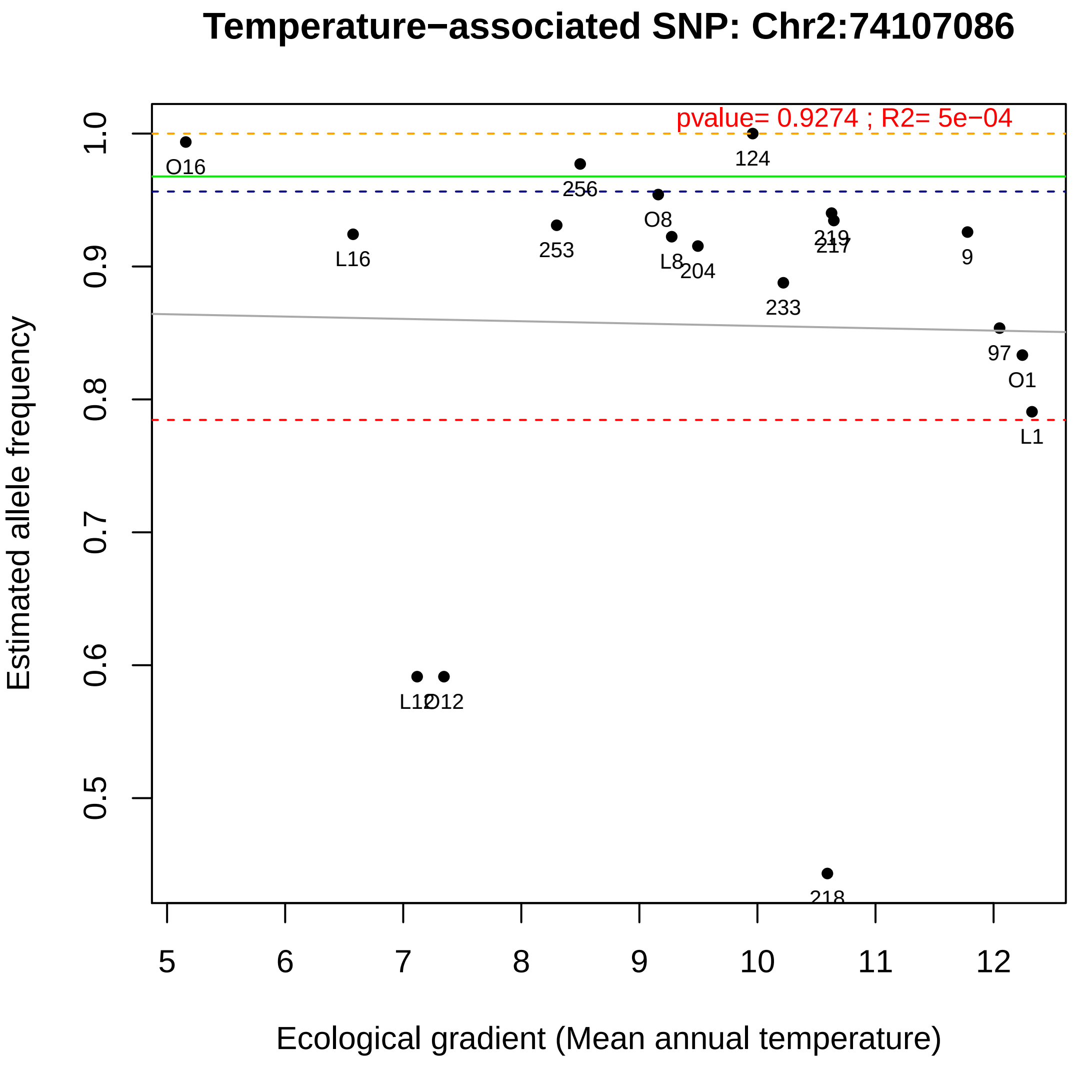

Temperature−associated SNP: Chr2:74107086
pvalue= 0.9274 ; R2= 5e−04
Estimated allele frequency
1.0
0.9
0.8
0.7
0.6
0.5
O16
256
124
O8
L16
253
L8
204
219
217
9
233
97
O1
L1
L12
O12
218
5
6
7
8
9
10
11
12
Ecological gradient (Mean annual temperature)

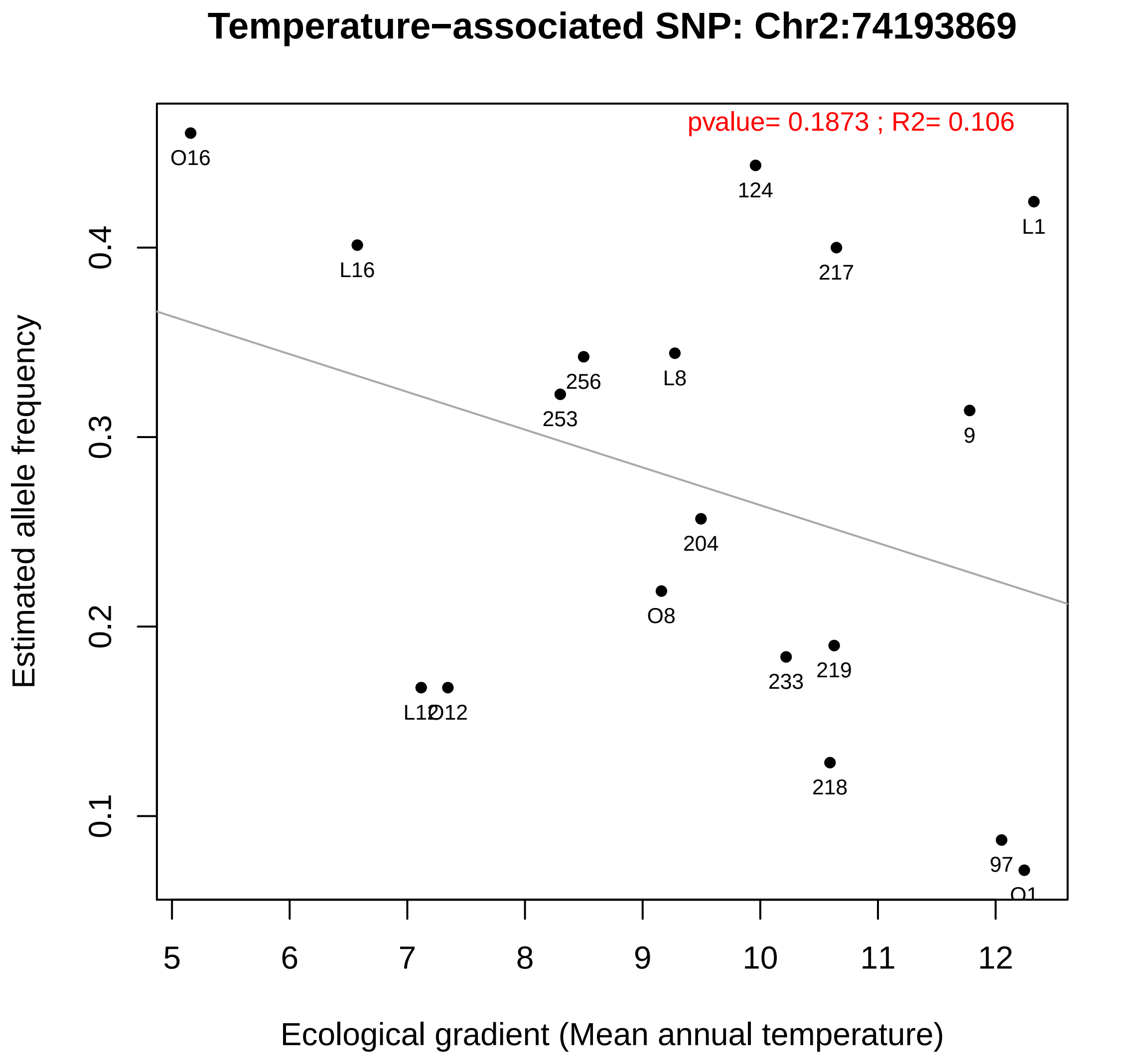

Temperature−associated SNP: Chr2:74193869
pvalue= 0.1873 ; R2= 0.106
Estimated allele frequency
0.1
0.2
0.3
0.4
5
6
7
8
9
10
11
12
Ecological gradient (Mean annual temperature)
O16
L16
124
L1
217
256
L8
253
9
204
O8
233
219
L12
O12
218
97

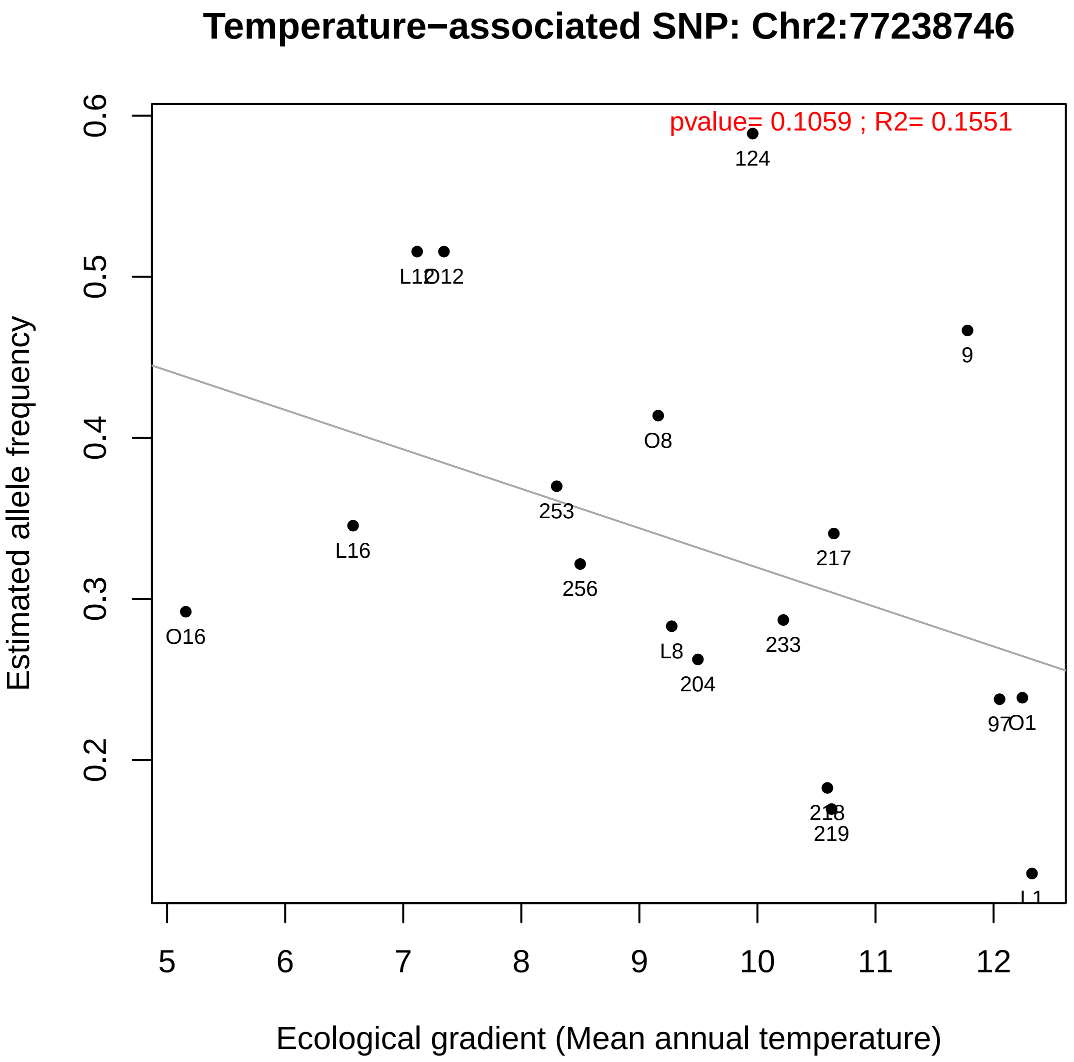
Temperature−associated SNP: Chr2:77238746
pvalue= 0.1059 ; R2= 0.1551
Estimated allele frequency
0.6
0.5
0.4
0.3
0.2
124
L12
O12
9
O8
253
L16
217
256
O16
233
L8
204
97
O1
218
219
L1
5
6
7
8
9
10
11
12
Ecological gradient (Mean annual temperature)

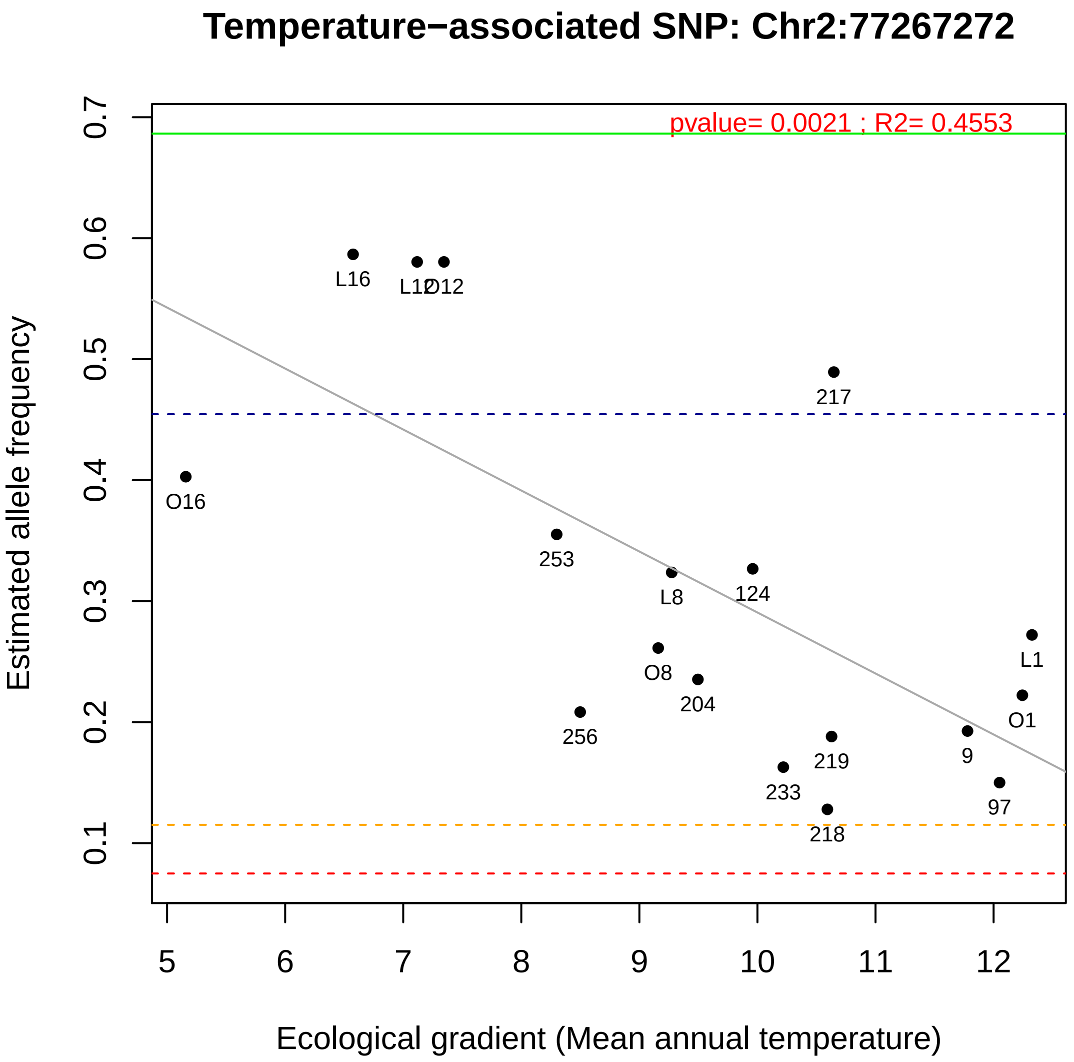
Temperature−associated SNP: Chr2:77267272
pvalue= 0.0021 ; R2= 0.4553
Estimated allele frequency
0.1
0.2
0.3
0.4
0.5
0.6
0.7
L16
L12
O12
217
O16
253
L8
124
O8
204
256
L1
O1
219
9
233
97
218
5
6
7
8
9
10
11
12
Ecological gradient (Mean annual temperature)

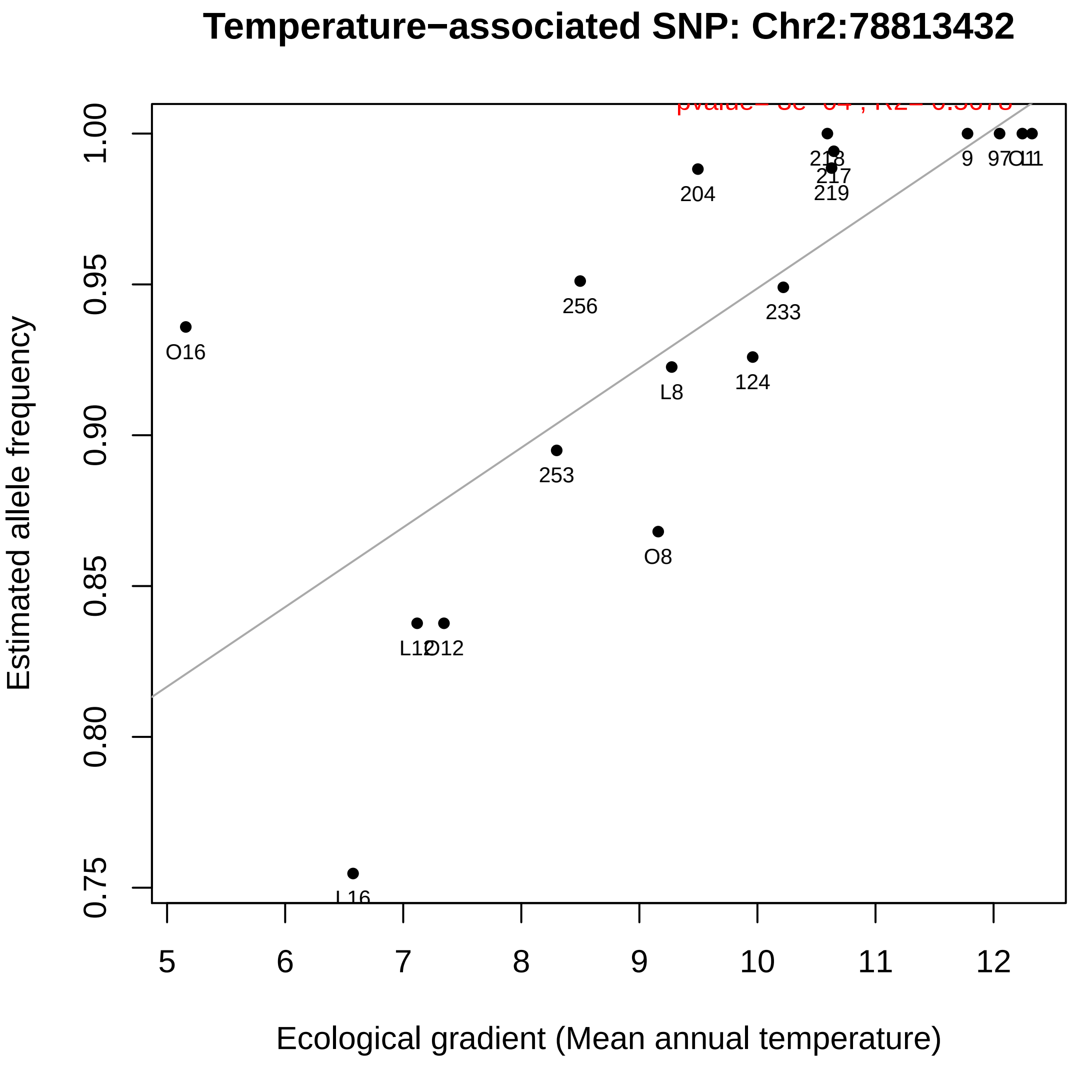
Temperature−associated SNP: Chr2:78813432
Estimated allele frequency
Ecological gradient (Mean annual temperature)
1.00
0.95
0.90
0.85
0.80
0.75
5
6
7
8
9
10
11
12
O16
256
253
204
218
217
219
233
124
L8
O8
L12
O12
L16
9
97
O1
1

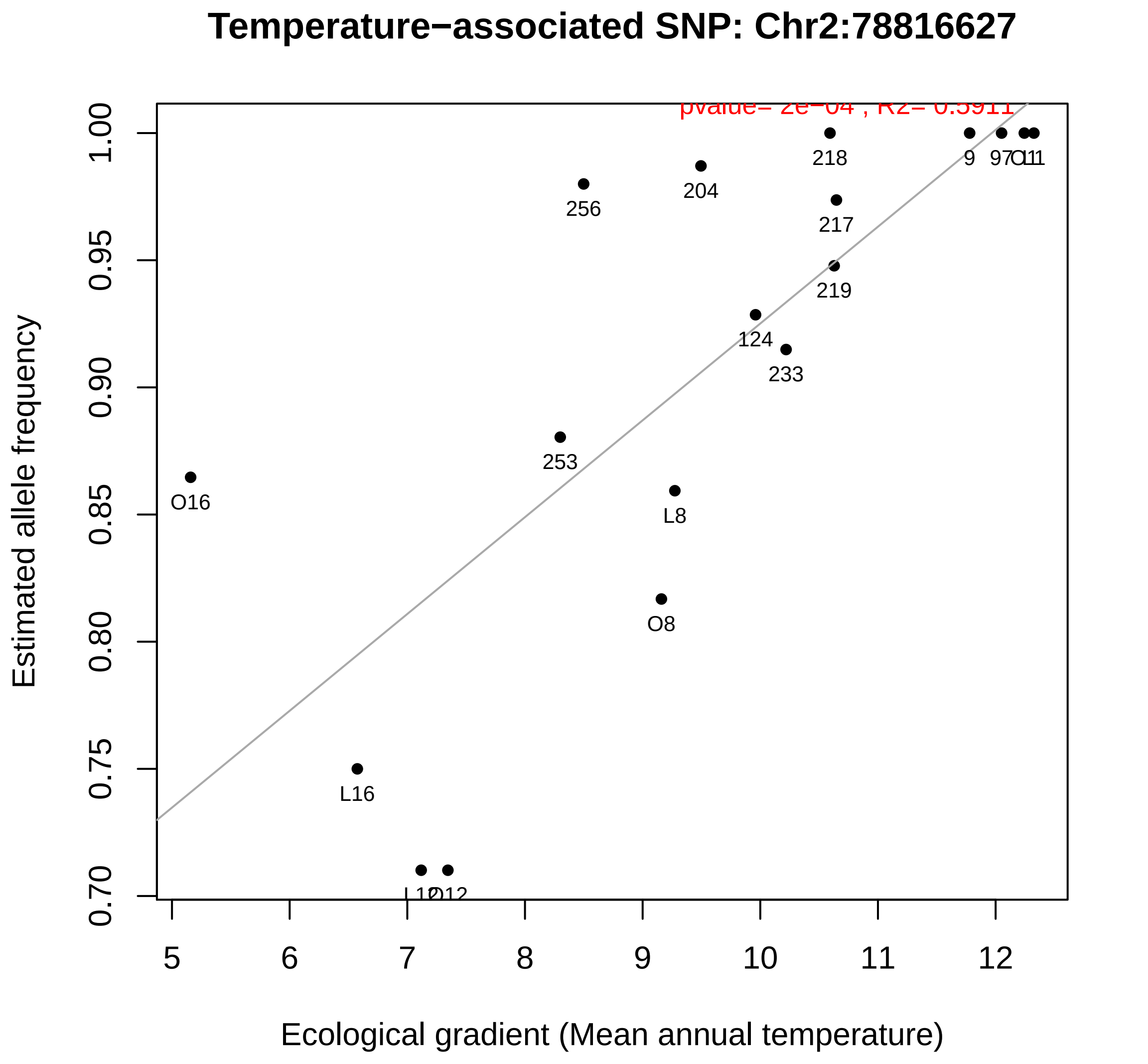

Temperature−associated SNP: Chr2:78816627
pvalue= 2e−04 , R2= 0.5911
Estimated allele frequency
Ecological gradient (Mean annual temperature)
1.00
0.95
0.90
0.85
0.80
0.75
0.70
5
6
7
8
9
10
11
12
218
9
204
256
217
219
124
233
253
O16
L8
O8
L16
O12

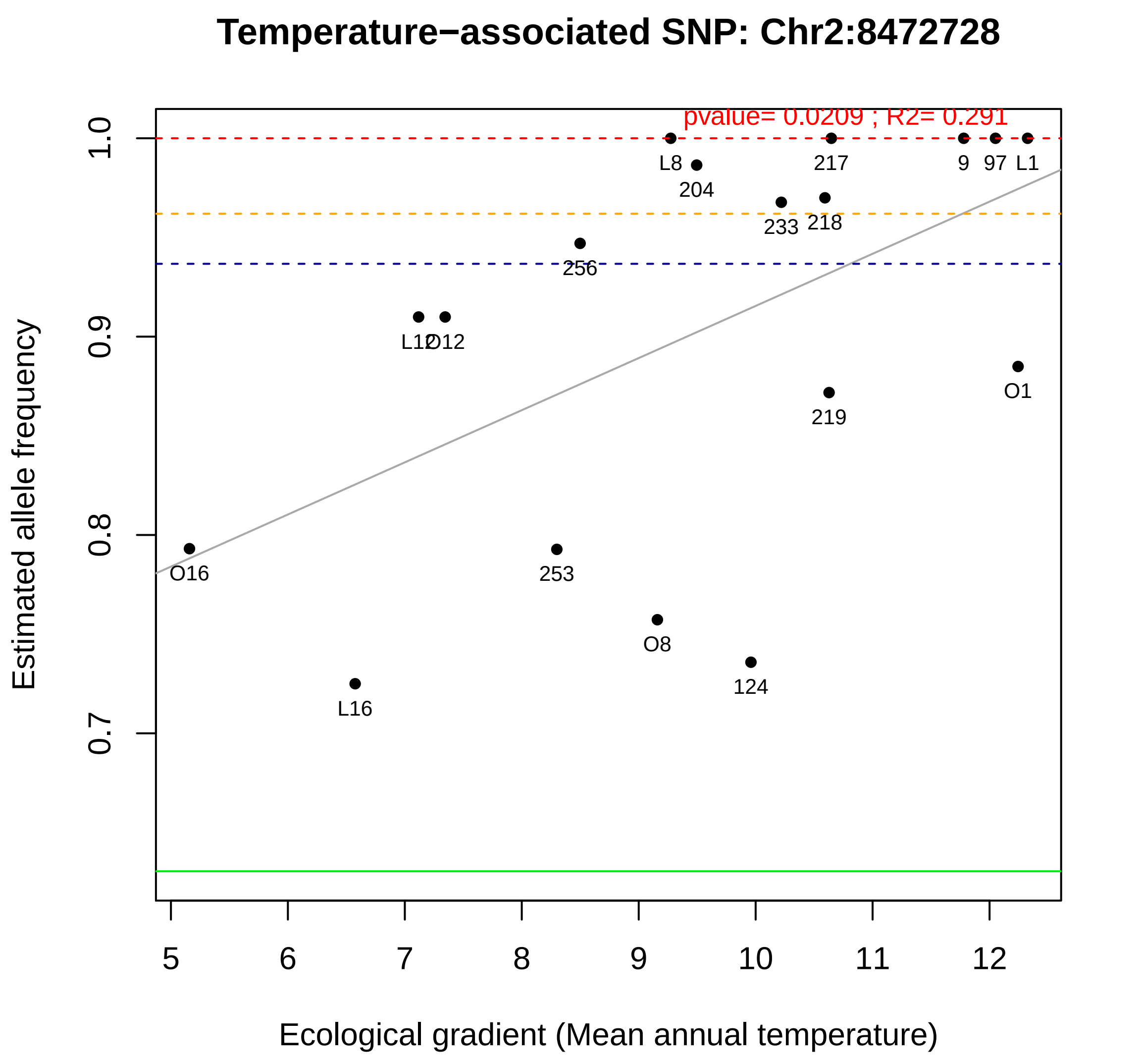

Temperature−associated SNP: Chr2:8472728
pvalue= 0.0209 ; R2= 0.291
Estimated allele frequency
Ecological gradient (Mean annual temperature)
1.0
0.9
0.8
0.7
5
6
7
8
9
10
11
12
L8
204
217
9
97
L1
233
218
256
L12
O12
O1
219
O16
253
O8
124
L16

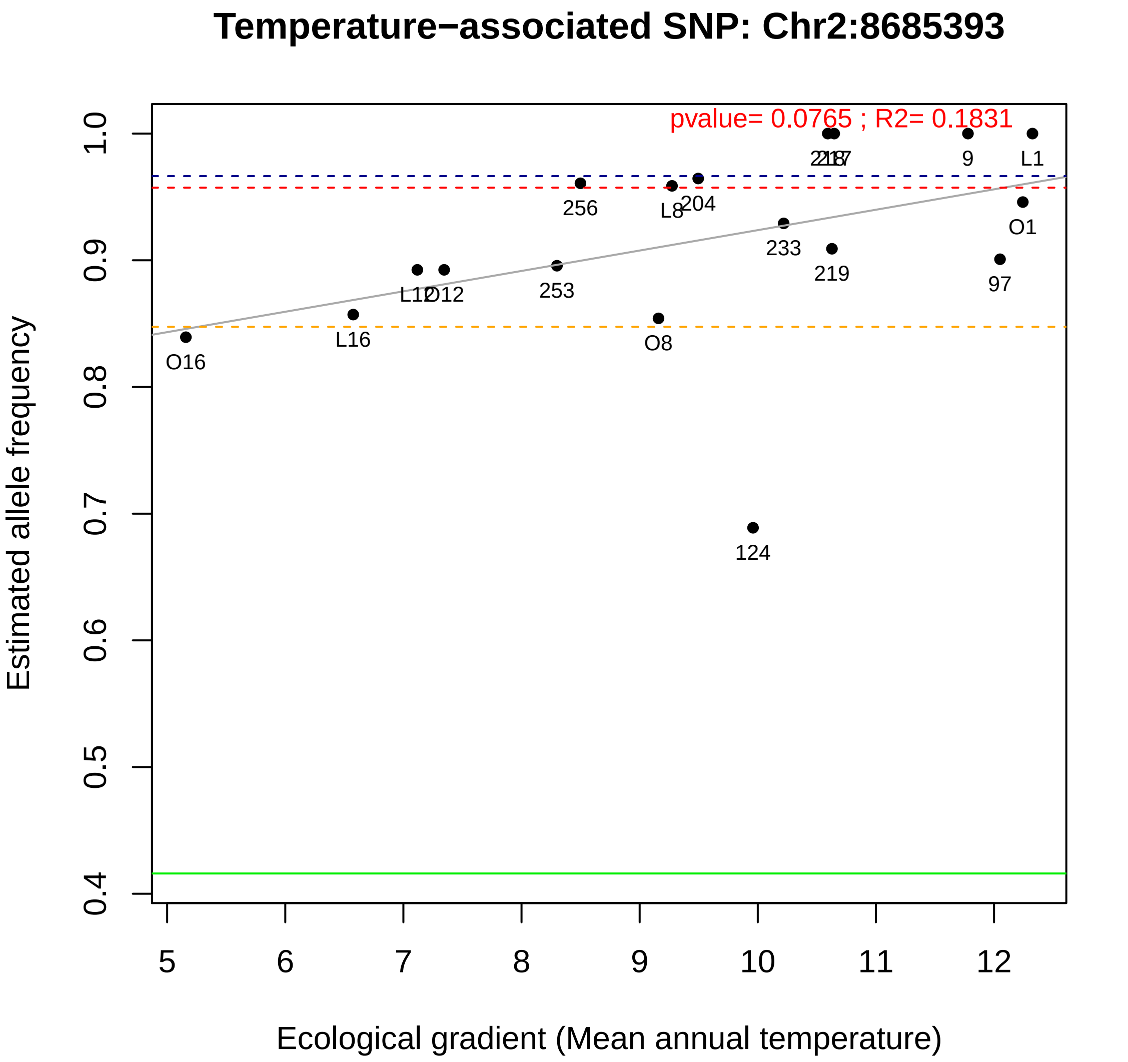
Temperature−associated SNP: Chr2:8685393
pvalue= 0.0765 ; R2= 0.1831
Estimated allele frequency
Ecological gradient (Mean annual temperature)
1.0
0.9
0.8
0.7
0.6
0.5
0.4
5
6
7
8
9
10
11
12
O16
L16
L12
O12
253
256
O8
L8
204
124
233
219
9
97
O1
L1

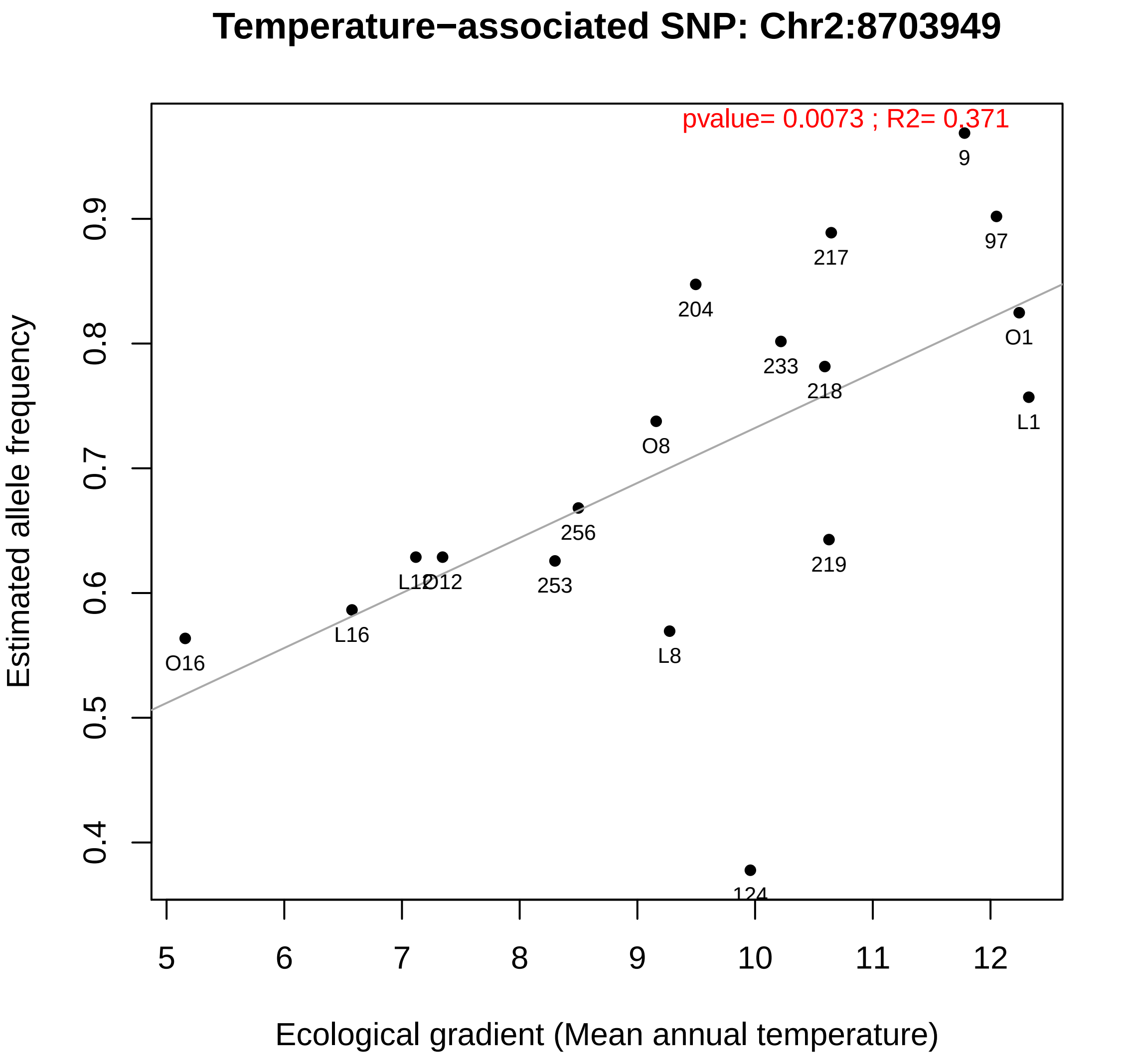

Temperature−associated SNP: Chr2:8703949
pvalue= 0.0073 ; R2= 0.371
Estimated allele frequency
Ecological gradient (Mean annual temperature)
0.4
0.5
0.6
0.7
0.8
0.9
5
6
7
8
9
10
11
12
9
97
217
204
O1
233
218
L1
O8
256
219
L12
O12
253
L16
L8
O16
124

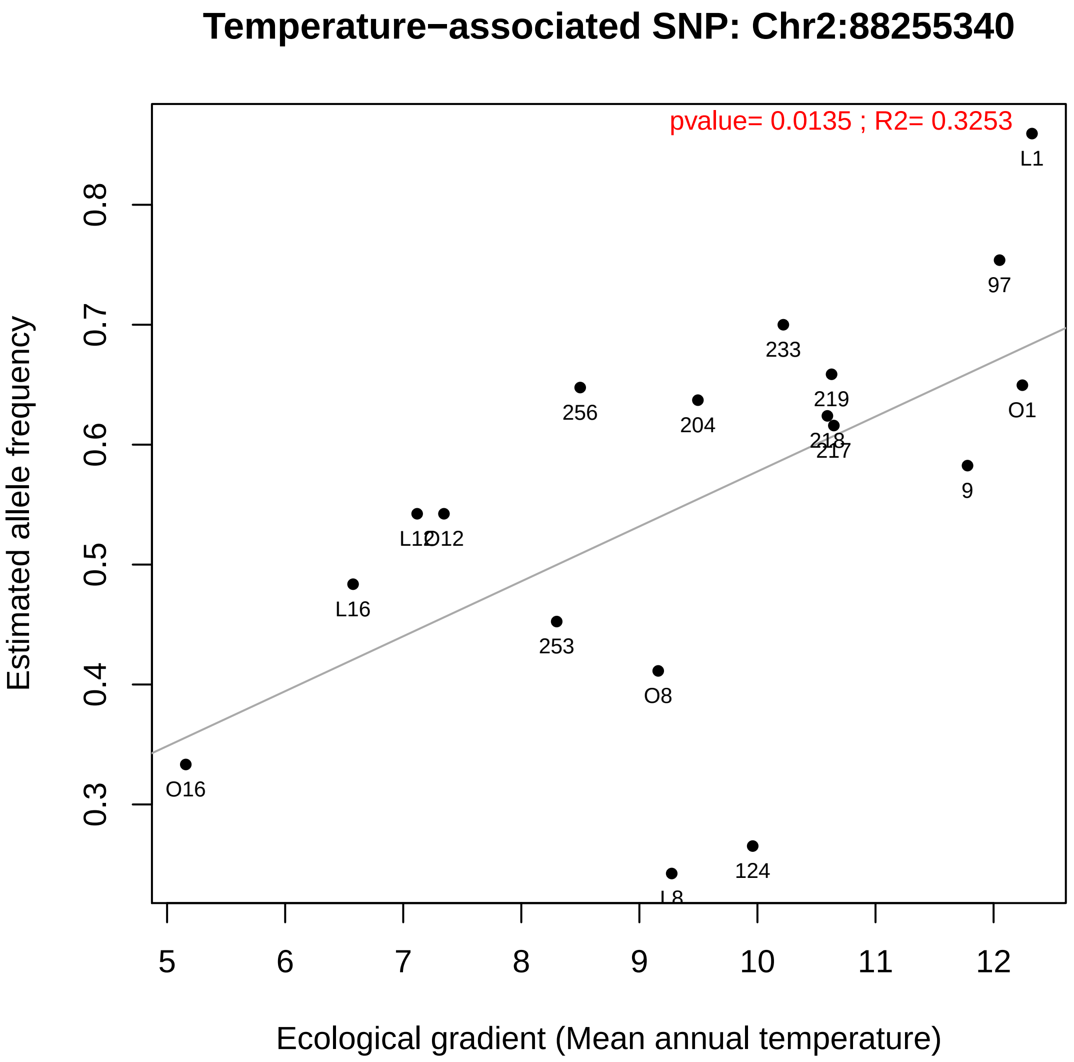
Temperature−associated SNP: Chr2:88255340
pvalue= 0.0135 ; R2= 0.3253
Estimated allele frequency
0.3
0.4
0.5
0.6
0.7
0.8
5
6
7
8
9
10
11
12
Ecological gradient (Mean annual temperature)
L1
97
233
219
O1
256
204
218
217
9
L12
O12
L16
253
O8
O16
124
L8

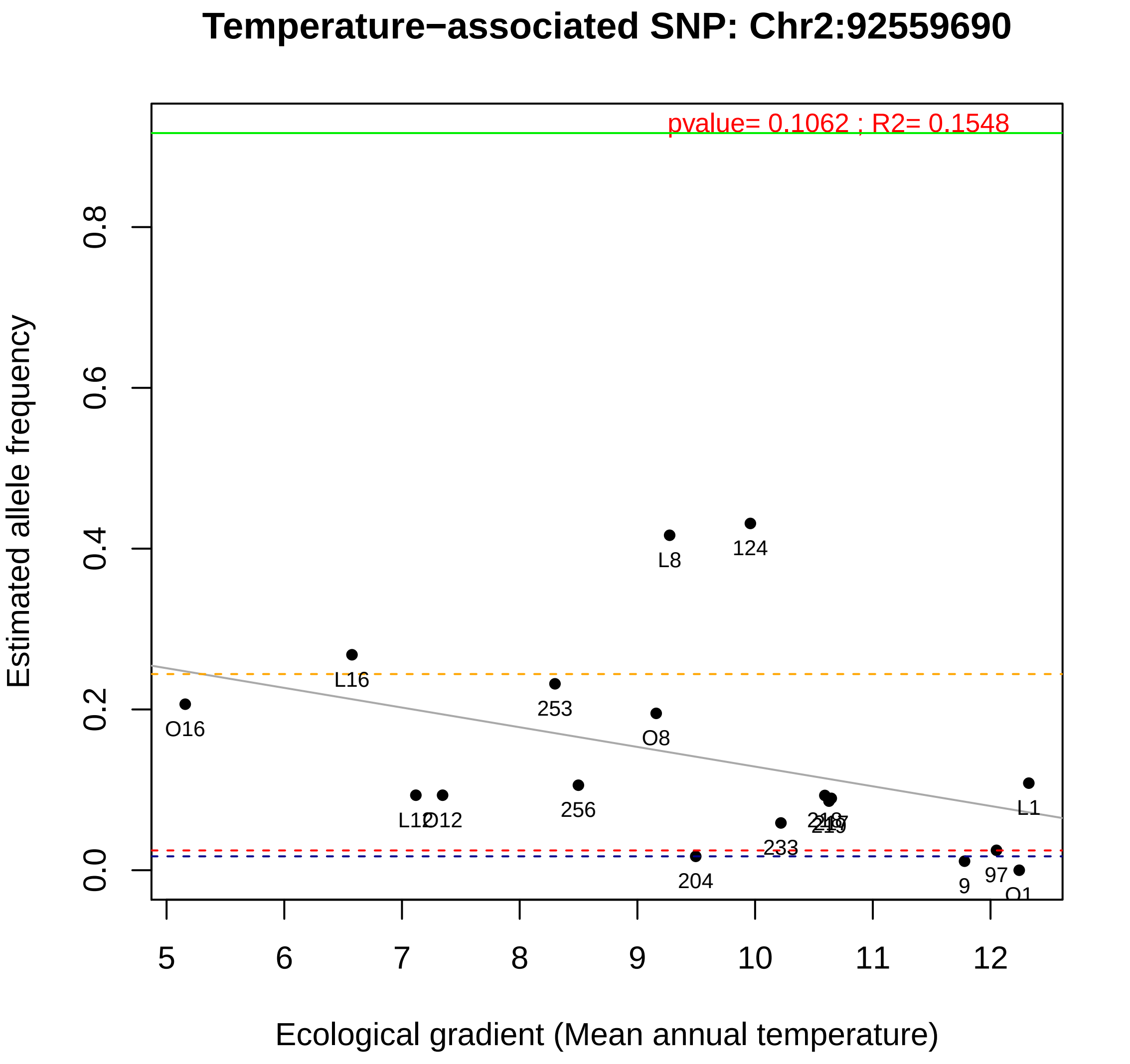

Temperature−associated SNP: Chr2:92559690
pvalue= 0.1062 ; R2= 0.1548
Estimated allele frequency
0.0
0.2
0.4
0.6
0.8
5
6
7
8
9
10
11
12
Ecological gradient (Mean annual temperature)
O16
L16
L12
O12
253
256
L8
124
O8
204
233
L1
9
97
O1

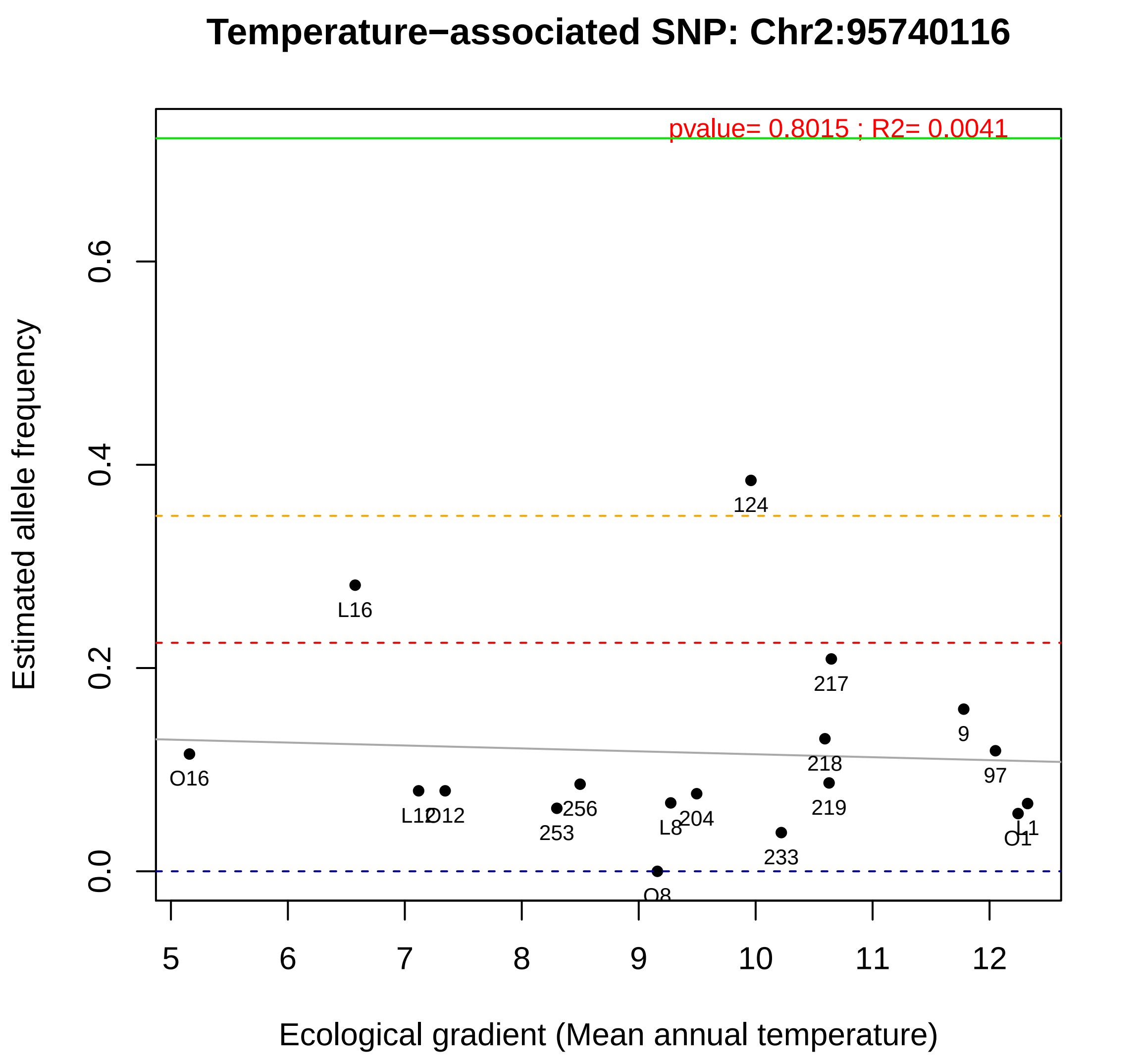
Temperature−associated SNP: Chr2:95740116
pvalue= 0.8015 ; R2= 0.0041
Estimated allele frequency
0.0
0.2
0.4
0.6
Ecological gradient (Mean annual temperature)
5
6
7
8
9
10
11
12
124
L16
217
9
218
97
219
O16
L12
O12
256
253
L8
204
233
L1
O1
O8

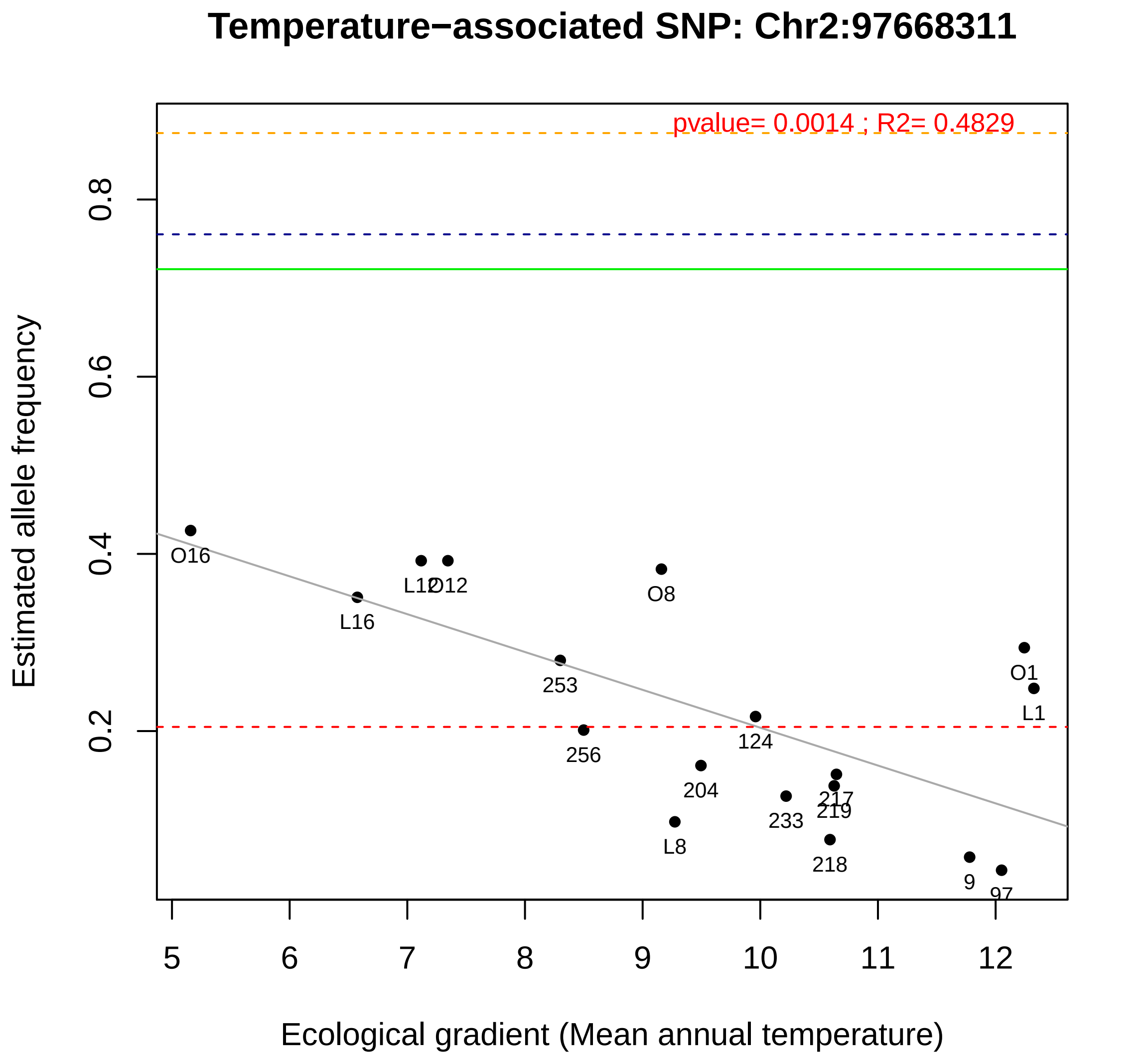

Temperature−associated SNP: Chr2:97668311
pvalue= 0.0014 ; R2= 0.4829
Estimated allele frequency
0.2
0.4
0.6
0.8
Ecological gradient (Mean annual temperature)
5
6
7
8
9
10
11
12
O16
L16
L12
O12
O8
253
256
124
204
L8
233
217
219
218
O1
L1
9
97

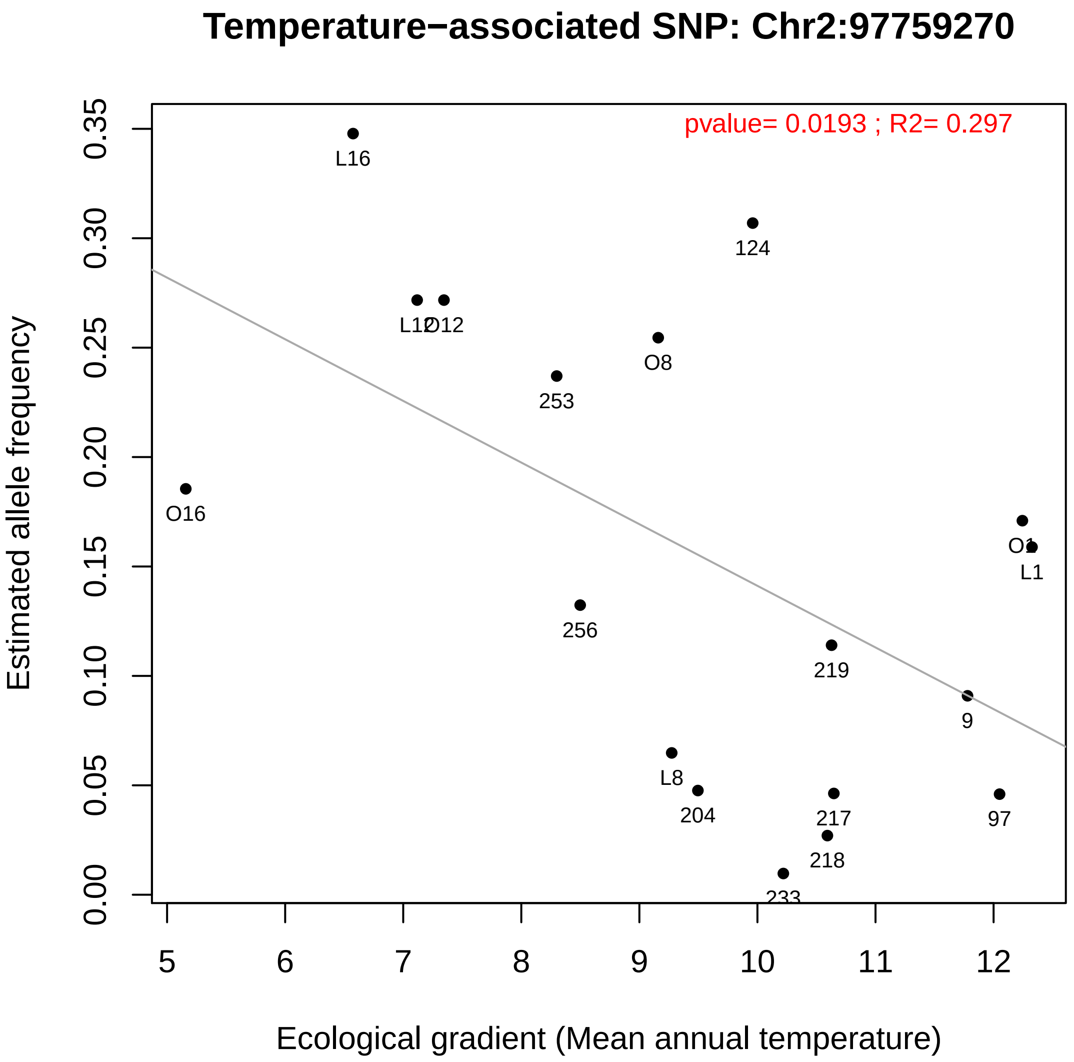
Temperature−associated SNP: Chr2:97759270
pvalue= 0.0193 ; R2= 0.297
Estimated allele frequency
0.00
0.05
0.10
0.15
0.20
0.25
0.30
0.35
5
6
7
8
9
10
11
12
Ecological gradient (Mean annual temperature)
L16
124
L12
O12
O8
253
O16
O1
L1
256
219
9
L8
204
217
97
218
233

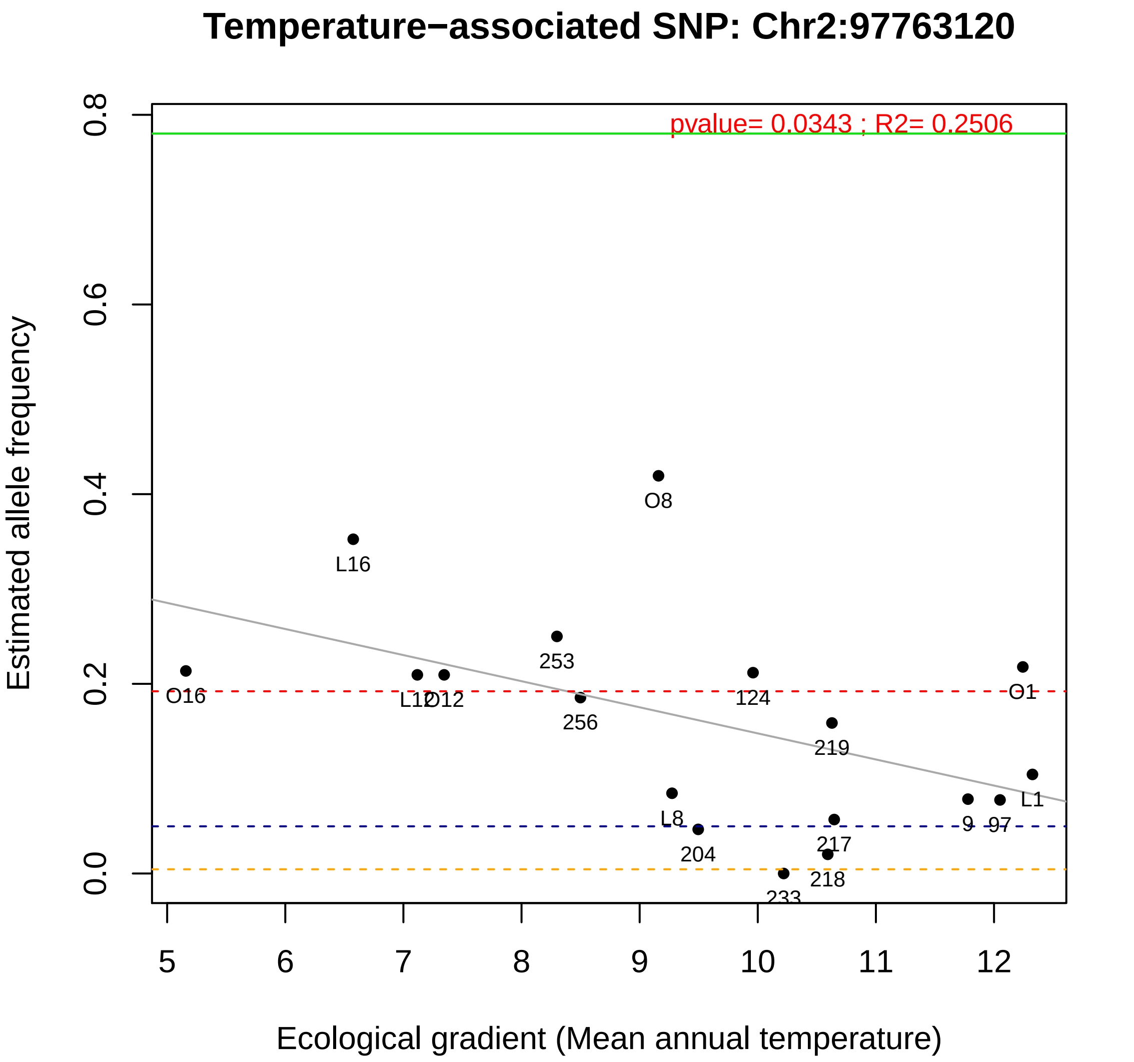
Temperature−associated SNP: Chr2:97763120
pvalue= 0.0343 ; R2= 0.2506
Estimated allele frequency
0.0
0.2
0.4
0.6
0.8
O8
L16
253
O16
L12
O12
256
124
O1
219
L1
L8
204
9
97
217
218
233
5
6
7
8
9
10
11
12
Ecological gradient (Mean annual temperature)

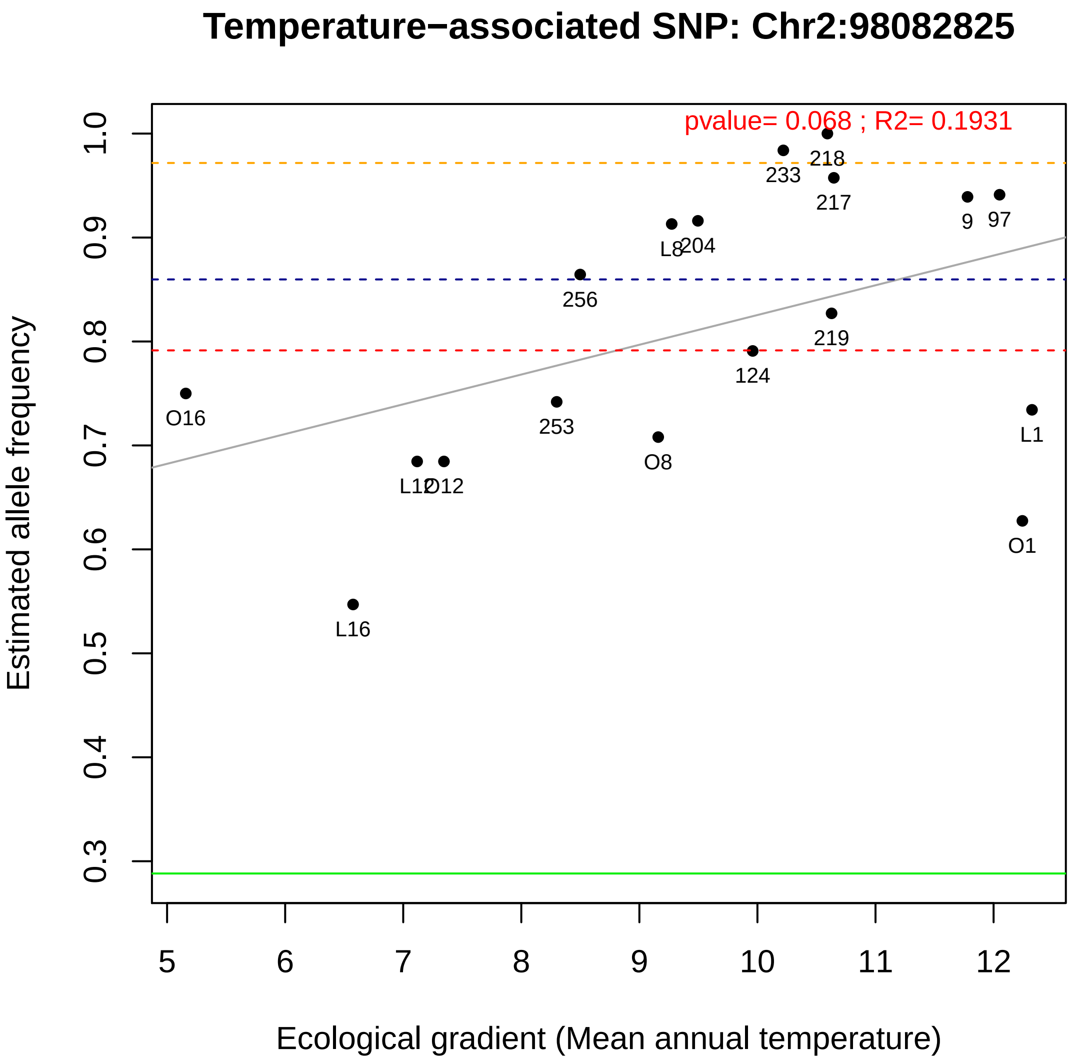
Temperature−associated SNP: Chr2:98082825
pvalue= 0.068 ; R2= 0.1931
Estimated allele frequency
Ecological gradient (Mean annual temperature)
1.0
0.9
0.8
0.7
0.6
0.5
0.4
0.3
5
6
7
8
9
10
11
12
O16
L16
L12
O12
253
256
L8
204
O8
124
233
218
217
219
9
97
L1
O1

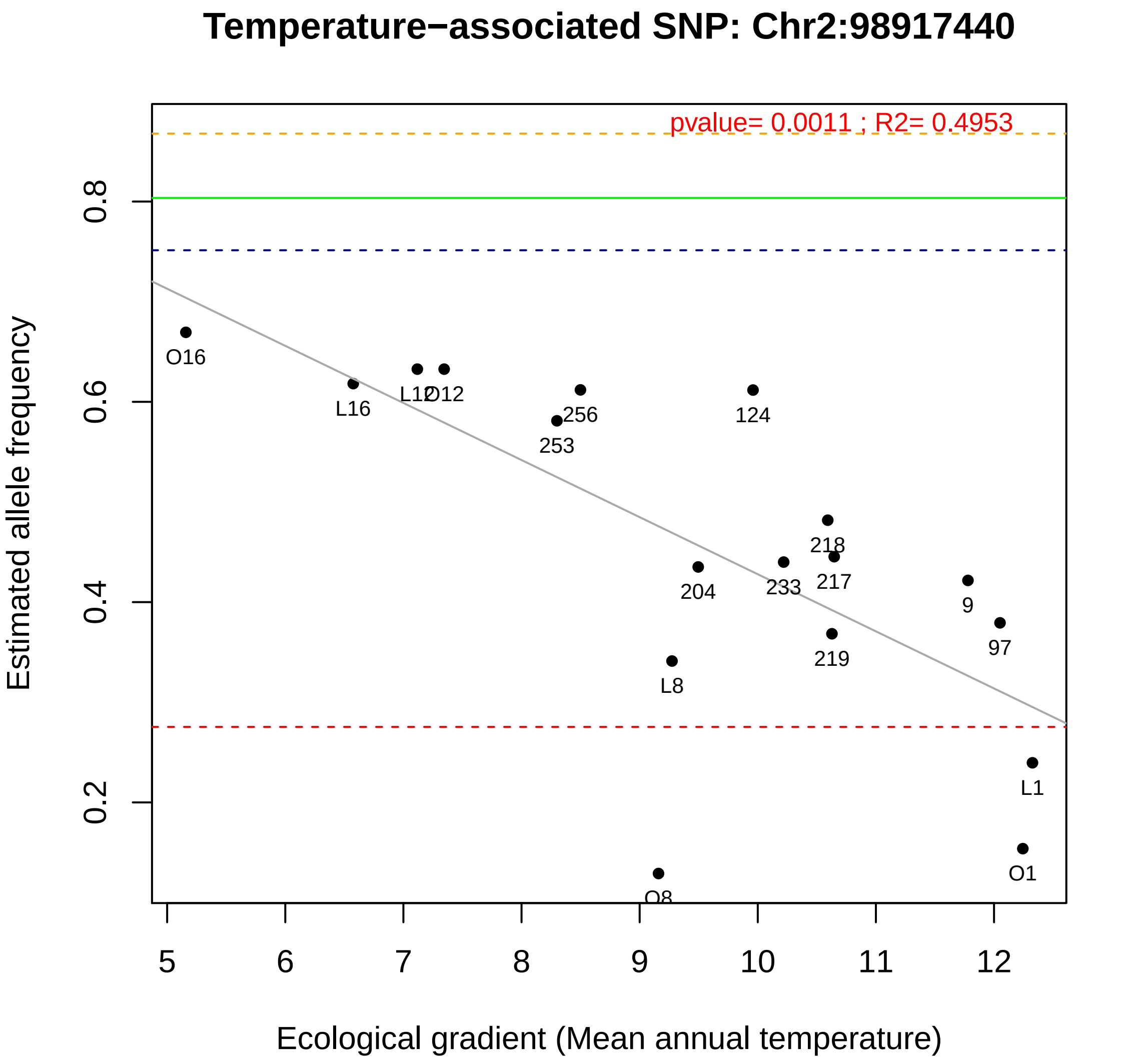

Temperature−associated SNP: Chr2:98917440
pvalue= 0.0011 ; R2= 0.4953
Estimated allele frequency
0.2
0.4
0.6
0.8
Ecological gradient (Mean annual temperature)
5
6
7
8
9
10
11
12
O16
L16
L12
O12
256
253
124
218
217
233
204
9
97
219
L8
L1
O1
O8

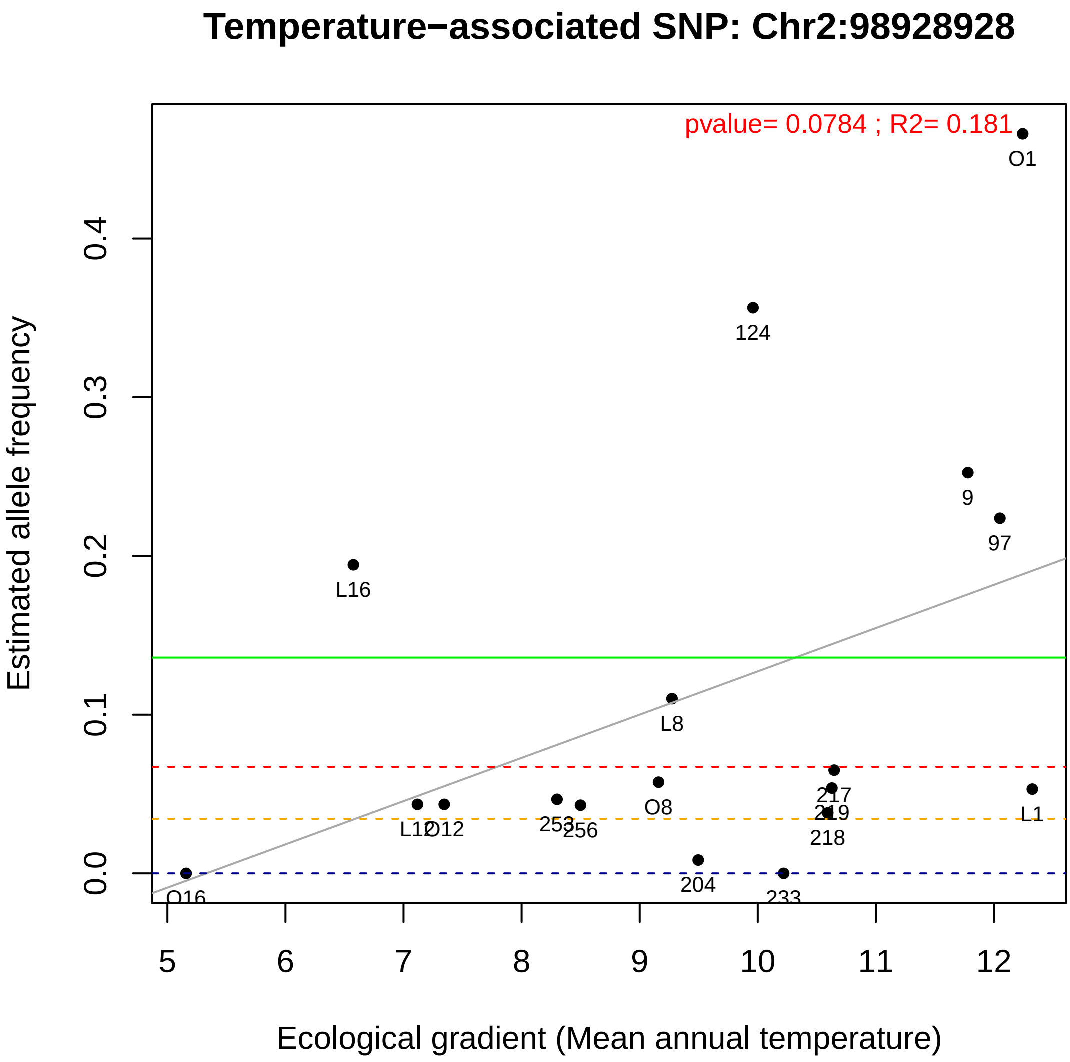
Temperature−associated SNP: Chr2:98928928
pvalue= 0.0784 ; R2= 0.181
Estimated allele frequency
Ecological gradient (Mean annual temperature)
0.0
0.1
0.2
0.3
0.4
5
6
7
8
9
10
11
12
O1
124
9
97
L16
L8
217
219
218
L1
O8
L12
O12
253
256
204
233
O16

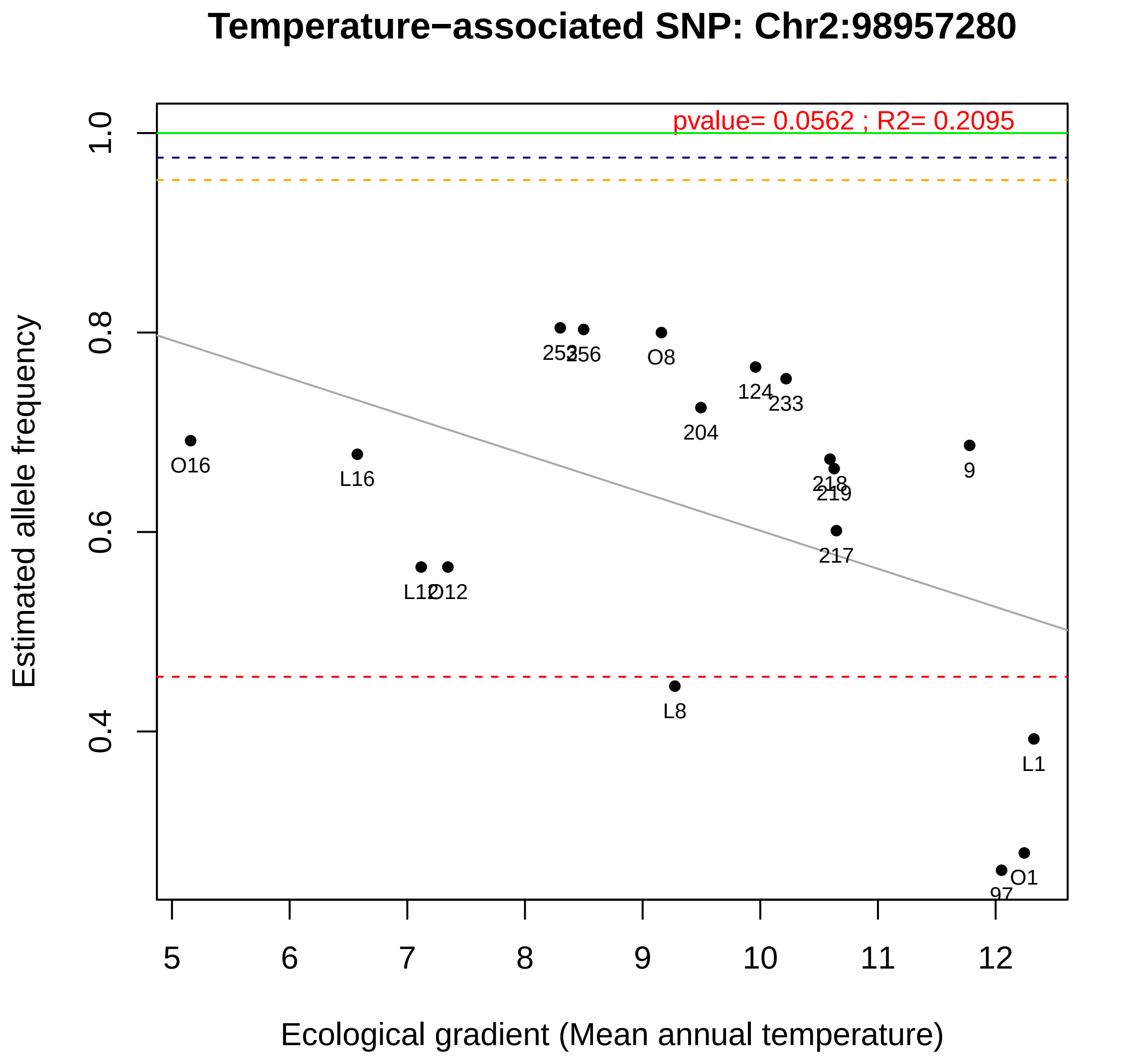
Temperature−associated SNP: Chr2:98957280
pvalue= 0.0562 ; R2= 0.2095
Estimated allele frequency
1.0
0.8
0.6
0.4
O16
L16
L12
O12
253
256
O8
204
124
233
218
219
217
9
L8
L1
O1
97
5
6
7
8
9
10
11
12
Ecological gradient (Mean annual temperature)

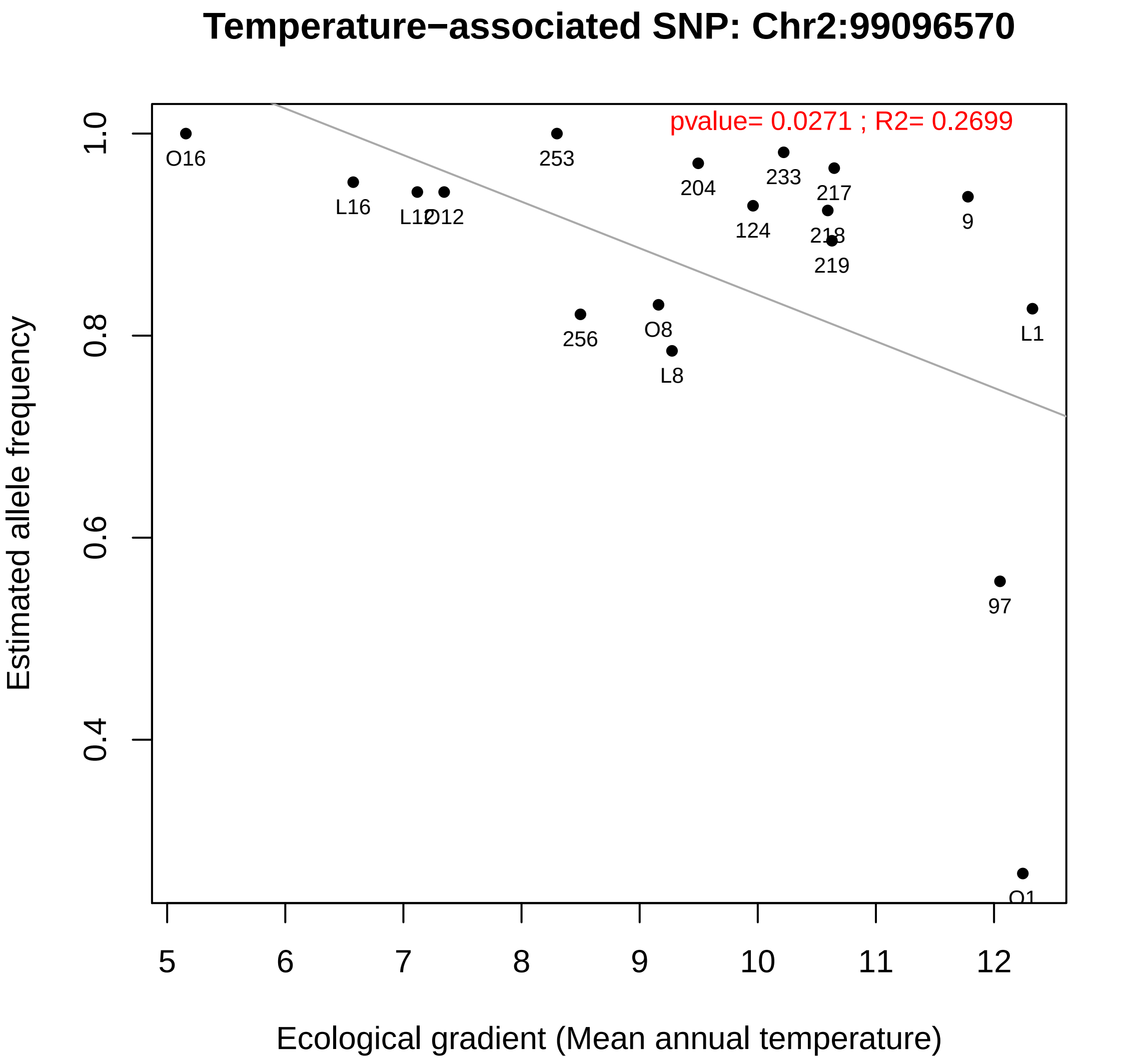
Temperature−associated SNP: Chr2:99096570
pvalue= 0.0271 ; R2= 0.2699
Estimated allele frequency
Ecological gradient (Mean annual temperature)
1.0
0.8
0.6
0.4
5
6
7
8
9
10
11
12
O16
L16
L12
O12
253
204
233
217
124
218
219
9
256
O8
L8
L1
97
O1

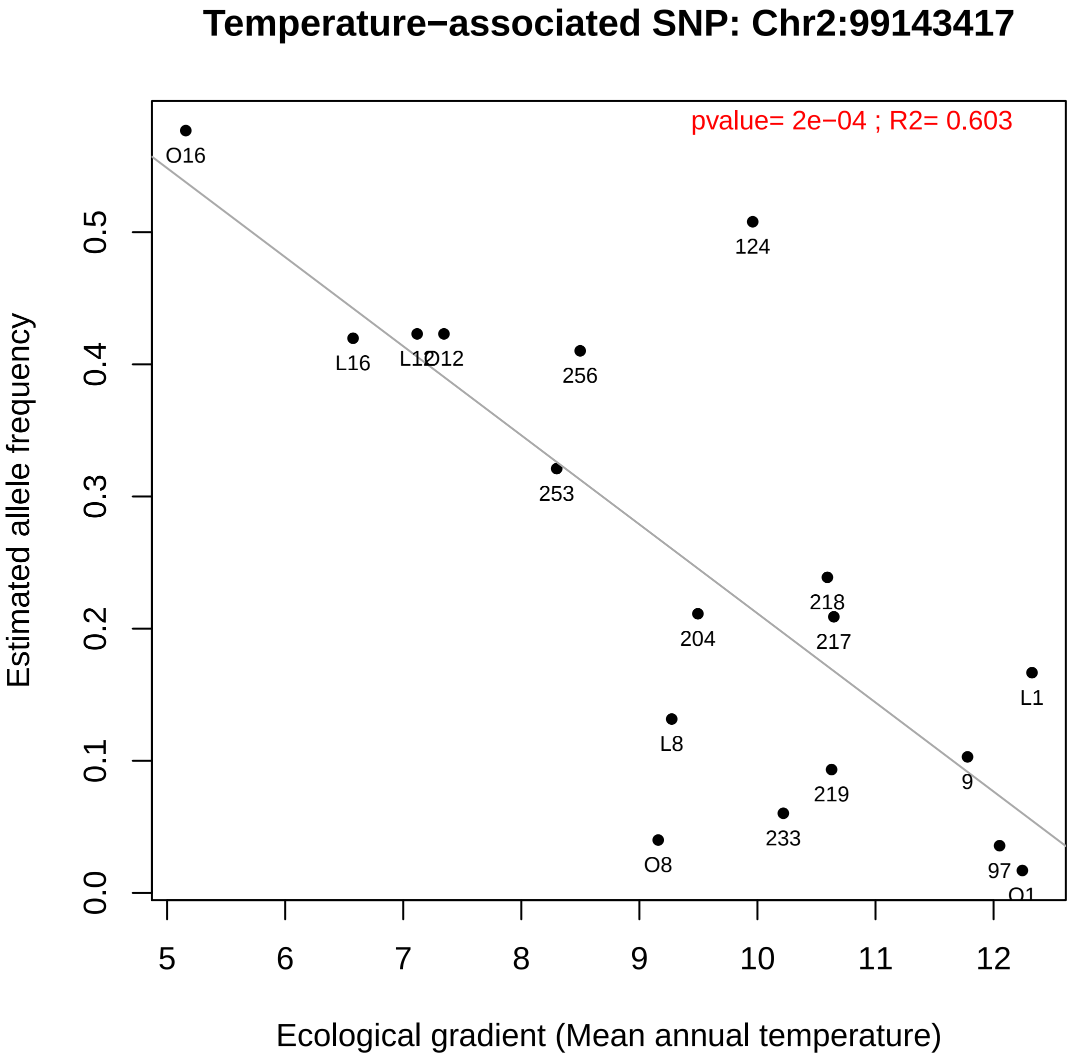
Temperature−associated SNP: Chr2:99143417
pvalue= 2e−04 ; R2= 0.603
O16
124
L16
L12
O12
256
253
218
204
217
L1
L8
9
219
233
O8
97
O1
Estimated allele frequency
0.0
0.1
0.2
0.3
0.4
0.5
5
6
7
8
9
10
11
12
Ecological gradient (Mean annual temperature)

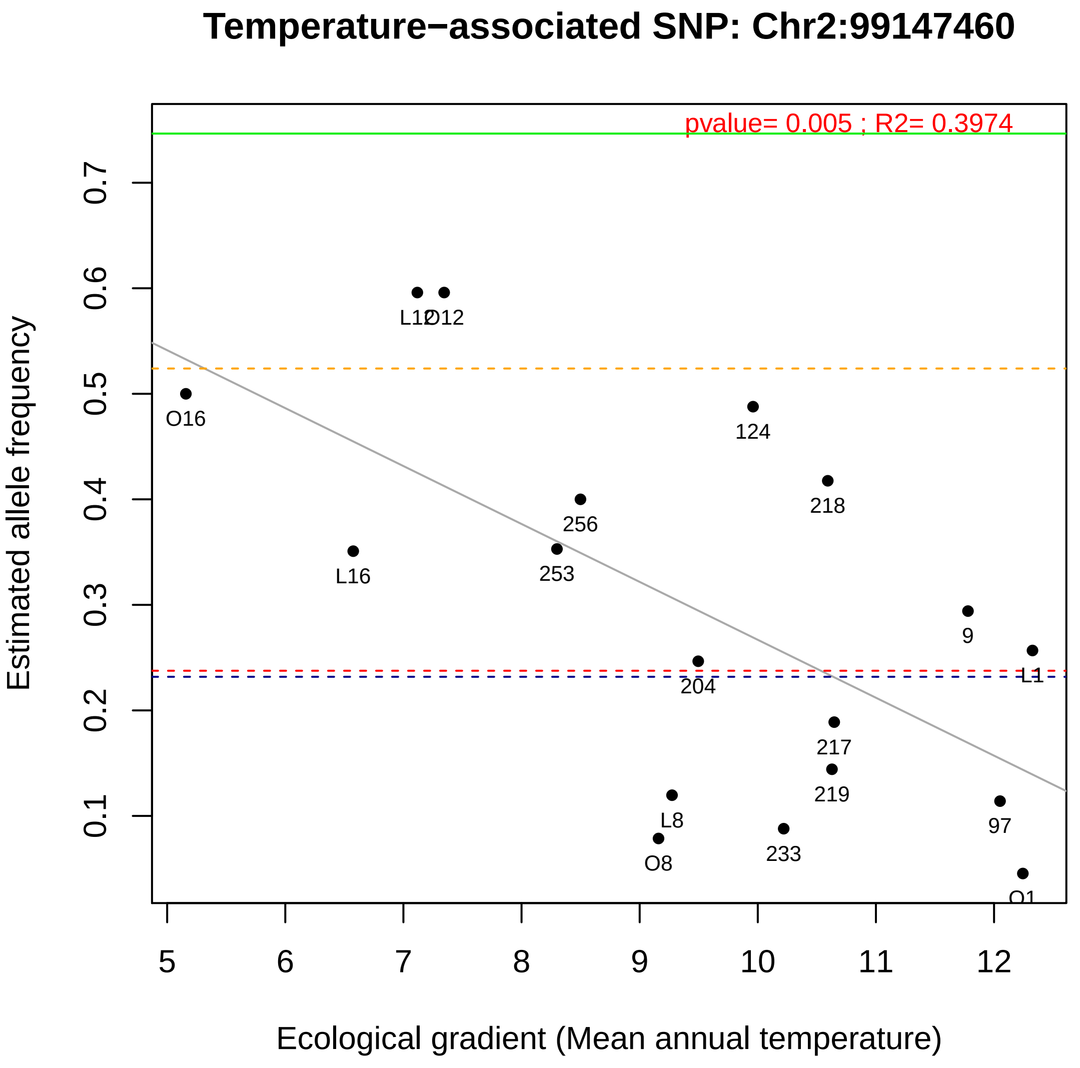

Temperature−associated SNP: Chr2:99147460
pvalue= 0.005 ; R2= 0.3974
Estimated allele frequency
Ecological gradient (Mean annual temperature)
0.1
0.2
0.3
0.4
0.5
0.6
0.7
5
6
7
8
9
10
11
12
L12
O12
O16
124
218
256
L16
253
9
L1
204
217
219
L8
233
97
O8
O1

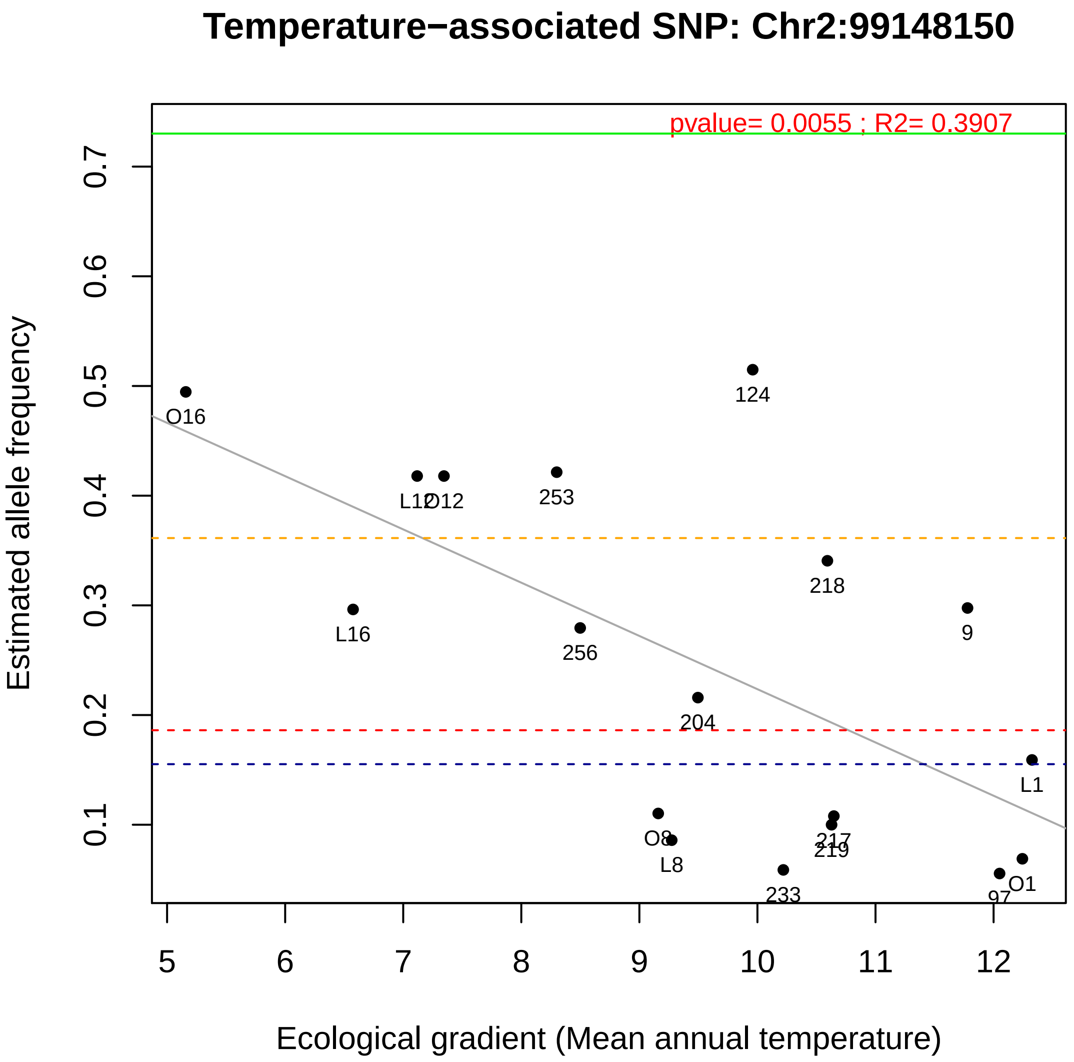

Temperature−associated SNP: Chr2:99148150
pvalue= 0.0055 ; R2= 0.3907
Estimated allele frequency
0.1
0.2
0.3
0.4
0.5
0.6
0.7
O16
L16
L12
O12
253
256
124
204
218
9
L1
O8
L8
233
217
219
97
O1
5
6
7
8
9
10
11
12
Ecological gradient (Mean annual temperature)

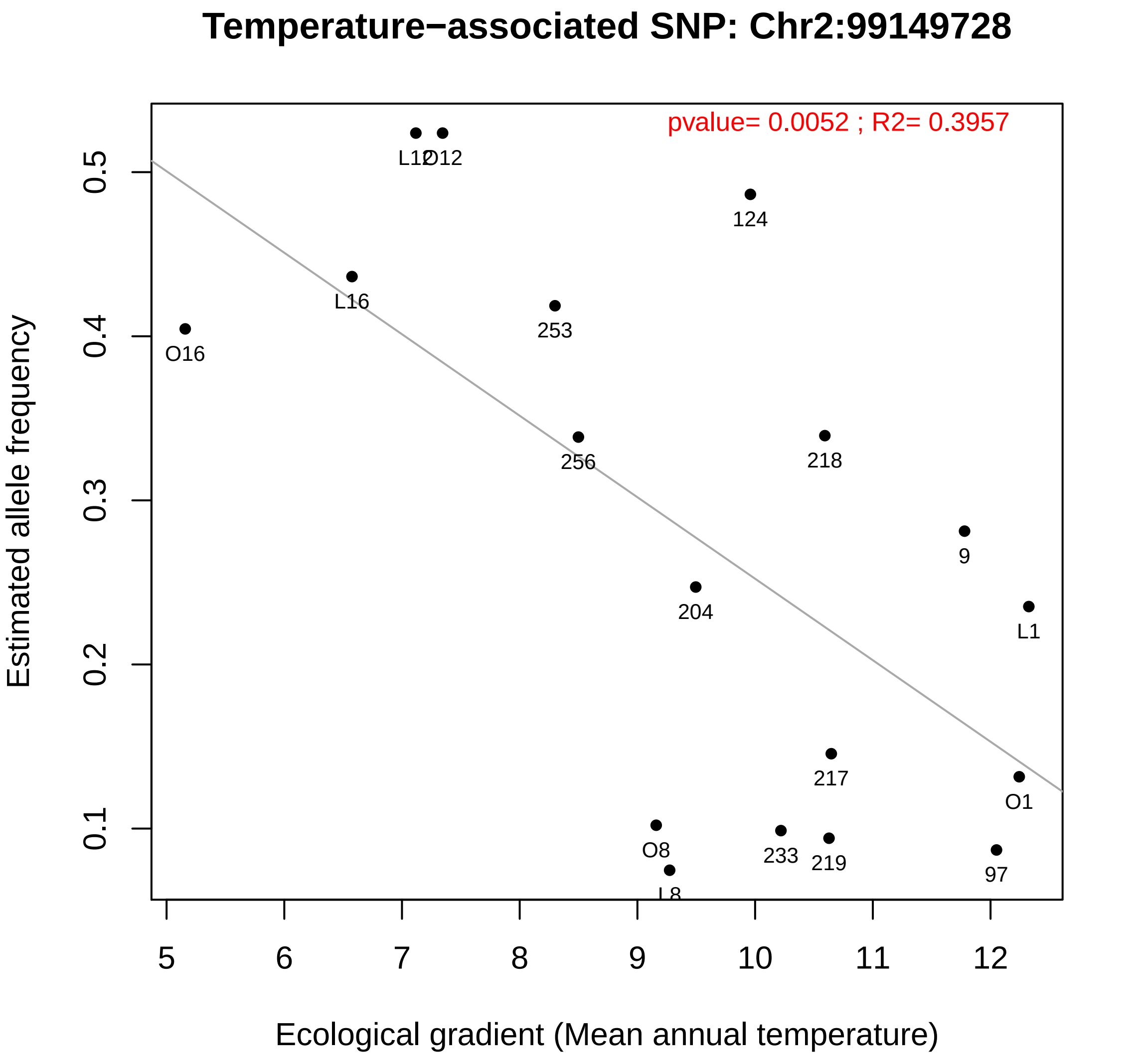
Temperature−associated SNP: Chr2:99149728
pvalue= 0.0052 ; R2= 0.3957
Estimated allele frequency
0.1
0.2
0.3
0.4
0.5
5
6
7
8
9
10
11
12
Ecological gradient (Mean annual temperature)
L12
O12
124
L16
253
O16
256
218
9
204
L1
217
O1
O8
233
219
97
L8

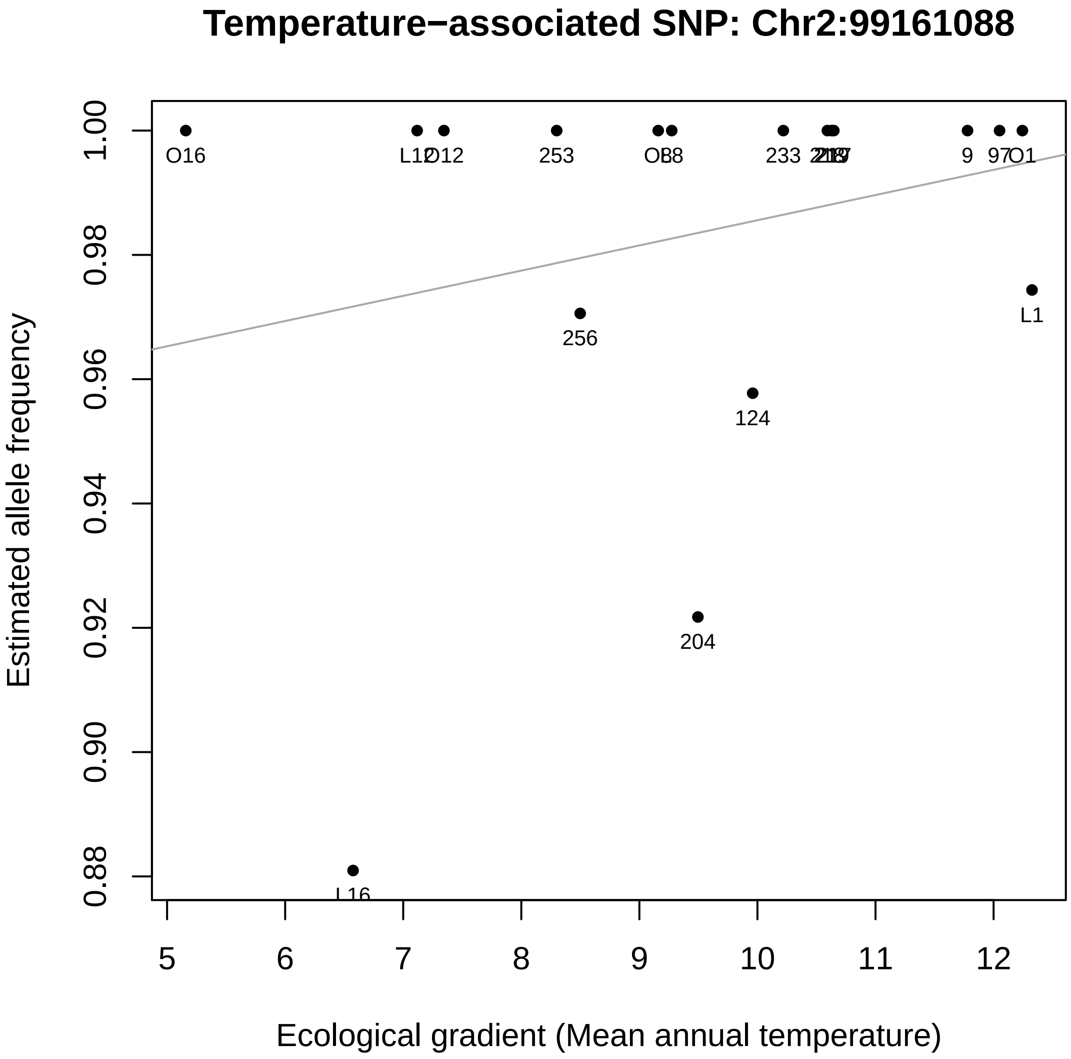

Temperature−associated SNP: Chr2:99161088
Estimated allele frequency
Ecological gradient (Mean annual temperature)
1.00
0.98
0.96
0.94
0.92
0.90
0.88
5
6
7
8
9
10
11
12
O16
253
233
9
256
L1
124
204

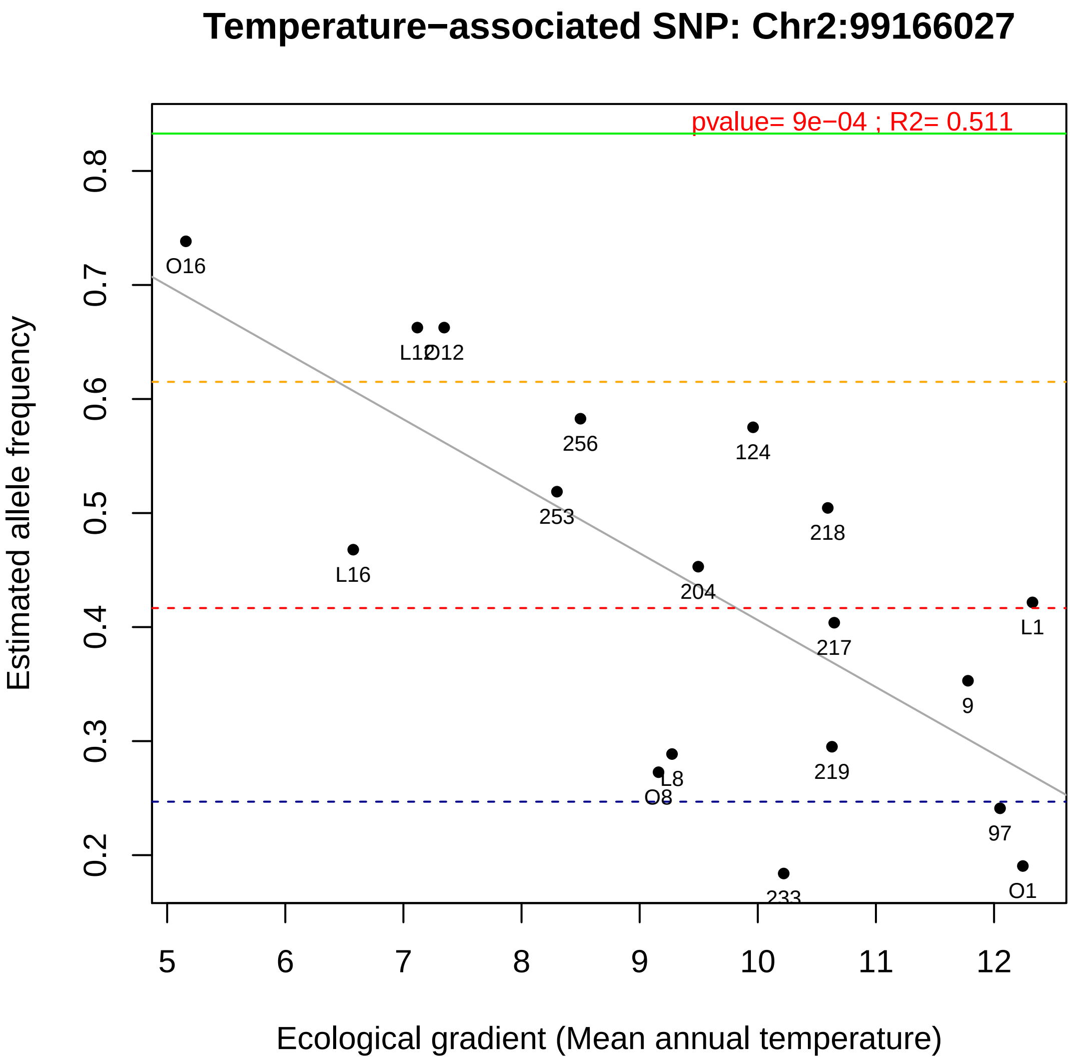

Temperature−associated SNP: Chr2:99166027
pvalue= 9e−04 ; R2= 0.511
Estimated allele frequency
0.2
0.3
0.4
0.5
0.6
0.7
0.8
Ecological gradient (Mean annual temperature)
5
6
7
8
9
10
11
12
O16
L12
O12
256
124
253
218
L16
204
L1
217
9
L8
O8
219
97
233
O1

**Temperature−associated SNP: Chr2:99167531**

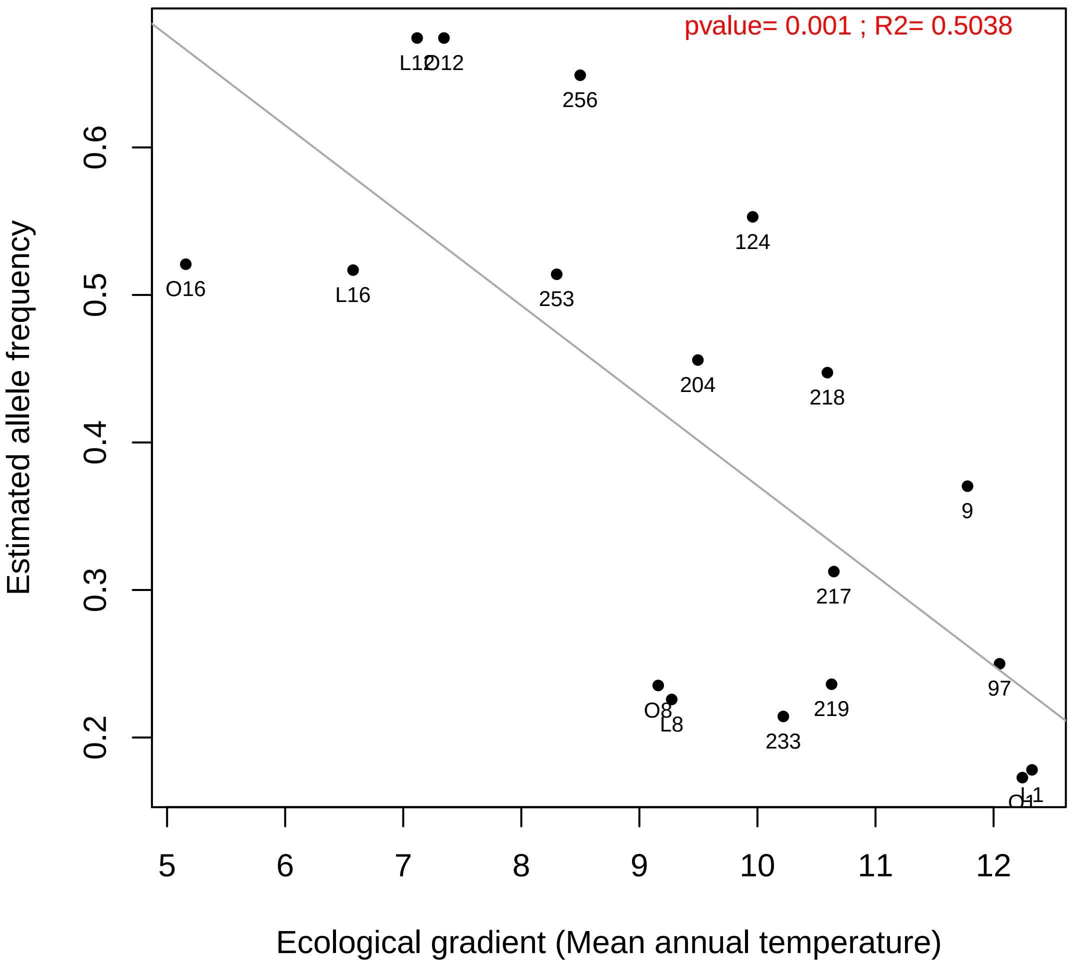

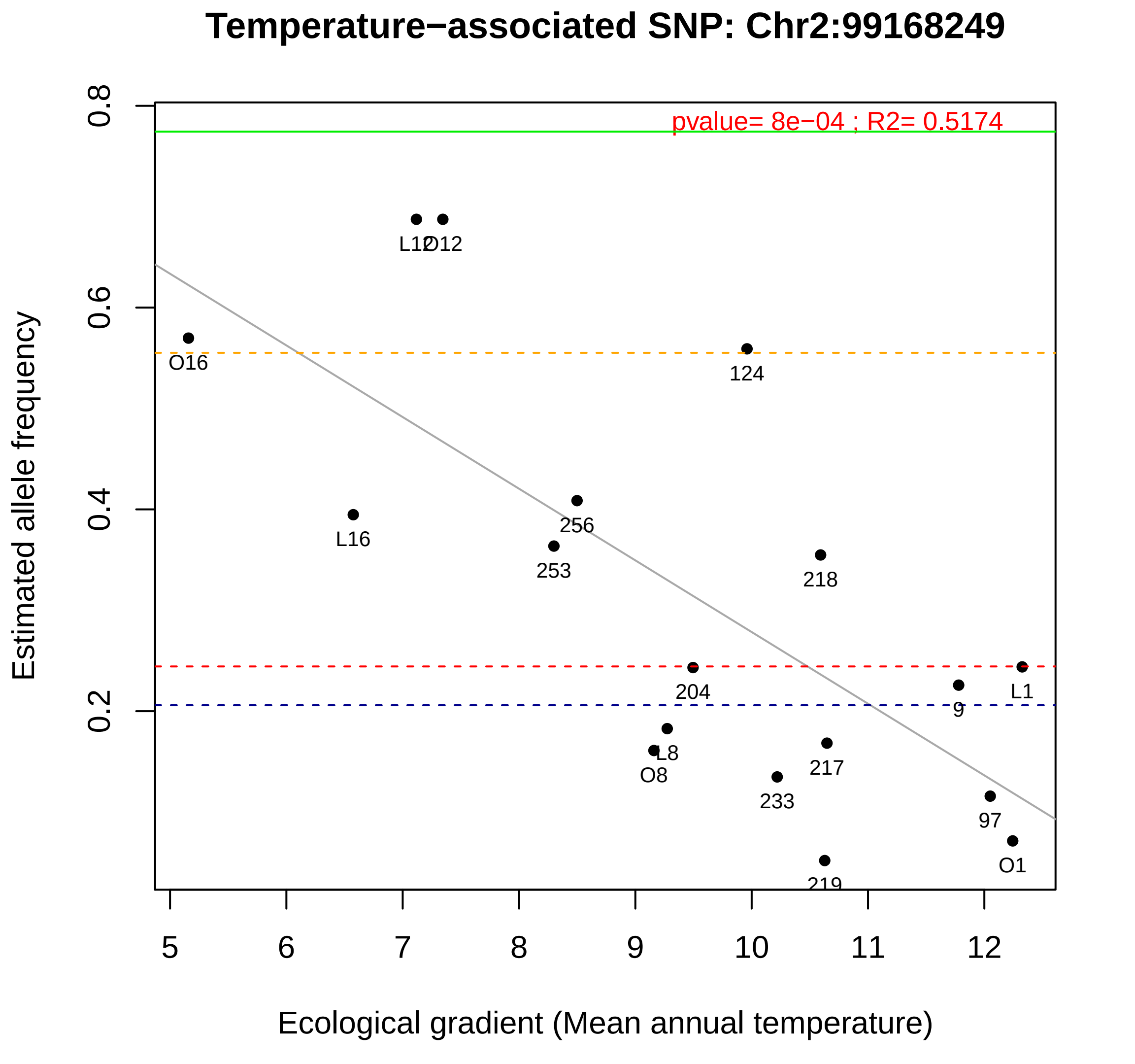

Temperature−associated SNP: Chr2:99168249
pvalue= 8e−04 ; R2= 0.5174
Estimated allele frequency
0.2
0.4
0.6
0.8
Ecological gradient (Mean annual temperature)
5
6
7
8
9
10
11
12
L12
O12
O16
124
L16
256
253
218
204
L1
9
L8
O8
217
233
97
O1
219

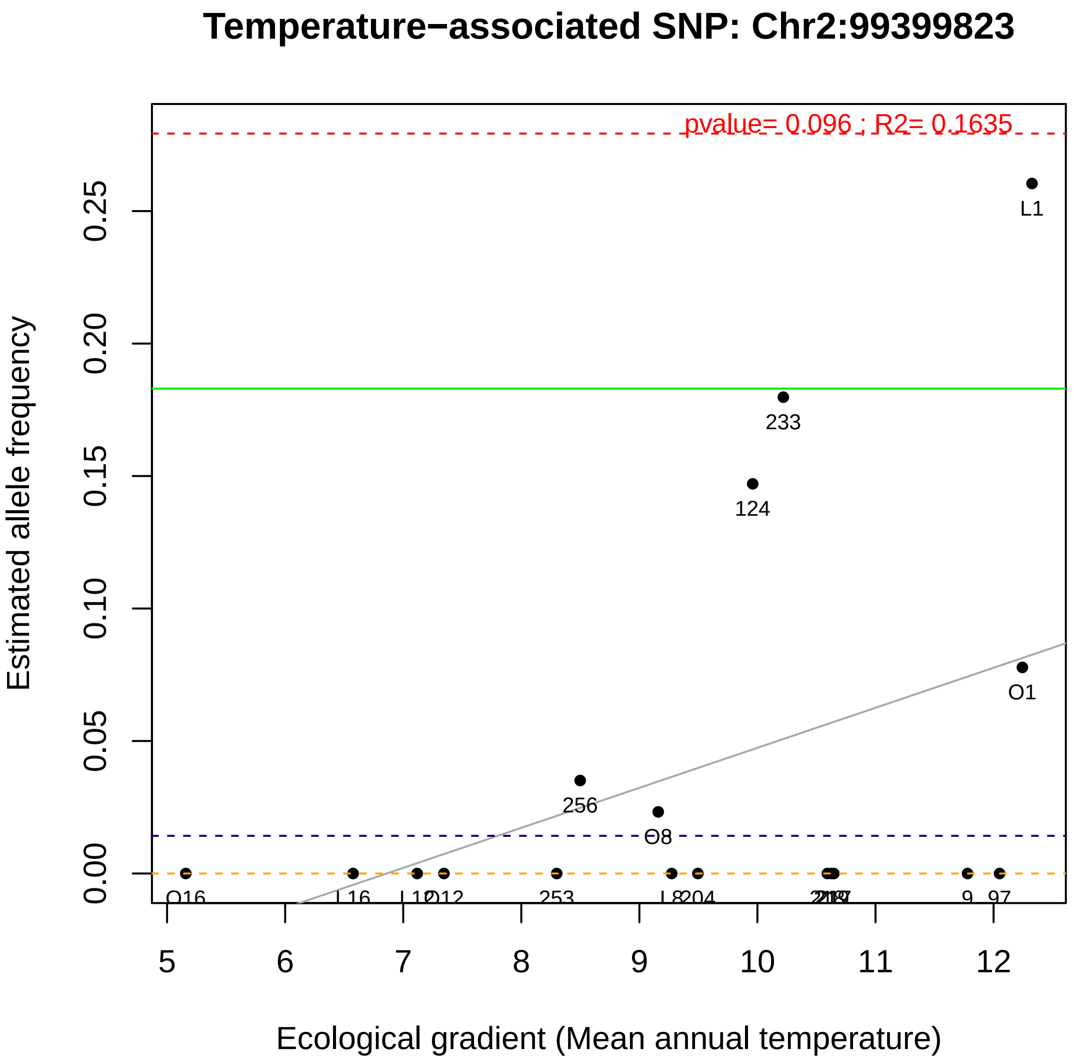
Temperature−associated SNP: Chr2:99399823
pvalue= 0.096 ; R2= 0.1635
Estimated allele frequency
0.00
0.05
0.10
0.15
0.20
0.25
Ecological gradient (Mean annual temperature)
5
6
7
8
9
10
11
12
L1
233
124
O1
256
O8
O16
L16
O12
253
L8
204
9
97

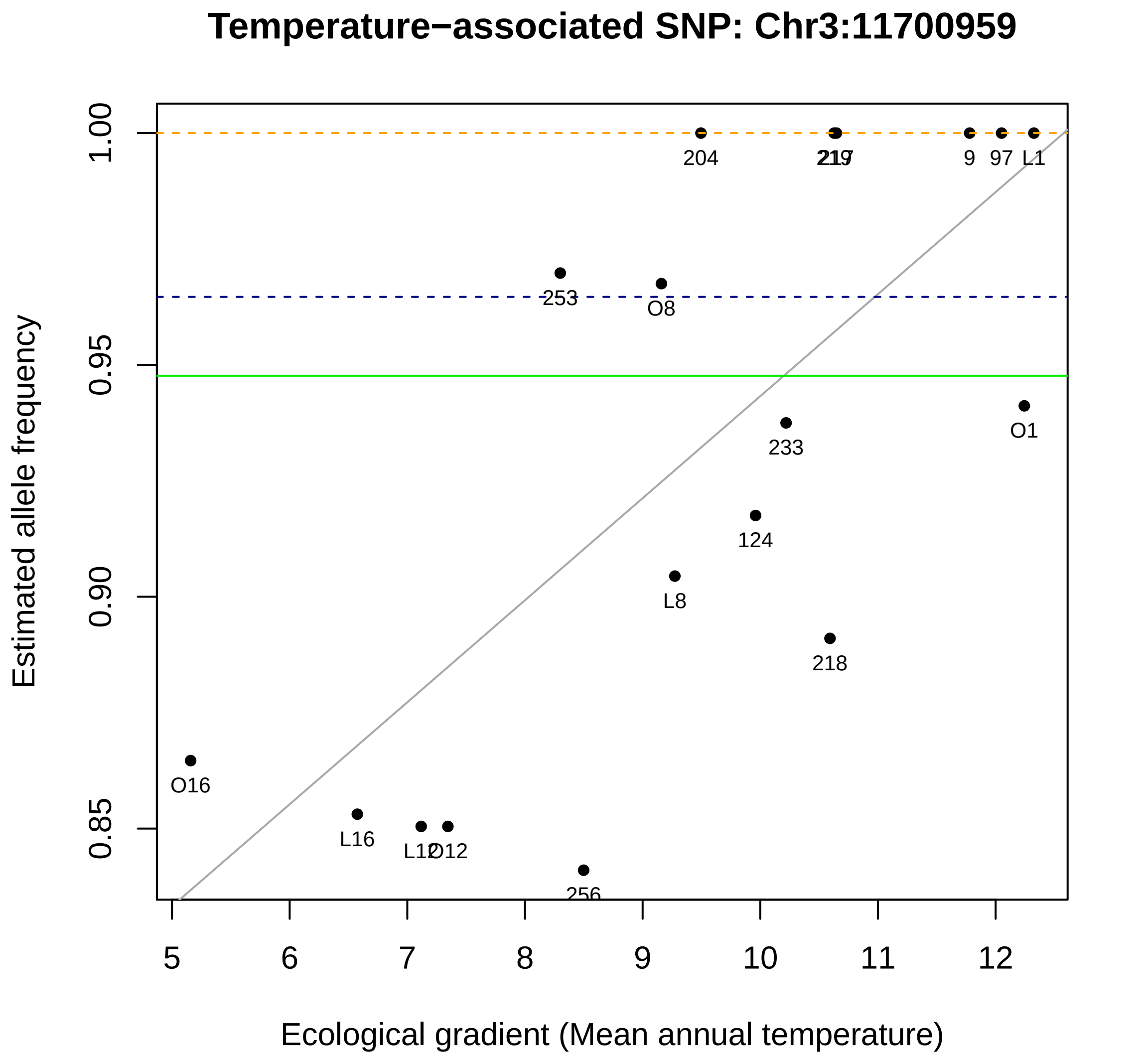
Temperature−associated SNP: Chr3:11700959
Estimated allele frequency
Ecological gradient (Mean annual temperature)
0.85
0.90
0.95
1.00
5
6
7
8
9
10
11
12
204
9
97
L1
253
O8
O1
233
124
L8
218
O16
L16
256

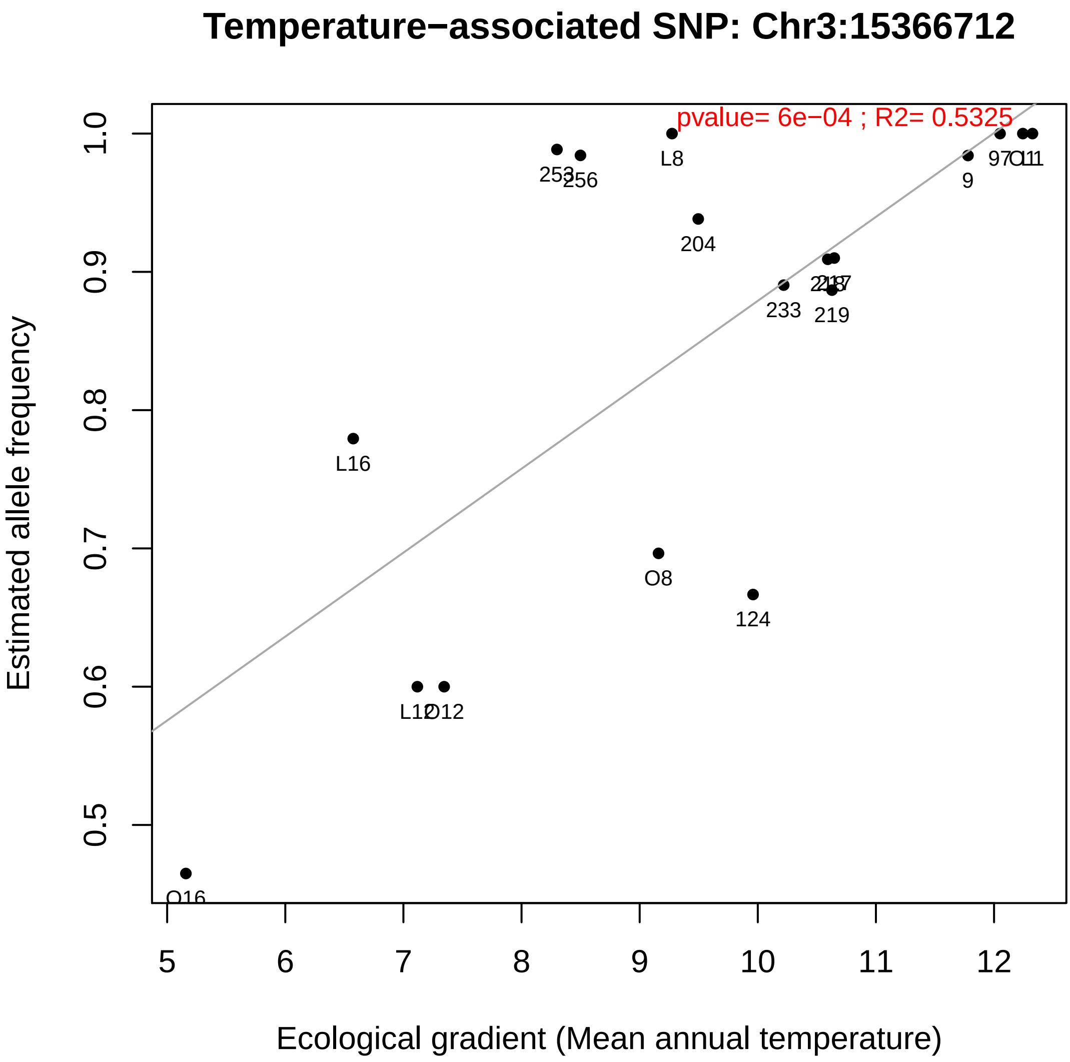
Temperature−associated SNP: Chr3:15366712
pvalue= 6e−04 ; R2= 0.5325
Estimated allele frequency
Ecological gradient (Mean annual temperature)
1.0
0.9
0.8
0.7
0.6
0.5
5
6
7
8
9
10
11
12
L8
253
256
204
233
219
9
L16
O8
124
L12
O12
O16

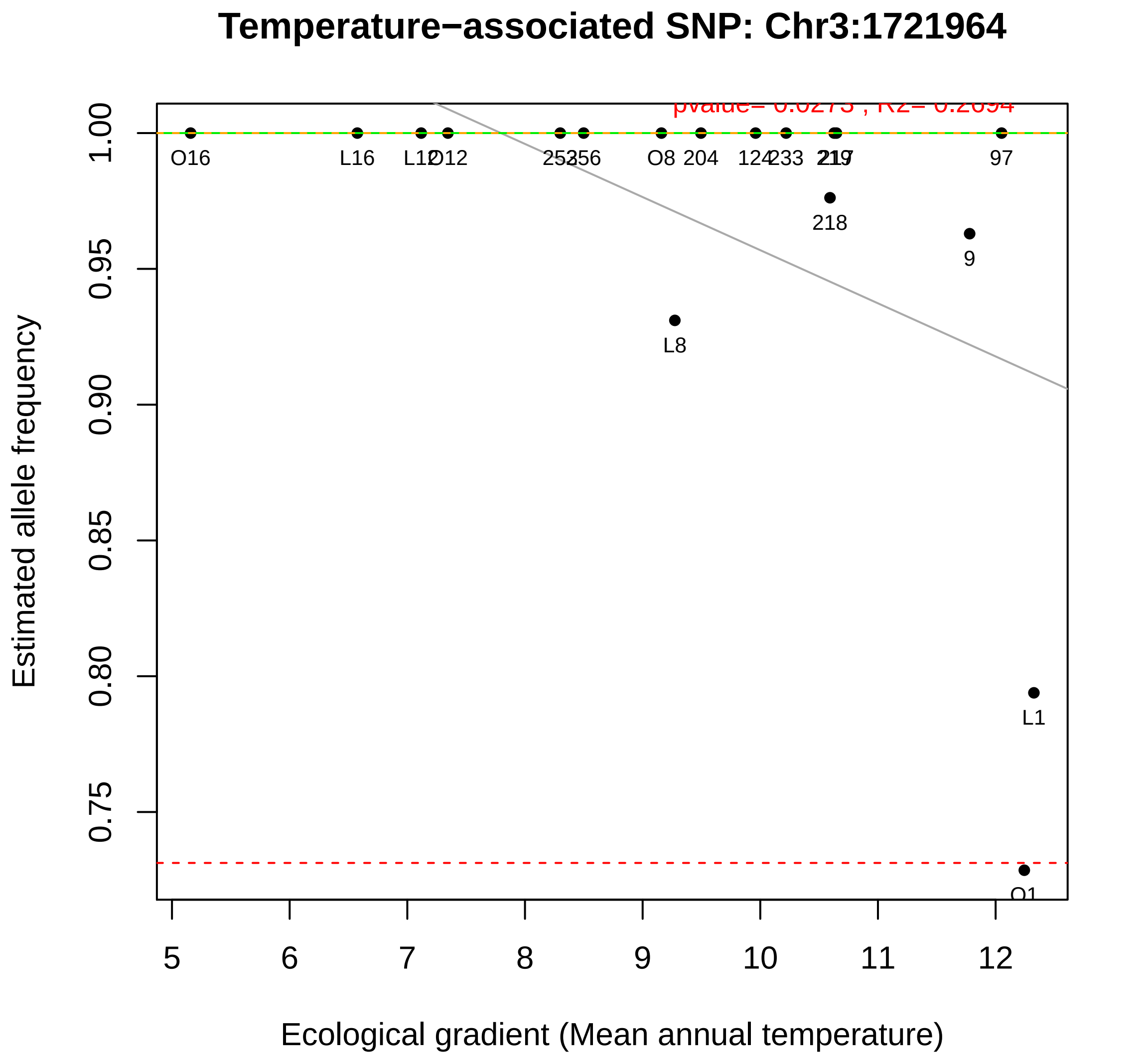

Temperature−associated SNP: Chr3:1721964
pvalue= 0.0273 , R2= 0.2094
Estimated allele frequency
1.00
0.95
0.90
0.85
0.80
0.75
O16
L16
L12
O12
253
256
O8
204
124
233
217
218
9
97
L8
L1
O1
5
6
7
8
9
10
11
12
Ecological gradient (Mean annual temperature)

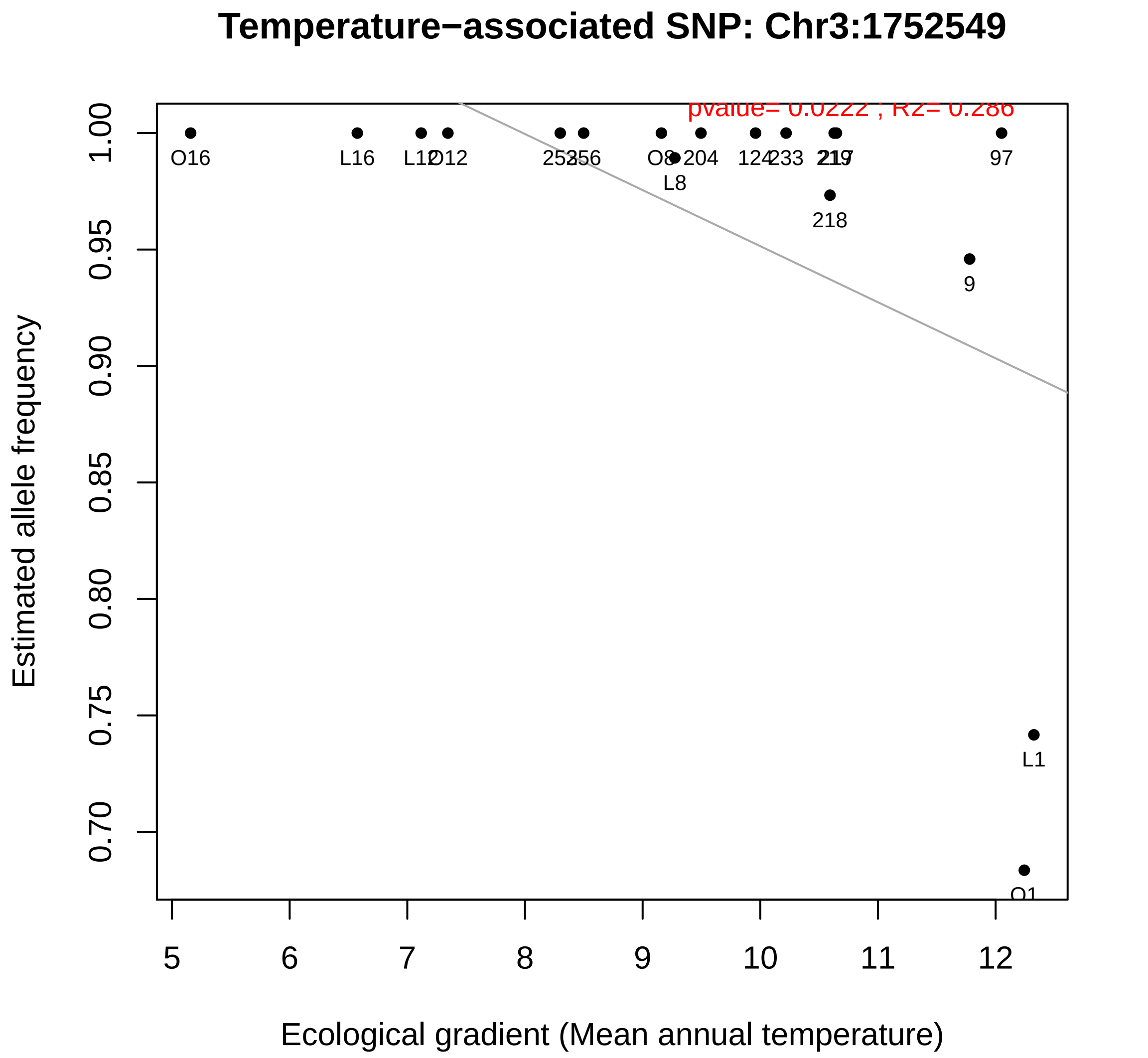

Temperature−associated SNP: Chr3:1752549
pvalue= 0.0222 ; R2= 0.286
Estimated allele frequency
Ecological gradient (Mean annual temperature)
1.00
0.95
0.90
0.85
0.80
0.75
0.70
5
6
7
8
9
10
11
12
O16
L16
L12
O12
O8
L8
204
124
233
218
9
97
L1
O1

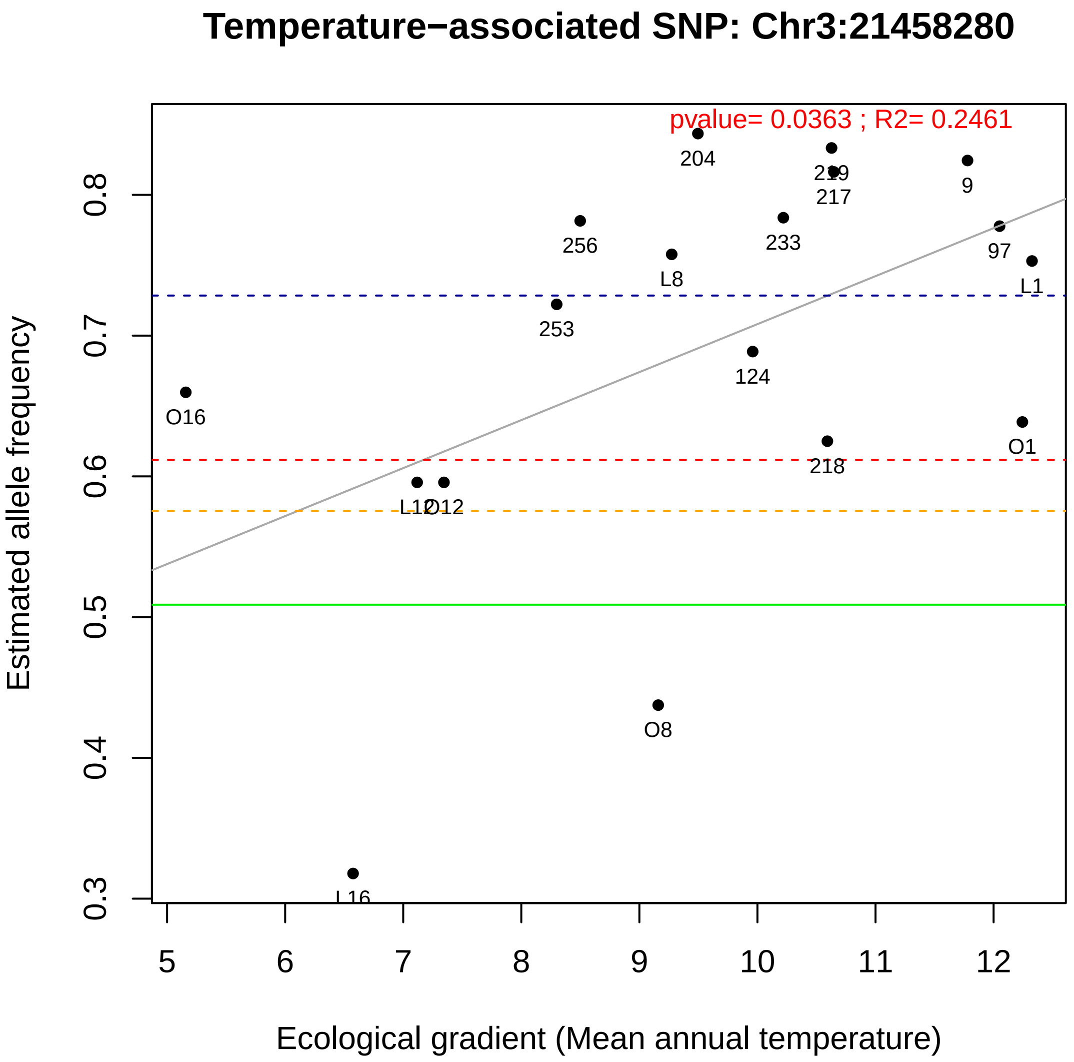

Temperature−associated SNP: Chr3:21458280
pvalue= 0.0363 ; R2= 0.2461
Estimated allele frequency
0.3
0.4
0.5
0.6
0.7
0.8
5
6
7
8
9
10
11
12
Ecological gradient (Mean annual temperature)
204
219
217
9
256
233
97
L8
L1
253
124
O16
O1
218
L12
O12
O8
L16

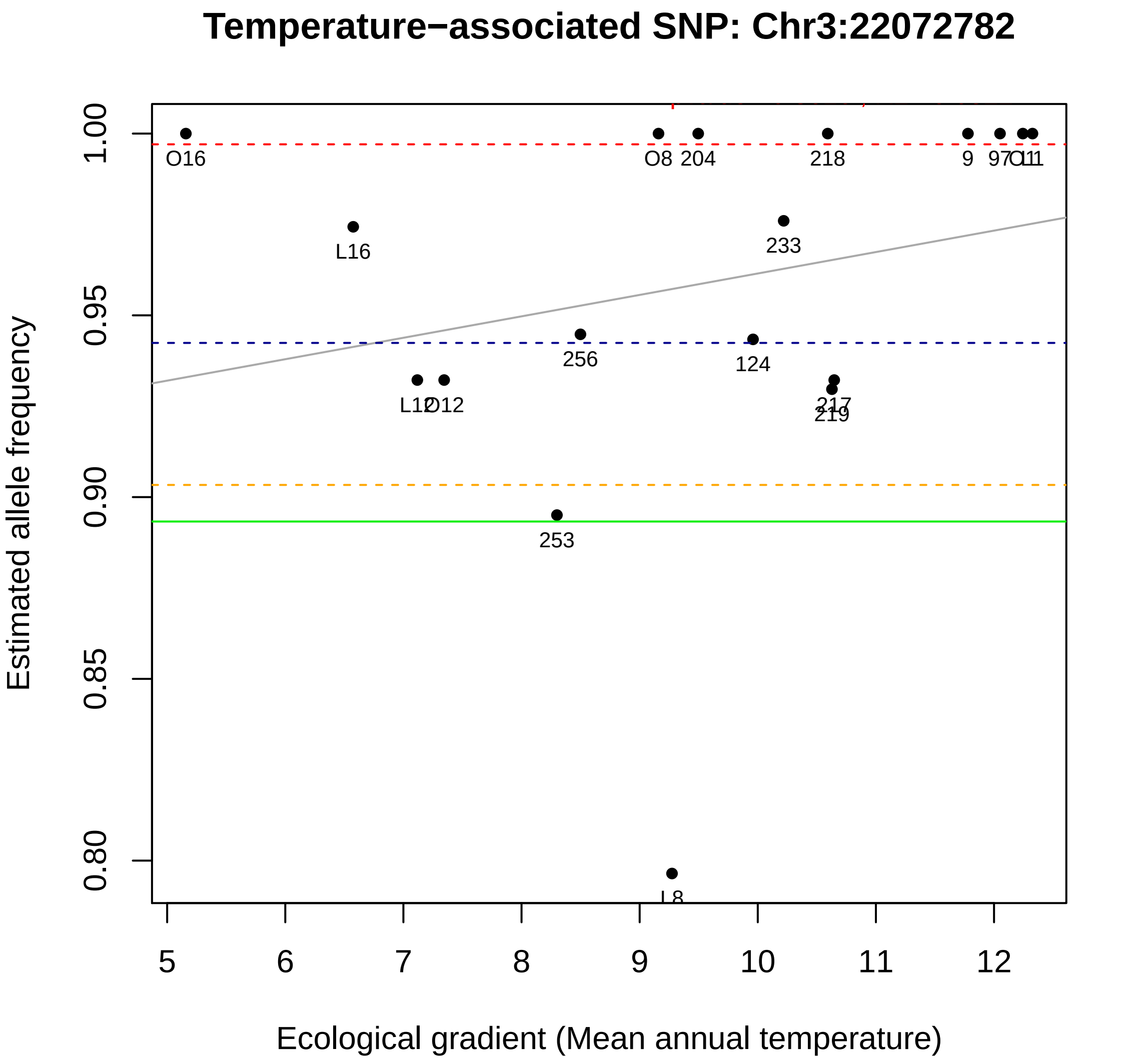

Temperature−associated SNP: Chr3:22072782
Estimated allele frequency
Ecological gradient (Mean annual temperature)
1.00
0.95
0.90
0.85
0.80
5
6
7
8
9
10
11
12
O16
O8
204
218
9
97
O11
L16
233
256
124
L12
O12
217
219
253
L8

**Temperature−associated SNP: Chr3:22630387**

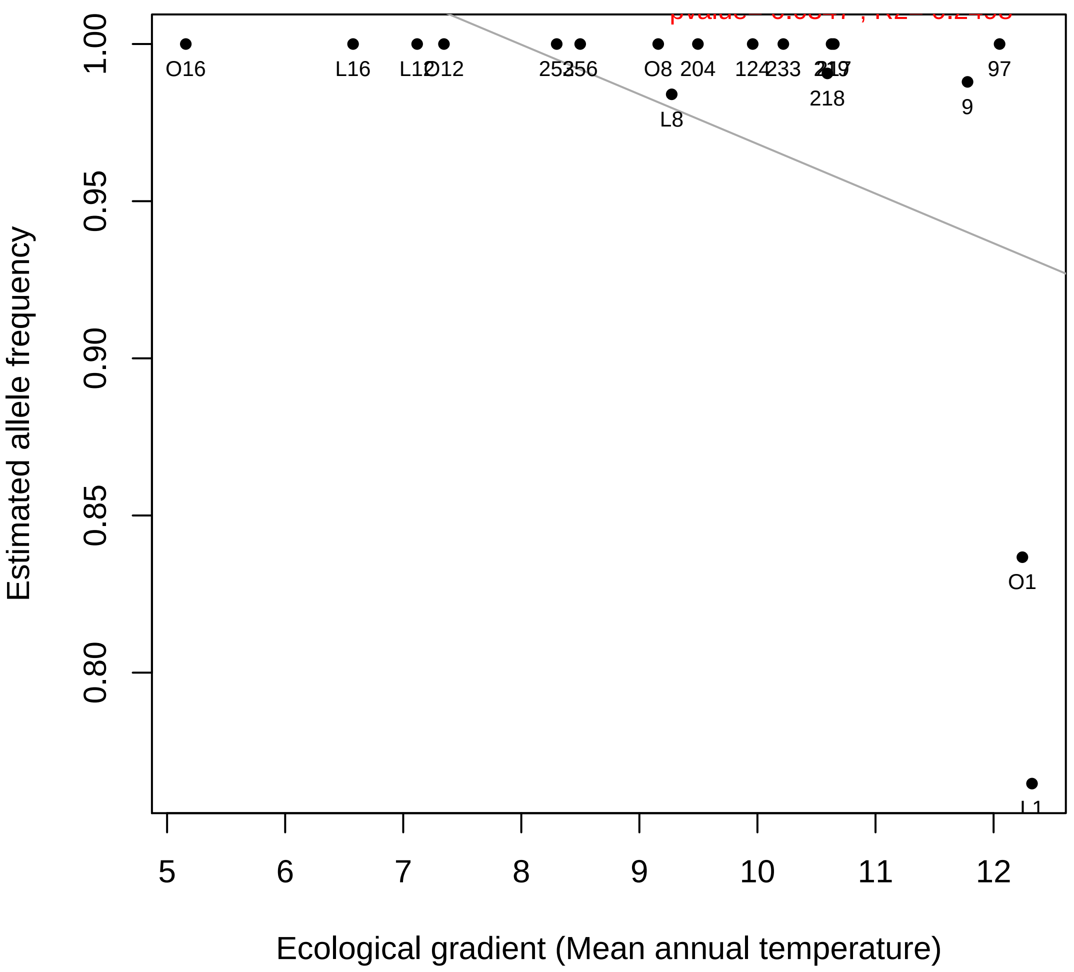

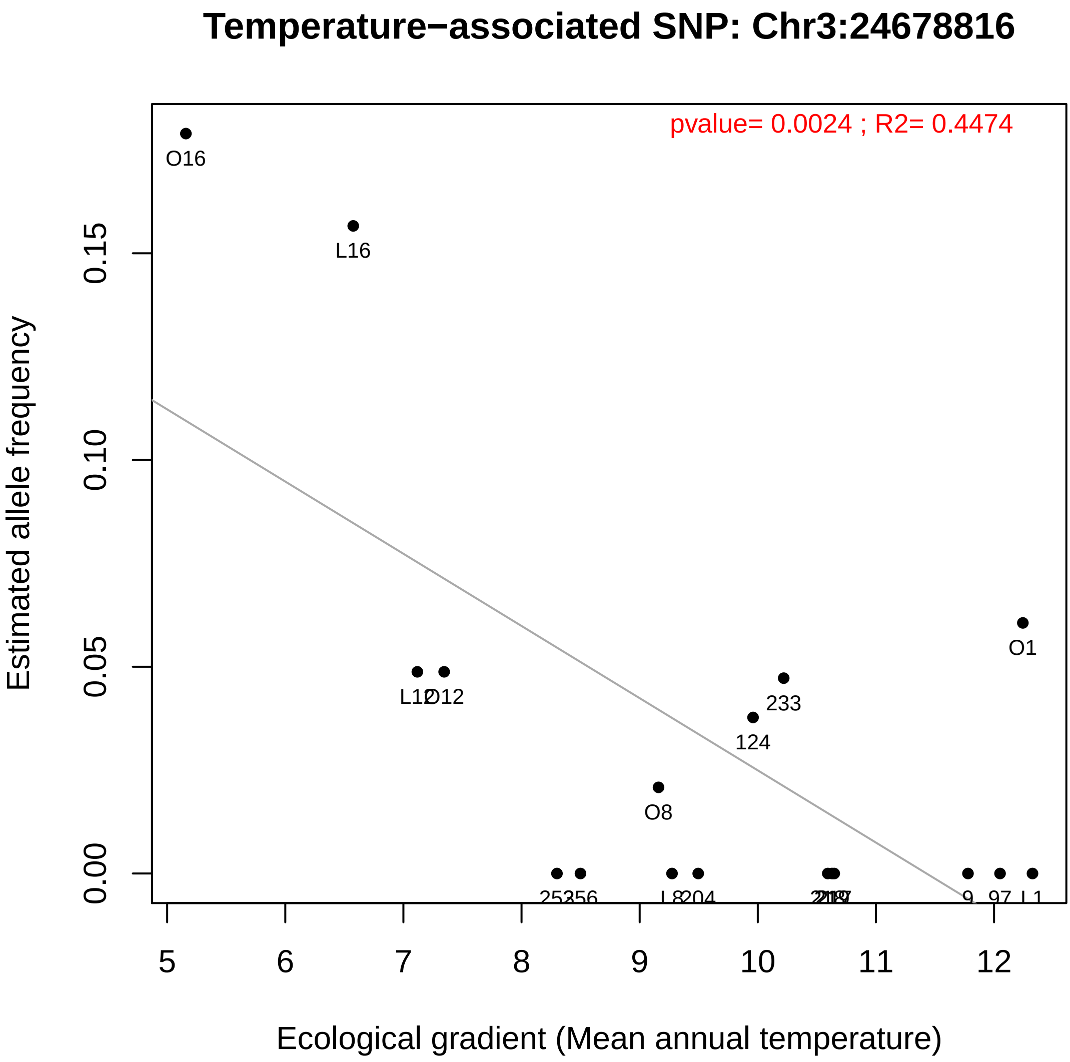

Temperature−associated SNP: Chr3:24678816
pvalue= 0.0024 ; R2= 0.4474
O16
L16
L12
O12
O1
233
124
O8
Estimated allele frequency
0.00
0.05
0.10
0.15
5
6
7
8
9
10
11
12
Ecological gradient (Mean annual temperature)

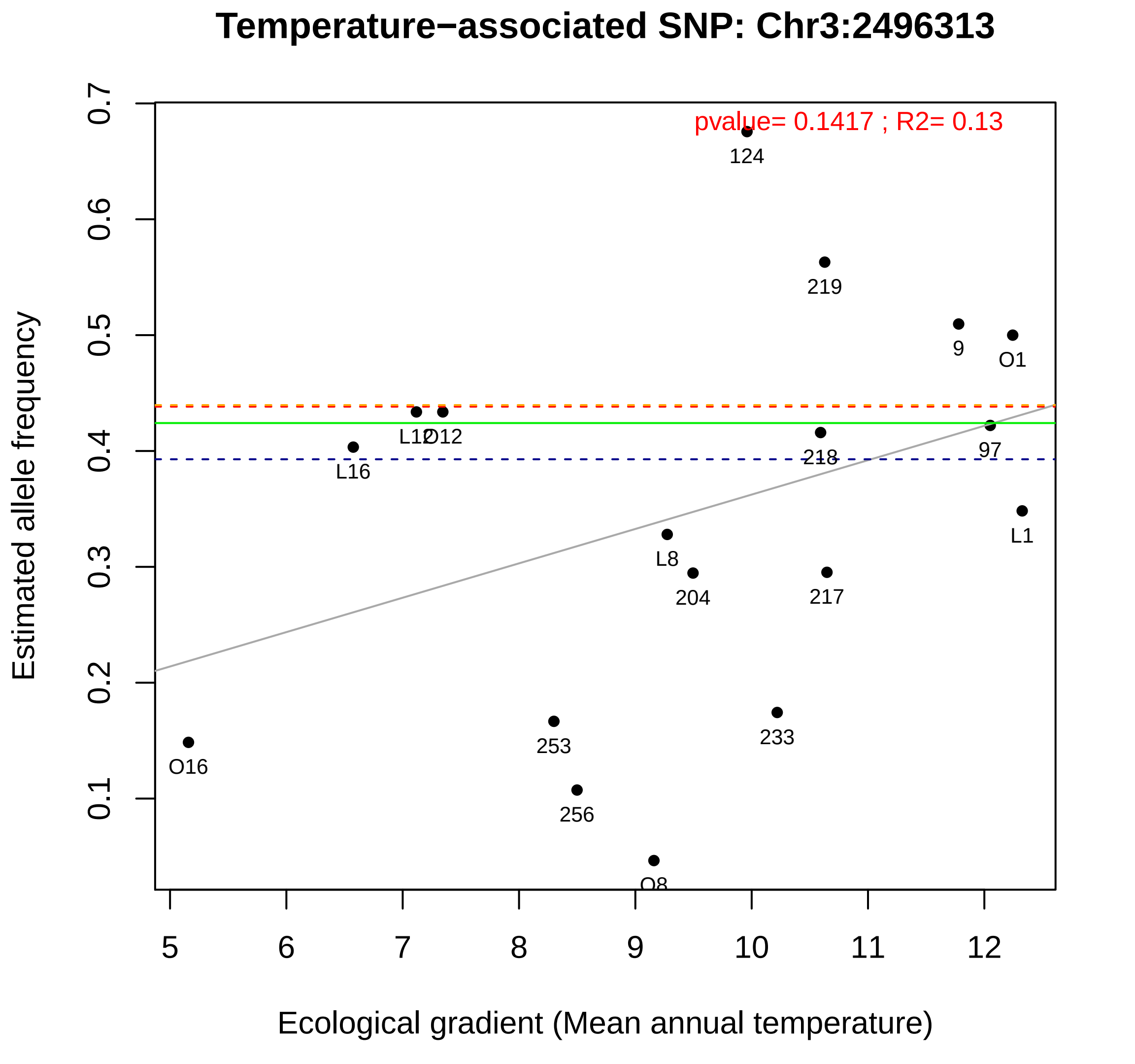
Temperature−associated SNP: Chr3:2496313
pvalue= 0.1417 ; R2= 0.13
Estimated allele frequency
Ecological gradient (Mean annual temperature)
0.1
0.2
0.3
0.4
0.5
0.6
0.7
5
6
7
8
9
10
11
12
124
219
9
O1
L12
O12
218
97
L16
L1
L8
204
217
253
233
O16
256
O8

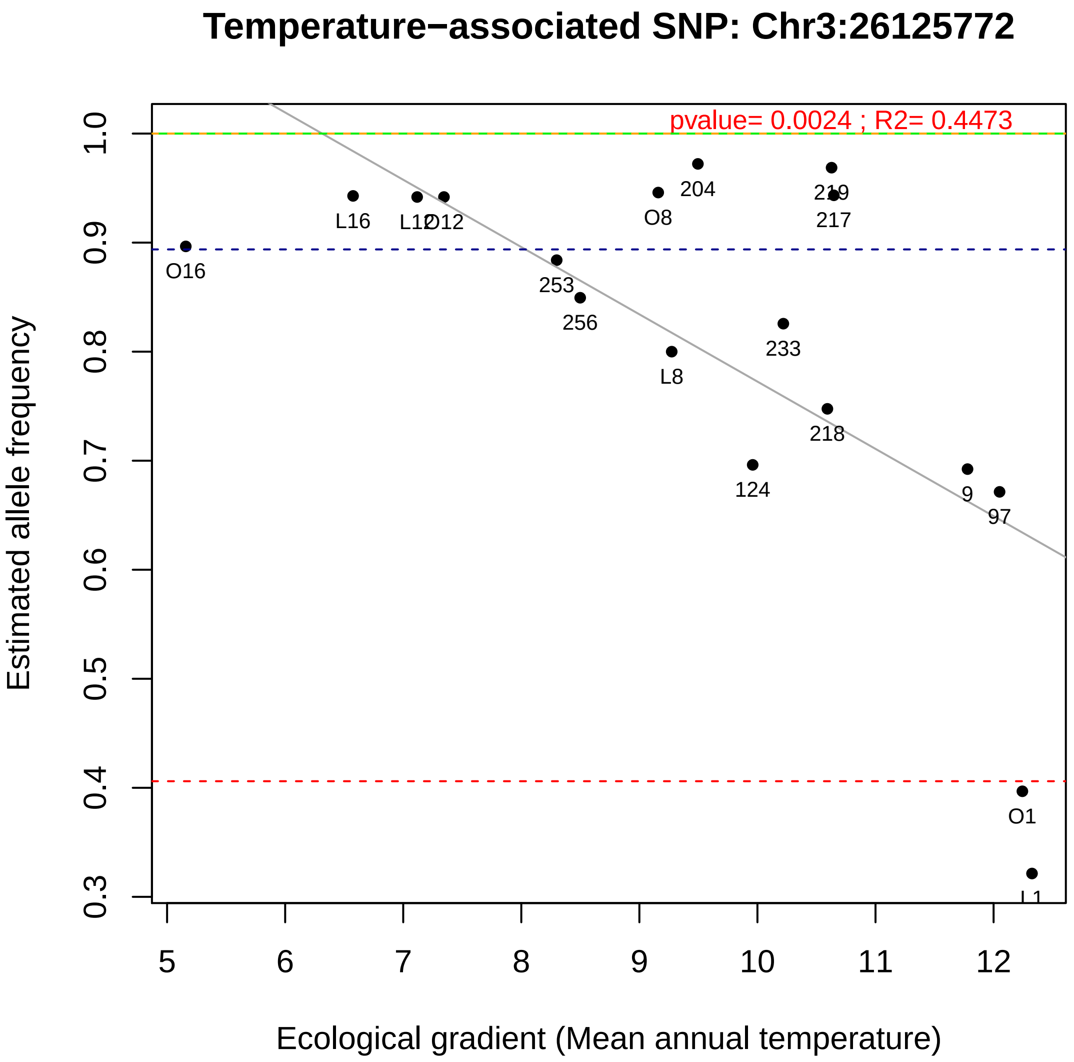
Temperature−associated SNP: Chr3:26125772
pvalue= 0.0024 ; R2= 0.4473
Estimated allele frequency
0.3
0.4
0.5
0.6
0.7
0.8
0.9
1.0
5
6
7
8
9
10
11
12
Ecological gradient (Mean annual temperature)
O16
L16
L12
O12
253
256
O8
204
L8
124
233
218
219
217
9
97
O1

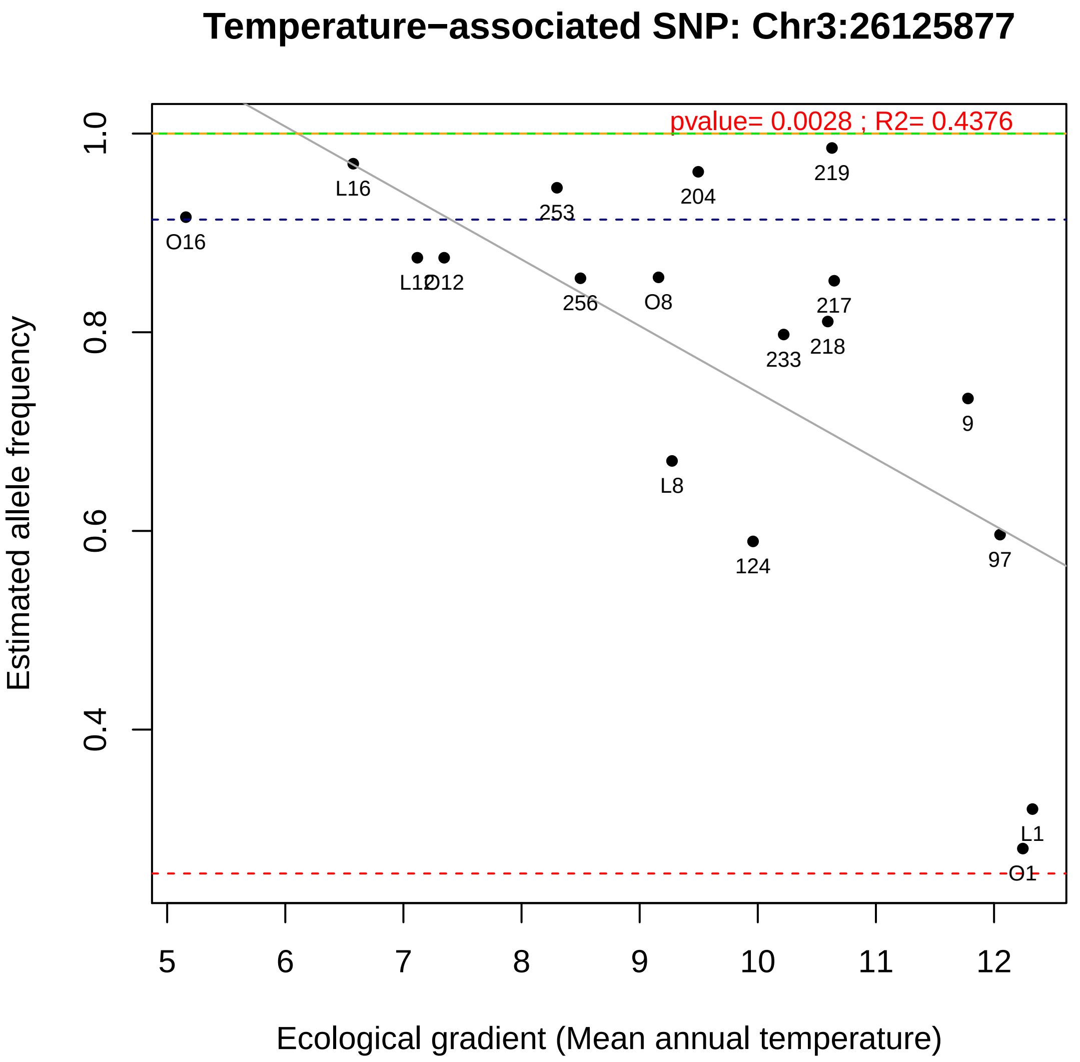

Temperature−associated SNP: Chr3:26125877
pvalue= 0.0028 ; R2= 0.4376
Estimated allele frequency
Ecological gradient (Mean annual temperature)
0.4
0.6
0.8
1.0
5
6
7
8
9
10
11
12
O16
L16
L12
O12
253
256
204
O8
L8
219
217
218
233
124
9
97
L1
O1

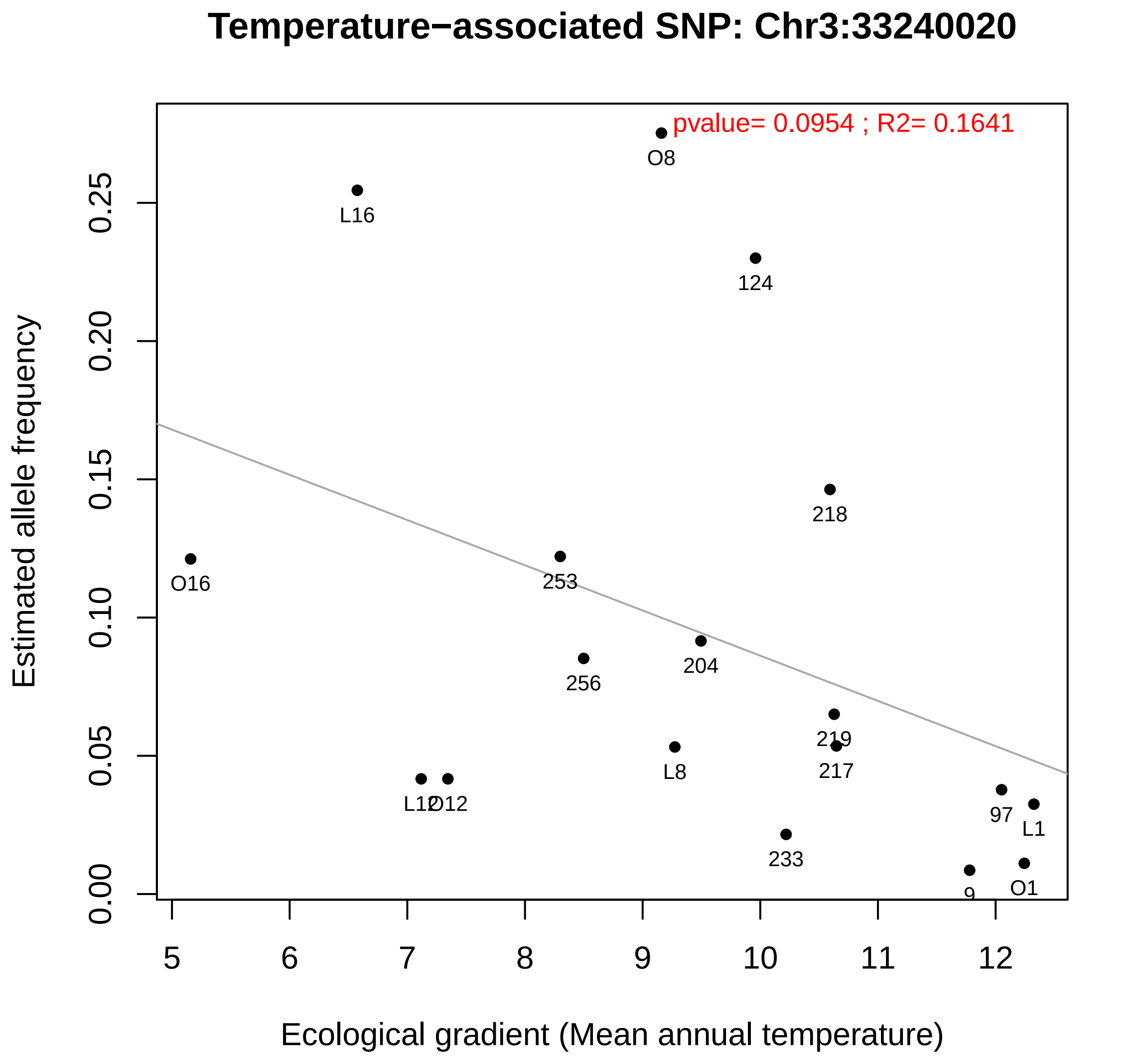
Temperature−associated SNP: Chr3:33240020
pvalue= 0.0954 ; R2= 0.1641
Estimated allele frequency
0.00
0.05
0.10
0.15
0.20
0.25
Ecological gradient (Mean annual temperature)
5
6
7
8
9
10
11
12
O8
L16
124
218
O16
253
204
256
219
217
L8
L12
O12
97
L1
233
9
O1

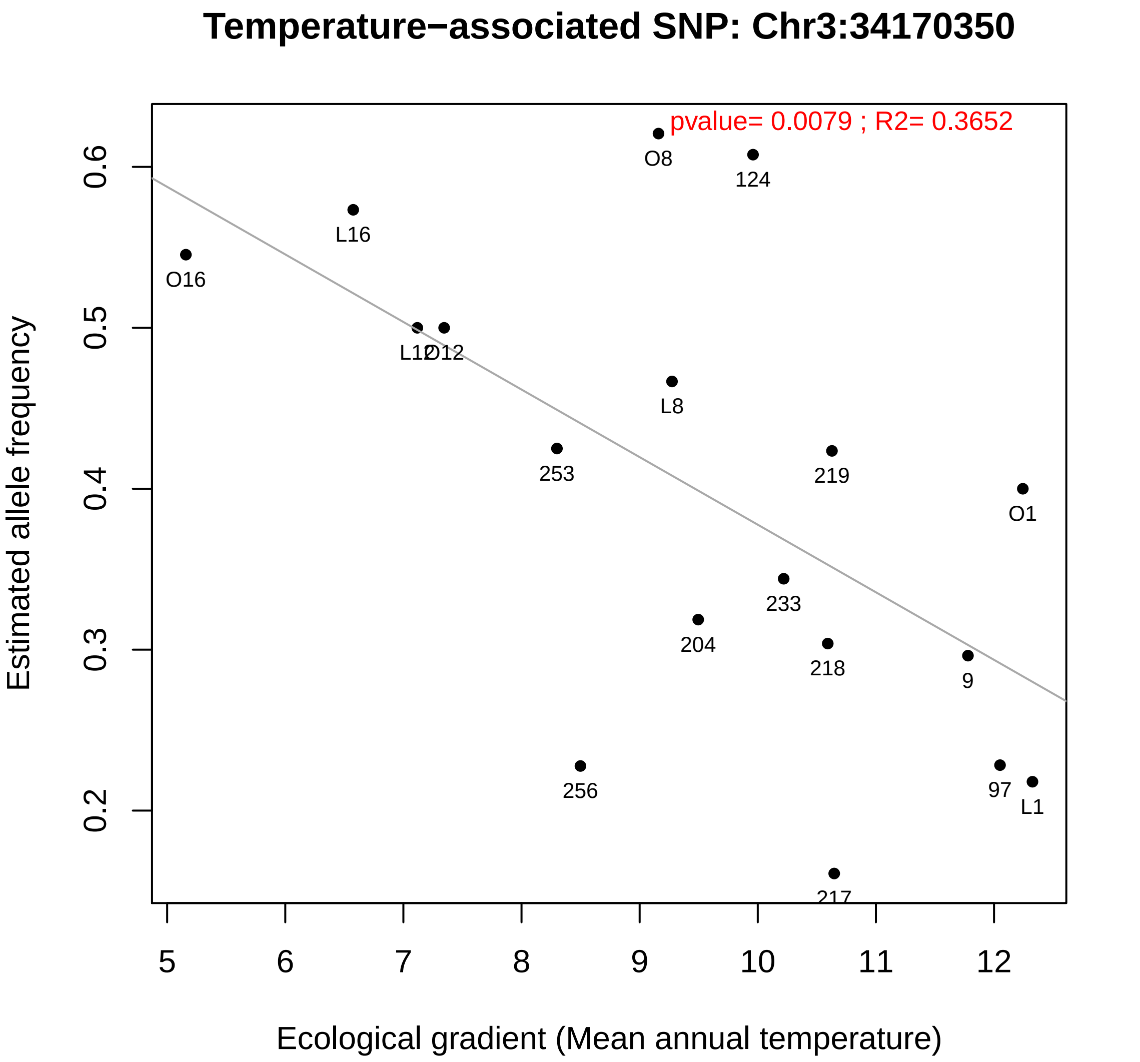
Temperature−associated SNP: Chr3:34170350
pvalue= 0.0079 ; R2= 0.3652
O8
124
L16
O16
L12
O12
L8
253
219
O1
233
204
218
9
256
97
L1
217
0.2
0.3
0.4
0.5
0.6
5
6
7
8
9
10
11
12
Estimated allele frequency
Ecological gradient (Mean annual temperature)

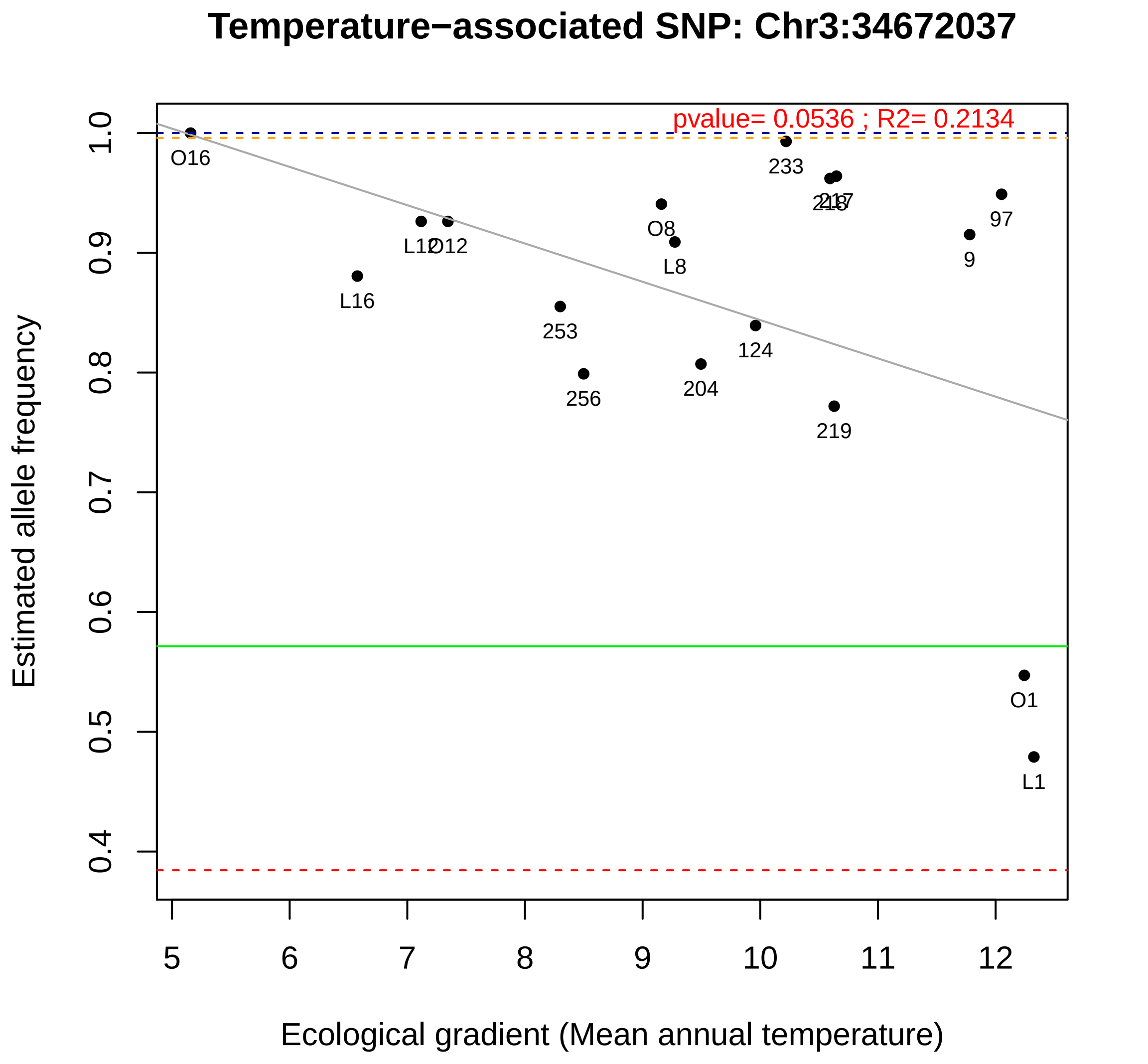

Temperature−associated SNP: Chr3:34672037
pvalue= 0.0536 ; R2= 0.2134
Estimated allele frequency
Ecological gradient (Mean annual temperature)
0.4
0.5
0.6
0.7
0.8
0.9
1.0
5
6
7
8
9
10
11
12
O16
233
217
218
97
O8
L12
O12
L8
9
L16
253
124
256
204
219
O1
L1

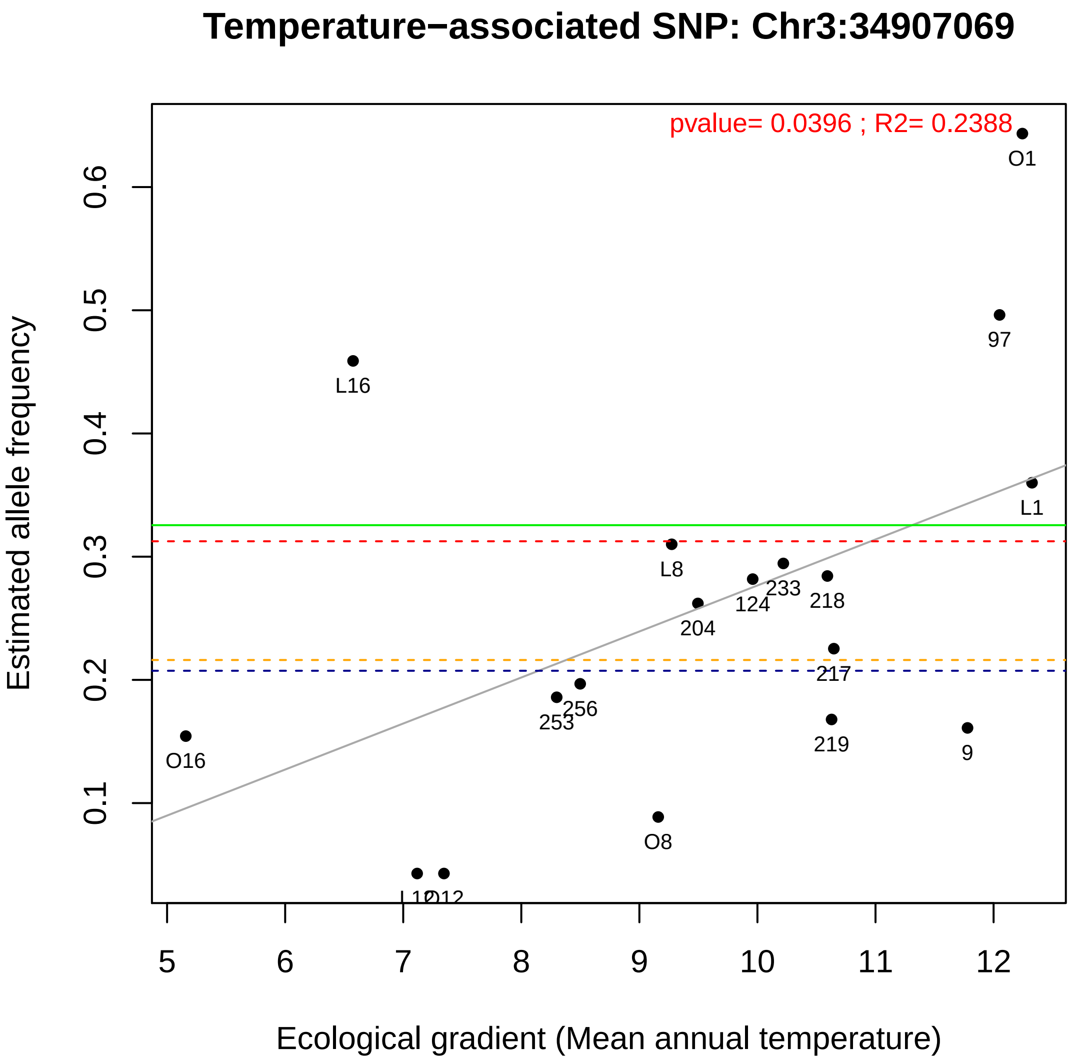

Temperature−associated SNP: Chr3:34907069
pvalue= 0.0396 ; R2= 0.2388
Estimated allele frequency
0.1
0.2
0.3
0.4
0.5
0.6
Ecological gradient (Mean annual temperature)
5
6
7
8
9
10
11
12
O1
97
L16
L1
L8
233
124
218
204
217
256
253
219
9
O16
O8
O12

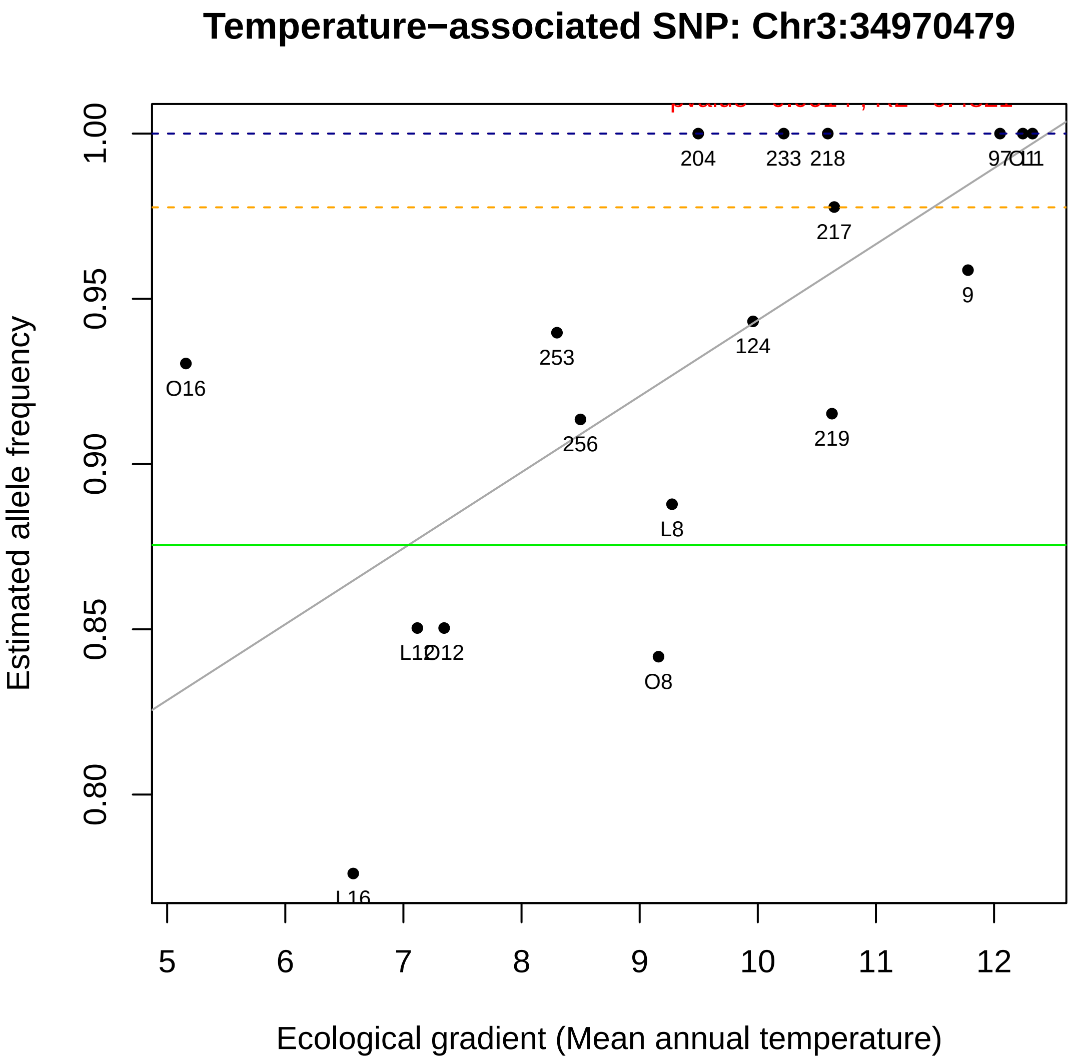
Temperature−associated SNP: Chr3:34970479
Estimated allele frequency
1.00
0.95
0.90
0.85
0.80
5
6
7
8
9
10
11
12
Ecological gradient (Mean annual temperature)
204
233
218
97
O11
217
9
253
124
O16
256
219
L8
L12
O12
O8
L16

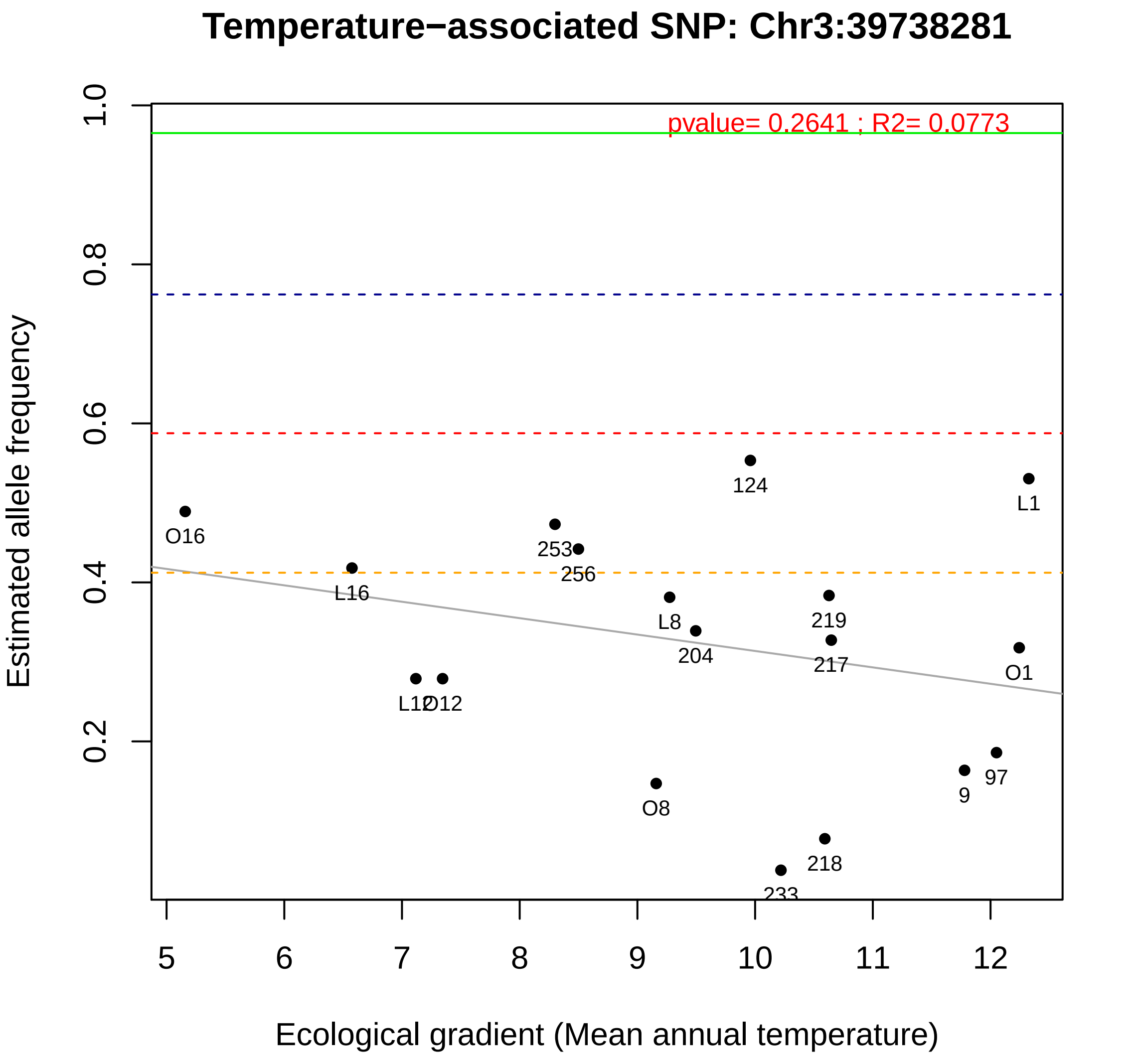

Temperature−associated SNP: Chr3:39738281
pvalue= 0.2641 ; R2= 0.0773
Estimated allele frequency
Ecological gradient (Mean annual temperature)
1.0
0.8
0.6
0.4
0.2
5
6
7
8
9
10
11
12
O16
L16
L12
O12
253
256
124
L8
204
O8
233
218
219
217
9
97
O1
L1

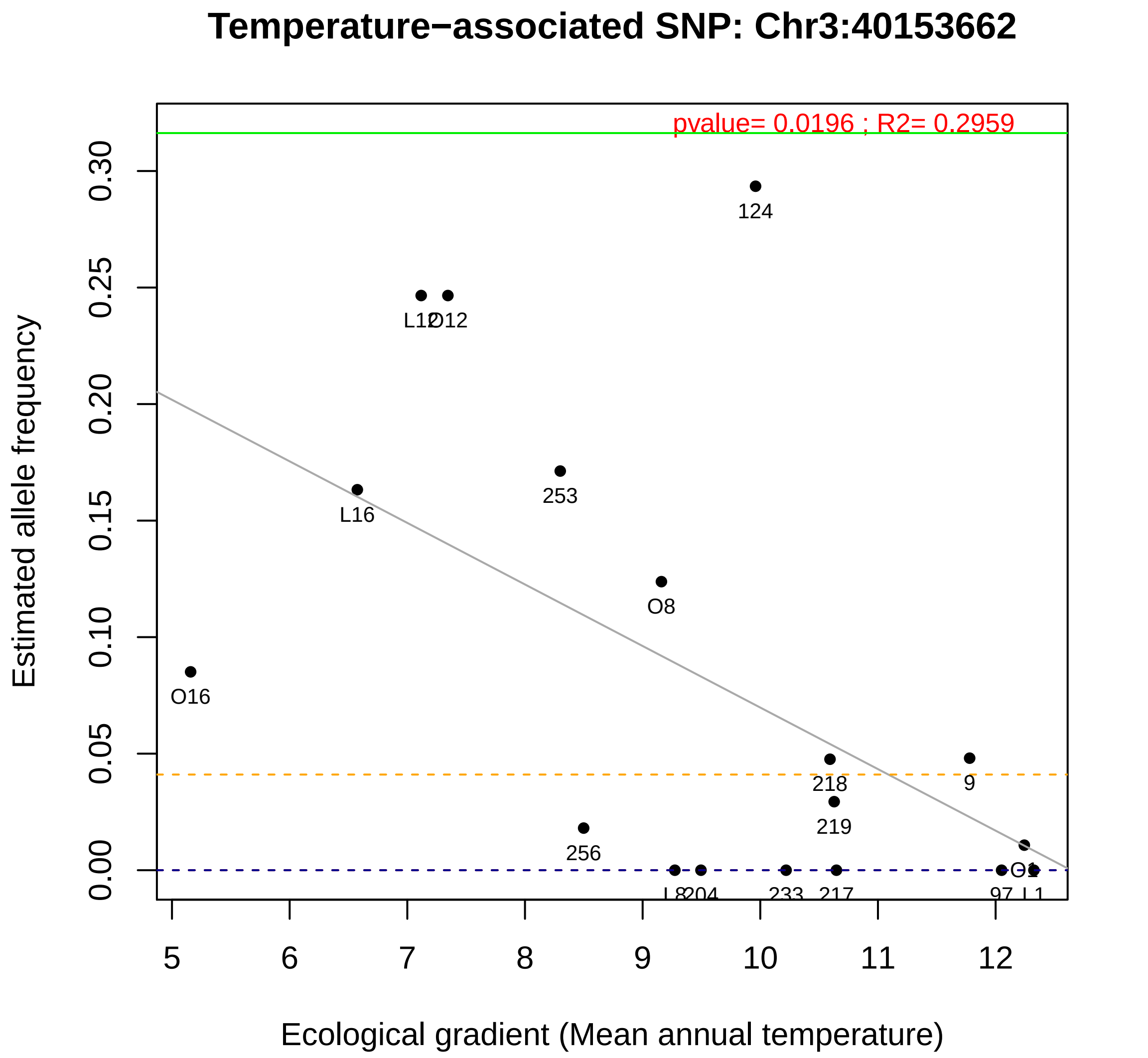

Temperature−associated SNP: Chr3:40153662
pvalue= 0.0196 ; R2= 0.2959
Estimated allele frequency
0.00
0.05
0.10
0.15
0.20
0.25
0.30
5
6
7
8
9
10
11
12
Ecological gradient (Mean annual temperature)
124
L12
O12
253
L16
O8
O16
218
9
219
256
L8
204
233
217
97
O1
L1

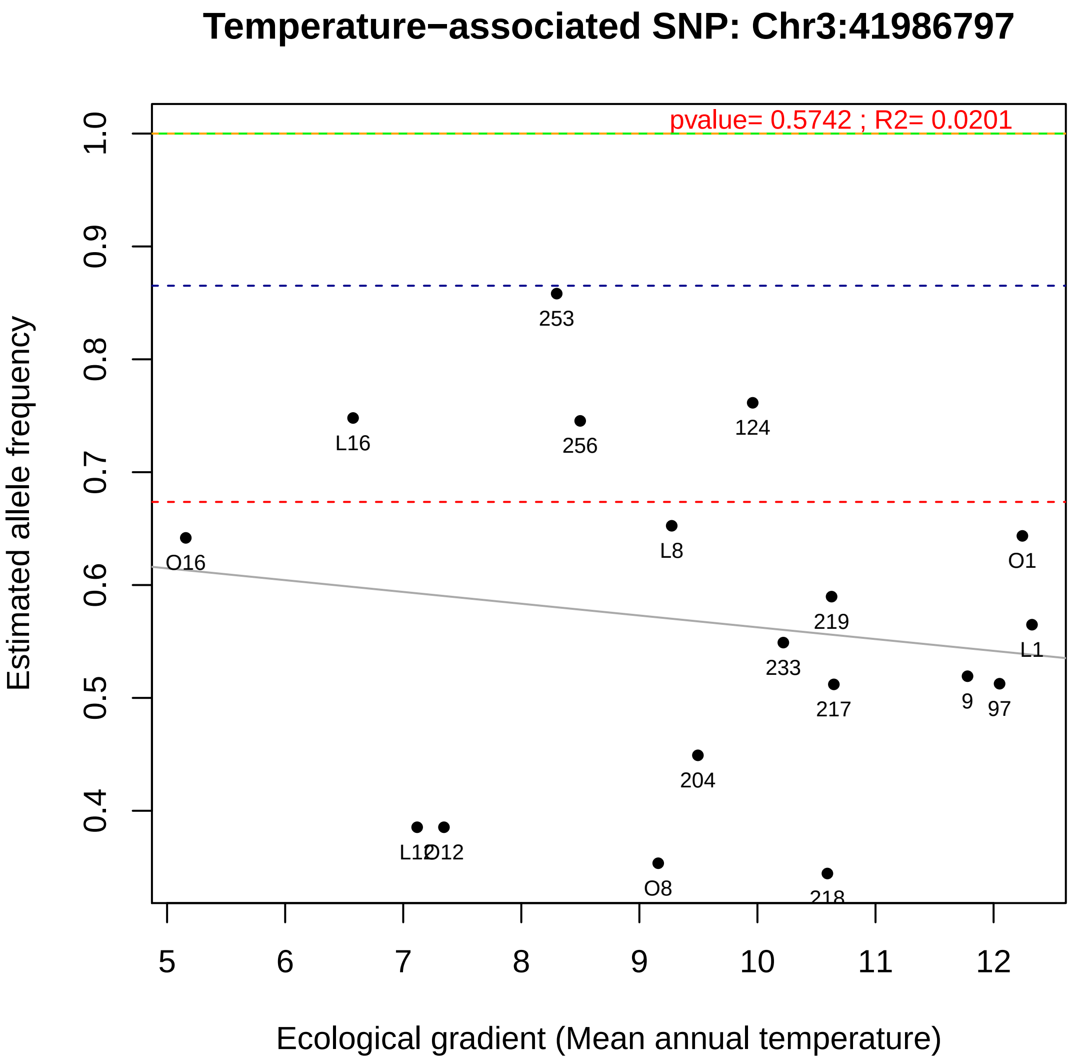

Temperature−associated SNP: Chr3:41986797
pvalue= 0.5742 ; R2= 0.0201
Estimated allele frequency
Ecological gradient (Mean annual temperature)
1.0
0.9
0.8
0.7
0.6
0.5
0.4
5
6
7
8
9
10
11
12
253
L16
256
124
O16
L8
O1
219
L1
233
217
9
97
204
L12
O12
O8

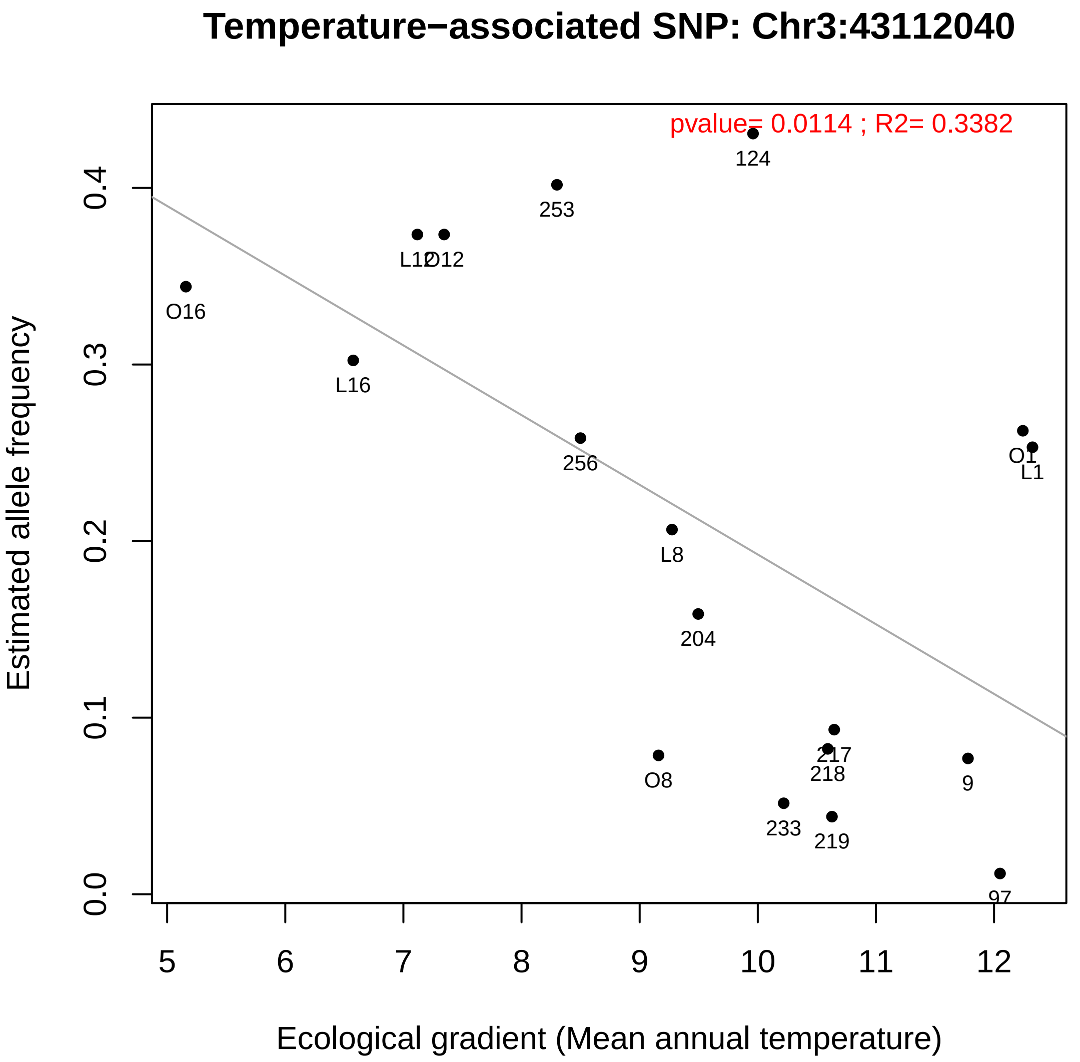
Temperature−associated SNP: Chr3:43112040
pvalue= 0.0114 ; R2= 0.3382
124
253
L12
O12
O16
L16
256
O1
L1
L8
204
217
218
O8
9
233
219
97
0.0
0.1
0.2
0.3
0.4
Estimated allele frequency
5
6
7
8
9
10
11
12
Ecological gradient (Mean annual temperature)

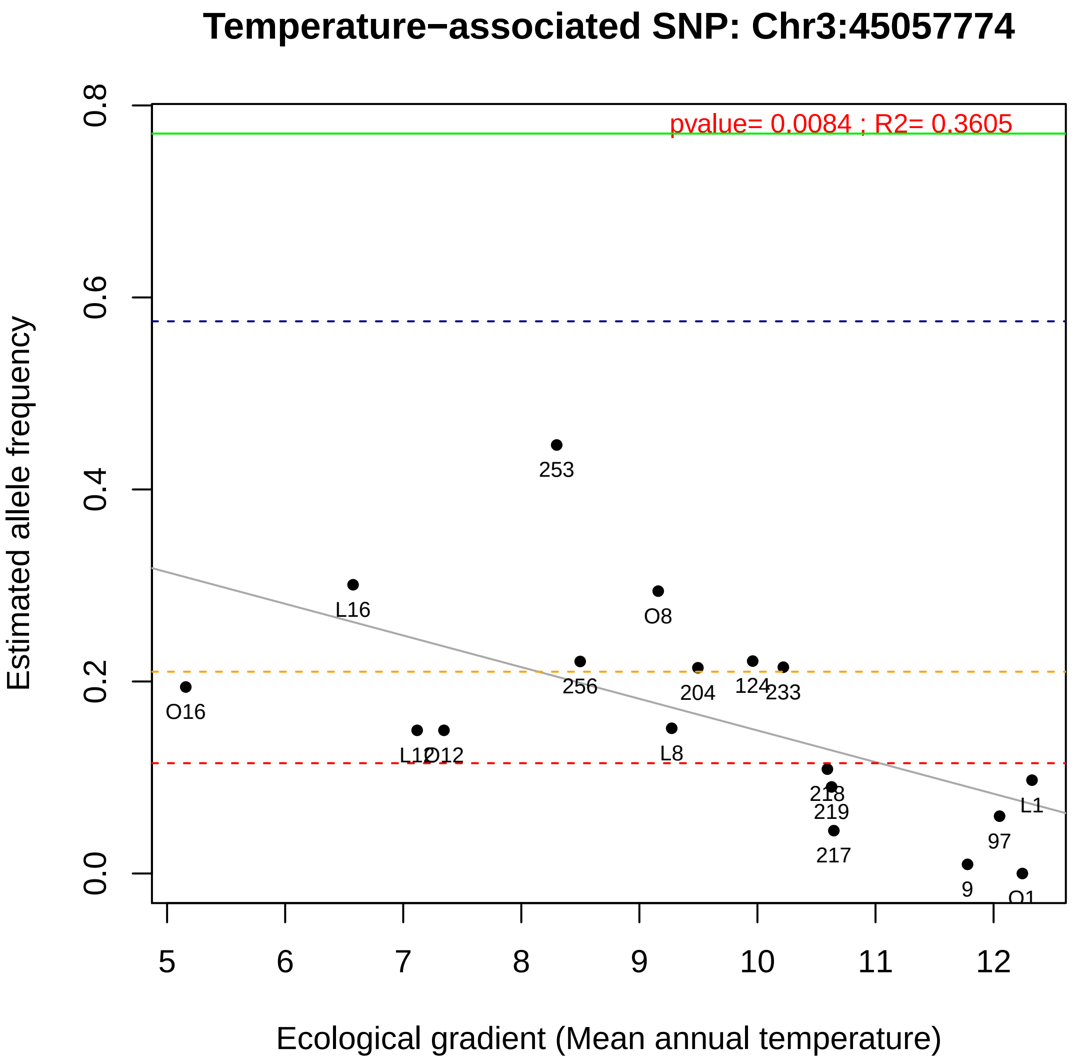
Temperature−associated SNP: Chr3:45057774
pvalue= 0.0084 ; R2= 0.3605
Estimated allele frequency
0.0
0.2
0.4
0.6
0.8
Ecological gradient (Mean annual temperature)
5
6
7
8
9
10
11
12
253
L16
O8
256
204
124
233
O16
L12
O12
L8
218
219
L1
97
217
9
O1

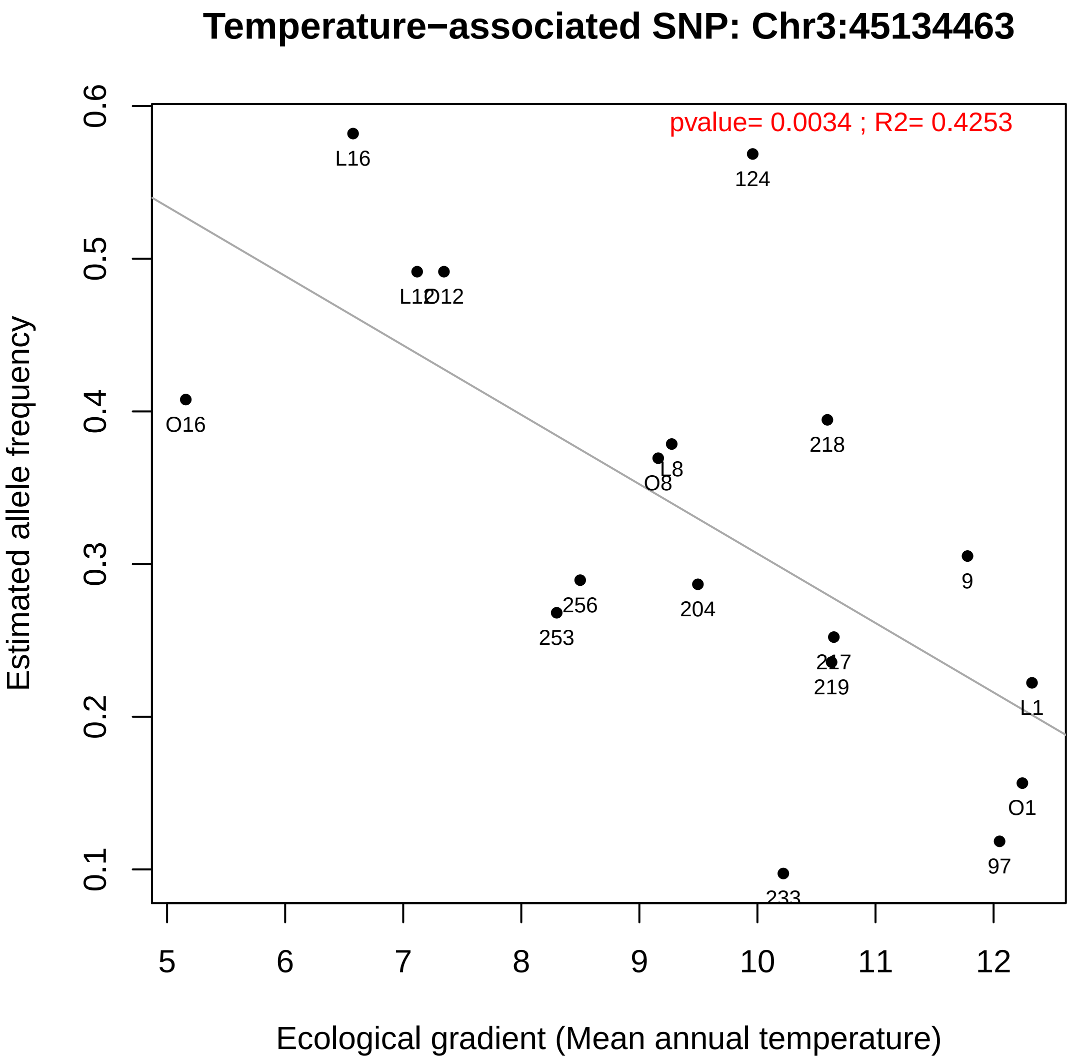
Temperature−associated SNP: Chr3:45134463
pvalue= 0.0034 ; R2= 0.4253
Estimated allele frequency
Ecological gradient (Mean annual temperature)
0.1
0.2
0.3
0.4
0.5
0.6
5
6
7
8
9
10
11
12
L16
124
L12
O12
O16
218
L8
O8
9
256
204
253
217
219
L1
O1
97
233

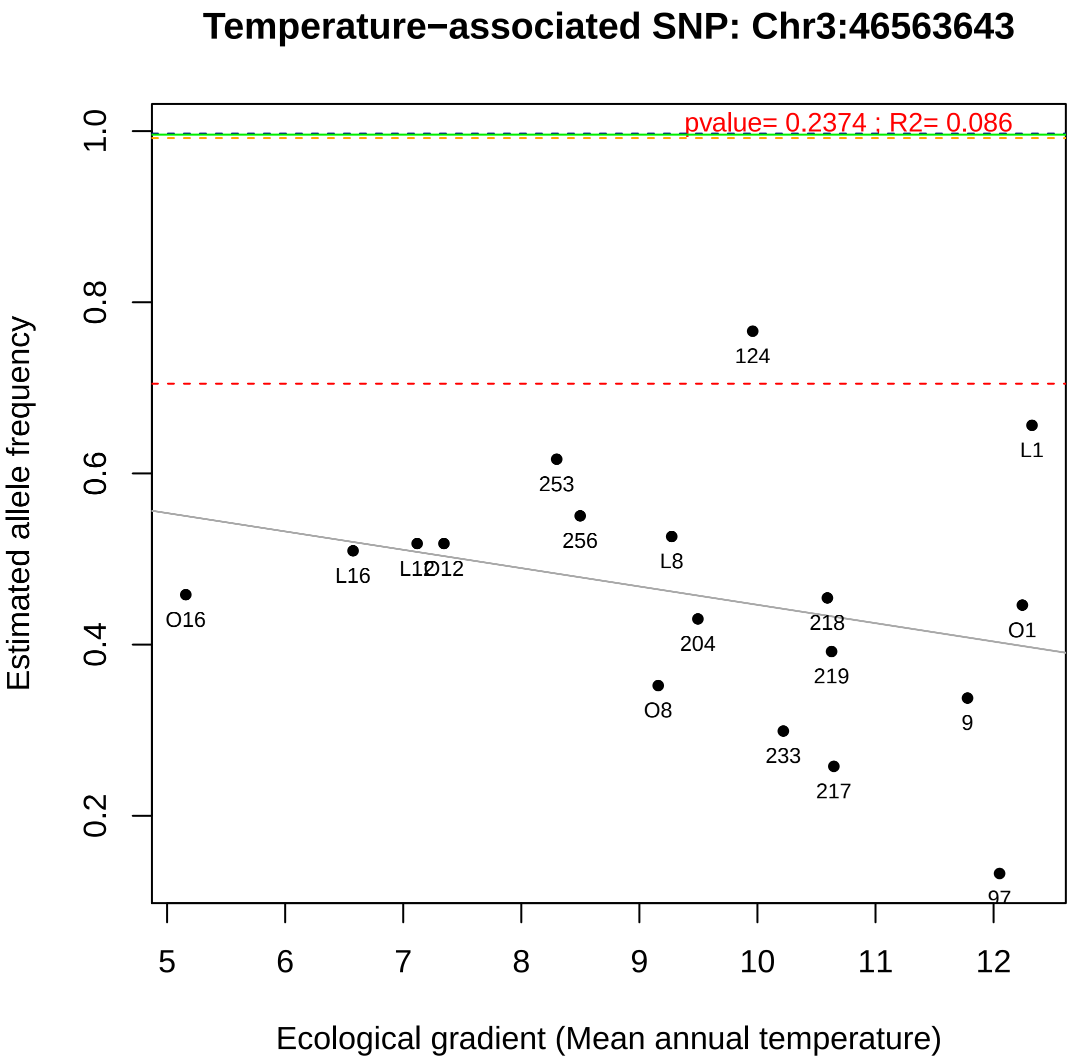

Temperature−associated SNP: Chr3:46563643
pvalue= 0.2374 ; R2= 0.086
Estimated allele frequency
0.2
0.4
0.6
0.8
1.0
124
L1
253
256
L8
L16
L12
O12
O16
218
O1
204
219
O8
9
233
217
97
5
6
7
8
9
10
11
12
Ecological gradient (Mean annual temperature)

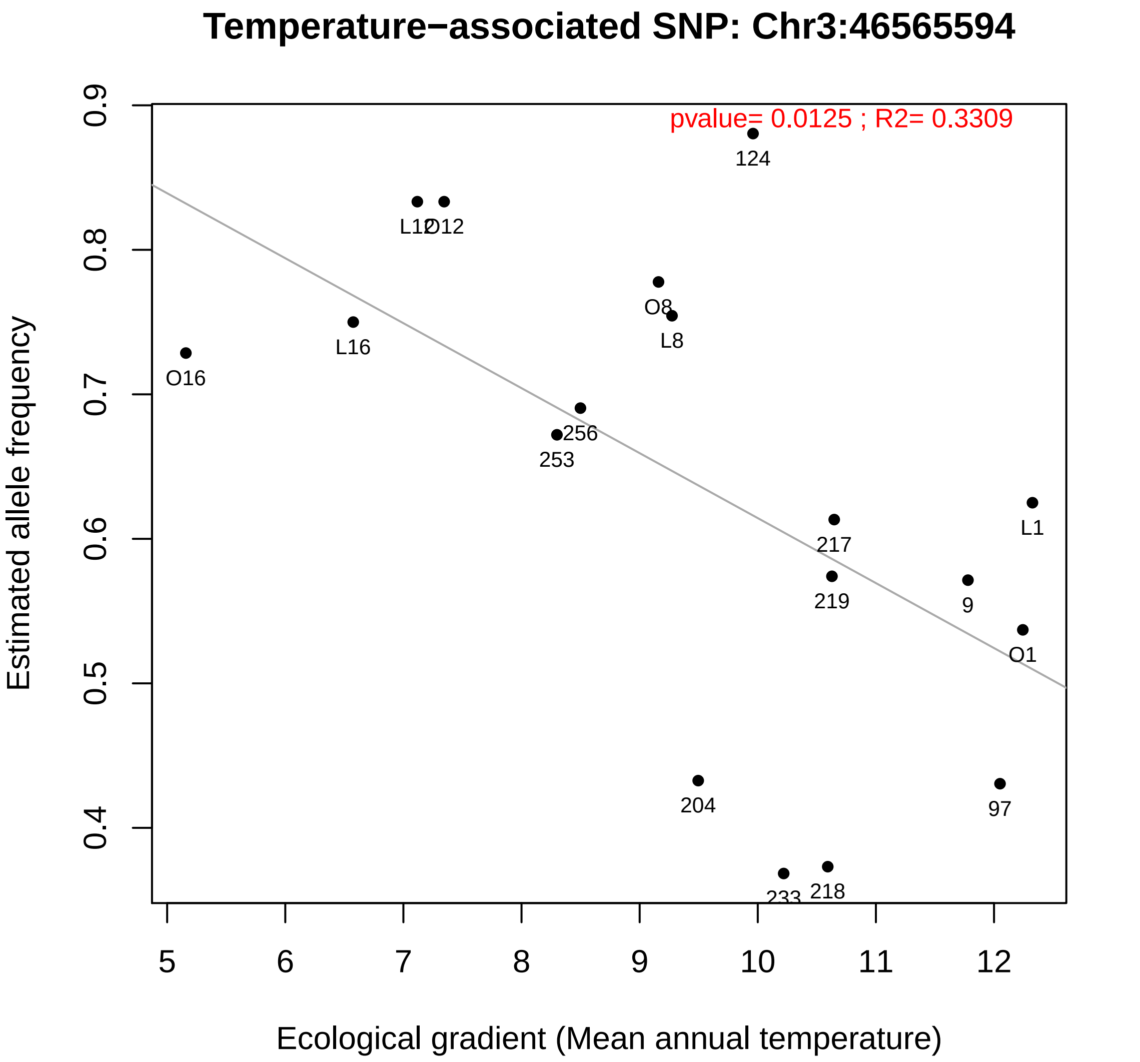

Temperature−associated SNP: Chr3:46565594
pvalue= 0.0125 ; R2= 0.3309
Estimated allele frequency
0.4
0.5
0.6
0.7
0.8
0.9
5
6
7
8
9
10
11
12
Ecological gradient (Mean annual temperature)
124
L12
O12
O8
L8
L16
O16
256
253
217
L1
219
9
O1
204
97
233
218

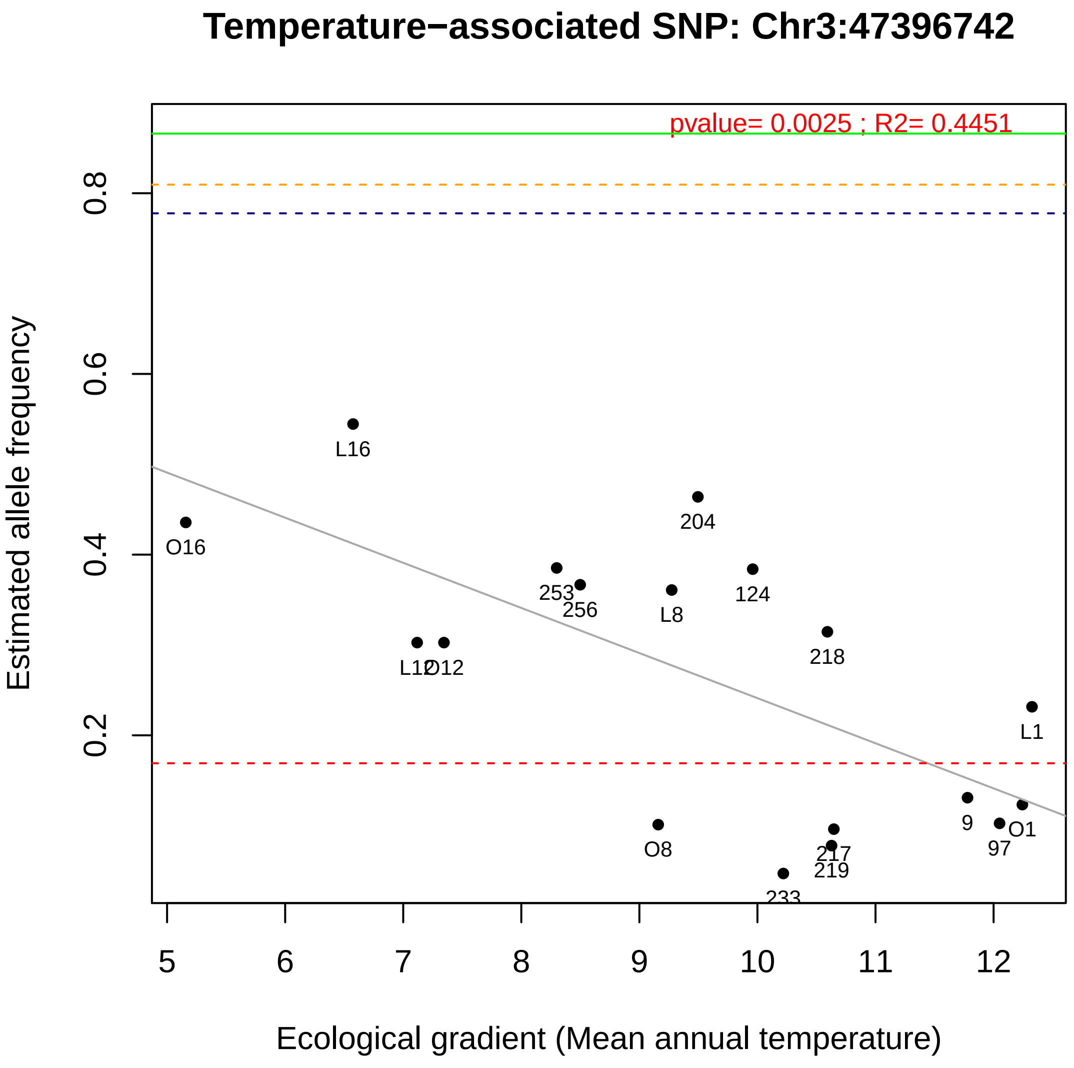
Temperature−associated SNP: Chr3:47396742
pvalue= 0.0025 ; R2= 0.4451
Estimated allele frequency
0.2
0.4
0.6
0.8
Ecological gradient (Mean annual temperature)
5
6
7
8
9
10
11
12
L16
O16
204
253
256
L8
124
L12
O12
218
L1
9
O1
97
O8
217
219
233

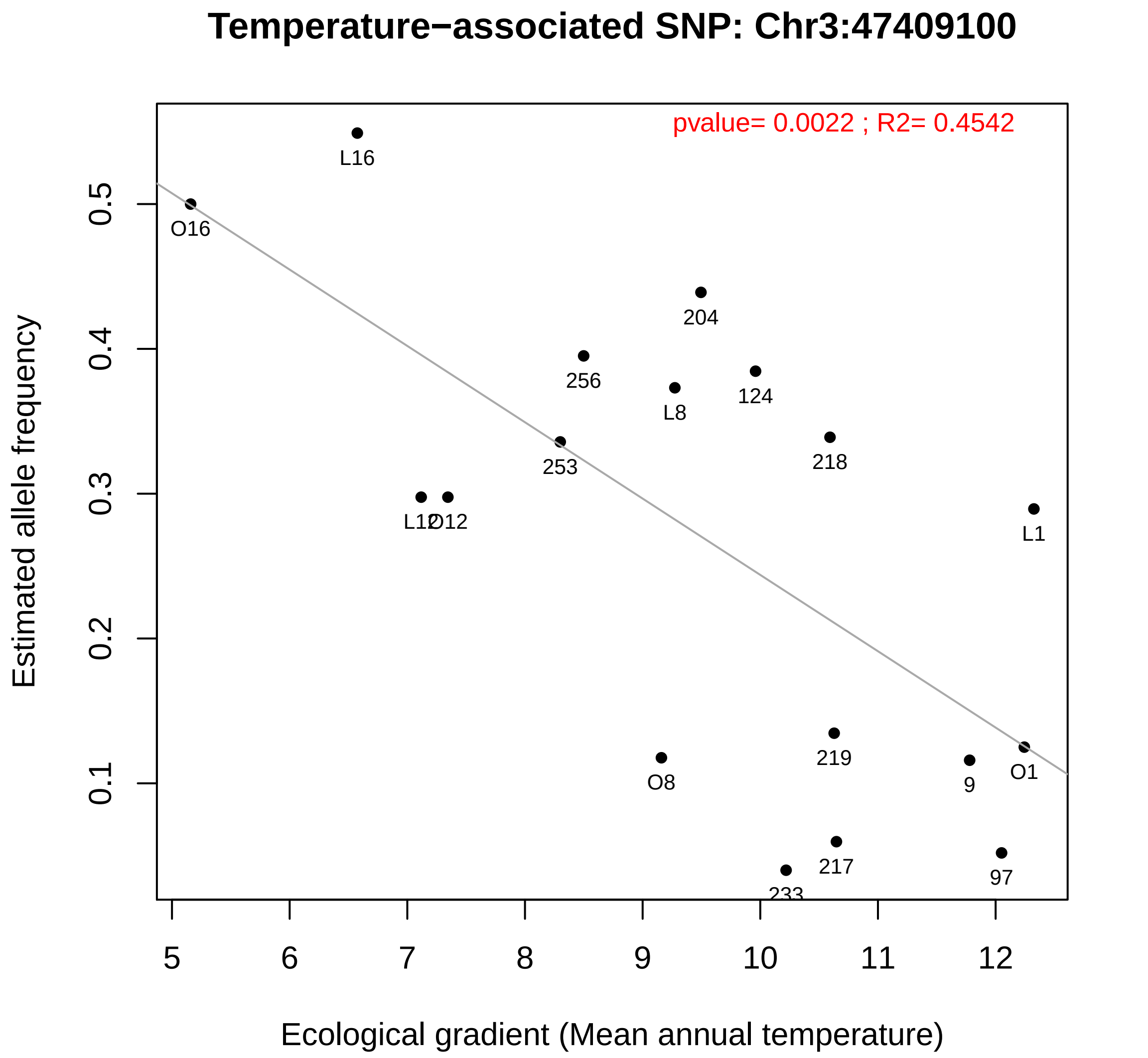
Temperature−associated SNP: Chr3:47409100
pvalue= 0.0022 ; R2= 0.4542
Estimated allele frequency
0.1
0.2
0.3
0.4
0.5
5
6
7
8
9
10
11
12
Ecological gradient (Mean annual temperature)
L16
O16
204
256
124
L8
218
253
L12
O12
L1
219
O1
O8
9
217
97
233

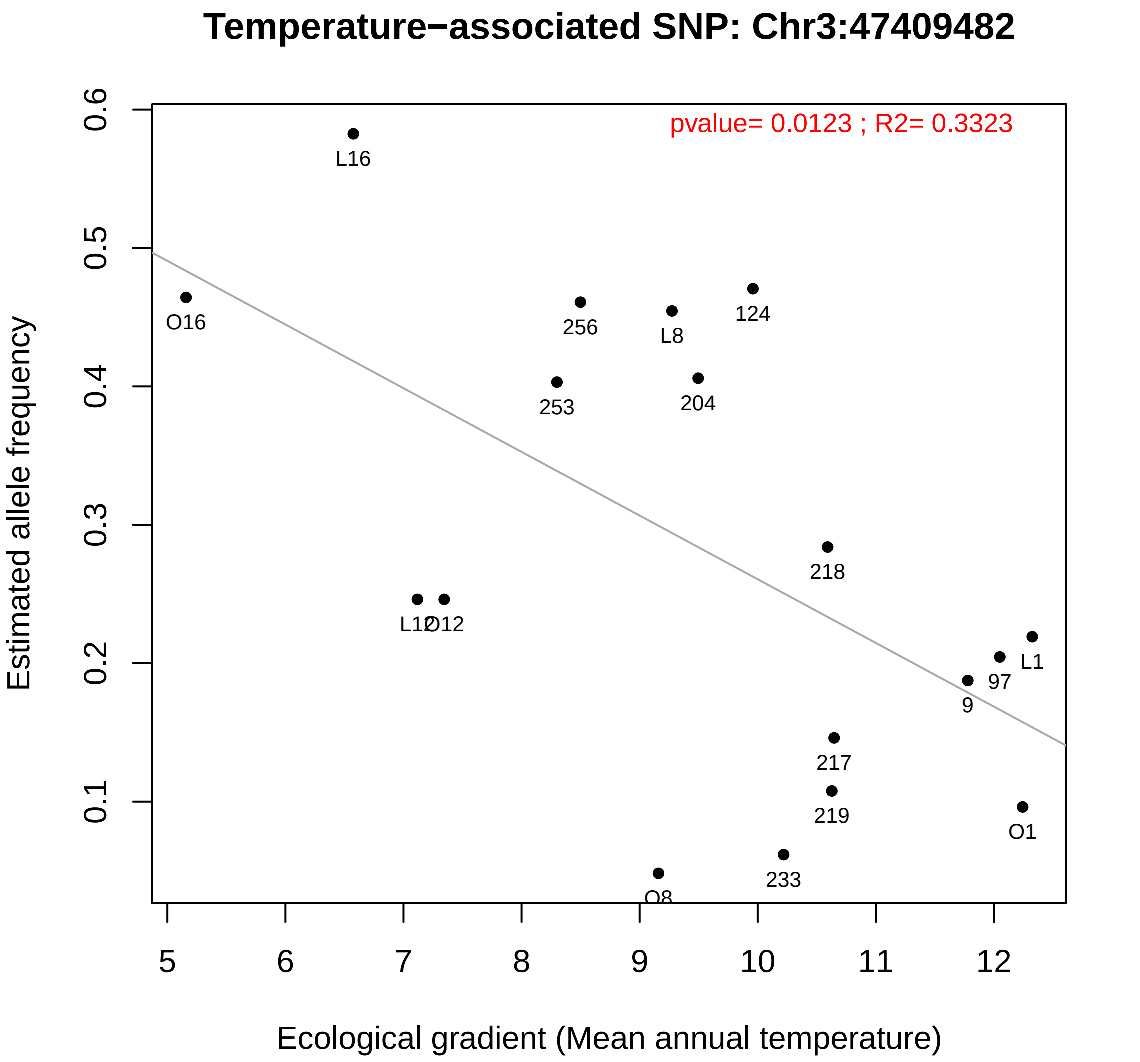
Temperature−associated SNP: Chr3:47409482
pvalue= 0.0123 ; R2= 0.3323
Estimated allele frequency
0.1
0.2
0.3
0.4
0.5
0.6
5
6
7
8
9
10
11
12
Ecological gradient (Mean annual temperature)
L16
O16
256
L8
124
253
204
218
L12
O12
L1
97
9
217
219
O1
233
O8

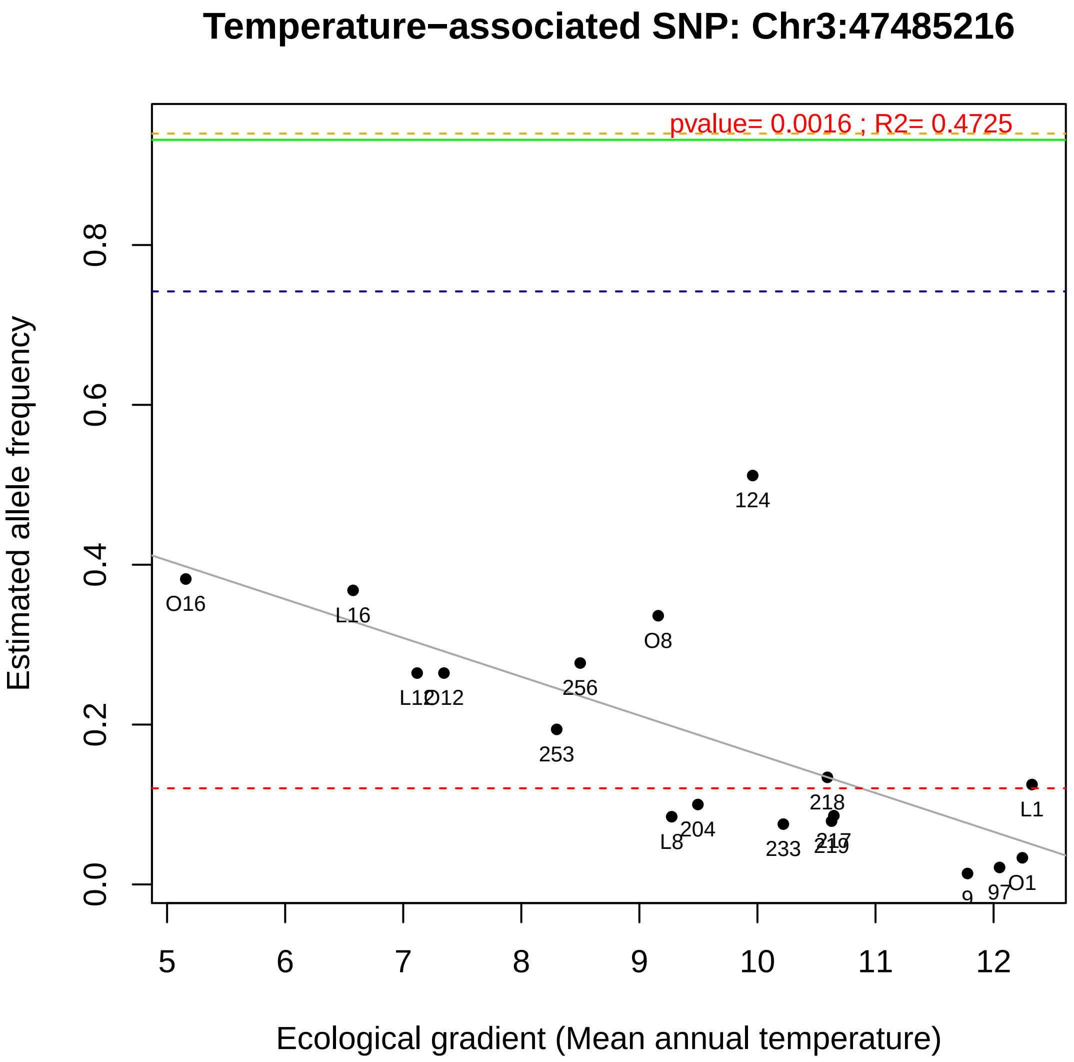

Temperature−associated SNP: Chr3:47485216
pvalue= 0.0016 ; R2= 0.4725
Estimated allele frequency
0.0
0.2
0.4
0.6
0.8
O16
L16
L12
O12
253
256
O8
124
L8
204
233
218
217
219
L1
9
97
O1
5
6
7
8
9
10
11
12
Ecological gradient (Mean annual temperature)

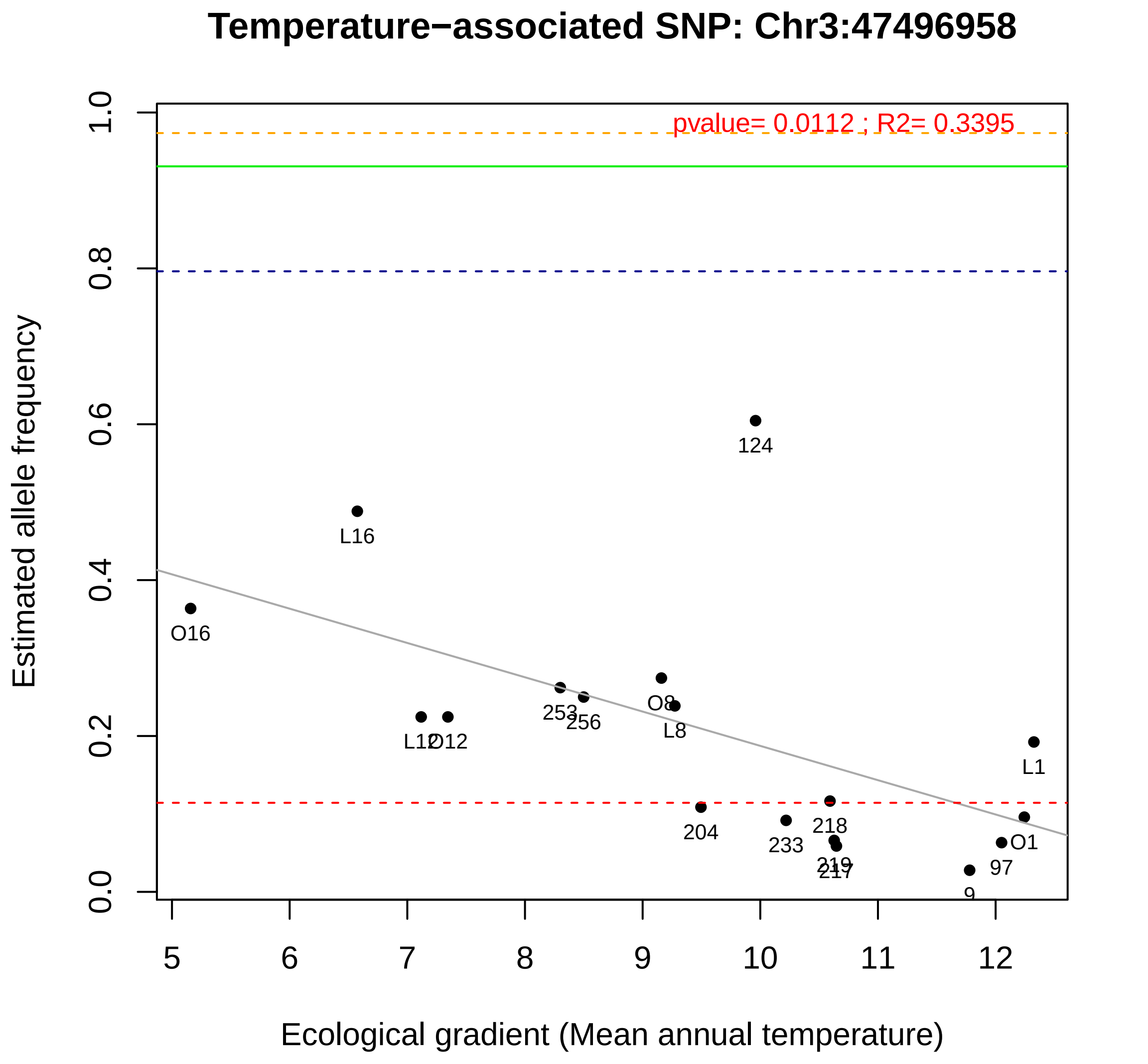
Temperature−associated SNP: Chr3:47496958
pvalue= 0.0112 ; R2= 0.3395
Estimated allele frequency
0.0
0.2
0.4
0.6
0.8
1.0
5
6
7
8
9
10
11
12
Ecological gradient (Mean annual temperature)
124
L16
O16
253
256
O8
L8
L12
O12
L1
204
233
218
219
217
O1
97
9

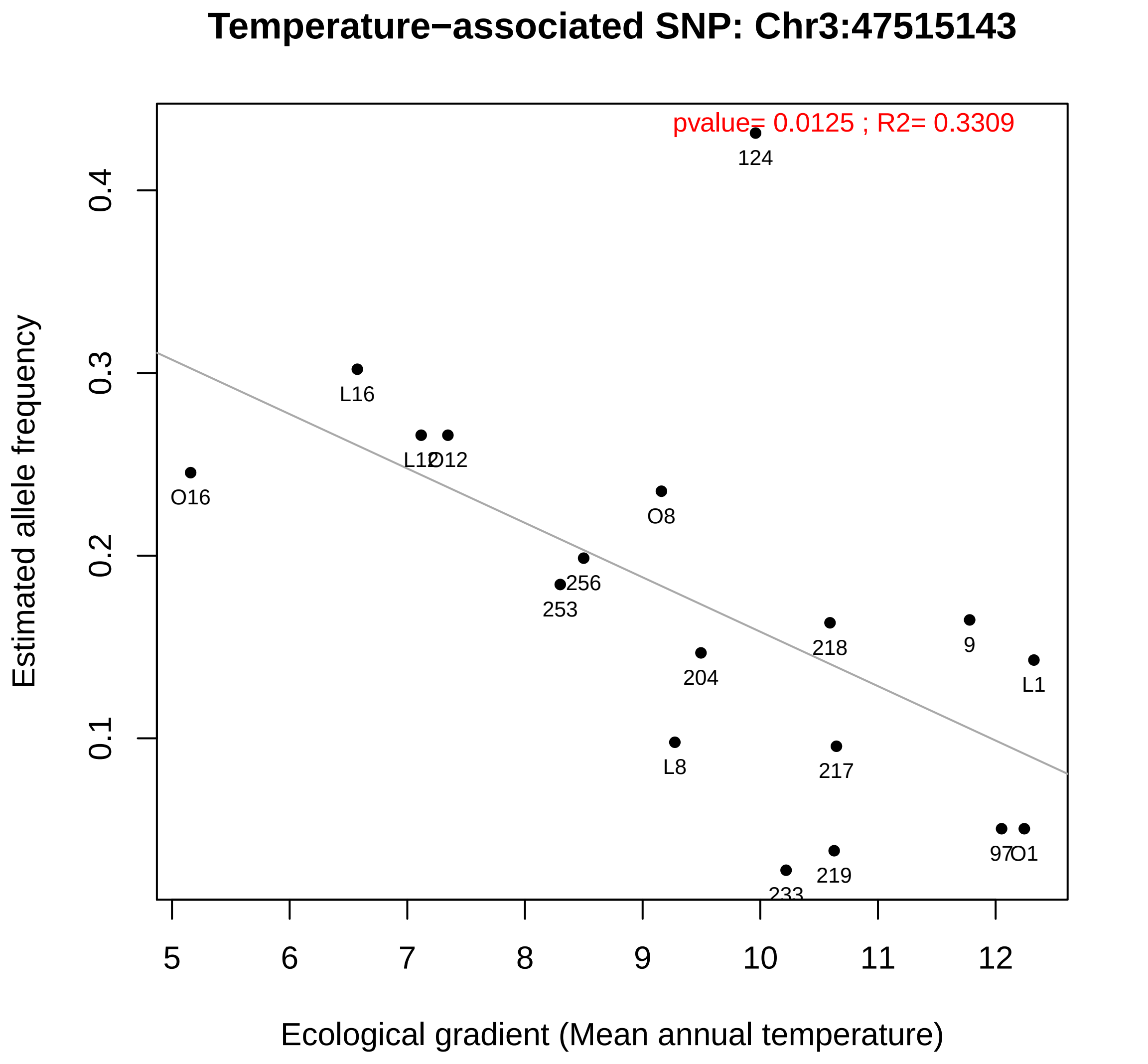
Temperature−associated SNP: Chr3:47515143
pvalue= 0.0125 ; R2= 0.3309
124
L16
L12
O12
O16
O8
256
253
218
9
204
L1
L8
217
97
O1
219
233
0.1
0.2
0.3
0.4
Estimated allele frequency
5
6
7
8
9
10
11
12
Ecological gradient (Mean annual temperature)

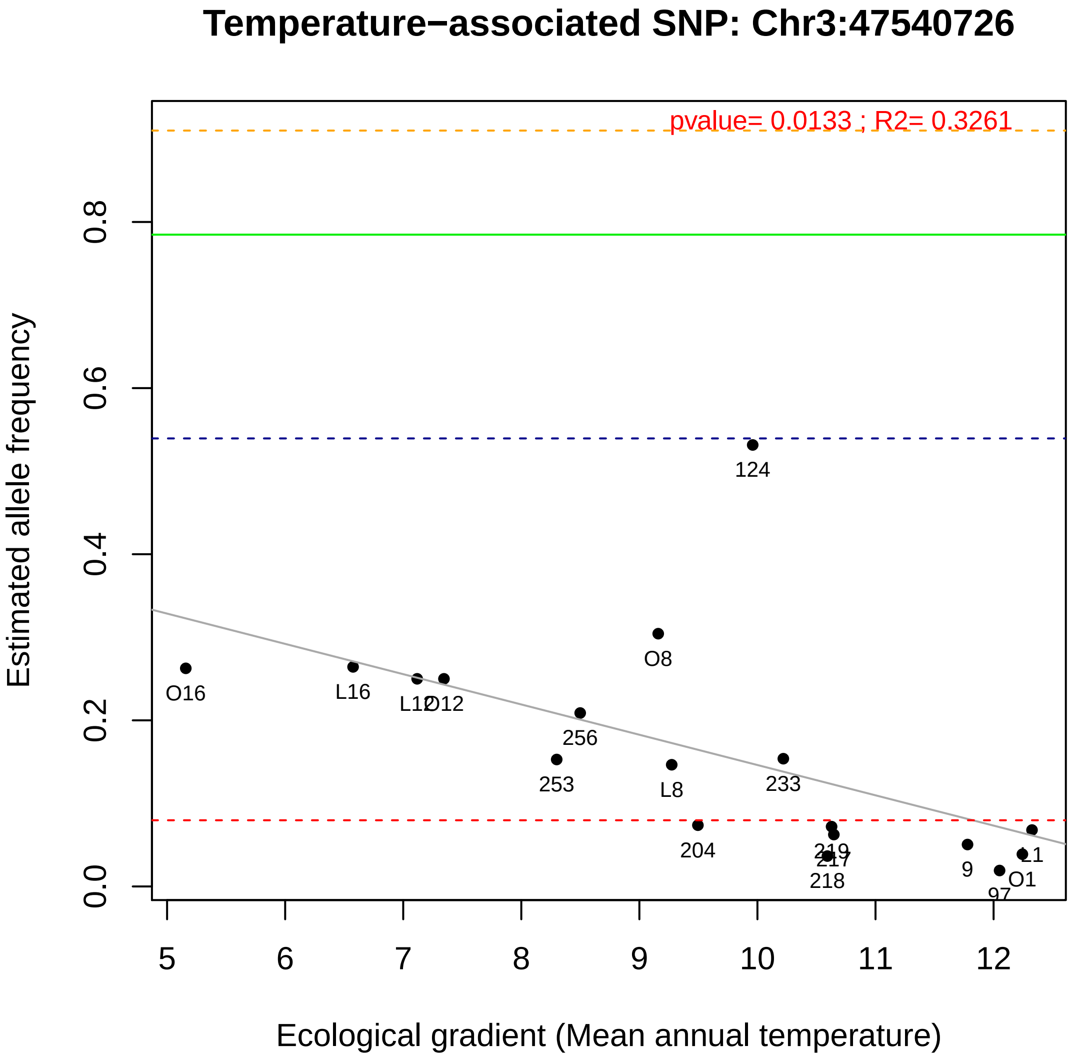

Temperature−associated SNP: Chr3:47540726
pvalue= 0.0133 ; R2= 0.3261
Estimated allele frequency
0.0
0.2
0.4
0.6
0.8
5
6
7
8
9
10
11
12
Ecological gradient (Mean annual temperature)
124
O8
O16
L16
L12
O12
256
253
L8
233
204
219
217
218
9
L1
O1
97

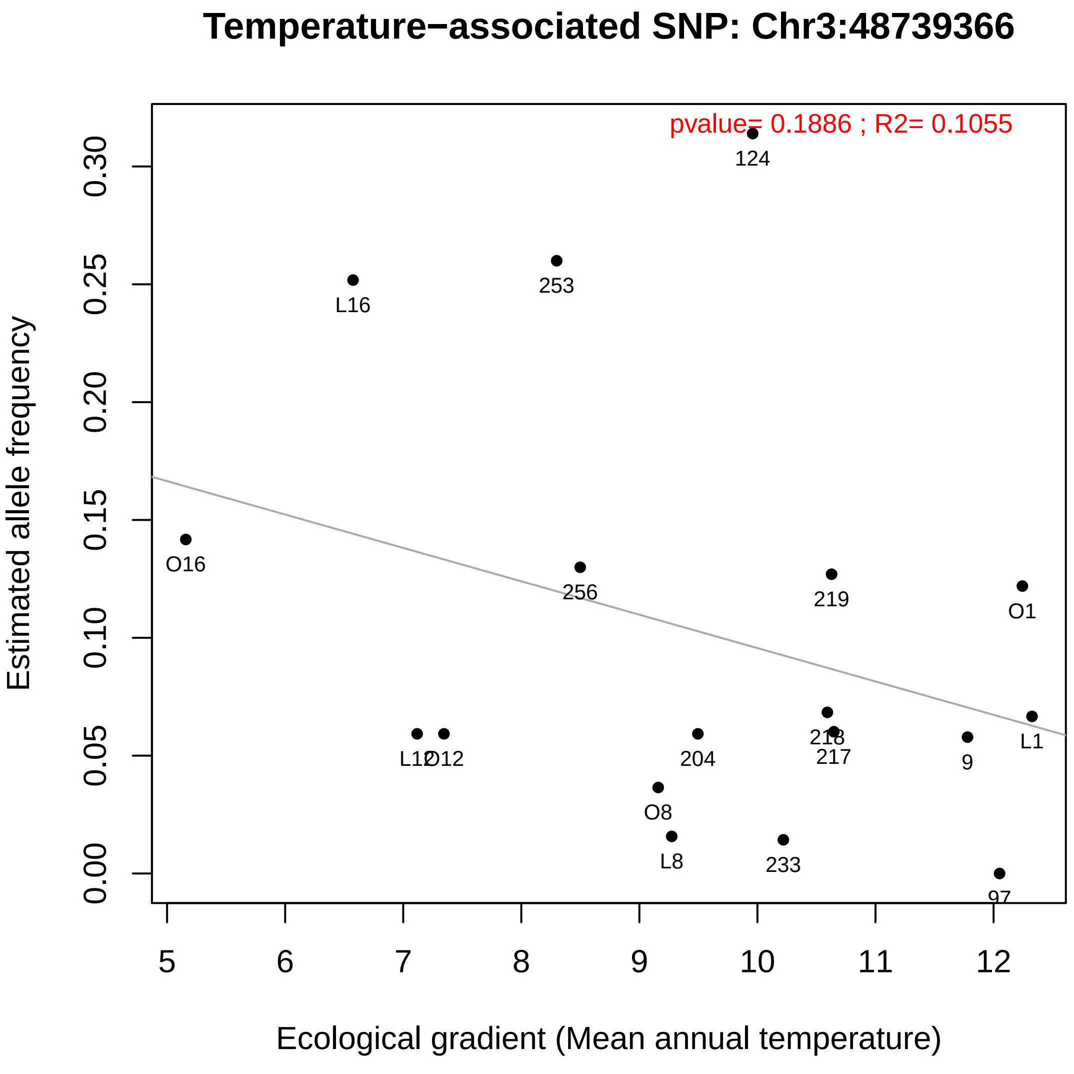

Temperature−associated SNP: Chr3:48739366
pvalue= 0.1886 ; R2= 0.1055
Estimated allele frequency
0.00
0.05
0.10
0.15
0.20
0.25
0.30
5
6
7
8
9
10
11
12
Ecological gradient (Mean annual temperature)
124
253
L16
O16
256
219
O1
L12
O12
204
218
217
L1
9
O8
L8
233
97

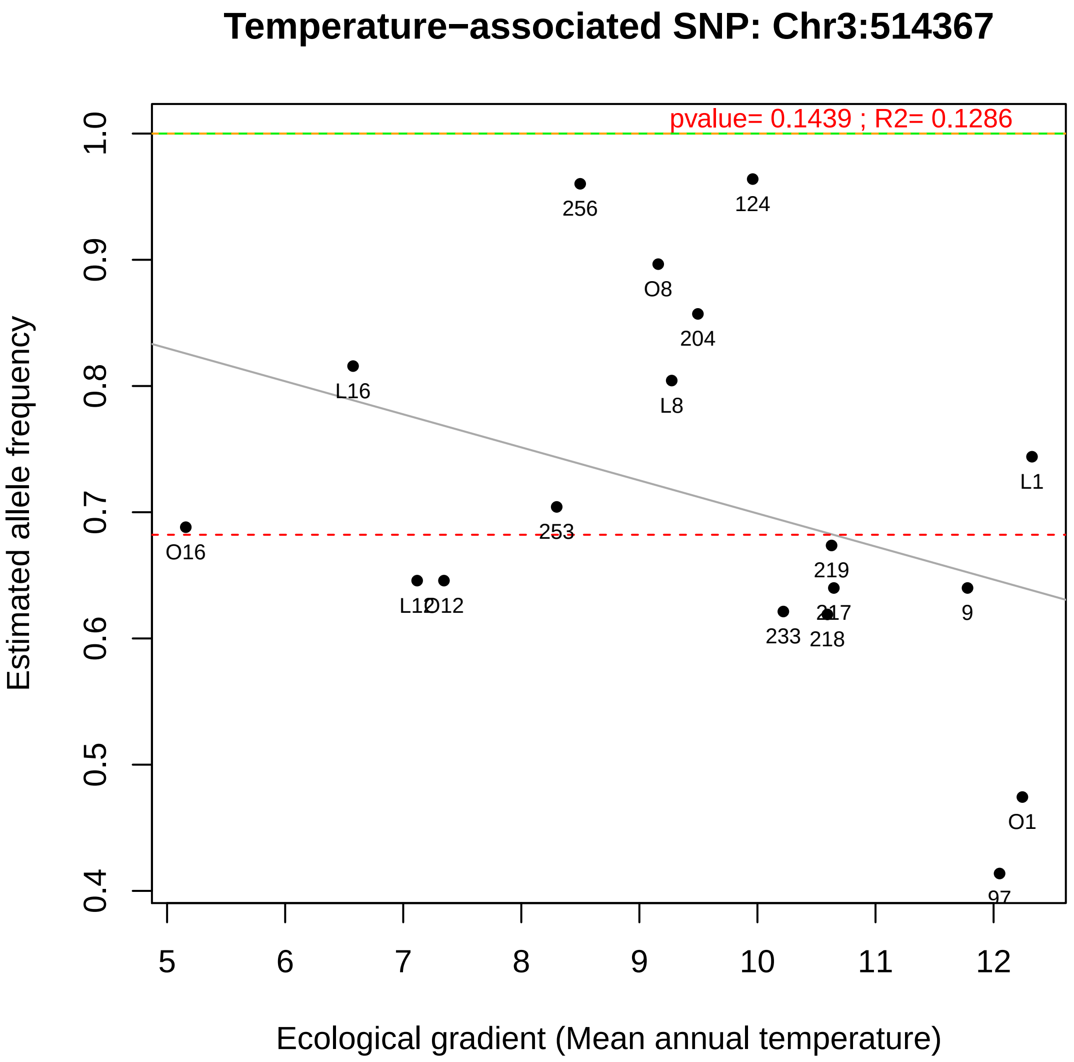
Temperature−associated SNP: Chr3:514367
pvalue= 0.1439 ; R2= 0.1286
Estimated allele frequency
Ecological gradient (Mean annual temperature)
0.4
0.5
0.6
0.7
0.8
0.9
1.0
5
6
7
8
9
10
11
12
256
124
O8
204
L16
L8
L1
253
O16
219
L12
O12
217
9
233
218
O1
97

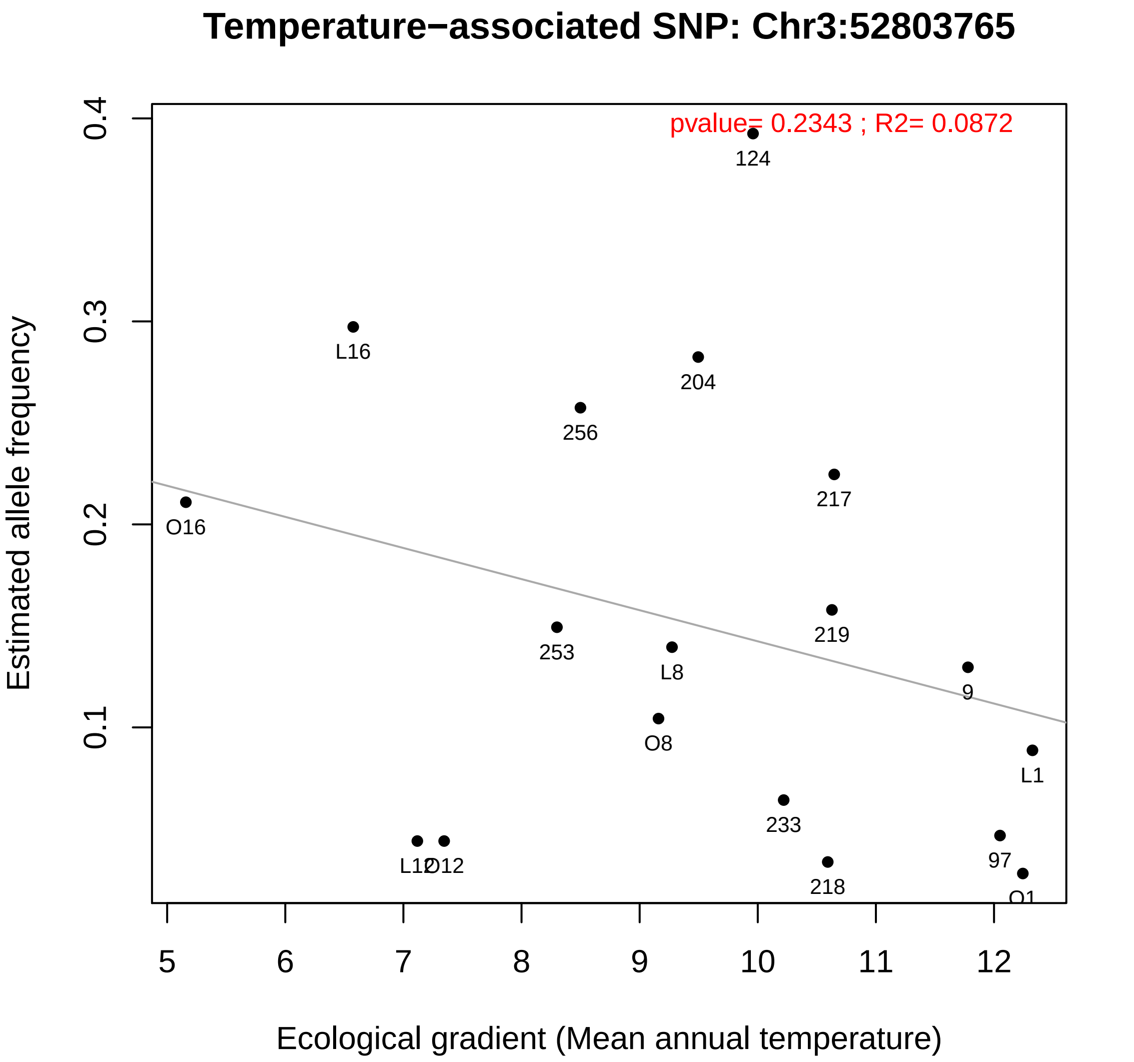
Temperature−associated SNP: Chr3:52803765
pvalue= 0.2343 ; R2= 0.0872
Estimated allele frequency
0.1
0.2
0.3
0.4
5
6
7
8
9
10
11
12
Ecological gradient (Mean annual temperature)
124
L16
204
256
217
O16
219
253
L8
9
O8
L1
233
L12
O12
97
218
O1

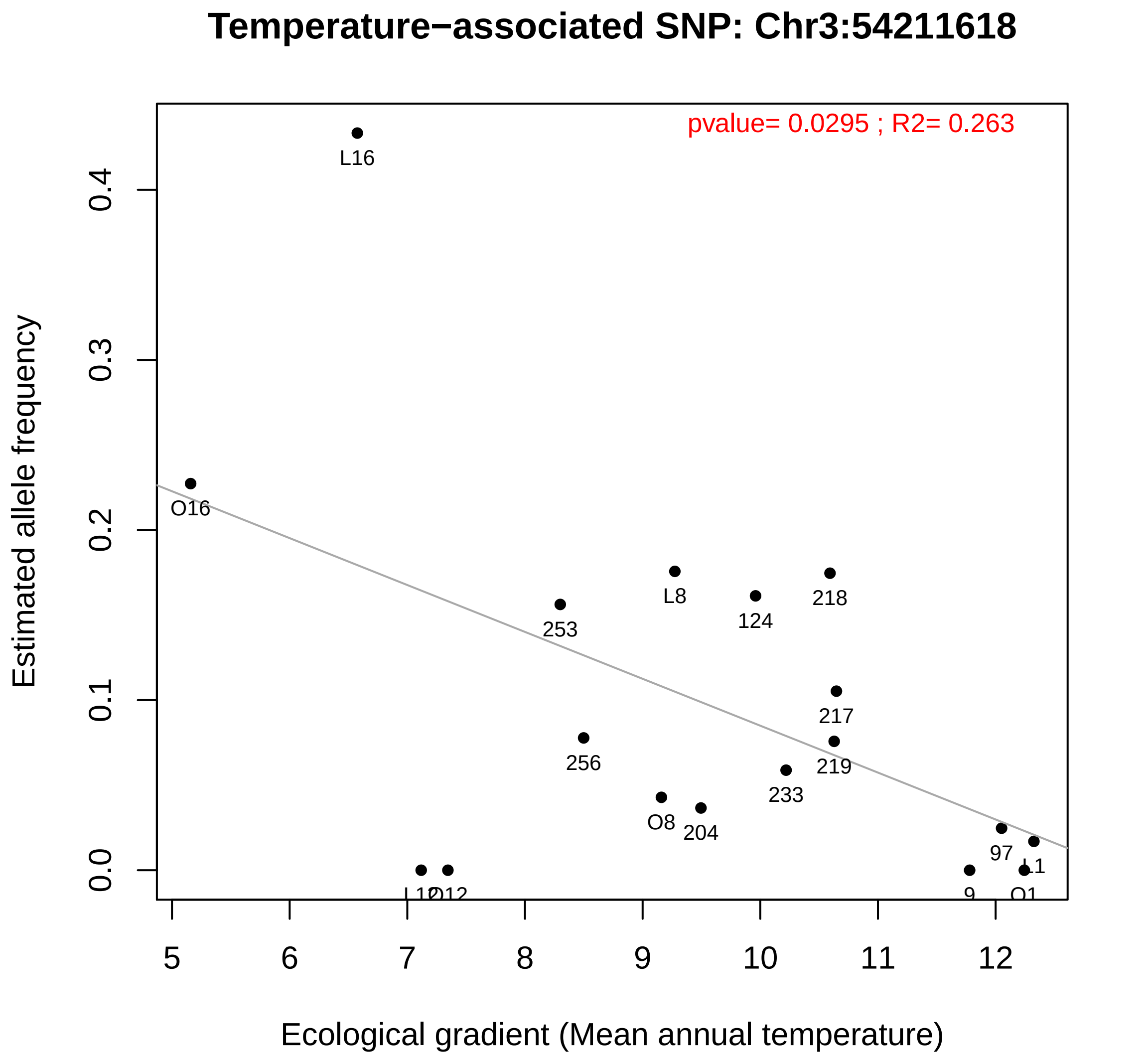
Temperature−associated SNP: Chr3:54211618
pvalue= 0.0295 ; R2= 0.263
Estimated allele frequency
0.0
0.1
0.2
0.3
0.4
Ecological gradient (Mean annual temperature)
5
6
7
8
9
10
11
12
L16
O16
253
L8
124
218
217
256
219
233
O8
204
97
L1
O12
9
O1

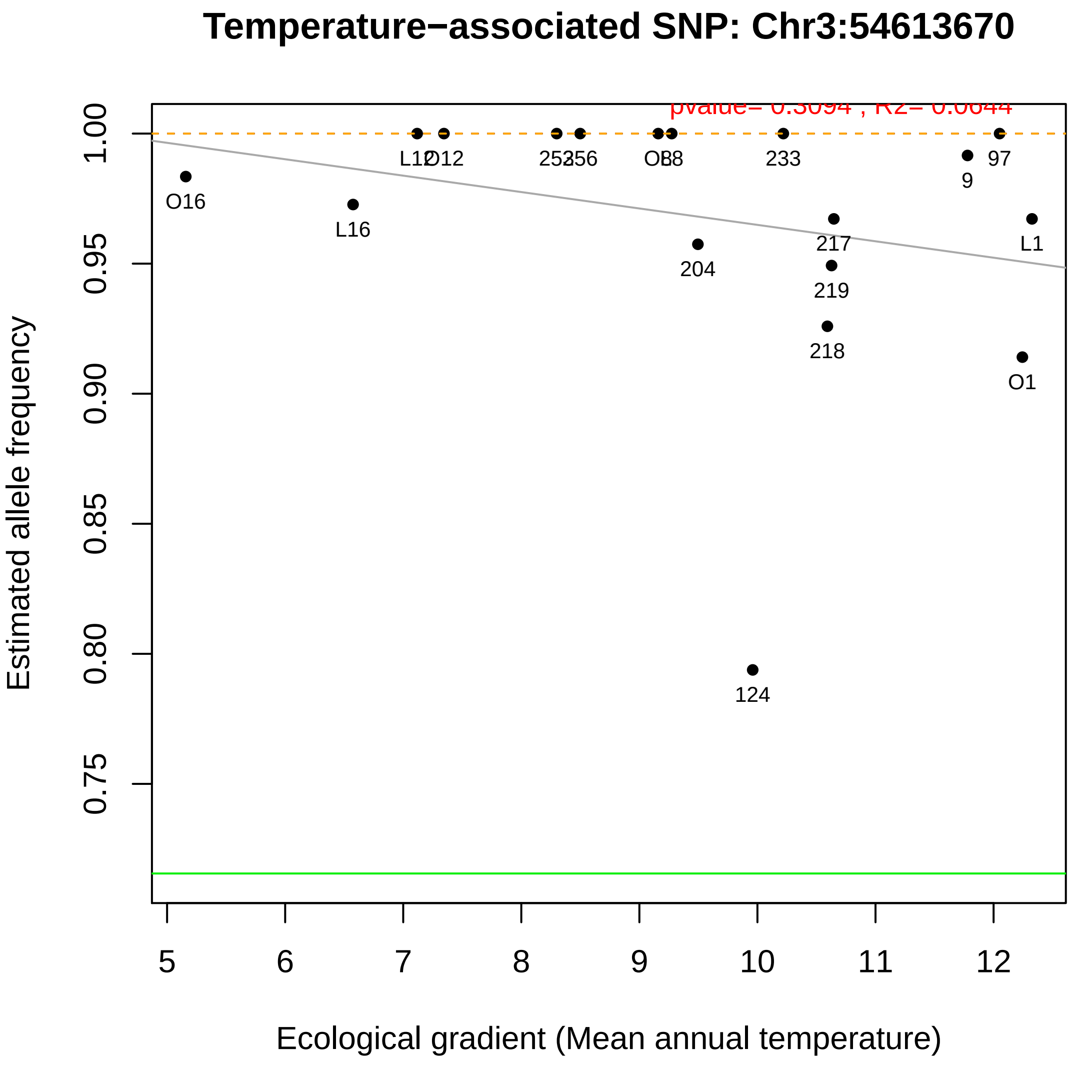

Temperature−associated SNP: Chr3:54613670
pvalue= 0.3094 , R2= 0.0644
Estimated allele frequency
Ecological gradient (Mean annual temperature)
0.75
0.80
0.85
0.90
0.95
1.00
5
6
7
8
9
10
11
12
O16
L16
L12
O12
256
233
97
9
204
217
L1
219
218
O1
124

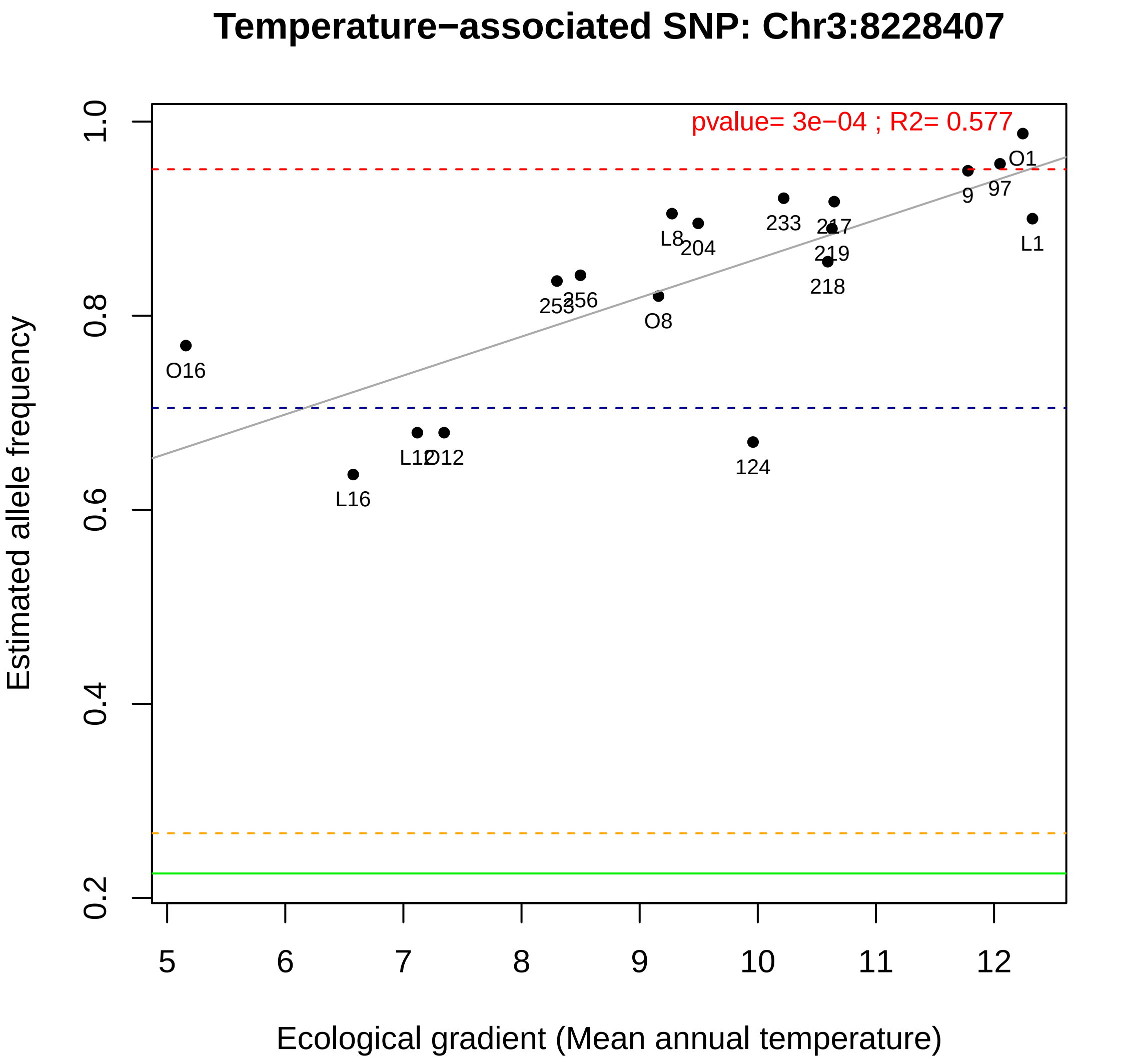
Temperature−associated SNP: Chr3:8228407
pvalue= 3e−04 ; R2= 0.577
Estimated allele frequency
1.0
0.8
0.6
0.4
0.2
5
6
7
8
9
10
11
12
Ecological gradient (Mean annual temperature)
O16
L16
L12
O12
253
256
O8
L8
204
124
233
217
219
218
9
97
O1
L1

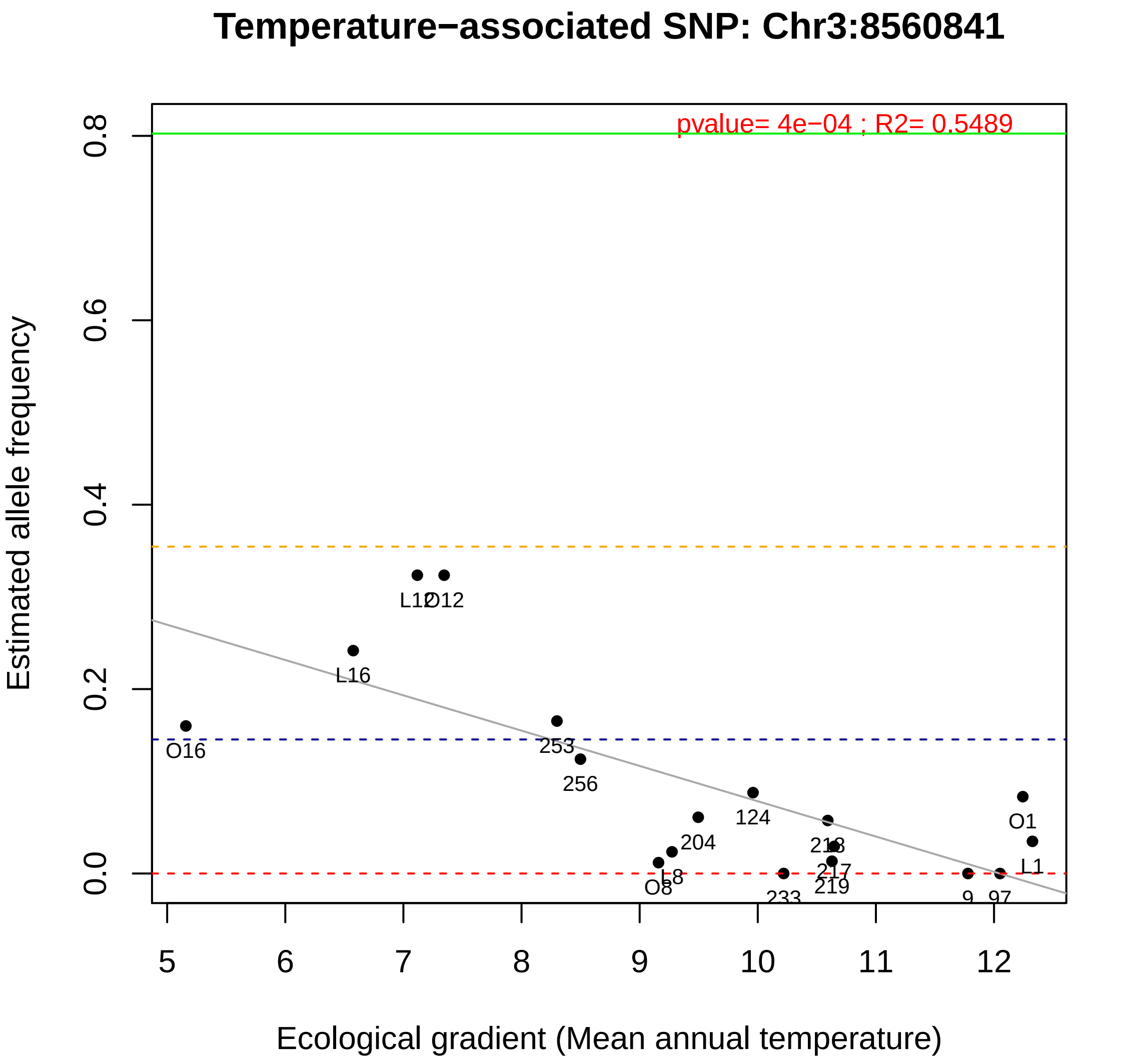
Temperature−associated SNP: Chr3:8560841
pvalue= 4e−04 ; R2= 0.5489
Estimated allele frequency
0.0
0.2
0.4
0.6
0.8
5
6
7
8
9
10
11
12
Ecological gradient (Mean annual temperature)
O16
L16
L12
O12
253
256
O8
L8
204
124
233
218
217
219
9
97
O1
L1

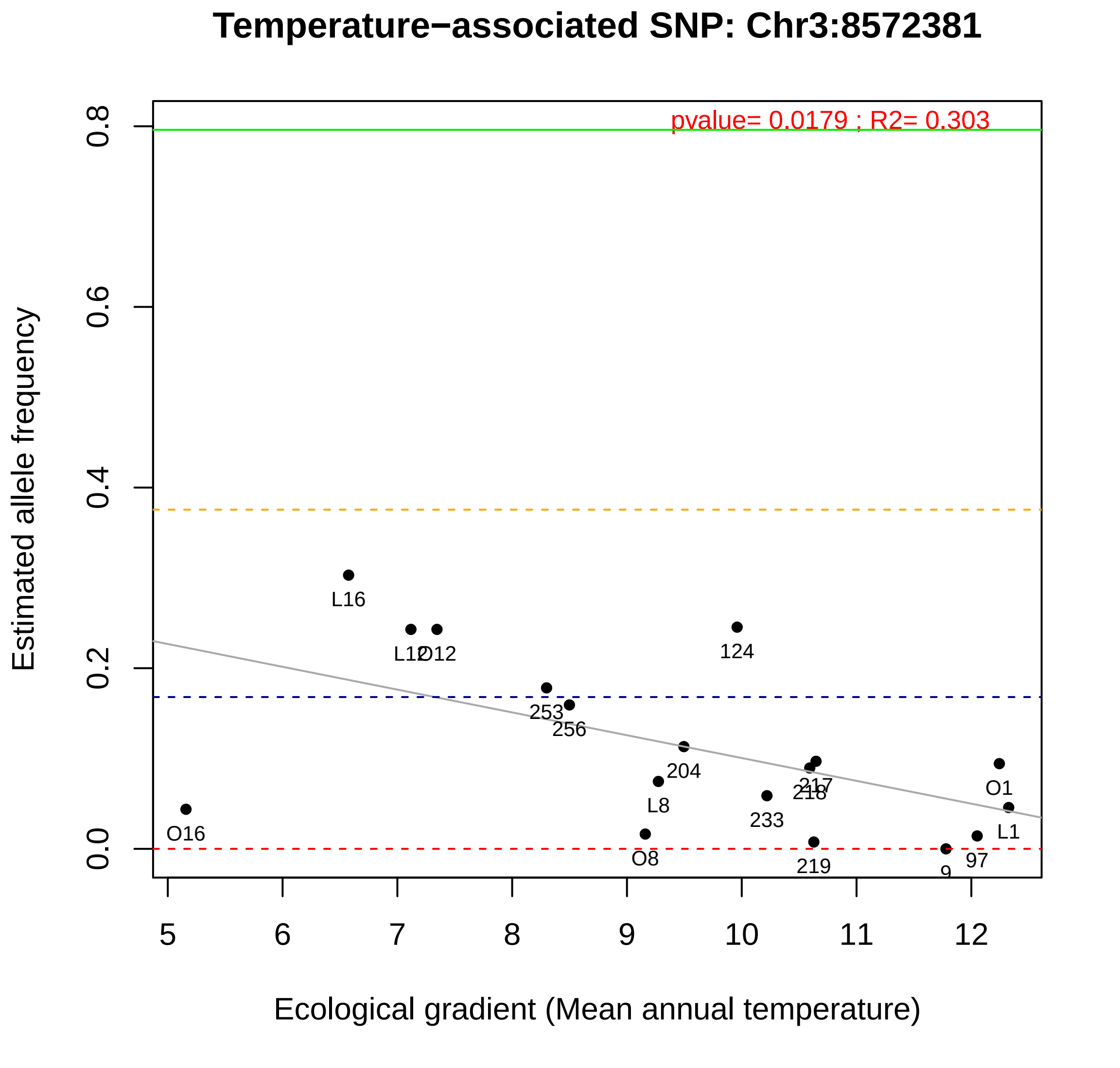
Temperature−associated SNP: Chr3:8572381
pvalue= 0.0179 ; R2= 0.303
Estimated allele frequency
0.0
0.2
0.4
0.6
0.8
5
6
7
8
9
10
11
12
Ecological gradient (Mean annual temperature)
L16
L12
O12
124
253
256
204
217
218
L8
O1
233
L1
O16
O8
219
97
9

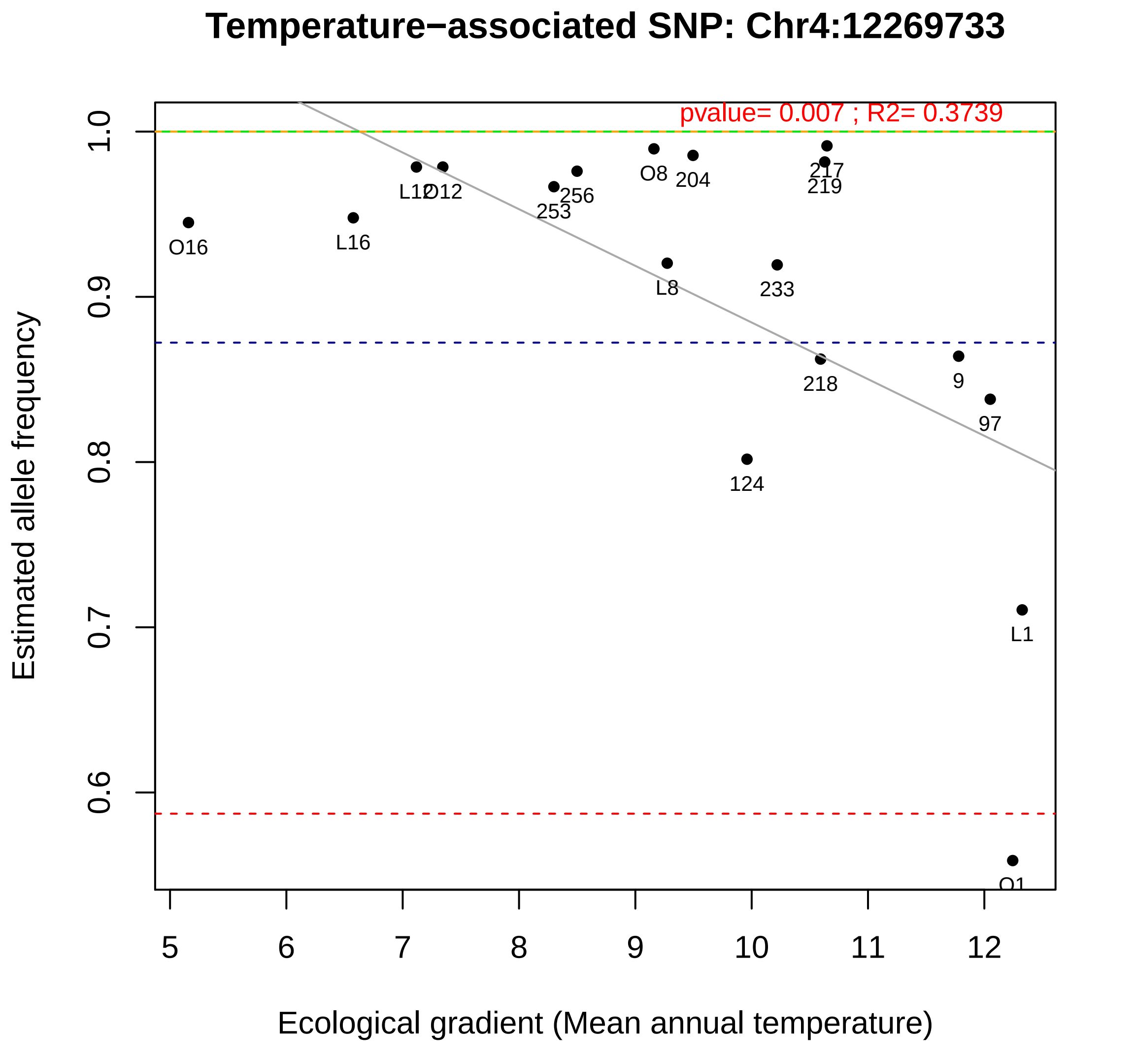
Temperature−associated SNP: Chr4:12269733
pvalue= 0.007 ; R2= 0.3739
Estimated allele frequency
1.0
0.9
0.8
0.7
0.6
O16
L16
L12
O12
253
256
O8
204
217
219
L8
233
218
9
97
124
L1
O1
5
6
7
8
9
10
11
12
Ecological gradient (Mean annual temperature)

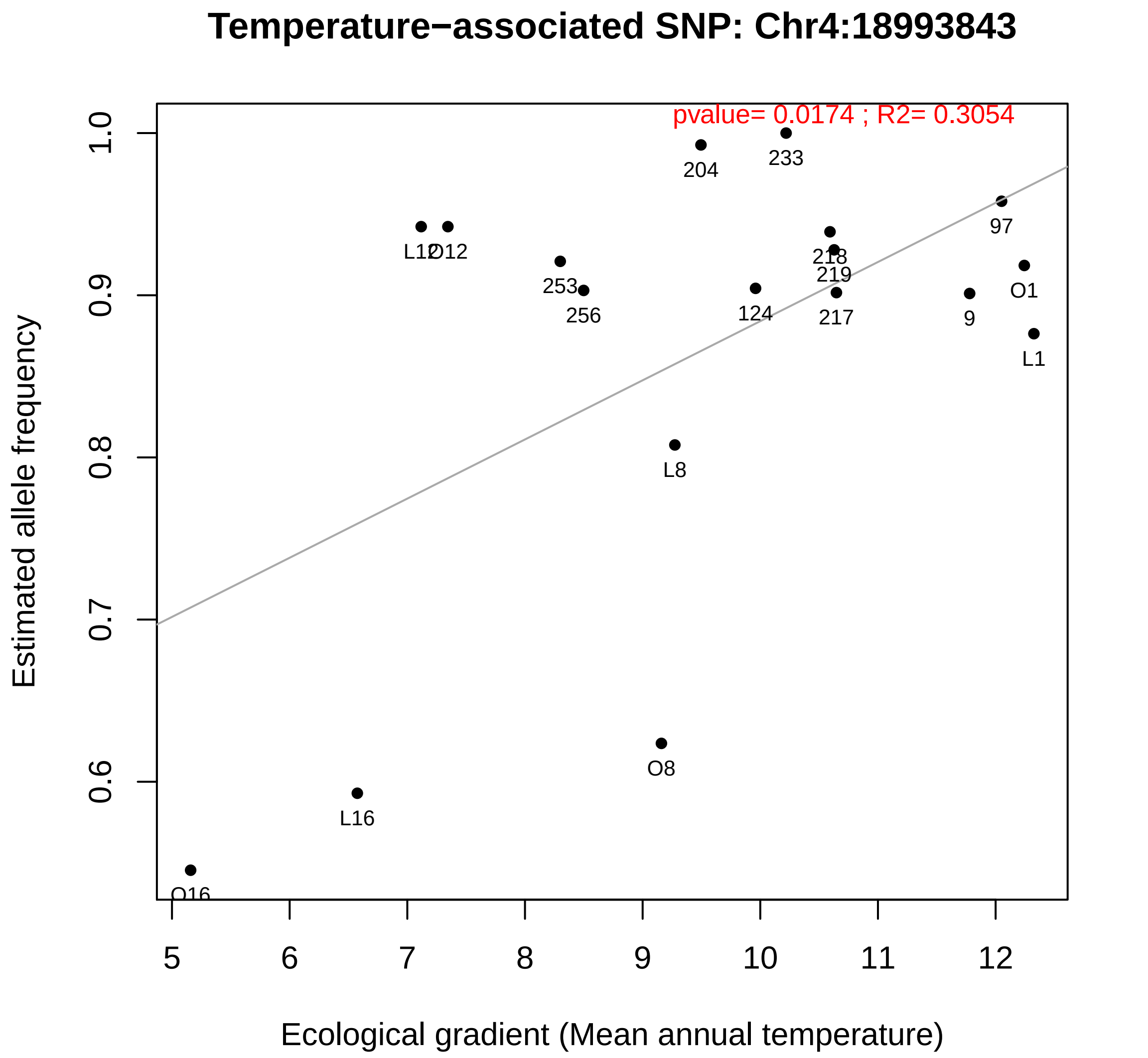

Temperature−associated SNP: Chr4:18993843
pvalue= 0.0174 ; R2= 0.3054
Estimated allele frequency
Ecological gradient (Mean annual temperature)
0.6
0.7
0.8
0.9
1.0
5
6
7
8
9
10
11
12
204
233
97
L12
O12
253
256
218
219
124
217
O1
9
L1
L8
O8
L16
O16

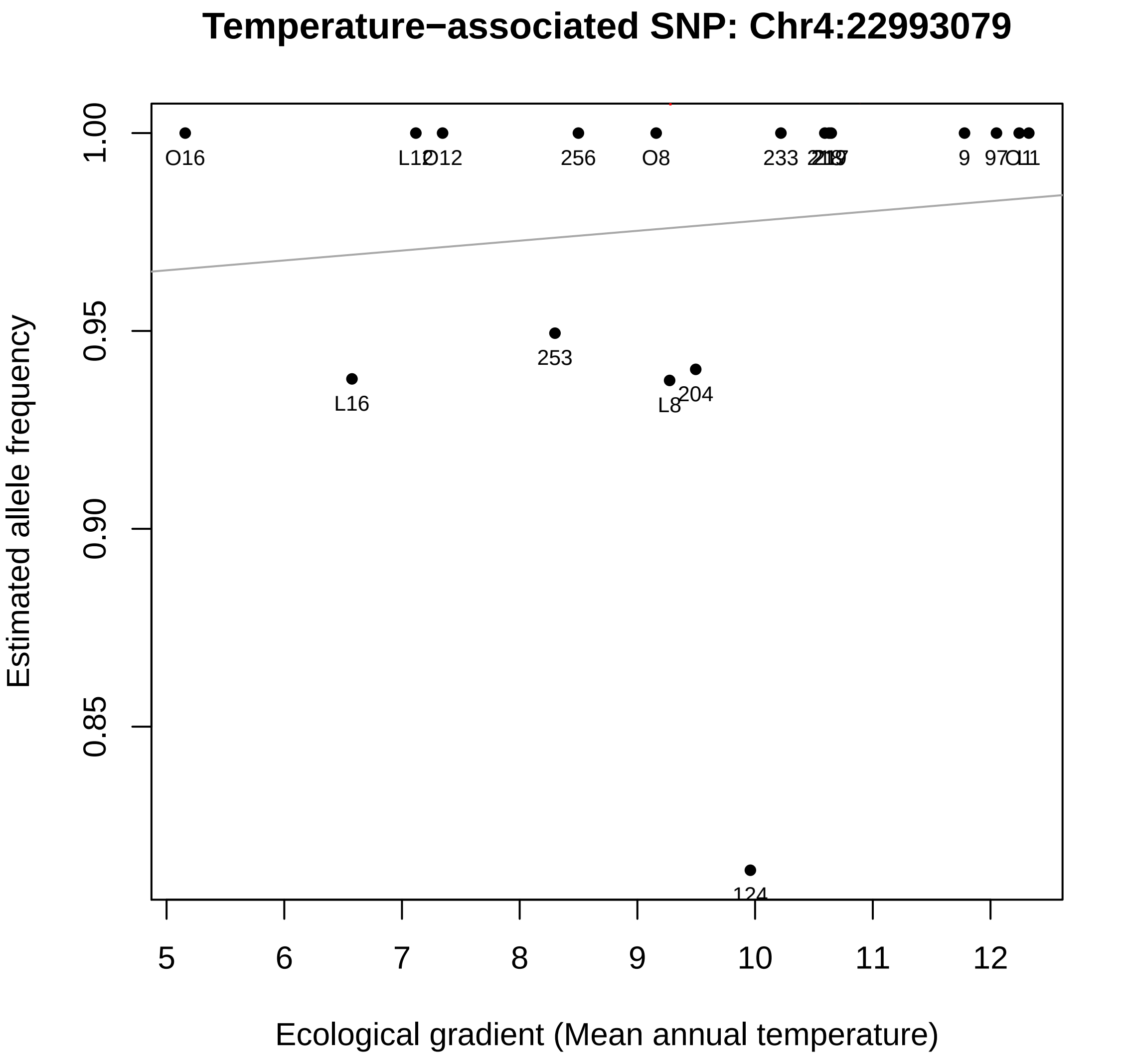

Temperature−associated SNP: Chr4:22993079
Estimated allele frequency
1.00
0.95
0.90
0.85
O16
L12
O12
256
O8
233
9
L16
253
L8
204
124
5
6
7
8
9
10
11
12
Ecological gradient (Mean annual temperature)

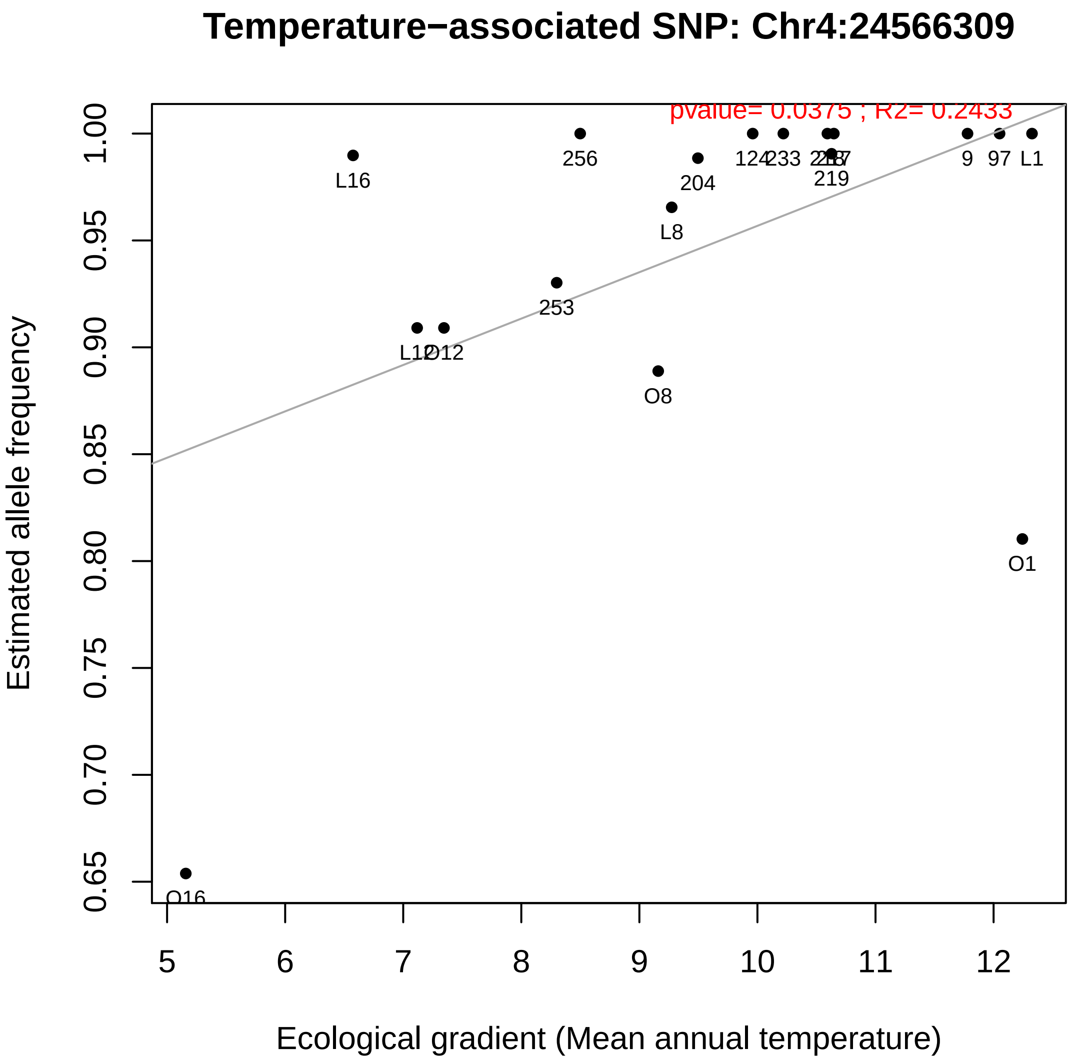
Temperature−associated SNP: Chr4:24566309
pvalue= 0.0375 ; R2= 0.2433
Estimated allele frequency
Ecological gradient (Mean annual temperature)
1.00
0.95
0.90
0.85
0.80
0.75
0.70
0.65
5
6
7
8
9
10
11
12
L16
256
204
124
233
219
9
97
L1
L8
253
L12
O12
O8
O1
O16

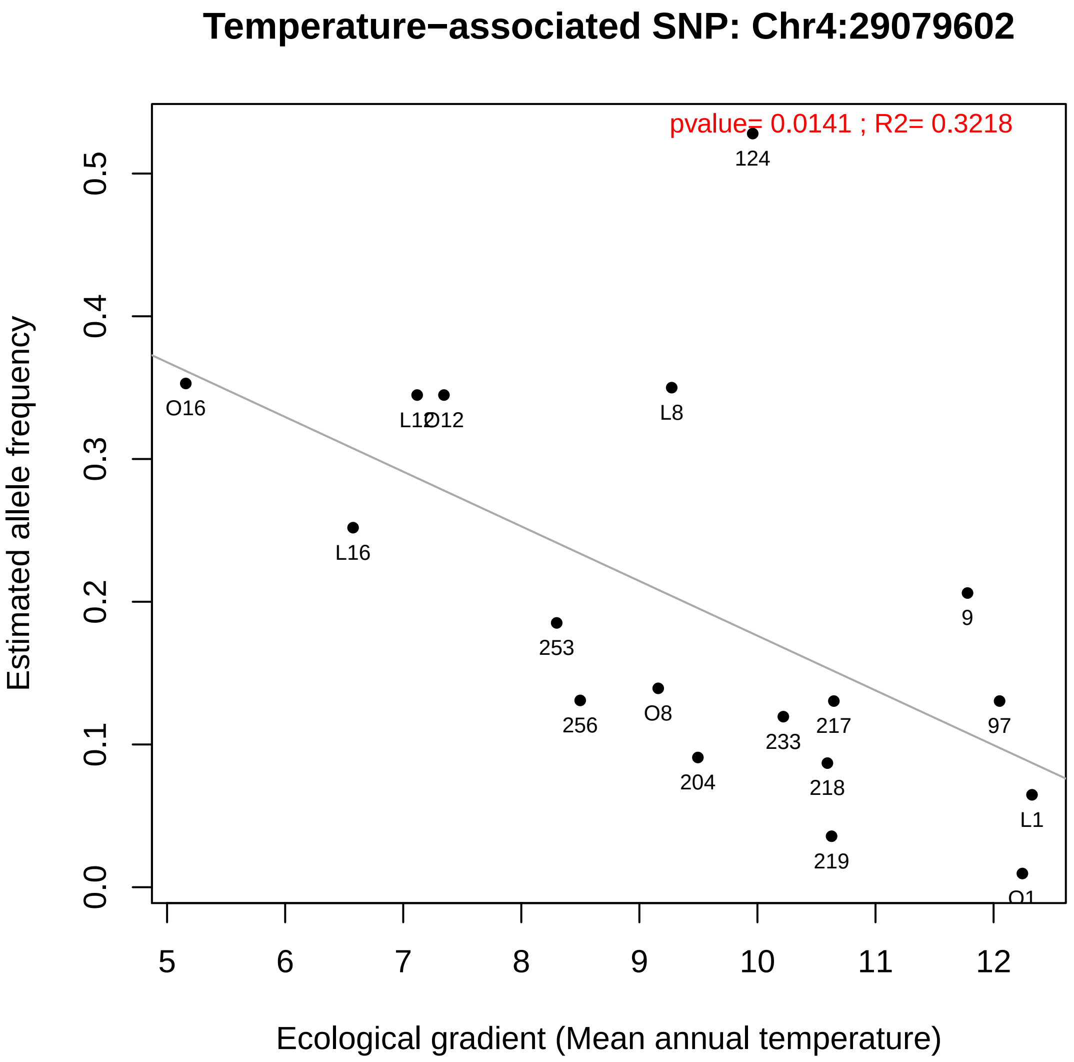

Temperature−associated SNP: Chr4:29079602
pvalue= 0.0141 ; R2= 0.3218
Estimated allele frequency
0.0
0.1
0.2
0.3
0.4
0.5
5
6
7
8
9
10
11
12
Ecological gradient (Mean annual temperature)
124
O16
L12
O12
L8
L16
9
253
256
O8
233
217
97
204
218
L1
219
O1

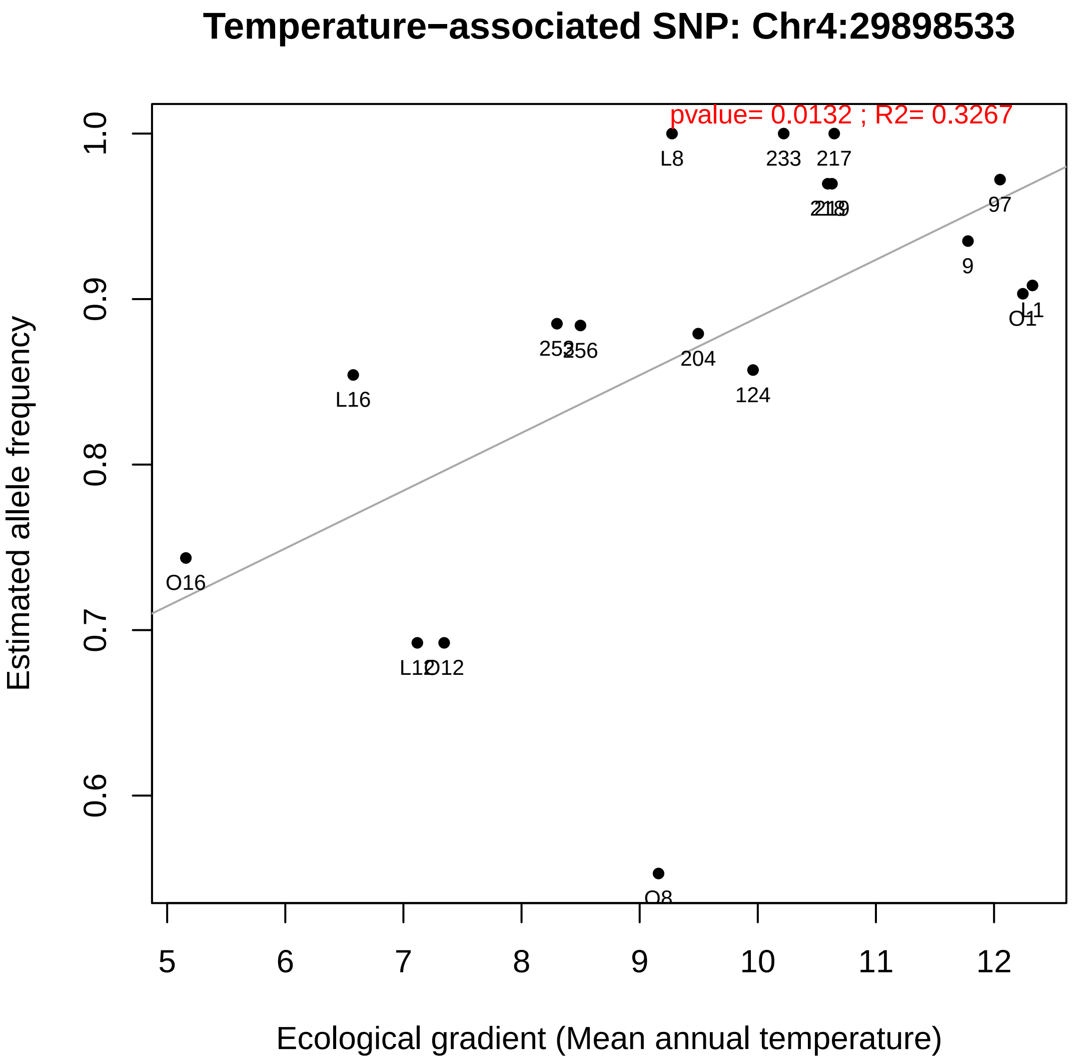

Temperature−associated SNP: Chr4:29898533
pvalue= 0.0132 ; R2= 0.3267
Estimated allele frequency
Ecological gradient (Mean annual temperature)
1.0
0.9
0.8
0.7
0.6
5
6
7
8
9
10
11
12
L8
233
217
97
9
O1
204
124
L16
O16
O12
O8

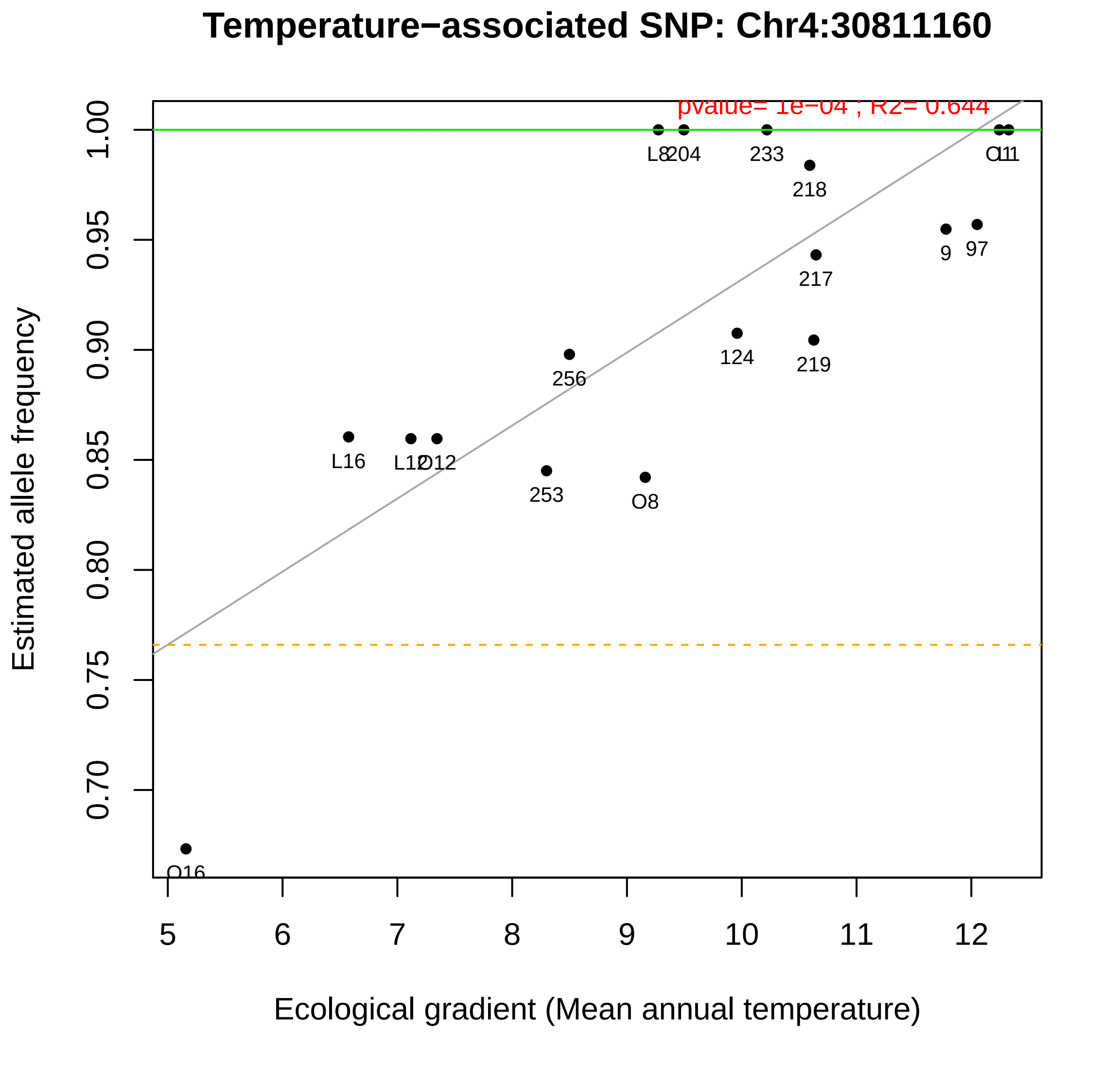

Temperature−associated SNP: Chr4:30811160
pvalue= 1e−04 ; R2= 0.644
Estimated allele frequency
1.00
0.95
0.90
0.85
0.80
0.75
0.70
L8
204
233
218
O11
217
9
97
124
219
256
L16
L12
O12
253
O8
O16
5
6
7
8
9
10
11
12
Ecological gradient (Mean annual temperature)

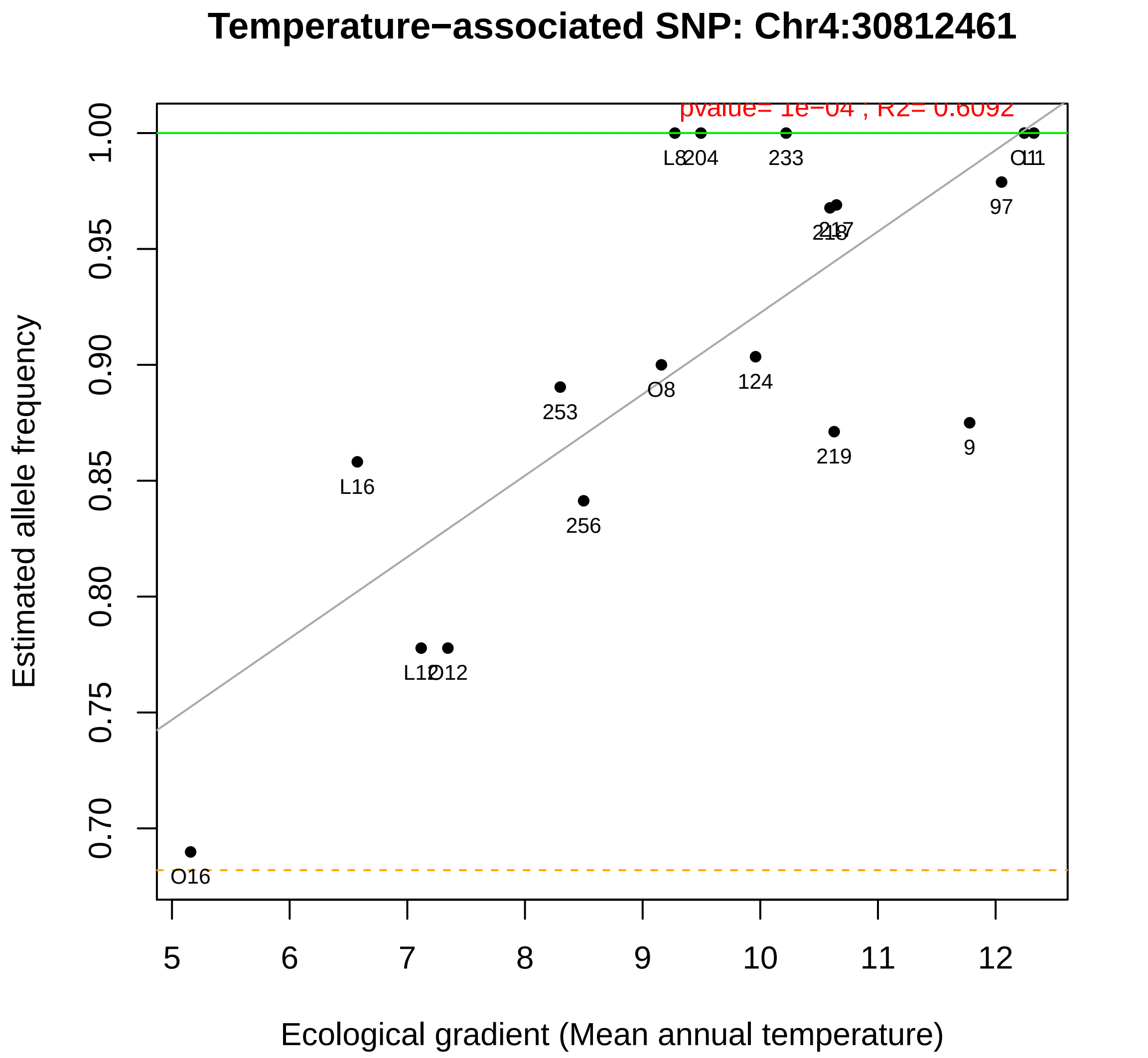

Temperature−associated SNP: Chr4:30812461
pvalue= 1e−04 ; R2= 0.6092
Estimated allele frequency
Ecological gradient (Mean annual temperature)
1.00
0.95
0.90
0.85
0.80
0.75
0.70
5
6
7
8
9
10
11
12
L8
204
233
97
253
O8
124
219
9
L16
256
L12
O12
O16

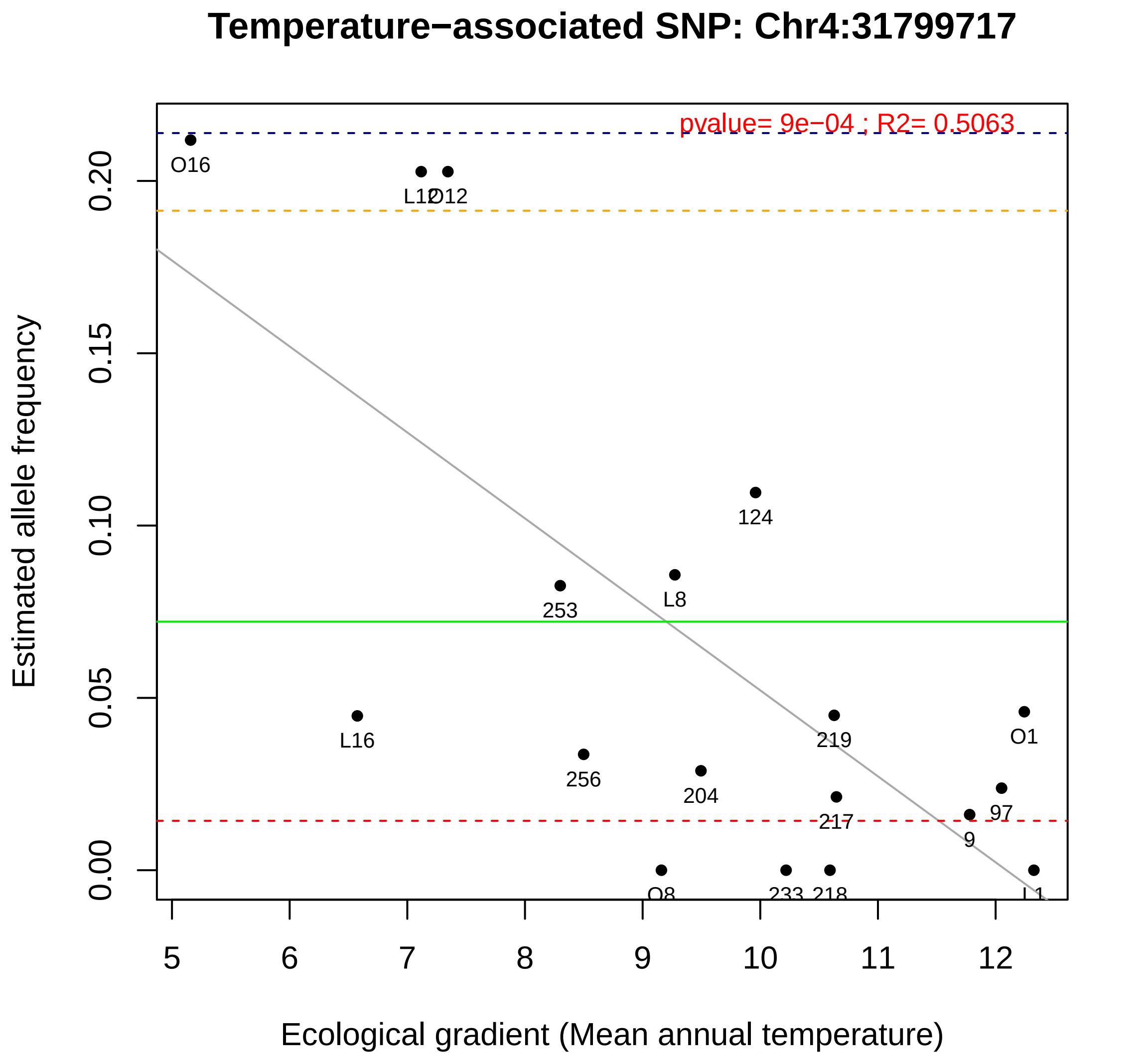
Temperature−associated SNP: Chr4:31799717
pvalue= 9e−04 ; R2= 0.5063
Estimated allele frequency
0.00
0.05
0.10
0.15
0.20
O16
L12
O12
124
253
L8
L16
219
O1
256
204
217
97
9
O8
233
218
L1
5
6
7
8
9
10
11
12
Ecological gradient (Mean annual temperature)

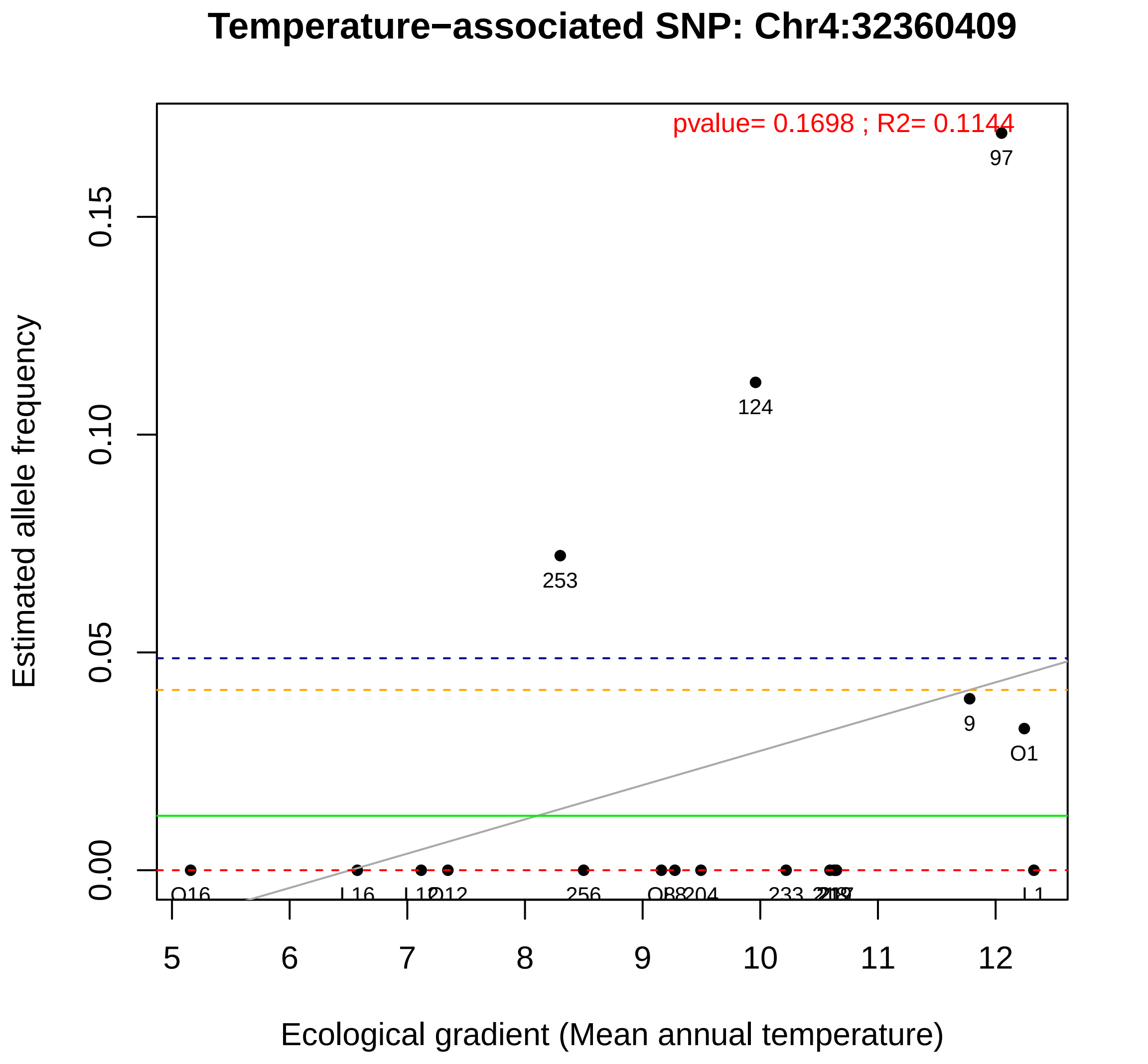

Temperature−associated SNP: Chr4:32360409
pvalue= 0.1698 ; R2= 0.1144
Estimated allele frequency
0.00
0.05
0.10
0.15
97
124
253
9
O1
O16
L16
L12
O12
256
O8
88
204
233
218
197
L1
5
6
7
8
9
10
11
12
Ecological gradient (Mean annual temperature)

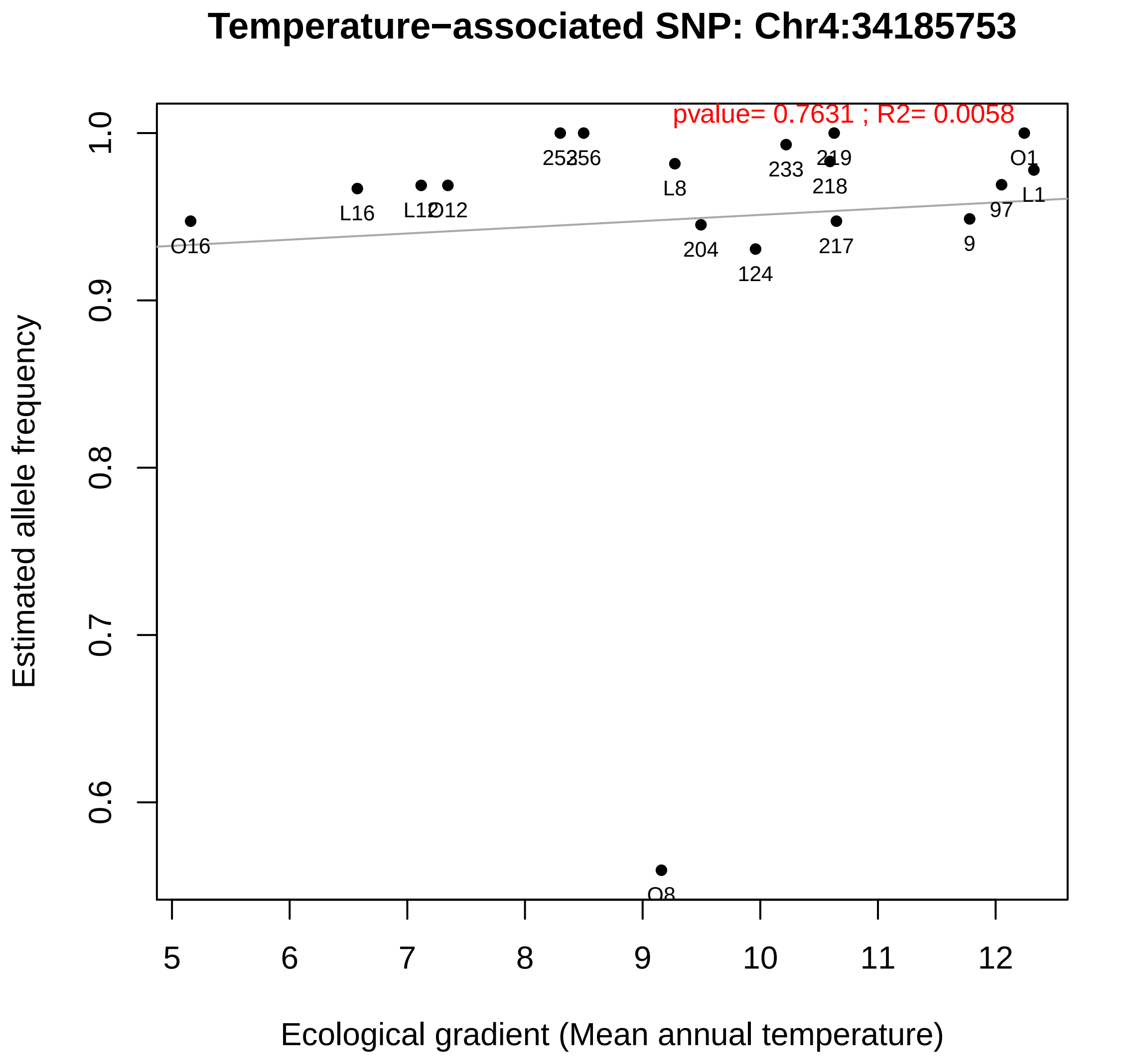

Temperature−associated SNP: Chr4:34185753
pvalue= 0.7631 ; R2= 0.0058
Estimated allele frequency
Ecological gradient (Mean annual temperature)
0.6
0.7
0.8
0.9
1.0
5
6
7
8
9
10
11
12
O16
L16
L12
O12
253
256
L8
204
124
233
219
218
217
9
97
O1
L1
O8

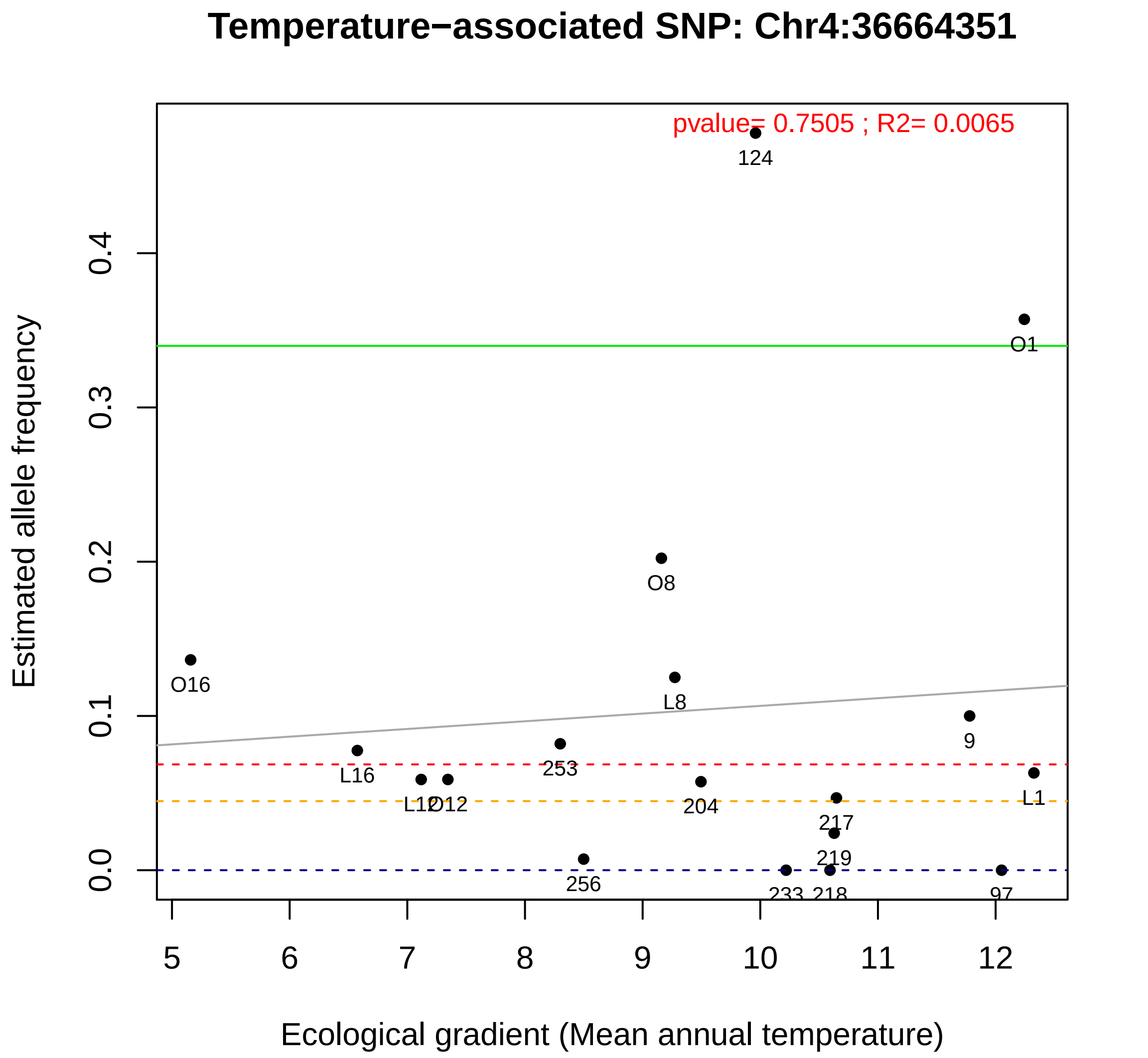
Temperature−associated SNP: Chr4:36664351
pvalue= 0.7505 ; R2= 0.0065
Estimated allele frequency
0.0
0.1
0.2
0.3
0.4
124
O1
O8
O16
L8
9
L16
253
L12
O12
204
L1
217
219
256
233
218
97
5
6
7
8
9
10
11
12
Ecological gradient (Mean annual temperature)

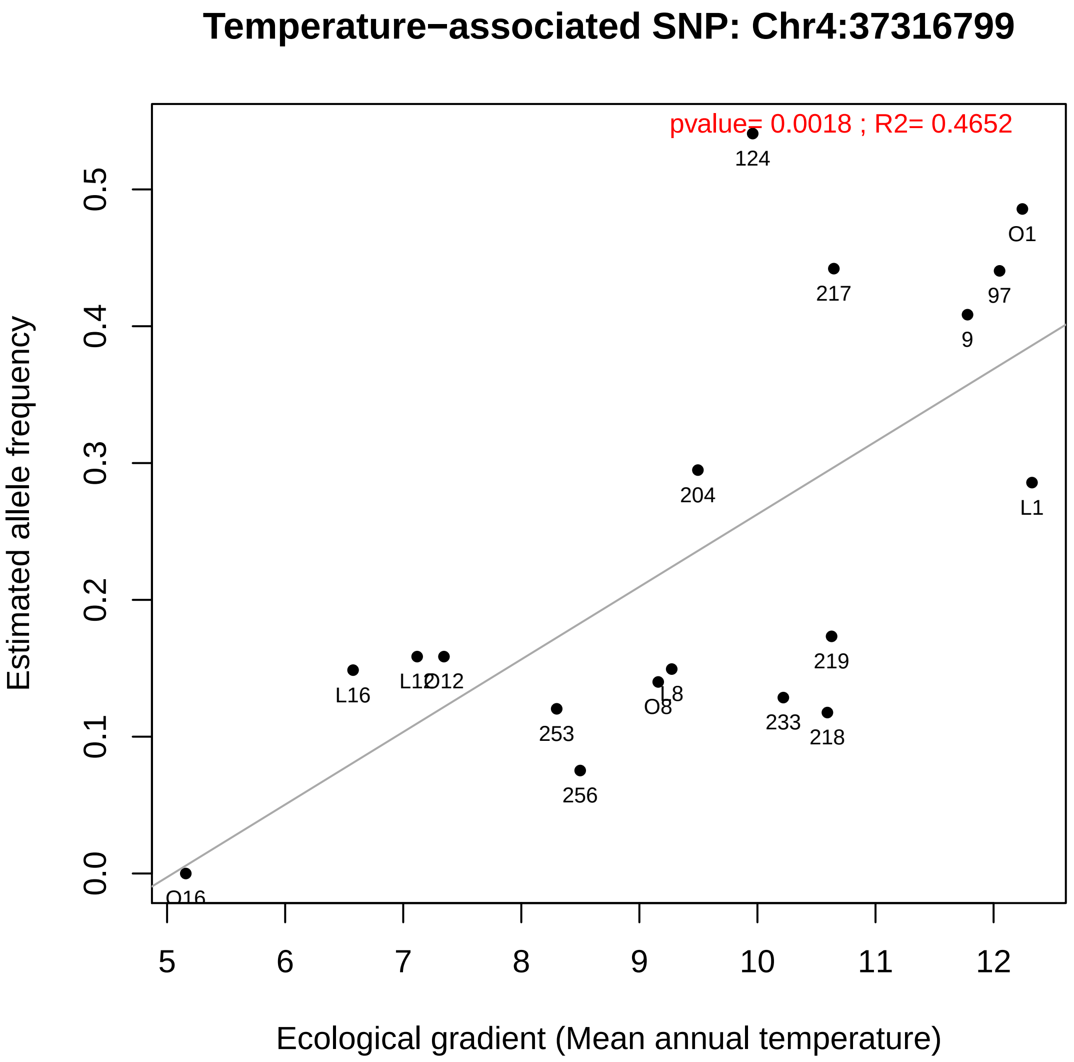

Temperature−associated SNP: Chr4:37316799
pvalue= 0.0018 ; R2= 0.4652
Estimated allele frequency
Ecological gradient (Mean annual temperature)
0.0
0.1
0.2
0.3
0.4
0.5
5
6
7
8
9
10
11
12
124
O1
217
97
9
204
L1
219
L16
L12
O12
L8
O8
233
218
253
256
O16

**Temperature−associated SNP: Chr4:40498101**

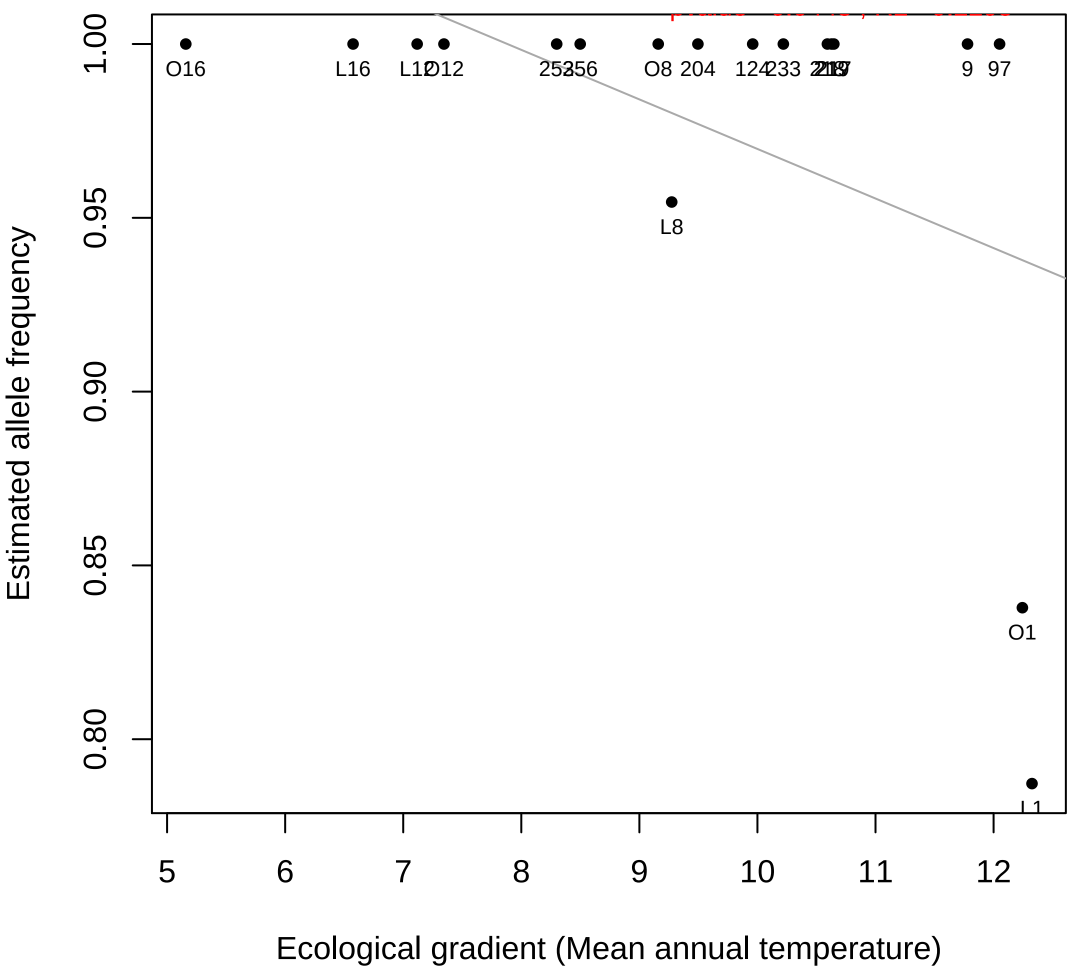

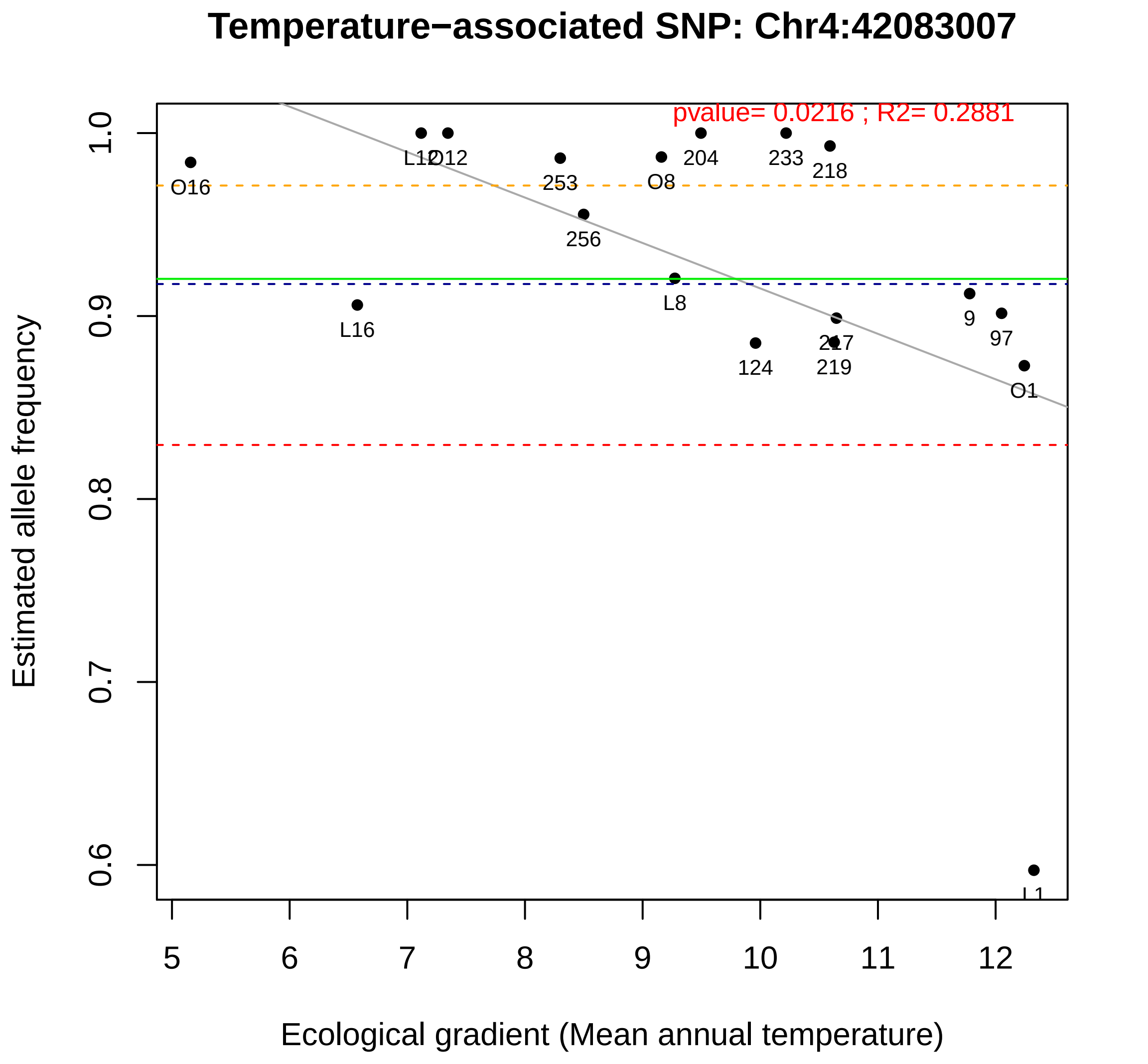

Temperature−associated SNP: Chr4:42083007
pvalue= 0.0216 ; R2= 0.2881
Estimated allele frequency
Ecological gradient (Mean annual temperature)
0.6
0.7
0.8
0.9
1.0
5
6
7
8
9
10
11
12
O16
L12
O12
253
O8
204
233
218
256
L8
L16
9
97
217
219
124
O1

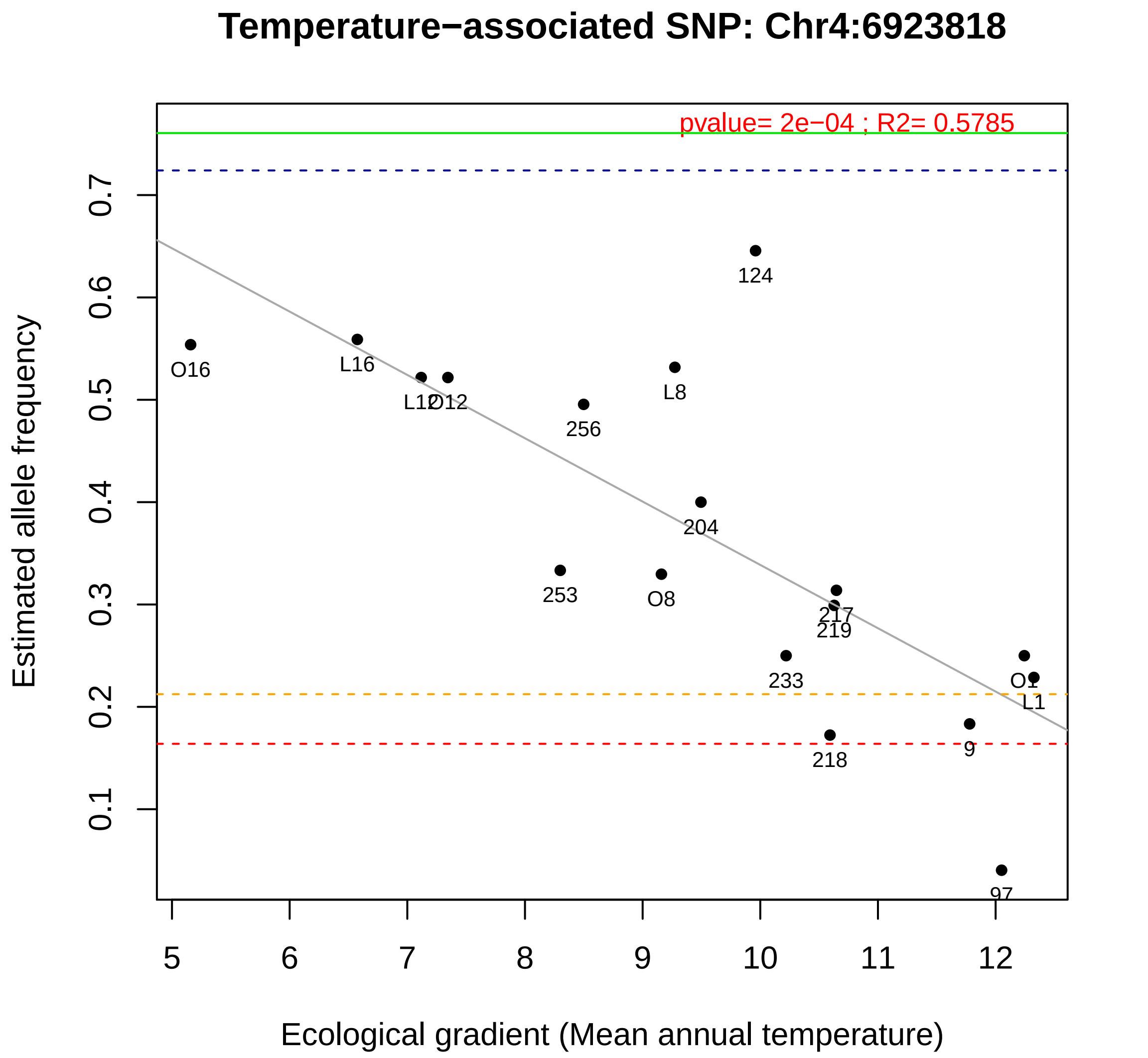

Temperature−associated SNP: Chr4:6923818
pvalue= 2e−04 ; R2= 0.5785
Estimated allele frequency
0.1
0.2
0.3
0.4
0.5
0.6
0.7
Ecological gradient (Mean annual temperature)
5
6
7
8
9
10
11
12
O16
L16
L12
O12
256
253
L8
O8
124
204
217
219
233
218
O1
L1
9
97

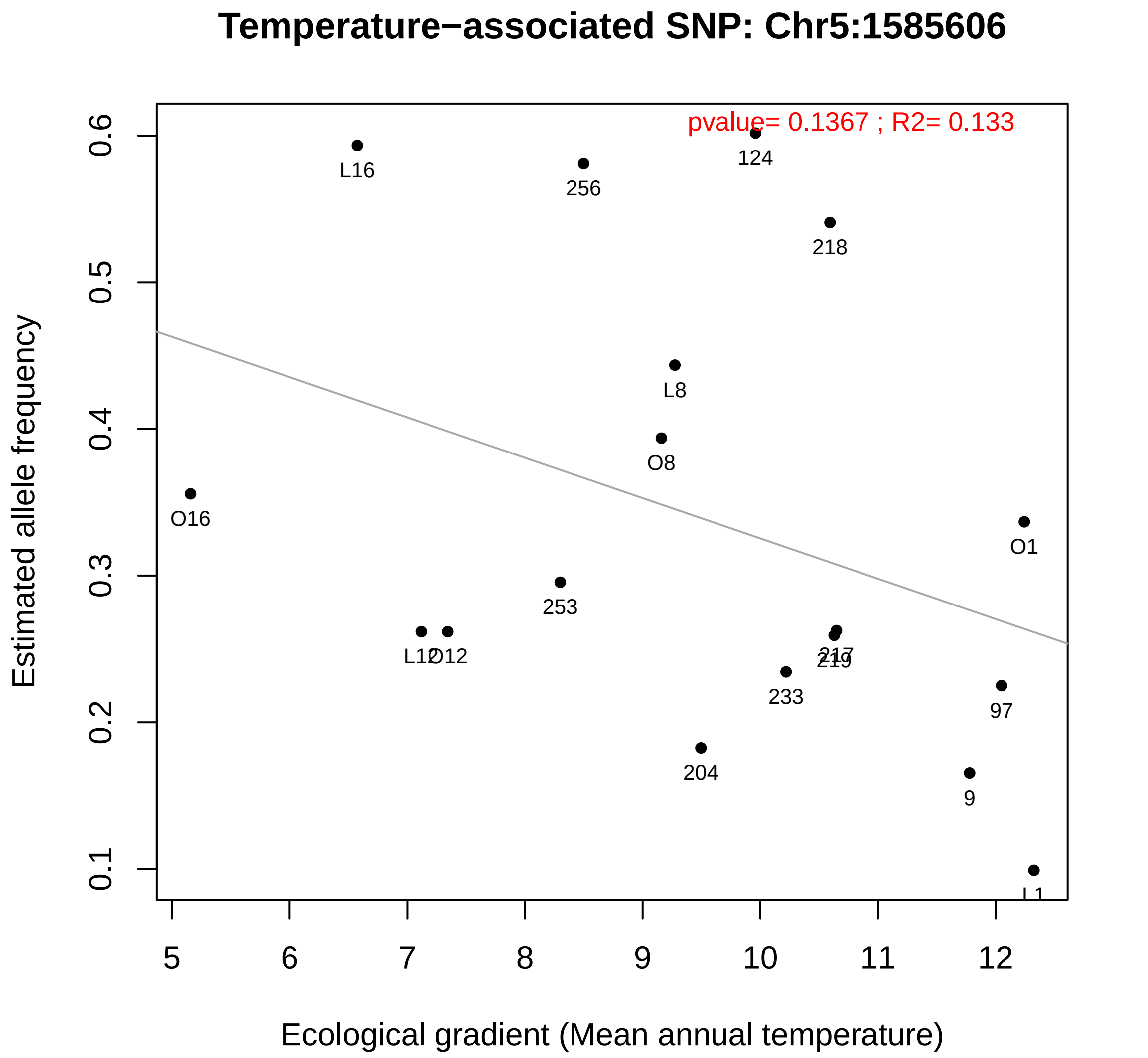
Temperature−associated SNP: Chr5:1585606
pvalue= 0.1367 ; R2= 0.133
Estimated allele frequency
Ecological gradient (Mean annual temperature)
0.1
0.2
0.3
0.4
0.5
0.6
5
6
7
8
9
10
11
12
L16
256
124
218
L8
O8
O16
O1
253
L12
O12
217
219
233
97
204
9
L1

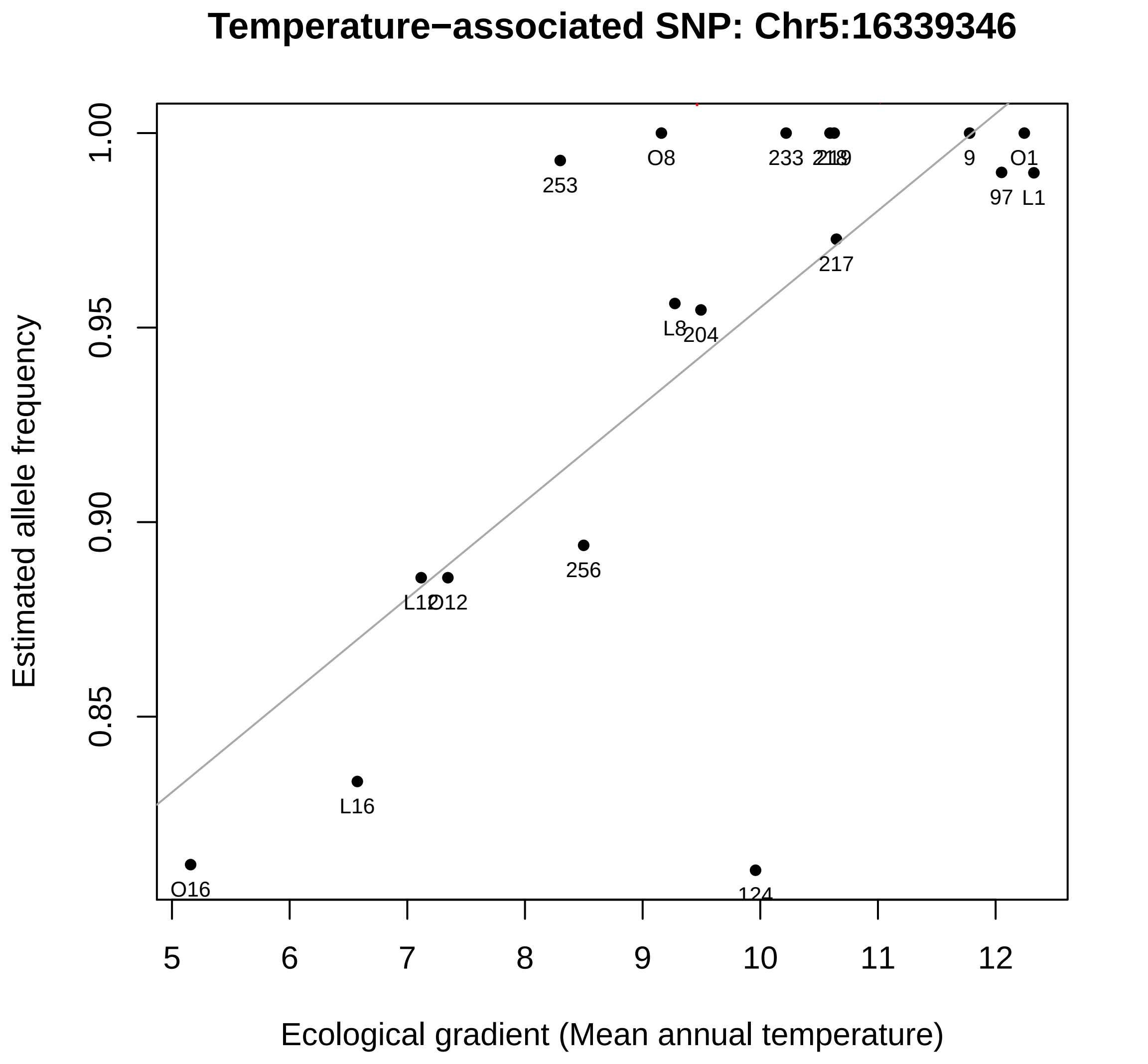

Temperature−associated SNP: Chr5:16339346
Estimated allele frequency
Ecological gradient (Mean annual temperature)
0.85
0.90
0.95
1.00
5
6
7
8
9
10
11
12
O16
L16
L12
O12
256
253
O8
L8
204
124
233
218
219
217
9
O1
97
L1

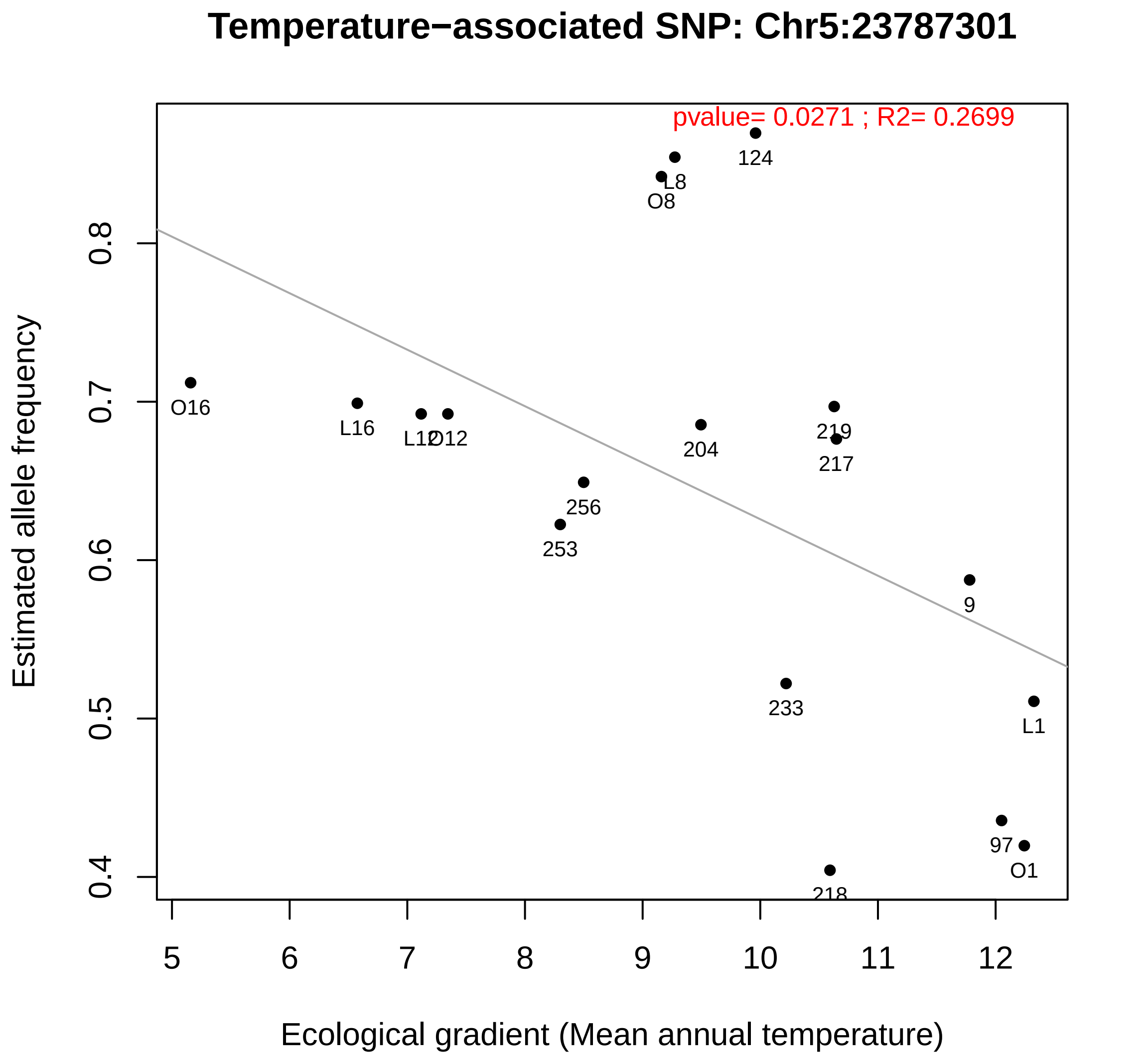
Temperature−associated SNP: Chr5:23787301
pvalue= 0.0271 ; R2= 0.2699
124
L8
O8
O16
L16
L12
O12
204
219
217
256
253
9
233
L1
97
O1
218
0.4
0.5
0.6
0.7
0.8
5
6
7
8
9
10
11
12
Estimated allele frequency
Ecological gradient (Mean annual temperature)

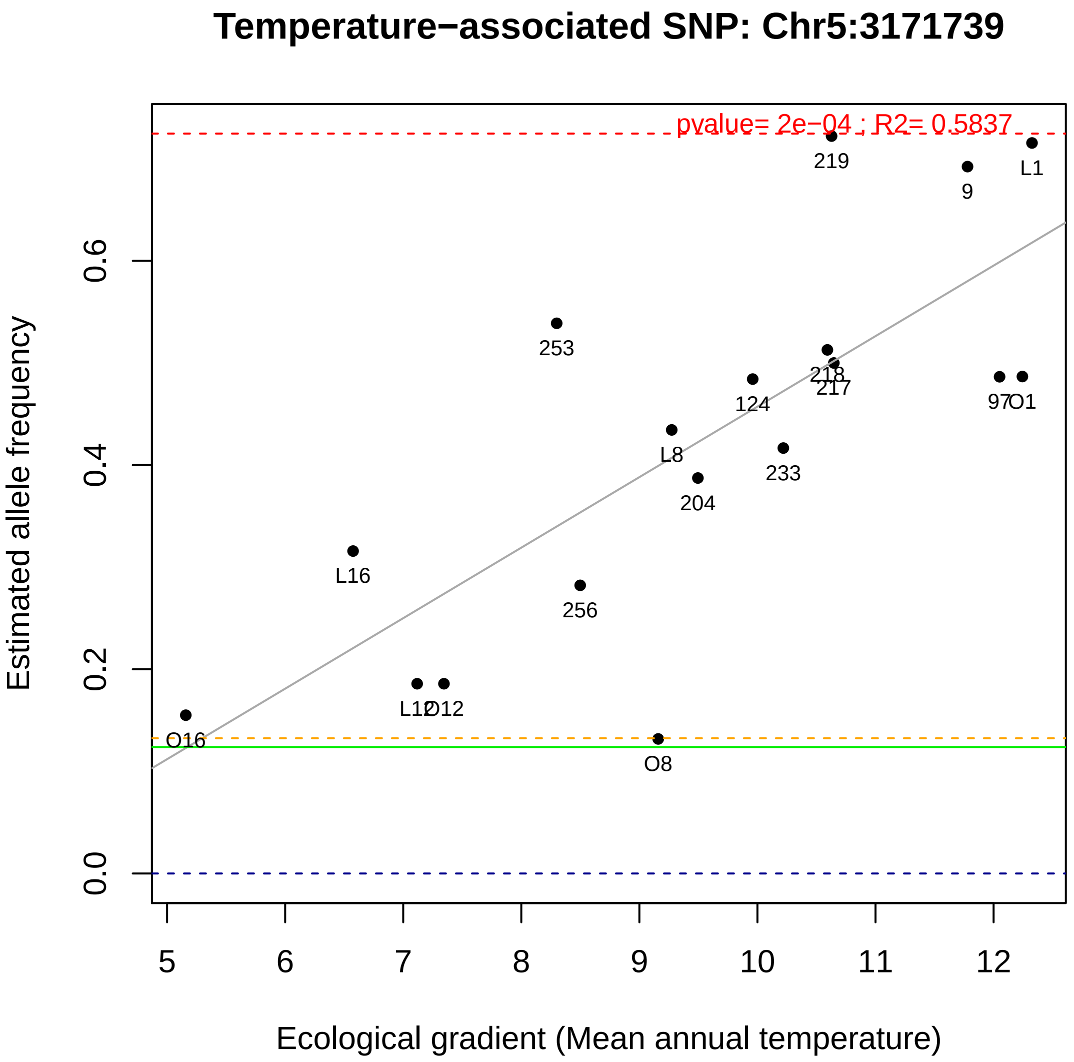

Temperature−associated SNP: Chr5:3171739
pvalue= 2e−04 ; R2= 0.5837
219
L1
9
253
218
217
124
97
O1
L8
233
204
L16
256
L12
O12
O16
O8
0.0
0.2
0.4
0.6
Estimated allele frequency
5
6
7
8
9
10
11
12
Ecological gradient (Mean annual temperature)

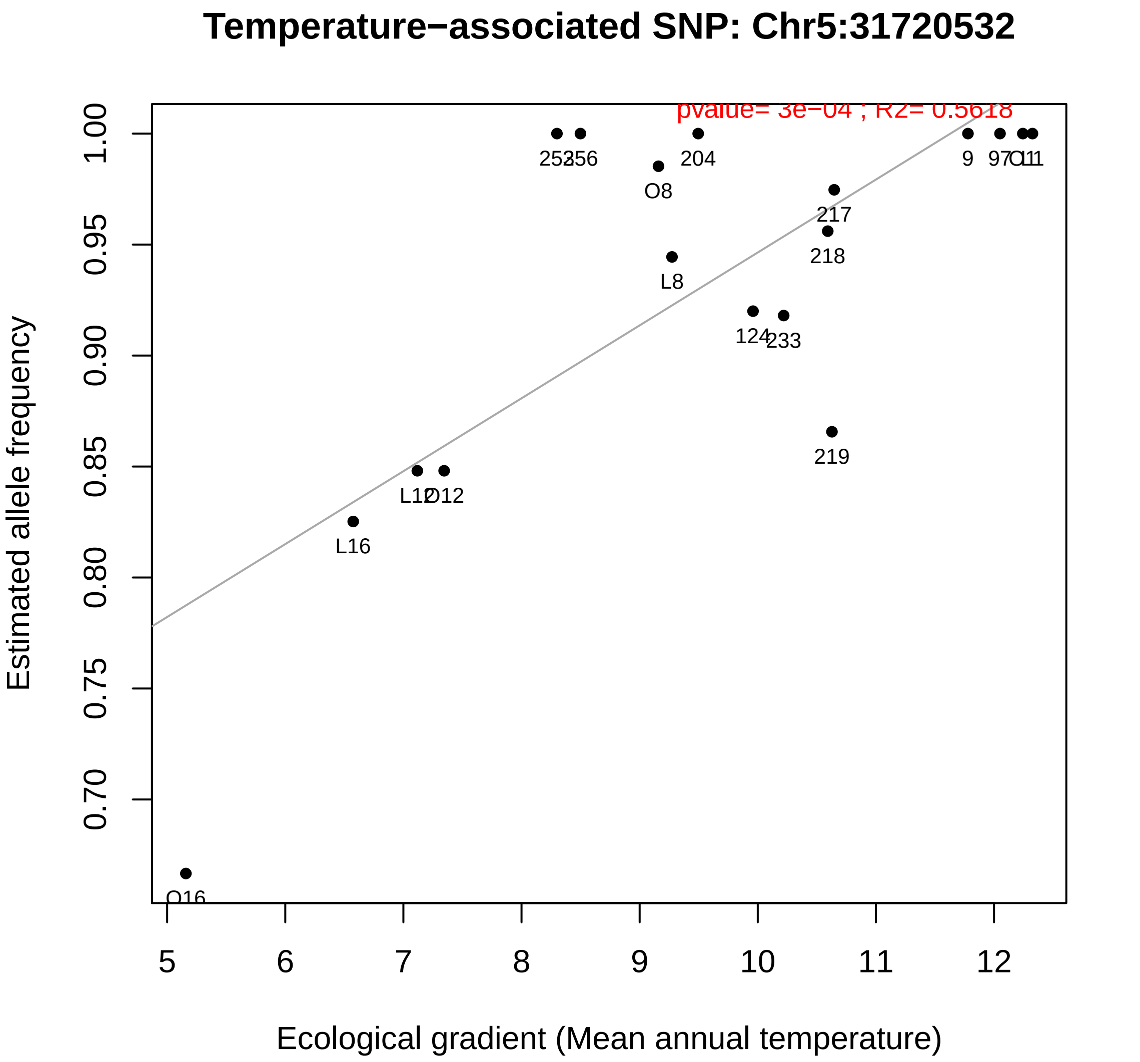
Temperature−associated SNP: Chr5:31720532
pvalue= 3e−04 ; R2= 0.5618
Estimated allele frequency
0.70
0.75
0.80
0.85
0.90
0.95
1.00
5
6
7
8
9
10
11
12
Ecological gradient (Mean annual temperature)
256
204
O8
217
218
L8
124
233
219
L16
O12
O16
9
97

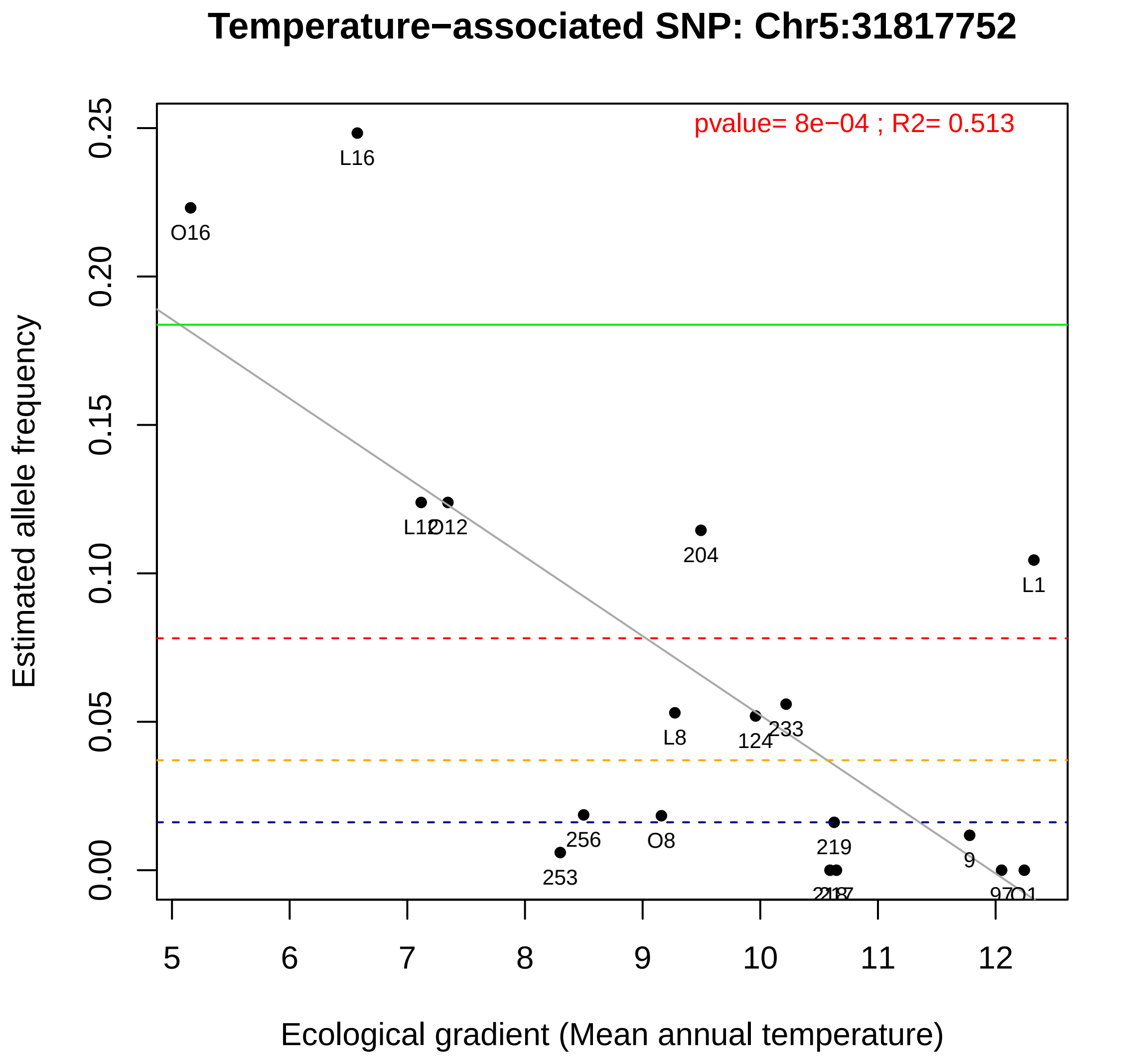
Temperature−associated SNP: Chr5:31817752
pvalue= 8e−04 ; R2= 0.513
Estimated allele frequency
0.00
0.05
0.10
0.15
0.20
0.25
5
6
7
8
9
10
11
12
Ecological gradient (Mean annual temperature)
L16
O16
L12
O12
204
L1
L8
124
233
256
253
O8
219
9
97
O1

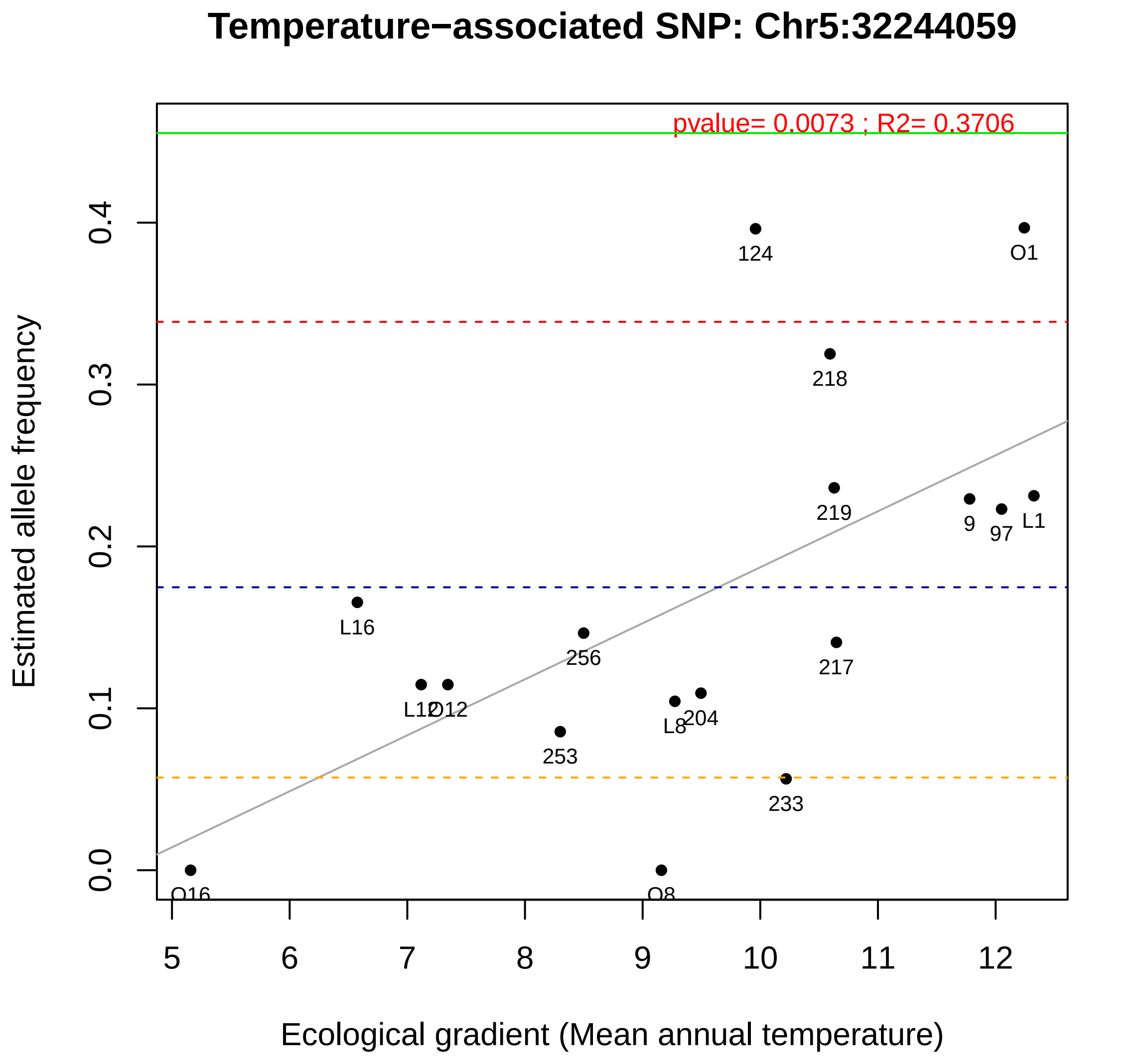
Temperature−associated SNP: Chr5:32244059
pvalue= 0.0073 ; R2= 0.3706
Estimated allele frequency
0.0
0.1
0.2
0.3
0.4
124
O1
218
219
9
97
L1
L16
256
217
L12
O12
253
L8
204
233
O16
O8
5
6
7
8
9
10
11
12
Ecological gradient (Mean annual temperature)

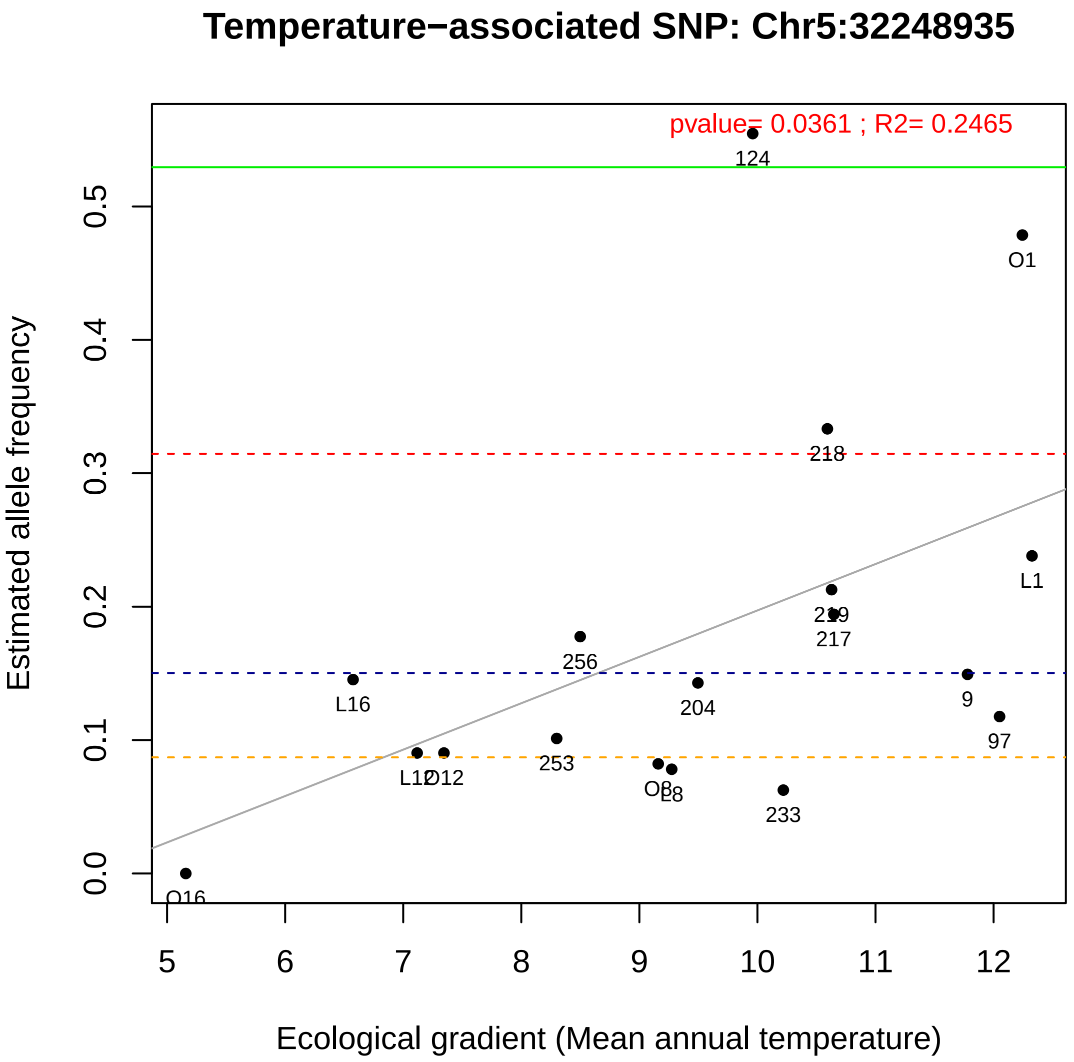

Temperature−associated SNP: Chr5:32248935
pvalue= 0.0361 ; R2= 0.2465
Estimated allele frequency
0.0
0.1
0.2
0.3
0.4
0.5
5
6
7
8
9
10
11
12
Ecological gradient (Mean annual temperature)
124
O1
218
L1
219
217
256
L16
204
9
97
253
L12
O12
O8
L8
233
O16

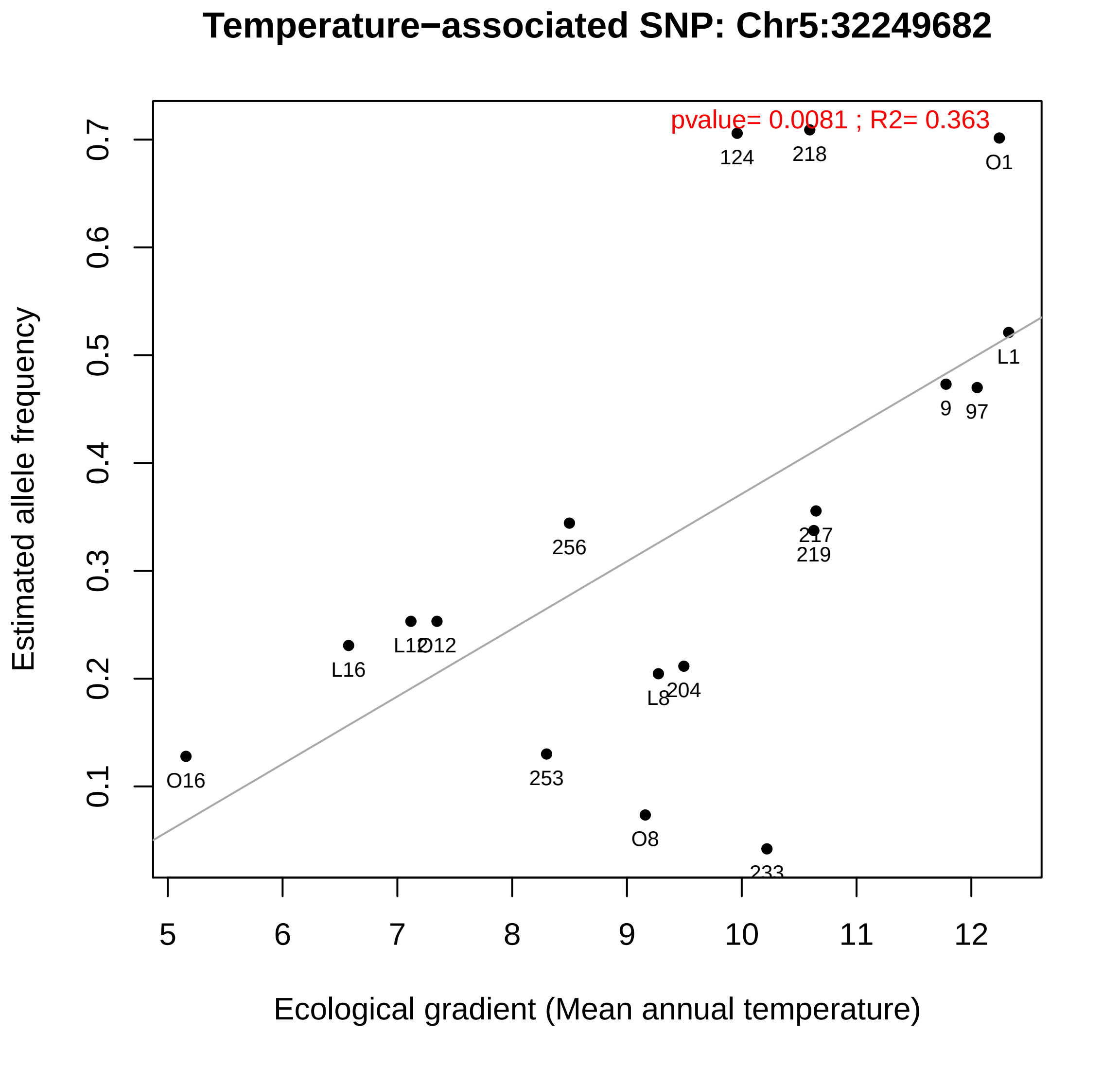

Temperature−associated SNP: Chr5:32249682
pvalue= 0.0081 ; R2= 0.363
Estimated allele frequency
Ecological gradient (Mean annual temperature)
0.1
0.2
0.3
0.4
0.5
0.6
0.7
5
6
7
8
9
10
11
12
124
218
O1
L1
9
97
217
219
256
L12
O12
L16
L8
204
O16
253
O8
233

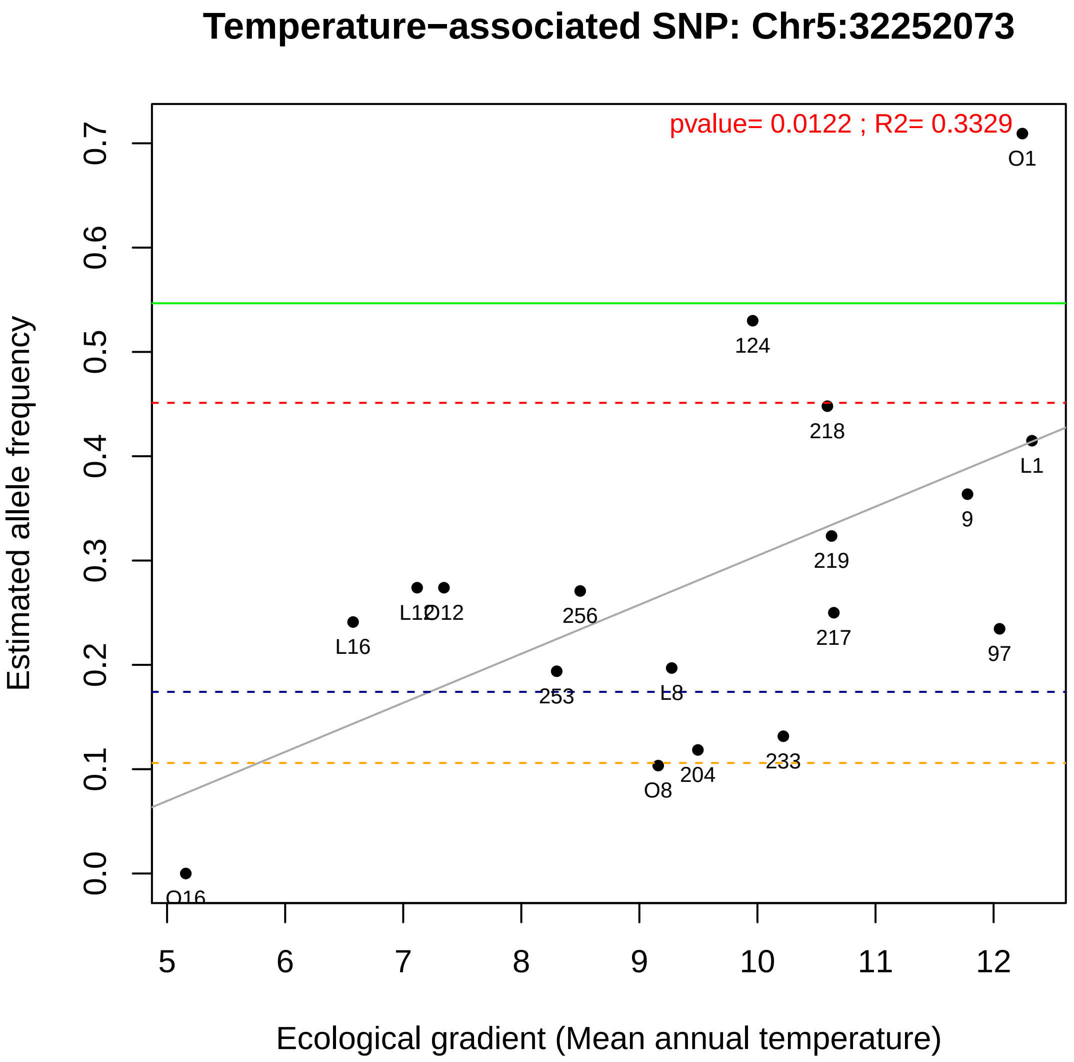
Temperature−associated SNP: Chr5:32252073
pvalue= 0.0122 ; R2= 0.3329
Estimated allele frequency
Ecological gradient (Mean annual temperature)
0.0
0.1
0.2
0.3
0.4
0.5
0.6
0.7
5
6
7
8
9
10
11
12
O1
124
218
L1
9
219
217
97
256
L12
O12
L16
253
L8
233
204
O8
O16

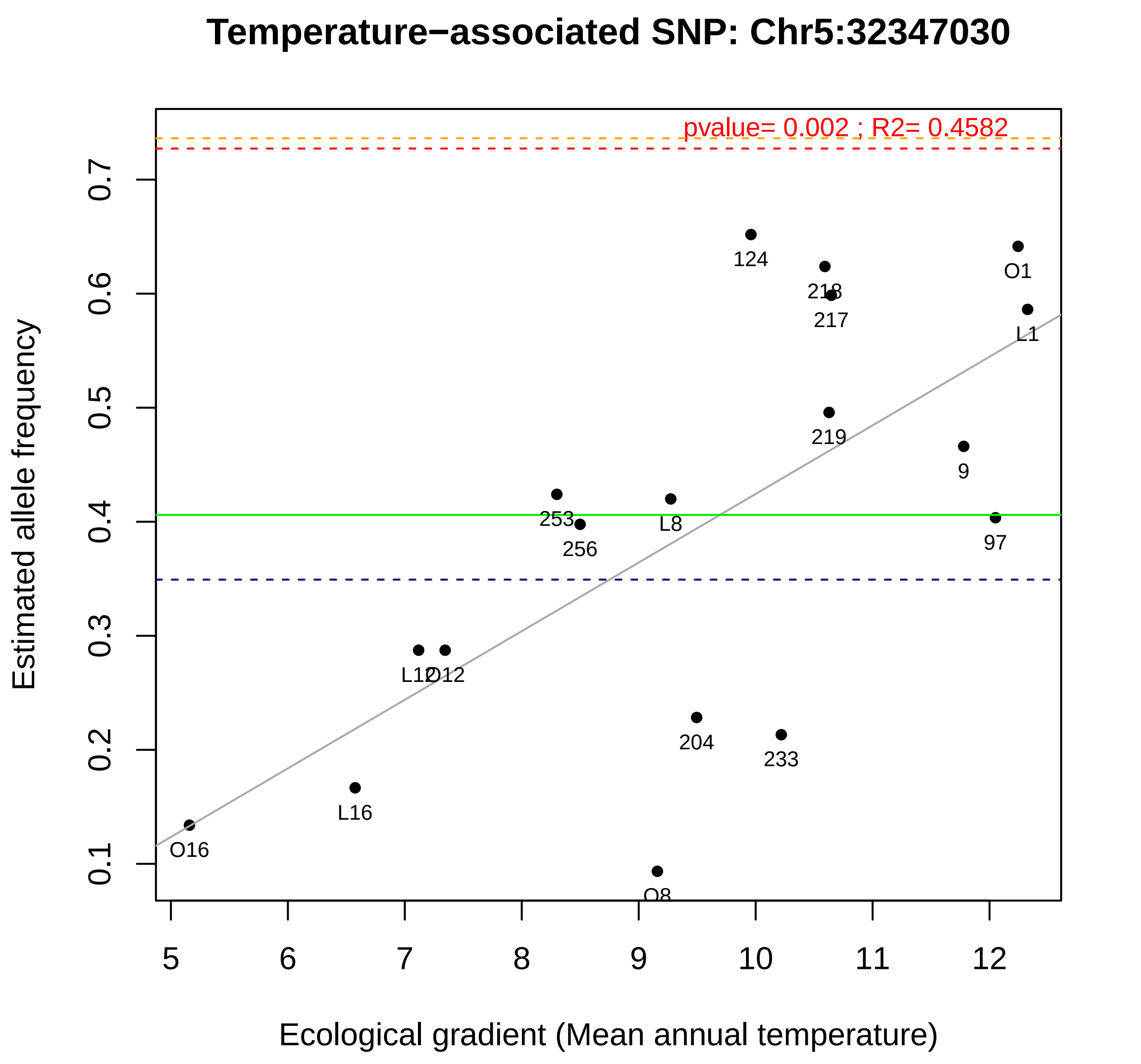

Temperature−associated SNP: Chr5:32347030
pvalue= 0.002 ; R2= 0.4582
Estimated allele frequency
0.1
0.2
0.3
0.4
0.5
0.6
0.7
5
6
7
8
9
10
11
12
Ecological gradient (Mean annual temperature)
124
218
217
O1
L1
219
9
253
256
L8
97
L12
O12
204
233
L16
O16
O8

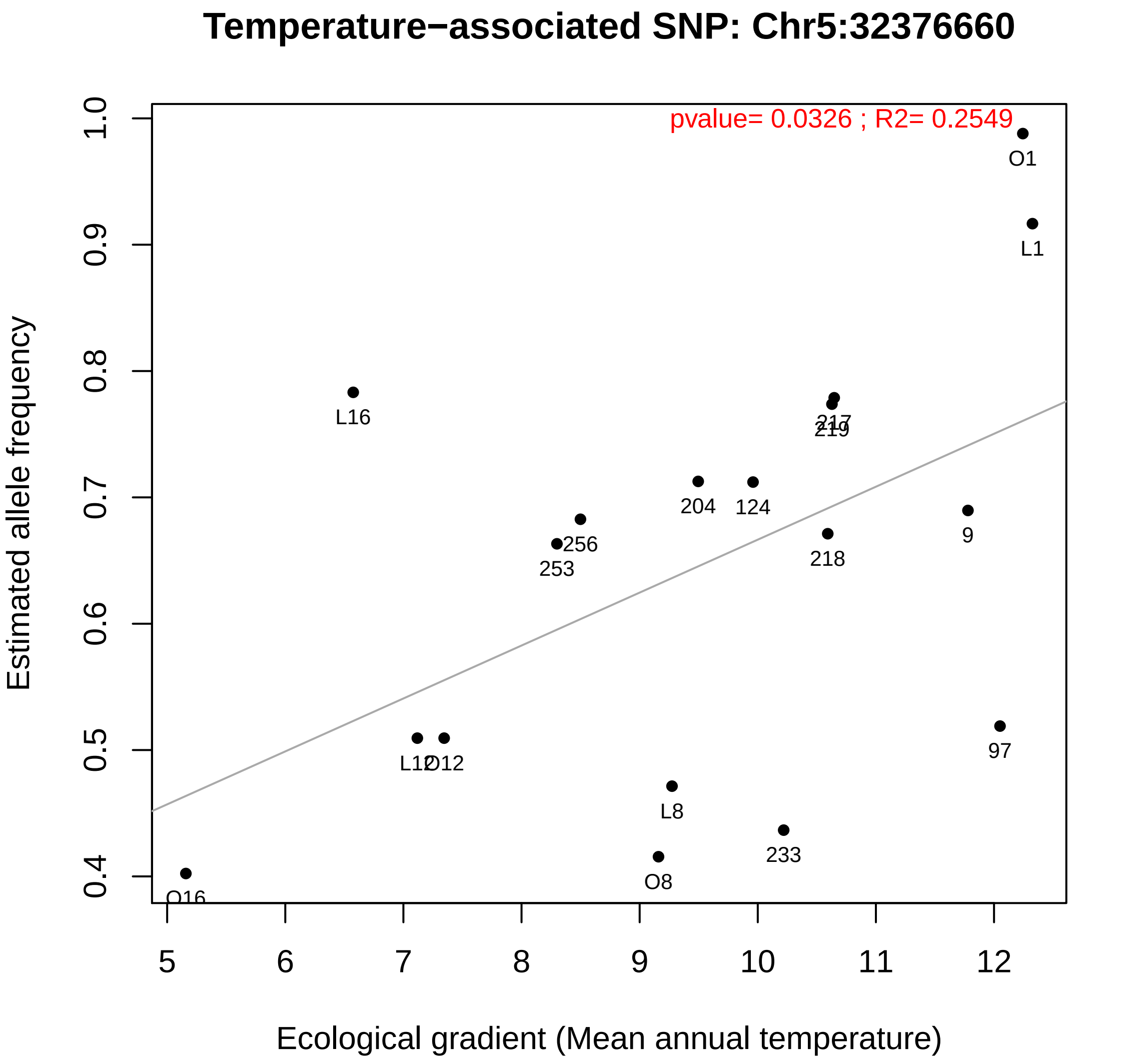
Temperature−associated SNP: Chr5:32376660
pvalue= 0.0326 ; R2= 0.2549
Estimated allele frequency
0.4
0.5
0.6
0.7
0.8
0.9
1.0
5
6
7
8
9
10
11
12
Ecological gradient (Mean annual temperature)
O1
L1
L16
217
219
204
124
256
253
218
9
L12
O12
L8
233
O8
97
O16

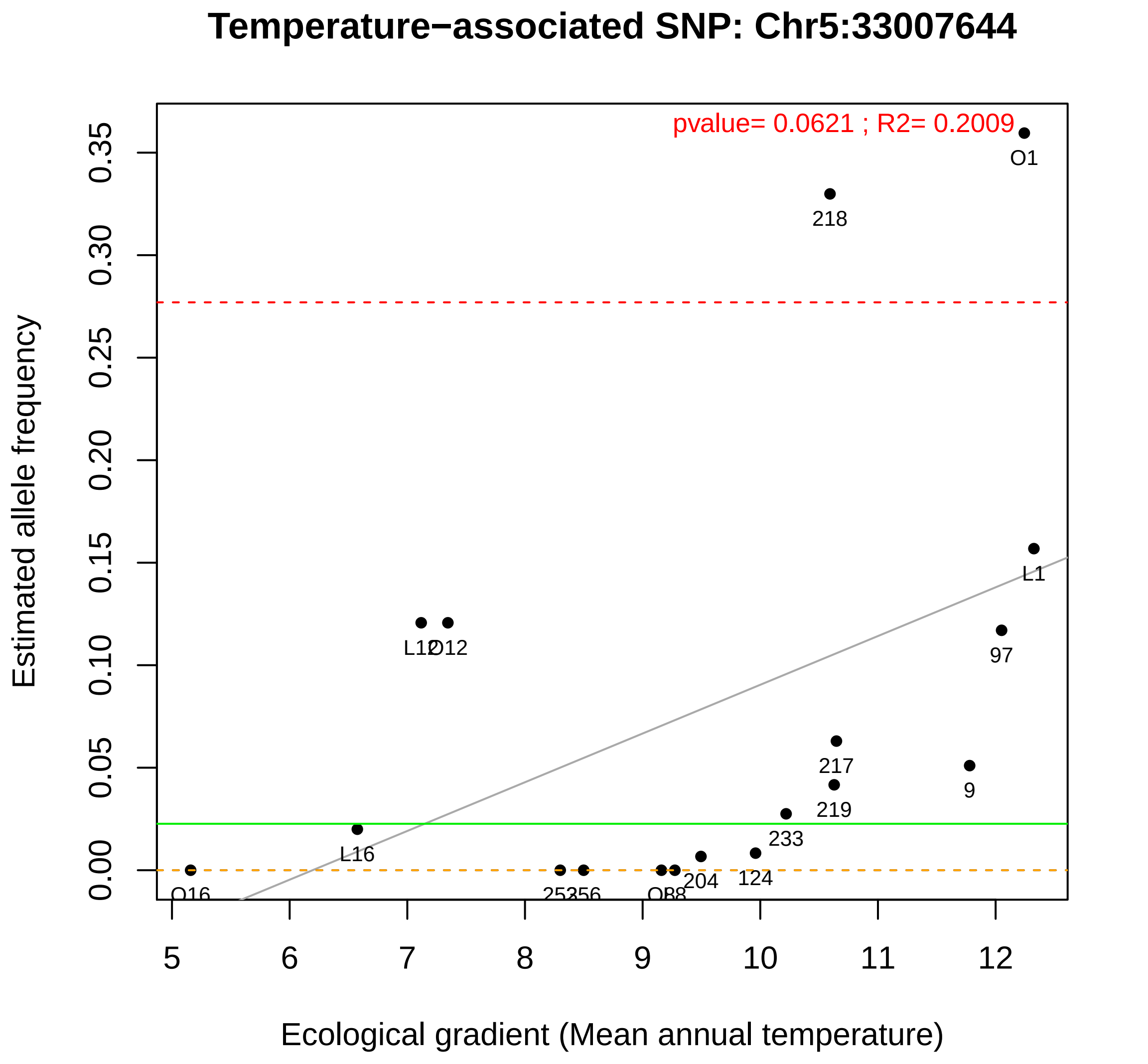

Temperature−associated SNP: Chr5:33007644
pvalue= 0.0621 ; R2= 0.2009
Estimated allele frequency
Ecological gradient (Mean annual temperature)
0.00
0.05
0.10
0.15
0.20
0.25
0.30
0.35
5
6
7
8
9
10
11
12
O1
218
L1
97
217
9
219
233
124
204
L16
O16

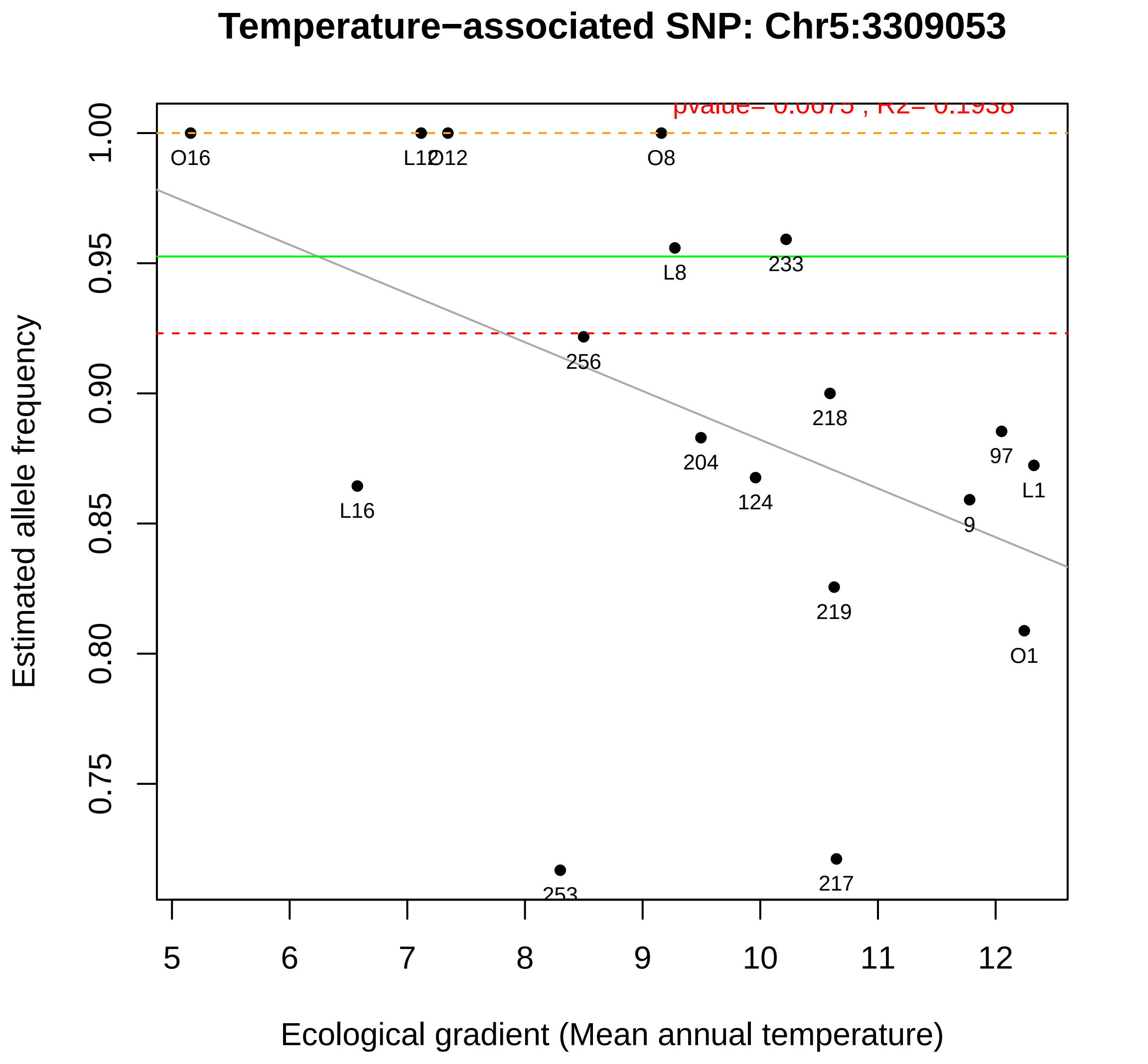
Temperature−associated SNP: Chr5:3309053
pvalue= 0.0675 , R2= 0.1938
Estimated allele frequency
1.00
0.95
0.90
0.85
0.80
0.75
O16
L12
O12
O8
L8
233
256
218
204
97
124
L1
L16
9
219
O1
217
253
5
6
7
8
9
10
11
12
Ecological gradient (Mean annual temperature)

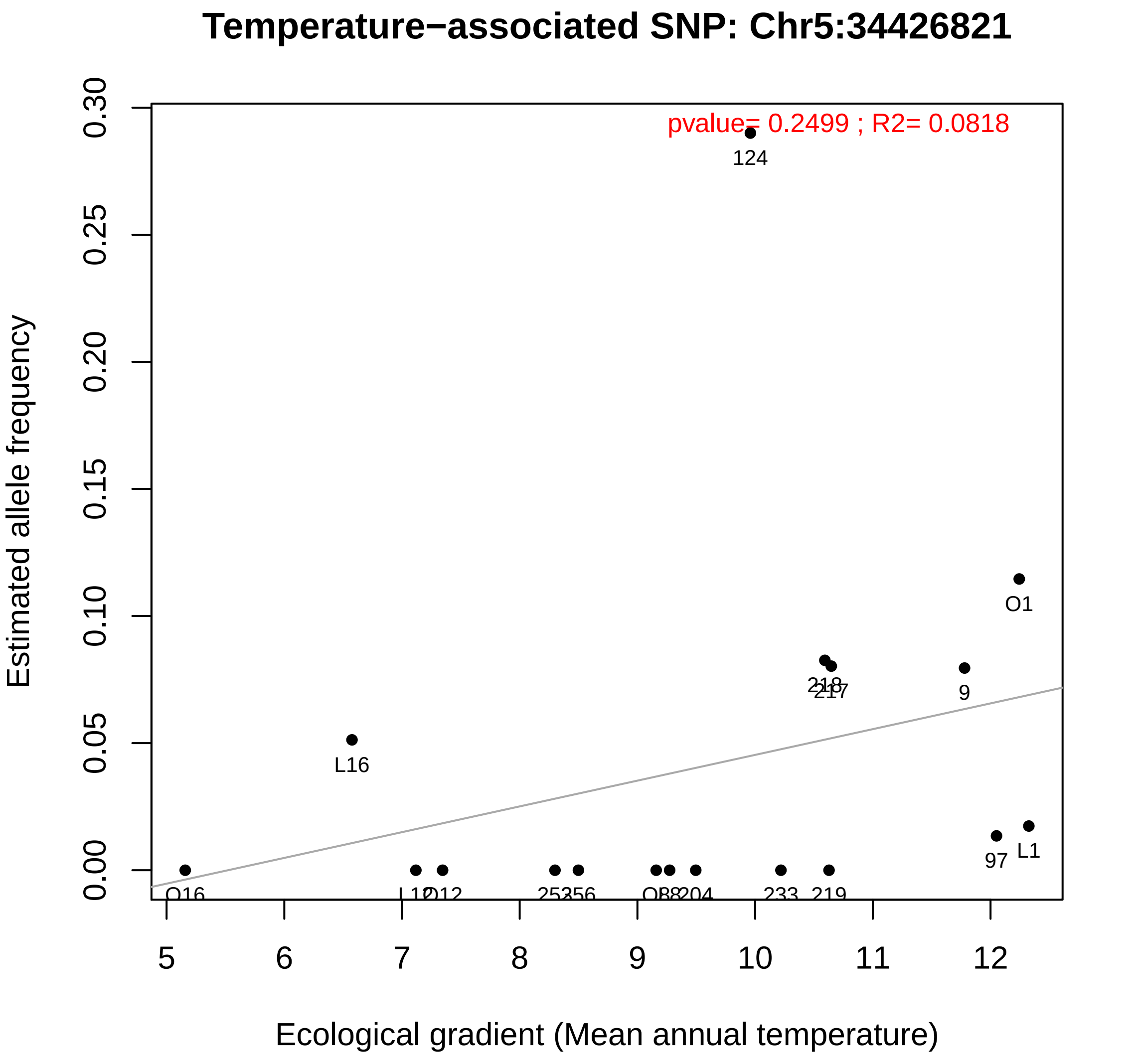
Temperature−associated SNP: Chr5:34426821
pvalue= 0.2499 ; R2= 0.0818
Estimated allele frequency
0.00
0.05
0.10
0.15
0.20
0.25
0.30
5
6
7
8
9
10
11
12
Ecological gradient (Mean annual temperature)
124
O1
218
217
9
L16
97
L1
O16
204
233
219

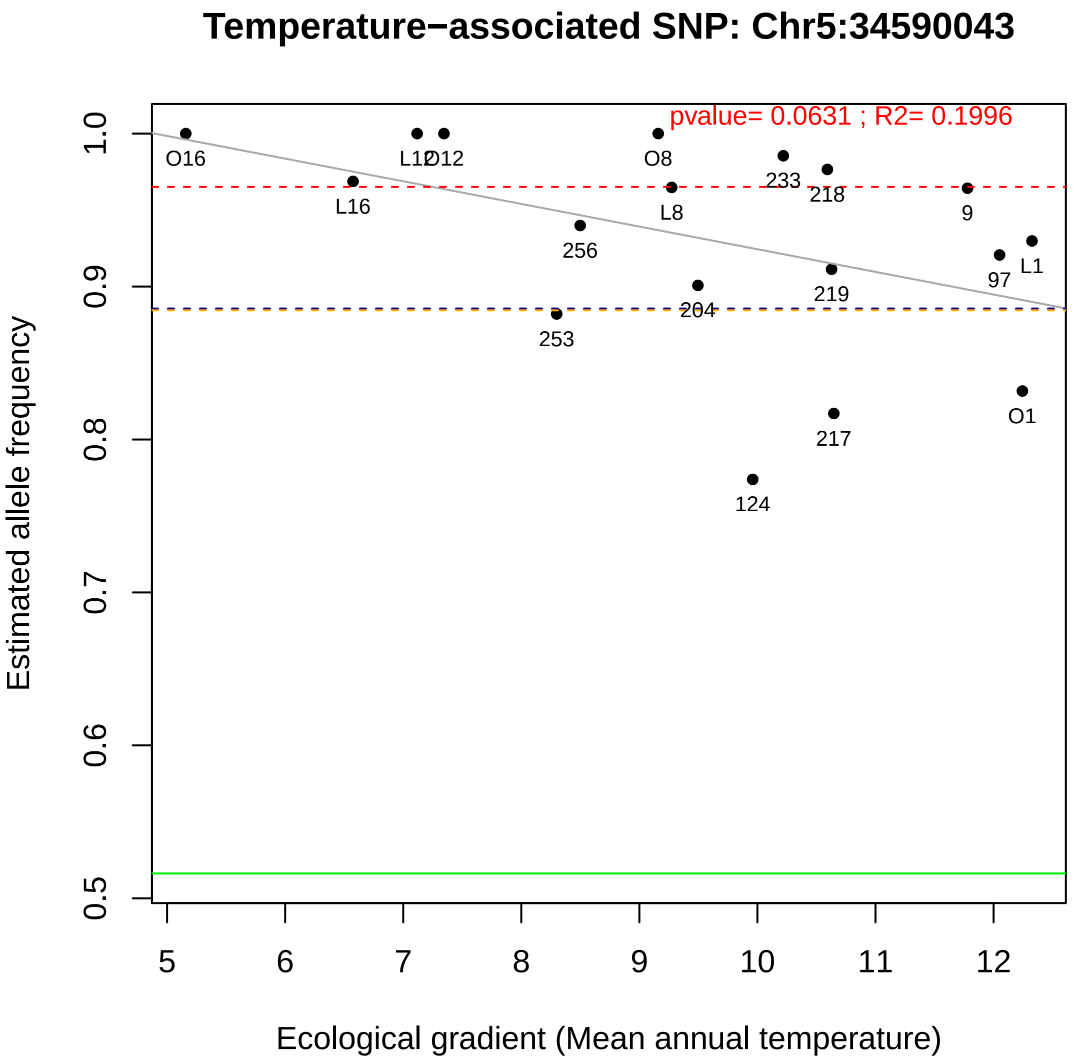
Temperature−associated SNP: Chr5:34590043
pvalue= 0.0631 ; R2= 0.1996
Estimated allele frequency
1.0
0.9
0.8
0.7
0.6
0.5
O16
L16
L12
O12
256
253
O8
L8
204
233
124
218
219
217
9
97
L1
O1
5
6
7
8
9
10
11
12
Ecological gradient (Mean annual temperature)

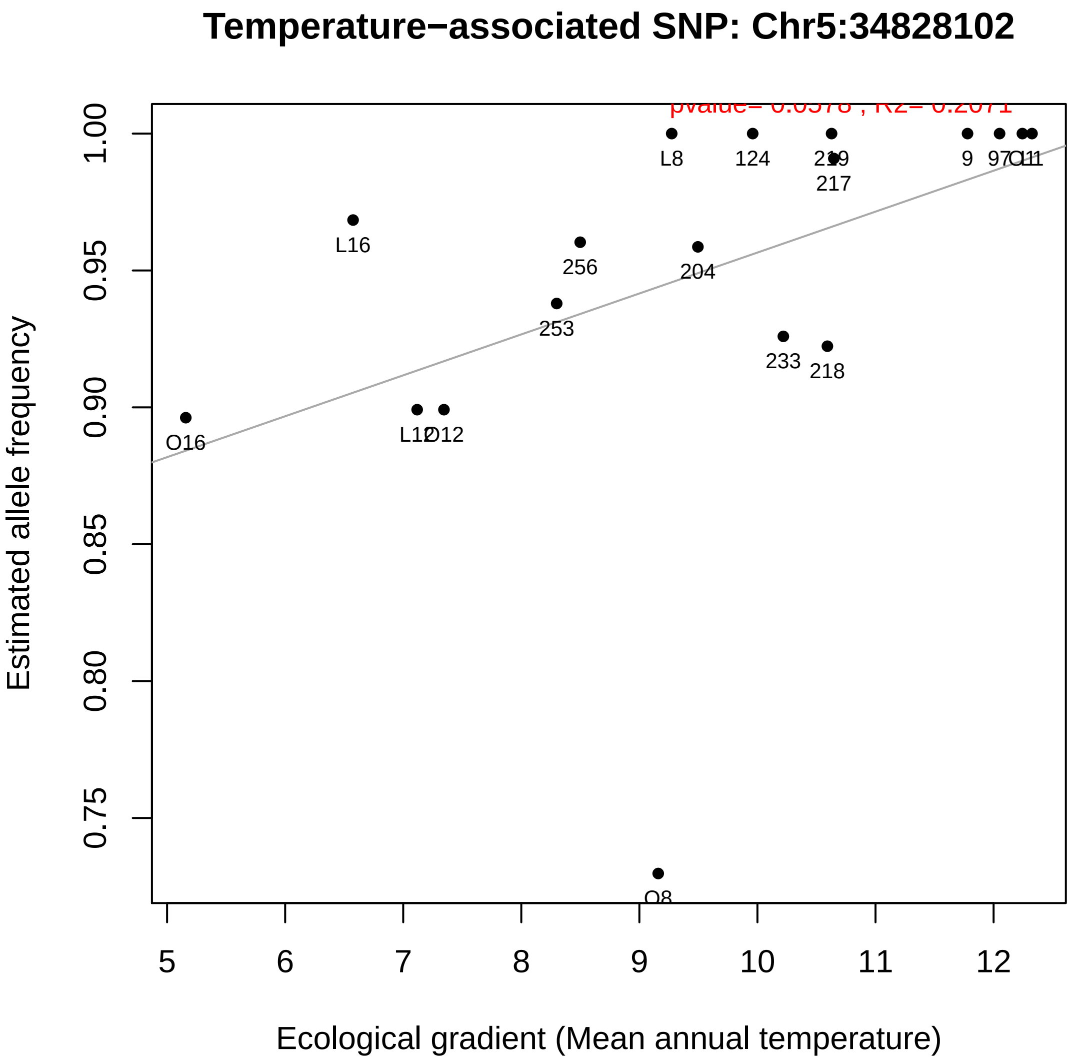
Temperature−associated SNP: Chr5:34828102
Estimated allele frequency
1.00
0.95
0.90
0.85
0.80
0.75
Ecological gradient (Mean annual temperature)
5
6
7
8
9
10
11
12
L8
124
219
217
9
97
O11
L16
256
204
253
233
218
O16
L12
O12
O8

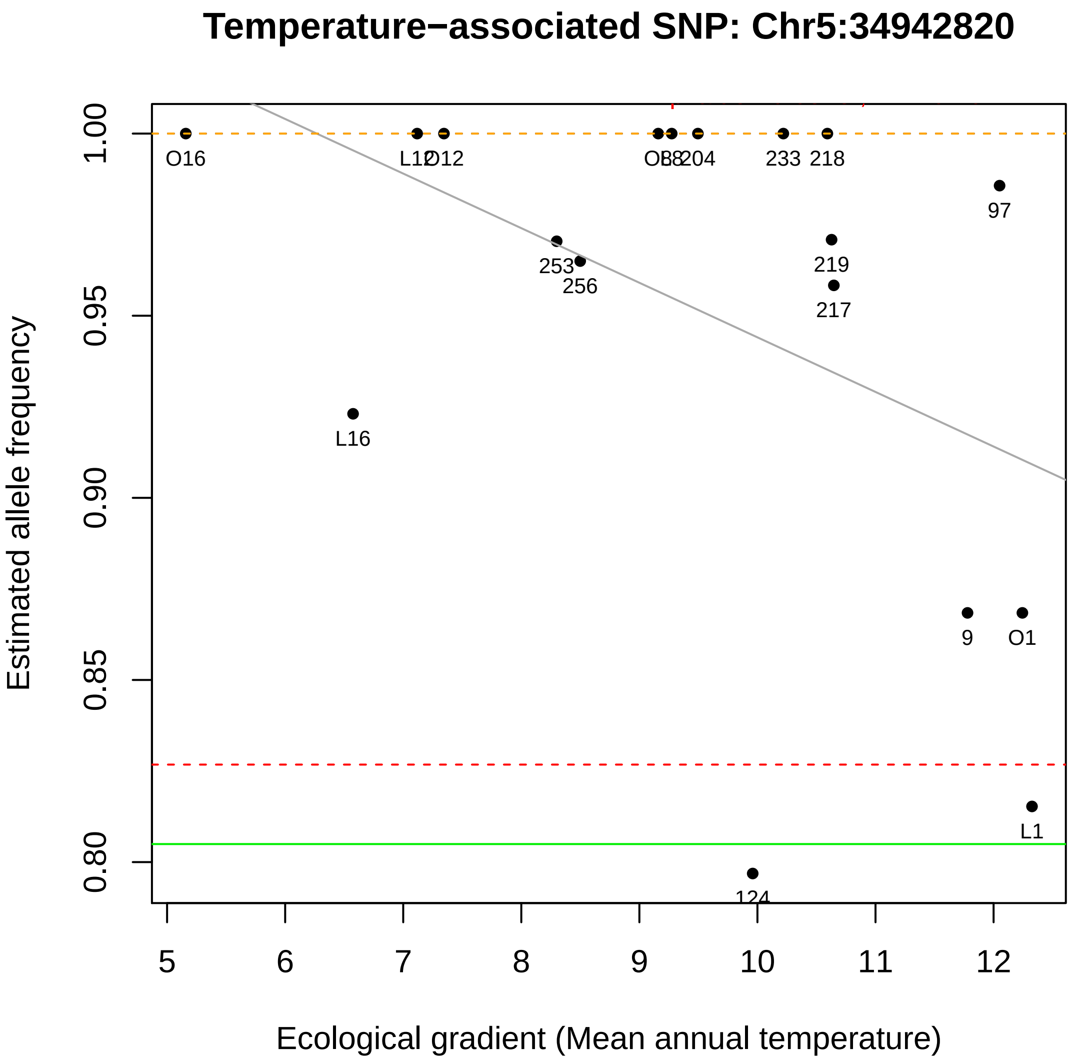
Temperature−associated SNP: Chr5:34942820
Estimated allele frequency
Ecological gradient (Mean annual temperature)
1.00
0.95
0.90
0.85
0.80
5
6
7
8
9
10
11
12
O16
L12
O12
204
233
218
97
253
256
219
217
L16
9
O1
L1
124

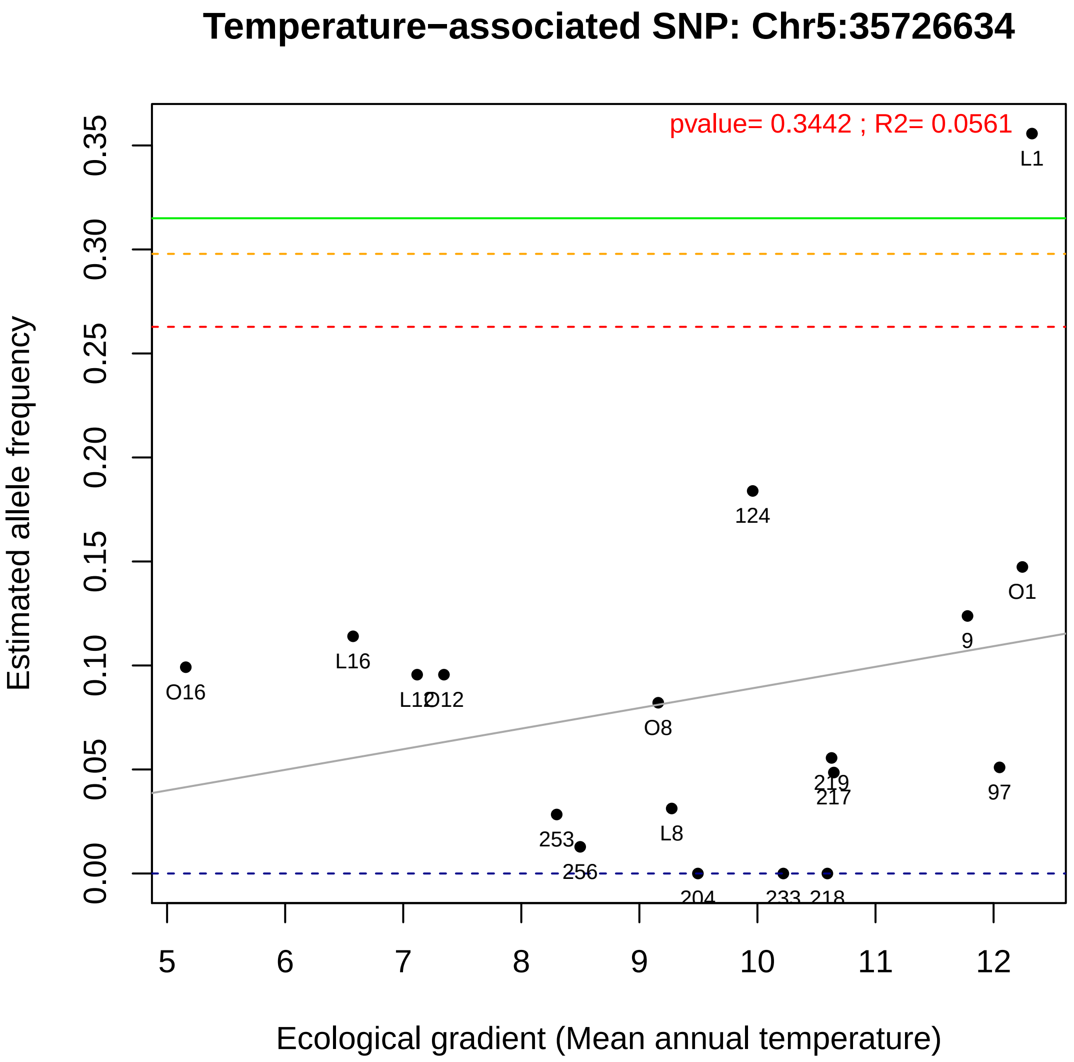

Temperature−associated SNP: Chr5:35726634
pvalue= 0.3442 ; R2= 0.0561
Estimated allele frequency
0.00
0.05
0.10
0.15
0.20
0.25
0.30
0.35
Ecological gradient (Mean annual temperature)
5
6
7
8
9
10
11
12
L1
124
O1
9
L16
O16
L12
O12
O8
219
217
97
253
256
L8
204
233
218

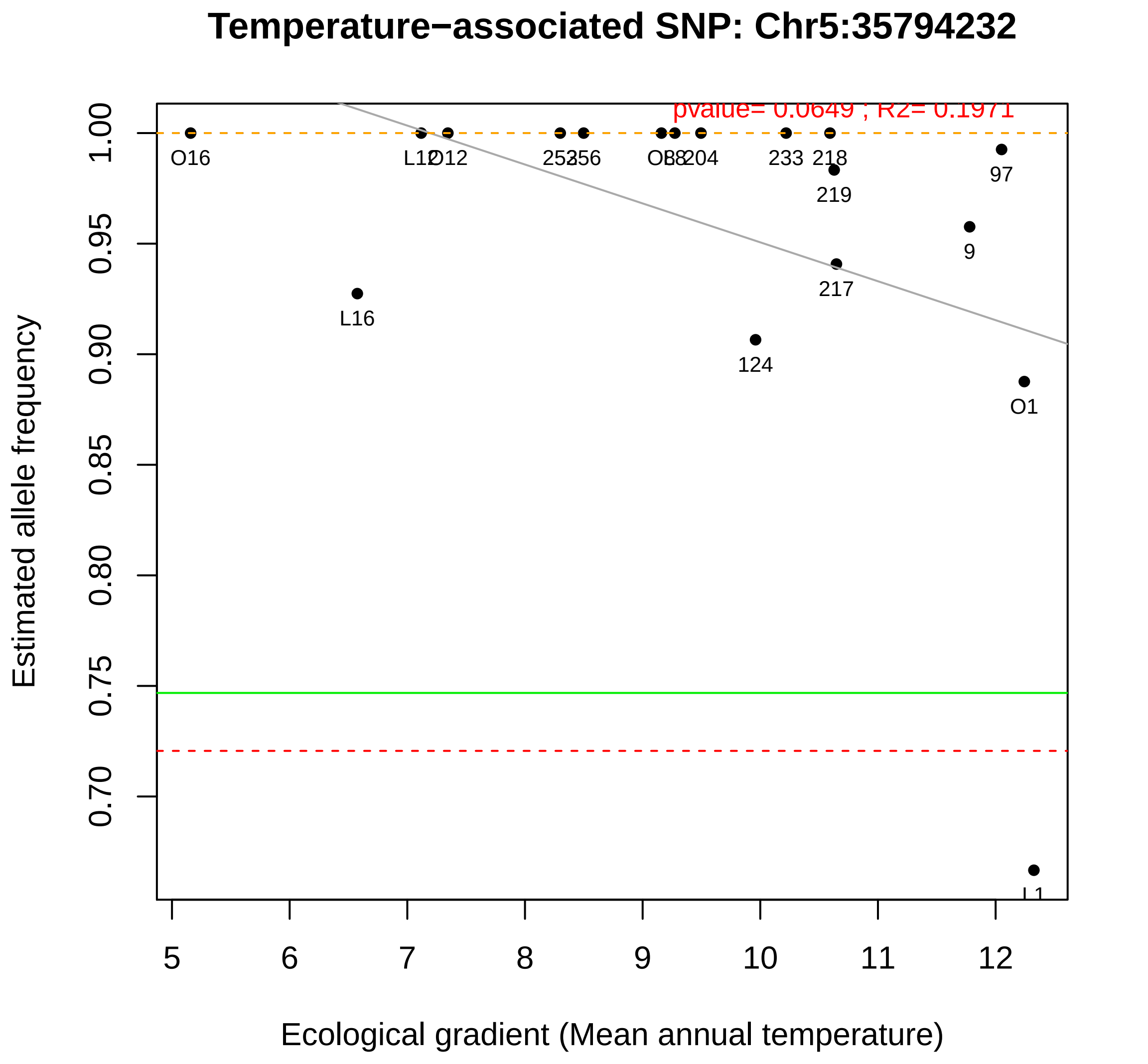

Temperature−associated SNP: Chr5:35794232
pvalue= 0.0649 ; R2= 0.1971
Estimated allele frequency
Ecological gradient (Mean annual temperature)
1.00
0.95
0.90
0.85
0.80
0.75
0.70
5
6
7
8
9
10
11
12
O16
L16
L12
O12
256
204
233
218
219
217
97
9
124
O1

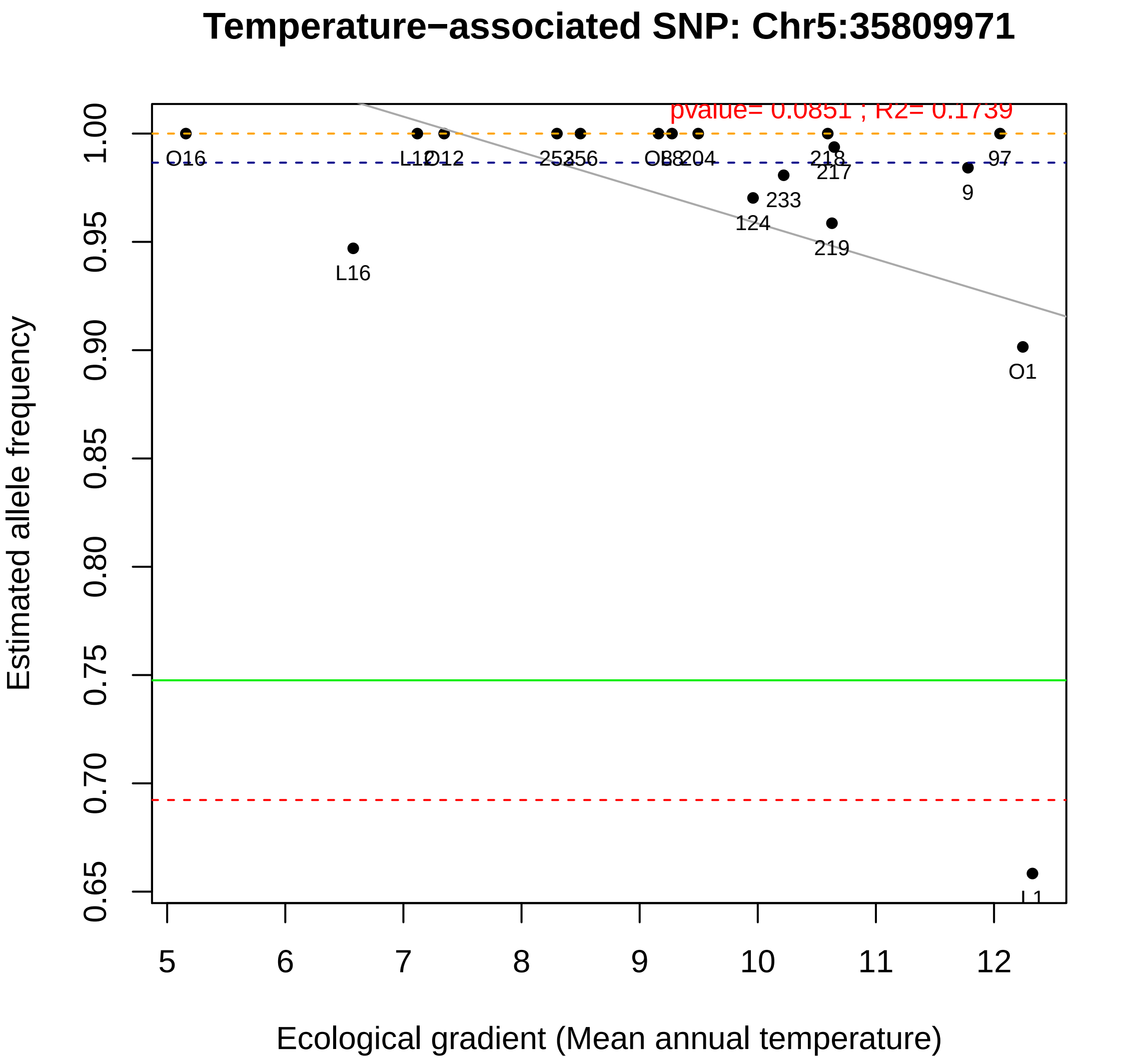
Temperature−associated SNP: Chr5:35809971
pvalue= 0.0851 ; R2= 0.1739
Estimated allele frequency
Ecological gradient (Mean annual temperature)
1.00
0.95
0.90
0.85
0.80
0.75
0.70
0.65
5
6
7
8
9
10
11
12
O16
L16
L12
O12
253
256
O3
L8
204
124
233
218
217
219
9
97
O1
L1

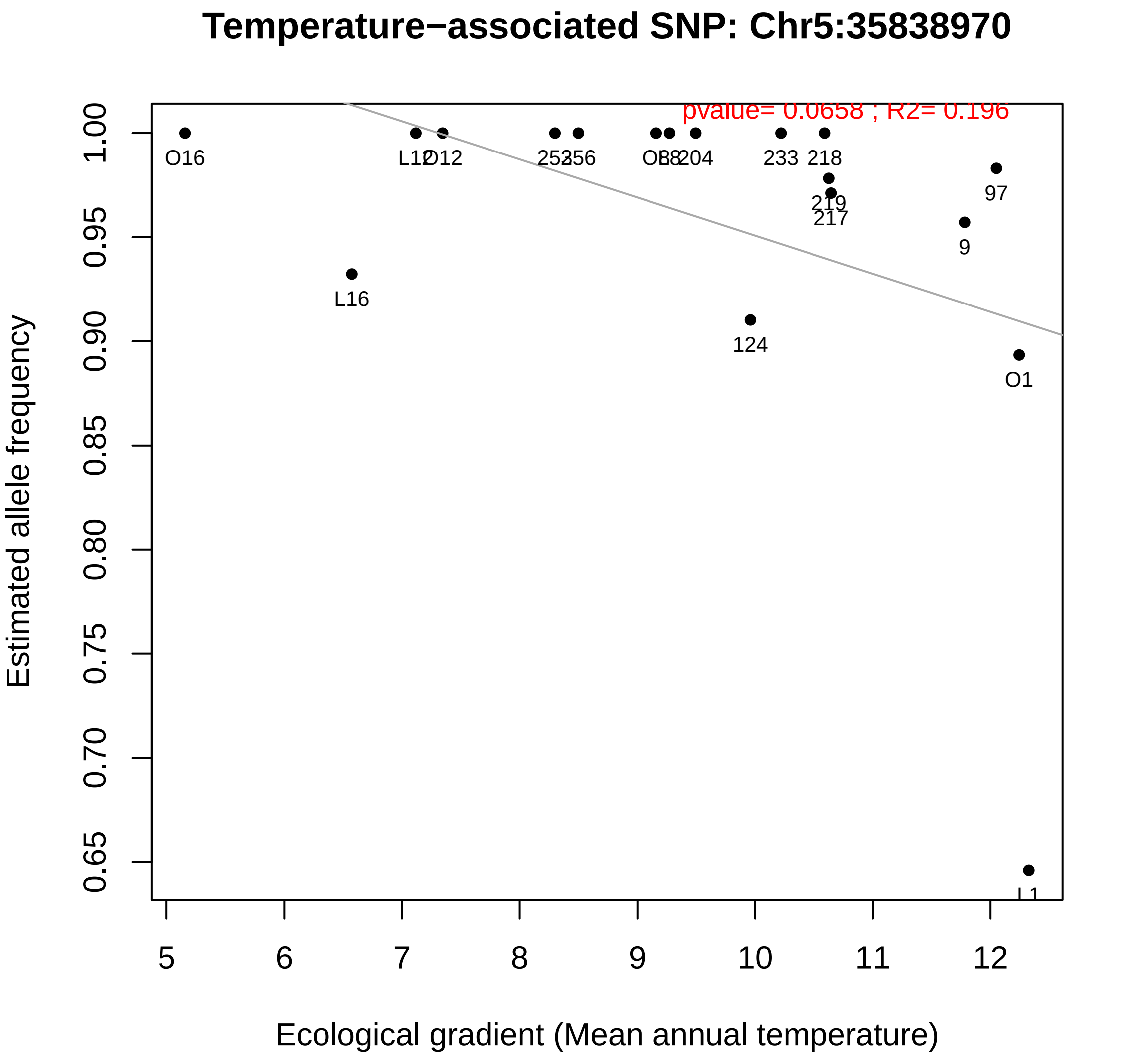

Temperature−associated SNP: Chr5:35838970
pvalue= 0.0658 ; R2= 0.196
Estimated allele frequency
1.00
0.95
0.90
0.85
0.80
0.75
0.70
0.65
O16
L12
O12
253
256
O3
18
204
233
218
219
217
97
9
L16
124
O1
5
6
7
8
9
10
11
12
Ecological gradient (Mean annual temperature)

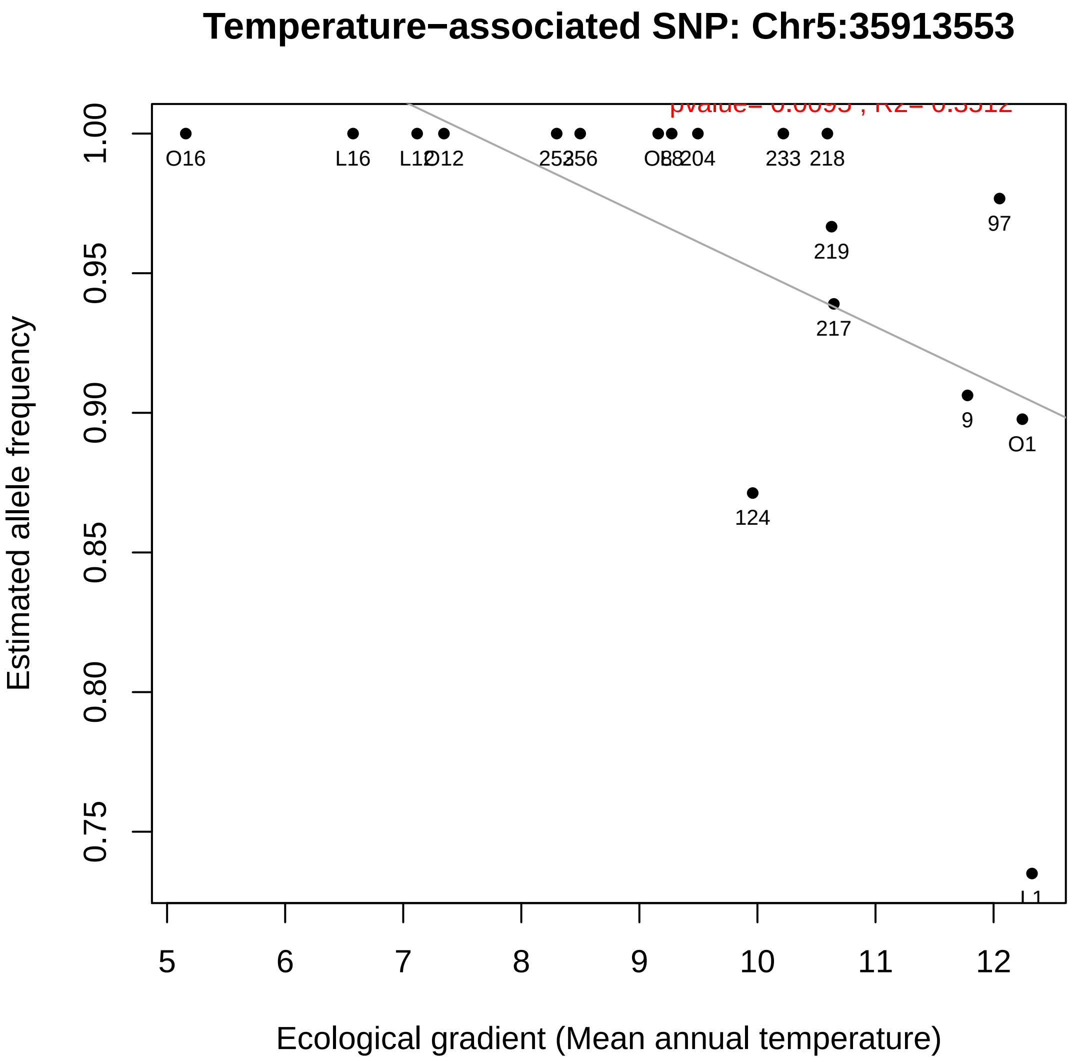
Temperature−associated SNP: Chr5:35913553
pvalue= 0.0093 ; R2= 0.3512
Estimated allele frequency
0.75
0.80
0.85
0.90
0.95
1.00
5
6
7
8
9
10
11
12
Ecological gradient (Mean annual temperature)
O16
L16
O12
233
218
97
219
217
9
O1
124

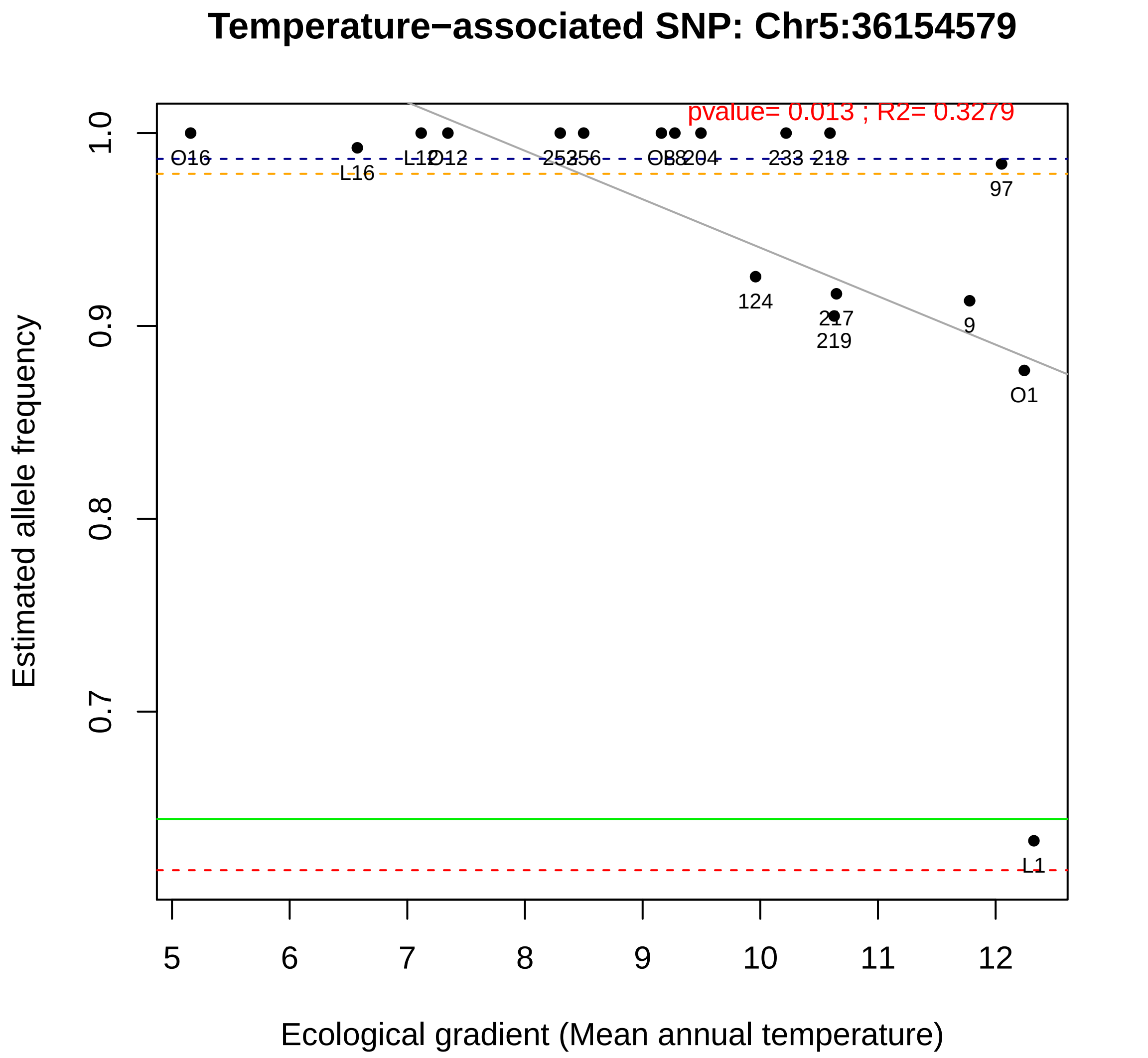

Temperature−associated SNP: Chr5:36154579
pvalue= 0.013 ; R2= 0.3279
O16
L16
L12
O12
253
256
O8
L8
204
233
218
97
124
217
219
9
O1
L1
Estimated allele frequency
1.0
0.9
0.8
0.7
5
6
7
8
9
10
11
12
Ecological gradient (Mean annual temperature)

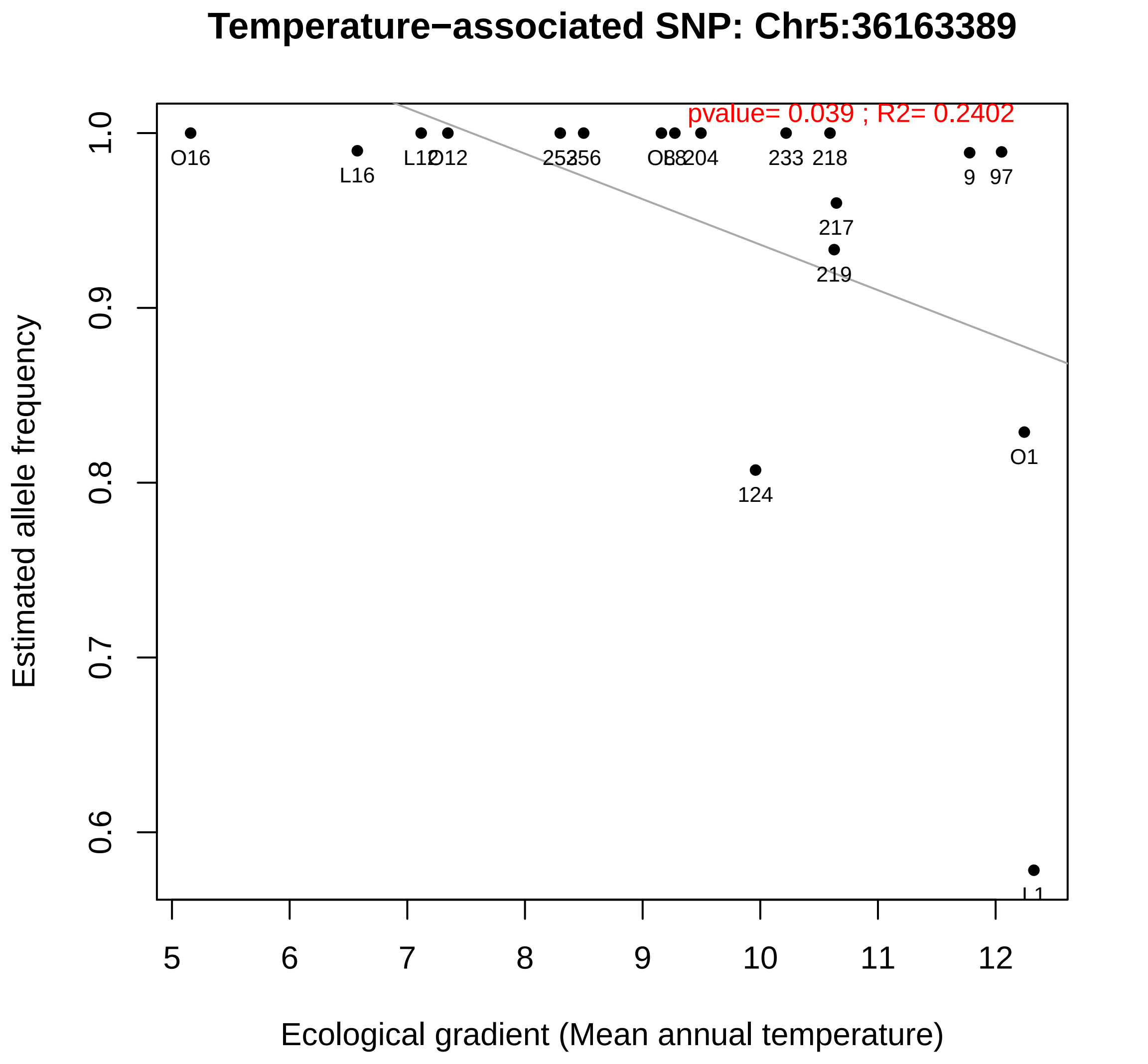

Temperature−associated SNP: Chr5:36163389
pvalue= 0.039 ; R2= 0.2402
Estimated allele frequency
0.6
0.7
0.8
0.9
1.0
5
6
7
8
9
10
11
12
Ecological gradient (Mean annual temperature)
O16
L16
233
218
9
97
217
219
124
O1

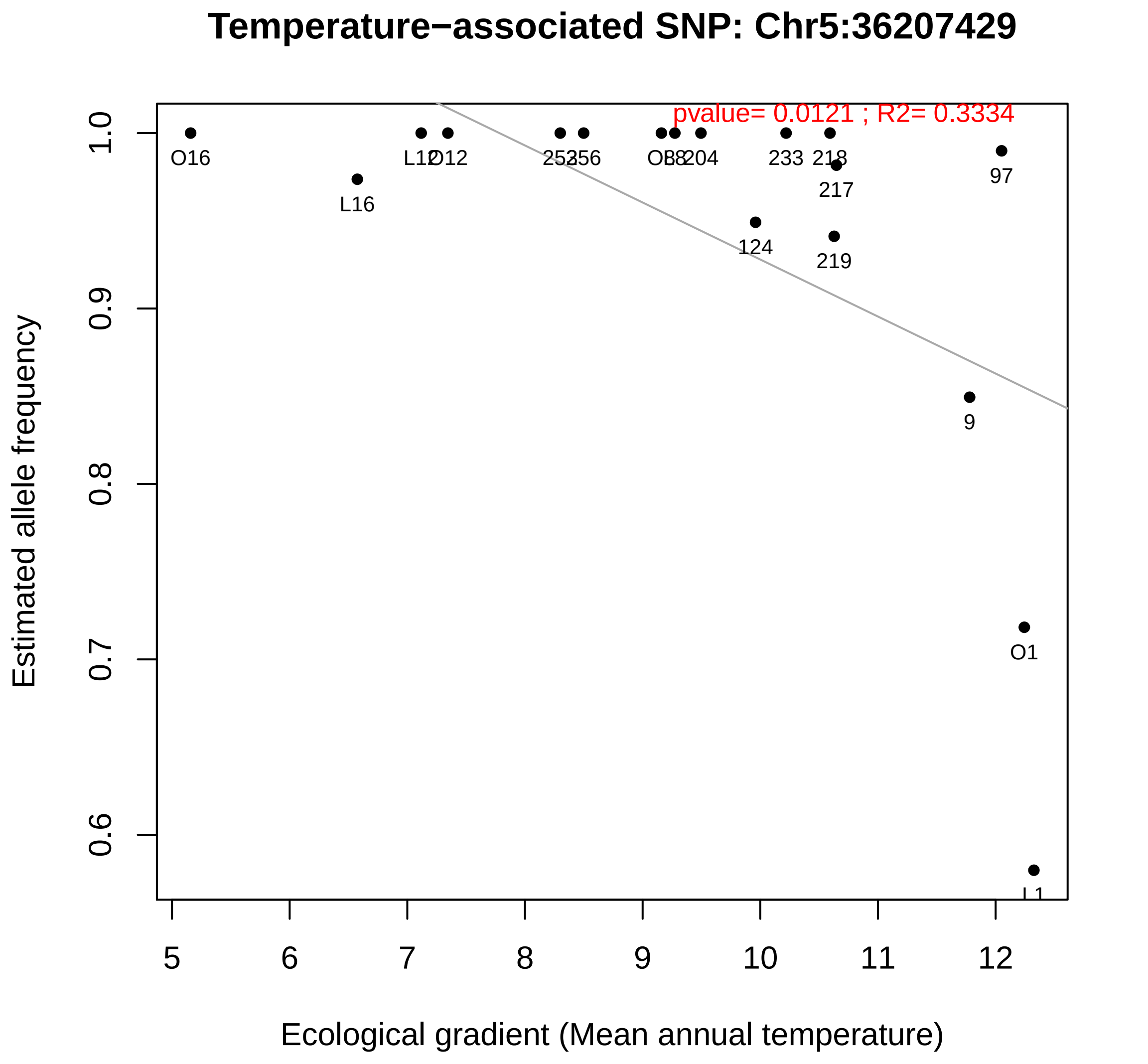

Temperature−associated SNP: Chr5:36207429
pvalue= 0.0121 ; R2= 0.3334
Estimated allele frequency
Ecological gradient (Mean annual temperature)
1.0
0.9
0.8
0.7
0.6
5
6
7
8
9
10
11
12
O16
L16
L12
O12
253
256
O8
L8
204
233
218
217
124
219
97
9
O1
L1

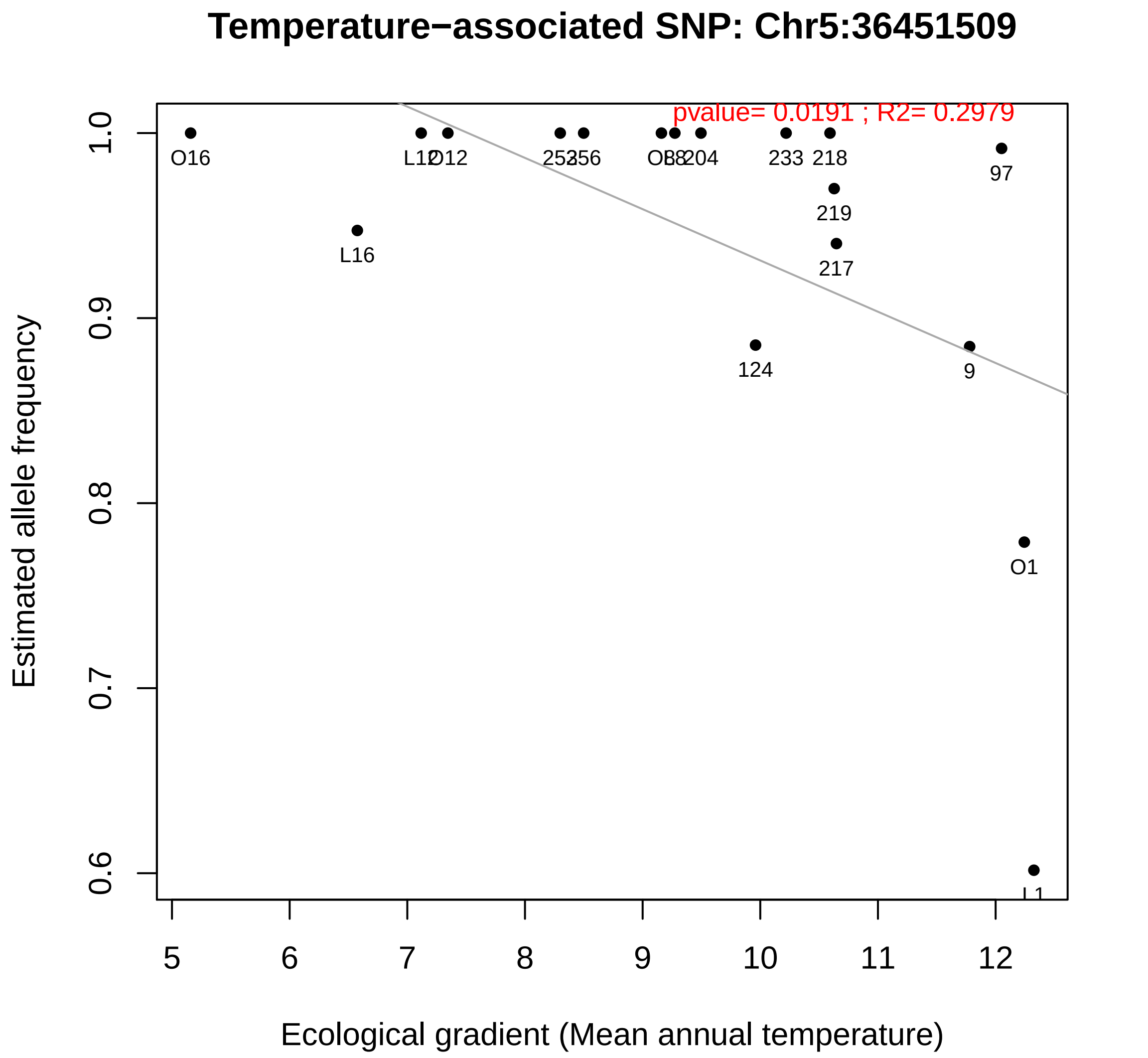

Temperature−associated SNP: Chr5:36451509
pvalue= 0.0191 ; R2= 0.2979
Estimated allele frequency
Ecological gradient (Mean annual temperature)
1.0
0.9
0.8
0.7
0.6
5
6
7
8
9
10
11
12
O16
L16
233
218
97
219
217
124
9
O1

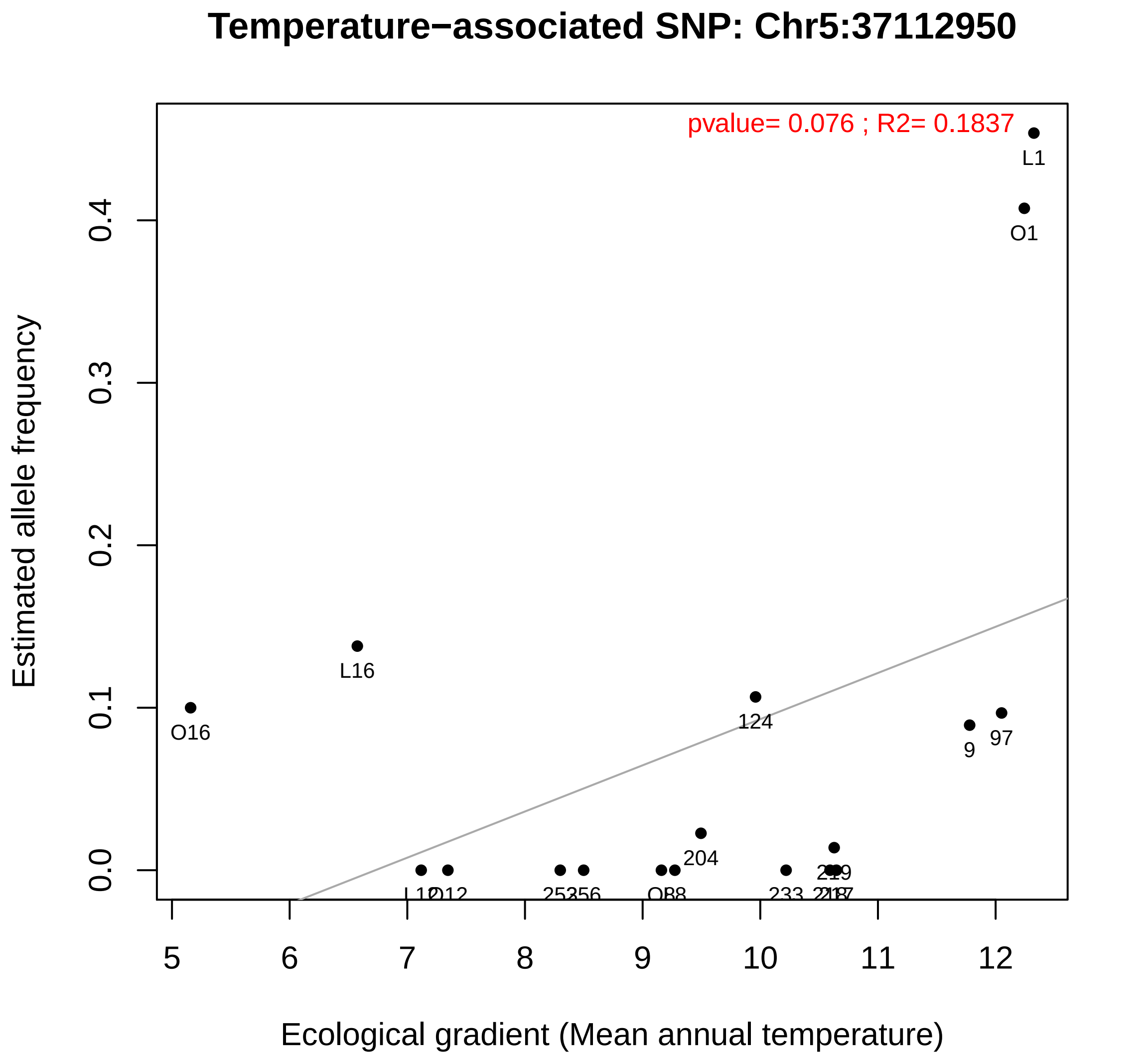

Temperature−associated SNP: Chr5:37112950
pvalue= 0.076 ; R2= 0.1837
Estimated allele frequency
Ecological gradient (Mean annual temperature)
0.0
0.1
0.2
0.3
0.4
5
6
7
8
9
10
11
12
L1
O1
L16
O16
124
9
97
204
233

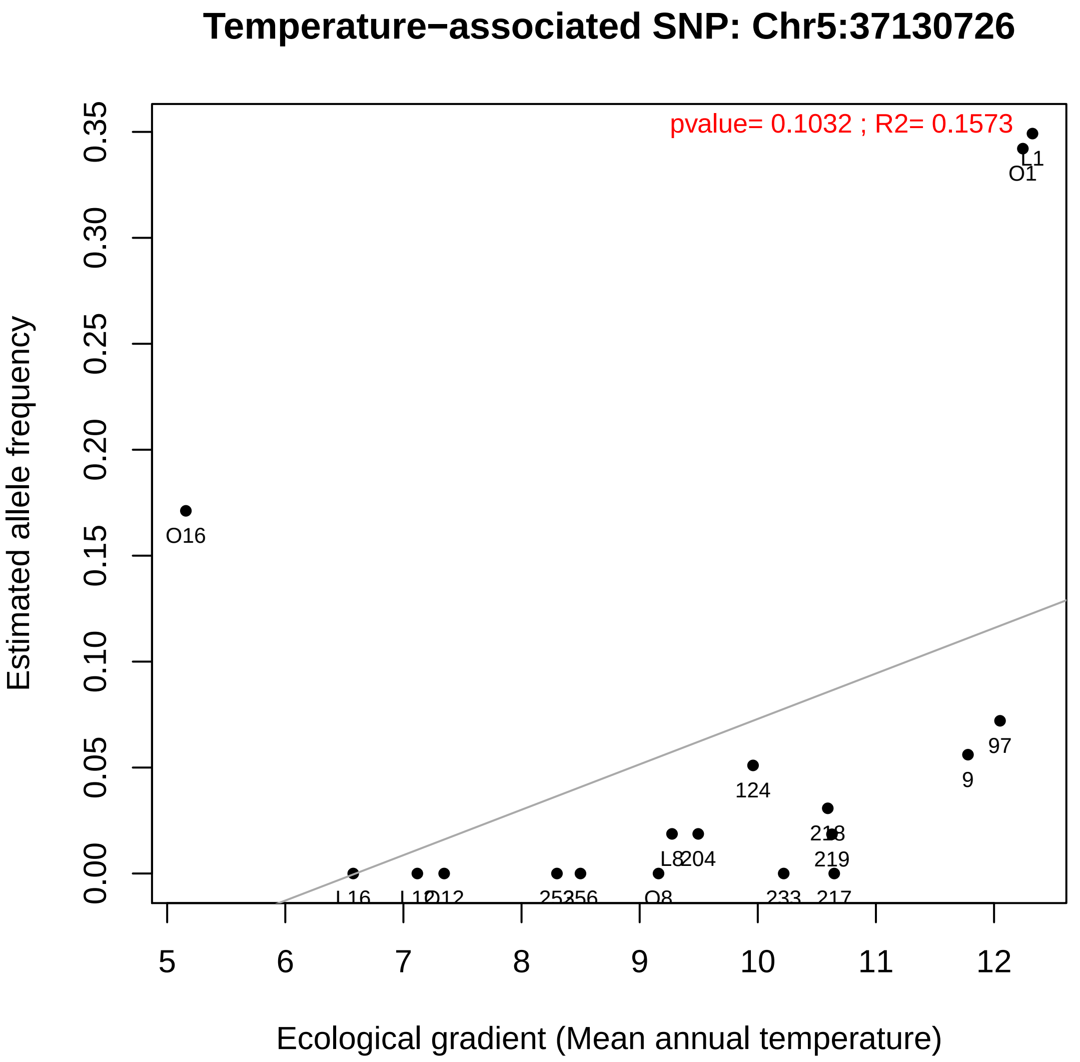

Temperature−associated SNP: Chr5:37130726
pvalue= 0.1032 ; R2= 0.1573
Estimated allele frequency
0.00
0.05
0.10
0.15
0.20
0.25
0.30
0.35
5
6
7
8
9
10
11
12
Ecological gradient (Mean annual temperature)
L1
O1
O16
97
9
124
218
219
L8
204
233
217
L16
O12
O8

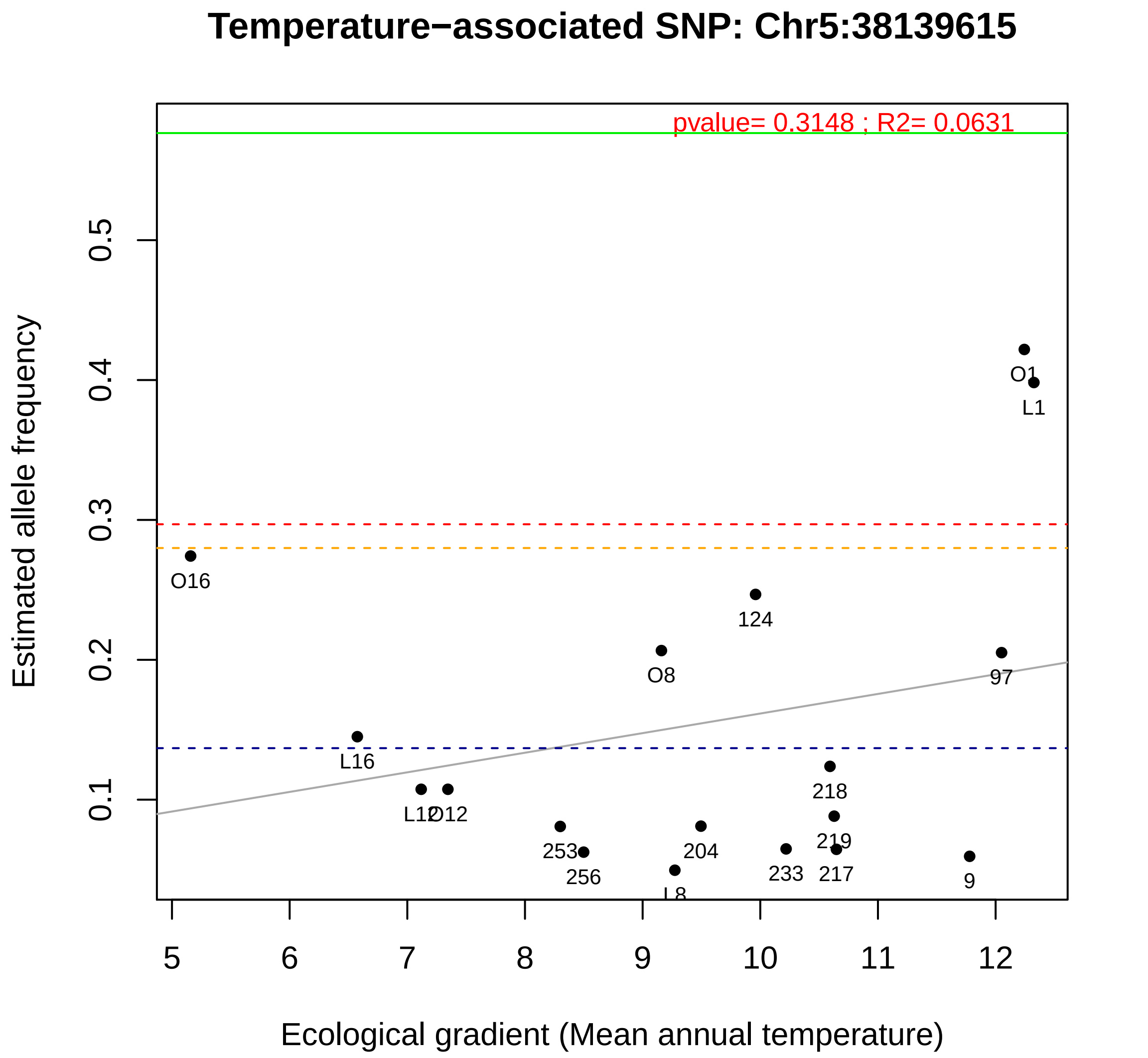
Temperature−associated SNP: Chr5:38139615
pvalue= 0.3148 ; R2= 0.0631
Estimated allele frequency
0.1
0.2
0.3
0.4
0.5
5
6
7
8
9
10
11
12
Ecological gradient (Mean annual temperature)
O16
L16
L12
O12
253
256
L8
O8
204
124
233
218
219
217
97
9
O1
L1

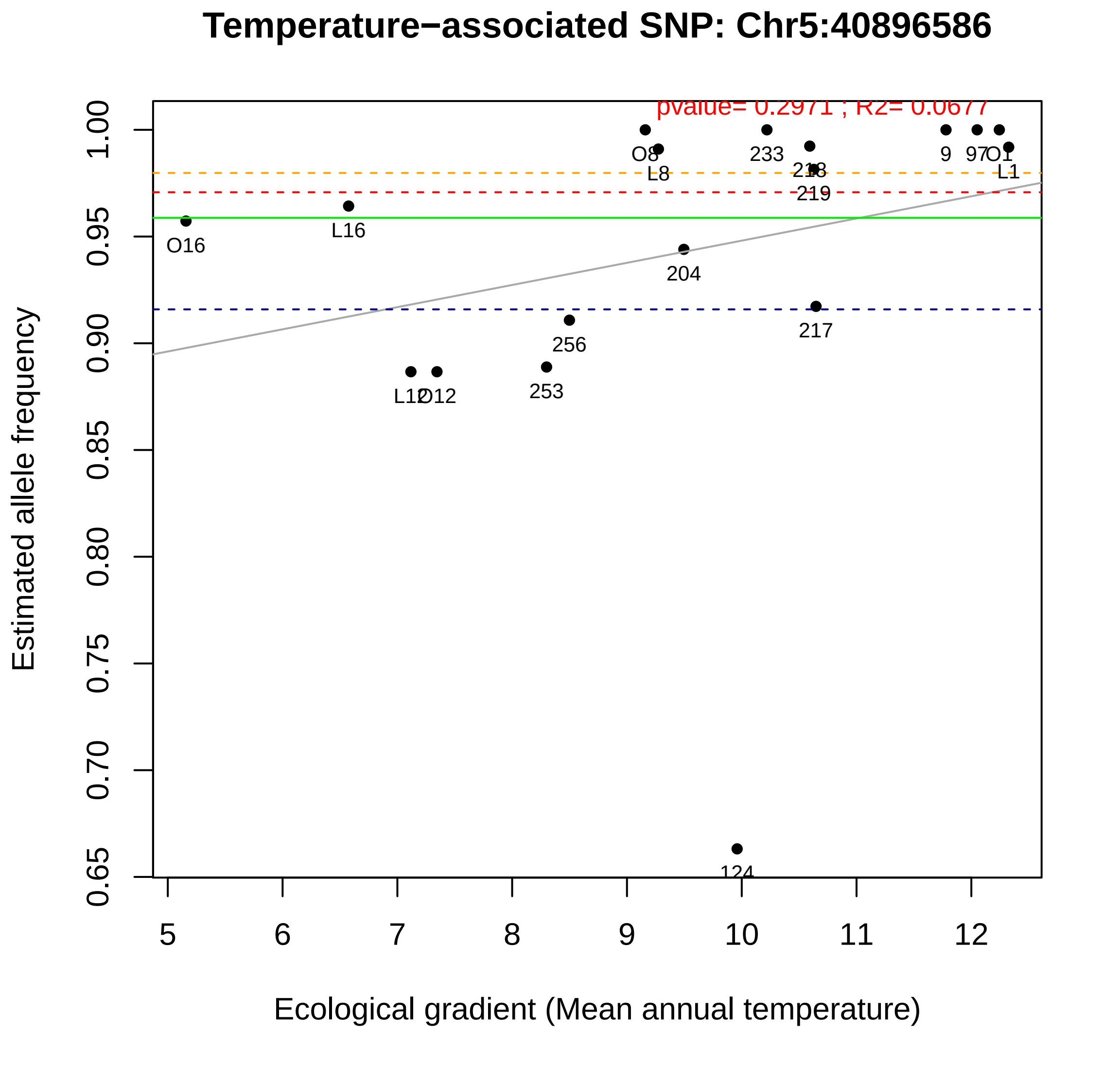
Temperature−associated SNP: Chr5:40896586
pvalue= 0.2971 ; R2= 0.0677
Estimated allele frequency
Ecological gradient (Mean annual temperature)
1.00
0.95
0.90
0.85
0.80
0.75
0.70
0.65
5
6
7
8
9
10
11
12
O16
L16
L12
O12
253
256
O8
L8
204
233
218
219
217
124
9
97
O1
L1

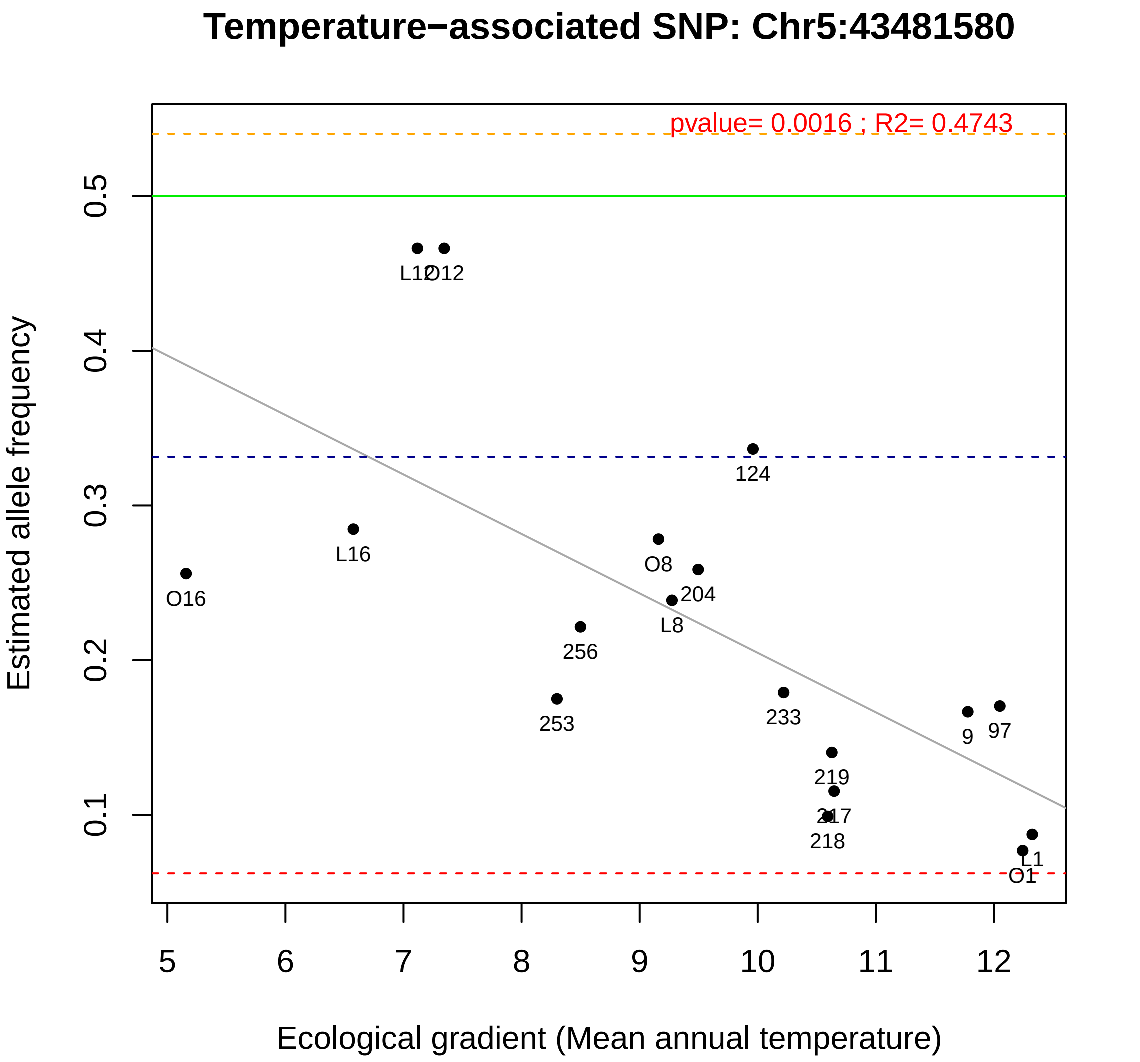

Temperature−associated SNP: Chr5:43481580
pvalue= 0.0016 ; R2= 0.4743
Estimated allele frequency
0.1
0.2
0.3
0.4
0.5
5
6
7
8
9
10
11
12
Ecological gradient (Mean annual temperature)
L12
O12
124
L16
O16
O8
204
L8
256
253
233
9
97
219
217
218
L1
O1

**Temperature−associated SNP: Chr5:44187515**

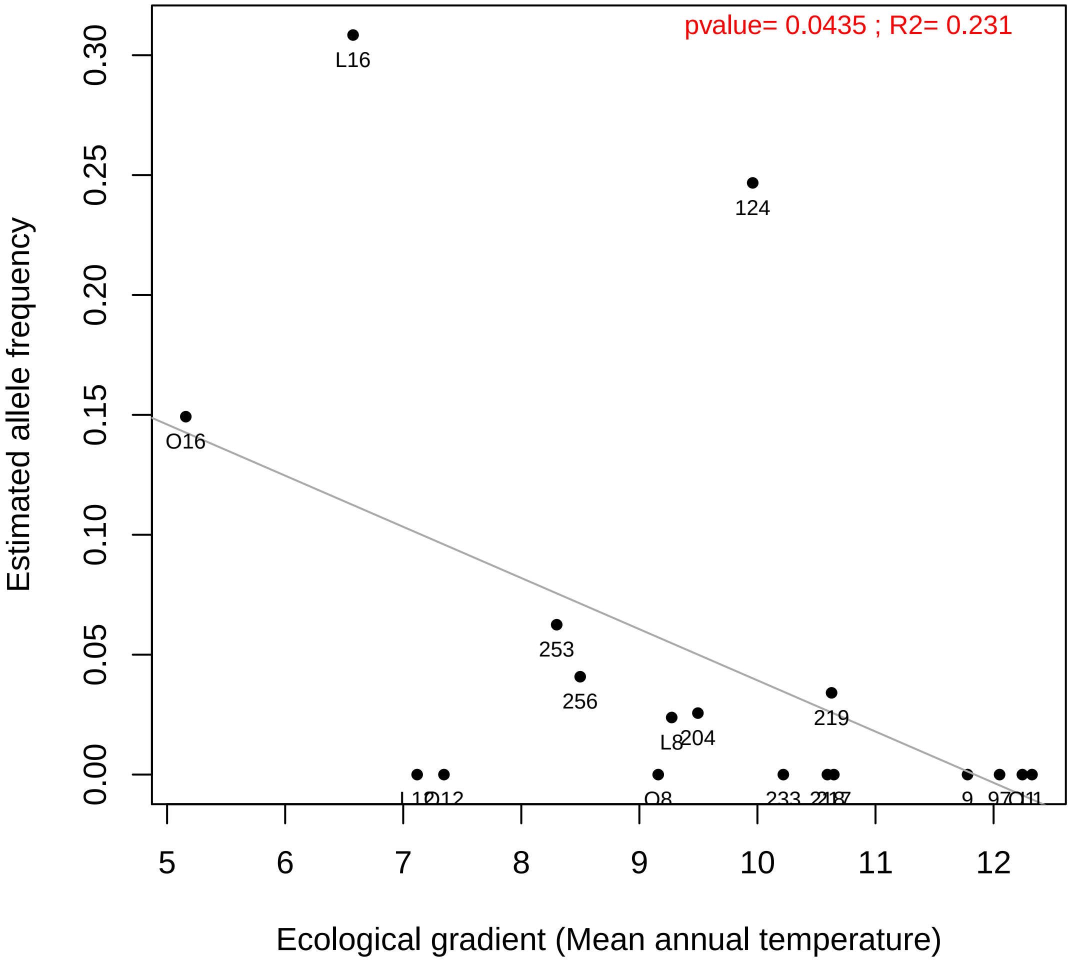

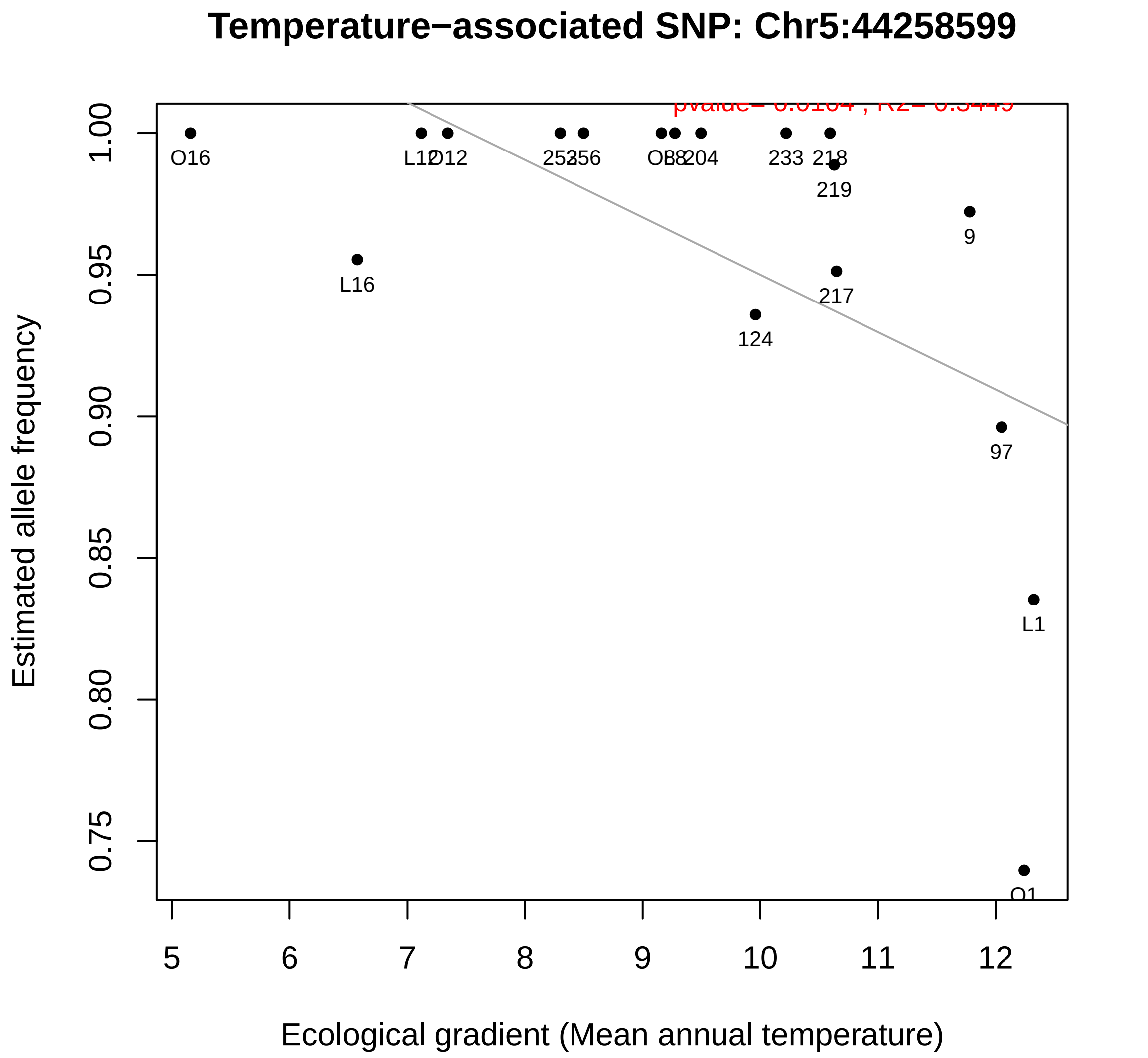

Temperature−associated SNP: Chr5:44258599
pvalue= 0.0104 ; R2= 0.3449
Estimated allele frequency
Ecological gradient (Mean annual temperature)
1.00
0.95
0.90
0.85
0.80
0.75
5
6
7
8
9
10
11
12
O16
L12
O12
253
256
O3
38
204
233
218
219
9
L16
217
124
97
L1
O1

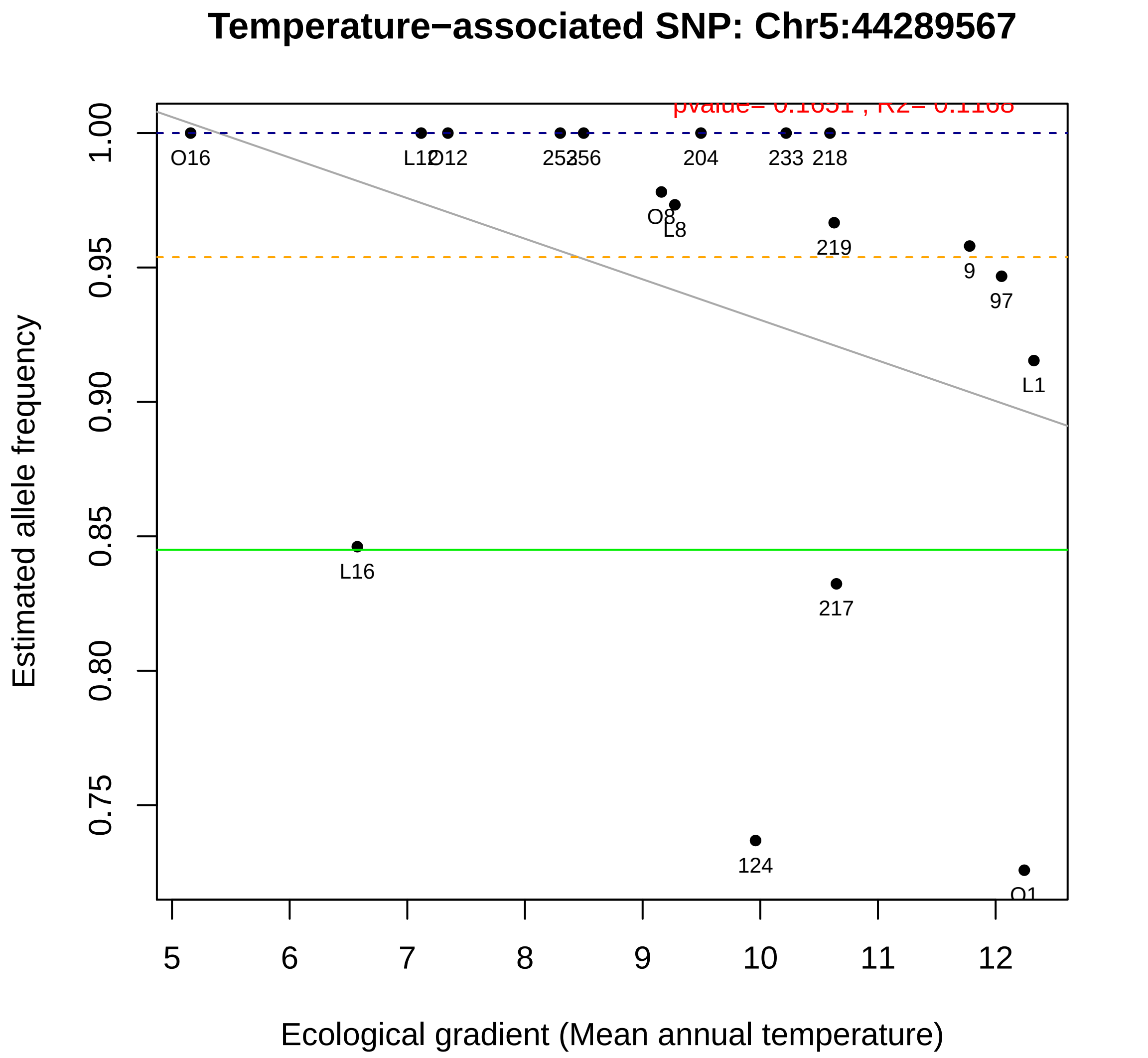

Temperature−associated SNP: Chr5:44289567
pvalue= 0.1651 , R2= 0.1168
Estimated allele frequency
0.75
0.80
0.85
0.90
0.95
1.00
5
6
7
8
9
10
11
12
Ecological gradient (Mean annual temperature)
O16
L12
O12
253
256
204
233
218
O8
L8
219
9
97
L1
L16
217
124
O1

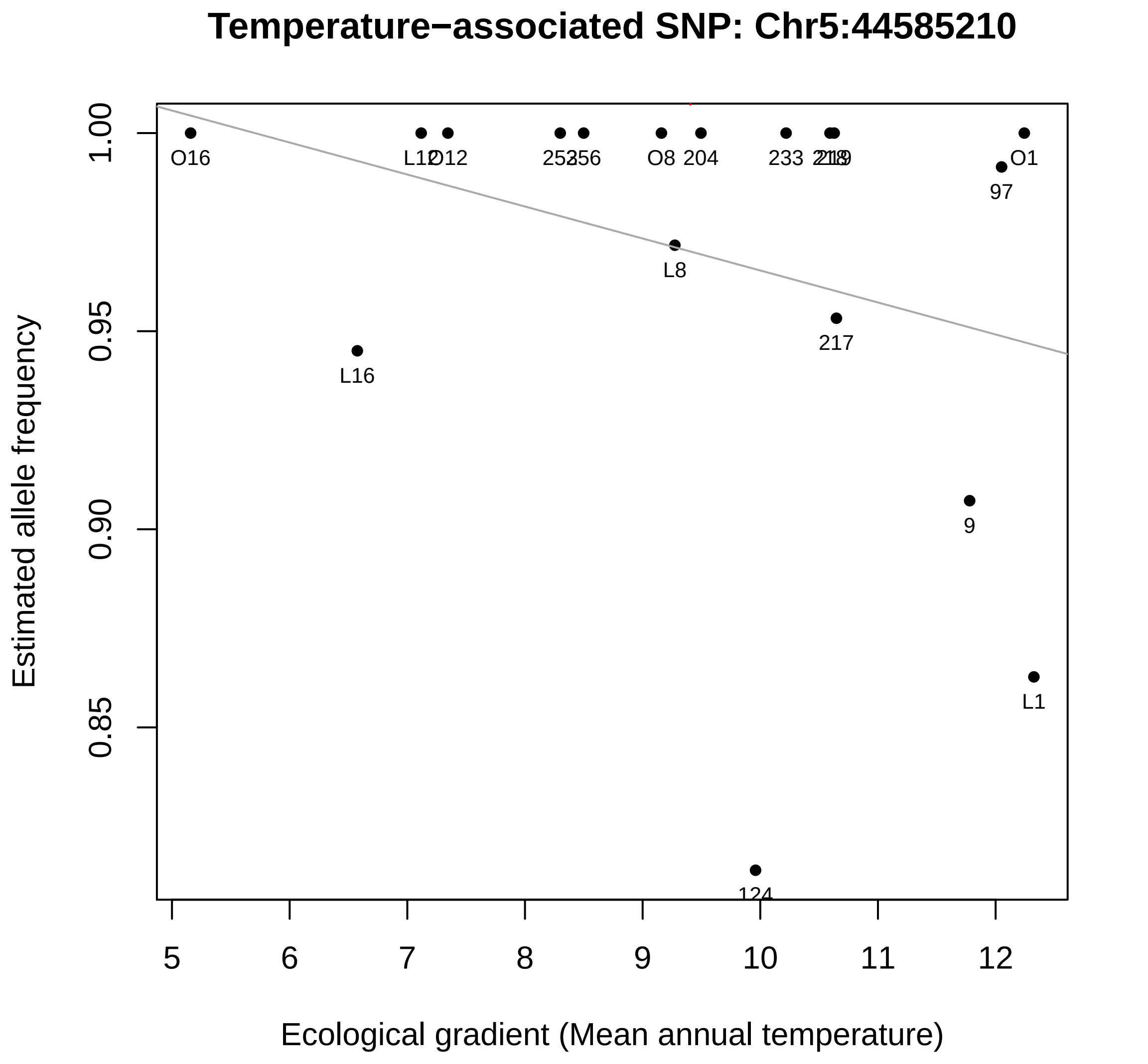
Temperature−associated SNP: Chr5:44585210
Estimated allele frequency
1.00
0.95
0.90
0.85
O16
L12
O12
253
256
O8
204
233
218
219
O1
97
L8
217
L16
9
L1
124
5
6
7
8
9
10
11
12
Ecological gradient (Mean annual temperature)

### Temperature−associated SNP: Chr5:4496394

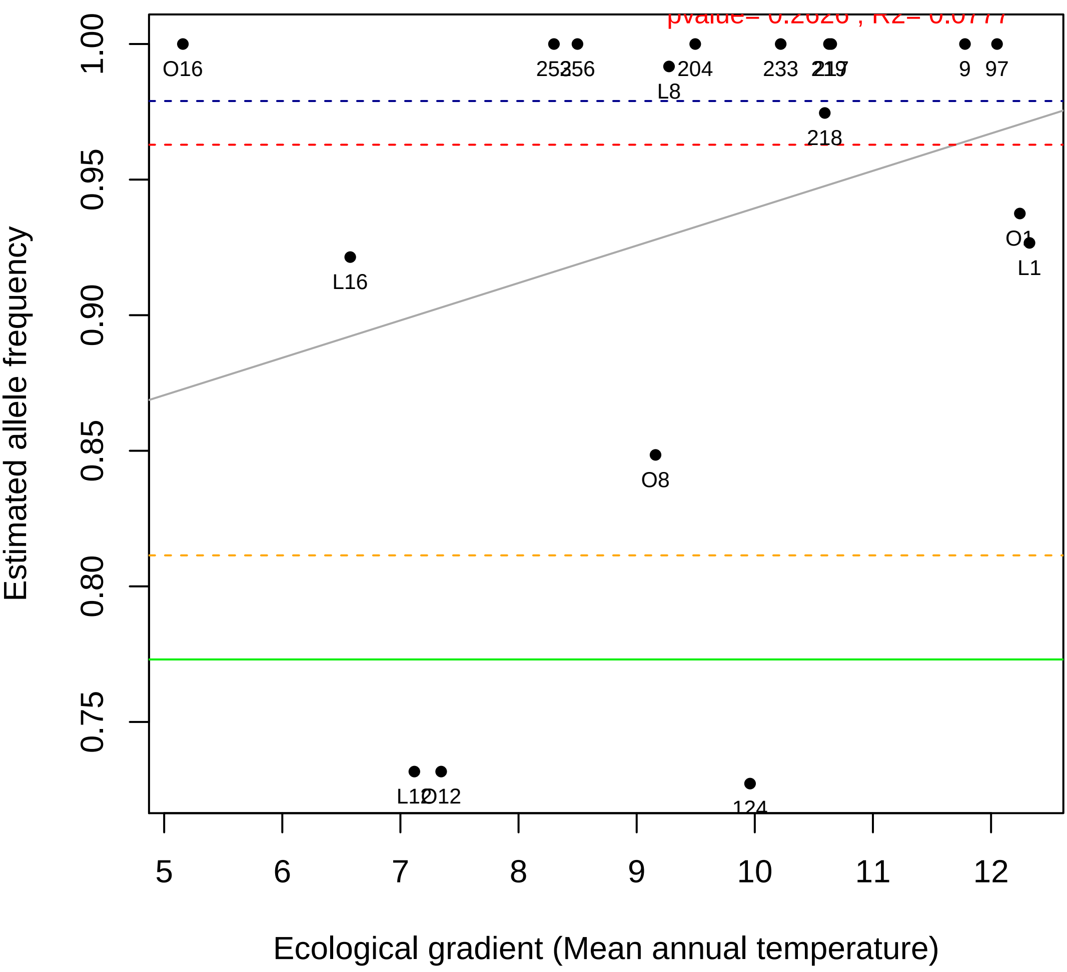

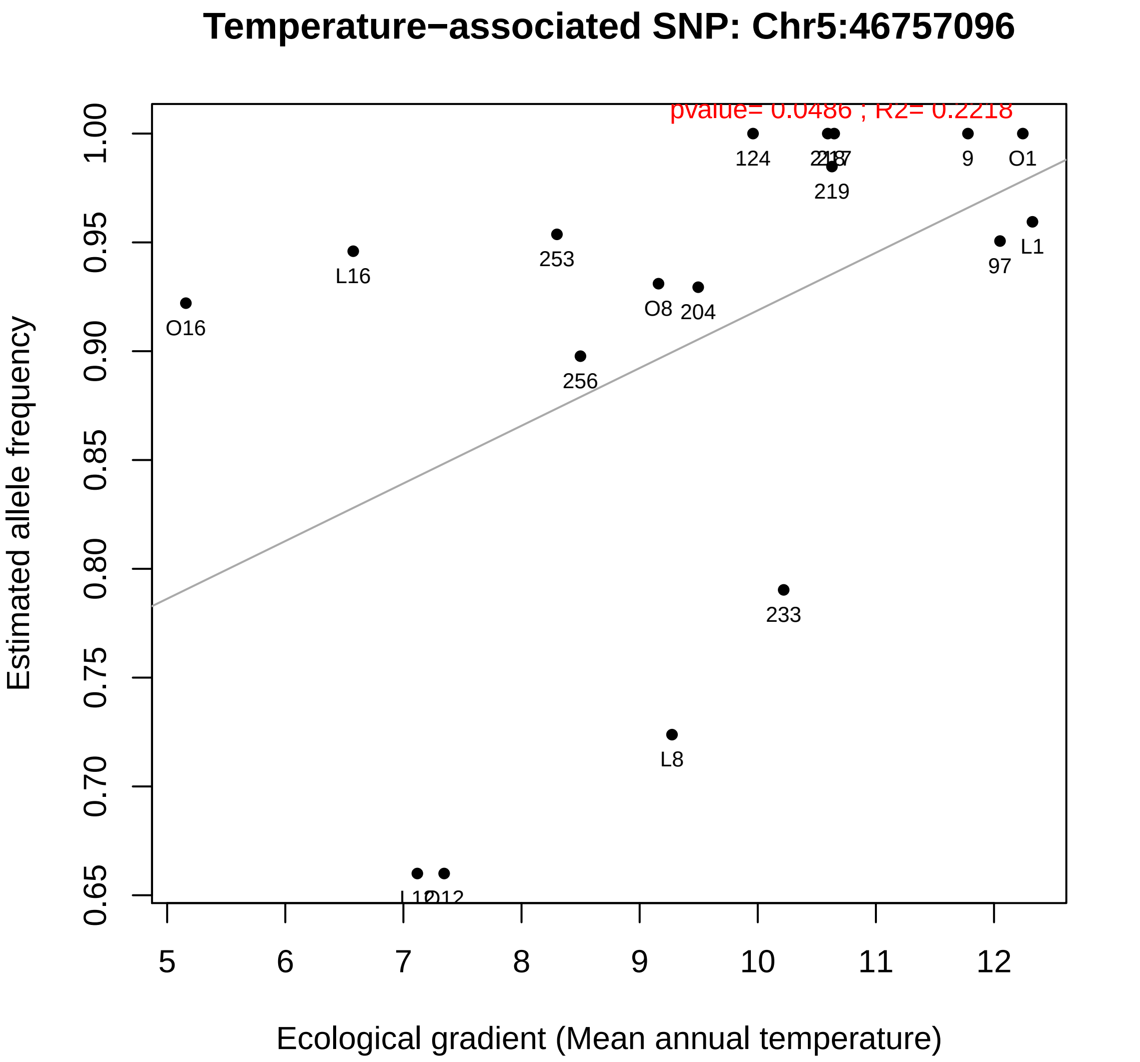
Temperature−associated SNP: Chr5:46757096
pvalue= 0.0486 ; R2= 0.2218
Estimated allele frequency
Ecological gradient (Mean annual temperature)
0.65
0.70
0.75
0.80
0.85
0.90
0.95
1.00
5
6
7
8
9
10
11
12
124
219
9
O1
L1
97
253
L16
O16
O8
204
256
233
L8
O12

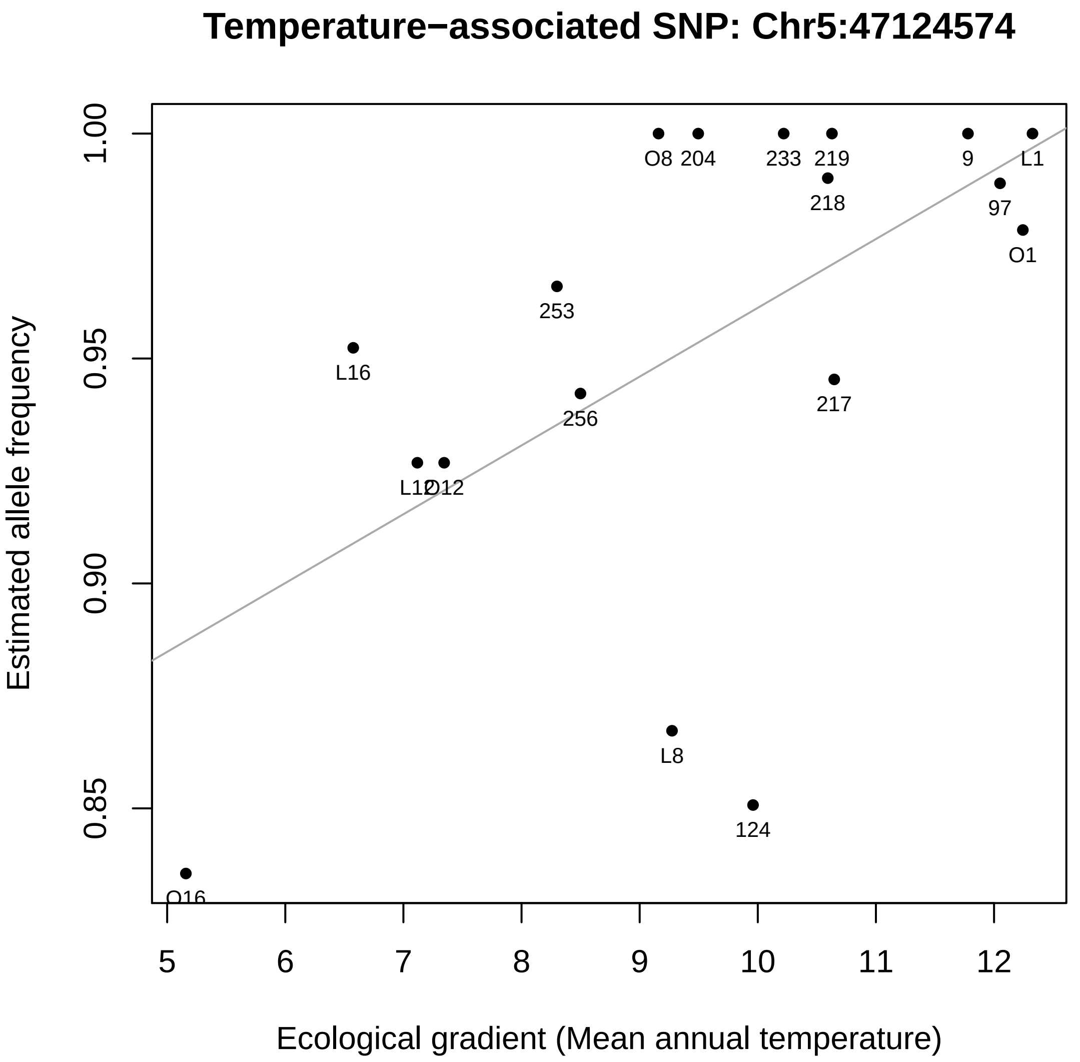

Temperature−associated SNP: Chr5:47124574
Estimated allele frequency
Ecological gradient (Mean annual temperature)
1.00
0.95
0.90
0.85
5
6
7
8
9
10
11
12
O8
204
233
219
9
L1
218
97
O1
253
L16
256
217
L12
O12
L8
124
O16

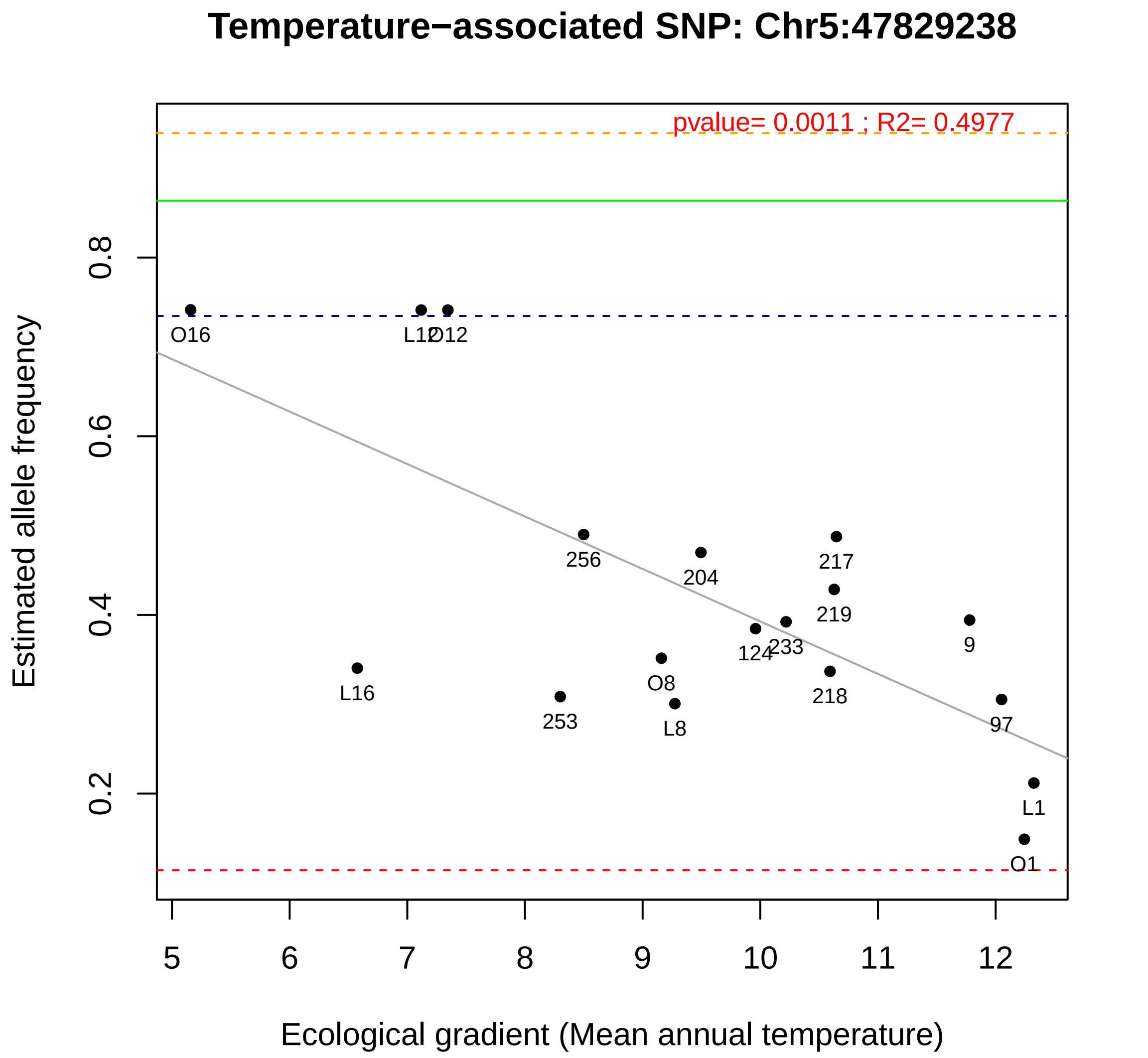

Temperature−associated SNP: Chr5:47829238
pvalue= 0.0011 ; R2= 0.4977
Estimated allele frequency
0.2
0.4
0.6
0.8
5
6
7
8
9
10
11
12
Ecological gradient (Mean annual temperature)
O16
L12
O12
256
204
217
219
9
124
233
218
L16
253
O8
L8
97
L1
O1

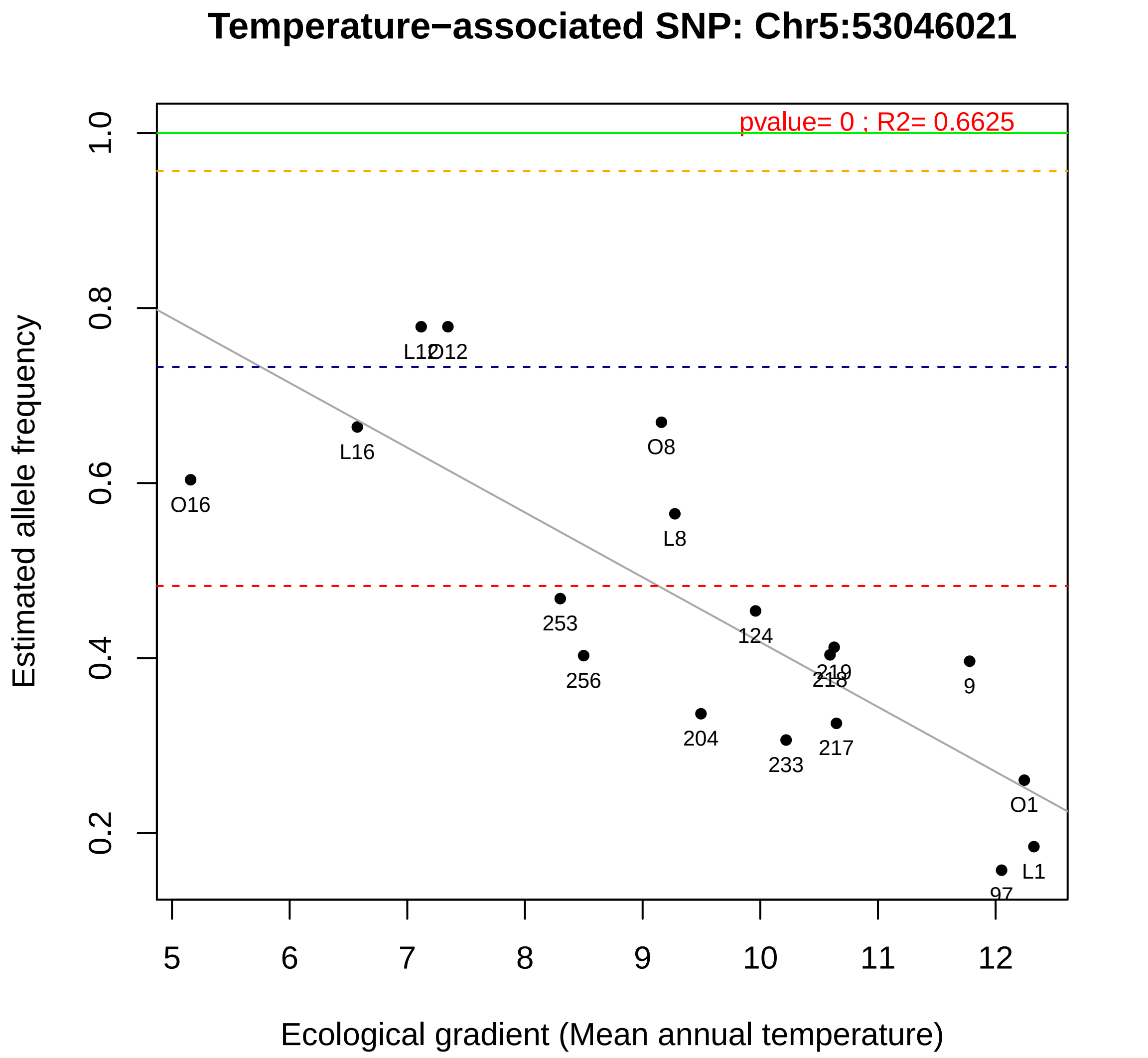
Temperature−associated SNP: Chr5:53046021
pvalue= 0 ; R2= 0.6625
Estimated allele frequency
Ecological gradient (Mean annual temperature)
0.2
0.4
0.6
0.8
1.0
5
6
7
8
9
10
11
12
O16
L16
L12
O12
O8
L8
253
256
204
124
233
219
218
217
9
O1
L1
97

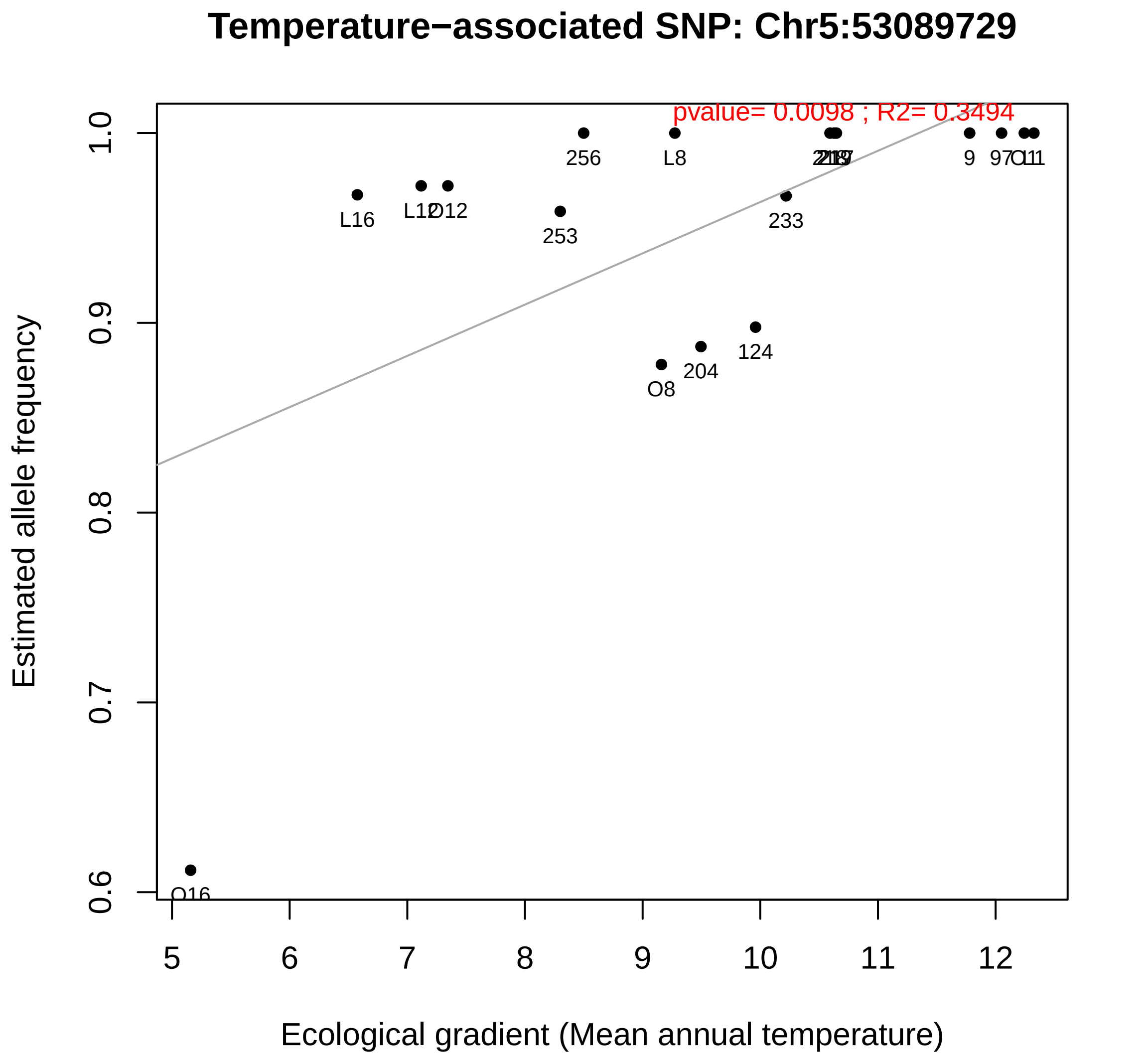

Temperature−associated SNP: Chr5:53089729
pvalue= 0.0098 ; R2= 0.3494
Estimated allele frequency
Ecological gradient (Mean annual temperature)
1.0
0.9
0.8
0.7
0.6
5
6
7
8
9
10
11
12
256
L8
9
L16
L12
O12
253
233
124
204
O8
O16

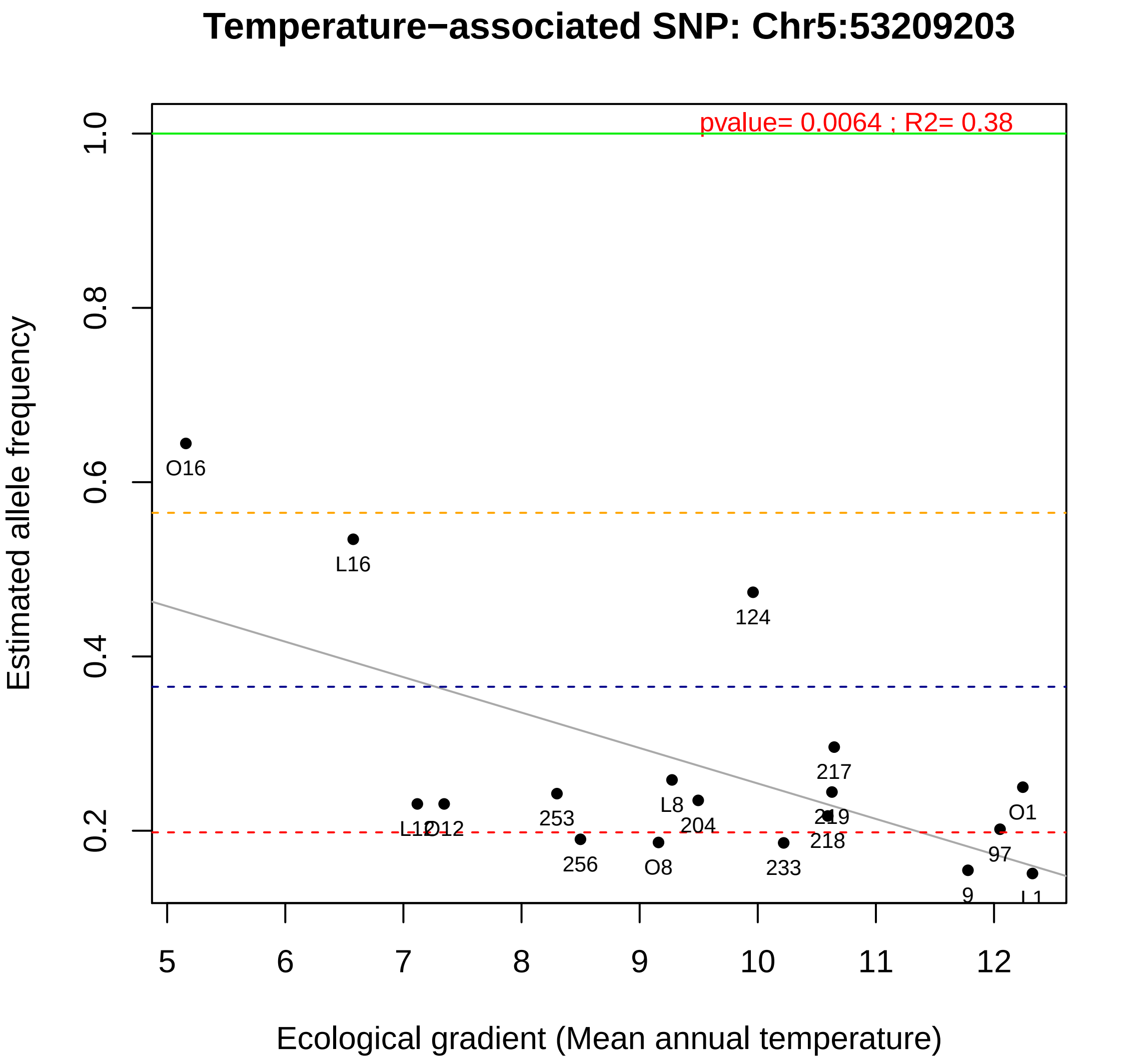
Temperature−associated SNP: Chr5:53209203
pvalue= 0.0064 ; R2= 0.38
Estimated allele frequency
1.0
0.8
0.6
0.4
0.2
O16
L16
124
217
L8
253
204
219
O1
L12
O12
218
97
256
O8
233
9
5
6
7
8
9
10
11
12
Ecological gradient (Mean annual temperature)

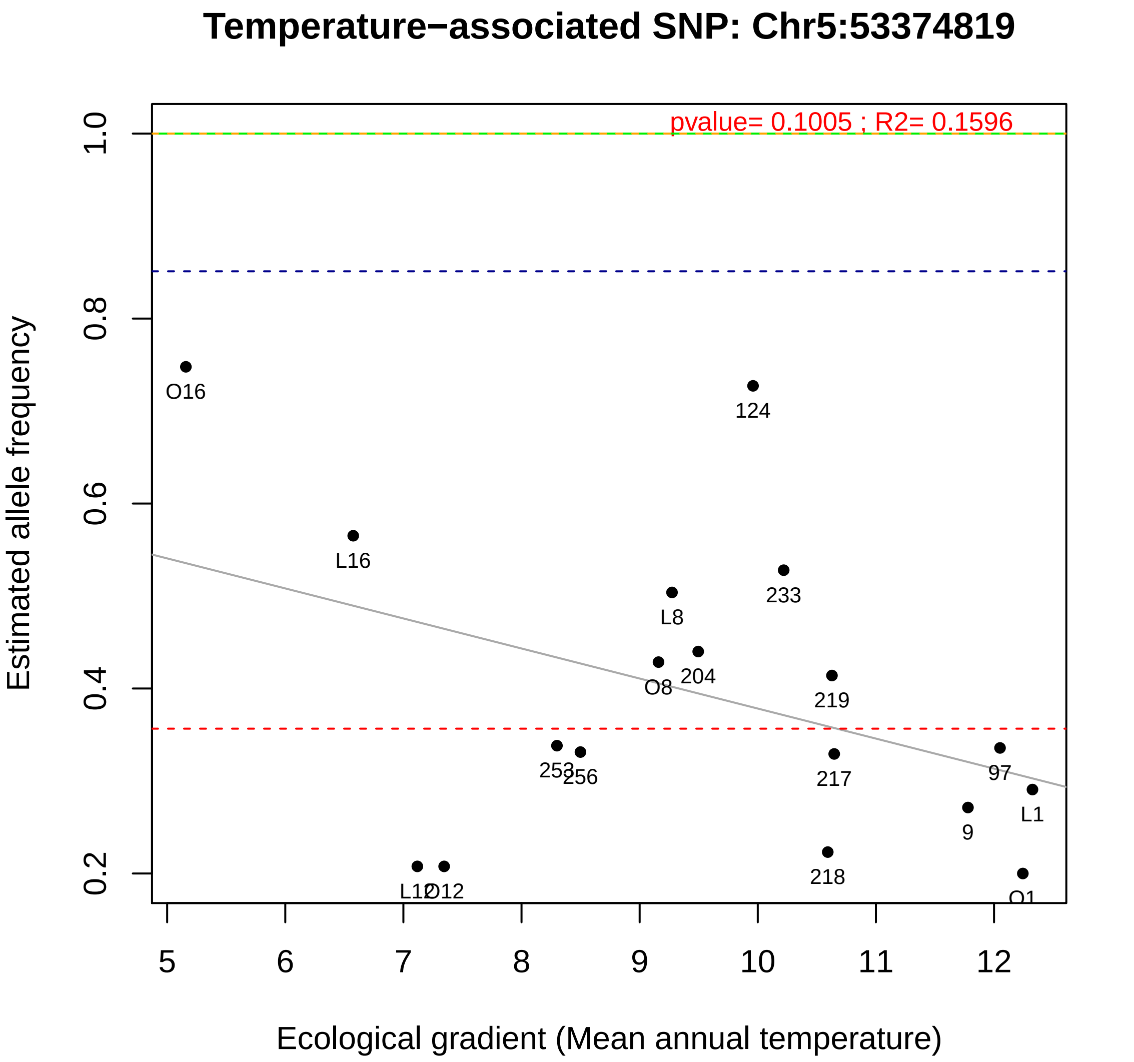
Temperature−associated SNP: Chr5:53374819
pvalue= 0.1005 ; R2= 0.1596
Estimated allele frequency
Ecological gradient (Mean annual temperature)
0.2
0.4
0.6
0.8
1.0
5
6
7
8
9
10
11
12
O16
124
L16
233
L8
204
O8
219
253
256
217
97
L1
9
218
L12
O12
O1

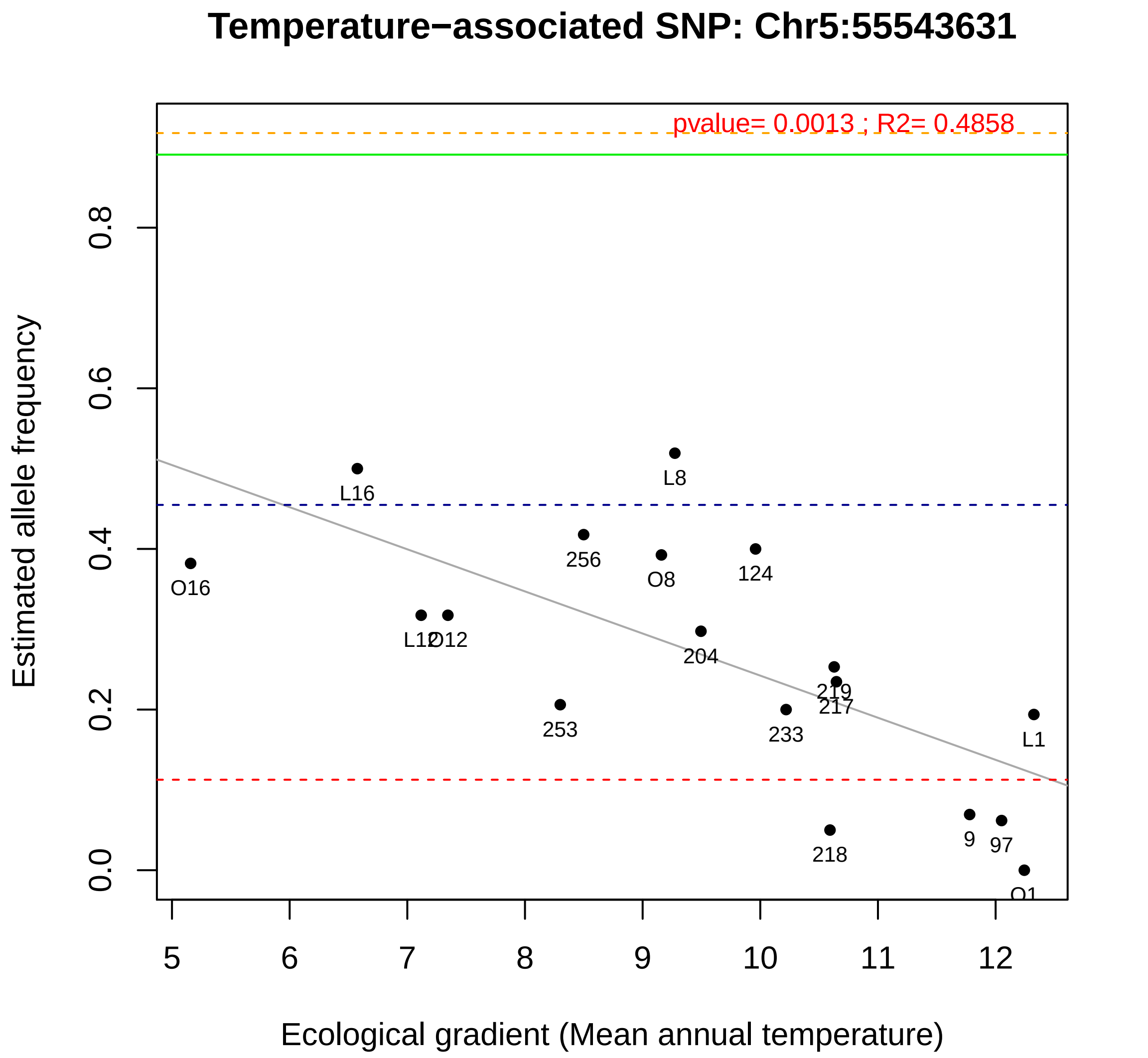

Temperature−associated SNP: Chr5:55543631
pvalue= 0.0013 ; R2= 0.4858
Estimated allele frequency
0.0
0.2
0.4
0.6
0.8
5
6
7
8
9
10
11
12
Ecological gradient (Mean annual temperature)
O16
L16
L12
O12
253
256
O8
L8
204
124
233
219
217
218
9
97
O1
L1

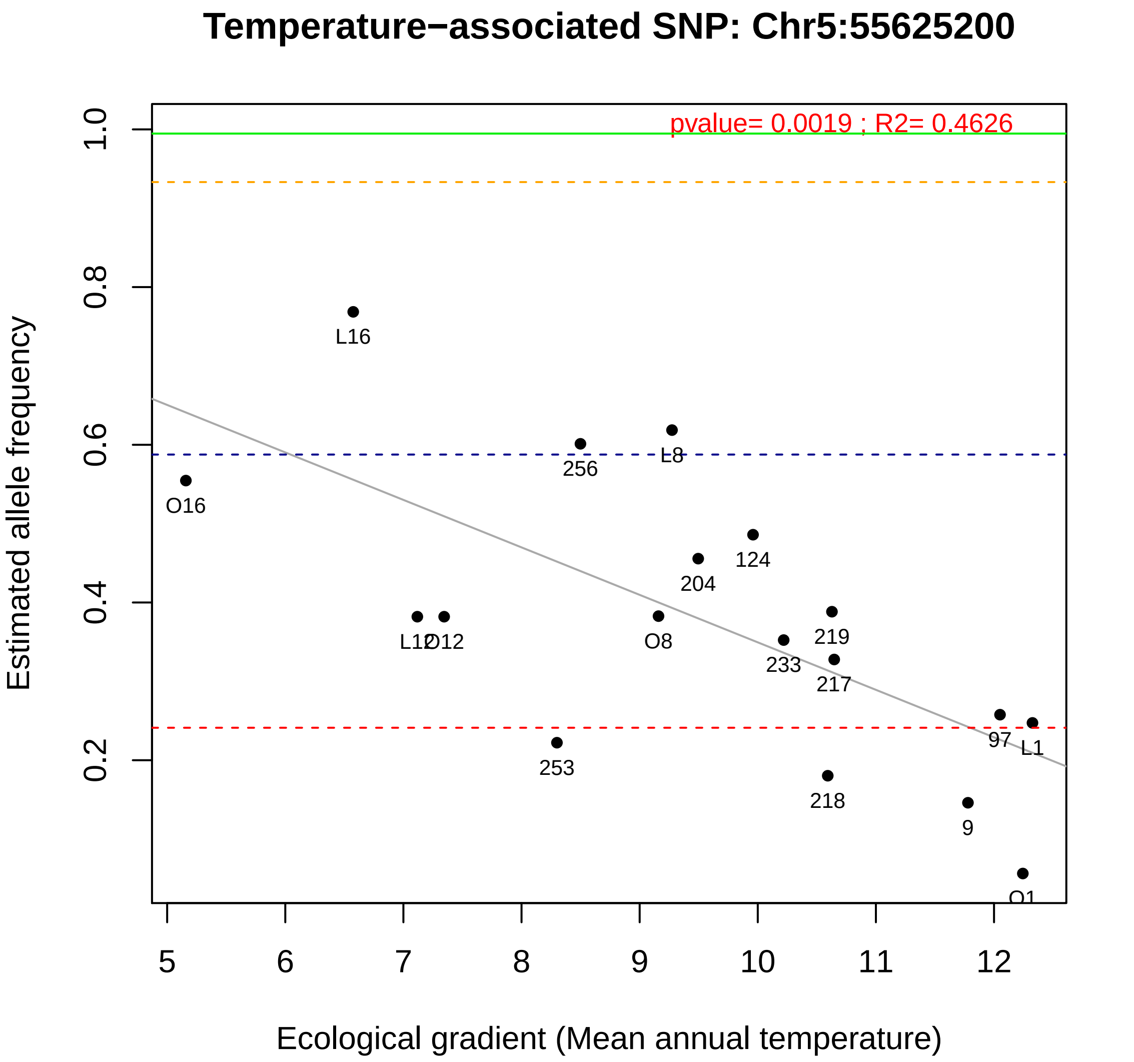
Temperature−associated SNP: Chr5:55625200
pvalue= 0.0019 ; R2= 0.4626
Estimated allele frequency
1.0
0.8
0.6
0.4
0.2
L16
256
L8
O16
124
204
L12
O12
O8
219
233
217
97
L1
253
218
9
O1
5
6
7
8
9
10
11
12
Ecological gradient (Mean annual temperature)

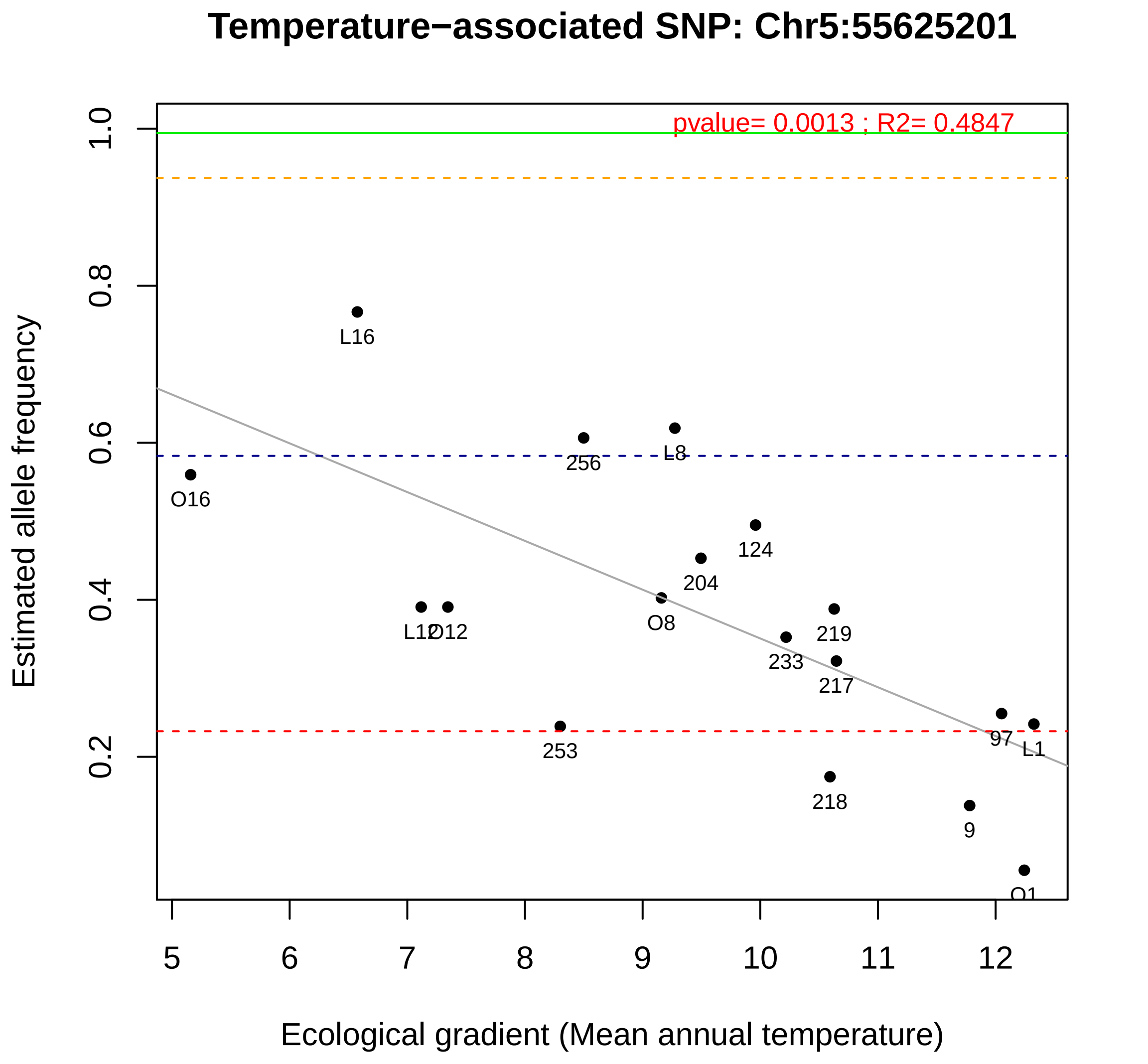

Temperature−associated SNP: Chr5:55625201
pvalue= 0.0013 ; R2= 0.4847
Estimated allele frequency
1.0
0.8
0.6
0.4
0.2
L16
256
L8
O16
124
204
O8
L12
O12
219
233
217
253
97
L1
218
9
O1
5
6
7
8
9
10
11
12
Ecological gradient (Mean annual temperature)

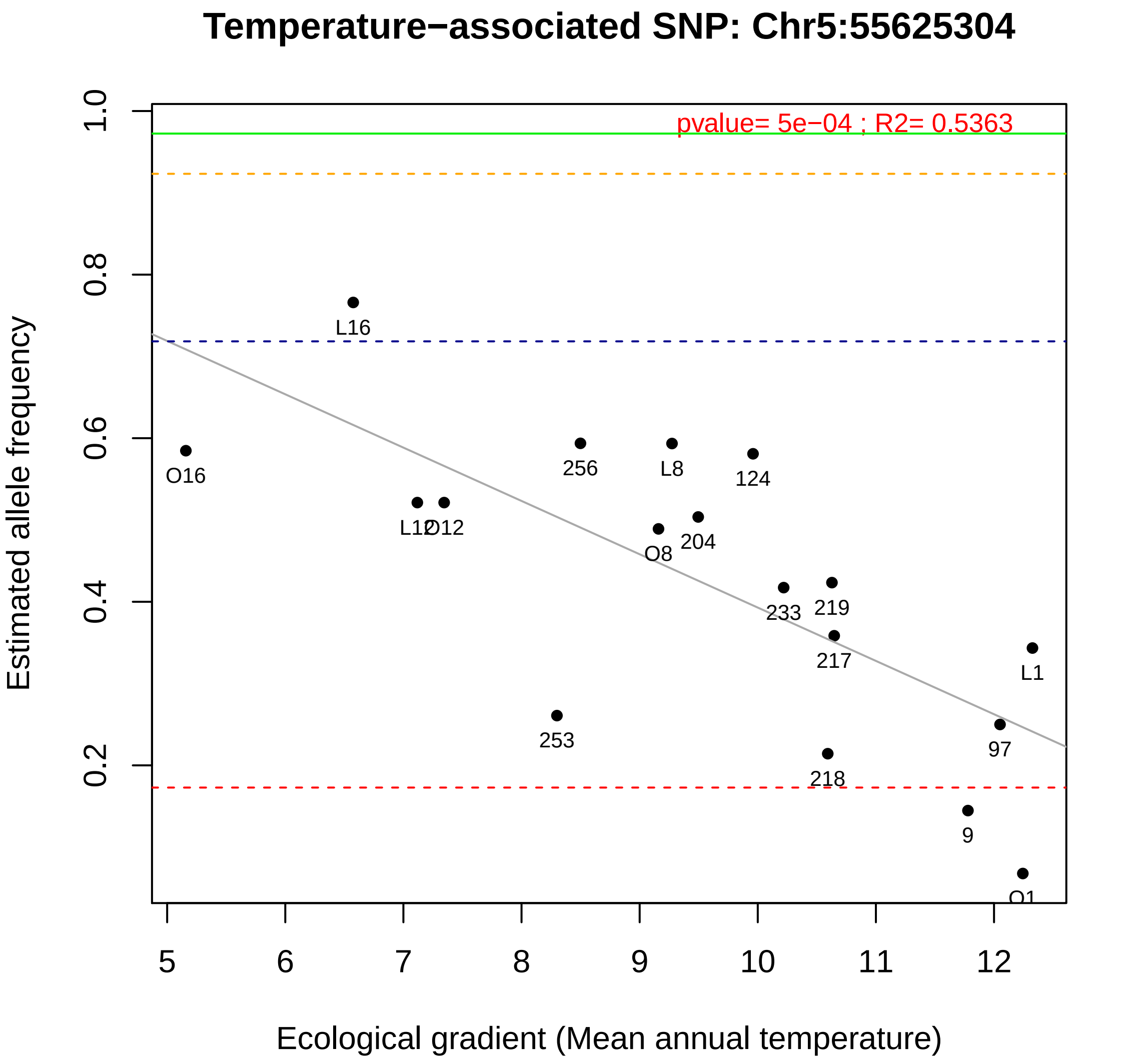

Temperature−associated SNP: Chr5:55625304
pvalue= 5e−04 ; R2= 0.5363
Estimated allele frequency
Ecological gradient (Mean annual temperature)
1.0
0.8
0.6
0.4
0.2
5
6
7
8
9
10
11
12
L16
O16
L12
O12
256
L8
124
O8
204
233
219
217
L1
253
97
218
9
O1

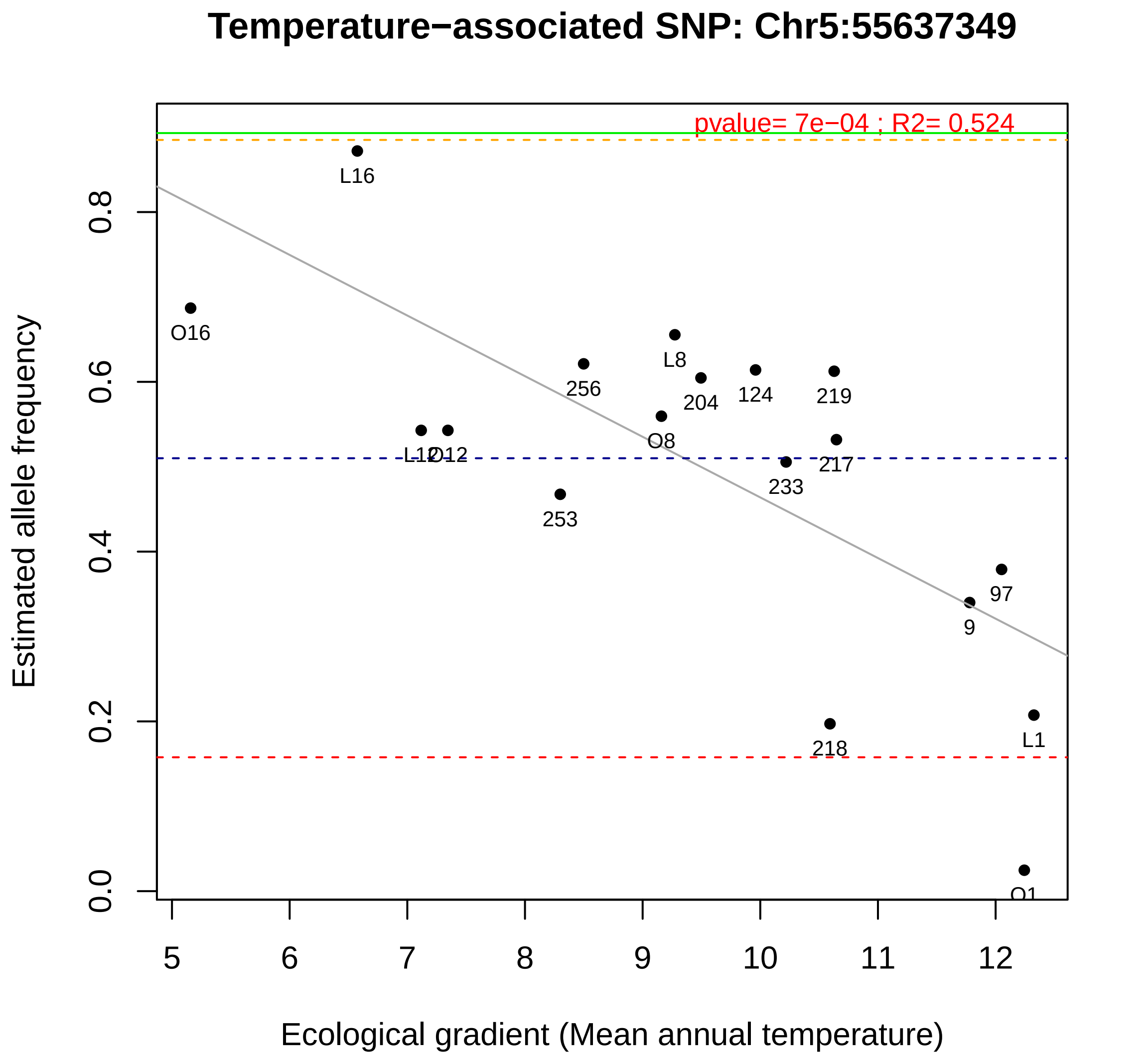
Temperature−associated SNP: Chr5:55637349
pvalue= 7e−04 ; R2= 0.524
Estimated allele frequency
0.0
0.2
0.4
0.6
0.8
Ecological gradient (Mean annual temperature)
5
6
7
8
9
10
11
12
L16
O16
L8
256
204
124
219
O8
L12
O12
233
217
253
97
9
218
L1
O1

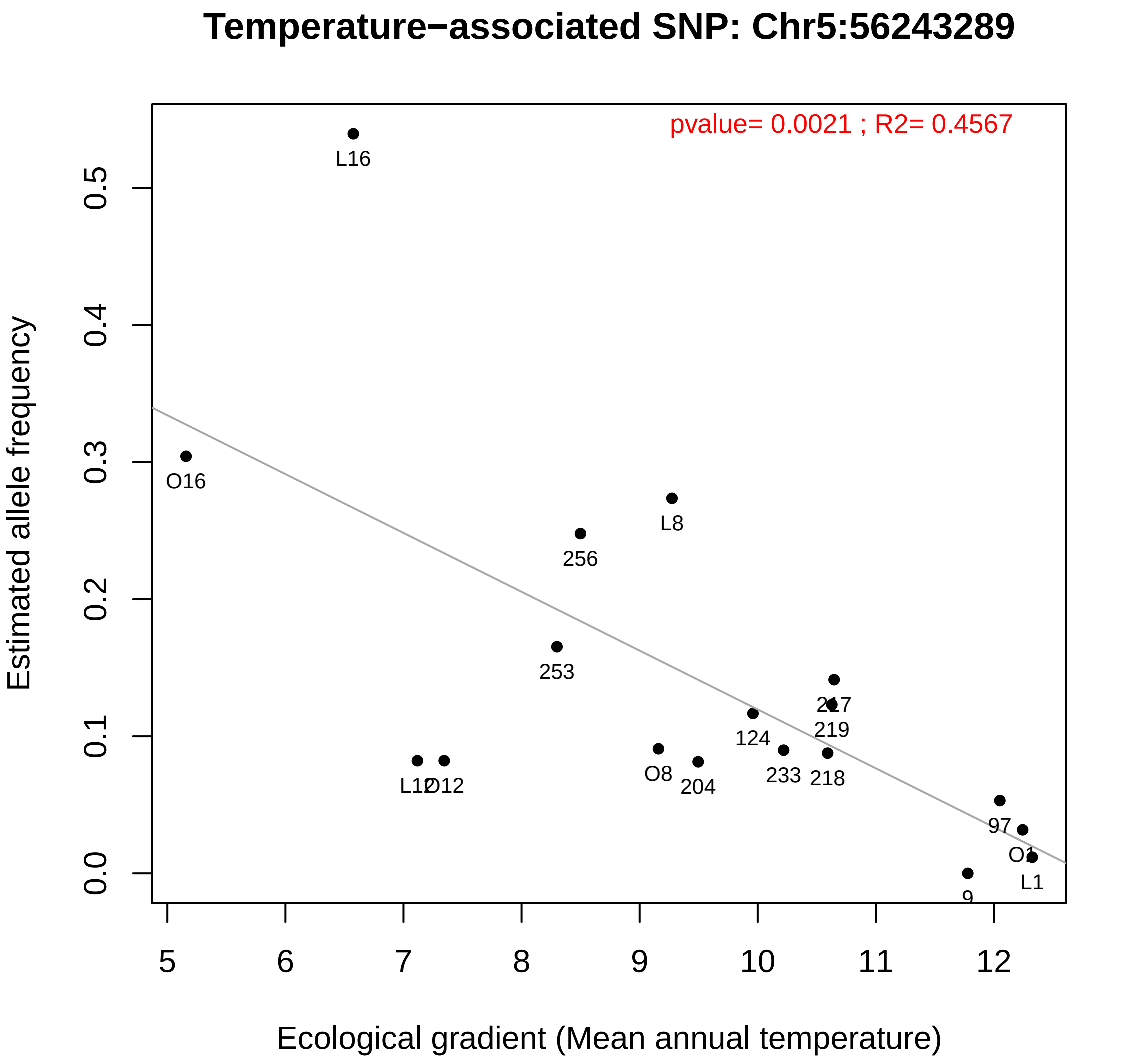
Temperature−associated SNP: Chr5:56243289
pvalue= 0.0021 ; R2= 0.4567
Estimated allele frequency
0.0
0.1
0.2
0.3
0.4
0.5
Ecological gradient (Mean annual temperature)
5
6
7
8
9
10
11
12
L16
O16
256
L8
253
124
219
233
218
O8
204
O12
97
L1
9

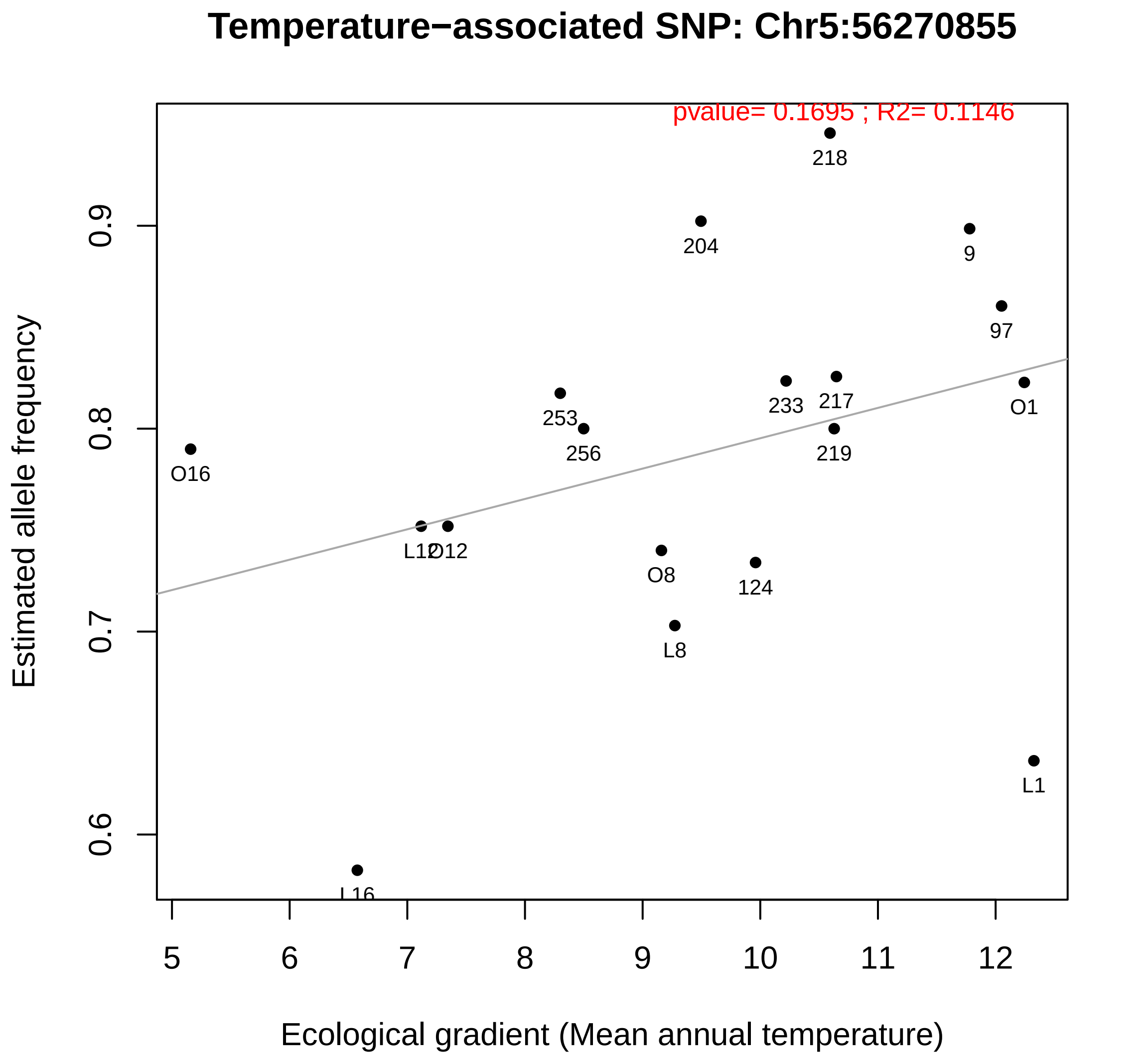
Temperature−associated SNP: Chr5:56270855
pvalue= 0.1695 ; R2= 0.1146
Estimated allele frequency
0.6
0.7
0.8
0.9
5
6
7
8
9
10
11
12
Ecological gradient (Mean annual temperature)
218
204
9
97
233
217
O1
253
256
219
O16
L12
O12
O8
124
L8
L1
L16

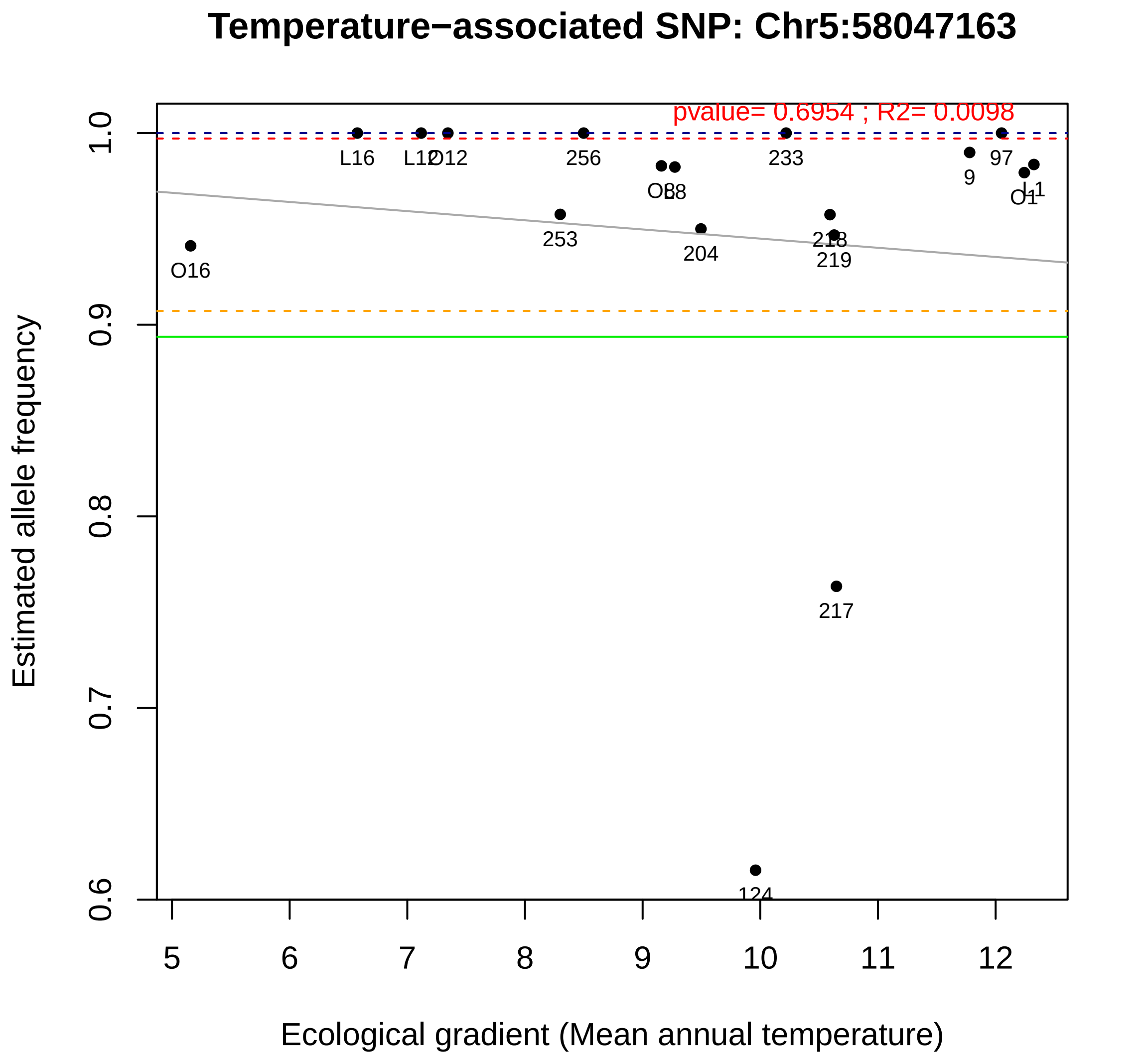
Temperature−associated SNP: Chr5:58047163
pvalue= 0.6954 ; R2= 0.0098
Estimated allele frequency
Ecological gradient (Mean annual temperature)
1.0
0.9
0.8
0.7
0.6
5
6
7
8
9
10
11
12
L16
L12
O12
256
233
97
9
253
204
218
219
217
124
O16
O1
L1

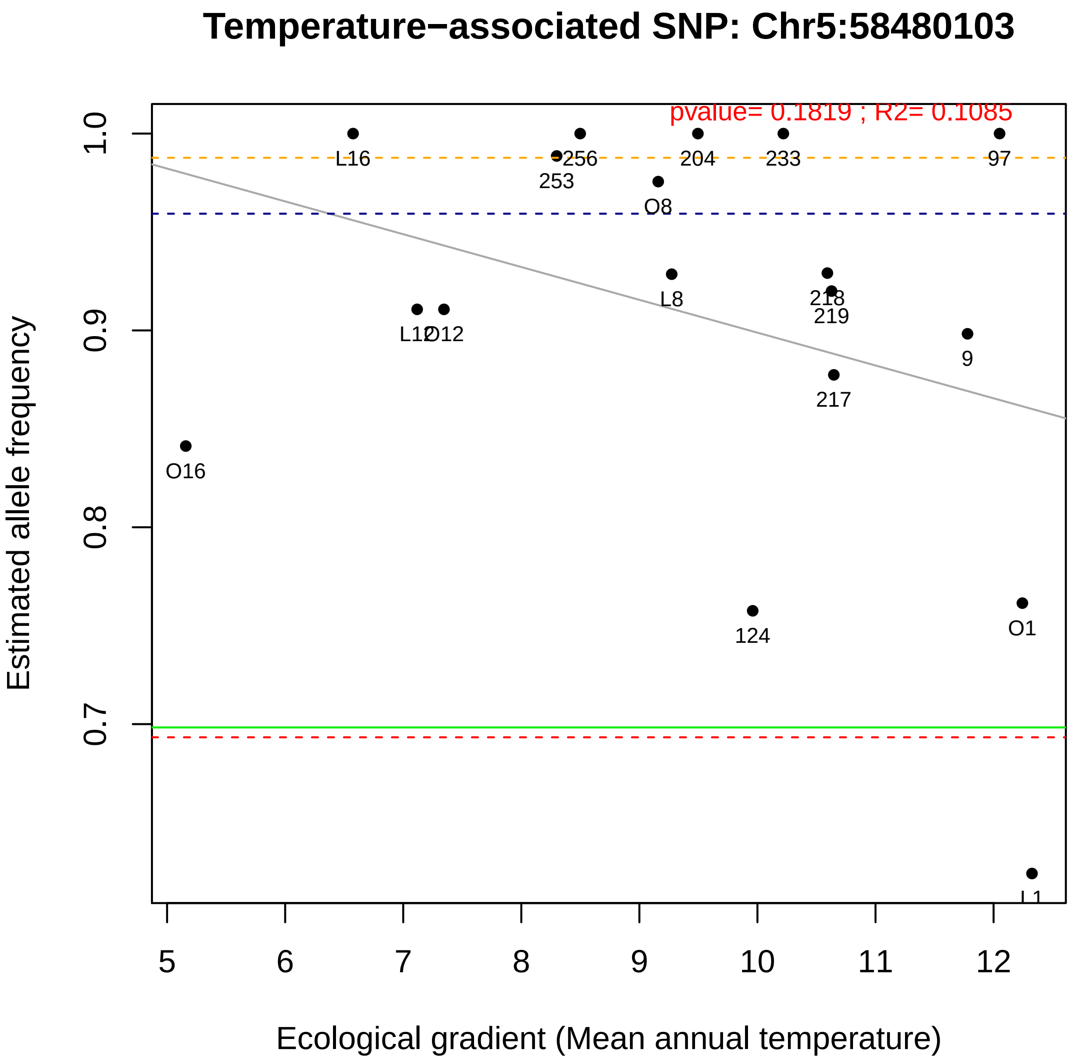
Temperature−associated SNP: Chr5:58480103
pvalue= 0.1819 ; R2= 0.1085
Estimated allele frequency
1.0
0.9
0.8
0.7
L16
256
253
204
233
97
O8
L8
218
219
L12
O12
9
217
O16
124
O1
L1
5
6
7
8
9
10
11
12
Ecological gradient (Mean annual temperature)

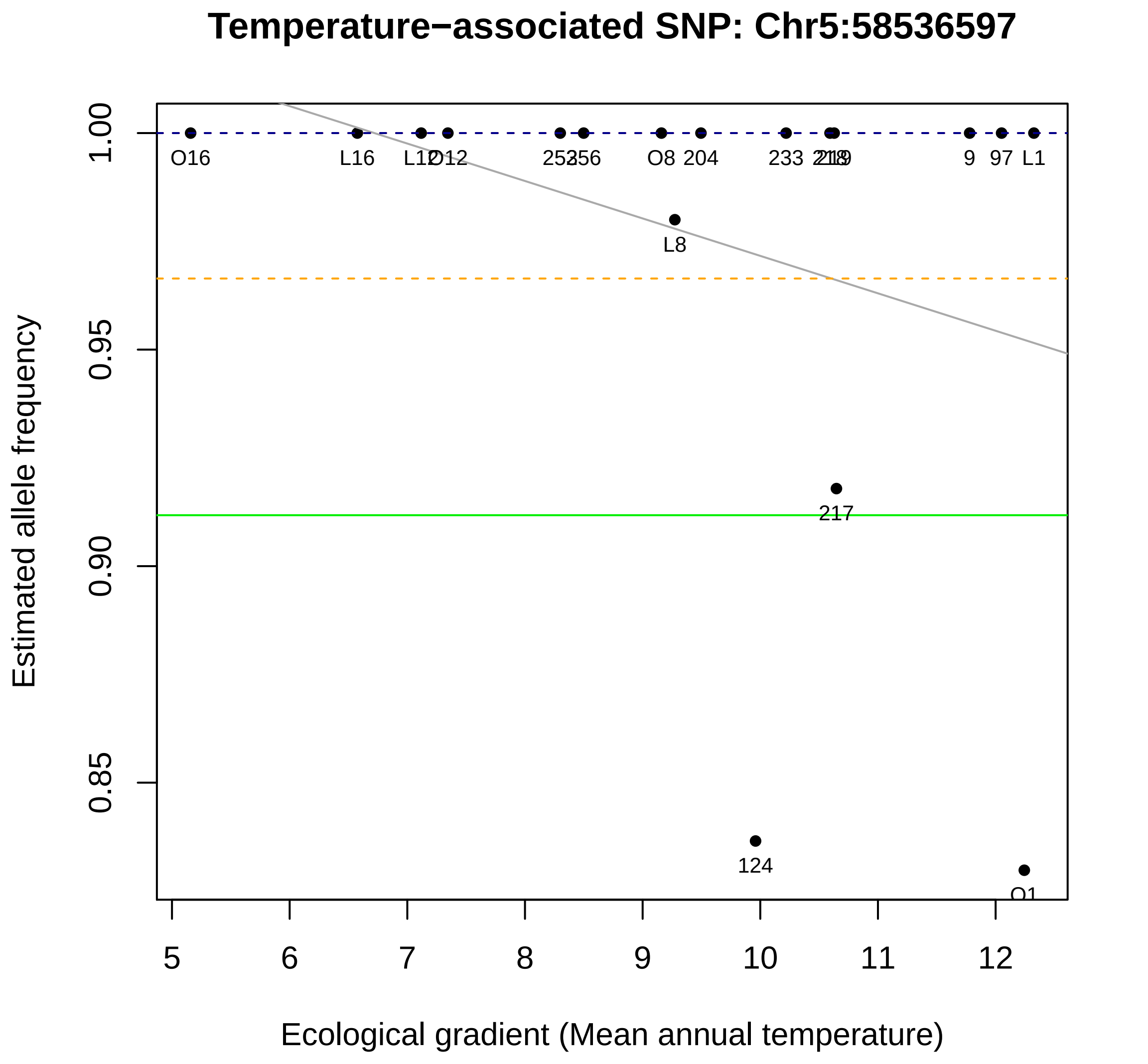
Temperature−associated SNP: Chr5:58536597
Estimated allele frequency
Ecological gradient (Mean annual temperature)
1.00
0.95
0.90
0.85
5
6
7
8
9
10
11
12
O16
L16
L12
O12
253
256
O8
204
233
218
219
9
97
L1
L8
217
124
O1

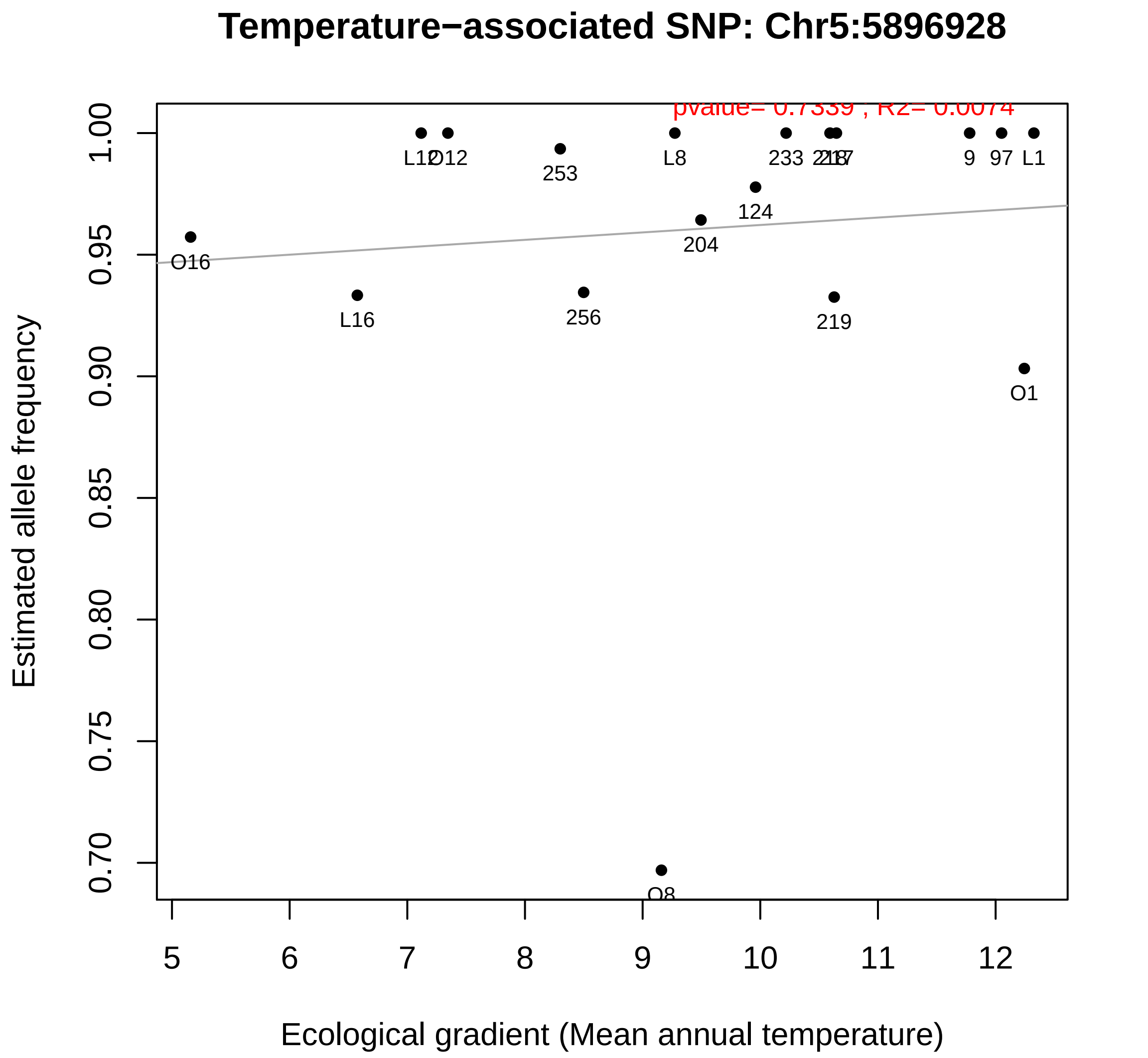

Temperature−associated SNP: Chr5:5896928
pvalue= 0.7339 ; R2= 0.0074
Estimated allele frequency
Ecological gradient (Mean annual temperature)
1.00
0.95
0.90
0.85
0.80
0.75
0.70
5
6
7
8
9
10
11
12
O16
L16
L12
O12
253
256
L8
204
124
233
219
9
97
L1
O1
O8

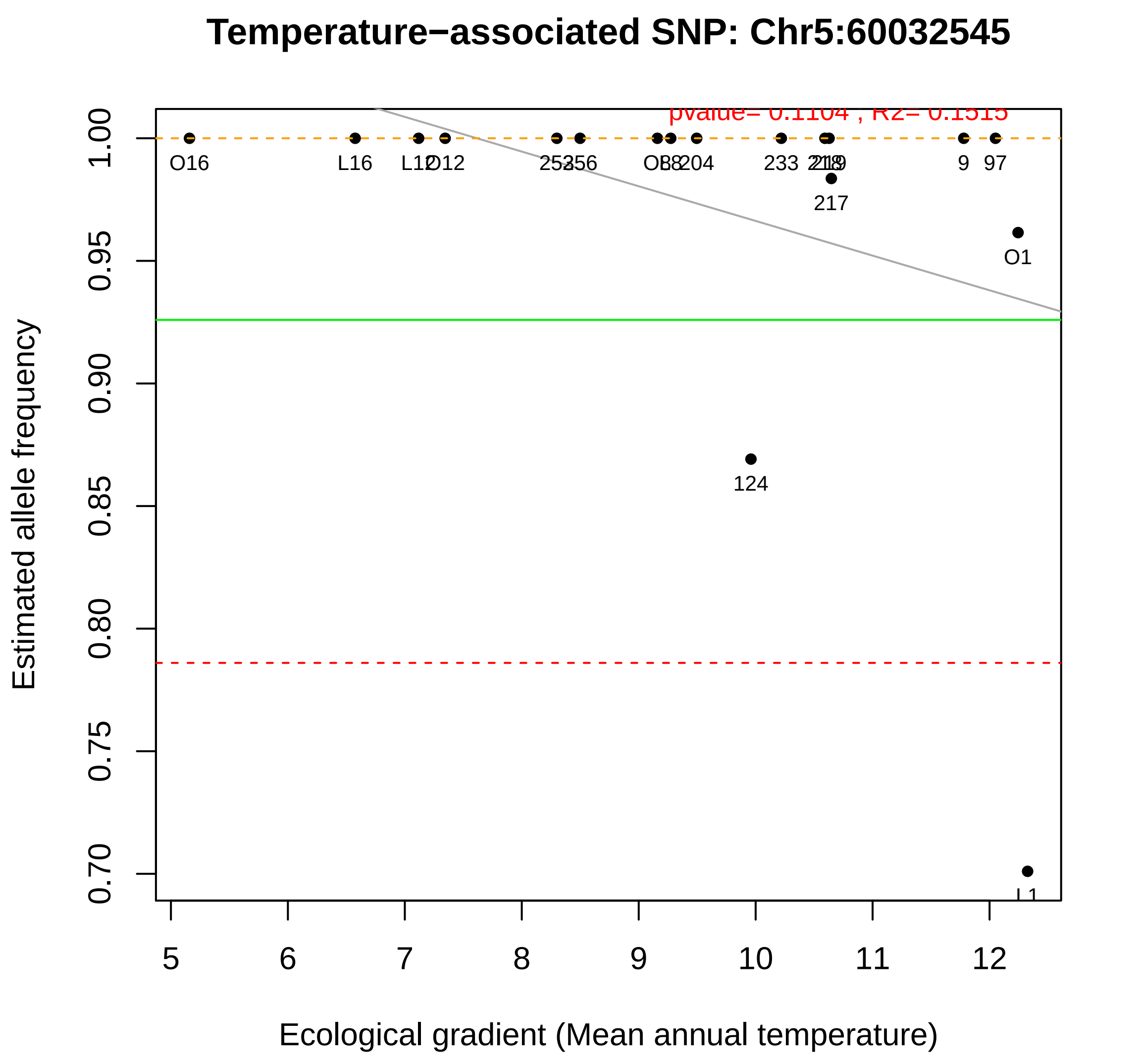

Temperature−associated SNP: Chr5:60032545
pvalue= 0.1104 , R2= 0.1515
Estimated allele frequency
Ecological gradient (Mean annual temperature)
1.00
0.95
0.90
0.85
0.80
0.75
0.70
5
6
7
8
9
10
11
12
O16
L16
O12
253
256
204
233
217
9
97
O1
124

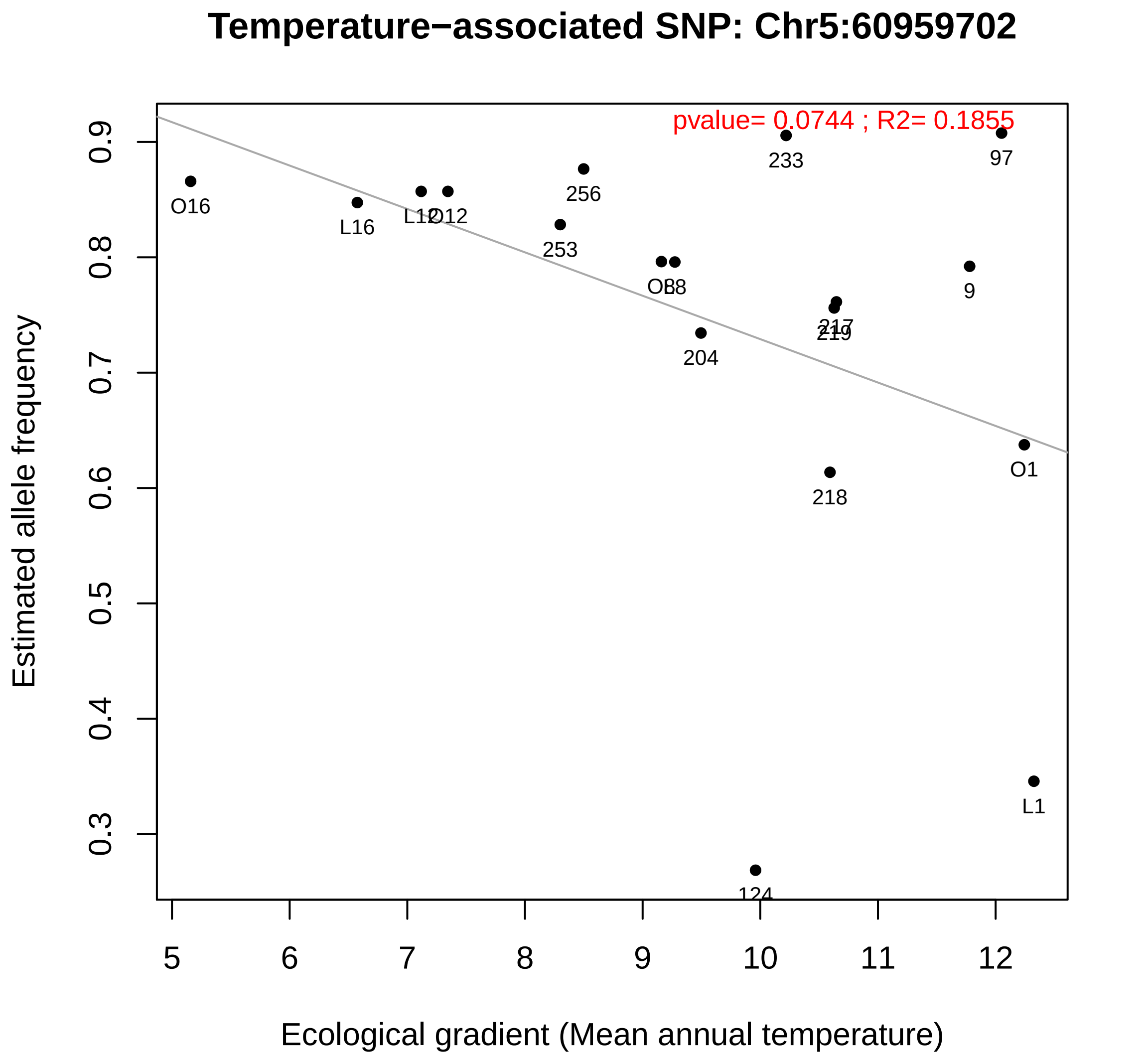

Temperature−associated SNP: Chr5:60959702
pvalue= 0.0744 ; R2= 0.1855
Estimated allele frequency
Ecological gradient (Mean annual temperature)
0.3
0.4
0.5
0.6
0.7
0.8
0.9
5
6
7
8
9
10
11
12
O16
L16
L12
O12
253
256
233
97
9
204
218
O1
L1
124

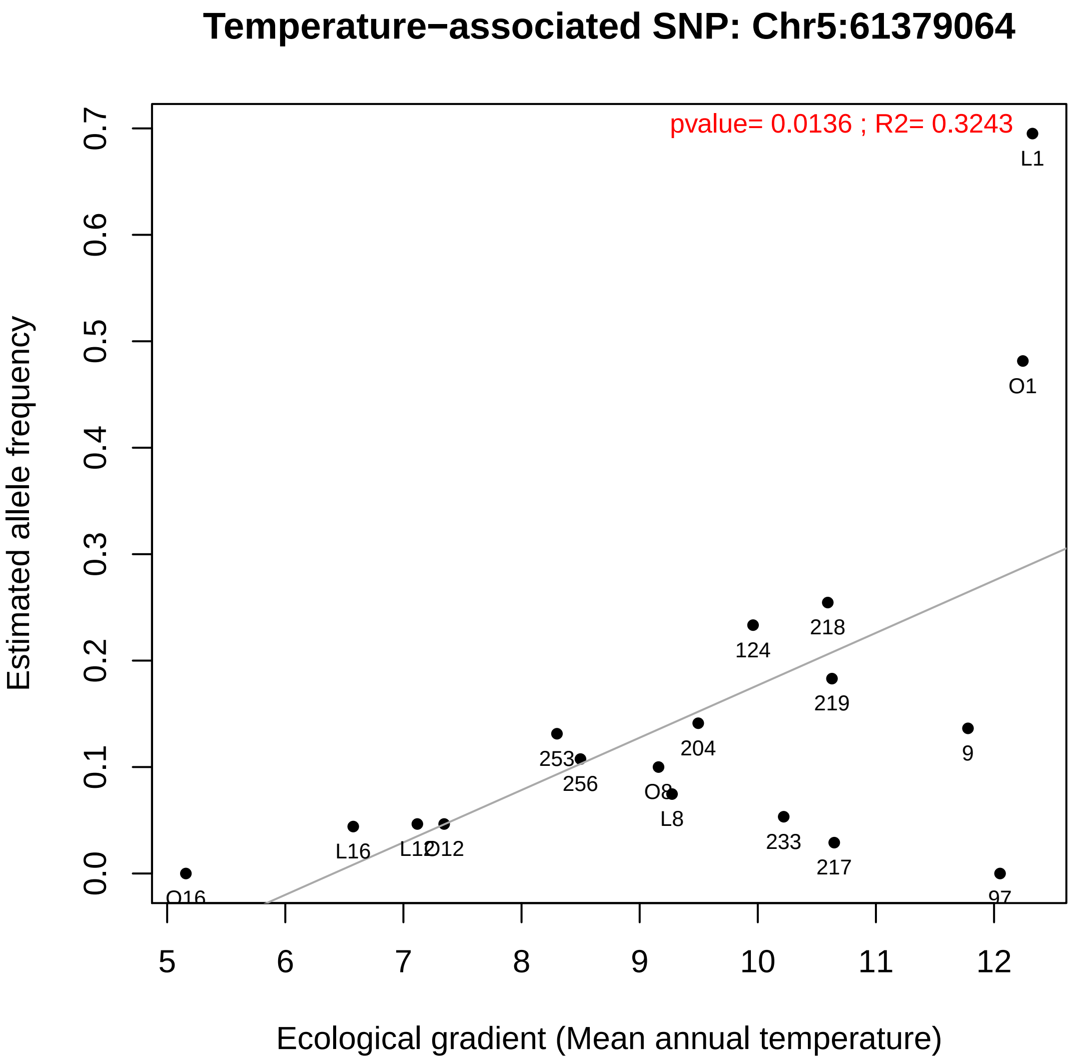

Temperature−associated SNP: Chr5:61379064
pvalue= 0.0136 ; R2= 0.3243
Estimated allele frequency
Ecological gradient (Mean annual temperature)
0.0
0.1
0.2
0.3
0.4
0.5
0.6
0.7
5
6
7
8
9
10
11
12
L1
O1
218
124
219
204
9
253
256
O8
L8
233
217
L16
L12
O12
O16
97

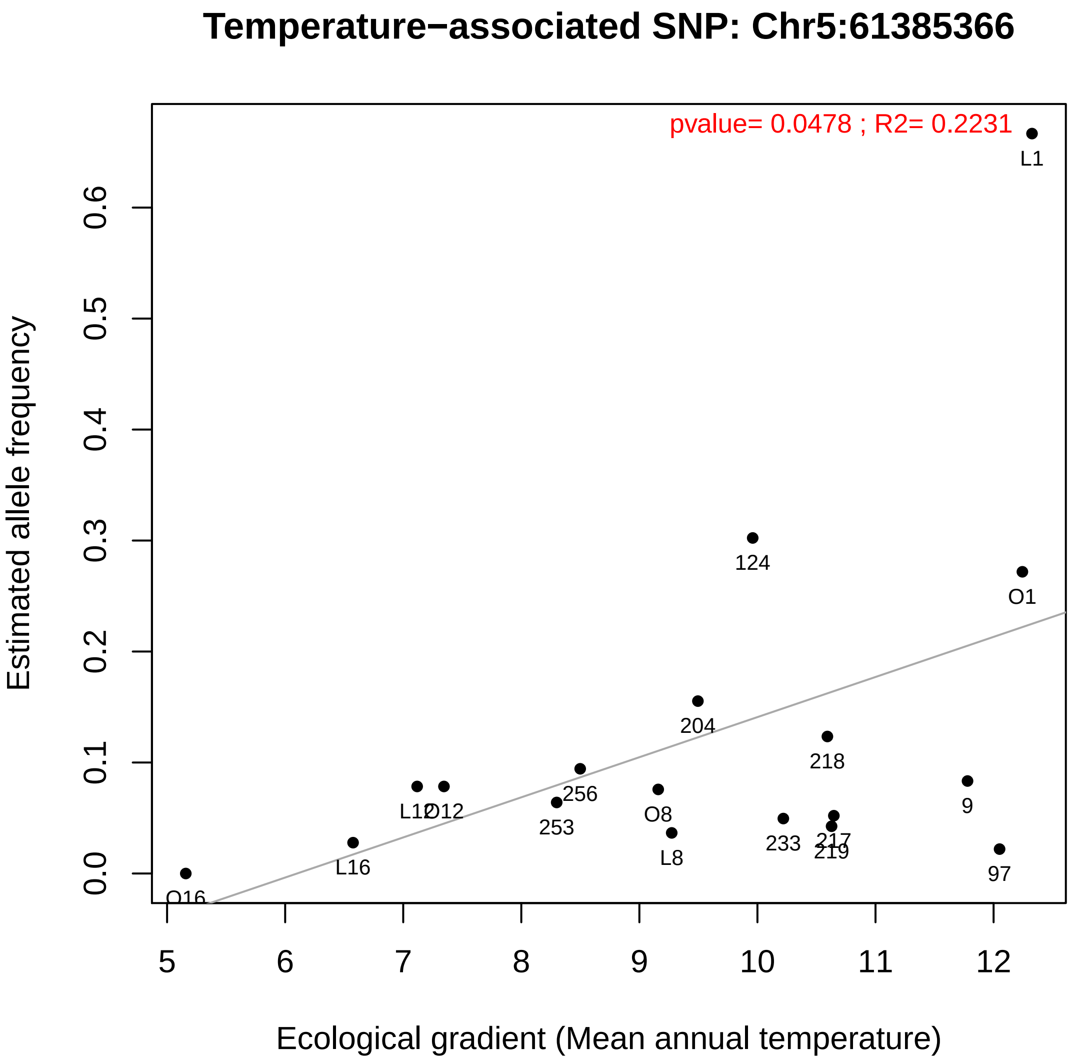
Temperature−associated SNP: Chr5:61385366
pvalue= 0.0478 ; R2= 0.2231
Estimated allele frequency
0.0
0.1
0.2
0.3
0.4
0.5
0.6
Ecological gradient (Mean annual temperature)
5
6
7
8
9
10
11
12
L1
124
O1
204
218
256
L12
O12
O8
253
9
L8
233
217
219
97
L16
O16

#### Temperature−associated SNP: Chr5:64837022

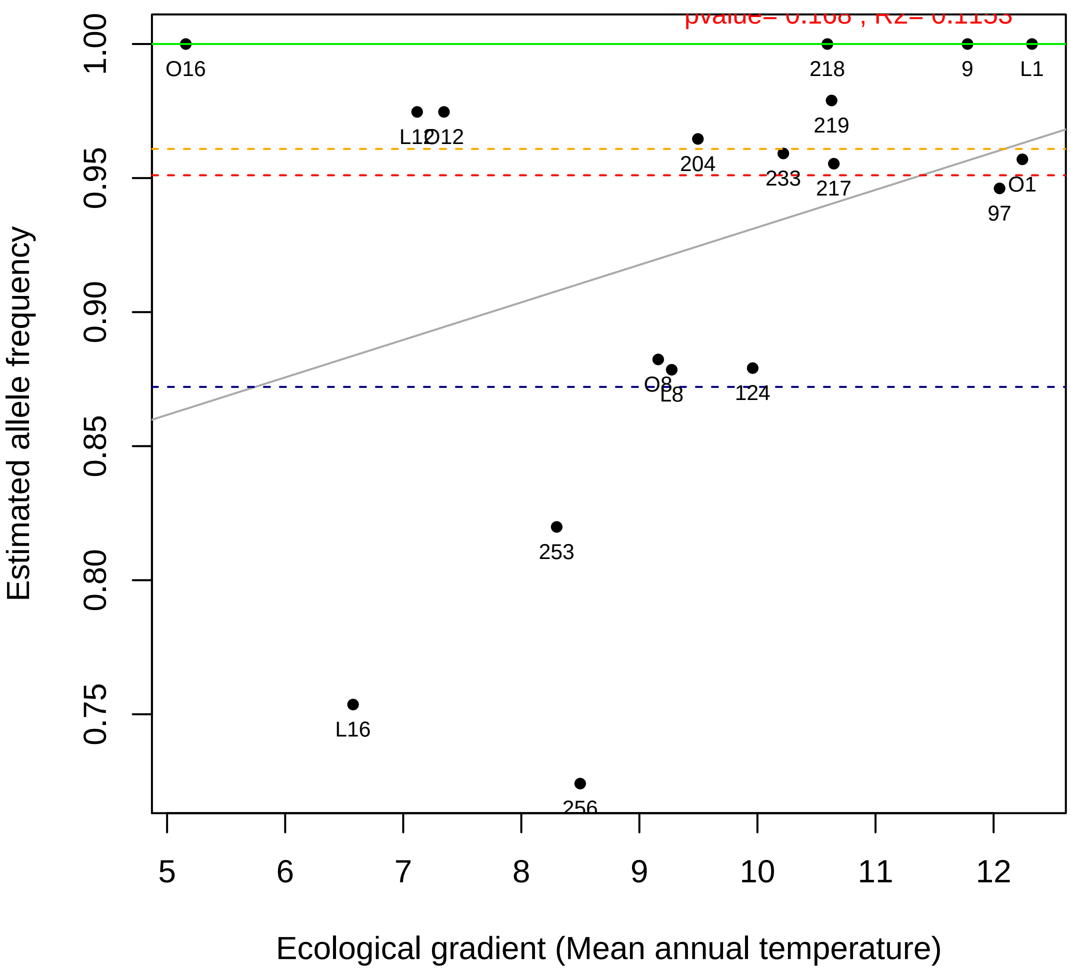

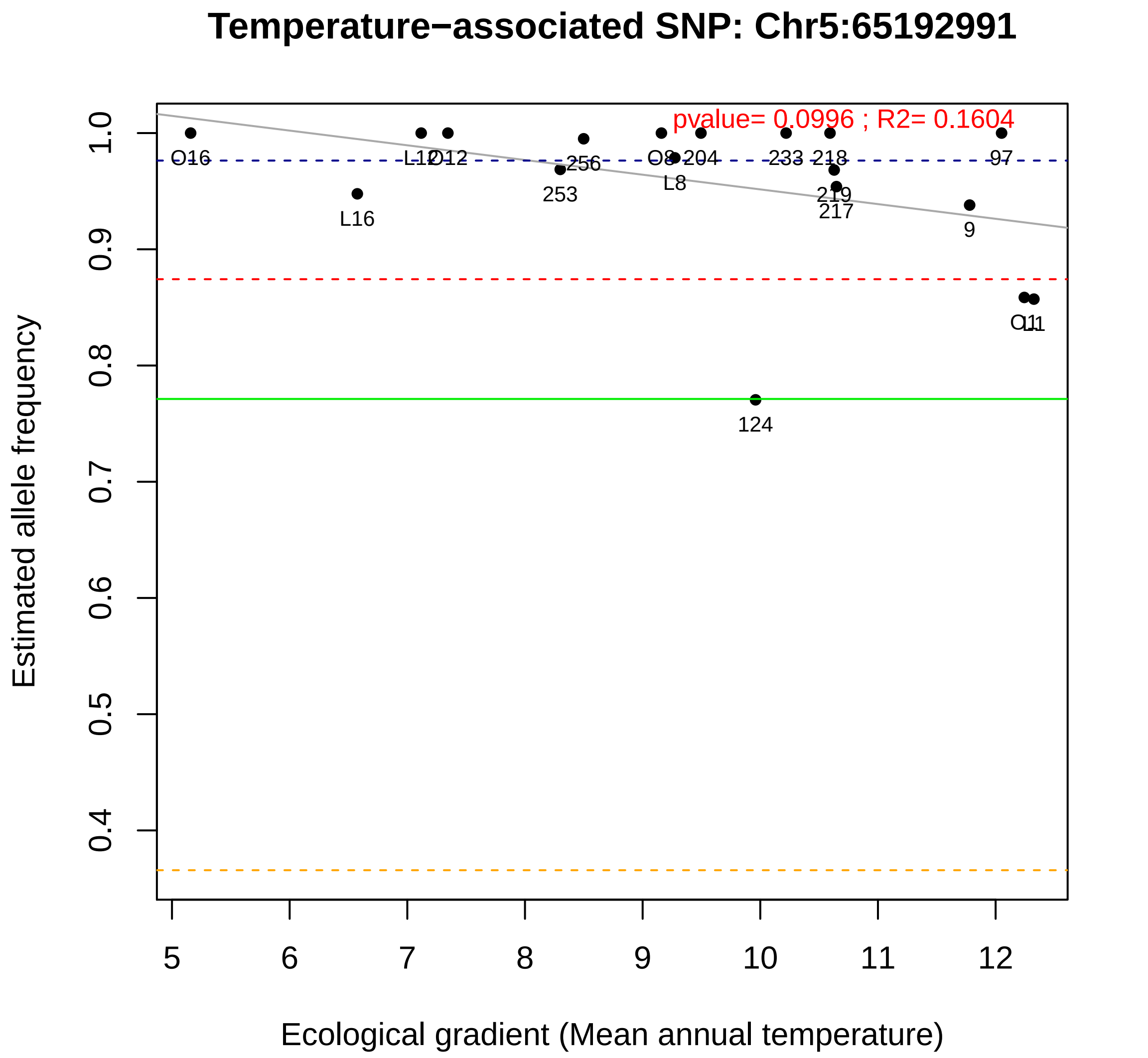

Temperature−associated SNP: Chr5:65192991
pvalue= 0.0996 ; R2= 0.1604
O16
L16
L12
O12
256
253
O8
L8
204
233
218
219
217
9
97
O1
L1
124
1.0
0.9
0.8
0.7
0.6
0.5
0.4
5
6
7
8
9
10
11
12
Estimated allele frequency
Ecological gradient (Mean annual temperature)

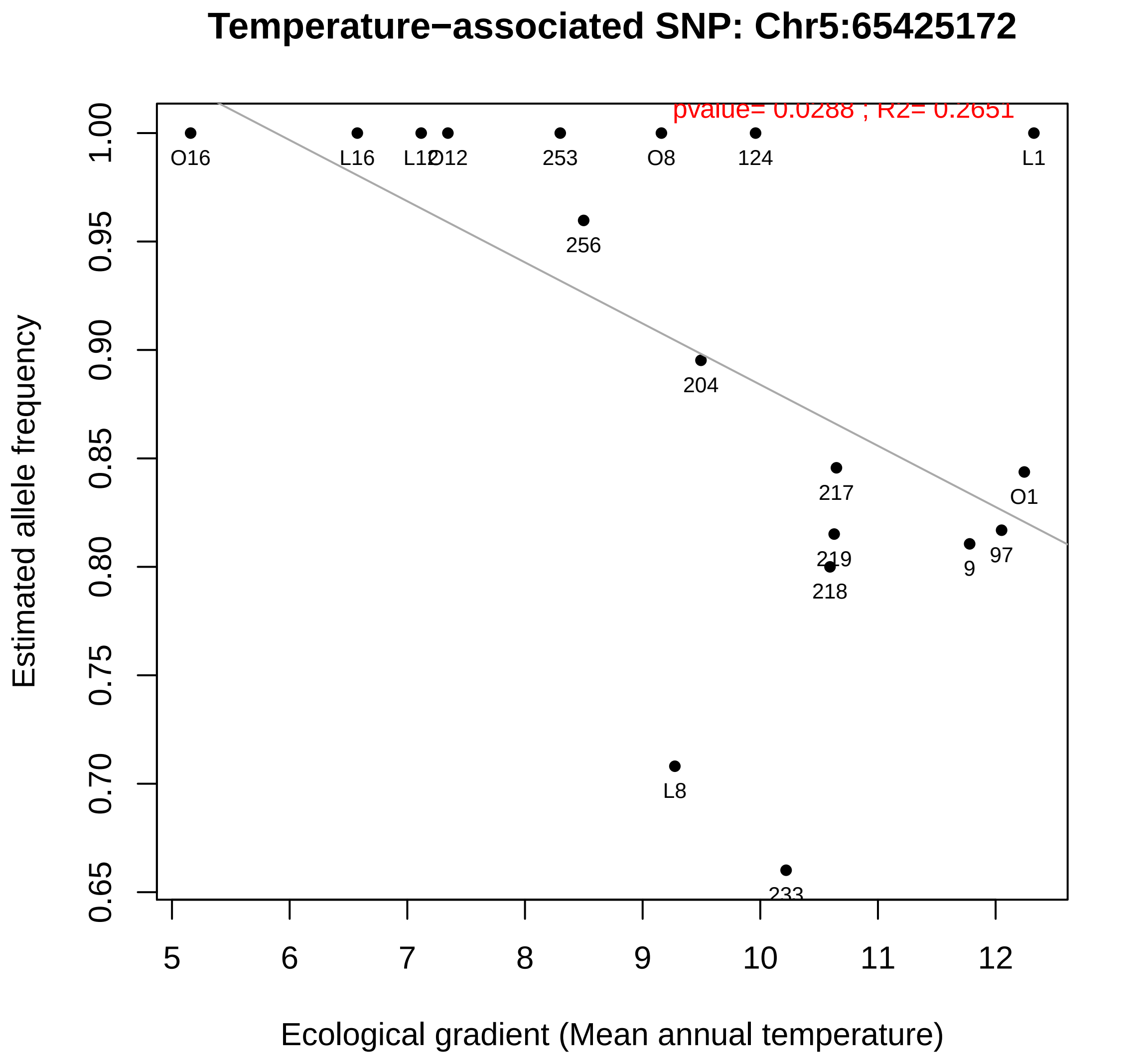
Temperature−associated SNP: Chr5:65425172
pvalue= 0.0288 ; R2= 0.2651
Estimated allele frequency
1.00
0.95
0.90
0.85
0.80
0.75
0.70
0.65
O16
L16
L12
O12
253
O8
124
L1
256
204
217
O1
219
218
9
97
L8
233
5
6
7
8
9
10
11
12
Ecological gradient (Mean annual temperature)

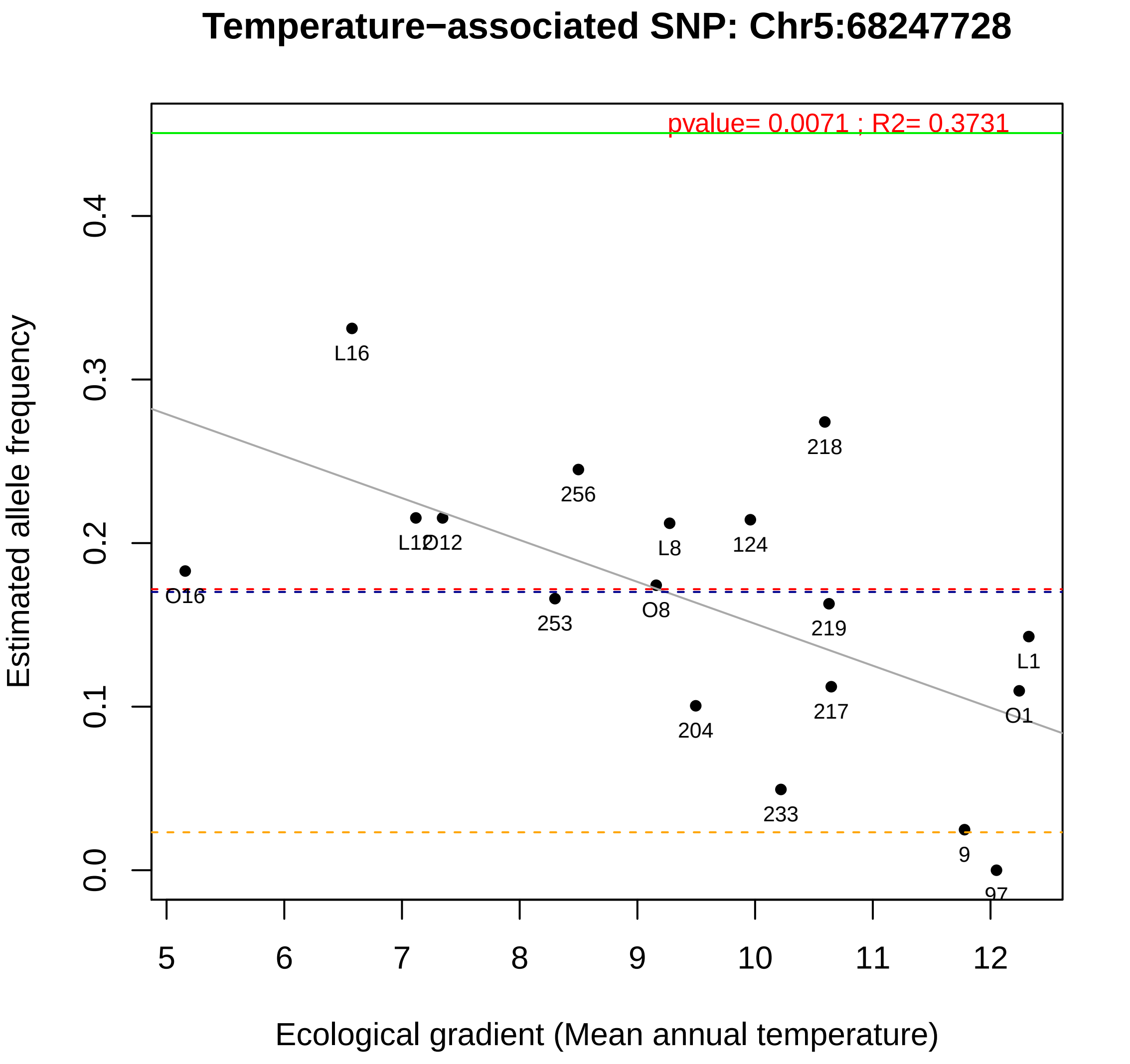
Temperature−associated SNP: Chr5:68247728
pvalue= 0.0071 ; R2= 0.3731
Estimated allele frequency
0.0
0.1
0.2
0.3
0.4
L16
218
256
L12
O12
L8
124
O16
253
O8
219
L1
217
O1
204
233
9
97
5
6
7
8
9
10
11
12
Ecological gradient (Mean annual temperature)

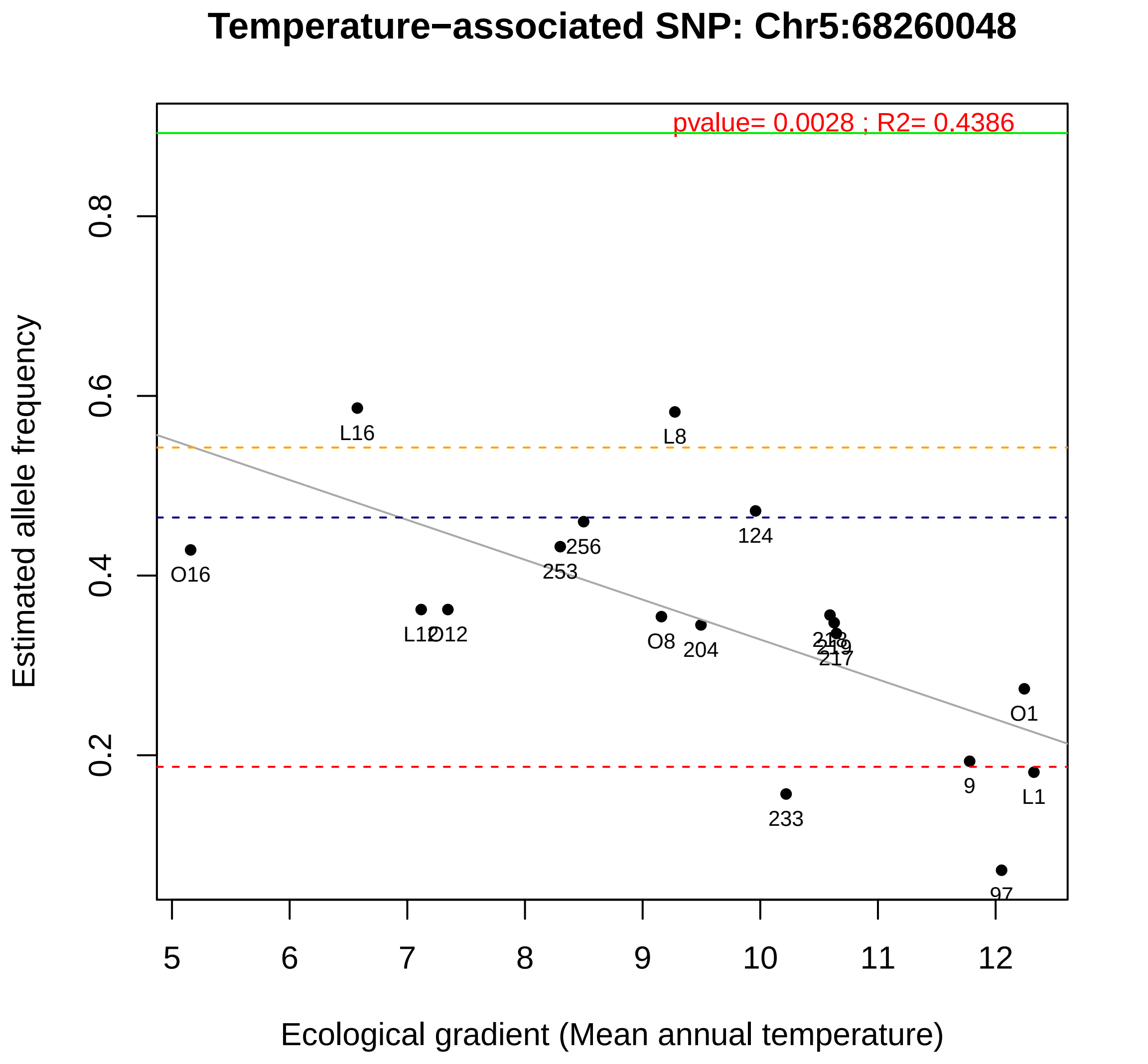

Temperature−associated SNP: Chr5:68260048
pvalue= 0.0028 ; R2= 0.4386
Estimated allele frequency
0.2
0.4
0.6
0.8
5
6
7
8
9
10
11
12
Ecological gradient (Mean annual temperature)
L16
L8
124
256
253
O16
L12
O12
O8
204
217
O1
9
L1
233
97

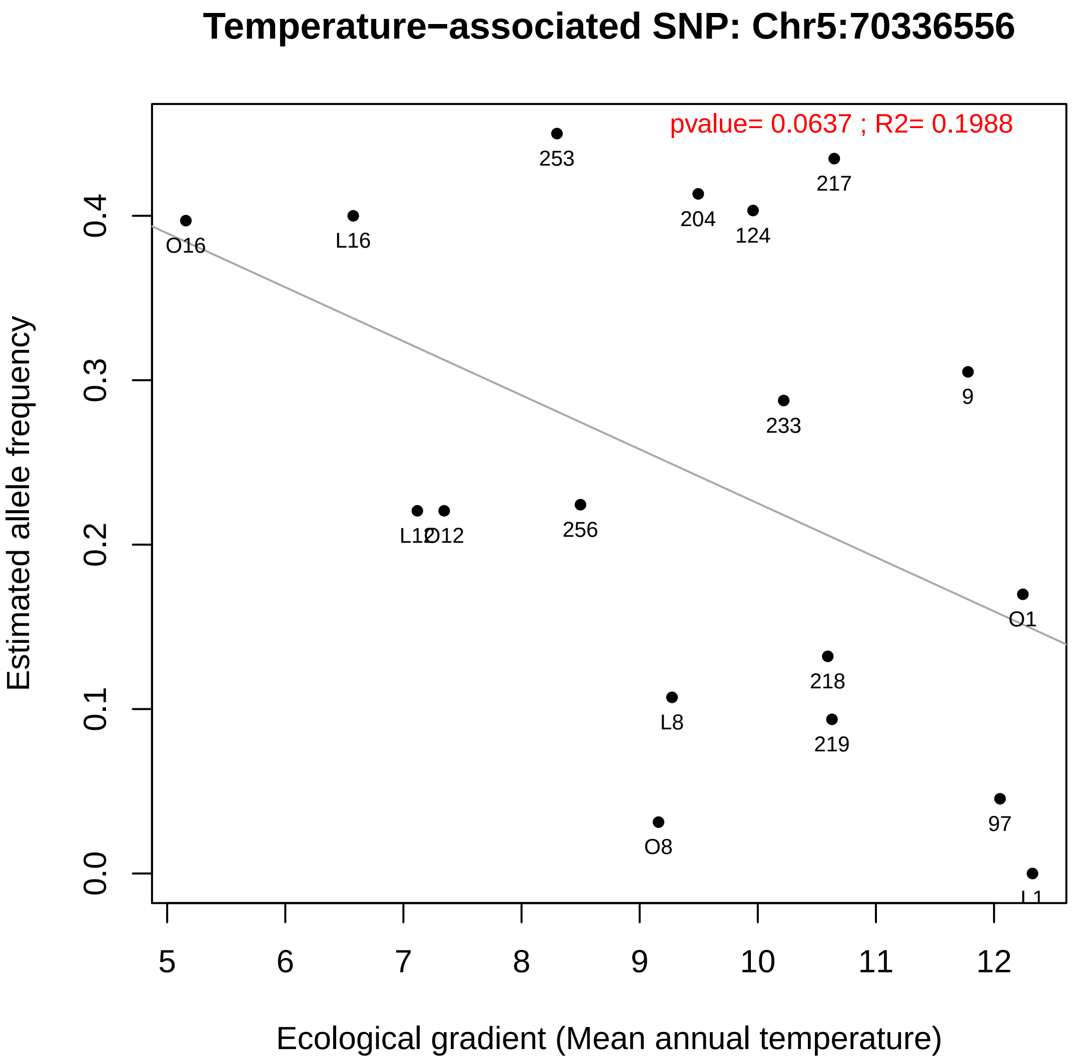
Temperature−associated SNP: Chr5:70336556
pvalue= 0.0637 ; R2= 0.1988
253
217
204
124
O16
L16
233
9
L12
O12
256
O1
218
L8
219
97
O8
L1
0.0
0.1
0.2
0.3
0.4
Estimated allele frequency
5
6
7
8
9
10
11
12
Ecological gradient (Mean annual temperature)

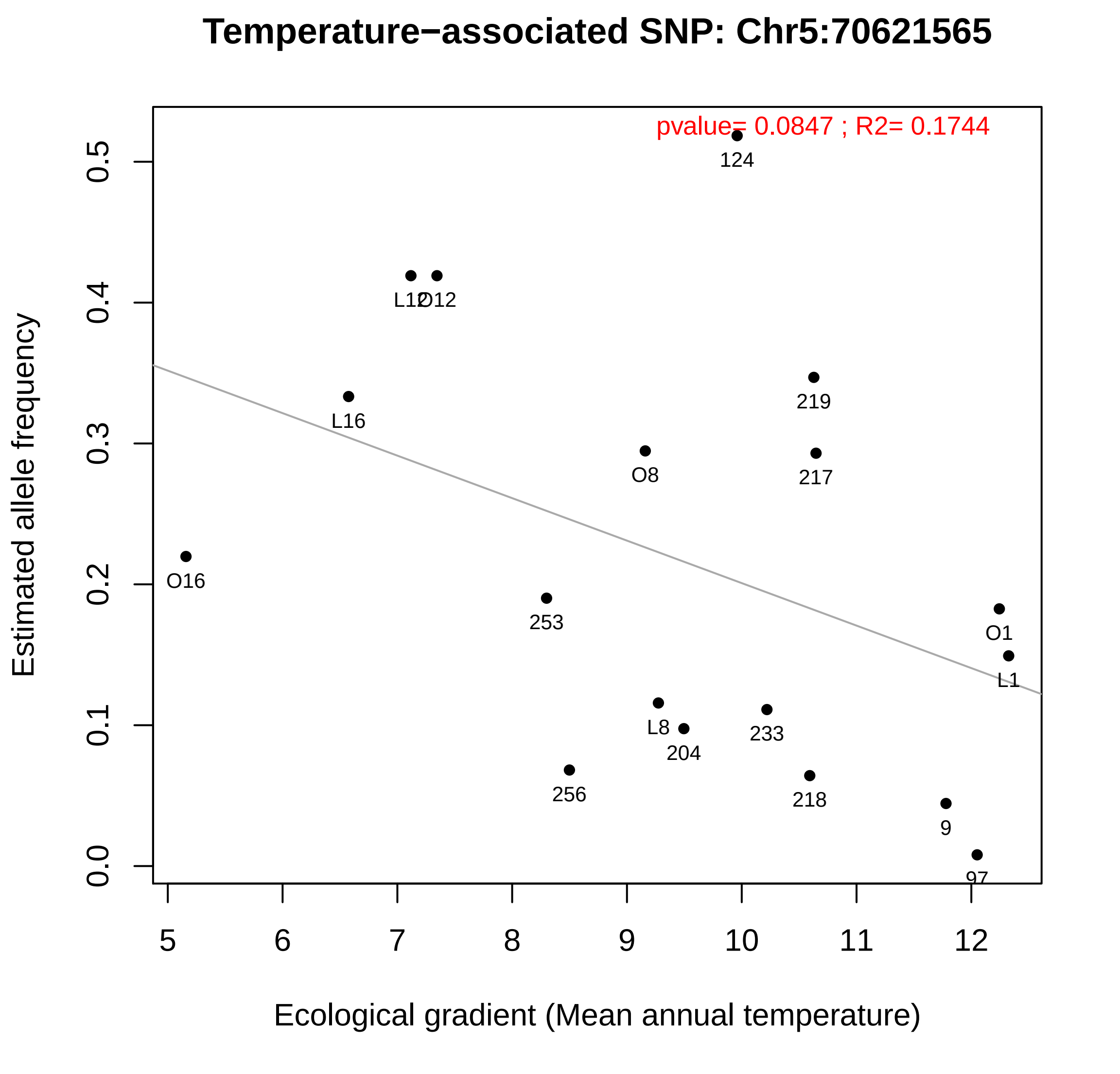

Temperature−associated SNP: Chr5:70621565
pvalue= 0.0847 ; R2= 0.1744
Estimated allele frequency
0.0
0.1
0.2
0.3
0.4
0.5
5
6
7
8
9
10
11
12
Ecological gradient (Mean annual temperature)
124
L12
O12
L16
219
O8
217
O16
253
O1
L1
L8
204
233
256
218
9
97

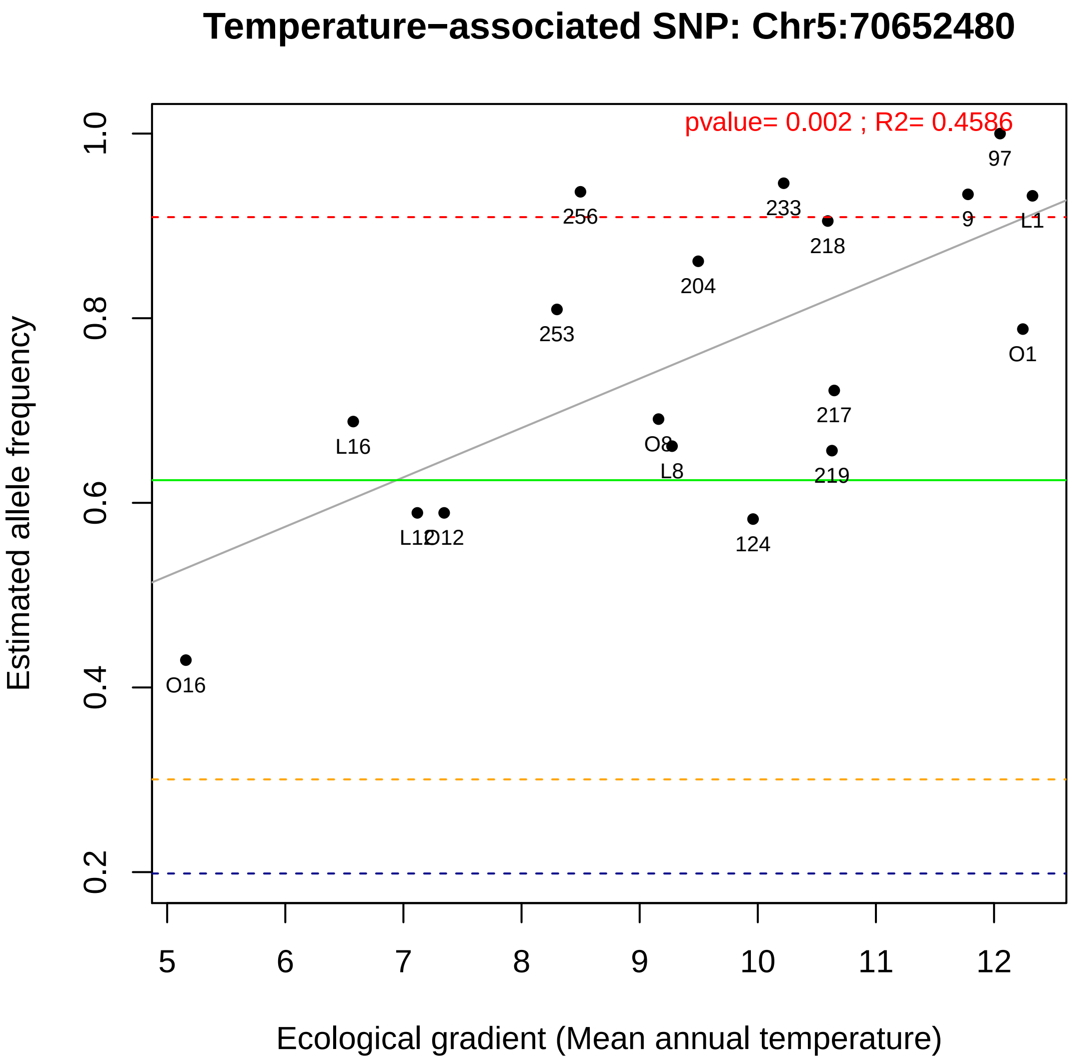

Temperature−associated SNP: Chr5:70652480
pvalue= 0.002 ; R2= 0.4586
Estimated allele frequency
0.2
0.4
0.6
0.8
1.0
5
6
7
8
9
10
11
12
Ecological gradient (Mean annual temperature)
97
256
233
9
L1
218
204
253
O1
217
L16
O8
L8
219
L12
O12
124
O16

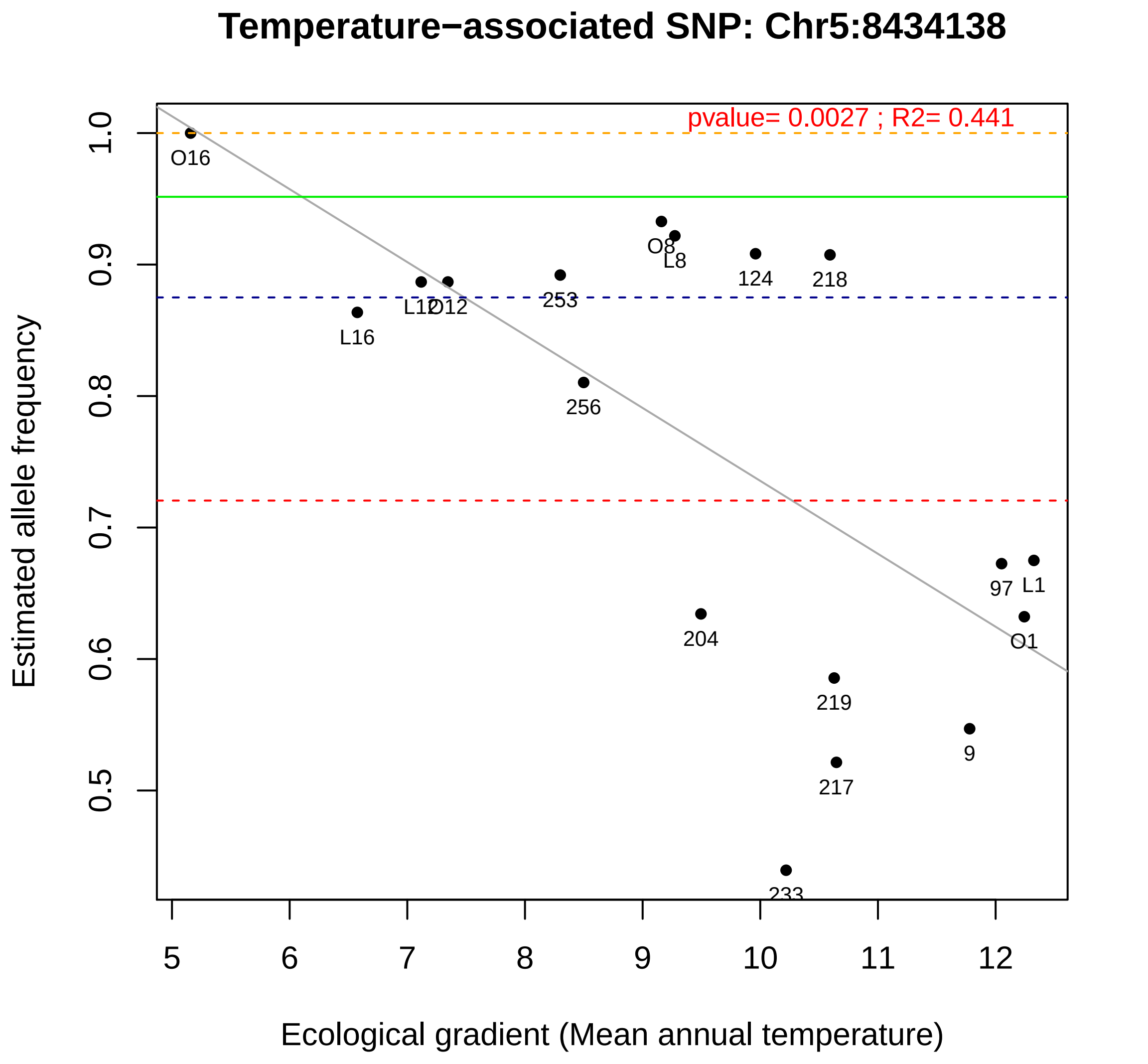

Temperature−associated SNP: Chr5:8434138
pvalue= 0.0027 ; R2= 0.441
Estimated allele frequency
Ecological gradient (Mean annual temperature)
1.0
0.9
0.8
0.7
0.6
0.5
5
6
7
8
9
10
11
12
O16
L16
L12
O12
253
256
O8
L8
124
218
204
219
217
233
9
97
L1
O1

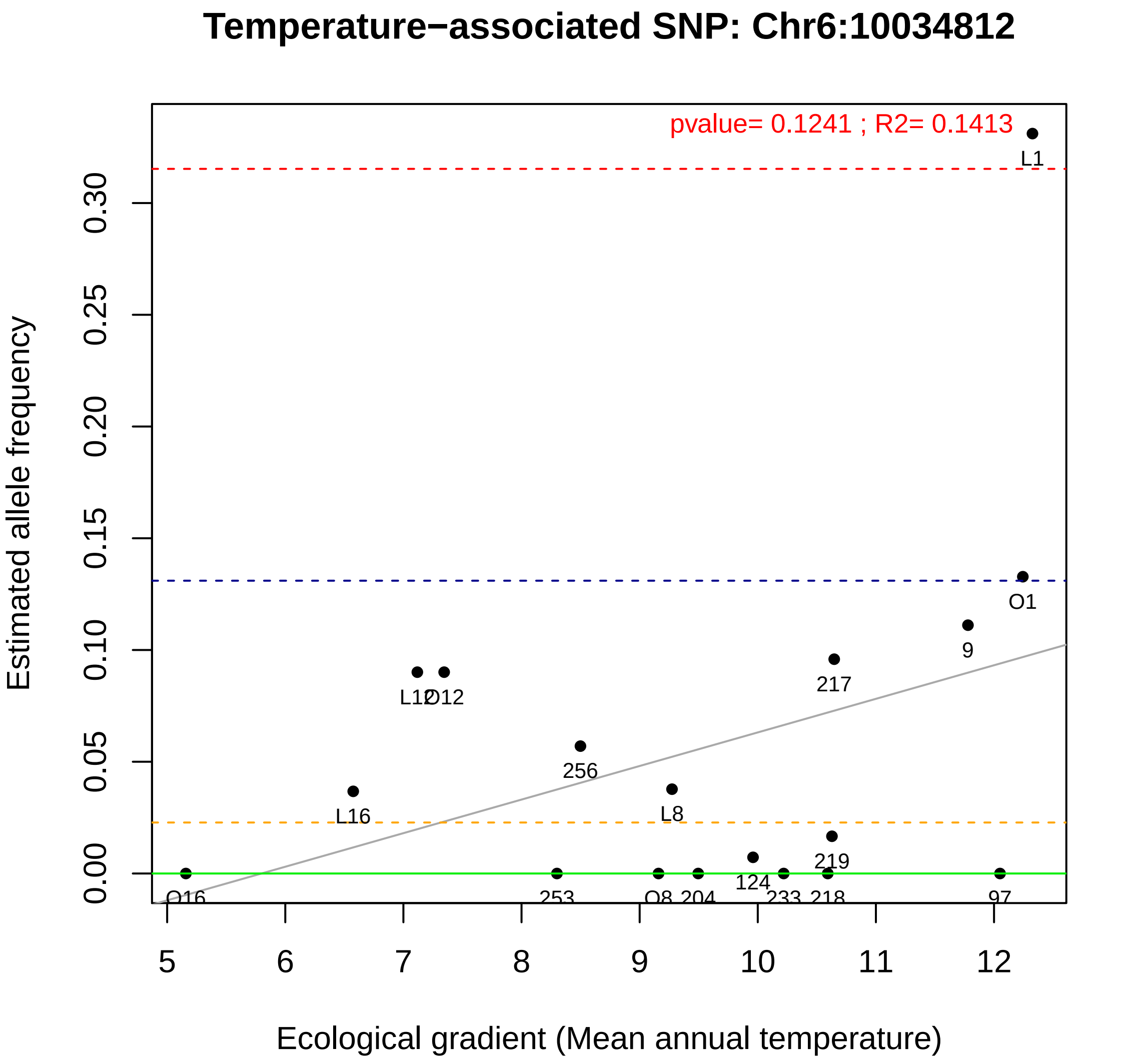

Temperature−associated SNP: Chr6:10034812
pvalue= 0.1241 ; R2= 0.1413
Estimated allele frequency
0.00
0.05
0.10
0.15
0.20
0.25
0.30
5
6
7
8
9
10
11
12
Ecological gradient (Mean annual temperature)
L1
O1
9
217
L12
O12
256
L16
L8
219
124
O16
253
O8
204
233
218
97

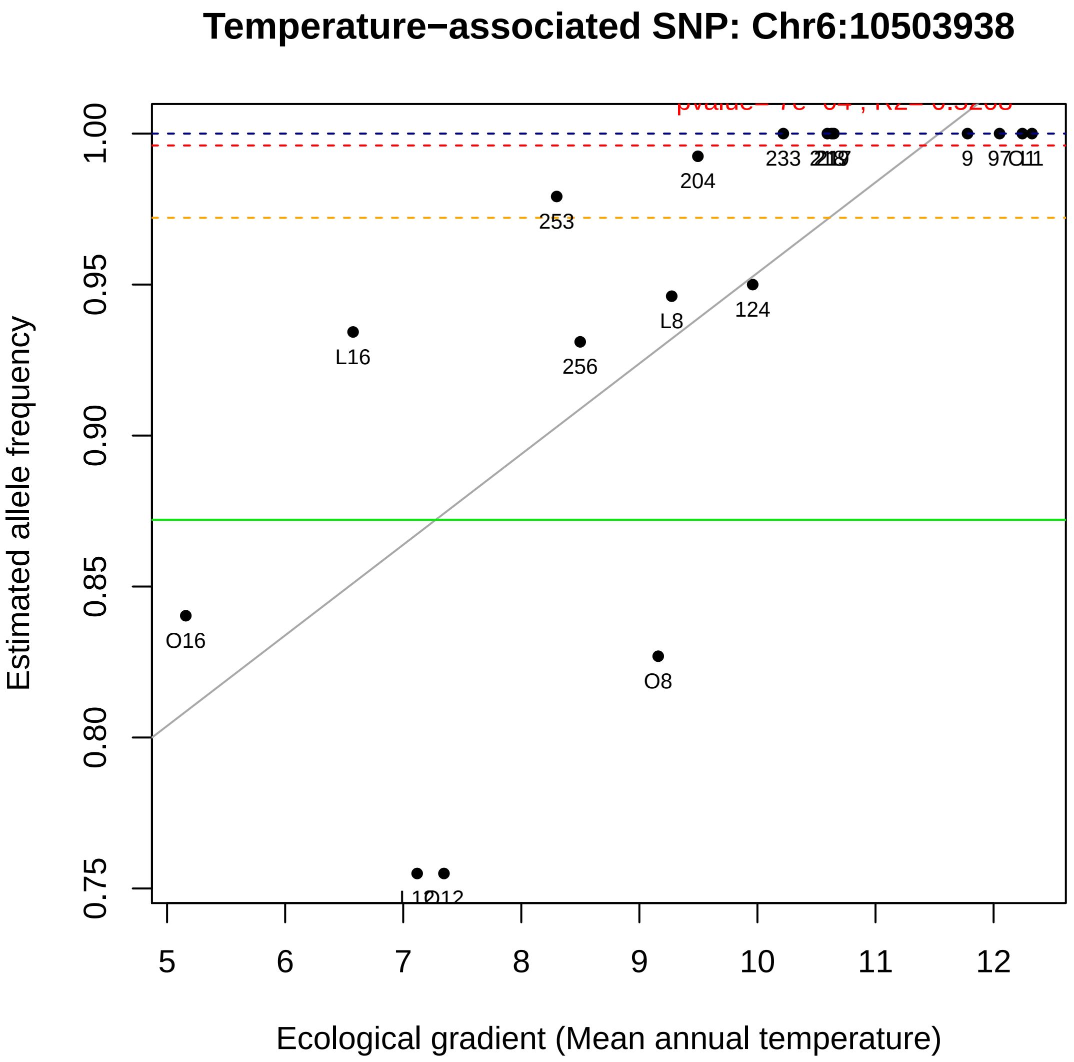

Temperature−associated SNP: Chr6:10503938
Estimated allele frequency
Ecological gradient (Mean annual temperature)
1.00
0.95
0.90
0.85
0.80
0.75
5
6
7
8
9
10
11
12
233
9
204
253
124
L8
L16
256
O16
O8

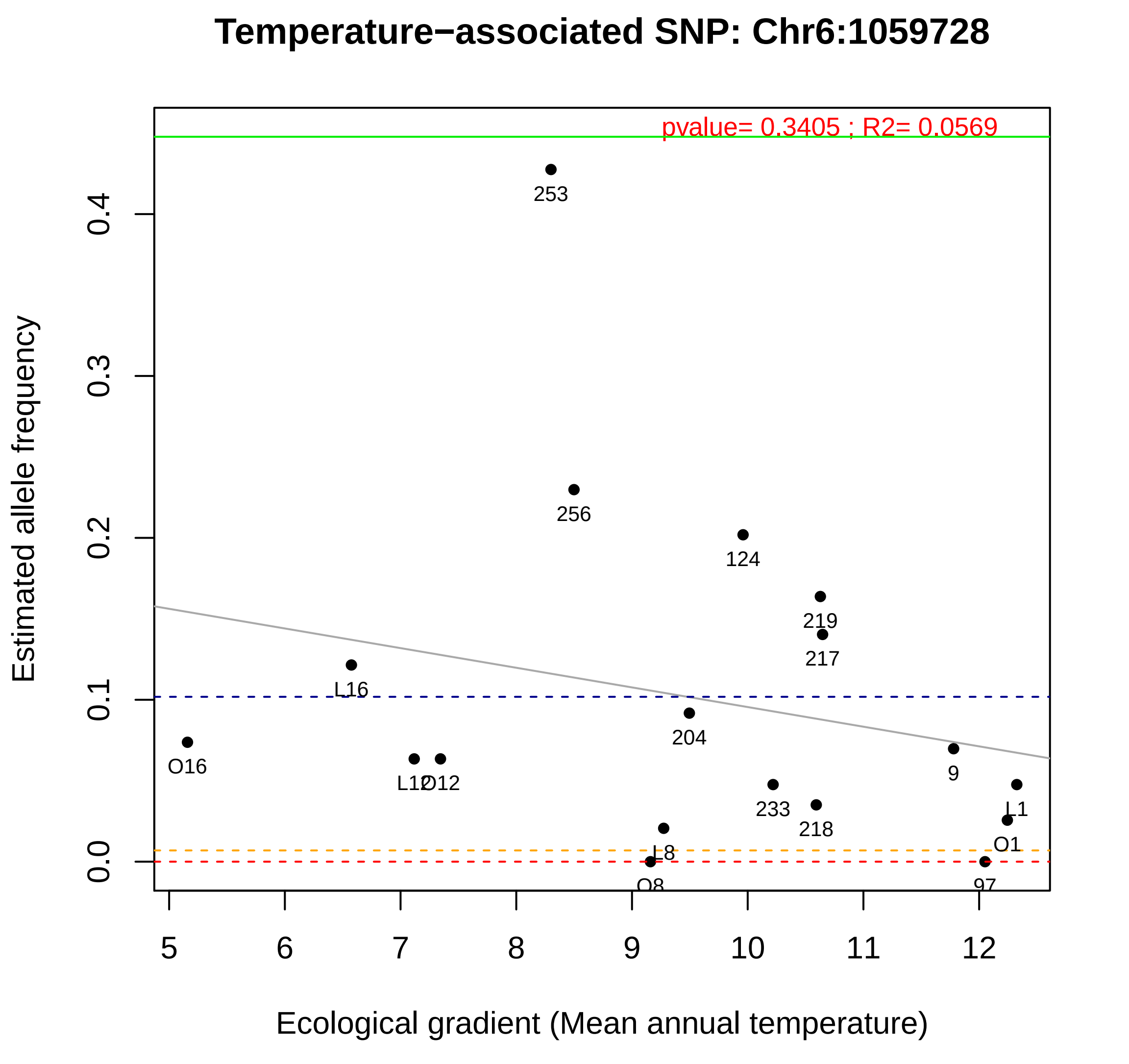

Temperature−associated SNP: Chr6:1059728
pvalue= 0.3405 ; R2= 0.0569
Estimated allele frequency
Ecological gradient (Mean annual temperature)
0.0
0.1
0.2
0.3
0.4
5
6
7
8
9
10
11
12
253
256
124
219
217
L16
204
O16
L12
O12
9
233
L1
218
O1
L8
O8
97

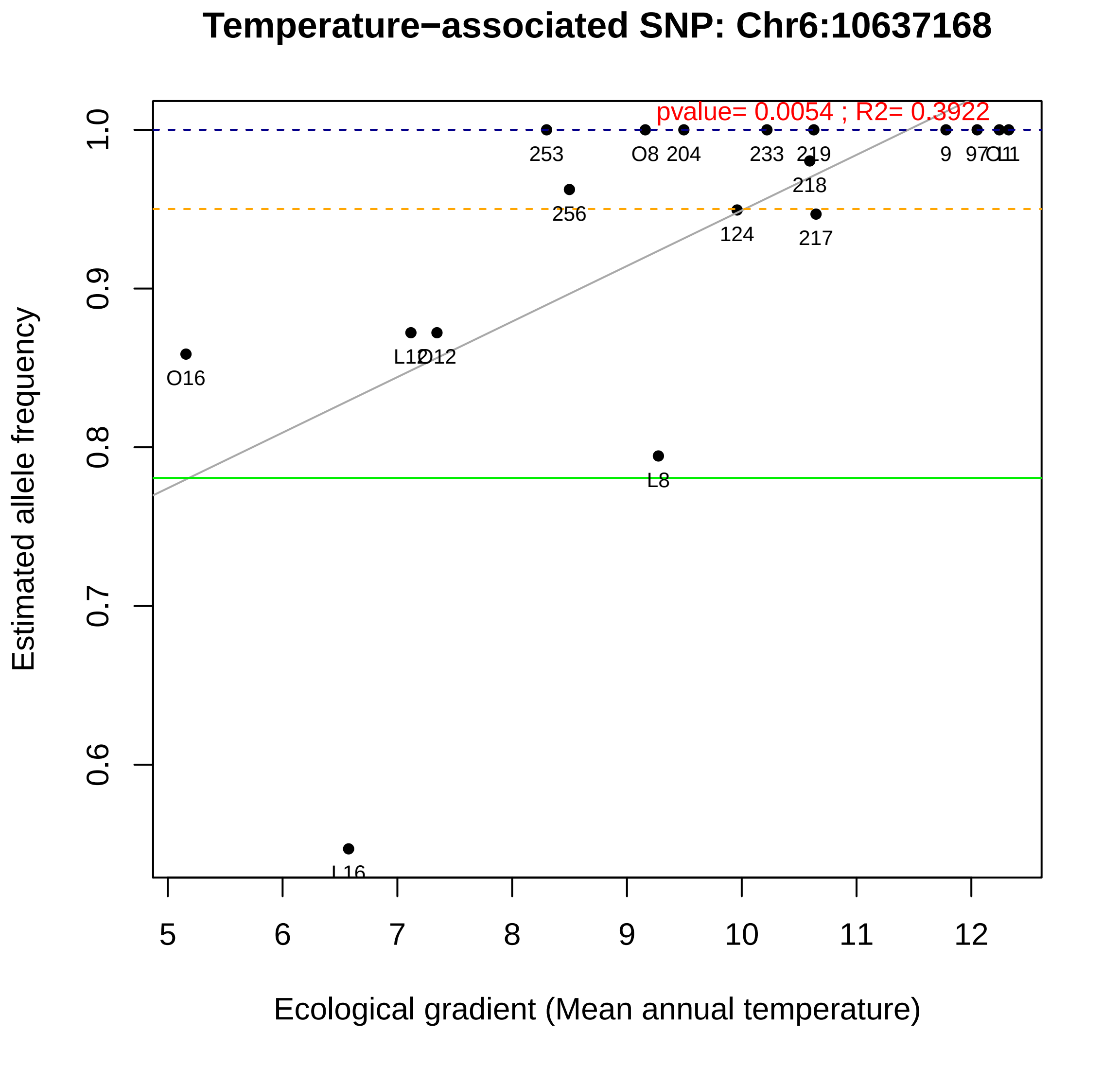

Temperature−associated SNP: Chr6:10637168
pvalue= 0.0054 ; R2= 0.3922
Estimated allele frequency
Ecological gradient (Mean annual temperature)
0.6
0.7
0.8
0.9
1.0
5
6
7
8
9
10
11
12
253
O8
204
233
219
9
218
256
124
217
O16
L12
O12
L8
L16

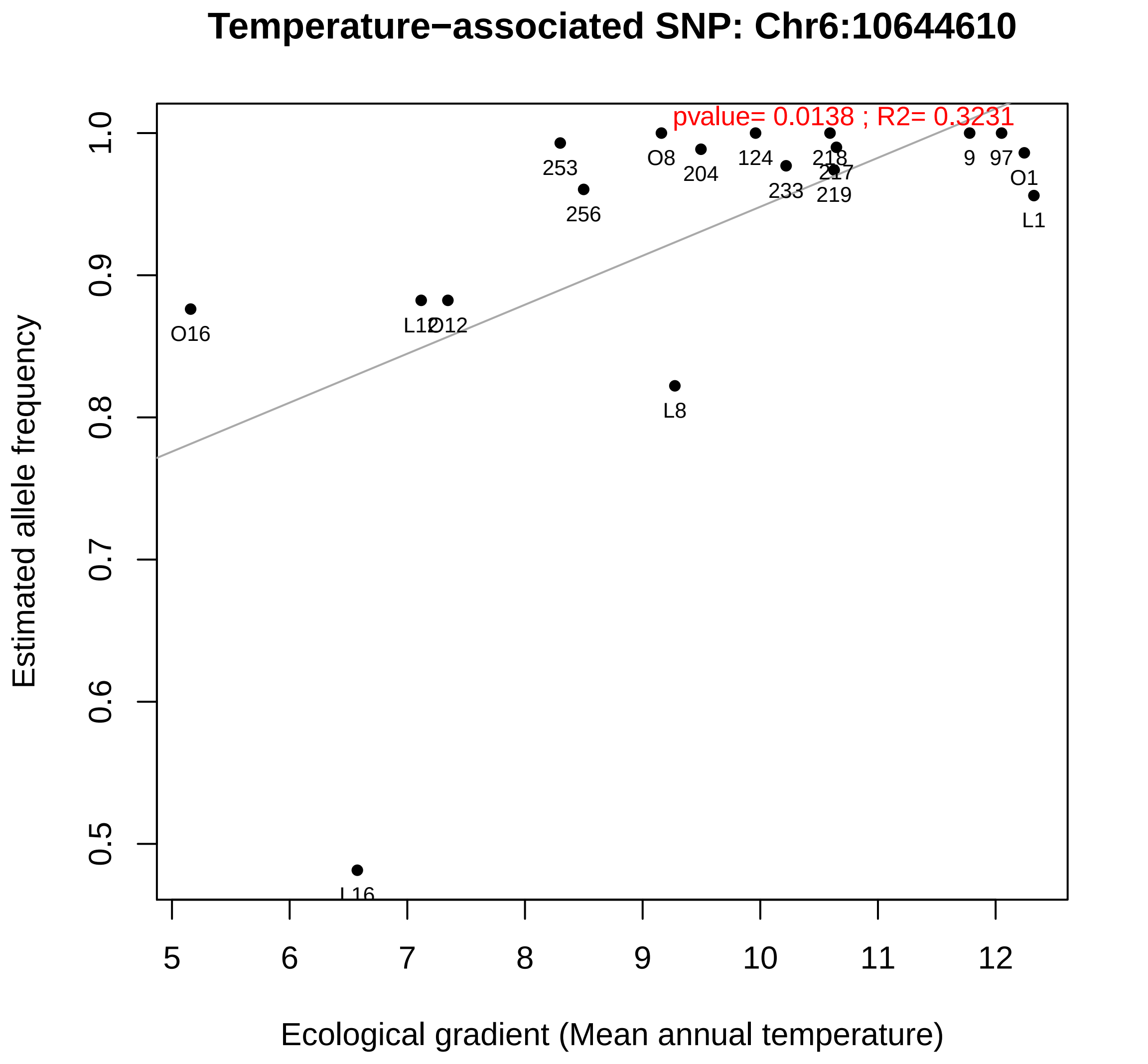

Temperature−associated SNP: Chr6:10644610
pvalue= 0.0138 ; R2= 0.3231
Estimated allele frequency
1.0
0.9
0.8
0.7
0.6
0.5
Ecological gradient (Mean annual temperature)
5
6
7
8
9
10
11
12
253
256
O8
204
124
233
218
217
219
9
97
O1
L1
O16
L12
O12
L8
L16

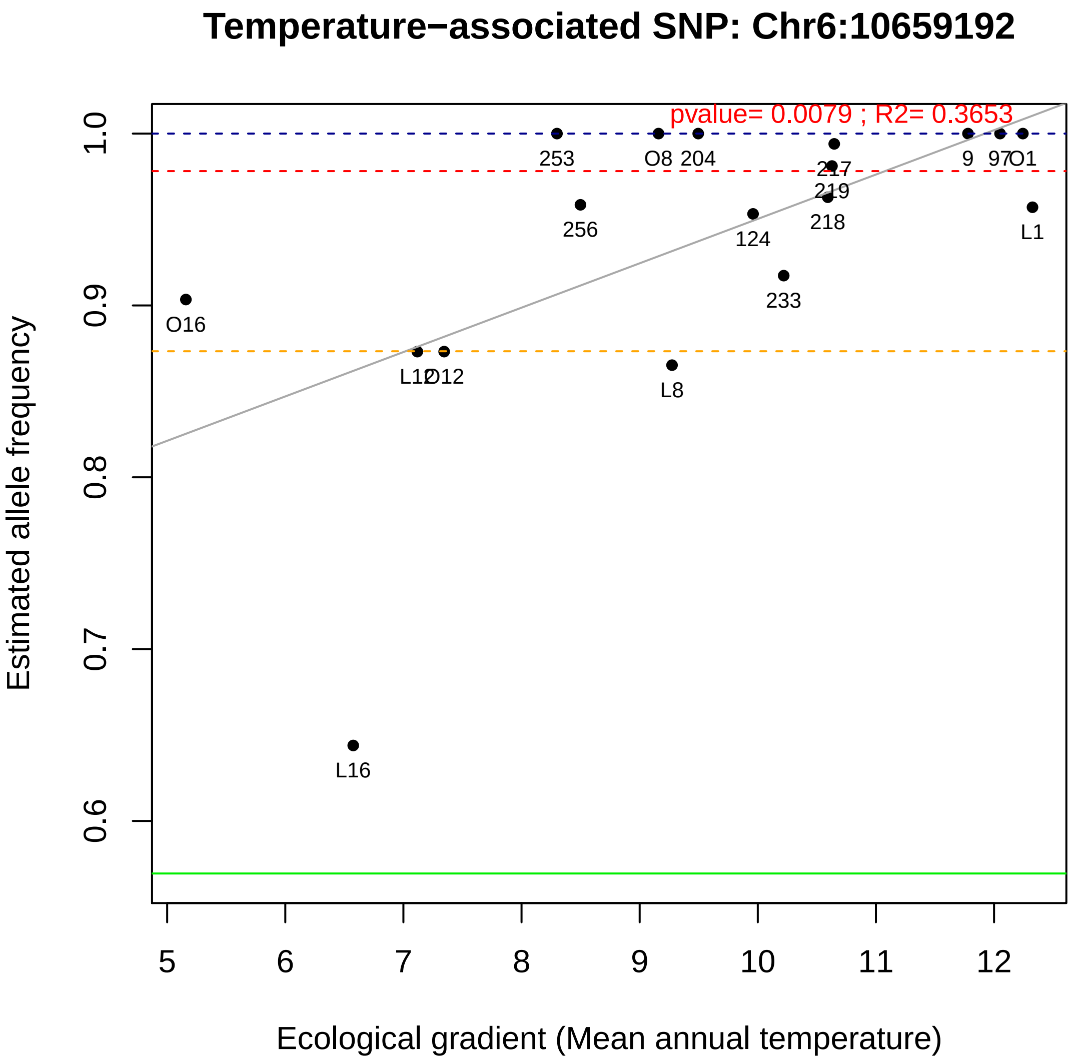

Temperature−associated SNP: Chr6:10659192
pvalue= 0.0079 ; R2= 0.3653
Estimated allele frequency
Ecological gradient (Mean annual temperature)
1.0
0.9
0.8
0.7
0.6
5
6
7
8
9
10
11
12
253
O8
204
217
219
218
9
97
O1
256
124
L1
233
O16
L12
O12
L8
L16

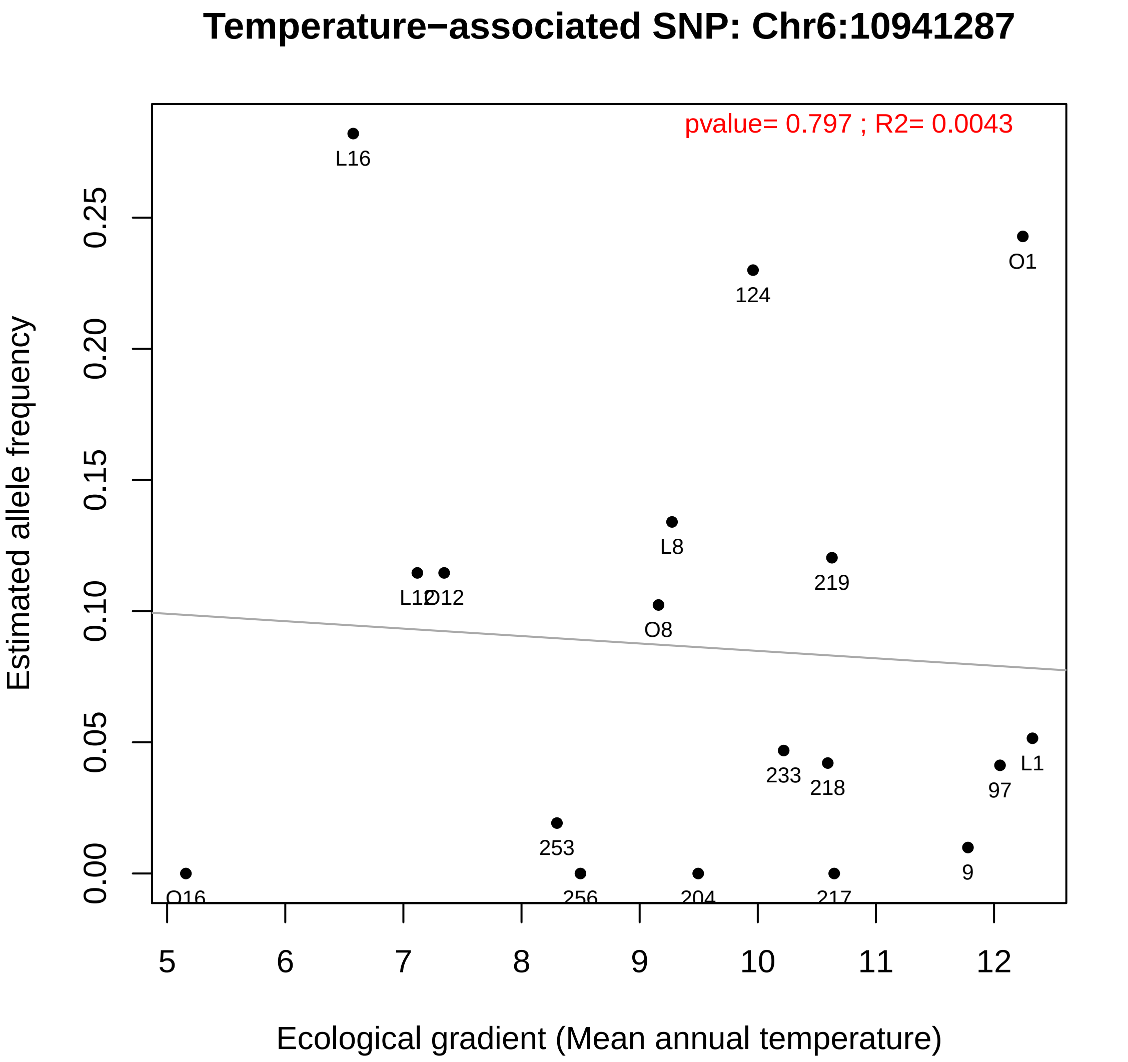
Temperature−associated SNP: Chr6:10941287
pvalue= 0.797 ; R2= 0.0043
Estimated allele frequency
0.00
0.05
0.10
0.15
0.20
0.25
5
6
7
8
9
10
11
12
Ecological gradient (Mean annual temperature)
L16
O1
124
L8
219
L12
O12
O8
233
218
L1
97
253
9
O16
256
204
217

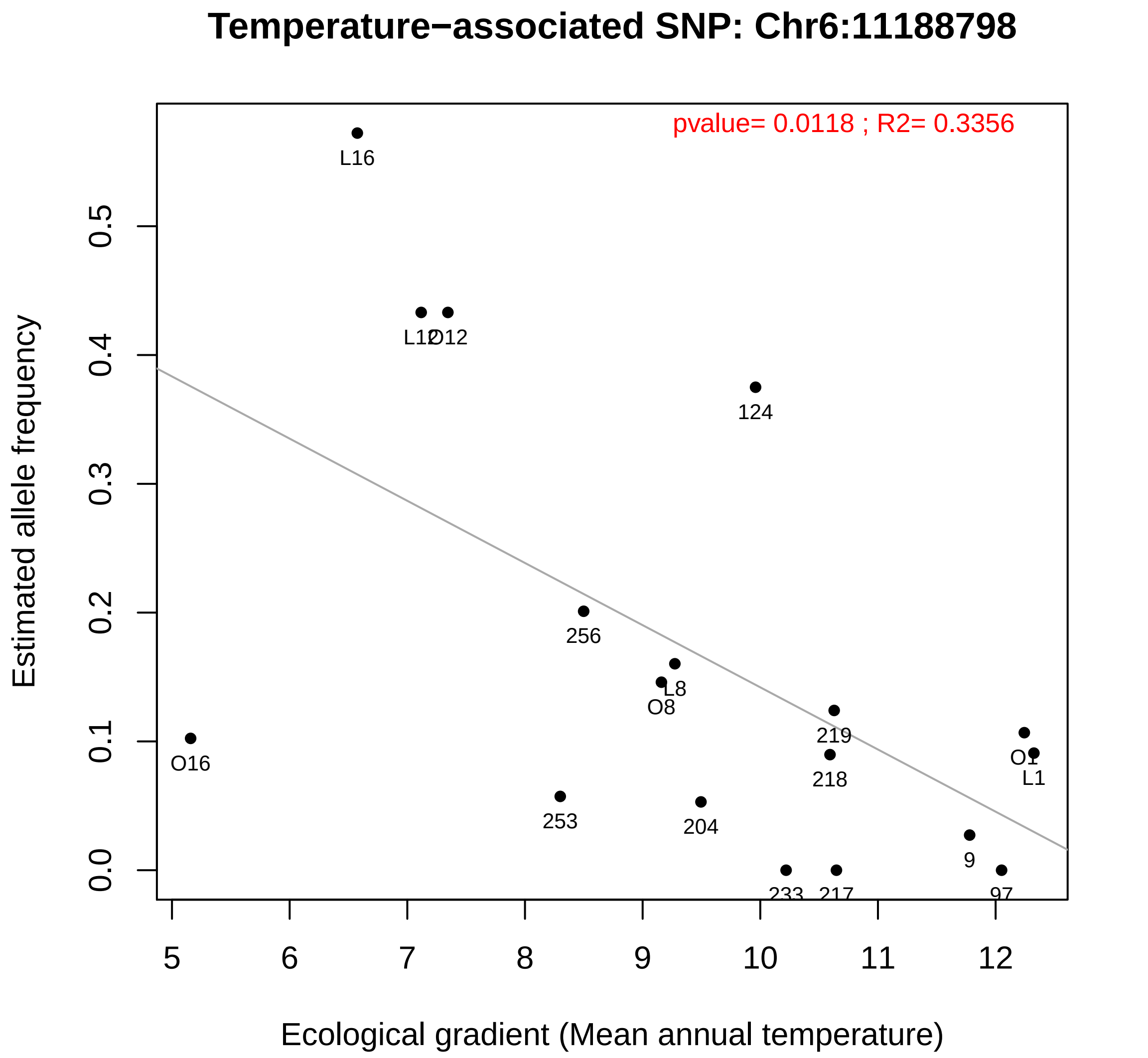
Temperature−associated SNP: Chr6:11188798
pvalue= 0.0118 ; R2= 0.3356
Estimated allele frequency
0.0
0.1
0.2
0.3
0.4
0.5
5
6
7
8
9
10
11
12
Ecological gradient (Mean annual temperature)
L16
L12
O12
124
256
L8
O8
219
218
O16
253
204
O1
L1
9
233
217
97

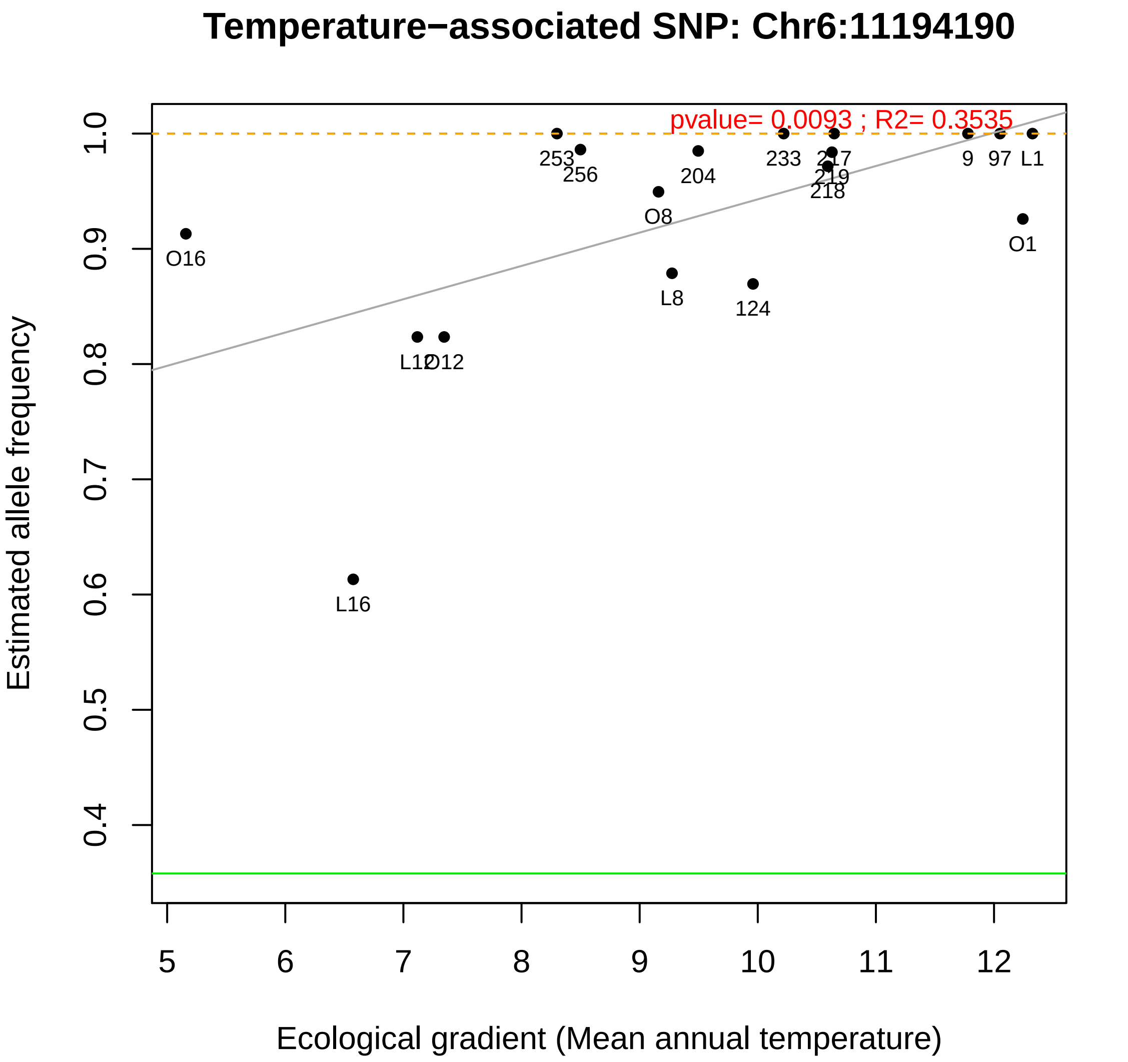

Temperature−associated SNP: Chr6:11194190
pvalue= 0.0093 ; R2= 0.3535
Estimated allele frequency
Ecological gradient (Mean annual temperature)
0.4
0.5
0.6
0.7
0.8
0.9
1.0
5
6
7
8
9
10
11
12
O16
L16
L12
O12
253
256
O8
L8
204
124
233
217
219
218
9
97
L1
O1

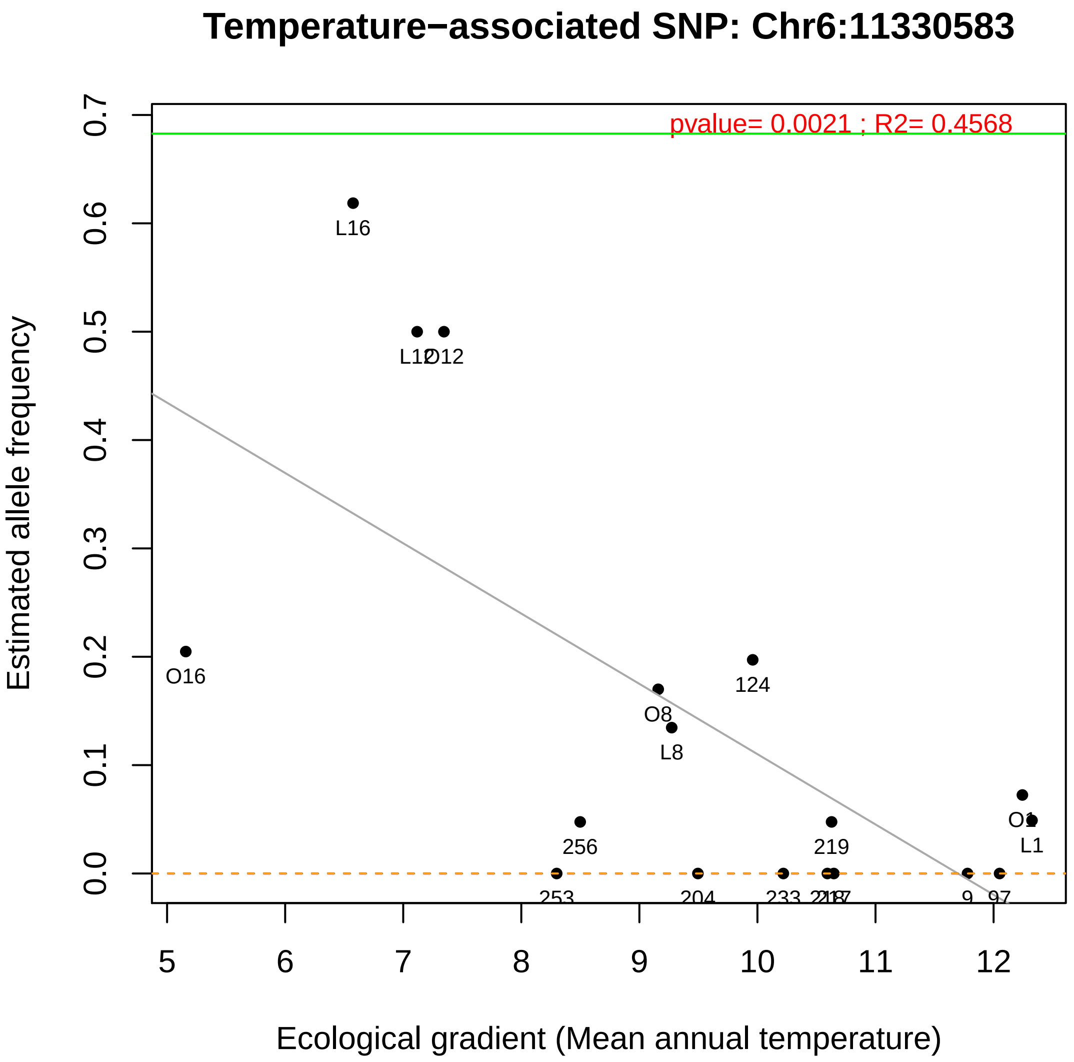

Temperature−associated SNP: Chr6:11330583
pvalue= 0.0021 ; R2= 0.4568
Estimated allele frequency
0.0
0.1
0.2
0.3
0.4
0.5
0.6
0.7
5
6
7
8
9
10
11
12
Ecological gradient (Mean annual temperature)
L16
L12
O12
O16
124
O8
L8
256
219
L1

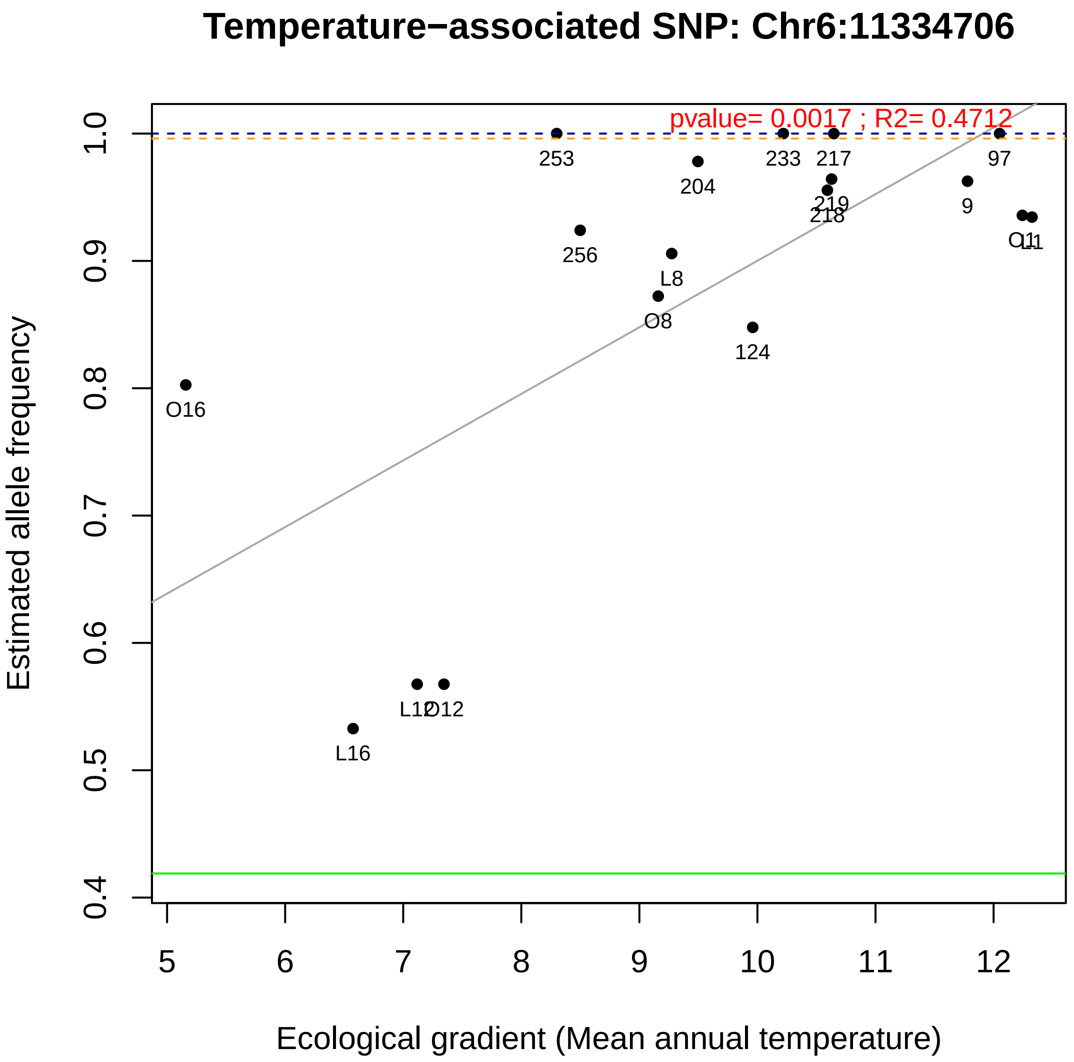

Temperature−associated SNP: Chr6:11334706
pvalue= 0.0017 ; R2= 0.4712
Estimated allele frequency
Ecological gradient (Mean annual temperature)
0.4
0.5
0.6
0.7
0.8
0.9
1.0
5
6
7
8
9
10
11
12
253
233
217
97
204
219
218
9
256
L8
O8
124
O16
L12
O12
L16
O11
L11

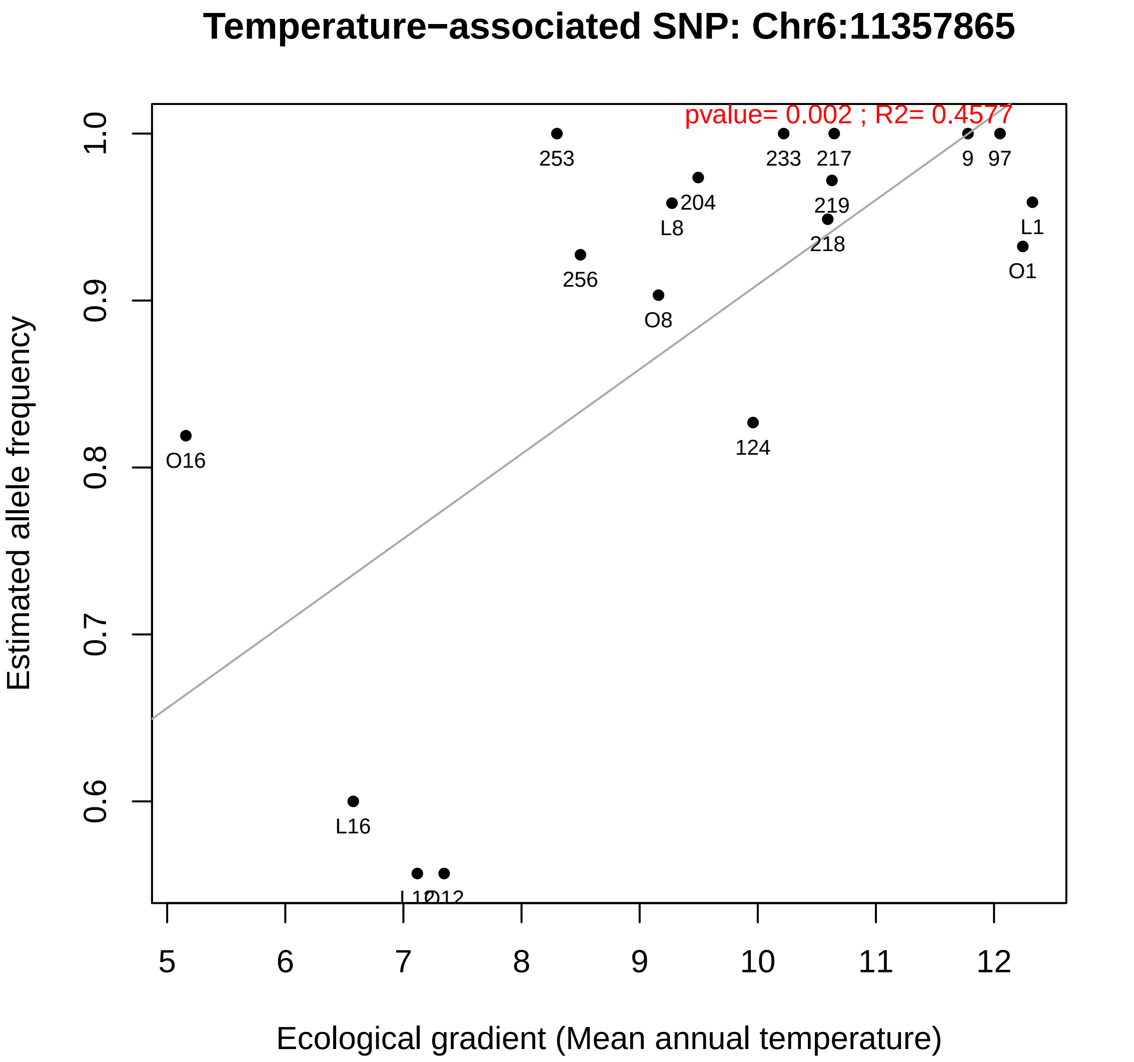
Temperature−associated SNP: Chr6:11357865
pvalue= 0.002 ; R2= 0.4577
Estimated allele frequency
0.6
0.7
0.8
0.9
1.0
5
6
7
8
9
10
11
12
Ecological gradient (Mean annual temperature)
253
233
217
9
97
204
L8
219
218
L1
O1
256
O8
124
O16
L16
O12

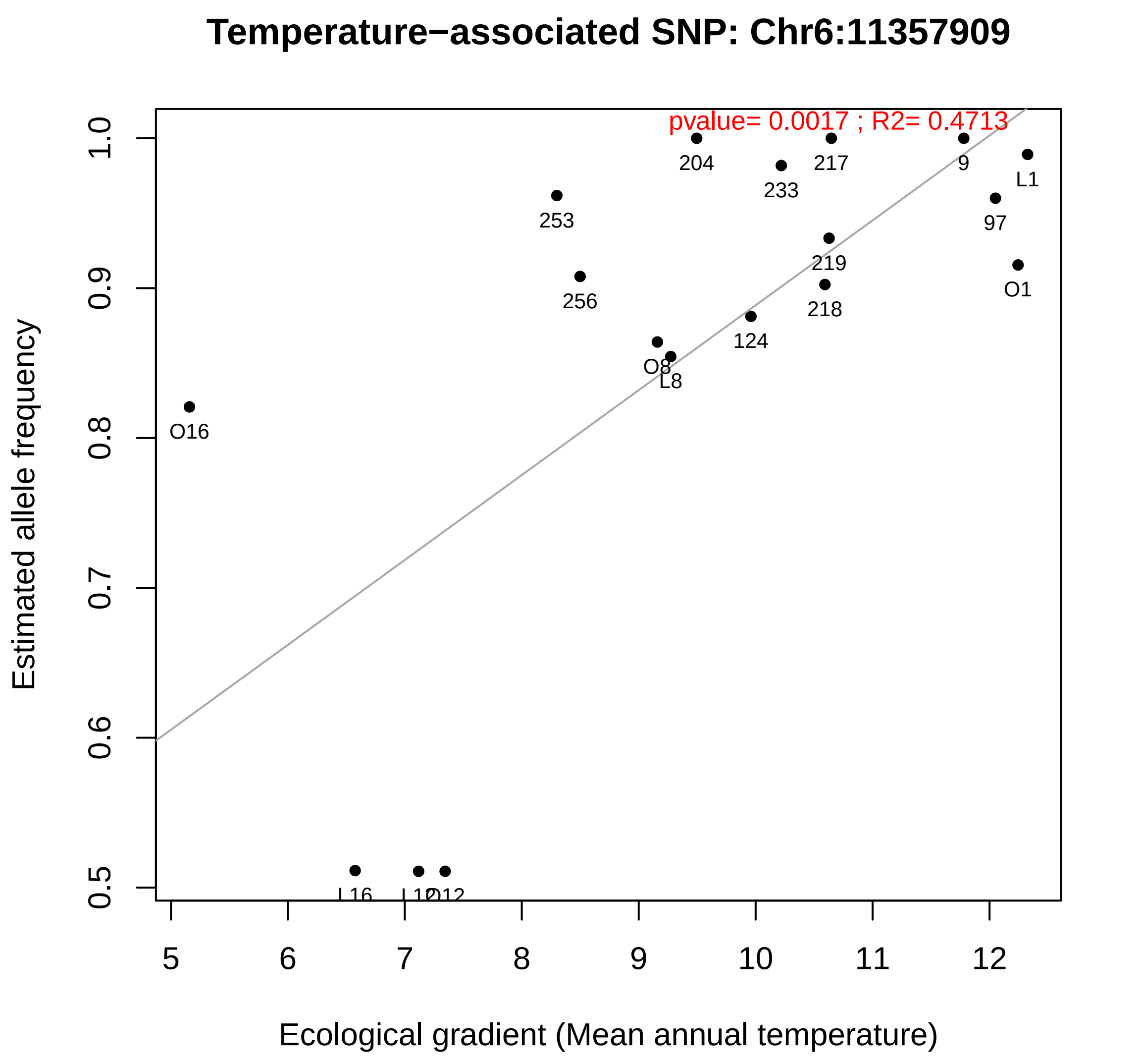

Temperature−associated SNP: Chr6:11357909
pvalue= 0.0017 ; R2= 0.4713
Estimated allele frequency
Ecological gradient (Mean annual temperature)
0.5
0.6
0.7
0.8
0.9
1.0
5
6
7
8
9
10
11
12
204
217
9
L1
233
253
97
219
O1
256
218
124
O8
L8
O16
L16

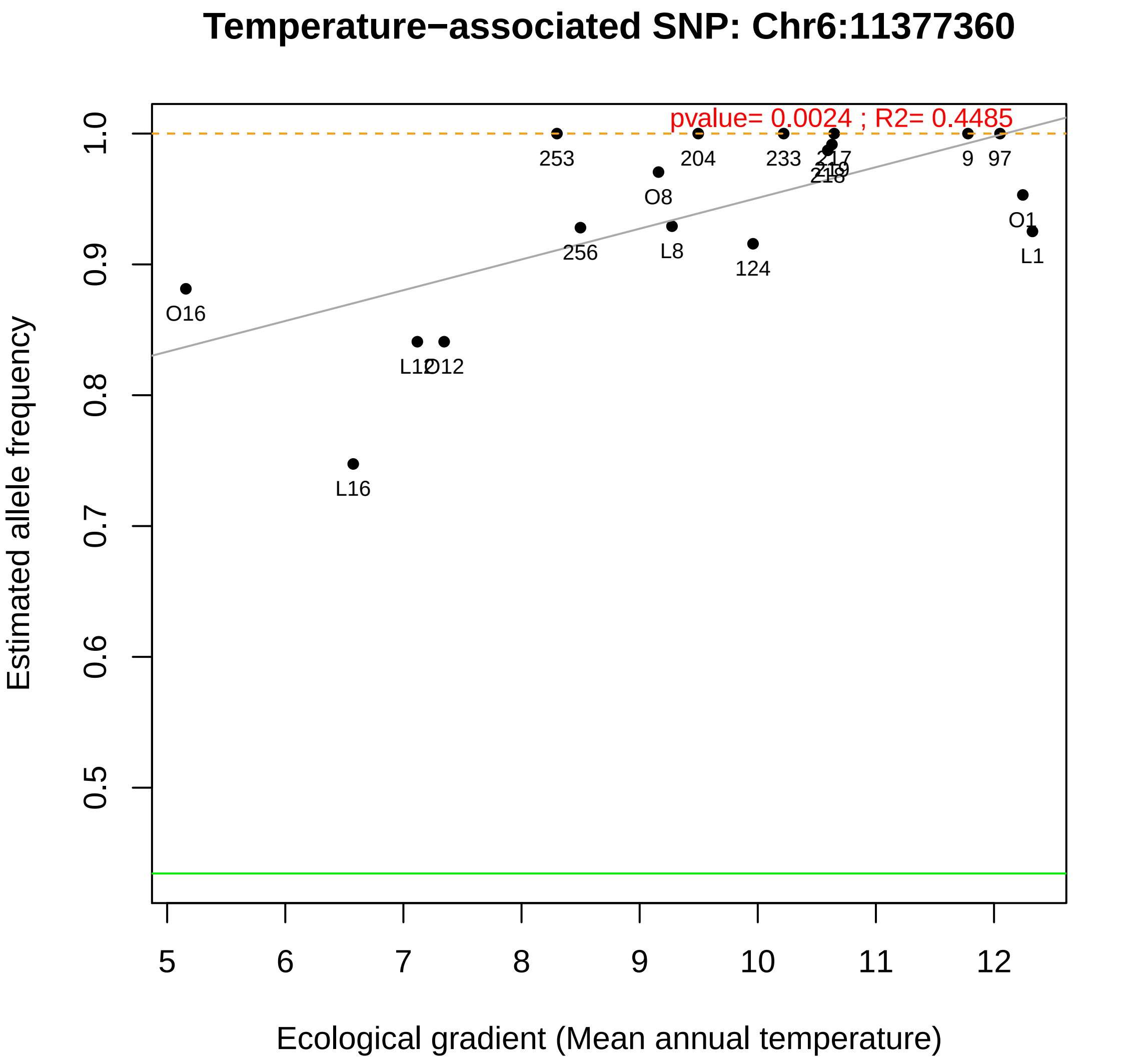

Temperature−associated SNP: Chr6:11377360
pvalue= 0.0024 ; R2= 0.4485
Estimated allele frequency
Ecological gradient (Mean annual temperature)
1.0
0.9
0.8
0.7
0.6
0.5
5
6
7
8
9
10
11
12
253
204
233
217
219
218
9
97
O8
O1
256
L8
L1
124
O16
L12
O12
L16

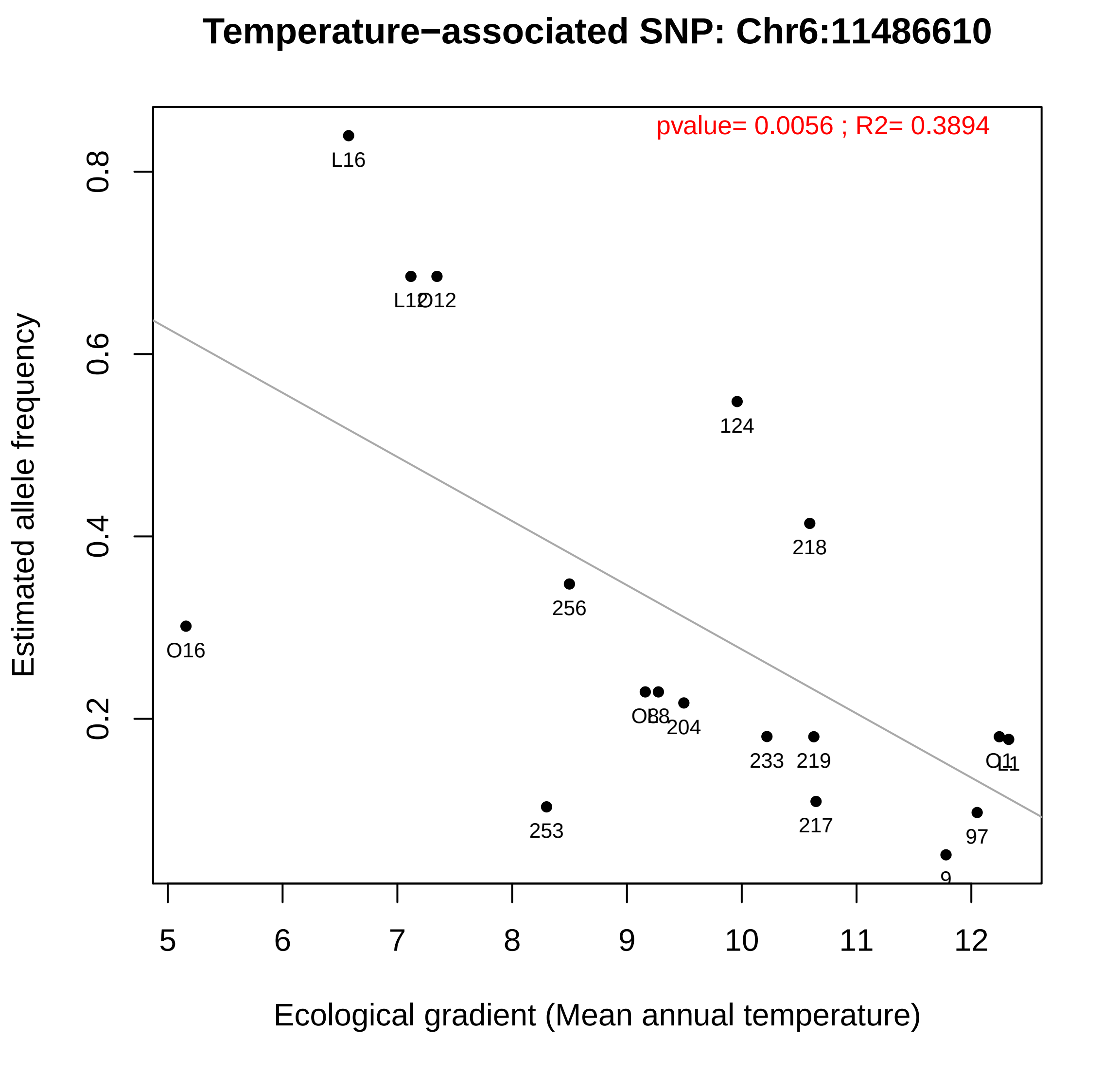
Temperature−associated SNP: Chr6:11486610
pvalue= 0.0056 ; R2= 0.3894
Estimated allele frequency
0.2
0.4
0.6
0.8
L16
L12
O12
124
218
256
O16
O8
38
204
233
219
O1
L1
253
217
97
9
5
6
7
8
9
10
11
12
Ecological gradient (Mean annual temperature)

**Temperature−associated SNP: Chr6:11486612**

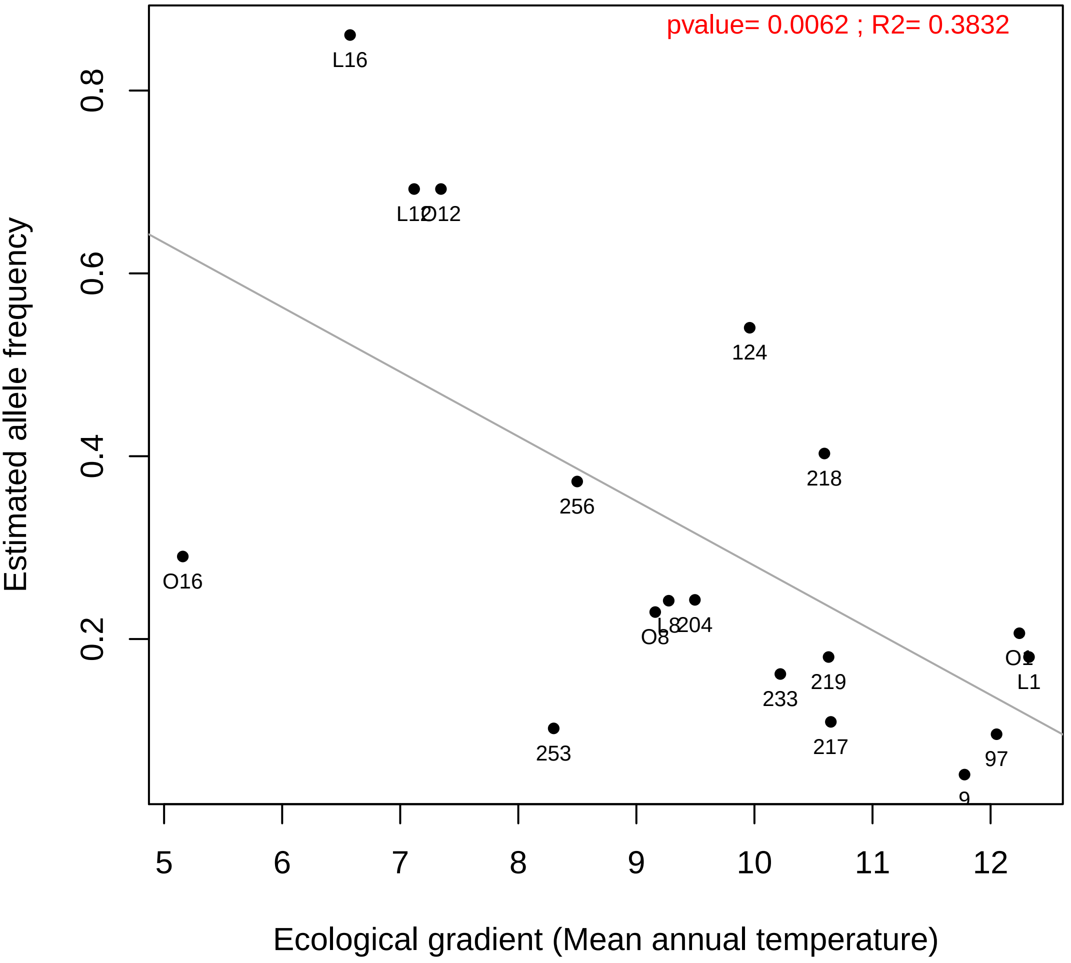

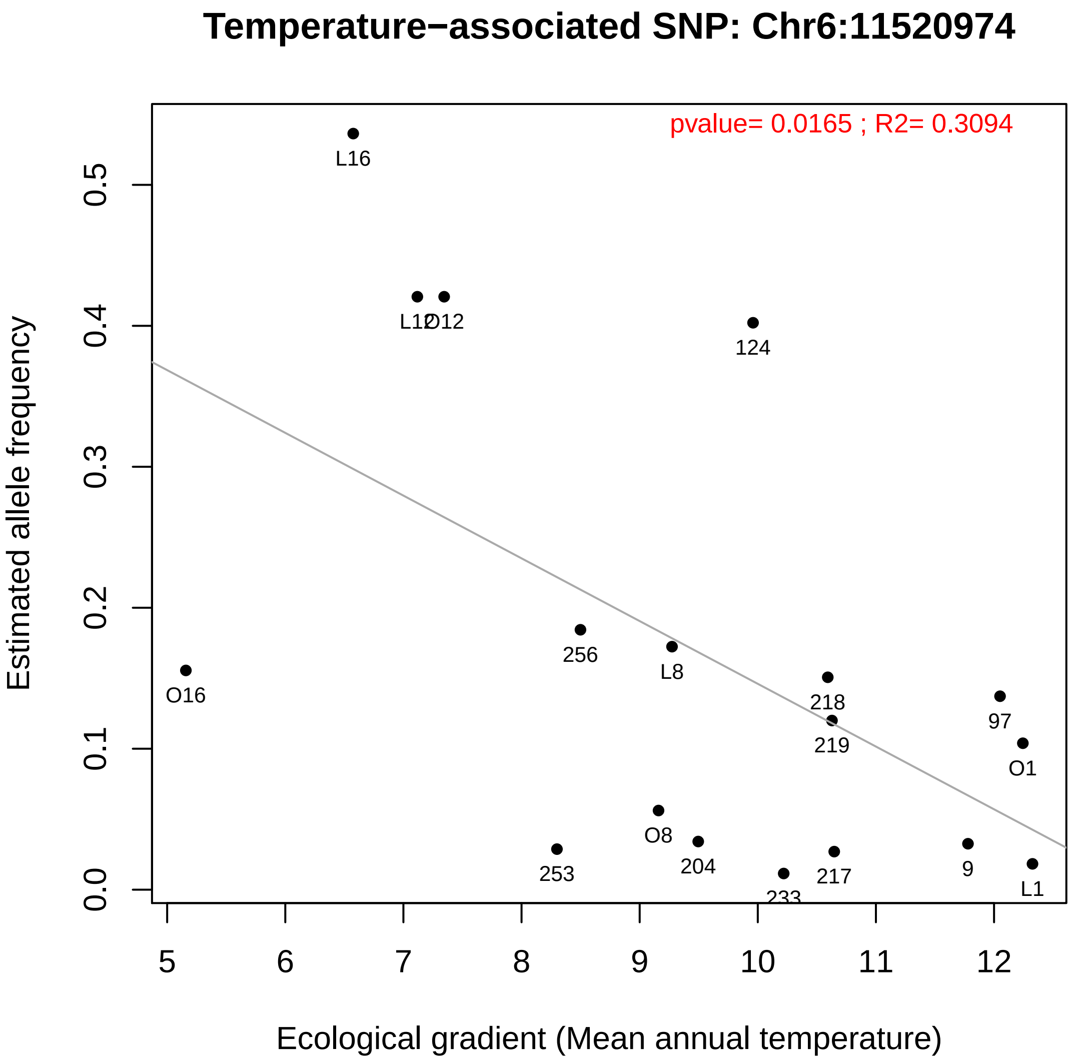

Temperature−associated SNP: Chr6:11520974
pvalue= 0.0165 ; R2= 0.3094
L16
L12
O12
124
256
L8
O16
218
97
219
O1
O8
204
253
233
217
9
L1
0.0
0.1
0.2
0.3
0.4
0.5
5
6
7
8
9
10
11
12
Estimated allele frequency
Ecological gradient (Mean annual temperature)

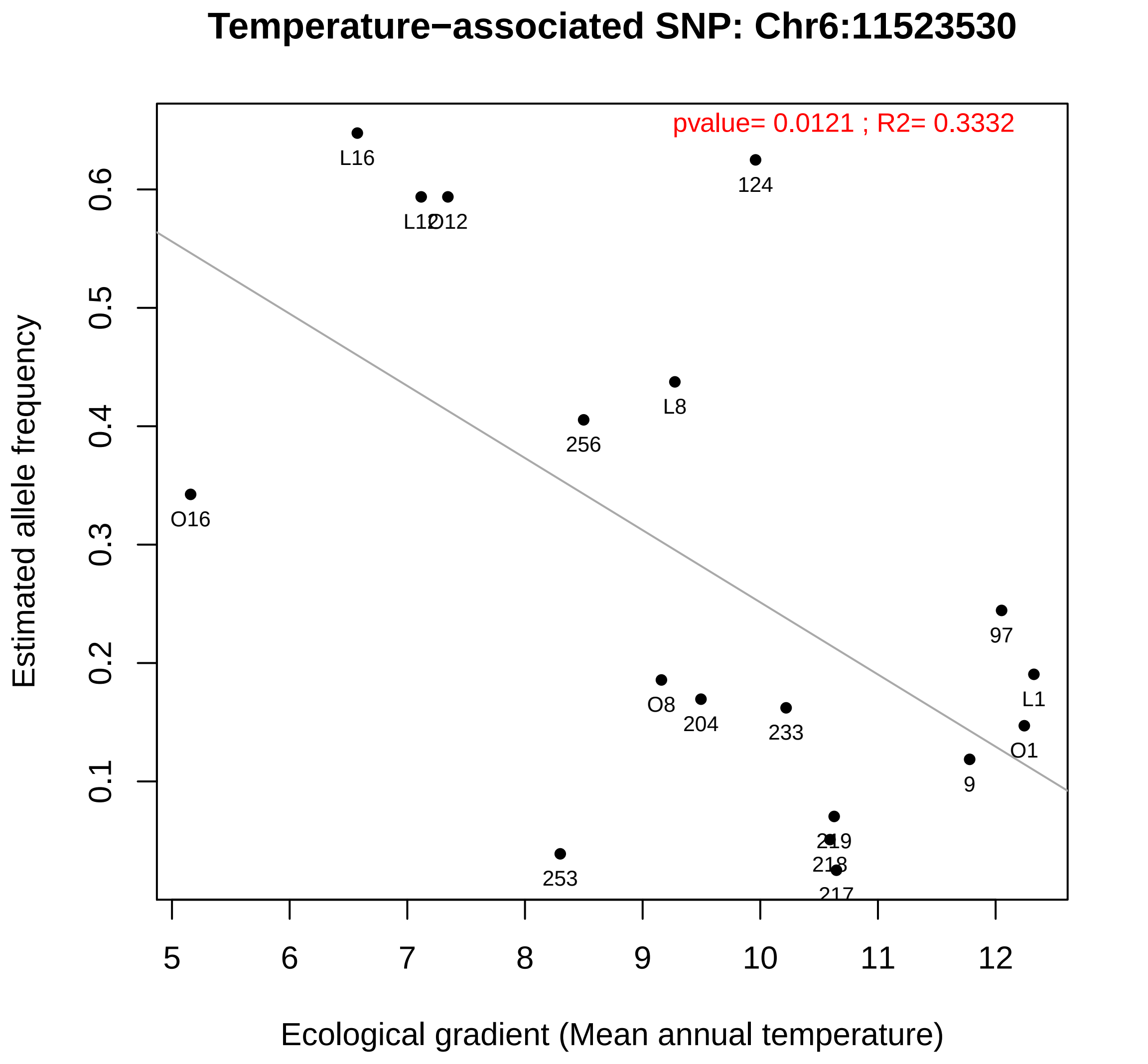

Temperature−associated SNP: Chr6:11523530
pvalue= 0.0121 ; R2= 0.3332
Estimated allele frequency
0.1
0.2
0.3
0.4
0.5
0.6
Ecological gradient (Mean annual temperature)
5
6
7
8
9
10
11
12
L16
124
L12
O12
L8
256
O16
97
L1
O8
204
233
O1
9
219
218
217
253

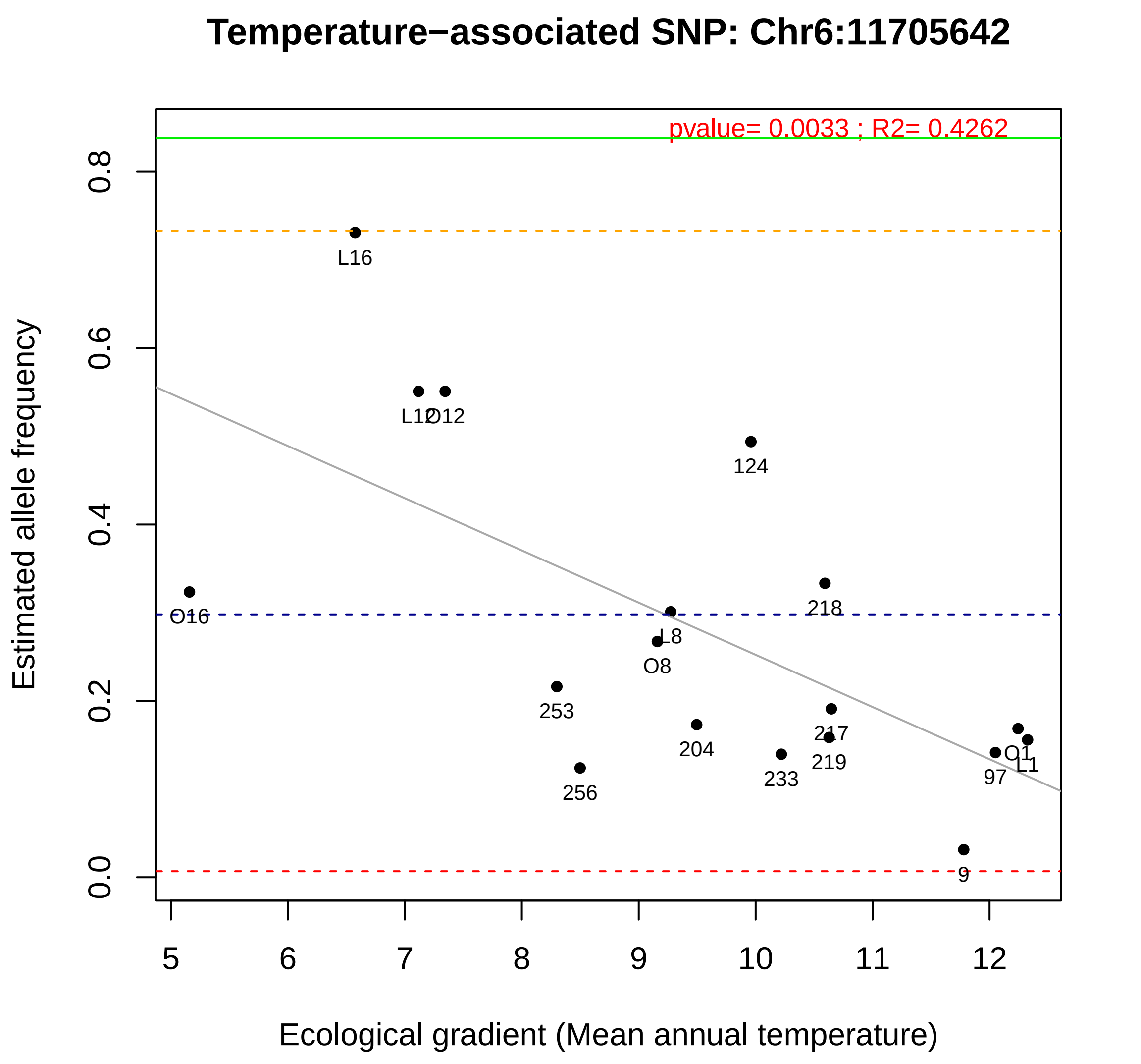

Temperature−associated SNP: Chr6:11705642
pvalue= 0.0033 ; R2= 0.4262
Estimated allele frequency
0.0
0.2
0.4
0.6
0.8
5
6
7
8
9
10
11
12
Ecological gradient (Mean annual temperature)
L16
L12
O12
124
O16
218
L8
O8
253
204
217
219
233
256
O1
L1
97
9

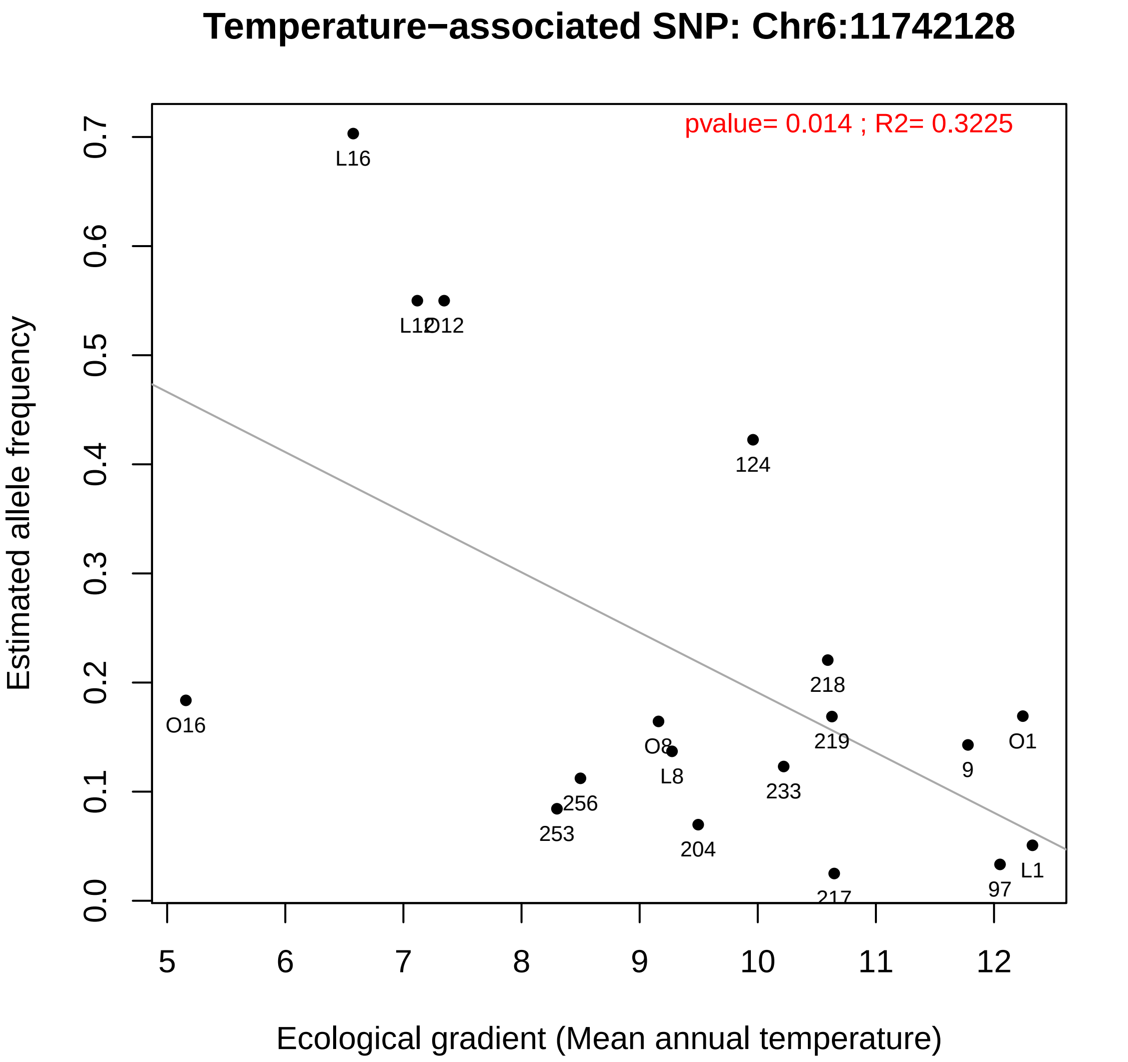

Temperature−associated SNP: Chr6:11742128
pvalue= 0.014 ; R2= 0.3225
Estimated allele frequency
0.0
0.1
0.2
0.3
0.4
0.5
0.6
0.7
5
6
7
8
9
10
11
12
Ecological gradient (Mean annual temperature)
L16
L12
O12
124
218
219
O1
O16
O8
L8
9
233
256
253
204
217
97
L1

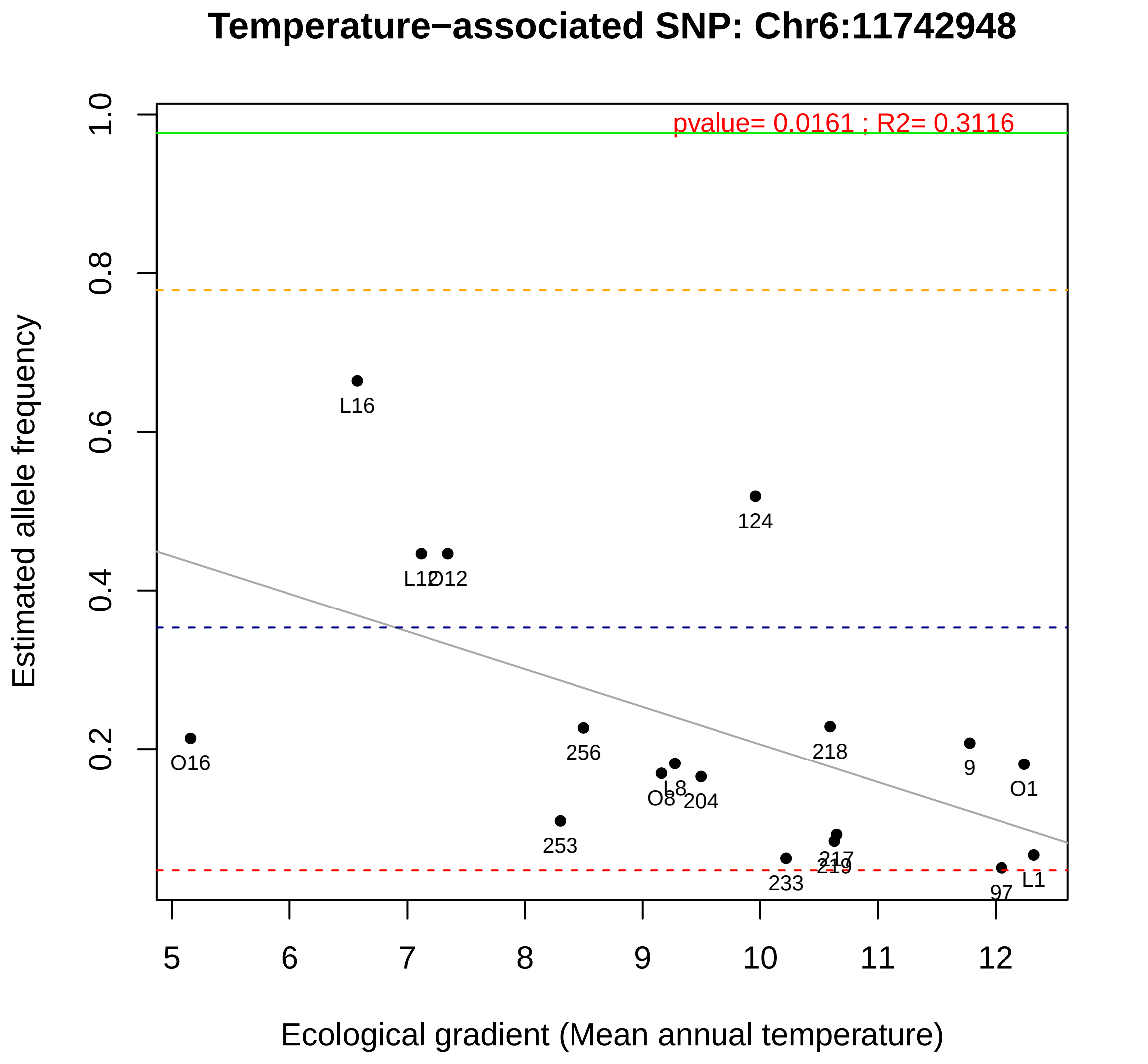

Temperature−associated SNP: Chr6:11742948
pvalue= 0.0161 ; R2= 0.3116
Estimated allele frequency
Ecological gradient (Mean annual temperature)
0.2
0.4
0.6
0.8
1.0
5
6
7
8
9
10
11
12
L16
124
L12
O12
O16
256
218
9
O1
L8
O8
204
253
217
219
233
97
L1

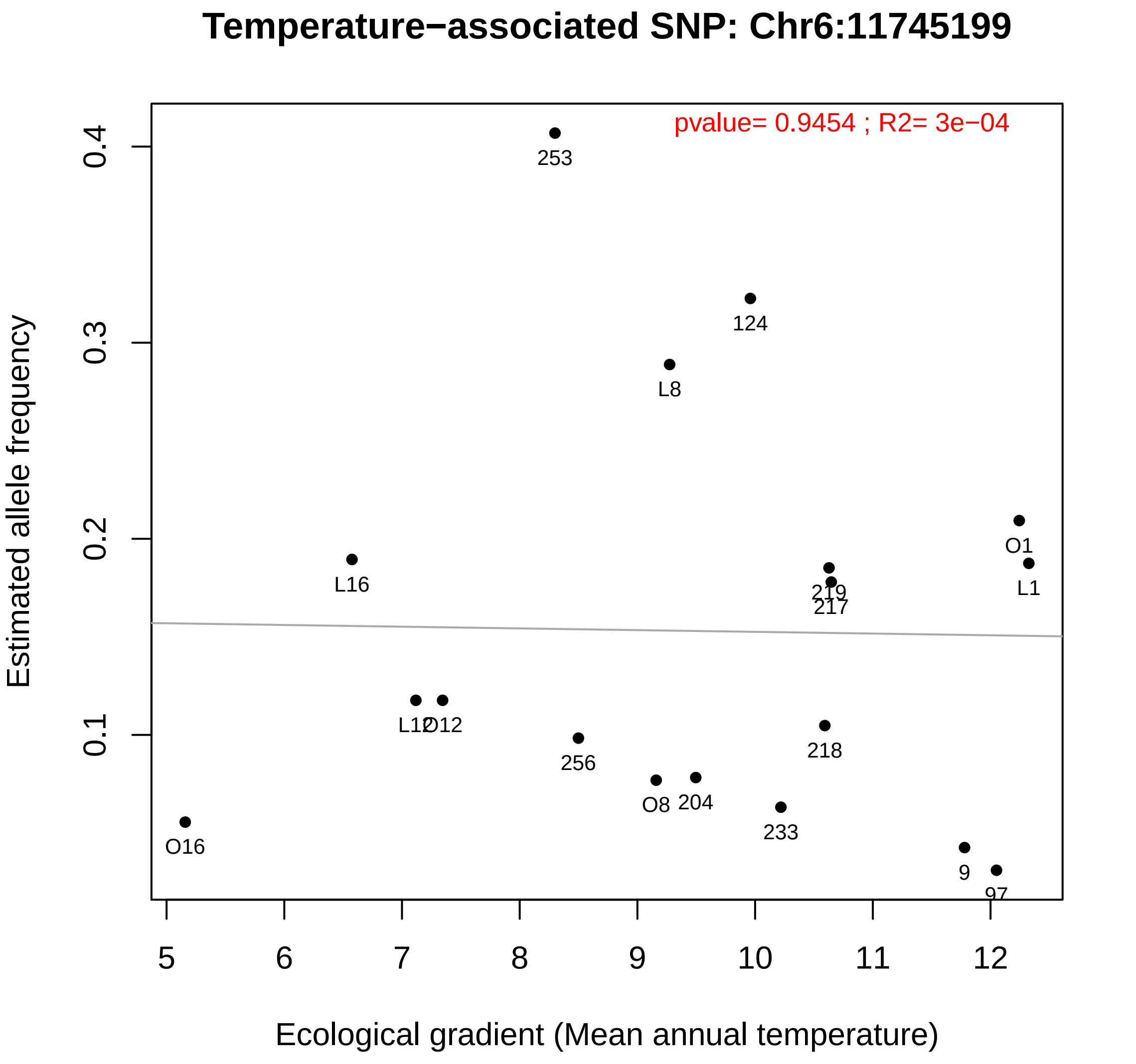
Temperature−associated SNP: Chr6:11745199
pvalue= 0.9454 ; R2= 3e−04
Estimated allele frequency
0.1
0.2
0.3
0.4
Ecological gradient (Mean annual temperature)
5
6
7
8
9
10
11
12
253
124
L8
L16
O1
L1
219
217
L12
O12
256
218
O8
204
233
O16
9
97

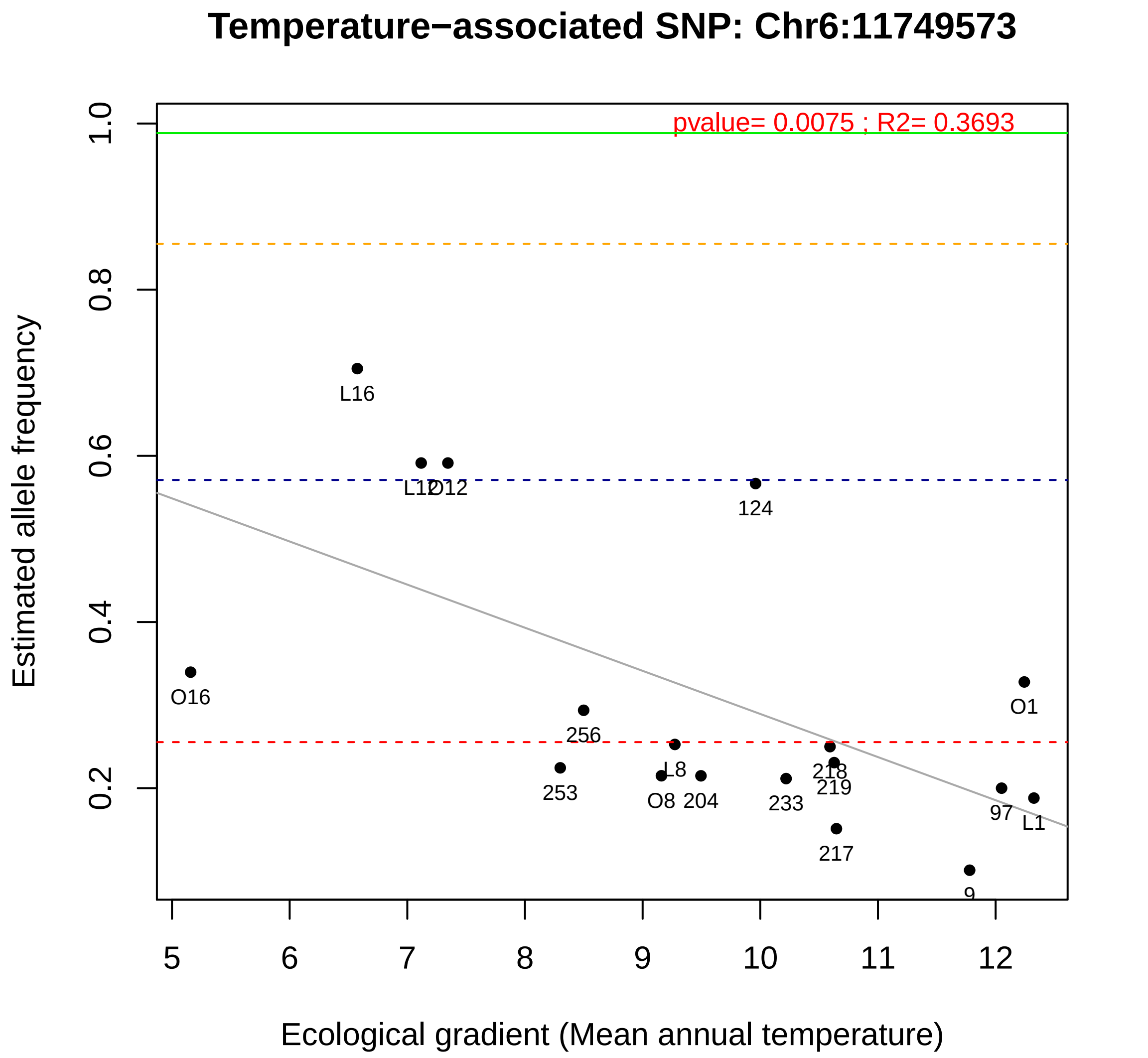
Temperature−associated SNP: Chr6:11749573
pvalue= 0.0075 ; R2= 0.3693
Estimated allele frequency
0.2
0.4
0.6
0.8
1.0
Ecological gradient (Mean annual temperature)
5
6
7
8
9
10
11
12
L16
L12
O12
124
O16
256
253
L8
O8
204
233
218
219
217
O1
97
L1
9

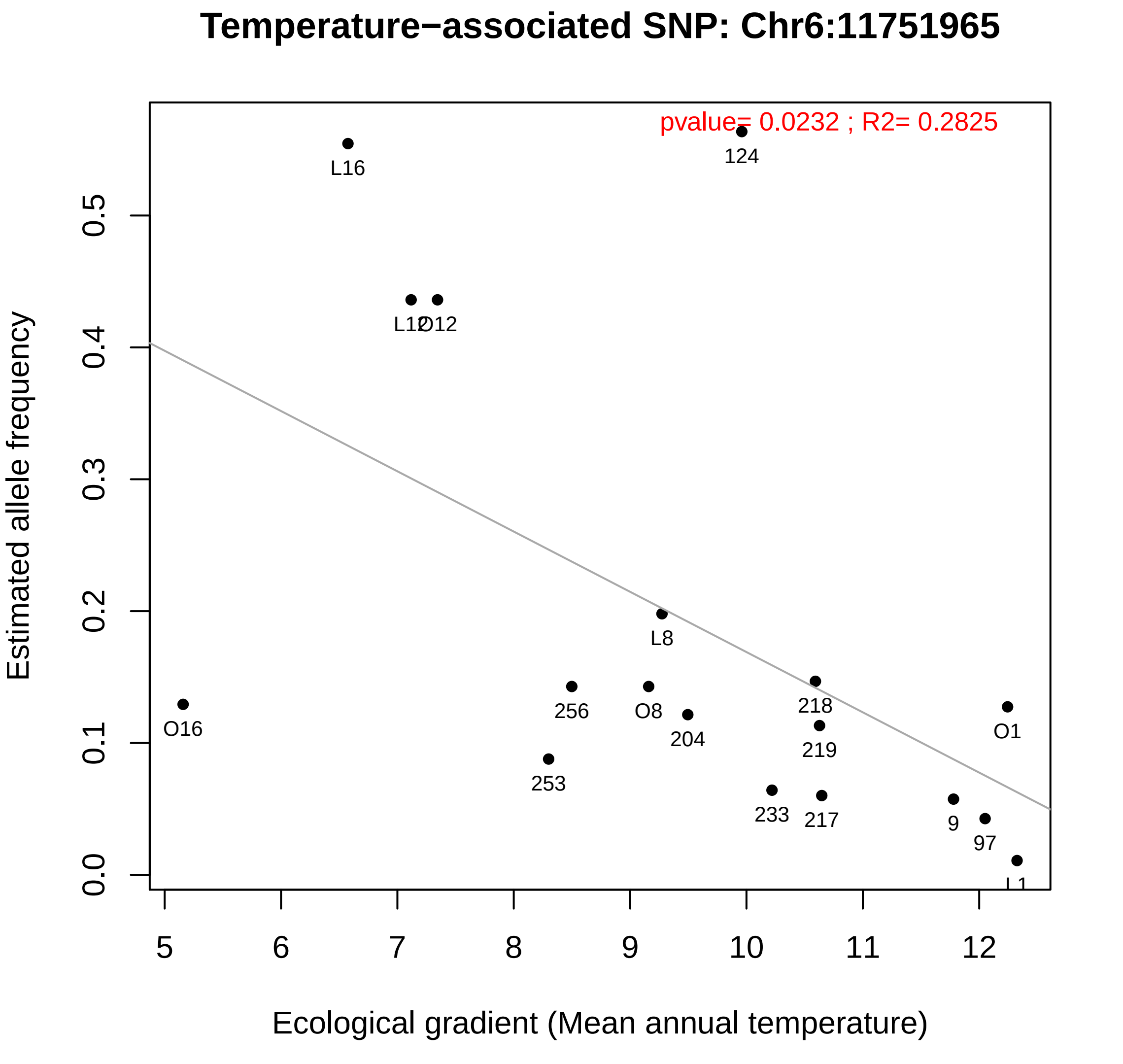

Temperature−associated SNP: Chr6:11751965
pvalue= 0.0232 ; R2= 0.2825
L16
124
L12
O12
L8
O16
256
O8
204
218
219
O1
253
233
217
9
97
L1
0.0
0.1
0.2
0.3
0.4
0.5
5
6
7
8
9
10
11
12
Estimated allele frequency
Ecological gradient (Mean annual temperature)

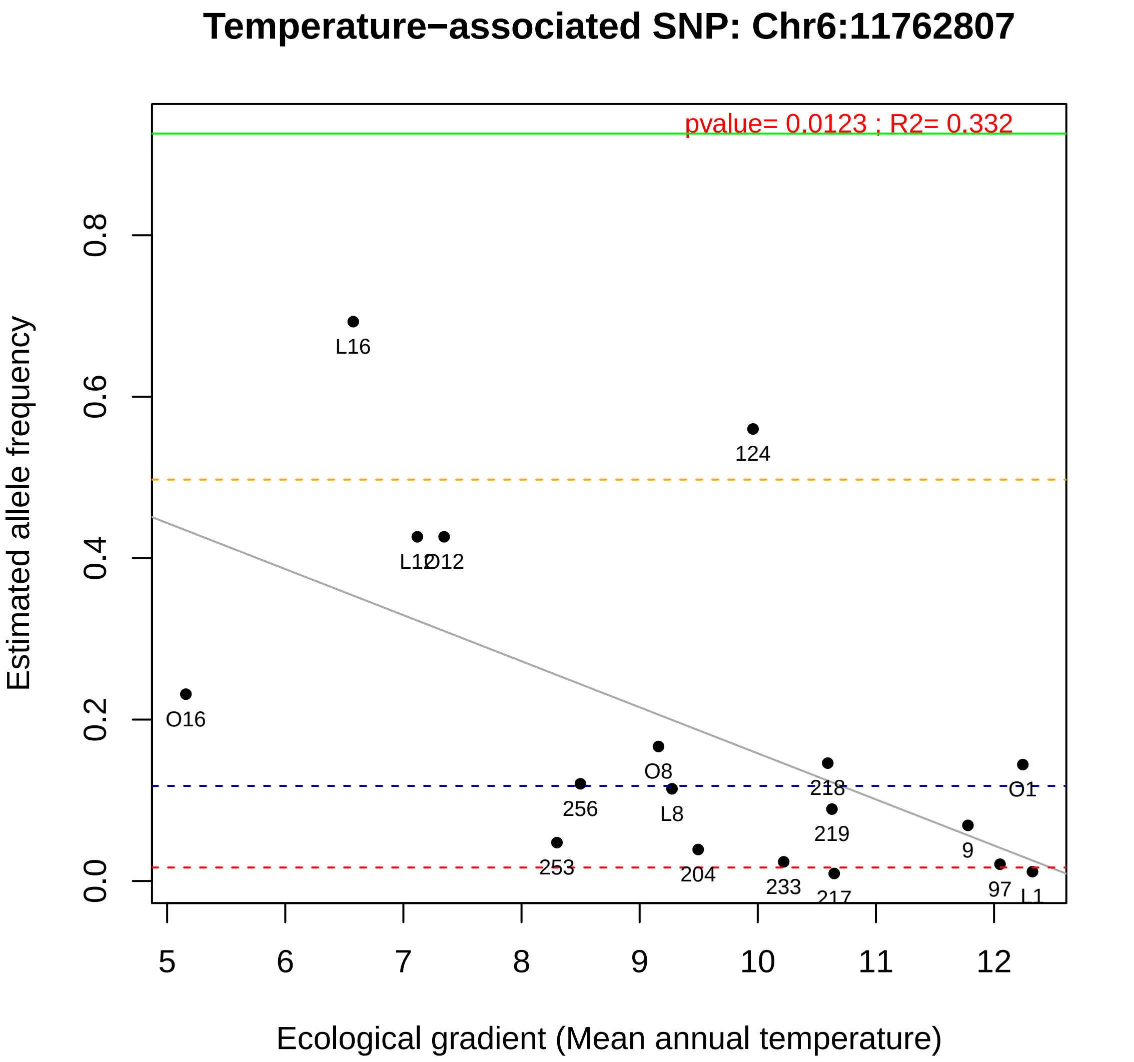
Temperature−associated SNP: Chr6:11762807
pvalue= 0.0123 ; R2= 0.332
Estimated allele frequency
0.0
0.2
0.4
0.6
0.8
Ecological gradient (Mean annual temperature)
5
6
7
8
9
10
11
12
L16
124
L12
O12
O16
O8
L8
256
218
219
O1
9
253
204
233
217
97
L1

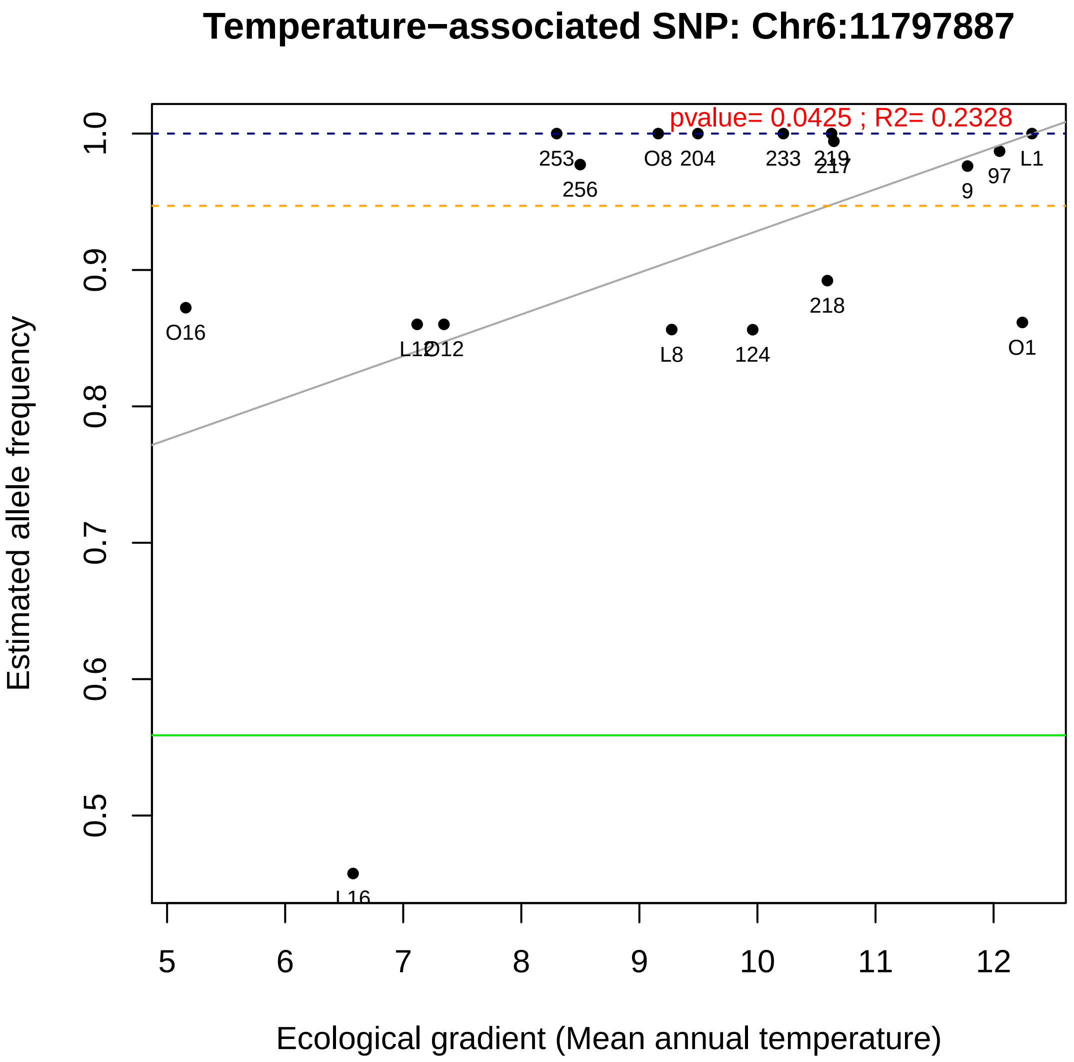

Temperature−associated SNP: Chr6:11797887
pvalue= 0.0425 ; R2= 0.2328
Estimated allele frequency
1.0
0.9
0.8
0.7
0.6
0.5
253
256
O8
204
233
219
217
L1
97
9
218
O16
L12
O12
L8
124
O1
L16
5
6
7
8
9
10
11
12
Ecological gradient (Mean annual temperature)

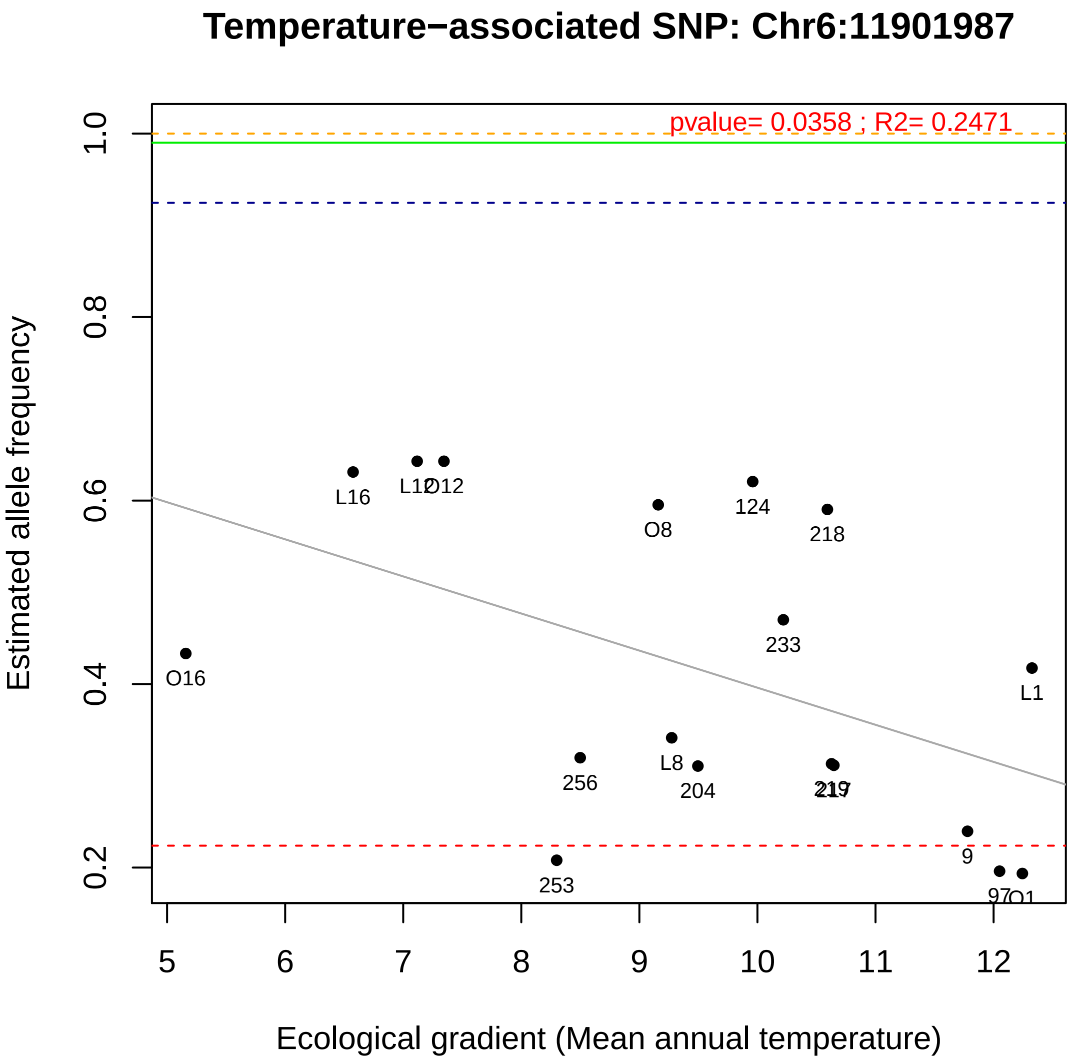

Temperature−associated SNP: Chr6:11901987
pvalue= 0.0358 ; R2= 0.2471
Estimated allele frequency
Ecological gradient (Mean annual temperature)
0.2
0.4
0.6
0.8
1.0
5
6
7
8
9
10
11
12
L16
L12
O12
O8
124
218
233
O16
L1
L8
256
204
253
9
97
O1

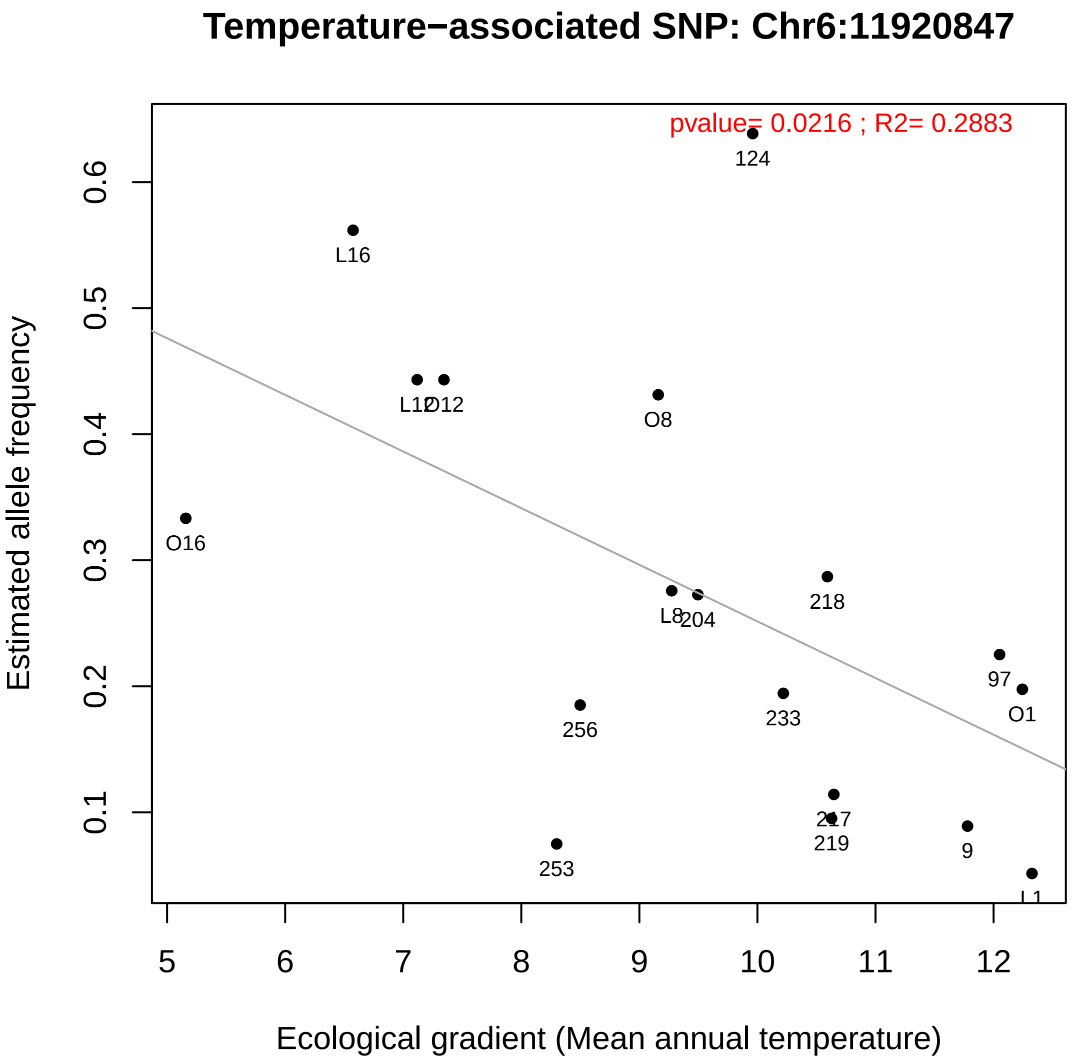
Temperature−associated SNP: Chr6:11920847
pvalue= 0.0216 ; R2= 0.2883
Estimated allele frequency
Ecological gradient (Mean annual temperature)
0.1
0.2
0.3
0.4
0.5
0.6
5
6
7
8
9
10
11
12
124
L16
L12
O12
O8
O16
L8
204
218
97
O1
256
233
217
219
9
253

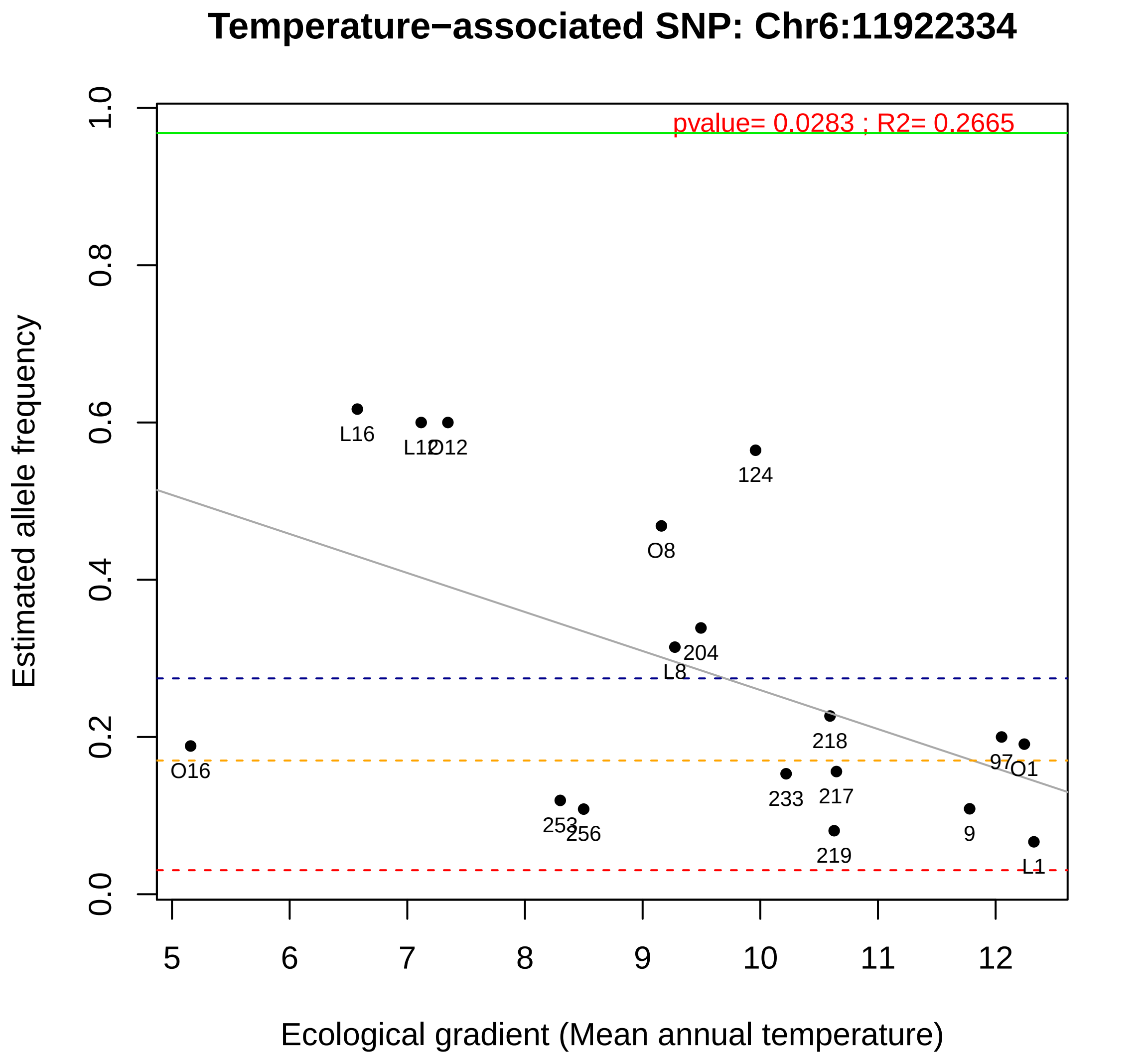
Temperature−associated SNP: Chr6:11922334
pvalue= 0.0283 ; R2= 0.2665
Estimated allele frequency
0.0
0.2
0.4
0.6
0.8
1.0
Ecological gradient (Mean annual temperature)
5
6
7
8
9
10
11
12
L16
L12
O12
124
O8
204
L8
218
O16
97
O1
233
217
253
256
9
219
L1

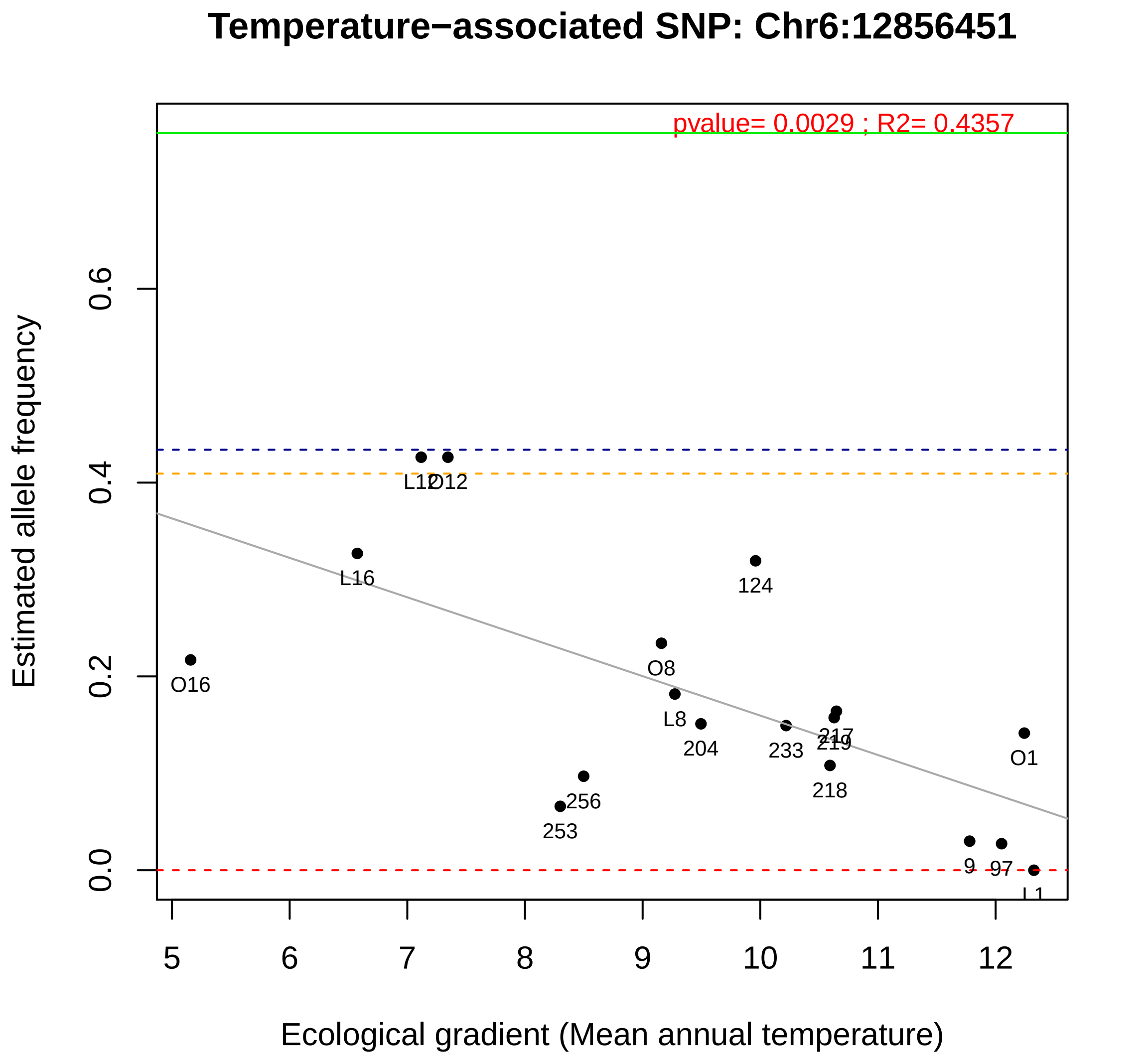
Temperature−associated SNP: Chr6:12856451
pvalue= 0.0029 ; R2= 0.4357
Estimated allele frequency
0.0
0.2
0.4
0.6
5
6
7
8
9
10
11
12
Ecological gradient (Mean annual temperature)
L12
O12
L16
124
O16
O8
L8
204
233
217
219
O1
218
256
253
9
97
L1

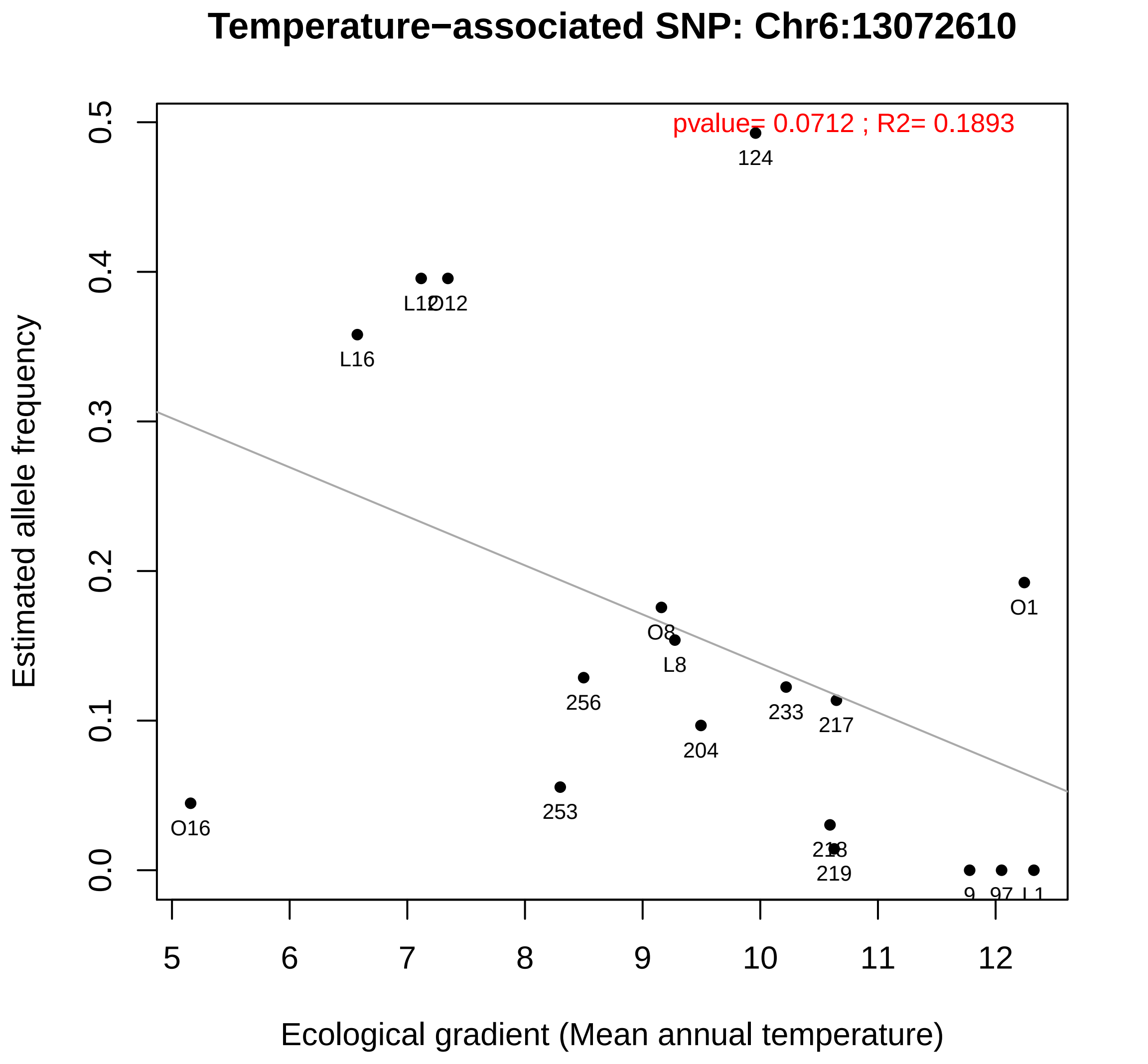
Temperature−associated SNP: Chr6:13072610
pvalue= 0.0712 ; R2= 0.1893
Estimated allele frequency
Ecological gradient (Mean annual temperature)
0.0
0.1
0.2
0.3
0.4
0.5
5
6
7
8
9
10
11
12
124
L12
O12
L16
O1
O8
L8
256
233
217
204
253
O16
218
219

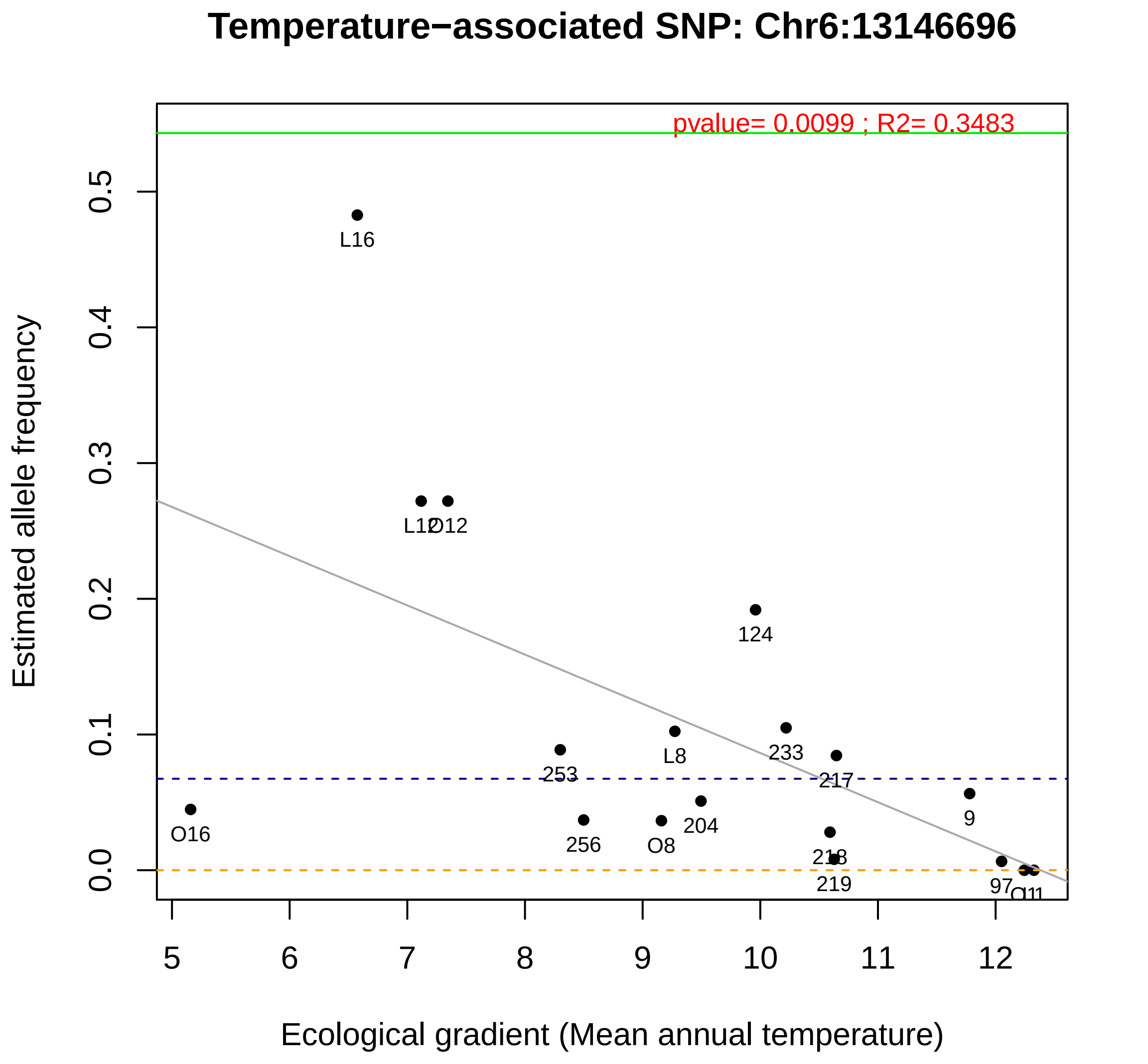
Temperature−associated SNP: Chr6:13146696
pvalue= 0.0099 ; R2= 0.3483
Estimated allele frequency
0.0
0.1
0.2
0.3
0.4
0.5
L16
L12
O12
124
253
L8
233
217
O16
256
O8
204
9
218
219
97
O11
5
6
7
8
9
10
11
12
Ecological gradient (Mean annual temperature)

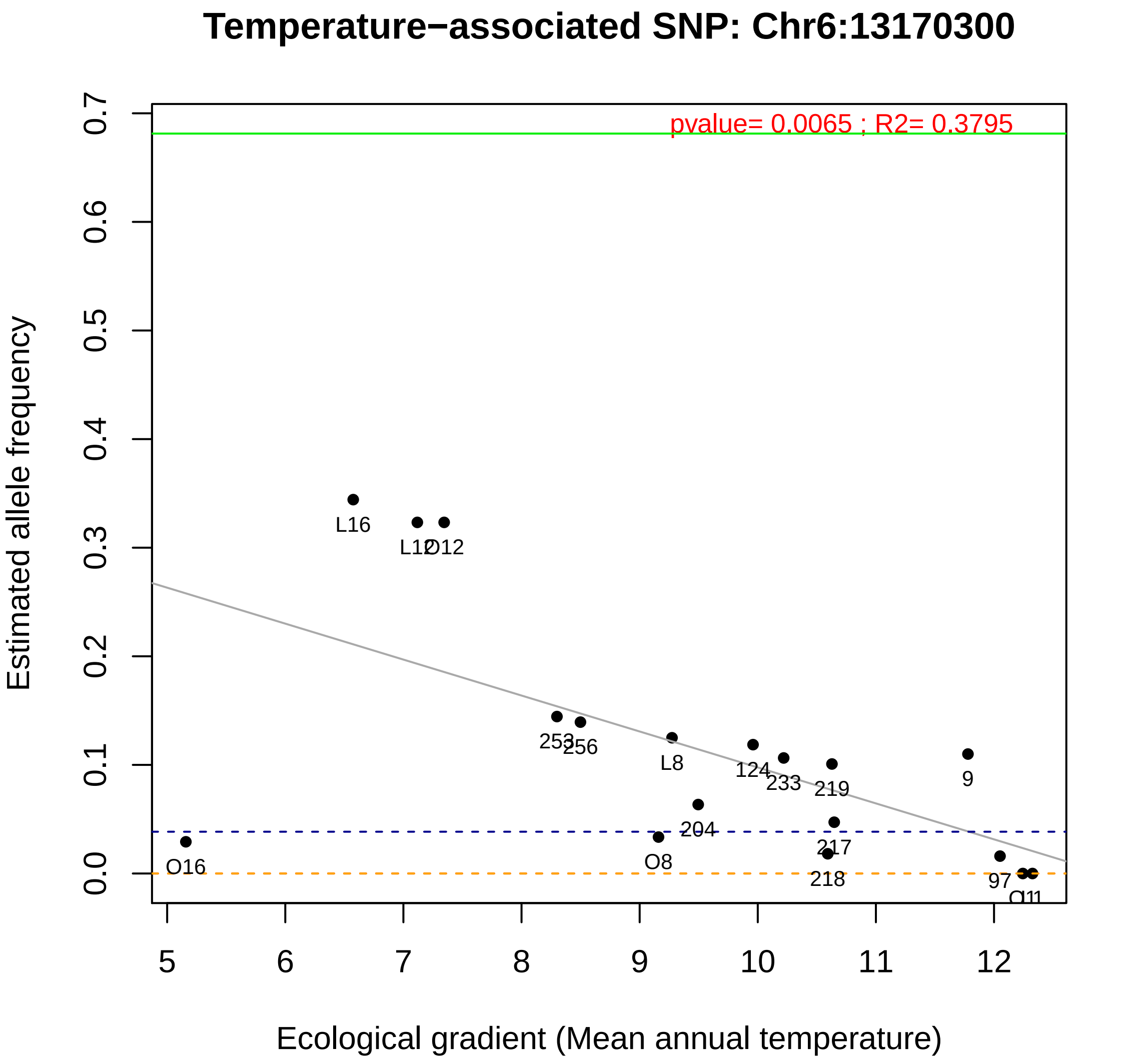

Temperature−associated SNP: Chr6:13170300
pvalue= 0.0065 ; R2= 0.3795
Estimated allele frequency
0.0
0.1
0.2
0.3
0.4
0.5
0.6
0.7
5
6
7
8
9
10
11
12
Ecological gradient (Mean annual temperature)
L16
L12
O12
253
256
L8
124
233
219
9
204
217
218
O16
O8
97
O11

**Temperature−associated SNP: Chr6:13459484**

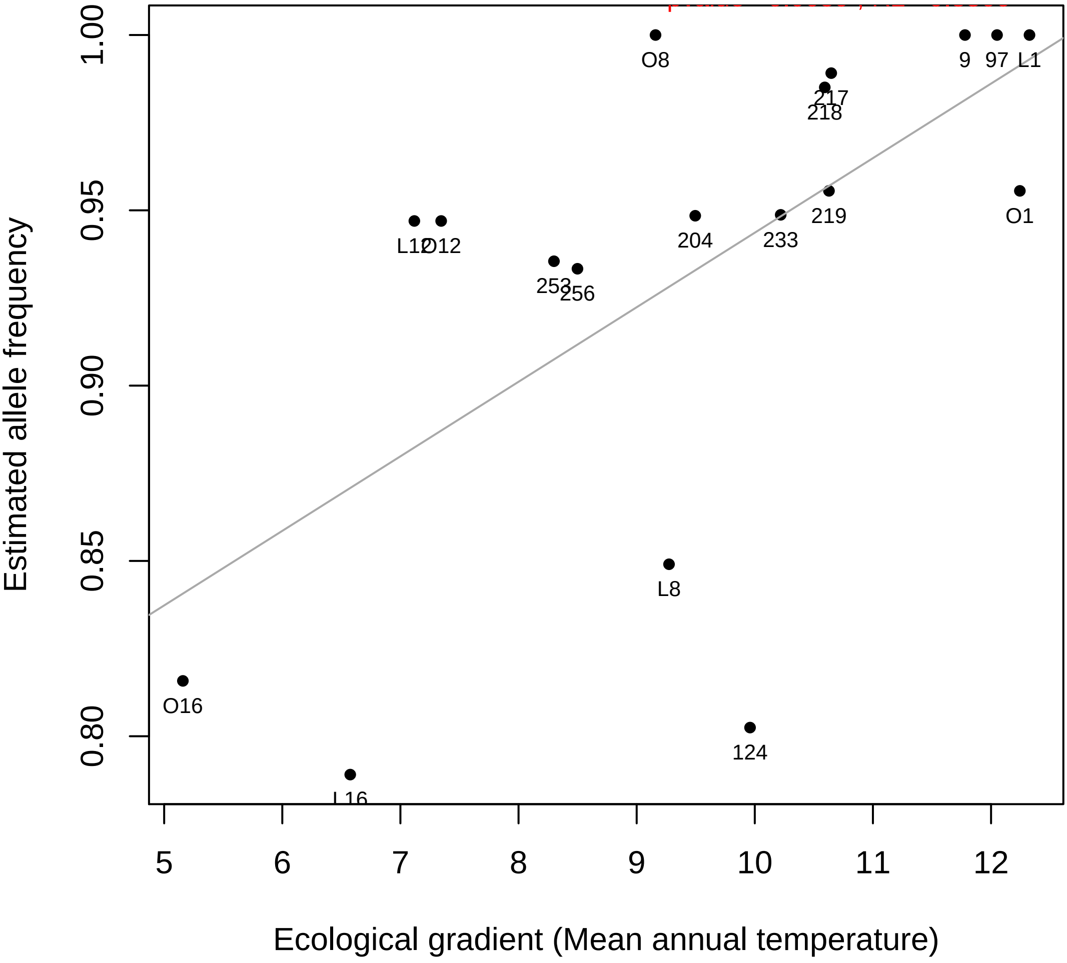

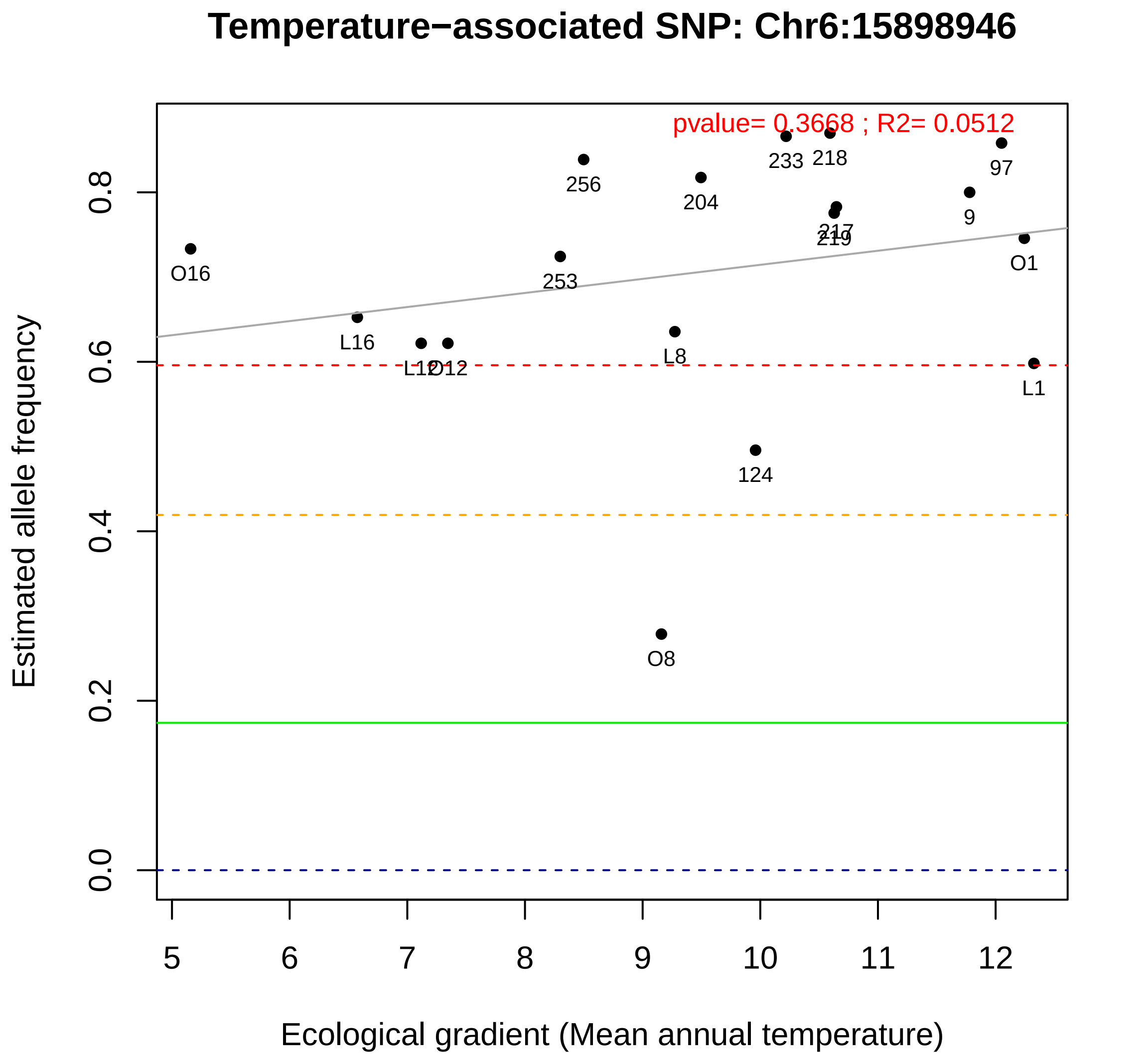

Temperature−associated SNP: Chr6:15898946
pvalue= 0.3668 ; R2= 0.0512
Estimated allele frequency
Ecological gradient (Mean annual temperature)
0.0
0.2
0.4
0.6
0.8
5
6
7
8
9
10
11
12
O16
L16
L12
O12
253
256
204
L8
O8
124
233
218
217
219
9
97
O1
L1

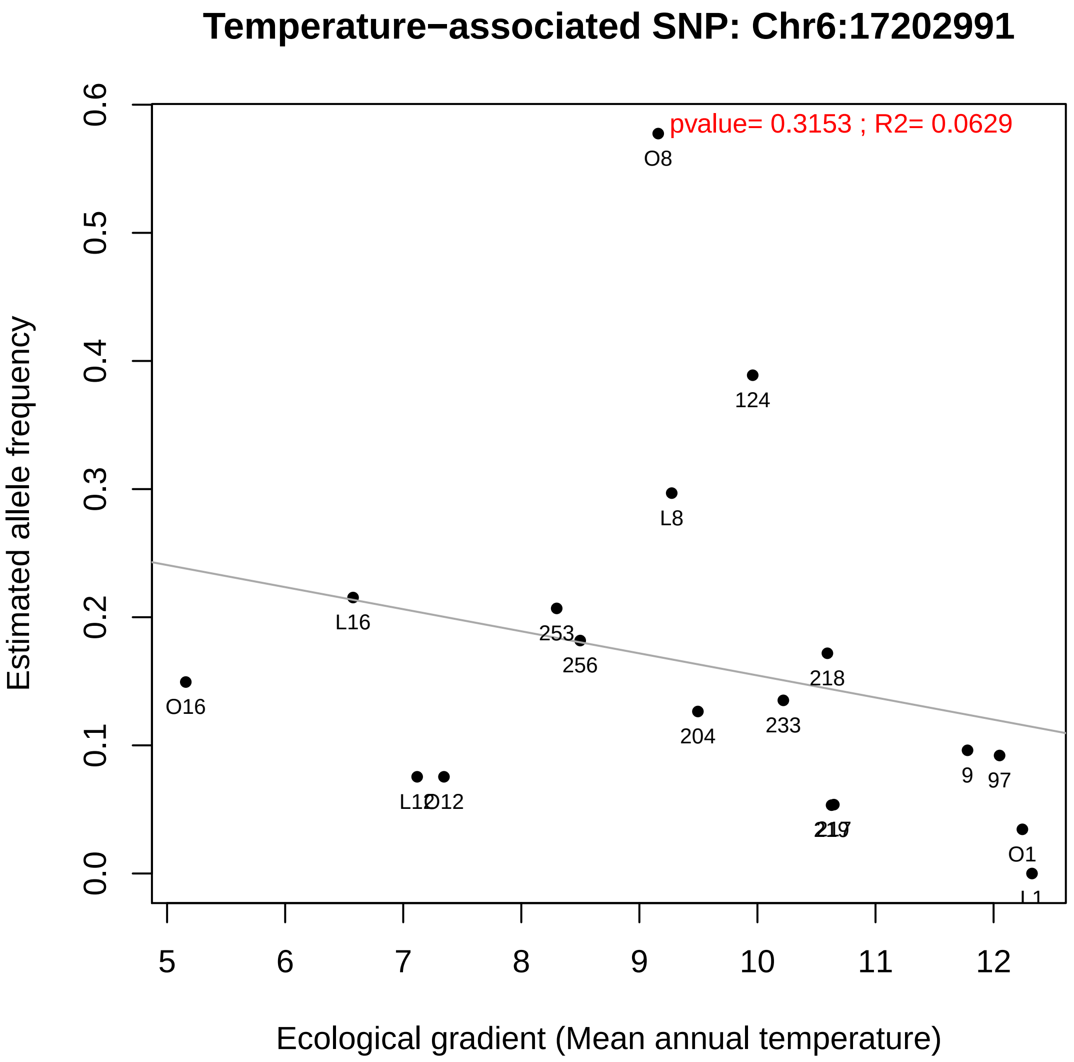
Temperature−associated SNP: Chr6:17202991
pvalue= 0.3153 ; R2= 0.0629
Estimated allele frequency
0.0
0.1
0.2
0.3
0.4
0.5
0.6
O8
124
L8
L16
253
256
O16
218
233
204
9
97
L12
O12
O1
L1
5
6
7
8
9
10
11
12
Ecological gradient (Mean annual temperature)

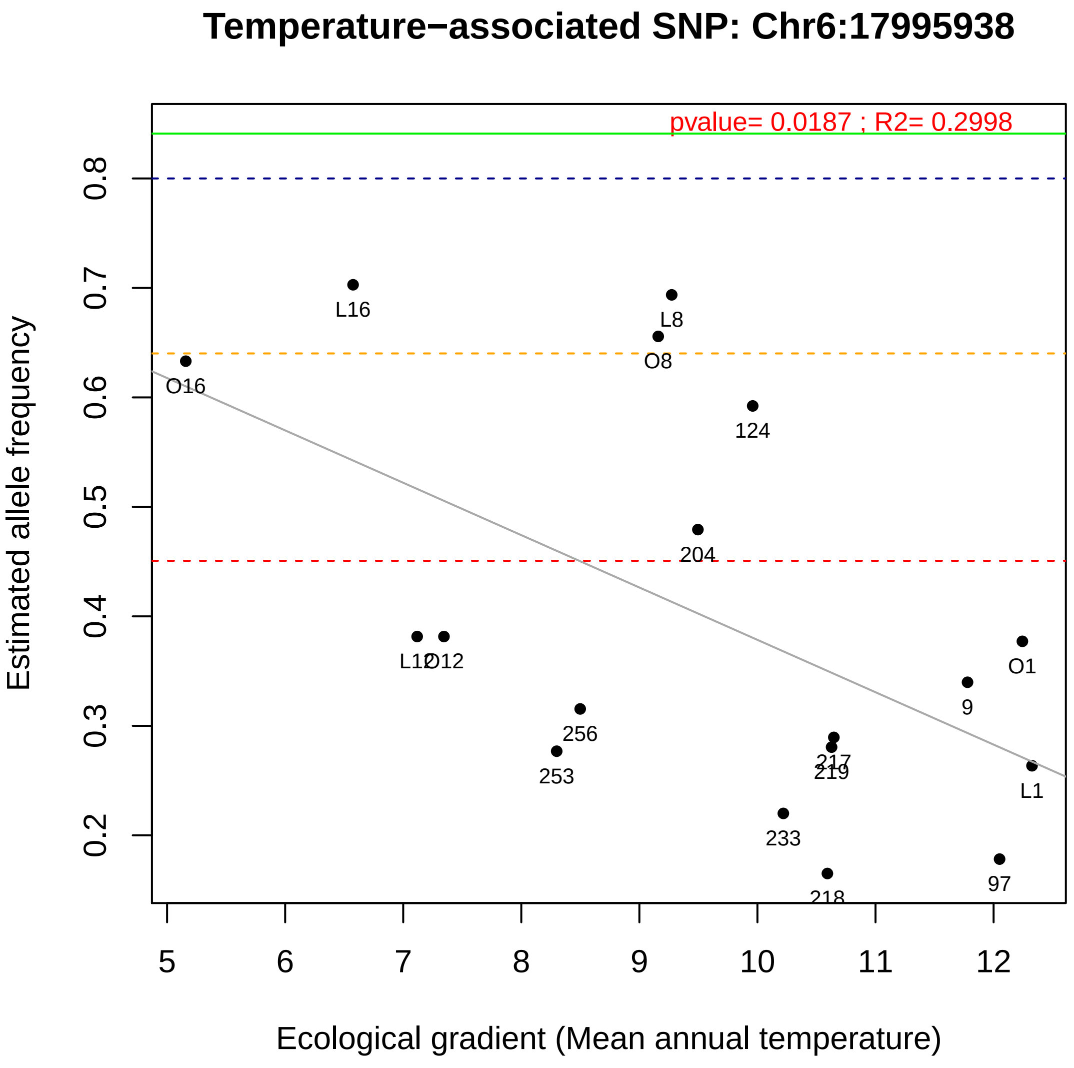

Temperature−associated SNP: Chr6:17995938
pvalue= 0.0187 ; R2= 0.2998
Estimated allele frequency
Ecological gradient (Mean annual temperature)
0.2
0.3
0.4
0.5
0.6
0.7
0.8
5
6
7
8
9
10
11
12
L16
L8
O8
O16
124
204
L12
O12
O1
9
256
253
217
219
L1
233
97
218

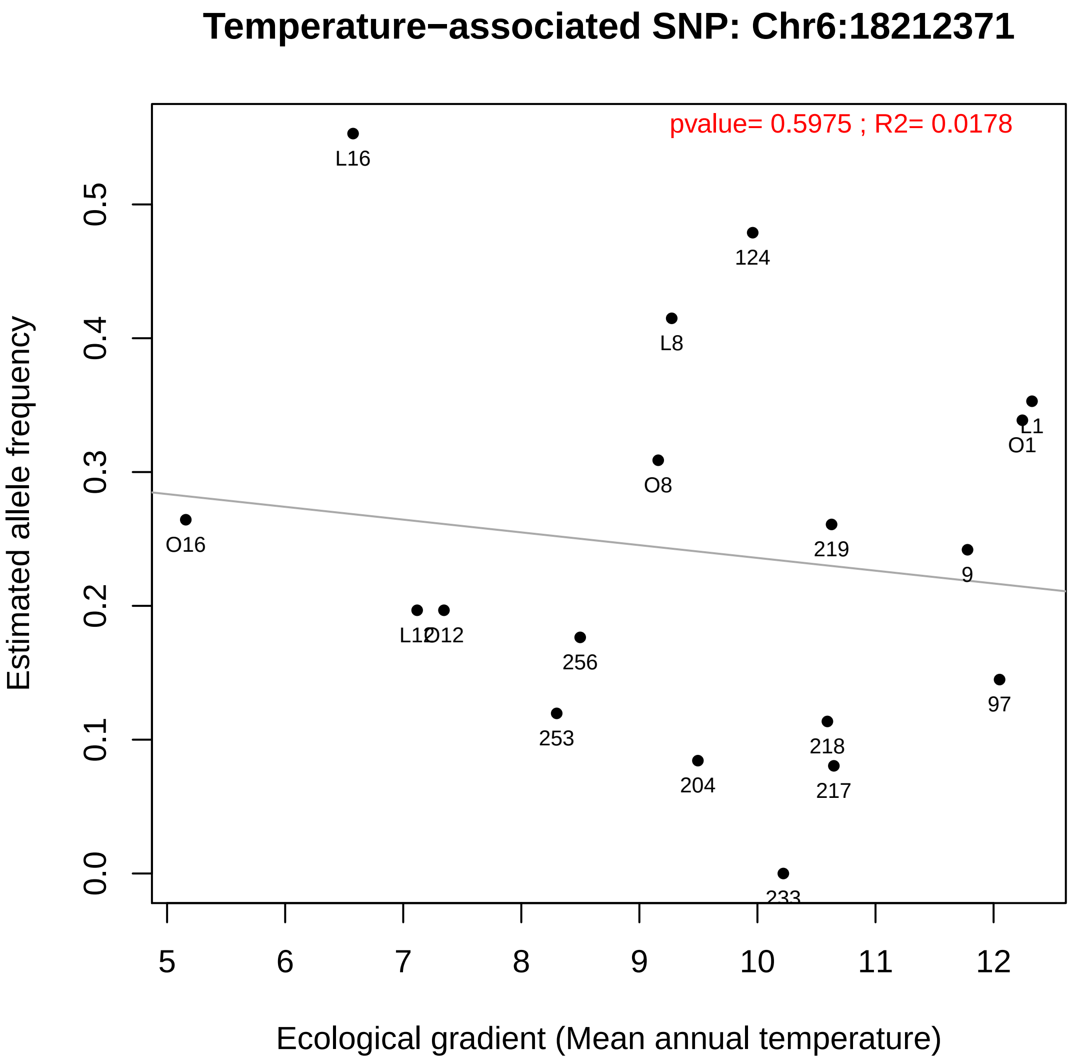

Temperature−associated SNP: Chr6:18212371
pvalue= 0.5975 ; R2= 0.0178
L16
124
L8
L1
O1
O8
O16
219
9
L12
O12
256
97
253
218
204
217
233
0.0
0.1
0.2
0.3
0.4
0.5
5
6
7
8
9
10
11
12
Estimated allele frequency
Ecological gradient (Mean annual temperature)

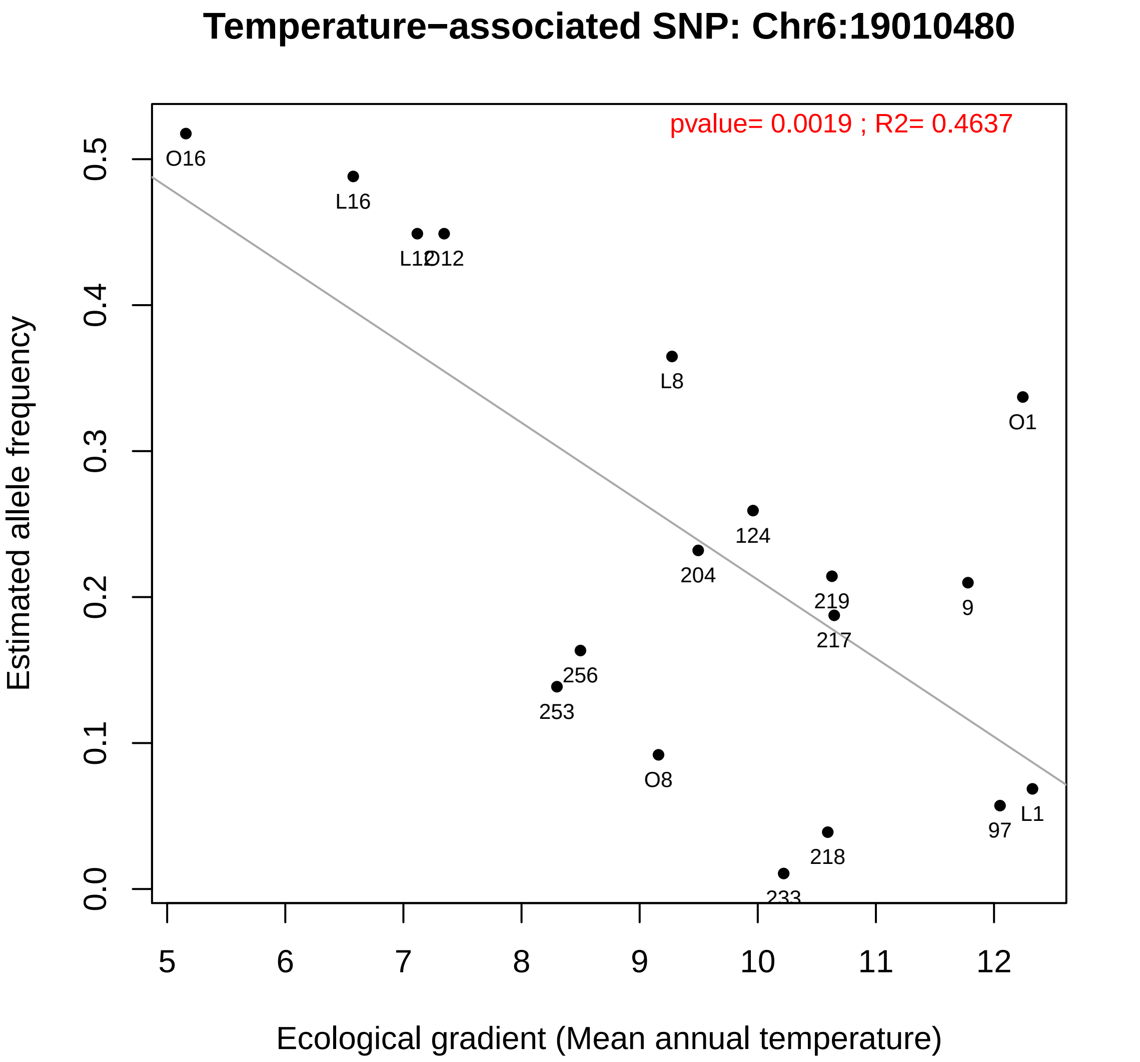

Temperature−associated SNP: Chr6:19010480
pvalue= 0.0019 ; R2= 0.4637
Estimated allele frequency
Ecological gradient (Mean annual temperature)
0.0
0.1
0.2
0.3
0.4
0.5
5
6
7
8
9
10
11
12
O16
L16
L12
O12
L8
O1
124
204
219
9
217
256
253
O8
L1
97
218
233

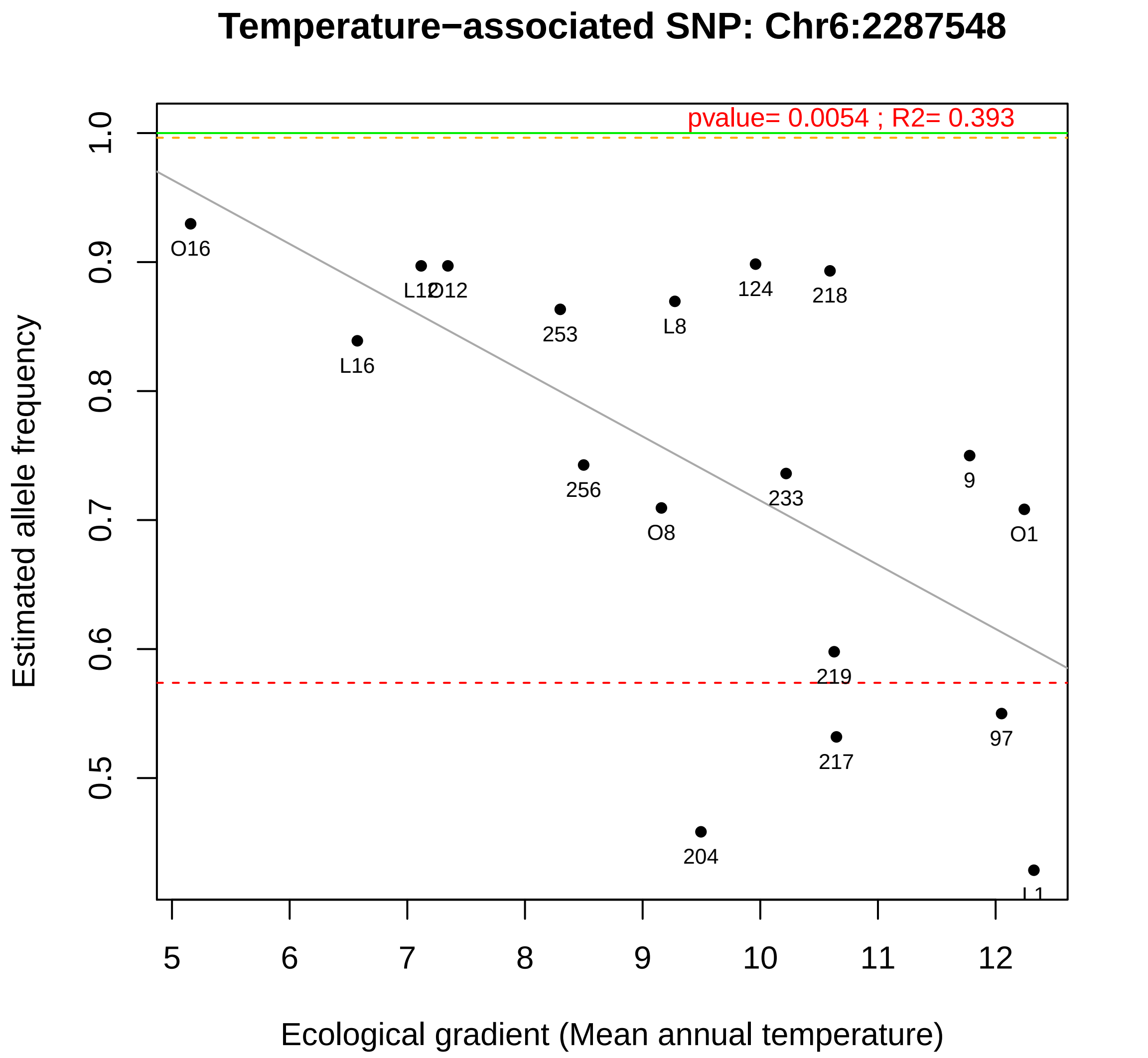
Temperature−associated SNP: Chr6:2287548
pvalue= 0.0054 ; R2= 0.393
Estimated allele frequency
1.0
0.9
0.8
0.7
0.6
0.5
O16
L12
O12
L16
253
L8
124
218
256
O8
233
9
O1
219
217
97
204
5
6
7
8
9
10
11
12
Ecological gradient (Mean annual temperature)

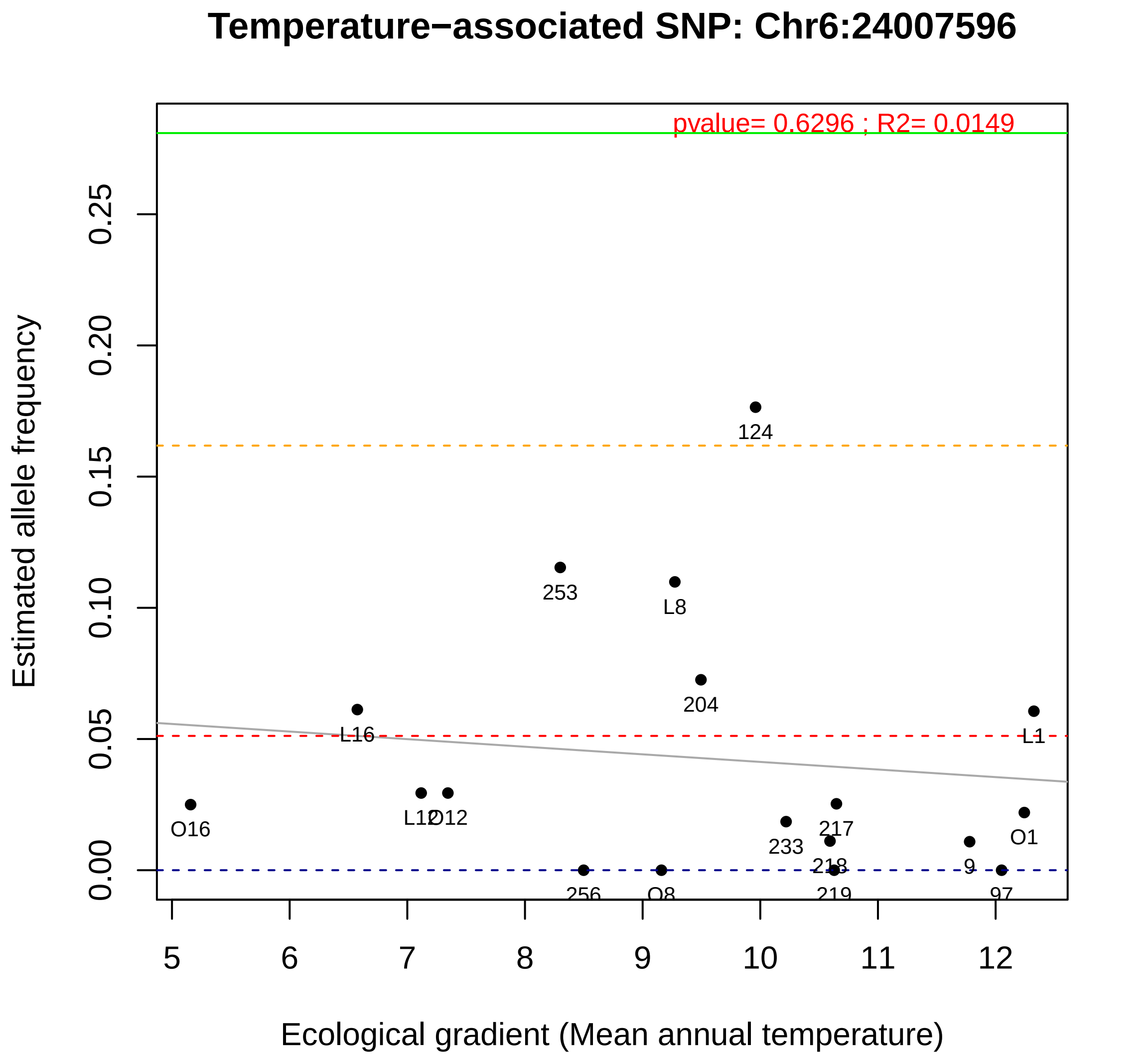

Temperature−associated SNP: Chr6:24007596
pvalue= 0.6296 ; R2= 0.0149
Estimated allele frequency
0.00
0.05
0.10
0.15
0.20
0.25
5
6
7
8
9
10
11
12
Ecological gradient (Mean annual temperature)
124
253
L8
204
L16
L1
O16
L12
O12
233
217
218
219
256
O8
9
97
O1

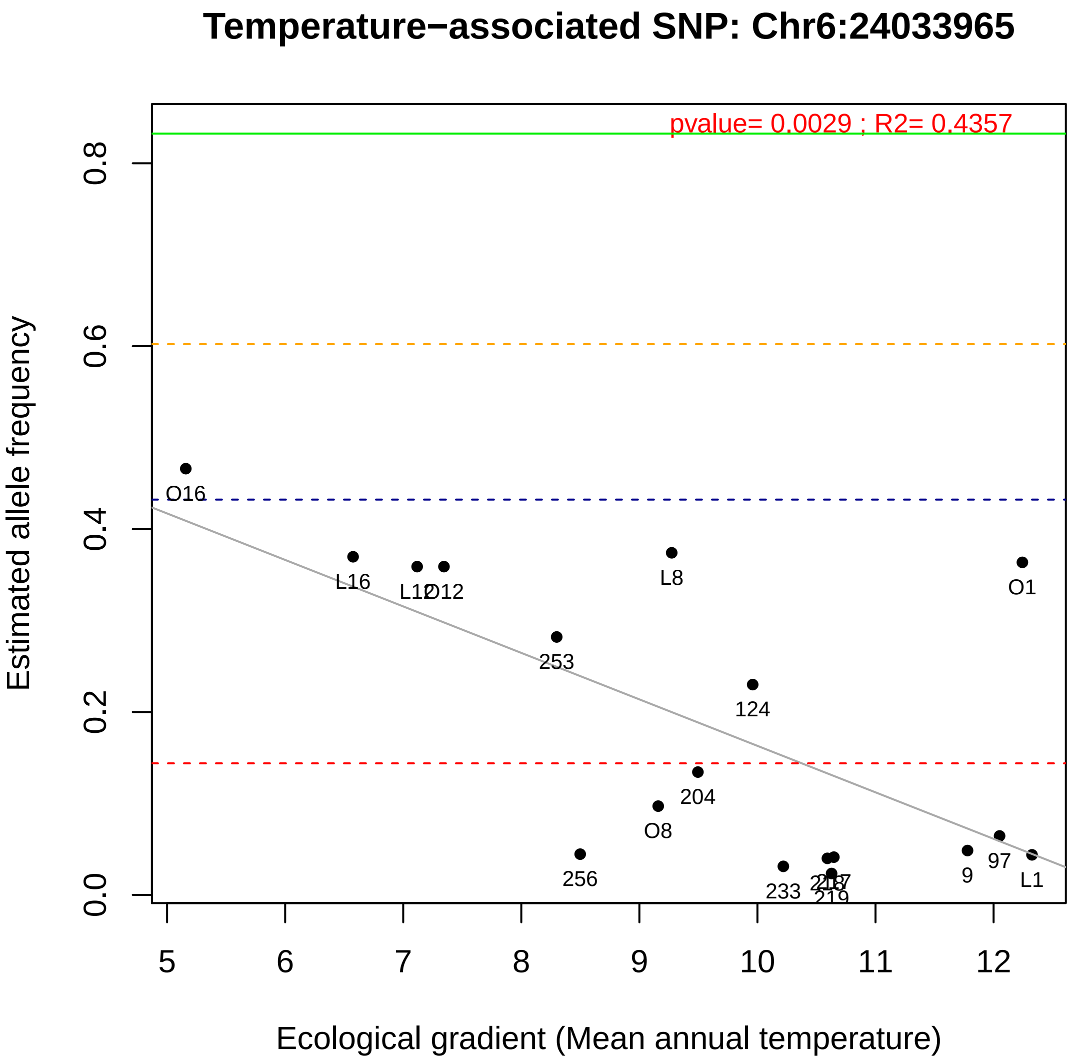
Temperature−associated SNP: Chr6:24033965
pvalue= 0.0029 ; R2= 0.4357
Estimated allele frequency
0.0
0.2
0.4
0.6
0.8
Ecological gradient (Mean annual temperature)
5
6
7
8
9
10
11
12
O16
L16
L12
O12
L8
O1
253
124
204
O8
256
233
219
9
97
L1

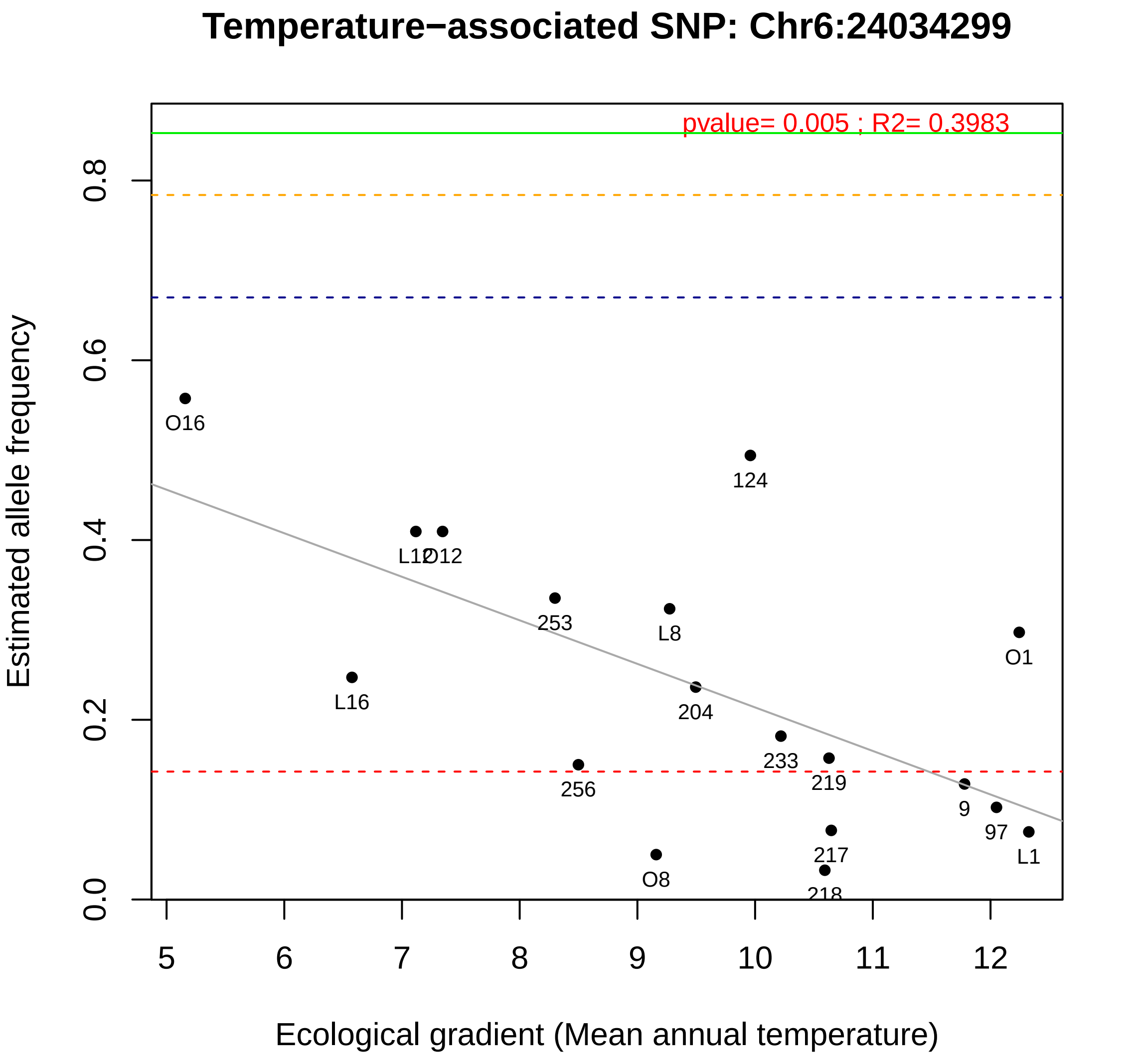

Temperature−associated SNP: Chr6:24034299
pvalue= 0.005 ; R2= 0.3983
Estimated allele frequency
0.0
0.2
0.4
0.6
0.8
5
6
7
8
9
10
11
12
Ecological gradient (Mean annual temperature)
O16
124
L12
O12
253
L8
O1
L16
204
233
219
256
9
97
217
L1
O8
218

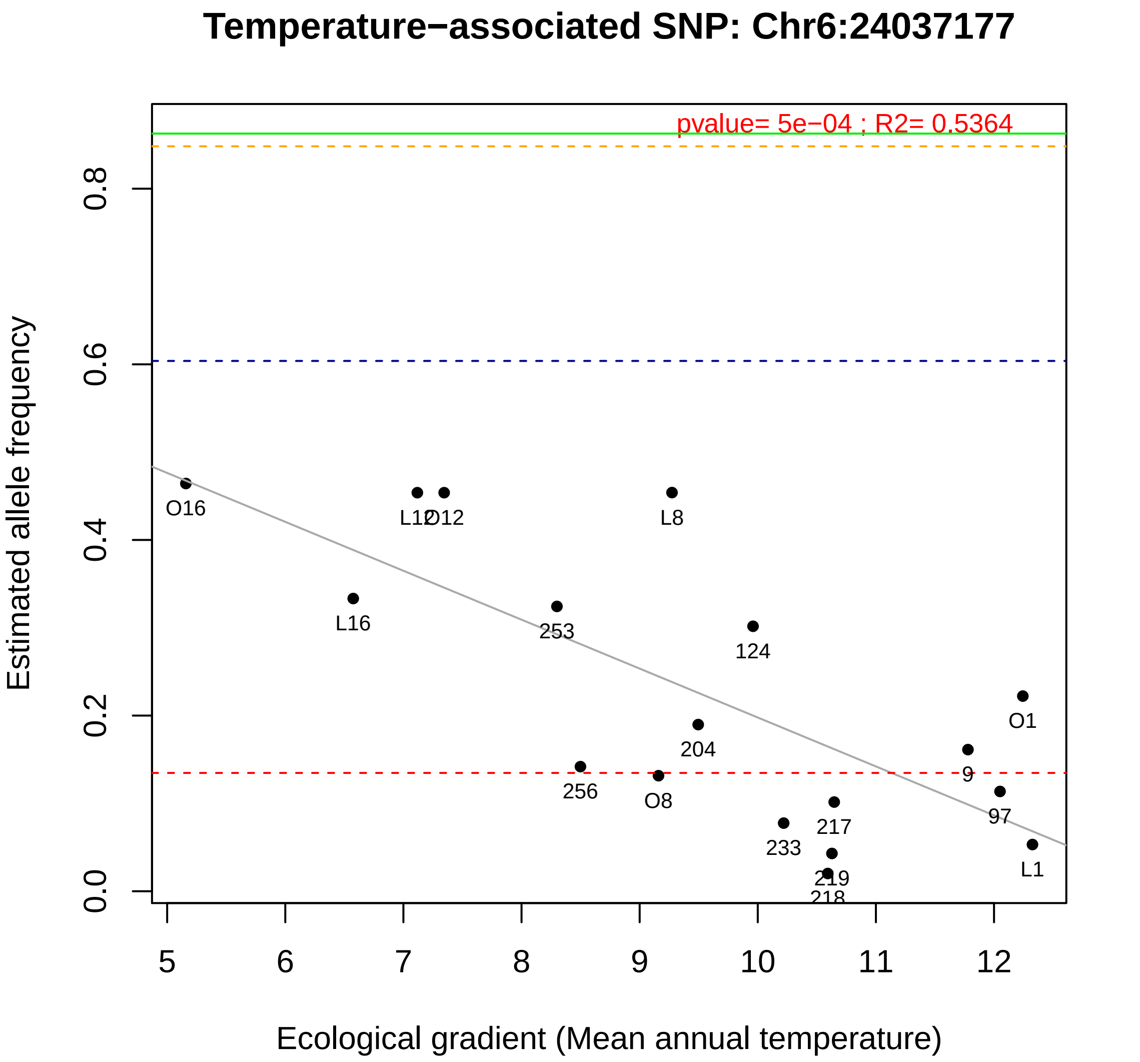

Temperature−associated SNP: Chr6:24037177
pvalue= 5e−04 ; R2= 0.5364
Estimated allele frequency
0.0
0.2
0.4
0.6
0.8
5
6
7
8
9
10
11
12
Ecological gradient (Mean annual temperature)
O16
L16
L12
O12
L8
253
124
256
O8
204
233
217
219
218
9
O1
97
L1

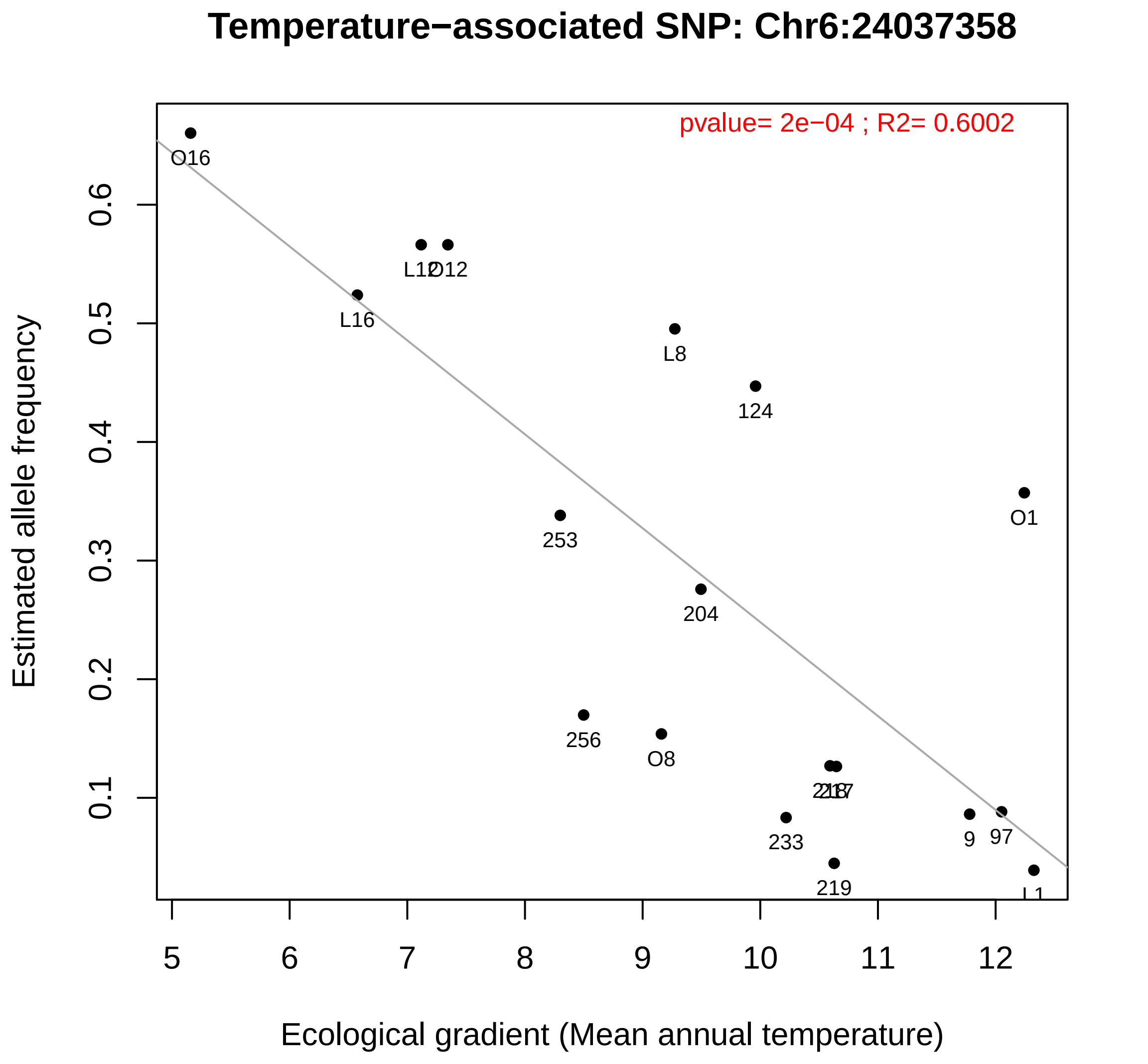

Temperature−associated SNP: Chr6:24037358
pvalue= 2e−04 ; R2= 0.6002
O16
L12
O12
L16
L8
124
O1
253
204
256
O8
218
217
233
219
9
97
L1
0.1
0.2
0.3
0.4
0.5
0.6
5
6
7
8
9
10
11
12
Estimated allele frequency
Ecological gradient (Mean annual temperature)

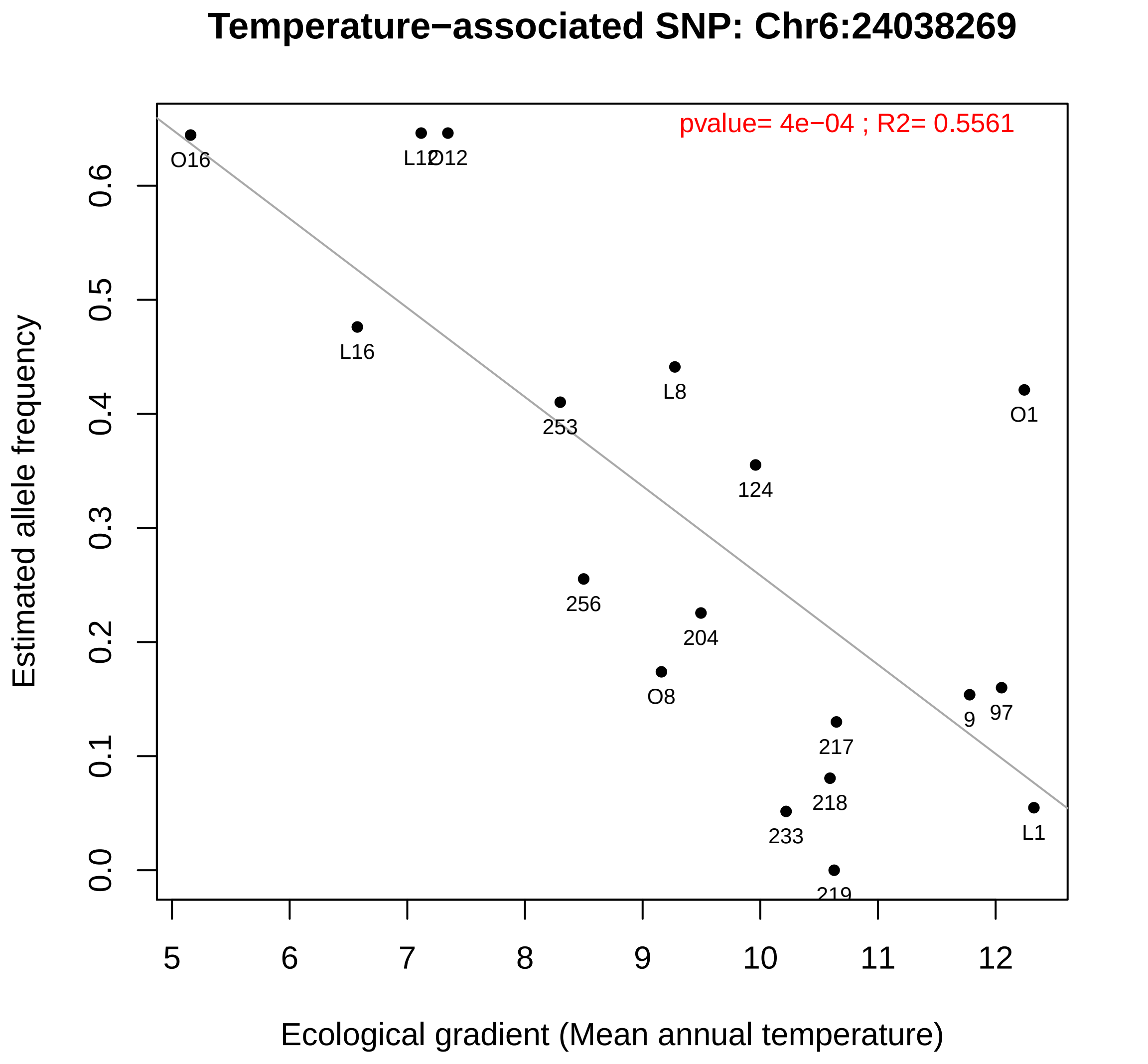

Temperature−associated SNP: Chr6:24038269
pvalue= 4e−04 ; R2= 0.5561
Estimated allele frequency
Ecological gradient (Mean annual temperature)
0.0
0.1
0.2
0.3
0.4
0.5
0.6
5
6
7
8
9
10
11
12
O16
L12
O12
L16
253
L8
O1
124
256
204
O8
9
97
217
218
233
L1
219

**Temperature−associated SNP: Chr6:24038322**

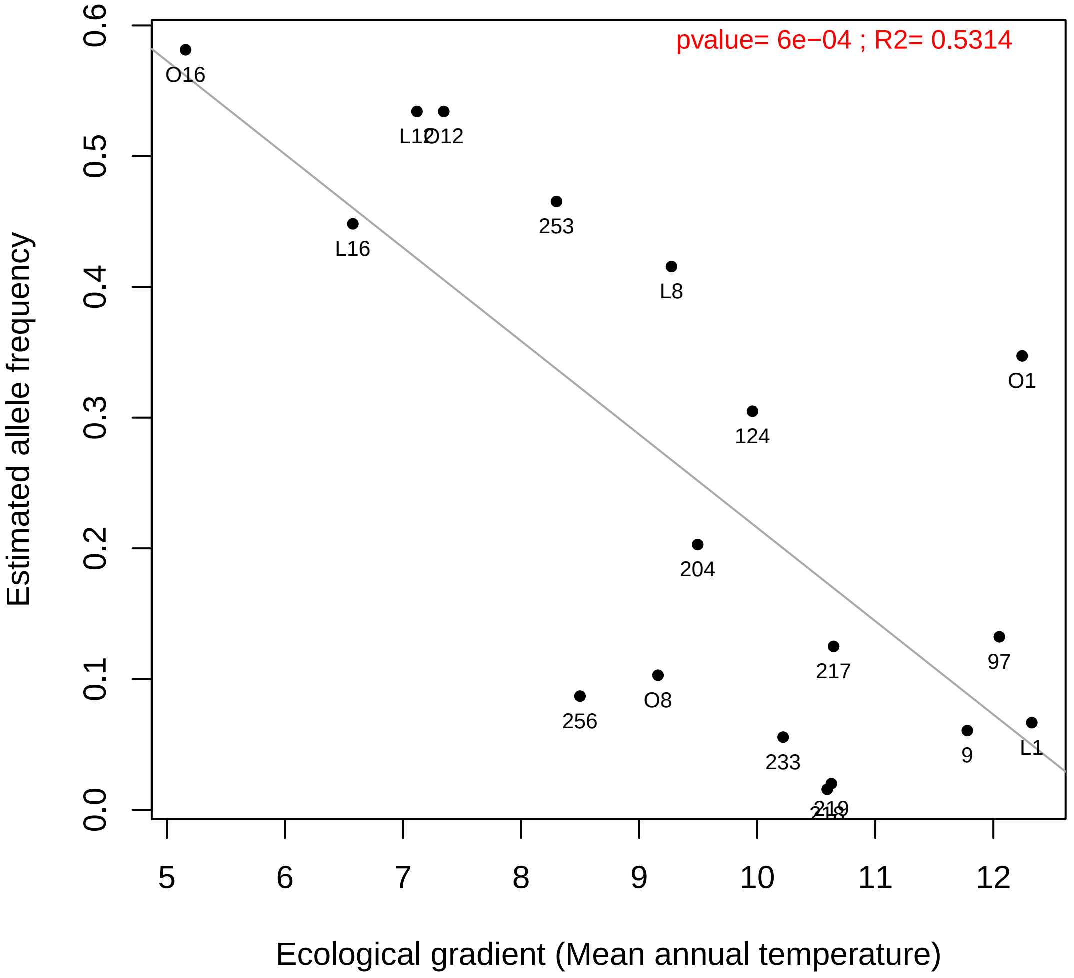

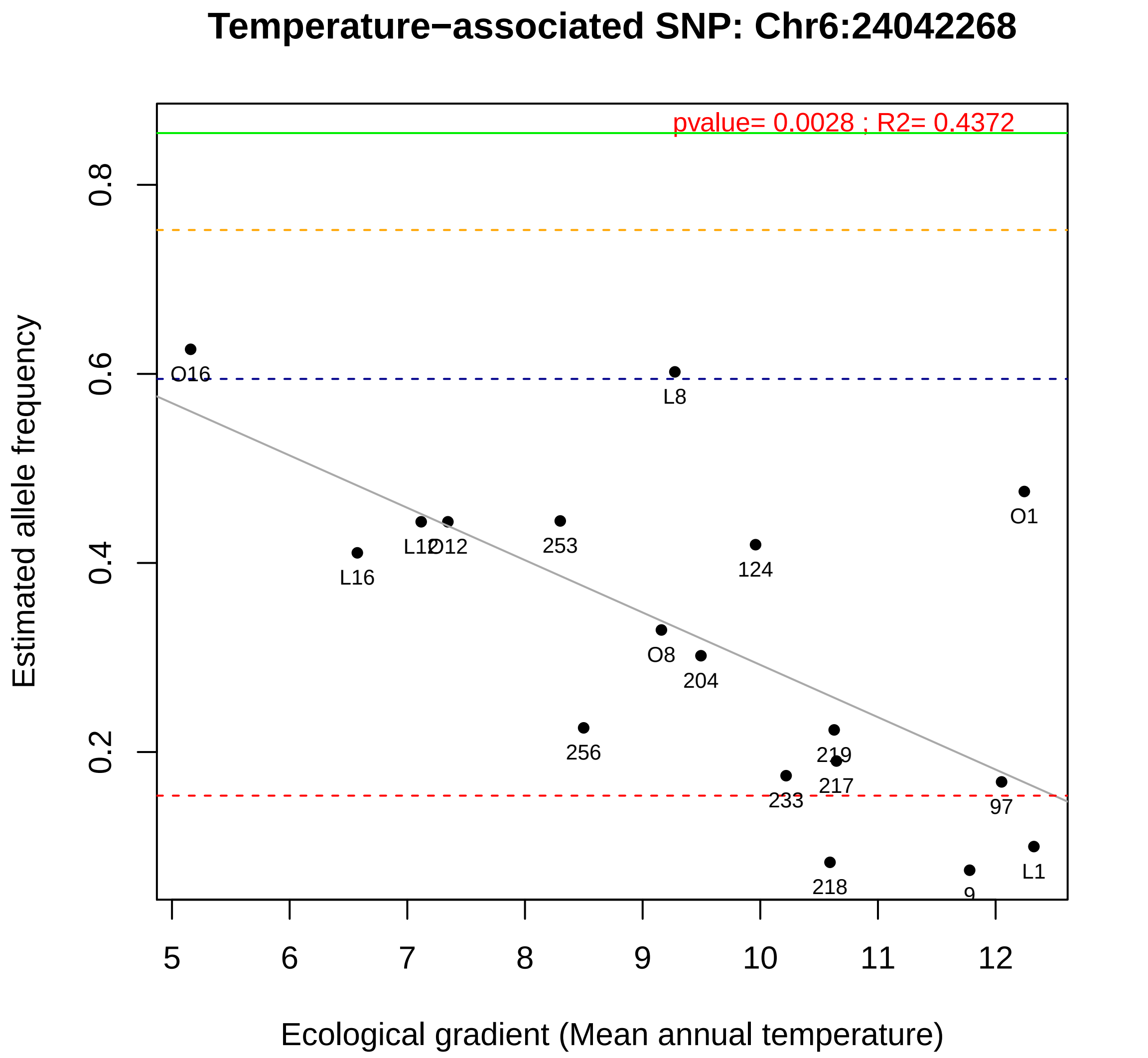
Temperature−associated SNP: Chr6:24042268
pvalue= 0.0028 ; R2= 0.4372
Estimated allele frequency
0.2
0.4
0.6
0.8
Ecological gradient (Mean annual temperature)
5
6
7
8
9
10
11
12
O16
L8
O1
L12
O12
253
L16
124
O8
204
256
219
217
233
97
218
9
L1

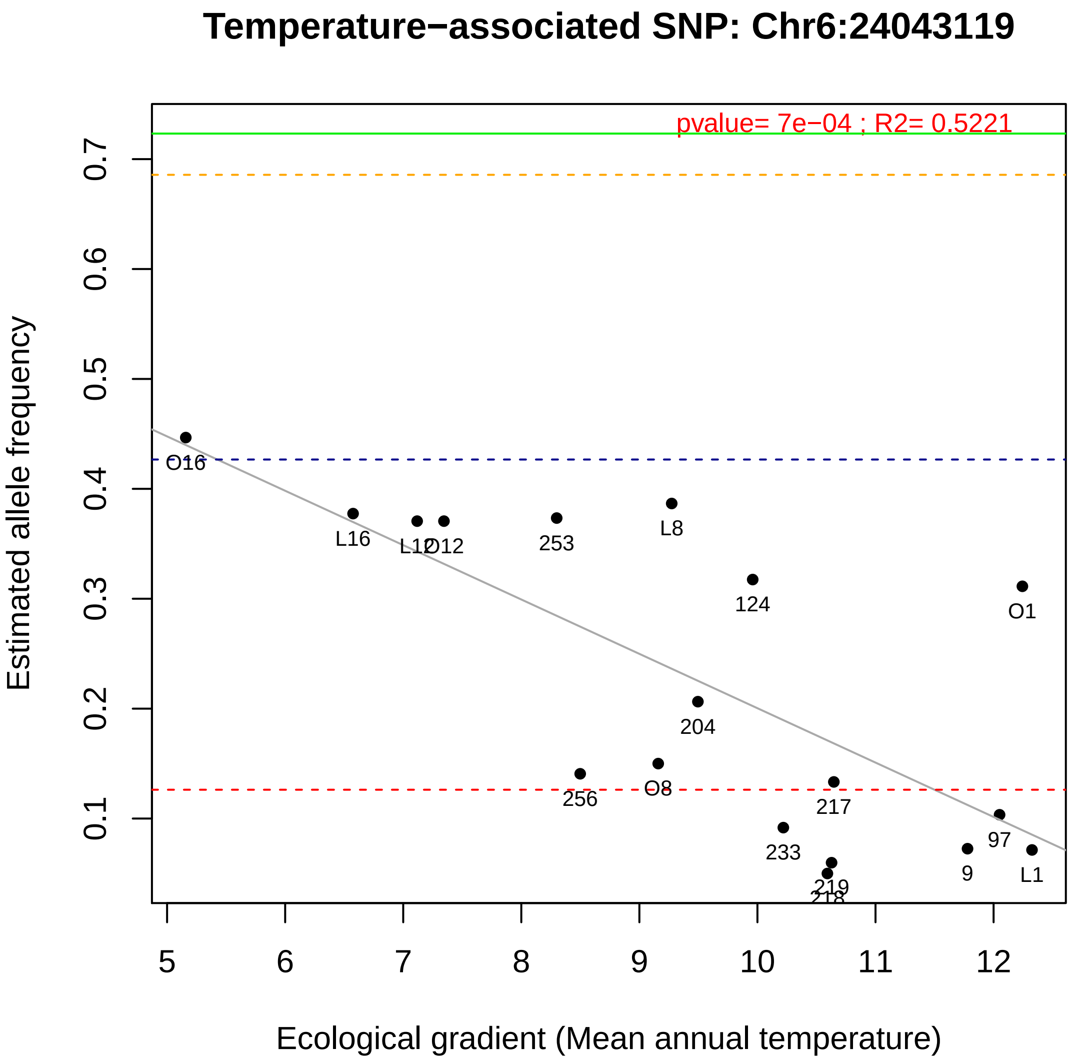
Temperature−associated SNP: Chr6:24043119
pvalue= 7e−04 ; R2= 0.5221
Estimated allele frequency
0.1
0.2
0.3
0.4
0.5
0.6
0.7
Ecological gradient (Mean annual temperature)
5
6
7
8
9
10
11
12
O16
L16
L12
O12
253
L8
124
O1
204
256
O8
217
233
97
9
L1
219
218

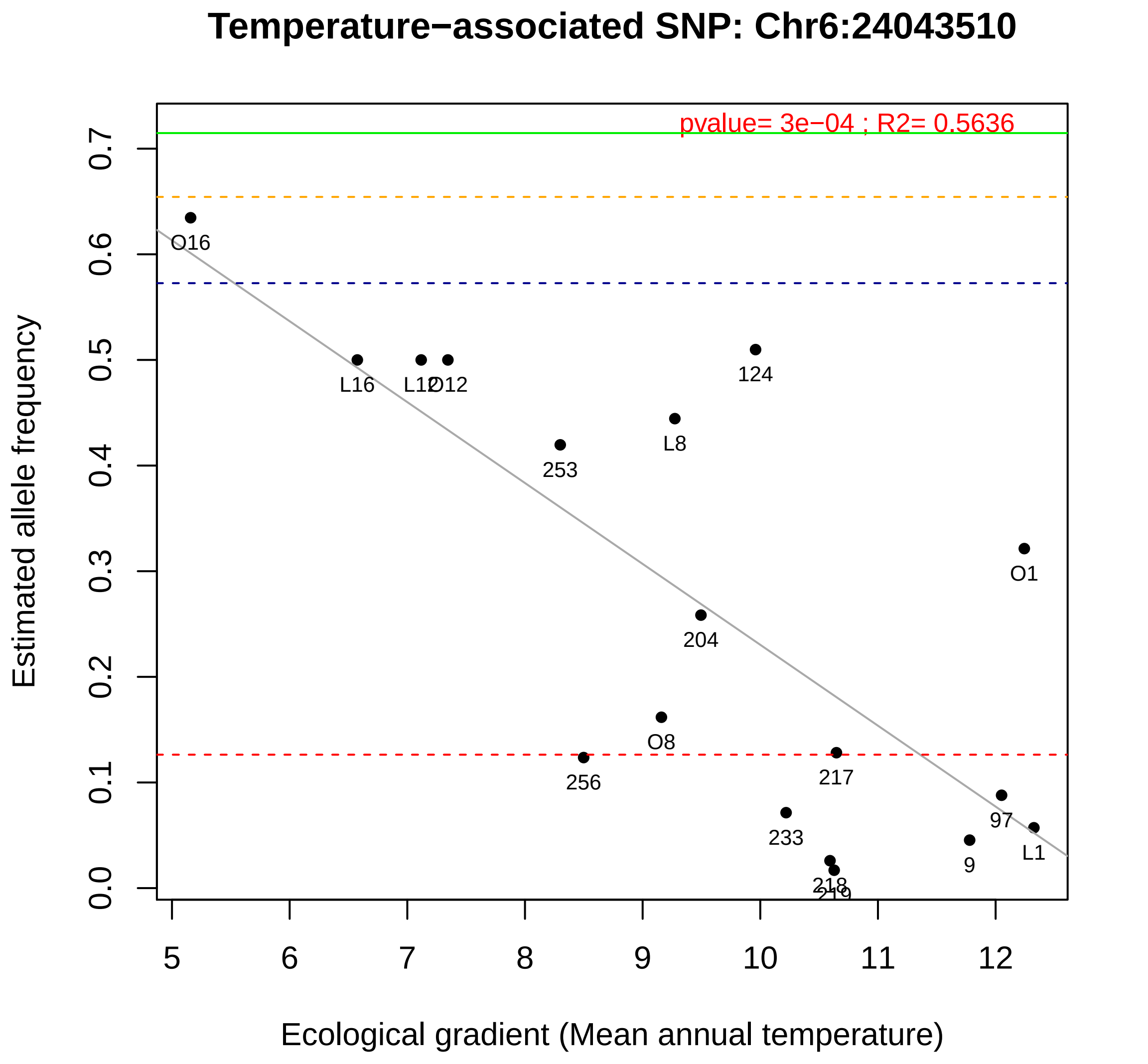

Temperature−associated SNP: Chr6:24043510
pvalue= 3e−04 ; R2= 0.5636
Estimated allele frequency
Ecological gradient (Mean annual temperature)
0.0
0.1
0.2
0.3
0.4
0.5
0.6
0.7
5
6
7
8
9
10
11
12
O16
L16
L12
O12
124
L8
253
O1
204
O8
256
217
233
97
L1
9
218
219

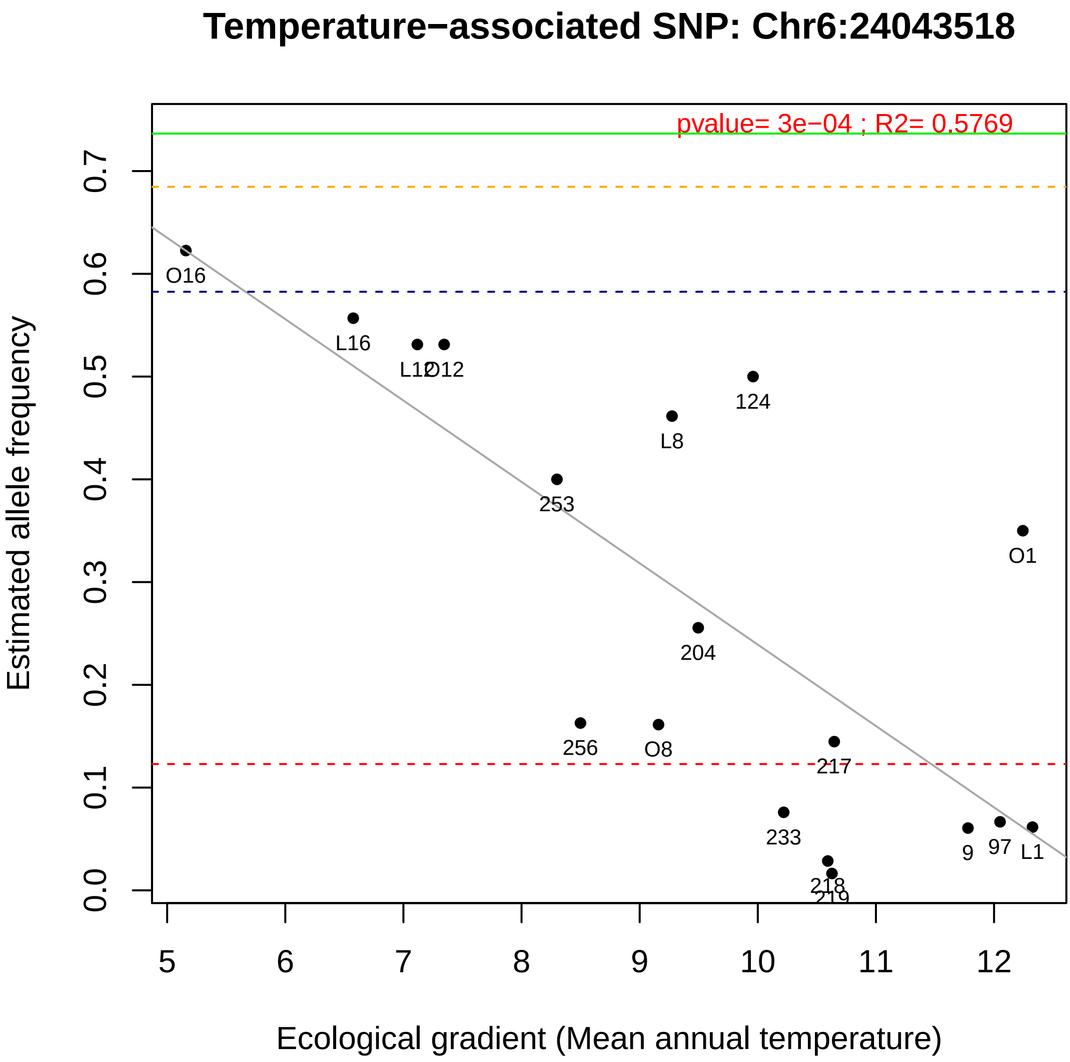
Temperature−associated SNP: Chr6:24043518
pvalue= 3e−04 ; R2= 0.5769
Estimated allele frequency
0.0
0.1
0.2
0.3
0.4
0.5
0.6
0.7
O16
L16
L12
O12
124
L8
253
O1
204
256
O8
217
233
218
219
9
97
L1
5
6
7
8
9
10
11
12
Ecological gradient (Mean annual temperature)

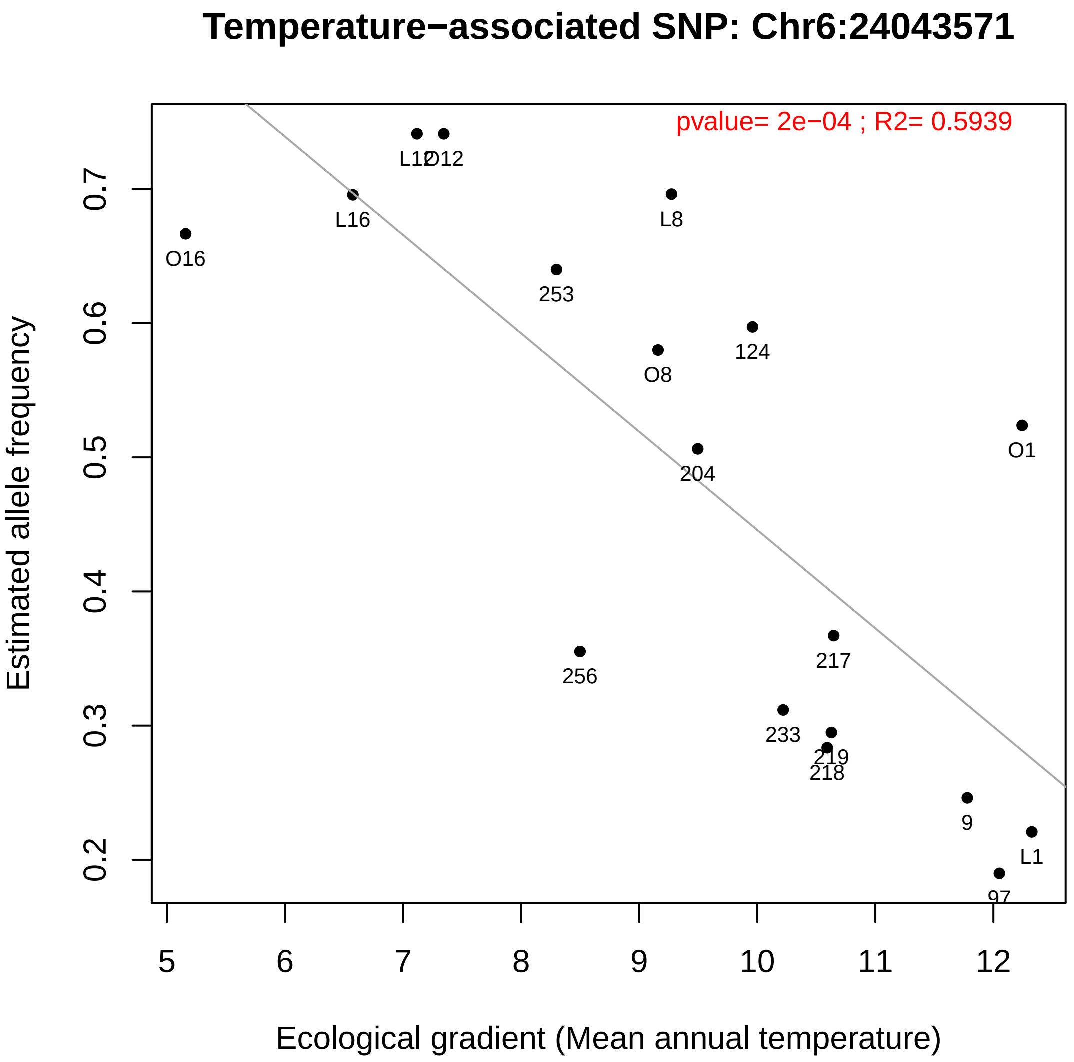
Temperature−associated SNP: Chr6:24043571
pvalue= 2e−04 ; R2= 0.5939
L12
O12
L16
L8
O16
253
124
O8
O1
204
256
217
233
219
218
9
L1
97
0.7
0.6
0.5
0.4
0.3
0.2
Estimated allele frequency
5
6
7
8
9
10
11
12
Ecological gradient (Mean annual temperature)

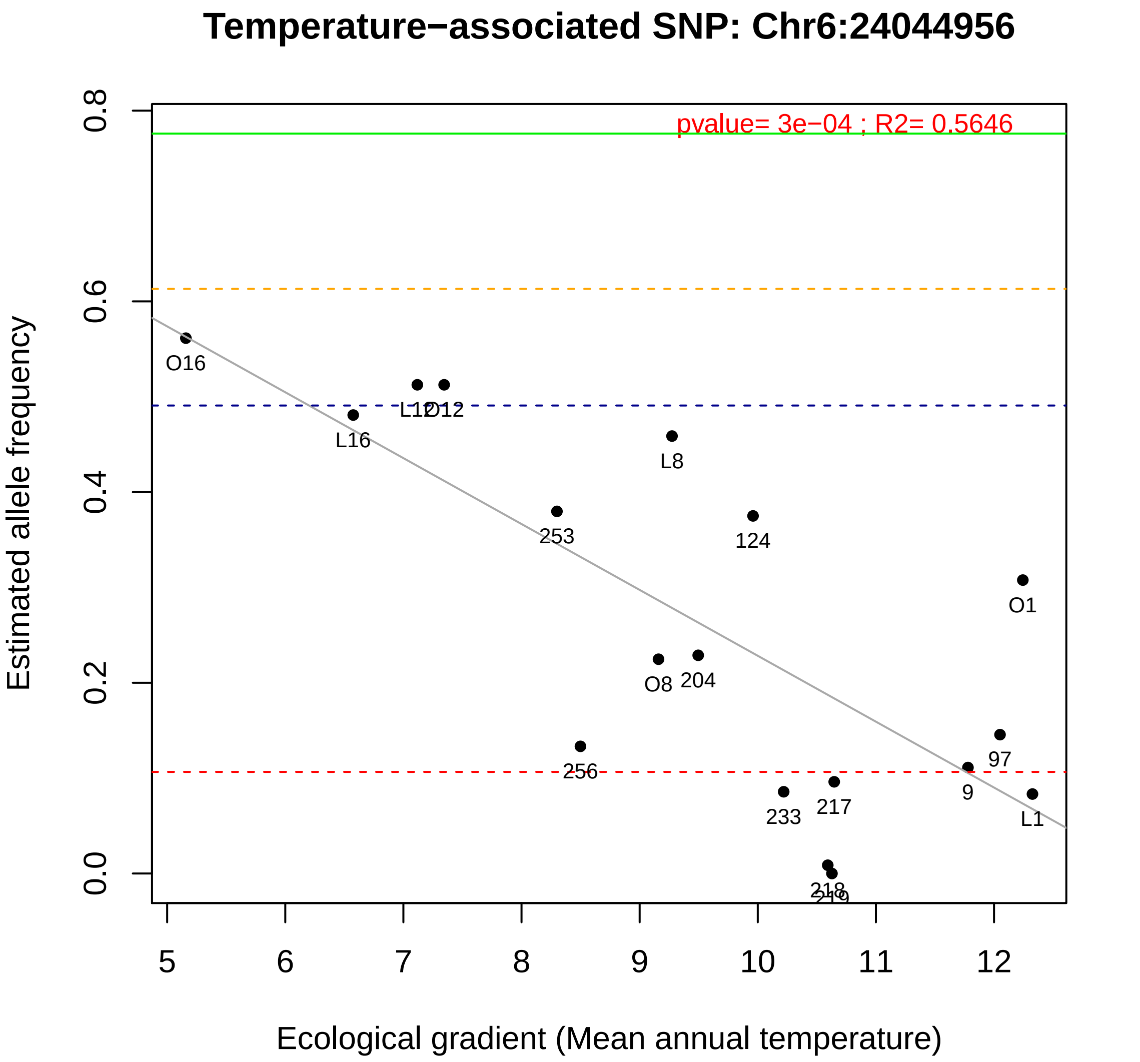

Temperature−associated SNP: Chr6:24044956
pvalue= 3e−04 ; R2= 0.5646
Estimated allele frequency
0.0
0.2
0.4
0.6
0.8
Ecological gradient (Mean annual temperature)
5
6
7
8
9
10
11
12
O16
L16
L12
O12
253
256
L8
O8
204
124
233
217
218
219
O1
97
9
L1

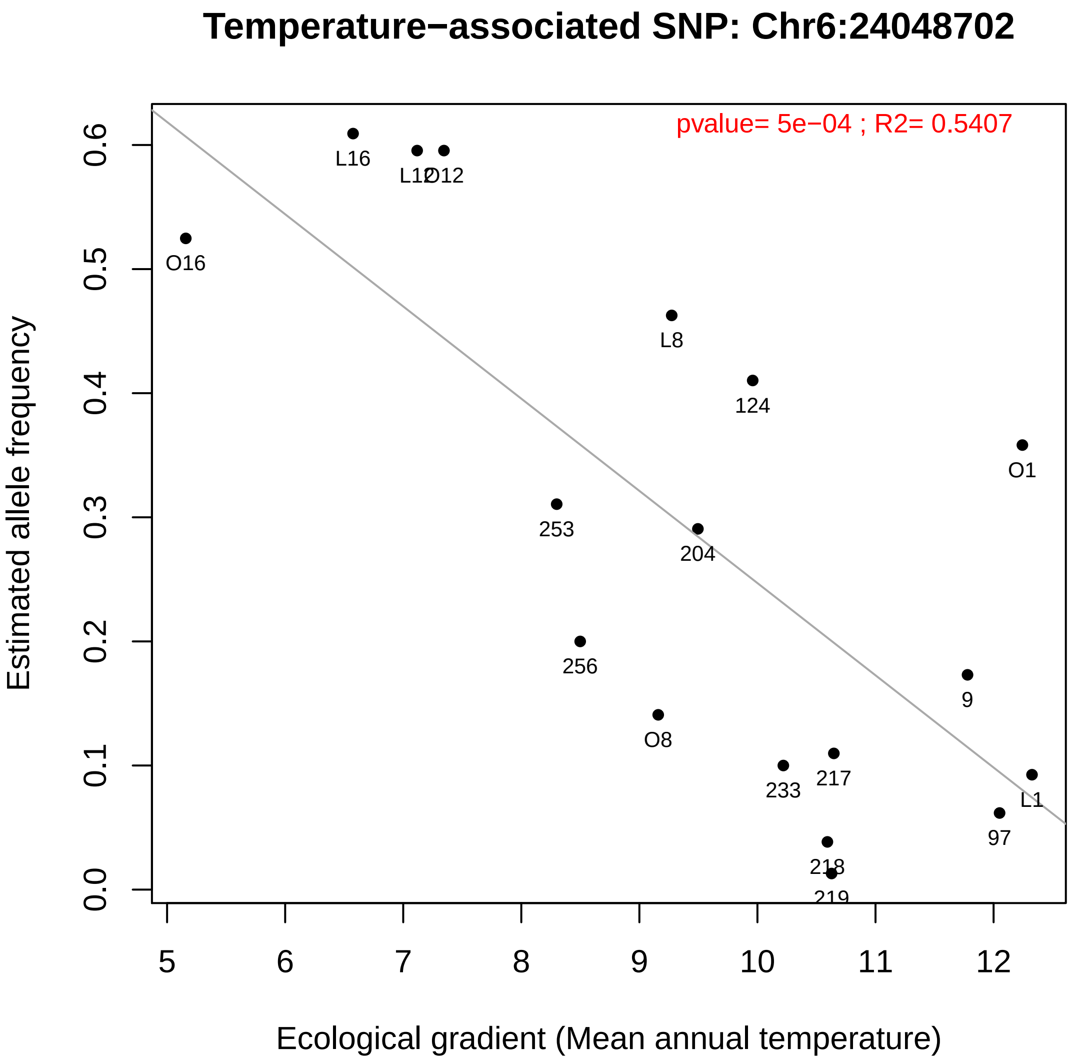
Temperature−associated SNP: Chr6:24048702
pvalue= 5e−04 ; R2= 0.5407
Estimated allele frequency
Ecological gradient (Mean annual temperature)
0.0
0.1
0.2
0.3
0.4
0.5
0.6
5
6
7
8
9
10
11
12
L16
L12
O12
O16
L8
124
O1
253
204
256
9
O8
233
217
L1
97
218
219

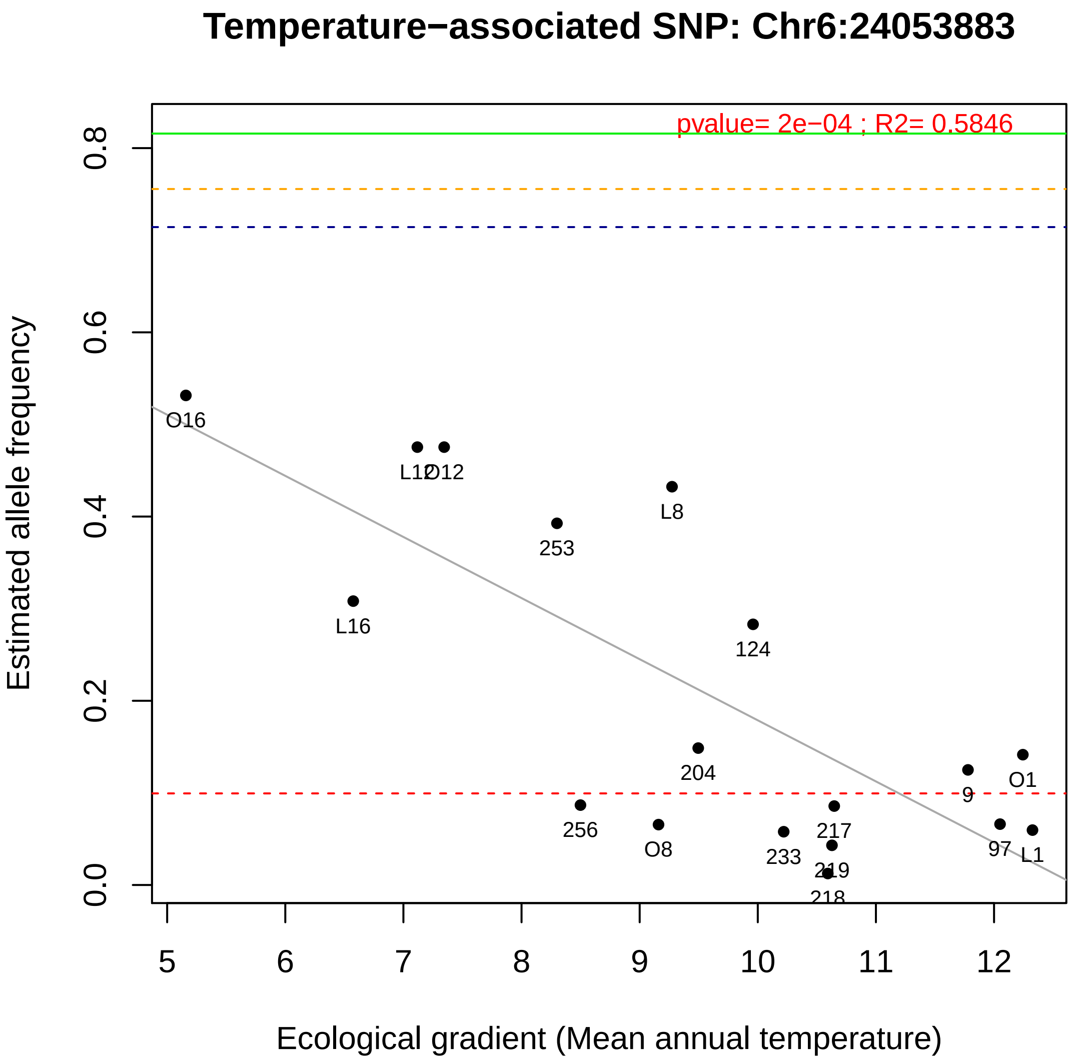
Temperature−associated SNP: Chr6:24053883
pvalue= 2e−04 ; R2= 0.5846
Estimated allele frequency
0.0
0.2
0.4
0.6
0.8
5
6
7
8
9
10
11
12
Ecological gradient (Mean annual temperature)
O16
L12
O12
L8
253
L16
124
204
256
O8
233
217
219
218
9
O1
97
L1

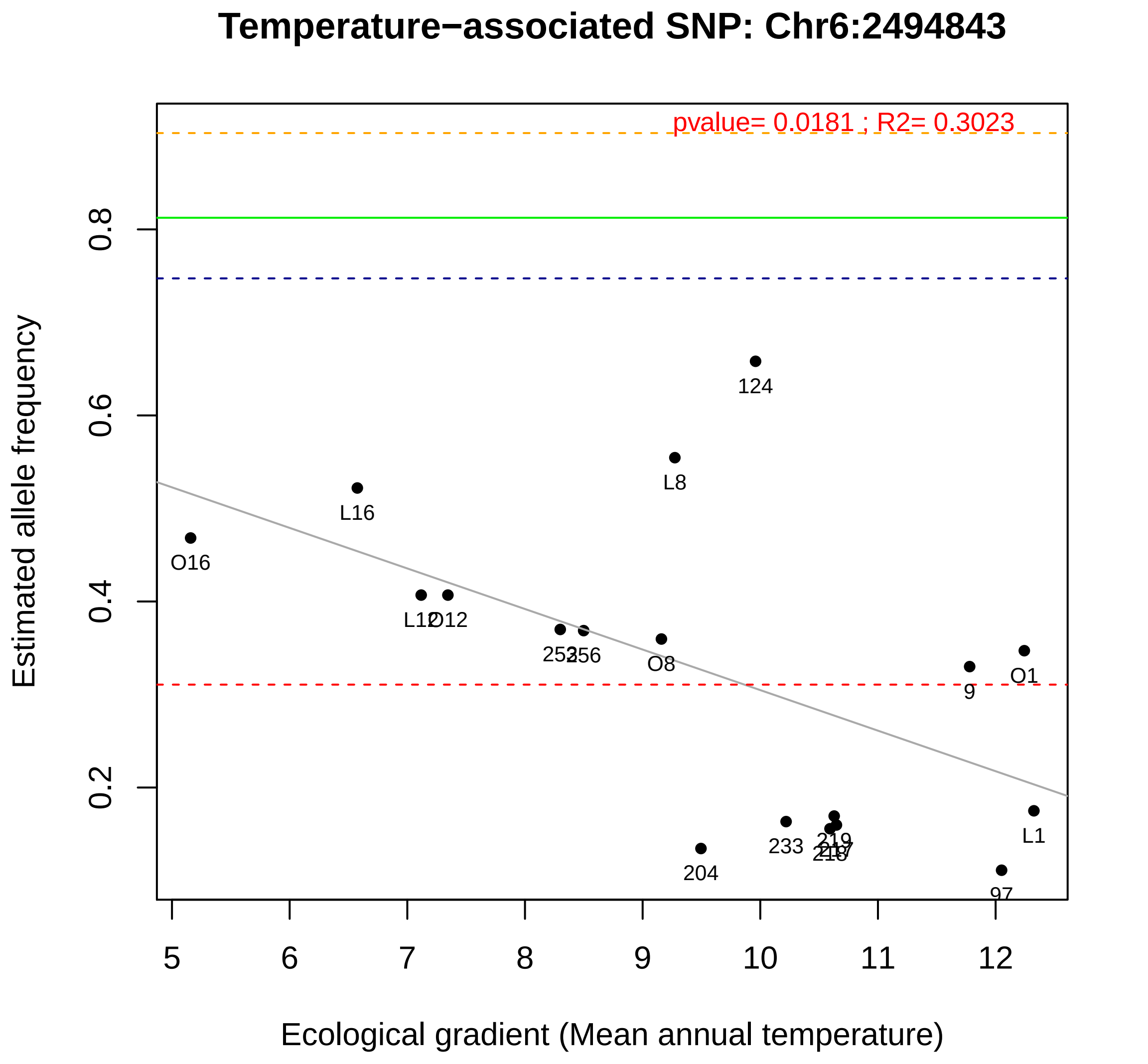

Temperature−associated SNP: Chr6:2494843
pvalue= 0.0181 ; R2= 0.3023
Estimated allele frequency
0.2
0.4
0.6
0.8
Ecological gradient (Mean annual temperature)
5
6
7
8
9
10
11
12
124
L8
L16
O16
O12
O8
O1
9
L1
204
233
219
97

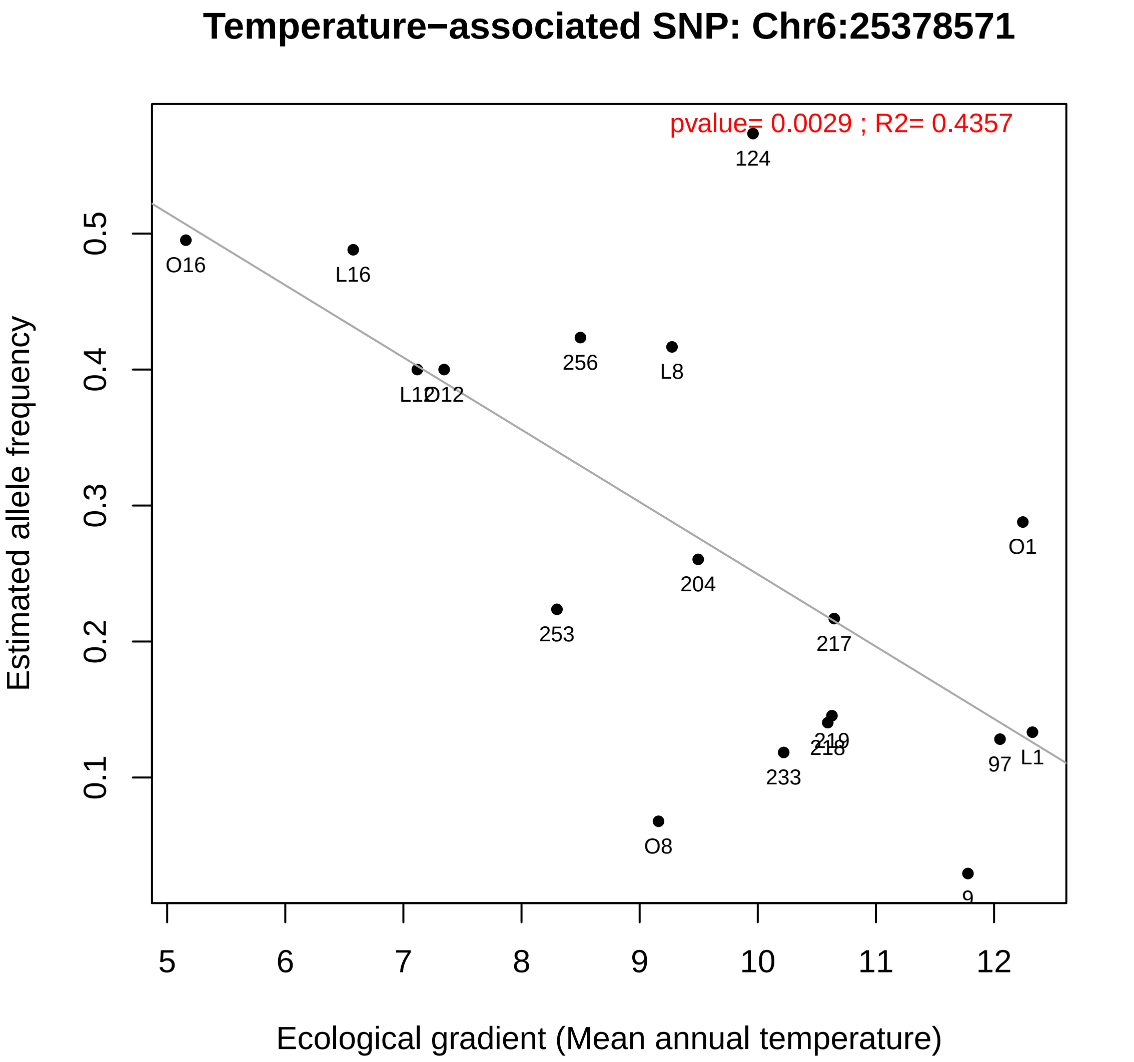
Temperature−associated SNP: Chr6:25378571
pvalue= 0.0029 ; R2= 0.4357
Estimated allele frequency
0.1
0.2
0.3
0.4
0.5
Ecological gradient (Mean annual temperature)
5
6
7
8
9
10
11
12
124
O16
L16
256
L8
L12
O12
O1
204
253
217
219
218
233
97
L1
O8
9

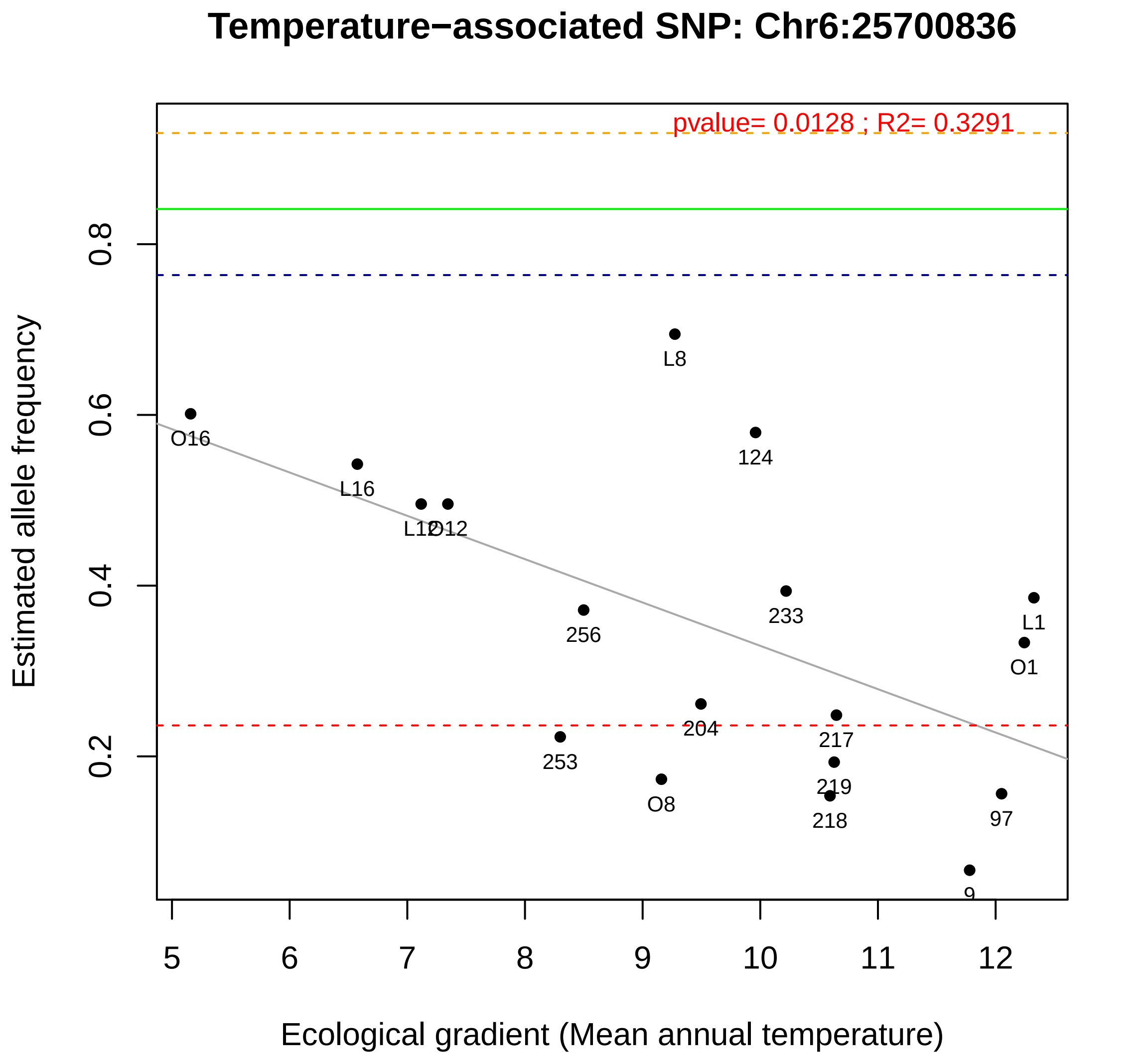
Temperature−associated SNP: Chr6:25700836
pvalue= 0.0128 ; R2= 0.3291
Estimated allele frequency
0.2
0.4
0.6
0.8
Ecological gradient (Mean annual temperature)
5
6
7
8
9
10
11
12
O16
L16
L12
O12
256
253
L8
O8
204
124
233
217
219
218
9
97
L1
O1

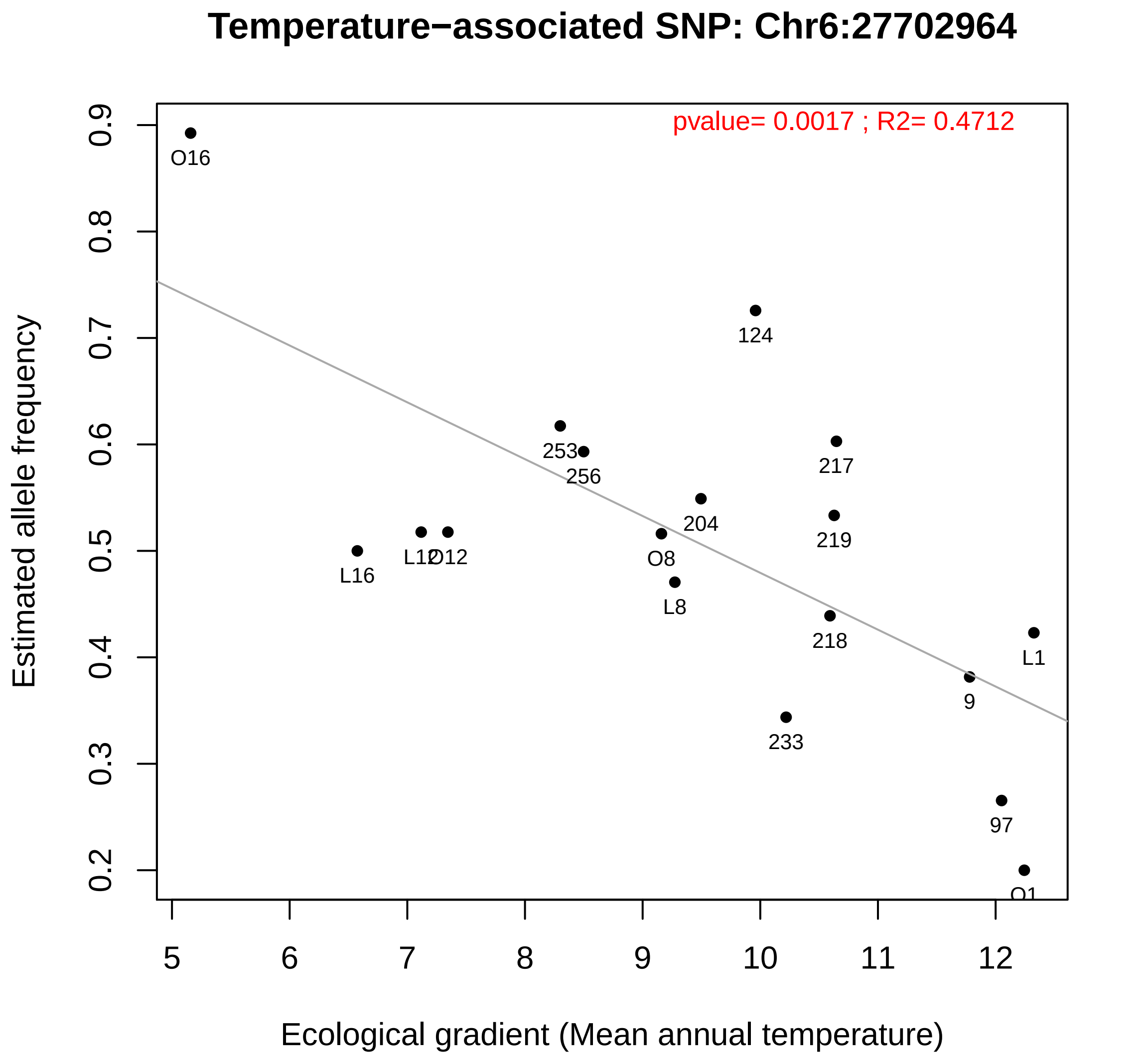
Temperature−associated SNP: Chr6:27702964
pvalue= 0.0017 ; R2= 0.4712
Estimated allele frequency
0.2
0.3
0.4
0.5
0.6
0.7
0.8
0.9
5
6
7
8
9
10
11
12
Ecological gradient (Mean annual temperature)
O16
124
253
256
217
204
219
L16
L12
O12
O8
L8
218
L1
9
233
97
O1

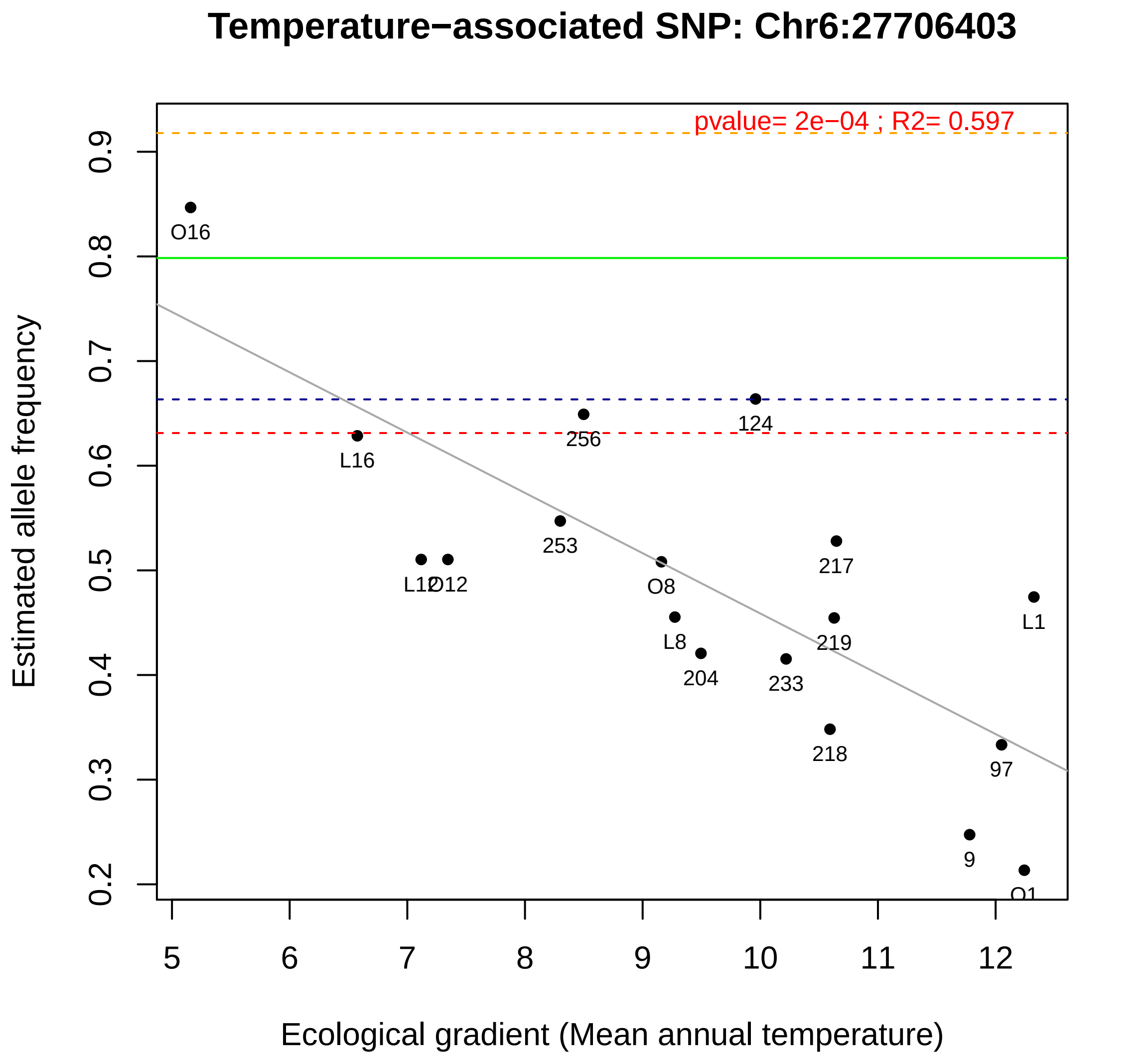
Temperature−associated SNP: Chr6:27706403
pvalue= 2e−04 ; R2= 0.597
Estimated allele frequency
0.2
0.3
0.4
0.5
0.6
0.7
0.8
0.9
5
6
7
8
9
10
11
12
Ecological gradient (Mean annual temperature)
O16
L16
L12
O12
253
256
124
O8
L8
204
233
217
219
218
L1
97
9
O1

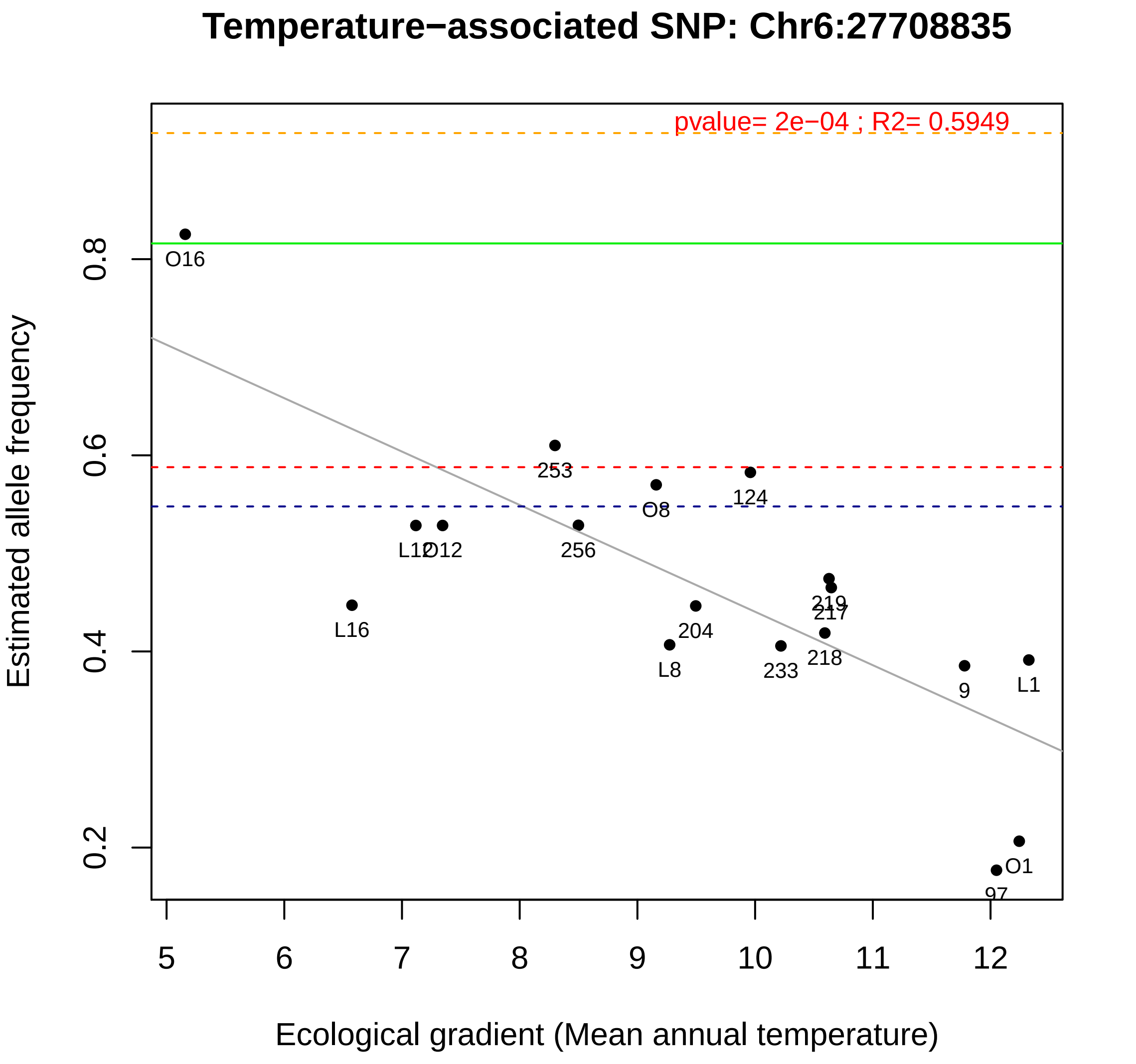
Temperature−associated SNP: Chr6:27708835
pvalue= 2e−04 ; R2= 0.5949
Estimated allele frequency
0.2
0.4
0.6
0.8
5
6
7
8
9
10
11
12
Ecological gradient (Mean annual temperature)
O16
253
124
O8
L12
O12
256
L16
204
L8
233
219
217
218
9
L1
O1
97

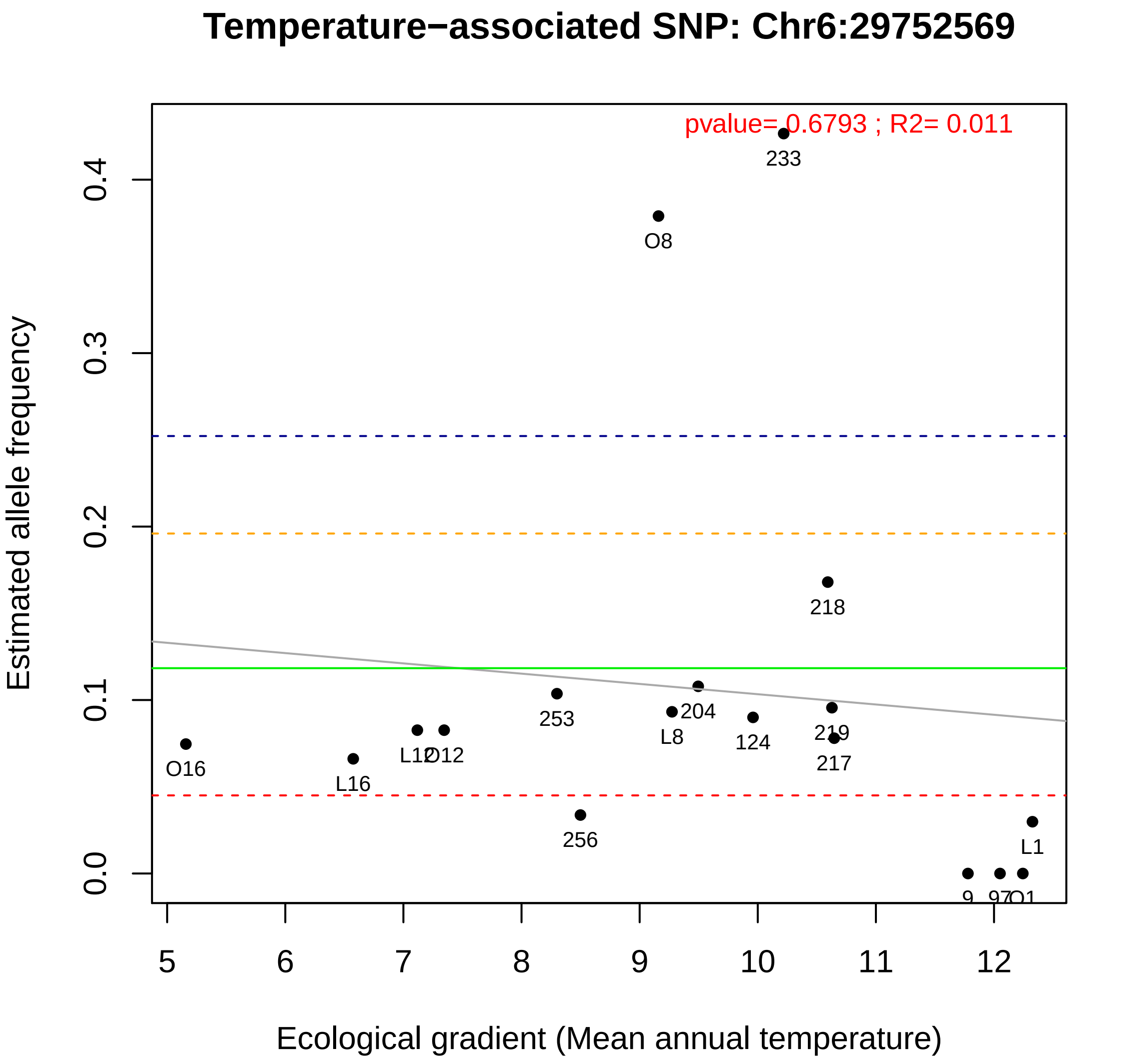
Temperature−associated SNP: Chr6:29752569
pvalue= 0.6793 ; R2= 0.011
Estimated allele frequency
0.0
0.1
0.2
0.3
0.4
5
6
7
8
9
10
11
12
Ecological gradient (Mean annual temperature)
233
O8
218
253
204
L8
124
219
217
O16
L16
L12
O12
256
L1
9
97
O1

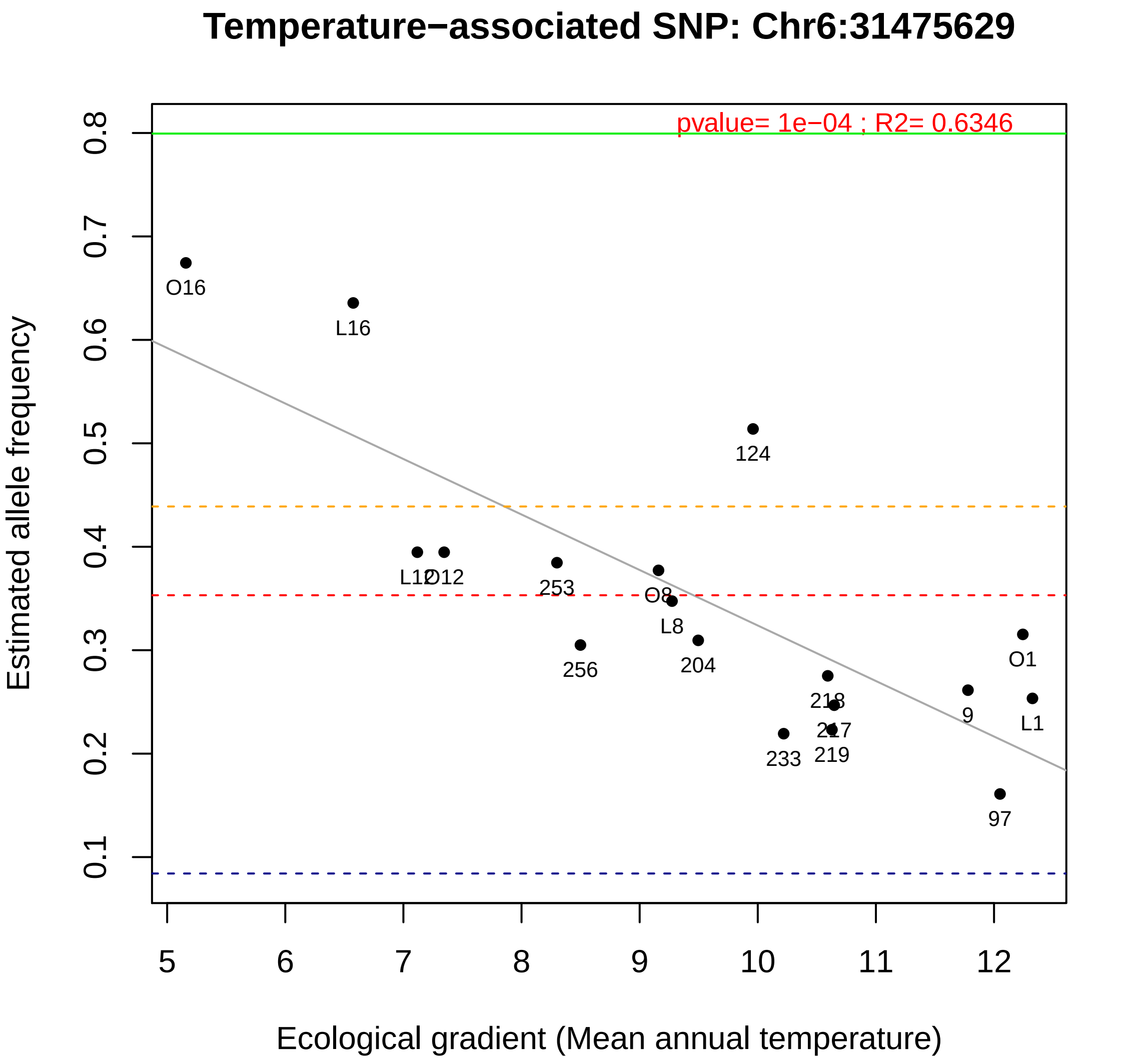

Temperature−associated SNP: Chr6:31475629
pvalue= 1e−04 ; R2= 0.6346
Estimated allele frequency
0.1
0.2
0.3
0.4
0.5
0.6
0.7
0.8
5
6
7
8
9
10
11
12
Ecological gradient (Mean annual temperature)
O16
L16
124
L12
O12
253
O8
L8
256
204
O1
218
217
219
233
9
L1
97

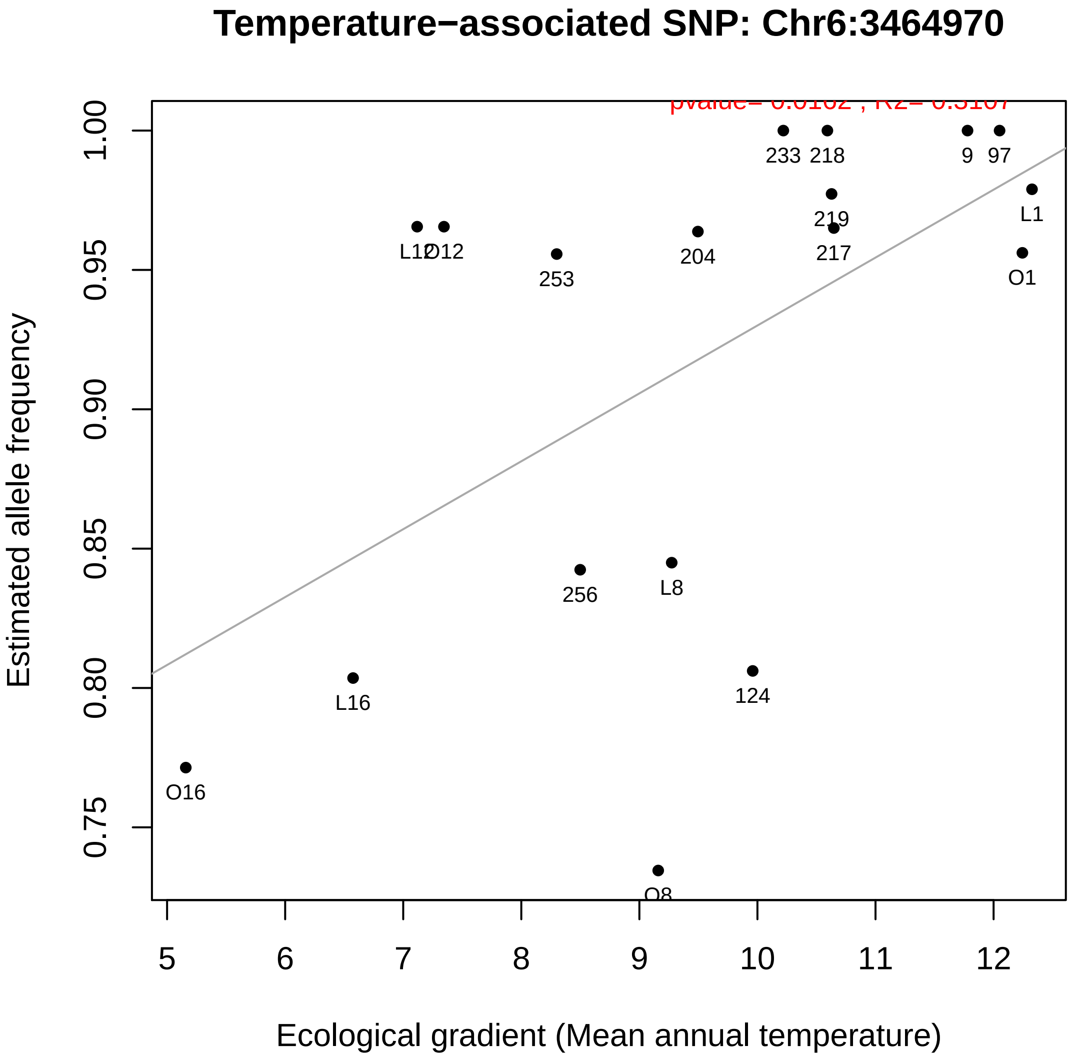

Temperature−associated SNP: Chr6:3464970
Estimated allele frequency
Ecological gradient (Mean annual temperature)
1.00
0.95
0.90
0.85
0.80
0.75
5
6
7
8
9
10
11
12
233
218
9
97
219
217
L1
O1
204
253
256
L8
L16
124
O16

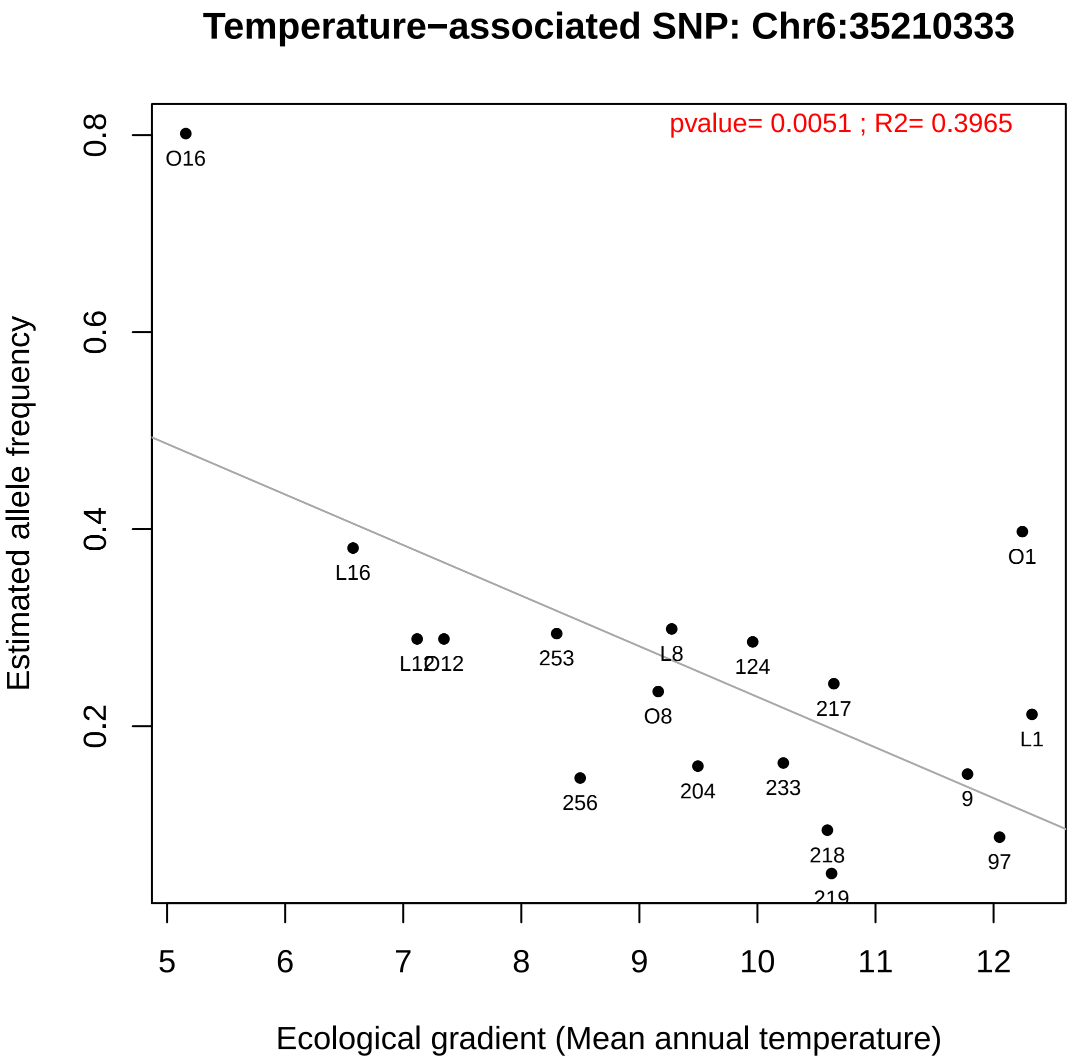

Temperature−associated SNP: Chr6:35210333
pvalue= 0.0051 ; R2= 0.3965
Estimated allele frequency
0.2
0.4
0.6
0.8
Ecological gradient (Mean annual temperature)
5
6
7
8
9
10
11
12
O16
L16
L12
O12
253
256
L8
O8
204
124
233
217
218
219
9
97
O1
L1

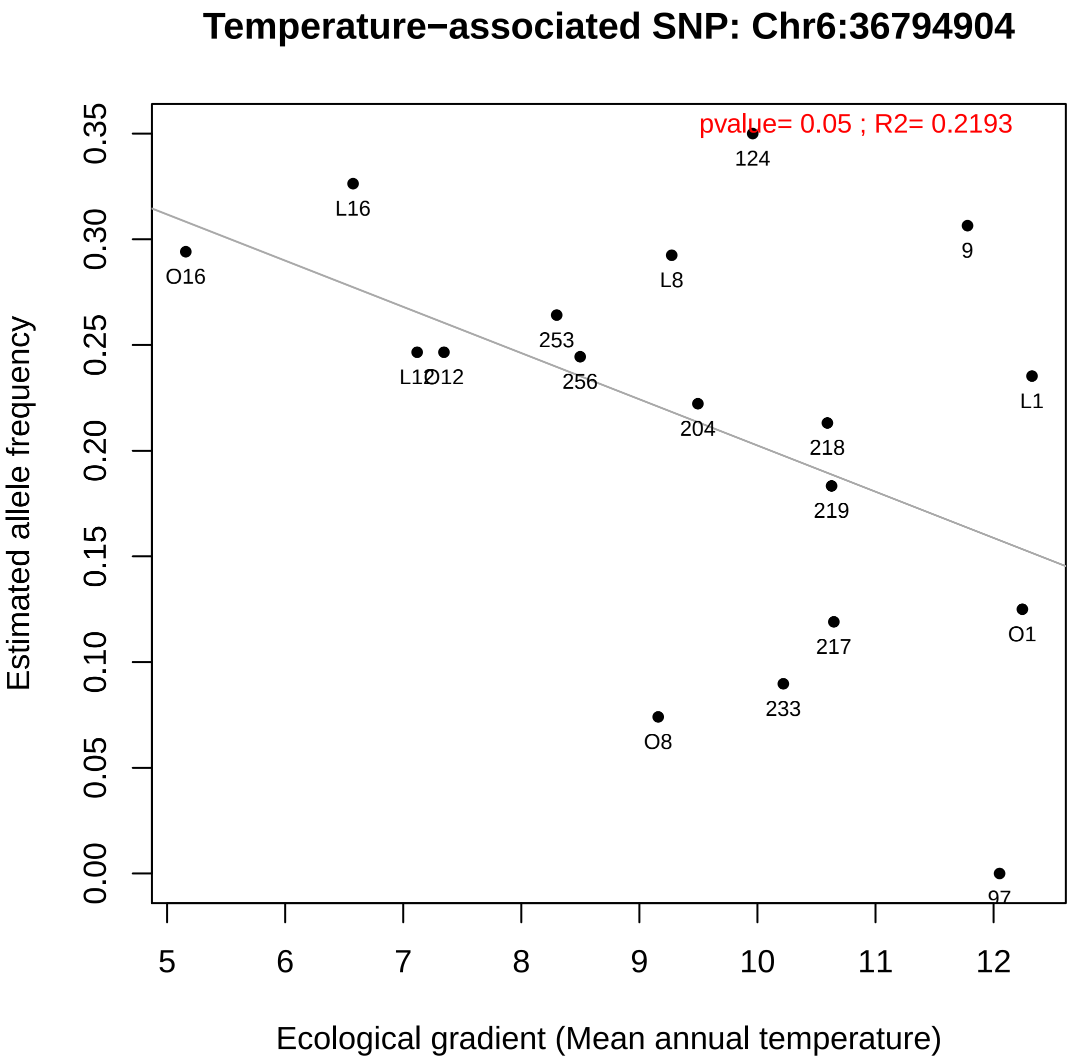

Temperature−associated SNP: Chr6:36794904
pvalue= 0.05 ; R2= 0.2193
Estimated allele frequency
0.00
0.05
0.10
0.15
0.20
0.25
0.30
0.35
5
6
7
8
9
10
11
12
Ecological gradient (Mean annual temperature)
124
L16
O16
9
L8
253
L12
O12
256
L1
204
218
219
O1
217
233
O8
97

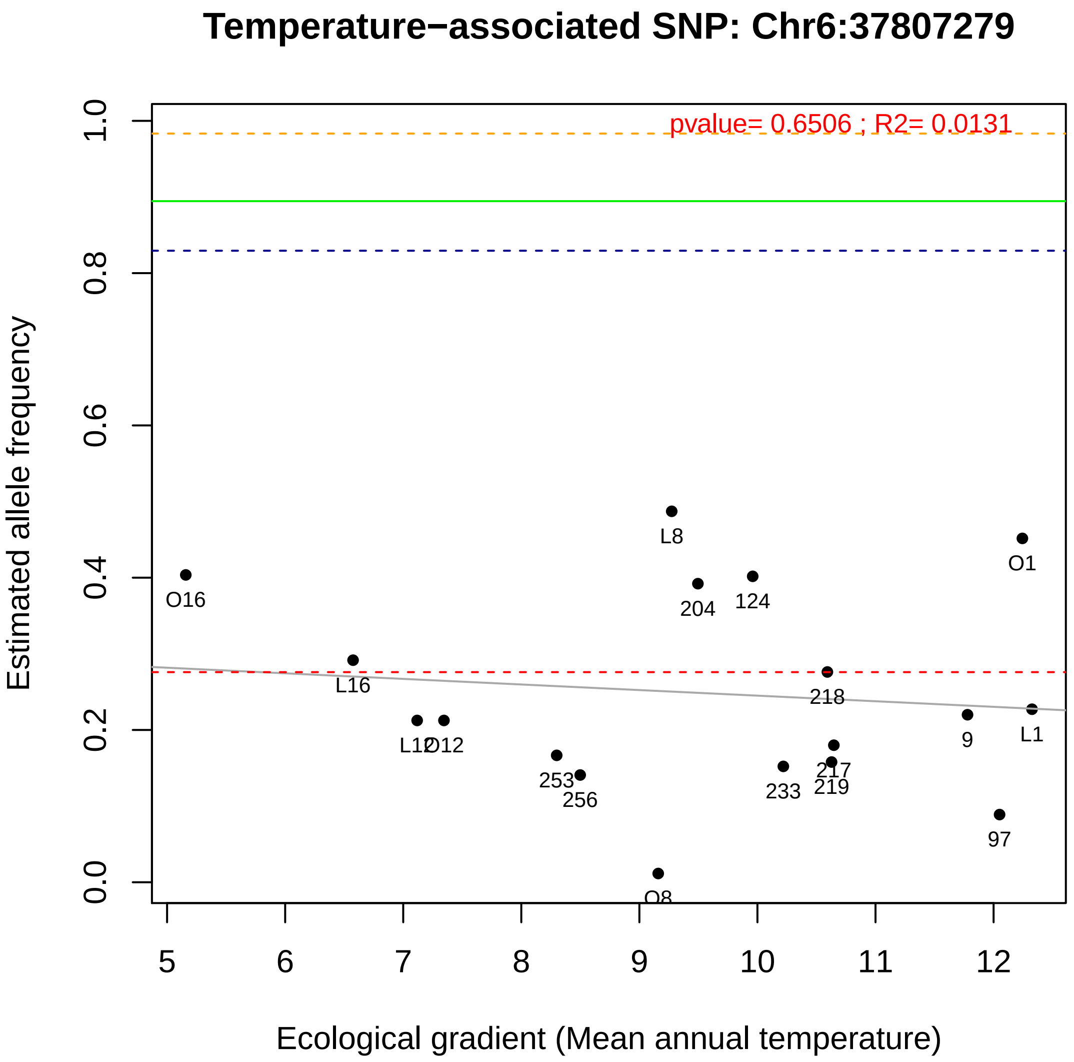

Temperature−associated SNP: Chr6:37807279
pvalue= 0.6506 ; R2= 0.0131
Estimated allele frequency
0.0
0.2
0.4
0.6
0.8
1.0
5
6
7
8
9
10
11
12
Ecological gradient (Mean annual temperature)
O16
L16
L12
O12
253
256
O8
L8
204
124
233
218
217
219
9
O1
L1
97

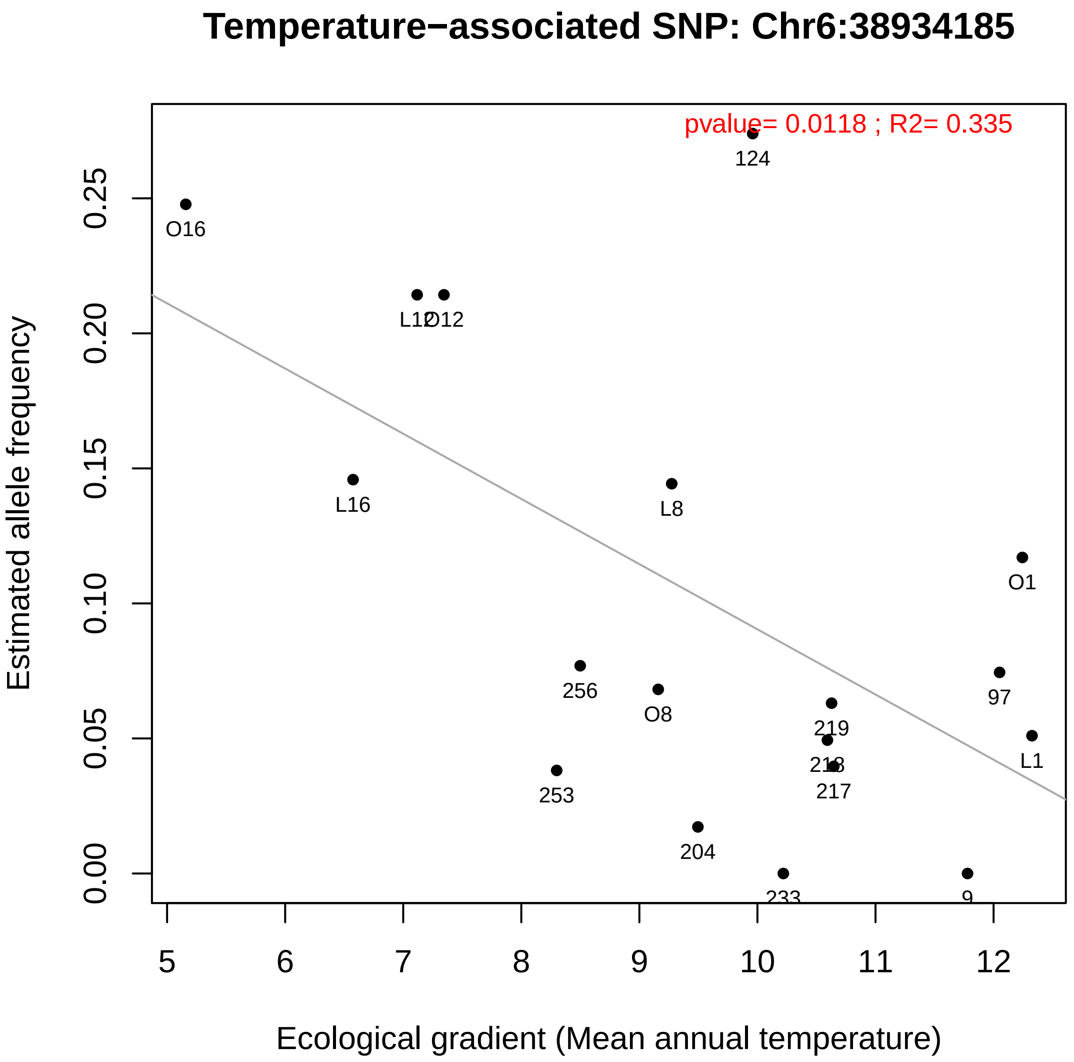
Temperature−associated SNP: Chr6:38934185
pvalue= 0.0118 ; R2= 0.335
Estimated allele frequency
0.00
0.05
0.10
0.15
0.20
0.25
5
6
7
8
9
10
11
12
Ecological gradient (Mean annual temperature)
O16
L12
O12
124
L16
L8
O1
256
O8
97
219
218
217
L1
253
204
233
9

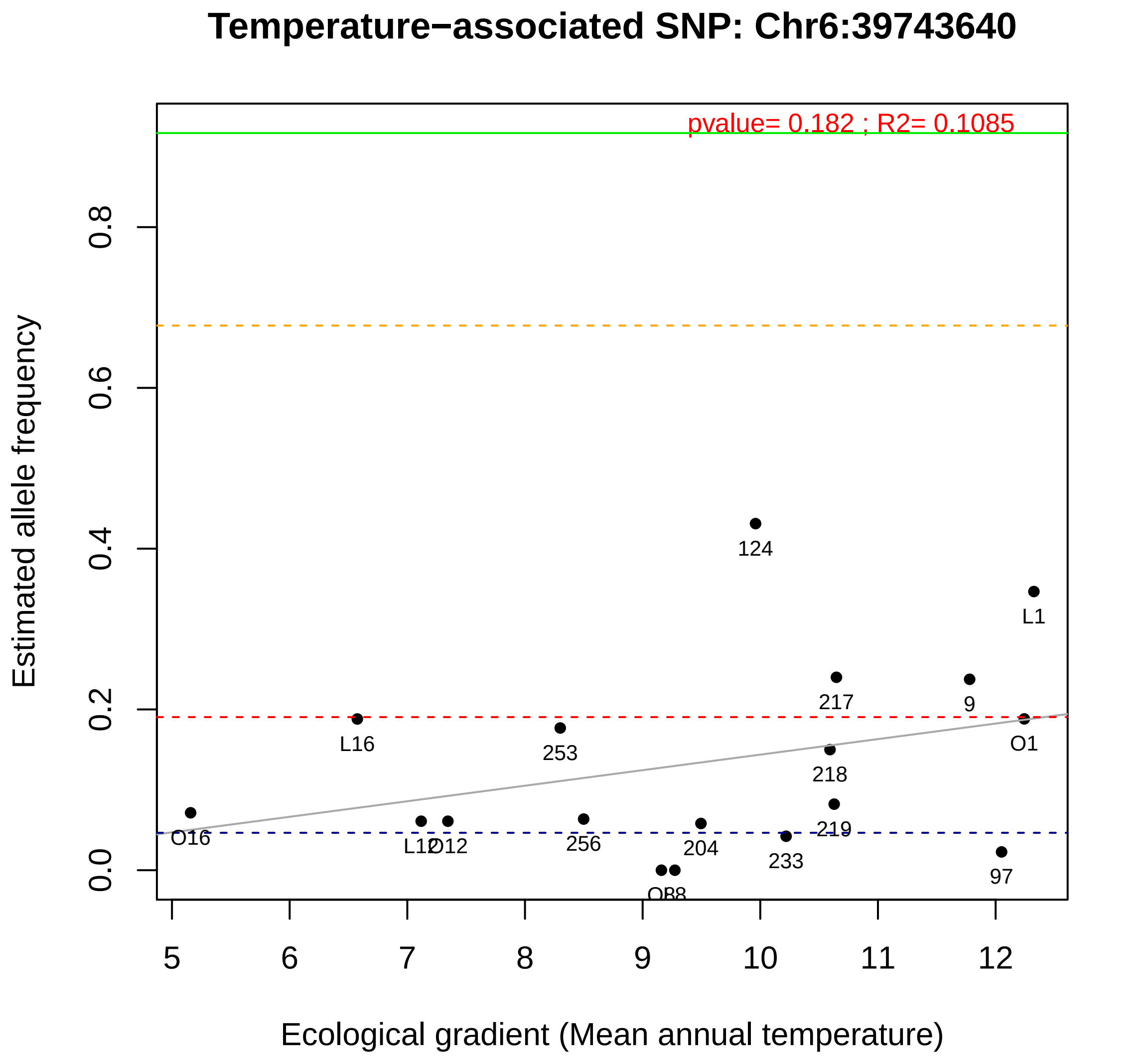

Temperature−associated SNP: Chr6:39743640
pvalue= 0.182 ; R2= 0.1085
Estimated allele frequency
0.0
0.2
0.4
0.6
0.8
5
6
7
8
9
10
11
12
Ecological gradient (Mean annual temperature)
O16
L16
L12
O12
253
256
204
233
124
217
218
219
9
L1
O1
97

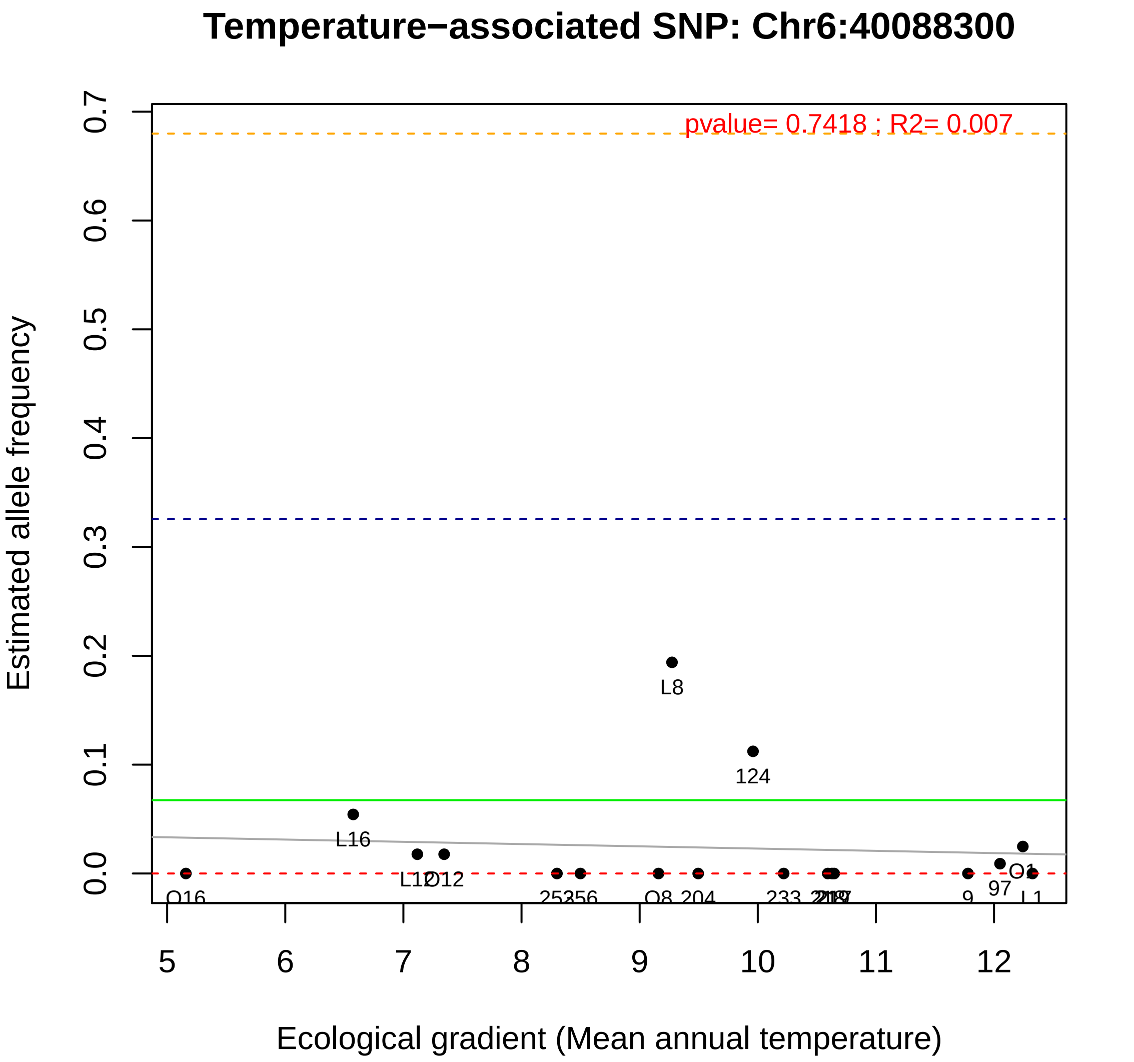

Temperature−associated SNP: Chr6:40088300
pvalue= 0.7418 ; R2= 0.007
Estimated allele frequency
0.0
0.1
0.2
0.3
0.4
0.5
0.6
0.7
5
6
7
8
9
10
11
12
Ecological gradient (Mean annual temperature)
L8
124
L16
L12
O12
O16
O8
204
233
9
97
O1
L1

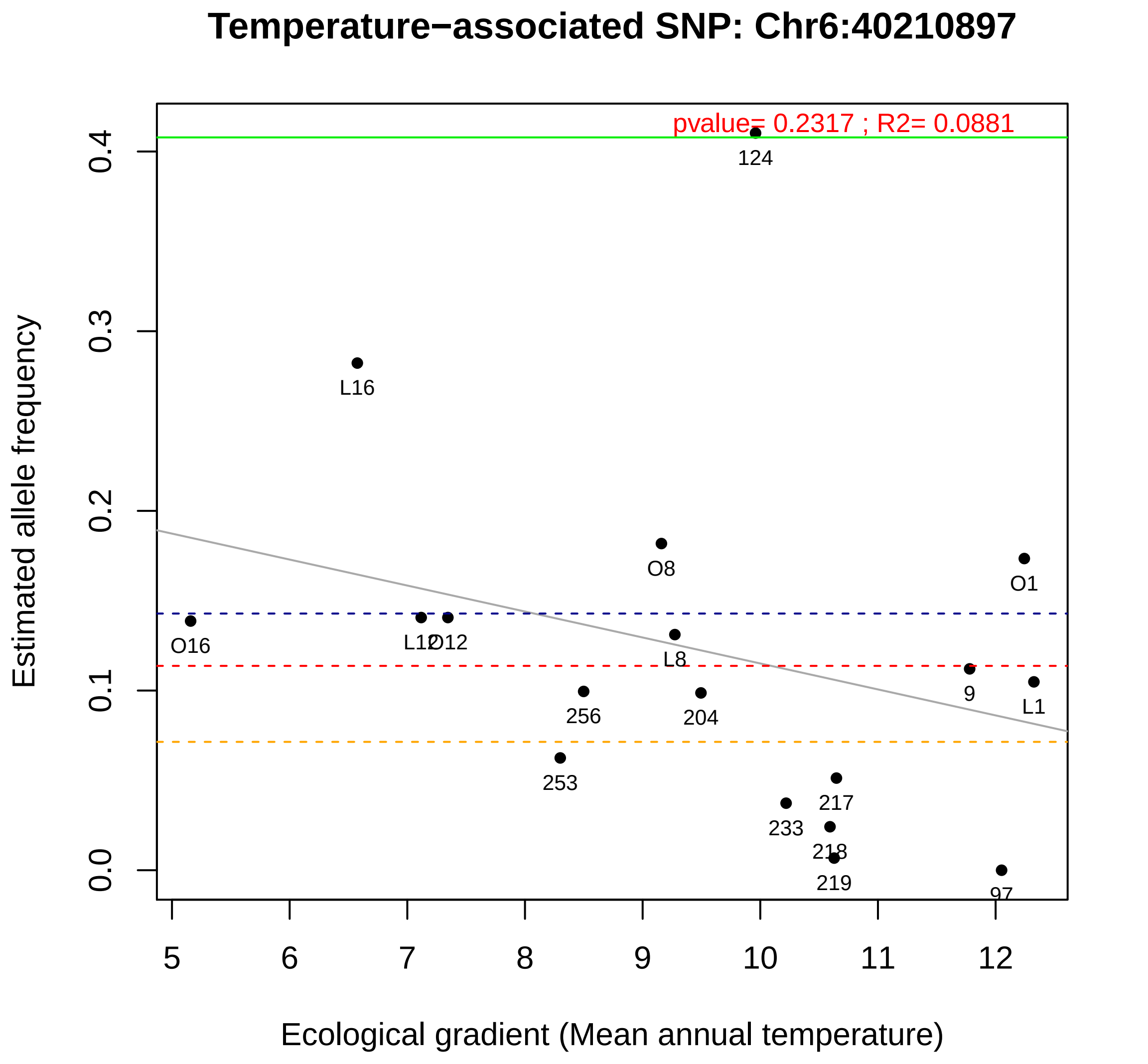

Temperature−associated SNP: Chr6:40210897
pvalue= 0.2317 ; R2= 0.0881
124
L16
O8
O1
O16
L12
O12
L8
9
L1
256
204
253
217
233
218
219
97
0.0
0.1
0.2
0.3
0.4
5
6
7
8
9
10
11
12
Estimated allele frequency
Ecological gradient (Mean annual temperature)

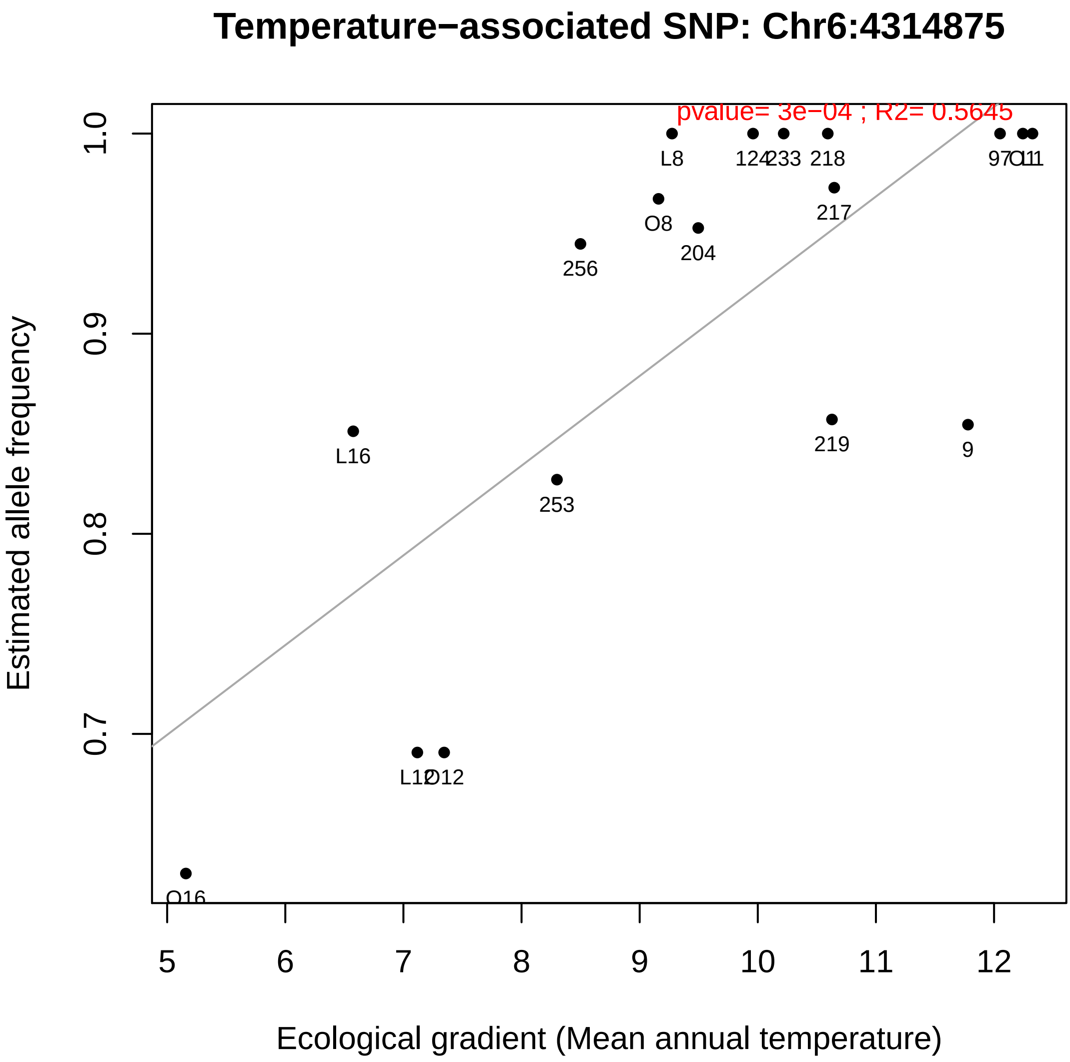

Temperature−associated SNP: Chr6:4314875
pvalue= 3e−04 ; R2= 0.5645
Estimated allele frequency
Ecological gradient (Mean annual temperature)
1.0
0.9
0.8
0.7
5
6
7
8
9
10
11
12
L8
124
233
218
97
O11
O8
217
204
256
L16
253
219
9
L12
O12
O16

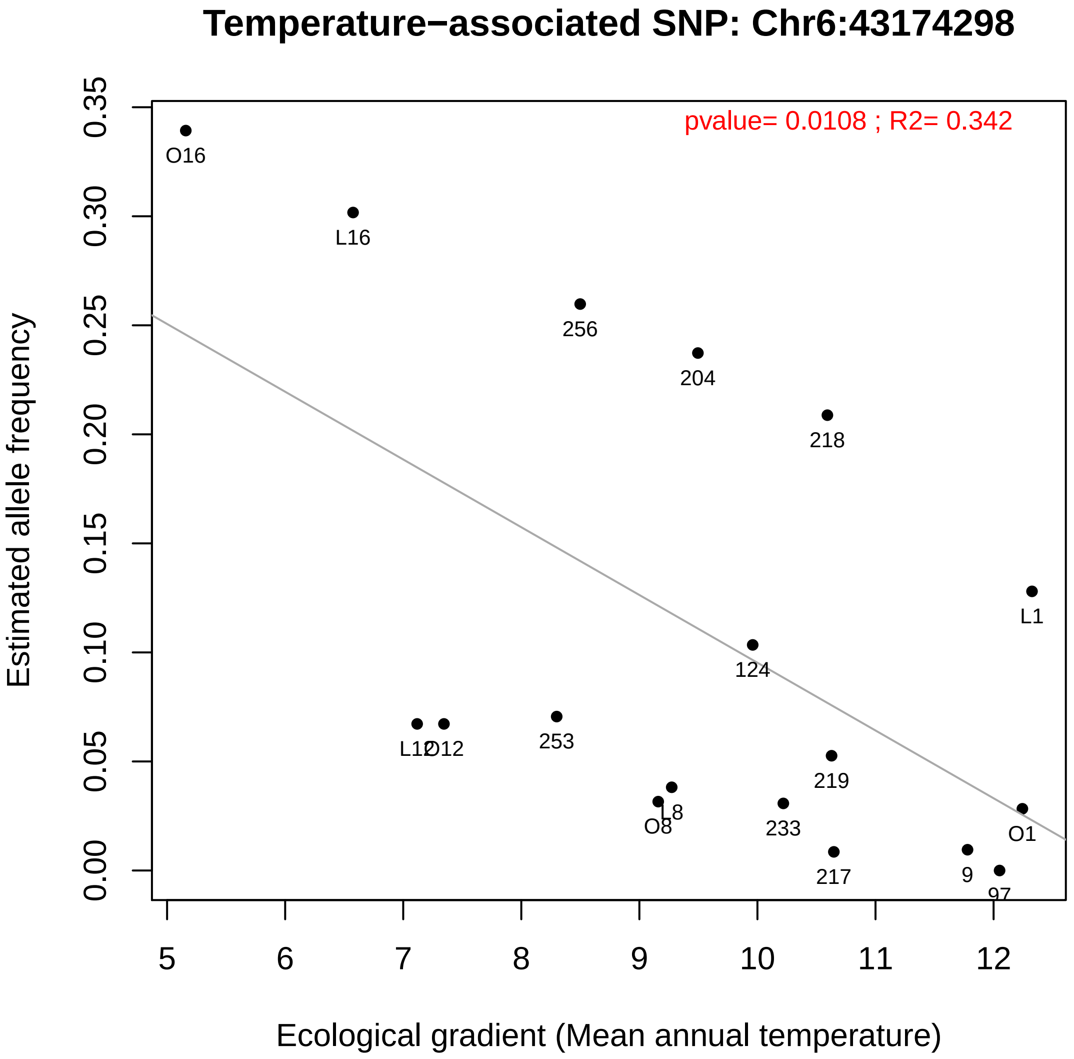
Temperature−associated SNP: Chr6:43174298
pvalue= 0.0108 ; R2= 0.342
Estimated allele frequency
0.00
0.05
0.10
0.15
0.20
0.25
0.30
0.35
Ecological gradient (Mean annual temperature)
5
6
7
8
9
10
11
12
O16
L16
256
204
218
L1
124
L12
O12
253
219
L8
O8
233
O1
217
9
97

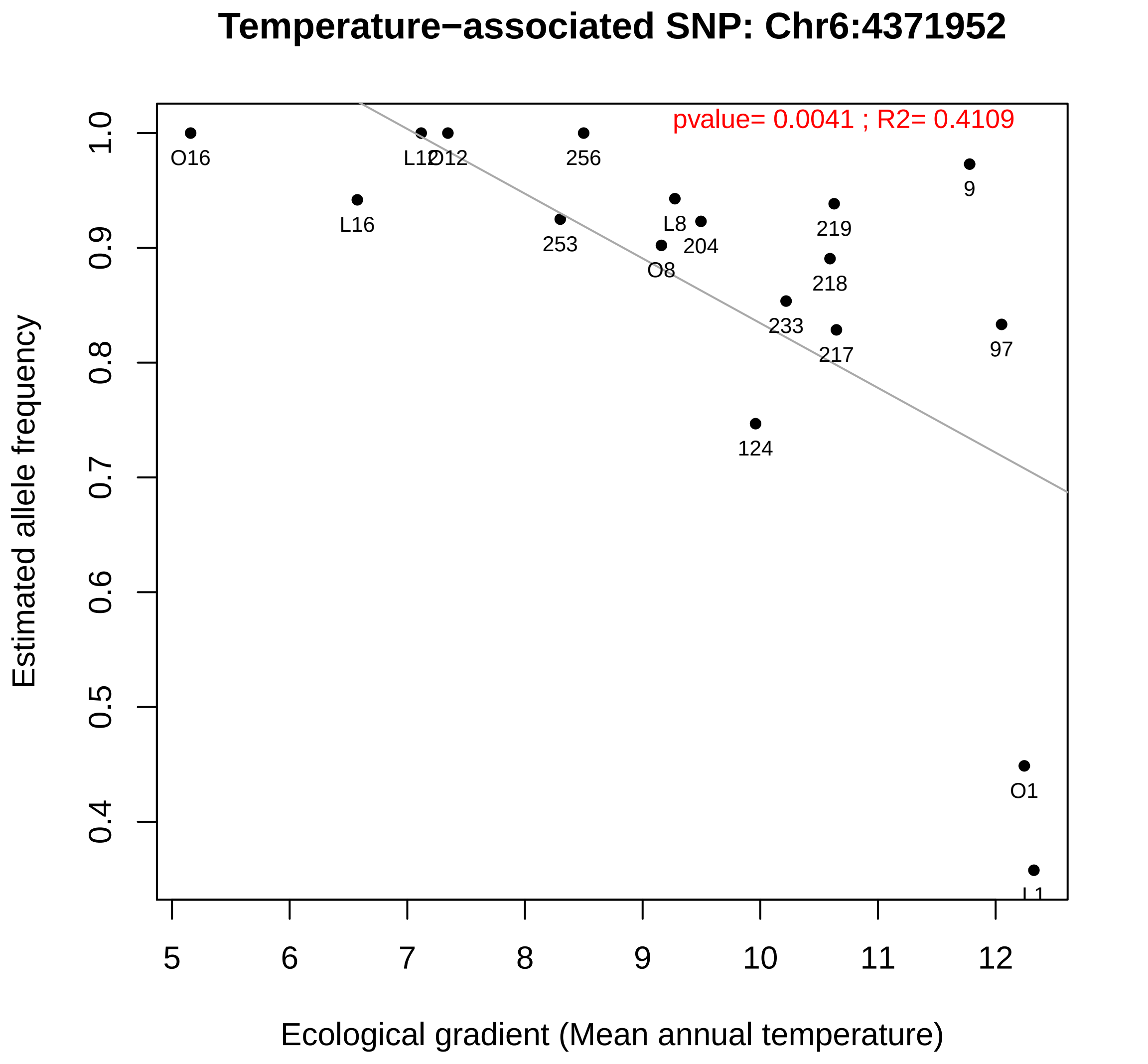
Temperature−associated SNP: Chr6:4371952
pvalue= 0.0041 ; R2= 0.4109
Estimated allele frequency
Ecological gradient (Mean annual temperature)
1.0
0.9
0.8
0.7
0.6
0.5
0.4
5
6
7
8
9
10
11
12
O16
L12
O12
256
9
L16
253
L8
204
219
O8
218
233
217
97
124
O1
L1

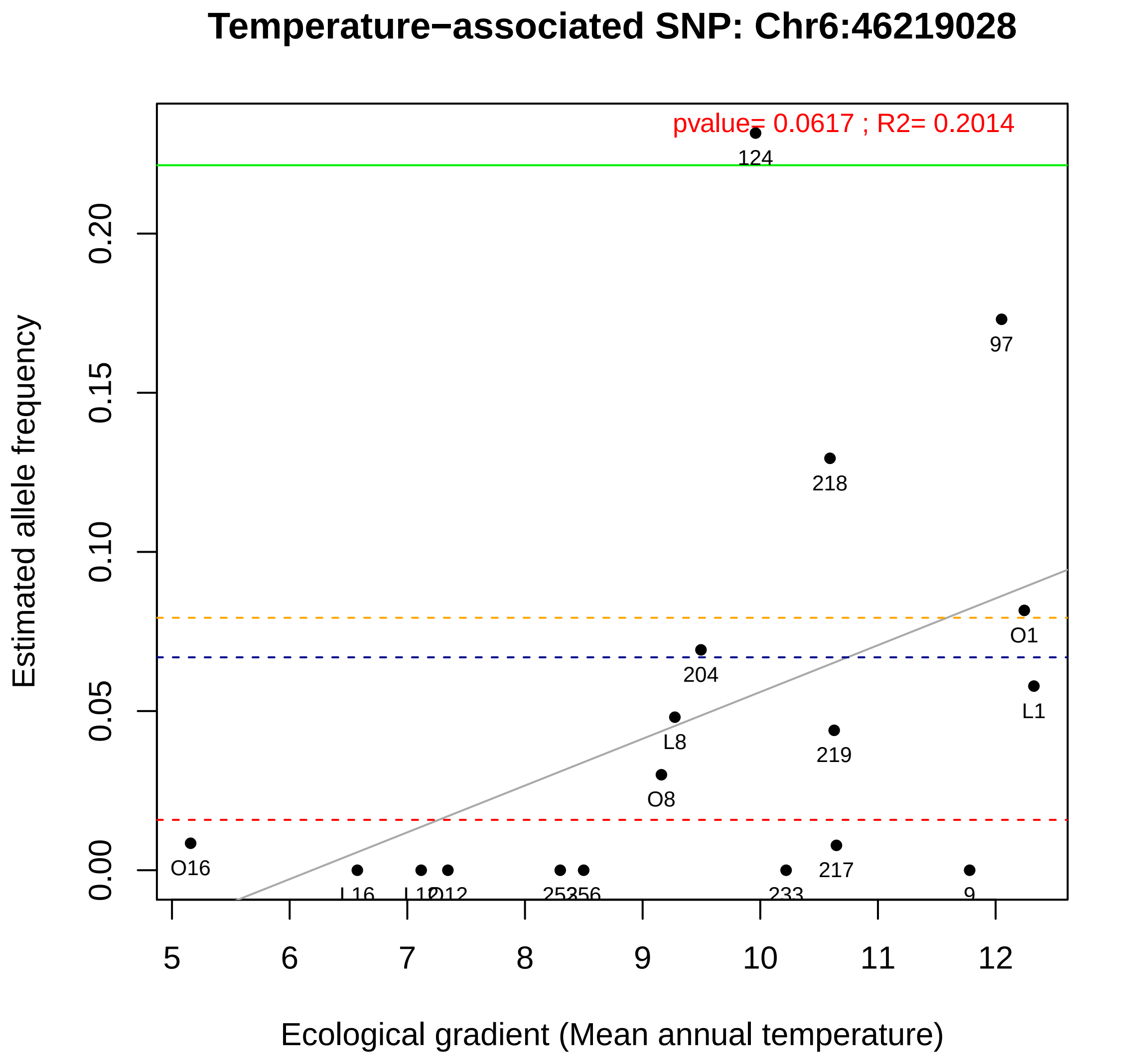

Temperature−associated SNP: Chr6:46219028
pvalue= 0.0617 ; R2= 0.2014
Estimated allele frequency
0.00
0.05
0.10
0.15
0.20
5
6
7
8
9
10
11
12
Ecological gradient (Mean annual temperature)
124
97
218
O1
204
L1
L8
219
O8
O16
217

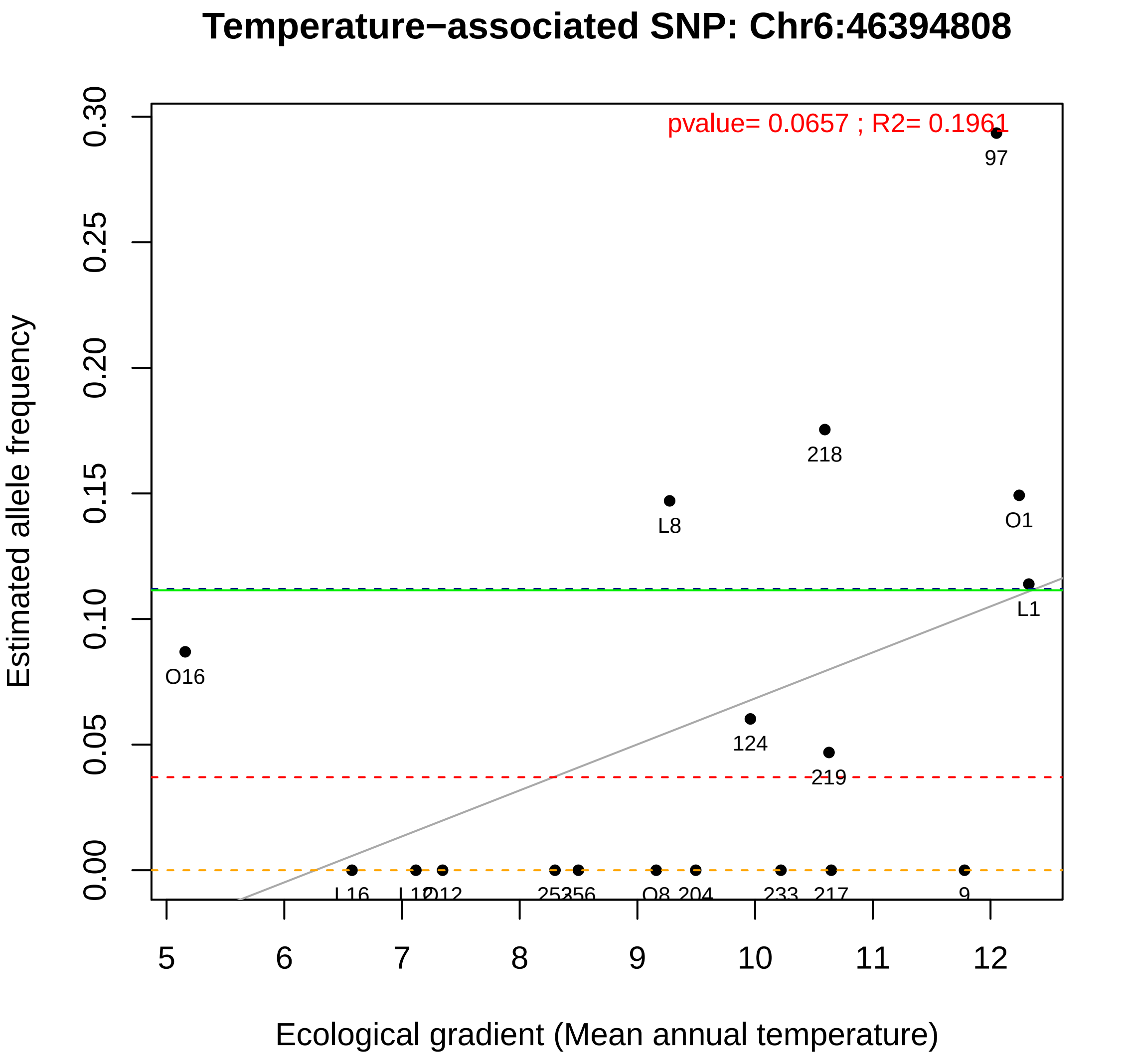
Temperature−associated SNP: Chr6:46394808
pvalue= 0.0657 ; R2= 0.1961
Estimated allele frequency
Ecological gradient (Mean annual temperature)
0.00
0.05
0.10
0.15
0.20
0.25
0.30
5
6
7
8
9
10
11
12
97
218
L8
O1
L1
O16
124
219
L16
O12
O8
204
233
217
9

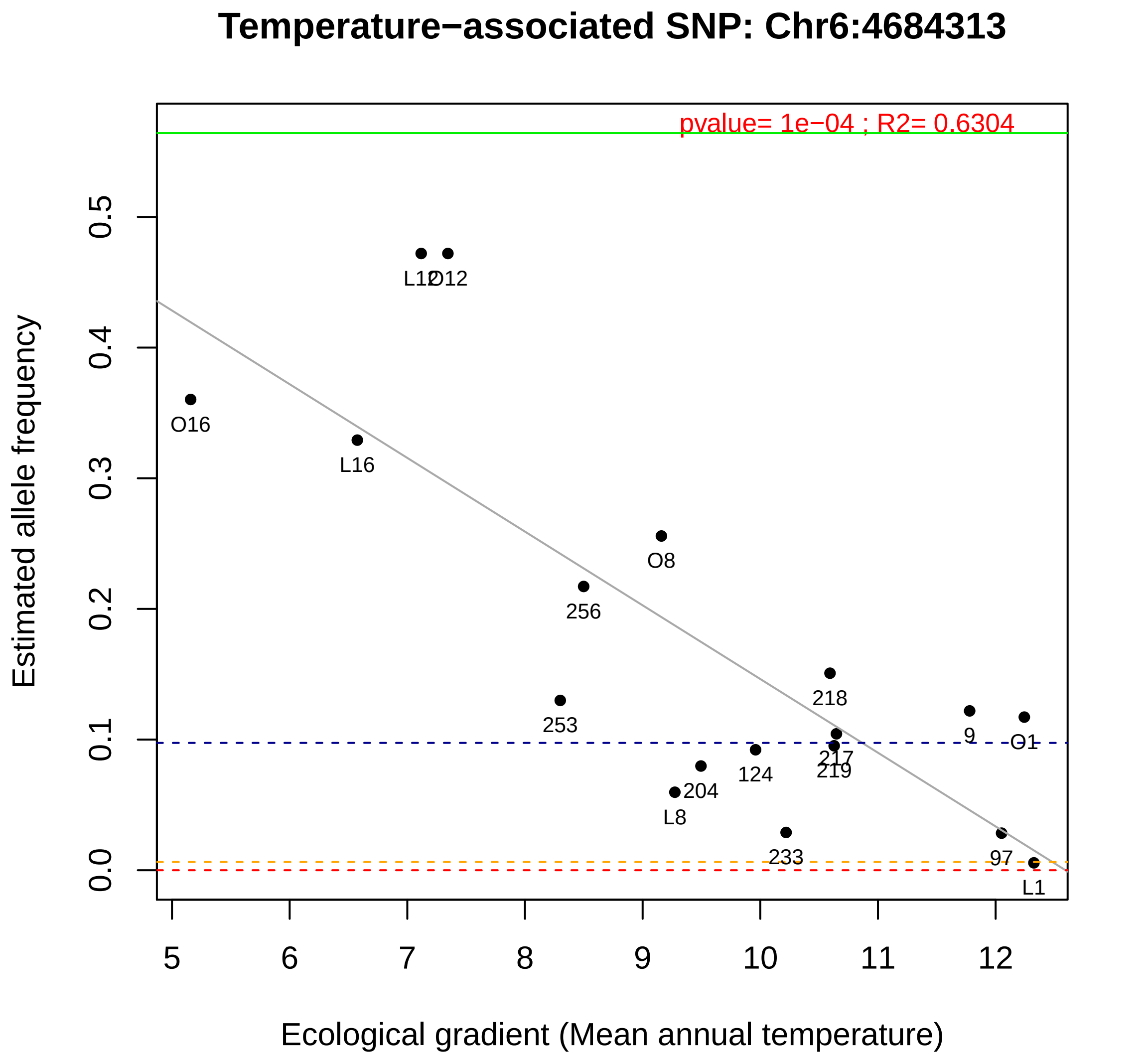

Temperature−associated SNP: Chr6:4684313
pvalue= 1e−04 ; R2= 0.6304
Estimated allele frequency
0.0
0.1
0.2
0.3
0.4
0.5
5
6
7
8
9
10
11
12
Ecological gradient (Mean annual temperature)
L12
O12
O16
L16
O8
256
218
253
9
O1
217
219
124
204
L8
233
97
L1

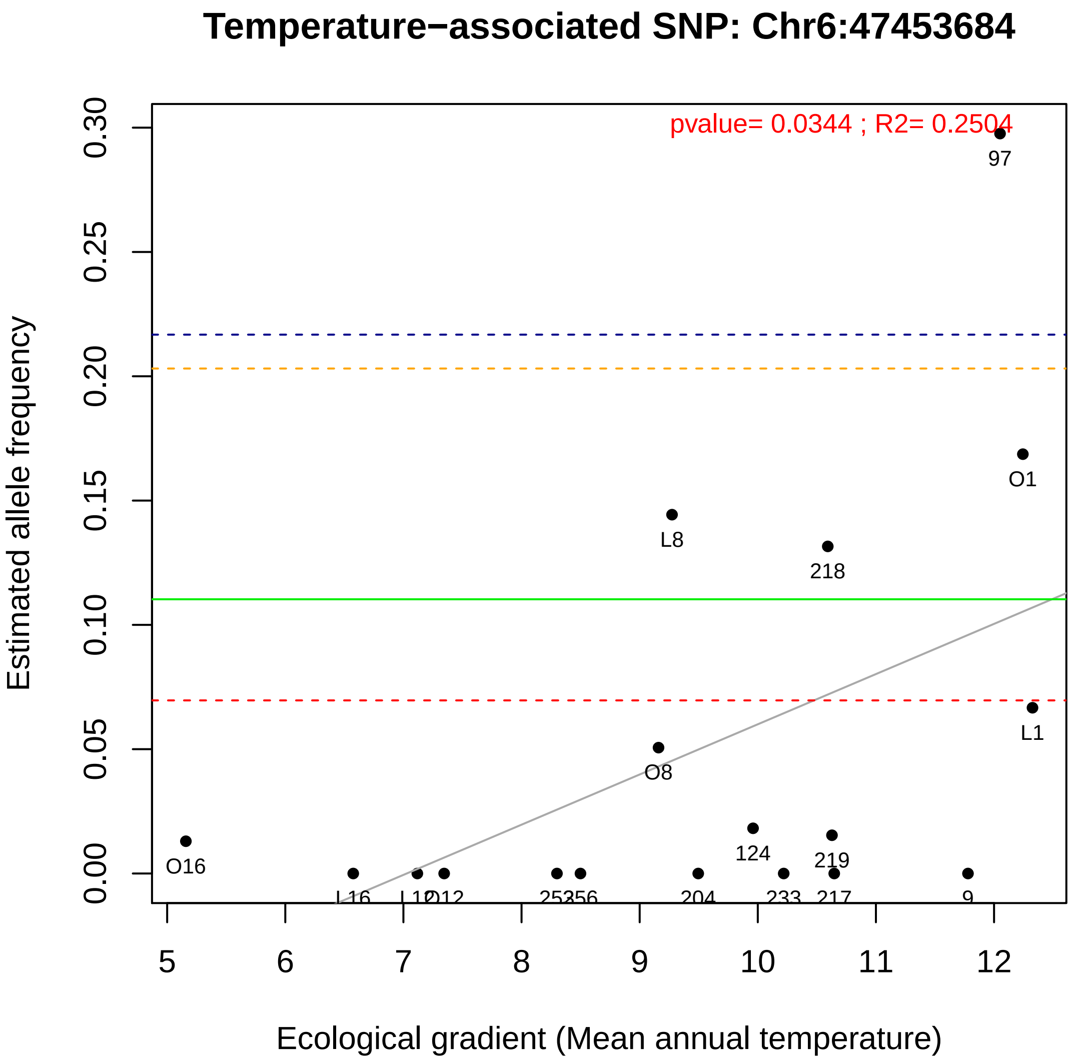

Temperature−associated SNP: Chr6:47453684
pvalue= 0.0344 ; R2= 0.2504
Estimated allele frequency
0.00
0.05
0.10
0.15
0.20
0.25
0.30
97
O1
L8
218
L1
O8
124
219
O16
L16
O12
204
233
217
9
5
6
7
8
9
10
11
12
Ecological gradient (Mean annual temperature)

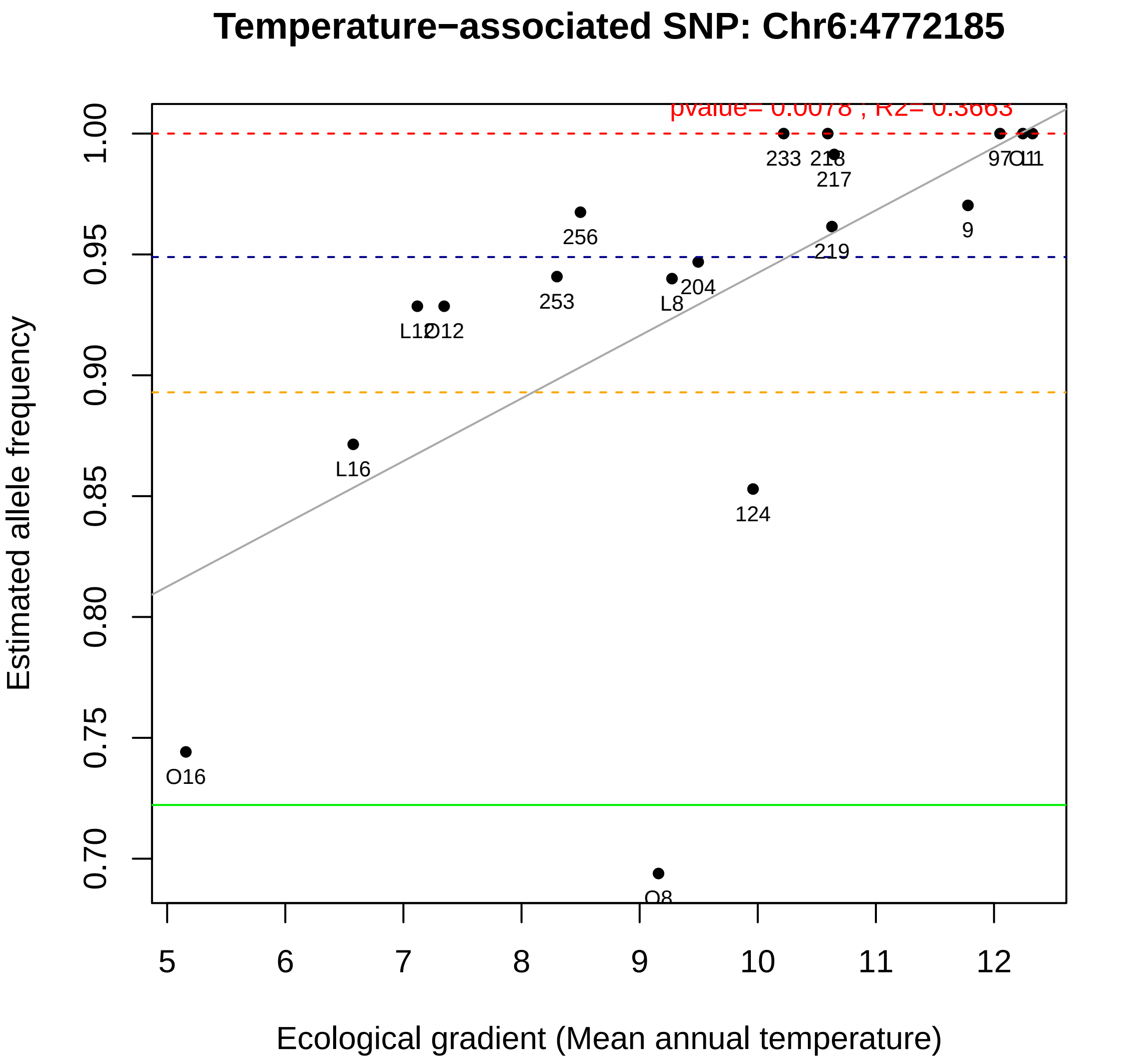
Temperature−associated SNP: Chr6:4772185
pvalue= 0.0078 ; R2= 0.3663
Estimated allele frequency
Ecological gradient (Mean annual temperature)
1.00
0.95
0.90
0.85
0.80
0.75
0.70
5
6
7
8
9
10
11
12
233
218
217
97
O11
256
9
219
253
L8
204
L12
O12
L16
124
O16
O8

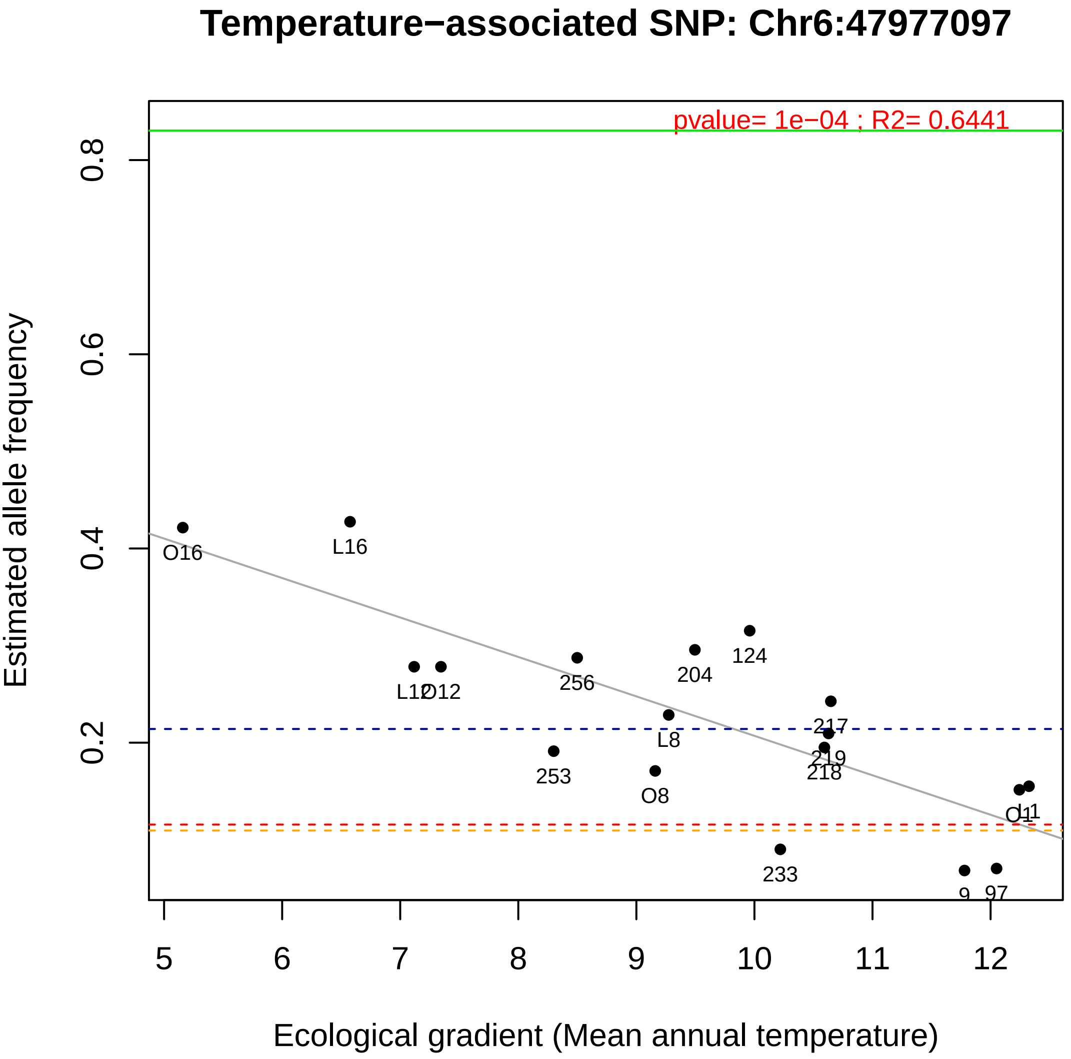
Temperature−associated SNP: Chr6:47977097
pvalue= 1e−04 ; R2= 0.6441
Estimated allele frequency
0.2
0.4
0.6
0.8
Ecological gradient (Mean annual temperature)
5
6
7
8
9
10
11
12
O16
L16
L12
O12
256
253
204
124
L8
O8
217
219
218
233
O1
L1
9
97

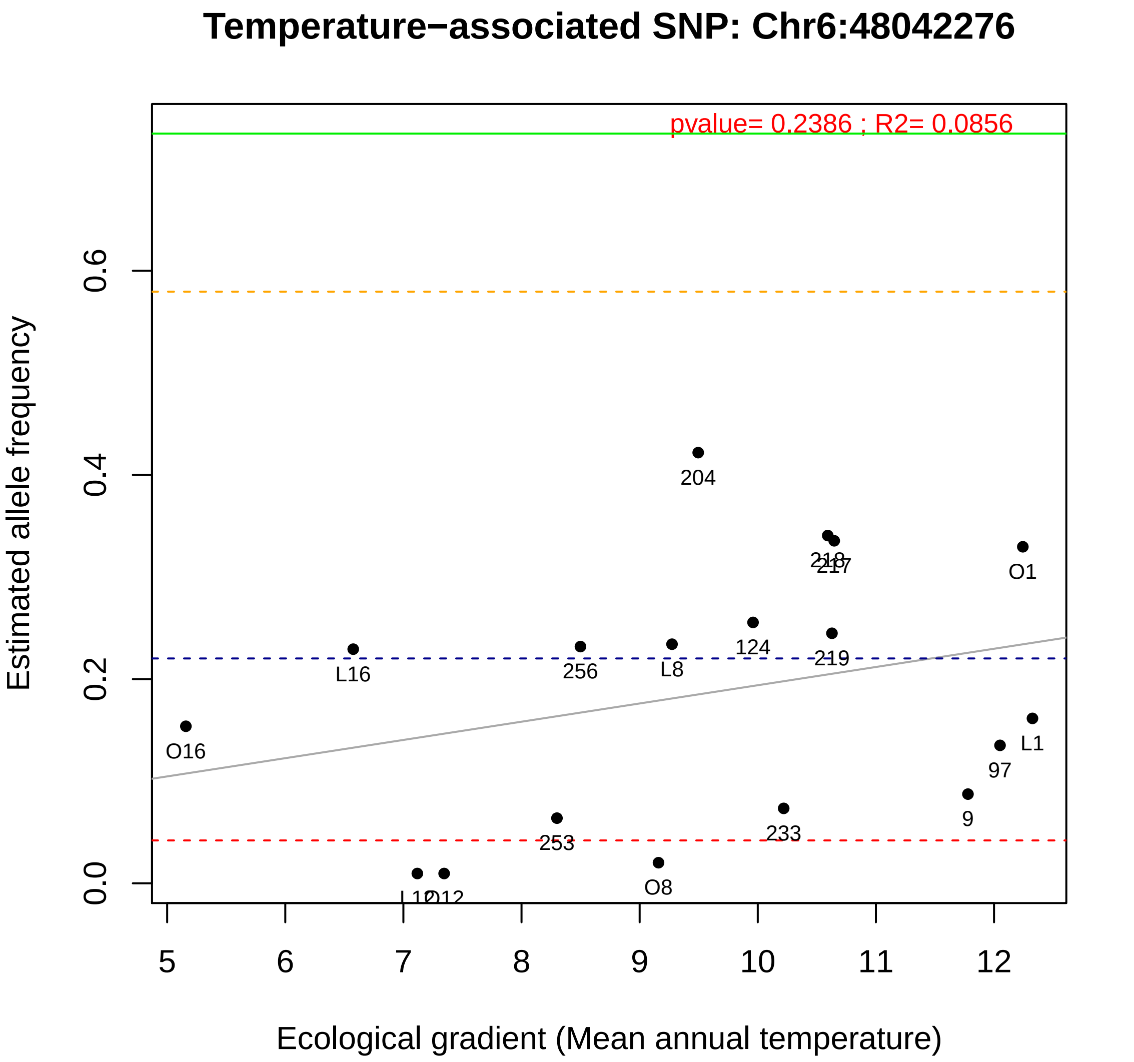
Temperature−associated SNP: Chr6:48042276
pvalue= 0.2386 ; R2= 0.0856
Estimated allele frequency
0.0
0.2
0.4
0.6
Ecological gradient (Mean annual temperature)
5
6
7
8
9
10
11
12
204
218
217
O1
124
219
L16
256
L8
O16
L1
97
9
233
253
O8
O12

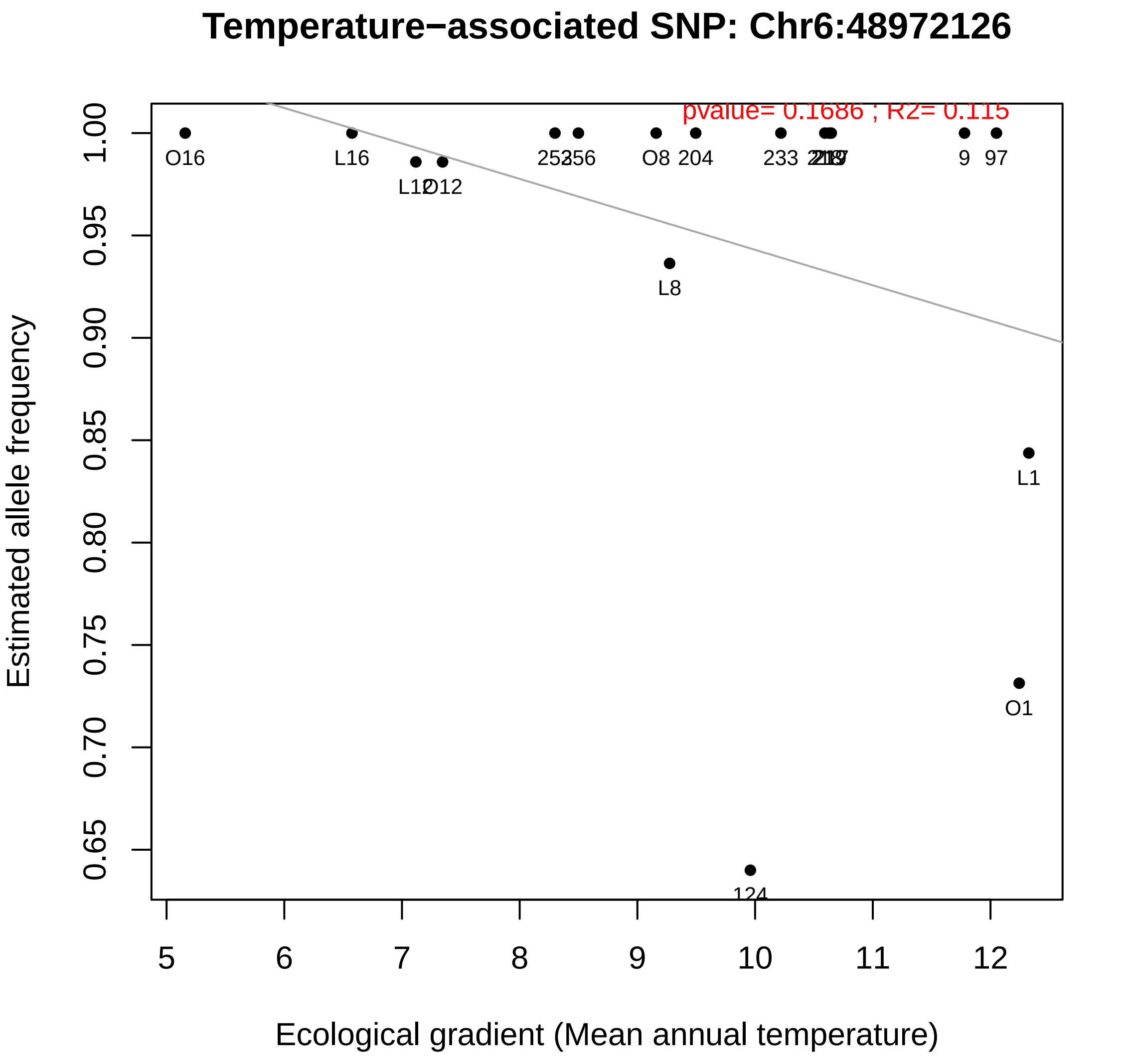
Temperature−associated SNP: Chr6:48972126
pvalue= 0.1686 ; R2= 0.115
Estimated allele frequency
Ecological gradient (Mean annual temperature)
0.65
0.70
0.75
0.80
0.85
0.90
0.95
1.00
5
6
7
8
9
10
11
12
O16
L16
L12
O12
O8
204
233
9
97
L8
L1
O1
124

### Temperature−associated SNP: Chr6:49019626

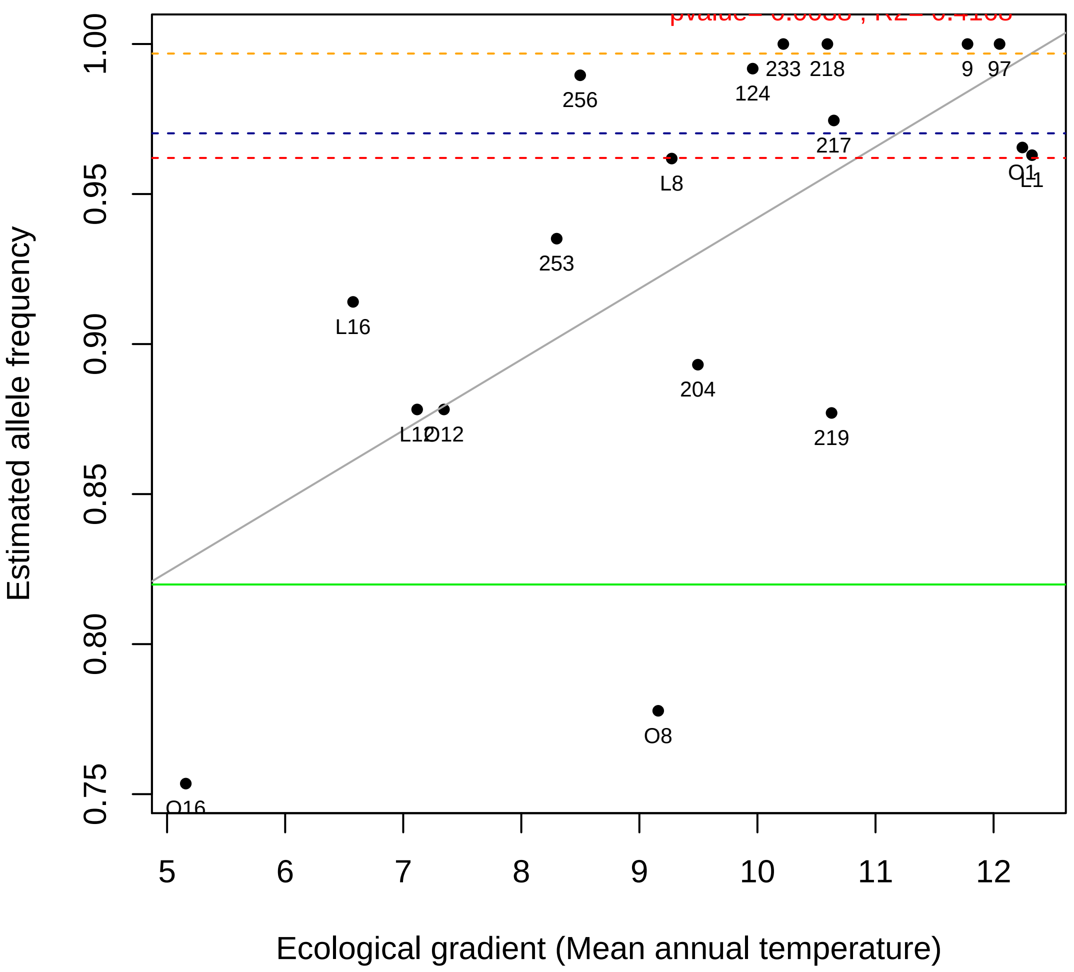

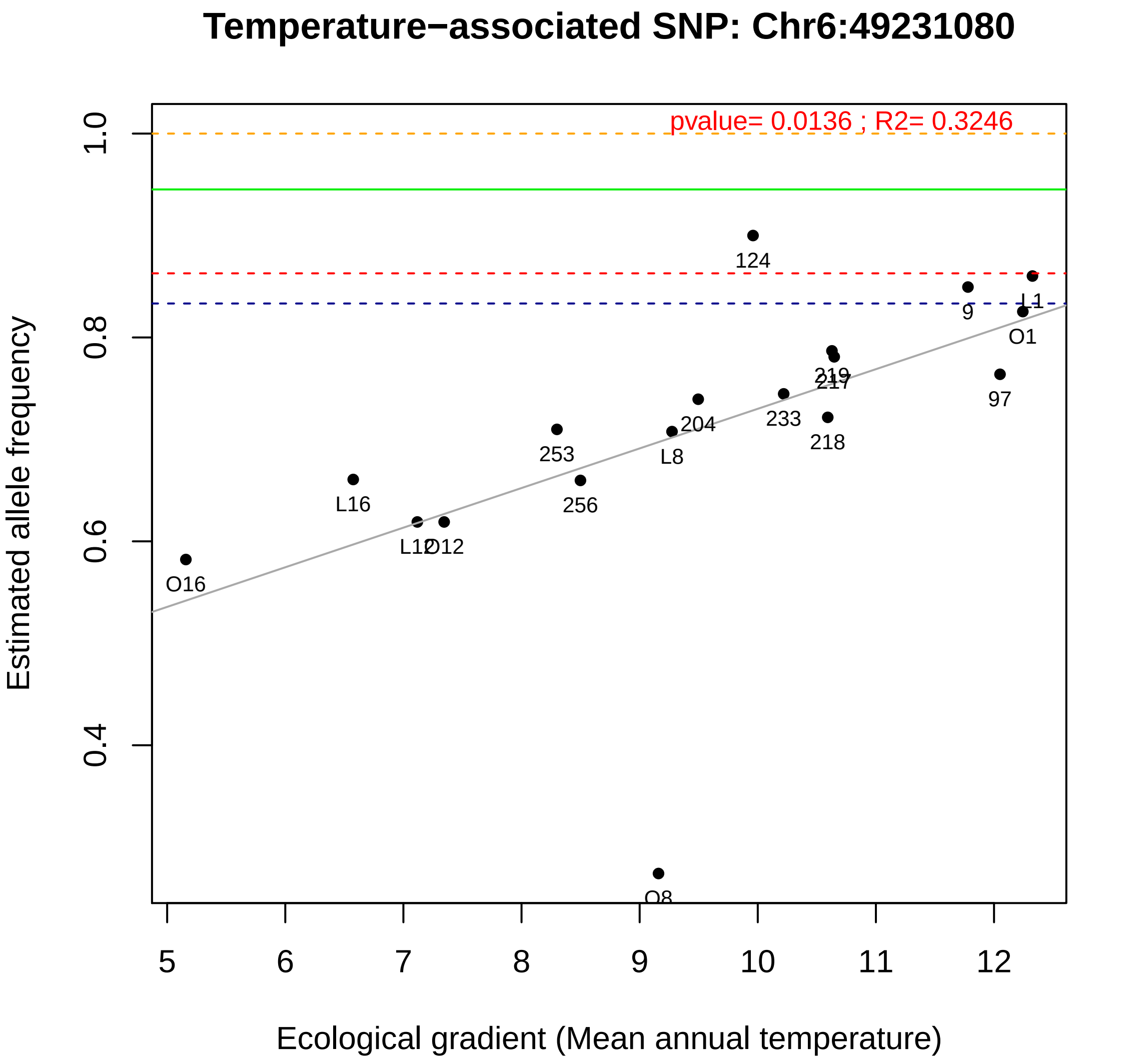

Temperature−associated SNP: Chr6:49231080
pvalue= 0.0136 ; R2= 0.3246
Estimated allele frequency
1.0
0.8
0.6
0.4
124
9
L1
O1
219
217
97
233
204
218
253
L8
L16
256
L12
O12
O16
O8
5
6
7
8
9
10
11
12
Ecological gradient (Mean annual temperature)

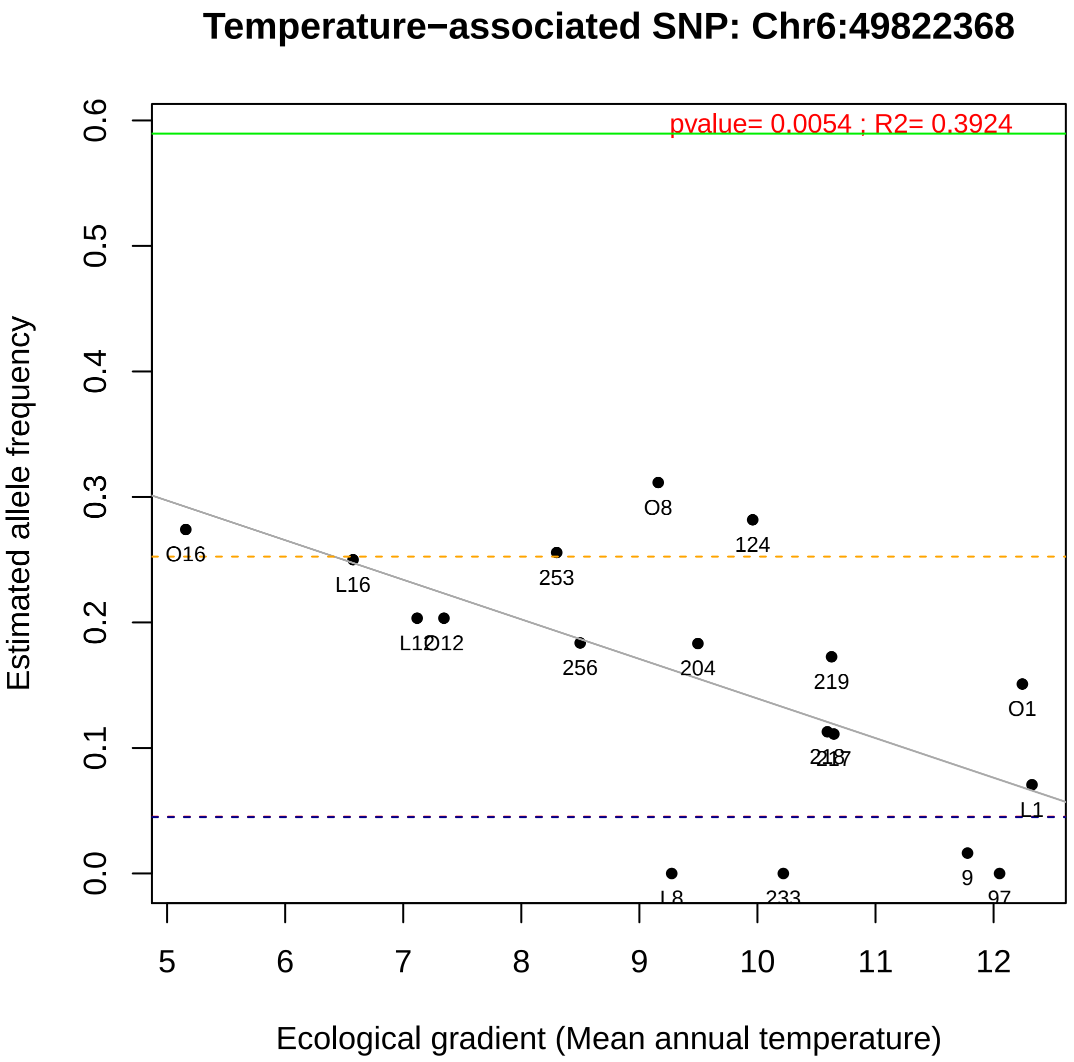

Temperature−associated SNP: Chr6:49822368
pvalue= 0.0054 ; R2= 0.3924
Estimated allele frequency
0.0
0.1
0.2
0.3
0.4
0.5
0.6
O16
L16
L12
O12
253
256
O8
204
124
219
218
217
O1
L1
L8
233
9
97
5
6
7
8
9
10
11
12
Ecological gradient (Mean annual temperature)

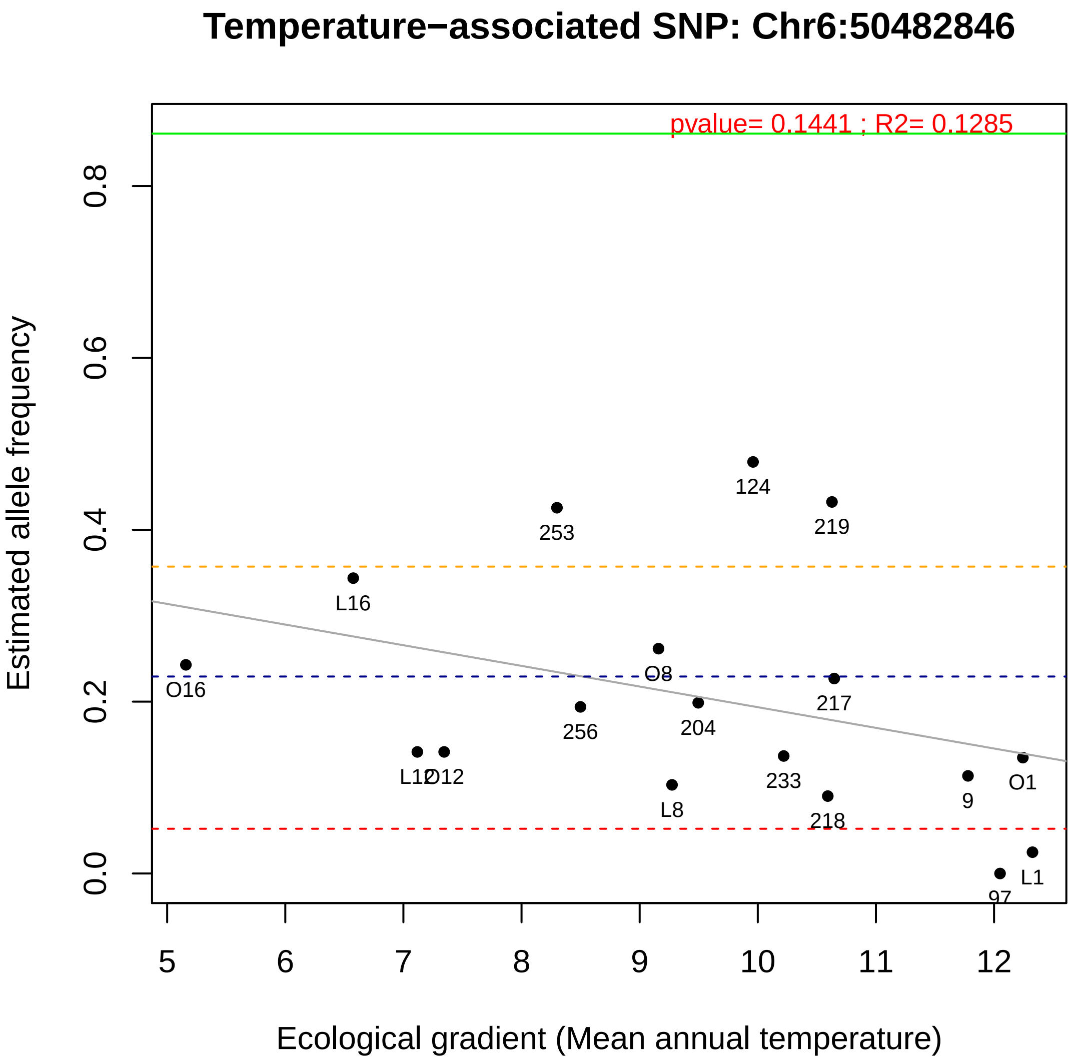

Temperature−associated SNP: Chr6:50482846
pvalue= 0.1441 ; R2= 0.1285
Estimated allele frequency
0.0
0.2
0.4
0.6
0.8
5
6
7
8
9
10
11
12
Ecological gradient (Mean annual temperature)
O16
L16
L12
O12
253
256
O8
L8
204
124
233
219
217
218
9
O1
97
L1

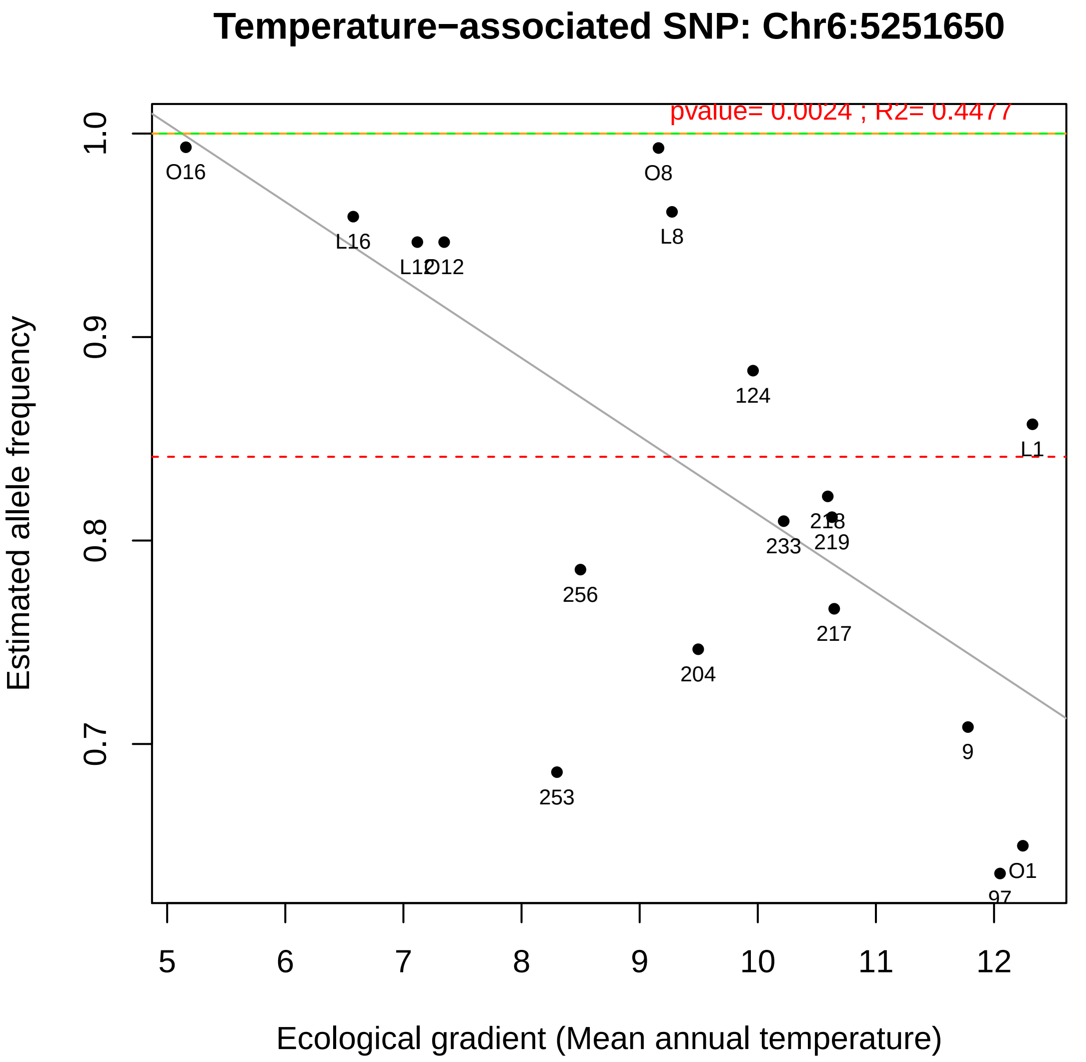

Temperature−associated SNP: Chr6:5251650
pvalue= 0.0024 ; R2= 0.4477
Estimated allele frequency
1.0
0.9
0.8
0.7
O16
O8
L16
L8
L12
O12
124
L1
218
219
233
256
217
204
9
253
O1
97
5
6
7
8
9
10
11
12
Ecological gradient (Mean annual temperature)

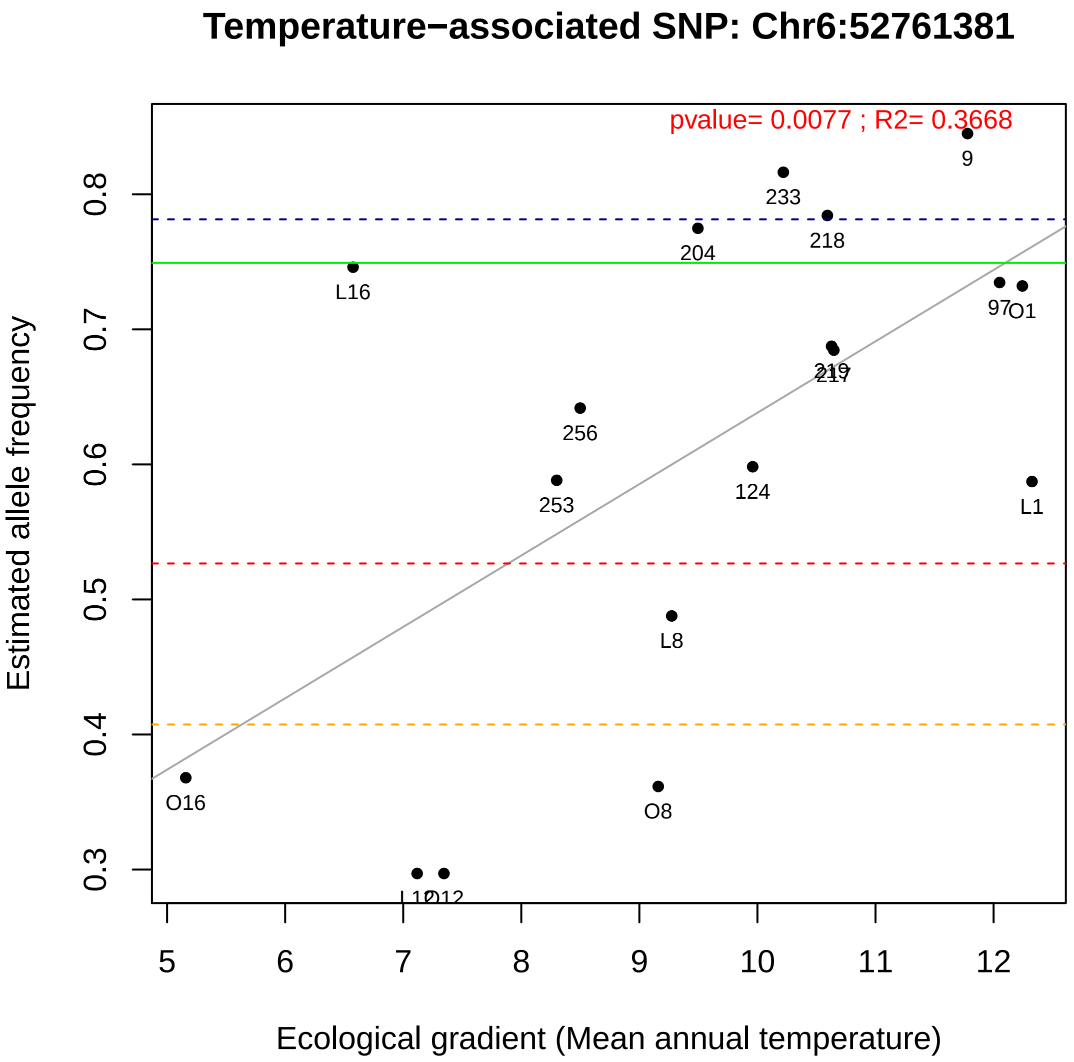
Temperature−associated SNP: Chr6:52761381
pvalue= 0.0077 ; R2= 0.3668
Estimated allele frequency
0.3
0.4
0.5
0.6
0.7
0.8
Ecological gradient (Mean annual temperature)
5
6
7
8
9
10
11
12
9
233
218
204
L16
97
O1
219
217
256
253
124
L1
L8
O16
O8
O12

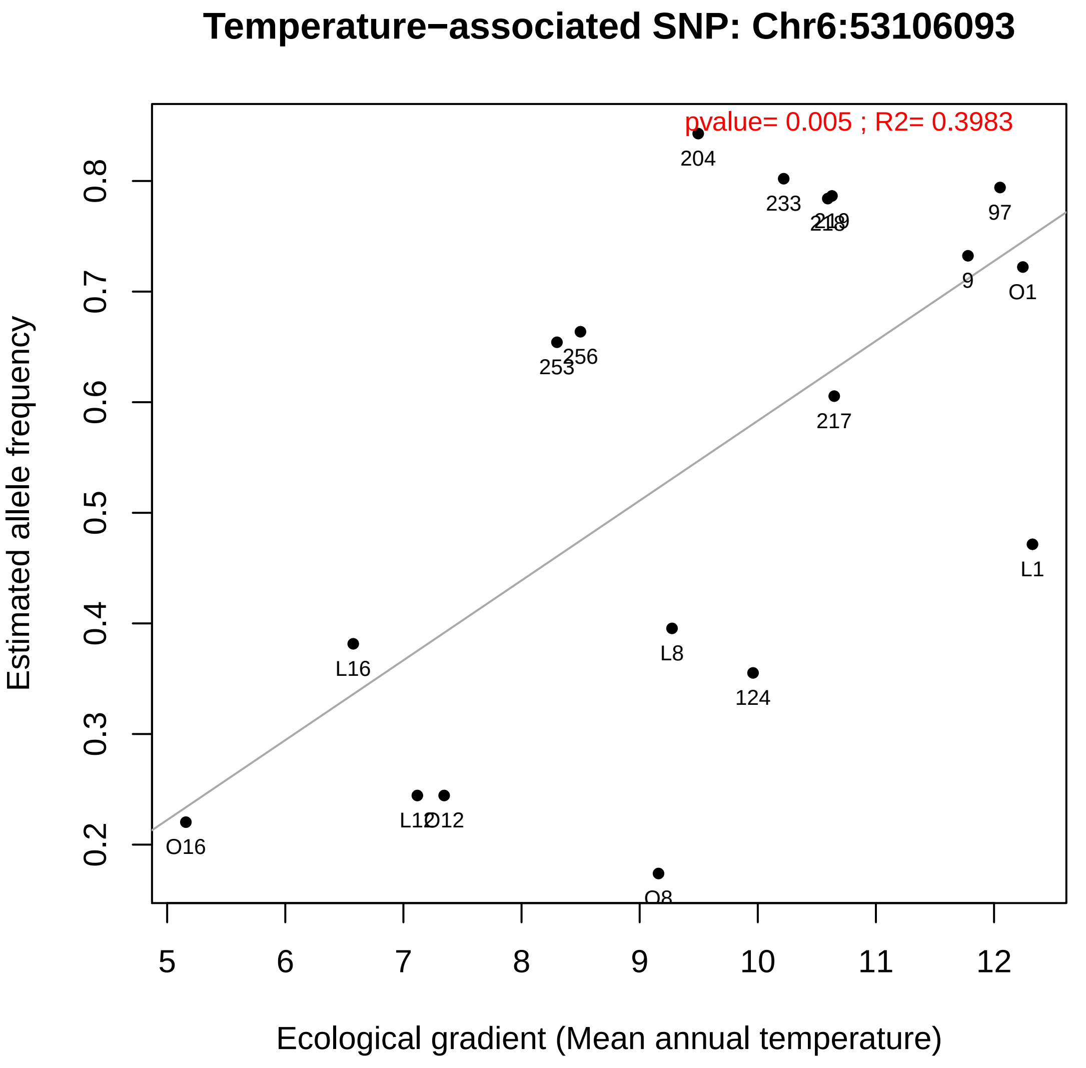
Temperature−associated SNP: Chr6:53106093
pvalue= 0.005 ; R2= 0.3983
Estimated allele frequency
Ecological gradient (Mean annual temperature)
0.2
0.3
0.4
0.5
0.6
0.7
0.8
5
6
7
8
9
10
11
12
204
233
219
218
97
9
O1
253
256
217
L1
L16
L8
124
L12
O12
O16
O8

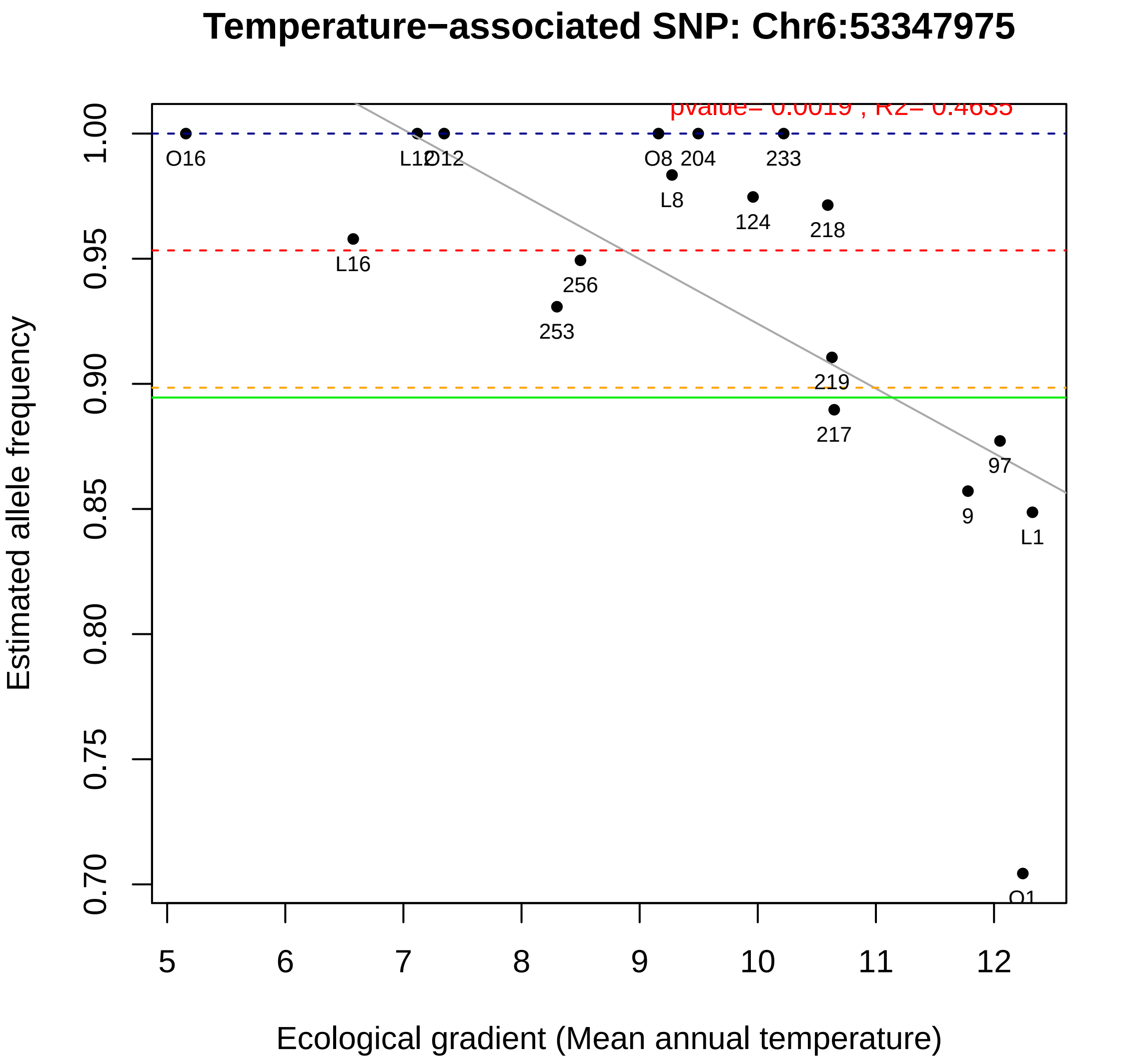

Temperature−associated SNP: Chr6:53347975
pvalue= 0.0019 , R2= 0.4635
Estimated allele frequency
Ecological gradient (Mean annual temperature)
1.00
0.95
0.90
0.85
0.80
0.75
0.70
5
6
7
8
9
10
11
12
O16
L12
O12
O8
204
233
L8
124
218
L16
256
253
219
217
97
9
L1
O1

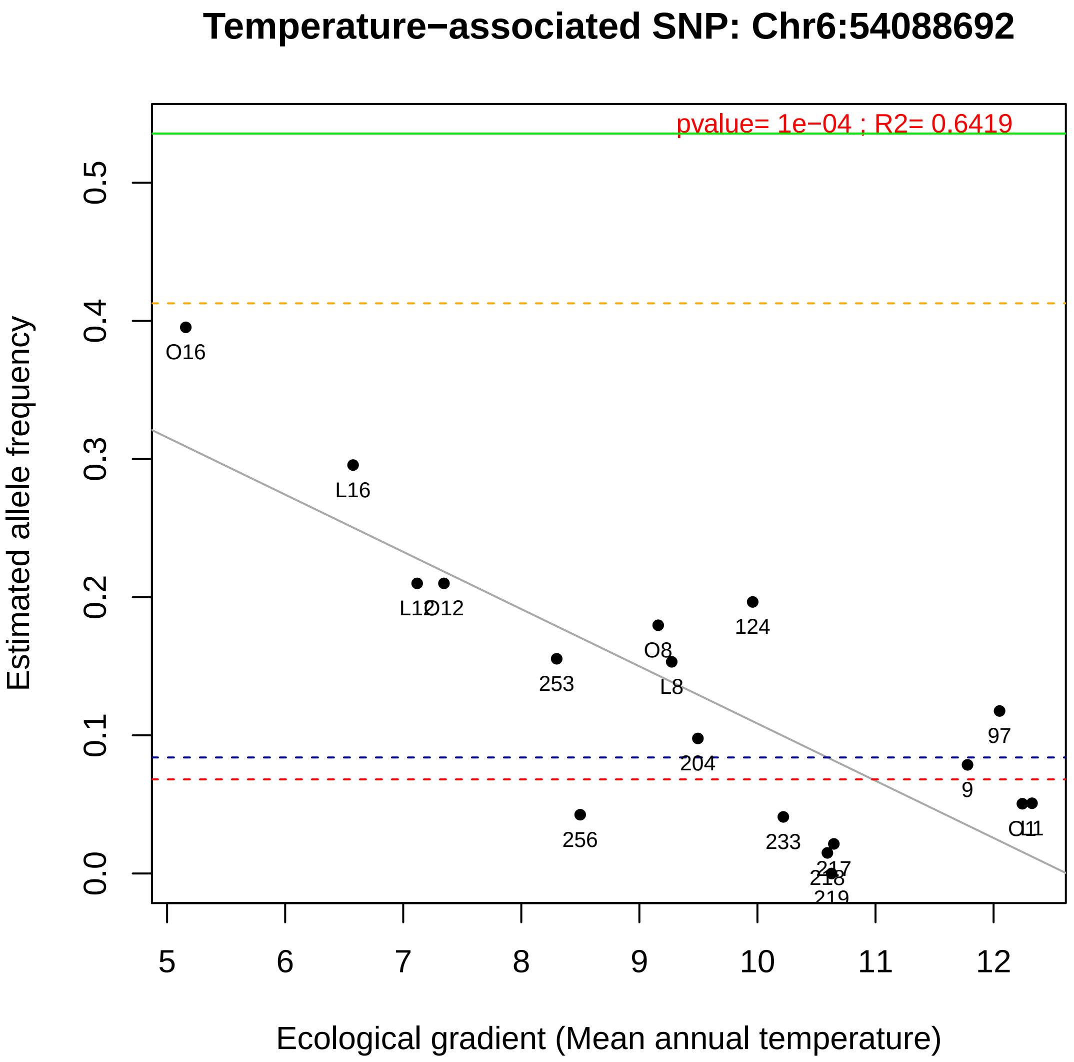

Temperature−associated SNP: Chr6:54088692
pvalue= 1e−04 ; R2= 0.6419
Estimated allele frequency
0.0
0.1
0.2
0.3
0.4
0.5
O16
L16
L12
O12
253
O8
L8
124
204
256
233
217
218
219
9
97
O11
L11
5
6
7
8
9
10
11
12
Ecological gradient (Mean annual temperature)

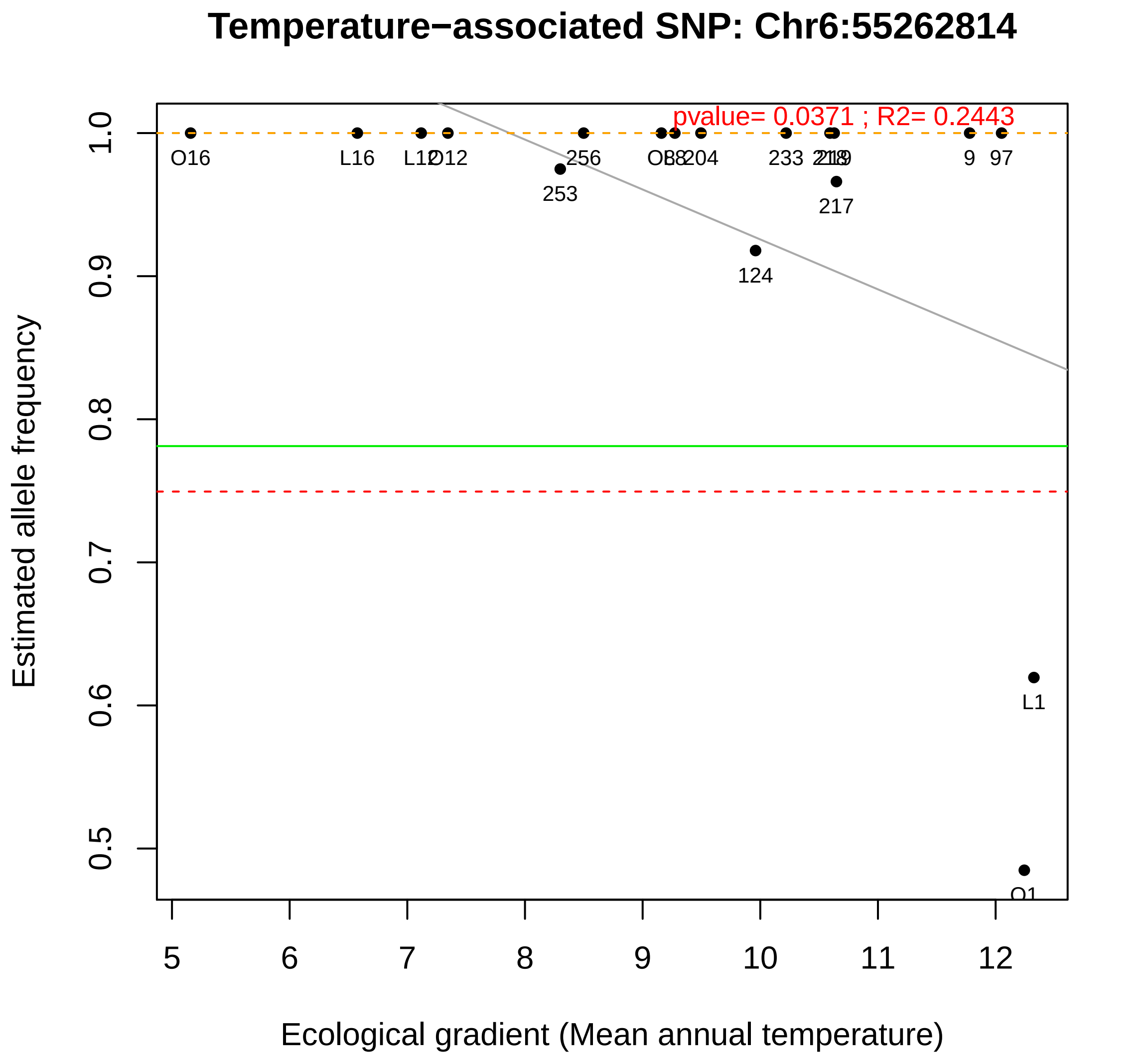

Temperature−associated SNP: Chr6:55262814
pvalue= 0.0371 ; R2= 0.2443
Estimated allele frequency
Ecological gradient (Mean annual temperature)
O16
L16
L12
O12
256
253
204
233
217
124
9
97
L1
O1
0.5
0.6
0.7
0.8
0.9
1.0
5
6
7
8
9
10
11
12

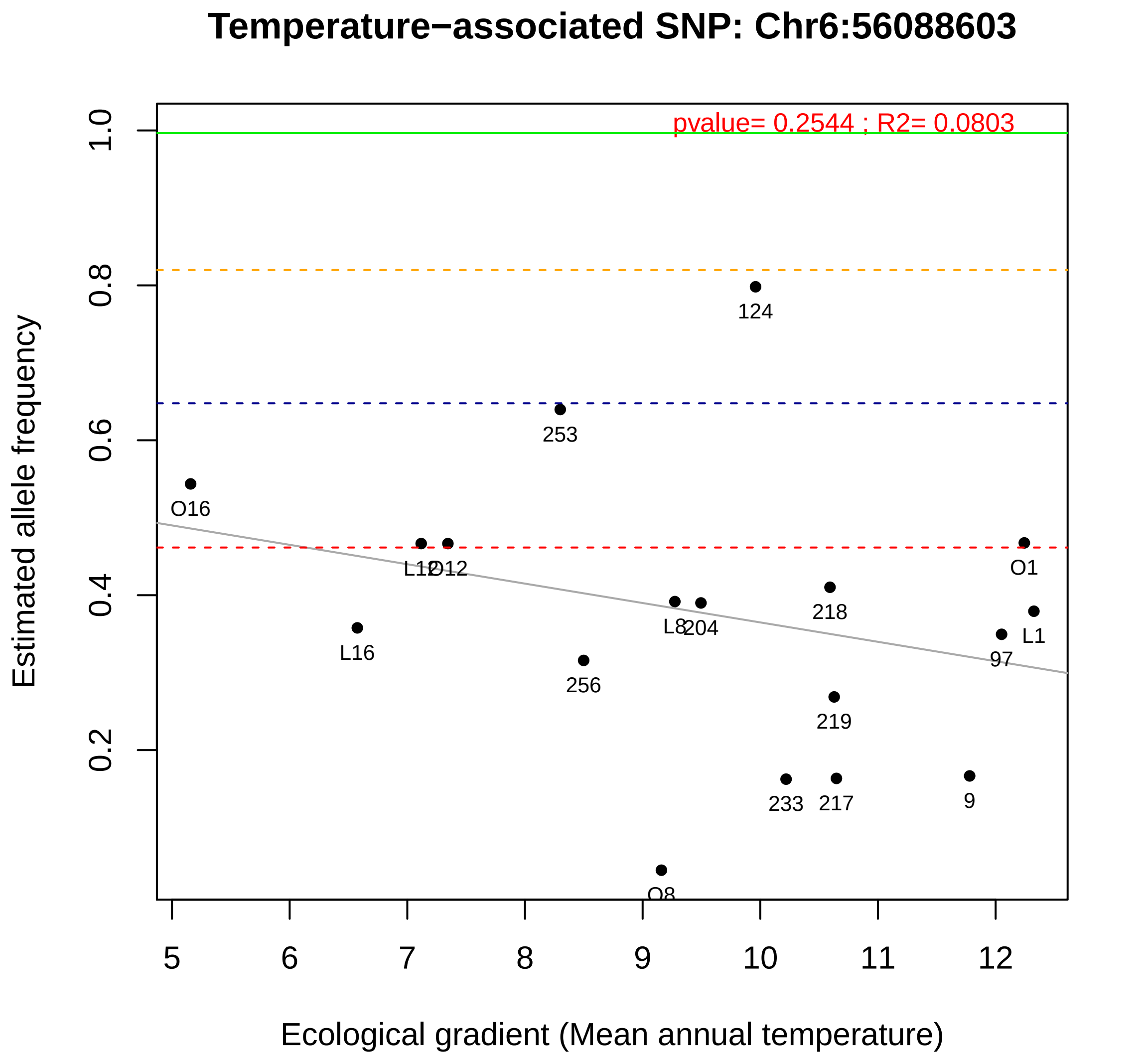

Temperature−associated SNP: Chr6:56088603
pvalue= 0.2544 ; R2= 0.0803
Estimated allele frequency
0.2
0.4
0.6
0.8
1.0
5
6
7
8
9
10
11
12
Ecological gradient (Mean annual temperature)
O16
L16
L12
O12
253
256
O8
L8
204
124
233
218
219
217
9
97
O1
L1

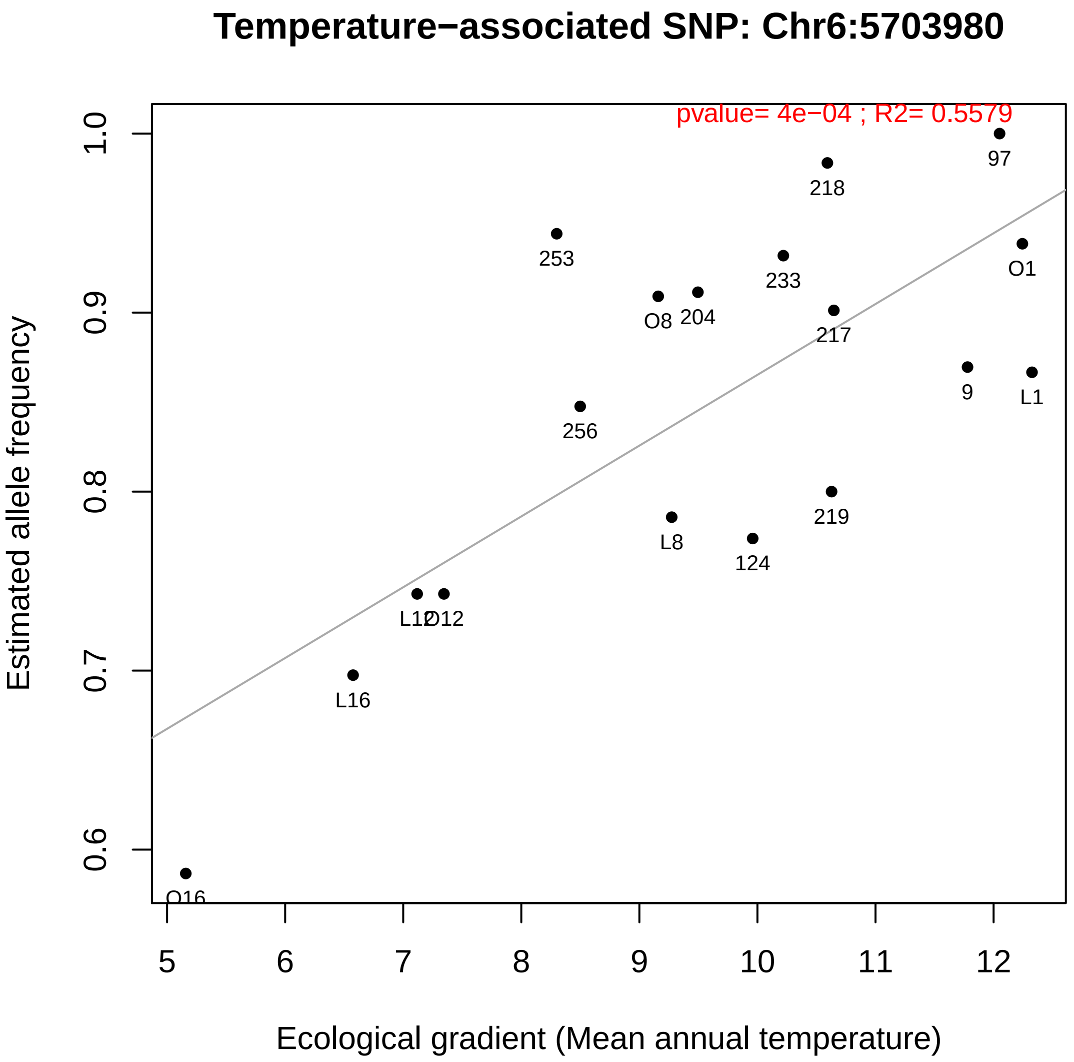
Temperature−associated SNP: Chr6:5703980
pvalue= 4e−04 ; R2= 0.5579
Estimated allele frequency
Ecological gradient (Mean annual temperature)
0.6
0.7
0.8
0.9
1.0
5
6
7
8
9
10
11
12
97
218
253
O1
233
O8
204
217
9
L1
256
219
L8
124
L12
O12
L16
O16

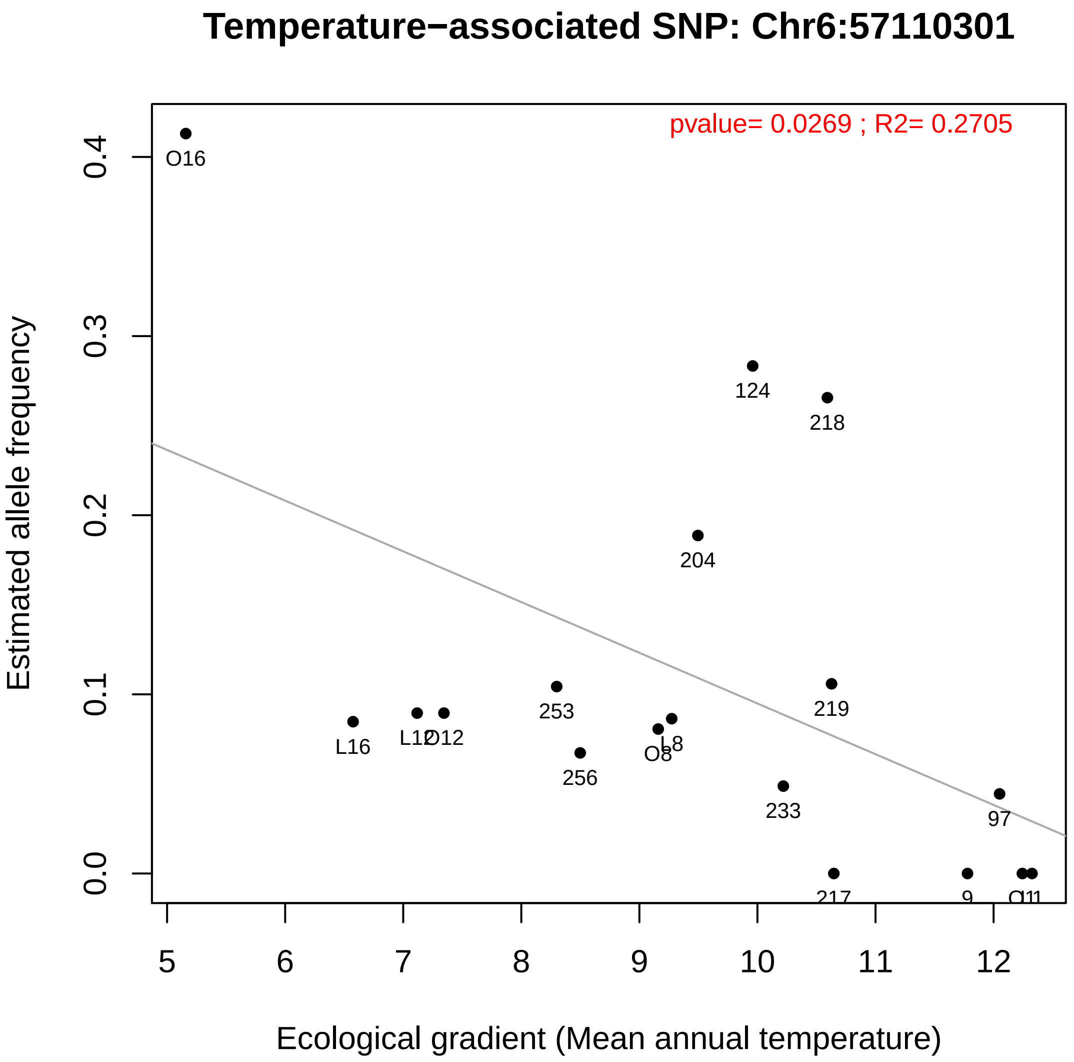

Temperature−associated SNP: Chr6:57110301
pvalue= 0.0269 ; R2= 0.2705
Estimated allele frequency
0.0
0.1
0.2
0.3
0.4
Ecological gradient (Mean annual temperature)
5
6
7
8
9
10
11
12
O16
124
218
204
253
219
L16
L12
O12
256
O8
L8
233
97
217
9
O11

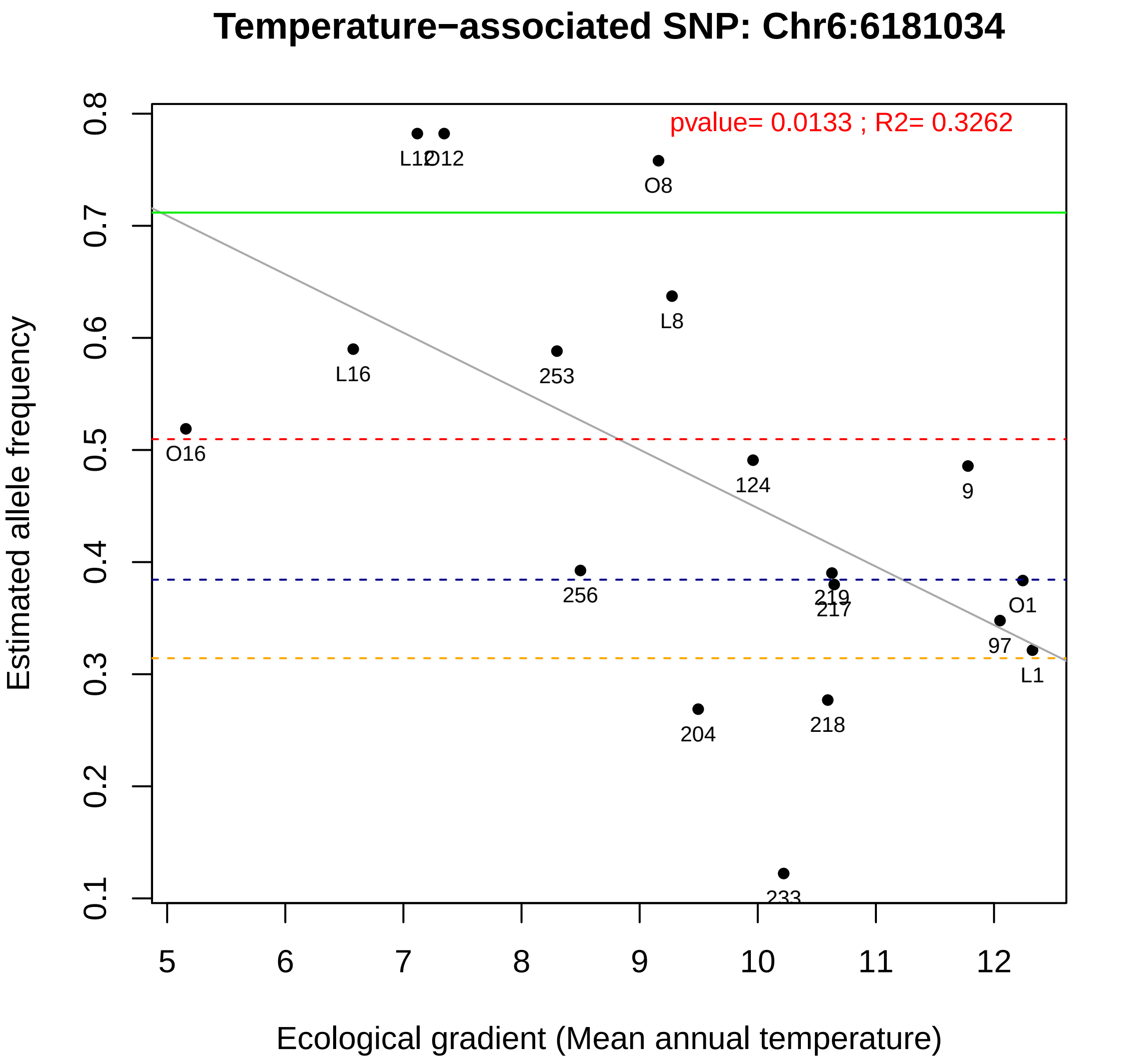

Temperature−associated SNP: Chr6:6181034
pvalue= 0.0133 ; R2= 0.3262
L12
O12
O8
L8
L16
253
O16
124
9
256
219
217
O1
97
L1
204
218
233
0.1
0.2
0.3
0.4
0.5
0.6
0.7
0.8
5
6
7
8
9
10
11
12
Estimated allele frequency
Ecological gradient (Mean annual temperature)

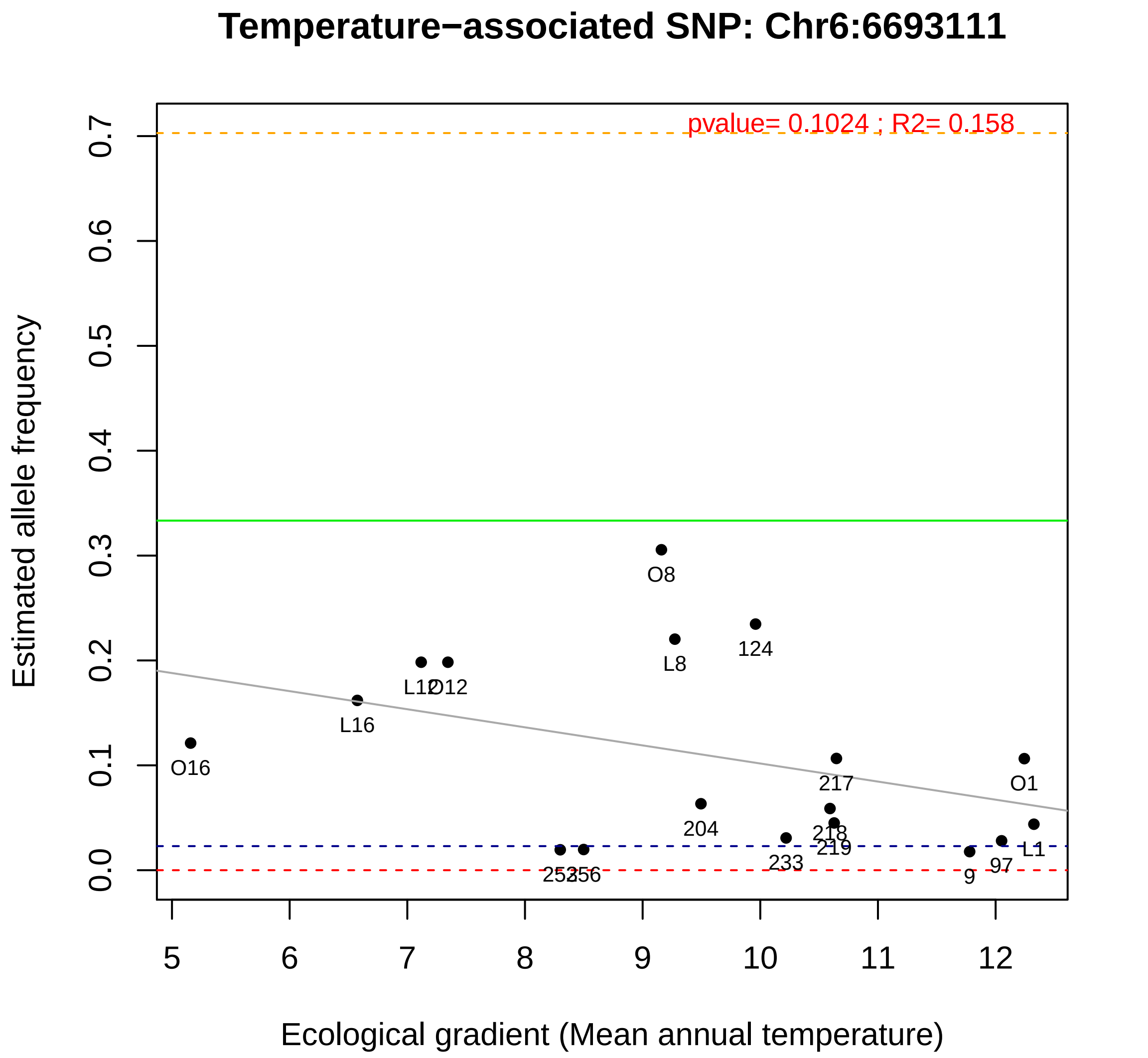

Temperature−associated SNP: Chr6:6693111
pvalue= 0.1024 ; R2= 0.158
Estimated allele frequency
Ecological gradient (Mean annual temperature)
0.0
0.1
0.2
0.3
0.4
0.5
0.6
0.7
5
6
7
8
9
10
11
12
O16
L16
L12
O12
O8
L8
124
204
217
O1
218
219
233
L1
97
9
253
256

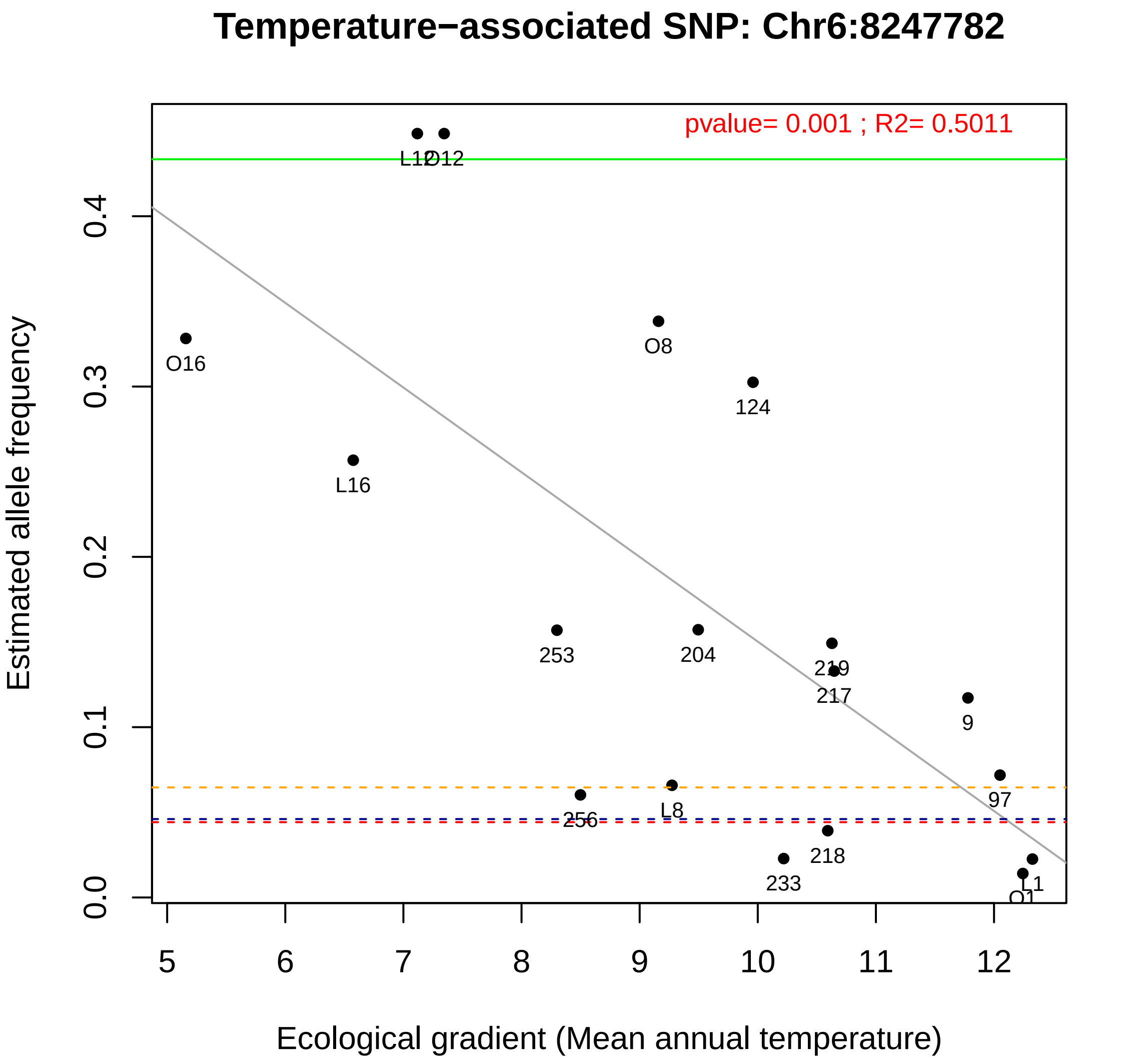
Temperature−associated SNP: Chr6:8247782
pvalue= 0.001 ; R2= 0.5011
Estimated allele frequency
0.0
0.1
0.2
0.3
0.4
5
6
7
8
9
10
11
12
Ecological gradient (Mean annual temperature)
L12
O12
O16
O8
124
L16
253
204
219
217
9
97
256
L8
218
233
L1
O1

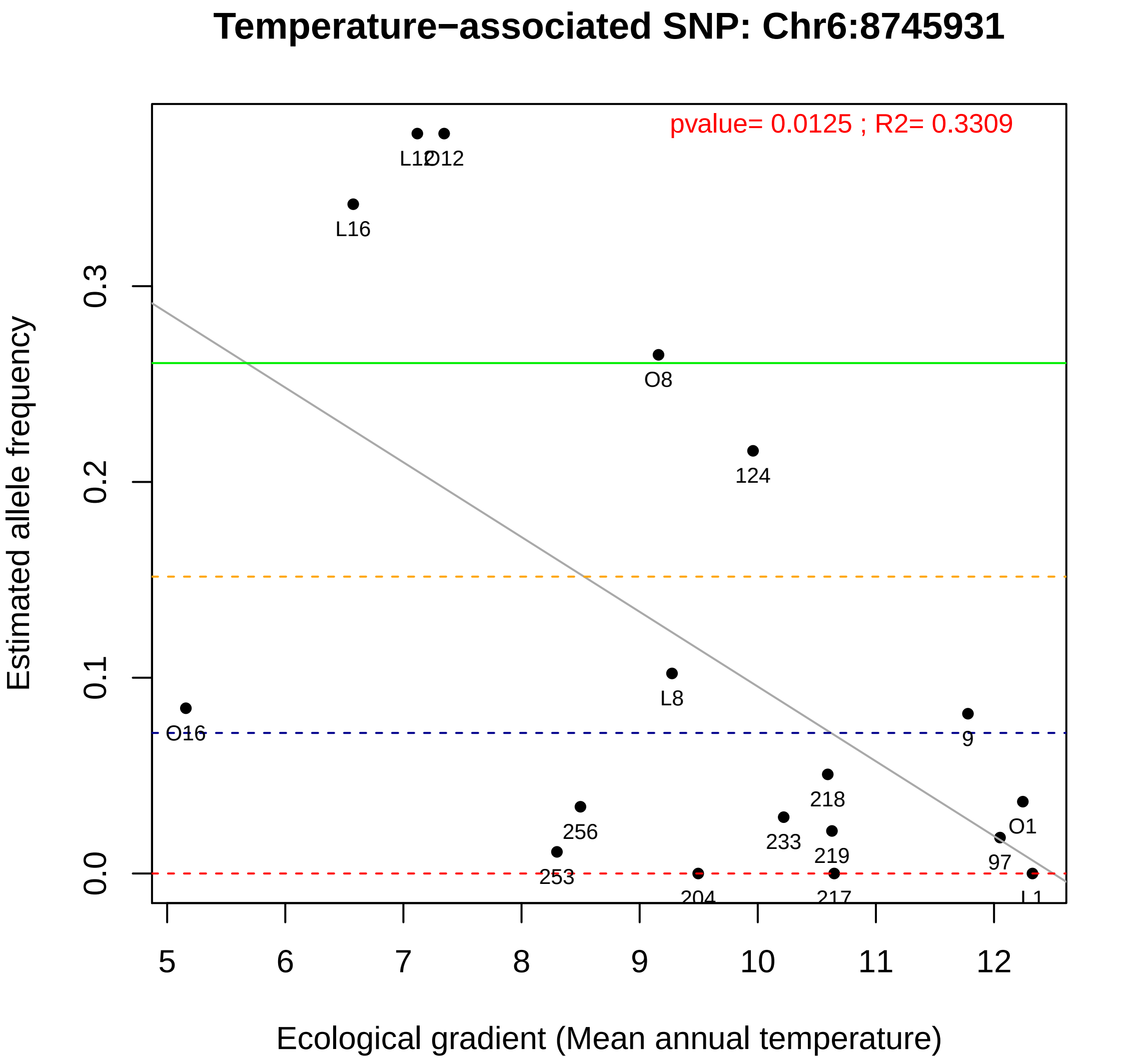
Temperature−associated SNP: Chr6:8745931
pvalue= 0.0125 ; R2= 0.3309
Estimated allele frequency
0.0
0.1
0.2
0.3
5
6
7
8
9
10
11
12
Ecological gradient (Mean annual temperature)
L12
O12
L16
O8
124
L8
O16
9
218
256
O1
233
219
97
253
204
217
L1

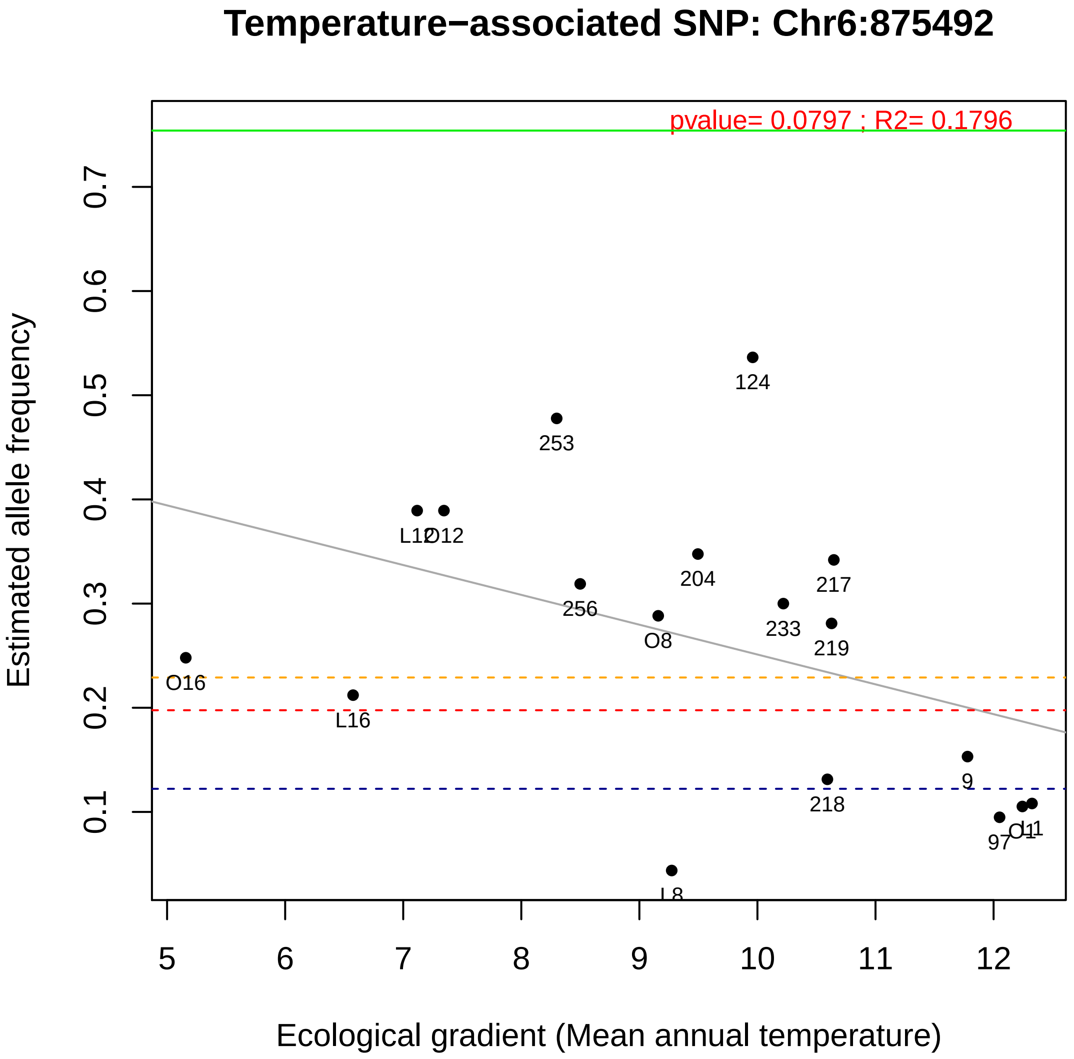

Temperature−associated SNP: Chr6:875492
pvalue= 0.0797 ; R2= 0.1796
Estimated allele frequency
0.1
0.2
0.3
0.4
0.5
0.6
0.7
O16
L16
L12
O12
253
256
124
204
O8
L8
233
217
219
218
9
97
O1
L1
5
6
7
8
9
10
11
12
Ecological gradient (Mean annual temperature)

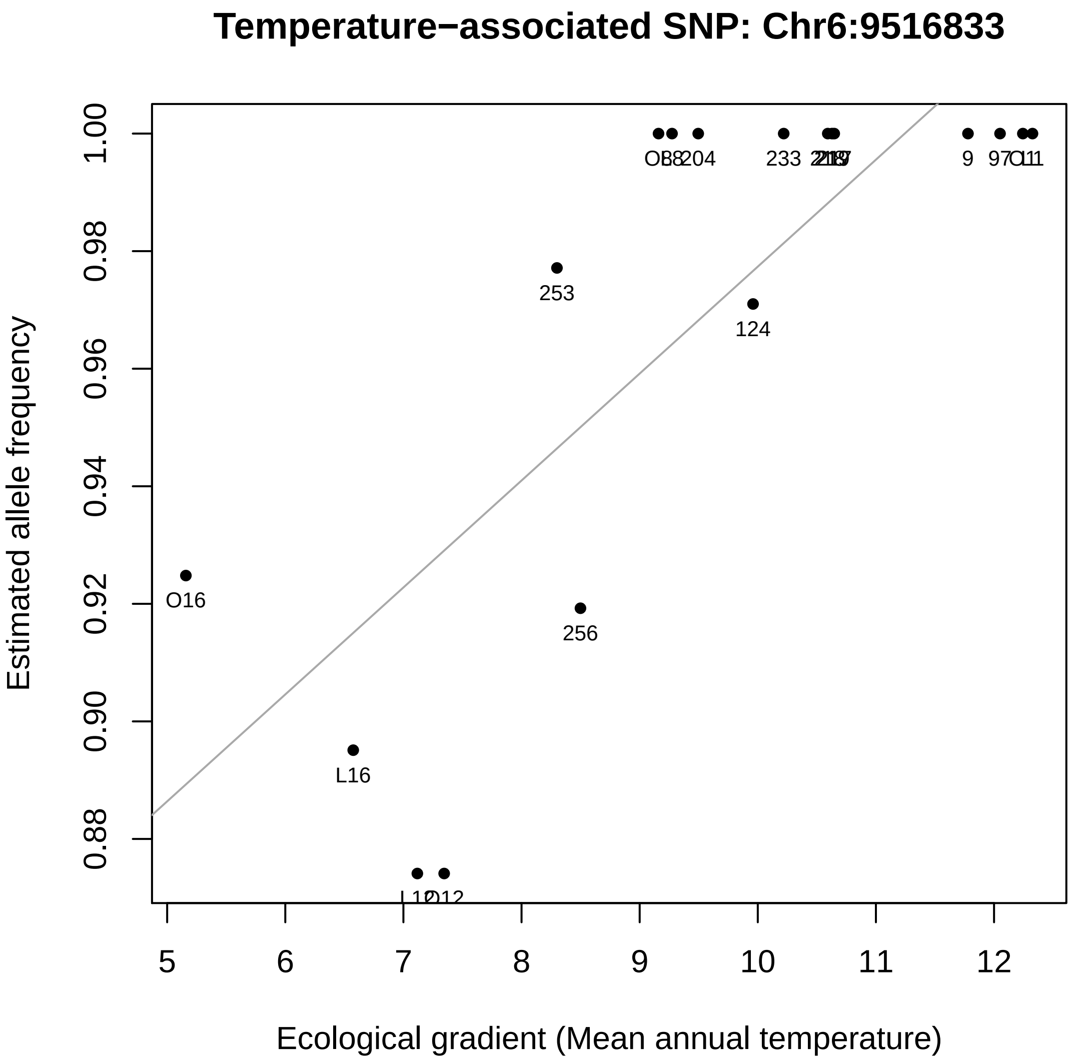
Temperature−associated SNP: Chr6:9516833
Estimated allele frequency
0.88
0.90
0.92
0.94
0.96
0.98
1.00
Ecological gradient (Mean annual temperature)
5
6
7
8
9
10
11
12
204
233
9
253
124
O16
256
L16

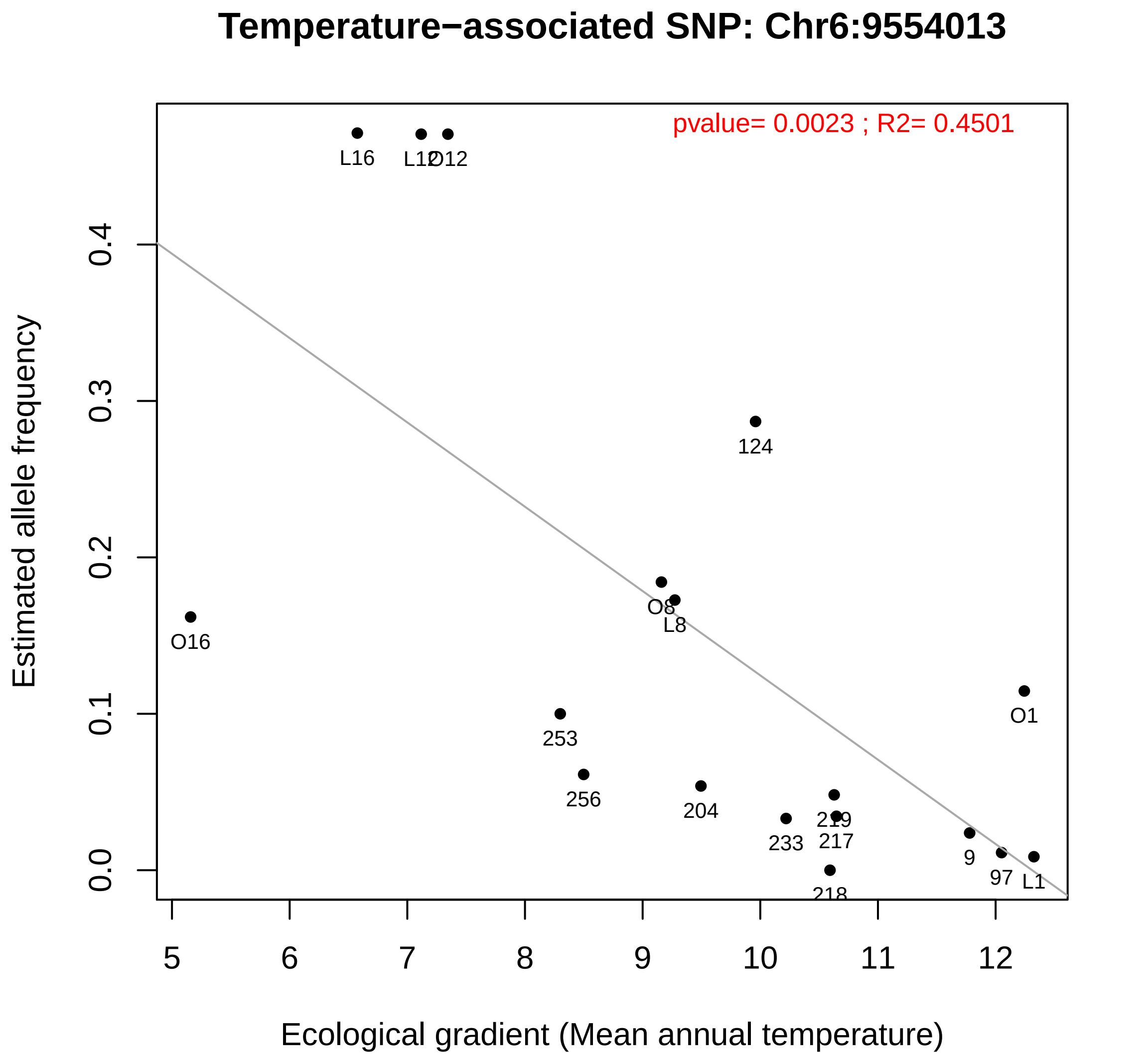

Temperature−associated SNP: Chr6:9554013
pvalue= 0.0023 ; R2= 0.4501
Estimated allele frequency
0.0
0.1
0.2
0.3
0.4
5
6
7
8
9
10
11
12
Ecological gradient (Mean annual temperature)
L16
L12
O12
124
O8
L8
O16
253
O1
256
204
219
217
233
9
97
L1
218

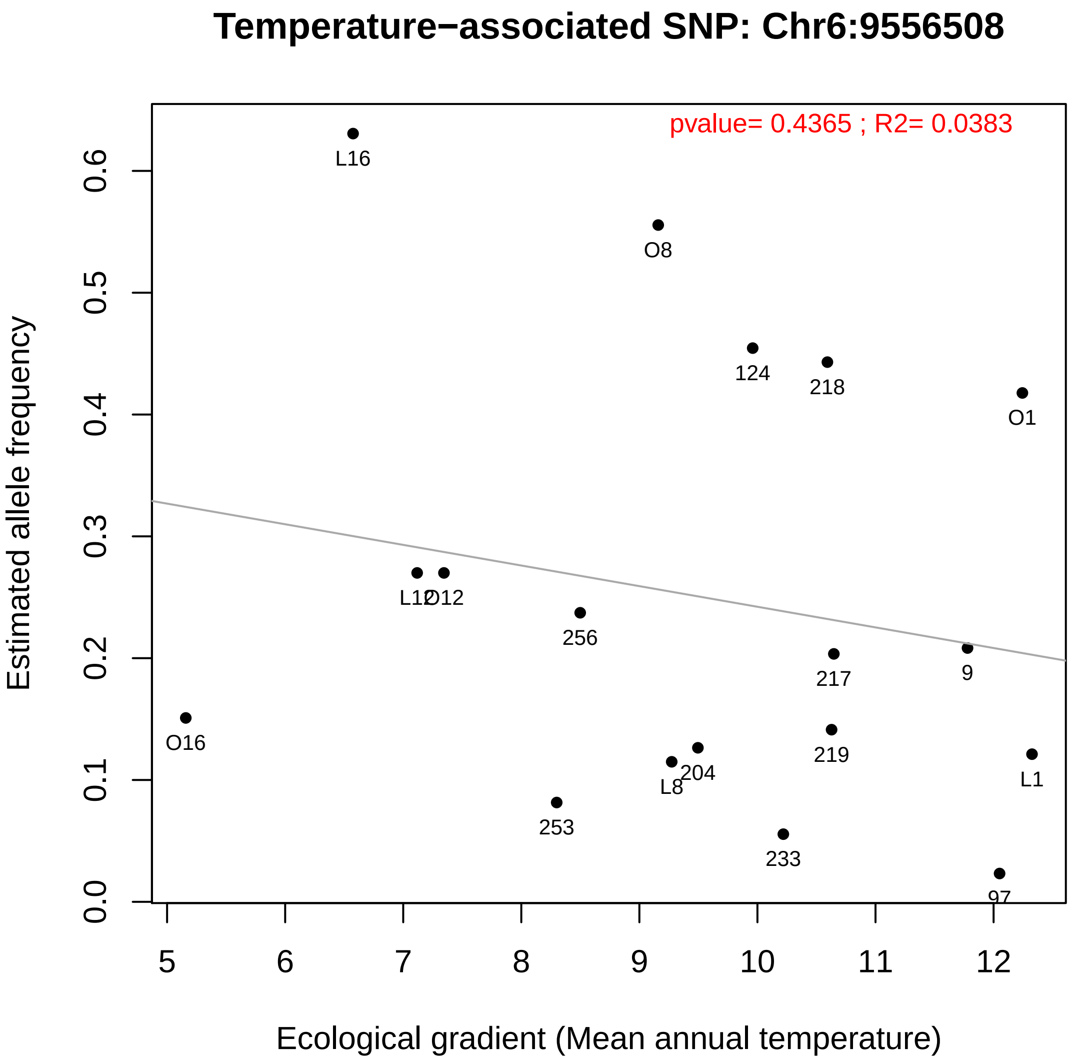

Temperature−associated SNP: Chr6:9556508
pvalue= 0.4365 ; R2= 0.0383
Estimated allele frequency
Ecological gradient (Mean annual temperature)
0.0
0.1
0.2
0.3
0.4
0.5
0.6
5
6
7
8
9
10
11
12
L16
O8
124
218
O1
O12
256
217
9
O16
219
204
L8
L1
253
233
97

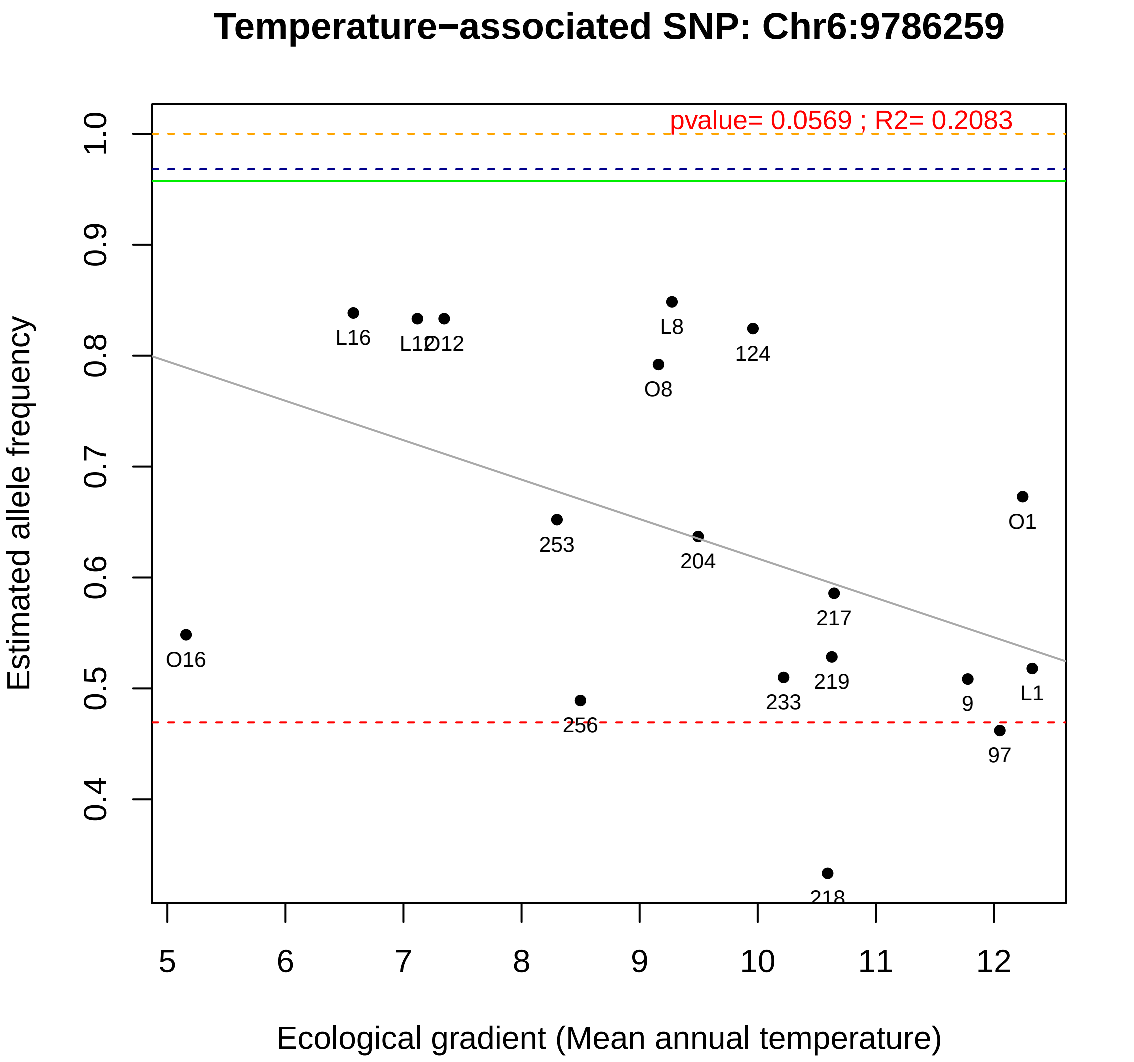

Temperature−associated SNP: Chr6:9786259
pvalue= 0.0569 ; R2= 0.2083
Estimated allele frequency
0.4
0.5
0.6
0.7
0.8
0.9
1.0
Ecological gradient (Mean annual temperature)
5
6
7
8
9
10
11
12
L16
L12
O12
L8
124
O8
253
204
O1
217
O16
219
233
9
L1
256
97
218

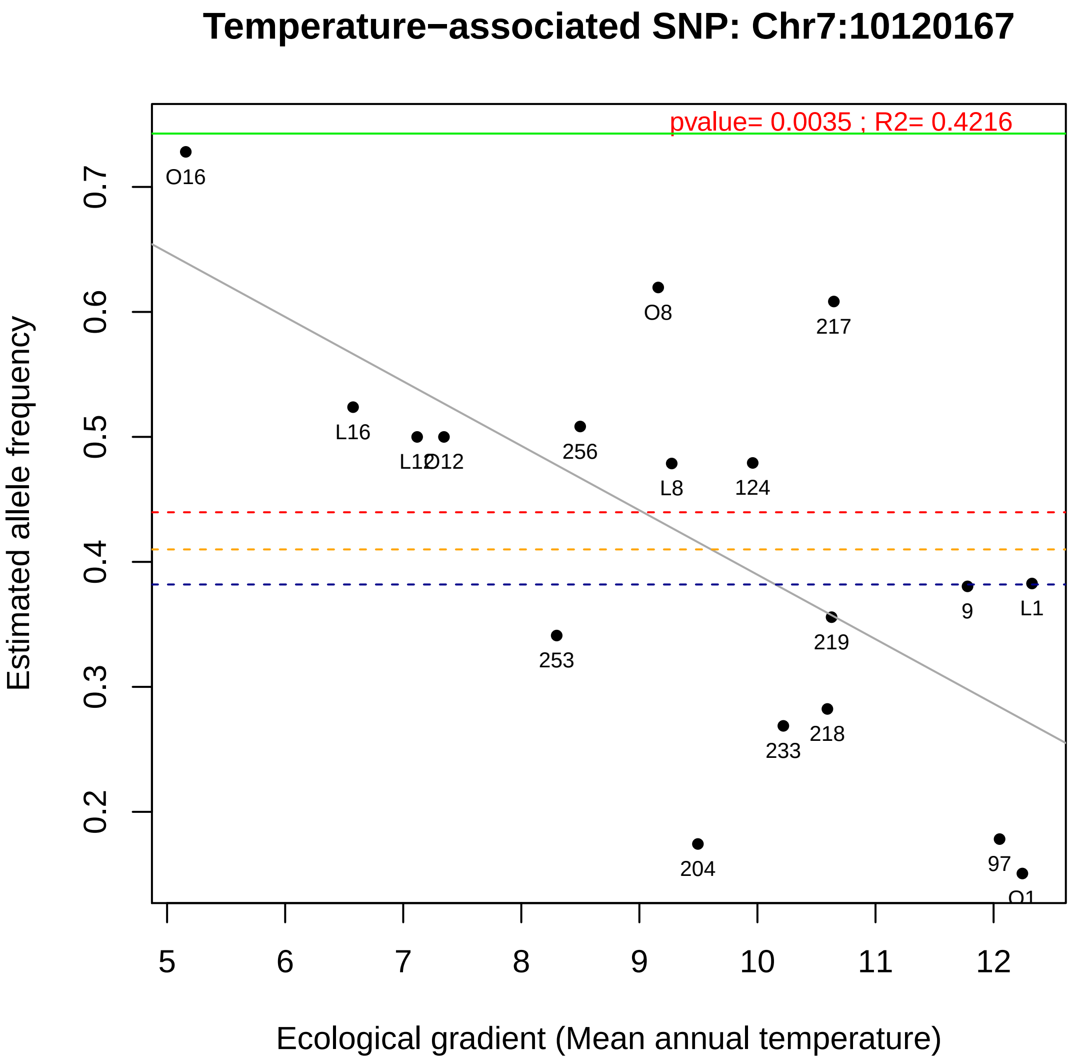

Temperature−associated SNP: Chr7:10120167
pvalue= 0.0035 ; R2= 0.4216
Estimated allele frequency
Ecological gradient (Mean annual temperature)
0.2
0.3
0.4
0.5
0.6
0.7
5
6
7
8
9
10
11
12
O16
O8
217
L16
L12
O12
256
L8
124
9
L1
219
253
218
233
204
97
O1

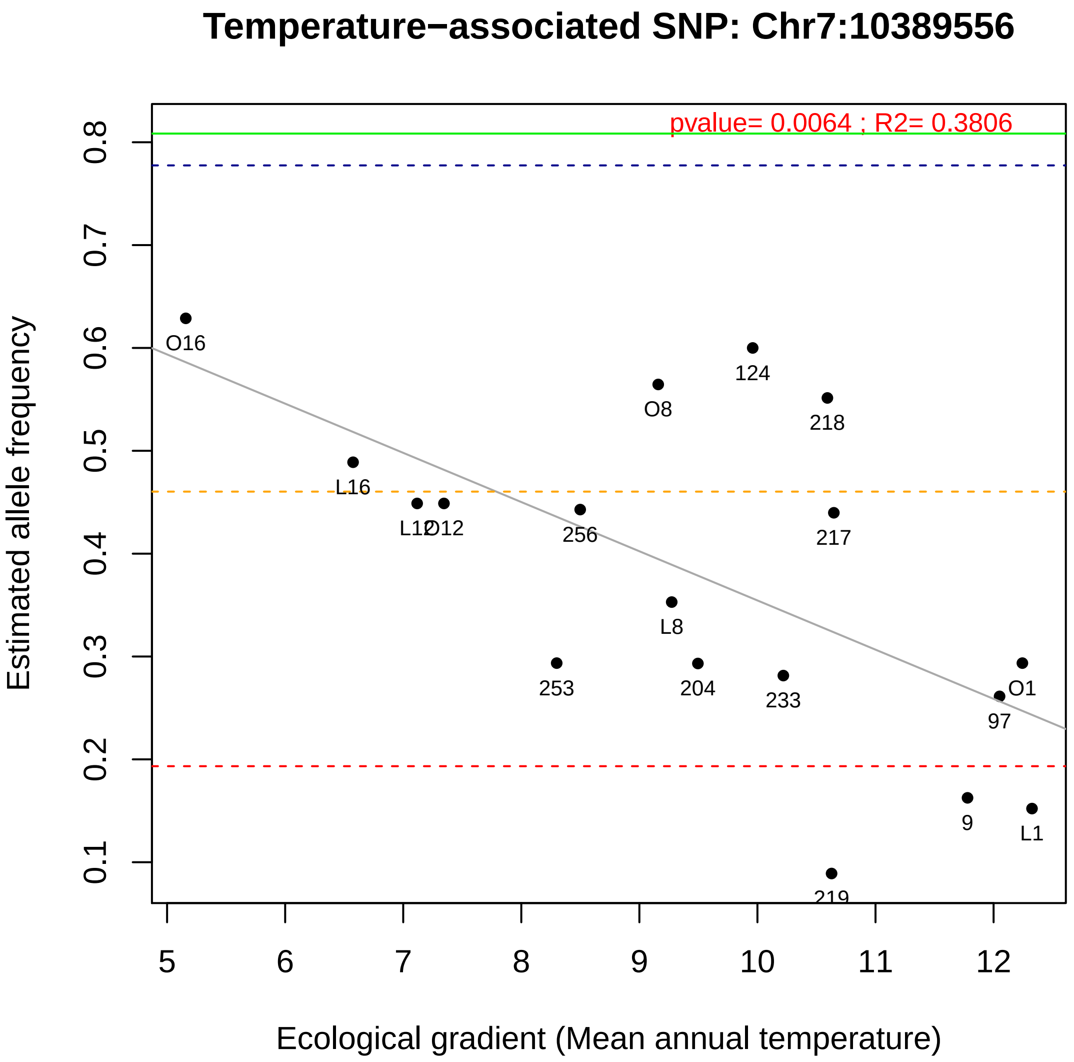

Temperature−associated SNP: Chr7:10389556
pvalue= 0.0064 ; R2= 0.3806
Estimated allele frequency
0.1
0.2
0.3
0.4
0.5
0.6
0.7
0.8
5
6
7
8
9
10
11
12
Ecological gradient (Mean annual temperature)
O16
L16
L12
O12
256
253
O8
L8
204
124
218
217
233
219
9
97
O1
L1

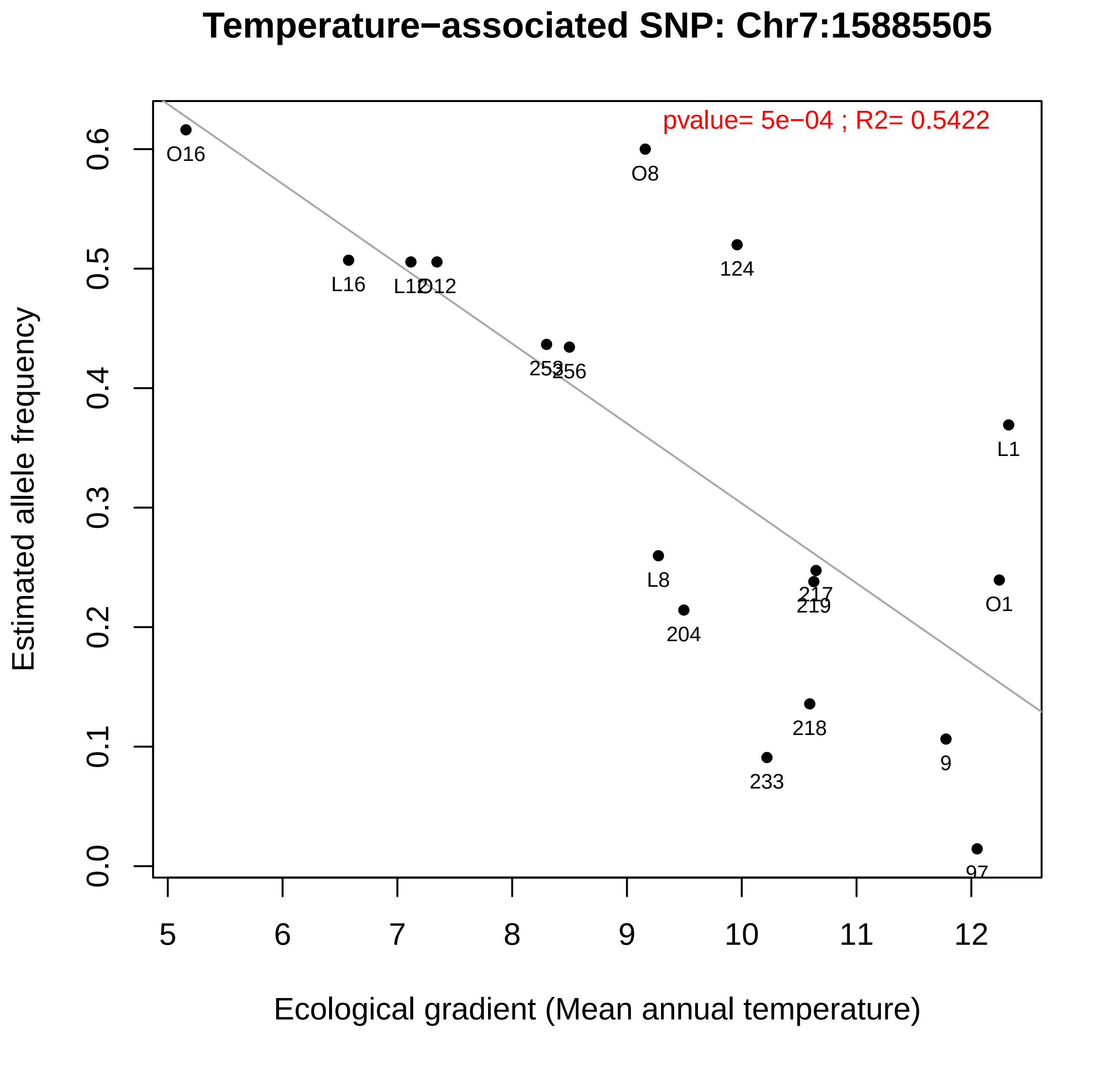

Temperature−associated SNP: Chr7:15885505
pvalue= 5e−04 ; R2= 0.5422
Estimated allele frequency
Ecological gradient (Mean annual temperature)
0.0
0.1
0.2
0.3
0.4
0.5
0.6
5
6
7
8
9
10
11
12
O16
O8
L16
L12
O12
124
253
256
L1
L8
217
219
O1
204
218
233
9
97

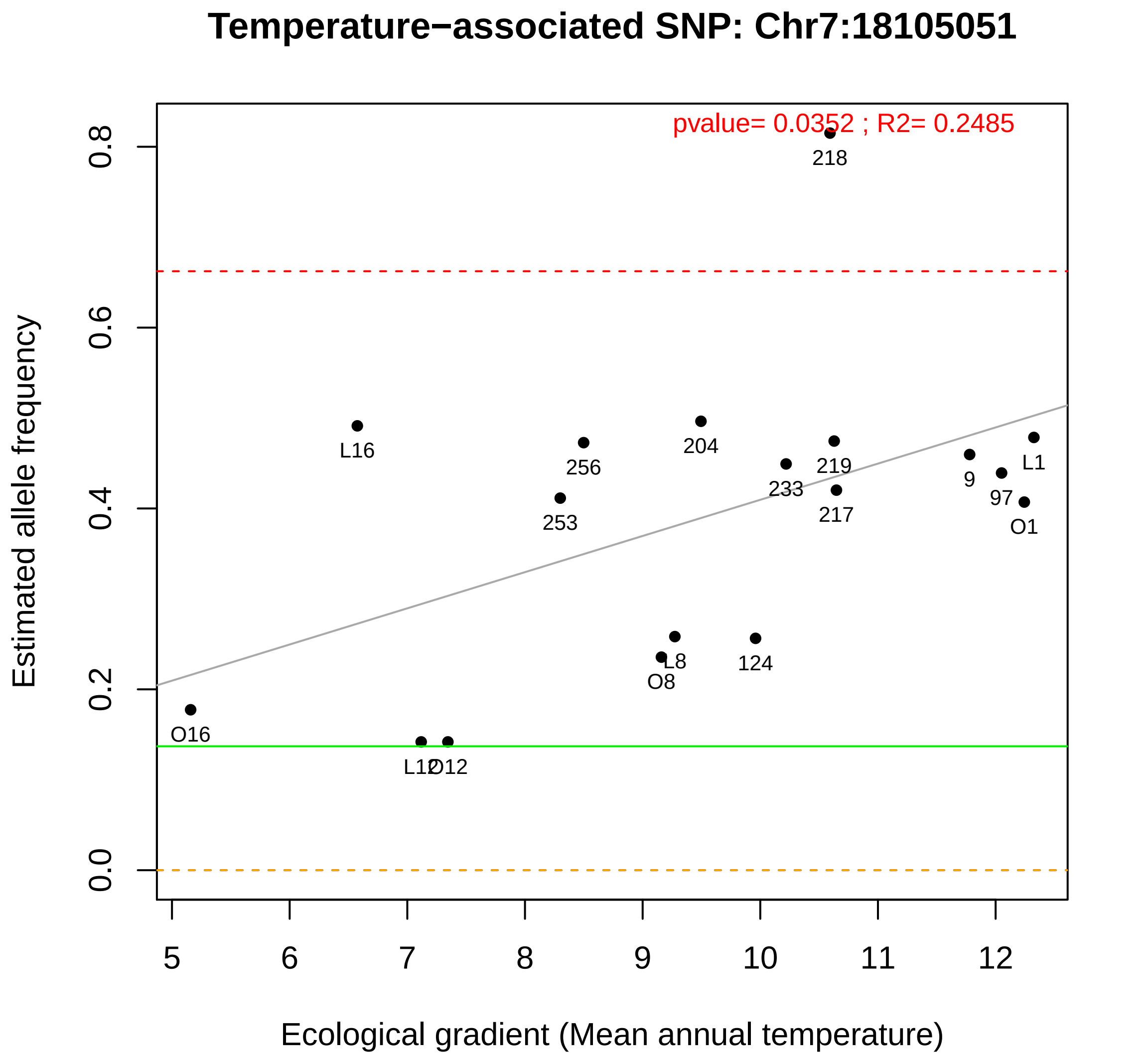
Temperature−associated SNP: Chr7:18105051
pvalue= 0.0352 ; R2= 0.2485
Estimated allele frequency
0.0
0.2
0.4
0.6
0.8
5
6
7
8
9
10
11
12
Ecological gradient (Mean annual temperature)
218
L16
256
204
219
233
253
217
9
L1
97
O1
L8
O8
124
O16
L12
O12

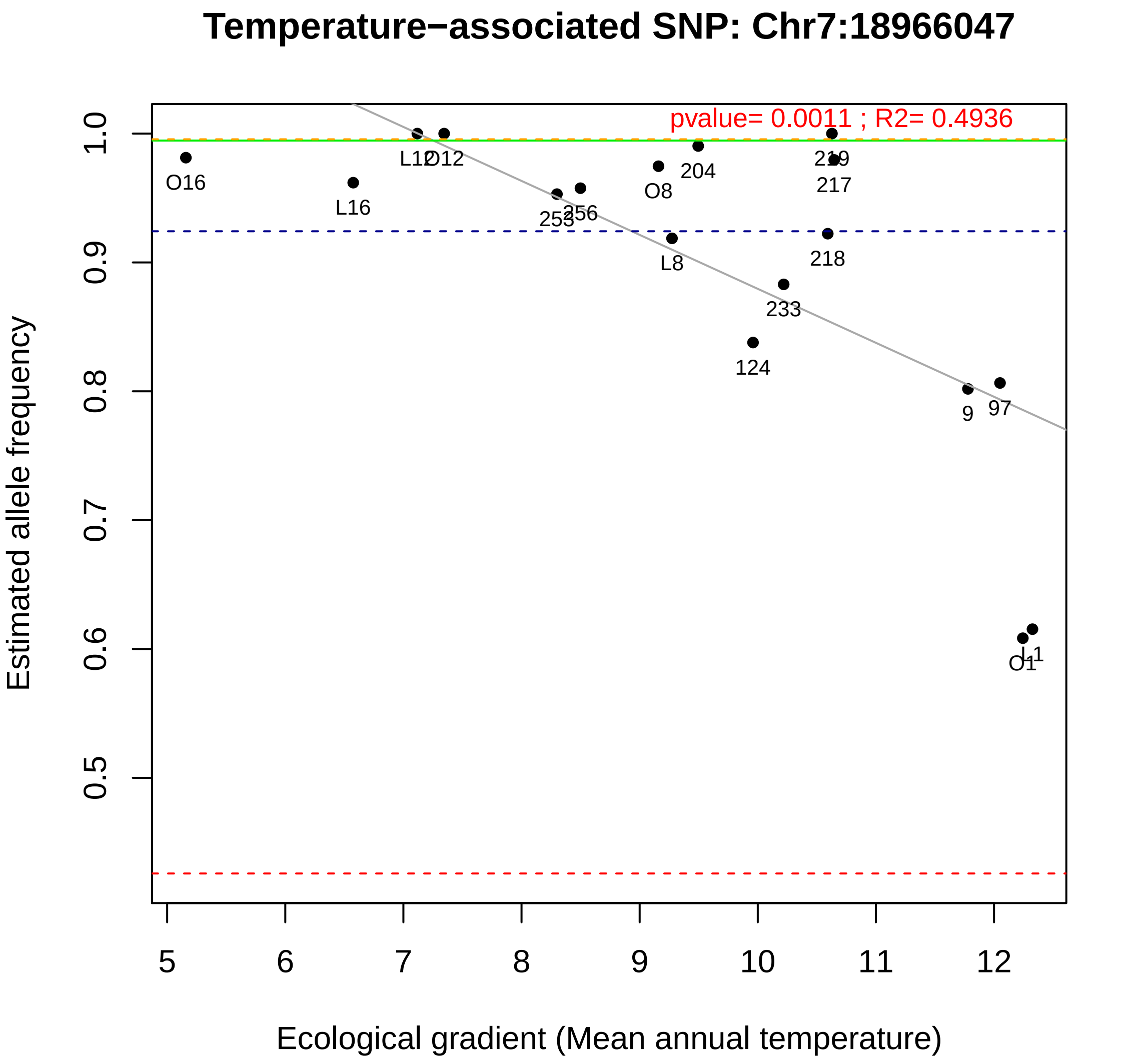
Temperature−associated SNP: Chr7:18966047
pvalue= 0.0011 ; R2= 0.4936
Estimated allele frequency
Ecological gradient (Mean annual temperature)
0.5
0.6
0.7
0.8
0.9
1.0
5
6
7
8
9
10
11
12
O16
L16
L12
O12
253
256
O8
204
L8
219
217
218
233
124
9
97
O1
L1

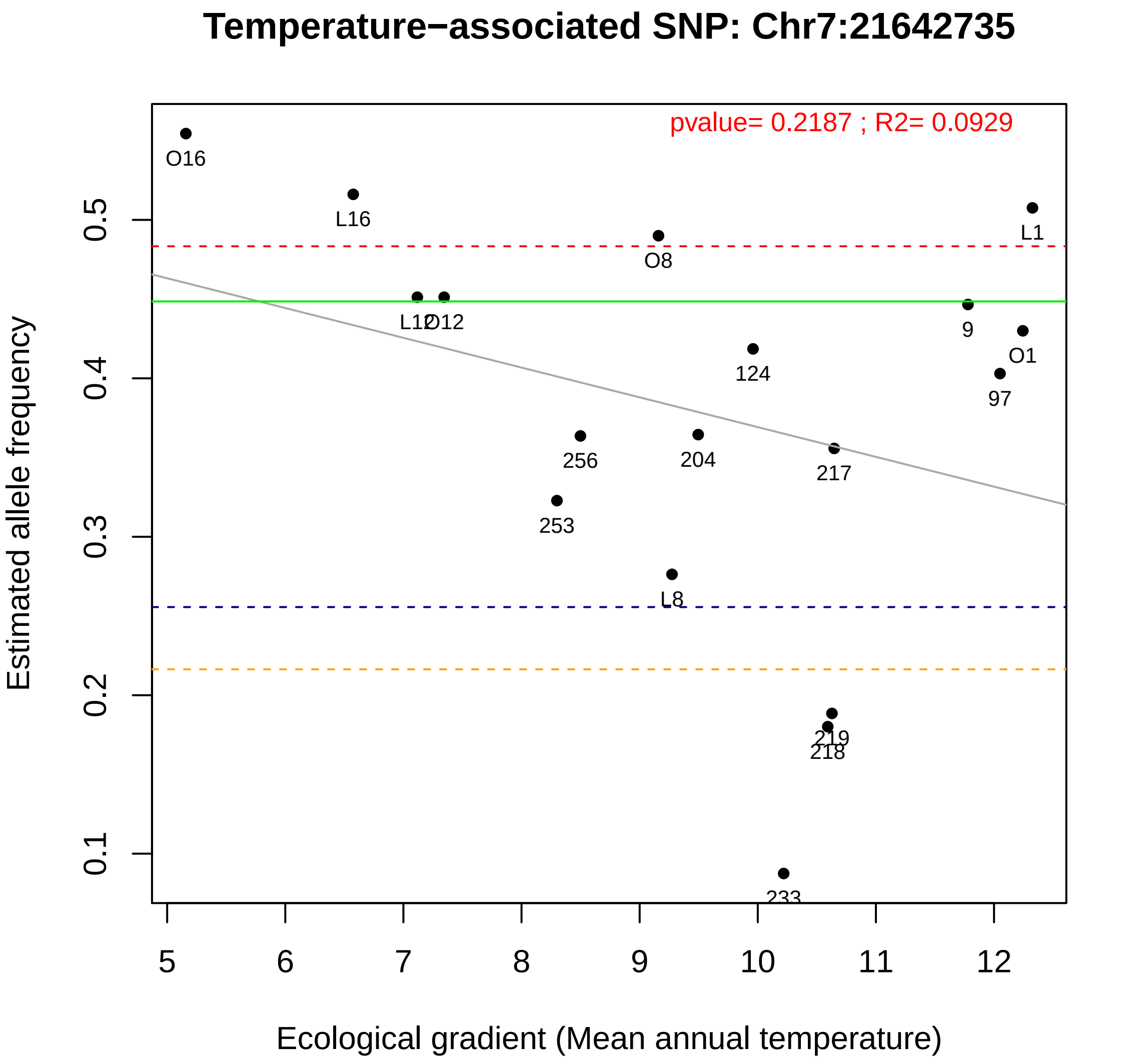
Temperature−associated SNP: Chr7:21642735
pvalue= 0.2187 ; R2= 0.0929
Estimated allele frequency
0.1
0.2
0.3
0.4
0.5
Ecological gradient (Mean annual temperature)
5
6
7
8
9
10
11
12
O16
L16
L12
O12
O8
L1
9
O1
124
97
256
204
217
253
L8
219
218
233

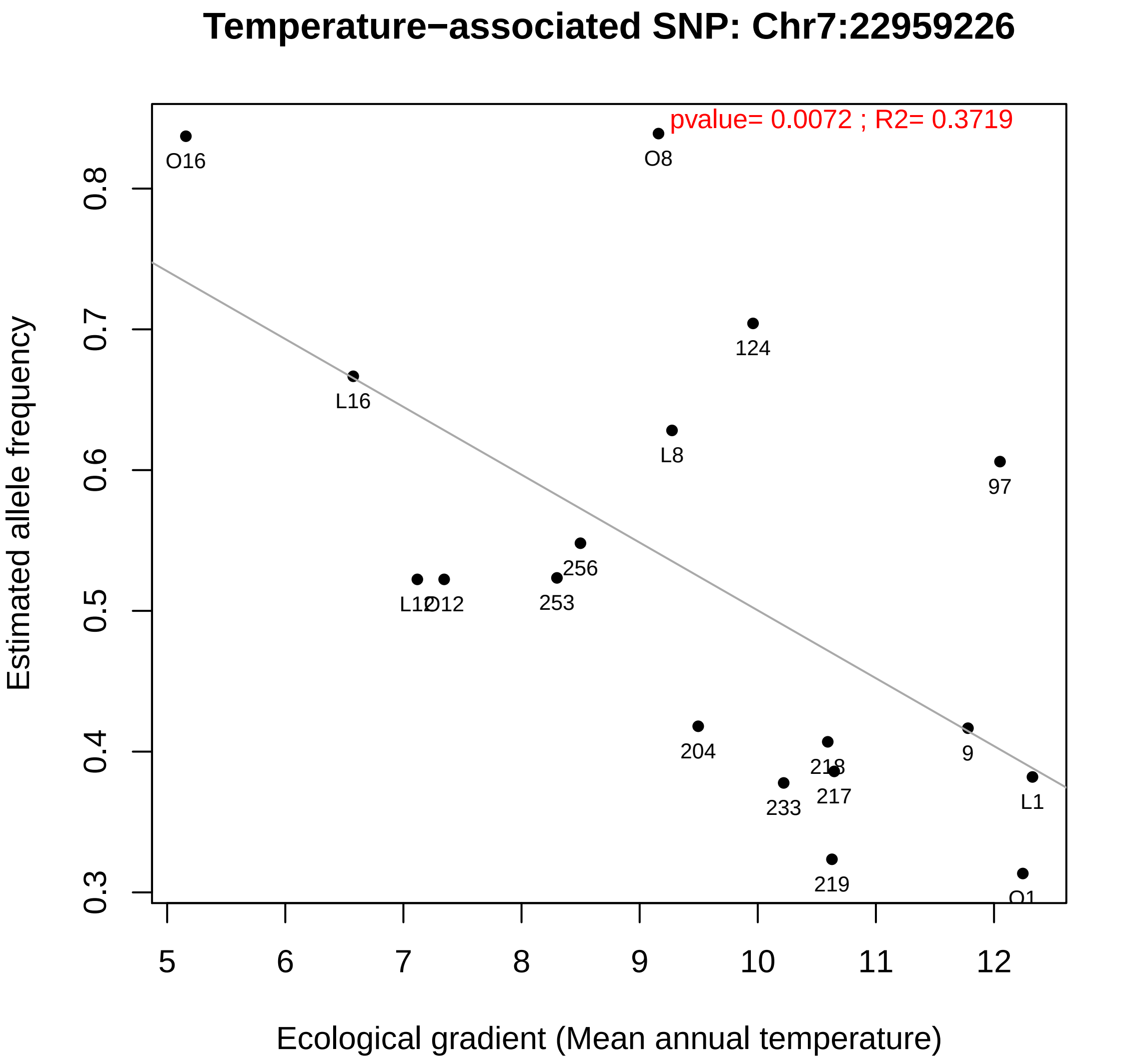
Temperature−associated SNP: Chr7:22959226
pvalue= 0.0072 ; R2= 0.3719
O16
O8
124
L16
L8
97
256
253
L12
O12
204
218
217
9
233
L1
219
O1
0.3
0.4
0.5
0.6
0.7
0.8
Estimated allele frequency
5
6
7
8
9
10
11
12
Ecological gradient (Mean annual temperature)

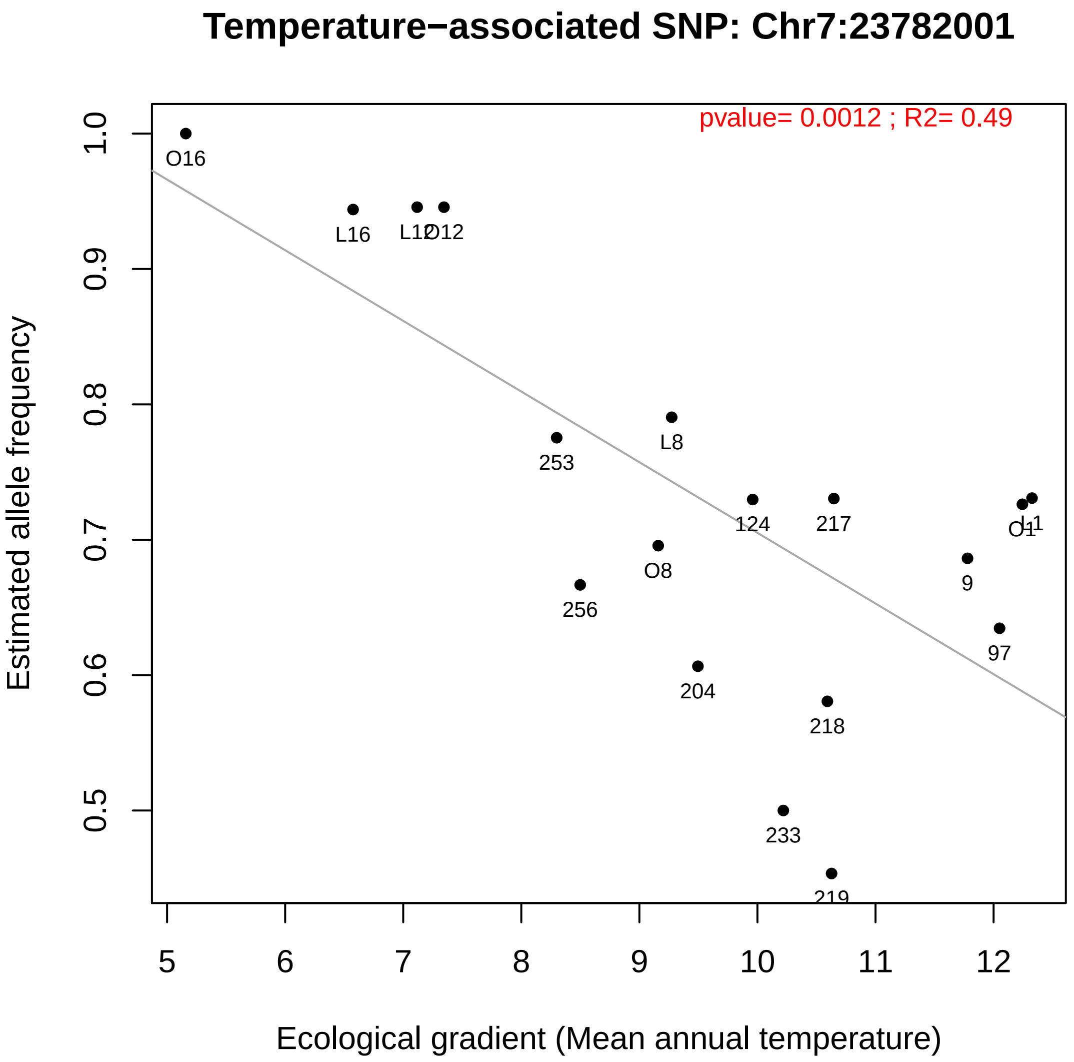

Temperature−associated SNP: Chr7:23782001
pvalue= 0.0012 ; R2= 0.49
Estimated allele frequency
Ecological gradient (Mean annual temperature)
1.0
0.9
0.8
0.7
0.6
0.5
5
6
7
8
9
10
11
12
O16
L16
L12
O12
253
L8
124
217
O1
L1
O8
256
9
97
204
218
233
219

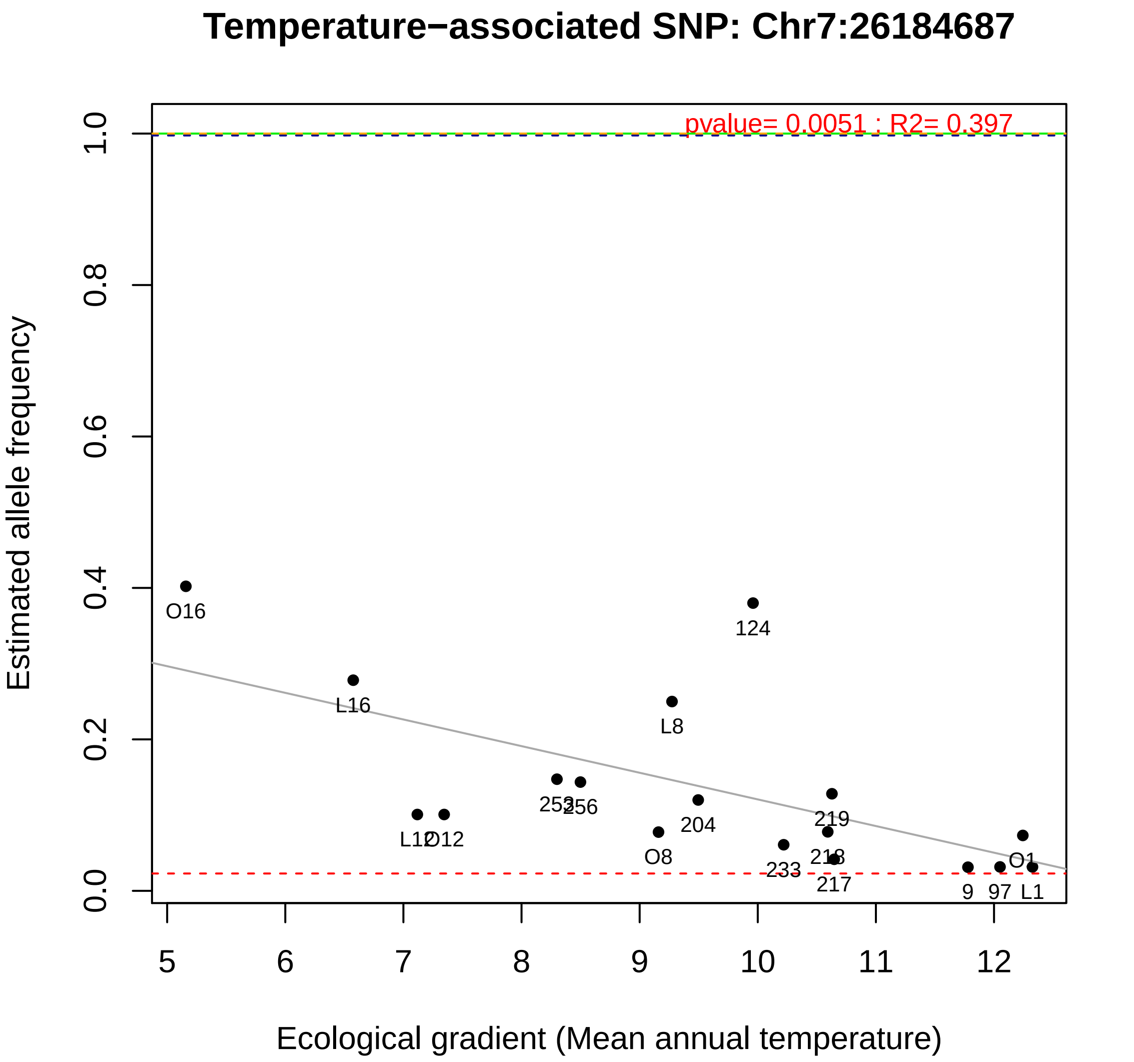
Temperature−associated SNP: Chr7:26184687
pvalue= 0.0051 ; R2= 0.397
Estimated allele frequency
0.0
0.2
0.4
0.6
0.8
1.0
5
6
7
8
9
10
11
12
Ecological gradient (Mean annual temperature)
O16
L16
124
L8
L12
O12
252
256
O8
204
233
219
218
217
9
97
O1
L1

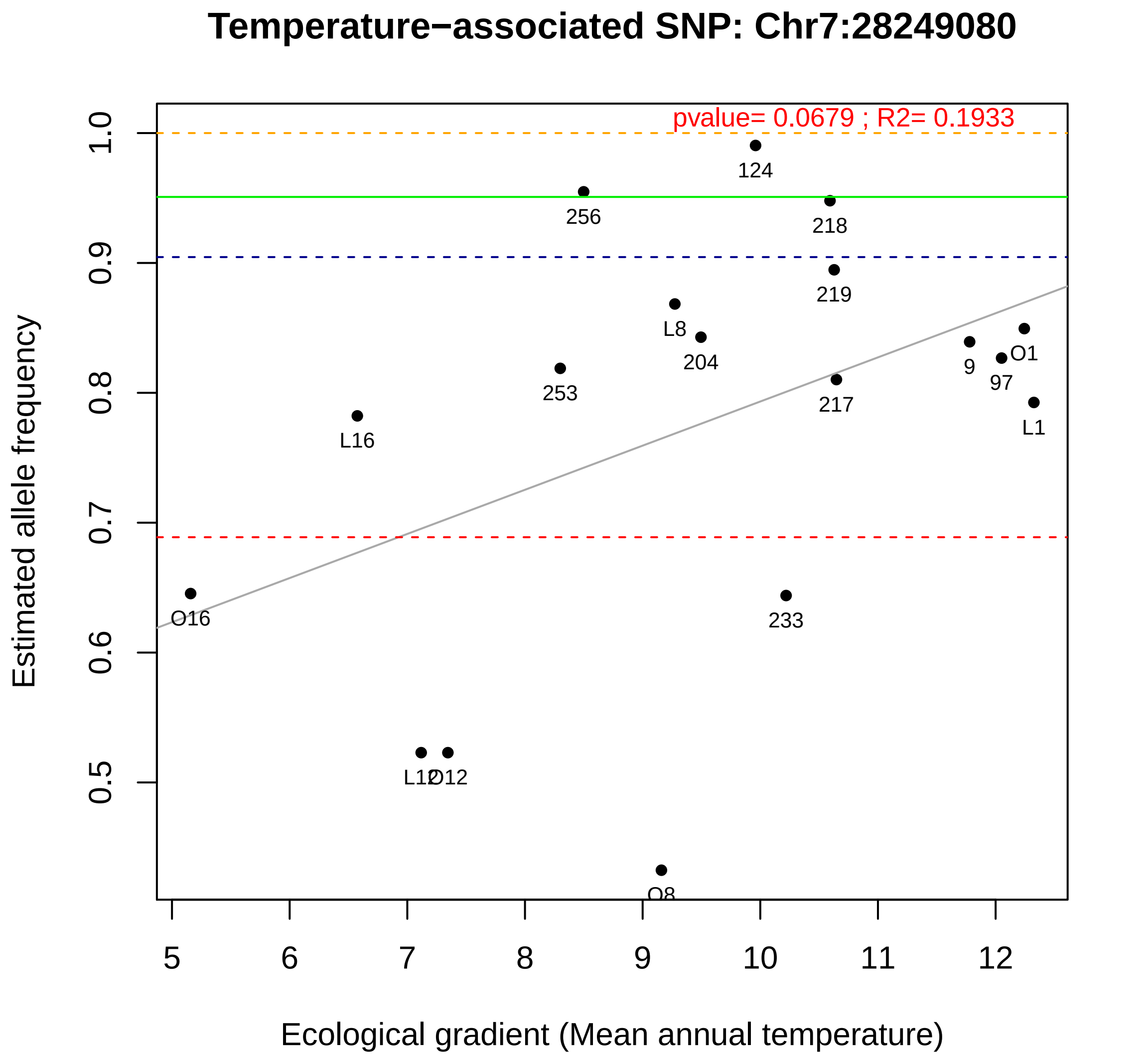

Temperature−associated SNP: Chr7:28249080
pvalue= 0.0679 ; R2= 0.1933
124
256
218
219
L8
204
253
L16
9
O1
97
217
L1
O16
233
L12
O12
O8
1.0
0.9
0.8
0.7
0.6
0.5
5
6
7
8
9
10
11
12
Estimated allele frequency
Ecological gradient (Mean annual temperature)

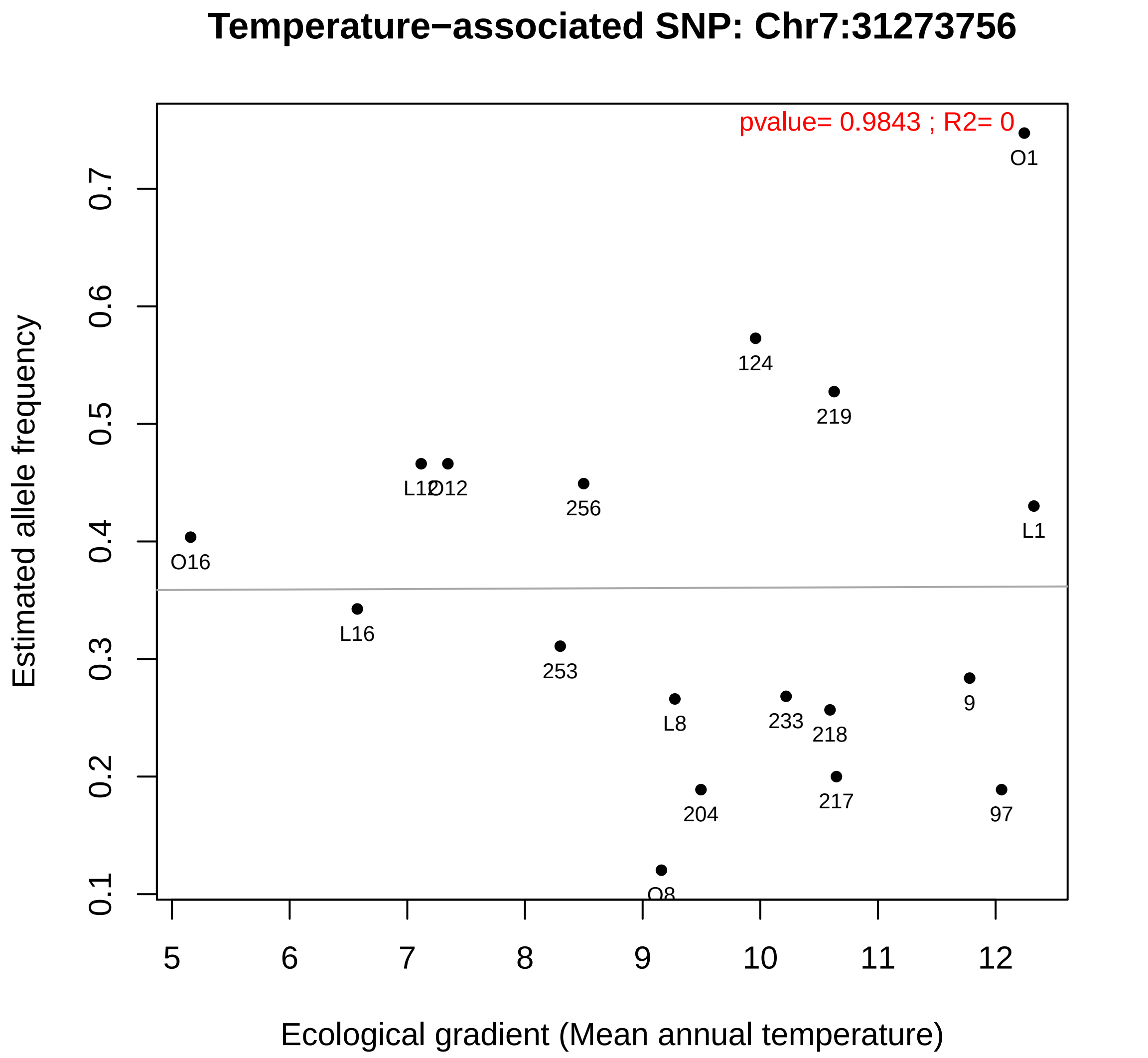

Temperature−associated SNP: Chr7:31273756
pvalue= 0.9843 ; R2= 0
O1
124
219
L12
O12
256
L1
O16
L16
253
L8
233
218
9
217
204
97
O8
0.1
0.2
0.3
0.4
0.5
0.6
0.7
5
6
7
8
9
10
11
12
Estimated allele frequency
Ecological gradient (Mean annual temperature)

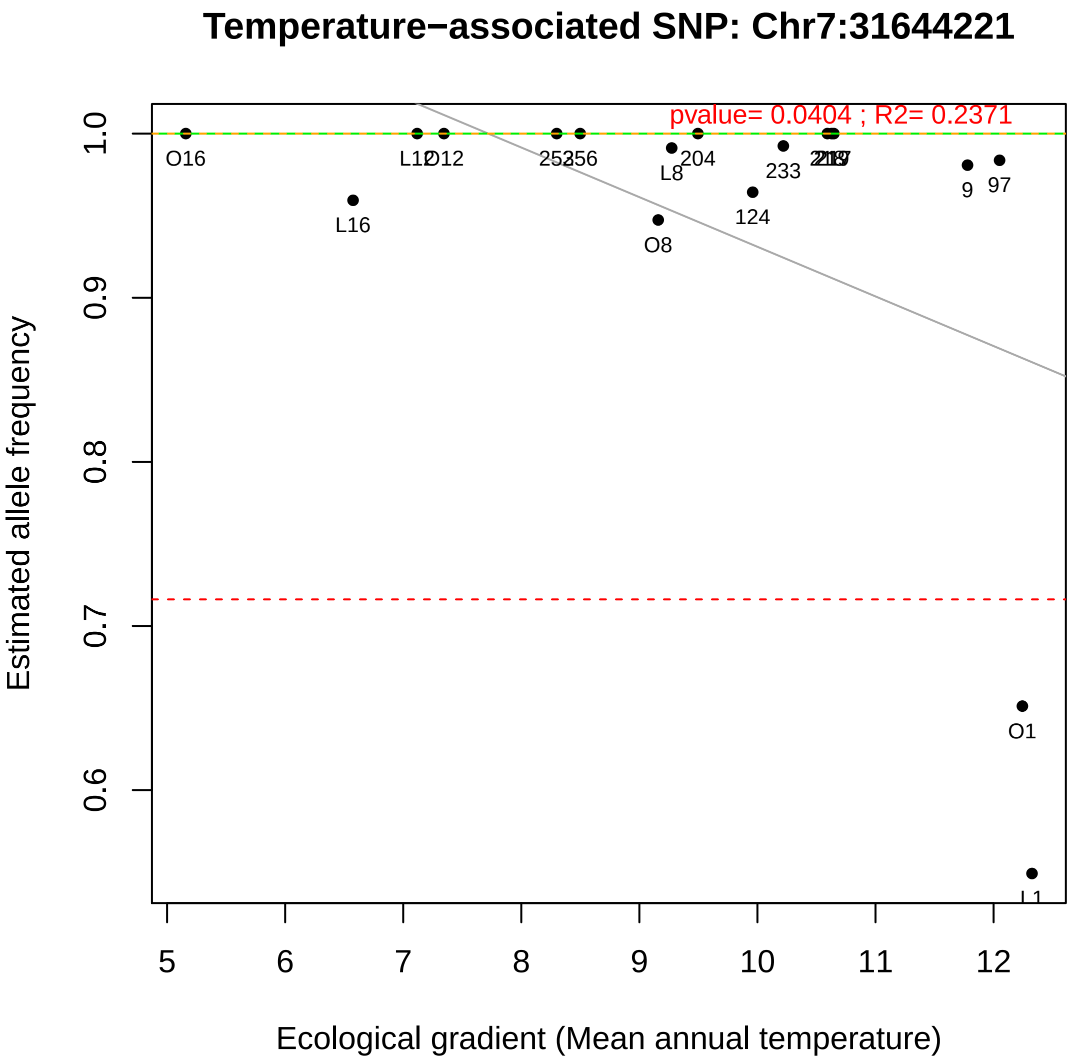

Temperature−associated SNP: Chr7:31644221
pvalue= 0.0404 ; R2= 0.2371
Estimated allele frequency
Ecological gradient (Mean annual temperature)
1.0
0.9
0.8
0.7
0.6
5
6
7
8
9
10
11
12
O16
L16
L12
O12
253
256
O8
L8
204
124
233
9
97
O1

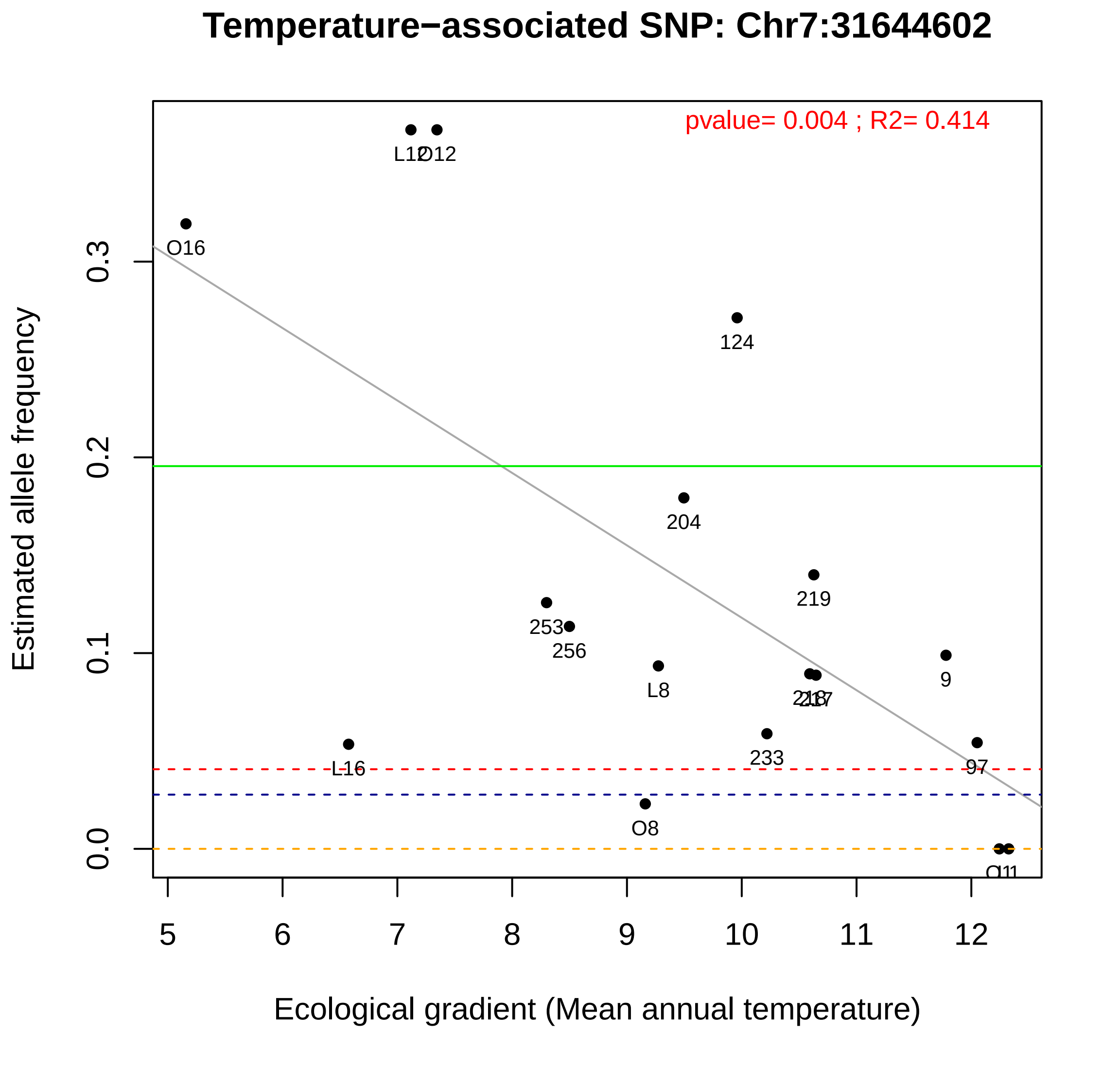
Temperature−associated SNP: Chr7:31644602
pvalue= 0.004 ; R2= 0.414
Estimated allele frequency
Ecological gradient (Mean annual temperature)
0.0
0.1
0.2
0.3
5
6
7
8
9
10
11
12
L12
O12
O16
124
204
219
253
256
L8
9
233
L16
97
O8

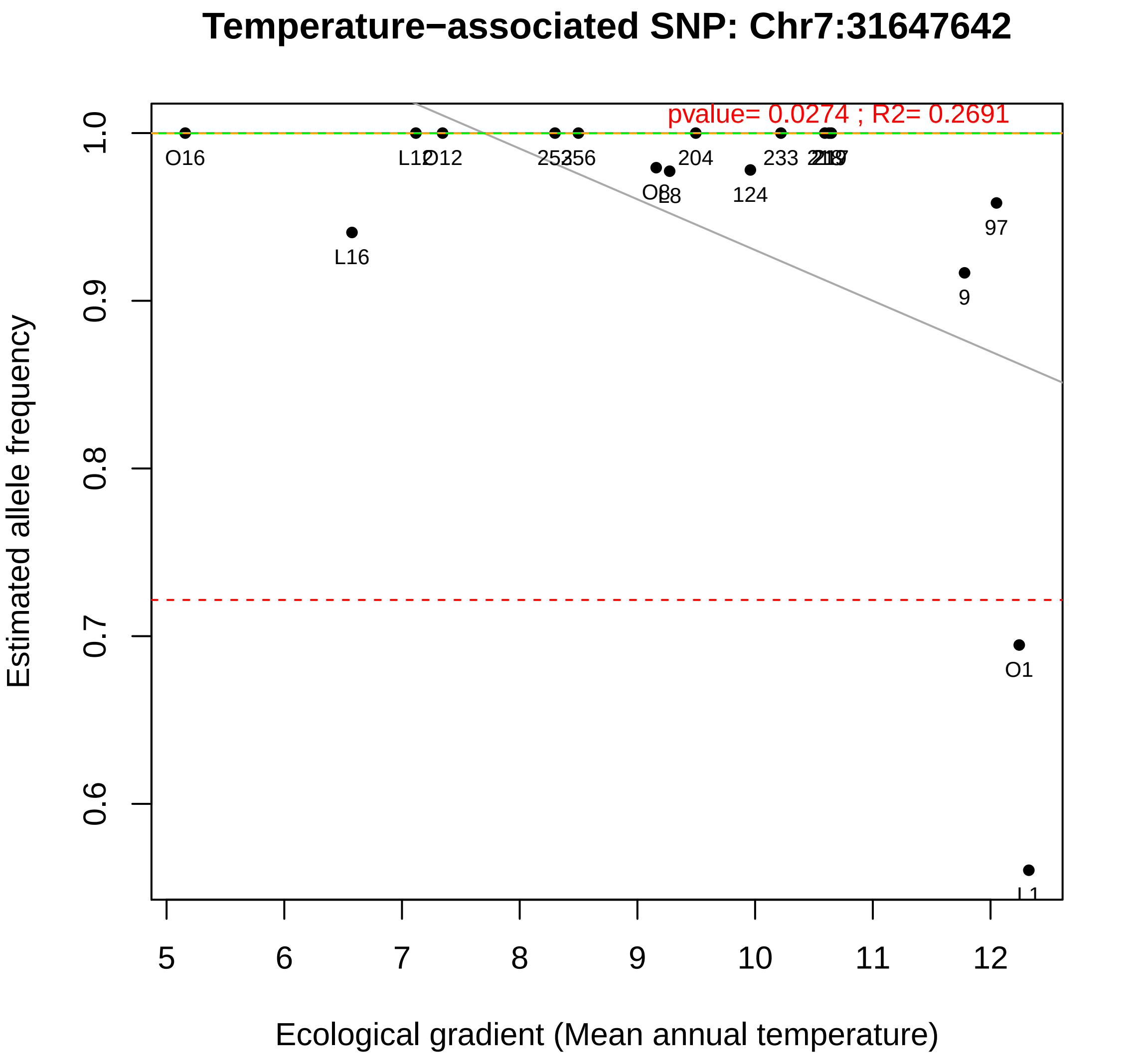

Temperature−associated SNP: Chr7:31647642
pvalue= 0.0274 ; R2= 0.2691
O16
L12
O12
L16
253
256
O8
L8
204
124
233
97
9
O1
L1
1.0
0.9
0.8
0.7
0.6
5
6
7
8
9
10
11
12
Estimated allele frequency
Ecological gradient (Mean annual temperature)

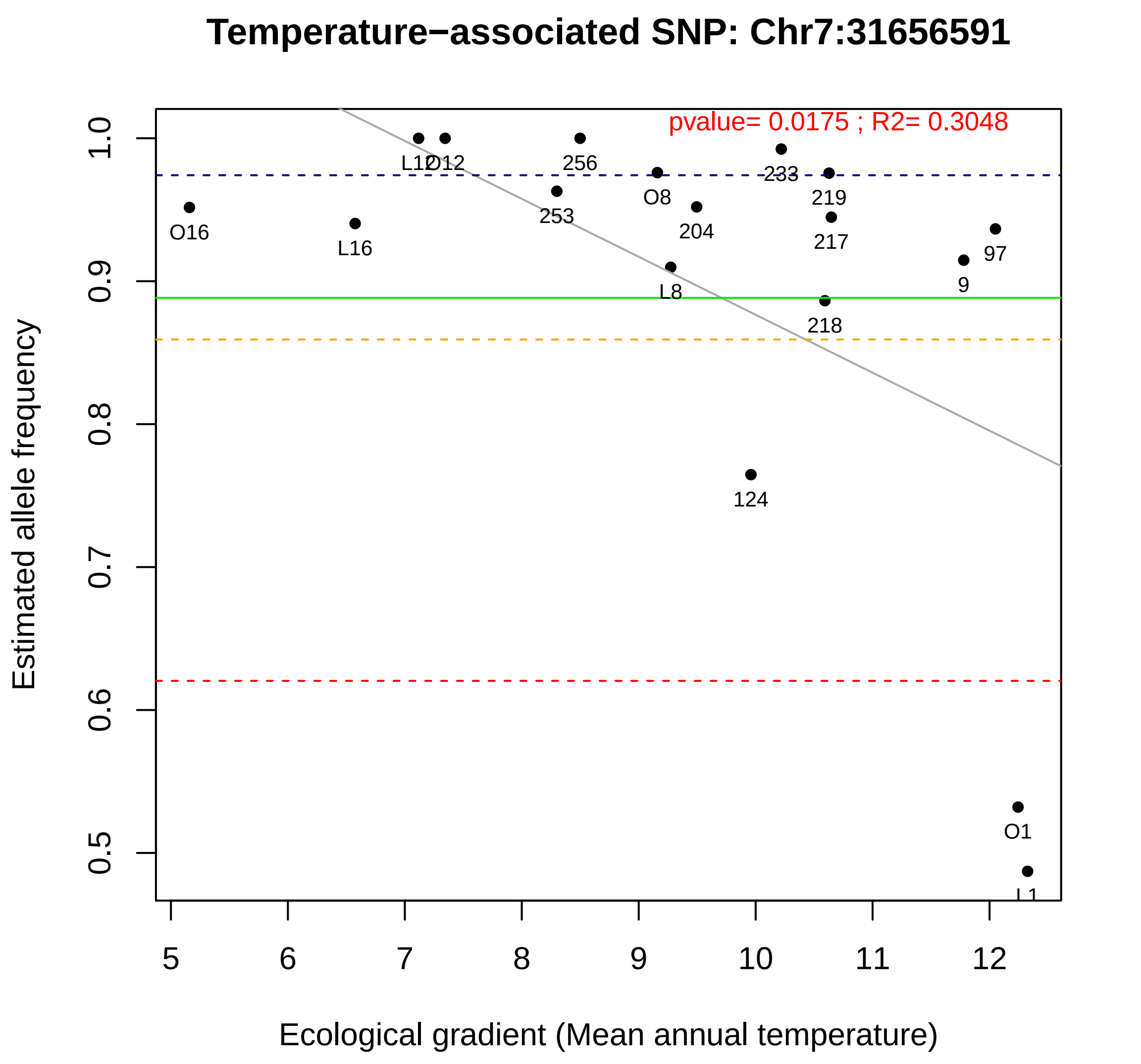

Temperature−associated SNP: Chr7:31656591
pvalue= 0.0175 ; R2= 0.3048
Estimated allele frequency
Ecological gradient (Mean annual temperature)
0.5
0.6
0.7
0.8
0.9
1.0
5
6
7
8
9
10
11
12
O16
L16
L12
O12
253
256
O8
204
L8
124
233
219
217
218
9
97
O1

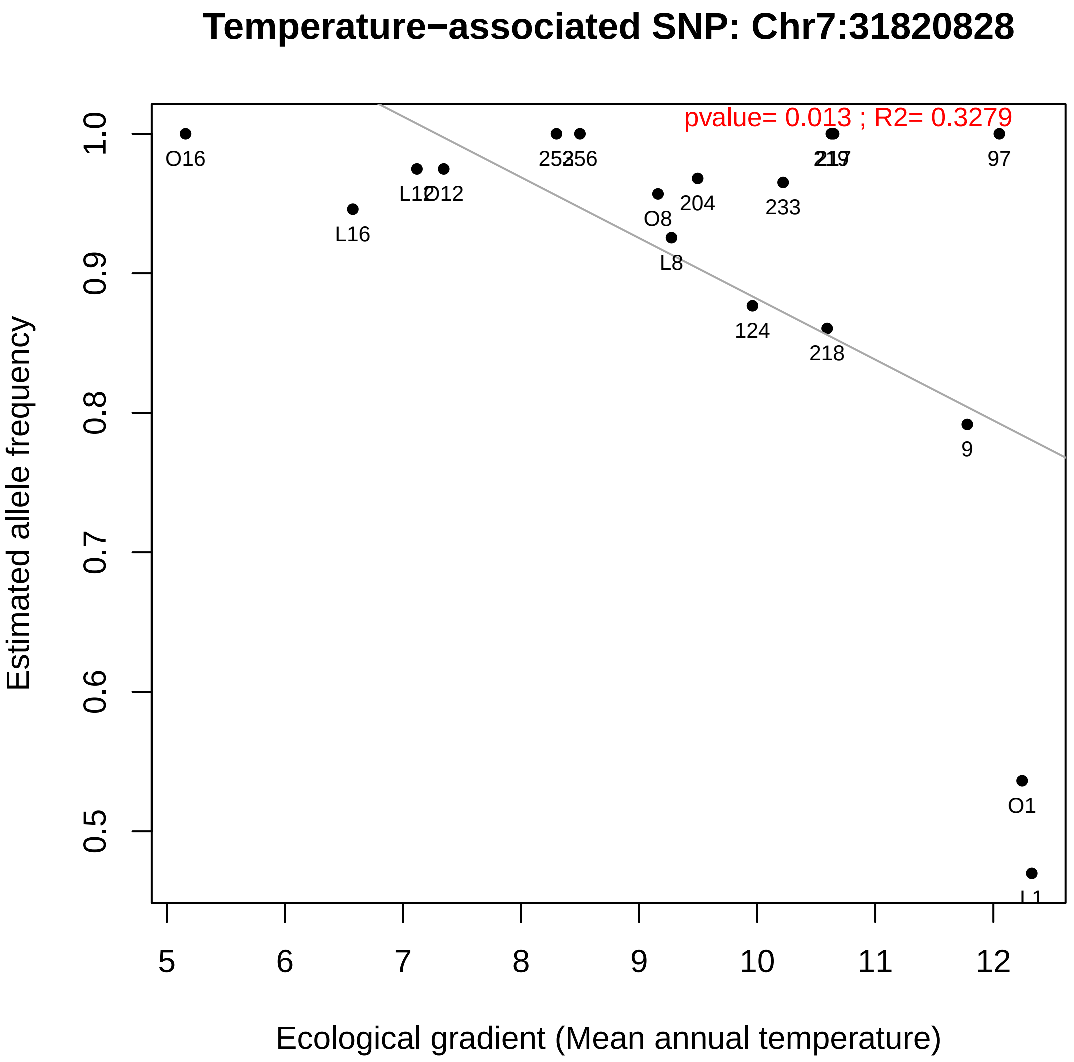

Temperature−associated SNP: Chr7:31820828
pvalue= 0.013 ; R2= 0.3279
Estimated allele frequency
Ecological gradient (Mean annual temperature)
1.0
0.9
0.8
0.7
0.6
0.5
5
6
7
8
9
10
11
12
O16
L16
L12
O12
O8
204
L8
233
124
218
97
9
O1

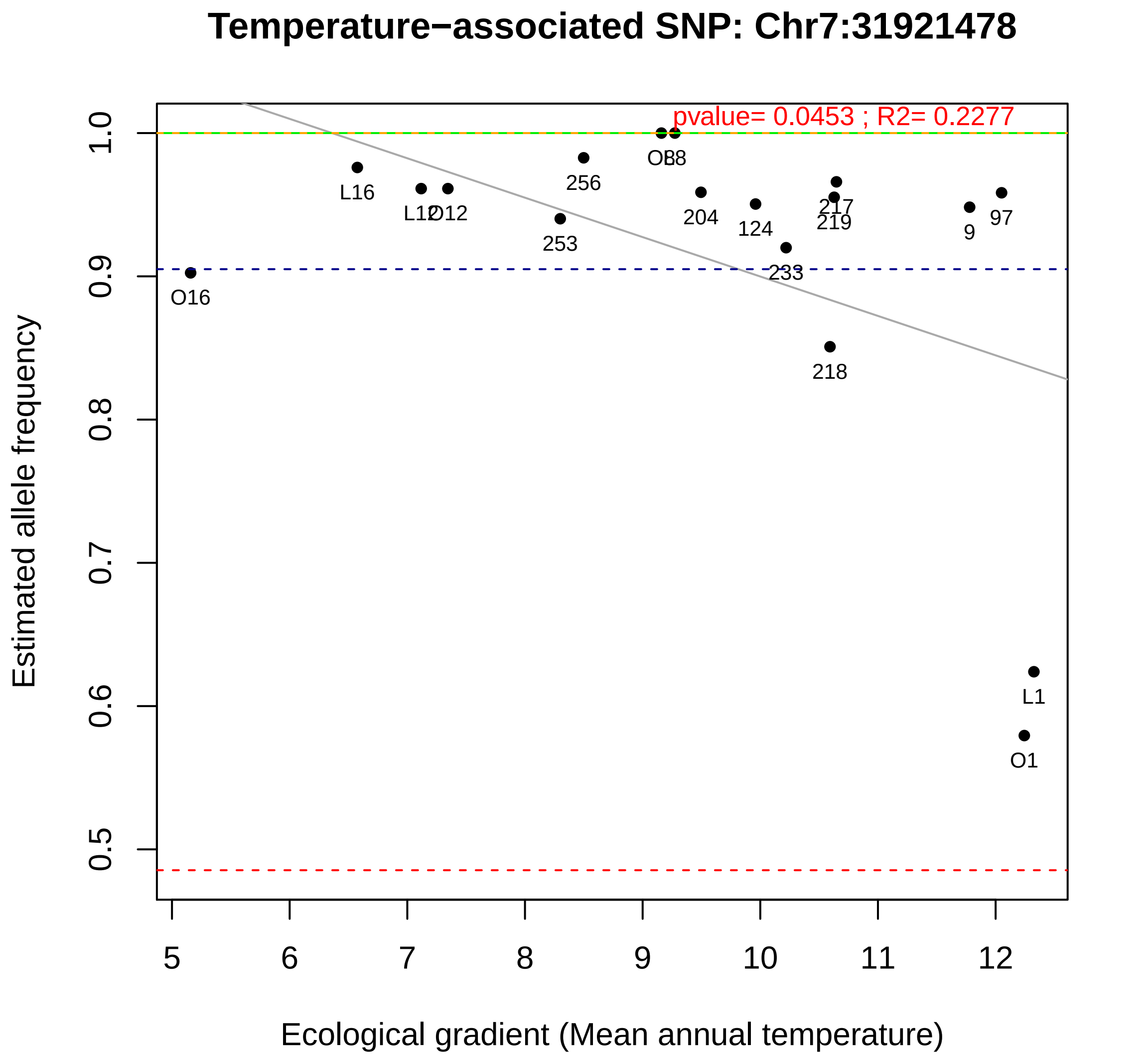

Temperature−associated SNP: Chr7:31921478
pvalue= 0.0453 ; R2= 0.2277
Estimated allele frequency
Ecological gradient (Mean annual temperature)
0.5
0.6
0.7
0.8
0.9
1.0
5
6
7
8
9
10
11
12
O16
L16
L12
O12
253
256
204
124
233
217
219
218
9
97
L1
O1

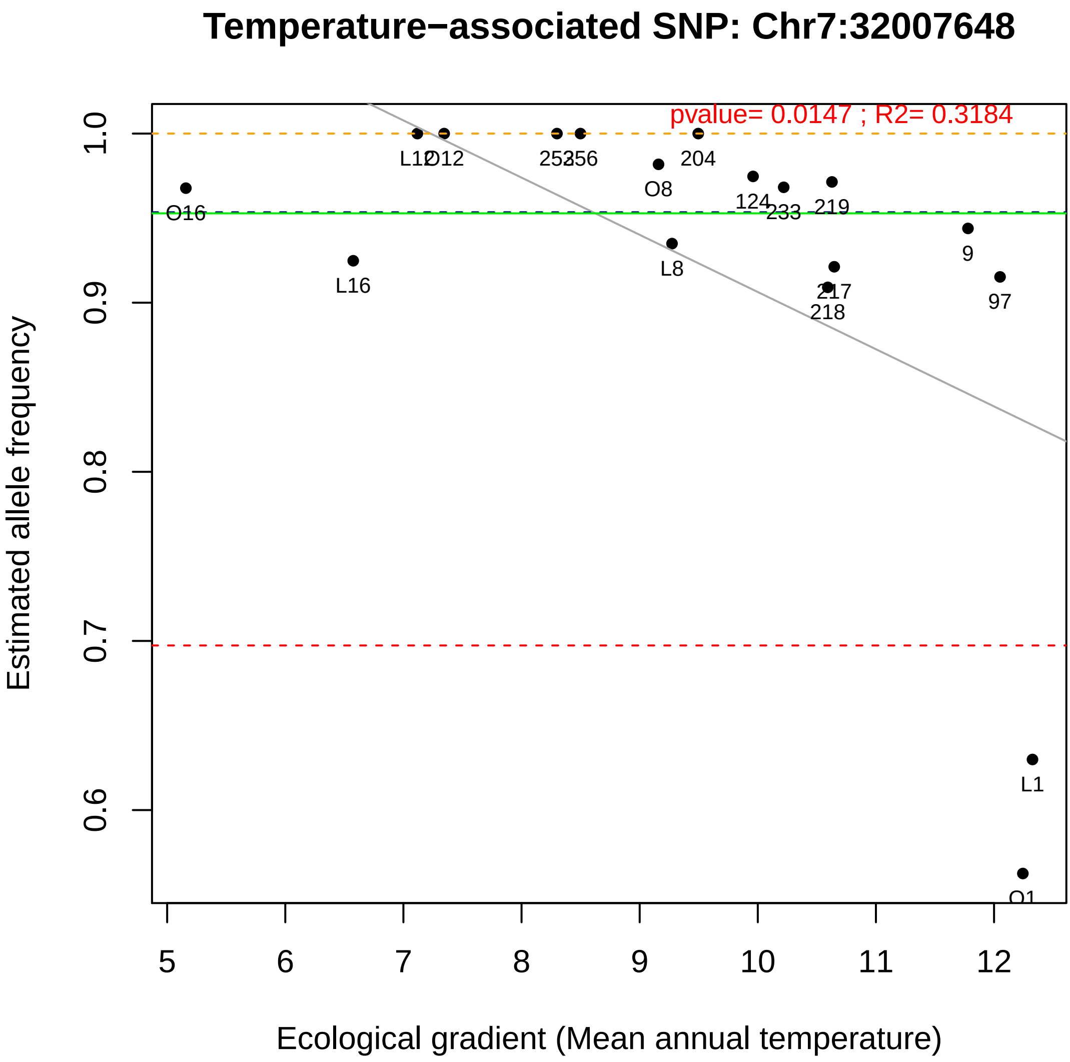
Temperature−associated SNP: Chr7:32007648
pvalue= 0.0147 ; R2= 0.3184
Estimated allele frequency
Ecological gradient (Mean annual temperature)
1.0
0.9
0.8
0.7
0.6
5
6
7
8
9
10
11
12
O16
L16
L12
O12
253
256
O8
204
L8
124
233
219
217
218
9
97
L1
O1

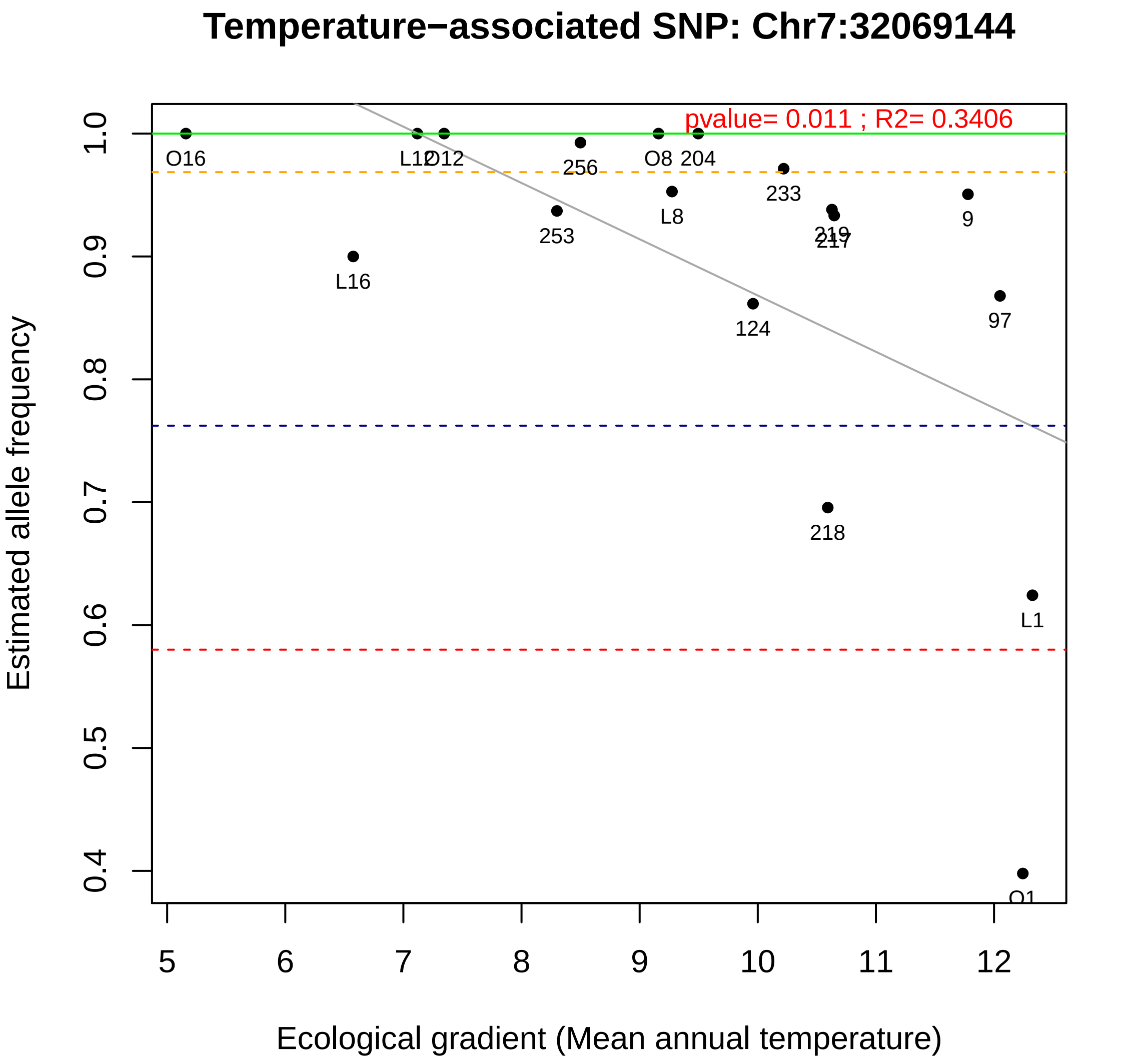

Temperature−associated SNP: Chr7:32069144
pvalue= 0.011 ; R2= 0.3406
Estimated allele frequency
Ecological gradient (Mean annual temperature)
1.0
0.9
0.8
0.7
0.6
0.5
0.4
5
6
7
8
9
10
11
12
O16
L12
O12
256
O8
204
233
L8
253
219
217
9
L16
124
97
218
L1
O1

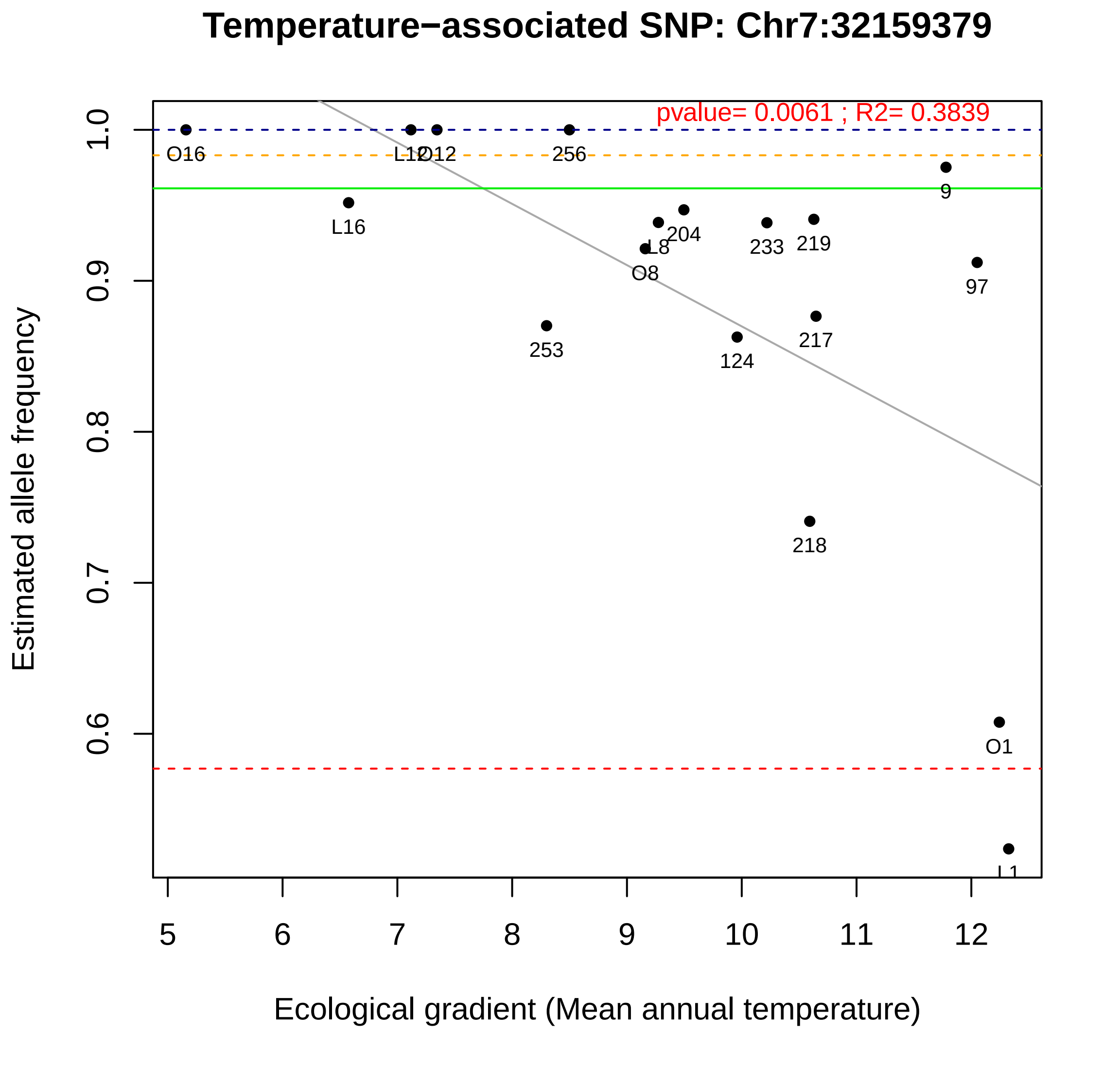
Temperature−associated SNP: Chr7:32159379
pvalue= 0.0061 ; R2= 0.3839
Estimated allele frequency
Ecological gradient (Mean annual temperature)
1.0
0.9
0.8
0.7
0.6
5
6
7
8
9
10
11
12
O16
L12
O12
256
9
L16
204
L8
233
219
O8
97
253
124
217
218
O1

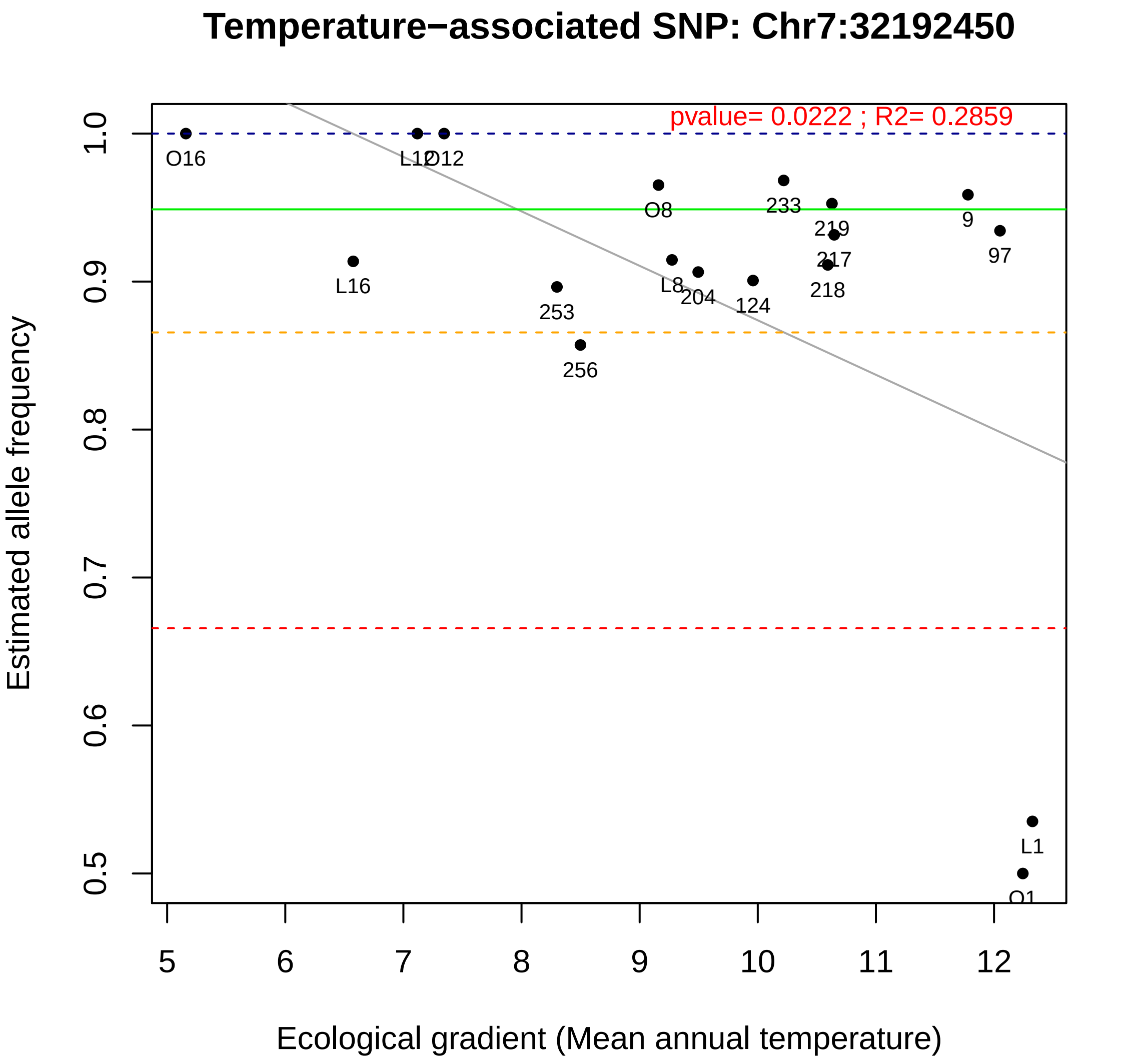

Temperature−associated SNP: Chr7:32192450
pvalue= 0.0222 ; R2= 0.2859
Estimated allele frequency
Ecological gradient (Mean annual temperature)
1.0
0.9
0.8
0.7
0.6
0.5
5
6
7
8
9
10
11
12
O16
L12
O12
O8
233
219
9
97
217
L16
253
L8
204
124
218
256
L1
O1

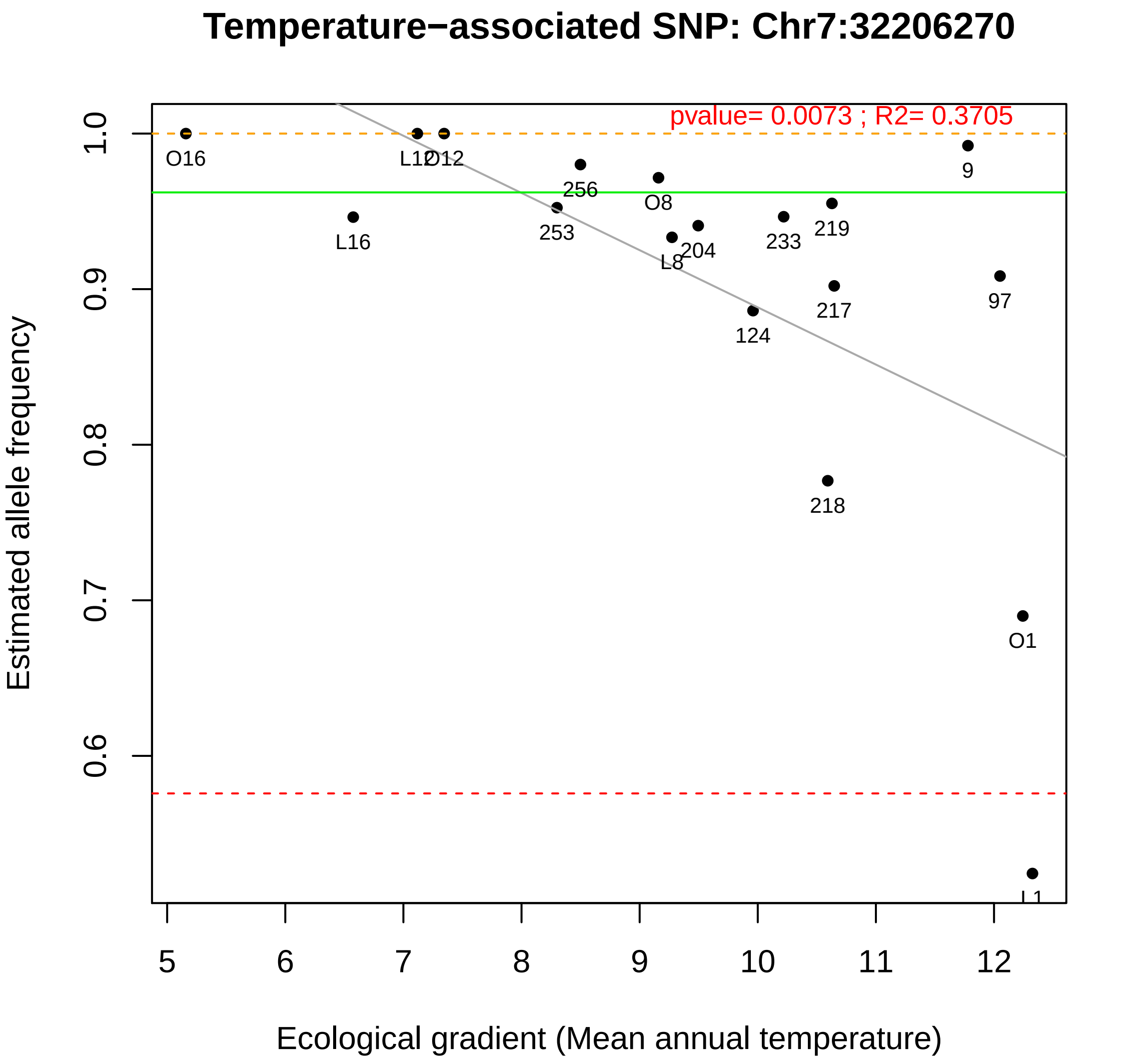
Temperature−associated SNP: Chr7:32206270
pvalue= 0.0073 ; R2= 0.3705
Estimated allele frequency
Ecological gradient (Mean annual temperature)
1.0
0.9
0.8
0.7
0.6
5
6
7
8
9
10
11
12
O16
L12
O12
256
O8
9
L16
253
204
L8
233
219
217
97
124
218
O1

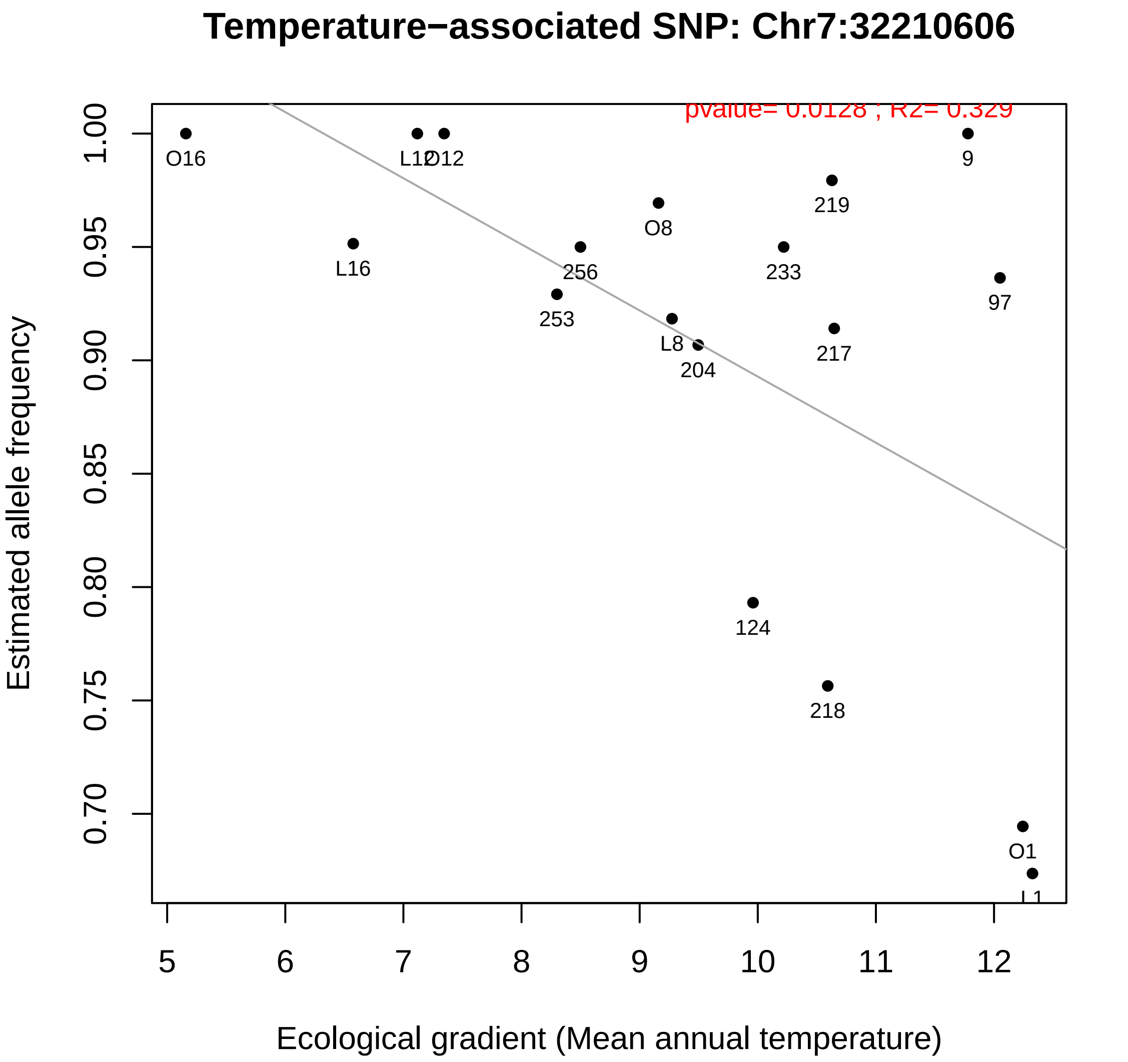

Temperature−associated SNP: Chr7:32210606
pvalue= 0.0128 ; R2= 0.329
O16
L12
O12
9
219
O8
L16
256
233
97
253
L8
204
217
124
218
O1
L1
1.00
0.95
0.90
0.85
0.80
0.75
0.70
5
6
7
8
9
10
11
12
Estimated allele frequency
Ecological gradient (Mean annual temperature)

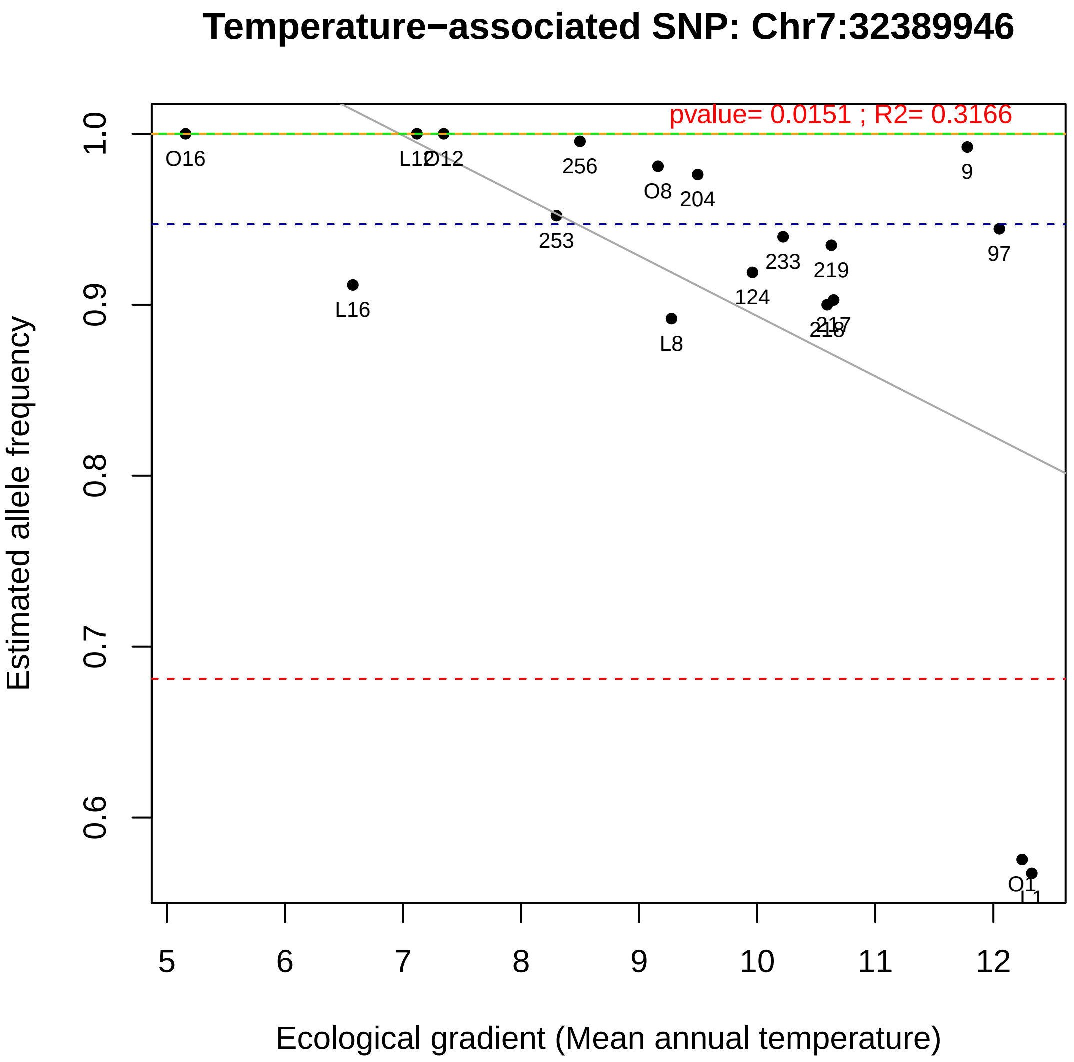
Temperature−associated SNP: Chr7:32389946
pvalue= 0.0151 ; R2= 0.3166
Estimated allele frequency
Ecological gradient (Mean annual temperature)
1.0
0.9
0.8
0.7
0.6
5
6
7
8
9
10
11
12
O16
L12
O12
256
O8
204
9
253
233
219
97
124
L16
L8
217
218
O1
L1

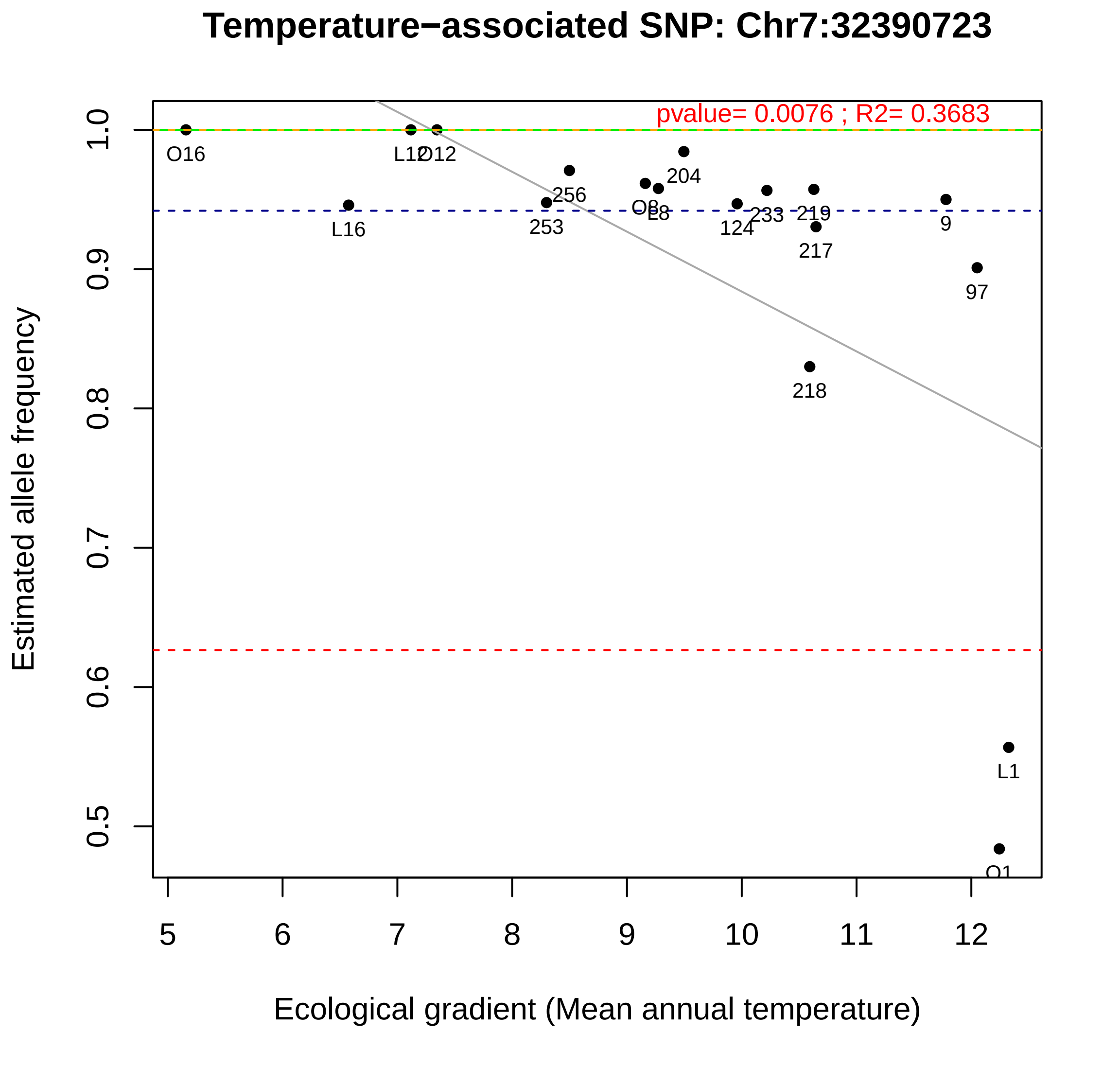

Temperature−associated SNP: Chr7:32390723
pvalue= 0.0076 ; R2= 0.3683
Estimated allele frequency
Ecological gradient (Mean annual temperature)
1.0
0.9
0.8
0.7
0.6
0.5
5
6
7
8
9
10
11
12
O16
L12
O12
L16
256
253
O8
L8
204
124
233
219
217
9
97
218
L1
O1

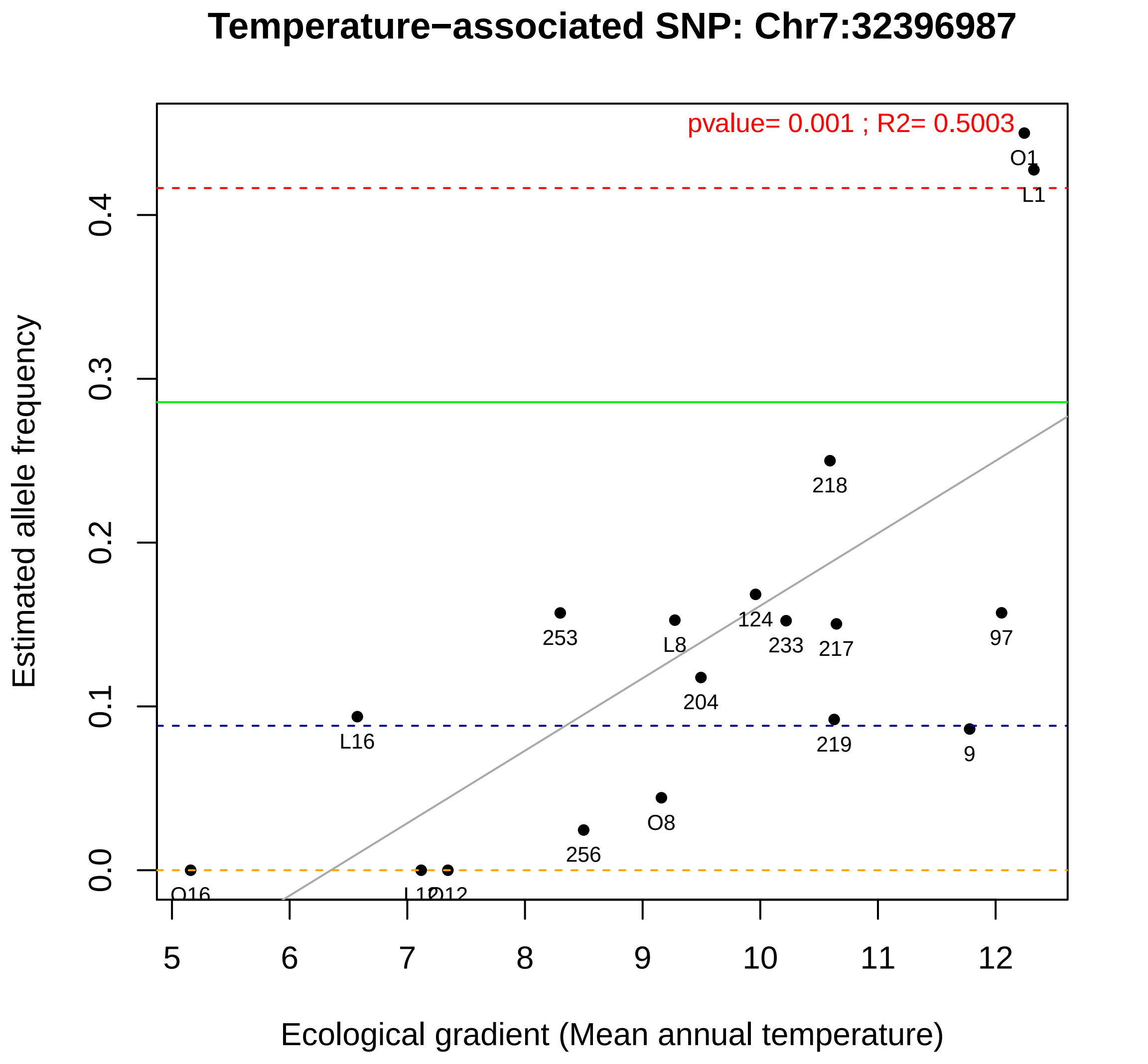
Temperature−associated SNP: Chr7:32396987
pvalue= 0.001 ; R2= 0.5003
Estimated allele frequency
0.0
0.1
0.2
0.3
0.4
Ecological gradient (Mean annual temperature)
5
6
7
8
9
10
11
12
O1
L1
218
124
253
L8
233
217
97
204
L16
219
9
O8
256
O16
O12

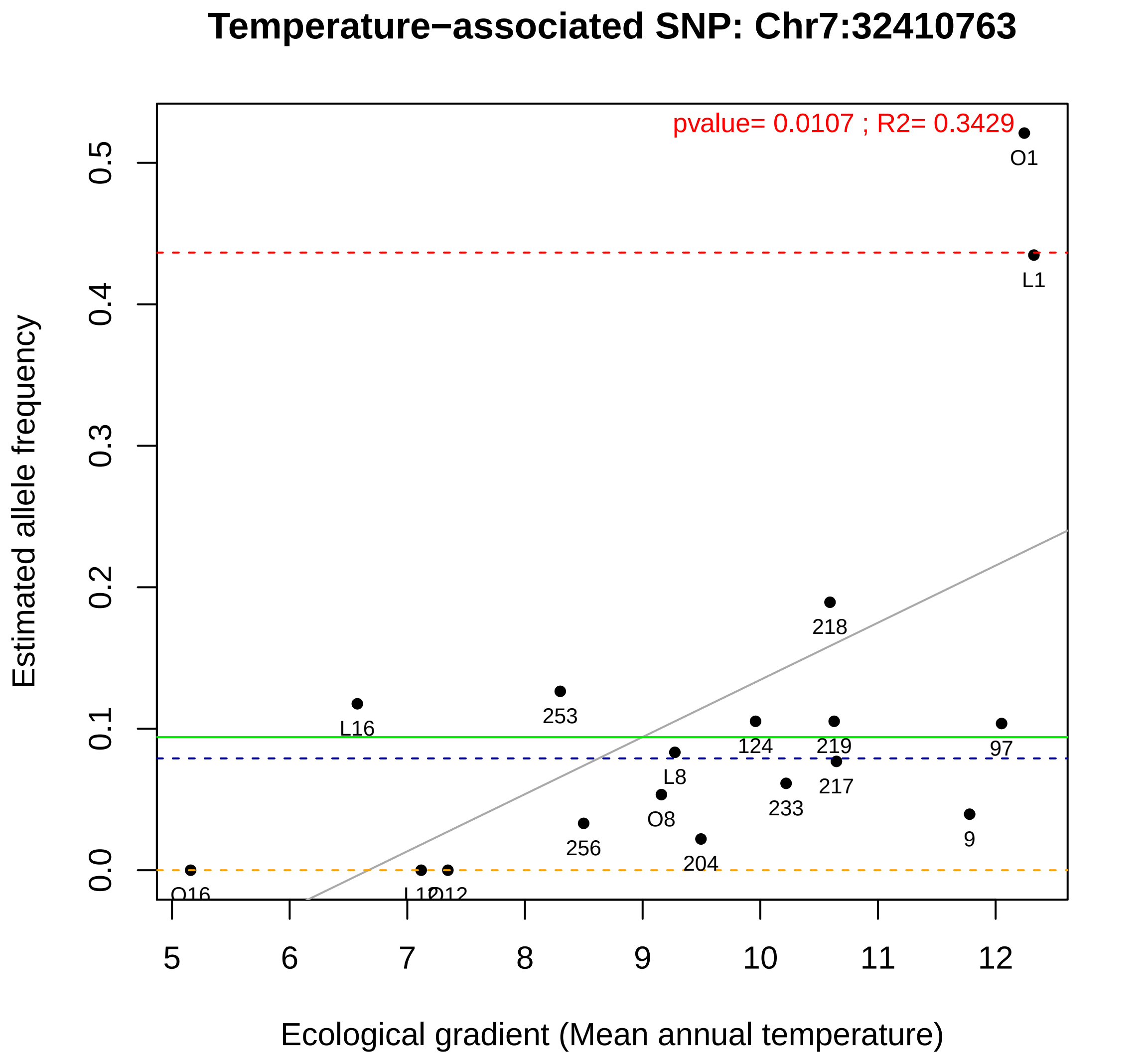

Temperature−associated SNP: Chr7:32410763
pvalue= 0.0107 ; R2= 0.3429
Estimated allele frequency
0.0
0.1
0.2
0.3
0.4
0.5
5
6
7
8
9
10
11
12
Ecological gradient (Mean annual temperature)
O1
L1
218
L16
253
124
219
97
L8
217
233
O8
256
204
9
O16
O12

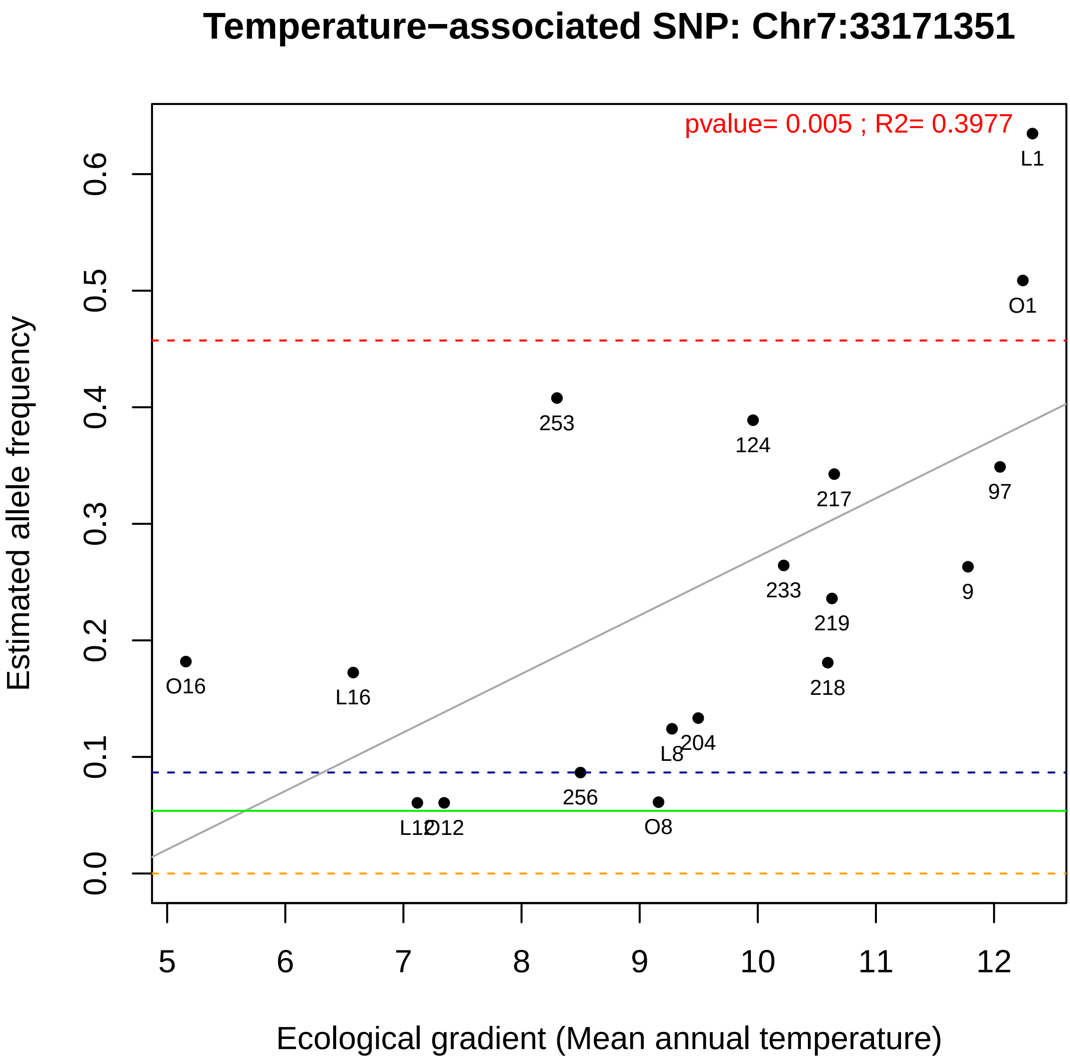
Temperature−associated SNP: Chr7:33171351
pvalue= 0.005 ; R2= 0.3977
Estimated allele frequency
0.0
0.1
0.2
0.3
0.4
0.5
0.6
5
6
7
8
9
10
11
12
Ecological gradient (Mean annual temperature)
L1
O1
253
124
217
97
233
9
219
O16
L16
218
204
L8
256
L12
O12
O8

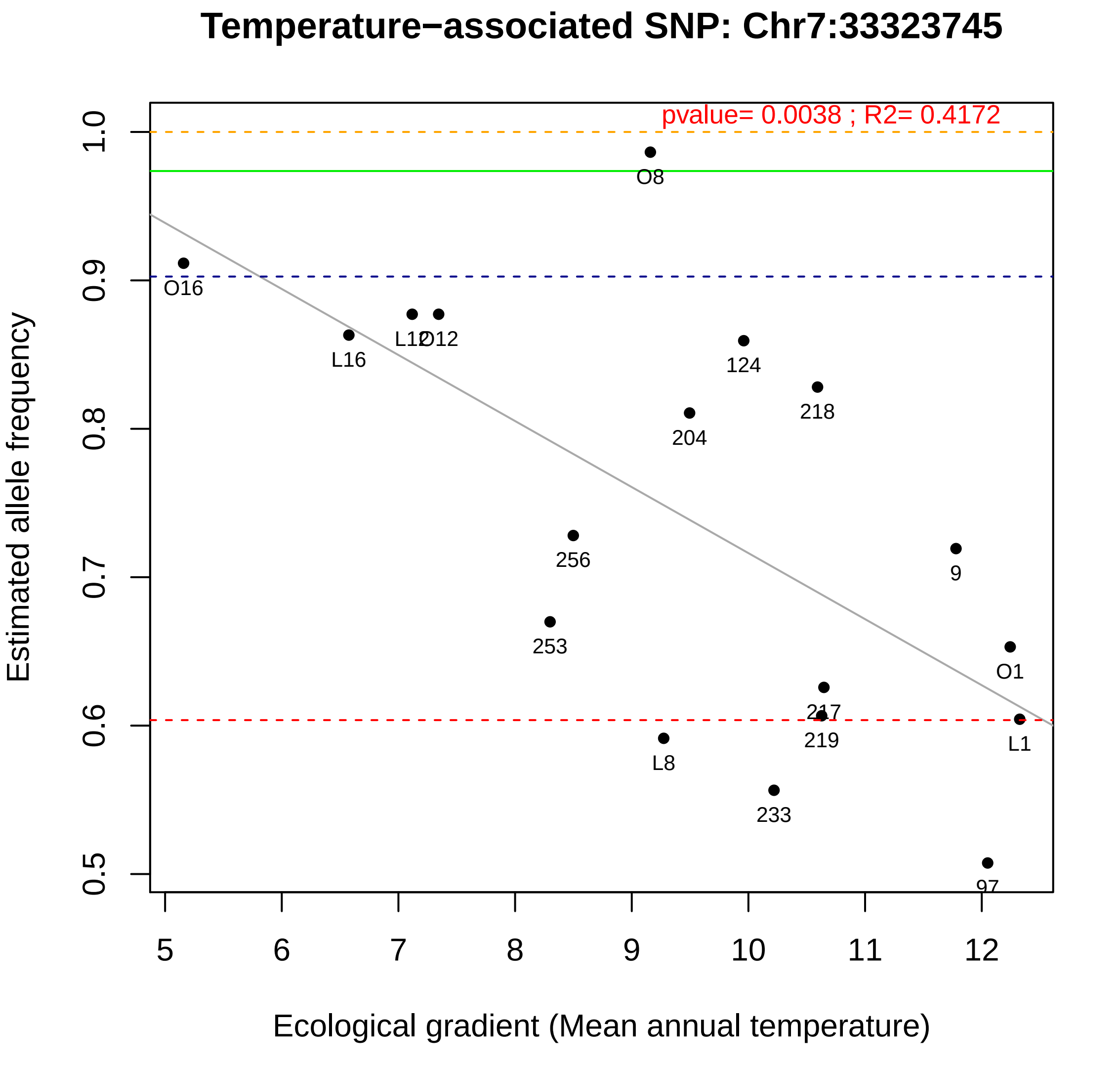

Temperature−associated SNP: Chr7:33323745
pvalue= 0.0038 ; R2= 0.4172
Estimated allele frequency
Ecological gradient (Mean annual temperature)
0.5
0.6
0.7
0.8
0.9
1.0
5
6
7
8
9
10
11
12
O8
O16
L16
L12
O12
124
218
204
256
253
9
O1
217
219
L1
L8
233
97

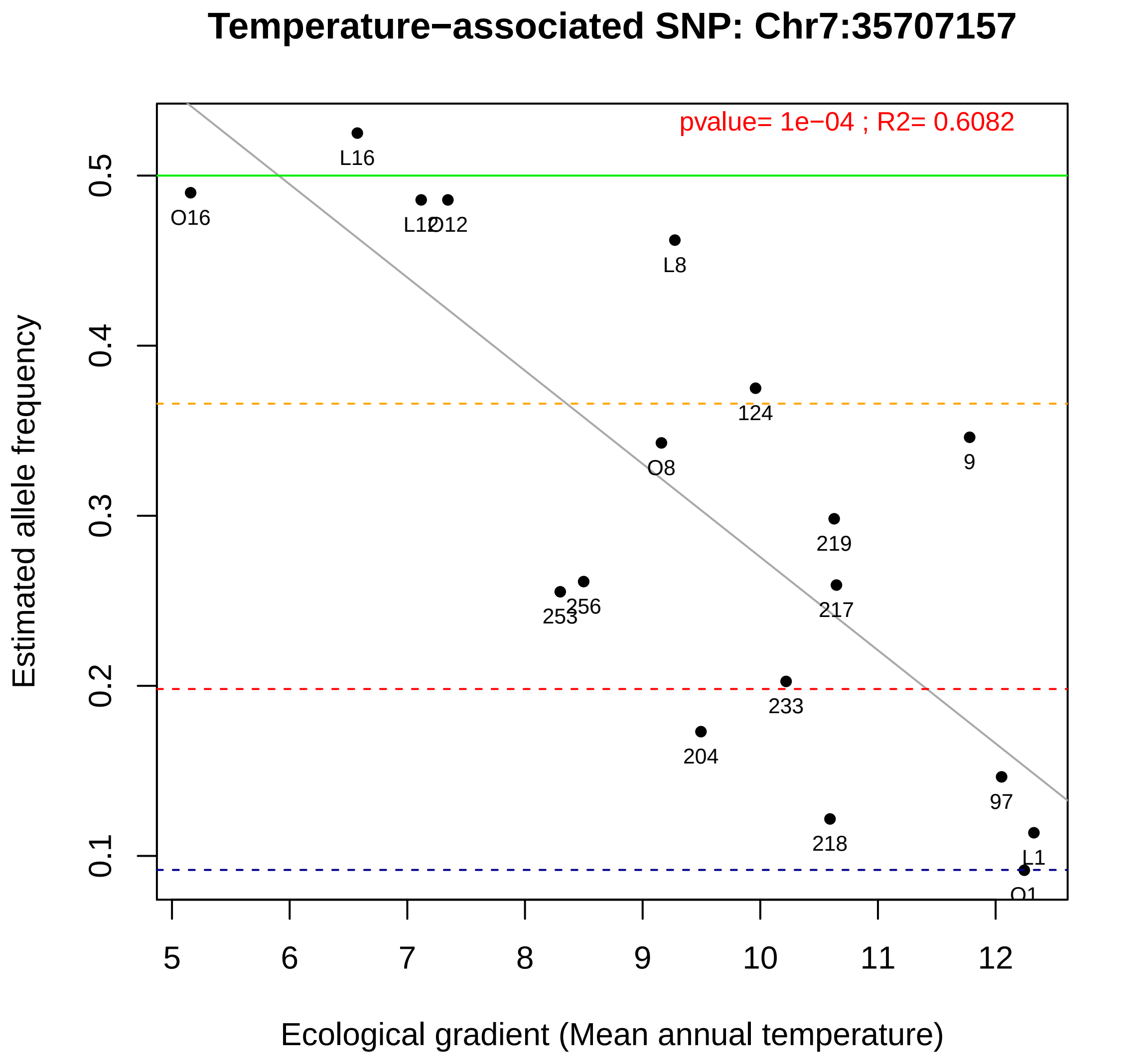

Temperature−associated SNP: Chr7:35707157
pvalue= 1e−04 ; R2= 0.6082
Estimated allele frequency
0.1
0.2
0.3
0.4
0.5
5
6
7
8
9
10
11
12
Ecological gradient (Mean annual temperature)
L16
O16
L12
O12
L8
124
O8
9
219
253
256
217
233
204
97
218
L1
O1

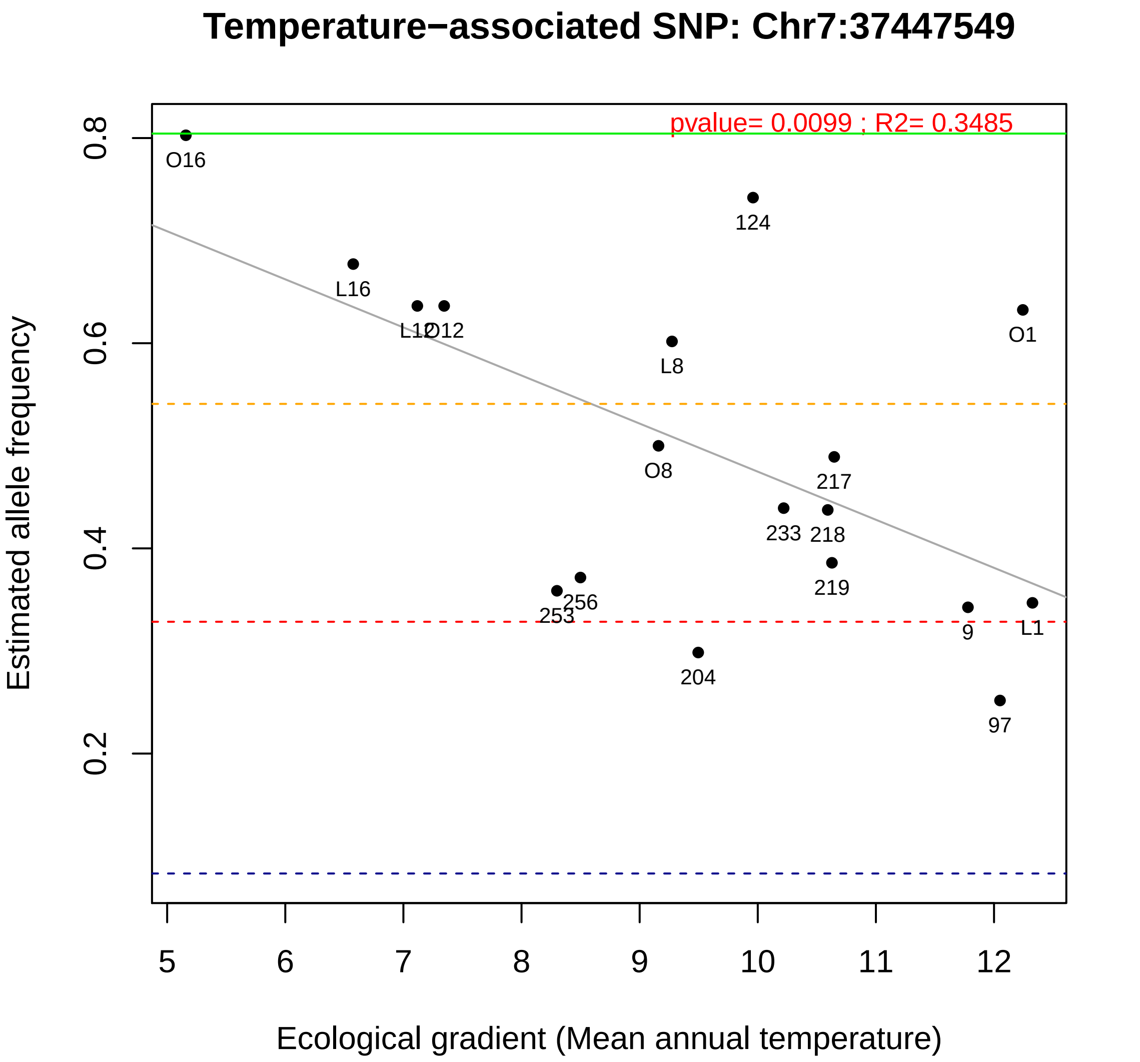

Temperature−associated SNP: Chr7:37447549
pvalue= 0.0099 ; R2= 0.3485
Estimated allele frequency
0.2
0.4
0.6
0.8
5
6
7
8
9
10
11
12
Ecological gradient (Mean annual temperature)
O16
124
L16
L12
O12
O1
L8
O8
217
233
218
219
256
253
9
L1
204
97

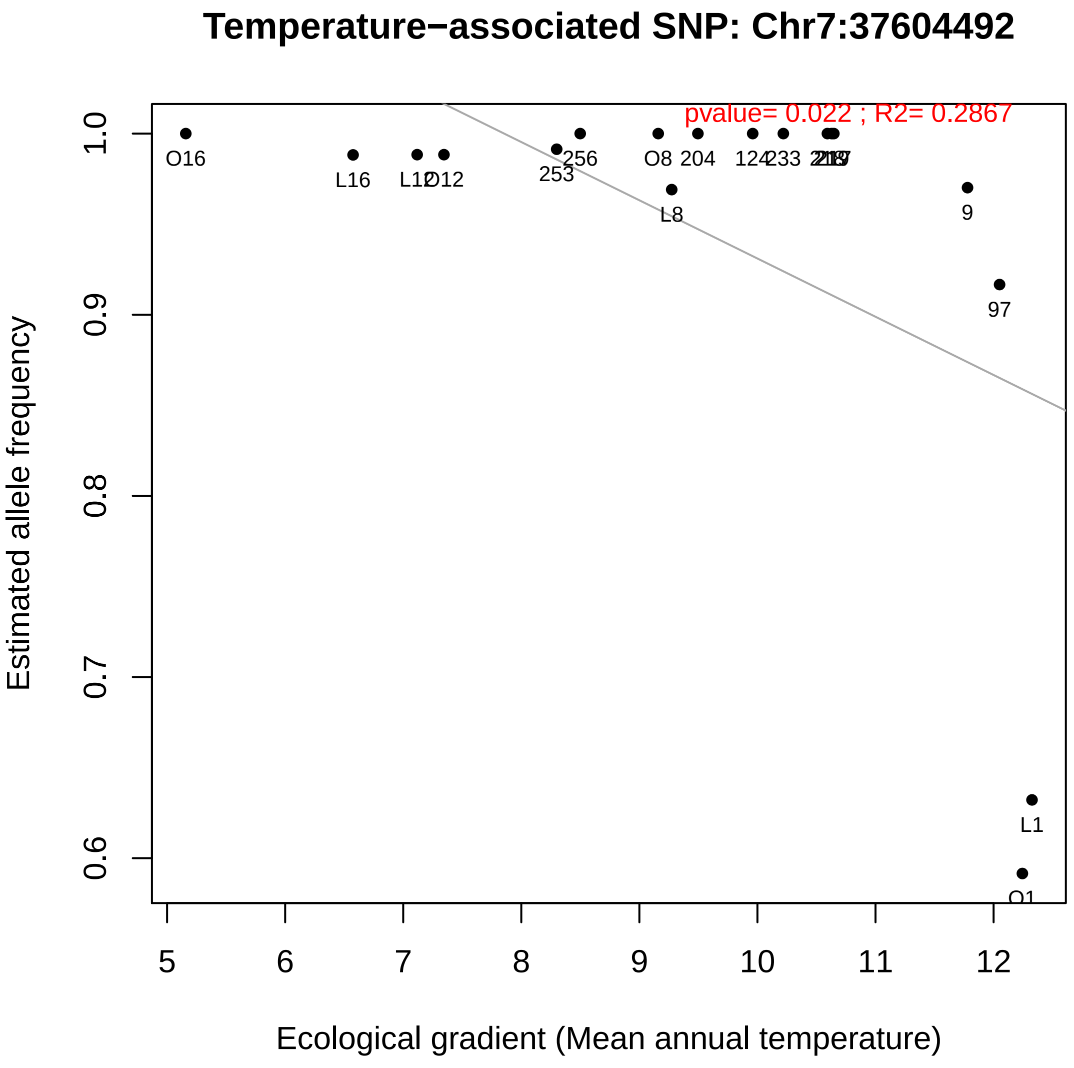
Temperature−associated SNP: Chr7:37604492
pvalue= 0.022 ; R2= 0.2867
Estimated allele frequency
Ecological gradient (Mean annual temperature)
O16
L16
L12
O12
253
256
O8
204
L8
124
233
9
97
L1
O1
1.0
0.9
0.8
0.7
0.6
5
6
7
8
9
10
11
12

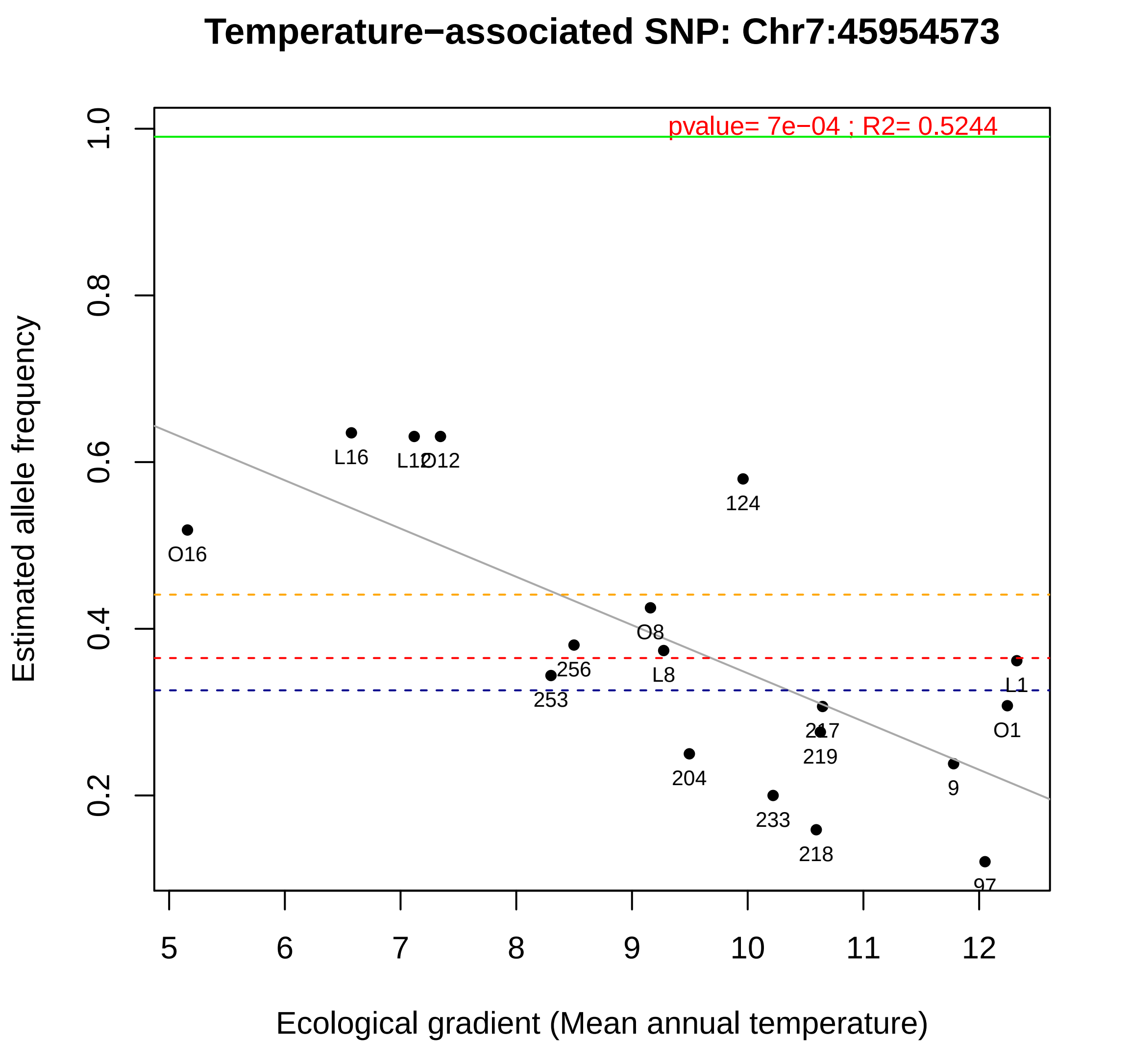

Temperature−associated SNP: Chr7:45954573
pvalue= 7e−04 ; R2= 0.5244
Estimated allele frequency
0.2
0.4
0.6
0.8
1.0
Ecological gradient (Mean annual temperature)
5
6
7
8
9
10
11
12
L16
L12
O12
124
O16
O8
256
L8
253
L1
217
O1
219
204
9
233
218
97

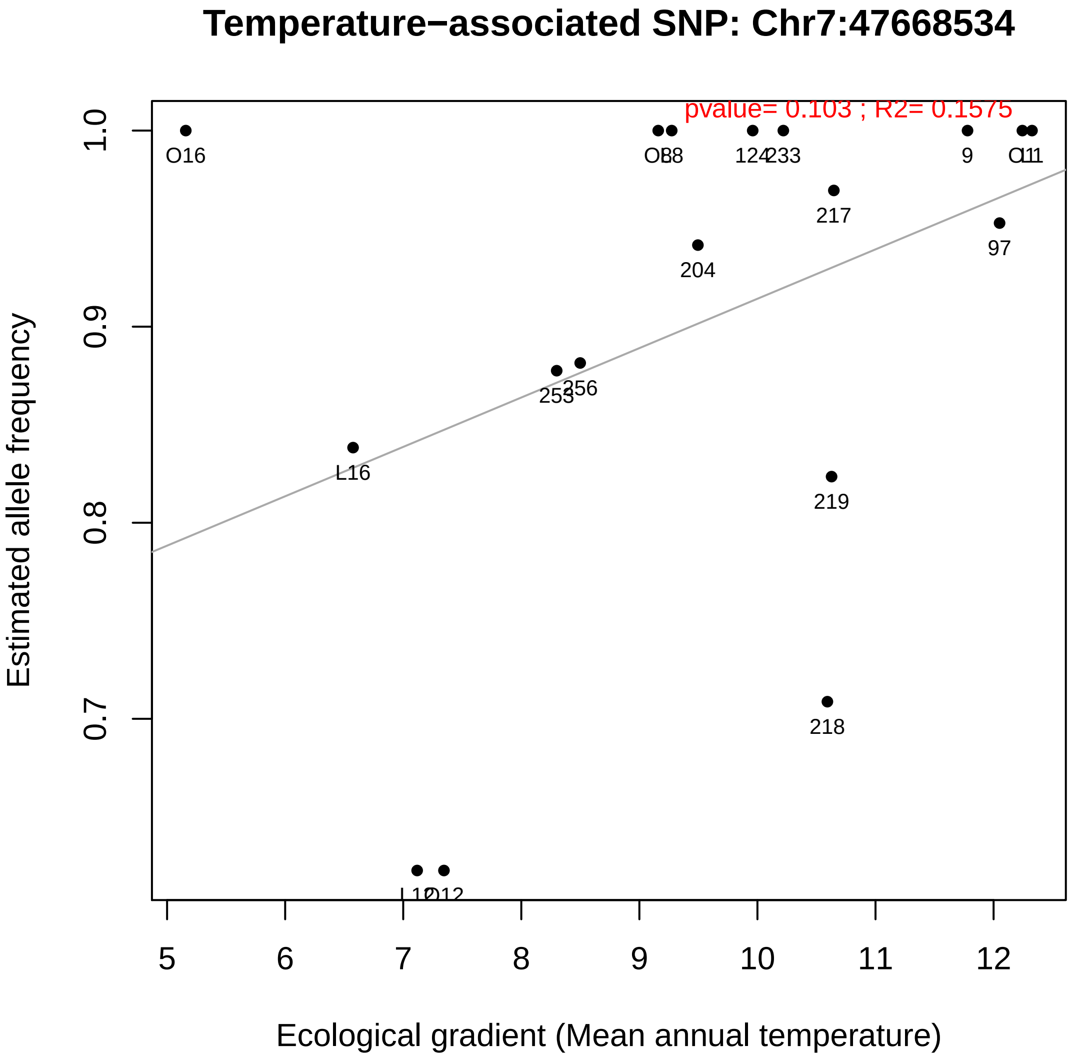

Temperature−associated SNP: Chr7:47668534
pvalue= 0.103 ; R2= 0.1575
O16
124
233
9
217
97
204
253
256
L16
219
218
1.0
0.9
0.8
0.7
5
6
7
8
9
10
11
12
Estimated allele frequency
Ecological gradient (Mean annual temperature)

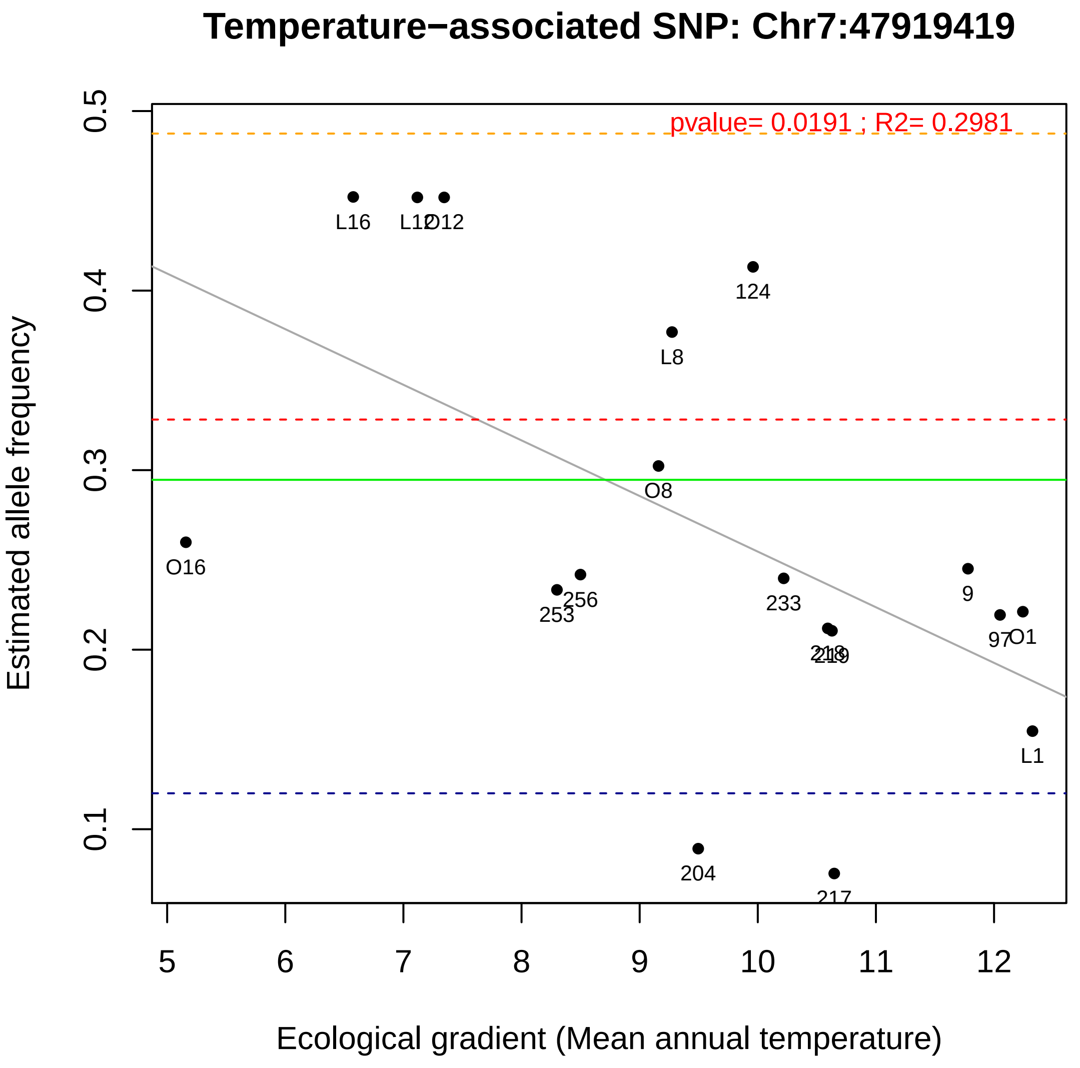

Temperature−associated SNP: Chr7:47919419
pvalue= 0.0191 ; R2= 0.2981
Estimated allele frequency
0.1
0.2
0.3
0.4
0.5
Ecological gradient (Mean annual temperature)
5
6
7
8
9
10
11
12
L16
O12
124
L8
O8
O16
256
253
233
9
97
O1
L1
204
217

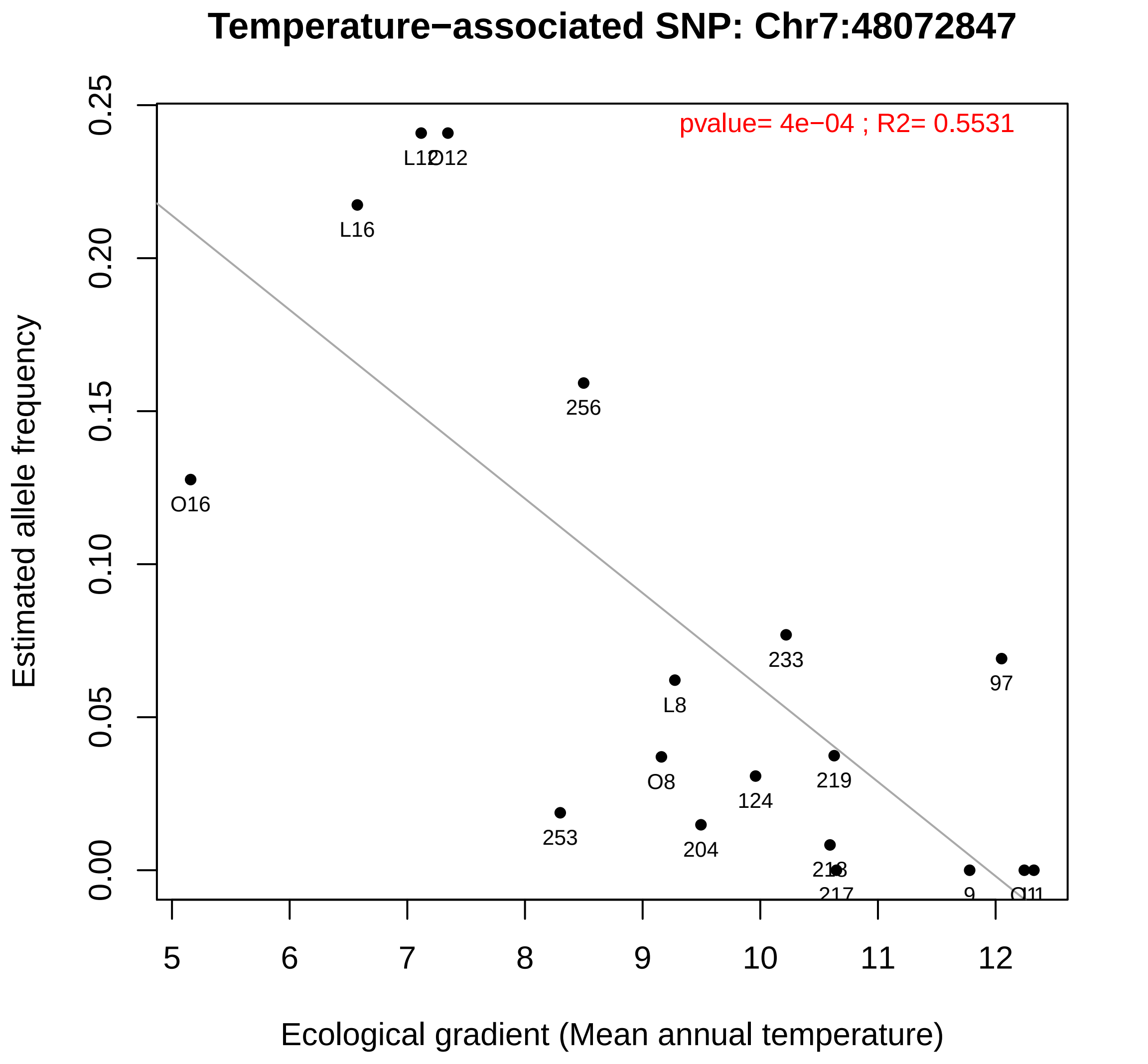

Temperature−associated SNP: Chr7:48072847
pvalue= 4e−04 ; R2= 0.5531
Estimated allele frequency
0.00
0.05
0.10
0.15
0.20
0.25
5
6
7
8
9
10
11
12
Ecological gradient (Mean annual temperature)
L16
O12
256
O16
233
97
L8
O8
219
124
253
204
9

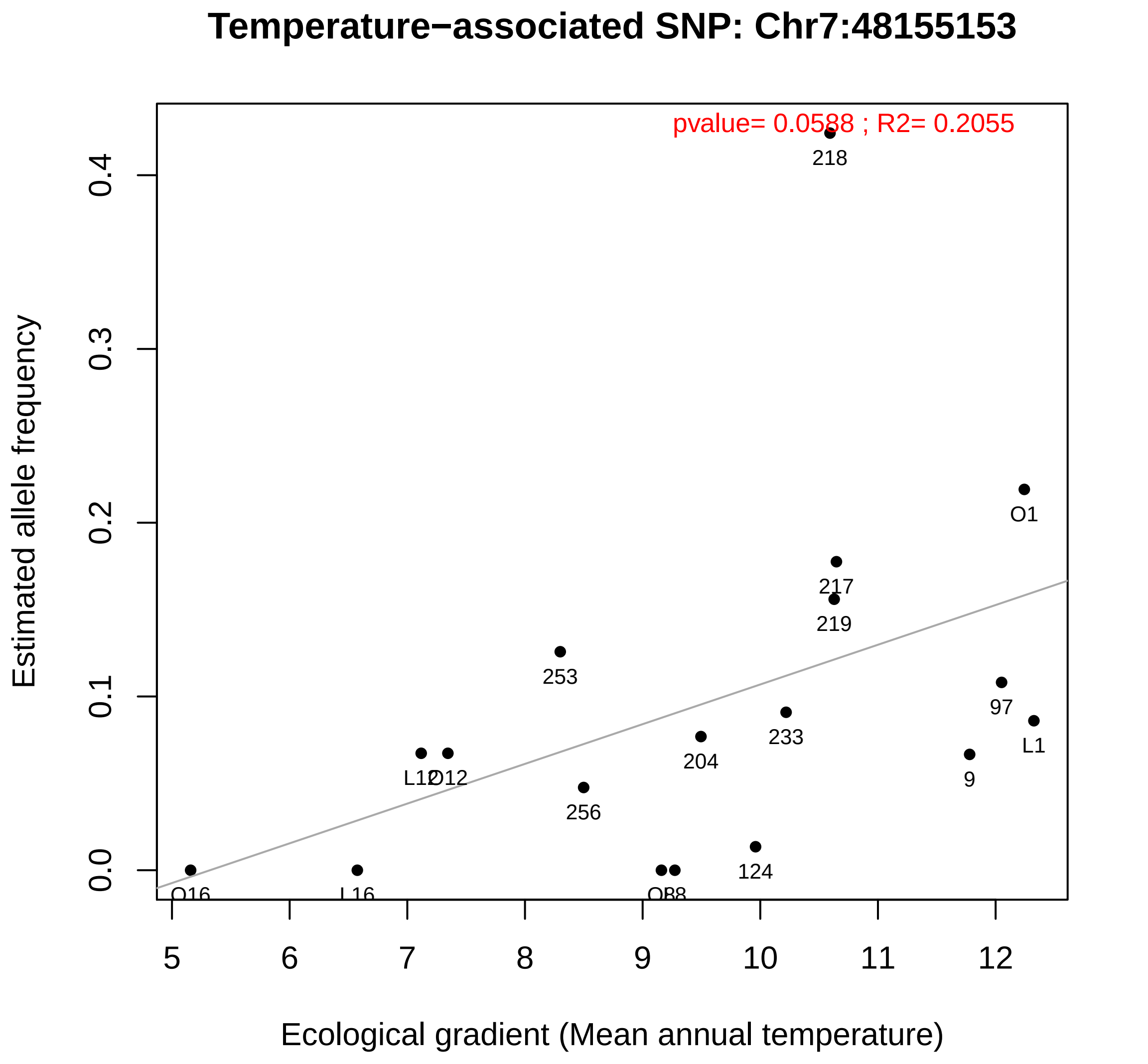
Temperature−associated SNP: Chr7:48155153
pvalue= 0.0588 ; R2= 0.2055
Estimated allele frequency
0.0
0.1
0.2
0.3
0.4
5
6
7
8
9
10
11
12
Ecological gradient (Mean annual temperature)
218
O1
217
219
253
97
233
L1
204
9
256
124
O16
L16

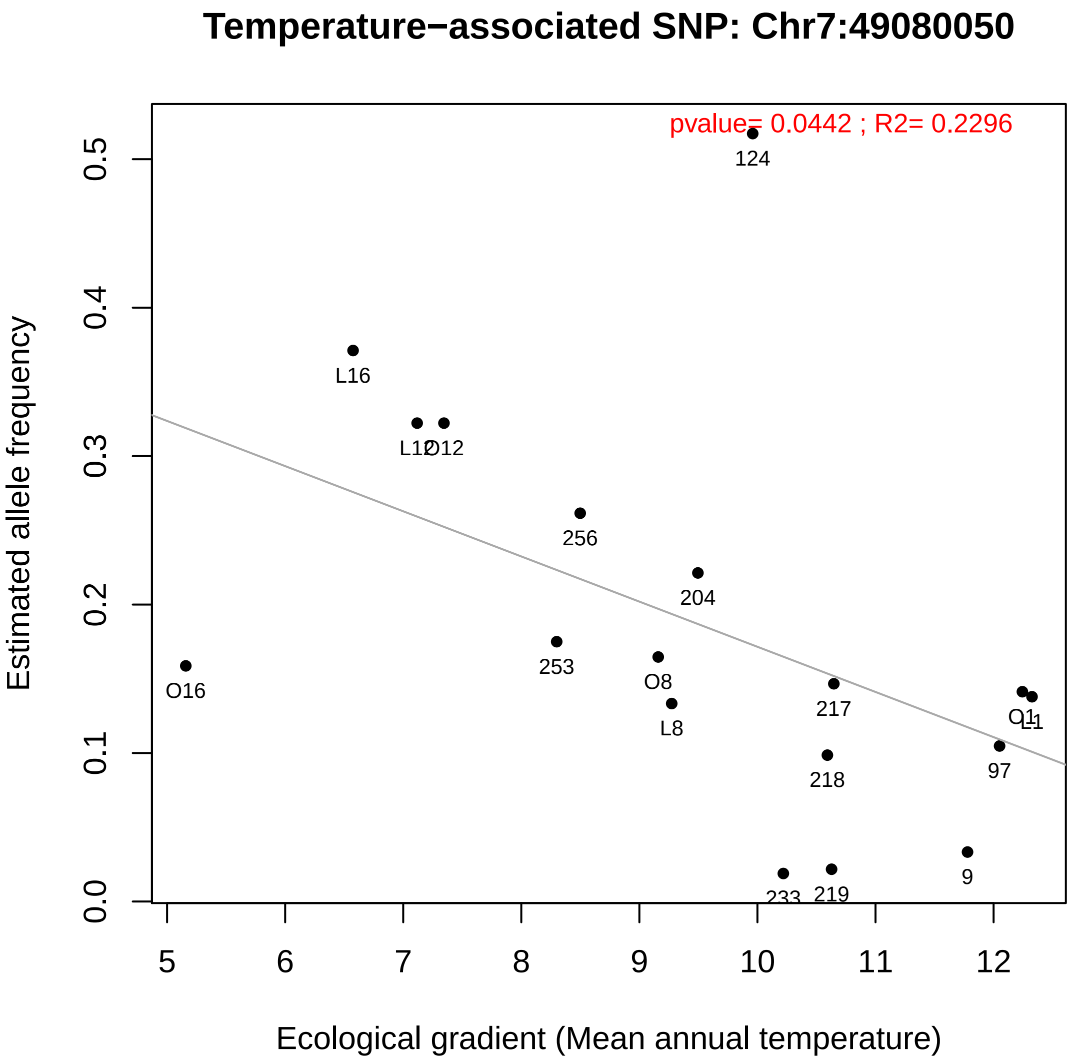
Temperature−associated SNP: Chr7:49080050
pvalue= 0.0442 ; R2= 0.2296
Estimated allele frequency
0.0
0.1
0.2
0.3
0.4
0.5
Ecological gradient (Mean annual temperature)
5
6
7
8
9
10
11
12
124
L16
L12
O12
256
204
253
O8
L8
O16
217
O1
L1
97
218
9
233
219

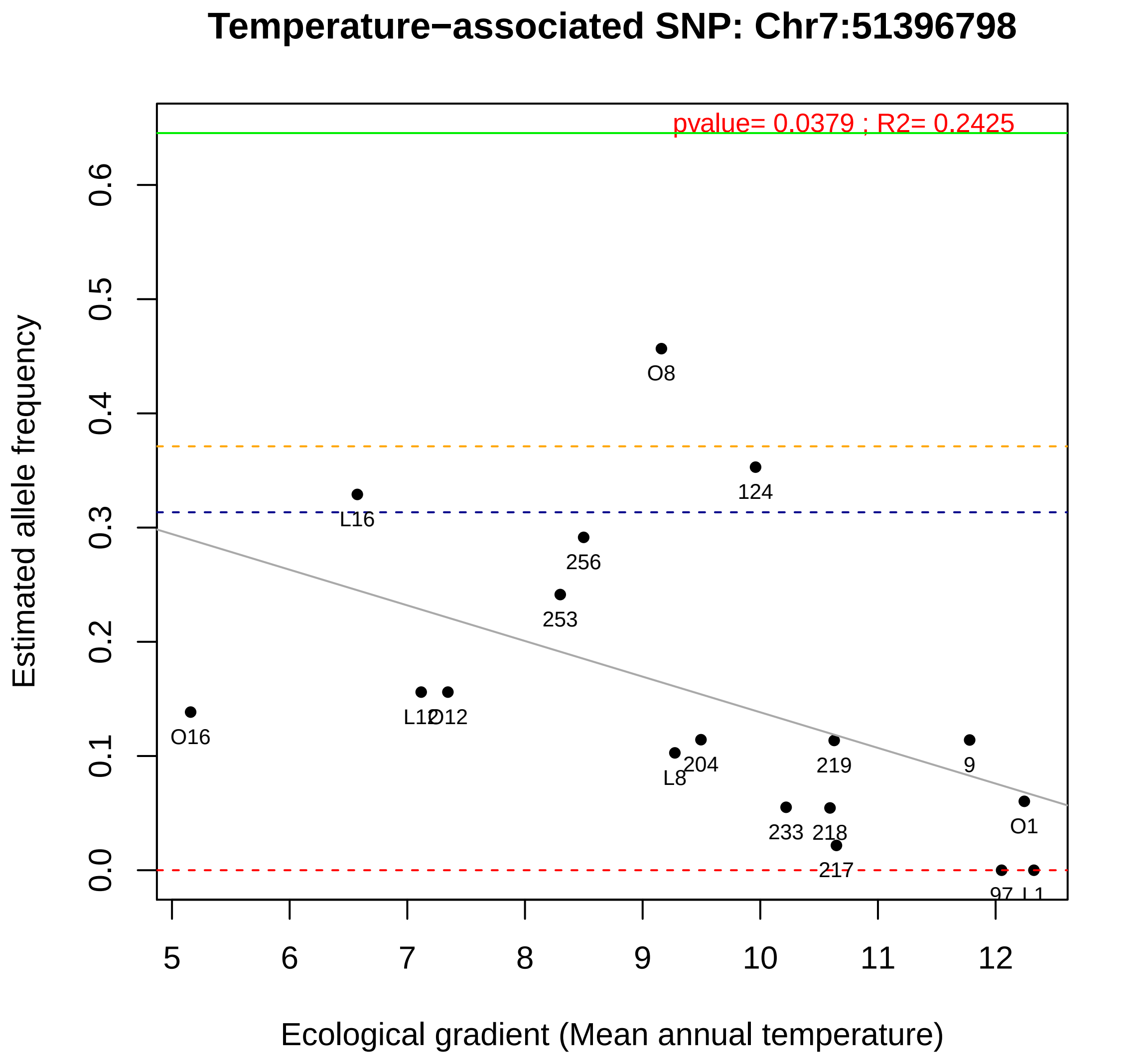
Temperature−associated SNP: Chr7:51396798
pvalue= 0.0379 ; R2= 0.2425
Estimated allele frequency
0.0
0.1
0.2
0.3
0.4
0.5
0.6
Ecological gradient (Mean annual temperature)
5
6
7
8
9
10
11
12
O8
124
L16
256
253
L12
O12
O16
204
L8
219
9
233
218
O1
217

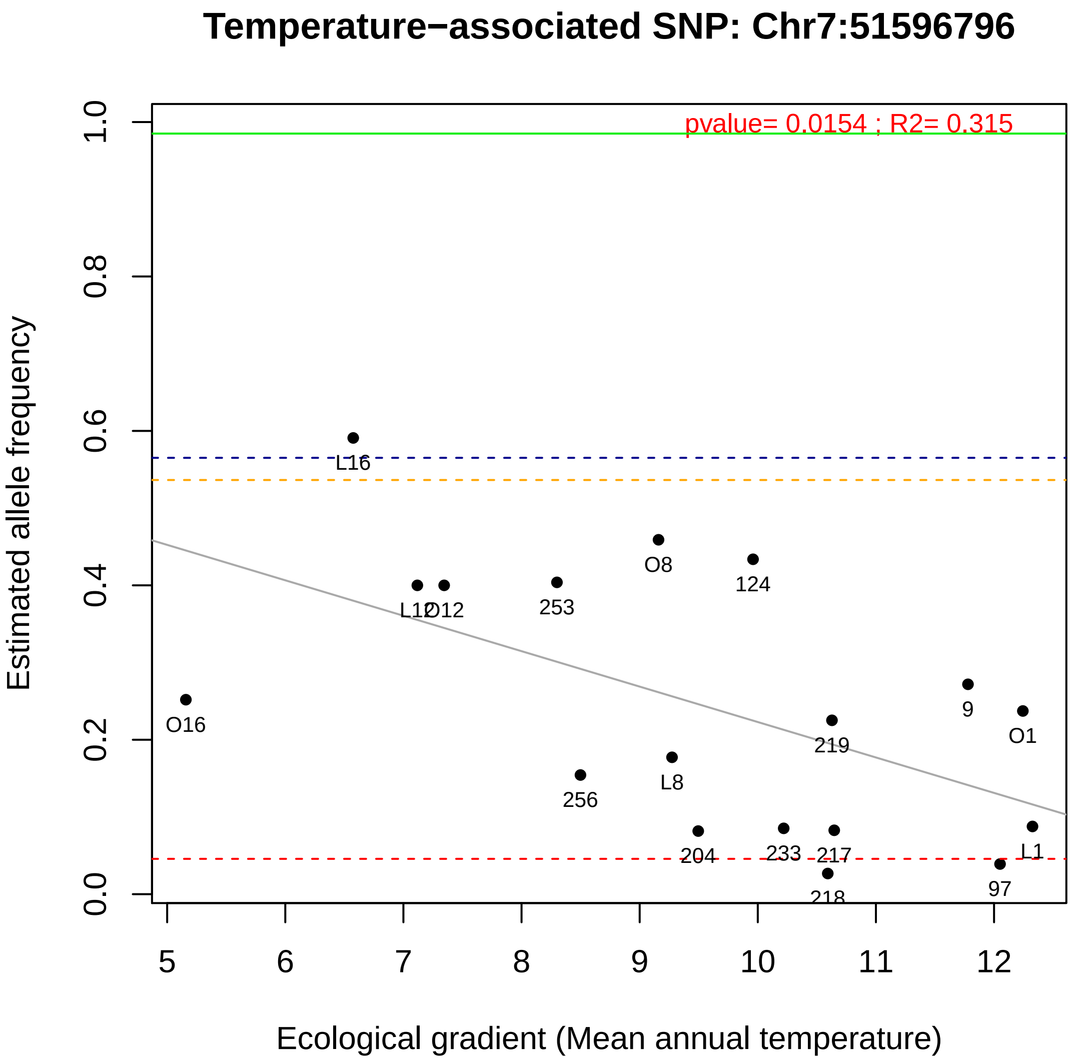

Temperature−associated SNP: Chr7:51596796
pvalue= 0.0154 ; R2= 0.315
Estimated allele frequency
0.0
0.2
0.4
0.6
0.8
1.0
L16
O8
124
L12
O12
253
O16
9
O1
219
L8
256
204
233
217
L1
218
97
5
6
7
8
9
10
11
12
Ecological gradient (Mean annual temperature)

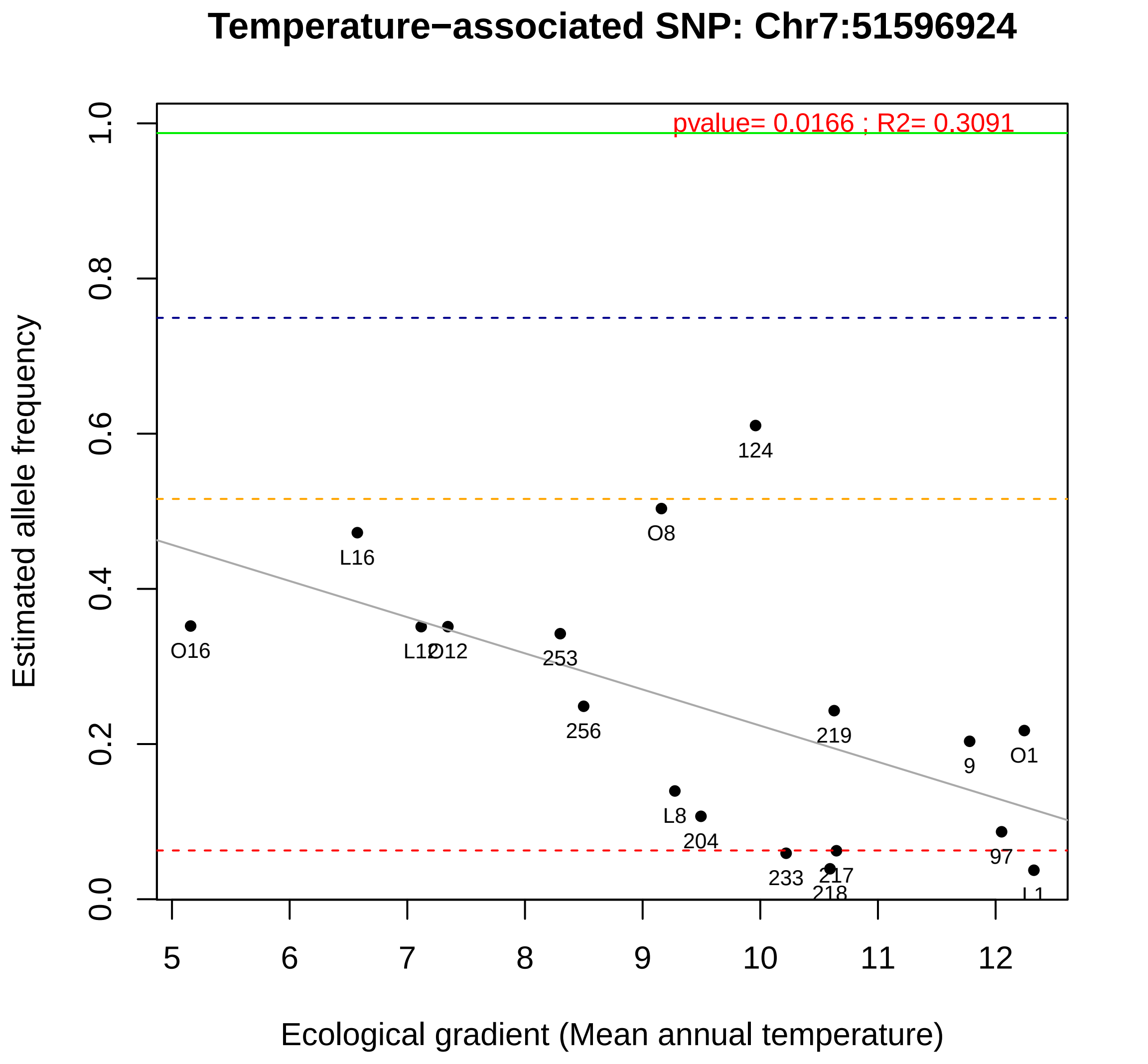

Temperature−associated SNP: Chr7:51596924
pvalue= 0.0166 ; R2= 0.3091
Estimated allele frequency
Ecological gradient (Mean annual temperature)
0.0
0.2
0.4
0.6
0.8
1.0
5
6
7
8
9
10
11
12
O16
L16
L12
O12
253
256
O8
124
L8
204
233
218
217
219
9
O1
97

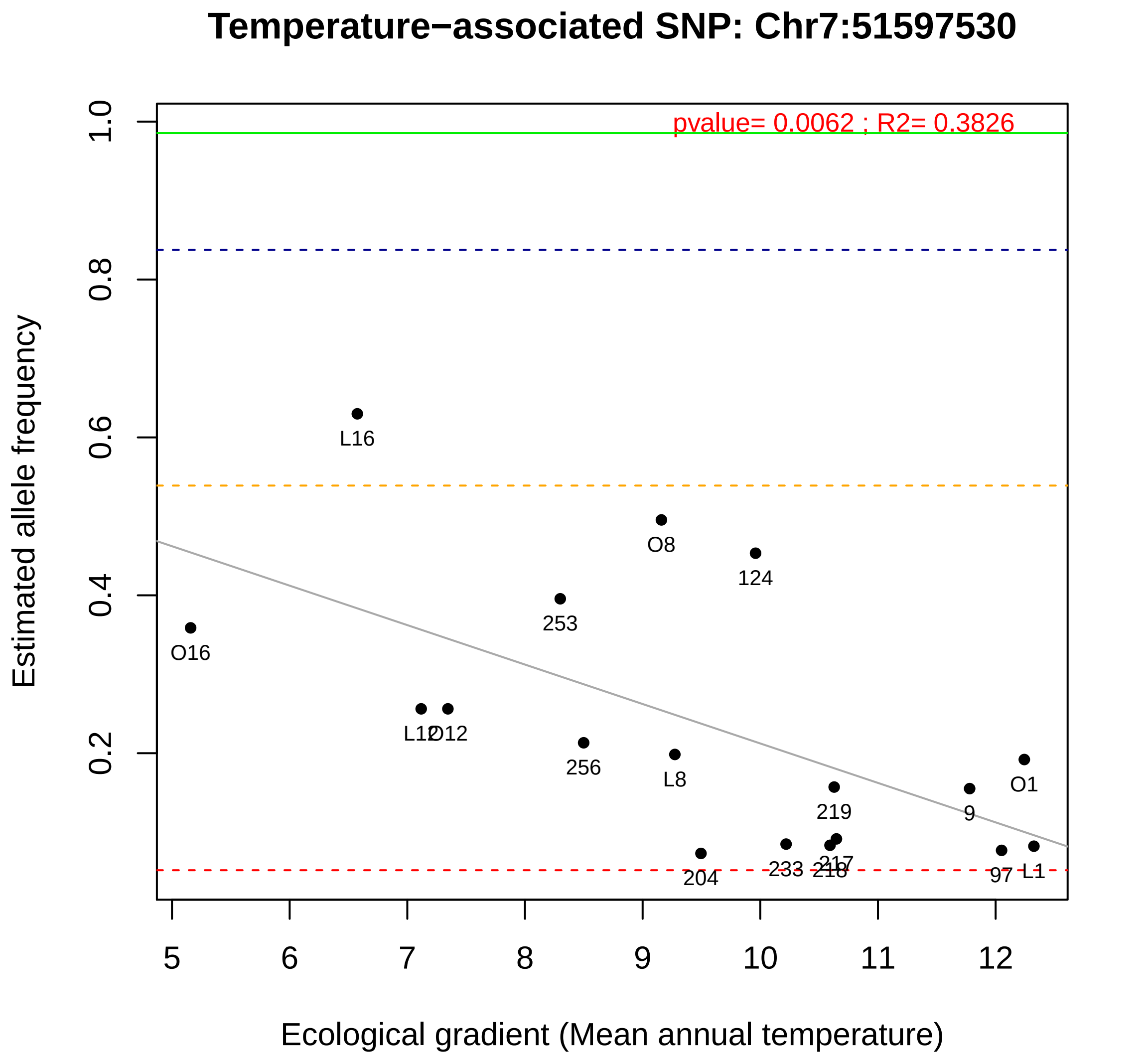
Temperature−associated SNP: Chr7:51597530
pvalue= 0.0062 ; R2= 0.3826
Estimated allele frequency
0.2
0.4
0.6
0.8
1.0
5
6
7
8
9
10
11
12
Ecological gradient (Mean annual temperature)
L16
O8
124
253
O16
L12
O12
256
L8
219
9
O1
204
233
217
218
97
L1

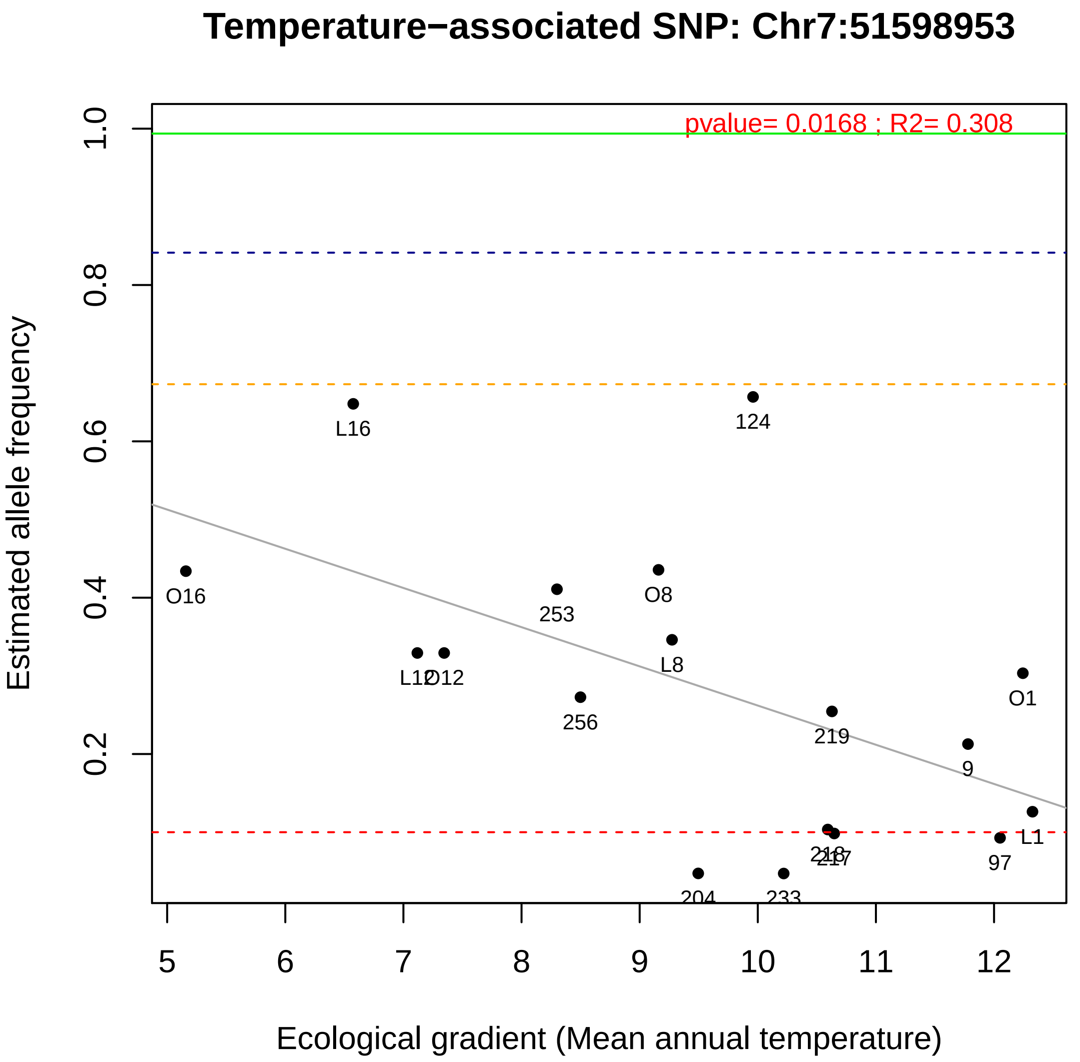

Temperature−associated SNP: Chr7:51598953
pvalue= 0.0168 ; R2= 0.308
Estimated allele frequency
0.2
0.4
0.6
0.8
1.0
5
6
7
8
9
10
11
12
Ecological gradient (Mean annual temperature)
L16
124
O16
253
O8
L8
L12
O12
256
O1
219
9
L1
97
218
217
204
233

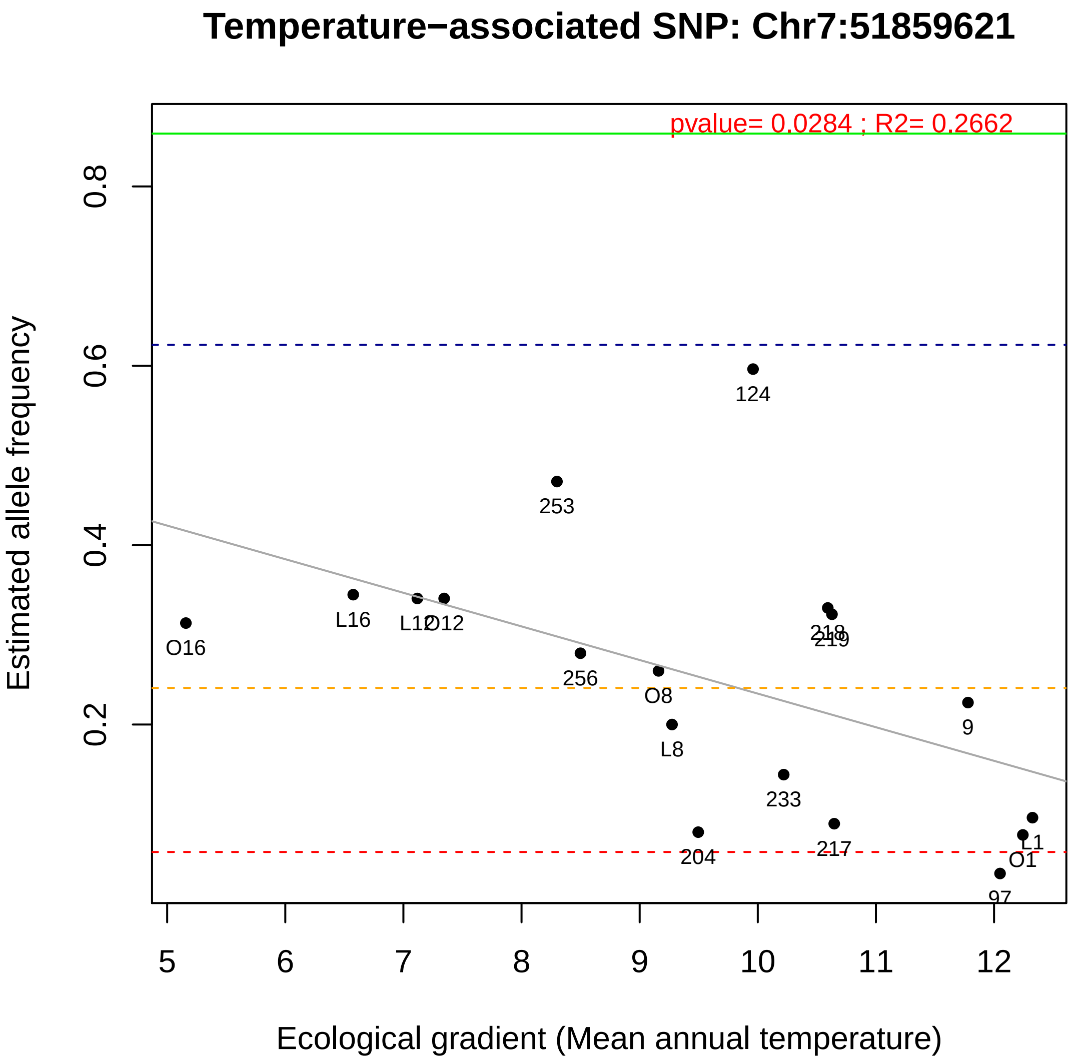

Temperature−associated SNP: Chr7:51859621
pvalue= 0.0284 ; R2= 0.2662
Estimated allele frequency
0.2
0.4
0.6
0.8
5
6
7
8
9
10
11
12
Ecological gradient (Mean annual temperature)
124
253
L16
L12
O12
O16
256
O8
L8
218
219
9
233
204
217
L1
O1
97

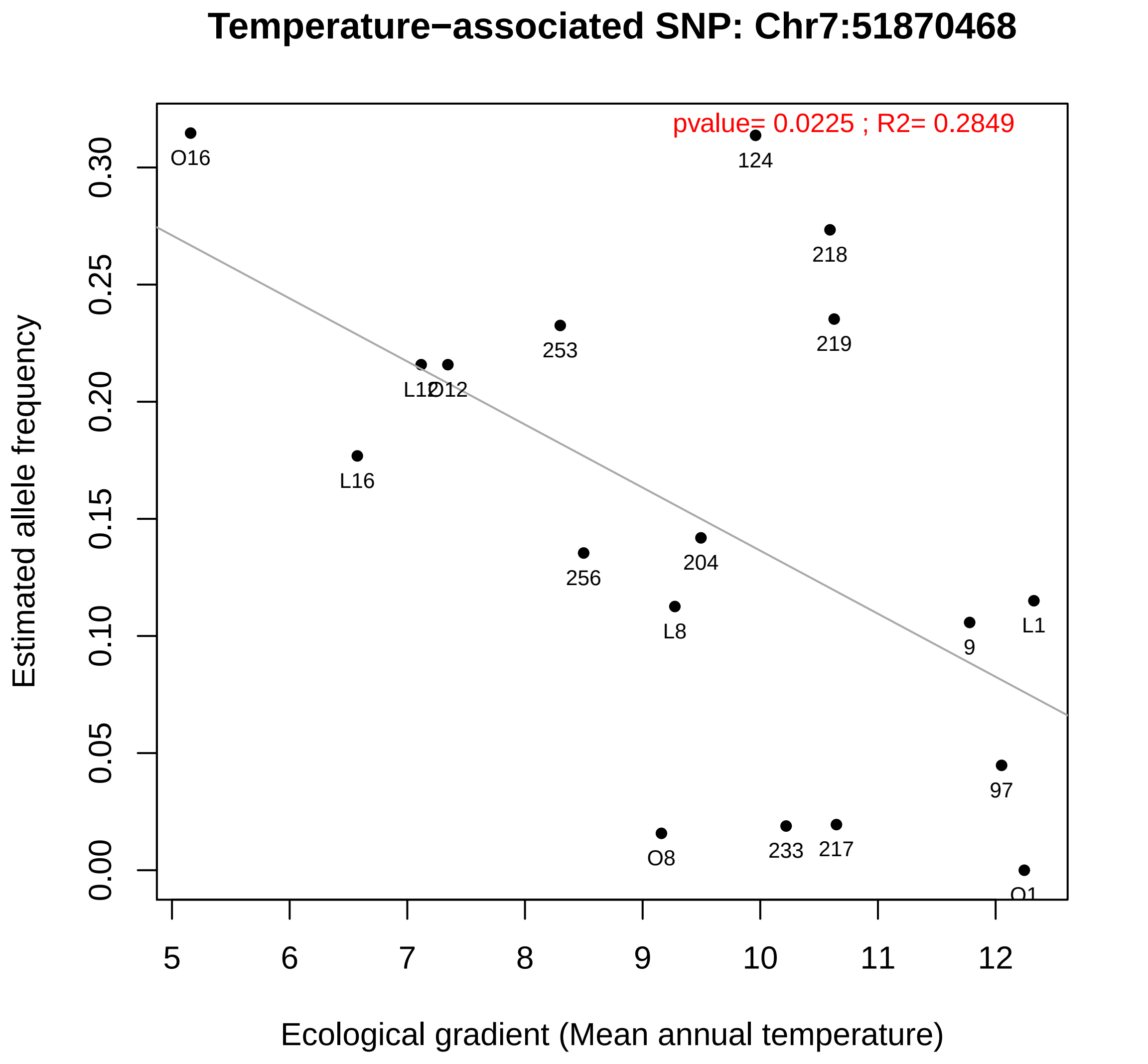

Temperature−associated SNP: Chr7:51870468
pvalue= 0.0225 ; R2= 0.2849
O16
124
218
219
253
L12
O12
L16
204
256
L8
9
L1
97
O8
233
217
O1
0.00
0.05
0.10
0.15
0.20
0.25
0.30
5
6
7
8
9
10
11
12
Estimated allele frequency
Ecological gradient (Mean annual temperature)

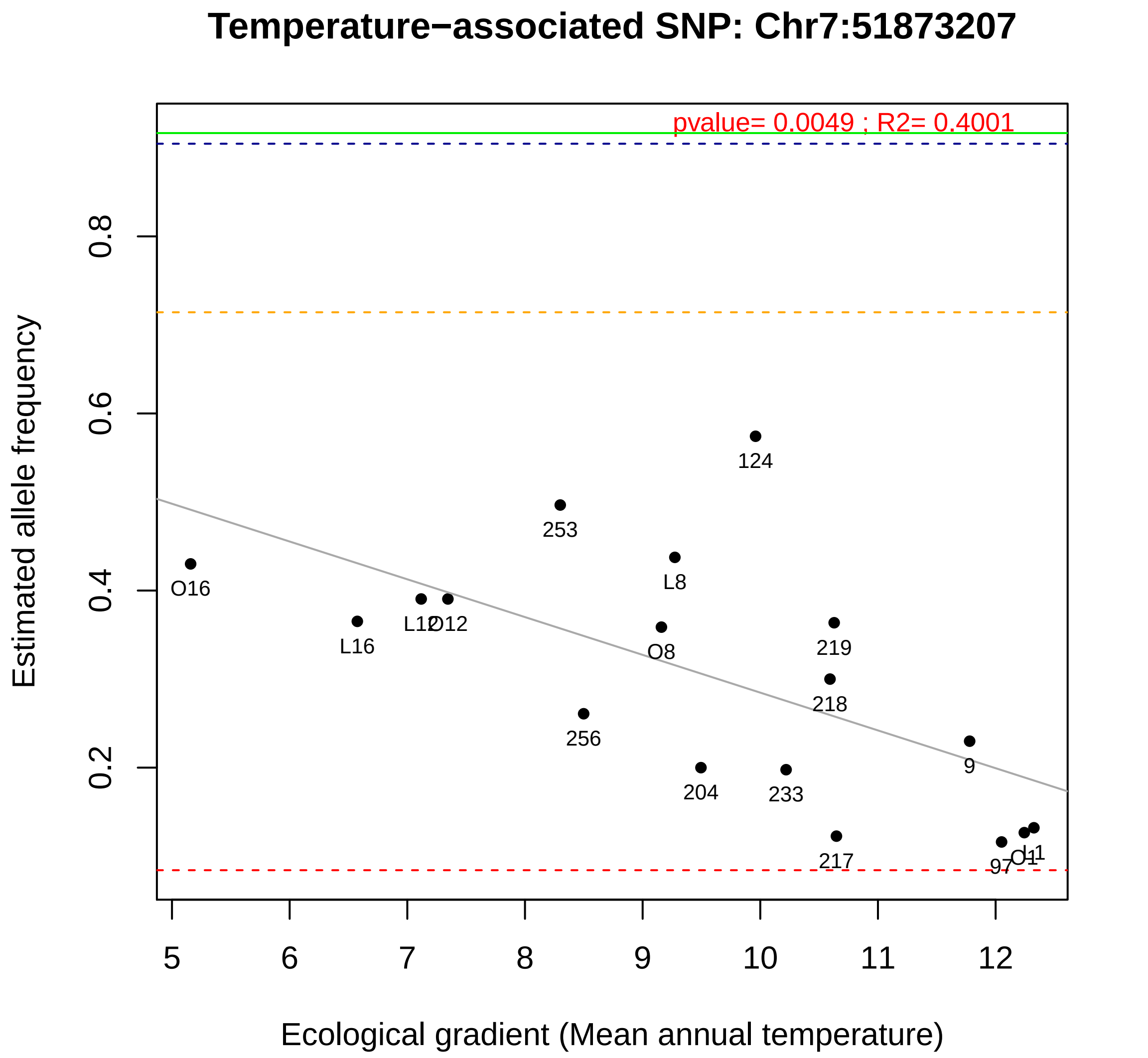
Temperature−associated SNP: Chr7:51873207
pvalue= 0.0049 ; R2= 0.4001
Estimated allele frequency
0.2
0.4
0.6
0.8
5
6
7
8
9
10
11
12
Ecological gradient (Mean annual temperature)
O16
L16
L12
O12
253
256
O8
L8
124
204
233
219
218
217
9
97
O1
L1

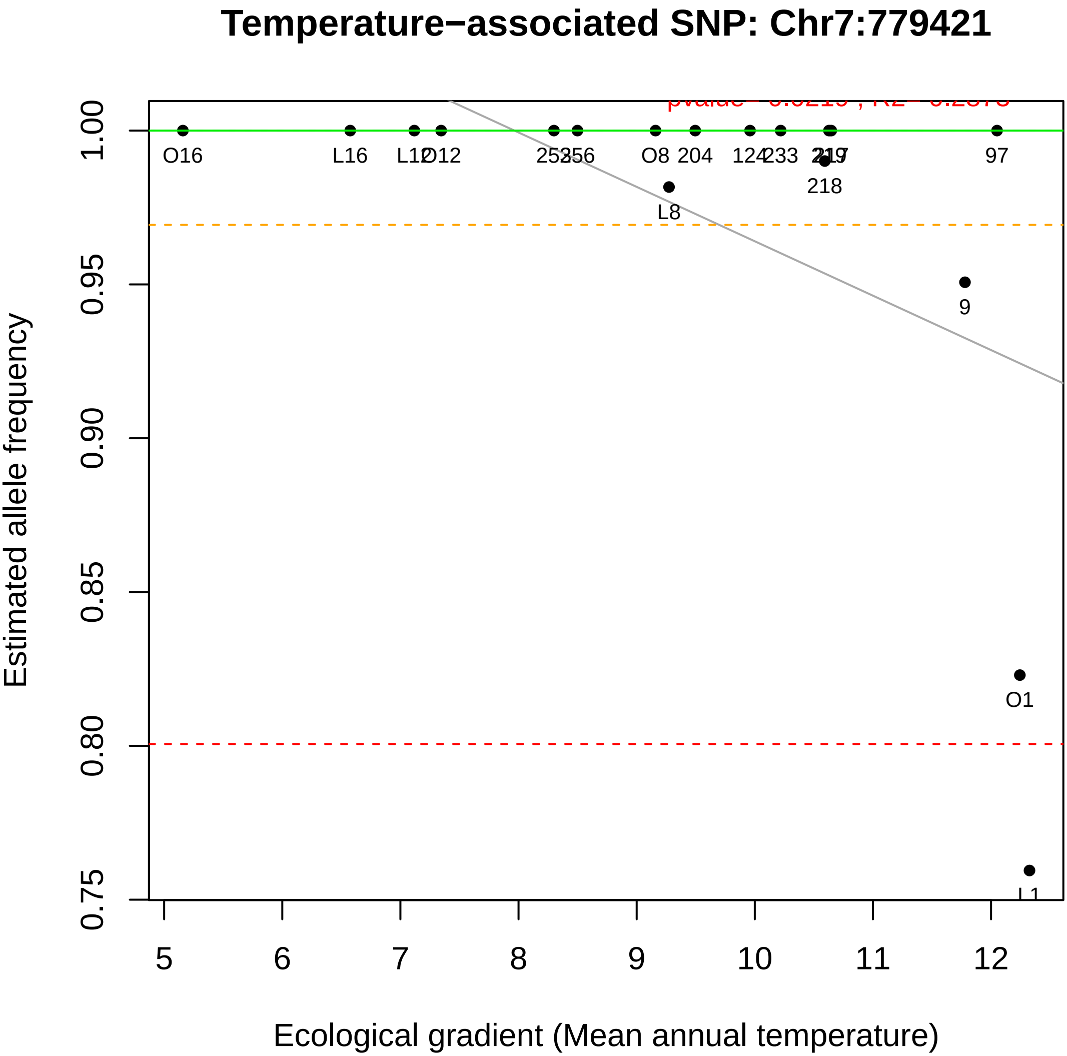
Temperature−associated SNP: Chr7:779421
Estimated allele frequency
Ecological gradient (Mean annual temperature)
0.75
0.80
0.85
0.90
0.95
1.00
5
6
7
8
9
10
11
12
O16
L16
O12
O8
204
97
L8
218
9
O1

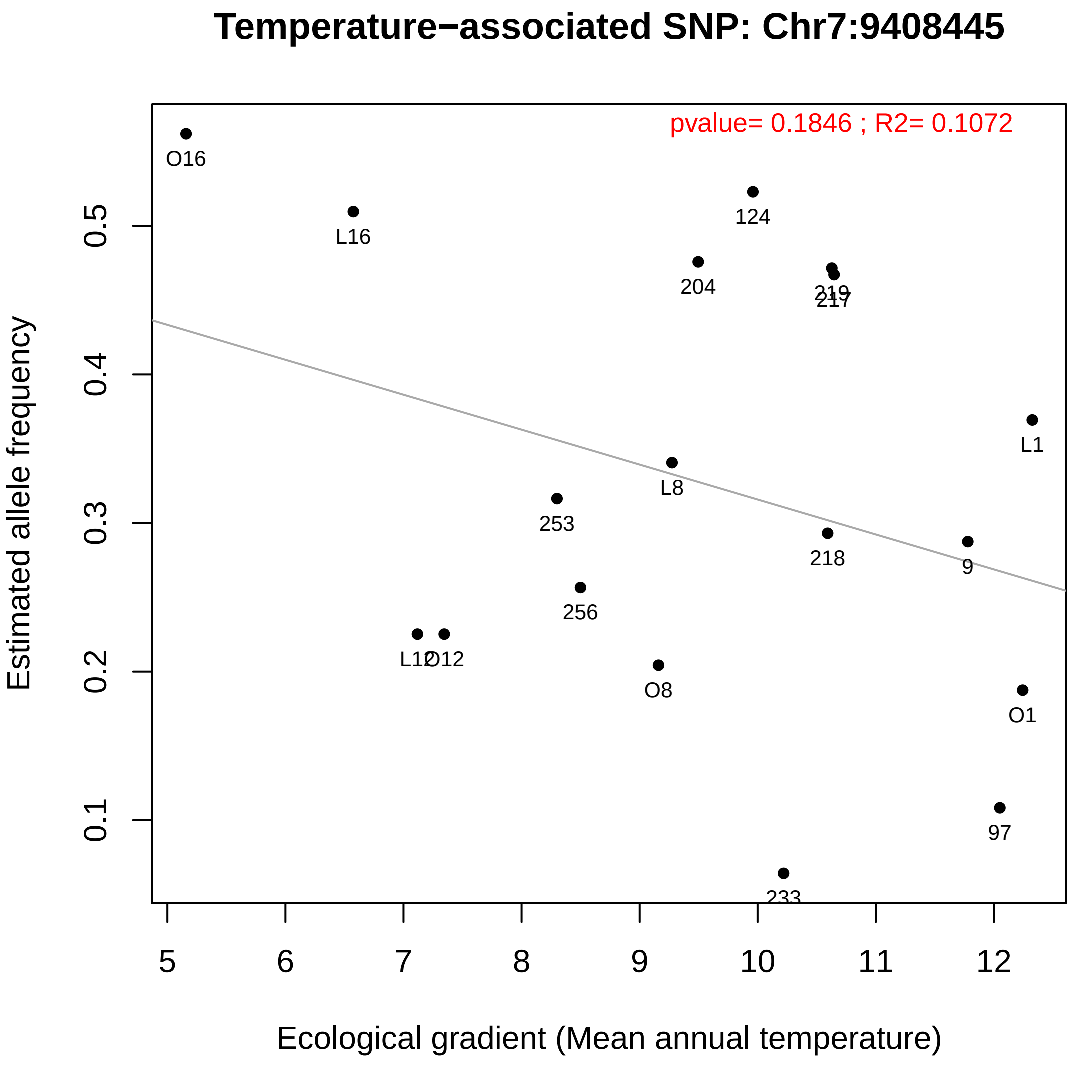
Temperature−associated SNP: Chr7:9408445
pvalue= 0.1846 ; R2= 0.1072
Estimated allele frequency
0.1
0.2
0.3
0.4
0.5
Ecological gradient (Mean annual temperature)
5
6
7
8
9
10
11
12
O16
L16
124
204
219
217
L1
L8
253
218
9
256
L12
O12
O8
O1
97
233

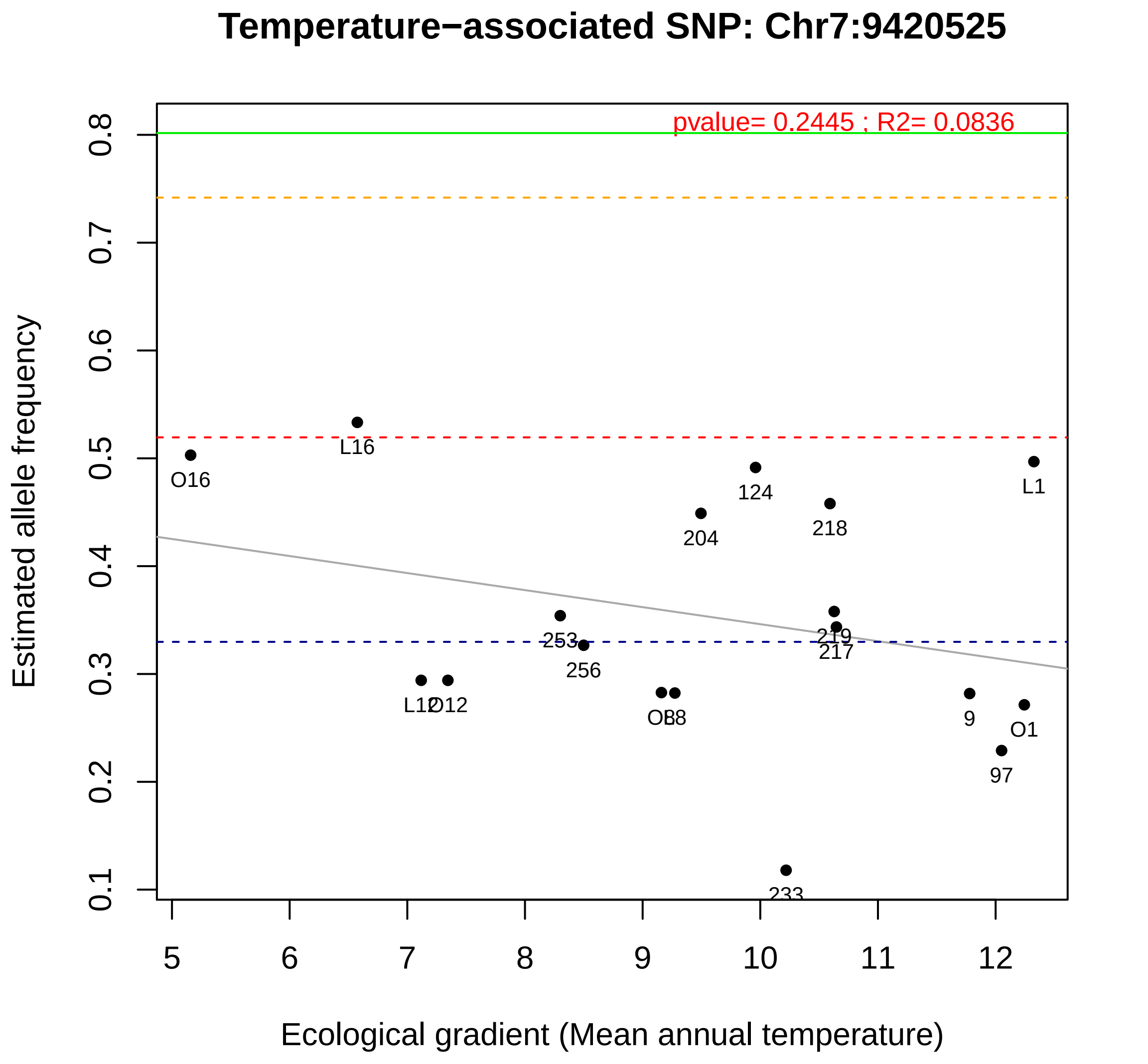

Temperature−associated SNP: Chr7:9420525
pvalue= 0.2445 ; R2= 0.0836
Estimated allele frequency
Ecological gradient (Mean annual temperature)
0.1
0.2
0.3
0.4
0.5
0.6
0.7
0.8
5
6
7
8
9
10
11
12
O16
L16
L12
O12
253
256
204
124
218
219
217
233
9
O1
97
L1

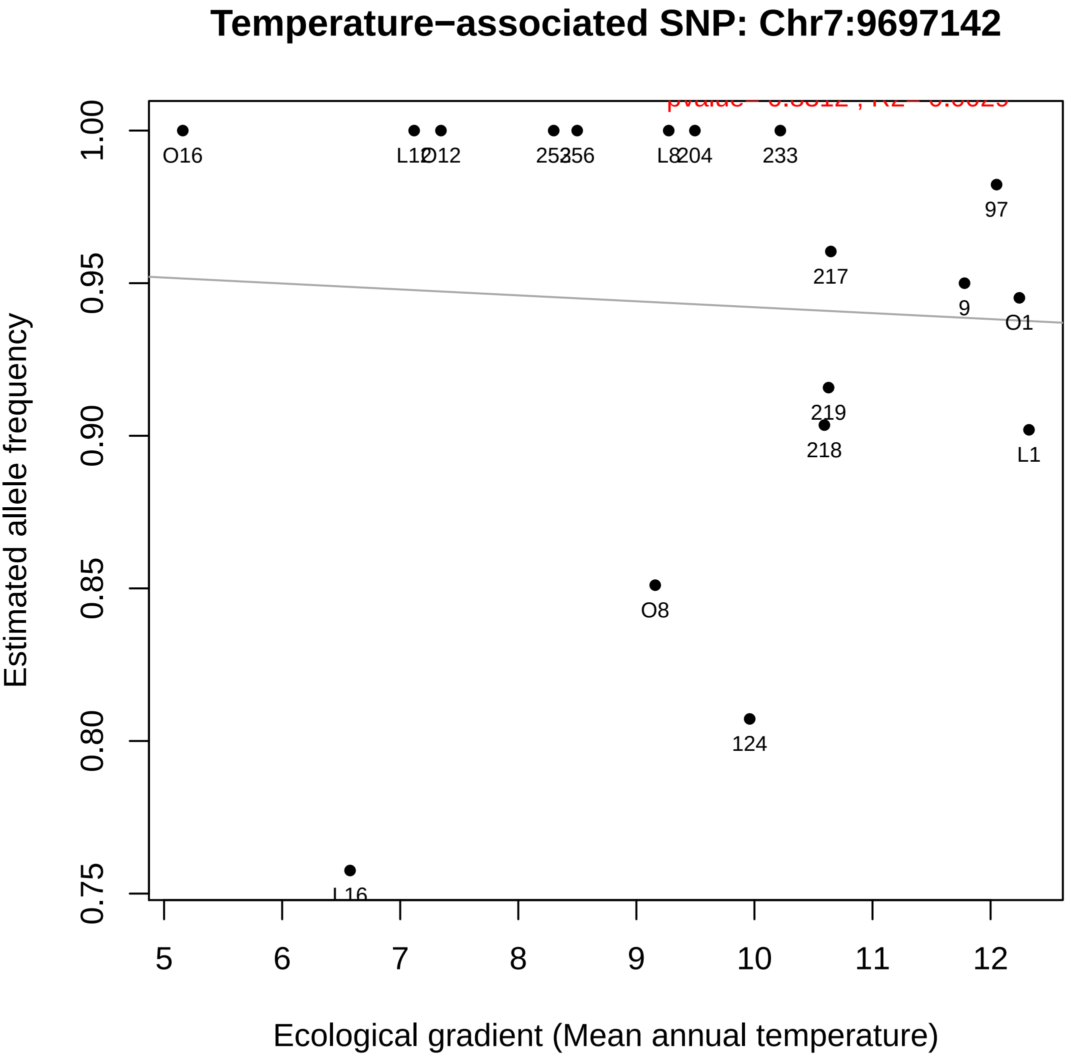

Temperature−associated SNP: Chr7:9697142
Estimated allele frequency
Ecological gradient (Mean annual temperature)
0.75
0.80
0.85
0.90
0.95
1.00
5
6
7
8
9
10
11
12
O16
L12
O12
256
L8
204
233
97
217
9
O1
219
218
L1
O8
124
L16

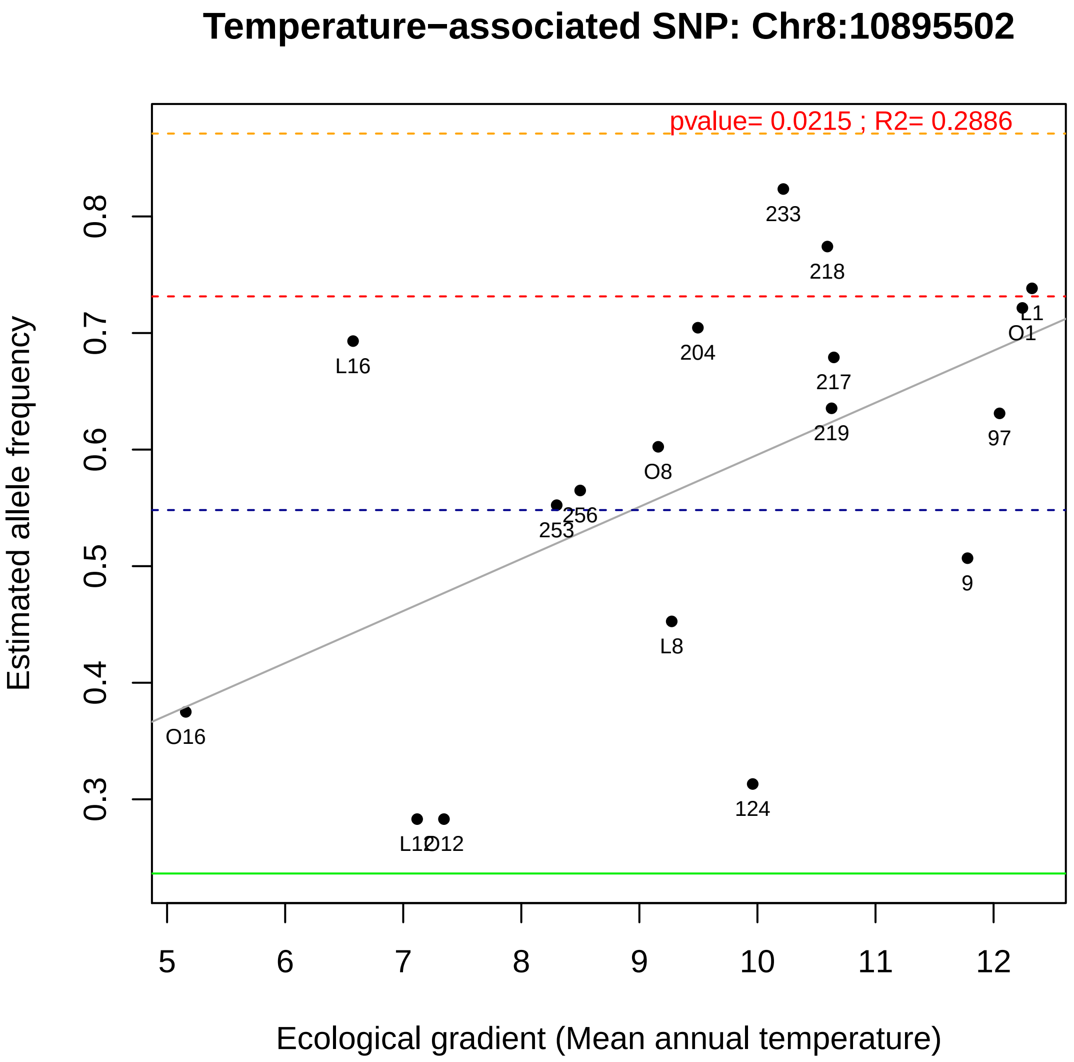

Temperature−associated SNP: Chr8:10895502
pvalue= 0.0215 ; R2= 0.2886
Estimated allele frequency
0.3
0.4
0.5
0.6
0.7
0.8
5
6
7
8
9
10
11
12
Ecological gradient (Mean annual temperature)
233
218
L1
O1
L16
204
217
219
97
O8
256
253
9
L8
O16
124
L12
O12

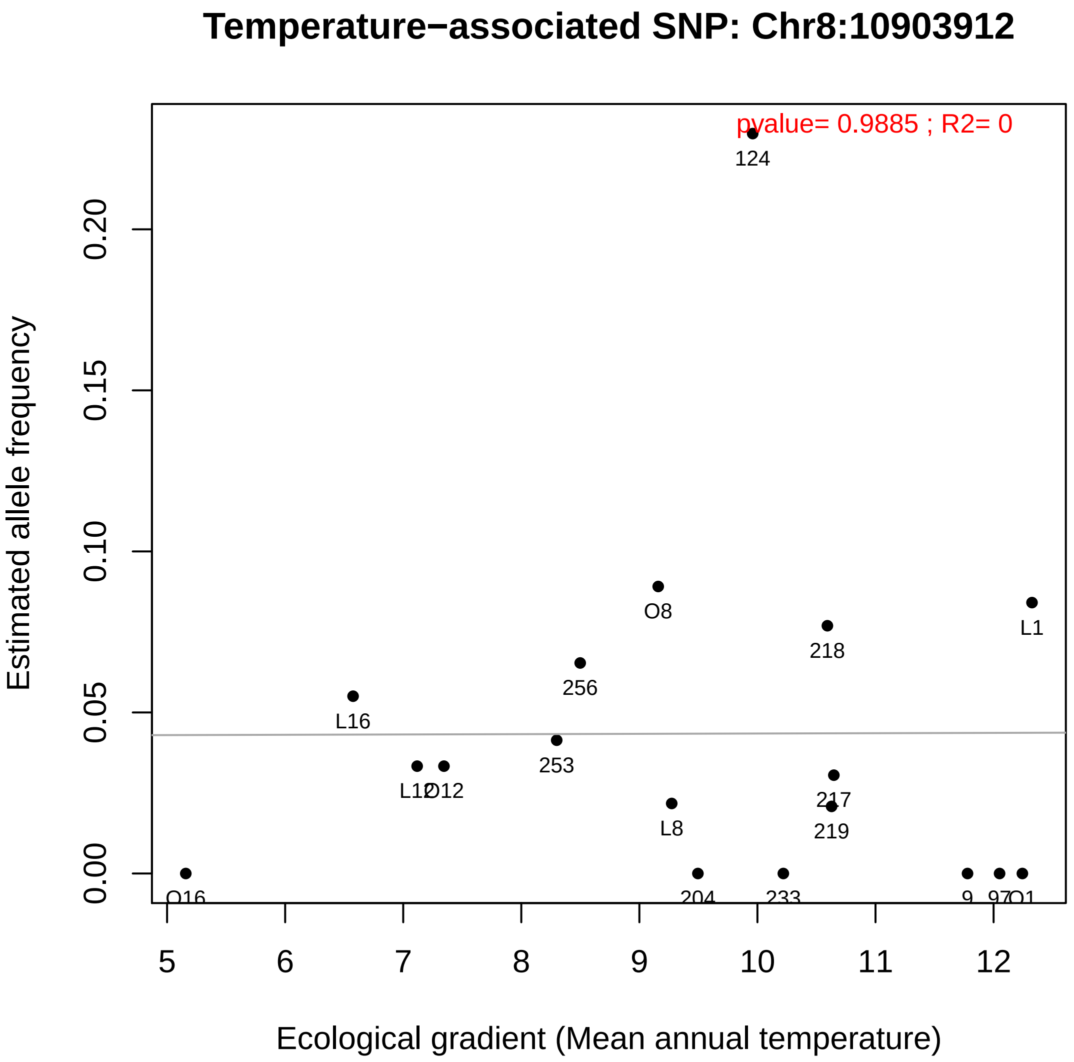

Temperature−associated SNP: Chr8:10903912
pvalue= 0.9885 ; R2= 0
124
O8
L1
218
256
L16
253
L12
O12
217
L8
219
O16
204
233
9
97
O1
0.00
0.05
0.10
0.15
0.20
5
6
7
8
9
10
11
12
Estimated allele frequency
Ecological gradient (Mean annual temperature)

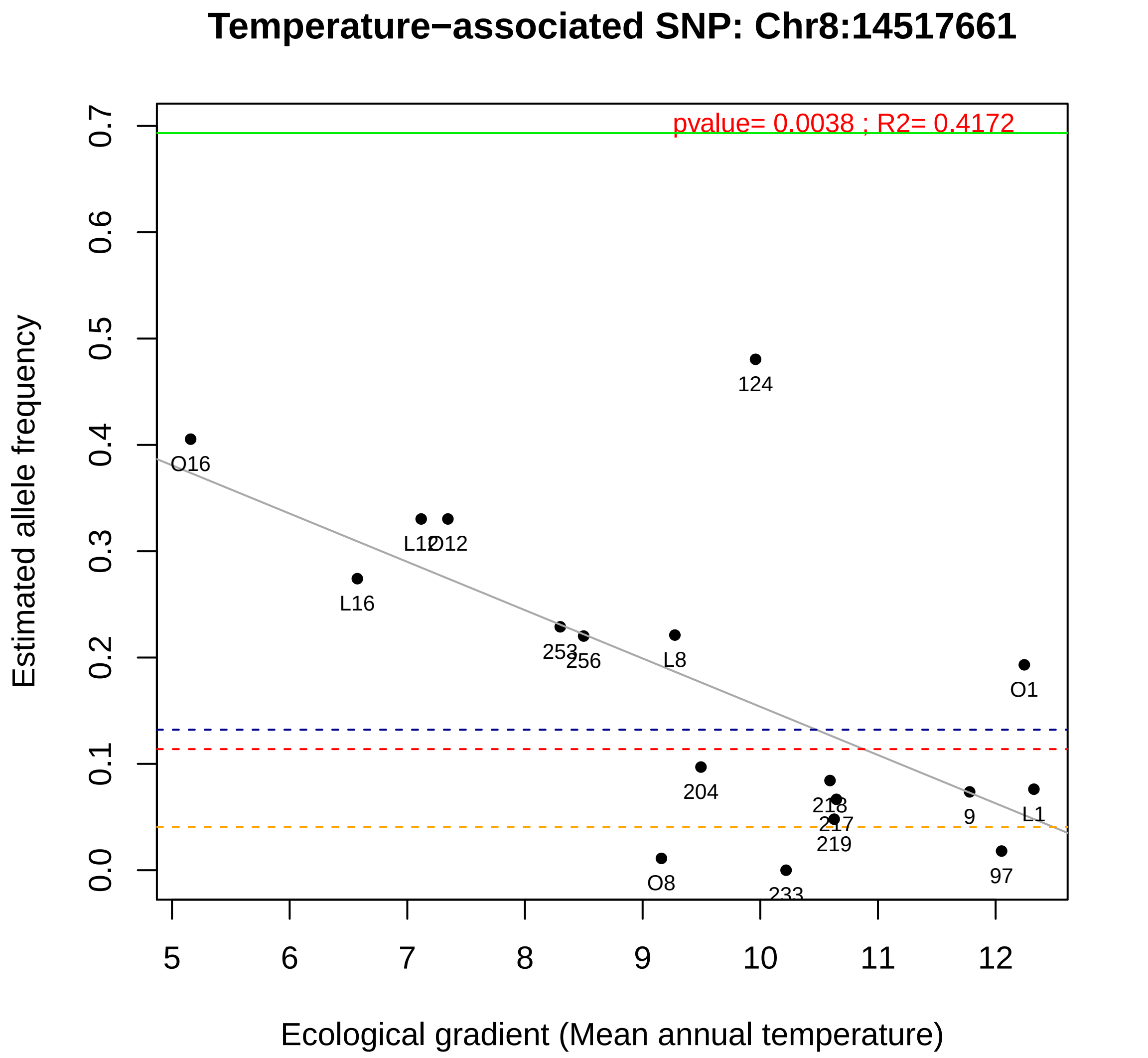
Temperature−associated SNP: Chr8:14517661
pvalue= 0.0038 ; R2= 0.4172
Estimated allele frequency
0.0
0.1
0.2
0.3
0.4
0.5
0.6
0.7
Ecological gradient (Mean annual temperature)
5
6
7
8
9
10
11
12
O16
L16
L12
O12
253
256
L8
124
204
O8
233
218
217
219
9
97
O1
L1

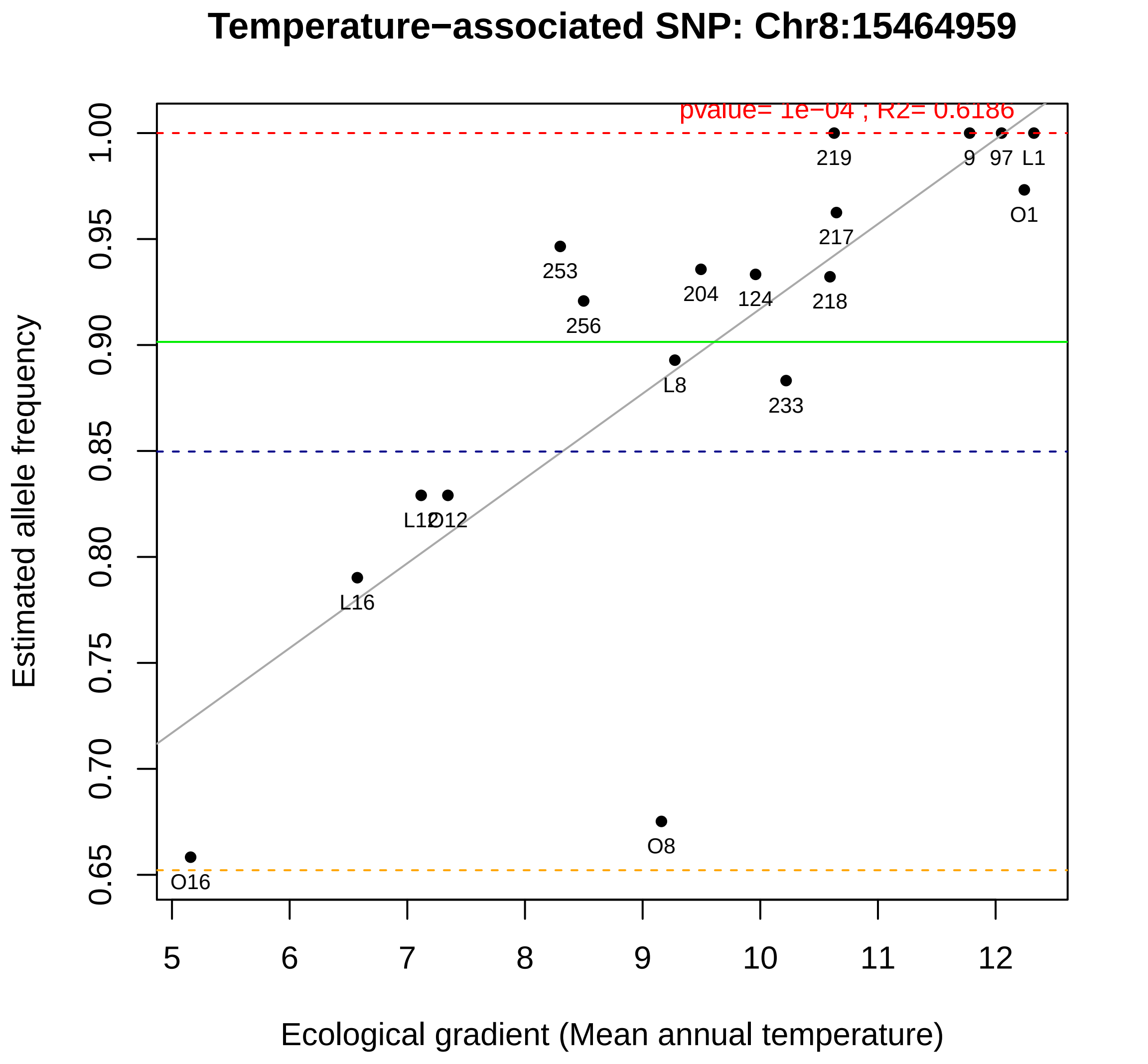
Temperature−associated SNP: Chr8:15464959
pvalue= 1e−04 ; R2= 0.6186
Estimated allele frequency
0.65
0.70
0.75
0.80
0.85
0.90
0.95
1.00
5
6
7
8
9
10
11
12
Ecological gradient (Mean annual temperature)
219
9
97
L1
O1
217
253
204
124
218
256
L8
233
L12
O12
L16
O8
O16

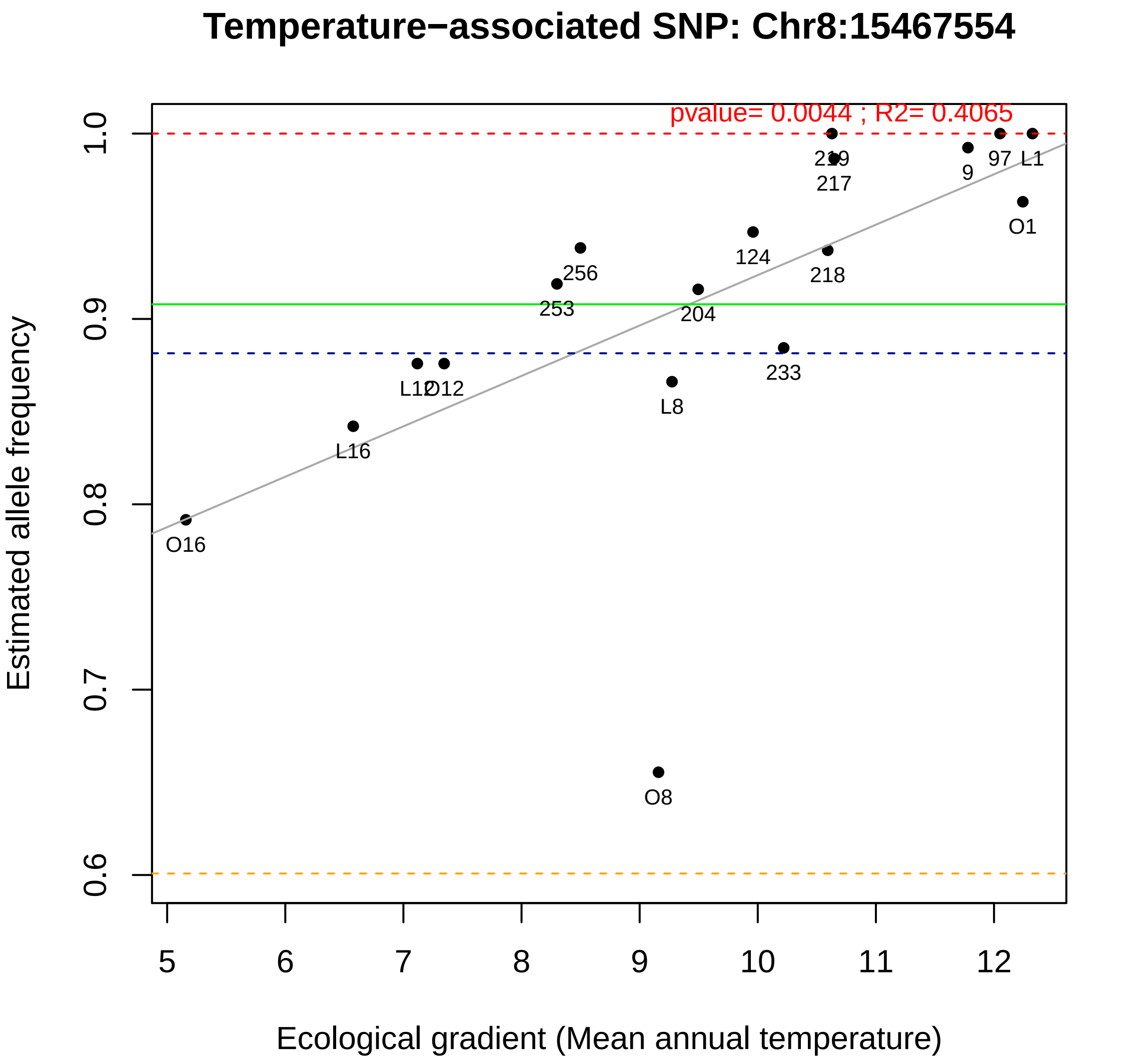
Temperature−associated SNP: Chr8:15467554
pvalue= 0.0044 ; R2= 0.4065
Estimated allele frequency
Ecological gradient (Mean annual temperature)
1.0
0.9
0.8
0.7
0.6
5
6
7
8
9
10
11
12
O16
L16
L12
O12
253
256
204
L8
O8
124
233
218
219
217
9
97
L1
O1

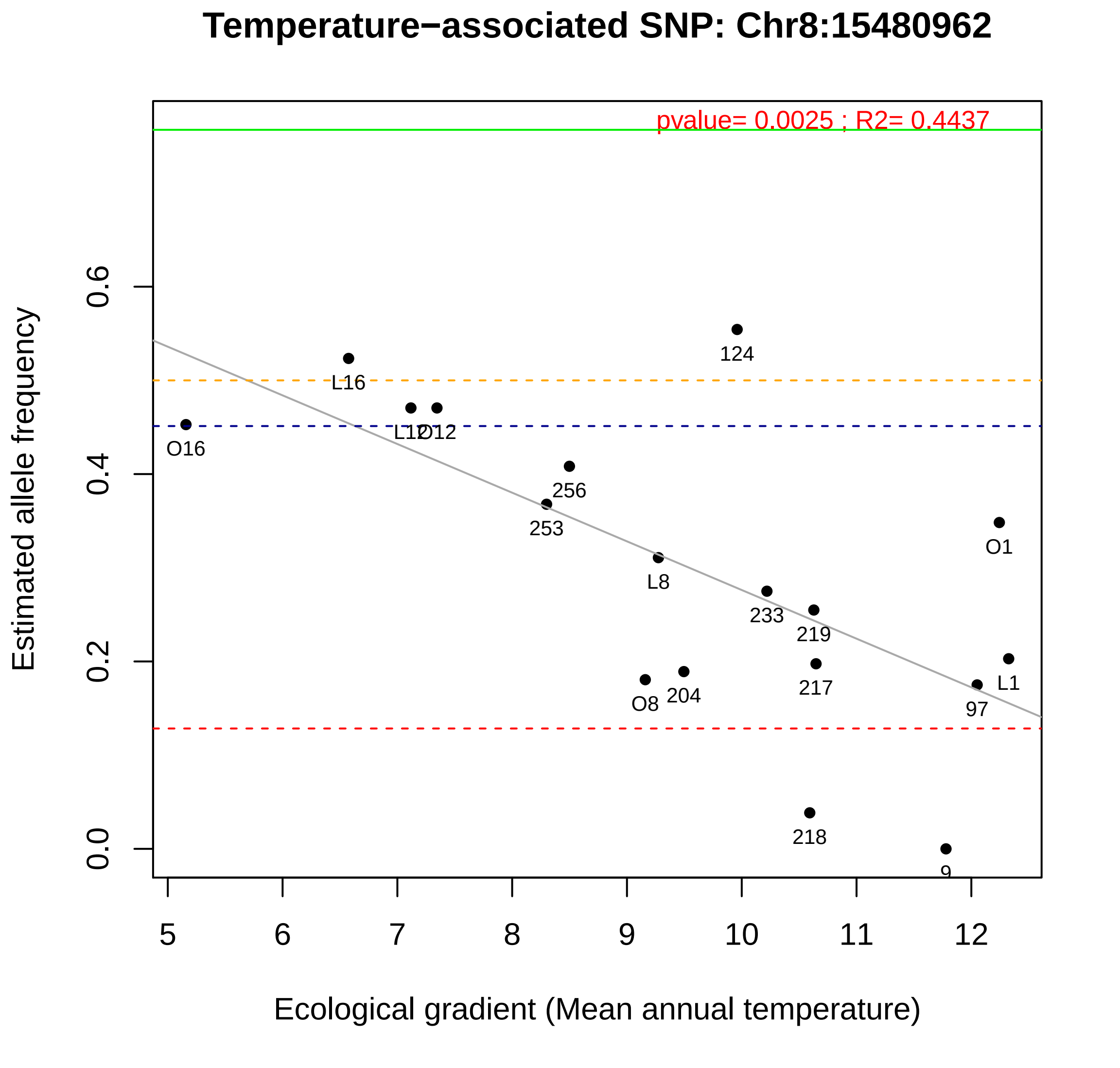

Temperature−associated SNP: Chr8:15480962
pvalue= 0.0025 ; R2= 0.4437
Estimated allele frequency
0.0
0.2
0.4
0.6
O16
L16
L12
O12
124
256
253
L8
233
219
O1
O8
204
217
L1
97
218
9
5
6
7
8
9
10
11
12
Ecological gradient (Mean annual temperature)

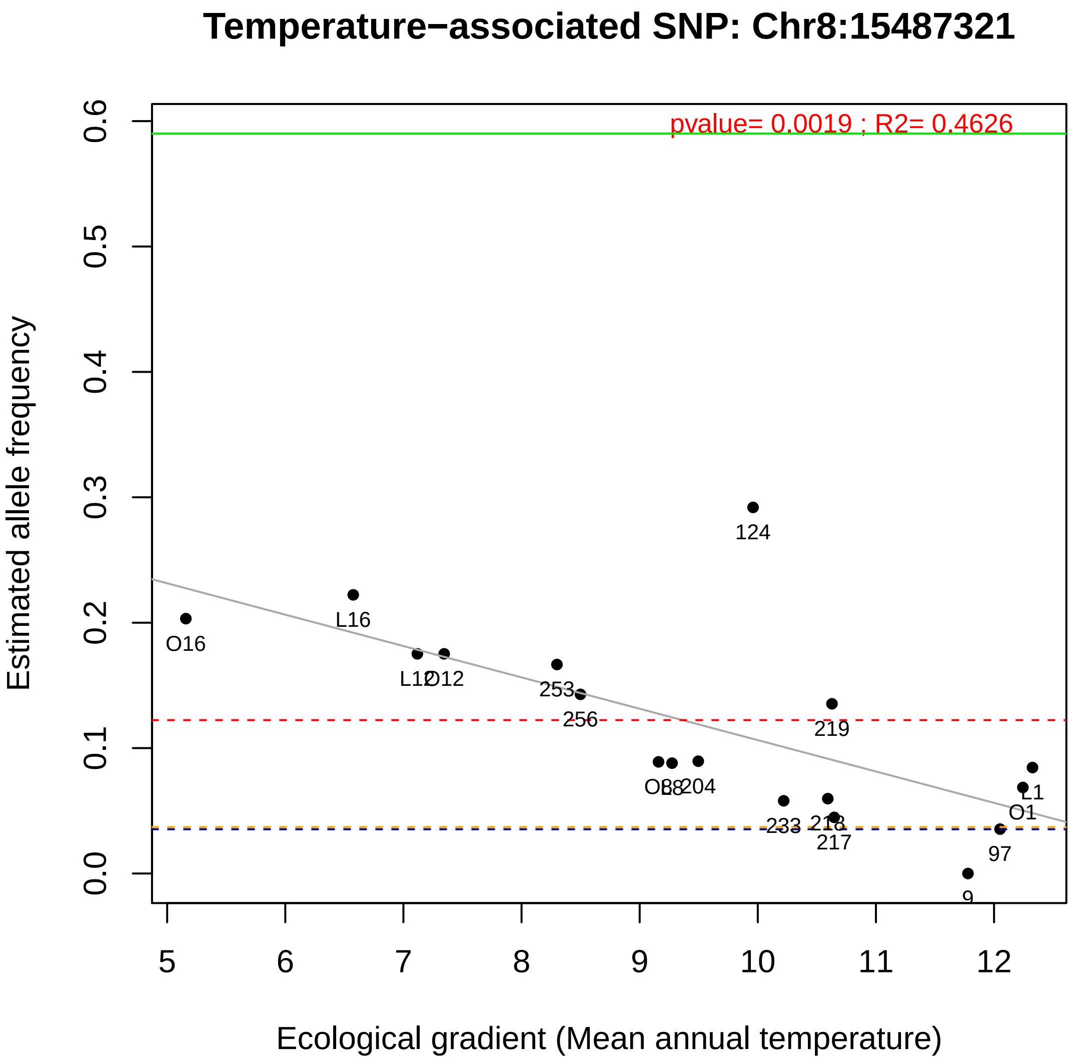

Temperature−associated SNP: Chr8:15487321
pvalue= 0.0019 ; R2= 0.4626
Estimated allele frequency
Ecological gradient (Mean annual temperature)
0.0
0.1
0.2
0.3
0.4
0.5
0.6
5
6
7
8
9
10
11
12
124
L16
O16
L12
O12
253
256
219
O8
L8
204
233
218
217
L1
O1
97
9

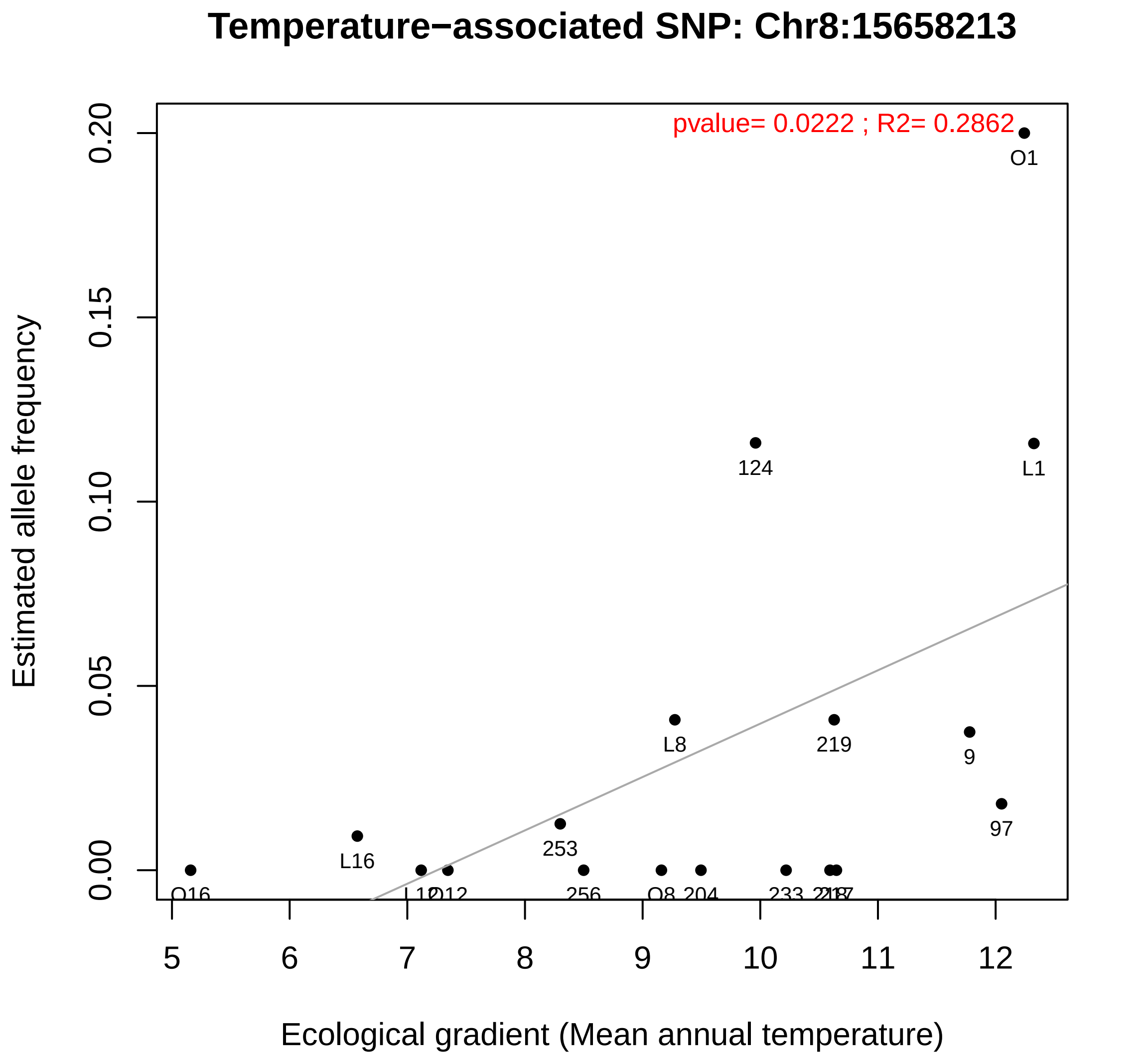
Temperature−associated SNP: Chr8:15658213
pvalue= 0.0222 ; R2= 0.2862
Estimated allele frequency
0.00
0.05
0.10
0.15
0.20
5
6
7
8
9
10
11
12
Ecological gradient (Mean annual temperature)
O1
124
L1
L8
219
9
97
253
L16
O16
256
O8
204
233

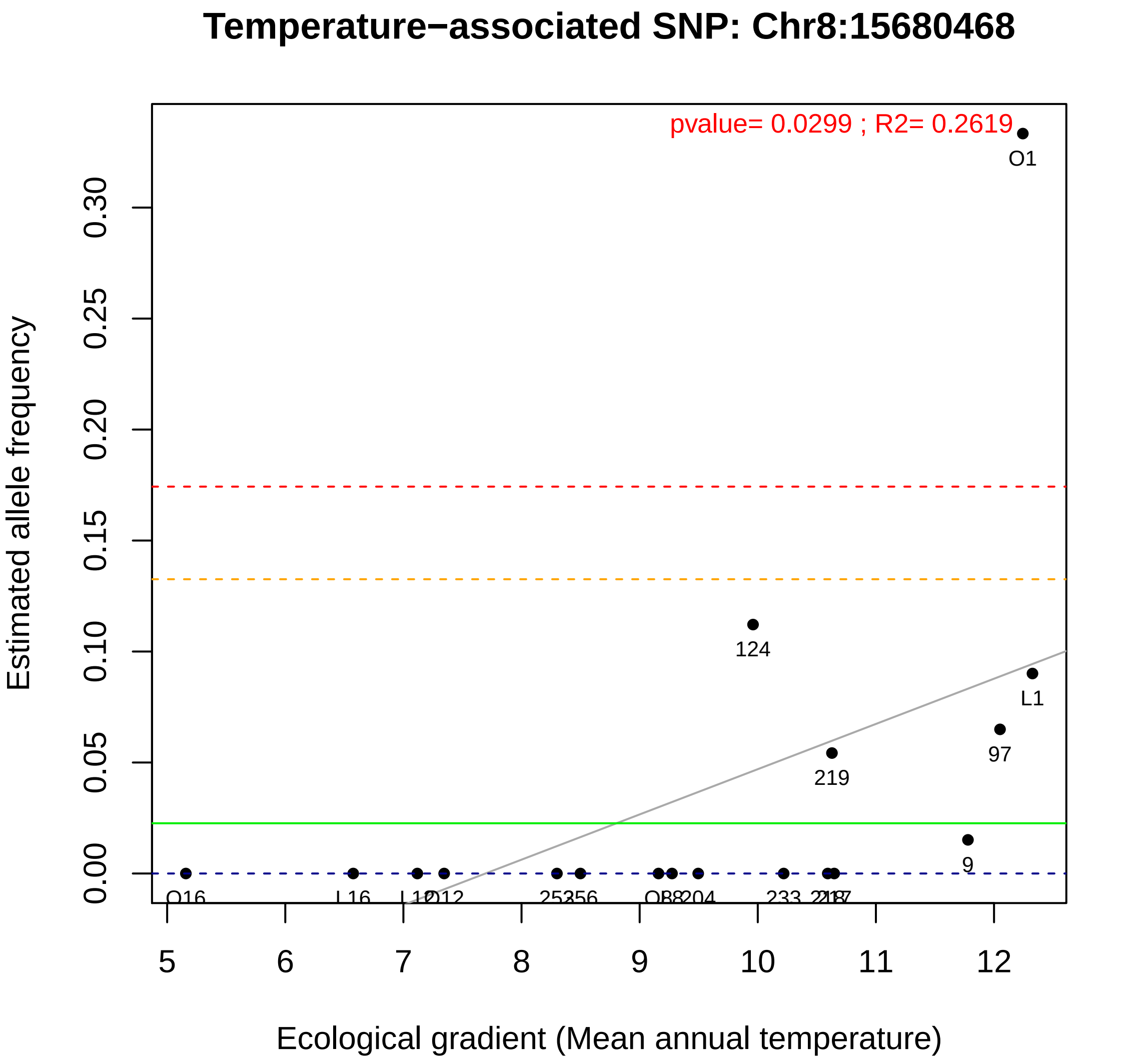

Temperature−associated SNP: Chr8:15680468
pvalue= 0.0299 ; R2= 0.2619
Estimated allele frequency
Ecological gradient (Mean annual temperature)
0.00
0.05
0.10
0.15
0.20
0.25
0.30
5
6
7
8
9
10
11
12
O1
124
L1
97
219
9
O16
L16
233
204

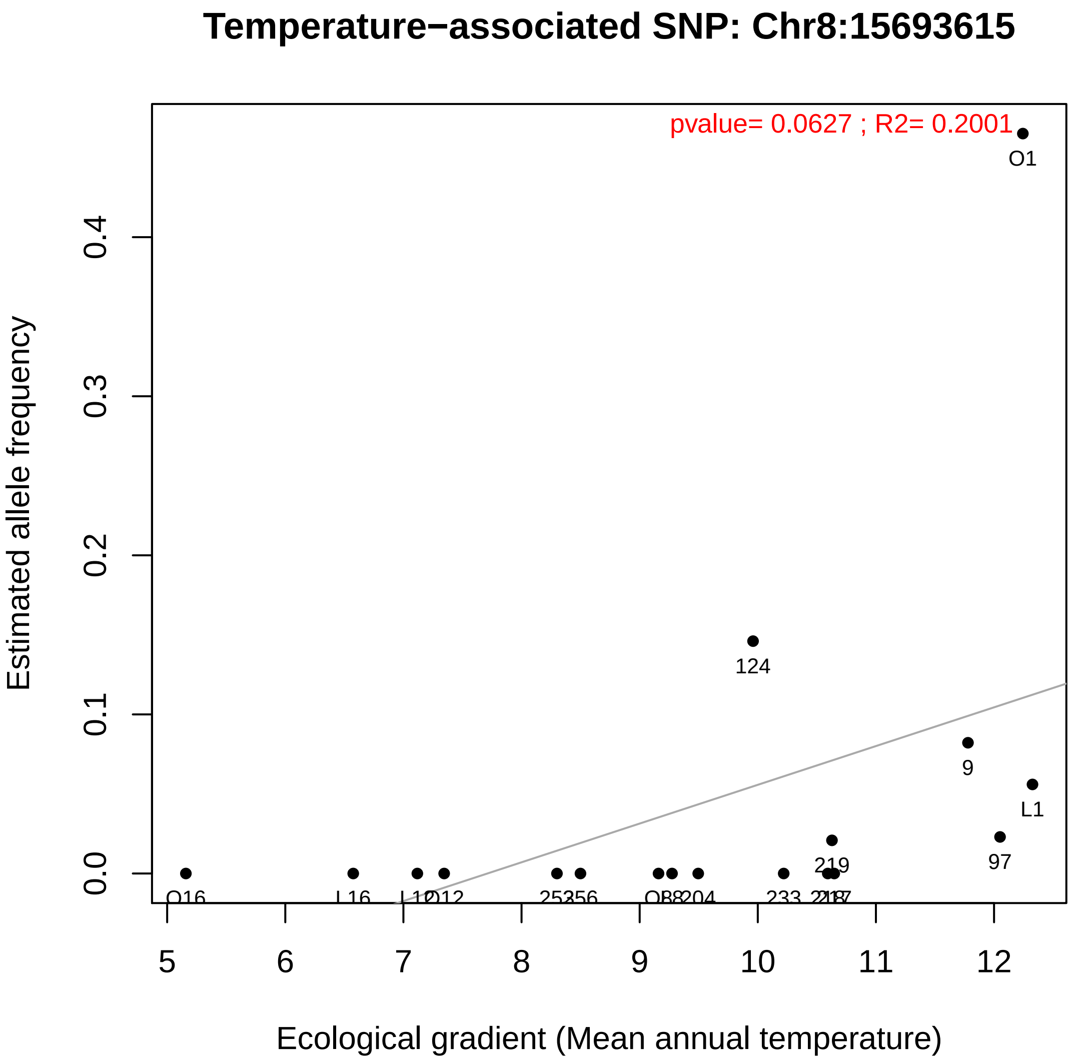

Temperature−associated SNP: Chr8:15693615
pvalue= 0.0627 ; R2= 0.2001
Estimated allele frequency
0.0
0.1
0.2
0.3
0.4
Ecological gradient (Mean annual temperature)
5
6
7
8
9
10
11
12
O1
124
9
L1
97
219
O16
L16
233
204

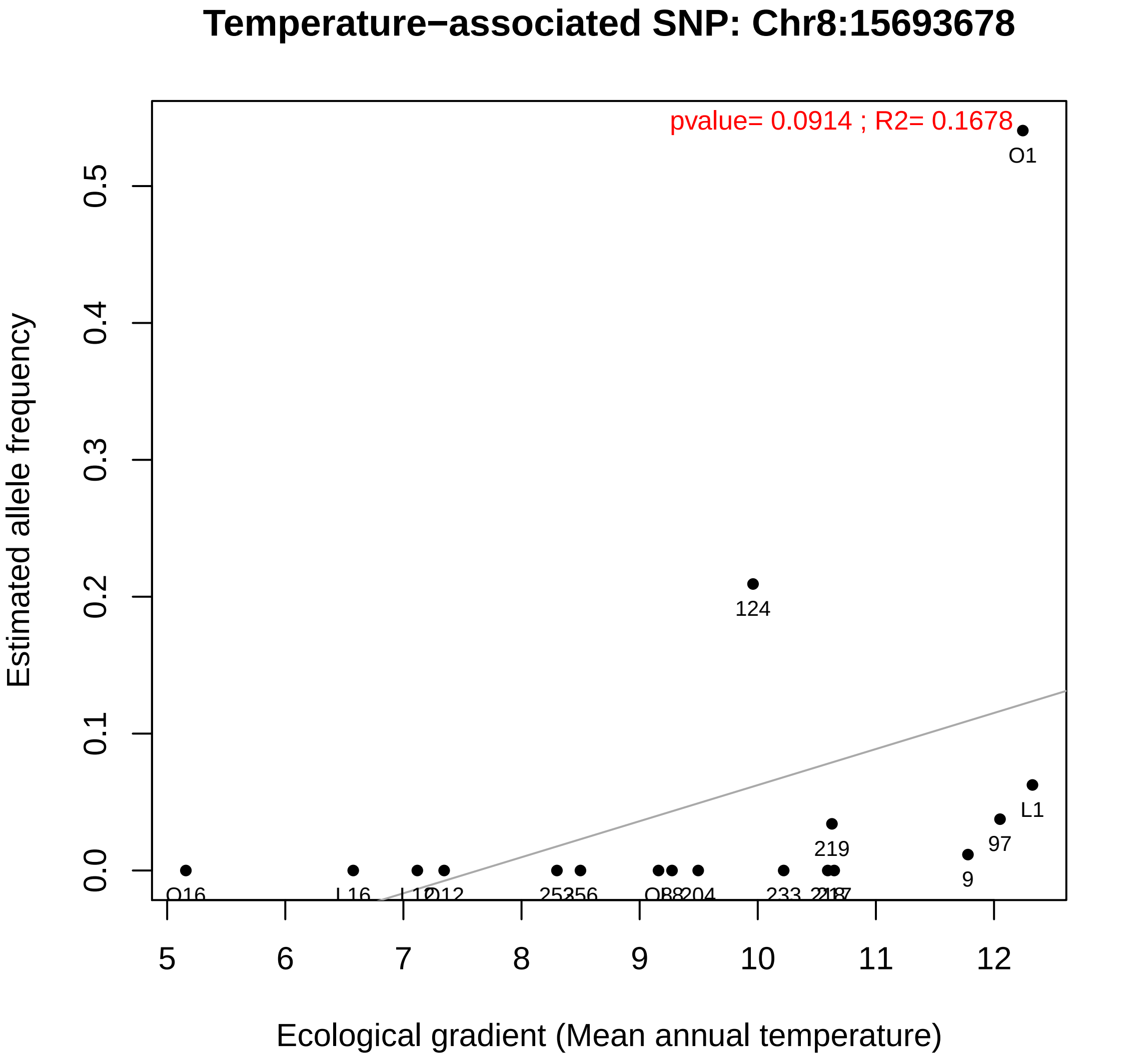

Temperature−associated SNP: Chr8:15693678
pvalue= 0.0914 ; R2= 0.1678
Estimated allele frequency
0.0
0.1
0.2
0.3
0.4
0.5
5
6
7
8
9
10
11
12
Ecological gradient (Mean annual temperature)
O1
124
219
97
L1
9
O16
L16
233

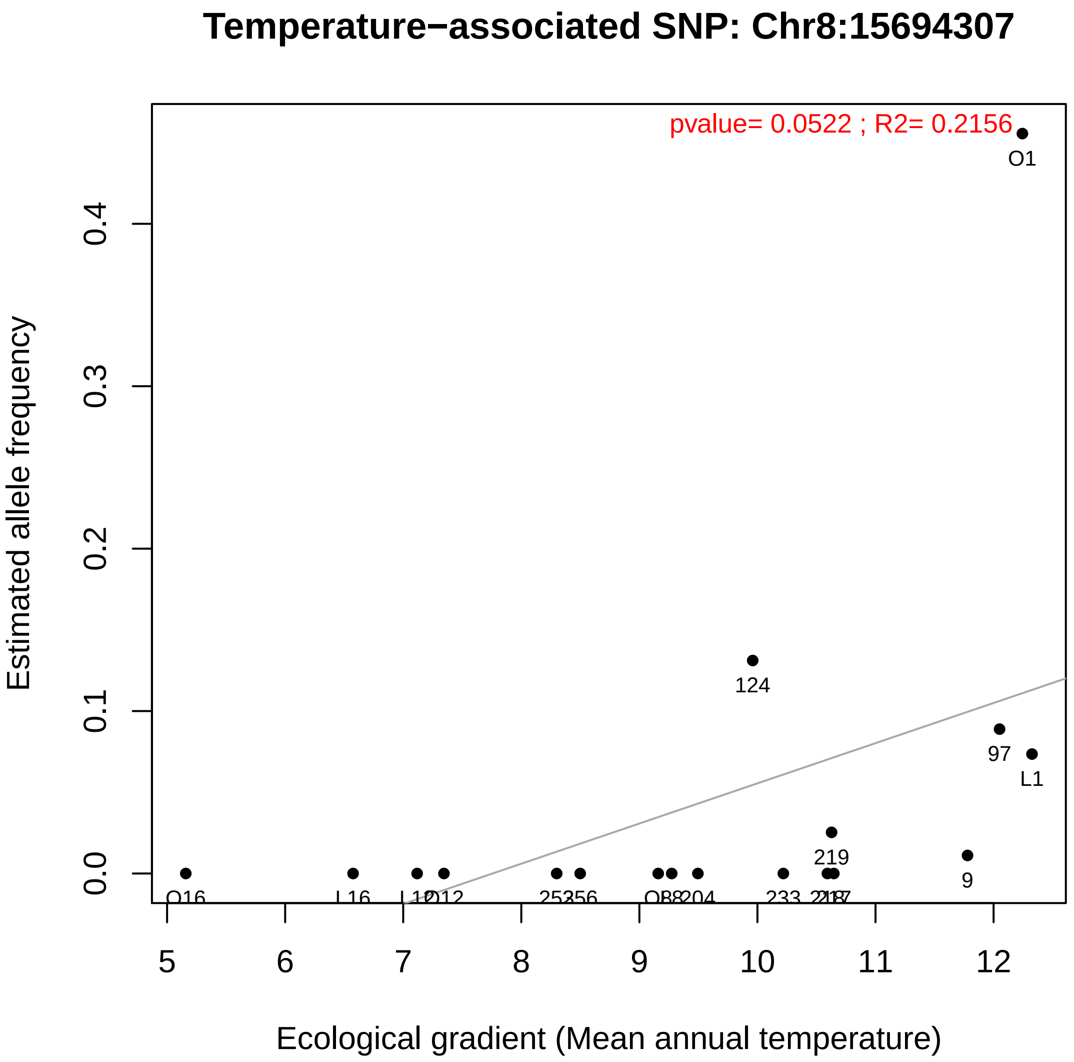
Temperature−associated SNP: Chr8:15694307
pvalue= 0.0522 ; R2= 0.2156
Estimated allele frequency
0.0
0.1
0.2
0.3
0.4
Ecological gradient (Mean annual temperature)
5
6
7
8
9
10
11
12
O1
124
97
L1
219
9
O16
L16
O12
204
233

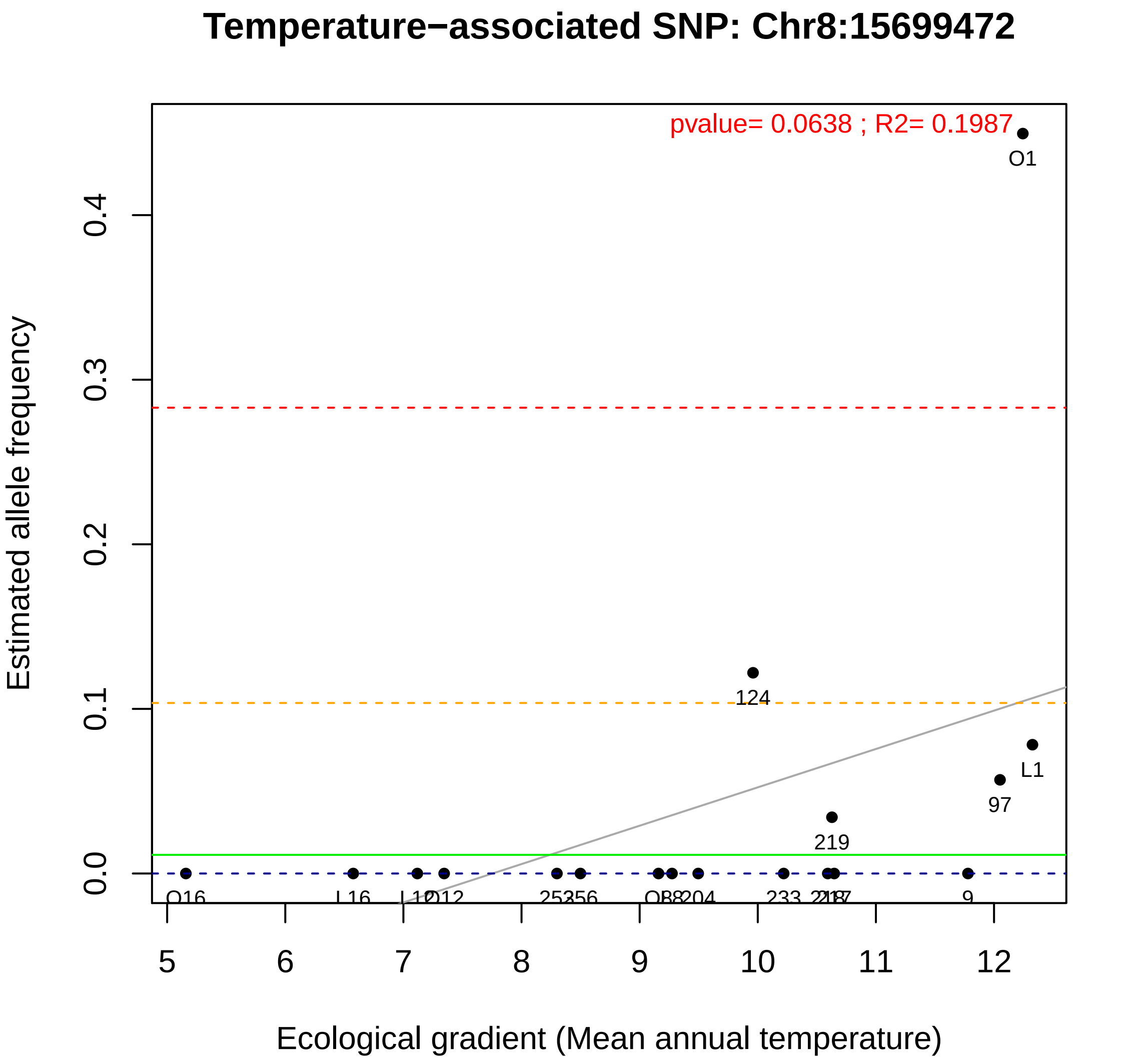

Temperature−associated SNP: Chr8:15699472
pvalue= 0.0638 ; R2= 0.1987
Estimated allele frequency
0.0
0.1
0.2
0.3
0.4
O1
124
L1
97
219
O16
L16
O12
204
233
9
5
6
7
8
9
10
11
12
Ecological gradient (Mean annual temperature)

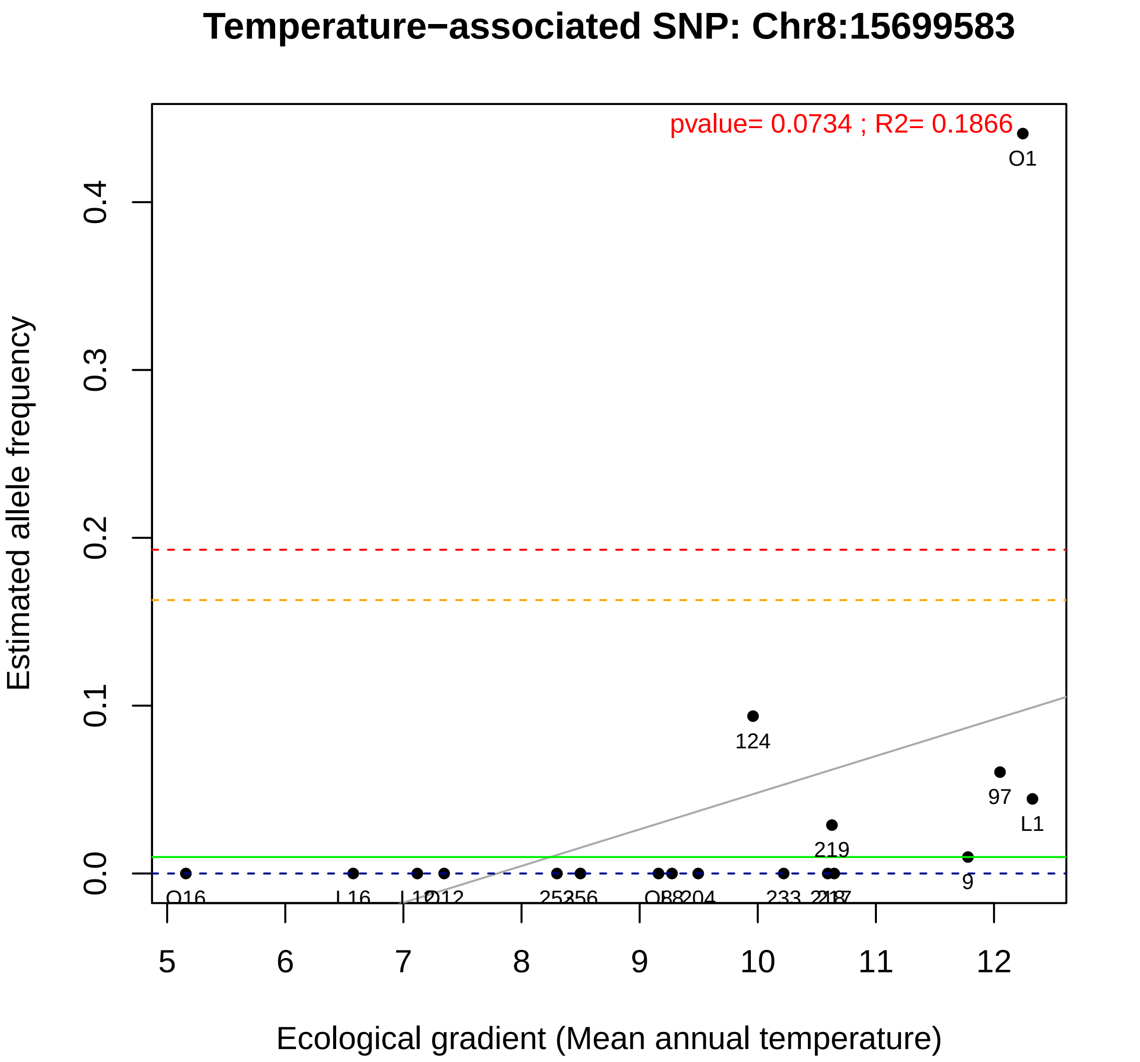
Temperature−associated SNP: Chr8:15699583
pvalue= 0.0734 ; R2= 0.1866
O1
124
97
L1
219
9
O16
L16
233
Estimated allele frequency
0.0
0.1
0.2
0.3
0.4
5
6
7
8
9
10
11
12
Ecological gradient (Mean annual temperature)

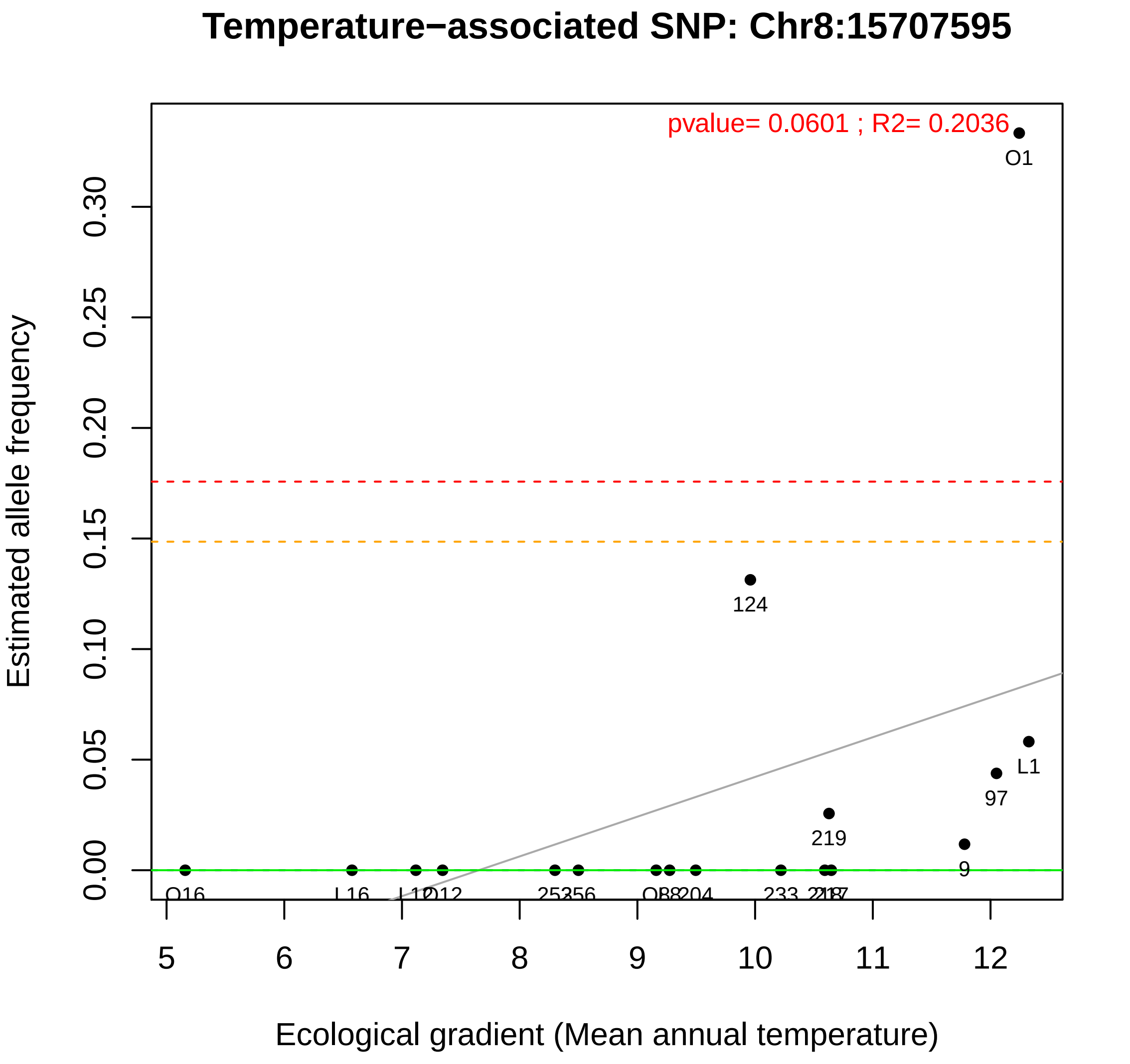
Temperature−associated SNP: Chr8:15707595
pvalue= 0.0601 ; R2= 0.2036
Estimated allele frequency
0.00
0.05
0.10
0.15
0.20
0.25
0.30
Ecological gradient (Mean annual temperature)
5
6
7
8
9
10
11
12
O1
124
219
97
L1
9
O16
L16
233
204

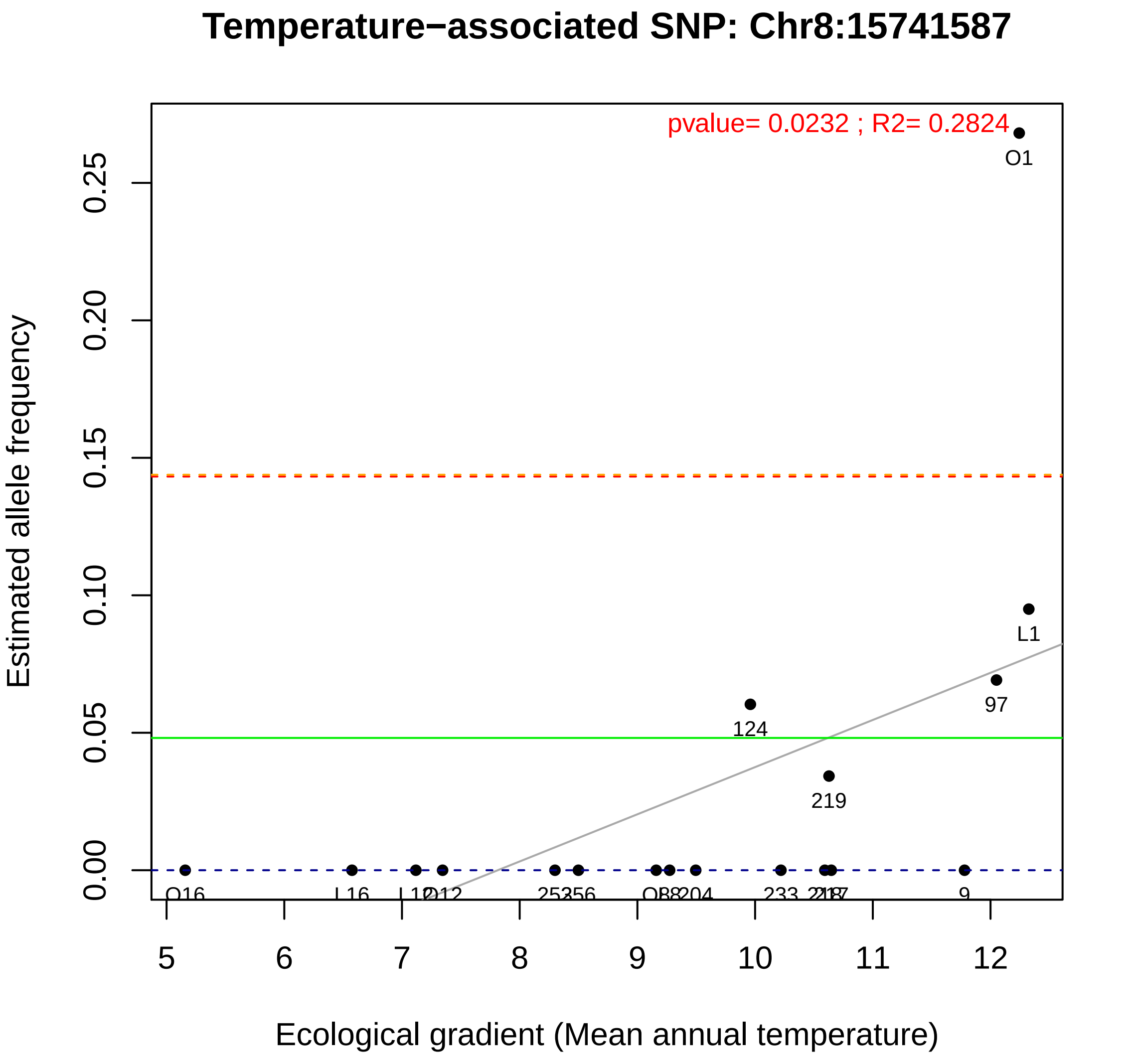
Temperature−associated SNP: Chr8:15741587
pvalue= 0.0232 ; R2= 0.2824
Estimated allele frequency
0.00
0.05
0.10
0.15
0.20
0.25
Ecological gradient (Mean annual temperature)
5
6
7
8
9
10
11
12
O1
L1
97
124
219
O16
L16
204
233
9

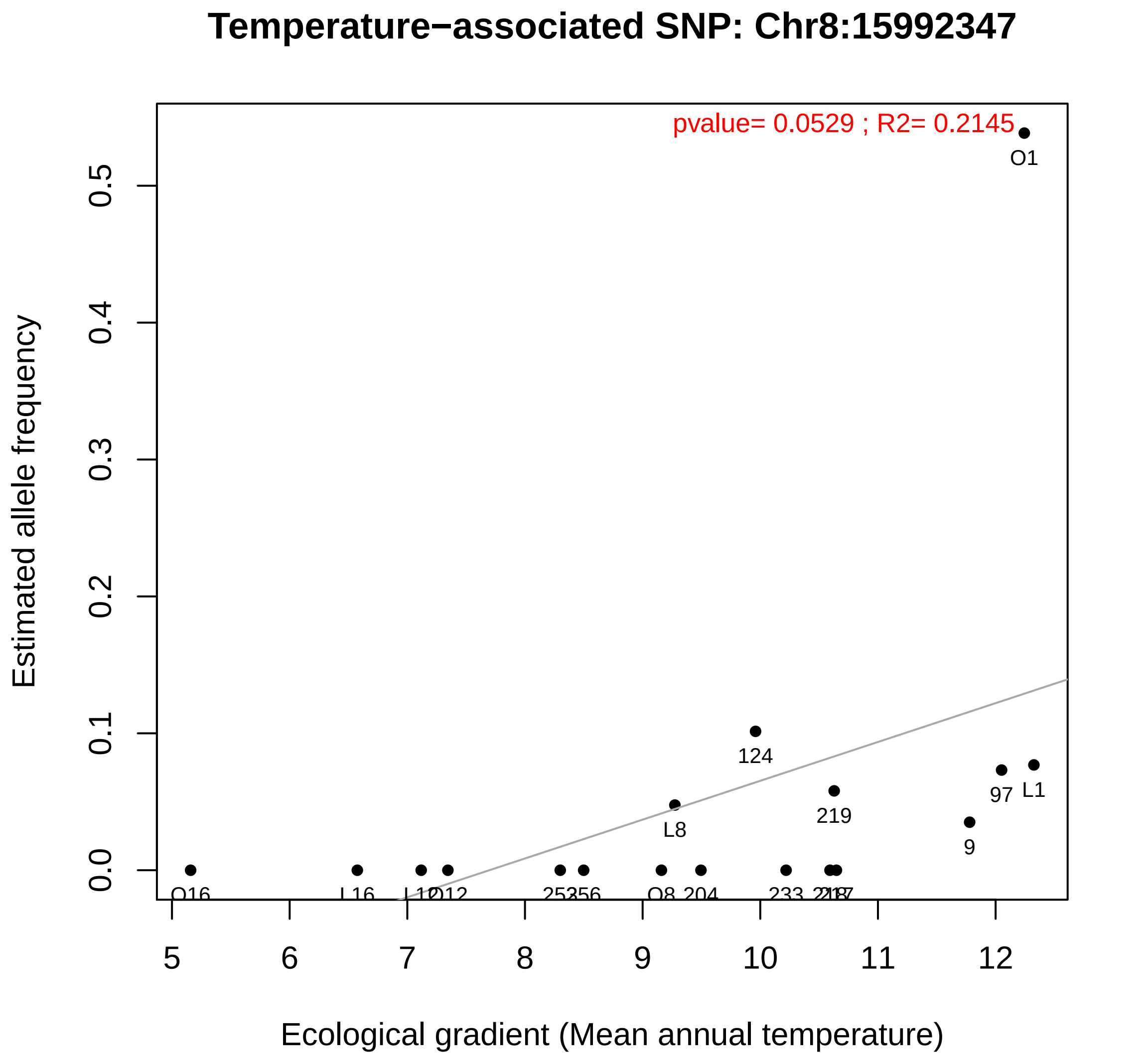

Temperature−associated SNP: Chr8:15992347
pvalue= 0.0529 ; R2= 0.2145
Estimated allele frequency
0.0
0.1
0.2
0.3
0.4
0.5
5
6
7
8
9
10
11
12
Ecological gradient (Mean annual temperature)
O1
124
97
L1
L8
219
9
O16
L16
O12
O8
204
233

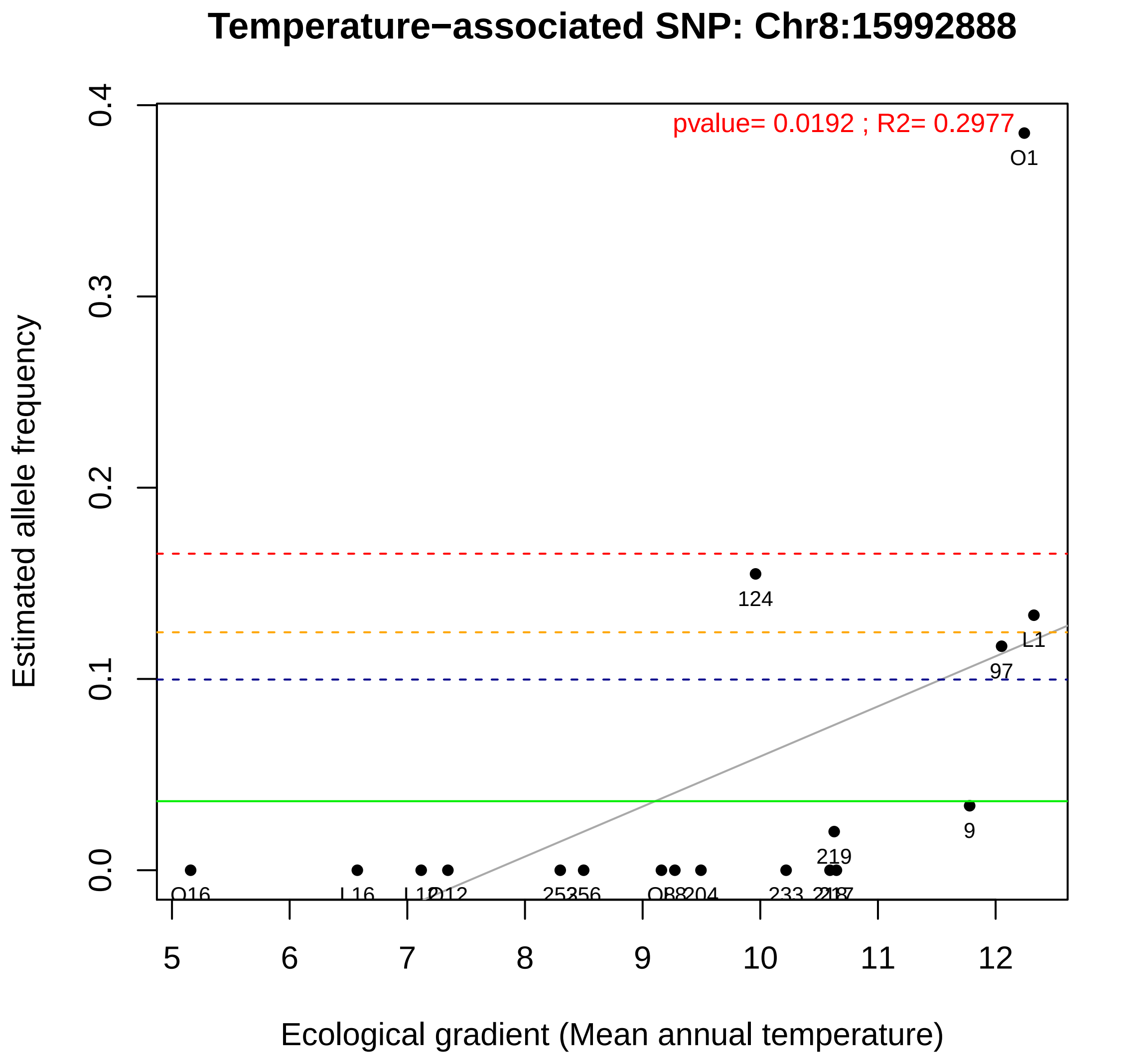

Temperature−associated SNP: Chr8:15992888
pvalue= 0.0192 ; R2= 0.2977
Estimated allele frequency
0.0
0.1
0.2
0.3
0.4
Ecological gradient (Mean annual temperature)
5
6
7
8
9
10
11
12
O1
124
L1
97
9
219
O16
L16
233
204

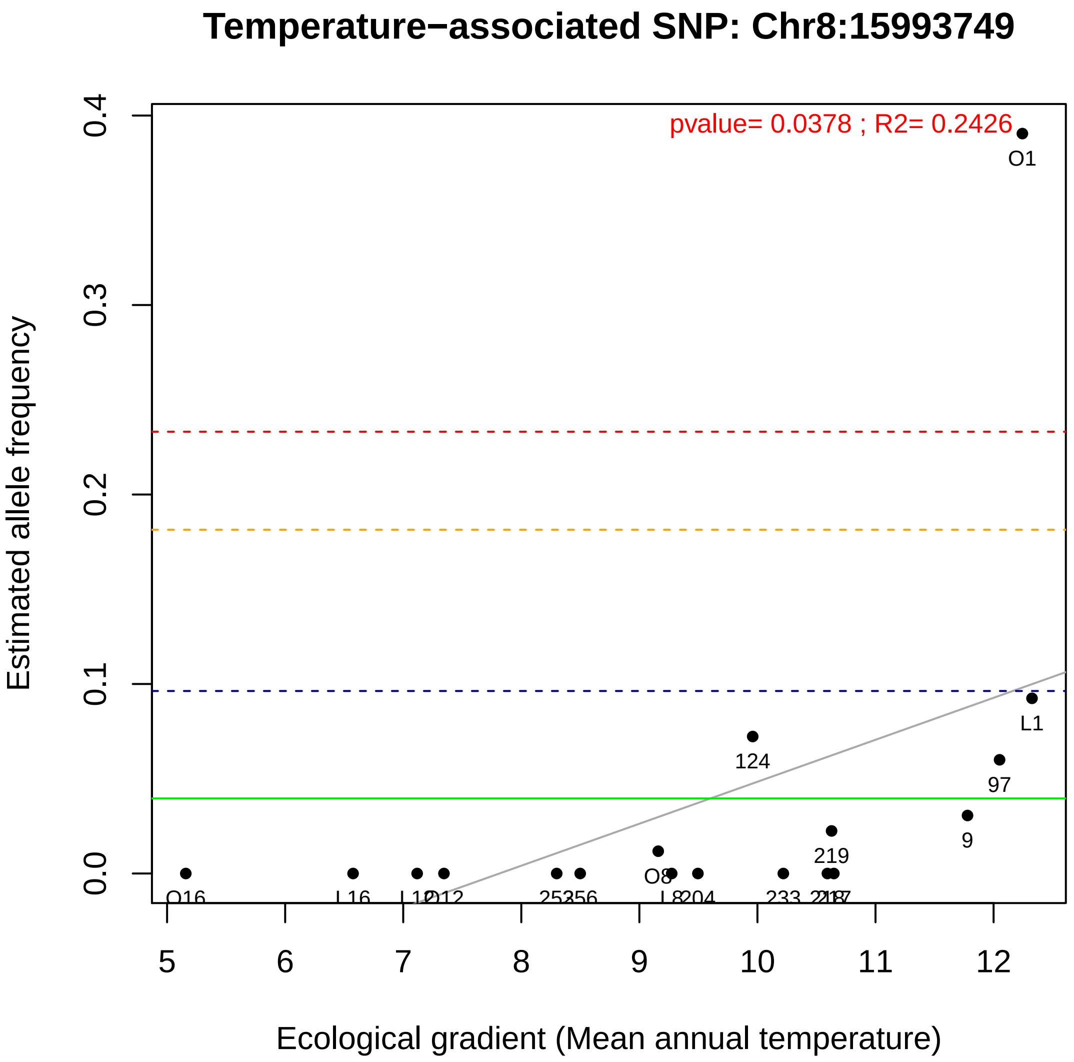
Temperature−associated SNP: Chr8:15993749
pvalue= 0.0378 ; R2= 0.2426
Estimated allele frequency
0.0
0.1
0.2
0.3
0.4
O1
L1
124
97
9
219
O8
O16
L16
L8
204
233
5
6
7
8
9
10
11
12
Ecological gradient (Mean annual temperature)

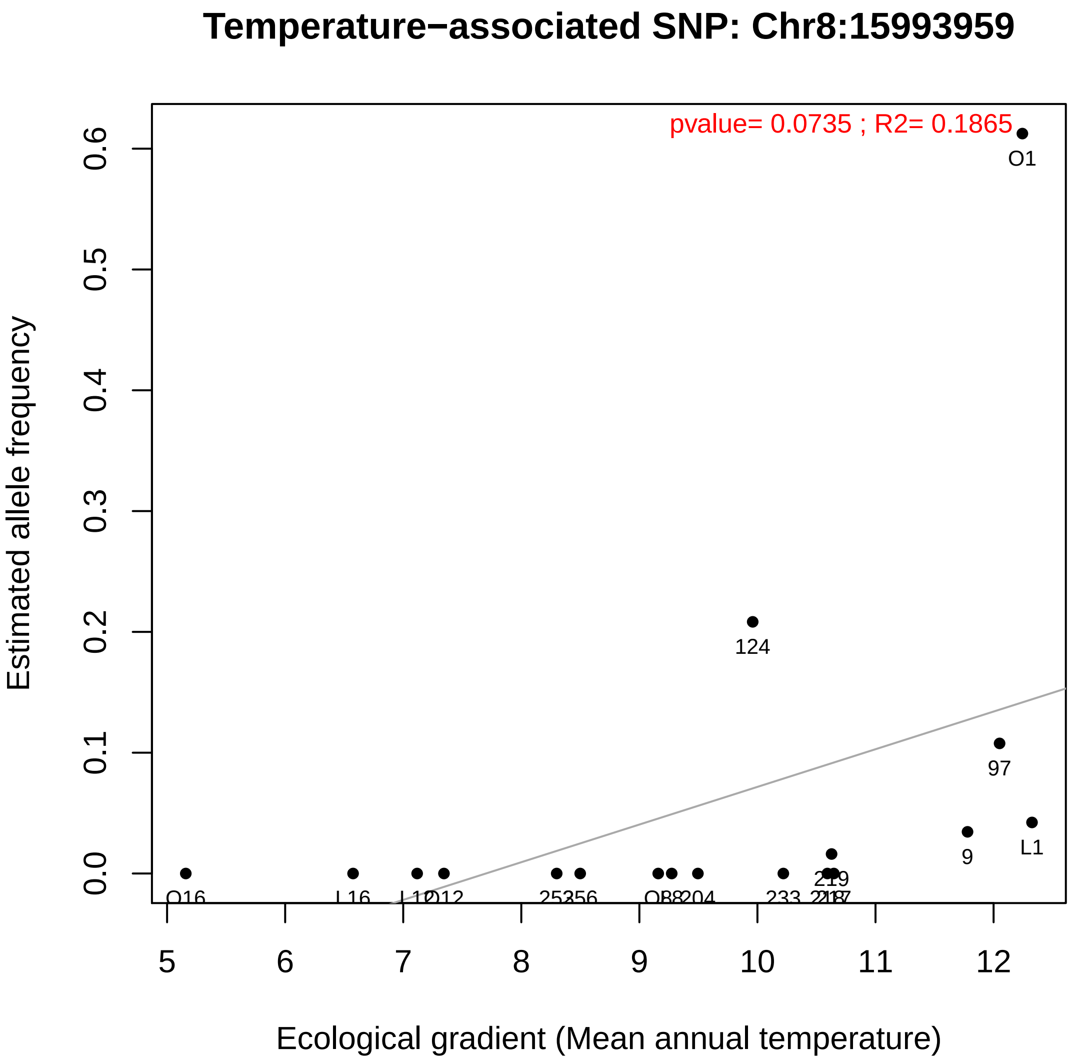
Temperature−associated SNP: Chr8:15993959
pvalue= 0.0735 ; R2= 0.1865
Estimated allele frequency
Ecological gradient (Mean annual temperature)
0.0
0.1
0.2
0.3
0.4
0.5
0.6
5
6
7
8
9
10
11
12
O1
124
97
9
L1
O16
L16
233

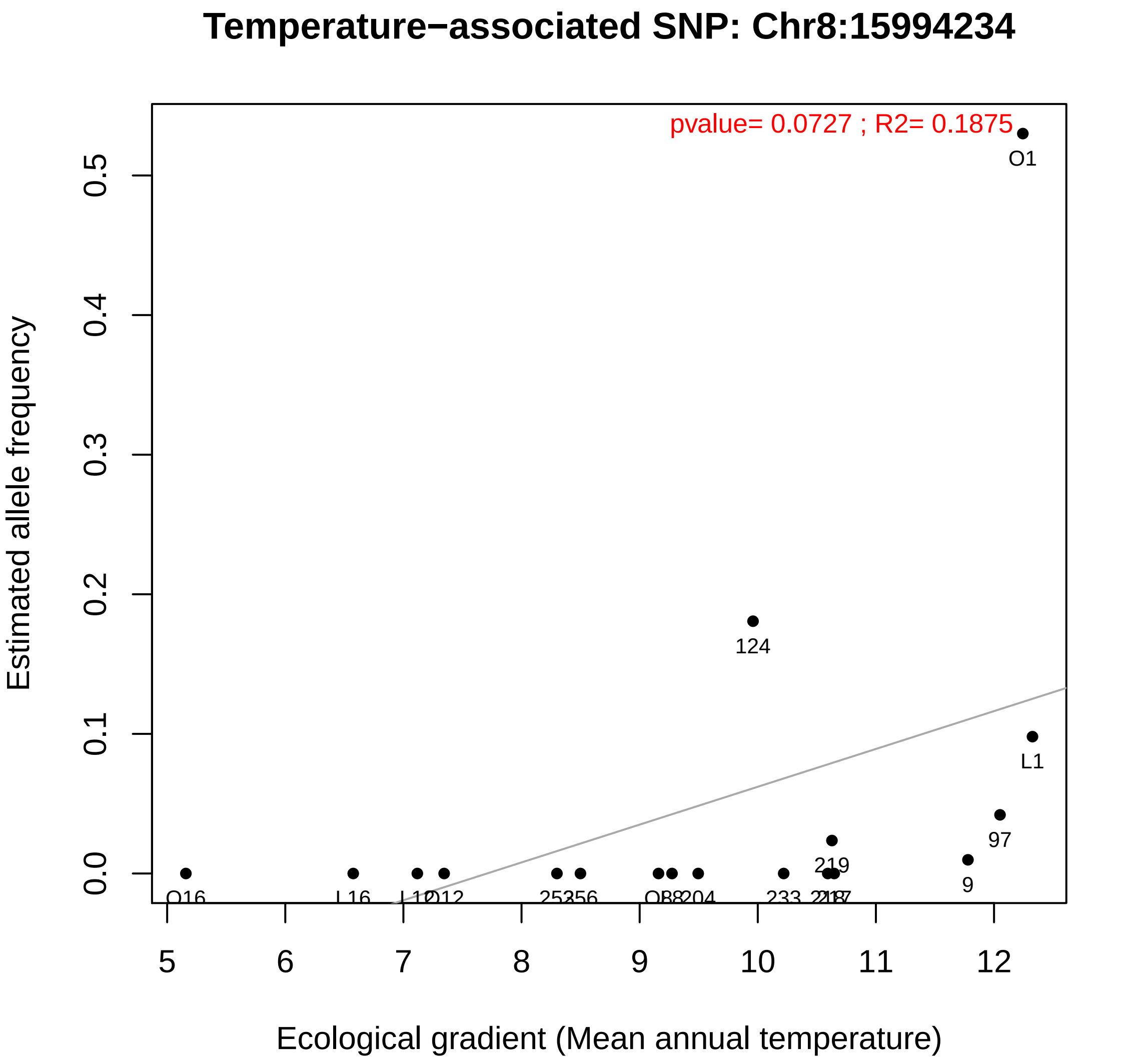

Temperature−associated SNP: Chr8:15994234
pvalue= 0.0727 ; R2= 0.1875
O1
124
L1
97
219
9
O16
L16
O12
233
0.0
0.1
0.2
0.3
0.4
0.5
5
6
7
8
9
10
11
12
Estimated allele frequency
Ecological gradient (Mean annual temperature)

**Temperature−associated SNP: Chr8:15994277**

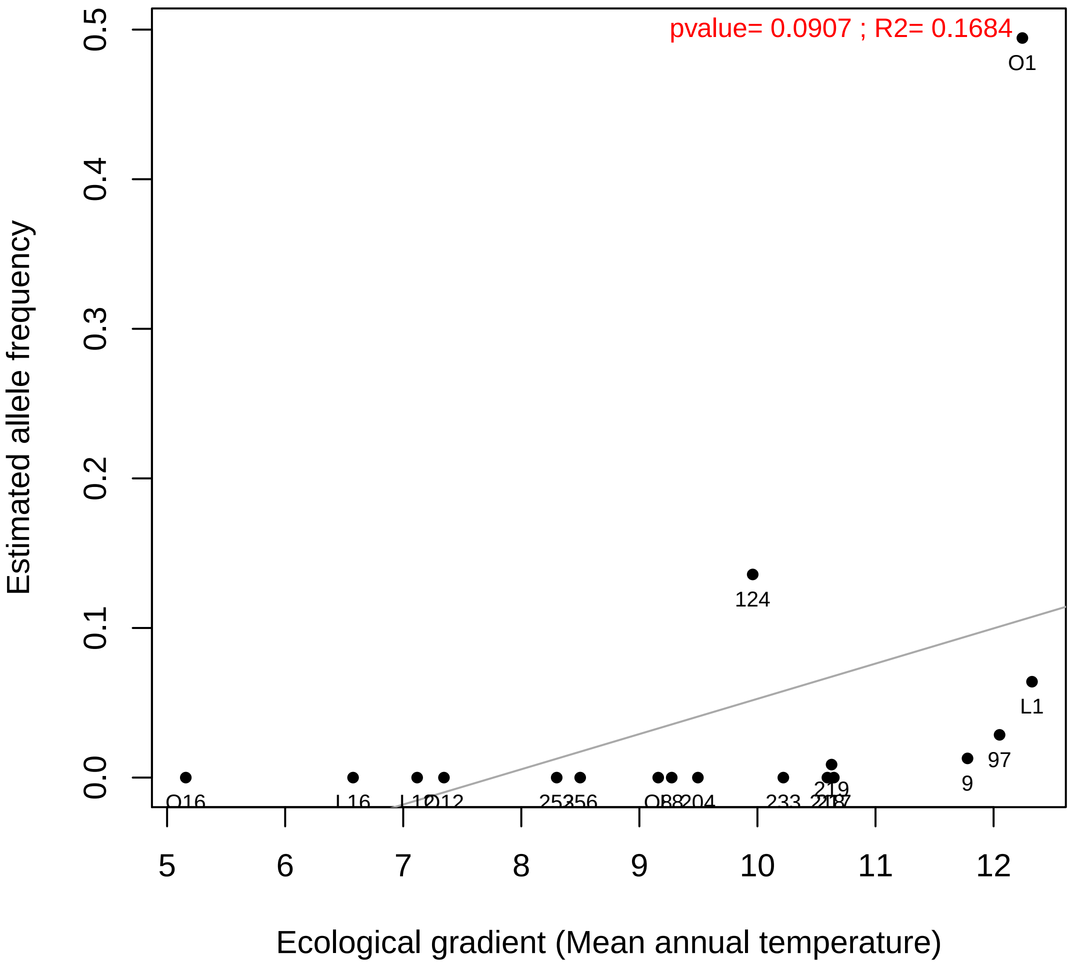

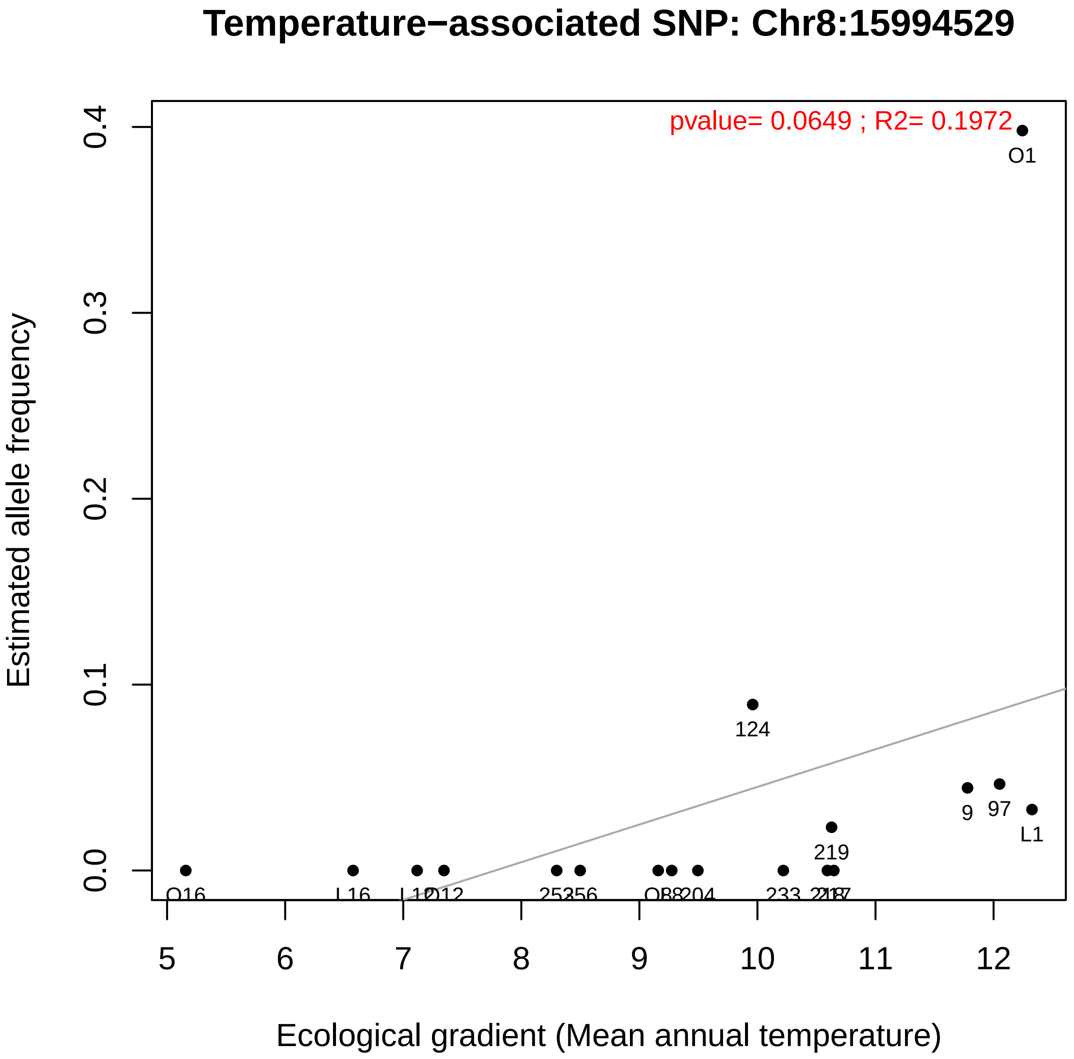
Temperature−associated SNP: Chr8:15994529
pvalue= 0.0649 ; R2= 0.1972
Estimated allele frequency
0.0
0.1
0.2
0.3
0.4
Ecological gradient (Mean annual temperature)
5
6
7
8
9
10
11
12
O1
124
219
9
97
L1
O16
L16
233
204

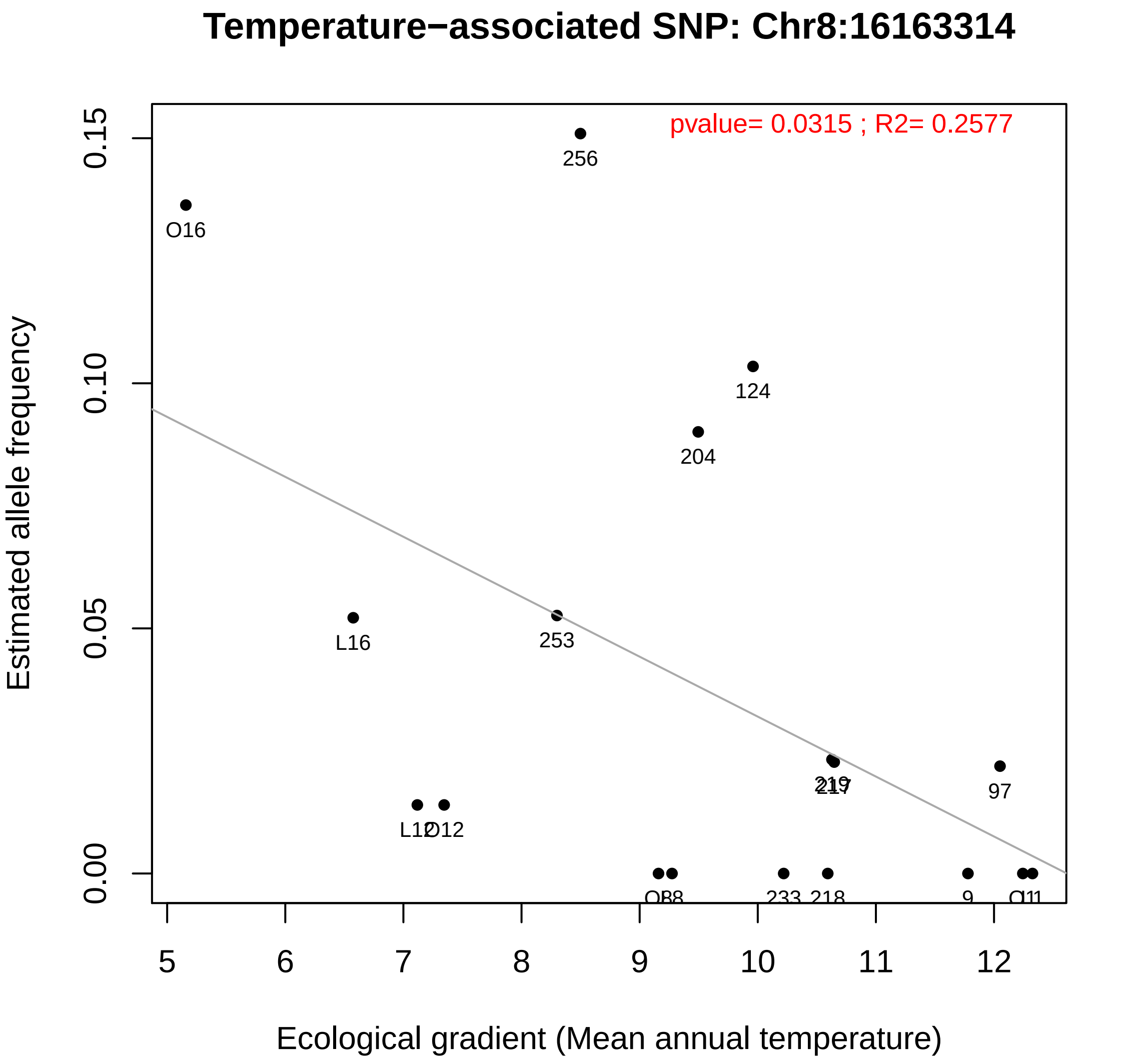

Temperature−associated SNP: Chr8:16163314
pvalue= 0.0315 ; R2= 0.2577
Estimated allele frequency
Ecological gradient (Mean annual temperature)
0.00
0.05
0.10
0.15
5
6
7
8
9
10
11
12
256
O16
124
204
L16
253
97
L12
O12
233
218
9

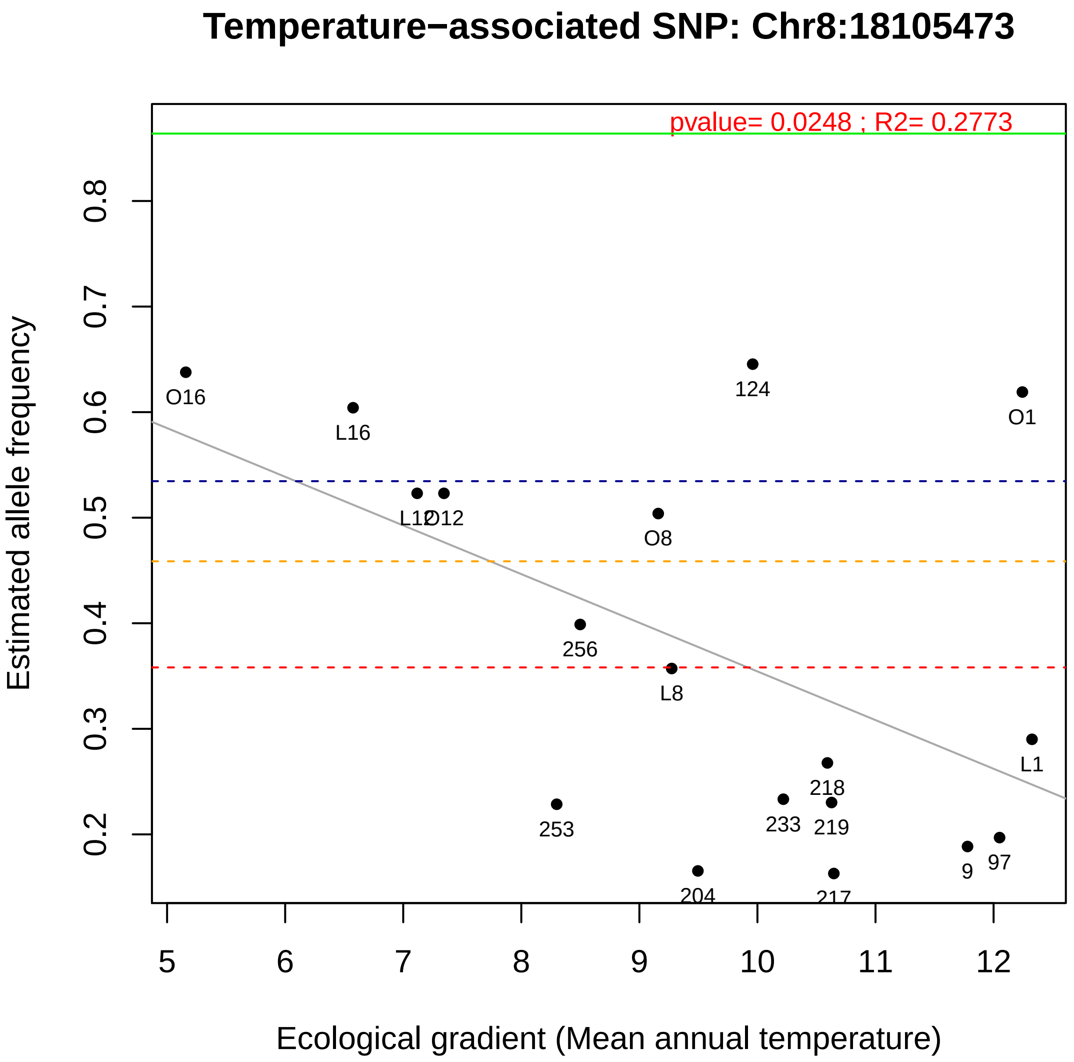
Temperature−associated SNP: Chr8:18105473
pvalue= 0.0248 ; R2= 0.2773
Estimated allele frequency
0.2
0.3
0.4
0.5
0.6
0.7
0.8
5
6
7
8
9
10
11
12
Ecological gradient (Mean annual temperature)
O16
L16
124
O1
L12
O12
O8
256
L8
L1
218
233
219
253
204
217
9
97

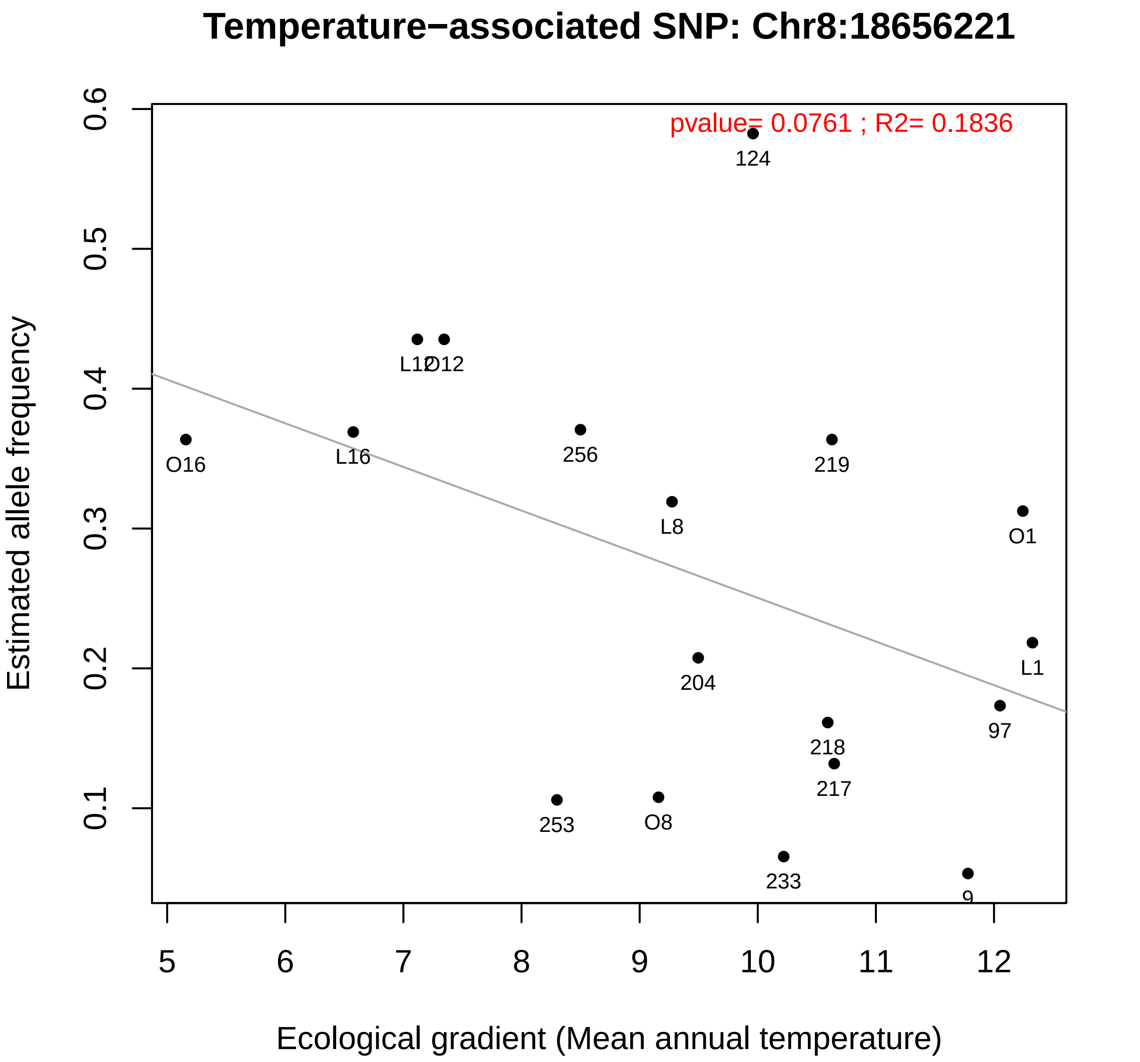
Temperature−associated SNP: Chr8:18656221
pvalue= 0.0761 ; R2= 0.1836
Estimated allele frequency
0.1
0.2
0.3
0.4
0.5
0.6
5
6
7
8
9
10
11
12
Ecological gradient (Mean annual temperature)
124
L12
O12
O16
L16
256
219
L8
O1
L1
204
218
97
217
253
O8
233
9

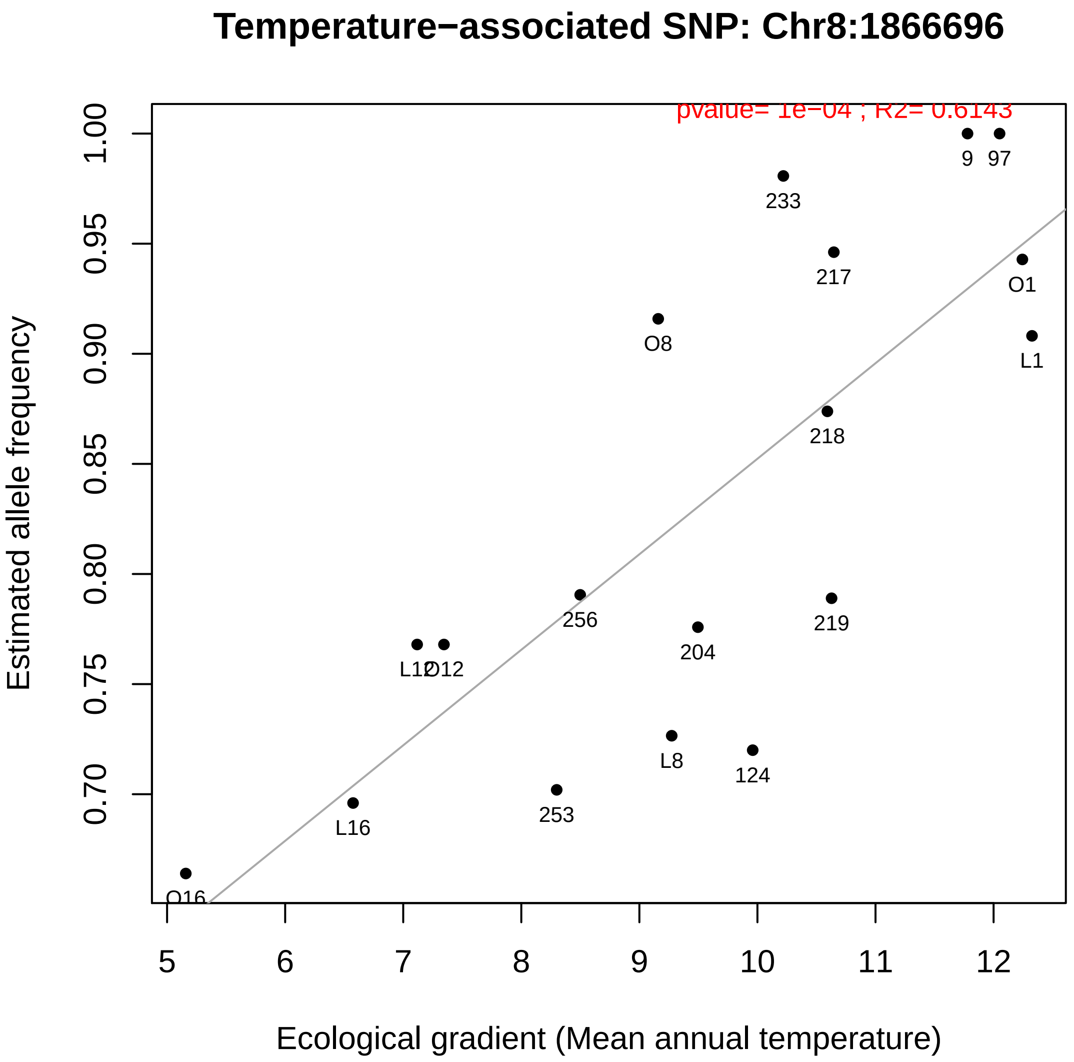
Temperature−associated SNP: Chr8:1866696
pvalue= 1e−04 ; R2= 0.6143
Estimated allele frequency
Ecological gradient (Mean annual temperature)
1.00
0.95
0.90
0.85
0.80
0.75
0.70
5
6
7
8
9
10
11
12
9
97
233
217
O1
O8
L1
218
256
219
204
L12
O12
L8
124
253
L16
O16

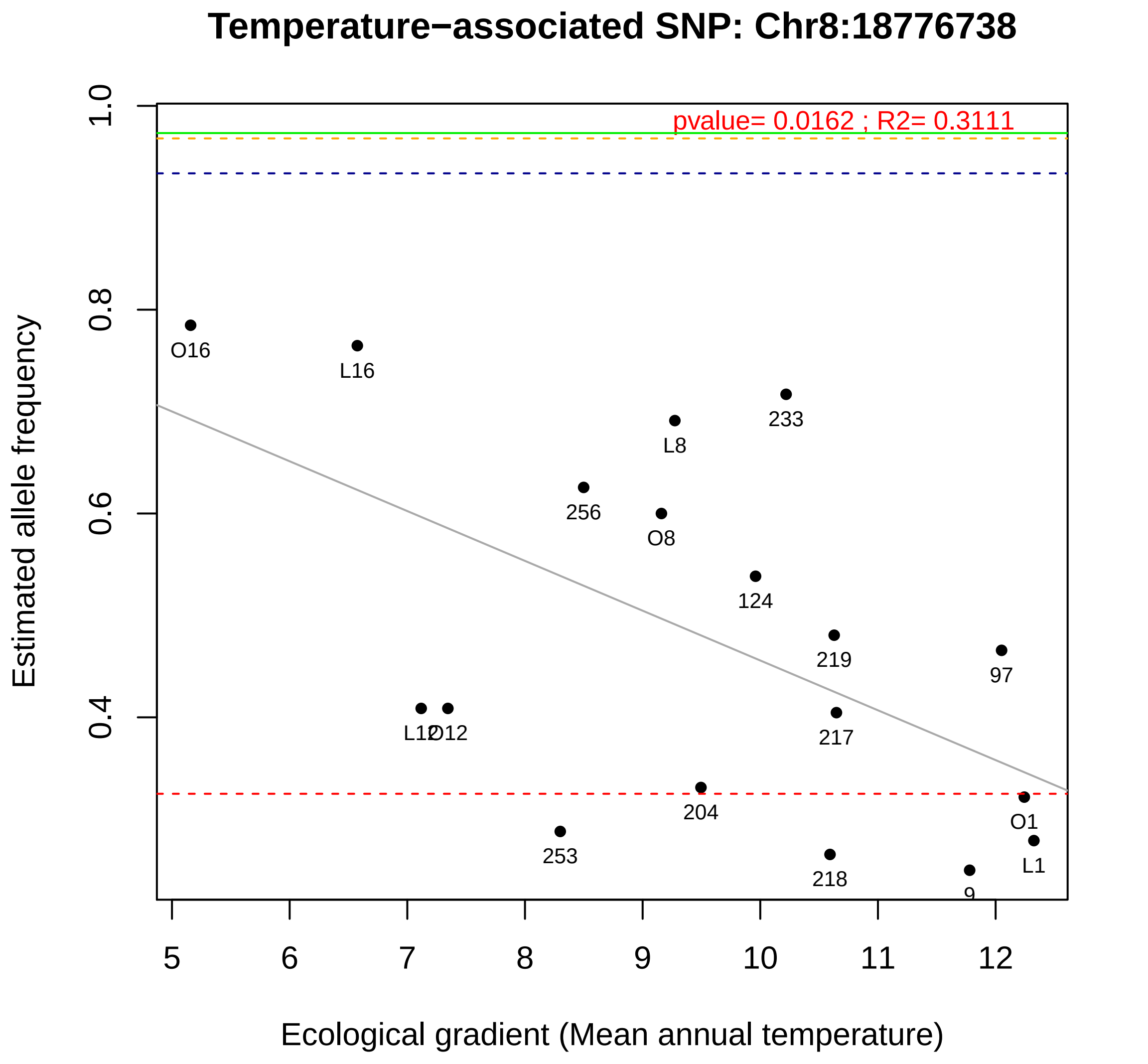

Temperature−associated SNP: Chr8:18776738
pvalue= 0.0162 ; R2= 0.3111
Estimated allele frequency
0.4
0.6
0.8
1.0
5
6
7
8
9
10
11
12
Ecological gradient (Mean annual temperature)
O16
L16
233
L8
256
O8
124
219
97
L12
O12
217
204
O1
253
L1
218
9

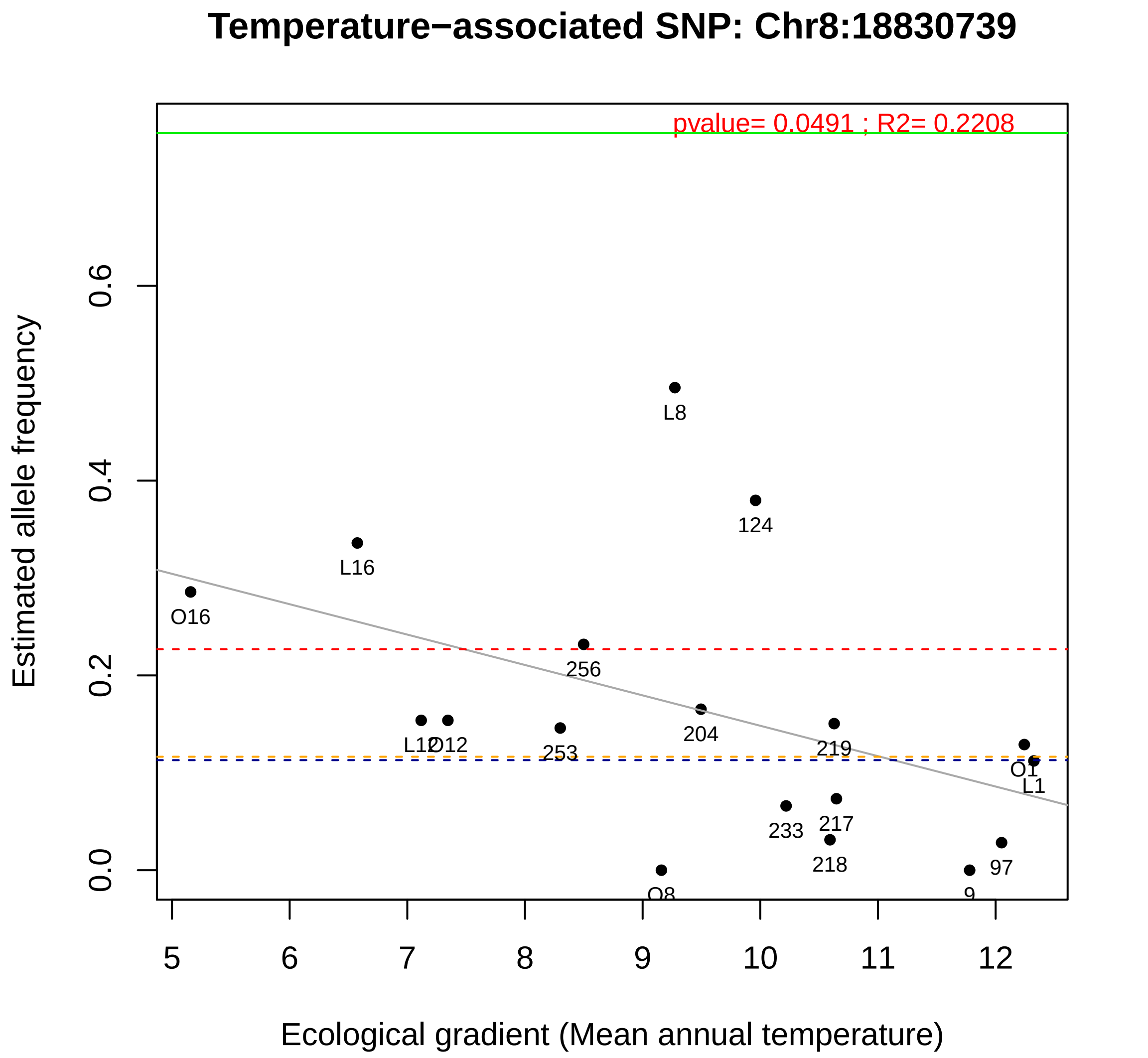
Temperature−associated SNP: Chr8:18830739
pvalue= 0.0491 ; R2= 0.2208
Estimated allele frequency
0.0
0.2
0.4
0.6
L8
124
L16
O16
256
204
L12
O12
253
219
O1
L1
233
217
218
97
O8
9
5
6
7
8
9
10
11
12
Ecological gradient (Mean annual temperature)

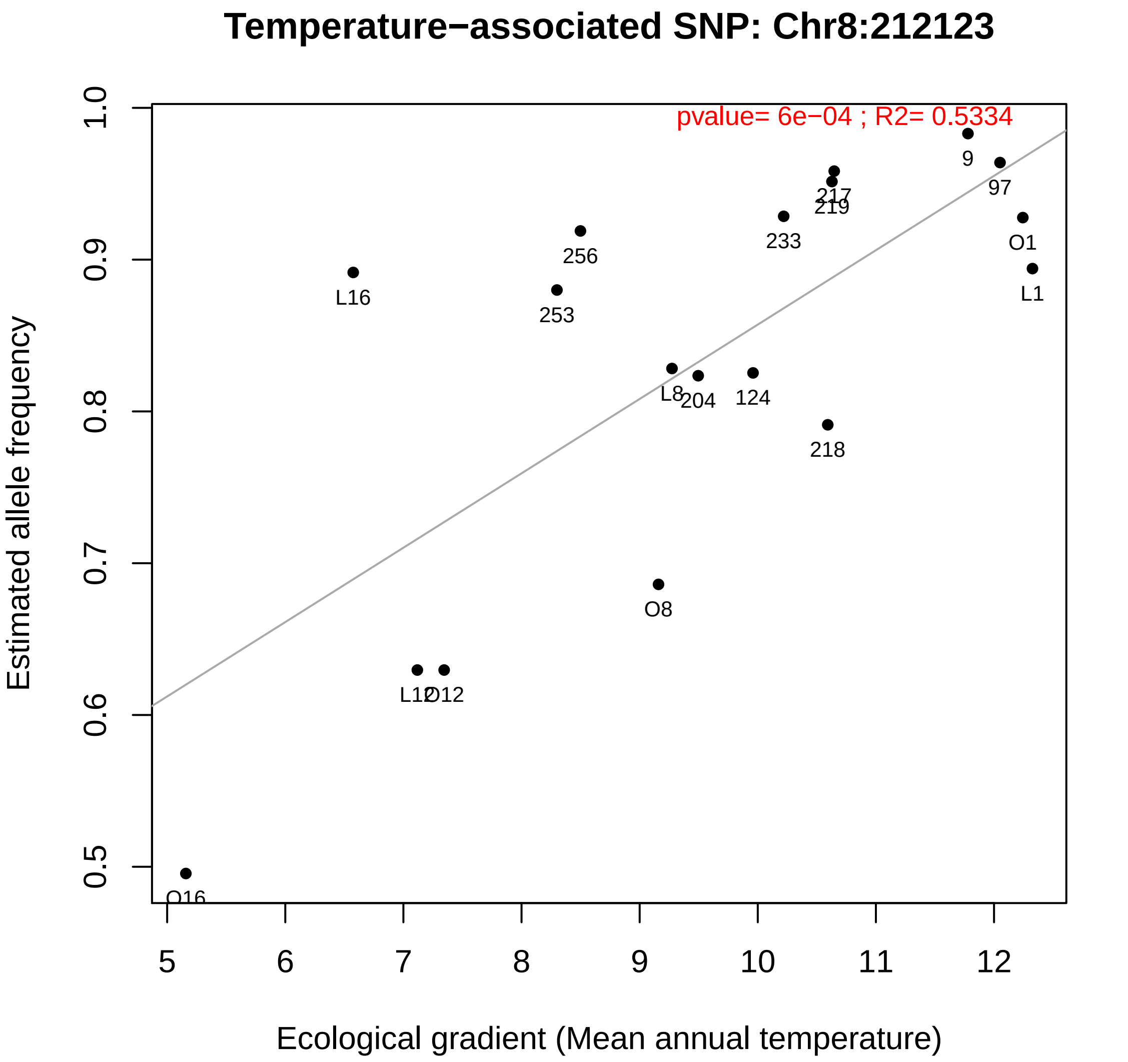

Temperature−associated SNP: Chr8:212123
pvalue= 6e−04 ; R2= 0.5334
Estimated allele frequency
Ecological gradient (Mean annual temperature)
0.5
0.6
0.7
0.8
0.9
1.0
5
6
7
8
9
10
11
12
9
97
217
219
233
O1
L1
256
L16
253
L8
204
124
218
O8
L12
O12
O16

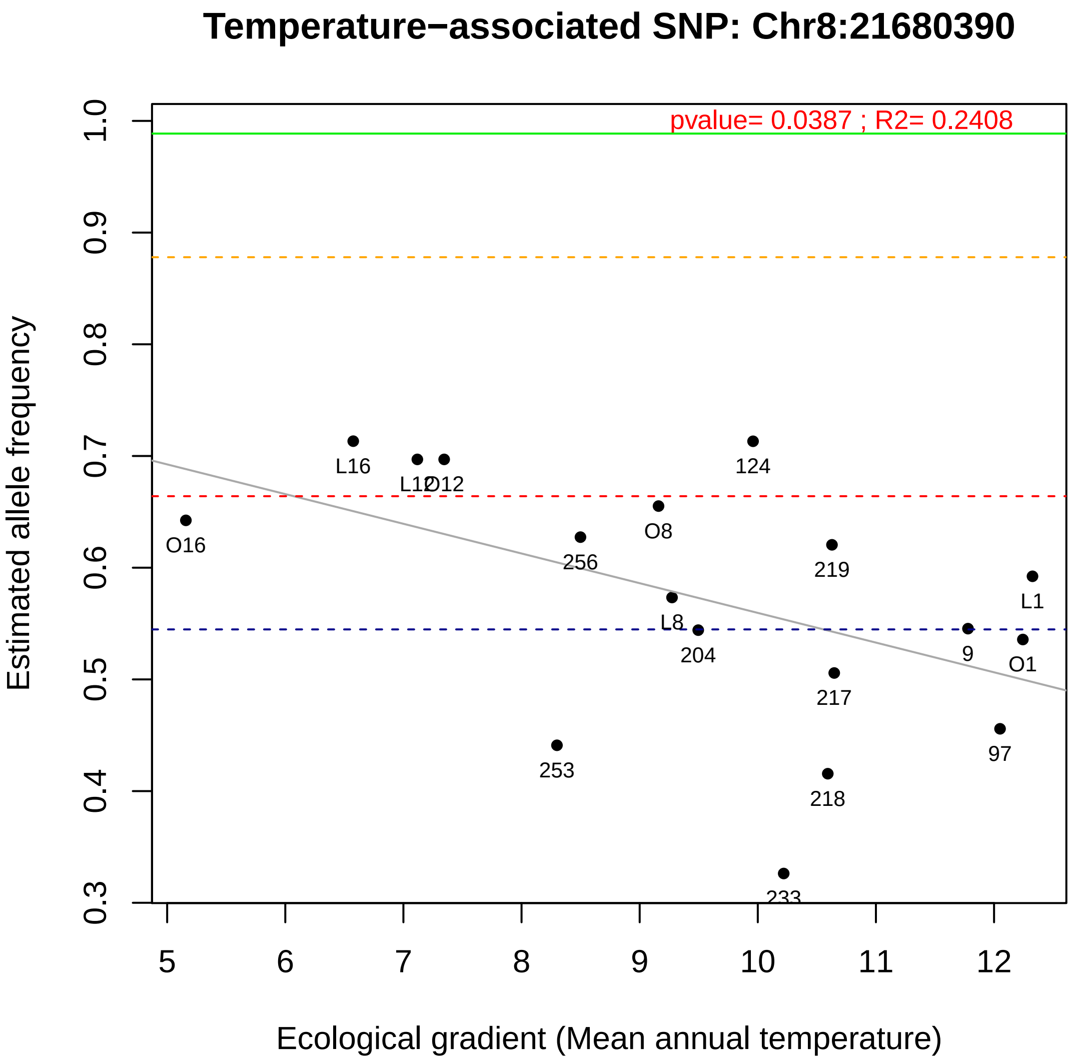
Temperature−associated SNP: Chr8:21680390
pvalue= 0.0387 ; R2= 0.2408
Estimated allele frequency
0.3
0.4
0.5
0.6
0.7
0.8
0.9
1.0
5
6
7
8
9
10
11
12
Ecological gradient (Mean annual temperature)
L16
L12
O12
124
O16
256
O8
219
L1
L8
204
9
O1
217
253
97
218
233

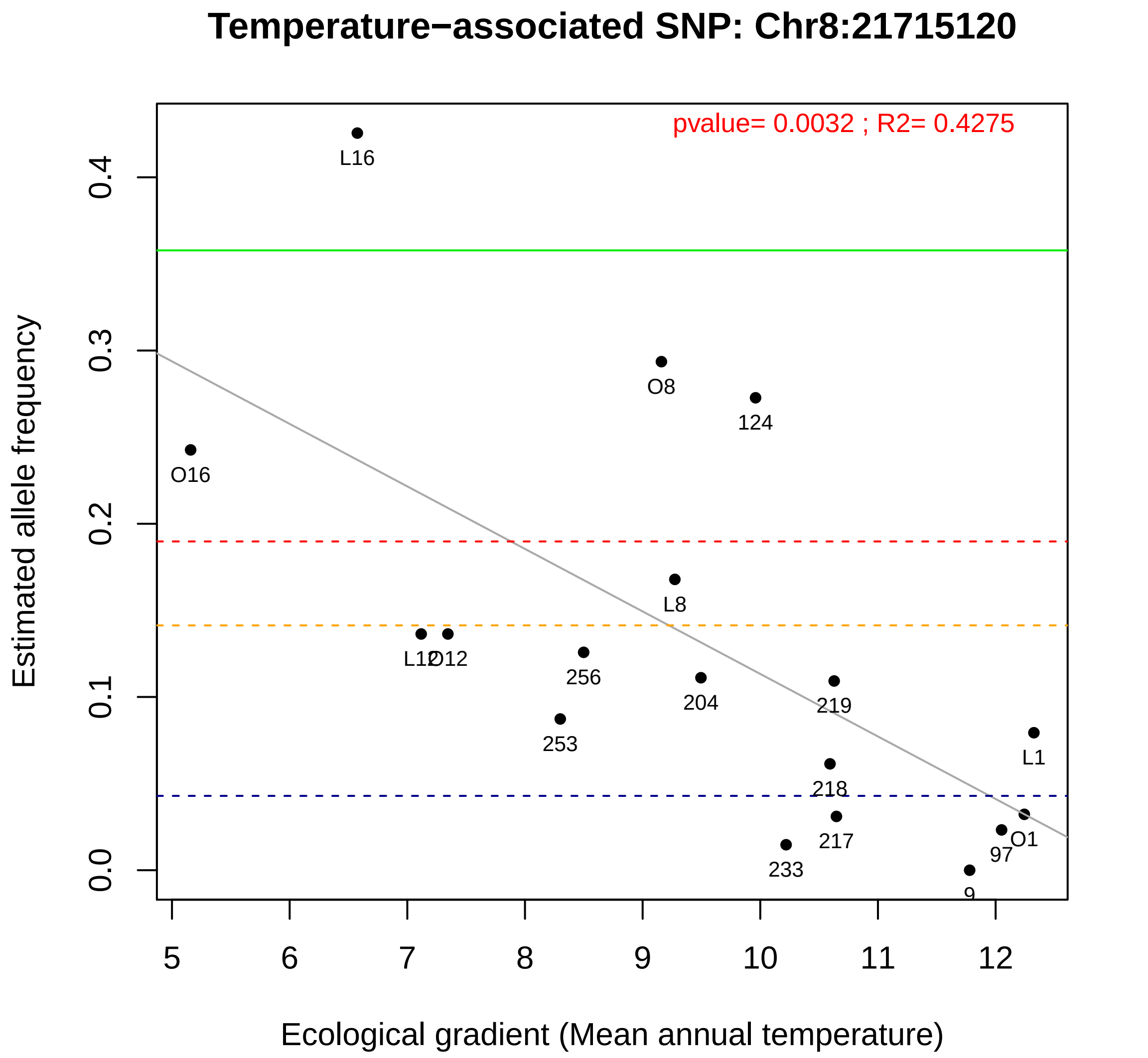
Temperature−associated SNP: Chr8:21715120
pvalue= 0.0032 ; R2= 0.4275
Estimated allele frequency
0.0
0.1
0.2
0.3
0.4
Ecological gradient (Mean annual temperature)
5
6
7
8
9
10
11
12
L16
O8
124
O16
L8
L12
O12
256
204
219
253
L1
218
217
O1
97
233
9

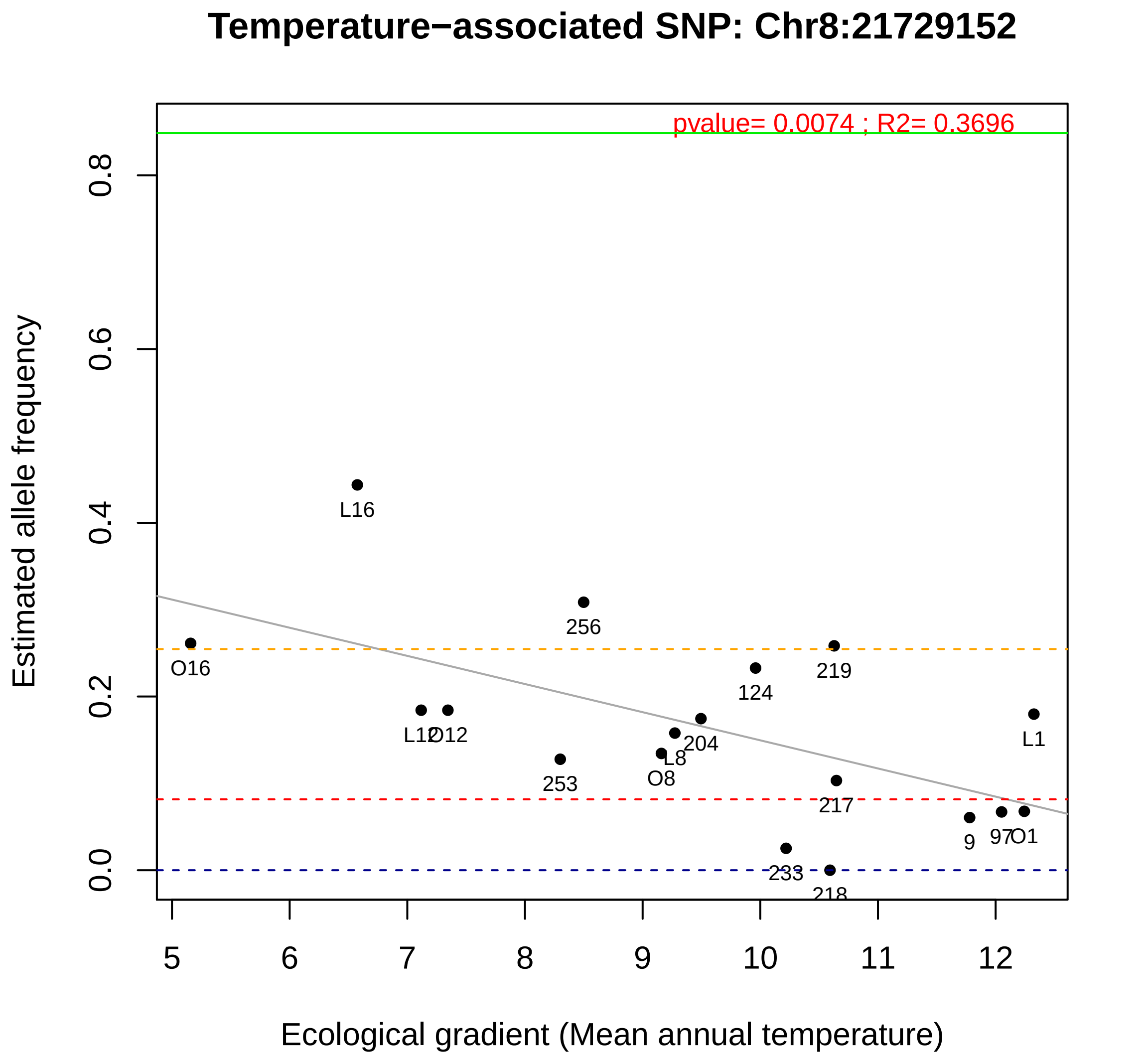

Temperature−associated SNP: Chr8:21729152
pvalue= 0.0074 ; R2= 0.3696
Estimated allele frequency
0.0
0.2
0.4
0.6
0.8
5
6
7
8
9
10
11
12
Ecological gradient (Mean annual temperature)
L16
256
O16
219
124
L12
O12
204
L1
L8
253
O8
217
9
97
O1
233
218

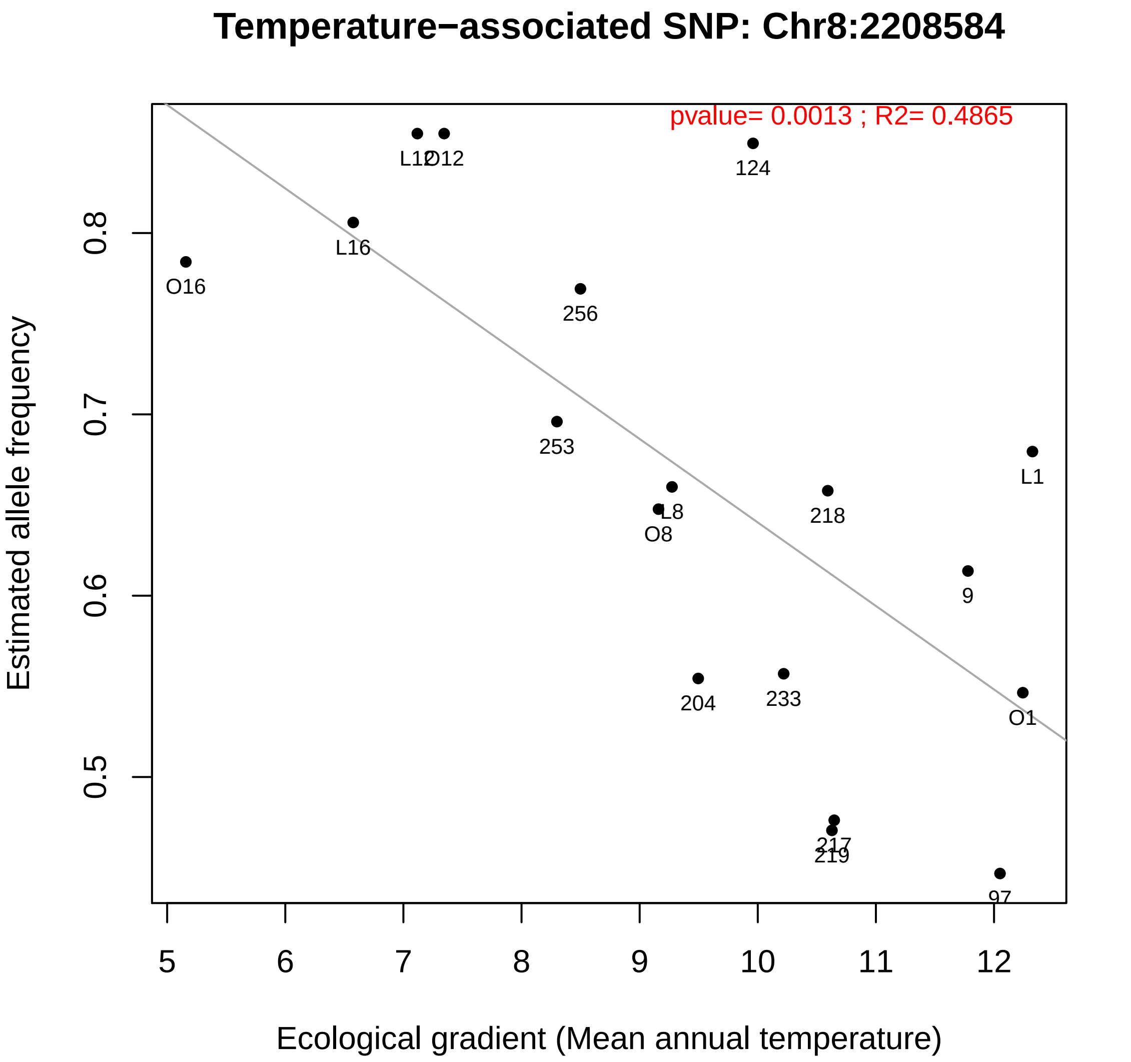
Temperature−associated SNP: Chr8:2208584
pvalue= 0.0013 ; R2= 0.4865
Estimated allele frequency
0.5
0.6
0.7
0.8
Ecological gradient (Mean annual temperature)
5
6
7
8
9
10
11
12
L12
O12
124
L16
O16
256
253
L1
L8
O8
218
9
204
233
O1
217
219
97

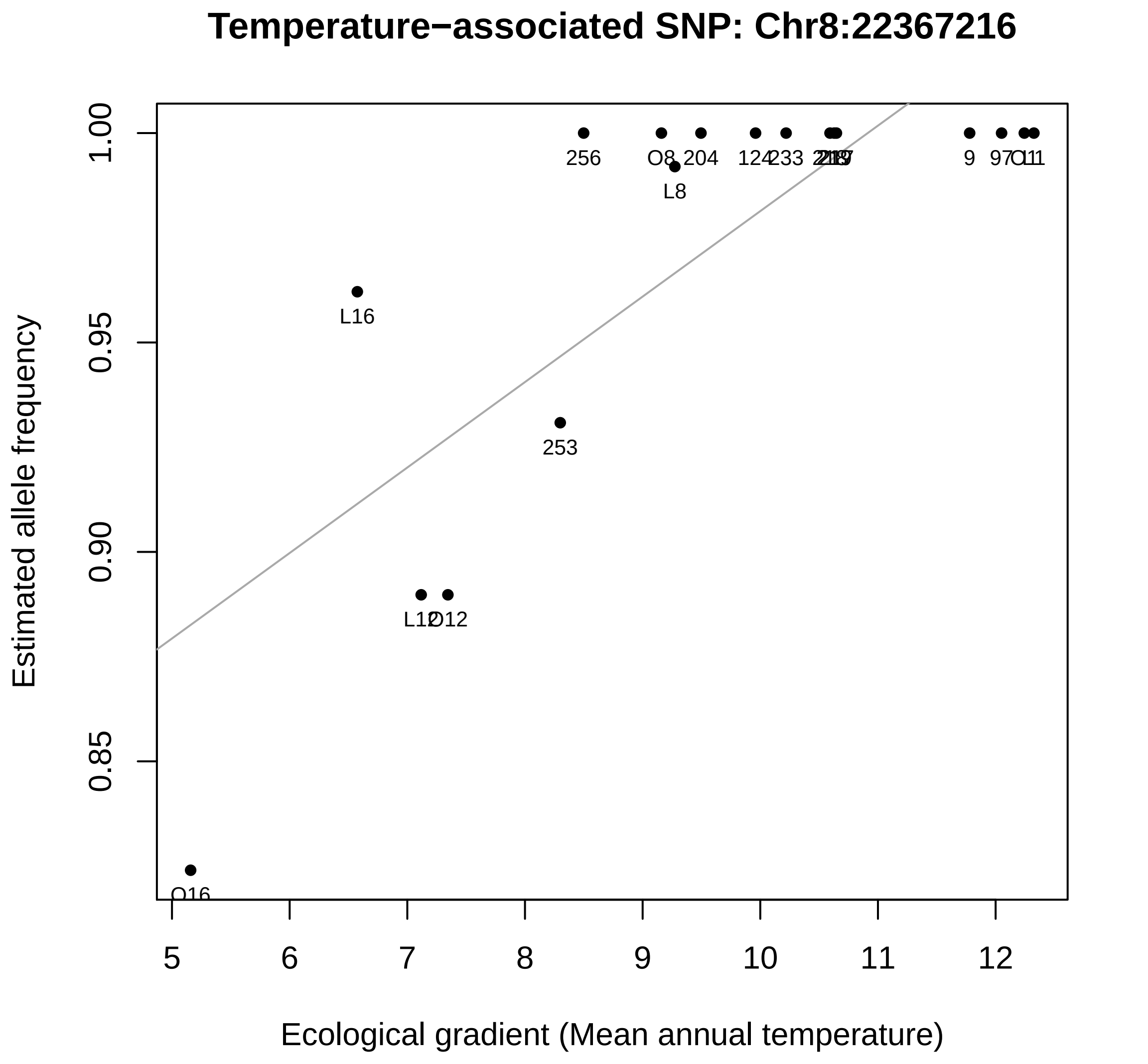

Temperature−associated SNP: Chr8:22367216
Estimated allele frequency
Ecological gradient (Mean annual temperature)
1.00
0.95
0.90
0.85
5
6
7
8
9
10
11
12
256
O8
204
124
233
L8
9
L16
253
L12
O12
O16

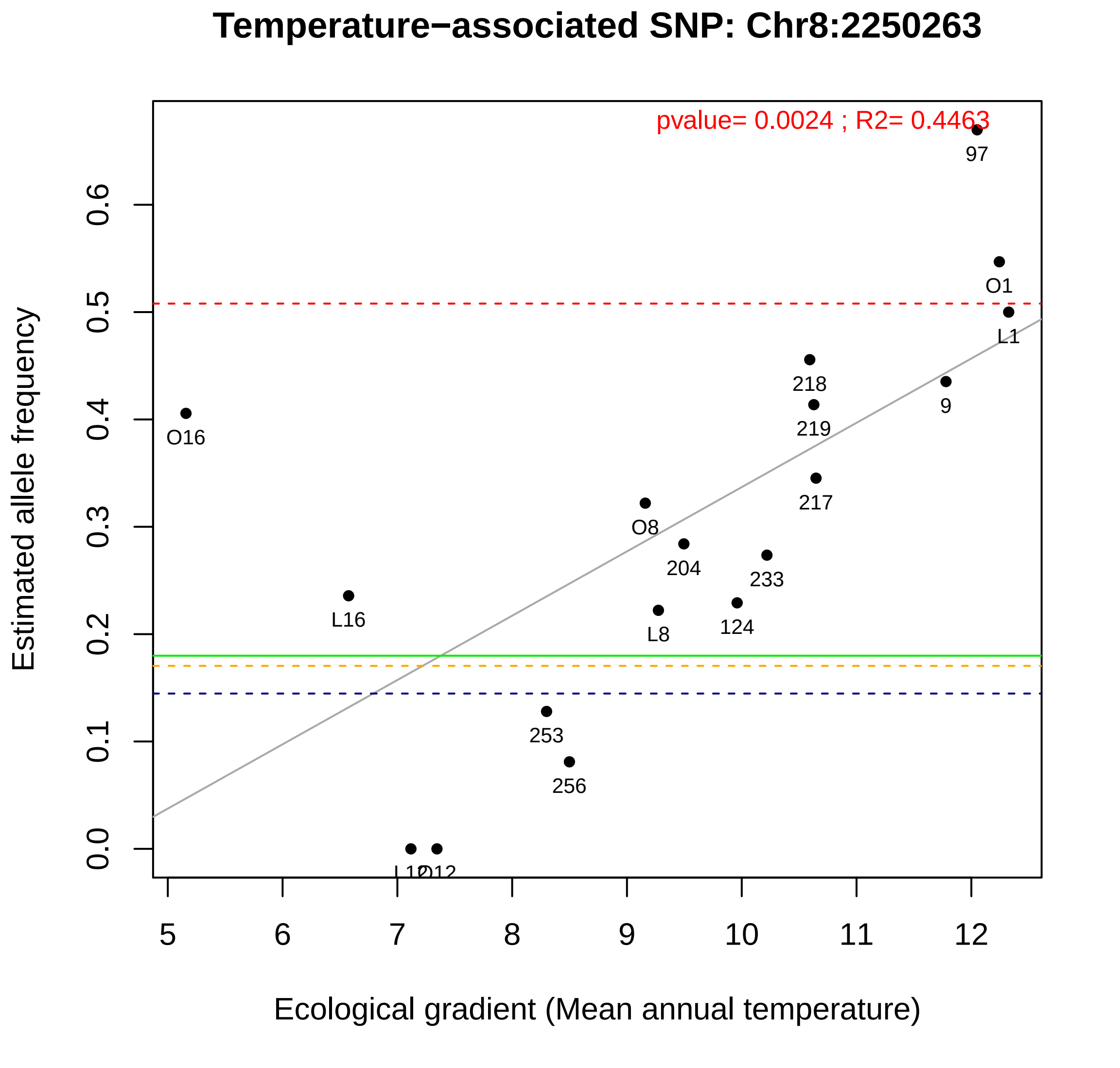
Temperature−associated SNP: Chr8:2250263
pvalue= 0.0024 ; R2= 0.4463
Estimated allele frequency
0.0
0.1
0.2
0.3
0.4
0.5
0.6
5
6
7
8
9
10
11
12
Ecological gradient (Mean annual temperature)
97
O1
L1
218
9
219
O16
217
O8
204
233
L16
L8
124
253
256
O12

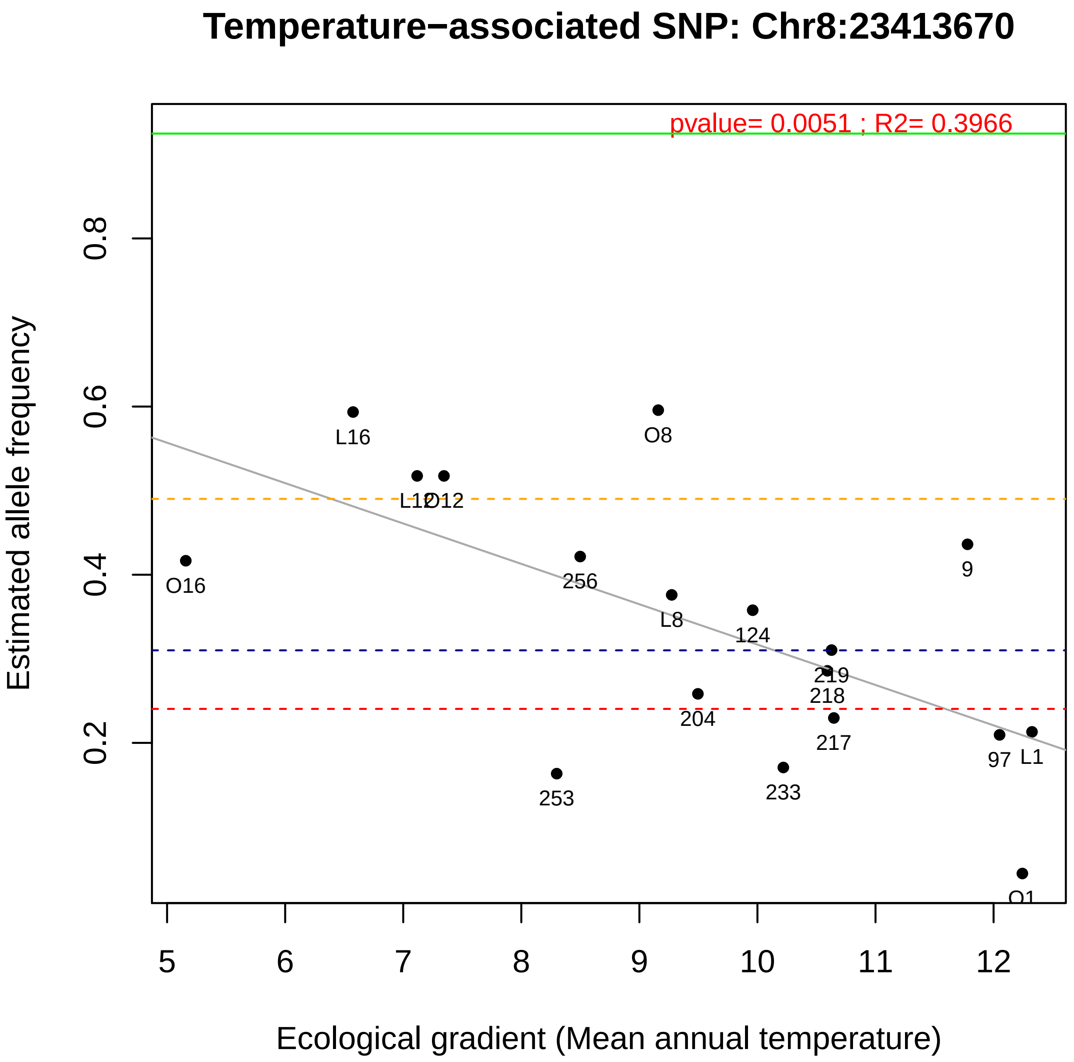

Temperature−associated SNP: Chr8:23413670
pvalue= 0.0051 ; R2= 0.3966
Estimated allele frequency
0.2
0.4
0.6
0.8
5
6
7
8
9
10
11
12
Ecological gradient (Mean annual temperature)
L16
O8
L12
O12
O16
256
9
L8
124
219
218
204
217
97
L1
253
233
O1

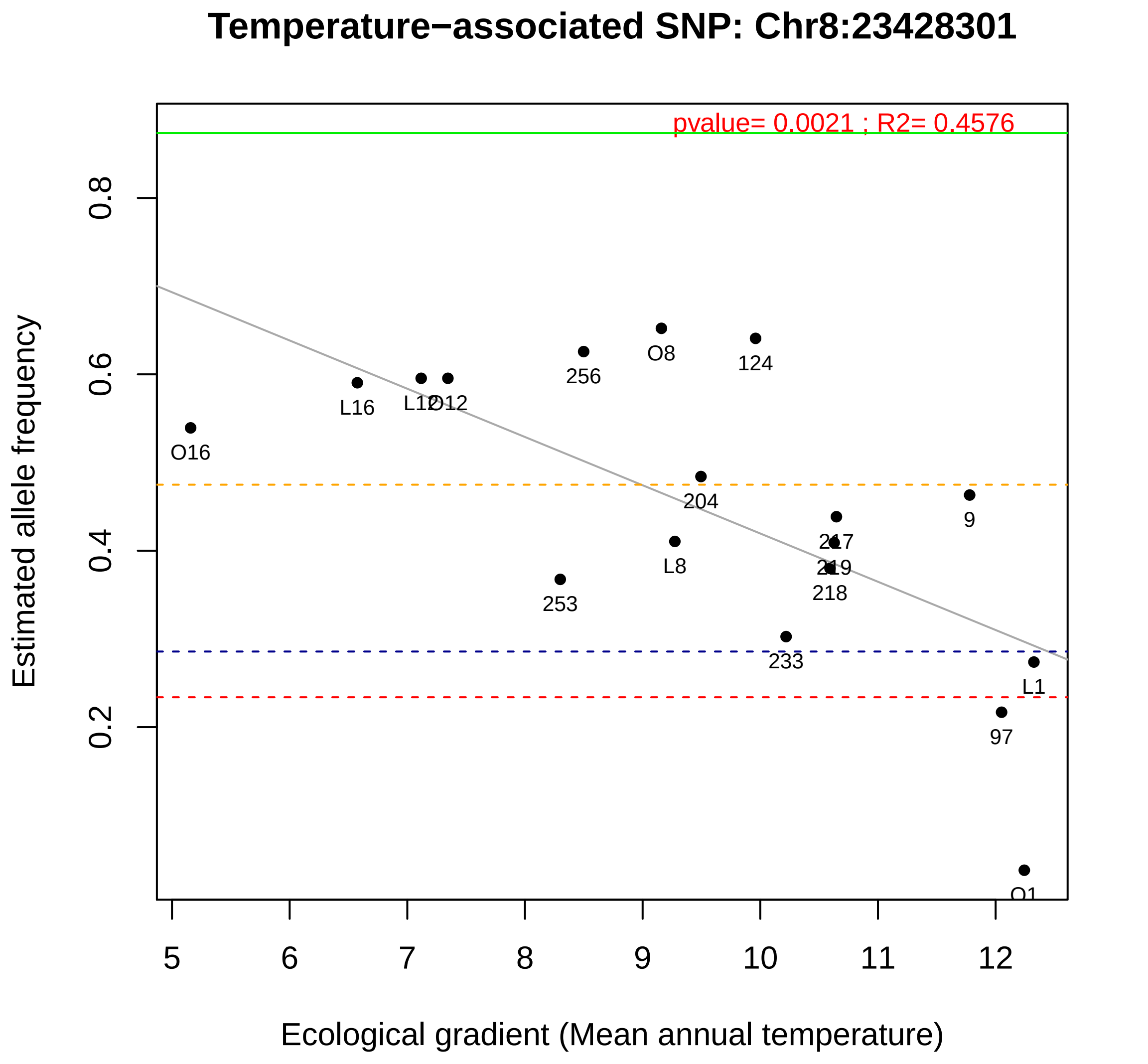

Temperature−associated SNP: Chr8:23428301
pvalue= 0.0021 ; R2= 0.4576
Estimated allele frequency
0.2
0.4
0.6
0.8
5
6
7
8
9
10
11
12
Ecological gradient (Mean annual temperature)
O16
L16
L12
O12
256
O8
124
204
9
217
219
218
L8
253
233
L1
97
O1

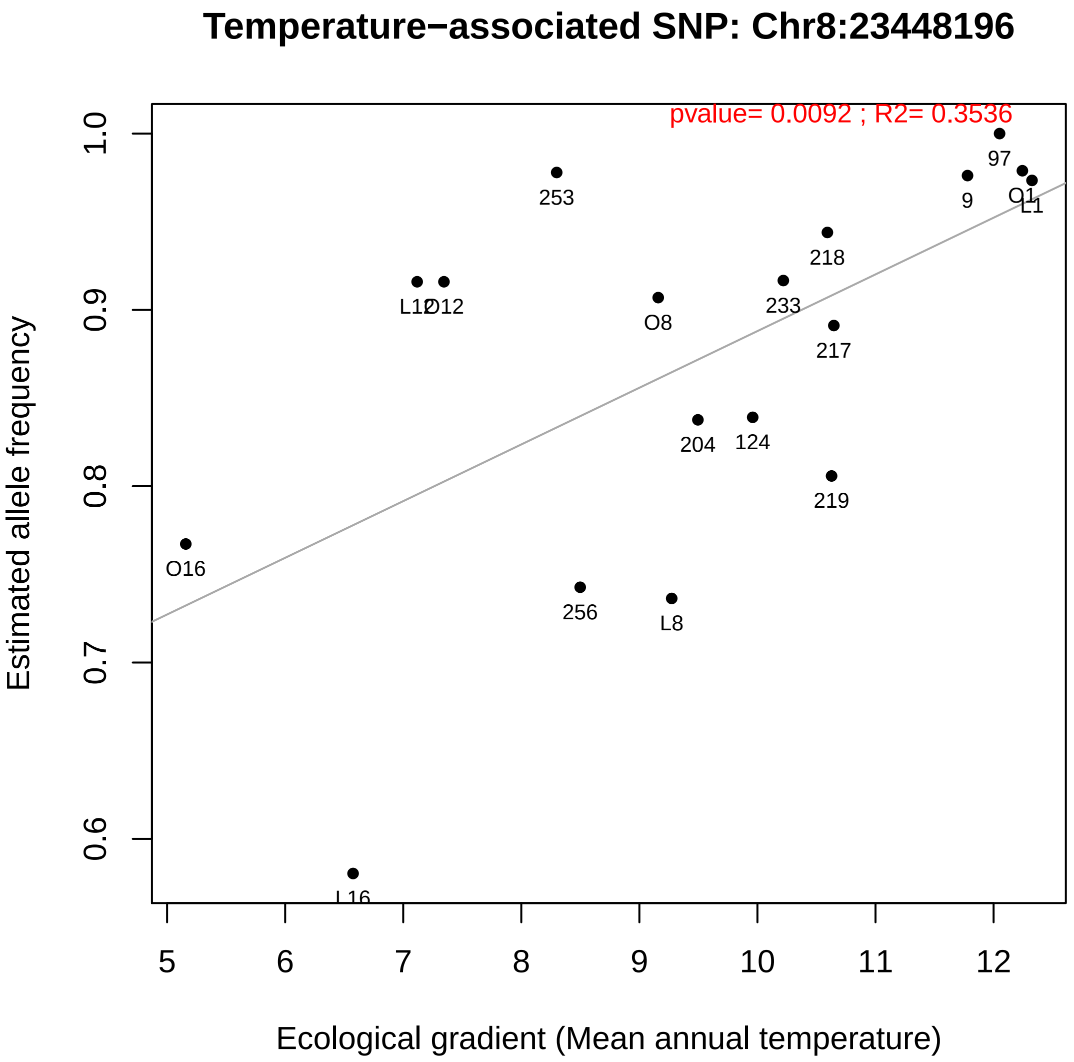

Temperature−associated SNP: Chr8:23448196
pvalue= 0.0092 ; R2= 0.3536
Estimated allele frequency
Ecological gradient (Mean annual temperature)
0.6
0.7
0.8
0.9
1.0
5
6
7
8
9
10
11
12
97
9
O1
L1
253
218
L12
O12
O8
233
217
204
124
219
O16
256
L8
L16

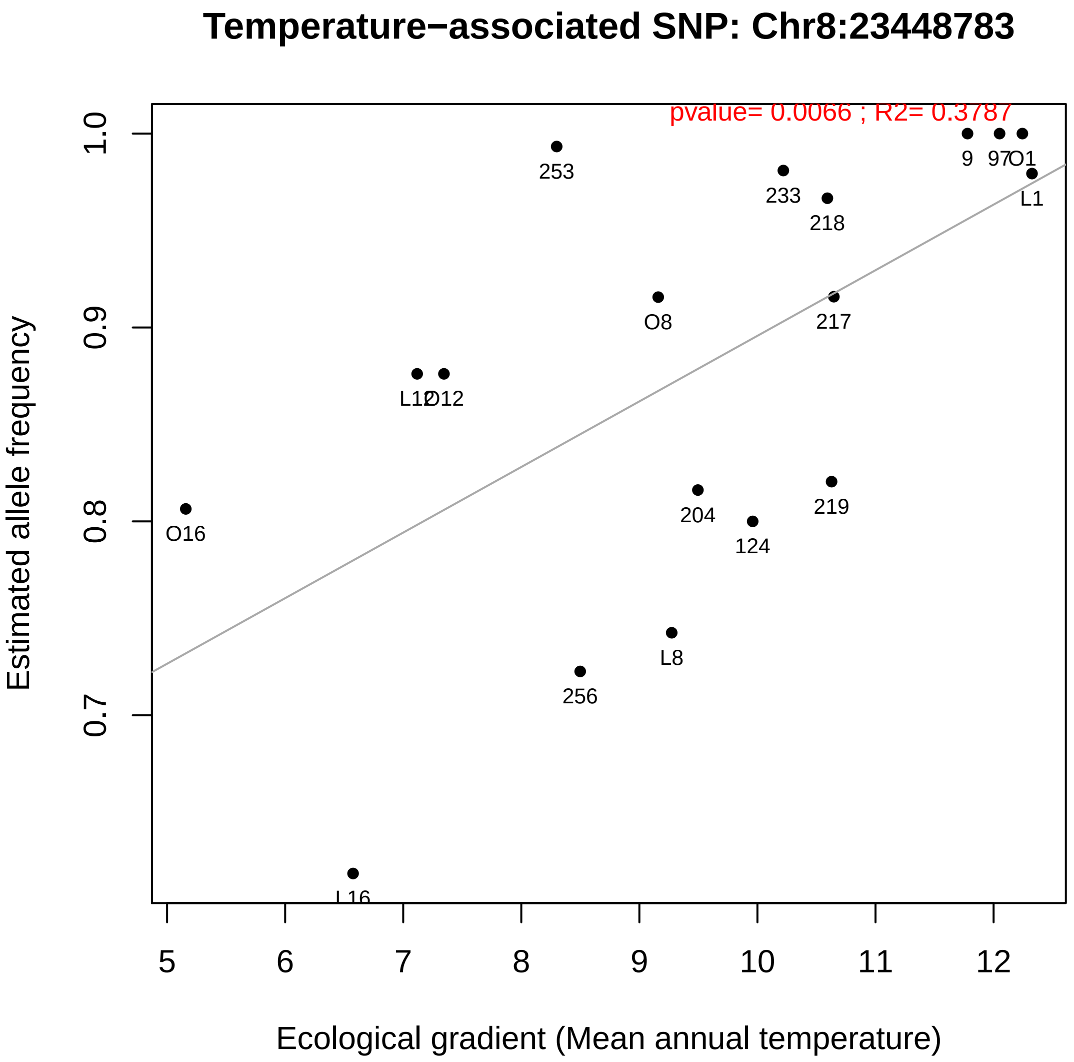
Temperature−associated SNP: Chr8:23448783
pvalue= 0.0066 ; R2= 0.3787
Estimated allele frequency
Ecological gradient (Mean annual temperature)
1.0
0.9
0.8
0.7
5
6
7
8
9
10
11
12
253
9
97
O1
L1
233
218
O8
217
L12
O12
O16
204
124
219
L8
256
L16

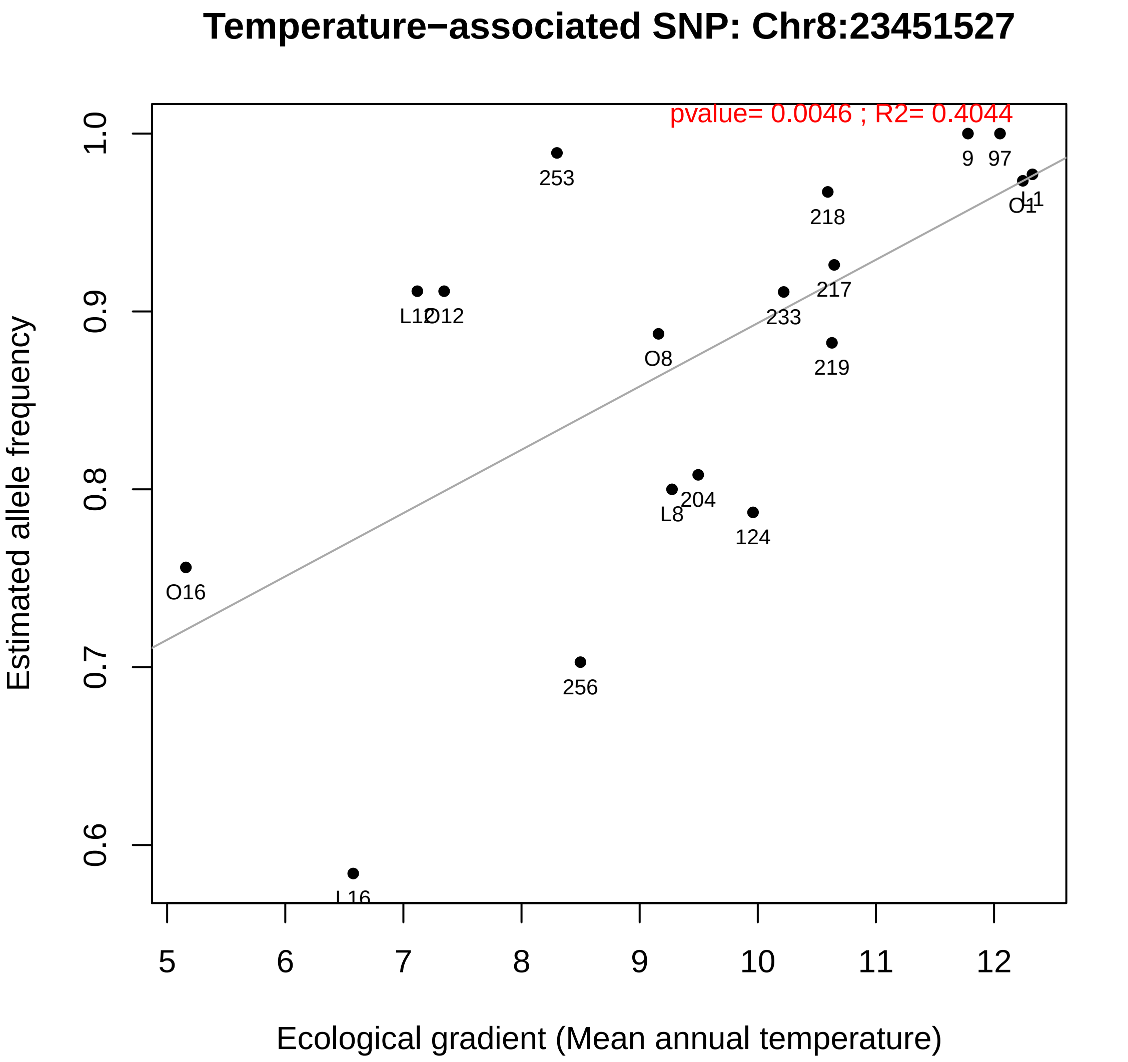

Temperature−associated SNP: Chr8:23451527
pvalue= 0.0046 ; R2= 0.4044
Estimated allele frequency
Ecological gradient (Mean annual temperature)
0.6
0.7
0.8
0.9
1.0
5
6
7
8
9
10
11
12
9
97
253
218
217
233
219
O8
L12
O12
204
L8
124
O16
256

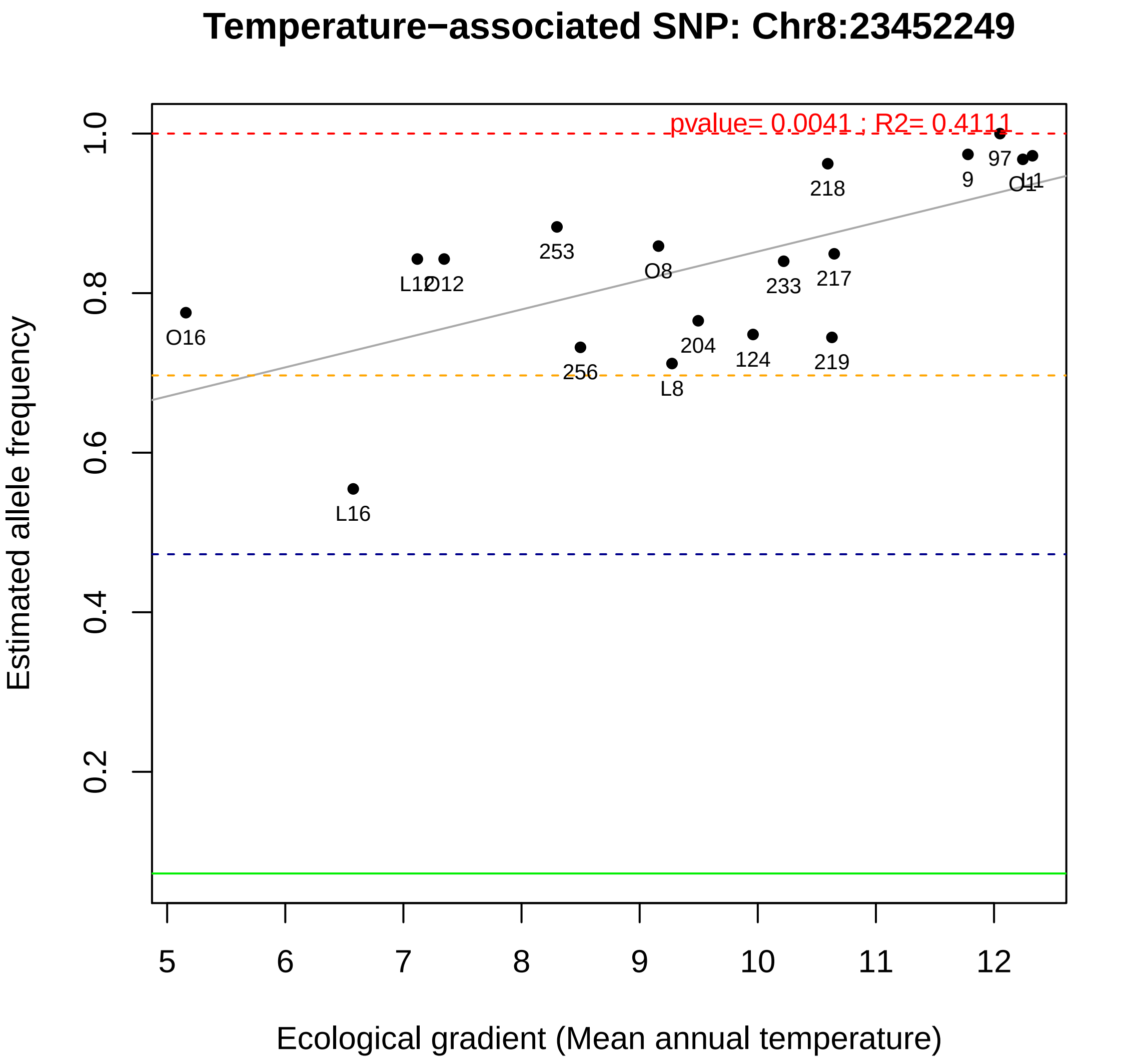

Temperature−associated SNP: Chr8:23452249
pvalue= 0.0041 ; R2= 0.4111
Estimated allele frequency
Ecological gradient (Mean annual temperature)
0.2
0.4
0.6
0.8
1.0
5
6
7
8
9
10
11
12
O16
L16
L12
O12
253
256
O8
L8
204
124
233
217
219
218
9
97

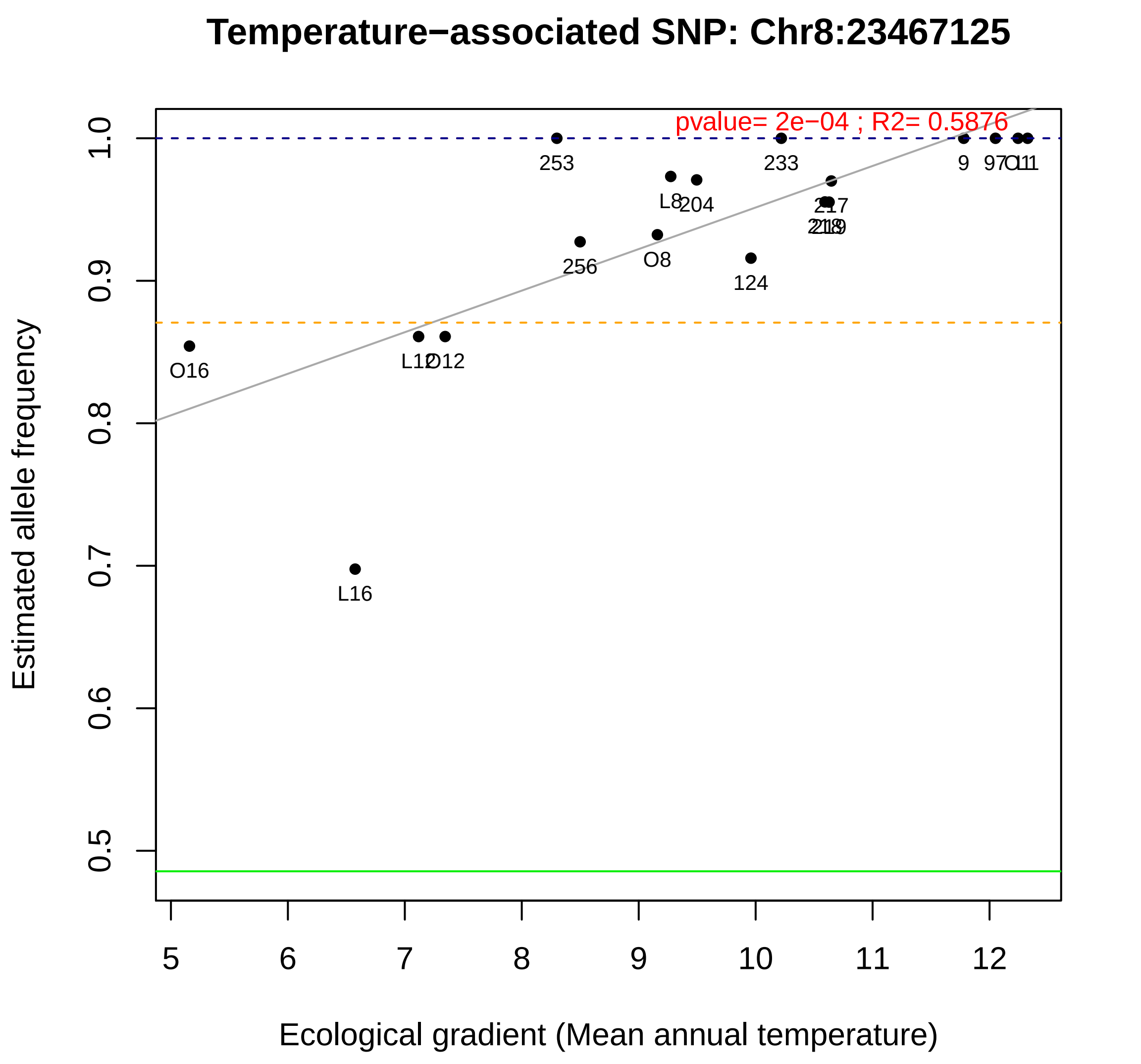

Temperature−associated SNP: Chr8:23467125
pvalue= 2e−04 ; R2= 0.5876
Estimated allele frequency
0.5
0.6
0.7
0.8
0.9
1.0
5
6
7
8
9
10
11
12
Ecological gradient (Mean annual temperature)
253
233
9
L8
204
217
256
O8
124
O16
L12
O12
L16

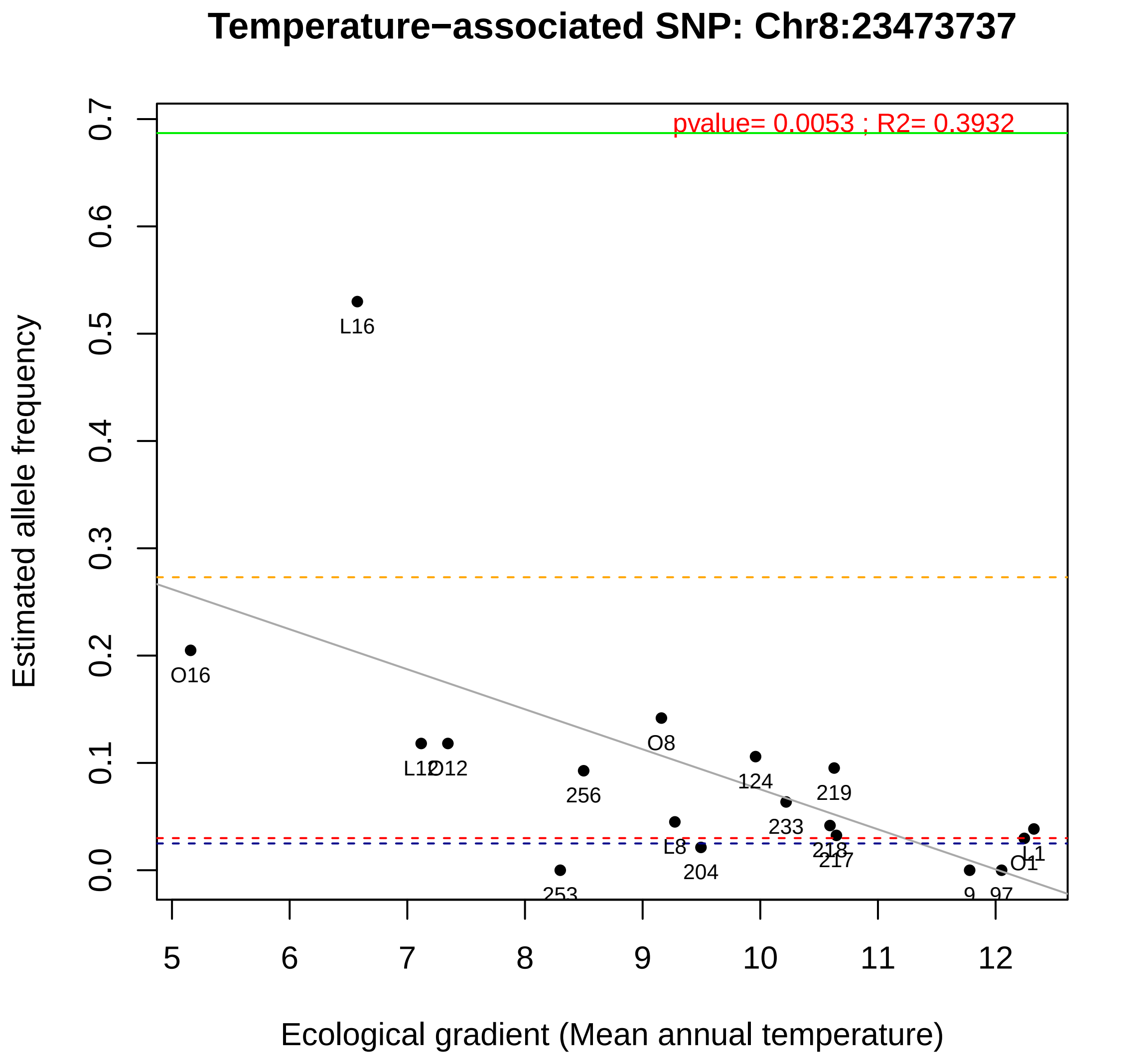

Temperature−associated SNP: Chr8:23473737
pvalue= 0.0053 ; R2= 0.3932
Estimated allele frequency
Ecological gradient (Mean annual temperature)
0.0
0.1
0.2
0.3
0.4
0.5
0.6
0.7
5
6
7
8
9
10
11
12
L16
O16
L12
O12
256
253
O8
E8
204
124
233
219
218
217
9
97
O1
L1

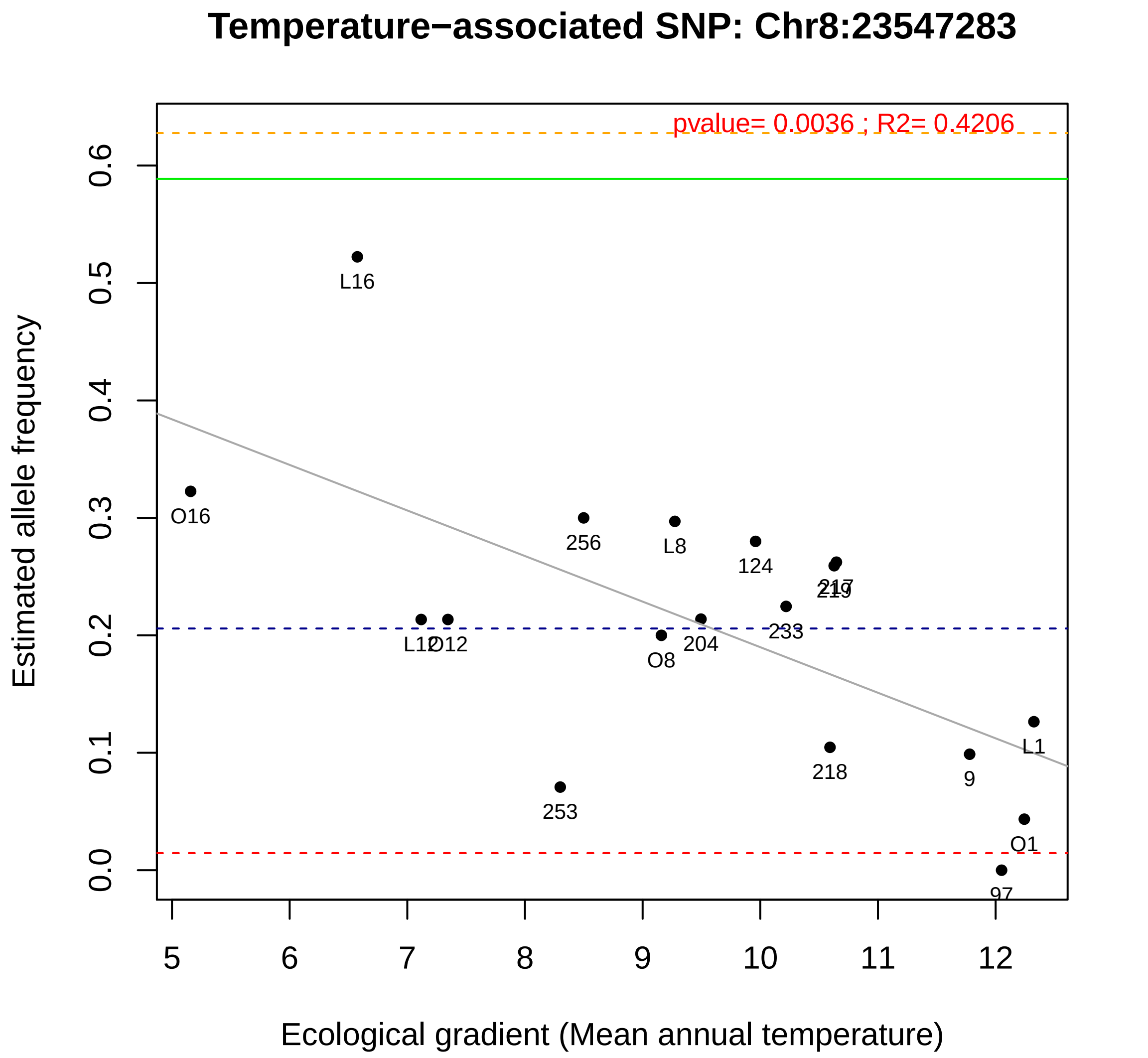

Temperature−associated SNP: Chr8:23547283
pvalue= 0.0036 ; R2= 0.4206
Estimated allele frequency
0.0
0.1
0.2
0.3
0.4
0.5
0.6
5
6
7
8
9
10
11
12
Ecological gradient (Mean annual temperature)
L16
O16
256
L8
124
217
219
233
204
O8
L12
O12
253
218
9
L1
O1
97

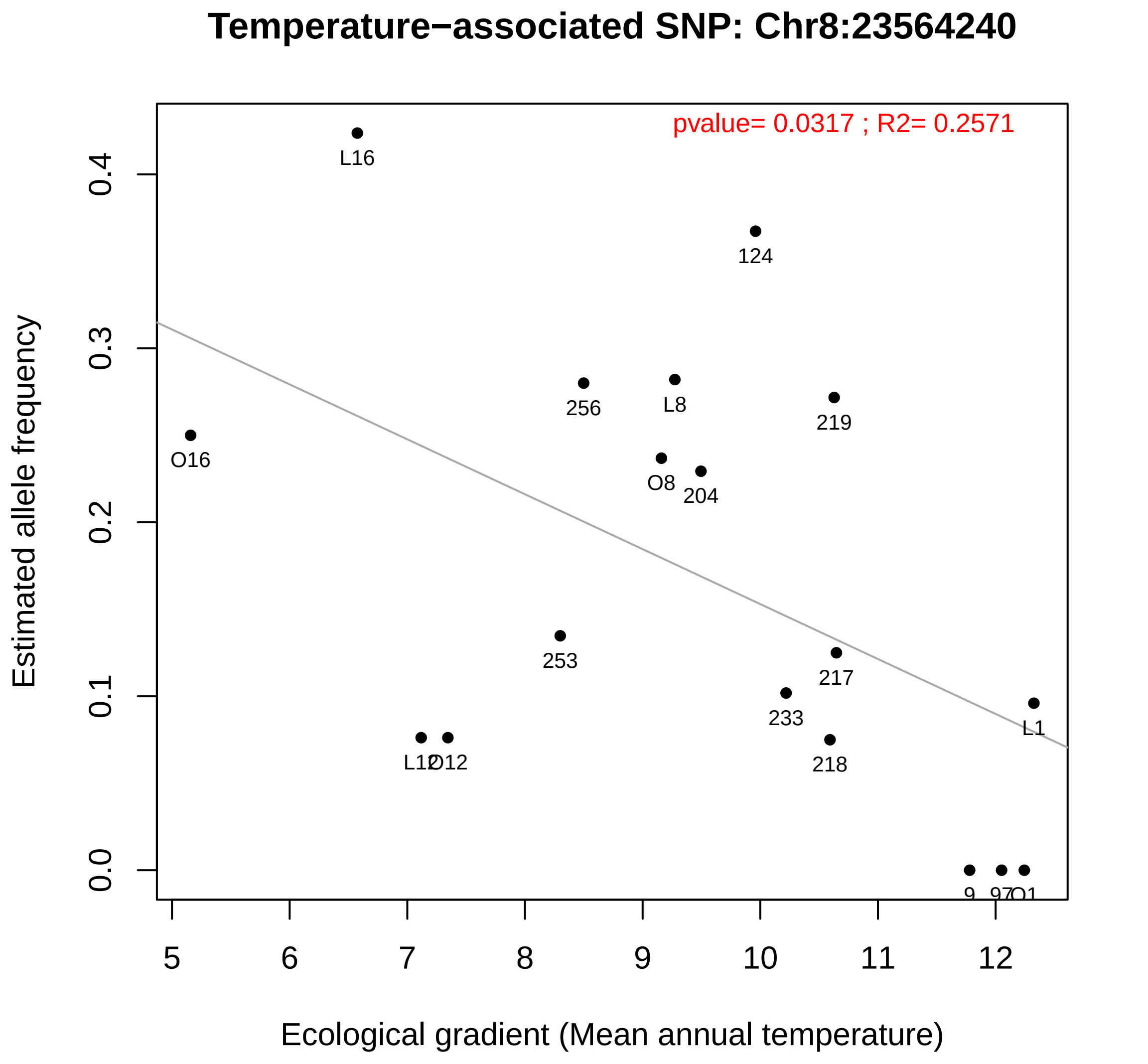
Temperature−associated SNP: Chr8:23564240
pvalue= 0.0317 ; R2= 0.2571
Estimated allele frequency
Ecological gradient (Mean annual temperature)
0.0
0.1
0.2
0.3
0.4
5
6
7
8
9
10
11
12
L16
124
256
L8
219
O16
O8
204
253
217
233
L1
L12
O12
218
9
97
O1

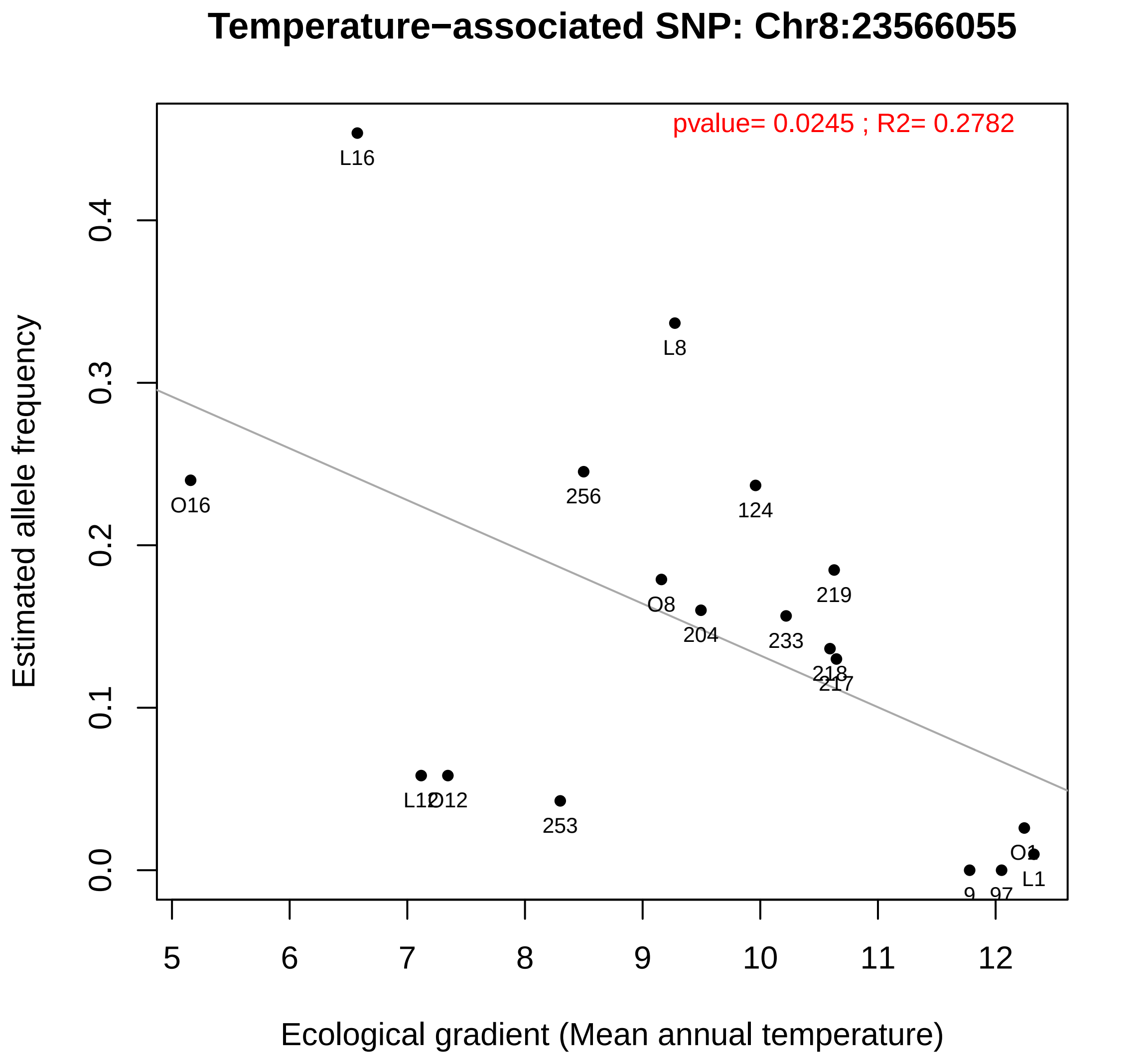

Temperature−associated SNP: Chr8:23566055
pvalue= 0.0245 ; R2= 0.2782
Estimated allele frequency
0.0
0.1
0.2
0.3
0.4
5
6
7
8
9
10
11
12
Ecological gradient (Mean annual temperature)
L16
L8
O16
256
124
O8
204
219
233
218
217
L12
O12
253
O1
L1
9
97

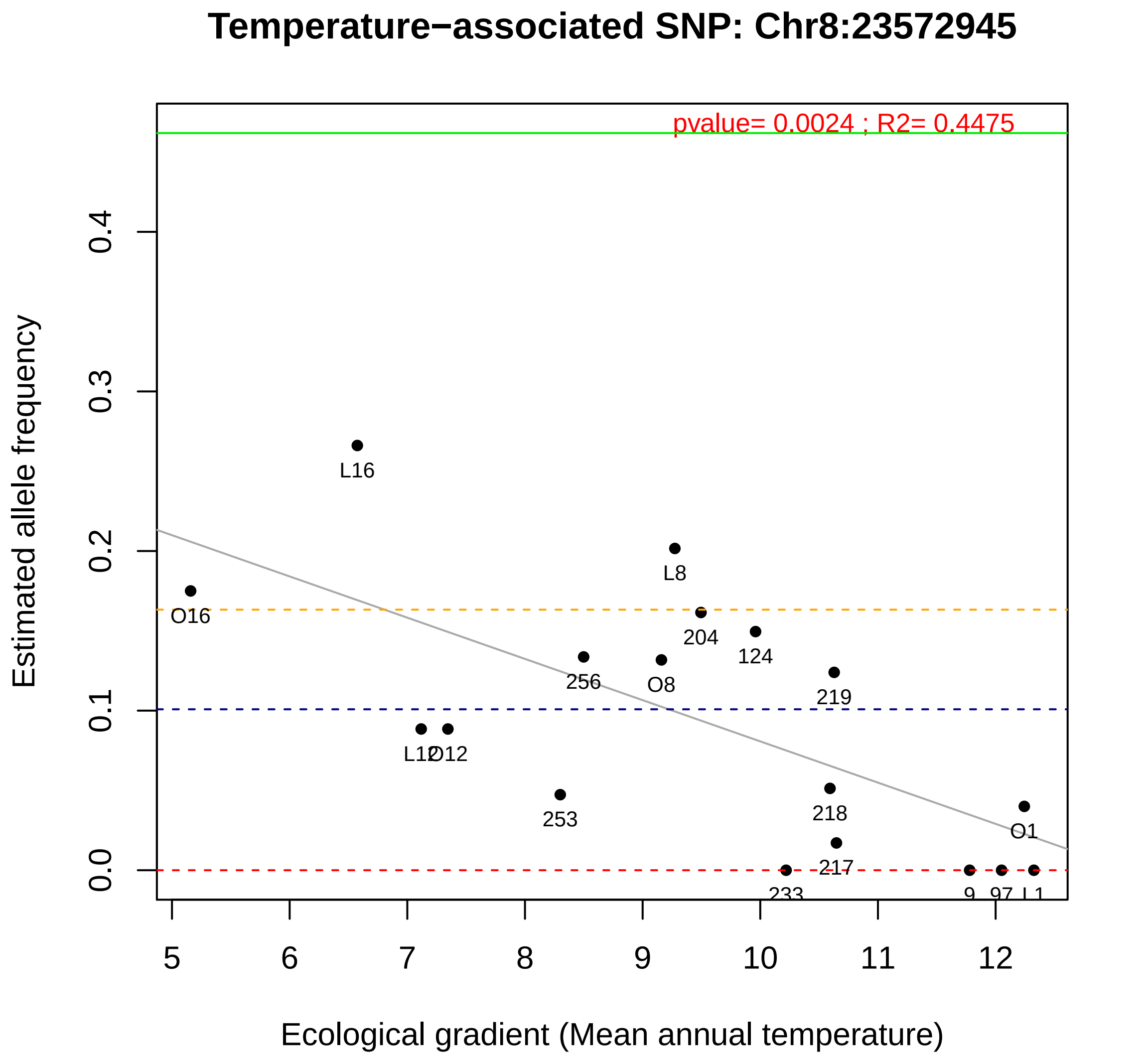
Temperature−associated SNP: Chr8:23572945
pvalue= 0.0024 ; R2= 0.4475
Estimated allele frequency
0.0
0.1
0.2
0.3
0.4
5
6
7
8
9
10
11
12
Ecological gradient (Mean annual temperature)
L16
O16
L8
204
124
256
O8
219
L12
O12
253
218
O1
217

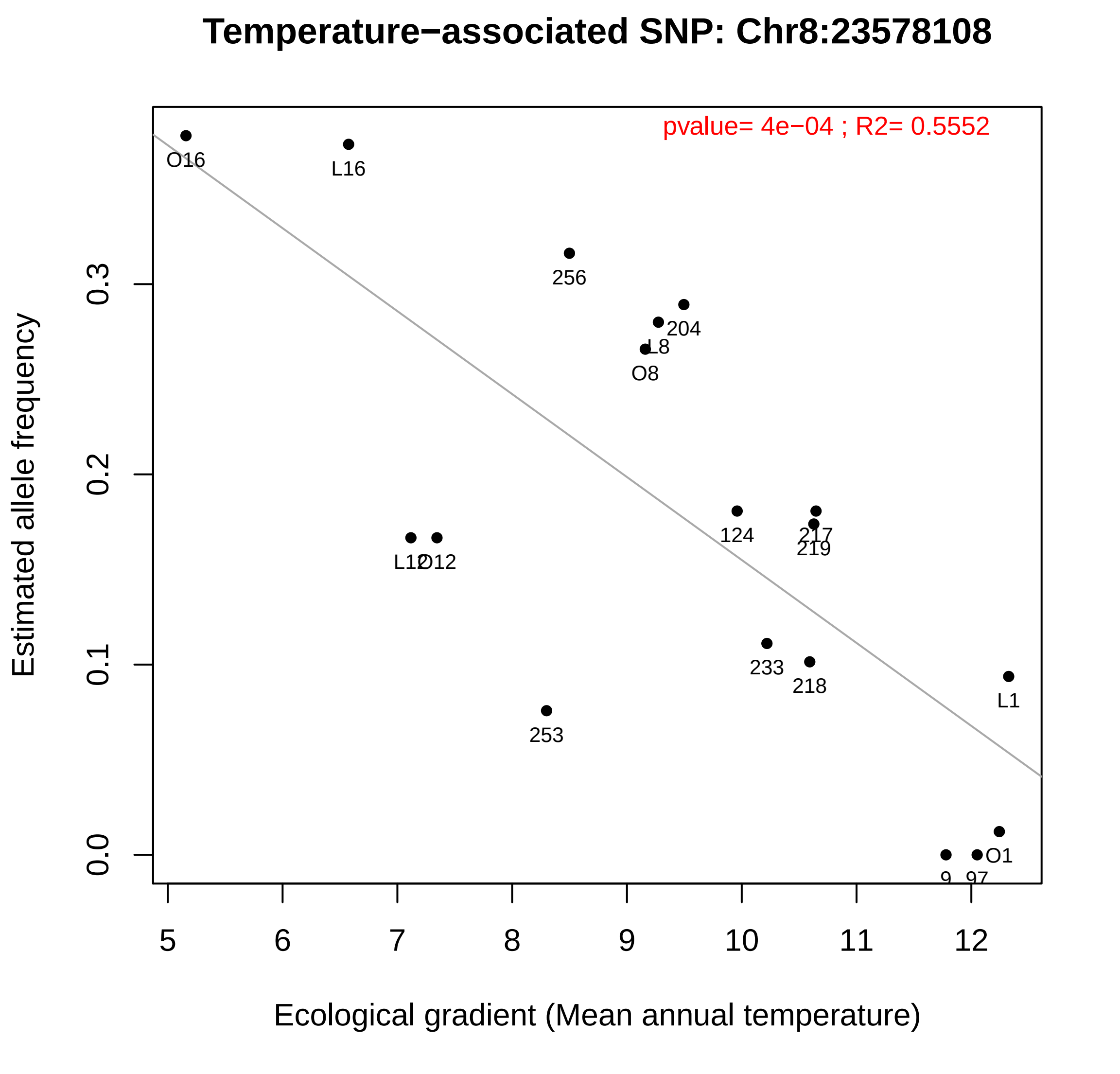

Temperature−associated SNP: Chr8:23578108
pvalue= 4e−04 ; R2= 0.5552
Estimated allele frequency
Ecological gradient (Mean annual temperature)
0.0
0.1
0.2
0.3
5
6
7
8
9
10
11
12
O16
L16
256
204
L8
O8
124
217
219
L12
O12
233
218
L1
253
O1
9
97

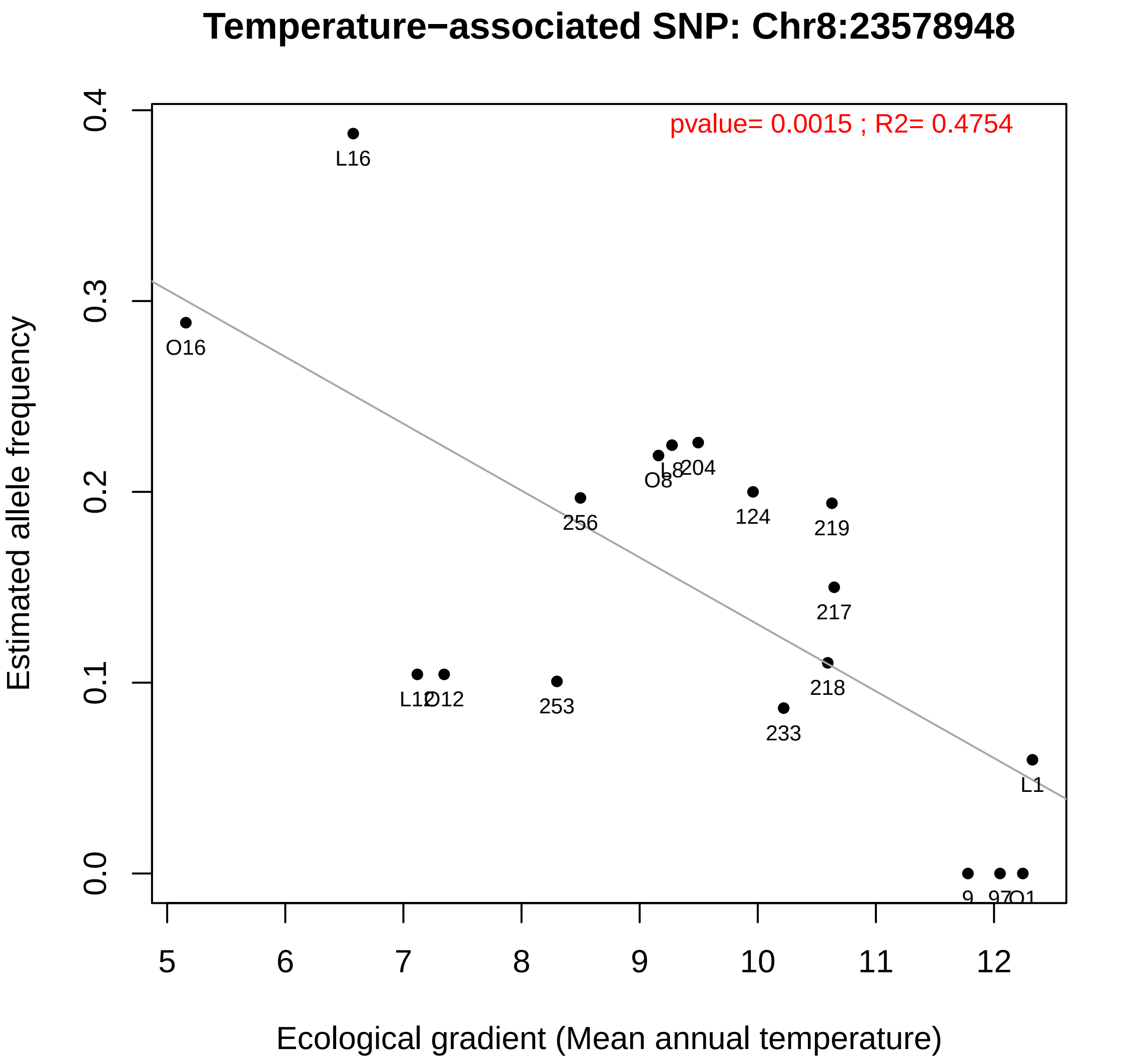
Temperature−associated SNP: Chr8:23578948
pvalue= 0.0015 ; R2= 0.4754
Estimated allele frequency
0.0
0.1
0.2
0.3
0.4
5
6
7
8
9
10
11
12
Ecological gradient (Mean annual temperature)
L16
O16
256
O8
L8
204
124
219
217
218
233
253
L12
O12
L1
9
97
O1

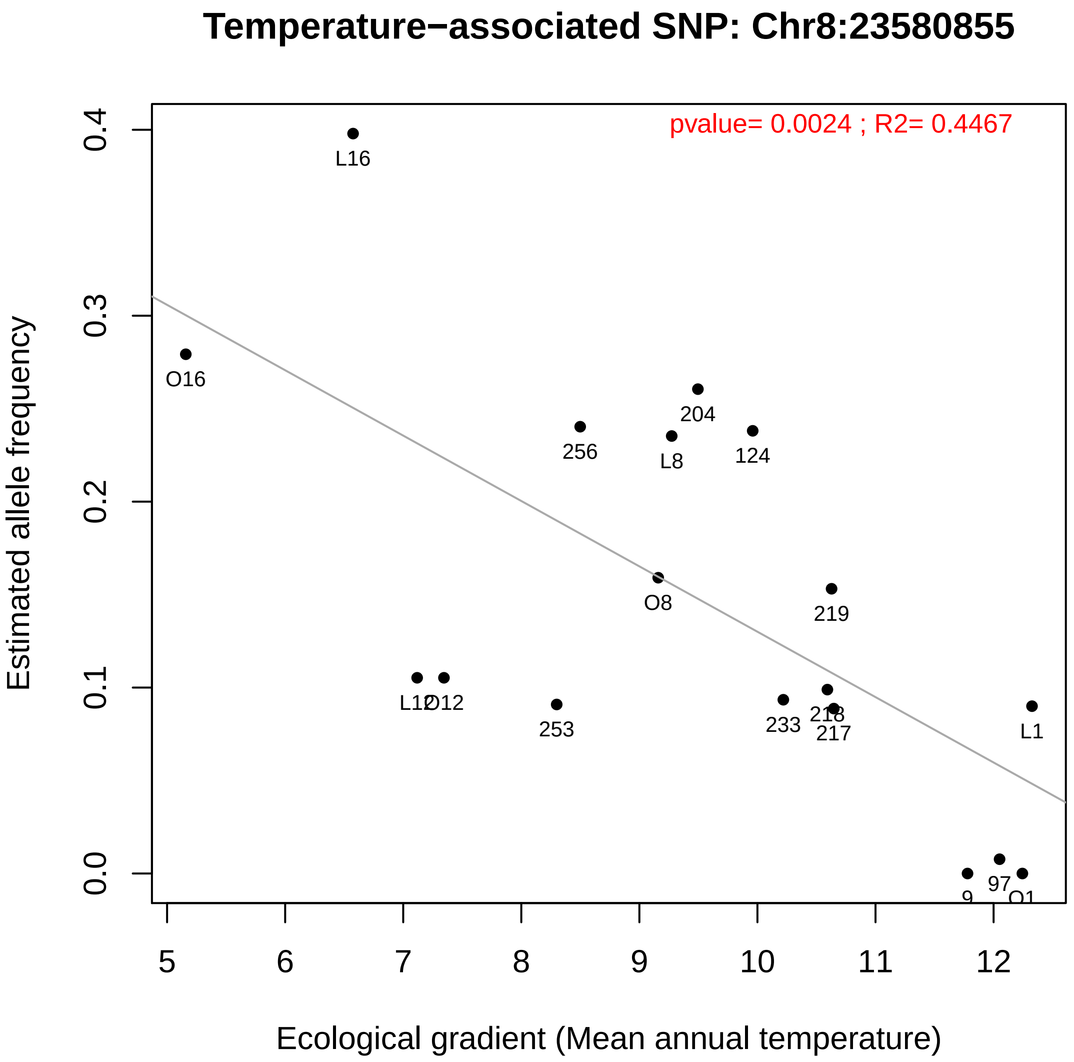
Temperature−associated SNP: Chr8:23580855
pvalue= 0.0024 ; R2= 0.4467
Estimated allele frequency
Ecological gradient (Mean annual temperature)
0.0
0.1
0.2
0.3
0.4
5
6
7
8
9
10
11
12
L16
O16
256
204
L8
124
O8
219
L12
O12
253
233
218
217
L1
97

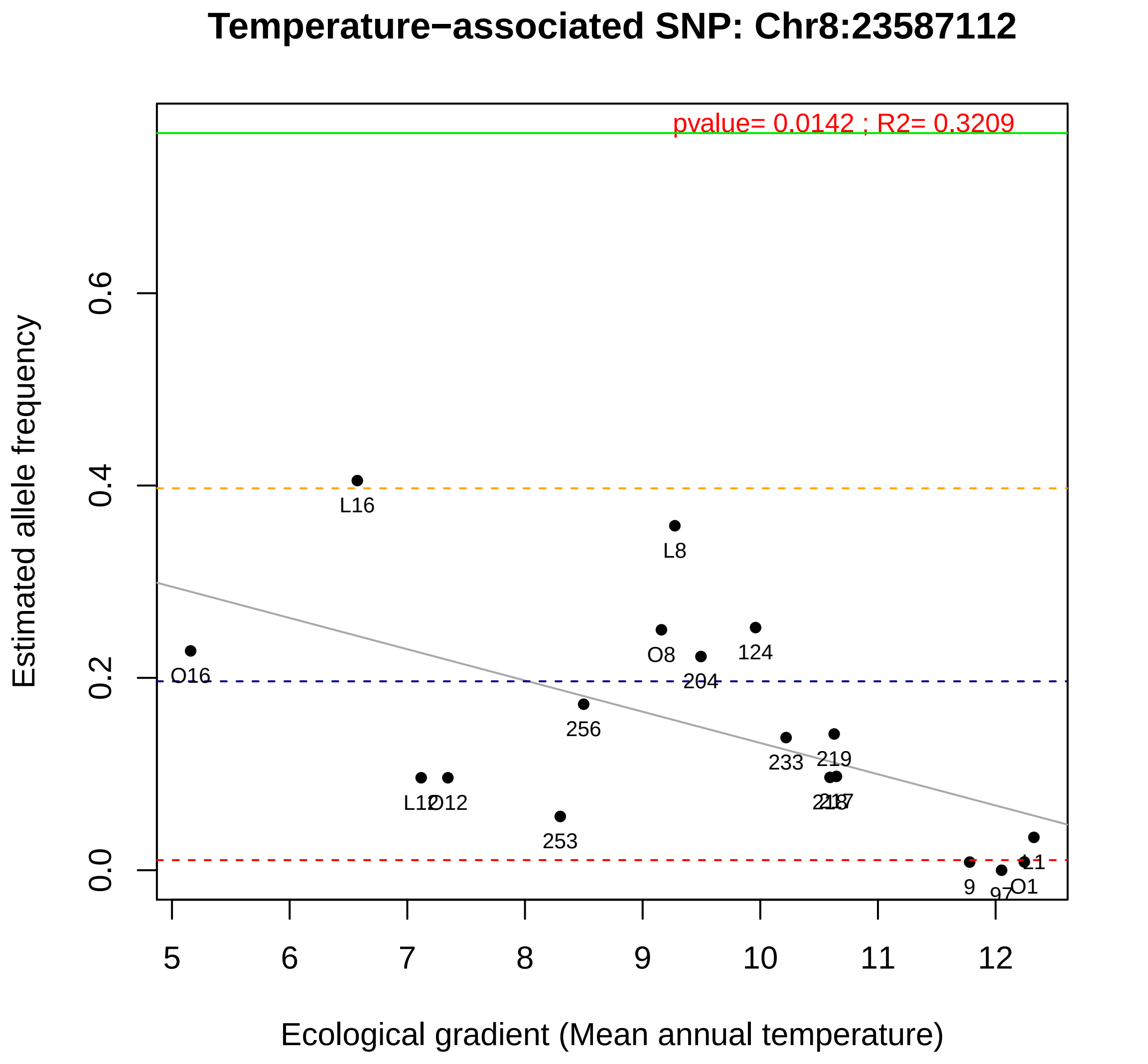
Temperature−associated SNP: Chr8:23587112
pvalue= 0.0142 ; R2= 0.3209
Estimated allele frequency
0.0
0.2
0.4
0.6
5
6
7
8
9
10
11
12
Ecological gradient (Mean annual temperature)
L16
L8
O8
204
124
O16
256
233
219
L12
O12
253
9
O1
L1

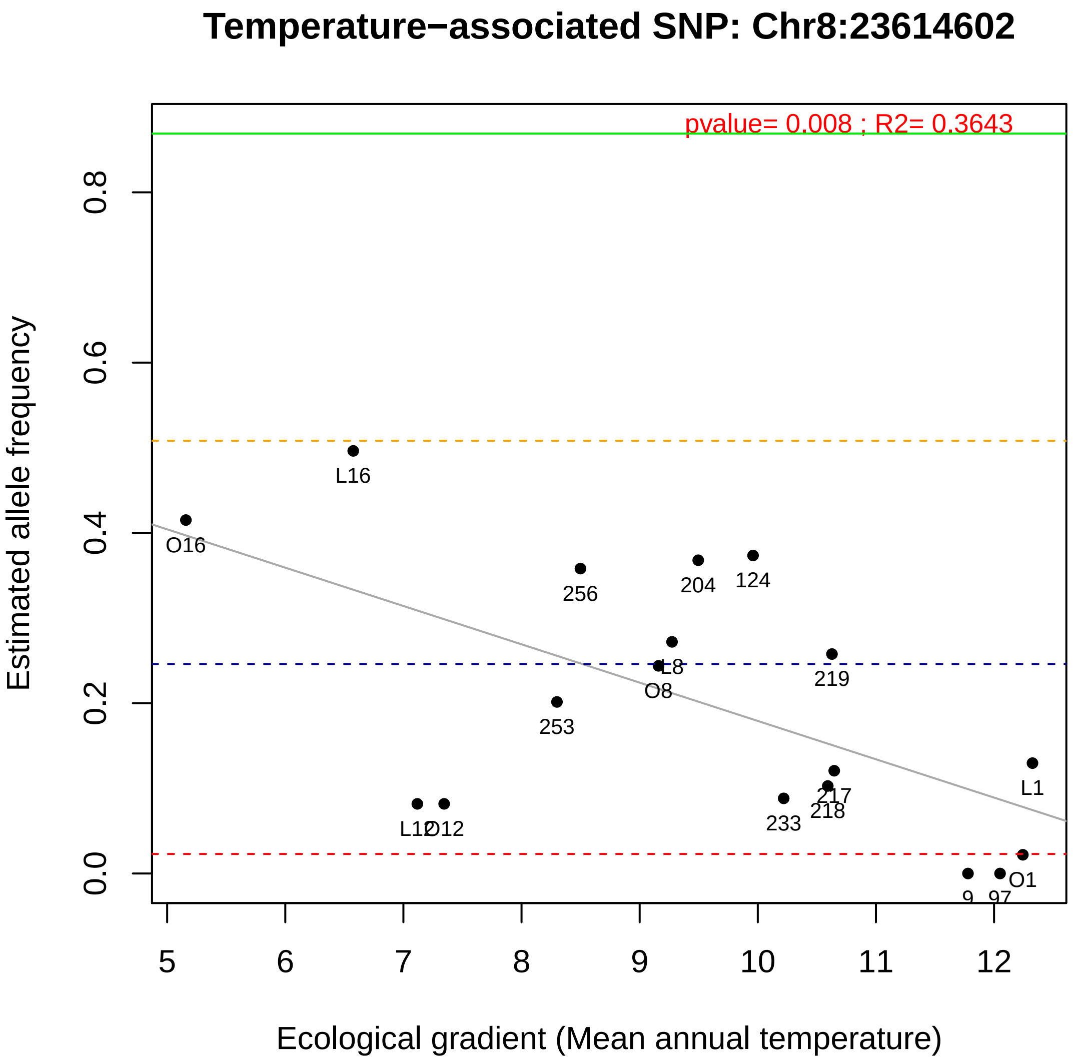

Temperature−associated SNP: Chr8:23614602
pvalue= 0.008 ; R2= 0.3643
Estimated allele frequency
0.0
0.2
0.4
0.6
0.8
5
6
7
8
9
10
11
12
Ecological gradient (Mean annual temperature)
L16
O16
256
204
124
L8
O8
219
253
L12
O12
233
217
218
L1
O1
9
97

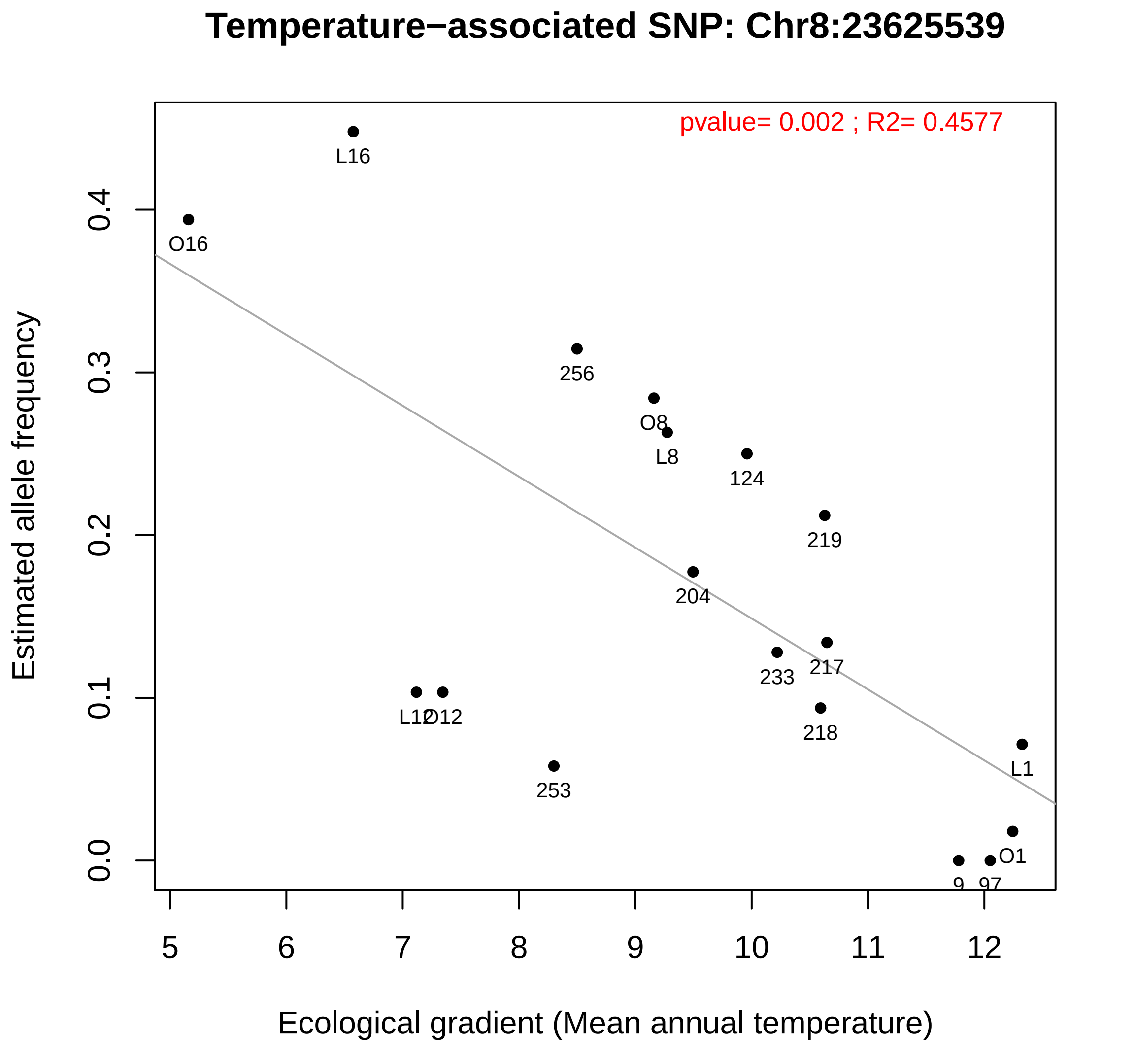

Temperature−associated SNP: Chr8:23625539
pvalue= 0.002 ; R2= 0.4577
Estimated allele frequency
0.0
0.1
0.2
0.3
0.4
5
6
7
8
9
10
11
12
Ecological gradient (Mean annual temperature)
L16
O16
256
O8
L8
124
219
204
233
217
L12
O12
218
253
L1
O1
9
97

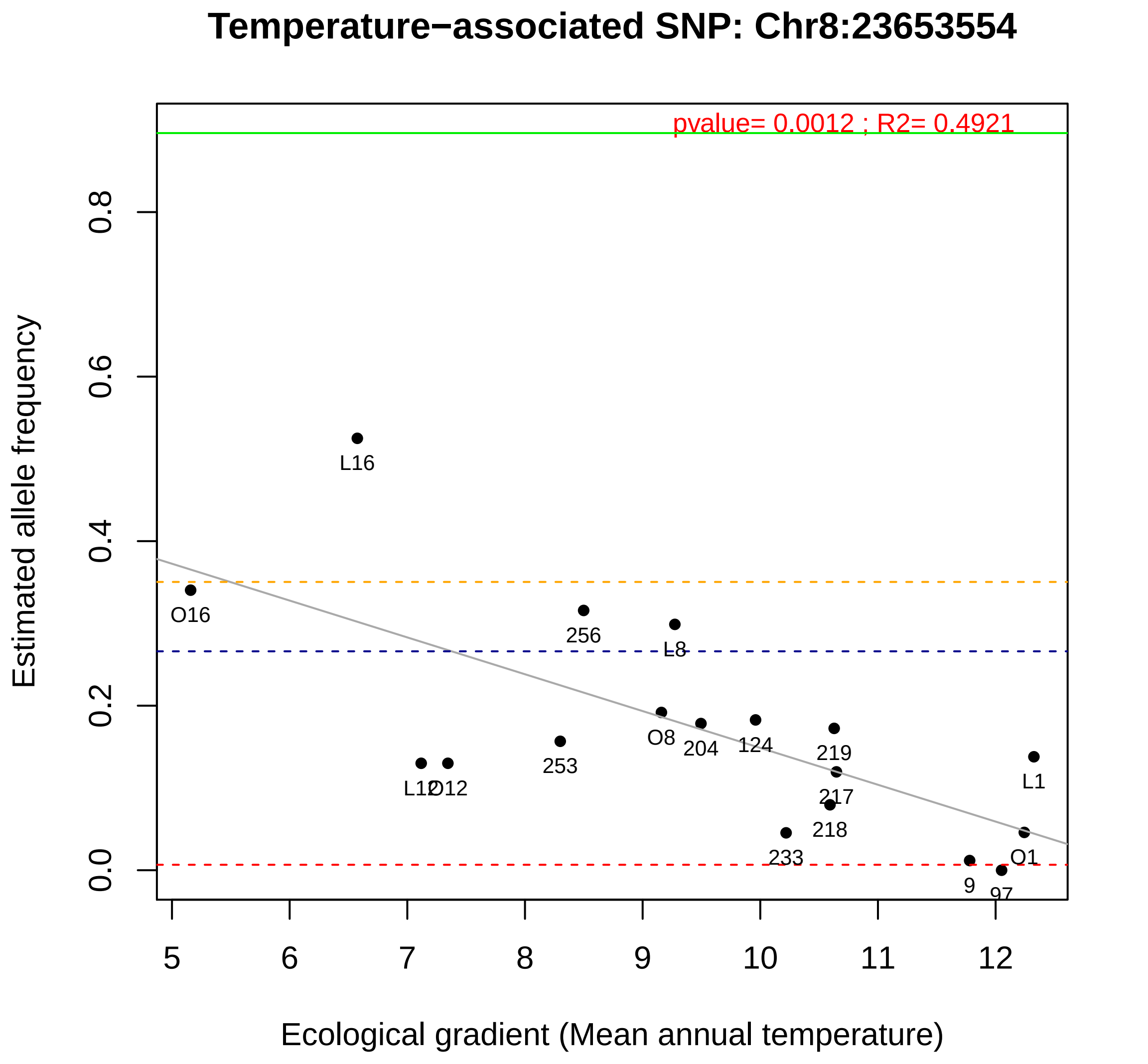

Temperature−associated SNP: Chr8:23653554
pvalue= 0.0012 ; R2= 0.4921
Estimated allele frequency
0.0
0.2
0.4
0.6
0.8
5
6
7
8
9
10
11
12
Ecological gradient (Mean annual temperature)
L16
O16
256
L8
253
O8
204
124
219
217
218
233
L1
O1
9
97
L12
O12

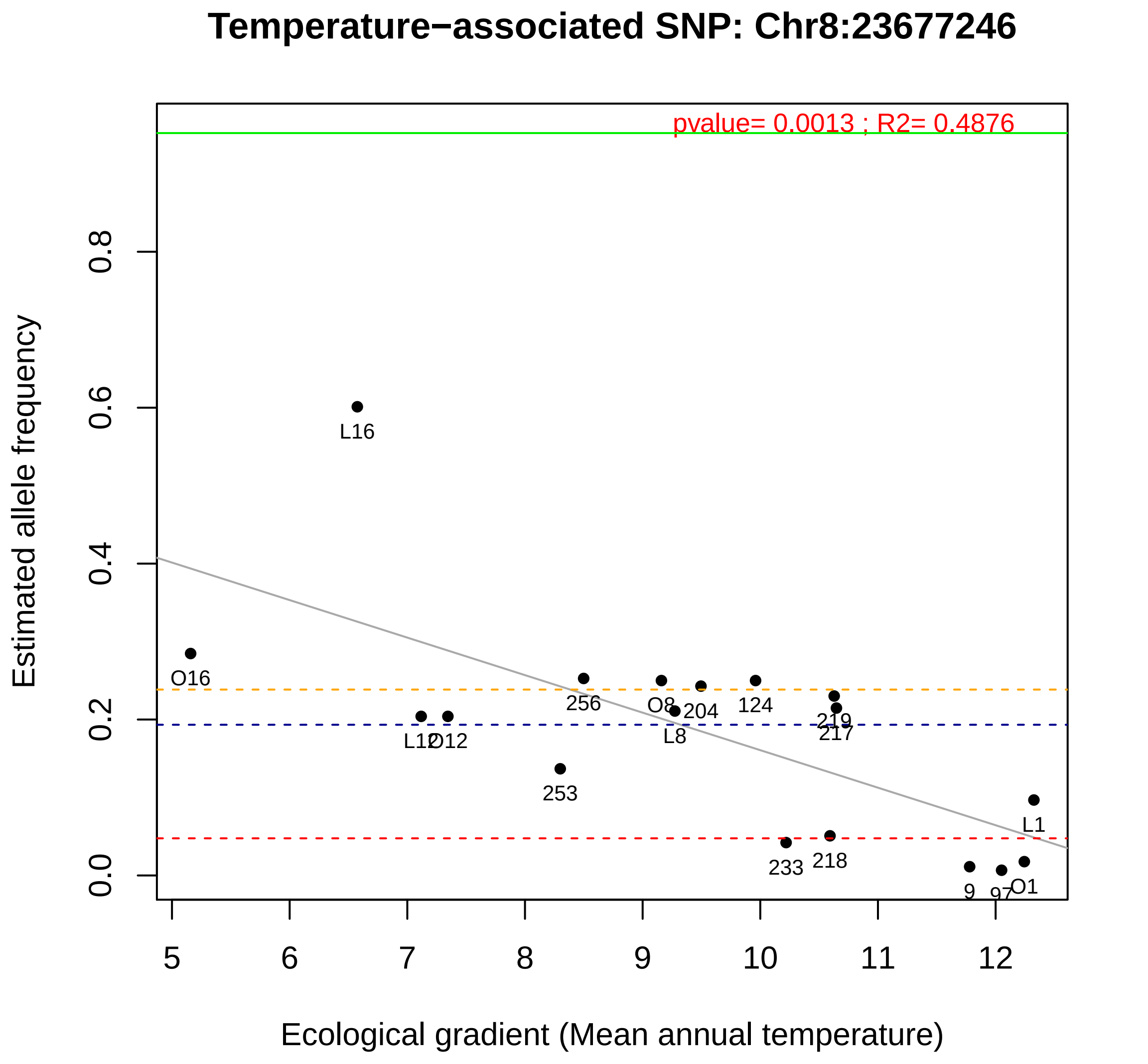
Temperature−associated SNP: Chr8:23677246
pvalue= 0.0013 ; R2= 0.4876
Estimated allele frequency
0.0
0.2
0.4
0.6
0.8
L16
O16
L12
O12
256
253
O8
L8
204
124
219
217
233
218
L1
9
97
O1
5
6
7
8
9
10
11
12
Ecological gradient (Mean annual temperature)

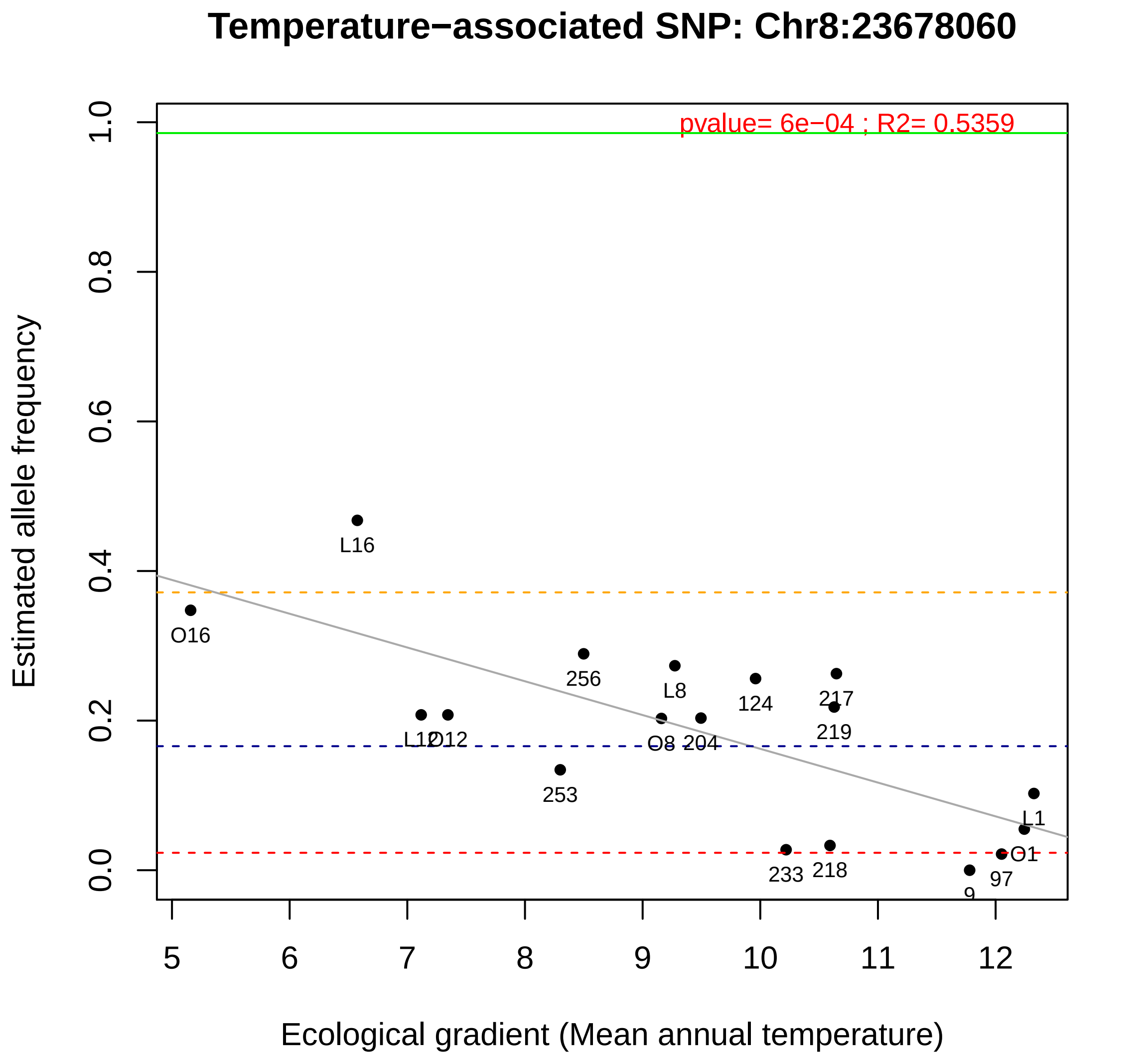

Temperature−associated SNP: Chr8:23678060
pvalue= 6e−04 ; R2= 0.5359
Estimated allele frequency
0.0
0.2
0.4
0.6
0.8
1.0
Ecological gradient (Mean annual temperature)
5
6
7
8
9
10
11
12
L16
O16
256
L8
124
217
219
L12
O12
O8
204
253
L1
O1
233
218
97
9

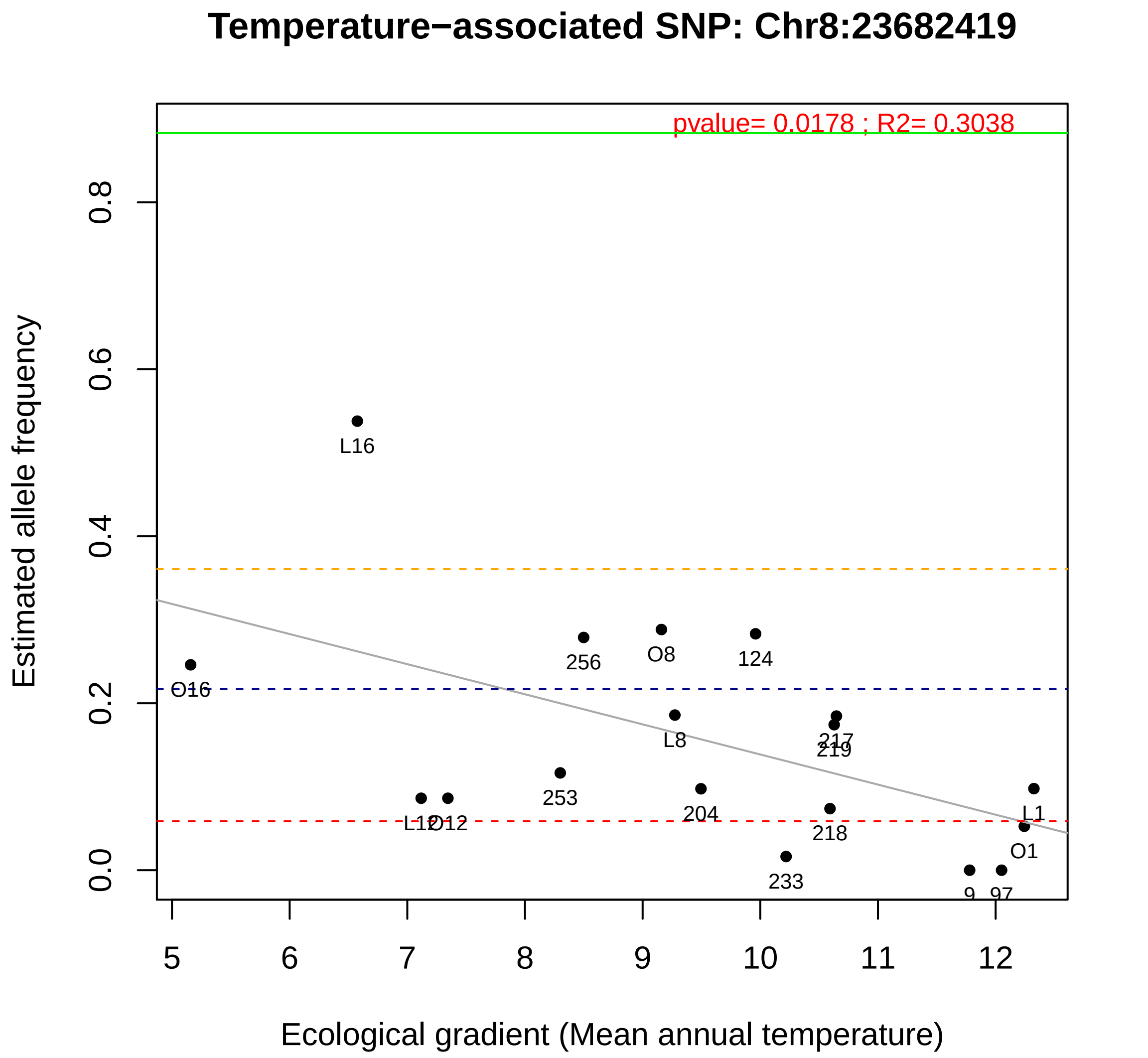
Temperature−associated SNP: Chr8:23682419
pvalue= 0.0178 ; R2= 0.3038
Estimated allele frequency
0.0
0.2
0.4
0.6
0.8
5
6
7
8
9
10
11
12
Ecological gradient (Mean annual temperature)
L16
O16
256
O8
124
L8
217
219
253
L12
O12
204
218
L1
O1
233
9
97

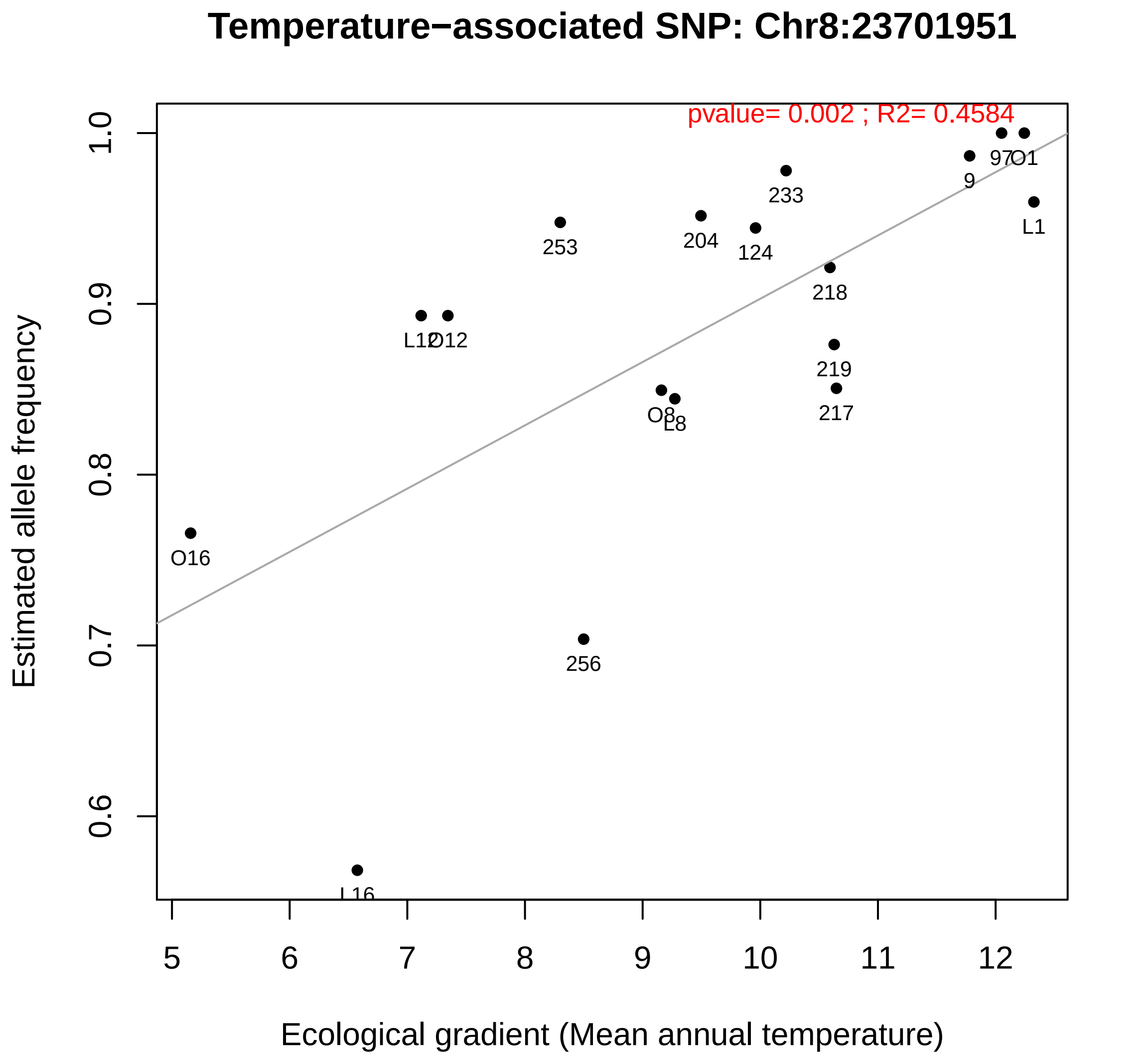

Temperature−associated SNP: Chr8:23701951
pvalue= 0.002 ; R2= 0.4584
Estimated allele frequency
0.6
0.7
0.8
0.9
1.0
5
6
7
8
9
10
11
12
Ecological gradient (Mean annual temperature)
97
O1
9
233
L1
204
124
253
218
L12
O12
219
217
O8
L8
O16
256
L16

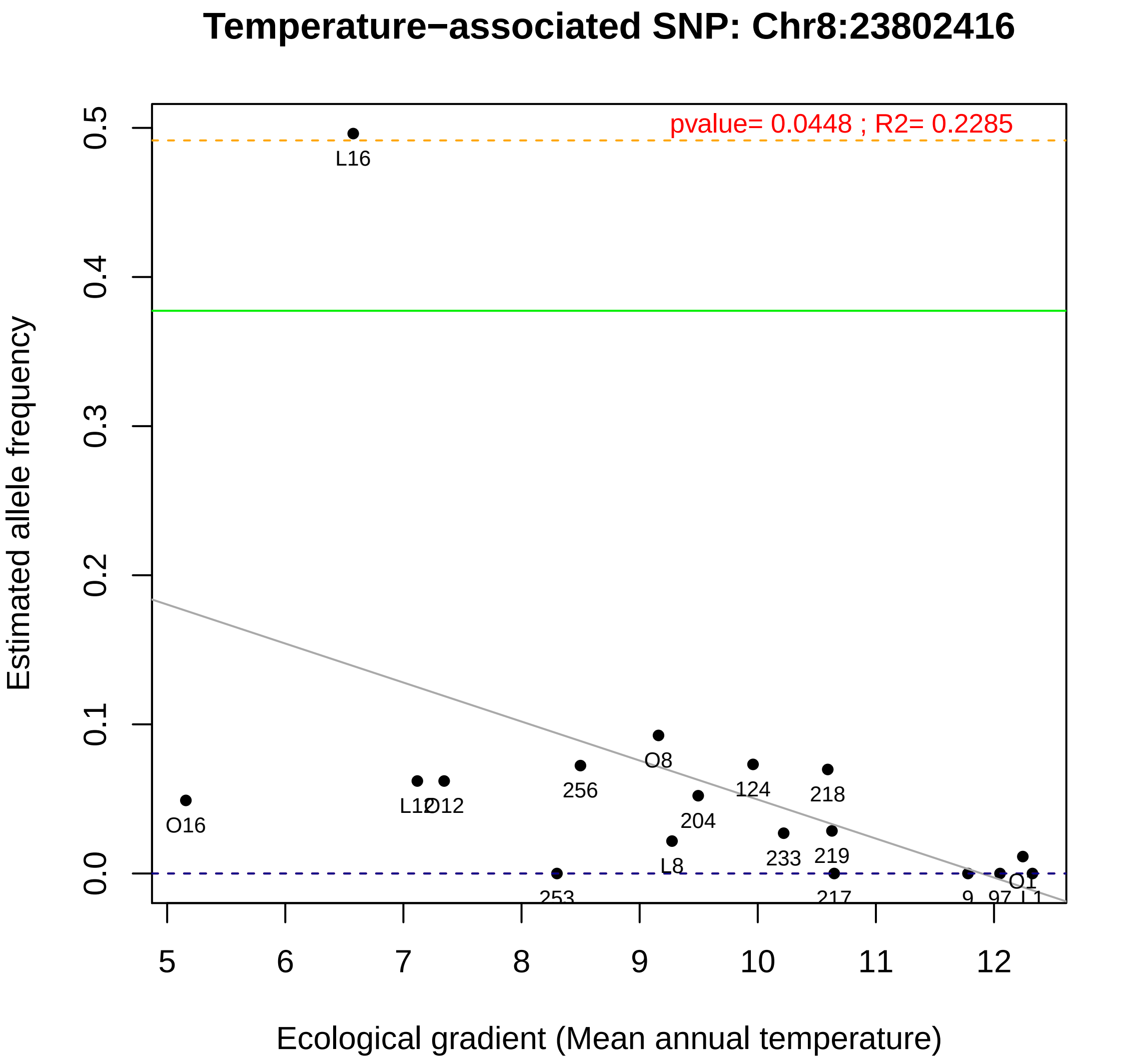

Temperature−associated SNP: Chr8:23802416
pvalue= 0.0448 ; R2= 0.2285
Estimated allele frequency
0.0
0.1
0.2
0.3
0.4
0.5
5
6
7
8
9
10
11
12
Ecological gradient (Mean annual temperature)
L16
O16
L12
O12
256
O8
204
L8
124
218
233
219
217
253
9
97
O1
L1

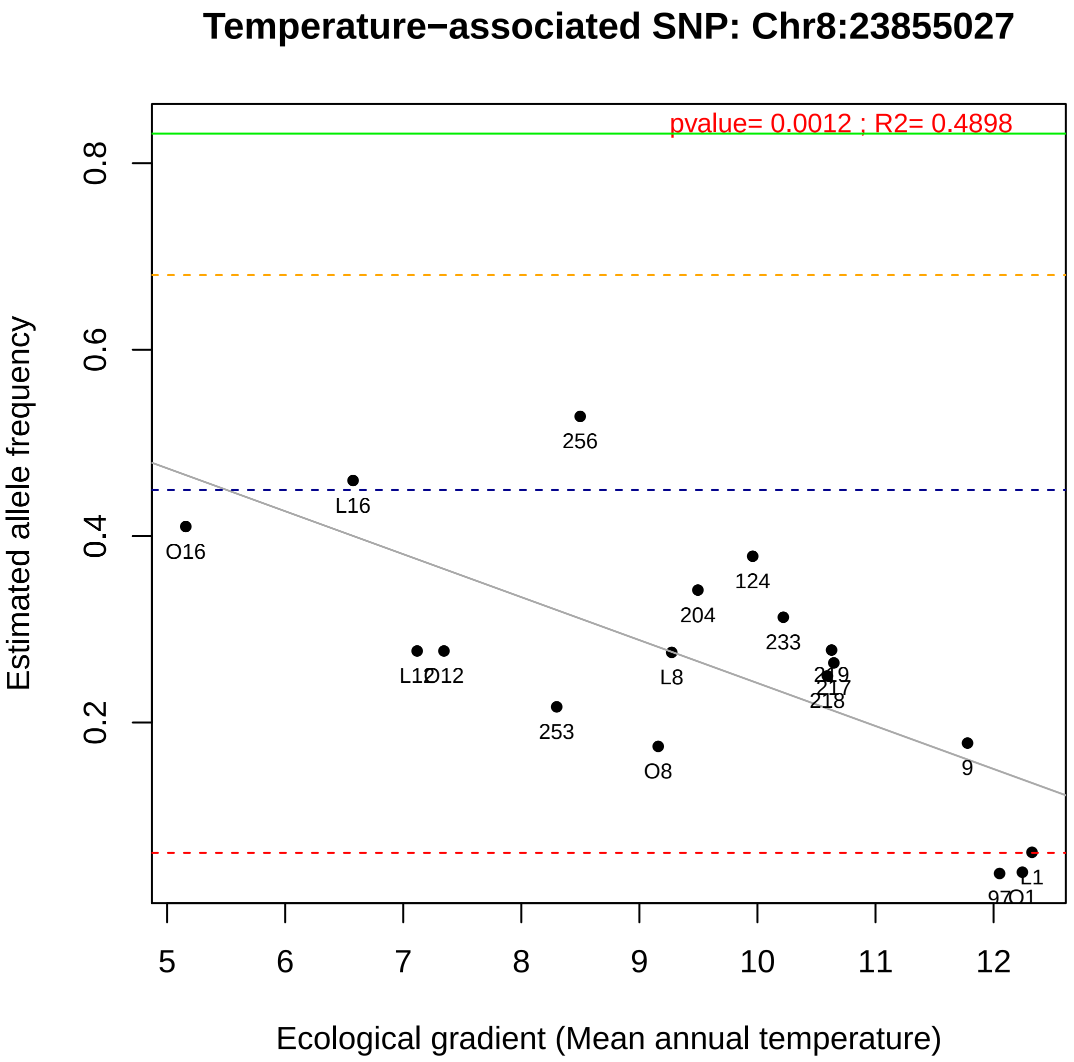

Temperature−associated SNP: Chr8:23855027
pvalue= 0.0012 ; R2= 0.4898
Estimated allele frequency
0.2
0.4
0.6
0.8
5
6
7
8
9
10
11
12
Ecological gradient (Mean annual temperature)
256
L16
O16
124
204
233
L12
O12
L8
219
217
218
253
O8
9
L1
97
O1

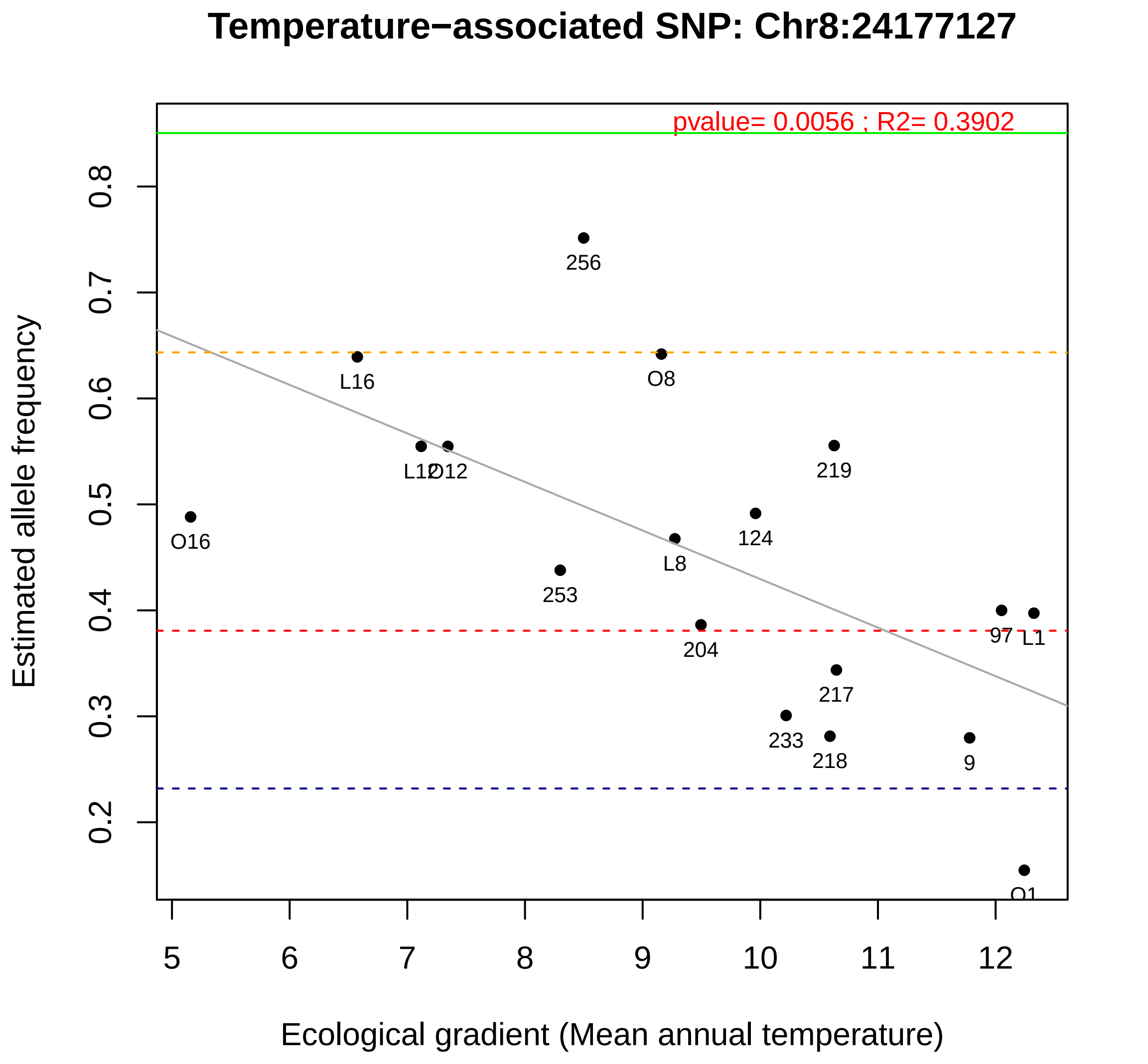
Temperature−associated SNP: Chr8:24177127
pvalue= 0.0056 ; R2= 0.3902
Estimated allele frequency
Ecological gradient (Mean annual temperature)
0.2
0.3
0.4
0.5
0.6
0.7
0.8
5
6
7
8
9
10
11
12
256
L16
O8
L12
O12
219
O16
124
L8
253
204
97
L1
217
233
218
9
O1

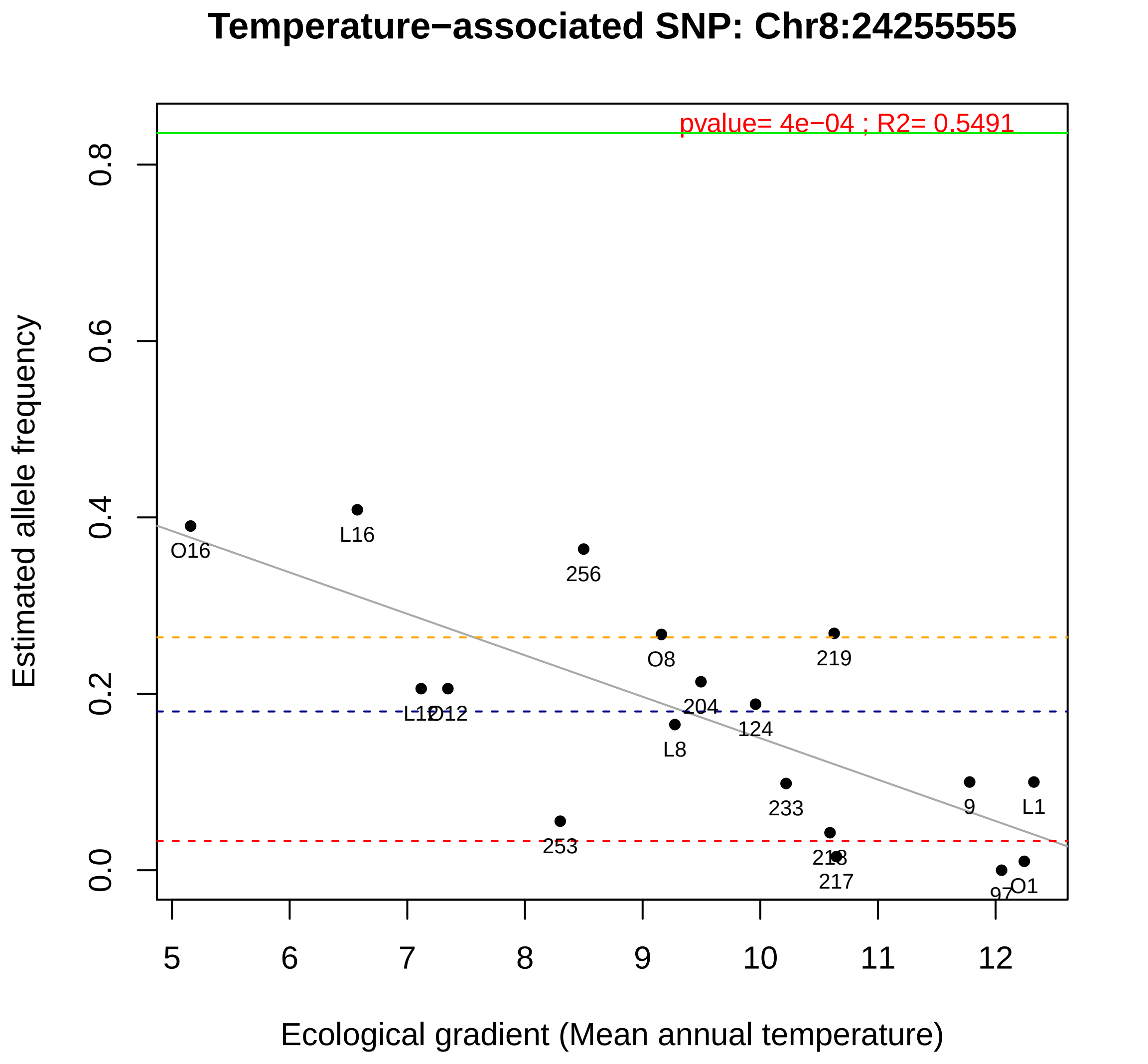

Temperature−associated SNP: Chr8:24255555
pvalue= 4e−04 ; R2= 0.5491
Estimated allele frequency
0.0
0.2
0.4
0.6
0.8
5
6
7
8
9
10
11
12
Ecological gradient (Mean annual temperature)
O16
L16
256
O8
219
204
L12
O12
124
L8
233
9
L1
253
218
217
97
O1

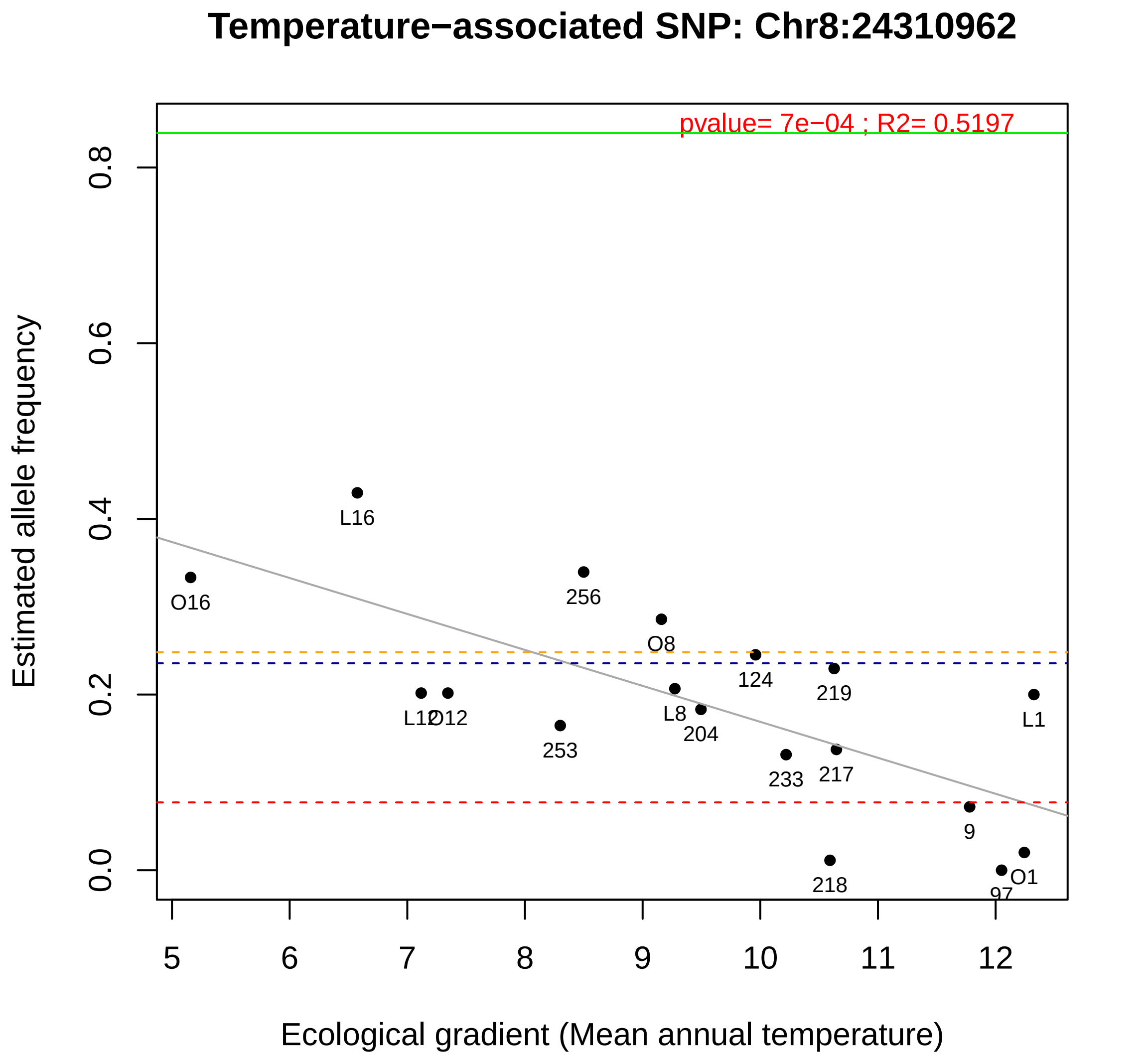

Temperature−associated SNP: Chr8:24310962
pvalue= 7e−04 ; R2= 0.5197
Estimated allele frequency
0.0
0.2
0.4
0.6
0.8
5
6
7
8
9
10
11
12
Ecological gradient (Mean annual temperature)
L16
O16
256
O8
124
219
L12
O12
253
L8
204
L1
233
217
9
218
97
O1

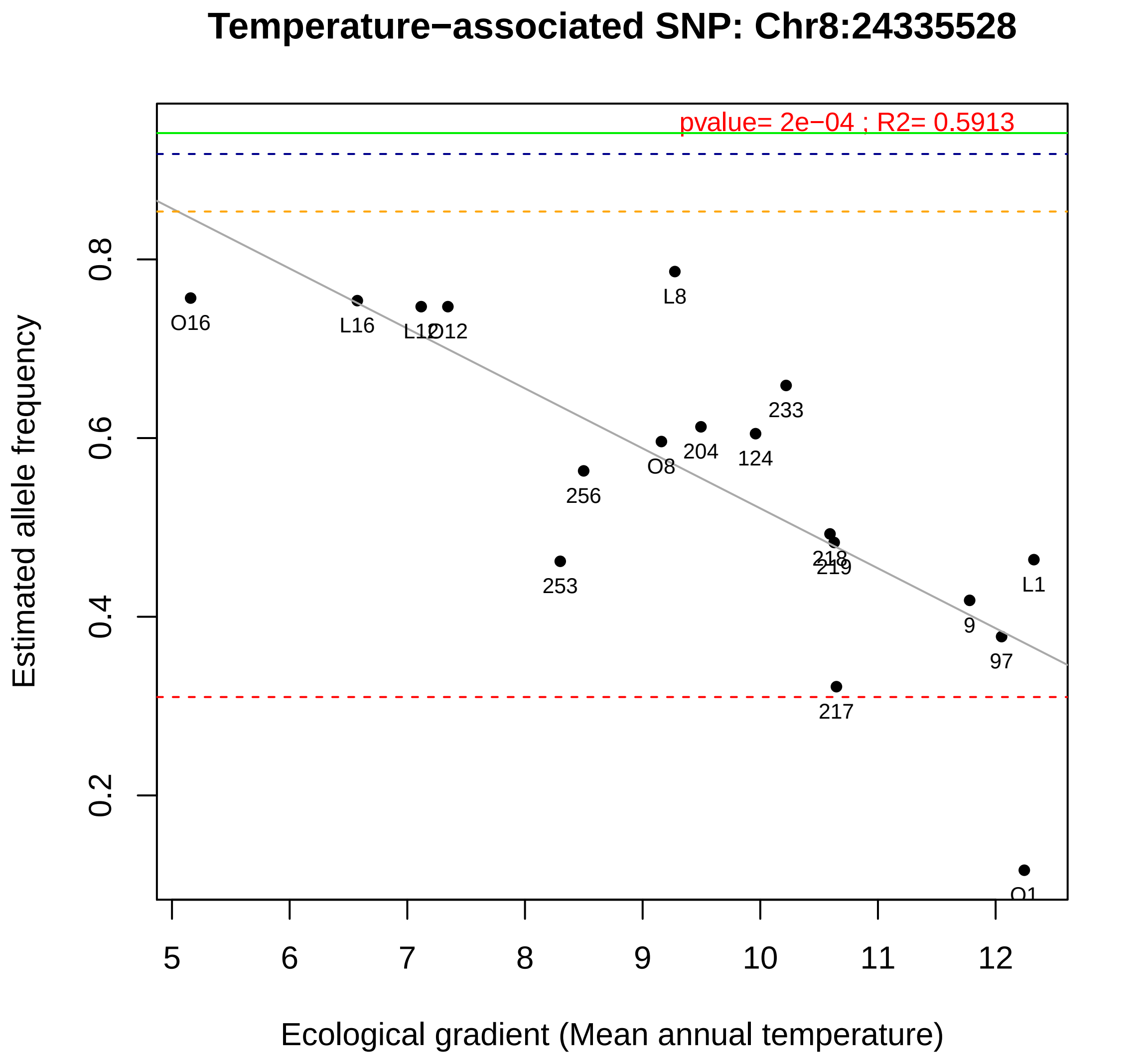
Temperature−associated SNP: Chr8:24335528
pvalue= 2e−04 ; R2= 0.5913
Estimated allele frequency
0.2
0.4
0.6
0.8
Ecological gradient (Mean annual temperature)
5
6
7
8
9
10
11
12
O16
L16
L12
O12
L8
233
204
124
O8
256
253
218
219
L1
9
97
217
O1

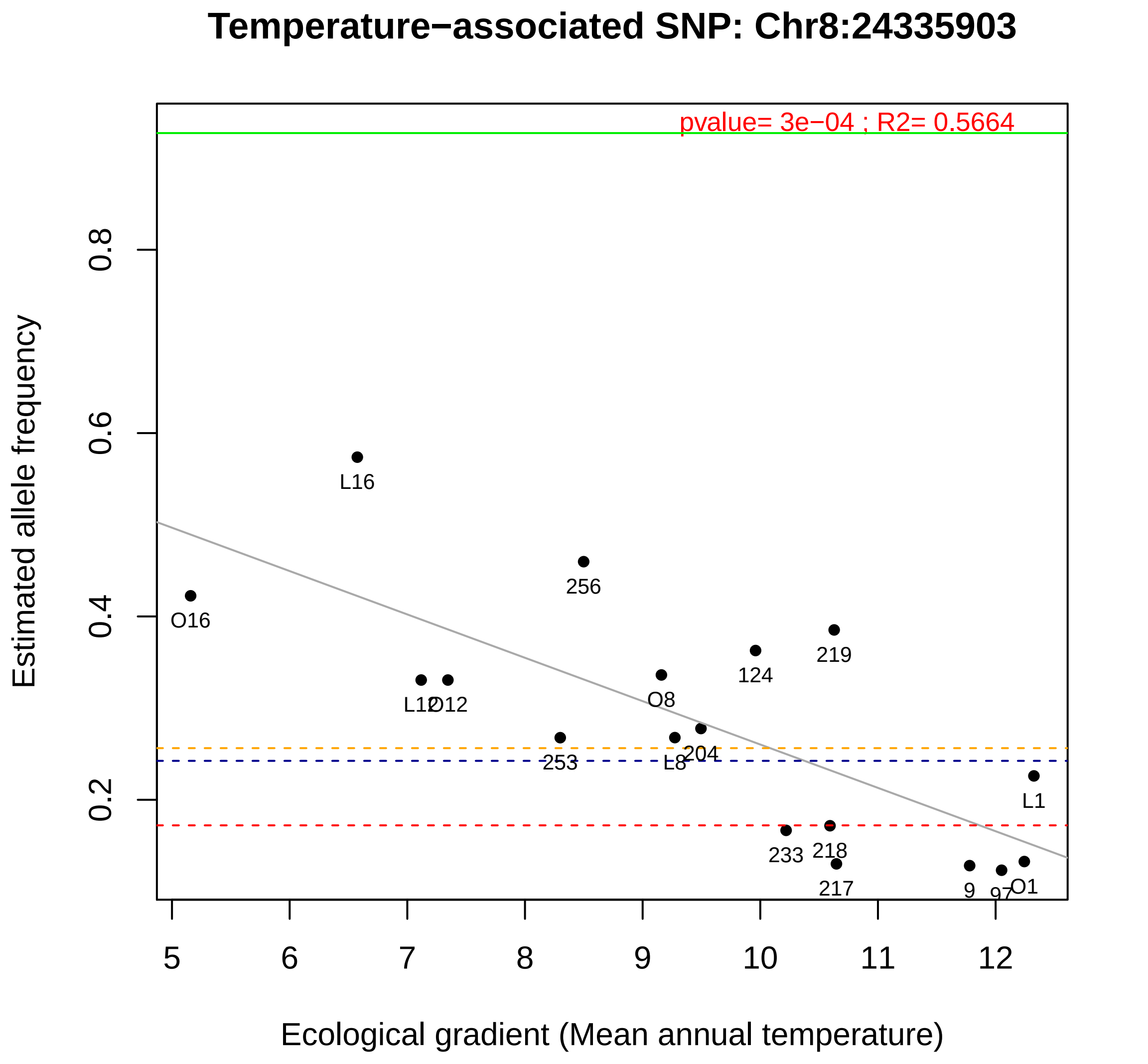

Temperature−associated SNP: Chr8:24335903
pvalue= 3e−04 ; R2= 0.5664
Estimated allele frequency
0.2
0.4
0.6
0.8
5
6
7
8
9
10
11
12
Ecological gradient (Mean annual temperature)
L16
256
O16
219
124
L12
O12
O8
253
L8
204
L1
233
218
217
9
97
O1

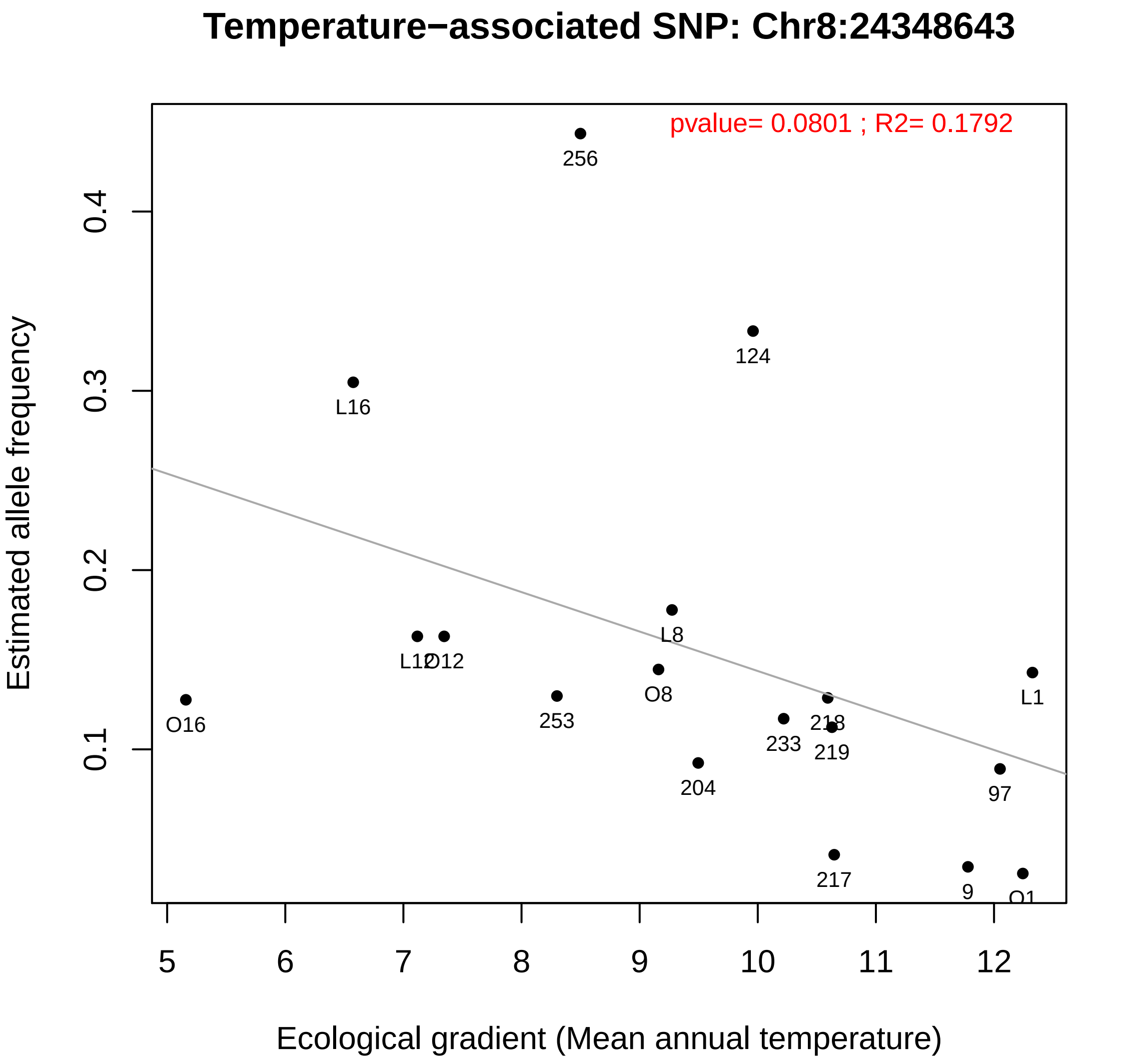
Temperature−associated SNP: Chr8:24348643
pvalue= 0.0801 ; R2= 0.1792
Estimated allele frequency
0.1
0.2
0.3
0.4
5
6
7
8
9
10
11
12
Ecological gradient (Mean annual temperature)
256
124
L16
L8
L12
O12
O8
L1
O16
253
218
233
219
204
97
217
9
O1

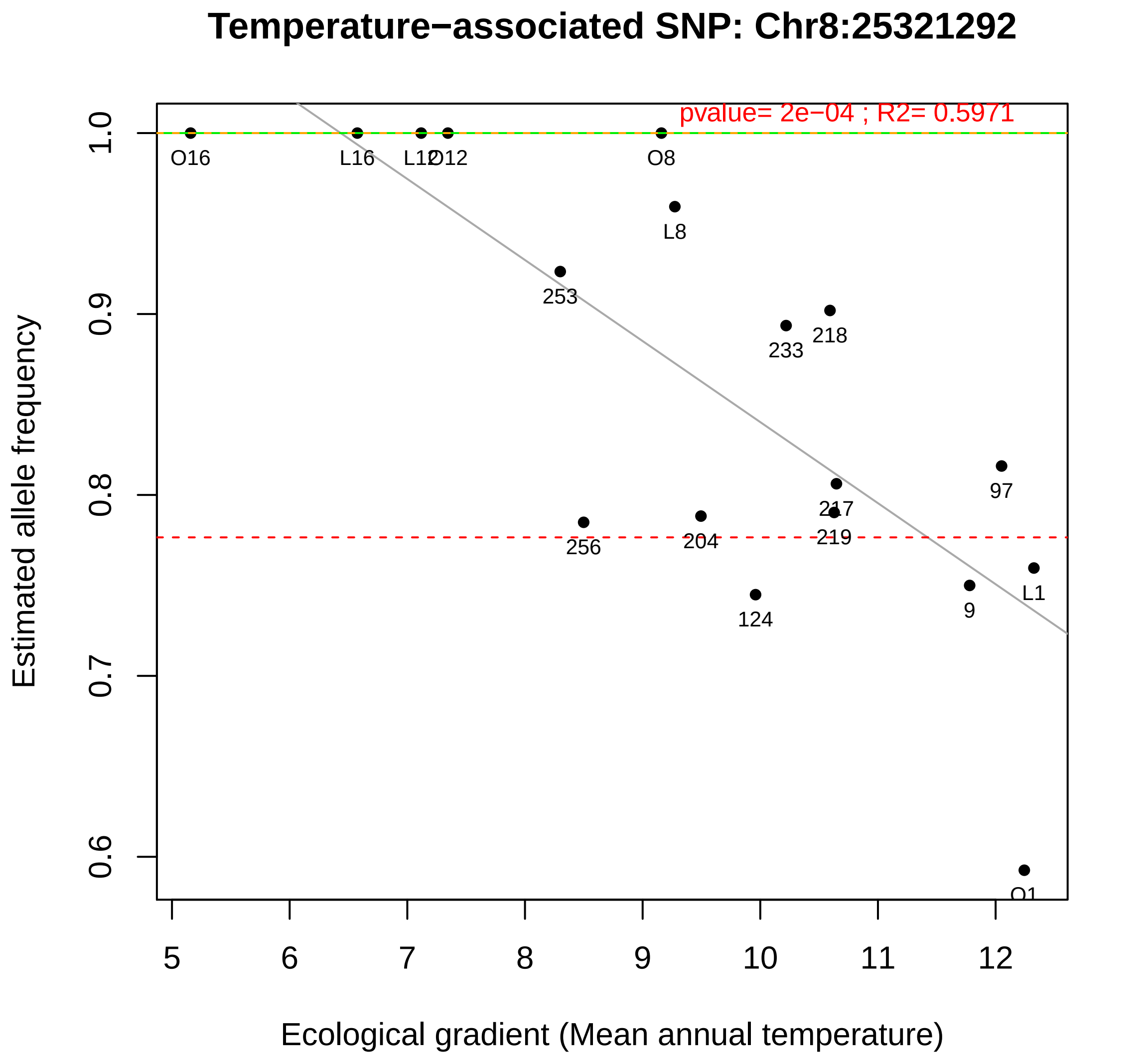
Temperature−associated SNP: Chr8:25321292
pvalue= 2e−04 ; R2= 0.5971
Estimated allele frequency
Ecological gradient (Mean annual temperature)
1.0
0.9
0.8
0.7
0.6
5
6
7
8
9
10
11
12
O16
L16
L12
O12
O8
L8
253
233
218
97
217
219
256
204
124
9
L1
O1

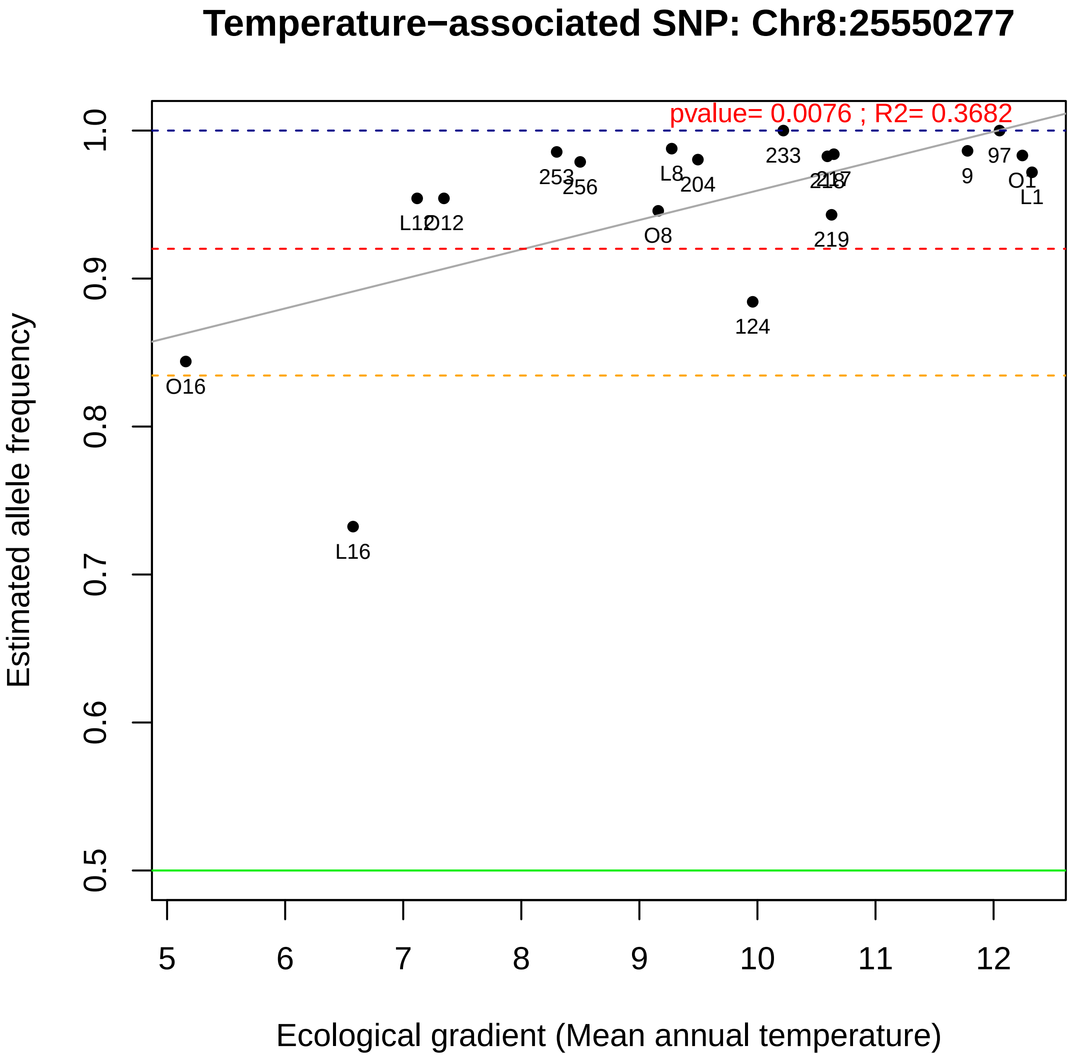

Temperature−associated SNP: Chr8:25550277
pvalue= 0.0076 ; R2= 0.3682
Estimated allele frequency
Ecological gradient (Mean annual temperature)
0.5
0.6
0.7
0.8
0.9
1.0
5
6
7
8
9
10
11
12
O16
L16
L12
O12
253
256
L8
O8
204
124
233
219
9
97
O1
L1

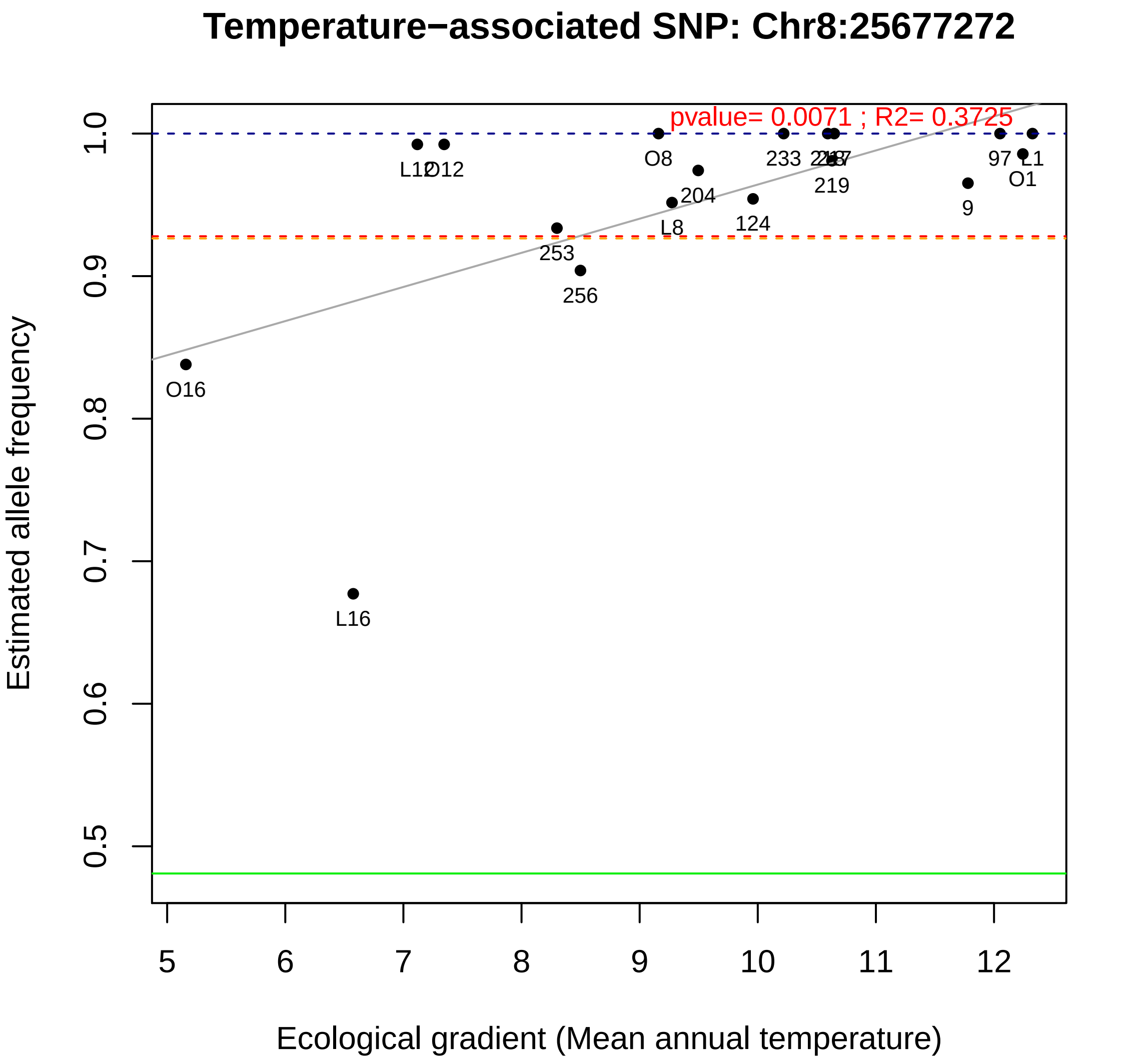
Temperature−associated SNP: Chr8:25677272
pvalue= 0.0071 ; R2= 0.3725
Estimated allele frequency
Ecological gradient (Mean annual temperature)
0.5
0.6
0.7
0.8
0.9
1.0
5
6
7
8
9
10
11
12
O16
L16
L12
O12
253
256
O8
L8
204
124
233
219
9
97
L1
O1

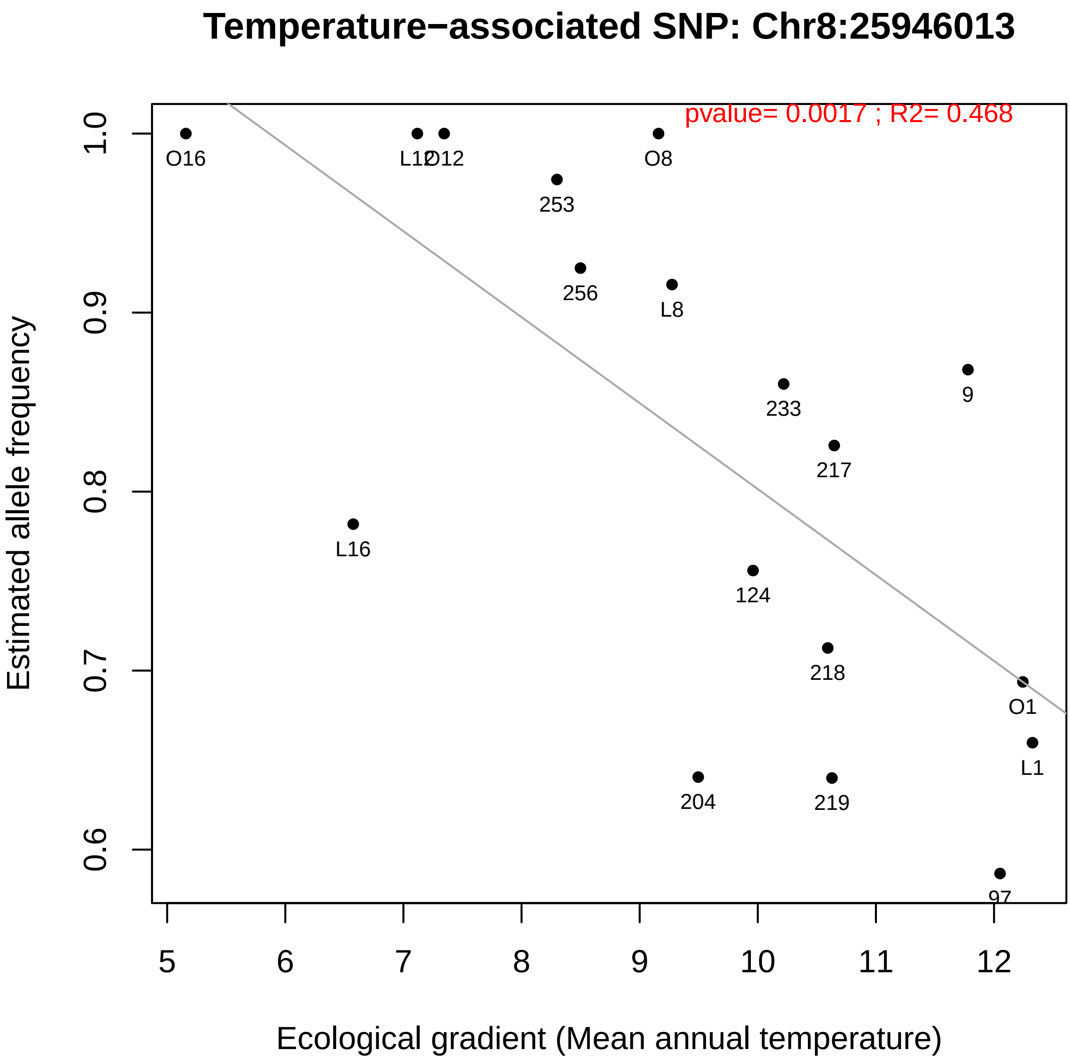

Temperature−associated SNP: Chr8:25946013
pvalue= 0.0017 ; R2= 0.468
Estimated allele frequency
Ecological gradient (Mean annual temperature)
1.0
0.9
0.8
0.7
0.6
5
6
7
8
9
10
11
12
O16
L12
O12
O8
253
256
L8
233
9
217
L16
124
218
O1
L1
204
219
97

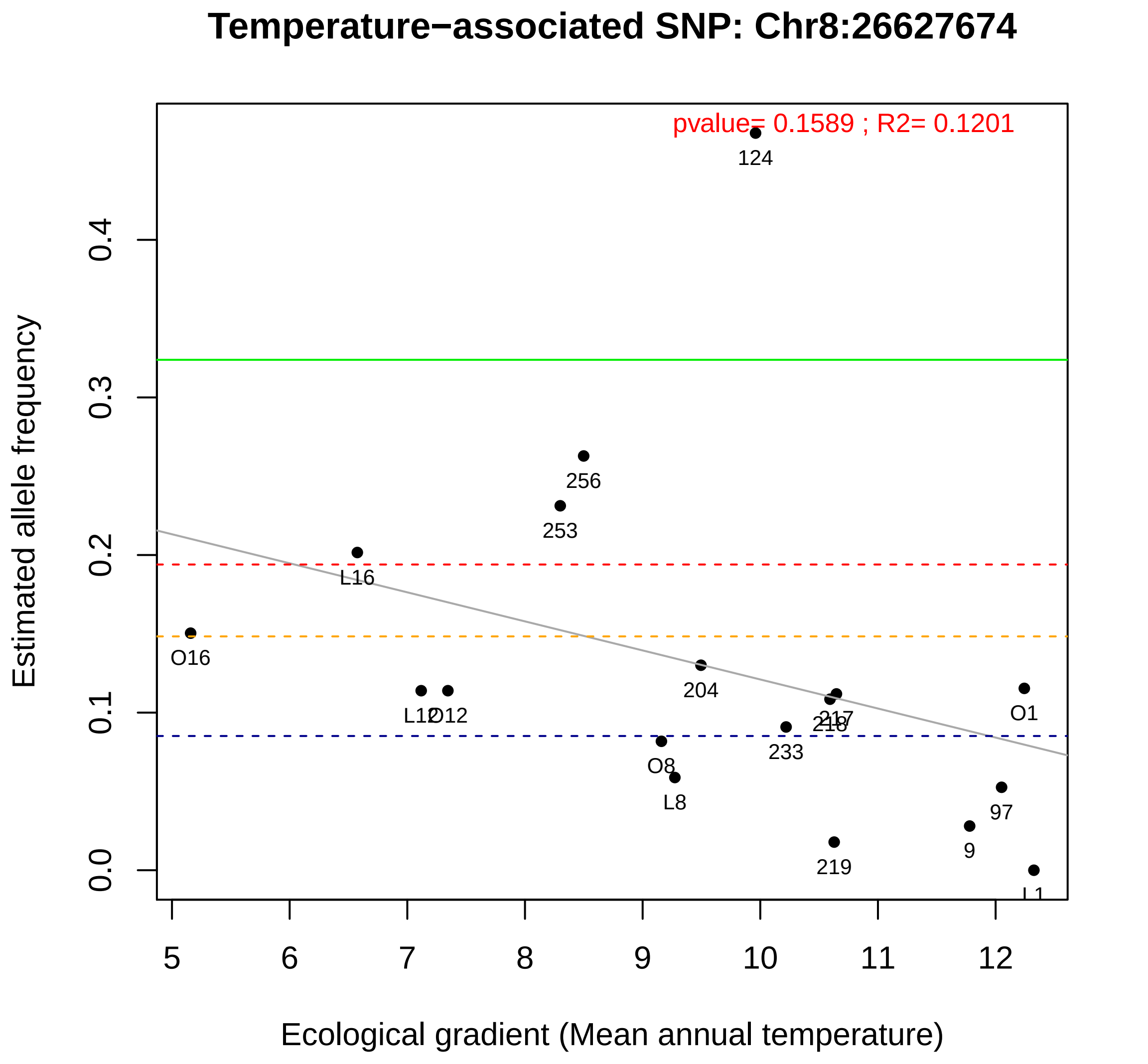
Temperature−associated SNP: Chr8:26627674
pvalue= 0.1589 ; R2= 0.1201
124
256
253
L16
O16
L12
O12
204
217
218
O1
233
O8
L8
97
9
219
L1
0.0
0.1
0.2
0.3
0.4
5
6
7
8
9
10
11
12
Estimated allele frequency
Ecological gradient (Mean annual temperature)

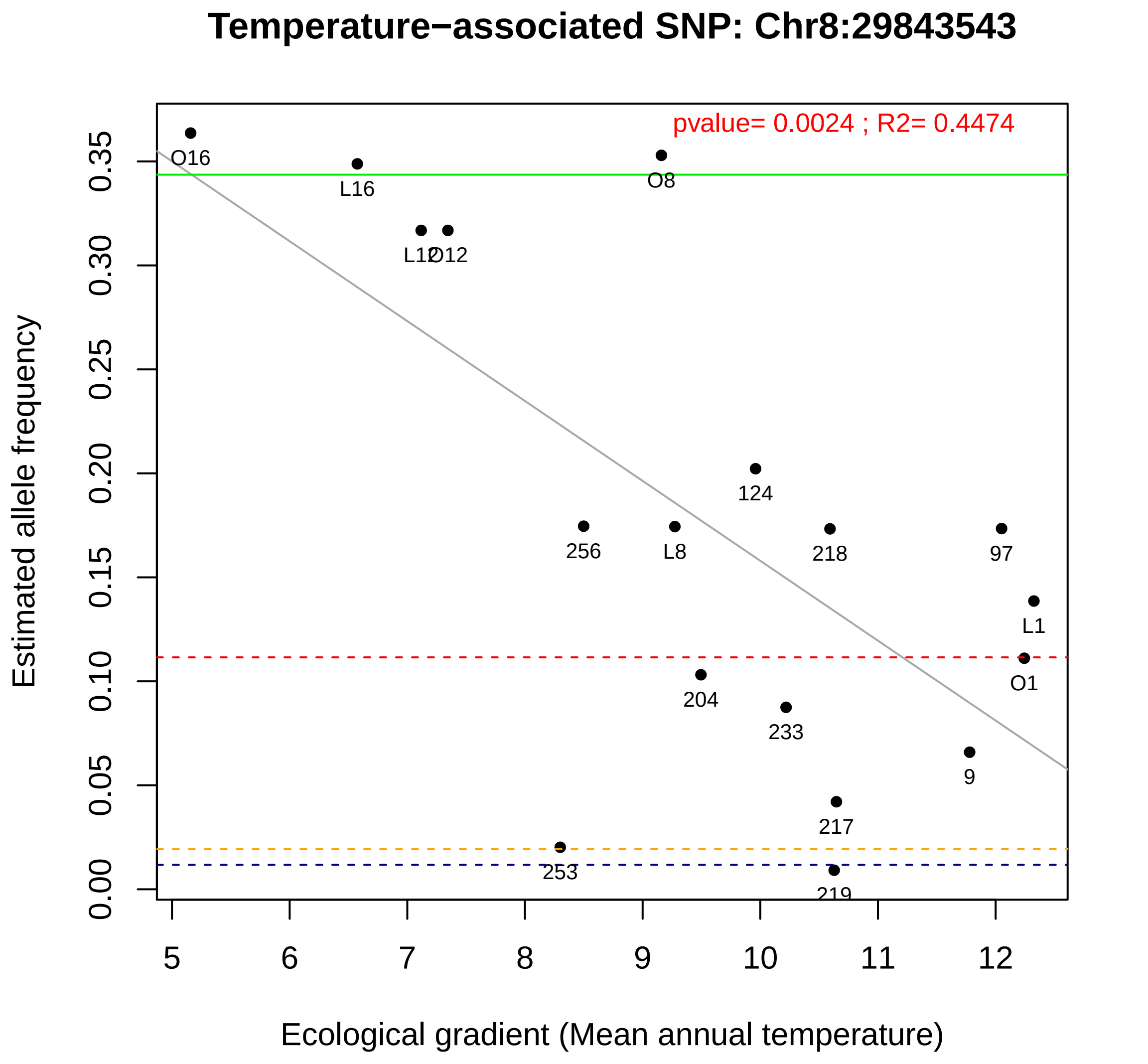
Temperature−associated SNP: Chr8:29843543
pvalue= 0.0024 ; R2= 0.4474
Estimated allele frequency
0.00
0.05
0.10
0.15
0.20
0.25
0.30
0.35
5
6
7
8
9
10
11
12
Ecological gradient (Mean annual temperature)
O16
L16
O8
L12
O12
124
256
L8
218
97
L1
O1
204
233
9
217
253
219

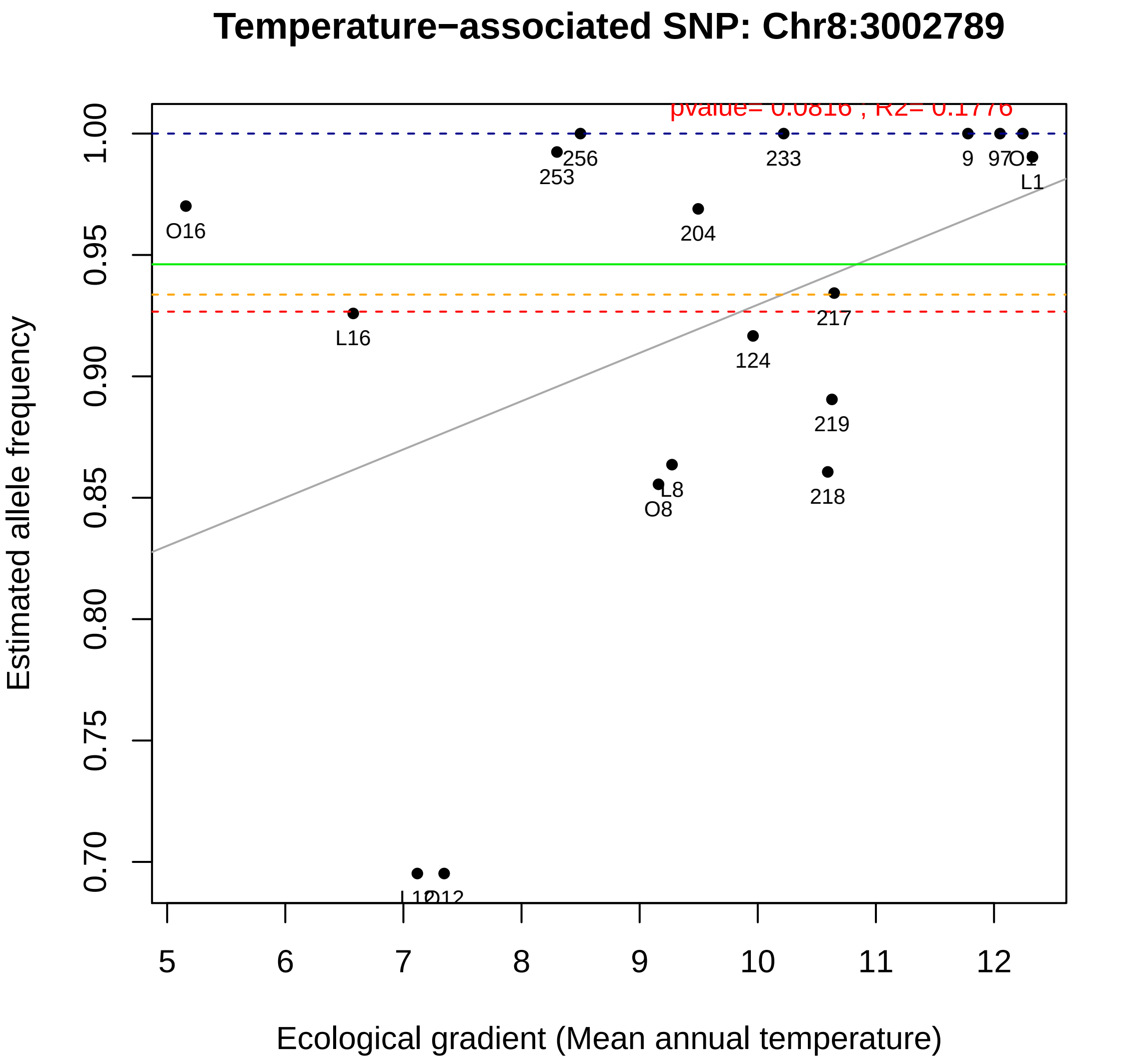

Temperature−associated SNP: Chr8:3002789
pvalue= 0.0816 ; R2= 0.1776
Estimated allele frequency
Ecological gradient (Mean annual temperature)
1.00
0.95
0.90
0.85
0.80
0.75
0.70
5
6
7
8
9
10
11
12
O16
L16
253
256
204
233
124
217
219
218
L8
O8
9
97
O1
L1
O12

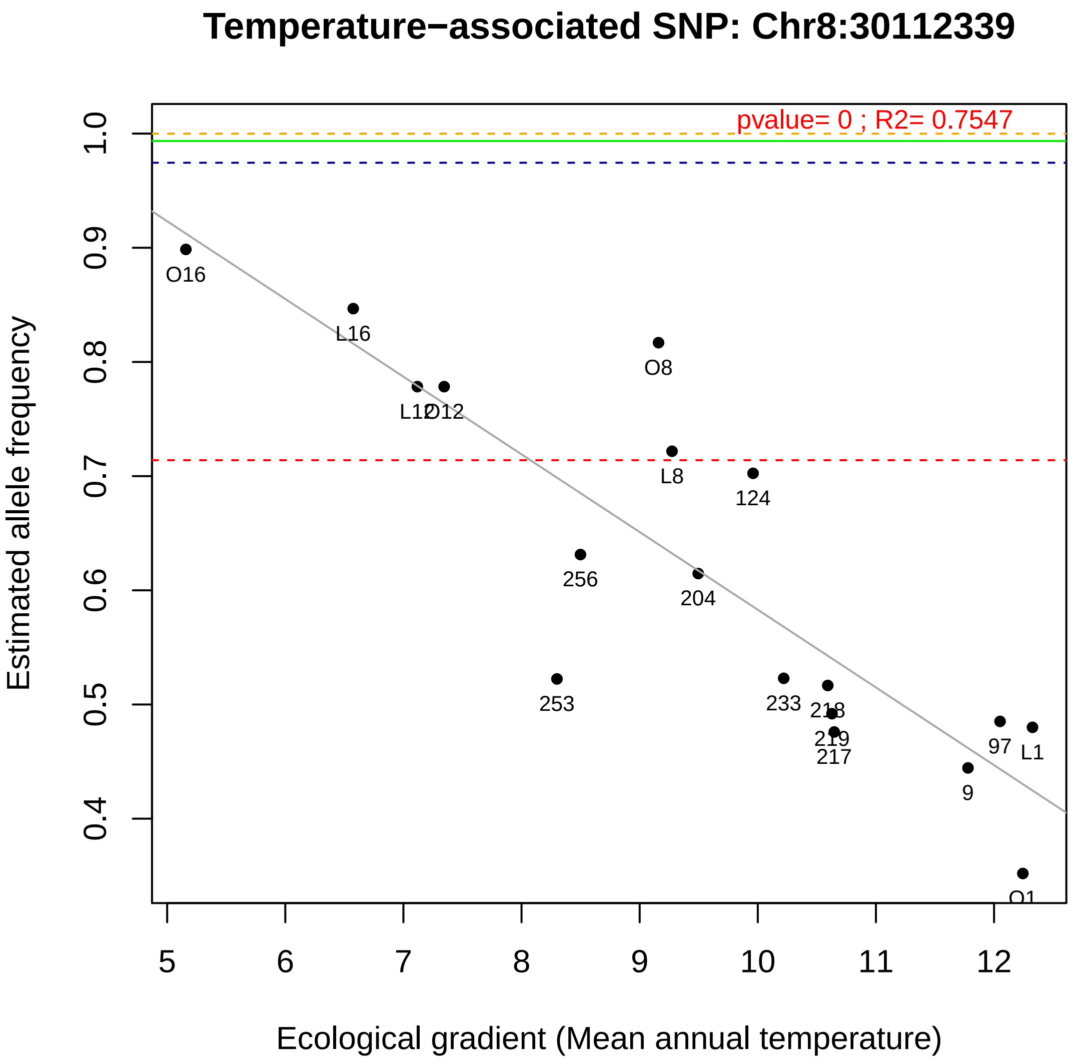

Temperature−associated SNP: Chr8:30112339
pvalue= 0 ; R2= 0.7547
Estimated allele frequency
Ecological gradient (Mean annual temperature)
0.4
0.5
0.6
0.7
0.8
0.9
1.0
5
6
7
8
9
10
11
12
O16
L16
L12
O12
O8
L8
124
256
204
253
233
218
219
217
9
97
L1
O1

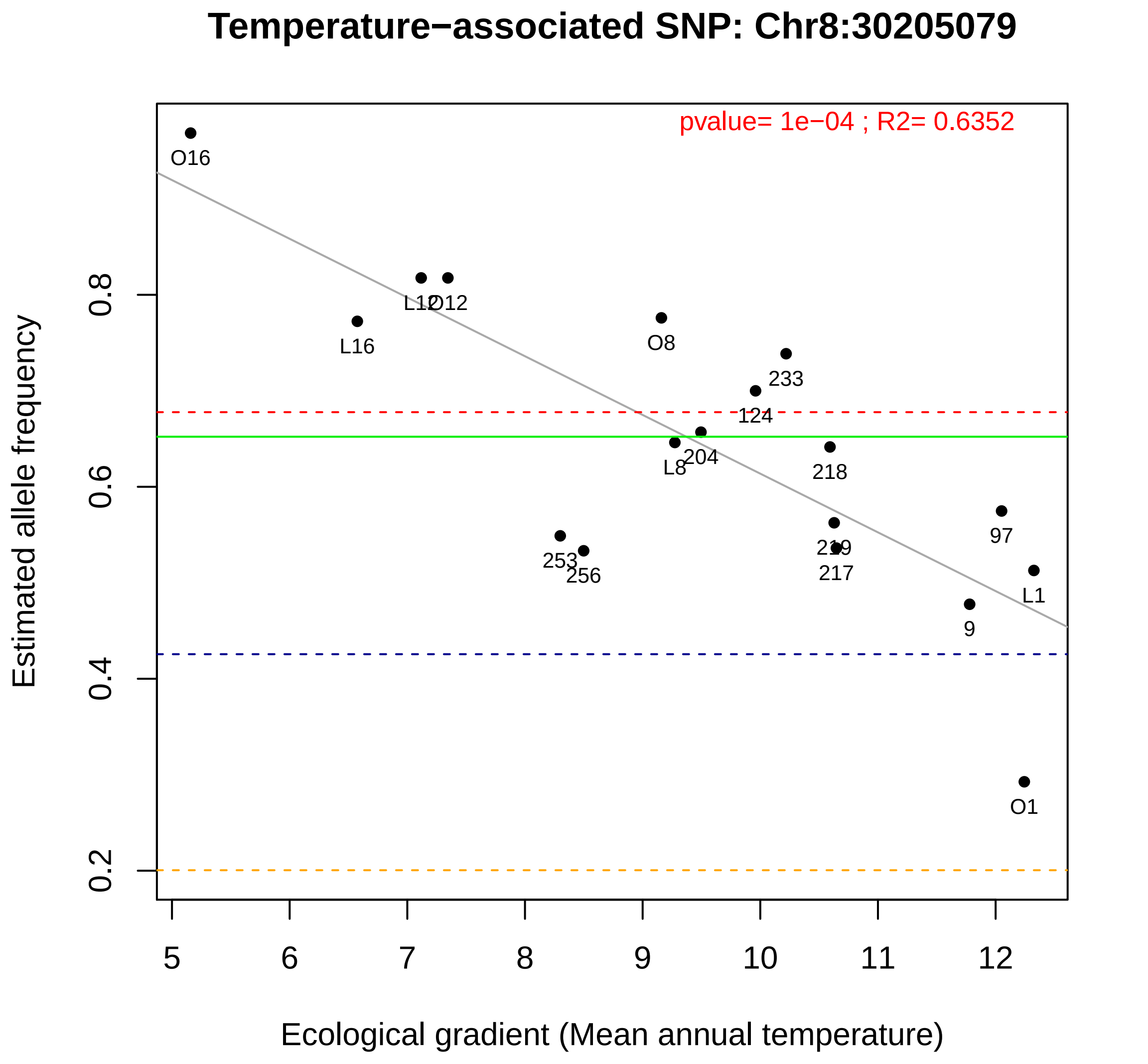

Temperature−associated SNP: Chr8:30205079
pvalue= 1e−04 ; R2= 0.6352
O16
L16
L12
O12
O8
233
124
204
L8
218
253
256
219
217
97
L1
9
O1
0.2
0.4
0.6
0.8
Estimated allele frequency
5
6
7
8
9
10
11
12
Ecological gradient (Mean annual temperature)

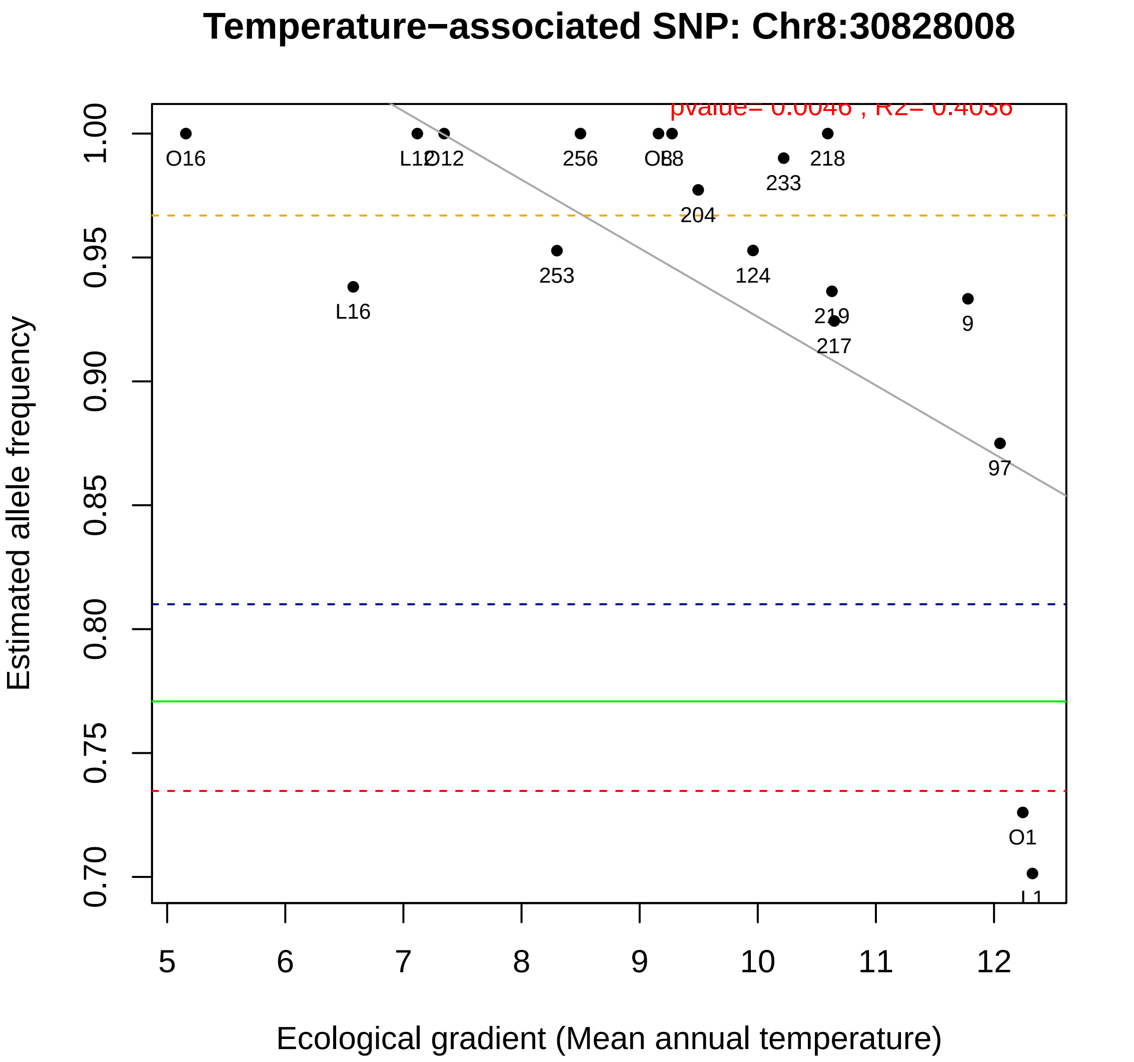
Temperature−associated SNP: Chr8:30828008
pvalue= 0.0046 , R2= 0.4036
Estimated allele frequency
Ecological gradient (Mean annual temperature)
O16
L16
L12
O12
253
256
O8
L8
204
124
233
218
219
217
9
97
O1
L1
1.00
0.95
0.90
0.85
0.80
0.75
0.70
5
6
7
8
9
10
11
12

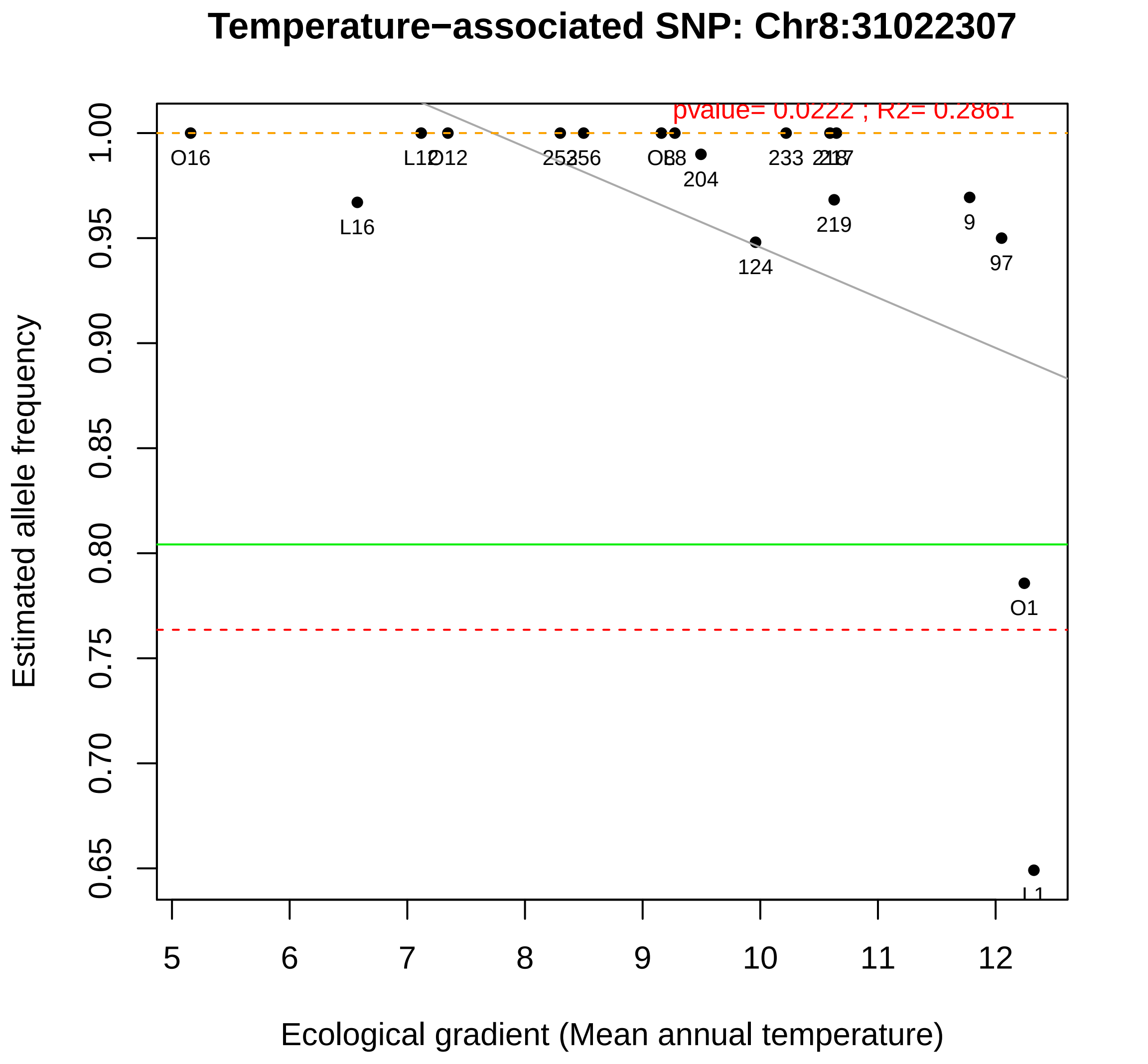

Temperature−associated SNP: Chr8:31022307
pvalue= 0.0222 ; R2= 0.2861
Estimated allele frequency
Ecological gradient (Mean annual temperature)
1.00
0.95
0.90
0.85
0.80
0.75
0.70
0.65
5
6
7
8
9
10
11
12
O16
L16
L12
O12
253
256
O3
18
204
233
218
7
124
219
9
97
O1

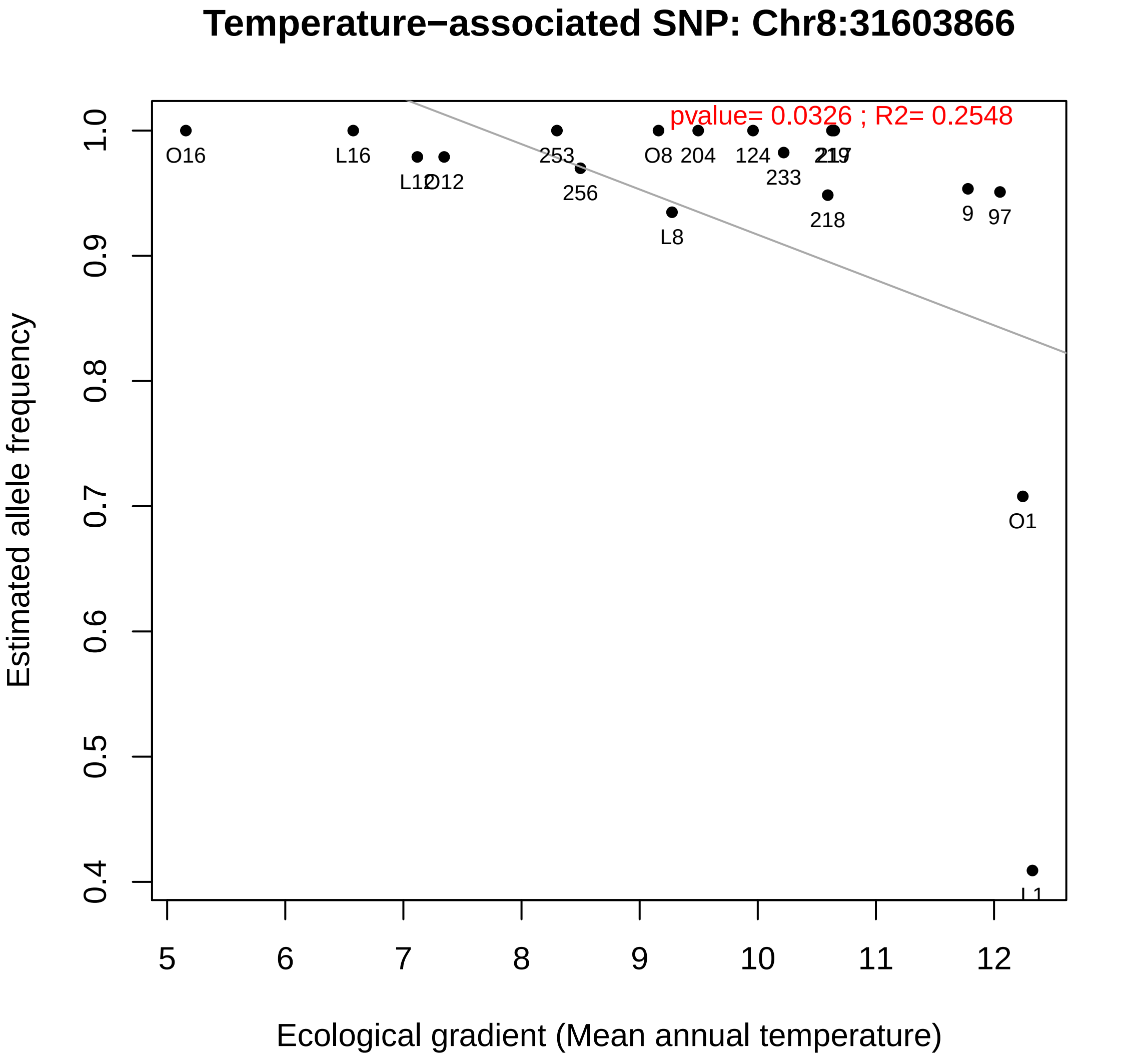

Temperature−associated SNP: Chr8:31603866
pvalue= 0.0326 ; R2= 0.2548
Estimated allele frequency
Ecological gradient (Mean annual temperature)
1.0
0.9
0.8
0.7
0.6
0.5
0.4
5
6
7
8
9
10
11
12
O16
L16
O12
253
256
O8
204
124
233
L8
218
9
97
O1

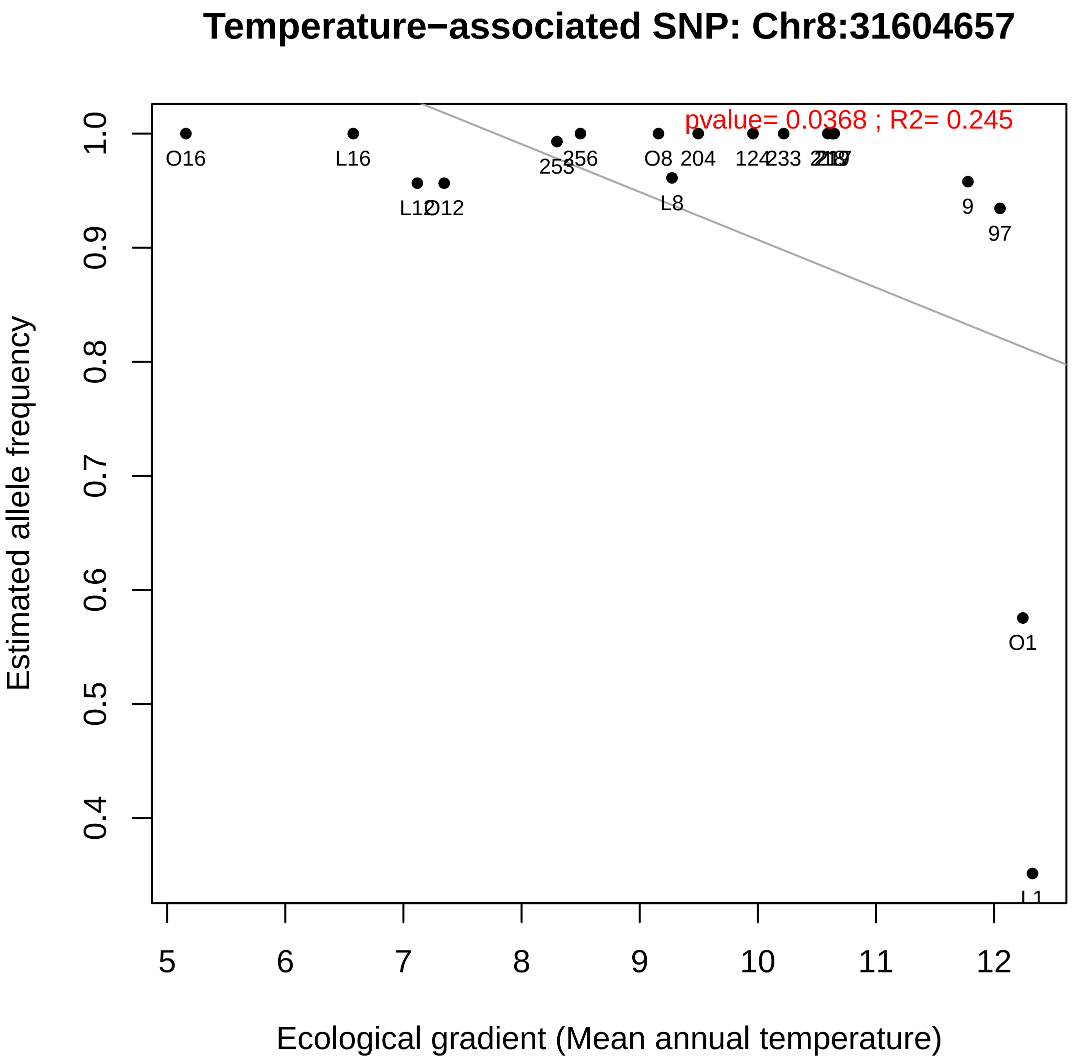

Temperature−associated SNP: Chr8:31604657
pvalue= 0.0368 ; R2= 0.245
Estimated allele frequency
Ecological gradient (Mean annual temperature)
O16
L16
L12
O12
253
256
O8
L8
204
124
233
9
97
O1

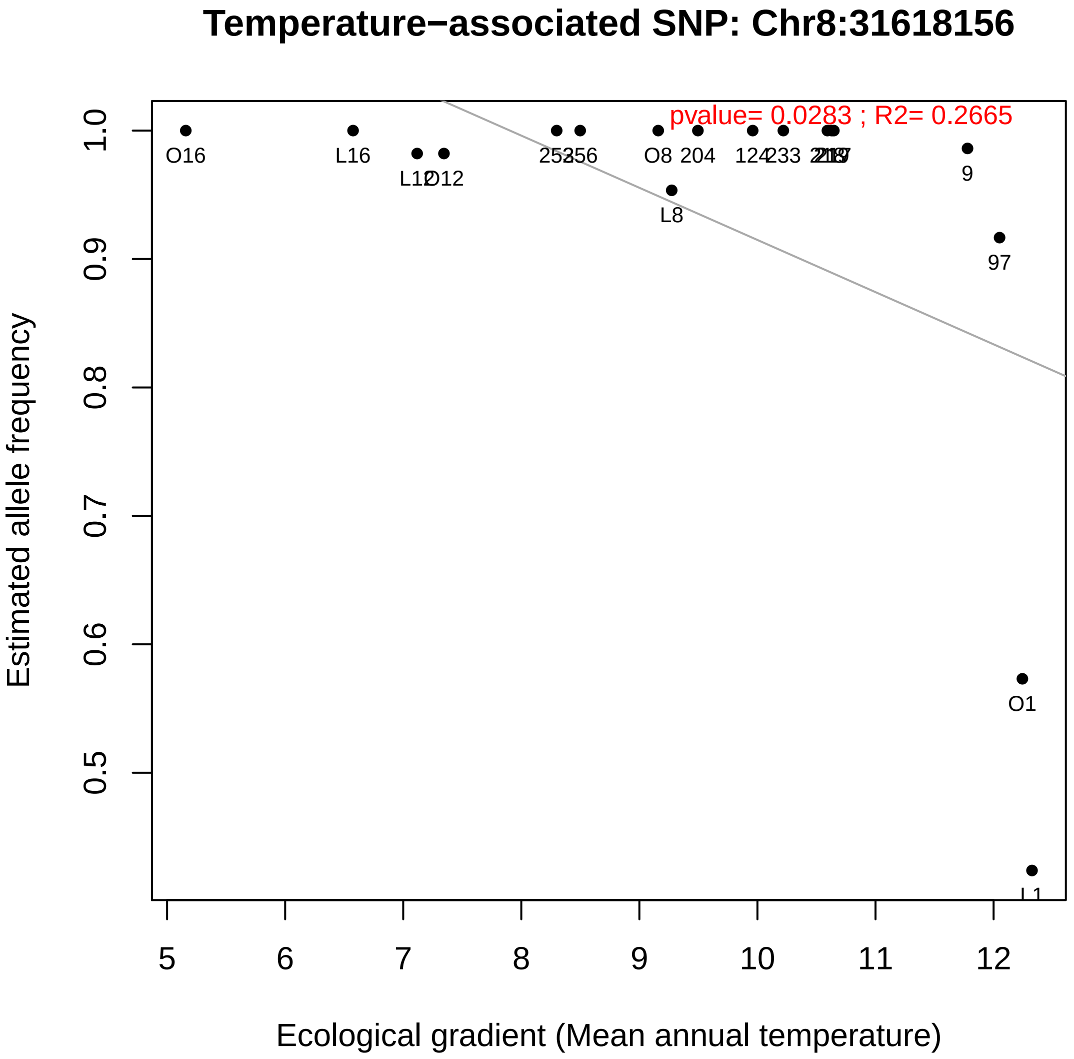
Temperature−associated SNP: Chr8:31618156
pvalue= 0.0283 ; R2= 0.2665
Estimated allele frequency
Ecological gradient (Mean annual temperature)
1.0
0.9
0.8
0.7
0.6
0.5
5
6
7
8
9
10
11
12
O16
L16
L12
O12
O8
204
L8
9
97
O1

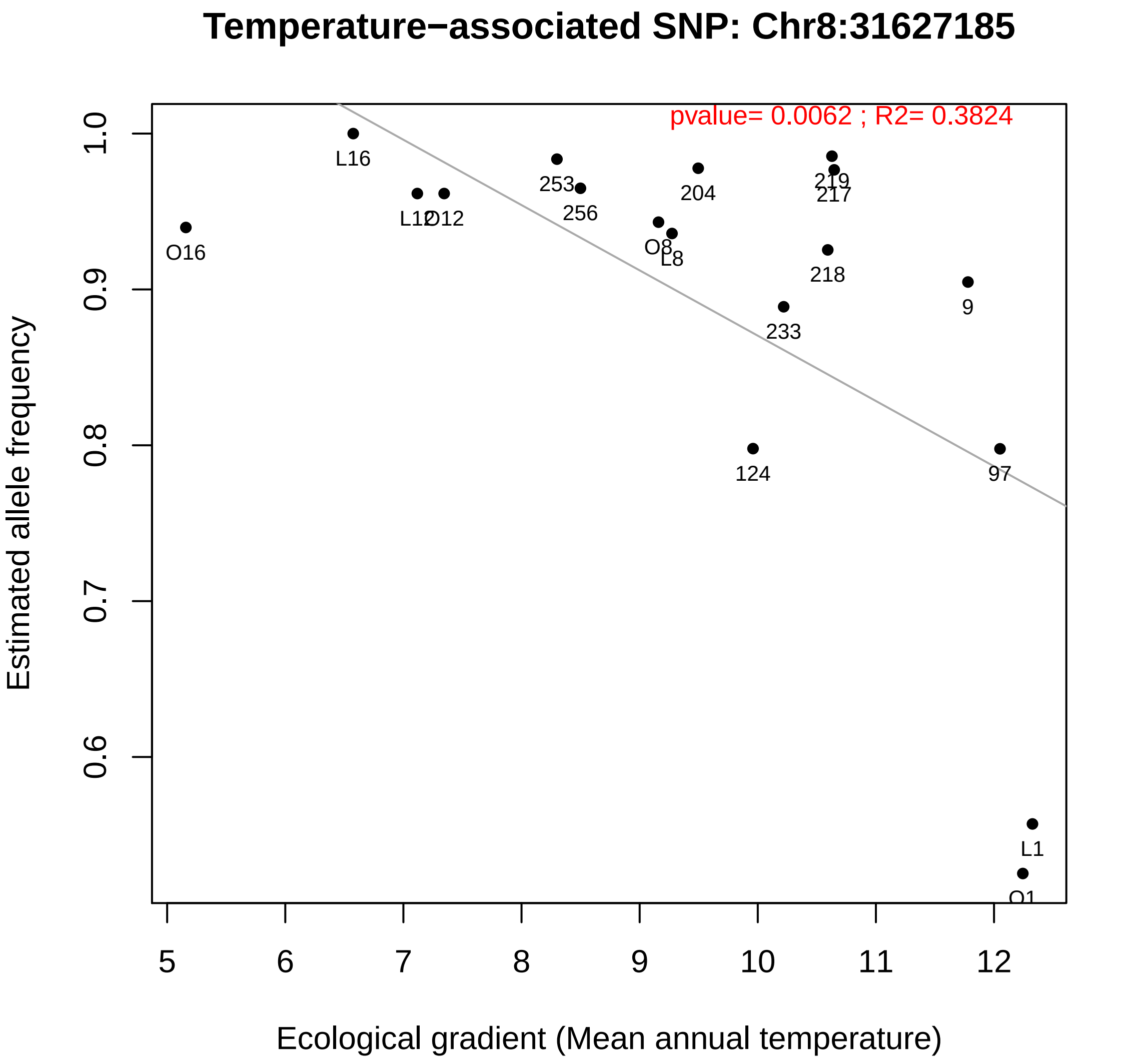

Temperature−associated SNP: Chr8:31627185
pvalue= 0.0062 ; R2= 0.3824
Estimated allele frequency
Ecological gradient (Mean annual temperature)
0.6
0.7
0.8
0.9
1.0
5
6
7
8
9
10
11
12
L16
L12
O12
O16
253
256
204
219
217
O8
L8
218
233
9
124
97
L1
O1

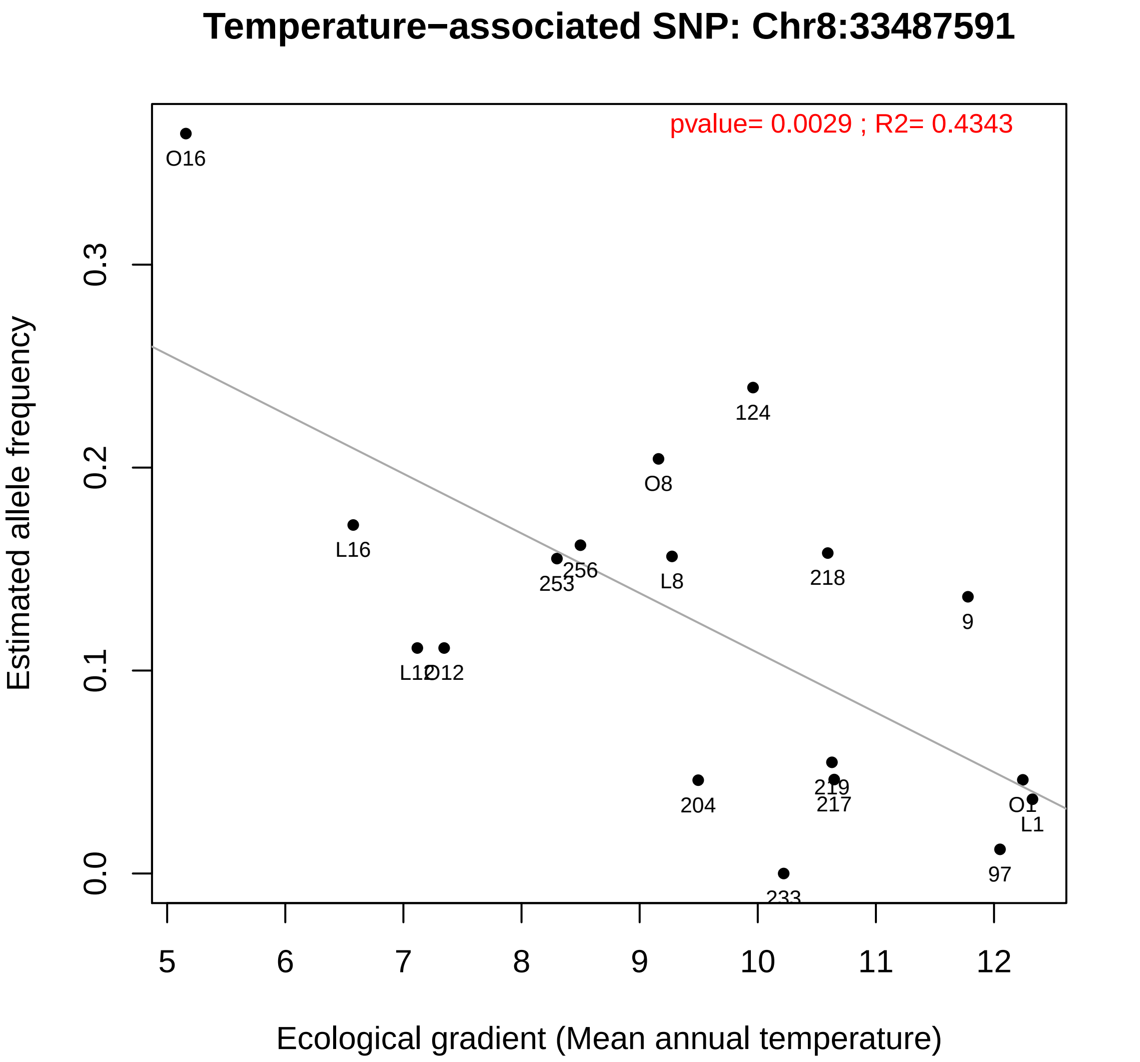

Temperature−associated SNP: Chr8:33487591
pvalue= 0.0029 ; R2= 0.4343
Estimated allele frequency
Ecological gradient (Mean annual temperature)
0.0
0.1
0.2
0.3
5
6
7
8
9
10
11
12
O16
L16
L12
O12
253
256
O8
124
L8
218
9
204
233
219
217
97
O1
L1

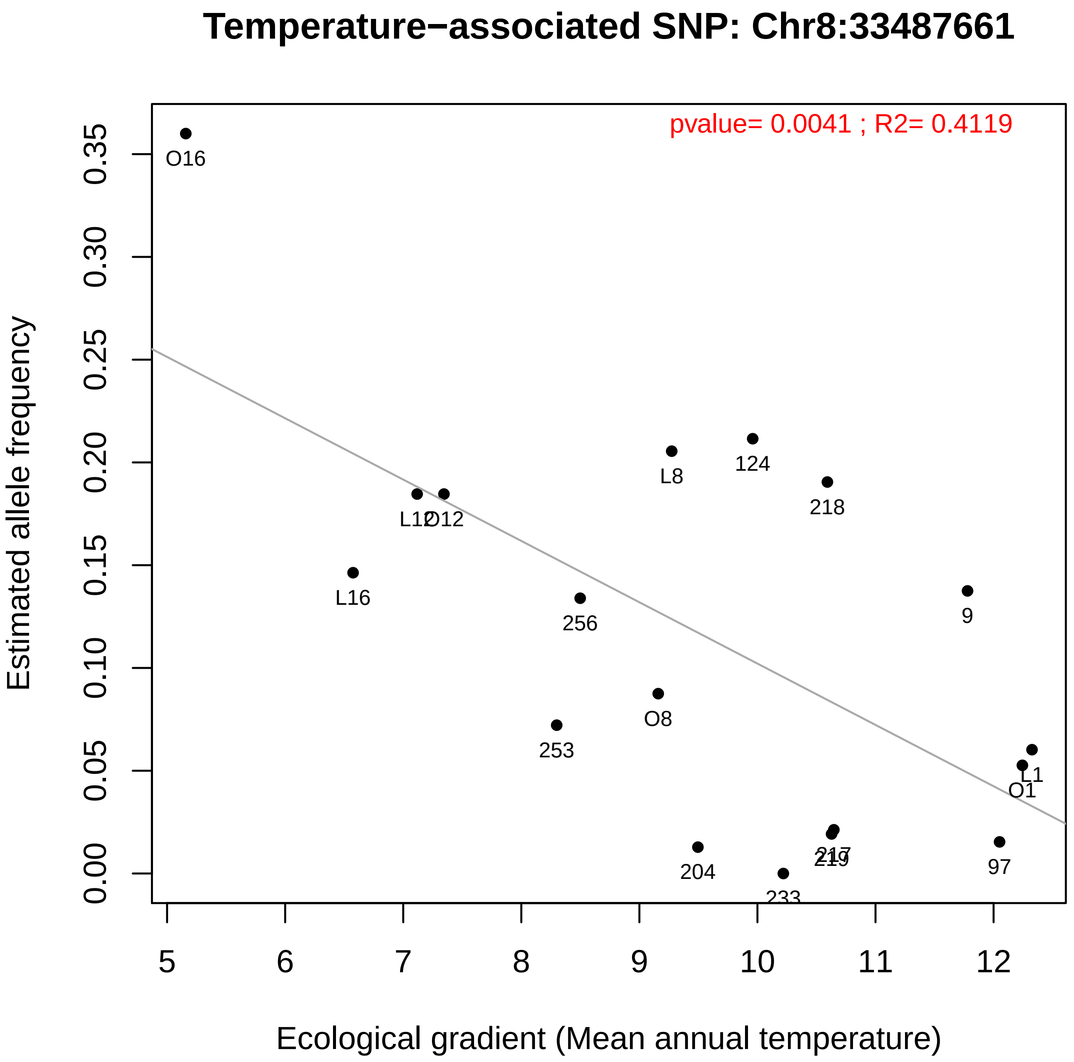
Temperature−associated SNP: Chr8:33487661
pvalue= 0.0041 ; R2= 0.4119
Estimated allele frequency
0.00
0.05
0.10
0.15
0.20
0.25
0.30
0.35
5
6
7
8
9
10
11
12
Ecological gradient (Mean annual temperature)
O16
L12
O12
L16
L8
124
218
256
9
253
O8
L1
O1
204
233
217
219
97

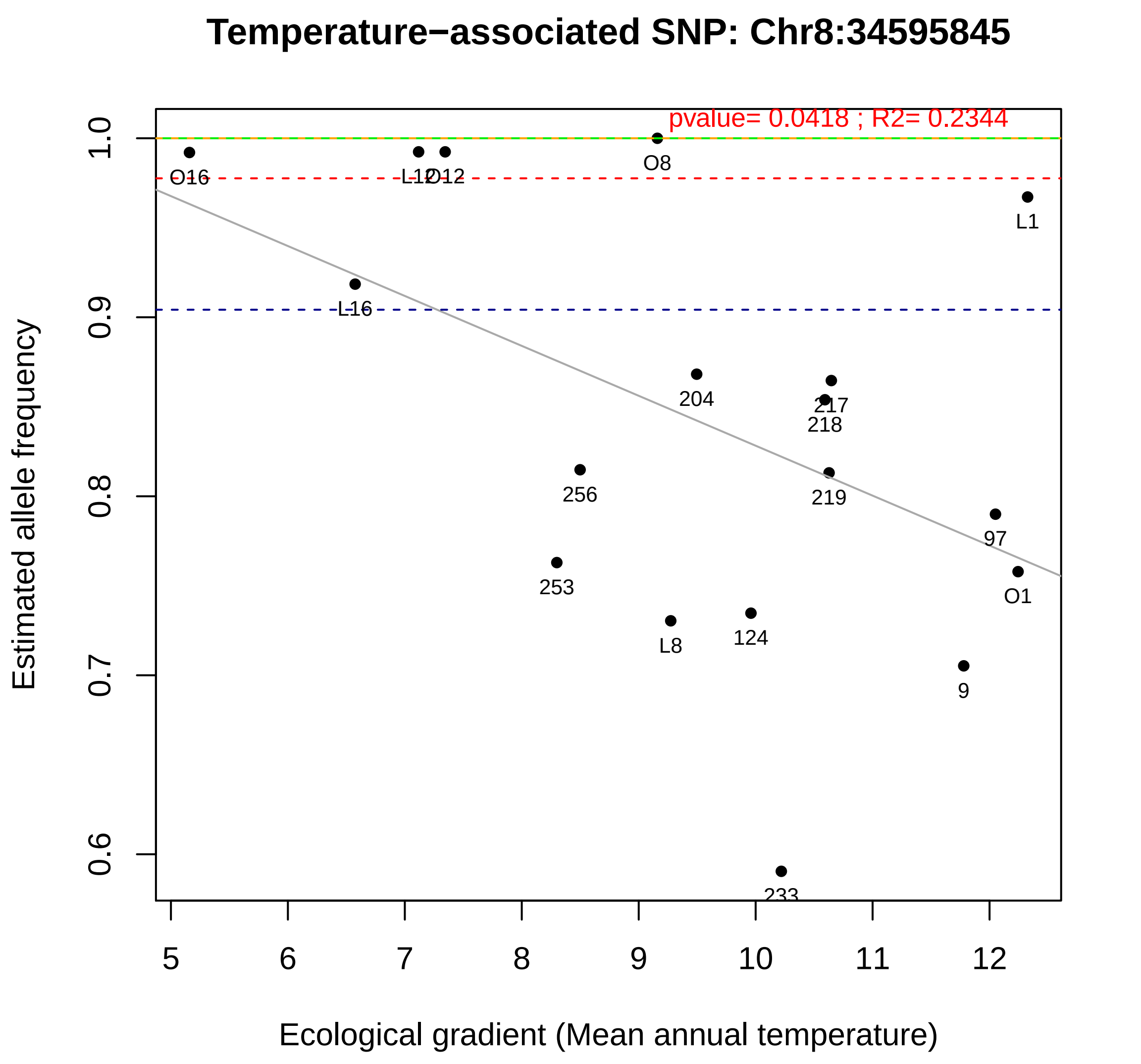

Temperature−associated SNP: Chr8:34595845
pvalue= 0.0418 ; R2= 0.2344
Estimated allele frequency
Ecological gradient (Mean annual temperature)
1.0
0.9
0.8
0.7
0.6
5
6
7
8
9
10
11
12
O16
L12
O12
O8
L1
L16
204
217
218
219
256
97
253
O1
L8
124
9
233

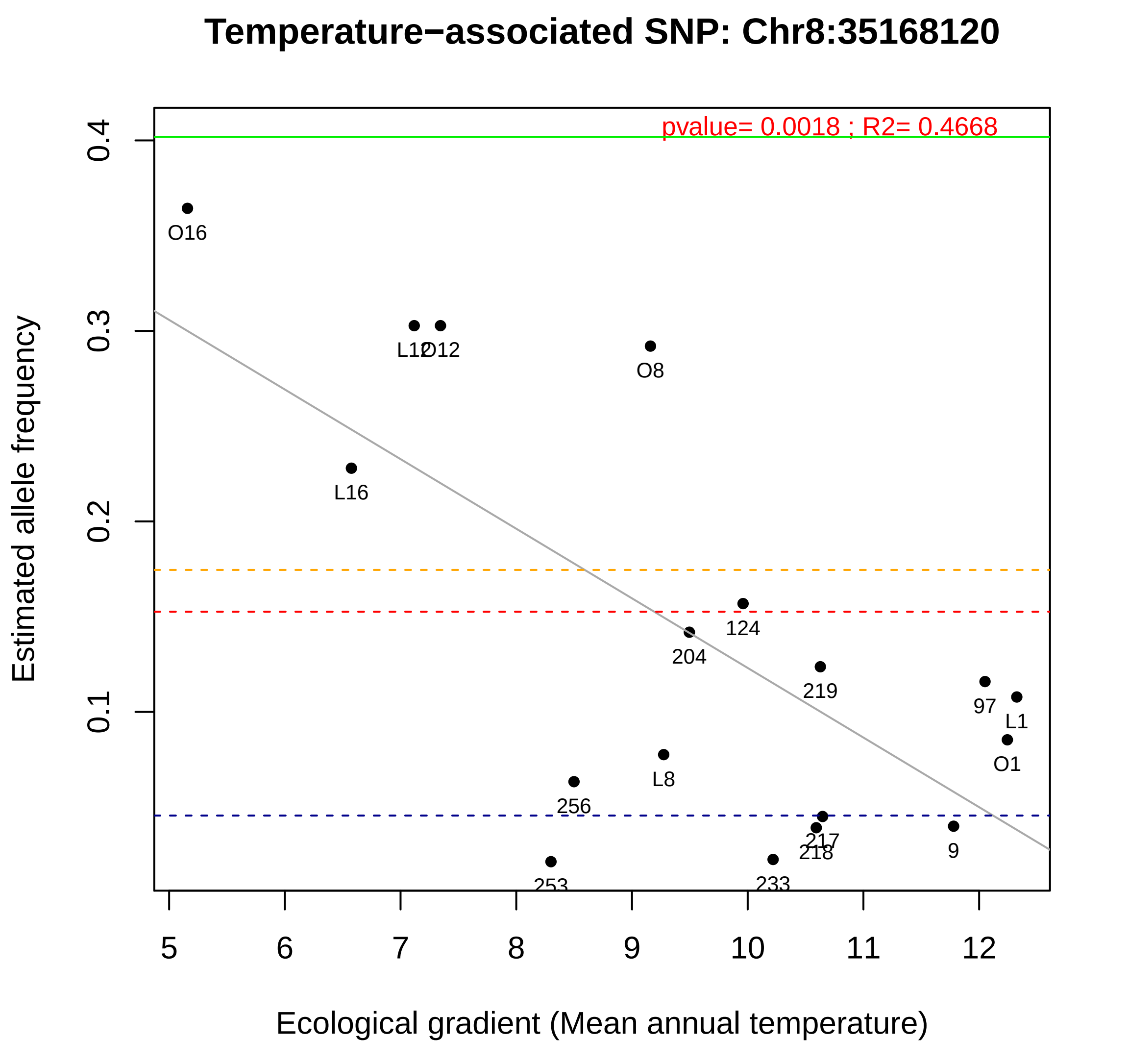

Temperature−associated SNP: Chr8:35168120
pvalue= 0.0018 ; R2= 0.4668
Estimated allele frequency
0.1
0.2
0.3
0.4
5
6
7
8
9
10
11
12
Ecological gradient (Mean annual temperature)
O16
L12
O12
O8
L16
124
204
219
97
L1
O1
L8
256
217
218
9
233
253

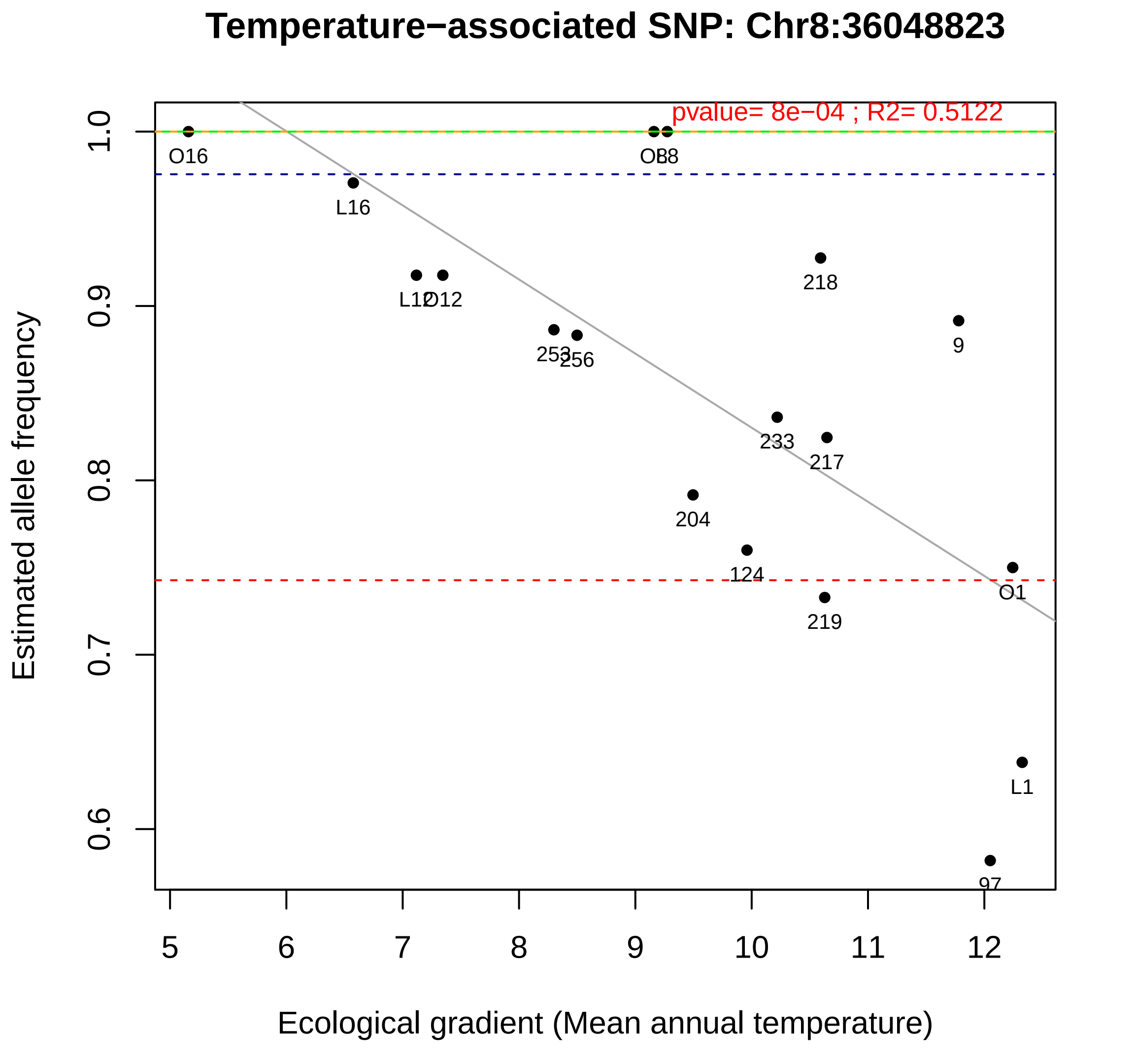

Temperature−associated SNP: Chr8:36048823
pvalue= 8e−04 ; R2= 0.5122
Estimated allele frequency
Ecological gradient (Mean annual temperature)
1.0
0.9
0.8
0.7
0.6
5
6
7
8
9
10
11
12
O16
L16
L12
O12
253
256
218
9
233
217
204
124
O1
219
L1
97

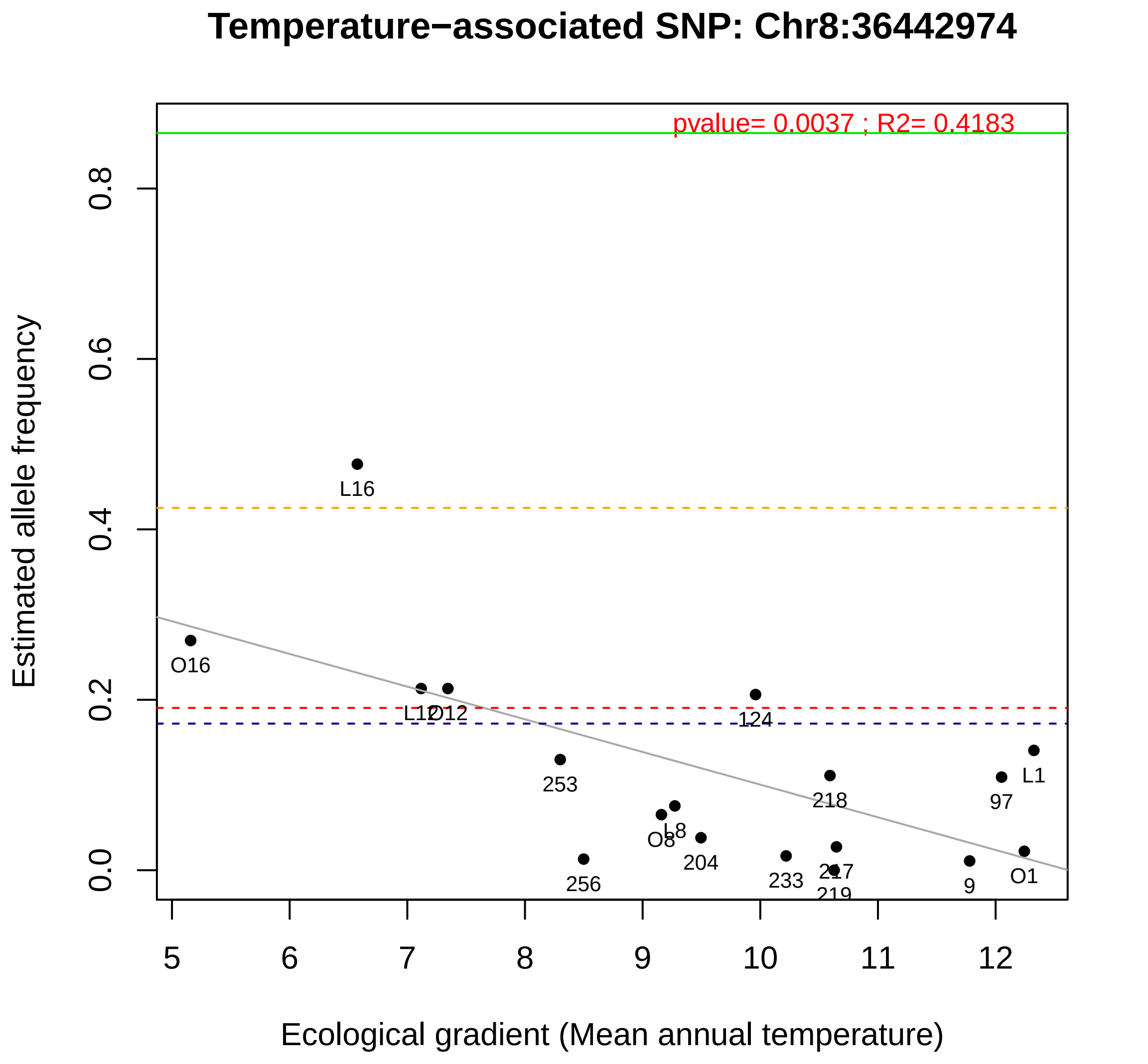
Temperature−associated SNP: Chr8:36442974
pvalue= 0.0037 ; R2= 0.4183
Estimated allele frequency
0.0
0.2
0.4
0.6
0.8
5
6
7
8
9
10
11
12
Ecological gradient (Mean annual temperature)
L16
O16
L12
O12
124
253
218
97
L1
L8
O8
204
256
233
217
219
9
O1

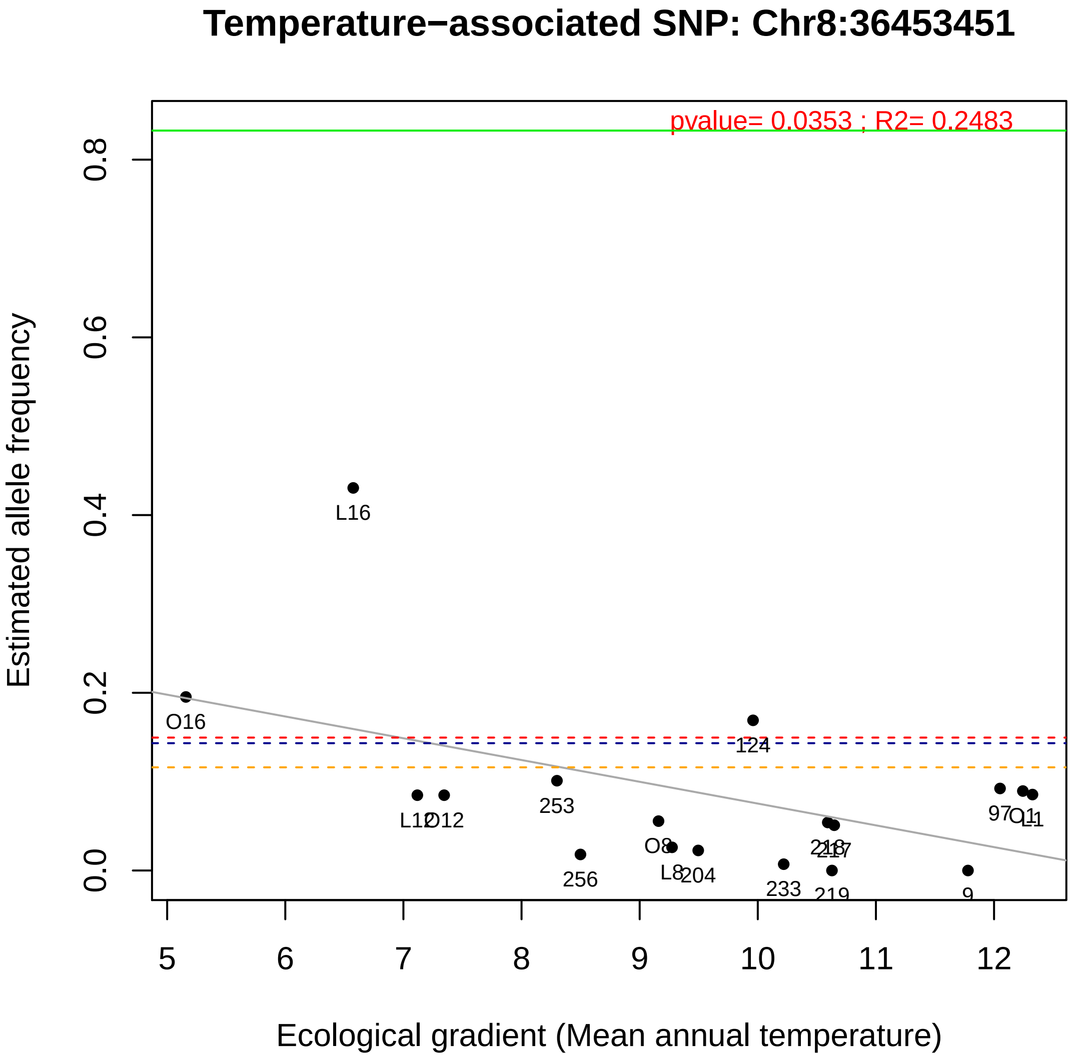
Temperature−associated SNP: Chr8:36453451
pvalue= 0.0353 ; R2= 0.2483
Estimated allele frequency
0.0
0.2
0.4
0.6
0.8
Ecological gradient (Mean annual temperature)
5
6
7
8
9
10
11
12
L16
O16
124
253
256
O8
L8
204
233
219
9
97
L1
O12

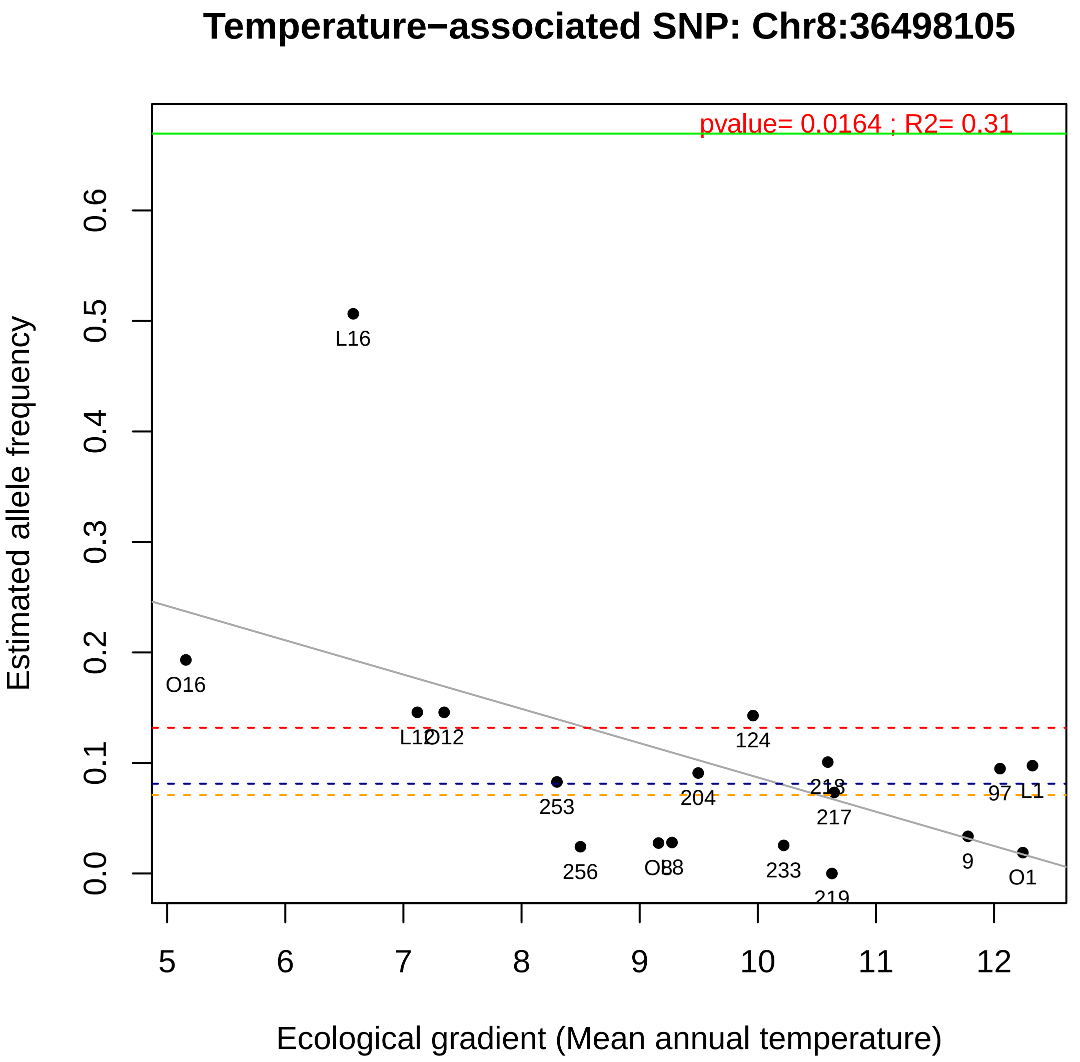
Temperature−associated SNP: Chr8:36498105
pvalue= 0.0164 ; R2= 0.31
Estimated allele frequency
0.0
0.1
0.2
0.3
0.4
0.5
0.6
5
6
7
8
9
10
11
12
Ecological gradient (Mean annual temperature)
L16
O16
L12
O12
124
253
204
218
217
97
L1
256
233
219
9
O1

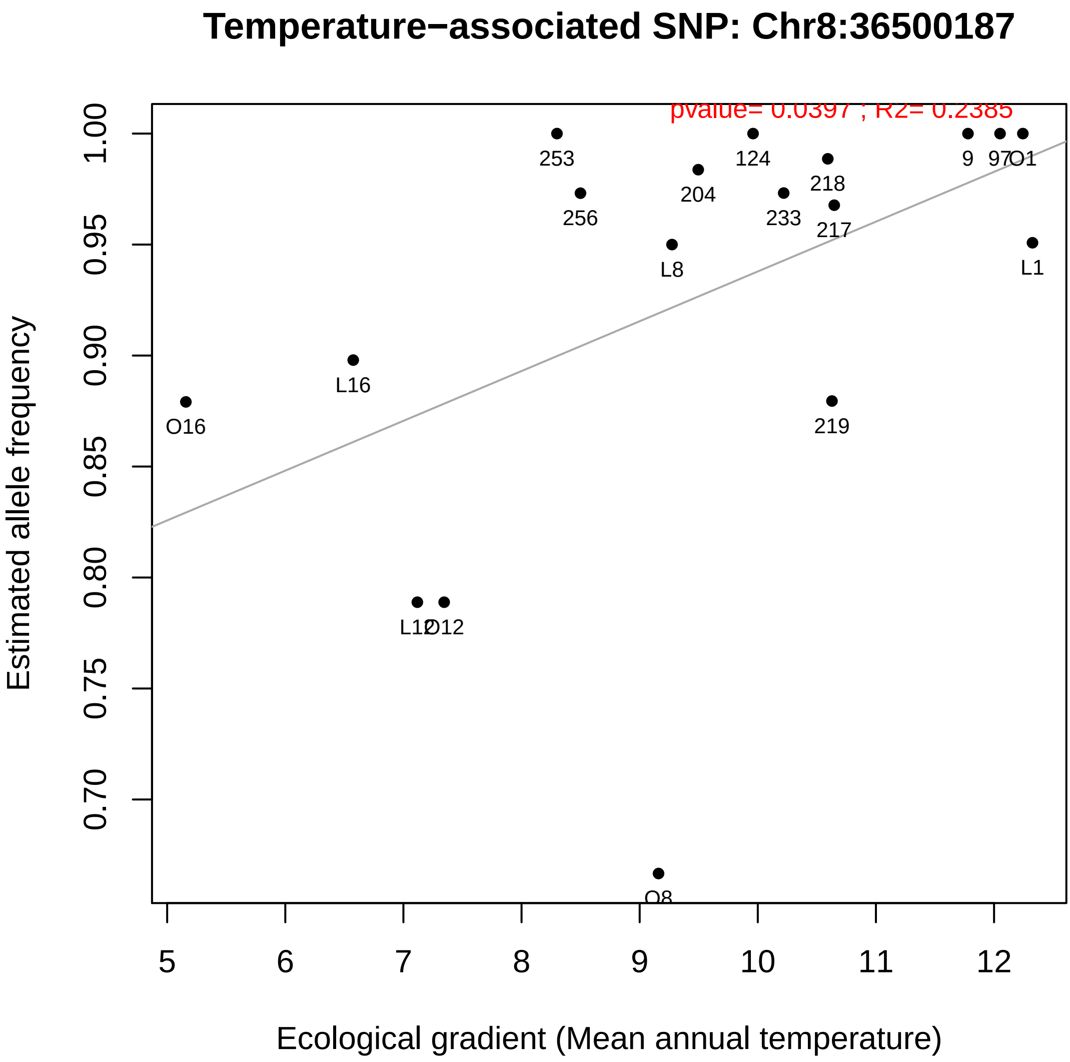

Temperature−associated SNP: Chr8:36500187
pvalue= 0.0397 ; R2= 0.2385
Estimated allele frequency
Ecological gradient (Mean annual temperature)
1.00
0.95
0.90
0.85
0.80
0.75
0.70
5
6
7
8
9
10
11
12
253
124
9
97
O1
204
218
256
233
217
L8
L1
L16
O16
219
L12
O12
O8

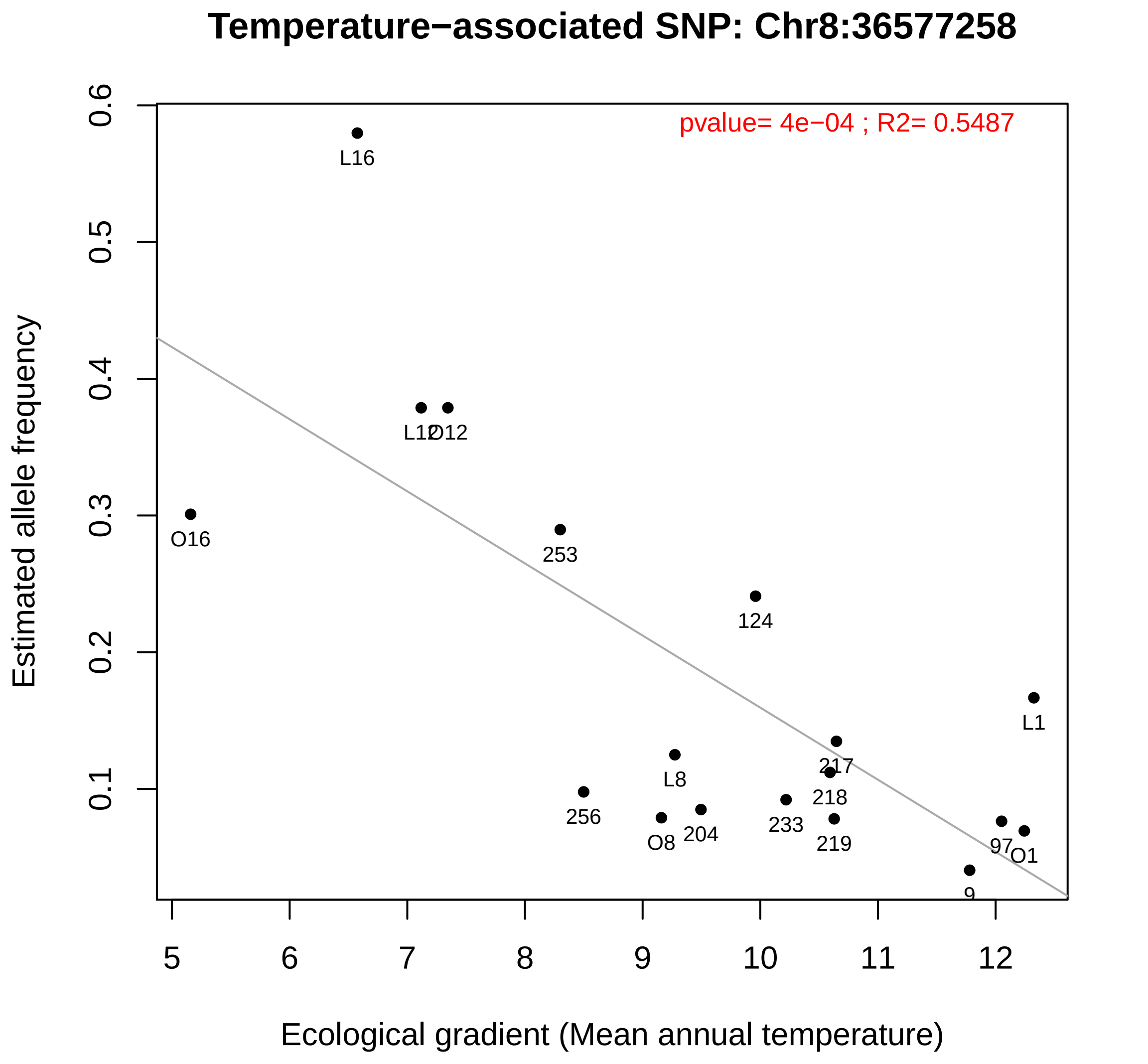
Temperature−associated SNP: Chr8:36577258
pvalue= 4e−04 ; R2= 0.5487
Estimated allele frequency
0.6
0.5
0.4
0.3
0.2
0.1
L16
L12
O12
O16
253
124
L1
217
218
L8
256
233
219
204
O8
97
O1
9
5
6
7
8
9
10
11
12
Ecological gradient (Mean annual temperature)

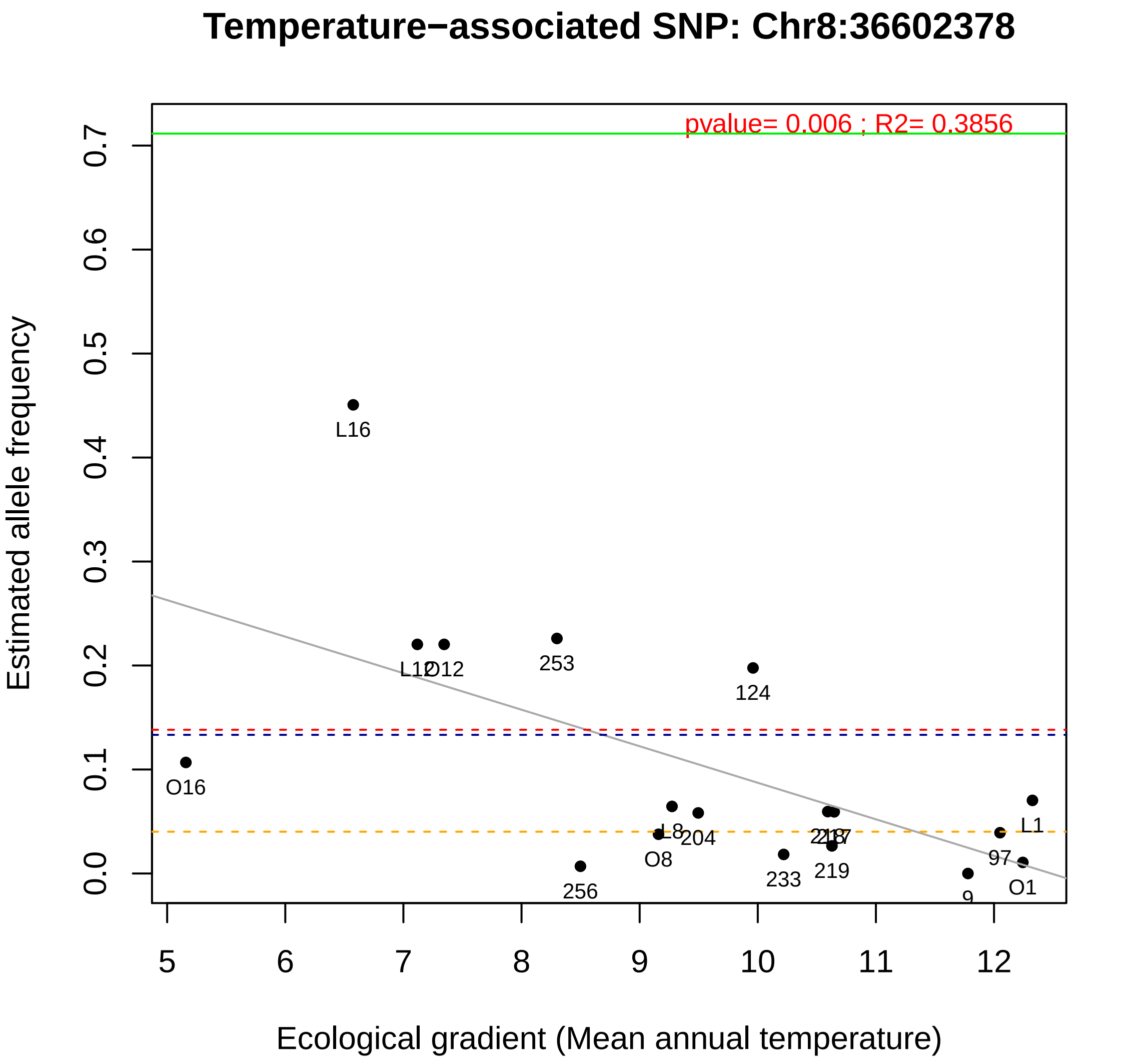
Temperature−associated SNP: Chr8:36602378
pvalue= 0.006 ; R2= 0.3856
Estimated allele frequency
0.0
0.1
0.2
0.3
0.4
0.5
0.6
0.7
Ecological gradient (Mean annual temperature)
5
6
7
8
9
10
11
12
L16
253
124
O16
L8
204
O8
256
233
219
L1
97
O1
9

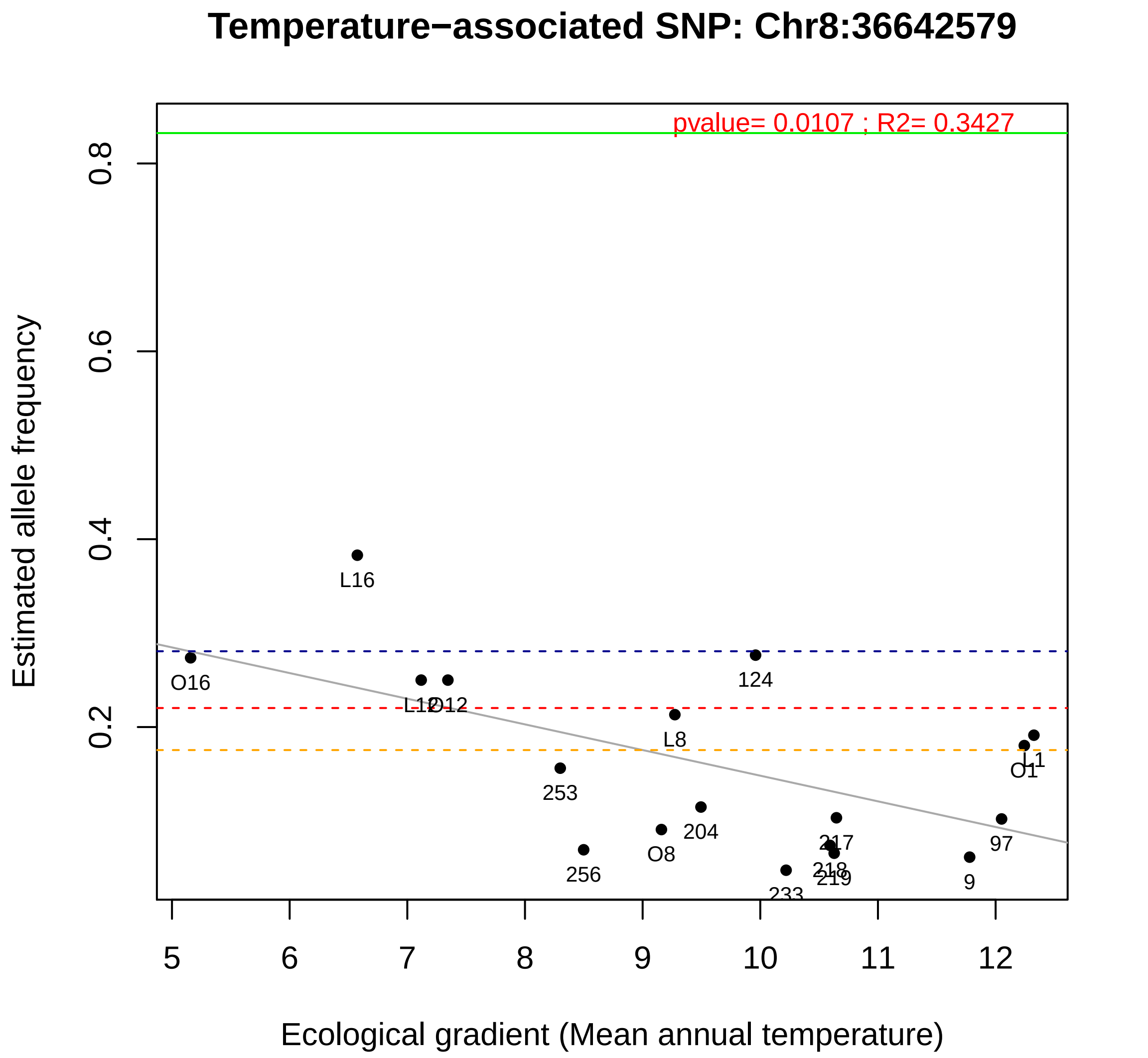
Temperature−associated SNP: Chr8:36642579
pvalue= 0.0107 ; R2= 0.3427
Estimated allele frequency
0.2
0.4
0.6
0.8
5
6
7
8
9
10
11
12
Ecological gradient (Mean annual temperature)
L16
O16
124
L12
O12
L8
253
204
O8
256
217
218
219
233
9
97
O1
L1

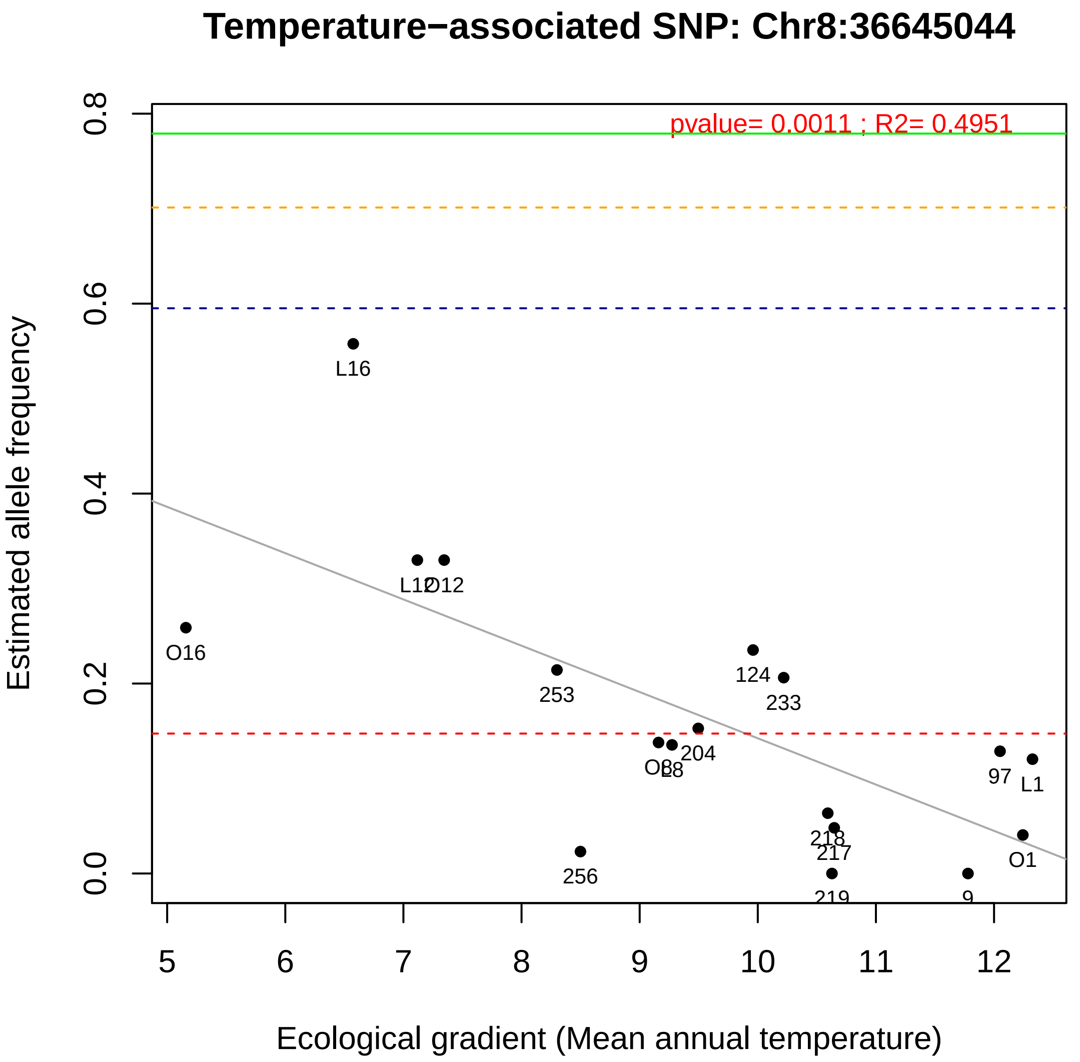

Temperature−associated SNP: Chr8:36645044
pvalue= 0.0011 ; R2= 0.4951
Estimated allele frequency
0.0
0.2
0.4
0.6
0.8
L16
L12
O12
O16
253
124
233
204
O8
L8
97
L1
218
217
O1
256
219
9
5
6
7
8
9
10
11
12
Ecological gradient (Mean annual temperature)

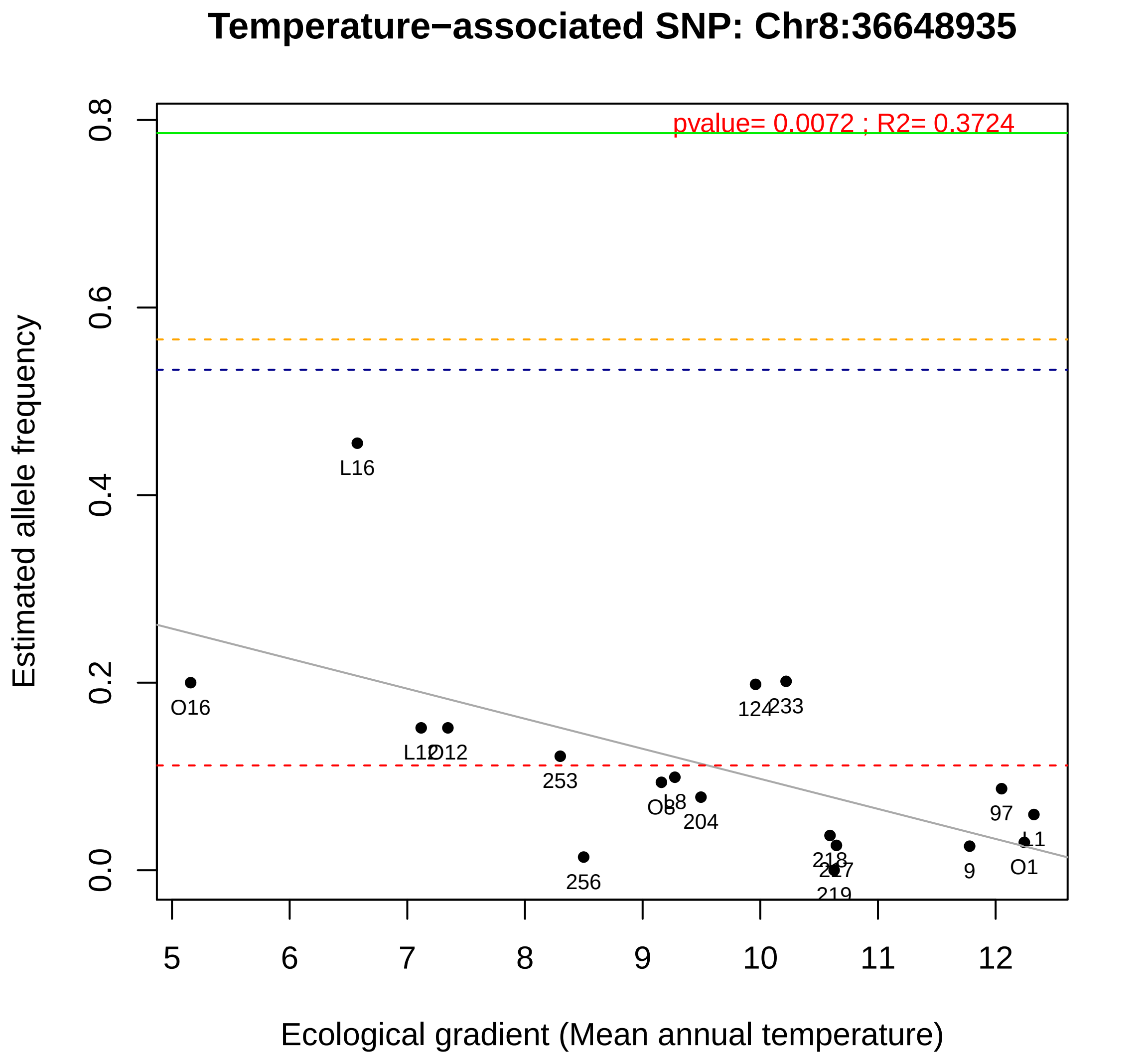
Temperature−associated SNP: Chr8:36648935
pvalue= 0.0072 ; R2= 0.3724
Estimated allele frequency
0.0
0.2
0.4
0.6
0.8
5
6
7
8
9
10
11
12
Ecological gradient (Mean annual temperature)
L16
O16
L12
O12
253
256
O8
L8
204
124
233
218
217
219
9
97
L1
O1

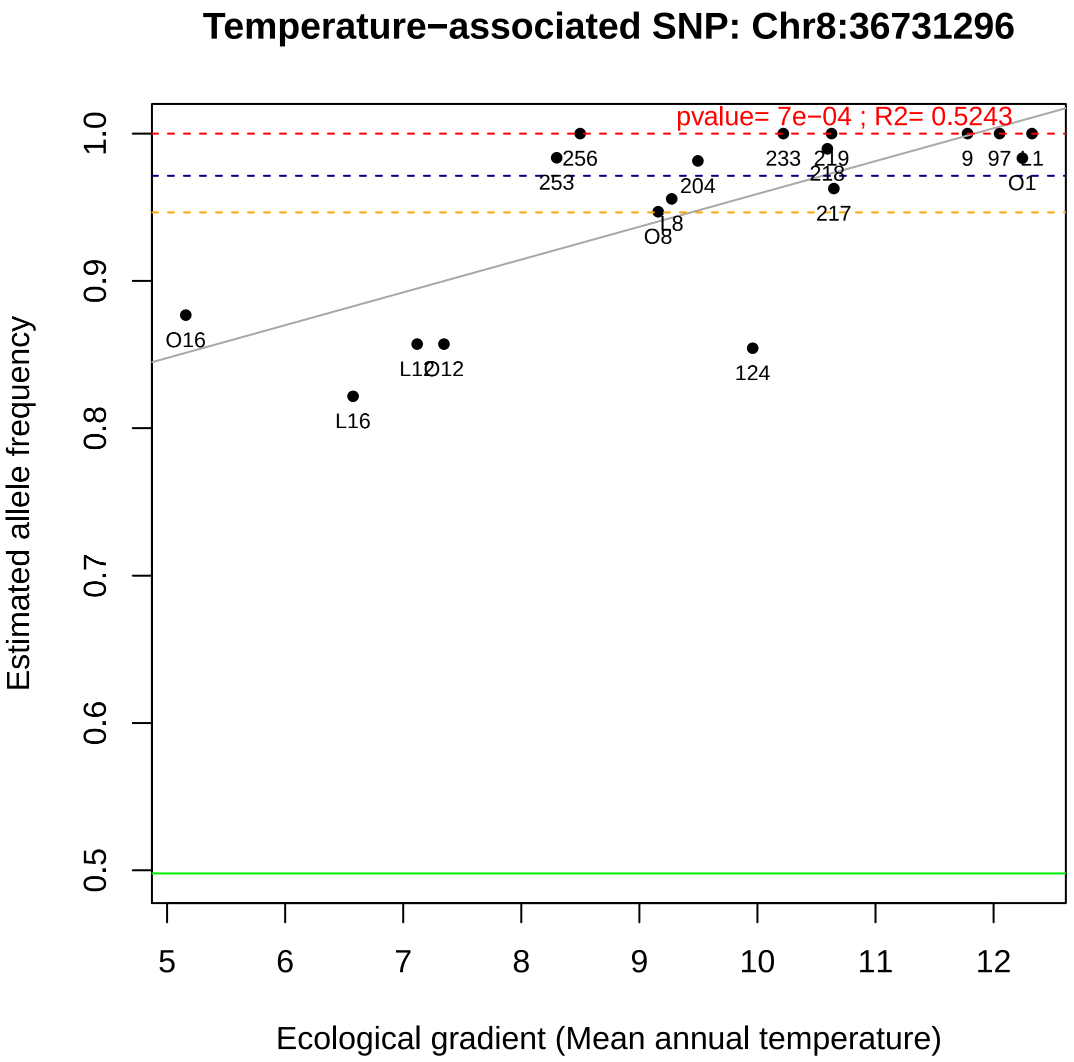

Temperature−associated SNP: Chr8:36731296
pvalue= 7e−04 ; R2= 0.5243
Estimated allele frequency
Ecological gradient (Mean annual temperature)
1.0
0.9
0.8
0.7
0.6
0.5
5
6
7
8
9
10
11
12
256
253
204
L8
O8
233
219
218
217
9
97
L1
O1
O16
L16
L12
O12
124

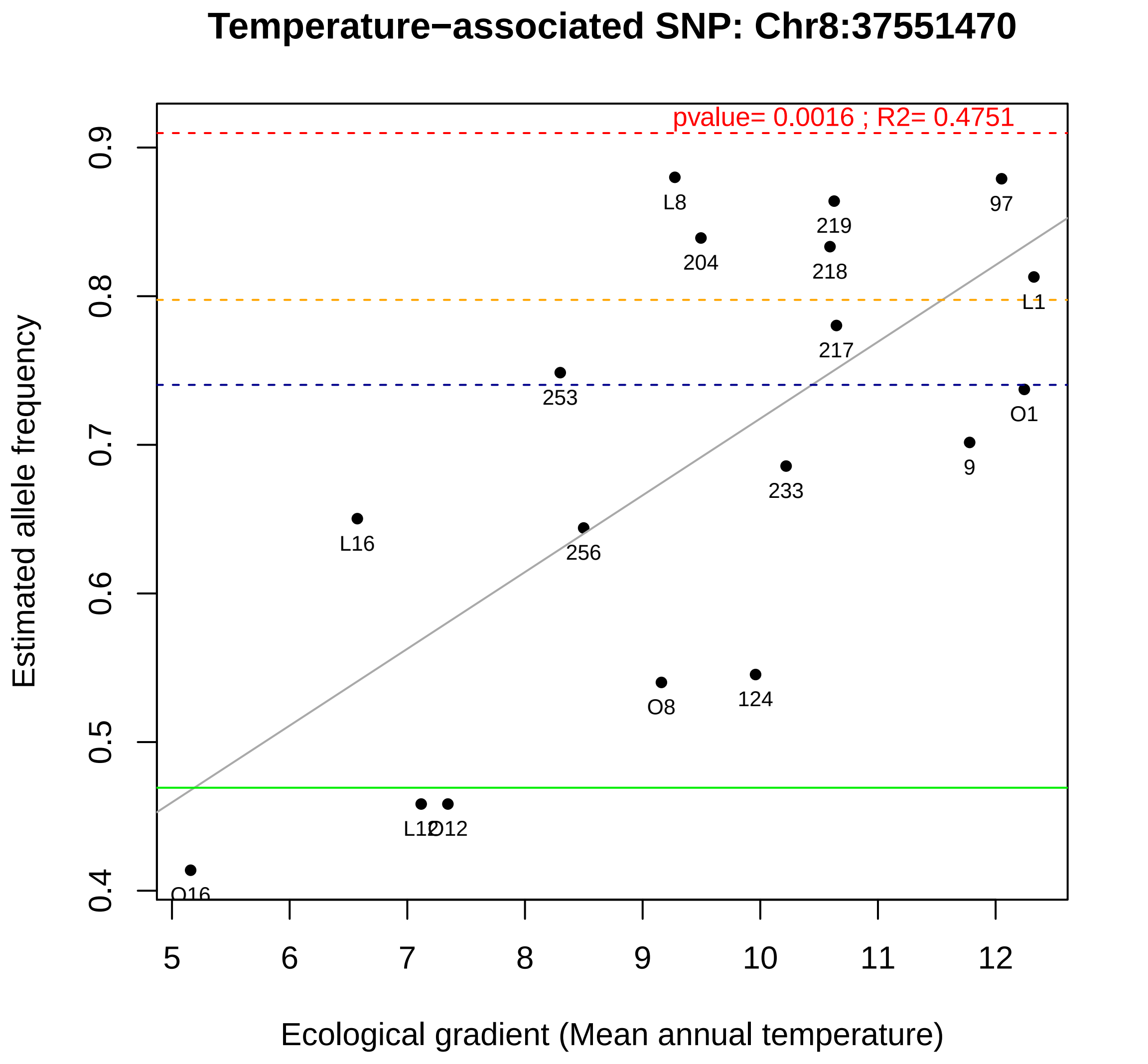
Temperature−associated SNP: Chr8:37551470
pvalue= 0.0016 ; R2= 0.4751
Estimated allele frequency
0.4
0.5
0.6
0.7
0.8
0.9
5
6
7
8
9
10
11
12
Ecological gradient (Mean annual temperature)
L8
97
219
204
218
L1
217
253
O1
9
233
L16
256
O8
124
L12
O12
O16

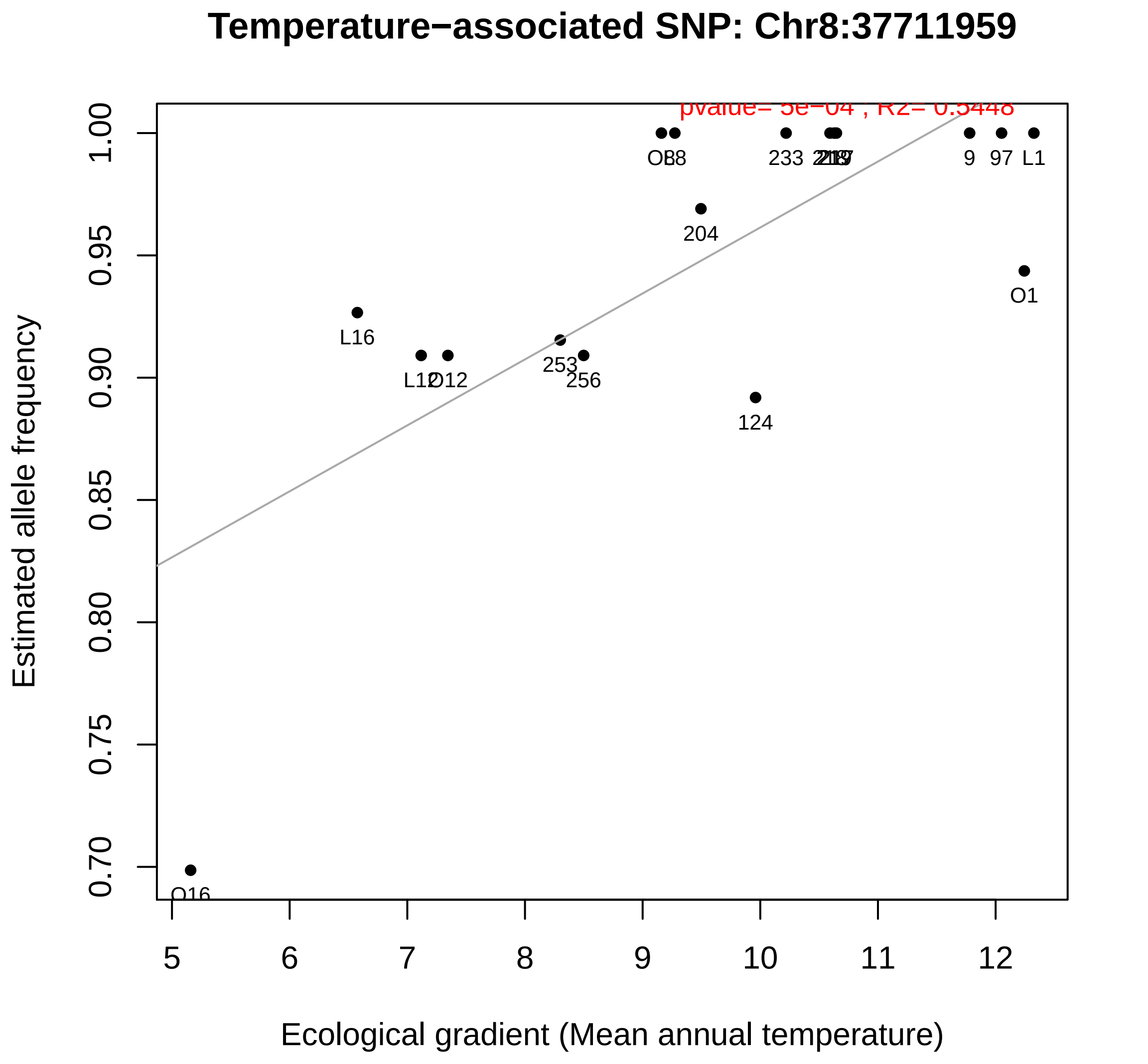

Temperature−associated SNP: Chr8:37711959
pvalue= 5e−04 , R2= 0.5448
Estimated allele frequency
Ecological gradient (Mean annual temperature)
0.70
0.75
0.80
0.85
0.90
0.95
1.00
5
6
7
8
9
10
11
12
O16
L16
L12
O12
253
256
204
124
233
9
97
L1
O1

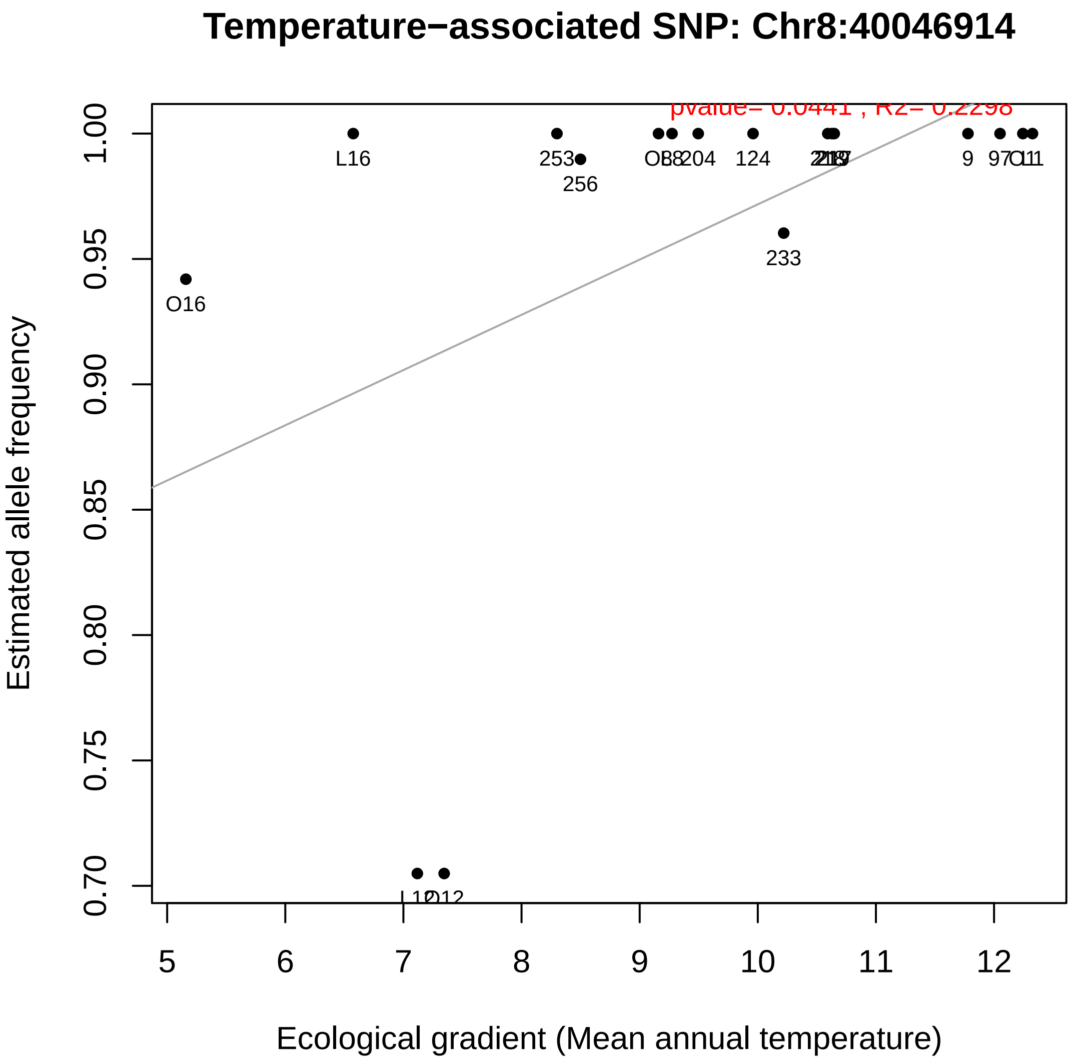

Temperature−associated SNP: Chr8:40046914
pvalue= 0.0441 , R2= 0.2298
Estimated allele frequency
Ecological gradient (Mean annual temperature)
1.00
0.95
0.90
0.85
0.80
0.75
0.70
5
6
7
8
9
10
11
12
L16
253
256
204
124
233
O16
9

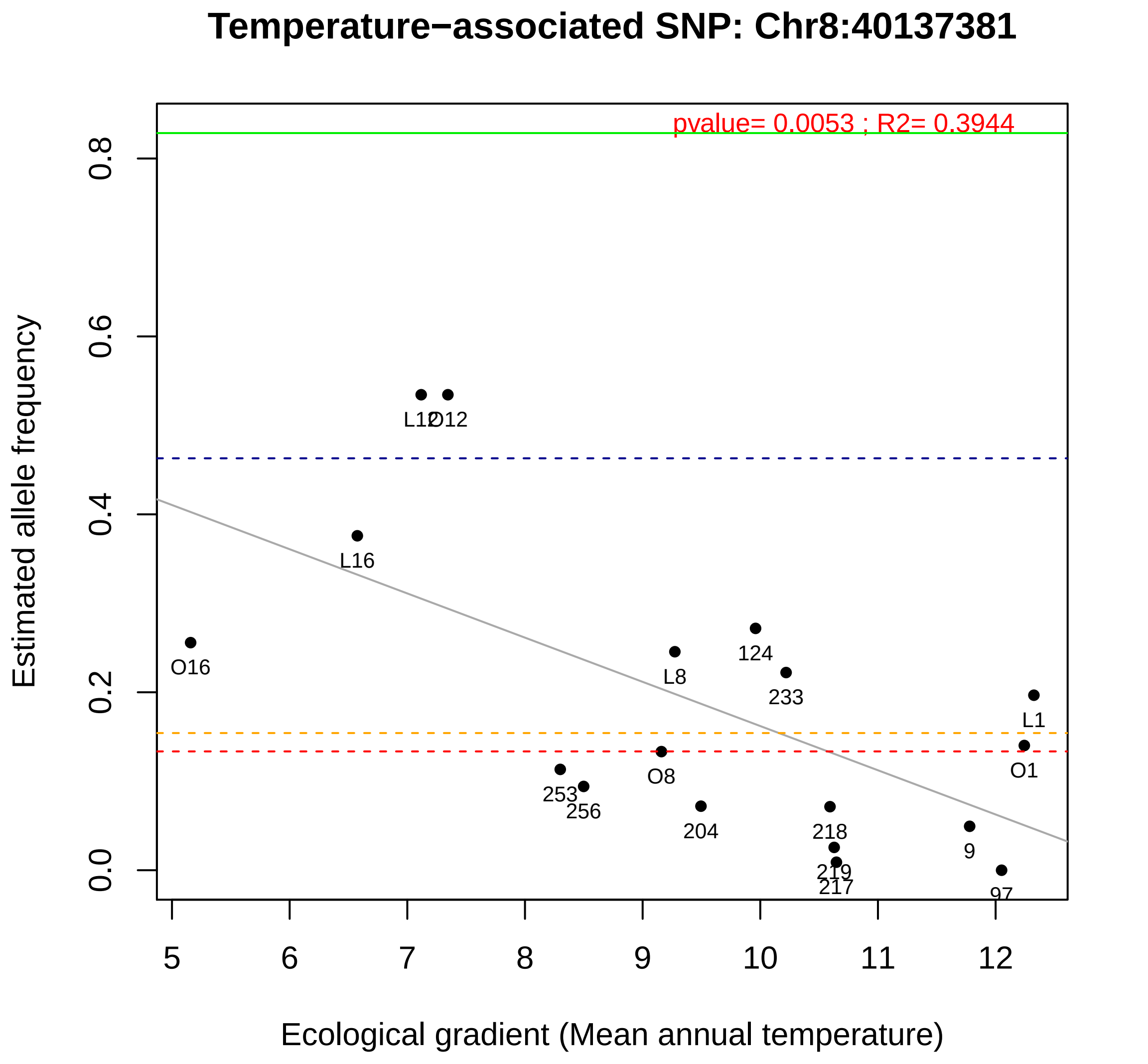

Temperature−associated SNP: Chr8:40137381
pvalue= 0.0053 ; R2= 0.3944
Estimated allele frequency
0.0
0.2
0.4
0.6
0.8
Ecological gradient (Mean annual temperature)
5
6
7
8
9
10
11
12
L12
O12
L16
O16
L8
124
233
L1
O1
O8
253
256
204
218
219
217
9
97

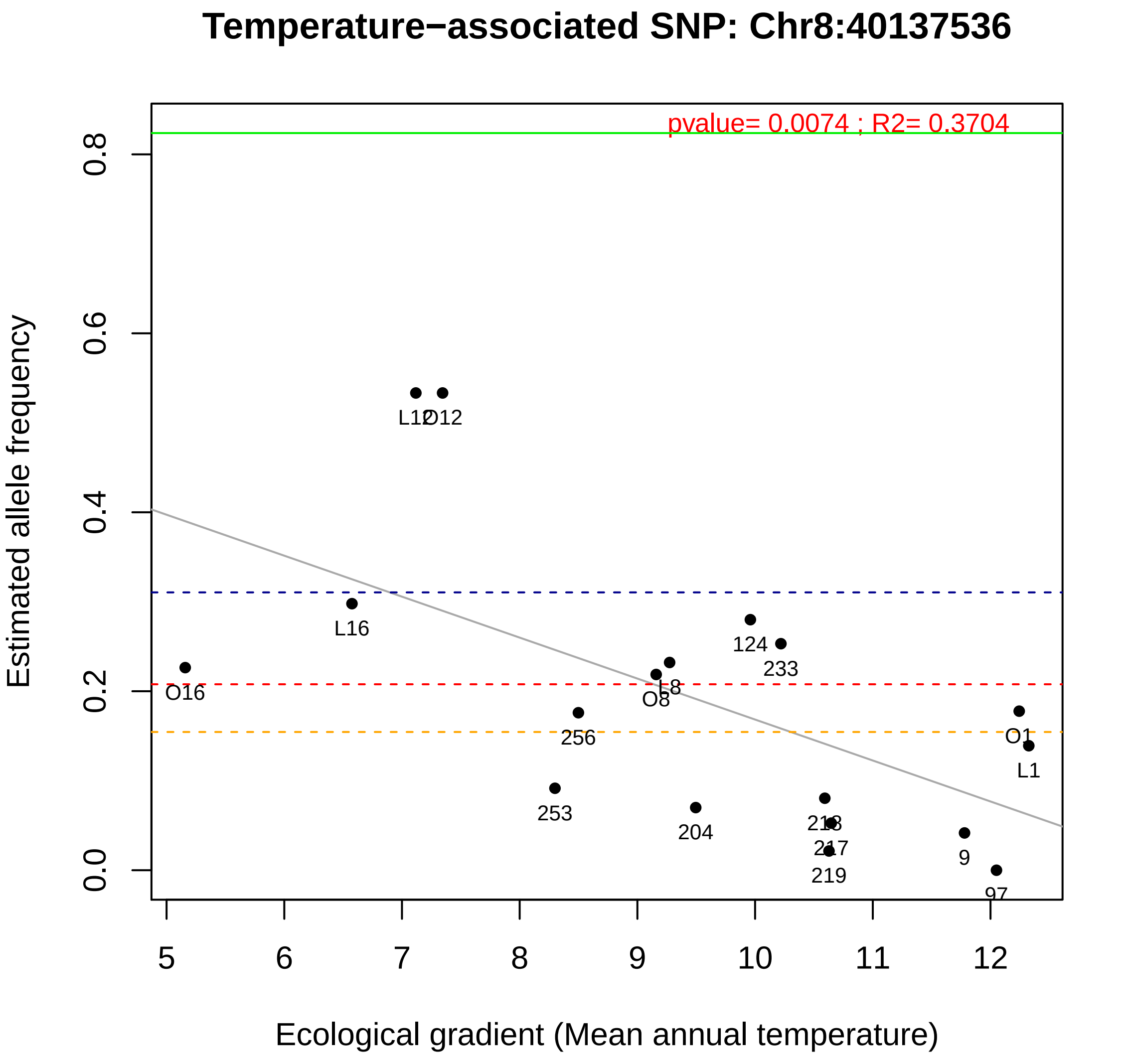
Temperature−associated SNP: Chr8:40137536
pvalue= 0.0074 ; R2= 0.3704
Estimated allele frequency
0.0
0.2
0.4
0.6
0.8
L12
O12
L16
O16
124
233
L8
O8
256
O1
L1
253
204
218
217
219
9
97
5
6
7
8
9
10
11
12
Ecological gradient (Mean annual temperature)

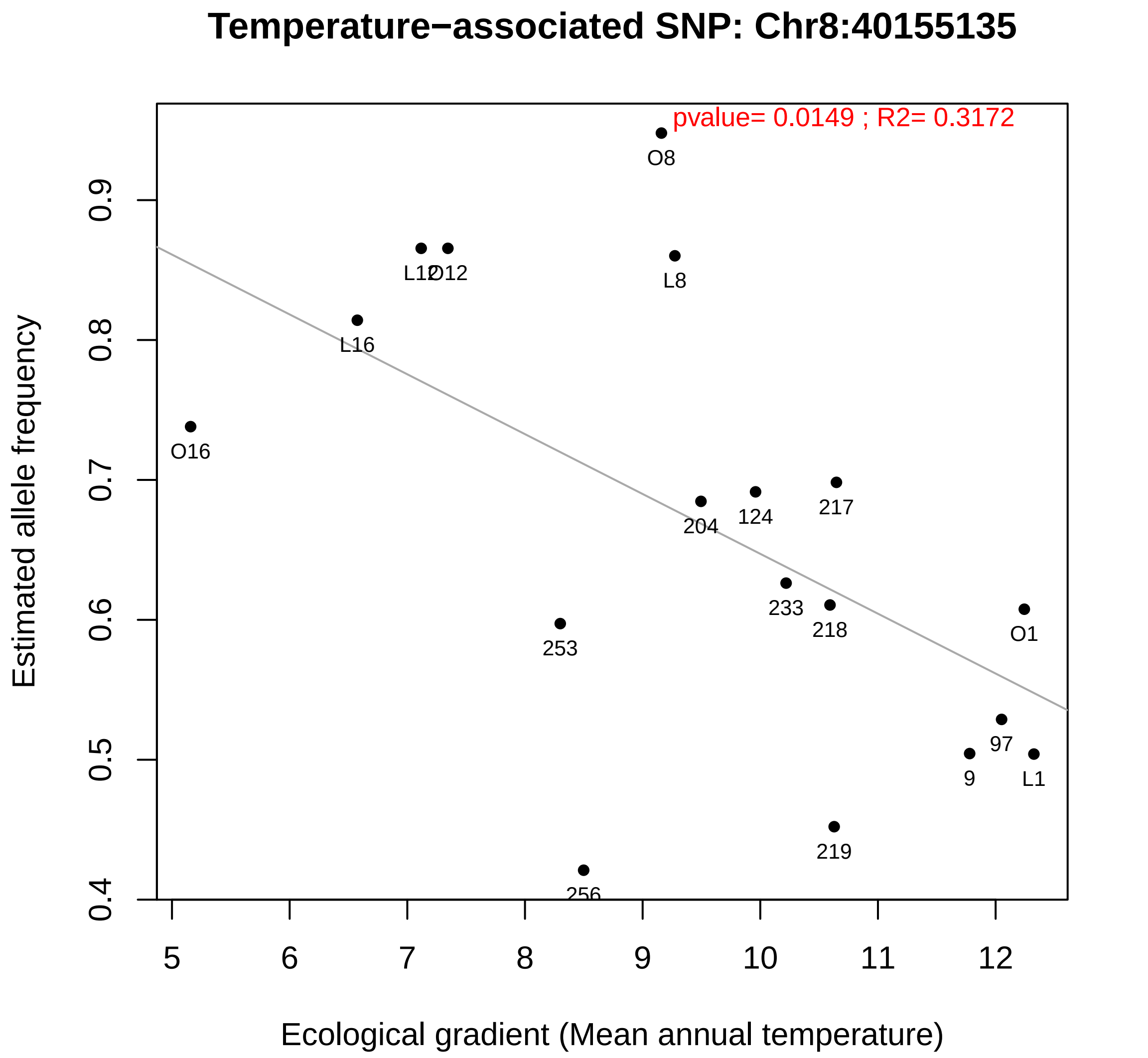
Temperature−associated SNP: Chr8:40155135
pvalue= 0.0149 ; R2= 0.3172
O8
L12
O12
L8
L16
O16
204
124
217
233
218
O1
253
97
9
L1
219
256
0.4
0.5
0.6
0.7
0.8
0.9
5
6
7
8
9
10
11
12
Estimated allele frequency
Ecological gradient (Mean annual temperature)

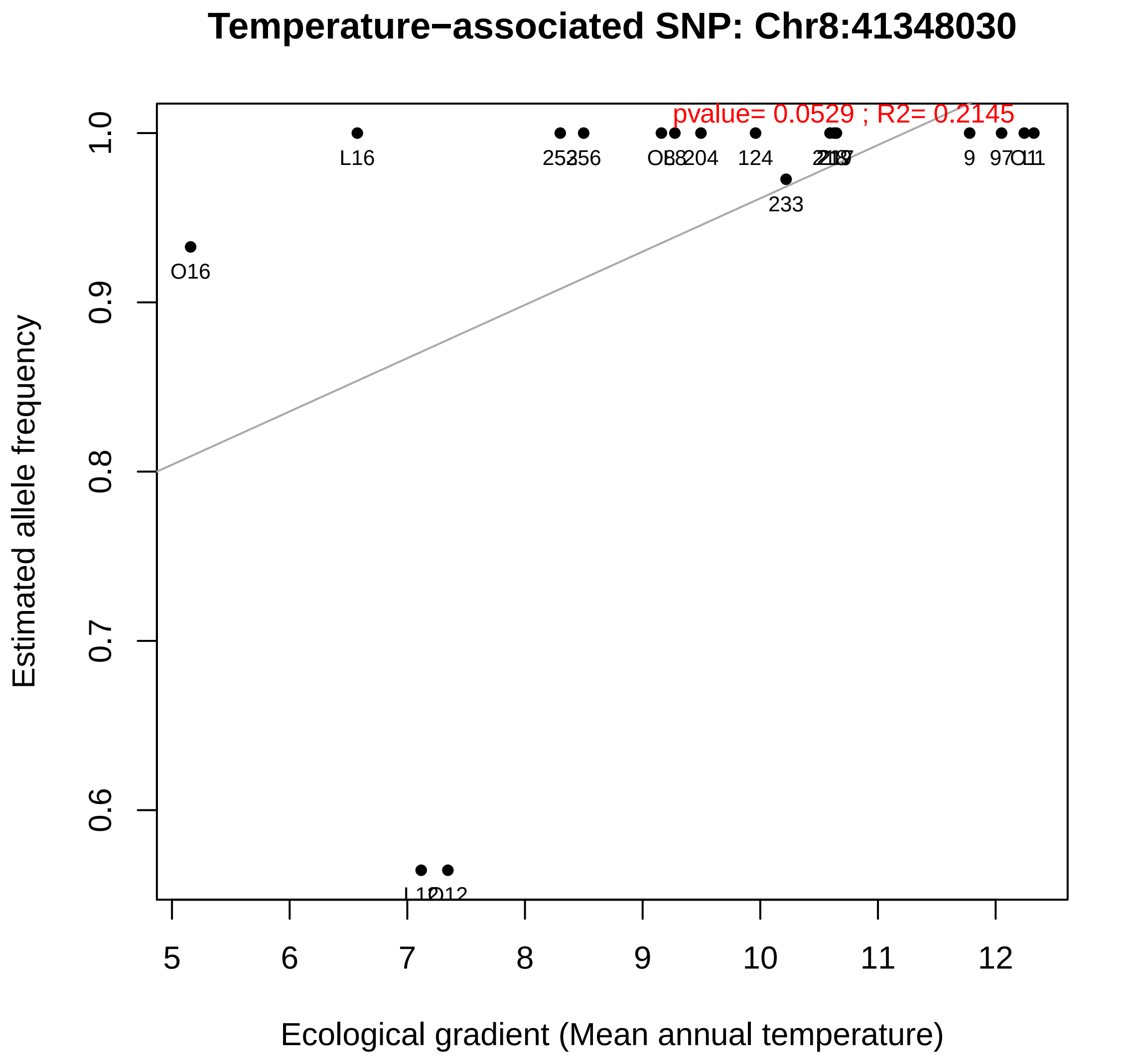
Temperature−associated SNP: Chr8:41348030
pvalue= 0.0529 ; R2= 0.2145
Estimated allele frequency
Ecological gradient (Mean annual temperature)
0.6
0.7
0.8
0.9
1.0
5
6
7
8
9
10
11
12
L16
O16
124
233
9
O12

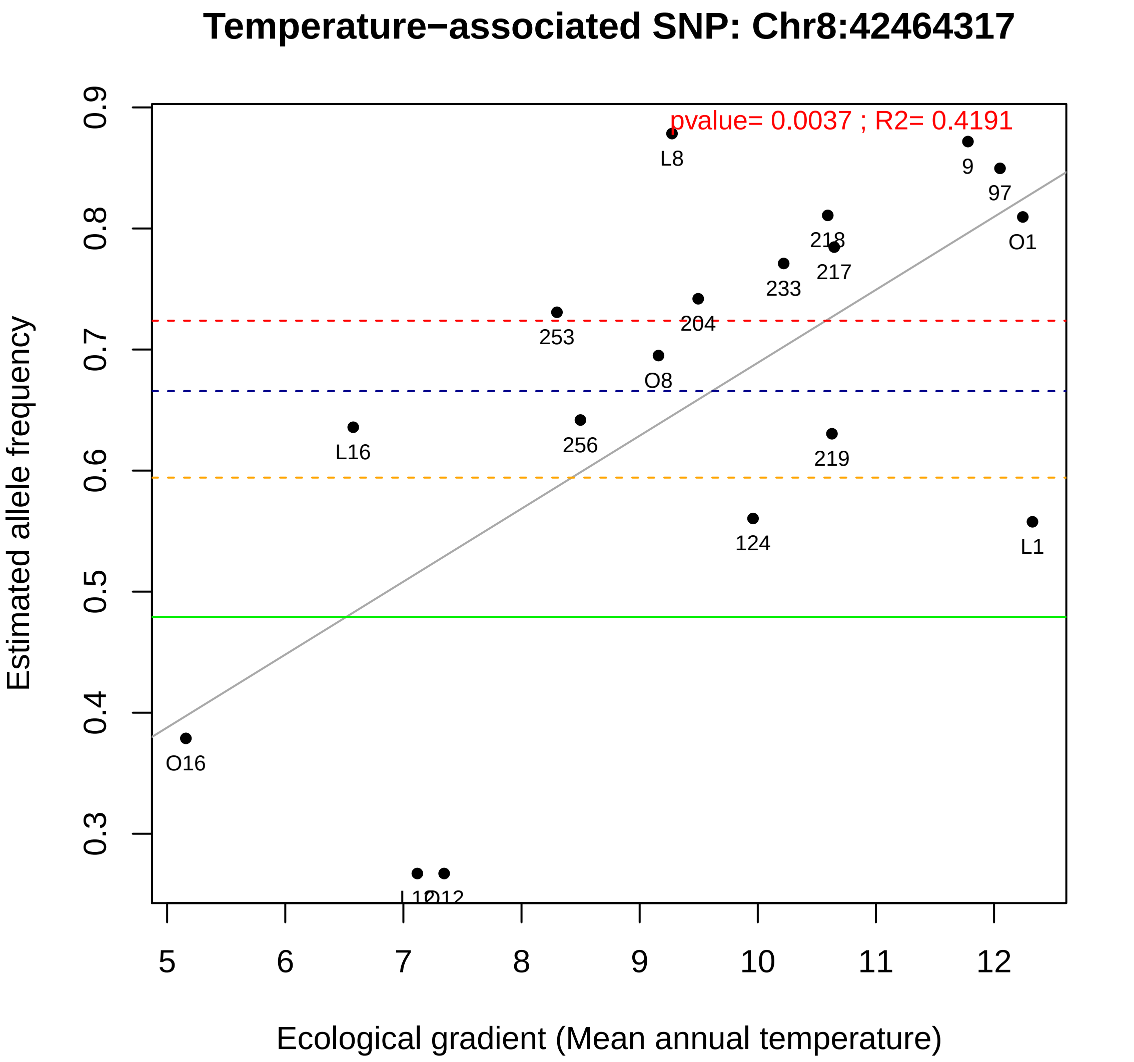
Temperature−associated SNP: Chr8:42464317
pvalue= 0.0037 ; R2= 0.4191
Estimated allele frequency
0.3
0.4
0.5
0.6
0.7
0.8
0.9
5
6
7
8
9
10
11
12
Ecological gradient (Mean annual temperature)
L8
9
97
O1
218
217
233
204
253
O8
L16
256
219
124
L1
O16
O12

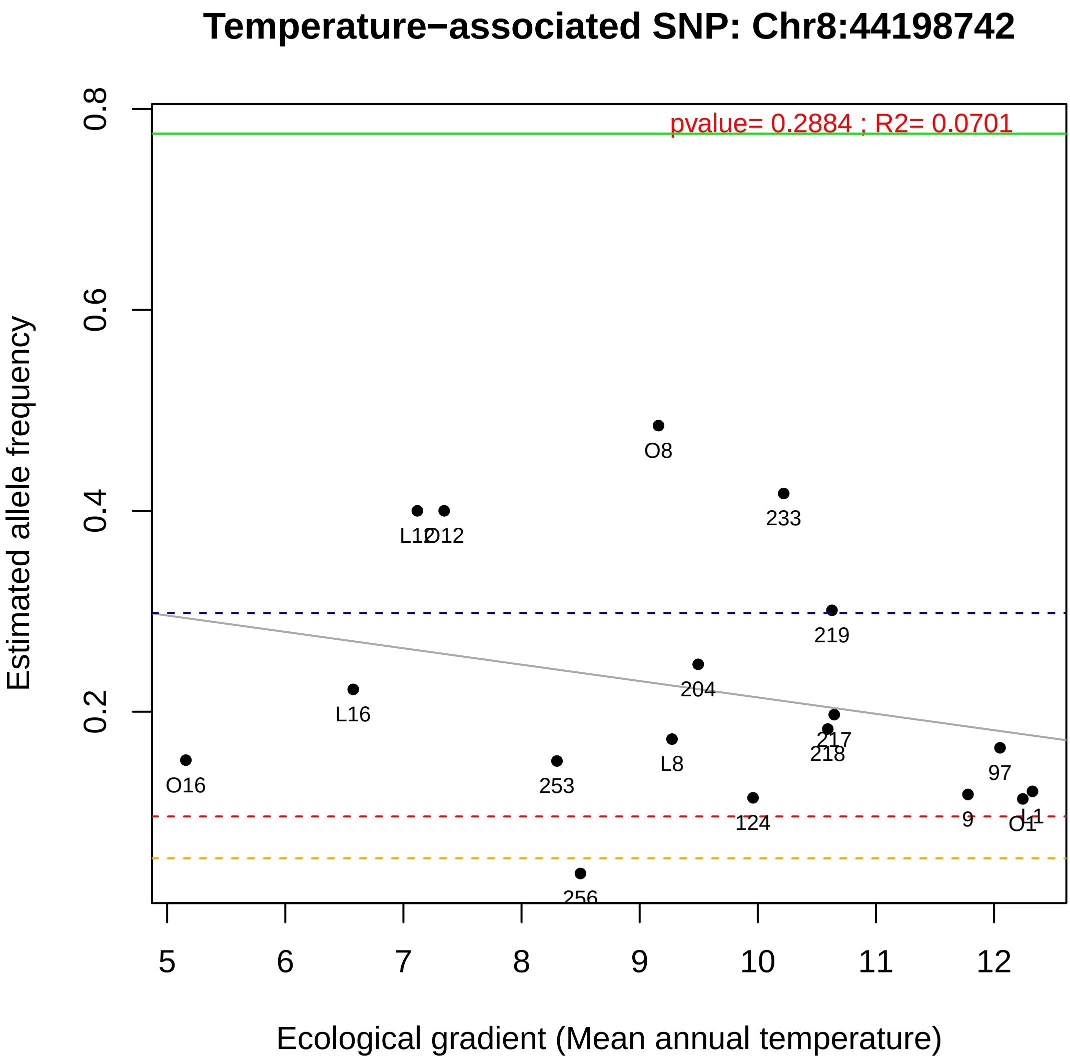

Temperature−associated SNP: Chr8:44198742
pvalue= 0.2884 ; R2= 0.0701
Estimated allele frequency
0.8
0.6
0.4
0.2
O8
233
L12
O12
219
204
L16
217
218
L8
O16
253
97
124
9
L1
O1
256
5
6
7
8
9
10
11
12
Ecological gradient (Mean annual temperature)

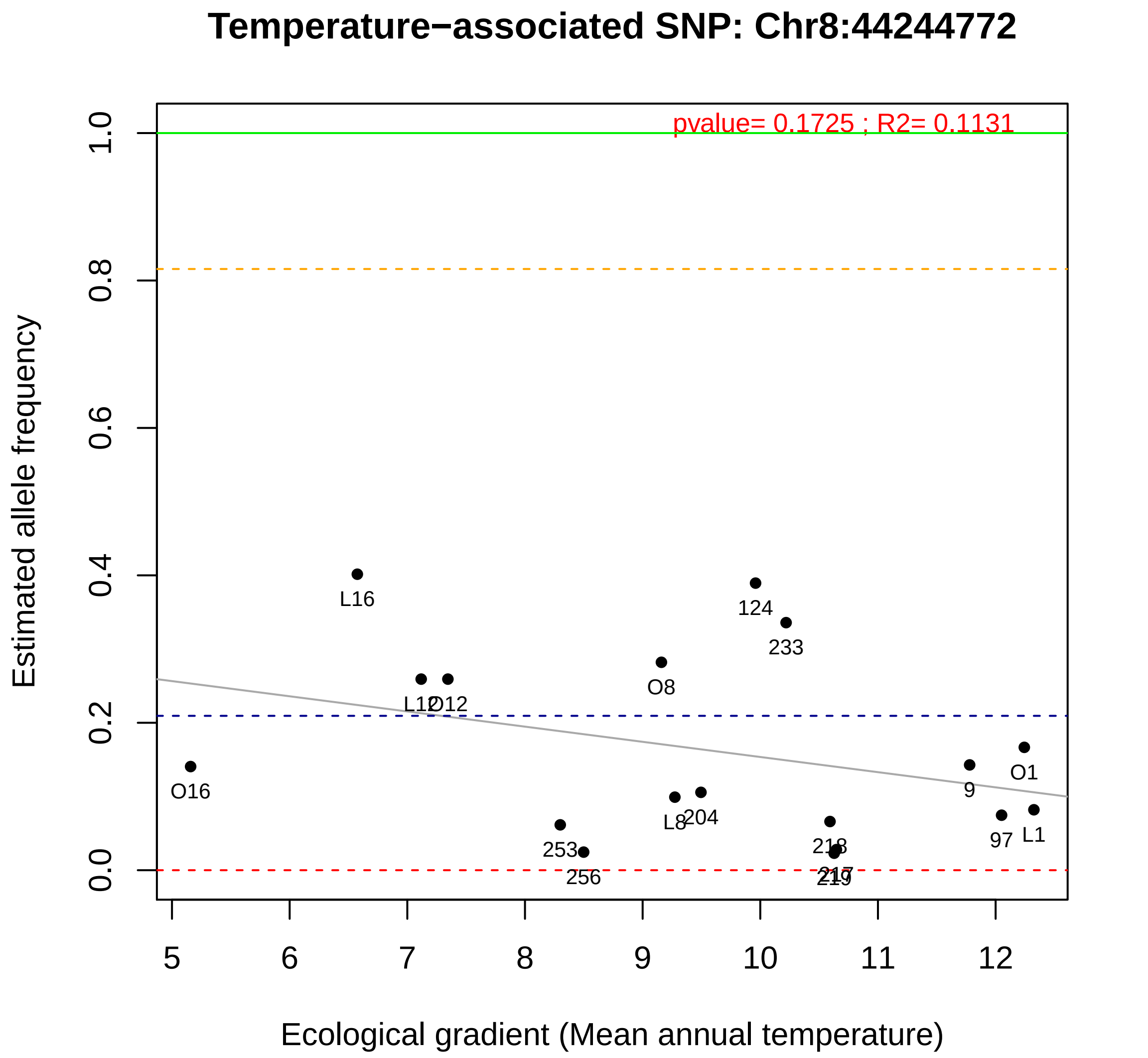

Temperature−associated SNP: Chr8:44244772
pvalue= 0.1725 ; R2= 0.1131
Estimated allele frequency
Ecological gradient (Mean annual temperature)
0.0
0.2
0.4
0.6
0.8
1.0
5
6
7
8
9
10
11
12
L16
124
233
O8
L12
O12
O16
O1
9
L8
204
253
256
218
217
219
97
L1

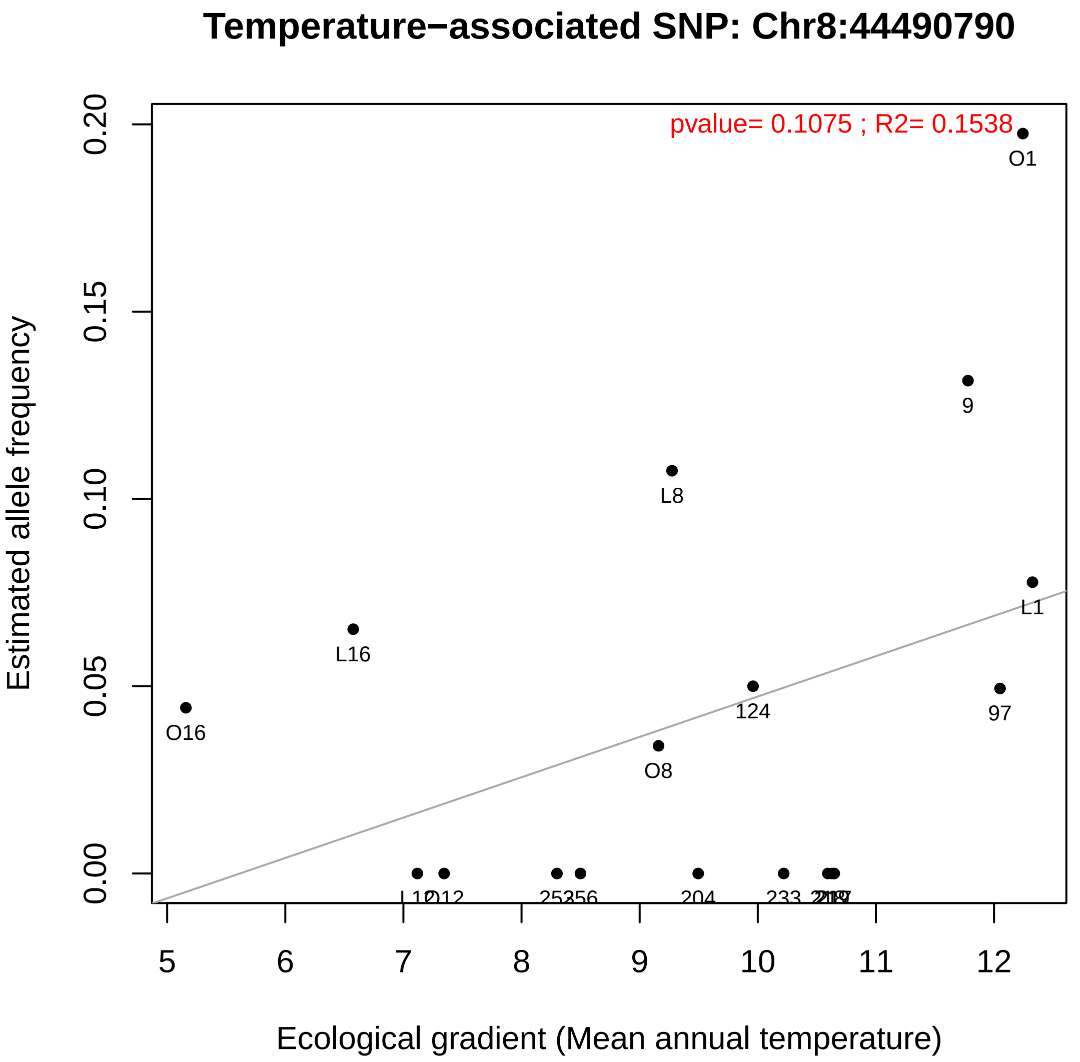
Temperature−associated SNP: Chr8:44490790
pvalue= 0.1075 ; R2= 0.1538
Estimated allele frequency
0.00
0.05
0.10
0.15
0.20
O1
9
L8
L1
L16
124
97
O16
O8
204
5
6
7
8
9
10
11
12
Ecological gradient (Mean annual temperature)

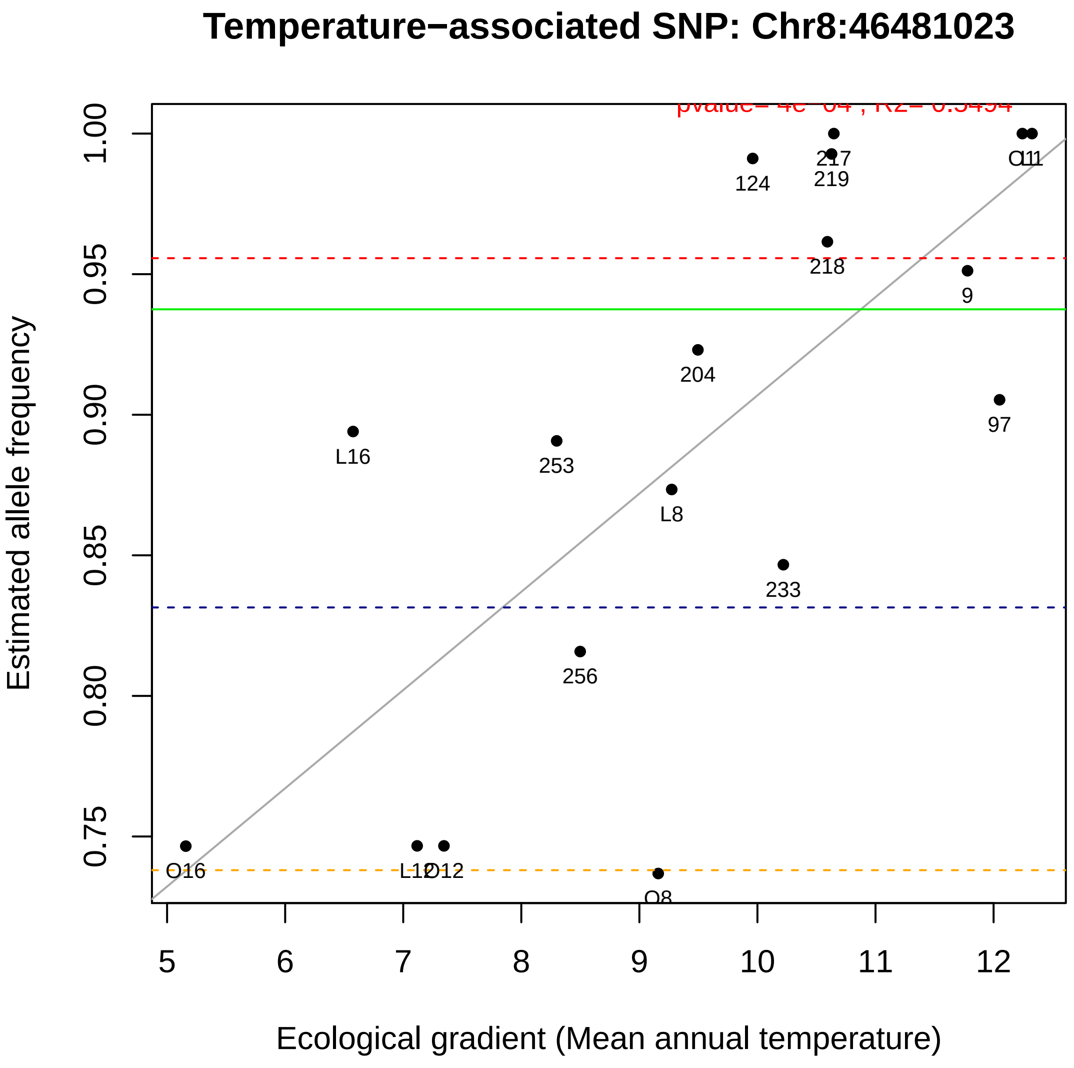
Temperature−associated SNP: Chr8:46481023
pvalue= 4e−04 ; R2= 0.3494
Estimated allele frequency
Ecological gradient (Mean annual temperature)
1.00
0.95
0.90
0.85
0.80
0.75
5
6
7
8
9
10
11
12
124
217
219
O11
218
9
204
97
L16
253
L8
233
256
O16
L12
O12
O8

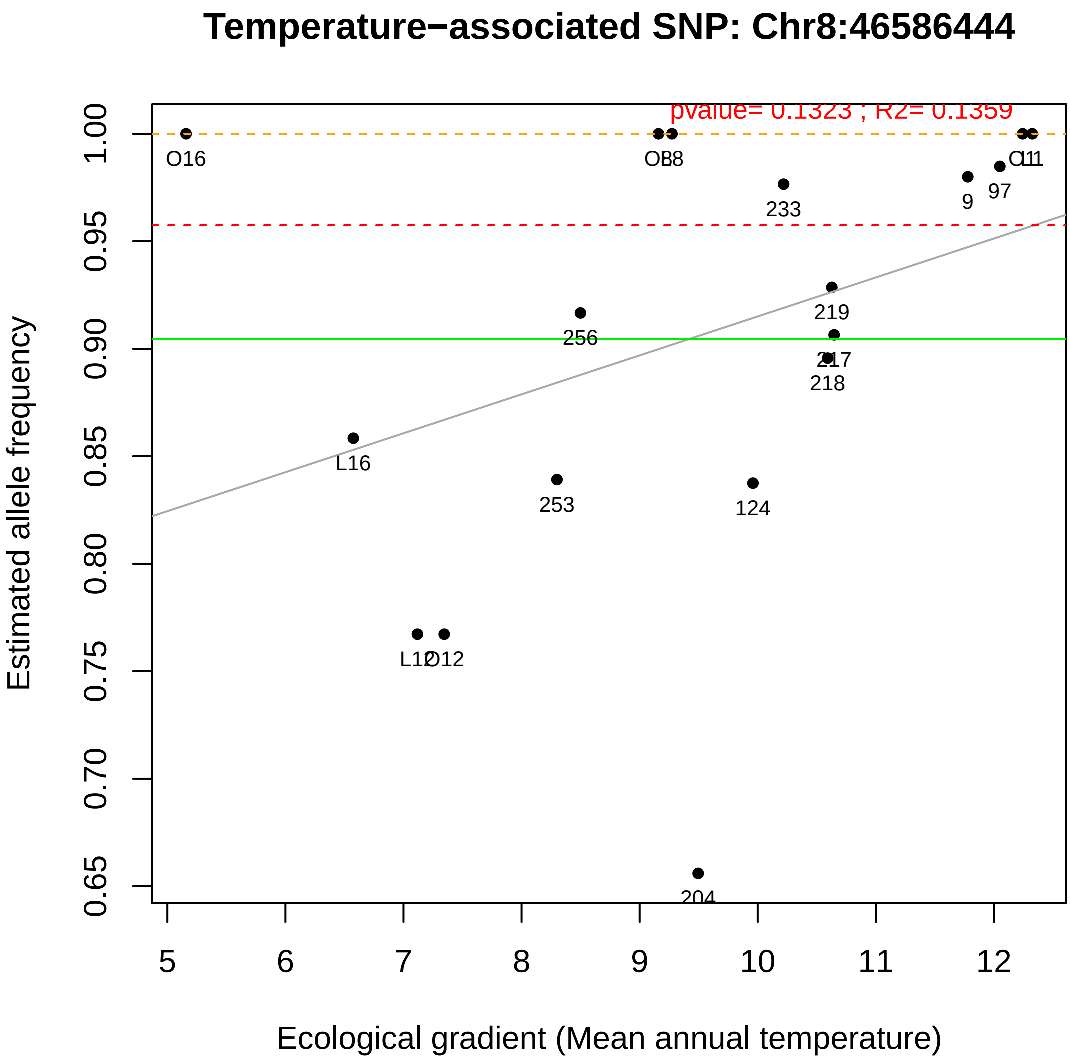

Temperature−associated SNP: Chr8:46586444
pvalue= 0.1323 ; R2= 0.1359
Estimated allele frequency
Ecological gradient (Mean annual temperature)
0.65
0.70
0.75
0.80
0.85
0.90
0.95
1.00
5
6
7
8
9
10
11
12
O16
L16
L12
O12
253
256
204
124
233
219
217
218
9
97

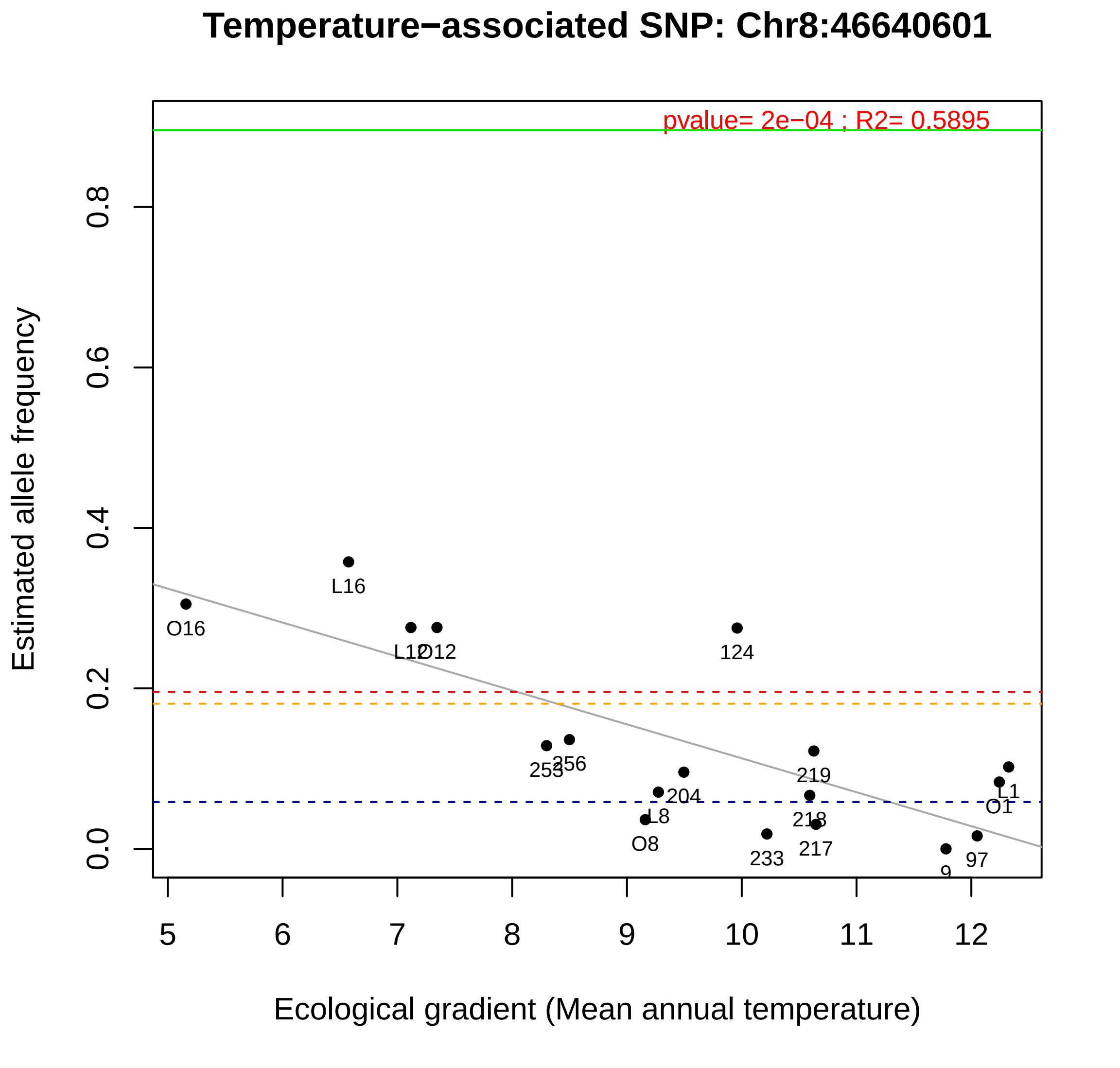

Temperature−associated SNP: Chr8:46640601
pvalue= 2e−04 ; R2= 0.5895
Estimated allele frequency
0.0
0.2
0.4
0.6
0.8
5
6
7
8
9
10
11
12
Ecological gradient (Mean annual temperature)
L16
O16
L12
O12
124
253
256
219
204
L8
O8
218
217
233
L1
O1
97
9

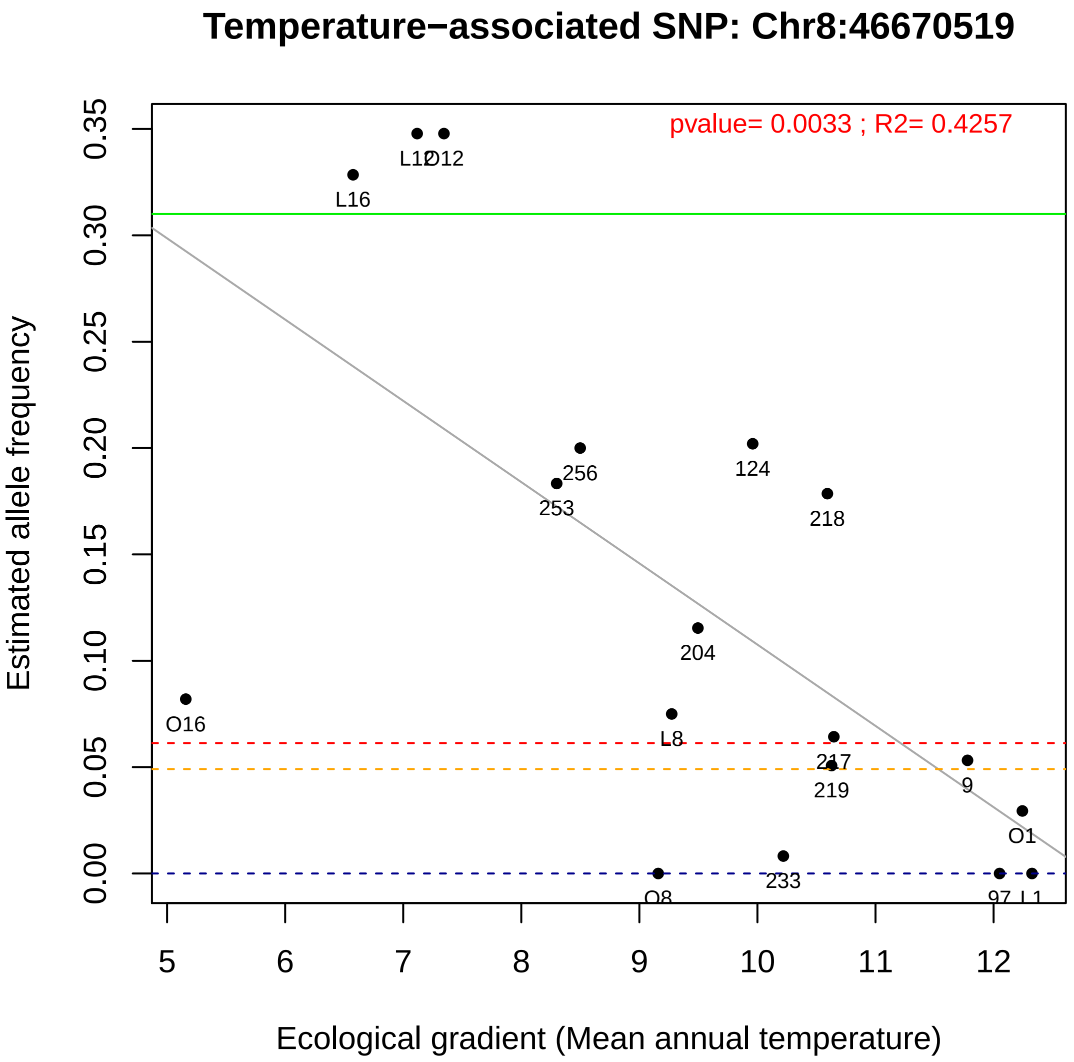

Temperature−associated SNP: Chr8:46670519
pvalue= 0.0033 ; R2= 0.4257
Estimated allele frequency
Ecological gradient (Mean annual temperature)
0.00
0.05
0.10
0.15
0.20
0.25
0.30
0.35
5
6
7
8
9
10
11
12
L16
L12
O12
256
253
124
218
204
O16
L8
217
219
9
O1
233

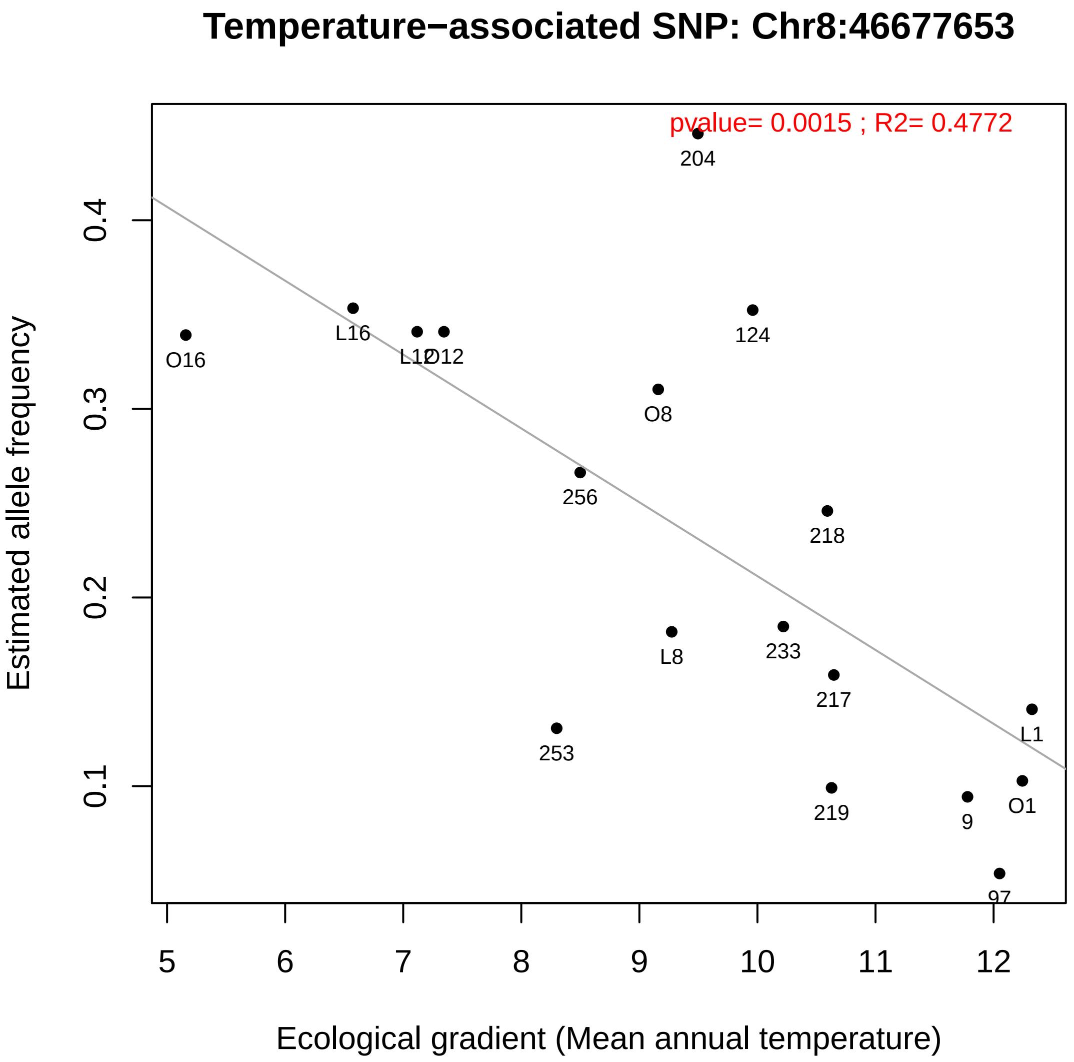

Temperature−associated SNP: Chr8:46677653
pvalue= 0.0015 ; R2= 0.4772
204
124
O16
L16
L12
O12
O8
256
218
L8
233
217
L1
253
219
O1
9
97
0.1
0.2
0.3
0.4
5
6
7
8
9
10
11
12
Estimated allele frequency
Ecological gradient (Mean annual temperature)

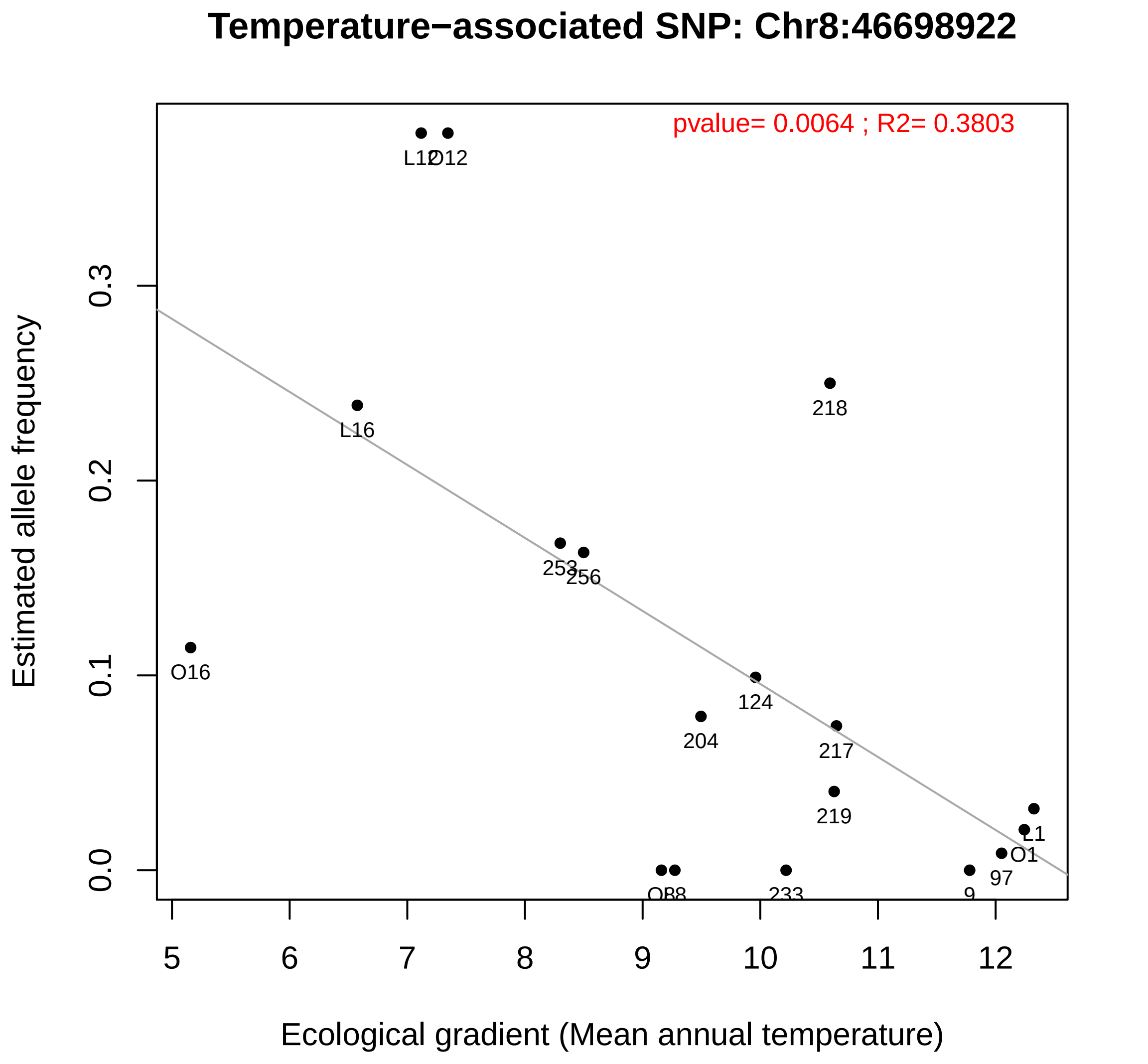

Temperature−associated SNP: Chr8:46698922
pvalue= 0.0064 ; R2= 0.3803
Estimated allele frequency
0.0
0.1
0.2
0.3
5
6
7
8
9
10
11
12
Ecological gradient (Mean annual temperature)
L12
O12
L16
218
253
256
O16
124
204
217
219
L1
O1
97
233

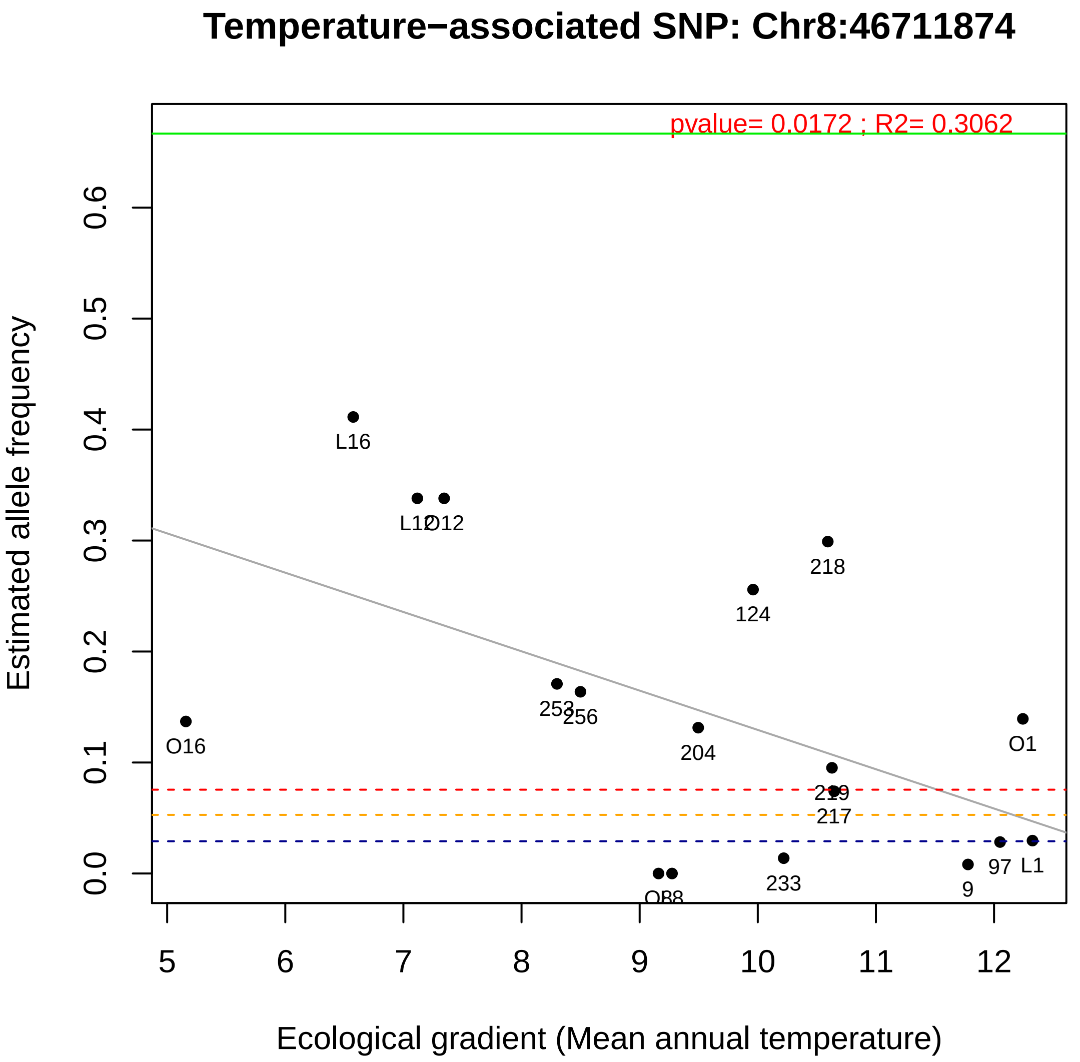
Temperature−associated SNP: Chr8:46711874
pvalue= 0.0172 ; R2= 0.3062
Estimated allele frequency
Ecological gradient (Mean annual temperature)
0.0
0.1
0.2
0.3
0.4
0.5
0.6
5
6
7
8
9
10
11
12
L16
L12
O12
218
124
253
256
O16
204
O1
219
217
233
97
L1
9

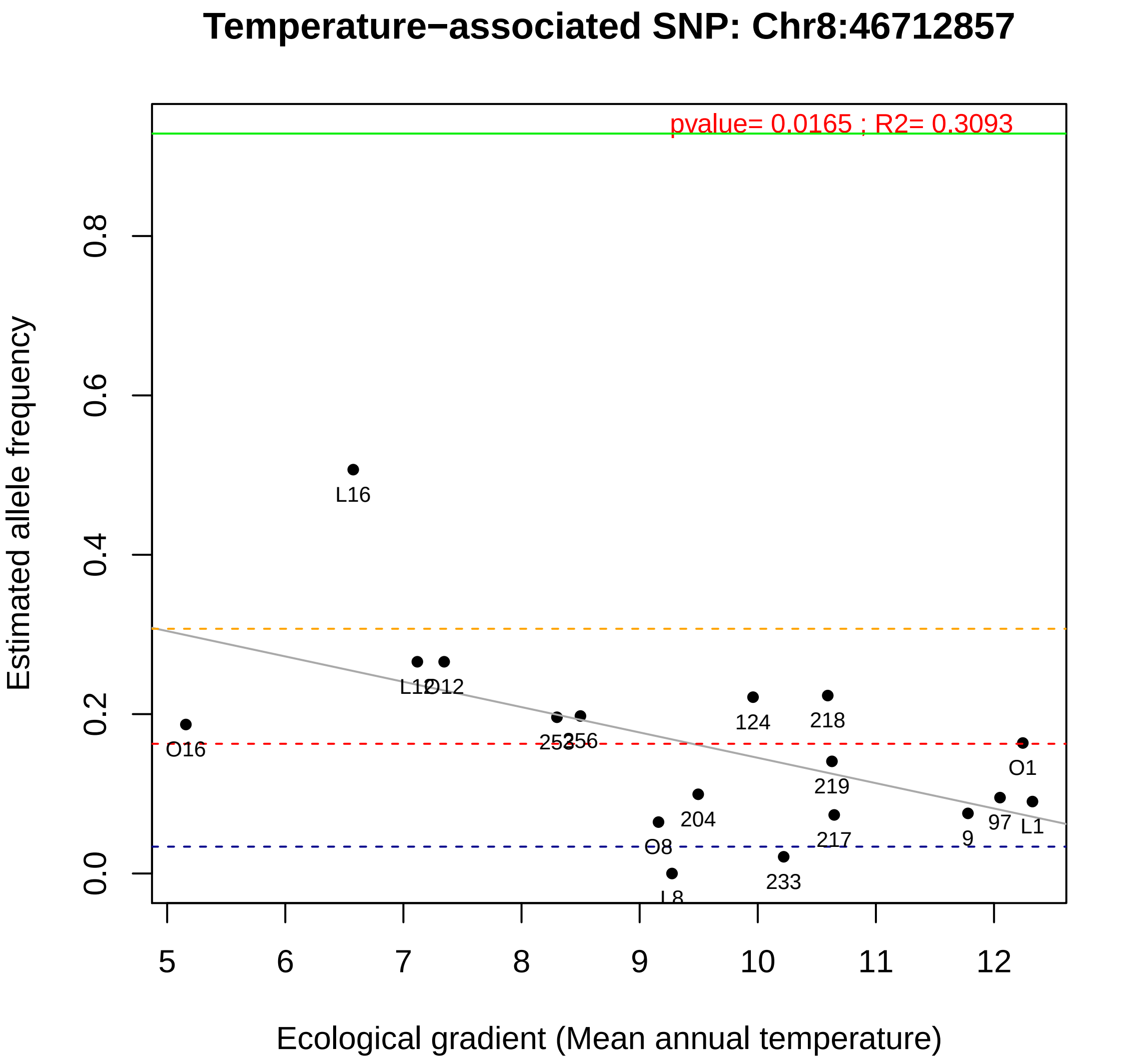
Temperature−associated SNP: Chr8:46712857
pvalue= 0.0165 ; R2= 0.3093
Estimated allele frequency
0.0
0.2
0.4
0.6
0.8
5
6
7
8
9
10
11
12
Ecological gradient (Mean annual temperature)
L16
L12
O12
O16
124
218
219
O1
204
O8
233
217
9
97
L1

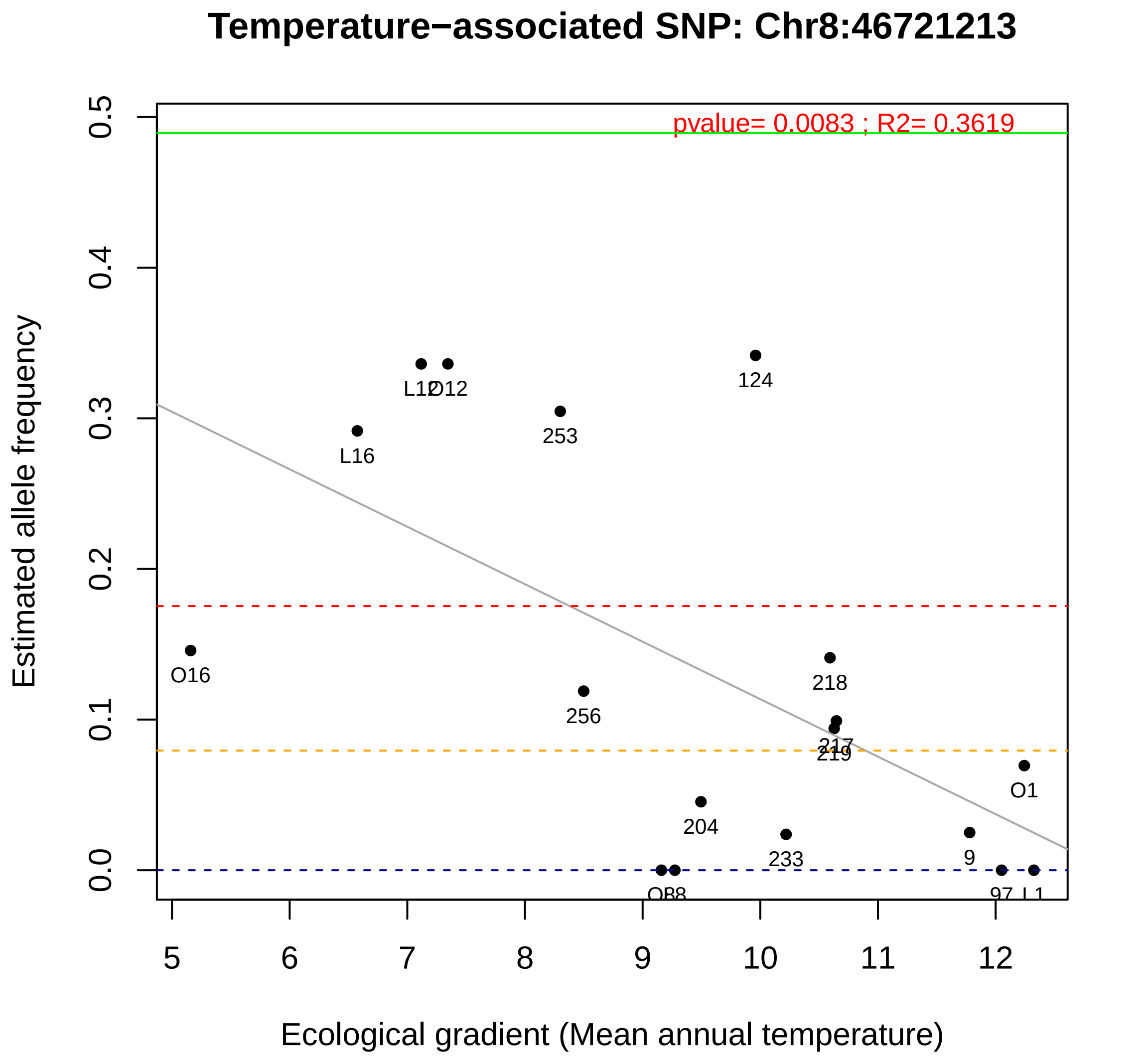

Temperature−associated SNP: Chr8:46721213
pvalue= 0.0083 ; R2= 0.3619
Estimated allele frequency
0.0
0.1
0.2
0.3
0.4
0.5
O16
L16
L12
O12
253
256
124
204
233
218
217
219
O1
9
97
L1
5
6
7
8
9
10
11
12
Ecological gradient (Mean annual temperature)

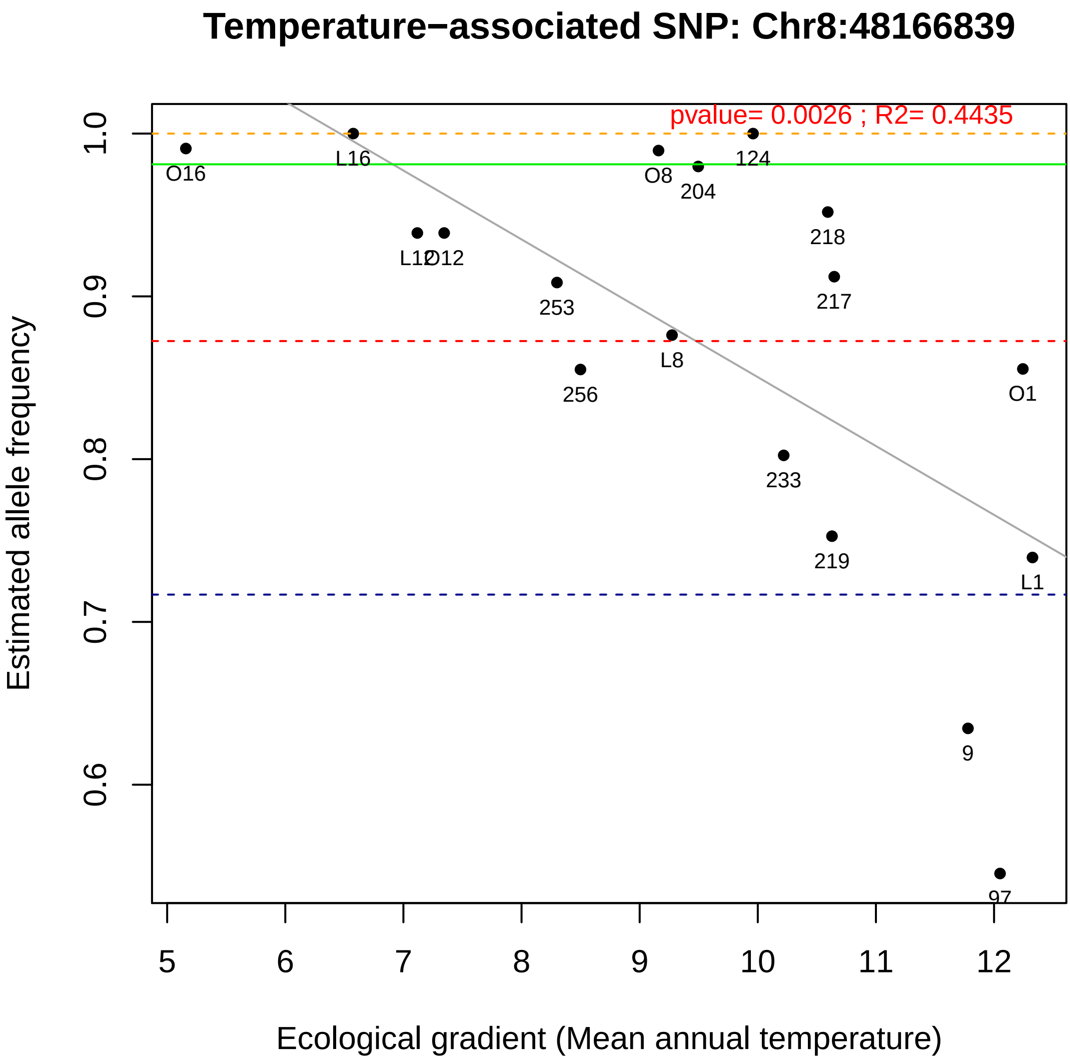

Temperature−associated SNP: Chr8:48166839
pvalue= 0.0026 ; R2= 0.4435
O16
L16
L12
O12
253
256
L8
O8
204
124
218
217
233
219
O1
L1
9
97
1.0
0.9
0.8
0.7
0.6
5
6
7
8
9
10
11
12
Estimated allele frequency
Ecological gradient (Mean annual temperature)

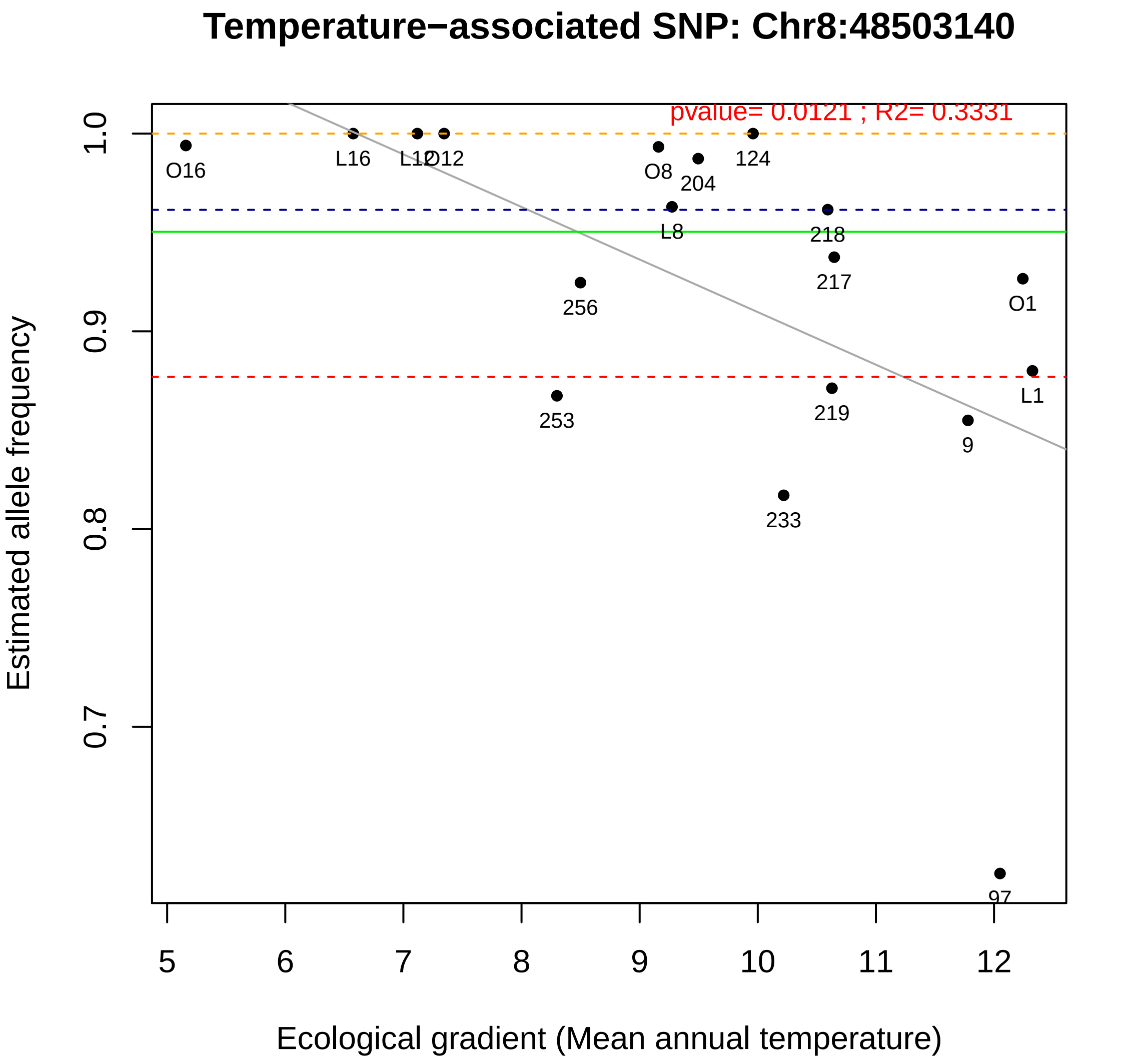

Temperature−associated SNP: Chr8:48503140
pvalue= 0.0121 ; R2= 0.3331
Estimated allele frequency
Ecological gradient (Mean annual temperature)
1.0
0.9
0.8
0.7
5
6
7
8
9
10
11
12
O16
L16
L12
O12
O8
204
124
L8
218
217
256
O1
L1
253
219
9
233
97

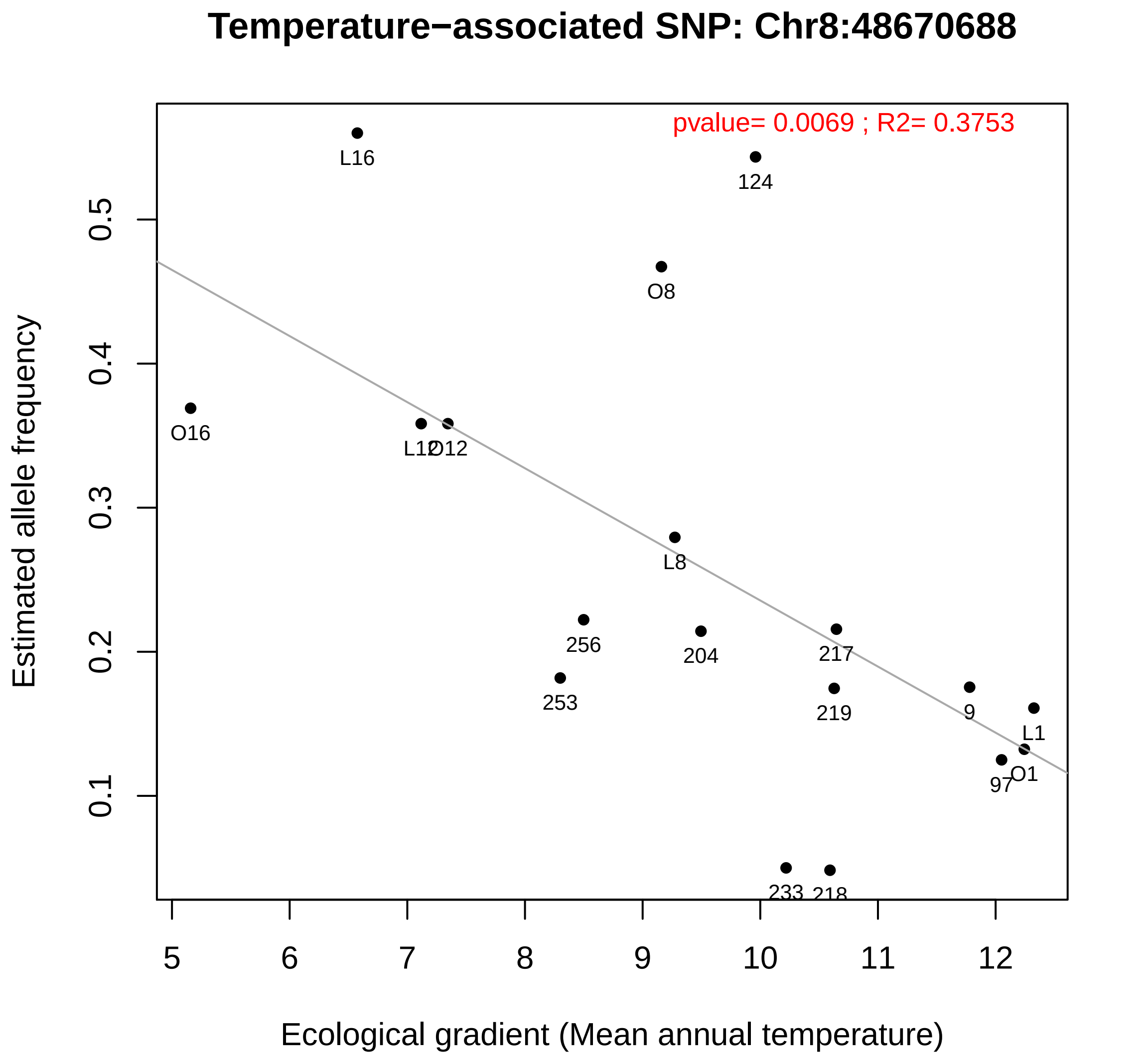

Temperature−associated SNP: Chr8:48670688
pvalue= 0.0069 ; R2= 0.3753
L16
124
O8
O16
L12
O12
L8
256
204
217
253
219
9
L1
O1
97
233
218
0.1
0.2
0.3
0.4
0.5
5
6
7
8
9
10
11
12
Estimated allele frequency
Ecological gradient (Mean annual temperature)

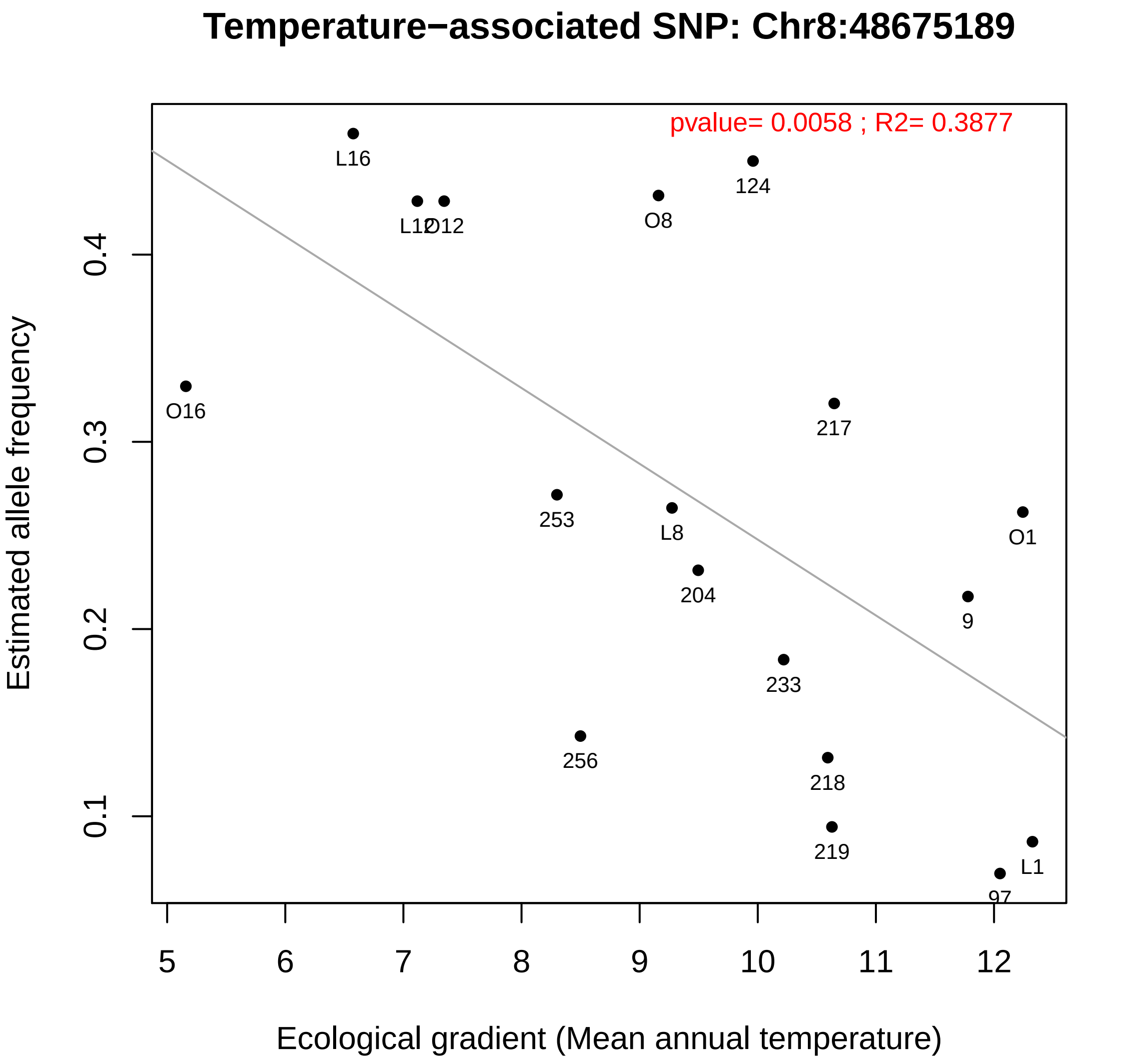

Temperature−associated SNP: Chr8:48675189
pvalue= 0.0058 ; R2= 0.3877
L16
124
L12
O12
O8
O16
217
253
L8
O1
204
9
233
256
218
219
L1
97
0.1
0.2
0.3
0.4
Estimated allele frequency
5
6
7
8
9
10
11
12
Ecological gradient (Mean annual temperature)

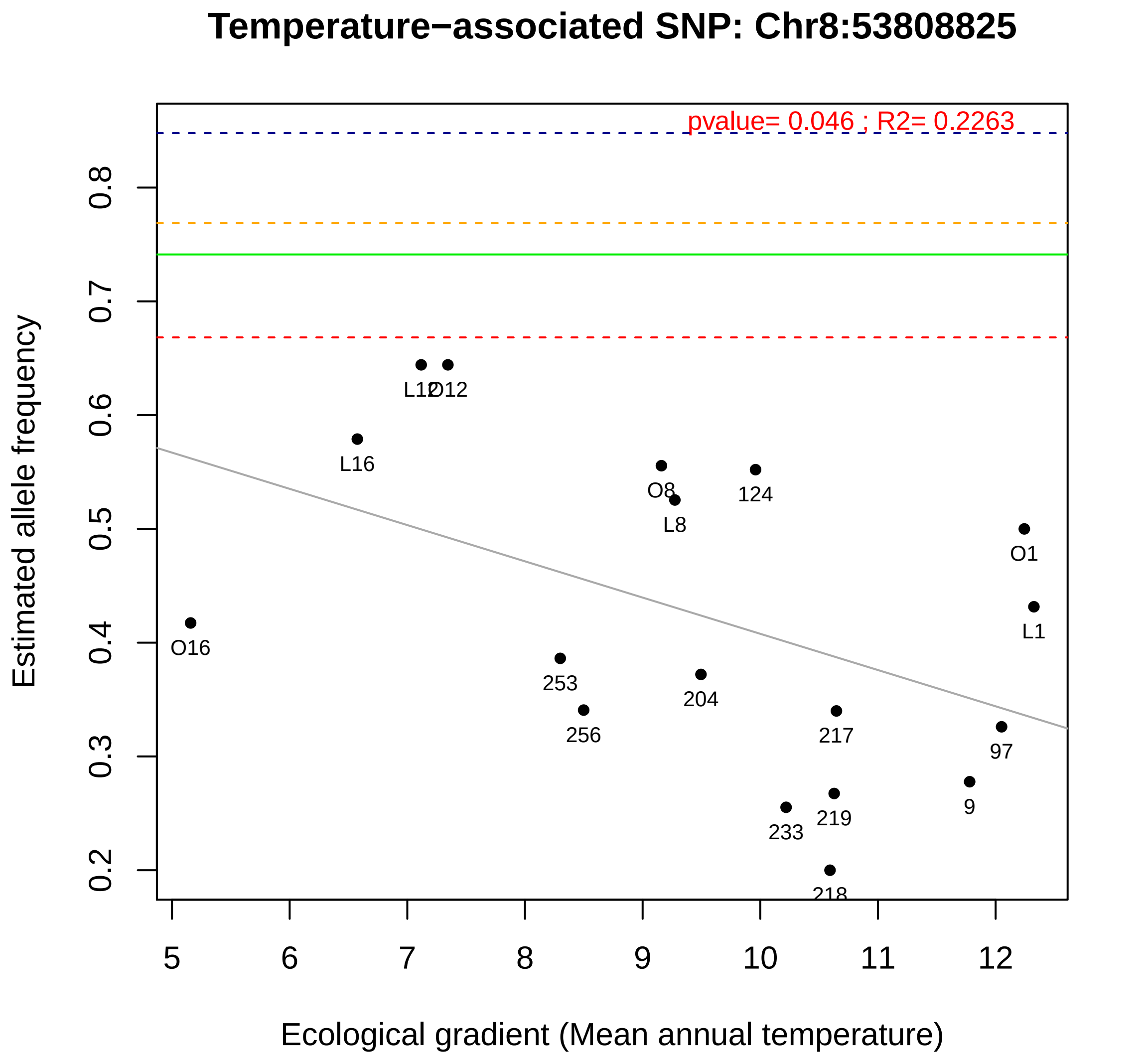
Temperature−associated SNP: Chr8:53808825
pvalue= 0.046 ; R2= 0.2263
Estimated allele frequency
Ecological gradient (Mean annual temperature)
0.2
0.3
0.4
0.5
0.6
0.7
0.8
5
6
7
8
9
10
11
12
L12
O12
L16
O8
L8
124
O1
L1
O16
253
256
204
217
97
9
219
233
218

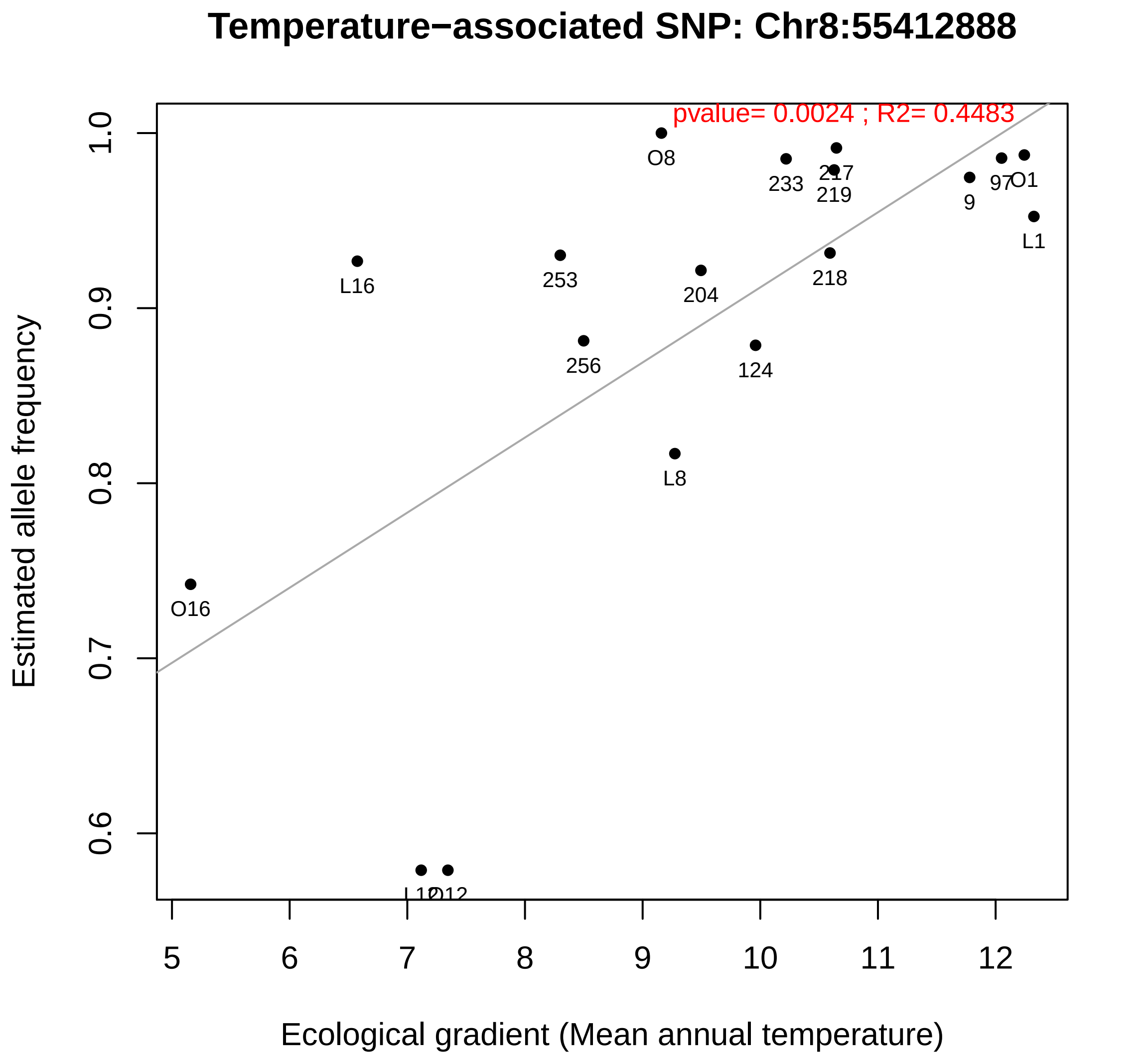

Temperature−associated SNP: Chr8:55412888
pvalue= 0.0024 ; R2= 0.4483
Estimated allele frequency
Ecological gradient (Mean annual temperature)
1.0
0.9
0.8
0.7
0.6
5
6
7
8
9
10
11
12
O8
233
217
219
9
97
O1
L1
L16
253
204
218
256
124
L8
O16
O12

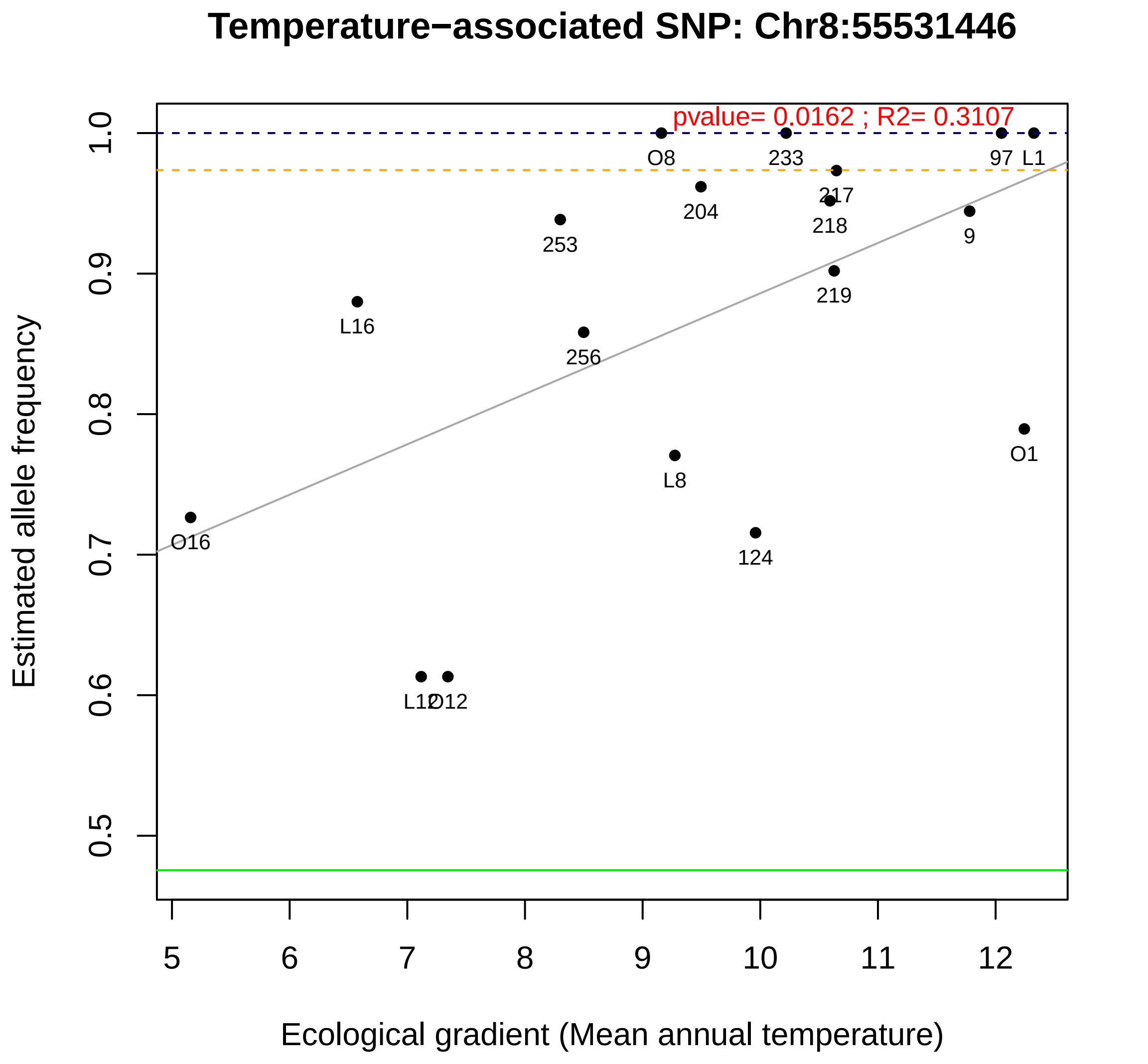

Temperature−associated SNP: Chr8:55531446
pvalue= 0.0162 ; R2= 0.3107
Estimated allele frequency
Ecological gradient (Mean annual temperature)
0.5
0.6
0.7
0.8
0.9
1.0
5
6
7
8
9
10
11
12
O8
233
97
L1
217
204
218
253
9
219
L16
256
O1
L8
O16
124
L12
O12

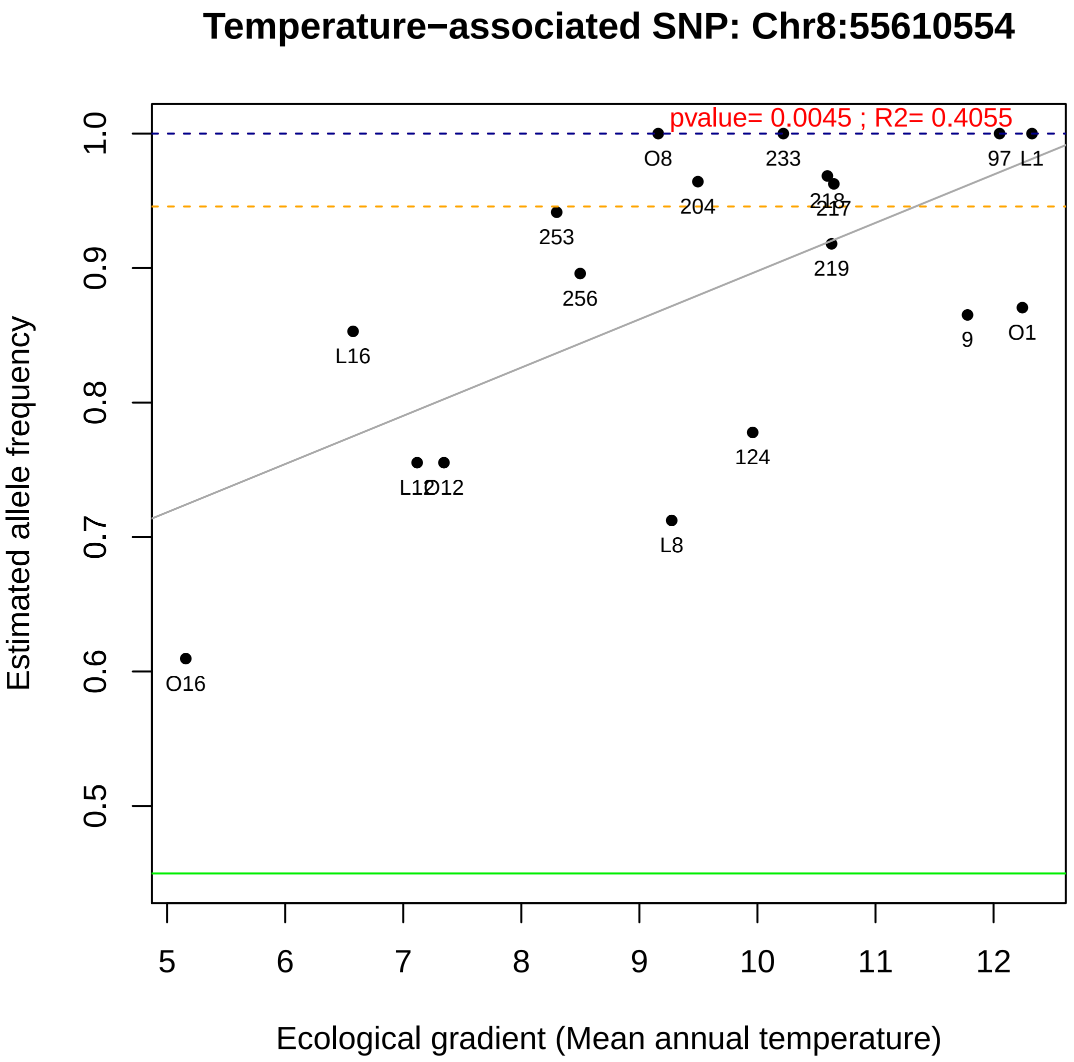

Temperature−associated SNP: Chr8:55610554
pvalue= 0.0045 ; R2= 0.4055
O8
233
97
L1
204
218
217
253
219
256
9
O1
L16
124
L12
O12
L8
O16
1.0
0.9
0.8
0.7
0.6
0.5
5
6
7
8
9
10
11
12
Estimated allele frequency
Ecological gradient (Mean annual temperature)

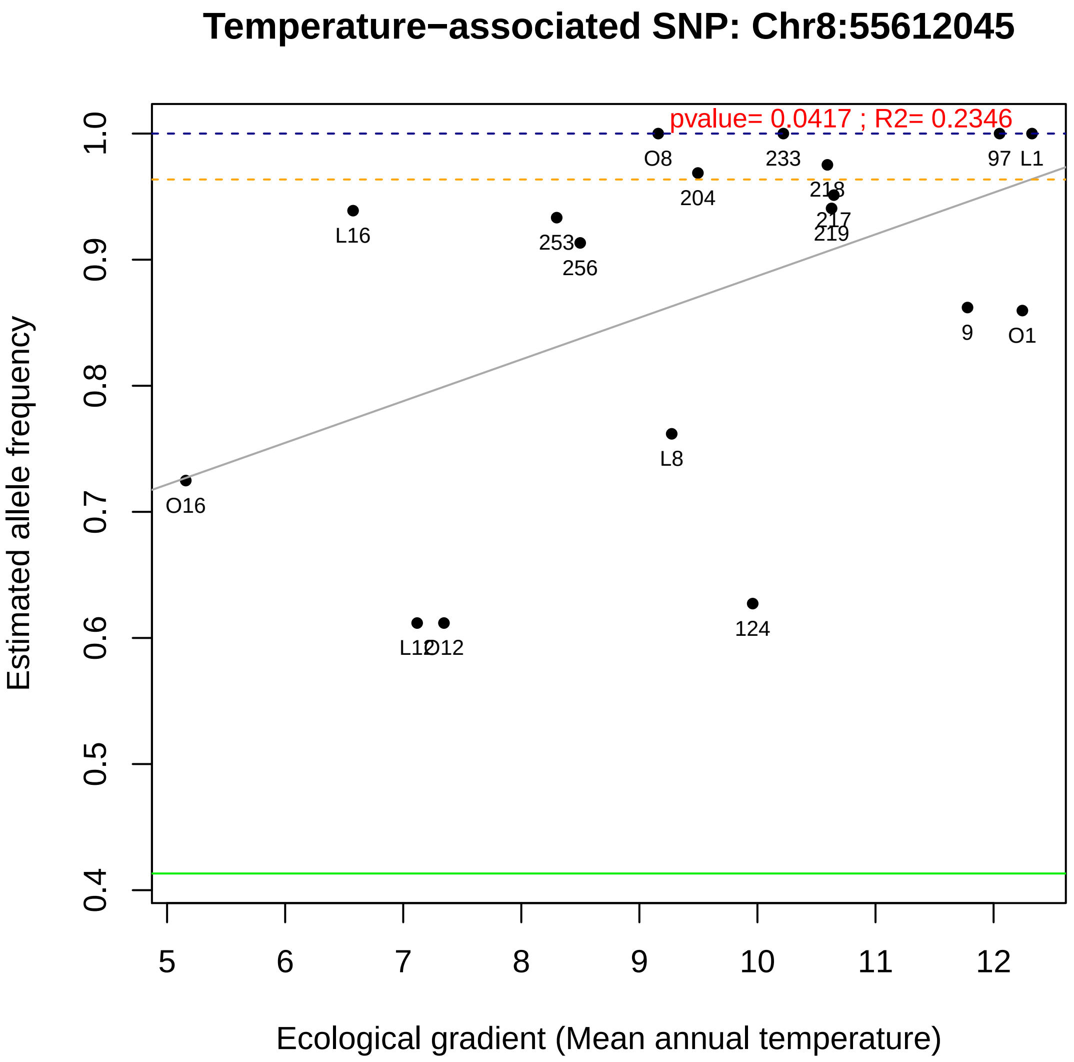

Temperature−associated SNP: Chr8:55612045
pvalue= 0.0417 ; R2= 0.2346
Estimated allele frequency
Ecological gradient (Mean annual temperature)
1.0
0.9
0.8
0.7
0.6
0.5
0.4
5
6
7
8
9
10
11
12
O8
233
97
L1
204
218
217
219
L16
253
256
9
O1
L8
O16
L12
O12
124

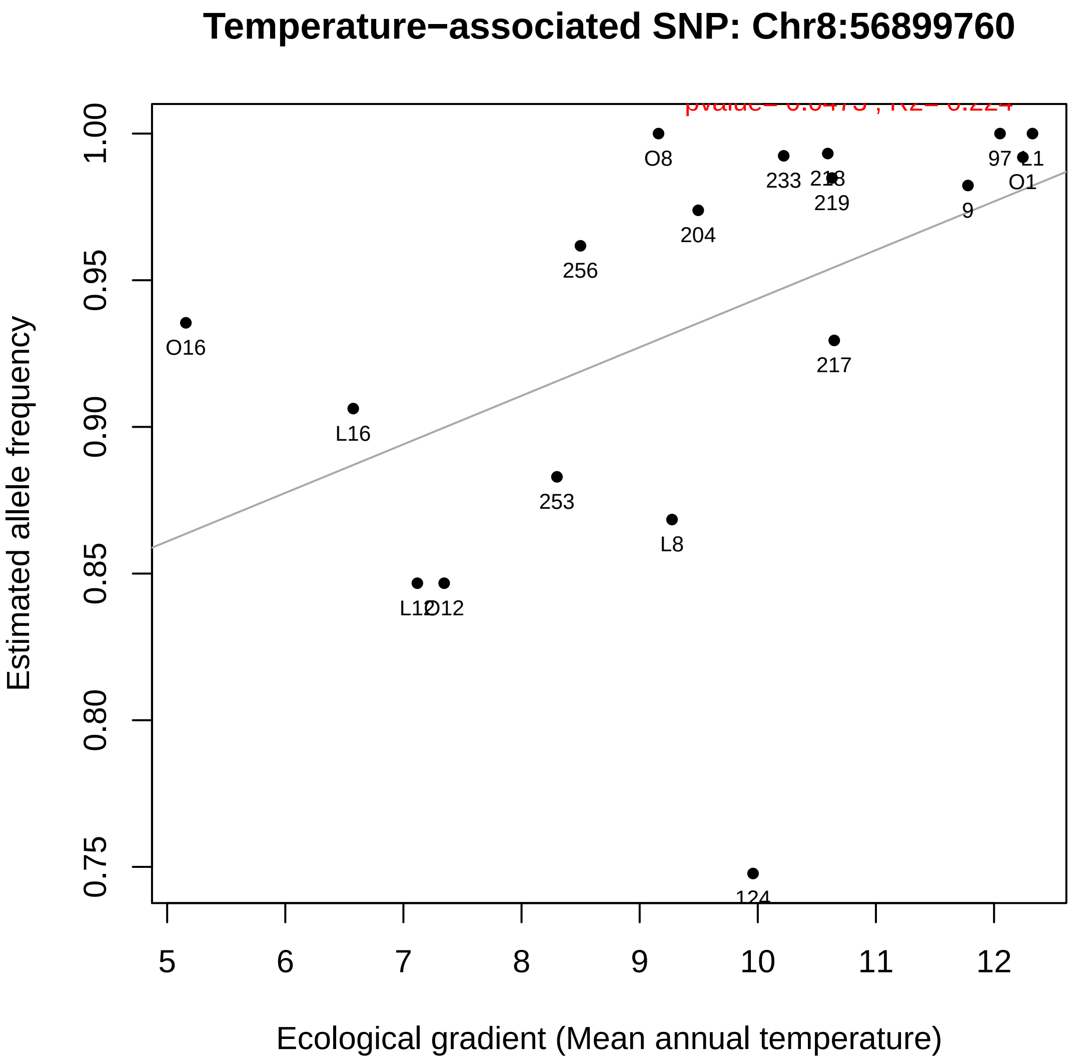

Temperature−associated SNP: Chr8:56899760
pvalue= 0.0473 ; R2= 0.224
Estimated allele frequency
Ecological gradient (Mean annual temperature)
1.00
0.95
0.90
0.85
0.80
0.75
5
6
7
8
9
10
11
12
O8
233
218
219
97
L1
O1
9
204
256
O16
217
L16
253
L8
L12
O12
124

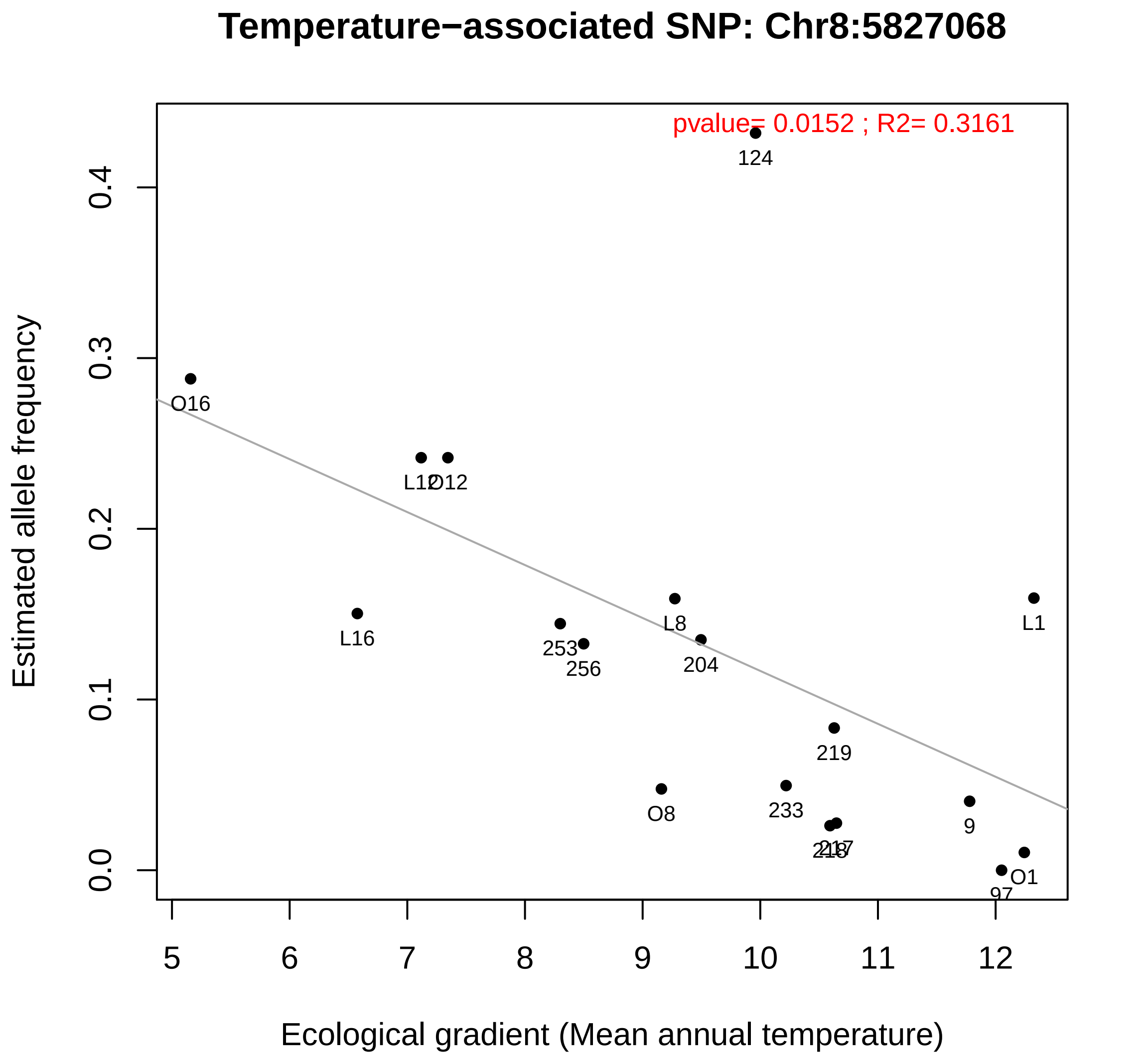

Temperature−associated SNP: Chr8:5827068
pvalue= 0.0152 ; R2= 0.3161
Estimated allele frequency
Ecological gradient (Mean annual temperature)
0.0
0.1
0.2
0.3
0.4
5
6
7
8
9
10
11
12
124
O16
L12
O12
L16
253
256
L8
204
L1
219
O8
233
9
O1
97

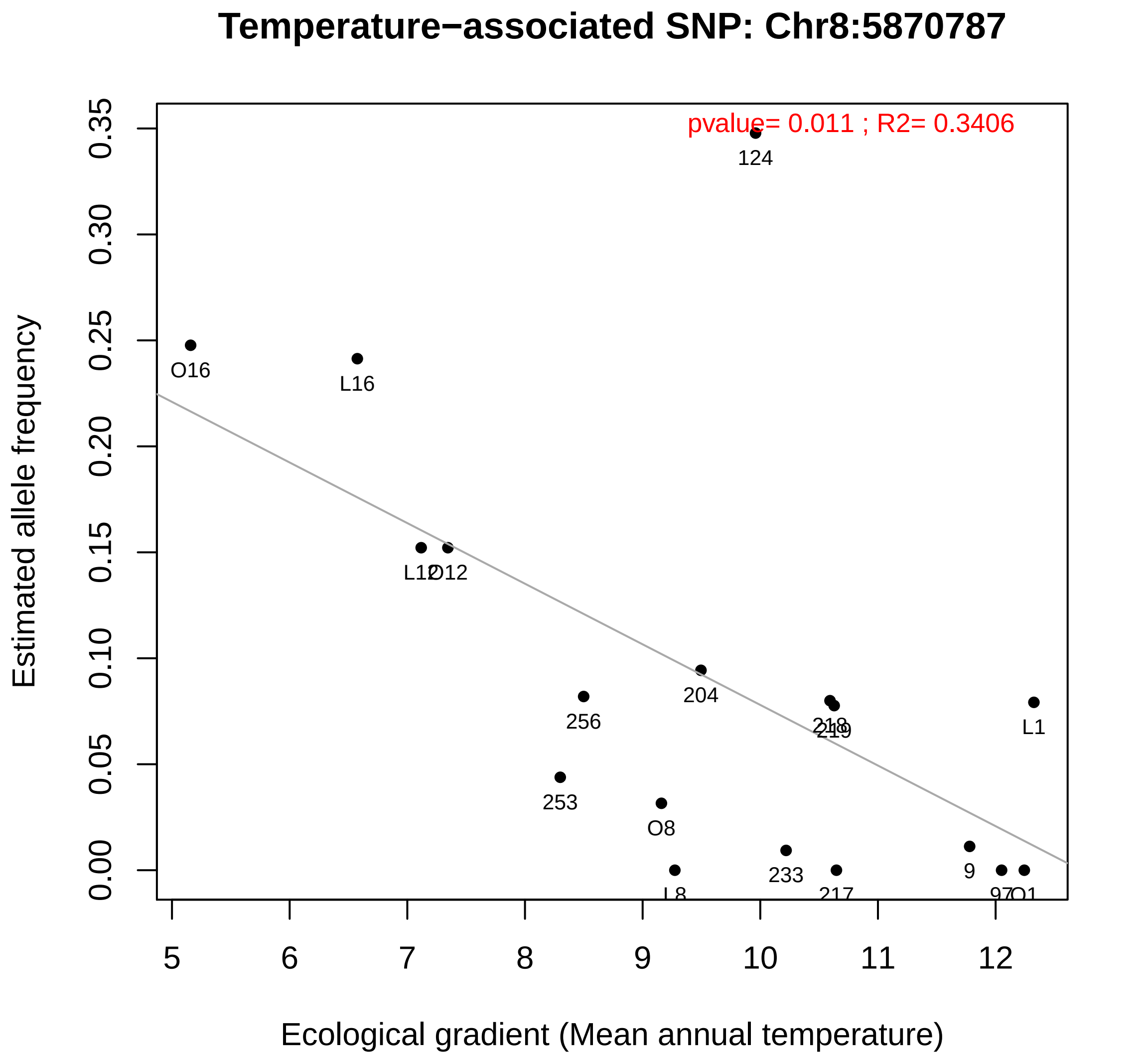

Temperature−associated SNP: Chr8:5870787
pvalue= 0.011 ; R2= 0.3406
Estimated allele frequency
Ecological gradient (Mean annual temperature)
0.00
0.05
0.10
0.15
0.20
0.25
0.30
0.35
5
6
7
8
9
10
11
12
124
O16
L16
L12
O12
204
256
218
219
L1
253
O8
233
9
L8
217
97
O1

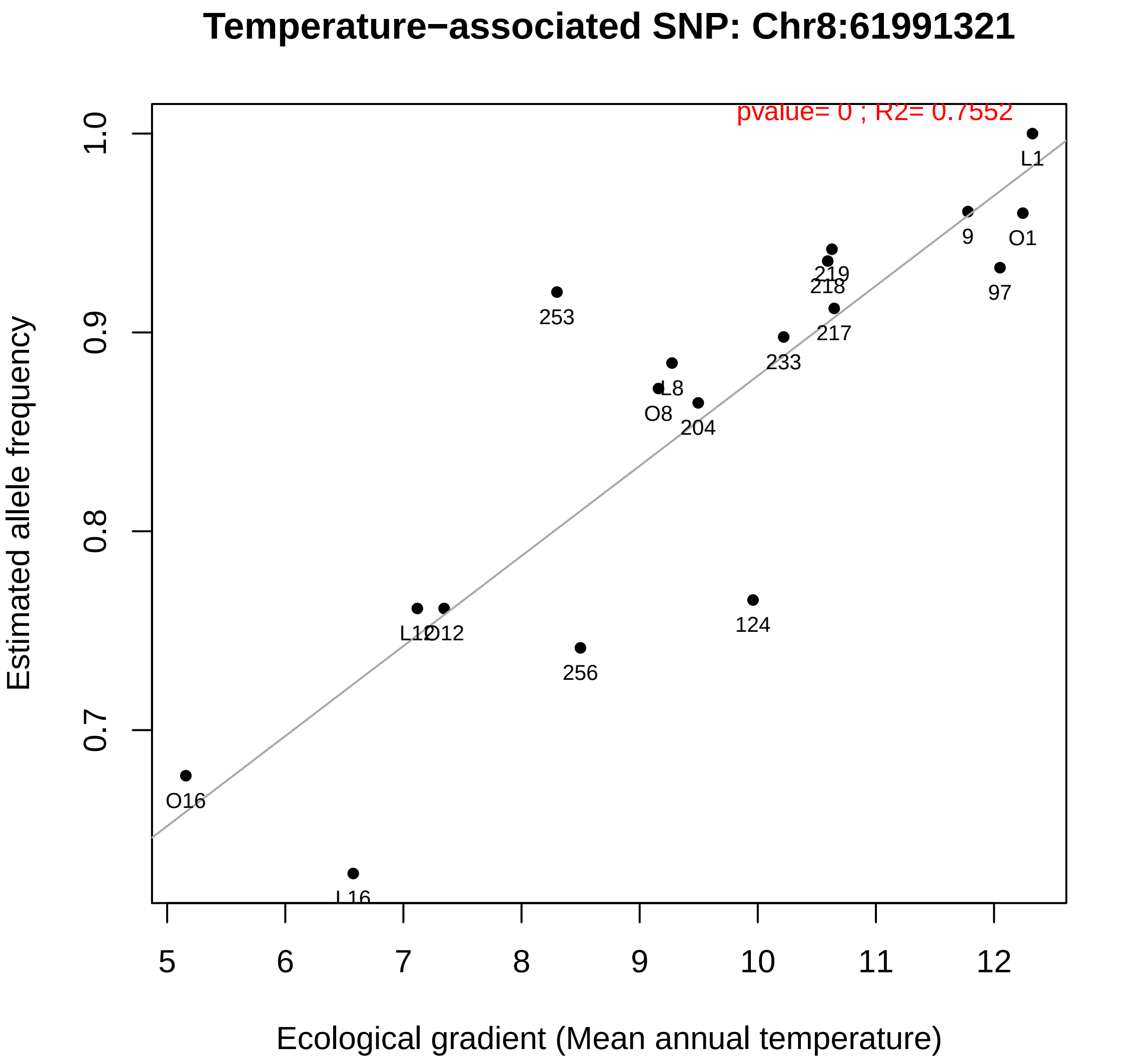

Temperature−associated SNP: Chr8:61991321
pvalue= 0 ; R2= 0.7552
L1
9
O1
219
218
97
253
217
233
L8
O8
204
L12
O12
124
256
O16
L16
1.0
0.9
0.8
0.7
Estimated allele frequency
5
6
7
8
9
10
11
12
Ecological gradient (Mean annual temperature)

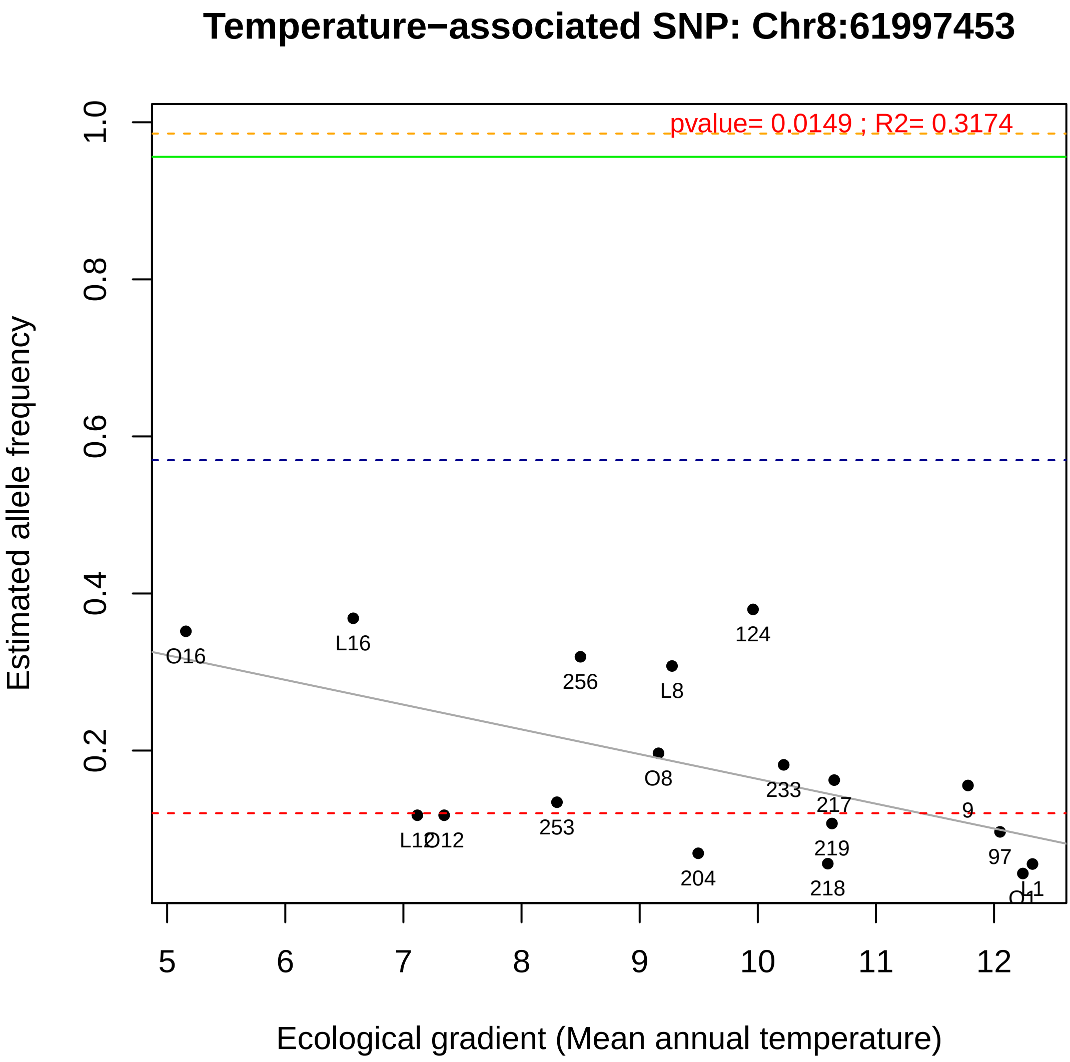

Temperature−associated SNP: Chr8:61997453
pvalue= 0.0149 ; R2= 0.3174
Estimated allele frequency
0.2
0.4
0.6
0.8
1.0
Ecological gradient (Mean annual temperature)
5
6
7
8
9
10
11
12
O16
L16
124
256
L8
O8
233
217
9
253
L12
O12
219
97
204
218
L1

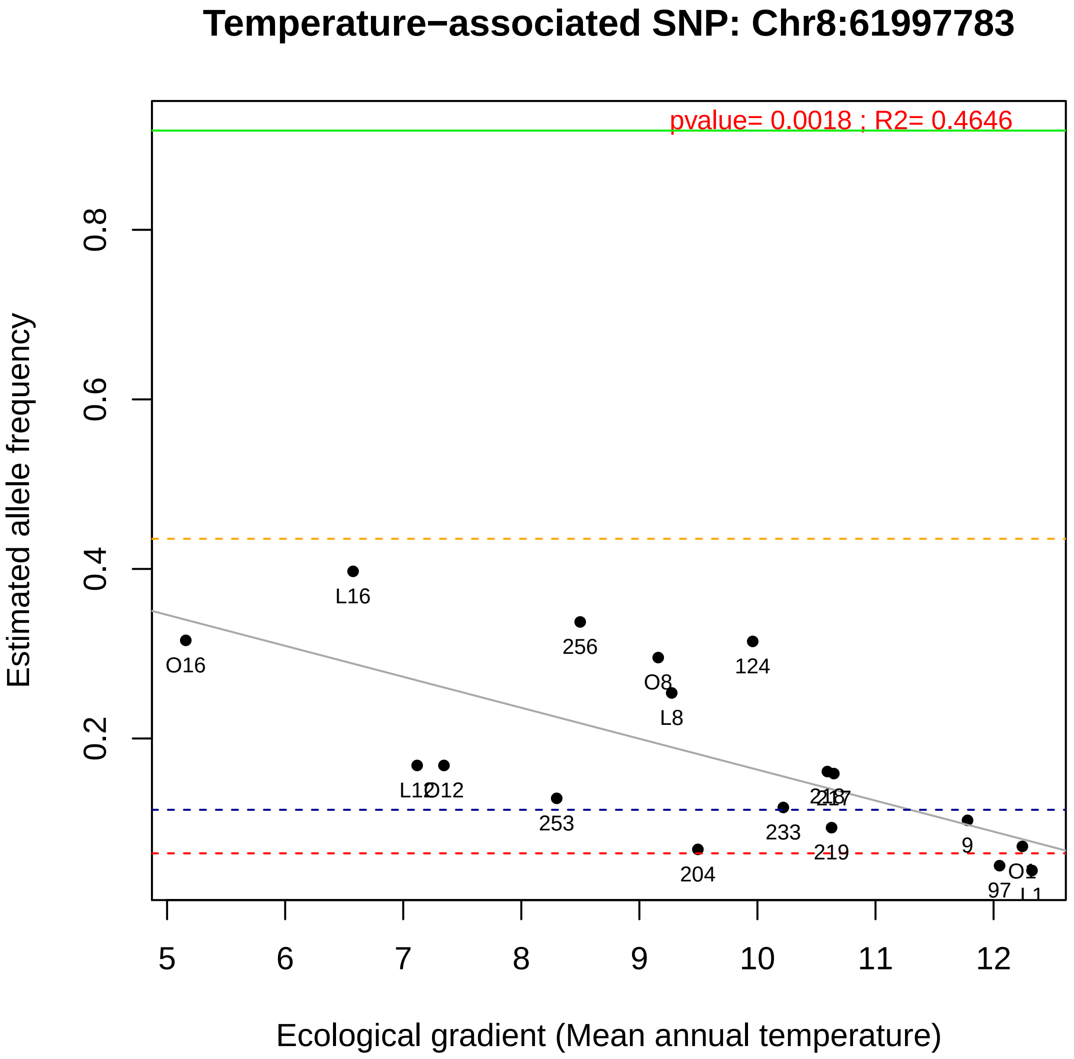
Temperature−associated SNP: Chr8:61997783
pvalue= 0.0018 ; R2= 0.4646
Estimated allele frequency
0.2
0.4
0.6
0.8
Ecological gradient (Mean annual temperature)
5
6
7
8
9
10
11
12
L16
O16
256
124
O8
L8
L12
O12
253
233
204
219
9
97

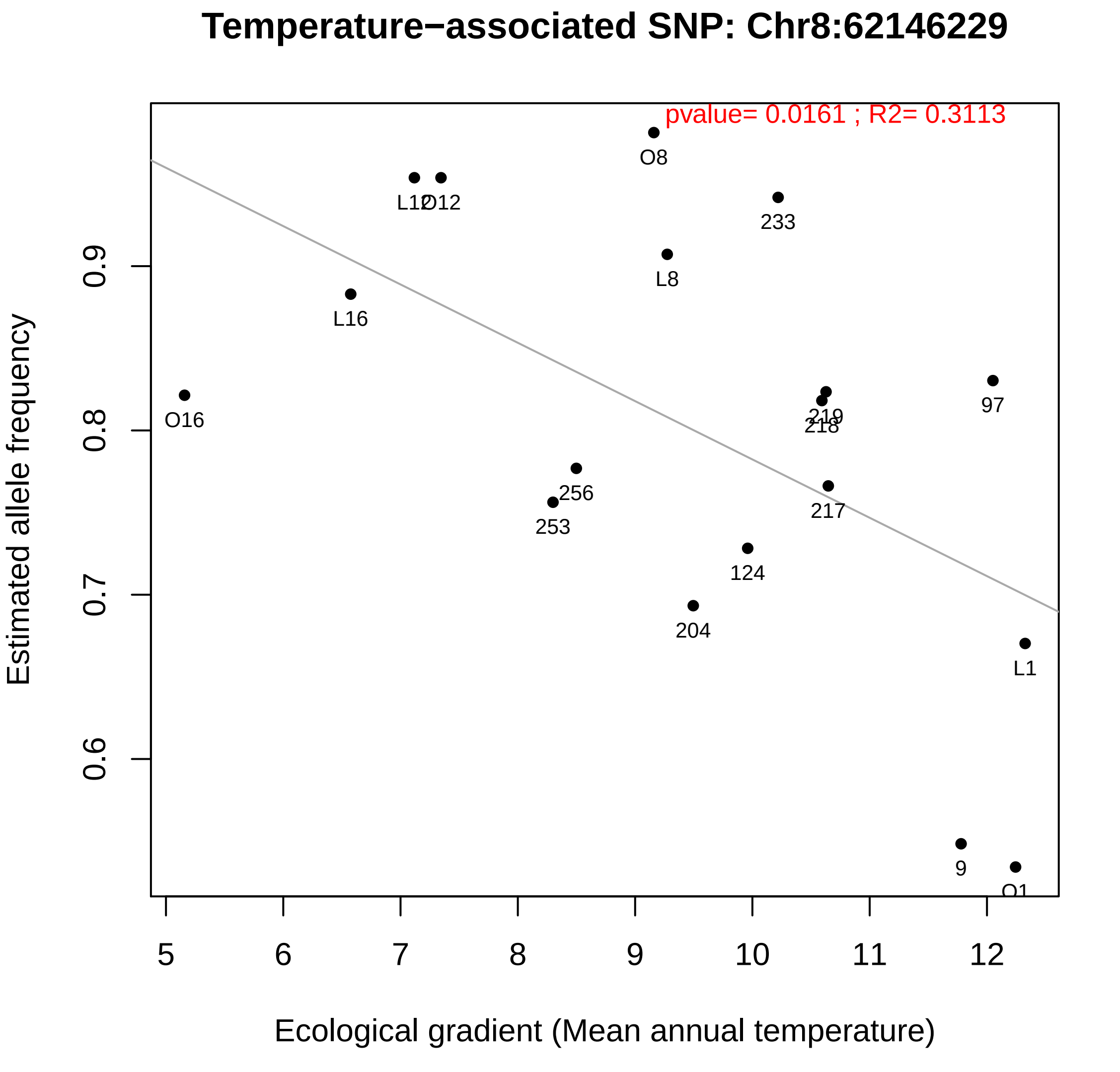
Temperature−associated SNP: Chr8:62146229
pvalue= 0.0161 ; R2= 0.3113
O8
L12
O12
233
L8
L16
O16
219
218
97
256
253
217
124
204
L1
9
O1
0.6
0.7
0.8
0.9
5
6
7
8
9
10
11
12
Estimated allele frequency
Ecological gradient (Mean annual temperature)

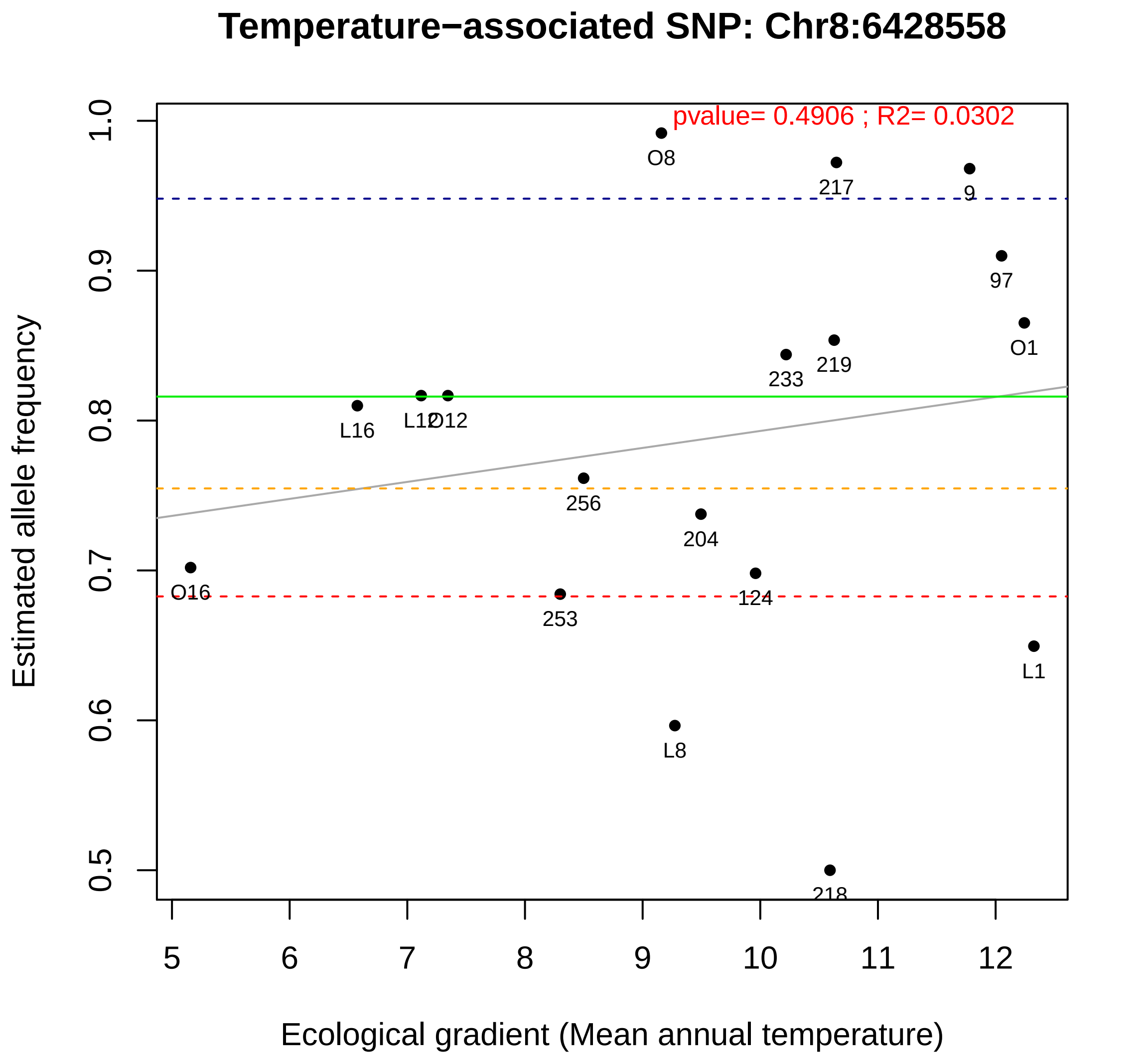
Temperature−associated SNP: Chr8:6428558
pvalue= 0.4906 ; R2= 0.0302
Estimated allele frequency
Ecological gradient (Mean annual temperature)
0.5
0.6
0.7
0.8
0.9
1.0
5
6
7
8
9
10
11
12
O8
217
9
97
O1
219
233
L16
L12
O12
256
204
O16
253
124
L1
L8
218

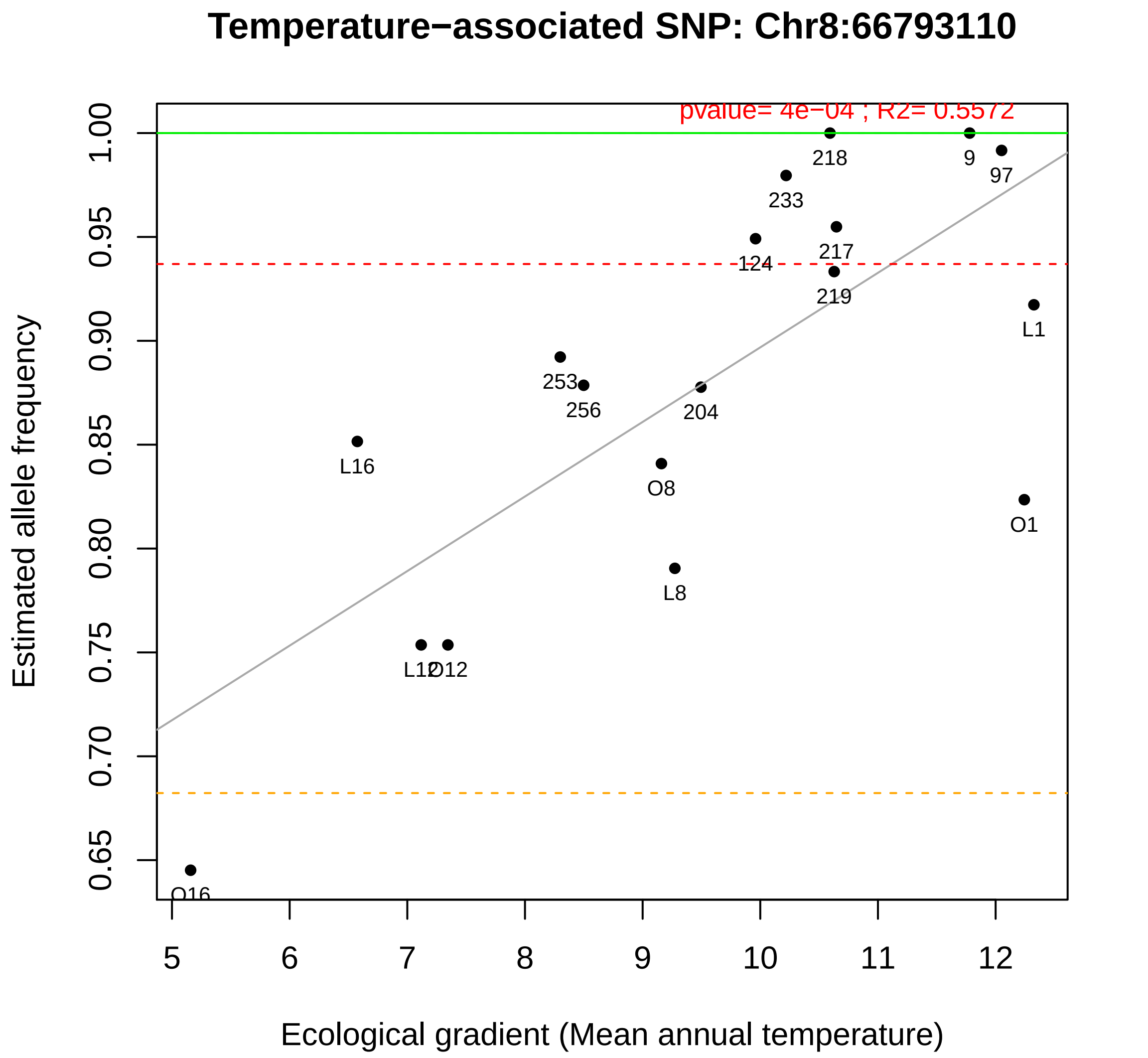
Temperature−associated SNP: Chr8:66793110
pvalue= 4e−04 ; R2= 0.5572
Estimated allele frequency
Ecological gradient (Mean annual temperature)
1.00
0.95
0.90
0.85
0.80
0.75
0.70
0.65
5
6
7
8
9
10
11
12
218
9
97
233
217
124
219
L1
253
256
204
L16
O8
O1
L8
L12
O12
O16

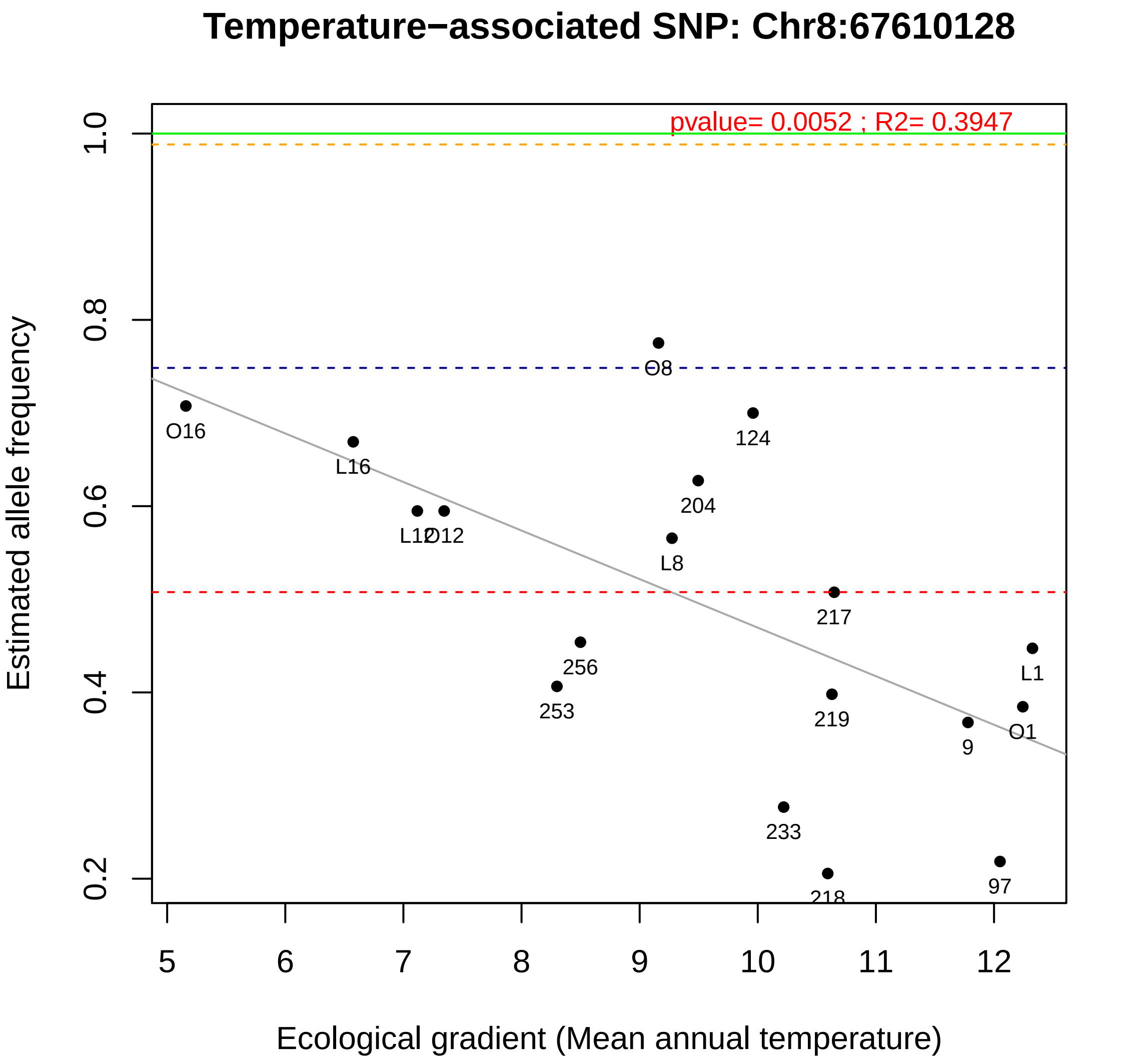

Temperature−associated SNP: Chr8:67610128
pvalue= 0.0052 ; R2= 0.3947
Estimated allele frequency
1.0
0.8
0.6
0.4
0.2
O8
O16
124
L16
204
L12
O12
L8
217
L1
256
253
219
O1
9
233
218
97
5
6
7
8
9
10
11
12
Ecological gradient (Mean annual temperature)

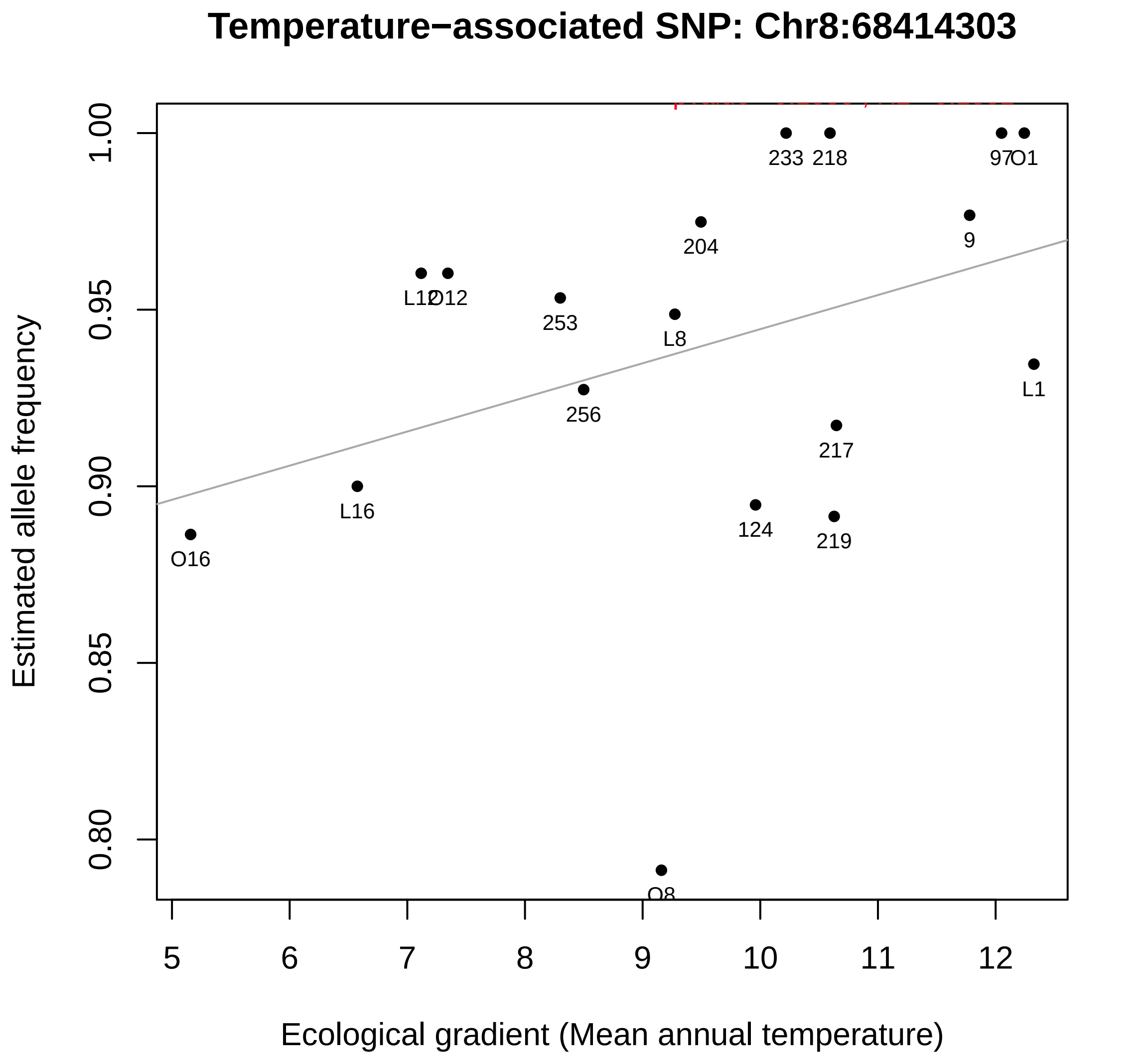
Temperature−associated SNP: Chr8:68414303
Estimated allele frequency
Ecological gradient (Mean annual temperature)
1.00
0.95
0.90
0.85
0.80
5
6
7
8
9
10
11
12
233
218
97
O1
204
9
L12
O12
253
L8
L1
256
217
L16
124
219
O16
O8

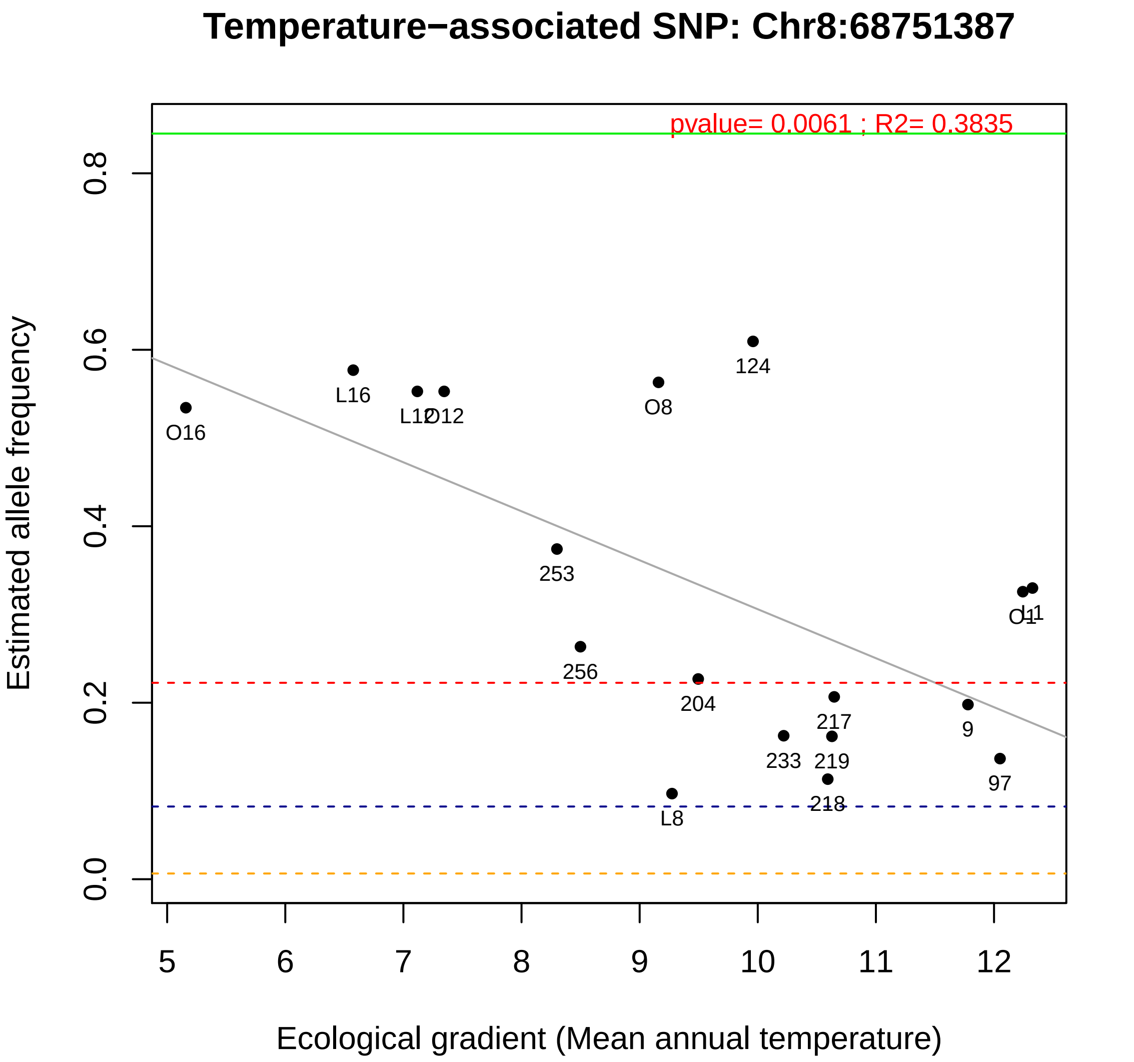
Temperature−associated SNP: Chr8:68751387
pvalue= 0.0061 ; R2= 0.3835
Estimated allele frequency
0.0
0.2
0.4
0.6
0.8
5
6
7
8
9
10
11
12
Ecological gradient (Mean annual temperature)
O16
L16
L12
O12
O8
124
253
256
204
217
233
219
218
L8
9
97
O1
L1

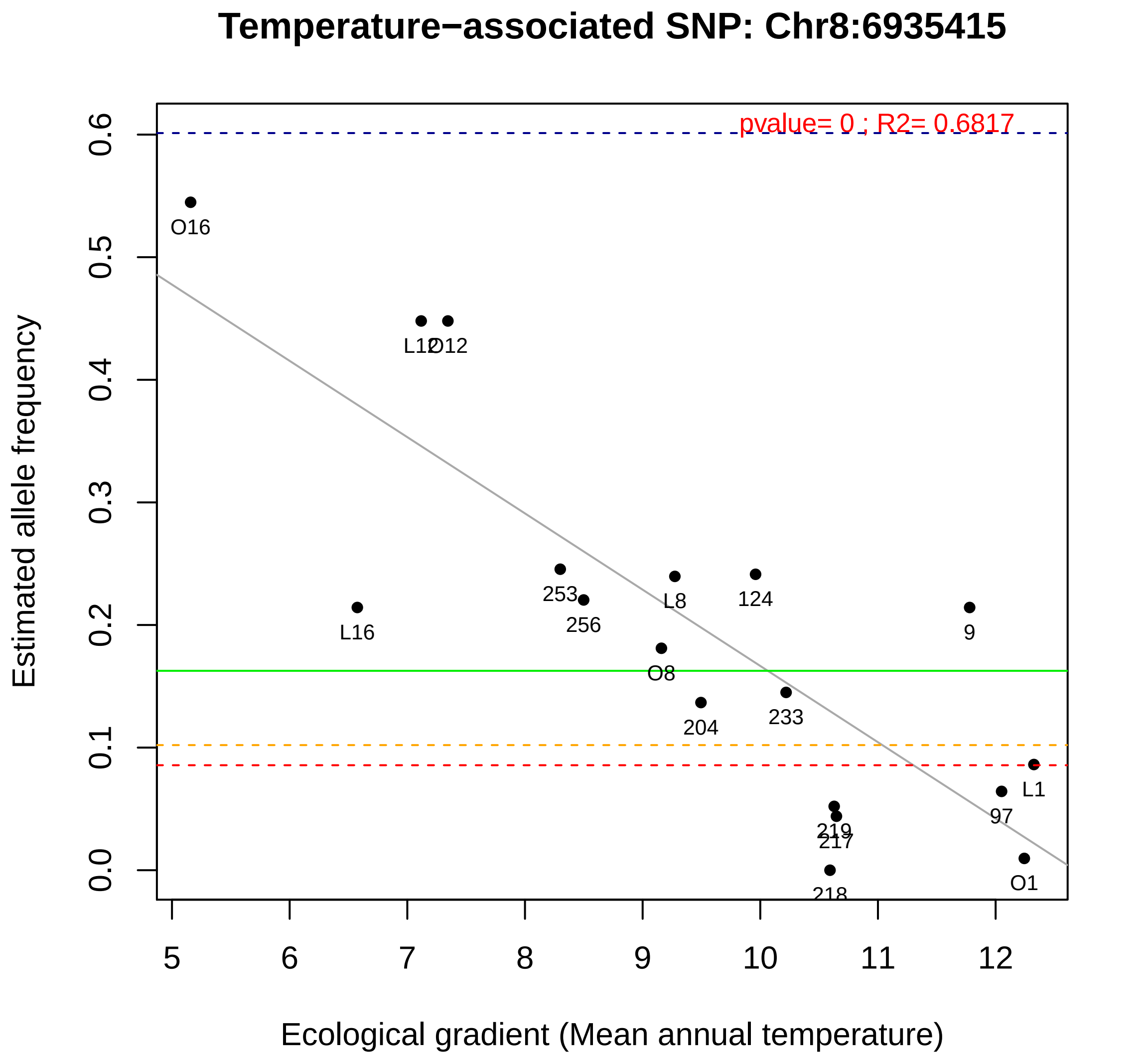

Temperature−associated SNP: Chr8:6935415
pvalue= 0 ; R2= 0.6817
Estimated allele frequency
0.0
0.1
0.2
0.3
0.4
0.5
0.6
Ecological gradient (Mean annual temperature)
5
6
7
8
9
10
11
12
O16
L12
O12
L16
253
256
L8
124
9
O8
204
233
219
217
218
L1
97
O1

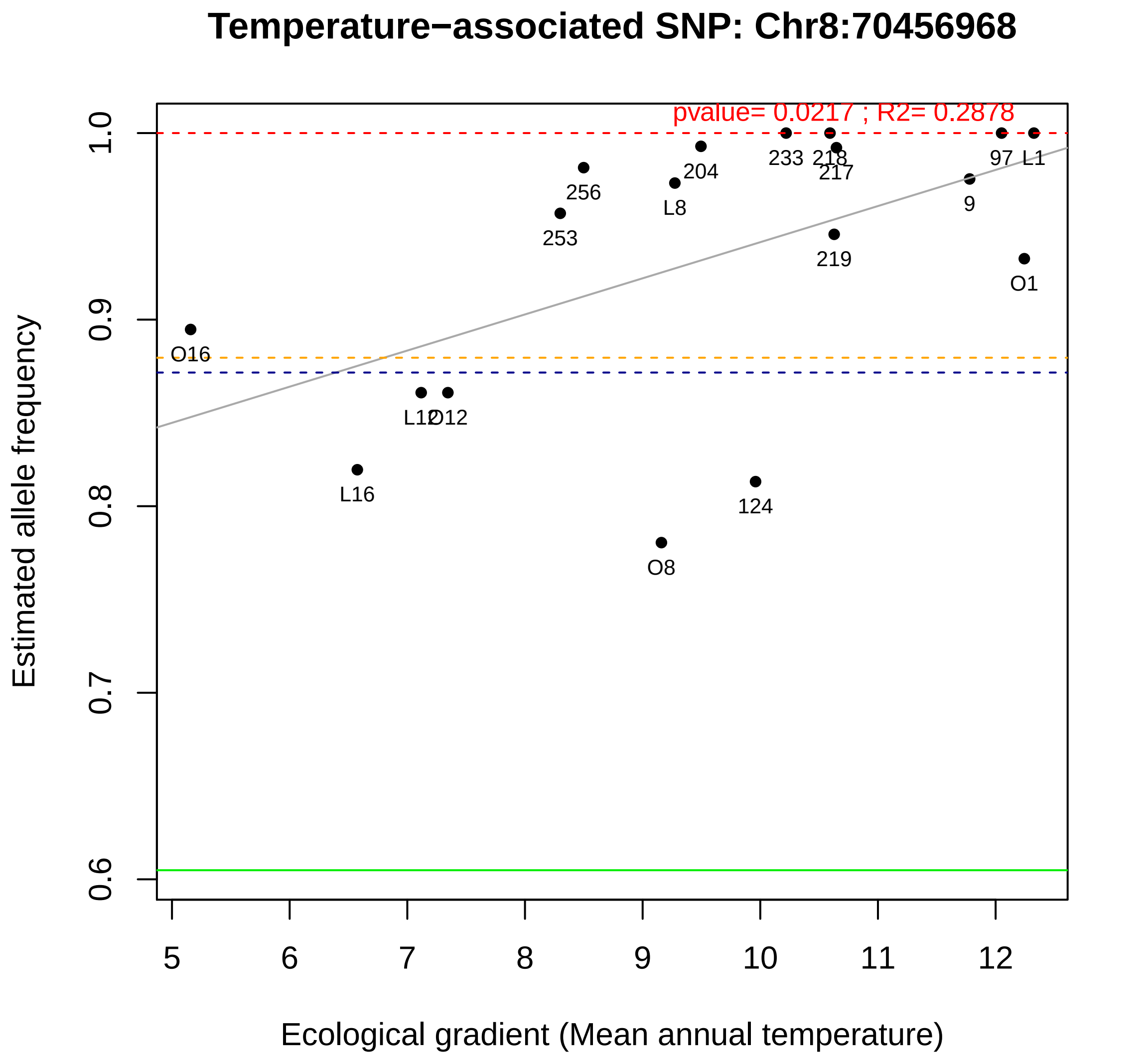
Temperature−associated SNP: Chr8:70456968
pvalue= 0.0217 ; R2= 0.2878
Estimated allele frequency
Ecological gradient (Mean annual temperature)
1.0
0.9
0.8
0.7
0.6
5
6
7
8
9
10
11
12
O16
L16
L12
O12
253
256
L8
204
O8
124
233
218
217
219
9
97
L1
O1

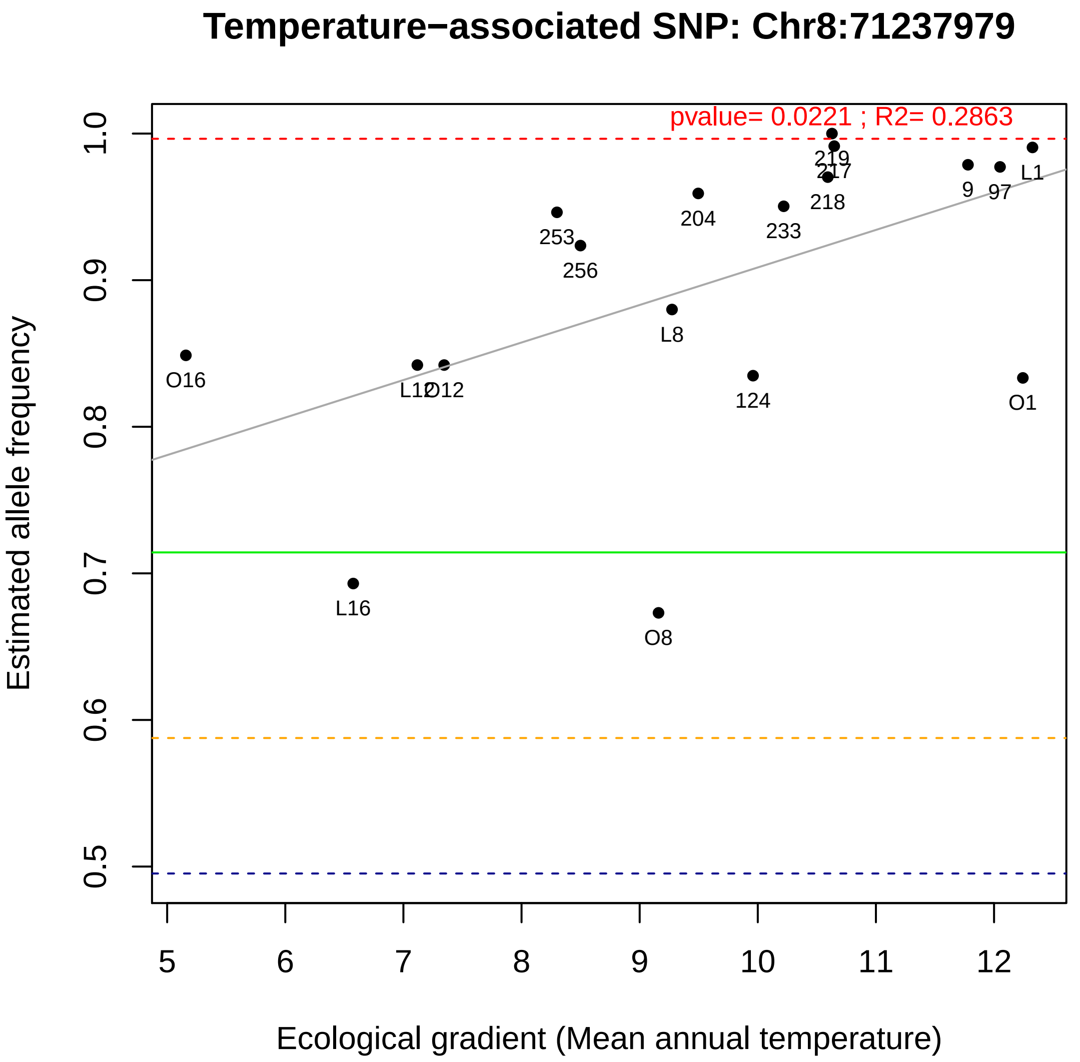

Temperature−associated SNP: Chr8:71237979
pvalue= 0.0221 ; R2= 0.2863
Estimated allele frequency
0.5
0.6
0.7
0.8
0.9
1.0
5
6
7
8
9
10
11
12
Ecological gradient (Mean annual temperature)
O16
L12
O12
L16
253
256
204
L8
O8
124
233
219
217
218
9
97
L1
O1

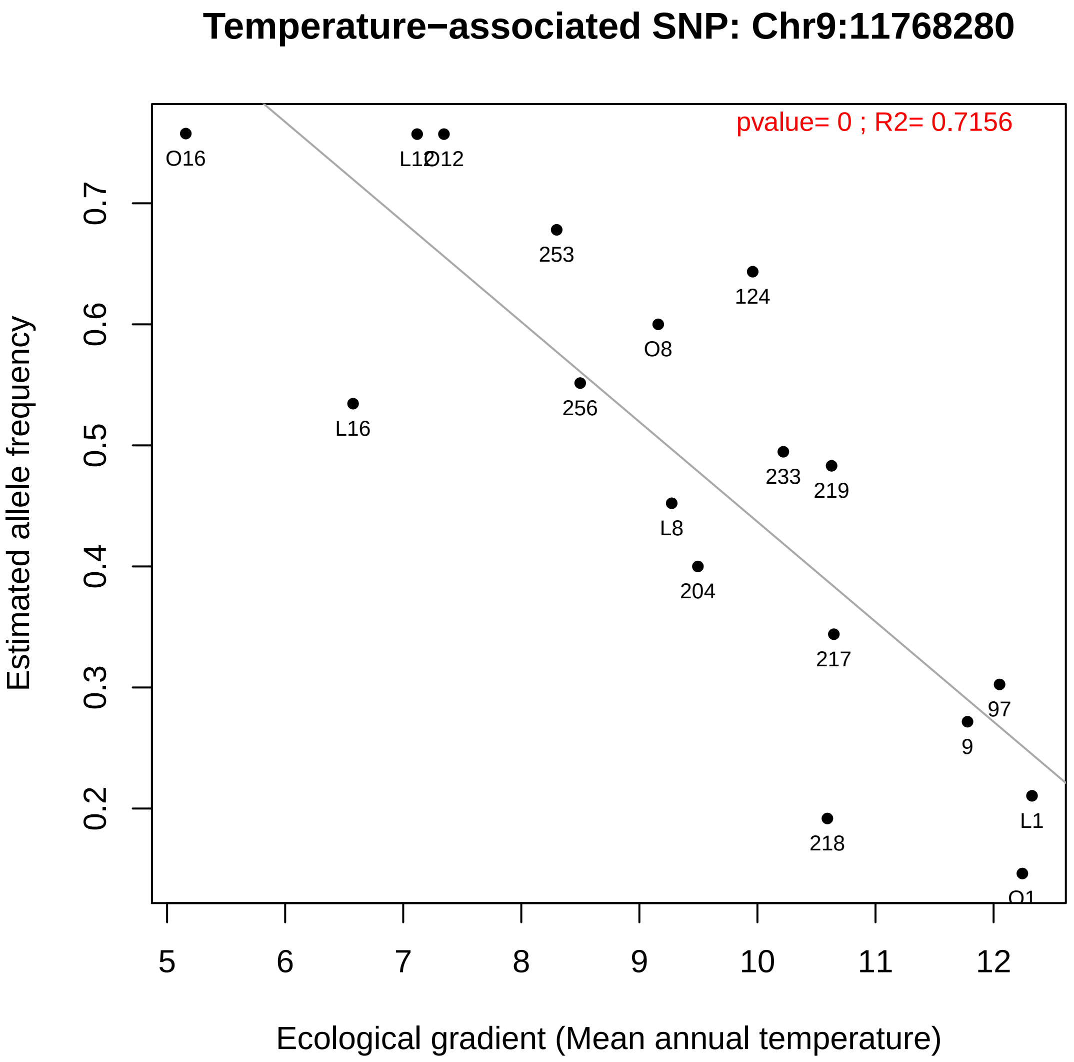
Temperature−associated SNP: Chr9:11768280
pvalue= 0 ; R2= 0.7156
O16
L12
O12
253
124
O8
256
L16
233
219
L8
204
217
97
9
L1
218
O1
0.2
0.3
0.4
0.5
0.6
0.7
5
6
7
8
9
10
11
12
Estimated allele frequency
Ecological gradient (Mean annual temperature)

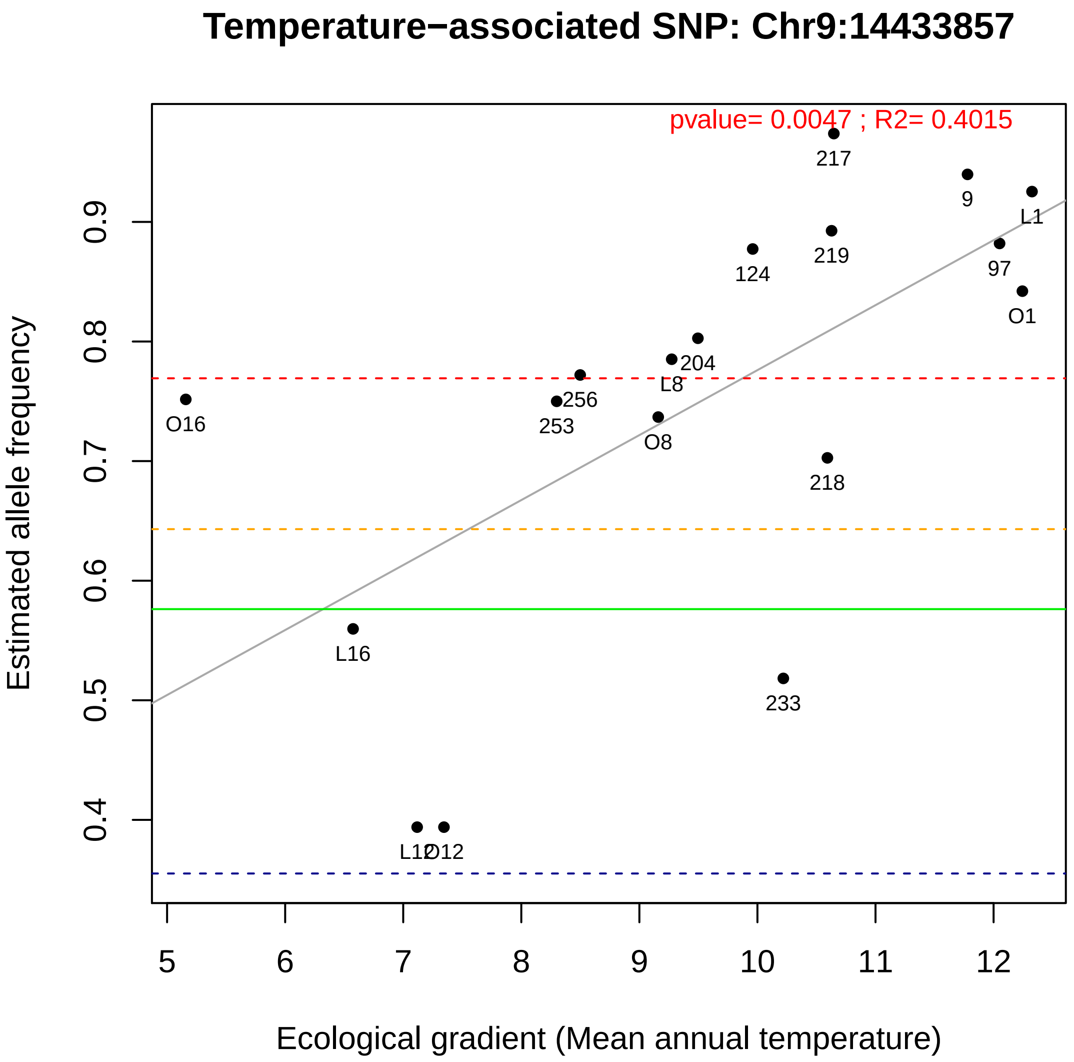

Temperature−associated SNP: Chr9:14433857
pvalue= 0.0047 ; R2= 0.4015
217
9
L1
219
124
97
O1
204
L8
256
253
O16
O8
218
L16
233
L12
O12
0.4
0.5
0.6
0.7
0.8
0.9
5
6
7
8
9
10
11
12
Estimated allele frequency
Ecological gradient (Mean annual temperature)

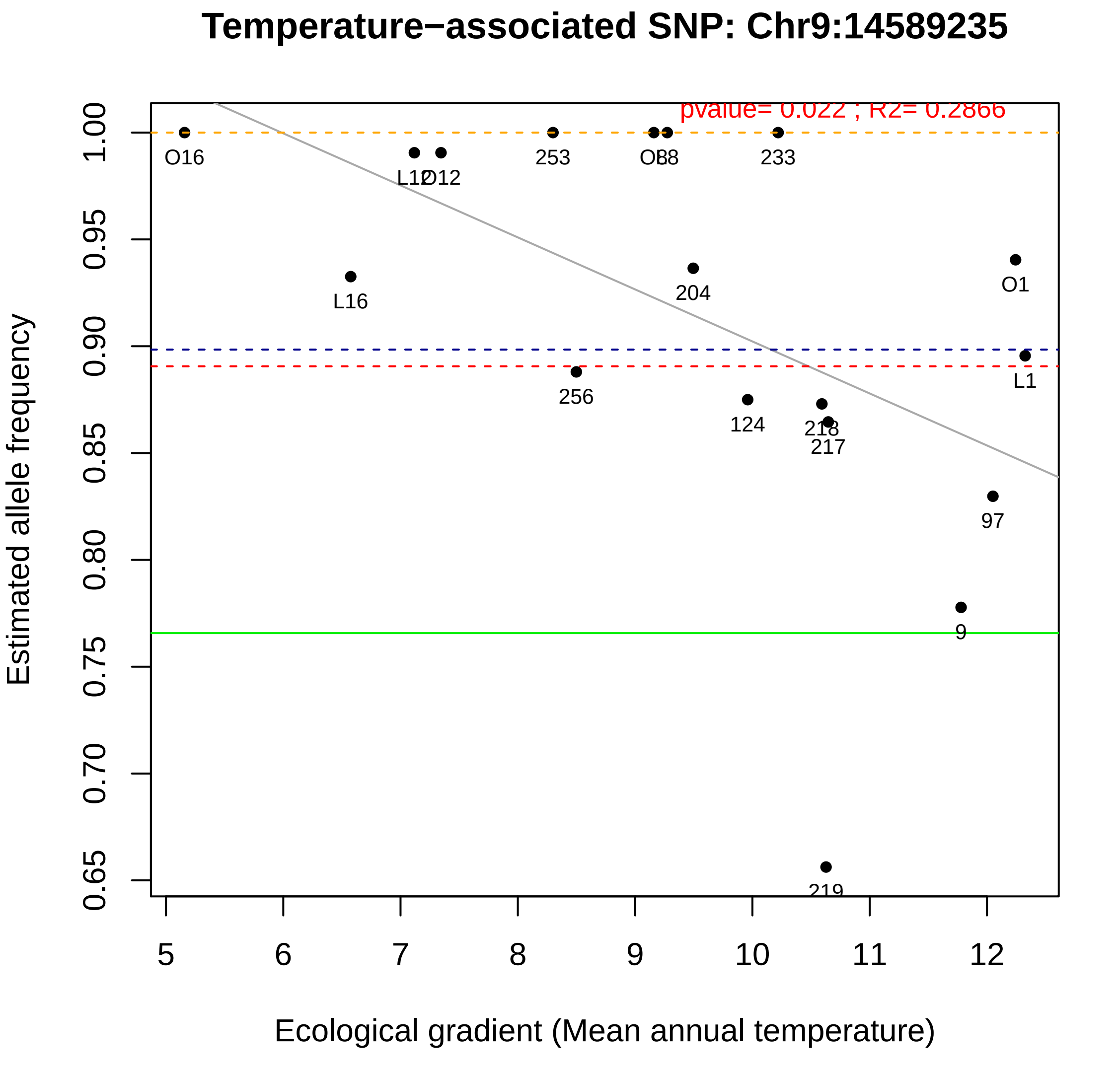
Temperature−associated SNP: Chr9:14589235
pvalue= 0.022 ; R2= 0.2866
Estimated allele frequency
Ecological gradient (Mean annual temperature)
1.00
0.95
0.90
0.85
0.80
0.75
0.70
0.65
5
6
7
8
9
10
11
12
O16
L12
O12
253
233
L16
204
O1
L1
256
124
218
217
97
9
219

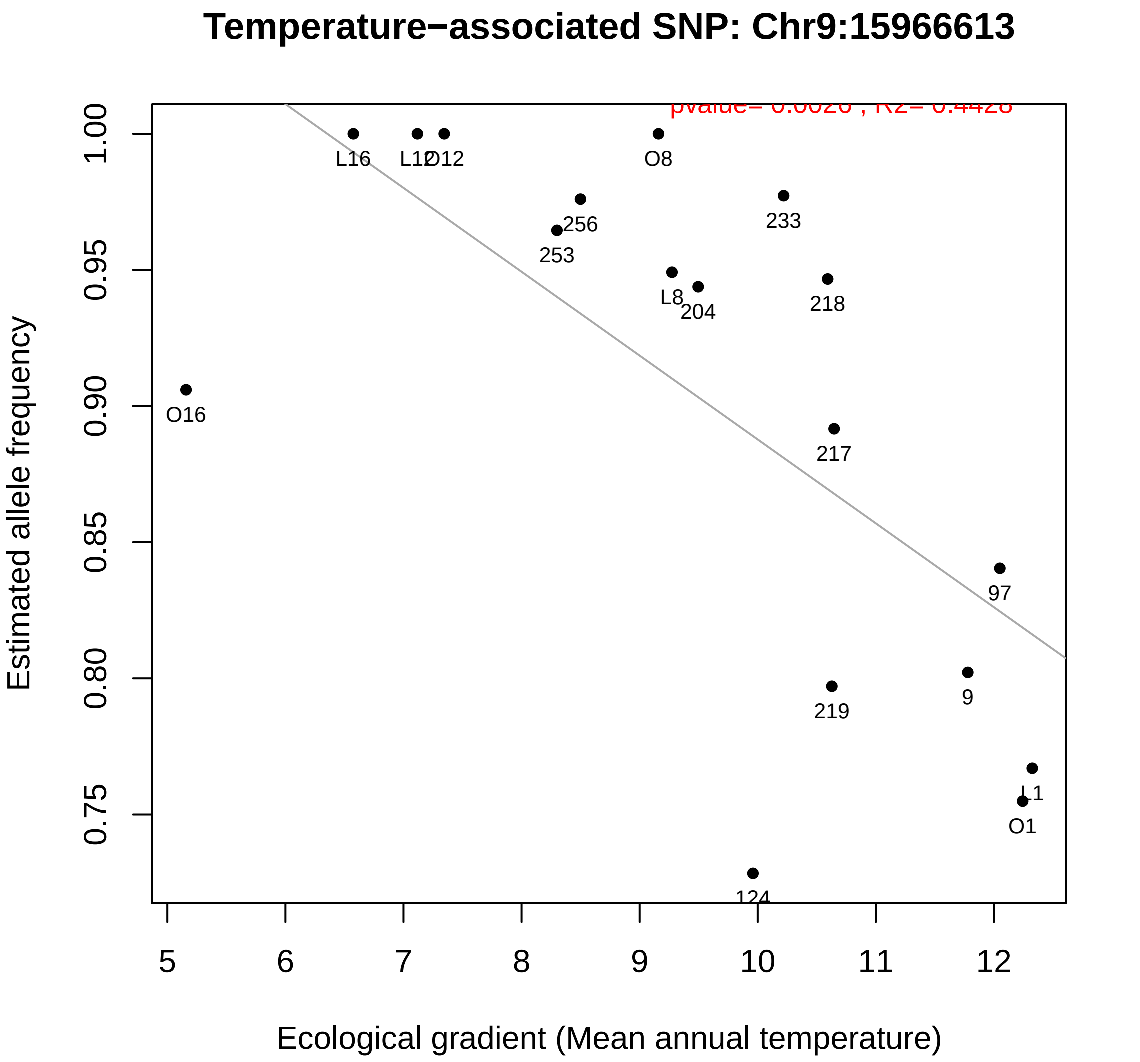

Temperature−associated SNP: Chr9:15966613
pvalue= 0.0026 ; R2= 0.4428
Estimated allele frequency
Ecological gradient (Mean annual temperature)
1.00
0.95
0.90
0.85
0.80
0.75
5
6
7
8
9
10
11
12
L16
L12
O12
O8
256
253
233
L8
204
218
O16
217
97
9
219
L1
O1
124

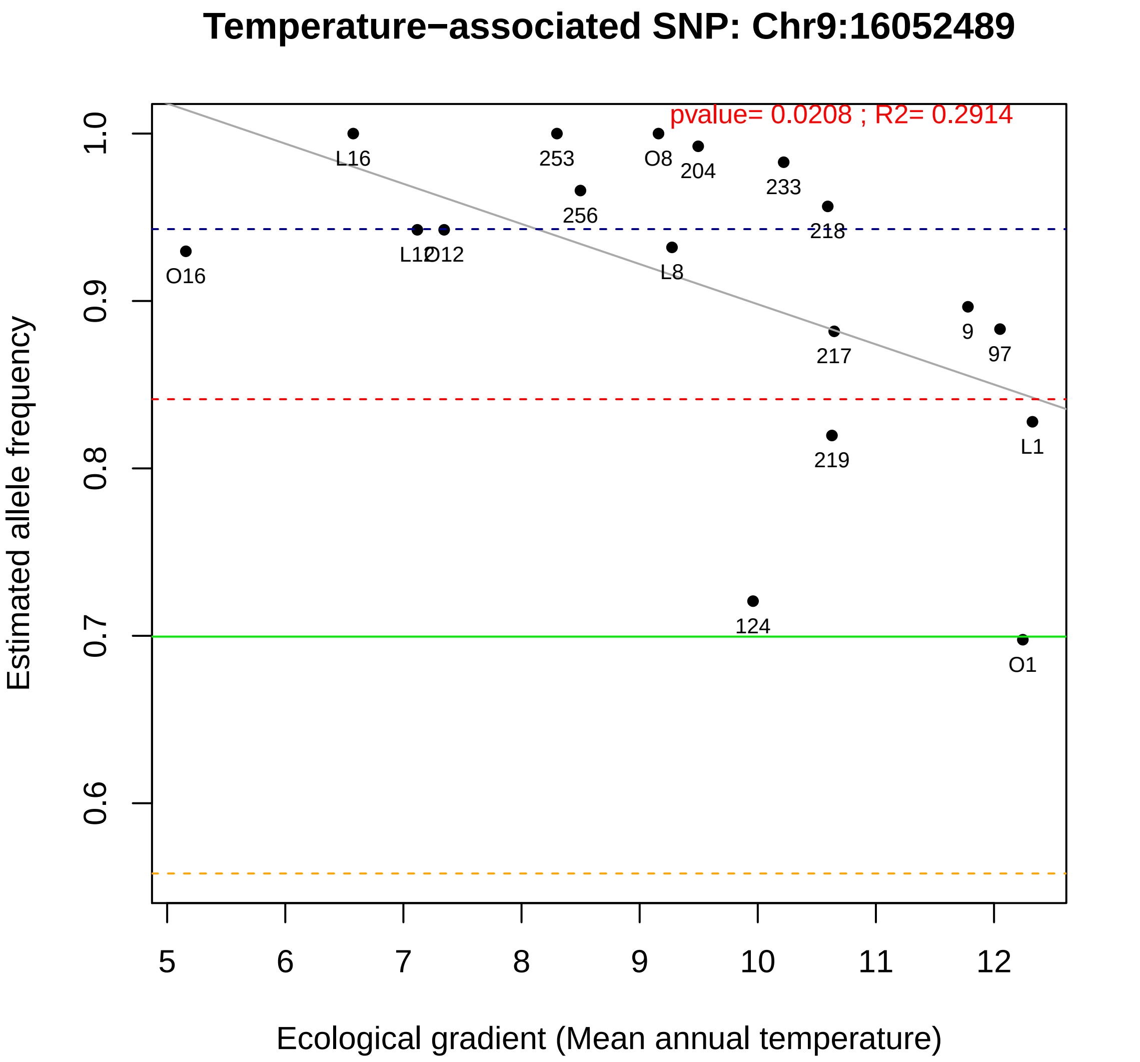
Temperature−associated SNP: Chr9:16052489
pvalue= 0.0208 ; R2= 0.2914
Estimated allele frequency
Ecological gradient (Mean annual temperature)
1.0
0.9
0.8
0.7
0.6
5
6
7
8
9
10
11
12
O16
L16
L12
O12
253
256
O8
204
L8
233
218
217
219
124
9
97
L1
O1

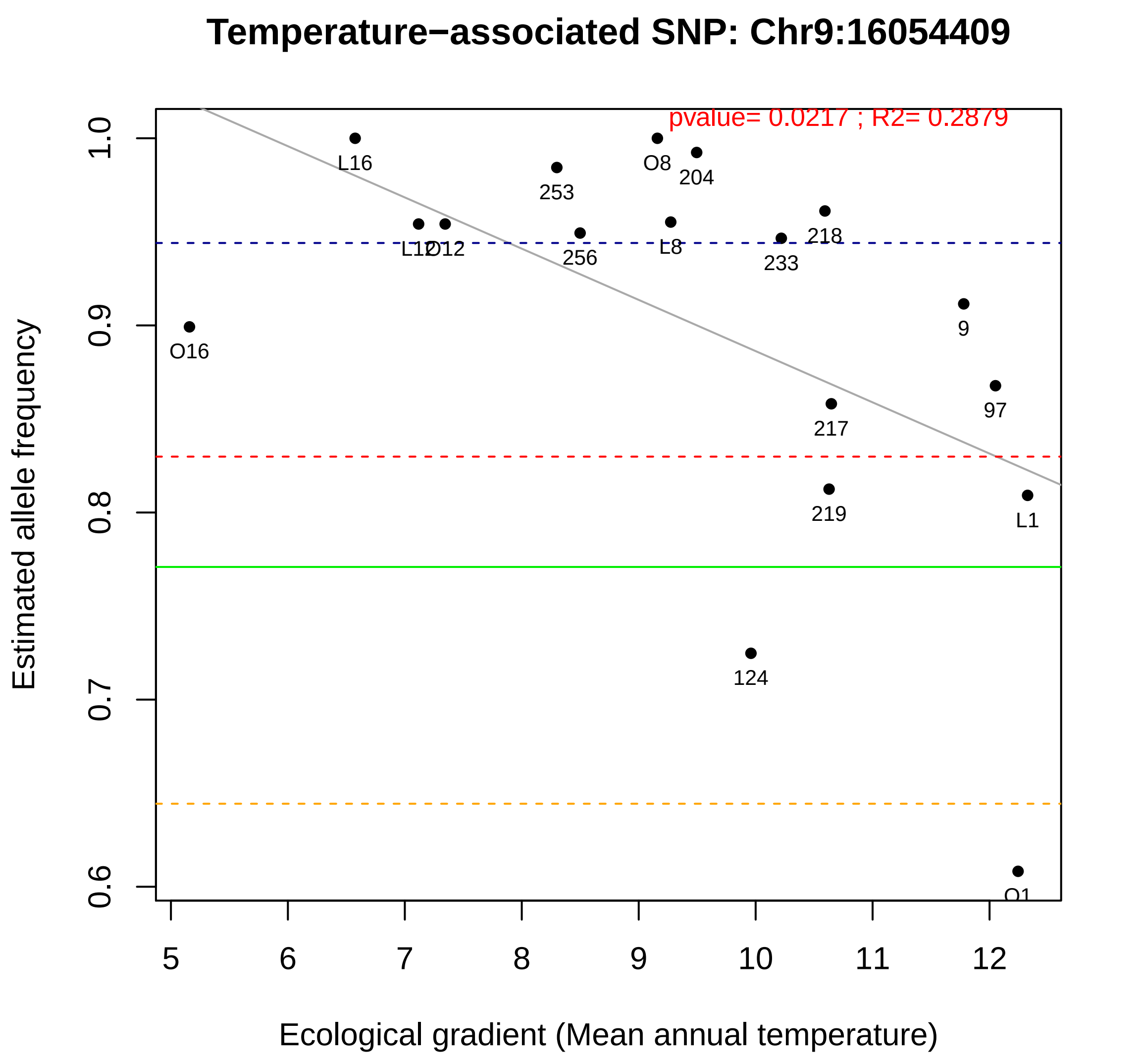

Temperature−associated SNP: Chr9:16054409
pvalue= 0.0217 ; R2= 0.2879
Estimated allele frequency
Ecological gradient (Mean annual temperature)
0.6
0.7
0.8
0.9
1.0
5
6
7
8
9
10
11
12
L16
O8
204
253
L12
O12
256
L8
218
233
O16
9
97
217
219
L1
124
O1

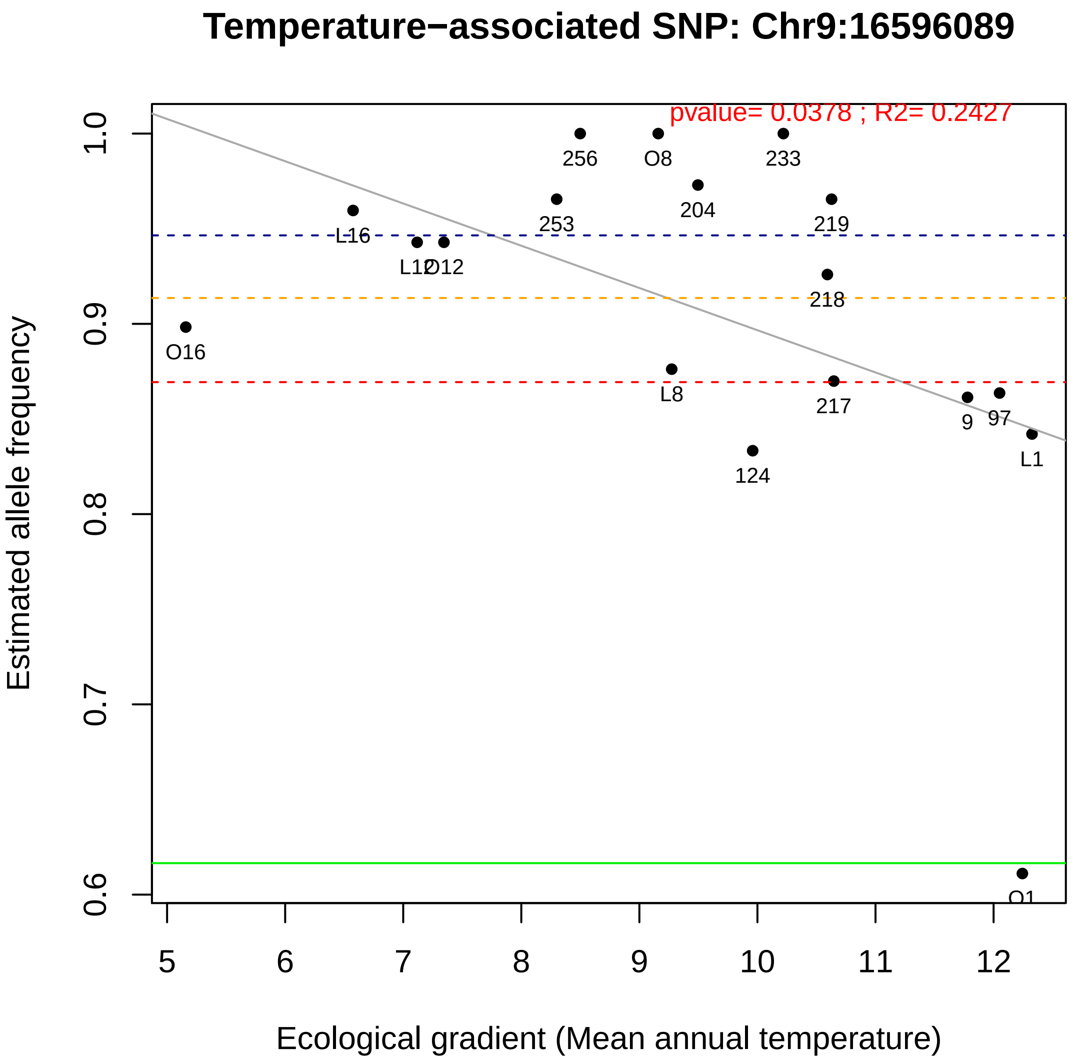
Temperature−associated SNP: Chr9:16596089
pvalue= 0.0378 ; R2= 0.2427
Estimated allele frequency
Ecological gradient (Mean annual temperature)
1.0
0.9
0.8
0.7
0.6
5
6
7
8
9
10
11
12
256
O8
233
204
253
219
L16
L12
O12
218
O16
L8
217
9
97
L1
124
O1

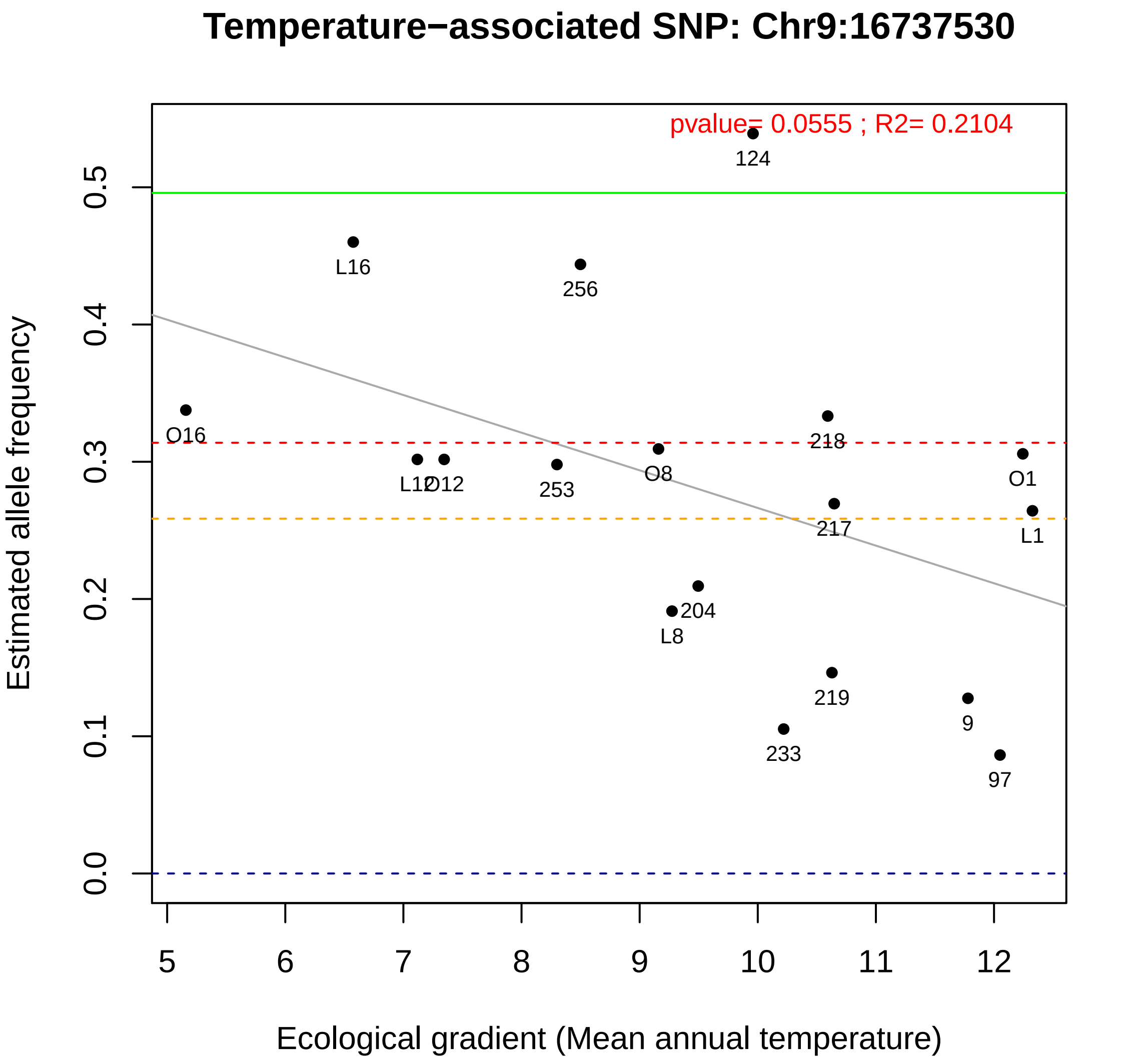
Temperature−associated SNP: Chr9:16737530
pvalue= 0.0555 ; R2= 0.2104
124
L16
256
O16
L12
O12
253
O8
218
O1
217
L1
204
L8
219
9
233
97
0.0
0.1
0.2
0.3
0.4
0.5
Estimated allele frequency
5
6
7
8
9
10
11
12
Ecological gradient (Mean annual temperature)

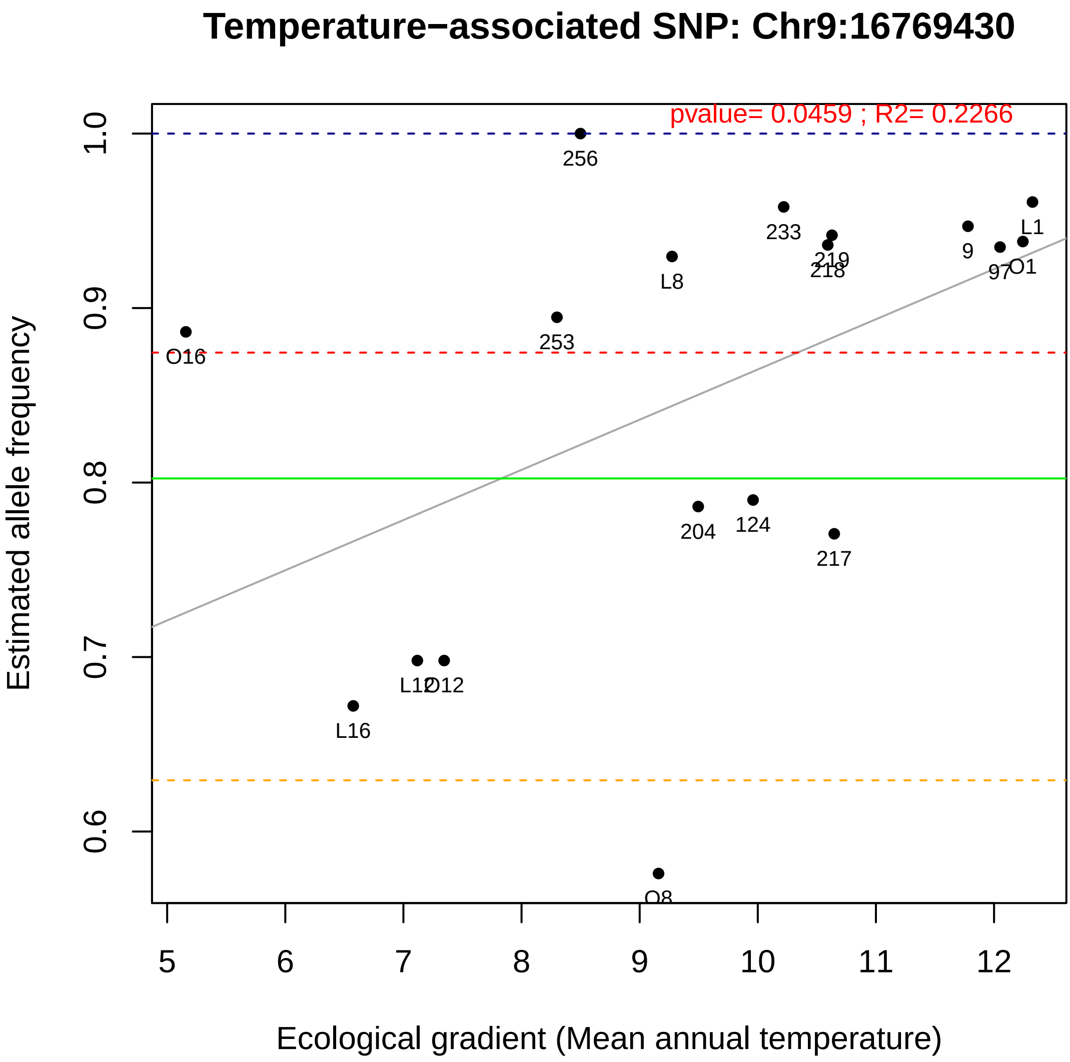

Temperature−associated SNP: Chr9:16769430
pvalue= 0.0459 ; R2= 0.2266
Estimated allele frequency
Ecological gradient (Mean annual temperature)
0.6
0.7
0.8
0.9
1.0
5
6
7
8
9
10
11
12
256
233
L1
9
219
218
97
O1
L8
253
O16
204
124
217
L12
O12
L16
O8

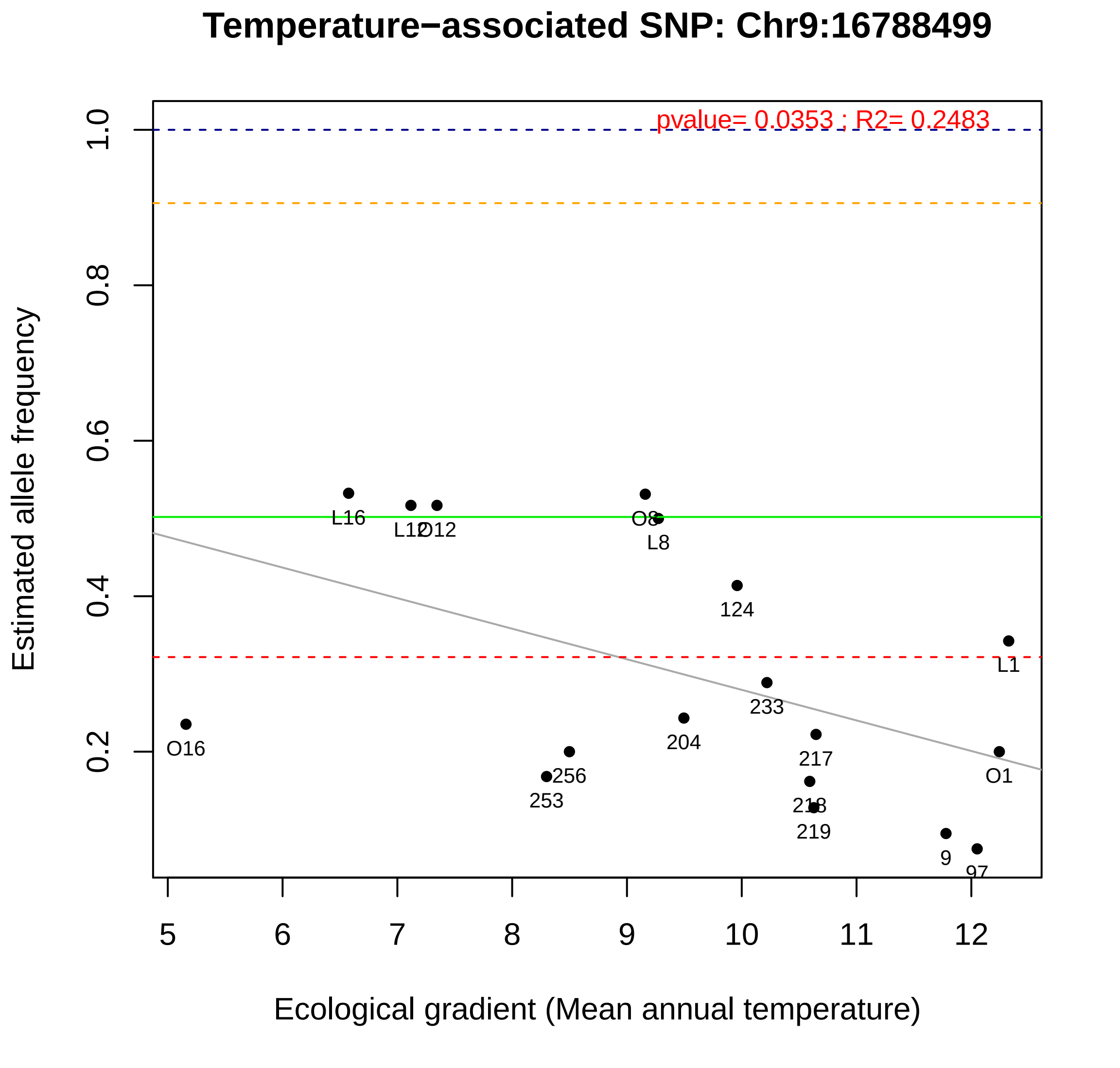
Temperature−associated SNP: Chr9:16788499
pvalue= 0.0353 ; R2= 0.2483
Estimated allele frequency
1.0
0.8
0.6
0.4
0.2
L16
L12
O12
O8
L8
124
L1
233
O16
204
217
256
O1
253
218
219
9
97
5
6
7
8
9
10
11
12
Ecological gradient (Mean annual temperature)

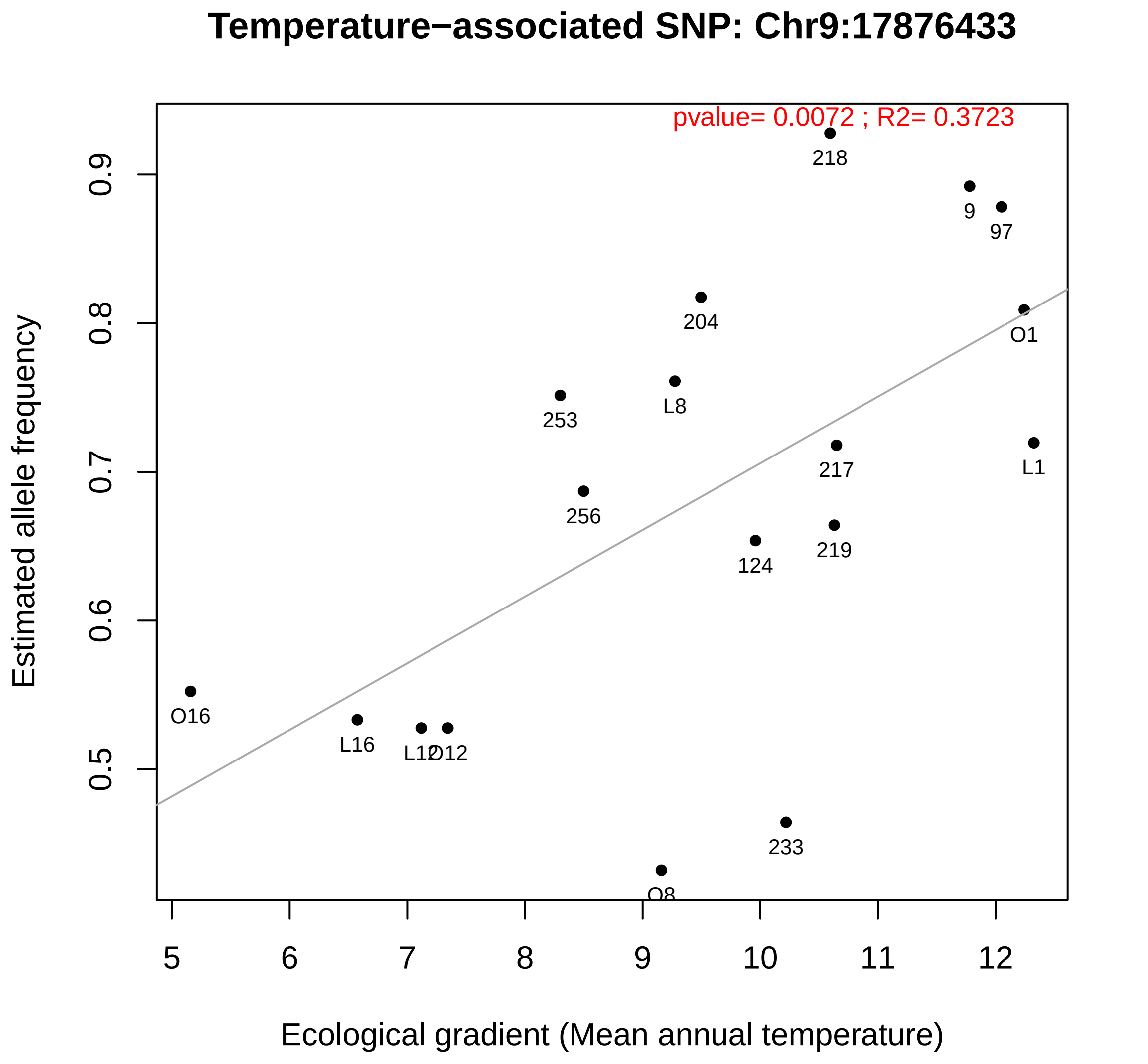
Temperature−associated SNP: Chr9:17876433
pvalue= 0.0072 ; R2= 0.3723
Estimated allele frequency
0.5
0.6
0.7
0.8
0.9
5
6
7
8
9
10
11
12
Ecological gradient (Mean annual temperature)
218
9
97
204
O1
L8
253
217
L1
256
124
219
O16
L16
L12
O12
233
O8

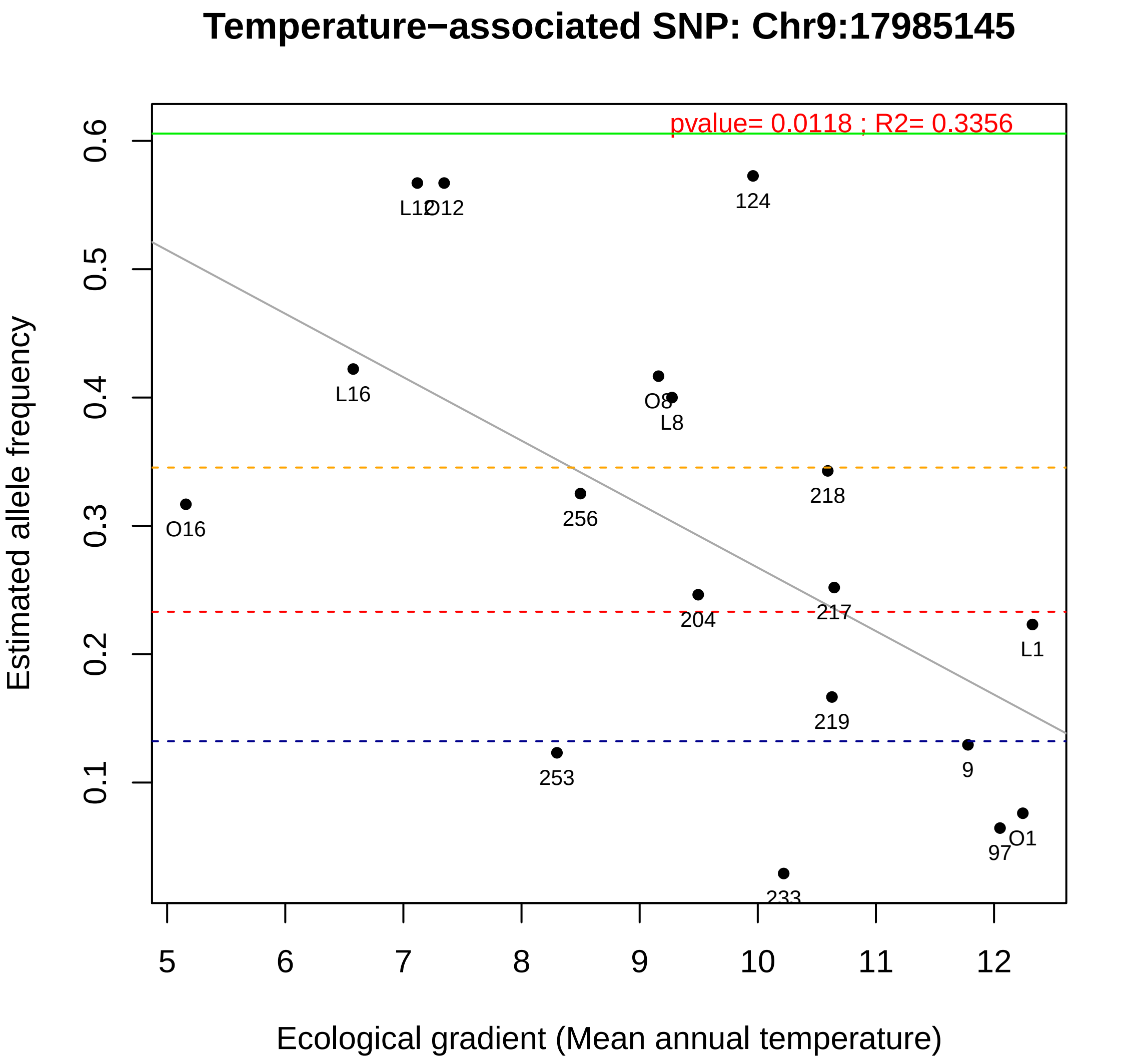

Temperature−associated SNP: Chr9:17985145
pvalue= 0.0118 ; R2= 0.3356
Estimated allele frequency
0.1
0.2
0.3
0.4
0.5
0.6
5
6
7
8
9
10
11
12
Ecological gradient (Mean annual temperature)
L12
O12
124
L16
O8
L8
O16
256
218
204
217
L1
219
253
9
O1
97
233

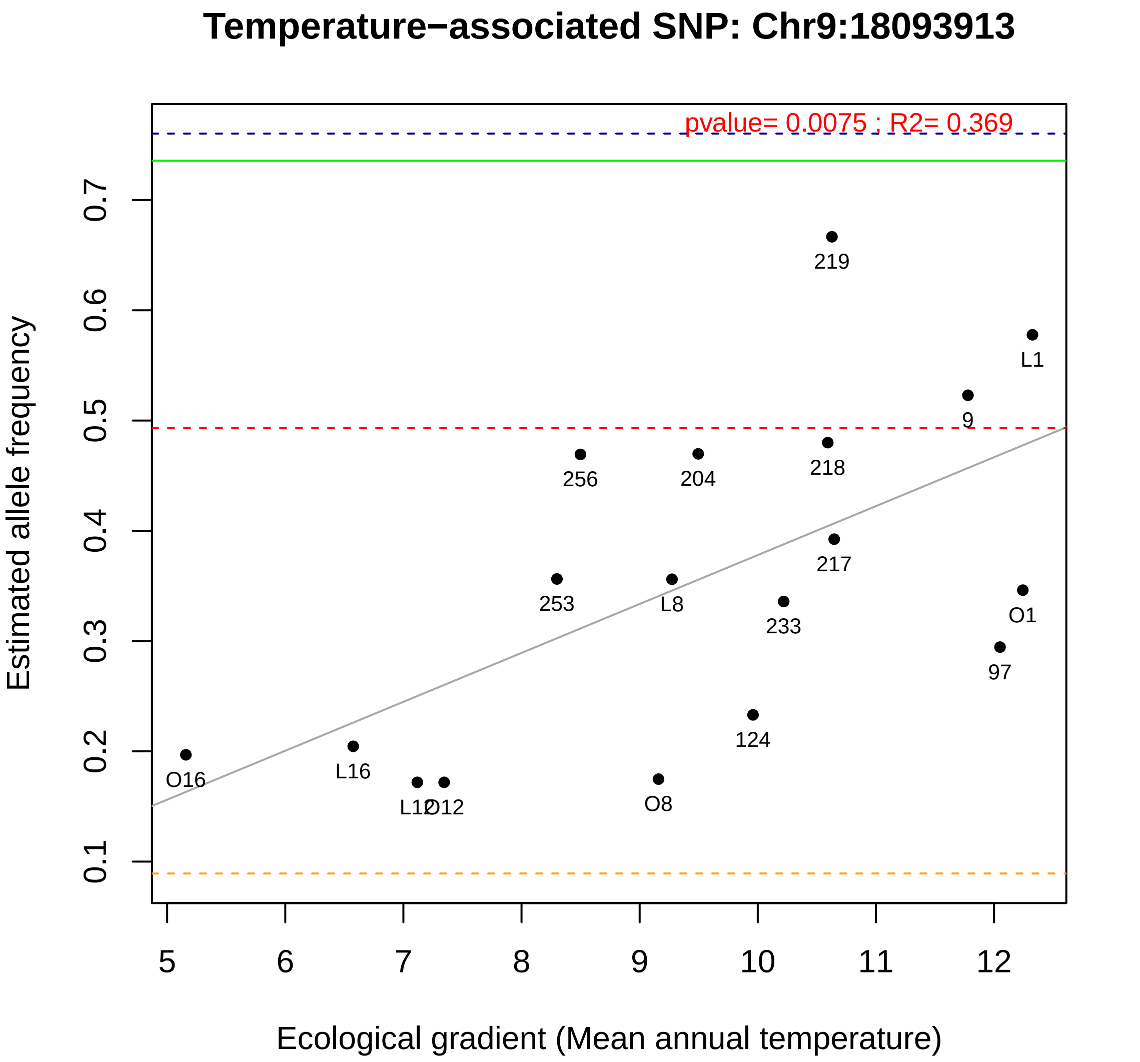
Temperature−associated SNP: Chr9:18093913
pvalue= 0.0075 ; R2= 0.369
Estimated allele frequency
0.1
0.2
0.3
0.4
0.5
0.6
0.7
5
6
7
8
9
10
11
12
Ecological gradient (Mean annual temperature)
219
L1
9
256
204
218
217
253
L8
233
O1
97
124
O16
L16
L12
O12
O8

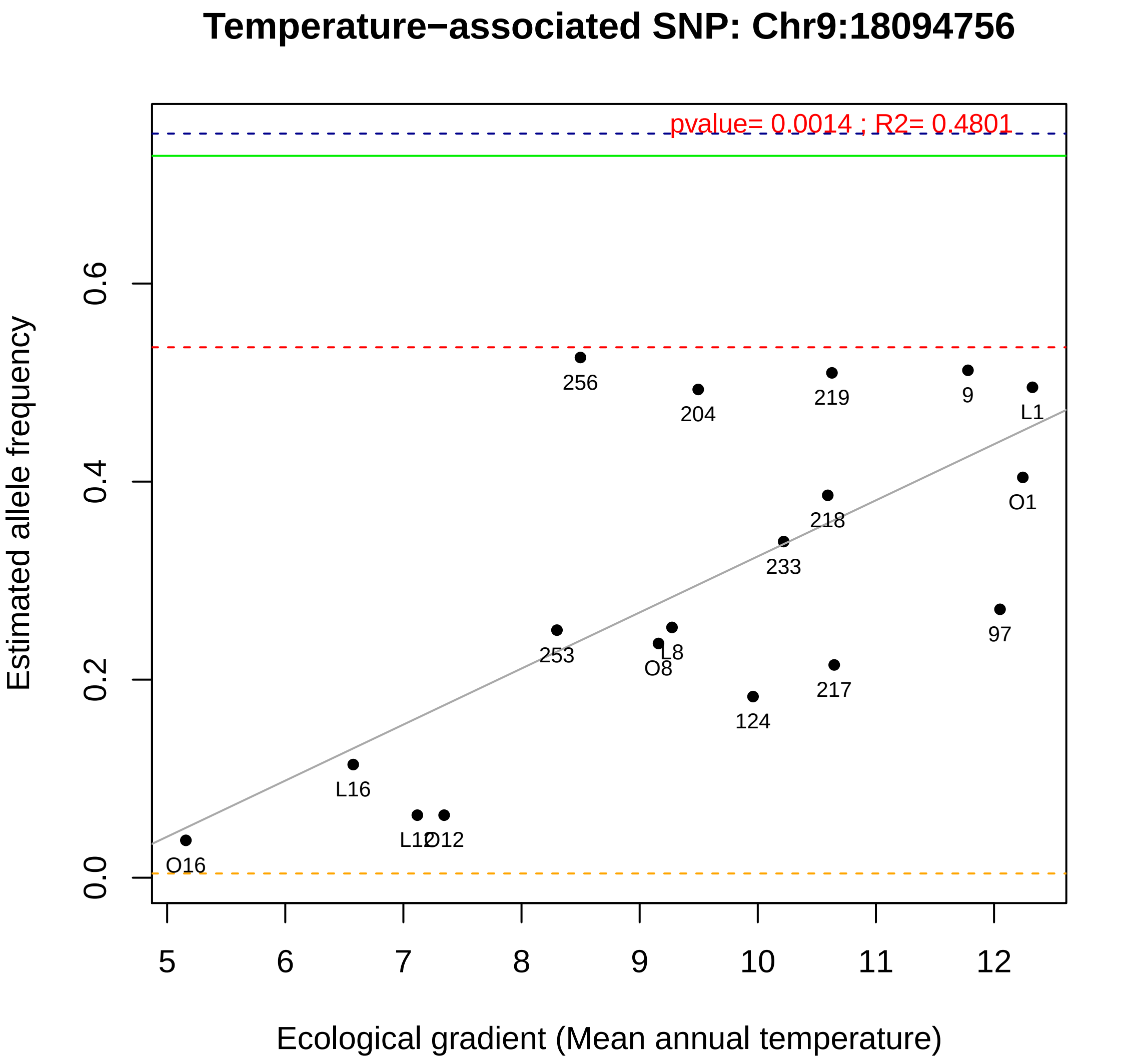
Temperature−associated SNP: Chr9:18094756
pvalue= 0.0014 ; R2= 0.4801
Estimated allele frequency
0.0
0.2
0.4
0.6
5
6
7
8
9
10
11
12
Ecological gradient (Mean annual temperature)
256
204
219
9
L1
O1
218
233
97
253
L8
O8
124
217
L16
L12
O12
O16

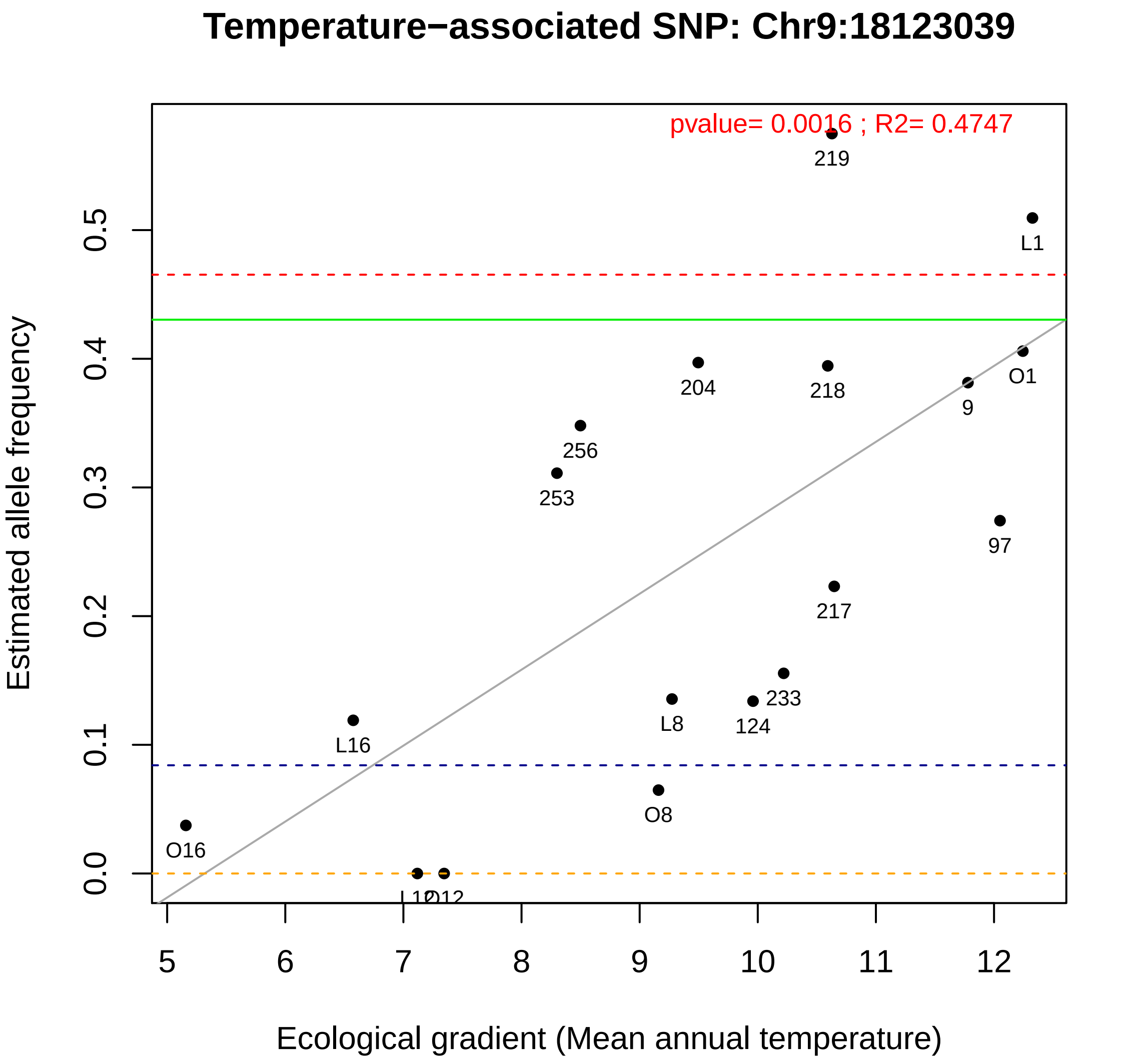
Temperature−associated SNP: Chr9:18123039
pvalue= 0.0016 ; R2= 0.4747
Estimated allele frequency
0.0
0.1
0.2
0.3
0.4
0.5
5
6
7
8
9
10
11
12
Ecological gradient (Mean annual temperature)
219
L1
O1
204
218
9
256
253
97
217
233
L8
124
L16
O8
O16
O12

**Temperature−associated SNP: Chr9:18145590**

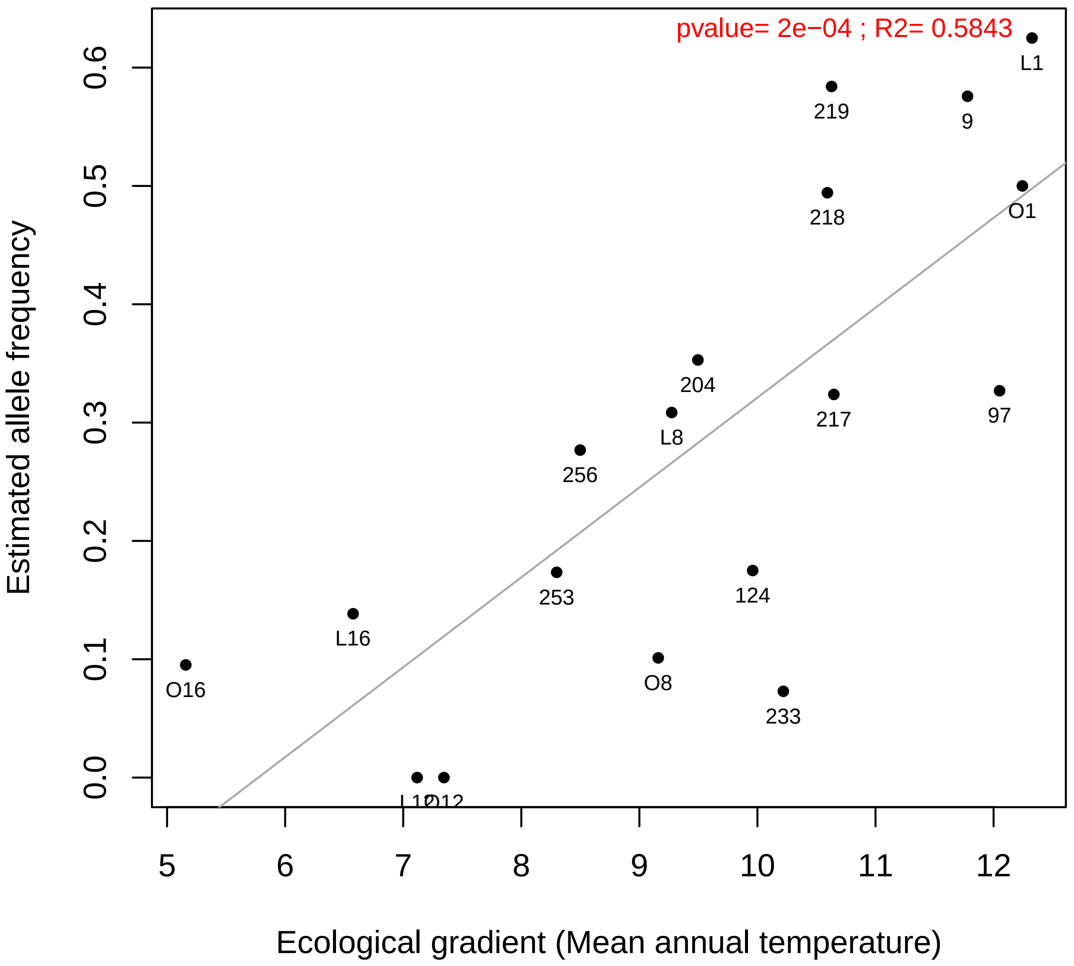

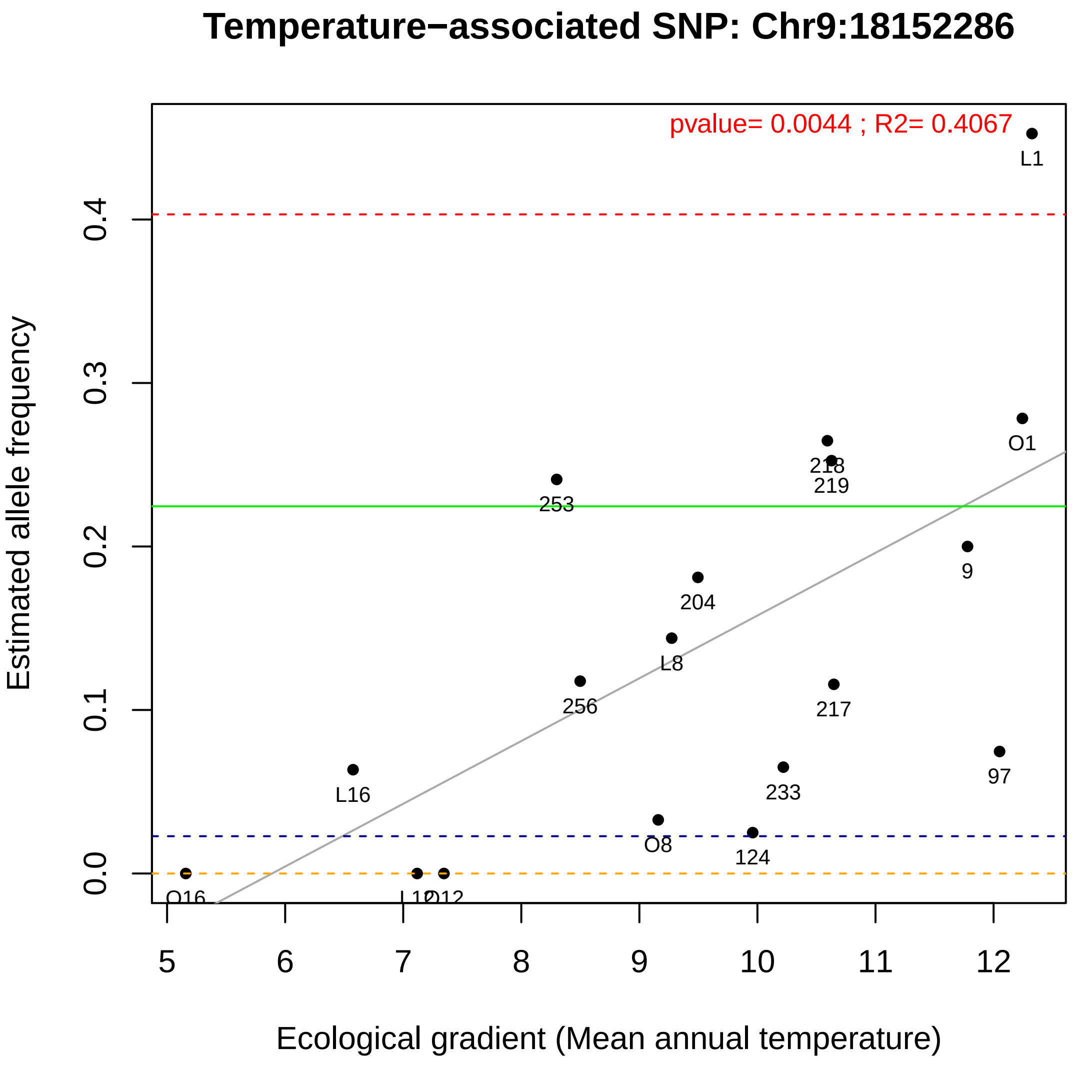

Temperature−associated SNP: Chr9:18152286
pvalue= 0.0044 ; R2= 0.4067
Estimated allele frequency
0.0
0.1
0.2
0.3
0.4
Ecological gradient (Mean annual temperature)
5
6
7
8
9
10
11
12
L1
O1
218
219
253
9
204
L8
256
217
97
L16
233
O8
124
O16
O12

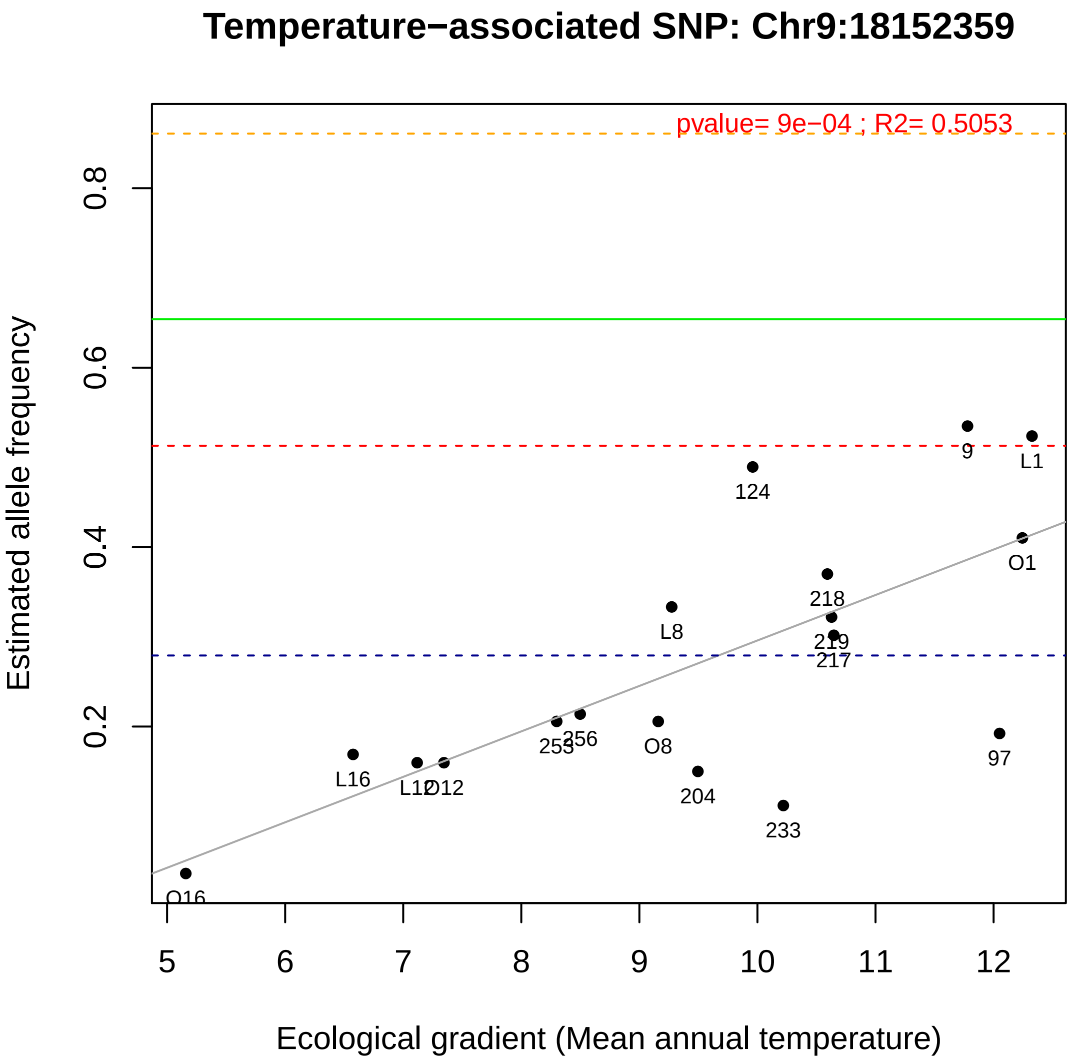
Temperature−associated SNP: Chr9:18152359
pvalue= 9e−04 ; R2= 0.5053
Estimated allele frequency
0.2
0.4
0.6
0.8
Ecological gradient (Mean annual temperature)
5
6
7
8
9
10
11
12
O16
L16
L12
O12
253
256
O8
204
233
L8
124
218
219
217
9
L1
O1
97

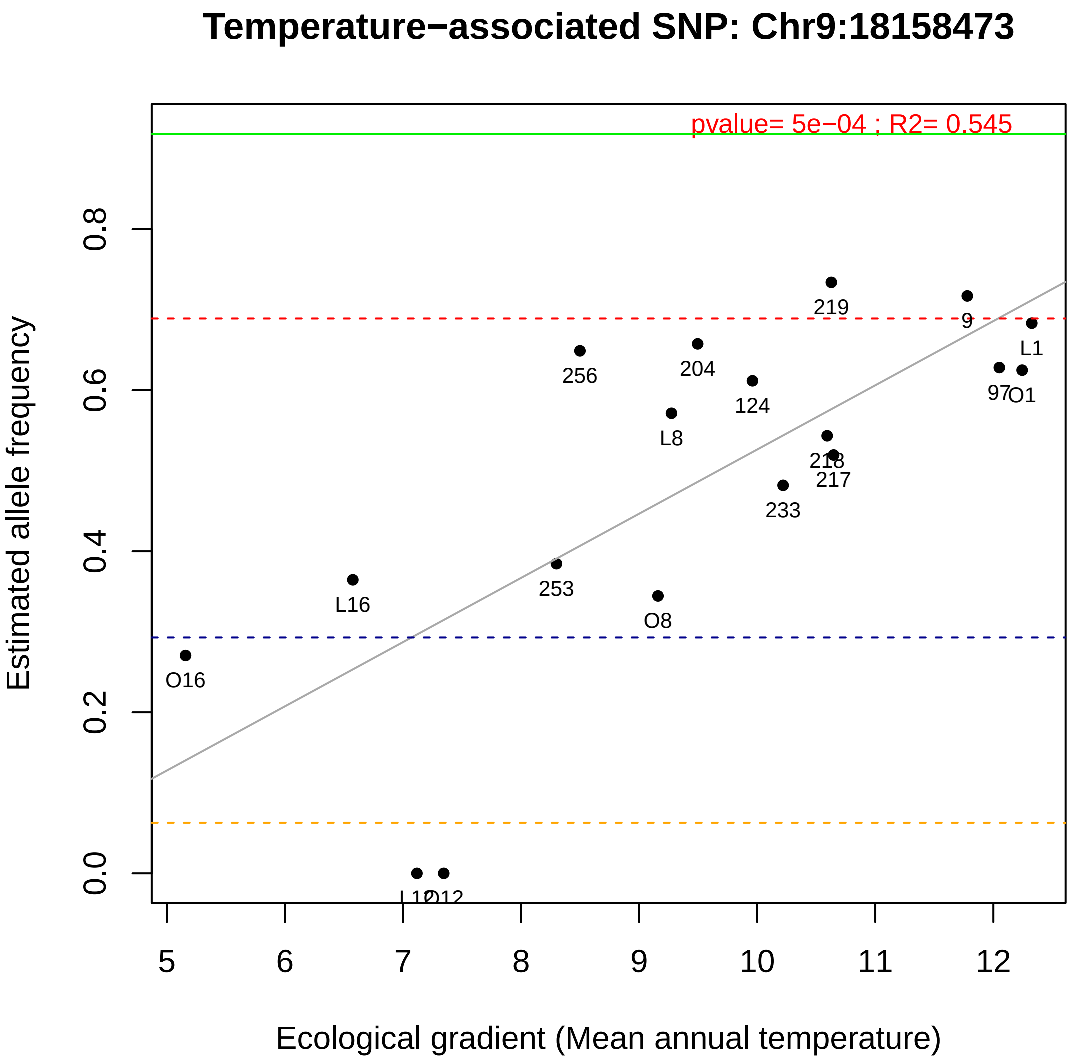

Temperature−associated SNP: Chr9:18158473
pvalue= 5e−04 ; R2= 0.545
Estimated allele frequency
0.0
0.2
0.4
0.6
0.8
5
6
7
8
9
10
11
12
Ecological gradient (Mean annual temperature)
O16
L16
253
256
L8
O8
204
124
233
218
217
219
9
L1
97
O1

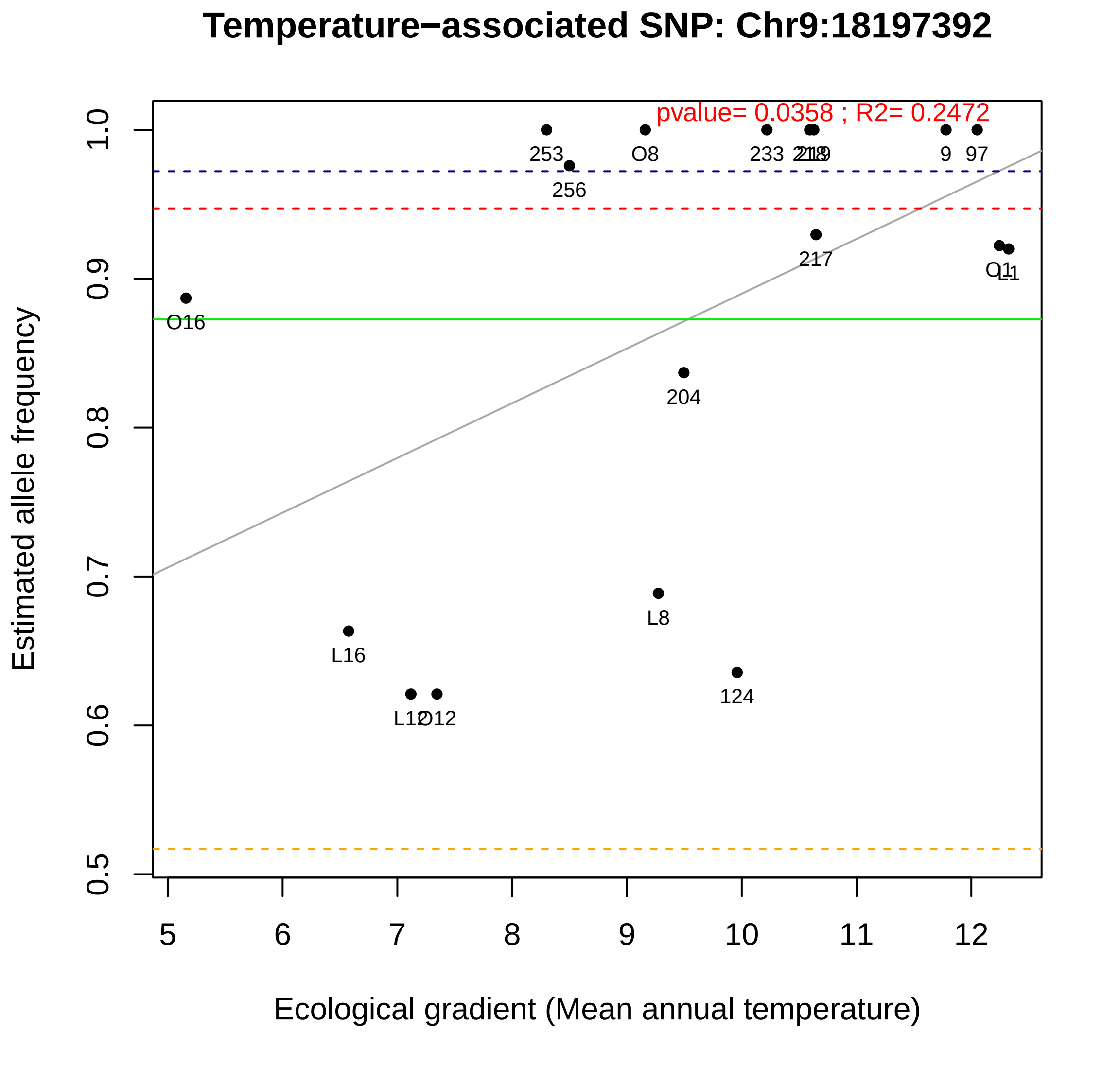

Temperature−associated SNP: Chr9:18197392
pvalue= 0.0358 ; R2= 0.2472
253
256
O8
233
9
97
217
O16
204
L8
L16
124
L12
O12
1.0
0.9
0.8
0.7
0.6
0.5
5
6
7
8
9
10
11
12
Estimated allele frequency
Ecological gradient (Mean annual temperature)

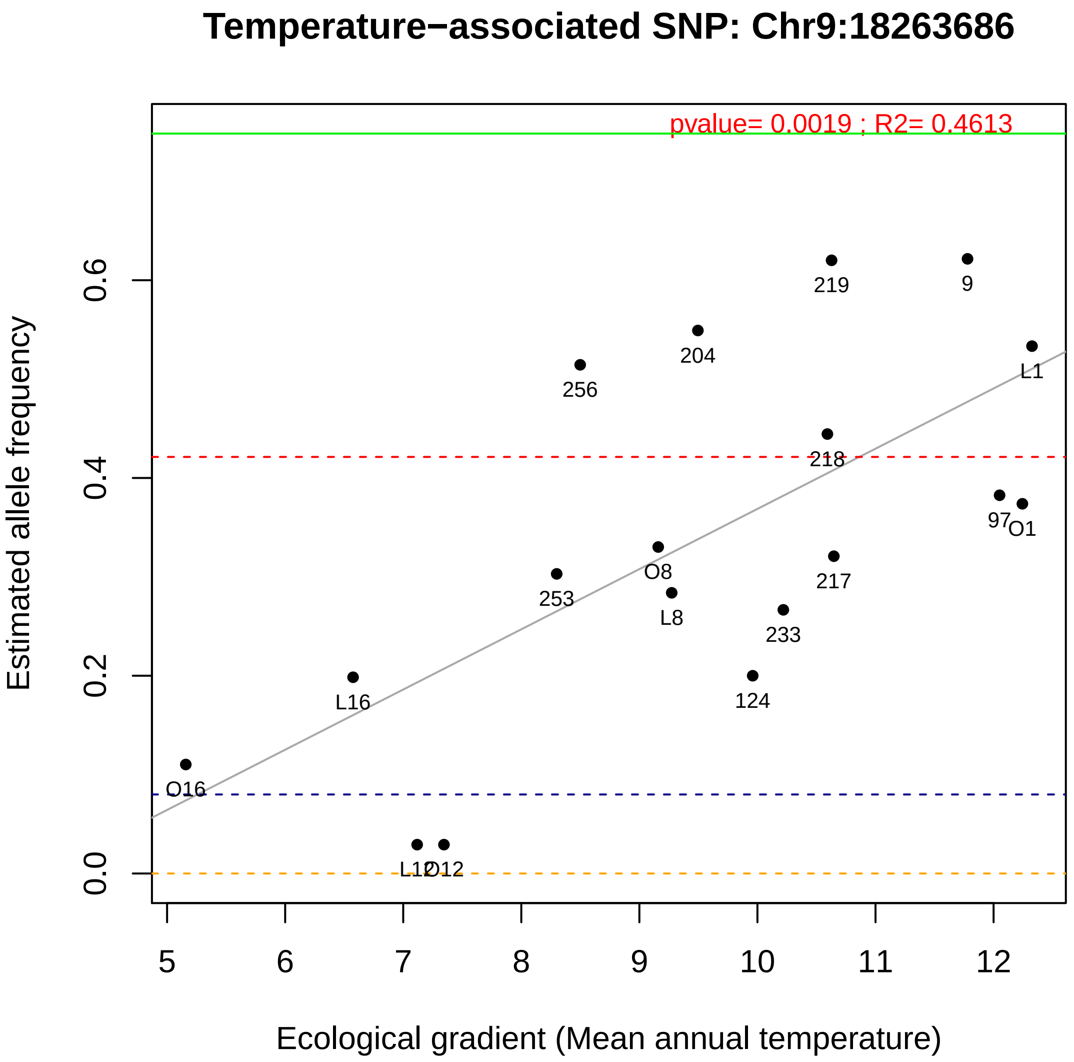

Temperature−associated SNP: Chr9:18263686
pvalue= 0.0019 ; R2= 0.4613
Estimated allele frequency
0.0
0.2
0.4
0.6
5
6
7
8
9
10
11
12
Ecological gradient (Mean annual temperature)
219
9
204
256
L1
218
97
O1
O8
217
253
L8
233
L16
124
O16
L12
O12

**Temperature−associated SNP: Chr9:18264591**

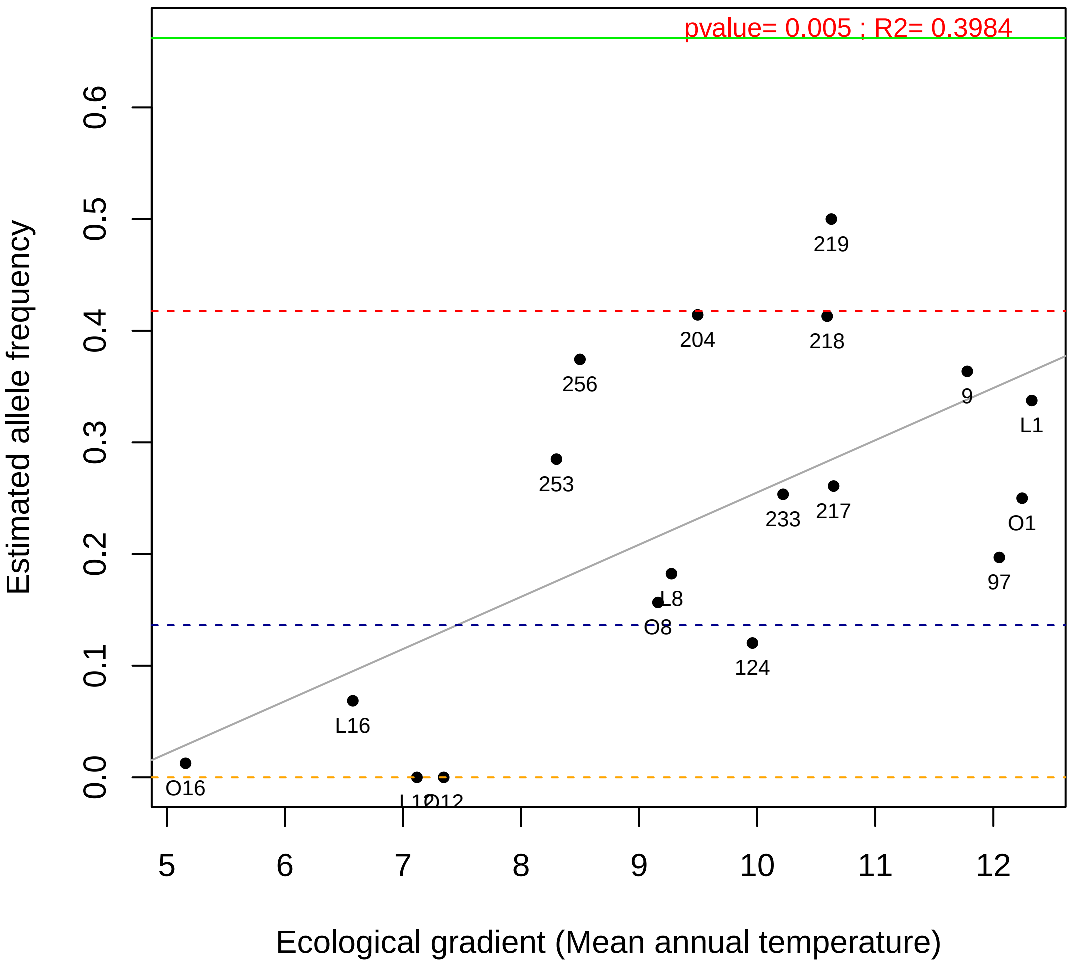

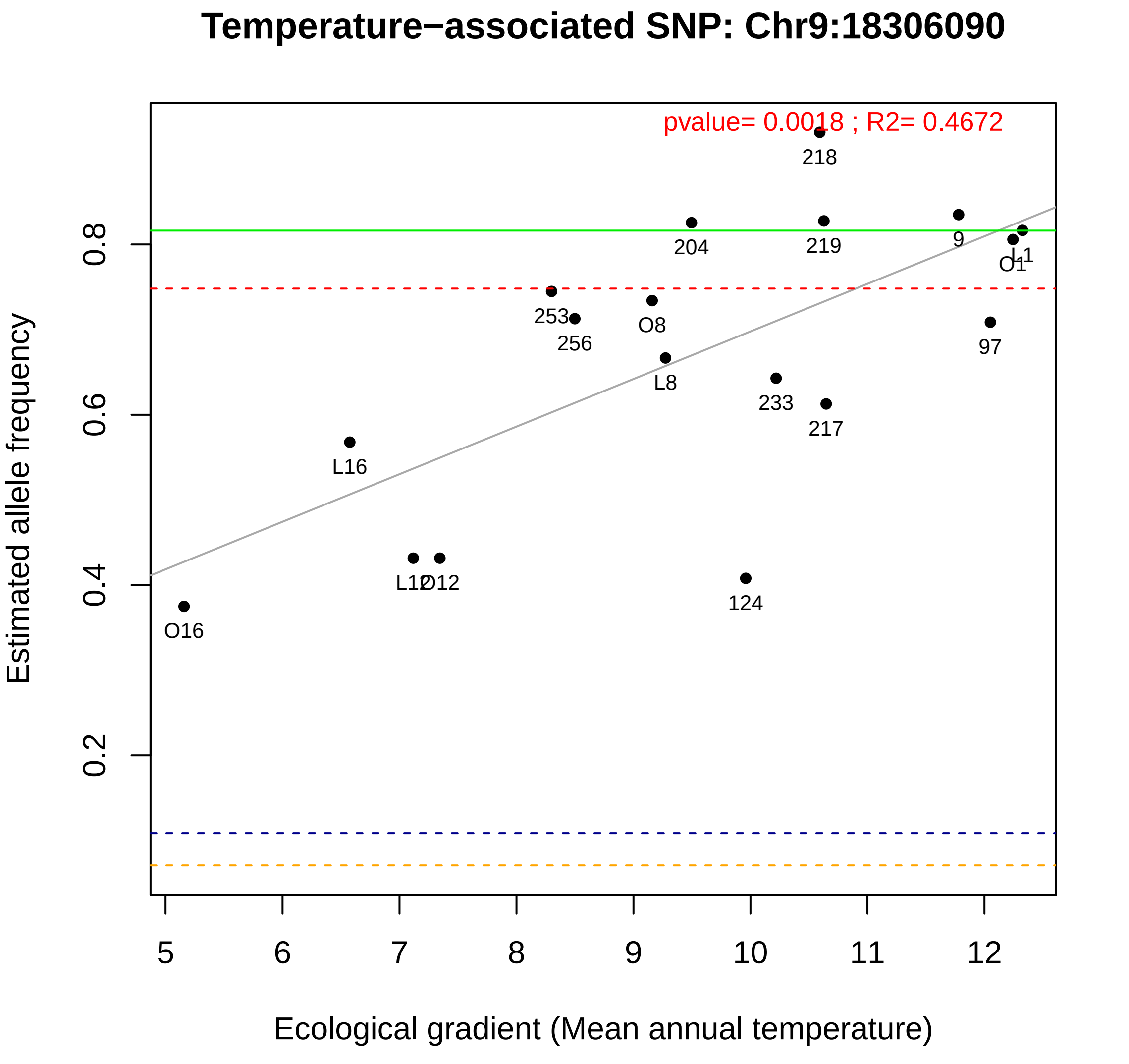
Temperature−associated SNP: Chr9:18306090
pvalue= 0.0018 ; R2= 0.4672
Estimated allele frequency
0.2
0.4
0.6
0.8
Ecological gradient (Mean annual temperature)
5
6
7
8
9
10
11
12
218
204
219
9
L1
O1
253
256
O8
L8
97
233
217
L16
L12
O12
124
O16

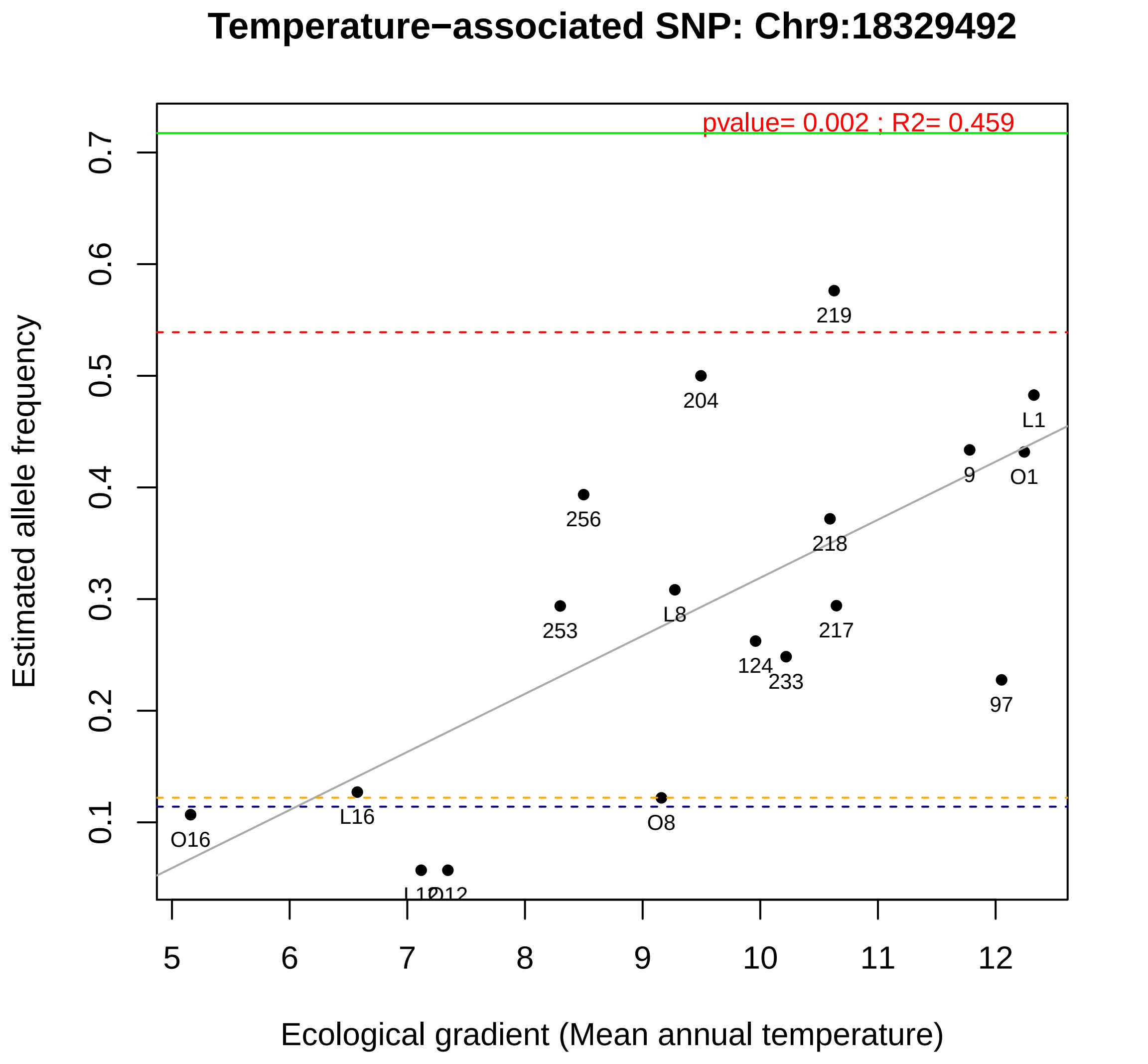

Temperature−associated SNP: Chr9:18329492
pvalue= 0.002 ; R2= 0.459
Estimated allele frequency
0.1
0.2
0.3
0.4
0.5
0.6
0.7
219
204
L1
9
O1
256
218
L8
253
217
124
233
97
L16
O8
O16
Ecological gradient (Mean annual temperature)
5
6
7
8
9
10
11
12

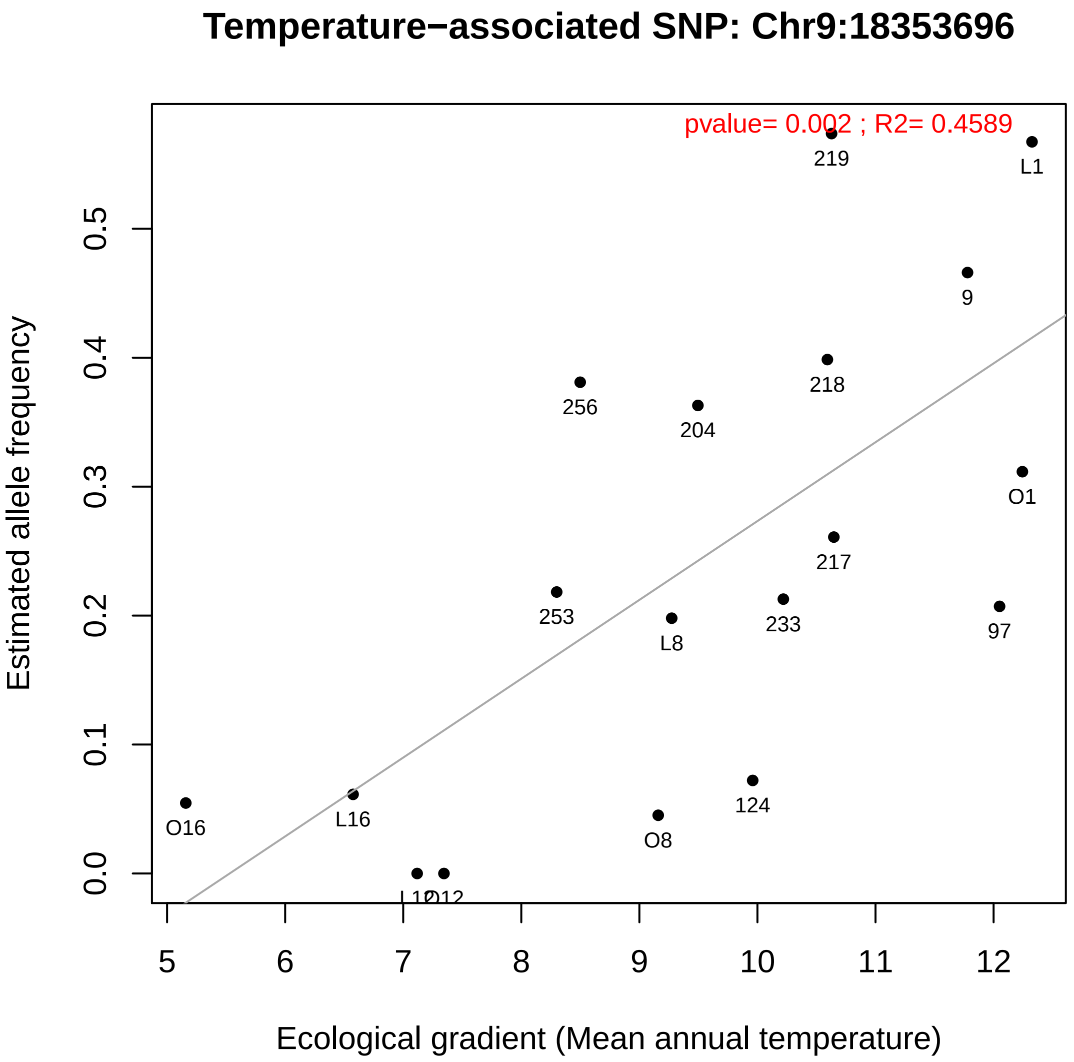
Temperature−associated SNP: Chr9:18353696
pvalue= 0.002 ; R2= 0.4589
Estimated allele frequency
0.0
0.1
0.2
0.3
0.4
0.5
Ecological gradient (Mean annual temperature)
5
6
7
8
9
10
11
12
219
L1
9
218
256
204
O1
217
253
L8
233
97
124
O16
L16
O8
O12

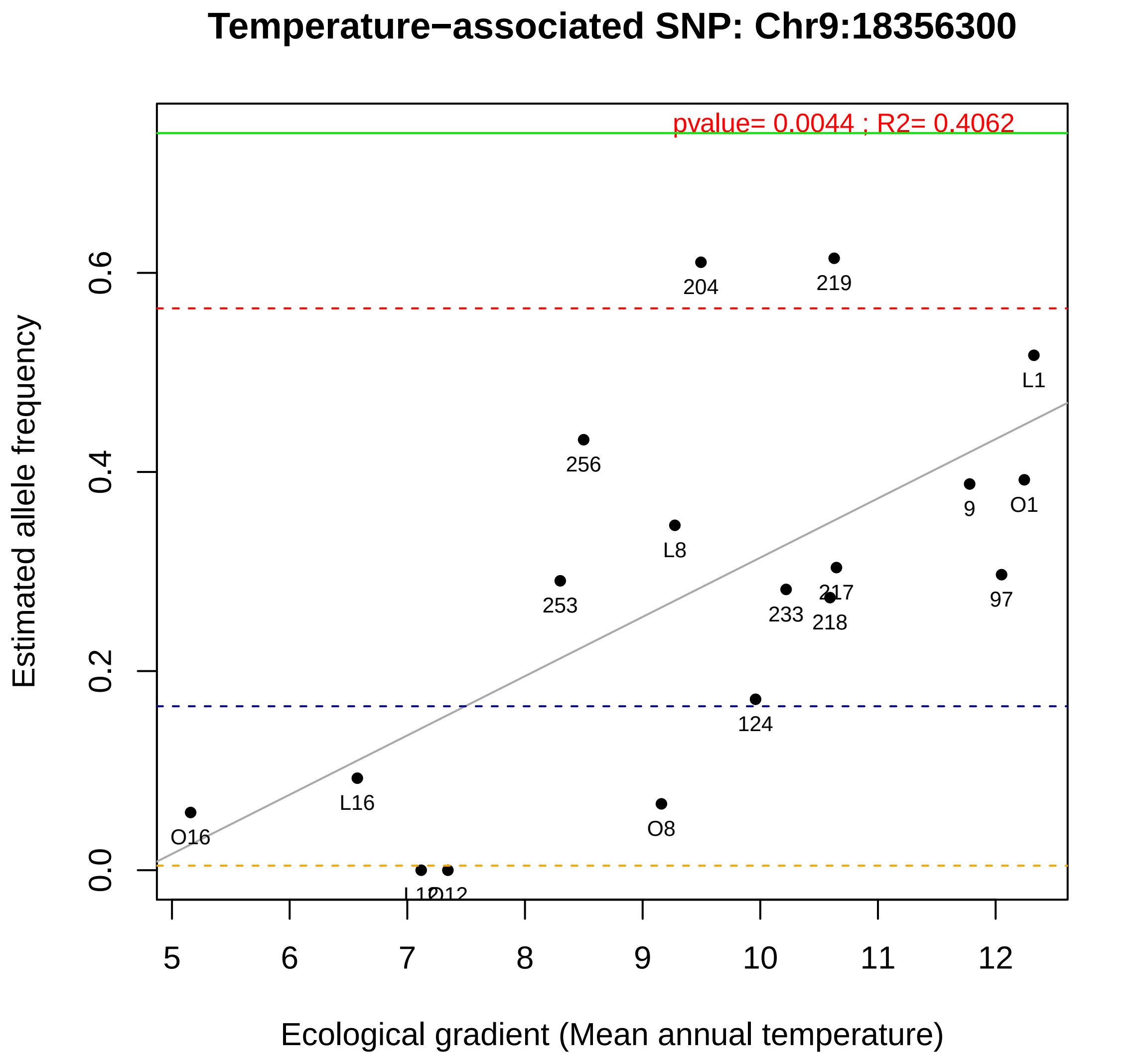
Temperature−associated SNP: Chr9:18356300
pvalue= 0.0044 ; R2= 0.4062
204
219
L1
256
9
O1
L8
217
97
253
233
218
124
L16
O16
O8
L12
O12
0.0
0.2
0.4
0.6
5
6
7
8
9
10
11
12
Estimated allele frequency
Ecological gradient (Mean annual temperature)

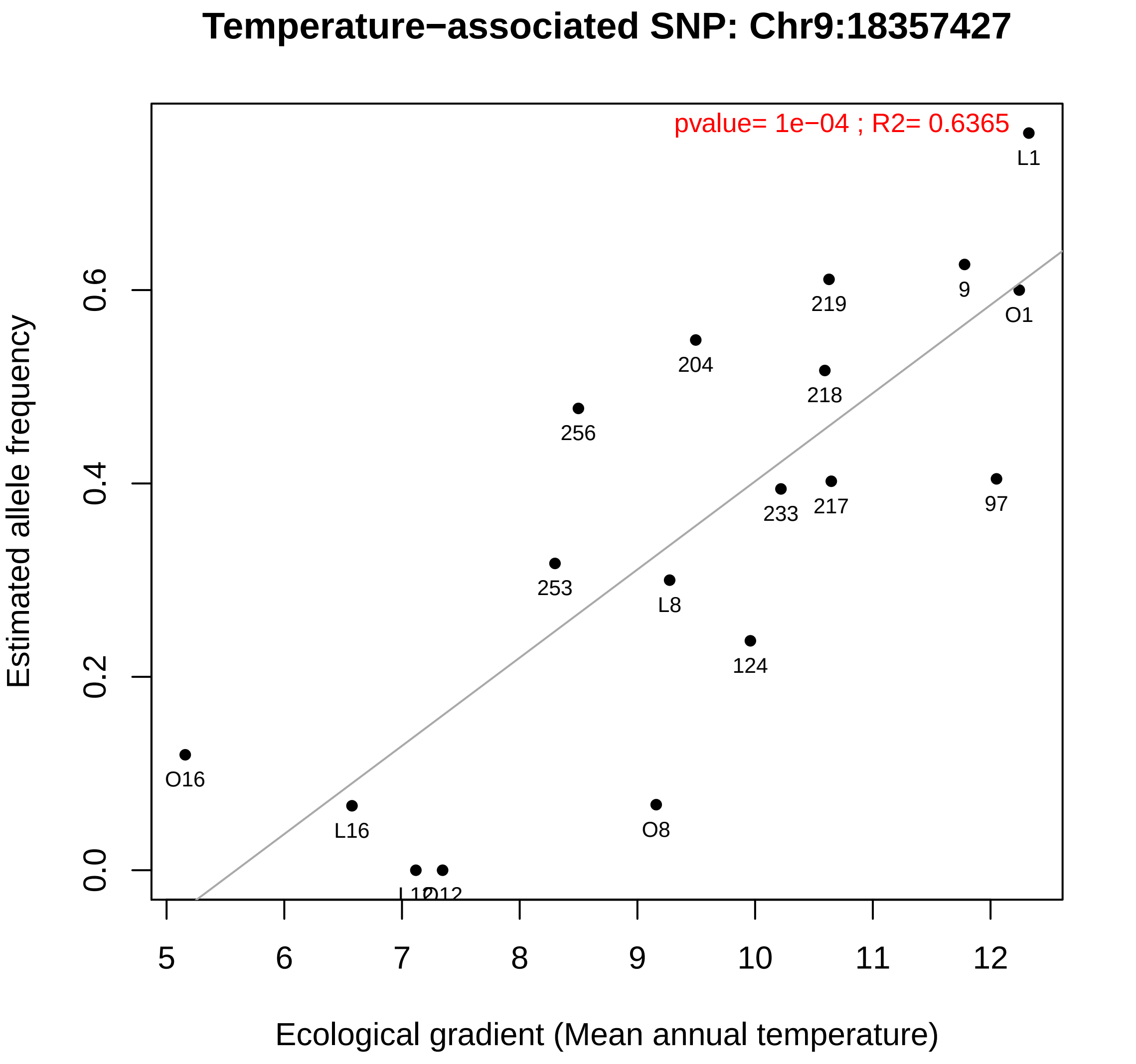

Temperature−associated SNP: Chr9:18357427
pvalue= 1e−04 ; R2= 0.6365
Estimated allele frequency
0.0
0.2
0.4
0.6
5
6
7
8
9
10
11
12
Ecological gradient (Mean annual temperature)
L1
9
O1
219
204
218
256
233
217
97
253
L8
124
O16
L16
O8
O12

**Temperature−associated SNP: Chr9:18357968**

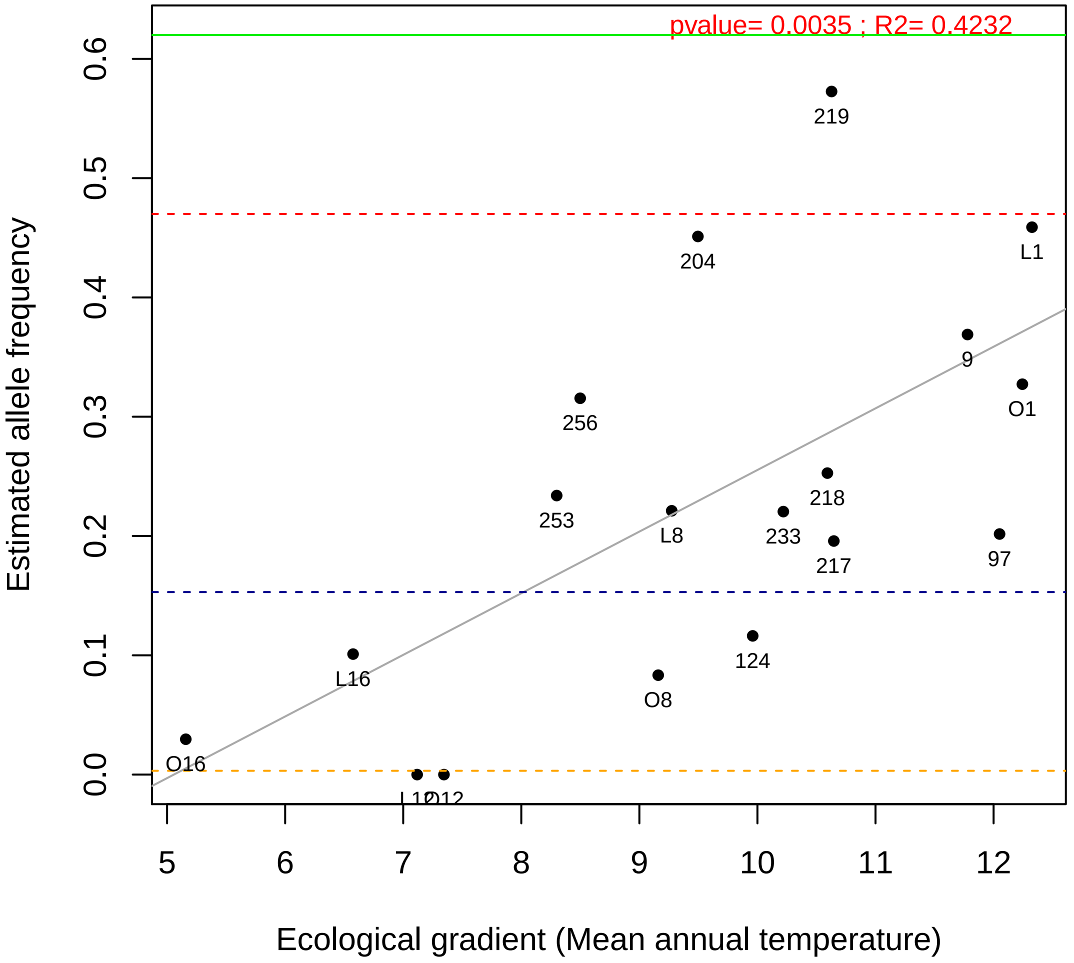

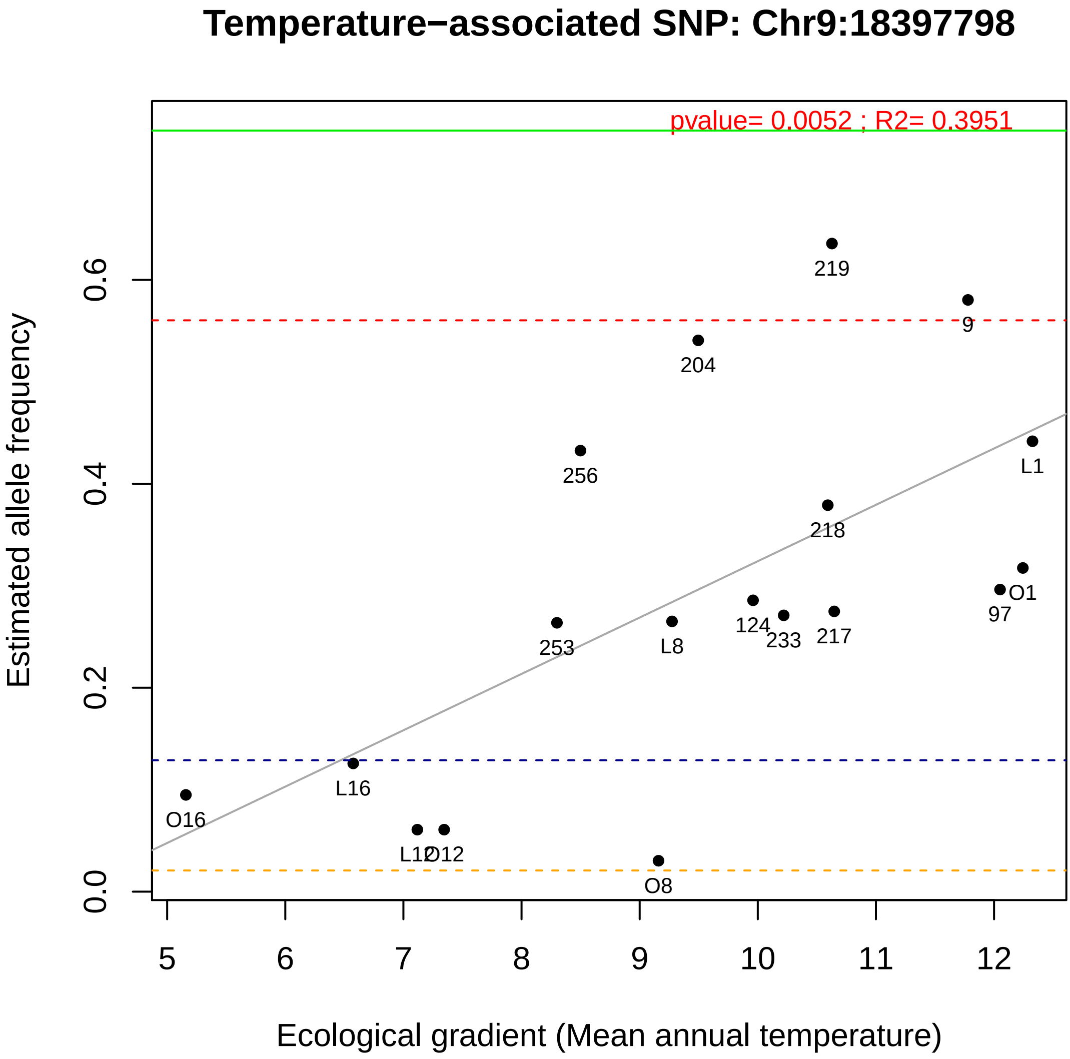
Temperature−associated SNP: Chr9:18397798
pvalue= 0.0052 ; R2= 0.3951
Estimated allele frequency
0.0
0.2
0.4
0.6
5
6
7
8
9
10
11
12
Ecological gradient (Mean annual temperature)
219
9
204
256
L1
218
O1
97
124
233
217
253
L8
L16
O16
L12
O12
O8

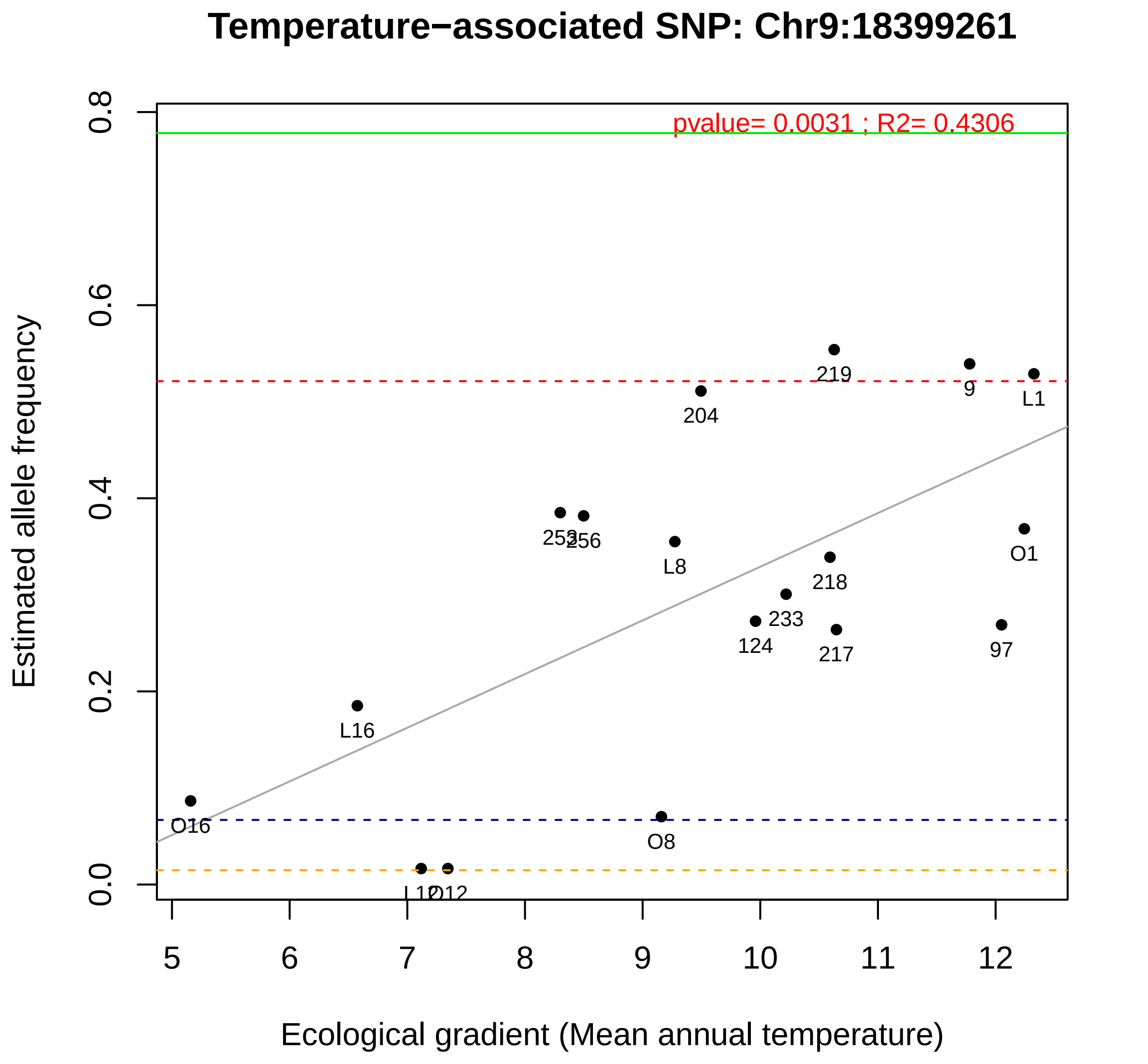
Temperature−associated SNP: Chr9:18399261
pvalue= 0.0031 ; R2= 0.4306
Estimated allele frequency
0.0
0.2
0.4
0.6
0.8
5
6
7
8
9
10
11
12
Ecological gradient (Mean annual temperature)
219
9
L1
204
253
256
L8
O1
218
233
124
217
97
L16
O16
O8
L12
O12

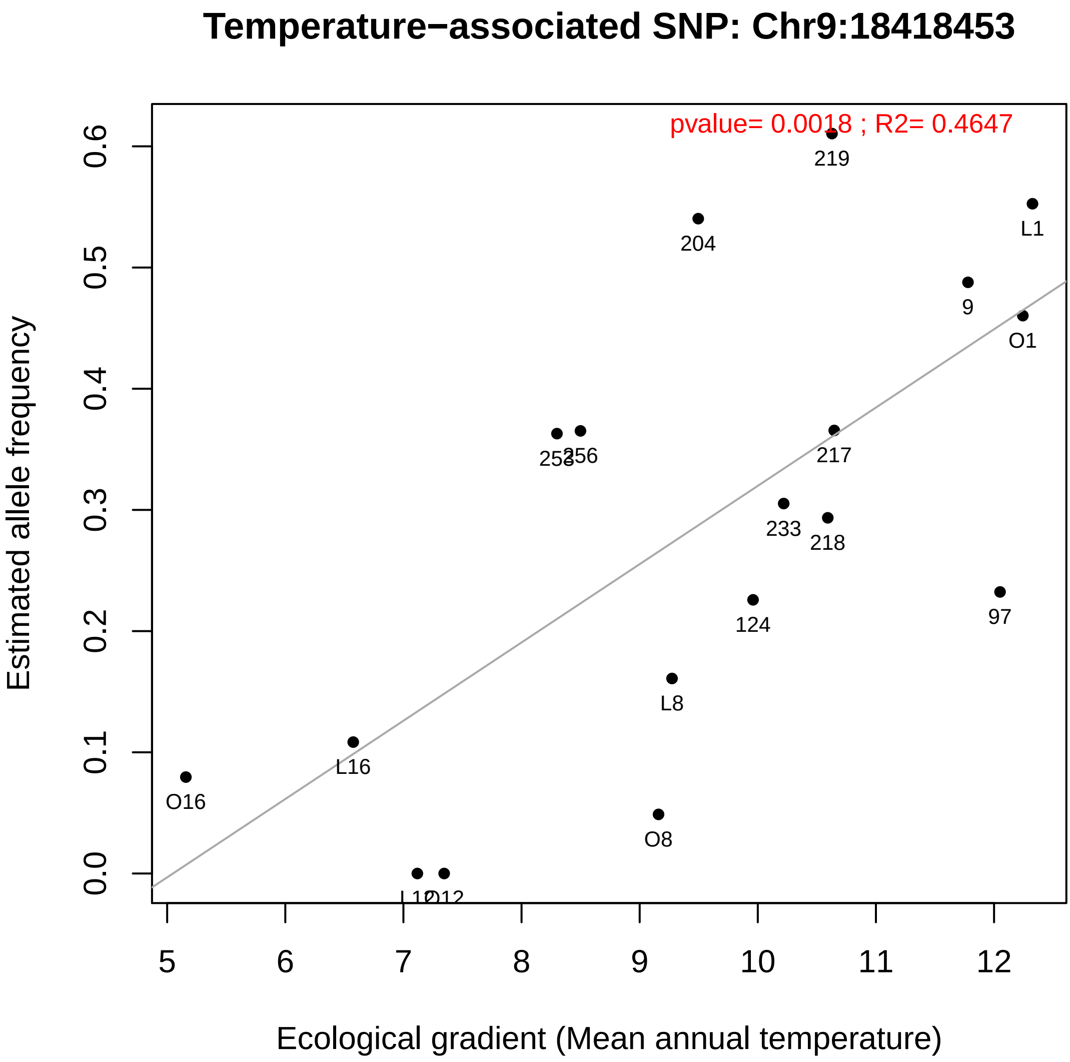

Temperature−associated SNP: Chr9:18418453
pvalue= 0.0018 ; R2= 0.4647
Estimated allele frequency
0.0
0.1
0.2
0.3
0.4
0.5
0.6
5
6
7
8
9
10
11
12
Ecological gradient (Mean annual temperature)
219
L1
204
9
O1
256
217
233
218
124
97
L8
L16
O16
O8

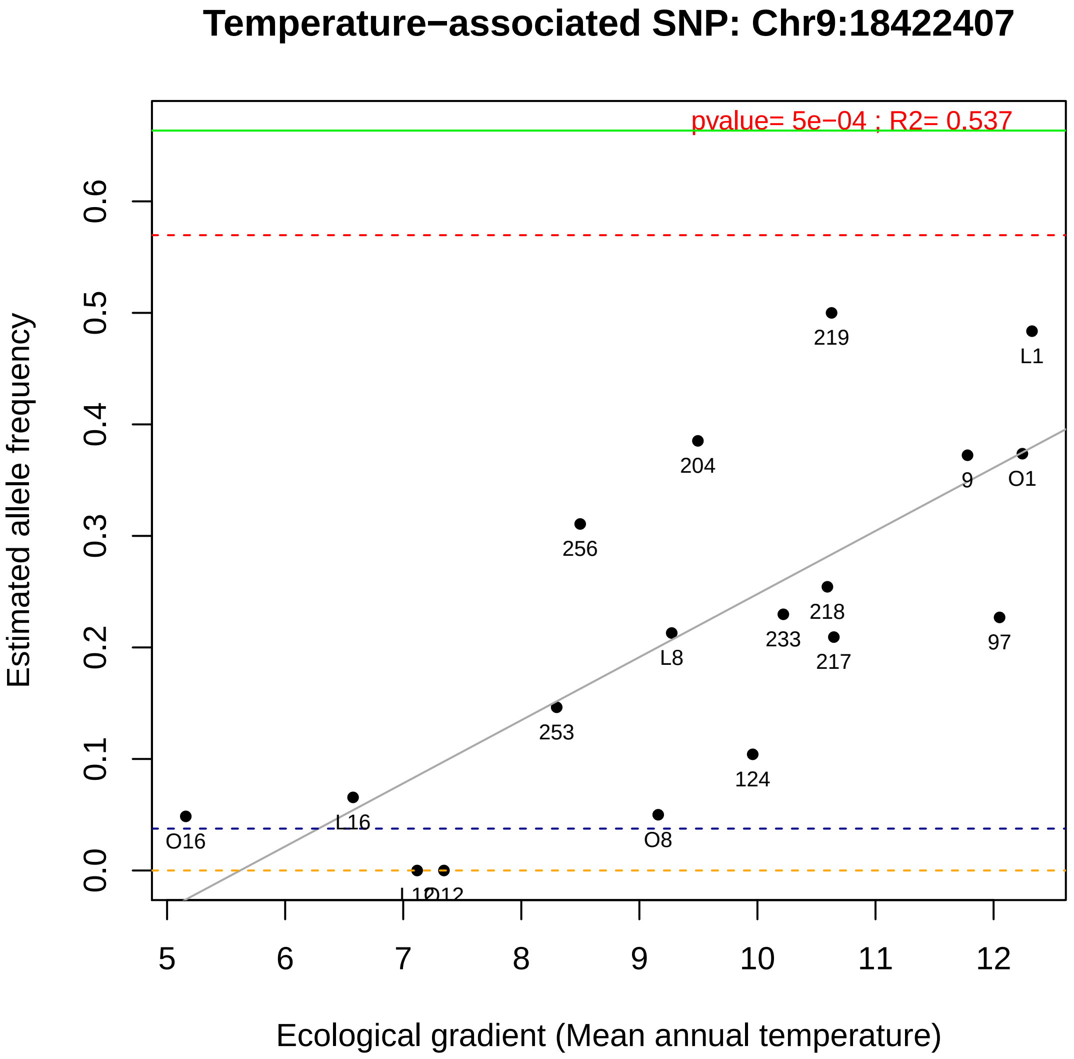
Temperature−associated SNP: Chr9:18422407
pvalue= 5e−04 ; R2= 0.537
Estimated allele frequency
0.0
0.1
0.2
0.3
0.4
0.5
0.6
5
6
7
8
9
10
11
12
Ecological gradient (Mean annual temperature)
219
L1
204
9
O1
256
218
233
217
97
L8
253
124
L16
O16
O8
O12

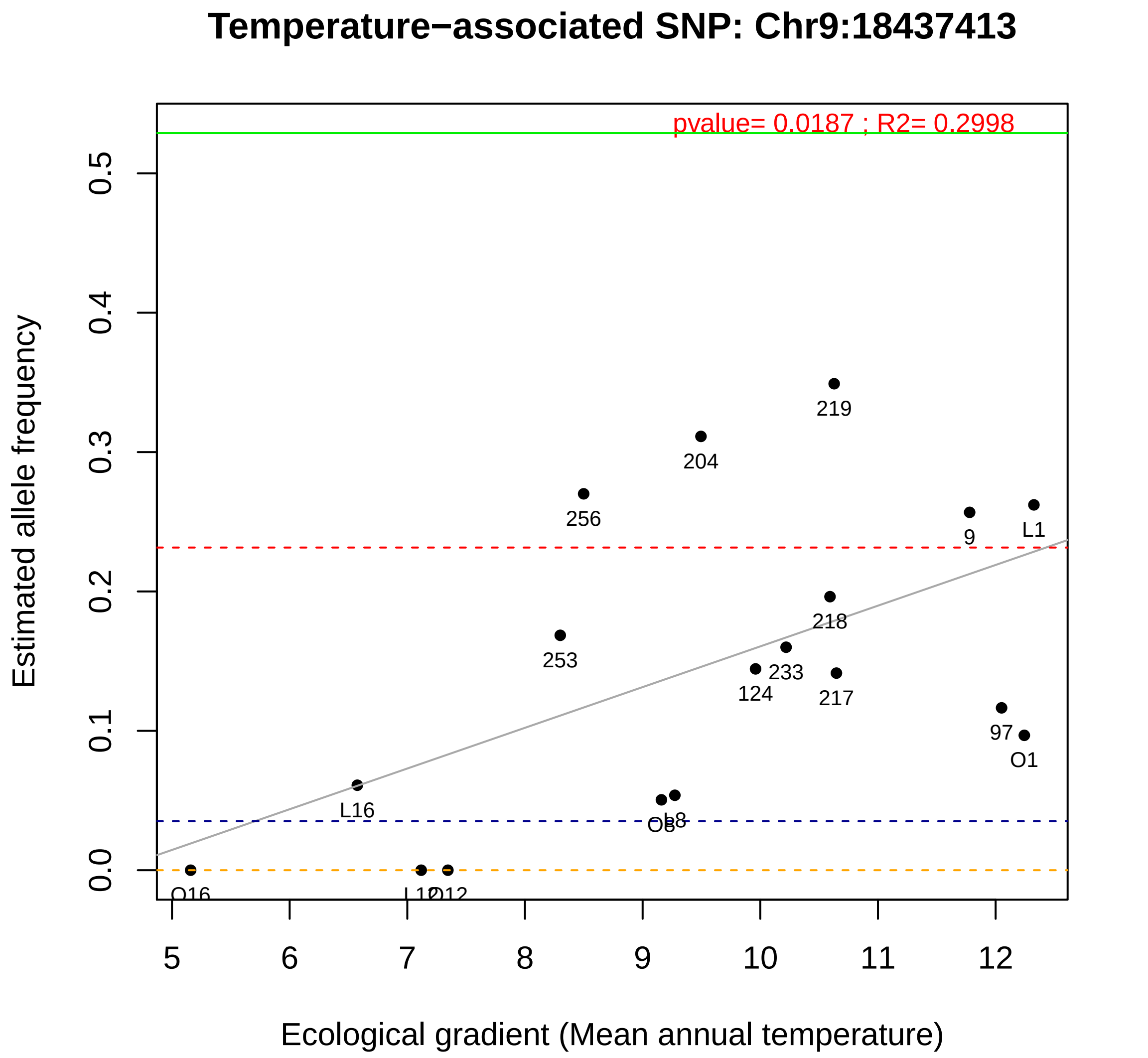
Temperature−associated SNP: Chr9:18437413
pvalue= 0.0187 ; R2= 0.2998
Estimated allele frequency
0.0
0.1
0.2
0.3
0.4
0.5
5
6
7
8
9
10
11
12
Ecological gradient (Mean annual temperature)
219
204
256
9
L1
218
253
233
124
217
97
O1
L16
O16
L8
O12

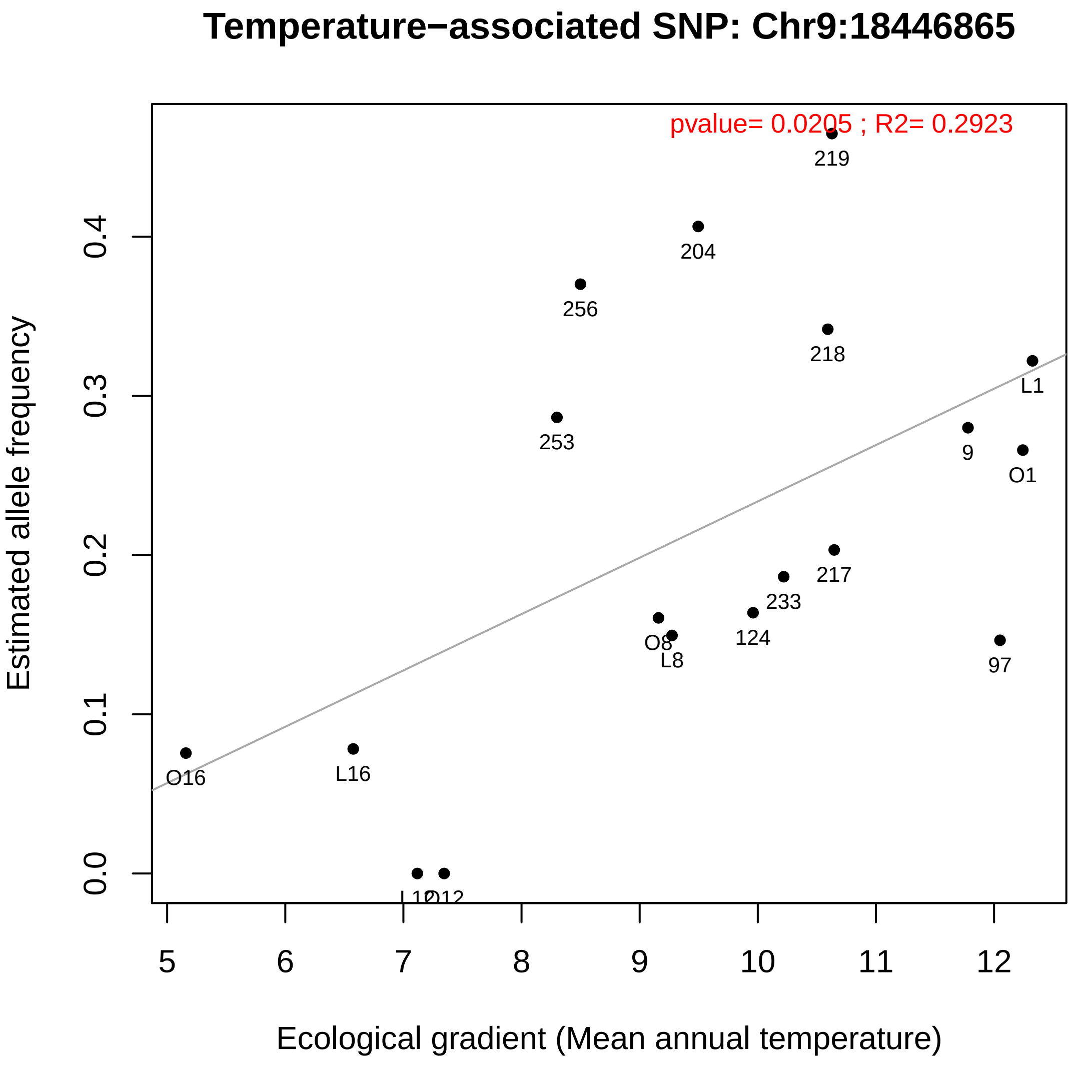
Temperature−associated SNP: Chr9:18446865
pvalue= 0.0205 ; R2= 0.2923
Estimated allele frequency
0.0
0.1
0.2
0.3
0.4
Ecological gradient (Mean annual temperature)
5
6
7
8
9
10
11
12
219
204
256
218
L1
253
9
O1
217
233
124
O8
L8
97
O16
L16
O12

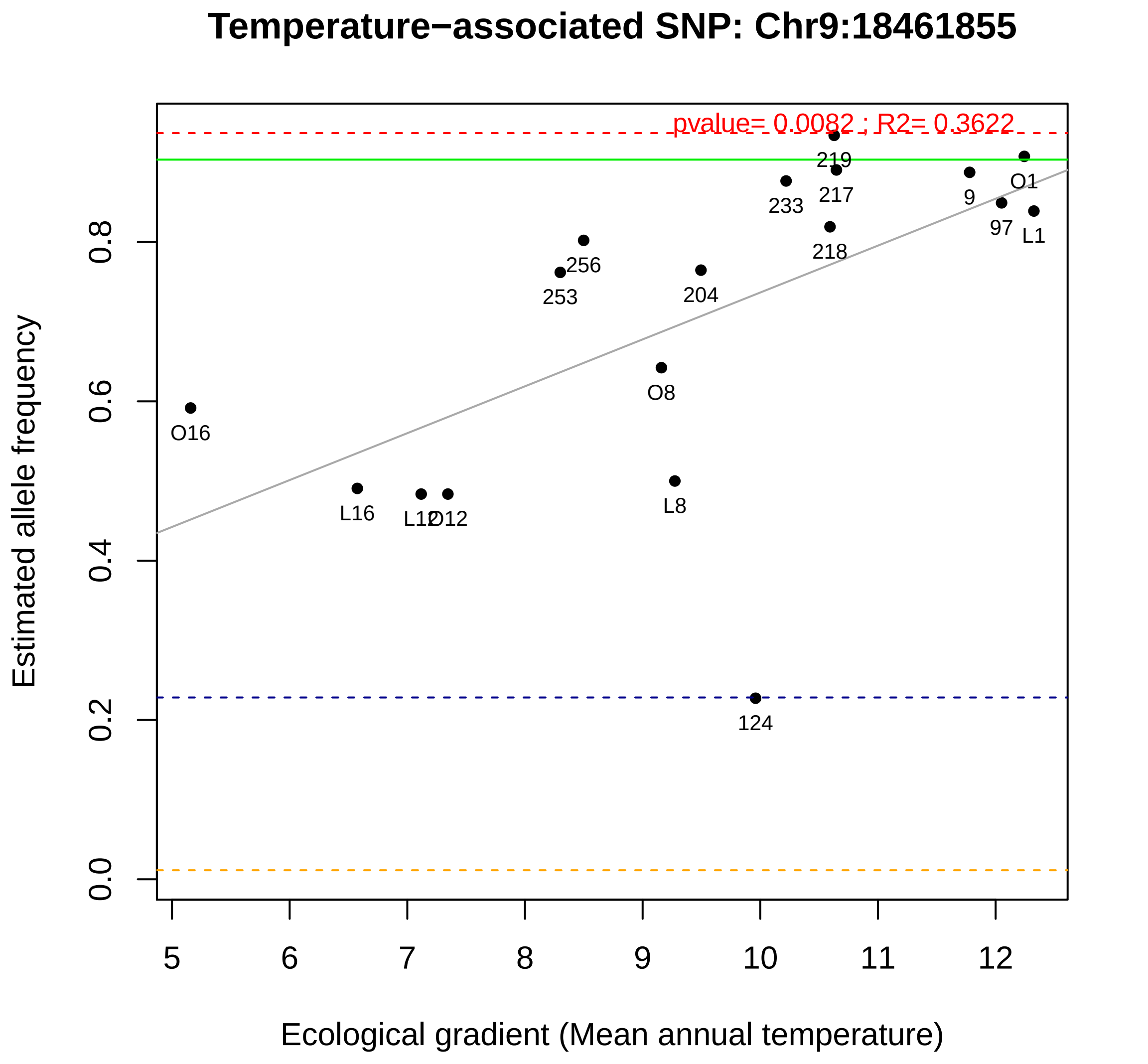
Temperature−associated SNP: Chr9:18461855
pvalue= 0.0082 ; R2= 0.3622
Estimated allele frequency
0.0
0.2
0.4
0.6
0.8
5
6
7
8
9
10
11
12
Ecological gradient (Mean annual temperature)
O16
L16
L12
O12
253
256
O8
L8
204
124
233
219
217
218
9
O1
97
L1

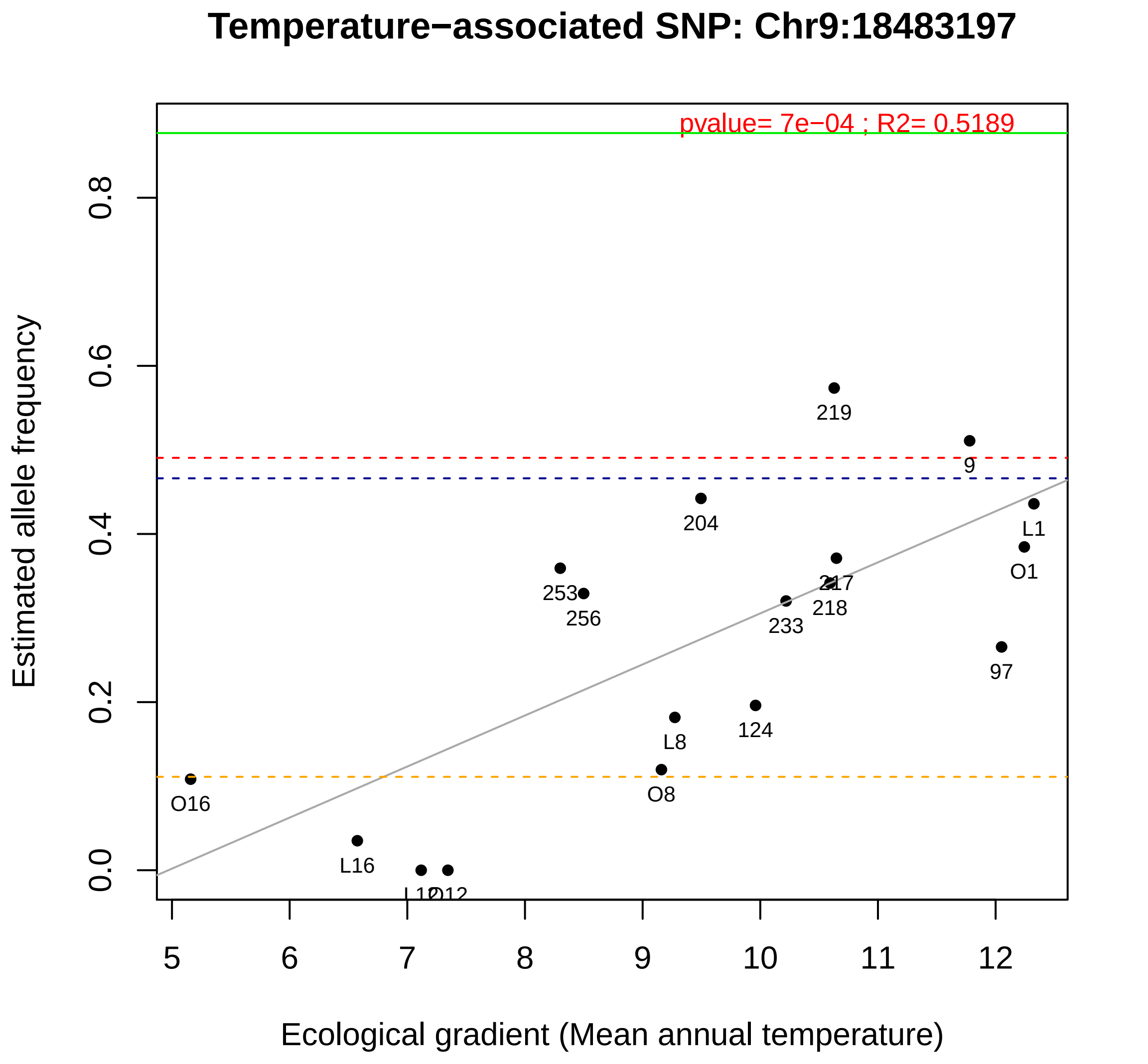

Temperature−associated SNP: Chr9:18483197
pvalue= 7e−04 ; R2= 0.5189
Estimated allele frequency
0.0
0.2
0.4
0.6
0.8
5
6
7
8
9
10
11
12
Ecological gradient (Mean annual temperature)
219
9
204
L1
O1
253
256
217
218
233
97
124
L8
O8
O16
L16
O12

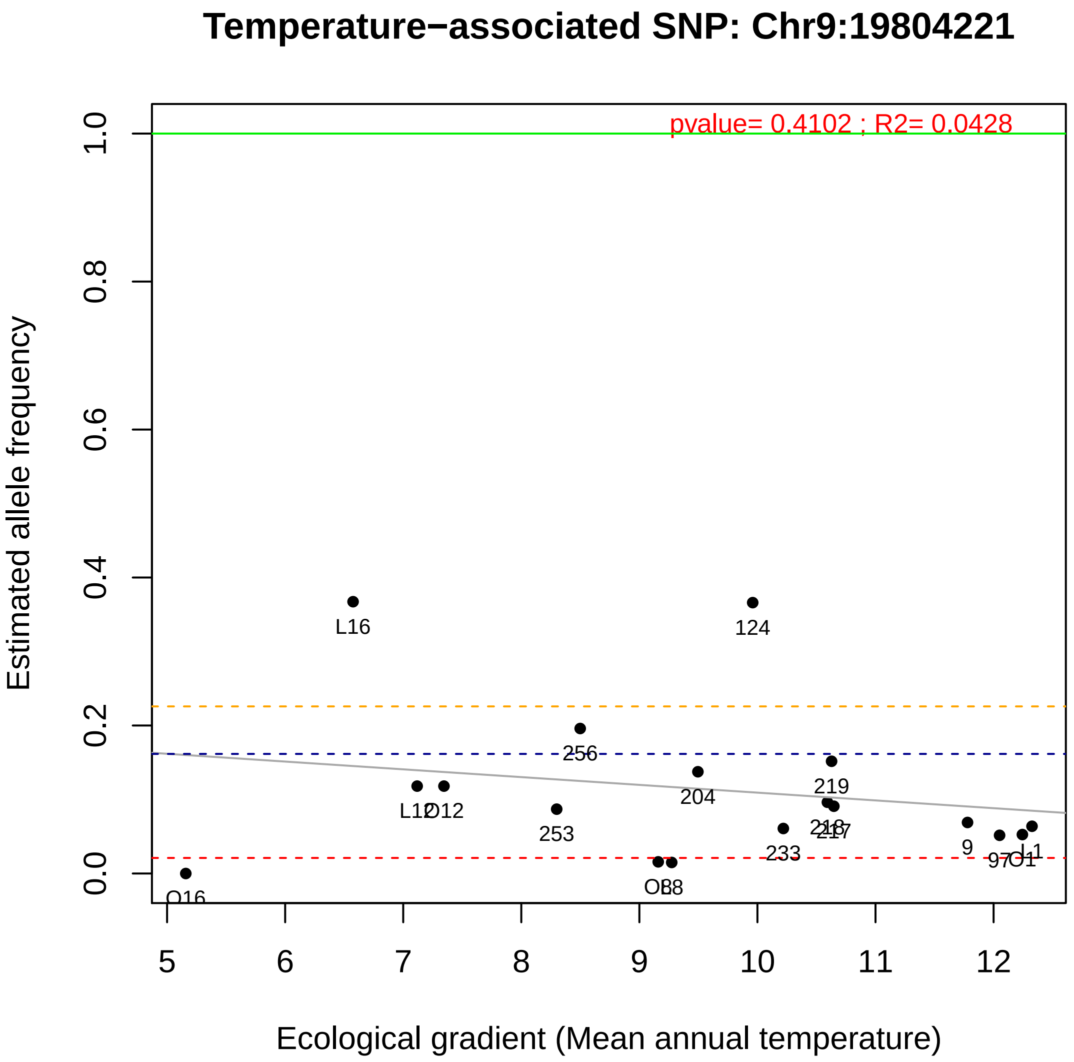
Temperature−associated SNP: Chr9:19804221
pvalue= 0.4102 ; R2= 0.0428
Estimated allele frequency
0.0
0.2
0.4
0.6
0.8
1.0
5
6
7
8
9
10
11
12
Ecological gradient (Mean annual temperature)
L16
124
256
204
219
253
233
9
O16

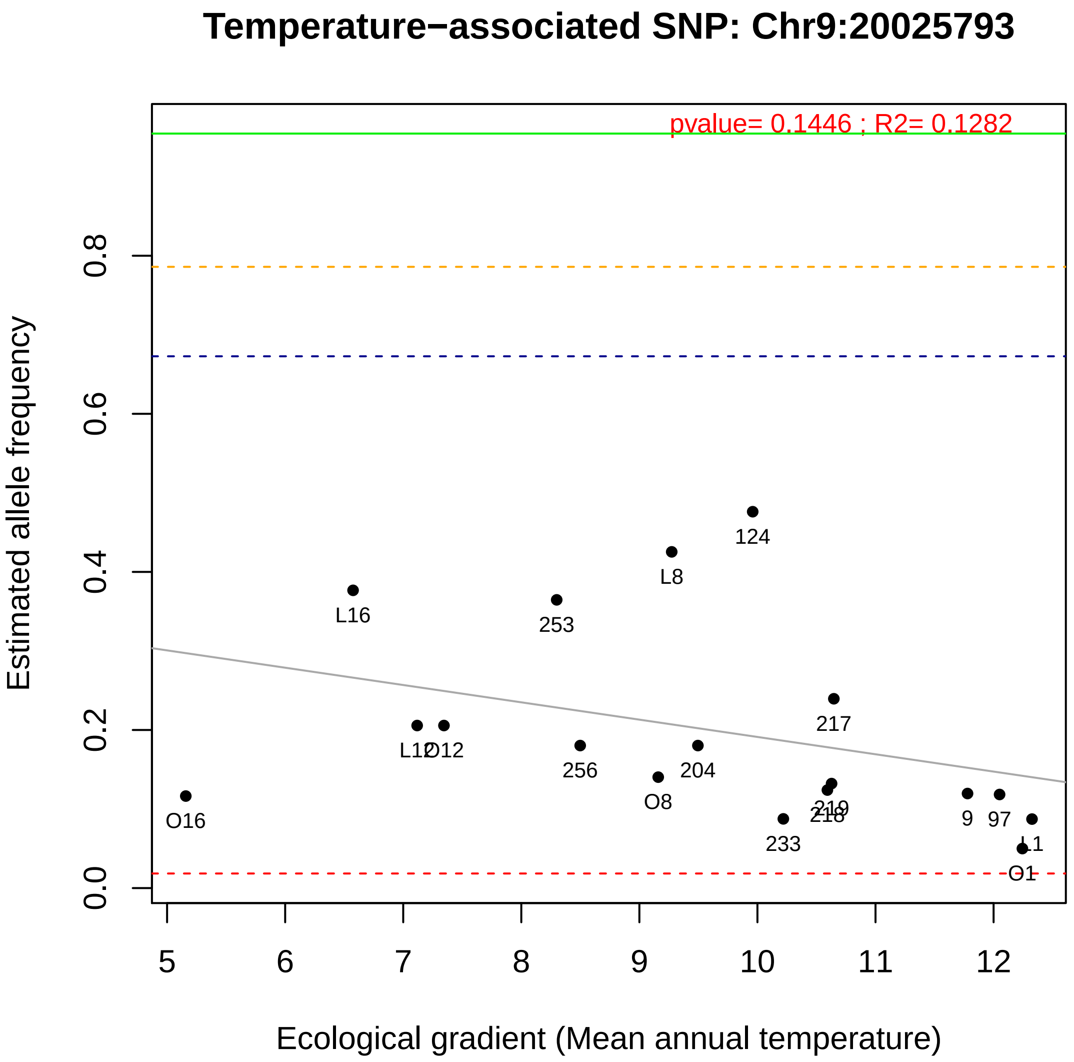

Temperature−associated SNP: Chr9:20025793
pvalue= 0.1446 ; R2= 0.1282
Estimated allele frequency
0.0
0.2
0.4
0.6
0.8
5
6
7
8
9
10
11
12
Ecological gradient (Mean annual temperature)
O16
L16
L12
O12
253
256
L8
O8
124
204
217
233
219
218
9
97
L1
O1

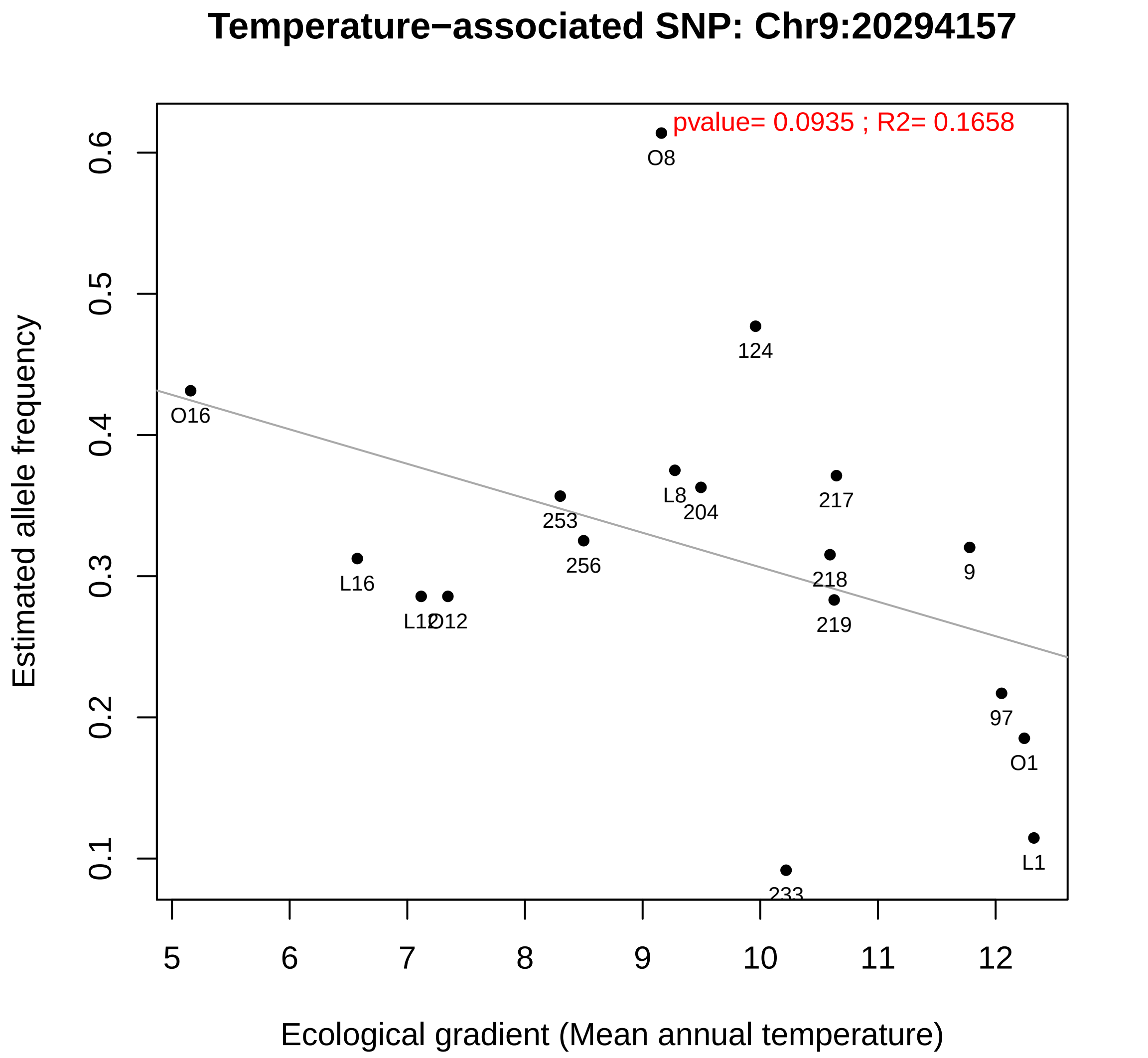

Temperature−associated SNP: Chr9:20294157
pvalue= 0.0935 ; R2= 0.1658
Estimated allele frequency
0.1
0.2
0.3
0.4
0.5
0.6
5
6
7
8
9
10
11
12
Ecological gradient (Mean annual temperature)
O8
124
O16
L8
204
217
253
256
218
9
L16
219
97
O1
L1
233

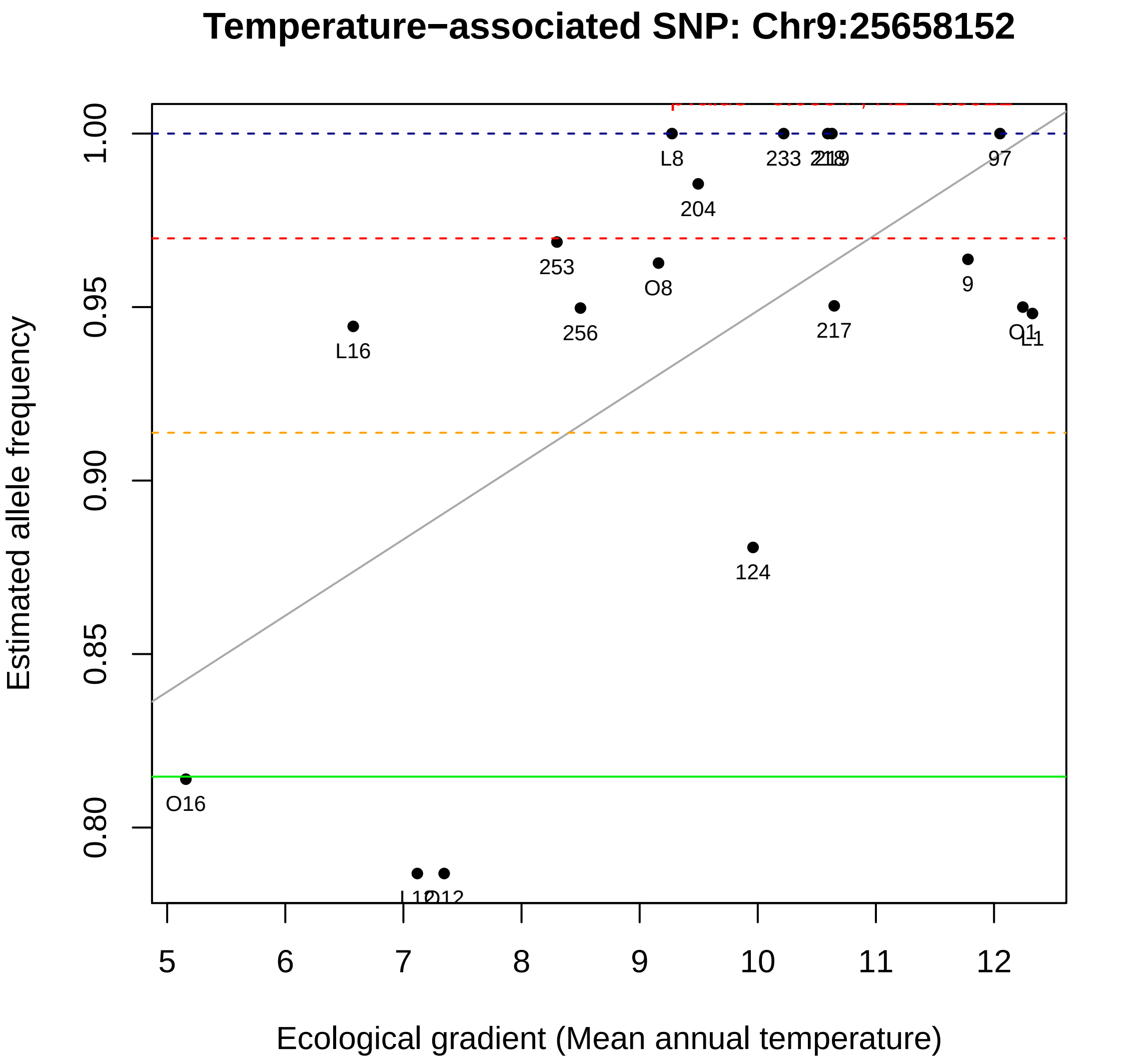
Temperature−associated SNP: Chr9:25658152
Estimated allele frequency
Ecological gradient (Mean annual temperature)
1.00
0.95
0.90
0.85
0.80
5
6
7
8
9
10
11
12
L8
233
97
204
253
O8
9
256
217
L16
124
O16

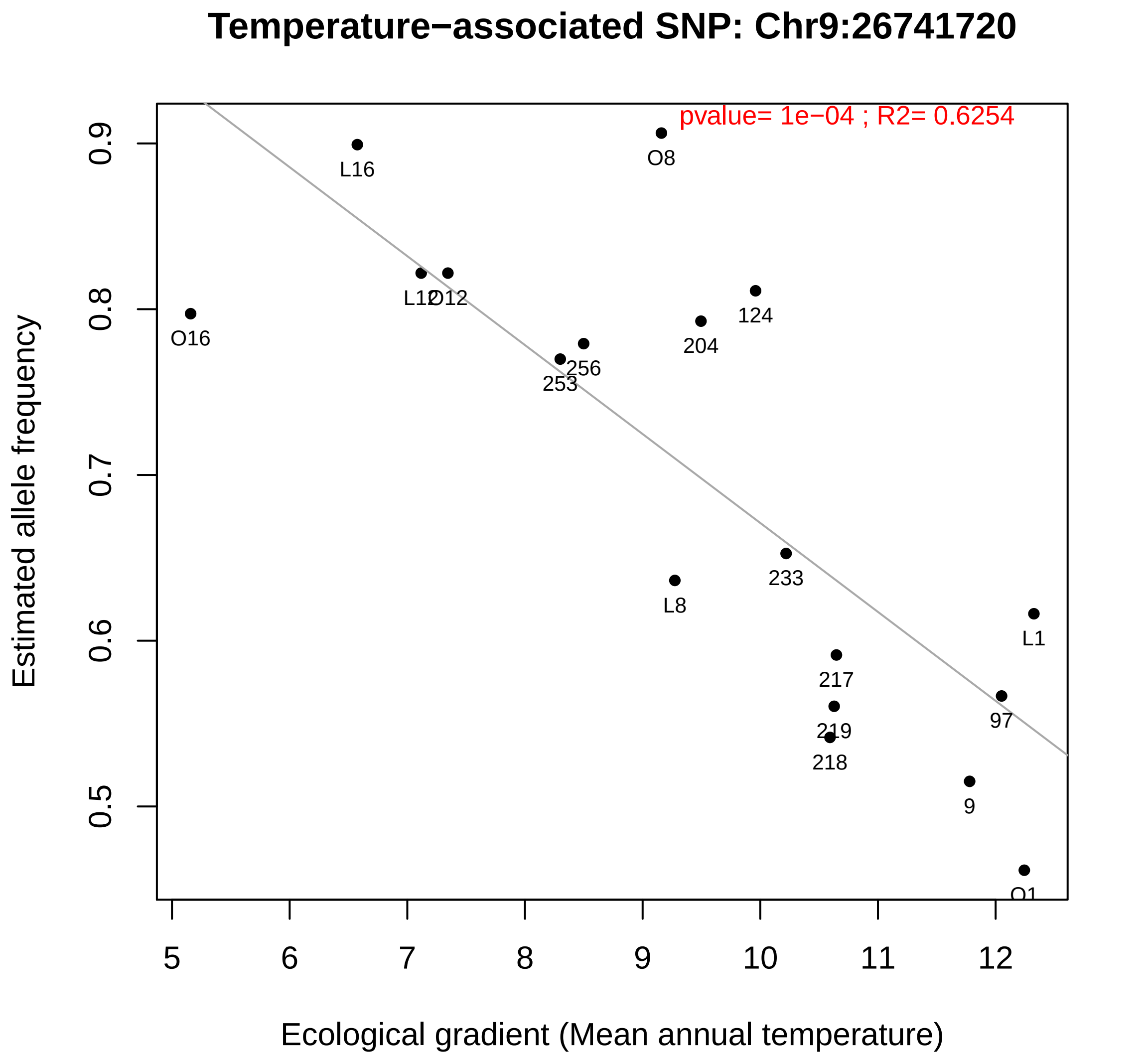
Temperature−associated SNP: Chr9:26741720
pvalue= 1e−04 ; R2= 0.6254
Estimated allele frequency
0.5
0.6
0.7
0.8
0.9
5
6
7
8
9
10
11
12
Ecological gradient (Mean annual temperature)
L16
O8
O16
L12
O12
124
204
253
256
233
L8
L1
217
219
218
97
9
O1

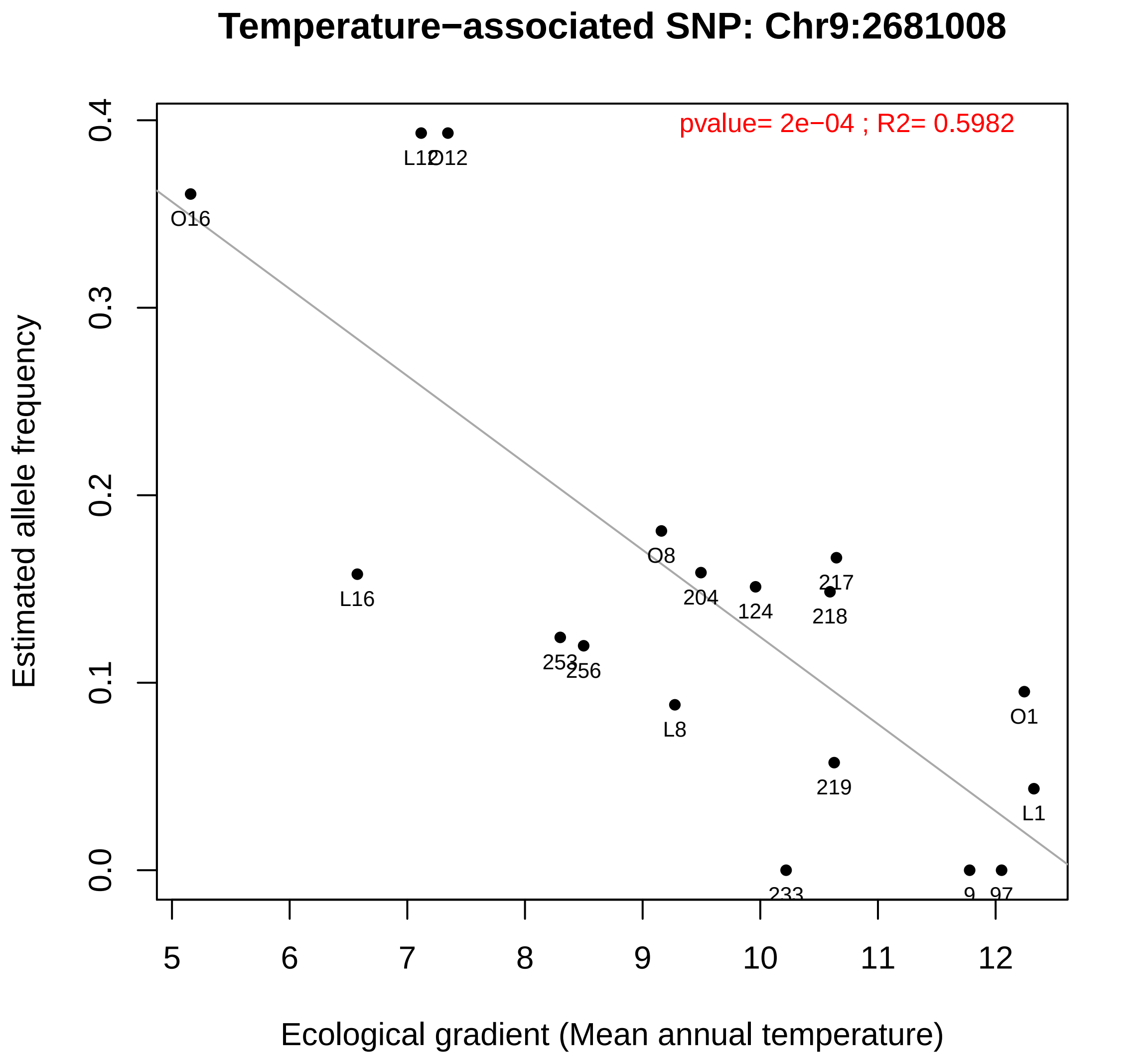

Temperature−associated SNP: Chr9:2681008
pvalue= 2e−04 ; R2= 0.5982
Estimated allele frequency
0.0
0.1
0.2
0.3
0.4
5
6
7
8
9
10
11
12
Ecological gradient (Mean annual temperature)
L12
O12
O16
O8
L16
204
124
217
218
253
256
L8
O1
219
L1
233
9
97

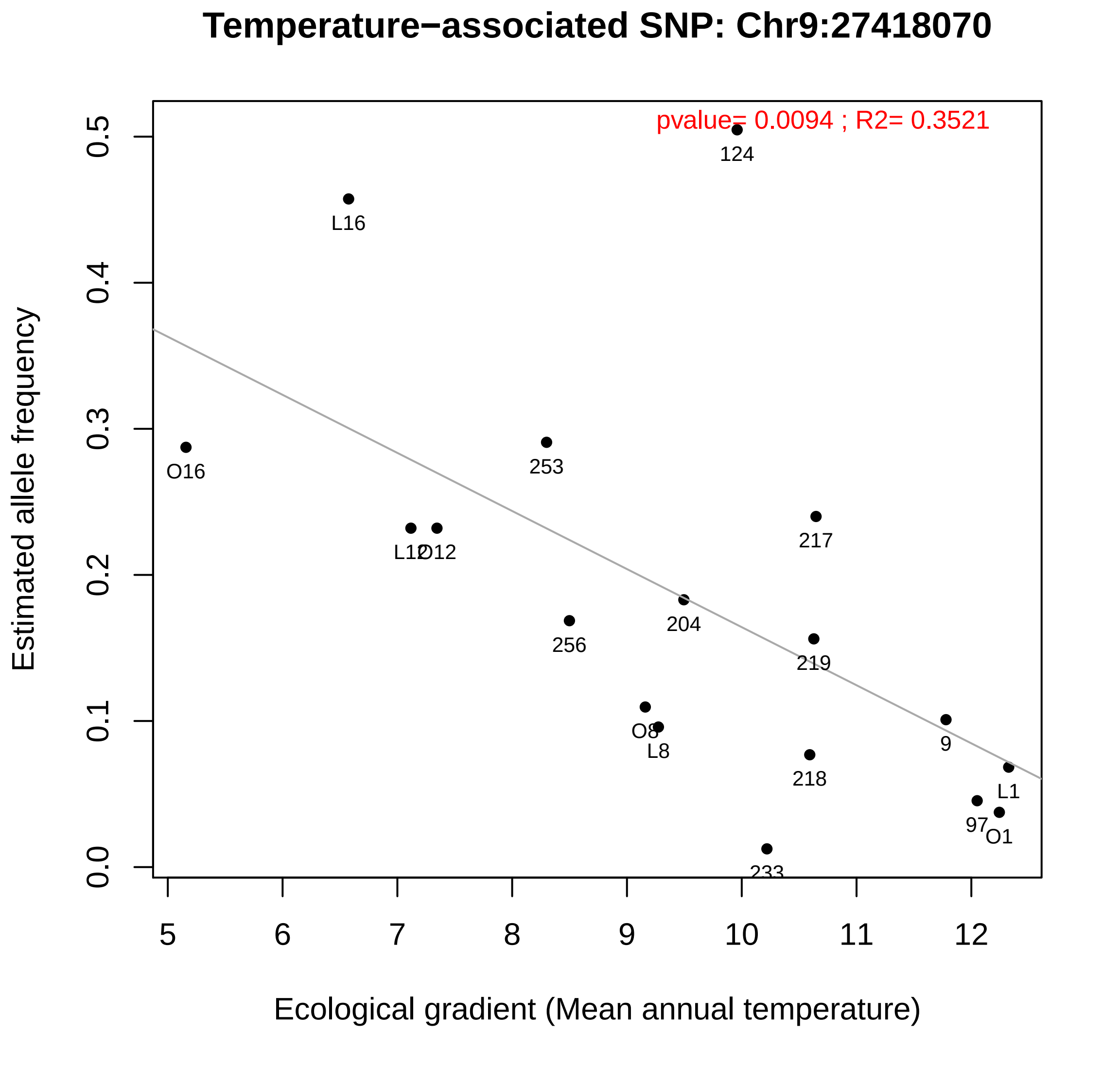
Temperature−associated SNP: Chr9:27418070
pvalue= 0.0094 ; R2= 0.3521
Estimated allele frequency
0.0
0.1
0.2
0.3
0.4
0.5
5
6
7
8
9
10
11
12
Ecological gradient (Mean annual temperature)
124
L16
O16
253
L12
O12
217
204
256
219
O8
L8
9
218
L1
97
O1
233

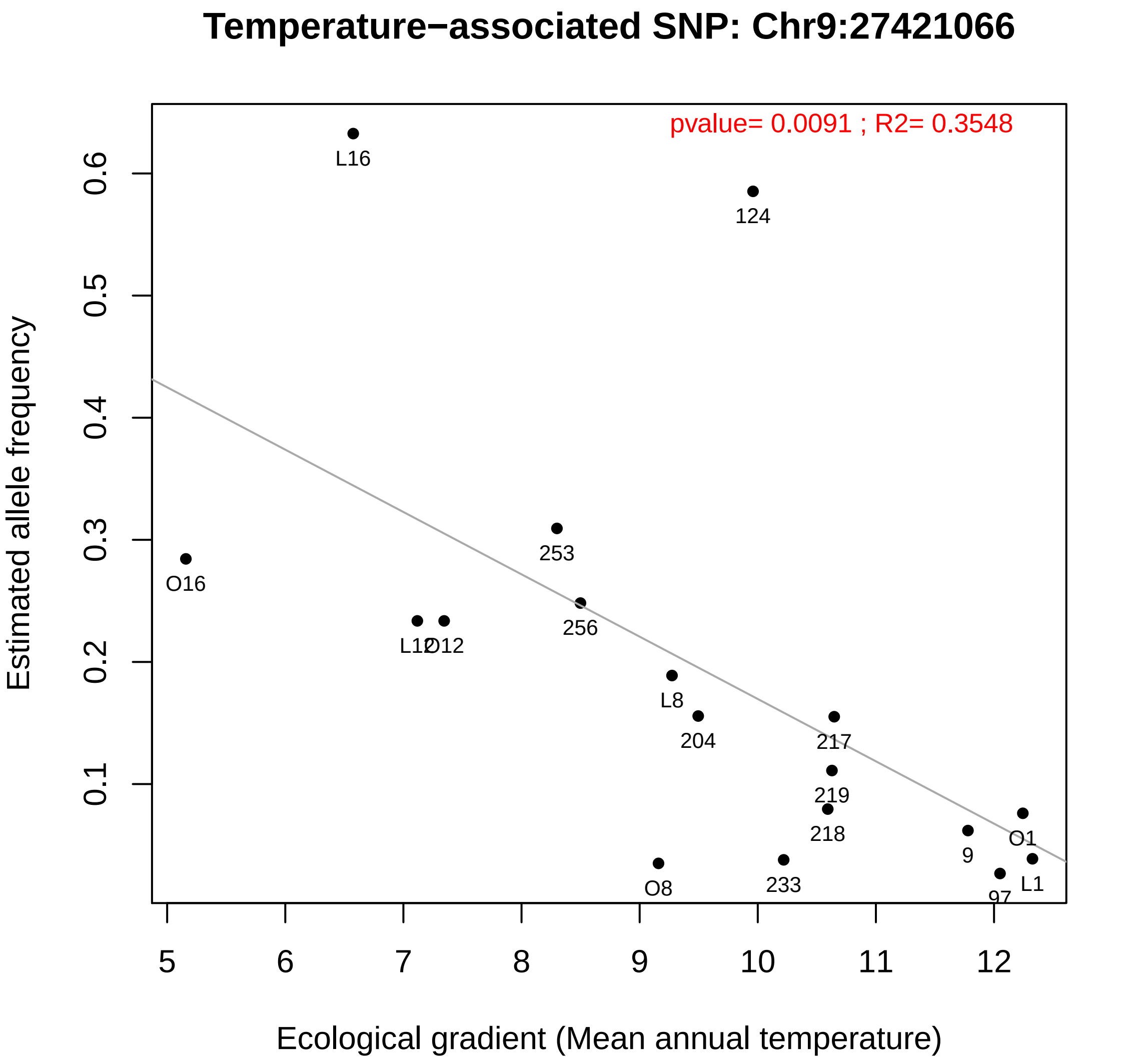

Temperature−associated SNP: Chr9:27421066
pvalue= 0.0091 ; R2= 0.3548
Estimated allele frequency
0.1
0.2
0.3
0.4
0.5
0.6
L16
124
253
O16
256
L12
O12
L8
204
217
219
218
O1
9
L1
97
O8
233
5
6
7
8
9
10
11
12
Ecological gradient (Mean annual temperature)

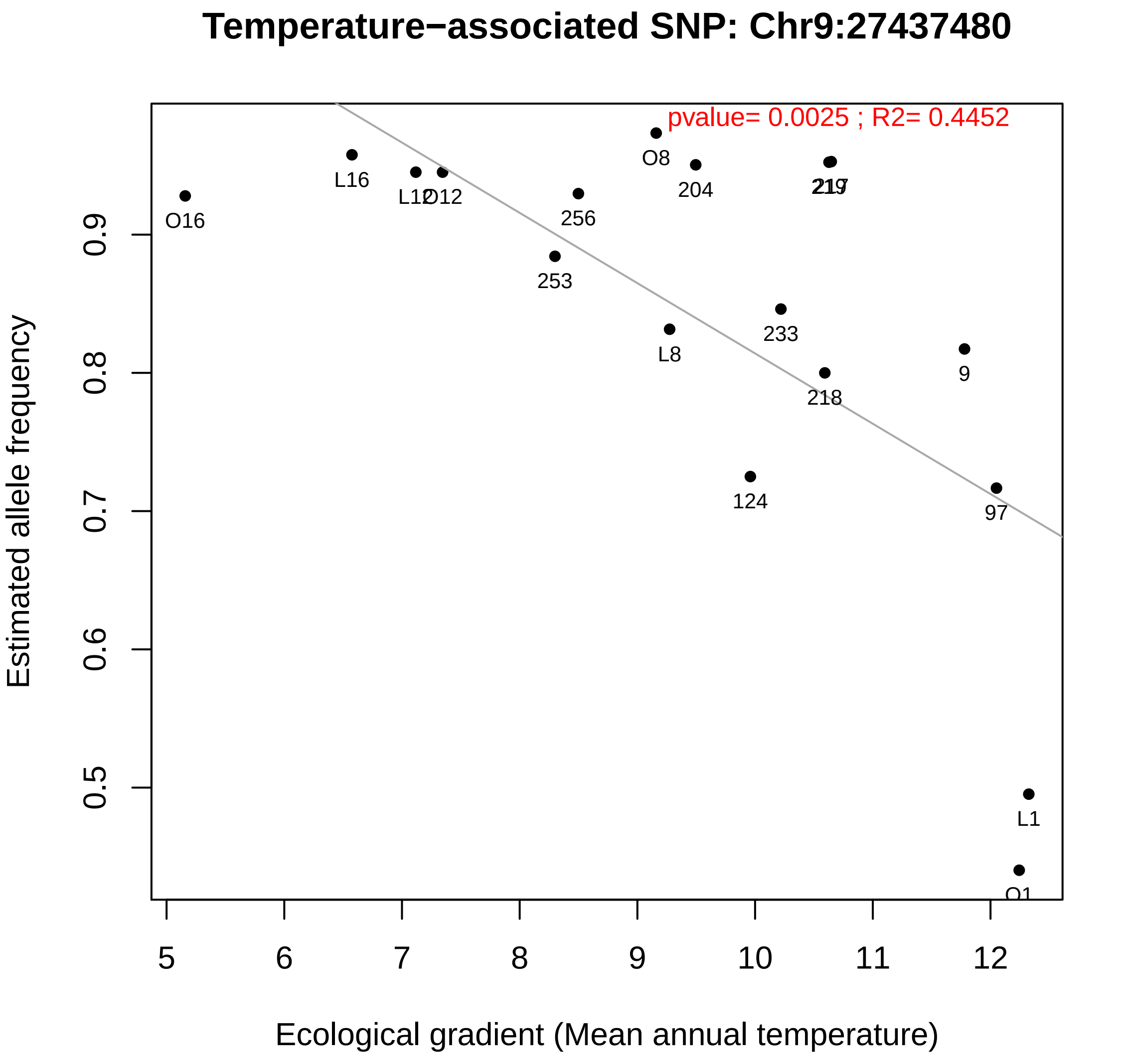

Temperature−associated SNP: Chr9:27437480
pvalue= 0.0025 ; R2= 0.4452
Estimated allele frequency
Ecological gradient (Mean annual temperature)
0.5
0.6
0.7
0.8
0.9
5
6
7
8
9
10
11
12
O16
L16
L12
O12
256
253
O8
204
L8
233
124
218
9
97
L1
O1

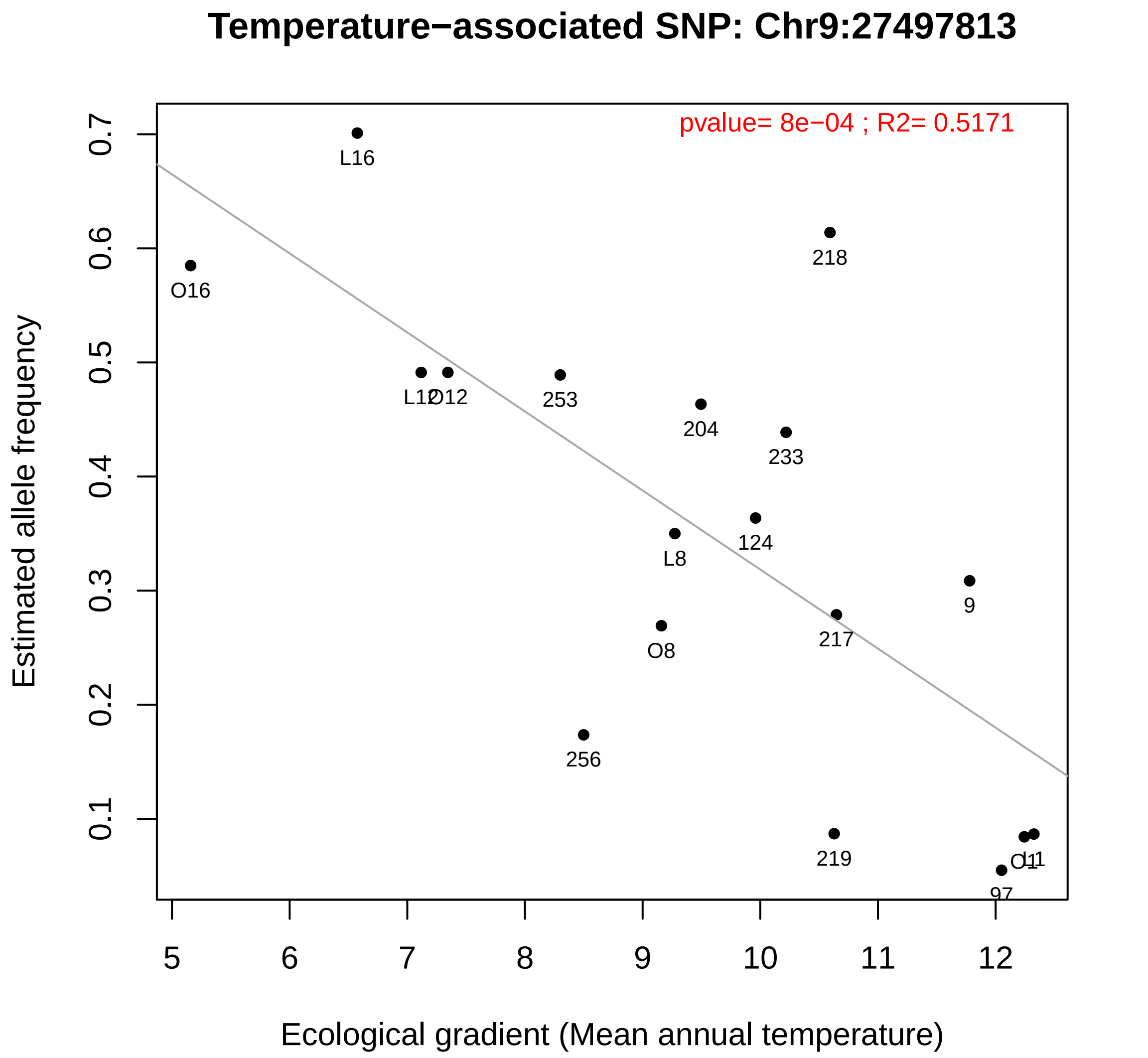
Temperature−associated SNP: Chr9:27497813
pvalue= 8e−04 ; R2= 0.5171
Estimated allele frequency
0.1
0.2
0.3
0.4
0.5
0.6
0.7
Ecological gradient (Mean annual temperature)
5
6
7
8
9
10
11
12
L16
O16
L12
O12
253
204
233
218
124
L8
9
217
O8
256
219
97

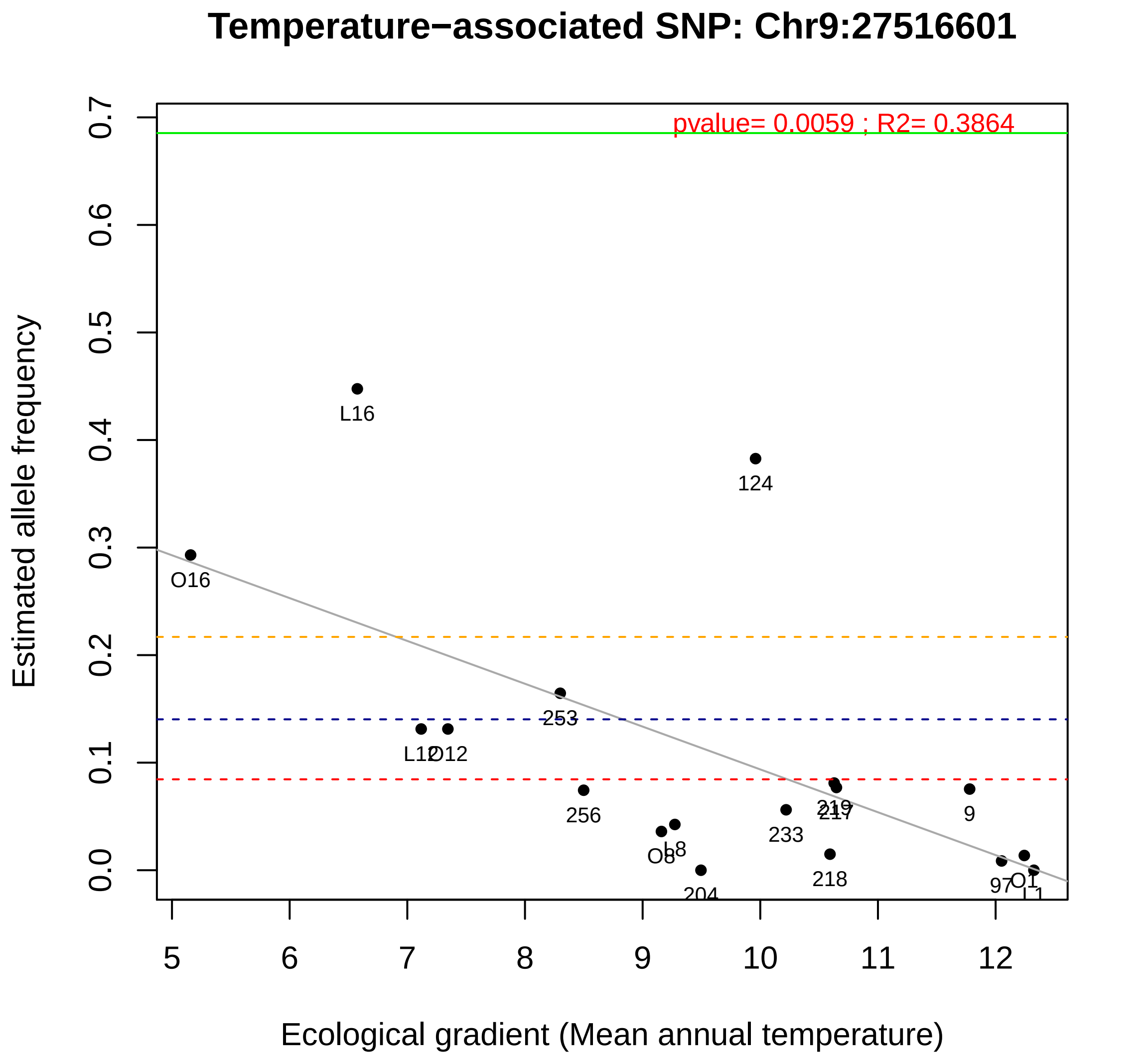
Temperature−associated SNP: Chr9:27516601
pvalue= 0.0059 ; R2= 0.3864
Estimated allele frequency
0.0
0.1
0.2
0.3
0.4
0.5
0.6
0.7
L16
124
O16
253
L12
O12
256
219
217
9
233
O8
L8
204
218
97
O1
L1
5
6
7
8
9
10
11
12
Ecological gradient (Mean annual temperature)

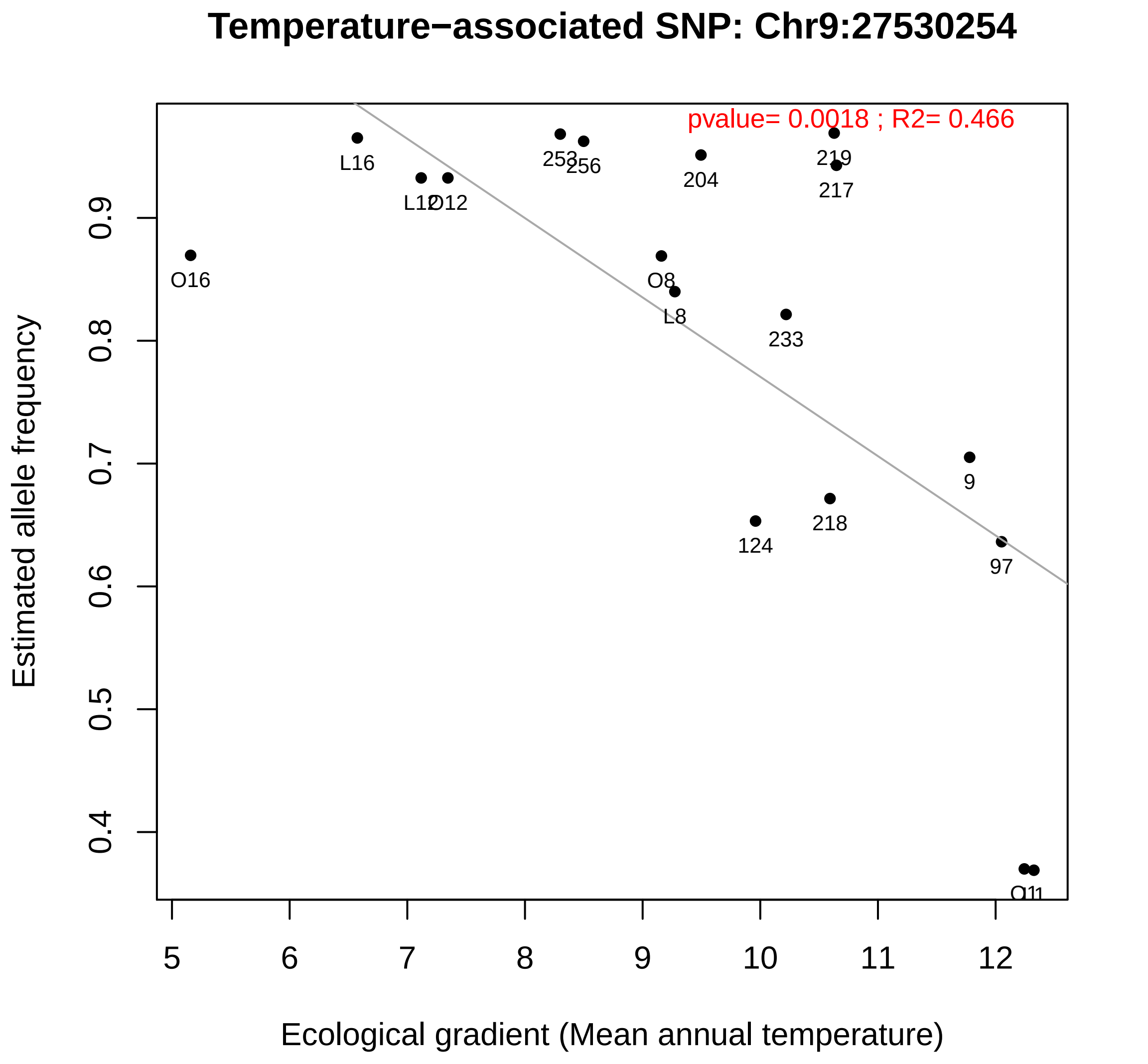
Temperature−associated SNP: Chr9:27530254
pvalue= 0.0018 ; R2= 0.466
Estimated allele frequency
Ecological gradient (Mean annual temperature)
0.4
0.5
0.6
0.7
0.8
0.9
5
6
7
8
9
10
11
12
L16
L12
O12
253
256
204
219
217
O16
O8
L8
233
9
218
124
97
O11

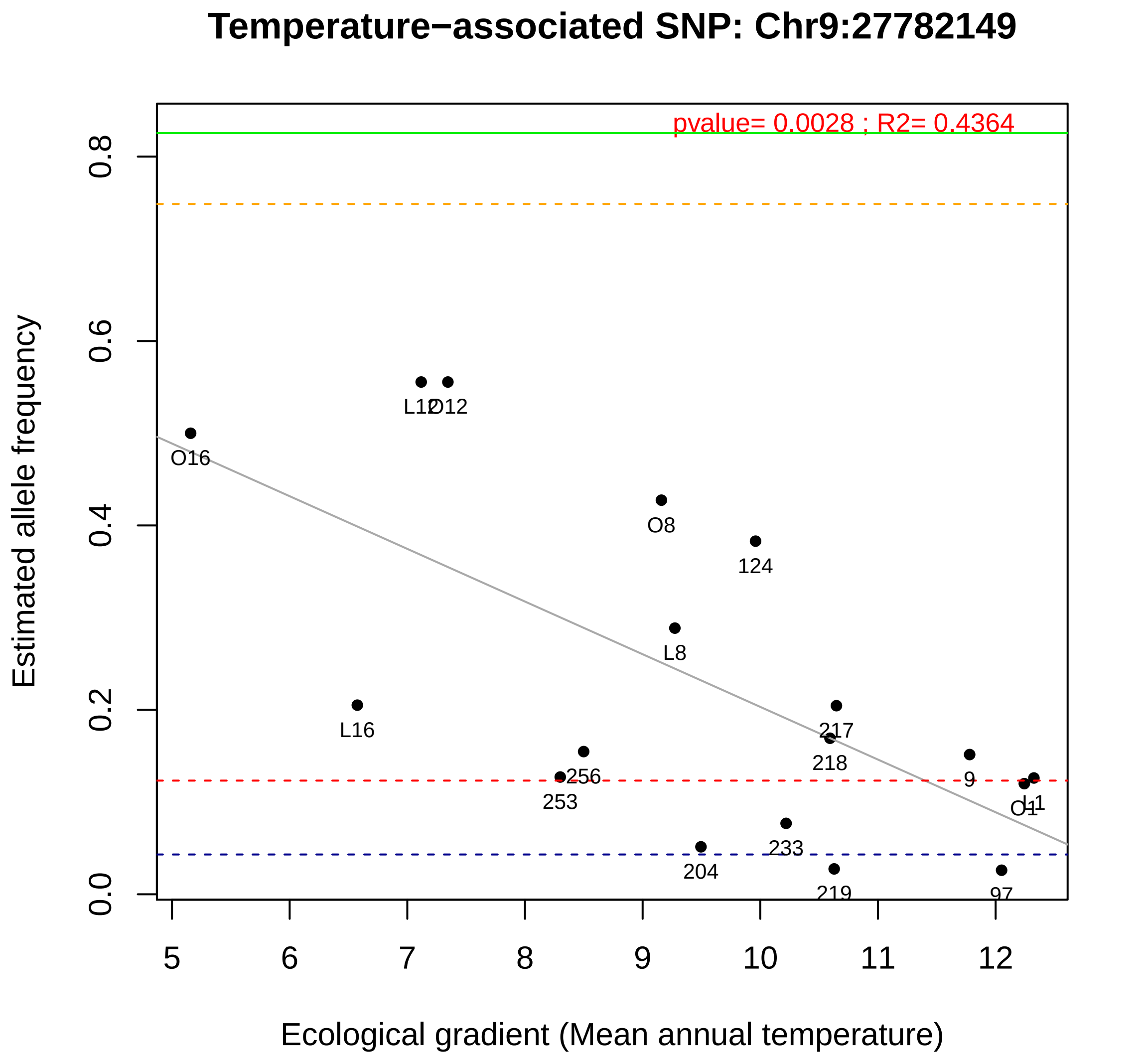

Temperature−associated SNP: Chr9:27782149
pvalue= 0.0028 ; R2= 0.4364
Estimated allele frequency
0.0
0.2
0.4
0.6
0.8
5
6
7
8
9
10
11
12
Ecological gradient (Mean annual temperature)
O16
L12
O12
O8
124
L8
L16
217
218
256
253
9
O1
L1
233
204
219
97

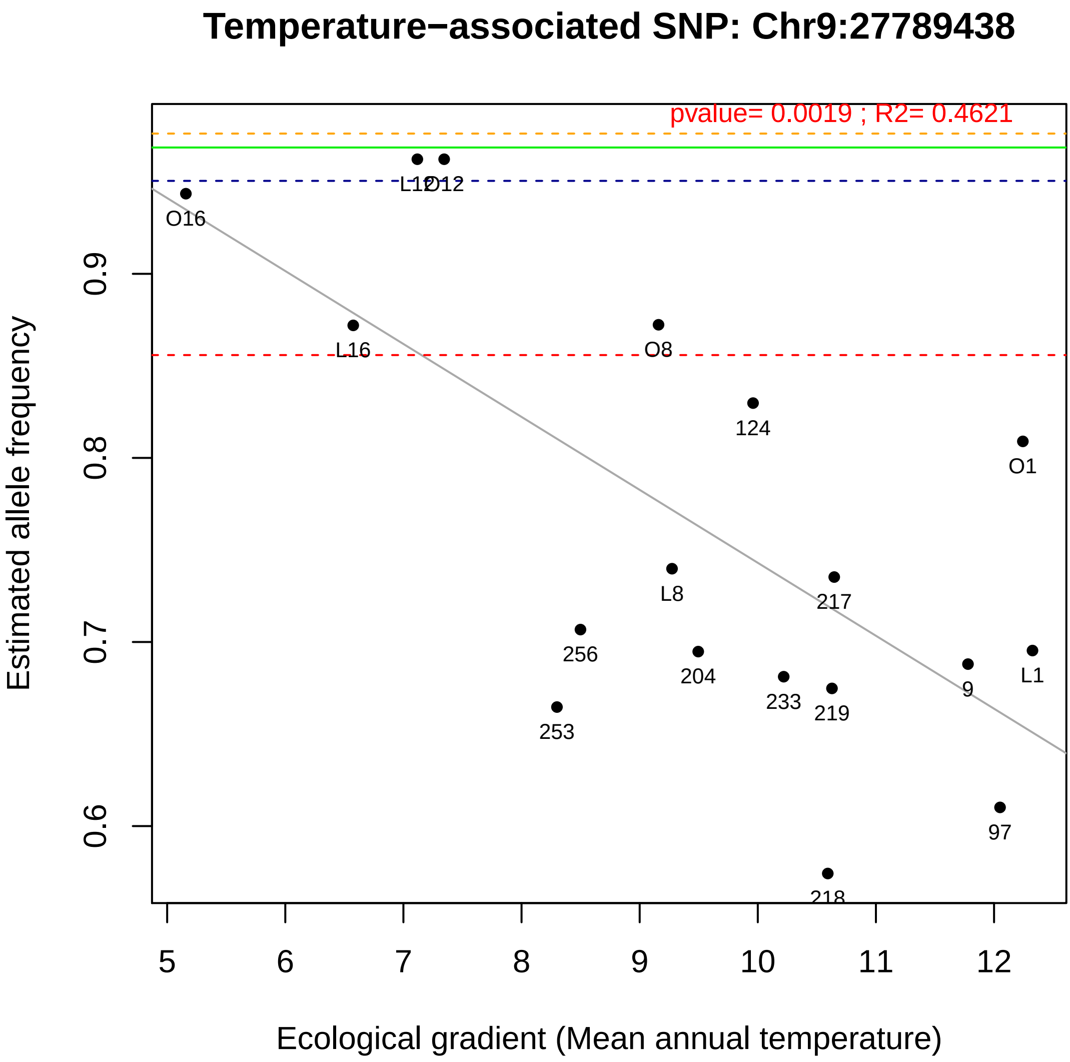

Temperature−associated SNP: Chr9:27789438
pvalue= 0.0019 ; R2= 0.4621
Estimated allele frequency
Ecological gradient (Mean annual temperature)
0.6
0.7
0.8
0.9
5
6
7
8
9
10
11
12
O16
L12
O12
L16
O8
124
O1
L8
217
256
204
L1
9
233
219
253
97
218

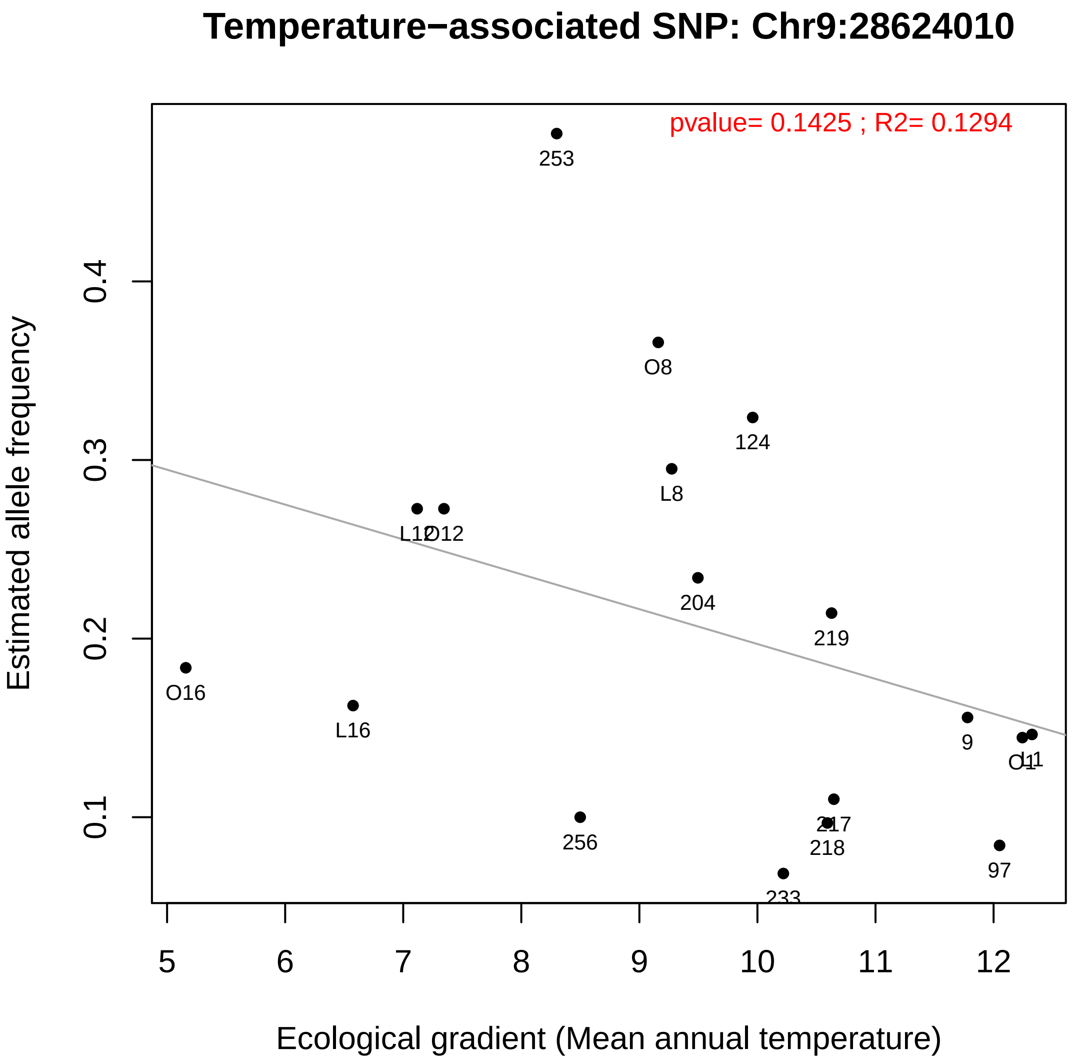
Temperature−associated SNP: Chr9:28624010
pvalue= 0.1425 ; R2= 0.1294
Estimated allele frequency
0.1
0.2
0.3
0.4
5
6
7
8
9
10
11
12
Ecological gradient (Mean annual temperature)
253
O8
124
L8
L12
O12
204
219
O16
L16
9
O1
L1
256
217
218
97
233

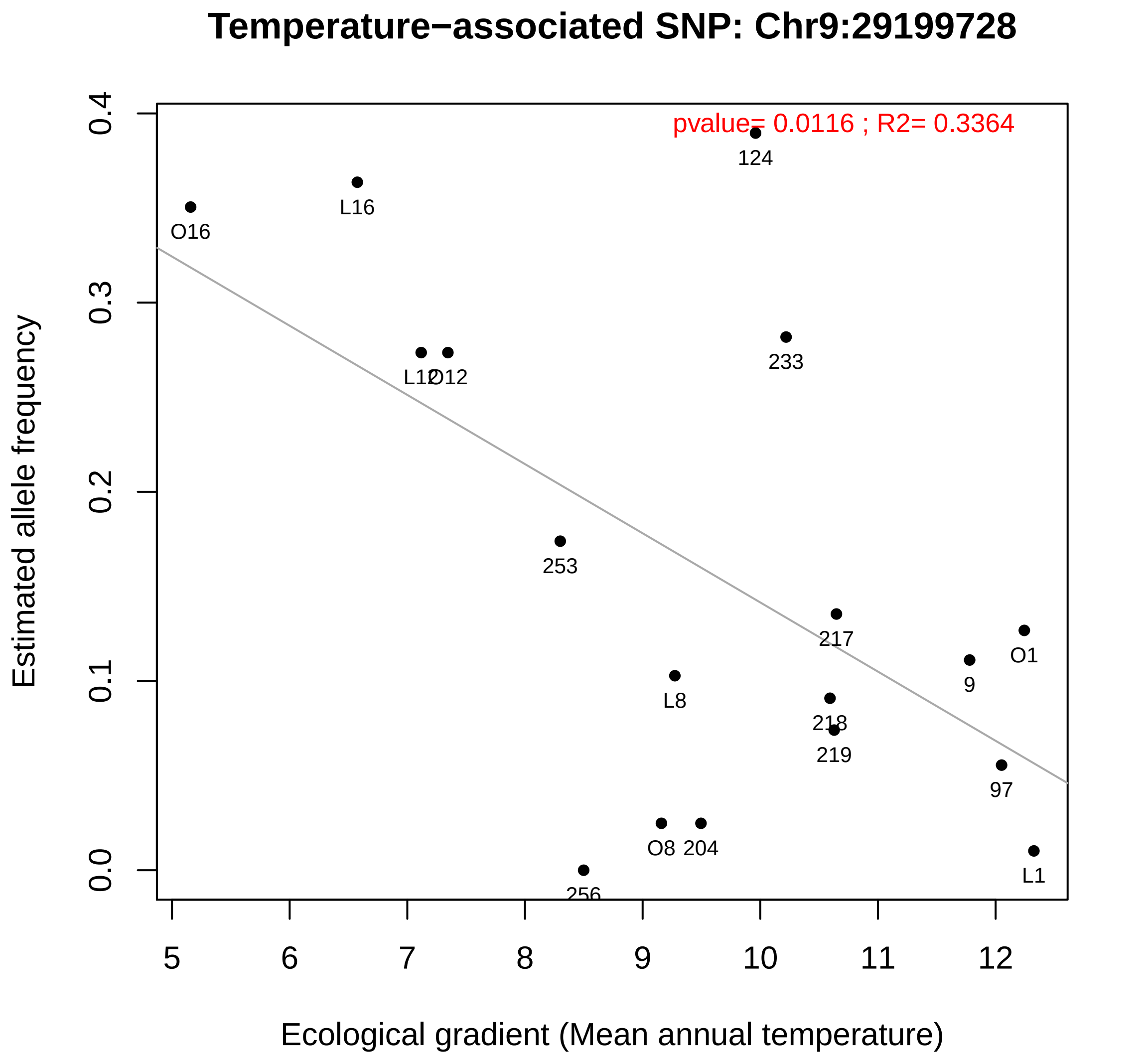
Temperature−associated SNP: Chr9:29199728
pvalue= 0.0116 ; R2= 0.3364
Estimated allele frequency
Ecological gradient (Mean annual temperature)
0.0
0.1
0.2
0.3
0.4
5
6
7
8
9
10
11
12
O16
L16
124
L12
O12
233
253
217
O1
9
L8
218
219
97
O8
204
256
L1

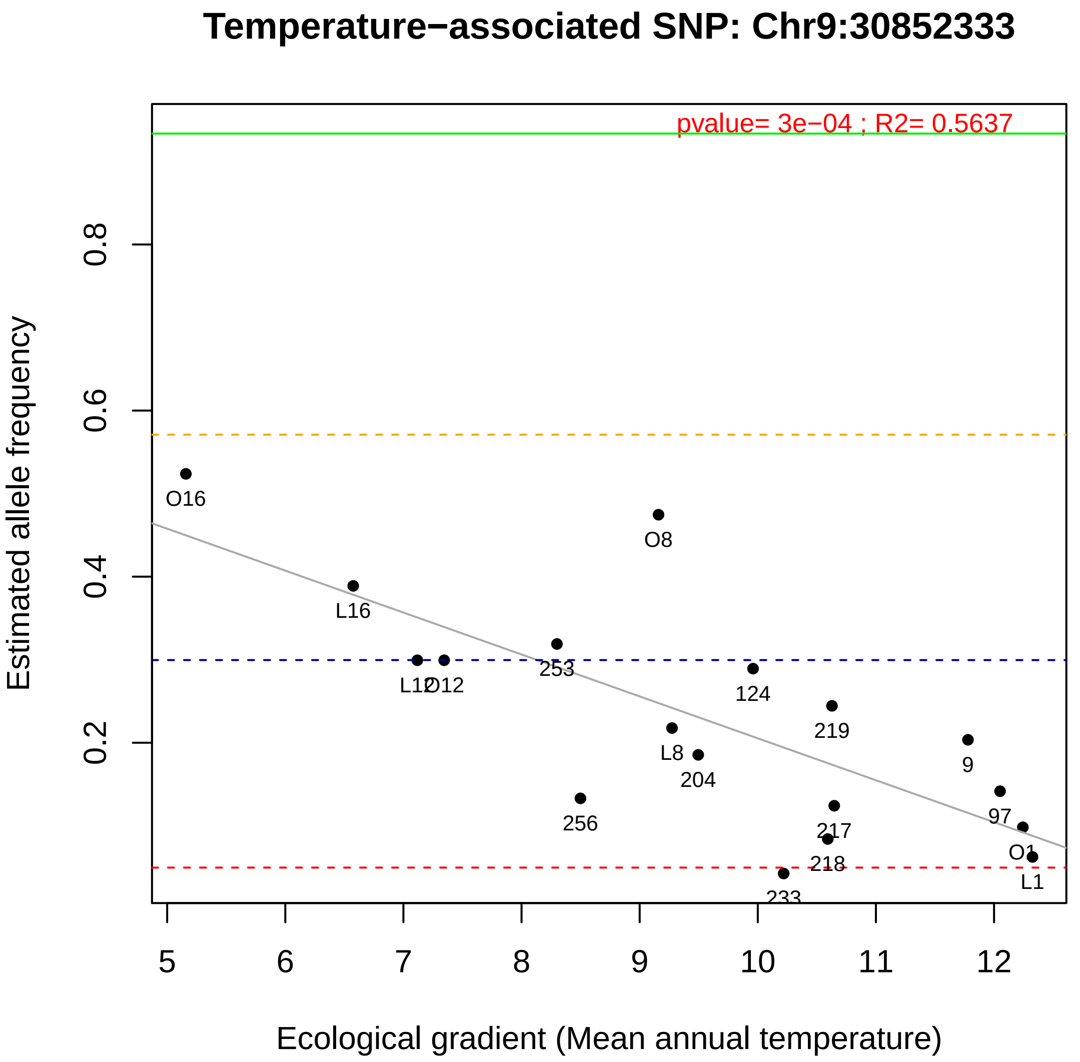

Temperature−associated SNP: Chr9:30852333
pvalue= 3e−04 ; R2= 0.5637
Estimated allele frequency
0.2
0.4
0.6
0.8
Ecological gradient (Mean annual temperature)
5
6
7
8
9
10
11
12
O16
L16
L12
O12
253
O8
124
219
L8
204
256
9
97
217
218
O1
L1
233

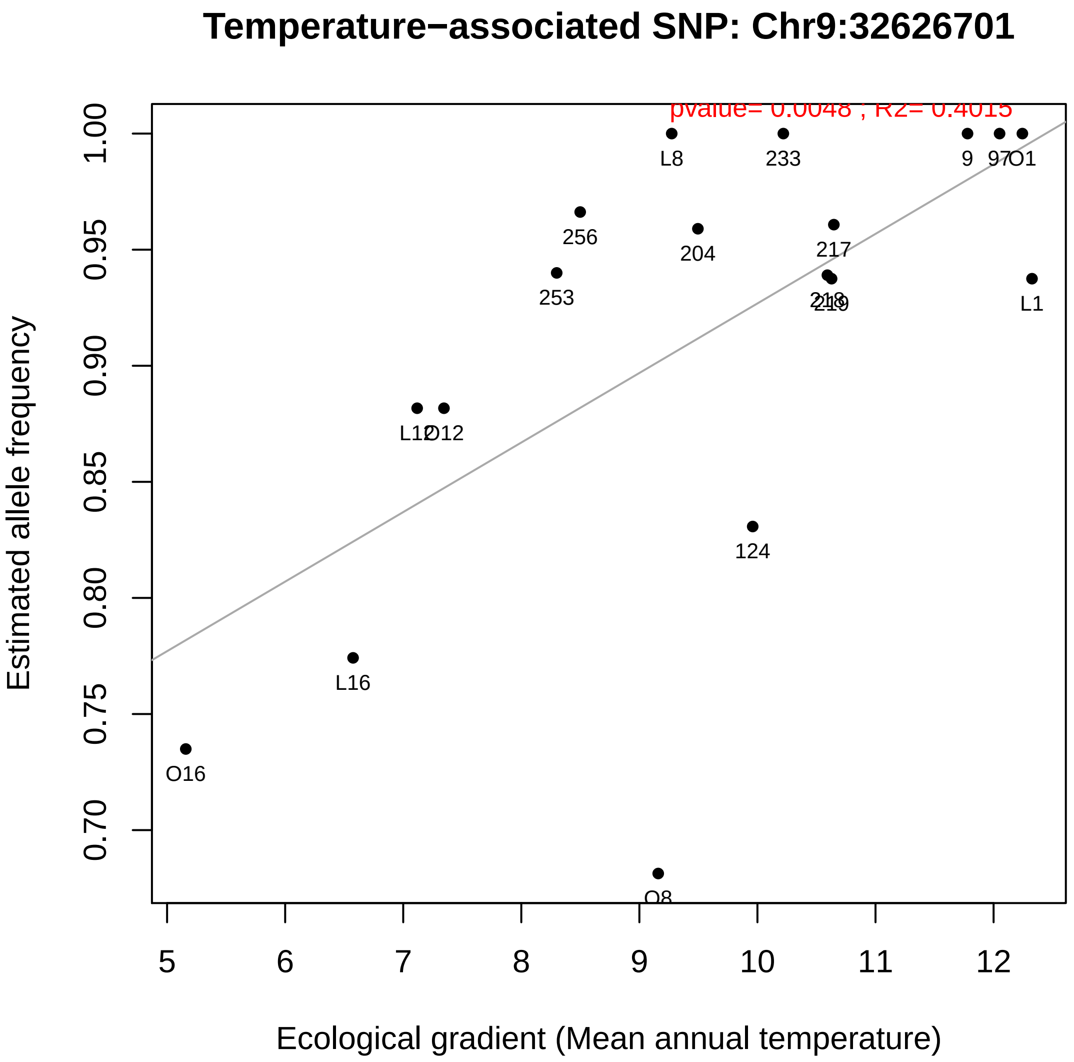
Temperature−associated SNP: Chr9:32626701
pvalue= 0.0048 ; R2= 0.4015
Estimated allele frequency
Ecological gradient (Mean annual temperature)
1.00
0.95
0.90
0.85
0.80
0.75
0.70
5
6
7
8
9
10
11
12
L8
233
9
97
O1
256
204
217
253
218
219
L1
L12
O12
124
L16
O16
O8

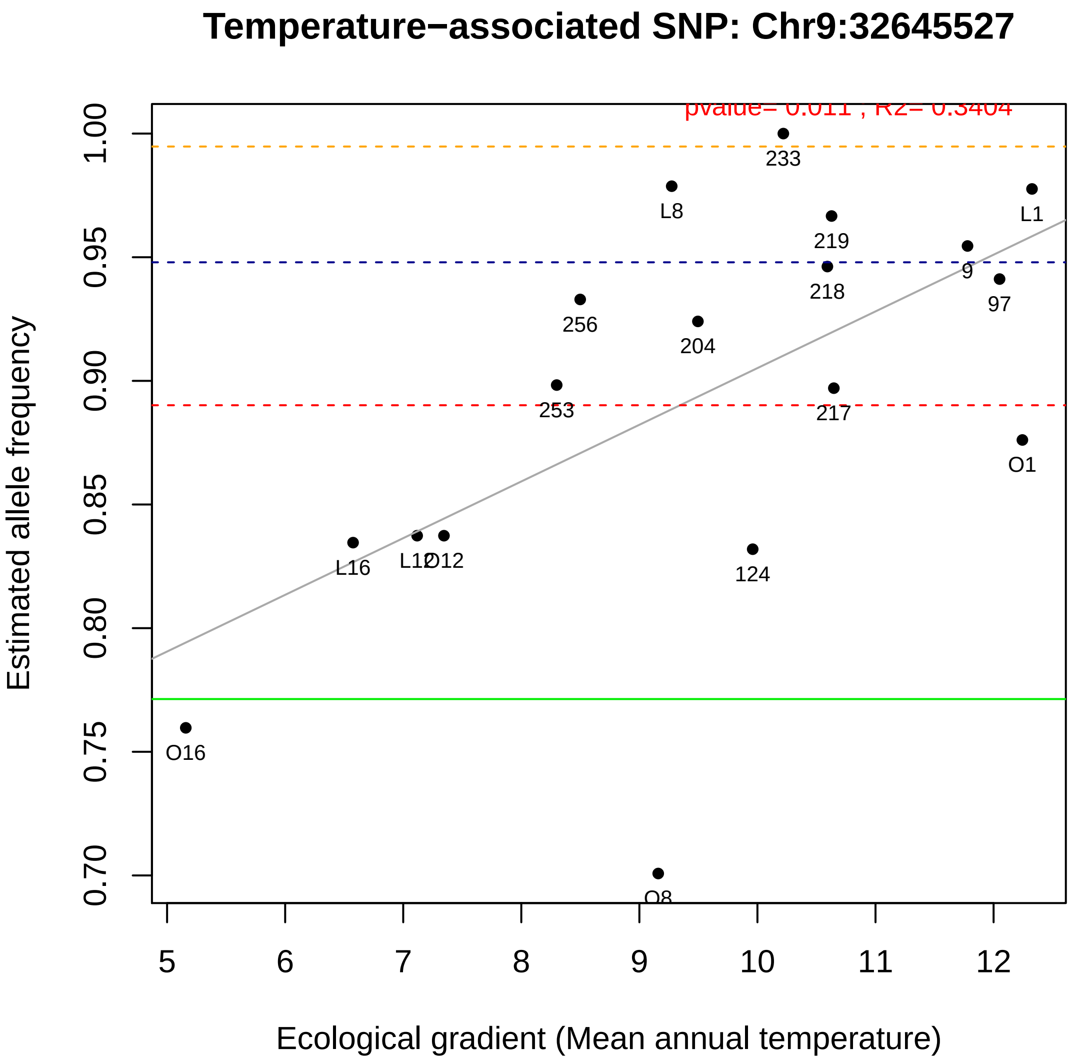

Temperature−associated SNP: Chr9:32645527
pvalue= 0.011 , R2= 0.3404
Estimated allele frequency
Ecological gradient (Mean annual temperature)
0.70
0.75
0.80
0.85
0.90
0.95
1.00
5
6
7
8
9
10
11
12
233
L8
L1
219
9
218
97
256
204
253
217
O1
L16
L12
O12
124
O16
O8

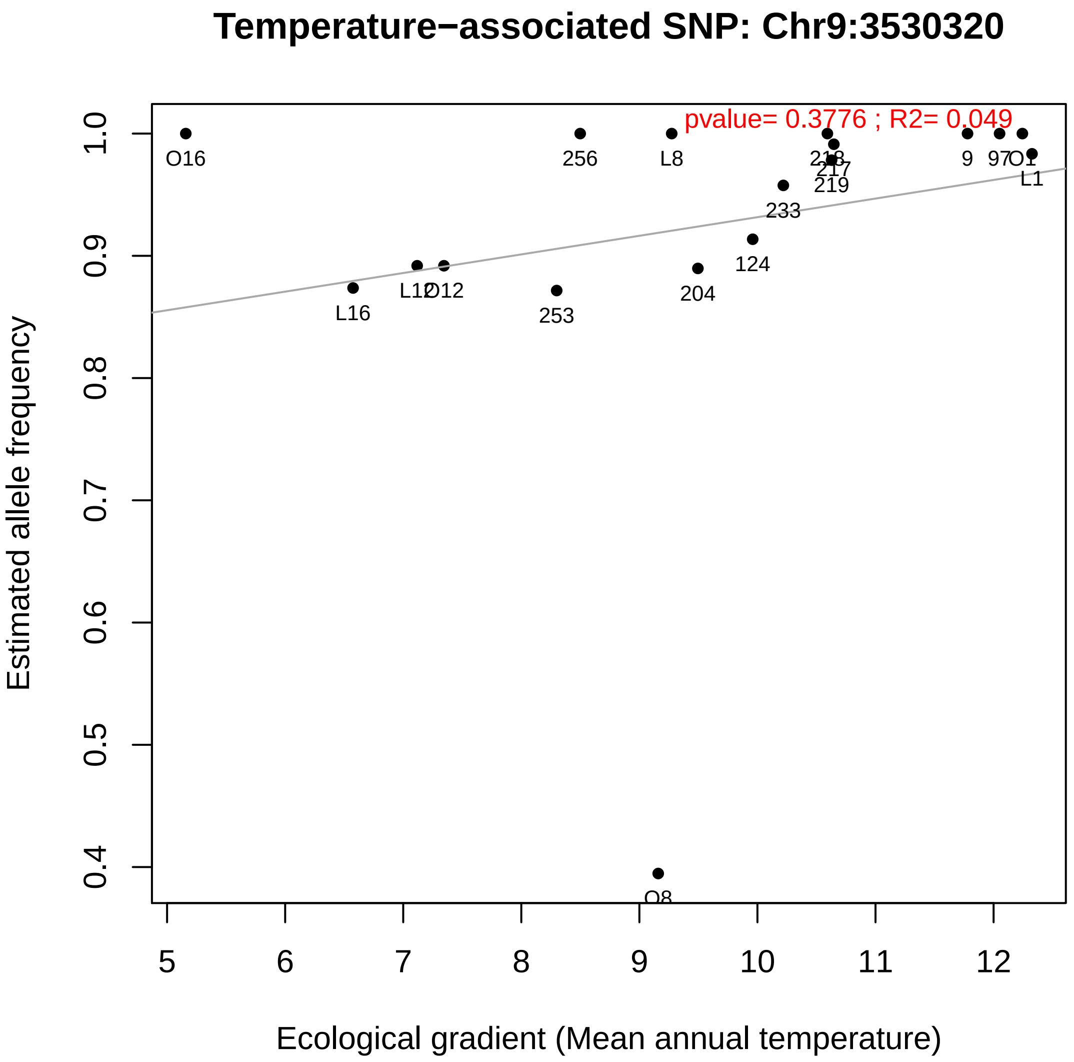
Temperature−associated SNP: Chr9:3530320
pvalue= 0.3776 ; R2= 0.049
Estimated allele frequency
Ecological gradient (Mean annual temperature)
O16
256
L8
218
217
219
9
97
O1
L1
233
124
L16
L12
O12
253
204
O8
1.0
0.9
0.8
0.7
0.6
0.5
0.4
5
6
7
8
9
10
11
12

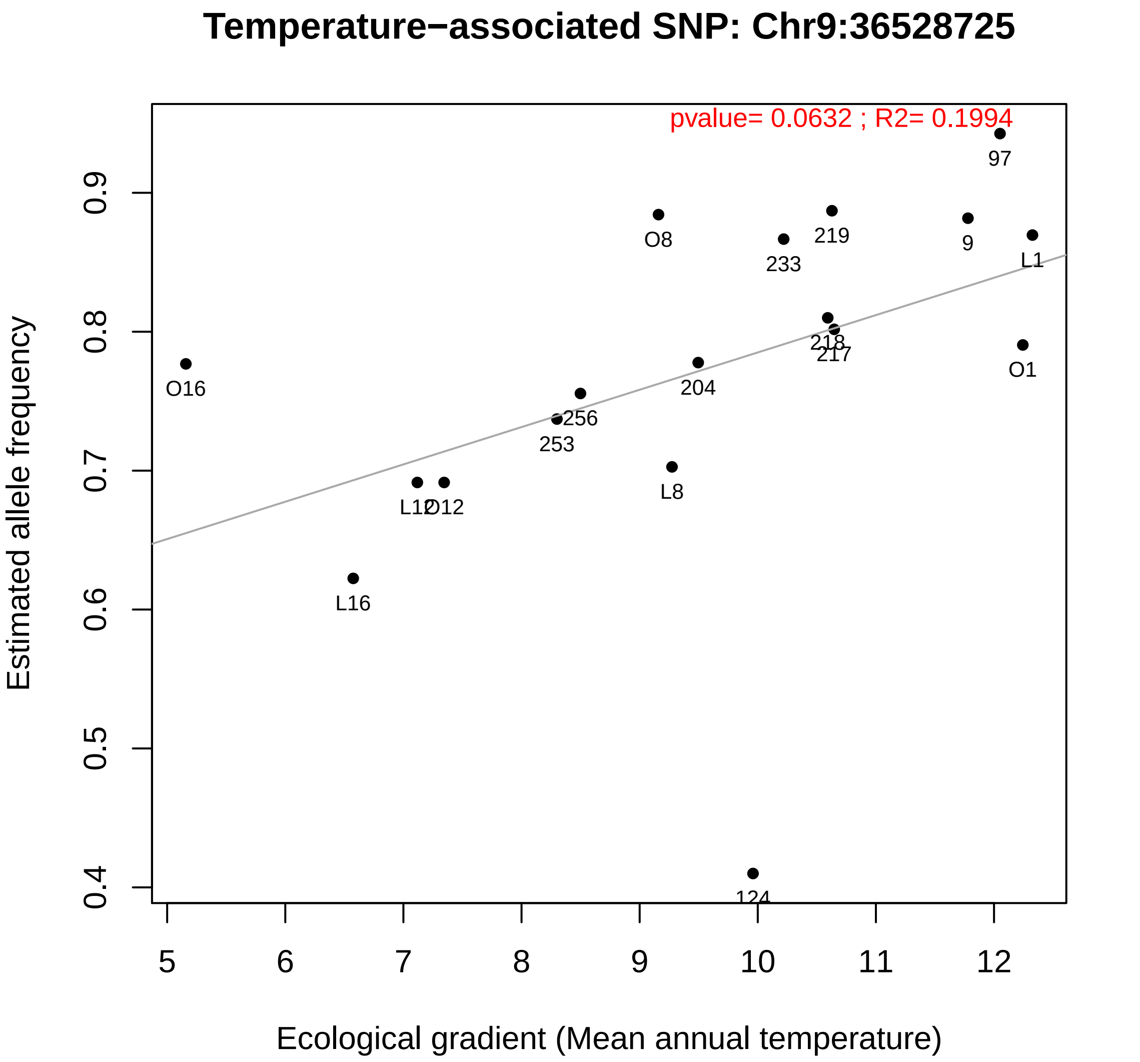
Temperature−associated SNP: Chr9:36528725
pvalue= 0.0632 ; R2= 0.1994
Estimated allele frequency
Ecological gradient (Mean annual temperature)
0.4
0.5
0.6
0.7
0.8
0.9
5
6
7
8
9
10
11
12
97
O8
219
9
233
L1
218
217
O1
O16
204
256
253
L8
L12
O12
L16
124

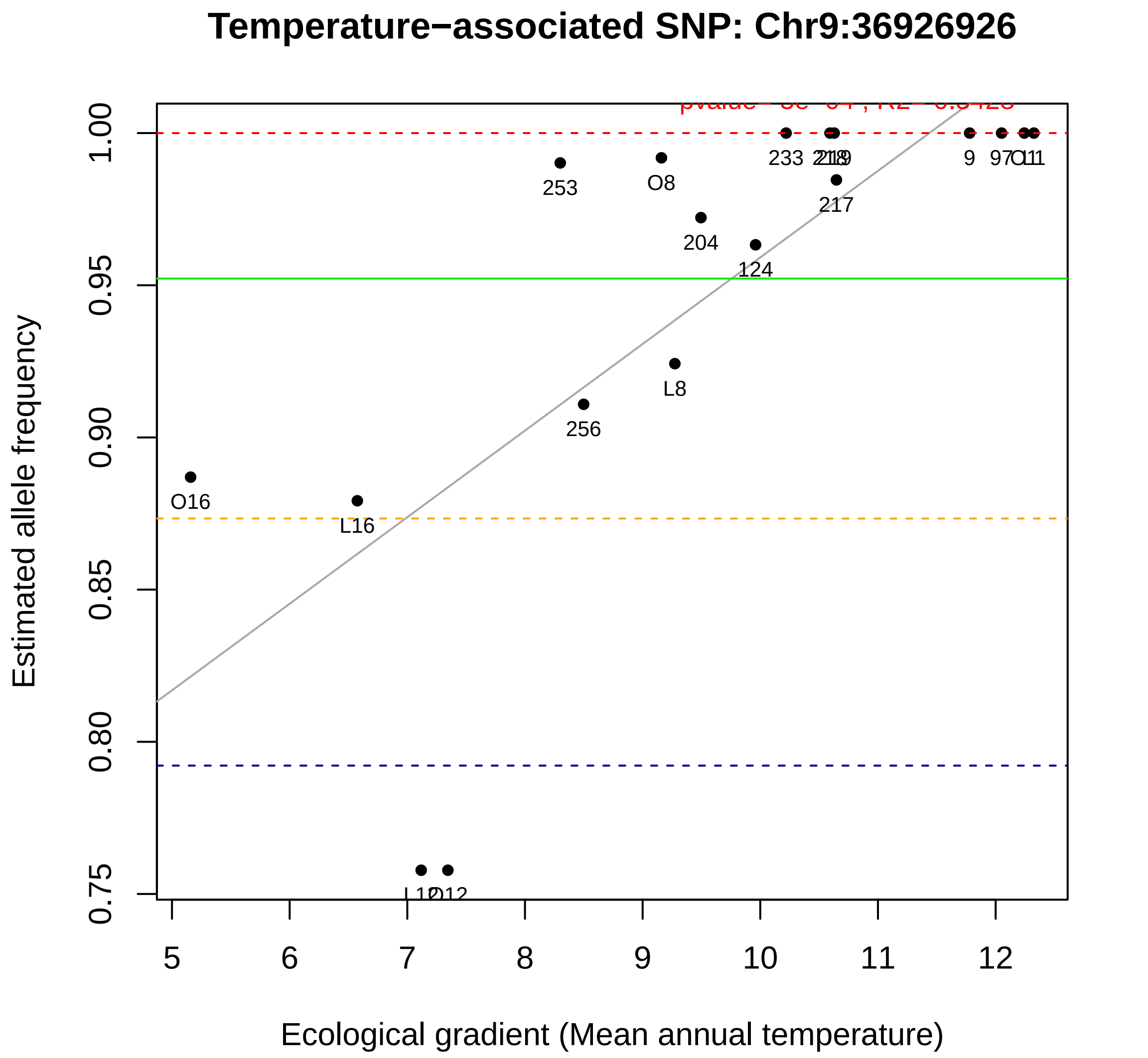

Temperature−associated SNP: Chr9:36926926
Estimated allele frequency
Ecological gradient (Mean annual temperature)
0.75
0.80
0.85
0.90
0.95
1.00
5
6
7
8
9
10
11
12
233
9
253
O8
217
204
124
L8
256
O16
L16

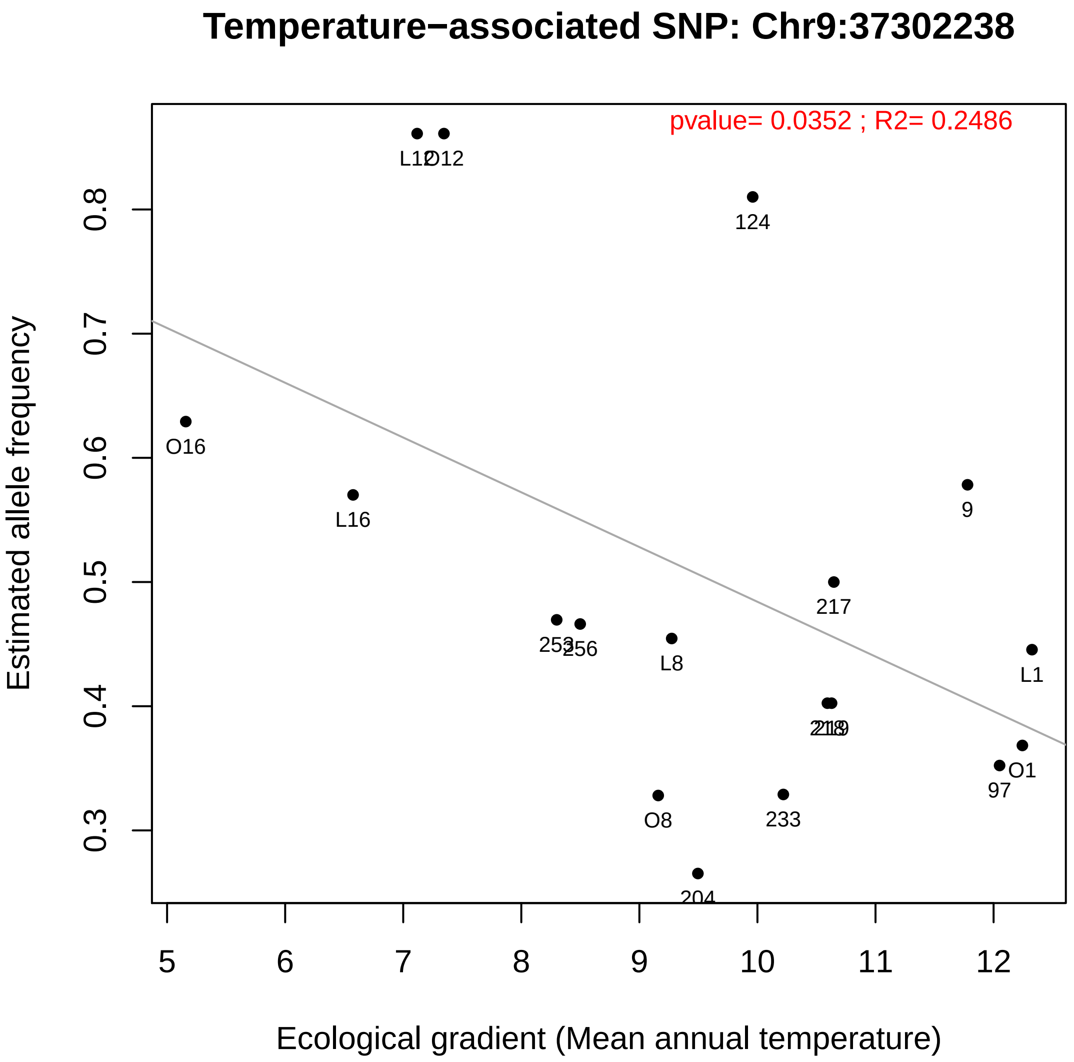

Temperature−associated SNP: Chr9:37302238
pvalue= 0.0352 ; R2= 0.2486
L12
O12
124
O16
L16
9
217
253
256
L8
L1
218
219
O1
97
O8
233
204
0.3
0.4
0.5
0.6
0.7
0.8
5
6
7
8
9
10
11
12
Estimated allele frequency
Ecological gradient (Mean annual temperature)

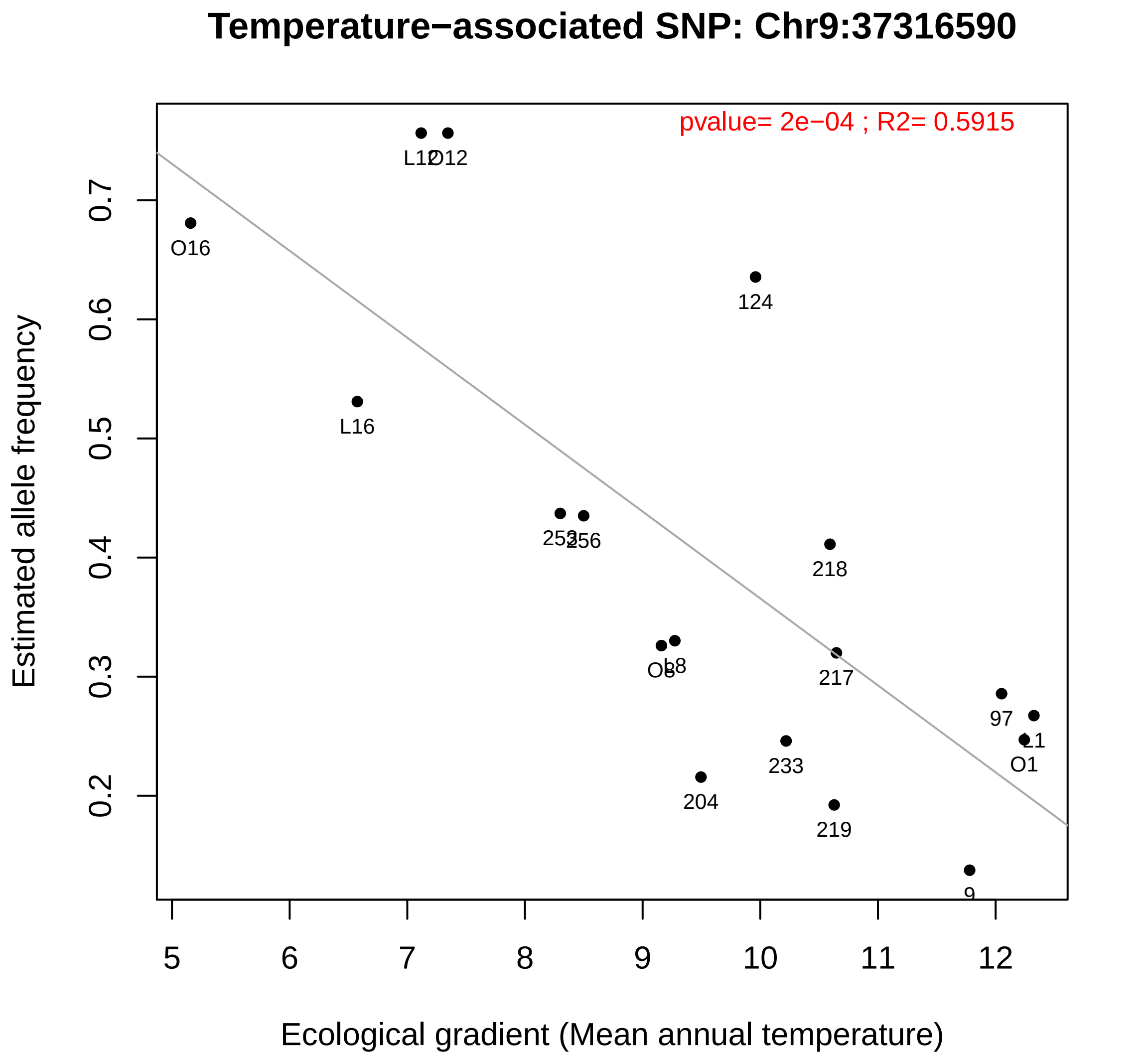

Temperature−associated SNP: Chr9:37316590
pvalue= 2e−04 ; R2= 0.5915
Estimated allele frequency
0.2
0.3
0.4
0.5
0.6
0.7
5
6
7
8
9
10
11
12
Ecological gradient (Mean annual temperature)
O16
L12
O12
124
L16
252
256
218
O8
L8
217
97
L1
O1
233
204
219
9

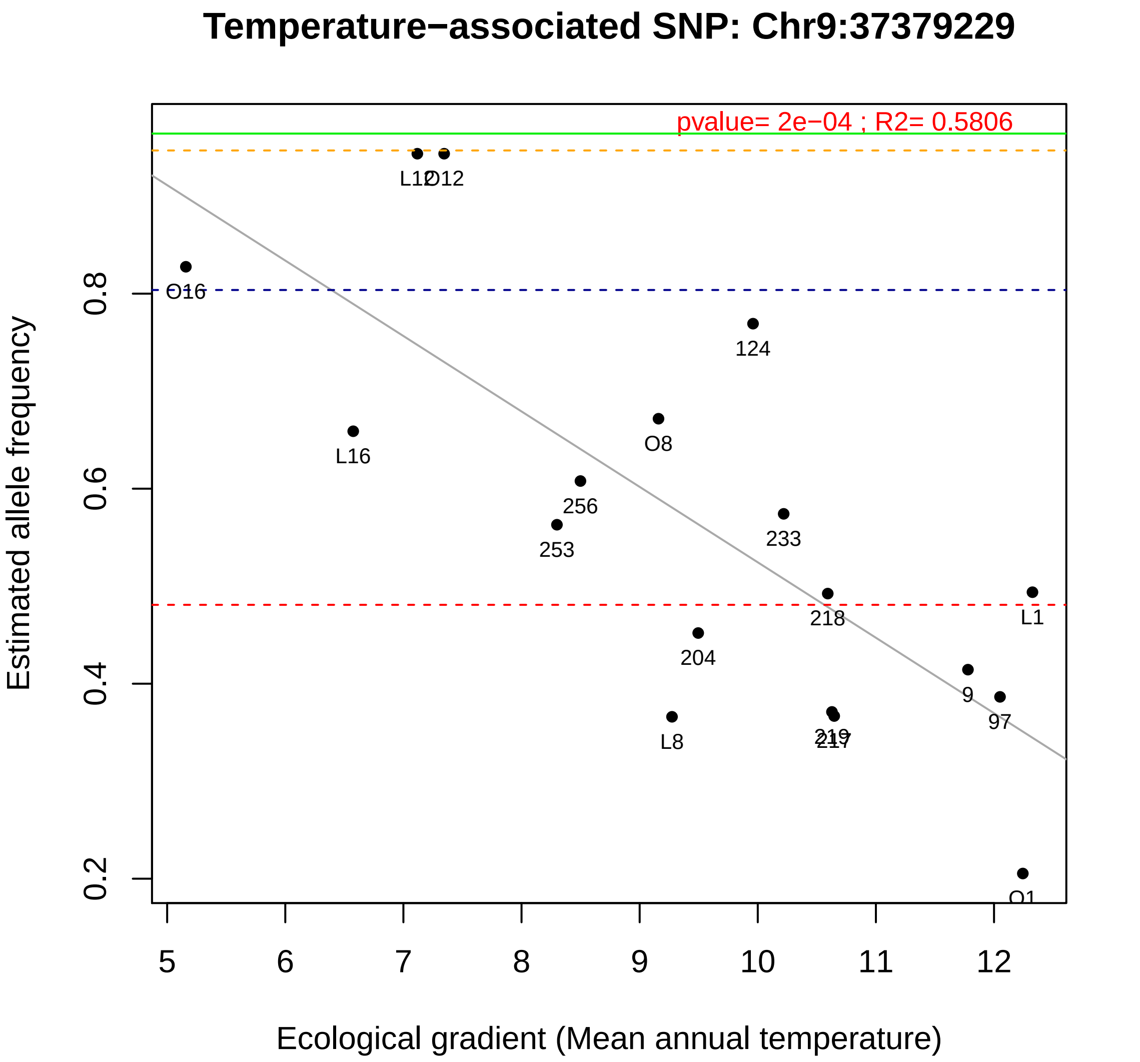

Temperature−associated SNP: Chr9:37379229
pvalue= 2e−04 ; R2= 0.5806
Estimated allele frequency
Ecological gradient (Mean annual temperature)
0.2
0.4
0.6
0.8
5
6
7
8
9
10
11
12
L12
O12
O16
124
L16
O8
256
253
233
218
L1
204
9
97
L8
219
217
O1

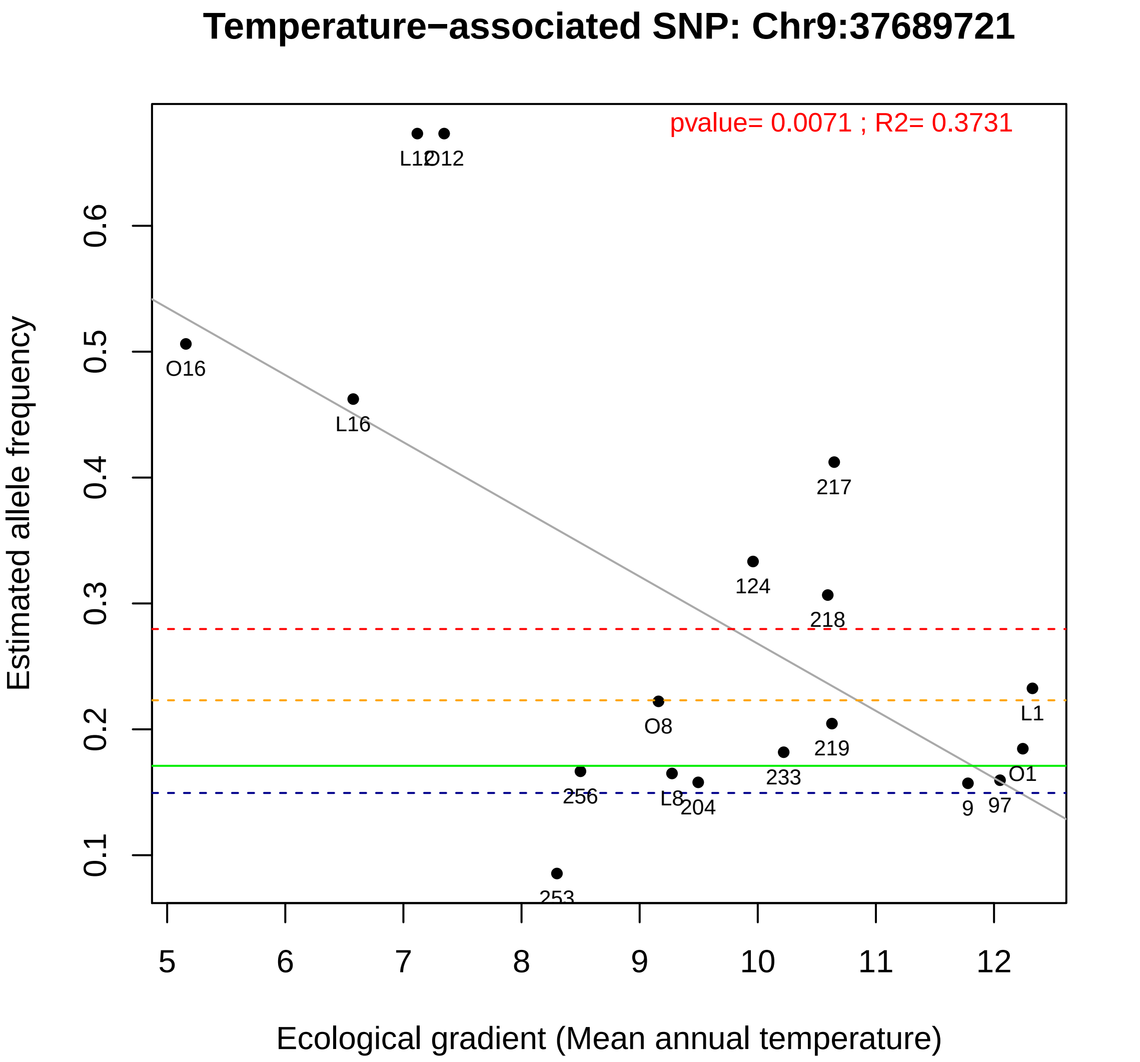

Temperature−associated SNP: Chr9:37689721
pvalue= 0.0071 ; R2= 0.3731
Estimated allele frequency
Ecological gradient (Mean annual temperature)
0.1
0.2
0.3
0.4
0.5
0.6
5
6
7
8
9
10
11
12
L12
O12
O16
L16
217
124
218
O8
L1
219
233
O1
256
L8
204
9
97
253

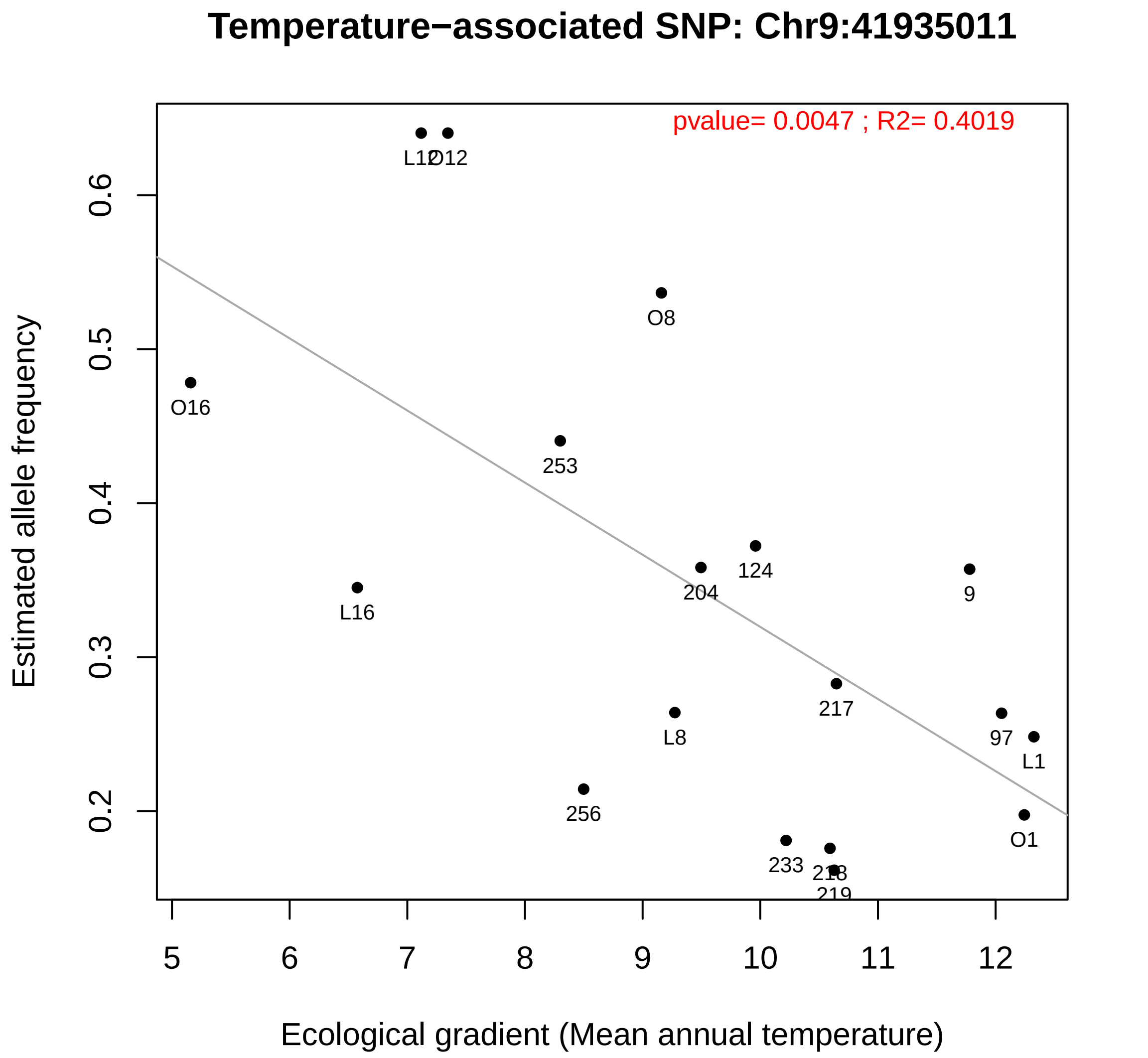
Temperature−associated SNP: Chr9:41935011
pvalue= 0.0047 ; R2= 0.4019
Estimated allele frequency
0.2
0.3
0.4
0.5
0.6
Ecological gradient (Mean annual temperature)
5
6
7
8
9
10
11
12
L12
O12
O8
O16
253
124
204
9
L16
217
L8
97
L1
256
O1
233
218
219

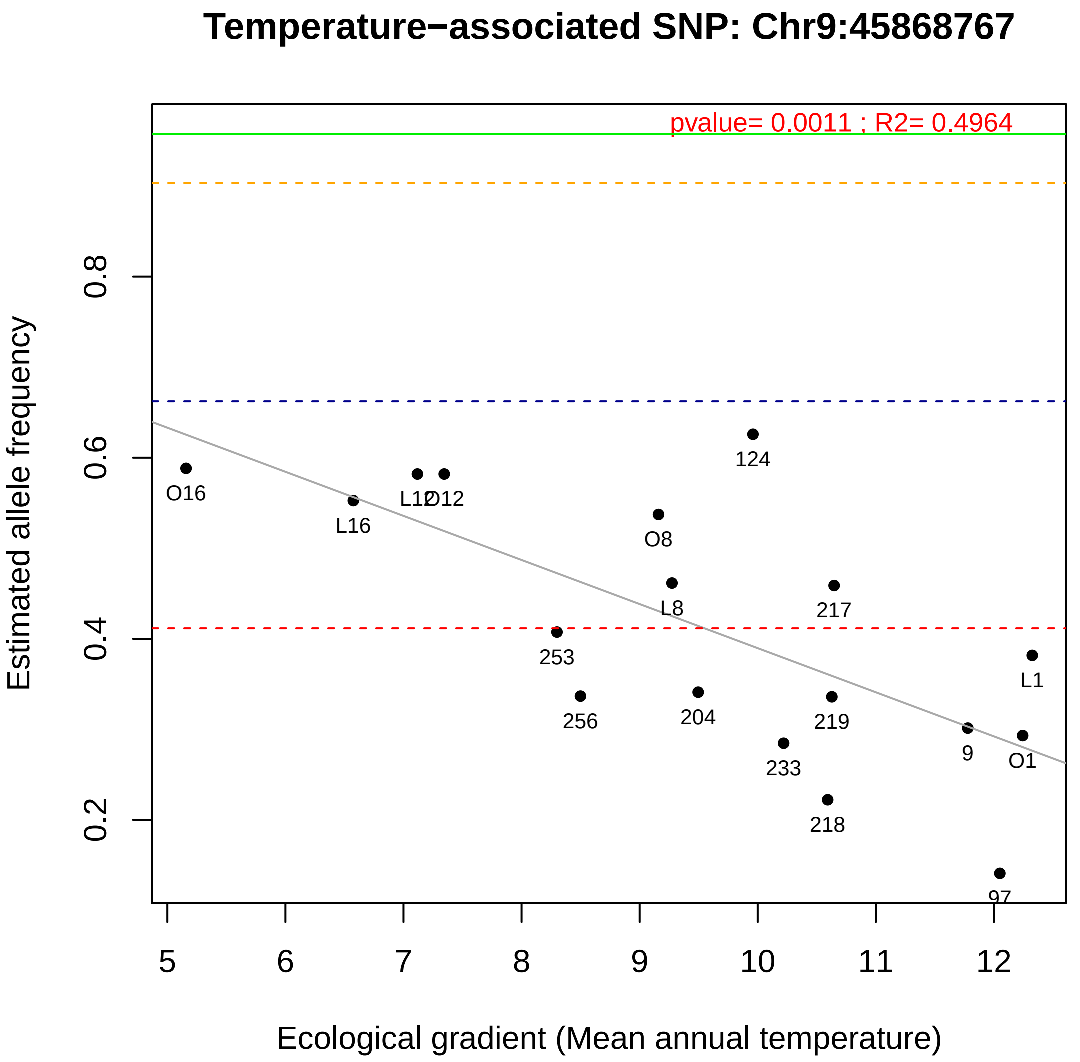

Temperature−associated SNP: Chr9:45868767
pvalue= 0.0011 ; R2= 0.4964
Estimated allele frequency
0.2
0.4
0.6
0.8
5
6
7
8
9
10
11
12
Ecological gradient (Mean annual temperature)
O16
L16
L12
O12
253
256
O8
L8
204
124
233
218
217
219
9
97
O1
L1

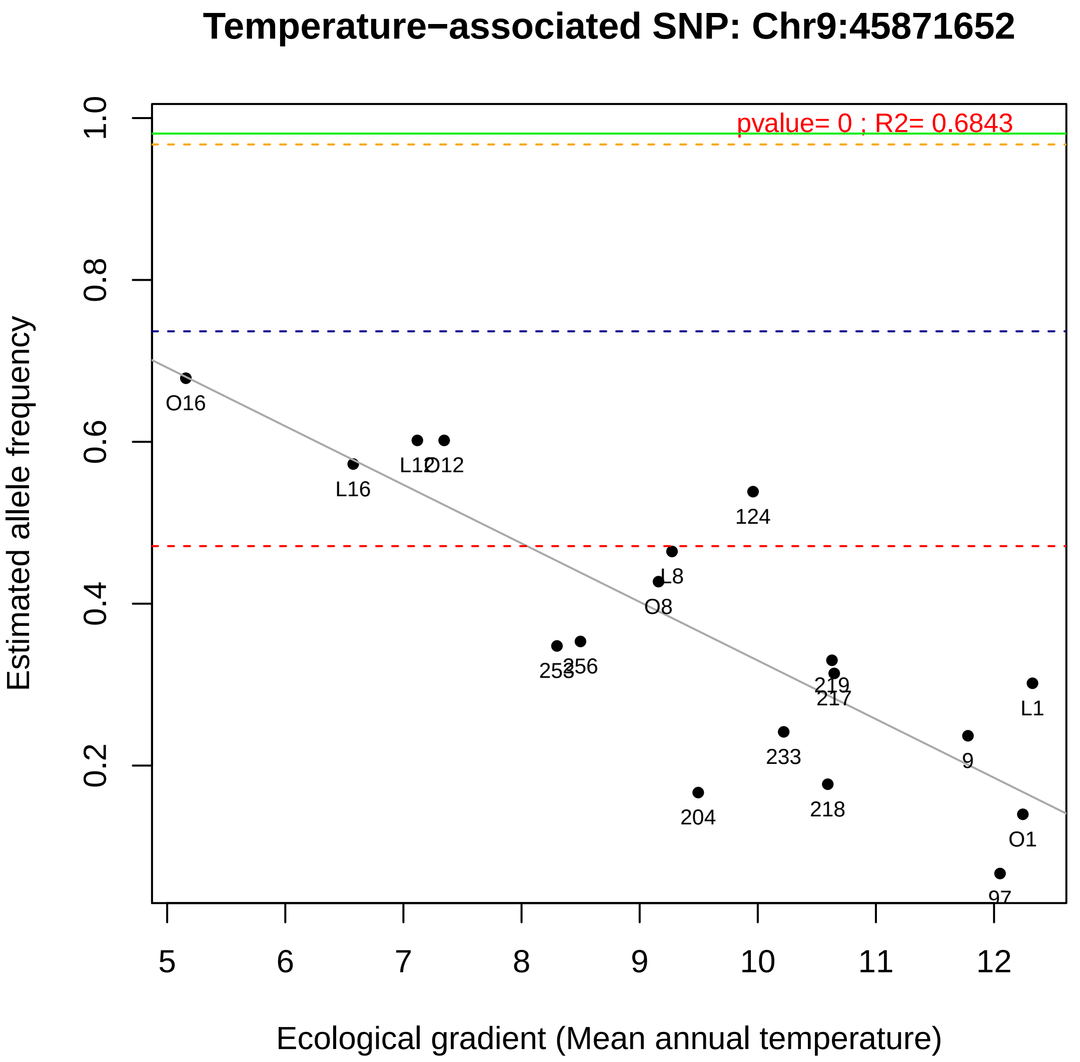

Temperature−associated SNP: Chr9:45871652
pvalue= 0 ; R2= 0.6843
Estimated allele frequency
Ecological gradient (Mean annual temperature)
1.0
0.8
0.6
0.4
0.2
5
6
7
8
9
10
11
12
O16
L16
L12
O12
124
L8
O8
253
256
219
217
L1
233
9
204
218
O1
97

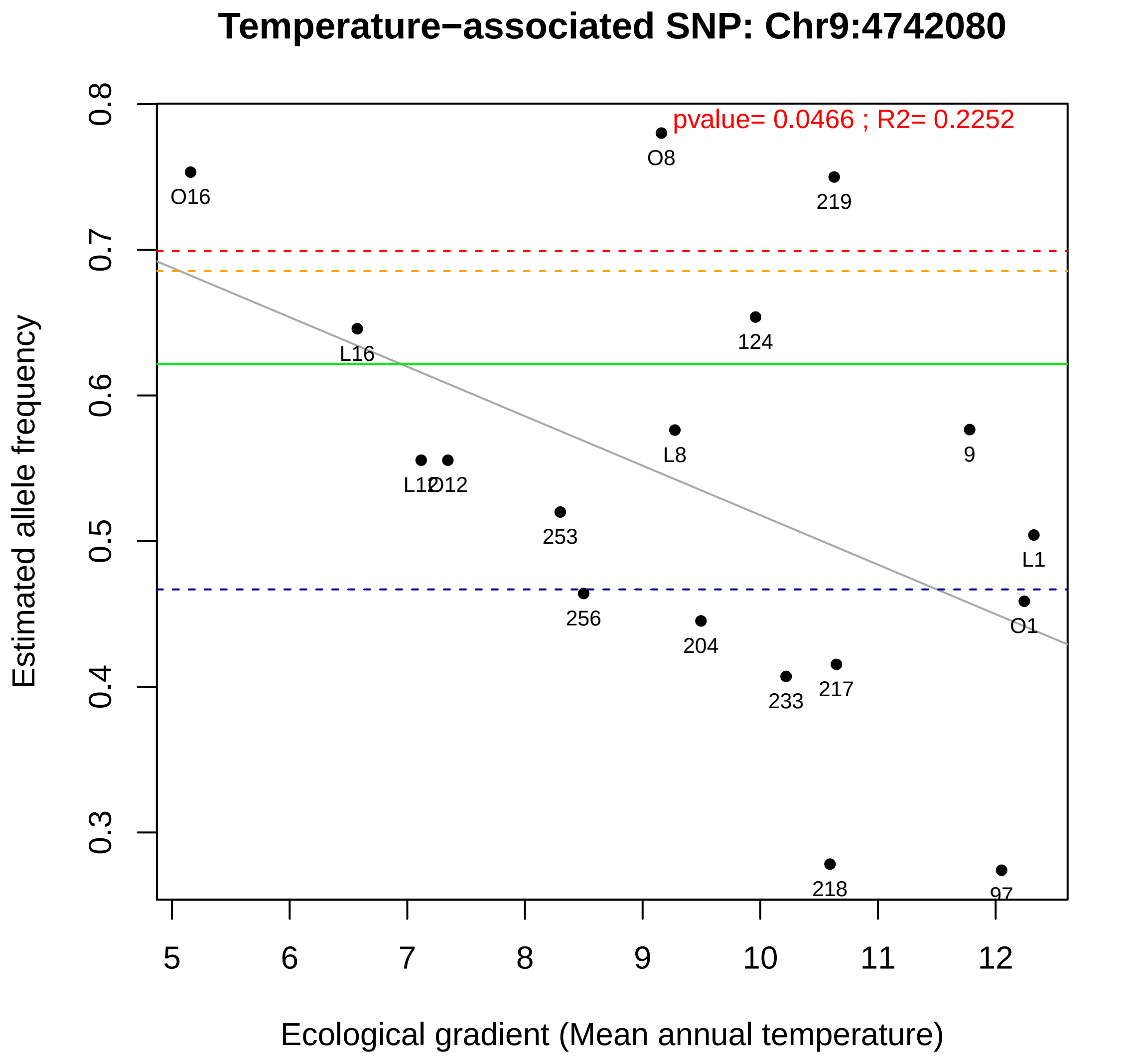
Temperature−associated SNP: Chr9:4742080
pvalue= 0.0466 ; R2= 0.2252
Estimated allele frequency
0.3
0.4
0.5
0.6
0.7
0.8
Ecological gradient (Mean annual temperature)
5
6
7
8
9
10
11
12
O16
O8
219
L16
124
L12
O12
L8
9
253
L1
256
O1
204
217
233
218
97

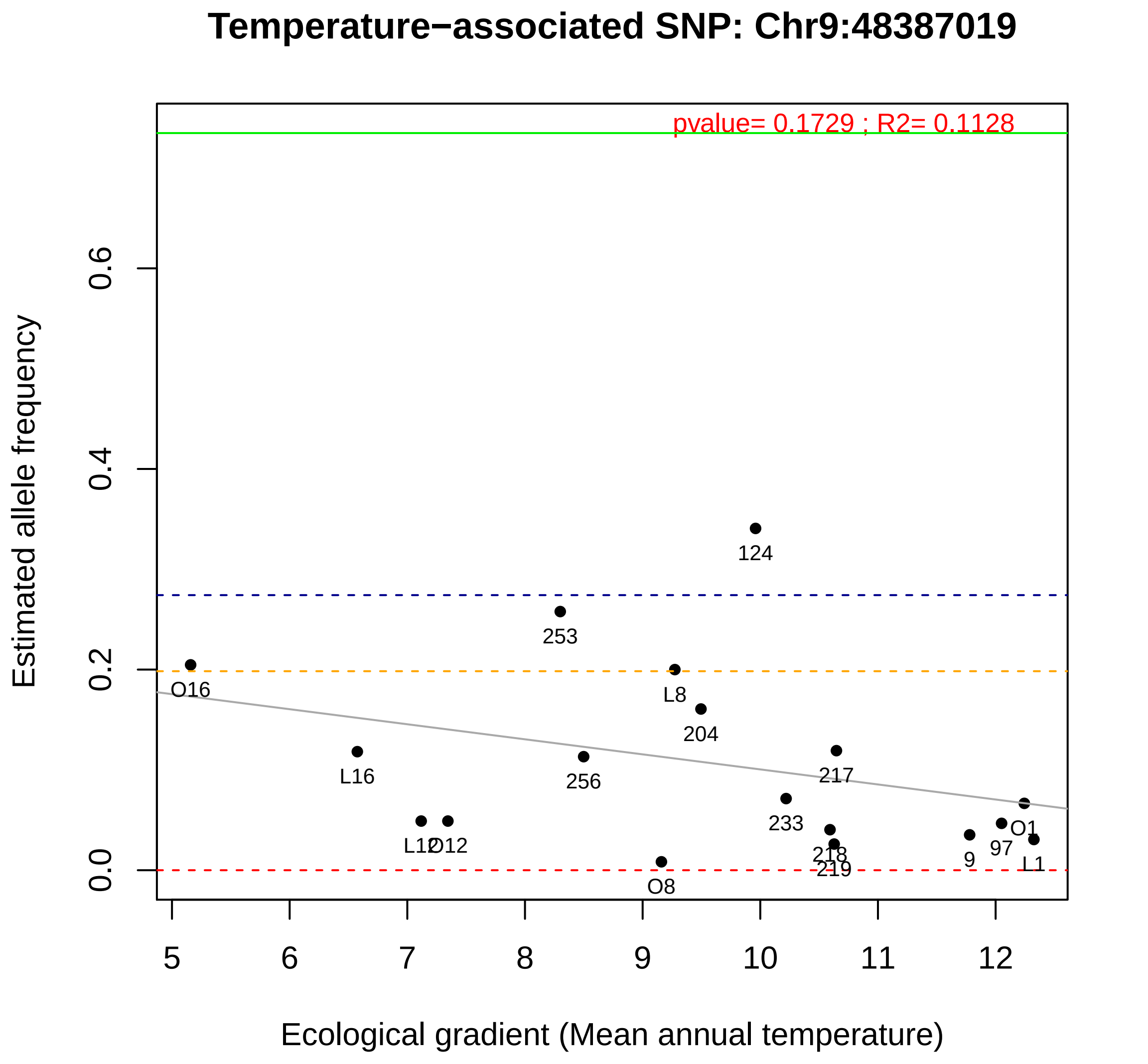

Temperature−associated SNP: Chr9:48387019
pvalue= 0.1729 ; R2= 0.1128
Estimated allele frequency
0.0
0.2
0.4
0.6
5
6
7
8
9
10
11
12
Ecological gradient (Mean annual temperature)
O16
L16
L12
O12
253
256
O8
L8
204
124
233
218
219
217
9
97
O1
L1

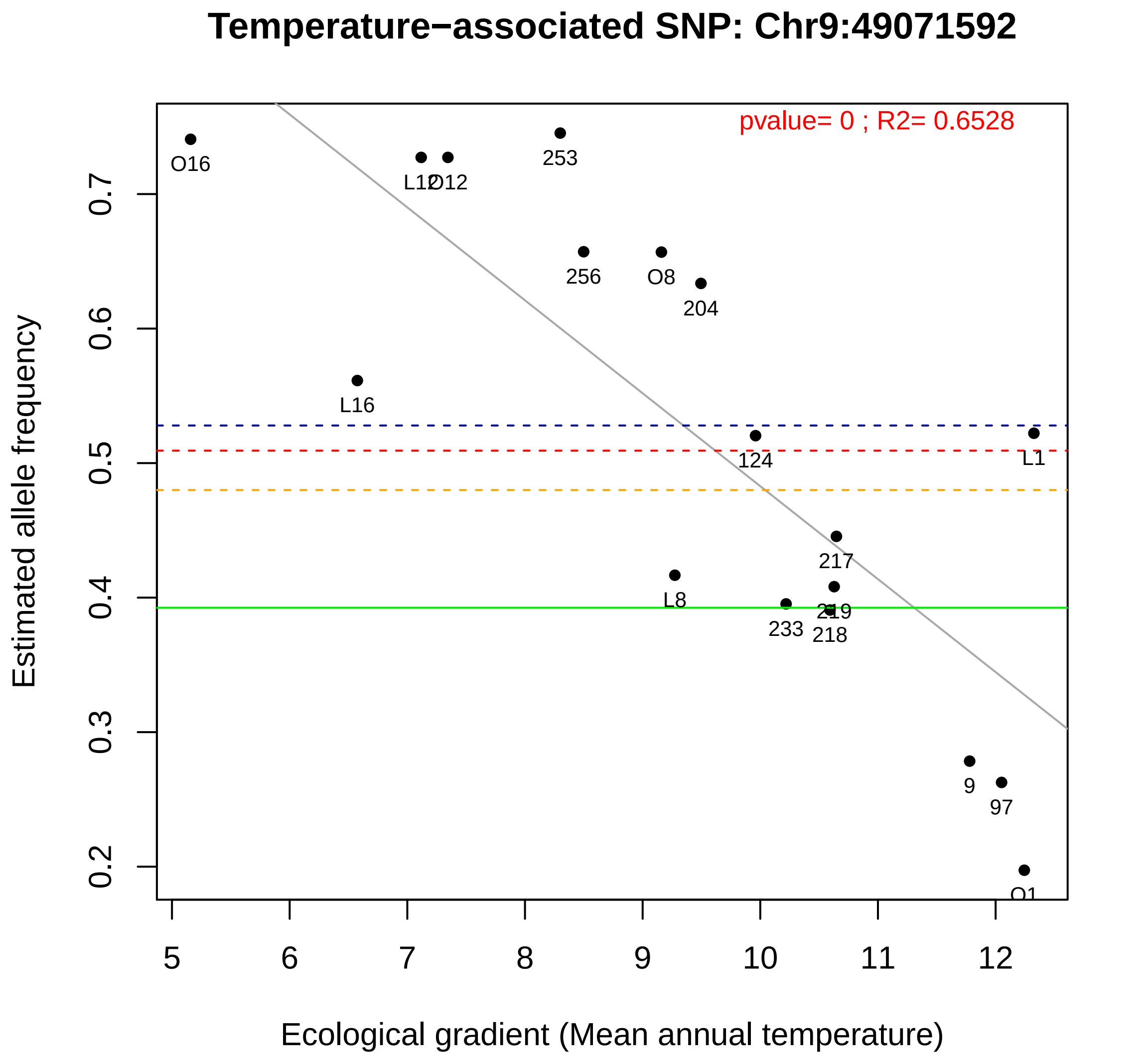

Temperature−associated SNP: Chr9:49071592
pvalue= 0 ; R2= 0.6528
O16
L12
O12
253
256
O8
204
L16
124
L1
217
L8
219
233
218
9
97
O1
0.2
0.3
0.4
0.5
0.6
0.7
Estimated allele frequency
5
6
7
8
9
10
11
12
Ecological gradient (Mean annual temperature)

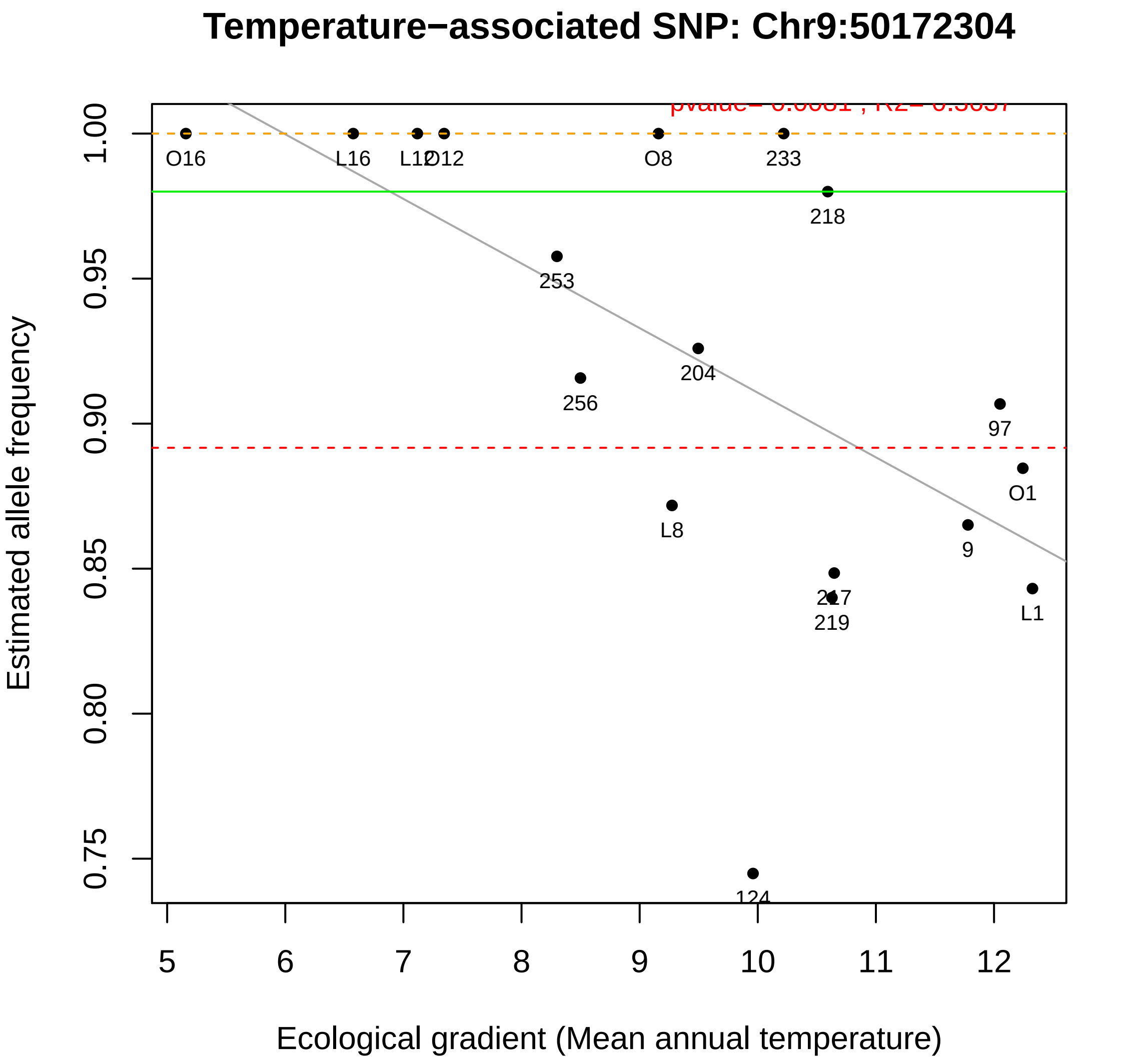

Temperature−associated SNP: Chr9:50172304
Estimated allele frequency
Ecological gradient (Mean annual temperature)
0.75
0.80
0.85
0.90
0.95
1.00
5
6
7
8
9
10
11
12
O16
L16
L12
O12
O8
233
218
253
204
256
97
O1
L8
9
217
219
L1
124

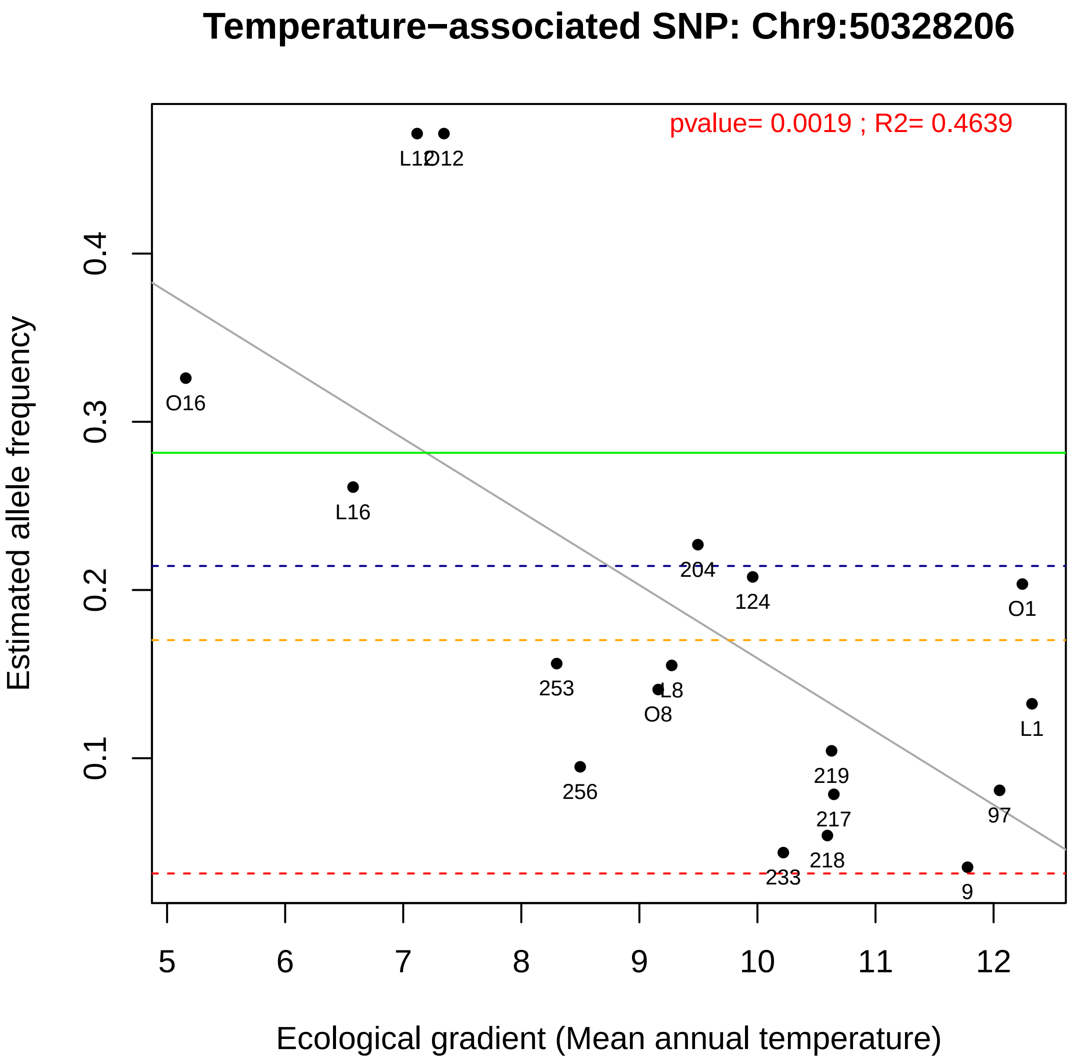
Temperature−associated SNP: Chr9:50328206
pvalue= 0.0019 ; R2= 0.4639
Estimated allele frequency
0.1
0.2
0.3
0.4
Ecological gradient (Mean annual temperature)
5
6
7
8
9
10
11
12
L12
O12
O16
L16
204
124
O1
253
L8
O8
L1
256
219
217
218
233
97
9

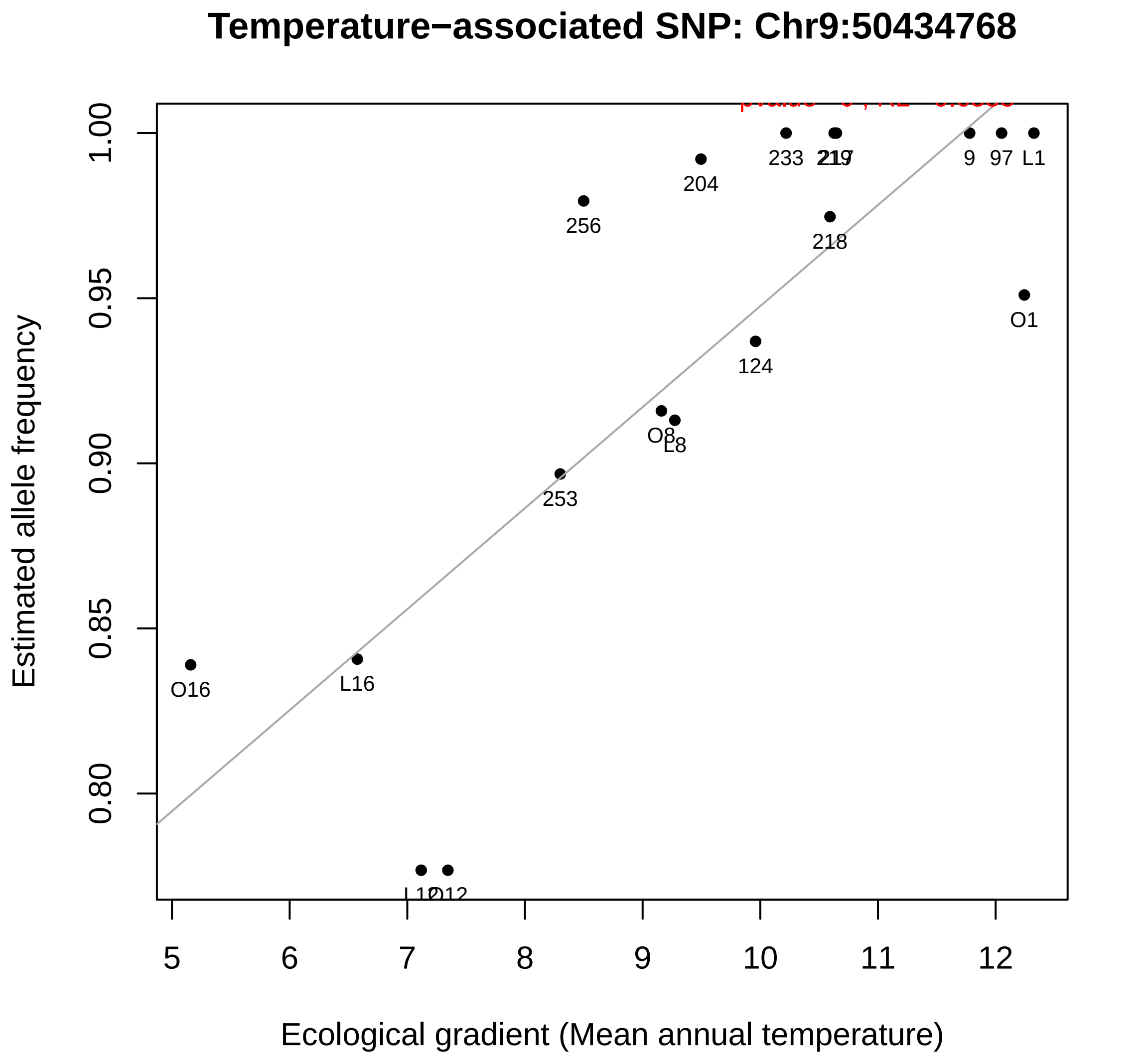
Temperature−associated SNP: Chr9:50434768
Estimated allele frequency
0.80
0.85
0.90
0.95
1.00
5
6
7
8
9
10
11
12
Ecological gradient (Mean annual temperature)
233
204
256
218
9
97
L1
O1
124
O8
L8
253
O16
L16
L12
O12

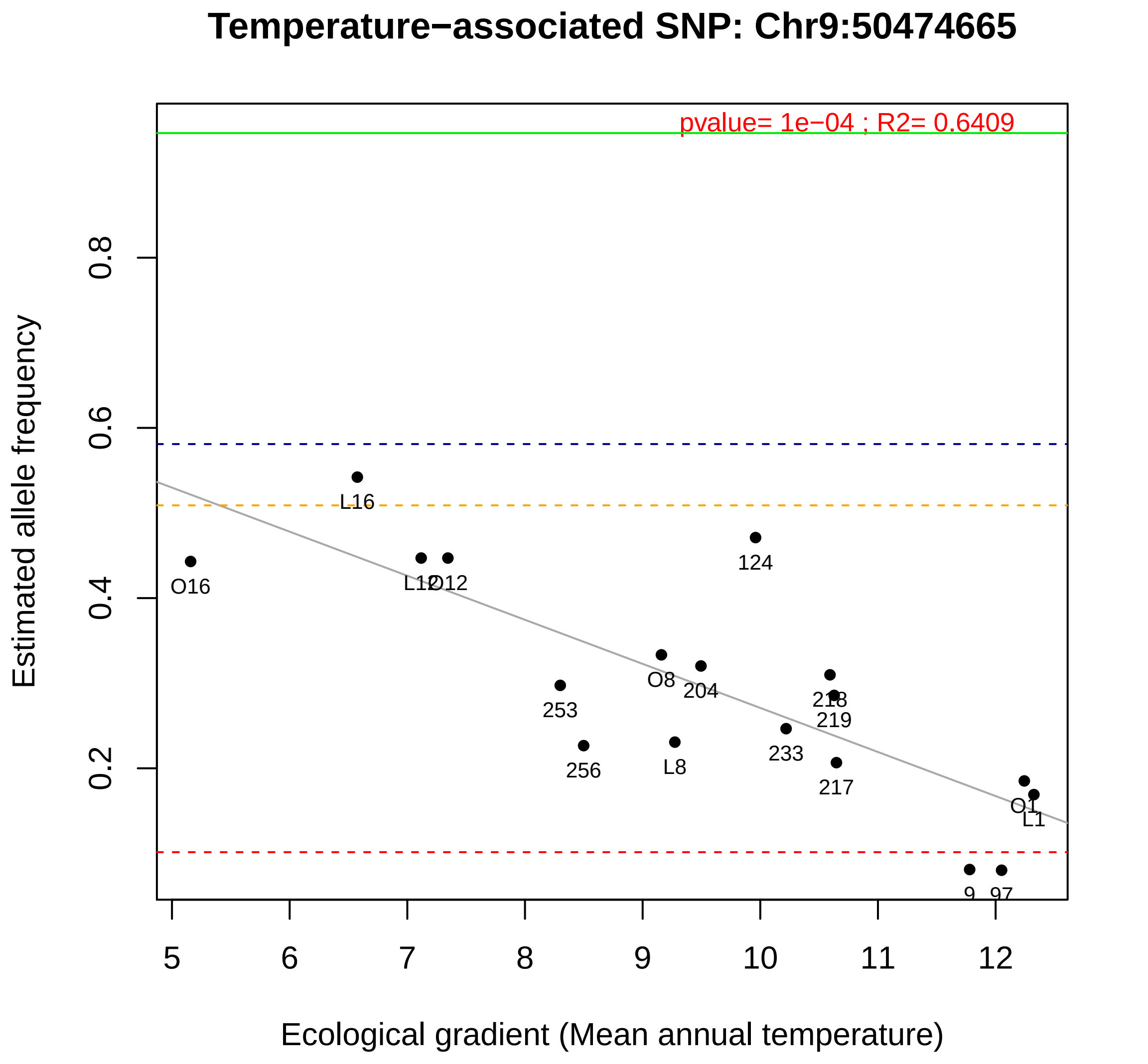
Temperature−associated SNP: Chr9:50474665
pvalue= 1e−04 ; R2= 0.6409
Estimated allele frequency
0.2
0.4
0.6
0.8
5
6
7
8
9
10
11
12
Ecological gradient (Mean annual temperature)
L16
O16
L12
O12
124
253
256
O8
204
L8
233
218
219
217
O1
L1
9
97

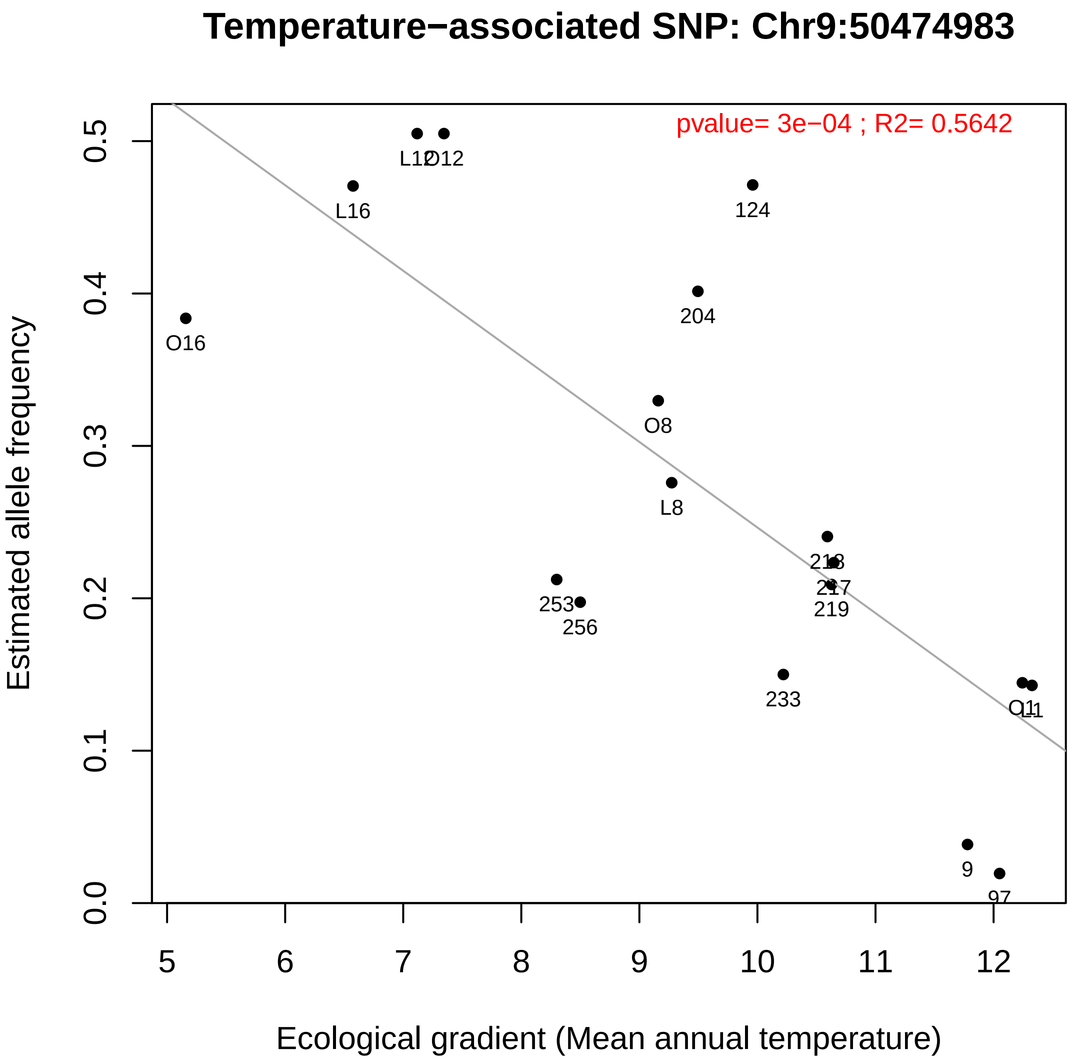

Temperature−associated SNP: Chr9:50474983
pvalue= 3e−04 ; R2= 0.5642
Estimated allele frequency
Ecological gradient (Mean annual temperature)
0.0
0.1
0.2
0.3
0.4
0.5
5
6
7
8
9
10
11
12
L12
O12
L16
124
204
O16
O8
L8
218
217
219
253
256
233
O1
L1
9
97

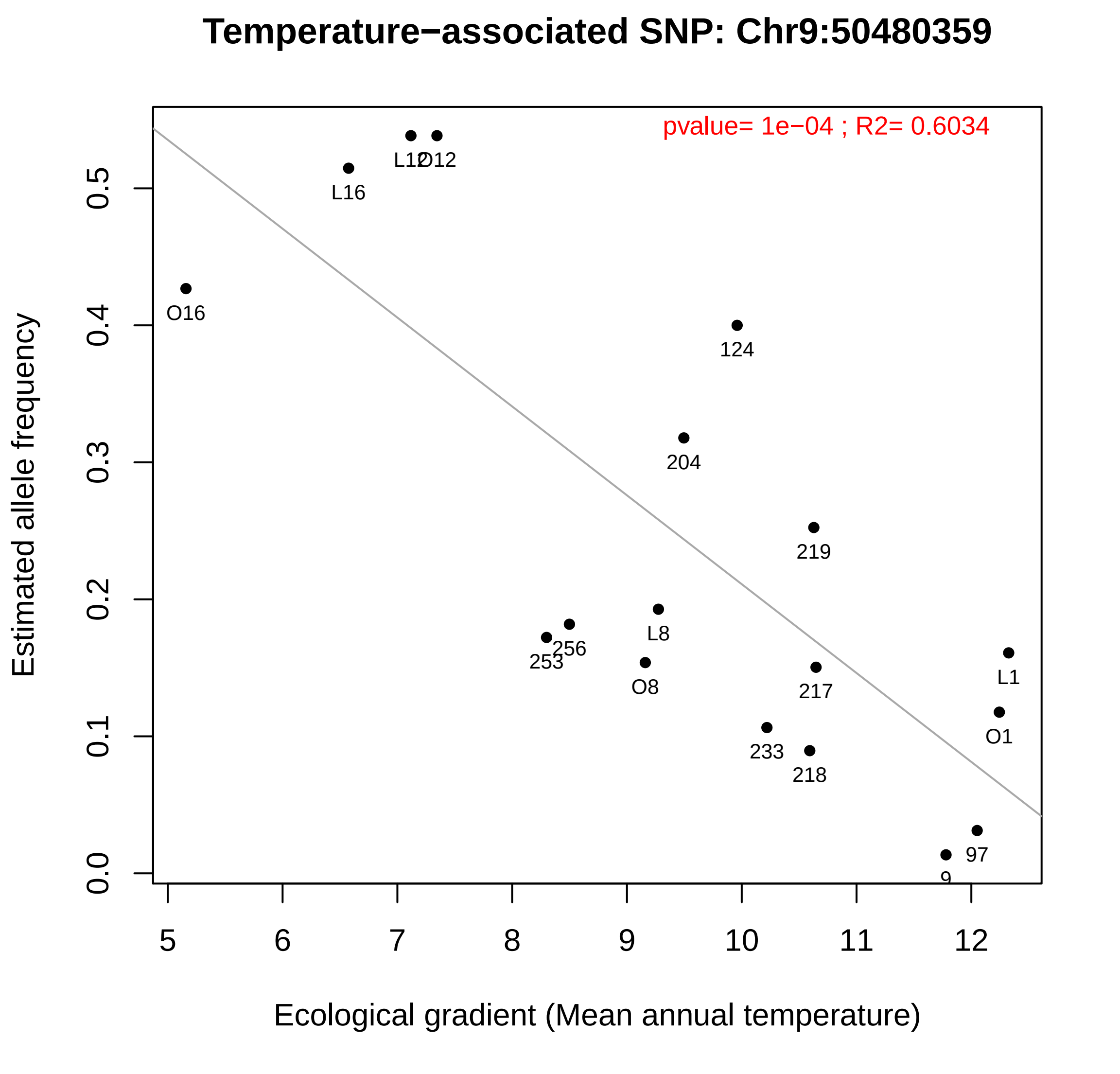
Temperature−associated SNP: Chr9:50480359
pvalue= 1e−04 ; R2= 0.6034
Estimated allele frequency
0.0
0.1
0.2
0.3
0.4
0.5
5
6
7
8
9
10
11
12
Ecological gradient (Mean annual temperature)
O16
L16
L12
O12
124
204
219
253
256
L8
O8
217
233
218
L1
O1
97
9

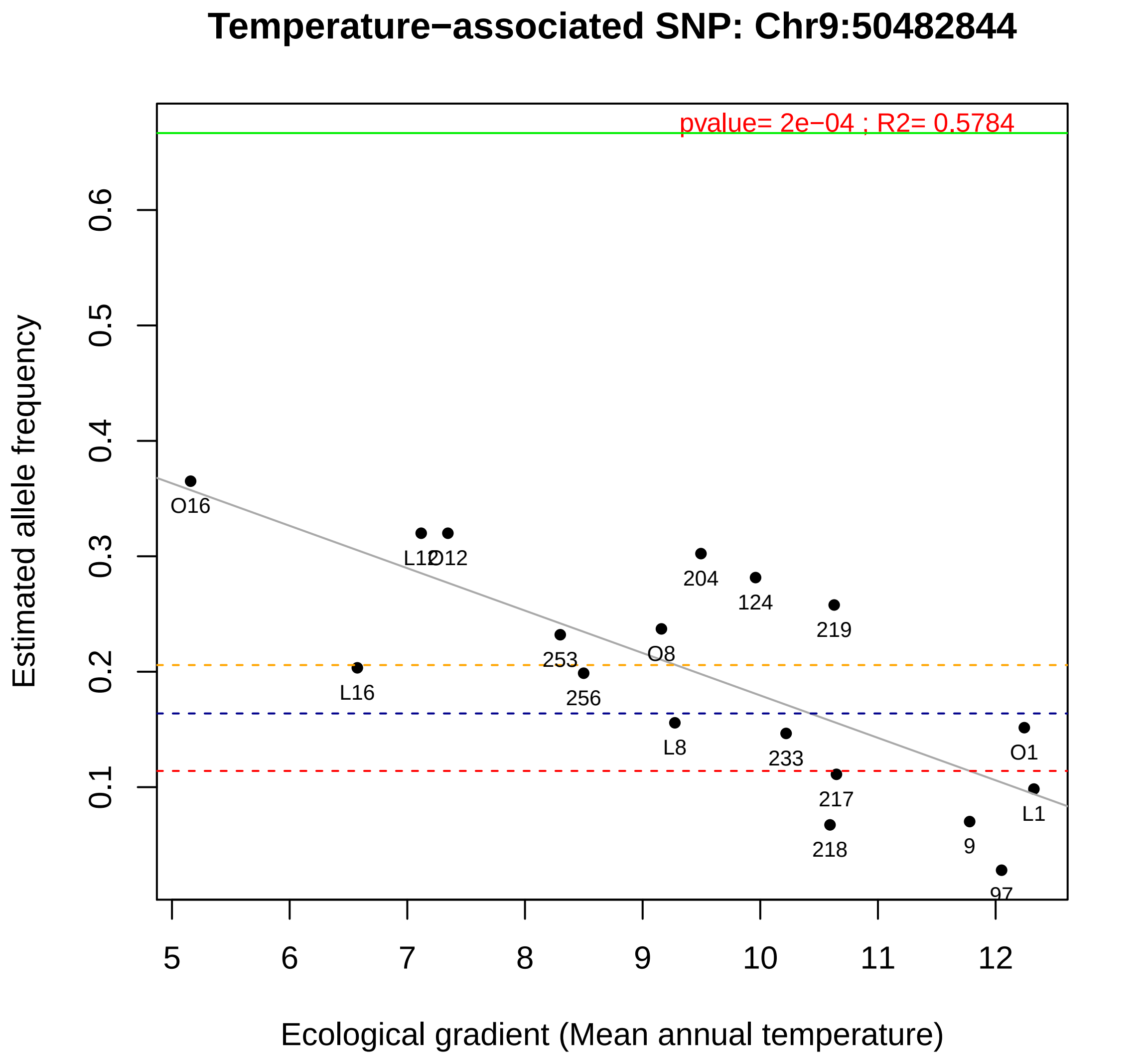

Temperature−associated SNP: Chr9:50482844
pvalue= 2e−04 ; R2= 0.5784
Estimated allele frequency
0.1
0.2
0.3
0.4
0.5
0.6
O16
L12
O12
204
124
219
O8
253
256
L16
L8
233
O1
217
L1
218
9
97
5
6
7
8
9
10
11
12
Ecological gradient (Mean annual temperature)

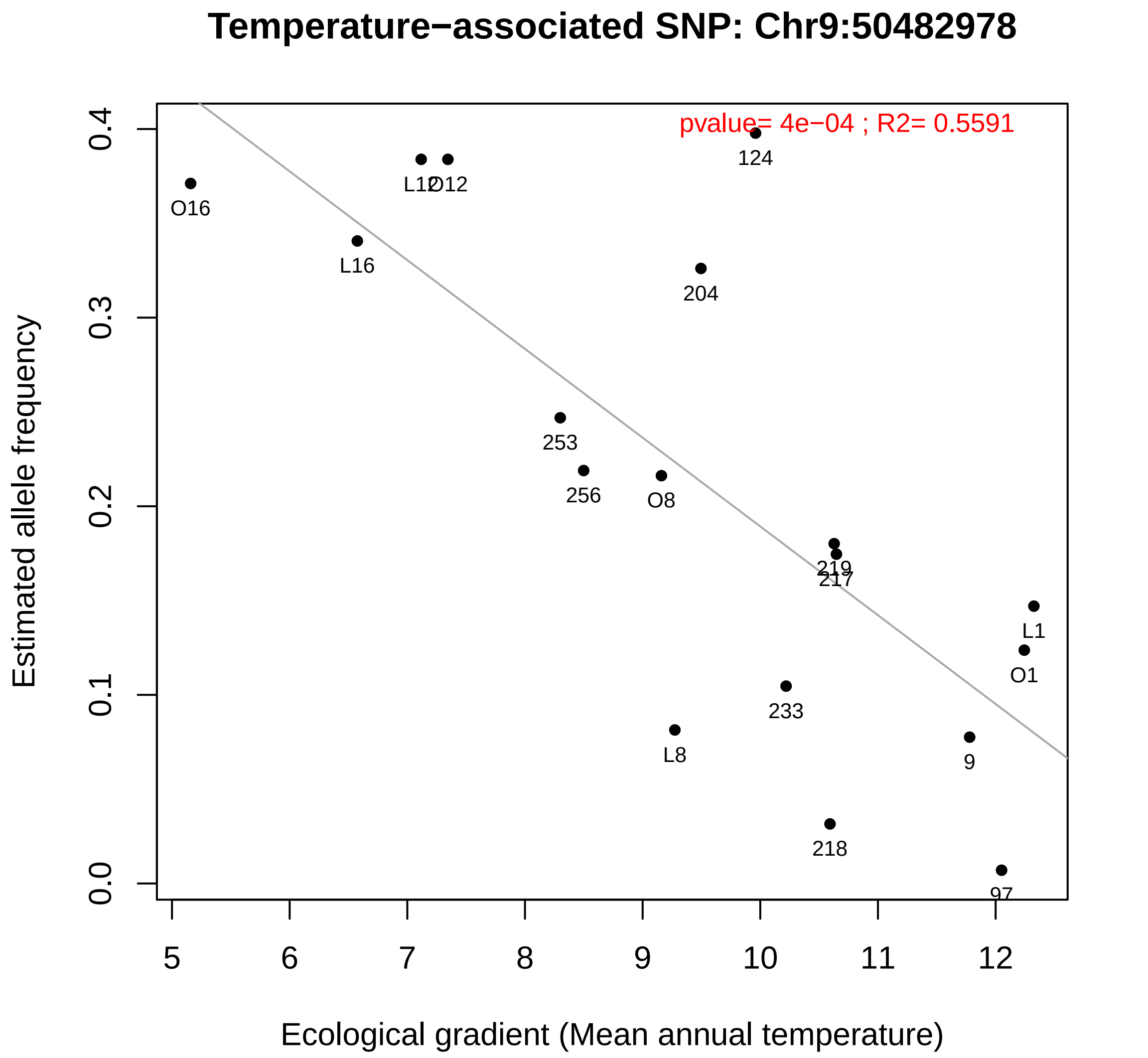
Temperature−associated SNP: Chr9:50482978
pvalue= 4e−04 ; R2= 0.5591
Estimated allele frequency
0.0
0.1
0.2
0.3
0.4
5
6
7
8
9
10
11
12
Ecological gradient (Mean annual temperature)
O16
L16
L12
O12
124
204
253
256
O8
219
217
L1
O1
233
L8
9
218
97

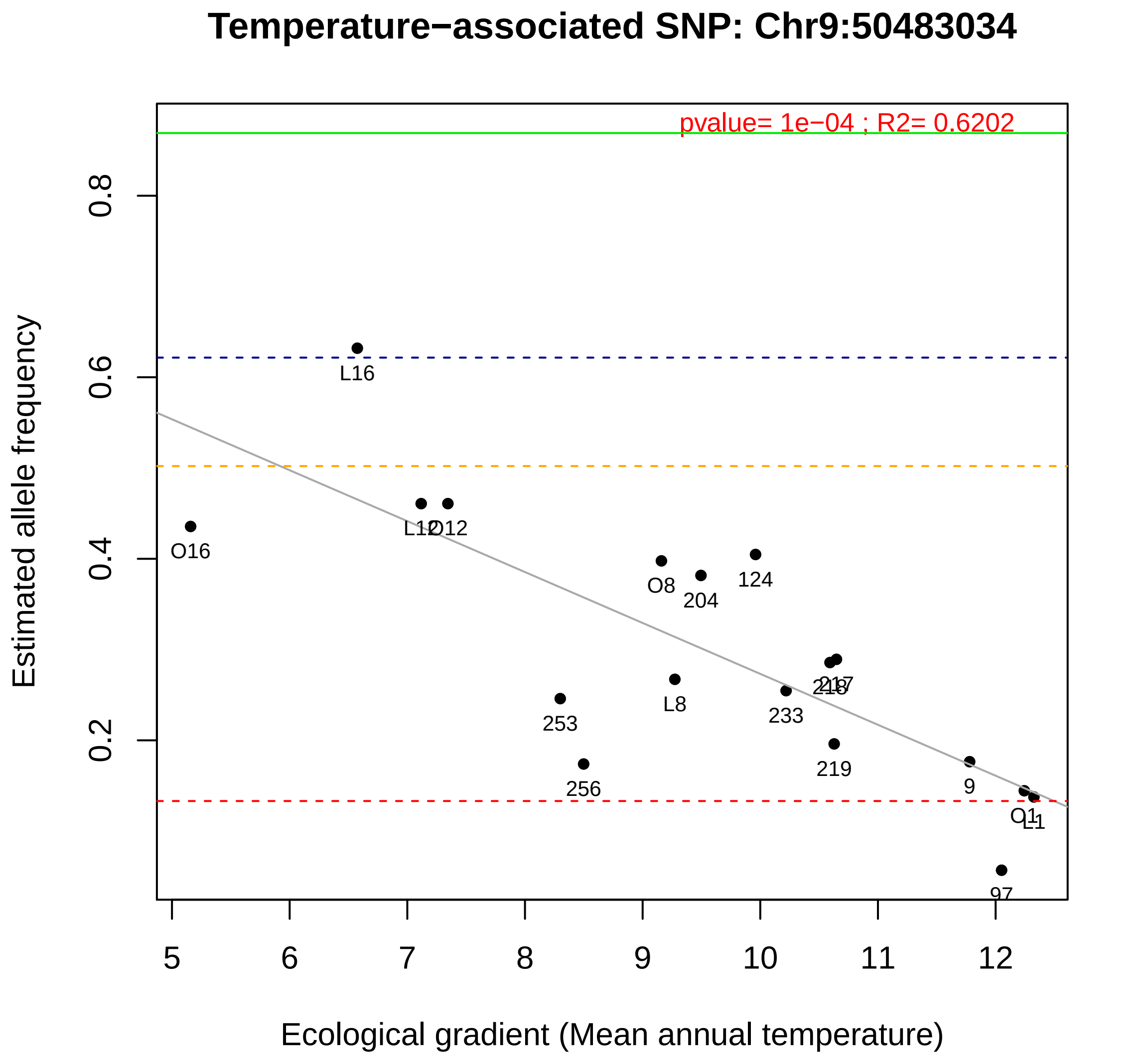

Temperature−associated SNP: Chr9:50483034
pvalue= 1e−04 ; R2= 0.6202
Estimated allele frequency
0.2
0.4
0.6
0.8
5
6
7
8
9
10
11
12
Ecological gradient (Mean annual temperature)
L16
O16
L12
O12
O8
204
124
L8
253
256
233
217
218
219
9
O1
L1
97

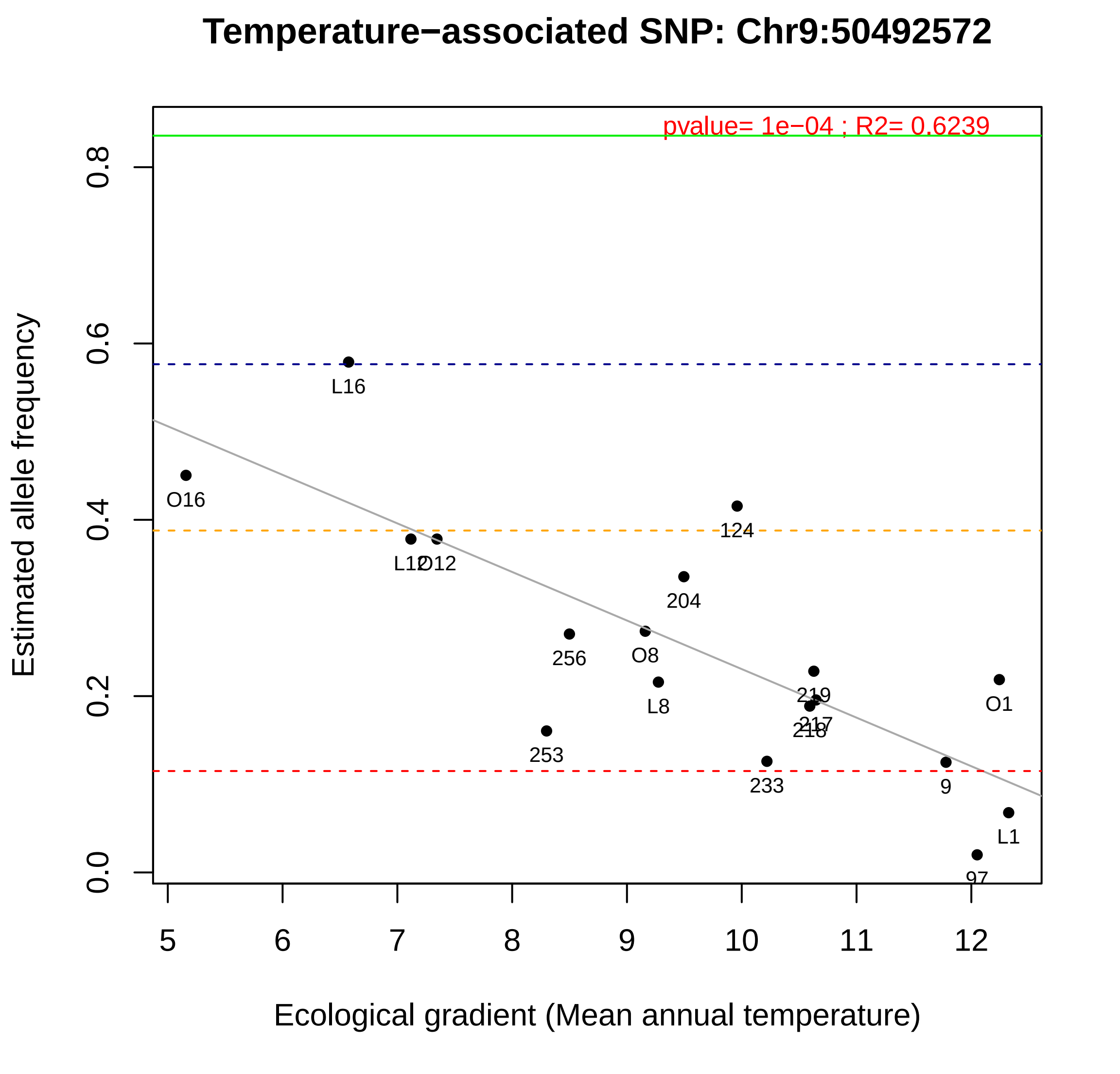

Temperature−associated SNP: Chr9:50492572
pvalue= 1e−04 ; R2= 0.6239
Estimated allele frequency
0.0
0.2
0.4
0.6
0.8
5
6
7
8
9
10
11
12
Ecological gradient (Mean annual temperature)
L16
O16
L12
O12
124
204
256
O8
L8
253
219
217
218
233
O1
9
L1
97

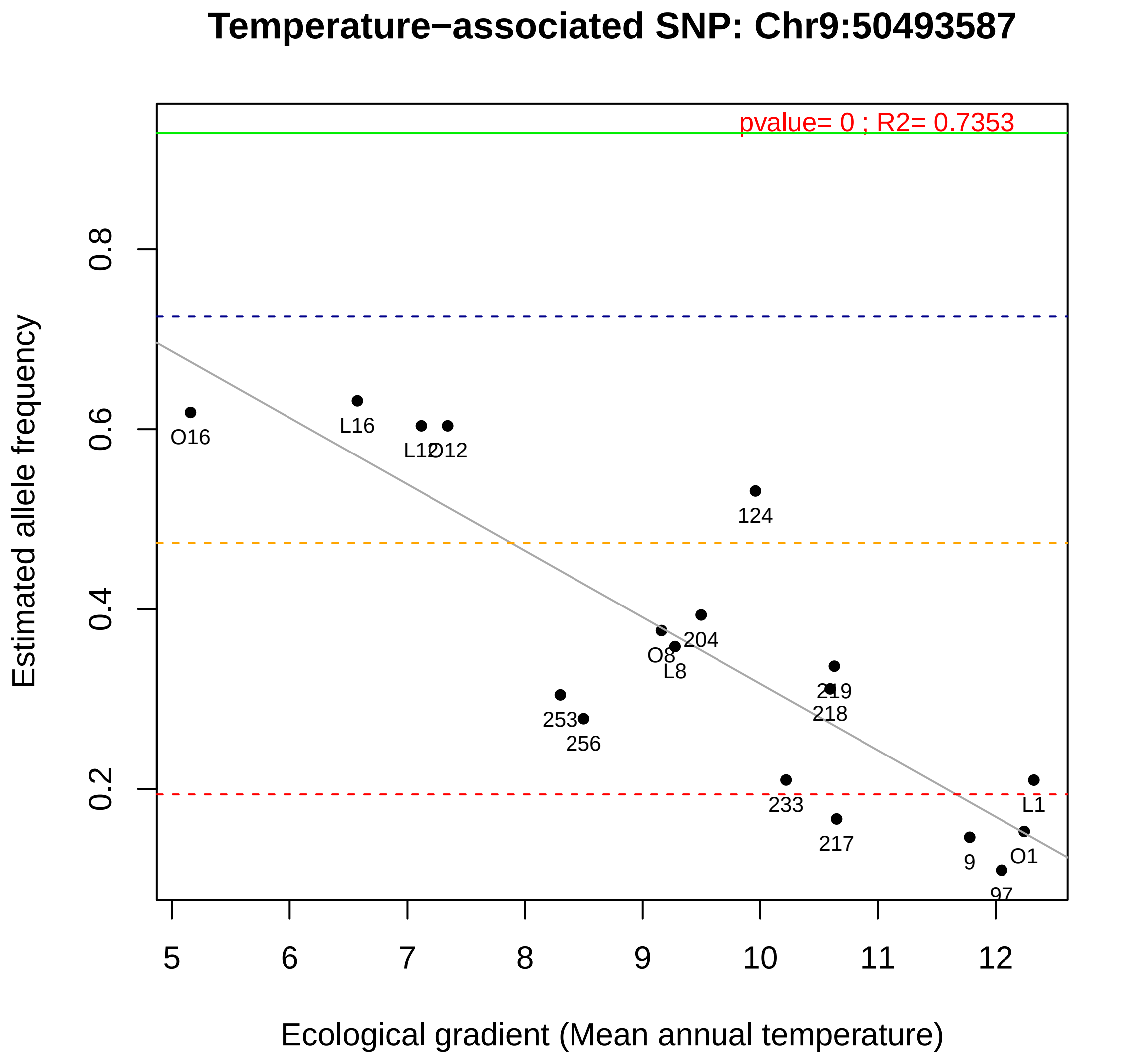

Temperature−associated SNP: Chr9:50493587
pvalue= 0 ; R2= 0.7353
Estimated allele frequency
0.2
0.4
0.6
0.8
O16
L16
L12
O12
124
204
O8
L8
219
218
253
256
233
217
L1
9
O1
97
5
6
7
8
9
10
11
12
Ecological gradient (Mean annual temperature)

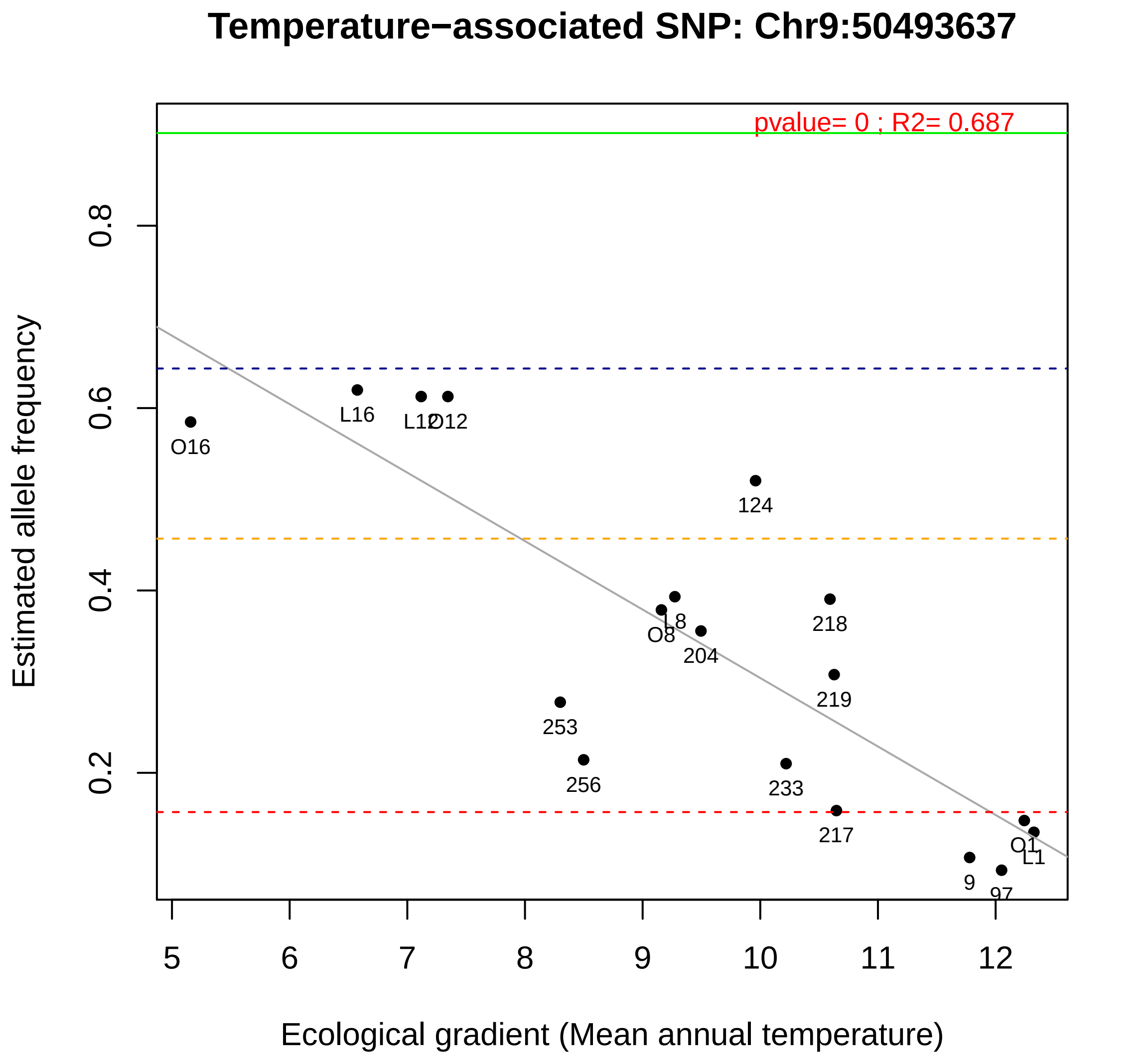

Temperature−associated SNP: Chr9:50493637
pvalue= 0 ; R2= 0.687
Estimated allele frequency
0.2
0.4
0.6
0.8
Ecological gradient (Mean annual temperature)
5
6
7
8
9
10
11
12
O16
L16
L12
O12
124
L8
O8
204
218
219
253
256
233
217
O1
L1
9
97

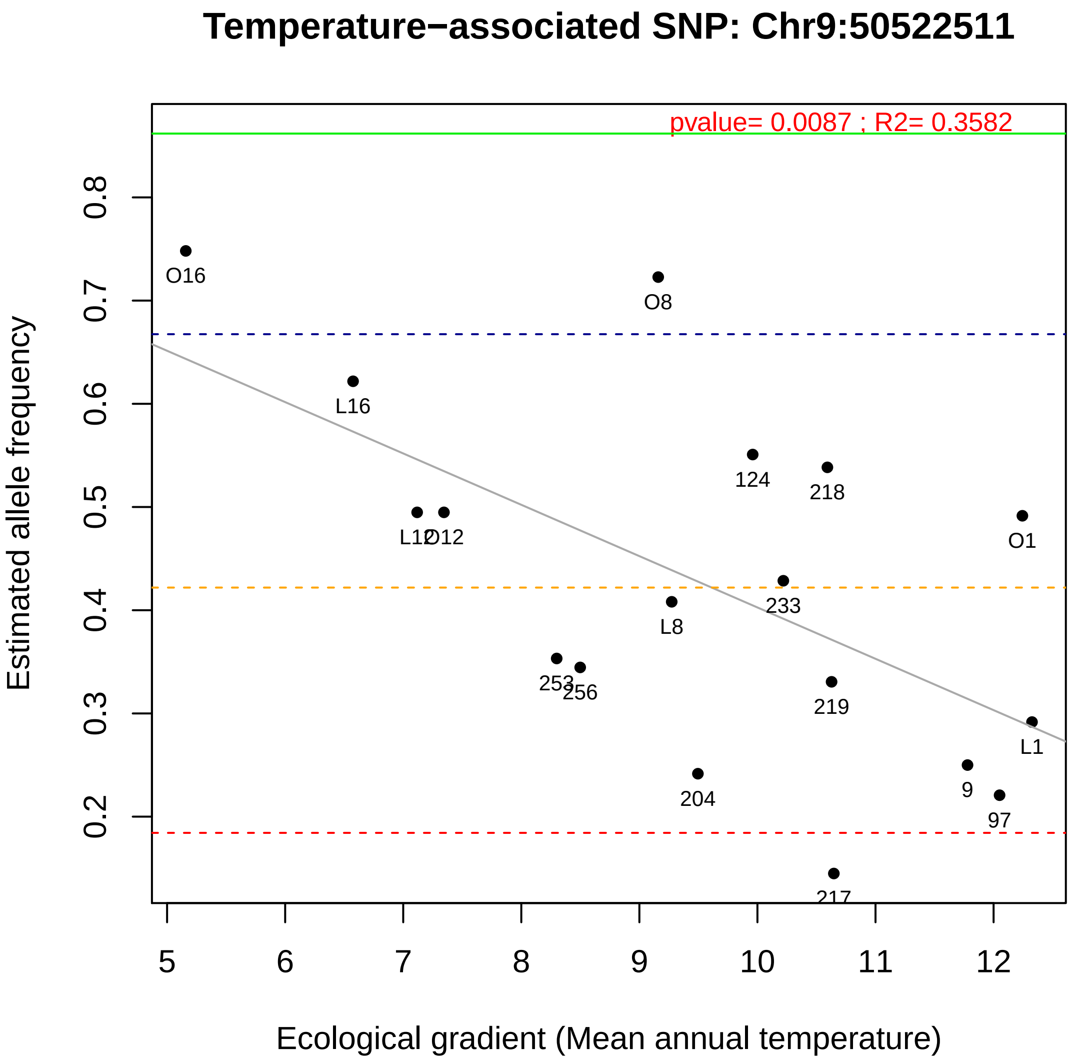

Temperature−associated SNP: Chr9:50522511
pvalue= 0.0087 ; R2= 0.3582
Estimated allele frequency
Ecological gradient (Mean annual temperature)
0.2
0.3
0.4
0.5
0.6
0.7
0.8
5
6
7
8
9
10
11
12
O16
O8
L16
124
218
L12
O12
O1
233
L8
253
256
219
L1
204
9
97
217

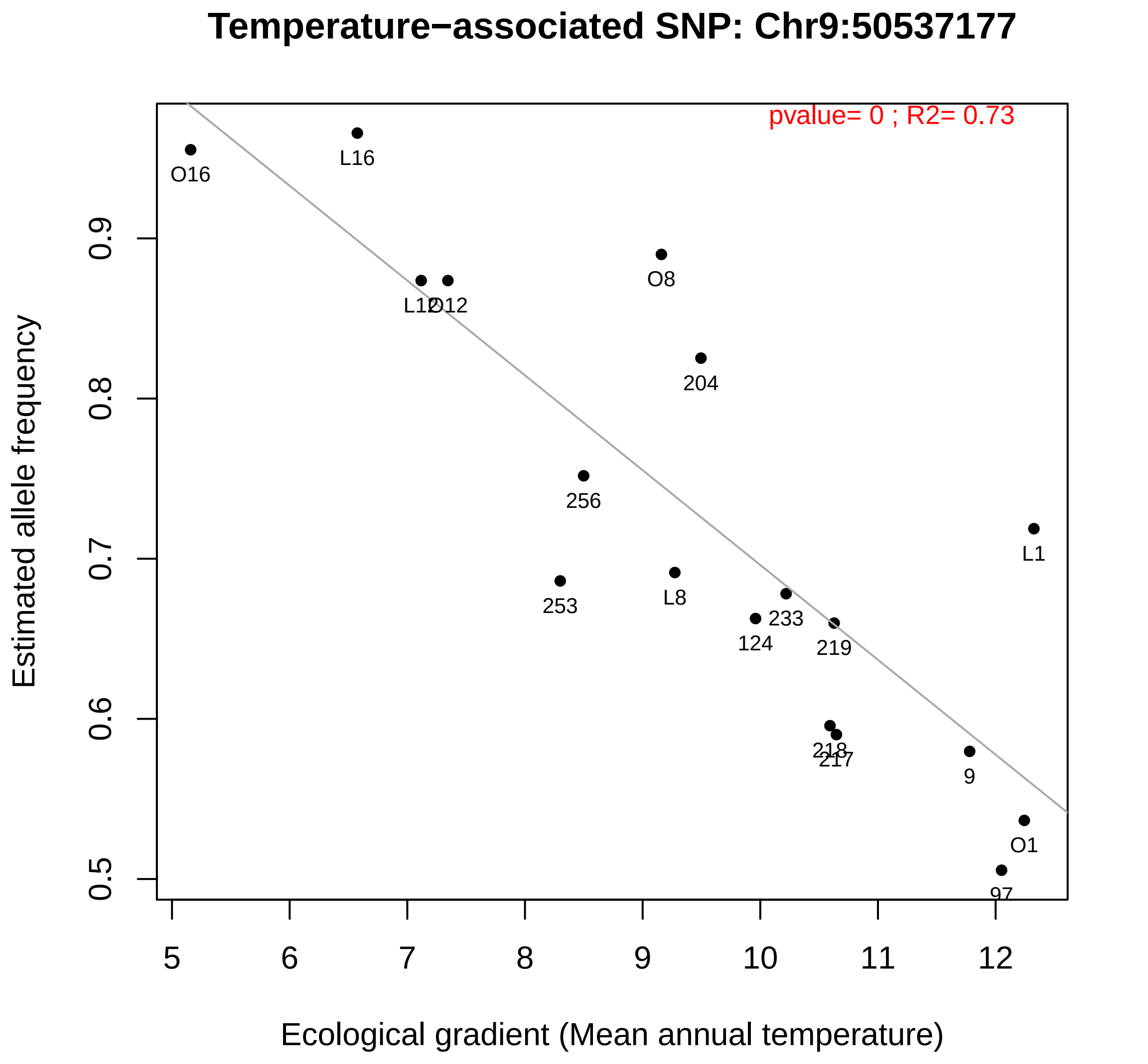
Temperature−associated SNP: Chr9:50537177
pvalue= 0 ; R2= 0.73
O16
L16
L12
O12
O8
204
256
L1
253
L8
233
124
219
218
217
9
O1
97
Estimated allele frequency
0.5
0.6
0.7
0.8
0.9
5
6
7
8
9
10
11
12
Ecological gradient (Mean annual temperature)

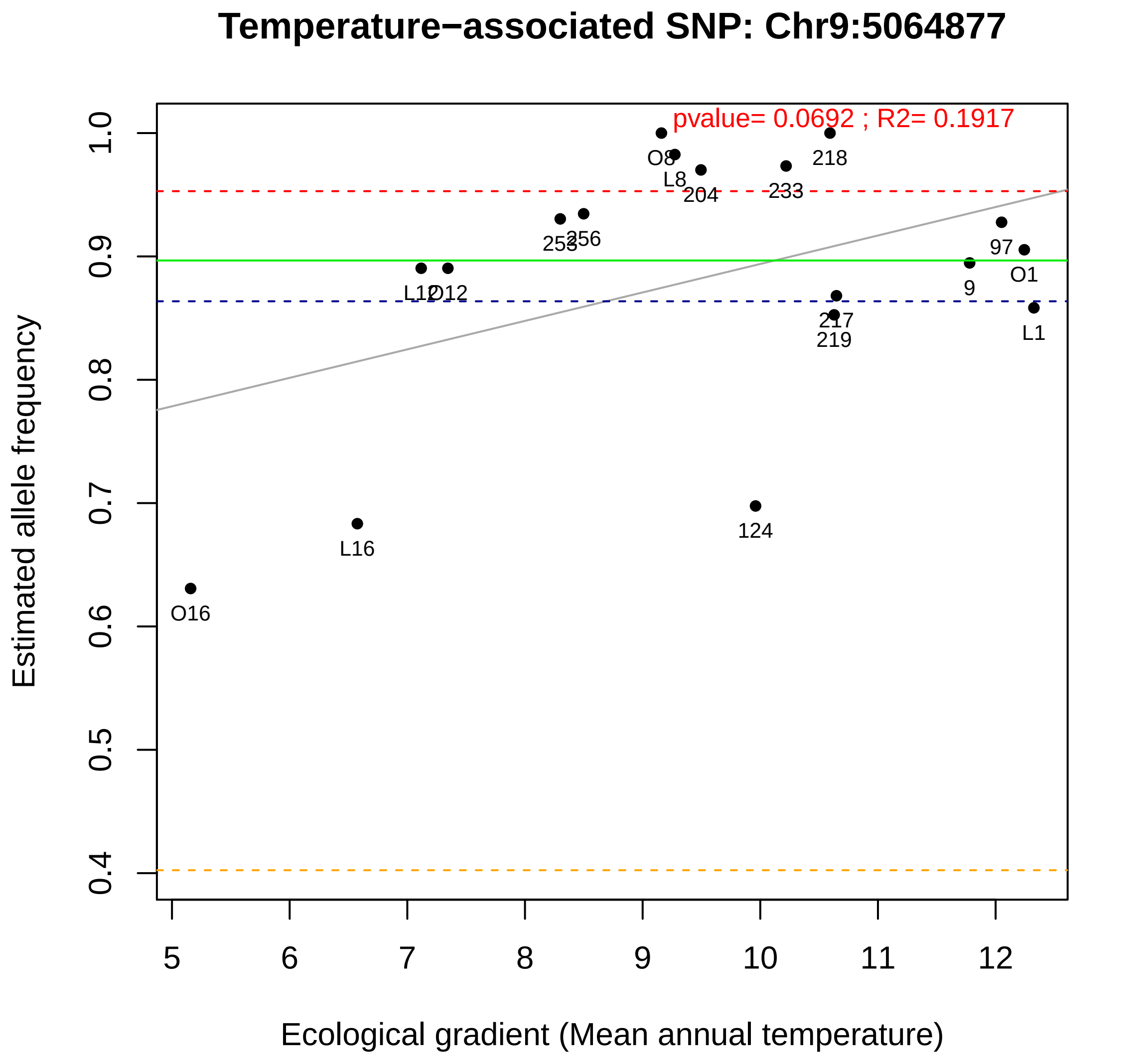
Temperature−associated SNP: Chr9:5064877
pvalue= 0.0692 ; R2= 0.1917
Estimated allele frequency
Ecological gradient (Mean annual temperature)
0.4
0.5
0.6
0.7
0.8
0.9
1.0
5
6
7
8
9
10
11
12
O8
L8
204
233
218
253
256
97
O1
9
L12
O12
217
219
L1
124
L16
O16

**Temperature−associated SNP: Chr9:5299934**

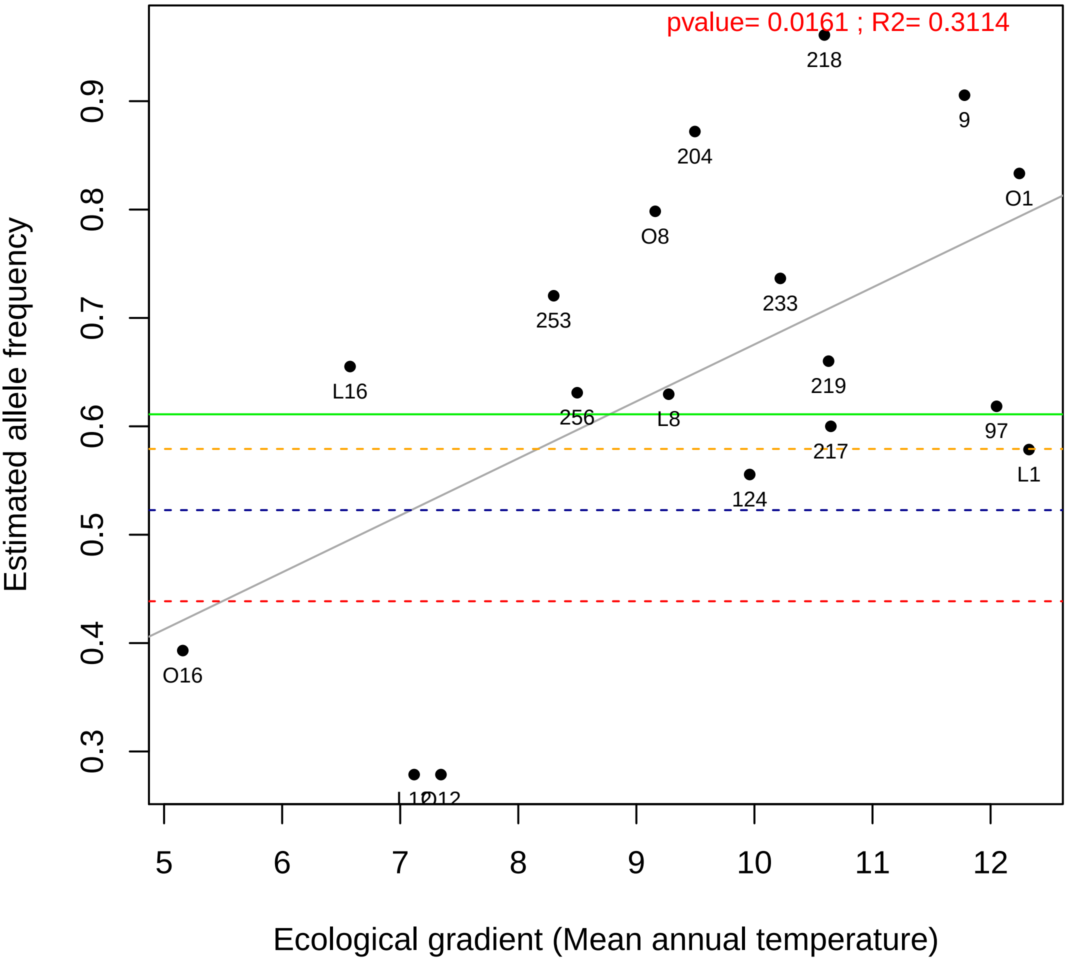

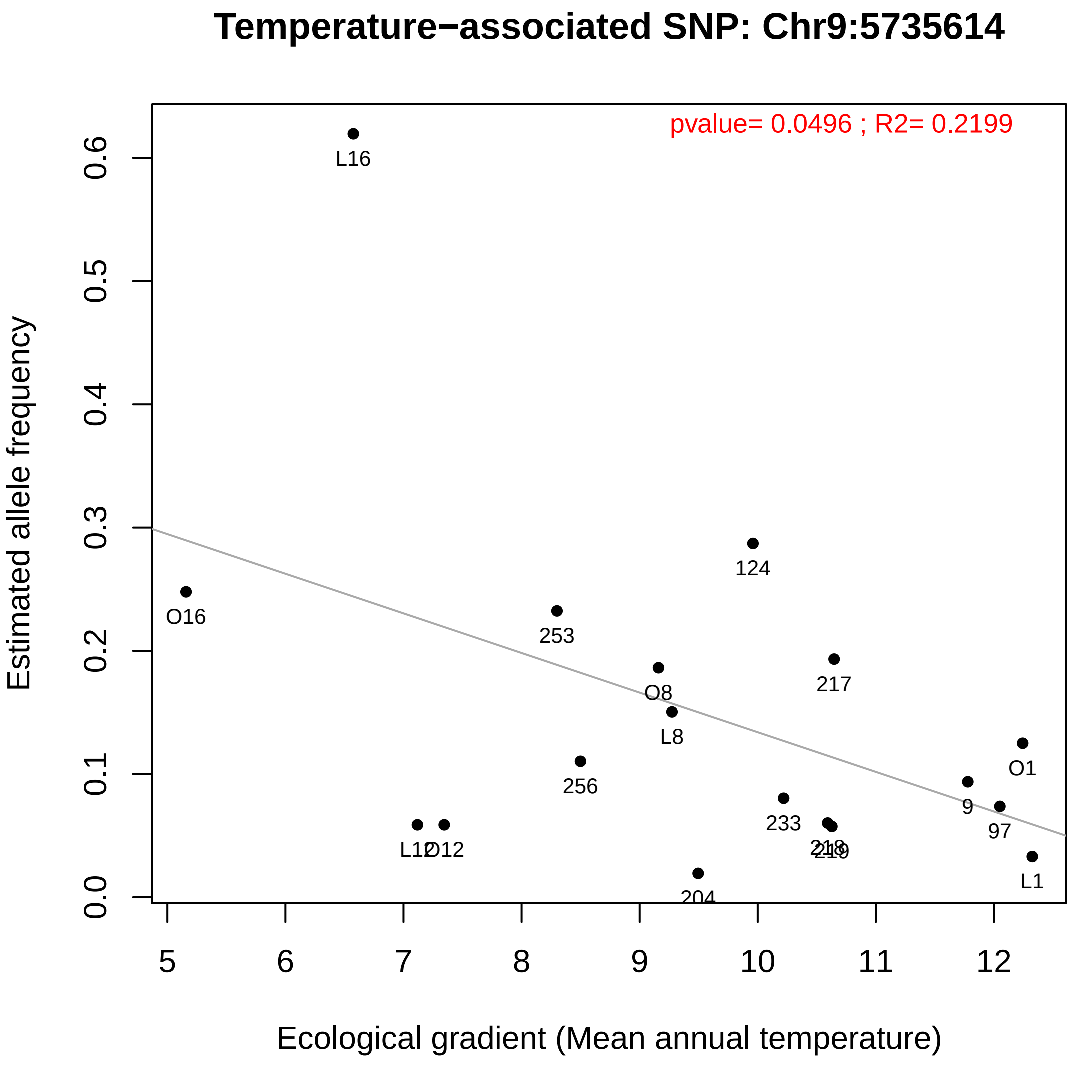

Temperature−associated SNP: Chr9:5735614
pvalue= 0.0496 ; R2= 0.2199
Estimated allele frequency
0.0
0.1
0.2
0.3
0.4
0.5
0.6
5
6
7
8
9
10
11
12
Ecological gradient (Mean annual temperature)
L16
O16
253
124
O8
217
L8
256
O1
9
233
97
L12
O12
204
L1

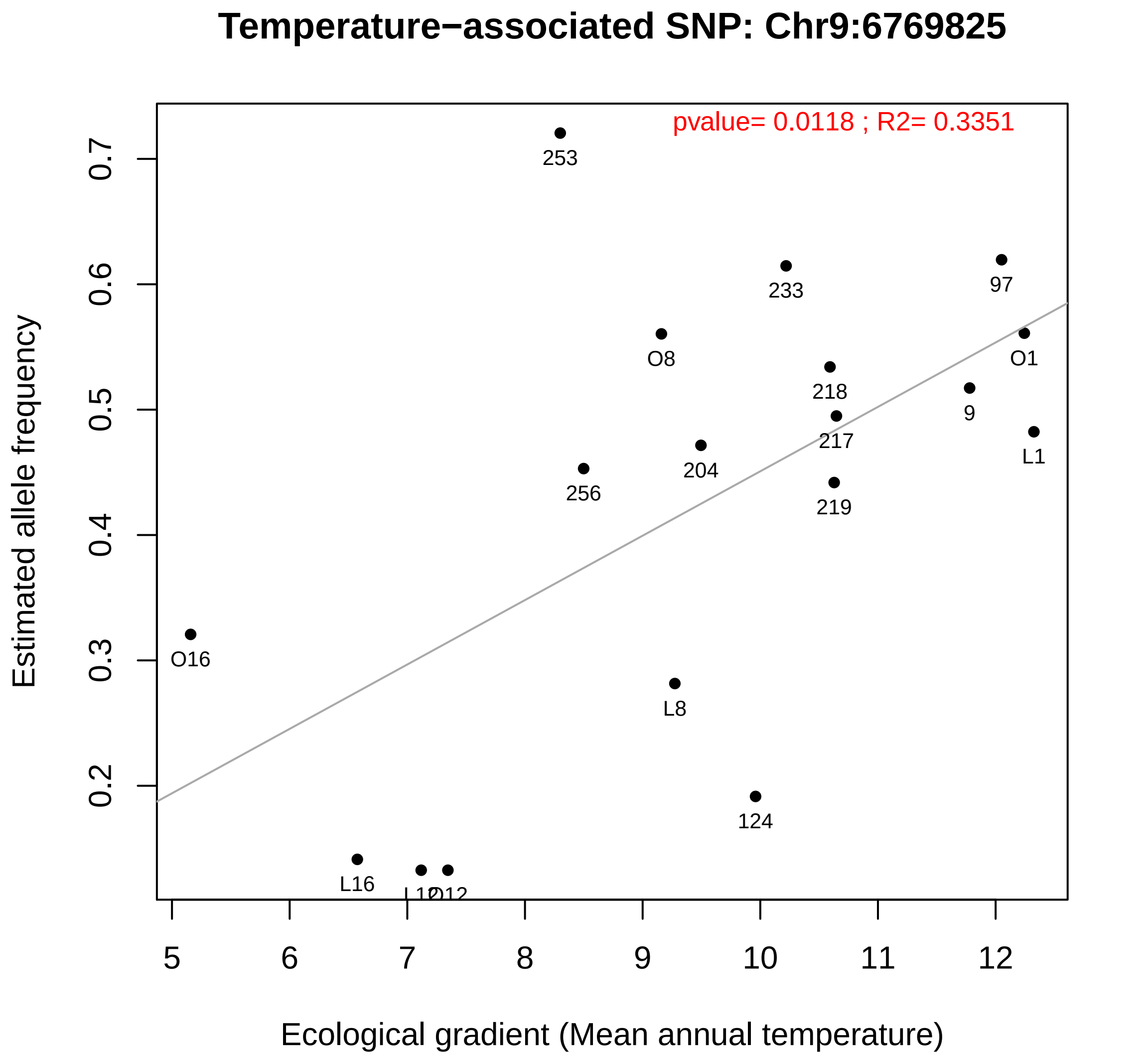
Temperature−associated SNP: Chr9:6769825
pvalue= 0.0118 ; R2= 0.3351
Estimated allele frequency
0.2
0.3
0.4
0.5
0.6
0.7
Ecological gradient (Mean annual temperature)
5
6
7
8
9
10
11
12
253
233
97
O8
O1
218
9
217
L1
204
256
219
O16
L8
124
L16
O12

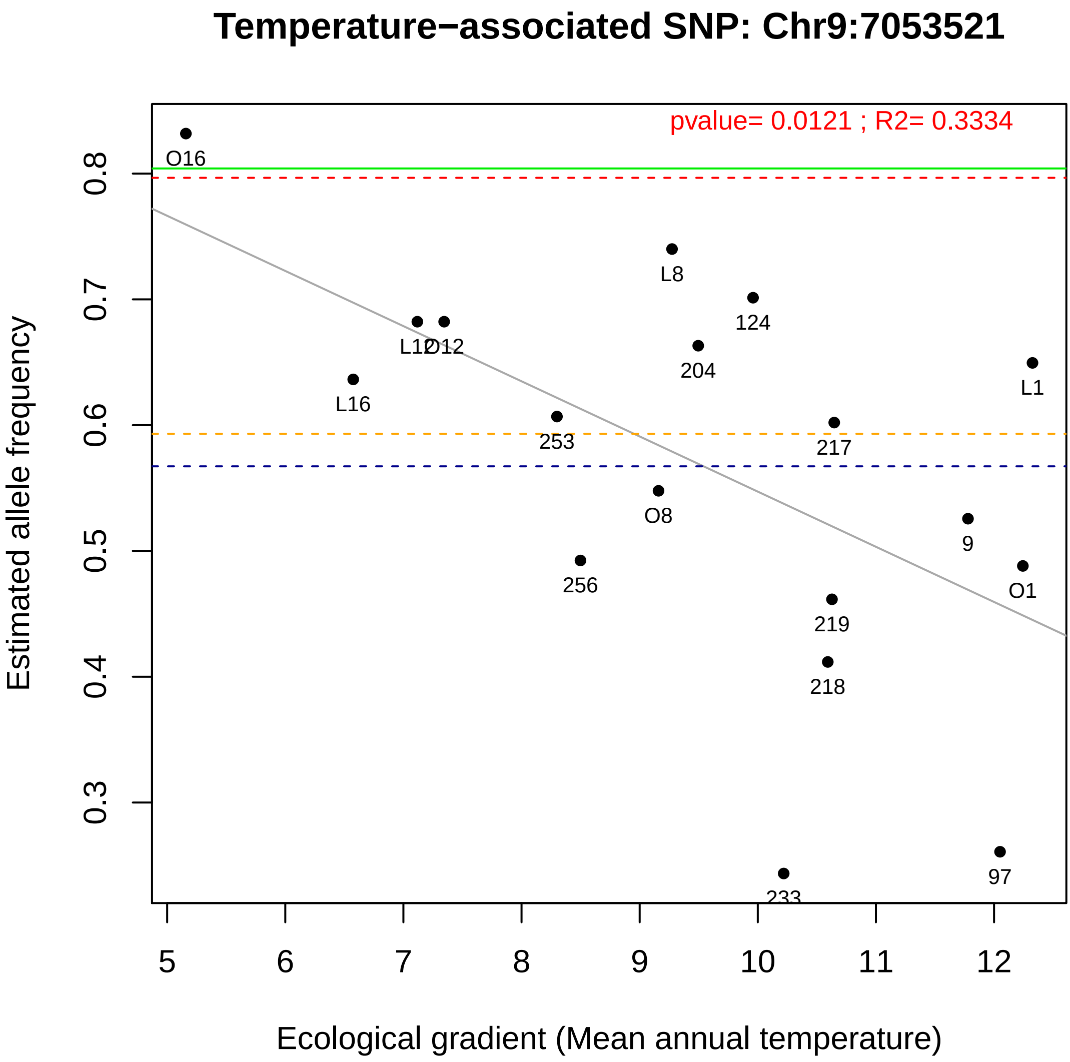

Temperature−associated SNP: Chr9:7053521
pvalue= 0.0121 ; R2= 0.3334
Estimated allele frequency
Ecological gradient (Mean annual temperature)
0.3
0.4
0.5
0.6
0.7
0.8
5
6
7
8
9
10
11
12
O16
L8
124
L12
O12
204
L1
L16
253
217
O8
9
256
O1
219
218
97
233

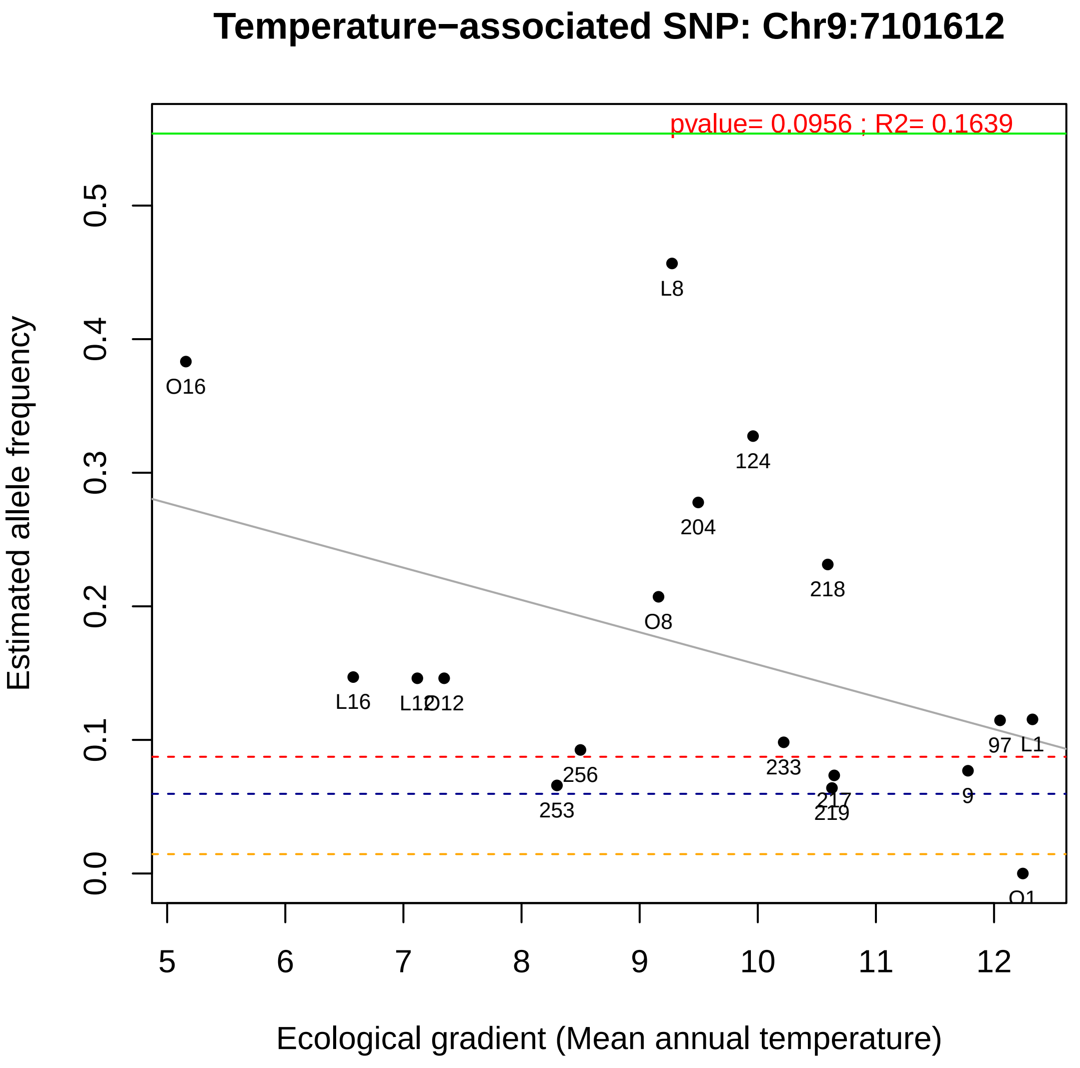

Temperature−associated SNP: Chr9:7101612
pvalue= 0.0956 ; R2= 0.1639
Estimated allele frequency
0.0
0.1
0.2
0.3
0.4
0.5
L8
O16
124
204
218
O8
L16
L12
O12
97
L1
233
256
253
217
219
9
O1
5
6
7
8
9
10
11
12
Ecological gradient (Mean annual temperature)

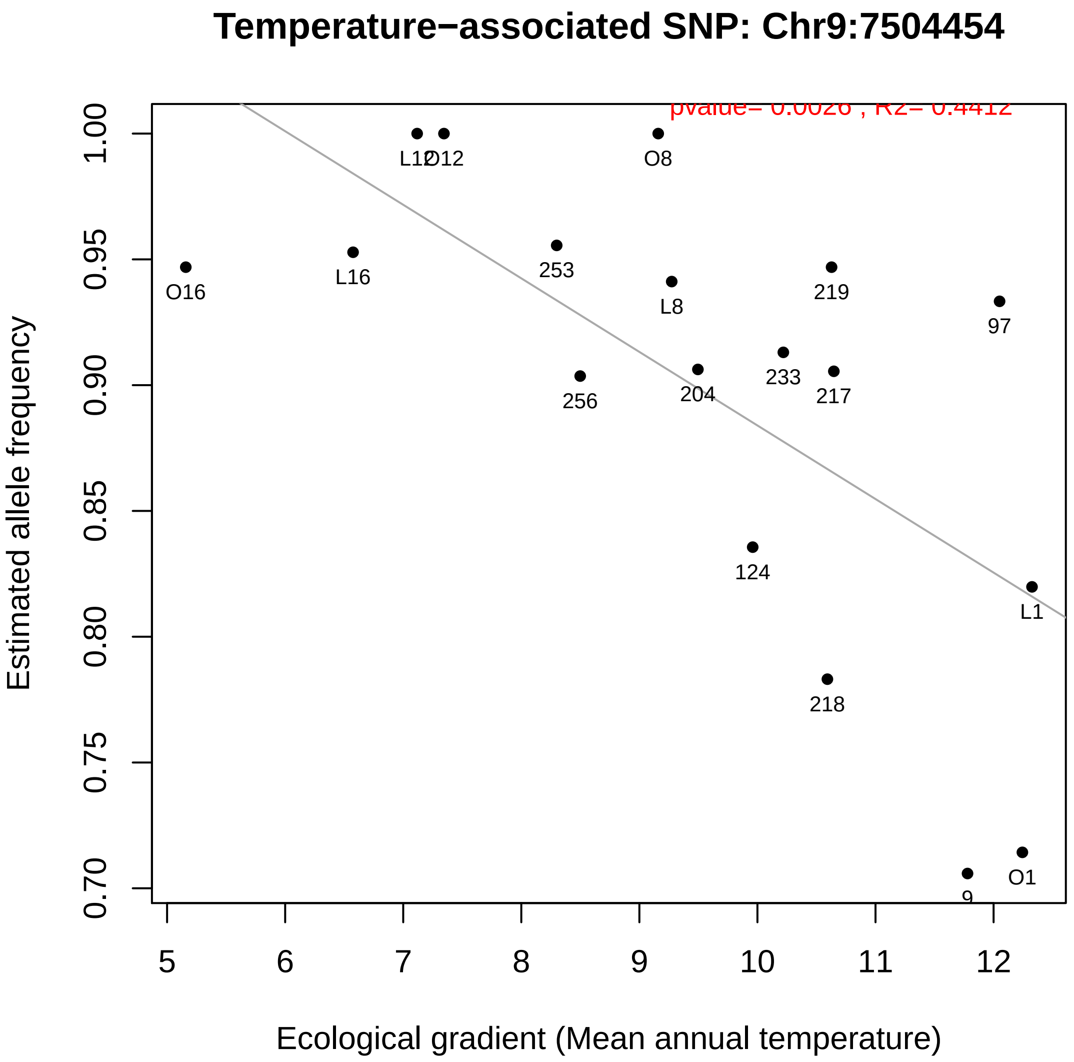

Temperature−associated SNP: Chr9:7504454
pvalue= 0.0026 , R2= 0.4412
Estimated allele frequency
1.00
0.95
0.90
0.85
0.80
0.75
0.70
5
6
7
8
9
10
11
12
Ecological gradient (Mean annual temperature)
L12
O12
O8
O16
L16
253
L8
219
97
256
204
233
217
124
L1
218
9
O1

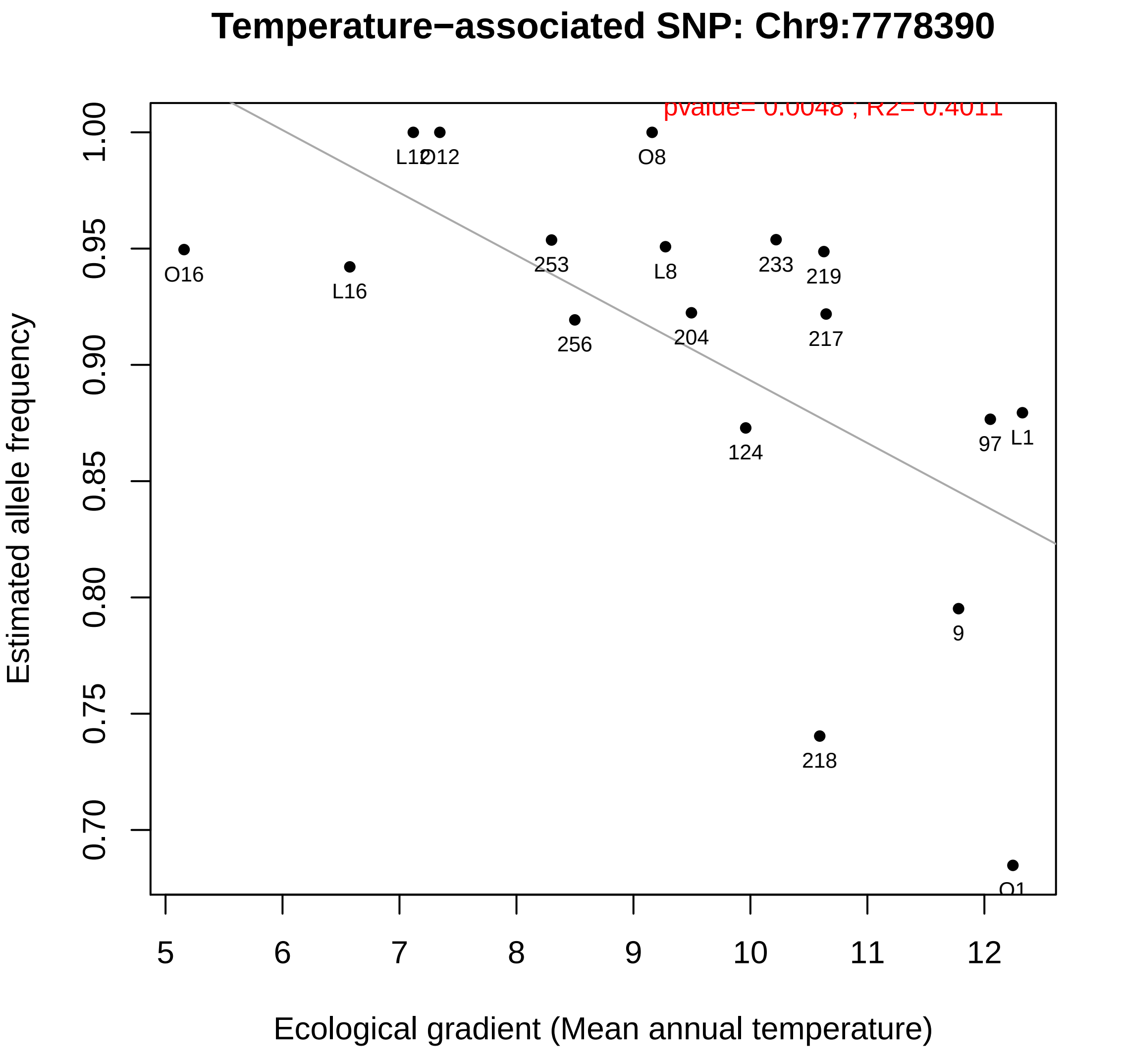
Temperature−associated SNP: Chr9:7778390
pvalue= 0.0048 ; R2= 0.4011
Estimated allele frequency
Ecological gradient (Mean annual temperature)
1.00
0.95
0.90
0.85
0.80
0.75
0.70
5
6
7
8
9
10
11
12
L12
O12
O8
O16
L16
253
256
L8
204
233
219
217
124
97
L1
9
218
O1

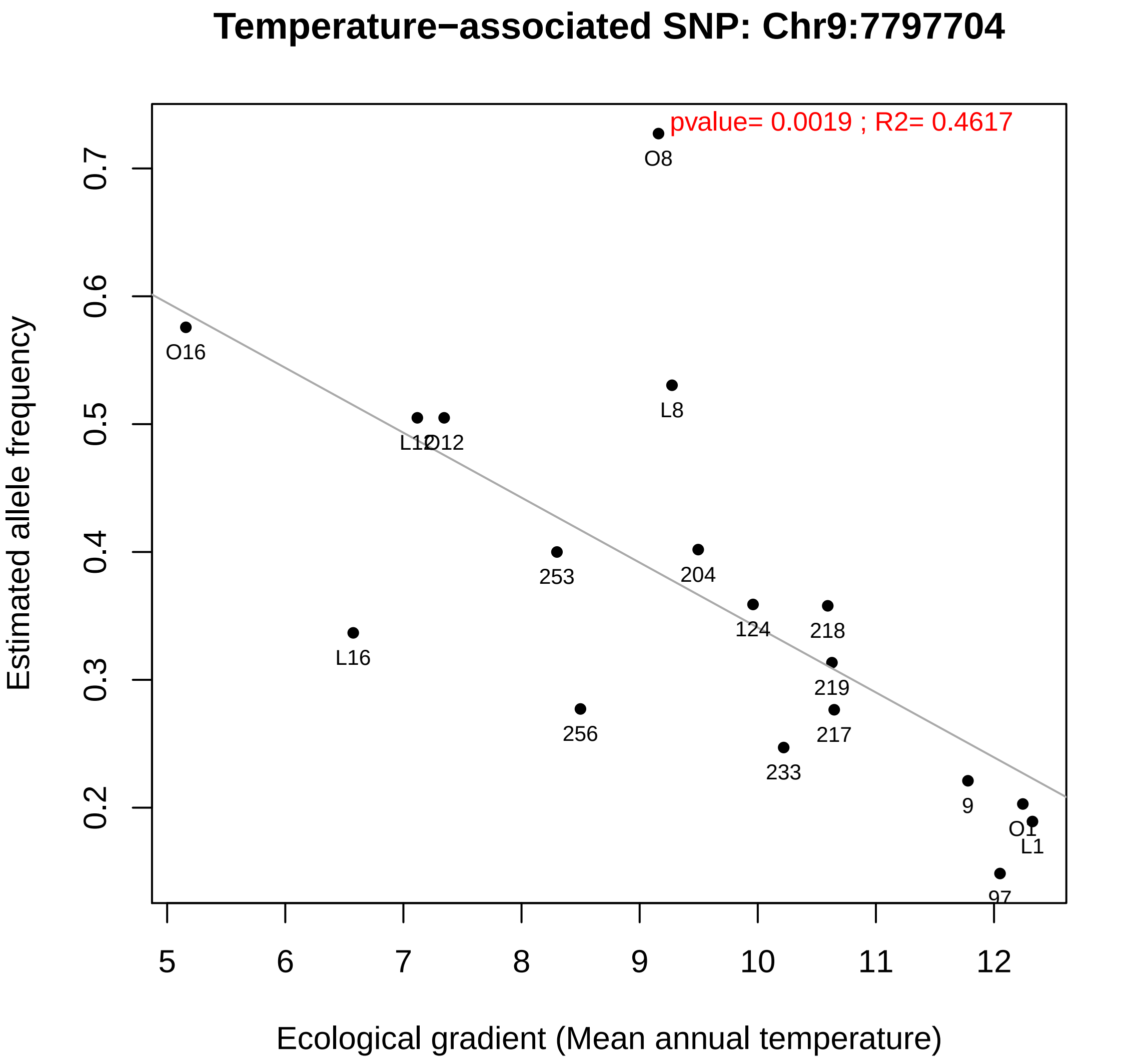
Temperature−associated SNP: Chr9:7797704
pvalue= 0.0019 ; R2= 0.4617
O8
O16
L8
L12
O12
253
204
124
218
L16
219
256
217
233
9
O1
L1
97
0.2
0.3
0.4
0.5
0.6
0.7
Estimated allele frequency
5
6
7
8
9
10
11
12
Ecological gradient (Mean annual temperature)

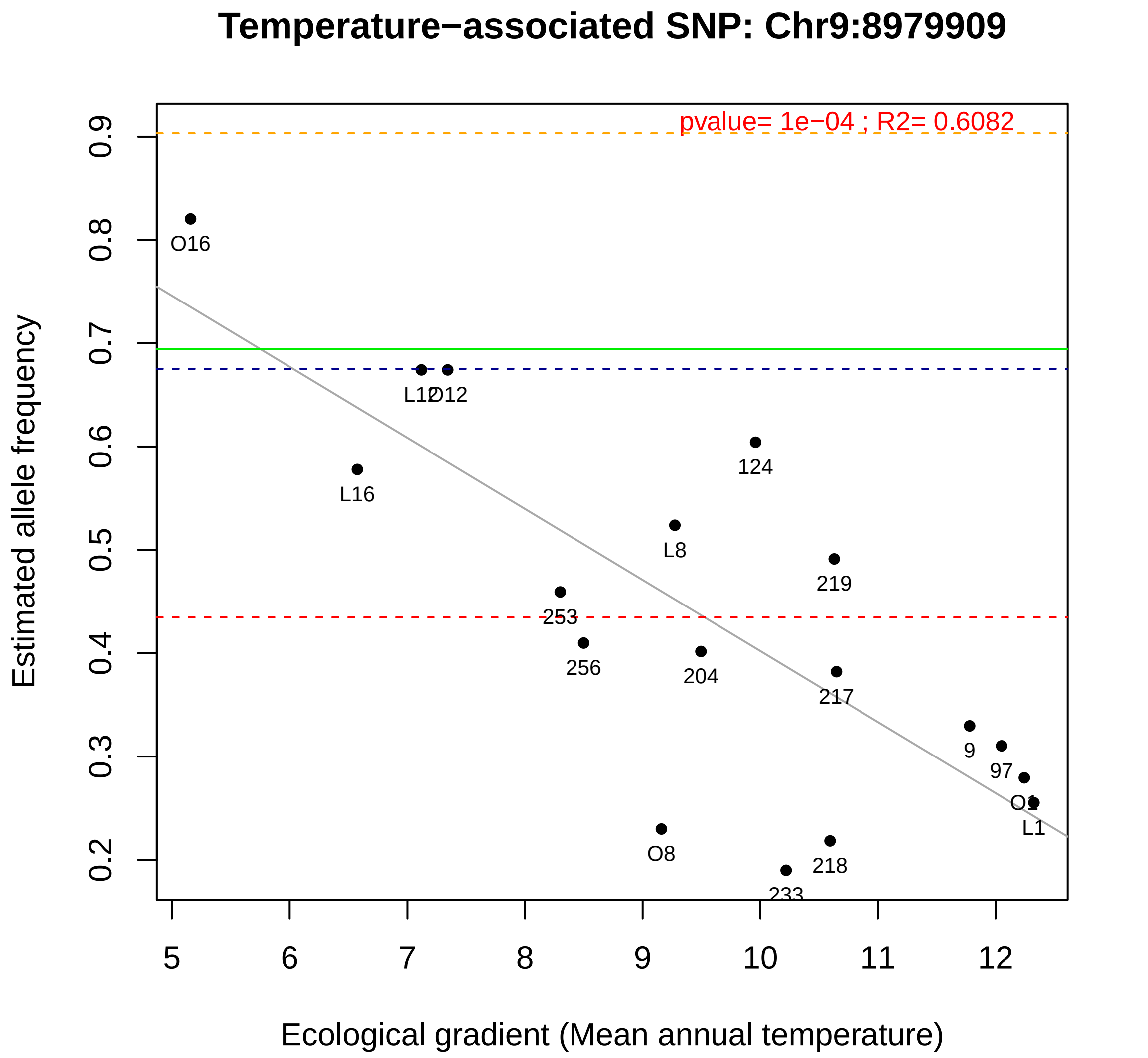
Temperature−associated SNP: Chr9:8979909
pvalue= 1e−04 ; R2= 0.6082
Estimated allele frequency
0.2
0.3
0.4
0.5
0.6
0.7
0.8
0.9
5
6
7
8
9
10
11
12
Ecological gradient (Mean annual temperature)
O16
L12
O12
L16
124
L8
219
253
256
204
217
9
97
O1
L1
O8
218
233

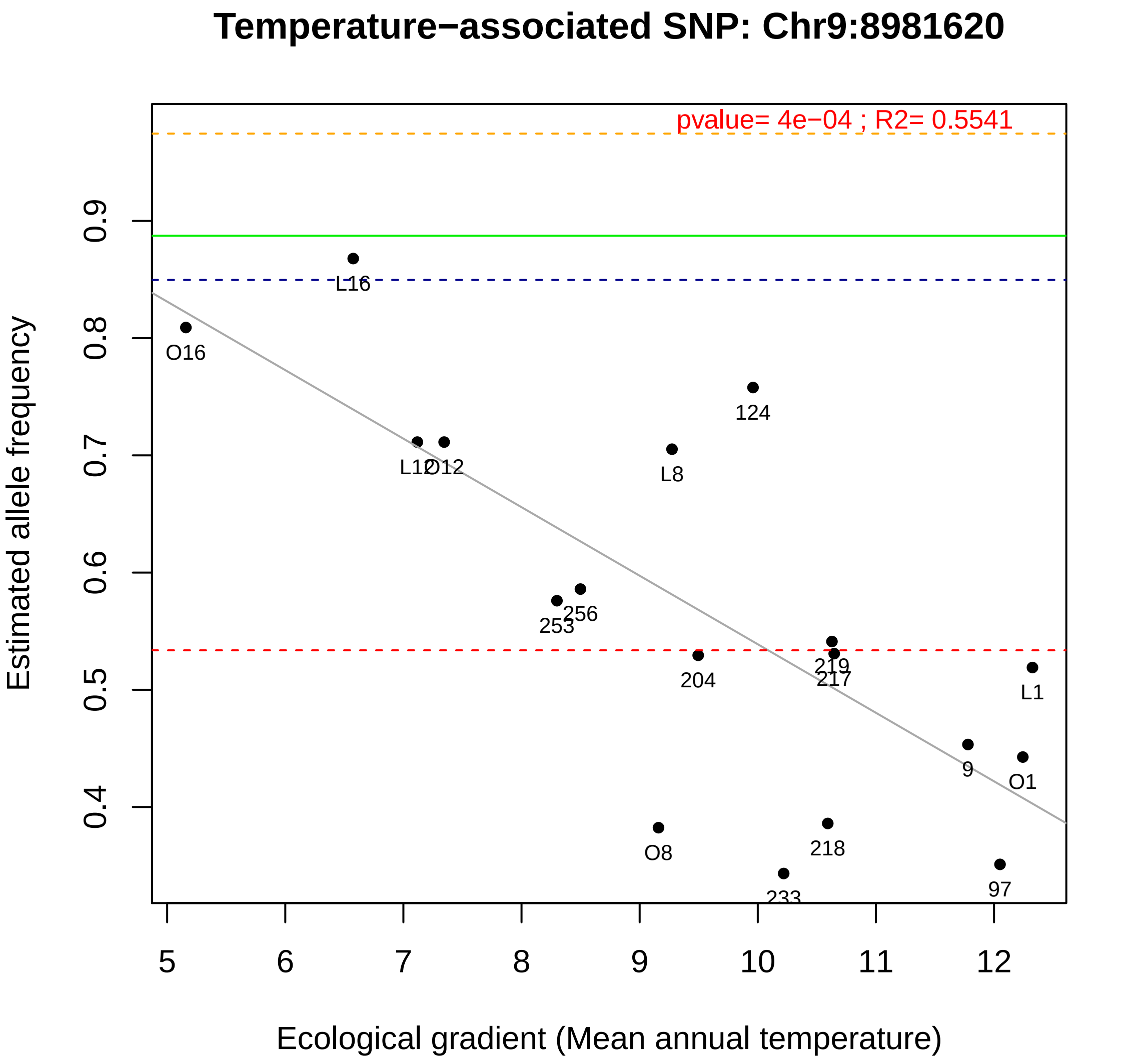
Temperature−associated SNP: Chr9:8981620
pvalue= 4e−04 ; R2= 0.5541
Estimated allele frequency
0.4
0.5
0.6
0.7
0.8
0.9
5
6
7
8
9
10
11
12
Ecological gradient (Mean annual temperature)
L16
O16
124
L12
O12
L8
253
256
204
219
217
L1
9
O1
O8
218
233
97

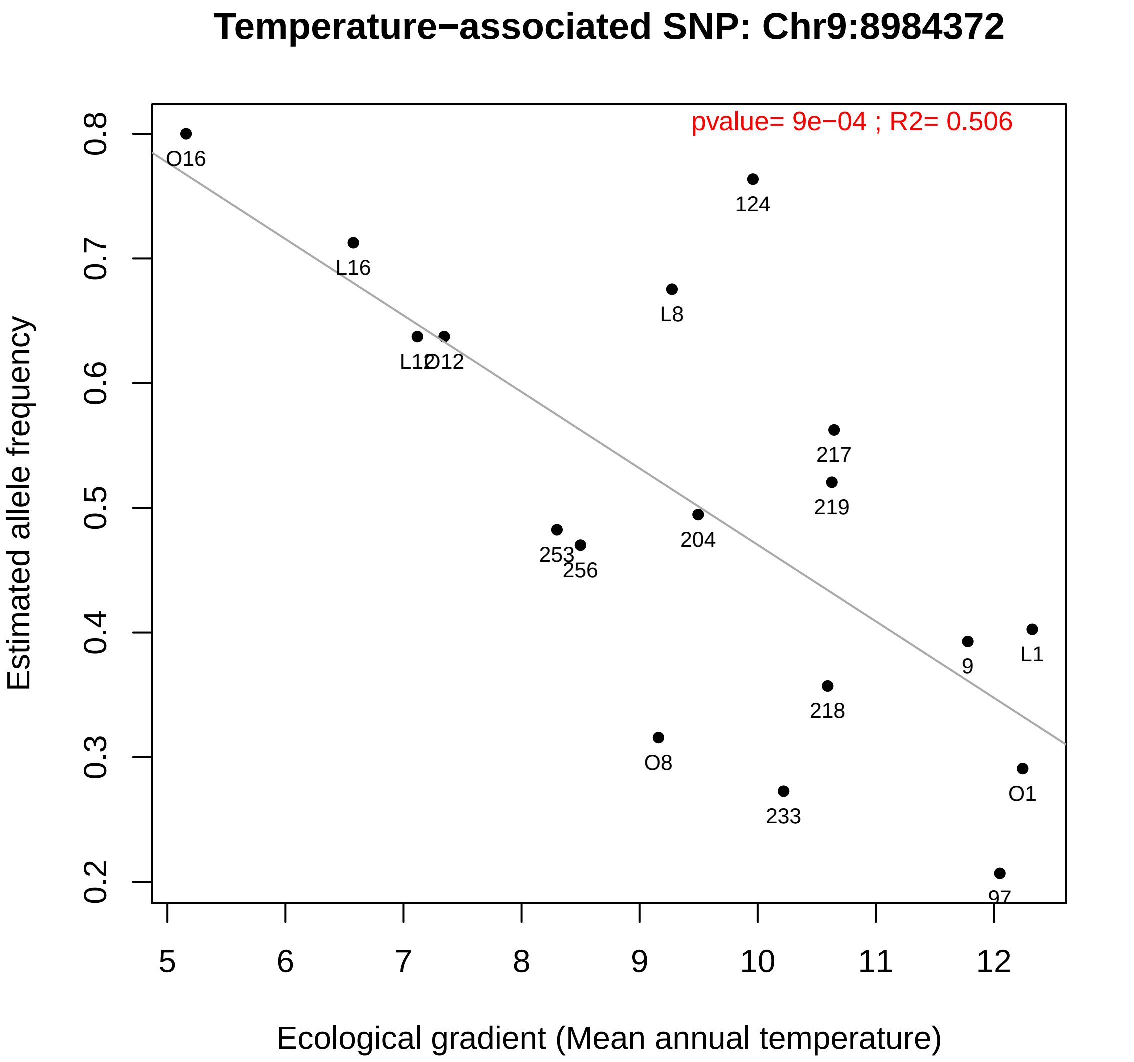

Temperature−associated SNP: Chr9:8984372
pvalue= 9e−04 ; R2= 0.506
Estimated allele frequency
Ecological gradient (Mean annual temperature)
0.2
0.3
0.4
0.5
0.6
0.7
0.8
5
6
7
8
9
10
11
12
O16
L16
L12
O12
124
L8
217
219
204
253
256
9
L1
218
O8
233
O1
97

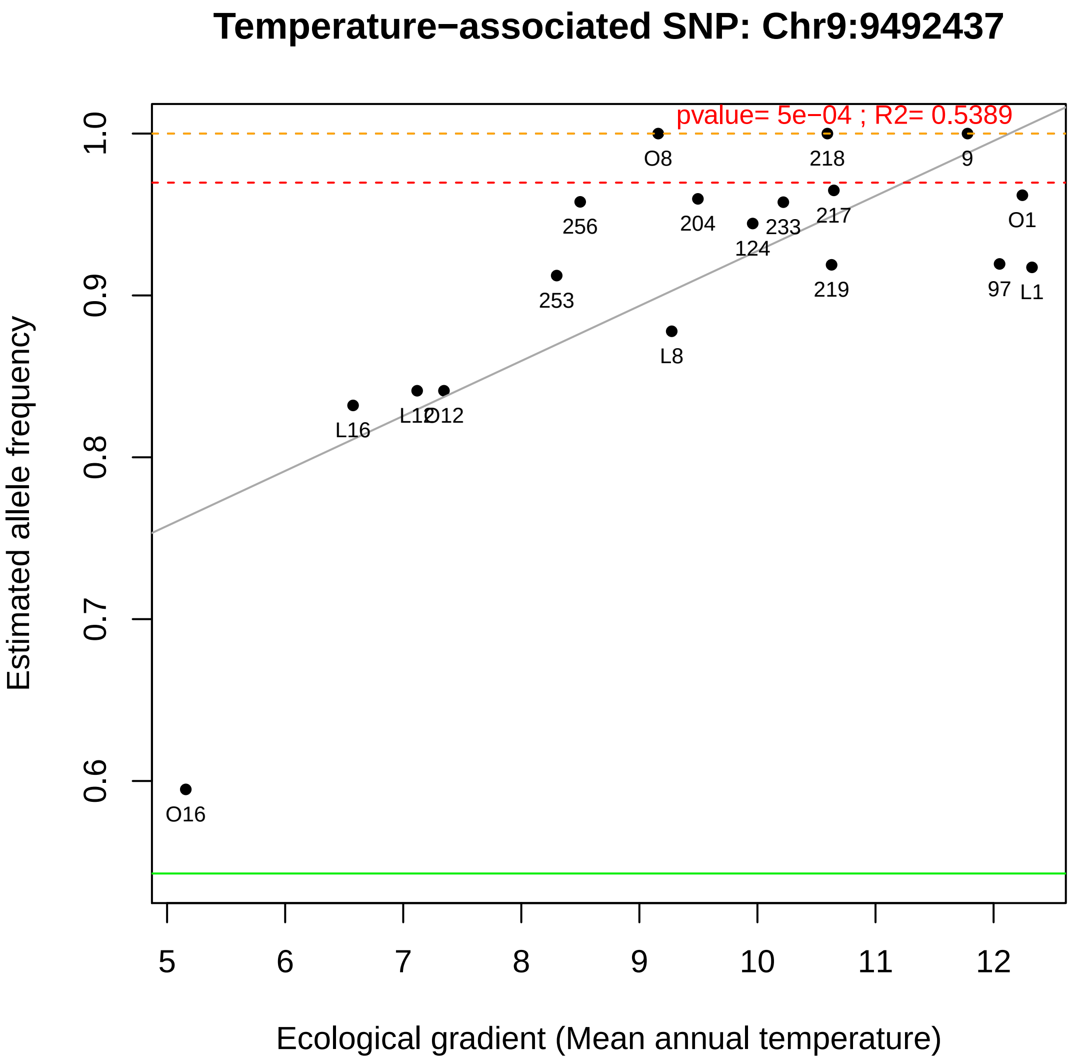
Temperature−associated SNP: Chr9:9492437
pvalue= 5e−04 ; R2= 0.5389
Estimated allele frequency
Ecological gradient (Mean annual temperature)
0.6
0.7
0.8
0.9
1.0
5
6
7
8
9
10
11
12
O16
L16
L12
O12
253
256
L8
O8
204
124
233
218
217
219
9
O1
97
L1

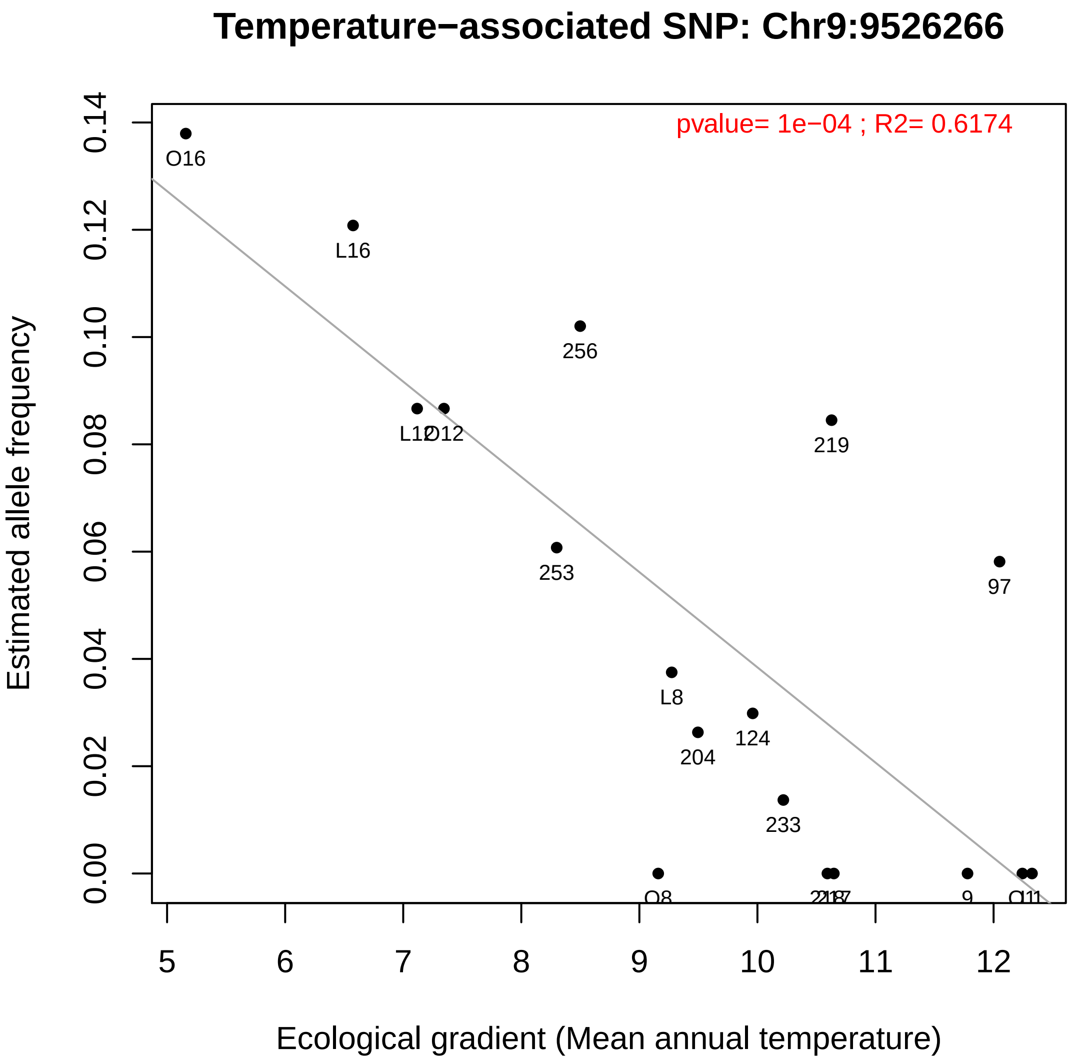
Temperature−associated SNP: Chr9:9526266
pvalue= 1e−04 ; R2= 0.6174
Estimated allele frequency
0.00
0.02
0.04
0.06
0.08
0.10
0.12
0.14
Ecological gradient (Mean annual temperature)
5
6
7
8
9
10
11
12
O16
L16
256
L12
O12
219
253
97
L8
124
204
233
O8
9

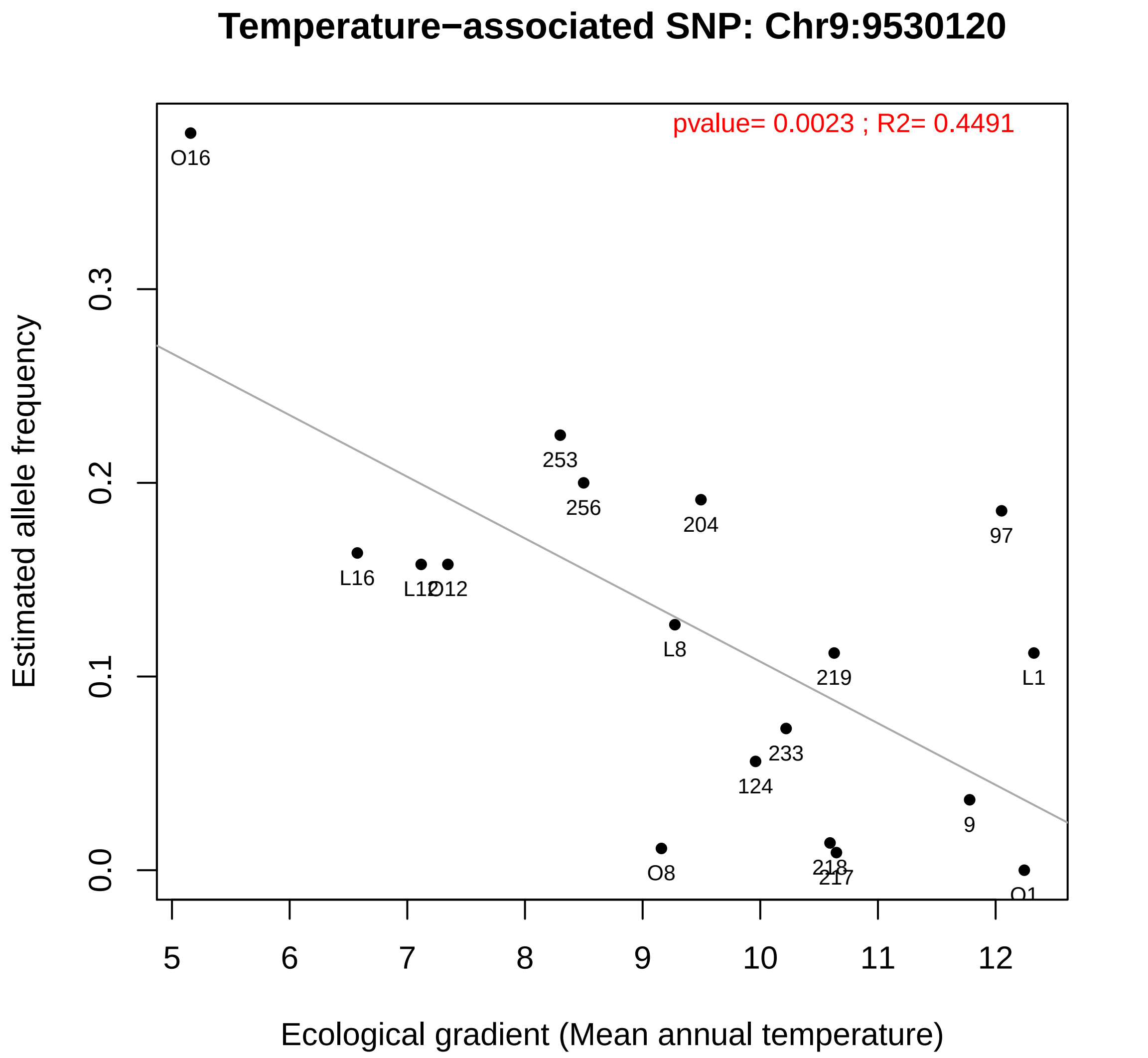

Temperature−associated SNP: Chr9:9530120
pvalue= 0.0023 ; R2= 0.4491
Estimated allele frequency
0.0
0.1
0.2
0.3
5
6
7
8
9
10
11
12
Ecological gradient (Mean annual temperature)
O16
253
256
204
97
L16
L12
O12
L8
219
L1
233
124
9
O8
218
217
O1

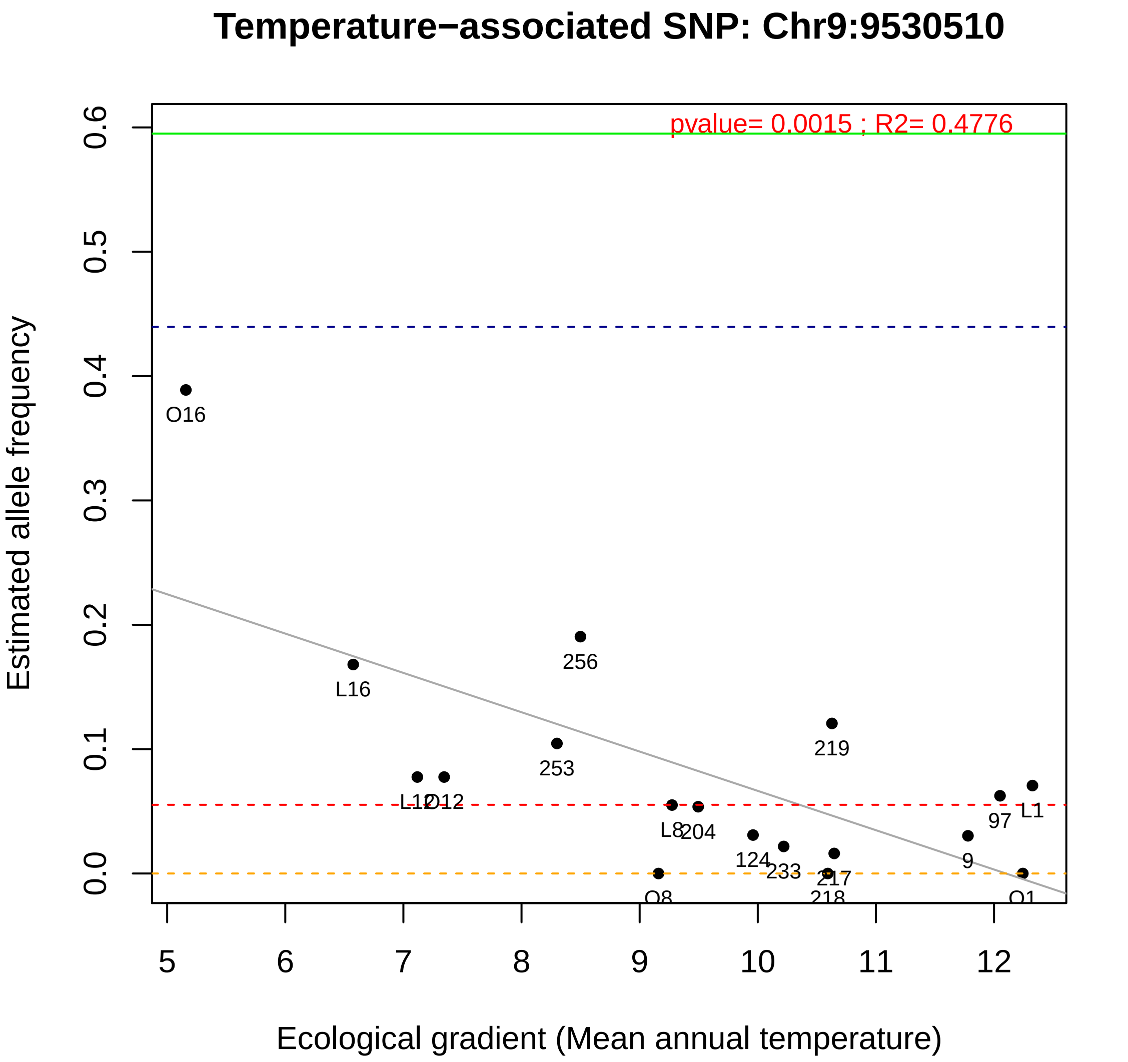
Temperature−associated SNP: Chr9:9530510
pvalue= 0.0015 ; R2= 0.4776
Estimated allele frequency
0.0
0.1
0.2
0.3
0.4
0.5
0.6
Ecological gradient (Mean annual temperature)
5
6
7
8
9
10
11
12
O16
L16
256
253
L12
O12
219
L8
204
124
233
217
218
O8
9
97
L1
O1

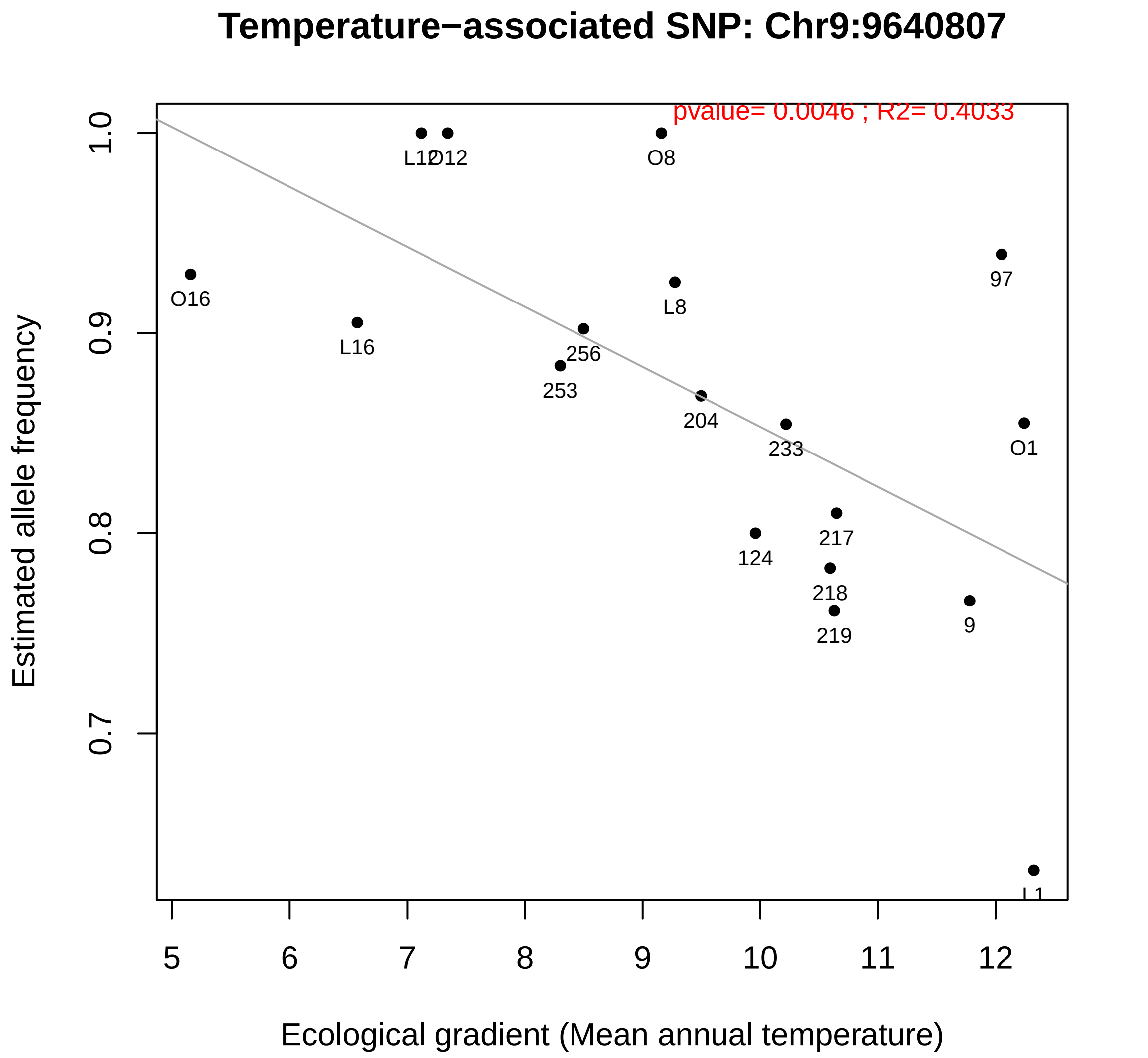

Temperature−associated SNP: Chr9:9640807
pvalue= 0.0046 ; R2= 0.4033
Estimated allele frequency
Ecological gradient (Mean annual temperature)
1.0
0.9
0.8
0.7
5
6
7
8
9
10
11
12
L12
O12
O8
O16
L16
97
L8
256
253
204
233
O1
217
124
218
219
9

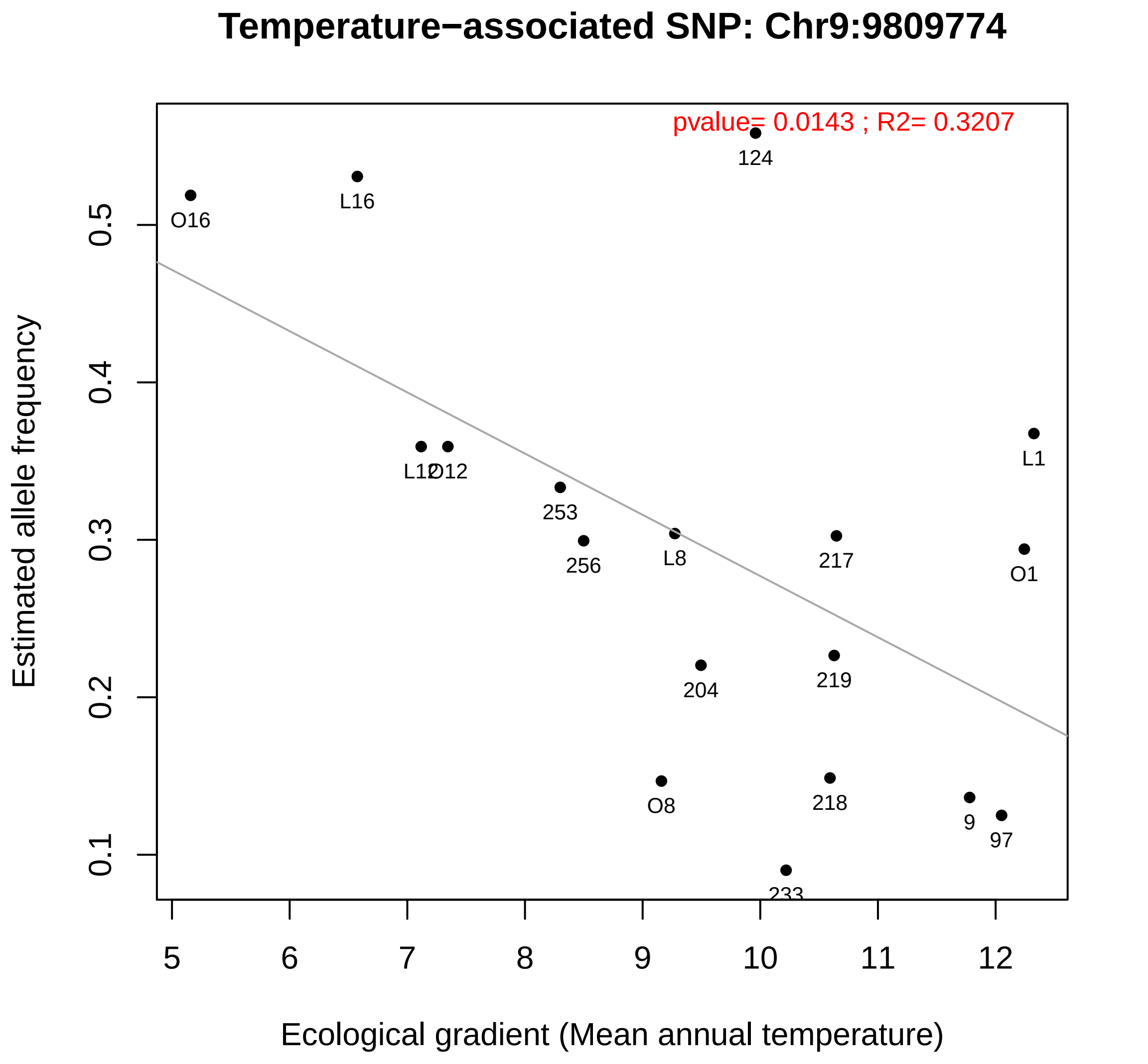
Temperature−associated SNP: Chr9:9809774
pvalue= 0.0143 ; R2= 0.3207
Estimated allele frequency
0.1
0.2
0.3
0.4
0.5
Ecological gradient (Mean annual temperature)
5
6
7
8
9
10
11
12
O16
L16
124
L12
O12
253
256
L8
217
L1
O1
204
219
O8
218
9
97
233

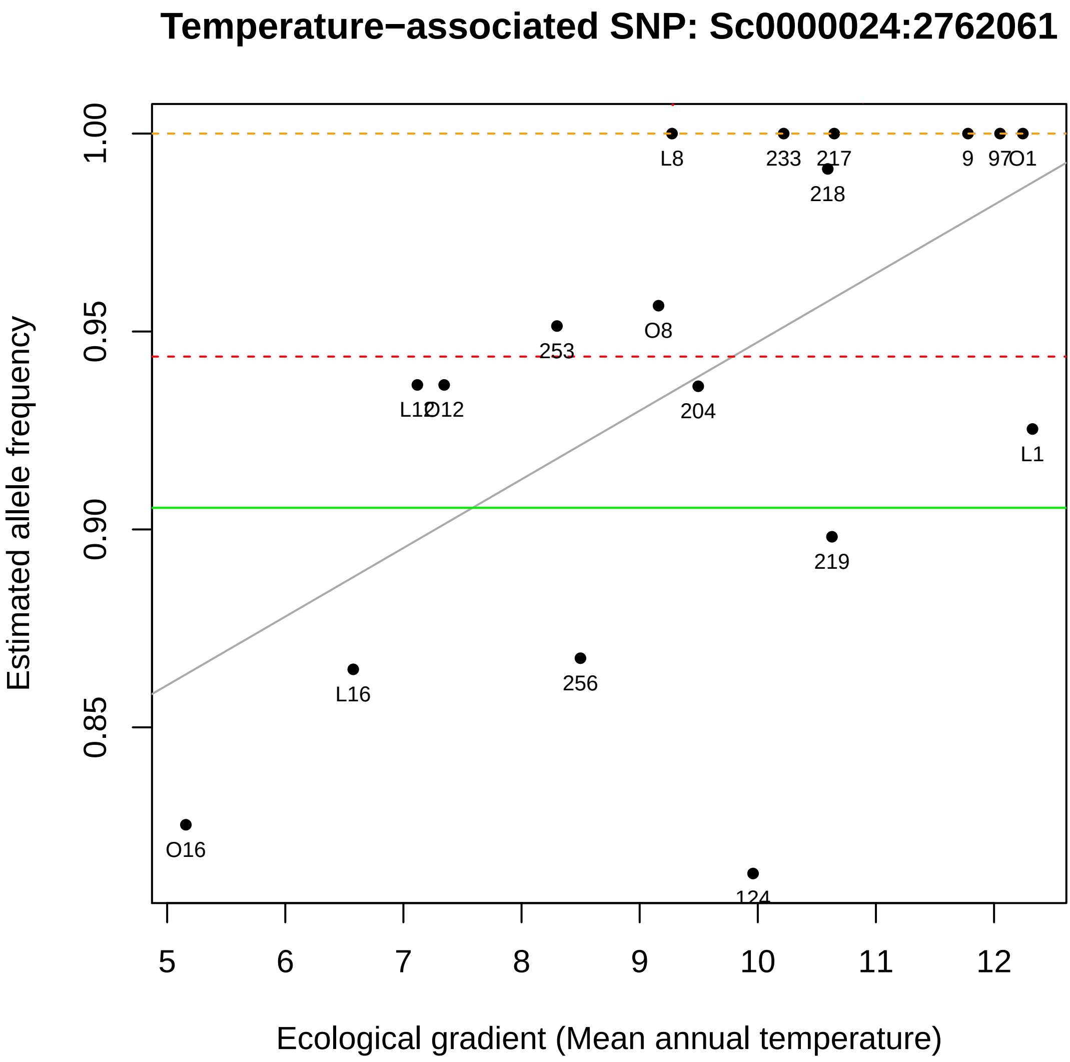

Temperature−associated SNP: Sc0000024:2762061
Estimated allele frequency
Ecological gradient (Mean annual temperature)
1.00
0.95
0.90
0.85
5
6
7
8
9
10
11
12
L8
233
217
218
9
97
O1
O8
253
L12
O12
204
L1
219
L16
256
O16
124

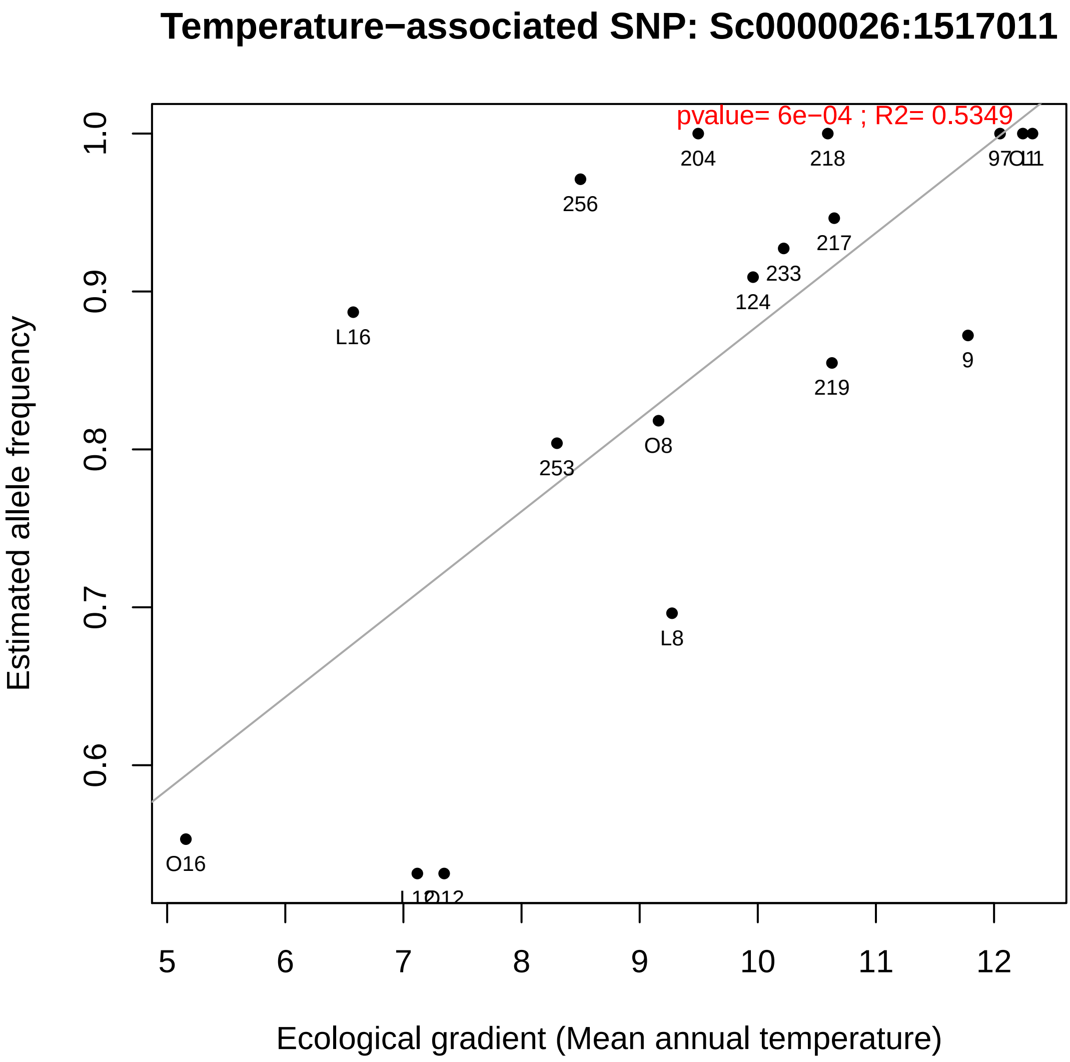
Temperature−associated SNP: Sc0000026:1517011
pvalue= 6e−04 ; R2= 0.5349
Estimated allele frequency
Ecological gradient (Mean annual temperature)
0.6
0.7
0.8
0.9
1.0
5
6
7
8
9
10
11
12
204
218
256
217
233
124
L16
9
219
O8
253
L8
O16
O12

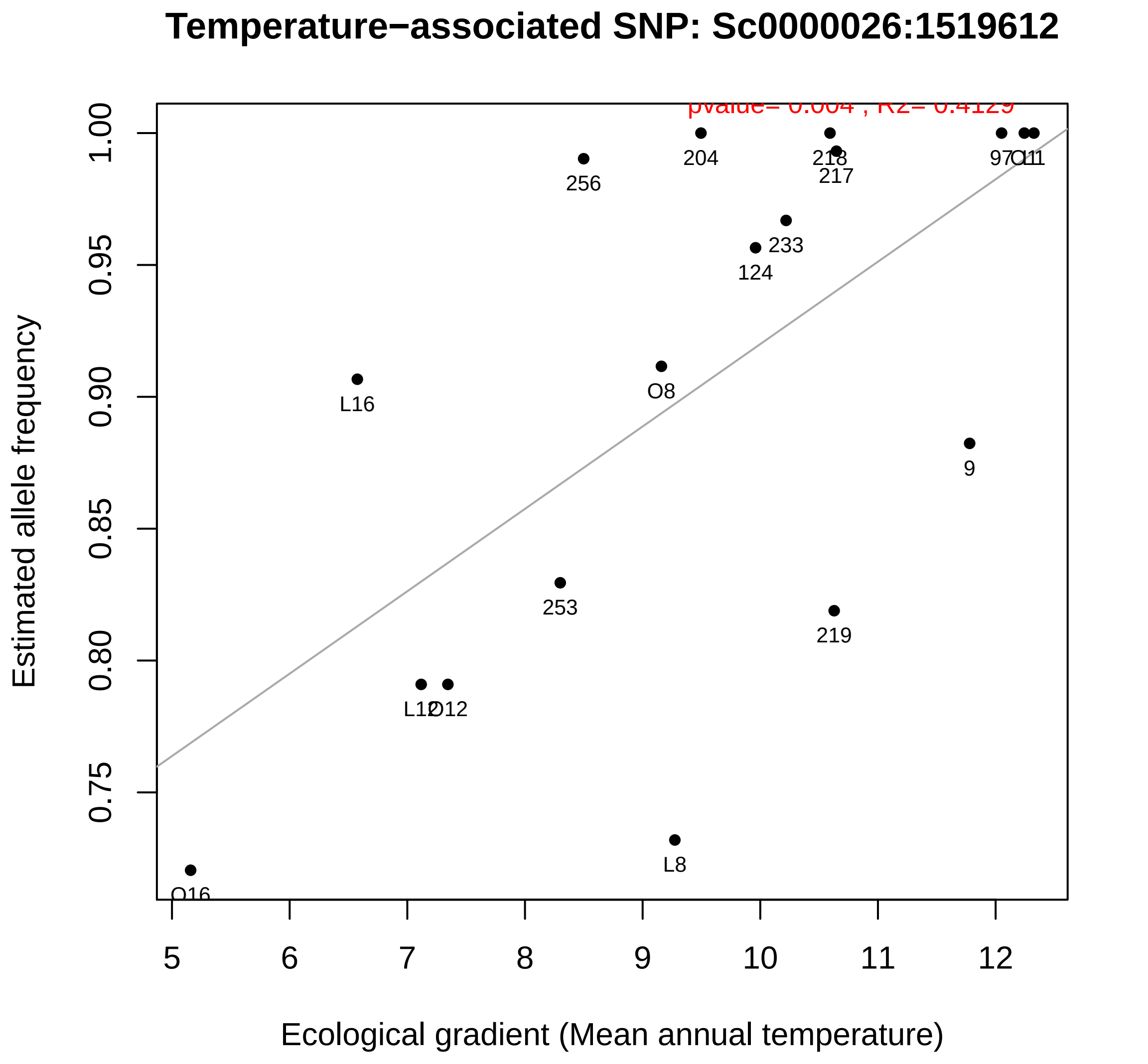
Temperature−associated SNP: Sc0000026:1519612
pvalue= 0.004 , R2= 0.4129
Estimated allele frequency
1.00
0.95
0.90
0.85
0.80
0.75
Ecological gradient (Mean annual temperature)
5
6
7
8
9
10
11
12
256
204
218
217
97
233
124
L16
O8
9
253
219
L12
O12
L8
O16

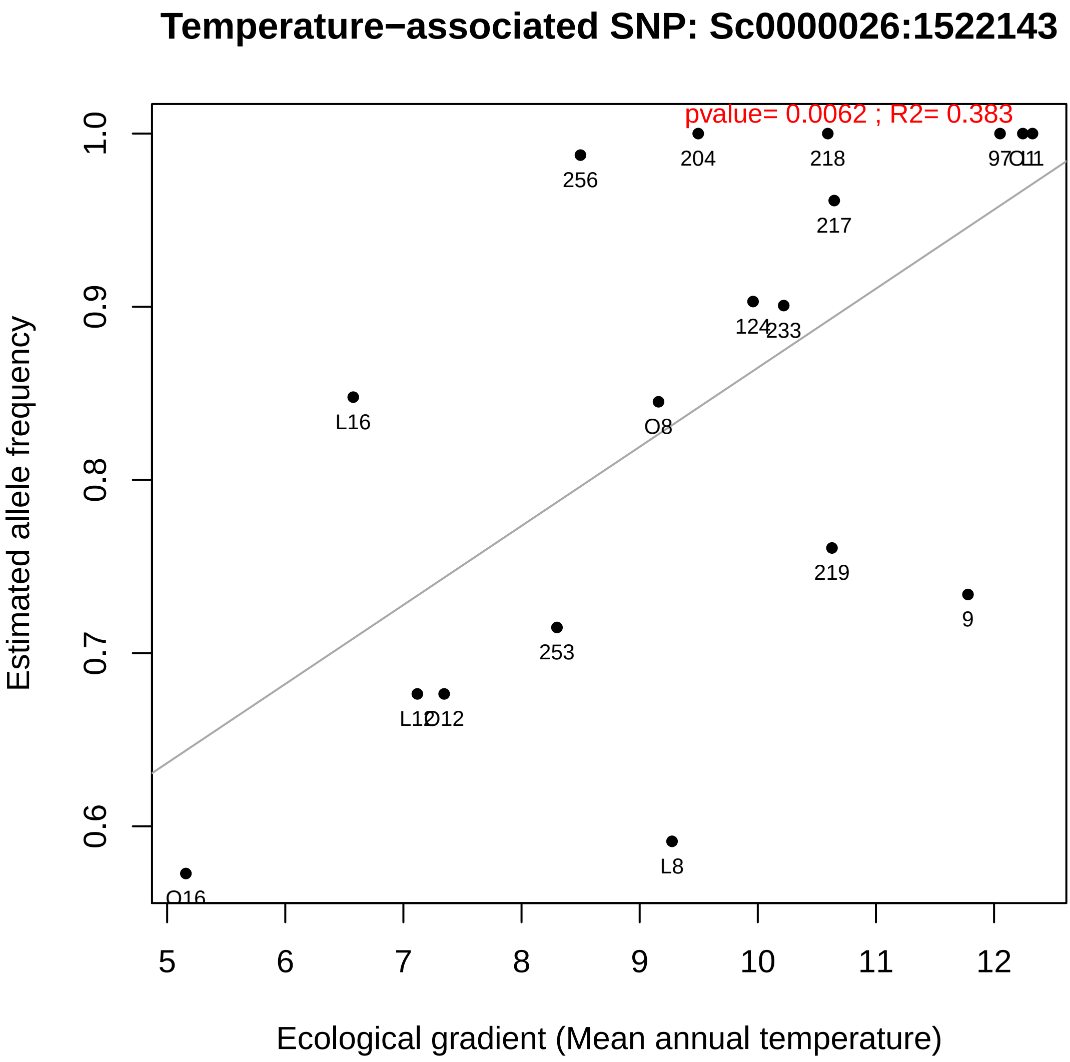

Temperature−associated SNP: Sc0000026:1522143
pvalue= 0.0062 ; R2= 0.383
Estimated allele frequency
Ecological gradient (Mean annual temperature)
1.0
0.9
0.8
0.7
0.6
5
6
7
8
9
10
11
12
256
204
218
97
O11
217
124
233
L16
O8
219
9
253
L12
O12
L8
O16

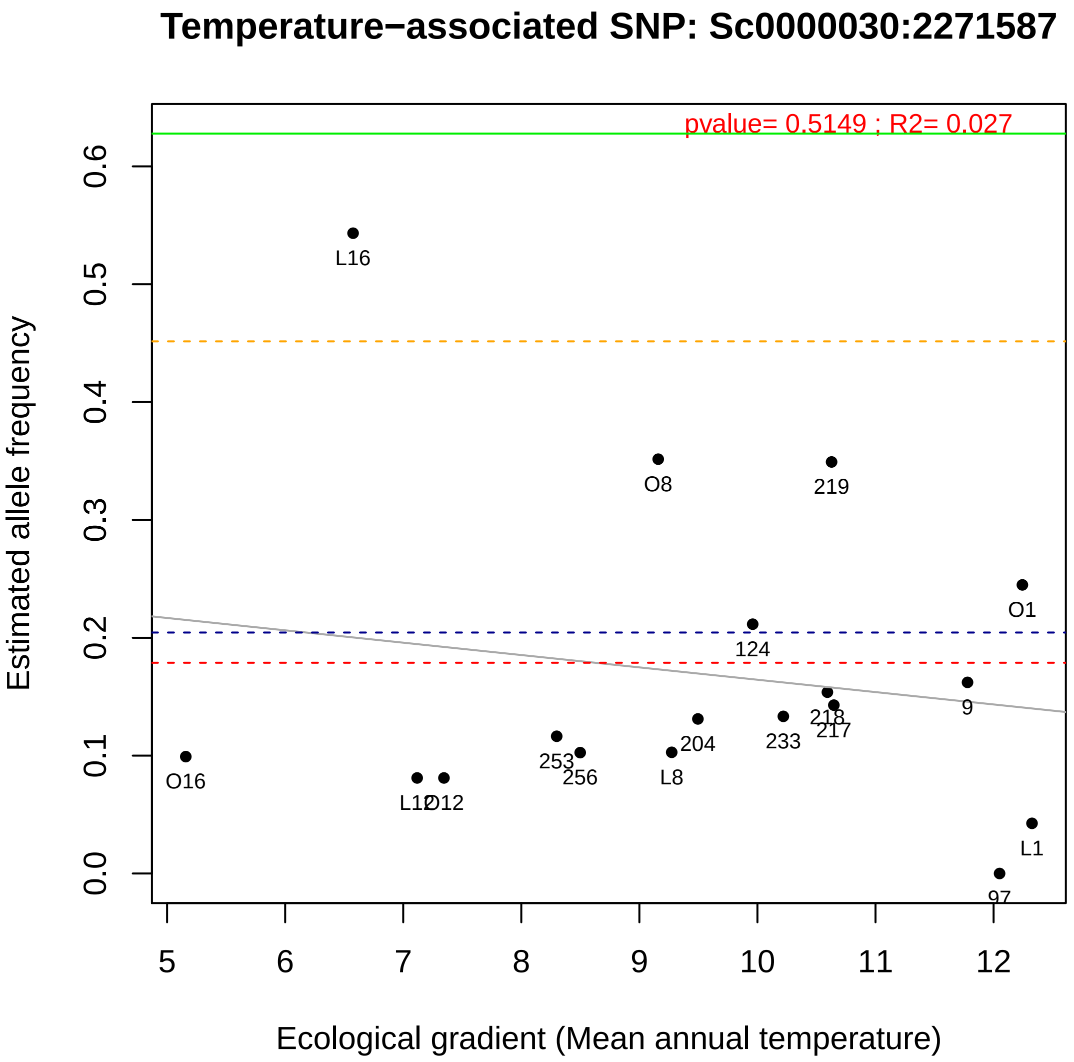

Temperature−associated SNP: Sc0000030:2271587
pvalue= 0.5149 ; R2= 0.027
Estimated allele frequency
0.0
0.1
0.2
0.3
0.4
0.5
0.6
L16
O8
219
O1
124
9
218
217
233
204
253
256
L8
O16
L12
O12
L1
97
5
6
7
8
9
10
11
12
Ecological gradient (Mean annual temperature)

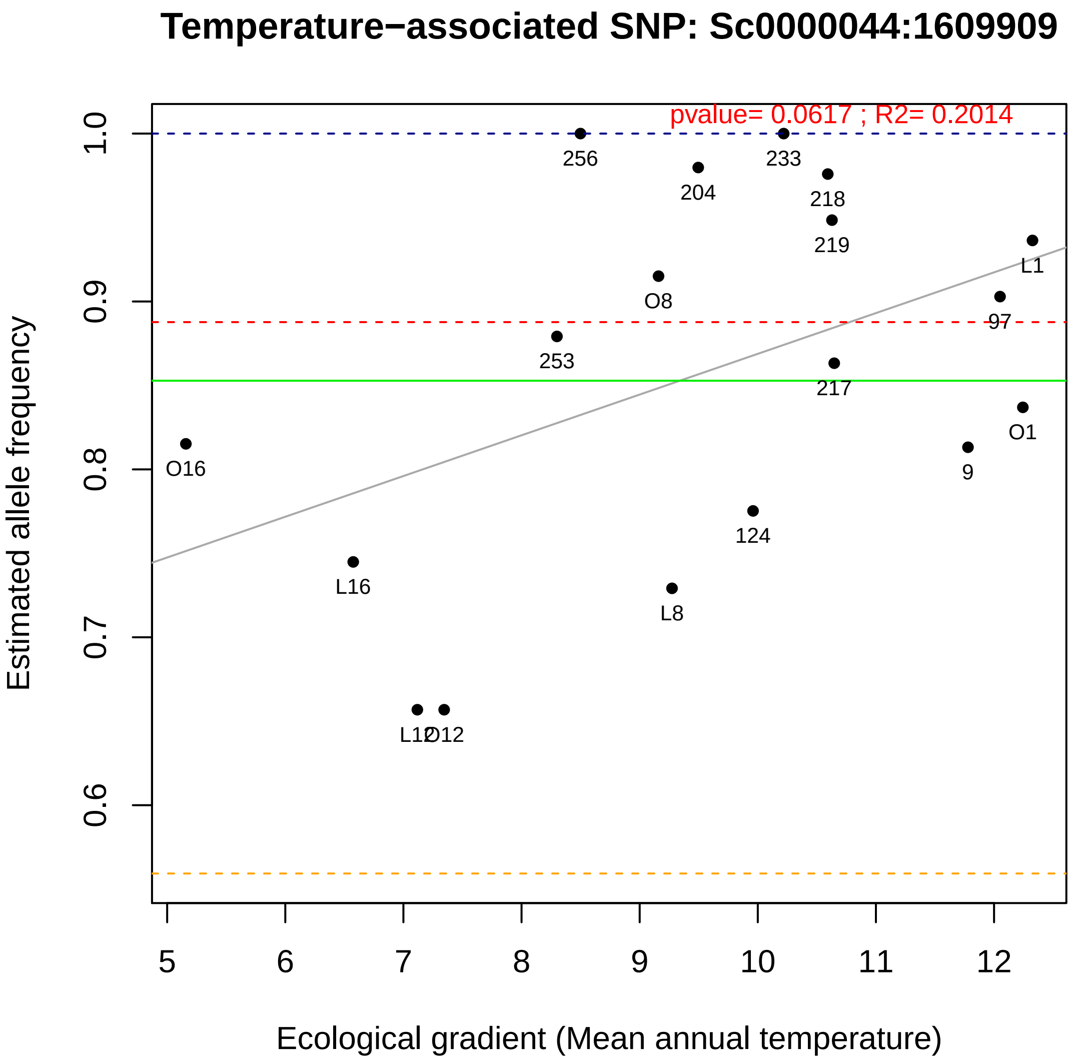

Temperature−associated SNP: Sc0000044:1609909
pvalue= 0.0617 ; R2= 0.2014
Estimated allele frequency
1.0
0.9
0.8
0.7
0.6
256
233
204
218
219
L1
O8
97
253
217
O1
O16
9
124
L16
L8
L12
O12
5
6
7
8
9
10
11
12
Ecological gradient (Mean annual temperature)

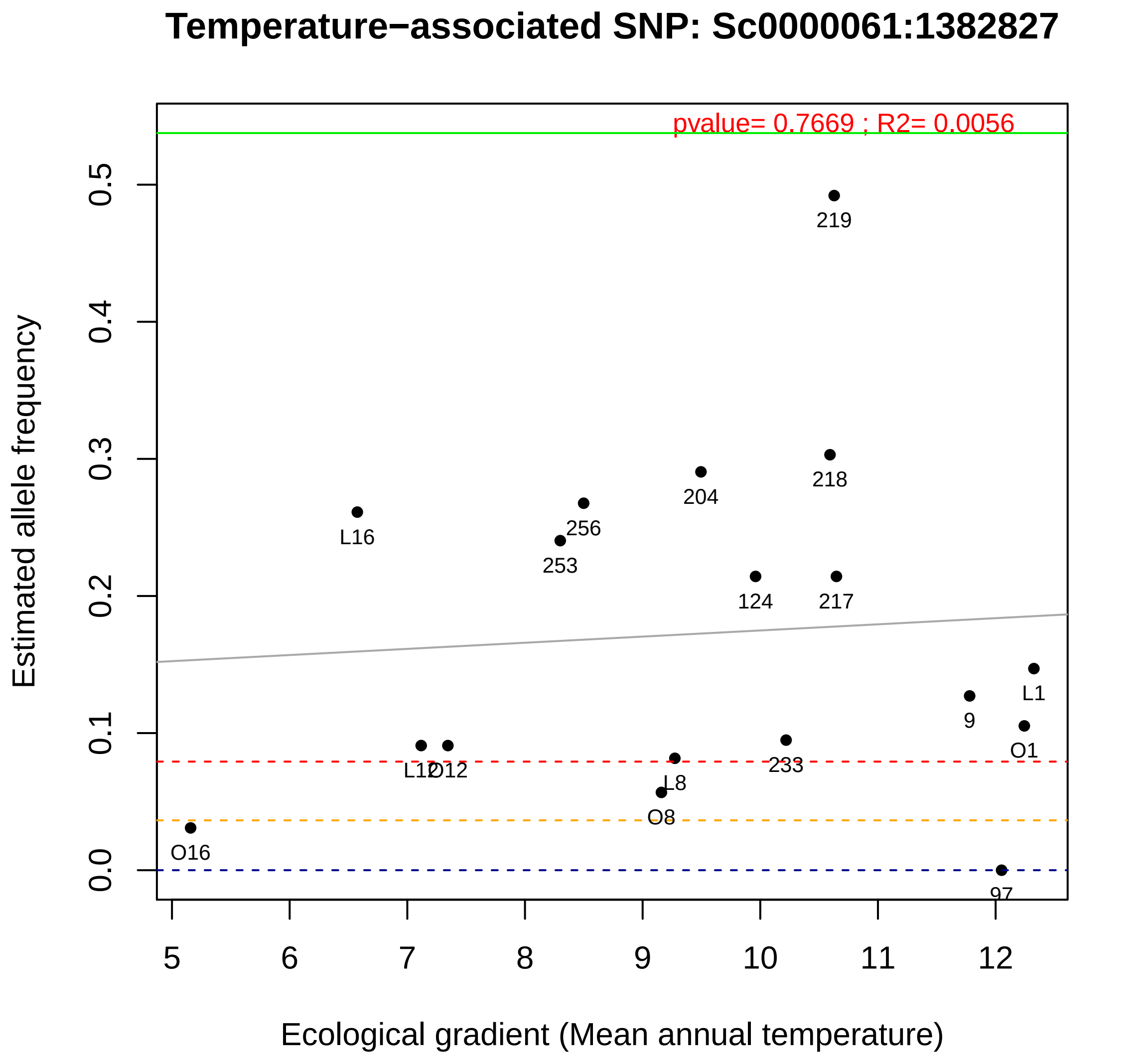
Temperature−associated SNP: Sc0000061:1382827
pvalue= 0.7669 ; R2= 0.0056
Estimated allele frequency
0.0
0.1
0.2
0.3
0.4
0.5
Ecological gradient (Mean annual temperature)
5
6
7
8
9
10
11
12
219
218
204
256
L16
253
124
217
L1
9
O1
233
L12
O12
L8
O8
O16
97

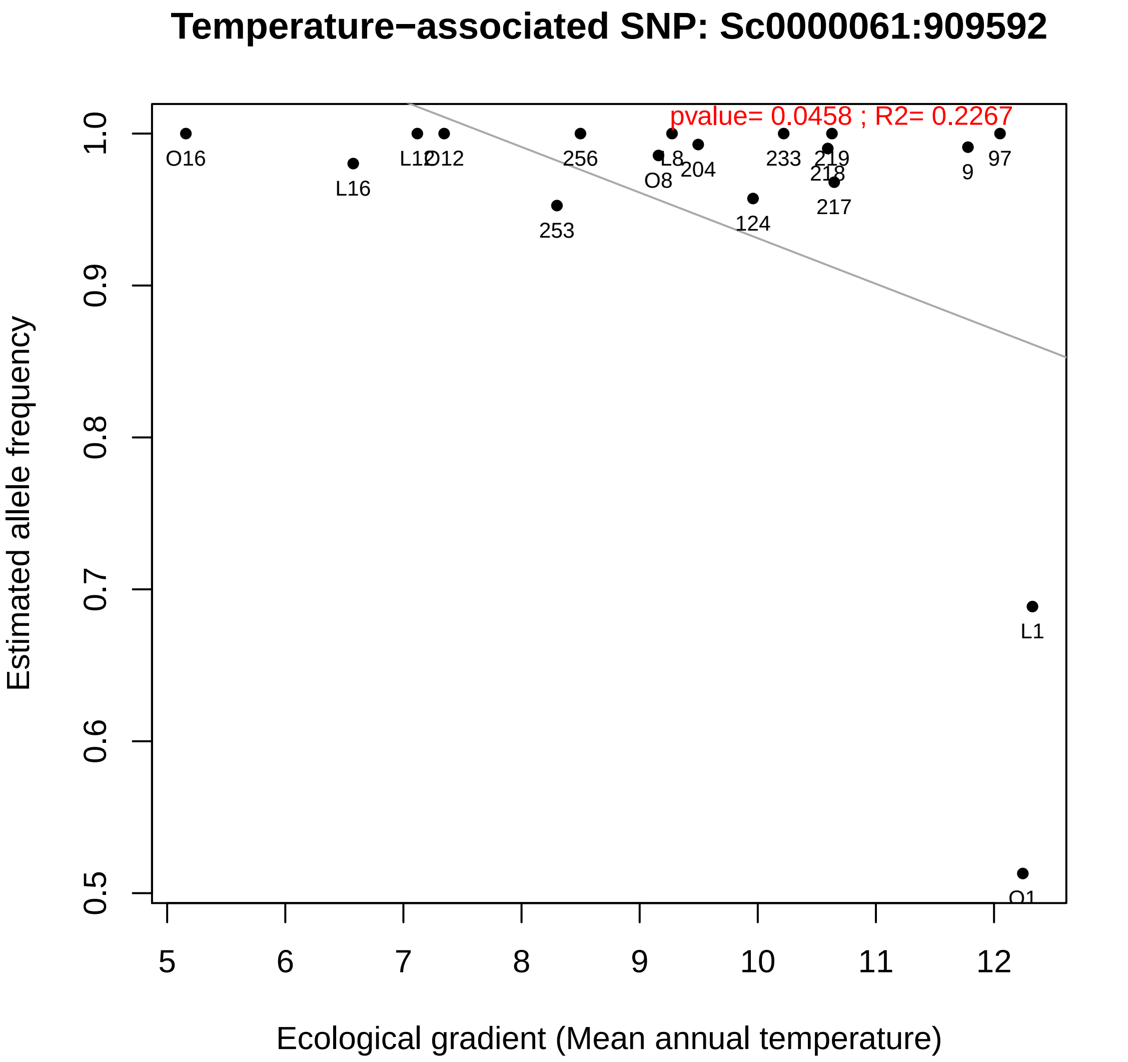

Temperature−associated SNP: Sc0000061:909592
pvalue= 0.0458 ; R2= 0.2267
Estimated allele frequency
Ecological gradient (Mean annual temperature)
0.5
0.6
0.7
0.8
0.9
1.0
5
6
7
8
9
10
11
12
O16
L16
L12
O12
256
253
O8
L8
204
124
233
219
218
217
9
97
L1
O1

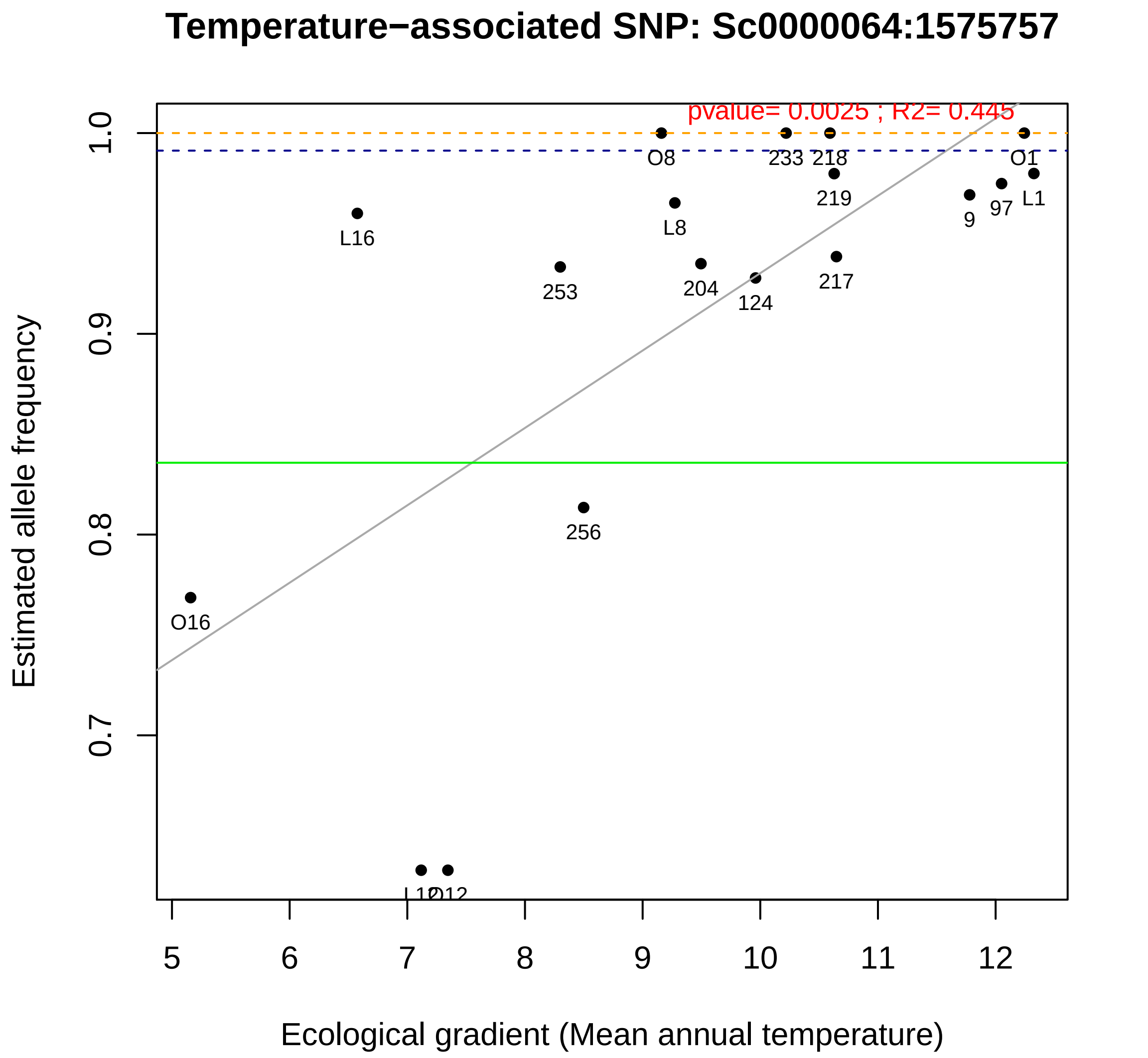
Temperature−associated SNP: Sc0000064:1575757
pvalue= 0.0025 ; R2= 0.445
Estimated allele frequency
0.7
0.8
0.9
1.0
Ecological gradient (Mean annual temperature)
5
6
7
8
9
10
11
12
O8
233
218
O1
219
L1
9
97
L16
L8
253
204
124
217
256
O16
O12

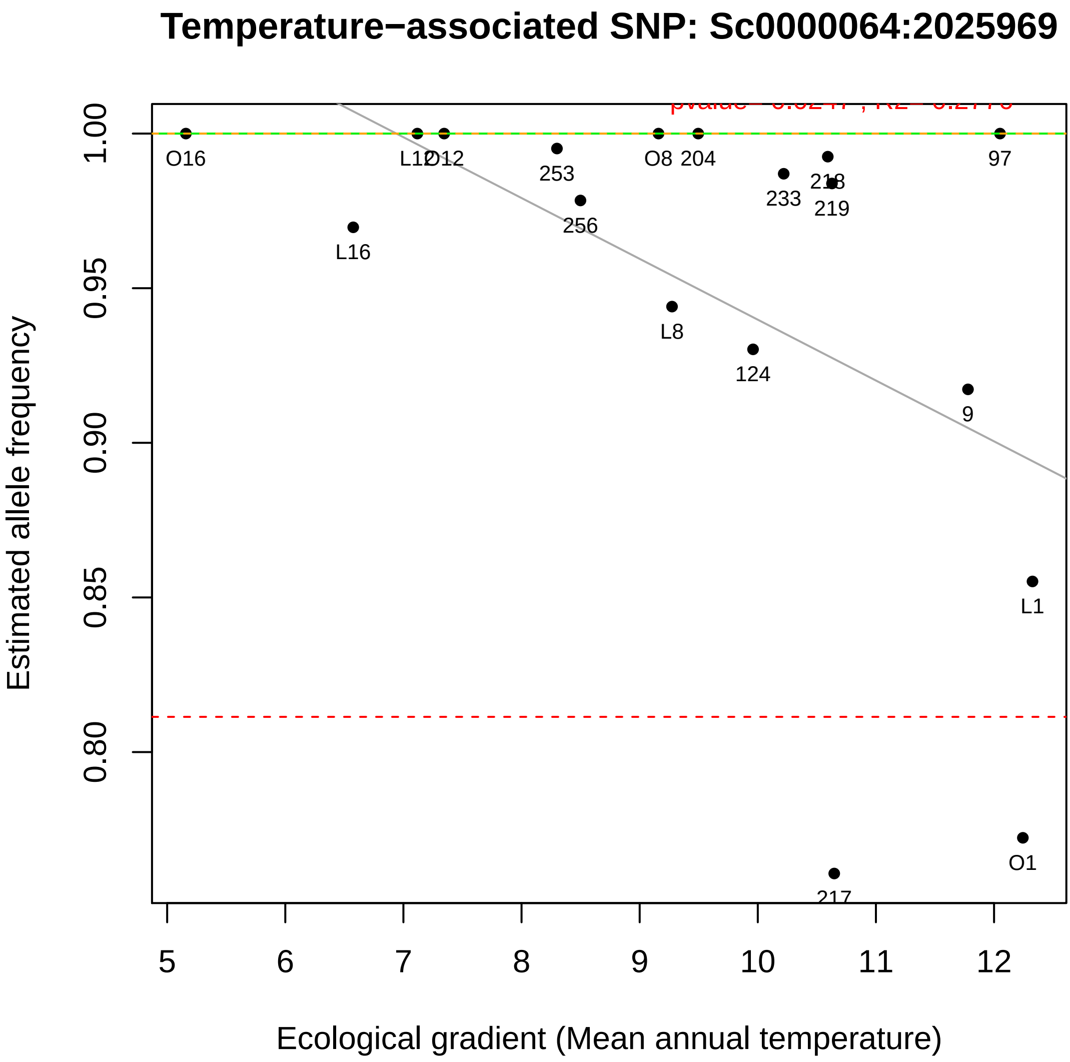

Temperature−associated SNP: Sc0000064:2025969
Estimated allele frequency
Ecological gradient (Mean annual temperature)
1.00
0.95
0.90
0.85
0.80
5
6
7
8
9
10
11
12
O16
L12
O12
253
256
O8
204
233
218
219
97
L16
L8
124
9
L1
O1
217

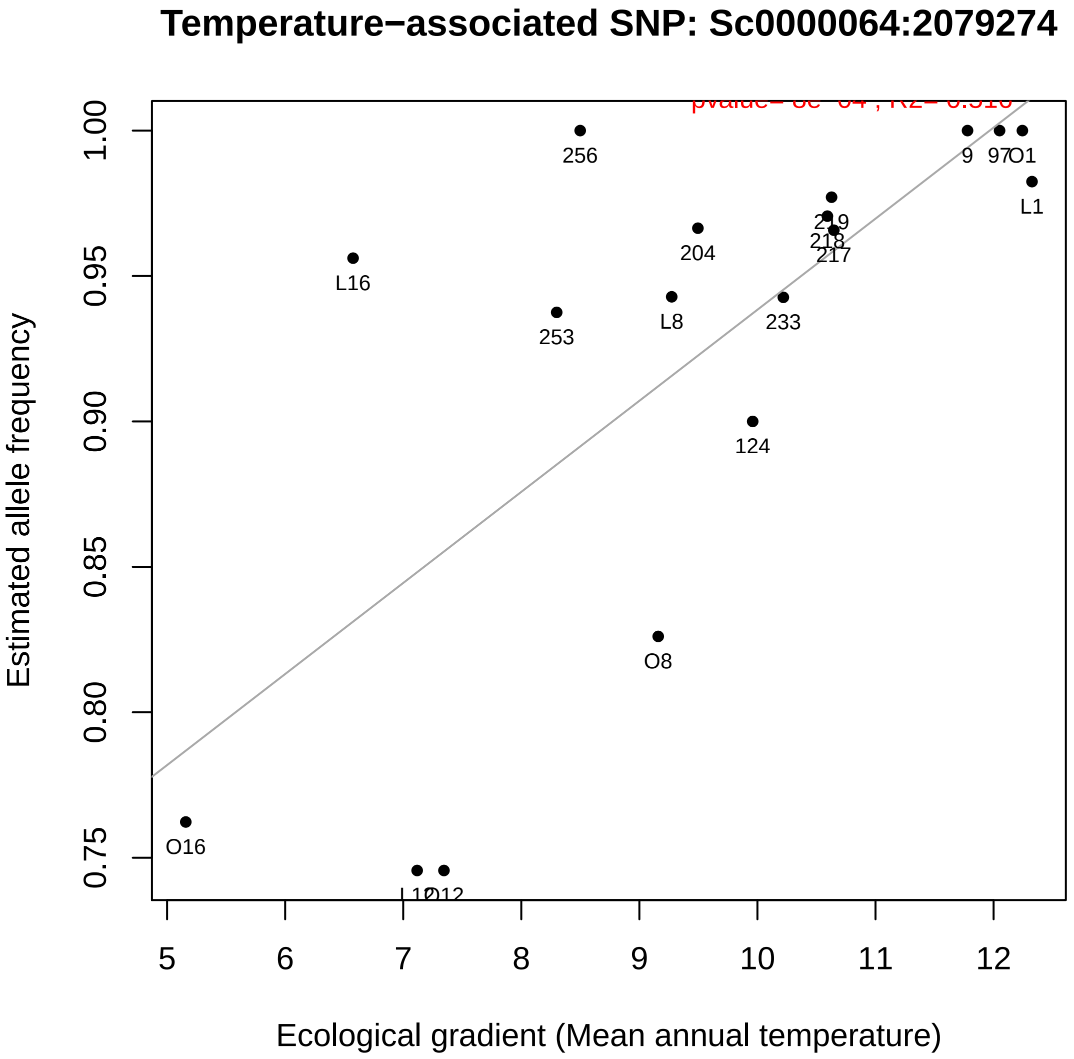
Temperature−associated SNP: Sc0000064:2079274
Estimated allele frequency
Ecological gradient (Mean annual temperature)
1.00
0.95
0.90
0.85
0.80
0.75
5
6
7
8
9
10
11
12
256
9
97
O1
L1
219
218
217
204
L16
253
L8
233
124
O8
O16
O12

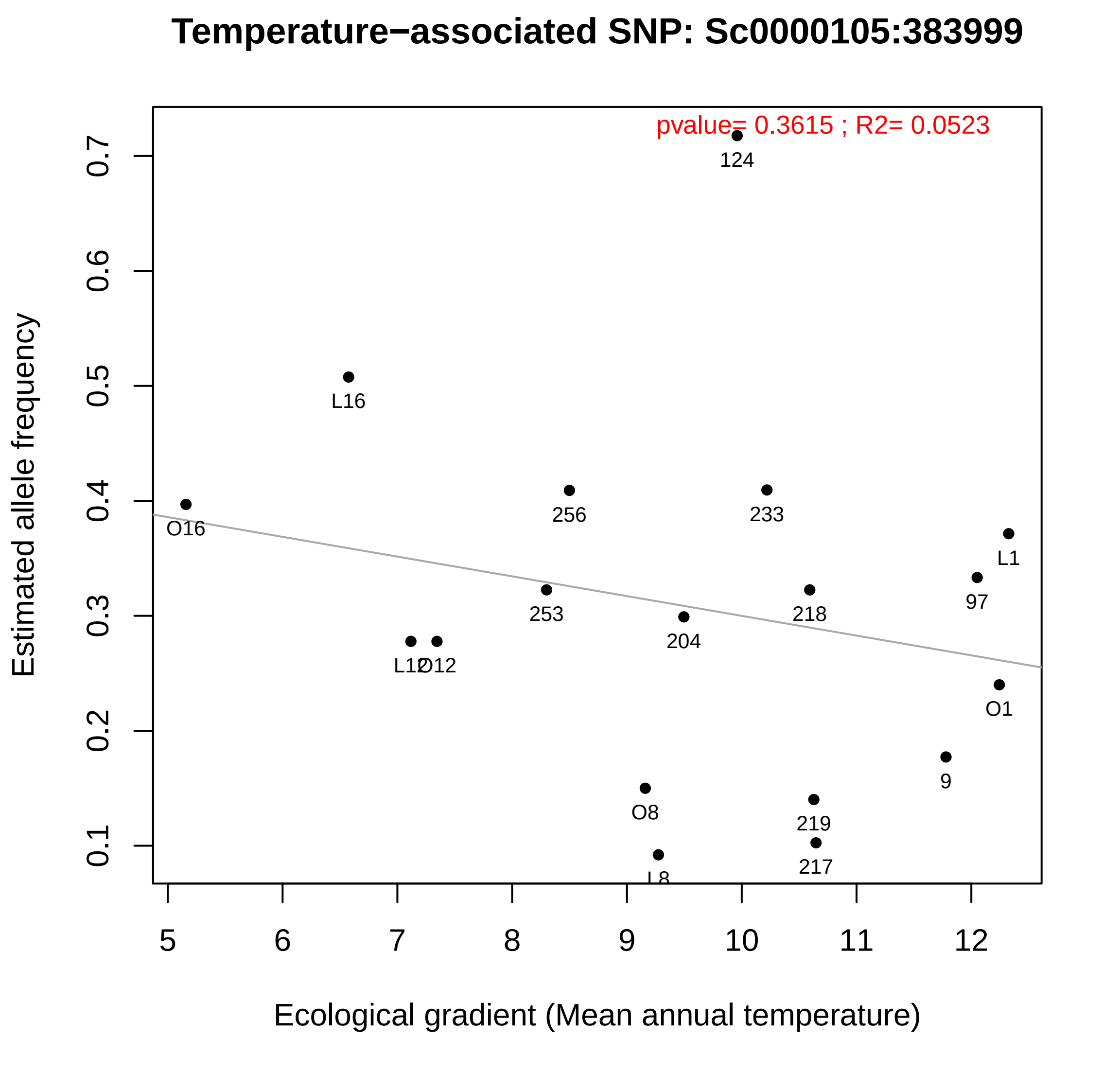
Temperature−associated SNP: Sc0000105:383999
pvalue= 0.3615 ; R2= 0.0523
Estimated allele frequency
0.1
0.2
0.3
0.4
0.5
0.6
0.7
Ecological gradient (Mean annual temperature)
5
6
7
8
9
10
11
12
124
L16
O16
256
233
L1
97
253
218
204
L12
O12
O1
9
O8
219
217
L8

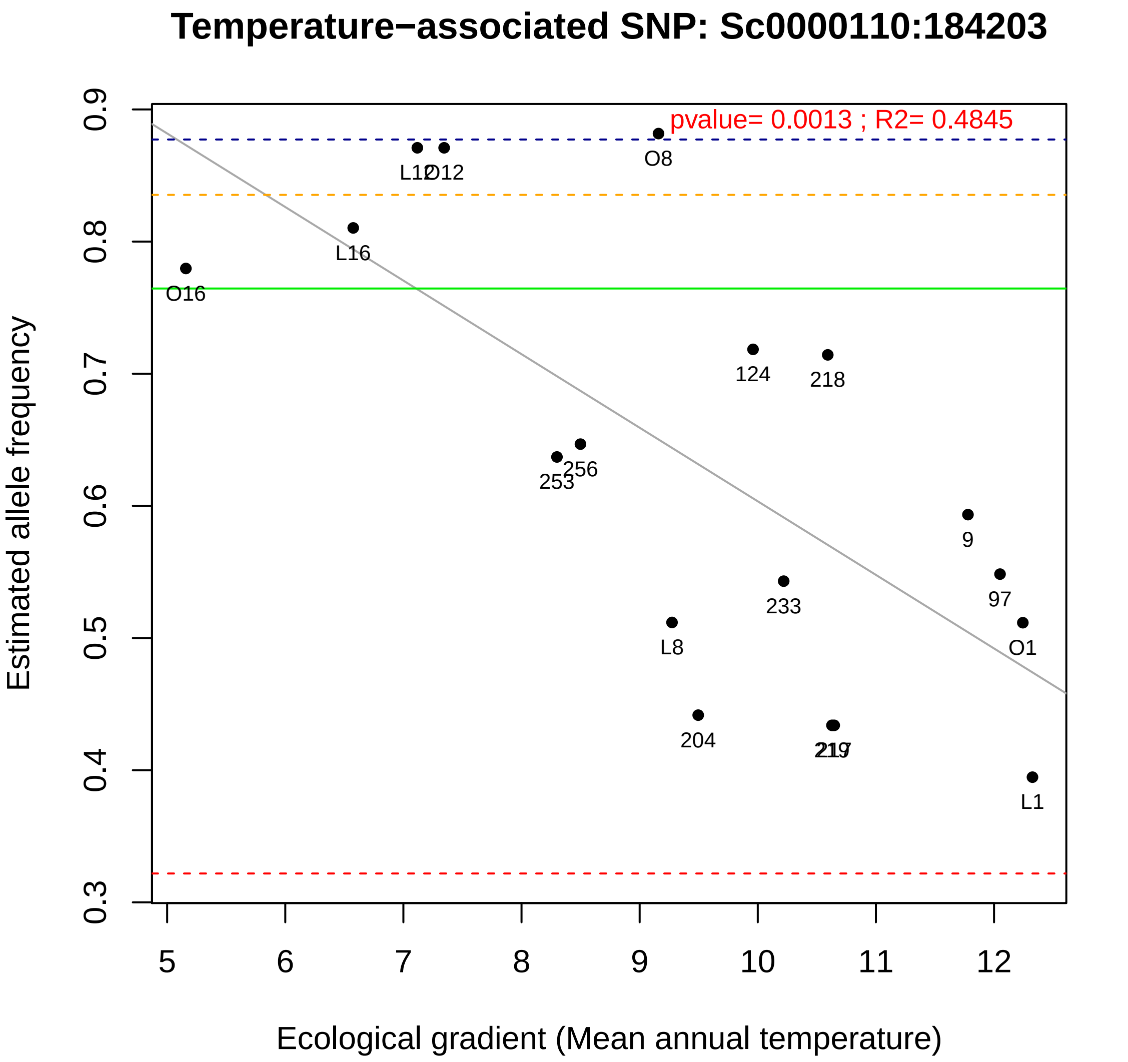
Temperature−associated SNP: Sc0000110:184203
pvalue= 0.0013 ; R2= 0.4845
Estimated allele frequency
Ecological gradient (Mean annual temperature)
0.3
0.4
0.5
0.6
0.7
0.8
0.9
5
6
7
8
9
10
11
12
O16
L16
L12
O12
O8
253
256
124
218
L8
204
233
9
97
O1
L1

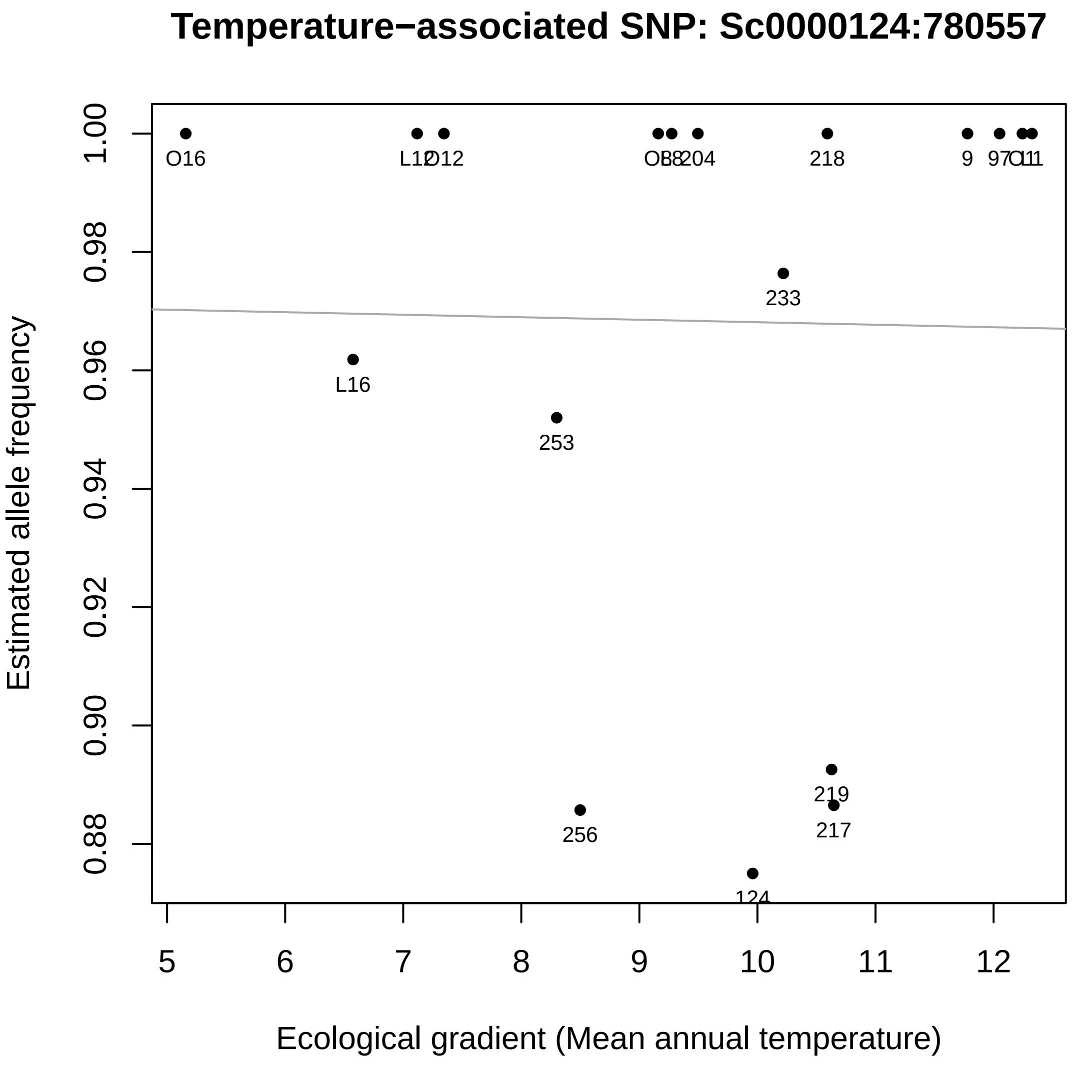

Temperature−associated SNP: Sc0000124:780557
Estimated allele frequency
Ecological gradient (Mean annual temperature)
0.88
0.90
0.92
0.94
0.96
0.98
1.00
5
6
7
8
9
10
11
12
O16
L12
O12
218
204
9
233
L16
253
219
256
217
124

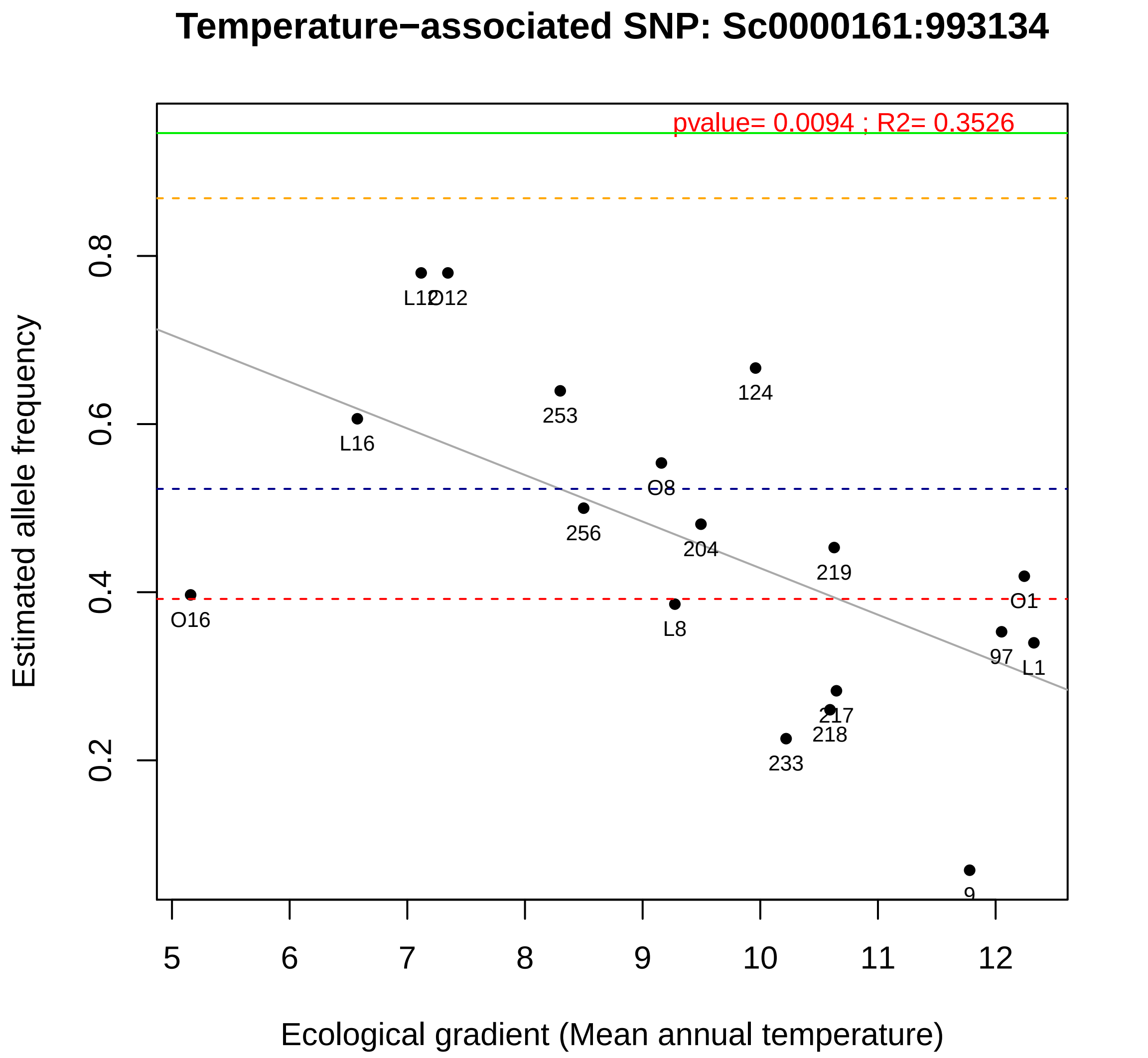

Temperature−associated SNP: Sc0000161:993134
pvalue= 0.0094 ; R2= 0.3526
Estimated allele frequency
0.2
0.4
0.6
0.8
5
6
7
8
9
10
11
12
Ecological gradient (Mean annual temperature)
L12
O12
253
124
L16
O8
256
204
219
O16
L8
O1
97
L1
217
218
233
9

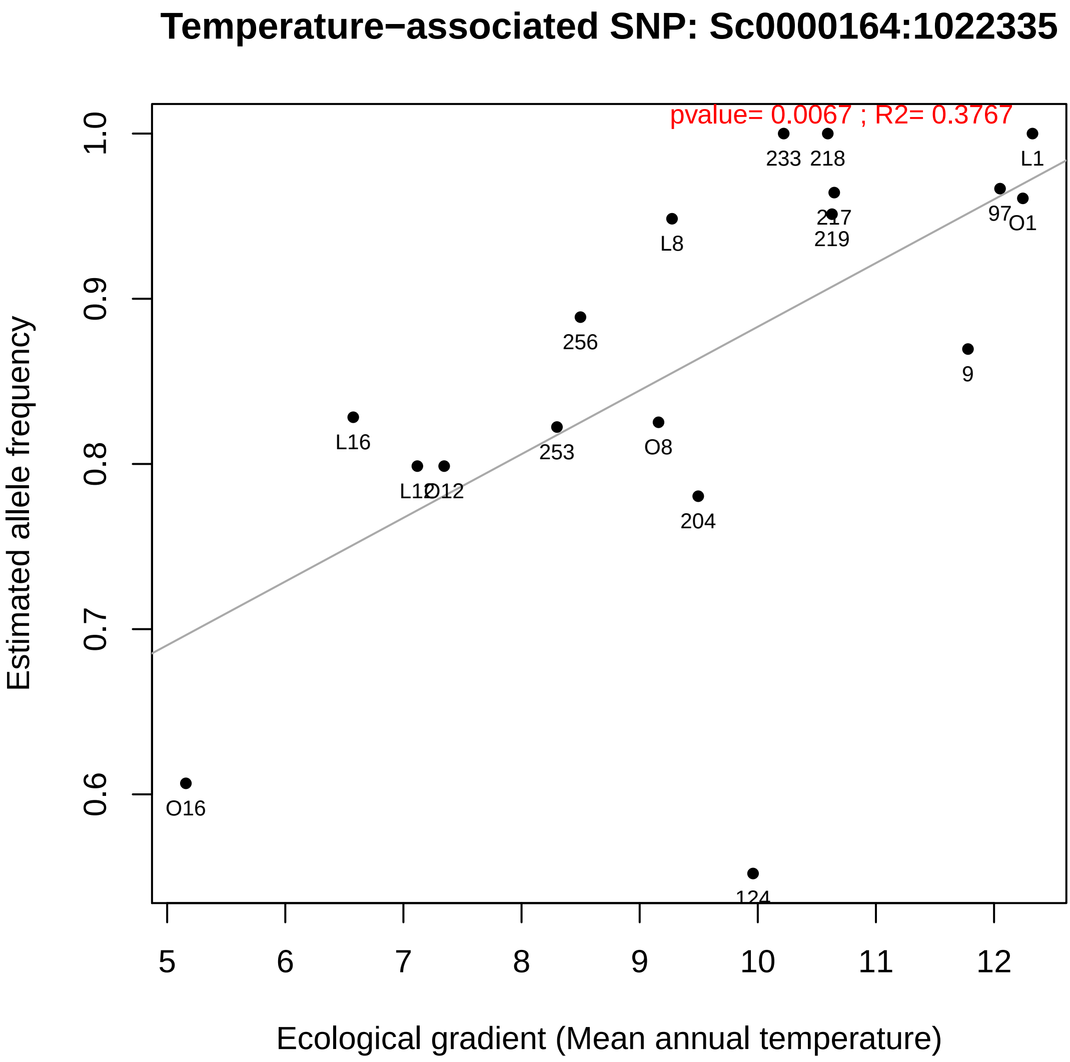
Temperature−associated SNP: Sc0000164:1022335
pvalue= 0.0067 ; R2= 0.3767
Estimated allele frequency
Ecological gradient (Mean annual temperature)
1.0
0.9
0.8
0.7
0.6
5
6
7
8
9
10
11
12
233
218
L1
217
219
97
O1
L8
256
9
L16
L12
O12
253
O8
204
O16
124

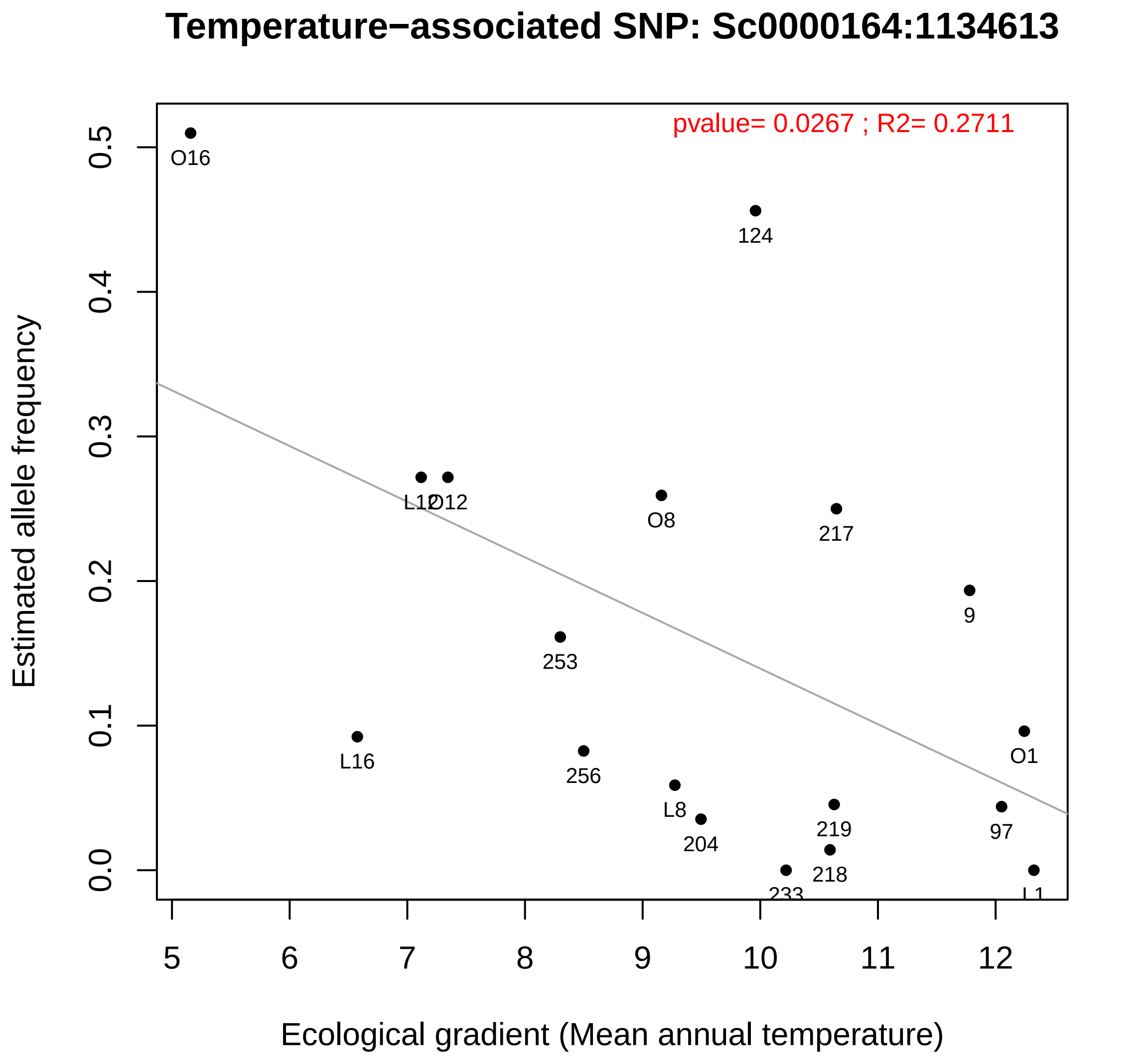

Temperature−associated SNP: Sc0000164:1134613
pvalue= 0.0267 ; R2= 0.2711
Estimated allele frequency
Ecological gradient (Mean annual temperature)
0.0
0.1
0.2
0.3
0.4
0.5
5
6
7
8
9
10
11
12
O16
124
L12
O12
O8
217
9
253
L16
256
O1
L8
204
219
97
218
233
L1

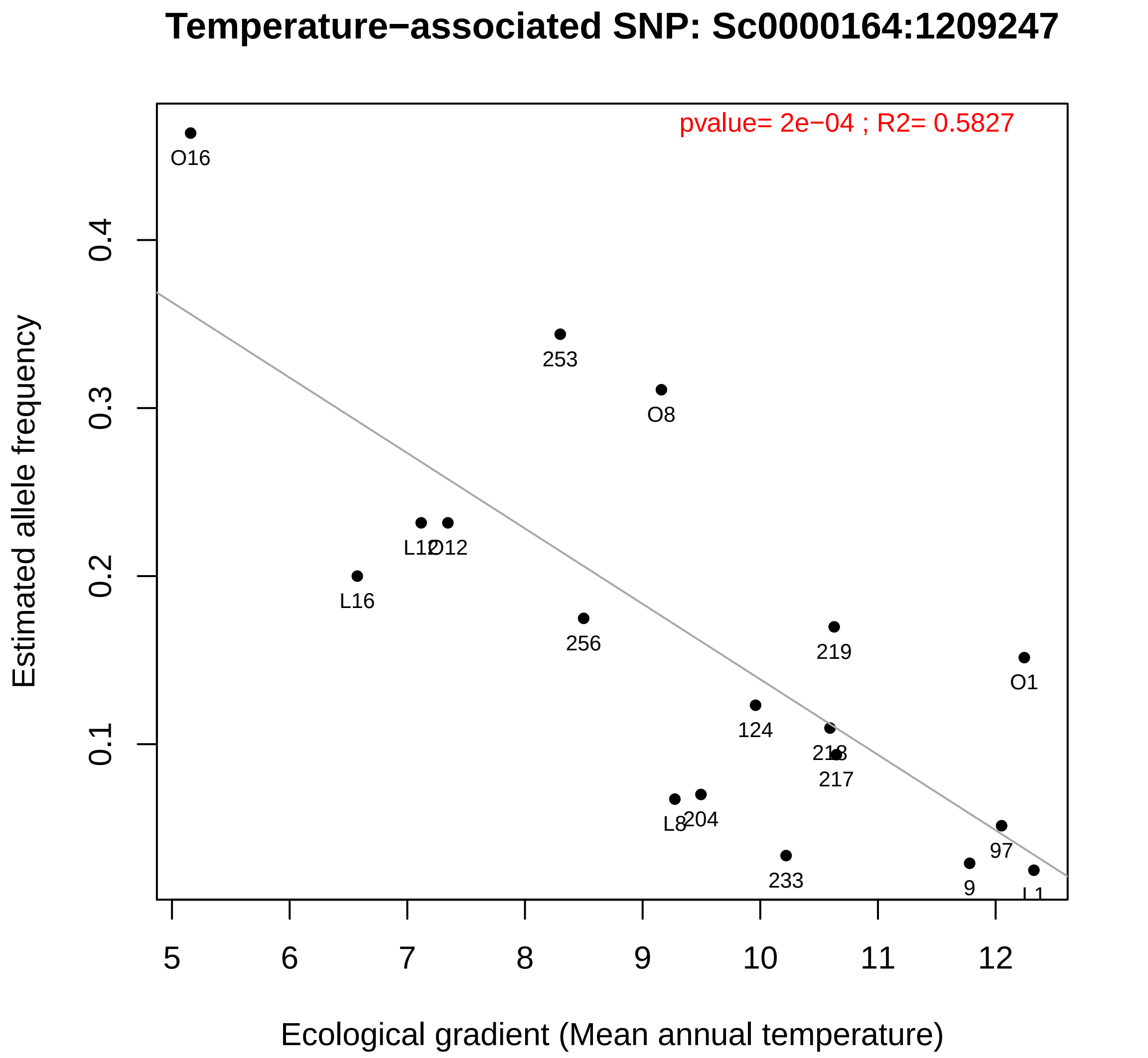

Temperature−associated SNP: Sc0000164:1209247
pvalue= 2e−04 ; R2= 0.5827
Estimated allele frequency
Ecological gradient (Mean annual temperature)
0.1
0.2
0.3
0.4
5
6
7
8
9
10
11
12
O16
253
O8
L12
O12
L16
256
219
O1
124
218
217
L8
204
97
233
9
L1

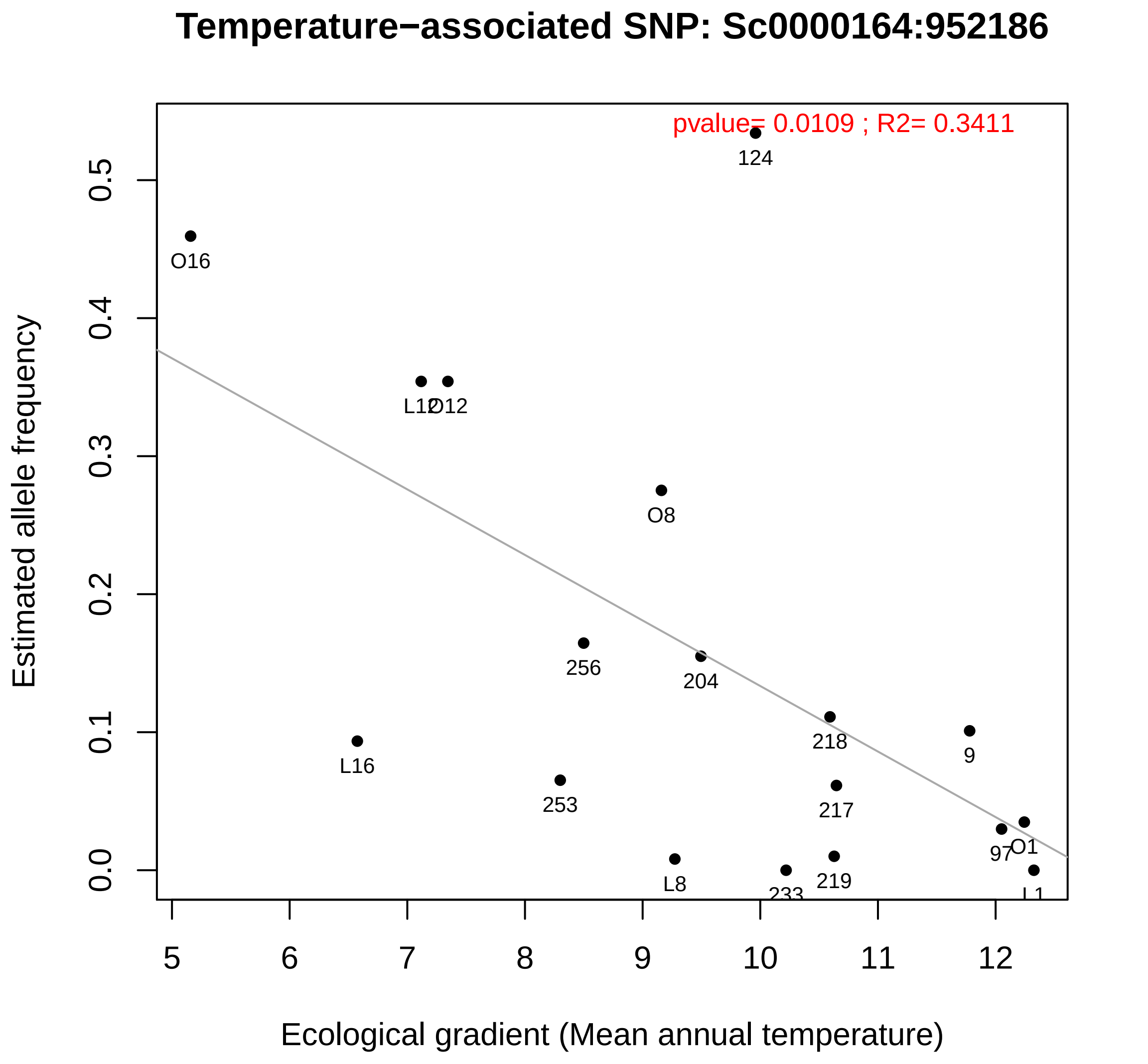

Temperature−associated SNP: Sc0000164:952186
pvalue= 0.0109 ; R2= 0.3411
Estimated allele frequency
0.0
0.1
0.2
0.3
0.4
0.5
5
6
7
8
9
10
11
12
Ecological gradient (Mean annual temperature)
124
O16
L12
O12
O8
256
204
218
9
L16
253
217
97
O1
L8
233
219
L1

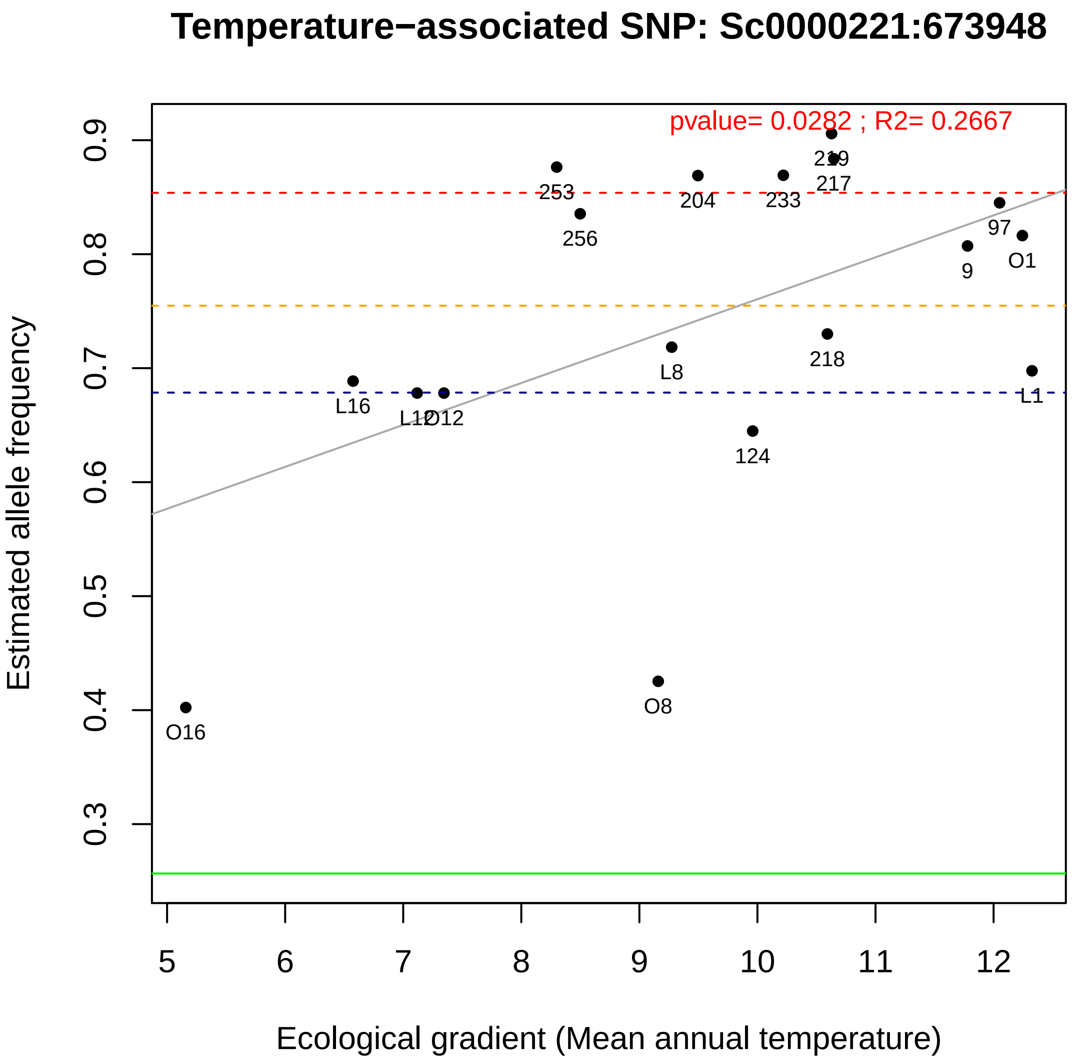
Temperature−associated SNP: Sc0000221:673948
pvalue= 0.0282 ; R2= 0.2667
Estimated allele frequency
0.3
0.4
0.5
0.6
0.7
0.8
0.9
Ecological gradient (Mean annual temperature)
5
6
7
8
9
10
11
12
253
256
204
233
219
217
97
O1
9
218
L8
L1
L16
L12
O12
124
O8
O16

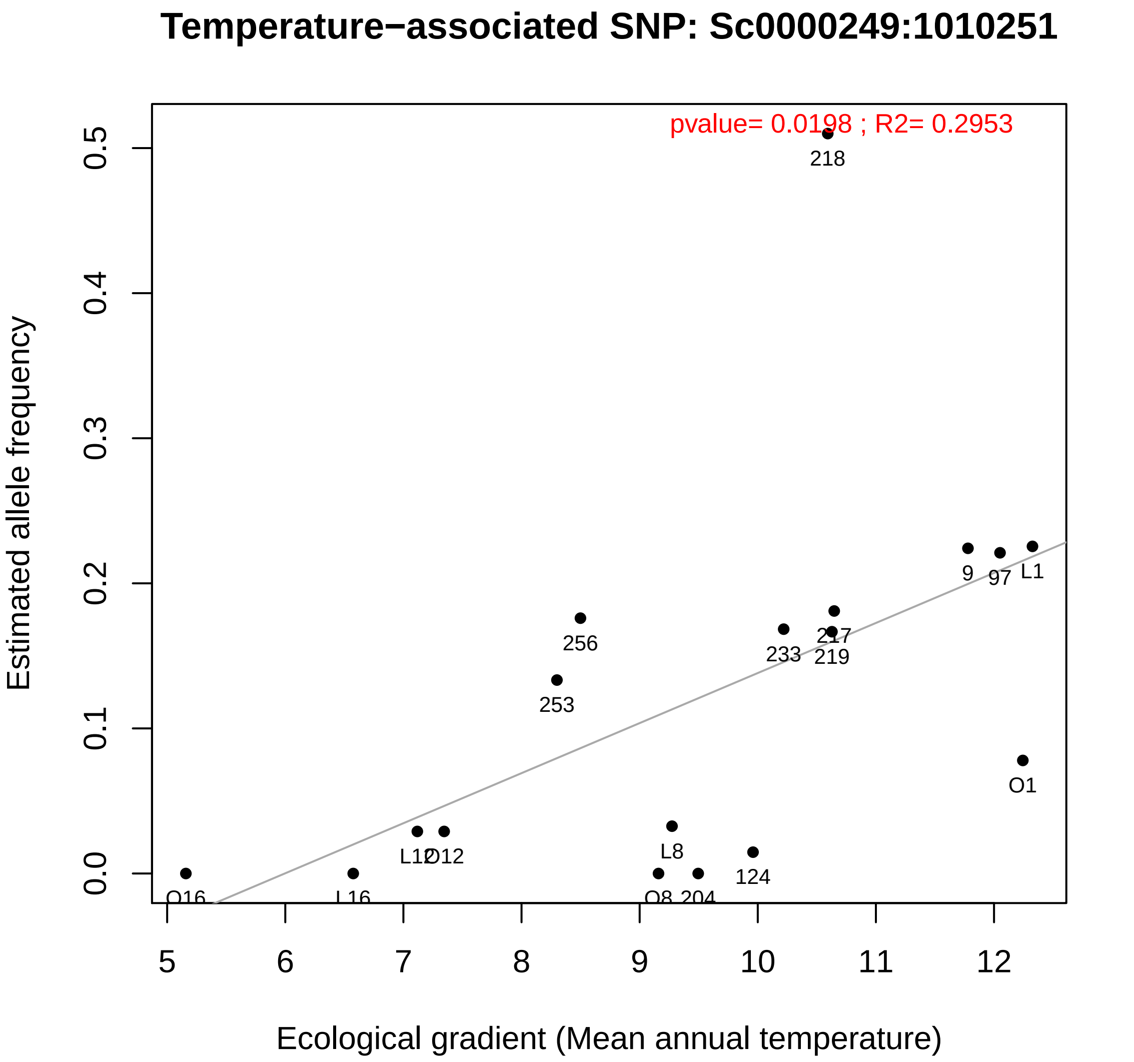
Temperature−associated SNP: Sc0000249:1010251
pvalue= 0.0198 ; R2= 0.2953
Estimated allele frequency
Ecological gradient (Mean annual temperature)
0.0
0.1
0.2
0.3
0.4
0.5
5
6
7
8
9
10
11
12
218
9
97
L1
256
233
217
219
253
O1
L8
L12
O12
124
O16
L16
O8
204

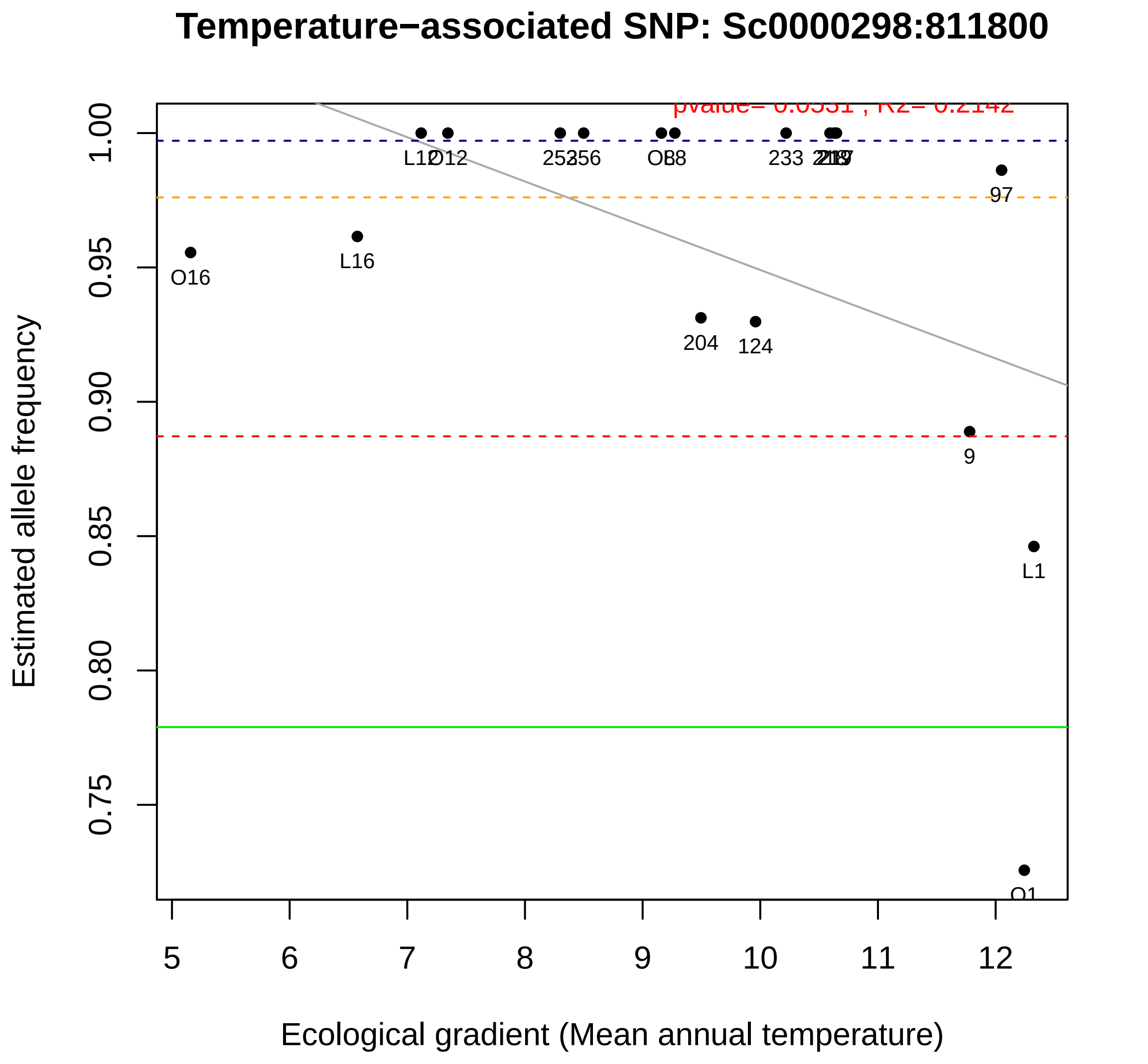

Temperature−associated SNP: Sc0000298:811800
pvalue= 0.0531 , R2= 0.2142
Estimated allele frequency
Ecological gradient (Mean annual temperature)
1.00
0.95
0.90
0.85
0.80
0.75
5
6
7
8
9
10
11
12
O16
L16
204
124
233
97
9
L1

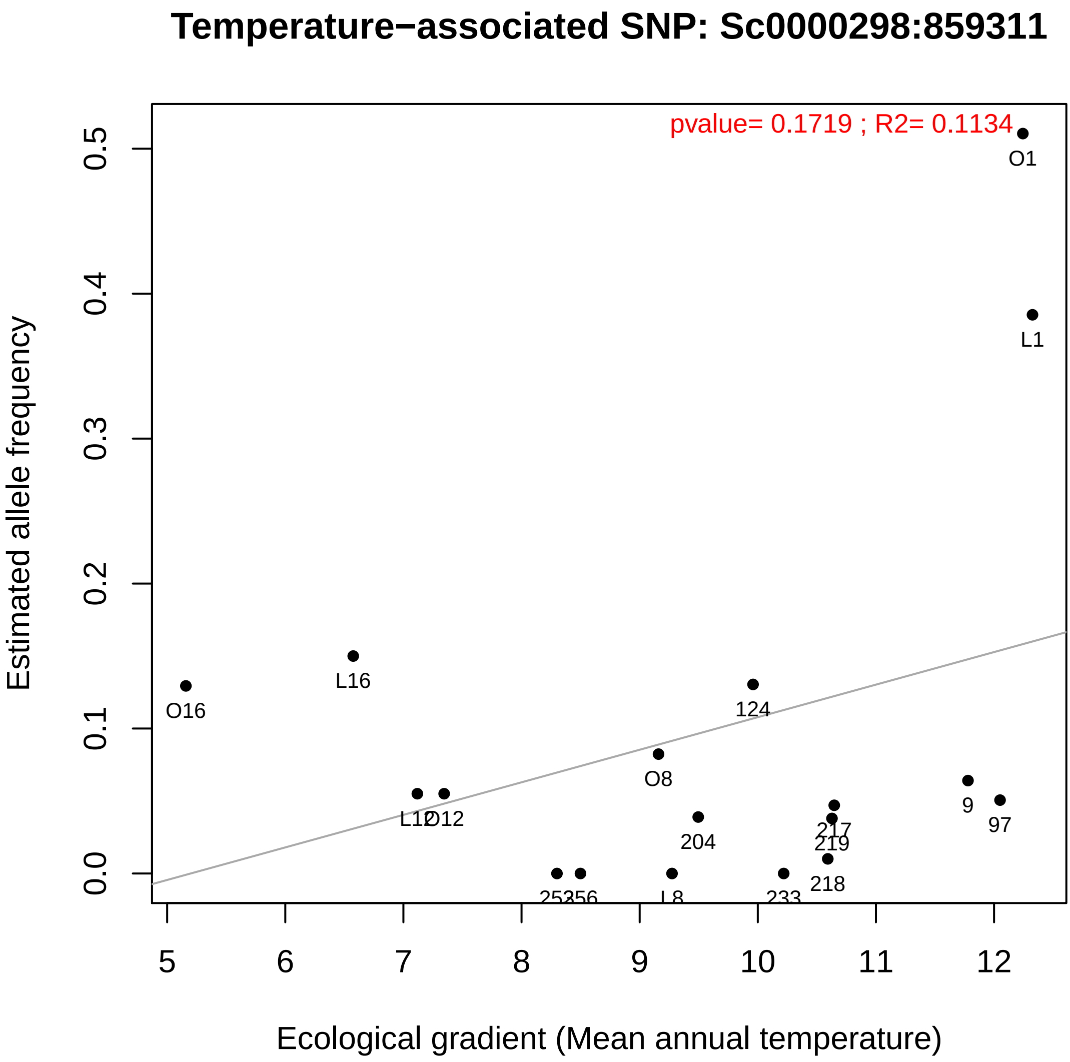

Temperature−associated SNP: Sc0000298:859311
pvalue= 0.1719 ; R2= 0.1134
Estimated allele frequency
0.0
0.1
0.2
0.3
0.4
0.5
5
6
7
8
9
10
11
12
Ecological gradient (Mean annual temperature)
O1
L1
O16
L16
124
O8
L12
O12
204
9
97
217
219
218
233
L8

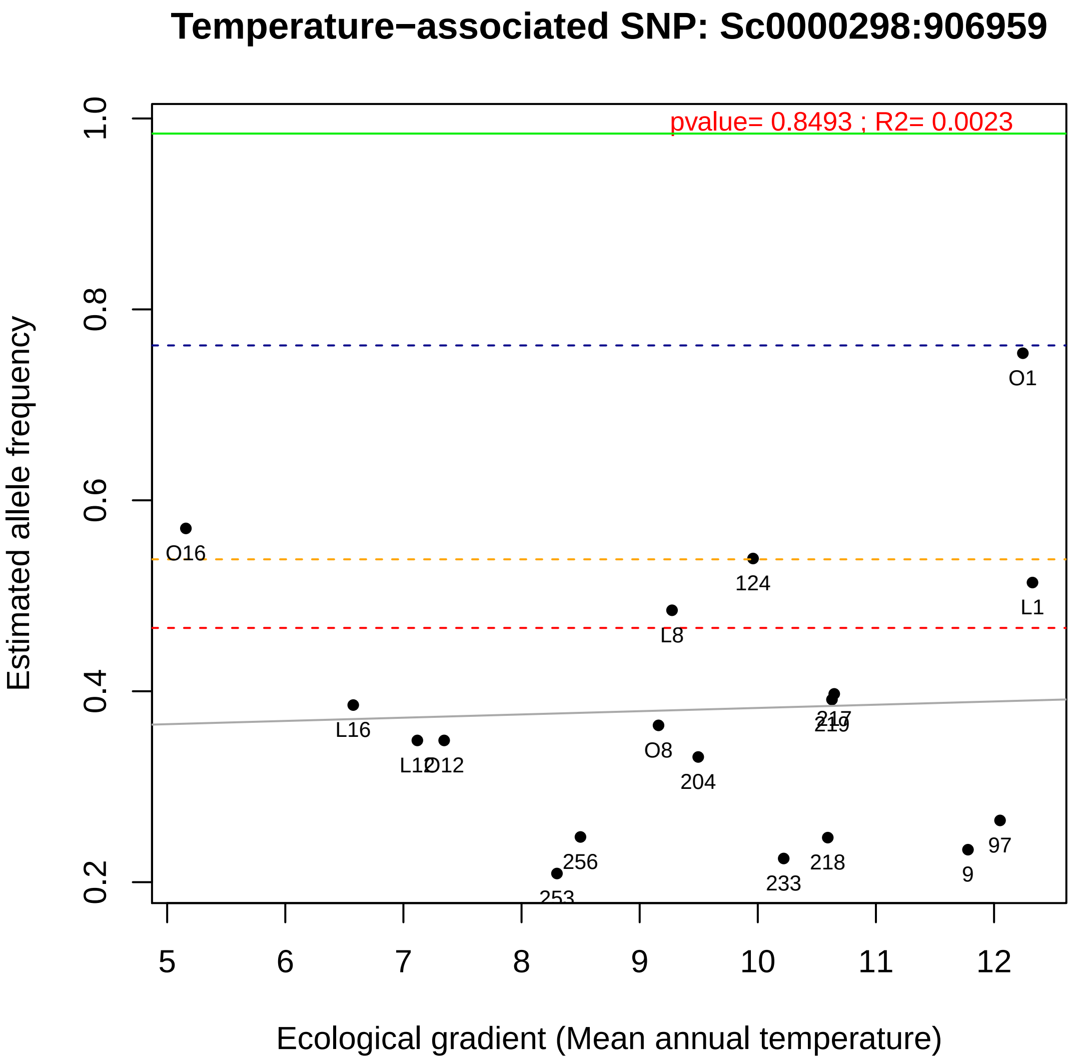

Temperature−associated SNP: Sc0000298:906959
pvalue= 0.8493 ; R2= 0.0023
Estimated allele frequency
Ecological gradient (Mean annual temperature)
0.2
0.4
0.6
0.8
1.0
5
6
7
8
9
10
11
12
O1
O16
124
L1
L8
L16
217
219
O8
L12
O12
204
256
253
218
233
97
9

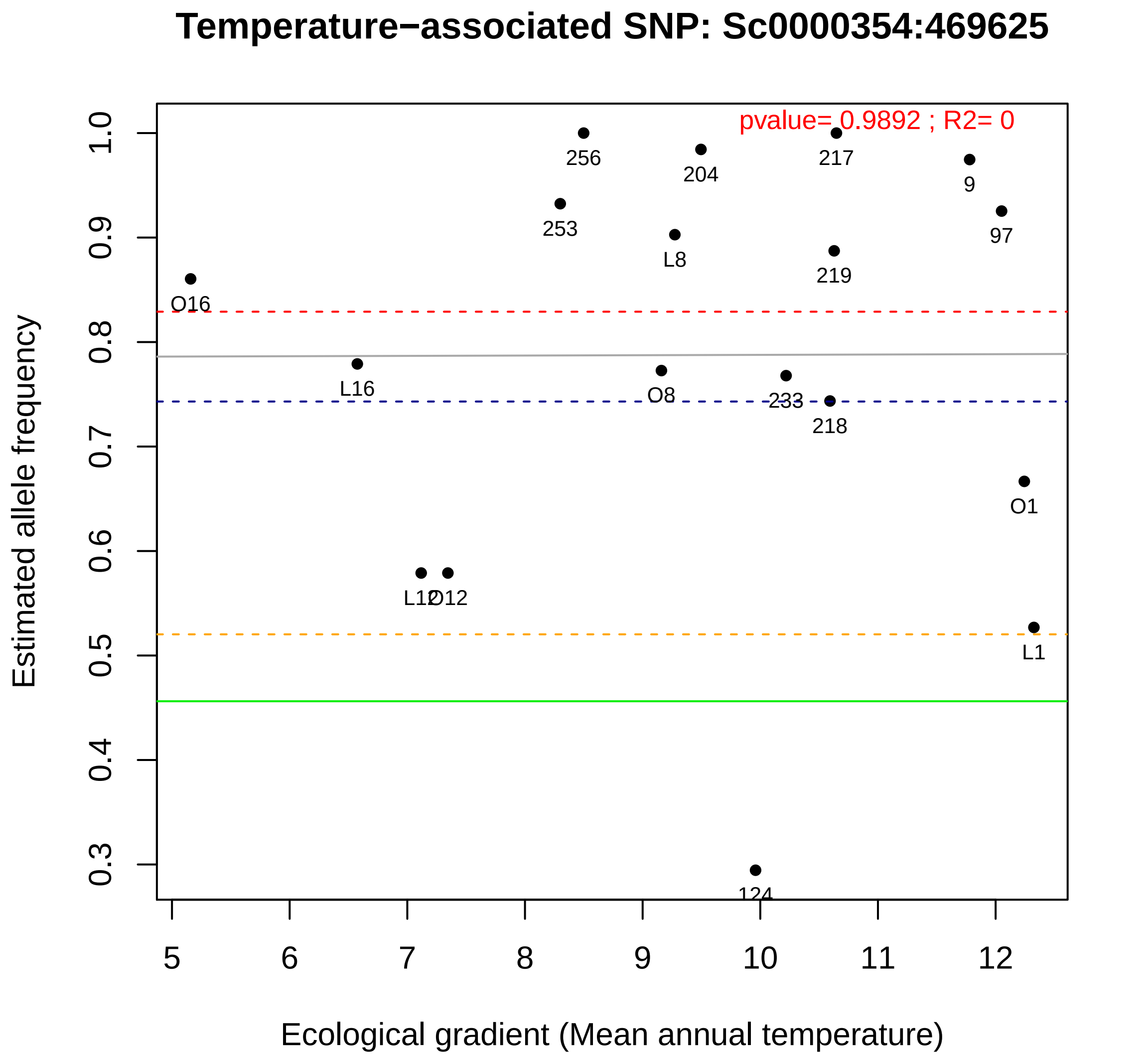
Temperature−associated SNP: Sc0000354:469625
pvalue= 0.9892 ; R2= 0
Estimated allele frequency
Ecological gradient (Mean annual temperature)
0.3
0.4
0.5
0.6
0.7
0.8
0.9
1.0
5
6
7
8
9
10
11
12
O16
L16
L12
O12
253
256
L8
O8
204
124
233
217
219
218
9
97
O1
L1

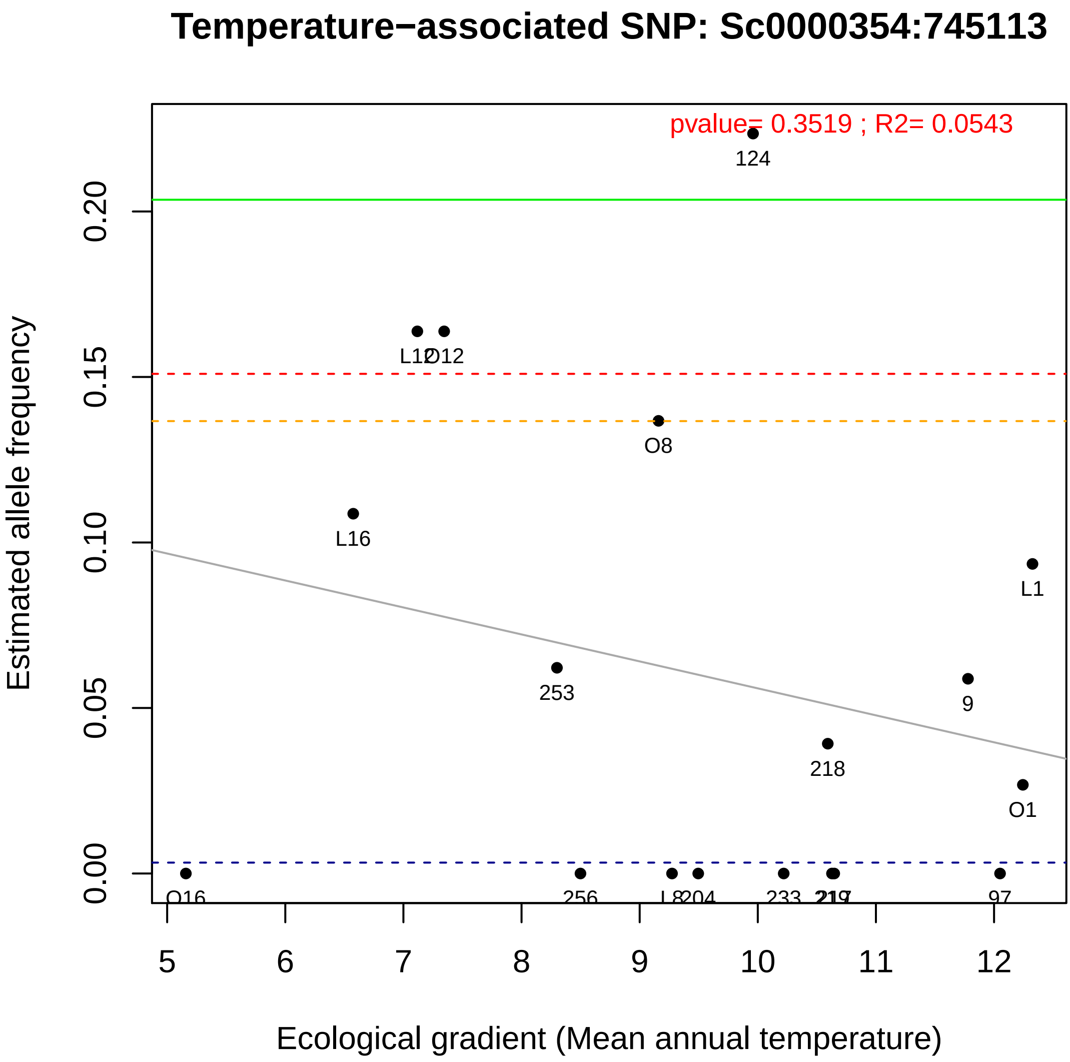
Temperature−associated SNP: Sc0000354:745113
pvalue= 0.3519 ; R2= 0.0543
Estimated allele frequency
0.00
0.05
0.10
0.15
0.20
5
6
7
8
9
10
11
12
Ecological gradient (Mean annual temperature)
124
O8
L16
253
9
L1
218
O1
O16
256
L8
204
233
97

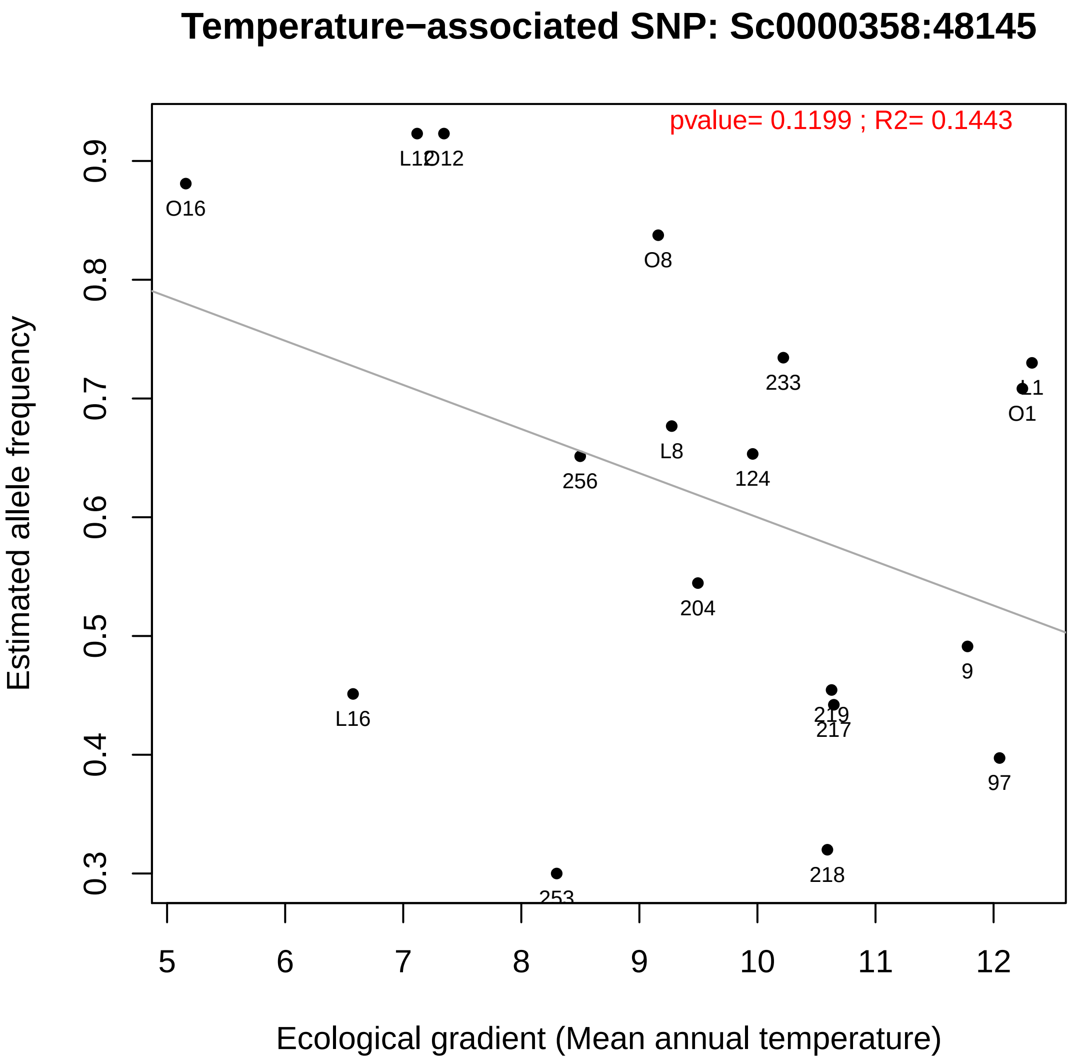
Temperature−associated SNP: Sc0000358:48145
pvalue= 0.1199 ; R2= 0.1443
Estimated allele frequency
0.3
0.4
0.5
0.6
0.7
0.8
0.9
5
6
7
8
9
10
11
12
Ecological gradient (Mean annual temperature)
O16
L12
O12
O8
233
L1
O1
L8
256
124
204
9
L16
219
217
97
218
253

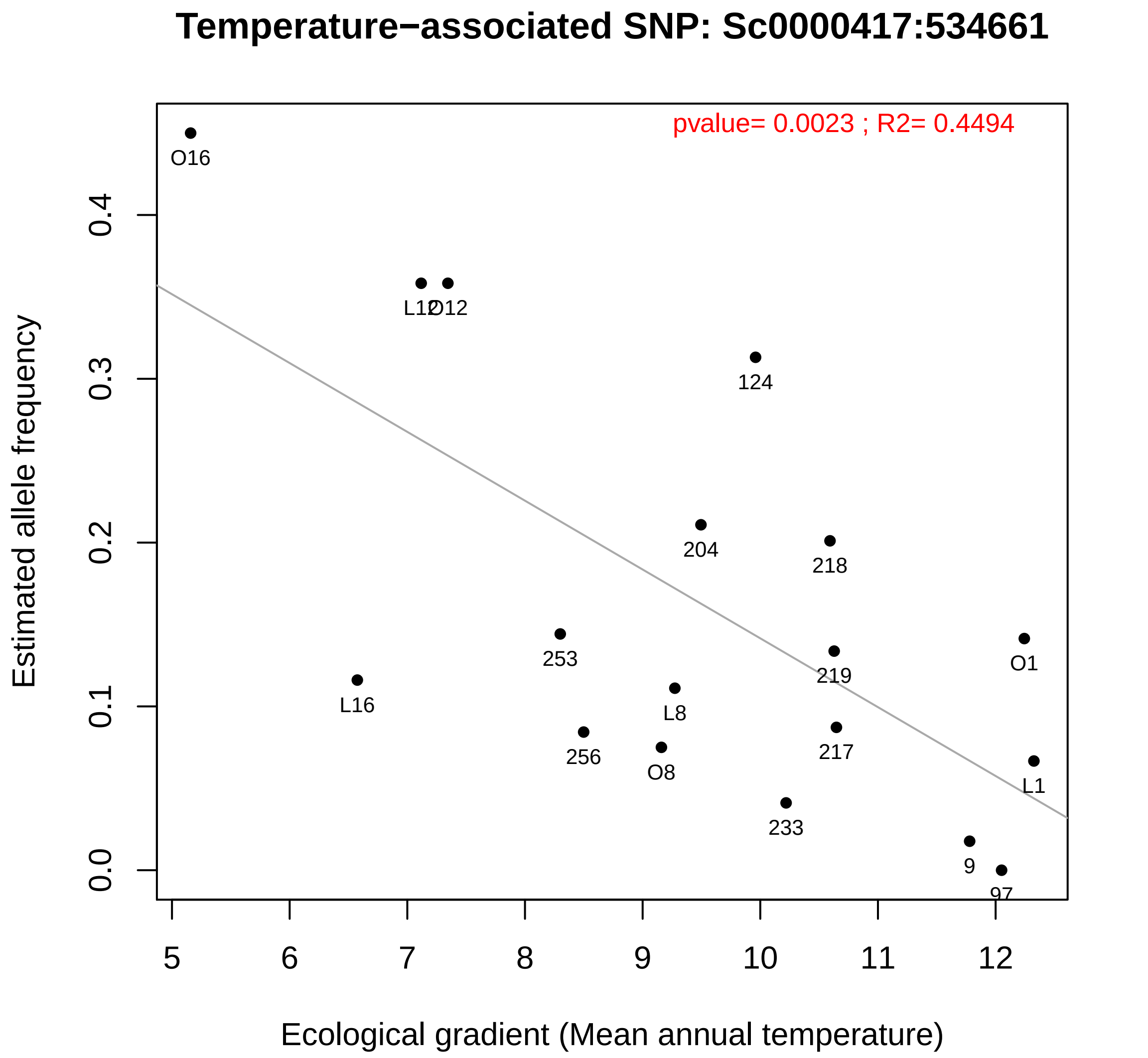

Temperature−associated SNP: Sc0000417:534661
pvalue= 0.0023 ; R2= 0.4494
Estimated allele frequency
0.0
0.1
0.2
0.3
0.4
5
6
7
8
9
10
11
12
Ecological gradient (Mean annual temperature)
O16
L12
O12
124
204
218
253
219
O1
L16
L8
256
217
O8
L1
233
9
97

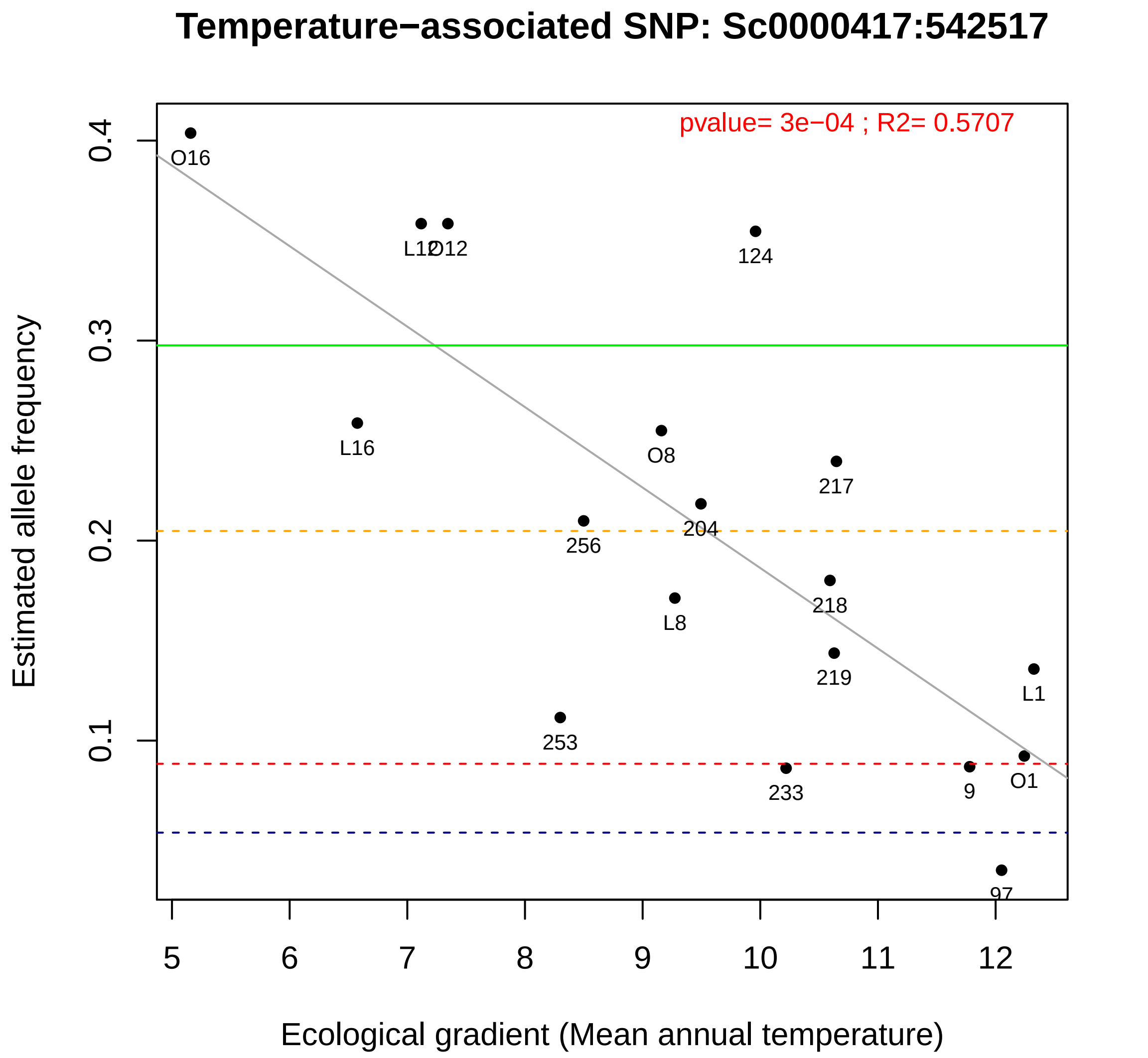
Temperature−associated SNP: Sc0000417:542517
pvalue= 3e−04 ; R2= 0.5707
O16
L12
O12
124
L16
O8
217
204
256
L8
218
219
L1
253
233
9
O1
97
0.1
0.2
0.3
0.4
Estimated allele frequency
5
6
7
8
9
10
11
12
Ecological gradient (Mean annual temperature)

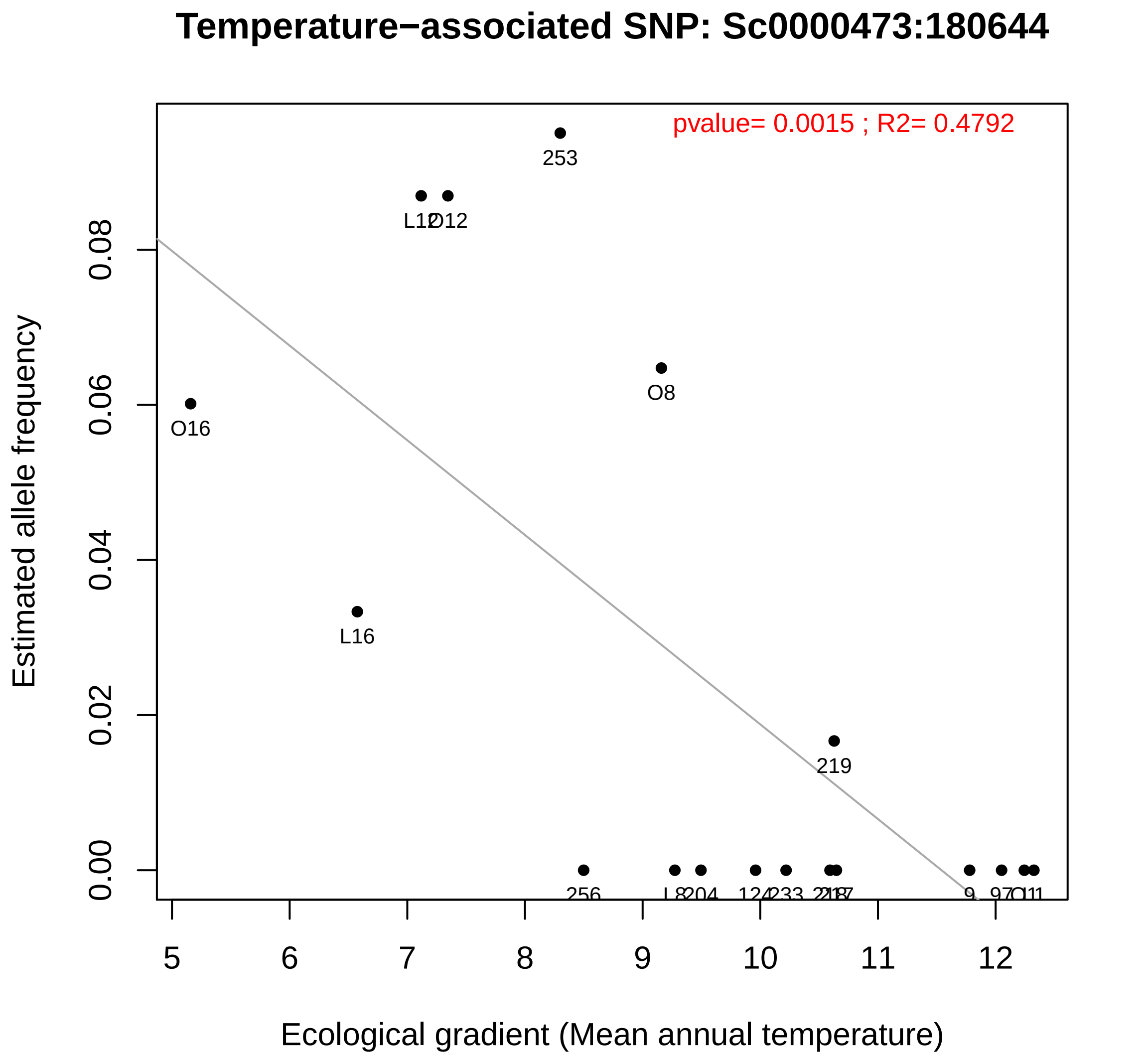
Temperature−associated SNP: Sc0000473:180644
pvalue= 0.0015 ; R2= 0.4792
Estimated allele frequency
Ecological gradient (Mean annual temperature)
0.00
0.02
0.04
0.06
0.08
5
6
7
8
9
10
11
12
253
O12
O8
O16
L16
219
256
204

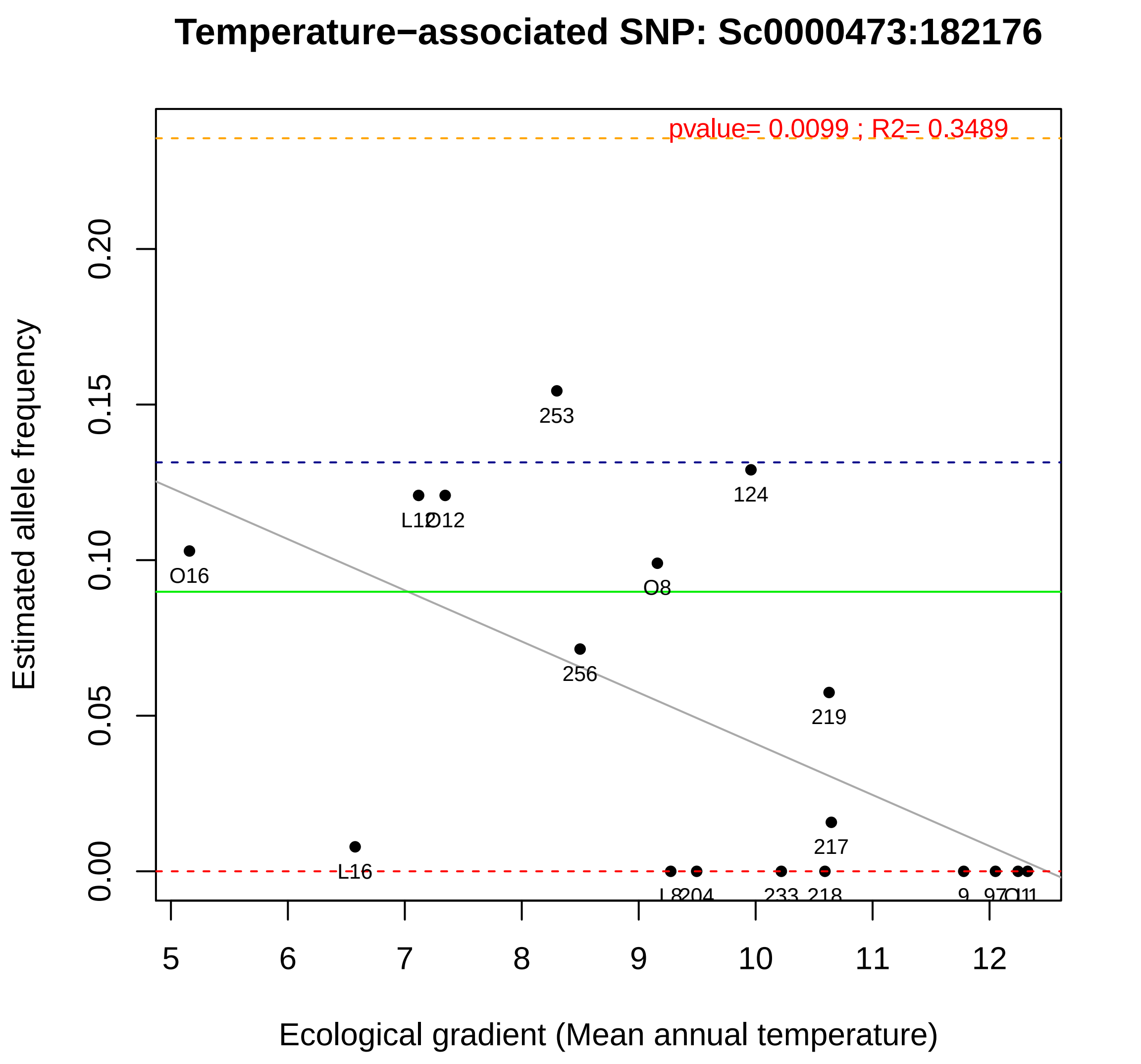
Temperature−associated SNP: Sc0000473:182176
pvalue= 0.0099 ; R2= 0.3489
Estimated allele frequency
0.00
0.05
0.10
0.15
0.20
5
6
7
8
9
10
11
12
Ecological gradient (Mean annual temperature)
O16
L16
L12
O12
253
256
O8
124
219
217
204
233
218

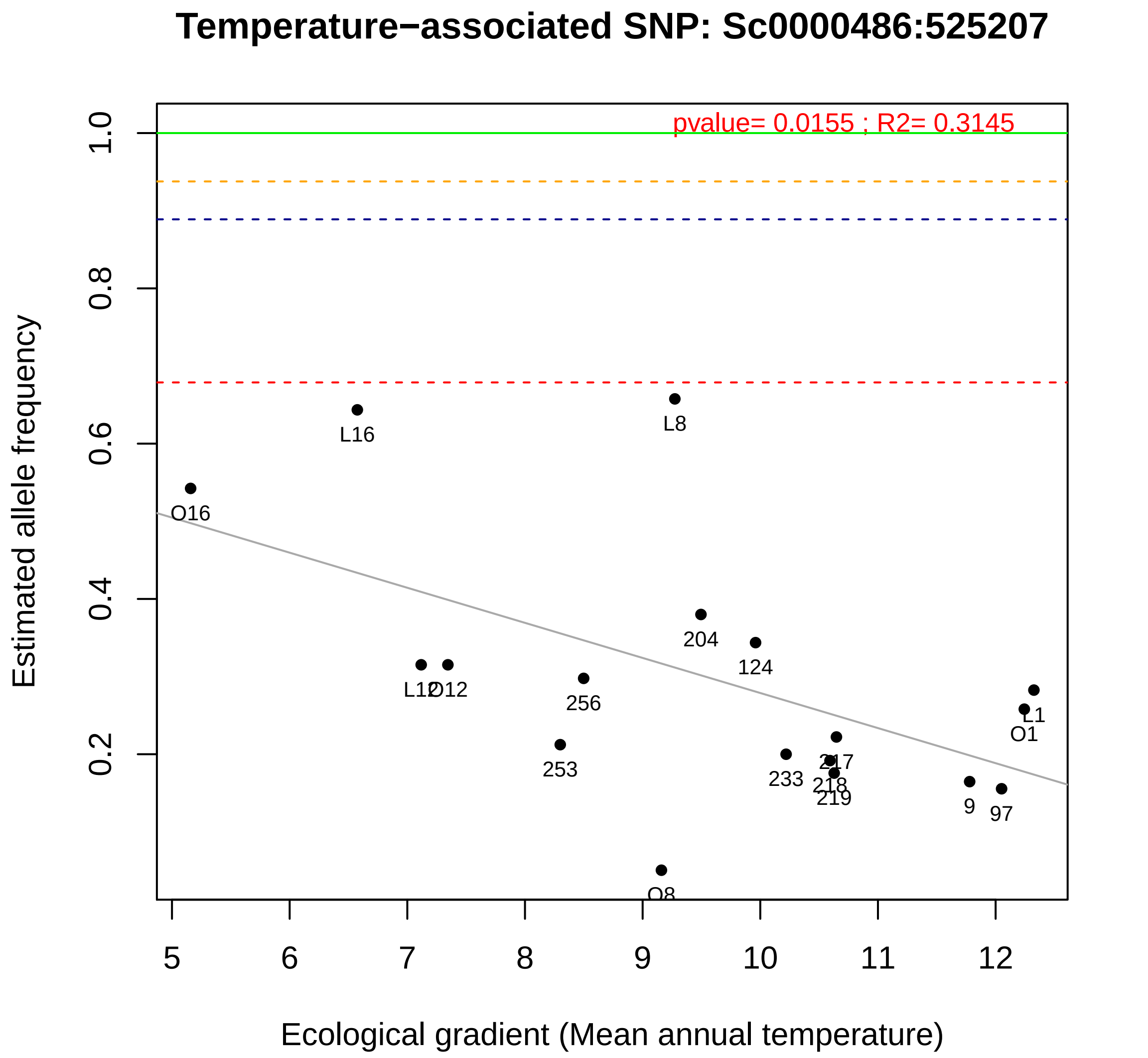

Temperature−associated SNP: Sc0000486:525207
pvalue= 0.0155 ; R2= 0.3145
Estimated allele frequency
0.2
0.4
0.6
0.8
1.0
5
6
7
8
9
10
11
12
Ecological gradient (Mean annual temperature)
O16
L16
L8
204
124
L12
O12
256
253
233
217
218
219
9
97
L1
O1
O8

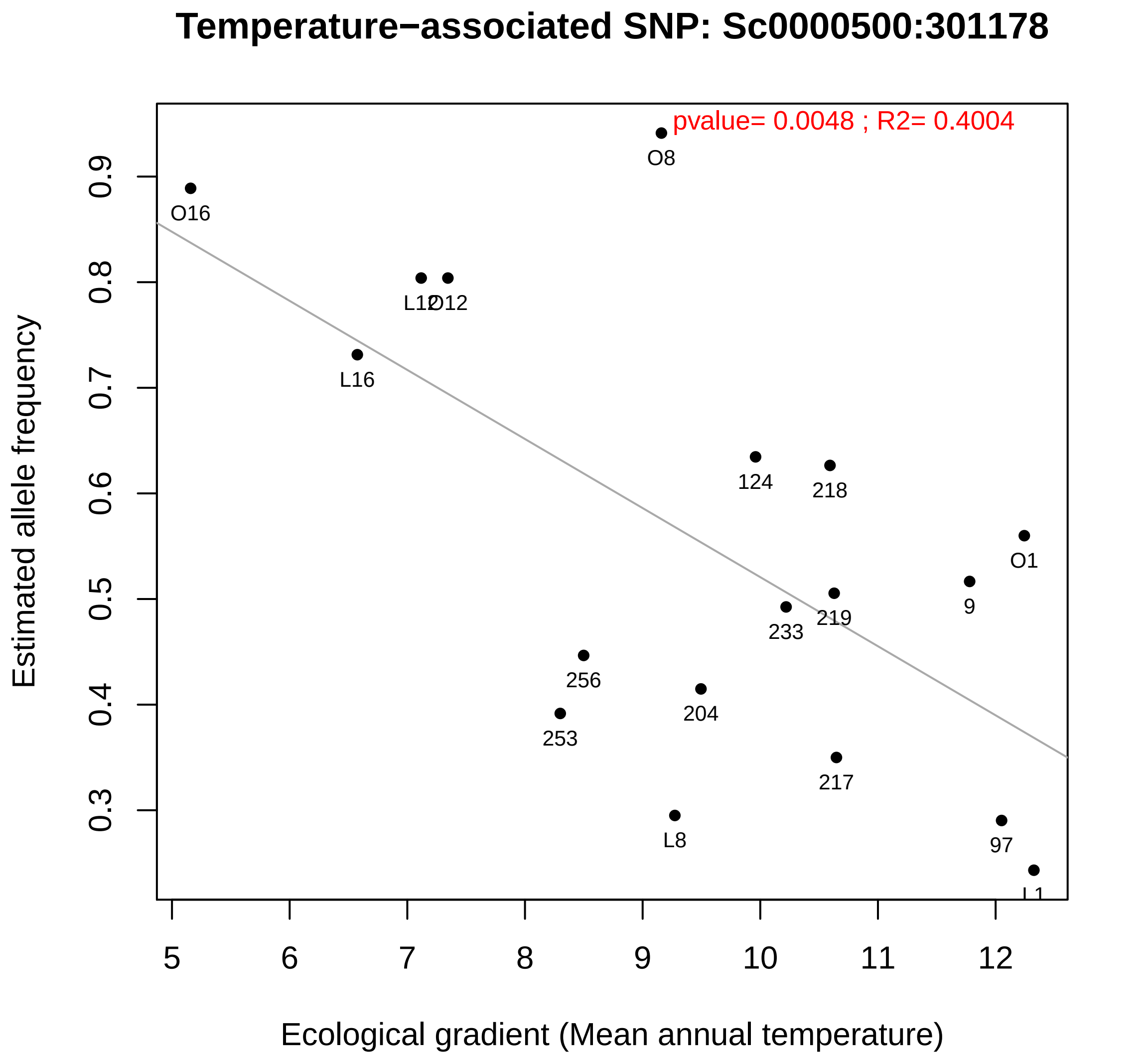

Temperature−associated SNP: Sc0000500:301178
pvalue= 0.0048 ; R2= 0.4004
Estimated allele frequency
Ecological gradient (Mean annual temperature)
0.3
0.4
0.5
0.6
0.7
0.8
0.9
5
6
7
8
9
10
11
12
O8
O16
L12
O12
L16
124
218
O1
9
219
233
256
204
253
217
L8
97

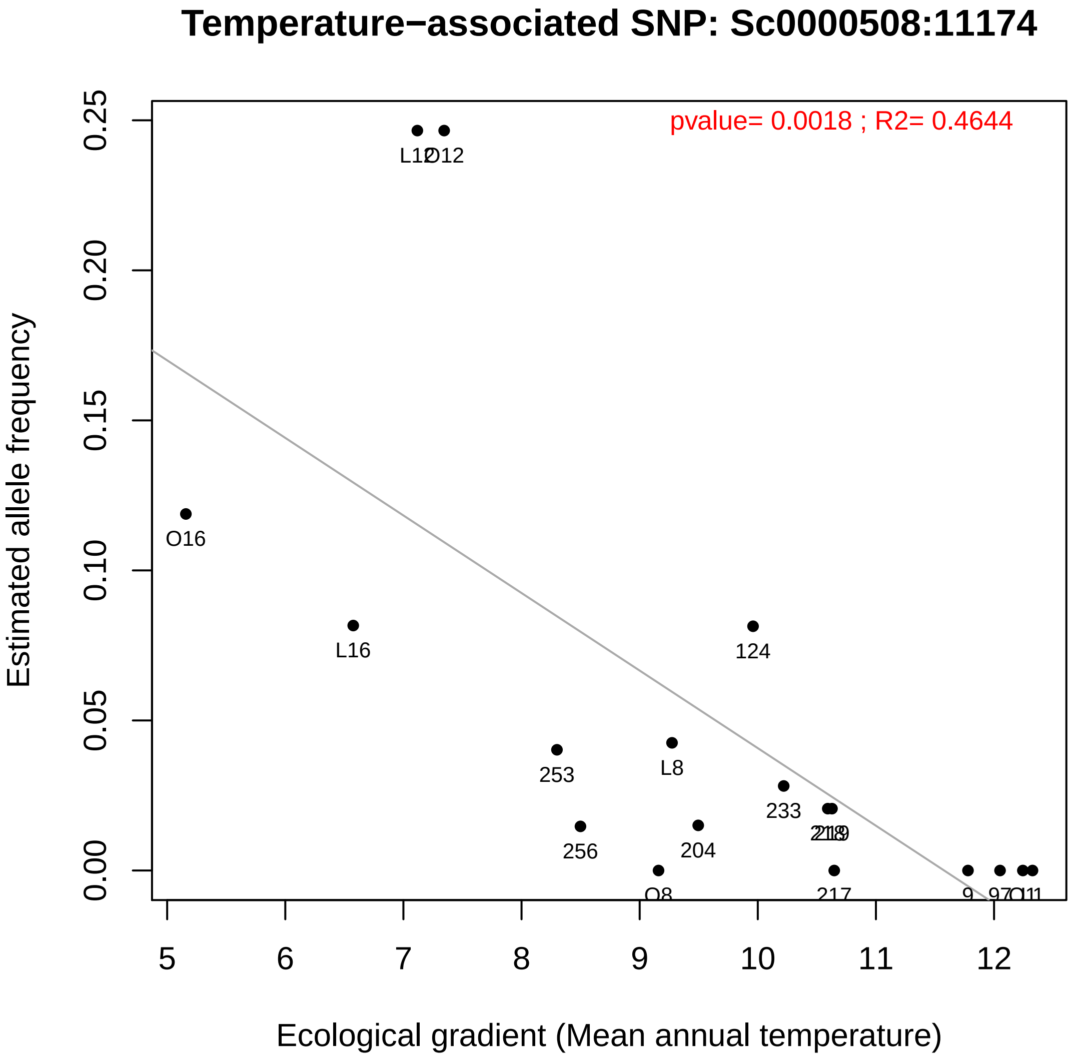
Temperature−associated SNP: Sc0000508:11174
pvalue= 0.0018 ; R2= 0.4644
Estimated allele frequency
Ecological gradient (Mean annual temperature)
0.00
0.05
0.10
0.15
0.20
0.25
5
6
7
8
9
10
11
12
L12
O12
O16
L16
124
253
L8
233
256
204
217
O8

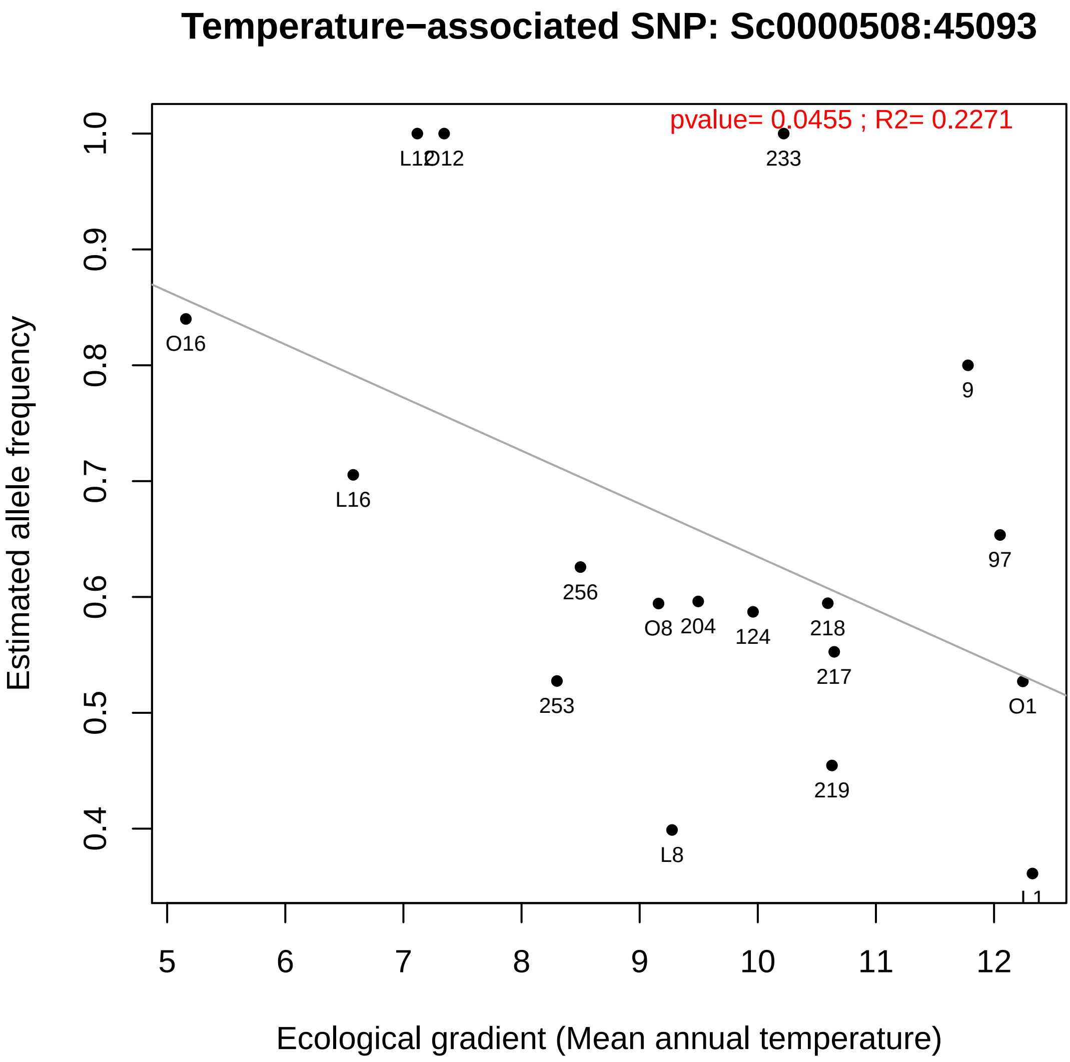

Temperature−associated SNP: Sc0000508:45093
pvalue= 0.0455 ; R2= 0.2271
Estimated allele frequency
Ecological gradient (Mean annual temperature)
0.4
0.5
0.6
0.7
0.8
0.9
1.0
5
6
7
8
9
10
11
12
L12
O12
233
O16
9
L16
97
256
O8
204
124
218
217
O1
253
219
L8

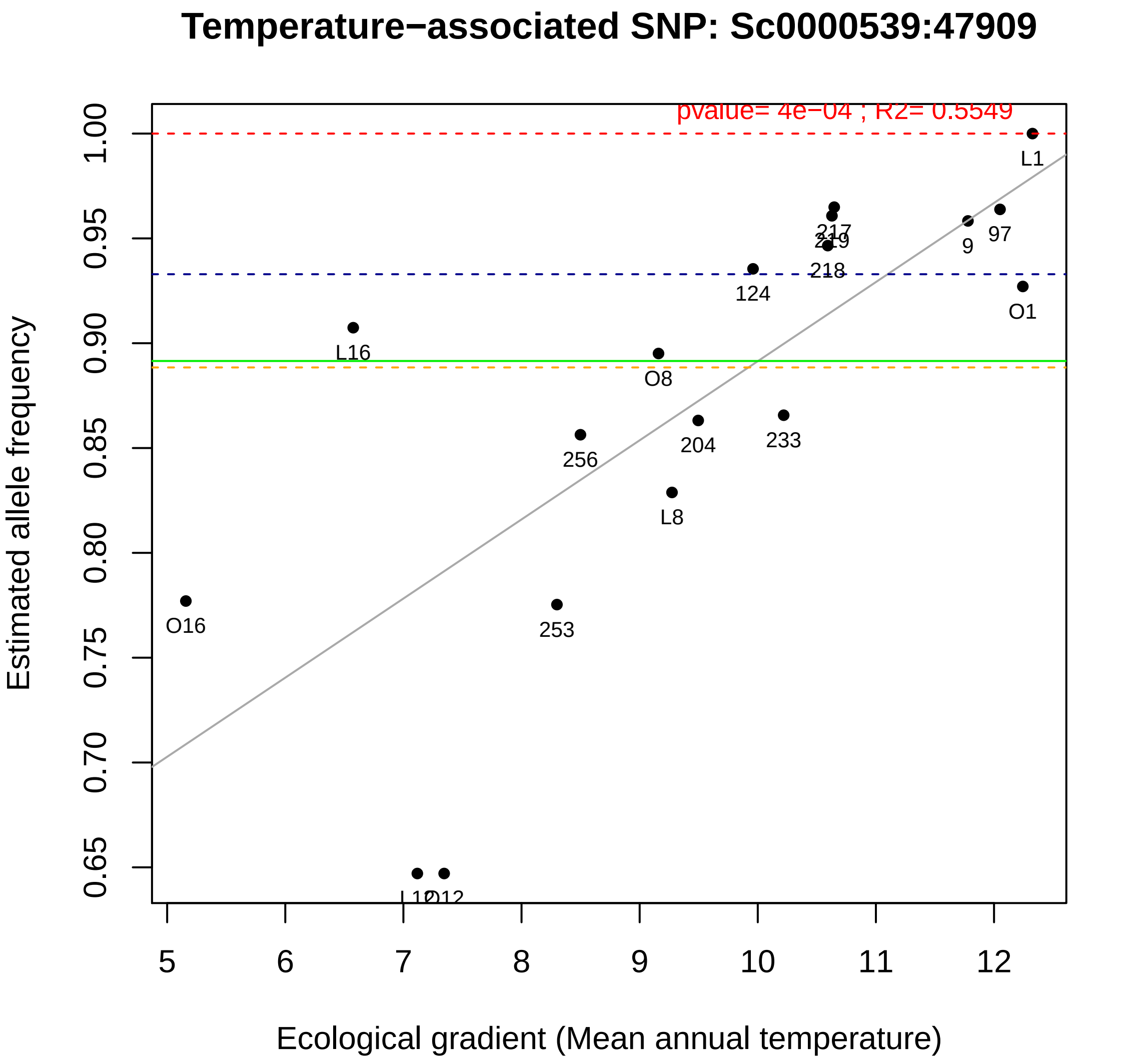

Temperature−associated SNP: Sc0000539:47909
pvalue= 4e−04 ; R2= 0.5549
Estimated allele frequency
Ecological gradient (Mean annual temperature)
1.00
0.95
0.90
0.85
0.80
0.75
0.70
0.65
5
6
7
8
9
10
11
12
L1
217
219
218
9
97
124
O1
L16
O8
256
204
233
L8
O16
253
O12

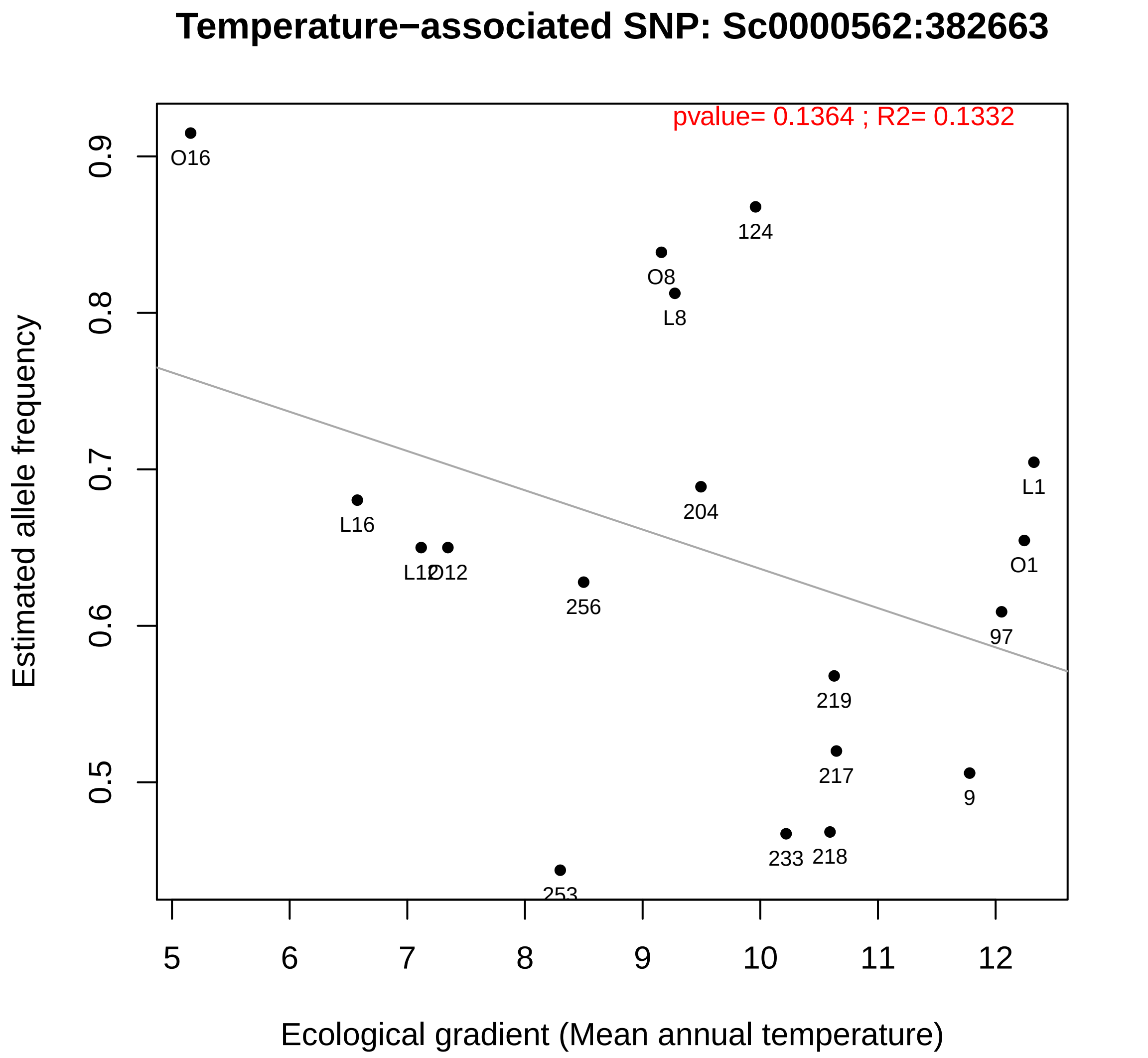

Temperature−associated SNP: Sc0000562:382663
pvalue= 0.1364 ; R2= 0.1332
Estimated allele frequency
0.5
0.6
0.7
0.8
0.9
Ecological gradient (Mean annual temperature)
5
6
7
8
9
10
11
12
O16
124
O8
L8
L1
204
L16
L12
O12
O1
256
97
219
217
9
233
218
253

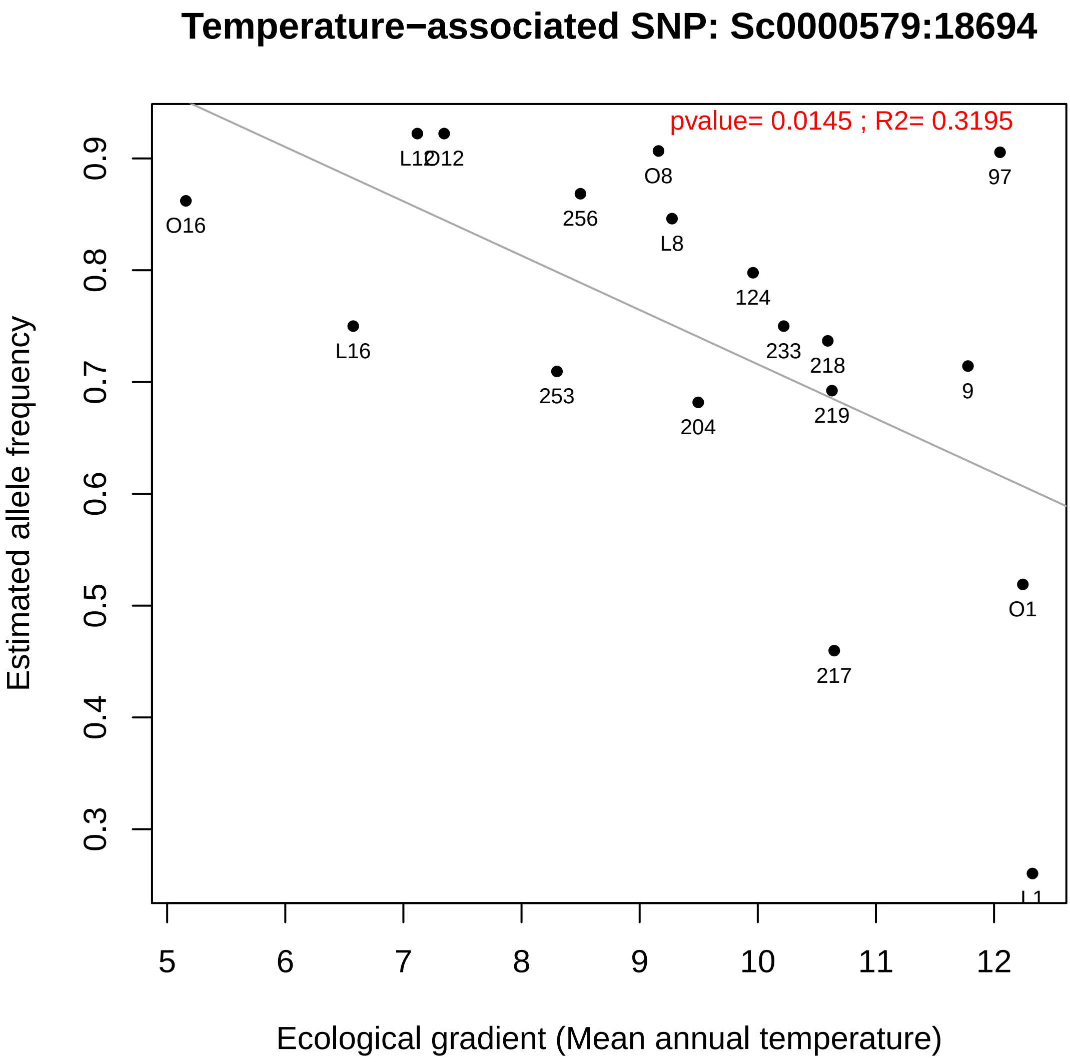
Temperature−associated SNP: Sc0000579:18694
pvalue= 0.0145 ; R2= 0.3195
Estimated allele frequency
0.3
0.4
0.5
0.6
0.7
0.8
0.9
5
6
7
8
9
10
11
12
Ecological gradient (Mean annual temperature)
L12
O12
O8
97
O16
256
L8
124
L16
233
218
9
253
219
204
O1
217

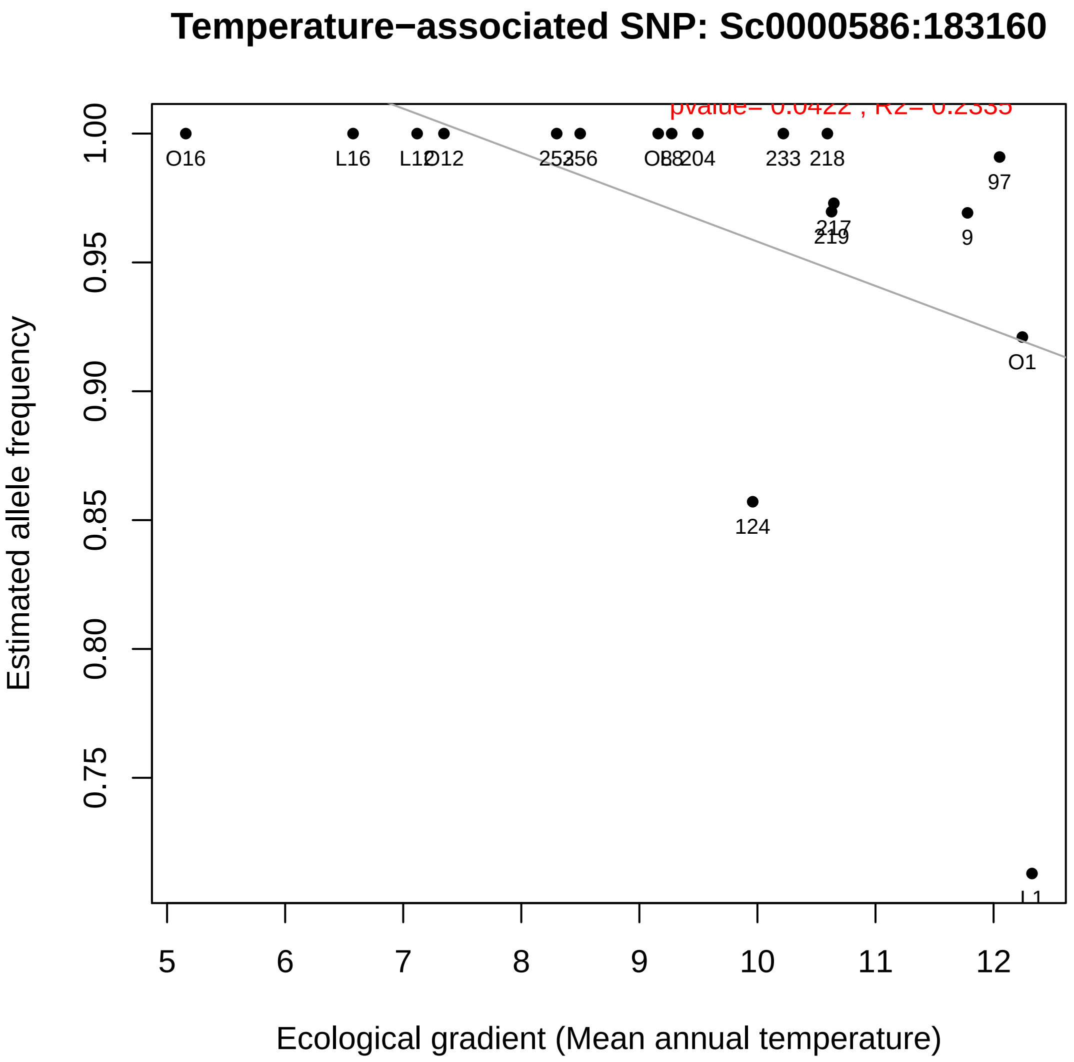
Temperature−associated SNP: Sc0000586:183160
pvalue= 0.0422 , R2= 0.2335
O16
L16
O12
256
204
233
218
97
9
124
O1
Estimated allele frequency
Ecological gradient (Mean annual temperature)
1.00
0.95
0.90
0.85
0.80
0.75
5
6
7
8
9
10
11
12

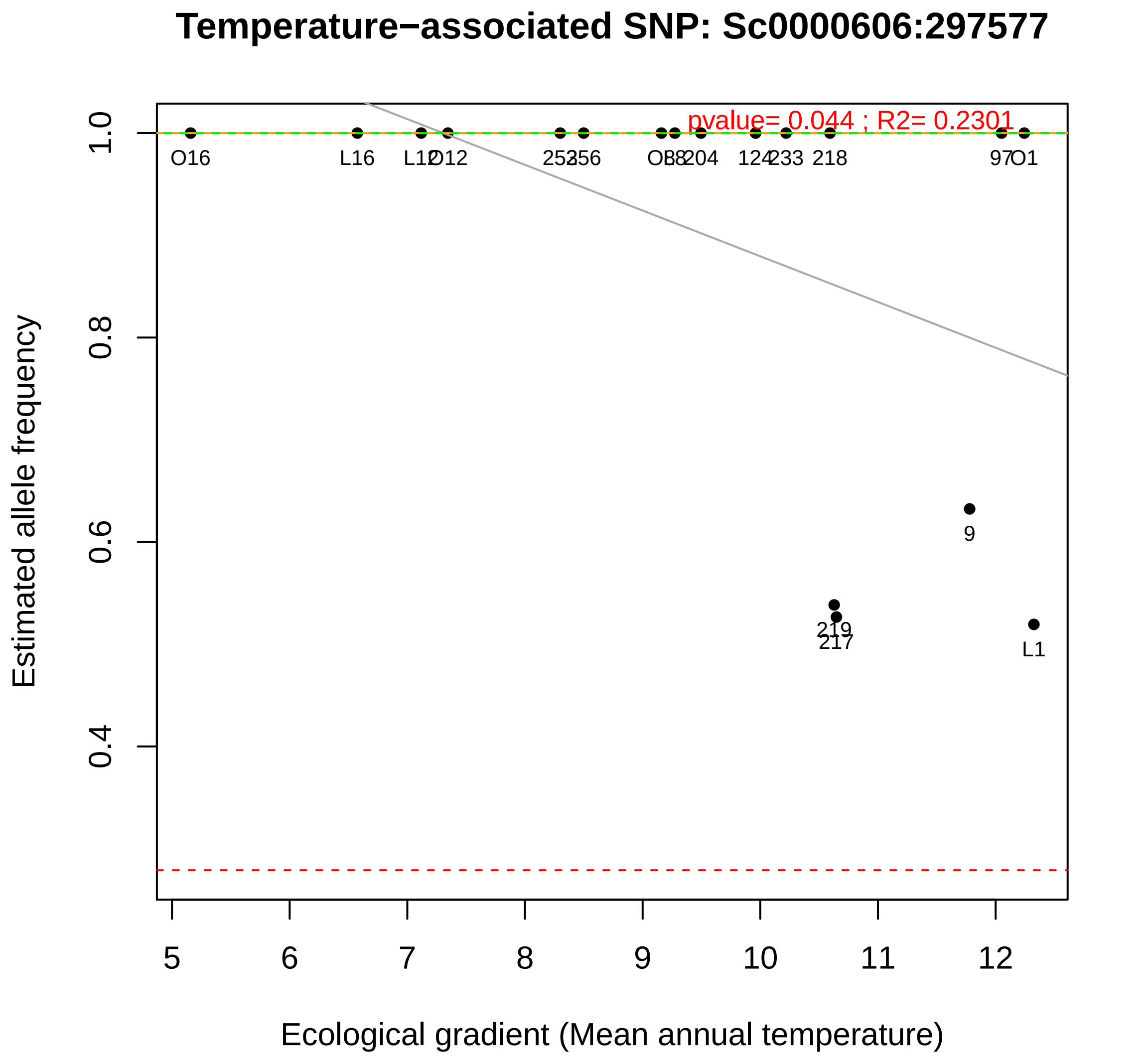

Temperature−associated SNP: Sc0000606:297577
pvalue= 0.044 ; R2= 0.2301
Estimated allele frequency
Ecological gradient (Mean annual temperature)
1.0
0.8
0.6
0.4
5
6
7
8
9
10
11
12
O16
L16
O12
218
9
219
217
L1

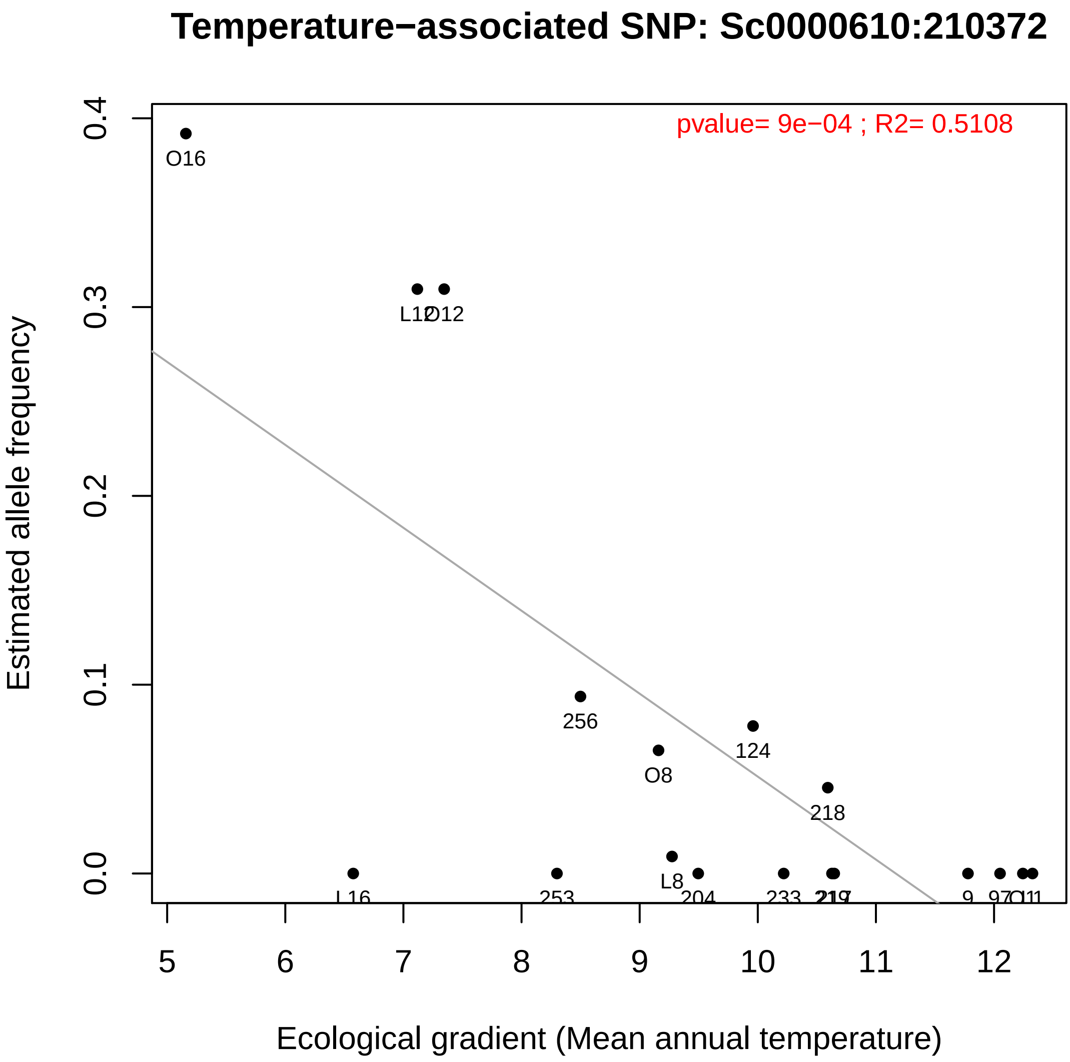

Temperature−associated SNP: Sc0000610:210372
pvalue= 9e−04 ; R2= 0.5108
Estimated allele frequency
0.0
0.1
0.2
0.3
0.4
Ecological gradient (Mean annual temperature)
5
6
7
8
9
10
11
12
O16
O12
256
124
O8
218
L8

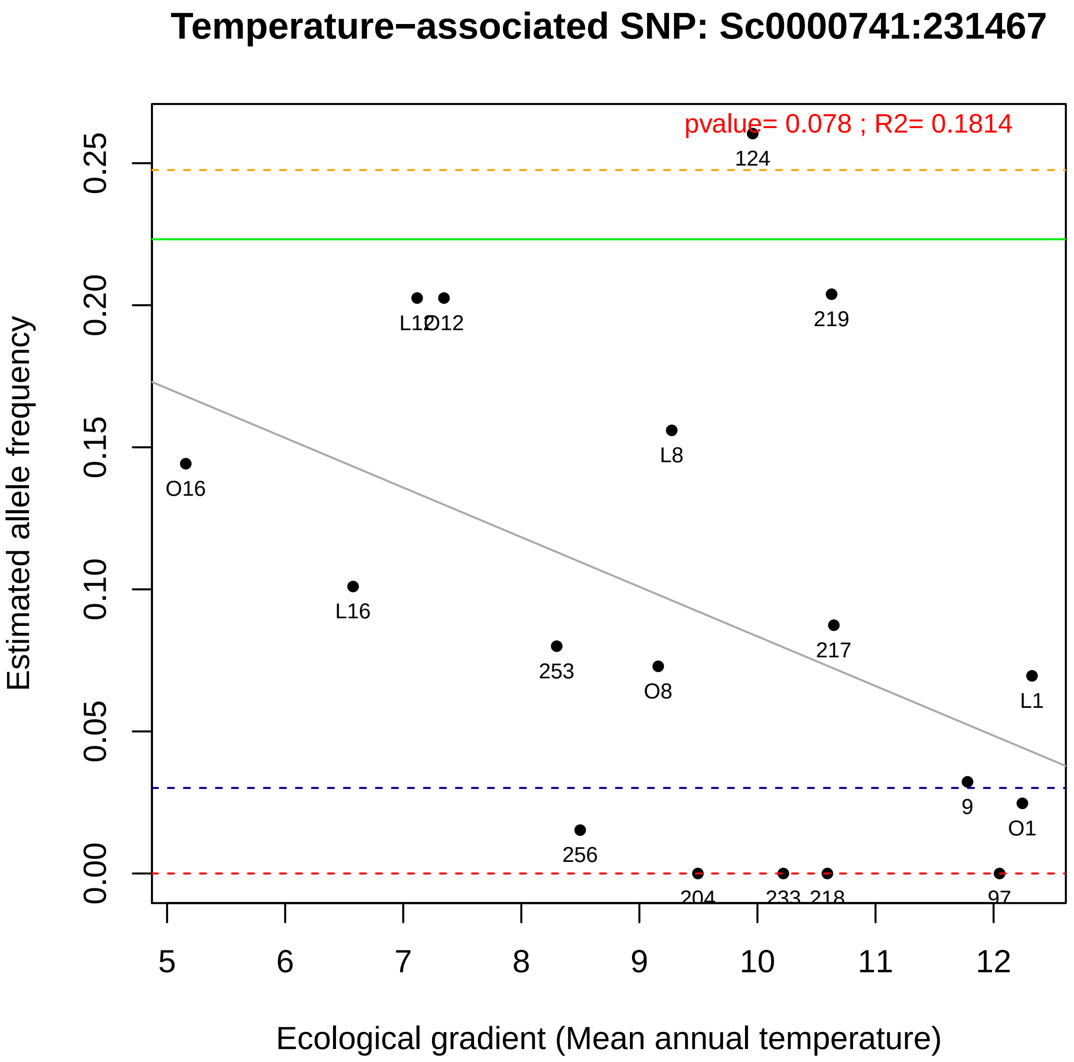

Temperature−associated SNP: Sc0000741:231467
pvalue= 0.078 ; R2= 0.1814
Estimated allele frequency
0.00
0.05
0.10
0.15
0.20
0.25
5
6
7
8
9
10
11
12
Ecological gradient (Mean annual temperature)
124
L12
O12
219
L8
O16
L16
253
O8
217
L1
9
O1
256
204
233
218
97

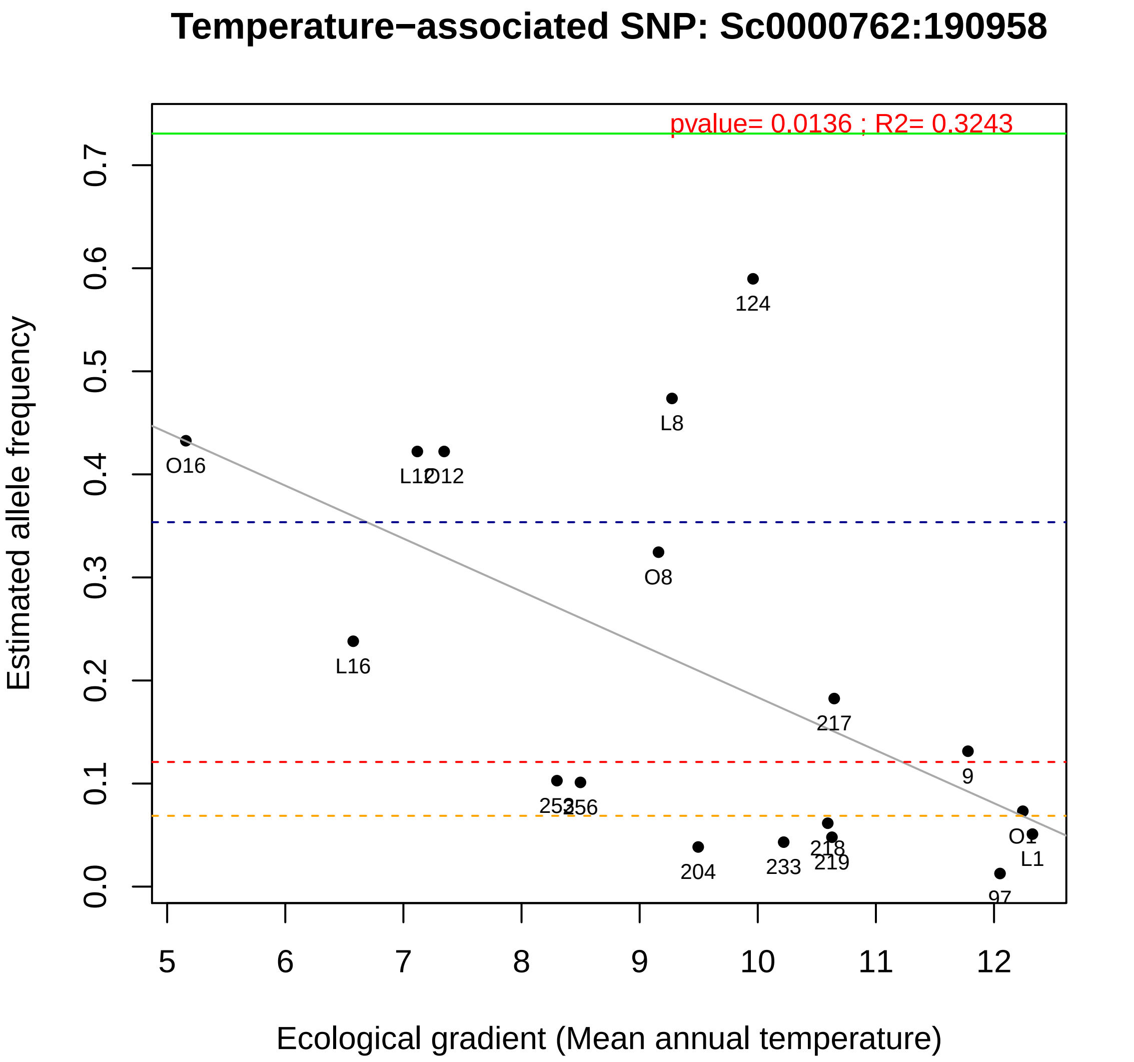
Temperature−associated SNP: Sc0000762:190958
pvalue= 0.0136 ; R2= 0.3243
Estimated allele frequency
0.0
0.1
0.2
0.3
0.4
0.5
0.6
0.7
5
6
7
8
9
10
11
12
Ecological gradient (Mean annual temperature)
O16
L12
O12
L16
252
256
L8
O8
124
204
233
217
218
219
9
97
O1
L1

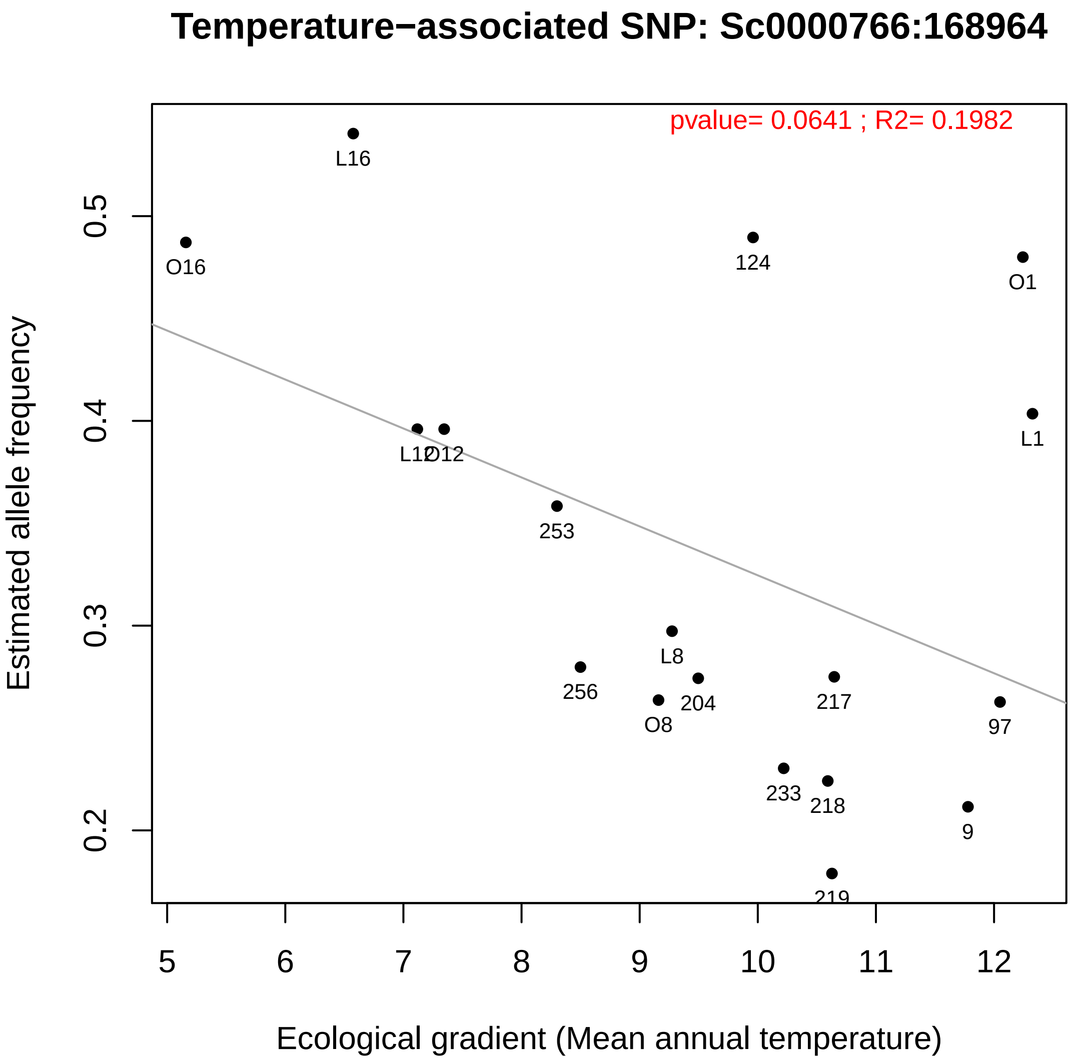

Temperature−associated SNP: Sc0000766:168964
pvalue= 0.0641 ; R2= 0.1982
L16
O16
124
O1
L12
O12
L1
253
L8
256
204
217
O8
97
233
218
9
219
0.5
0.4
0.3
0.2
5
6
7
8
9
10
11
12
Estimated allele frequency
Ecological gradient (Mean annual temperature)

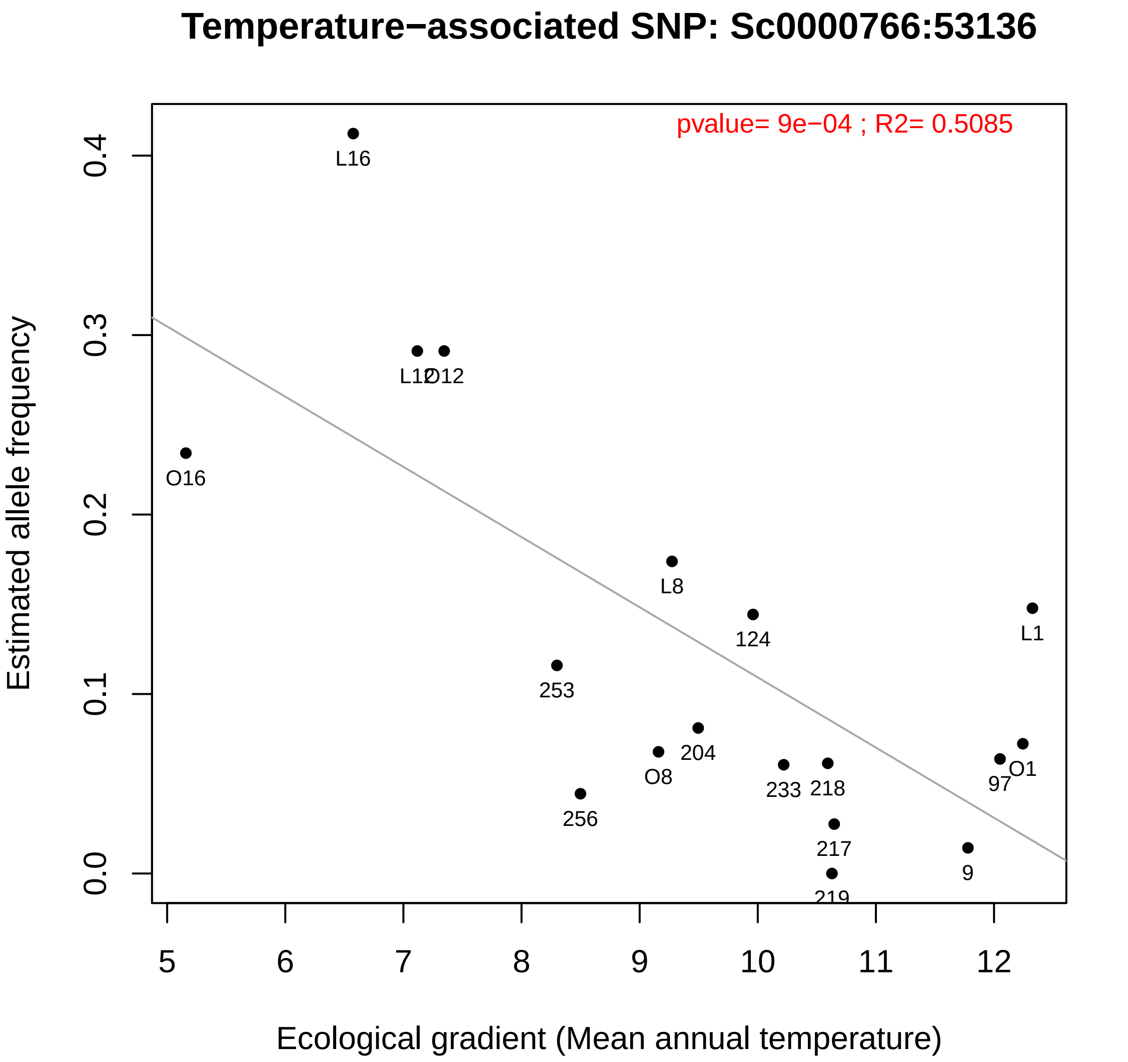
Temperature−associated SNP: Sc0000766:53136
pvalue= 9e−04 ; R2= 0.5085
Estimated allele frequency
0.0
0.1
0.2
0.3
0.4
Ecological gradient (Mean annual temperature)
5
6
7
8
9
10
11
12
L16
L12
O12
O16
L8
124
L1
253
204
O8
233
218
97
O1
256
217
9
219

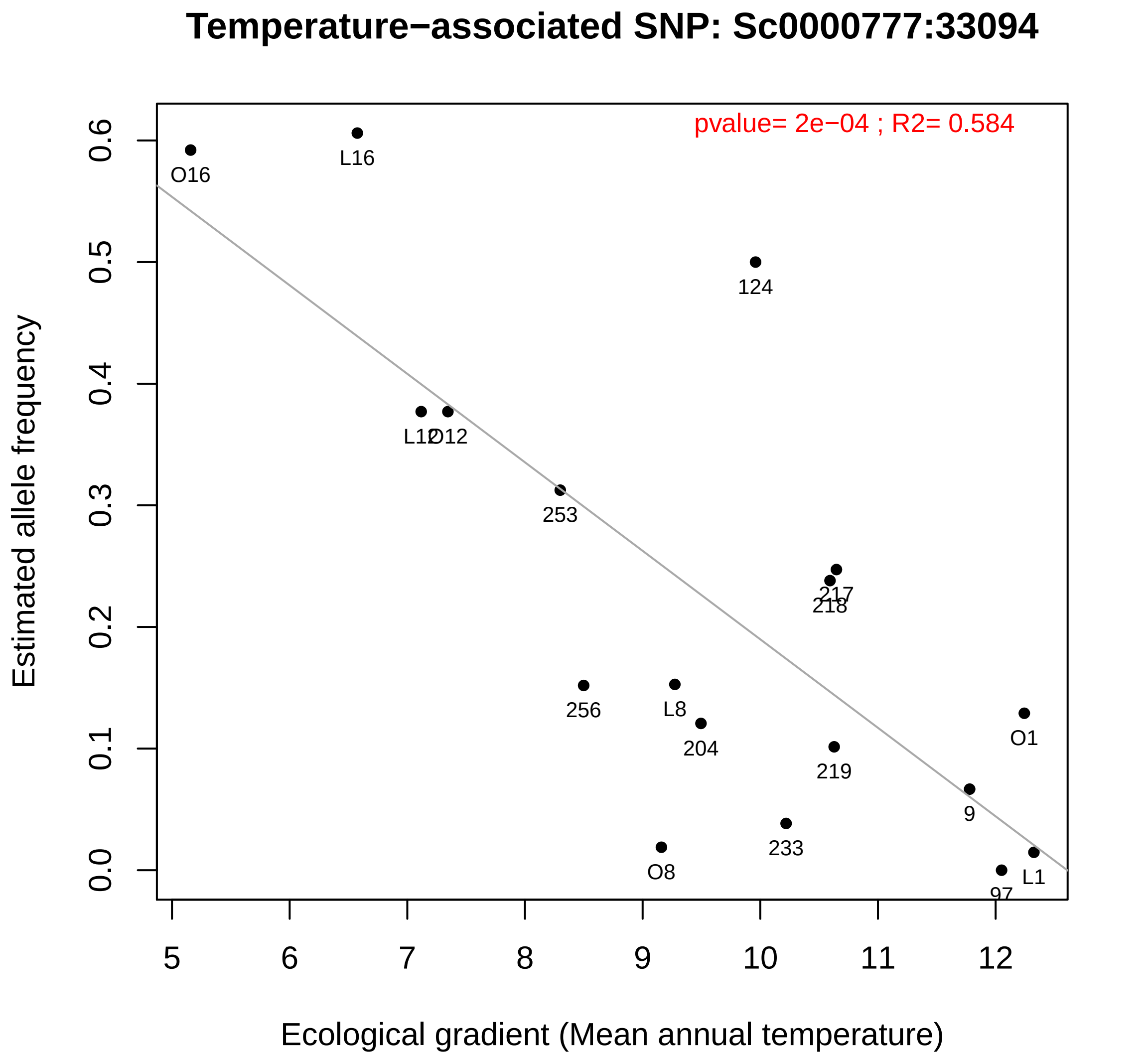
Temperature−associated SNP: Sc0000777:33094
pvalue= 2e−04 ; R2= 0.584
Estimated allele frequency
Ecological gradient (Mean annual temperature)
0.0
0.1
0.2
0.3
0.4
0.5
0.6
5
6
7
8
9
10
11
12
O16
L16
124
L12
O12
253
217
218
256
L8
204
219
O1
9
233
O8
L1
97

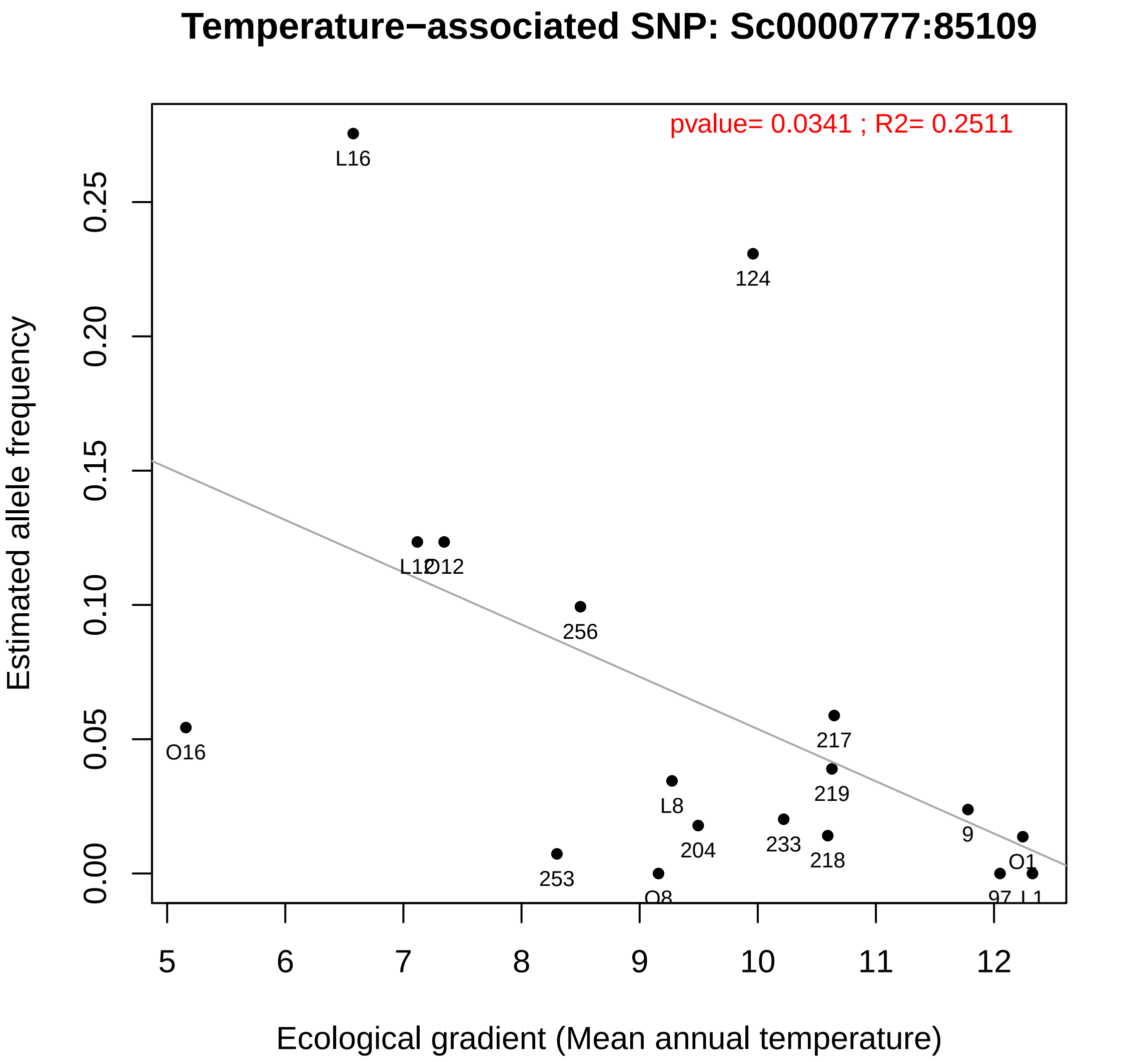
Temperature−associated SNP: Sc0000777:85109
pvalue= 0.0341 ; R2= 0.2511
Estimated allele frequency
0.00
0.05
0.10
0.15
0.20
0.25
5
6
7
8
9
10
11
12
Ecological gradient (Mean annual temperature)
L16
124
L12
O12
256
O16
217
219
L8
204
233
218
9
O1
253
O8
97
L1

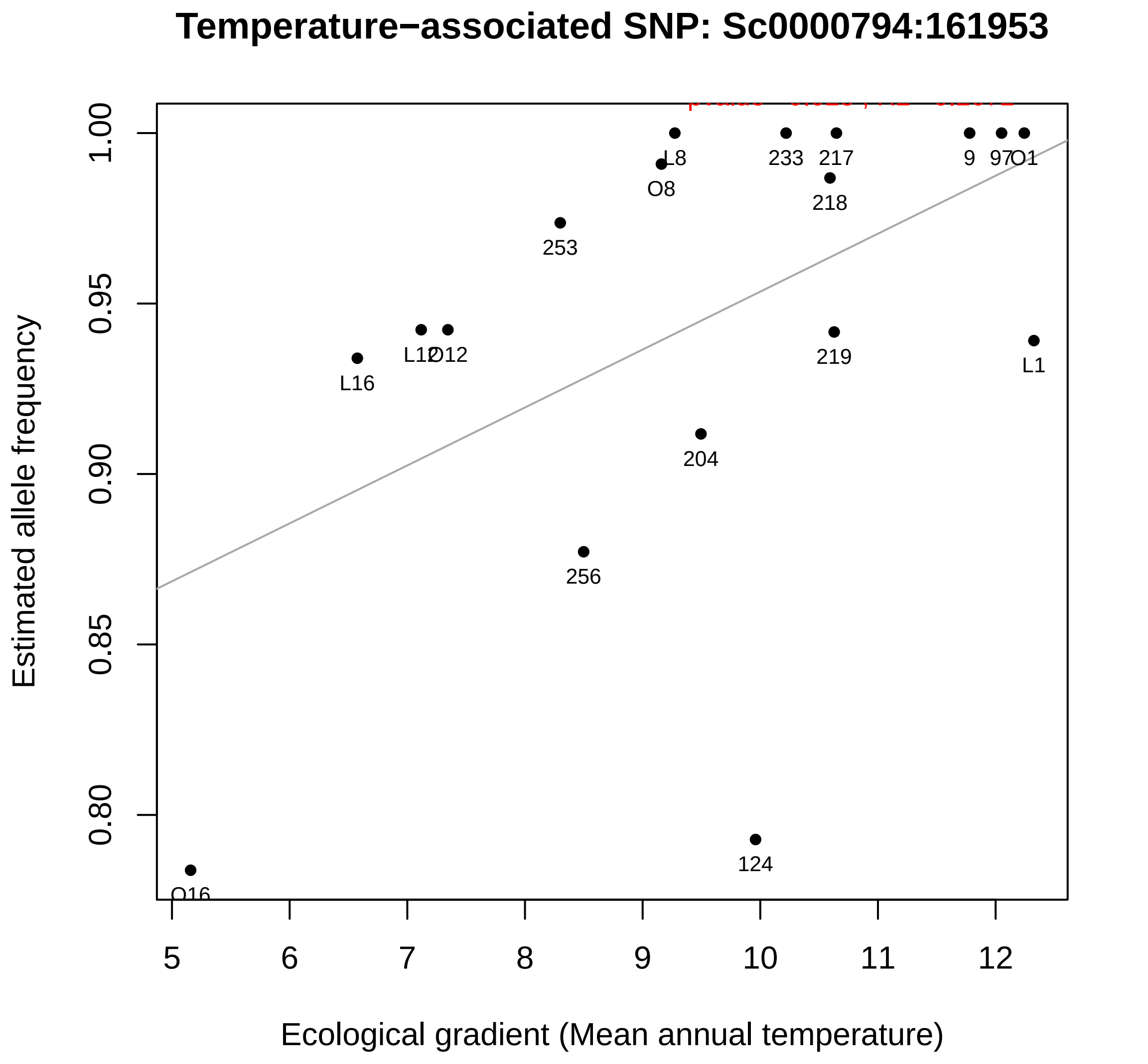

Temperature−associated SNP: Sc0000794:161953
Estimated allele frequency
Ecological gradient (Mean annual temperature)
1.00
0.95
0.90
0.85
0.80
5
6
7
8
9
10
11
12
L8
O8
233
217
218
9
97
O1
253
L12
O12
L16
219
L1
204
256
124
O16

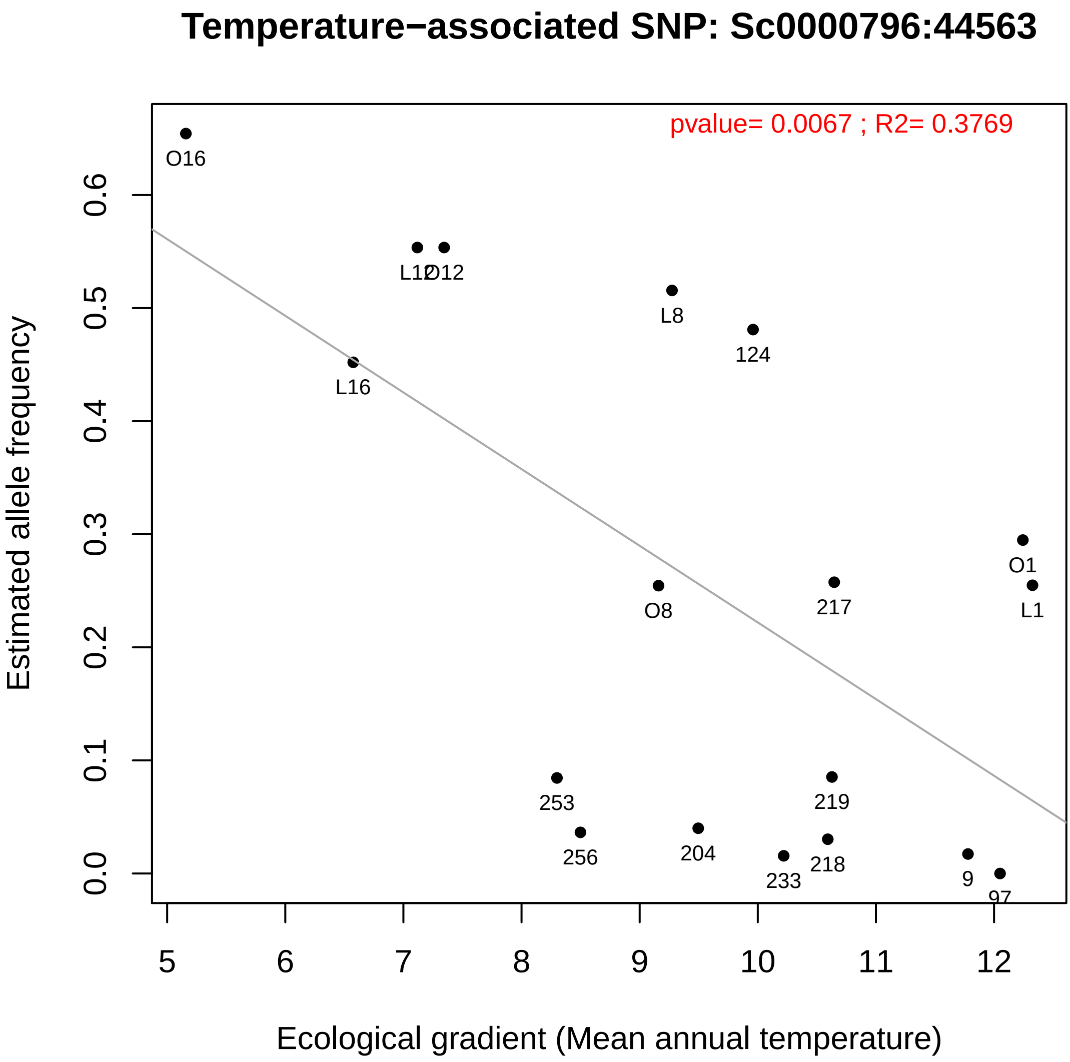

Temperature−associated SNP: Sc0000796:44563
pvalue= 0.0067 ; R2= 0.3769
Estimated allele frequency
0.0
0.1
0.2
0.3
0.4
0.5
0.6
5
6
7
8
9
10
11
12
Ecological gradient (Mean annual temperature)
O16
L12
O12
L8
124
L16
O1
L1
O8
217
253
219
256
204
218
233
9
97

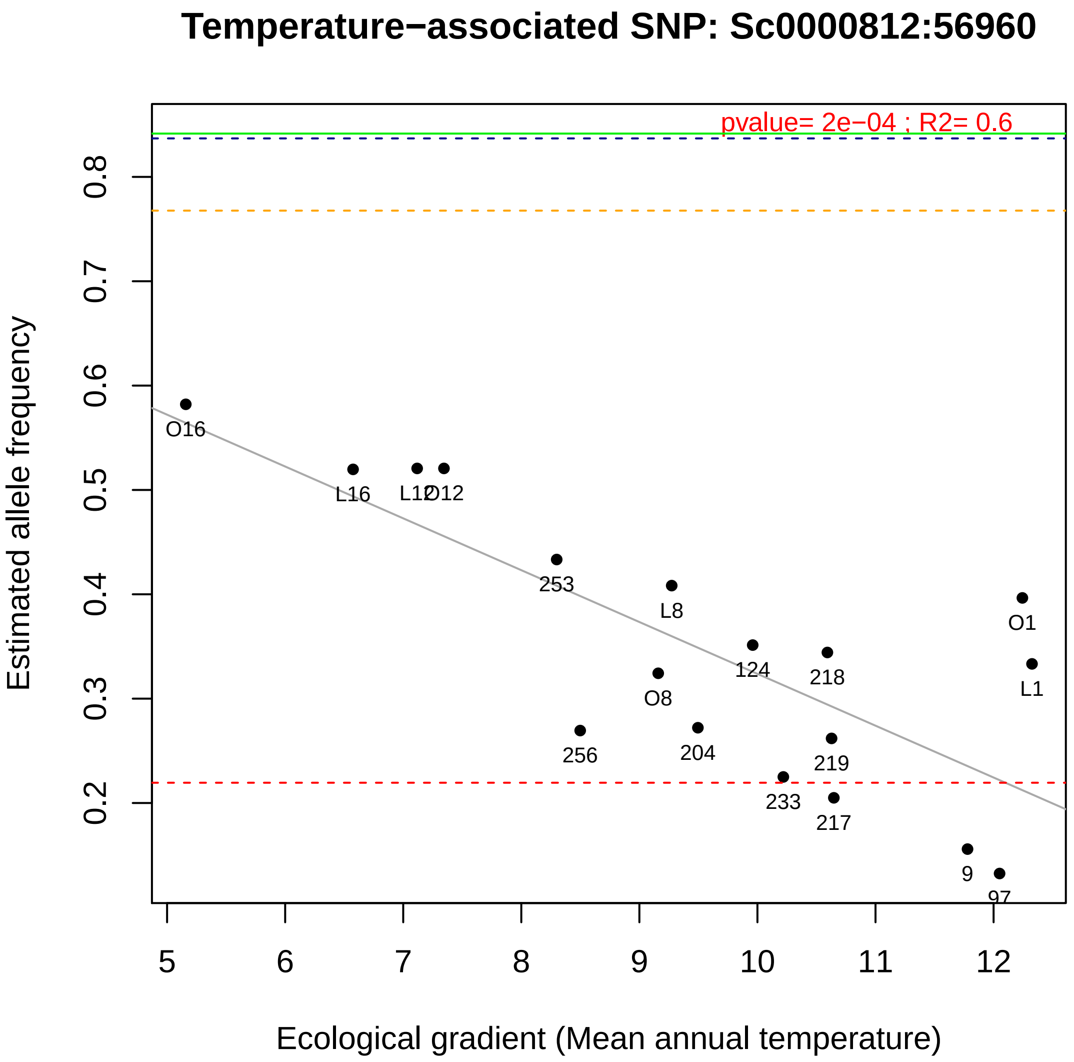
Temperature−associated SNP: Sc0000812:56960
pvalue= 2e−04 ; R2= 0.6
Estimated allele frequency
0.2
0.3
0.4
0.5
0.6
0.7
0.8
O16
L16
L12
O12
253
L8
O1
124
218
L1
O8
256
204
219
233
217
9
97
5
6
7
8
9
10
11
12
Ecological gradient (Mean annual temperature)

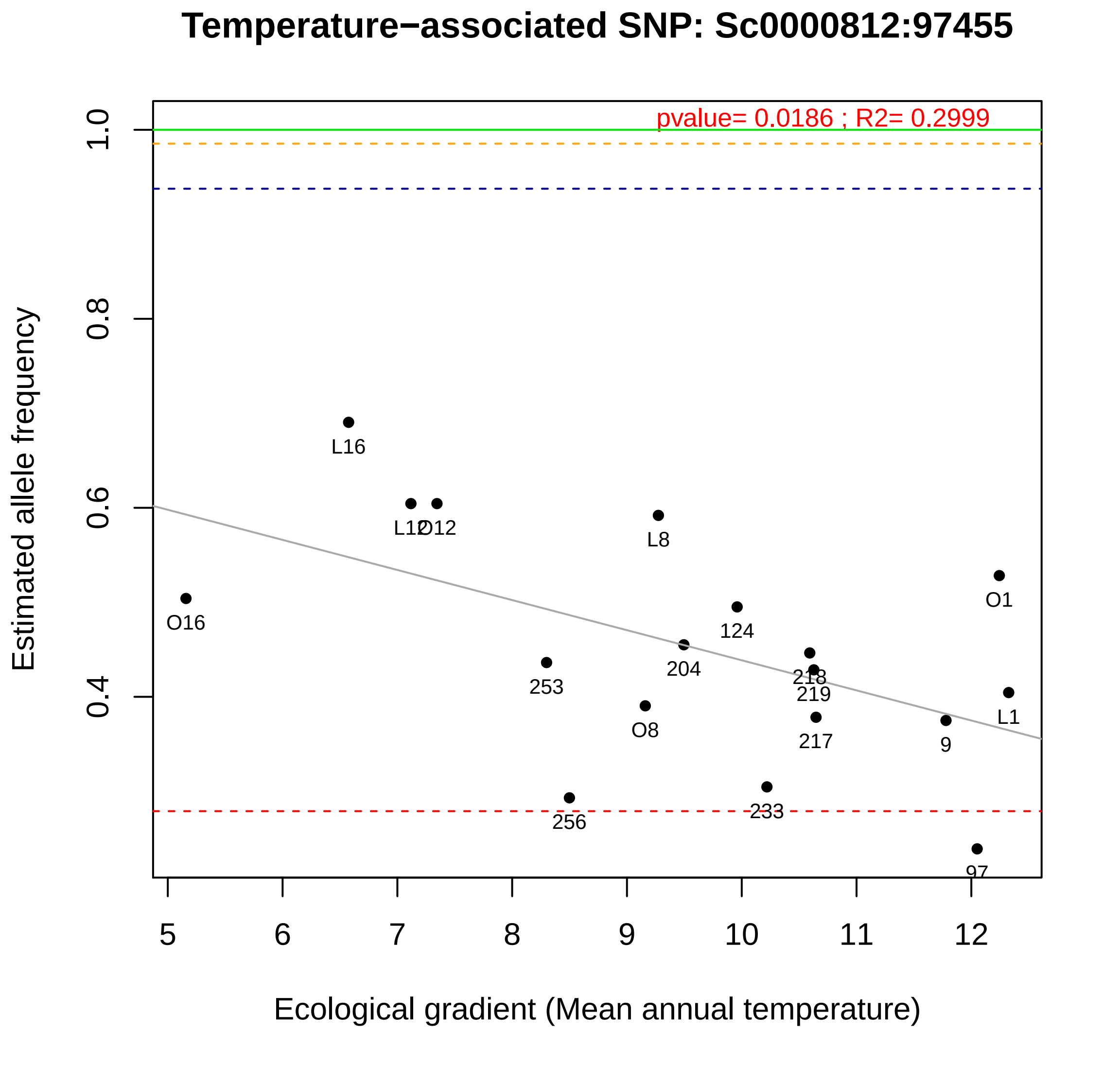
Temperature−associated SNP: Sc0000812:97455
pvalue= 0.0186 ; R2= 0.2999
Estimated allele frequency
0.4
0.6
0.8
1.0
L16
L12
O12
L8
O16
O1
124
204
253
218
219
L1
O8
217
9
256
233
97
5
6
7
8
9
10
11
12
Ecological gradient (Mean annual temperature)

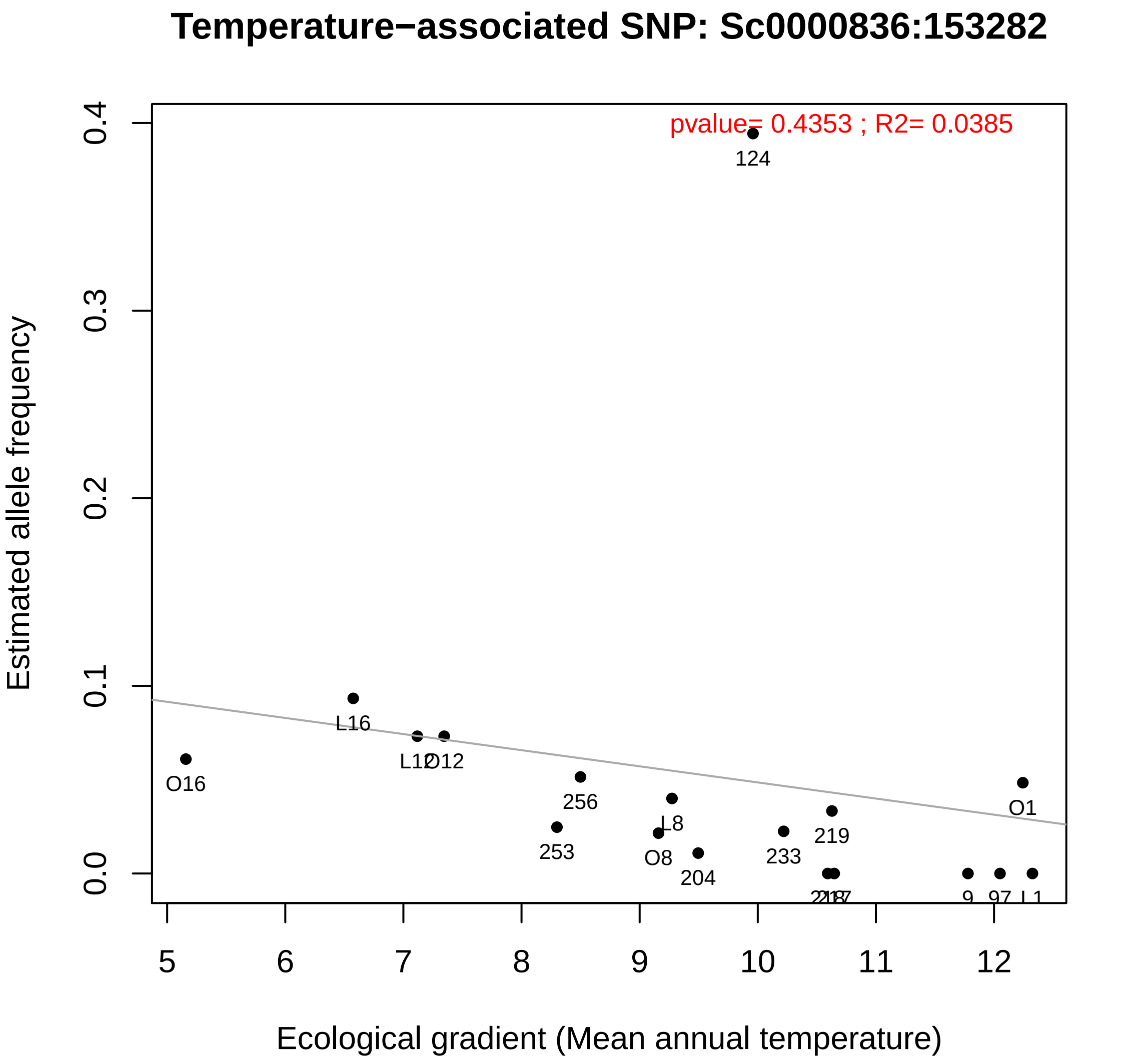
Temperature−associated SNP: Sc0000836:153282
pvalue= 0.4353 ; R2= 0.0385
Estimated allele frequency
0.0
0.1
0.2
0.3
0.4
5
6
7
8
9
10
11
12
Ecological gradient (Mean annual temperature)
124
L16
L12
O12
O16
256
253
L8
O8
204
233
219
O1
9
97
L1

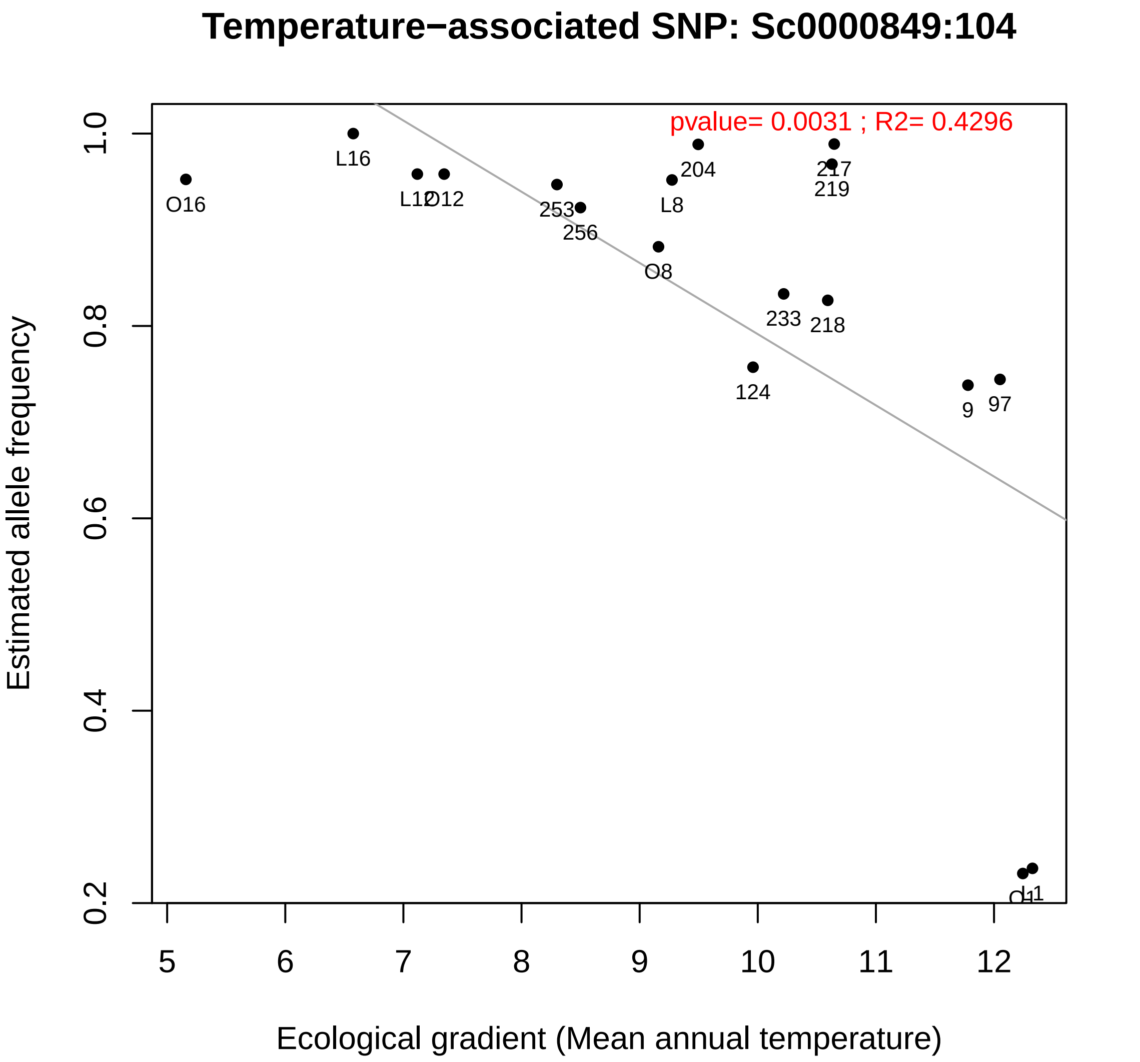
Temperature−associated SNP: Sc0000849:104
pvalue= 0.0031 ; R2= 0.4296
Estimated allele frequency
Ecological gradient (Mean annual temperature)
0.2
0.4
0.6
0.8
1.0
5
6
7
8
9
10
11
12
O16
L16
L12
O12
253
256
O8
L8
204
217
219
233
218
124
9
97

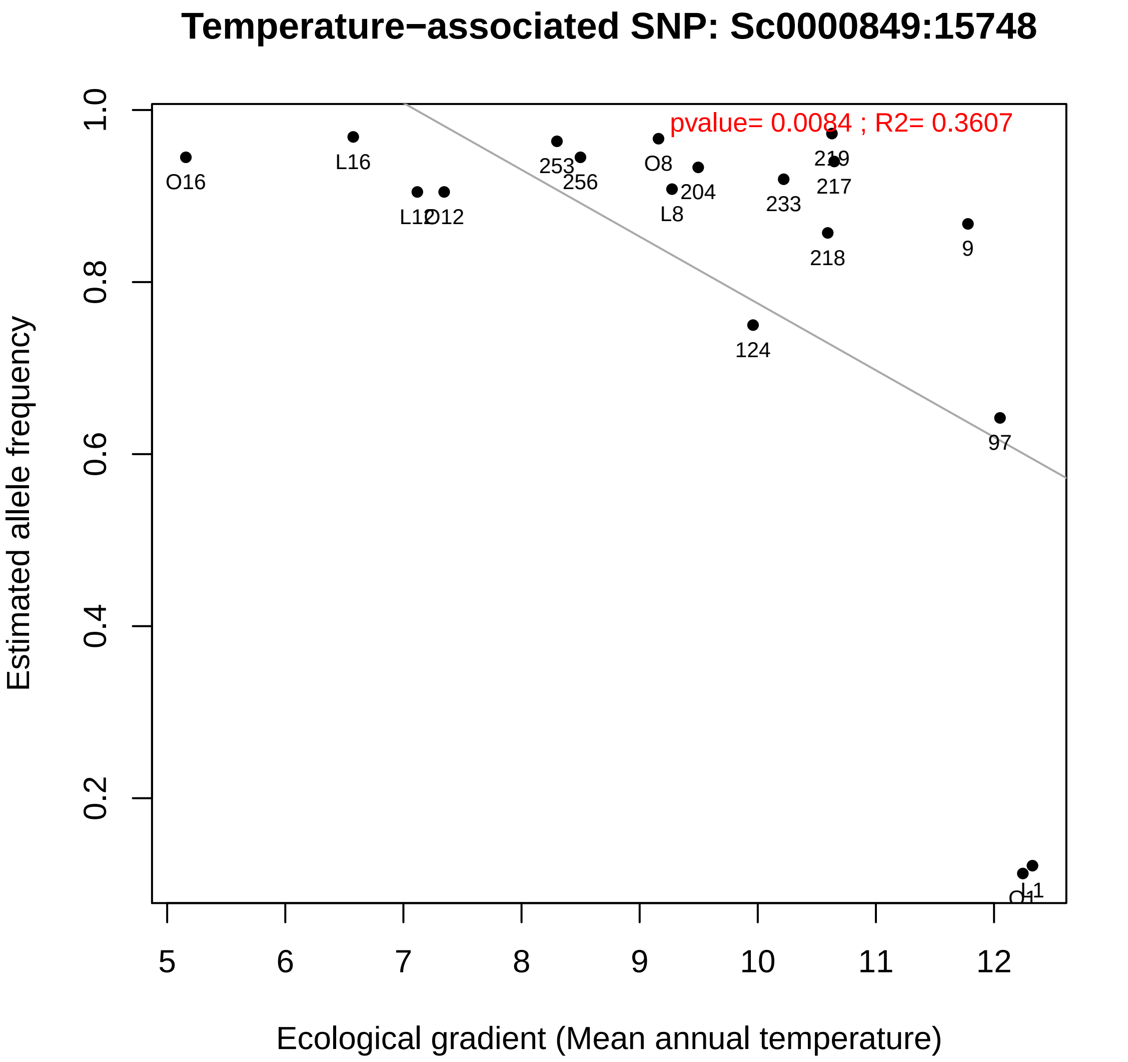
Temperature−associated SNP: Sc0000849:15748
pvalue= 0.0084 ; R2= 0.3607
Estimated allele frequency
0.2
0.4
0.6
0.8
1.0
Ecological gradient (Mean annual temperature)
5
6
7
8
9
10
11
12
O16
L16
L12
O12
253
256
O8
L8
204
233
219
217
218
124
9
97
O1
L1

#### Temperature−associated SNP: Sc0000849:15755

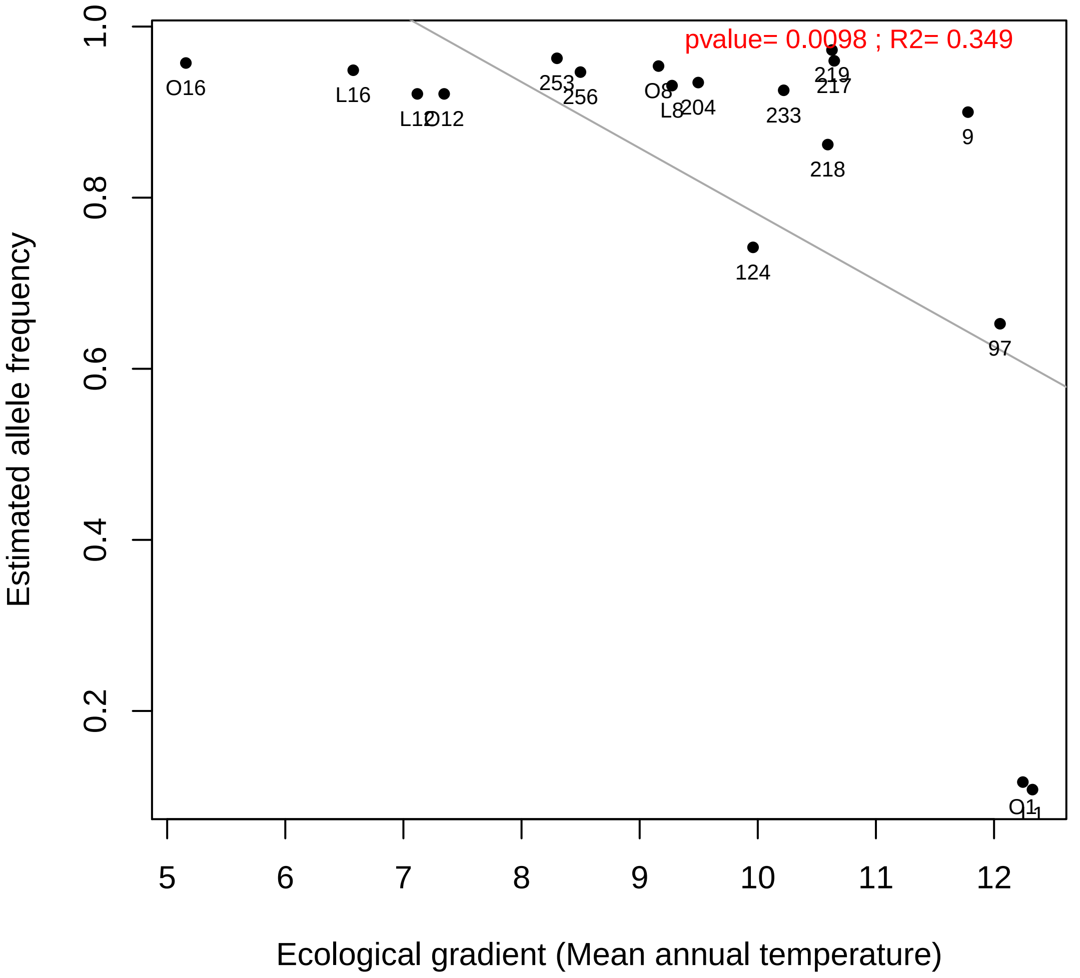

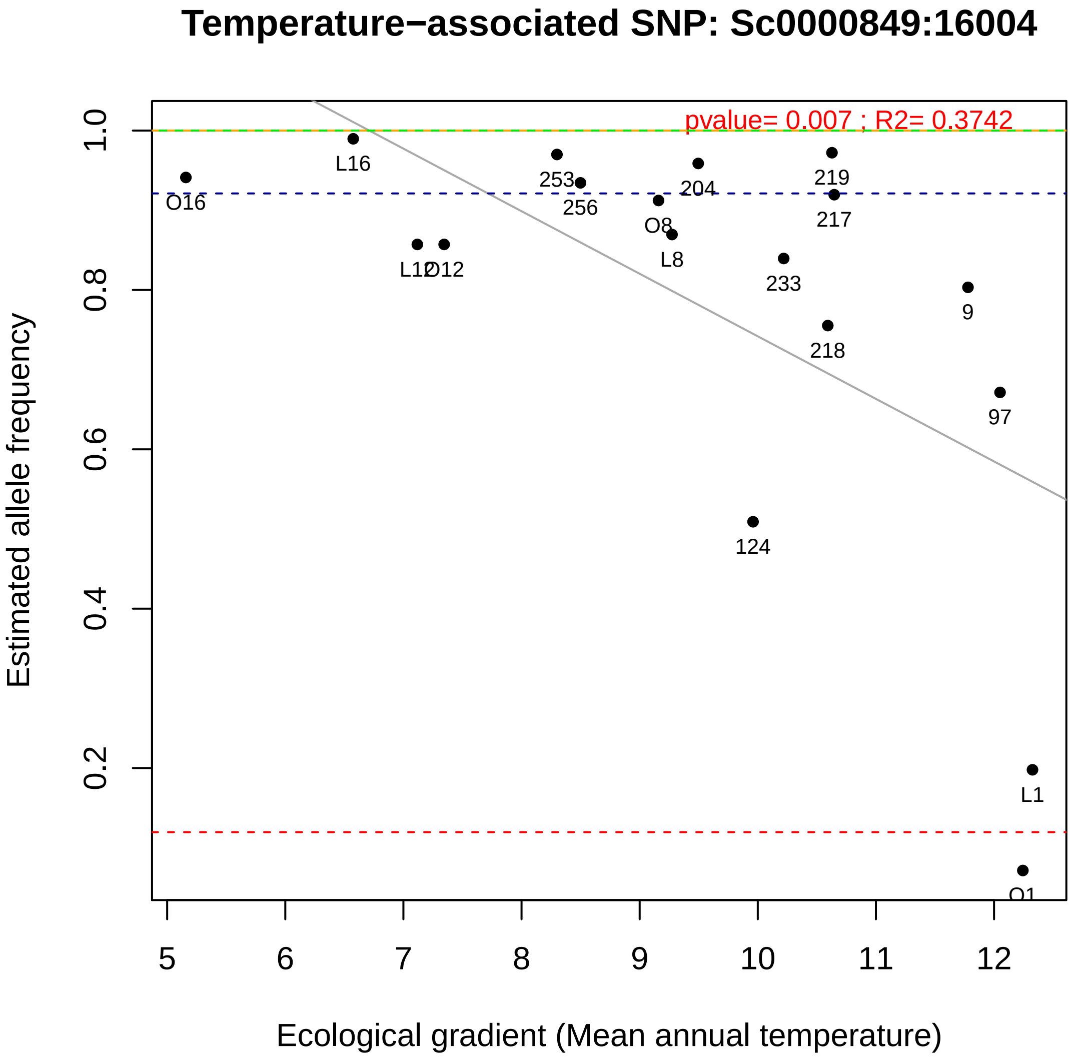

Temperature−associated SNP: Sc0000849:16004
pvalue= 0.007 ; R2= 0.3742
Estimated allele frequency
0.2
0.4
0.6
0.8
1.0
5
6
7
8
9
10
11
12
Ecological gradient (Mean annual temperature)
O16
L16
L12
O12
253
256
O8
L8
204
124
233
219
217
218
9
97
L1
O1

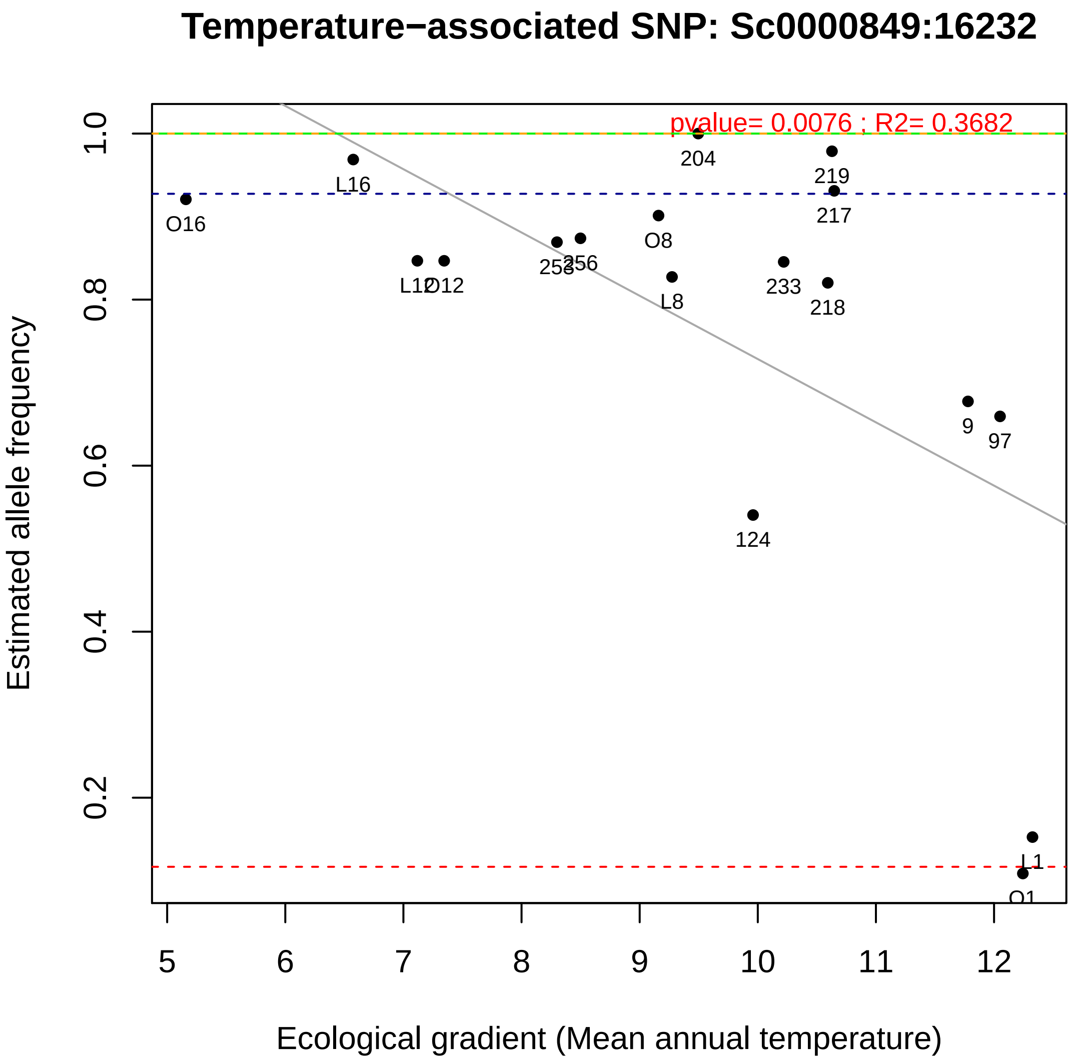

Temperature−associated SNP: Sc0000849:16232
pvalue= 0.0076 ; R2= 0.3682
Estimated allele frequency
Ecological gradient (Mean annual temperature)
0.2
0.4
0.6
0.8
1.0
5
6
7
8
9
10
11
12
O16
L16
L12
O12
253
256
O8
L8
204
233
219
217
218
124
9
97
L1
O1

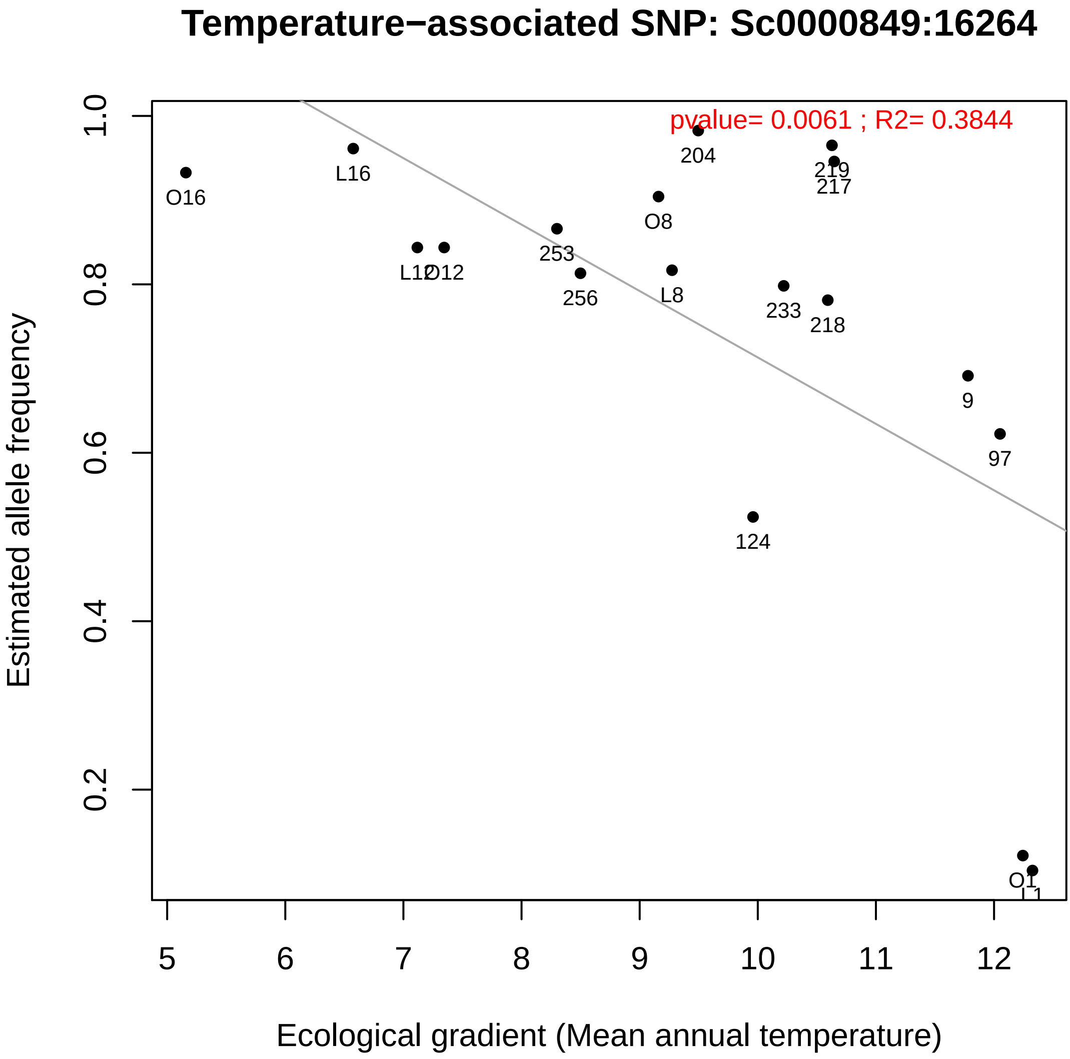

Temperature−associated SNP: Sc0000849:16264
pvalue= 0.0061 ; R2= 0.3844
Estimated allele frequency
Ecological gradient (Mean annual temperature)
0.2
0.4
0.6
0.8
1.0
5
6
7
8
9
10
11
12
O16
L16
L12
O12
253
256
O8
L8
204
219
217
233
218
124
9
97
O1

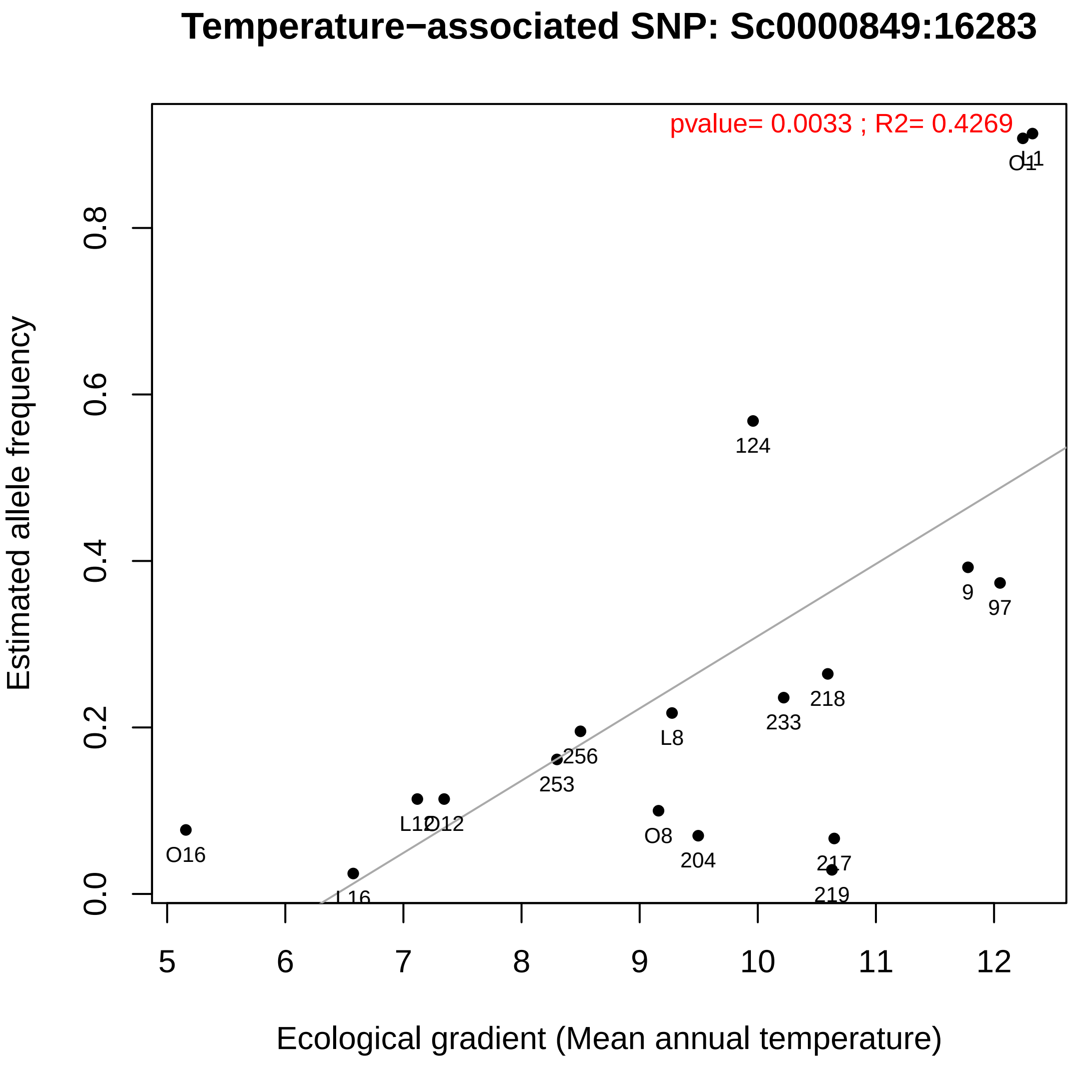
Temperature−associated SNP: Sc0000849:16283
pvalue= 0.0033 ; R2= 0.4269
Estimated allele frequency
0.0
0.2
0.4
0.6
0.8
5
6
7
8
9
10
11
12
Ecological gradient (Mean annual temperature)
O1
L1
124
9
97
218
233
L8
256
253
L12
O12
O8
204
217
219
O16
L16

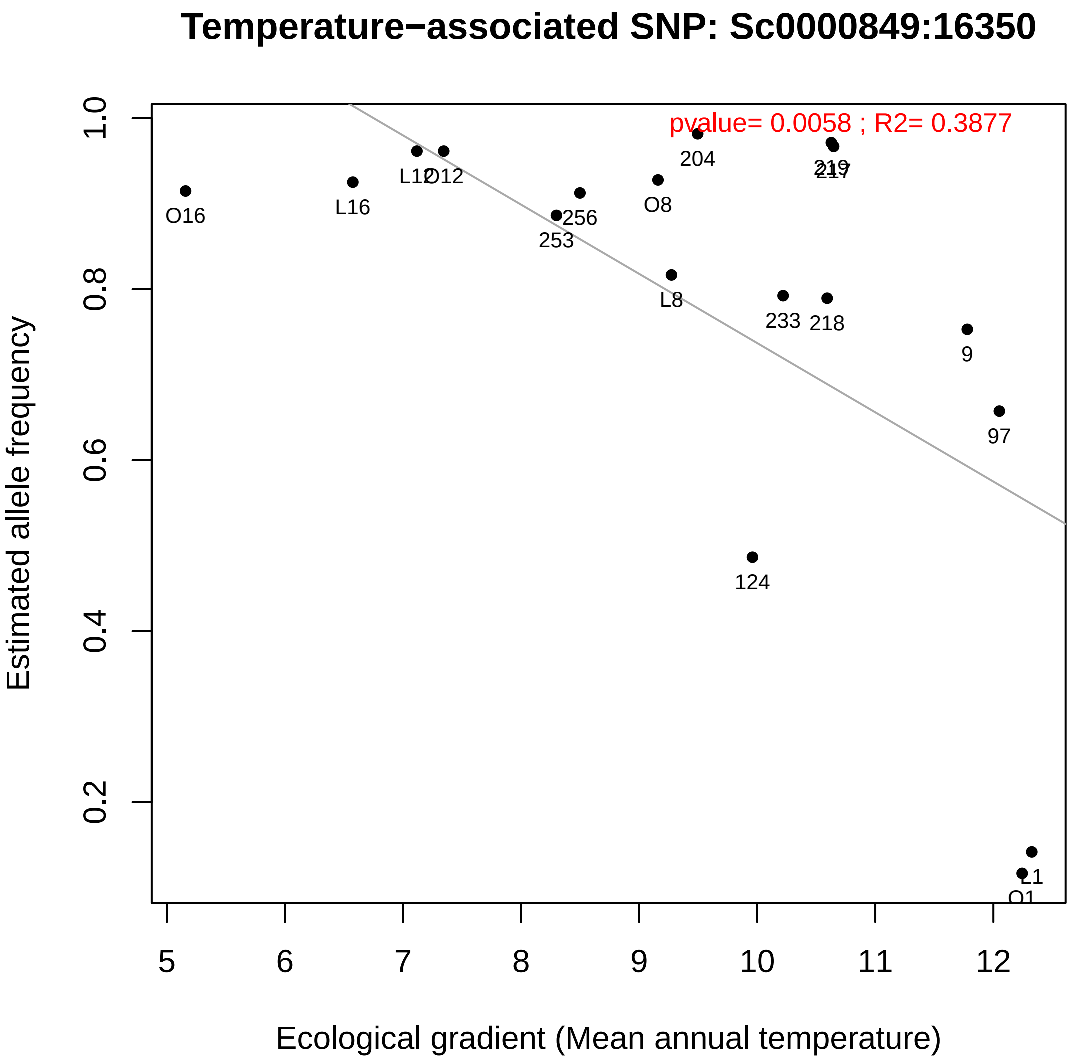
Temperature−associated SNP: Sc0000849:16350
pvalue= 0.0058 ; R2= 0.3877
Estimated allele frequency
Ecological gradient (Mean annual temperature)
O16
L16
L12
O12
253
256
O8
204
219
217
L8
233
218
9
97
124
L1
O1
0.2
0.4
0.6
0.8
1.0
5
6
7
8
9
10
11
12

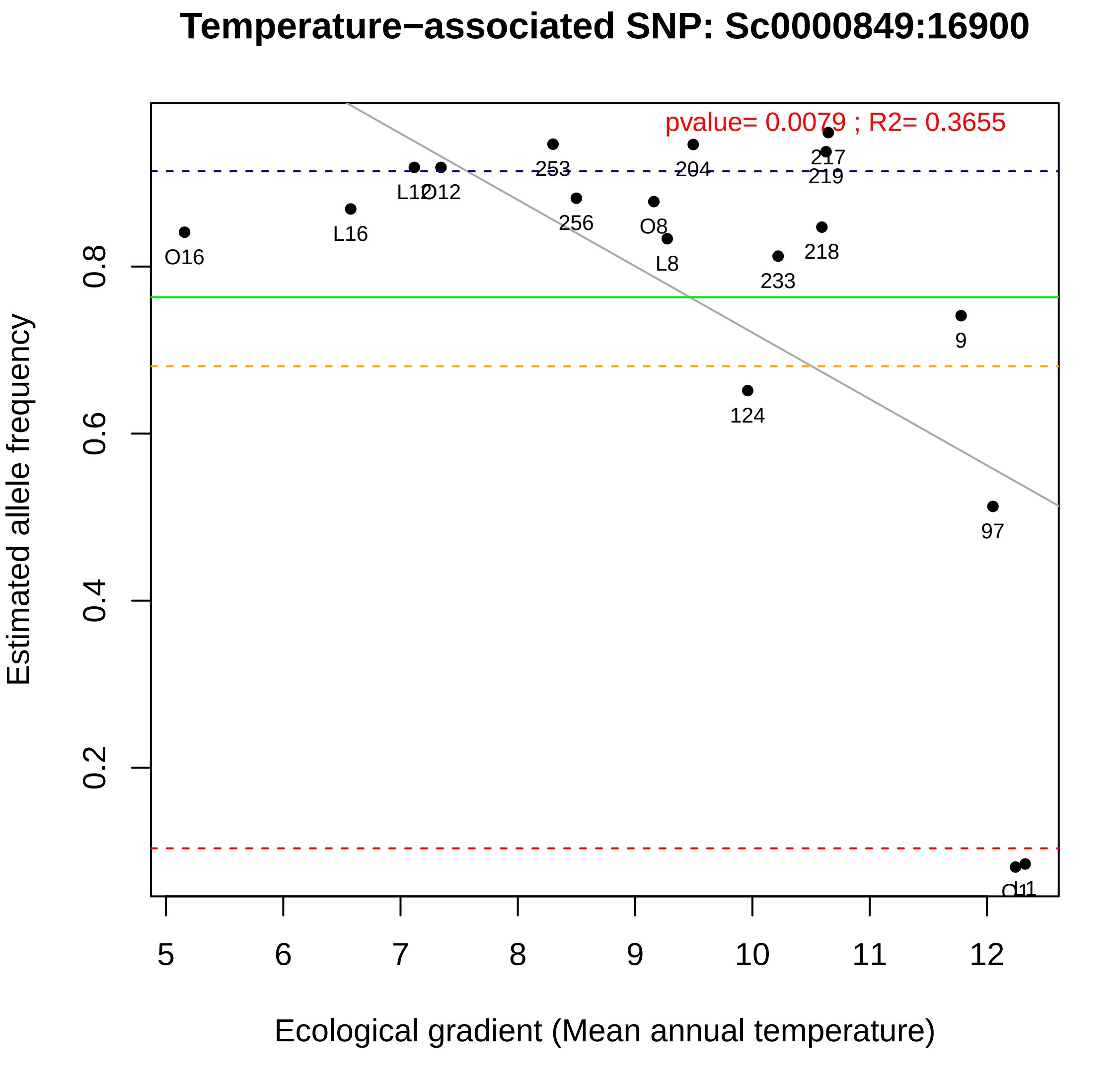

Temperature−associated SNP: Sc0000849:16900
pvalue= 0.0079 ; R2= 0.3655
Estimated allele frequency
0.2
0.4
0.6
0.8
5
6
7
8
9
10
11
12
Ecological gradient (Mean annual temperature)
O16
L16
L12
O12
253
256
204
O8
L8
233
217
219
218
9
124
97

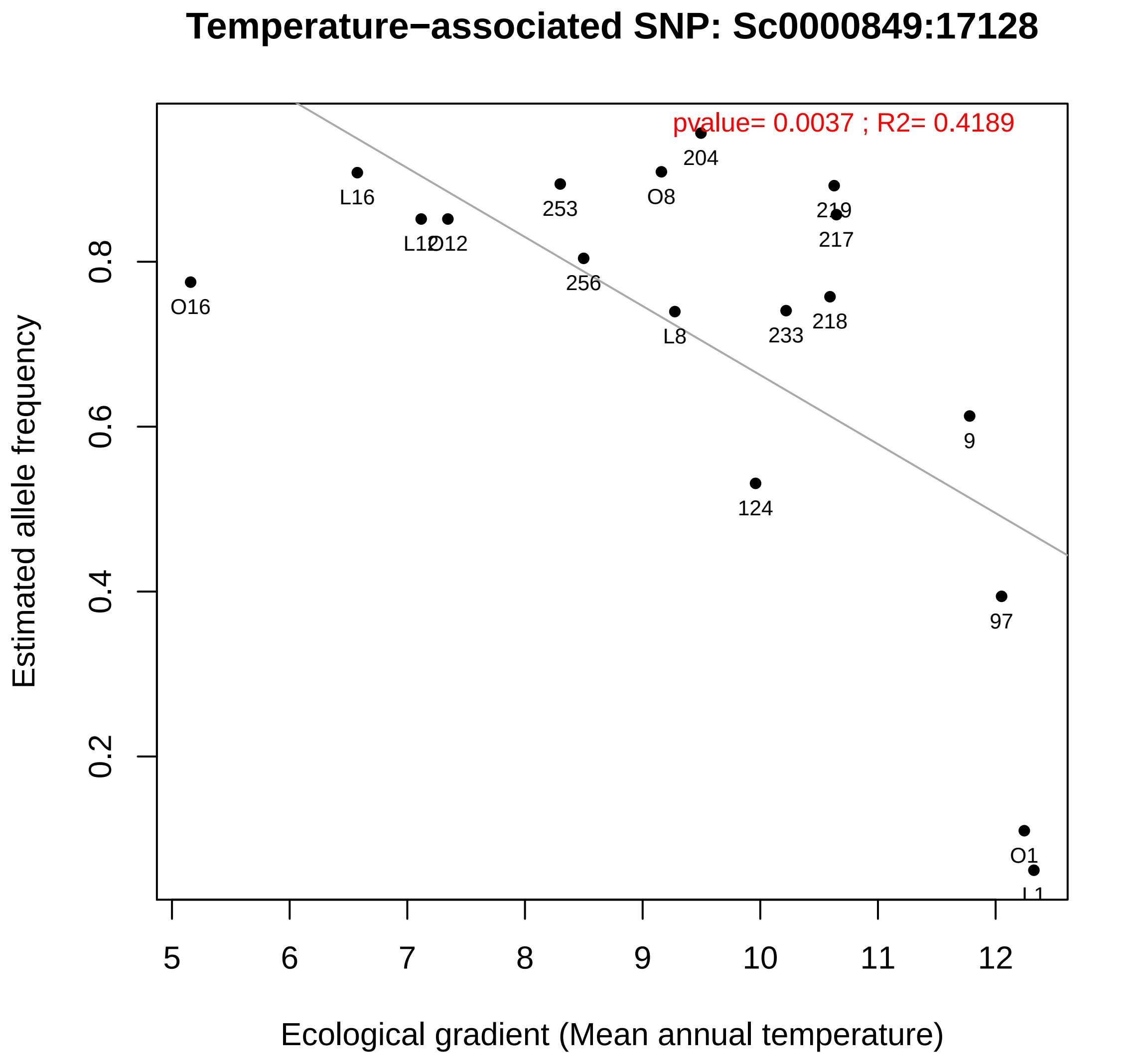

Temperature−associated SNP: Sc0000849:17128
pvalue= 0.0037 ; R2= 0.4189
O16
L16
L12
O12
253
256
O8
204
L8
124
233
219
217
218
9
97
O1
0.2
0.4
0.6
0.8
Estimated allele frequency
5
6
7
8
9
10
11
12
Ecological gradient (Mean annual temperature)

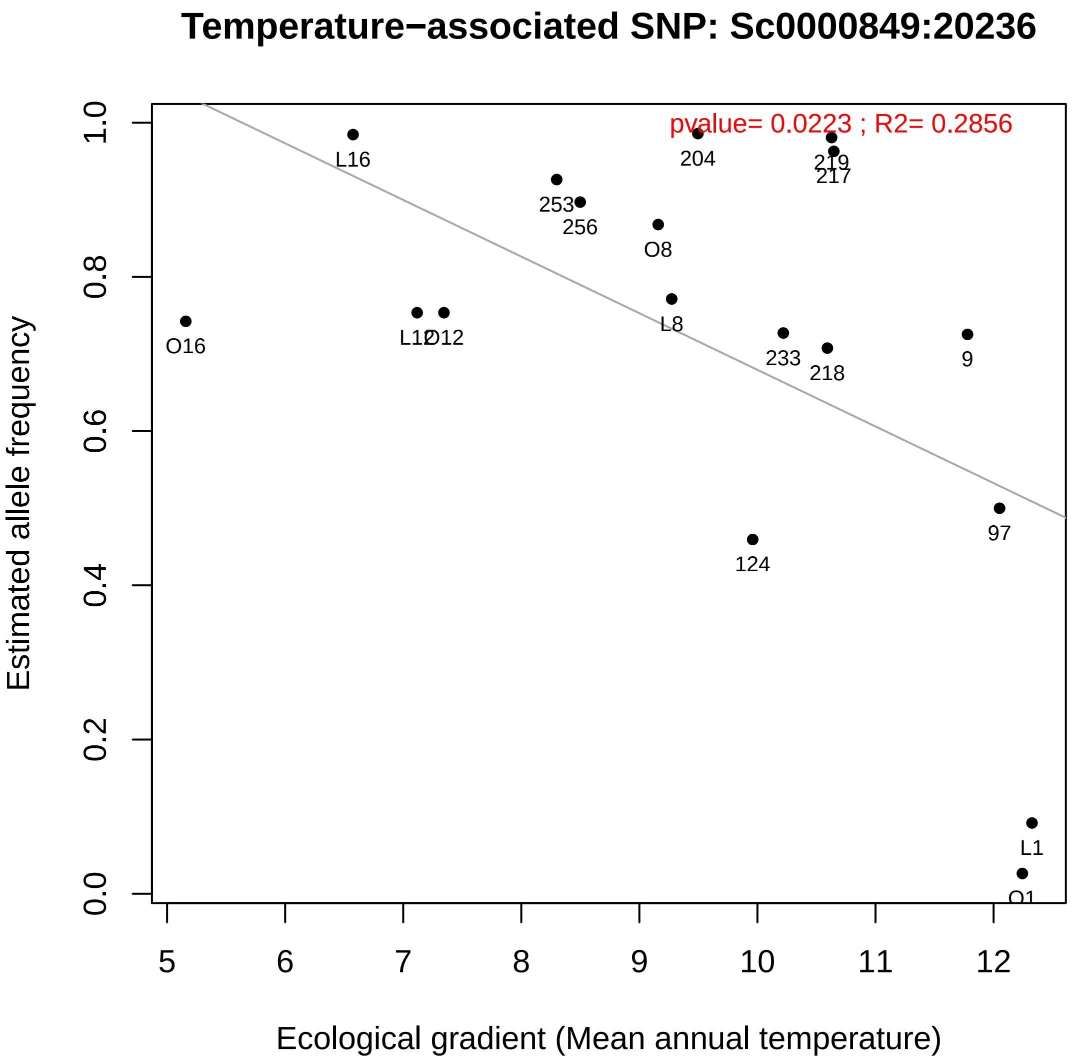

Temperature−associated SNP: Sc0000849:20236
pvalue= 0.0223 ; R2= 0.2856
Estimated allele frequency
0.0
0.2
0.4
0.6
0.8
1.0
Ecological gradient (Mean annual temperature)
5
6
7
8
9
10
11
12
O16
L16
L12
O12
253
256
O8
L8
204
124
233
219
217
218
9
97
L1
O1

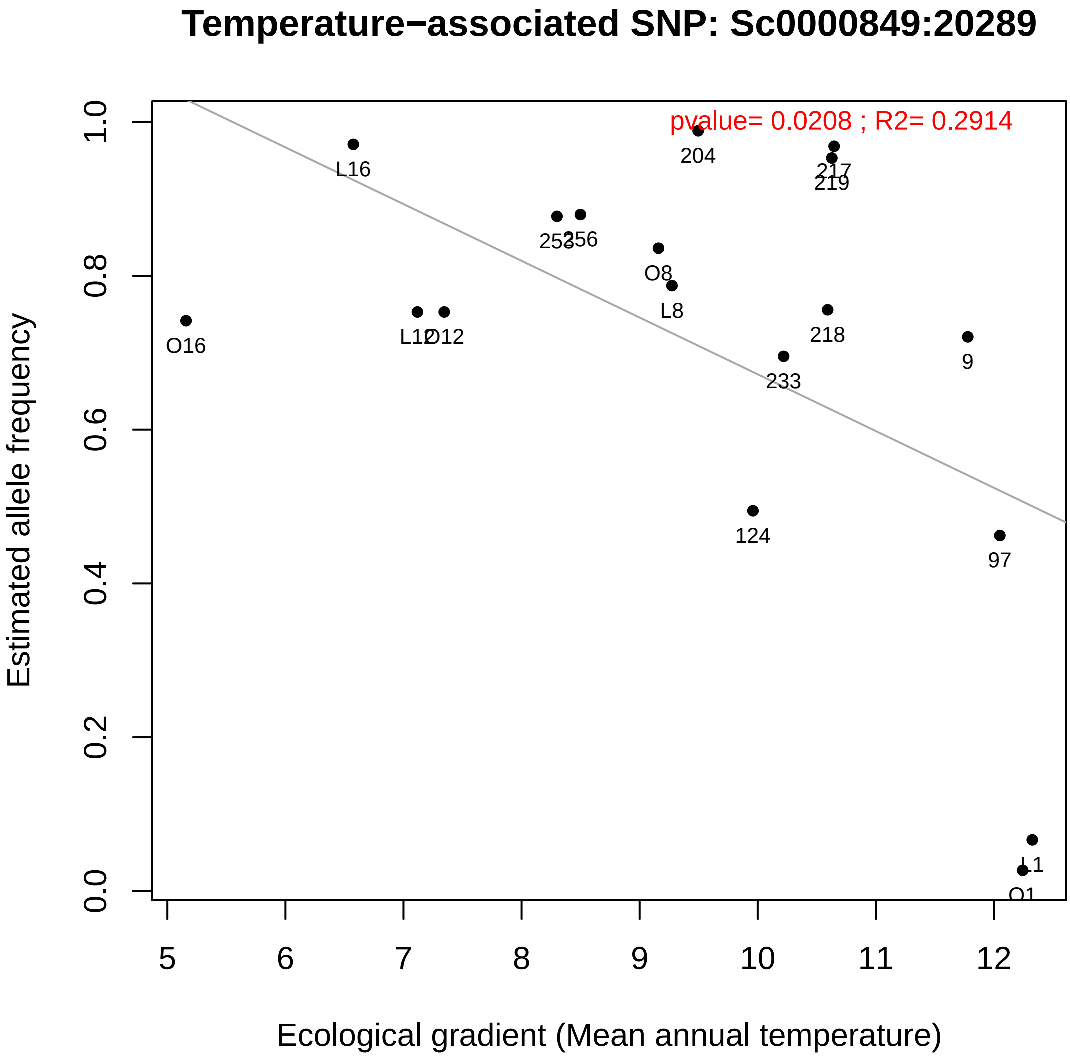
Temperature−associated SNP: Sc0000849:20289
pvalue= 0.0208 ; R2= 0.2914
Estimated allele frequency
Ecological gradient (Mean annual temperature)
0.0
0.2
0.4
0.6
0.8
1.0
5
6
7
8
9
10
11
12
L16
204
217
219
256
O8
L8
O16
O12
218
233
9
124
97
L1
O1

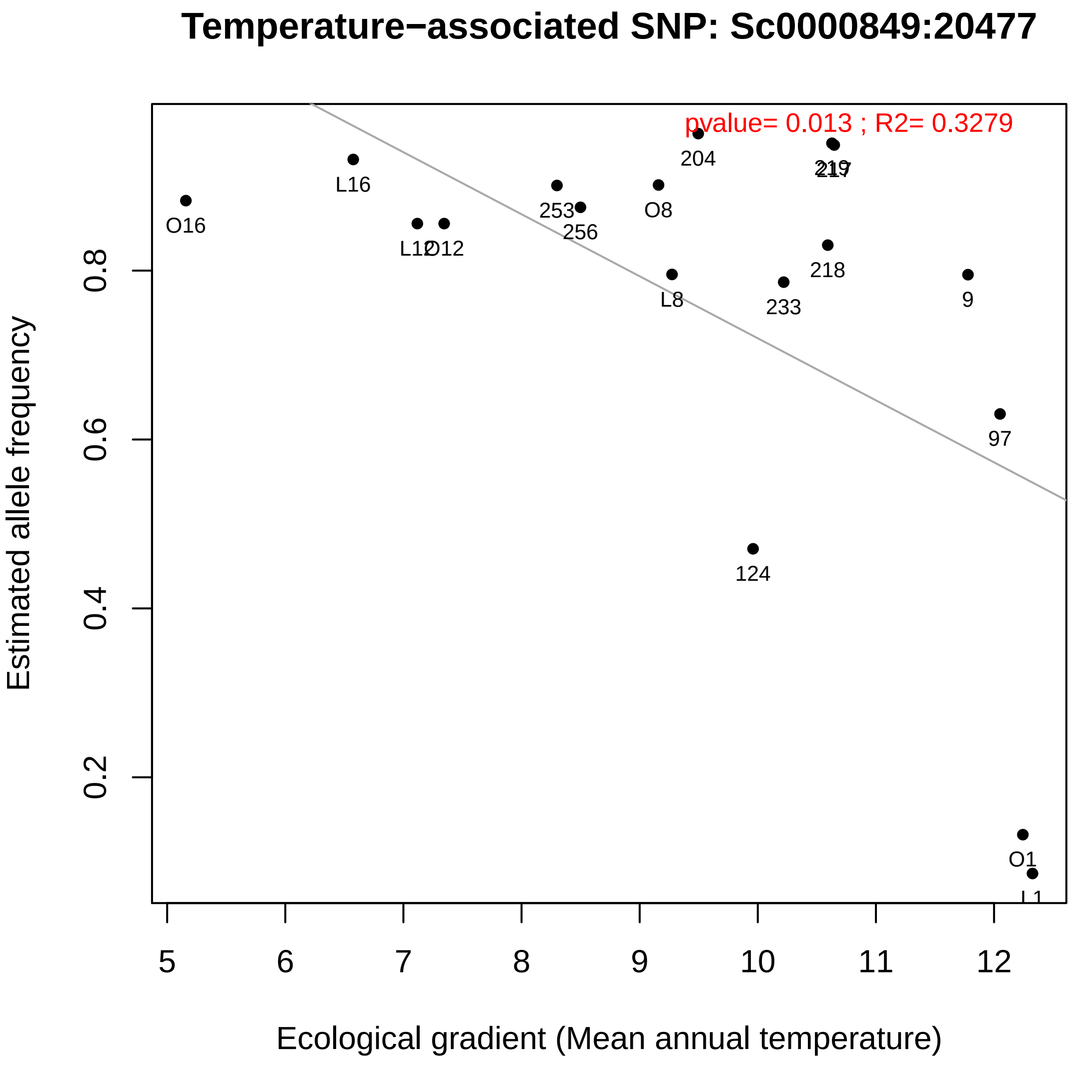
Temperature−associated SNP: Sc0000849:20477
pvalue= 0.013 ; R2= 0.3279
Estimated allele frequency
0.2
0.4
0.6
0.8
5
6
7
8
9
10
11
12
Ecological gradient (Mean annual temperature)
O16
L16
L12
O12
253
256
O8
204
L8
233
218
9
97
124
O1
L1

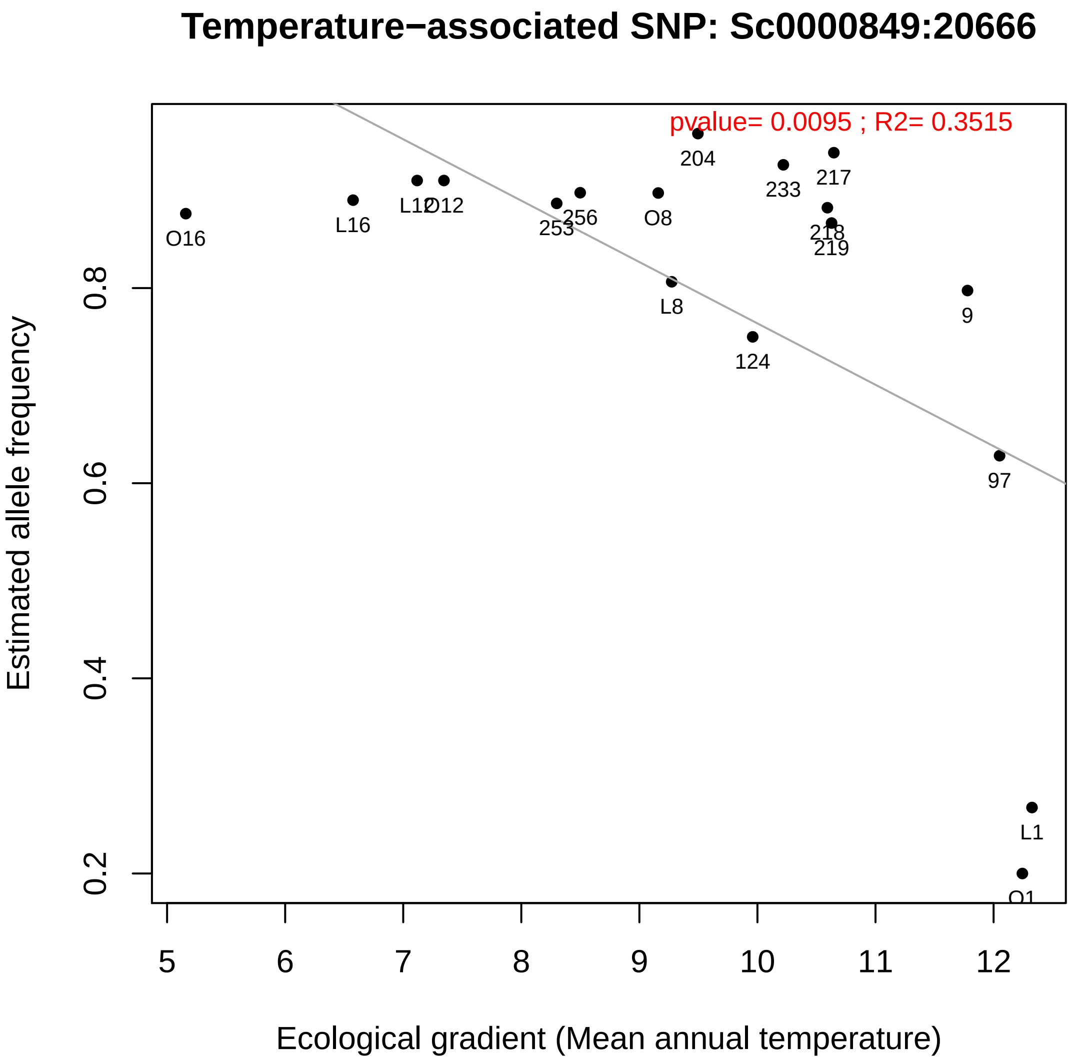
Temperature−associated SNP: Sc0000849:20666
pvalue= 0.0095 ; R2= 0.3515
Estimated allele frequency
0.2
0.4
0.6
0.8
5
6
7
8
9
10
11
12
Ecological gradient (Mean annual temperature)
O16
L16
L12
O12
253
256
O8
204
233
217
218
219
L8
124
9
97
L1
O1

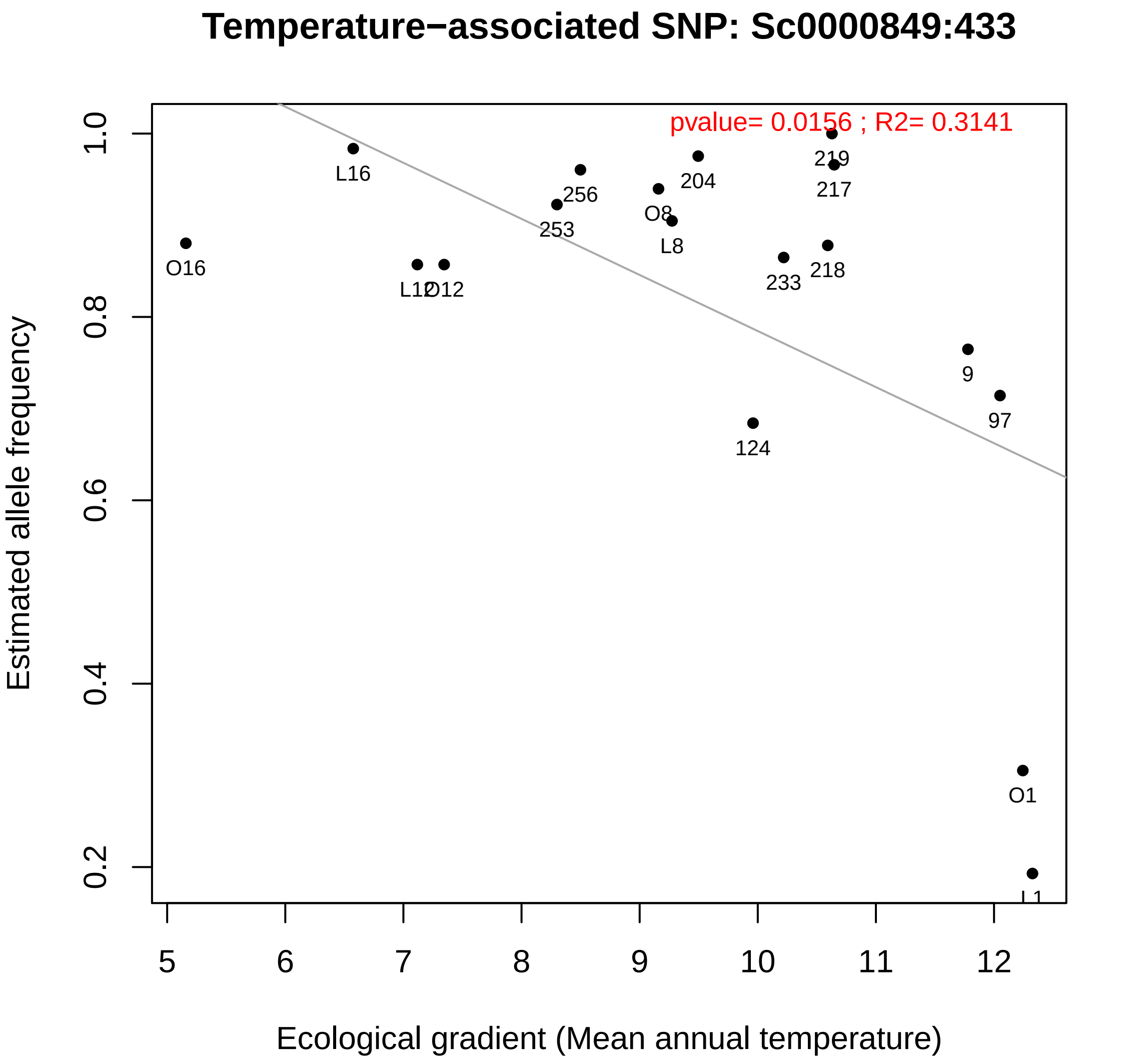

Temperature−associated SNP: Sc0000849:433
pvalue= 0.0156 ; R2= 0.3141
Estimated allele frequency
Ecological gradient (Mean annual temperature)
0.2
0.4
0.6
0.8
1.0
5
6
7
8
9
10
11
12
O16
L16
L12
O12
253
256
O8
L8
204
233
218
219
217
124
9
97
O1

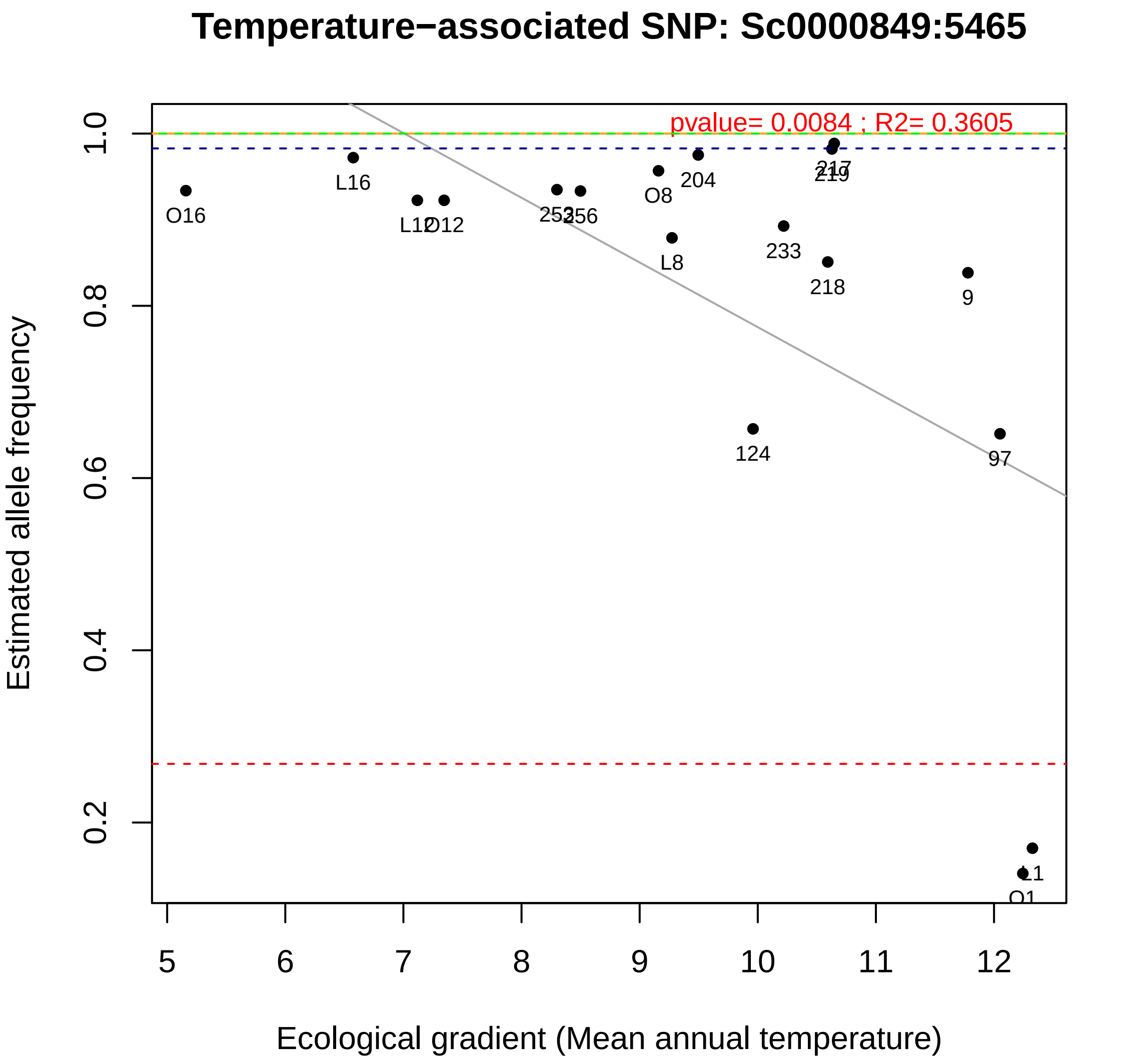
Temperature−associated SNP: Sc0000849:5465
pvalue= 0.0084 ; R2= 0.3605
Estimated allele frequency
1.0
0.8
0.6
0.4
0.2
5
6
7
8
9
10
11
12
Ecological gradient (Mean annual temperature)
O16
L16
L12
O12
256
204
O8
217
219
233
L8
218
9
124
97
L1
O1

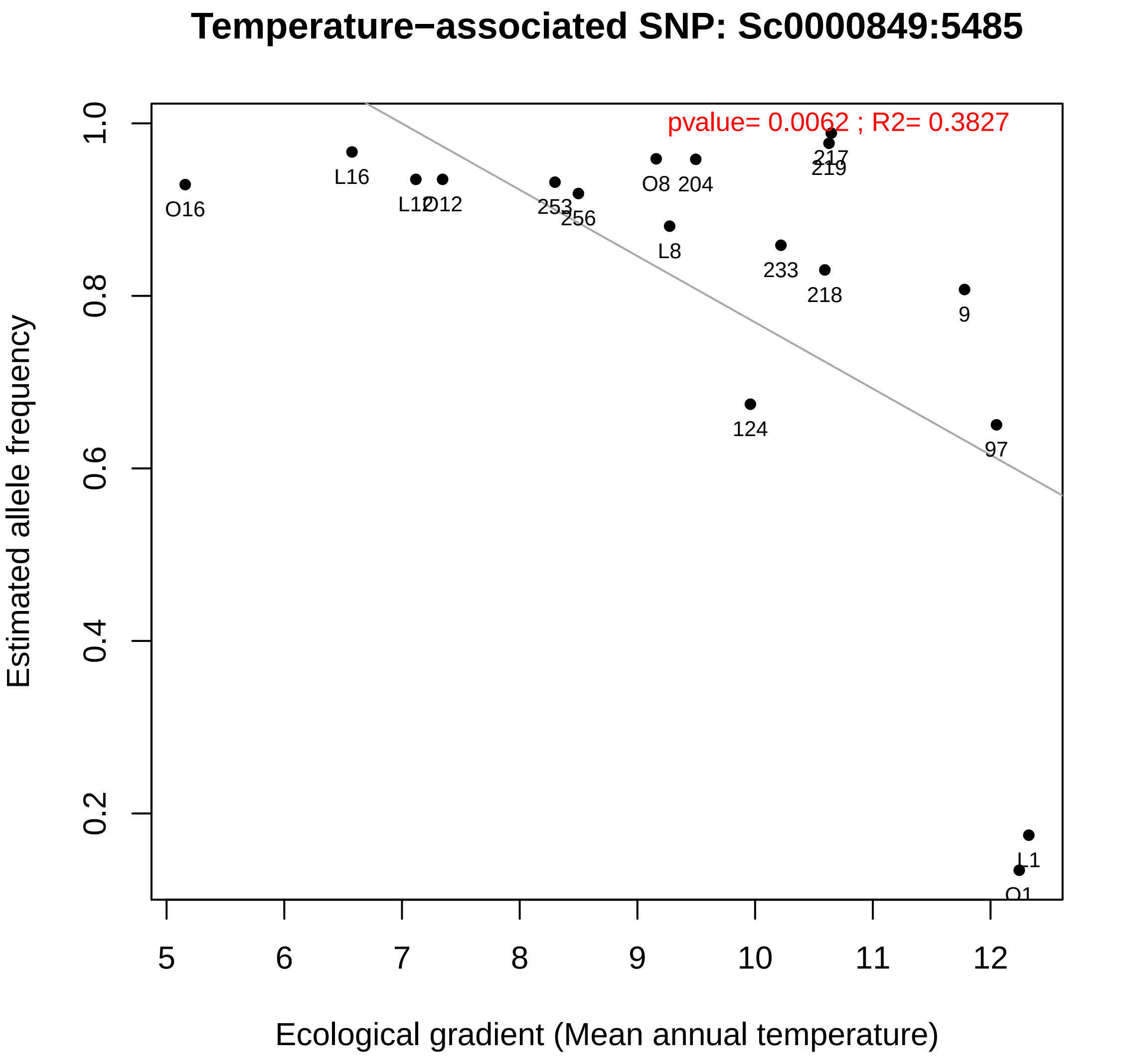
Temperature−associated SNP: Sc0000849:5485
pvalue= 0.0062 ; R2= 0.3827
Estimated allele frequency
Ecological gradient (Mean annual temperature)
1.0
0.8
0.6
0.4
0.2
5
6
7
8
9
10
11
12
O16
L16
L12
O12
253
256
O8
204
L8
217
219
233
218
124
9
97
L1
O1

**Temperature−associated SNP: Sc0000849:5617**

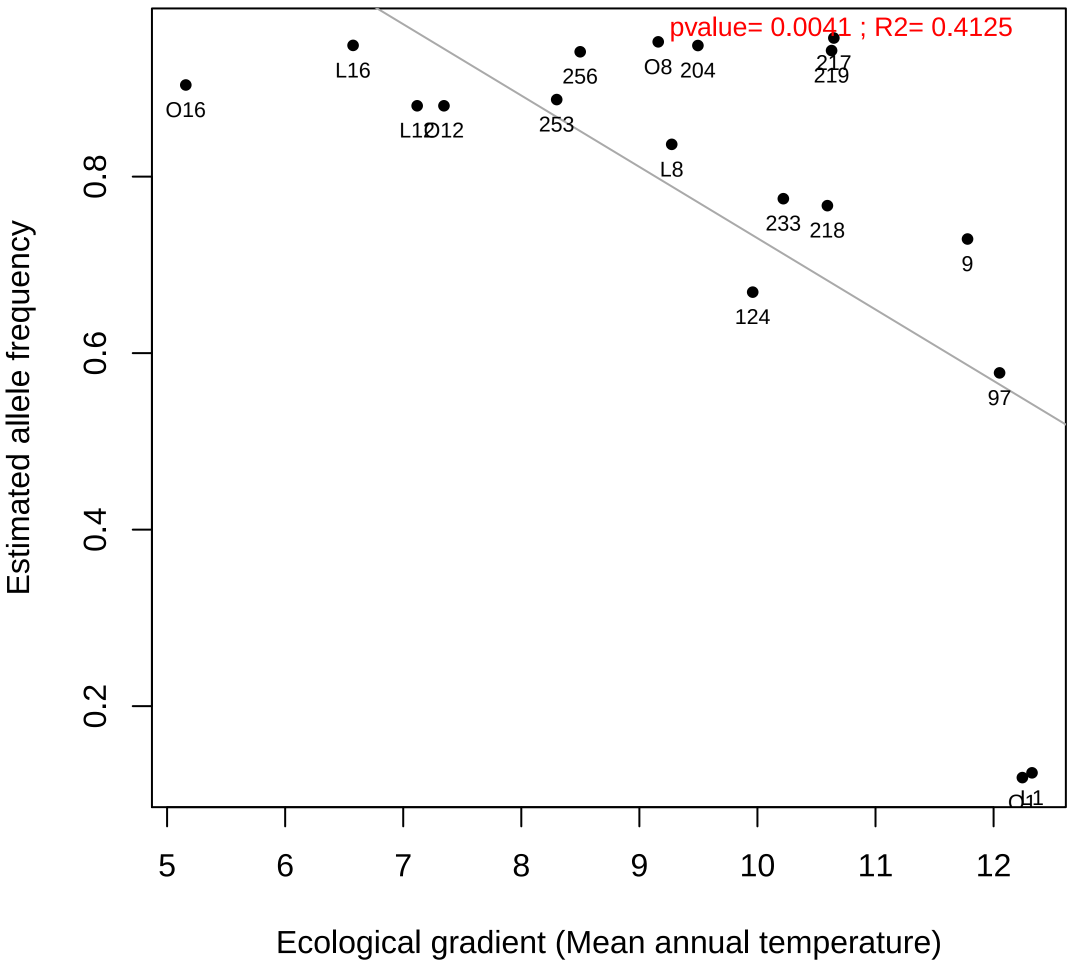

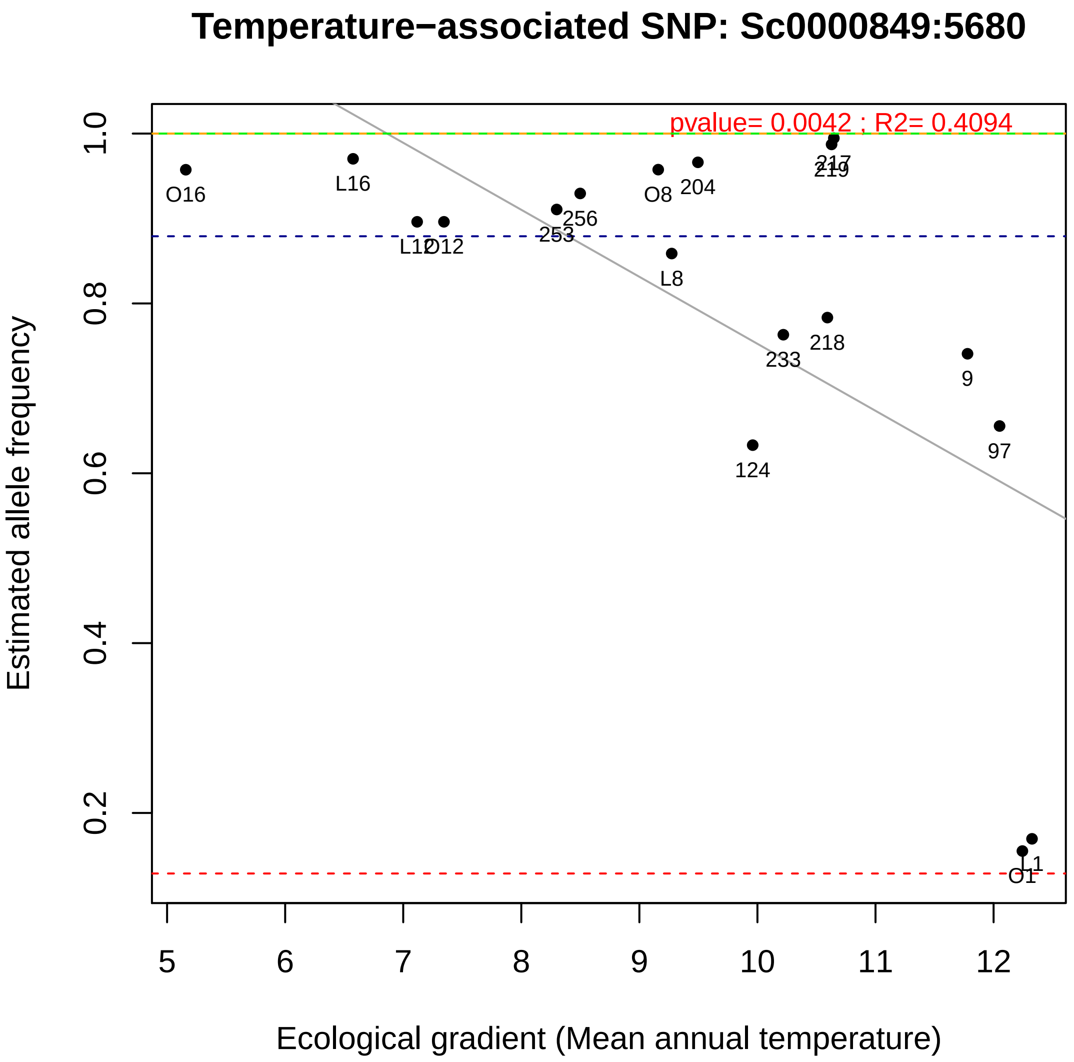
Temperature−associated SNP: Sc0000849:5680
pvalue= 0.0042 ; R2= 0.4094
Estimated allele frequency
Ecological gradient (Mean annual temperature)
0.2
0.4
0.6
0.8
1.0
5
6
7
8
9
10
11
12
O16
L16
L12
O12
253
256
O8
204
L8
217
219
233
218
124
9
97
L1
O1

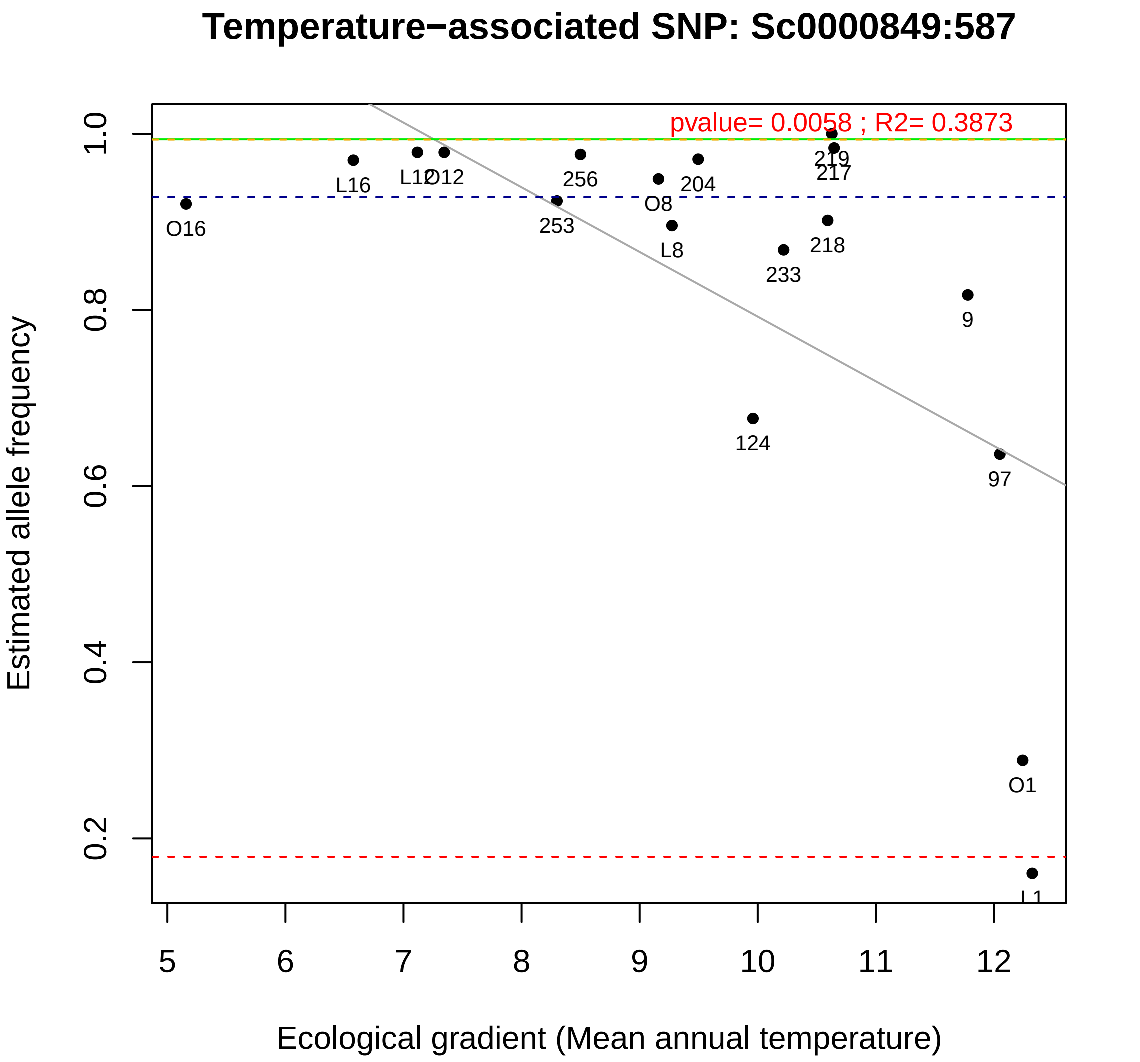

Temperature−associated SNP: Sc0000849:587
pvalue= 0.0058 ; R2= 0.3873
Estimated allele frequency
0.2
0.4
0.6
0.8
1.0
5
6
7
8
9
10
11
12
Ecological gradient (Mean annual temperature)
O16
L16
L12
O12
256
253
O8
204
L8
219
217
218
233
9
124
97
O1

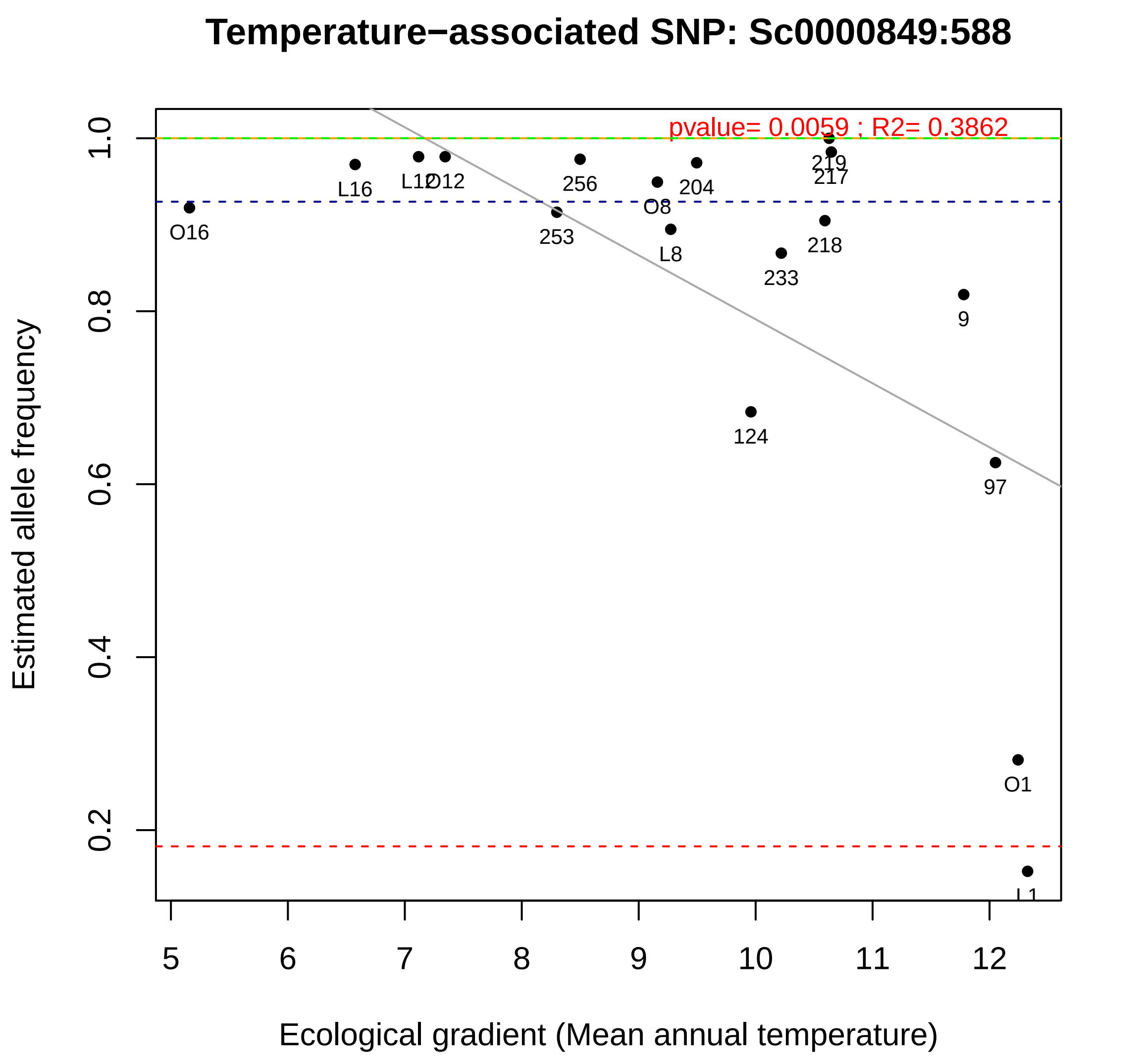

Temperature−associated SNP: Sc0000849:588
pvalue= 0.0059 ; R2= 0.3862
Estimated allele frequency
Ecological gradient (Mean annual temperature)
1.0
0.8
0.6
0.4
0.2
5
6
7
8
9
10
11
12
O16
L16
L12
O12
256
253
O8
204
L8
219
217
218
233
9
124
97
O1

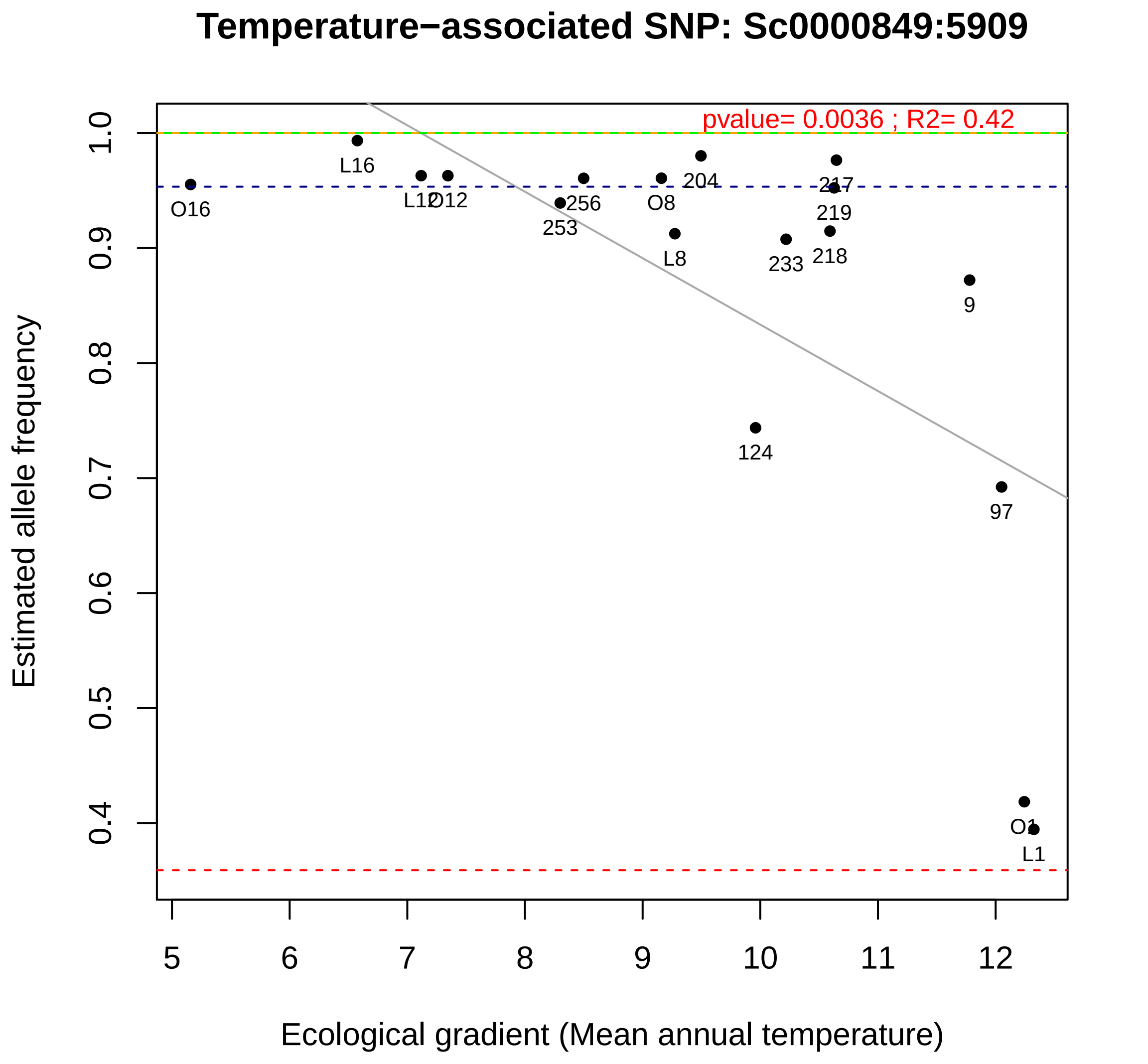
Temperature−associated SNP: Sc0000849:5909
pvalue= 0.0036 ; R2= 0.42
Estimated allele frequency
Ecological gradient (Mean annual temperature)
1.0
0.9
0.8
0.7
0.6
0.5
0.4
5
6
7
8
9
10
11
12
O16
L16
L12
O12
253
256
O8
204
L8
233
217
219
218
9
124
97
O1
L1

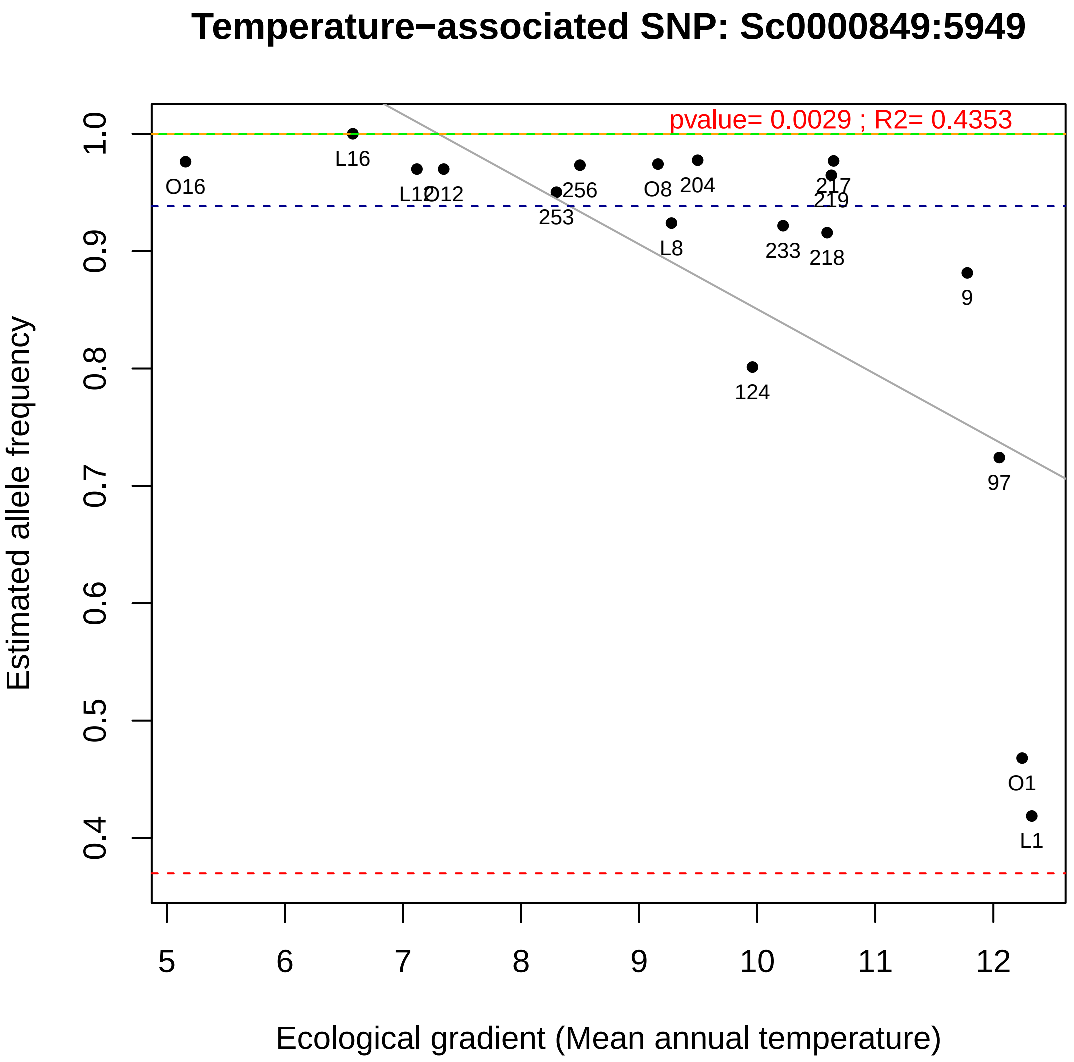

Temperature−associated SNP: Sc0000849:5949
pvalue= 0.0029 ; R2= 0.4353
Estimated allele frequency
1.0
0.9
0.8
0.7
0.6
0.5
0.4
5
6
7
8
9
10
11
12
Ecological gradient (Mean annual temperature)
O16
L16
L12
O12
253
256
O8
204
L8
217
219
233
218
124
9
97
O1
L1

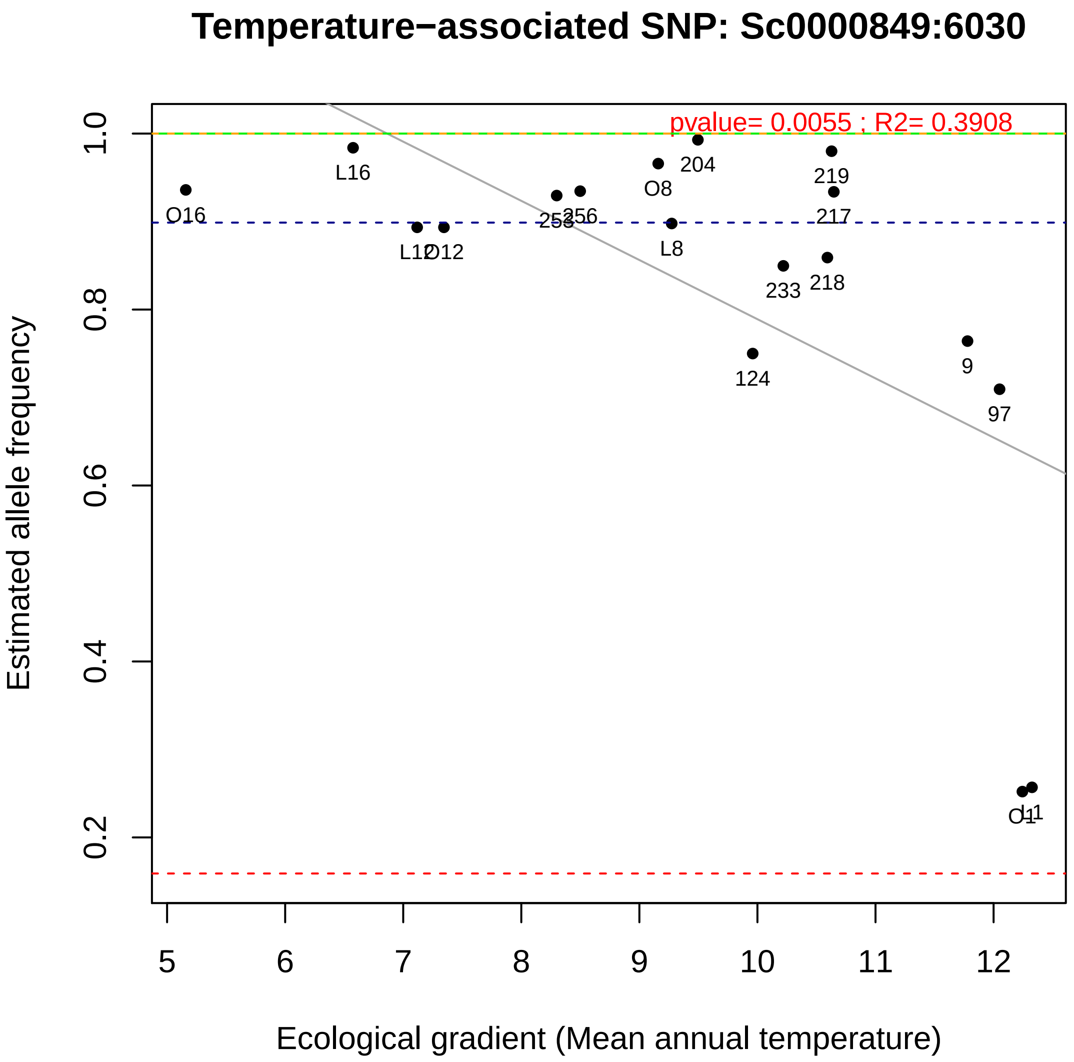

Temperature−associated SNP: Sc0000849:6030
pvalue= 0.0055 ; R2= 0.3908
Estimated allele frequency
Ecological gradient (Mean annual temperature)
0.2
0.4
0.6
0.8
1.0
5
6
7
8
9
10
11
12
O16
L16
L12
O12
253
256
O8
204
L8
124
233
219
217
218
9
97
O1
L1

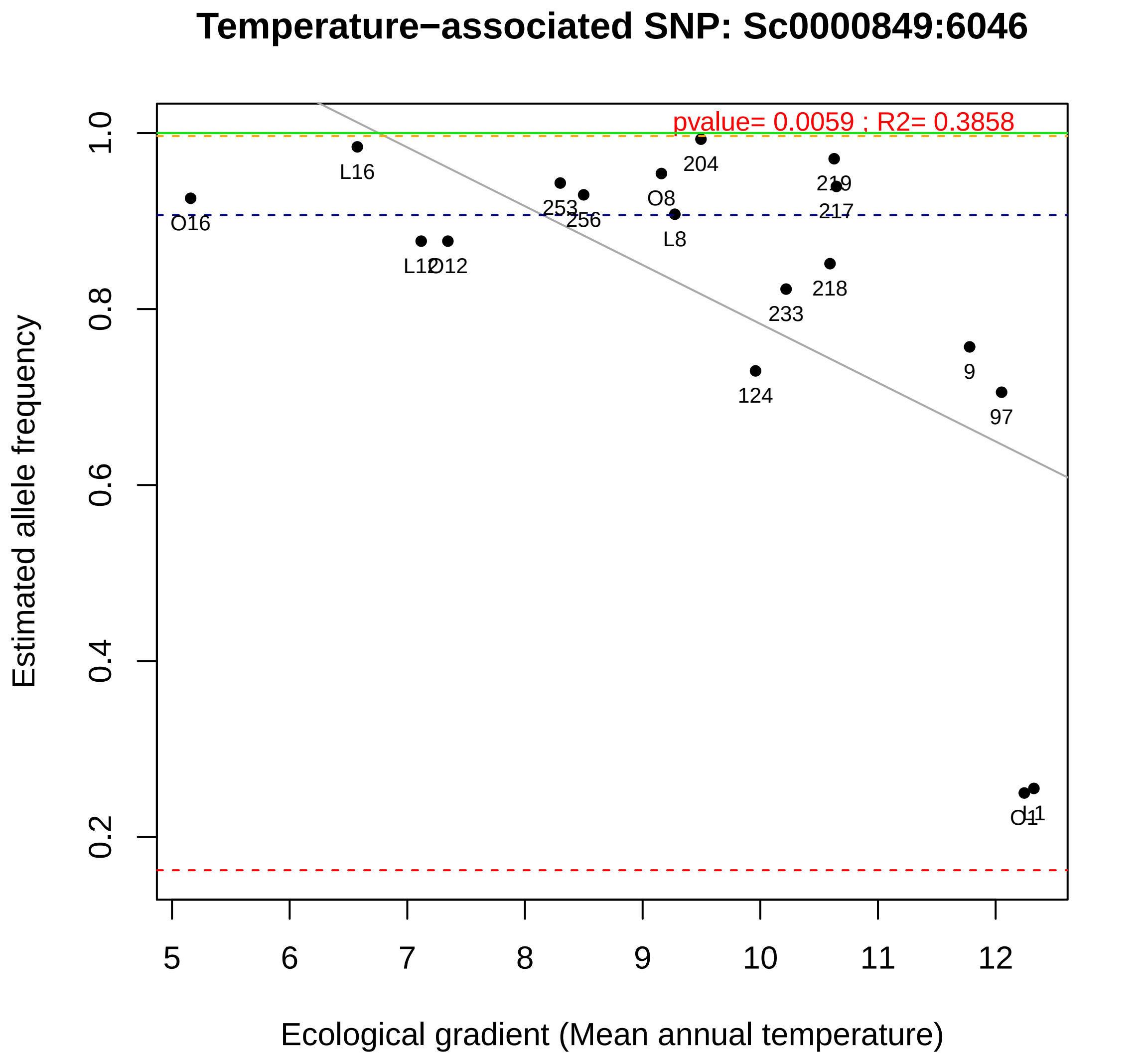
Temperature−associated SNP: Sc0000849:6046
pvalue= 0.0059 ; R2= 0.3858
Estimated allele frequency
0.2
0.4
0.6
0.8
1.0
Ecological gradient (Mean annual temperature)
5
6
7
8
9
10
11
12
O16
L16
L12
O12
253
256
O8
L8
204
219
217
218
233
124
9
97
O1
L1

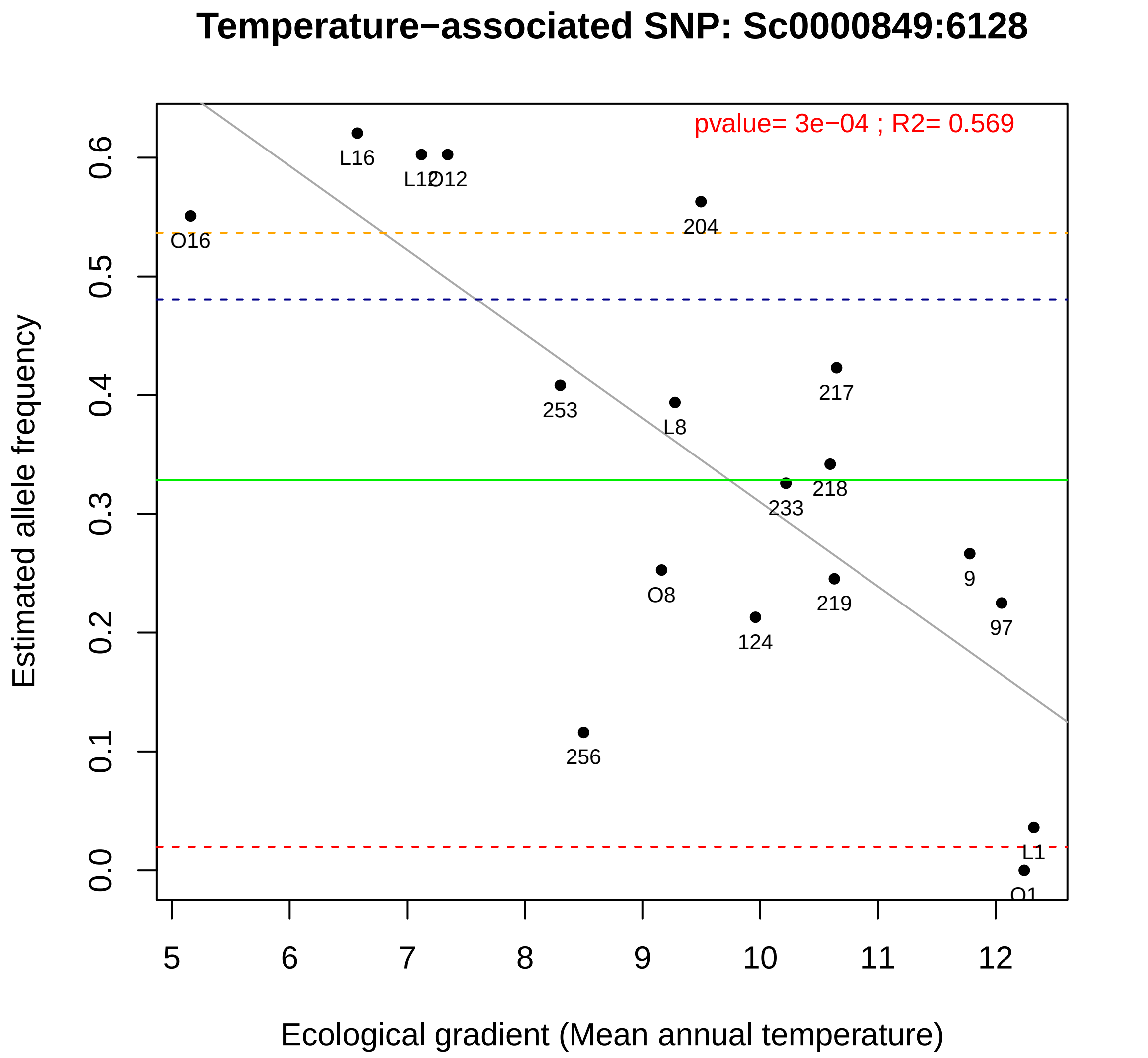

Temperature−associated SNP: Sc0000849:6128
pvalue= 3e−04 ; R2= 0.569
Estimated allele frequency
Ecological gradient (Mean annual temperature)
0.0
0.1
0.2
0.3
0.4
0.5
0.6
5
6
7
8
9
10
11
12
L16
L12
O12
O16
204
253
L8
217
218
233
O8
124
219
9
97
256
L1
O1

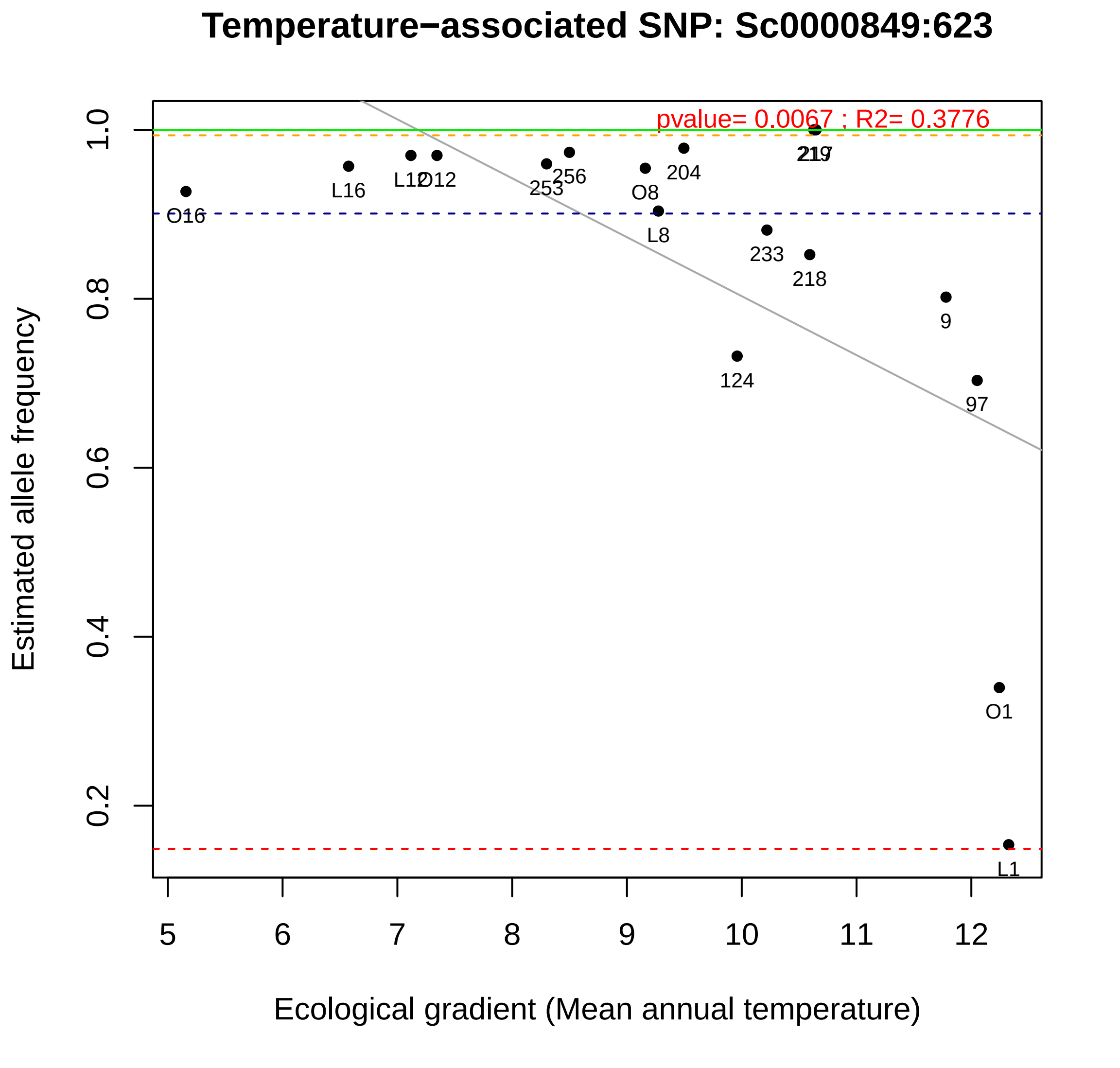
Temperature−associated SNP: Sc0000849:623
pvalue= 0.0067 ; R2= 0.3776
Estimated allele frequency
Ecological gradient (Mean annual temperature)
1.0
0.8
0.6
0.4
0.2
5
6
7
8
9
10
11
12
O16
L16
L12
O12
253
256
O8
204
219
217
L8
233
218
9
124
97
O1
L1

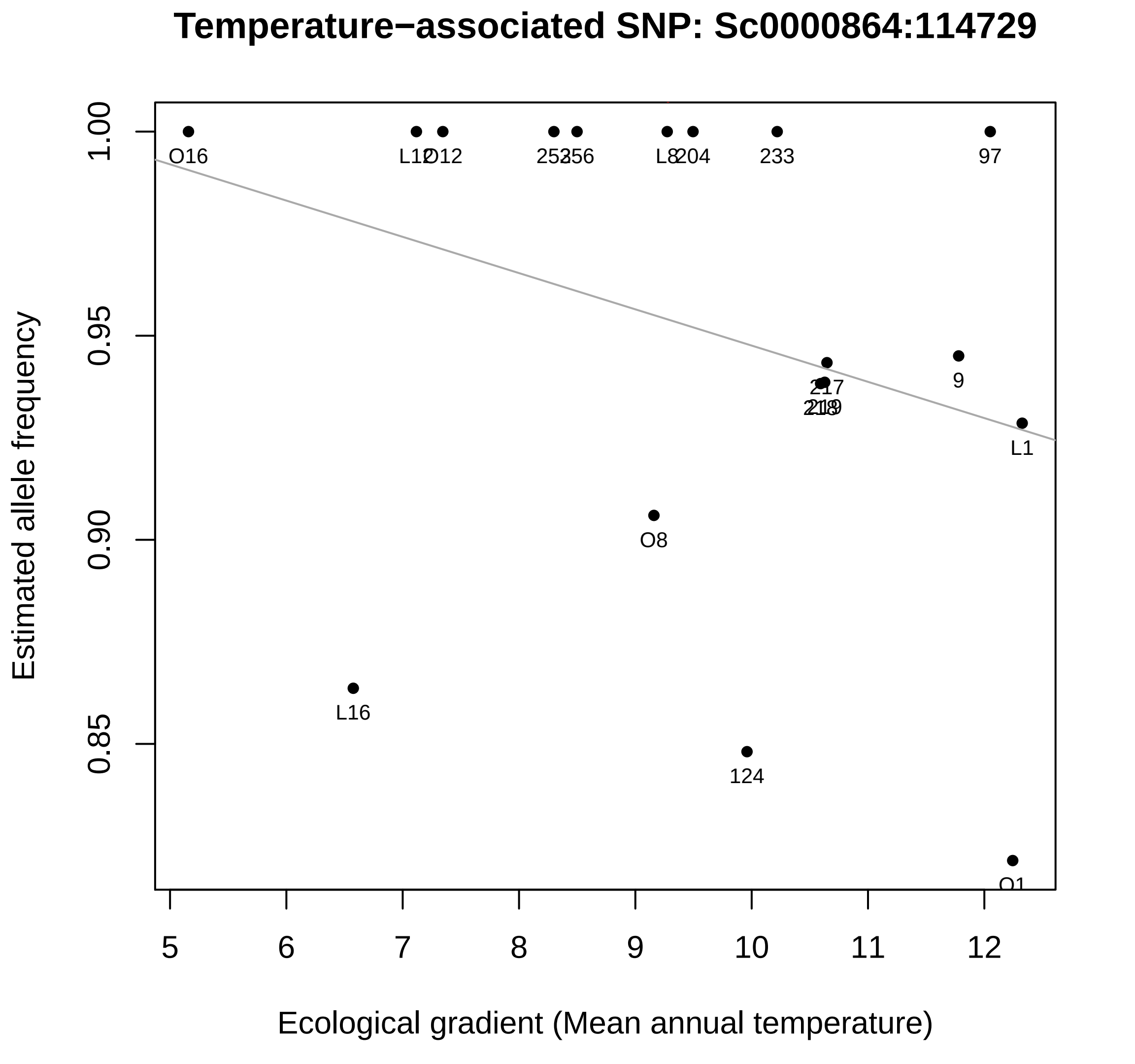

Temperature−associated SNP: Sc0000864:114729
Estimated allele frequency
Ecological gradient (Mean annual temperature)
1.00
0.95
0.90
0.85
5
6
7
8
9
10
11
12
O16
L12
O12
253
256
L8
204
233
97
217
218
219
9
L1
O8
L16
124
O1

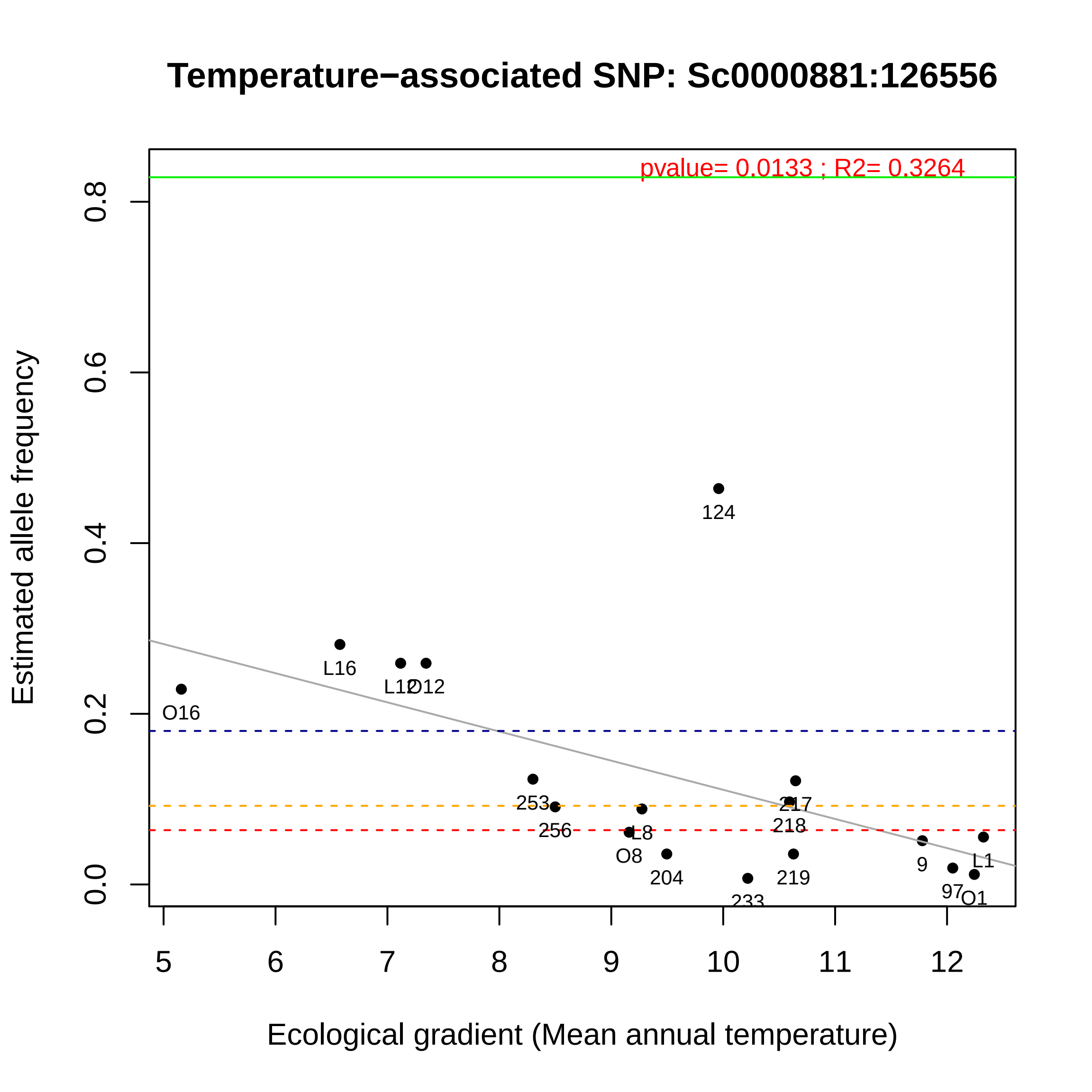
Temperature−associated SNP: Sc0000881:126556
pvalue= 0.0133 ; R2= 0.3264
Estimated allele frequency
0.0
0.2
0.4
0.6
0.8
Ecological gradient (Mean annual temperature)
5
6
7
8
9
10
11
12
O16
L16
L12
O12
124
253
256
L8
O8
204
233
217
218
219
9
97
O1
L1

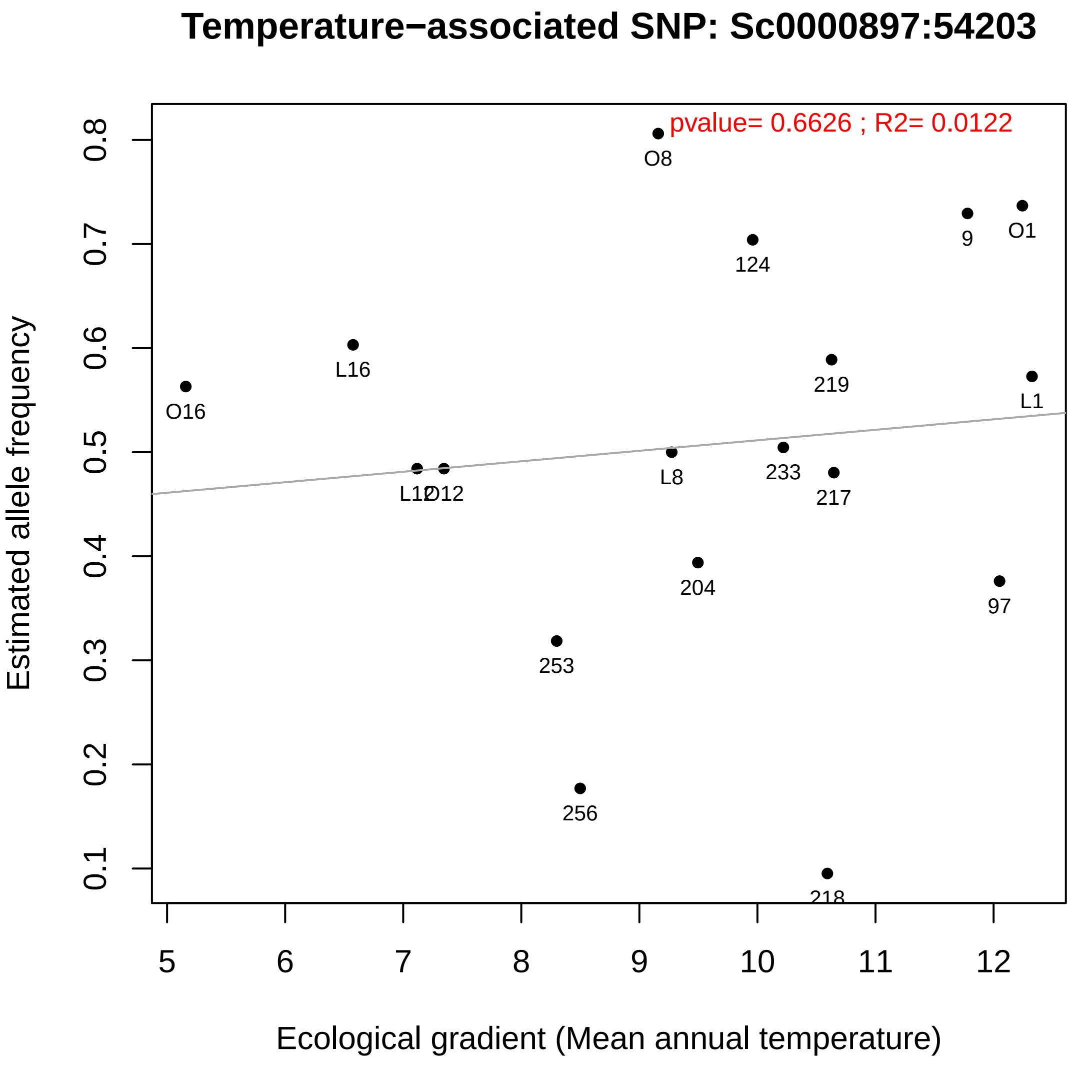
Temperature−associated SNP: Sc0000897:54203
pvalue= 0.6626 ; R2= 0.0122
Estimated allele frequency
0.1
0.2
0.3
0.4
0.5
0.6
0.7
0.8
5
6
7
8
9
10
11
12
Ecological gradient (Mean annual temperature)
O8
9
O1
124
L16
O16
219
L1
L12
O12
L8
233
217
204
97
253
256
218

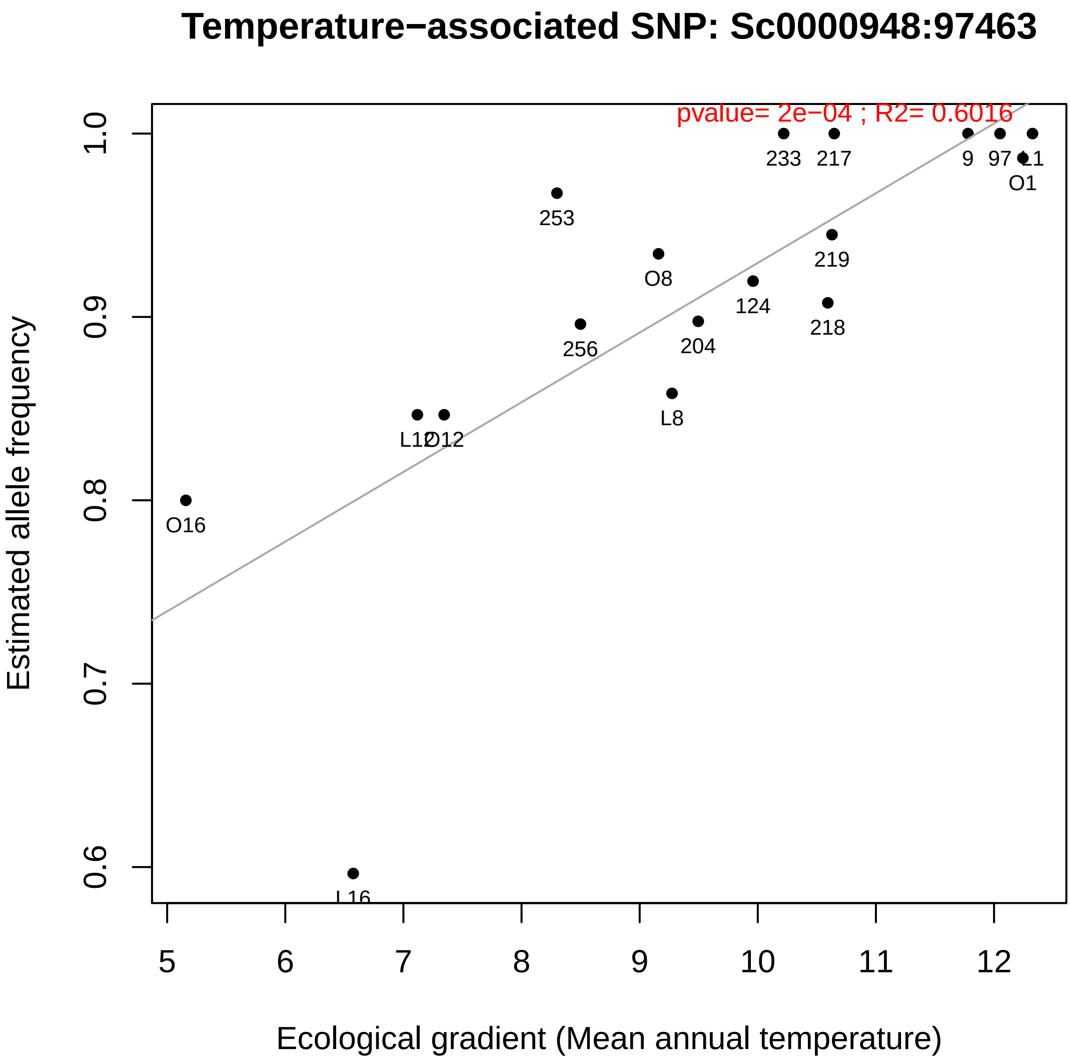
Temperature−associated SNP: Sc0000948:97463
pvalue= 2e−04 ; R2= 0.6016
Estimated allele frequency
Ecological gradient (Mean annual temperature)
0.6
0.7
0.8
0.9
1.0
5
6
7
8
9
10
11
12
233
217
9
97
L1
O1
253
O8
219
124
218
256
204
L8
L12
O12
O16
L16

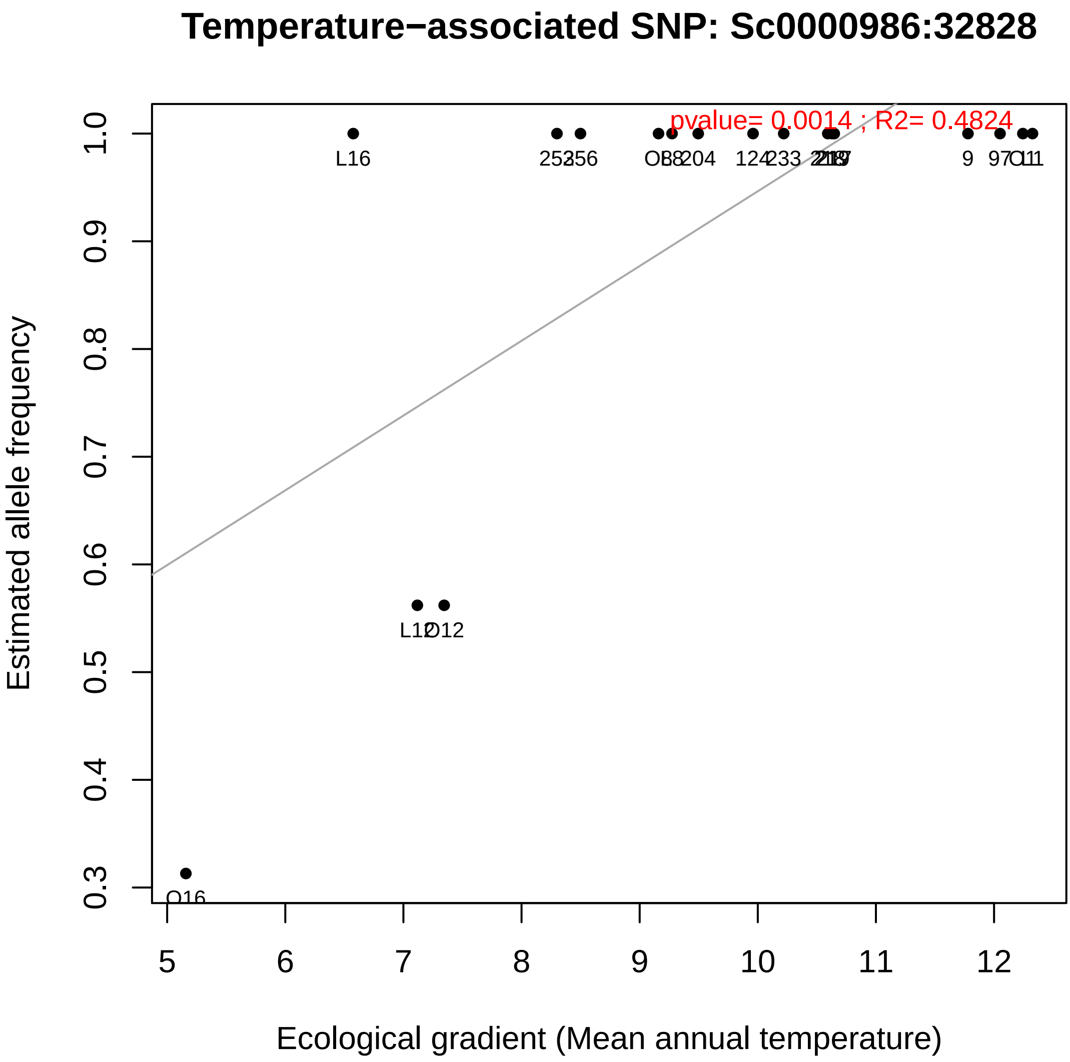

Temperature−associated SNP: Sc0000986:32828
pvalue= 0.0014 ; R2= 0.4824
Estimated allele frequency
Ecological gradient (Mean annual temperature)
L16
L12
O12
O16
9

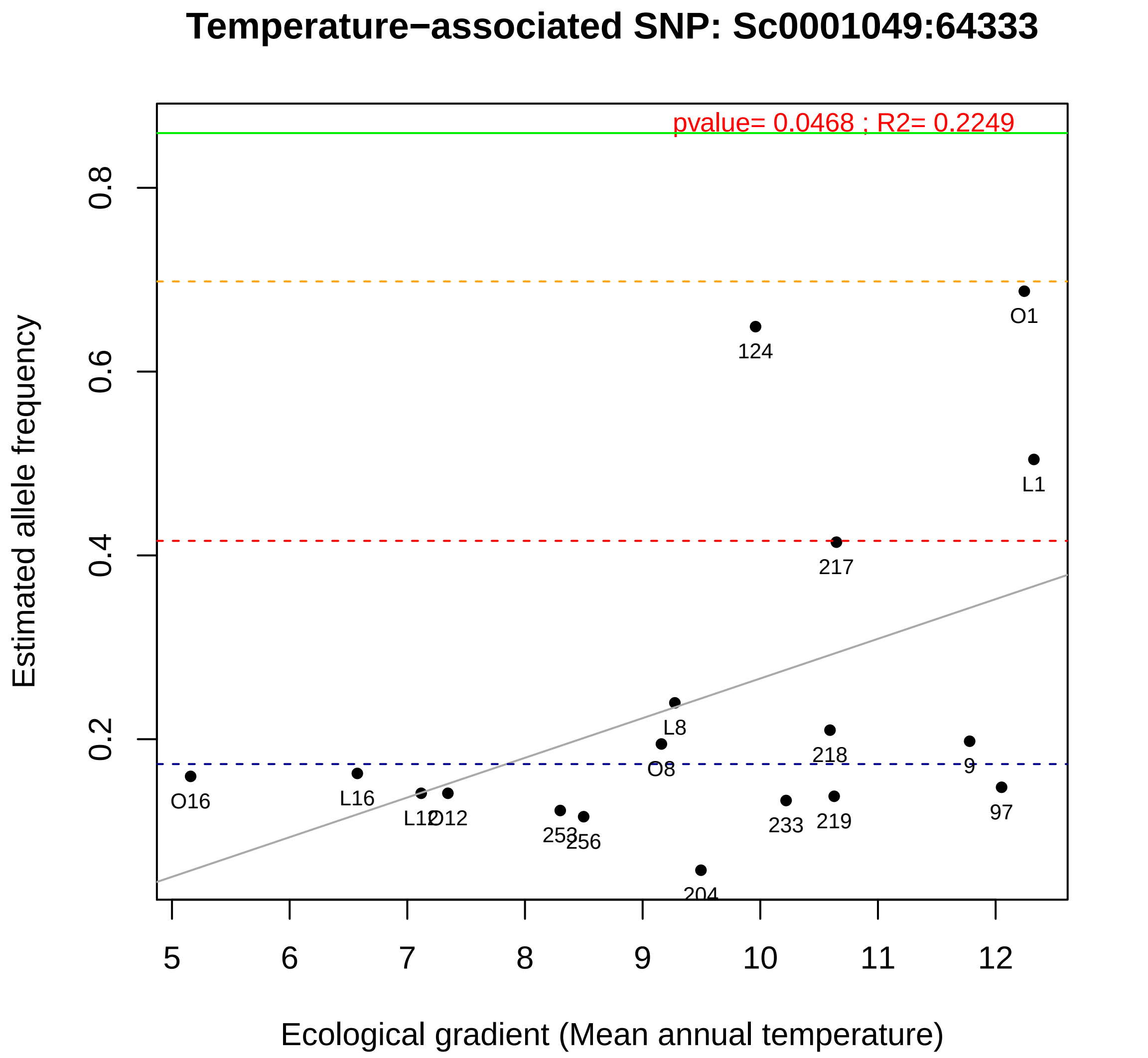

Temperature−associated SNP: Sc0001049:64333
pvalue= 0.0468 ; R2= 0.2249
Estimated allele frequency
0.2
0.4
0.6
0.8
5
6
7
8
9
10
11
12
Ecological gradient (Mean annual temperature)
O1
124
L1
217
L8
218
O8
9
O16
L16
L12
O12
253
256
233
219
97
204

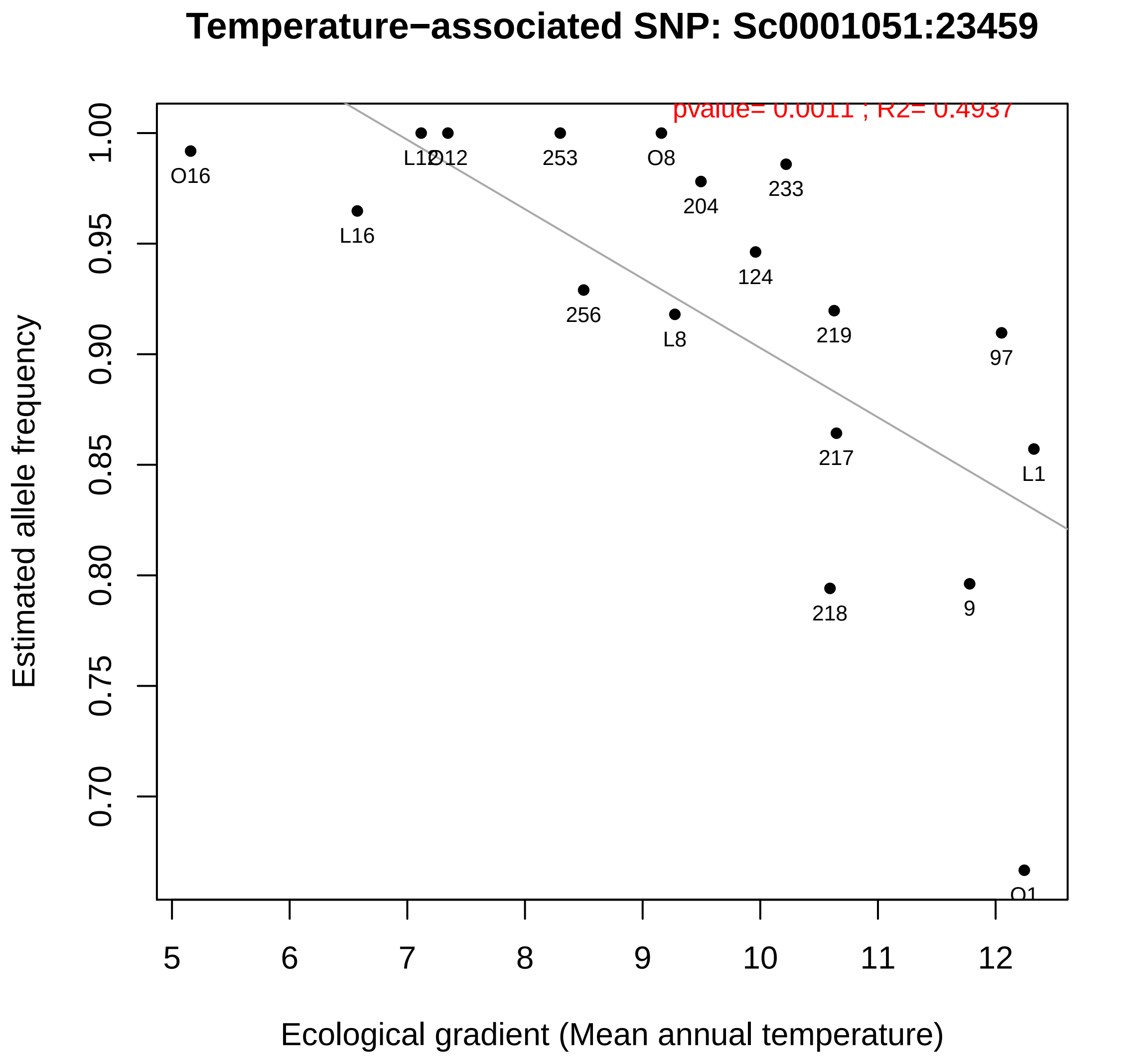

Temperature−associated SNP: Sc0001051:23459
pvalue= 0.0011 ; R2= 0.4937
Estimated allele frequency
1.00
0.95
0.90
0.85
0.80
0.75
0.70
O16
L16
L12
O12
253
256
O8
L8
204
233
124
219
217
218
97
9
L1
O1
5
6
7
8
9
10
11
12
Ecological gradient (Mean annual temperature)

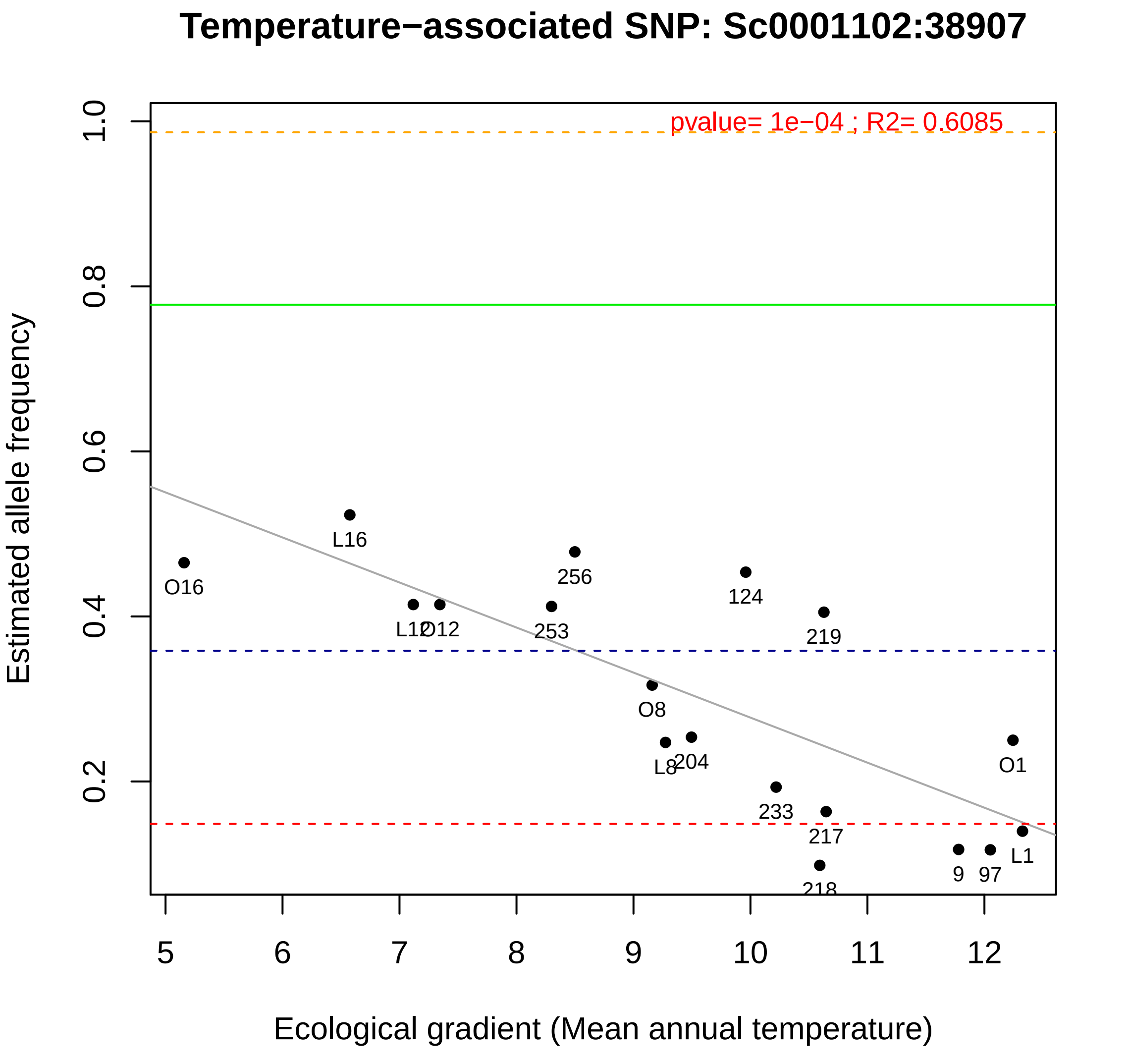

Temperature−associated SNP: Sc0001102:38907
pvalue= 1e−04 ; R2= 0.6085
Estimated allele frequency
0.2
0.4
0.6
0.8
1.0
5
6
7
8
9
10
11
12
Ecological gradient (Mean annual temperature)
O16
L16
L12
O12
253
256
124
219
O8
L8
204
233
217
218
9
97
L1
O1
